## Supplementary material for "CDK12 loss in cancer cells affects DNA damage response genes through premature cleavage and polyadenylation"

**Supplementary Table 1: fold-change expression microarray data**

| Name | Description | Kelly_400nM_THZ531_vs_DMSO | IMR32_400nM_THZ531_vs_DMSO |
|---|---|---|---|
| A2M | NM_000014 | -0.2405 | 0.2345 |
| NAT2 | NM_000015 | 0.1555 | -0.0255 |
| ACADM | NM_000016 | -0.8865 | -0.9625 |
| ACADS | NM_000017 | -0.1445 | -0.7225 |
| ACADVL | NM_000018 | 0.2025 | -0.2045 |
| ACAT1 | NM_000019 | -0.626 | -0.48 |
| ACVRL1 | NM_000020 | -0.4475 | -0.5065 |
| PSEN1 | NM_000021 | -1.579 | -2.5275 |
| ADA | NM_000022 | 0.4335 | -0.2465 |
| SGCA | NM_000023 | 0.0825 | -0.5015 |
| ADRB2 | NM_000024 | -0.8365 | -0.3815 |
| ADRB3 | NM_000025 | 0.1165 | -0.1475 |
| ADSL | NM_000026 | 0.1215 | 0.096 |
| AGA | NM_000027 | -0.795 | -0.6705 |
| AGL | NM_000028 | -0.9155 | -1.669 |
| AGT | NM_000029 | 0.0345 | -0.402 |
| AGXT | NM_000030 | -0.0275 | -0.4955 |
| ALAD | NM_000031 | -0.6495 | -2.1865 |
| ALAS2 | NM_000032 | -0.359 | -0.387 |
| ABCD1 | NM_000033 | -0.0915 | -0.487 |
| ALDOA | NM_000034 | 0.305 | -0.157 |
| ALDOB | NM_000035 | -0.181 | 0.087 |
| AMPD1 | NM_000036 | -0.0935 | -0.156 |
| ANK1 | NM_000037 | -0.1485 | -0.816 |
| APC | NM_000038 | -0.032 | -1.032 |
| APOA1 | NM_000039 | 0.1095 | 0.059 |
| APOC3 | NM_000040 | 0.198 | -0.0975 |
| APOE | NM_000041 | -0.0005 | -0.2705 |
| APOH | NM_000042 | -0.4405 | -0.5365 |
| FAS | NM_000043 | -0.544 | -1.3825 |
| AR | NM_000044 | 0.607 | 0.2115 |
| ARG1 | NM_000045 | 0.1215 | -0.424 |
| ARSB | NM_000046 | -0.3585 | -0.6745 |
| ARSE | NM_000047 | 0.2 | -0.433 |
| ASL | NM_000048 | 0.125 | -0.4295 |
| ASPA | NM_000049 | -0.087 | -0.202 |
| ASS1 | NM_000050 | 0.267 | -0.304 |
| ATM | NM_000051 | 0.2805 | -0.0285 |
| ATP7A | NM_000052 | -1.199 | -2.7875 |
| ATP7B | NM_000053 | -0.3195 | -1.4865 |
| AVPR2 | NM_000054 | 0.0455 | -0.3515 |

| | | | |
|---|---|---|---|
| BCHE | NM_000055 | -1.271 | -1.2165 |
| BCKDHB | NM_000056 | -0.408 | -0.5885 |
| BLM | NM_000057 | -1.3735 | -3.418 |
| BRCA2 | NM_000059 | -0.671 | -1.874 |
| BTD | NM_000060 | -0.233 | -0.593 |
| BTK | NM_000061 | -0.2495 | -0.368 |
| SERPING1 | NM_000062 | -0.0875 | -0.3745 |
| C2 | NM_000063 | -0.1915 | -0.1815 |
| C3 | NM_000064 | -0.467 | -0.163 |
| C8B | NM_000066 | -0.0515 | -0.0385 |
| CA2 | NM_000067 | 0.094 | 0.0875 |
| CACNA1A | NM_000068 | 0.246 | -0.356 |
| CACNA1S | NM_000069 | 0.149 | -0.1375 |
| CAPN3 | NM_000070 | -0.009 | -0.367 |
| CBS | NM_000071 | -0.478 | -0.8445 |
| CD36 | NM_000072 | 0.484 | -0.838 |
| CD3G | NM_000073 | -0.2765 | -0.2375 |
| CD40LG | NM_000074 | -0.045 | -0.8605 |
| CDK4 | NM_000075 | -0.252 | -1.1015 |
| CDKN1C | NM_000076 | 0.4475 | 0.2755 |
| CDKN2A | NM_000077 | 0.204 | 0.138 |
| CETP | NM_000078 | -0.1255 | -0.108 |
| CHRNA1 | NM_000079 | -0.018 | -0.4185 |
| CHRNE | NM_000080 | 0.0365 | -0.2905 |
| LYST | NM_000081 | -0.127 | 0.491 |
| ERCC8 | NM_000082 | -1.0925 | -1.4615 |
| CLCN1 | NM_000083 | -0.4725 | -0.419 |
| CLCN5 | NM_000084 | -0.076 | -0.404 |
| CLCNKB | NM_000085 | 0.2105 | -0.2405 |
| CLN3 | NM_000086 | -0.39 | -1.5685 |
| CNGA1 | NM_000087 | -0.128 | 0.114 |
| COL1A1 | NM_000088 | -0.114 | -0.519 |
| COL1A2 | NM_000089 | 0.196 | 0.135 |
| COL3A1 | NM_000090 | 0.1385 | -0.728 |
| COL4A3 | NM_000091 | -0.0035 | -0.3135 |
| COL4A4 | NM_000092 | 0.645 | 0.2865 |
| COL5A1 | NM_000093 | -0.14 | -0.734 |
| COL7A1 | NM_000094 | 0.1535 | 0.1365 |
| COMP | NM_000095 | 0.115 | -0.2015 |
| CP | NM_000096 | -0.272 | -0.0465 |
| CPOX | NM_000097 | -0.867 | -2.9025 |
| CPT2 | NM_000098 | -0.5325 | -2.738 |
| CST3 | NM_000099 | 0.1825 | -0.3885 |

| | | | |
|---|---|---|---|
| CSTB | NM_000100 | 0.073 | -0.6965 |
| CYBA | NM_000101 | 0.3465 | -0.155 |
| CYP17A1 | NM_000102 | -0.062 | -0.528 |
| CYP19A1 | NM_000103 | 0.0665 | 0.0925 |
| CYP1B1 | NM_000104 | -0.4945 | -0.796 |
| CYP2D6 | NM_000106 | -0.237 | -0.605 |
| DDB2 | NM_000107 | -0.9235 | -1.489 |
| DLD | NM_000108 | -0.558 | -1.562 |
| DMD | NM_000109 | -0.1335 | -0.38 |
| DPYD | NM_000110 | -0.4415 | -0.968 |
| SLC26A3 | NM_000111 | -0.0035 | 0.2905 |
| SLC26A2 | NM_000112 | -1.891 | -2.965 |
| TOR1A | NM_000113 | -1.4235 | -3.1935 |
| EDN3 | NM_000114 | 0.004 | -0.3385 |
| EDNRB | NM_000115 | -0.4215 | 0.038 |
| TAZ | NM_000116 | -0.391 | -1.057 |
| EMD | NM_000117 | 0.0795 | -0.421 |
| ENG | NM_000118 | -0.126 | -0.2855 |
| EPB42 | NM_000119 | -0.426 | -0.2975 |
| EPOR | NM_000121 | -0.3795 | -0.8455 |
| ERCC3 | NM_000122 | -0.696 | -1.9215 |
| ERCC5 | NM_000123 | -0.939 | -2.501 |
| ESR1 | NM_000125 | -0.121 | -0.142 |
| ETFA | NM_000126 | -0.816 | -0.856 |
| EXT1 | NM_000127 | -0.323 | -2.684 |
| F11 | NM_000128 | -0.4305 | 0.2235 |
| F13A1 | NM_000129 | -0.1185 | -0.2465 |
| F5 | NM_000130 | -0.41 | 0.05 |
| F7 | NM_000131 | -0.0465 | -0.5435 |
| F8 | NM_000132 | 0.245 | -0.3385 |
| F9 | NM_000133 | -0.295 | -0.1845 |
| FABP2 | NM_000134 | -0.2325 | -1.0605 |
| FANCA | NM_000135 | -0.397 | -1.195 |
| FANCC | NM_000136 | -0.3885 | -1.2935 |
| FAH | NM_000137 | 0.223 | -0.0505 |
| FBN1 | NM_000138 | -0.8585 | -1.0585 |
| MS4A2 | NM_000139 | -0.25 | -0.081 |
| FECH | NM_000140 | -0.3075 | -1.092 |
| FGFR2 | NM_000141 | -1.4165 | -3.3415 |
| FGFR3 | NM_000142 | 0.069 | 0.864 |
| FH | NM_000143 | -0.528 | -1.1565 |
| FXN | NM_000144 | 0.019 | -1.496 |
| FTL | NM_000146 | 0.2285 | -0.1005 |

| | | | |
|---|---|---|---|
| FUCA1 | NM_000147 | 0.031 | -0.477 |
| FUT1 | NM_000148 | -0.0885 | -0.3855 |
| FUT3 | NM_000149 | 0.1225 | -0.0305 |
| FUT6 | NM_000150 | -0.058 | -0.0995 |
| G6PC | NM_000151 | -0.2325 | -0.4555 |
| GAA | NM_000152 | -0.333 | -0.7525 |
| GALC | NM_000153 | -0.8875 | -0.897 |
| GALK1 | NM_000154 | -0.1165 | -0.8265 |
| GALT | NM_000155 | -0.2685 | -1.0215 |
| GAMT | NM_000156 | 0.0595 | -0.2325 |
| GBA | NM_000157 | -0.6095 | -1.2035 |
| GBE1 | NM_000158 | -0.972 | -0.7425 |
| GCDH | NM_000159 | -0.0065 | -0.6735 |
| GCGR | NM_000160 | 0.076 | -0.2715 |
| GCH1 | NM_000161 | -1.115 | -1.5515 |
| GCK | NM_000162 | -0.3445 | -0.586 |
| GHR | NM_000163 | -0.4875 | 0.1065 |
| GIPR | NM_000164 | 0.0005 | -0.9265 |
| GJA1 | NM_000165 | -0.0545 | -0.052 |
| GJB1 | NM_000166 | 0.1495 | -0.3985 |
| GK | NM_000167 | -1.097 | -1.8855 |
| GLI3 | NM_000168 | 0.0875 | -1.2275 |
| GLA | NM_000169 | -0.8215 | -1.722 |
| GLDC | NM_000170 | -0.489 | -0.9455 |
| GLRA1 | NM_000171 | -0.1185 | -0.2605 |
| GNAT1 | NM_000172 | -0.057 | -0.5365 |
| GP1BA | NM_000173 | 0.6505 | -0.0945 |
| GP9 | NM_000174 | -0.0535 | -0.2765 |
| GPI | NM_000175 | 0.324 | 0.001 |
| NR3C1 | NM_000176 | -0.2885 | -0.689 |
| GSS | NM_000178 | -0.146 | -0.758 |
| MSH6 | NM_000179 | -1.202 | -2.4675 |
| GUCY2D | NM_000180 | 0.032 | -0.409 |
| GUSB | NM_000181 | -0.4875 | -1.068 |
| HADHA | NM_000182 | -0.159 | -0.4635 |
| HADHB | NM_000183 | -0.1925 | -1.079 |
| HBG2 | NM_000184 | -0.056 | -0.1715 |
| SERPIND1 | NM_000185 | -0.49 | -0.761 |
| CFH | NM_000186 | -0.2185 | -0.651 |
| HGD | NM_000187 | 0.4845 | 0.0435 |
| HK1 | NM_000188 | -0.054 | -0.65 |
| HK2 | NM_000189 | -0.761 | -1.309 |
| HMGCL | NM_000191 | -0.481 | -1.2395 |

| | | | |
|---|---|---|---|
| TBX5 | NM_000192 | 0.241 | -0.285 |
| SHH | NM_000193 | -0.016 | -0.2215 |
| HPRT1 | NM_000194 | -0.838 | -0.686 |
| HPS1 | NM_000195 | 0.005 | -0.6805 |
| HSD11B2 | NM_000196 | 0.0595 | -0.3435 |
| HSD17B3 | NM_000197 | -0.1735 | -0.0625 |
| HSD3B2 | NM_000198 | 0.465 | -0.124 |
| SGSH | NM_000199 | 0.0885 | -0.2245 |
| HTN3 | NM_000200 | 0.061 | 0.0195 |
| ICAM1 | NM_000201 | 0.087 | -0.549 |
| IDS | NM_000202 | -1.307 | -0.9705 |
| IDUA | NM_000203 | 0.4745 | -0.078 |
| CFI | NM_000204 | 0.5585 | 0.06 |
| IL2RG | NM_000206 | 0.0245 | -0.3015 |
| INS | NM_000207 | 0.237 | -0.374 |
| INSR | NM_000208 | 0.539 | -2.069 |
| PDX1 | NM_000209 | 0.056 | 0.053 |
| ITGA6 | NM_000210 | -0.445 | 0.3145 |
| ITGB2 | NM_000211 | 0.0965 | 0.032 |
| ITGB3 | NM_000212 | 0.1955 | 0.085 |
| ITGB4 | NM_000213 | -0.194 | -0.5105 |
| JAG1 | NM_000214 | -0.3 | 0.118 |
| JAK3 | NM_000215 | 0.059 | -0.369 |
| KAL1 | NM_000216 | -0.131 | -1.5895 |
| KCNA1 | NM_000217 | 0.0225 | -0.305 |
| KCNQ1 | NM_000218 | 0.1635 | -0.2765 |
| KCNE1 | NM_000219 | -0.046 | -0.102 |
| KCNJ1 | NM_000220 | -0.357 | -0.0905 |
| KHK | NM_000221 | -0.3475 | -0.618 |
| KIT | NM_000222 | -0.1835 | -2.558 |
| KRT12 | NM_000223 | 0.079 | -0.2025 |
| KRT18 | NM_000224 | 0.1865 | -0.2155 |
| KRT9 | NM_000226 | -0.007 | 0.1985 |
| LAMB3 | NM_000228 | -0.129 | -0.0855 |
| LCAT | NM_000229 | -0.297 | -0.347 |
| LEP | NM_000230 | -0.1565 | -0.43 |
| SGCG | NM_000231 | 0.091 | 0.252 |
| SGCB | NM_000232 | -0.0585 | -1.1165 |
| LHCGR | NM_000233 | -0.395 | -0.3245 |
| LIG1 | NM_000234 | 0.07 | -0.855 |
| LIPA | NM_000235 | -1.19 | -1.315 |
| LIPC | NM_000236 | 0.1965 | 0.097 |
| LPL | NM_000237 | -0.067 | -0.2905 |

| | | | |
|---|---|---|---|
| KCNH2 | NM_000238 | -0.4365 | -1.796 |
| LYZ | NM_000239 | 0.0595 | -0.029 |
| MAOA | NM_000240 | -1.023 | -0.6985 |
| MBL2 | NM_000242 | -0.041 | -0.1 |
| MEFV | NM_000243 | 0.0875 | -0.229 |
| MET | NM_000245 | -0.731 | -0.075 |
| CIITA | NM_000246 | 0.238 | -0.325 |
| MICA | NM_000247 | -0.09 | -0.0155 |
| MITF | NM_000248 | -0.596 | -0.749 |
| MLH1 | NM_000249 | -1.0075 | -1.4605 |
| MPO | NM_000250 | -0.0595 | -0.49 |
| MSH2 | NM_000251 | -0.945 | -1.3655 |
| MTM1 | NM_000252 | -0.698 | -1.9895 |
| MTTP | NM_000253 | -0.0205 | 0.087 |
| MTR | NM_000254 | -1.5325 | -1.5395 |
| MUT | NM_000255 | -0.8775 | -1.8925 |
| MYBPC3 | NM_000256 | 0.099 | -0.221 |
| MYH7 | NM_000257 | -0.132 | -0.389 |
| MYL3 | NM_000258 | 0.2155 | -0.327 |
| MYO5A | NM_000259 | -0.588 | -0.993 |
| MYO7A | NM_000260 | -0.1115 | -0.541 |
| MYOC | NM_000261 | 0.228 | 0.1285 |
| NAGA | NM_000262 | -0.369 | -1.2885 |
| NAGLU | NM_000263 | -0.836 | -1.9 |
| PTCH1 | NM_000264 | -1.717 | -5.325 |
| NDP | NM_000266 | -0.013 | -0.0315 |
| NF1 | NM_000267 | -0.6645 | -1.328 |
| NF2 | NM_000268 | -0.0255 | -0.984 |
| PNP | NM_000270 | -0.6435 | -1.9975 |
| NPC1 | NM_000271 | -0.511 | -1.8825 |
| NPHP1 | NM_000272 | -0.3955 | -0.1105 |
| GPR143 | NM_000273 | -0.44 | -0.0635 |
| OAT | NM_000274 | -0.907 | -0.0905 |
| OCA2 | NM_000275 | -0.371 | -0.5445 |
| OCRL | NM_000276 | -0.9435 | -0.8775 |
| PAH | NM_000277 | -0.0065 | -0.2105 |
| PAX2 | NM_000278 | 0.2985 | -0.22 |
| PAX6 | NM_000280 | 0.165 | -0.474 |
| PCBD1 | NM_000281 | 0.2225 | -0.2735 |
| PCCA | NM_000282 | -0.787 | -1.8335 |
| PDE6B | NM_000283 | -0.055 | -0.7385 |
| PDHA1 | NM_000284 | -0.099 | -0.538 |
| PEPD | NM_000285 | 0.256 | -0.468 |

| | | | |
|---|---|---|---|
| PEX12 | NM_000286 | -1.4965 | -2.614 |
| PEX6 | NM_000287 | -0.2795 | -0.7525 |
| PEX7 | NM_000288 | -0.5145 | -1.7055 |
| PFKM | NM_000289 | 0.0145 | -0.251 |
| PGAM2 | NM_000290 | 0.1 | 0.1145 |
| PGK1 | NM_000291 | 0.1695 | -0.2665 |
| PHKA2 | NM_000292 | -0.2945 | -0.8945 |
| PHKB | NM_000293 | -0.8825 | -0.503 |
| PHKG2 | NM_000294 | -0.378 | -1.344 |
| SERPINA1 | NM_000295 | -0.127 | -0.2305 |
| PKD2 | NM_000297 | -2.388 | -3.067 |
| PKLR | NM_000298 | 0.0545 | -0.254 |
| PKP1 | NM_000299 | -0.137 | -0.3875 |
| PLA2G2A | NM_000300 | -0.135 | 0.0185 |
| PLG | NM_000301 | 0.01 | -0.478 |
| PLOD1 | NM_000302 | -0.532 | -0.945 |
| PMM2 | NM_000303 | -1.245 | -2.4295 |
| PMP22 | NM_000304 | -0.822 | -1.9125 |
| PON2 | NM_000305 | -0.8165 | -1.068 |
| POU1F1 | NM_000306 | 0.0815 | 0.1225 |
| POU3F4 | NM_000307 | -0.1565 | -0.3655 |
| CTSA | NM_000308 | -0.249 | -1.347 |
| PPOX | NM_000309 | -0.1875 | -0.3395 |
| PPT1 | NM_000310 | -0.6435 | -0.6965 |
| PRNP | NM_000311 | -1.2005 | -1.22 |
| PROC | NM_000312 | -0.5295 | -0.3825 |
| PROS1 | NM_000313 | -0.7335 | -0.5515 |
| PTEN | NM_000314 | -1.5115 | -3.465 |
| PTH | NM_000315 | 0.1695 | -0.163 |
| PTH1R | NM_000316 | -0.034 | -0.324 |
| PTS | NM_000317 | -0.8535 | -2.2475 |
| PEX2 | NM_000318 | -0.915 | -1.3515 |
| PEX5 | NM_000319 | -0.3075 | -1.087 |
| QDPR | NM_000320 | -0.042 | -0.5465 |
| RB1 | NM_000321 | -1.094 | -1.3235 |
| PRPH2 | NM_000322 | -0.69 | -1.3025 |
| RHAG | NM_000324 | -0.34 | -0.3575 |
| PITX2 | NM_000325 | -0.084 | -0.4735 |
| RLBP1 | NM_000326 | 0.0315 | -0.071 |
| ROM1 | NM_000327 | -0.014 | -0.502 |
| RPGR | NM_000328 | -1.108 | -1.5155 |
| RPE65 | NM_000329 | -0.257 | -0.1695 |
| RS1 | NM_000330 | -0.2015 | -0.121 |

| | | | |
|---|---|---|---|
| SAA1 | NM_000331 | 0.341 | 0.388 |
| ATXN1 | NM_000332 | -0.821 | -1.2505 |
| ATXN7 | NM_000333 | -1.679 | -2.367 |
| SCN4A | NM_000334 | -0.1235 | -0.4755 |
| SCN5A | NM_000335 | 0.021 | -0.709 |
| SCNN1B | NM_000336 | 0.1485 | -0.4205 |
| SGCD | NM_000337 | 0.104 | -0.1815 |
| SLC12A1 | NM_000338 | 0.1415 | 0.0065 |
| SLC12A3 | NM_000339 | -0.0575 | -0.321 |
| SLC2A2 | NM_000340 | 0.225 | -0.3295 |
| SLC3A1 | NM_000341 | -0.435 | -0.3465 |
| SLC4A1 | NM_000342 | -0.033 | -0.3225 |
| SLC5A1 | NM_000343 | -0.121 | -0.1215 |
| SMN1 | NM_000344 | -1.2465 | -2.9415 |
| SNCA | NM_000345 | -0.884 | -1.472 |
| SOX9 | NM_000346 | 0.336 | 0.492 |
| SPTB | NM_000347 | -0.0655 | -0.693 |
| SRD5A2 | NM_000348 | -0.0415 | 0.151 |
| STAR | NM_000349 | 0.268 | -0.2715 |
| ABCA4 | NM_000350 | 0.246 | -0.1455 |
| STS | NM_000351 | -0.833 | -0.988 |
| ABCC8 | NM_000352 | -0.2925 | -1.0085 |
| TAT | NM_000353 | 0.009 | -0.2515 |
| SERPINA7 | NM_000354 | -0.068 | 0.004 |
| TCN2 | NM_000355 | -0.11 | -0.502 |
| TCOF1 | NM_000356 | -0.2495 | -0.907 |
| TGFBI | NM_000358 | -0.1045 | -0.078 |
| TGM1 | NM_000359 | -0.077 | -0.3475 |
| TH | NM_000360 | 0.162 | -0.2025 |
| THBD | NM_000361 | -0.322 | -0.302 |
| TIMP3 | NM_000362 | -0.2295 | -1.415 |
| TNNI3 | NM_000363 | 0.001 | -0.125 |
| TNNT2 | NM_000364 | 0.049 | -0.0645 |
| TPI1 | NM_000365 | 0.128 | -0.2305 |
| TPM1 | NM_000366 | 0.047 | -0.349 |
| TPMT | NM_000367 | -0.9655 | -1.4915 |
| TSC1 | NM_000368 | -2.043 | -2.7515 |
| TSHR | NM_000369 | -0.3615 | -1.4765 |
| TTPA | NM_000370 | -0.0755 | -0.59 |
| TTR | NM_000371 | -0.2525 | -0.531 |
| TYR | NM_000372 | 0.0085 | -0.4235 |
| UMPS | NM_000373 | -1.5695 | -5.0975 |
| UROD | NM_000374 | 0.1625 | -0.6445 |

| | | | |
|---|---|---|---|
| UROS | NM_000375 | -0.186 | -0.5865 |
| VDR | NM_000376 | 0.0385 | -0.6 |
| WAS | NM_000377 | -0.04 | 0.089 |
| WT1 | NM_000378 | 0.373 | -0.2905 |
| XDH | NM_000379 | 0.267 | 0.2705 |
| XPA | NM_000380 | -0.5255 | -1.216 |
| MID1 | NM_000381 | -0.5825 | -1.5365 |
| ALDH3A2 | NM_000382 | -0.9925 | -1.6615 |
| AIRE | NM_000383 | -0.1245 | -0.4725 |
| APOB | NM_000384 | 0.208 | -0.0665 |
| BLMH | NM_000386 | -0.368 | -1.006 |
| SLC25A20 | NM_000387 | -0.358 | -1.19 |
| CASR | NM_000388 | 0.16 | 0.001 |
| CHM | NM_000390 | -0.972 | -2.3085 |
| TPP1 | NM_000391 | -0.168 | -0.6845 |
| ABCC2 | NM_000392 | -0.261 | -0.3835 |
| COL5A2 | NM_000393 | -0.846 | -0.586 |
| CRYAA | NM_000394 | 0.175 | 0.08 |
| CSF2RB | NM_000395 | 0.0525 | 0.0675 |
| CTSK | NM_000396 | -0.8745 | -0.82 |
| CYBB | NM_000397 | 0.1385 | -0.1255 |
| CYB5R3 | NM_000398 | 0.0845 | -0.5685 |
| EGR2 | NM_000399 | -0.005 | -0.105 |
| ERCC2 | NM_000400 | 0.0625 | -0.9705 |
| EXT2 | NM_000401 | -0.1755 | -1.341 |
| G6PD | NM_000402 | 0.1555 | -0.416 |
| GALE | NM_000403 | -0.1105 | -1.0535 |
| GLB1 | NM_000404 | -0.364 | -1.3295 |
| GM2A | NM_000405 | 0.162 | -0.357 |
| GNRHR | NM_000406 | -0.2185 | -0.093 |
| GP1BB | NM_000407 | -0.0585 | -0.138 |
| GPD2 | NM_000408 | -0.9915 | -0.978 |
| GUCA1A | NM_000409 | -0.3025 | -0.375 |
| HFE | NM_000410 | 0.087 | -0.1325 |
| HLCS | NM_000411 | -0.572 | -1.0345 |
| HRG | NM_000412 | -0.204 | -0.6115 |
| HSD17B1 | NM_000413 | 0.4115 | -0.5445 |
| HSD17B4 | NM_000414 | -0.8495 | -0.8125 |
| IAPP | NM_000415 | -0.277 | -0.0945 |
| IFNGR1 | NM_000416 | -1.0445 | -2.3245 |
| IL2RA | NM_000417 | 0.144 | -0.581 |
| IL4R | NM_000418 | -0.1765 | -1.718 |
| ITGA2B | NM_000419 | -0.1175 | -0.1865 |

| | | | |
|---|---|---|---|
| KEL | NM_000420 | -0.012 | -0.0885 |
| KRT10 | NM_000421 | -0.2615 | -0.8565 |
| KRT17 | NM_000422 | 0.465 | 0.438 |
| KRT2 | NM_000423 | 0.43 | -0.1665 |
| KRT5 | NM_000424 | -0.029 | -0.133 |
| L1CAM | NM_000425 | 0.135 | -0.474 |
| LAMA2 | NM_000426 | 0.2085 | -0.102 |
| LOR | NM_000427 | -0.022 | -0.033 |
| LTBP2 | NM_000428 | 0.055 | -0.6 |
| MAT1A | NM_000429 | 0.197 | -0.2815 |
| PAFAH1B1 | NM_000430 | -1.7695 | -2.873 |
| MVK | NM_000431 | -0.6345 | -1.7265 |
| MYL2 | NM_000432 | 0.1405 | -0.117 |
| NCF2 | NM_000433 | -0.082 | -0.836 |
| NEU1 | NM_000434 | -0.596 | -1.2715 |
| NOTCH3 | NM_000435 | 0.0805 | -0.4685 |
| OXCT1 | NM_000436 | -0.8235 | -0.87 |
| PAFAH2 | NM_000437 | -1.131 | -1.121 |
| PAX3 | NM_000438 | 0.264 | -0.16 |
| PCSK1 | NM_000439 | 0.0825 | -0.6175 |
| PDE6A | NM_000440 | -0.019 | -0.335 |
| SLC26A4 | NM_000441 | 0.0005 | 0.126 |
| PECAM1 | NM_000442 | -0.327 | -0.3655 |
| ABCB4 | NM_000443 | -0.2445 | -0.103 |
| PHEX | NM_000444 | -0.1235 | -0.5695 |
| PON1 | NM_000446 | -0.1045 | 0.212 |
| PSEN2 | NM_000447 | -0.302 | -1.2705 |
| RAG1 | NM_000448 | 0.2735 | -0.2235 |
| RFX5 | NM_000449 | -1.658 | -2.875 |
| SELE | NM_000450 | 0.1925 | -0.406 |
| SHOX | NM_000451 | 0.3105 | -0.4925 |
| SLC10A2 | NM_000452 | -0.343 | -0.1665 |
| SLC5A5 | NM_000453 | -0.0955 | -0.159 |
| SOD1 | NM_000454 | -0.115 | -0.328 |
| STK11 | NM_000455 | -0.524 | -1.2585 |
| SUOX | NM_000456 | -0.973 | -2.0965 |
| HNF4A | NM_000457 | -0.024 | 0.0685 |
| HNF1B | NM_000458 | -0.2325 | -0.226 |
| TEK | NM_000459 | -0.6045 | 0.0455 |
| THPO | NM_000460 | -0.1595 | -0.2275 |
| THRB | NM_000461 | 0.3855 | -0.679 |
| UBE3A | NM_000462 | -1.2775 | -2.2785 |
| UGT1A1 | NM_000463 | 0.021 | 0.0035 |

| | | | |
|---|---|---|---|
| BARD1 | NM_000465 | -1.6665 | -2.8415 |
| PEX1 | NM_000466 | -1.745 | -2.777 |
| TWIST1 | NM_000474 | -0.976 | -2.949 |
| NR0B1 | NM_000475 | 0.0865 | -0.0155 |
| AK1 | NM_000476 | 0.2035 | -0.549 |
| ALB | NM_000477 | 0.1805 | -0.3135 |
| ALPL | NM_000478 | 0.0015 | -0.5595 |
| AMH | NM_000479 | 0.1735 | 0.733 |
| AMPD3 | NM_000480 | -0.631 | -0.786 |
| AMT | NM_000481 | -0.056 | -0.721 |
| APOA4 | NM_000482 | -0.2 | 0.2665 |
| APOC2 | NM_000483 | 0.0045 | -0.219 |
| APP | NM_000484 | -0.3025 | -0.3875 |
| APRT | NM_000485 | 0.28 | -0.254 |
| AQP2 | NM_000486 | -0.1245 | -0.3475 |
| ARSA | NM_000487 | -0.097 | -0.137 |
| SERPINC1 | NM_000488 | -0.0255 | -0.19 |
| ATRX | NM_000489 | 0.3615 | -1.5905 |
| AVP | NM_000490 | 0.176 | -0.5195 |
| C1QB | NM_000491 | -0.091 | -0.62 |
| CFTR | NM_000492 | 0.2015 | -0.4285 |
| COL10A1 | NM_000493 | -0.1075 | -0.256 |
| COL17A1 | NM_000494 | 0.131 | -0.2235 |
| COL4A5 | NM_000495 | 0.0465 | -0.197 |
| CRYBB2 | NM_000496 | -0.1125 | -0.654 |
| CYP11B1 | NM_000497 | -0.1315 | -0.43 |
| CYP11B2 | NM_000498 | -0.272 | -0.0755 |
| CYP1A1 | NM_000499 | 0.127 | -0.2105 |
| CYP21A2 | NM_000500 | -0.2815 | -0.1725 |
| ELN | NM_000501 | -0.008 | -0.4465 |
| EPX | NM_000502 | -0.046 | -0.2745 |
| EYA1 | NM_000503 | -1.4595 | -2.6105 |
| F10 | NM_000504 | -0.1955 | -0.1645 |
| F12 | NM_000505 | 0.007 | -0.6085 |
| F2 | NM_000506 | 0.087 | -0.507 |
| FBP1 | NM_000507 | -0.0905 | -0.2685 |
| FGA | NM_000508 | -0.1405 | -0.1215 |
| FGG | NM_000509 | -0.077 | -0.1765 |
| FSHB | NM_000510 | 0.032 | -0.2385 |
| FUT2 | NM_000511 | -0.256 | -0.049 |
| GALNS | NM_000512 | -0.663 | -1.168 |
| OPN1MW | NM_000513 | 0.226 | 0.0605 |
| GDNF | NM_000514 | -0.214 | -0.568 |

| | | | |
|---|---|---|---|
| GH1 | NM_000515 | 0.11 | -0.1405 |
| GNAS | NM_000516 | 0.0835 | -0.2445 |
| HBB | NM_000518 | 0.0535 | -0.399 |
| HBD | NM_000519 | -0.1185 | -0.209 |
| HEXA | NM_000520 | 0.0555 | -0.6475 |
| HEXB | NM_000521 | -1.2175 | -1.627 |
| HOXA13 | NM_000522 | 0.003 | -0.4095 |
| HOXD13 | NM_000523 | -0.4845 | -0.087 |
| HTR1A | NM_000524 | 0.359 | -0.357 |
| KCNJ11 | NM_000525 | -0.1825 | -0.8775 |
| KRT14 | NM_000526 | 0.085 | 0.081 |
| LDLR | NM_000527 | -1.2675 | -3.441 |
| MAN2B1 | NM_000528 | 0.117 | -0.2625 |
| MC2R | NM_000529 | -0.005 | -0.5425 |
| MPZ | NM_000530 | -0.287 | -0.726 |
| OTC | NM_000531 | -0.2325 | -0.682 |
| PCCB | NM_000532 | 0.146 | -0.722 |
| PLP1 | NM_000533 | 0.0055 | -0.3675 |
| PMS1 | NM_000534 | -1.4985 | -3.287 |
| RAG2 | NM_000536 | -0.0205 | -0.069 |
| REN | NM_000537 | 0.3545 | -0.079 |
| RFXAP | NM_000538 | -1.1255 | -2.747 |
| RHO | NM_000539 | -0.0685 | -0.256 |
| RYR1 | NM_000540 | -0.1625 | -0.191 |
| SAG | NM_000541 | 0.1685 | -0.4275 |
| SFTPB | NM_000542 | 0.1175 | -0.316 |
| SMPD1 | NM_000543 | -0.4185 | -1.018 |
| TAP2 | NM_000544 | -0.2035 | -0.459 |
| HNF1A | NM_000545 | 0.037 | -0.369 |
| TP53 | NM_000546 | -0.5945 | -0.8345 |
| TPO | NM_000547 | 0.244 | -0.0135 |
| TSC2 | NM_000548 | 0.0595 | -0.401 |
| TSHB | NM_000549 | 0.1205 | -0.1115 |
| TYRP1 | NM_000550 | -0.2 | -0.371 |
| VHL | NM_000551 | -0.342 | -0.81 |
| VWF | NM_000552 | 0.055 | -0.3055 |
| WRN | NM_000553 | -0.903 | -1.727 |
| CRX | NM_000554 | -0.0965 | -0.151 |
| DCX | NM_000555 | -0.96 | -0.4395 |
| GDF5 | NM_000557 | 0.18 | -0.3305 |
| HBG1 | NM_000559 | 0.2185 | 0.542 |
| CD53 | NM_000560 | 0 | -0.4635 |
| C8A | NM_000562 | 0.0465 | 0.1155 |

| | | | |
|---|---|---|---|
| IL5RA | NM_000564 | 0.056 | -0.797 |
| IL6R | NM_000565 | -0.468 | -0.3015 |
| FCGR1A | NM_000566 | 0.1395 | -0.1645 |
| CRP | NM_000567 | 0.052 | -0.0035 |
| FCGR3A | NM_000569 | 0.4365 | -0.566 |
| FCGR3B | NM_000570 | -0.361 | -0.2015 |
| IL10 | NM_000572 | 0.842 | 0.157 |
| CR1 | NM_000573 | 0.1535 | 0.026 |
| CD55 | NM_000574 | -1.036 | 0.004 |
| IL1A | NM_000575 | -0.08 | -0.625 |
| IL1B | NM_000576 | -0.0805 | -0.5055 |
| IL1RN | NM_000577 | -0.094 | -0.1175 |
| SLC11A1 | NM_000578 | -0.123 | -0.352 |
| CCR5 | NM_000579 | 0.0005 | -0.04 |
| GPX1 | NM_000581 | 0.216 | -0.2285 |
| SPP1 | NM_000582 | 0.3405 | 0.0675 |
| GC | NM_000583 | -0.1785 | 0.3455 |
| IL8 | NM_000584 | 0.3165 | -0.1375 |
| IL15 | NM_000585 | 0.0515 | -0.575 |
| IL2 | NM_000586 | 0.138 | 0.341 |
| C7 | NM_000587 | -0.1435 | -0.2415 |
| IL3 | NM_000588 | 0.038 | -0.05 |
| IL4 | NM_000589 | -0.089 | -0.429 |
| IL9 | NM_000590 | 0.396 | -0.2505 |
| TAP1 | NM_000593 | 0.136 | -0.403 |
| TNF | NM_000594 | -0.012 | -0.041 |
| LTA | NM_000595 | 0.209 | -0.024 |
| IGFBP1 | NM_000596 | -0.0425 | 0.041 |
| IGFBP2 | NM_000597 | 0.2715 | -0.372 |
| IGFBP3 | NM_000598 | -0.1475 | -0.8595 |
| IGFBP5 | NM_000599 | -1.0025 | -1.358 |
| IL6 | NM_000600 | -0.1465 | -0.5765 |
| HGF | NM_000601 | 0.329 | 0.368 |
| SERPINE1 | NM_000602 | 0.2535 | -0.0945 |
| NOS3 | NM_000603 | 0.2665 | -0.11 |
| IFNA2 | NM_000605 | 0.2785 | 0.332 |
| C8G | NM_000606 | 0.111 | -0.624 |
| ORM1 | NM_000607 | -0.0245 | 0.01 |
| ORM2 | NM_000608 | -0.2245 | -0.334 |
| CXCL12 | NM_000609 | -0.2575 | -1.2595 |
| CD44 | NM_000610 | -1.315 | -1.637 |
| CD59 | NM_000611 | -0.806 | -0.8585 |
| HPX | NM_000613 | 0.4875 | 0.0305 |

| | | | |
|---|---|---|---|
| CNTF | NM_000614 | -0.1845 | -0.4025 |
| NCAM1 | NM_000615 | -0.7595 | -0.572 |
| CD4 | NM_000616 | 0.0055 | -0.527 |
| SLC11A2 | NM_000617 | -0.876 | -1.3705 |
| IGF1 | NM_000618 | 0.1515 | -0.487 |
| IFNG | NM_000619 | -0.176 | 0.194 |
| NOS1 | NM_000620 | 0.011 | -0.471 |
| HTR2A | NM_000621 | -0.1865 | 0.1495 |
| BDKRB2 | NM_000623 | -0.531 | -0.2465 |
| SERPINA5 | NM_000624 | -0.583 | 0.4035 |
| NOS2 | NM_000625 | 0.199 | -0.609 |
| CD79B | NM_000626 | -0.1475 | -0.3155 |
| LTBP1 | NM_000627 | -0.0975 | -0.986 |
| IL10RB | NM_000628 | -1.0005 | -1.7715 |
| IFNAR1 | NM_000629 | -1.49 | -2.707 |
| ITGAM | NM_000632 | -0.0195 | -0.1615 |
| BCL2 | NM_000633 | -1.7365 | -3.4365 |
| CXCR1 | NM_000634 | -0.222 | -0.633 |
| RFX2 | NM_000635 | -0.166 | -0.346 |
| SOD2 | NM_000636 | 0.1 | -0.1 |
| GSR | NM_000637 | -0.3685 | -1.5555 |
| VTN | NM_000638 | 0.176 | -0.3125 |
| FASLG | NM_000639 | -0.015 | -0.266 |
| IL13RA2 | NM_000640 | 0.084 | 0.344 |
| IL11 | NM_000641 | -0.0445 | -0.457 |
| AGL | NM_000642 | 0.551 | -0.7535 |
| AGL | NM_000643 | 0.551 | -0.7535 |
| AGL | NM_000644 | 0.551 | -0.7535 |
| AGL | NM_000645 | 0.551 | -0.7535 |
| AGL | NM_000646 | 0.551 | -0.7535 |
| CR1 | NM_000651 | 0.1535 | 0.026 |
| SELL | NM_000655 | 0.016 | 0.0455 |
| BCL2 | NM_000657 | 0.227 | -0.322 |
| AIRE | NM_000658 | -0.1245 | -0.4725 |
| TGFB1 | NM_000660 | 0.1855 | -0.1205 |
| RPL9 | NM_000661 | -0.0175 | -0.3365 |
| NAT1 | NM_000662 | -1.268 | -0.7705 |
| ABAT | NM_000663 | 0.065 | -0.351 |
| ACHE | NM_000665 | 0.2575 | -0.317 |
| ACY1 | NM_000666 | -0.225 | -0.602 |
| ADH1A | NM_000667 | -0.044 | -0.091 |
| ADH1B | NM_000668 | -0.3255 | -0.7485 |
| ADH1C | NM_000669 | -0.1415 | -0.3805 |

| | | | |
|---|---|---|---|
| ADH4 | NM_000670 | 0.512 | -0.4745 |
| ADH5 | NM_000671 | -0.6865 | -1.3275 |
| ADH6 | NM_000672 | 0.1775 | -0.358 |
| ADH7 | NM_000673 | 0.147 | -0.218 |
| ADORA1 | NM_000674 | -0.0715 | -0.722 |
| ADORA2A | NM_000675 | -0.3705 | -1.009 |
| ADORA2B | NM_000676 | -1.2775 | -2.83 |
| ADORA3 | NM_000677 | 0.0815 | -0.046 |
| ADRA1D | NM_000678 | -0.086 | -0.2645 |
| ADRA1B | NM_000679 | 0.5305 | 0.0275 |
| ADRA1A | NM_000680 | -0.2555 | -0.1675 |
| ADRA2A | NM_000681 | -0.776 | -1.815 |
| ADRA2B | NM_000682 | 0.0525 | -0.182 |
| ADRA2C | NM_000683 | -0.639 | -1.596 |
| ADRB1 | NM_000684 | 0.018 | -0.4715 |
| AGTR1 | NM_000685 | -0.793 | -0.37 |
| AGTR2 | NM_000686 | 0.3515 | -0.387 |
| AHCY | NM_000687 | 0.316 | -0.219 |
| ALAS1 | NM_000688 | -0.5745 | -2.806 |
| ALDH1A1 | NM_000689 | 0.1825 | -0.2555 |
| ALDH2 | NM_000690 | 0.194 | -0.382 |
| ALDH1B1 | NM_000692 | -0.8505 | -2.2995 |
| ALDH1A3 | NM_000693 | 0.0315 | -1.349 |
| ALDH3B1 | NM_000694 | -0.188 | -0.3895 |
| ALDH3B2 | NM_000695 | 0.112 | -0.3485 |
| ALDH9A1 | NM_000696 | -0.286 | -0.9605 |
| ALOX12 | NM_000697 | 0.0675 | 0.053 |
| ALOX5 | NM_000698 | -0.0625 | -0.1795 |
| AMY2A | NM_000699 | -0.847 | -0.5775 |
| ANXA1 | NM_000700 | -0.232 | -0.335 |
| ATP1A1 | NM_000701 | -0.086 | -0.716 |
| ATP1A2 | NM_000702 | 0.238 | -0.0225 |
| ATP4A | NM_000704 | -0.0005 | -0.193 |
| ATP4B | NM_000705 | 0.143 | -0.318 |
| AVPR1A | NM_000706 | -0.0605 | -0.129 |
| AVPR1B | NM_000707 | -0.045 | -0.099 |
| BCKDHA | NM_000709 | -0.1865 | -0.732 |
| BDKRB1 | NM_000710 | -0.222 | -0.415 |
| BLVRA | NM_000712 | -0.373 | -1.14 |
| BLVRB | NM_000713 | -0.345 | -0.28 |
| TSPO | NM_000714 | 0.219 | -0.3105 |
| C4BPA | NM_000715 | -0.457 | 0.7375 |
| C4BPB | NM_000716 | -0.2635 | -0.591 |

| | | | |
|---|---|---|---|
| CA4 | NM_000717 | -0.0775 | -0.1145 |
| CACNA1B | NM_000718 | -1.21 | -1.025 |
| CACNA1C | NM_000719 | -0.2715 | -0.3655 |
| CACNA1D | NM_000720 | -0.3465 | 0.065 |
| CACNA1E | NM_000721 | -0.149 | -0.064 |
| CACNA2D1 | NM_000722 | -0.065 | -0.7315 |
| CACNB1 | NM_000723 | -0.268 | -1.1035 |
| CACNB2 | NM_000724 | 0.037 | -0.4365 |
| CACNB3 | NM_000725 | -0.331 | -0.988 |
| CACNB4 | NM_000726 | -0.508 | -0.9685 |
| CACNG1 | NM_000727 | 0.1375 | -0.4635 |
| CALCB | NM_000728 | -0.1445 | -0.6105 |
| CCK | NM_000729 | -0.147 | -0.0285 |
| CCKAR | NM_000730 | 0.063 | -0.282 |
| CD3D | NM_000732 | 0.624 | -0.0825 |
| CD3E | NM_000733 | -0.058 | -0.0865 |
| CGA | NM_000735 | -0.136 | -0.4805 |
| CHRM1 | NM_000738 | -0.685 | -1.475 |
| CHRM2 | NM_000739 | 0.095 | -0.344 |
| CHRM3 | NM_000740 | -0.8795 | -0.8805 |
| CHRM4 | NM_000741 | -0.1055 | -0.5125 |
| CHRNA2 | NM_000742 | -0.0765 | -0.1495 |
| CHRNA3 | NM_000743 | -1.3335 | -2.343 |
| CHRNA4 | NM_000744 | 0.047 | -0.189 |
| CHRNA5 | NM_000745 | -0.57 | -1.558 |
| CHRNB1 | NM_000747 | -0.4265 | -0.7255 |
| CHRNB2 | NM_000748 | -1.397 | -2.979 |
| CHRNB3 | NM_000749 | -0.091 | -0.1985 |
| CHRNB4 | NM_000750 | -0.5465 | -1.894 |
| CHRND | NM_000751 | 0.0335 | -0.286 |
| COMT | NM_000754 | -0.6255 | -0.8935 |
| CRAT | NM_000755 | -0.026 | -0.777 |
| CRH | NM_000756 | -0.11 | -0.2545 |
| CSF1 | NM_000757 | 0.068 | -0.4195 |
| CSF2 | NM_000758 | -0.071 | -0.646 |
| CSF3 | NM_000759 | 0.093 | -0.3545 |
| CSF3R | NM_000760 | 0.041 | 0.0825 |
| CYP1A2 | NM_000761 | 0.089 | -0.439 |
| CYP2A7 | NM_000764 | 0.121 | -0.241 |
| CYP3A7 | NM_000765 | -0.1895 | -0.2635 |
| CYP2A13 | NM_000766 | 0.2945 | -0.3705 |
| CYP2B6 | NM_000767 | -0.209 | -0.0955 |
| CYP2C19 | NM_000769 | -0.0295 | -0.115 |

| | | | |
|---|---|---|---|
| CYP2C8 | NM_000770 | -0.4925 | -0.3515 |
| CYP2C9 | NM_000771 | -0.09 | -0.3035 |
| CYP2C18 | NM_000772 | -0.048 | -0.335 |
| CYP2E1 | NM_000773 | -0.084 | -0.318 |
| CYP2F1 | NM_000774 | -0.096 | -0.448 |
| CYP2J2 | NM_000775 | -0.5235 | -1.2255 |
| CYP3A5 | NM_000777 | -0.6525 | -0.515 |
| CYP4A11 | NM_000778 | 0.002 | -0.319 |
| CYP4B1 | NM_000779 | -0.083 | -0.0915 |
| CYP7A1 | NM_000780 | 0.3445 | -0.1165 |
| CYP11A1 | NM_000781 | 0.135 | -0.059 |
| CYP24A1 | NM_000782 | -0.3545 | -0.228 |
| CYP26A1 | NM_000783 | 0.175 | -0.312 |
| CYP27A1 | NM_000784 | 0.019 | -0.6385 |
| CYP27B1 | NM_000785 | -0.1145 | -1.0625 |
| CYP51A1 | NM_000786 | -1.0375 | -1.3685 |
| DBH | NM_000787 | -0.21 | -0.577 |
| DCK | NM_000788 | -1.331 | -2.9735 |
| ACE | NM_000789 | -0.234 | -0.243 |
| DDC | NM_000790 | 0.0235 | -0.3285 |
| DHFR | NM_000791 | -1.0545 | -0.824 |
| DIO1 | NM_000792 | -0.1625 | -0.1355 |
| DIO2 | NM_000793 | 0.0605 | -0.2345 |
| DRD1 | NM_000794 | 0.0415 | -0.199 |
| DRD2 | NM_000795 | -0.4845 | -1.0025 |
| DRD3 | NM_000796 | 0.596 | 0.507 |
| DRD4 | NM_000797 | -0.0425 | -0.904 |
| DRD5 | NM_000798 | -0.1425 | -0.3075 |
| EPO | NM_000799 | -0.0715 | 0.035 |
| FGF1 | NM_000800 | 0.2095 | -0.095 |
| FKBP1A | NM_000801 | 0.2695 | -0.1105 |
| FOLR2 | NM_000803 | -0.0485 | 0.096 |
| FOLR3 | NM_000804 | -0.002 | -0.747 |
| GAST | NM_000805 | -0.1905 | 0.275 |
| GABRA1 | NM_000806 | -0.4125 | -0.7615 |
| GABRA2 | NM_000807 | -0.223 | -0.2635 |
| GABRA3 | NM_000808 | 0.0035 | -0.3305 |
| GABRA4 | NM_000809 | 0.055 | 0.162 |
| GABRA5 | NM_000810 | 0.038 | -0.9535 |
| GABRA6 | NM_000811 | -0.1345 | -0.237 |
| GABRB1 | NM_000812 | -0.1355 | -0.726 |
| GABRB2 | NM_000813 | -0.188 | -0.324 |
| GABRB3 | NM_000814 | -1.149 | -1.38 |

| | | | |
|---|---|---|---|
| GABRD | NM_000815 | -0.407 | -0.138 |
| GABRG2 | NM_000816 | 0.011 | -0.577 |
| GAD1 | NM_000817 | 0.131 | -0.098 |
| GAD2 | NM_000818 | 0.1075 | -0.389 |
| GART | NM_000819 | -0.557 | -1.4935 |
| GAS6 | NM_000820 | -0.203 | -0.3345 |
| GGCX | NM_000821 | -0.1275 | -0.799 |
| GHRHR | NM_000823 | 0.1405 | -0.352 |
| GLRB | NM_000824 | -1.6365 | -1.7595 |
| GNRH1 | NM_000825 | -0.257 | -0.711 |
| GRIA2 | NM_000826 | -0.6775 | -1.264 |
| GRIA1 | NM_000827 | -0.7495 | -0.244 |
| GRIA3 | NM_000828 | 0.146 | 0.326 |
| GRIA4 | NM_000829 | -0.663 | -0.1215 |
| GRIK1 | NM_000830 | 0.1 | -0.0175 |
| GRIK3 | NM_000831 | -1.0635 | -1.434 |
| GRIN1 | NM_000832 | 0.2565 | 0.227 |
| GRIN2A | NM_000833 | 0.2525 | -0.051 |
| GRIN2B | NM_000834 | 0.237 | -0.3905 |
| GRIN2C | NM_000835 | 0.026 | 0.315 |
| GRIN2D | NM_000836 | -0.0585 | -0.783 |
| GRINA | NM_000837 | -0.0775 | -0.431 |
| GRM1 | NM_000838 | 0.23 | 0.0755 |
| GRM2 | NM_000839 | -0.4 | -0.7465 |
| GRM3 | NM_000840 | 0.408 | -0.466 |
| GRM4 | NM_000841 | -0.3345 | -0.808 |
| GRM5 | NM_000842 | -0.0525 | -0.4285 |
| GRM6 | NM_000843 | -0.142 | -0.1245 |
| GRM7 | NM_000844 | 0.146 | -0.7625 |
| GRM8 | NM_000845 | -1.312 | -1.6935 |
| GSTA2 | NM_000846 | 0.179 | -0.1935 |
| GSTA3 | NM_000847 | 0.1875 | -0.143 |
| GSTM3 | NM_000849 | -0.0165 | -0.132 |
| GSTM4 | NM_000850 | 0.1655 | -0.5215 |
| GSTM5 | NM_000851 | -0.2305 | -0.9055 |
| GSTP1 | NM_000852 | 0.2485 | -0.227 |
| GSTT1 | NM_000853 | 0.0695 | -0.5425 |
| GUCY1A2 | NM_000855 | 0.282 | -0.59 |
| GUCY1A3 | NM_000856 | -1.0835 | -2.0525 |
| GUCY1B3 | NM_000857 | -1.9595 | -2.76 |
| GUK1 | NM_000858 | 0.4125 | -0.165 |
| HMGCR | NM_000859 | -1.3985 | -2.906 |
| HPGD | NM_000860 | -0.3415 | -0.8265 |

| | | | |
|---|---|---|---|
| HRH1 | NM_000861 | -0.043 | -0.601 |
| HSD3B1 | NM_000862 | 0.086 | -0.224 |
| HTR1B | NM_000863 | -0.09 | -0.432 |
| HTR1D | NM_000864 | -0.0395 | -0.15 |
| HTR1E | NM_000865 | -0.312 | -1.168 |
| HTR1F | NM_000866 | -0.0955 | 0.08 |
| HTR2B | NM_000867 | 0.074 | 0.3975 |
| HTR2C | NM_000868 | -0.486 | -0.2735 |
| HTR3A | NM_000869 | -0.1285 | -0.6555 |
| HTR6 | NM_000871 | -0.0925 | -0.23 |
| HTR7 | NM_000872 | -0.0655 | -0.482 |
| ICAM2 | NM_000873 | -1.088 | -0.6605 |
| IFNAR2 | NM_000874 | -0.7095 | -0.8435 |
| IGF1R | NM_000875 | -0.3475 | -0.8765 |
| IGF2R | NM_000876 | -0.437 | -1.634 |
| IL1R1 | NM_000877 | 0.4215 | -0.4995 |
| IL2RB | NM_000878 | -0.361 | -0.077 |
| IL5 | NM_000879 | 0.074 | -0.341 |
| IL7 | NM_000880 | -0.1255 | -0.126 |
| IL12A | NM_000882 | -0.9195 | -0.6115 |
| IMPDH1 | NM_000883 | -0.211 | -1.6375 |
| IMPDH2 | NM_000884 | 0.1835 | -0.2625 |
| ITGA4 | NM_000885 | -0.713 | -1.871 |
| ITGAX | NM_000887 | -0.202 | -0.3785 |
| ITGB6 | NM_000888 | 0.284 | 0.0585 |
| ITGB7 | NM_000889 | -0.022 | -0.0895 |
| KCNJ5 | NM_000890 | 0.0295 | 0.164 |
| KCNJ2 | NM_000891 | -0.2535 | 0.112 |
| KLKB1 | NM_000892 | -0.129 | -0.446 |
| KNG1 | NM_000893 | 0.2715 | 0.046 |
| LHB | NM_000894 | 0.0085 | -0.586 |
| LTA4H | NM_000895 | -0.771 | -0.447 |
| CYP4F3 | NM_000896 | 0.4855 | -0.774 |
| MAOB | NM_000898 | -0.108 | -0.2525 |
| KITLG | NM_000899 | -0.1425 | -1.928 |
| MGP | NM_000900 | -0.256 | -0.3085 |
| NR3C2 | NM_000901 | -1.2945 | -0.8685 |
| MME | NM_000902 | -0.403 | -0.471 |
| NQO1 | NM_000903 | 0.231 | -0.5815 |
| NQO2 | NM_000904 | -0.5545 | -1.1465 |
| NPY | NM_000905 | -0.1745 | -0.528 |
| NPR1 | NM_000906 | -0.0095 | -0.538 |
| NPR3 | NM_000908 | 0.214 | -0.245 |

| | | | |
|---|---|---|---|
| NPY1R | NM_000909 | 0.17 | -0.239 |
| NPY2R | NM_000910 | 0.024 | -1.3065 |
| OPRD1 | NM_000911 | 0.0355 | -0.459 |
| OPRK1 | NM_000912 | 0.407 | -0.666 |
| OPRL1 | NM_000913 | -1.03 | -1.4635 |
| OPRM1 | NM_000914 | -0.147 | -0.562 |
| OXT | NM_000915 | 0.008 | -0.353 |
| OXTR | NM_000916 | -0.756 | -0.6035 |
| P4HB | NM_000918 | -0.02 | -0.453 |
| PAM | NM_000919 | -0.739 | -0.8865 |
| PC | NM_000920 | 0.171 | -0.143 |
| PDE3A | NM_000921 | 0.092 | 0.1 |
| PDE3B | NM_000922 | -1.0255 | -2.343 |
| PDE4C | NM_000923 | -0.183 | -0.1575 |
| PDE1B | NM_000924 | 0.0265 | -0.2955 |
| PDHB | NM_000925 | -1.0295 | -2.649 |
| PGR | NM_000926 | 0.006 | -0.2725 |
| ABCB1 | NM_000927 | -0.175 | -0.566 |
| PLA2G1B | NM_000928 | 0.168 | -0.3105 |
| PLA2G5 | NM_000929 | 0.137 | -0.0945 |
| PLAT | NM_000930 | -0.6655 | -1.295 |
| PLCB3 | NM_000932 | 0.049 | -0.422 |
| PLCB4 | NM_000933 | -0.5485 | -1.781 |
| SERPINF2 | NM_000934 | -0.152 | -0.4985 |
| PLOD2 | NM_000935 | -0.396 | -1.107 |
| PNLIP | NM_000936 | -0.1455 | 0.235 |
| POLR2A | NM_000937 | 0.286 | -0.637 |
| POLR2B | NM_000938 | -1.0405 | -1.344 |
| POMC | NM_000939 | 0.0485 | -0.093 |
| PON3 | NM_000940 | -0.487 | -0.3565 |
| POR | NM_000941 | -0.1365 | -0.7675 |
| PPIB | NM_000942 | -0.0885 | -0.264 |
| PPIC | NM_000943 | 0.196 | -0.532 |
| PPP3CA | NM_000944 | -1.0855 | -1.8885 |
| PPP3R1 | NM_000945 | -1.026 | -2.164 |
| PRIM1 | NM_000946 | -1.6305 | -2.508 |
| PRIM2 | NM_000947 | -0.889 | -1.2565 |
| PRL | NM_000948 | -0.4175 | -0.0695 |
| PRLR | NM_000949 | -0.028 | 0.0215 |
| PRRG1 | NM_000950 | -1.3485 | -2.7765 |
| PRRG2 | NM_000951 | 0.845 | 0.4425 |
| PTAFR | NM_000952 | -0.3325 | -0.208 |
| PTGDR | NM_000953 | 0.5725 | -0.117 |

| | | | |
|---|---|---|---|
| PTGDS | NM_000954 | -0.181 | -0.0135 |
| PTGER1 | NM_000955 | -0.0685 | -0.1505 |
| PTGER2 | NM_000956 | -0.269 | -0.213 |
| PTGER4 | NM_000958 | 0.1745 | 0.1895 |
| PTGFR | NM_000959 | -0.3595 | -1.171 |
| PTGIR | NM_000960 | -0.38 | -0.3355 |
| PTGIS | NM_000961 | -0.125 | -0.4605 |
| PTGS1 | NM_000962 | 0.3395 | -0.4795 |
| PTGS2 | NM_000963 | -0.004 | -0.363 |
| RARA | NM_000964 | -0.49 | -1.943 |
| RARB | NM_000965 | 0.0095 | -1.128 |
| RARG | NM_000966 | -0.129 | -0.906 |
| RPL3 | NM_000967 | 0.0645 | -0.235 |
| RPL4 | NM_000968 | -0.3075 | -0.309 |
| RPL5 | NM_000969 | -0.271 | -0.2035 |
| RPL6 | NM_000970 | -0.259 | -0.331 |
| RPL7 | NM_000971 | -0.343 | -0.102 |
| RPL7A | NM_000972 | 0.0625 | -0.313 |
| RPL8 | NM_000973 | 0.2675 | -0.1125 |
| RPL11 | NM_000975 | -0.081 | -0.1535 |
| RPL12 | NM_000976 | 0.1935 | -0.2525 |
| RPL13 | NM_000977 | 0.0625 | -0.329 |
| RPL23 | NM_000978 | 0.0935 | -0.2 |
| RPL18 | NM_000979 | 0.1935 | -0.236 |
| RPL18A | NM_000980 | 0.111 | -0.3455 |
| RPL19 | NM_000981 | -0.033 | -0.209 |
| RPL21 | NM_000982 | -0.1095 | 0.002 |
| RPL22 | NM_000983 | -0.0545 | -0.259 |
| RPL23A | NM_000984 | 0.0125 | -0.2295 |
| RPL17 | NM_000985 | -0.1785 | -0.184 |
| RPL24 | NM_000986 | -0.0115 | -0.2285 |
| RPL26 | NM_000987 | -0.0315 | -0.1815 |
| RPL27 | NM_000988 | 0.0475 | -0.3385 |
| RPL30 | NM_000989 | 0.1595 | -0.227 |
| RPL27A | NM_000990 | -0.1445 | -0.456 |
| RPL28 | NM_000991 | 0.1075 | -0.274 |
| RPL29 | NM_000992 | 0.1315 | -0.313 |
| RPL31 | NM_000993 | -0.0905 | -0.3225 |
| RPL32 | NM_000994 | 0.057 | -0.362 |
| RPL34 | NM_000995 | -0.0295 | -0.214 |
| RPL35A | NM_000996 | -0.6345 | -0.7155 |
| RPL37 | NM_000997 | 0.1405 | -0.3225 |
| RPL37A | NM_000998 | 0.017 | -0.3955 |

| | | | |
|---|---|---|---|
| RPL38 | NM_000999 | 0.0725 | -0.151 |
| RPL39 | NM_001000 | 0.014 | -0.3055 |
| RPL36AL | NM_001001 | -0.6265 | -1.2485 |
| ITSN1 | NM_001001132 | -1.046 | -1.7135 |
| PRSS58 | NM_001001317 | 0.1435 | -0.027 |
| ATP2B1 | NM_001001323 | -0.169 | -1.4015 |
| SPINK14 | NM_001001325 | 0.1045 | -0.1035 |
| PRKCSH | NM_001001329 | 0.0595 | -0.344 |
| REEP3 | NM_001001330 | -0.5485 | -0.754 |
| ATP2B2 | NM_001001331 | -1.334 | -2.4225 |
| BLOC1S2 | NM_001001342 | -1.1145 | -0.969 |
| ATP2B3 | NM_001001344 | 0.0715 | -0.5615 |
| CLDN20 | NM_001001346 | -0.1125 | 0.182 |
| NKIRAS2 | NM_001001349 | -0.569 | -1.22 |
| CD44 | NM_001001389 | 0.1785 | -0.749 |
| CD44 | NM_001001390 | 0.1785 | -0.749 |
| CD44 | NM_001001391 | 0.1785 | -0.749 |
| CD44 | NM_001001392 | 0.1785 | -0.749 |
| DNAJB3 | NM_001001394 | 0.191 | 0.3525 |
| LMO3 | NM_001001395 | -1.304 | -1.7735 |
| ATP2B4 | NM_001001396 | -1.222 | -1.6475 |
| C16orf42 | NM_001001410 | -0.3085 | -1.3595 |
| ZNF676 | NM_001001411 | -0.4295 | -0.5445 |
| CALHM1 | NM_001001412 | -0.2535 | 0.1835 |
| NCCRP1 | NM_001001414 | -0.0735 | -0.103 |
| ZNF429 | NM_001001415 | -0.5915 | -2.609 |
| SMAD5 | NM_001001419 | -1.112 | -1.9855 |
| SMAD5 | NM_001001420 | -1.112 | -1.9855 |
| STX16 | NM_001001433 | -1.417 | -1.407 |
| C16orf87 | NM_001001436 | -0.6905 | -2.488 |
| LSS | NM_001001438 | -0.157 | -0.532 |
| SLC35E4 | NM_001001479 | -0.1465 | -0.739 |
| KRTAP5-5 | NM_001001480 | -0.1215 | 0.057 |
| UBE2W | NM_001001481 | -1.268 | -3.192 |
| ZDHHC13 | NM_001001483 | -1.217 | -2.068 |
| PTER | NM_001001484 | -0.155 | -0.582 |
| ATP2C1 | NM_001001485 | -0.972 | -1.569 |
| ATP2C1 | NM_001001486 | -0.972 | -1.569 |
| ATP2C1 | NM_001001487 | -0.972 | -1.569 |
| SNCB | NM_001001502 | 0.0735 | -0.398 |
| NDUFV3 | NM_001001503 | -0.3885 | -1.2635 |
| HDGFRP2 | NM_001001520 | -0.215 | -1.342 |
| TAGLN | NM_001001522 | 0.149 | -0.3335 |

| | | | |
|---|---|---|---|
| RORC | NM_001001523 | 0.021 | -0.4155 |
| TM6SF2 | NM_001001524 | -0.428 | -0.7495 |
| CD36 | NM_001001547 | 0.1265 | -0.518 |
| CD36 | NM_001001548 | 0.1265 | -0.518 |
| GRB10 | NM_001001549 | 0.1195 | -0.5235 |
| GRB10 | NM_001001550 | 0.1195 | -0.5235 |
| C9orf103 | NM_001001551 | 0.2525 | -0.4835 |
| LEMD1 | NM_001001552 | -0.236 | -0.6165 |
| GRB10 | NM_001001555 | -0.9305 | -2.207 |
| GALK2 | NM_001001556 | -1.106 | -2.3835 |
| GDF6 | NM_001001557 | -1.197 | -0.6035 |
| GGA1 | NM_001001560 | -0.682 | -1.404 |
| GGA1 | NM_001001561 | -0.682 | -1.404 |
| TIMM50 | NM_001001563 | -0.038 | -0.821 |
| PDE9A | NM_001001571 | -0.9145 | -0.971 |
| ALKBH2 | NM_001001655 | -0.805 | -2.2345 |
| OR9A4 | NM_001001656 | -0.139 | -0.2805 |
| OR2Y1 | NM_001001657 | -0.1095 | -0.019 |
| OR9A2 | NM_001001658 | 0.3265 | 0.452 |
| OR2A14 | NM_001001659 | 0.043 | -0.5115 |
| LYRM5 | NM_001001660 | -1.499 | -1.892 |
| ZNF425 | NM_001001661 | -0.7925 | -1.742 |
| ZNF782 | NM_001001662 | -0.094 | -0.632 |
| TMEM211 | NM_001001663 | -0.0705 | -0.5275 |
| SPOPL | NM_001001664 | -0.525 | -1.7255 |
| CYP27C1 | NM_001001665 | -0.206 | -1.435 |
| ANO7 | NM_001001666 | -0.1645 | -0.5885 |
| OR6V1 | NM_001001667 | 0.016 | 0.089 |
| ZNF470 | NM_001001668 | 0.0225 | -1.551 |
| ARHGEF37 | NM_001001669 | -0.0495 | 0.165 |
| FAM75D1 | NM_001001670 | 0.108 | -0.463 |
| MAP3K15 | NM_001001671 | -0.1695 | -0.7745 |
| RNASE9 | NM_001001673 | -0.1285 | -0.148 |
| OR4F15 | NM_001001674 | 0.0035 | -0.295 |
| LCN9 | NM_001001676 | -0.1 | 0.308 |
| MED11 | NM_001001683 | -1.0425 | -2.037 |
| IL17REL | NM_001001694 | -0.017 | -0.298 |
| C4orf3 | NM_001001701 | -0.366 | -0.53 |
| C9orf170 | NM_001001709 | -0.053 | -0.259 |
| FAM166A | NM_001001710 | -0.103 | 0.0355 |
| DDI1 | NM_001001711 | 0.0745 | -0.0935 |
| LCN10 | NM_001001712 | 0.0675 | -0.294 |
| FARP1 | NM_001001715 | -0.2055 | -0.598 |

| | | | |
|---|---|---|---|
| CC2D2B | NM_001001732 | -0.083 | -0.0005 |
| ATP1A4 | NM_001001734 | 0.128 | -0.1845 |
| RFWD2 | NM_001001740 | -1.255 | -2.924 |
| BLID | NM_001001786 | 0.1455 | -0.0605 |
| RAET1G | NM_001001788 | 0.0035 | -0.4595 |
| TOMM5 | NM_001001790 | -0.0055 | -0.544 |
| C10orf55 | NM_001001791 | 0.105 | -0.4375 |
| FAM116B | NM_001001794 | -0.055 | -0.325 |
| C8orf82 | NM_001001795 | -0.408 | -1.4845 |
| OR2A42 | NM_001001802 | -0.067 | -0.159 |
| OR2T34 | NM_001001821 | 0.7265 | 0.23 |
| OR2T27 | NM_001001824 | -0.0045 | -0.474 |
| STX19 | NM_001001850 | 0.0585 | -0.1125 |
| ITIH5 | NM_001001851 | -0.036 | -0.487 |
| PIM3 | NM_001001852 | -0.664 | -2.059 |
| C14orf37 | NM_001001872 | -0.303 | -0.685 |
| TPD52L3 | NM_001001874 | 0.0165 | -0.4795 |
| TPD52L3 | NM_001001875 | -0.0025 | -0.232 |
| RUNX1 | NM_001001890 | 0.0375 | -0.054 |
| ANO7 | NM_001001891 | -0.1645 | -0.5885 |
| TTC3 | NM_001001894 | 0.0705 | -0.3775 |
| UBASH3A | NM_001001895 | -0.044 | -0.2395 |
| OR4E2 | NM_001001912 | 0.271 | 0.106 |
| OR52N1 | NM_001001913 | -0.126 | 0.1585 |
| OR2G3 | NM_001001914 | 0.011 | -0.356 |
| OR2G2 | NM_001001915 | -0.161 | -0.483 |
| OR52J3 | NM_001001916 | 0.172 | 0.012 |
| OR56A1 | NM_001001917 | -0.243 | 0.1225 |
| OR14C36 | NM_001001918 | -0.1485 | 0.2705 |
| OR13C4 | NM_001001919 | 0.4725 | -0.3875 |
| OR4C15 | NM_001001920 | 0.001 | -0.1935 |
| OR5AS1 | NM_001001921 | -0.1185 | -0.29 |
| OR52N5 | NM_001001922 | -0.1315 | -0.307 |
| OR5C1 | NM_001001923 | -0.1535 | -0.2925 |
| MTUS1 | NM_001001924 | -1.1065 | -3.0205 |
| MTUS1 | NM_001001925 | -0.8945 | -2.8875 |
| PPARA | NM_001001928 | 0.0355 | -0.1175 |
| MTUS1 | NM_001001931 | -0.8945 | -2.8875 |
| LHX8 | NM_001001933 | -1.1155 | -1.855 |
| AFAP1L2 | NM_001001936 | 0.0205 | -0.0525 |
| ATP5A1 | NM_001001937 | -0.1785 | -0.2575 |
| C9orf47 | NM_001001938 | 0.33 | 0.3055 |
| OR5D18 | NM_001001952 | 0.062 | -0.061 |

| | | | |
|---|---|---|---|
| OR10G9 | NM_001001953 | 0.331 | -0.3415 |
| OR5A2 | NM_001001954 | 0.0225 | -0.3115 |
| OR4C13 | NM_001001955 | -0.1025 | 0.0545 |
| OR13C9 | NM_001001956 | 0.195 | -0.0125 |
| OR2W3 | NM_001001957 | 0.1665 | -0.209 |
| OR7G3 | NM_001001958 | 0.094 | 0.206 |
| OR11L1 | NM_001001959 | 0.228 | -0.1975 |
| OR5W2 | NM_001001960 | 0.054 | 0.168 |
| OR13C3 | NM_001001961 | -0.568 | -0.4025 |
| OR2T11 | NM_001001964 | 0.0715 | -0.4365 |
| OR4D5 | NM_001001965 | -0.142 | -0.351 |
| OR14A16 | NM_001001966 | -0.0585 | -0.4095 |
| OR5D13 | NM_001001967 | 0.1015 | -0.016 |
| OR6S1 | NM_001001968 | -0.023 | -0.443 |
| FAM13C | NM_001001971 | -0.277 | -1.1595 |
| ATP5C1 | NM_001001973 | -0.432 | -0.5025 |
| PLEKHA1 | NM_001001974 | -1.204 | -1.5215 |
| ATP5D | NM_001001975 | 0.2845 | -0.324 |
| ATE1 | NM_001001976 | -0.5925 | -1.1715 |
| ATP5E | NM_001001977 | 0.103 | -0.0615 |
| PAMR1 | NM_001001991 | 0.061 | -0.1 |
| USP16 | NM_001001992 | -0.374 | -1.7405 |
| GPM6B | NM_001001994 | -1.02 | -0.3295 |
| GPM6B | NM_001001995 | -1.02 | -0.3295 |
| GPM6B | NM_001001996 | -1.02 | -0.3295 |
| EXOSC10 | NM_001001998 | -1.096 | -1.952 |
| RPLP0 | NM_001002 | 0.125 | -0.2855 |
| GMPR2 | NM_001002001 | -0.4185 | -1.231 |
| NT5C1B | NM_001002006 | -0.0925 | -0.479 |
| NT5C3 | NM_001002010 | -0.746 | -1.2545 |
| HCFC1R1 | NM_001002017 | 0.0785 | -0.683 |
| HCFC1R1 | NM_001002018 | 0.016 | -0.87 |
| PUS1 | NM_001002019 | -1.322 | -2.637 |
| PUS1 | NM_001002020 | -1.048 | -2.034 |
| CLDN18 | NM_001002026 | 0.037 | -0.2065 |
| ATP5G1 | NM_001002027 | 0.284 | -0.066 |
| ECHDC1 | NM_001002030 | -0.8545 | -1.684 |
| HN1 | NM_001002032 | -0.35 | -1.5975 |
| HN1 | NM_001002033 | -0.35 | -1.5975 |
| FAM109B | NM_001002034 | -0.103 | -0.3315 |
| DEFB108B | NM_001002035 | -0.143 | -0.7315 |
| ASTL | NM_001002036 | -0.125 | -0.279 |
| KLK2 | NM_001002231 | 0.1735 | -0.465 |

| | | | |
|---|---|---|---|
| SERPINA1 | NM_001002235 | -0.127 | -0.2305 |
| SERPINA1 | NM_001002236 | -0.127 | -0.2305 |
| AFTPH | NM_001002243 | -1.2 | -2.3835 |
| ANAPC11 | NM_001002244 | 0.2395 | -0.369 |
| ANAPC11 | NM_001002245 | 0.2395 | -0.369 |
| ANAPC11 | NM_001002246 | 0.2395 | -0.369 |
| ANAPC11 | NM_001002247 | 0.2395 | -0.369 |
| ANAPC11 | NM_001002248 | 0.2395 | -0.369 |
| ANAPC11 | NM_001002249 | 0.2395 | -0.369 |
| ARL6IP4 | NM_001002251 | 0.269 | -0.3275 |
| ARL6IP4 | NM_001002252 | 0.269 | -0.3275 |
| AWAT2 | NM_001002254 | -0.041 | -0.1075 |
| SUMO4 | NM_001002255 | 0.433 | -0.3015 |
| LCLAT1 | NM_001002257 | -0.567 | -1.0845 |
| ATP5G3 | NM_001002258 | 0.0935 | -0.4055 |
| CAPRIN2 | NM_001002259 | -1.5535 | -2.0835 |
| ZFYVE27 | NM_001002261 | -0.2705 | -1.3395 |
| ZFYVE27 | NM_001002262 | -0.2705 | -1.3395 |
| EPSTI1 | NM_001002264 | -0.266 | -0.101 |
| EXOSC3 | NM_001002269 | -1.556 | -3.747 |
| FCGR2B | NM_001002273 | -0.2175 | -0.407 |
| FCGR2B | NM_001002274 | -0.2175 | -0.407 |
| FCGR2B | NM_001002275 | -0.2175 | -0.407 |
| WLS | NM_001002292 | -0.092 | -0.74 |
| FMO3 | NM_001002294 | -0.108 | 0.1175 |
| GATA3 | NM_001002295 | -0.7485 | -1.9475 |
| GOLGA7 | NM_001002296 | -1.293 | -1.831 |
| NFU1 | NM_001002755 | -1.074 | -1.8545 |
| PRY2 | NM_001002758 | -0.173 | -0.2565 |
| DNAJB12 | NM_001002762 | -0.7045 | -2.1445 |
| MCTP1 | NM_001002796 | -0.0905 | -0.036 |
| SMC4 | NM_001002800 | -0.278 | -1.8885 |
| PDE4DIP | NM_001002810 | -0.9925 | -0.862 |
| PDE4DIP | NM_001002811 | -0.9925 | -0.862 |
| PDE4DIP | NM_001002812 | -0.9925 | -0.862 |
| RAB11FIP1 | NM_001002814 | 0.026 | -0.933 |
| ZNF787 | NM_001002836 | -0.4 | -1.5605 |
| INPP5J | NM_001002837 | -0.1895 | -0.535 |
| MYL4 | NM_001002841 | -0.2895 | 0.0125 |
| ZNF280D | NM_001002843 | -0.5185 | -0.6775 |
| ZNF280D | NM_001002844 | -0.5185 | -0.6775 |
| ANXA2 | NM_001002857 | -0.2605 | -0.2505 |
| ANXA2 | NM_001002858 | -0.2605 | -0.2505 |

| | | | |
|---|---|---|---|
| BTBD7 | NM_001002860 | -1.9965 | -4.493 |
| DERL3 | NM_001002862 | 0.165 | -0.181 |
| THOC5 | NM_001002878 | 0.0445 | -0.883 |
| CBY1 | NM_001002880 | -1.5265 | -2.169 |
| FCRLB | NM_001002901 | 0.194 | -0.1855 |
| OR8G1 | NM_001002905 | 0.1485 | -0.7475 |
| OR8K1 | NM_001002907 | -0.2265 | 0.2605 |
| GPATCH8 | NM_001002909 | -1.026 | -2.1815 |
| GPR139 | NM_001002911 | -0.0965 | -0.576 |
| C1orf173 | NM_001002912 | -0.6765 | 0.045 |
| PTRH1 | NM_001002913 | 0.111 | -0.9905 |
| KCTD11 | NM_001002914 | -0.5445 | -0.7345 |
| IGFL2 | NM_001002915 | 0.0055 | -0.2705 |
| H2BFWT | NM_001002916 | 0.2545 | -0.234 |
| OR8D1 | NM_001002917 | -0.086 | -0.0235 |
| OR8D2 | NM_001002918 | -0.29 | -0.2515 |
| FAM150B | NM_001002919 | 0.0915 | -1.2905 |
| IGFL4 | NM_001002923 | -0.1215 | -0.561 |
| OR5AP2 | NM_001002925 | -0.1895 | -0.153 |
| TWISTNB | NM_001002926 | -0.314 | -0.3925 |
| RPLP1 | NM_001003 | 0.0785 | -0.3505 |
| TPD52L1 | NM_001003395 | -0.23 | -0.633 |
| BICD1 | NM_001003398 | -0.8615 | -1.9545 |
| CACNA1I | NM_001003406 | -0.0065 | -0.2005 |
| ABLIM1 | NM_001003407 | -0.6685 | -1.1735 |
| ABLIM1 | NM_001003408 | -1.0475 | -1.0985 |
| OR56A3 | NM_001003443 | -0.324 | -0.3525 |
| SMAD2 | NM_001003652 | -1.341 | -2.2695 |
| C1orf95 | NM_001003665 | -0.3035 | -0.8595 |
| C18orf1 | NM_001003674 | -0.1455 | -2.338 |
| C18orf1 | NM_001003675 | -0.1455 | -2.338 |
| C11orf49 | NM_001003676 | -0.4635 | -1.062 |
| C11orf49 | NM_001003677 | -0.31 | -0.7105 |
| C11orf49 | NM_001003678 | -0.31 | -0.7105 |
| LEPR | NM_001003679 | -0.63 | -0.3545 |
| LEPR | NM_001003680 | -0.362 | -0.5775 |
| HMGXB4 | NM_001003681 | -1.501 | -2.162 |
| TMEM200B | NM_001003682 | -0.15 | -0.6185 |
| PDE1A | NM_001003683 | 0.0345 | -0.835 |
| UQCR10 | NM_001003684 | 0.0625 | -0.4215 |
| SMAD1 | NM_001003688 | -1.6175 | -2.971 |
| MAD2L1BP | NM_001003690 | -0.8955 | -2.6965 |
| LY6G6F | NM_001003693 | -0.1005 | -0.028 |

| | | | |
|---|---|---|---|
| BRPF1 | NM_001003694 | -0.693 | -2.1945 |
| RREB1 | NM_001003698 | 0.0685 | -0.8065 |
| RREB1 | NM_001003699 | 0.0685 | -0.8065 |
| RREB1 | NM_001003700 | 0.0685 | -0.8065 |
| ATP5J | NM_001003703 | -0.084 | -0.0935 |
| OSBPL8 | NM_001003712 | -0.0895 | -0.896 |
| ATP5J2 | NM_001003713 | 0.2605 | -0.2165 |
| ATP5J2 | NM_001003714 | 0.2425 | -0.187 |
| RECQL5 | NM_001003715 | -0.791 | -1.203 |
| RECQL5 | NM_001003716 | -0.791 | -1.203 |
| GLE1 | NM_001003722 | -0.827 | -1.7725 |
| OR10A3 | NM_001003745 | 0.0235 | -0.411 |
| OR8I2 | NM_001003750 | -0.1575 | -0.249 |
| KLHL31 | NM_001003760 | 0.2405 | -0.3005 |
| STRADA | NM_001003786 | -0.9125 | -1.7355 |
| STRADA | NM_001003787 | -0.9125 | -1.7355 |
| STRADA | NM_001003788 | -0.9125 | -1.7355 |
| ERLIN2 | NM_001003790 | -1.133 | -1.754 |
| ERLIN2 | NM_001003791 | -1.133 | -1.754 |
| RBMS3 | NM_001003792 | -0.315 | -0.709 |
| RBMS3 | NM_001003793 | -0.449 | -0.8745 |
| MGLL | NM_001003794 | -0.246 | -0.8355 |
| TARP | NM_001003799 | 0.0195 | 0.0605 |
| BICD2 | NM_001003800 | -0.4905 | -0.841 |
| SMARCD3 | NM_001003801 | 0.0975 | -0.4005 |
| SMARCD3 | NM_001003802 | 0.0975 | -0.4005 |
| ATP5S | NM_001003803 | -0.2015 | -0.9055 |
| ATP5S | NM_001003805 | -0.2015 | -0.9055 |
| DLGAP1 | NM_001003809 | -0.322 | -0.227 |
| HNRNPD | NM_001003810 | -0.4805 | -1.078 |
| TEX11 | NM_001003811 | 0.508 | 0.42 |
| TRIM6 | NM_001003818 | -0.17 | -0.341 |
| TRIM6-TRIM34 | NM_001003819 | 0.1225 | -0.3925 |
| PARVB | NM_001003828 | 0.0445 | 0.1985 |
| SLC6A19 | NM_001003841 | 0.0225 | -0.3055 |
| SP5 | NM_001003845 | 0.2295 | -0.971 |
| MED15 | NM_001003891 | -0.4365 | -1.3195 |
| DUPD1 | NM_001003892 | 0.071 | -0.159 |
| CDY1B | NM_001003894 | 0.063 | -0.747 |
| MANBAL | NM_001003897 | -0.3005 | -1.377 |
| PARP3 | NM_001003931 | -0.193 | -0.867 |
| TXNDC8 | NM_001003936 | -0.52 | -0.0265 |
| TSPYL6 | NM_001003937 | 0.056 | -0.286 |

| | | | |
|---|---|---|---|
| HBM | NM_001003938 | -0.1125 | -0.273 |
| BMF | NM_001003940 | 0.206 | -0.8675 |
| OGDH | NM_001003941 | -0.7535 | -1.7415 |
| BMF | NM_001003942 | 0.206 | -0.8675 |
| BMF | NM_001003943 | 0.206 | -0.8675 |
| ANXA13 | NM_001003954 | 0.172 | -0.287 |
| CAPNS1 | NM_001003962 | 0.2915 | -0.159 |
| RPLP2 | NM_001004 | 0.161 | -0.31 |
| FBLN2 | NM_001004019 | 0.09 | -0.6295 |
| DYRK3 | NM_001004023 | -1.1895 | -2.4955 |
| GPRASP2 | NM_001004051 | -1.109 | -1.6075 |
| OR52B2 | NM_001004052 | -0.0845 | -0.228 |
| LRRC29 | NM_001004055 | 0.0185 | -0.2165 |
| GRK4 | NM_001004056 | -0.2485 | -0.5915 |
| GRK4 | NM_001004057 | -0.9455 | -1.2855 |
| OR8K5 | NM_001004058 | -0.2595 | -0.6935 |
| OR4S2 | NM_001004059 | -0.044 | -0.3775 |
| NOMO2 | NM_001004060 | -0.2645 | -0.9555 |
| OR4K1 | NM_001004063 | 0.0265 | 0.0495 |
| OR8J3 | NM_001004064 | -0.3645 | -0.795 |
| GRK6 | NM_001004105 | -0.6775 | -1.1955 |
| GRK6 | NM_001004106 | -0.6775 | -1.1955 |
| OR4P4 | NM_001004124 | 0.2065 | -0.1845 |
| TUSC1 | NM_001004125 | -0.0865 | -1.0495 |
| ALG11 | NM_001004127 | -0.6205 | -1.2115 |
| OR10AD1 | NM_001004134 | -0.286 | -0.4605 |
| OR2A12 | NM_001004135 | 0.183 | -0.3145 |
| OR2T2 | NM_001004136 | 0.0685 | -0.225 |
| OR52M1 | NM_001004137 | 0.2125 | -0.182 |
| CD200 | NM_001004196 | -1.6905 | -2.4175 |
| OR13A1 | NM_001004297 | -0.012 | -0.351 |
| C10orf90 | NM_001004298 | -0.1055 | -0.36 |
| ZNF813 | NM_001004301 | -0.9045 | -0.952 |
| C1orf168 | NM_001004303 | 0.132 | -0.1685 |
| ZNF740 | NM_001004304 | -1.4555 | -2.8335 |
| CCDC144NL | NM_001004306 | -0.1555 | -0.0255 |
| ZCCHC16 | NM_001004308 | -0.324 | -0.321 |
| ZNF774 | NM_001004309 | -0.5285 | -1.053 |
| FCRL6 | NM_001004310 | -0.1195 | -0.452 |
| FIGLA | NM_001004311 | 0.3625 | -0.0125 |
| RTP2 | NM_001004312 | -0.055 | -0.536 |
| TMEM220 | NM_001004313 | -0.265 | -0.5025 |
| LEKR1 | NM_001004316 | 0.2065 | -0.5935 |

| | | | |
|---|---|---|---|
| LIN28B | NM_001004317 | -1.5245 | -4.224 |
| PAPL | NM_001004318 | 0.1 | -0.2605 |
| VHLL | NM_001004319 | 0.382 | -0.138 |
| AGMO | NM_001004320 | -0.383 | -0.003 |
| C7orf61 | NM_001004323 | 0.1125 | -0.237 |
| KRTAP5-2 | NM_001004325 | 0.4245 | -0.2605 |
| SLC22A20 | NM_001004326 | -0.032 | -0.4975 |
| ZNF705A | NM_001004328 | 0.068 | 0.0995 |
| DBX2 | NM_001004329 | -0.4645 | -0.165 |
| PLEKHG7 | NM_001004330 | 0.062 | -0.224 |
| RNASEK | NM_001004333 | 0.035 | -0.358 |
| GPR179 | NM_001004334 | -0.113 | -0.2385 |
| ZYG11A | NM_001004339 | -0.3585 | -0.4495 |
| FCGR1B | NM_001004340 | -0.1445 | -0.729 |
| ETV3L | NM_001004341 | -0.1925 | -0.452 |
| TRIM67 | NM_001004342 | -1.5615 | -2.2955 |
| MAP1LC3C | NM_001004343 | 0.065 | -0.227 |
| SPDYE3 | NM_001004351 | 0.161 | -0.048 |
| C9orf173 | NM_001004353 | 0.1305 | -0.281 |
| NRARP | NM_001004354 | -0.2215 | 1.1215 |
| FGFRL1 | NM_001004356 | -0.1025 | -0.5225 |
| FGFRL1 | NM_001004358 | 0.019 | -0.8995 |
| DPP10 | NM_001004360 | -0.0165 | -0.1135 |
| UMODL1 | NM_001004416 | 0.2685 | -0.2605 |
| CLEC2D | NM_001004419 | 0.0845 | -0.721 |
| PLA2G6 | NM_001004426 | -0.287 | -0.6265 |
| METRNL | NM_001004431 | -0.5565 | -1.6165 |
| LINGO4 | NM_001004432 | -0.345 | -0.0945 |
| SLC30A2 | NM_001004434 | -0.0905 | 0.188 |
| ITGA11 | NM_001004439 | -0.319 | -0.598 |
| FAM19A3 | NM_001004440 | -0.332 | -0.478 |
| ANKRD34B | NM_001004441 | 0.126 | 0.0285 |
| OR1B1 | NM_001004450 | -0.041 | 0.059 |
| OR1J1 | NM_001004451 | -0.3015 | -0.3325 |
| OR1J4 | NM_001004452 | -0.2805 | -0.2445 |
| OR1L8 | NM_001004454 | -0.3415 | -0.0575 |
| OR1M1 | NM_001004456 | 0.2375 | -0.1145 |
| OR1N2 | NM_001004457 | -0.004 | -0.2 |
| OR1S2 | NM_001004459 | -0.1085 | -0.2355 |
| OR10A2 | NM_001004460 | -0.243 | 0.121 |
| OR10A6 | NM_001004461 | -0.071 | -0.4205 |
| OR10G4 | NM_001004462 | -0.1045 | -0.19 |
| OR10G7 | NM_001004463 | 0.065 | -0.1775 |

| | | | |
|---|---|---|---|
| OR10G8 | NM_001004464 | -0.2335 | 0.176 |
| OR10H4 | NM_001004465 | -0.1505 | 0.2745 |
| OR10H5 | NM_001004466 | -0.3295 | -0.157 |
| OR10J3 | NM_001004467 | 0.2995 | -0.3265 |
| OR10J5 | NM_001004469 | -0.322 | -0.3105 |
| ST8SIA6 | NM_001004470 | 0.0265 | -0.029 |
| OR10Q1 | NM_001004471 | 0.206 | 0.075 |
| OR10R2 | NM_001004472 | 0.2205 | -0.3255 |
| OR10K1 | NM_001004473 | -0.175 | 0.3245 |
| OR10S1 | NM_001004474 | 0.1895 | -0.0295 |
| OR10T2 | NM_001004475 | -0.3015 | -0.36 |
| OR10K2 | NM_001004476 | 0.181 | -0.4355 |
| OR10X1 | NM_001004477 | 0.2995 | -0.2845 |
| OR10Z1 | NM_001004478 | -0.3605 | -0.309 |
| OR11H4 | NM_001004479 | -0.3585 | -0.0985 |
| OR11H6 | NM_001004480 | -0.0305 | -0.126 |
| OR13C2 | NM_001004481 | -0.1695 | -0.62 |
| OR13C5 | NM_001004482 | -0.2285 | -0.452 |
| OR13C8 | NM_001004483 | 0.3955 | -0.275 |
| OR13D1 | NM_001004484 | -0.216 | -0.354 |
| OR13F1 | NM_001004485 | -0.1225 | -0.377 |
| OR13H1 | NM_001004486 | -0.1255 | -0.5385 |
| OR13J1 | NM_001004487 | 0.1785 | -0.3275 |
| OR2A25 | NM_001004488 | -0.2065 | -0.1065 |
| OR2AG1 | NM_001004489 | -0.003 | -0.022 |
| OR2AG2 | NM_001004490 | -0.2365 | -0.9165 |
| OR2AK2 | NM_001004491 | -0.036 | -0.3175 |
| OR2B11 | NM_001004492 | -0.1465 | 0.336 |
| OR2D3 | NM_001004684 | -0.201 | -0.179 |
| OR2F2 | NM_001004685 | -0.1685 | -0.287 |
| OR2L2 | NM_001004686 | 0.696 | -0.146 |
| OR2L3 | NM_001004687 | 0.095 | -0.1425 |
| OR2M2 | NM_001004688 | -0.2315 | 0.5955 |
| OR2M3 | NM_001004689 | -0.0685 | -0.262 |
| OR2M5 | NM_001004690 | -0.144 | -0.5805 |
| OR2M7 | NM_001004691 | 0.1315 | -0.559 |
| OR2T10 | NM_001004693 | 0.618 | 0.0085 |
| OR2T4 | NM_001004696 | -0.1485 | -0.08 |
| OR2W5 | NM_001004698 | 0.0265 | -0.16 |
| OR2Z1 | NM_001004699 | 0.0065 | -0.2035 |
| OR4C11 | NM_001004700 | -0.191 | -0.5015 |
| OR4C16 | NM_001004701 | -0.273 | 0.379 |
| OR4C3 | NM_001004702 | -0.0105 | -0.1905 |

| | | | |
|---|---|---|---|
| OR4C46 | NM_001004703 | 0.2515 | -0.4115 |
| OR4C6 | NM_001004704 | -0.1535 | -0.031 |
| OR4D10 | NM_001004705 | -0.137 | -0.5725 |
| OR4D11 | NM_001004706 | -0.1635 | -0.288 |
| OR4D2 | NM_001004707 | 0.029 | -0.06 |
| OR4D6 | NM_001004708 | 0.0315 | -0.11 |
| OR4D9 | NM_001004711 | -0.0315 | -0.0395 |
| OR4K14 | NM_001004712 | -0.159 | -0.2465 |
| OR1I1 | NM_001004713 | 0.0085 | -0.1145 |
| OR4K13 | NM_001004714 | -0.01 | -0.171 |
| OR4K17 | NM_001004715 | -0.022 | -0.0295 |
| OR4L1 | NM_001004717 | 0.254 | 0.6535 |
| NCK2 | NM_001004720 | -0.2155 | -1.735 |
| NCK2 | NM_001004722 | -0.2155 | -1.735 |
| OR4N2 | NM_001004723 | 0.1855 | -0.5335 |
| OR4N5 | NM_001004724 | -0.3 | -0.0075 |
| OR4S1 | NM_001004725 | -0.244 | -0.489 |
| OR4X1 | NM_001004726 | -0.165 | -0.4175 |
| OR4X2 | NM_001004727 | -0.344 | -0.1825 |
| OR5A1 | NM_001004728 | -0.134 | 0.2375 |
| OR5AN1 | NM_001004729 | -0.08 | -0.3995 |
| OR5AR1 | NM_001004730 | -0.1015 | -0.5125 |
| OR5AU1 | NM_001004731 | 0.161 | -0.3125 |
| OR5B12 | NM_001004733 | -0.009 | -0.314 |
| OR14I1 | NM_001004734 | -0.0735 | -0.472 |
| OR5D14 | NM_001004735 | 0.1465 | -0.0195 |
| OR5K1 | NM_001004736 | 0.247 | -0.9905 |
| OR5K2 | NM_001004737 | 0.7155 | 0.1965 |
| OR5L1 | NM_001004738 | 0.014 | 0.092 |
| OR5L2 | NM_001004739 | 0.4205 | 0.2815 |
| OR5M1 | NM_001004740 | -0.1625 | -0.1965 |
| OR5M10 | NM_001004741 | 0.061 | 0.3685 |
| OR5M3 | NM_001004742 | 0.0745 | -0.1425 |
| OR5M9 | NM_001004743 | -0.0985 | -0.6845 |
| OR5R1 | NM_001004744 | -0.314 | -0.0825 |
| OR5T1 | NM_001004745 | 0.256 | 0.1855 |
| OR5T2 | NM_001004746 | -0.0895 | -0.286 |
| OR5T3 | NM_001004747 | 0.113 | 0.324 |
| OR51A7 | NM_001004749 | 0.1765 | -0.3285 |
| OR51B6 | NM_001004750 | -0.0905 | 0.2075 |
| OR51D1 | NM_001004751 | 0.078 | -0.467 |
| OR51F1 | NM_001004752 | 0.2005 | 0.117 |
| OR51F2 | NM_001004753 | -0.2555 | 0.0195 |

| | | | |
|---|---|---|---|
| OR51I2 | NM_001004754 | -0.0025 | 0.2555 |
| OR51L1 | NM_001004755 | -0.257 | -0.0325 |
| OR51M1 | NM_001004756 | -0.1245 | -0.335 |
| OR51Q1 | NM_001004757 | -0.005 | -0.2245 |
| OR51S1 | NM_001004758 | -0.2965 | -0.4255 |
| OR51T1 | NM_001004759 | 0.0065 | 0.078 |
| OR51V1 | NM_001004760 | 0.0295 | -0.3615 |
| RPS3 | NM_001005 | 0.1885 | -0.2855 |
| SFMBT1 | NM_001005158 | -0.647 | -1.472 |
| SFMBT1 | NM_001005159 | -0.647 | -1.472 |
| OR52A5 | NM_001005160 | -0.23 | -0.2625 |
| OR52B4 | NM_001005161 | 0.4745 | 0.035 |
| OR52B6 | NM_001005162 | -0.2915 | -0.217 |
| OR52D1 | NM_001005163 | -0.042 | -0.1425 |
| OR52E2 | NM_001005164 | 0.0845 | -0.074 |
| OR52E4 | NM_001005165 | 0.083 | 0.0355 |
| OR52E6 | NM_001005167 | -0.0765 | 0.228 |
| OR52E8 | NM_001005168 | -0.138 | 0.9025 |
| OR52I1 | NM_001005169 | -0.018 | -0.132 |
| OR52K1 | NM_001005171 | -0.0075 | 0.054 |
| OR52K2 | NM_001005172 | -0.036 | -0.2705 |
| OR52L1 | NM_001005173 | -0.19 | -0.269 |
| OR52N2 | NM_001005174 | -0.3245 | -0.057 |
| OR52N4 | NM_001005175 | -0.1345 | -0.079 |
| SP140 | NM_001005176 | 0.2105 | 0.0325 |
| OR52R1 | NM_001005177 | -0.138 | -0.4325 |
| OR52W1 | NM_001005178 | -0.0075 | 0.103 |
| OR56A4 | NM_001005179 | 0.12 | 0.0475 |
| OR56B1 | NM_001005180 | 0.281 | 0.1465 |
| OR56B4 | NM_001005181 | 0.02 | -0.3075 |
| OR6C1 | NM_001005182 | -0.0895 | 0.0245 |
| OR6C76 | NM_001005183 | -0.033 | -0.329 |
| OR6K6 | NM_001005184 | 0.122 | -0.381 |
| OR6N1 | NM_001005185 | 0.1845 | -0.2415 |
| OR6Q1 | NM_001005186 | -0.161 | -0.365 |
| OR6T1 | NM_001005187 | 0.08 | -0.427 |
| OR6X1 | NM_001005188 | -0.332 | -0.3295 |
| OR6Y1 | NM_001005189 | 0.1455 | 0.221 |
| OR7A10 | NM_001005190 | -0.267 | 0.0395 |
| OR7D4 | NM_001005191 | -0.099 | -0.5035 |
| OR7G1 | NM_001005192 | -0.114 | 0.2545 |
| OR7G2 | NM_001005193 | -0.142 | -0.0835 |
| OR8A1 | NM_001005194 | -0.2485 | -0.1285 |

| | | | |
|---|---|---|---|
| OR8B12 | NM_001005195 | -0.265 | -0.333 |
| OR8B4 | NM_001005196 | -0.1815 | -0.0545 |
| OR8D4 | NM_001005197 | -0.1945 | 0.0445 |
| OR8G5 | NM_001005198 | 0.6575 | -0.53 |
| OR8H1 | NM_001005199 | -0.08 | -0.05 |
| OR8H2 | NM_001005200 | 0.1025 | -0.406 |
| OR8H3 | NM_001005201 | 0.051 | -0.542 |
| OR8K3 | NM_001005202 | -0.15 | 0.2095 |
| OR8S1 | NM_001005203 | 0.029 | -0.16 |
| OR8J1 | NM_001005205 | 0.3395 | 0.121 |
| TRIM37 | NM_001005207 | -1.2675 | -1.0395 |
| TMEM198 | NM_001005209 | -0.587 | -1.7525 |
| LRRC55 | NM_001005210 | -0.3875 | -0.472 |
| OR9I1 | NM_001005211 | 0.1745 | -0.3765 |
| OR9Q1 | NM_001005212 | 0.1765 | 0.0435 |
| OR9G1 | NM_001005213 | 0.1 | -0.031 |
| LRRC52 | NM_001005214 | -0.264 | -0.4455 |
| OR2J3 | NM_001005216 | -0.5015 | 0.1035 |
| OR5B21 | NM_001005218 | 0.2235 | 0.07 |
| OR2B3 | NM_001005226 | -0.165 | 0.3445 |
| OR1L3 | NM_001005234 | 0.0885 | -0.392 |
| OR1L4 | NM_001005235 | 0.015 | 0.0355 |
| OR1L1 | NM_001005236 | -0.058 | -0.0875 |
| OR51G1 | NM_001005237 | -0.21 | -0.385 |
| OR51G2 | NM_001005238 | 0.269 | -0.1555 |
| OR4N4 | NM_001005241 | -0.025 | -0.1195 |
| PKP2 | NM_001005242 | -1.0725 | -0.488 |
| OR9K2 | NM_001005243 | -0.304 | -0.303 |
| OR5M11 | NM_001005245 | 0.055 | -0.223 |
| OR4C12 | NM_001005270 | -0.2125 | -0.195 |
| CHD3 | NM_001005271 | 0.1465 | -0.3305 |
| OR4A5 | NM_001005272 | -0.1775 | -0.2945 |
| CHD3 | NM_001005273 | 0.1465 | -0.3305 |
| OR4A16 | NM_001005274 | -0.1215 | -0.405 |
| OR4A15 | NM_001005275 | -0.2115 | -0.055 |
| OR2AE1 | NM_001005276 | -0.1235 | 0.125 |
| OR6N2 | NM_001005278 | 0.042 | -0.0585 |
| OR6K2 | NM_001005279 | -0.129 | 0.1005 |
| OR10A7 | NM_001005280 | -0.1165 | -0.482 |
| OR6B1 | NM_001005281 | 0 | 0.029 |
| OR5M8 | NM_001005282 | 0.148 | 0.128 |
| OR9Q2 | NM_001005283 | 0.048 | -0.238 |
| OR9G4 | NM_001005284 | 0.198 | -0.05 |

| | | | |
|---|---|---|---|
| OR2AT4 | NM_001005285 | -0.1795 | -0.016 |
| OR6F1 | NM_001005286 | -0.019 | -0.279 |
| OR51I1 | NM_001005288 | 0.0565 | -0.446 |
| OR52H1 | NM_001005289 | 0.3425 | 0.283 |
| SREBF1 | NM_001005291 | -0.514 | -1.0535 |
| OR5AK2 | NM_001005323 | -0.1005 | -0.687 |
| OR10V1 | NM_001005324 | 0.2645 | -0.21 |
| OR6M1 | NM_001005325 | -0.2315 | -0.2985 |
| OR4F6 | NM_001005326 | -0.1745 | -0.254 |
| OR6K3 | NM_001005327 | 0.2765 | -0.682 |
| MAGED1 | NM_001005332 | 0.138 | -0.3165 |
| MAGED1 | NM_001005333 | 0.0995 | -0.539 |
| HNRNPL | NM_001005335 | -0.005 | -0.5555 |
| DNM1 | NM_001005336 | -0.2845 | -0.342 |
| PKP1 | NM_001005337 | -0.137 | -0.3875 |
| OR5H1 | NM_001005338 | 0.1945 | -0.05 |
| GPNMB | NM_001005340 | -0.148 | -0.0895 |
| AK4 | NM_001005353 | -0.494 | -0.979 |
| PRR13 | NM_001005354 | -0.3535 | -1.1185 |
| POTEG | NM_001005356 | -0.008 | -0.311 |
| DNM2 | NM_001005360 | -0.215 | -1.3345 |
| DNM2 | NM_001005361 | -0.1715 | -0.9265 |
| DNM2 | NM_001005362 | -0.1715 | -0.9265 |
| POTEA | NM_001005365 | -0.077 | -0.203 |
| KDM2B | NM_001005366 | -0.952 | -2.0065 |
| TTYH1 | NM_001005367 | 0.0605 | -0.2985 |
| MTIF2 | NM_001005369 | -1.2525 | -2.2685 |
| LRSAM1 | NM_001005373 | -0.2465 | -0.847 |
| LRSAM1 | NM_001005374 | -0.2465 | -0.847 |
| PLAUR | NM_001005376 | -0.253 | -1.3375 |
| PLAUR | NM_001005377 | -0.437 | -1.4495 |
| ACTR2 | NM_001005386 | -0.667 | -0.681 |
| NFASC | NM_001005388 | 0.3565 | -0.278 |
| NFASC | NM_001005389 | 0.3565 | -0.278 |
| YPEL2 | NM_001005404 | -1.046 | -1.264 |
| KRTAP5-11 | NM_001005405 | -0.018 | -0.2845 |
| SF3A1 | NM_001005409 | 0.1675 | -1.004 |
| ZWINT | NM_001005413 | -0.5895 | -1.3875 |
| 2-Mar | NM_001005415 | -0.032 | -0.758 |
| B4GALT2 | NM_001005417 | -0.223 | -0.958 |
| EBF3 | NM_001005463 | 0.0755 | -0.588 |
| OR10G3 | NM_001005465 | 0.2575 | -0.3215 |
| OR10G2 | NM_001005466 | -0.3025 | -0.251 |

| | | | |
|---|---|---|---|
| OR8B3 | NM_001005467 | -0.1695 | -0.4075 |
| OR5B3 | NM_001005469 | 0.4405 | -0.5655 |
| OR4B1 | NM_001005470 | -0.191 | -0.185 |
| OR2T6 | NM_001005471 | -0.1105 | -0.027 |
| PLCXD3 | NM_001005473 | 0.187 | -0.3255 |
| NFKBIZ | NM_001005474 | -0.336 | 0.606 |
| PKP4 | NM_001005476 | -1.0155 | -2.478 |
| OR5H6 | NM_001005479 | 0.144 | -0.104 |
| OR2A2 | NM_001005480 | 0.0695 | -0.15 |
| OR5H2 | NM_001005482 | -0.095 | -0.4445 |
| OR4K5 | NM_001005483 | -0.0635 | -0.0365 |
| OR4K15 | NM_001005486 | -0.211 | 0.1325 |
| OR13G1 | NM_001005487 | 0.25 | -0.116 |
| OR5B17 | NM_001005489 | -0.1225 | -0.479 |
| OR6C74 | NM_001005490 | 0.092 | -0.604 |
| OR10AG1 | NM_001005491 | 0.138 | -0.0475 |
| OR5J2 | NM_001005492 | 0.2595 | -0.11 |
| OR6C6 | NM_001005493 | -0.033 | -0.4925 |
| OR6C4 | NM_001005494 | -0.678 | -0.1535 |
| OR5D16 | NM_001005496 | -0.0565 | -0.2285 |
| OR6C75 | NM_001005497 | -0.454 | -0.4605 |
| RHBDF2 | NM_001005498 | -0.4875 | -0.7025 |
| OR6C70 | NM_001005499 | -0.184 | -0.0695 |
| OR4K2 | NM_001005501 | 0.0225 | -0.084 |
| CPM | NM_001005502 | 0.222 | -0.293 |
| OR11G2 | NM_001005503 | 0.006 | -0.0705 |
| CACNA2D2 | NM_001005505 | -0.297 | -0.7505 |
| OR4A47 | NM_001005512 | -0.0905 | -0.203 |
| OR4C45 | NM_001005513 | 0.3175 | -0.2105 |
| OR5H14 | NM_001005514 | 0.254 | -0.1345 |
| OR5K3 | NM_001005516 | 0.035 | -0.1895 |
| OR5K4 | NM_001005517 | 0 | -0.1535 |
| OR6C65 | NM_001005518 | -0.284 | -0.281 |
| OR6C68 | NM_001005519 | 0.0705 | 0.017 |
| SF3B1 | NM_001005526 | -0.352 | -1.205 |
| OR5B2 | NM_001005566 | 0.0715 | -0.0625 |
| OR51B5 | NM_001005567 | 0.2095 | -0.6415 |
| EDA | NM_001005609 | -0.0235 | 0.383 |
| EDA | NM_001005610 | -0.0235 | 0.383 |
| EDA | NM_001005612 | -0.0235 | 0.383 |
| EDA | NM_001005613 | -0.0235 | 0.383 |
| ITGB4 | NM_001005619 | -0.194 | -0.5105 |
| ITGB4 | NM_001005731 | -0.194 | -0.5105 |

| | | | |
|---|---|---|---|
| CHEK2 | NM_001005735 | -1.0785 | -2.1535 |
| FPR2 | NM_001005738 | -0.0935 | -0.3865 |
| VPS54 | NM_001005739 | -1.312 | -3.535 |
| GBA | NM_001005741 | -0.6095 | -1.2035 |
| GBA | NM_001005742 | -0.6095 | -1.2035 |
| NUMB | NM_001005743 | -0.986 | -2.4315 |
| NUMB | NM_001005744 | -0.986 | -2.4315 |
| NUMB | NM_001005745 | -0.986 | -2.4315 |
| CACNB4 | NM_001005746 | -0.508 | -0.9685 |
| CACNB4 | NM_001005747 | -0.136 | -1.5415 |
| SUMO1 | NM_001005781 | -0.9765 | -0.777 |
| SUMO1 | NM_001005782 | -0.9765 | -0.777 |
| HAO2 | NM_001005783 | -0.293 | -0.407 |
| SUMO2 | NM_001005849 | -0.0805 | -0.3705 |
| ZNF835 | NM_001005850 | 0.0975 | -0.5315 |
| ZNF780B | NM_001005851 | -0.502 | -0.7155 |
| TXLNG2P | NM_001005852 | 0.3575 | -0.806 |
| OR6B2 | NM_001005853 | -0.166 | -0.1775 |
| ATP8B2 | NM_001005855 | -0.1625 | -0.925 |
| ERBB2 | NM_001005862 | -0.0025 | 0.099 |
| IP6K2 | NM_001005909 | -0.747 | -2.991 |
| IP6K2 | NM_001005910 | 0.385 | -0.656 |
| IP6K2 | NM_001005911 | 0.385 | -0.656 |
| SEMA3B | NM_001005914 | -0.146 | -0.431 |
| ERBB3 | NM_001005915 | 0.225 | 0.0825 |
| ATP7B | NM_001005918 | 0.299 | -0.5255 |
| JMJD8 | NM_001005920 | -0.2965 | -0.7735 |
| KRTAP5-1 | NM_001005922 | -0.13 | -0.2315 |
| RPS3A | NM_001006 | -0.247 | -0.1785 |
| C3orf37 | NM_001006109 | -0.6235 | -1.7985 |
| YBEY | NM_001006114 | -0.4835 | -0.5375 |
| IP6K1 | NM_001006115 | -1.048 | -2.4815 |
| ERBB2IP | NM_001006600 | -0.84 | -3.27 |
| FAM69A | NM_001006605 | -0.9505 | -3.5175 |
| LRRC37A2 | NM_001006607 | -0.534 | -2.0295 |
| SIAH1 | NM_001006610 | -1.064 | -2.284 |
| WBP5 | NM_001006612 | -1.3325 | -2.078 |
| WBP5 | NM_001006613 | -0.9305 | -1.434 |
| WBP5 | NM_001006614 | -1.3325 | -2.078 |
| TSPAN17 | NM_001006616 | -0.0675 | -0.8525 |
| MAPKAP1 | NM_001006617 | -0.387 | -1.259 |
| MAPKAP1 | NM_001006618 | -0.387 | -1.259 |
| MAPKAP1 | NM_001006619 | -0.387 | -1.259 |

| | | | |
|---|---|---|---|
| MAPKAP1 | NM_001006620 | -0.387 | -1.259 |
| MAPKAP1 | NM_001006621 | -0.387 | -1.259 |
| WDR33 | NM_001006622 | -1.522 | -3.302 |
| WDR33 | NM_001006623 | -1.522 | -3.302 |
| PDPN | NM_001006624 | 0.369 | -0.079 |
| PDPN | NM_001006625 | 0.369 | -0.079 |
| CHRM2 | NM_001006626 | 0.095 | -0.344 |
| CHRM2 | NM_001006627 | 0.095 | -0.344 |
| CHRM2 | NM_001006628 | 0.095 | -0.344 |
| CHRM2 | NM_001006629 | 0.095 | -0.344 |
| CHRM2 | NM_001006630 | 0.095 | -0.344 |
| CHRM2 | NM_001006631 | 0.095 | -0.344 |
| CHRM2 | NM_001006632 | 0.095 | -0.344 |
| ARHGAP17 | NM_001006634 | -0.505 | -1.715 |
| GTDC1 | NM_001006636 | 0.021 | -0.2115 |
| RAB37 | NM_001006638 | -0.155 | -0.2165 |
| TCEAL1 | NM_001006639 | -0.9445 | -1.035 |
| TCEAL1 | NM_001006640 | -0.9445 | -1.035 |
| SLC25A25 | NM_001006641 | -0.6215 | -2.116 |
| SLC25A25 | NM_001006642 | -0.6215 | -2.116 |
| SLC25A25 | NM_001006643 | -0.6215 | -2.116 |
| FAM149A | NM_001006655 | 0.214 | -0.4905 |
| ZNF473 | NM_001006656 | -1.374 | -2.3005 |
| WDR35 | NM_001006657 | -1.0685 | -2.47 |
| CR2 | NM_001006658 | 0.1755 | 0.042 |
| RPS6KA1 | NM_001006665 | -0.2155 | -1.1745 |
| APOBEC3F | NM_001006666 | -0.714 | -0.301 |
| RPS6KA2 | NM_001006932 | -0.6265 | -1.588 |
| TCEAL3 | NM_001006933 | -0.6985 | -1.2315 |
| TCEAL4 | NM_001006935 | -0.824 | -1.113 |
| TCEAL4 | NM_001006937 | -0.824 | -1.113 |
| TCEAL6 | NM_001006938 | -0.509 | -1.1865 |
| LRRC18 | NM_001006939 | 0.0225 | 0.046 |
| ALG3 | NM_001006941 | -0.2825 | -1.162 |
| EPHA8 | NM_001006943 | -0.591 | -0.5 |
| RPS6KA4 | NM_001006944 | -0.4875 | -1.7495 |
| LRIF1 | NM_001006945 | -1.772 | -4.346 |
| SDC1 | NM_001006946 | -0.717 | -1.498 |
| UHRF1BP1L | NM_001006947 | 0.346 | -0.087 |
| RPS4X | NM_001007 | -0.1225 | -0.2505 |
| ODF2L | NM_001007022 | -0.362 | -0.7635 |
| DIO2 | NM_001007023 | 0.0605 | -0.2345 |
| GOSR1 | NM_001007024 | -1.479 | -2.7135 |

| | | | |
|---|---|---|---|
| GOSR1 | NM_001007025 | -1.479 | -2.7135 |
| ATN1 | NM_001007026 | -0.5645 | -0.325 |
| ALG8 | NM_001007027 | -1.277 | -1.198 |
| CLEC6A | NM_001007033 | -0.4335 | 0.2975 |
| SDCBP | NM_001007067 | -0.3655 | -0.421 |
| SDCBP | NM_001007068 | -0.3655 | -0.421 |
| SDCBP | NM_001007069 | -0.3655 | -0.421 |
| SDCBP | NM_001007070 | -0.3655 | -0.421 |
| ZSCAN2 | NM_001007072 | 0.173 | -0.643 |
| RPL32 | NM_001007073 | 0.057 | -0.362 |
| RPL32 | NM_001007074 | 0.057 | -0.362 |
| KLHL5 | NM_001007075 | -0.8775 | -1.448 |
| ZNF182 | NM_001007088 | -0.49 | -1.236 |
| RESP18 | NM_001007089 | 0.051 | -0.1755 |
| C8orf48 | NM_001007090 | -0.928 | -1.7325 |
| ZNF37A | NM_001007094 | -0.73 | -1.6925 |
| NTRK2 | NM_001007097 | -0.042 | -0.1995 |
| SCP2 | NM_001007098 | -0.5505 | -0.4715 |
| SCP2 | NM_001007099 | -0.5505 | -0.4715 |
| SCP2 | NM_001007100 | -0.5505 | -0.4715 |
| ZNF484 | NM_001007101 | -0.17 | -0.7185 |
| L3MBTL3 | NM_001007102 | -0.453 | -1.039 |
| FSD2 | NM_001007122 | -0.1145 | -0.4095 |
| C20orf201 | NM_001007125 | -0.2015 | -0.3005 |
| IGF2 | NM_001007139 | -0.029 | -0.401 |
| NTRK3 | NM_001007156 | -0.2265 | -0.6735 |
| ZNF483 | NM_001007169 | 0.0615 | 0.1535 |
| C8orf22 | NM_001007176 | 0.04 | 0.0215 |
| DEPDC5 | NM_001007188 | -1.6815 | -3.1645 |
| IGIP | NM_001007189 | -0.245 | -1.49 |
| CACYBP | NM_001007214 | -0.6125 | -1.4635 |
| IGF2BP2 | NM_001007225 | 0.0105 | -2.1195 |
| SPOP | NM_001007226 | -0.491 | -2.0825 |
| SPOP | NM_001007227 | -0.491 | -2.0825 |
| SPOP | NM_001007228 | -0.491 | -2.0825 |
| SPOP | NM_001007229 | -0.491 | -2.0825 |
| SPOP | NM_001007230 | -0.491 | -2.0825 |
| ARHGAP25 | NM_001007231 | 0.145 | -0.1345 |
| CARD17 | NM_001007232 | -0.343 | -0.2365 |
| IGSF3 | NM_001007237 | -0.3675 | -1.1225 |
| METTL13 | NM_001007239 | -0.7815 | -3.855 |
| GP2 | NM_001007240 | 0.1395 | -0.1095 |
| GP2 | NM_001007241 | 0.1395 | -0.1095 |

| | | | |
|---|---|---|---|
| GP2 | NM_001007242 | 0.1395 | -0.1095 |
| IFRD1 | NM_001007245 | -1.735 | -3.0135 |
| BRWD1 | NM_001007246 | -0.9065 | -1.164 |
| ZNF599 | NM_001007248 | -0.844 | -1.607 |
| OR8G2 | NM_001007249 | -0.157 | -0.121 |
| SCP2 | NM_001007250 | -0.5505 | -0.4715 |
| ERV3-1 | NM_001007253 | 0.1305 | -0.085 |
| KLHDC9 | NM_001007255 | -0.163 | -0.1045 |
| PRDM2 | NM_001007257 | -0.841 | -0.9575 |
| SETD4 | NM_001007259 | 0.041 | -0.8355 |
| PLA2R1 | NM_001007267 | 0.0735 | 0.058 |
| GEMIN7 | NM_001007269 | -1.229 | -2.997 |
| GEMIN7 | NM_001007270 | -1.229 | -2.997 |
| DUSP13 | NM_001007271 | -0.0295 | -0.3005 |
| DUSP13 | NM_001007272 | -0.0005 | -0.2145 |
| DUSP13 | NM_001007273 | -0.0005 | -0.2145 |
| EI24 | NM_001007277 | 0.1685 | -0.3725 |
| TRIM13 | NM_001007278 | -1.2885 | -3.7325 |
| RASL10A | NM_001007279 | 0.234 | -0.3255 |
| TULP4 | NM_001007466 | -1.237 | -2.0945 |
| SFI1 | NM_001007467 | -0.214 | -0.554 |
| SMARCB1 | NM_001007468 | -0.373 | -1.1505 |
| TRPM3 | NM_001007470 | 0.08 | 0.028 |
| TRPM3 | NM_001007471 | 0.08 | 0.028 |
| NWD1 | NM_001007525 | 0.028 | -0.3455 |
| LMBRD2 | NM_001007527 | -0.5985 | -0.903 |
| NKAPL | NM_001007531 | 0.046 | -0.562 |
| STH | NM_001007532 | -0.0905 | -0.557 |
| C1QTNF9B | NM_001007537 | -0.1725 | -0.1735 |
| SHISA2 | NM_001007538 | -0.3555 | 0.071 |
| CDHR4 | NM_001007540 | -0.1785 | -0.406 |
| C1orf186 | NM_001007544 | -0.0885 | -0.1805 |
| CSDE1 | NM_001007553 | -0.4585 | -0.404 |
| SS18 | NM_001007559 | -1.7305 | -2.509 |
| IRGQ | NM_001007561 | -0.908 | -1.5515 |
| IGFBPL1 | NM_001007563 | 0.0455 | -0.269 |
| C2CD4B | NM_001007595 | -0.9465 | -0.223 |
| NTRK1 | NM_001007792 | -0.088 | -0.389 |
| BUB3 | NM_001007793 | -0.662 | -0.772 |
| CEPT1 | NM_001007794 | -1.63 | -3.414 |
| RPS4Y1 | NM_001008 | -0.689 | -0.293 |
| OPTN | NM_001008211 | -0.034 | -0.4645 |
| OPTN | NM_001008212 | -0.034 | -0.4645 |

| | | | |
|---|---|---|---|
| OPTN | NM_001008213 | -0.034 | -0.4645 |
| COA5 | NM_001008215 | -1.391 | -1.85 |
| GALE | NM_001008216 | -0.1105 | -1.0535 |
| CPLX2 | NM_001008220 | -0.204 | -0.964 |
| ZDHHC9 | NM_001008222 | -0.174 | -1.033 |
| C1QL4 | NM_001008223 | -0.406 | -0.9445 |
| UACA | NM_001008224 | -0.0525 | -0.8955 |
| CNOT4 | NM_001008225 | 0.61 | -0.9195 |
| FAM154B | NM_001008226 | -0.0835 | -0.176 |
| MOG | NM_001008228 | -0.501 | 0.329 |
| MOG | NM_001008229 | -0.501 | 0.329 |
| TTC32 | NM_001008237 | -1.1815 | -1.6625 |
| C18orf25 | NM_001008239 | -1.4265 | -2.5865 |
| TMEM89 | NM_001008269 | 0.0855 | -0.431 |
| PRSS37 | NM_001008270 | -0.193 | -0.1805 |
| SCXA | NM_001008271 | -0.241 | -0.5685 |
| TRIM72 | NM_001008274 | -0.0855 | -0.116 |
| REG3G | NM_001008387 | 0.1515 | -0.042 |
| CISD2 | NM_001008388 | -0.9265 | -1.681 |
| CGGBP1 | NM_001008390 | -1.2975 | -1.9045 |
| CCDC73 | NM_001008391 | -0.2395 | -0.7615 |
| CTDSPL | NM_001008392 | -0.992 | -2.586 |
| C4orf46 | NM_001008393 | -1.5275 | -2.212 |
| EID3 | NM_001008394 | 1.4985 | 1.756 |
| C7orf59 | NM_001008395 | 0.1075 | -0.6595 |
| WDHD1 | NM_001008396 | -0.756 | -2.1825 |
| GPX8 | NM_001008397 | -0.6175 | -1.971 |
| ZNF761 | NM_001008401 | -0.8555 | -1.6785 |
| C14orf180 | NM_001008404 | -0.1855 | -0.484 |
| BCAP29 | NM_001008405 | -1.4795 | -1.146 |
| TTLL9 | NM_001008409 | 0.0765 | -0.1245 |
| STEAP3 | NM_001008410 | -0.179 | -0.492 |
| SLC41A3 | NM_001008485 | 0.056 | -0.745 |
| SLC41A3 | NM_001008486 | 0.056 | -0.745 |
| SLC41A3 | NM_001008487 | 0.056 | -0.745 |
| PHOSPHO2 | NM_001008489 | -1.7395 | -2.025 |
| 2-Sep | NM_001008491 | -0.811 | -0.436 |
| ENAH | NM_001008493 | -0.8325 | -0.2935 |
| ISX | NM_001008494 | 0.1625 | -0.043 |
| TMEM64 | NM_001008495 | -1.0095 | -1.795 |
| PIWIL3 | NM_001008496 | 0.006 | -0.333 |
| OPRM1 | NM_001008503 | -0.147 | -0.562 |
| OPRM1 | NM_001008504 | -0.147 | -0.562 |

| | | | |
|---|---|---|---|
| OPRM1 | NM_001008505 | -0.147 | -0.562 |
| MXRA7 | NM_001008528 | 0.173 | -0.265 |
| MXRA7 | NM_001008529 | 0.173 | -0.265 |
| LGMN | NM_001008530 | 0.082 | -1.197 |
| AKAP14 | NM_001008535 | 0.742 | -0.3805 |
| TCHHL1 | NM_001008536 | -0.0895 | -0.324 |
| KIAA2022 | NM_001008537 | -0.1005 | -1.609 |
| SLC7A2 | NM_001008539 | -1.211 | -2.2195 |
| CXCR4 | NM_001008540 | 0.2665 | -0.1165 |
| MXI1 | NM_001008541 | -0.881 | -1.754 |
| NUPL1 | NM_001008564 | -1.767 | -2.8835 |
| TPST2 | NM_001008566 | -0.7835 | -3.0595 |
| TCOF1 | NM_001008657 | -0.2495 | -0.907 |
| PICALM | NM_001008660 | -0.851 | -1.871 |
| CCBL2 | NM_001008662 | -0.7525 | -1.1565 |
| CST9 | NM_001008693 | 0.195 | -0.181 |
| THAP7 | NM_001008695 | -1.2625 | -2.6985 |
| TFIP11 | NM_001008697 | -0.8425 | -2.262 |
| IL4R | NM_001008699 | -0.343 | -1.372 |
| LPHN1 | NM_001008701 | 0.0045 | -1.145 |
| C6orf1 | NM_001008703 | -0.592 | -1.6475 |
| C6orf1 | NM_001008704 | -0.592 | -1.6475 |
| EML1 | NM_001008707 | -0.534 | -1.474 |
| CHAC2 | NM_001008708 | -2.0915 | -3.342 |
| PPP1CA | NM_001008709 | 0.144 | -0.4505 |
| RBPMS | NM_001008710 | -0.1975 | -1.048 |
| RBPMS | NM_001008711 | -0.1975 | -1.048 |
| RBPMS | NM_001008712 | -0.2585 | -1.1885 |
| CCDC147 | NM_001008723 | 0.4125 | -0.251 |
| WDR89 | NM_001008726 | -0.7005 | -1.7335 |
| ZNF121 | NM_001008727 | -0.4385 | -1.6395 |
| LOC401052 | NM_001008737 | 0.027 | -0.3805 |
| FNIP1 | NM_001008738 | 0.5005 | 1.0215 |
| C6orf226 | NM_001008739 | 0.2075 | 0.0865 |
| TMCO2 | NM_001008740 | 0.1145 | -0.6345 |
| SULT1C3 | NM_001008743 | 0.373 | -0.2335 |
| TDP1 | NM_001008744 | -1.678 | -3.915 |
| RAB19 | NM_001008749 | 0.1825 | -0.3315 |
| FBXO47 | NM_001008777 | 0.4015 | 0.009 |
| SPDYC | NM_001008778 | 0.1135 | -0.518 |
| SPDYA | NM_001008779 | 0.0155 | 0.325 |
| FAT3 | NM_001008781 | -0.95 | -1.7485 |
| SLC35D3 | NM_001008783 | 0.0115 | -0.095 |

| | | | |
|---|---|---|---|
| CD200R1L | NM_001008784 | -0.1725 | -0.327 |
| CCT3 | NM_001008800 | -0.166 | -0.5775 |
| ZNF468 | NM_001008801 | -0.6015 | -1.45 |
| DSP | NM_001008844 | -0.057 | 0.0875 |
| NIPA2 | NM_001008860 | -1.056 | -2.0195 |
| NIPA2 | NM_001008892 | -1.056 | -2.0195 |
| NIPA2 | NM_001008894 | -1.056 | -2.0195 |
| CUL4A | NM_001008895 | -1.5185 | -1.7745 |
| TCP1 | NM_001008897 | -0.597 | -0.7485 |
| GTSF1L | NM_001008901 | -0.1805 | 0.5135 |
| STK16 | NM_001008910 | -0.356 | -1.018 |
| RCHY1 | NM_001008925 | -0.779 | -2.1975 |
| CKAP5 | NM_001008938 | -0.548 | -1.541 |
| RPS5 | NM_001009 | 0.1365 | -0.257 |
| GRINA | NM_001009184 | -0.206 | -0.574 |
| ACSL6 | NM_001009185 | -0.249 | -0.309 |
| CCT6A | NM_001009186 | -1.4135 | -0.945 |
| PPP2CB | NM_001009552 | -0.7745 | -1.786 |
| MFAP3L | NM_001009554 | -0.2535 | -0.274 |
| SH3D19 | NM_001009555 | -1.105 | -1.9275 |
| SLCO1B7 | NM_001009562 | -0.103 | -0.263 |
| CDKL4 | NM_001009565 | -0.236 | -0.353 |
| CLSTN1 | NM_001009566 | -0.0065 | -0.791 |
| SMPDL3B | NM_001009568 | 0.354 | -0.428 |
| CCT7 | NM_001009570 | 0.0705 | -0.455 |
| CADPS2 | NM_001009571 | 0.137 | -0.251 |
| ARMCX6 | NM_001009584 | -0.5045 | -1.0455 |
| HS3ST6 | NM_001009606 | 0.1145 | 0.011 |
| C20orf94 | NM_001009608 | -0.616 | -0.816 |
| SPANXN3 | NM_001009609 | 0.4405 | -0.4425 |
| FAM22D | NM_001009610 | -0.2925 | -0.9755 |
| C20orf202 | NM_001009612 | 0.0315 | -0.694 |
| SPANXN4 | NM_001009613 | 0.1315 | -0.0055 |
| SPANXN1 | NM_001009614 | 0.166 | 0.0325 |
| SPANXN2 | NM_001009615 | 0.0285 | -0.042 |
| SPANXN5 | NM_001009616 | 0.078 | -0.056 |
| AZI1 | NM_001009811 | -0.2505 | -1.0285 |
| LBX2 | NM_001009812 | -0.2135 | -1.073 |
| MEIS3 | NM_001009813 | 0.037 | -0.27 |
| KIAA0564 | NM_001009814 | -0.501 | -1.349 |
| BRD9 | NM_001009877 | -0.3105 | -1.066 |
| ZCCHC11 | NM_001009881 | -1.3105 | -2.9285 |
| C12orf29 | NM_001009894 | -1.343 | -2.702 |

| | | | |
|---|---|---|---|
| KIAA2018 | NM_001009899 | -0.566 | -2.5465 |
| B3GNTL1 | NM_001009905 | -0.4665 | -0.8095 |
| LUZP2 | NM_001009909 | -0.2375 | -0.123 |
| VPS8 | NM_001009921 | -0.574 | -0.9475 |
| RCHY1 | NM_001009922 | -0.779 | -2.1975 |
| C20orf30 | NM_001009925 | -0.278 | -0.312 |
| HRNR | NM_001009931 | -0.339 | -0.2355 |
| DNASE1L1 | NM_001009932 | -0.4735 | -1.1035 |
| DNASE1L1 | NM_001009933 | -0.4735 | -1.1035 |
| DNASE1L1 | NM_001009934 | -0.4735 | -1.1035 |
| PHF19 | NM_001009936 | 0.068 | -0.7825 |
| ANKRD16 | NM_001009941 | -0.906 | -1.9465 |
| ANKRD16 | NM_001009943 | -0.5055 | -1.1495 |
| PKD1 | NM_001009944 | -0.695 | -1.0915 |
| SSBP3 | NM_001009955 | -0.017 | -0.824 |
| ZNF655 | NM_001009958 | -0.9175 | -1.7805 |
| ERMN | NM_001009959 | -0.109 | -0.1205 |
| ZNF655 | NM_001009960 | -0.9175 | -1.7805 |
| C20orf194 | NM_001009984 | -0.299 | -1.4005 |
| SYTL3 | NM_001009991 | 0.007 | -1.0465 |
| ZNF648 | NM_001009992 | -0.102 | 0.081 |
| FAM168B | NM_001009993 | -0.658 | -0.901 |
| RIPPLY2 | NM_001009994 | -0.306 | -0.478 |
| DALRD3 | NM_001009996 | 0.1585 | -0.5235 |
| C10orf62 | NM_001009997 | -0.0495 | -0.1925 |
| SSBP4 | NM_001009998 | 0.106 | -0.328 |
| RPS6 | NM_001010 | -0.2005 | -0.222 |
| ARHGAP28 | NM_001010000 | -0.8285 | -2.7505 |
| IRAK1BP1 | NM_001010844 | -1.5165 | -2.219 |
| ACSM2A | NM_001010845 | -0.193 | -0.148 |
| SHE | NM_001010846 | -0.0815 | 0.1435 |
| NRG3 | NM_001010848 | -0.1615 | -0.614 |
| ZNF766 | NM_001010851 | -1.6405 | -3.763 |
| CLVS2 | NM_001010852 | -0.1835 | -2.3745 |
| PM20D2 | NM_001010853 | -1.064 | -1.465 |
| TTC7B | NM_001010854 | -0.4695 | -1.7895 |
| PIK3R6 | NM_001010855 | -0.001 | -0.028 |
| LELP1 | NM_001010857 | 0.005 | -0.087 |
| RNF187 | NM_001010858 | -0.2965 | -0.816 |
| C22orf42 | NM_001010859 | 0.2255 | 0.1385 |
| SAMD15 | NM_001010860 | 0.037 | -0.1205 |
| LPPR5 | NM_001010861 | -1.132 | -1.9615 |
| SPIN3 | NM_001010862 | -0.8325 | -0.9125 |

| | | | |
|---|---|---|---|
| C10orf128 | NM_001010863 | -0.36 | -0.61 |
| EME2 | NM_001010865 | -0.528 | -0.584 |
| TMEM201 | NM_001010866 | 0.0115 | -0.51 |
| IBA57 | NM_001010867 | -0.2455 | -1.151 |
| TDRD6 | NM_001010870 | -0.054 | -0.792 |
| FAM83B | NM_001010872 | -0.114 | 0.0265 |
| TSPO2 | NM_001010873 | 0.0115 | -0.551 |
| TECRL | NM_001010874 | 0.1005 | -0.148 |
| SLC25A30 | NM_001010875 | -1.0055 | -1.896 |
| C6orf191 | NM_001010876 | -0.0785 | -0.481 |
| ZNF311 | NM_001010877 | -0.102 | -0.145 |
| C16orf91 | NM_001010878 | -0.6965 | -1.25 |
| ZIK1 | NM_001010879 | -1.2515 | -2.5195 |
| ZNF780A | NM_001010880 | -0.344 | -0.0595 |
| FAM102B | NM_001010883 | -1.2325 | -1.866 |
| C6orf127 | NM_001010886 | 0.1245 | -0.6395 |
| ACER2 | NM_001010887 | -0.391 | -0.063 |
| ZC3H12B | NM_001010888 | -0.1405 | 0.3595 |
| MTX3 | NM_001010891 | -0.937 | -1.653 |
| RSPH4A | NM_001010892 | -0.3035 | -0.5905 |
| SLC10A5 | NM_001010893 | -0.2475 | 0.3255 |
| C9orf102 | NM_001010895 | -0.411 | -1.8795 |
| C10orf113 | NM_001010896 | -0.2955 | -0.594 |
| SERP2 | NM_001010897 | -0.393 | -0.9725 |
| SLC6A17 | NM_001010898 | -0.564 | -1.056 |
| C6orf222 | NM_001010903 | 0.1055 | -0.401 |
| GLYATL3 | NM_001010904 | -0.143 | -0.4905 |
| C6orf58 | NM_001010905 | -0.2025 | -0.1595 |
| C8orf80 | NM_001010906 | -0.881 | -0.2205 |
| C9orf153 | NM_001010907 | 0.369 | -0.077 |
| C1QL3 | NM_001010908 | -0.7005 | -1.4005 |
| MUC21 | NM_001010909 | -0.192 | -0.4025 |
| C10orf114 | NM_001010911 | -0.8415 | -1.6325 |
| C10orf120 | NM_001010912 | -0.356 | -0.325 |
| PTPLAD2 | NM_001010915 | -0.39 | -0.336 |
| GOLGA7B | NM_001010917 | -0.1295 | -0.853 |
| FAM26F | NM_001010919 | 0.182 | -0.517 |
| BCL2L15 | NM_001010922 | -0.2755 | -0.5515 |
| THEMIS | NM_001010923 | 0.157 | -0.1605 |
| FAM171A1 | NM_001010924 | -0.4015 | -1.4695 |
| HES5 | NM_001010926 | 0.212 | -0.0445 |
| TIAM2 | NM_001010927 | -0.8985 | -2.3525 |
| HGF | NM_001010931 | 0.329 | 0.368 |

| | | | |
|---|---|---|---|
| HGF | NM_001010932 | 0.329 | 0.368 |
| HGF | NM_001010933 | 0.329 | 0.368 |
| HGF | NM_001010934 | 0.329 | 0.368 |
| RAP1A | NM_001010935 | -1.231 | -2.913 |
| TNK2 | NM_001010938 | -0.0645 | -0.7045 |
| LIPJ | NM_001010939 | 0.2645 | -0.0285 |
| C9orf135 | NM_001010940 | -0.191 | -0.1395 |
| RAP1B | NM_001010942 | -0.98 | -1.353 |
| CYP4A22 | NM_001010969 | 0.1155 | -0.2825 |
| SAMD13 | NM_001010971 | -0.367 | -1.358 |
| BCAS4 | NM_001010974 | -0.137 | -0.6095 |
| METTL21C | NM_001010977 | 0.336 | 0.243 |
| LDLRAD1 | NM_001010978 | 0.097 | -0.1835 |
| C1orf189 | NM_001010979 | 0.187 | -0.3435 |
| C1orf130 | NM_001010980 | 0.226 | 0.0505 |
| AFMID | NM_001010982 | -0.3455 | -1.381 |
| GLT8D1 | NM_001010983 | -1.635 | -2.303 |
| C1orf124 | NM_001010984 | -0.6125 | -1.391 |
| MYBPHL | NM_001010985 | -0.083 | 0.017 |
| IFIT1B | NM_001010987 | -0.251 | 0.056 |
| HERPUD1 | NM_001010989 | -1.1025 | -2.921 |
| HERPUD1 | NM_001010990 | -1.1025 | -2.921 |
| RPS7 | NM_001011 | -0.1 | -0.3165 |
| PDLIM5 | NM_001011513 | 1.1995 | 2.0065 |
| PDLIM5 | NM_001011515 | 1.1995 | 2.0065 |
| PDLIM5 | NM_001011516 | 1.1995 | 2.0065 |
| FYTTD1 | NM_001011537 | -0.7315 | -1.058 |
| MAGEA11 | NM_001011544 | -0.0115 | 0.2275 |
| DSTN | NM_001011546 | -0.27 | -0.4695 |
| SLC5A9 | NM_001011547 | -0.021 | -0.094 |
| MAGEA4 | NM_001011548 | -0.2955 | -0.294 |
| MAGEA4 | NM_001011549 | -0.2955 | -0.294 |
| MAGEA4 | NM_001011550 | -0.2955 | -0.294 |
| C1GALT1C1 | NM_001011551 | -1.4495 | -2.3095 |
| SLC13A3 | NM_001011554 | 0.38 | -0.2435 |
| AR | NM_001011645 | 0.607 | 0.2115 |
| CDK5RAP2 | NM_001011649 | -0.822 | -1.2085 |
| TMEM44 | NM_001011655 | -1.114 | -1.2125 |
| ZMAT1 | NM_001011657 | 0.1925 | -0.226 |
| TRAPPC2 | NM_001011658 | -0.6405 | -1.1545 |
| PCGF6 | NM_001011663 | -1.4975 | -3.0625 |
| CREB5 | NM_001011666 | -0.219 | -1.026 |
| CHCHD7 | NM_001011667 | -0.5065 | -0.8085 |

| | | | |
|---|---|---|---|
| CHCHD7 | NM_001011668 | -0.5065 | -0.8085 |
| CHCHD7 | NM_001011669 | -0.5065 | -0.8085 |
| CHCHD7 | NM_001011670 | -0.5065 | -0.8085 |
| CHCHD7 | NM_001011671 | -0.5065 | -0.8085 |
| MCCD1 | NM_001011700 | -0.2855 | -0.3685 |
| FAM125B | NM_001011703 | -0.775 | -2.4845 |
| LECT1 | NM_001011705 | 0.3485 | -0.4305 |
| OLA1 | NM_001011708 | -0.701 | -0.7115 |
| PNLIPRP3 | NM_001011709 | 0.121 | -0.205 |
| NAA30 | NM_001011713 | -1.248 | -2.889 |
| NMS | NM_001011717 | 0.2025 | -0.277 |
| XKR7 | NM_001011718 | -0.3585 | -0.3585 |
| ARSH | NM_001011719 | 0.2305 | -0.4095 |
| XKR9 | NM_001011720 | -0.388 | -0.57 |
| ARHGEF10L | NM_001011722 | -0.5445 | -1.896 |
| HNRNPA1L2 | NM_001011724 | 0.23 | -0.441 |
| HNRNPA1L2 | NM_001011725 | 0.23 | -0.441 |
| DEFB121 | NM_001011878 | -0.15 | -0.2465 |
| CLEC18B | NM_001011880 | -0.1175 | -0.335 |
| BTBD1 | NM_001011885 | -0.8315 | -1.1125 |
| RPS8 | NM_001012 | 0.022 | -0.3055 |
| MSL1 | NM_001012241 | -1.2945 | -1.457 |
| RNASE13 | NM_001012264 | -0.0135 | -0.1515 |
| CENPP | NM_001012267 | -0.399 | -1.821 |
| BIRC5 | NM_001012270 | -0.116 | -0.84 |
| BIRC5 | NM_001012271 | -0.116 | -0.84 |
| C6orf174 | NM_001012279 | -1.317 | -1.87 |
| CRLF2 | NM_001012288 | 0.139 | -0.4215 |
| MCRS1 | NM_001012300 | -0.2075 | -1.287 |
| ARSI | NM_001012301 | 0.3785 | -0.4625 |
| ANO9 | NM_001012302 | -0.2845 | -0.216 |
| ZNF302 | NM_001012320 | -1.4865 | -2.7615 |
| CTNNBIP1 | NM_001012329 | 0.0815 | -0.328 |
| MDK | NM_001012333 | 0.314 | -0.1 |
| MDK | NM_001012334 | 0.314 | -0.1 |
| ROPN1B | NM_001012337 | 0.1675 | -0.62 |
| NTRK3 | NM_001012338 | -0.273 | -0.8245 |
| DNAJC21 | NM_001012339 | -1.107 | -2.3615 |
| WDR31 | NM_001012361 | -0.167 | -0.2095 |
| OPCML | NM_001012393 | -0.0225 | -0.0685 |
| AKTIP | NM_001012398 | -1.0405 | -1.765 |
| SGOL1 | NM_001012409 | -0.9065 | -2.11 |
| SGOL1 | NM_001012410 | -0.38 | -0.871 |

| | | | |
|---|---|---|---|
| SGOL1 | NM_001012411 | -0.38 | -0.871 |
| SGOL1 | NM_001012412 | -0.38 | -0.871 |
| SGOL1 | NM_001012413 | -0.38 | -0.871 |
| TRIM61 | NM_001012414 | -0.091 | -0.165 |
| SOHLH1 | NM_001012415 | 0.0805 | -0.0405 |
| KRTAP5-6 | NM_001012416 | 0.054 | -0.3375 |
| MYLK4 | NM_001012418 | -0.1175 | -0.6705 |
| C1orf146 | NM_001012425 | 0.3525 | 0.294 |
| FOXP4 | NM_001012426 | -0.104 | -0.826 |
| FOXP4 | NM_001012427 | -0.104 | -0.826 |
| ASB11 | NM_001012428 | -0.091 | 0.024 |
| C9orf128 | NM_001012446 | -0.1535 | -0.197 |
| FAM71F2 | NM_001012454 | -0.3065 | -0.157 |
| ZSCAN23 | NM_001012455 | -0.5495 | -0.527 |
| U2AF2 | NM_001012478 | 0.216 | 0.0175 |
| C9orf117 | NM_001012502 | -0.0045 | -0.179 |
| KRTAP5-7 | NM_001012503 | -0.1745 | -0.2285 |
| FOXP1 | NM_001012505 | -1.1875 | -1.91 |
| CENPW | NM_001012507 | -0.473 | -1.154 |
| SPRN | NM_001012508 | 0.3325 | -0.2555 |
| SLC45A2 | NM_001012509 | -0.5245 | -0.3865 |
| GOSR2 | NM_001012511 | -0.48 | -1.608 |
| GRP | NM_001012512 | -0.3465 | -0.1805 |
| GRP | NM_001012513 | -0.3465 | -0.1805 |
| ITM2C | NM_001012514 | -0.206 | -0.9595 |
| FECH | NM_001012515 | -0.3075 | -1.092 |
| ITM2C | NM_001012516 | -0.206 | -0.9595 |
| CTBP1 | NM_001012614 | -0.4425 | -0.764 |
| IL32 | NM_001012631 | 0.0195 | -0.0945 |
| IL32 | NM_001012632 | 0.0195 | -0.0945 |
| IL32 | NM_001012633 | 0.0195 | -0.0945 |
| IL32 | NM_001012634 | 0.0195 | -0.0945 |
| IL32 | NM_001012635 | 0.0195 | -0.0945 |
| IL32 | NM_001012636 | 0.0195 | -0.0945 |
| GRAMD2 | NM_001012642 | 0.0395 | 0.0205 |
| MYPOP | NM_001012643 | -0.218 | -1.021 |
| ARGFX | NM_001012659 | 0.1565 | 0.529 |
| SLC3A2 | NM_001012662 | -0.5325 | -1.3455 |
| SLC3A2 | NM_001012664 | -0.5325 | -1.3455 |
| C14orf182 | NM_001012706 | 0.08 | 0.0785 |
| KRTAP5-3 | NM_001012708 | 0.058 | 0.163 |
| KRTAP5-4 | NM_001012709 | -0.1055 | -0.4435 |
| KRTAP5-10 | NM_001012710 | 0.313 | -0.1485 |

| | | | |
|---|---|---|---|
| C9orf106 | NM_001012715 | 0.2015 | 0.051 |
| C18orf56 | NM_001012716 | 0.1195 | -0.2925 |
| IL32 | NM_001012718 | 0.0195 | -0.0945 |
| RGR | NM_001012720 | -0.239 | -0.106 |
| RGR | NM_001012722 | 0.15 | -0.2545 |
| AGPAT2 | NM_001012727 | -0.75 | -1.357 |
| DPRX | NM_001012728 | 0.0885 | 0.054 |
| DUXA | NM_001012729 | 0.1145 | -0.5395 |
| DCTD | NM_001012732 | 0.0235 | -0.55 |
| ABI1 | NM_001012750 | -0.7035 | -1.5425 |
| ABI1 | NM_001012751 | -0.7035 | -1.5425 |
| ABI1 | NM_001012752 | -0.7035 | -1.5425 |
| ZNF763 | NM_001012753 | -0.567 | -1.085 |
| NHLRC3 | NM_001012754 | -1.942 | -1.754 |
| MCART6 | NM_001012755 | -0.2175 | -0.643 |
| ZNF260 | NM_001012756 | -1.0635 | -2.8335 |
| NUDT17 | NM_001012758 | 0.112 | -0.588 |
| CTU2 | NM_001012759 | 0.2275 | -0.829 |
| RGMB | NM_001012761 | -0.759 | -1.4405 |
| CTU2 | NM_001012762 | 0.2275 | -0.829 |
| GNRHR | NM_001012763 | -0.2185 | -0.093 |
| DISC1 | NM_001012957 | -0.2995 | -0.02 |
| DISC1 | NM_001012958 | -0.2995 | -0.02 |
| DISC1 | NM_001012959 | -0.2995 | -0.02 |
| KLK6 | NM_001012964 | -0.04 | -0.6725 |
| KLK6 | NM_001012965 | -0.04 | -0.6725 |
| DDX60L | NM_001012967 | 0.0575 | -0.4475 |
| SPIN4 | NM_001012968 | -1.4235 | -1.8135 |
| ADAL | NM_001012969 | -0.487 | -0.6275 |
| C1orf100 | NM_001012970 | -0.498 | -0.321 |
| C20orf106 | NM_001012971 | -0.0665 | -0.1805 |
| PLAC9 | NM_001012973 | 0.4335 | -0.456 |
| C6orf154 | NM_001012974 | 0.182 | -0.184 |
| RNASE10 | NM_001012975 | -0.051 | -0.8695 |
| PABPC1L2A | NM_001012977 | -0.0985 | -0.2085 |
| BEX5 | NM_001012978 | -0.088 | -0.0135 |
| TCEAL5 | NM_001012979 | -1.2615 | -1.8185 |
| SATL1 | NM_001012980 | 0.047 | -0.3915 |
| ZKSCAN2 | NM_001012981 | -1.4455 | -3.32 |
| C16orf86 | NM_001012984 | -0.103 | -0.5 |
| C1orf31 | NM_001012985 | -1.0805 | -3.1495 |
| UBE2NL | NM_001012989 | -0.1725 | -0.153 |
| ARL13A | NM_001012990 | 0.2345 | -0.4215 |

| | | | |
|---|---|---|---|
| C16orf88 | NM_001012991 | -0.5475 | -1.76 |
| C9orf152 | NM_001012993 | 0.2715 | -0.332 |
| SNX30 | NM_001012994 | -0.736 | -1.318 |
| RPS9 | NM_001013 | 0.344 | -0.071 |
| SORCS1 | NM_001013031 | -0.958 | -0.506 |
| SLC3A2 | NM_001013251 | -0.5325 | -1.3455 |
| LSP1 | NM_001013253 | 0.2225 | -0.3575 |
| LSP1 | NM_001013254 | 0.2225 | -0.3575 |
| LSP1 | NM_001013255 | 0.2225 | -0.3575 |
| BCAM | NM_001013257 | 0.147 | -0.4 |
| ZNF789 | NM_001013258 | -0.4935 | -0.631 |
| OR2G6 | NM_001013355 | -0.0235 | -0.359 |
| OR8U8 | NM_001013356 | -0.1485 | -0.2005 |
| IGFBP3 | NM_001013398 | -0.1475 | -0.8595 |
| CCNC | NM_001013399 | -1.5155 | -3.291 |
| CXorf66 | NM_001013403 | -0.1825 | 0.048 |
| KRIT1 | NM_001013406 | -1.236 | -1.9075 |
| FBXW7 | NM_001013415 | -0.993 | -1.8715 |
| PAGE5 | NM_001013435 | -0.265 | 0.3075 |
| MPST | NM_001013436 | -0.618 | -1.257 |
| SEH1L | NM_001013437 | -1.047 | -1.7625 |
| FXR1 | NM_001013438 | -0.8035 | -1.643 |
| FXR1 | NM_001013439 | -0.8035 | -1.643 |
| EPGN | NM_001013442 | 0.123 | -0.202 |
| AWAT1 | NM_001013579 | 0.1185 | -0.2945 |
| C1orf190 | NM_001013615 | -0.174 | -0.414 |
| AGPHD1 | NM_001013619 | -0.066 | -0.036 |
| ALG10B | NM_001013620 | -1.3235 | -2.2185 |
| FAM53A | NM_001013622 | -0.0575 | -0.1585 |
| C6orf94 | NM_001013623 | -0.0325 | -0.2765 |
| C1orf192 | NM_001013625 | 0.5885 | -0.1405 |
| PPP1R42 | NM_001013626 | 0.148 | 0.5525 |
| NHSL2 | NM_001013627 | -0.0445 | -0.18 |
| DCAF12L2 | NM_001013628 | 0.1125 | -0.9515 |
| AADACL4 | NM_001013630 | -0.097 | -0.3765 |
| HNRNPCL1 | NM_001013631 | -0.3115 | -0.039 |
| TCTEX1D4 | NM_001013632 | 0.083 | 0.14 |
| C12orf68 | NM_001013635 | -0.344 | -0.6625 |
| PRR25 | NM_001013638 | 0.158 | -0.1585 |
| TMEM82 | NM_001013641 | 0.197 | -0.334 |
| TRNP1 | NM_001013642 | -0.2745 | -0.357 |
| LOC646851 | NM_001013647 | -0.433 | -0.1805 |
| C2orf68 | NM_001013649 | -0.499 | -1.0955 |

| | | | |
|---|---|---|---|
| PRR23B | NM_001013650 | -0.0725 | -0.1305 |
| LRRC26 | NM_001013653 | 0.008 | -0.4385 |
| C15orf58 | NM_001013657 | -0.258 | -0.684 |
| PTX4 | NM_001013658 | -0.104 | -0.3435 |
| ZNF793 | NM_001013659 | -0.335 | -1.616 |
| FRRS1 | NM_001013660 | -0.1635 | -0.1225 |
| VSIG8 | NM_001013661 | -0.1925 | -0.679 |
| PTRHD1 | NM_001013663 | 0.208 | -0.352 |
| C17orf97 | NM_001013672 | 0.098 | -0.8565 |
| C1orf141 | NM_001013674 | -0.1885 | 0.032 |
| FIGNL2 | NM_001013690 | -0.6935 | -0.937 |
| LDLRAD2 | NM_001013693 | -0.181 | -0.3655 |
| SRRD | NM_001013694 | -1.2385 | -3.0575 |
| C12orf69 | NM_001013698 | 0.0625 | 0.181 |
| H3F3C | NM_001013699 | 0.241 | -0.595 |
| EIF2AK4 | NM_001013703 | -1.782 | -1.78 |
| PLIN5 | NM_001013706 | -0.0545 | -0.0205 |
| C5orf56 | NM_001013717 | -0.0585 | -0.602 |
| C6orf138 | NM_001013732 | -0.345 | -0.487 |
| RFPL4B | NM_001013734 | 0.0425 | 0.1145 |
| FOXB2 | NM_001013735 | 0.091 | -0.203 |
| FAM47C | NM_001013736 | 0.507 | 0.043 |
| DGKK | NM_001013742 | -0.007 | -0.2935 |
| TMEM225 | NM_001013743 | -0.3235 | -0.1965 |
| ZNF107 | NM_001013746 | -1.044 | -2.142 |
| MAD1L1 | NM_001013836 | -0.536 | -1.669 |
| RLTPR | NM_001013838 | -0.236 | -0.8055 |
| EXOC7 | NM_001013839 | -0.638 | -1.009 |
| STAP2 | NM_001013841 | -0.1985 | -0.5965 |
| C8orf58 | NM_001013842 | -0.011 | -0.624 |
| SLTM | NM_001013843 | -0.6445 | -1.596 |
| EXOC6 | NM_001013848 | -0.7935 | -1.8155 |
| RPS10 | NM_001014 | 0.1155 | -0.3135 |
| C5orf39 | NM_001014279 | -0.0335 | -0.3255 |
| DCUN1D2 | NM_001014283 | -0.9845 | -1.584 |
| FAM48A | NM_001014286 | -1.402 | -2.491 |
| SPRR2G | NM_001014291 | -0.191 | -0.5075 |
| IL31 | NM_001014336 | -0.0625 | -0.155 |
| FLG2 | NM_001014342 | 0.418 | -0.37 |
| KATNAL1 | NM_001014380 | -0.55 | -1.105 |
| AKT1 | NM_001014431 | -0.054 | -0.615 |
| AKT1 | NM_001014432 | -0.2855 | -0.678 |
| CUTA | NM_001014433 | 0.1705 | -0.2855 |

| | | | |
|---|---|---|---|
| LHX9 | NM_001014434 | -1.053 | -0.939 |
| DBNL | NM_001014436 | -0.088 | -0.6955 |
| CARS | NM_001014437 | -0.4275 | -1.4015 |
| ODF3B | NM_001014440 | 0.0665 | -0.1855 |
| C1QTNF9B-AS1 | NM_001014442 | -0.1515 | -0.1645 |
| USP21 | NM_001014443 | -0.396 | -1.078 |
| NLE1 | NM_001014445 | -0.5005 | -1.1845 |
| OCIAD2 | NM_001014446 | -0.1455 | -0.2765 |
| CPZ | NM_001014447 | -0.1255 | -0.1695 |
| DDX19B | NM_001014449 | -0.939 | -1.457 |
| SPRR2F | NM_001014450 | -0.272 | -0.3135 |
| DDX19B | NM_001014451 | -0.939 | -1.457 |
| TMEM93 | NM_001014764 | -0.7365 | -2.2115 |
| FBXO44 | NM_001014765 | -0.2985 | -0.644 |
| DDR2 | NM_001014796 | -1.2105 | -1.327 |
| KCNMA1 | NM_001014797 | 0.253 | -0.204 |
| CRMP1 | NM_001014809 | -0.006 | -0.489 |
| ME3 | NM_001014811 | -0.046 | -0.678 |
| FAM96A | NM_001014812 | -0.712 | -1.636 |
| C14orf183 | NM_001014830 | -0.424 | -0.28 |
| PAK4 | NM_001014831 | 0.006 | -1.046 |
| PAK4 | NM_001014832 | -0.203 | -0.8765 |
| PAK4 | NM_001014834 | -0.203 | -0.8765 |
| PAK4 | NM_001014835 | -0.203 | -0.8765 |
| CUTA | NM_001014837 | 0.1705 | -0.2855 |
| CUTA | NM_001014838 | 0.1705 | -0.2855 |
| NCDN | NM_001014839 | -0.191 | -0.6295 |
| CUTA | NM_001014840 | 0.1705 | -0.2855 |
| NCDN | NM_001014841 | -0.191 | -0.6295 |
| TM9SF1 | NM_001014842 | -0.3175 | -1.8215 |
| ZNF638 | NM_001014972 | -1.0405 | -2.5595 |
| CFH | NM_001014975 | -0.0225 | -0.261 |
| C16orf93 | NM_001014979 | -0.209 | -0.1215 |
| FAM132A | NM_001014980 | -0.1215 | -0.3235 |
| GLTPD2 | NM_001014985 | 0.3455 | -0.1155 |
| FOLH1 | NM_001014986 | -0.0475 | -0.4265 |
| LAT | NM_001014987 | -0.49 | -0.5675 |
| MRPL22 | NM_001014990 | -0.6005 | -2.3645 |
| RPS11 | NM_001015 | 0.113 | -0.263 |
| CKMT1A | NM_001015001 | 0.2115 | -0.4215 |
| LLGL2 | NM_001015002 | 0.161 | -0.42 |
| PAGE2B | NM_001015038 | 0.334 | -0.2205 |
| FAM13A | NM_001015045 | -1.3375 | -2.468 |

| | | | |
|---|---|---|---|
| BAG5 | NM_001015048 | -0.861 | -0.89 |
| BAG5 | NM_001015049 | -0.861 | -0.89 |
| ALG1L | NM_001015050 | 0.0515 | 0.008 |
| RUNX2 | NM_001015051 | 0.37 | -0.2655 |
| HDAC5 | NM_001015053 | -0.104 | -0.696 |
| RTKN | NM_001015055 | -0.792 | -1.192 |
| RTKN | NM_001015056 | -0.7365 | -1.2965 |
| UFSP1 | NM_001015072 | 0.1245 | -0.845 |
| PURG | NM_001015508 | -0.8295 | -1.563 |
| PHF6 | NM_001015877 | -0.8655 | -2.384 |
| AURKC | NM_001015878 | -0.348 | -0.578 |
| AURKC | NM_001015879 | -0.348 | -0.578 |
| PAPSS2 | NM_001015880 | -0.3295 | -1.385 |
| TSC22D3 | NM_001015881 | -0.063 | 0.017 |
| DNAJC25 | NM_001015882 | -0.6925 | -1.133 |
| RAE1 | NM_001015885 | -0.486 | -1.8385 |
| IGSF11 | NM_001015887 | 0.147 | -0.4015 |
| TAF9 | NM_001015891 | -0.354 | 0.077 |
| TAF9 | NM_001015892 | -0.354 | 0.077 |
| RPS13 | NM_001017 | 0.0555 | -0.2925 |
| C6orf221 | NM_001017361 | -0.2165 | -0.0085 |
| ARID3C | NM_001017363 | -0.1325 | -0.2515 |
| C4BPB | NM_001017364 | -0.2635 | -0.591 |
| C4BPB | NM_001017365 | -0.2635 | -0.591 |
| C4BPB | NM_001017366 | -0.2635 | -0.591 |
| C4BPB | NM_001017367 | -0.2635 | -0.591 |
| NHLRC3 | NM_001017370 | -1.0535 | -1.154 |
| SP3 | NM_001017371 | -0.703 | -2.25 |
| SLC27A6 | NM_001017372 | 0.8085 | 0.19 |
| SAMD3 | NM_001017373 | 0.04 | 0.0225 |
| TLR10 | NM_001017388 | 0.3965 | 0.028 |
| SUGP2 | NM_001017392 | -0.5725 | -0.0645 |
| TMCC1 | NM_001017395 | -1.7795 | -3.536 |
| ZNF2 | NM_001017396 | -0.4865 | -1.942 |
| TRIM36 | NM_001017397 | -0.2695 | -0.513 |
| TRIM36 | NM_001017398 | -0.12 | -0.751 |
| LAMB3 | NM_001017402 | -0.129 | -0.0855 |
| LGR6 | NM_001017403 | -0.0305 | -0.0485 |
| LGR6 | NM_001017404 | -0.0305 | -0.0485 |
| MAEA | NM_001017405 | -0.641 | -1.7785 |
| S100PBP | NM_001017406 | -1.4765 | -1.686 |
| GOPC | NM_001017408 | -2.2635 | -4.148 |
| USP1 | NM_001017415 | -0.87 | -2.201 |

| | | | |
|---|---|---|---|
| USP1 | NM_001017416 | -0.701 | -2.4895 |
| SPRR2B | NM_001017418 | -0.4435 | 0.0995 |
| ESCO2 | NM_001017420 | -1.202 | -2.583 |
| ALDH18A1 | NM_001017423 | -0.313 | -0.8985 |
| KCNK2 | NM_001017424 | 0.1875 | -1.1915 |
| KCNK2 | NM_001017425 | 0.1875 | -1.1915 |
| PGBD2 | NM_001017434 | -0.836 | -2.242 |
| CT45A3 | NM_001017435 | 0.0595 | -0.2615 |
| CCDC157 | NM_001017437 | -0.0365 | -0.374 |
| CALN1 | NM_001017440 | -0.072 | -1.047 |
| BTBD11 | NM_001017523 | -0.9125 | -2.1475 |
| SIRT3 | NM_001017524 | -0.296 | -1.67 |
| ARHGAP8 | NM_001017526 | 0.261 | -0.4215 |
| PRR5 | NM_001017528 | -0.5815 | -0.556 |
| PRR5 | NM_001017529 | -0.5815 | -0.556 |
| PRR5 | NM_001017530 | -0.5815 | -0.556 |
| CARD16 | NM_001017534 | 0.0185 | -0.255 |
| VDR | NM_001017535 | 0.0385 | -0.6 |
| VDR | NM_001017536 | 0.0385 | -0.6 |
| INPP5D | NM_001017915 | 0.219 | -0.327 |
| CYB561 | NM_001017916 | -0.9105 | -1.0105 |
| CYB561 | NM_001017917 | -0.8475 | -0.8595 |
| RCCD1 | NM_001017919 | -0.805 | -2.1675 |
| DAPL1 | NM_001017920 | 0.071 | -0.0475 |
| VMAC | NM_001017921 | -0.032 | -0.2905 |
| ERMAP | NM_001017922 | -0.1405 | -0.2345 |
| C14orf28 | NM_001017923 | -1.0065 | -1.52 |
| LRIT2 | NM_001017924 | -0.1 | -0.2375 |
| ZHX1 | NM_001017926 | -0.5085 | -2.0115 |
| C2orf76 | NM_001017927 | -0.681 | -2.1055 |
| CCDC58 | NM_001017928 | -1.3615 | -1.5605 |
| DCAF8L1 | NM_001017930 | -0.3045 | -0.141 |
| OS9 | NM_001017956 | -0.1515 | -0.6785 |
| OS9 | NM_001017957 | -0.1515 | -0.6785 |
| OS9 | NM_001017958 | -0.1515 | -0.6785 |
| FAM78B | NM_001017961 | -0.2285 | -1.514 |
| P4HA1 | NM_001017962 | -1.5875 | -1.2865 |
| HSP90AA1 | NM_001017963 | 0.0215 | -0.1525 |
| YDJC | NM_001017964 | -0.7635 | -1.9135 |
| MARVELD3 | NM_001017967 | 0.176 | 0.4905 |
| KIAA2026 | NM_001017969 | -0.0605 | -0.596 |
| TMEM30B | NM_001017970 | 0.0025 | 0.0145 |
| ATP6AP1L | NM_001017971 | -0.676 | -0.874 |

| | | | |
|---|---|---|---|
| HFM1 | NM_001017975 | -0.211 | -1.8395 |
| DCAF6 | NM_001017977 | -1.115 | -1.723 |
| CXorf61 | NM_001017978 | 0.2595 | 0.474 |
| RAB28 | NM_001017979 | -0.961 | -2.559 |
| VMA21 | NM_001017980 | -1.0465 | -0.7685 |
| RNF215 | NM_001017981 | -0.254 | -0.8505 |
| FCGR1B | NM_001017986 | -0.1445 | -0.729 |
| C5orf45 | NM_001017987 | -0.613 | -0.845 |
| OPA3 | NM_001017989 | -0.462 | -1.596 |
| H2AFB1 | NM_001017990 | 0.4955 | 0.194 |
| ACTBL2 | NM_001017992 | 0.0105 | 0.4615 |
| SH3PXD2B | NM_001017995 | -1.23 | -2.3555 |
| GNG10 | NM_001017998 | -1.13 | -0.8855 |
| RPS15 | NM_001018 | 0.12 | -0.1925 |
| SORBS3 | NM_001018003 | -0.02 | -0.284 |
| TPM1 | NM_001018004 | -1.03 | -0.866 |
| TPM1 | NM_001018005 | -0.0355 | -0.3935 |
| TPM1 | NM_001018006 | -0.0355 | -0.3935 |
| TPM1 | NM_001018007 | -0.0355 | -0.3935 |
| TPM1 | NM_001018008 | -0.0355 | -0.3935 |
| SH3BP5 | NM_001018009 | -0.358 | -0.844 |
| MUC1 | NM_001018016 | -0.1105 | -0.321 |
| MUC1 | NM_001018017 | -0.1105 | -0.321 |
| TPM1 | NM_001018020 | -0.0355 | -0.3935 |
| MTCP1NB | NM_001018024 | -0.105 | -0.8505 |
| MTCP1 | NM_001018025 | -0.533 | -0.799 |
| TSHR | NM_001018036 | 0.1545 | -0.2765 |
| VPS13A | NM_001018037 | -0.206 | -1.2185 |
| VPS13A | NM_001018038 | -0.1695 | -1.3065 |
| SFMBT2 | NM_001018039 | -0.0795 | -0.4315 |
| EPM2A | NM_001018041 | -0.5975 | -1.5745 |
| YSK4 | NM_001018046 | -0.136 | -0.2335 |
| PAEP | NM_001018049 | -0.2495 | -0.487 |
| POLR3H | NM_001018050 | -0.629 | -2.1645 |
| POLR3H | NM_001018052 | -0.629 | -2.1645 |
| PFKFB2 | NM_001018053 | -0.753 | -0.1705 |
| LRP8 | NM_001018054 | -1.334 | -2.2045 |
| BRCC3 | NM_001018055 | -0.2775 | -1.067 |
| DKK3 | NM_001018057 | -0.871 | -1.1315 |
| TFEC | NM_001018058 | -0.0955 | -0.0655 |
| AIFM3 | NM_001018060 | -0.0965 | 0.04 |
| NTRK2 | NM_001018064 | -0.042 | -0.1995 |
| NTRK2 | NM_001018065 | 0.0285 | -0.106 |

| | | | |
|---|---|---|---|
| NTRK2 | NM_001018066 | -0.042 | -0.1995 |
| SERBP1 | NM_001018067 | -1.056 | -1.278 |
| SERBP1 | NM_001018068 | -1.056 | -1.278 |
| SERBP1 | NM_001018069 | -1.056 | -1.278 |
| CORO1B | NM_001018070 | 0.083 | -0.4095 |
| FRMPD2 | NM_001018071 | -0.069 | 0.044 |
| BTBD11 | NM_001018072 | -1.1435 | -2.33 |
| PCK2 | NM_001018073 | -0.3455 | -1.109 |
| NR3C1 | NM_001018074 | -0.2885 | -0.689 |
| NR3C1 | NM_001018075 | -0.2885 | -0.689 |
| NR3C1 | NM_001018076 | -0.2885 | -0.689 |
| NR3C1 | NM_001018077 | -0.2885 | -0.689 |
| FPGS | NM_001018078 | 0.1 | -0.784 |
| FSHB | NM_001018080 | 0.032 | -0.2385 |
| ADIG | NM_001018082 | -0.0285 | -0.2965 |
| VPS13C | NM_001018088 | 0.0215 | -0.285 |
| NARG2 | NM_001018089 | -0.979 | -2.575 |
| GCOM1 | NM_001018090 | -1.346 | -2.307 |
| GCOM1 | NM_001018091 | -0.3855 | -0.511 |
| GCOM1 | NM_001018100 | -0.3855 | -0.511 |
| GRINL1A | NM_001018102 | -0.1435 | -0.441 |
| FAHD1 | NM_001018104 | -0.6825 | -1.601 |
| SERF2 | NM_001018108 | 0.172 | -0.2695 |
| PODXL | NM_001018111 | -0.3125 | -0.906 |
| FANCA | NM_001018112 | -0.397 | -1.195 |
| FANCB | NM_001018113 | -0.284 | -1.2735 |
| FANCD2 | NM_001018115 | -0.816 | -0.939 |
| MURC | NM_001018116 | -0.9575 | -0.8385 |
| NME1-NME2 | NM_001018136 | 0.3215 | 0.004 |
| PON2 | NM_001018161 | -0.8165 | -1.068 |
| HAX1 | NM_001018837 | -0.456 | -1.0295 |
| RPS15A | NM_001019 | 0.0655 | -0.254 |
| RPS16 | NM_001020 | 0.101 | -0.321 |
| PUM1 | NM_001020658 | -1.262 | -2.858 |
| MYADM | NM_001020818 | -0.7145 | -2.059 |
| MYADM | NM_001020819 | -0.7145 | -2.059 |
| MYADM | NM_001020820 | -0.7145 | -2.059 |
| MYADM | NM_001020821 | -0.7145 | -2.059 |
| NR3C1 | NM_001020825 | -0.2885 | -0.689 |
| RPS17 | NM_001021 | 0.1245 | -0.2485 |
| RPS19 | NM_001022 | 0.135 | -0.211 |
| RPS20 | NM_001023 | -0.139 | -0.1875 |
| ZNF187 | NM_001023560 | -1.4035 | -2.8105 |

| | | | |
|---|---|---|---|
| ZNF749 | NM_001023561 | -0.7905 | -0.7885 |
| ZNF805 | NM_001023563 | -1.693 | -3.1645 |
| IQCB1 | NM_001023570 | -0.9345 | -1.4145 |
| IQCB1 | NM_001023571 | -0.9345 | -1.4145 |
| ABCC5 | NM_001023587 | 0.3215 | -0.1295 |
| RPS21 | NM_001024 | 0.053 | -0.3 |
| HNMT | NM_001024074 | 0.0555 | -0.3355 |
| HNMT | NM_001024075 | 0.0555 | -0.3355 |
| NR3C1 | NM_001024094 | -0.2885 | -0.689 |
| SPRR2E | NM_001024209 | -0.022 | -0.3095 |
| S100A13 | NM_001024210 | 0.248 | -0.5225 |
| S100A13 | NM_001024211 | 0.248 | -0.5225 |
| S100A13 | NM_001024212 | 0.248 | -0.5225 |
| S100A13 | NM_001024213 | 0.248 | -0.5225 |
| FBLIM1 | NM_001024215 | -0.0575 | -0.515 |
| FBLIM1 | NM_001024216 | -0.0575 | -0.515 |
| GPHN | NM_001024218 | -1.026 | -2.481 |
| ARF1 | NM_001024226 | 0.118 | -0.4555 |
| ARF1 | NM_001024227 | 0.118 | -0.4555 |
| ARF1 | NM_001024228 | 0.118 | -0.4555 |
| BAALC | NM_001024372 | 0.117 | -0.706 |
| TM2D2 | NM_001024380 | -0.8485 | -2.6585 |
| TM2D2 | NM_001024381 | -0.8485 | -2.6585 |
| HMBS | NM_001024382 | -0.8255 | -2.0835 |
| SBK1 | NM_001024401 | -0.3735 | -1.045 |
| GLRA4 | NM_001024452 | -0.153 | -0.1645 |
| RGAG4 | NM_001024455 | 0.012 | -0.1985 |
| RGPD1 | NM_001024457 | 0.147 | -0.6125 |
| SOD2 | NM_001024465 | 0.1 | -0.1 |
| SOD2 | NM_001024466 | 0.1 | -0.1 |
| ZMYND17 | NM_001024593 | -1.168 | -0.9725 |
| C1orf53 | NM_001024594 | 0.4305 | -0.085 |
| ZNF772 | NM_001024596 | -2.0015 | -4.228 |
| HES3 | NM_001024598 | 0.078 | -0.329 |
| HIST2H2BF | NM_001024599 | 0.0245 | -0.5435 |
| C1orf227 | NM_001024601 | -0.2675 | -0.284 |
| C7orf66 | NM_001024607 | 0.0005 | -0.0775 |
| LRRC66 | NM_001024611 | -0.058 | 0.247 |
| FEZF1 | NM_001024613 | 0.1555 | 0.4565 |
| NRP1 | NM_001024628 | -2.015 | -3.5045 |
| NRP1 | NM_001024629 | -2.015 | -3.5045 |
| RUNX2 | NM_001024630 | 0.37 | -0.2655 |
| XCR1 | NM_001024644 | 0.0405 | -0.209 |

| | | | |
|---|---|---|---|
| RAB3IP | NM_001024647 | -0.321 | -0.6525 |
| CANX | NM_001024649 | -0.8695 | -0.3845 |
| KALRN | NM_001024660 | -0.1745 | -0.7935 |
| PRAMEF13 | NM_001024661 | -0.104 | -0.7215 |
| RPL6 | NM_001024662 | -0.3325 | -0.2975 |
| SH3KBP1 | NM_001024666 | -0.134 | -0.166 |
| LIN52 | NM_001024674 | -1.338 | -1.5785 |
| C20orf134 | NM_001024675 | 0.1885 | -0.4525 |
| LRRC24 | NM_001024678 | -0.391 | -0.941 |
| C1orf68 | NM_001024679 | -0.034 | -0.1405 |
| FBXO48 | NM_001024680 | 0.029 | -0.7325 |
| ZNF688 | NM_001024683 | -0.097 | -0.5025 |
| MECR | NM_001024732 | -0.2225 | -0.797 |
| CD276 | NM_001024736 | 0.0785 | -0.8745 |
| APLP1 | NM_001024807 | 0.286 | -0.285 |
| BCL7A | NM_001024808 | -0.735 | -1.6145 |
| RARA | NM_001024809 | -0.49 | -1.943 |
| RNASE12 | NM_001024822 | 0.226 | 0.0745 |
| TNRC6B | NM_001024843 | -2.032 | -3.354 |
| CD82 | NM_001024844 | 0.1735 | -0.383 |
| SLC6A9 | NM_001024845 | 0.164 | -0.447 |
| TGFBR2 | NM_001024847 | 0.172 | -0.011 |
| ZNF197 | NM_001024855 | -0.77 | -2.079 |
| SPTB | NM_001024858 | -0.0005 | -0.1755 |
| CEACAM1 | NM_001024912 | 0.085 | -0.347 |
| CBWD5 | NM_001024916 | -0.732 | -0.9245 |
| RPL9 | NM_001024921 | -0.0175 | -0.3365 |
| EXOC1 | NM_001024924 | -1.285 | -2.101 |
| WASF1 | NM_001024934 | -0.6535 | -1.242 |
| WASF1 | NM_001024935 | -0.6535 | -1.242 |
| WASF1 | NM_001024936 | -0.6815 | -1.272 |
| MINK1 | NM_001024937 | -0.192 | -0.971 |
| SLC2A11 | NM_001024938 | -0.092 | -0.506 |
| SLC2A11 | NM_001024939 | -0.092 | -0.506 |
| TRIM17 | NM_001024940 | -0.3515 | -0.479 |
| ASL | NM_001024943 | 0.125 | -0.4295 |
| ASL | NM_001024944 | 0.125 | -0.4295 |
| ASL | NM_001024946 | 0.125 | -0.4295 |
| YIPF5 | NM_001024947 | -1.051 | -1.5445 |
| FNBP1L | NM_001024948 | -0.844 | -1.5075 |
| SC5DL | NM_001024956 | -1.3535 | -1.9715 |
| BRMS1 | NM_001024957 | -0.3835 | -1.673 |
| ARPC4 | NM_001024959 | -0.0055 | -0.569 |

| | | | |
|---|---|---|---|
| ARPC4 | NM_001024960 | -0.0805 | -0.7065 |
| RPS23 | NM_001025 | -0.013 | -0.279 |
| ARPP21 | NM_001025068 | -0.4355 | -0.2805 |
| ARPP21 | NM_001025069 | -0.757 | 0.091 |
| RPS14 | NM_001025071 | -0.227 | -0.8075 |
| CELF2 | NM_001025076 | -0.901 | -0.5255 |
| CELF2 | NM_001025077 | -0.901 | -0.5255 |
| MBP | NM_001025081 | 0.0715 | -0.0095 |
| CELF4 | NM_001025087 | -1.4965 | -1.4715 |
| CELF4 | NM_001025088 | -1.4965 | -1.4715 |
| CELF4 | NM_001025089 | -1.4965 | -1.4715 |
| MBP | NM_001025090 | 0.0715 | -0.0095 |
| ABCF1 | NM_001025091 | -0.845 | -1.981 |
| MBP | NM_001025092 | 0.0715 | -0.0095 |
| MBP | NM_001025100 | 0.0715 | -0.0095 |
| MBP | NM_001025101 | 0.0715 | -0.0095 |
| CSN1S1 | NM_001025104 | 0.015 | -0.3585 |
| CSNK1A1 | NM_001025105 | -1.33 | -2.16 |
| ADAR | NM_001025107 | -0.567 | -1.21 |
| AFF3 | NM_001025108 | -1.2675 | -2.733 |
| CD34 | NM_001025109 | 0.366 | -0.309 |
| CD74 | NM_001025158 | -0.1105 | -0.6615 |
| CD74 | NM_001025159 | -0.1105 | -0.6615 |
| CD97 | NM_001025160 | -0.193 | 0.051 |
| CYP2D6 | NM_001025161 | -0.237 | -0.605 |
| MSLNL | NM_001025190 | -0.175 | -0.24 |
| CES1 | NM_001025195 | 0.2275 | 0.013 |
| CHI3L2 | NM_001025197 | -0.1455 | -0.1445 |
| CHI3L2 | NM_001025199 | -0.1455 | -0.1445 |
| CTRB2 | NM_001025200 | -0.098 | 0.6445 |
| CHN1 | NM_001025201 | -0.84 | -1.1335 |
| U2AF1 | NM_001025204 | -0.3195 | -1.0315 |
| AP2M1 | NM_001025205 | 0.182 | -0.3565 |
| KPRP | NM_001025231 | -0.2055 | -0.207 |
| CLLU1OS | NM_001025232 | -0.043 | -0.424 |
| CLLU1 | NM_001025233 | 0.135 | -0.304 |
| TSPAN4 | NM_001025234 | 0.2605 | -0.4945 |
| TSPAN4 | NM_001025235 | 0.2605 | -0.4945 |
| TSPAN4 | NM_001025236 | 0.188 | -0.432 |
| TSPAN4 | NM_001025237 | 0.2605 | -0.4945 |
| TSPAN4 | NM_001025238 | 0.2605 | -0.4945 |
| TSPAN4 | NM_001025239 | 0.2605 | -0.4945 |
| IRAK1 | NM_001025243 | -0.425 | -0.8285 |

| | | | |
|---|---|---|---|
| TAF5L | NM_001025247 | 0.6585 | 1.487 |
| DUT | NM_001025248 | -0.175 | -0.6775 |
| DUT | NM_001025249 | -0.175 | -0.6775 |
| TPD52 | NM_001025252 | -0.9245 | -0.882 |
| TPD52 | NM_001025253 | -0.9245 | -0.882 |
| CXorf65 | NM_001025265 | 0.221 | -0.5435 |
| C3orf70 | NM_001025266 | -1.0815 | -1.6 |
| DPPA5 | NM_001025290 | -0.025 | -0.3605 |
| IFITM5 | NM_001025295 | 0.0005 | 0.0735 |
| RAB12 | NM_001025300 | -0.443 | -0.1385 |
| ANO6 | NM_001025356 | -1.406 | -1.2875 |
| C8orf85 | NM_001025357 | -0.0905 | 0.054 |
| VEGFA | NM_001025366 | -0.661 | -0.7585 |
| VEGFA | NM_001025367 | -0.674 | -0.6745 |
| VEGFA | NM_001025368 | -0.674 | -0.6745 |
| VEGFA | NM_001025369 | -0.674 | -0.6745 |
| VEGFA | NM_001025370 | -0.674 | -0.6745 |
| AMPD3 | NM_001025389 | -0.631 | -0.786 |
| AMPD3 | NM_001025390 | -0.631 | -0.786 |
| NQO1 | NM_001025433 | 0.231 | -0.5815 |
| NQO1 | NM_001025434 | 0.231 | -0.5815 |
| SPAG16 | NM_001025436 | -0.8395 | -1.181 |
| NDEL1 | NM_001025579 | -0.7215 | -2.3775 |
| AMMECR1 | NM_001025580 | -1.402 | -2.528 |
| SCGBL | NM_001025591 | 0.0945 | -0.272 |
| ARFIP1 | NM_001025593 | -1.3235 | -3.1465 |
| ARFIP1 | NM_001025595 | -1.7455 | -3.6355 |
| CELF1 | NM_001025596 | -0.732 | -1.313 |
| ARHGAP30 | NM_001025598 | 0.1775 | -0.4315 |
| RFX5 | NM_001025603 | -1.658 | -2.875 |
| ARRDC2 | NM_001025604 | 0.0185 | -0.4235 |
| ARHGAP24 | NM_001025616 | -0.2485 | -0.394 |
| VRK3 | NM_001025778 | -0.092 | -1.031 |
| FAM108B1 | NM_001025780 | -0.917 | -2.666 |
| TTLL3 | NM_001025930 | -0.09 | -0.7 |
| RPS24 | NM_001026 | -0.179 | -0.225 |
| CYP11B1 | NM_001026213 | -0.1315 | -0.43 |
| CCDC7 | NM_001026383 | -0.374 | -0.083 |
| RPS25 | NM_001028 | -0.028 | -0.253 |
| RPS26 | NM_001029 | 0.1145 | -0.282 |
| CCM2 | NM_001029835 | -1.528 | -2.997 |
| C3orf23 | NM_001029839 | -0.394 | -0.4515 |
| C3orf23 | NM_001029840 | -1.564 | -3.481 |

| | | | |
|---|---|---|---|
| PDE8B | NM_001029851 | 0.1905 | -1.272 |
| PDE8B | NM_001029852 | 0.316 | -1.1125 |
| PDE8B | NM_001029853 | 0.316 | -1.1125 |
| PDE8B | NM_001029854 | 0.316 | -1.1125 |
| SLC35F1 | NM_001029858 | -0.2005 | -0.724 |
| KCTD21 | NM_001029859 | -0.712 | -1.346 |
| FBXO43 | NM_001029860 | -0.0815 | -1.798 |
| NANOS2 | NM_001029861 | 0.0675 | -0.1625 |
| C6orf120 | NM_001029863 | -1.719 | -0.748 |
| KIAA1755 | NM_001029864 | -0.0465 | -0.459 |
| DBX1 | NM_001029865 | 0.1615 | -0.4375 |
| PLAC8L1 | NM_001029869 | -0.5505 | -0.4515 |
| ANKRD56 | NM_001029870 | -0.5335 | -0.5085 |
| RSPO4 | NM_001029871 | -0.0195 | -0.3275 |
| REP15 | NM_001029874 | 0.0185 | -0.842 |
| RGS7BP | NM_001029875 | -0.5285 | -0.074 |
| SFMBT2 | NM_001029880 | -0.0795 | -0.4315 |
| CIB4 | NM_001029881 | -0.01 | -0.295 |
| AHDC1 | NM_001029882 | -0.6205 | -0.7935 |
| C2orf71 | NM_001029883 | 0.0075 | -0.2495 |
| PLEKHG1 | NM_001029884 | -0.2525 | -0.1975 |
| GLTPD1 | NM_001029885 | -0.687 | -1.53 |
| PFN3 | NM_001029886 | 0.1335 | -0.266 |
| HELT | NM_001029887 | -0.0415 | -0.205 |
| FAM24A | NM_001029888 | 0.0625 | 0.0225 |
| PGAM4 | NM_001029891 | 0.2375 | -0.215 |
| WDR45 | NM_001029896 | 0.257 | -0.379 |
| ZFAT | NM_001029939 | -0.164 | -0.7475 |
| CDNF | NM_001029954 | -0.0135 | -0.594 |
| DCAF4L1 | NM_001029955 | -0.353 | -0.3125 |
| TTLL13 | NM_001029964 | -0.0505 | -0.323 |
| ZNF16 | NM_001029976 | -1.0275 | -2.6235 |
| KIAA0101 | NM_001029989 | -0.8535 | -0.4175 |
| METTL17 | NM_001029991 | -0.1385 | -0.7995 |
| PCDP1 | NM_001029996 | 0.0365 | 0.018 |
| SLC10A7 | NM_001029998 | -0.6675 | -0.9285 |
| RPS27 | NM_001030 | 0.107 | -0.282 |
| RPS29 | NM_001030001 | 0.1375 | -0.2625 |
| GRB7 | NM_001030002 | 0.312 | -0.2765 |
| HNF4A | NM_001030003 | -0.024 | 0.0685 |
| HNF4A | NM_001030004 | -0.024 | 0.0685 |
| AP2B1 | NM_001030006 | -0.333 | -0.748 |
| AP1G1 | NM_001030007 | -0.51 | -1.1665 |

| | | | |
|---|---|---|---|
| RPS15A | NM_001030009 | 0.0655 | -0.254 |
| ALDH3B1 | NM_001030010 | -0.188 | -0.3895 |
| NPS | NM_001030013 | -0.1745 | 0.0745 |
| OPN4 | NM_001030015 | -0.226 | -0.2685 |
| APRT | NM_001030018 | 0.28 | -0.254 |
| SUN3 | NM_001030019 | -0.274 | -0.2025 |
| KLK3 | NM_001030047 | -0.024 | -0.0285 |
| KLK3 | NM_001030048 | -0.024 | -0.0285 |
| KLK3 | NM_001030050 | -0.024 | -0.0285 |
| ARHGAP5 | NM_001030055 | -0.89 | -2.686 |
| PPAPDC1A | NM_001030059 | -0.232 | -1.142 |
| SAMD5 | NM_001030060 | -0.207 | -0.202 |
| ARNTL | NM_001030272 | -0.2265 | -1.008 |
| ARNTL | NM_001030273 | -0.2265 | -1.008 |
| ATF3 | NM_001030287 | -0.3345 | 1.001 |
| SPN | NM_001030288 | 0.1035 | -0.475 |
| CERKL | NM_001030311 | -0.0855 | -1.3575 |
| CERKL | NM_001030312 | -0.0855 | -1.3575 |
| CERKL | NM_001030313 | -0.0855 | -1.3575 |
| RPS28 | NM_001031 | 0.124 | -0.2275 |
| ALDH3B2 | NM_001031615 | 0.112 | -0.3485 |
| COX19 | NM_001031617 | -0.815 | -1.3135 |
| SPDYE2 | NM_001031618 | -1.3675 | -2.0585 |
| ZNF451 | NM_001031623 | -1.3515 | -3.1765 |
| SMAGP | NM_001031628 | -0.103 | -1.298 |
| ZNF816 | NM_001031665 | 0.0585 | -1.928 |
| CYB5RL | NM_001031672 | -0.7655 | -1.019 |
| RAB24 | NM_001031677 | -0.8445 | -1.505 |
| MSRB3 | NM_001031679 | -0.2365 | -0.701 |
| RUNX3 | NM_001031680 | -0.0435 | -0.659 |
| CTNS | NM_001031681 | -1.33 | -2.5025 |
| IFIT3 | NM_001031683 | -0.1885 | -0.458 |
| SRSF7 | NM_001031684 | -0.889 | -0.46 |
| TP53BP2 | NM_001031685 | -0.9605 | -1.718 |
| PLAA | NM_001031689 | -1.0395 | -2.6895 |
| FAM131B | NM_001031690 | 0.1315 | -0.118 |
| LRRC17 | NM_001031692 | 0.322 | -1.2345 |
| HHLA3 | NM_001031693 | -0.1015 | 0.0395 |
| SCMH1 | NM_001031694 | -0.253 | -1.367 |
| RBFOX2 | NM_001031695 | -0.862 | -1.1125 |
| PRPF40B | NM_001031698 | -0.0935 | -0.595 |
| FAM198B | NM_001031700 | -0.33 | -0.2505 |
| NT5DC3 | NM_001031701 | -0.344 | -1.656 |

| | | | |
|---|---|---|---|
| SEMA5B | NM_001031702 | 0.022 | -0.274 |
| C3orf75 | NM_001031703 | -0.583 | -1.7865 |
| RNLS | NM_001031709 | 0.3085 | -0.048 |
| KLHL7 | NM_001031710 | 0.4635 | -0.0885 |
| ERGIC1 | NM_001031711 | -0.057 | -0.7155 |
| TRMT11 | NM_001031712 | -0.7395 | -1.6915 |
| CCDC90A | NM_001031713 | -0.0685 | -0.7505 |
| INF2 | NM_001031714 | -0.3 | -0.759 |
| IQCH | NM_001031715 | 0.229 | -0.1345 |
| OBFC2A | NM_001031716 | -1.992 | -1.8 |
| CRELD1 | NM_001031717 | -0.554 | -0.848 |
| DHRS12 | NM_001031719 | 0.052 | -0.6175 |
| GSTCD | NM_001031720 | -0.6875 | -1.435 |
| ZNF613 | NM_001031721 | -0.7815 | -2.612 |
| ATAT1 | NM_001031722 | -0.7065 | -1.499 |
| DNAJB14 | NM_001031723 | -0.724 | -1.077 |
| DDX59 | NM_001031725 | -1.4815 | -2.7005 |
| C19orf12 | NM_001031726 | -0.531 | -1.53 |
| MRI1 | NM_001031727 | -0.2705 | -0.745 |
| YTHDC1 | NM_001031732 | -1.3355 | -2.657 |
| CALML4 | NM_001031733 | -0.737 | -1.1135 |
| FDX1L | NM_001031734 | -0.794 | -1.6335 |
| IZUMO4 | NM_001031735 | 0.294 | -0.044 |
| CCDC78 | NM_001031737 | -0.1395 | -0.314 |
| TMEM150A | NM_001031738 | -0.119 | -0.5455 |
| ASB9 | NM_001031739 | 0.199 | -0.333 |
| MANEAL | NM_001031740 | -0.876 | -1.801 |
| C6orf165 | NM_001031743 | -0.0255 | -0.5765 |
| RIBC1 | NM_001031745 | 0.272 | -0.1385 |
| C12orf40 | NM_001031748 | 0.1505 | 0.256 |
| LYPD5 | NM_001031749 | 0.2225 | -0.2255 |
| TIPRL | NM_001031800 | -0.6605 | -1.4925 |
| LIMK2 | NM_001031801 | -0.9735 | -2.8255 |
| LLGL2 | NM_001031803 | 0.2995 | -0.694 |
| MAF | NM_001031804 | -0.238 | -0.441 |
| ALDH3A2 | NM_001031806 | -0.9925 | -1.6615 |
| SKOR1 | NM_001031807 | 0.0455 | -0.2615 |
| CSNK1G3 | NM_001031812 | -0.839 | -2.3445 |
| BOLA2 | NM_001031827 | -0.504 | -1.034 |
| RAB40AL | NM_001031834 | 0.181 | -0.2055 |
| PHKB | NM_001031835 | -0.8825 | -0.503 |
| KCNU1 | NM_001031836 | -0.3135 | -0.6195 |
| CPT1A | NM_001031847 | -0.816 | -1.0275 |

| | | | |
|---|---|---|---|
| SERPINB8 | NM_001031848 | -0.041 | -0.4225 |
| MASP1 | NM_001031849 | 0.167 | -0.0785 |
| PSG6 | NM_001031850 | 0.225 | 0.348 |
| INSC | NM_001031853 | 0.0705 | -0.524 |
| ACCSL | NM_001031854 | -0.073 | -0.01 |
| LONRF3 | NM_001031855 | 0.081 | -0.4965 |
| RPS29 | NM_001032 | 0.1375 | -0.2625 |
| STXBP1 | NM_001032221 | -1.2315 | -1.146 |
| MMP28 | NM_001032278 | -0.276 | -0.392 |
| TFAP2A | NM_001032280 | 0.0025 | -0.092 |
| TFPI | NM_001032281 | -0.269 | -0.344 |
| KLF10 | NM_001032282 | -0.221 | 0.0325 |
| TMPO | NM_001032283 | -1.5825 | -1.827 |
| TMPO | NM_001032284 | -1.5825 | -1.827 |
| NR2C1 | NM_001032287 | -0.899 | -2.0125 |
| UBE2V1 | NM_001032288 | -0.2425 | -1.1055 |
| SLC35A2 | NM_001032289 | -0.6345 | -1.8085 |
| ZNF174 | NM_001032292 | -0.8265 | -1.5565 |
| ZNF207 | NM_001032293 | -0.662 | -1.4985 |
| SERPING1 | NM_001032295 | -0.0875 | -0.3745 |
| STK24 | NM_001032296 | -0.0975 | -1.0485 |
| MINOS1 | NM_001032363 | 0.0625 | -0.5 |
| SPINT1 | NM_001032367 | 0.127 | -0.468 |
| ZNF226 | NM_001032372 | -0.596 | -0.881 |
| ZNF226 | NM_001032373 | -0.596 | -0.881 |
| ZNF226 | NM_001032374 | -0.596 | -0.881 |
| SULT6B1 | NM_001032377 | 0.0725 | -0.382 |
| PQBP1 | NM_001032381 | -0.1635 | -0.5965 |
| PQBP1 | NM_001032382 | -0.1635 | -0.5965 |
| PQBP1 | NM_001032383 | -0.1635 | -0.5965 |
| PQBP1 | NM_001032384 | -0.183 | -0.81 |
| SUOX | NM_001032386 | -0.8505 | -1.3955 |
| SUOX | NM_001032387 | -0.8505 | -1.3955 |
| HNRNPH2 | NM_001032393 | -0.79 | -0.9495 |
| GPR126 | NM_001032394 | 0.149 | 0.0315 |
| GPR126 | NM_001032395 | 0.149 | 0.0315 |
| PJA1 | NM_001032396 | -0.633 | -1.645 |
| OAS1 | NM_001032409 | -0.025 | -0.293 |
| USP16 | NM_001032410 | -0.374 | -1.7405 |
| RAB41 | NM_001032726 | -0.253 | -0.4515 |
| USP48 | NM_001032730 | -0.76 | -1.467 |
| OAS2 | NM_001032731 | -0.0745 | -0.4055 |
| KYNU | NM_001032998 | 0.233 | -0.398 |

| | | | |
|---|---|---|---|
| CBFA2T2 | NM_001032999 | -0.9095 | -2.0895 |
| RRM1 | NM_001033 | -0.73 | -0.6475 |
| RPAIN | NM_001033002 | -1.303 | -1.4555 |
| DEFB135 | NM_001033017 | 0.543 | -0.066 |
| DEFB136 | NM_001033018 | -0.4205 | -0.266 |
| DEFB134 | NM_001033019 | -0.1105 | -0.346 |
| FBXO7 | NM_001033024 | -0.237 | -1.433 |
| EXTL2 | NM_001033025 | -1.1575 | -1.1395 |
| C19orf6 | NM_001033026 | -0.242 | -0.7965 |
| CYFIP1 | NM_001033028 | -0.013 | -0.2075 |
| FAIM | NM_001033031 | -0.9405 | -2.512 |
| FAIM | NM_001033032 | -0.9405 | -2.512 |
| GLUL | NM_001033044 | -0.3065 | -1.226 |
| GPR155 | NM_001033045 | -0.756 | -0.979 |
| C17orf62 | NM_001033046 | -0.531 | -1.6595 |
| NPNT | NM_001033047 | -0.2665 | -0.1805 |
| MTERFD3 | NM_001033050 | -1.8725 | -3.0075 |
| NLRP1 | NM_001033053 | -0.5045 | -0.3005 |
| AIPL1 | NM_001033054 | -0.0765 | -0.6315 |
| AIPL1 | NM_001033055 | -0.0765 | -0.6315 |
| GLUL | NM_001033056 | -0.3065 | -1.226 |
| MAGI1 | NM_001033057 | -1.3115 | -3.687 |
| AMD1 | NM_001033059 | -0.98 | -2.807 |
| TAAR2 | NM_001033080 | 0.351 | -0.5435 |
| MYCL1 | NM_001033081 | 0.082 | -0.437 |
| MYCL1 | NM_001033082 | 0.082 | -0.437 |
| PBLD | NM_001033083 | -0.8705 | -0.753 |
| MACROD2 | NM_001033087 | -0.328 | -0.334 |
| NGRN | NM_001033088 | 0.24 | -0.0685 |
| PAIP2 | NM_001033112 | -0.556 | -0.3605 |
| ENTPD8 | NM_001033113 | -0.091 | -0.5355 |
| SRGAP3 | NM_001033117 | -0.8975 | -1.18 |
| SAR1B | NM_001033503 | -0.672 | -1.69 |
| TMEM98 | NM_001033504 | 0.109 | -0.293 |
| CSTF3 | NM_001033505 | -0.897 | -1.512 |
| CSTF3 | NM_001033506 | -0.5865 | -1.4935 |
| SERINC4 | NM_001033517 | -0.155 | -0.5705 |
| WIPI2 | NM_001033518 | -0.1835 | -1.579 |
| WIPI2 | NM_001033519 | -0.1835 | -1.579 |
| WIPI2 | NM_001033520 | -0.1835 | -1.579 |
| CSTF1 | NM_001033521 | -1.246 | -3.079 |
| CSTF1 | NM_001033522 | -1.685 | -3.3255 |
| TOM1L2 | NM_001033551 | 0.091 | -0.5285 |

| | | | |
|---|---|---|---|
| SPECC1 | NM_001033553 | -0.3345 | -0.4385 |
| SPECC1 | NM_001033554 | 0.016 | -0.4035 |
| SPECC1 | NM_001033555 | -0.3345 | -0.4385 |
| PPM1B | NM_001033556 | -1.071 | -2.7885 |
| PPM1B | NM_001033557 | -1.071 | -2.7885 |
| DYX1C1 | NM_001033559 | -0.0905 | -0.2295 |
| DYX1C1 | NM_001033560 | -0.0905 | -0.2295 |
| PHF12 | NM_001033561 | -0.732 | -2.664 |
| C6orf225 | NM_001033564 | 0.1595 | -0.3205 |
| RHOT1 | NM_001033566 | -1.606 | -2.0195 |
| RHOT1 | NM_001033568 | -1.606 | -2.0195 |
| DUSP28 | NM_001033575 | -0.403 | -0.9635 |
| UNC45B | NM_001033576 | 0.1015 | 0.0325 |
| SGK3 | NM_001033578 | -0.0005 | -0.778 |
| MYO19 | NM_001033580 | -1.2365 | -2.001 |
| PRKCZ | NM_001033581 | -0.0375 | -0.907 |
| PRKCZ | NM_001033582 | -0.0375 | -0.907 |
| ACOT9 | NM_001033583 | 0.013 | -0.547 |
| MTUS2 | NM_001033602 | -0.7245 | -0.3505 |
| BBS9 | NM_001033604 | -0.615 | -1.0105 |
| BBS9 | NM_001033605 | -0.615 | -1.0105 |
| LY9 | NM_001033667 | 0.1435 | -0.6675 |
| CABP1 | NM_001033677 | -0.022 | -0.203 |
| TRPT1 | NM_001033678 | -0.433 | -1.0655 |
| NOP2 | NM_001033714 | -0.476 | -1.5105 |
| ZNF404 | NM_001033719 | -1.216 | -1.4195 |
| ZNF704 | NM_001033723 | -0.257 | -0.3895 |
| VEGFA | NM_001033756 | -0.674 | -0.6745 |
| RPL3 | NM_001033853 | 0.1215 | -0.275 |
| DCLRE1C | NM_001033855 | -1.3045 | -1.667 |
| DCLRE1C | NM_001033857 | -1.3045 | -1.667 |
| DCLRE1C | NM_001033858 | -1.3045 | -1.667 |
| ACADVL | NM_001033859 | 0.2025 | -0.2045 |
| SMAGP | NM_001033873 | 0.0265 | -0.7455 |
| CXCL12 | NM_001033886 | -0.2575 | -1.2595 |
| TRAF5 | NM_001033910 | -0.905 | -1.337 |
| TIAL1 | NM_001033925 | -0.8975 | -1.4415 |
| UBA52 | NM_001033930 | 0.2265 | -0.1225 |
| CALCA | NM_001033952 | 0.0585 | -0.5015 |
| CALCA | NM_001033953 | 0.0585 | -0.5015 |
| RRM2 | NM_001034 | -0.8335 | -0.8185 |
| DMAP1 | NM_001034023 | 0.1275 | -0.9345 |
| DMAP1 | NM_001034024 | 0.154 | -0.8035 |

| | | | |
|---|---|---|---|
| ERP29 | NM_001034025 | 0.269 | -0.6125 |
| HIST2H4B | NM_001034077 | -0.1045 | -0.088 |
| EIF2B4 | NM_001034116 | -0.8805 | -2.379 |
| MCART2 | NM_001034172 | -0.1645 | -0.8525 |
| ALDH1L2 | NM_001034173 | -0.29 | -0.4185 |
| EXOSC9 | NM_001034194 | -1.131 | -2.6995 |
| NHP2 | NM_001034833 | 0.164 | -0.3505 |
| RDM1 | NM_001034836 | -0.173 | -0.392 |
| KCNIP1 | NM_001034837 | -0.2155 | -0.5205 |
| KCNIP1 | NM_001034838 | -0.239 | -0.458 |
| ITPRIPL2 | NM_001034841 | -1.7105 | -3.177 |
| PTCHD3 | NM_001034842 | -0.053 | 0.2775 |
| GALNTL6 | NM_001034845 | 0.2285 | 0.58 |
| FAM134B | NM_001034850 | -0.3635 | -0.695 |
| SMOC1 | NM_001034852 | -0.058 | -0.889 |
| RPGR | NM_001034853 | -1.108 | -1.5155 |
| KCNIP3 | NM_001034914 | -0.201 | -0.671 |
| ESRP1 | NM_001034915 | 0.333 | -0.0835 |
| SORBS1 | NM_001034954 | -0.6285 | -1.6185 |
| SORBS1 | NM_001034955 | -0.3905 | -0.84 |
| SORBS1 | NM_001034956 | -0.3905 | -0.84 |
| SORBS1 | NM_001034957 | -0.3905 | -0.84 |
| RPL14 | NM_001034996 | -0.0915 | -0.2095 |
| RYR2 | NM_001035 | 0.019 | -0.718 |
| KCNIP4 | NM_001035003 | -0.668 | 0.1505 |
| KCNIP4 | NM_001035004 | -0.668 | 0.1505 |
| C18orf32 | NM_001035005 | -1.0075 | -1.261 |
| RPL17 | NM_001035006 | -0.1785 | -0.184 |
| RGL3 | NM_001035223 | 0.191 | -0.346 |
| RABGAP1L | NM_001035230 | -0.5975 | -0.655 |
| SRA1 | NM_001035235 | -0.064 | -1.308 |
| UFD1L | NM_001035247 | 0.0395 | -0.415 |
| FAM102A | NM_001035254 | -0.4385 | -0.9435 |
| POMC | NM_001035256 | 0.0485 | -0.093 |
| RPL38 | NM_001035258 | 0.155 | -0.237 |
| VPS26A | NM_001035260 | -0.4455 | -1.8035 |
| RPL41 | NM_001035267 | 0.038 | -0.227 |
| BOLA3 | NM_001035505 | -0.485 | -2.5875 |
| AGBL5 | NM_001035507 | 0.2305 | -0.4205 |
| SDHC | NM_001035511 | 0.1615 | -0.5105 |
| SDHC | NM_001035512 | 0.1615 | -0.5105 |
| SDHC | NM_001035513 | 0.1615 | -0.5105 |
| DMKN | NM_001035516 | 0.229 | -0.1485 |

| | | | |
|---|---|---|---|
| GTF3C2 | NM_001035521 | -1.144 | -2.2895 |
| FAM129B | NM_001035534 | -0.5435 | -1.9905 |
| RYR3 | NM_001036 | -0.0145 | -1.067 |
| HHLA3 | NM_001036645 | 0.422 | 0.051 |
| SCN1B | NM_001037 | -0.114 | -0.5 |
| UNKL | NM_001037125 | -0.868 | -1.4625 |
| EXOC4 | NM_001037126 | -0.55 | -0.6115 |
| AGAP1 | NM_001037131 | -0.997 | -1.5385 |
| NRCAM | NM_001037132 | -0.8265 | -0.685 |
| CNTROB | NM_001037144 | -0.3825 | -1.755 |
| CYS1 | NM_001037160 | 0.239 | -0.0525 |
| ACOT6 | NM_001037162 | 0.1315 | -0.0445 |
| C7orf70 | NM_001037163 | -1.2385 | -2.9965 |
| ARL4A | NM_001037164 | -0.918 | -2.4755 |
| FOXK1 | NM_001037165 | -1.146 | -2.5085 |
| ACOT9 | NM_001037171 | 0.013 | -0.547 |
| ACPL2 | NM_001037172 | -1.268 | -2.4135 |
| ARL5A | NM_001037174 | -0.956 | -1.7965 |
| SUSD4 | NM_001037175 | 0.037 | -0.27 |
| C11orf85 | NM_001037225 | -0.378 | -0.005 |
| LOC285033 | NM_001037228 | -0.3065 | -1.0905 |
| ZNF829 | NM_001037232 | 0.0095 | -1.1425 |
| GGPS1 | NM_001037277 | -1.767 | -2.246 |
| PARD6A | NM_001037281 | -0.847 | -1.216 |
| EIF3B | NM_001037283 | -0.047 | -0.533 |
| CMTM5 | NM_001037288 | -0.152 | -0.7815 |
| BCO2 | NM_001037290 | 0.077 | 0.07 |
| PALM2 | NM_001037293 | -0.0065 | -0.687 |
| LPPR5 | NM_001037317 | -1.132 | -1.9615 |
| ECE2 | NM_001037324 | 0.059 | -0.2655 |
| STAU1 | NM_001037328 | -0.7415 | -1.708 |
| CNGA4 | NM_001037329 | -0.1655 | 0.108 |
| C18orf62 | NM_001037331 | -0.274 | -0.2005 |
| CYFIP2 | NM_001037332 | -0.0385 | -0.562 |
| CYFIP2 | NM_001037333 | -0.0385 | -0.562 |
| USP14 | NM_001037334 | -0.734 | -1.408 |
| PRIC285 | NM_001037335 | -0.15 | -0.5925 |
| PDE4B | NM_001037339 | -0.9285 | -1.947 |
| PDE4B | NM_001037340 | -0.9285 | -1.947 |
| PDE4B | NM_001037341 | -0.9285 | -1.947 |
| RUFY3 | NM_001037442 | -1.1765 | -1.148 |
| DYNLL1 | NM_001037494 | -0.274 | -0.403 |
| DYNLL1 | NM_001037495 | -0.274 | -0.403 |

| | | | |
|---|---|---|---|
| DEFB110 | NM_001037497 | -0.5735 | -0.33 |
| DEFB112 | NM_001037498 | 0.444 | -0.1695 |
| DEFB114 | NM_001037499 | 0.345 | -0.2015 |
| DEFB124 | NM_001037500 | 0.0525 | -0.0145 |
| GON4L | NM_001037533 | -1.2575 | -2.5065 |
| SCML1 | NM_001037535 | -0.4205 | -0.1625 |
| SCML1 | NM_001037536 | -0.428 | -0.3135 |
| SCML1 | NM_001037540 | -0.428 | -0.3135 |
| AGPAT3 | NM_001037553 | -0.856 | -1.2755 |
| HEPN1 | NM_001037558 | -0.1285 | -0.079 |
| SCD5 | NM_001037582 | -0.49 | -0.5125 |
| CTLA4 | NM_001037631 | 0.019 | -0.2365 |
| SIL1 | NM_001037633 | -0.2665 | -1.1565 |
| BTF3 | NM_001037637 | -0.2145 | -0.619 |
| PARL | NM_001037639 | -0.4265 | -1.456 |
| GATSL3 | NM_001037666 | -0.0695 | -0.339 |
| DEFB107A | NM_001037668 | -0.128 | 0.083 |
| C12orf74 | NM_001037671 | -0.2655 | -0.393 |
| DEFB113 | NM_001037729 | -0.1565 | -0.561 |
| DEFB115 | NM_001037730 | -0.065 | -0.5205 |
| DEFB116 | NM_001037731 | -0.0195 | -0.8535 |
| DEFB128 | NM_001037732 | -0.3405 | -0.0985 |
| ZNF630 | NM_001037735 | -0.1555 | -0.8255 |
| NPM1 | NM_001037738 | -0.3505 | -0.38 |
| COL28A1 | NM_001037763 | -0.202 | -0.1195 |
| DEFB130 | NM_001037804 | -0.2485 | 0.168 |
| NCKAP5L | NM_001037806 | -0.3345 | -1.4095 |
| EIF3C | NM_001037808 | 0.0845 | -0.68 |
| ZNF284 | NM_001037813 | -1.5445 | -2.31 |
| GAB4 | NM_001037814 | 0.1395 | -0.285 |
| REV1 | NM_001037872 | -0.8115 | -2.483 |
| DIXDC1 | NM_001037954 | -2.3785 | -4.033 |
| ALAS2 | NM_001037967 | -0.359 | -0.387 |
| ALAS2 | NM_001037968 | -0.359 | -0.387 |
| SLC38A10 | NM_001037984 | -0.3355 | -0.5775 |
| SCNN1A | NM_001038 | -0.116 | -0.194 |
| DLX1 | NM_001038493 | -0.032 | -0.0315 |
| MARVELD2 | NM_001038603 | 0.0625 | 0.2025 |
| NARF | NM_001038618 | -0.3915 | -1.5865 |
| B3GALNT1 | NM_001038628 | 0.0155 | -0.871 |
| RSPO1 | NM_001038633 | 0.044 | -0.102 |
| CDC42SE2 | NM_001038702 | -1.5315 | -1.9045 |
| C17orf47 | NM_001038704 | -0.0145 | 0.4645 |

| | | | |
|---|---|---|---|
| GPR149 | NM_001038705 | -0.026 | -0.0635 |
| CDC42SE1 | NM_001038707 | -0.811 | -1.568 |
| SCNN1G | NM_001039 | 0.2915 | -0.1335 |
| LRTM2 | NM_001039029 | -0.0535 | -0.401 |
| PRPS2 | NM_001039091 | -1.203 | -1.6035 |
| TRIM71 | NM_001039111 | -0.543 | -1.15 |
| FER1L6 | NM_001039112 | -0.01 | -0.755 |
| ZNF718 | NM_001039127 | -0.397 | 0.146 |
| ALOX15B | NM_001039130 | 0.24 | -0.1405 |
| ALOX15B | NM_001039131 | 0.24 | -0.1405 |
| ICAM4 | NM_001039132 | -0.2025 | -0.241 |
| C20orf27 | NM_001039140 | 0.13 | -0.64 |
| TRIOBP | NM_001039141 | -0.5405 | -0.9545 |
| RGS21 | NM_001039152 | -0.1875 | -0.194 |
| MRGPRE | NM_001039165 | 0.058 | -0.3465 |
| ATP5J2 | NM_001039178 | 0.2425 | -0.187 |
| TTPAL | NM_001039199 | -0.902 | -1.741 |
| ALG13 | NM_001039210 | -0.6645 | -1.719 |
| ATAD3C | NM_001039211 | 0.032 | -0.6045 |
| CEACAM16 | NM_001039213 | -0.0185 | -0.42 |
| EFEMP1 | NM_001039348 | 0.2345 | 0.3475 |
| EFEMP1 | NM_001039349 | 0.2345 | 0.3475 |
| DPP6 | NM_001039350 | -0.537 | -0.9885 |
| SLC25A29 | NM_001039355 | -0.895 | -2.0295 |
| ZBTB7C | NM_001039360 | -0.044 | -0.4055 |
| PRAMEF10 | NM_001039361 | -0.214 | -0.3155 |
| ATP6V1C2 | NM_001039362 | 0.134 | -0.2445 |
| ATP6V1E1 | NM_001039366 | -1.1765 | -1.0575 |
| ATP6V1E1 | NM_001039367 | -1.1765 | -1.0575 |
| HEPACAM2 | NM_001039372 | 0.185 | 0.0755 |
| KIAA1984 | NM_001039374 | 0.5275 | 0.118 |
| C20orf7 | NM_001039375 | -0.965 | -2.578 |
| C10orf25 | NM_001039380 | 0.168 | -0.587 |
| C9orf68 | NM_001039395 | -0.1 | -0.021 |
| MPEG1 | NM_001039396 | 0.2565 | -0.5025 |
| TBC1D28 | NM_001039397 | 0.101 | 0.0145 |
| ATP6V0B | NM_001039457 | 0.324 | 0.007 |
| ELMO1 | NM_001039459 | -0.1695 | -0.73 |
| HEATR8 | NM_001039464 | 0.1145 | -0.4725 |
| SRSF5 | NM_001039465 | -0.813 | -0.776 |
| MARK2 | NM_001039469 | -0.758 | -2.485 |
| NPRL3 | NM_001039476 | -0.0855 | -0.605 |
| C1orf38 | NM_001039477 | -0.032 | -0.253 |

| | | | |
|---|---|---|---|
| KIAA0317 | NM_001039479 | -0.592 | -2.5525 |
| ETNK1 | NM_001039481 | -0.969 | -1.2885 |
| CD151 | NM_001039490 | -0.2565 | -1.2915 |
| FHL2 | NM_001039492 | -0.5065 | -2.9275 |
| C11orf20 | NM_001039496 | 0.0435 | -0.4955 |
| VWA5B1 | NM_001039500 | -0.34 | -0.2675 |
| PRSS53 | NM_001039503 | -0.1745 | -0.068 |
| SIRPG | NM_001039508 | -0.05 | -0.39 |
| RUSC1-AS1 | NM_001039517 | -0.058 | -0.1945 |
| CHRNA1 | NM_001039523 | -0.018 | -0.4185 |
| SKA1 | NM_001039535 | -1.0645 | -3.4865 |
| MAP2 | NM_001039538 | -1.005 | -1.082 |
| GK5 | NM_001039547 | -1.5575 | -2.0395 |
| KLHL35 | NM_001039548 | -0.324 | -0.6825 |
| DNAJB2 | NM_001039550 | -0.4895 | -0.996 |
| RPS4Y2 | NM_001039567 | 0.084 | -0.096 |
| AP1S3 | NM_001039569 | 0.0405 | -0.7145 |
| KREMEN1 | NM_001039570 | -0.426 | -1.38 |
| SEC14L1 | NM_001039573 | 0.111 | -0.986 |
| KCNC4 | NM_001039574 | -0.0495 | -0.3955 |
| CCNL2 | NM_001039577 | -0.676 | -1.05 |
| MAP9 | NM_001039580 | -0.845 | -3.1585 |
| PNCK | NM_001039582 | 0.171 | -0.203 |
| PICK1 | NM_001039583 | -0.6855 | -0.5985 |
| PICK1 | NM_001039584 | -0.6855 | -0.5985 |
| PTGFR | NM_001039585 | -0.3595 | -1.171 |
| DPH2 | NM_001039589 | -0.5445 | -1.6 |
| USP9X | NM_001039590 | -0.292 | -1.5735 |
| USP9X | NM_001039591 | -0.292 | -1.5735 |
| SPAG8 | NM_001039592 | -0.1515 | -0.1645 |
| IAH1 | NM_001039613 | -0.6465 | -1.473 |
| C15orf59 | NM_001039614 | -0.5045 | -1.5125 |
| SPRED3 | NM_001039616 | -0.309 | -0.675 |
| ZDHHC19 | NM_001039617 | 0.087 | -0.5095 |
| CREBZF | NM_001039618 | -1.9725 | -2.7395 |
| PRMT5 | NM_001039619 | -0.319 | -1.413 |
| ZMYM5 | NM_001039649 | -0.51 | -0.9615 |
| ZMYM5 | NM_001039650 | -0.44 | -1.673 |
| ZNF550 | NM_001039654 | -0.5625 | -1.016 |
| MTL5 | NM_001039656 | -0.734 | -0.2215 |
| IL18BP | NM_001039659 | 0.238 | -0.157 |
| IL18BP | NM_001039660 | -0.38 | -0.2835 |
| TIRAP | NM_001039661 | -0.676 | -0.7195 |

| | | | |
|---|---|---|---|
| TNFRSF25 | NM_001039664 | -0.183 | -0.233 |
| IFFO1 | NM_001039670 | -0.8575 | -1.7445 |
| YIF1B | NM_001039671 | 0.239 | -0.397 |
| YIF1B | NM_001039672 | 0.239 | -0.397 |
| YIF1B | NM_001039673 | 0.239 | -0.397 |
| UNC45A | NM_001039675 | 0.0065 | -0.698 |
| CHTF8 | NM_001039690 | -1.003 | -3.6565 |
| TYW5 | NM_001039693 | -1.0885 | -1.575 |
| SNAPC3 | NM_001039697 | -0.681 | -1.145 |
| OLAH | NM_001039702 | -0.186 | -0.3725 |
| NBPF10 | NM_001039703 | -0.337 | -0.5095 |
| TRO | NM_001039705 | -0.4355 | -1.2345 |
| C7orf63 | NM_001039706 | 0.103 | 0.016 |
| SDCCAG3 | NM_001039707 | -0.7325 | -2.1505 |
| SDCCAG3 | NM_001039708 | -0.705 | -1.3425 |
| CBFA2T2 | NM_001039709 | -0.9095 | -2.0895 |
| DEDD | NM_001039711 | -1.01 | -2.161 |
| DEDD | NM_001039712 | -1.01 | -2.161 |
| DIO1 | NM_001039715 | -0.1625 | -0.1355 |
| DIO1 | NM_001039716 | -0.1625 | -0.1355 |
| C4orf29 | NM_001039717 | -1.359 | -1.9195 |
| NOSTRIN | NM_001039724 | -0.3015 | -0.4185 |
| SLC22A10 | NM_001039752 | 0.042 | -0.8585 |
| EML6 | NM_001039753 | -1.295 | -0.8665 |
| FAM196A | NM_001039762 | -0.0345 | -0.535 |
| TMEM232 | NM_001039763 | -0.4865 | 0.107 |
| TMPPE | NM_001039770 | 0.137 | 0.3945 |
| CBLN3 | NM_001039771 | -0.003 | -0.327 |
| AIM1L | NM_001039775 | 0.264 | -0.095 |
| CCNI2 | NM_001039780 | -0.065 | -0.1245 |
| HRCT1 | NM_001039792 | -0.1775 | -0.4245 |
| CDC42 | NM_001039802 | -0.3145 | -0.3855 |
| CDK20 | NM_001039803 | -0.449 | -1.2395 |
| CHIC1 | NM_001039840 | -0.1435 | -0.212 |
| ARHGAP11B | NM_001039841 | -0.227 | -0.712 |
| C17orf90 | NM_001039842 | -0.862 | -1.3795 |
| ACBD7 | NM_001039844 | -0.3275 | -0.2765 |
| MDH1B | NM_001039845 | -0.233 | -0.945 |
| IZUMO4 | NM_001039846 | 0.294 | -0.044 |
| GPX4 | NM_001039847 | 0.345 | -0.2735 |
| GPX4 | NM_001039848 | 0.345 | -0.2735 |
| C19orf46 | NM_001039876 | 0.008 | -0.286 |
| STRN4 | NM_001039877 | -0.066 | -0.72 |

| | | | |
|---|---|---|---|
| FKRP | NM_001039885 | -1.1795 | -1.837 |
| ZNF808 | NM_001039886 | -0.026 | -0.732 |
| C19orf55 | NM_001039887 | -0.162 | -0.716 |
| ANKRD34A | NM_001039888 | -0.3445 | -0.6895 |
| ZNF674 | NM_001039891 | -0.7745 | -1.489 |
| C15orf56 | NM_001039905 | -0.175 | -0.303 |
| ZNF384 | NM_001039920 | -0.8115 | -2.357 |
| CD79B | NM_001039933 | 0.0665 | -0.1825 |
| CHN2 | NM_001039936 | 0.2065 | 0.1045 |
| INTS6 | NM_001039937 | -2.319 | -3.86 |
| INTS6 | NM_001039938 | -2.319 | -3.86 |
| SGSM1 | NM_001039948 | -0.293 | -0.5015 |
| MESP2 | NM_001039958 | 0.29 | 0.0415 |
| SLC4A8 | NM_001039960 | 1.0945 | 0.3425 |
| GPER | NM_001039966 | 0.005 | -0.5815 |
| PTPN5 | NM_001039970 | -0.1585 | -0.496 |
| FAM83G | NM_001039999 | 0.0485 | -0.2295 |
| SHBG | NM_001040 | -0.0225 | -0.5525 |
| MLLT4 | NM_001040000 | -0.349 | -0.9015 |
| MEOX1 | NM_001040002 | 0.0855 | 0.1905 |
| TCERG1 | NM_001040006 | -1.516 | -2.5565 |
| RSPO4 | NM_001040007 | -0.028 | 0.012 |
| SWI5 | NM_001040011 | -0.938 | -1.99 |
| FAM3C | NM_001040020 | -1.714 | -2.532 |
| CD14 | NM_001040021 | -0.166 | 0.092 |
| SIRPA | NM_001040022 | -0.4105 | -0.587 |
| SIRPA | NM_001040023 | -0.4105 | -0.587 |
| ARL16 | NM_001040025 | -0.363 | -1.3 |
| CD37 | NM_001040031 | 0.5415 | 0.209 |
| C8orf74 | NM_001040032 | -0.0105 | -0.0245 |
| CD53 | NM_001040033 | 0 | -0.4635 |
| CD63 | NM_001040034 | 0.1715 | -0.2165 |
| SPP1 | NM_001040058 | 0.3405 | 0.0675 |
| CD68 | NM_001040059 | -0.088 | 0.0115 |
| SPP1 | NM_001040060 | 0.3405 | 0.0675 |
| C3orf72 | NM_001040061 | -0.041 | 0.102 |
| LOC650293 | NM_001040071 | 0.036 | 0.2295 |
| ENPP2 | NM_001040092 | 0.134 | -0.556 |
| MOSPD3 | NM_001040097 | 0.1455 | -0.4725 |
| SPTSSB | NM_001040100 | 0.1065 | -0.4135 |
| D4S234E | NM_001040101 | -0.266 | -1.21 |
| OPALIN | NM_001040102 | -0.561 | 0.612 |
| OPALIN | NM_001040103 | -0.561 | 0.612 |

| | | | |
|---|---|---|---|
| MUC17 | NM_001040105 | 0.253 | -0.2815 |
| HVCN1 | NM_001040107 | 0.058 | -0.405 |
| MLH3 | NM_001040108 | -0.898 | -0.9805 |
| MLN | NM_001040109 | 0.1215 | -0.576 |
| NRF1 | NM_001040110 | -0.898 | -2.977 |
| MYH11 | NM_001040113 | -0.05 | -0.205 |
| MYH11 | NM_001040114 | -0.05 | -0.205 |
| ARAP1 | NM_001040118 | -0.3265 | -0.8935 |
| PQLC2 | NM_001040125 | -0.583 | -0.741 |
| PQLC2 | NM_001040126 | -0.583 | -0.741 |
| SPINK13 | NM_001040129 | 0 | -0.428 |
| PALM | NM_001040134 | 0.3305 | 0.3745 |
| SCN2A | NM_001040142 | 0.1875 | 0.159 |
| SCN2A | NM_001040143 | 0.1875 | 0.159 |
| SERPINB7 | NM_001040147 | -0.385 | 0.3165 |
| SCN3B | NM_001040151 | -0.656 | -0.9335 |
| PEG10 | NM_001040152 | -0.4375 | 0.29 |
| SLAIN1 | NM_001040153 | -1.978 | -4.0495 |
| H2AFY | NM_001040158 | 0.04 | -0.4995 |
| SPOCK3 | NM_001040159 | -0.615 | -0.4285 |
| C16orf13 | NM_001040160 | 0.23 | -0.2425 |
| C16orf13 | NM_001040161 | 0.23 | -0.2425 |
| C16orf13 | NM_001040162 | 0.23 | -0.2425 |
| C16orf13 | NM_001040165 | 0.23 | -0.2425 |
| LFNG | NM_001040167 | -0.3805 | -1.4655 |
| LFNG | NM_001040168 | -0.3805 | -1.4655 |
| HTR4 | NM_001040172 | 0.001 | -0.051 |
| AKR1E2 | NM_001040177 | -0.378 | -1.065 |
| MCHR2 | NM_001040179 | 0.2015 | -0.241 |
| CLDND1 | NM_001040181 | -1.2655 | -1.8485 |
| CLDND1 | NM_001040182 | -1.2655 | -1.8485 |
| CLDND1 | NM_001040183 | -1.2655 | -1.8485 |
| ZNF765 | NM_001040185 | -0.3815 | -1.209 |
| DNAJC28 | NM_001040192 | -0.789 | -0.0755 |
| AGTRAP | NM_001040194 | -0.3905 | -0.525 |
| AGTRAP | NM_001040195 | -0.3905 | -0.525 |
| AGTRAP | NM_001040196 | -0.3905 | -0.525 |
| AGTRAP | NM_001040197 | -0.3905 | -0.525 |
| CLDND1 | NM_001040199 | -1.2575 | -1.902 |
| CLDND1 | NM_001040200 | -1.2655 | -1.8485 |
| PAQR3 | NM_001040202 | -1.5195 | -1.3645 |
| NKAIN2 | NM_001040214 | -0.2975 | -0.4165 |
| FAM63A | NM_001040217 | -0.6285 | -0.816 |

| | | | |
|---|---|---|---|
| DCLK2 | NM_001040260 | -0.185 | -1.0295 |
| DCLK2 | NM_001040261 | -0.185 | -1.0295 |
| ADAMTSL1 | NM_001040272 | -0.2905 | -1.0215 |
| TYSND1 | NM_001040273 | -1.1575 | -1.819 |
| SYCP2L | NM_001040274 | -1.066 | -0.5265 |
| ESR2 | NM_001040275 | -0.0185 | -0.086 |
| ESR2 | NM_001040276 | -0.0185 | -0.086 |
| CD83 | NM_001040280 | -0.552 | -1.466 |
| DMRTC2 | NM_001040283 | -0.2535 | -0.537 |
| PAPD5 | NM_001040284 | -1.4275 | -3.5945 |
| PAPD5 | NM_001040285 | -1.4275 | -3.5945 |
| DCUN1D4 | NM_001040402 | -1.0695 | -1.9695 |
| C9orf131 | NM_001040410 | 0.3055 | -0.0195 |
| PRDM15 | NM_001040424 | -1.0505 | -2.1465 |
| U2AF1L4 | NM_001040425 | -0.3245 | -0.8565 |
| HAGH | NM_001040427 | -0.167 | -0.805 |
| SPATA7 | NM_001040428 | -0.373 | -0.3375 |
| PCDH17 | NM_001040429 | -0.7445 | -0.9225 |
| CCDC56 | NM_001040431 | -0.6805 | -1.077 |
| ZCWPW2 | NM_001040432 | -0.4455 | -0.369 |
| SPINK9 | NM_001040433 | -0.2395 | 0.041 |
| YARS2 | NM_001040436 | -0.785 | -2.6345 |
| C6orf48 | NM_001040437 | -0.2465 | -0.8445 |
| C6orf48 | NM_001040438 | -0.3275 | -0.8595 |
| MAPK8IP3 | NM_001040439 | -0.16 | -0.589 |
| CCDC112 | NM_001040440 | -0.766 | -2.538 |
| ZBTB8A | NM_001040441 | -1.176 | -2.454 |
| PHF11 | NM_001040443 | -0.001 | -0.717 |
| PHF11 | NM_001040444 | -0.001 | -0.717 |
| ASB1 | NM_001040445 | -0.7085 | -1.781 |
| MTMR12 | NM_001040446 | -1.623 | -4.0795 |
| DEFB131 | NM_001040448 | 0.3035 | 0.1615 |
| FAM63B | NM_001040450 | -1.2095 | -2.0045 |
| RUFY1 | NM_001040451 | -0.7755 | -1.9895 |
| RUFY1 | NM_001040452 | -0.7755 | -1.9895 |
| FAM63B | NM_001040453 | -1.2095 | -2.0045 |
| SLC26A6 | NM_001040454 | -0.228 | -0.785 |
| SIDT2 | NM_001040455 | -0.0285 | -0.288 |
| RHBDD2 | NM_001040457 | -0.2615 | -1.2195 |
| ERAP1 | NM_001040458 | -0.4265 | -0.664 |
| BTNL8 | NM_001040462 | -0.0105 | 0.1015 |
| TMEM70 | NM_001040613 | -1.2235 | -4.072 |
| LINS | NM_001040616 | -1.548 | -3.044 |

| | | | |
|---|---|---|---|
| ATF3 | NM_001040619 | -0.5615 | 1.166 |
| NCALD | NM_001040624 | -0.857 | -2.6 |
| NCALD | NM_001040625 | -1.0055 | -1.9685 |
| NCALD | NM_001040626 | -1.0055 | -1.9685 |
| NCALD | NM_001040627 | -1.0055 | -1.9685 |
| NCALD | NM_001040628 | -1.0055 | -1.9685 |
| NCALD | NM_001040629 | -1.0055 | -1.9685 |
| NCALD | NM_001040630 | -1.0055 | -1.9685 |
| PRKAG2 | NM_001040633 | -0.789 | -1.3805 |
| CRCP | NM_001040647 | -0.735 | -1.8625 |
| CRCP | NM_001040648 | -0.735 | -1.8625 |
| ACP1 | NM_001040649 | -0.5725 | -0.618 |
| CD3D | NM_001040651 | -0.091 | -0.1825 |
| ZXDC | NM_001040653 | -1.6455 | -2.1275 |
| TTC23 | NM_001040655 | -1.3155 | -1.6855 |
| TTC23 | NM_001040656 | -0.5295 | -1.3105 |
| TTC23 | NM_001040657 | -0.5295 | -1.3105 |
| TTC23 | NM_001040658 | -0.5295 | -1.3105 |
| TTC23 | NM_001040659 | -0.5295 | -1.3105 |
| TTC23 | NM_001040660 | -0.5295 | -1.3105 |
| SLC29A4 | NM_001040661 | -0.444 | -1.0065 |
| STEAP2 | NM_001040665 | 0.2585 | -0.0985 |
| STEAP2 | NM_001040666 | 0.2585 | -0.0985 |
| HSF4 | NM_001040667 | -0.302 | -0.6445 |
| INCENP | NM_001040694 | -1.0225 | -1.862 |
| UEVLD | NM_001040697 | -0.1565 | -1.233 |
| FUT6 | NM_001040701 | -0.058 | -0.0995 |
| DEFB106B | NM_001040704 | 0.2255 | 0.0725 |
| HEY1 | NM_001040708 | -0.309 | 0.649 |
| SYPL2 | NM_001040709 | 0.1525 | -0.201 |
| C2orf84 | NM_001040710 | -0.398 | 0.0665 |
| PTPRD | NM_001040712 | -0.291 | -1.5925 |
| KIAA0895L | NM_001040715 | -0.5375 | -0.853 |
| PC | NM_001040716 | 0.171 | -0.143 |
| HIST2H2AA4 | NM_001040874 | 0.5485 | 0.231 |
| GDAP1 | NM_001040875 | -0.695 | -0.7505 |
| ABCE1 | NM_001040876 | -0.9235 | -2.7145 |
| SI | NM_001041 | -0.022 | -0.144 |
| SLC2A4 | NM_001042 | -0.0015 | -0.5365 |
| G6PD | NM_001042351 | 0.1535 | -0.269 |
| FAM110A | NM_001042353 | -0.599 | -2.4875 |
| CYLD | NM_001042355 | -0.745 | -1.417 |
| C15orf60 | NM_001042367 | 0.206 | 0.055 |

| | | | |
|---|---|---|---|
| RALGDS | NM_001042368 | -0.5245 | -1.1245 |
| TROVE2 | NM_001042369 | -0.8535 | -1.218 |
| TROVE2 | NM_001042370 | -0.8535 | -1.218 |
| PGP | NM_001042371 | -0.635 | -1.225 |
| INS-IGF2 | NM_001042376 | -0.1345 | -0.5825 |
| CEP63 | NM_001042383 | -0.6295 | -1.7645 |
| CEP63 | NM_001042384 | -0.6295 | -1.7645 |
| PREPL | NM_001042385 | -0.8635 | -0.928 |
| PREPL | NM_001042386 | -0.8635 | -0.928 |
| PPP4R1 | NM_001042388 | -1.1755 | -1.829 |
| PTPN20A | NM_001042389 | -0.114 | -0.0495 |
| CCDC41 | NM_001042399 | -0.5175 | -2.417 |
| CEP63 | NM_001042400 | -0.6295 | -1.7645 |
| NAAA | NM_001042402 | -0.23 | -0.5155 |
| USP44 | NM_001042403 | -0.121 | -2.6125 |
| CEP170 | NM_001042404 | 0.4865 | -0.928 |
| CEP170 | NM_001042405 | 0.4865 | -0.928 |
| HMGCLL1 | NM_001042406 | 0.2295 | -0.454 |
| ANKZF1 | NM_001042410 | -0.7855 | -1.479 |
| CYLD | NM_001042412 | -0.745 | -1.417 |
| GLIS3 | NM_001042413 | 0.036 | -0.011 |
| PSPC1 | NM_001042414 | 0.0525 | -0.791 |
| RUFY2 | NM_001042417 | -0.706 | -2.015 |
| SLC16A3 | NM_001042422 | 0.023 | -0.6925 |
| SLC16A3 | NM_001042423 | 0.023 | -0.6925 |
| WHSC1 | NM_001042424 | -0.668 | -1.475 |
| TFAP2A | NM_001042425 | 0.0025 | -0.092 |
| CENPA | NM_001042426 | -1.5115 | -4.3695 |
| C10orf53 | NM_001042427 | -0.053 | -0.2365 |
| FAM98B | NM_001042429 | -0.579 | -0.465 |
| CLN3 | NM_001042432 | -0.39 | -1.5685 |
| ST3GAL5 | NM_001042437 | -0.6065 | -2.274 |
| CAST | NM_001042440 | -0.609 | -1.1605 |
| MST4 | NM_001042452 | -1.014 | -1.011 |
| MST4 | NM_001042453 | -0.958 | -0.739 |
| TGFB1I1 | NM_001042454 | -0.545 | -1.7215 |
| FILIP1L | NM_001042459 | 0.005 | -0.454 |
| TRAPPC5 | NM_001042461 | -0.4215 | -0.8795 |
| TRAPPC5 | NM_001042462 | -0.4215 | -0.8795 |
| TMEM80 | NM_001042463 | -0.3835 | -1.0875 |
| PSAP | NM_001042465 | 0.31 | -0.2785 |
| PSAP | NM_001042466 | 0.31 | -0.2785 |
| MLPH | NM_001042467 | 0.2095 | -0.456 |

| | | | |
|---|---|---|---|
| SUMF2 | NM_001042469 | 0.2845 | -0.377 |
| SUMF2 | NM_001042470 | 0.3035 | -0.3715 |
| ABHD12 | NM_001042472 | -0.1675 | -0.8155 |
| ACBD5 | NM_001042473 | -1.221 | -1.9905 |
| C6orf204 | NM_001042475 | -0.733 | -2.4905 |
| CARHSP1 | NM_001042476 | 0.125 | -0.8365 |
| AJAP1 | NM_001042478 | 0.2065 | -0.2095 |
| GEMIN8 | NM_001042479 | -0.436 | -0.917 |
| GEMIN8 | NM_001042480 | -0.436 | -0.917 |
| FRMD6 | NM_001042481 | -1.082 | -4.15 |
| TPK1 | NM_001042482 | 0.3035 | -0.326 |
| NUPR1 | NM_001042483 | 0.222 | -0.0375 |
| DLGAP4 | NM_001042486 | -0.685 | -1.695 |
| NF1 | NM_001042492 | -0.6645 | -1.328 |
| C6orf162 | NM_001042493 | -0.46 | -1.2885 |
| SLC12A6 | NM_001042494 | -0.356 | -0.688 |
| SLC12A6 | NM_001042495 | -0.356 | -0.688 |
| SLC12A6 | NM_001042496 | -0.946 | -1.067 |
| SLC12A6 | NM_001042497 | -0.356 | -0.688 |
| SLC35A2 | NM_001042498 | -1.1725 | -2.372 |
| DEFA1B | NM_001042500 | 0.0085 | -0.283 |
| PABPC1L2B | NM_001042506 | 0.066 | -0.083 |
| LGALS7B | NM_001042507 | 0.0545 | 0.3975 |
| ZNF706 | NM_001042510 | -0.6945 | -2.1645 |
| ZNF706 | NM_001042511 | -0.6945 | -2.1645 |
| DIAPH3 | NM_001042517 | -1.074 | -2.7205 |
| C2orf88 | NM_001042519 | -0.243 | -0.498 |
| C2orf88 | NM_001042520 | 0.3055 | -0.476 |
| C2orf88 | NM_001042521 | 0.3055 | -0.476 |
| SPRED3 | NM_001042522 | 0.1735 | -0.1635 |
| COASY | NM_001042529 | -0.0625 | -0.8425 |
| COASY | NM_001042530 | -0.0625 | -0.8425 |
| COASY | NM_001042531 | -0.0625 | -0.8425 |
| COASY | NM_001042532 | -0.0625 | -0.8425 |
| MINA | NM_001042533 | -0.971 | -2.2405 |
| AGAP3 | NM_001042535 | 0.1235 | -0.532 |
| INSC | NM_001042536 | 0.0705 | -0.524 |
| SLC9A6 | NM_001042537 | -1.075 | -2.2795 |
| MAZ | NM_001042539 | 0.0975 | -0.1115 |
| NCBP2 | NM_001042540 | -0.7165 | -0.402 |
| GLRA3 | NM_001042543 | 0.2565 | -0.1075 |
| LTBP4 | NM_001042544 | -0.3845 | -0.861 |
| LTBP4 | NM_001042545 | -0.3845 | -0.861 |

| | | | |
|---|---|---|---|
| ATPAF1 | NM_001042546 | -0.403 | -0.375 |
| FEZ2 | NM_001042548 | -0.769 | -1.257 |
| NSL1 | NM_001042549 | -1.1745 | -2.4495 |
| SMC2 | NM_001042550 | 0.068 | -1.8885 |
| SMC2 | NM_001042551 | 0.068 | -1.8885 |
| TATDN3 | NM_001042552 | -1.1835 | -1.194 |
| TATDN3 | NM_001042553 | -1.1835 | -1.194 |
| FRS2 | NM_001042555 | -1.1355 | -2.8635 |
| EIF4G2 | NM_001042559 | -0.829 | -0.506 |
| CHD2 | NM_001042572 | -1.271 | -3.384 |
| ENGASE | NM_001042573 | -0.578 | -1.2825 |
| CRTC3 | NM_001042574 | -0.2145 | -0.6955 |
| TMPRSS7 | NM_001042575 | -0.056 | -0.2755 |
| RRBP1 | NM_001042576 | -0.296 | -0.8495 |
| SNUPN | NM_001042581 | -0.6075 | -1.559 |
| CD1E | NM_001042583 | 0.0335 | -0.3035 |
| CD1E | NM_001042584 | 0.0335 | -0.3035 |
| CD1E | NM_001042585 | 0.0335 | -0.3035 |
| CD1E | NM_001042586 | 0.0335 | -0.3035 |
| CD1E | NM_001042587 | 0.0335 | -0.3035 |
| SNUPN | NM_001042588 | -0.6075 | -1.559 |
| TMEM8B | NM_001042589 | 0.2775 | -0.7995 |
| TMEM8B | NM_001042590 | 0.2775 | -0.7995 |
| POT1 | NM_001042594 | -1.327 | -2.0685 |
| TMEM91 | NM_001042595 | 0.2265 | -0.65 |
| ERBB4 | NM_001042599 | -0.1035 | 0.122 |
| MAP4K1 | NM_001042600 | 0.19 | -0.238 |
| TTC14 | NM_001042601 | -0.826 | -1.667 |
| KDM5A | NM_001042603 | -1.061 | -1.8615 |
| XRN1 | NM_001042604 | -1.3255 | -2.6375 |
| DBNDD1 | NM_001042610 | -0.6185 | -1.1715 |
| PIGY | NM_001042616 | -0.6355 | -1.19 |
| PARP2 | NM_001042618 | -0.9395 | -1.605 |
| CAPSL | NM_001042625 | 0.09 | -0.561 |
| SDHAF1 | NM_001042631 | -1.208 | -3.2225 |
| SNX21 | NM_001042632 | -0.511 | -0.79 |
| SNX21 | NM_001042633 | -0.511 | -0.79 |
| NGDN | NM_001042635 | -1.4705 | -2.6565 |
| TRAK1 | NM_001042646 | -0.093 | -0.963 |
| PLEKHG5 | NM_001042663 | -0.023 | -0.397 |
| PLEKHG5 | NM_001042664 | -0.055 | -0.5065 |
| PLEKHG5 | NM_001042665 | -0.055 | -0.5065 |
| ARHGAP24 | NM_001042669 | -0.2485 | -0.394 |

| | | | |
|---|---|---|---|
| RHOC | NM_001042678 | 0.094 | -0.803 |
| RHOC | NM_001042679 | 0.094 | -0.803 |
| C19orf28 | NM_001042680 | -0.6325 | -0.999 |
| RERE | NM_001042681 | -0.1985 | -1.225 |
| RERE | NM_001042682 | -0.1985 | -1.225 |
| SHPRH | NM_001042683 | -1.0245 | -2.5705 |
| LGALS9B | NM_001042685 | -0.062 | -0.094 |
| C4orf44 | NM_001042690 | -0.194 | -0.4915 |
| FAM159A | NM_001042693 | 0.03 | -0.4695 |
| ZSWIM7 | NM_001042697 | -0.6415 | -0.7395 |
| ZSWIM7 | NM_001042698 | -0.6415 | -0.7395 |
| DFNB59 | NM_001042702 | -1.0285 | -0.5055 |
| CLCNKA | NM_001042704 | -0.1665 | -0.2825 |
| IQCJ | NM_001042705 | -0.0535 | -0.0445 |
| IQCJ | NM_001042706 | -0.15 | -0.0035 |
| GTPBP10 | NM_001042717 | -0.6835 | -1.3085 |
| RYR1 | NM_001042723 | -0.1625 | -0.191 |
| PVRL2 | NM_001042724 | -0.4665 | -0.3275 |
| RARG | NM_001042728 | -0.129 | -0.906 |
| FGR | NM_001042729 | -0.091 | -0.2515 |
| SEC24B | NM_001042734 | -1.174 | -2.9535 |
| FGR | NM_001042747 | -0.0205 | 0.0015 |
| STAG2 | NM_001042749 | -0.396 | -2.1605 |
| STAG2 | NM_001042750 | -0.396 | -2.1605 |
| STAG2 | NM_001042751 | -0.396 | -2.1605 |
| SRGAP2 | NM_001042758 | -0.146 | -0.501 |
| LCK | NM_001042771 | -0.47 | -0.194 |
| TNNT3 | NM_001042780 | -0.0795 | 0.0055 |
| TNNT3 | NM_001042781 | -0.0795 | 0.0055 |
| TNNT3 | NM_001042782 | -0.0795 | 0.0055 |
| CCDC158 | NM_001042784 | 0.513 | -0.1335 |
| SLC6A2 | NM_001043 | -0.9595 | -0.931 |
| METTL12 | NM_001043229 | 0.963 | 1.652 |
| TPM3 | NM_001043351 | -0.2085 | -0.625 |
| TPM3 | NM_001043352 | -0.2085 | -0.625 |
| TPM3 | NM_001043353 | -0.2085 | -0.625 |
| SLC6A3 | NM_001044 | -0.4005 | -0.11 |
| MSMP | NM_001044264 | -0.066 | -0.4665 |
| SMAP1 | NM_001044305 | -0.7115 | -1.1265 |
| FAM69C | NM_001044369 | 0.0515 | -0.4825 |
| MPPED1 | NM_001044370 | -0.0595 | -0.8455 |
| ZNF557 | NM_001044387 | -1.3795 | -1.586 |
| ZNF557 | NM_001044388 | -1.3795 | -1.586 |

| | | | |
|---|---|---|---|
| MUC1 | NM_001044390 | -0.1105 | -0.321 |
| MUC1 | NM_001044391 | -0.1105 | -0.321 |
| MUC1 | NM_001044392 | -0.1105 | -0.321 |
| MUC1 | NM_001044393 | -0.1105 | -0.321 |
| CSNK1G3 | NM_001044722 | -0.839 | -2.3445 |
| CSNK1G3 | NM_001044723 | -0.839 | -2.3445 |
| SLC6A4 | NM_001045 | 0.075 | -0.536 |
| WDR38 | NM_001045476 | -0.029 | -0.357 |
| FAM22G | NM_001045477 | -0.2065 | -0.462 |
| S100A7L2 | NM_001045479 | -0.3835 | -0.1695 |
| SLA | NM_001045556 | 0.084 | -0.2425 |
| SLA | NM_001045557 | 0.084 | -0.2425 |
| SLC12A2 | NM_001046 | -1.4425 | -1.9465 |
| SRD5A1 | NM_001047 | -0.1115 | -0.787 |
| NET1 | NM_001047160 | -1.103 | -1.8655 |
| DPH3 | NM_001047434 | -0.619 | -1.8035 |
| NBPF7 | NM_001047980 | -0.059 | -0.3005 |
| SST | NM_001048 | 0.521 | -0.078 |
| STIL | NM_001048166 | -1.6015 | -3.456 |
| MUTYH | NM_001048173 | -0.603 | -1.328 |
| PHACTR4 | NM_001048183 | -0.907 | -1.9745 |
| RCC1 | NM_001048194 | -0.3655 | -1.5915 |
| RCC1 | NM_001048195 | -0.3655 | -1.5915 |
| RCC1 | NM_001048199 | -0.3655 | -1.5915 |
| HIPK3 | NM_001048200 | -0.1805 | -0.3865 |
| UHRF1 | NM_001048201 | -0.588 | -1.676 |
| REC8 | NM_001048205 | -0.3225 | -0.742 |
| NTM | NM_001048209 | -0.022 | -0.5335 |
| CLCC1 | NM_001048210 | -0.8185 | -1.2665 |
| CEMP1 | NM_001048212 | -0.083 | -0.225 |
| SCYL1 | NM_001048218 | -0.2065 | -1.1025 |
| DBNDD2 | NM_001048223 | -0.3305 | -1.387 |
| ADORA1 | NM_001048230 | -0.0715 | -0.722 |
| UBL5 | NM_001048241 | 0.2965 | -0.218 |
| C5orf43 | NM_001048249 | -1.481 | -3.3185 |
| CTXN3 | NM_001048252 | -0.155 | -0.1465 |
| C9orf116 | NM_001048265 | -0.1445 | -0.5975 |
| SSTR1 | NM_001049 | 0.012 | -0.0645 |
| SSTR2 | NM_001050 | -0.493 | -2.7395 |
| SSTR3 | NM_001051 | 0.017 | -0.314 |
| SSTR4 | NM_001052 | -0.3765 | -0.751 |
| SSTR5 | NM_001053 | 0.738 | -0.2825 |
| SULT1C2 | NM_001056 | 0.141 | -0.333 |

| | | | |
|---|---|---|---|
| TACR2 | NM_001057 | 0.0595 | -0.1545 |
| TACR1 | NM_001058 | -0.1275 | -0.196 |
| TACR3 | NM_001059 | -0.033 | -0.533 |
| TBXA2R | NM_001060 | 0.088 | -0.4375 |
| TBXAS1 | NM_001061 | 0.099 | -0.327 |
| TCN1 | NM_001062 | 0.131 | -0.182 |
| TF | NM_001063 | 0.202 | -0.1305 |
| TKT | NM_001064 | 0.2215 | -0.3755 |
| TNFRSF1A | NM_001065 | -0.8165 | -1.6265 |
| TNFRSF1B | NM_001066 | -0.2635 | -0.1685 |
| TOP2A | NM_001067 | -0.0925 | -1.4645 |
| TOP2B | NM_001068 | -0.087 | -1.006 |
| TUBB2A | NM_001069 | 0.389 | 0.0965 |
| TUBG1 | NM_001070 | -0.5485 | -2.245 |
| TYMS | NM_001071 | -0.144 | -0.4415 |
| MZT1 | NM_001071775 | -0.7765 | -0.9955 |
| UGT1A6 | NM_001072 | 0.4505 | 0.043 |
| UGT2B11 | NM_001073 | 0.528 | 0.4605 |
| UGT2B7 | NM_001074 | -0.081 | -0.616 |
| UGT2B10 | NM_001075 | -0.319 | -0.1645 |
| MOCS1 | NM_001075098 | -0.2925 | -1.621 |
| SKAP1 | NM_001075099 | 0.0145 | -0.096 |
| UGT2B15 | NM_001076 | 0.166 | -0.1115 |
| ACSS2 | NM_001076552 | 0.1255 | -0.623 |
| ZNF626 | NM_001076675 | -0.4565 | -0.362 |
| CLTA | NM_001076677 | -0.0495 | -0.5025 |
| ZNF493 | NM_001076678 | -1.129 | -2.08 |
| C17orf108 | NM_001076680 | -0.615 | -0.6195 |
| NCAM1 | NM_001076682 | -0.7595 | -0.572 |
| UBTF | NM_001076683 | -0.6445 | -1.406 |
| UBTF | NM_001076684 | -0.4465 | -1.622 |
| FAM107A | NM_001076778 | 0.159 | -0.139 |
| PKD1L2 | NM_001076780 | 0.2125 | 0.014 |
| ZNF391 | NM_001076781 | -0.279 | -0.621 |
| SLC7A6 | NM_001076785 | -0.861 | -1.1865 |
| QSER1 | NM_001076786 | -0.922 | -3.322 |
| TP53I11 | NM_001076787 | -0.6055 | -1.23 |
| UGT2B17 | NM_001077 | 0.104 | 0.2495 |
| METTL9 | NM_001077180 | -0.547 | -0.784 |
| CDC14B | NM_001077181 | -0.3795 | -1.1515 |
| FSCN2 | NM_001077182 | 0.0055 | -0.263 |
| TSC2 | NM_001077183 | 0.0595 | -0.401 |
| MYH14 | NM_001077186 | -0.1625 | 0.0935 |

| | | | |
|---|---|---|---|
| HS6ST2 | NM_001077188 | -0.872 | -0.7075 |
| GPBAR1 | NM_001077191 | 0.1155 | -0.4315 |
| ZNF436 | NM_001077195 | -1.4275 | -1.886 |
| PDE11A | NM_001077196 | 0.1995 | -0.4175 |
| PDE11A | NM_001077197 | 0.1995 | -0.4175 |
| ATG9A | NM_001077198 | -0.385 | -1.16 |
| SREK1 | NM_001077199 | -0.4285 | -1.9685 |
| SENP7 | NM_001077203 | -0.981 | -1.1655 |
| SEC31A | NM_001077206 | -0.6105 | -0.4885 |
| SEC31A | NM_001077207 | -0.8125 | -0.954 |
| SEC31A | NM_001077208 | -0.6105 | -0.4885 |
| SPPL2B | NM_001077238 | -0.3725 | -0.6545 |
| RNF214 | NM_001077239 | -1.114 | -2.7785 |
| SLC25A45 | NM_001077241 | -0.099 | -0.9425 |
| DEPDC7 | NM_001077242 | -0.401 | -2.366 |
| GRIA4 | NM_001077243 | -0.843 | -0.705 |
| GRIA4 | NM_001077244 | -0.663 | -0.1215 |
| NCOR2 | NM_001077261 | 0.005 | -0.558 |
| UBXN11 | NM_001077262 | 0.113 | -0.131 |
| TMPRSS13 | NM_001077263 | -0.0495 | 0.0885 |
| ZFYVE19 | NM_001077268 | -0.3105 | -1.12 |
| WIPF1 | NM_001077269 | -1.691 | -2.723 |
| ZNF682 | NM_001077349 | -0.16 | -1.1405 |
| NPRL3 | NM_001077350 | -0.013 | -0.3335 |
| RBM23 | NM_001077351 | -1.154 | -2.968 |
| RBM23 | NM_001077352 | -1.154 | -2.968 |
| PDE11A | NM_001077358 | 0.1995 | -0.4175 |
| POMT1 | NM_001077365 | -0.875 | -2.2155 |
| POMT1 | NM_001077366 | -0.4295 | -0.8615 |
| DPH5 | NM_001077394 | -1.2555 | -1.7365 |
| DPH5 | NM_001077395 | -1.2555 | -1.7365 |
| IRF2BP2 | NM_001077397 | -0.8925 | -1.7395 |
| PNKD | NM_001077399 | 0.074 | -0.7585 |
| ACVRL1 | NM_001077401 | -0.417 | -0.5285 |
| CRELD1 | NM_001077415 | -0.554 | -0.848 |
| TMEM231 | NM_001077416 | -0.303 | -1.359 |
| TMEM231 | NM_001077418 | -0.303 | -1.359 |
| TMEM231 | NM_001077419 | -0.303 | -1.359 |
| ZNF586 | NM_001077426 | -1.2695 | -1.7975 |
| LYPD1 | NM_001077427 | -0.1915 | -1.6205 |
| BCLAF1 | NM_001077440 | 0.361 | -0.795 |
| BCLAF1 | NM_001077441 | 0.361 | -0.795 |
| HNRNPC | NM_001077442 | -0.1685 | -0.3245 |

| | | | |
|---|---|---|---|
| HNRNPC | NM_001077443 | -0.1685 | -0.3245 |
| PHF16 | NM_001077445 | -1.2335 | -2.48 |
| TSEN34 | NM_001077446 | -0.2055 | -1.2655 |
| PPCS | NM_001077447 | -1.1855 | -2.6475 |
| NR1I3 | NM_001077469 | 0.0515 | 0.029 |
| NR1I3 | NM_001077470 | 0.0515 | 0.029 |
| NR1I3 | NM_001077471 | 0.0515 | 0.029 |
| NR1I3 | NM_001077472 | 0.0515 | 0.029 |
| NR1I3 | NM_001077473 | 0.0515 | 0.029 |
| NR1I3 | NM_001077474 | 0.0515 | 0.029 |
| NR1I3 | NM_001077475 | 0.0515 | 0.029 |
| NR1I3 | NM_001077476 | 0.0515 | 0.029 |
| NR1I3 | NM_001077477 | 0.0515 | 0.029 |
| NR1I3 | NM_001077478 | 0.0515 | 0.029 |
| NR1I3 | NM_001077479 | 0.0515 | 0.029 |
| NR1I3 | NM_001077480 | 0.0515 | 0.029 |
| NR1I3 | NM_001077481 | 0.0515 | 0.029 |
| NR1I3 | NM_001077482 | 0.0515 | 0.029 |
| SLC38A1 | NM_001077484 | -1.147 | -0.9795 |
| GNAS | NM_001077488 | 0.0835 | -0.2445 |
| GNAS | NM_001077489 | 0.0835 | -0.2445 |
| GNAS | NM_001077490 | 0.0835 | -0.2445 |
| NFKB2 | NM_001077493 | -0.201 | -0.837 |
| PRR3 | NM_001077497 | -1.292 | -3.2065 |
| C17orf63 | NM_001077498 | -0.8405 | -2.8985 |
| KLK10 | NM_001077500 | 0.0135 | -0.1875 |
| TCF19 | NM_001077511 | -0.791 | -1.8185 |
| AP3D1 | NM_001077523 | -0.206 | -1.224 |
| MTMR14 | NM_001077525 | -0.1115 | -1.78 |
| MTMR14 | NM_001077526 | -0.1115 | -1.78 |
| JRK | NM_001077527 | -0.3155 | -1.278 |
| EXOC3L4 | NM_001077594 | 0.1175 | -0.218 |
| UBXN2B | NM_001077619 | -1.33 | -1.279 |
| PRCD | NM_001077620 | -0.012 | -0.3815 |
| VPS37D | NM_001077621 | -0.254 | -0.926 |
| ZNF846 | NM_001077624 | -0.034 | -0.4855 |
| APH1A | NM_001077628 | -0.174 | -0.71 |
| WTH3DI | NM_001077637 | 0.01 | -0.384 |
| FAM55D | NM_001077639 | 0.0295 | -0.083 |
| TBX20 | NM_001077653 | 0.322 | -0.0875 |
| TNFAIP8 | NM_001077654 | -1.058 | -2.854 |
| C3orf43 | NM_001077657 | -0.347 | -0.1245 |
| URGCP | NM_001077663 | -0.426 | -2.246 |

| | | | |
|---|---|---|---|
| URGCP | NM_001077664 | -0.709 | -2.1165 |
| AGAP7 | NM_001077685 | -1.1195 | -2.1465 |
| ALG9 | NM_001077690 | -0.122 | -0.8265 |
| ALG9 | NM_001077691 | -1.1955 | -1.534 |
| ALG9 | NM_001077692 | -0.122 | -0.8265 |
| ECSCR | NM_001077693 | -0.1845 | -0.1695 |
| MIER1 | NM_001077700 | -0.8485 | -3.324 |
| MIER1 | NM_001077701 | -0.8485 | -3.324 |
| MIER1 | NM_001077702 | -0.8485 | -3.324 |
| MIER1 | NM_001077703 | -0.8485 | -3.324 |
| MIER1 | NM_001077704 | -0.8485 | -3.324 |
| ECT2L | NM_001077706 | -0.1245 | -0.4175 |
| FAM110C | NM_001077710 | 0.052 | -0.1025 |
| KRTAP27-1 | NM_001077711 | -0.2625 | -0.0385 |
| VCAM1 | NM_001078 | -0.458 | -0.438 |
| SRSF1 | NM_001078166 | -1.407 | -1.605 |
| FAM127A | NM_001078171 | -0.2135 | -0.656 |
| FAM127C | NM_001078173 | -0.9215 | -1.769 |
| SLC29A1 | NM_001078174 | 0.1325 | -0.6855 |
| SLC29A1 | NM_001078175 | 0.1325 | -0.6855 |
| SLC29A1 | NM_001078176 | 0.1325 | -0.6855 |
| SLC29A1 | NM_001078177 | -0.5735 | -1.837 |
| CDC16 | NM_001078645 | -1.309 | -2.5375 |
| TMEM134 | NM_001078650 | 0.124 | -0.9555 |
| TMEM134 | NM_001078651 | 0.124 | -0.9555 |
| ZAP70 | NM_001079 | -0.0875 | -0.316 |
| FAM18A | NM_001079512 | -0.5545 | -1.299 |
| UBN1 | NM_001079514 | -0.268 | -1.5145 |
| TBCE | NM_001079515 | -1.139 | -1.7905 |
| MED24 | NM_001079518 | -0.0585 | -0.808 |
| FAM177A1 | NM_001079519 | -1.005 | -1.0975 |
| DACT1 | NM_001079520 | -0.099 | -1.0915 |
| KTN1 | NM_001079521 | 0.2755 | -1.327 |
| KTN1 | NM_001079522 | 0.2755 | -1.327 |
| PAICS | NM_001079524 | -0.696 | -0.5225 |
| PAICS | NM_001079525 | -0.696 | -0.5225 |
| IKZF2 | NM_001079526 | -0.0565 | 0.004 |
| ABCC6 | NM_001079528 | -0.2955 | -0.488 |
| FAM153B | NM_001079529 | -0.059 | 0.2075 |
| CFC1B | NM_001079530 | -0.02 | -0.569 |
| CPEB1 | NM_001079533 | -0.2155 | -1.676 |
| CPEB1 | NM_001079534 | -0.2155 | -1.676 |
| CPEB1 | NM_001079535 | -0.2155 | -1.676 |

| | | | |
|---|---|---|---|
| ART5 | NM_001079536 | -0.121 | -0.4685 |
| TRAPPC6B | NM_001079537 | -1.113 | -1.809 |
| XBP1 | NM_001079539 | -1.467 | -3.776 |
| HNRNPUL2 | NM_001079559 | -0.241 | -0.7755 |
| NKX2-1 | NM_001079668 | 0.0405 | -0.429 |
| TMTC4 | NM_001079669 | -0.7565 | -1.9185 |
| CAB39L | NM_001079670 | -0.5415 | -1.0815 |
| FNDC3A | NM_001079673 | -1.2875 | -2.941 |
| ETV4 | NM_001079675 | -0.0325 | -0.5565 |
| N4BP2L1 | NM_001079691 | -1.521 | -2.131 |
| MS4A14 | NM_001079692 | 0.1455 | -0.2355 |
| FKTN | NM_001079802 | -0.868 | -2.1205 |
| GAA | NM_001079803 | 0.304 | -0.368 |
| GAA | NM_001079804 | 0.304 | -0.368 |
| PGA3 | NM_001079807 | 0.0425 | -0.077 |
| TMEM183B | NM_001079809 | -0.1115 | -0.8455 |
| XIRP2 | NM_001079810 | 0.0105 | -0.206 |
| GLB1 | NM_001079811 | -0.2935 | -1.355 |
| DIAPH1 | NM_001079812 | -0.243 | -0.9065 |
| INSR | NM_001079817 | 0.539 | -2.069 |
| ITGA6 | NM_001079818 | -1.159 | -1.0485 |
| NLRP3 | NM_001079821 | 0.214 | -0.318 |
| LAMA2 | NM_001079823 | 0.2085 | -0.102 |
| CLRN2 | NM_001079827 | 0.082 | -0.261 |
| OCIAD1 | NM_001079839 | -1.113 | -1.1115 |
| OCIAD1 | NM_001079840 | -1.113 | -1.1115 |
| OCIAD1 | NM_001079841 | -1.113 | -1.1115 |
| OCIAD1 | NM_001079842 | -1.113 | -1.1115 |
| CASZ1 | NM_001079843 | 0.257 | -0.387 |
| CREBBP | NM_001079846 | -0.55 | -2.652 |
| GYG2 | NM_001079855 | -0.1645 | -0.707 |
| GPR64 | NM_001079858 | 0.0395 | -0.498 |
| GPR64 | NM_001079859 | 0.0395 | -0.498 |
| GPR64 | NM_001079860 | 0.0395 | -0.498 |
| DBI | NM_001079862 | -0.241 | -0.2425 |
| DBI | NM_001079863 | -0.241 | -0.2425 |
| TAX1BP1 | NM_001079864 | -0.0635 | -0.6615 |
| BCS1L | NM_001079866 | -0.31 | -1.3055 |
| PEX2 | NM_001079867 | -0.915 | -1.3515 |
| HAP1 | NM_001079870 | -0.5305 | -0.9255 |
| HAP1 | NM_001079871 | -0.4085 | 0.046 |
| CUL4B | NM_001079872 | -0.5495 | -0.806 |
| VAV3 | NM_001079874 | -0.2875 | -1.226 |

| | | | |
|---|---|---|---|
| RASA4 | NM_001079877 | -1.054 | -1.5585 |
| CNGA3 | NM_001079878 | -0.1505 | -1.043 |
| PRKD2 | NM_001079880 | -0.3955 | -1.4515 |
| PRKD2 | NM_001079881 | -0.3955 | -1.4515 |
| PRKD2 | NM_001079882 | -0.3955 | -1.4515 |
| SPACA5B | NM_001079900 | -0.1685 | -0.6165 |
| ZNF331 | NM_001079906 | -1.416 | -1.737 |
| ZNF331 | NM_001079907 | -1.416 | -1.737 |
| LRRIQ1 | NM_001079910 | -0.027 | -0.045 |
| DMP1 | NM_001079911 | 0.206 | -0.102 |
| OR7E24 | NM_001079935 | -0.0815 | -0.0075 |
| ALDH5A1 | NM_001080 | 0.112 | -0.777 |
| LDB3 | NM_001080114 | -0.218 | -0.4 |
| LDB3 | NM_001080115 | -0.218 | -0.4 |
| LDB3 | NM_001080116 | -0.218 | -0.4 |
| PRNP | NM_001080121 | -1.532 | -1.462 |
| PRNP | NM_001080122 | -1.532 | -1.462 |
| PRNP | NM_001080123 | -1.532 | -1.462 |
| CASP8 | NM_001080124 | 0.105 | 0.061 |
| CASP8 | NM_001080125 | 0.105 | 0.061 |
| ARHGAP9 | NM_001080156 | -0.1585 | -0.6555 |
| ARHGAP9 | NM_001080157 | -0.1585 | -0.6555 |
| TMEM200C | NM_001080209 | -0.3105 | -2.5965 |
| PACRG | NM_001080378 | -0.186 | -0.394 |
| GJC1 | NM_001080383 | -1.5865 | -2.9855 |
| SP100 | NM_001080391 | 0.134 | -0.2535 |
| KIAA1147 | NM_001080392 | -1.4355 | -2.3105 |
| GXYLT2 | NM_001080393 | -0.2855 | -0.382 |
| KIAA0146 | NM_001080394 | -0.2295 | -0.743 |
| AATK | NM_001080395 | -0.5425 | -1.185 |
| FAM155A | NM_001080396 | -0.562 | -0.702 |
| SLC45A1 | NM_001080397 | -0.0165 | -0.3355 |
| KIAA0368 | NM_001080398 | -0.323 | -1.218 |
| OC90 | NM_001080399 | 0.225 | -0.566 |
| PLIN4 | NM_001080400 | -0.251 | -0.492 |
| PPM1N | NM_001080401 | 0.026 | -0.2465 |
| CCDC61 | NM_001080402 | -0.663 | -1.0625 |
| ZNF878 | NM_001080404 | -0.3545 | -0.8465 |
| CEACAM18 | NM_001080405 | 0.009 | -0.5055 |
| C12orf71 | NM_001080406 | 0.1605 | 0.2135 |
| GLB1L3 | NM_001080407 | 0.0525 | -0.7055 |
| PROX2 | NM_001080408 | 0.2715 | -0.435 |
| ZNF99 | NM_001080409 | -0.223 | 0.196 |

| | | | |
|---|---|---|---|
| FBXO41 | NM_001080410 | -0.641 | -1.473 |
| ZNF433 | NM_001080411 | -0.015 | -1.177 |
| ZBTB38 | NM_001080412 | -1.3775 | -2.8695 |
| NOBOX | NM_001080413 | -0.115 | 0.231 |
| CCDC88C | NM_001080414 | -0.6955 | -2.514 |
| MYBL1 | NM_001080416 | -1.568 | -2.6565 |
| ZNF629 | NM_001080417 | -1.212 | -2.6025 |
| DLGAP3 | NM_001080418 | -0.1805 | -0.692 |
| UNK | NM_001080419 | -0.854 | -2.2825 |
| SHANK3 | NM_001080420 | -0.4005 | -1.0685 |
| UNC13A | NM_001080421 | -1.136 | -1.385 |
| GRIP2 | NM_001080423 | -0.3425 | -0.697 |
| KDM6B | NM_001080424 | -1.2725 | -2.3135 |
| BEX4 | NM_001080425 | -0.9835 | -1.339 |
| DUSP27 | NM_001080426 | 0.14 | 0.475 |
| THSD7B | NM_001080427 | -0.799 | -1.4405 |
| TOX3 | NM_001080430 | -1.866 | -3.5335 |
| SLC45A4 | NM_001080431 | -0.8485 | -3.1485 |
| FTO | NM_001080432 | -0.3365 | -0.7245 |
| CCDC85A | NM_001080433 | 0.073 | -1.591 |
| LMTK3 | NM_001080434 | -0.3545 | -1.3415 |
| WHAMM | NM_001080435 | -0.567 | -1.924 |
| WTIP | NM_001080436 | 0.35 | -0.317 |
| SNED1 | NM_001080437 | -0.257 | -0.6835 |
| HSF5 | NM_001080439 | -0.103 | -0.285 |
| OTOL1 | NM_001080440 | -0.162 | 0.051 |
| TTC36 | NM_001080441 | 0.157 | -0.0065 |
| SLC38A8 | NM_001080442 | -0.297 | 0.005 |
| KIF18B | NM_001080443 | -0.716 | -2.2295 |
| IGSF5 | NM_001080444 | -0.0235 | -0.418 |
| C11orf94 | NM_001080446 | -0.1505 | -0.0215 |
| EPHA6 | NM_001080448 | 0.1895 | -0.7605 |
| DNA2 | NM_001080449 | -1.592 | -2.4805 |
| BEND3 | NM_001080450 | -0.734 | -2.2455 |
| SERPINA11 | NM_001080451 | 0.246 | -0.049 |
| GPR108 | NM_001080452 | -0.2655 | -0.2865 |
| INTS1 | NM_001080453 | 0.1195 | -0.5845 |
| ACSM4 | NM_001080454 | 0.0355 | 0.1585 |
| ZSCAN5B | NM_001080456 | 0.0965 | -0.206 |
| LRRC4B | NM_001080457 | -1.0995 | -1.9595 |
| EVX2 | NM_001080458 | -0.0915 | -0.698 |
| LRRIQ4 | NM_001080460 | -0.0525 | -0.143 |
| UNCX | NM_001080461 | 0.2325 | -0.104 |

| | | | |
|---|---|---|---|
| TMEM202 | NM_001080462 | -0.1875 | -0.1935 |
| DYNC2H1 | NM_001080463 | 0.123 | -1.0755 |
| ASPG | NM_001080464 | -0.096 | -0.8185 |
| C17orf98 | NM_001080465 | 0.088 | -0.3725 |
| BTBD17 | NM_001080466 | -0.142 | -2.0705 |
| MYO5B | NM_001080467 | 0.666 | -0.8845 |
| SYCN | NM_001080468 | -0.515 | -0.1765 |
| FBXO46 | NM_001080469 | -0.766 | -1.092 |
| ZNF697 | NM_001080470 | -1.148 | -2.6065 |
| PEAR1 | NM_001080471 | -0.6955 | -0.565 |
| FITM2 | NM_001080472 | -0.019 | -0.795 |
| MFSD2B | NM_001080473 | 0.1005 | -0.515 |
| C2orf78 | NM_001080474 | 0.116 | -0.2815 |
| PLEKHM3 | NM_001080475 | -1.31 | -2.917 |
| GRXCR1 | NM_001080476 | -0.426 | 0.019 |
| ODZ3 | NM_001080477 | -1.165 | -1.9255 |
| LRRC14B | NM_001080478 | -0.1535 | -0.2305 |
| RGNEF | NM_001080479 | -0.874 | -1.9285 |
| MBOAT1 | NM_001080480 | -0.0025 | -1.0165 |
| USP45 | NM_001080481 | -0.065 | 0.491 |
| C9orf172 | NM_001080482 | 0.08 | -0.31 |
| TMEM8C | NM_001080483 | -0.197 | -0.309 |
| KIAA1751 | NM_001080484 | 0.232 | -0.0895 |
| ZNF275 | NM_001080485 | -0.9955 | -1.796 |
| PABPN1L | NM_001080487 | -0.3265 | -0.4095 |
| ONECUT3 | NM_001080488 | 0.045 | -0.207 |
| GLOD5 | NM_001080489 | 0.224 | 0.364 |
| USP6NL | NM_001080491 | -0.8545 | -1.573 |
| PRSS54 | NM_001080492 | 0.1425 | -0.268 |
| ZNF823 | NM_001080493 | -1.7065 | -3.4225 |
| TTC39A | NM_001080494 | -0.463 | -0.7525 |
| TNRC18 | NM_001080495 | -1.1845 | -2.2035 |
| RGP1 | NM_001080496 | -0.9985 | -1.752 |
| MEGF9 | NM_001080497 | -0.6995 | -2.63 |
| VWC2L | NM_001080500 | 0.069 | -0.3115 |
| TMEM223 | NM_001080501 | -0.5465 | -1.0125 |
| CCDC159 | NM_001080503 | -0.188 | -0.7475 |
| RBM44 | NM_001080504 | -0.003 | -0.038 |
| SHISA3 | NM_001080505 | 0.365 | -0.547 |
| TMEM150C | NM_001080506 | 0.094 | -0.0035 |
| OOEP | NM_001080507 | 0.181 | -0.311 |
| TBX18 | NM_001080508 | -0.569 | -0.389 |
| TSPAN11 | NM_001080509 | -0.4925 | -1.288 |

| | | | |
|---|---|---|---|
| METTL23 | NM_001080510 | -1.803 | -3.388 |
| CLEC2L | NM_001080511 | 0.1175 | -0.476 |
| BICC1 | NM_001080512 | -0.816 | -2.9655 |
| CPN2 | NM_001080513 | -0.2515 | -0.407 |
| GRXCR2 | NM_001080516 | 0.324 | -0.7235 |
| SETD5 | NM_001080517 | -1.214 | -2.767 |
| LIPK | NM_001080518 | 0.0155 | -0.3255 |
| BAHCC1 | NM_001080519 | -1.135 | -1.2815 |
| DRGX | NM_001080520 | -0.165 | -0.097 |
| RASSF10 | NM_001080521 | -0.048 | 0.0645 |
| CC2D2A | NM_001080522 | -1.302 | -2.023 |
| ARRDC5 | NM_001080523 | 0.0045 | -0.413 |
| C16orf90 | NM_001080524 | -0.308 | -0.1945 |
| SPINK8 | NM_001080525 | -0.1435 | -0.0215 |
| FABP9 | NM_001080526 | -0.1335 | 0.021 |
| MYO7B | NM_001080527 | 0.1935 | -0.3695 |
| LOC646498 | NM_001080528 | 0.0695 | -0.53 |
| WIPF3 | NM_001080529 | -0.1045 | 0.0415 |
| C4orf51 | NM_001080531 | 0.401 | -0.204 |
| TMC3 | NM_001080532 | -0.368 | -0.48 |
| UNC119B | NM_001080533 | -0.6245 | -1.32 |
| UNC13C | NM_001080534 | -0.248 | -0.081 |
| SNTN | NM_001080537 | -0.3465 | -0.216 |
| AKR1B15 | NM_001080538 | 0.222 | 0.1005 |
| CCDC150 | NM_001080539 | -0.5215 | -1.7715 |
| MGA | NM_001080541 | -0.6075 | -2.6415 |
| FBF1 | NM_001080542 | 0.109 | -0.6205 |
| C19orf29 | NM_001080543 | -0.6355 | -1.629 |
| PPP1R1C | NM_001080545 | 0.1305 | -0.2945 |
| TMEM218 | NM_001080546 | -0.6325 | -2.025 |
| SPI1 | NM_001080547 | 0.2625 | -0.652 |
| C9orf84 | NM_001080551 | 0.2575 | -0.4255 |
| GSG1 | NM_001080554 | 0.109 | 0.452 |
| GSG1 | NM_001080555 | 0.109 | 0.452 |
| DGKG | NM_001080744 | 0.0795 | -0.2595 |
| DGKG | NM_001080745 | 0.0795 | -0.2595 |
| KIR2DL4 | NM_001080770 | 0.013 | -0.415 |
| KIR2DL4 | NM_001080772 | 0.013 | -0.415 |
| MYO1C | NM_001080779 | -0.737 | -1.396 |
| MAGEA9B | NM_001080790 | -0.2595 | -0.2745 |
| C15orf57 | NM_001080791 | -0.812 | -2.1395 |
| C15orf57 | NM_001080792 | -0.812 | -2.1395 |
| C2orf89 | NM_001080824 | -0.104 | -0.123 |

| | | | |
|---|---|---|---|
| TMEM120B | NM_001080825 | -0.504 | -0.7055 |
| SGK223 | NM_001080826 | -0.8325 | -2.903 |
| PRAMEF12 | NM_001080830 | 0.323 | -0.558 |
| PRR21 | NM_001080835 | 0.2375 | -0.218 |
| MEIG1 | NM_001080836 | 0.0765 | -0.3385 |
| GSTT2B | NM_001080843 | -0.23 | -0.6435 |
| CSAG2 | NM_001080848 | 0.2865 | -0.104 |
| DNLZ | NM_001080849 | 0.032 | -0.312 |
| CCDC30 | NM_001080850 | 0.0665 | 0.128 |
| DMRTC1B | NM_001080851 | 0.064 | -0.3425 |
| PXN | NM_001080855 | -0.1285 | -0.966 |
| MYO1C | NM_001080950 | -0.007 | -0.775 |
| PLAGL1 | NM_001080951 | -0.0175 | -0.963 |
| PLAGL1 | NM_001080952 | -0.0175 | -0.963 |
| PLAGL1 | NM_001080953 | -0.327 | -0.991 |
| PLAGL1 | NM_001080954 | -0.0175 | -0.963 |
| PLAGL1 | NM_001080955 | -0.0175 | -0.963 |
| PLAGL1 | NM_001080956 | -0.0175 | -0.963 |
| REPS2 | NM_001080975 | 0.017 | -0.4125 |
| DSE | NM_001080976 | -0.966 | -2.887 |
| LILRB2 | NM_001080978 | 0.013 | 0.206 |
| FRG2B | NM_001080998 | -0.078 | 0.7785 |
| CUBN | NM_001081 | -0.1595 | -0.194 |
| COMMD5 | NM_001081003 | -0.6445 | -2.1015 |
| COMMD5 | NM_001081004 | -0.6445 | -2.1015 |
| LILRB4 | NM_001081438 | 0.0845 | -0.528 |
| LILRB5 | NM_001081442 | 0.386 | -0.259 |
| LILRB5 | NM_001081443 | 0.386 | -0.259 |
| LILRB3 | NM_001081450 | 0.0225 | -0.585 |
| P2RY14 | NM_001081455 | 0.01 | -0.1365 |
| JMJD6 | NM_001081461 | -1.127 | -2.3835 |
| NXF1 | NM_001081491 | 0.33 | -0.0235 |
| THOC2 | NM_001081550 | -0.4445 | -1.845 |
| DEFB103A | NM_001081551 | 0.4465 | -0.0305 |
| SPAG11A | NM_001081552 | -0.5055 | -0.2915 |
| CPSF4 | NM_001081559 | -0.5865 | -1.1025 |
| DMPK | NM_001081560 | -0.673 | -0.582 |
| DMPK | NM_001081562 | -0.673 | -0.582 |
| DMPK | NM_001081563 | -0.673 | -0.582 |
| GAB3 | NM_001081573 | 0.1865 | -0.74 |
| LILRB1 | NM_001081637 | 0.454 | -0.641 |
| LILRB1 | NM_001081638 | 0.454 | -0.641 |
| LILRB1 | NM_001081639 | 0.454 | -0.641 |

| | | | |
|---|---|---|---|
| PRKDC | NM_001081640 | -0.496 | -0.521 |
| KLHL38 | NM_001081675 | 0.223 | -0.332 |
| SCN3A | NM_001081676 | 0.1245 | 0.005 |
| SCN3A | NM_001081677 | 0.1245 | 0.005 |
| G6PC2 | NM_001081686 | -0.3455 | -0.6325 |
| ELN | NM_001081752 | -0.008 | -0.4465 |
| ELN | NM_001081753 | -0.008 | -0.4465 |
| ELN | NM_001081754 | -0.008 | -0.4465 |
| ELN | NM_001081755 | -0.008 | -0.4465 |
| RGS9 | NM_001081955 | -0.0445 | -0.1075 |
| CYP4F2 | NM_001082 | 0.2575 | -0.5815 |
| ZNF623 | NM_001082480 | -1.27 | -2.375 |
| ACD | NM_001082486 | -0.6515 | -1.4305 |
| ACD | NM_001082487 | -0.4235 | -1.2055 |
| CA10 | NM_001082533 | -0.378 | -0.3395 |
| CA10 | NM_001082534 | -0.378 | -0.3395 |
| TCTN1 | NM_001082537 | -0.269 | -0.5095 |
| TCTN1 | NM_001082538 | -0.269 | -0.5095 |
| RBFOX3 | NM_001082575 | 0.2705 | -0.261 |
| RBFOX2 | NM_001082576 | -0.681 | -1.2645 |
| RBFOX2 | NM_001082577 | -0.681 | -1.2645 |
| RBFOX2 | NM_001082578 | -0.681 | -1.2645 |
| RBFOX2 | NM_001082579 | -0.681 | -1.2645 |
| CD33 | NM_001082618 | 0.26 | -0.237 |
| PIH1D2 | NM_001082619 | -0.0735 | -0.122 |
| SCARB1 | NM_001082959 | -0.3435 | -0.969 |
| FAM19A5 | NM_001082967 | -0.477 | -1.6575 |
| TOM1L2 | NM_001082968 | -0.9045 | -1.753 |
| C11orf57 | NM_001082969 | -1.7485 | -2.789 |
| C11orf57 | NM_001082970 | -1.7485 | -2.789 |
| DDC | NM_001082971 | -0.2835 | -0.81 |
| CASC1 | NM_001082972 | -0.017 | -0.1445 |
| CASC1 | NM_001082973 | -0.017 | -0.1445 |
| PDE5A | NM_001083 | -0.417 | -2.186 |
| GNRH1 | NM_001083111 | 0.06 | -0.7765 |
| GPD2 | NM_001083112 | -0.9915 | -0.978 |
| PRF1 | NM_001083116 | 0.172 | -0.109 |
| FAM75A3 | NM_001083124 | 0.05 | 0.061 |
| PYDC2 | NM_001083308 | -0.0715 | -0.604 |
| CHMP1A | NM_001083314 | -0.289 | -1.2075 |
| ZFP112 | NM_001083335 | -0.5425 | -2.2045 |
| CEP57L1 | NM_001083535 | -1.0785 | -1.9845 |
| FGD3 | NM_001083536 | 0.0665 | -0.804 |

| | | | |
|---|---|---|---|
| POTEE | NM_001083538 | 0.104 | -0.1925 |
| KIR3DS1 | NM_001083539 | 0.1095 | -0.011 |
| RABEP1 | NM_001083585 | -0.5515 | -1.61 |
| TMEM214 | NM_001083590 | -0.4525 | -1.1425 |
| CELF2 | NM_001083591 | -0.901 | -0.5255 |
| ROR1 | NM_001083592 | -0.561 | -2.197 |
| NAA60 | NM_001083601 | -0.331 | -1.7705 |
| PTCH1 | NM_001083602 | -1.717 | -5.325 |
| PTCH1 | NM_001083603 | -1.717 | -5.325 |
| PTCH1 | NM_001083604 | -1.717 | -5.325 |
| PTCH1 | NM_001083605 | -1.717 | -5.325 |
| PTCH1 | NM_001083606 | -1.717 | -5.325 |
| PTCH1 | NM_001083607 | -1.717 | -5.325 |
| NARF | NM_001083608 | -0.3915 | -1.5865 |
| AGPHD1 | NM_001083612 | -0.0245 | -0.3435 |
| TMEM219 | NM_001083613 | 0.2185 | -0.4965 |
| EARS2 | NM_001083614 | -0.7045 | -2.0425 |
| MYO3B | NM_001083615 | -0.0665 | 0.1395 |
| RB1CC1 | NM_001083617 | -0.525 | -2.3125 |
| GRIA2 | NM_001083619 | -0.6775 | -1.264 |
| GRIA2 | NM_001083620 | -0.6775 | -1.264 |
| ZBTB40 | NM_001083621 | -1.173 | -2.494 |
| ANKRD12 | NM_001083625 | 0.2205 | -0.6865 |
| STRN3 | NM_001083893 | -1.472 | -3.2245 |
| GP6 | NM_001083899 | -0.1205 | -0.242 |
| BANK1 | NM_001083907 | 0.4275 | -0.518 |
| GPR123 | NM_001083909 | 0.0725 | 0.2545 |
| SIRPB1 | NM_001083910 | 0.046 | -0.012 |
| C10orf26 | NM_001083913 | -1.1695 | -3.171 |
| CTBP2 | NM_001083914 | -0.9365 | -1.7355 |
| C1orf116 | NM_001083924 | -0.096 | -0.247 |
| ASRGL1 | NM_001083926 | -0.61 | -1.0175 |
| C2orf56 | NM_001083946 | -1.2545 | -1.942 |
| TMPRSS4 | NM_001083947 | -0.072 | -0.4105 |
| THADA | NM_001083953 | -1.246 | -1.931 |
| ZNF655 | NM_001083956 | -1.9335 | -2.7445 |
| WDR62 | NM_001083961 | -0.31 | -1.095 |
| TCF4 | NM_001083962 | -1.0505 | -2.2855 |
| TDRKH | NM_001083963 | -1.0575 | -0.972 |
| TDRKH | NM_001083964 | -1.0575 | -0.972 |
| TDRKH | NM_001083965 | -1.0575 | -0.972 |
| PLOD3 | NM_001084 | -0.1255 | -1.2535 |
| DDTL | NM_001084393 | -0.5435 | -0.6715 |

| | | | |
|---|---|---|---|
| SERPINA3 | NM_001085 | -0.1615 | 0.125 |
| MRAS | NM_001085049 | -1.0805 | -1.3675 |
| TOR2A | NM_001085347 | -0.2205 | -0.4615 |
| MORC4 | NM_001085354 | -0.482 | -0.7485 |
| BTLA | NM_001085357 | -0.0195 | 0.135 |
| MZT2A | NM_001085365 | 0.3295 | -0.0205 |
| ZNF655 | NM_001085366 | -0.9175 | -1.7805 |
| ZNF655 | NM_001085367 | -0.9175 | -1.7805 |
| ZNF655 | NM_001085368 | -0.9175 | -1.7805 |
| C11orf83 | NM_001085372 | -0.7995 | -2.1545 |
| C1orf226 | NM_001085375 | -0.191 | -1.031 |
| MCC | NM_001085377 | -1.3125 | -2.6785 |
| PSAPL1 | NM_001085382 | 0.212 | -0.539 |
| ZNF154 | NM_001085384 | 0.0945 | -0.542 |
| TLX1NB | NM_001085398 | -0.323 | 0.0065 |
| RELL1 | NM_001085399 | -0.512 | -0.333 |
| RELL1 | NM_001085400 | -0.896 | -1.668 |
| C6orf201 | NM_001085401 | -0.079 | -0.043 |
| PLSCR5 | NM_001085420 | -0.138 | -1.502 |
| ARSA | NM_001085425 | -0.097 | -0.137 |
| ARSA | NM_001085426 | -0.097 | -0.137 |
| ARSA | NM_001085427 | -0.097 | -0.137 |
| ARSA | NM_001085428 | -0.022 | -0.4915 |
| TMEM213 | NM_001085429 | -0.0445 | 0.3125 |
| C17orf67 | NM_001085430 | -0.106 | -0.2365 |
| C2orf54 | NM_001085437 | 0.0125 | -0.4925 |
| C2orf77 | NM_001085447 | -0.007 | -0.174 |
| LNP1 | NM_001085451 | -1.446 | -2.342 |
| GIT1 | NM_001085454 | -0.018 | -0.6135 |
| KRTAP24-1 | NM_001085455 | -0.131 | -0.3795 |
| CTNND1 | NM_001085458 | -0.546 | -1.48 |
| CTNND1 | NM_001085459 | -0.546 | -1.48 |
| CTNND1 | NM_001085460 | -0.546 | -1.48 |
| CTNND1 | NM_001085461 | -0.546 | -1.48 |
| CTNND1 | NM_001085462 | -0.546 | -1.48 |
| CTNND1 | NM_001085463 | -0.546 | -1.48 |
| CTNND1 | NM_001085464 | -0.546 | -1.48 |
| CTNND1 | NM_001085465 | -0.546 | -1.48 |
| CTNND1 | NM_001085466 | -0.546 | -1.48 |
| CTNND1 | NM_001085467 | -0.546 | -1.48 |
| CTNND1 | NM_001085468 | -0.546 | -1.48 |
| CTNND1 | NM_001085469 | -0.546 | -1.48 |
| FOXN3 | NM_001085471 | -1.488 | -2.1125 |

| | | | |
|---|---|---|---|
| LOC646627 | NM_001085474 | 0.3405 | -0.135 |
| IQCF3 | NM_001085479 | 0.28 | 0.086 |
| FAM162B | NM_001085480 | -0.7255 | -0.8845 |
| MAP1LC3B2 | NM_001085481 | 0.137 | -0.1605 |
| SEPP1 | NM_001085486 | -0.3935 | -0.502 |
| MYSM1 | NM_001085487 | -1.677 | -2.5265 |
| TMEM150B | NM_001085488 | 0.0465 | -0.691 |
| AADAC | NM_001086 | 0.076 | -0.534 |
| C17orf89 | NM_001086521 | 0.0595 | -0.653 |
| AAMP | NM_001087 | 0.1695 | -1.09 |
| AANAT | NM_001088 | -0.3885 | 0.124 |
| ABCA3 | NM_001089 | -0.3805 | -0.726 |
| C5orf49 | NM_001089584 | 0.0155 | -0.4305 |
| UQCRHL | NM_001089591 | 0.1955 | 0.064 |
| ABCF1 | NM_001090 | -0.761 | -1.957 |
| ABP1 | NM_001091 | 0.0135 | -0.1565 |
| ABR | NM_001092 | -0.7775 | -0.998 |
| ACACB | NM_001093 | -0.615 | -1.029 |
| MEX3A | NM_001093725 | -1.195 | -2.418 |
| CCDC102B | NM_001093729 | -0.788 | -0.183 |
| DYTN | NM_001093730 | 0.1435 | -0.3475 |
| C5orf44 | NM_001093755 | -1.5265 | -2.229 |
| C5orf44 | NM_001093756 | -1.5265 | -2.229 |
| FAM195B | NM_001093767 | 0.155 | -0.4665 |
| SFTPA1 | NM_001093770 | 0.117 | 0.288 |
| TXNRD1 | NM_001093771 | -1.177 | -1.714 |
| KIT | NM_001093772 | -0.129 | -0.187 |
| ACCN1 | NM_001094 | -0.571 | -0.3355 |
| ACCN2 | NM_001095 | 0.152 | -0.2345 |
| ACLY | NM_001096 | -0.2915 | -0.77 |
| ACR | NM_001097 | -0.059 | -0.262 |
| GPR34 | NM_001097579 | 0.3895 | -0.223 |
| TMEM22 | NM_001097599 | -1.4185 | -3.657 |
| TMEM22 | NM_001097600 | -1.4185 | -3.657 |
| KNCN | NM_001097611 | -0.393 | -0.186 |
| GPR89A | NM_001097612 | -0.846 | -1.2885 |
| GPR89A | NM_001097613 | -0.846 | -1.2885 |
| POLR2J3 | NM_001097615 | 0.2595 | -0.3075 |
| GPR89C | NM_001097616 | -1.436 | -1.8385 |
| TMEM184A | NM_001097620 | -0.0365 | -0.3375 |
| OCM | NM_001097622 | 0.4325 | -0.287 |
| GCNT1 | NM_001097633 | -0.308 | -0.466 |
| GCNT1 | NM_001097634 | -0.308 | -0.466 |

| | | | |
|---|---|---|---|
| GCNT1 | NM_001097635 | -0.308 | -0.466 |
| GCNT1 | NM_001097636 | -0.308 | -0.466 |
| FUT2 | NM_001097638 | -0.256 | -0.049 |
| FUT3 | NM_001097639 | 0.1225 | -0.0305 |
| FUT3 | NM_001097640 | 0.1225 | -0.0305 |
| FUT3 | NM_001097641 | 0.1225 | -0.0305 |
| TAS2R30 | NM_001097643 | 0.181 | -0.246 |
| ACO2 | NM_001098 | 0.0985 | -0.378 |
| BSX | NM_001098169 | 0.3425 | 0.1445 |
| PRDM7 | NM_001098173 | -0.0855 | 0.176 |
| ENTPD1 | NM_001098175 | -0.2325 | -0.583 |
| GPR1 | NM_001098199 | -0.39 | -0.3715 |
| GPR18 | NM_001098200 | 0.473 | 0.5005 |
| GPER | NM_001098201 | 0.005 | -0.5815 |
| HIC1 | NM_001098202 | -0.1405 | -0.188 |
| HNRNPF | NM_001098204 | -0.7185 | -2.124 |
| HNRNPF | NM_001098205 | -0.7185 | -2.124 |
| HNRNPF | NM_001098206 | -0.7185 | -2.124 |
| HNRNPF | NM_001098207 | -0.7185 | -2.124 |
| HNRNPF | NM_001098208 | -0.7185 | -2.124 |
| CTNNB1 | NM_001098209 | -0.298 | -0.466 |
| CTNNB1 | NM_001098210 | -0.298 | -0.466 |
| HRH1 | NM_001098211 | -0.043 | -0.601 |
| HRH1 | NM_001098212 | -0.043 | -0.601 |
| HRH1 | NM_001098213 | -0.043 | -0.601 |
| MGAT3 | NM_001098270 | -0.05 | -0.6035 |
| HMGCS1 | NM_001098272 | -1.2025 | -1.4315 |
| COPA | NM_001098398 | -0.594 | -0.618 |
| ZNF295 | NM_001098402 | -1.4495 | -3.176 |
| ZNF295 | NM_001098403 | -1.4495 | -3.176 |
| GAGE2D | NM_001098407 | 0.3635 | -0.304 |
| GAGE12H | NM_001098410 | 0.2055 | -0.0315 |
| GAGE10 | NM_001098413 | 0.444 | 0.2855 |
| ZNF621 | NM_001098414 | -0.616 | -1.577 |
| HDAC7 | NM_001098416 | -0.093 | -0.681 |
| DLG1 | NM_001098424 | -1.5135 | -3.089 |
| SMARCD2 | NM_001098426 | -0.568 | -1.8545 |
| TDRD10 | NM_001098475 | -0.539 | -0.113 |
| GRSF1 | NM_001098477 | -1.0765 | -1.905 |
| HLA-F | NM_001098478 | 0.083 | -0.4835 |
| HLA-F | NM_001098479 | 0.083 | -0.4835 |
| CRTC1 | NM_001098482 | -0.4185 | -1.0555 |
| C10orf125 | NM_001098483 | -0.145 | -0.476 |

| | | | |
|---|---|---|---|
| SLC4A4 | NM_001098484 | 0.0425 | -0.0635 |
| SLC17A3 | NM_001098486 | -0.089 | -0.248 |
| ZNF419 | NM_001098491 | -1.2995 | -2.445 |
| ZNF419 | NM_001098492 | -1.2995 | -2.445 |
| ZNF419 | NM_001098493 | -1.2995 | -2.445 |
| ZNF419 | NM_001098494 | -1.2995 | -2.445 |
| ZNF419 | NM_001098495 | -0.9345 | -2.193 |
| ZNF419 | NM_001098496 | -1.2995 | -2.445 |
| SGSM1 | NM_001098497 | -0.293 | -0.5015 |
| SGSM1 | NM_001098498 | -0.293 | -0.5015 |
| KIAA1217 | NM_001098500 | 0.0705 | -0.1695 |
| KIAA1217 | NM_001098501 | 0.0705 | -0.1695 |
| CHCHD4 | NM_001098502 | -1.061 | -3.238 |
| PTPRJ | NM_001098503 | 0.667 | 0.2955 |
| DDX17 | NM_001098504 | -1.0625 | -2.2805 |
| CEACAM21 | NM_001098506 | 0.077 | -0.1175 |
| ZNF207 | NM_001098507 | -0.662 | -1.4985 |
| SGSM2 | NM_001098509 | -0.461 | -0.858 |
| PELI3 | NM_001098510 | 0.012 | -0.4545 |
| KIF2A | NM_001098511 | -1.127 | -2.455 |
| PRKG1 | NM_001098512 | 0.092 | -0.6315 |
| MRGPRF | NM_001098515 | -0.2185 | -0.2895 |
| MUC20 | NM_001098516 | 0.0055 | -0.557 |
| CADM1 | NM_001098517 | -0.1275 | -0.554 |
| GPR116 | NM_001098518 | -0.0625 | -0.218 |
| HTATIP2 | NM_001098520 | 0.04 | -0.531 |
| HTATIP2 | NM_001098521 | 0.04 | -0.531 |
| HTATIP2 | NM_001098522 | 0.04 | -0.531 |
| HTATIP2 | NM_001098523 | 0.242 | -0.036 |
| FAM129C | NM_001098524 | 0.0795 | -0.346 |
| CKAP2 | NM_001098525 | -1.05 | -3.19 |
| AMICA1 | NM_001098526 | 0.1805 | 0.0645 |
| RFPL2 | NM_001098527 | 0.159 | -0.118 |
| TXNDC2 | NM_001098529 | 0.188 | 0.1015 |
| RAPGEF3 | NM_001098531 | -0.2535 | -0.278 |
| RAPGEF3 | NM_001098532 | -0.4965 | -0.2995 |
| CDK10 | NM_001098533 | -0.3685 | -0.763 |
| BAG6 | NM_001098534 | 0.166 | -0.382 |
| USP5 | NM_001098536 | 0.1605 | -0.43 |
| PNPLA7 | NM_001098537 | -0.0645 | -0.3575 |
| PRR4 | NM_001098538 | -0.1965 | -0.8115 |
| HPSE | NM_001098540 | -0.8815 | -0.394 |
| TMBIM6 | NM_001098576 | 0.1265 | -0.491 |

| | | | |
|---|---|---|---|
| RPL31 | NM_001098577 | -0.0905 | -0.3225 |
| MRVI1 | NM_001098579 | 0.1675 | -0.759 |
| SIGLEC14 | NM_001098612 | 0.067 | -0.073 |
| PUS7L | NM_001098614 | -0.454 | -0.847 |
| PUS7L | NM_001098615 | -0.454 | -0.847 |
| C1orf43 | NM_001098616 | -0.665 | -1.5265 |
| PCBP2 | NM_001098620 | 0.152 | -0.2755 |
| C14orf109 | NM_001098621 | -1.533 | -3.3375 |
| NANOS3 | NM_001098622 | -0.0225 | -0.3125 |
| OBSCN | NM_001098623 | 0.063 | -0.0625 |
| MID1 | NM_001098624 | -0.5825 | -1.5365 |
| JKAMP | NM_001098625 | -1.635 | -2.8215 |
| IRF5 | NM_001098627 | -0.1315 | -0.227 |
| IRF5 | NM_001098629 | -0.1315 | -0.227 |
| IRF5 | NM_001098630 | -0.1315 | -0.227 |
| AKT1S1 | NM_001098632 | -0.1365 | -0.59 |
| AKT1S1 | NM_001098633 | -0.1365 | -0.59 |
| RBM47 | NM_001098634 | 0.1795 | 0.1535 |
| SEZ6 | NM_001098635 | -0.35 | -0.602 |
| RNF169 | NM_001098638 | -1.1625 | -2.2205 |
| RASGRP2 | NM_001098671 | -0.378 | -0.756 |
| HEPHL1 | NM_001098672 | 0.0015 | -0.1915 |
| GNG4 | NM_001098721 | 0.1385 | -0.3365 |
| GNG4 | NM_001098722 | 0.1385 | -0.3365 |
| TCL1A | NM_001098725 | -0.0035 | 0.0105 |
| RPP14 | NM_001098783 | -1.4495 | -3.006 |
| FAM89B | NM_001098784 | -0.1805 | -1.0865 |
| C7orf68 | NM_001098786 | -0.896 | -2.7265 |
| BET1L | NM_001098787 | -0.779 | -2.2775 |
| MID1IP1 | NM_001098790 | -0.5775 | -1.1735 |
| MID1IP1 | NM_001098791 | -0.5775 | -1.1735 |
| FAM160A2 | NM_001098794 | -0.2845 | -0.2965 |
| TOX2 | NM_001098796 | -0.1435 | -0.454 |
| TOX2 | NM_001098797 | -0.1435 | -0.454 |
| TOX2 | NM_001098798 | -0.1435 | -0.454 |
| MAGED4 | NM_001098800 | -0.271 | -0.6755 |
| C18orf19 | NM_001098801 | -1.275 | -2.711 |
| CEP78 | NM_001098802 | -0.441 | -1.242 |
| C9orf129 | NM_001098808 | 0.143 | -0.7325 |
| 8-Sep | NM_001098811 | -0.6105 | -1.3525 |
| SRL | NM_001098814 | -0.15 | -0.527 |
| KIAA0748 | NM_001098815 | -0.0115 | -0.113 |
| ODZ4 | NM_001098816 | -0.9665 | -2.263 |

| | | | |
|---|---|---|---|
| INO80C | NM_001098817 | -0.067 | -1.0965 |
| PDE4C | NM_001098818 | -0.183 | -0.1575 |
| PDE4C | NM_001098819 | -0.183 | -0.1575 |
| TMEM91 | NM_001098821 | 0.2265 | -0.65 |
| TMEM91 | NM_001098822 | 0.186 | -0.3295 |
| TMEM91 | NM_001098823 | 0.2265 | -0.65 |
| TMEM91 | NM_001098824 | 0.2265 | -0.65 |
| TMEM91 | NM_001098825 | 0.2265 | -0.65 |
| MORN4 | NM_001098831 | -0.435 | -0.611 |
| FAM104A | NM_001098832 | -1.5285 | -3.527 |
| ATXN7L3 | NM_001098833 | -0.452 | -1.0835 |
| GBX1 | NM_001098834 | -0.142 | -0.065 |
| MS4A15 | NM_001098835 | 0.11 | -0.317 |
| CXorf30 | NM_001098843 | 0.076 | 0.195 |
| ANXA8L1 | NM_001098845 | 0.085 | 0.1205 |
| ACPP | NM_001099 | -0.3445 | -0.5955 |
| RAD51AP2 | NM_001099218 | 0.2065 | 0.4195 |
| KRTAP19-8 | NM_001099219 | -0.383 | -0.2825 |
| ZNF862 | NM_001099220 | -0.556 | -0.592 |
| TIFAB | NM_001099221 | -0.031 | -0.5255 |
| IFT74 | NM_001099222 | -0.8065 | -1.8385 |
| IFT74 | NM_001099224 | 0.0245 | -0.7565 |
| ZNF506 | NM_001099269 | -0.883 | -1.969 |
| ZBTB34 | NM_001099270 | -1.023 | -1.466 |
| POC5 | NM_001099271 | -1.519 | -2.5115 |
| BTBD9 | NM_001099272 | -0.2065 | -0.124 |
| TINF2 | NM_001099274 | -1.467 | -3.3055 |
| HEATR7A | NM_001099280 | -0.095 | -0.565 |
| HEATR7A | NM_001099281 | -0.095 | -0.565 |
| PTMA | NM_001099285 | 0.0515 | -0.2865 |
| FAM54A | NM_001099286 | -1.8985 | -3.635 |
| NIPAL4 | NM_001099287 | -0.086 | -0.439 |
| SH3RF3 | NM_001099289 | -0.794 | -1.646 |
| KIF4B | NM_001099293 | -1.1615 | -1.361 |
| KIAA1644 | NM_001099294 | -0.6715 | -0.13 |
| C2orf80 | NM_001099334 | -0.7175 | 0.0425 |
| PHYHIP | NM_001099335 | -0.493 | -0.5165 |
| C12orf42 | NM_001099336 | 0.295 | 0.327 |
| FAM22A | NM_001099338 | 0.031 | -0.6295 |
| SGCE | NM_001099400 | -0.92 | -1.5595 |
| SGCE | NM_001099401 | -1.5125 | -1.894 |
| CCNK | NM_001099402 | -0.2185 | -1.532 |
| PRDM8 | NM_001099403 | -0.266 | -1.5015 |

| | | | |
|---|---|---|---|
| SCN5A | NM_001099404 | 0.021 | -0.709 |
| SCN5A | NM_001099405 | -0.6195 | -1.359 |
| SLC39A6 | NM_001099406 | -0.9015 | -0.6785 |
| DZANK1 | NM_001099407 | -1.2705 | -2.0765 |
| EIF4E1B | NM_001099408 | -0.176 | 0.02 |
| EHBP1L1 | NM_001099409 | 0.014 | -0.1115 |
| GPRASP1 | NM_001099410 | -0.836 | -0.7705 |
| GPRASP1 | NM_001099411 | -0.836 | -0.7705 |
| KAT6A | NM_001099412 | -1.43 | -3.683 |
| HAVCR1 | NM_001099414 | -0.21 | -0.3655 |
| POM121C | NM_001099415 | -0.633 | -2.4665 |
| BCAS3 | NM_001099432 | -0.0395 | -0.6885 |
| JAKMIP1 | NM_001099433 | -1.466 | -2.789 |
| DCDC2B | NM_001099434 | -0.2065 | -0.3535 |
| ULK3 | NM_001099436 | 0.0845 | -0.7865 |
| ZNF30 | NM_001099437 | -1.384 | -2.6945 |
| ZNF30 | NM_001099438 | -1.384 | -2.6945 |
| EPHA10 | NM_001099439 | -0.626 | -0.726 |
| CPPED1 | NM_001099455 | -0.7375 | -1.844 |
| NPW | NM_001099456 | 0.2265 | -0.171 |
| FAM54B | NM_001099625 | -0.4945 | -1.672 |
| FAM54B | NM_001099626 | -0.4595 | -1.327 |
| FAM54B | NM_001099627 | -0.4945 | -1.672 |
| ZNF146 | NM_001099638 | -1.133 | -3.801 |
| ZNF146 | NM_001099639 | -1.161 | -3.777 |
| TMEM100 | NM_001099640 | -0.7665 | -0.6835 |
| FTSJD1 | NM_001099642 | -2.079 | -3.6435 |
| RPL22L1 | NM_001099645 | -0.3215 | -0.035 |
| SLC47A2 | NM_001099646 | 0.3 | -0.191 |
| GXYLT1 | NM_001099650 | -0.128 | -1.3775 |
| GPR137C | NM_001099652 | -1.1065 | -2.064 |
| LRRN3 | NM_001099658 | -0.572 | -4.192 |
| LRRN3 | NM_001099660 | -0.572 | -4.192 |
| PTAR1 | NM_001099666 | -1.1365 | -2.152 |
| ARMS2 | NM_001099667 | -0.18 | -0.309 |
| HIGD1A | NM_001099668 | -1.236 | -1.123 |
| C8orf59 | NM_001099670 | -0.7235 | -0.5105 |
| C8orf59 | NM_001099671 | -0.7235 | -0.5105 |
| C8orf59 | NM_001099672 | -0.7235 | -0.5105 |
| C8orf59 | NM_001099673 | -0.7235 | -0.5105 |
| C12orf56 | NM_001099676 | 0.4515 | -0.1175 |
| LRRC58 | NM_001099678 | -1.172 | -1.8605 |
| TRIM32 | NM_001099679 | -0.808 | -3.729 |

| | | | |
|---|---|---|---|
| MAGIX | NM_001099680 | -0.2885 | -0.31 |
| MAGIX | NM_001099681 | -0.096 | -0.555 |
| MAGIX | NM_001099682 | -0.2885 | -0.31 |
| RHOXF2B | NM_001099685 | -0.4835 | -0.3755 |
| TGFA | NM_001099691 | -0.284 | -0.276 |
| EIF5AL1 | NM_001099692 | -0.065 | -0.583 |
| RPL31 | NM_001099693 | -0.0905 | -0.3225 |
| ZNF578 | NM_001099694 | -0.175 | -0.8705 |
| REPIN1 | NM_001099695 | -0.3255 | -0.718 |
| REPIN1 | NM_001099696 | -0.3255 | -0.718 |
| RSPH10B2 | NM_001099697 | -0.6425 | -1.138 |
| ADCYAP1 | NM_001099733 | -0.6705 | -1.418 |
| WDR83 | NM_001099737 | -0.299 | -0.8255 |
| SYBU | NM_001099743 | -0.7135 | -2.029 |
| SYBU | NM_001099744 | -0.8675 | -2.754 |
| SYBU | NM_001099745 | -0.7135 | -2.029 |
| SYBU | NM_001099746 | -0.7135 | -2.029 |
| SYBU | NM_001099747 | -0.7135 | -2.029 |
| SYBU | NM_001099748 | -0.7135 | -2.029 |
| SYBU | NM_001099749 | -0.7135 | -2.029 |
| SYBU | NM_001099750 | -0.7135 | -2.029 |
| SYBU | NM_001099751 | -0.7135 | -2.029 |
| SYBU | NM_001099752 | -0.7135 | -2.029 |
| SYBU | NM_001099753 | -0.7135 | -2.029 |
| SYBU | NM_001099754 | -0.7135 | -2.029 |
| SYBU | NM_001099755 | -0.7135 | -2.029 |
| SYBU | NM_001099756 | -0.7135 | -2.029 |
| POTEF | NM_001099771 | 0.102 | 0.081 |
| CYP4B1 | NM_001099772 | -0.083 | -0.0915 |
| CYP11A1 | NM_001099773 | 0.2835 | -0.344 |
| C3orf55 | NM_001099777 | -0.011 | -0.2205 |
| PSMB11 | NM_001099780 | 0.23 | -0.42 |
| GGT5 | NM_001099781 | -0.1495 | -0.3845 |
| GGT5 | NM_001099782 | -0.1495 | -0.3845 |
| FBXL19 | NM_001099784 | -0.529 | -1.4715 |
| ICAM2 | NM_001099786 | -1.088 | -0.6605 |
| ICAM2 | NM_001099787 | -1.088 | -0.6605 |
| ICAM2 | NM_001099788 | -1.088 | -0.6605 |
| ICAM2 | NM_001099789 | -1.088 | -0.6605 |
| PRAMEF19 | NM_001099790 | 0.175 | -0.6385 |
| SYS1 | NM_001099791 | -0.6285 | -1.7545 |
| PRAMEF17 | NM_001099851 | 0.225 | -0.544 |
| PRAMEF20 | NM_001099852 | -0.489 | -0.182 |

| | | | |
|---|---|---|---|
| MCF2 | NM_001099855 | 0.004 | -0.1765 |
| IKBKG | NM_001099856 | -0.315 | -0.9625 |
| IKBKG | NM_001099857 | -0.315 | -0.9625 |
| C7orf25 | NM_001099858 | -0.8455 | -1.381 |
| MAGEB16 | NM_001099921 | -0.305 | -0.1315 |
| ALG13 | NM_001099922 | -0.6645 | -1.719 |
| FAM166B | NM_001099951 | 0.09 | -0.0755 |
| ITPR1 | NM_001099952 | -1.355 | -2.454 |
| ACTA1 | NM_001100 | 0.172 | -0.0055 |
| LOC286238 | NM_001100111 | -0.5155 | 0.0285 |
| MYH2 | NM_001100112 | -0.2985 | 0.202 |
| RIMS2 | NM_001100117 | -1.074 | -2.011 |
| XRCC3 | NM_001100118 | -0.0735 | -0.912 |
| ECE2 | NM_001100120 | 0.059 | -0.2655 |
| ECE2 | NM_001100121 | 0.059 | -0.2655 |
| SBNO2 | NM_001100122 | 0.1355 | -0.512 |
| C7orf57 | NM_001100159 | 0.429 | 0.5605 |
| MRVI1 | NM_001100163 | 0.1675 | -0.759 |
| PHACTR2 | NM_001100164 | -0.047 | -1.1355 |
| PHACTR2 | NM_001100165 | -0.047 | -1.1355 |
| PHACTR2 | NM_001100166 | -0.047 | -1.1355 |
| MRVI1 | NM_001100167 | 0.1675 | -0.759 |
| CCR5 | NM_001100168 | 0.0005 | -0.04 |
| IMMT | NM_001100169 | -0.7905 | -1.062 |
| IMMT | NM_001100170 | -0.7905 | -1.062 |
| TMEM192 | NM_001100389 | -1.213 | -1.587 |
| IQCE | NM_001100390 | -0.1225 | -0.487 |
| RALYL | NM_001100391 | 0.284 | -1.265 |
| RALYL | NM_001100392 | 0.284 | -1.265 |
| RALYL | NM_001100393 | 0.284 | -1.265 |
| C2orf73 | NM_001100396 | -0.098 | -0.248 |
| RAPGEF4 | NM_001100397 | -0.226 | -0.721 |
| RAP1GAP2 | NM_001100398 | 0.069 | -0.3735 |
| PDS5A | NM_001100399 | 0.8015 | 0.317 |
| PDS5A | NM_001100400 | 0.8015 | 0.317 |
| C17orf62 | NM_001100407 | -0.531 | -1.6595 |
| C17orf62 | NM_001100408 | -0.531 | -1.6595 |
| SENP6 | NM_001100409 | -1.6285 | -2.9675 |
| FAM184A | NM_001100411 | -0.605 | -2.9825 |
| DROSHA | NM_001100412 | -0.919 | -1.884 |
| C19orf60 | NM_001100418 | 0.088 | -0.72 |
| C19orf60 | NM_001100419 | 0.1565 | -0.6095 |
| C21orf91 | NM_001100420 | -1.9175 | -4.3995 |

| | | | |
|---|---|---|---|
| C21orf91 | NM_001100421 | -0.4885 | -1.794 |
| SPATS2L | NM_001100422 | -0.8245 | -2.3115 |
| SPATS2L | NM_001100423 | -0.8245 | -2.3115 |
| SPATS2L | NM_001100424 | -0.8245 | -2.3115 |
| KIAA0895 | NM_001100425 | -2.1685 | -3.382 |
| RAP1GDS1 | NM_001100426 | -0.9505 | -0.795 |
| RAP1GDS1 | NM_001100427 | -0.9505 | -0.795 |
| RAP1GDS1 | NM_001100428 | -0.9505 | -0.795 |
| RAP1GDS1 | NM_001100429 | -0.9505 | -0.795 |
| RAP1GDS1 | NM_001100430 | -0.9505 | -0.795 |
| VSIG4 | NM_001100431 | -0.1545 | 0.0275 |
| RC3H2 | NM_001100588 | -0.4275 | -1.5575 |
| KIAA0232 | NM_001100590 | -1.492 | -3.0915 |
| ATP6V0E2 | NM_001100592 | 0.172 | -0.544 |
| SNRK | NM_001100594 | -0.6325 | -1.627 |
| SKA2 | NM_001100595 | -1.0405 | -1.059 |
| ZNF707 | NM_001100598 | -0.0455 | -0.405 |
| ZNF707 | NM_001100599 | -0.0455 | -0.405 |
| MMD2 | NM_001100600 | -0.222 | -0.371 |
| KDELR2 | NM_001100603 | -0.2355 | -0.555 |
| SERPINA10 | NM_001100607 | -0.1485 | 0.1235 |
| CABLES1 | NM_001100619 | -0.5665 | -1.7955 |
| TROAP | NM_001100620 | -0.026 | -0.516 |
| C17orf80 | NM_001100621 | -1.7265 | -3.598 |
| C17orf80 | NM_001100622 | -1.7265 | -3.598 |
| PLEKHB2 | NM_001100623 | -0.6275 | -0.928 |
| CENPN | NM_001100624 | -1.6045 | -3.747 |
| CENPN | NM_001100625 | -1.6045 | -3.747 |
| ERLIN1 | NM_001100626 | -0.7365 | -1.7955 |
| PRAMEF22 | NM_001100631 | 0.226 | -0.499 |
| CXCL16 | NM_001100812 | -0.004 | -0.244 |
| TMEM55B | NM_001100814 | -1.153 | -2.83 |
| MED7 | NM_001100816 | -2.1275 | -3.3335 |
| TCEB3CL | NM_001100817 | 0.3235 | -0.285 |
| PID1 | NM_001100818 | -0.0435 | -0.535 |
| MOB4 | NM_001100819 | -1.1345 | -2.377 |
| TMEM170B | NM_001100829 | -1.906 | -3.687 |
| C16orf46 | NM_001100873 | -0.9645 | -1.314 |
| SLC9B1 | NM_001100874 | -0.3115 | -1.484 |
| 10-Mar | NM_001100875 | 0.151 | -0.053 |
| PHYHD1 | NM_001100876 | 0.0375 | -0.1295 |
| PHYHD1 | NM_001100877 | 0.0375 | -0.1295 |
| C8orf73 | NM_001100878 | -0.3 | -0.426 |

| | | | |
|---|---|---|---|
| ST20 | NM_001100879 | -0.3235 | -1.081 |
| ST20 | NM_001100880 | -0.3235 | -1.081 |
| MS4A13 | NM_001100909 | -0.0655 | 0.5855 |
| BEND7 | NM_001100912 | -0.178 | -2.358 |
| PACS2 | NM_001100913 | -0.2195 | -0.367 |
| KCTD19 | NM_001100915 | -0.12 | -0.017 |
| MBOAT4 | NM_001100916 | -0.292 | -0.1525 |
| TSPAN19 | NM_001100917 | -0.4035 | -0.7125 |
| ACTB | NM_001101 | 0.154 | -0.2855 |
| SERPINE3 | NM_001101320 | -0.1205 | -0.338 |
| LOC728819 | NM_001101330 | -1.0865 | -0.753 |
| C3orf79 | NM_001101337 | 0.3245 | 0.0445 |
| ZNF883 | NM_001101338 | -1.0815 | -1.8775 |
| C12orf77 | NM_001101339 | 0.11 | 0.2545 |
| C19orf76 | NM_001101340 | -0.5005 | -0.118 |
| SFTA3 | NM_001101341 | -0.2155 | 0.1935 |
| CCDC160 | NM_001101357 | 0.2325 | -0.0425 |
| KBTBD13 | NM_001101362 | 0.2955 | -0.7995 |
| IGLON5 | NM_001101372 | 0.1335 | -0.3635 |
| FAM183A | NM_001101376 | -0.0655 | -0.087 |
| PIRT | NM_001101387 | 0.0055 | 0.234 |
| CLDN25 | NM_001101389 | 0.0975 | 0.0065 |
| LINGO3 | NM_001101391 | 0.027 | -0.0455 |
| SBK2 | NM_001101401 | 0.128 | -0.2335 |
| SH2D7 | NM_001101404 | 0.1605 | 0.0425 |
| ZNF541 | NM_001101419 | -0.1575 | -0.4895 |
| MYO1H | NM_001101421 | 0.29 | -0.028 |
| ISPD | NM_001101426 | -0.839 | -2.5275 |
| NPC1L1 | NM_001101648 | -0.0785 | -0.547 |
| COMMD9 | NM_001101653 | 0.121 | -0.64 |
| CXXC1 | NM_001101654 | 0.0585 | -0.5545 |
| NRD1 | NM_001101662 | -1.1065 | -0.7305 |
| ACOX3 | NM_001101667 | -0.232 | -1.5875 |
| INPP4B | NM_001101669 | -0.044 | -0.407 |
| ZNF219 | NM_001101672 | -0.5185 | -1.2055 |
| SAMD12 | NM_001101676 | -0.088 | -0.6785 |
| FAM13B | NM_001101800 | -1.296 | -2.7485 |
| FAM13B | NM_001101801 | -0.7795 | -1.69 |
| PHF21A | NM_001101802 | -1.179 | -2.4045 |
| ACTN1 | NM_001102 | -0.259 | -1.6145 |
| C14orf159 | NM_001102366 | 0.069 | -0.2425 |
| C14orf159 | NM_001102367 | 0.069 | -0.2425 |
| C14orf159 | NM_001102368 | 0.069 | -0.2425 |

| | | | |
|---|---|---|---|
| C14orf159 | NM_001102369 | 0.069 | -0.2425 |
| FOXRED2 | NM_001102371 | -0.468 | -0.9155 |
| GNAT3 | NM_001102386 | -0.1235 | 0.197 |
| SIKE1 | NM_001102396 | -1.312 | -1.59 |
| HNRNPR | NM_001102397 | -0.601 | -0.736 |
| HNRNPR | NM_001102398 | -0.601 | -0.736 |
| HNRNPR | NM_001102399 | -0.5085 | -0.7005 |
| PCTP | NM_001102402 | -0.8715 | -2.726 |
| ALPK1 | NM_001102406 | -0.3575 | -0.2435 |
| KNG1 | NM_001102416 | 0.342 | 0.365 |
| ZFAND5 | NM_001102420 | -1.4895 | -2.4895 |
| ZFAND5 | NM_001102421 | -1.4895 | -2.4895 |
| TBC1D8 | NM_001102426 | -0.561 | -1.7605 |
| RGS4 | NM_001102445 | -1.5255 | -3.067 |
| RGS8 | NM_001102450 | 0.169 | -0.3095 |
| ZNF219 | NM_001102454 | -0.598 | -2.1515 |
| LIPN | NM_001102469 | -0.339 | -0.2235 |
| ADH6 | NM_001102470 | 0.1775 | -0.358 |
| PPAPDC1B | NM_001102559 | -0.4715 | -0.549 |
| PPAPDC1B | NM_001102560 | 0.05 | -0.311 |
| PCP4L1 | NM_001102566 | -0.1735 | -0.116 |
| SNX18 | NM_001102575 | -0.994 | -1.925 |
| DTX2 | NM_001102594 | -0.5255 | -1.3885 |
| DTX2 | NM_001102595 | -0.5255 | -1.3885 |
| DTX2 | NM_001102596 | -0.5255 | -1.3885 |
| CEACAM20 | NM_001102600 | 0.146 | -0.46 |
| ZNF160 | NM_001102603 | -1.411 | -2.7915 |
| IMPDH1 | NM_001102605 | -0.0955 | -1.069 |
| COL6A6 | NM_001102608 | -0.7045 | -0.2475 |
| C5orf58 | NM_001102609 | -0.3965 | -0.341 |
| TUBGCP5 | NM_001102610 | -1.499 | -2.3035 |
| PGPEP1L | NM_001102612 | 0.1385 | -0.463 |
| SLC35G6 | NM_001102614 | 0.0725 | -0.382 |
| ZNF554 | NM_001102651 | -0.1305 | -1.3405 |
| OTUD4 | NM_001102653 | -1.223 | -2.577 |
| NTF3 | NM_001102654 | -0.5465 | -0.176 |
| ZNF836 | NM_001102657 | 0.228 | -0.266 |
| CT62 | NM_001102658 | 0.307 | 0.065 |
| LOC200726 | NM_001102659 | 0.3725 | -0.246 |
| EPN2 | NM_001102664 | -0.2555 | -0.8275 |
| PSMA4 | NM_001102667 | -0.9245 | -0.6685 |
| PSMA4 | NM_001102668 | -0.908 | -0.769 |
| RRAS2 | NM_001102669 | -0.9475 | -2.199 |

| | | | |
|---|---|---|---|
| ACTN2 | NM_001103 | -0.6525 | -1.4755 |
| GIGYF2 | NM_001103146 | -1.4825 | -2.957 |
| GIGYF2 | NM_001103147 | -1.4825 | -2.957 |
| GIGYF2 | NM_001103148 | -1.4825 | -2.957 |
| PNMAL1 | NM_001103149 | -1.3745 | -1.5295 |
| PNMA5 | NM_001103150 | -0.1525 | -0.2185 |
| PNMA5 | NM_001103151 | -0.111 | -0.5855 |
| ARL17B | NM_001103154 | -0.106 | 0.3675 |
| SH2D5 | NM_001103160 | -0.1305 | -0.6545 |
| SH2D5 | NM_001103161 | -0.7285 | -0.698 |
| ZGLP1 | NM_001103167 | -0.1925 | -0.7575 |
| AADACL3 | NM_001103170 | -0.063 | -0.0625 |
| CCDC64B | NM_001103175 | -0.5625 | -0.2505 |
| CNPY1 | NM_001103176 | -0.112 | -0.01 |
| FMN1 | NM_001103184 | -0.9705 | -2.137 |
| ACTN3 | NM_001104 | 0.0695 | 0.186 |
| FAM70A | NM_001104544 | -0.3695 | -0.6185 |
| FAM70A | NM_001104545 | -0.3695 | -0.6185 |
| RPP30 | NM_001104546 | -0.88 | -2.174 |
| LOC642587 | NM_001104548 | -0.327 | 0.0755 |
| PAQR5 | NM_001104554 | -0.6285 | -0.4595 |
| ELOVL7 | NM_001104558 | 0.237 | -1.5305 |
| GPR172B | NM_001104577 | 0.0285 | -0.614 |
| ALS2CR8 | NM_001104586 | -0.28 | -0.1025 |
| SLFN11 | NM_001104587 | -0.0355 | 0.0725 |
| SLFN11 | NM_001104588 | -0.0355 | 0.0725 |
| SLFN11 | NM_001104589 | -0.0355 | 0.0725 |
| SLFN11 | NM_001104590 | -0.0355 | 0.0725 |
| FAM118A | NM_001104595 | -1.1545 | -2.2575 |
| C4orf19 | NM_001104629 | -0.2505 | -0.039 |
| PDE4D | NM_001104631 | -1.413 | -2.114 |
| SLC25A36 | NM_001104647 | -0.7045 | -1.165 |
| PCMTD2 | NM_001104925 | -1.248 | -0.864 |
| ACVR1 | NM_001105 | -0.925 | -2.6475 |
| ACSM2B | NM_001105069 | -0.0065 | -0.023 |
| MECOM | NM_001105077 | -0.401 | -0.379 |
| MECOM | NM_001105078 | -0.9225 | -0.1355 |
| FBRS | NM_001105079 | -0.1445 | -0.8585 |
| TLE3 | NM_001105192 | -0.616 | -2.0775 |
| FAM123C | NM_001105193 | -0.124 | -0.984 |
| FAM123C | NM_001105194 | -0.124 | -0.984 |
| FAM123C | NM_001105195 | -0.124 | -0.984 |
| TMEM177 | NM_001105198 | -1.0385 | -3.4295 |

| | | | |
|---|---|---|---|
| TMEM177 | NM_001105199 | -1.05 | -3.27 |
| RUSC1 | NM_001105203 | -0.186 | -0.759 |
| RUSC1 | NM_001105204 | -0.186 | -0.759 |
| RUSC1 | NM_001105205 | -0.186 | -0.759 |
| LAMA4 | NM_001105206 | 0.188 | -0.2665 |
| LAMA4 | NM_001105207 | 0.188 | -0.2665 |
| LAMA4 | NM_001105208 | 0.188 | -0.2665 |
| LAMA4 | NM_001105209 | 0.188 | -0.2665 |
| ASH2L | NM_001105214 | -0.946 | -1.8355 |
| PCDH19 | NM_001105243 | -1.295 | -2.411 |
| PTPRM | NM_001105244 | -0.8185 | -1.0735 |
| ARMC5 | NM_001105247 | 0.003 | -0.978 |
| TMC5 | NM_001105248 | 0.147 | 0.37 |
| TMC5 | NM_001105249 | 0.147 | 0.37 |
| NRXN3 | NM_001105250 | -0.261 | -0.418 |
| ZFYVE16 | NM_001105251 | -0.7555 | -2.052 |
| FABP12 | NM_001105281 | 0.026 | -0.506 |
| TTC21A | NM_001105513 | -0.0765 | -0.3015 |
| ABCC4 | NM_001105515 | -1.143 | -1.335 |
| FAM58BP | NM_001105517 | 0.211 | -0.442 |
| C2orf70 | NM_001105519 | 0.014 | -0.3555 |
| C17orf100 | NM_001105520 | -0.5565 | -2.1455 |
| JAKMIP3 | NM_001105521 | -1.0135 | -0.778 |
| C18orf34 | NM_001105528 | 0.258 | 0.106 |
| ATP8A1 | NM_001105529 | -0.7385 | -0.2995 |
| CAP1 | NM_001105530 | -0.699 | -0.8875 |
| FAM135A | NM_001105531 | -1.5035 | -3.084 |
| C7orf58 | NM_001105533 | -0.0785 | -0.0015 |
| ZNF142 | NM_001105537 | -0.943 | -2.974 |
| MYBBP1A | NM_001105538 | -0.534 | -0.937 |
| ZBTB10 | NM_001105539 | -0.486 | -1.5785 |
| DGKZ | NM_001105540 | -0.2985 | -0.85 |
| NEFM | NM_001105541 | -0.097 | 0.7965 |
| PLXNA4 | NM_001105543 | 0.4225 | -0.5845 |
| ECHDC1 | NM_001105544 | -1.0715 | -1.5455 |
| ECHDC1 | NM_001105545 | -1.0715 | -1.5455 |
| ZNF83 | NM_001105549 | -0.8165 | -1.541 |
| C1orf38 | NM_001105556 | -0.032 | -0.253 |
| WEE2 | NM_001105558 | 0.138 | 0.0455 |
| UBE4B | NM_001105562 | -0.5295 | -1.183 |
| CCHCR1 | NM_001105563 | -0.454 | -1.013 |
| CCHCR1 | NM_001105564 | -0.454 | -1.013 |
| SMTNL1 | NM_001105565 | -0.1335 | -0.337 |

| | | | |
|---|---|---|---|
| KIF13A | NM_001105566 | -0.256 | -0.6685 |
| KIF13A | NM_001105567 | -0.256 | -0.6685 |
| KIF13A | NM_001105568 | -0.256 | -0.6685 |
| MSGN1 | NM_001105569 | 0.11 | -0.2975 |
| NUDT19 | NM_001105570 | -1.2345 | -3.4875 |
| DHRS7C | NM_001105571 | -0.1685 | -0.254 |
| PLA2G2C | NM_001105572 | -0.023 | -0.252 |
| FBXO45 | NM_001105573 | -1.556 | -2.611 |
| HMX3 | NM_001105574 | -0.196 | -0.65 |
| ANKRD58 | NM_001105576 | -0.1825 | -0.47 |
| PRHOXNB | NM_001105577 | 0.1395 | -0.259 |
| SYCE2 | NM_001105578 | 0.304 | -0.2555 |
| SYNDIG1L | NM_001105579 | 0.0395 | -0.482 |
| GABRR3 | NM_001105580 | 0.2905 | -0.847 |
| LRRC30 | NM_001105581 | 0.1715 | -0.0105 |
| LRRIQ3 | NM_001105659 | -0.527 | -0.179 |
| USP17 | NM_001105662 | 0.07 | 0.047 |
| NUDT7 | NM_001105663 | -0.7985 | -0.7065 |
| TTC24 | NM_001105669 | 0.156 | -0.2435 |
| ACVR2B | NM_001106 | -1.2895 | -1.7825 |
| ADAM8 | NM_001109 | -0.0615 | -0.2615 |
| HIGD1C | NM_001109619 | -0.3965 | 0.051 |
| C16orf92 | NM_001109659 | -0.081 | 0.07 |
| C16orf92 | NM_001109660 | -0.081 | 0.07 |
| C12orf51 | NM_001109662 | -0.223 | -0.9085 |
| ZNF250 | NM_001109689 | -1.084 | -2.694 |
| PTPRB | NM_001109754 | 0.138 | -0.102 |
| GSG1L | NM_001109763 | -0.044 | -0.5835 |
| ZFP57 | NM_001109809 | 0.005 | -0.3955 |
| TBX22 | NM_001109878 | 0.1475 | 0.218 |
| TBX22 | NM_001109879 | 0.1475 | 0.218 |
| RNFT2 | NM_001109903 | -0.295 | -0.9045 |
| C6orf136 | NM_001109938 | -0.2985 | -1.448 |
| SYNPO | NM_001109974 | -0.5025 | -0.46 |
| FAM160A1 | NM_001109977 | 0.0425 | -0.019 |
| KLHL33 | NM_001109997 | 0.5925 | -0.122 |
| ADAM10 | NM_001110 | -0.526 | -1.6225 |
| SRRM3 | NM_001110199 | -1.1425 | -1.3885 |
| PRH2 | NM_001110213 | -0.1825 | -0.4995 |
| CENPM | NM_001110215 | -0.239 | -1.397 |
| GJB6 | NM_001110219 | -0.523 | -0.47 |
| GJB6 | NM_001110220 | -0.523 | -0.47 |
| GJB6 | NM_001110221 | -0.523 | -0.47 |

| | | | |
|---|---|---|---|
| ZP3 | NM_001110354 | -0.325 | -0.7655 |
| RNASE9 | NM_001110356 | -0.0975 | -0.1575 |
| RNASE9 | NM_001110357 | -0.1285 | -0.148 |
| RNASE9 | NM_001110358 | -0.1285 | -0.148 |
| RNASE9 | NM_001110359 | -0.1285 | -0.148 |
| RNASE9 | NM_001110360 | -0.1285 | -0.148 |
| RNASE9 | NM_001110361 | -0.1285 | -0.148 |
| TMEM87A | NM_001110503 | -1.5265 | -2.5045 |
| EBF4 | NM_001110514 | 0.1705 | -0.6095 |
| FLNA | NM_001110556 | 0.1165 | -0.328 |
| SLC35E2B | NM_001110781 | -0.7245 | -1.293 |
| MECP2 | NM_001110792 | -0.846 | -1.771 |
| NAA16 | NM_001110798 | -1.0985 | -2.0525 |
| TDRD12 | NM_001110822 | 0.2865 | -0.171 |
| ADAR | NM_001111 | -0.567 | -1.21 |
| NAV2 | NM_001111018 | -0.2205 | -1.109 |
| NAV2 | NM_001111019 | -0.2205 | -1.109 |
| SUPT5H | NM_001111020 | 0.0555 | -0.7165 |
| ACVR1C | NM_001111031 | -0.0915 | -0.6225 |
| ACVR1C | NM_001111032 | -0.0915 | -0.6225 |
| ACVR1C | NM_001111033 | -0.0915 | -0.6225 |
| FBXL13 | NM_001111038 | -0.0325 | 0.042 |
| ZNF187 | NM_001111039 | -1.4035 | -2.8105 |
| NHLH2 | NM_001111061 | -0.489 | -0.589 |
| ACVR1 | NM_001111067 | -0.3035 | -2.0435 |
| EZR | NM_001111077 | -0.026 | -1.0215 |
| LYN | NM_001111097 | -0.4895 | -1.253 |
| CNRIP1 | NM_001111101 | -0.7045 | -1.154 |
| UBE2K | NM_001111112 | -0.7565 | -1.5355 |
| UBE2K | NM_001111113 | -0.7565 | -1.5355 |
| IQSEC2 | NM_001111125 | 0.006 | -0.2885 |
| IGF1 | NM_001111283 | 0.1515 | -0.487 |
| IGF1 | NM_001111284 | 0.1515 | -0.487 |
| IGF1 | NM_001111285 | 0.1515 | -0.487 |
| PPIL6 | NM_001111298 | -0.089 | 0.242 |
| PDE4A | NM_001111307 | -0.6795 | -1.967 |
| PDE4A | NM_001111308 | -0.085 | -0.6775 |
| PDE4A | NM_001111309 | -0.085 | -0.6775 |
| CLDN22 | NM_001111319 | -0.0015 | -0.395 |
| DDX54 | NM_001111322 | 0.012 | -0.959 |
| ADARB1 | NM_001112 | -0.8415 | -1.3015 |
| VAX1 | NM_001112704 | 0.106 | -0.047 |
| SCIN | NM_001112706 | 0.2455 | -0.4325 |

| | | | |
|---|---|---|---|
| TLK2 | NM_001112707 | -1.519 | -3.608 |
| LIMCH1 | NM_001112717 | -0.6805 | -0.9945 |
| LIMCH1 | NM_001112718 | -0.6805 | -0.9945 |
| LIMCH1 | NM_001112719 | -0.6805 | -0.9945 |
| LIMCH1 | NM_001112720 | -0.6805 | -0.9945 |
| STK32A | NM_001112724 | 0.191 | 0.166 |
| KIAA0284 | NM_001112726 | -0.3325 | -1.1155 |
| MCF2L | NM_001112732 | -0.772 | -0.633 |
| ZSCAN30 | NM_001112734 | -0.899 | -1.633 |
| KCNC1 | NM_001112741 | -0.4945 | -0.9015 |
| SLC8A1 | NM_001112800 | -0.3575 | -0.271 |
| SLC8A1 | NM_001112801 | -0.3575 | -0.271 |
| SLC8A1 | NM_001112802 | -0.3575 | -0.271 |
| FPGT-TNNI3K | NM_001112808 | 0.381 | 0.051 |
| GRIA4 | NM_001112812 | -0.663 | -0.1215 |
| NFAT5 | NM_001113178 | -1.0085 | -2.852 |
| BRD2 | NM_001113182 | 0.032 | -1.0205 |
| NACA | NM_001113201 | -0.099 | -0.188 |
| NACA | NM_001113202 | -0.099 | -0.188 |
| NACA | NM_001113203 | -0.099 | -0.188 |
| TSTD1 | NM_001113207 | 0.4055 | 0.1365 |
| NTNG1 | NM_001113226 | -0.0105 | -1.3755 |
| NTNG1 | NM_001113228 | -0.0105 | -1.3755 |
| HIPK2 | NM_001113239 | -0.8355 | -1.335 |
| TEN1 | NM_001113324 | 0.429 | -0.457 |
| ECE1 | NM_001113347 | -0.1565 | -0.8735 |
| ECE1 | NM_001113348 | -0.1565 | -0.8735 |
| ECE1 | NM_001113349 | -0.1565 | -0.8735 |
| TBC1D14 | NM_001113361 | -0.682 | -1.4365 |
| TBC1D14 | NM_001113363 | 0.0385 | -0.9445 |
| FANCI | NM_001113378 | -0.91 | -1.1085 |
| RGS4 | NM_001113380 | -1.2155 | -2.4785 |
| RGS4 | NM_001113381 | -1.2155 | -2.4785 |
| FER1L5 | NM_001113382 | -0.186 | -0.02 |
| ZNF385B | NM_001113397 | 0.574 | -0.417 |
| ZNF385B | NM_001113398 | 0.574 | -0.417 |
| AMN1 | NM_001113402 | -1.468 | -2.112 |
| LDB1 | NM_001113407 | -0.446 | -1.63 |
| FGGY | NM_001113411 | -0.036 | -0.356 |
| C17orf51 | NM_001113434 | -0.892 | -1.1965 |
| NOXRED1 | NM_001113475 | -0.131 | -0.567 |
| MANEAL | NM_001113482 | -0.876 | -1.801 |
| AMOT | NM_001113490 | -0.921 | -0.796 |

| | | | |
|---|---|---|---|
| 9-Sep | NM_001113491 | 0.113 | -0.631 |
| MDGA2 | NM_001113498 | -0.4705 | 0.1505 |
| ARHGEF7 | NM_001113511 | -0.5705 | -1.762 |
| ARHGEF7 | NM_001113512 | -0.5705 | -1.762 |
| ARHGEF7 | NM_001113513 | -0.5705 | -1.762 |
| PARP15 | NM_001113523 | -0.1725 | -0.109 |
| ZNF276 | NM_001113525 | -0.412 | -0.8895 |
| FOLR2 | NM_001113534 | -0.0485 | 0.096 |
| FOLR2 | NM_001113535 | -0.1865 | -0.125 |
| FOLR2 | NM_001113536 | -0.0485 | 0.096 |
| LIMA1 | NM_001113546 | -1.0115 | -2.3605 |
| LIMA1 | NM_001113547 | -1.0115 | -2.3605 |
| RNF180 | NM_001113561 | -0.1955 | 0.3595 |
| C17orf76 | NM_001113567 | -0.2425 | -0.3605 |
| CDKL3 | NM_001113575 | -0.8285 | -1.061 |
| ARL17A | NM_001113738 | -0.431 | -0.133 |
| TYMP | NM_001113755 | -0.071 | -0.255 |
| TYMP | NM_001113756 | 0.1555 | -0.1235 |
| ADCY7 | NM_001114 | -0.101 | -0.323 |
| CLIC5 | NM_001114086 | -0.1505 | -0.399 |
| ENTPD6 | NM_001114089 | -0.53 | -1.749 |
| NGEF | NM_001114090 | 0.016 | -0.085 |
| CDC27 | NM_001114091 | -0.7895 | -2.749 |
| THUMPD3 | NM_001114092 | -1.6515 | -3.024 |
| LSM14A | NM_001114093 | -1.281 | -1.746 |
| SEZ6L2 | NM_001114099 | -0.1135 | -0.5305 |
| SEZ6L2 | NM_001114100 | -0.1095 | -0.6625 |
| SLC44A3 | NM_001114106 | 0.137 | -0.4 |
| PDLIM3 | NM_001114107 | -0.334 | -1.205 |
| TTC22 | NM_001114108 | -0.1495 | -0.453 |
| C17orf85 | NM_001114118 | -0.443 | -1.603 |
| DEPDC1 | NM_001114120 | -1.5305 | -4.08 |
| CHEK1 | NM_001114121 | -0.789 | -1.633 |
| CHEK1 | NM_001114122 | -1.066 | -1.552 |
| ELK1 | NM_001114123 | -0.3155 | -0.972 |
| NBEAL1 | NM_001114132 | -0.05 | -0.413 |
| SYNPO2L | NM_001114133 | -0.0025 | 0.267 |
| EPB49 | NM_001114135 | -0.2505 | -0.205 |
| EPB49 | NM_001114136 | -0.023 | -0.579 |
| EPB49 | NM_001114137 | -0.088 | -0.5745 |
| EPB49 | NM_001114138 | -0.2505 | -0.205 |
| EPB49 | NM_001114139 | -0.2505 | -0.205 |
| FOSB | NM_001114171 | 0.038 | -0.2555 |

| | | | |
|---|---|---|---|
| PIK3R3 | NM_001114172 | -1.1455 | -3.346 |
| CTSC | NM_001114173 | -0.4375 | -2.279 |
| GABRA2 | NM_001114175 | -0.0315 | -0.273 |
| AEBP2 | NM_001114176 | -1.5545 | -3.7785 |
| IRAK4 | NM_001114182 | -0.4565 | -0.8125 |
| GRIA1 | NM_001114183 | -0.7495 | -0.244 |
| MTRF1L | NM_001114184 | -0.924 | -1.3715 |
| MVK | NM_001114185 | -0.8125 | -2.01 |
| ELF3 | NM_001114309 | 0.209 | -0.283 |
| GRM1 | NM_001114329 | 0.23 | 0.0755 |
| CLCN7 | NM_001114331 | -0.386 | -0.9145 |
| C4orf47 | NM_001114357 | -0.1965 | 0.212 |
| FOXP3 | NM_001114377 | -0.14 | -0.278 |
| ITGAL | NM_001114380 | -0.0375 | -0.0335 |
| TSC2 | NM_001114382 | 0.0595 | -0.401 |
| TMPRSS11A | NM_001114387 | -0.013 | 0.0625 |
| PAPD4 | NM_001114393 | -1.3575 | -2.8305 |
| PAPD4 | NM_001114394 | -1.3575 | -2.8305 |
| CNTLN | NM_001114395 | -0.5065 | -1.2525 |
| KLRD1 | NM_001114396 | 0.099 | -0.2005 |
| DHX36 | NM_001114397 | -1.176 | -2.453 |
| UPK3BL | NM_001114403 | 0.008 | -0.368 |
| ASPDH | NM_001114598 | 0.121 | -0.232 |
| C1orf144 | NM_001114600 | 0.1575 | -0.953 |
| MFGE8 | NM_001114614 | -0.1305 | -0.1175 |
| MGAT1 | NM_001114617 | -0.157 | -1 |
| MGAT1 | NM_001114618 | -0.446 | -1.8305 |
| MGAT1 | NM_001114619 | -0.446 | -1.8305 |
| MGAT1 | NM_001114620 | -0.446 | -1.8305 |
| JMJD7 | NM_001114632 | -0.336 | -1.4605 |
| PLAG1 | NM_001114634 | 0.097 | -0.1135 |
| PLAG1 | NM_001114635 | 0.097 | -0.1135 |
| FANCL | NM_001114636 | -0.9345 | -1.58 |
| PRRT4 | NM_001114726 | -0.823 | -0.802 |
| PABPC4L | NM_001114734 | -0.177 | -0.0455 |
| BCL2A1 | NM_001114735 | 0.3805 | -0.1965 |
| C1orf70 | NM_001114748 | 0.0385 | -0.098 |
| CD55 | NM_001114752 | -1.036 | 0.004 |
| ENG | NM_001114753 | -0.173 | -0.52 |
| ZNF683 | NM_001114759 | 0.188 | 0.039 |
| SH2D1A | NM_001114937 | 0.481 | -0.218 |
| CCDC17 | NM_001114938 | 0.0095 | -0.0055 |
| SMTNL2 | NM_001114974 | -0.311 | -0.5095 |

| | | | |
|---|---|---|---|
| TP63 | NM_001114978 | -0.1155 | 0.2695 |
| TP63 | NM_001114979 | -0.1155 | 0.2695 |
| TP63 | NM_001114980 | -0.1155 | 0.2695 |
| TP63 | NM_001114981 | -0.1155 | 0.2695 |
| TP63 | NM_001114982 | -0.1155 | 0.2695 |
| ADCY8 | NM_001115 | -0.0915 | -0.2435 |
| LIN54 | NM_001115007 | -0.743 | -2.6445 |
| LIN54 | NM_001115008 | -0.743 | -2.6445 |
| KIAA1310 | NM_001115016 | -0.2375 | -0.688 |
| WDR26 | NM_001115113 | -1.4135 | -3.47 |
| ANKRD53 | NM_001115116 | -0.315 | -0.2785 |
| C6 | NM_001115131 | -0.02 | 0.378 |
| CD300LD | NM_001115152 | -0.3095 | -0.178 |
| GDI2 | NM_001115156 | -0.9145 | -0.7325 |
| ADCY9 | NM_001116 | -0.7475 | -2.6095 |
| ADCYAP1 | NM_001117 | -0.6705 | -1.418 |
| ADCYAP1R1 | NM_001118 | -0.4205 | -2.2315 |
| GLRA2 | NM_001118885 | -0.0015 | -0.655 |
| GLRA2 | NM_001118886 | -0.0015 | -0.655 |
| ANGPT2 | NM_001118887 | -0.2975 | -0.1125 |
| ANGPT2 | NM_001118888 | -0.2975 | -0.1125 |
| GLRX | NM_001118890 | -0.7235 | -0.853 |
| ADD1 | NM_001119 | 0.0445 | -0.8975 |
| MFSD10 | NM_001120 | -0.053 | -0.7115 |
| ADD3 | NM_001121 | -0.794 | -1.0915 |
| PLIN2 | NM_001122 | -0.946 | -1.7575 |
| LAMP2 | NM_001122606 | -0.9845 | -0.8595 |
| RUNX1 | NM_001122607 | 0.0375 | -0.054 |
| HCCS | NM_001122608 | -0.662 | -2.824 |
| CDKN1C | NM_001122630 | 0.4475 | 0.2755 |
| CDKN1C | NM_001122631 | 0.4475 | 0.2755 |
| CPS1 | NM_001122633 | -0.208 | 0.025 |
| CPS1 | NM_001122634 | -0.208 | 0.025 |
| GALNT9 | NM_001122636 | 0.0435 | -0.297 |
| FAM178B | NM_001122646 | 0.196 | 0.1295 |
| EDNRB | NM_001122659 | -0.4215 | 0.038 |
| DDX3Y | NM_001122665 | -0.423 | -0.929 |
| PHKA1 | NM_001122670 | -0.603 | -0.5625 |
| CCBL1 | NM_001122671 | -0.0575 | -0.472 |
| ABCD3 | NM_001122674 | -0.8785 | -1.197 |
| ODZ2 | NM_001122679 | -0.4585 | -0.61 |
| SH3BP2 | NM_001122681 | -0.299 | -1.011 |
| CXorf64 | NM_001122716 | -0.65 | -0.5205 |

| | | | |
|---|---|---|---|
| MICAL3 | NM_001122731 | -0.21 | -0.3985 |
| ESR1 | NM_001122740 | -0.121 | -0.142 |
| ESR1 | NM_001122741 | -0.121 | -0.142 |
| ESR1 | NM_001122742 | -0.121 | -0.142 |
| POU1F1 | NM_001122757 | 0.116 | -0.547 |
| PPOX | NM_001122764 | -0.2605 | -1.1285 |
| LCA5 | NM_001122769 | -1.0105 | -1.56 |
| ZBTB37 | NM_001122770 | -0.079 | -0.437 |
| AGAP2 | NM_001122772 | 0.3625 | -0.276 |
| KIF17 | NM_001122819 | -0.4085 | -0.335 |
| SET | NM_001122821 | -0.3805 | -0.4045 |
| GTF3C5 | NM_001122823 | -0.9325 | -2.521 |
| TACC1 | NM_001122824 | -0.8015 | -1.557 |
| ESRP1 | NM_001122825 | 0.333 | -0.0835 |
| ESRP1 | NM_001122826 | 0.333 | -0.0835 |
| ESRP1 | NM_001122827 | 0.333 | -0.0835 |
| STK31 | NM_001122833 | -0.3935 | -0.1425 |
| HHAT | NM_001122834 | -0.2445 | -0.559 |
| NAPEPLD | NM_001122838 | -1.3605 | -2.712 |
| SLC50A1 | NM_001122839 | -0.179 | -0.8575 |
| FRMD1 | NM_001122841 | 0.17 | -0.1935 |
| NCOA7 | NM_001122842 | -1.2485 | -2.957 |
| SLC6A12 | NM_001122847 | 0.0565 | -0.349 |
| SLC6A12 | NM_001122848 | 0.0565 | -0.349 |
| MYOZ3 | NM_001122853 | -0.077 | -0.1185 |
| PPM1M | NM_001122870 | -0.252 | -0.72 |
| GGT6 | NM_001122890 | -0.2525 | -0.139 |
| CD99 | NM_001122898 | -0.1205 | -0.304 |
| DARC | NM_001122951 | -0.175 | -0.4375 |
| BSCL2 | NM_001122955 | -0.107 | -0.9375 |
| DBNL | NM_001122956 | -0.088 | -0.6955 |
| BCKDK | NM_001122957 | -0.2665 | -0.9875 |
| C1orf194 | NM_001122961 | -0.2485 | -0.529 |
| SIRPB2 | NM_001122962 | 0.005 | -0.438 |
| SMEK2 | NM_001122964 | -1.7515 | -2.847 |
| RPTN | NM_001122965 | 0.32 | -0.5675 |
| ADK | NM_001123 | -0.902 | -0.845 |
| C3orf71 | NM_001123040 | 1.41 | 1.7145 |
| CCR2 | NM_001123041 | -0.0365 | -0.128 |
| C7orf65 | NM_001123065 | -0.07 | 0.334 |
| MAPT | NM_001123066 | 0.0845 | -0.5055 |
| MAPT | NM_001123067 | 0.0845 | -0.5055 |
| SYCE3 | NM_001123225 | 0.3195 | -0.1725 |

| | | | |
|---|---|---|---|
| MTO1 | NM_001123226 | -1.785 | -3.24 |
| TMEM14E | NM_001123228 | -0.1705 | 0.0265 |
| ZBTB1 | NM_001123329 | -1.133 | -2.026 |
| PPP6C | NM_001123355 | -0.4145 | -1.4785 |
| C6orf186 | NM_001123364 | 0.0035 | -0.293 |
| HMSD | NM_001123366 | 0.547 | -0.586 |
| PPP6C | NM_001123369 | -0.4145 | -1.4785 |
| HIST2H3D | NM_001123375 | 2.404 | 1.464 |
| TMEM72 | NM_001123376 | -0.179 | -0.3735 |
| PARK7 | NM_001123377 | -0.0085 | -0.1105 |
| ADSL | NM_001123378 | -0.3455 | -1.097 |
| BCOR | NM_001123383 | -1.034 | -3.5405 |
| BCOR | NM_001123384 | -1.034 | -3.5405 |
| BCOR | NM_001123385 | -0.975 | -4.032 |
| KRTAP2-1 | NM_001123387 | -0.3465 | -0.256 |
| CLDN19 | NM_001123395 | 0.065 | -0.5455 |
| CCR2 | NM_001123396 | 0.107 | -0.159 |
| ADM | NM_001124 | 0.1095 | -0.8985 |
| PABPC1L | NM_001124756 | -0.893 | -1.4565 |
| SPNS2 | NM_001124758 | -0.2245 | -1.4445 |
| FRG2C | NM_001124759 | -0.6385 | 0.7415 |
| C3orf78 | NM_001124767 | 0.109 | -0.3625 |
| ADPRH | NM_001125 | 0.3775 | -0.1205 |
| ADSS | NM_001126 | -1.7405 | -3.4825 |
| PTGER3 | NM_001126044 | -0.2095 | -0.2735 |
| KLLN | NM_001126049 | -0.8385 | -0.5055 |
| HDGF | NM_001126050 | 0.248 | -0.3115 |
| HDGF | NM_001126051 | 0.248 | -0.3115 |
| CASK | NM_001126054 | -1.1375 | -1.4375 |
| CASK | NM_001126055 | -1.1375 | -1.4375 |
| DMKN | NM_001126056 | 0.229 | -0.1485 |
| DMKN | NM_001126057 | 0.229 | -0.1485 |
| DMKN | NM_001126058 | 0.229 | -0.1485 |
| DMKN | NM_001126059 | 0.229 | -0.1485 |
| NOS1AP | NM_001126060 | 0.02 | -0.5025 |
| KHDC1L | NM_001126063 | -0.389 | -0.23 |
| HP | NM_001126102 | 0.243 | -0.0175 |
| RACGAP1 | NM_001126103 | -1.4795 | -1.8495 |
| RACGAP1 | NM_001126104 | -1.4795 | -1.8495 |
| SLC7A7 | NM_001126106 | -0.0995 | 0.0345 |
| SLC12A3 | NM_001126107 | -0.0575 | -0.321 |
| SLC12A3 | NM_001126108 | -0.0575 | -0.321 |
| OSGIN2 | NM_001126111 | -0.321 | -1.4185 |

| | | | |
|---|---|---|---|
| TP53 | NM_001126112 | -0.5945 | -0.8345 |
| TP53 | NM_001126113 | -0.5945 | -0.8345 |
| TP53 | NM_001126114 | -0.5945 | -0.8345 |
| TP53 | NM_001126115 | -0.5945 | -0.8345 |
| TP53 | NM_001126116 | -0.166 | -0.501 |
| TP53 | NM_001126117 | -0.5945 | -0.8345 |
| SLC25A19 | NM_001126121 | -0.869 | -2.739 |
| SLC25A19 | NM_001126122 | -0.869 | -2.739 |
| ENOSF1 | NM_001126123 | -0.779 | -0.6855 |
| PROK2 | NM_001126128 | 0.253 | -0.2675 |
| GINS3 | NM_001126130 | -0.775 | -2.2675 |
| TNNT1 | NM_001126132 | 0.266 | 0.288 |
| TNNT1 | NM_001126133 | 0.29 | 0.1125 |
| NRGN | NM_001126181 | 0.129 | -0.064 |
| TP73 | NM_001126240 | -0.0585 | -0.73 |
| TP73 | NM_001126241 | -0.0585 | -0.73 |
| TP73 | NM_001126242 | -0.0585 | -0.73 |
| LNX1 | NM_001126328 | -0.3485 | -0.7315 |
| VCAN | NM_001126336 | -0.48 | -1.449 |
| TUFT1 | NM_001126337 | 0.0535 | -0.3995 |
| THTPA | NM_001126339 | -0.4225 | -0.666 |
| ORAI2 | NM_001126340 | -0.5505 | -1.0075 |
| AP1B1 | NM_001127 | 0.0475 | -0.406 |
| CADM3 | NM_001127173 | -0.0515 | -0.0635 |
| MRO | NM_001127174 | 0.0155 | 0.159 |
| MRO | NM_001127175 | 0.0155 | 0.159 |
| MRO | NM_001127176 | 0.0155 | 0.159 |
| PIGG | NM_001127178 | -0.8 | -1.669 |
| MYO7A | NM_001127179 | 0.4125 | -0.2555 |
| MYO7A | NM_001127180 | 0.4125 | -0.2555 |
| CENPL | NM_001127181 | -1.4525 | -3.2925 |
| CEP55 | NM_001127182 | -1.8455 | -3.662 |
| CFLAR | NM_001127183 | -0.587 | -0.769 |
| CFLAR | NM_001127184 | -0.651 | -0.44 |
| CSK | NM_001127190 | -0.28 | -1.5995 |
| CNBP | NM_001127192 | -0.682 | -0.669 |
| CNBP | NM_001127193 | -0.682 | -0.669 |
| CNBP | NM_001127194 | -0.682 | -0.669 |
| CNBP | NM_001127195 | -0.682 | -0.669 |
| CNBP | NM_001127196 | -0.682 | -0.669 |
| TMC6 | NM_001127198 | -0.0285 | -0.358 |
| PCID2 | NM_001127202 | -1.3035 | -2.211 |
| PCID2 | NM_001127203 | -1.5115 | -2.244 |

| | | | |
|---|---|---|---|
| HMOX2 | NM_001127204 | -0.2595 | -1.9745 |
| HMOX2 | NM_001127205 | -0.2595 | -1.9745 |
| HMOX2 | NM_001127206 | -0.2595 | -1.9745 |
| SMARCAL1 | NM_001127207 | -1.7225 | -3.108 |
| TET2 | NM_001127208 | -1.049 | -3.293 |
| KIAA1598 | NM_001127211 | 0.117 | -0.1755 |
| ACSF3 | NM_001127214 | -0.204 | -0.9045 |
| GFI1 | NM_001127215 | 0.225 | -0.352 |
| GFI1 | NM_001127216 | 0.225 | -0.352 |
| SMAD9 | NM_001127217 | -0.631 | -1.7815 |
| POLD2 | NM_001127218 | 0.156 | -0.415 |
| ACCS | NM_001127219 | -0.0495 | -0.109 |
| CACNA1A | NM_001127221 | 0.246 | -0.356 |
| CACNA1A | NM_001127222 | 0.246 | -0.356 |
| CD59 | NM_001127223 | -0.806 | -0.8585 |
| CD59 | NM_001127225 | -0.806 | -0.8585 |
| CD59 | NM_001127226 | -0.806 | -0.8585 |
| CD59 | NM_001127227 | -0.806 | -0.8585 |
| AURKAIP1 | NM_001127229 | -0.1445 | -0.823 |
| AURKAIP1 | NM_001127230 | -0.1445 | -0.823 |
| AUTS2 | NM_001127231 | -0.6005 | -2.613 |
| AUTS2 | NM_001127232 | -0.148 | -2.2845 |
| GPBP1 | NM_001127235 | -0.808 | -2.45 |
| GPBP1 | NM_001127236 | -0.808 | -2.45 |
| BBC3 | NM_001127240 | -0.1295 | 0.108 |
| BBC3 | NM_001127241 | -0.1295 | 0.108 |
| BBC3 | NM_001127242 | -0.1295 | 0.108 |
| LRRC8A | NM_001127244 | -0.714 | -1.6635 |
| LRRC8A | NM_001127245 | -0.8835 | -2.146 |
| NLRP7 | NM_001127255 | 0.7245 | -0.7405 |
| SLC39A10 | NM_001127257 | -1.1375 | -1.6265 |
| HHIPL1 | NM_001127258 | 0.01 | -0.2275 |
| TMEM129 | NM_001127266 | -0.466 | -0.71 |
| UBAP2L | NM_001127320 | -0.3455 | -1.631 |
| CBX5 | NM_001127321 | -1.341 | -1.0305 |
| CBX5 | NM_001127322 | -1.341 | -1.0305 |
| GRM8 | NM_001127323 | -1.5425 | -2.1155 |
| ACADM | NM_001127328 | -0.8865 | -0.9625 |
| PHTF2 | NM_001127357 | -1.417 | -2.733 |
| PHTF2 | NM_001127358 | -1.417 | -2.733 |
| PHTF2 | NM_001127359 | -1.417 | -2.733 |
| PHTF2 | NM_001127360 | -1.417 | -2.733 |
| RNF19B | NM_001127361 | -0.081 | -1.442 |

| | | | |
|---|---|---|---|
| NR2C1 | NM_001127362 | -0.899 | -2.0125 |
| C7orf46 | NM_001127364 | -1.001 | -1.393 |
| C7orf46 | NM_001127365 | -1.001 | -1.393 |
| PAX3 | NM_001127366 | 0.1345 | 0.0185 |
| CDCA7L | NM_001127370 | -0.28 | -1.7325 |
| CDCA7L | NM_001127371 | -0.28 | -1.7325 |
| ZNF84 | NM_001127372 | -1.0375 | -2.924 |
| SAA2 | NM_001127380 | -0.1515 | -0.3145 |
| PPP2R2B | NM_001127381 | -1.281 | -1.527 |
| CYBRD1 | NM_001127383 | -0.2815 | -0.442 |
| CTNNA3 | NM_001127384 | -0.1425 | -0.549 |
| CTXN3 | NM_001127385 | -0.155 | -0.1465 |
| ALS2CR12 | NM_001127391 | -0.3915 | -0.2665 |
| C11orf9 | NM_001127392 | -0.282 | -0.9 |
| C14orf2 | NM_001127393 | 0.2335 | -0.119 |
| TSEN15 | NM_001127394 | -1.017 | -1.3245 |
| ERLEC1 | NM_001127397 | -1.4385 | -3.3535 |
| ERLEC1 | NM_001127398 | -1.4385 | -3.3535 |
| YPEL5 | NM_001127399 | -0.6555 | -1.0835 |
| YPEL5 | NM_001127400 | -0.6555 | -1.0835 |
| YPEL5 | NM_001127401 | -0.6555 | -1.0835 |
| CPA5 | NM_001127441 | -0.335 | -0.2305 |
| CD36 | NM_001127443 | 0.1265 | -0.518 |
| CD36 | NM_001127444 | 0.1265 | -0.518 |
| ABAT | NM_001127448 | 0.065 | -0.351 |
| DFNA5 | NM_001127453 | -0.833 | -0.781 |
| CRY2 | NM_001127457 | -0.408 | -1.4755 |
| CRLS1 | NM_001127458 | -0.914 | -2.4385 |
| PAN2 | NM_001127460 | -0.653 | -0.872 |
| NLRP3 | NM_001127461 | -0.0565 | 0.0615 |
| NLRP3 | NM_001127462 | -0.0565 | 0.0615 |
| ZNF469 | NM_001127464 | -0.092 | -0.801 |
| ANK2 | NM_001127493 | -0.69 | -1.0435 |
| SPRY4 | NM_001127496 | -0.172 | -0.609 |
| MET | NM_001127500 | -2.003 | -0.7975 |
| ALPL | NM_001127501 | 0.1715 | -0.4185 |
| ASAH1 | NM_001127505 | -0.843 | -0.7025 |
| BANK1 | NM_001127507 | 0.4275 | -0.518 |
| APC | NM_001127510 | -0.032 | -1.032 |
| APC | NM_001127511 | -0.032 | -1.032 |
| ING4 | NM_001127582 | -0.7445 | -2.024 |
| ING4 | NM_001127583 | -0.7445 | -2.024 |
| ING4 | NM_001127584 | -0.7445 | -2.024 |

| | | | |
|---|---|---|---|
| ING4 | NM_001127585 | -0.7445 | -2.024 |
| ING4 | NM_001127586 | -0.7445 | -2.024 |
| FCGR3A | NM_001127592 | 0.4365 | -0.566 |
| FCGR3A | NM_001127593 | 0.4365 | -0.566 |
| FCGR3A | NM_001127595 | 0.4365 | -0.566 |
| FCGR3A | NM_001127596 | 0.4365 | -0.566 |
| LIPA | NM_001127605 | -1.102 | -1.2775 |
| BAAT | NM_001127610 | -0.0015 | -0.465 |
| PAX6 | NM_001127612 | 0.165 | -0.474 |
| ENOX1 | NM_001127615 | -1.115 | -1.3835 |
| ALDOA | NM_001127617 | 0.305 | -0.157 |
| GALE | NM_001127621 | -0.1105 | -1.0535 |
| LAMB3 | NM_001127641 | -0.129 | -0.0855 |
| GABRA1 | NM_001127643 | -0.4125 | -0.7615 |
| GABRA1 | NM_001127644 | -0.4125 | -0.7615 |
| GABRA1 | NM_001127645 | -0.4125 | -0.7615 |
| GABRA1 | NM_001127646 | -0.4125 | -0.7615 |
| GABRA1 | NM_001127647 | -0.4125 | -0.7615 |
| GABRA1 | NM_001127648 | -0.4125 | -0.7615 |
| PEX26 | NM_001127649 | 0.079 | -0.819 |
| NCF2 | NM_001127651 | -0.082 | -0.836 |
| MFN2 | NM_001127660 | -0.5245 | -1.6125 |
| GSN | NM_001127662 | -0.04 | -0.0725 |
| KCNE1 | NM_001127668 | -0.046 | -0.102 |
| KCNE1 | NM_001127669 | -0.046 | -0.102 |
| KCNE1 | NM_001127670 | -0.046 | -0.102 |
| LIFR | NM_001127671 | -0.5505 | -1.532 |
| ADIPOR1 | NM_001127687 | -0.1255 | -2.337 |
| RHD | NM_001127691 | 0.1115 | -0.5965 |
| PCCA | NM_001127692 | -0.8495 | -1.0835 |
| CTSA | NM_001127695 | -0.484 | -1.676 |
| ATXN3 | NM_001127696 | -0.521 | -2.1695 |
| ATXN3 | NM_001127697 | -0.521 | -2.1695 |
| SPINK5 | NM_001127698 | -0.4955 | -0.411 |
| SPINK5 | NM_001127699 | -0.4955 | -0.411 |
| SERPINA1 | NM_001127700 | -0.127 | -0.2305 |
| SERPINA1 | NM_001127701 | 0.0925 | -0.1035 |
| SERPINA1 | NM_001127702 | -0.127 | -0.2305 |
| SERPINA1 | NM_001127703 | -0.127 | -0.2305 |
| SERPINA1 | NM_001127704 | -0.127 | -0.2305 |
| SERPINA1 | NM_001127705 | -0.127 | -0.2305 |
| SERPINA1 | NM_001127706 | -0.127 | -0.2305 |
| SERPINA1 | NM_001127707 | -0.127 | -0.2305 |

| | | | |
|---|---|---|---|
| PRG4 | NM_001127708 | -0.2445 | -0.2815 |
| PRG4 | NM_001127709 | -0.2445 | -0.2815 |
| PRG4 | NM_001127710 | -0.2445 | -0.2815 |
| TMEM14B | NM_001127711 | -0.9575 | -0.698 |
| ATL1 | NM_001127713 | -1.3525 | -2.599 |
| HIVEP3 | NM_001127714 | 0.0365 | -0.5415 |
| STXBP5 | NM_001127715 | -0.8715 | -1.8025 |
| ETFA | NM_001127716 | -0.8085 | -0.7995 |
| LAMA3 | NM_001127717 | -0.045 | -0.357 |
| MMP2 | NM_001127891 | 0.2795 | -0.31 |
| SALL1 | NM_001127892 | -0.299 | -0.479 |
| CEACAM19 | NM_001127893 | 0.1165 | -0.6515 |
| RPGRIP1L | NM_001127897 | -1.289 | -2.3945 |
| CLCN5 | NM_001127898 | -0.076 | -0.404 |
| CLCN5 | NM_001127899 | -1.489 | -1.783 |
| AP1G1 | NM_001128 | -0.51 | -1.1665 |
| SPDYE4 | NM_001128076 | 0.0295 | -0.5475 |
| KRTAP20-3 | NM_001128077 | 0.451 | -0.127 |
| AP4S1 | NM_001128126 | -1.2495 | -1.2885 |
| GK | NM_001128127 | -1.097 | -1.8855 |
| ZEB1 | NM_001128128 | -0.308 | -2.162 |
| DPEP1 | NM_001128141 | -0.1255 | -0.213 |
| RWDD3 | NM_001128142 | -1.2525 | -2.6935 |
| PMS1 | NM_001128143 | 1.231 | 1.245 |
| PMS1 | NM_001128144 | 1.231 | 1.245 |
| NF1 | NM_001128147 | -0.6645 | -1.328 |
| TFRC | NM_001128148 | -0.9985 | -0.5185 |
| ATXN7 | NM_001128149 | -1.679 | -2.367 |
| ORMDL1 | NM_001128150 | -0.831 | -1.5085 |
| VPS53 | NM_001128159 | -0.7235 | -0.988 |
| UBP1 | NM_001128160 | -0.825 | -1.241 |
| UBP1 | NM_001128161 | -0.966 | -0.97 |
| ATXN1 | NM_001128164 | 0.0255 | -0.4575 |
| FBLN7 | NM_001128165 | -0.184 | -0.0455 |
| PAK3 | NM_001128166 | -0.9495 | -1.583 |
| PAK3 | NM_001128167 | -0.9495 | -1.583 |
| PAK3 | NM_001128168 | -0.9495 | -1.583 |
| PAK3 | NM_001128172 | -0.9495 | -1.583 |
| PAK3 | NM_001128173 | -0.9495 | -1.583 |
| UGT8 | NM_001128174 | -0.725 | -2.499 |
| DTNA | NM_001128175 | -0.5815 | -1.7065 |
| THRB | NM_001128176 | 0.3855 | -0.679 |
| THRB | NM_001128177 | 0.3855 | -0.679 |

| | | | |
|---|---|---|---|
| NPHP1 | NM_001128178 | -0.3955 | -0.1105 |
| NPHP1 | NM_001128179 | -0.3955 | -0.1105 |
| C10orf122 | NM_001128202 | -0.0645 | -0.6895 |
| PLA2G16 | NM_001128203 | -0.3085 | -0.512 |
| SULF1 | NM_001128204 | 0.0655 | -0.37 |
| SULF1 | NM_001128205 | 0.0655 | -0.37 |
| SULF1 | NM_001128206 | 0.0655 | -0.37 |
| LEPROTL1 | NM_001128208 | -1.173 | -1.911 |
| SGCD | NM_001128209 | 0.104 | -0.1815 |
| SPRED2 | NM_001128210 | -0.6615 | -2.3885 |
| NUDCD1 | NM_001128211 | -1.199 | -2.8695 |
| WDSUB1 | NM_001128212 | -0.8975 | -1.4785 |
| WDSUB1 | NM_001128213 | -1.1275 | -2.6255 |
| LIPM | NM_001128215 | -0.329 | -0.3405 |
| PSIP1 | NM_001128217 | -0.356 | -0.6365 |
| VGLL4 | NM_001128219 | -0.5375 | -1.5645 |
| VGLL4 | NM_001128220 | -0.198 | -1.1195 |
| VGLL4 | NM_001128221 | -0.198 | -1.1195 |
| ZNF717 | NM_001128223 | 0.042 | -0.553 |
| TMCC1 | NM_001128224 | -0.9735 | -1.5765 |
| SLC39A13 | NM_001128225 | -0.418 | -1.019 |
| DIS3 | NM_001128226 | -0.638 | -1.895 |
| GNE | NM_001128227 | -1.128 | -2.1295 |
| TPRN | NM_001128228 | -0.0255 | -0.2165 |
| LYRM1 | NM_001128301 | -0.7275 | -0.7225 |
| LYRM1 | NM_001128302 | -0.7275 | -0.7225 |
| LACC1 | NM_001128303 | -0.272 | -0.5065 |
| PLSCR4 | NM_001128306 | -0.1515 | -0.1005 |
| TSPAN14 | NM_001128309 | 0.418 | -1.1895 |
| SPARCL1 | NM_001128310 | -0.2195 | 0.1805 |
| ANUBL1 | NM_001128324 | -0.8995 | -2.39 |
| SPON2 | NM_001128325 | 0.019 | -0.038 |
| BSPH1 | NM_001128326 | 0.4775 | 0.173 |
| MPV17L | NM_001128423 | 0.1715 | -1.206 |
| FAM198B | NM_001128424 | -0.33 | -0.2505 |
| AP1AR | NM_001128426 | -1.1245 | -2.4445 |
| FSTL5 | NM_001128427 | -0.3675 | -0.3765 |
| FSTL5 | NM_001128428 | -0.3675 | -0.3765 |
| SMARCAD1 | NM_001128429 | -1.1235 | -4.0625 |
| SMARCAD1 | NM_001128430 | -1.1235 | -4.0625 |
| SLC39A14 | NM_001128431 | -0.6365 | -1.573 |
| SLC14A1 | NM_001128588 | -0.054 | 0.3315 |
| CYP21A2 | NM_001128590 | -0.2815 | -0.1725 |

| | | | |
|---|---|---|---|
| PSMG4 | NM_001128591 | -0.4755 | -1.3075 |
| PSMG4 | NM_001128592 | -0.4755 | -1.3075 |
| TC2N | NM_001128595 | -0.216 | -0.501 |
| KRTAP25-1 | NM_001128598 | -0.1325 | -0.1235 |
| LCE6A | NM_001128600 | -0.3455 | -0.322 |
| RASGRP1 | NM_001128602 | 0.0795 | 0.168 |
| MAPKBP1 | NM_001128608 | -0.2935 | -1.0835 |
| USP8 | NM_001128610 | -0.683 | -2.3825 |
| USP8 | NM_001128611 | -0.683 | -2.3825 |
| NUDT22 | NM_001128613 | -0.3095 | -0.7485 |
| ARHGEF3 | NM_001128615 | -0.9255 | -2.32 |
| ARHGEF3 | NM_001128616 | -0.9255 | -2.32 |
| REPS1 | NM_001128617 | -0.734 | -2.017 |
| C9orf57 | NM_001128618 | 0.4505 | 0.259 |
| LUZP6 | NM_001128619 | -0.844 | -0.433 |
| PAK1 | NM_001128620 | -0.3045 | -1.6395 |
| SPIRE1 | NM_001128626 | -1.6745 | -2.523 |
| SPIRE1 | NM_001128627 | -1.6745 | -2.523 |
| PAK6 | NM_001128628 | -0.0785 | -0.326 |
| PAK6 | NM_001128629 | -0.0785 | -0.326 |
| DCAKD | NM_001128631 | -0.2775 | -0.7405 |
| RIMBP3C | NM_001128633 | -0.056 | -0.937 |
| ELFN1 | NM_001128636 | -0.09 | -0.799 |
| NCS1 | NM_001128826 | -0.2275 | -0.618 |
| DLG4 | NM_001128827 | -0.229 | -0.5625 |
| CA1 | NM_001128829 | 0.052 | -0.7995 |
| CA1 | NM_001128830 | 0.141 | -0.382 |
| CA1 | NM_001128831 | 0.052 | -0.7995 |
| ZBTB4 | NM_001128833 | -1.529 | -2.598 |
| PLP1 | NM_001128834 | 0.0055 | -0.3675 |
| TRAPPC2 | NM_001128835 | -0.239 | -0.6835 |
| CACNA1D | NM_001128839 | -0.3465 | 0.065 |
| CACNA1D | NM_001128840 | -0.3465 | 0.065 |
| TNIP3 | NM_001128843 | 0.3175 | -0.134 |
| SMARCA4 | NM_001128844 | -0.2165 | -0.7055 |
| SMARCA4 | NM_001128845 | -0.2165 | -0.7055 |
| SMARCA4 | NM_001128846 | -0.2165 | -0.7055 |
| SMARCA4 | NM_001128847 | -0.2165 | -0.7055 |
| SMARCA4 | NM_001128848 | -0.2165 | -0.7055 |
| SMARCA4 | NM_001128849 | -0.2165 | -0.7055 |
| RRAD | NM_001128850 | 0.1065 | 0.256 |
| SRRT | NM_001128852 | 0.2585 | 0.405 |
| SRRT | NM_001128853 | 0.2585 | 0.405 |

| | | | |
|---|---|---|---|
| SRRT | NM_001128854 | 0.2585 | 0.405 |
| GTPBP3 | NM_001128855 | 0.1175 | -0.088 |
| PCBP2 | NM_001128911 | 0.152 | -0.2755 |
| PCBP2 | NM_001128912 | 0.152 | -0.2755 |
| PCBP2 | NM_001128913 | 0.152 | -0.2755 |
| PCBP2 | NM_001128914 | 0.152 | -0.2755 |
| CYP24A1 | NM_001128915 | -0.3545 | -0.228 |
| TOMM40 | NM_001128916 | -0.439 | -1.1905 |
| TOMM40 | NM_001128917 | -0.439 | -1.1905 |
| MARK3 | NM_001128918 | -1.1385 | -2.5945 |
| MARK3 | NM_001128919 | -1.1385 | -2.5945 |
| MARK3 | NM_001128920 | -1.1385 | -2.5945 |
| MARK3 | NM_001128921 | -1.1385 | -2.5945 |
| LRRC32 | NM_001128922 | -0.333 | -0.5815 |
| SH3D19 | NM_001128923 | -1.2825 | -1.9435 |
| SH3D19 | NM_001128924 | -1.2825 | -1.9435 |
| CYP2C18 | NM_001128925 | -0.048 | -0.335 |
| FAM71F2 | NM_001128926 | -0.43 | -0.3645 |
| INPP1 | NM_001128928 | -0.6145 | -2.4895 |
| ENTPD4 | NM_001128930 | -1.1605 | -2.0365 |
| NAF1 | NM_001128931 | -0.3115 | -1.2255 |
| CYP4F11 | NM_001128932 | 0.001 | -0.2155 |
| SYNPO2 | NM_001128933 | -0.0725 | -1.4085 |
| SYNPO2 | NM_001128934 | -0.0725 | -1.4085 |
| AEBP1 | NM_001129 | -0.003 | -0.6755 |
| ZNF837 | NM_001129730 | -0.2295 | -1.207 |
| PLEKHG4 | NM_001129731 | -0.2885 | -1.1345 |
| CALHM3 | NM_001129742 | -0.207 | -0.293 |
| NSDHL | NM_001129765 | -0.562 | -2.284 |
| GRAPL | NM_001129778 | 0.2015 | 0.1945 |
| CYB5R3 | NM_001129819 | 0.241 | -0.4135 |
| SLFN14 | NM_001129820 | 0.0285 | -0.256 |
| CACNA1C | NM_001129827 | -0.2715 | -0.3655 |
| CACNA1C | NM_001129829 | -0.2715 | -0.3655 |
| CACNA1C | NM_001129830 | -0.2715 | -0.3655 |
| CACNA1C | NM_001129831 | -0.2715 | -0.3655 |
| CACNA1C | NM_001129832 | -0.2715 | -0.3655 |
| CACNA1C | NM_001129833 | -0.2715 | -0.3655 |
| CACNA1C | NM_001129834 | -0.2715 | -0.3655 |
| CACNA1C | NM_001129835 | -0.2715 | -0.3655 |
| CACNA1C | NM_001129836 | -0.2715 | -0.3655 |
| CACNA1C | NM_001129837 | -0.2715 | -0.3655 |
| CACNA1C | NM_001129838 | -0.2715 | -0.3655 |

| | | | |
|---|---|---|---|
| CACNA1C | NM_001129839 | -0.2715 | -0.3655 |
| CACNA1C | NM_001129840 | -0.2715 | -0.3655 |
| CACNA1C | NM_001129841 | -0.2715 | -0.3655 |
| CACNA1C | NM_001129842 | -0.2715 | -0.3655 |
| CACNA1C | NM_001129843 | -0.2715 | -0.3655 |
| CACNA1C | NM_001129844 | -0.2715 | -0.3655 |
| CACNA1C | NM_001129846 | -0.2715 | -0.3655 |
| PATE3 | NM_001129883 | -0.232 | 0.1245 |
| CPSF4L | NM_001129885 | -0.11 | -0.1955 |
| DCT | NM_001129889 | 0.051 | -0.506 |
| LRRC69 | NM_001129890 | -0.1035 | -0.286 |
| FAM196B | NM_001129891 | 0.1585 | -0.4 |
| HGC6.3 | NM_001129895 | 0.031 | -0.234 |
| SYTL4 | NM_001129896 | -0.1455 | -1.6585 |
| ZNF673 | NM_001129899 | -1.433 | -1.5525 |
| TMPRSS11BNL | NM_001129907 | 0.056 | -0.567 |
| FAM198A | NM_001129908 | -0.253 | 0.009 |
| SYCE1L | NM_001129979 | 0.5875 | 0.294 |
| KIAA1841 | NM_001129993 | -1.062 | -1.1745 |
| KCTD15 | NM_001129994 | -0.9515 | -1.9305 |
| KCTD15 | NM_001129995 | -0.9515 | -1.9305 |
| ZNF222 | NM_001129996 | -1.366 | -2.197 |
| CLEC12B | NM_001129998 | -0.0965 | 0.042 |
| AES | NM_001130 | 0.245 | -0.215 |
| C3orf55 | NM_001130001 | -0.011 | -0.2205 |
| C3orf55 | NM_001130002 | -0.011 | -0.2205 |
| SYNPR | NM_001130003 | -0.62 | 0.132 |
| ACTN1 | NM_001130004 | -0.259 | -1.6145 |
| ACTN1 | NM_001130005 | -0.259 | -1.6145 |
| GSPT1 | NM_001130006 | -0.6225 | -1.308 |
| GSPT1 | NM_001130007 | -0.6225 | -1.308 |
| GEN1 | NM_001130009 | -0.458 | -2.017 |
| TEX101 | NM_001130011 | 0.254 | -0.3075 |
| SLC9A3R2 | NM_001130012 | 0.049 | -0.5035 |
| PSG5 | NM_001130014 | -0.32 | 0.4065 |
| ANKRD33 | NM_001130015 | 0.058 | -0.303 |
| ART3 | NM_001130016 | -0.047 | -0.3845 |
| ART3 | NM_001130017 | -0.047 | -0.3845 |
| ATP6V0A1 | NM_001130020 | -0.608 | -1.662 |
| ATP6V0A1 | NM_001130021 | -0.52 | -1.1825 |
| ZNF680 | NM_001130022 | -1.107 | -1.6155 |
| FAM115C | NM_001130025 | 0.0335 | -0.7205 |
| FAM115C | NM_001130026 | 0.0335 | -0.7205 |

| | | | |
|---|---|---|---|
| CLK3 | NM_001130028 | -0.3935 | -2.1085 |
| RELL2 | NM_001130029 | -0.307 | -0.804 |
| ZNF562 | NM_001130031 | -1.5165 | -3.2635 |
| ZNF562 | NM_001130032 | -1.0175 | -2.2645 |
| PLEKHB1 | NM_001130033 | 0.339 | -0.3015 |
| PLEKHB1 | NM_001130034 | 0.339 | -0.3015 |
| PLEKHB1 | NM_001130035 | 0.339 | -0.3015 |
| PLEKHB1 | NM_001130036 | 0.339 | -0.3015 |
| ELMOD1 | NM_001130037 | -0.701 | -0.3935 |
| SHC1 | NM_001130040 | -0.1715 | -1.0885 |
| SHC1 | NM_001130041 | -0.1715 | -1.0885 |
| CRYZ | NM_001130042 | -0.6095 | -0.7745 |
| TTLL10 | NM_001130045 | 0.7405 | 0.399 |
| TBCEL | NM_001130047 | -0.4105 | -1.07 |
| DOCK9 | NM_001130048 | -0.9655 | -1.1575 |
| DOCK9 | NM_001130049 | -0.676 | -1.6925 |
| DOCK9 | NM_001130050 | 0.121 | -0.086 |
| EEF1D | NM_001130053 | 0.3765 | -0.1945 |
| EEF1D | NM_001130055 | 0.3765 | -0.1945 |
| EEF1D | NM_001130056 | 0.3765 | -0.1945 |
| EEF1D | NM_001130057 | 0.3765 | -0.1945 |
| SLC44A5 | NM_001130058 | -0.913 | -0.834 |
| ATF7 | NM_001130059 | -0.6065 | -0.2775 |
| ATF7 | NM_001130060 | -0.6065 | -0.2775 |
| GRM2 | NM_001130063 | -0.2915 | -0.241 |
| GAP43 | NM_001130064 | 0.1145 | -0.4145 |
| MYO9B | NM_001130065 | -0.645 | -0.4465 |
| TRIM2 | NM_001130067 | -1.1615 | -1.5725 |
| SUMF2 | NM_001130069 | 0.214 | -0.3995 |
| EPN1 | NM_001130071 | 0.0635 | -0.6425 |
| EPN1 | NM_001130072 | 0.0565 | -0.4545 |
| MOV10 | NM_001130079 | -0.039 | -0.6235 |
| IFI27 | NM_001130080 | -0.0045 | -0.358 |
| PLD1 | NM_001130081 | -0.092 | 0.1985 |
| PLXNB1 | NM_001130082 | -0.3655 | -0.81 |
| ABLIM2 | NM_001130083 | 0.0865 | 0.1585 |
| ABLIM2 | NM_001130084 | 0.0865 | 0.1585 |
| ABLIM2 | NM_001130085 | 0.0865 | 0.1585 |
| ABLIM2 | NM_001130086 | 0.0865 | 0.1585 |
| ABLIM2 | NM_001130087 | 0.0865 | 0.1585 |
| ABLIM2 | NM_001130088 | 0.0865 | 0.1585 |
| KARS | NM_001130089 | -0.63 | -0.588 |
| LRRC48 | NM_001130090 | 0.0285 | -0.669 |

| | | | |
|---|---|---|---|
| LRRC48 | NM_001130091 | 0.0285 | -0.669 |
| LRRC48 | NM_001130092 | 0.0285 | -0.669 |
| KIFC3 | NM_001130099 | -0.1945 | -0.42 |
| KIFC3 | NM_001130100 | -0.6955 | -0.8625 |
| COL13A1 | NM_001130103 | 0.2635 | -0.061 |
| COL4A3BP | NM_001130105 | -0.8425 | -1.933 |
| HSPBP1 | NM_001130106 | -0.09 | -0.873 |
| KLC1 | NM_001130107 | -0.539 | -0.653 |
| SETBP1 | NM_001130110 | -0.6415 | -1.959 |
| FAM108A1 | NM_001130111 | 0.019 | -0.9785 |
| RFC5 | NM_001130112 | -0.9315 | -1.808 |
| ISLR2 | NM_001130136 | -0.135 | -0.6735 |
| ISLR2 | NM_001130137 | -0.113 | -0.8215 |
| ISLR2 | NM_001130138 | -0.113 | -0.8215 |
| ERAP2 | NM_001130140 | -0.0005 | -0.449 |
| PCBP3 | NM_001130141 | -0.4945 | -1.0485 |
| VWA5A | NM_001130142 | -0.1645 | -0.2525 |
| PPP1R14D | NM_001130143 | 0.033 | -0.189 |
| LTBP3 | NM_001130144 | -0.3685 | -0.559 |
| YAP1 | NM_001130145 | -0.309 | -1.111 |
| CCDC77 | NM_001130146 | -1.3375 | -2.9705 |
| CCDC77 | NM_001130147 | -1.3375 | -2.9705 |
| CCDC77 | NM_001130148 | -1.3375 | -2.9705 |
| MLF1 | NM_001130156 | -0.534 | -0.676 |
| MLF1 | NM_001130157 | -0.534 | -0.676 |
| MYO1B | NM_001130158 | -0.744 | -1.4695 |
| STOX1 | NM_001130159 | -0.2655 | -0.51 |
| STOX1 | NM_001130160 | -1.181 | -1.8705 |
| STOX1 | NM_001130161 | -0.382 | -1.028 |
| MYB | NM_001130172 | -0.286 | -1.0575 |
| MYB | NM_001130173 | -0.286 | -1.0575 |
| DNAJA4 | NM_001130182 | 0.96 | -0.0175 |
| DNAJA4 | NM_001130183 | 0.96 | -0.0175 |
| PRR20C | NM_001130405 | -0.188 | 0.2025 |
| ACAA1 | NM_001130410 | 0.064 | -0.877 |
| SCNN1D | NM_001130413 | -0.3945 | -0.488 |
| APBA2 | NM_001130414 | -0.2215 | -0.696 |
| APOLD1 | NM_001130415 | -1.3115 | -1.416 |
| SMARCC2 | NM_001130420 | 0.0015 | -0.5455 |
| SPTAN1 | NM_001130438 | -0.1575 | -0.6805 |
| SRP9 | NM_001130440 | -0.7285 | -0.369 |
| HRAS | NM_001130442 | -0.062 | -0.789 |
| SSFA2 | NM_001130445 | -1.161 | -3.3495 |

| | | | |
|---|---|---|---|
| C10orf131 | NM_001130446 | -0.25 | 0.1705 |
| HAUS2 | NM_001130447 | -0.9595 | -1.693 |
| C15orf62 | NM_001130448 | 0.0585 | -0.7865 |
| DYSF | NM_001130455 | -0.1995 | -0.4835 |
| THAP5 | NM_001130475 | -1.464 | -2.5725 |
| LY6H | NM_001130478 | 0.0305 | -0.7285 |
| VRK2 | NM_001130480 | -0.494 | -1.714 |
| VRK2 | NM_001130481 | -0.494 | -1.714 |
| VRK2 | NM_001130482 | -0.494 | -1.714 |
| VRK2 | NM_001130483 | -0.494 | -1.714 |
| NDRG4 | NM_001130487 | -0.529 | -0.8035 |
| C19orf69 | NM_001130514 | -0.198 | -0.029 |
| MPST | NM_001130517 | -0.618 | -1.257 |
| CSGALNACT1 | NM_001130518 | 0.0685 | 0.0865 |
| ZNF195 | NM_001130519 | -1.2155 | -2.4815 |
| ZNF195 | NM_001130520 | -1.2155 | -2.4815 |
| CSDE1 | NM_001130523 | -0.4585 | -0.404 |
| AP1M1 | NM_001130524 | -0.029 | -0.902 |
| SPAG9 | NM_001130527 | -1.214 | -1.697 |
| SPAG9 | NM_001130528 | -1.214 | -1.697 |
| CALU | NM_001130674 | -0.828 | -0.878 |
| CLGN | NM_001130675 | -1.112 | -1.686 |
| C17orf96 | NM_001130677 | -0.5055 | 0.367 |
| EIF4E | NM_001130678 | -1.155 | -1.9695 |
| EIF4E | NM_001130679 | -1.155 | -1.9695 |
| GUCY1A3 | NM_001130682 | -1.0835 | -2.0525 |
| GUCY1A3 | NM_001130683 | -1.0835 | -2.0525 |
| GUCY1A3 | NM_001130684 | -1.0835 | -2.0525 |
| GUCY1A3 | NM_001130685 | -1.0835 | -2.0525 |
| GUCY1A3 | NM_001130686 | -0.617 | -1.884 |
| GUCY1A3 | NM_001130687 | -1.0835 | -2.0525 |
| HMGB2 | NM_001130688 | -1.223 | -0.61 |
| HMGB2 | NM_001130689 | -1.223 | -0.61 |
| PDE10A | NM_001130690 | -1.016 | -2.6285 |
| PPP3CA | NM_001130691 | -0.6765 | -1.326 |
| PPP3CA | NM_001130692 | -0.6765 | -1.326 |
| TRPC3 | NM_001130698 | -0.2595 | -0.5005 |
| IPCEF1 | NM_001130699 | 0.046 | 0.296 |
| IPCEF1 | NM_001130700 | 0.046 | 0.296 |
| STYX | NM_001130701 | -1.3385 | -2.7085 |
| BSCL2 | NM_001130702 | -0.107 | -0.9375 |
| C14orf93 | NM_001130706 | -0.785 | -1.9555 |
| C14orf93 | NM_001130708 | -0.785 | -1.9555 |

| | | | |
|---|---|---|---|
| RUFY3 | NM_001130709 | -1.1765 | -1.148 |
| LSM5 | NM_001130710 | -0.4875 | -0.5105 |
| CLEC2A | NM_001130711 | 0.472 | 0.065 |
| LEF1 | NM_001130713 | -0.1305 | -0.2025 |
| LEF1 | NM_001130714 | -0.1305 | -0.2025 |
| PLAC8 | NM_001130715 | 0.002 | -0.258 |
| PLAC8 | NM_001130716 | 0.002 | -0.258 |
| AHCYL2 | NM_001130720 | -0.0475 | -0.544 |
| ELOVL6 | NM_001130721 | -1.554 | -2.3615 |
| AHCYL2 | NM_001130722 | -0.0475 | -0.544 |
| AHCYL2 | NM_001130723 | -0.0475 | -0.544 |
| PSMB5 | NM_001130725 | -0.234 | -1.0895 |
| CCDC149 | NM_001130726 | -0.173 | -0.4095 |
| PACRGL | NM_001130727 | -1.421 | -2.629 |
| RABL5 | NM_001130820 | -0.4325 | -0.978 |
| RABL5 | NM_001130821 | -0.4325 | -0.978 |
| RABL5 | NM_001130822 | -0.4325 | -0.978 |
| DNMT1 | NM_001130823 | -0.1225 | -0.8115 |
| SUPT5H | NM_001130824 | -0.0065 | -0.121 |
| SUPT5H | NM_001130825 | -0.0065 | -0.121 |
| GAS7 | NM_001130831 | -0.1265 | -0.5135 |
| LDB2 | NM_001130834 | -1.461 | -1.3235 |
| RTCD1 | NM_001130841 | -1.0415 | -2.6275 |
| BCL6 | NM_001130845 | -0.725 | -1.059 |
| CAB39 | NM_001130849 | -1.6915 | -2.837 |
| CAB39 | NM_001130850 | -1.1265 | -2.199 |
| CDKN3 | NM_001130851 | -1.4765 | -2.2735 |
| PPIP5K1 | NM_001130858 | -0.2065 | -0.879 |
| PPIP5K1 | NM_001130859 | -0.2065 | -0.879 |
| HEPH | NM_001130860 | 0.1965 | 0.139 |
| CLDN5 | NM_001130861 | -0.1605 | -0.4165 |
| RAD51AP1 | NM_001130862 | -0.7315 | -1.125 |
| ENPP2 | NM_001130863 | -0.1515 | -0.7605 |
| PWWP2A | NM_001130864 | -1.501 | -1.749 |
| DMRT2 | NM_001130865 | 0.0885 | -0.145 |
| ERCC2 | NM_001130867 | 0.0625 | -0.9705 |
| CCRL2 | NM_001130910 | 0.082 | 0.054 |
| SUGT1 | NM_001130912 | -0.8635 | -1.433 |
| ZNF720 | NM_001130913 | -1.6785 | -2.916 |
| MAMSTR | NM_001130915 | -0.119 | -0.156 |
| TGFBR1 | NM_001130916 | -0.846 | -1.0815 |
| LILRA2 | NM_001130917 | -0.151 | -0.7585 |
| TTLL6 | NM_001130918 | -0.0095 | -0.0815 |

| | | | |
|---|---|---|---|
| RABL2B | NM_001130921 | -0.6655 | -0.7335 |
| TMEM201 | NM_001130924 | -0.067 | -0.142 |
| MEF2A | NM_001130926 | -0.93 | -2.3655 |
| MEF2A | NM_001130927 | -0.93 | -2.3655 |
| MEF2A | NM_001130928 | -0.93 | -2.3655 |
| C7orf73 | NM_001130929 | -0.4005 | -0.357 |
| ARHGEF18 | NM_001130955 | -0.7155 | -2.2705 |
| C1orf101 | NM_001130957 | -0.1035 | -0.8605 |
| FABP6 | NM_001130958 | 0.245 | -0.234 |
| PLCH1 | NM_001130960 | 0.017 | -0.4845 |
| PLCH1 | NM_001130961 | -0.078 | -1.174 |
| TMEM194A | NM_001130963 | -1.743 | -1.707 |
| PLCD1 | NM_001130964 | -0.03 | -0.284 |
| SUN1 | NM_001130965 | -0.5 | -1.0515 |
| TBXAS1 | NM_001130966 | 0.099 | -0.327 |
| ZNF385A | NM_001130967 | 0.02 | -0.49 |
| ZNF385A | NM_001130968 | -0.3215 | -0.6205 |
| NELF | NM_001130969 | -0.0835 | -0.7205 |
| NELF | NM_001130970 | -0.0835 | -0.7205 |
| NELF | NM_001130971 | -0.0835 | -0.7205 |
| DYSF | NM_001130976 | -0.1995 | -0.4835 |
| DYSF | NM_001130977 | -0.1995 | -0.4835 |
| DYSF | NM_001130978 | -0.1995 | -0.4835 |
| DYSF | NM_001130979 | -0.1995 | -0.4835 |
| DYSF | NM_001130980 | -0.1995 | -0.4835 |
| DYSF | NM_001130981 | -0.1995 | -0.4835 |
| DYSF | NM_001130982 | -0.1995 | -0.4835 |
| DYSF | NM_001130983 | -0.1995 | -0.4835 |
| DYSF | NM_001130984 | -0.1995 | -0.4835 |
| DYSF | NM_001130985 | -0.1995 | -0.4835 |
| DYSF | NM_001130986 | -0.1995 | -0.4835 |
| DYSF | NM_001130987 | -0.1995 | -0.4835 |
| HYOU1 | NM_001130991 | 0.1975 | -0.405 |
| RBP1 | NM_001130992 | 0.0575 | -0.2345 |
| RBP1 | NM_001130993 | 0.0575 | -0.2345 |
| HMGCR | NM_001130996 | -1.3985 | -2.906 |
| FAM58A | NM_001130997 | -0.494 | -2.5665 |
| CRISP1 | NM_001131 | -0.225 | -0.141 |
| MEF2C | NM_001131005 | -1.922 | -0.404 |
| KIAA0922 | NM_001131007 | -0.758 | -2.888 |
| PTPN12 | NM_001131008 | -1.3705 | -2.3275 |
| PTPN12 | NM_001131009 | -1.3705 | -2.3275 |
| SATB1 | NM_001131010 | -1.747 | -2.552 |

| | | | |
|---|---|---|---|
| CIZ1 | NM_001131015 | 0.1275 | -0.617 |
| CIZ1 | NM_001131016 | 0.1205 | -0.4265 |
| CIZ1 | NM_001131017 | 0.1205 | -0.4265 |
| CIZ1 | NM_001131018 | 0.1205 | -0.4265 |
| GFAP | NM_001131019 | 0.0515 | -0.47 |
| PEX5 | NM_001131023 | -0.3075 | -1.087 |
| PEX5 | NM_001131024 | -0.3075 | -1.087 |
| PEX5 | NM_001131025 | -0.2065 | -1.256 |
| PEX5 | NM_001131026 | -0.3075 | -1.087 |
| PDLIM4 | NM_001131027 | -0.476 | -1.322 |
| ATG10 | NM_001131028 | -0.8715 | -1.924 |
| RNF212 | NM_001131034 | -0.092 | -0.003 |
| ZCCHC9 | NM_001131036 | -1.7915 | -3.0825 |
| HRH2 | NM_001131055 | 0.1455 | -0.2045 |
| SAP30L | NM_001131062 | -0.9355 | -2.2765 |
| SAP30L | NM_001131063 | -0.9355 | -2.2765 |
| ACP2 | NM_001131064 | -0.4 | -1.225 |
| RFESD | NM_001131065 | -1.7005 | -2.2875 |
| RFESD | NM_001131066 | -1.7005 | -2.2875 |
| AFM | NM_001133 | 0.196 | 0.3555 |
| AFP | NM_001134 | -0.572 | -0.1935 |
| ACPP | NM_001134194 | 0.3385 | -0.218 |
| IRX2 | NM_001134222 | 0.0735 | -0.2405 |
| USP46 | NM_001134223 | -0.482 | -1.186 |
| INPP4A | NM_001134224 | -0.6765 | -1.123 |
| INPP4A | NM_001134225 | -0.6765 | -1.123 |
| NT5DC2 | NM_001134231 | 0.2165 | -0.149 |
| TMEM106B | NM_001134232 | -0.9485 | -1.4925 |
| C1orf204 | NM_001134233 | 0.3855 | -0.411 |
| ESRRG | NM_001134285 | -1.2545 | -4.612 |
| AP3M2 | NM_001134296 | -1.3865 | -1.628 |
| FAM127B | NM_001134321 | -0.4405 | -1.0605 |
| EIF2AK1 | NM_001134335 | -0.508 | -0.8085 |
| FAM40B | NM_001134336 | -0.407 | -0.353 |
| RNF24 | NM_001134337 | -0.8565 | -2.1095 |
| RNF24 | NM_001134338 | -0.8565 | -2.1095 |
| RBM20 | NM_001134363 | -0.856 | -1.9835 |
| MAP4 | NM_001134364 | -0.397 | -1.094 |
| GAD2 | NM_001134366 | 0.1075 | -0.389 |
| SLC6A6 | NM_001134367 | 0.6295 | -0.3535 |
| SLC6A6 | NM_001134368 | 0.6295 | -0.3535 |
| LRRFIP2 | NM_001134369 | -1.1075 | -2.1585 |
| NT5C2 | NM_001134373 | -1.418 | -1.362 |

| | | | |
|---|---|---|---|
| SPATC1 | NM_001134374 | -0.0015 | -0.236 |
| CCNJ | NM_001134375 | -1.128 | -3.4635 |
| CCNJ | NM_001134376 | -0.6185 | -1.9185 |
| TBC1D5 | NM_001134380 | -0.458 | -0.404 |
| TBC1D5 | NM_001134381 | -0.458 | -0.404 |
| IQSEC1 | NM_001134382 | -0.8365 | -1.756 |
| ZDHHC4 | NM_001134387 | 0.0195 | -0.739 |
| ZDHHC4 | NM_001134388 | 0.0195 | -0.739 |
| ZDHHC4 | NM_001134389 | 0.0195 | -0.739 |
| VAV2 | NM_001134398 | -0.723 | -1.7655 |
| CYB561D1 | NM_001134400 | 0.0715 | -0.298 |
| CYB561D1 | NM_001134402 | 0.0715 | -0.298 |
| CYB561D1 | NM_001134403 | 0.0715 | -0.298 |
| CYB561D1 | NM_001134404 | -0.6875 | -1.8605 |
| RUNDC3B | NM_001134405 | -0.717 | -1.447 |
| RUNDC3B | NM_001134406 | -0.717 | -1.447 |
| GRIN2A | NM_001134407 | 0.2525 | -0.051 |
| GRIN2A | NM_001134408 | 0.2525 | -0.051 |
| TBCCD1 | NM_001134415 | -1.582 | -3.1835 |
| LEPREL1 | NM_001134418 | 0.0385 | -0.356 |
| CDC7 | NM_001134419 | -1.24 | -2.8305 |
| CDC7 | NM_001134420 | -1.24 | -2.8305 |
| CDV3 | NM_001134422 | -0.9405 | -1.1705 |
| CDV3 | NM_001134423 | -0.5875 | -1.1235 |
| TOR2A | NM_001134430 | -0.2205 | -0.4615 |
| TOR2A | NM_001134431 | -0.2205 | -0.4615 |
| AZI2 | NM_001134432 | -0.797 | -1.6665 |
| AZI2 | NM_001134433 | -0.797 | -1.6665 |
| SPOCK2 | NM_001134434 | 0.0785 | -0.178 |
| PHLDB2 | NM_001134437 | -0.3555 | -2.6365 |
| PHLDB2 | NM_001134438 | -0.3555 | -2.6365 |
| PHLDB2 | NM_001134439 | -1.498 | -2.934 |
| ZNF502 | NM_001134440 | -0.9835 | -2.178 |
| ZNF502 | NM_001134441 | -0.9835 | -2.178 |
| ZNF502 | NM_001134442 | -0.9835 | -2.178 |
| DDAH1 | NM_001134445 | -0.714 | -0.511 |
| TMEM130 | NM_001134450 | -1.053 | -1.3895 |
| TMEM130 | NM_001134451 | -1.053 | -1.3895 |
| FAM55C | NM_001134456 | -1.3355 | -1.7835 |
| NOTO | NM_001134462 | -0.217 | -0.2535 |
| C3orf58 | NM_001134470 | -0.7575 | -0.043 |
| KIAA0182 | NM_001134473 | -0.799 | -1.3995 |
| LRRC8B | NM_001134476 | -1.1245 | -2.2545 |

| | | | |
|---|---|---|---|
| PARN | NM_001134477 | -1.042 | -1.239 |
| LRRC8D | NM_001134479 | -1.705 | -3.6305 |
| TOMM5 | NM_001134484 | -0.0055 | -0.544 |
| TOMM5 | NM_001134485 | -0.0055 | -0.544 |
| GBP5 | NM_001134486 | 0.171 | -0.0565 |
| HS2ST1 | NM_001134492 | -0.987 | -2.4845 |
| TOMM6 | NM_001134493 | -0.1835 | -0.566 |
| AFAP1 | NM_001134647 | -0.6875 | -2.016 |
| EIF4E3 | NM_001134649 | -0.939 | -0.9835 |
| EIF4E3 | NM_001134650 | -0.939 | -0.9835 |
| EIF4E3 | NM_001134651 | -0.939 | -0.9835 |
| ZNF213 | NM_001134655 | -0.112 | -0.505 |
| ZNF662 | NM_001134656 | -0.145 | 0.0635 |
| PRR23C | NM_001134657 | -0.486 | -0.1365 |
| PRR23A | NM_001134659 | -0.01 | -0.3215 |
| RG9MTD2 | NM_001134665 | -0.4635 | -0.608 |
| RG9MTD2 | NM_001134666 | -0.4635 | -0.608 |
| DERL1 | NM_001134671 | 0.183 | -0.776 |
| NFIA | NM_001134673 | -0.9245 | -2.006 |
| OST4 | NM_001134693 | 0.221 | -0.267 |
| SARDH | NM_001134707 | -0.15 | -0.133 |
| DEK | NM_001134709 | -0.6915 | -1.241 |
| C1orf94 | NM_001134734 | 0.159 | -0.625 |
| BCL6 | NM_001134738 | 0.303 | -0.525 |
| SLC4A4 | NM_001134742 | 0.0425 | -0.0635 |
| LRRTM4 | NM_001134745 | 0.357 | -0.345 |
| ARFRP1 | NM_001134758 | -0.7575 | -1.4965 |
| SLC12A5 | NM_001134771 | 0.072 | -0.159 |
| STX16 | NM_001134772 | -1.417 | -1.407 |
| STX16 | NM_001134773 | -1.3845 | -1.4995 |
| KLC2 | NM_001134774 | -0.104 | -0.639 |
| KLC2 | NM_001134775 | -0.104 | -0.639 |
| KLC2 | NM_001134776 | -0.104 | -0.639 |
| PBX3 | NM_001134778 | -1.205 | -1.776 |
| IPO11 | NM_001134779 | -0.531 | -1.8255 |
| HYLS1 | NM_001134793 | -1.126 | -3.6475 |
| AHI1 | NM_001134830 | -0.9195 | -1.136 |
| AHI1 | NM_001134831 | -0.9195 | -1.136 |
| AHI1 | NM_001134832 | -0.9195 | -1.136 |
| SIRPB2 | NM_001134836 | 0.005 | -0.438 |
| CCDC152 | NM_001134848 | -0.3255 | -0.839 |
| TCF7 | NM_001134851 | -0.874 | -1.8875 |
| TRIM17 | NM_001134855 | -0.503 | -0.4715 |

| | | | |
|---|---|---|---|
| PPP1R18 | NM_001134870 | -0.3375 | -0.6055 |
| C14orf80 | NM_001134875 | -0.534 | -1.2305 |
| KIF9 | NM_001134878 | -0.163 | -0.91 |
| RTL1 | NM_001134888 | -0.1305 | -0.3115 |
| FIP1L1 | NM_001134937 | -1.338 | -2.24 |
| FIP1L1 | NM_001134938 | -1.338 | -2.24 |
| GHRL | NM_001134941 | -0.155 | 0.1135 |
| GHRL | NM_001134944 | -0.155 | 0.1135 |
| GHRL | NM_001134945 | -0.155 | 0.1135 |
| GHRL | NM_001134946 | -0.155 | 0.1135 |
| FERMT2 | NM_001134999 | -1.0665 | -2.2655 |
| ACAN | NM_001135 | -0.2335 | -0.147 |
| FERMT2 | NM_001135000 | -0.7075 | -1.6395 |
| DNAJB5 | NM_001135004 | -0.865 | -2.141 |
| DNAJB5 | NM_001135005 | -1.2885 | -2.6135 |
| ELMOD3 | NM_001135021 | -0.6425 | -1.4745 |
| ELMOD3 | NM_001135022 | -0.422 | -1.257 |
| ELMOD3 | NM_001135023 | -0.6425 | -1.4745 |
| PDHX | NM_001135024 | -0.9945 | -2.4695 |
| MFAP3 | NM_001135037 | -1.5585 | -2.1475 |
| MAPK9 | NM_001135044 | -0.5785 | -1.1935 |
| JDP2 | NM_001135047 | -0.9605 | -2.184 |
| JDP2 | NM_001135048 | -0.9605 | -2.184 |
| JDP2 | NM_001135049 | -0.9605 | -2.184 |
| FAM160B1 | NM_001135051 | 0.5845 | -1.0385 |
| SYK | NM_001135052 | 0.0805 | -0.3045 |
| SIGIRR | NM_001135053 | 0.1875 | -0.411 |
| SIGIRR | NM_001135054 | 0.337 | -0.445 |
| TKT | NM_001135055 | 0.2215 | -0.3755 |
| LRRC15 | NM_001135057 | -0.0675 | -0.0795 |
| COCH | NM_001135058 | -1.0935 | -1.3035 |
| PRSS41 | NM_001135086 | 0.086 | -0.1885 |
| MUC15 | NM_001135091 | -0.3955 | 0.0725 |
| MUC15 | NM_001135092 | 0.2045 | -0.1605 |
| FNDC3B | NM_001135095 | -0.866 | -2.2945 |
| UPP2 | NM_001135098 | 0.0085 | -0.036 |
| TMPRSS2 | NM_001135099 | 0.0215 | -0.5965 |
| CRELD2 | NM_001135101 | -0.156 | -0.212 |
| KCNK16 | NM_001135105 | -0.1535 | -0.362 |
| KCNK16 | NM_001135106 | 0.0945 | -0.499 |
| RIC3 | NM_001135109 | -0.736 | -1.2185 |
| DNAJA3 | NM_001135110 | -1.437 | -3.122 |
| KCNK17 | NM_001135111 | -0.6405 | -0.1825 |

| | | | |
|---|---|---|---|
| SLC39A8 | NM_001135146 | -0.3845 | -1.0445 |
| SLC39A8 | NM_001135147 | -0.067 | -0.998 |
| SLC39A8 | NM_001135148 | -0.067 | -0.998 |
| SLC39A14 | NM_001135153 | -0.6365 | -1.573 |
| SLC39A14 | NM_001135154 | -0.6365 | -1.573 |
| DPF1 | NM_001135155 | -0.418 | -1.191 |
| DPF1 | NM_001135156 | -0.6885 | -2.1595 |
| COMT | NM_001135161 | -0.6255 | -0.8935 |
| COMT | NM_001135162 | -0.6255 | -0.8935 |
| SIGLEC11 | NM_001135163 | 0.1865 | -0.342 |
| ALDH3A1 | NM_001135168 | 0.053 | -0.1305 |
| C1QTNF7 | NM_001135170 | 0.1575 | 0.0505 |
| C1QTNF7 | NM_001135171 | 0.363 | -0.433 |
| NEDD1 | NM_001135175 | -1.072 | -2.8115 |
| NEDD1 | NM_001135176 | -1.072 | -2.8115 |
| NEDD1 | NM_001135177 | -1.072 | -2.8115 |
| ZNF397 | NM_001135178 | -1.106 | -1.745 |
| ZDHHC3 | NM_001135179 | -0.351 | -1.815 |
| SLC5A9 | NM_001135181 | 0.01 | -0.0695 |
| AGFG1 | NM_001135187 | -0.5755 | -1.4755 |
| AGFG1 | NM_001135188 | -0.5755 | -1.4755 |
| AGFG1 | NM_001135189 | -0.5755 | -1.4755 |
| ARAP1 | NM_001135190 | -0.278 | -0.885 |
| ASAP2 | NM_001135191 | -1.6535 | -1.854 |
| SLC39A5 | NM_001135195 | -0.1445 | -0.2075 |
| C10orf71 | NM_001135196 | 0.4625 | -0.345 |
| CCDC36 | NM_001135197 | -0.2905 | -0.3135 |
| FKBP7 | NM_001135212 | -1.426 | -1.331 |
| GIT2 | NM_001135213 | -0.6875 | -2.828 |
| GIT2 | NM_001135214 | -0.6875 | -2.828 |
| LRRC23 | NM_001135217 | 0.153 | -0.44 |
| TAF12 | NM_001135218 | -1.0015 | -1.56 |
| PIP5KL1 | NM_001135219 | -0.1435 | -0.3725 |
| LDHA | NM_001135239 | -0.2955 | -0.612 |
| C1orf226 | NM_001135240 | 0.022 | -0.2365 |
| AKR1C2 | NM_001135241 | 0.161 | -0.448 |
| NDRG1 | NM_001135242 | 0.1135 | -0.508 |
| TCOF1 | NM_001135243 | -0.2455 | -1.0685 |
| TCOF1 | NM_001135244 | -0.2495 | -0.907 |
| TCOF1 | NM_001135245 | -0.2495 | -0.907 |
| MFAP2 | NM_001135247 | 0.051 | -0.4275 |
| MFAP2 | NM_001135248 | 0.051 | -0.4275 |
| PAX7 | NM_001135254 | 0.2385 | -0.4755 |

| | | | |
|---|---|---|---|
| RBBP4 | NM_001135255 | -0.5985 | -0.552 |
| RBBP4 | NM_001135256 | -0.5985 | -0.552 |
| MKNK1 | NM_001135553 | -0.7305 | -0.8235 |
| EPB41L2 | NM_001135554 | -0.624 | -0.5595 |
| EPB41L2 | NM_001135555 | -0.624 | -0.5595 |
| DYNC1I1 | NM_001135556 | 0.1855 | 0.5 |
| DYNC1I1 | NM_001135557 | 0.1855 | 0.5 |
| HSF2 | NM_001135564 | -1.4485 | -3.69 |
| HDHD1 | NM_001135565 | -1.1245 | -2.6055 |
| C12orf73 | NM_001135570 | -0.661 | -0.8425 |
| C6orf228 | NM_001135575 | -1.0725 | -1.1645 |
| C19orf71 | NM_001135580 | -0.327 | -0.168 |
| SLC2A5 | NM_001135585 | -0.306 | -0.32 |
| C5orf24 | NM_001135586 | -1.716 | -1.8755 |
| GDAP2 | NM_001135589 | 0.181 | -0.4085 |
| ZNF577 | NM_001135590 | -1.0785 | -1.5895 |
| RPS27A | NM_001135592 | -0.1275 | -0.04 |
| CCDC88A | NM_001135597 | -0.1135 | -2.4695 |
| C2orf63 | NM_001135598 | -1.5685 | -1.0725 |
| TGFB2 | NM_001135599 | 0.1295 | -0.414 |
| GLB1 | NM_001135602 | -0.364 | -1.3295 |
| ESM1 | NM_001135604 | -0.14 | -0.407 |
| ARHGAP26 | NM_001135608 | -1.27 | -2.303 |
| PRDM2 | NM_001135610 | -0.841 | -0.9575 |
| C11orf68 | NM_001135635 | -0.2885 | -1.123 |
| PIP5K1A | NM_001135636 | -0.1475 | -1.0495 |
| PIP5K1A | NM_001135637 | -0.1475 | -1.0495 |
| PIP5K1A | NM_001135638 | -0.714 | -2.138 |
| CNGB1 | NM_001135639 | -0.416 | -0.6195 |
| INPP5K | NM_001135642 | -0.279 | -1.1715 |
| DCTN4 | NM_001135643 | -0.835 | -0.785 |
| DCTN4 | NM_001135644 | -0.8215 | -0.6185 |
| FAM53C | NM_001135647 | -0.144 | -0.8355 |
| PTPRK | NM_001135648 | -0.9365 | -1.6405 |
| FOXI3 | NM_001135649 | 0.02 | -0.6785 |
| EEF1E1 | NM_001135650 | -1.621 | -2.9335 |
| EIF2AK2 | NM_001135651 | -0.702 | -1.022 |
| EIF2AK2 | NM_001135652 | -0.702 | -1.022 |
| PABPC4 | NM_001135653 | 0.16 | -0.396 |
| PABPC4 | NM_001135654 | 0.0305 | -0.3275 |
| NRXN1 | NM_001135659 | -0.3315 | -0.859 |
| RAB7L1 | NM_001135662 | -1.158 | -0.9295 |
| RAB7L1 | NM_001135663 | -1.5845 | -1.2965 |

| | | | |
|---|---|---|---|
| RAB7L1 | NM_001135664 | -1.158 | -0.9295 |
| XPR1 | NM_001135669 | -0.751 | -0.929 |
| MSRA | NM_001135670 | 0.296 | -0.4945 |
| MSRA | NM_001135671 | 0.296 | -0.4945 |
| ATL2 | NM_001135673 | -1.434 | -2.358 |
| C8orf40 | NM_001135675 | -1.285 | -1.905 |
| LTK | NM_001135685 | -0.3495 | -0.2165 |
| TARM1 | NM_001135686 | 0.1325 | -0.0105 |
| PENK | NM_001135690 | -0.0405 | -0.248 |
| VDAC3 | NM_001135694 | -0.731 | -1.1245 |
| SGCA | NM_001135697 | 0.0825 | -0.5015 |
| YWHAZ | NM_001135699 | -0.584 | -0.5065 |
| YWHAZ | NM_001135700 | -0.584 | -0.5065 |
| YWHAZ | NM_001135701 | -0.584 | -0.5065 |
| YWHAZ | NM_001135702 | -0.584 | -0.5065 |
| LRP12 | NM_001135703 | -0.736 | -1.767 |
| ACBD4 | NM_001135704 | -0.031 | -0.6915 |
| ACBD4 | NM_001135705 | -0.031 | -0.6915 |
| ACBD4 | NM_001135706 | -0.031 | -0.6915 |
| ACBD4 | NM_001135707 | -0.031 | -0.6915 |
| PIWIL2 | NM_001135721 | 0.1355 | -0.133 |
| HMBOX1 | NM_001135726 | -0.2745 | -1.3475 |
| TOM1 | NM_001135729 | -0.1625 | -0.9325 |
| TOM1 | NM_001135730 | -0.1625 | -0.9325 |
| TOM1 | NM_001135732 | -0.1625 | -0.9325 |
| TP53INP1 | NM_001135733 | -1.2965 | -2.076 |
| ZNF384 | NM_001135734 | -0.7905 | -2.308 |
| PNCK | NM_001135740 | -0.4305 | -0.049 |
| ALS2 | NM_001135745 | -1.659 | -2.0115 |
| NEIL2 | NM_001135746 | -0.31 | -1.0285 |
| NEIL2 | NM_001135747 | -0.31 | -1.0285 |
| NEIL2 | NM_001135748 | -0.31 | -1.0285 |
| PSMG4 | NM_001135750 | -0.387 | -1.4645 |
| DERL3 | NM_001135751 | 0.165 | -0.181 |
| ECD | NM_001135752 | -0.819 | -2.3385 |
| ECD | NM_001135753 | -1.0745 | -2.271 |
| PVR | NM_001135768 | -1.2145 | -1.957 |
| PVR | NM_001135769 | -1.2145 | -1.957 |
| PVR | NM_001135770 | -1.2145 | -1.957 |
| RPN2 | NM_001135771 | -0.1615 | -0.345 |
| LRP5L | NM_001135772 | -0.1795 | -1.1655 |
| SPATA2 | NM_001135773 | -0.232 | -1.0245 |
| SYN3 | NM_001135774 | -0.103 | -0.304 |

| | | | |
|---|---|---|---|
| C9orf7 | NM_001135775 | -0.2285 | -0.8915 |
| ZBTB43 | NM_001135776 | -0.8695 | -1.6645 |
| SYT1 | NM_001135805 | -1.046 | -2.128 |
| SYT1 | NM_001135806 | -0.7285 | -1.8615 |
| FAM60A | NM_001135811 | -1.344 | -1.1145 |
| FAM60A | NM_001135812 | -0.9635 | -0.9235 |
| TMEM2 | NM_001135820 | -0.1435 | -1.1785 |
| FDPS | NM_001135821 | -0.284 | -0.5885 |
| FDPS | NM_001135822 | -0.284 | -0.5885 |
| SSPN | NM_001135823 | -1.489 | -1.289 |
| RNF185 | NM_001135825 | -0.576 | -1.27 |
| SIRPB1 | NM_001135844 | 0.1595 | -0.2875 |
| PHPT1 | NM_001135861 | 0.434 | 0.001 |
| PHF21B | NM_001135862 | -0.3435 | -0.999 |
| METTL20 | NM_001135863 | -0.627 | -1.4225 |
| PIK3IP1 | NM_001135911 | -0.592 | -0.71 |
| KCP | NM_001135914 | -0.0625 | -0.301 |
| DOLPP1 | NM_001135917 | -0.833 | -2.2675 |
| SLC46A3 | NM_001135919 | -0.134 | -1.1445 |
| KIAA1432 | NM_001135920 | -0.562 | -1.129 |
| VWDE | NM_001135924 | -0.222 | -0.1575 |
| POSTN | NM_001135934 | 0.2475 | -0.192 |
| POSTN | NM_001135935 | 0.2475 | -0.192 |
| POSTN | NM_001135936 | 0.2475 | -0.192 |
| SMAD2 | NM_001135937 | -1.341 | -2.2695 |
| MYOT | NM_001135940 | 0.1605 | -0.3045 |
| MADD | NM_001135943 | -0.668 | -1.477 |
| MADD | NM_001135944 | -0.668 | -1.477 |
| URM1 | NM_001135947 | -0.0755 | -0.9225 |
| GKAP1 | NM_001135953 | -0.6645 | -1.772 |
| UCK1 | NM_001135954 | -1.2175 | -2.1285 |
| TRPC4 | NM_001135955 | 0.2865 | -0.5715 |
| TRPC4 | NM_001135956 | 0.2865 | -0.5715 |
| TRPC4 | NM_001135957 | 0.2865 | -0.5715 |
| TRPC4 | NM_001135958 | 0.2865 | -0.5715 |
| TTC39C | NM_001135993 | -1.234 | -2.3885 |
| ATXN3L | NM_001135995 | 0.0565 | -0.1245 |
| RUSC2 | NM_001135999 | -0.534 | -2.126 |
| AGER | NM_001136 | -0.0285 | -0.2605 |
| ABL2 | NM_001136000 | -1.382 | -2.9035 |
| ABL2 | NM_001136001 | -1.382 | -2.9035 |
| TMEM229A | NM_001136002 | -1.3885 | -1.2845 |
| C2CD4D | NM_001136003 | 0.215 | -0.344 |

| | | | |
|---|---|---|---|
| MICAL3 | NM_001136004 | 0.852 | 0.0615 |
| GART | NM_001136005 | -0.557 | -1.4935 |
| GART | NM_001136006 | -0.557 | -1.4935 |
| FXYD3 | NM_001136007 | 0.4865 | -0.123 |
| FXYD3 | NM_001136008 | 0.4865 | -0.123 |
| FXYD3 | NM_001136009 | 0.4865 | -0.123 |
| FXYD3 | NM_001136010 | 0.4865 | -0.123 |
| FXYD3 | NM_001136011 | 0.4865 | -0.123 |
| FXYD3 | NM_001136012 | 0.4865 | -0.123 |
| ANXA2 | NM_001136015 | -0.2605 | -0.2505 |
| APP | NM_001136016 | -0.3025 | -0.3875 |
| CCND3 | NM_001136017 | -0.6295 | -1.572 |
| EPHX1 | NM_001136018 | 0.0295 | -0.6485 |
| FCGRT | NM_001136019 | -0.3565 | -0.9975 |
| ICA1 | NM_001136020 | -0.608 | -1.057 |
| NFATC2 | NM_001136021 | -0.2475 | -0.6905 |
| NFATC4 | NM_001136022 | -0.1935 | -0.7425 |
| NFE2 | NM_001136023 | 0.1205 | -0.0555 |
| NHS | NM_001136024 | -0.2435 | -0.3225 |
| PLS3 | NM_001136025 | -0.468 | -0.3215 |
| DEPDC5 | NM_001136029 | -1.6815 | -3.1645 |
| KIAA0748 | NM_001136030 | -0.0115 | -0.113 |
| ATG7 | NM_001136031 | -0.2245 | -0.7755 |
| PUF60 | NM_001136033 | 0.0035 | -0.874 |
| TRMT1 | NM_001136035 | -0.1185 | -0.694 |
| ZNF692 | NM_001136036 | -0.6755 | -1.8195 |
| LIMS2 | NM_001136037 | -0.007 | -0.366 |
| NIF3L1 | NM_001136039 | -1.756 | -3.9685 |
| CPSF7 | NM_001136040 | -1.14 | -2.4285 |
| HDAC11 | NM_001136041 | 0.1065 | -0.638 |
| AARSD1 | NM_001136042 | -0.0025 | -0.647 |
| TMUB1 | NM_001136044 | -0.1405 | -1.0925 |
| ZMYND15 | NM_001136046 | -0.349 | -0.3105 |
| LMLN | NM_001136049 | -0.6 | -0.8235 |
| DHRS1 | NM_001136050 | -0.0005 | -0.633 |
| LRIG3 | NM_001136051 | -0.4595 | -1.3465 |
| CPT1C | NM_001136052 | -0.4325 | -0.8675 |
| TMEM132C | NM_001136103 | -0.067 | -0.4035 |
| C11orf93 | NM_001136105 | -0.082 | -0.3175 |
| BEAN1 | NM_001136106 | 0.0135 | -0.285 |
| METTL11B | NM_001136107 | -0.0595 | -0.229 |
| R3HCC1 | NM_001136108 | -0.4 | -1.3835 |
| CASP5 | NM_001136109 | -0.085 | -0.228 |

| | | | |
|---|---|---|---|
| CASP5 | NM_001136110 | -0.085 | -0.228 |
| CASP5 | NM_001136112 | -0.085 | -0.228 |
| POMT1 | NM_001136113 | -0.4295 | -0.8615 |
| POMT1 | NM_001136114 | -0.4295 | -0.8615 |
| FBXO28 | NM_001136115 | -2.225 | -4.7375 |
| ZNF879 | NM_001136116 | -0.755 | -1.5325 |
| FAM178A | NM_001136123 | -1.141 | -2.2115 |
| CCND3 | NM_001136125 | -0.6295 | -1.572 |
| CCND3 | NM_001136126 | -0.6295 | -1.572 |
| DNM3 | NM_001136127 | -0.598 | -1.1115 |
| APP | NM_001136129 | -0.3025 | -0.3875 |
| APP | NM_001136130 | -0.3025 | -0.3875 |
| APP | NM_001136131 | -0.3025 | -0.3875 |
| RPL28 | NM_001136134 | 0.1075 | -0.274 |
| RPL28 | NM_001136135 | 0.1075 | -0.274 |
| RPL28 | NM_001136136 | 0.1075 | -0.274 |
| RPL28 | NM_001136137 | 0.1075 | -0.274 |
| RPS6KC1 | NM_001136138 | -0.9745 | -2.086 |
| TCF3 | NM_001136139 | -0.076 | -1.1245 |
| CMPK1 | NM_001136140 | -1.017 | -1.233 |
| ALG1L2 | NM_001136152 | -0.4025 | -0.4345 |
| ATF6B | NM_001136153 | -0.2215 | -0.791 |
| ERG | NM_001136154 | -0.1435 | -0.526 |
| ERG | NM_001136155 | -0.1435 | -0.526 |
| ZNF507 | NM_001136156 | -1.163 | -3.222 |
| OTUD5 | NM_001136157 | -0.415 | -1.06 |
| OTUD5 | NM_001136158 | -0.415 | -1.06 |
| OTUD5 | NM_001136159 | -0.415 | -1.06 |
| C8orf45 | NM_001136160 | 0.395 | -0.1675 |
| C8orf45 | NM_001136161 | 0.395 | -0.1675 |
| EGR2 | NM_001136177 | -0.005 | -0.105 |
| EGR2 | NM_001136178 | -0.005 | -0.105 |
| EGR2 | NM_001136179 | -0.005 | -0.105 |
| HSBP1L1 | NM_001136180 | -0.048 | -0.5105 |
| LAX1 | NM_001136190 | 0.0505 | -0.408 |
| KANK2 | NM_001136191 | -0.14 | -0.694 |
| FASTKD2 | NM_001136193 | -1.006 | -2.7225 |
| FASTKD2 | NM_001136194 | -1.006 | -2.7225 |
| TNPO2 | NM_001136195 | -0.709 | -1.032 |
| TNPO2 | NM_001136196 | -0.709 | -1.032 |
| FZR1 | NM_001136197 | -0.7205 | -1.2115 |
| FZR1 | NM_001136198 | -0.7205 | -1.2115 |
| GRAMD1A | NM_001136199 | -0.774 | -1.7275 |

| | | | |
|---|---|---|---|
| C10orf32 | NM_001136200 | -0.799 | -0.813 |
| ISOC2 | NM_001136202 | -0.186 | -1.0835 |
| RCC2 | NM_001136204 | -0.254 | -1.216 |
| KCTD1 | NM_001136205 | -1.2195 | -3.109 |
| POTEH | NM_001136213 | -0.045 | -0.2295 |
| ARTN | NM_001136215 | -0.0085 | -0.564 |
| TMEM51 | NM_001136216 | -0.569 | -2.0175 |
| TMEM51 | NM_001136217 | -0.569 | -2.0175 |
| TMEM51 | NM_001136218 | -0.569 | -2.0175 |
| FCGR2A | NM_001136219 | 0.0625 | -0.108 |
| RCOR3 | NM_001136223 | -1.8 | -2.731 |
| RCOR3 | NM_001136224 | -1.8 | -2.731 |
| RCOR3 | NM_001136225 | -1.8 | -2.731 |
| HSD17B13 | NM_001136230 | 0.304 | -0.367 |
| SEC13 | NM_001136232 | -0.008 | -0.7355 |
| FAM48B2 | NM_001136233 | -0.3215 | -0.408 |
| FAM48B1 | NM_001136234 | -0.0655 | -0.1045 |
| TMPPE | NM_001136238 | 0.22 | 0.2075 |
| PRDM6 | NM_001136239 | 0.336 | -0.548 |
| SGMS2 | NM_001136257 | 0.002 | 0.184 |
| SGMS2 | NM_001136258 | 0.002 | 0.184 |
| ATXN7L3B | NM_001136262 | -0.627 | -1.8085 |
| C2CD4C | NM_001136263 | -0.3925 | -1.1605 |
| DAZAP2 | NM_001136264 | -0.3285 | -0.667 |
| IFFO2 | NM_001136265 | -0.5665 | -0.952 |
| DAZAP2 | NM_001136266 | -0.3285 | -0.667 |
| DAZAP2 | NM_001136267 | -0.3285 | -0.667 |
| DAZAP2 | NM_001136268 | -0.3285 | -0.667 |
| DAZAP2 | NM_001136269 | -0.3285 | -0.667 |
| NKX2-6 | NM_001136271 | 0.172 | -0.415 |
| ZFP92 | NM_001136273 | -0.1005 | -0.5055 |
| AKIRIN1 | NM_001136275 | -0.918 | -1.868 |
| TMEM108 | NM_001136469 | -0.148 | -2.2635 |
| LITAF | NM_001136472 | -0.104 | -1.0455 |
| LITAF | NM_001136473 | -0.104 | -1.0455 |
| VASH2 | NM_001136474 | -1.1995 | -1.716 |
| VASH2 | NM_001136475 | -1.1995 | -1.716 |
| ANP32E | NM_001136478 | -0.689 | -0.6605 |
| ANP32E | NM_001136479 | -0.6 | -0.5525 |
| C19orf38 | NM_001136482 | -0.132 | -0.184 |
| C17orf105 | NM_001136483 | -0.1675 | -0.169 |
| C11orf86 | NM_001136485 | 0.0825 | -0.11 |
| TRIM64 | NM_001136486 | -0.2735 | -0.4265 |

| | | | |
|---|---|---|---|
| MFSD2A | NM_001136493 | -0.4865 | -1.5115 |
| C1orf198 | NM_001136494 | 0.021 | -0.698 |
| C1orf198 | NM_001136495 | 0.021 | -0.698 |
| BTF3L4 | NM_001136497 | -1.462 | -2.232 |
| CISD3 | NM_001136498 | -0.2515 | -0.6785 |
| ZNF841 | NM_001136499 | -0.3205 | -0.817 |
| ZNF844 | NM_001136501 | -0.0345 | -0.5475 |
| C19orf77 | NM_001136503 | 0.0075 | -0.0125 |
| SYT2 | NM_001136504 | -0.559 | -0.4935 |
| CCDC79 | NM_001136505 | -0.158 | -0.1555 |
| SLC22A24 | NM_001136506 | -0.4205 | -0.3605 |
| MBD3L5 | NM_001136507 | 0.014 | -0.2555 |
| C1orf185 | NM_001136508 | 0.5875 | -0.8555 |
| ZNF843 | NM_001136509 | -0.083 | -0.434 |
| SERPINE2 | NM_001136528 | -0.353 | -0.7755 |
| SERPINE2 | NM_001136530 | -0.353 | -0.7755 |
| DCAF8L2 | NM_001136533 | 0.1455 | 0.0115 |
| TMEM233 | NM_001136534 | -0.1535 | -0.297 |
| DTHD1 | NM_001136536 | 0.155 | 0.0545 |
| BTBD19 | NM_001136537 | 0.3455 | -0.3045 |
| ACAD10 | NM_001136538 | -0.416 | -0.9865 |
| APOL1 | NM_001136540 | 0.1205 | -0.137 |
| APOL1 | NM_001136541 | 0.193 | -0.3855 |
| LYSMD1 | NM_001136543 | -0.434 | -0.9025 |
| TLK1 | NM_001136554 | -1.191 | -2.977 |
| TLK1 | NM_001136555 | -1.191 | -2.977 |
| GPR107 | NM_001136557 | -1.4385 | -1.2465 |
| GPR107 | NM_001136558 | -0.1445 | -0.608 |
| LOC440563 | NM_001136561 | -0.329 | -0.071 |
| RAD21L1 | NM_001136566 | -0.2215 | -0.164 |
| FAM47E | NM_001136570 | -0.53 | -0.8185 |
| ZAR1L | NM_001136571 | 0.01 | -0.163 |
| LANCL1 | NM_001136574 | -0.8505 | -1.104 |
| LANCL1 | NM_001136575 | -0.8505 | -1.104 |
| LRRFIP1 | NM_001137550 | -1.425 | -2.078 |
| LRRFIP1 | NM_001137551 | -1.425 | -2.078 |
| LRRFIP1 | NM_001137552 | -1.425 | -2.078 |
| LRRFIP1 | NM_001137553 | -1.425 | -2.078 |
| MCTS1 | NM_001137554 | -0.325 | -0.86 |
| ANAPC5 | NM_001137559 | -0.1575 | -0.765 |
| TMEM151B | NM_001137560 | -0.405 | -1.0195 |
| ZBTB42 | NM_001137601 | -0.515 | -1.145 |
| POLR1B | NM_001137604 | -1.5995 | -3.9215 |

| | | | |
|---|---|---|---|
| PARVG | NM_001137605 | 0.088 | -0.186 |
| PARVG | NM_001137606 | 0.088 | -0.186 |
| ZNF732 | NM_001137608 | 0.245 | -0.334 |
| FAM86B2 | NM_001137610 | -0.245 | -0.3095 |
| ANAPC7 | NM_001137664 | -1.09 | -1.641 |
| CASP8AP2 | NM_001137667 | -1.7405 | -3.8585 |
| CASP8AP2 | NM_001137668 | -1.7405 | -3.8585 |
| RGSL1 | NM_001137669 | 0.1635 | -0.3735 |
| POTEC | NM_001137671 | -0.439 | -0.757 |
| ILF3 | NM_001137673 | 0.135 | -0.262 |
| ZNF860 | NM_001137674 | 0.216 | -0.208 |
| ATXN1L | NM_001137675 | -0.9495 | -1.768 |
| AGRP | NM_001138 | 0.3155 | -0.168 |
| ALOX12B | NM_001139 | 0.099 | -0.263 |
| TTLL11 | NM_001139442 | -0.1205 | -0.5465 |
| BEST1 | NM_001139443 | 0.126 | -0.161 |
| BET3L | NM_001139444 | 0.109 | 0.2875 |
| SVOPL | NM_001139456 | -0.282 | -0.3475 |
| BCAP31 | NM_001139457 | 0.154 | -0.236 |
| CNST | NM_001139459 | -1.806 | -3.636 |
| TBL1X | NM_001139466 | -0.9075 | -2.2995 |
| TBL1X | NM_001139467 | -0.5815 | -1.7555 |
| TBL1X | NM_001139468 | -0.5815 | -1.7555 |
| RASGRP3 | NM_001139488 | -0.1455 | 0.0325 |
| TRAF3IP1 | NM_001139490 | -1.008 | -1.422 |
| FGF13 | NM_001139498 | -0.0635 | -0.424 |
| LSM5 | NM_001139499 | -0.4875 | -0.5105 |
| FGF13 | NM_001139500 | -0.0635 | -0.424 |
| FGF13 | NM_001139501 | -0.0635 | -0.424 |
| FGF13 | NM_001139502 | -0.0635 | -0.424 |
| ECHDC1 | NM_001139510 | -1.0715 | -1.5455 |
| DACH2 | NM_001139514 | -0.0405 | -0.161 |
| DACH2 | NM_001139515 | -0.0405 | -0.161 |
| PRKRA | NM_001139517 | -0.8245 | -1.3795 |
| PRKRA | NM_001139518 | -0.8245 | -1.3795 |
| ALOX15 | NM_001140 | -0.016 | -0.2015 |
| ALOX15B | NM_001141 | 0.24 | -0.1405 |
| FAM205A | NM_001141917 | 0.1685 | -0.329 |
| XG | NM_001141919 | 0.1985 | -0.0995 |
| XG | NM_001141920 | 0.1985 | -0.0995 |
| C4orf48 | NM_001141936 | 0.3275 | -0.357 |
| ACTA2 | NM_001141945 | -0.22 | -0.5805 |
| CCDC66 | NM_001141947 | -0.9165 | -2.2135 |

| | | | |
|---|---|---|---|
| TPTE2 | NM_001141968 | -0.1185 | -0.1605 |
| DAXX | NM_001141969 | -0.2725 | -0.882 |
| DAXX | NM_001141970 | -0.2815 | -0.934 |
| ATPBD4 | NM_001141972 | -0.9095 | -1.098 |
| ATP13A2 | NM_001141973 | -0.1365 | -0.628 |
| ATP13A2 | NM_001141974 | -0.1365 | -0.628 |
| TP53BP1 | NM_001141979 | -0.5995 | -1.662 |
| TP53BP1 | NM_001141980 | -0.9105 | -1.82 |
| AMELX | NM_001142 | -0.0465 | -0.4 |
| GGCX | NM_001142269 | -0.1275 | -0.799 |
| RAB11FIP3 | NM_001142272 | -0.814 | -1.623 |
| CLASP1 | NM_001142273 | -0.155 | -0.906 |
| CLASP1 | NM_001142274 | -0.155 | -0.906 |
| NIPA1 | NM_001142275 | -0.081 | -0.3915 |
| APLP2 | NM_001142276 | -0.0115 | -0.5195 |
| APLP2 | NM_001142277 | -0.0215 | -0.5775 |
| APLP2 | NM_001142278 | -0.3565 | -0.861 |
| RNASEH2B | NM_001142279 | -0.8505 | -1.7145 |
| ARCN1 | NM_001142281 | -1.5615 | -3.4985 |
| RPS24 | NM_001142282 | -0.179 | -0.225 |
| RPS24 | NM_001142283 | -0.179 | -0.225 |
| RPS24 | NM_001142284 | -0.179 | -0.225 |
| RPS24 | NM_001142285 | -0.179 | -0.225 |
| SMC6 | NM_001142286 | -0.2945 | -2.4585 |
| SEMA4D | NM_001142287 | -0.301 | -1.2555 |
| COX4NB | NM_001142288 | -0.415 | -2.2255 |
| MGRN1 | NM_001142290 | -0.5735 | -1.226 |
| LMAN2L | NM_001142292 | -1.639 | -3.117 |
| ARFGAP3 | NM_001142293 | -0.4595 | -1.513 |
| SPG20 | NM_001142294 | -1.3075 | -2.834 |
| SPG20 | NM_001142295 | -1.3075 | -2.834 |
| SPG20 | NM_001142296 | -1.3075 | -2.834 |
| SQSTM1 | NM_001142298 | -0.709 | -1.4875 |
| SQSTM1 | NM_001142299 | -0.709 | -1.4875 |
| CCNYL1 | NM_001142300 | -1.425 | -2.709 |
| TMEM67 | NM_001142301 | -1.1445 | -1.783 |
| CCDC113 | NM_001142302 | -0.1935 | -0.5415 |
| ZNF771 | NM_001142305 | -0.264 | -0.311 |
| C5orf60 | NM_001142306 | 0.1875 | -0.335 |
| GTF2H1 | NM_001142307 | -1.641 | -3.4835 |
| TMEM169 | NM_001142310 | -1.8375 | -3.2545 |
| TMEM169 | NM_001142311 | -1.5175 | -2.4055 |
| TMEM169 | NM_001142312 | -1.5175 | -2.4055 |

| | | | |
|---|---|---|---|
| DUSP19 | NM_001142314 | -0.4655 | -1.05 |
| LMO2 | NM_001142315 | 0.0105 | -0.5865 |
| LMO2 | NM_001142316 | 0.0105 | -0.5865 |
| TXNL4B | NM_001142317 | -0.7305 | -1.514 |
| TXNL4B | NM_001142318 | -0.7305 | -1.514 |
| C2orf44 | NM_001142319 | -1.976 | -4.4465 |
| DMTF1 | NM_001142326 | -1.036 | -1.9595 |
| DMTF1 | NM_001142327 | -1.036 | -1.9595 |
| RBFOX1 | NM_001142333 | -0.464 | -0.8375 |
| RBFOX1 | NM_001142334 | -0.464 | -0.8375 |
| GNAL | NM_001142339 | 0.2605 | -0.6505 |
| CMKLR1 | NM_001142343 | -0.142 | -0.557 |
| CMKLR1 | NM_001142344 | 0.234 | 0.0755 |
| CMKLR1 | NM_001142345 | 0.234 | 0.0755 |
| SCN4B | NM_001142348 | 0.382 | -0.2585 |
| SCN4B | NM_001142349 | 0.382 | -0.2585 |
| ST6GAL2 | NM_001142351 | 0.0225 | -0.685 |
| ST6GAL2 | NM_001142352 | 0.0225 | -0.685 |
| PPP3CB | NM_001142353 | -1.9175 | -2.2115 |
| PPP3CB | NM_001142354 | -1.485 | -2.206 |
| NIF3L1 | NM_001142355 | -1.756 | -3.9685 |
| NIF3L1 | NM_001142356 | -1.756 | -3.9685 |
| ALG5 | NM_001142364 | -1.1745 | -1.4165 |
| GSTM2 | NM_001142368 | 0.3265 | -0.6315 |
| PTPN18 | NM_001142370 | -0.2045 | -0.5775 |
| PDK3 | NM_001142386 | 0.3405 | -0.721 |
| PNPLA4 | NM_001142389 | -0.3905 | -0.8045 |
| SLC10A3 | NM_001142391 | -0.4345 | -0.6675 |
| NEDD9 | NM_001142393 | -0.455 | -1.4445 |
| FAHD1 | NM_001142398 | -0.6825 | -1.601 |
| CD164 | NM_001142401 | -0.8875 | -1.047 |
| CD164 | NM_001142402 | -0.8875 | -1.047 |
| CD164 | NM_001142403 | -0.8875 | -1.047 |
| CD164 | NM_001142404 | -0.8875 | -1.047 |
| SLMO1 | NM_001142405 | -0.7705 | -1.0715 |
| CRISP2 | NM_001142407 | 0.2505 | 0.003 |
| CRISP2 | NM_001142408 | 0.2505 | 0.003 |
| C9orf47 | NM_001142413 | 0.33 | 0.3055 |
| CRCP | NM_001142414 | -0.735 | -1.8625 |
| AIMP1 | NM_001142415 | -0.4575 | -0.4495 |
| AIMP1 | NM_001142416 | -0.4575 | -0.4495 |
| CRISP2 | NM_001142417 | 0.2505 | 0.003 |
| MORF4L2 | NM_001142418 | -1.1285 | -0.881 |

| | | | |
|---|---|---|---|
| MORF4L2 | NM_001142419 | -1.2745 | -0.9635 |
| MORF4L2 | NM_001142420 | -1.2745 | -0.9635 |
| MORF4L2 | NM_001142421 | -1.2745 | -0.9635 |
| MORF4L2 | NM_001142422 | -1.2745 | -0.9635 |
| MORF4L2 | NM_001142423 | -1.2745 | -0.9635 |
| MORF4L2 | NM_001142424 | -1.2745 | -0.9635 |
| MORF4L2 | NM_001142425 | -1.2745 | -0.9635 |
| MORF4L2 | NM_001142426 | -1.2745 | -0.9635 |
| MORF4L2 | NM_001142427 | -1.2745 | -0.9635 |
| MORF4L2 | NM_001142428 | -1.2745 | -0.9635 |
| MORF4L2 | NM_001142429 | -1.2745 | -0.9635 |
| MORF4L2 | NM_001142430 | -1.2745 | -0.9635 |
| MORF4L2 | NM_001142431 | -1.2745 | -0.9635 |
| MORF4L2 | NM_001142432 | -1.2745 | -0.9635 |
| MGEA5 | NM_001142434 | -0.893 | -1.4035 |
| CRISP2 | NM_001142435 | 0.2505 | 0.003 |
| EDC3 | NM_001142443 | -0.6485 | -2.071 |
| EDC3 | NM_001142444 | -0.6485 | -2.21 |
| ANK1 | NM_001142445 | -0.1485 | -0.816 |
| ANK1 | NM_001142446 | -0.1485 | -0.816 |
| ATP1B4 | NM_001142447 | 0.03 | -0.2615 |
| SPNS1 | NM_001142448 | -0.065 | -0.543 |
| SPNS1 | NM_001142449 | -0.065 | -0.543 |
| SPNS1 | NM_001142450 | -0.065 | -0.543 |
| SPNS1 | NM_001142451 | -0.065 | -0.543 |
| ASB10 | NM_001142459 | 0.2975 | -0.385 |
| ASB10 | NM_001142460 | 0.2975 | -0.385 |
| OSR2 | NM_001142462 | -0.4925 | -0.084 |
| DKC1 | NM_001142463 | -0.7235 | -0.959 |
| GPT2 | NM_001142466 | 0.186 | -1.051 |
| C16orf45 | NM_001142469 | -0.404 | -1.225 |
| C6orf203 | NM_001142470 | -1.8855 | -4.4665 |
| FAIM3 | NM_001142473 | -0.066 | -0.257 |
| C5orf13 | NM_001142474 | -0.34 | -0.3645 |
| C5orf13 | NM_001142475 | -0.34 | -0.3645 |
| C5orf13 | NM_001142476 | -0.34 | -0.3645 |
| C5orf13 | NM_001142477 | -0.34 | -0.3645 |
| C5orf13 | NM_001142478 | -0.34 | -0.3645 |
| C5orf13 | NM_001142479 | -0.34 | -0.3645 |
| C5orf13 | NM_001142480 | -0.34 | -0.3645 |
| C5orf13 | NM_001142481 | -0.34 | -0.3645 |
| C5orf13 | NM_001142482 | -0.34 | -0.3645 |
| C5orf13 | NM_001142483 | -0.34 | -0.3645 |

| | | | |
|---|---|---|---|
| MYO5A | NM_001142495 | -0.251 | -0.545 |
| MLKL | NM_001142497 | 0.202 | 0.2045 |
| SIRT1 | NM_001142498 | -1.502 | -3.002 |
| PPP1R13L | NM_001142502 | -0.085 | 0.076 |
| STARD8 | NM_001142503 | 0.104 | -0.245 |
| STARD8 | NM_001142504 | 0.104 | -0.245 |
| ABCG4 | NM_001142505 | 0.021 | -0.364 |
| FAM111A | NM_001142519 | -0.195 | -0.1145 |
| FAM111A | NM_001142520 | -0.5785 | 0 |
| FAM111A | NM_001142521 | -0.195 | -0.1145 |
| FBXO5 | NM_001142522 | -1.378 | -2.408 |
| IRAK3 | NM_001142523 | -0.0905 | -0.294 |
| BHLHB9 | NM_001142524 | -0.7825 | -1.764 |
| SH2D3C | NM_001142531 | -0.452 | -1.414 |
| SH2D3C | NM_001142532 | -0.452 | -1.414 |
| SH2D3C | NM_001142533 | -0.452 | -1.414 |
| SH2D3C | NM_001142534 | -0.452 | -1.414 |
| LGALS12 | NM_001142535 | -0.091 | 0.1045 |
| SLC35B3 | NM_001142540 | -2.031 | -3.208 |
| SLC35B3 | NM_001142541 | -2.031 | -3.208 |
| ADCK1 | NM_001142545 | -0.288 | -1.438 |
| LUZP1 | NM_001142546 | -0.5905 | -1.251 |
| RPUSD3 | NM_001142547 | -0.944 | -1.7425 |
| RAD54L | NM_001142548 | -0.519 | -1.697 |
| DDX4 | NM_001142549 | -0.1035 | -0.1605 |
| WDR47 | NM_001142550 | -0.4135 | -2.223 |
| WDR47 | NM_001142551 | -0.4135 | -2.223 |
| DCHS2 | NM_001142552 | -0.188 | 0.388 |
| DCHS2 | NM_001142553 | -0.0375 | -0.1165 |
| TRMT1 | NM_001142554 | -0.1185 | -0.694 |
| ADCK4 | NM_001142555 | -0.433 | -1.817 |
| HMMR | NM_001142556 | -1.0135 | -2.5545 |
| HMMR | NM_001142557 | -1.0135 | -2.5545 |
| CNGA1 | NM_001142564 | -0.128 | 0.114 |
| CPSF7 | NM_001142565 | -1.14 | -2.4285 |
| BBX | NM_001142568 | -0.131 | -1.1405 |
| C1orf106 | NM_001142569 | -0.053 | -0.225 |
| RAD51D | NM_001142571 | 0.2145 | -0.3015 |
| ZNF669 | NM_001142572 | -0.5315 | -1.281 |
| IMPDH1 | NM_001142573 | -0.0955 | -1.069 |
| IMPDH1 | NM_001142574 | -0.0955 | -1.069 |
| IMPDH1 | NM_001142575 | -0.0955 | -1.069 |
| IMPDH1 | NM_001142576 | -0.251 | -1.6415 |

| | | | |
|---|---|---|---|
| ZNF780A | NM_001142577 | -0.344 | -0.0595 |
| ZNF780A | NM_001142578 | -0.344 | -0.0595 |
| ZNF780A | NM_001142579 | -1.268 | -3.0165 |
| NFYC | NM_001142587 | -0.3825 | -1.432 |
| NFYC | NM_001142588 | -0.3825 | -1.432 |
| NFYC | NM_001142589 | -0.3825 | -1.432 |
| NFYC | NM_001142590 | -0.3825 | -1.432 |
| ITPK1 | NM_001142593 | 0.0235 | -0.9185 |
| ITPK1 | NM_001142594 | 0.0235 | -0.9185 |
| CLP1 | NM_001142597 | -0.594 | -1.5375 |
| SPHK1 | NM_001142601 | -0.1685 | -0.8105 |
| SPHK1 | NM_001142602 | -0.1685 | -0.8105 |
| KIAA0141 | NM_001142603 | -0.4585 | -0.8325 |
| PPT1 | NM_001142604 | -0.6435 | -0.6965 |
| EFTUD2 | NM_001142605 | -0.1155 | -0.8155 |
| ULK2 | NM_001142610 | -1.286 | -1.7435 |
| EHBP1 | NM_001142614 | -0.7665 | -1.437 |
| EHBP1 | NM_001142615 | -0.7665 | -1.437 |
| EHBP1 | NM_001142616 | -0.7665 | -1.437 |
| STRA6 | NM_001142617 | -0.683 | -1.1345 |
| STRA6 | NM_001142618 | -0.683 | -1.1345 |
| STRA6 | NM_001142619 | -0.4695 | -0.8435 |
| STRA6 | NM_001142620 | -0.683 | -1.1345 |
| GHDC | NM_001142623 | -0.2425 | -0.6435 |
| RAB34 | NM_001142625 | 0.0325 | -1.7095 |
| TSHR | NM_001142626 | 0.1545 | -0.2765 |
| SEC61A2 | NM_001142627 | -0.119 | -0.3935 |
| SEC61A2 | NM_001142628 | -0.119 | -0.3935 |
| PIK3R5 | NM_001142633 | 0.015 | -0.1775 |
| TNRC6C | NM_001142640 | -1.0965 | -3.3925 |
| FBRSL1 | NM_001142641 | -0.6845 | -1.321 |
| CASKIN2 | NM_001142643 | -0.5995 | -1.6095 |
| SPHKAP | NM_001142644 | -0.1105 | -0.4535 |
| TMEM194B | NM_001142645 | 0.083 | -1.6595 |
| SAR1A | NM_001142648 | -0.899 | -0.331 |
| ANO5 | NM_001142649 | -0.182 | -0.879 |
| HNRPLL | NM_001142650 | -1.6025 | -2.7035 |
| NEURL1B | NM_001142651 | -1.1335 | -3.699 |
| AARSD1 | NM_001142653 | -0.5425 | -0.8315 |
| AARSD1 | NM_001142654 | -0.415 | -0.5785 |
| ATG13 | NM_001142673 | -0.4675 | -1.8515 |
| CHID1 | NM_001142674 | 0.211 | -0.2895 |
| CHID1 | NM_001142675 | 0.211 | -0.2895 |

| | | | |
|---|---|---|---|
| CHID1 | NM_001142676 | 0.211 | -0.2895 |
| CHID1 | NM_001142677 | 0.211 | -0.2895 |
| ANO6 | NM_001142678 | -1.454 | -1.2275 |
| ANO6 | NM_001142679 | -1.454 | -1.2275 |
| ANO6 | NM_001142680 | -1.454 | -1.2275 |
| ZMYM5 | NM_001142684 | -1.318 | -1.8505 |
| ARHGAP32 | NM_001142685 | -1.033 | -2.2805 |
| DLG2 | NM_001142699 | -0.104 | -0.1645 |
| DLG2 | NM_001142700 | -0.104 | -0.1645 |
| DLG2 | NM_001142702 | -0.104 | -0.1645 |
| FAM111B | NM_001142703 | -0.364 | -0.31 |
| FAM111B | NM_001142704 | -0.364 | -0.31 |
| ERI2 | NM_001142725 | -1.446 | -2.213 |
| KCTD1 | NM_001142730 | -1.2195 | -3.109 |
| ASB14 | NM_001142733 | 0.049 | -0.1455 |
| C15orf23 | NM_001142761 | -2.1465 | -3.481 |
| C15orf23 | NM_001142762 | -2.1465 | -3.481 |
| PCDH15 | NM_001142763 | 0.3695 | 0.4005 |
| PCDH15 | NM_001142764 | 0.3695 | 0.4005 |
| PCDH15 | NM_001142765 | 0.3695 | 0.4005 |
| PCDH15 | NM_001142766 | 0.3695 | 0.4005 |
| PCDH15 | NM_001142767 | 0.3695 | 0.4005 |
| PCDH15 | NM_001142768 | 0.3695 | 0.4005 |
| PCDH15 | NM_001142769 | 0.3695 | 0.4005 |
| PCDH15 | NM_001142770 | 0.0015 | -0.7555 |
| PCDH15 | NM_001142771 | 0.1375 | 0.377 |
| PCDH15 | NM_001142772 | 0.3695 | 0.4005 |
| PCDH15 | NM_001142773 | 0.3695 | 0.4005 |
| CHAC1 | NM_001142776 | -0.6065 | -0.544 |
| HSD3B7 | NM_001142777 | 0.085 | -0.3355 |
| HSD3B7 | NM_001142778 | 0.085 | -0.3355 |
| MAGI3 | NM_001142782 | -0.9535 | -2.801 |
| IL11RA | NM_001142784 | 0.022 | -0.6485 |
| CXCR3 | NM_001142797 | 0.031 | -0.051 |
| C17orf49 | NM_001142798 | 0.1705 | -0.3875 |
| EYS | NM_001142800 | 0.3525 | -0.5735 |
| EYS | NM_001142801 | 0.3525 | -0.5735 |
| SLC6A8 | NM_001142805 | -0.3295 | -0.8945 |
| SLC6A8 | NM_001142806 | -0.3295 | -0.8945 |
| ACOXL | NM_001142807 | -0.0835 | -0.2795 |
| C21orf56 | NM_001142854 | -0.3795 | -0.951 |
| EFCAB1 | NM_001142857 | -0.043 | -0.168 |
| SMAD6 | NM_001142861 | -0.6655 | -1.897 |

| | | | |
|---|---|---|---|
| IP6K3 | NM_001142883 | 0.0835 | -0.271 |
| TMOD2 | NM_001142885 | -0.376 | -1.073 |
| EPHX3 | NM_001142886 | -0.474 | -0.5045 |
| LRRC61 | NM_001142928 | -0.9145 | -1.967 |
| CHAT | NM_001142929 | 0.0335 | -0.886 |
| API5 | NM_001142930 | -1.603 | -1.6575 |
| API5 | NM_001142931 | -1.267 | -1.653 |
| CHAT | NM_001142933 | 0.0335 | -0.886 |
| CHAT | NM_001142934 | 0.0335 | -0.886 |
| MXD3 | NM_001142935 | 0.044 | -0.3425 |
| DAGLB | NM_001142936 | -0.874 | -2.324 |
| C11orf21 | NM_001142946 | -0.0635 | -0.2615 |
| LPAR5 | NM_001142961 | -0.143 | -0.3575 |
| C22orf46 | NM_001142964 | -1.317 | -1.2565 |
| GREB1L | NM_001142966 | -0.468 | -2.017 |
| AMELY | NM_001143 | -0.1455 | -0.398 |
| ZBED5 | NM_001143667 | -1.885 | -4.337 |
| AMIGO2 | NM_001143668 | 0.3685 | -1.7235 |
| BRP44 | NM_001143674 | -0.0225 | -0.882 |
| SGK1 | NM_001143676 | 1.832 | 2.4965 |
| SGK1 | NM_001143677 | 0.649 | 1.4025 |
| SGK1 | NM_001143678 | 1.359 | 2.539 |
| GSTK1 | NM_001143679 | -0.256 | -0.5815 |
| GSTK1 | NM_001143680 | -0.256 | -0.5815 |
| GSTK1 | NM_001143681 | -0.256 | -0.5815 |
| CALCOCO1 | NM_001143682 | -0.1165 | -0.447 |
| CES5A | NM_001143685 | -0.0455 | -0.294 |
| DCST1 | NM_001143687 | -0.0625 | -0.238 |
| DIS3L | NM_001143688 | -1.497 | -1.8245 |
| LRRC27 | NM_001143757 | -0.3155 | -1.0585 |
| LRRC27 | NM_001143758 | 0.1985 | -0.558 |
| LRRC27 | NM_001143759 | -0.3155 | -1.0585 |
| EIF5A | NM_001143760 | 0.2005 | -0.1815 |
| EIF5A | NM_001143761 | 0.2005 | -0.1815 |
| EIF5A | NM_001143762 | 0.2005 | -0.1815 |
| SYCE1 | NM_001143763 | -0.0115 | 0.16 |
| SYCE1 | NM_001143764 | -0.0115 | 0.16 |
| ZNF438 | NM_001143770 | -0.828 | -1.152 |
| FAM13C | NM_001143773 | -1.1515 | -2.573 |
| PHYHIPL | NM_001143774 | -0.879 | -2.4345 |
| CTDNEP1 | NM_001143775 | -0.337 | -1.6245 |
| ASAP3 | NM_001143778 | 0.02 | -0.2315 |
| IFT81 | NM_001143779 | -1.009 | -2.189 |

| | | | |
|---|---|---|---|
| SLC25A39 | NM_001143780 | 0.265 | -0.2935 |
| FKBP11 | NM_001143781 | -0.572 | -0.5555 |
| FKBP11 | NM_001143782 | -0.521 | -0.6705 |
| FES | NM_001143784 | -0.186 | -0.4545 |
| PPHLN1 | NM_001143787 | -1.1515 | -2.0805 |
| PPHLN1 | NM_001143788 | -1.1515 | -2.0805 |
| PPHLN1 | NM_001143789 | -1.1515 | -2.0805 |
| PHOSPHO1 | NM_001143804 | 0.083 | -0.3285 |
| BDNF | NM_001143805 | -0.605 | -1.907 |
| BDNF | NM_001143806 | -0.605 | -1.907 |
| BDNF | NM_001143807 | -0.605 | -1.907 |
| BDNF | NM_001143808 | -0.605 | -1.907 |
| BDNF | NM_001143809 | -0.605 | -1.907 |
| BDNF | NM_001143810 | -0.605 | -1.907 |
| BDNF | NM_001143811 | -0.605 | -1.907 |
| BDNF | NM_001143812 | -0.605 | -1.907 |
| BDNF | NM_001143813 | -0.605 | -1.907 |
| BDNF | NM_001143814 | -0.605 | -1.907 |
| BDNF | NM_001143816 | -0.605 | -1.907 |
| SERPINB2 | NM_001143818 | 0.3125 | -0.5285 |
| TPCN1 | NM_001143819 | -0.53 | -1.053 |
| ETS1 | NM_001143820 | -0.345 | -1.2305 |
| PLEKHA5 | NM_001143821 | -1.354 | -3.347 |
| ZFP161 | NM_001143823 | -1.4405 | -2.7725 |
| SLC38A4 | NM_001143824 | 0.3245 | 0.3095 |
| MAPRE2 | NM_001143826 | -0.7635 | -1.343 |
| MAPRE2 | NM_001143827 | -0.7635 | -1.343 |
| HRH4 | NM_001143828 | 0.2355 | -0.062 |
| CCDC68 | NM_001143829 | -0.815 | -0.938 |
| GAS2 | NM_001143830 | -0.19 | -0.1115 |
| GRM5 | NM_001143831 | -0.0525 | -0.4285 |
| LEUTX | NM_001143832 | 0.0765 | -0.11 |
| IQCF6 | NM_001143833 | 0.0985 | -0.3145 |
| NFRKB | NM_001143835 | -0.6905 | -2.1255 |
| NOX4 | NM_001143836 | 0.0195 | -0.5715 |
| NOX4 | NM_001143837 | 0.0195 | -0.5715 |
| SLC13A5 | NM_001143838 | 0.199 | -0.514 |
| PDE2A | NM_001143839 | -0.4595 | -1.1105 |
| TMEM106C | NM_001143841 | -0.1905 | -0.1445 |
| TMEM106C | NM_001143842 | -0.1905 | -0.1445 |
| TMEM106C | NM_001143843 | -0.1905 | -0.1445 |
| TCTN2 | NM_001143850 | -0.57 | -0.842 |
| TCHP | NM_001143852 | -0.906 | -1.507 |

| | | | |
|---|---|---|---|
| RPH3A | NM_001143854 | 0.0845 | -0.2505 |
| PPIAL4B | NM_001143883 | -0.3885 | -0.214 |
| PPP1R12A | NM_001143885 | -0.3175 | -1.7575 |
| PPP1R12A | NM_001143886 | -0.3175 | -1.7575 |
| COPS2 | NM_001143887 | -0.4095 | -1.847 |
| BSDC1 | NM_001143888 | -0.2225 | -0.8095 |
| BSDC1 | NM_001143889 | -0.2225 | -0.8095 |
| BSDC1 | NM_001143890 | -0.2225 | -0.8095 |
| C12orf65 | NM_001143905 | -1.623 | -3.1635 |
| TRAFD1 | NM_001143906 | -0.581 | -1.643 |
| LOC729020 | NM_001143909 | -0.118 | -0.489 |
| FAM76A | NM_001143912 | -0.3435 | -0.4015 |
| FAM76A | NM_001143913 | -0.3435 | -0.4015 |
| FAM76A | NM_001143914 | -0.3435 | -0.4015 |
| FAM76A | NM_001143915 | -0.3435 | -0.4015 |
| LTB4R | NM_001143919 | -0.785 | -0.6295 |
| CROT | NM_001143935 | -1.1135 | -1.595 |
| C15orf61 | NM_001143936 | -0.454 | -1.704 |
| PSMA1 | NM_001143937 | -0.564 | -0.096 |
| ZNF534 | NM_001143939 | -0.221 | -0.4545 |
| LGSN | NM_001143940 | -0.26 | -0.779 |
| RBM24 | NM_001143941 | -0.1465 | -0.9645 |
| RBM24 | NM_001143942 | -0.4675 | -0.5245 |
| EFCAB2 | NM_001143943 | 0.225 | -0.224 |
| LEMD2 | NM_001143944 | -1.1345 | -2.5345 |
| GAS6 | NM_001143945 | -0.203 | -0.3345 |
| GAS6 | NM_001143946 | -0.203 | -0.3345 |
| C6orf105 | NM_001143948 | -0.1875 | -0.553 |
| GPR63 | NM_001143957 | -0.7555 | -2.0085 |
| TMEM30A | NM_001143958 | -0.827 | -1.3535 |
| C2orf74 | NM_001143959 | -1.6525 | -2.2885 |
| CAPN8 | NM_001143962 | 0.03 | -0.024 |
| L1CAM | NM_001143963 | 0.457 | -0.229 |
| ARL5C | NM_001143968 | 0.189 | -0.2905 |
| CDYL | NM_001143970 | -1.21 | -1.1755 |
| CDYL | NM_001143971 | -1.21 | -1.1755 |
| TCTN3 | NM_001143973 | -0.857 | -2.2265 |
| UBTFL1 | NM_001143975 | -0.018 | -0.1355 |
| ZCCHC18 | NM_001143978 | -0.4715 | -0.8005 |
| NDE1 | NM_001143979 | -1.4135 | -2.1175 |
| CCDC154 | NM_001143980 | 0.0185 | -0.404 |
| CHRDL1 | NM_001143981 | -0.23 | -0.1735 |
| CHRDL1 | NM_001143982 | 0.032 | 0.1005 |

| | | | |
|---|---|---|---|
| CHRDL1 | NM_001143983 | 0.032 | 0.1005 |
| PTPMT1 | NM_001143984 | -0.5465 | -1.355 |
| BANF1 | NM_001143985 | 0.2465 | -0.493 |
| TLE6 | NM_001143986 | -0.188 | -0.5995 |
| NBPF4 | NM_001143989 | -0.3085 | -1.3915 |
| WRAP53 | NM_001143990 | -0.332 | -0.602 |
| WRAP53 | NM_001143991 | -0.332 | -0.602 |
| WRAP53 | NM_001143992 | -0.332 | -0.602 |
| RASSF7 | NM_001143993 | -0.4195 | -1.298 |
| RASSF7 | NM_001143994 | -0.4195 | -1.298 |
| LPXN | NM_001143995 | -0.78 | -1.0555 |
| OGDHL | NM_001143996 | -0.3015 | -0.8615 |
| OGDHL | NM_001143997 | -0.3015 | -0.8615 |
| SEC14L1 | NM_001143998 | -0.4435 | -0.6915 |
| SEC14L1 | NM_001143999 | 0.111 | -0.986 |
| AMFR | NM_001144 | 0.246 | -0.398 |
| AGAP5 | NM_001144000 | -0.7305 | -1.6 |
| SEC14L1 | NM_001144001 | 0.111 | -0.986 |
| CTPS2 | NM_001144002 | -1.0275 | -0.89 |
| SLITRK2 | NM_001144003 | -0.1435 | -0.487 |
| SLITRK2 | NM_001144004 | -0.1435 | -0.487 |
| SLITRK2 | NM_001144005 | -0.1435 | -0.487 |
| SLITRK2 | NM_001144006 | -0.1435 | -0.487 |
| SLITRK2 | NM_001144008 | -0.1435 | -0.487 |
| SLITRK2 | NM_001144009 | -0.1435 | -0.487 |
| SLITRK2 | NM_001144010 | -0.1435 | -0.487 |
| TMX2 | NM_001144012 | -0.195 | -0.7195 |
| RGPD3 | NM_001144013 | 0.239 | 0.1165 |
| NDOR1 | NM_001144026 | -1.0745 | -1.4655 |
| NDOR1 | NM_001144027 | -0.3475 | -0.622 |
| NDOR1 | NM_001144028 | -0.3475 | -0.622 |
| NAT10 | NM_001144030 | -0.5435 | -2.465 |
| STOML3 | NM_001144033 | -0.122 | 0.043 |
| TMEM25 | NM_001144034 | -0.588 | -2.037 |
| TMEM25 | NM_001144035 | -0.588 | -2.037 |
| TMEM25 | NM_001144036 | -0.588 | -2.037 |
| TMEM25 | NM_001144037 | -0.588 | -2.037 |
| TMEM25 | NM_001144038 | -0.588 | -2.037 |
| NTM | NM_001144058 | -0.022 | -0.5335 |
| NTM | NM_001144059 | -0.022 | -0.5335 |
| NHSL1 | NM_001144060 | 0.085 | -0.484 |
| COPB1 | NM_001144061 | -0.7555 | -1.998 |
| COPB1 | NM_001144062 | -0.7555 | -1.998 |

| | | | |
|---|---|---|---|
| OSBPL5 | NM_001144063 | -0.3035 | -0.723 |
| ZNF772 | NM_001144068 | -1.1105 | -1.8145 |
| ABCC3 | NM_001144070 | -0.3115 | -0.1385 |
| LDHAL6A | NM_001144071 | -0.3455 | 0.027 |
| UBAC2 | NM_001144072 | -0.317 | -1.719 |
| CHORDC1 | NM_001144073 | -1.0505 | -2.5735 |
| DET1 | NM_001144074 | -1.097 | -3.0075 |
| PLCL2 | NM_001144382 | -1.0225 | -2.5125 |
| CDH17 | NM_001144663 | -0.21 | -0.1885 |
| NPFFR2 | NM_001144756 | 0.664 | -0.5265 |
| SCG5 | NM_001144757 | -0.294 | -0.6975 |
| PHLDB1 | NM_001144758 | -0.1505 | -1.0515 |
| PHLDB1 | NM_001144759 | -0.1505 | -1.0515 |
| TLE2 | NM_001144761 | 0.052 | -0.3565 |
| TLE2 | NM_001144762 | 0.052 | -0.3565 |
| DNAJC7 | NM_001144766 | -0.7385 | -1.0455 |
| UGT2B10 | NM_001144767 | -0.319 | -0.1645 |
| NSMAF | NM_001144772 | -0.968 | -1.4405 |
| ELAVL4 | NM_001144774 | -1.8505 | -0.916 |
| ELAVL4 | NM_001144775 | -1.8505 | -0.916 |
| ELAVL4 | NM_001144776 | -1.8505 | -0.916 |
| ELAVL4 | NM_001144777 | -1.8505 | -0.916 |
| CD58 | NM_001144822 | -1.438 | -2.471 |
| DENND4A | NM_001144823 | -0.6575 | -1.5325 |
| ZNF234 | NM_001144824 | -1.3605 | -2.1535 |
| RUNDC3A | NM_001144825 | -0.1255 | -0.548 |
| RUNDC3A | NM_001144826 | -0.1315 | -0.6425 |
| RPUSD4 | NM_001144827 | -0.904 | -2.3975 |
| FMO5 | NM_001144829 | -0.0655 | -0.1195 |
| PHB2 | NM_001144831 | 0.081 | -0.4365 |
| TTC39A | NM_001144832 | -0.463 | -0.7525 |
| PLEKHG6 | NM_001144856 | 0.168 | -0.7195 |
| PLEKHG6 | NM_001144857 | 0.168 | -0.7195 |
| LIPT2 | NM_001144869 | -0.505 | -1.0635 |
| VSTM5 | NM_001144871 | -0.0165 | -0.146 |
| CCDC42B | NM_001144872 | -0.142 | -0.422 |
| PATE4 | NM_001144874 | -0.284 | -0.173 |
| DOK3 | NM_001144876 | -0.391 | -0.7645 |
| SCAI | NM_001144877 | -0.5355 | -0.128 |
| IMPA1 | NM_001144878 | -0.9455 | -1.2825 |
| IMPA1 | NM_001144879 | -0.9455 | -1.2825 |
| PRICKLE1 | NM_001144881 | -0.785 | -2.75 |
| PRICKLE1 | NM_001144882 | -0.785 | -2.75 |

| | | | |
|---|---|---|---|
| PRICKLE1 | NM_001144883 | -0.785 | -2.75 |
| SLC30A7 | NM_001144884 | -1.33 | -2.6405 |
| CITED1 | NM_001144887 | -0.12 | -0.893 |
| BAIAP2 | NM_001144888 | -0.5985 | -1.2375 |
| SLC23A3 | NM_001144890 | -0.2 | -0.056 |
| MBIP | NM_001144891 | -1.276 | -2.536 |
| FGF1 | NM_001144892 | 0.2095 | -0.095 |
| CD209 | NM_001144893 | 0.0665 | -0.056 |
| CD209 | NM_001144894 | 0.0665 | -0.056 |
| CD209 | NM_001144895 | 0.0665 | -0.056 |
| CD209 | NM_001144896 | 0.0665 | -0.056 |
| CD209 | NM_001144897 | 0.0665 | -0.056 |
| CD209 | NM_001144899 | 0.0665 | -0.056 |
| SMCR7 | NM_001144900 | -0.5355 | -1.235 |
| TM6SF1 | NM_001144903 | -0.2825 | -0.3745 |
| CLEC4M | NM_001144904 | -0.3085 | -0.2745 |
| CLEC4M | NM_001144905 | -0.3085 | -0.2745 |
| CLEC4M | NM_001144906 | -0.3085 | -0.2745 |
| CLEC4M | NM_001144907 | -0.3085 | -0.2745 |
| CLEC4M | NM_001144908 | -0.3085 | -0.2745 |
| CLEC4M | NM_001144909 | -0.3085 | -0.2745 |
| CLEC4M | NM_001144910 | -0.3085 | -0.2745 |
| CLEC4M | NM_001144911 | -0.3085 | -0.2745 |
| ATG7 | NM_001144912 | -0.2245 | -0.7755 |
| FGFR2 | NM_001144913 | -1.4165 | -3.3415 |
| FGFR2 | NM_001144914 | -1.4165 | -3.3415 |
| FGFR2 | NM_001144915 | -1.4165 | -3.3415 |
| FGFR2 | NM_001144916 | -1.4165 | -3.3415 |
| FGFR2 | NM_001144917 | -1.4165 | -3.3415 |
| FGFR2 | NM_001144918 | -1.2 | -2.651 |
| FGFR2 | NM_001144919 | -1.4165 | -3.3415 |
| TTC26 | NM_001144920 | 0.077 | -0.6115 |
| TTC26 | NM_001144923 | 0.077 | -0.6115 |
| MX1 | NM_001144925 | 0.138 | 0.0445 |
| NKIRAS2 | NM_001144927 | -0.569 | -1.22 |
| NKIRAS2 | NM_001144928 | -0.569 | -1.22 |
| NKIRAS2 | NM_001144929 | -0.569 | -1.22 |
| LOC391322 | NM_001144931 | 0.872 | 0.8155 |
| PSMB5 | NM_001144932 | -0.298 | -1.16 |
| EFCAB3 | NM_001144933 | -0.874 | -0.955 |
| FGF1 | NM_001144934 | 0.2095 | -0.095 |
| FGF1 | NM_001144935 | 0.2095 | -0.095 |
| C11orf95 | NM_001144936 | -1.0685 | -1.309 |

| | | | |
|---|---|---|---|
| FNDC7 | NM_001144937 | -0.154 | -0.788 |
| RAB34 | NM_001144943 | 0.0315 | -1.905 |
| MYL12B | NM_001144944 | -1.3565 | -0.325 |
| MYL12B | NM_001144945 | -1.3565 | -0.325 |
| SSC5D | NM_001144950 | 0.1065 | -0.5325 |
| GLYCTK | NM_001144951 | 0.35 | -0.0325 |
| SDK2 | NM_001144952 | -1.2395 | -1.173 |
| C5orf47 | NM_001144954 | -0.6105 | 0.167 |
| DTWD1 | NM_001144955 | -1.87 | -2.782 |
| RIIAD1 | NM_001144956 | 0.0675 | -0.4225 |
| EFCAB4B | NM_001144958 | 0.1915 | -0.324 |
| ANKUB1 | NM_001144960 | -0.045 | -0.589 |
| NFKBIL1 | NM_001144961 | -0.7705 | -1.652 |
| NFKBIL1 | NM_001144962 | -0.7705 | -1.652 |
| NFKBIL1 | NM_001144963 | -0.7705 | -1.652 |
| NEDD4L | NM_001144964 | -0.9225 | -2.059 |
| NEDD4L | NM_001144965 | -0.9225 | -2.059 |
| NEDD4L | NM_001144966 | -0.9225 | -2.059 |
| NEDD4L | NM_001144967 | -0.9225 | -2.059 |
| NEDD4L | NM_001144968 | -0.9225 | -2.059 |
| NEDD4L | NM_001144969 | -0.9225 | -2.059 |
| NEDD4L | NM_001144970 | -0.9225 | -2.059 |
| NEDD4L | NM_001144971 | -0.9225 | -2.059 |
| SNX20 | NM_001144972 | 0.034 | -0.208 |
| MTHFD2L | NM_001144978 | -1.3875 | -1.545 |
| CCDC169 | NM_001144985 | 0.003 | -0.819 |
| ZNF814 | NM_001144989 | -0.395 | -1.047 |
| KIAA1239 | NM_001144990 | -0.005 | -0.908 |
| C2orf72 | NM_001144994 | 0.126 | -0.775 |
| CCDC85C | NM_001144995 | -0.517 | -1.6715 |
| ITGAV | NM_001144999 | -1.023 | -1.5975 |
| ANG | NM_001145 | -0.2205 | 0.0415 |
| ITGAV | NM_001145000 | -1.023 | -1.5975 |
| NEK6 | NM_001145001 | -0.1015 | -0.482 |
| MUC7 | NM_001145006 | -0.4325 | -0.7875 |
| BTN3A1 | NM_001145008 | -0.839 | -0.7685 |
| BTN3A1 | NM_001145009 | -0.839 | -0.7685 |
| C12orf70 | NM_001145010 | -0.216 | -0.4185 |
| C16orf96 | NM_001145011 | 0.165 | -0.367 |
| RFPL4A | NM_001145014 | -0.353 | 0.0135 |
| SLC36A3 | NM_001145017 | -0.198 | -0.1625 |
| CCDC153 | NM_001145018 | 0.127 | -0.152 |
| C6orf52 | NM_001145020 | 0.019 | -0.386 |

| | | | |
|---|---|---|---|
| SCRN2 | NM_001145023 | -0.2485 | -1.397 |
| C11orf34 | NM_001145024 | -0.0595 | -0.0115 |
| EDEM2 | NM_001145025 | -0.12 | 0.143 |
| PTPRQ | NM_001145026 | 0.003 | -0.2345 |
| PALM3 | NM_001145028 | -0.2535 | -0.5585 |
| ANKRD30B | NM_001145029 | -0.696 | -0.237 |
| C3orf77 | NM_001145030 | 0.211 | -0.103 |
| PLAU | NM_001145031 | -0.252 | 0.5425 |
| C11orf96 | NM_001145033 | 0.52 | 1.252 |
| COG2 | NM_001145036 | -1.412 | -3.549 |
| SLCO3A1 | NM_001145044 | -0.267 | -0.953 |
| LOC388946 | NM_001145051 | -0.066 | -0.0715 |
| EFCAB5 | NM_001145053 | 0.022 | -0.161 |
| C2orf81 | NM_001145054 | -0.031 | -0.555 |
| IL18BP | NM_001145055 | -0.38 | -0.2835 |
| SLC44A2 | NM_001145056 | -0.108 | -0.9795 |
| IL18BP | NM_001145057 | -0.38 | -0.2835 |
| IQCF5 | NM_001145059 | 0.198 | -0.1985 |
| GATSL2 | NM_001145064 | 0.1955 | -0.689 |
| FAM190A | NM_001145065 | -0.1495 | -0.544 |
| USP27X | NM_001145073 | -1.2075 | -1.6515 |
| EML4 | NM_001145076 | -0.646 | -1.529 |
| LRRC10B | NM_001145077 | 0.424 | -0.764 |
| ZNF805 | NM_001145078 | -1.693 | -3.1645 |
| COG6 | NM_001145079 | -1.2695 | -2.292 |
| C17orf104 | NM_001145080 | -0.0055 | -1.4675 |
| ZNF619 | NM_001145082 | -0.387 | -1.166 |
| ZNF619 | NM_001145083 | -0.4985 | -1.1255 |
| WDR67 | NM_001145088 | -0.791 | -1.8465 |
| ZNF619 | NM_001145093 | -0.387 | -1.166 |
| HHLA1 | NM_001145095 | -0.0035 | -0.301 |
| SKIL | NM_001145097 | -0.6595 | -0.461 |
| SKIL | NM_001145098 | -0.6595 | -0.461 |
| SLC2A6 | NM_001145099 | -0.682 | -1.334 |
| BTBD18 | NM_001145101 | 0.0505 | 0.103 |
| SMAD3 | NM_001145102 | -0.0265 | -1.451 |
| SMAD3 | NM_001145103 | -0.0265 | -1.451 |
| SMAD3 | NM_001145104 | -0.0265 | -1.451 |
| FIBCD1 | NM_001145106 | -0.067 | -0.416 |
| NELL2 | NM_001145107 | -0.61 | -0.706 |
| NELL2 | NM_001145108 | -0.6475 | -0.7445 |
| NELL2 | NM_001145109 | -0.61 | -0.706 |
| NELL2 | NM_001145110 | -0.61 | -0.706 |

| | | | |
|---|---|---|---|
| PATL2 | NM_001145112 | -0.1805 | -0.813 |
| MYADML2 | NM_001145113 | 0.0105 | -0.1285 |
| RRP12 | NM_001145114 | -0.0985 | -1.081 |
| PPP1R3G | NM_001145115 | -0.1155 | -0.324 |
| GRID2IP | NM_001145118 | 0.031 | 0.0255 |
| CAPN14 | NM_001145122 | -0.2725 | -0.1945 |
| MKRN1 | NM_001145125 | -0.3455 | -0.87 |
| EVPLL | NM_001145127 | -0.145 | -0.224 |
| AKD1 | NM_001145128 | -0.2595 | -0.167 |
| C5orf52 | NM_001145132 | -0.673 | -0.107 |
| RELA | NM_001145138 | -0.7155 | -2.538 |
| CXorf49B | NM_001145139 | 0.1455 | -0.3245 |
| HTR3D | NM_001145143 | -0.136 | -0.3185 |
| SLC1A5 | NM_001145144 | -0.7095 | -2.018 |
| SLC1A5 | NM_001145145 | -0.7095 | -2.018 |
| CRHR1 | NM_001145146 | 0.2285 | -1.308 |
| VAMP7 | NM_001145149 | -0.849 | -0.845 |
| VPS37A | NM_001145152 | -1.5415 | -3.6465 |
| DNAH14 | NM_001145154 | 0.163 | -0.345 |
| NR2F2 | NM_001145155 | -0.3325 | -0.513 |
| NR2F2 | NM_001145156 | -0.3325 | -0.513 |
| NR2F2 | NM_001145157 | -0.3325 | -0.513 |
| HBS1L | NM_001145158 | -0.5595 | -0.957 |
| INTS9 | NM_001145159 | -0.9445 | -2.658 |
| TPM4 | NM_001145160 | -0.74 | -0.729 |
| UBE2QL1 | NM_001145161 | -1.0185 | -3.1075 |
| DOHH | NM_001145165 | -0.1555 | -1.9745 |
| PRMT3 | NM_001145166 | -0.884 | -1.36 |
| PRMT3 | NM_001145167 | -0.884 | -1.36 |
| GPR113 | NM_001145168 | -0.1235 | -0.5355 |
| GPR113 | NM_001145169 | -0.1235 | -0.5355 |
| SHISA7 | NM_001145176 | -0.027 | -0.2845 |
| FAM200B | NM_001145191 | -0.376 | -1.1565 |
| C18orf42 | NM_001145194 | -0.82 | -0.21 |
| SLC39A12 | NM_001145195 | -0.3685 | -0.402 |
| C12orf75 | NM_001145199 | 0.021 | -0.6115 |
| PRAP1 | NM_001145201 | -0.0205 | -0.1765 |
| SHISA9 | NM_001145204 | -0.174 | -0.301 |
| SHISA9 | NM_001145205 | -0.174 | -0.301 |
| KIAA1671 | NM_001145206 | -0.8645 | -1.9985 |
| HBS1L | NM_001145207 | -0.5595 | -0.957 |
| DEPDC1B | NM_001145208 | -1.4085 | -1.461 |
| ANKRD65 | NM_001145210 | -0.0085 | -0.3625 |

| | | | |
|---|---|---|---|
| SLCO2B1 | NM_001145211 | 0.073 | -0.4525 |
| SLCO2B1 | NM_001145212 | 0.073 | -0.4525 |
| GOLGA6D | NM_001145224 | -0.1425 | -0.089 |
| SP9 | NM_001145250 | -0.084 | -0.387 |
| IKBKG | NM_001145255 | -0.315 | -0.9625 |
| IRAK4 | NM_001145256 | -0.4565 | -0.8125 |
| IRAK4 | NM_001145257 | -0.4565 | -0.8125 |
| IRAK4 | NM_001145258 | -0.4565 | -0.8125 |
| NCOA4 | NM_001145260 | -0.9855 | -1.4745 |
| NCOA4 | NM_001145261 | -0.9855 | -1.4745 |
| NCOA4 | NM_001145262 | -0.9855 | -1.4745 |
| NCOA4 | NM_001145263 | -0.9855 | -1.4745 |
| SLC35C1 | NM_001145265 | -0.5575 | -1.1465 |
| SLC35C1 | NM_001145266 | -0.5575 | -1.1465 |
| FAM185A | NM_001145268 | -0.402 | -1.9555 |
| FAM185A | NM_001145269 | -0.621 | -2.041 |
| ADAMDEC1 | NM_001145271 | 0.052 | 0.2435 |
| ADAMDEC1 | NM_001145272 | 0.052 | 0.2435 |
| ZFY | NM_001145275 | 0.373 | -1.1505 |
| ZFY | NM_001145276 | 0.373 | -1.1505 |
| NECAP2 | NM_001145277 | -0.6085 | -1.8115 |
| NECAP2 | NM_001145278 | -0.6085 | -1.8115 |
| OPRM1 | NM_001145279 | -0.147 | -0.562 |
| OPRM1 | NM_001145280 | -0.147 | -0.562 |
| OPRM1 | NM_001145281 | -0.147 | -0.562 |
| OPRM1 | NM_001145282 | -0.147 | -0.562 |
| OPRM1 | NM_001145283 | -0.147 | -0.562 |
| OPRM1 | NM_001145284 | -0.147 | -0.562 |
| OPRM1 | NM_001145285 | -0.147 | -0.562 |
| OPRM1 | NM_001145286 | -0.147 | -0.562 |
| OPRM1 | NM_001145287 | -0.147 | -0.562 |
| SLC17A8 | NM_001145288 | -0.2775 | -0.166 |
| SLC37A2 | NM_001145290 | 0.4695 | 0.1785 |
| PDE6B | NM_001145291 | -0.055 | -0.7385 |
| PDE6B | NM_001145292 | -0.055 | -0.7385 |
| NGLY1 | NM_001145293 | -1.0265 | -1.912 |
| NGLY1 | NM_001145294 | -1.0265 | -1.912 |
| NGLY1 | NM_001145295 | -1.0265 | -1.912 |
| EXOC7 | NM_001145297 | -0.1865 | -0.535 |
| EXOC7 | NM_001145298 | -0.1865 | -0.535 |
| EXOC7 | NM_001145299 | -0.1865 | -0.535 |
| RARA | NM_001145301 | -0.49 | -1.943 |
| RARA | NM_001145302 | -0.49 | -1.943 |

| | | | |
|---|---|---|---|
| TMC4 | NM_001145303 | -0.156 | -0.095 |
| KIAA1683 | NM_001145304 | -0.1375 | -0.7415 |
| CDK6 | NM_001145306 | -0.7675 | -1.011 |
| LRTOMT | NM_001145307 | -0.1105 | -0.392 |
| LRTOMT | NM_001145308 | -0.0875 | -0.4045 |
| PLIN1 | NM_001145311 | -0.0525 | -0.5135 |
| ETV3 | NM_001145312 | -0.823 | -1.1555 |
| FSD1L | NM_001145313 | -0.061 | -1.284 |
| CEP44 | NM_001145314 | -0.7585 | -1.4275 |
| DSN1 | NM_001145315 | -1.969 | -2.419 |
| DSN1 | NM_001145316 | -1.969 | -2.419 |
| DSN1 | NM_001145317 | -1.969 | -2.419 |
| DSN1 | NM_001145318 | -1.969 | -2.419 |
| PLS1 | NM_001145319 | -1.0865 | -1.9235 |
| ADAMTSL2 | NM_001145320 | -0.482 | -1.021 |
| UBE2Q2 | NM_001145335 | -0.8905 | -1.397 |
| ZNF566 | NM_001145343 | -1.2225 | -2.2965 |
| ZNF566 | NM_001145344 | -1.2225 | -2.2965 |
| ZNF566 | NM_001145345 | -2.0015 | -3.1065 |
| RBMXL3 | NM_001145346 | 0.0135 | -0.1485 |
| ZNF576 | NM_001145347 | -0.7 | -2.022 |
| JMJD5 | NM_001145348 | -0.4175 | -0.9105 |
| IPP | NM_001145349 | -0.6225 | -0.883 |
| C20orf173 | NM_001145350 | 0.2795 | -0.133 |
| ELF1 | NM_001145353 | -0.088 | -1.1125 |
| MKLN1 | NM_001145354 | -0.4855 | -2.427 |
| SIN3A | NM_001145357 | -1.285 | -3.2455 |
| SIN3A | NM_001145358 | -1.244 | -2.3705 |
| ABHD11 | NM_001145364 | -0.2895 | -0.9 |
| ZNF652 | NM_001145365 | -1.5065 | -2.7635 |
| PTPN3 | NM_001145368 | -0.329 | -1.728 |
| PTPN3 | NM_001145369 | -0.329 | -1.728 |
| PTPN3 | NM_001145370 | -0.329 | -1.728 |
| PTPN3 | NM_001145371 | -0.329 | -1.728 |
| PTPN3 | NM_001145372 | -0.329 | -1.728 |
| OTUD1 | NM_001145373 | -0.181 | -0.817 |
| ALKBH2 | NM_001145374 | -0.952 | -2.8075 |
| ALKBH2 | NM_001145375 | -0.805 | -2.2345 |
| OXSM | NM_001145391 | -1.5345 | -2.6195 |
| TSEN2 | NM_001145392 | -1.9225 | -3.4725 |
| TSEN2 | NM_001145393 | -1.9225 | -3.4725 |
| TSEN2 | NM_001145394 | -1.9225 | -3.4725 |
| TSEN2 | NM_001145395 | -1.9225 | -3.4725 |

| | | | |
|---|---|---|---|
| ALDH16A1 | NM_001145396 | -0.082 | -0.603 |
| TEF | NM_001145398 | -0.3755 | -0.684 |
| MPPED2 | NM_001145399 | -1.0105 | -2.2935 |
| FAM71E2 | NM_001145402 | -0.1545 | -0.0695 |
| ZNF506 | NM_001145404 | -0.687 | -0.1635 |
| NONO | NM_001145408 | -0.189 | -0.1935 |
| NONO | NM_001145409 | -0.183 | -0.3515 |
| NONO | NM_001145410 | -0.183 | -0.3515 |
| CCDC110 | NM_001145411 | 0.6145 | -0.5245 |
| NFE2L2 | NM_001145412 | -1.476 | -3.3855 |
| NFE2L2 | NM_001145413 | -1.476 | -3.3855 |
| CHM | NM_001145414 | -0.15 | -0.3585 |
| SETDB1 | NM_001145415 | -1.5125 | -3.0715 |
| TMEM205 | NM_001145416 | -0.047 | -0.5585 |
| MSL2 | NM_001145417 | -0.771 | -1.2225 |
| TTC28 | NM_001145418 | -0.6155 | -1.5495 |
| NR1D2 | NM_001145425 | -0.5465 | -1.259 |
| CSDA | NM_001145426 | 0.1425 | -0.201 |
| SNX18 | NM_001145427 | -0.994 | -1.925 |
| C4orf52 | NM_001145432 | -0.8105 | -1.498 |
| ZNF880 | NM_001145434 | 0.1855 | -0.5385 |
| LSS | NM_001145437 | -0.521 | -0.956 |
| PGAP2 | NM_001145438 | -0.393 | -1.112 |
| TYW1B | NM_001145440 | 0.03 | -2.112 |
| TYW1B | NM_001145441 | -0.006 | -1.156 |
| POTEM | NM_001145442 | -1.108 | 0.025 |
| PFKFB3 | NM_001145443 | -0.076 | -0.8195 |
| SCAF4 | NM_001145444 | -1.237 | -3.9705 |
| ZNF200 | NM_001145446 | -1.8995 | -3.8065 |
| ZNF200 | NM_001145447 | -1.8995 | -3.8065 |
| ZNF200 | NM_001145448 | -1.8995 | -3.8065 |
| MORN2 | NM_001145450 | -0.177 | -0.2735 |
| ARHGEF33 | NM_001145451 | -0.7055 | -0.289 |
| GFRA1 | NM_001145453 | 0.0155 | -0.6755 |
| STMN1 | NM_001145454 | -0.072 | -0.3225 |
| NCR1 | NM_001145457 | 0.136 | 0.169 |
| NCR1 | NM_001145458 | 0.136 | 0.169 |
| HOPX | NM_001145459 | -0.0895 | -0.5305 |
| HOPX | NM_001145460 | -0.0895 | -0.5305 |
| YIF1B | NM_001145461 | 0.239 | -0.397 |
| YIF1B | NM_001145462 | 0.239 | -0.397 |
| YIF1B | NM_001145463 | 0.239 | -0.397 |
| EXOG | NM_001145464 | -1.0555 | -2.0335 |

| | | | |
|---|---|---|---|
| NANOGNB | NM_001145465 | 0.333 | -0.254 |
| NCR3 | NM_001145466 | -0.35 | -0.3055 |
| SPECC1L | NM_001145468 | -1.5055 | -2.427 |
| LOXHD1 | NM_001145472 | 0.357 | 0.1095 |
| LOXHD1 | NM_001145473 | 0.0045 | -0.3325 |
| ATPAF1-AS1 | NM_001145474 | -0.139 | -0.222 |
| FAM186A | NM_001145475 | -0.1465 | -0.245 |
| NFIA | NM_001145511 | -0.9245 | -2.006 |
| NFIA | NM_001145512 | -0.9245 | -2.006 |
| SCRN1 | NM_001145513 | -1.1465 | -0.6265 |
| SCRN1 | NM_001145514 | -0.981 | -0.5245 |
| SCRN1 | NM_001145515 | -1.1465 | -0.6265 |
| RAI14 | NM_001145520 | -0.8165 | -1.194 |
| RAI14 | NM_001145521 | -0.8165 | -1.194 |
| RAI14 | NM_001145522 | -0.8165 | -1.194 |
| RAI14 | NM_001145523 | -0.8165 | -1.194 |
| YPEL3 | NM_001145524 | -0.1035 | -0.3585 |
| RAI14 | NM_001145525 | -0.7985 | -0.7875 |
| AFMID | NM_001145526 | -0.3455 | -1.381 |
| EHMT1 | NM_001145527 | -0.5765 | -2.0305 |
| C7orf71 | NM_001145531 | -0.3895 | -0.275 |
| C17orf107 | NM_001145536 | 0.435 | -0.176 |
| TCP11L1 | NM_001145541 | -0.449 | -1.4305 |
| ZSCAN18 | NM_001145542 | -0.326 | -0.8385 |
| ZSCAN18 | NM_001145543 | -0.326 | -0.8385 |
| ZSCAN18 | NM_001145544 | -0.326 | -0.8385 |
| C16orf82 | NM_001145545 | 0.0535 | -0.022 |
| RBM17 | NM_001145547 | -0.789 | -1.2275 |
| ZDHHC7 | NM_001145548 | -0.434 | -1.5585 |
| FLJ25363 | NM_001145553 | 0.12 | 0.012 |
| TRIM45 | NM_001145635 | -0.448 | -1.0545 |
| C1orf228 | NM_001145636 | 0.041 | -0.1835 |
| GPSM1 | NM_001145638 | -0.581 | -1.1565 |
| ZFR2 | NM_001145640 | -0.771 | -0.4115 |
| SRRM5 | NM_001145641 | 0.16 | 0.0605 |
| KIAA0226 | NM_001145642 | -0.9985 | -1.733 |
| PHGR1 | NM_001145643 | -0.0365 | 0.122 |
| TNFSF13B | NM_001145645 | -0.0015 | -0.0165 |
| APH1B | NM_001145646 | -1.1155 | -2.003 |
| RASGRF1 | NM_001145648 | -0.0115 | 0.137 |
| ZNF529 | NM_001145649 | -1.4525 | -2.2325 |
| ZNF529 | NM_001145650 | -1.4525 | -2.2325 |
| C6orf141 | NM_001145652 | 0.1735 | -0.6035 |

| | | | |
|---|---|---|---|
| RAP1GAP | NM_001145657 | -0.398 | -1.0045 |
| CTAGE9 | NM_001145659 | 0.5965 | 0.1145 |
| GATA2 | NM_001145661 | -0.6295 | -1.9175 |
| GATA2 | NM_001145662 | -0.829 | -1.959 |
| RFX8 | NM_001145664 | -0.2395 | -0.3495 |
| ZNF181 | NM_001145665 | -1.8585 | -2.8545 |
| GLG1 | NM_001145666 | -0.6955 | -1.193 |
| GLG1 | NM_001145667 | -0.6955 | -1.193 |
| CTXN2 | NM_001145668 | 0.088 | -0.1725 |
| SORBS2 | NM_001145670 | -0.5915 | -1.332 |
| SORBS2 | NM_001145671 | -0.5915 | -1.332 |
| SORBS2 | NM_001145672 | -0.5915 | -1.332 |
| SORBS2 | NM_001145673 | -0.5915 | -1.332 |
| SORBS2 | NM_001145674 | -0.5915 | -1.332 |
| SORBS2 | NM_001145675 | -0.5915 | -1.332 |
| KIAA0825 | NM_001145678 | -0.412 | -0.09 |
| LOC375190 | NM_001145710 | -0.754 | -1.02 |
| LOC389493 | NM_001145712 | 0.0865 | -0.3525 |
| SAE1 | NM_001145713 | -0.134 | -0.7595 |
| SAE1 | NM_001145714 | -0.134 | -0.7595 |
| KPNA7 | NM_001145715 | 0.0745 | -0.5315 |
| PNPLA1 | NM_001145716 | -0.1375 | -0.1595 |
| PNPLA1 | NM_001145717 | -0.1375 | -0.1595 |
| CT47B1 | NM_001145718 | 0.2695 | 0.218 |
| ZBTB8B | NM_001145720 | -0.114 | -0.686 |
| HOMER3 | NM_001145721 | -0.1185 | -0.629 |
| HOMER3 | NM_001145722 | -0.1185 | -0.629 |
| HOMER3 | NM_001145724 | -0.1185 | -0.629 |
| LYAR | NM_001145725 | -0.9265 | -1.915 |
| IFLTD1 | NM_001145727 | -0.7185 | -0.891 |
| IFLTD1 | NM_001145728 | -0.7185 | -0.891 |
| IFLTD1 | NM_001145729 | -0.7185 | -0.891 |
| CDC20B | NM_001145734 | -0.0955 | -0.157 |
| XPC | NM_001145769 | -1.0635 | -2.6125 |
| GPR56 | NM_001145770 | -0.1105 | -0.4785 |
| GPR56 | NM_001145771 | -0.1105 | -0.4785 |
| GPR56 | NM_001145772 | -0.1105 | -0.4785 |
| GPR56 | NM_001145773 | -0.1105 | -0.4785 |
| GPR56 | NM_001145774 | -0.6515 | -1.437 |
| FKBP5 | NM_001145775 | -0.789 | -1.4185 |
| FKBP5 | NM_001145776 | -0.789 | -1.4185 |
| FKBP5 | NM_001145777 | -0.789 | -1.4185 |
| MEF2BNB | NM_001145783 | -1.138 | -2.3915 |

| | | | |
|---|---|---|---|
| MEF2BNB | NM_001145784 | -0.649 | -0.863 |
| MEF2B | NM_001145785 | 0.024 | -0.4255 |
| ANTXR2 | NM_001145794 | -1.623 | -2.2605 |
| SH2B1 | NM_001145795 | -0.333 | -0.928 |
| SH2B1 | NM_001145796 | -0.333 | -0.928 |
| SH2B1 | NM_001145797 | -0.533 | -1.5105 |
| CACNB4 | NM_001145798 | -0.508 | -0.9685 |
| IRGM | NM_001145805 | -0.0125 | 0.3875 |
| ITGAM | NM_001145808 | -0.0195 | -0.1615 |
| MYH14 | NM_001145809 | -0.1625 | 0.0935 |
| SOX6 | NM_001145811 | -0.325 | 0.2215 |
| SH2B1 | NM_001145812 | -0.333 | -0.928 |
| AMDHD2 | NM_001145815 | -0.0785 | -0.3915 |
| HPGD | NM_001145816 | -0.3415 | -0.8265 |
| SOX6 | NM_001145819 | -0.325 | 0.2215 |
| TNNI2 | NM_001145829 | 0.3555 | -0.17 |
| MTFR1 | NM_001145838 | -1.025 | -2.0125 |
| MTFR1 | NM_001145839 | -1.025 | -2.0125 |
| TNNI2 | NM_001145841 | 0.3555 | -0.17 |
| ROBO1 | NM_001145845 | -0.903 | -1.004 |
| PROM1 | NM_001145847 | -0.308 | -0.5795 |
| PROM1 | NM_001145848 | -0.308 | -0.5795 |
| PROM1 | NM_001145849 | -0.308 | -0.5795 |
| PROM1 | NM_001145850 | -0.308 | -0.5795 |
| PROM1 | NM_001145851 | -0.308 | -0.5795 |
| PROM1 | NM_001145852 | -0.308 | -0.5795 |
| WFS1 | NM_001145853 | -0.1785 | -1.0815 |
| SH3BP2 | NM_001145855 | -0.299 | -1.011 |
| SH3BP2 | NM_001145856 | -0.299 | -1.011 |
| POP1 | NM_001145860 | -1.3625 | -2.3805 |
| POP1 | NM_001145861 | -0.8435 | -2.6675 |
| MTMR11 | NM_001145862 | -0.2175 | -0.113 |
| CD8A | NM_001145873 | 0.212 | -0.582 |
| SARS2 | NM_001145901 | -0.3055 | -0.755 |
| C2 | NM_001145903 | -0.241 | 0.3285 |
| TANC1 | NM_001145909 | -1.0355 | -2.089 |
| SCAPER | NM_001145923 | -0.6495 | -1.4675 |
| SAP130 | NM_001145928 | -0.8345 | -3.2985 |
| TKTL1 | NM_001145933 | 0.0995 | -0.0545 |
| TKTL1 | NM_001145934 | 0.1615 | -0.022 |
| MMP1 | NM_001145938 | 0.2665 | -0.448 |
| PAFAH1B3 | NM_001145939 | 0.1065 | -0.5855 |
| SLCO1C1 | NM_001145944 | 0.281 | -0.1675 |

| | | | |
|---|---|---|---|
| SLCO1C1 | NM_001145945 | 0.281 | -0.1675 |
| SLCO1C1 | NM_001145946 | -0.1745 | 0.015 |
| TIMM8A | NM_001145951 | -0.1585 | -0.1855 |
| SLC12A4 | NM_001145961 | 0.111 | -0.6285 |
| SLC12A4 | NM_001145962 | 0.111 | -0.6285 |
| SLC12A4 | NM_001145963 | 0.111 | -0.6285 |
| SLC12A4 | NM_001145964 | 0.111 | -0.6285 |
| MKI67 | NM_001145966 | 0.0225 | -1.609 |
| RDH13 | NM_001145971 | -0.7765 | -1.8235 |
| SLC13A2 | NM_001145975 | 0.1895 | -0.123 |
| SLC13A2 | NM_001145976 | 0.1895 | -0.123 |
| SLC15A2 | NM_001145998 | -0.1275 | -0.373 |
| ANGPT1 | NM_001146 | -0.203 | -0.3305 |
| NEK11 | NM_001146003 | -0.1855 | -0.549 |
| IGFALS | NM_001146006 | -0.024 | -0.2015 |
| SLCO5A1 | NM_001146008 | -0.1355 | -1.0045 |
| SLCO5A1 | NM_001146009 | -0.1355 | -1.0045 |
| DLGAP5 | NM_001146015 | -0.8455 | -2.7575 |
| TJAP1 | NM_001146016 | -1.0215 | -2.418 |
| TJAP1 | NM_001146017 | -1.0215 | -2.418 |
| TJAP1 | NM_001146018 | -1.0215 | -2.418 |
| TJAP1 | NM_001146019 | -1.0215 | -2.418 |
| TJAP1 | NM_001146020 | -1.0215 | -2.418 |
| JPH4 | NM_001146028 | -0.361 | -0.7505 |
| SEMA7A | NM_001146029 | -0.0805 | -0.4105 |
| SEMA7A | NM_001146030 | -0.0805 | -0.4105 |
| FCHO2 | NM_001146032 | 0.0785 | -0.866 |
| OR56A5 | NM_001146033 | 0.244 | -0.2385 |
| SLC14A1 | NM_001146036 | -0.054 | 0.3315 |
| SLC14A1 | NM_001146037 | -0.054 | 0.3315 |
| GORAB | NM_001146039 | -2.007 | -4.367 |
| GLRA1 | NM_001146040 | -0.1185 | -0.2605 |
| KRTAP4-9 | NM_001146041 | -0.1165 | 0.1585 |
| HSDL1 | NM_001146051 | -1.411 | -1.7925 |
| SNCA | NM_001146054 | -0.884 | -1.472 |
| SNCA | NM_001146055 | -0.884 | -1.472 |
| MBOAT7 | NM_001146056 | -1.4555 | -1.2885 |
| CAPN2 | NM_001146068 | -0.709 | -0.3605 |
| MFSD10 | NM_001146069 | -0.053 | -0.7115 |
| TDRD3 | NM_001146071 | -1.082 | -2.2515 |
| RPAP3 | NM_001146075 | -0.424 | -2.4425 |
| RPAP3 | NM_001146076 | -0.424 | -2.4425 |
| MBOAT7 | NM_001146082 | -1.4555 | -1.2885 |

| | | | |
|---|---|---|---|
| MBOAT7 | NM_001146083 | -0.491 | -1.177 |
| FAN1 | NM_001146094 | -1.914 | -2.345 |
| FAN1 | NM_001146095 | -1.914 | -2.345 |
| FAN1 | NM_001146096 | -1.914 | -2.345 |
| NEK3 | NM_001146099 | -0.764 | -1.376 |
| PARP9 | NM_001146102 | -0.0065 | -1.1455 |
| PARP9 | NM_001146103 | -0.0065 | -1.1455 |
| PARP9 | NM_001146104 | -0.0065 | -1.1455 |
| PARP9 | NM_001146105 | -0.0065 | -1.1455 |
| PARP9 | NM_001146106 | -0.0065 | -1.1455 |
| MIER1 | NM_001146110 | -0.8485 | -3.324 |
| MIER1 | NM_001146111 | -0.8485 | -3.324 |
| MIER1 | NM_001146112 | -0.8485 | -3.324 |
| MIER1 | NM_001146113 | -0.8485 | -3.324 |
| DIP2A | NM_001146114 | -1.243 | -1.7675 |
| DIP2A | NM_001146115 | -1.243 | -1.7675 |
| DIP2A | NM_001146116 | -1.243 | -1.7675 |
| AVPR2 | NM_001146151 | 0.0455 | -0.3515 |
| CYP51A1 | NM_001146152 | -1.1295 | -1.392 |
| PTGR2 | NM_001146154 | -1.001 | -0.6575 |
| PTGR2 | NM_001146155 | -1.001 | -0.6575 |
| GSK3B | NM_001146156 | -1.4495 | -3.4265 |
| FAM25A | NM_001146157 | 0.082 | -0.154 |
| MOGS | NM_001146158 | -0.453 | -1.262 |
| TATDN1 | NM_001146160 | -0.833 | -1.039 |
| TRIM77P | NM_001146162 | 0.0345 | 0.0125 |
| TATDN3 | NM_001146169 | -1.1835 | -1.194 |
| TATDN3 | NM_001146170 | -1.1835 | -1.194 |
| TATDN3 | NM_001146171 | -1.1835 | -1.194 |
| IP6K2 | NM_001146178 | 0.385 | -0.656 |
| IP6K2 | NM_001146179 | 0.385 | -0.656 |
| PEG3 | NM_001146184 | -0.1245 | 0.076 |
| PEG3 | NM_001146185 | -0.1245 | 0.076 |
| PEG3 | NM_001146186 | -0.1245 | 0.076 |
| PEG3 | NM_001146187 | -0.1245 | 0.076 |
| ZNF407 | NM_001146189 | 0.342 | -0.5495 |
| ZNF407 | NM_001146190 | 0.342 | -0.5495 |
| MPZL1 | NM_001146191 | -0.7115 | -1.592 |
| ZMYND12 | NM_001146192 | 0.2445 | -0.3005 |
| CCDC168 | NM_001146197 | -0.125 | -0.112 |
| RASGRP4 | NM_001146202 | -0.2685 | -0.173 |
| RASGRP4 | NM_001146203 | -0.2685 | -0.173 |
| RASGRP4 | NM_001146204 | -0.2685 | -0.173 |

| | | | |
|---|---|---|---|
| RASGRP4 | NM_001146205 | -0.2685 | -0.173 |
| RASGRP4 | NM_001146206 | -0.2685 | -0.173 |
| RASGRP4 | NM_001146207 | -0.2685 | -0.173 |
| PDE2A | NM_001146209 | -0.4595 | -1.1105 |
| SPDYE6 | NM_001146210 | -0.464 | -0.4915 |
| TBC1D15 | NM_001146213 | -0.7245 | -2.5675 |
| TBC1D15 | NM_001146214 | -0.7245 | -2.5675 |
| TACC1 | NM_001146216 | -0.8015 | -1.557 |
| WRB | NM_001146218 | -1.1845 | -1.306 |
| ZNF226 | NM_001146220 | -0.596 | -0.881 |
| KRT72 | NM_001146225 | -0.041 | -0.643 |
| KRT72 | NM_001146226 | -0.041 | -0.643 |
| RPS20 | NM_001146227 | -0.139 | -0.1875 |
| PODNL1 | NM_001146254 | 0.1085 | -0.0735 |
| PODNL1 | NM_001146255 | 0.1085 | -0.0735 |
| ZDHHC15 | NM_001146256 | 0.114 | -0.7405 |
| ZDHHC15 | NM_001146257 | 0.114 | -0.7405 |
| PIP4K2C | NM_001146259 | -0.1515 | -1.1625 |
| SYT14 | NM_001146261 | -0.4435 | -1.3615 |
| SYT14 | NM_001146262 | -0.4435 | -1.3615 |
| SYT14 | NM_001146264 | -0.4435 | -1.3615 |
| GPR85 | NM_001146265 | -0.6355 | -1.333 |
| GPR85 | NM_001146266 | -0.6355 | -1.333 |
| GPR85 | NM_001146267 | -0.6355 | -1.333 |
| TMEM64 | NM_001146273 | -1.0095 | -1.795 |
| TCF7L2 | NM_001146274 | -0.4665 | -1.848 |
| NCEH1 | NM_001146276 | -0.841 | -1.0105 |
| NCEH1 | NM_001146277 | -0.841 | -1.0105 |
| NCEH1 | NM_001146278 | -0.841 | -1.0105 |
| SHBG | NM_001146279 | -0.0225 | -0.5525 |
| SHBG | NM_001146280 | -0.0225 | -0.5525 |
| SHBG | NM_001146281 | -0.0225 | -0.5525 |
| SRP54 | NM_001146282 | -1.039 | -1.845 |
| TCF7L2 | NM_001146283 | -0.4665 | -1.848 |
| TCF7L2 | NM_001146284 | -0.4665 | -1.848 |
| TCF7L2 | NM_001146285 | -0.4665 | -1.848 |
| TCF7L2 | NM_001146286 | -0.4665 | -1.848 |
| AIFM3 | NM_001146288 | -0.0965 | 0.04 |
| LEPRE1 | NM_001146289 | -0.778 | -1.6275 |
| ZNF674 | NM_001146291 | -0.7745 | -1.489 |
| C1orf86 | NM_001146310 | 0.1065 | -0.4025 |
| MYOCD | NM_001146312 | 0.2005 | -0.0665 |
| MYOCD | NM_001146313 | 0.2005 | -0.0665 |

| | | | |
|---|---|---|---|
| ABHD14B | NM_001146314 | 0.21 | -0.294 |
| SPSB2 | NM_001146316 | -0.332 | -0.7905 |
| GRAMD3 | NM_001146319 | -0.033 | -0.2755 |
| GRAMD3 | NM_001146320 | -0.033 | -0.2755 |
| GRAMD3 | NM_001146321 | -0.033 | -0.2755 |
| GRAMD3 | NM_001146322 | -0.033 | -0.2755 |
| SUMF2 | NM_001146333 | 0.3035 | -0.3715 |
| NACAD | NM_001146334 | -1.144 | -2.3665 |
| SLC6A15 | NM_001146335 | -0.975 | -1.2145 |
| TMEM114 | NM_001146336 | 0.359 | -0.6945 |
| AFAP1L1 | NM_001146337 | -0.4675 | -0.085 |
| VSTM2B | NM_001146339 | -0.917 | -1.749 |
| NKX1-2 | NM_001146340 | 0.297 | -0.2265 |
| ANKRD34C | NM_001146341 | 0.36 | -0.293 |
| PQLC1 | NM_001146343 | -0.204 | -0.913 |
| PQLC1 | NM_001146345 | -0.204 | -0.913 |
| AGXT2L1 | NM_001146590 | 0.017 | -0.49 |
| AGXT2L1 | NM_001146627 | 0.017 | -0.49 |
| C14orf176 | NM_001146683 | -0.1215 | -0.582 |
| RNF222 | NM_001146684 | 0.118 | -0.1015 |
| TMEM88B | NM_001146685 | 0.1125 | -0.4055 |
| KDM3A | NM_001146688 | -0.9725 | -2.629 |
| KDM4C | NM_001146694 | -0.8685 | -1.857 |
| KDM4C | NM_001146695 | -0.8685 | -1.857 |
| KDM4C | NM_001146696 | -0.8685 | -1.857 |
| RBM19 | NM_001146698 | -0.588 | -1.405 |
| RBM19 | NM_001146699 | -0.588 | -1.405 |
| KDM5C | NM_001146702 | 0.0145 | -0.985 |
| KDM5D | NM_001146705 | 0.0245 | -1.352 |
| KDM5D | NM_001146706 | 0.0245 | -1.352 |
| TIMD4 | NM_001146726 | 0.191 | 0.006 |
| HRASLS5 | NM_001146728 | 0.0455 | 0.006 |
| HRASLS5 | NM_001146729 | 0.0455 | 0.006 |
| ANGPT2 | NM_001147 | -0.2975 | -0.1125 |
| ANK2 | NM_001148 | -0.69 | -1.0435 |
| ANK3 | NM_001149 | -0.1175 | -1.2675 |
| ANPEP | NM_001150 | -0.3545 | -0.202 |
| SLC25A4 | NM_001151 | -0.3875 | -1.3445 |
| SLC25A5 | NM_001152 | 0.1225 | -0.4045 |
| ANXA4 | NM_001153 | -0.6415 | -1.0265 |
| SCOC | NM_001153446 | -0.5325 | -1.0695 |
| SCOC | NM_001153484 | -0.5325 | -1.0695 |
| SCOC | NM_001153552 | -0.5325 | -1.0695 |

| | | | |
|---|---|---|---|
| SCOC | NM_001153585 | -0.5325 | -1.0695 |
| SCOC | NM_001153635 | -0.5325 | -1.0695 |
| SCOC | NM_001153663 | -0.5325 | -1.0695 |
| SCOC | NM_001153690 | -0.5325 | -1.0695 |
| ANXA5 | NM_001154 | -0.037 | -0.322 |
| ANXA6 | NM_001155 | 0.1015 | -0.61 |
| ANXA7 | NM_001156 | -0.5935 | -1.062 |
| CCDC81 | NM_001156474 | -0.467 | -0.517 |
| ANXA11 | NM_001157 | 0.1595 | -0.83 |
| AOX1 | NM_001159 | -0.3245 | -0.787 |
| ZNF716 | NM_001159279 | -0.0875 | -0.177 |
| ADAL | NM_001159280 | -0.7105 | -0.587 |
| HDAC10 | NM_001159286 | -0.0905 | -0.133 |
| TPI1 | NM_001159287 | 0.128 | -0.2305 |
| ZNF737 | NM_001159293 | -0.532 | -0.7375 |
| KLF8 | NM_001159296 | -0.096 | -0.2575 |
| PLA2G4C | NM_001159322 | -0.087 | -0.472 |
| PLA2G4C | NM_001159323 | -0.087 | -0.472 |
| REG4 | NM_001159352 | -0.144 | -0.446 |
| REG4 | NM_001159353 | -0.144 | -0.446 |
| MTHFSD | NM_001159377 | 0.244 | -0.499 |
| MTHFSD | NM_001159378 | 0.244 | -0.499 |
| MTHFSD | NM_001159379 | 0.244 | -0.499 |
| MTHFSD | NM_001159380 | 0.244 | -0.499 |
| B4GALNT2 | NM_001159387 | 0.1105 | -0.324 |
| B4GALNT2 | NM_001159388 | 0.1105 | -0.324 |
| GUK1 | NM_001159390 | 0.352 | -0.18 |
| GUK1 | NM_001159391 | 0.352 | -0.18 |
| FCRL1 | NM_001159397 | -0.391 | -0.3955 |
| FCRL1 | NM_001159398 | -0.391 | -0.3955 |
| BRI3 | NM_001159491 | -0.4865 | 0.2345 |
| IVD | NM_001159508 | -0.4105 | -1.1085 |
| ZNF735 | NM_001159524 | 0.2405 | -0.2955 |
| BEGAIN | NM_001159531 | -1.0055 | -1.2105 |
| RFPL2 | NM_001159545 | 0.159 | -0.118 |
| RFPL2 | NM_001159546 | 0.159 | -0.118 |
| BEND4 | NM_001159547 | -1.4905 | -2.1415 |
| P2RX6 | NM_001159554 | -0.1535 | -0.2975 |
| SCNN1A | NM_001159575 | -0.116 | -0.194 |
| SCNN1A | NM_001159576 | -0.116 | -0.194 |
| CD109 | NM_001159587 | -0.1945 | 0.07 |
| CD109 | NM_001159588 | -0.1945 | 0.07 |
| BIVM | NM_001159596 | -1.053 | -1.2625 |

| | | | |
|---|---|---|---|
| SH3YL1 | NM_001159597 | -1.0815 | -1.397 |
| RAB28 | NM_001159601 | -0.961 | -2.559 |
| BNIPL | NM_001159642 | 0.2465 | 0.062 |
| MCTP2 | NM_001159643 | 0.006 | -0.7005 |
| MCTP2 | NM_001159644 | 0.006 | -0.7005 |
| BOD1 | NM_001159651 | -0.4935 | -1.3225 |
| SYNCRIP | NM_001159673 | -0.617 | -0.922 |
| SYNCRIP | NM_001159674 | -0.617 | -0.922 |
| SYNCRIP | NM_001159675 | -0.617 | -0.922 |
| SYNCRIP | NM_001159676 | -0.617 | -0.922 |
| SYNCRIP | NM_001159677 | -0.617 | -0.922 |
| EMCN | NM_001159694 | 0.0025 | -0.5065 |
| FHL1 | NM_001159699 | -0.136 | -0.286 |
| FHL1 | NM_001159700 | -0.136 | -0.286 |
| FHL1 | NM_001159701 | -0.136 | -0.286 |
| FHL1 | NM_001159702 | -0.136 | -0.286 |
| FHL1 | NM_001159703 | -0.136 | -0.286 |
| FHL1 | NM_001159704 | -0.136 | -0.286 |
| BTNL8 | NM_001159707 | -0.0105 | 0.1015 |
| BTNL8 | NM_001159708 | -0.0105 | 0.1015 |
| BTNL8 | NM_001159709 | -0.0105 | 0.1015 |
| BTNL8 | NM_001159710 | -0.0105 | 0.1015 |
| TSPO2 | NM_001159726 | 0.0115 | -0.551 |
| PLBD2 | NM_001159727 | 0.132 | -0.3055 |
| LTA | NM_001159740 | 0.209 | -0.024 |
| ABR | NM_001159746 | -0.7775 | -0.998 |
| RIOK2 | NM_001159749 | -1.0185 | -2.7605 |
| BZW2 | NM_001159767 | -0.725 | -0.9785 |
| SLC39A11 | NM_001159770 | 0.2905 | -0.1365 |
| CANT1 | NM_001159772 | -0.1745 | -1.4215 |
| CANT1 | NM_001159773 | -0.1745 | -1.4215 |
| ZNF583 | NM_001159860 | -0.8585 | -1.5245 |
| ZNF583 | NM_001159861 | -0.8585 | -1.5245 |
| FLT1 | NM_001159920 | -0.074 | 0.037 |
| FBXW12 | NM_001159927 | 0.1725 | -0.029 |
| FBXW12 | NM_001159929 | 0.1725 | -0.029 |
| EBNA1BP2 | NM_001159936 | -0.383 | -1.1075 |
| EVI5L | NM_001159944 | 0.1055 | -0.58 |
| EPS15 | NM_001159969 | -0.9405 | -1.0495 |
| NRG1 | NM_001159995 | -0.1445 | -0.887 |
| NRG1 | NM_001159996 | -0.1445 | -0.887 |
| NRG1 | NM_001159999 | -0.1445 | -0.887 |
| APAF1 | NM_001160 | -0.295 | -1.123 |

| | | | |
|---|---|---|---|
| NRG1 | NM_001160001 | -0.1445 | -0.887 |
| NRG1 | NM_001160002 | -0.1445 | -0.887 |
| NRG1 | NM_001160004 | -0.1445 | -0.887 |
| NRG1 | NM_001160005 | -0.1445 | -0.887 |
| NRG1 | NM_001160007 | -0.1445 | -0.887 |
| NRG1 | NM_001160008 | -0.1445 | -0.887 |
| ST6GALNAC3 | NM_001160011 | -1.064 | -1.891 |
| FLT1 | NM_001160030 | -0.074 | 0.037 |
| FLT1 | NM_001160031 | -0.074 | 0.037 |
| SGOL2 | NM_001160033 | 0.0415 | -1.0965 |
| RHOBTB2 | NM_001160036 | -0.084 | -0.6705 |
| RHOBTB2 | NM_001160037 | -0.084 | -0.6705 |
| IQCC | NM_001160042 | -1.032 | -0.9575 |
| SGOL2 | NM_001160046 | 0.0415 | -1.0965 |
| TXNDC16 | NM_001160047 | -0.4725 | -0.5905 |
| CLDN10 | NM_001160100 | 0.126 | -1.3265 |
| LPO | NM_001160102 | 0.0965 | -0.295 |
| ZC3H14 | NM_001160103 | -0.9785 | -2.06 |
| ZC3H14 | NM_001160104 | -0.9785 | -2.06 |
| NOS3 | NM_001160109 | 0.2665 | -0.11 |
| NOS3 | NM_001160110 | 0.2665 | -0.11 |
| NOS3 | NM_001160111 | 0.2665 | -0.11 |
| C15orf40 | NM_001160113 | 0.7175 | 0.2555 |
| C15orf40 | NM_001160114 | 0.7175 | 0.2555 |
| C15orf40 | NM_001160115 | 0.7175 | 0.2555 |
| C15orf40 | NM_001160116 | 0.7175 | 0.2555 |
| KLF6 | NM_001160124 | -0.1815 | 0.227 |
| KLF6 | NM_001160125 | -0.1815 | 0.227 |
| KCNQ5 | NM_001160130 | 0.5015 | -0.1945 |
| KCNQ5 | NM_001160132 | 0.5015 | -0.1945 |
| KCNQ5 | NM_001160133 | 0.5015 | -0.1945 |
| KCNQ5 | NM_001160134 | 0.5015 | -0.1945 |
| C7orf63 | NM_001160138 | 0.103 | 0.016 |
| DDHD1 | NM_001160147 | -0.76 | -1.2145 |
| DDHD1 | NM_001160148 | -0.76 | -1.2145 |
| MGAT4A | NM_001160154 | -0.1515 | -0.028 |
| SCN5A | NM_001160160 | 0.021 | -0.709 |
| SCN5A | NM_001160161 | 0.021 | -0.709 |
| HRH4 | NM_001160166 | 0.2355 | -0.062 |
| PRR5L | NM_001160167 | 0.1745 | -0.223 |
| PRR5L | NM_001160168 | 0.1745 | -0.223 |
| PRR5L | NM_001160169 | 0.1745 | -0.223 |
| NAT1 | NM_001160170 | -1.268 | -0.7705 |

| | | | |
|---|---|---|---|
| NAT1 | NM_001160171 | -1.268 | -0.7705 |
| NAT1 | NM_001160172 | -1.268 | -0.7705 |
| NAT1 | NM_001160173 | -1.268 | -0.7705 |
| NAT1 | NM_001160174 | -1.268 | -0.7705 |
| NAT1 | NM_001160175 | -1.268 | -0.7705 |
| NAT1 | NM_001160176 | -1.268 | -0.7705 |
| NAT1 | NM_001160179 | -1.268 | -0.7705 |
| ZNF138 | NM_001160183 | -0.8825 | -1.2765 |
| PLEKHN1 | NM_001160184 | -0.0885 | -0.185 |
| SLC25A13 | NM_001160210 | -0.4575 | -1.218 |
| PMFBP1 | NM_001160213 | -0.027 | -0.218 |
| RNF170 | NM_001160223 | -0.4265 | -1.0105 |
| RNF170 | NM_001160224 | -0.4265 | -1.0105 |
| RNF170 | NM_001160225 | -0.4265 | -1.0105 |
| CNR1 | NM_001160226 | 0.0025 | -0.3535 |
| ADARB1 | NM_001160230 | 0.249 | -0.6755 |
| ATP1A1 | NM_001160233 | 0.0115 | -0.621 |
| ATP1A1 | NM_001160234 | 0.0115 | -0.621 |
| RPAIN | NM_001160243 | -1.303 | -1.4555 |
| RPAIN | NM_001160244 | -1.303 | -1.4555 |
| RPAIN | NM_001160246 | -1.303 | -1.4555 |
| CNR1 | NM_001160258 | 0.0025 | -0.3535 |
| CNR1 | NM_001160259 | 0.0025 | -0.3535 |
| FEZF1 | NM_001160264 | 0.1555 | 0.4565 |
| RPAIN | NM_001160266 | -1.303 | -1.4555 |
| RPAIN | NM_001160267 | -1.303 | -1.4555 |
| CERKL | NM_001160277 | -0.0855 | -1.3575 |
| PANX2 | NM_001160300 | 0.004 | -0.707 |
| DPYD | NM_001160301 | -0.4415 | -0.968 |
| SYNJ1 | NM_001160302 | -0.59 | -1.34 |
| SETD6 | NM_001160305 | -0.143 | -0.7975 |
| SYNJ1 | NM_001160306 | -0.59 | -1.34 |
| SETDB2 | NM_001160308 | -0.928 | -2.114 |
| OR6P1 | NM_001160325 | 0.2565 | -0.5775 |
| NFASC | NM_001160331 | 0.3565 | -0.278 |
| NFASC | NM_001160332 | 0.3565 | -0.278 |
| NFASC | NM_001160333 | 0.3565 | -0.278 |
| LY6K | NM_001160354 | -0.0195 | -0.6825 |
| LY6K | NM_001160355 | -0.0195 | -0.6825 |
| TMC7 | NM_001160364 | -0.2535 | -0.4015 |
| CDK10 | NM_001160367 | -0.3685 | -0.763 |
| TRPT1 | NM_001160389 | -0.1225 | -0.352 |
| TRPT1 | NM_001160390 | -0.1225 | -0.352 |

| | | | |
|---|---|---|---|
| TRPT1 | NM_001160392 | -0.1225 | -0.352 |
| TRPT1 | NM_001160393 | -0.1225 | -0.352 |
| TLCD1 | NM_001160407 | -0.528 | -0.706 |
| TULP3 | NM_001160408 | -1.0875 | -1.925 |
| ZBP1 | NM_001160417 | -0.198 | 0.0605 |
| ZBP1 | NM_001160418 | -0.198 | 0.0605 |
| ZBP1 | NM_001160419 | -0.198 | 0.0605 |
| IGF2BP1 | NM_001160423 | -0.893 | -3.278 |
| SCEL | NM_001160706 | -0.0095 | -0.0995 |
| POLR3B | NM_001160708 | -1.205 | -3.336 |
| NUDT2 | NM_001161 | -0.6 | -1.3805 |
| ARMC10 | NM_001161009 | -0.149 | -0.8295 |
| ARMC10 | NM_001161010 | -0.149 | -0.8295 |
| ARMC10 | NM_001161011 | -0.149 | -0.8295 |
| ARMC10 | NM_001161012 | -0.149 | -0.8295 |
| ARMC10 | NM_001161013 | -0.149 | -0.8295 |
| SSH1 | NM_001161330 | -1.212 | -2.3545 |
| SSH1 | NM_001161331 | -1.212 | -2.3545 |
| HIST2H2BF | NM_001161334 | 0.0245 | -0.5435 |
| CHFR | NM_001161344 | -0.195 | -1.4025 |
| CHFR | NM_001161345 | -0.195 | -1.4025 |
| CHFR | NM_001161346 | -0.195 | -1.4025 |
| CHFR | NM_001161347 | -0.195 | -1.4025 |
| KCNMA1 | NM_001161352 | 0.253 | -0.204 |
| KCNMA1 | NM_001161353 | 0.253 | -0.204 |
| NPTN | NM_001161363 | -0.6125 | -1.858 |
| NPTN | NM_001161364 | -0.6125 | -1.858 |
| SEC14L4 | NM_001161368 | 0.0305 | -0.5145 |
| C6orf136 | NM_001161376 | -0.2985 | -1.448 |
| LIMS2 | NM_001161403 | -0.007 | -0.366 |
| LIMS2 | NM_001161404 | -0.007 | -0.366 |
| NDFIP2 | NM_001161407 | -0.695 | -0.857 |
| GPR17 | NM_001161415 | 0.1635 | -0.439 |
| GPR17 | NM_001161416 | 0.1635 | -0.439 |
| GPR17 | NM_001161417 | 0.1635 | -0.439 |
| ZNF610 | NM_001161425 | -0.783 | -2.4 |
| ZNF610 | NM_001161426 | -0.783 | -2.4 |
| ZNF610 | NM_001161427 | -0.783 | -2.4 |
| RANBP3L | NM_001161429 | -0.434 | -0.025 |
| PTPRH | NM_001161440 | -0.338 | -0.687 |
| SH2D2A | NM_001161441 | 0.388 | -1.1415 |
| SH2D2A | NM_001161442 | 0.388 | -1.1415 |
| SH2D2A | NM_001161443 | 0.388 | -1.1415 |

| | | | |
|---|---|---|---|
| SH2D2A | NM_001161444 | 0.388 | -1.1415 |
| CYBASC3 | NM_001161452 | -0.5905 | -1.257 |
| CYBASC3 | NM_001161454 | -0.5905 | -1.257 |
| JMJD4 | NM_001161465 | -1.207 | -1.8605 |
| ALDH3B1 | NM_001161473 | -0.188 | -0.3895 |
| WDR25 | NM_001161476 | -0.0745 | -0.562 |
| ZNF611 | NM_001161499 | -0.9895 | -1.4905 |
| ZNF611 | NM_001161500 | -0.9895 | -1.4905 |
| ZNF611 | NM_001161501 | -0.9895 | -1.4905 |
| ALDH4A1 | NM_001161504 | -0.1265 | -0.537 |
| COG5 | NM_001161520 | -0.827 | -1.1225 |
| KLHL2 | NM_001161521 | -0.98 | -1.7285 |
| KLHL2 | NM_001161522 | -0.98 | -1.7285 |
| CSF2RA | NM_001161529 | -0.4185 | -0.0625 |
| CSF2RA | NM_001161530 | -0.4185 | -0.0625 |
| CSF2RA | NM_001161531 | -0.4185 | -0.0625 |
| CSF2RA | NM_001161532 | -0.4185 | -0.0625 |
| RBM34 | NM_001161533 | 0.392 | 1.2725 |
| TNIK | NM_001161560 | -0.5465 | -1.6965 |
| TNIK | NM_001161561 | -0.5465 | -1.6965 |
| TNIK | NM_001161562 | -0.5465 | -1.6965 |
| TNIK | NM_001161563 | -0.5465 | -1.6965 |
| TNIK | NM_001161564 | -0.5465 | -1.6965 |
| TNIK | NM_001161565 | -0.5465 | -1.6965 |
| TNIK | NM_001161566 | -0.5465 | -1.6965 |
| MAFF | NM_001161572 | 0.0575 | -0.5275 |
| MAFF | NM_001161573 | 0.0575 | -0.5275 |
| MAFF | NM_001161574 | 0.0575 | -0.5275 |
| SAMD4A | NM_001161576 | -0.2185 | -0.4035 |
| SAMD4A | NM_001161577 | -0.2185 | -0.4035 |
| POC1A | NM_001161580 | -0.5565 | -2.072 |
| POC1A | NM_001161581 | -0.5565 | -2.072 |
| TEX10 | NM_001161584 | -1.65 | -3.9535 |
| C9orf11 | NM_001161585 | -0.0335 | -0.1285 |
| ME3 | NM_001161586 | -0.046 | -0.678 |
| GYS1 | NM_001161587 | -0.446 | -0.8565 |
| RGL3 | NM_001161616 | 0.191 | -0.346 |
| DUS3L | NM_001161619 | -0.094 | -1.0905 |
| NXNL2 | NM_001161625 | 0.245 | -0.148 |
| KDM4DL | NM_001161630 | 0.2135 | -0.1765 |
| MYB | NM_001161656 | -0.286 | -1.0575 |
| MYB | NM_001161657 | -0.286 | -1.0575 |
| MYB | NM_001161658 | -0.286 | -1.0575 |

| | | | |
|---|---|---|---|
| MYB | NM_001161659 | -0.286 | -1.0575 |
| MYB | NM_001161660 | -0.286 | -1.0575 |
| WWC1 | NM_001161661 | -0.3225 | -1.103 |
| WWC1 | NM_001161662 | -0.3225 | -1.103 |
| RSPH4A | NM_001161664 | -0.3035 | -0.5905 |
| FXN | NM_001161706 | 0.019 | -1.496 |
| KIRREL3 | NM_001161707 | -0.179 | -0.3365 |
| SYNC | NM_001161708 | -0.5325 | -0.3575 |
| PPP2R5C | NM_001161725 | -2.0235 | -3.1815 |
| PPP2R5C | NM_001161726 | -2.0235 | -3.1815 |
| PLA2G2A | NM_001161727 | -0.135 | 0.0185 |
| PLA2G2A | NM_001161728 | -0.135 | 0.0185 |
| PLA2G2A | NM_001161729 | -0.135 | 0.0185 |
| AHCY | NM_001161766 | 0.316 | -0.219 |
| HTR3A | NM_001161772 | -0.1285 | -0.6555 |
| PDP1 | NM_001161778 | -0.826 | -1.52 |
| PDP1 | NM_001161779 | -0.826 | -1.52 |
| PDP1 | NM_001161780 | -0.826 | -1.52 |
| PDP1 | NM_001161781 | -0.826 | -1.52 |
| GPR144 | NM_001161808 | -0.0245 | -0.4965 |
| DAOA | NM_001161812 | -0.1605 | -0.088 |
| DAOA | NM_001161814 | -0.1605 | -0.088 |
| MYO1B | NM_001161819 | -0.744 | -1.4695 |
| C7orf72 | NM_001161834 | 0.1585 | -0.693 |
| SULF2 | NM_001161841 | -0.5715 | -1.1015 |
| .OC10012854[illegible] | NM_001162367 | -0.1985 | -0.019 |
| LOC728392 | NM_001162371 | 0.102 | -0.414 |
| ARHGEF2 | NM_001162383 | -0.355 | -2.111 |
| ARHGEF2 | NM_001162384 | -0.355 | -2.111 |
| CLK1 | NM_001162407 | -0.908 | -1.031 |
| ETS1 | NM_001162422 | -0.345 | -1.2305 |
| TSC1 | NM_001162426 | 0.5605 | -0.125 |
| TSC1 | NM_001162427 | 0.5605 | -0.125 |
| PDCD6IP | NM_001162429 | -0.661 | -1.2085 |
| .OC10028771[illegible] | NM_001162435 | -0.4025 | -0.3225 |
| C2orf83 | NM_001162483 | 0.1715 | 0.206 |
| ARL13A | NM_001162490 | 0.47 | -0.085 |
| ARL13A | NM_001162491 | 0.47 | -0.085 |
| C21orf62 | NM_001162495 | -0.321 | -0.094 |
| C21orf62 | NM_001162496 | -0.321 | -0.094 |
| LPAR6 | NM_001162497 | 0.064 | 0.3105 |
| LPAR6 | NM_001162498 | 0.064 | 0.3105 |
| CAND2 | NM_001162499 | -0.7965 | -1.762 |

| | | | |
|---|---|---|---|
| TNRC6B | NM_001162501 | -2.032 | -3.354 |
| TMEM189 | NM_001162505 | -0.8755 | -2.279 |
| FAM135A | NM_001162529 | -1.5035 | -3.084 |
| RBMXL1 | NM_001162536 | -0.711 | -2.3305 |
| COX11 | NM_001162861 | -1.2915 | -2.066 |
| COX11 | NM_001162862 | -1.2915 | -2.066 |
| KIAA0040 | NM_001162893 | -0.591 | 0.2585 |
| KIAA0040 | NM_001162894 | -0.591 | 0.2585 |
| KIAA0040 | NM_001162895 | -0.591 | 0.2585 |
| TMEM217 | NM_001162900 | -0.034 | -0.026 |
| TYW3 | NM_001162916 | -0.172 | -0.754 |
| SYTL2 | NM_001162951 | -0.046 | -0.2065 |
| SYTL2 | NM_001162952 | -0.046 | -0.2065 |
| SYTL2 | NM_001162953 | -0.046 | -0.2065 |
| APBA1 | NM_001163 | -0.378 | -2.549 |
| PLK1S1 | NM_001163022 | -1.326 | -1.6115 |
| PLK1S1 | NM_001163023 | -1.326 | -1.6115 |
| RPTOR | NM_001163034 | -0.723 | -1.185 |
| C17orf99 | NM_001163075 | -0.0595 | -0.3995 |
| RDM1 | NM_001163120 | -0.173 | -0.392 |
| RDM1 | NM_001163121 | -0.173 | -0.392 |
| RDM1 | NM_001163122 | -0.173 | -0.392 |
| RDM1 | NM_001163124 | -0.173 | -0.392 |
| RDM1 | NM_001163125 | -0.173 | -0.392 |
| RDM1 | NM_001163130 | -0.173 | -0.392 |
| ETV1 | NM_001163147 | -1.24 | -2.281 |
| ETV1 | NM_001163148 | -1.24 | -2.281 |
| ETV1 | NM_001163149 | -1.24 | -2.281 |
| ETV1 | NM_001163150 | -1.24 | -2.281 |
| ETV1 | NM_001163151 | -1.24 | -2.281 |
| ETV1 | NM_001163152 | -1.24 | -2.281 |
| FGFR3 | NM_001163213 | 0.069 | 0.864 |
| PLXNB3 | NM_001163257 | -0.03 | -0.178 |
| FAM63A | NM_001163258 | -0.6285 | -0.816 |
| FAM63A | NM_001163259 | -0.6285 | -0.816 |
| FAM63A | NM_001163260 | -0.6285 | -0.816 |
| PCYT1B | NM_001163264 | -0.1205 | -0.516 |
| PCYT1B | NM_001163265 | -0.1205 | -0.516 |
| ODZ1 | NM_001163278 | -0.0075 | -0.5005 |
| ODZ1 | NM_001163279 | -0.0075 | -0.5005 |
| HTATSF1 | NM_001163280 | -0.78 | -0.327 |
| EWSR1 | NM_001163285 | -0.296 | -0.8785 |
| EWSR1 | NM_001163286 | -0.296 | -0.8785 |

| | | | |
|---|---|---|---|
| EWSR1 | NM_001163287 | -0.296 | -0.8785 |
| MARK2 | NM_001163296 | -0.758 | -2.485 |
| MARK2 | NM_001163297 | -0.758 | -2.485 |
| FBXL17 | NM_001163315 | -0.7295 | -1.8625 |
| CCDC120 | NM_001163321 | -0.284 | -0.137 |
| CCDC120 | NM_001163322 | -0.284 | -0.137 |
| CCDC120 | NM_001163323 | -0.284 | -0.137 |
| SYTL5 | NM_001163334 | -0.244 | -0.192 |
| SYTL5 | NM_001163335 | -0.244 | -0.192 |
| RAB2B | NM_001163380 | -0.589 | -0.8845 |
| ZSCAN12 | NM_001163391 | 0.159 | -0.6535 |
| FAM172A | NM_001163417 | -0.6135 | -0.6715 |
| FAM172A | NM_001163418 | -0.6135 | -0.6715 |
| MYEOV2 | NM_001163424 | 0.2875 | -0.5235 |
| CPA4 | NM_001163446 | 0.1735 | -0.166 |
| ZNF746 | NM_001163474 | -0.307 | -1.6505 |
| DCAF11 | NM_001163484 | -0.0445 | -0.6475 |
| FBXO24 | NM_001163499 | 0.3045 | -0.2115 |
| DCAF4 | NM_001163508 | -0.5925 | -1.5045 |
| DCAF4 | NM_001163509 | -0.5925 | -1.5045 |
| SYNRG | NM_001163544 | -0.907 | -2.1575 |
| SYNRG | NM_001163545 | -0.907 | -2.1575 |
| SYNRG | NM_001163546 | -0.907 | -2.1575 |
| SYNRG | NM_001163547 | -0.907 | -2.1575 |
| C2orf61 | NM_001163561 | -0.086 | -0.235 |
| C1orf129 | NM_001163629 | 0.465 | 0.2465 |
| GTF2I | NM_001163636 | -0.1905 | -0.6505 |
| HTR3D | NM_001163646 | -0.136 | -0.3185 |
| WDR81 | NM_001163673 | -0.9245 | -2.243 |
| KCNMB3 | NM_001163677 | -0.5575 | -0.341 |
| SHOX2 | NM_001163678 | -0.6445 | -0.635 |
| MYO19 | NM_001163735 | -1.2365 | -2.001 |
| COL11A2 | NM_001163771 | 0.2385 | -0.5075 |
| ROD1 | NM_001163788 | -1.0745 | -1.801 |
| ROD1 | NM_001163790 | -1.0745 | -1.801 |
| WDR81 | NM_001163809 | -0.9245 | -2.243 |
| WDR81 | NM_001163811 | -0.9245 | -2.243 |
| C10orf2 | NM_001163812 | -1.032 | -2.465 |
| C10orf2 | NM_001163813 | -1.032 | -2.465 |
| C10orf2 | NM_001163814 | -1.032 | -2.465 |
| DHCR7 | NM_001163817 | -0.636 | -1.5385 |
| LRTM2 | NM_001163925 | -0.0535 | -0.401 |
| LRTM2 | NM_001163926 | -0.0535 | -0.401 |

| | | | |
|---|---|---|---|
| ABCB5 | NM_001163941 | -0.39 | -0.031 |
| ABCB5 | NM_001163942 | -0.39 | -0.031 |
| RAB37 | NM_001163989 | -0.2305 | -0.536 |
| RAB37 | NM_001163990 | -0.2305 | -0.536 |
| FAM170A | NM_001163991 | -0.2355 | -0.1995 |
| ABCB5 | NM_001163993 | -0.39 | -0.031 |
| MECOM | NM_001163999 | -0.401 | -0.379 |
| APBB1 | NM_001164 | -0.201 | -1.038 |
| MECOM | NM_001164000 | -0.401 | -0.379 |
| COPS7A | NM_001164093 | -0.3265 | -1.836 |
| COPS7A | NM_001164094 | -0.3265 | -1.836 |
| COPS7A | NM_001164095 | -0.3265 | -1.836 |
| VCAN | NM_001164097 | -0.48 | -1.449 |
| VCAN | NM_001164098 | -0.48 | -1.449 |
| MOV10L1 | NM_001164104 | -0.022 | -0.046 |
| MOV10L1 | NM_001164105 | -0.022 | -0.046 |
| MOV10L1 | NM_001164106 | -0.022 | -0.046 |
| CASS4 | NM_001164114 | 0.2985 | -0.203 |
| CASS4 | NM_001164115 | 0.2985 | -0.203 |
| CASS4 | NM_001164116 | 0.2985 | -0.203 |
| PPP6R3 | NM_001164160 | -0.8895 | -1.973 |
| PPP6R3 | NM_001164161 | -0.8895 | -1.973 |
| PPP6R3 | NM_001164162 | -0.8895 | -1.973 |
| PPP6R3 | NM_001164163 | -0.8895 | -1.973 |
| PPP6R3 | NM_001164164 | -0.8895 | -1.973 |
| ENTPD1 | NM_001164178 | -0.2325 | -0.583 |
| ENTPD1 | NM_001164179 | -0.2325 | -0.583 |
| ENTPD1 | NM_001164181 | -0.2325 | -0.583 |
| ENTPD1 | NM_001164182 | -0.2325 | -0.583 |
| ENTPD1 | NM_001164183 | -0.2325 | -0.583 |
| PLIN3 | NM_001164189 | -0.1495 | -0.7095 |
| PLIN3 | NM_001164194 | -0.1495 | -0.7095 |
| LRCH1 | NM_001164211 | -0.158 | -1.2395 |
| LRCH1 | NM_001164213 | -0.158 | -1.2395 |
| DDHD2 | NM_001164232 | -1.3185 | -1.3035 |
| DDHD2 | NM_001164234 | -1.3185 | -1.3035 |
| NLGN4Y | NM_001164238 | 0.0205 | -0.5245 |
| DHX16 | NM_001164239 | -1.009 | -2.9925 |
| C1orf27 | NM_001164245 | -0.6855 | -1.2995 |
| C1orf27 | NM_001164246 | -0.6855 | -1.2995 |
| C17orf72 | NM_001164257 | 0.046 | -0.262 |
| LTBP3 | NM_001164266 | -0.3685 | -0.559 |
| WDR46 | NM_001164267 | -0.2155 | -0.726 |

| | | | |
|---|---|---|---|
| RAD51 | NM_001164269 | -0.7715 | -2.366 |
| RAD51 | NM_001164270 | -0.7715 | -2.366 |
| DLC1 | NM_001164271 | -1.246 | -2.5865 |
| MGA | NM_001164273 | -0.6075 | -2.6415 |
| SLC37A4 | NM_001164277 | -0.202 | -1.2205 |
| SLC37A4 | NM_001164278 | -0.202 | -1.2205 |
| SLC37A4 | NM_001164279 | -0.202 | -1.2205 |
| SLC37A4 | NM_001164280 | -0.202 | -1.2205 |
| TRAF3IP2 | NM_001164281 | 0.058 | -0.0505 |
| TRAF3IP2 | NM_001164282 | 0.058 | -0.0505 |
| TRAF3IP2 | NM_001164283 | 0.058 | -0.0505 |
| FAM166B | NM_001164310 | 0.09 | -0.0755 |
| ZBTB24 | NM_001164313 | -1.7475 | -3.846 |
| ANKRD36 | NM_001164315 | -0.156 | -0.757 |
| FLNB | NM_001164317 | -0.094 | -0.873 |
| FLNB | NM_001164318 | -0.094 | -0.873 |
| FLNB | NM_001164319 | -0.094 | -0.873 |
| BRD8 | NM_001164326 | -1.2755 | -2.346 |
| ZBTB20 | NM_001164342 | -0.405 | -0.2505 |
| ZBTB20 | NM_001164343 | -0.405 | -0.2505 |
| ZBTB20 | NM_001164344 | -0.405 | -0.2505 |
| ZBTB20 | NM_001164345 | -0.405 | -0.2505 |
| ZBTB20 | NM_001164346 | -0.405 | -0.2505 |
| ZBTB20 | NM_001164347 | -0.405 | -0.2505 |
| THAP4 | NM_001164356 | -0.718 | -2.4365 |
| GPN3 | NM_001164372 | -0.998 | -1.9035 |
| GPN3 | NM_001164373 | -0.998 | -1.9035 |
| STAU2 | NM_001164380 | -0.786 | -0.834 |
| STAU2 | NM_001164381 | -0.786 | -0.834 |
| STAU2 | NM_001164382 | -0.786 | -0.834 |
| STAU2 | NM_001164383 | -0.786 | -0.834 |
| STAU2 | NM_001164384 | -0.786 | -0.834 |
| STAU2 | NM_001164385 | -0.786 | -0.834 |
| RAPGEF6 | NM_001164386 | -0.88 | -3.2735 |
| RAPGEF6 | NM_001164387 | -0.88 | -3.2735 |
| RAPGEF6 | NM_001164388 | -0.88 | -3.2735 |
| RAPGEF6 | NM_001164389 | -0.88 | -3.2735 |
| RAPGEF6 | NM_001164390 | -0.88 | -3.2735 |
| MTF2 | NM_001164391 | -1.0445 | -2.919 |
| MTF2 | NM_001164392 | -1.0445 | -2.919 |
| MTF2 | NM_001164393 | -1.0445 | -2.919 |
| CDK5 | NM_001164410 | -0.0365 | -0.6165 |
| KRTAP22-2 | NM_001164434 | 0.523 | -0.43 |

| | | | |
|---|---|---|---|
| KRTAP21-3 | NM_001164435 | -0.12 | 0.3095 |
| ACTR3C | NM_001164458 | 0.0665 | 0.1085 |
| ACTR3C | NM_001164459 | 0.0665 | 0.1085 |
| MUC12 | NM_001164462 | -0.203 | 0.0175 |
| FNBP1L | NM_001164473 | -0.464 | -1.51 |
| SLC41A3 | NM_001164475 | 0.056 | -0.745 |
| ADAM8 | NM_001164489 | -0.0615 | -0.2615 |
| ADAM8 | NM_001164490 | -0.0615 | -0.2615 |
| WDR52 | NM_001164496 | -0.3865 | -0.9105 |
| EIF4ENIF1 | NM_001164501 | -0.713 | -1.327 |
| EIF4ENIF1 | NM_001164502 | -0.713 | -1.327 |
| NEB | NM_001164507 | -0.594 | -0.371 |
| NEB | NM_001164508 | -0.594 | -0.371 |
| ZNF135 | NM_001164527 | -0.054 | -0.975 |
| ZNF135 | NM_001164529 | -0.054 | -0.975 |
| ZNF135 | NM_001164530 | -0.054 | -0.975 |
| DISC1 | NM_001164537 | -0.2995 | -0.02 |
| DISC1 | NM_001164538 | -0.2995 | -0.02 |
| DISC1 | NM_001164539 | -0.2995 | -0.02 |
| DISC1 | NM_001164540 | -0.2995 | -0.02 |
| DISC1 | NM_001164541 | -0.2995 | -0.02 |
| DISC1 | NM_001164542 | -0.2995 | -0.02 |
| DISC1 | NM_001164544 | -0.2995 | -0.02 |
| DISC1 | NM_001164545 | -0.2995 | -0.02 |
| DISC1 | NM_001164546 | -0.2995 | -0.02 |
| DISC1 | NM_001164547 | -0.2995 | -0.02 |
| DISC1 | NM_001164548 | -0.2995 | -0.02 |
| DISC1 | NM_001164549 | -0.2995 | -0.02 |
| DISC1 | NM_001164550 | -0.2995 | -0.02 |
| DISC1 | NM_001164551 | -0.2995 | -0.02 |
| DISC1 | NM_001164552 | -0.2995 | -0.02 |
| DISC1 | NM_001164553 | -0.2995 | -0.02 |
| DISC1 | NM_001164554 | -0.2995 | -0.02 |
| DISC1 | NM_001164555 | -0.2995 | -0.02 |
| DISC1 | NM_001164556 | -0.2995 | -0.02 |
| PDZRN4 | NM_001164595 | 0.2225 | 0.181 |
| ASXL1 | NM_001164603 | -1.0175 | -3.4285 |
| FXYD5 | NM_001164605 | -0.0945 | 0.022 |
| FZD6 | NM_001164615 | -1.104 | -0.792 |
| FZD6 | NM_001164616 | -1.104 | -0.792 |
| GPC3 | NM_001164617 | -0.0175 | -0.2355 |
| GPC3 | NM_001164618 | -0.0175 | -0.2355 |
| GPC3 | NM_001164619 | -0.0175 | -0.2355 |

| | | | |
|---|---|---|---|
| EIF2C2 | NM_001164623 | 0.2285 | -0.883 |
| GTDC1 | NM_001164629 | 0.021 | -0.2115 |
| MAST4 | NM_001164664 | 0.227 | -0.2985 |
| KIAA1549 | NM_001164665 | -0.8425 | -3.1455 |
| SUMF1 | NM_001164674 | -0.4965 | -0.8625 |
| SUMF1 | NM_001164675 | -0.4965 | -0.8625 |
| CCR3 | NM_001164680 | -0.0235 | -0.3235 |
| THEMIS | NM_001164685 | 0.157 | -0.1605 |
| THEMIS | NM_001164687 | 0.157 | -0.1605 |
| RD3 | NM_001164688 | -0.0705 | -0.56 |
| AMHR2 | NM_001164690 | -0.179 | -0.147 |
| AMHR2 | NM_001164691 | -0.179 | -0.147 |
| LTB4R2 | NM_001164692 | -0.164 | -0.52 |
| IYD | NM_001164694 | -0.101 | 0.0085 |
| IYD | NM_001164695 | -0.101 | 0.0085 |
| AMT | NM_001164710 | -0.056 | -0.721 |
| AMT | NM_001164711 | -0.056 | -0.721 |
| AMT | NM_001164712 | -0.056 | -0.721 |
| ZNF605 | NM_001164715 | -0.468 | -1.842 |
| TGFB1I1 | NM_001164719 | -0.545 | -1.7215 |
| PTAFR | NM_001164721 | -0.3325 | -0.208 |
| PTAFR | NM_001164722 | -0.3325 | -0.208 |
| PTAFR | NM_001164723 | -0.3325 | -0.208 |
| REEP1 | NM_001164730 | -1.119 | -1.0765 |
| REEP1 | NM_001164731 | -1.119 | -1.0765 |
| REEP1 | NM_001164732 | -1.119 | -1.0765 |
| CALCR | NM_001164737 | 0.01 | 0.0765 |
| CALCR | NM_001164738 | 0.01 | 0.0765 |
| ARHGAP4 | NM_001164741 | -0.0045 | -0.3475 |
| RASSF8 | NM_001164746 | -0.5525 | -1.803 |
| RASSF8 | NM_001164747 | -0.5525 | -1.803 |
| RASSF8 | NM_001164748 | -0.5525 | -1.803 |
| NPAS3 | NM_001164749 | 0.0345 | -0.345 |
| ASPH | NM_001164750 | -1.0635 | -0.5415 |
| ASPH | NM_001164751 | -1.0635 | -0.5415 |
| ASPH | NM_001164752 | -1.0635 | -0.5415 |
| ASPH | NM_001164753 | -1.0635 | -0.5415 |
| ASPH | NM_001164754 | -1.0635 | -0.5415 |
| ASPH | NM_001164755 | -1.0635 | -0.5415 |
| ASPH | NM_001164756 | -1.0635 | -0.5415 |
| NOS1AP | NM_001164757 | 0.02 | -0.5025 |
| PRKAR1B | NM_001164758 | -0.343 | -1.096 |
| PRKAR1B | NM_001164759 | -0.343 | -1.096 |

| | | | |
|---|---|---|---|
| PRKAR1B | NM_001164760 | -0.343 | -1.096 |
| PRKAR1B | NM_001164761 | -0.343 | -1.096 |
| PRKAR1B | NM_001164762 | -0.343 | -1.096 |
| ZFHX3 | NM_001164766 | -0.4935 | -1.687 |
| SLC7A2 | NM_001164771 | -1.211 | -2.2195 |
| ATXN3 | NM_001164774 | -0.521 | -2.1695 |
| ATXN3 | NM_001164776 | -0.521 | -2.1695 |
| ATXN3 | NM_001164777 | -0.521 | -2.1695 |
| ATXN3 | NM_001164778 | -0.521 | -2.1695 |
| ATXN3 | NM_001164779 | -0.521 | -2.1695 |
| ATXN3 | NM_001164780 | -0.521 | -2.1695 |
| ATXN3 | NM_001164781 | -0.521 | -2.1695 |
| ATXN3 | NM_001164782 | -0.521 | -2.1695 |
| SLC25A26 | NM_001164796 | -0.112 | -1.016 |
| RBMX | NM_001164803 | -0.7635 | -1.229 |
| ACIN1 | NM_001164814 | -0.559 | -1.4105 |
| ACIN1 | NM_001164815 | -0.559 | -1.4105 |
| ACIN1 | NM_001164816 | -0.559 | -1.4105 |
| ACIN1 | NM_001164817 | -0.559 | -1.4105 |
| DCAF17 | NM_001164821 | -0.496 | -0.617 |
| C1orf212 | NM_001164824 | -0.8555 | -2.974 |
| C1orf212 | NM_001164825 | -0.8555 | -2.974 |
| CA1 | NM_001164830 | 0.052 | -0.7995 |
| FXYD6 | NM_001164831 | 0.187 | -0.2205 |
| FXYD6 | NM_001164832 | 0.187 | -0.2205 |
| L1TD1 | NM_001164835 | -0.078 | 0.206 |
| FXYD6 | NM_001164836 | 0.187 | -0.2205 |
| FXYD6 | NM_001164837 | 0.187 | -0.2205 |
| LYRM4 | NM_001164840 | -0.0965 | -0.564 |
| LYRM4 | NM_001164841 | -0.0965 | -0.564 |
| CTNNA2 | NM_001164883 | -0.539 | -0.1275 |
| BIRC3 | NM_001165 | -0.24 | -0.0535 |
| TMEM41B | NM_001165030 | -1.2185 | -1.4945 |
| FBLN2 | NM_001165035 | 0.09 | -0.6295 |
| OGDH | NM_001165036 | -0.7535 | -1.7415 |
| GABRA5 | NM_001165037 | 0.038 | -0.9535 |
| GFRA2 | NM_001165038 | -0.613 | -2.2025 |
| GFRA2 | NM_001165039 | -0.613 | -2.2025 |
| LOC730755 | NM_001165252 | 0.1155 | 0.1995 |
| MNX1 | NM_001165255 | 0.6805 | 1.474 |
| SCG3 | NM_001165257 | -0.2465 | -0.4465 |
| NFKB1 | NM_001165412 | -0.6035 | -2.378 |
| SERPINE1 | NM_001165413 | 0.2535 | -0.0945 |

| | | | |
|---|---|---|---|
| LDHA | NM_001165414 | -0.2955 | -0.612 |
| LDHA | NM_001165415 | -0.2955 | -0.612 |
| LDHA | NM_001165416 | -0.2955 | -0.612 |
| SLC25A11 | NM_001165417 | -0.9435 | -1.366 |
| SLC25A11 | NM_001165418 | -0.9435 | -1.366 |
| ATP5L2 | NM_001165877 | -0.4325 | -0.6675 |
| ZNF268 | NM_001165887 | -1.119 | -1.924 |
| NPAS3 | NM_001165893 | 0.0345 | -0.345 |
| PDE4D | NM_001165899 | -1.413 | -2.114 |
| STX1A | NM_001165903 | -0.0585 | -0.746 |
| SERPINF2 | NM_001165920 | -0.152 | -0.4985 |
| SERPINF2 | NM_001165921 | -0.152 | -0.4985 |
| HNF1B | NM_001165923 | 0.0275 | -0.069 |
| MKS1 | NM_001165927 | -0.4345 | -1.096 |
| DAG1 | NM_001165928 | -0.4615 | -1.0555 |
| RRM2 | NM_001165931 | -0.8335 | -0.8185 |
| RGS9 | NM_001165933 | -0.0445 | -0.1075 |
| STARD3 | NM_001165937 | -0.309 | -1.516 |
| STARD3 | NM_001165938 | -0.309 | -1.516 |
| CLCNKB | NM_001165945 | 0.2105 | -0.2405 |
| IDE | NM_001165946 | -0.728 | -1.5025 |
| HTR2A | NM_001165947 | -0.1865 | 0.1495 |
| ALOXE3 | NM_001165960 | 0.1215 | -0.1975 |
| ELAC2 | NM_001165962 | -0.345 | -1.142 |
| SCN1A | NM_001165963 | -0.315 | -0.3145 |
| SCN1A | NM_001165964 | -0.315 | -0.3145 |
| PITPNM3 | NM_001165966 | -0.115 | -0.208 |
| STRADA | NM_001165969 | -0.9125 | -1.7355 |
| STRADA | NM_001165970 | -0.9125 | -1.7355 |
| NRG3 | NM_001165972 | -0.1615 | -0.614 |
| NRG3 | NM_001165973 | -0.1615 | -0.614 |
| UROC1 | NM_001165974 | -0.0645 | -0.48 |
| PDE1B | NM_001165975 | 0.0265 | -0.2955 |
| PROM2 | NM_001165977 | 0.0475 | -0.427 |
| PROM2 | NM_001165978 | 0.0475 | -0.427 |
| PLCE1 | NM_001165979 | 0.0295 | -0.3815 |
| GDPD1 | NM_001165993 | -2.4975 | -3.392 |
| GDPD1 | NM_001165994 | -2.4975 | -3.392 |
| BIRC2 | NM_001166 | -1.4025 | -2.2095 |
| APOBEC3H | NM_001166002 | -0.27 | 0.0105 |
| APOBEC3H | NM_001166003 | -0.27 | 0.0105 |
| APOBEC3H | NM_001166004 | -0.27 | 0.0105 |
| EPB41 | NM_001166005 | -0.672 | -0.825 |

| | | | |
|---|---|---|---|
| EPB41 | NM_001166006 | -0.672 | -0.825 |
| EPB41 | NM_001166007 | -0.672 | -0.825 |
| ZSCAN30 | NM_001166012 | -0.899 | -1.633 |
| SKA3 | NM_001166017 | -2.0545 | -2.899 |
| AP1B1 | NM_001166019 | -0.03 | -0.202 |
| SBSN | NM_001166034 | -0.3535 | -0.274 |
| SBSN | NM_001166035 | -0.3535 | -0.274 |
| ZNF260 | NM_001166036 | -0.5865 | -1.8515 |
| ZNF260 | NM_001166037 | -0.5865 | -1.8515 |
| ZNF260 | NM_001166038 | -0.5865 | -1.8515 |
| APBB2 | NM_001166050 | -0.1085 | -0.418 |
| APBB2 | NM_001166051 | -0.1085 | -0.418 |
| APBB2 | NM_001166052 | -0.1085 | -0.418 |
| APBB2 | NM_001166053 | -0.1085 | -0.418 |
| APBB2 | NM_001166054 | -0.1085 | -0.418 |
| EDNRA | NM_001166055 | -0.9585 | -0.9835 |
| RXFP2 | NM_001166058 | -0.215 | -0.3625 |
| HMGCL | NM_001166059 | -0.481 | -1.2395 |
| GLRB | NM_001166060 | -1.6365 | -1.7595 |
| GLRB | NM_001166061 | -1.6365 | -1.7595 |
| SPINT2 | NM_001166103 | -0.0695 | -0.1925 |
| NR3C2 | NM_001166104 | -0.812 | -0.0415 |
| TADA2A | NM_001166105 | -1.5365 | -2.296 |
| PALLD | NM_001166108 | -0.998 | -2.4165 |
| PALLD | NM_001166109 | -0.998 | -2.4165 |
| PALLD | NM_001166110 | -0.998 | -2.4165 |
| PNPLA6 | NM_001166111 | -0.166 | -0.6985 |
| PNPLA6 | NM_001166112 | -0.166 | -0.6985 |
| PNPLA6 | NM_001166113 | -0.166 | -0.6985 |
| PNPLA6 | NM_001166114 | -0.166 | -0.6985 |
| TMOD1 | NM_001166116 | -0.4135 | -0.9585 |
| LEF1 | NM_001166119 | -0.1305 | -0.2025 |
| FRAS1 | NM_001166133 | -0.3195 | -0.8905 |
| RBM28 | NM_001166135 | -0.7105 | -1.5725 |
| EVC2 | NM_001166136 | -0.6345 | -0.0795 |
| FAM75C2 | NM_001166137 | -0.0945 | -0.191 |
| LCORL | NM_001166139 | -0.7695 | -1.135 |
| PDHX | NM_001166158 | -0.9945 | -2.4695 |
| NDUFS2 | NM_001166159 | -0.1405 | -0.7055 |
| PPP1R9A | NM_001166160 | -0.331 | -0.5415 |
| PPP1R9A | NM_001166161 | -0.331 | -0.5415 |
| PPP1R9A | NM_001166162 | -0.331 | -0.5415 |
| PPP1R9A | NM_001166163 | -0.331 | -0.5415 |

| | | | |
|---|---|---|---|
| NEK6 | NM_001166167 | -0.1015 | -0.482 |
| NEK6 | NM_001166168 | -0.1015 | -0.482 |
| NEK6 | NM_001166169 | -0.1015 | -0.482 |
| NEK6 | NM_001166170 | -0.1015 | -0.482 |
| NEK6 | NM_001166171 | -0.1015 | -0.482 |
| NKX2-5 | NM_001166175 | 0.061 | -0.2805 |
| NKX2-5 | NM_001166176 | 0.061 | -0.2805 |
| SYNPO | NM_001166208 | -0.5025 | -0.46 |
| SYNPO | NM_001166209 | -0.5025 | -0.46 |
| CLCF1 | NM_001166212 | -0.0295 | -0.1525 |
| S1PR5 | NM_001166215 | 0.3555 | -0.0875 |
| UTP14A | NM_001166221 | -0.661 | -1.016 |
| CARNS1 | NM_001166222 | -0.1435 | -0.3585 |
| CEP120 | NM_001166226 | -1.6895 | -3.209 |
| GSDMD | NM_001166237 | -0.0245 | -0.3735 |
| CGREF1 | NM_001166239 | -0.1165 | -0.707 |
| CGREF1 | NM_001166240 | -0.1165 | -0.707 |
| CGREF1 | NM_001166241 | -0.1165 | -0.707 |
| C22orf39 | NM_001166242 | -0.0905 | -0.945 |
| FHIT | NM_001166243 | -0.2035 | -0.466 |
| HPSE2 | NM_001166244 | -0.2705 | -0.563 |
| HPSE2 | NM_001166245 | -0.2705 | -0.563 |
| HPSE2 | NM_001166246 | -0.2705 | -0.563 |
| GRIK2 | NM_001166247 | -0.8425 | -1.477 |
| LPPR4 | NM_001166252 | -1.1065 | -0.724 |
| TBXAS1 | NM_001166253 | 0.099 | -0.327 |
| TBXAS1 | NM_001166254 | 0.099 | -0.327 |
| TRIP13 | NM_001166260 | -0.694 | -0.936 |
| MTIF3 | NM_001166261 | -1.0275 | -1.766 |
| MTIF3 | NM_001166262 | -1.0275 | -1.766 |
| MTIF3 | NM_001166263 | -1.0275 | -1.766 |
| LTBP1 | NM_001166264 | -0.0975 | -0.986 |
| LTBP1 | NM_001166265 | -0.0975 | -0.986 |
| LTBP1 | NM_001166266 | -0.0975 | -0.986 |
| SPATA13 | NM_001166271 | -0.197 | -0.9725 |
| ARHGAP25 | NM_001166276 | 0.145 | -0.1345 |
| ARHGAP25 | NM_001166277 | 0.145 | -0.1345 |
| DLG3 | NM_001166278 | -0.8345 | -2.8375 |
| RGMA | NM_001166283 | -0.8385 | -2.211 |
| RGMA | NM_001166286 | -0.8385 | -2.211 |
| RGMA | NM_001166287 | -0.8385 | -2.211 |
| RGMA | NM_001166288 | -0.8385 | -2.211 |
| RGMA | NM_001166289 | -0.8385 | -2.211 |

| | | | |
|---|---|---|---|
| KCNJ11 | NM_001166290 | -0.1825 | -0.8775 |
| PTCH2 | NM_001166292 | -0.2615 | -0.5365 |
| SSX2IP | NM_001166293 | -0.6025 | -1.314 |
| SSX2IP | NM_001166294 | -0.6025 | -1.314 |
| SSX2IP | NM_001166295 | -0.6025 | -1.314 |
| DHX40 | NM_001166301 | -0.9805 | -0.88 |
| PIGX | NM_001166304 | -1.06 | -1.315 |
| TMEM44 | NM_001166305 | -1.114 | -1.2125 |
| TMEM44 | NM_001166306 | -1.114 | -1.2125 |
| MFNG | NM_001166343 | -0.0455 | -0.46 |
| MDFIC | NM_001166345 | 0.2375 | -0.2935 |
| MDFIC | NM_001166346 | 0.2375 | -0.2935 |
| LFNG | NM_001166355 | -0.3805 | -1.4655 |
| SHMT2 | NM_001166356 | 0.348 | -0.3655 |
| SHMT2 | NM_001166357 | 0.348 | -0.3655 |
| SHMT2 | NM_001166358 | 0.348 | -0.3655 |
| SHMT2 | NM_001166359 | 0.348 | -0.3655 |
| MAGEA12 | NM_001166386 | -0.0605 | -0.1465 |
| MAGEA12 | NM_001166387 | -0.0605 | -0.1465 |
| MTUS1 | NM_001166393 | -0.8945 | -2.8875 |
| CHST4 | NM_001166395 | -0.3985 | -0.394 |
| SMOC2 | NM_001166412 | -0.2995 | -0.7415 |
| EHHADH | NM_001166415 | -0.193 | -0.6 |
| SSX2IP | NM_001166417 | -0.6025 | -1.314 |
| HDAC8 | NM_001166418 | 0.4635 | -0.028 |
| HDAC8 | NM_001166419 | 0.4635 | -0.028 |
| HDAC8 | NM_001166420 | 0.4635 | -0.028 |
| HDAC8 | NM_001166422 | 0.4635 | -0.028 |
| PGC | NM_001166424 | 0.2665 | -0.182 |
| GNAI2 | NM_001166425 | 0.096 | -0.5715 |
| WDR13 | NM_001166426 | -0.0425 | -0.8285 |
| ITIH1 | NM_001166434 | -0.223 | -0.309 |
| ITIH1 | NM_001166435 | -0.223 | -0.309 |
| ITIH1 | NM_001166436 | -0.223 | -0.309 |
| HDAC8 | NM_001166448 | 0.4635 | -0.028 |
| KNG1 | NM_001166451 | 0.342 | 0.365 |
| MPP1 | NM_001166460 | 0.061 | -0.6075 |
| MPP1 | NM_001166461 | 0.061 | -0.6075 |
| MPP1 | NM_001166462 | 0.061 | -0.6075 |
| DEFB133 | NM_001166478 | -0.701 | -0.3825 |
| SLC16A1 | NM_001166496 | -1.225 | -1.0865 |
| HPSE | NM_001166498 | -0.8815 | -0.394 |
| ADH7 | NM_001166504 | 0.147 | -0.218 |

| | | | |
|---|---|---|---|
| DDX4 | NM_001166533 | -0.1035 | -0.1605 |
| DDX4 | NM_001166534 | -0.1035 | -0.1605 |
| LST1 | NM_001166538 | -0.04 | -0.218 |
| IDS | NM_001166550 | -0.7255 | -0.5545 |
| AANAT | NM_001166579 | -0.3885 | 0.124 |
| EIF2B3 | NM_001166588 | -1.4105 | -1.2655 |
| APOA5 | NM_001166598 | 0.058 | -0.0475 |
| FAM122B | NM_001166599 | -0.416 | -1.2235 |
| FAM122B | NM_001166600 | -0.416 | -1.2235 |
| CFHR3 | NM_001166624 | -0.5635 | -0.217 |
| NLGN3 | NM_001166660 | -0.1905 | -0.3365 |
| CD244 | NM_001166663 | -0.2845 | -0.0245 |
| CD244 | NM_001166664 | -0.2845 | -0.0245 |
| AFF1 | NM_001166693 | -0.826 | -1.6925 |
| CHRNA3 | NM_001166694 | -1.153 | -1.99 |
| SLC1A3 | NM_001166695 | -0.2025 | -0.363 |
| SLC1A3 | NM_001166696 | -0.2025 | -0.363 |
| FAM13C | NM_001166698 | -0.277 | -1.1595 |
| FAM104B | NM_001166699 | -0.595 | -0.7705 |
| FAM104B | NM_001166700 | -0.595 | -0.7705 |
| FAM104B | NM_001166701 | -0.595 | -0.7705 |
| FAM104B | NM_001166702 | -0.595 | -0.7705 |
| FAM104B | NM_001166703 | -0.595 | -0.7705 |
| FAM104B | NM_001166704 | -0.595 | -0.7705 |
| XIAP | NM_001167 | -2.2095 | -3.0085 |
| C9orf82 | NM_001167575 | -1.624 | -3.2525 |
| TRPC7 | NM_001167576 | -0.1875 | 0.019 |
| TRPC7 | NM_001167577 | -0.1875 | 0.019 |
| SLC34A1 | NM_001167579 | 0.189 | -0.0565 |
| SV2B | NM_001167580 | -0.1735 | -0.309 |
| RBM6 | NM_001167582 | -1.173 | -1.2265 |
| ZFAT | NM_001167583 | -0.164 | -0.7475 |
| CTSA | NM_001167594 | -0.249 | -1.347 |
| AMACR | NM_001167595 | -0.4105 | -1.661 |
| XPNPEP1 | NM_001167604 | -0.154 | -0.6625 |
| RAB35 | NM_001167606 | -0.469 | -1.728 |
| GM2A | NM_001167607 | 0.162 | -0.357 |
| RHBDD1 | NM_001167608 | -1.1225 | -1.4295 |
| LEMD3 | NM_001167614 | -1.6875 | -3.657 |
| MLH1 | NM_001167617 | -1.0075 | -1.4605 |
| MLH1 | NM_001167618 | -1.0075 | -1.4605 |
| MLH1 | NM_001167619 | -1.0075 | -1.4605 |
| ATXN10 | NM_001167621 | -0.5975 | -0.7965 |

| | | | |
|---|---|---|---|
| CACNA1C | NM_001167623 | -0.2715 | -0.3655 |
| CACNA1C | NM_001167624 | -0.2715 | -0.3655 |
| CACNA1C | NM_001167625 | -0.2715 | -0.3655 |
| CDH18 | NM_001167667 | -0.238 | -0.5555 |
| LPP | NM_001167671 | -0.087 | -1.078 |
| LPP | NM_001167672 | -0.087 | -1.078 |
| CADM2 | NM_001167674 | -0.113 | -0.487 |
| CADM2 | NM_001167675 | -0.113 | -0.487 |
| FLI1 | NM_001167681 | 0.058 | 0.076 |
| VARS2 | NM_001167733 | 0.05 | -0.762 |
| VARS2 | NM_001167734 | 0.05 | -0.762 |
| NAV1 | NM_001167738 | -1.2845 | -1.6955 |
| SMYD3 | NM_001167740 | 0.9405 | 2.0725 |
| MTX3 | NM_001167741 | -0.5855 | -0.6855 |
| ADCY10 | NM_001167749 | -0.397 | -0.2165 |
| C20orf26 | NM_001167816 | 0.1555 | -0.5125 |
| FHL1 | NM_001167819 | -0.136 | -0.286 |
| BLCAP | NM_001167820 | -0.487 | -1.549 |
| BLCAP | NM_001167821 | -0.487 | -1.549 |
| BLCAP | NM_001167822 | -0.487 | -1.549 |
| BLCAP | NM_001167823 | -0.487 | -1.549 |
| TFEB | NM_001167827 | -0.2265 | -0.874 |
| TMED5 | NM_001167830 | -0.9455 | -1.8555 |
| SBNO1 | NM_001167856 | -0.5575 | -1.7525 |
| PPP1R12B | NM_001167857 | 0.8205 | 0.5085 |
| PPP1R12B | NM_001167858 | 0.8205 | 0.5085 |
| NSMCE4A | NM_001167865 | -1.3165 | -2.2585 |
| ATP5SL | NM_001167867 | -0.2885 | -1.115 |
| ATP5SL | NM_001167868 | -0.2885 | -1.115 |
| ATP5SL | NM_001167869 | -0.2885 | -1.115 |
| ATP5SL | NM_001167870 | -0.2885 | -1.115 |
| ATP5SL | NM_001167871 | -0.2885 | -1.115 |
| LHPP | NM_001167880 | -0.0585 | -0.051 |
| ANKRD50 | NM_001167882 | -1.129 | -3.358 |
| PGPEP1L | NM_001167902 | 0.1385 | -0.463 |
| MFSD1 | NM_001167903 | -0.865 | -0.943 |
| VEPH1 | NM_001167911 | 0.139 | -0.02 |
| VEPH1 | NM_001167912 | 0.139 | -0.02 |
| VEPH1 | NM_001167915 | 0.139 | -0.02 |
| VEPH1 | NM_001167916 | 0.139 | -0.02 |
| VEPH1 | NM_001167917 | 0.139 | -0.02 |
| IL1RAP | NM_001167928 | 0.0075 | -0.354 |
| IL1RAP | NM_001167929 | 0.0075 | -0.354 |

| | | | |
|---|---|---|---|
| IL1RAP | NM_001167930 | 0.0075 | -0.354 |
| IL1RAP | NM_001167931 | 0.0075 | -0.354 |
| CCM2 | NM_001167934 | -1.528 | -2.997 |
| CCM2 | NM_001167935 | -1.528 | -2.997 |
| CADPS2 | NM_001167940 | 0.137 | -0.251 |
| WDR76 | NM_001167941 | -0.3815 | -2.414 |
| TNFAIP8L1 | NM_001167942 | -0.725 | -1.731 |
| CACNB2 | NM_001167945 | 0.037 | -0.4365 |
| TMEM178 | NM_001167959 | -0.4395 | -1.705 |
| KCTD7 | NM_001167961 | 0.297 | -0.3965 |
| SLC26A5 | NM_001167962 | -0.0105 | -0.4415 |
| WDR65 | NM_001167965 | -0.287 | -0.379 |
| TRMT2B | NM_001167970 | 0.3785 | -0.6125 |
| TRMT2B | NM_001167971 | 0.3785 | -0.6125 |
| TRMT2B | NM_001167972 | 0.3785 | -0.6125 |
| PQBP1 | NM_001167989 | -0.1635 | -0.5965 |
| PQBP1 | NM_001167990 | -0.1635 | -0.5965 |
| PQBP1 | NM_001167992 | -0.1635 | -0.5965 |
| BIRC5 | NM_001168 | -0.116 | -0.84 |
| TMEM92 | NM_001168215 | -0.41 | -0.4895 |
| TBC1D17 | NM_001168222 | -0.288 | -0.636 |
| ABL2 | NM_001168236 | -1.382 | -2.9035 |
| ABL2 | NM_001168237 | -1.382 | -2.9035 |
| ABL2 | NM_001168238 | -1.382 | -2.9035 |
| ABL2 | NM_001168239 | -1.382 | -2.9035 |
| OCIAD1 | NM_001168254 | -1.113 | -1.1115 |
| GPR156 | NM_001168271 | -0.1445 | -0.1305 |
| ITPR1 | NM_001168272 | -1.355 | -2.454 |
| WWTR1 | NM_001168278 | -0.4415 | -0.417 |
| WWTR1 | NM_001168280 | -0.4415 | -0.417 |
| CXCR2 | NM_001168298 | -0.132 | -0.1315 |
| KLHL13 | NM_001168299 | -1.0605 | -1.8235 |
| KLHL13 | NM_001168300 | -1.0605 | -1.8235 |
| KLHL13 | NM_001168301 | -1.0605 | -1.8235 |
| KLHL13 | NM_001168302 | -1.0605 | -1.8235 |
| KLHL13 | NM_001168303 | -1.0605 | -1.8235 |
| EDN1 | NM_001168319 | 0.329 | -0.3645 |
| TSPAN9 | NM_001168320 | -0.4695 | -0.8035 |
| CD300LG | NM_001168322 | -0.1235 | -0.406 |
| CD300LG | NM_001168323 | -0.1235 | -0.406 |
| CD300LG | NM_001168324 | -0.1235 | -0.406 |
| NDUFB4 | NM_001168331 | -0.1255 | -0.642 |
| ME2 | NM_001168335 | -1.2725 | -2.0435 |

| | | | |
|---|---|---|---|
| PLG | NM_001168338 | 0.01 | -0.478 |
| TTC39B | NM_001168339 | 0.2655 | -0.91 |
| TTC39B | NM_001168340 | 0.2655 | -0.91 |
| TTC39B | NM_001168341 | 0.2655 | -0.91 |
| TTC39B | NM_001168342 | 0.2655 | -0.91 |
| RREB1 | NM_001168344 | 0.0685 | -0.8065 |
| NSUN5 | NM_001168347 | -0.653 | -2.114 |
| NSUN5 | NM_001168348 | -0.653 | -2.114 |
| OSMR | NM_001168355 | -0.006 | -0.5335 |
| PLA2G7 | NM_001168357 | -0.4225 | 0.139 |
| TDRD6 | NM_001168359 | -0.054 | -0.792 |
| GALNTL1 | NM_001168368 | -0.1395 | -0.6665 |
| CUL7 | NM_001168370 | -0.3485 | -0.556 |
| KIAA0319 | NM_001168374 | -0.3735 | -0.3995 |
| KIAA0319 | NM_001168375 | -0.3735 | -0.3995 |
| KIAA0319 | NM_001168376 | -0.3735 | -0.3995 |
| KIAA0319 | NM_001168377 | -0.3735 | -0.3995 |
| ZIC4 | NM_001168378 | 0.116 | -0.0595 |
| ZIC4 | NM_001168379 | 0.116 | -0.0595 |
| ALG13 | NM_001168385 | -0.6645 | -1.719 |
| CITED2 | NM_001168388 | 0.203 | 0.9055 |
| CITED2 | NM_001168389 | 0.203 | 0.9055 |
| C5orf41 | NM_001168393 | -0.2265 | -0.516 |
| C5orf41 | NM_001168394 | -0.2265 | -0.516 |
| SLC35A1 | NM_001168398 | -1.678 | -1.95 |
| RIMS1 | NM_001168407 | -0.1775 | -0.633 |
| RIMS1 | NM_001168408 | -0.1775 | -0.633 |
| RIMS1 | NM_001168409 | -0.1775 | -0.633 |
| RIMS1 | NM_001168410 | -0.1775 | -0.633 |
| RIMS1 | NM_001168411 | -0.1775 | -0.633 |
| TSPAN3 | NM_001168412 | -0.7005 | -0.5225 |
| MAP7D2 | NM_001168465 | -0.6265 | -1.088 |
| MAP7D2 | NM_001168466 | -0.6265 | -1.088 |
| MAP7D2 | NM_001168467 | -0.6265 | -1.088 |
| PDZD3 | NM_001168468 | -0.073 | -0.4995 |
| TAF7L | NM_001168474 | -0.556 | -0.1925 |
| CNDP2 | NM_001168499 | -0.3655 | -0.9445 |
| ASB9 | NM_001168530 | 0.199 | -0.333 |
| ASB9 | NM_001168531 | 0.199 | -0.333 |
| TMEM48 | NM_001168551 | -1.147 | -1.555 |
| IFT46 | NM_001168618 | -1.0425 | -2.1835 |
| CNKSR2 | NM_001168647 | -1.3135 | -2.7785 |
| CNKSR2 | NM_001168648 | -1.3135 | -2.7785 |

| | | | |
|---|---|---|---|
| CNKSR2 | NM_001168649 | -1.3135 | -2.7785 |
| TXLNG | NM_001168683 | -1.8695 | -3.742 |
| TMEM135 | NM_001168724 | -0.9945 | -2.259 |
| AQP8 | NM_001169 | 0.1545 | -0.44 |
| STIM2 | NM_001169117 | -0.5705 | -1.546 |
| STIM2 | NM_001169118 | -0.5705 | -1.546 |
| AFF2 | NM_001169122 | 0.0525 | -0.377 |
| AFF2 | NM_001169123 | 0.0525 | -0.377 |
| AFF2 | NM_001169124 | 0.0525 | -0.377 |
| AFF2 | NM_001169125 | 0.0525 | -0.377 |
| CXorf41 | NM_001169154 | -0.192 | -0.3135 |
| CXorf58 | NM_001169574 | 0.275 | -0.0525 |
| AQP7 | NM_001170 | 0.2705 | 0.1885 |
| C4orf3 | NM_001170330 | -0.366 | -0.53 |
| LANCL3 | NM_001170331 | -0.3095 | -0.303 |
| CDC20B | NM_001170402 | -0.0955 | -0.157 |
| CDK1 | NM_001170406 | -1.2655 | -2.246 |
| CDK1 | NM_001170407 | -1.2655 | -2.246 |
| TJP2 | NM_001170414 | -0.8935 | -1.5035 |
| TJP2 | NM_001170415 | -0.8935 | -1.5035 |
| TJP2 | NM_001170416 | -0.8935 | -1.5035 |
| MOG | NM_001170418 | -0.501 | 0.329 |
| PRSS35 | NM_001170423 | -0.1805 | -0.6685 |
| CDK16 | NM_001170460 | -0.188 | -1.0355 |
| CDK17 | NM_001170464 | -1.3335 | -2.9845 |
| ATAD3A | NM_001170535 | -0.3505 | -1.0495 |
| ATAD3A | NM_001170536 | -0.3505 | -1.0495 |
| PGAM5 | NM_001170543 | -0.229 | -1.028 |
| PGAM5 | NM_001170544 | -0.229 | -1.028 |
| VSIG1 | NM_001170553 | 0.1535 | -0.3755 |
| HHAT | NM_001170564 | -0.2445 | -0.559 |
| FAM46D | NM_001170574 | 0.169 | -0.1685 |
| HHAT | NM_001170580 | -0.2445 | -0.559 |
| PLB1 | NM_001170585 | -0.139 | -0.082 |
| HHAT | NM_001170587 | -0.2445 | -0.559 |
| HHAT | NM_001170588 | -0.2445 | -0.559 |
| AFF2 | NM_001170628 | 0.0525 | -0.377 |
| CHD8 | NM_001170629 | -0.9515 | -2.3895 |
| TJP2 | NM_001170630 | -0.8935 | -1.5035 |
| FUS | NM_001170634 | 0.078 | -0.447 |
| SRGAP2 | NM_001170637 | -0.146 | -0.501 |
| CDK20 | NM_001170639 | -0.936 | -1.6585 |
| CDK20 | NM_001170640 | -0.936 | -1.6585 |

| | | | |
|---|---|---|---|
| LAS1L | NM_001170649 | -0.053 | -0.292 |
| LAS1L | NM_001170650 | -0.053 | -0.292 |
| ZNHIT6 | NM_001170670 | -0.7825 | -2.087 |
| MIB2 | NM_001170686 | -0.306 | -0.029 |
| MIB2 | NM_001170687 | -0.306 | -0.029 |
| MIB2 | NM_001170688 | -0.306 | -0.029 |
| MIB2 | NM_001170689 | -0.306 | -0.029 |
| SCUBE2 | NM_001170690 | -0.1885 | -0.153 |
| CAGE1 | NM_001170692 | 0.414 | 0.325 |
| CAGE1 | NM_001170693 | 0.414 | 0.325 |
| SPATA22 | NM_001170695 | 0.5785 | 0.1205 |
| SPATA22 | NM_001170696 | 0.5785 | 0.1205 |
| SPATA22 | NM_001170697 | 0.5785 | 0.1205 |
| SPATA22 | NM_001170698 | 0.5785 | 0.1205 |
| SPATA22 | NM_001170699 | 0.5785 | 0.1205 |
| DTHD1 | NM_001170700 | 0.155 | 0.0545 |
| MBNL3 | NM_001170701 | 0.1225 | -0.466 |
| MBNL3 | NM_001170702 | 0.1225 | -0.466 |
| MBNL3 | NM_001170703 | 0.1225 | -0.466 |
| MBNL3 | NM_001170704 | 0.1225 | -0.466 |
| BCAR1 | NM_001170714 | -0.474 | -1.64 |
| BCAR1 | NM_001170715 | -0.474 | -1.64 |
| BCAR1 | NM_001170716 | -0.474 | -1.64 |
| BCAR1 | NM_001170717 | -0.474 | -1.64 |
| BCAR1 | NM_001170718 | -0.474 | -1.64 |
| BCAR1 | NM_001170719 | -0.474 | -1.64 |
| BCAR1 | NM_001170720 | -0.474 | -1.64 |
| BCAR1 | NM_001170721 | -0.474 | -1.64 |
| C1orf49 | NM_001170722 | -0.119 | -0.0755 |
| C1orf49 | NM_001170723 | -0.119 | -0.0755 |
| C1orf49 | NM_001170724 | -0.119 | -0.0755 |
| GPR137 | NM_001170726 | 0.158 | -0.7635 |
| IQSEC3 | NM_001170738 | 0.5215 | 0.376 |
| PIN4 | NM_001170747 | -0.824 | -1.142 |
| C1orf127 | NM_001170754 | 0.1505 | -0.254 |
| FAM122B | NM_001170756 | -0.416 | -1.2235 |
| FAM122B | NM_001170757 | -0.416 | -1.2235 |
| FAM122C | NM_001170779 | -0.3205 | -0.2315 |
| FAM122C | NM_001170780 | -0.3205 | -0.2315 |
| FAM122C | NM_001170781 | -0.3205 | -0.2315 |
| FAM122C | NM_001170782 | -0.3205 | -0.2315 |
| FAM122C | NM_001170783 | -0.3205 | -0.2315 |
| FAM122C | NM_001170784 | -0.3205 | -0.2315 |

| | | | |
|---|---|---|---|
| FAM82A1 | NM_001170791 | -0.477 | -1.316 |
| FAM82A1 | NM_001170792 | -0.477 | -1.316 |
| FAM82A1 | NM_001170793 | -0.477 | -1.316 |
| BACH2 | NM_001170794 | -0.419 | -0.6775 |
| C2orf28 | NM_001170795 | -0.192 | -0.8825 |
| ZFAND1 | NM_001170796 | -1.4555 | -1.211 |
| ZFAND1 | NM_001170797 | -1.4555 | -1.211 |
| LARP4 | NM_001170803 | -0.0595 | -1.7255 |
| LARP4 | NM_001170804 | -0.0595 | -1.7255 |
| FHL5 | NM_001170807 | 0.146 | -0.066 |
| LARP4 | NM_001170808 | -0.0595 | -1.7255 |
| GPR137 | NM_001170880 | 0.158 | -0.7635 |
| GPR137 | NM_001170881 | 0.158 | -0.7635 |
| ZNF736 | NM_001170905 | 0.1235 | -0.378 |
| FOXO4 | NM_001170931 | -0.0255 | -0.5755 |
| FUS | NM_001170937 | 0.078 | -0.447 |
| ISYNA1 | NM_001170938 | 0.3265 | -0.31 |
| ISYNA1 | NM_001170939 | 0.3265 | -0.31 |
| SMG6 | NM_001170957 | -0.1935 | -0.4215 |
| ABCC6 | NM_001171 | -0.2955 | -0.488 |
| MUM1L1 | NM_001171020 | 0.2975 | -0.1205 |
| PTPRD | NM_001171025 | -0.291 | -1.5925 |
| SMPD4 | NM_001171083 | -0.4145 | -1.509 |
| CLCN2 | NM_001171087 | -0.2475 | -0.5305 |
| CLCN2 | NM_001171088 | -0.2475 | -0.5305 |
| CLCN2 | NM_001171089 | -0.2475 | -0.5305 |
| UBXN6 | NM_001171091 | 0.2825 | -0.2915 |
| CLDN2 | NM_001171092 | -0.2865 | -0.2975 |
| CLDN2 | NM_001171095 | -0.2865 | -0.2975 |
| FAM133A | NM_001171109 | 0.0405 | -0.0425 |
| FAM133A | NM_001171110 | 0.0405 | -0.0425 |
| FAM133A | NM_001171111 | 0.0405 | -0.0425 |
| FAM3A | NM_001171132 | -0.1035 | -0.785 |
| FAM3A | NM_001171133 | -0.1035 | -0.785 |
| FAM3A | NM_001171134 | -0.1035 | -0.785 |
| ZBED1 | NM_001171135 | -0.09 | -1.0735 |
| ZBED1 | NM_001171136 | -0.09 | -1.0735 |
| STRBP | NM_001171137 | -0.697 | -1.6225 |
| ZMYM3 | NM_001171162 | -0.313 | -0.944 |
| ZMYM3 | NM_001171163 | -0.313 | -0.944 |
| CAMTA2 | NM_001171166 | -0.282 | -0.4655 |
| CAMTA2 | NM_001171167 | -0.282 | -0.4655 |
| CAMTA2 | NM_001171168 | -0.282 | -0.4655 |

| | | | |
|---|---|---|---|
| SLC25A21 | NM_001171170 | -0.031 | -0.247 |
| CX3CR1 | NM_001171171 | -0.0255 | -0.0575 |
| CX3CR1 | NM_001171172 | -0.0255 | -0.0575 |
| CX3CR1 | NM_001171174 | -0.0255 | -0.0575 |
| CENPL | NM_001171182 | -1.4525 | -3.2925 |
| DRP2 | NM_001171184 | 0.095 | -0.083 |
| FAM9A | NM_001171186 | -0.1935 | -0.031 |
| UBAP1 | NM_001171201 | -0.7235 | -2.371 |
| UBAP1 | NM_001171202 | -0.7235 | -2.371 |
| UBAP1 | NM_001171203 | -0.7235 | -2.371 |
| UBAP1 | NM_001171204 | -0.7235 | -2.371 |
| C17orf53 | NM_001171251 | -0.793 | -2.348 |
| MCFD2 | NM_001171506 | -0.6425 | -1.091 |
| MCFD2 | NM_001171507 | -0.6425 | -1.091 |
| MCFD2 | NM_001171508 | -0.6425 | -1.091 |
| MCFD2 | NM_001171509 | -0.6425 | -1.091 |
| MCFD2 | NM_001171510 | -0.6425 | -1.091 |
| MCFD2 | NM_001171511 | -0.6425 | -1.091 |
| ITM2A | NM_001171581 | 0.0705 | 0.0155 |
| PREPL | NM_001171603 | -0.8635 | -0.928 |
| PREPL | NM_001171606 | -0.8635 | -0.928 |
| LDB3 | NM_001171610 | -0.218 | -0.4 |
| LDB3 | NM_001171611 | -0.218 | -0.4 |
| PREPL | NM_001171613 | -0.8635 | -0.928 |
| PREPL | NM_001171617 | -0.8635 | -0.928 |
| VEGFA | NM_001171622 | -0.674 | -0.6745 |
| VEGFA | NM_001171623 | -0.674 | -0.6745 |
| VEGFA | NM_001171624 | -0.674 | -0.6745 |
| VEGFA | NM_001171625 | -0.674 | -0.6745 |
| VEGFA | NM_001171626 | -0.674 | -0.6745 |
| VEGFA | NM_001171627 | -0.674 | -0.6745 |
| VEGFA | NM_001171628 | -0.674 | -0.6745 |
| VEGFA | NM_001171629 | -0.674 | -0.6745 |
| VEGFA | NM_001171630 | -0.674 | -0.6745 |
| CCDC148 | NM_001171637 | 0.315 | 0.084 |
| MYO3B | NM_001171642 | -0.0665 | 0.1395 |
| ZEB2 | NM_001171653 | -1.3735 | -2.6525 |
| KLHL5 | NM_001171654 | -0.8775 | -1.448 |
| CYB5R3 | NM_001171660 | 0.0845 | -0.5685 |
| CYB5R3 | NM_001171661 | 0.0845 | -0.5685 |
| AMMECR1 | NM_001171689 | -1.402 | -2.528 |
| FBXL2 | NM_001171713 | -0.983 | -1.5135 |
| C3orf52 | NM_001171747 | -0.2535 | -0.4715 |

| | | | |
|---|---|---|---|
| C8orf83 | NM_001171795 | -0.9265 | -1.4765 |
| C8orf83 | NM_001171796 | -0.9265 | -1.4765 |
| C8orf83 | NM_001171797 | -0.9265 | -1.4765 |
| C8orf83 | NM_001171798 | -0.9265 | -1.4765 |
| C8orf83 | NM_001171799 | -0.9265 | -1.4765 |
| PAPD7 | NM_001171805 | -0.591 | -1.569 |
| PAPD7 | NM_001171806 | -0.591 | -1.569 |
| GBA | NM_001171811 | -0.6095 | -1.2035 |
| GBA | NM_001171812 | -0.6095 | -1.2035 |
| OAT | NM_001171814 | -0.907 | -0.0905 |
| RNF166 | NM_001171815 | -0.7085 | -0.3725 |
| RNF166 | NM_001171816 | -0.7085 | -0.3725 |
| PPARD | NM_001171818 | -0.119 | -0.264 |
| PPARD | NM_001171819 | -0.119 | -0.264 |
| PPARD | NM_001171820 | -0.119 | -0.264 |
| DEFB121 | NM_001171832 | -0.298 | 0.1585 |
| TUBAL3 | NM_001171864 | -0.028 | -0.454 |
| UGT3A1 | NM_001171873 | 0.2045 | -0.8015 |
| MCF2 | NM_001171876 | 0.004 | -0.1765 |
| MCF2 | NM_001171877 | 0.004 | -0.1765 |
| MCF2 | NM_001171878 | 0.004 | -0.1765 |
| MCF2 | NM_001171879 | 0.004 | -0.1765 |
| FGFR1OP2 | NM_001171887 | -1.003 | -1.799 |
| FGFR1OP2 | NM_001171888 | -1.003 | -1.799 |
| MEF2A | NM_001171894 | -0.93 | -2.3655 |
| VPRBP | NM_001171904 | -1.292 | -3.776 |
| NUDT16 | NM_001171905 | -1.109 | -1.566 |
| NUDT16 | NM_001171906 | -1.109 | -1.566 |
| CDH23 | NM_001171930 | 0.637 | 0.184 |
| CDH23 | NM_001171931 | 0.637 | 0.184 |
| CDH23 | NM_001171932 | 0.637 | 0.184 |
| CDH23 | NM_001171933 | 0.637 | 0.184 |
| CDH23 | NM_001171934 | 0.637 | 0.184 |
| CDH23 | NM_001171935 | 0.637 | 0.184 |
| CDH23 | NM_001171936 | 0.637 | 0.184 |
| FUZ | NM_001171937 | -0.147 | -0.6085 |
| FNDC5 | NM_001171940 | -0.1825 | -0.6595 |
| FNDC5 | NM_001171941 | -0.1825 | -0.6595 |
| GLRA2 | NM_001171942 | -0.0015 | -0.655 |
| SUN1 | NM_001171944 | -0.5265 | -1.293 |
| SUN1 | NM_001171945 | -0.5265 | -1.293 |
| SUN1 | NM_001171946 | -0.5265 | -1.293 |
| C19orf50 | NM_001171948 | -0.479 | -1.207 |

| | | | |
|---|---|---|---|
| C19orf50 | NM_001171949 | -0.479 | -1.207 |
| PRSS37 | NM_001171951 | -0.193 | -0.1805 |
| CDHR5 | NM_001171968 | -0.0115 | -0.41 |
| CDHR1 | NM_001171971 | -0.003 | -0.004 |
| AGA | NM_001171988 | -0.795 | -0.6705 |
| HCCS | NM_001171991 | -0.662 | -2.824 |
| C6orf223 | NM_001171992 | -0.2375 | -0.5085 |
| ARG2 | NM_001172 | -0.7945 | -1.6415 |
| TBP | NM_001172085 | -1.5125 | -3.825 |
| PEX2 | NM_001172086 | -0.915 | -1.3515 |
| PEX2 | NM_001172087 | -0.915 | -1.3515 |
| GRAMD1C | NM_001172105 | 0.271 | -0.3075 |
| IL16 | NM_001172128 | -0.113 | -0.7195 |
| SCMH1 | NM_001172218 | -0.3695 | -0.8695 |
| SCMH1 | NM_001172219 | -0.3695 | -0.8695 |
| SCMH1 | NM_001172220 | -0.3695 | -0.8695 |
| SCMH1 | NM_001172221 | -0.3695 | -0.8695 |
| SCMH1 | NM_001172222 | -0.3695 | -0.8695 |
| MOB2 | NM_001172223 | -0.4325 | -1.0765 |
| ZNF540 | NM_001172225 | -0.205 | -1.361 |
| ZNF540 | NM_001172226 | -0.205 | -1.361 |
| NIPAL4 | NM_001172292 | 0.0955 | 0.49 |
| MASTL | NM_001172303 | -1.912 | -4.0075 |
| MASTL | NM_001172304 | -1.912 | -4.0075 |
| NEXN | NM_001172309 | -0.2235 | 0.029 |
| PLS1 | NM_001172312 | -1.0865 | -1.9235 |
| PLS3 | NM_001172335 | -0.468 | -0.3215 |
| SFTPC | NM_001172357 | 0.128 | -0.5065 |
| SFTPC | NM_001172410 | 0.128 | -0.5065 |
| VANGL1 | NM_001172411 | -0.4445 | -2.0235 |
| VANGL1 | NM_001172412 | -0.4445 | -2.0235 |
| BAG1 | NM_001172415 | -0.2585 | -1.336 |
| KCNJ13 | NM_001172416 | -0.021 | -0.533 |
| KCNJ13 | NM_001172417 | -0.021 | -0.533 |
| BTBD9 | NM_001172418 | -0.2065 | -0.124 |
| EFHC1 | NM_001172420 | -1.005 | -0.877 |
| LEFTY2 | NM_001172425 | -0.145 | 0.346 |
| KLHL7 | NM_001172428 | 0.4635 | -0.0885 |
| AMPD3 | NM_001172430 | -0.631 | -0.786 |
| AMPD3 | NM_001172431 | -0.631 | -0.786 |
| PHKG2 | NM_001172432 | -0.378 | -1.344 |
| RAB3GAP1 | NM_001172435 | -0.928 | -1.5545 |
| PHKA1 | NM_001172436 | -0.603 | -0.5625 |

| | | | |
|---|---|---|---|
| PEG10 | NM_001172437 | -0.4375 | 0.29 |
| PEG10 | NM_001172438 | -0.4375 | 0.29 |
| ENDOU | NM_001172439 | 0.2245 | 0.3265 |
| ENDOU | NM_001172440 | 0.2245 | 0.3265 |
| RRM2B | NM_001172477 | -1.2105 | -2.8885 |
| RRM2B | NM_001172478 | -1.2105 | -2.8885 |
| SLC6A2 | NM_001172501 | -0.9595 | -0.931 |
| SLC6A2 | NM_001172502 | -0.9595 | -0.931 |
| SLC6A2 | NM_001172504 | -0.9595 | -0.931 |
| SATB2 | NM_001172509 | -0.5765 | -0.742 |
| SATB2 | NM_001172517 | -0.5765 | -0.742 |
| GOLGA3 | NM_001172557 | -0.584 | -1.377 |
| INTS9 | NM_001172562 | -0.9445 | -2.658 |
| MYD88 | NM_001172566 | -0.3095 | -1.104 |
| MYD88 | NM_001172567 | -0.3095 | -1.104 |
| MYD88 | NM_001172568 | -0.3095 | -1.104 |
| MYD88 | NM_001172569 | -0.3095 | -1.104 |
| MCPH1 | NM_001172574 | -0.097 | -1.6855 |
| MCPH1 | NM_001172575 | -0.097 | -1.6855 |
| NEO1 | NM_001172623 | -0.089 | -0.86 |
| NEO1 | NM_001172624 | -0.089 | -0.86 |
| ARHGAP33 | NM_001172630 | -0.0135 | -0.299 |
| TECPR2 | NM_001172631 | 0.061 | -1.837 |
| OLR1 | NM_001172632 | -0.316 | -0.3935 |
| OLR1 | NM_001172633 | -0.316 | -0.3935 |
| ZFP62 | NM_001172638 | -1.4165 | -2.934 |
| CELF1 | NM_001172639 | -0.732 | -1.313 |
| CELF1 | NM_001172640 | -0.732 | -1.313 |
| TK2 | NM_001172643 | -0.4315 | -1.081 |
| TK2 | NM_001172644 | -0.4315 | -1.081 |
| TK2 | NM_001172645 | -0.4315 | -1.081 |
| PLCB4 | NM_001172646 | -0.5485 | -1.781 |
| CELF3 | NM_001172648 | -0.2615 | -0.9645 |
| CELF3 | NM_001172649 | -0.2615 | -0.9645 |
| ZNF701 | NM_001172655 | -0.749 | -1.0985 |
| ZFYVE28 | NM_001172656 | -0.178 | -0.5695 |
| ZFYVE28 | NM_001172657 | -0.178 | -0.5695 |
| ZFYVE28 | NM_001172658 | -0.178 | -0.5695 |
| ZFYVE28 | NM_001172659 | -0.178 | -0.5695 |
| ZFYVE28 | NM_001172660 | -0.178 | -0.5695 |
| RAB40C | NM_001172663 | -0.3865 | -1.1185 |
| RAB40C | NM_001172664 | -0.3865 | -1.1185 |
| RAB40C | NM_001172665 | -0.3865 | -1.1185 |

| | | | |
|---|---|---|---|
| RAB40C | NM_001172666 | -0.3865 | -1.1185 |
| ZNF668 | NM_001172668 | 0.0395 | -0.2525 |
| ZNF668 | NM_001172669 | 0.0395 | -0.2525 |
| ZNF668 | NM_001172670 | 0.0395 | -0.2525 |
| ZNF430 | NM_001172671 | -1.0255 | -1.9455 |
| PNPLA4 | NM_001172672 | -0.3905 | -0.8045 |
| ZNF347 | NM_001172674 | -0.207 | -1.622 |
| ZNF347 | NM_001172675 | -0.207 | -1.622 |
| ZNF641 | NM_001172681 | -1.107 | -2.528 |
| ZNF641 | NM_001172682 | -1.107 | -2.528 |
| CELF6 | NM_001172684 | 0.277 | -0.693 |
| CELF6 | NM_001172685 | 0.277 | -0.693 |
| GGA1 | NM_001172687 | -0.682 | -1.404 |
| GGA1 | NM_001172688 | -0.682 | -1.404 |
| ZNF573 | NM_001172689 | -1.6785 | -1.9385 |
| ZNF573 | NM_001172690 | -1.6785 | -1.9385 |
| ZNF573 | NM_001172691 | -1.6785 | -1.9385 |
| ZNF573 | NM_001172692 | -1.6785 | -1.9385 |
| TSFM | NM_001172695 | 0.118 | -1.6475 |
| TSFM | NM_001172696 | 0.118 | -1.6475 |
| TSFM | NM_001172697 | 0.118 | -1.6475 |
| PPARGC1B | NM_001172698 | -0.3485 | -0.471 |
| PPARGC1B | NM_001172699 | -0.3485 | -0.471 |
| SHROOM1 | NM_001172700 | -0.0935 | -0.1175 |
| SLC38A6 | NM_001172702 | -0.9925 | -1.0935 |
| GGA3 | NM_001172703 | -0.9555 | -2.739 |
| GGA3 | NM_001172704 | -0.9555 | -2.739 |
| EIF4G2 | NM_001172705 | -0.829 | -0.506 |
| SP3 | NM_001172712 | -0.703 | -2.25 |
| GOLGA4 | NM_001172713 | 0.349 | -1.046 |
| RAI2 | NM_001172732 | -0.295 | -0.8365 |
| PPP1R16B | NM_001172735 | -0.1695 | -0.905 |
| RAI2 | NM_001172739 | -0.295 | -0.8365 |
| RAI2 | NM_001172743 | -0.295 | -0.8365 |
| SEC23B | NM_001172745 | -0.6955 | -1.268 |
| SEC23B | NM_001172746 | -0.6955 | -1.268 |
| FOXP2 | NM_001172766 | 0.8085 | -0.3895 |
| FOXP2 | NM_001172767 | 0.8085 | -0.3895 |
| ZNF548 | NM_001172773 | -1.348 | -2.2395 |
| DPY19L3 | NM_001172774 | 1.985 | 2.916 |
| LRRC34 | NM_001172779 | -1.203 | -0.87 |
| LRRC34 | NM_001172780 | -1.203 | -0.87 |
| DNAI2 | NM_001172810 | 0.384 | 0.2545 |

| | | | |
|---|---|---|---|
| SLC30A8 | NM_001172811 | -0.0215 | 0.104 |
| SLC30A8 | NM_001172813 | -0.0215 | 0.104 |
| SLC30A8 | NM_001172814 | -0.0215 | 0.104 |
| SLC30A8 | NM_001172815 | -0.0215 | 0.104 |
| PGM1 | NM_001172818 | 0.017 | -0.4035 |
| PGM1 | NM_001172819 | 0.017 | -0.4035 |
| YTHDF2 | NM_001172828 | -1.191 | -3.014 |
| ZNF300 | NM_001172831 | -1.809 | -2.724 |
| ZNF300 | NM_001172832 | -1.809 | -2.724 |
| CAV1 | NM_001172895 | 0.087 | -0.168 |
| CAV1 | NM_001172896 | 0.087 | -0.168 |
| CAV1 | NM_001172897 | 0.087 | -0.168 |
| ARHGAP5 | NM_001173 | -0.89 | -2.686 |
| YTHDF2 | NM_001173128 | -1.191 | -3.014 |
| LOXHD1 | NM_001173129 | 0.357 | 0.1095 |
| HAVCR1 | NM_001173393 | -0.21 | -0.3655 |
| OBSL1 | NM_001173408 | -0.39 | -0.521 |
| OBSL1 | NM_001173431 | -0.39 | -0.521 |
| TFCP2 | NM_001173452 | -1.779 | -1.78 |
| TFCP2 | NM_001173453 | -1.779 | -1.78 |
| PDHA1 | NM_001173454 | -0.099 | -0.538 |
| PDHA1 | NM_001173455 | -0.099 | -0.538 |
| PDHA1 | NM_001173456 | -0.099 | -0.538 |
| SHISA6 | NM_001173461 | 0.3635 | 0.0285 |
| SHISA6 | NM_001173462 | 0.3635 | 0.0285 |
| KIF21A | NM_001173463 | -0.5345 | -2.2535 |
| KIF21A | NM_001173464 | -0.5345 | -2.2535 |
| KIF21A | NM_001173465 | -0.5345 | -2.2535 |
| SP7 | NM_001173467 | -0.002 | -0.1105 |
| PDHB | NM_001173468 | -1.0295 | -2.649 |
| SPEG | NM_001173476 | -0.5125 | -1.422 |
| ARHGEF9 | NM_001173479 | 0.1965 | -0.1395 |
| ARHGEF9 | NM_001173480 | 0.1965 | -0.1395 |
| CRBN | NM_001173482 | -1.2105 | -2.0545 |
| NEBL | NM_001173484 | 0.1015 | 0.1055 |
| NKRF | NM_001173487 | -1.094 | -2.1835 |
| NKRF | NM_001173488 | -1.094 | -2.1835 |
| MAN2B1 | NM_001173498 | 0.117 | -0.2625 |
| MRRF | NM_001173512 | -0.8035 | -1.159 |
| TXNRD3 | NM_001173513 | -0.673 | -1.183 |
| TYROBP | NM_001173514 | 0.058 | -0.44 |
| TYROBP | NM_001173515 | 0.058 | -0.44 |
| MAP7D3 | NM_001173516 | -0.876 | -1.4865 |

| | | | |
|---|---|---|---|
| MAP7D3 | NM_001173517 | -0.876 | -1.4865 |
| PCDH7 | NM_001173523 | -0.437 | -0.099 |
| TROVE2 | NM_001173524 | -0.8535 | -1.218 |
| TROVE2 | NM_001173525 | -0.8535 | -1.218 |
| POU5F1 | NM_001173531 | -0.0935 | 0.0605 |
| DGCR2 | NM_001173533 | -0.5775 | -1.2855 |
| DGCR2 | NM_001173534 | -0.5775 | -1.2855 |
| BANP | NM_001173539 | -0.715 | -1.8985 |
| BANP | NM_001173540 | -0.715 | -1.8985 |
| BANP | NM_001173541 | -0.715 | -1.8985 |
| BANP | NM_001173542 | -0.715 | -1.8985 |
| BANP | NM_001173543 | -0.715 | -1.8985 |
| TMPRSS4 | NM_001173551 | -0.072 | -0.4105 |
| TMPRSS4 | NM_001173552 | -0.072 | -0.4105 |
| TCTN1 | NM_001173975 | -0.269 | -0.5095 |
| TCTN1 | NM_001173976 | -0.269 | -0.5095 |
| CHST11 | NM_001173982 | -0.643 | -1.778 |
| BRD7 | NM_001173984 | -0.96 | -1.752 |
| C9orf86 | NM_001173988 | -0.3275 | -1.032 |
| C9orf86 | NM_001173989 | -0.3275 | -1.032 |
| CACNA2D2 | NM_001174051 | -0.3505 | -0.4495 |
| OSBPL10 | NM_001174060 | 0.0155 | -1.174 |
| SMG7 | NM_001174061 | -0.2385 | -1.0785 |
| FGFR1 | NM_001174063 | -0.0765 | -1.193 |
| FGFR1 | NM_001174064 | -0.0765 | -1.193 |
| FGFR1 | NM_001174065 | -0.0765 | -1.193 |
| FGFR1 | NM_001174066 | -0.0765 | -1.193 |
| FGFR1 | NM_001174067 | -0.0765 | -1.193 |
| SYTL4 | NM_001174068 | 0.148 | -1.633 |
| LMX1A | NM_001174069 | 0.307 | -0.0985 |
| FAM53A | NM_001174070 | -1.0635 | -1.7765 |
| SERINC5 | NM_001174071 | -0.4275 | -0.101 |
| SERINC5 | NM_001174072 | -0.4275 | -0.101 |
| CACNA1A | NM_001174080 | 0.246 | -0.356 |
| NLRP2 | NM_001174081 | 0.1685 | -0.2315 |
| NLRP2 | NM_001174082 | 0.1685 | -0.2315 |
| NLRP2 | NM_001174083 | 0.1685 | -0.2315 |
| POLL | NM_001174084 | -0.2525 | -0.865 |
| POLL | NM_001174085 | -0.2525 | -0.865 |
| NCOA3 | NM_001174087 | -1.302 | -1.833 |
| NCOA3 | NM_001174088 | -1.302 | -1.833 |
| SLC4A11 | NM_001174089 | 0.1205 | -0.5975 |
| SLC4A11 | NM_001174090 | 0.1205 | -0.5975 |

| | | | |
|---|---|---|---|
| TMEM185A | NM_001174092 | 0.696 | -0.1825 |
| ZEB1 | NM_001174093 | 0.001 | -1.4665 |
| ZEB1 | NM_001174094 | 0.001 | -1.4665 |
| ZEB1 | NM_001174095 | 0.001 | -1.4665 |
| ZEB1 | NM_001174096 | 0.001 | -1.4665 |
| LDHB | NM_001174097 | -0.503 | -0.293 |
| SLC29A3 | NM_001174098 | -0.593 | -1.936 |
| SGK494 | NM_001174103 | -1.0105 | -0.787 |
| DMXL2 | NM_001174116 | -0.469 | -1.567 |
| DMXL2 | NM_001174117 | -0.469 | -1.567 |
| MEX3D | NM_001174118 | -0.7905 | -2.4225 |
| ZFYVE27 | NM_001174119 | -0.2705 | -1.3395 |
| ZFYVE27 | NM_001174120 | -0.2705 | -1.3395 |
| ZFYVE27 | NM_001174121 | -0.2705 | -1.3395 |
| ZFYVE27 | NM_001174122 | -0.2705 | -1.3395 |
| GOLGA7 | NM_001174124 | -1.293 | -1.831 |
| SLC11A2 | NM_001174125 | -0.876 | -1.3705 |
| SLC11A2 | NM_001174126 | -0.876 | -1.3705 |
| SLC11A2 | NM_001174127 | -0.876 | -1.3705 |
| SLC11A2 | NM_001174128 | -0.876 | -1.3705 |
| SLC11A2 | NM_001174129 | -0.876 | -1.3705 |
| SLC11A2 | NM_001174130 | -0.876 | -1.3705 |
| CCK | NM_001174138 | 0.2685 | -0.1495 |
| LMX1B | NM_001174146 | -0.278 | -0.1 |
| LMX1B | NM_001174147 | -0.278 | -0.1 |
| ARL13B | NM_001174150 | -1.172 | -2.9525 |
| ARL13B | NM_001174151 | -1.172 | -2.9525 |
| RABEPK | NM_001174152 | -0.829 | -1.7325 |
| RABEPK | NM_001174153 | -0.829 | -1.7325 |
| ZFAT | NM_001174157 | -0.164 | -0.7475 |
| ZFAT | NM_001174158 | -0.164 | -0.7475 |
| SH2D4A | NM_001174159 | -0.1745 | -0.106 |
| SH2D4A | NM_001174160 | -0.1745 | -0.106 |
| PRRT4 | NM_001174164 | -0.823 | -0.802 |
| SLC16A6 | NM_001174166 | -1.038 | -1.1295 |
| SYK | NM_001174167 | 0.0805 | -0.3045 |
| SYK | NM_001174168 | 0.0805 | -0.3045 |
| ARHGDIB | NM_001175 | -0.3645 | -0.435 |
| ARHGDIG | NM_001176 | 0.2375 | -0.613 |
| ARL1 | NM_001177 | -1.0635 | -2.377 |
| PLSCR4 | NM_001177304 | -0.1515 | -0.1005 |
| PAM | NM_001177306 | -0.739 | -0.8865 |
| TRPM6 | NM_001177310 | 0.2155 | -0.3425 |

| | | | |
|---|---|---|---|
| TRPM6 | NM_001177311 | 0.2155 | -0.3425 |
| RRAS2 | NM_001177314 | -0.9475 | -2.199 |
| RRAS2 | NM_001177315 | -0.9475 | -2.199 |
| GPR137 | NM_001177358 | 0.158 | -0.7635 |
| CPEB2 | NM_001177381 | 0.569 | -0.0055 |
| CPEB2 | NM_001177382 | 0.569 | -0.0055 |
| CPEB2 | NM_001177383 | 0.569 | -0.0055 |
| CPEB2 | NM_001177384 | 0.569 | -0.0055 |
| ATXN7 | NM_001177387 | -1.679 | -2.367 |
| LGALS3 | NM_001177388 | 0.069 | -0.728 |
| RPS27A | NM_001177413 | -0.1275 | -0.04 |
| TRPV4 | NM_001177428 | 0.2725 | -0.2715 |
| TRPV4 | NM_001177431 | 0.2725 | -0.2715 |
| TRPV4 | NM_001177433 | 0.2725 | -0.2715 |
| MAMLD1 | NM_001177465 | -0.061 | -0.4465 |
| MAMLD1 | NM_001177466 | -0.061 | -0.4465 |
| MOSPD2 | NM_001177475 | -1.6895 | -2.4475 |
| HDX | NM_001177478 | -0.3865 | -1.7905 |
| HDX | NM_001177479 | -0.3865 | -1.7905 |
| C10orf113 | NM_001177483 | -0.2955 | -0.594 |
| AOAH | NM_001177506 | -0.0155 | -0.3145 |
| AOAH | NM_001177507 | -0.0155 | -0.3145 |
| MICA | NM_001177519 | -0.0455 | -0.166 |
| ALPL | NM_001177520 | 0.0015 | -0.5595 |
| SIGLEC6 | NM_001177547 | 0.088 | 0.198 |
| SIGLEC6 | NM_001177548 | 0.088 | 0.198 |
| SIGLEC6 | NM_001177549 | 0.088 | 0.198 |
| PPP2R1B | NM_001177562 | -0.8865 | -1.919 |
| PPP2R1B | NM_001177563 | -0.8865 | -1.919 |
| PPP2R2A | NM_001177591 | -1.072 | -2.6155 |
| THPO | NM_001177597 | -0.1595 | -0.2275 |
| THPO | NM_001177598 | -0.1595 | -0.2275 |
| SUCLG2 | NM_001177599 | -0.7795 | -0.8435 |
| CD33 | NM_001177608 | 0.26 | -0.237 |
| CINP | NM_001177611 | 0.2125 | -0.6645 |
| CINP | NM_001177612 | 0.2125 | -0.6645 |
| DAG1 | NM_001177634 | -0.4615 | -1.0555 |
| DAG1 | NM_001177635 | -0.4615 | -1.0555 |
| DAG1 | NM_001177636 | -0.4615 | -1.0555 |
| DAG1 | NM_001177637 | -0.4615 | -1.0555 |
| DAG1 | NM_001177638 | -0.4615 | -1.0555 |
| DAG1 | NM_001177639 | -0.4615 | -1.0555 |
| DAG1 | NM_001177640 | -0.4615 | -1.0555 |

| | | | |
|---|---|---|---|
| DAG1 | NM_001177641 | -0.4615 | -1.0555 |
| DAG1 | NM_001177642 | -0.4615 | -1.0555 |
| DAG1 | NM_001177643 | -0.4615 | -1.0555 |
| DAG1 | NM_001177644 | -0.4615 | -1.0555 |
| SLC9A6 | NM_001177651 | -1.075 | -2.2795 |
| HABP2 | NM_001177660 | 0.177 | 0.3795 |
| RIF1 | NM_001177663 | -0.0155 | -2.036 |
| RIF1 | NM_001177664 | -0.0155 | -2.036 |
| RIF1 | NM_001177665 | -0.0155 | -2.036 |
| PNPLA5 | NM_001177675 | -0.0895 | -0.7535 |
| GPR68 | NM_001177676 | 0.087 | -0.4795 |
| RGNEF | NM_001177693 | -0.874 | -1.9285 |
| RBMS3 | NM_001177711 | -0.315 | -0.709 |
| RBMS3 | NM_001177712 | -0.315 | -0.709 |
| KLF11 | NM_001177716 | -0.193 | -0.739 |
| KLF11 | NM_001177718 | -0.193 | -0.739 |
| ADIPOQ | NM_001177800 | 0.2835 | -0.412 |
| RANGRF | NM_001177801 | -0.052 | -0.961 |
| RANGRF | NM_001177802 | -0.052 | -0.961 |
| SRPR | NM_001177842 | -0.577 | -1.9565 |
| PCSK1 | NM_001177875 | 0.0825 | -0.6175 |
| PCSK1 | NM_001177876 | 0.0825 | -0.6175 |
| SYT12 | NM_001177880 | 0.1 | -0.4725 |
| VIT | NM_001177969 | 0.118 | 0.1785 |
| VIT | NM_001177970 | 0.118 | 0.1785 |
| VIT | NM_001177971 | 0.118 | 0.1785 |
| VIT | NM_001177972 | 0.118 | 0.1785 |
| SCN8A | NM_001177984 | -0.232 | -0.931 |
| FAM109A | NM_001177996 | -0.196 | -1.0595 |
| FAM109A | NM_001177997 | -0.196 | -1.0595 |
| SLC34A2 | NM_001177998 | 0.213 | -0.1625 |
| SLC34A2 | NM_001177999 | 0.213 | -0.1625 |
| ARNTL | NM_001178 | -0.2265 | -1.008 |
| PIKFYVE | NM_001178000 | -0.0555 | -0.526 |
| ATP8B3 | NM_001178002 | -0.1395 | -0.1355 |
| WDR66 | NM_001178003 | 0.2105 | -0.496 |
| PCCA | NM_001178004 | -0.8495 | -1.0835 |
| BHMT2 | NM_001178005 | 0.2245 | -0.583 |
| BBS12 | NM_001178007 | -0.312 | -1.1115 |
| CBS | NM_001178008 | -0.478 | -0.8445 |
| CBS | NM_001178009 | -0.478 | -0.8445 |
| CDC45 | NM_001178010 | -0.455 | -1.2005 |
| CDC45 | NM_001178011 | -0.455 | -1.2005 |

| | | | |
|---|---|---|---|
| PCCB | NM_001178014 | 0.146 | -0.722 |
| SLC4A10 | NM_001178015 | 0.189 | 0.0315 |
| SLC4A10 | NM_001178016 | 0.189 | 0.0315 |
| DBI | NM_001178017 | -0.241 | -0.2425 |
| SF1 | NM_001178030 | -0.0435 | -0.7715 |
| SF1 | NM_001178031 | -0.0435 | -0.7715 |
| ZC4H2 | NM_001178032 | -0.3345 | -0.7665 |
| ZC4H2 | NM_001178033 | -0.3345 | -0.7665 |
| DPP10 | NM_001178034 | -0.0165 | -0.1135 |
| C6orf204 | NM_001178035 | -0.733 | -2.4905 |
| DPP10 | NM_001178036 | -0.0165 | -0.1135 |
| DPP10 | NM_001178037 | -0.0165 | -0.1135 |
| STX3 | NM_001178040 | -0.2125 | -1.331 |
| DBI | NM_001178041 | -0.241 | -0.2425 |
| DBI | NM_001178042 | -0.241 | -0.2425 |
| DBI | NM_001178043 | -0.241 | -0.2425 |
| SLC44A4 | NM_001178044 | -0.0065 | -0.3865 |
| SLC44A4 | NM_001178045 | -0.0065 | -0.3865 |
| TAC3 | NM_001178054 | 0.1885 | -0.1635 |
| ACE | NM_001178057 | -0.234 | -0.243 |
| SEMA6C | NM_001178061 | 0.038 | -0.625 |
| SEMA6C | NM_001178062 | 0.038 | -0.625 |
| C2 | NM_001178063 | -0.241 | 0.3285 |
| NELF | NM_001178064 | -0.0835 | -0.7205 |
| CASR | NM_001178065 | -0.093 | -0.282 |
| ASNS | NM_001178075 | -0.3555 | -0.499 |
| ASNS | NM_001178076 | -0.3555 | -0.499 |
| ASNS | NM_001178077 | -0.3555 | -0.499 |
| STAT6 | NM_001178078 | -0.142 | 0.0855 |
| STAT6 | NM_001178079 | -0.142 | 0.0855 |
| STAT6 | NM_001178080 | -0.142 | 0.0855 |
| STAT6 | NM_001178081 | -0.142 | 0.0855 |
| EXT2 | NM_001178083 | -0.1755 | -1.341 |
| ZFX | NM_001178084 | -0.271 | -0.653 |
| ZFX | NM_001178085 | -0.271 | -0.653 |
| ZFX | NM_001178086 | -0.271 | -0.653 |
| SYNJ2 | NM_001178088 | -0.825 | -1.2495 |
| ZNF454 | NM_001178089 | -0.4255 | -0.699 |
| ZNF454 | NM_001178090 | -0.4255 | -0.699 |
| BCAT1 | NM_001178091 | -0.724 | -0.6965 |
| BCAT1 | NM_001178092 | -0.724 | -0.6965 |
| BCAT1 | NM_001178093 | -0.724 | -0.6965 |
| BCAT1 | NM_001178094 | -0.724 | -0.6965 |

| | | | |
|---|---|---|---|
| ZFX | NM_001178095 | -0.271 | -0.653 |
| F3 | NM_001178096 | -0.706 | -1.129 |
| C12orf74 | NM_001178097 | -0.2655 | -0.393 |
| CD19 | NM_001178098 | -0.148 | -0.193 |
| ZNF182 | NM_001178099 | -0.49 | -1.236 |
| CD8B | NM_001178100 | 0.0235 | 0.1305 |
| ZNF37A | NM_001178101 | -0.73 | -1.6925 |
| LOX | NM_001178102 | 0.059 | -0.4155 |
| ZNF185 | NM_001178106 | 0.0615 | -0.333 |
| ZNF185 | NM_001178107 | 0.0615 | -0.333 |
| ZNF185 | NM_001178108 | 0.0615 | -0.333 |
| ZNF185 | NM_001178109 | 0.0615 | -0.333 |
| ZNF185 | NM_001178110 | 0.0615 | -0.333 |
| SCAMP5 | NM_001178111 | -0.65 | -1.723 |
| SCAMP5 | NM_001178112 | -0.65 | -1.723 |
| ZNF185 | NM_001178113 | 0.0615 | -0.333 |
| ZNF185 | NM_001178115 | 0.0615 | -0.333 |
| ABI1 | NM_001178116 | -0.7035 | -1.5425 |
| ABI1 | NM_001178119 | -0.7035 | -1.5425 |
| ABI1 | NM_001178120 | -0.7035 | -1.5425 |
| ABI1 | NM_001178121 | -0.7035 | -1.5425 |
| ABI1 | NM_001178122 | -0.7035 | -1.5425 |
| ABI1 | NM_001178123 | -0.7035 | -1.5425 |
| ABI1 | NM_001178124 | -0.7035 | -1.5425 |
| ABI1 | NM_001178125 | -0.7035 | -1.5425 |
| IGLL5 | NM_001178126 | 0.017 | -0.0175 |
| SEMA3E | NM_001178129 | -0.013 | 0.157 |
| EGF | NM_001178130 | 0.5775 | 0.32 |
| EGF | NM_001178131 | 0.5775 | 0.32 |
| SUFU | NM_001178133 | 0.273 | -0.1215 |
| CXCL12 | NM_001178134 | -0.2575 | -1.2595 |
| HDHD1 | NM_001178135 | -1.5915 | -2.954 |
| HDHD1 | NM_001178136 | -1.5915 | -2.954 |
| CPEB3 | NM_001178137 | -1.214 | -2.07 |
| TFDP2 | NM_001178138 | -0.717 | -1.247 |
| TFDP2 | NM_001178139 | -0.717 | -1.247 |
| TFDP2 | NM_001178140 | -0.717 | -1.247 |
| TFDP2 | NM_001178141 | -0.717 | -1.247 |
| TFDP2 | NM_001178142 | -0.717 | -1.247 |
| IGSF10 | NM_001178145 | -0.2155 | -0.419 |
| IGSF10 | NM_001178146 | -0.2155 | -0.419 |
| CSF3 | NM_001178147 | 0.093 | -0.3545 |
| ART3 | NM_001179 | -0.047 | -0.3845 |

| | | | |
|---|---|---|---|
| ASGR2 | NM_001181 | 0.198 | -0.1445 |
| ALDH7A1 | NM_001182 | -0.467 | -0.5525 |
| ATP6AP1 | NM_001183 | 0.1285 | -0.3175 |
| ATR | NM_001184 | -1.0255 | -2.898 |
| NPNT | NM_001184690 | -0.2665 | -0.1805 |
| NPNT | NM_001184691 | -0.2665 | -0.1805 |
| NPNT | NM_001184692 | -0.2665 | -0.1805 |
| NPNT | NM_001184693 | -0.2665 | -0.1805 |
| MEPE | NM_001184694 | -0.2565 | -0.5515 |
| MEPE | NM_001184695 | -0.2565 | -0.5515 |
| MEPE | NM_001184696 | -0.2565 | -0.5515 |
| MEPE | NM_001184697 | -0.2565 | -0.5515 |
| MLN | NM_001184698 | 0.1215 | -0.576 |
| UGDH | NM_001184700 | -1.1075 | -1.923 |
| UGDH | NM_001184701 | -1.1075 | -1.923 |
| GYG2 | NM_001184702 | -0.1645 | -0.707 |
| GYG2 | NM_001184703 | -0.1645 | -0.707 |
| GYG2 | NM_001184704 | -0.1645 | -0.707 |
| HADH | NM_001184705 | -0.8285 | -1.3545 |
| SLAMF6 | NM_001184714 | 0.203 | -0.102 |
| SLAMF6 | NM_001184715 | 0.203 | -0.102 |
| SLAMF6 | NM_001184716 | 0.203 | -0.102 |
| TIPARP | NM_001184717 | -0.491 | -0.315 |
| TIPARP | NM_001184718 | -0.491 | -0.315 |
| GYG1 | NM_001184720 | -0.8115 | -1.111 |
| GYG1 | NM_001184721 | -0.8115 | -1.111 |
| GPI | NM_001184722 | 0.324 | 0.001 |
| TM4SF18 | NM_001184723 | 0.1375 | 0.6385 |
| TCN2 | NM_001184726 | -0.148 | -0.5935 |
| GPRASP1 | NM_001184727 | -0.836 | -0.7705 |
| PIGT | NM_001184728 | 0.2525 | -0.284 |
| PIGT | NM_001184729 | 0.2525 | -0.284 |
| PIGT | NM_001184730 | 0.2525 | -0.284 |
| SLC22A8 | NM_001184732 | -0.0395 | -0.388 |
| SLC22A8 | NM_001184733 | -0.0395 | -0.388 |
| ASTN2 | NM_001184734 | 0.029 | -1.152 |
| ASTN2 | NM_001184735 | 0.029 | -1.152 |
| SLC22A8 | NM_001184736 | -0.0395 | -0.388 |
| CTTN | NM_001184740 | -0.321 | -0.5175 |
| FGB | NM_001184741 | -0.06 | -0.553 |
| ZBTB33 | NM_001184742 | -1.261 | -2.149 |
| PGBD1 | NM_001184743 | -1.4745 | -2.4205 |
| PPP1R3F | NM_001184745 | -0.6525 | -1.2665 |

| | | | |
|---|---|---|---|
| PAFAH1B2 | NM_001184746 | -1.1985 | -1.8985 |
| PAFAH1B2 | NM_001184747 | -1.1985 | -1.8985 |
| PAFAH1B2 | NM_001184748 | -1.1985 | -1.8985 |
| CLTA | NM_001184760 | -0.0495 | -0.5025 |
| CLTA | NM_001184761 | -0.0495 | -0.5025 |
| CLTA | NM_001184762 | -0.0495 | -0.5025 |
| UHMK1 | NM_001184763 | -0.505 | -1.07 |
| ODF2L | NM_001184765 | -0.362 | -0.7635 |
| ODF2L | NM_001184766 | -0.362 | -0.7635 |
| BEND2 | NM_001184767 | -0.1025 | -0.1145 |
| ARMCX6 | NM_001184768 | -0.5045 | -1.0455 |
| SEZ6L | NM_001184773 | -0.135 | -0.801 |
| SEZ6L | NM_001184774 | -0.135 | -0.801 |
| SEZ6L | NM_001184775 | -0.135 | -0.801 |
| SEZ6L | NM_001184776 | -0.135 | -0.801 |
| SEZ6L | NM_001184777 | -0.135 | -0.801 |
| NOX5 | NM_001184779 | -0.027 | -0.1445 |
| NOX5 | NM_001184780 | -0.027 | -0.1445 |
| DGCR2 | NM_001184781 | -0.5775 | -1.2855 |
| CXorf57 | NM_001184782 | 0.0195 | -0.1775 |
| VDAC2 | NM_001184783 | -0.0035 | -0.4675 |
| PARD3 | NM_001184785 | -0.282 | -0.83 |
| PARD3 | NM_001184786 | -0.282 | -0.83 |
| PARD3 | NM_001184787 | -0.282 | -0.83 |
| PARD3 | NM_001184788 | -0.282 | -0.83 |
| PARD3 | NM_001184789 | -0.282 | -0.83 |
| PARD3 | NM_001184790 | -0.282 | -0.83 |
| PARD3 | NM_001184791 | -0.282 | -0.83 |
| PARD3 | NM_001184792 | -0.282 | -0.83 |
| PARD3 | NM_001184793 | -0.282 | -0.83 |
| PARD3 | NM_001184794 | -0.282 | -0.83 |
| PEX11B | NM_001184795 | -0.329 | -1.5255 |
| TMLHE | NM_001184797 | -0.5795 | -1.2935 |
| UBR2 | NM_001184801 | -0.696 | -1.729 |
| CD99L2 | NM_001184808 | -0.5455 | -0.894 |
| CEACAM1 | NM_001184813 | 0.085 | -0.347 |
| CEACAM1 | NM_001184815 | 0.085 | -0.347 |
| CEACAM1 | NM_001184816 | 0.085 | -0.347 |
| GNL3L | NM_001184819 | -0.0475 | -0.7615 |
| VDAC2 | NM_001184823 | -0.0035 | -0.4675 |
| PRMT7 | NM_001184824 | -0.21 | -0.9525 |
| PSG1 | NM_001184825 | -0.1935 | -0.236 |
| PSG1 | NM_001184826 | -0.1935 | -0.236 |

| | | | |
|---|---|---|---|
| VSIG4 | NM_001184830 | -0.1545 | 0.0275 |
| VSIG4 | NM_001184831 | -0.1545 | 0.0275 |
| SLC12A1 | NM_001184832 | 0.1415 | 0.0065 |
| GPR64 | NM_001184833 | 0.0395 | -0.498 |
| GPR64 | NM_001184834 | 0.0395 | -0.498 |
| GPR64 | NM_001184835 | 0.0395 | -0.498 |
| GPR64 | NM_001184836 | 0.0395 | -0.498 |
| GPR64 | NM_001184837 | 0.0395 | -0.498 |
| FCRLA | NM_001184866 | -0.2045 | -0.5575 |
| FCRLA | NM_001184867 | -0.2045 | -0.5575 |
| FCRLA | NM_001184870 | -0.2045 | -0.5575 |
| FCRLA | NM_001184871 | -0.2045 | -0.5575 |
| FCRLA | NM_001184872 | -0.2045 | -0.5575 |
| FCRLA | NM_001184873 | -0.2045 | -0.5575 |
| CD84 | NM_001184879 | -0.316 | -0.0675 |
| PCDH19 | NM_001184880 | -1.295 | -2.411 |
| CD84 | NM_001184881 | -0.316 | -0.0675 |
| CD84 | NM_001184882 | -0.316 | -0.0675 |
| PLCB3 | NM_001184883 | 0.049 | -0.422 |
| NIPA2 | NM_001184888 | -1.056 | -2.0195 |
| NIPA2 | NM_001184889 | -1.056 | -2.0195 |
| FLAD1 | NM_001184891 | -0.7995 | -1.8245 |
| FLAD1 | NM_001184892 | -0.7995 | -1.8245 |
| PHF8 | NM_001184896 | -0.405 | -1.2095 |
| PHF8 | NM_001184897 | -0.405 | -1.2095 |
| PHF8 | NM_001184898 | -0.405 | -1.2095 |
| CARD8 | NM_001184900 | -0.39 | -1.377 |
| CARD8 | NM_001184901 | -0.39 | -1.377 |
| CARD8 | NM_001184902 | -0.39 | -1.377 |
| CARD8 | NM_001184903 | -0.39 | -1.377 |
| CARD8 | NM_001184904 | -0.39 | -1.377 |
| FBXL20 | NM_001184906 | -0.9725 | -1.695 |
| PCYT2 | NM_001184917 | -0.4005 | -0.8435 |
| PNMA5 | NM_001184924 | -0.1525 | -0.2185 |
| NRG2 | NM_001184935 | -0.0085 | -0.3145 |
| EPB41L5 | NM_001184937 | -0.123 | -0.4955 |
| EPB41L5 | NM_001184938 | -0.123 | -0.4955 |
| EPB41L5 | NM_001184939 | -0.123 | -0.4955 |
| C9orf25 | NM_001184940 | -0.236 | -0.622 |
| C9orf25 | NM_001184941 | -0.236 | -0.622 |
| C9orf25 | NM_001184942 | -0.236 | -0.622 |
| C9orf25 | NM_001184943 | -0.236 | -0.622 |
| C9orf25 | NM_001184945 | -0.236 | -0.622 |

| | | | |
|---|---|---|---|
| SH3KBP1 | NM_001184960 | -0.134 | -0.166 |
| PEG10 | NM_001184961 | -0.4375 | 0.29 |
| PEG10 | NM_001184962 | -0.4375 | 0.29 |
| WDR44 | NM_001184965 | -0.9015 | -2.158 |
| WDR44 | NM_001184966 | -0.9015 | -2.158 |
| MITF | NM_001184967 | -0.596 | -0.749 |
| MITF | NM_001184968 | -0.596 | -0.749 |
| PACSIN2 | NM_001184970 | -0.074 | -1.3545 |
| PACSIN2 | NM_001184971 | -0.074 | -1.3545 |
| PACSIN3 | NM_001184974 | -0.4965 | -1.181 |
| PACSIN3 | NM_001184975 | -0.4965 | -1.181 |
| NCKAP1L | NM_001184976 | -0.004 | -0.252 |
| UQCC | NM_001184977 | -1.13 | -1.8685 |
| NDUFS5 | NM_001184979 | 0.116 | -0.216 |
| WNK1 | NM_001184985 | -0.809 | -1.332 |
| RNF135 | NM_001184992 | -0.099 | -0.3995 |
| RNF17 | NM_001184993 | 0.1105 | -0.457 |
| RNF32 | NM_001184996 | -0.0975 | 0.048 |
| RNF32 | NM_001184997 | -0.0975 | 0.048 |
| KIAA0430 | NM_001184998 | -0.9565 | -2.1035 |
| KIAA0430 | NM_001184999 | -0.9565 | -2.1035 |
| AZGP1 | NM_001185 | -0.0755 | -0.1025 |
| RNF4 | NM_001185009 | -0.83 | -3.0035 |
| RNF4 | NM_001185010 | -0.83 | -3.0035 |
| NDUFA2 | NM_001185012 | 0.2915 | -0.197 |
| SP110 | NM_001185015 | -0.213 | -0.626 |
| CLDN7 | NM_001185022 | 0.1675 | -0.329 |
| CLDN7 | NM_001185023 | 0.1675 | -0.329 |
| ZDHHC8 | NM_001185024 | 0.1035 | -0.4645 |
| ACOX1 | NM_001185039 | -1.148 | -2.459 |
| ADD2 | NM_001185054 | -0.0145 | -0.5655 |
| ADD2 | NM_001185055 | -0.0145 | -0.5655 |
| CLDN11 | NM_001185056 | 0.3 | -0.482 |
| NOL3 | NM_001185057 | -0.1915 | -0.91 |
| NOL3 | NM_001185058 | -0.1915 | -0.91 |
| ZCCHC6 | NM_001185059 | 0.1915 | -0.011 |
| AQP1 | NM_001185060 | -0.4675 | -0.332 |
| AQP1 | NM_001185061 | -0.4675 | -0.332 |
| AQP1 | NM_001185062 | -0.4675 | -0.332 |
| ZNF75D | NM_001185063 | -1.483 | -1.221 |
| CLDN12 | NM_001185072 | -0.4135 | -1.025 |
| CLDN12 | NM_001185073 | -0.4135 | -1.025 |
| ZCCHC6 | NM_001185074 | 0.1915 | -0.011 |

| | | | |
|---|---|---|---|
| FMR1 | NM_001185075 | -1.4225 | -1.6075 |
| FMR1 | NM_001185076 | -1.4225 | -1.6075 |
| ARHGDIA | NM_001185077 | -0.1785 | -1.075 |
| ARHGDIA | NM_001185078 | -0.1785 | -1.075 |
| CLDN15 | NM_001185080 | -0.387 | -1.104 |
| FMR1 | NM_001185081 | -1.4225 | -1.6075 |
| FMR1 | NM_001185082 | -1.4225 | -1.6075 |
| GRIN1 | NM_001185090 | 0.2565 | 0.227 |
| GRIN1 | NM_001185091 | 0.2565 | 0.227 |
| NIT1 | NM_001185092 | -0.992 | -2.1745 |
| NIT1 | NM_001185093 | -0.992 | -2.1745 |
| NIT1 | NM_001185094 | -0.992 | -2.1745 |
| AIF1L | NM_001185095 | 0.253 | -0.097 |
| AIF1L | NM_001185096 | 0.253 | -0.097 |
| CD22 | NM_001185099 | 0.2145 | 0.0145 |
| CD22 | NM_001185100 | 0.2145 | 0.0145 |
| CD22 | NM_001185101 | 0.2145 | 0.0145 |
| PLCXD2 | NM_001185106 | -1.049 | -2.5005 |
| CD1E | NM_001185107 | 0.0335 | -0.3035 |
| CD1E | NM_001185108 | 0.0335 | -0.3035 |
| CD1E | NM_001185110 | 0.0335 | -0.3035 |
| CD1E | NM_001185112 | 0.0335 | -0.3035 |
| CD1E | NM_001185113 | 0.0335 | -0.3035 |
| CD1E | NM_001185114 | 0.0335 | -0.3035 |
| CD1E | NM_001185115 | 0.0335 | -0.3035 |
| CLDN19 | NM_001185117 | 0.065 | -0.5455 |
| MYNN | NM_001185118 | -1.564 | -3.3895 |
| MYNN | NM_001185119 | -1.564 | -3.3895 |
| VAMP4 | NM_001185127 | -1.2335 | -1.6575 |
| IL24 | NM_001185156 | -0.124 | -0.286 |
| IL24 | NM_001185157 | -0.124 | -0.286 |
| IL24 | NM_001185158 | -0.124 | -0.286 |
| CES3 | NM_001185176 | -0.159 | -0.316 |
| CES3 | NM_001185177 | -0.159 | -0.316 |
| VAMP7 | NM_001185183 | -0.849 | -0.845 |
| BACH1 | NM_001186 | -0.896 | -2.286 |
| BAGE | NM_001187 | -0.8285 | -0.019 |
| BAK1 | NM_001188 | -0.2335 | -1.0645 |
| NKX3-2 | NM_001189 | 0.105 | -0.279 |
| BCAT2 | NM_001190 | -0.236 | -0.792 |
| CES5A | NM_001190158 | -0.0455 | -0.294 |
| CCDC17 | NM_001190182 | 0.0095 | -0.0055 |
| CES4A | NM_001190201 | -0.0045 | -0.428 |

| | | | |
|---|---|---|---|
| CES4A | NM_001190202 | -0.0045 | -0.428 |
| PPIP5K1 | NM_001190214 | -0.2065 | -0.879 |
| DNAJC19 | NM_001190233 | -0.699 | -1.694 |
| IFT80 | NM_001190241 | -0.642 | -1.0205 |
| IFT80 | NM_001190242 | -0.642 | -1.0205 |
| ZNF630 | NM_001190255 | -0.4205 | -1.2405 |
| GCET2 | NM_001190259 | 0.21 | 0.0825 |
| GCET2 | NM_001190260 | 0.21 | 0.0825 |
| C1D | NM_001190263 | -0.698 | -1.6015 |
| C1D | NM_001190265 | -0.698 | -1.6015 |
| ATG16L1 | NM_001190266 | -0.4715 | -2.0815 |
| ATG16L1 | NM_001190267 | -0.4715 | -2.0815 |
| FBXO11 | NM_001190274 | -1.3205 | -2.855 |
| IP6K2 | NM_001190316 | 0.385 | -0.656 |
| IP6K2 | NM_001190317 | 0.385 | -0.656 |
| DGCR8 | NM_001190326 | -0.218 | -0.0705 |
| YJEFN3 | NM_001190328 | -0.031 | -0.746 |
| ATP5G3 | NM_001190329 | 0.0935 | -0.4055 |
| DMKN | NM_001190347 | 0.229 | -0.1485 |
| DMKN | NM_001190348 | 0.229 | -0.1485 |
| DMKN | NM_001190349 | 0.229 | -0.1485 |
| GNE | NM_001190383 | -1.128 | -2.1295 |
| GNE | NM_001190384 | -1.128 | -2.1295 |
| GNE | NM_001190388 | -1.128 | -2.1295 |
| RPH3AL | NM_001190411 | -0.2765 | -0.5205 |
| RPH3AL | NM_001190412 | -0.2765 | -0.5205 |
| RPH3AL | NM_001190413 | -0.2765 | -0.5205 |
| TAF6 | NM_001190415 | -0.073 | -0.646 |
| ZNF674 | NM_001190417 | -0.7745 | -1.489 |
| NCOR1 | NM_001190438 | -0.6605 | -1.2695 |
| NCOR1 | NM_001190440 | -0.6605 | -1.2695 |
| CAST | NM_001190442 | -0.609 | -1.1605 |
| PPP2R3A | NM_001190447 | -0.734 | -1.83 |
| CORO2B | NM_001190456 | 0.014 | -0.493 |
| CORO2B | NM_001190457 | 0.014 | -0.493 |
| DOCK8 | NM_001190458 | 0.0285 | 0.1265 |
| GDNF | NM_001190468 | -0.214 | -0.568 |
| GDNF | NM_001190469 | -0.214 | -0.568 |
| CLSPN | NM_001190481 | -0.8865 | -1.5745 |
| PCSK5 | NM_001190482 | -0.7435 | -1.0985 |
| CYP3A5 | NM_001190484 | -0.131 | 0.34 |
| ALS2CL | NM_001190707 | 0.067 | -0.5855 |
| COL11A1 | NM_001190709 | 0.313 | -0.504 |

| | | | |
|---|---|---|---|
| DNM2 | NM_001190716 | -0.1715 | -0.9265 |
| IKBKB | NM_001190720 | -0.9915 | -2.568 |
| LRSAM1 | NM_001190723 | -0.2465 | -0.847 |
| ATAT1 | NM_001190724 | -0.7065 | -1.499 |
| RERG | NM_001190726 | -1.388 | -0.306 |
| RALGPS1 | NM_001190728 | -0.989 | -0.613 |
| RALGPS1 | NM_001190729 | -0.989 | -0.613 |
| RALGPS1 | NM_001190730 | -0.989 | -0.613 |
| NFIB | NM_001190737 | -0.775 | -1.879 |
| NFIB | NM_001190738 | -0.775 | -1.879 |
| NCF2 | NM_001190789 | -0.082 | -0.836 |
| ZNF317 | NM_001190791 | -0.5875 | -1.525 |
| NCF2 | NM_001190794 | -0.082 | -0.836 |
| NCK1 | NM_001190796 | -1.093 | -2.476 |
| PLK4 | NM_001190799 | -1.158 | -3.2425 |
| PLK4 | NM_001190801 | -1.158 | -3.2425 |
| CYB5A | NM_001190807 | -0.0135 | -0.7865 |
| DHX35 | NM_001190809 | -0.844 | -1.921 |
| DAZL | NM_001190811 | 0.037 | -0.12 |
| SMAD7 | NM_001190821 | -0.3085 | -0.4 |
| SMAD7 | NM_001190822 | -0.3085 | -0.4 |
| SMAD7 | NM_001190823 | -0.3085 | -0.4 |
| MAFIP | NM_001190825 | -1.091 | -1.2535 |
| C20orf177 | NM_001190826 | -0.304 | -0.4915 |
| C20orf177 | NM_001190827 | -0.304 | -0.4915 |
| FCGR2B | NM_001190828 | -0.2175 | -0.407 |
| ZNF726 | NM_001190829 | -0.2105 | -0.393 |
| MGP | NM_001190839 | -0.6225 | 0.2675 |
| CNOT4 | NM_001190847 | 0.61 | -0.9195 |
| CNOT4 | NM_001190848 | 0.61 | -0.9195 |
| CNOT4 | NM_001190849 | 0.61 | -0.9195 |
| CNOT4 | NM_001190850 | 0.61 | -0.9195 |
| PLEKHA5 | NM_001190860 | -1.354 | -3.347 |
| MRPS35 | NM_001190864 | -1.125 | -2.825 |
| ORC4 | NM_001190879 | -0.643 | -2.021 |
| HYI | NM_001190880 | 0.2225 | -0.255 |
| ORC4 | NM_001190881 | -0.643 | -2.021 |
| ORC4 | NM_001190882 | -0.643 | -2.021 |
| PDYN | NM_001190892 | -0.3215 | -0.2845 |
| PDYN | NM_001190898 | -0.3215 | -0.2845 |
| PDYN | NM_001190899 | -0.3215 | -0.2845 |
| PDYN | NM_001190900 | -0.3215 | -0.2845 |
| AKR1D1 | NM_001190906 | -0.1975 | -0.5885 |

| | | | |
|---|---|---|---|
| AKR1D1 | NM_001190907 | -0.1975 | -0.5885 |
| THRA | NM_001190918 | -0.483 | -3.231 |
| THRA | NM_001190919 | -0.483 | -3.231 |
| TNFSF10 | NM_001190942 | -0.391 | -0.174 |
| TNFSF10 | NM_001190943 | -0.391 | -0.174 |
| TRAF1 | NM_001190945 | -0.021 | -0.542 |
| FAM193B | NM_001190946 | -0.862 | -1.8655 |
| TRAF1 | NM_001190947 | -0.021 | -0.542 |
| ADAM18 | NM_001190956 | 0.1245 | -0.213 |
| ZMYM2 | NM_001190964 | -0.9155 | -2.786 |
| ZMYM2 | NM_001190965 | -0.9155 | -2.786 |
| YAF2 | NM_001190977 | -1.6075 | -2.688 |
| YAF2 | NM_001190979 | -1.6075 | -2.688 |
| YAF2 | NM_001190980 | -1.6075 | -2.688 |
| IL6ST | NM_001190981 | -0.292 | -1.308 |
| COQ7 | NM_001190983 | -0.613 | -1.8345 |
| CRISP3 | NM_001190986 | 0.2405 | -0.536 |
| SRSF11 | NM_001190987 | -1.044 | -1.3515 |
| CNPY2 | NM_001190991 | -0.4495 | -1.0025 |
| SLC33A1 | NM_001190992 | -1.543 | -2.122 |
| IPO8 | NM_001190995 | -0.949 | -1.62 |
| SLC6A13 | NM_001190997 | 0.1085 | -0.356 |
| BCL2L1 | NM_001191 | -0.325 | -1.5325 |
| SRSF10 | NM_001191005 | -1.748 | -1.245 |
| SRSF10 | NM_001191006 | -1.748 | -1.245 |
| SRSF10 | NM_001191007 | -1.748 | -1.245 |
| SRSF10 | NM_001191009 | -1.748 | -1.245 |
| GSTO2 | NM_001191013 | -0.043 | -0.7895 |
| GSTO2 | NM_001191014 | -0.043 | -0.7895 |
| GSTO2 | NM_001191015 | -0.043 | -0.7895 |
| CASP12 | NM_001191016 | -0.2185 | -0.359 |
| CTCF | NM_001191022 | -1.0065 | -3.079 |
| TNPO3 | NM_001191028 | -0.7495 | -1.4865 |
| C17orf72 | NM_001191029 | 0.046 | -0.262 |
| C17orf72 | NM_001191030 | 0.046 | -0.262 |
| C17orf72 | NM_001191031 | 0.046 | -0.262 |
| C8orf83 | NM_001191035 | -0.9265 | -1.4765 |
| C8orf83 | NM_001191036 | -0.9265 | -1.4765 |
| PSMD1 | NM_001191037 | -0.6925 | -1.134 |
| SEC31A | NM_001191049 | -0.6105 | -0.4885 |
| PDE1C | NM_001191056 | 0.0755 | -0.223 |
| PDE1C | NM_001191057 | 0.0755 | -0.223 |
| PDE1C | NM_001191058 | 0.0755 | -0.223 |

| | | | |
|---|---|---|---|
| PDE1C | NM_001191059 | 0.0755 | -0.223 |
| SLC25A22 | NM_001191060 | -0.3655 | -0.2805 |
| SLC25A22 | NM_001191061 | -0.3655 | -0.2805 |
| GABRB3 | NM_001191320 | -0.9845 | -1.441 |
| GABRB3 | NM_001191321 | -0.9845 | -1.441 |
| GREM1 | NM_001191322 | 0.0545 | 0.0425 |
| GREM1 | NM_001191323 | 0.0545 | 0.0425 |
| RNF138 | NM_001191324 | -1.3675 | -2.6415 |
| TNFRSF17 | NM_001192 | -0.2375 | 0.082 |
| SIRT5 | NM_001193267 | -0.7815 | -1.5085 |
| EML2 | NM_001193268 | -0.154 | -0.4775 |
| EML2 | NM_001193269 | -0.154 | -0.4775 |
| MSL3 | NM_001193270 | -1.0535 | -0.7935 |
| RBBP5 | NM_001193272 | -0.7855 | -1.6135 |
| RBBP5 | NM_001193273 | -0.7855 | -1.6135 |
| MID1 | NM_001193277 | -0.5825 | -1.5365 |
| MID1 | NM_001193278 | -0.5825 | -1.5365 |
| MID1 | NM_001193279 | -0.5825 | -1.5365 |
| MID1 | NM_001193280 | -0.5825 | -1.5365 |
| MID1 | NM_001193281 | -0.5825 | -1.5365 |
| SIRT6 | NM_001193285 | 0.287 | -0.2465 |
| SIRT2 | NM_001193286 | 0.186 | -0.3835 |
| SLC24A2 | NM_001193288 | 0.0105 | -0.2615 |
| APOBEC3A | NM_001193289 | -0.159 | -0.53 |
| SEMA4A | NM_001193300 | -0.0655 | -0.5805 |
| SEMA4A | NM_001193301 | -0.0655 | -0.5805 |
| SEMA4A | NM_001193302 | -0.0655 | -0.5805 |
| TMEM127 | NM_001193304 | -0.0225 | -1.0105 |
| FPR1 | NM_001193306 | 0.088 | -0.5575 |
| SYTL1 | NM_001193308 | -0.042 | -0.6325 |
| C7orf10 | NM_001193311 | -0.0205 | -0.8055 |
| C7orf10 | NM_001193312 | -0.0205 | -0.8055 |
| C7orf10 | NM_001193313 | -0.0205 | -0.8055 |
| RNF212 | NM_001193318 | -0.092 | -0.003 |
| IKBKE | NM_001193321 | -0.4 | 0.2085 |
| IKBKE | NM_001193322 | -0.4 | 0.2085 |
| ZNF692 | NM_001193328 | -0.6755 | -1.8195 |
| C9orf3 | NM_001193329 | -0.689 | -2.123 |
| C9orf3 | NM_001193331 | -0.689 | -2.123 |
| CORO1A | NM_001193333 | -0.0025 | -0.735 |
| WLS | NM_001193334 | -0.092 | -0.74 |
| ASPN | NM_001193335 | -0.352 | 0.1105 |
| FAIM3 | NM_001193338 | -0.066 | -0.257 |

| | | | |
|---|---|---|---|
| SLC13A3 | NM_001193339 | 0.38 | -0.2435 |
| SLC13A3 | NM_001193340 | 0.38 | -0.2435 |
| SLC13A3 | NM_001193342 | 0.38 | -0.2435 |
| MRPS18A | NM_001193343 | -0.711 | -2.2555 |
| MEF2C | NM_001193347 | -1.922 | -0.404 |
| MEF2C | NM_001193348 | -1.922 | -0.404 |
| MEF2C | NM_001193349 | -1.922 | -0.404 |
| MEF2C | NM_001193350 | -1.922 | -0.404 |
| NUP62 | NM_001193357 | -0.971 | -2.442 |
| EXD2 | NM_001193360 | -0.9125 | -1.425 |
| EXD2 | NM_001193361 | -0.9125 | -1.425 |
| EXD2 | NM_001193362 | -0.9125 | -1.425 |
| EXD2 | NM_001193363 | -0.9125 | -1.425 |
| DIDO1 | NM_001193369 | 0.457 | -1.4845 |
| DIDO1 | NM_001193370 | 0.457 | -1.4845 |
| TERT | NM_001193376 | 0.013 | -0.3745 |
| UCKL1 | NM_001193379 | 0.086 | -0.4115 |
| IL17RE | NM_001193380 | 0.0985 | -0.376 |
| UNKL | NM_001193388 | -0.868 | -1.4625 |
| UNKL | NM_001193389 | -0.868 | -1.4625 |
| PPP2R4 | NM_001193397 | -0.1025 | -0.9615 |
| TUBA8 | NM_001193414 | -0.343 | -1.4635 |
| DDX3X | NM_001193416 | -1.372 | -2.2 |
| DDX3X | NM_001193417 | -1.372 | -2.2 |
| TSHZ2 | NM_001193421 | -0.311 | -0.505 |
| SUV39H2 | NM_001193424 | -1.187 | -2.8455 |
| SUV39H2 | NM_001193425 | -1.187 | -2.8455 |
| SUV39H2 | NM_001193426 | -1.187 | -2.8455 |
| SUV39H2 | NM_001193427 | -1.187 | -2.8455 |
| PTPN22 | NM_001193431 | 0.167 | -0.009 |
| C10orf81 | NM_001193434 | 0.1905 | -0.272 |
| C10orf81 | NM_001193435 | 0.1905 | -0.272 |
| TMTC1 | NM_001193451 | -0.9035 | -1.299 |
| NUDT16L1 | NM_001193452 | -0.3915 | -0.98 |
| NSUN2 | NM_001193455 | -0.8135 | -1.825 |
| IFFO1 | NM_001193457 | -0.913 | -1.502 |
| MSRB3 | NM_001193460 | -0.2365 | -0.701 |
| MSRB3 | NM_001193461 | -0.2365 | -0.701 |
| KIAA1267 | NM_001193465 | -0.9175 | -3.09 |
| KIAA1267 | NM_001193466 | -0.9175 | -3.09 |
| FOLH1 | NM_001193471 | -0.4335 | -0.138 |
| FOLH1 | NM_001193472 | -0.4335 | -0.138 |
| FOLH1 | NM_001193473 | -0.4335 | -0.138 |

| | | | |
|---|---|---|---|
| CDADC1 | NM_001193478 | -0.5785 | -1.921 |
| ALDH8A1 | NM_001193480 | -0.583 | -0.1985 |
| LIMS1 | NM_001193482 | -0.657 | -0.715 |
| LIMS1 | NM_001193483 | -0.657 | -0.715 |
| LIMS1 | NM_001193484 | -0.657 | -0.715 |
| LIMS1 | NM_001193485 | -0.657 | -0.715 |
| LIMS1 | NM_001193488 | -0.657 | -0.715 |
| SECISBP2L | NM_001193489 | -0.7085 | -2.452 |
| SLC1A4 | NM_001193493 | -0.0625 | -0.888 |
| ADAR | NM_001193495 | -0.567 | -1.21 |
| ENO3 | NM_001193503 | 0.181 | -0.1135 |
| REST | NM_001193508 | -0.0355 | -0.42 |
| MAP3K12 | NM_001193511 | -1.0245 | -1.4335 |
| SLC30A6 | NM_001193513 | -1.377 | -2.531 |
| SLC30A6 | NM_001193514 | -1.377 | -2.531 |
| SLC30A6 | NM_001193515 | -1.377 | -2.531 |
| RASAL1 | NM_001193520 | -0.0615 | 0.079 |
| RASAL1 | NM_001193521 | -0.0615 | 0.079 |
| FAM65A | NM_001193522 | -0.3885 | -1.086 |
| FAM65A | NM_001193523 | -0.3885 | -1.086 |
| FAM65A | NM_001193524 | -0.3885 | -1.086 |
| SCRN3 | NM_001193528 | -1.4825 | -2.216 |
| CCT6B | NM_001193529 | -0.016 | -0.853 |
| CCT6B | NM_001193530 | -0.016 | -0.853 |
| NEK4 | NM_001193533 | -0.765 | -1.789 |
| FBXL5 | NM_001193534 | -1.564 | -3.956 |
| FBXL5 | NM_001193535 | -1.564 | -3.956 |
| DOCK8 | NM_001193536 | 0.0285 | 0.1265 |
| TMEM126B | NM_001193537 | -1.144 | -2.1265 |
| TMEM126B | NM_001193538 | -1.144 | -2.1265 |
| ANXA6 | NM_001193544 | 0.1015 | -0.61 |
| ENY2 | NM_001193557 | -0.716 | -0.688 |
| CSRP1 | NM_001193570 | 0.3115 | -0.303 |
| CSRP1 | NM_001193571 | 0.3115 | -0.303 |
| CSRP1 | NM_001193572 | 0.3115 | -0.303 |
| NRCAM | NM_001193582 | -1.005 | -0.5705 |
| NRCAM | NM_001193583 | -1.005 | -0.5705 |
| NRCAM | NM_001193584 | -1.005 | -0.5705 |
| SCP2 | NM_001193599 | -0.5505 | -0.4715 |
| SCP2 | NM_001193600 | -0.5505 | -0.4715 |
| TUBD1 | NM_001193609 | -2.037 | -2.406 |
| TUBD1 | NM_001193610 | -2.037 | -2.406 |
| TUBD1 | NM_001193611 | -2.037 | -2.406 |

| | | | |
|---|---|---|---|
| TUBD1 | NM_001193612 | -2.037 | -2.406 |
| TUBD1 | NM_001193613 | -2.037 | -2.406 |
| SCP2 | NM_001193617 | -0.5505 | -0.4715 |
| LOC390940 | NM_001193621 | -0.197 | -0.162 |
| LOC390940 | NM_001193622 | -0.197 | -0.162 |
| DHRS4L2 | NM_001193635 | -0.148 | -0.4855 |
| DHRS4L2 | NM_001193636 | -0.148 | -0.4855 |
| DHRS4L2 | NM_001193637 | -0.148 | -0.4855 |
| CRB1 | NM_001193640 | -0.674 | -0.994 |
| PEX19 | NM_001193644 | -0.245 | -0.8145 |
| ATF5 | NM_001193646 | 0.176 | -0.3215 |
| C17orf62 | NM_001193653 | -0.531 | -1.6595 |
| C17orf62 | NM_001193654 | -0.531 | -1.6595 |
| C17orf62 | NM_001193655 | -0.531 | -1.6595 |
| C17orf62 | NM_001193657 | -0.531 | -1.6595 |
| HCN2 | NM_001194 | -0.212 | -0.705 |
| POGZ | NM_001194937 | -1.1695 | -2.5125 |
| POGZ | NM_001194938 | -1.1695 | -2.5125 |
| EIF4G1 | NM_001194946 | -0.1565 | -0.948 |
| EIF4G1 | NM_001194947 | -0.1565 | -0.948 |
| MATR3 | NM_001194954 | -0.217 | -0.35 |
| MATR3 | NM_001194955 | -0.217 | -0.35 |
| MATR3 | NM_001194956 | -0.217 | -0.35 |
| MEPCE | NM_001194990 | -0.7145 | -3.606 |
| MEPCE | NM_001194991 | -0.7145 | -3.606 |
| MEPCE | NM_001194992 | -0.7145 | -3.606 |
| C12orf65 | NM_001194995 | -1.623 | -3.1635 |
| CARS | NM_001194997 | -0.4275 | -1.4015 |
| CEP152 | NM_001194998 | -0.4725 | -2.5485 |
| MR1 | NM_001194999 | -0.28 | -0.164 |
| BFSP1 | NM_001195 | -0.202 | -0.499 |
| MR1 | NM_001195000 | -0.28 | -0.164 |
| C11orf70 | NM_001195005 | -1.025 | -1.08 |
| PPIE | NM_001195007 | -0.108 | -0.9975 |
| CD4 | NM_001195014 | 0.0055 | -0.527 |
| CD4 | NM_001195015 | 0.0055 | -0.527 |
| CD4 | NM_001195016 | 0.0055 | -0.527 |
| CD4 | NM_001195017 | 0.0055 | -0.527 |
| GPCRLTM7 | NM_001195021 | 0.0715 | -0.372 |
| MR1 | NM_001195035 | -0.28 | -0.164 |
| ECT2L | NM_001195037 | -0.092 | -0.01 |
| YAP1 | NM_001195044 | -0.309 | -1.111 |
| YAP1 | NM_001195045 | -0.309 | -1.111 |

| | | | |
|---|---|---|---|
| DDIT3 | NM_001195053 | -0.677 | -1.316 |
| DDIT3 | NM_001195054 | -0.677 | -1.316 |
| DDIT3 | NM_001195055 | -0.677 | -1.316 |
| DDIT3 | NM_001195056 | -0.677 | -1.316 |
| DDIT3 | NM_001195057 | -0.677 | -1.316 |
| TMEM184B | NM_001195071 | -0.3755 | -0.9525 |
| TMEM184B | NM_001195072 | -0.3755 | -0.9525 |
| A2LD1 | NM_001195087 | -0.3365 | -0.382 |
| ANKRD28 | NM_001195098 | -1.155 | -2.4335 |
| ANKRD28 | NM_001195099 | -1.155 | -2.4335 |
| PTP4A2 | NM_001195100 | -0.817 | -0.334 |
| PTP4A2 | NM_001195101 | -0.817 | -0.334 |
| GSR | NM_001195102 | -0.3685 | -1.5555 |
| GSR | NM_001195103 | -0.3685 | -1.5555 |
| GSR | NM_001195104 | -0.3685 | -1.5555 |
| TLR10 | NM_001195106 | 0.3965 | 0.028 |
| TLR10 | NM_001195107 | 0.3965 | 0.028 |
| TLR10 | NM_001195108 | 0.3965 | 0.028 |
| CDKN2A | NM_001195132 | 0.204 | 0.138 |
| COG4 | NM_001195139 | -0.2175 | -0.8875 |
| TCOF1 | NM_001195141 | -0.2495 | -0.907 |
| ANKRD44 | NM_001195144 | 0.0395 | -0.3575 |
| C17orf57 | NM_001195192 | -0.3525 | 0.363 |
| NASP | NM_001195193 | -0.7075 | -1.004 |
| TP53AIP1 | NM_001195194 | 0.029 | -0.2835 |
| TP53AIP1 | NM_001195195 | 0.029 | -0.2835 |
| CCDC107 | NM_001195200 | -0.345 | -1.11 |
| CCDC107 | NM_001195201 | -0.345 | -1.11 |
| EEF1D | NM_001195203 | 0.3765 | -0.1945 |
| YIPF6 | NM_001195214 | -1.451 | -1.335 |
| DENND1B | NM_001195215 | 0.216 | -1.094 |
| DENND1B | NM_001195216 | 0.216 | -1.094 |
| CCDC107 | NM_001195217 | -0.345 | -1.11 |
| ALG1L | NM_001195223 | 0.0515 | 0.008 |
| PRODH | NM_001195226 | -0.132 | -0.128 |
| FAM64A | NM_001195228 | -0.8745 | -2.1875 |
| APTX | NM_001195248 | -0.647 | -1.9035 |
| APTX | NM_001195249 | -0.647 | -1.9035 |
| APTX | NM_001195250 | -0.647 | -1.9035 |
| APTX | NM_001195251 | -0.647 | -1.9035 |
| APTX | NM_001195252 | -0.647 | -1.9035 |
| APTX | NM_001195254 | -0.647 | -1.9035 |
| PDE4DIP | NM_001195260 | -0.9925 | -0.862 |

| | | | |
|---|---|---|---|
| PDE4DIP | NM_001195261 | -0.9925 | -0.862 |
| PDZD7 | NM_001195263 | -0.036 | -0.842 |
| FLVCR2 | NM_001195283 | 0.026 | -0.3215 |
| IRF4 | NM_001195286 | -0.5695 | -0.3175 |
| SERPINB6 | NM_001195291 | -0.41 | -0.8315 |
| MIPOL1 | NM_001195296 | -0.0125 | -0.2125 |
| MIPOL1 | NM_001195297 | -0.0125 | -0.2125 |
| C16orf57 | NM_001195302 | 0.0165 | -0.329 |
| RGS5 | NM_001195303 | -0.1725 | -0.09 |
| CGGBP1 | NM_001195308 | -1.2975 | -1.9045 |
| GPR35 | NM_001195381 | 0.0475 | -0.621 |
| GPR35 | NM_001195382 | 0.0475 | -0.621 |
| TMEM99 | NM_001195386 | -0.0735 | -1.925 |
| TMEM99 | NM_001195387 | -0.0735 | -1.925 |
| FCRL5 | NM_001195388 | -0.185 | -0.4305 |
| ARL4A | NM_001195396 | -0.918 | -2.4755 |
| DCLK1 | NM_001195415 | -1.1205 | -1.1045 |
| DCLK1 | NM_001195416 | -1.1205 | -1.1045 |
| GTPBP3 | NM_001195422 | 0.1175 | -0.088 |
| SRSF2 | NM_001195427 | -0.693 | -0.8275 |
| DCLK1 | NM_001195430 | -1.1205 | -1.1045 |
| MLF1 | NM_001195432 | -0.534 | -0.676 |
| MLF1 | NM_001195433 | -0.534 | -0.676 |
| MLF1 | NM_001195434 | -0.534 | -0.676 |
| SRSF7 | NM_001195446 | -0.889 | -0.46 |
| SATB1 | NM_001195470 | -1.235 | -2.3285 |
| SLC12A8 | NM_001195483 | 0.012 | -0.3145 |
| TLX1 | NM_001195517 | -0.356 | -0.2525 |
| SPTAN1 | NM_001195532 | -0.1575 | -0.6805 |
| CHMP5 | NM_001195536 | -0.3815 | -0.0455 |
| DCX | NM_001195553 | -0.96 | -0.4395 |
| CLINT1 | NM_001195555 | -1.0615 | -1.9445 |
| CLINT1 | NM_001195556 | -1.0615 | -1.9445 |
| CAMTA1 | NM_001195563 | 0.0475 | -0.016 |
| DICER1 | NM_001195573 | 0.0765 | -1.67 |
| SEPHS1 | NM_001195602 | -0.965 | -0.7845 |
| SEPHS1 | NM_001195604 | -0.965 | -0.7845 |
| PLEKHA1 | NM_001195608 | -1.204 | -1.5215 |
| DCDC2 | NM_001195610 | 0.201 | -0.089 |
| C16orf93 | NM_001195620 | -0.209 | -0.1215 |
| TOPORS | NM_001195622 | -0.643 | -1.386 |
| MLLT10 | NM_001195626 | 0.167 | -0.071 |
| MLLT10 | NM_001195627 | 0.167 | -0.071 |

| | | | |
|---|---|---|---|
| MLLT10 | NM_001195628 | 0.167 | -0.071 |
| MLLT10 | NM_001195630 | 0.167 | -0.071 |
| C8orf34 | NM_001195639 | 0.157 | -0.2925 |
| PLA2R1 | NM_001195641 | 0.0735 | 0.058 |
| DHFRL1 | NM_001195643 | -0.342 | -0.103 |
| VAPB | NM_001195677 | -0.3075 | -1.7435 |
| TGFBR3 | NM_001195683 | -0.535 | -0.8875 |
| TGFBR3 | NM_001195684 | -0.535 | -0.8875 |
| LYPD6 | NM_001195685 | -0.122 | -0.588 |
| SLC1A2 | NM_001195728 | 0.0675 | -0.4915 |
| CHPF | NM_001195731 | -0.043 | -0.6745 |
| PIP5K1C | NM_001195733 | 0.0385 | -0.6505 |
| C1orf93 | NM_001195736 | 0.09 | -0.5055 |
| C1orf93 | NM_001195737 | 0.09 | -0.5055 |
| C1orf93 | NM_001195738 | 0.09 | -0.5055 |
| C1orf93 | NM_001195740 | 0.09 | -0.5055 |
| C1orf93 | NM_001195741 | 0.09 | -0.5055 |
| THAP3 | NM_001195752 | -0.4755 | -0.9105 |
| THAP3 | NM_001195753 | -0.4755 | -0.9105 |
| CLRN1 | NM_001195794 | -0.0205 | -0.34 |
| LDLR | NM_001195798 | -1.0065 | -2.926 |
| LDLR | NM_001195799 | -1.0065 | -2.926 |
| LDLR | NM_001195800 | -1.0065 | -2.926 |
| LDLR | NM_001195802 | -1.0065 | -2.926 |
| LDLR | NM_001195803 | -1.0065 | -2.926 |
| PLD5 | NM_001195811 | -0.466 | -1.648 |
| PLD5 | NM_001195812 | -0.466 | -1.648 |
| HSDL2 | NM_001195822 | -0.789 | -0.6675 |
| WDR65 | NM_001195831 | -0.287 | -0.379 |
| RINL | NM_001195833 | 0.0075 | -0.277 |
| BID | NM_001196 | -0.028 | -1.151 |
| BIK | NM_001197 | -1.0355 | -0.3475 |
| PLEKHA8 | NM_001197026 | -0.9615 | -1.962 |
| PLEKHA8 | NM_001197027 | -0.9615 | -1.962 |
| ANKHD1 | NM_001197030 | 0.2395 | -0.272 |
| ZFP91 | NM_001197051 | -1.0505 | -2.0155 |
| IFRD1 | NM_001197079 | -1.735 | -3.0135 |
| IFRD1 | NM_001197080 | -1.735 | -3.0135 |
| PRSS3 | NM_001197097 | 0.118 | -0.569 |
| PRSS3 | NM_001197098 | 0.118 | -0.569 |
| IQCJ | NM_001197100 | -0.0535 | -0.0445 |
| MLL | NM_001197104 | -0.375 | -1.726 |
| GCLC | NM_001197115 | -1.575 | -2.98 |

| | | | |
|---|---|---|---|
| IRF3 | NM_001197122 | -0.355 | -1.256 |
| IRF3 | NM_001197123 | -0.355 | -1.256 |
| IRF3 | NM_001197124 | -0.355 | -1.256 |
| IRF3 | NM_001197125 | -0.355 | -1.256 |
| IRF3 | NM_001197126 | -0.355 | -1.256 |
| IRF3 | NM_001197127 | -0.355 | -1.256 |
| IRF3 | NM_001197128 | -0.355 | -1.256 |
| SYS1 | NM_001197129 | -0.6285 | -1.7545 |
| PPP1R12B | NM_001197131 | 0.8205 | 0.5085 |
| TUBB3 | NM_001197181 | -0.147 | -1.0135 |
| ASGR1 | NM_001197216 | -0.2435 | -0.2325 |
| PDE4D | NM_001197218 | -1.413 | -2.114 |
| PDE4D | NM_001197219 | -1.413 | -2.114 |
| PDE4D | NM_001197220 | -1.413 | -2.114 |
| PDE4D | NM_001197221 | -1.413 | -2.114 |
| PDE4D | NM_001197222 | -1.413 | -2.114 |
| PDE4D | NM_001197223 | -1.413 | -2.114 |
| BTN2A1 | NM_001197233 | -0.878 | -2.342 |
| BTN2A1 | NM_001197234 | -0.878 | -2.342 |
| BTN2A2 | NM_001197237 | -1.0235 | -1.553 |
| BTN2A2 | NM_001197238 | -1.0235 | -1.553 |
| BTN2A2 | NM_001197239 | -1.0235 | -1.553 |
| BTN2A2 | NM_001197240 | -1.0235 | -1.553 |
| BCL7B | NM_001197244 | -0.3765 | -1.8145 |
| BTN3A2 | NM_001197246 | -0.5525 | -0.435 |
| BTN3A2 | NM_001197247 | -0.5525 | -0.435 |
| BTN3A2 | NM_001197248 | -0.5525 | -0.435 |
| BTN3A2 | NM_001197249 | -0.5525 | -0.435 |
| ORC3 | NM_001197259 | -1.2905 | -1.8755 |
| DOK1 | NM_001197260 | -0.5415 | -2.156 |
| DPYSL2 | NM_001197293 | -0.1565 | -0.526 |
| DPYSL3 | NM_001197294 | 0.2595 | -0.4435 |
| ECM2 | NM_001197295 | 0.2135 | -0.0515 |
| ECM2 | NM_001197296 | 0.2135 | -0.0515 |
| GABPA | NM_001197297 | -0.8585 | -1.919 |
| DKKL1 | NM_001197301 | 0.1385 | -0.281 |
| DKKL1 | NM_001197302 | 0.1385 | -0.281 |
| CLEC2D | NM_001197317 | 0.0845 | -0.721 |
| CLEC2D | NM_001197318 | 0.0845 | -0.721 |
| CLEC2D | NM_001197319 | 0.0845 | -0.721 |
| MCL1 | NM_001197320 | 0.1675 | -0.209 |
| ARNT | NM_001197325 | -0.8185 | -2.4545 |
| HSPA12B | NM_001197327 | -0.0255 | -0.1845 |

| | | | |
|---|---|---|---|
| POLE2 | NM_001197330 | -1.604 | -3.319 |
| POLE2 | NM_001197331 | -1.604 | -3.319 |
| PRDM1 | NM_001198 | 0 | -0.304 |
| TCF7L2 | NM_001198525 | -0.4665 | -1.848 |
| TCF7L2 | NM_001198526 | -0.4665 | -1.848 |
| TCF7L2 | NM_001198527 | -0.4665 | -1.848 |
| TCF7L2 | NM_001198528 | -0.4665 | -1.848 |
| TCF7L2 | NM_001198529 | -0.4665 | -1.848 |
| TCF7L2 | NM_001198530 | -0.4665 | -1.848 |
| TCF7L2 | NM_001198531 | -0.4665 | -1.848 |
| OXR1 | NM_001198532 | -0.4415 | -0.855 |
| OXR1 | NM_001198533 | -0.4415 | -0.855 |
| OXR1 | NM_001198534 | -0.4415 | -0.855 |
| OXR1 | NM_001198535 | -0.4415 | -0.855 |
| MEFV | NM_001198536 | 0.0875 | -0.229 |
| ERAP1 | NM_001198541 | -0.239 | -0.649 |
| HYDIN | NM_001198542 | 0.1905 | 0.063 |
| HYDIN | NM_001198543 | 0.1905 | 0.063 |
| NOL4 | NM_001198546 | -1.113 | -1.8675 |
| NOL4 | NM_001198547 | -1.113 | -1.8675 |
| NOL4 | NM_001198548 | -1.113 | -1.8675 |
| NOL4 | NM_001198549 | -1.113 | -1.8675 |
| WT1 | NM_001198551 | 0.373 | -0.2905 |
| WT1 | NM_001198552 | 0.373 | -0.2905 |
| LMNB1 | NM_001198557 | -0.7785 | -2.475 |
| SIGLEC9 | NM_001198558 | -0.375 | -0.288 |
| DNAJC27 | NM_001198559 | -1.066 | -1.499 |
| ADCY4 | NM_001198568 | 0.1855 | -0.08 |
| ADCY4 | NM_001198592 | 0.1855 | -0.08 |
| STON1 | NM_001198595 | -1.346 | -2.0295 |
| MAP7 | NM_001198608 | -0.406 | -0.8655 |
| MAP7 | NM_001198609 | -0.406 | -0.8655 |
| MAP7 | NM_001198611 | -0.406 | -0.8655 |
| MAP7 | NM_001198614 | -0.406 | -0.8655 |
| MAP7 | NM_001198615 | -0.406 | -0.8655 |
| MAP7 | NM_001198616 | -0.406 | -0.8655 |
| MAP7 | NM_001198617 | -0.406 | -0.8655 |
| MAP7 | NM_001198618 | -0.406 | -0.8655 |
| MAP7 | NM_001198619 | -0.406 | -0.8655 |
| XIRP1 | NM_001198621 | -0.0875 | -0.3295 |
| RUNX1T1 | NM_001198625 | -1.1215 | -2.571 |
| RUNX1T1 | NM_001198626 | -1.1215 | -2.571 |
| RUNX1T1 | NM_001198627 | -1.1215 | -2.571 |

| | | | |
|---|---|---|---|
| RUNX1T1 | NM_001198628 | -1.1215 | -2.571 |
| RUNX1T1 | NM_001198629 | -1.1215 | -2.571 |
| RUNX1T1 | NM_001198630 | -1.1215 | -2.571 |
| RUNX1T1 | NM_001198631 | -1.1215 | -2.571 |
| RUNX1T1 | NM_001198632 | -1.1215 | -2.571 |
| RUNX1T1 | NM_001198633 | -1.1215 | -2.571 |
| RUNX1T1 | NM_001198634 | -1.1215 | -2.571 |
| TRIM6 | NM_001198644 | -0.17 | -0.341 |
| TRIM6 | NM_001198645 | -0.17 | -0.341 |
| ARHGEF12 | NM_001198665 | -1.0745 | -1.5105 |
| TMEM136 | NM_001198670 | -1.694 | -2.9355 |
| TMEM136 | NM_001198671 | -1.694 | -2.9355 |
| TMEM136 | NM_001198672 | -1.694 | -2.9355 |
| TMEM136 | NM_001198673 | -1.694 | -2.9355 |
| TMEM136 | NM_001198674 | -1.694 | -2.9355 |
| TMEM136 | NM_001198675 | -1.694 | -2.9355 |
| RUNX1T1 | NM_001198679 | -1.1215 | -2.571 |
| MFAP4 | NM_001198695 | -0.052 | -0.436 |
| AIFM2 | NM_001198696 | -0.165 | -0.909 |
| PRR5 | NM_001198721 | -0.5815 | -0.556 |
| ARHGAP8 | NM_001198726 | 0.261 | -0.4215 |
| GNGT2 | NM_001198754 | -0.4605 | -0.2865 |
| GNGT2 | NM_001198755 | -0.4605 | -0.2865 |
| GNGT2 | NM_001198756 | -0.4605 | -0.2865 |
| CD302 | NM_001198763 | -0.3835 | -1.126 |
| CD302 | NM_001198764 | -0.3835 | -1.126 |
| PI4KB | NM_001198773 | -0.2755 | -1.6825 |
| PI4KB | NM_001198774 | -0.2755 | -1.6825 |
| PI4KB | NM_001198775 | -0.2755 | -1.6825 |
| CUL2 | NM_001198777 | -1.0925 | -2.887 |
| CUL2 | NM_001198778 | -1.0925 | -2.887 |
| CUL2 | NM_001198779 | -1.0925 | -2.887 |
| ARPC4 | NM_001198780 | -0.0805 | -0.7065 |
| POU2F1 | NM_001198783 | -0.422 | -1.2635 |
| POU2F1 | NM_001198786 | -0.422 | -1.2635 |
| ASCC1 | NM_001198798 | -1.1025 | -1.866 |
| ASCC1 | NM_001198799 | -1.1025 | -1.866 |
| ASCC1 | NM_001198800 | -1.1025 | -1.866 |
| EIF4G3 | NM_001198801 | -0.836 | -1.756 |
| EIF4G3 | NM_001198802 | -0.836 | -1.756 |
| EIF4G3 | NM_001198803 | -0.836 | -1.756 |
| SLC43A1 | NM_001198810 | -0.5665 | -1.581 |
| A1CF | NM_001198818 | 0.0105 | -0.246 |

| | | | |
|---|---|---|---|
| A1CF | NM_001198819 | 0.0105 | -0.246 |
| A1CF | NM_001198820 | 0.0105 | -0.246 |
| C8orf58 | NM_001198827 | -0.011 | -0.624 |
| LIPF | NM_001198828 | 0.173 | 0.088 |
| LIPF | NM_001198829 | 0.173 | 0.088 |
| LIPF | NM_001198830 | 0.173 | 0.088 |
| PDE4DIP | NM_001198832 | -0.9925 | -0.862 |
| PDE4DIP | NM_001198834 | -0.9925 | -0.862 |
| RBM14 | NM_001198836 | 0.1865 | -1.0305 |
| RBM14 | NM_001198837 | 0.1865 | -1.0305 |
| RBM12 | NM_001198838 | -0.381 | -1.116 |
| RBM12 | NM_001198840 | -0.381 | -1.116 |
| CCT2 | NM_001198842 | -1.396 | -1.365 |
| RBM4 | NM_001198843 | -0.0515 | -0.2005 |
| RBM4 | NM_001198844 | -0.0515 | -0.2005 |
| FOXJ3 | NM_001198850 | -0.9695 | -1.7435 |
| FOXJ3 | NM_001198851 | -0.9695 | -1.7435 |
| FOXJ3 | NM_001198852 | -0.9695 | -1.7435 |
| CYP2C8 | NM_001198853 | -0.4925 | -0.3515 |
| CYP2C8 | NM_001198854 | -0.4925 | -0.3515 |
| CYP2C8 | NM_001198855 | -0.4925 | -0.3515 |
| TMEM206 | NM_001198862 | -0.7105 | -1.981 |
| CAPN1 | NM_001198868 | 0.0425 | -0.4865 |
| CAPN1 | NM_001198869 | 0.0425 | -0.4865 |
| ACY1 | NM_001198895 | 0.2045 | -1.045 |
| ACY1 | NM_001198896 | 0.2045 | -1.045 |
| ACY1 | NM_001198897 | 0.2045 | -1.045 |
| ACY1 | NM_001198898 | 0.2045 | -1.045 |
| YY1AP1 | NM_001198899 | -0.6905 | -2.053 |
| YY1AP1 | NM_001198900 | -0.6905 | -2.053 |
| YY1AP1 | NM_001198901 | -0.6905 | -2.053 |
| YY1AP1 | NM_001198902 | -0.6905 | -2.053 |
| YY1AP1 | NM_001198903 | -0.6905 | -2.053 |
| YY1AP1 | NM_001198904 | -0.6905 | -2.053 |
| YY1AP1 | NM_001198905 | -0.6905 | -2.053 |
| YY1AP1 | NM_001198906 | -0.6905 | -2.053 |
| PPFIBP1 | NM_001198915 | -0.7885 | -1.0285 |
| PPFIBP1 | NM_001198916 | -0.7885 | -1.0285 |
| DTNA | NM_001198938 | -0.5815 | -1.7065 |
| DTNA | NM_001198939 | -0.5815 | -1.7065 |
| DTNA | NM_001198940 | -0.5815 | -1.7065 |
| DTNA | NM_001198941 | -0.5815 | -1.7065 |
| DTNA | NM_001198942 | -0.5815 | -1.7065 |

| | | | |
|---|---|---|---|
| DTNA | NM_001198943 | -0.5815 | -1.7065 |
| DTNA | NM_001198944 | -0.5815 | -1.7065 |
| DTNA | NM_001198945 | -0.5815 | -1.7065 |
| MYO16 | NM_001198950 | -0.159 | -0.375 |
| ZFYVE21 | NM_001198953 | -0.025 | -0.635 |
| DCAF6 | NM_001198956 | -1.115 | -1.723 |
| DCAF6 | NM_001198957 | -1.115 | -1.723 |
| ECHDC2 | NM_001198961 | -0.0225 | -0.5385 |
| ECHDC2 | NM_001198962 | -0.0225 | -0.5385 |
| FKTN | NM_001198963 | -0.97 | -1.501 |
| NFATC4 | NM_001198965 | -0.1935 | -0.7425 |
| NFATC4 | NM_001198966 | -0.1935 | -0.7425 |
| NFATC4 | NM_001198967 | -0.1935 | -0.7425 |
| KIAA1522 | NM_001198972 | -0.495 | -1.898 |
| KIAA1522 | NM_001198973 | -0.495 | -1.898 |
| SMAP2 | NM_001198978 | -0.457 | -1.3915 |
| SMAP2 | NM_001198979 | -0.457 | -1.3915 |
| SMAP2 | NM_001198980 | -0.457 | -1.3915 |
| NFS1 | NM_001198989 | -0.4815 | -1.573 |
| NADK | NM_001198993 | 0.262 | -0.6915 |
| NADK | NM_001198994 | 0.262 | -0.6915 |
| NADK | NM_001198995 | 0.262 | -0.6915 |
| SEMA6D | NM_001198999 | -0.2595 | -0.628 |
| BMP1 | NM_001199 | 0.233 | -0.124 |
| DCTN5 | NM_001199011 | -0.685 | -1.5425 |
| C1orf201 | NM_001199012 | -0.407 | -0.1115 |
| C1orf201 | NM_001199013 | -0.407 | -0.1115 |
| C1orf201 | NM_001199014 | -0.407 | -0.1115 |
| LRRC49 | NM_001199017 | -0.469 | -1.252 |
| LRRC49 | NM_001199018 | -0.469 | -1.252 |
| SPON2 | NM_001199021 | 0.019 | -0.038 |
| SERINC2 | NM_001199037 | -0.1555 | -0.2945 |
| SERINC2 | NM_001199038 | -0.1555 | -0.2945 |
| SERINC2 | NM_001199039 | -0.1555 | -0.2945 |
| STRA6 | NM_001199040 | -0.683 | -1.1345 |
| STRA6 | NM_001199041 | -0.683 | -1.1345 |
| STRA6 | NM_001199042 | -0.683 | -1.1345 |
| C16orf5 | NM_001199054 | -0.7265 | -2.1975 |
| C16orf5 | NM_001199055 | -0.7265 | -2.1975 |
| C16orf5 | NM_001199056 | -0.7265 | -2.1975 |
| PODN | NM_001199080 | 0.2825 | -0.503 |
| PODN | NM_001199081 | 0.2825 | -0.503 |
| PODN | NM_001199082 | 0.2825 | -0.503 |

| | | | |
|---|---|---|---|
| TDRD5 | NM_001199085 | -0.375 | -0.2865 |
| NT5C1B | NM_001199086 | -0.0925 | -0.479 |
| NT5C1B | NM_001199087 | -0.0925 | -0.479 |
| NT5C1B | NM_001199088 | -0.0925 | -0.479 |
| TDRD5 | NM_001199089 | -0.375 | -0.2865 |
| TDRD5 | NM_001199091 | -0.375 | -0.2865 |
| TDRD5 | NM_001199092 | -0.375 | -0.2865 |
| BAIAP3 | NM_001199096 | 0.01 | 0.3965 |
| BAIAP3 | NM_001199097 | 0.01 | 0.3965 |
| BAIAP3 | NM_001199098 | 0.01 | 0.3965 |
| BAIAP3 | NM_001199099 | 0.01 | 0.3965 |
| TBC1D24 | NM_001199107 | 0.1475 | -0.4845 |
| MDH1 | NM_001199111 | -0.556 | -0.6175 |
| MDH1 | NM_001199112 | -0.556 | -0.6175 |
| COPS3 | NM_001199125 | -0.8075 | -1.259 |
| TANK | NM_001199135 | -1.848 | -3.981 |
| NLRC4 | NM_001199138 | 0.028 | -0.119 |
| NLRC4 | NM_001199139 | 0.028 | -0.119 |
| AMMECR1L | NM_001199140 | -0.889 | -2.873 |
| XIRP2 | NM_001199143 | 0.0105 | -0.206 |
| XIRP2 | NM_001199144 | 0.0105 | -0.206 |
| XIRP2 | NM_001199145 | 0.0105 | -0.206 |
| SLC38A11 | NM_001199148 | 0.096 | -0.3225 |
| LTF | NM_001199149 | 0.4125 | 0.1915 |
| PI16 | NM_001199159 | 0.0345 | -0.139 |
| USP19 | NM_001199160 | -0.294 | -1.7655 |
| USP19 | NM_001199161 | -0.294 | -1.7655 |
| USP19 | NM_001199162 | -0.294 | -1.7655 |
| PSMC5 | NM_001199163 | -0.3105 | -0.5585 |
| MGAT5B | NM_001199172 | -0.0295 | -0.273 |
| MLST8 | NM_001199173 | -0.014 | -0.842 |
| MLST8 | NM_001199174 | -0.014 | -0.842 |
| MLST8 | NM_001199175 | -0.014 | -0.842 |
| ATP2C1 | NM_001199179 | -0.972 | -1.569 |
| ATP2C1 | NM_001199180 | -0.972 | -1.569 |
| ATP2C1 | NM_001199181 | -0.972 | -1.569 |
| ATP2C1 | NM_001199182 | -0.972 | -1.569 |
| ATP2C1 | NM_001199183 | -0.972 | -1.569 |
| ATP2C1 | NM_001199184 | -0.972 | -1.569 |
| ATP2C1 | NM_001199185 | -0.972 | -1.569 |
| ARMC6 | NM_001199196 | -0.691 | -1.6735 |
| TBC1D23 | NM_001199198 | -0.9835 | -2.1075 |
| AK2 | NM_001199199 | -0.2575 | -1.04 |

| | | | |
|---|---|---|---|
| ZBBX | NM_001199201 | -0.29 | 0.098 |
| ZBBX | NM_001199202 | -0.29 | 0.098 |
| GTF2IRD1 | NM_001199207 | -0.802 | -2.0405 |
| CYP4F3 | NM_001199208 | 0.4855 | -0.774 |
| CYP4F3 | NM_001199209 | 0.4855 | -0.774 |
| STMN2 | NM_001199214 | -0.8015 | -0.3535 |
| INMT | NM_001199219 | 0.2125 | -0.1835 |
| SGOL1 | NM_001199251 | -0.38 | -0.871 |
| SGOL1 | NM_001199252 | -0.38 | -0.871 |
| SGOL1 | NM_001199253 | -0.38 | -0.871 |
| SGOL1 | NM_001199254 | -0.38 | -0.871 |
| SGOL1 | NM_001199255 | -0.38 | -0.871 |
| SGOL1 | NM_001199256 | -0.38 | -0.871 |
| SGOL1 | NM_001199257 | -0.38 | -0.871 |
| TOR1AIP2 | NM_001199260 | -0.5465 | -1.3555 |
| UCHL5 | NM_001199261 | -0.881 | -1.905 |
| UCHL5 | NM_001199262 | -0.881 | -1.905 |
| UCHL5 | NM_001199263 | -0.881 | -1.905 |
| SGK2 | NM_001199264 | -0.0075 | 0.1375 |
| DGKZ | NM_001199266 | -0.2985 | -0.85 |
| DGKZ | NM_001199267 | -0.2985 | -0.85 |
| DGKZ | NM_001199268 | -0.2985 | -0.85 |
| CHRNA6 | NM_001199279 | -0.0865 | -0.4445 |
| HAS3 | NM_001199280 | -0.0425 | -0.237 |
| CABIN1 | NM_001199281 | 0.189 | -0.578 |
| LRBA | NM_001199282 | -0.5395 | -0.953 |
| HSD17B4 | NM_001199291 | -1 | -1.197 |
| HSD17B4 | NM_001199292 | -1 | -1.197 |
| ZNF549 | NM_001199295 | -1.4575 | -3.6205 |
| UIMC1 | NM_001199297 | -1.672 | -2.1365 |
| UIMC1 | NM_001199298 | -1.672 | -2.1365 |
| CNOT2 | NM_001199302 | -1.895 | -3.0975 |
| CNOT2 | NM_001199303 | -1.895 | -3.0975 |
| PEX26 | NM_001199319 | 0.079 | -0.819 |
| MUTED | NM_001199322 | -1.377 | -2.427 |
| MUTED | NM_001199323 | -1.377 | -2.427 |
| ZNF615 | NM_001199324 | -0.449 | -2.3075 |
| FPGT | NM_001199328 | -1.2485 | -2.245 |
| FPGT | NM_001199329 | -1.2485 | -2.245 |
| RPL17 | NM_001199340 | -0.1785 | -0.184 |
| RPL17 | NM_001199341 | -0.1785 | -0.184 |
| RPL17 | NM_001199342 | -0.1785 | -0.184 |
| RPL17 | NM_001199343 | -0.1785 | -0.184 |

| | | | |
|---|---|---|---|
| RPL17 | NM_001199344 | -0.1785 | -0.184 |
| RPL17 | NM_001199345 | -0.1785 | -0.184 |
| C18orf32 | NM_001199346 | -1.0075 | -1.261 |
| ACSBG1 | NM_001199377 | -0.2995 | -0.4895 |
| GBF1 | NM_001199378 | -0.209 | -0.722 |
| GBF1 | NM_001199379 | -0.209 | -0.722 |
| RNF145 | NM_001199380 | -1.6055 | -2.5945 |
| RNF145 | NM_001199381 | -1.6055 | -2.5945 |
| RNF145 | NM_001199382 | -1.6055 | -2.5945 |
| RNF145 | NM_001199383 | -1.6055 | -2.5945 |
| EPB41L2 | NM_001199388 | -0.624 | -0.5595 |
| EPB41L2 | NM_001199389 | -0.624 | -0.5595 |
| NEK1 | NM_001199397 | -0.1285 | -0.993 |
| NEK1 | NM_001199398 | -0.1285 | -0.993 |
| NEK1 | NM_001199399 | -0.1285 | -0.993 |
| NEK1 | NM_001199400 | -0.1285 | -0.993 |
| ARHGAP23 | NM_001199417 | -0.136 | -0.343 |
| TRAF3 | NM_001199427 | -0.419 | -1.483 |
| NIP7 | NM_001199434 | -1.349 | -2.368 |
| BRD2 | NM_001199455 | 0.032 | -1.0205 |
| BRD2 | NM_001199456 | 0.032 | -1.0205 |
| PDCD2 | NM_001199461 | -0.431 | -1.326 |
| PDCD2 | NM_001199462 | -0.431 | -1.326 |
| PDCD2 | NM_001199463 | -0.431 | -1.326 |
| PDCD2 | NM_001199464 | -0.431 | -1.326 |
| ISY1 | NM_001199469 | -0.5875 | -1.8275 |
| ZNF193 | NM_001199479 | -0.5055 | -2.217 |
| ZNF193 | NM_001199480 | -0.5055 | -2.217 |
| PDCD4 | NM_001199492 | -1.3635 | -3.0195 |
| PHACTR3 | NM_001199505 | -0.8465 | -1.5995 |
| PHACTR3 | NM_001199506 | -0.8465 | -1.5995 |
| TGIF2 | NM_001199513 | -1.421 | -3.2615 |
| TGIF2 | NM_001199514 | -1.421 | -3.2615 |
| TGIF2 | NM_001199515 | -1.421 | -3.2615 |
| UMODL1 | NM_001199527 | 0.2685 | -0.2605 |
| UMODL1 | NM_001199528 | 0.2685 | -0.2605 |
| C20orf24 | NM_001199534 | -0.1675 | -1.47 |
| CIDEC | NM_001199551 | 0.0575 | -0.2065 |
| CIDEC | NM_001199552 | 0.0575 | -0.2065 |
| PLA2G6 | NM_001199562 | -0.287 | -0.6265 |
| BVES | NM_001199563 | -0.112 | -1.426 |
| CLIP3 | NM_001199570 | 0.1355 | -0.3365 |
| TRIM22 | NM_001199573 | -0.1175 | -0.168 |

| | | | |
|---|---|---|---|
| SUN2 | NM_001199579 | -0.535 | -1.1555 |
| SUN2 | NM_001199580 | -0.535 | -1.1555 |
| PACSIN1 | NM_001199583 | -0.0815 | -0.4885 |
| NCOA7 | NM_001199619 | -1.2485 | -2.957 |
| NCOA7 | NM_001199620 | -1.2485 | -2.957 |
| NCOA7 | NM_001199621 | -1.2485 | -2.957 |
| NCOA7 | NM_001199622 | -1.2485 | -2.957 |
| CIDEC | NM_001199623 | 0.0575 | -0.2065 |
| MYL6B | NM_001199629 | 0.374 | -0.2435 |
| SLC28A3 | NM_001199633 | 0.148 | -0.155 |
| ADCYAP1R1 | NM_001199635 | -0.4205 | -2.2315 |
| ADCYAP1R1 | NM_001199636 | -0.4205 | -2.2315 |
| ADCYAP1R1 | NM_001199637 | -0.4205 | -2.2315 |
| IL33 | NM_001199640 | -0.0065 | 0.086 |
| IL33 | NM_001199641 | -0.0065 | 0.086 |
| PTK2 | NM_001199649 | -1.0095 | -1.4185 |
| LETM2 | NM_001199659 | -0.39 | -1.135 |
| LETM2 | NM_001199660 | -0.39 | -1.135 |
| CALU | NM_001199671 | -0.828 | -0.878 |
| CALU | NM_001199672 | -0.828 | -0.878 |
| CALU | NM_001199673 | -0.828 | -0.878 |
| CALU | NM_001199674 | -0.828 | -0.878 |
| TMEM56 | NM_001199679 | -0.4725 | -1.1695 |
| RWDD3 | NM_001199682 | -1.2525 | -2.6935 |
| EDA2R | NM_001199687 | -0.285 | -0.6325 |
| SLC4A2 | NM_001199692 | -0.085 | -0.4575 |
| SLC4A2 | NM_001199693 | -0.085 | -0.4575 |
| SLC4A2 | NM_001199694 | -0.085 | -0.4575 |
| DHX33 | NM_001199699 | 0.1735 | -0.86 |
| KIAA0895 | NM_001199706 | -2.1685 | -3.382 |
| KIAA0895 | NM_001199707 | -2.1685 | -3.382 |
| KIAA0895 | NM_001199708 | -2.1685 | -3.382 |
| CRABP2 | NM_001199723 | -0.0495 | -0.416 |
| MSRA | NM_001199729 | 0.296 | -0.4945 |
| CTSS | NM_001199739 | 0.427 | -0.496 |
| DCTN5 | NM_001199743 | -0.685 | -1.5425 |
| ARL2 | NM_001199745 | -0.589 | -0.6905 |
| HOXD8 | NM_001199746 | -0.9985 | -1.8955 |
| HOXD8 | NM_001199747 | -0.9985 | -1.8955 |
| CPT1C | NM_001199752 | -0.537 | -0.947 |
| CPT1C | NM_001199753 | -0.537 | -0.947 |
| PPP2R5A | NM_001199756 | -0.7905 | -1.555 |
| ST20 | NM_001199757 | -0.3235 | -1.081 |

| | | | |
|---|---|---|---|
| MTHFS | NM_001199758 | -0.7945 | -3.214 |
| PTPRN | NM_001199763 | 0.0625 | -0.4285 |
| PTPRN | NM_001199764 | 0.0625 | -0.4285 |
| OTX1 | NM_001199770 | 0.053 | -0.16 |
| RDH5 | NM_001199771 | -0.2815 | -0.639 |
| CPD | NM_001199775 | -0.4815 | -1.2235 |
| POC1B | NM_001199777 | -0.9035 | -1.528 |
| PSMB2 | NM_001199779 | -0.0105 | -0.851 |
| PSMB2 | NM_001199780 | -0.0105 | -0.851 |
| SDCBP2 | NM_001199784 | 0.0295 | -0.399 |
| FKBP1A | NM_001199786 | 0.2695 | -0.1105 |
| SLC35E2 | NM_001199787 | -0.1305 | -0.6825 |
| PTPN7 | NM_001199797 | -0.427 | 0.095 |
| ILDR1 | NM_001199799 | 0.4515 | 0.0865 |
| ILDR1 | NM_001199800 | 0.4515 | 0.0865 |
| INTS7 | NM_001199809 | -1.678 | -2.6125 |
| INTS7 | NM_001199811 | -1.678 | -2.6125 |
| INTS7 | NM_001199812 | -1.678 | -2.6125 |
| ZNF812 | NM_001199814 | -0.151 | 0.005 |
| MDP1 | NM_001199821 | -0.2505 | -0.926 |
| MDP1 | NM_001199822 | -0.2505 | -0.926 |
| SPRR1A | NM_001199828 | 0.0425 | -0.465 |
| HORMAD1 | NM_001199829 | 0.0315 | -0.6115 |
| HPSE | NM_001199830 | -0.8815 | -0.394 |
| VCAM1 | NM_001199834 | -0.458 | -0.438 |
| SNX10 | NM_001199835 | 0.488 | -0.142 |
| SNX10 | NM_001199837 | 0.488 | -0.142 |
| SNX10 | NM_001199838 | 0.488 | -0.142 |
| BCL2L2 | NM_001199839 | -0.197 | -0.6445 |
| SMURF1 | NM_001199847 | -0.018 | -0.859 |
| DAP3 | NM_001199849 | -0.766 | -1.349 |
| DAP3 | NM_001199850 | -0.766 | -1.349 |
| DAP3 | NM_001199851 | -0.766 | -1.349 |
| AK3 | NM_001199852 | -1.0535 | -0.948 |
| AK3 | NM_001199853 | -1.0535 | -0.948 |
| AK3 | NM_001199854 | -1.0535 | -0.948 |
| AK3 | NM_001199855 | -1.0535 | -0.948 |
| AK3 | NM_001199856 | -1.0535 | -0.948 |
| ANGPT1 | NM_001199859 | 0.113 | 0.099 |
| KCNAB2 | NM_001199860 | -0.5345 | -0.7975 |
| KCNAB2 | NM_001199861 | -0.5345 | -0.7975 |
| KCNAB2 | NM_001199862 | -0.5345 | -0.7975 |
| KCNAB2 | NM_001199863 | -0.5345 | -0.7975 |

| | | | |
|---|---|---|---|
| KIF16B | NM_001199865 | 0.0925 | -0.7335 |
| KIF16B | NM_001199866 | 0.0925 | -0.7335 |
| MARK4 | NM_001199867 | -0.1 | -0.61 |
| GLRX3 | NM_001199868 | -0.5545 | -1.143 |
| SERF2 | NM_001199875 | 0.172 | -0.2695 |
| SERF2 | NM_001199876 | 0.172 | -0.2695 |
| SERF2 | NM_001199877 | 0.172 | -0.2695 |
| SERF2 | NM_001199878 | 0.172 | -0.2695 |
| EGR3 | NM_001199880 | -0.1135 | -0.415 |
| EGR3 | NM_001199881 | -0.1135 | -0.415 |
| IL7 | NM_001199886 | -0.1255 | -0.126 |
| IL7 | NM_001199887 | -0.1255 | -0.126 |
| IL7 | NM_001199888 | -0.1255 | -0.126 |
| ACTG2 | NM_001199893 | 0.1455 | 0.2195 |
| C20orf103 | NM_001199897 | -0.273 | -0.3755 |
| PDK2 | NM_001199898 | -0.2915 | -0.6255 |
| PDK2 | NM_001199899 | -0.2915 | -0.6255 |
| PDK2 | NM_001199900 | -0.2915 | -0.6255 |
| PGM3 | NM_001199917 | -0.843 | -1.77 |
| PGM3 | NM_001199918 | -0.843 | -1.77 |
| PGM3 | NM_001199919 | -0.843 | -1.77 |
| SIAE | NM_001199922 | -0.0585 | -0.3865 |
| SESN1 | NM_001199933 | -1.3455 | -2.732 |
| SESN1 | NM_001199934 | -1.3455 | -2.732 |
| DOPEY1 | NM_001199942 | -0.2565 | -1.544 |
| SH3GL1 | NM_001199943 | -0.3395 | -0.972 |
| SH3GL1 | NM_001199944 | -0.3395 | -0.972 |
| ACTG1 | NM_001199954 | 0.067 | -0.427 |
| NDUFB5 | NM_001199957 | -0.591 | -0.991 |
| NDUFB5 | NM_001199958 | -0.591 | -0.991 |
| RPL36A | NM_001199972 | -0.0595 | -0.52 |
| UQCRB | NM_001199975 | -0.2305 | -0.266 |
| PLSCR2 | NM_001199978 | -0.1205 | -0.294 |
| PLSCR2 | NM_001199979 | -0.1205 | -0.294 |
| NDUFS1 | NM_001199981 | -0.957 | -1.692 |
| NDUFS1 | NM_001199982 | -0.957 | -1.692 |
| NDUFS1 | NM_001199983 | -0.957 | -1.692 |
| NDUFS1 | NM_001199984 | -0.957 | -1.692 |
| NDUFB6 | NM_001199987 | -0.3485 | -0.7025 |
| BMP2 | NM_001200 | 0.053 | -0.7165 |
| NOTCH2 | NM_001200001 | -0.0975 | -0.29 |
| OPRL1 | NM_001200019 | -1.03 | -1.4635 |
| NMNAT3 | NM_001200047 | -0.9115 | -1.598 |

| | | | |
|---|---|---|---|
| NPL | NM_001200050 | -1.021 | -0.68 |
| NPL | NM_001200051 | -1.021 | -0.68 |
| NPL | NM_001200052 | -1.021 | -0.68 |
| NPL | NM_001200056 | -1.021 | -0.68 |
| BMP3 | NM_001201 | 0.28 | 0.256 |
| PPP1R3B | NM_001201329 | 0.03 | -0.4425 |
| C2orf3 | NM_001201334 | 0.155 | -1.125 |
| C2orf3 | NM_001201335 | 0.155 | -1.125 |
| SAFB | NM_001201338 | -0.7525 | -1.85 |
| SAFB | NM_001201339 | -0.7525 | -1.85 |
| SAFB | NM_001201340 | -0.7525 | -1.85 |
| ASGR2 | NM_001201352 | 0.198 | -0.1445 |
| CCL25 | NM_001201359 | 0.214 | -0.3075 |
| RNF14 | NM_001201365 | -1.5105 | -3.03 |
| DNAL1 | NM_001201366 | -0.487 | -0.7585 |
| RNF7 | NM_001201370 | -0.175 | -0.8935 |
| ALDH7A1 | NM_001201377 | -0.467 | -0.5525 |
| EDNRB | NM_001201397 | -0.4215 | 0.038 |
| GALC | NM_001201401 | -0.8875 | -0.897 |
| GALC | NM_001201402 | -0.8875 | -0.897 |
| WASF2 | NM_001201404 | -0.047 | -0.7055 |
| ZNF778 | NM_001201407 | -0.847 | -1.2845 |
| DAAM2 | NM_001201427 | 0.3375 | -0.422 |
| GORASP2 | NM_001201428 | -0.741 | -2.1635 |
| CABIN1 | NM_001201429 | 0.189 | -0.578 |
| FLJ42280 | NM_001201450 | -0.124 | -0.6025 |
| FLJ42280 | NM_001201451 | -0.124 | -0.6025 |
| ZNF513 | NM_001201459 | -0.829 | -1.818 |
| TTYH1 | NM_001201461 | 0.0605 | -0.2985 |
| NETO1 | NM_001201465 | -0.866 | -1.0225 |
| CORO7 | NM_001201472 | -0.336 | -0.551 |
| CORO7 | NM_001201473 | -0.336 | -0.551 |
| C18orf21 | NM_001201474 | -1.18 | -2.666 |
| C18orf21 | NM_001201475 | -1.18 | -2.666 |
| C18orf21 | NM_001201476 | -1.18 | -2.666 |
| NETO2 | NM_001201477 | -1.36 | -2.516 |
| OSBPL6 | NM_001201480 | -0.6335 | -1.7275 |
| OSBPL6 | NM_001201481 | -0.6335 | -1.7275 |
| OSBPL6 | NM_001201482 | -0.6335 | -1.7275 |
| ENO1 | NM_001201483 | 0.1355 | -0.367 |
| PCSK2 | NM_001201528 | -0.133 | -0.3435 |
| PCSK2 | NM_001201529 | -0.133 | -0.3435 |
| TAF1A | NM_001201536 | -0.956 | -1.7995 |

| | | | |
|---|---|---|---|
| FAM161A | NM_001201543 | -2.0005 | -1.9825 |
| RBM15 | NM_001201545 | -0.9735 | -2.0685 |
| SLC16A4 | NM_001201546 | 0.437 | -0.4225 |
| SLC16A4 | NM_001201547 | 0.437 | -0.4225 |
| SLC16A4 | NM_001201548 | 0.437 | -0.4225 |
| SLC16A4 | NM_001201549 | 0.437 | -0.4225 |
| CFHR4 | NM_001201550 | 0.1145 | -0.1675 |
| CFHR4 | NM_001201551 | 0.1145 | -0.1675 |
| ZNF821 | NM_001201552 | -0.663 | -1.671 |
| ZNF821 | NM_001201553 | -0.663 | -1.671 |
| ZNF821 | NM_001201554 | -0.663 | -1.671 |
| ZNF821 | NM_001201556 | -0.663 | -1.671 |
| PLRG1 | NM_001201564 | -1.224 | -2.0575 |
| SEC22C | NM_001201572 | -0.698 | -1.946 |
| NUBPL | NM_001201573 | -0.715 | -0.376 |
| NUBPL | NM_001201574 | -0.715 | -0.376 |
| CTSL2 | NM_001201575 | -0.622 | -1.111 |
| PLSCR3 | NM_001201576 | -0.332 | -0.6095 |
| ASB11 | NM_001201583 | -0.091 | 0.024 |
| SEC22C | NM_001201584 | -0.698 | -1.946 |
| BMP4 | NM_001202 | 0.2185 | -0.3545 |
| NR4A1 | NM_001202233 | -0.594 | -1.4755 |
| ASB6 | NM_001202403 | -0.696 | -2.7405 |
| ALDH7A1 | NM_001202404 | -0.467 | -0.5525 |
| ZNF559 | NM_001202406 | -1.098 | -3.5055 |
| ZNF559 | NM_001202407 | -1.098 | -3.5055 |
| ZNF559 | NM_001202408 | -1.098 | -3.5055 |
| ZNF559 | NM_001202409 | -1.098 | -3.5055 |
| ZNF559 | NM_001202410 | -1.098 | -3.5055 |
| ZNF559 | NM_001202411 | -1.098 | -3.5055 |
| ZNF559 | NM_001202412 | -1.098 | -3.5055 |
| PRDX1 | NM_001202431 | -0.138 | 0.063 |
| SCN1A | NM_001202435 | -0.315 | -0.3145 |
| C10orf137 | NM_001202438 | -1.547 | -3.0605 |
| ADK | NM_001202449 | -0.902 | -0.845 |
| ADK | NM_001202450 | -0.902 | -0.845 |
| ZMYND11 | NM_001202464 | -1.3235 | -2.104 |
| ZMYND11 | NM_001202465 | -1.3235 | -2.104 |
| ZMYND11 | NM_001202466 | -1.3235 | -2.104 |
| ZMYND11 | NM_001202467 | -1.3235 | -2.104 |
| ZMYND11 | NM_001202468 | -1.3235 | -2.104 |
| GBAS | NM_001202469 | -1.23 | -1.315 |
| PGR | NM_001202474 | 0.006 | -0.2725 |

| | | | |
|---|---|---|---|
| CRHR2 | NM_001202475 | -0.442 | -0.621 |
| UBE2E1 | NM_001202476 | -0.9535 | -0.9985 |
| CRHR2 | NM_001202481 | -0.442 | -0.621 |
| CRHR2 | NM_001202482 | -0.442 | -0.621 |
| CRHR2 | NM_001202483 | -0.442 | -0.621 |
| UBE2G2 | NM_001202489 | -0.591 | -0.8965 |
| UBE2H | NM_001202498 | -1.0205 | -1.6345 |
| NIPSNAP1 | NM_001202502 | -0.1395 | -0.6835 |
| CTDP1 | NM_001202504 | -0.689 | -2.1235 |
| MXD1 | NM_001202513 | 0.0775 | -0.305 |
| MXD1 | NM_001202514 | 0.0775 | -0.305 |
| CFLAR | NM_001202515 | -0.587 | -0.769 |
| CFLAR | NM_001202516 | -0.587 | -0.769 |
| CFLAR | NM_001202517 | -0.587 | -0.769 |
| CFLAR | NM_001202518 | -0.587 | -0.769 |
| CFLAR | NM_001202519 | -0.587 | -0.769 |
| DDR1 | NM_001202521 | -0.0885 | -0.7975 |
| DDR1 | NM_001202522 | -0.0885 | -0.7975 |
| DDR1 | NM_001202523 | -0.0885 | -0.7975 |
| CUX1 | NM_001202543 | 0.11 | -0.0355 |
| CUX1 | NM_001202544 | 0.11 | -0.0355 |
| CUX1 | NM_001202545 | 0.11 | -0.0355 |
| CUX1 | NM_001202546 | 0.11 | -0.0355 |
| WDR27 | NM_001202550 | -0.7935 | -0.7385 |
| PIGV | NM_001202554 | -0.641 | -0.985 |
| CD44 | NM_001202555 | 0.1785 | -0.749 |
| CD44 | NM_001202556 | 0.1785 | -0.749 |
| CD44 | NM_001202557 | 0.1785 | -0.749 |
| WBSCR22 | NM_001202560 | 0.266 | -0.35 |
| CYP3A4 | NM_001202855 | -0.127 | 0.2815 |
| ECM1 | NM_001202858 | -0.205 | -0.624 |
| SHC1 | NM_001202859 | -0.1715 | -1.0885 |
| BMPR1B | NM_001203 | -0.9485 | -2.009 |
| SEMA4G | NM_001203244 | -0.042 | -0.667 |
| RPS10 | NM_001203245 | 0.1155 | -0.3135 |
| GPBP1 | NM_001203246 | -0.808 | -2.45 |
| EZH2 | NM_001203247 | -1.045 | -1.709 |
| EZH2 | NM_001203248 | -1.045 | -1.709 |
| EZH2 | NM_001203249 | -1.045 | -1.709 |
| MAPT | NM_001203251 | 0.0845 | -0.5055 |
| MAPT | NM_001203252 | 0.0845 | -0.5055 |
| MTA1 | NM_001203258 | 0.022 | -0.6295 |
| IL17RC | NM_001203263 | -0.0215 | -0.42 |

| | | | |
|---|---|---|---|
| IL17RC | NM_001203264 | -0.0215 | -0.42 |
| IL17RC | NM_001203265 | -0.0215 | -0.42 |
| BMPR2 | NM_001204 | -1.4635 | -3.3875 |
| SLC25A27 | NM_001204051 | -0.4455 | -0.4175 |
| SLC25A27 | NM_001204052 | -0.4455 | -0.4175 |
| NDUFC2 | NM_001204054 | 0.1625 | -0.2805 |
| NDUFC2 | NM_001204055 | 0.1625 | -0.2805 |
| ANKRD12 | NM_001204056 | 0.2205 | -0.6865 |
| CHURC1 | NM_001204063 | -1.208 | -0.718 |
| CHURC1 | NM_001204064 | -1.208 | -0.718 |
| ANKS1B | NM_001204065 | -0.3535 | -0.4295 |
| ANKS1B | NM_001204066 | -0.3535 | -0.4295 |
| ANKS1B | NM_001204067 | -0.3535 | -0.4295 |
| ANKS1B | NM_001204068 | -0.3535 | -0.4295 |
| ANKS1B | NM_001204069 | -0.3535 | -0.4295 |
| ANKS1B | NM_001204070 | -0.3535 | -0.4295 |
| KATNA1 | NM_001204076 | -1.009 | -3.057 |
| UBE4A | NM_001204077 | -0.78 | -1.4065 |
| ATP6V1G2 | NM_001204078 | 0.0255 | -0.322 |
| ANKS1B | NM_001204079 | -0.3535 | -0.4295 |
| ANKS1B | NM_001204080 | -0.3535 | -0.4295 |
| ANKS1B | NM_001204081 | -0.3535 | -0.4295 |
| NBL1 | NM_001204084 | -0.0625 | -0.5305 |
| NBL1 | NM_001204085 | -0.0625 | -0.5305 |
| NBL1 | NM_001204086 | -0.0625 | -0.5305 |
| KCNN3 | NM_001204087 | -0.193 | 0.091 |
| COX16 | NM_001204090 | -0.1235 | -0.2095 |
| RPS10 | NM_001204091 | 0.1155 | -0.3135 |
| CASC1 | NM_001204101 | -0.017 | -0.1445 |
| CASC1 | NM_001204102 | -0.017 | -0.1445 |
| BCL2L11 | NM_001204106 | -0.4175 | -1.9595 |
| BCL2L11 | NM_001204107 | -0.4175 | -1.9595 |
| BCL2L11 | NM_001204108 | -0.4175 | -1.9595 |
| BCL2L11 | NM_001204109 | -0.4175 | -1.9595 |
| BCL2L11 | NM_001204110 | -0.4175 | -1.9595 |
| BCL2L11 | NM_001204111 | -0.4175 | -1.9595 |
| BCL2L11 | NM_001204112 | -0.4175 | -1.9595 |
| BCL2L11 | NM_001204113 | -0.4175 | -1.9595 |
| CLEC17A | NM_001204118 | -0.0825 | -0.149 |
| LRMP | NM_001204126 | -0.1405 | -0.466 |
| LRMP | NM_001204127 | -0.1405 | -0.466 |
| MBD1 | NM_001204136 | -0.503 | -2.472 |
| MBD1 | NM_001204137 | -0.503 | -2.472 |

| | | | |
|---|---|---|---|
| MBD1 | NM_001204138 | -0.503 | -2.472 |
| MBD1 | NM_001204139 | -0.503 | -2.472 |
| MBD1 | NM_001204140 | -0.503 | -2.472 |
| MBD1 | NM_001204141 | -0.503 | -2.472 |
| MBD1 | NM_001204142 | -0.503 | -2.472 |
| MBD1 | NM_001204143 | -0.503 | -2.472 |
| HDAC9 | NM_001204144 | -0.862 | -2.3525 |
| HDAC9 | NM_001204145 | -0.862 | -2.3525 |
| HDAC9 | NM_001204146 | -0.862 | -2.3525 |
| HDAC9 | NM_001204147 | -0.862 | -2.3525 |
| HDAC9 | NM_001204148 | -0.862 | -2.3525 |
| MBD1 | NM_001204151 | -0.503 | -2.472 |
| SPHK2 | NM_001204158 | 0.0025 | -0.1975 |
| SPHK2 | NM_001204159 | 0.0025 | -0.1975 |
| SPHK2 | NM_001204160 | 0.0025 | -0.1975 |
| MDM4 | NM_001204171 | -1.3115 | -1.2645 |
| MDM4 | NM_001204172 | -1.3115 | -1.2645 |
| CHODL | NM_001204174 | -0.395 | -0.7455 |
| CHODL | NM_001204175 | -0.395 | -0.7455 |
| CHODL | NM_001204176 | -0.395 | -0.7455 |
| CHODL | NM_001204177 | -0.395 | -0.7455 |
| CHODL | NM_001204178 | -0.395 | -0.7455 |
| NEK2 | NM_001204182 | -1.202 | -1.6875 |
| NEK2 | NM_001204183 | -1.202 | -1.6875 |
| TP73 | NM_001204184 | -0.0585 | -0.73 |
| TP73 | NM_001204185 | -0.0585 | -0.73 |
| TP73 | NM_001204186 | -0.0585 | -0.73 |
| TP73 | NM_001204187 | -0.0585 | -0.73 |
| TP73 | NM_001204188 | -0.0585 | -0.73 |
| TP73 | NM_001204189 | -0.0585 | -0.73 |
| TP73 | NM_001204190 | -0.0585 | -0.73 |
| TP73 | NM_001204191 | -0.0585 | -0.73 |
| TP73 | NM_001204192 | -0.0585 | -0.73 |
| SRP19 | NM_001204193 | -1.294 | -3.035 |
| SRP19 | NM_001204194 | -1.294 | -3.035 |
| SRP19 | NM_001204196 | -1.294 | -3.035 |
| NBEA | NM_001204197 | -1.4285 | -1.5005 |
| SRP19 | NM_001204199 | -1.294 | -3.035 |
| SEC14L2 | NM_001204204 | -0.0205 | -0.3355 |
| NOS1 | NM_001204213 | -0.1035 | -0.42 |
| NOS1 | NM_001204214 | -0.1035 | -0.42 |
| NOS1 | NM_001204218 | -0.1035 | -0.42 |
| ZNF695 | NM_001204221 | -0.0785 | -0.464 |

| | | | |
|---|---|---|---|
| TBC1D10A | NM_001204240 | -0.7015 | -1.9335 |
| SCARB2 | NM_001204255 | 0.284 | -0.1625 |
| NR3C1 | NM_001204258 | -0.2885 | -0.689 |
| NR3C1 | NM_001204259 | -0.2885 | -0.689 |
| NR3C1 | NM_001204260 | -0.2885 | -0.689 |
| NR3C1 | NM_001204261 | -0.2885 | -0.689 |
| NR3C1 | NM_001204262 | -0.2885 | -0.689 |
| NR3C1 | NM_001204263 | -0.2885 | -0.689 |
| NR3C1 | NM_001204264 | -0.2885 | -0.689 |
| NR3C1 | NM_001204265 | -0.2885 | -0.689 |
| GABRA4 | NM_001204266 | 0.055 | 0.162 |
| GABRA4 | NM_001204267 | 0.055 | 0.162 |
| KCNA2 | NM_001204269 | -0.035 | 0.022 |
| PPP5C | NM_001204284 | -0.004 | -0.82 |
| MUC1 | NM_001204285 | -0.1105 | -0.321 |
| MUC1 | NM_001204286 | -0.1105 | -0.321 |
| MUC1 | NM_001204287 | -0.1105 | -0.321 |
| MUC1 | NM_001204288 | -0.1105 | -0.321 |
| MUC1 | NM_001204289 | -0.1105 | -0.321 |
| MUC1 | NM_001204290 | -0.1105 | -0.321 |
| MUC1 | NM_001204291 | -0.1105 | -0.321 |
| MUC1 | NM_001204292 | -0.1105 | -0.321 |
| MUC1 | NM_001204293 | -0.1105 | -0.321 |
| MUC1 | NM_001204294 | -0.1105 | -0.321 |
| MUC1 | NM_001204295 | -0.1105 | -0.321 |
| MUC1 | NM_001204296 | -0.1105 | -0.321 |
| MUC1 | NM_001204297 | -0.1105 | -0.321 |
| ZNF664 | NM_001204298 | -0.9385 | -0.859 |
| ARHGAP19 | NM_001204300 | -1.024 | -1.638 |
| APP | NM_001204301 | -0.3025 | -0.3875 |
| APP | NM_001204302 | -0.3025 | -0.3875 |
| APP | NM_001204303 | -0.3025 | -0.3875 |
| PRLR | NM_001204314 | -0.028 | 0.0215 |
| PRLR | NM_001204315 | -0.028 | 0.0215 |
| PRLR | NM_001204316 | -0.028 | 0.0215 |
| PRLR | NM_001204317 | -0.028 | 0.0215 |
| PRLR | NM_001204318 | -0.028 | 0.0215 |
| TNFSF15 | NM_001204344 | -0.2215 | -0.1185 |
| SPOCK3 | NM_001204352 | -0.615 | -0.4285 |
| SPOCK3 | NM_001204353 | -0.615 | -0.4285 |
| SPOCK3 | NM_001204354 | -0.615 | -0.4285 |
| SPOCK3 | NM_001204355 | -0.615 | -0.4285 |
| SPOCK3 | NM_001204356 | -0.615 | -0.4285 |

| | | | |
|---|---|---|---|
| SPOCK3 | NM_001204358 | -0.615 | -0.4285 |
| SPOCK3 | NM_001204359 | -0.615 | -0.4285 |
| FAM24B | NM_001204364 | 0.246 | -0.287 |
| MGST2 | NM_001204366 | 0.0645 | -0.026 |
| MGST2 | NM_001204367 | 0.0645 | -0.026 |
| MGST2 | NM_001204368 | 0.0645 | -0.026 |
| NPR3 | NM_001204375 | 0.214 | -0.245 |
| NPR3 | NM_001204376 | 0.214 | -0.245 |
| VEGFA | NM_001204384 | -0.674 | -0.6745 |
| VEGFA | NM_001204385 | -0.674 | -0.6745 |
| DLG1 | NM_001204386 | -1.5135 | -3.089 |
| DLG1 | NM_001204387 | -1.5135 | -3.089 |
| DLG1 | NM_001204388 | -1.5135 | -3.089 |
| XIAP | NM_001204401 | -1.9625 | -2.4475 |
| PRPS1 | NM_001204402 | -0.487 | -0.536 |
| ANK3 | NM_001204403 | -0.1175 | -1.2675 |
| ANK3 | NM_001204404 | -0.1175 | -1.2675 |
| SEC14L1 | NM_001204408 | 0.111 | -0.986 |
| SEC14L1 | NM_001204410 | 0.111 | -0.986 |
| TINAGL1 | NM_001204414 | -0.337 | -0.2875 |
| TINAGL1 | NM_001204415 | -0.337 | -0.2875 |
| RGS6 | NM_001204416 | 0.1935 | -0.2925 |
| RGS6 | NM_001204417 | 0.1935 | -0.2925 |
| RGS6 | NM_001204418 | 0.1935 | -0.2925 |
| RGS6 | NM_001204419 | 0.1935 | -0.2925 |
| RGS6 | NM_001204420 | 0.1935 | -0.2925 |
| RGS6 | NM_001204421 | 0.1935 | -0.2925 |
| RGS6 | NM_001204422 | 0.1935 | -0.2925 |
| RGS6 | NM_001204423 | 0.1935 | -0.2925 |
| RGS6 | NM_001204424 | 0.1935 | -0.2925 |
| LIMK1 | NM_001204426 | -0.4925 | -0.764 |
| C20orf29 | NM_001204446 | -0.8105 | -1.6785 |
| C20orf29 | NM_001204447 | -0.8105 | -1.6785 |
| CCPG1 | NM_001204450 | -0.52 | -1.423 |
| CCPG1 | NM_001204451 | -0.52 | -1.423 |
| PSMC2 | NM_001204453 | -1.085 | -1.7615 |
| RBAK | NM_001204456 | -0.753 | -3.574 |
| TNFRSF19 | NM_001204458 | -0.231 | -0.731 |
| TNFRSF19 | NM_001204459 | -0.231 | -0.731 |
| RBM10 | NM_001204466 | -0.748 | -1.5345 |
| RBM10 | NM_001204467 | -0.748 | -1.5345 |
| RBM10 | NM_001204468 | -0.748 | -1.5345 |
| CAMK2G | NM_001204492 | -0.476 | -0.1155 |

| | | | |
|---|---|---|---|
| FLT3LG | NM_001204502 | -0.402 | -0.41 |
| FLT3LG | NM_001204503 | -0.402 | -0.41 |
| DGKH | NM_001204504 | -0.112 | -0.3965 |
| DGKH | NM_001204505 | -0.112 | -0.3965 |
| DGKH | NM_001204506 | -0.112 | -0.3965 |
| EIF4A1 | NM_001204510 | -0.018 | -0.2655 |
| KIFAP3 | NM_001204514 | -0.737 | -1.367 |
| KIFAP3 | NM_001204516 | -0.737 | -1.367 |
| KIFAP3 | NM_001204517 | -0.737 | -1.367 |
| P2RX5 | NM_001204519 | -0.3315 | -1.107 |
| P2RX5 | NM_001204520 | -0.3315 | -1.107 |
| SSR4 | NM_001204526 | 0.271 | -0.15 |
| SSR4 | NM_001204527 | 0.271 | -0.15 |
| CDH16 | NM_001204744 | 0.394 | -0.267 |
| CDH16 | NM_001204745 | 0.394 | -0.267 |
| CDH16 | NM_001204746 | 0.394 | -0.267 |
| RFC1 | NM_001204747 | -0.7545 | -2.2455 |
| TLL1 | NM_001204760 | -0.2835 | -0.729 |
| KCNH2 | NM_001204798 | -0.4365 | -1.796 |
| ANO6 | NM_001204803 | -1.454 | -1.2275 |
| NT5E | NM_001204813 | -0.5125 | -0.52 |
| ZNF586 | NM_001204814 | -1.2695 | -1.7975 |
| ZNF587 | NM_001204817 | -0.8765 | -0.9715 |
| XPNPEP3 | NM_001204827 | -1.33 | -2.076 |
| ZNF568 | NM_001204835 | -0.113 | -0.859 |
| ZNF568 | NM_001204836 | -0.113 | -0.859 |
| ZNF568 | NM_001204837 | -0.113 | -0.859 |
| ZNF568 | NM_001204838 | -0.113 | -0.859 |
| ZNF568 | NM_001204839 | -0.113 | -0.859 |
| SCNM1 | NM_001204856 | -0.1915 | -0.9175 |
| TCEB1 | NM_001204857 | -0.623 | -1.14 |
| TCEB1 | NM_001204858 | -0.623 | -1.14 |
| TCEB1 | NM_001204859 | -0.623 | -1.14 |
| TCEB1 | NM_001204860 | -0.623 | -1.14 |
| TCEB1 | NM_001204861 | -0.623 | -1.14 |
| TCEB1 | NM_001204862 | -0.623 | -1.14 |
| TCEB1 | NM_001204863 | -0.623 | -1.14 |
| TCEB1 | NM_001204864 | -0.623 | -1.14 |
| STX16 | NM_001204868 | -1.417 | -1.407 |
| WISP1 | NM_001204869 | 0.096 | -0.1165 |
| WISP1 | NM_001204870 | 0.096 | -0.1165 |
| BAG4 | NM_001204878 | -1.3245 | -2.9755 |
| UBE2D1 | NM_001204880 | -1.0135 | -1.645 |

| | | | |
|---|---|---|---|
| ALDH2 | NM_001204889 | 0.114 | -0.371 |
| C16orf57 | NM_001204911 | 0.0165 | -0.329 |
| PBX1 | NM_001204961 | -0.9275 | -0.778 |
| PBX1 | NM_001204963 | -0.9275 | -0.778 |
| BNIP1 | NM_001205 | -0.6045 | -1.8855 |
| GK | NM_001205019 | -1.097 | -1.8855 |
| MDM1 | NM_001205028 | -1.0145 | -0.964 |
| MDM1 | NM_001205029 | -1.0145 | -0.964 |
| ATG13 | NM_001205119 | -0.4675 | -1.8515 |
| ATG13 | NM_001205120 | -0.4675 | -1.8515 |
| ATG13 | NM_001205121 | -0.4675 | -1.8515 |
| ATG13 | NM_001205122 | -0.4675 | -1.8515 |
| LRTOMT | NM_001205138 | -0.0875 | -0.4045 |
| ALKBH2 | NM_001205179 | -0.805 | -2.2345 |
| ALKBH2 | NM_001205180 | -0.805 | -2.2345 |
| MECOM | NM_001205194 | -0.401 | -0.379 |
| SLC19A1 | NM_001205206 | -0.8135 | -1.8015 |
| SLC19A1 | NM_001205207 | -0.8135 | -1.8015 |
| CRISP1 | NM_001205220 | -0.225 | -0.141 |
| SORT1 | NM_001205228 | -1.048 | -1.288 |
| ACSL6 | NM_001205247 | -0.249 | -0.309 |
| ACSL6 | NM_001205248 | -0.249 | -0.309 |
| ACSL6 | NM_001205250 | -0.249 | -0.309 |
| ACSL6 | NM_001205251 | -0.249 | -0.309 |
| RAD54B | NM_001205262 | 0.2995 | -1.086 |
| RAD54B | NM_001205263 | 0.2995 | -1.086 |
| CACNA1E | NM_001205293 | -0.149 | -0.064 |
| CACNA1E | NM_001205294 | -0.149 | -0.064 |
| TTF1 | NM_001205296 | -0.692 | -1.0905 |
| EGFLAM | NM_001205301 | -0.277 | -0.894 |
| STEAP4 | NM_001205315 | 0.0535 | -0.2615 |
| STEAP4 | NM_001205316 | 0.0535 | -0.2615 |
| RIMS4 | NM_001205317 | -0.0065 | -0.483 |
| NXN | NM_001205319 | -0.008 | -0.6505 |
| JAM3 | NM_001205329 | -0.324 | -1.404 |
| CEACAM1 | NM_001205344 | 0.085 | -0.347 |
| KLF9 | NM_001206 | -0.995 | -2.3565 |
| USMG5 | NM_001206426 | 0.12 | 0.041 |
| USMG5 | NM_001206427 | 0.12 | 0.041 |
| ELMO1 | NM_001206480 | -0.1695 | -0.73 |
| ELMO1 | NM_001206482 | -0.1695 | -0.73 |
| ATF3 | NM_001206484 | -0.3345 | 1.001 |
| ATF3 | NM_001206485 | -0.3345 | 1.001 |

| | | | |
|---|---|---|---|
| ATF3 | NM_001206486 | -0.3345 | 1.001 |
| ATF3 | NM_001206488 | -0.3345 | 1.001 |
| MAVS | NM_001206491 | -1.15 | -1.6835 |
| CASP10 | NM_001206524 | 0.2235 | 0.0155 |
| CAPZB | NM_001206540 | 0.034 | -0.666 |
| CASP10 | NM_001206542 | 0.2235 | 0.0155 |
| KIAA1432 | NM_001206557 | -0.562 | -1.129 |
| POLR1D | NM_001206559 | -0.681 | -1.811 |
| IFI16 | NM_001206567 | 0.262 | -1.8705 |
| SORCS1 | NM_001206569 | -0.958 | -0.506 |
| SORCS1 | NM_001206570 | -0.958 | -0.506 |
| SORCS1 | NM_001206571 | -0.958 | -0.506 |
| SORCS1 | NM_001206572 | -0.958 | -0.506 |
| CHN1 | NM_001206602 | -0.9215 | -1.0645 |
| SELPLG | NM_001206609 | 0.181 | -0.117 |
| EHF | NM_001206615 | -0.2185 | -0.4535 |
| EHF | NM_001206616 | -0.2185 | -0.4535 |
| SH3GLB1 | NM_001206651 | -0.6525 | -2.0345 |
| SH3GLB1 | NM_001206652 | -0.6525 | -2.0345 |
| SH3GLB1 | NM_001206653 | -0.6525 | -2.0345 |
| NCOR2 | NM_001206654 | 0.005 | -0.558 |
| IGSF8 | NM_001206665 | 0.093 | -0.45 |
| PLA2G4E | NM_001206670 | 0.0675 | -0.083 |
| RIC3 | NM_001206671 | -0.736 | -1.2185 |
| RIC3 | NM_001206672 | -0.736 | -1.2185 |
| ABHD12B | NM_001206673 | -0.282 | -0.1365 |
| ATF7 | NM_001206682 | -0.6065 | -0.2775 |
| ATF7 | NM_001206683 | -0.6065 | -0.2775 |
| RFX4 | NM_001206691 | 0.2 | -0.0625 |
| IRF6 | NM_001206696 | 0.3485 | -0.2805 |
| SP100 | NM_001206701 | 0.134 | -0.2535 |
| SP100 | NM_001206702 | 0.134 | -0.2535 |
| SP100 | NM_001206703 | 0.134 | -0.2535 |
| SP100 | NM_001206704 | 0.134 | -0.2535 |
| PRKAG1 | NM_001206709 | -0.668 | -1.343 |
| PRKAG1 | NM_001206710 | -0.668 | -1.343 |
| AKT3 | NM_001206729 | -0.707 | -1.202 |
| NSFL1C | NM_001206736 | 0.013 | -0.898 |
| ITGB3BP | NM_001206739 | -0.905 | -1.0665 |
| HSD11B1 | NM_001206741 | 0.0395 | -0.0545 |
| TPO | NM_001206744 | 0.244 | -0.0135 |
| TPO | NM_001206745 | 0.244 | -0.0135 |
| CAV2 | NM_001206747 | 0.348 | -0.08 |

| | | | |
|---|---|---|---|
| CAV2 | NM_001206748 | 0.348 | -0.08 |
| DLG2 | NM_001206769 | -0.104 | -0.1645 |
| GTF3C3 | NM_001206774 | -1.034 | -2.012 |
| TMPRSS13 | NM_001206789 | 0.0195 | -0.0005 |
| TMPRSS13 | NM_001206790 | 0.0195 | -0.0005 |
| ARID4B | NM_001206794 | -0.3485 | -1.9785 |
| PKM2 | NM_001206796 | 0.2085 | -0.32 |
| PKM2 | NM_001206797 | 0.2085 | -0.32 |
| PKM2 | NM_001206798 | 0.2085 | -0.32 |
| PKM2 | NM_001206799 | 0.2085 | -0.32 |
| RFC5 | NM_001206801 | -0.6595 | -1.6615 |
| TP53I3 | NM_001206802 | -0.4655 | -1.214 |
| MAP2K5 | NM_001206804 | -0.421 | -0.4285 |
| RAB11A | NM_001206836 | -0.3755 | -0.5395 |
| PTPRZ1 | NM_001206838 | 0.0475 | -0.2985 |
| PTPRZ1 | NM_001206839 | 0.0475 | -0.2985 |
| TGOLN2 | NM_001206840 | -1.2205 | -1.5165 |
| TGOLN2 | NM_001206841 | -1.2205 | -1.5165 |
| GSG1 | NM_001206842 | 0.109 | 0.452 |
| GSG1 | NM_001206843 | 0.109 | 0.452 |
| TGOLN2 | NM_001206844 | -1.2205 | -1.5165 |
| GSG1 | NM_001206845 | 0.109 | 0.452 |
| ASPM | NM_001206846 | -0.921 | -3.1225 |
| NLGN4Y | NM_001206850 | 0.0205 | -0.5245 |
| CAPRIN2 | NM_001206856 | -1.5535 | -2.0835 |
| STRADB | NM_001206864 | -0.715 | -1.189 |
| IL6R | NM_001206866 | -0.468 | -0.3015 |
| CTDSP1 | NM_001206878 | -0.375 | -0.399 |
| CTDSP1 | NM_001206879 | -0.375 | -0.399 |
| MRVI1 | NM_001206880 | 0.1675 | -0.759 |
| MRVI1 | NM_001206881 | 0.1675 | -0.759 |
| CTNND1 | NM_001206883 | -0.546 | -1.48 |
| CTNND1 | NM_001206884 | -0.546 | -1.48 |
| CTNND1 | NM_001206885 | -0.546 | -1.48 |
| CTNND1 | NM_001206886 | -0.546 | -1.48 |
| CTNND1 | NM_001206887 | -0.546 | -1.48 |
| CTNND1 | NM_001206888 | -0.546 | -1.48 |
| CTNND1 | NM_001206889 | -0.546 | -1.48 |
| CTNND1 | NM_001206890 | -0.546 | -1.48 |
| CTNND1 | NM_001206891 | -0.546 | -1.48 |
| ALDH1A2 | NM_001206897 | -0.0655 | 0.0445 |
| TAX1BP1 | NM_001206901 | -0.0635 | -0.6615 |
| TAX1BP1 | NM_001206902 | -0.0635 | -0.6615 |

| | | | |
|---|---|---|---|
| CACNB3 | NM_001206915 | -0.331 | -0.988 |
| CACNB3 | NM_001206916 | -0.331 | -0.988 |
| CACNB3 | NM_001206917 | -0.331 | -0.988 |
| CD86 | NM_001206924 | -0.29 | -0.3095 |
| CD86 | NM_001206925 | -0.29 | -0.3095 |
| DNAH8 | NM_001206927 | -0.2485 | -0.6205 |
| AGER | NM_001206929 | -0.0285 | -0.2605 |
| SLC6A12 | NM_001206931 | 0.0565 | -0.349 |
| AGER | NM_001206932 | -0.0285 | -0.2605 |
| AGER | NM_001206934 | -0.0285 | -0.2605 |
| AGER | NM_001206936 | -0.0285 | -0.2605 |
| FAM115A | NM_001206938 | -0.791 | -1.2205 |
| AGER | NM_001206940 | -0.0285 | -0.2605 |
| FAM115A | NM_001206941 | -0.791 | -1.2205 |
| CSPG5 | NM_001206942 | 0.172 | -0.3795 |
| CSPG5 | NM_001206943 | 0.172 | -0.3795 |
| CSPG5 | NM_001206944 | 0.172 | -0.3795 |
| CSPG5 | NM_001206945 | 0.172 | -0.3795 |
| PICALM | NM_001206946 | -0.851 | -1.871 |
| PICALM | NM_001206947 | -0.851 | -1.871 |
| SLC16A3 | NM_001206950 | 0.023 | -0.6925 |
| SLC16A3 | NM_001206951 | 0.023 | -0.6925 |
| SLC16A3 | NM_001206952 | 0.023 | -0.6925 |
| AGER | NM_001206954 | -0.0285 | -0.2605 |
| RASSF1 | NM_001206957 | 0.833 | 0.5995 |
| AGER | NM_001206966 | -0.0285 | -0.2605 |
| PTPRB | NM_001206971 | 0.138 | -0.102 |
| PTPRB | NM_001206972 | 0.138 | -0.102 |
| DHPS | NM_001206974 | -0.027 | -0.3955 |
| NR1H4 | NM_001206977 | -0.443 | -0.311 |
| NR1H4 | NM_001206978 | -0.443 | -0.311 |
| NR1H4 | NM_001206979 | -0.443 | -0.311 |
| C4orf26 | NM_001206981 | -0.085 | 0.0265 |
| NR1H4 | NM_001206992 | -0.443 | -0.311 |
| NR1H4 | NM_001206993 | -0.443 | -0.311 |
| PPP2R2C | NM_001206994 | 0.0455 | -0.444 |
| PPP2R2C | NM_001206995 | 0.0455 | -0.444 |
| PPP2R2C | NM_001206996 | 0.0455 | -0.444 |
| C4orf22 | NM_001206997 | 0.5225 | 0.017 |
| CIT | NM_001206999 | -0.8815 | -1.848 |
| BTF3 | NM_001207 | -0.2145 | -0.619 |
| HNRPDL | NM_001207000 | -1.353 | -0.9925 |
| C5orf56 | NM_001207001 | -0.0585 | -0.602 |

| | | | |
|---|---|---|---|
| C5orf56 | NM_001207002 | -0.0585 | -0.602 |
| C5orf56 | NM_001207003 | -0.0585 | -0.602 |
| UGT2B28 | NM_001207004 | -0.4495 | -0.511 |
| ZNF233 | NM_001207005 | -1.236 | -0.688 |
| IL21 | NM_001207006 | -0.084 | -0.094 |
| MLLT4 | NM_001207008 | -0.349 | -0.9015 |
| ZNF497 | NM_001207009 | -0.279 | -0.3395 |
| CLEC12A | NM_001207010 | -0.1225 | 0.2765 |
| PGF | NM_001207012 | -0.8635 | -3.248 |
| PTPN2 | NM_001207013 | 0.016 | -1.1395 |
| SERPINH1 | NM_001207014 | -0.1115 | -1.661 |
| PTPRR | NM_001207015 | -0.95 | -1.8585 |
| PTPRR | NM_001207016 | -0.95 | -1.8585 |
| SMTN | NM_001207017 | -0.232 | -0.64 |
| SMTN | NM_001207018 | -0.232 | -0.64 |
| FCER2 | NM_001207019 | 0.1375 | -0.3115 |
| IQCF3 | NM_001207023 | 0.28 | 0.086 |
| M6PR | NM_001207024 | -0.5595 | -1.9085 |
| POU2F2 | NM_001207025 | -0.15 | -0.759 |
| POU2F2 | NM_001207026 | -0.15 | -0.759 |
| RNF40 | NM_001207033 | -0.2625 | -1.1055 |
| RNF40 | NM_001207034 | -0.2625 | -1.1055 |
| ETV7 | NM_001207035 | 0.2655 | -0.104 |
| ETV7 | NM_001207036 | 0.2655 | -0.104 |
| ETV7 | NM_001207037 | 0.2655 | -0.104 |
| ETV7 | NM_001207038 | 0.2655 | -0.104 |
| ETV7 | NM_001207039 | 0.2655 | -0.104 |
| ETV7 | NM_001207040 | 0.2655 | -0.104 |
| ETV7 | NM_001207041 | 0.2655 | -0.104 |
| PRMT1 | NM_001207042 | -0.1365 | -0.383 |
| LOC554223 | NM_001207043 | -0.529 | 0.273 |
| CLASP2 | NM_001207044 | -0.98 | -1.246 |
| ATRN | NM_001207047 | 0.1085 | -0.3095 |
| BACE1 | NM_001207048 | -0.613 | -1.0805 |
| BACE1 | NM_001207049 | -0.613 | -1.0805 |
| CLASP1 | NM_001207051 | -0.155 | -0.906 |
| KLK7 | NM_001207053 | 0.075 | -0.4105 |
| DNMT3B | NM_001207055 | -0.217 | -0.679 |
| DNMT3B | NM_001207056 | -0.217 | -0.679 |
| C15orf44 | NM_001207058 | -1.312 | -3.044 |
| C15orf44 | NM_001207059 | -1.312 | -3.044 |
| C22orf23 | NM_001207062 | -0.267 | -0.3445 |
| CXADR | NM_001207063 | -0.8815 | -1.0315 |

| | | | |
|---|---|---|---|
| CXADR | NM_001207064 | -0.8815 | -1.0315 |
| CXADR | NM_001207065 | -0.8815 | -1.0315 |
| CXADR | NM_001207066 | -0.8815 | -1.0315 |
| BZW1 | NM_001207067 | -0.8115 | -0.797 |
| BZW1 | NM_001207068 | -0.8115 | -0.797 |
| BZW1 | NM_001207069 | -0.8115 | -0.797 |
| C20orf152 | NM_001207076 | -0.0985 | -0.4025 |
| BUB1B | NM_001211 | -1.441 | -2.9435 |
| C1QBP | NM_001212 | -0.02 | -0.4515 |
| C16orf3 | NM_001214 | 0.115 | -0.174 |
| ESR2 | NM_001214902 | -0.0185 | -0.086 |
| ESR2 | NM_001214903 | -0.0185 | -0.086 |
| ZNF48 | NM_001214906 | -0.438 | -0.9335 |
| ZNF48 | NM_001214907 | -0.438 | -0.9335 |
| ZNF48 | NM_001214909 | -0.438 | -0.9335 |
| CA6 | NM_001215 | -0.206 | -0.271 |
| CA9 | NM_001216 | 0.07 | -0.4255 |
| CA11 | NM_001217 | -0.067 | -0.51 |
| CA12 | NM_001218 | -0.858 | -1.8465 |
| CALU | NM_001219 | -0.828 | -0.878 |
| CAMK2B | NM_001220 | -0.105 | -0.256 |
| PPFIA2 | NM_001220473 | -0.055 | -0.2715 |
| PPFIA2 | NM_001220474 | -0.055 | -0.2715 |
| PPFIA2 | NM_001220475 | -0.055 | -0.2715 |
| PPFIA2 | NM_001220476 | -0.055 | -0.2715 |
| PPFIA2 | NM_001220477 | -0.055 | -0.2715 |
| PPFIA2 | NM_001220478 | -0.055 | -0.2715 |
| PPFIA2 | NM_001220479 | -0.055 | -0.2715 |
| PPFIA2 | NM_001220480 | -0.055 | -0.2715 |
| MEIS2 | NM_001220482 | -0.9845 | -2.763 |
| CDH13 | NM_001220488 | -0.5675 | -0.28 |
| CDH13 | NM_001220489 | -0.5675 | -0.28 |
| CDH13 | NM_001220490 | -0.5675 | -0.28 |
| CDH13 | NM_001220491 | -0.5675 | -0.28 |
| CDH13 | NM_001220492 | -0.5675 | -0.28 |
| GLYATL1 | NM_001220494 | -0.0535 | -0.276 |
| GLYATL1 | NM_001220496 | -0.0535 | -0.276 |
| FCER2 | NM_001220500 | 0.1375 | -0.3115 |
| IKZF1 | NM_001220765 | -0.056 | -0.385 |
| IKZF1 | NM_001220766 | -0.056 | -0.385 |
| IKZF1 | NM_001220767 | -0.056 | -0.385 |
| IKZF1 | NM_001220768 | -0.056 | -0.385 |
| IKZF1 | NM_001220769 | -0.056 | -0.385 |

| | | | |
|---|---|---|---|
| IKZF1 | NM_001220770 | -0.056 | -0.385 |
| IKZF1 | NM_001220771 | -0.056 | -0.385 |
| IKZF1 | NM_001220772 | -0.056 | -0.385 |
| IKZF1 | NM_001220773 | -0.056 | -0.385 |
| IKZF1 | NM_001220774 | -0.056 | -0.385 |
| IKZF1 | NM_001220775 | -0.056 | -0.385 |
| IKZF1 | NM_001220776 | -0.056 | -0.385 |
| CAMK2D | NM_001221 | -0.8105 | -1.0155 |
| CAMK2G | NM_001222 | -0.476 | -0.1155 |
| CASP1 | NM_001223 | 0.024 | 0.318 |
| CASP2 | NM_001224 | -0.7065 | -1.2805 |
| CASP4 | NM_001225 | -0.092 | -0.2085 |
| CASP6 | NM_001226 | -0.571 | -1.925 |
| CASP7 | NM_001227 | -1.071 | -1.6805 |
| CASP8 | NM_001228 | 0.105 | 0.061 |
| CASP9 | NM_001229 | -0.281 | -0.646 |
| CASP10 | NM_001230 | 0.2235 | 0.0155 |
| CASQ1 | NM_001231 | 0.4935 | -0.025 |
| CASQ2 | NM_001232 | -0.1765 | -0.0485 |
| CAV2 | NM_001233 | 0.498 | 0.241 |
| CAV3 | NM_001234 | -0.012 | -0.465 |
| SERPINH1 | NM_001235 | -0.1115 | -1.661 |
| CBR3 | NM_001236 | 0.127 | -1.1155 |
| CCNA2 | NM_001237 | -1.279 | -2.102 |
| CCNE1 | NM_001238 | -0.7685 | -2.3855 |
| CCNH | NM_001239 | -0.8415 | -2.3375 |
| CCNT1 | NM_001240 | -0.432 | -1.0395 |
| CCNT2 | NM_001241 | -0.1945 | -1.091 |
| CD27 | NM_001242 | -0.125 | -0.943 |
| PITRM1 | NM_001242307 | -0.245 | -0.7995 |
| PITRM1 | NM_001242309 | -0.245 | -0.7995 |
| EDA2R | NM_001242310 | -0.285 | -0.6325 |
| FAM124A | NM_001242312 | -1.583 | -2.2085 |
| MAP3K13 | NM_001242314 | -0.6495 | -1.591 |
| MAP3K13 | NM_001242317 | -0.6495 | -1.591 |
| PDE7A | NM_001242318 | -0.941 | -2.5165 |
| LHX6 | NM_001242333 | -0.398 | -0.4705 |
| LHX6 | NM_001242334 | -0.398 | -0.4705 |
| LHX6 | NM_001242335 | -0.398 | -0.4705 |
| MFAP3 | NM_001242336 | -1.5585 | -2.1475 |
| C1orf101 | NM_001242340 | -0.1035 | -0.8605 |
| VNN2 | NM_001242350 | -0.0095 | 0.0735 |
| ODF2 | NM_001242352 | -0.4545 | -0.8675 |

| | | | |
|---|---|---|---|
| ODF2 | NM_001242353 | -0.4545 | -0.8675 |
| ODF2 | NM_001242354 | -0.4545 | -0.8675 |
| NCR1 | NM_001242356 | 0.136 | 0.169 |
| NCR1 | NM_001242357 | 0.136 | 0.169 |
| RHOBTB1 | NM_001242359 | -0.933 | -1.701 |
| NRD1 | NM_001242361 | -1.1065 | -0.7305 |
| SLC46A1 | NM_001242366 | -1.1485 | -1.6215 |
| C9orf7 | NM_001242369 | -0.2285 | -0.8915 |
| C9orf7 | NM_001242370 | -0.2285 | -0.8915 |
| ANAPC13 | NM_001242374 | -1.2855 | -1.5425 |
| ANAPC13 | NM_001242375 | -1.2855 | -1.5425 |
| GFAP | NM_001242376 | 0.0515 | -0.47 |
| DCP2 | NM_001242377 | -0.864 | -1.3175 |
| SYTL3 | NM_001242384 | 0.007 | -1.0465 |
| MAK | NM_001242385 | -0.356 | -0.1935 |
| ACSS2 | NM_001242393 | -0.1015 | -0.494 |
| SYTL3 | NM_001242394 | 0.007 | -1.0465 |
| SYTL3 | NM_001242395 | 0.007 | -1.0465 |
| TWF1 | NM_001242397 | -0.8495 | -1.426 |
| GHR | NM_001242399 | -0.4875 | 0.1065 |
| GHR | NM_001242400 | -0.4875 | 0.1065 |
| GHR | NM_001242401 | -0.4875 | 0.1065 |
| GHR | NM_001242402 | -0.4875 | 0.1065 |
| GHR | NM_001242403 | -0.4875 | 0.1065 |
| GHR | NM_001242404 | -0.4875 | 0.1065 |
| GHR | NM_001242405 | -0.4875 | 0.1065 |
| GHR | NM_001242406 | -0.4875 | 0.1065 |
| FAM59A | NM_001242409 | -0.3745 | -1.9465 |
| AHRR | NM_001242412 | 0.1635 | -0.353 |
| PRKCQ | NM_001242413 | 0.083 | -0.2895 |
| WDR20 | NM_001242414 | -1.131 | -2.623 |
| WDR20 | NM_001242415 | -1.131 | -2.623 |
| WDR20 | NM_001242416 | -1.131 | -2.623 |
| WDR20 | NM_001242417 | -1.131 | -2.623 |
| WDR20 | NM_001242418 | -1.131 | -2.623 |
| GRB7 | NM_001242442 | 0.312 | -0.2765 |
| GRB7 | NM_001242443 | 0.312 | -0.2765 |
| PHF21B | NM_001242450 | -0.3435 | -0.999 |
| IRF5 | NM_001242452 | -0.1315 | -0.227 |
| GHR | NM_001242460 | -0.4875 | 0.1065 |
| GHR | NM_001242461 | -0.4875 | 0.1065 |
| GHR | NM_001242462 | -0.4875 | 0.1065 |
| PIK3R1 | NM_001242466 | -0.308 | -1.1495 |

| | | | |
|---|---|---|---|
| ZNF345 | NM_001242472 | -0.843 | -1.283 |
| ZNF345 | NM_001242474 | -0.843 | -1.283 |
| ZNF345 | NM_001242475 | -0.843 | -1.283 |
| ZNF345 | NM_001242476 | -0.843 | -1.283 |
| EIF1AD | NM_001242481 | -0.597 | -1.5905 |
| EIF1AD | NM_001242482 | -0.597 | -1.5905 |
| EIF1AD | NM_001242483 | -0.597 | -1.5905 |
| EIF1AD | NM_001242484 | -0.597 | -1.5905 |
| EIF1AD | NM_001242485 | -0.597 | -1.5905 |
| EIF1AD | NM_001242486 | -0.597 | -1.5905 |
| KIAA0913 | NM_001242487 | -0.0525 | -0.028 |
| KIAA0913 | NM_001242488 | -0.0525 | -0.028 |
| MIF4GD | NM_001242498 | -0.5145 | -1.258 |
| MIF4GD | NM_001242500 | -0.5145 | -1.258 |
| MIF4GD | NM_001242501 | -0.5145 | -1.258 |
| DIO2 | NM_001242502 | 0.0605 | -0.2345 |
| DIO2 | NM_001242503 | 0.0605 | -0.2345 |
| GDA | NM_001242505 | -0.1385 | -0.5655 |
| GDA | NM_001242506 | -0.1385 | -0.5655 |
| GDA | NM_001242507 | -0.1385 | -0.5655 |
| OSBPL1A | NM_001242508 | -0.34 | -1.1105 |
| HLA-DPA1 | NM_001242524 | -0.19 | -0.095 |
| HLA-DPA1 | NM_001242525 | -0.19 | -0.095 |
| MFSD11 | NM_001242532 | -0.679 | -1.3765 |
| MFSD11 | NM_001242533 | -0.679 | -1.3765 |
| MFSD11 | NM_001242534 | -0.679 | -1.3765 |
| MFSD11 | NM_001242535 | -0.679 | -1.3765 |
| MFSD11 | NM_001242536 | -0.679 | -1.3765 |
| MFSD11 | NM_001242537 | -0.679 | -1.3765 |
| NCOA6 | NM_001242539 | -1.147 | -2.5645 |
| ANAPC16 | NM_001242546 | -0.5375 | -0.5575 |
| ANAPC16 | NM_001242547 | -0.5375 | -0.5575 |
| ANAPC16 | NM_001242548 | -0.5375 | -0.5575 |
| MAP4K4 | NM_001242559 | -0.341 | -0.495 |
| MAP4K4 | NM_001242560 | -0.341 | -0.495 |
| RIN2 | NM_001242581 | 0.1275 | -0.4685 |
| TREM1 | NM_001242589 | 0.0775 | -0.248 |
| TREM1 | NM_001242590 | 0.0775 | -0.248 |
| RBM39 | NM_001242599 | -1.0695 | -2.1315 |
| RBM39 | NM_001242600 | -1.0695 | -2.1315 |
| NCAM1 | NM_001242607 | -0.7595 | -0.572 |
| NCAM1 | NM_001242608 | -0.7595 | -0.572 |
| CD99L2 | NM_001242614 | -0.5455 | -0.894 |

| | | | |
|---|---|---|---|
| NXT2 | NM_001242617 | -0.5415 | -0.357 |
| NXT2 | NM_001242618 | -0.5415 | -0.357 |
| GFOD1 | NM_001242628 | -0.571 | -1.5355 |
| GFOD1 | NM_001242629 | -0.571 | -1.5355 |
| GFOD1 | NM_001242630 | -0.571 | -1.5355 |
| IL31RA | NM_001242636 | 0.0715 | -0.4265 |
| IL31RA | NM_001242637 | 0.0715 | -0.4265 |
| IL31RA | NM_001242638 | 0.0715 | -0.4265 |
| IL31RA | NM_001242639 | 0.0715 | -0.4265 |
| BRCC3 | NM_001242640 | -0.2775 | -1.067 |
| RAB2A | NM_001242644 | -1.148 | -2.2375 |
| PHACTR1 | NM_001242648 | -0.2605 | -0.5105 |
| AHCYL1 | NM_001242673 | -0.5435 | -1.022 |
| AHCYL1 | NM_001242674 | -0.5435 | -1.022 |
| AHCYL1 | NM_001242675 | -0.5435 | -1.022 |
| AHCYL1 | NM_001242676 | -0.5435 | -1.022 |
| SLC14A2 | NM_001242692 | 0.0035 | -0.3915 |
| CAMTA1 | NM_001242701 | 0.0475 | -0.016 |
| ZNF385C | NM_001242704 | -0.1125 | -0.336 |
| ARHGEF38 | NM_001242729 | -0.2565 | -0.236 |
| ZNF691 | NM_001242739 | -1.513 | -2.6305 |
| HLA-A | NM_001242758 | 0.184 | -0.362 |
| DLGAP1 | NM_001242761 | -0.322 | -0.227 |
| DLGAP1 | NM_001242762 | -0.322 | -0.227 |
| DLGAP1 | NM_001242763 | -0.322 | -0.227 |
| DLGAP1 | NM_001242764 | -0.322 | -0.227 |
| DLGAP1 | NM_001242765 | -0.322 | -0.227 |
| DLGAP1 | NM_001242766 | -0.322 | -0.227 |
| MTHFD1L | NM_001242767 | 0.879 | 0.4915 |
| MTHFD1L | NM_001242768 | 0.879 | 0.4915 |
| MTHFD1L | NM_001242769 | 0.879 | 0.4915 |
| TMEM139 | NM_001242773 | 0.2435 | -0.3775 |
| TMEM139 | NM_001242774 | 0.2435 | -0.3775 |
| TMEM139 | NM_001242775 | 0.2435 | -0.3775 |
| TMEM139 | NM_001242776 | 0.2435 | -0.3775 |
| TMEM139 | NM_001242777 | 0.2435 | -0.3775 |
| IKBKB | NM_001242778 | -0.9915 | -2.568 |
| TRIM26 | NM_001242783 | -0.9725 | -1.8285 |
| HLCS | NM_001242784 | -0.572 | -1.0345 |
| HLCS | NM_001242785 | -0.572 | -1.0345 |
| BRF1 | NM_001242786 | -0.009 | -0.1075 |
| BRF1 | NM_001242787 | -0.009 | -0.1075 |
| BRF1 | NM_001242788 | -0.009 | -0.1075 |

| | | | |
|---|---|---|---|
| BRF1 | NM_001242789 | -0.009 | -0.1075 |
| BRF1 | NM_001242790 | -0.009 | -0.1075 |
| SNAP91 | NM_001242792 | -1.1855 | -1.6045 |
| SNAP91 | NM_001242793 | -1.1855 | -1.6045 |
| SNAP91 | NM_001242794 | -1.1855 | -1.6045 |
| NUP93 | NM_001242795 | 0.009 | -1.033 |
| NUP93 | NM_001242796 | 0.009 | -1.033 |
| ZNF790 | NM_001242800 | -0.3995 | -0.951 |
| ZNF790 | NM_001242801 | -0.3995 | -0.951 |
| ZNF790 | NM_001242802 | -0.3995 | -0.951 |
| BTN3A3 | NM_001242803 | -0.139 | -0.7845 |
| ANKRD6 | NM_001242809 | -0.4595 | -0.767 |
| ANKRD6 | NM_001242811 | -0.4595 | -0.767 |
| ANKRD6 | NM_001242813 | -0.4595 | -0.767 |
| ANKRD6 | NM_001242814 | -0.4595 | -0.767 |
| DEF8 | NM_001242816 | -0.0555 | -0.379 |
| DEF8 | NM_001242817 | -0.0555 | -0.379 |
| DEF8 | NM_001242818 | -0.0555 | -0.379 |
| DEF8 | NM_001242819 | -0.0555 | -0.379 |
| DEF8 | NM_001242820 | -0.0555 | -0.379 |
| DEF8 | NM_001242821 | -0.0555 | -0.379 |
| DEF8 | NM_001242822 | -0.0555 | -0.379 |
| FDPS | NM_001242824 | -0.284 | -0.5885 |
| FDPS | NM_001242825 | -0.284 | -0.5885 |
| RNF41 | NM_001242826 | -0.776 | -2.4925 |
| SIRT5 | NM_001242827 | -0.7815 | -1.5085 |
| ELOVL5 | NM_001242828 | -1.283 | -1.5085 |
| IP6K1 | NM_001242829 | -0.3725 | -1.5875 |
| ELOVL5 | NM_001242830 | -1.283 | -1.5085 |
| ELOVL5 | NM_001242831 | -1.283 | -1.5085 |
| ARFGAP2 | NM_001242832 | -0.3625 | -0.8795 |
| NDRG4 | NM_001242833 | -0.529 | -0.8035 |
| NDRG4 | NM_001242834 | -0.529 | -0.8035 |
| NDRG4 | NM_001242835 | -0.529 | -0.8035 |
| NDRG4 | NM_001242836 | -0.529 | -0.8035 |
| AP2A2 | NM_001242837 | -0.423 | -1.131 |
| GUK1 | NM_001242839 | 0.352 | -0.18 |
| GUK1 | NM_001242840 | 0.352 | -0.18 |
| ZNF195 | NM_001242841 | -1.2155 | -2.4815 |
| ZNF195 | NM_001242842 | -1.2155 | -2.4815 |
| ZNF195 | NM_001242843 | -1.2155 | -2.4815 |
| RNF146 | NM_001242844 | -2.1675 | -4.566 |
| RNF146 | NM_001242845 | -2.1675 | -4.566 |

| | | | |
|---|---|---|---|
| RNF146 | NM_001242846 | -2.1675 | -4.566 |
| RNF146 | NM_001242849 | -2.1675 | -4.566 |
| RNF146 | NM_001242850 | -2.1675 | -4.566 |
| RNF146 | NM_001242851 | -2.1675 | -4.566 |
| RNF146 | NM_001242852 | -2.1675 | -4.566 |
| ARFIP2 | NM_001242854 | -0.4445 | -1.55 |
| ARFIP2 | NM_001242855 | -0.4445 | -1.55 |
| ARFIP2 | NM_001242856 | -0.4445 | -1.55 |
| PRKACB | NM_001242857 | -1.4275 | -1.8175 |
| PRKACB | NM_001242858 | -1.4275 | -1.8175 |
| PRKACB | NM_001242859 | -1.4275 | -1.8175 |
| PRKACB | NM_001242860 | -1.4275 | -1.8175 |
| PRKACB | NM_001242861 | -1.4275 | -1.8175 |
| PRKACB | NM_001242862 | -1.4275 | -1.8175 |
| ENOX1 | NM_001242863 | -1.115 | -1.3835 |
| PRMT2 | NM_001242864 | -0.777 | -0.847 |
| PRMT2 | NM_001242865 | -0.777 | -0.847 |
| PRMT2 | NM_001242866 | -0.777 | -0.847 |
| SLAIN1 | NM_001242868 | -1.978 | -4.0495 |
| SLAIN1 | NM_001242869 | -1.978 | -4.0495 |
| SLAIN1 | NM_001242870 | -1.978 | -4.0495 |
| SLAIN1 | NM_001242871 | -1.978 | -4.0495 |
| KLHDC3 | NM_001242872 | -0.0095 | -0.8585 |
| PRKCZ | NM_001242874 | -0.0375 | -0.907 |
| CARKD | NM_001242881 | -0.5245 | -1.336 |
| CARKD | NM_001242882 | -0.5245 | -1.336 |
| CARKD | NM_001242883 | -0.5245 | -1.336 |
| ZBTB17 | NM_001242884 | -0.1065 | -0.385 |
| .OC10028703( | NM_001242885 | 0.18 | -0.575 |
| DDC | NM_001242886 | 0.0235 | -0.3285 |
| DDC | NM_001242887 | 0.0235 | -0.3285 |
| DDC | NM_001242888 | 0.0235 | -0.3285 |
| DDC | NM_001242889 | 0.0235 | -0.3285 |
| DDC | NM_001242890 | 0.0235 | -0.3285 |
| CSDE1 | NM_001242891 | -0.4585 | -0.404 |
| CSDE1 | NM_001242892 | -0.4585 | -0.404 |
| CSDE1 | NM_001242893 | -0.4585 | -0.404 |
| DEPDC5 | NM_001242896 | -1.6815 | -3.1645 |
| DEPDC5 | NM_001242897 | -1.6815 | -3.1645 |
| PPP6R2 | NM_001242898 | -0.1485 | -0.811 |
| PPP6R2 | NM_001242899 | -0.1485 | -0.811 |
| PPP6R2 | NM_001242900 | -0.1485 | -0.811 |
| AKAP1 | NM_001242902 | -0.1795 | -1.1685 |

| | | | |
|---|---|---|---|
| AKAP1 | NM_001242903 | -0.1795 | -1.1685 |
| MPPE1 | NM_001242904 | -1.091 | -1.281 |
| RSPO1 | NM_001242908 | 0.044 | -0.102 |
| RSPO1 | NM_001242909 | 0.044 | -0.102 |
| RSPO1 | NM_001242910 | 0.044 | -0.102 |
| ZFAND6 | NM_001242911 | -0.9975 | -0.972 |
| ZFAND6 | NM_001242912 | -0.9975 | -0.972 |
| ZFAND6 | NM_001242913 | -0.9975 | -0.972 |
| ZFAND6 | NM_001242914 | -0.9975 | -0.972 |
| ZFAND6 | NM_001242915 | -0.9975 | -0.972 |
| ZFAND6 | NM_001242916 | -0.9975 | -0.972 |
| ZFAND6 | NM_001242917 | -0.9975 | -0.972 |
| ZFAND6 | NM_001242918 | -0.9975 | -0.972 |
| ZFAND6 | NM_001242919 | -0.9975 | -0.972 |
| EIF3L | NM_001242923 | -0.3145 | -0.186 |
| ZNF410 | NM_001242924 | -1.314 | -2.961 |
| TMCC2 | NM_001242925 | -0.5715 | -0.8105 |
| ZNF410 | NM_001242926 | -1.314 | -2.961 |
| ZNF410 | NM_001242927 | -1.314 | -2.961 |
| ZNF410 | NM_001242928 | -1.314 | -2.961 |
| ANKS3 | NM_001242929 | -0.58 | -1.421 |
| LSP1 | NM_001242932 | 0.2225 | -0.3575 |
| SNX1 | NM_001242933 | -0.4635 | -1.3525 |
| SNCAIP | NM_001242935 | -0.0295 | -0.8295 |
| FAM47E | NM_001242936 | 0.095 | 0.3505 |
| EPDR1 | NM_001242946 | -0.183 | -2.312 |
| EPDR1 | NM_001242948 | -0.183 | -2.312 |
| MAK | NM_001242957 | -0.356 | -0.1935 |
| PARN | NM_001242992 | -1.042 | -1.239 |
| TNFRSF8 | NM_001243 | 0.054 | -0.171 |
| ABCB9 | NM_001243013 | 0.0745 | -0.486 |
| ABCB9 | NM_001243014 | 0.0745 | -0.486 |
| AKT2 | NM_001243027 | -0.1 | -0.836 |
| AKT2 | NM_001243028 | -0.1 | -0.836 |
| HLA-C | NM_001243042 | 0.372 | -0.3075 |
| E2F3 | NM_001243076 | -1.7195 | -3.6495 |
| CD28 | NM_001243077 | -0.539 | -1.0495 |
| CD28 | NM_001243078 | -0.539 | -1.0495 |
| ELF5 | NM_001243080 | -0.3485 | -0.1935 |
| ELF5 | NM_001243081 | -0.3485 | -0.1935 |
| HIF1A | NM_001243084 | -0.736 | -1.572 |
| FOXM1 | NM_001243088 | -0.183 | -0.867 |
| FOXM1 | NM_001243089 | -0.183 | -0.867 |

| | | | |
|---|---|---|---|
| FYB | NM_001243093 | 0.1725 | -0.4775 |
| HSF2 | NM_001243094 | -1.4485 | -3.69 |
| IL5RA | NM_001243099 | 0.056 | -0.797 |
| NTRK3 | NM_001243101 | -0.273 | -0.8245 |
| PLD2 | NM_001243108 | -0.2145 | -0.564 |
| NFKBIB | NM_001243116 | -0.0265 | -0.657 |
| SKP2 | NM_001243120 | -0.9155 | -1.594 |
| PDE4A | NM_001243121 | -0.085 | -0.6775 |
| KLK7 | NM_001243126 | 0.075 | -0.4105 |
| PACS2 | NM_001243127 | -0.003 | -0.6055 |
| RPL13 | NM_001243130 | 0.0625 | -0.329 |
| RPL13 | NM_001243131 | 0.0625 | -0.329 |
| NLRP3 | NM_001243133 | -0.0565 | 0.0615 |
| PELI3 | NM_001243135 | 0.012 | -0.4545 |
| PELI3 | NM_001243136 | 0.012 | -0.4545 |
| PDE8A | NM_001243137 | -0.804 | -2.083 |
| NUSAP1 | NM_001243142 | -1.4285 | -1.266 |
| NUSAP1 | NM_001243143 | -1.4285 | -1.266 |
| NUSAP1 | NM_001243144 | -1.4285 | -1.266 |
| NVL | NM_001243146 | -1.339 | -1.9675 |
| NVL | NM_001243147 | -1.339 | -1.9675 |
| TAF1C | NM_001243156 | -0.0525 | -0.5505 |
| TAF1C | NM_001243157 | -0.0525 | -0.5505 |
| TAF1C | NM_001243158 | -0.0525 | -0.5505 |
| TAF1C | NM_001243159 | -0.0525 | -0.5505 |
| TAF1C | NM_001243160 | -0.0525 | -0.5505 |
| PTCRA | NM_001243168 | -0.0825 | -0.39 |
| PTCRA | NM_001243169 | -0.0825 | -0.39 |
| PTCRA | NM_001243170 | -0.0825 | -0.39 |
| ALDOA | NM_001243177 | 0.305 | -0.157 |
| ATP7B | NM_001243182 | 0.299 | -0.5255 |
| PIM1 | NM_001243186 | -0.252 | -0.9345 |
| INPP5F | NM_001243194 | -0.915 | -1.909 |
| INPP5F | NM_001243195 | -0.915 | -1.909 |
| HIP1 | NM_001243198 | -0.733 | -1.1245 |
| RANBP6 | NM_001243202 | -0.942 | -2.37 |
| RANBP6 | NM_001243203 | -0.942 | -2.37 |
| TCF4 | NM_001243226 | -1.0505 | -2.2855 |
| TCF4 | NM_001243227 | -1.0505 | -2.2855 |
| TCF4 | NM_001243228 | -1.0505 | -2.2855 |
| TCF4 | NM_001243230 | -1.0505 | -2.2855 |
| TCF4 | NM_001243231 | -1.0505 | -2.2855 |
| TCF4 | NM_001243232 | -1.0505 | -2.2855 |

| | | | |
|---|---|---|---|
| TCF4 | NM_001243233 | -1.0505 | -2.2855 |
| TCF4 | NM_001243234 | -1.0505 | -2.2855 |
| TCF4 | NM_001243235 | -1.0505 | -2.2855 |
| TCF4 | NM_001243236 | -1.0505 | -2.2855 |
| LEPRE1 | NM_001243246 | -0.778 | -1.6275 |
| NPRL3 | NM_001243247 | -0.013 | -0.3335 |
| NPRL3 | NM_001243248 | -0.013 | -0.3335 |
| NPRL3 | NM_001243249 | -0.013 | -0.3335 |
| NARS2 | NM_001243251 | -1.12 | -1.875 |
| EFHD1 | NM_001243252 | -0.305 | 0.015 |
| TIGD6 | NM_001243253 | -0.9915 | -1.5385 |
| DENND5A | NM_001243254 | -0.9235 | -2.7135 |
| ZIC4 | NM_001243256 | 0.116 | -0.0595 |
| HINFP | NM_001243259 | -0.5445 | -2.3425 |
| MS4A4A | NM_001243266 | -0.252 | -0.338 |
| CNTN5 | NM_001243270 | -0.29 | -0.2995 |
| CNTN5 | NM_001243271 | -0.29 | -0.2995 |
| ACSF3 | NM_001243279 | -0.204 | -0.9045 |
| ALCAM | NM_001243280 | -0.4825 | -0.9185 |
| ALCAM | NM_001243281 | -0.4825 | -0.9185 |
| ALCAM | NM_001243283 | -0.4825 | -0.9185 |
| PVRL3 | NM_001243286 | -0.6825 | -1.524 |
| PVRL3 | NM_001243288 | -0.6825 | -1.524 |
| APLP2 | NM_001243299 | -0.3565 | -0.861 |
| STK36 | NM_001243313 | -0.5865 | -0.712 |
| RAET1E | NM_001243325 | 0.065 | -0.2615 |
| RAET1E | NM_001243327 | 0.065 | -0.2615 |
| RAET1E | NM_001243328 | 0.065 | -0.2615 |
| SEZ6L2 | NM_001243332 | -0.099 | -0.6895 |
| SEZ6L2 | NM_001243333 | -0.099 | -0.6895 |
| CCDC40 | NM_001243342 | 0.022 | -0.0475 |
| SH3D19 | NM_001243349 | -1.2825 | -1.9435 |
| NUB1 | NM_001243351 | -0.6535 | -0.837 |
| CLCN3 | NM_001243372 | -1.6635 | -2.8755 |
| CLCN3 | NM_001243374 | -1.6635 | -2.8755 |
| PARVB | NM_001243385 | 0.0445 | 0.1985 |
| PARVB | NM_001243386 | 0.0445 | 0.1985 |
| SLC6A13 | NM_001243392 | 0.1085 | -0.356 |
| CLEC16A | NM_001243403 | -0.6945 | -1.2335 |
| KIF13A | NM_001243423 | -0.256 | -0.6685 |
| TTC39C | NM_001243425 | -1.234 | -2.3885 |
| ERG | NM_001243428 | -0.1435 | -0.526 |
| ERG | NM_001243429 | -0.1435 | -0.526 |

| | | | |
|---|---|---|---|
| ERG | NM_001243432 | -0.1435 | -0.526 |
| SPECC1 | NM_001243438 | -0.3345 | -0.4385 |
| SPECC1 | NM_001243439 | -0.3345 | -0.4385 |
| STARD13 | NM_001243466 | -1.199 | -0.698 |
| UBASH3A | NM_001243467 | -0.044 | -0.2395 |
| B9D1 | NM_001243473 | -0.1795 | -0.535 |
| STARD13 | NM_001243474 | -1.199 | -0.698 |
| B9D1 | NM_001243475 | -0.1795 | -0.535 |
| STARD13 | NM_001243476 | -1.199 | -0.698 |
| SETDB1 | NM_001243491 | -0.321 | -0.8105 |
| ESRRG | NM_001243505 | -1.2545 | -4.612 |
| ESRRG | NM_001243506 | -1.2545 | -4.612 |
| ESRRG | NM_001243507 | -1.2545 | -4.612 |
| ESRRG | NM_001243509 | -1.2545 | -4.612 |
| ESRRG | NM_001243510 | -1.2545 | -4.612 |
| ESRRG | NM_001243511 | -1.2545 | -4.612 |
| ESRRG | NM_001243512 | -1.2545 | -4.612 |
| ESRRG | NM_001243513 | -1.2545 | -4.612 |
| ESRRG | NM_001243514 | -1.2545 | -4.612 |
| ESRRG | NM_001243515 | -1.2545 | -4.612 |
| ESRRG | NM_001243518 | -1.2545 | -4.612 |
| ESRRG | NM_001243519 | -1.2545 | -4.612 |
| HYI | NM_001243526 | 0.2225 | -0.255 |
| IL15RA | NM_001243539 | -0.133 | -0.2235 |
| DHDDS | NM_001243564 | -0.5955 | -1.0075 |
| DHDDS | NM_001243565 | -0.5955 | -1.0075 |
| MTMR2 | NM_001243571 | -0.759 | -1.315 |
| CDON | NM_001243597 | -1.268 | -2.232 |
| LMO3 | NM_001243609 | -1.304 | -1.7735 |
| LMO3 | NM_001243610 | -1.304 | -1.7735 |
| LMO3 | NM_001243611 | -1.304 | -1.7735 |
| LMO3 | NM_001243612 | -1.304 | -1.7735 |
| LMO3 | NM_001243613 | -1.304 | -1.7735 |
| CFL2 | NM_001243645 | -0.8905 | -2.2675 |
| CD2BP2 | NM_001243646 | -0.538 | -1.4955 |
| GFOD2 | NM_001243650 | 0.353 | -0.8485 |
| GLRX | NM_001243658 | -0.7235 | -0.853 |
| GLRX | NM_001243659 | -0.7235 | -0.853 |
| LETMD1 | NM_001243689 | -0.3545 | -0.817 |
| ZFP161 | NM_001243702 | -1.4405 | -2.7725 |
| ZFP161 | NM_001243704 | -1.4405 | -2.7725 |
| RAB6A | NM_001243718 | -1.152 | -2.154 |
| RAB6A | NM_001243719 | -1.152 | -2.154 |

| | | | |
|---|---|---|---|
| SETMAR | NM_001243723 | -1.5105 | -3.311 |
| ATPAF1 | NM_001243728 | -0.403 | -0.375 |
| C3orf55 | NM_001243729 | -0.011 | -0.2205 |
| RARG | NM_001243730 | -0.129 | -0.906 |
| RARG | NM_001243731 | -0.129 | -0.906 |
| RARG | NM_001243732 | -0.129 | -0.906 |
| C5orf44 | NM_001243737 | -1.5265 | -2.229 |
| RGL2 | NM_001243738 | -0.023 | -0.6925 |
| ZNF124 | NM_001243740 | -0.9225 | -1.566 |
| FANCC | NM_001243743 | -0.2255 | -0.3485 |
| FANCC | NM_001243744 | -0.2255 | -0.3485 |
| CROT | NM_001243745 | -0.014 | 0.263 |
| FAM20A | NM_001243746 | 0.0275 | -0.2675 |
| API5 | NM_001243747 | -1.267 | -1.653 |
| C7orf49 | NM_001243749 | -1.075 | -2.6955 |
| NUDT8 | NM_001243750 | 0.027 | -0.4335 |
| C7orf49 | NM_001243751 | -1.075 | -2.6955 |
| C7orf49 | NM_001243752 | -1.075 | -2.6955 |
| C7orf49 | NM_001243753 | -1.075 | -2.6955 |
| C7orf49 | NM_001243754 | -1.075 | -2.6955 |
| C7orf49 | NM_001243755 | -1.075 | -2.6955 |
| PXN | NM_001243756 | -0.252 | -0.7805 |
| USP2 | NM_001243759 | 0.16 | 0.0275 |
| RABGAP1L | NM_001243763 | -0.5975 | -0.655 |
| RABGAP1L | NM_001243764 | -0.5975 | -0.655 |
| RABGAP1L | NM_001243765 | -0.5975 | -0.655 |
| POMGNT1 | NM_001243766 | -0.079 | -0.597 |
| BTF3L4 | NM_001243767 | -1.462 | -2.232 |
| FAM178A | NM_001243770 | -1.141 | -2.2115 |
| APH1A | NM_001243771 | -0.174 | -0.71 |
| APH1A | NM_001243772 | -0.174 | -0.71 |
| GNG2 | NM_001243773 | 0.0335 | -0.392 |
| GNG2 | NM_001243774 | 0.0335 | -0.392 |
| LIMA1 | NM_001243775 | -1.0115 | -2.3605 |
| CEP57 | NM_001243776 | -1.329 | -2.3185 |
| CEP57 | NM_001243777 | -1.329 | -2.3185 |
| C10orf58 | NM_001243778 | -0.401 | -0.456 |
| C10orf58 | NM_001243779 | -0.401 | -0.456 |
| C10orf58 | NM_001243780 | -0.401 | -0.456 |
| C10orf58 | NM_001243781 | -0.401 | -0.456 |
| C10orf58 | NM_001243782 | -0.401 | -0.456 |
| PDE2A | NM_001243784 | -0.4595 | -1.1105 |
| RAG2 | NM_001243785 | -0.0205 | -0.069 |

| | | | |
|---|---|---|---|
| RAG2 | NM_001243786 | -0.0205 | -0.069 |
| SMUG1 | NM_001243787 | 0.0505 | -0.914 |
| SMUG1 | NM_001243788 | 0.0505 | -0.914 |
| SMUG1 | NM_001243789 | 0.0505 | -0.914 |
| SMUG1 | NM_001243790 | 0.0505 | -0.914 |
| SMUG1 | NM_001243791 | 0.0505 | -0.914 |
| CHST12 | NM_001243794 | -1.021 | -2.4235 |
| CHST12 | NM_001243795 | -1.021 | -2.4235 |
| TSC22D1 | NM_001243797 | -0.259 | 0.5975 |
| TSC22D1 | NM_001243798 | -0.259 | 0.5975 |
| TSC22D1 | NM_001243799 | -0.259 | 0.5975 |
| ZC4H2 | NM_001243804 | -0.3345 | -0.7665 |
| CACNA1B | NM_001243812 | -1.21 | -1.025 |
| STAT4 | NM_001243835 | 0.17 | -0.4555 |
| CHST9 | NM_001243848 | -0.443 | -0.03 |
| SPHK2 | NM_001243876 | 0.0025 | -0.1975 |
| TOB1 | NM_001243877 | -0.637 | 0.3105 |
| FHL3 | NM_001243878 | -0.7535 | -1.1315 |
| TRA2B | NM_001243879 | -0.809 | -1.2385 |
| TOB1 | NM_001243885 | -0.637 | 0.3105 |
| TPD52L2 | NM_001243891 | -0.269 | -0.5175 |
| TPD52L2 | NM_001243892 | -0.269 | -0.5175 |
| TPD52L2 | NM_001243894 | -0.269 | -0.5175 |
| TPD52L2 | NM_001243895 | -0.269 | -0.5175 |
| HDLBP | NM_001243900 | -0.0575 | -0.3395 |
| MAPKAPK3 | NM_001243925 | 0.1425 | -0.6935 |
| MAPKAPK3 | NM_001243926 | 0.1425 | -0.6935 |
| PRPSAP2 | NM_001243936 | -1.0105 | -1.3285 |
| PRPSAP2 | NM_001243940 | -1.0105 | -1.3285 |
| PRPSAP2 | NM_001243941 | -1.0105 | -1.3285 |
| PRPSAP2 | NM_001243942 | -1.0105 | -1.3285 |
| PPP1R14A | NM_001243947 | -0.3045 | -1.1185 |
| KIF2A | NM_001243952 | -0.9335 | -2.404 |
| KIF2A | NM_001243953 | -0.9335 | -2.404 |
| NEDD4L | NM_001243960 | -0.9225 | -2.059 |
| HLA-DQB1 | NM_001243961 | 0.0105 | -0.2455 |
| HLA-DQB1 | NM_001243962 | 0.0105 | -0.2455 |
| LRCH2 | NM_001243963 | -0.531 | -1.3595 |
| HLA-DRB1 | NM_001243965 | -0.23 | -0.436 |
| RELA | NM_001243984 | -0.7155 | -2.538 |
| RELA | NM_001243985 | -0.7155 | -2.538 |
| SPIB | NM_001243998 | -0.0165 | 0.049 |
| SPIB | NM_001243999 | -0.0165 | 0.049 |

| | | | |
|---|---|---|---|
| TNFSF8 | NM_001244 | 0.062 | 0.006 |
| SPIB | NM_001244000 | -0.0165 | 0.049 |
| KIF1A | NM_001244008 | -0.346 | -0.6845 |
| C12orf53 | NM_001244014 | 0.0975 | -0.7195 |
| C12orf53 | NM_001244015 | 0.0975 | -0.7195 |
| ZNF726 | NM_001244038 | -0.2105 | -0.393 |
| GIMAP6 | NM_001244071 | 0.054 | 0.0995 |
| GIMAP6 | NM_001244072 | 0.054 | 0.0995 |
| MAP3K8 | NM_001244134 | -0.464 | -0.2965 |
| KIAA0586 | NM_001244189 | -0.7505 | -1.783 |
| KIAA0586 | NM_001244190 | -0.7505 | -1.783 |
| KIAA0586 | NM_001244191 | -0.7505 | -1.783 |
| KIAA0586 | NM_001244192 | -0.7505 | -1.783 |
| KIAA0586 | NM_001244193 | -0.7505 | -1.783 |
| HBP1 | NM_001244262 | -0.5595 | -2.4045 |
| RGNEF | NM_001244364 | -0.874 | -1.9285 |
| ARG1 | NM_001244438 | 0.1215 | -0.424 |
| BID | NM_001244567 | -0.028 | -1.151 |
| BID | NM_001244569 | -0.028 | -1.151 |
| BID | NM_001244570 | -0.028 | -1.151 |
| BID | NM_001244572 | -0.028 | -1.151 |
| TFEC | NM_001244583 | -0.0955 | -0.0655 |
| LUC7L2 | NM_001244584 | -0.965 | -1.8725 |
| LUC7L2 | NM_001244585 | -0.965 | -1.8725 |
| DPYSL2 | NM_001244604 | -0.1565 | -0.526 |
| IMMP2L | NM_001244606 | -0.346 | -0.2075 |
| ARID5B | NM_001244638 | -0.204 | -0.3765 |
| CHMP2B | NM_001244644 | -0.8055 | -1.9105 |
| THNSL2 | NM_001244676 | -0.258 | -0.7225 |
| THNSL2 | NM_001244678 | -0.258 | -0.7225 |
| POU2F3 | NM_001244682 | 0.138 | 0.3435 |
| PDE3A | NM_001244683 | 0.092 | 0.1 |
| ZFP36L1 | NM_001244698 | -0.349 | -0.4295 |
| ZFP36L1 | NM_001244701 | -0.349 | -0.4295 |
| CSAD | NM_001244705 | -0.7915 | -1.114 |
| CSAD | NM_001244706 | -0.7915 | -1.114 |
| GFPT1 | NM_001244710 | -0.9315 | -2.363 |
| RAD23B | NM_001244713 | -0.688 | -1.143 |
| RAD23B | NM_001244724 | -0.688 | -1.143 |
| MARVELD2 | NM_001244734 | 0.0625 | 0.2025 |
| FCGR3B | NM_001244753 | -0.361 | -0.2015 |
| TNIP3 | NM_001244764 | 0.3175 | -0.134 |
| FOXP1 | NM_001244808 | -1.1875 | -1.91 |

| | | | |
|---|---|---|---|
| FOXP1 | NM_001244810 | -1.1875 | -1.91 |
| FOXP1 | NM_001244812 | -1.1875 | -1.91 |
| FOXP1 | NM_001244813 | -1.1875 | -1.91 |
| FOXP1 | NM_001244814 | -1.1875 | -1.91 |
| FOXP1 | NM_001244815 | -1.1875 | -1.91 |
| FOXP1 | NM_001244816 | -1.1875 | -1.91 |
| CHEK1 | NM_001244846 | -1.066 | -1.552 |
| CLEC4G | NM_001244856 | 0.141 | -0.4265 |
| DAB2 | NM_001244871 | 0.1555 | 0.2915 |
| DLST | NM_001244883 | -0.164 | -1.708 |
| AGAP1 | NM_001244888 | -0.997 | -1.5385 |
| UNC5B | NM_001244889 | -0.3675 | -1.5 |
| ROD1 | NM_001244896 | -1.0745 | -1.801 |
| ROD1 | NM_001244897 | -1.0745 | -1.801 |
| ROD1 | NM_001244898 | -1.0745 | -1.801 |
| FCGR1B | NM_001244910 | -0.1445 | -0.729 |
| PRPF4 | NM_001244926 | -1.297 | -3.679 |
| MST1R | NM_001244937 | -0.0595 | -0.449 |
| TXN | NM_001244938 | -0.182 | -0.1135 |
| STEAP2 | NM_001244944 | 0.2585 | -0.0985 |
| STEAP2 | NM_001244945 | 0.2585 | -0.0985 |
| STEAP2 | NM_001244946 | 0.2585 | -0.0985 |
| GPAM | NM_001244949 | -1.475 | -1.881 |
| SPOCK2 | NM_001244950 | 0.0785 | -0.178 |
| FRMD3 | NM_001244959 | -0.405 | -1.2715 |
| FRMD3 | NM_001244960 | -0.405 | -1.2715 |
| FRMD3 | NM_001244961 | -0.405 | -1.2715 |
| FRMD3 | NM_001244962 | -0.405 | -1.2715 |
| NAV2 | NM_001244963 | -0.2205 | -1.109 |
| NRP1 | NM_001244972 | -2.015 | -3.5045 |
| NRP1 | NM_001244973 | -2.015 | -3.5045 |
| PPP1CC | NM_001244974 | -0.986 | -1.684 |
| SEL1L | NM_001244984 | -0.372 | -1.0265 |
| PPP1R12A | NM_001244990 | -0.3175 | -1.7575 |
| PPP1R12A | NM_001244992 | -0.3175 | -1.7575 |
| TMPRSS13 | NM_001244995 | 0.0195 | -0.0005 |
| SIGLEC6 | NM_001245 | 0.088 | 0.198 |
| NFIC | NM_001245002 | -0.9405 | -2.6905 |
| NFIC | NM_001245004 | -0.9405 | -2.6905 |
| NFIC | NM_001245005 | -0.9405 | -2.6905 |
| ENTPD2 | NM_001246 | 0.242 | -0.3105 |
| ENTPD6 | NM_001247 | -0.2145 | -1.0375 |
| SYT13 | NM_001247987 | 0.329 | -0.2555 |

| | | | |
|---|---|---|---|
| CTAGE5 | NM_001247988 | -0.6255 | -1.4575 |
| CTAGE5 | NM_001247989 | -0.6255 | -1.4575 |
| CTAGE5 | NM_001247990 | -0.6255 | -1.4575 |
| POU2F2 | NM_001247994 | -0.15 | -0.759 |
| ASAP1 | NM_001247996 | -0.5945 | -1.2915 |
| CLIP1 | NM_001247997 | -0.823 | -2.401 |
| MS4A6A | NM_001247999 | -0.1165 | 0.223 |
| ENTPD3 | NM_001248 | 0.0775 | -0.4 |
| SET | NM_001248000 | -0.3805 | -0.4045 |
| SET | NM_001248001 | -0.3805 | -0.4045 |
| ARNTL2 | NM_001248002 | -0.255 | -0.831 |
| ARNTL2 | NM_001248003 | -0.255 | -0.831 |
| ARNTL2 | NM_001248004 | -0.255 | -0.831 |
| ARNTL2 | NM_001248005 | -0.255 | -0.831 |
| TRIM3 | NM_001248006 | -0.428 | -1.2885 |
| TRIM3 | NM_001248007 | -0.428 | -1.2885 |
| SKIL | NM_001248008 | -0.6595 | -0.461 |
| ENTPD5 | NM_001249 | 0.068 | -0.3015 |
| CD40 | NM_001250 | -0.7055 | 0.427 |
| CD68 | NM_001251 | -0.088 | 0.0115 |
| SP1 | NM_001251825 | -1.318 | -2.7675 |
| SPP1 | NM_001251829 | 0.3405 | 0.0675 |
| SPP1 | NM_001251830 | 0.3405 | 0.0675 |
| PIK3R5 | NM_001251851 | 0.015 | -0.1775 |
| PIK3R5 | NM_001251852 | 0.015 | -0.1775 |
| PIK3R5 | NM_001251853 | 0.015 | -0.1775 |
| PIK3R5 | NM_001251855 | 0.015 | -0.1775 |
| USP4 | NM_001251877 | -0.3965 | -1.4 |
| VIPR1 | NM_001251882 | 0.117 | -0.0895 |
| VIPR1 | NM_001251883 | 0.117 | -0.0895 |
| VIPR1 | NM_001251884 | 0.117 | -0.0895 |
| VIPR1 | NM_001251885 | 0.117 | -0.0895 |
| ASPSCR1 | NM_001251888 | -0.623 | -1.1805 |
| CD83 | NM_001251901 | -0.552 | -1.466 |
| APPL2 | NM_001251904 | -1.113 | -1.472 |
| APPL2 | NM_001251905 | -1.113 | -1.472 |
| RAP1B | NM_001251917 | -0.98 | -1.353 |
| RAP1B | NM_001251918 | -0.98 | -1.353 |
| RAP1B | NM_001251921 | -0.98 | -1.353 |
| RAP1B | NM_001251922 | -0.98 | -1.353 |
| ARHGEF26 | NM_001251962 | 0.133 | -0.135 |
| ARHGEF26 | NM_001251963 | 0.133 | -0.135 |
| TP53AIP1 | NM_001251964 | 0.029 | -0.2835 |

| | | | |
|---|---|---|---|
| SPOCK3 | NM_001251967 | -0.615 | -0.4285 |
| SLC30A5 | NM_001251969 | -1.8635 | -2.8875 |
| SPAG9 | NM_001251971 | -1.214 | -1.697 |
| RCAN2 | NM_001251973 | 0.1425 | -0.3635 |
| RCAN2 | NM_001251974 | 0.1425 | -0.3635 |
| RCAN3 | NM_001251977 | -0.1435 | 0.322 |
| RCAN3 | NM_001251978 | -0.1435 | 0.322 |
| RCAN3 | NM_001251979 | -0.1435 | 0.322 |
| RCAN3 | NM_001251980 | -0.1435 | 0.322 |
| RCAN3 | NM_001251981 | -0.1435 | 0.322 |
| RCAN3 | NM_001251982 | -0.1435 | 0.322 |
| RCAN3 | NM_001251983 | -0.1435 | 0.322 |
| RCAN3 | NM_001251984 | -0.1435 | 0.322 |
| RCAN3 | NM_001251985 | -0.1435 | 0.322 |
| TRDN | NM_001251987 | -0.1255 | -0.53 |
| GMNN | NM_001251989 | -1.24 | -2.416 |
| GMNN | NM_001251990 | -1.24 | -2.416 |
| GMNN | NM_001251991 | -1.24 | -2.416 |
| CD70 | NM_001252 | 0.111 | -0.315 |
| PAPOLA | NM_001252006 | 0.144 | -0.3525 |
| PAPOLA | NM_001252007 | 0.144 | -0.3525 |
| LUZP2 | NM_001252008 | -0.2375 | -0.123 |
| LUZP2 | NM_001252010 | -0.2375 | -0.123 |
| TOR2A | NM_001252018 | -0.2205 | -0.4615 |
| TRPM1 | NM_001252020 | -0.016 | 0.0465 |
| TOR2A | NM_001252021 | -0.2205 | -0.4615 |
| TOR2A | NM_001252023 | -0.2205 | -0.4615 |
| TRPM1 | NM_001252024 | -0.016 | 0.0465 |
| TRPM1 | NM_001252030 | -0.016 | 0.0465 |
| RAB5B | NM_001252036 | -0.567 | -1.245 |
| RAB5B | NM_001252037 | -0.567 | -1.245 |
| RAB5C | NM_001252039 | 0.0955 | -0.6495 |
| TAOK2 | NM_001252043 | -0.1305 | -0.474 |
| PCMT1 | NM_001252049 | -0.662 | -0.839 |
| PCMT1 | NM_001252050 | -0.662 | -0.839 |
| PCMT1 | NM_001252051 | -0.662 | -0.839 |
| PCMT1 | NM_001252052 | -0.662 | -0.839 |
| PCMT1 | NM_001252053 | -0.662 | -0.839 |
| SYT7 | NM_001252065 | -0.215 | -0.5635 |
| CDC5L | NM_001253 | -0.534 | -1.5975 |
| CDC6 | NM_001254 | -1.746 | -3.934 |
| CDC20 | NM_001255 | -0.5365 | -1.2 |
| CDC27 | NM_001256 | -0.7895 | -2.749 |

| | | | |
|---|---|---|---|
| CDH13 | NM_001257 | -0.5675 | -0.28 |
| CDK3 | NM_001258 | -0.4905 | -1.116 |
| CDK6 | NM_001259 | -1.035 | -1.124 |
| CDK8 | NM_001260 | -1.292 | -2.581 |
| CDK9 | NM_001261 | -0.352 | -1.425 |
| CDKN2C | NM_001262 | -0.597 | -2.3365 |
| CDS1 | NM_001263 | -0.3555 | -0.8025 |
| CDSN | NM_001264 | 0.0085 | 0.0395 |
| CDX2 | NM_001265 | 0.1755 | -0.423 |
| CHAD | NM_001267 | 0.202 | -0.2705 |
| RCBTB2 | NM_001268 | -0.9045 | -0.7995 |
| RCC1 | NM_001269 | -0.3655 | -1.5915 |
| CHD1 | NM_001270 | -0.635 | -3.7085 |
| CHD2 | NM_001271 | -1.271 | -3.384 |
| CHD4 | NM_001273 | -0.2635 | -0.8005 |
| CHEK1 | NM_001274 | -1.066 | -1.552 |
| CHGA | NM_001275 | -0.0745 | -0.802 |
| CHI3L1 | NM_001276 | -0.295 | -0.1605 |
| CHKA | NM_001277 | -1.089 | -2.491 |
| CHUK | NM_001278 | -1.2005 | -1.2375 |
| CIDEA | NM_001279 | -0.2375 | -0.5555 |
| CIRBP | NM_001280 | 0.125 | -0.3975 |
| TBCB | NM_001281 | 0.2235 | -0.271 |
| AP2B1 | NM_001282 | -0.422 | -0.6825 |
| AP1S1 | NM_001283 | 0.2155 | -0.5485 |
| AP3S1 | NM_001284 | -0.2785 | -0.4085 |
| CLCA1 | NM_001285 | 0.1495 | -0.162 |
| CLCN6 | NM_001286 | -0.8425 | -1.713 |
| CLCN7 | NM_001287 | -0.386 | -0.9145 |
| CLIC1 | NM_001288 | -0.243 | -1.3405 |
| CLIC2 | NM_001289 | 0.2245 | -0.565 |
| LDB2 | NM_001290 | -1.461 | -1.3235 |
| CLNS1A | NM_001293 | -0.5285 | -0.262 |
| CLPTM1 | NM_001294 | -0.173 | -1.5285 |
| CCR1 | NM_001295 | 0.058 | -0.28 |
| CCBP2 | NM_001296 | 0.0165 | -0.109 |
| CNGB1 | NM_001297 | -0.416 | -0.6195 |
| CNGA3 | NM_001298 | -0.1505 | -1.043 |
| CNN1 | NM_001299 | -0.1815 | -0.277 |
| KLF6 | NM_001300 | -0.167 | 0.532 |
| CORT | NM_001302 | -0.0315 | -0.3635 |
| COX10 | NM_001303 | -0.3505 | -2.8545 |
| CPD | NM_001304 | -0.695 | -1.7215 |

| | | | |
|---|---|---|---|
| CLDN4 | NM_001305 | 0.1315 | -0.3835 |
| CLDN3 | NM_001306 | 0.0275 | 0.2245 |
| CLDN7 | NM_001307 | 0.1675 | -0.329 |
| CPN1 | NM_001308 | 0.225 | -0.087 |
| CREBL2 | NM_001310 | -2.0165 | -2.1665 |
| CRIP1 | NM_001311 | 0.242 | -0.3285 |
| CRIP2 | NM_001312 | 0.0305 | -0.3665 |
| CRMP1 | NM_001313 | 0.0265 | -0.3865 |
| MAPK14 | NM_001315 | -0.9775 | -2.637 |
| CSE1L | NM_001316 | -0.814 | -1.8675 |
| CSH1 | NM_001317 | -0.1945 | -0.0325 |
| CSHL1 | NM_001318 | -0.2255 | -0.2515 |
| CSNK1G2 | NM_001319 | -0.3445 | -0.898 |
| CSNK2B | NM_001320 | 0.1285 | -0.5685 |
| CSRP2 | NM_001321 | -0.018 | -0.9935 |
| CST2 | NM_001322 | 0.0365 | -0.321 |
| CST6 | NM_001323 | -0.2215 | -0.186 |
| CSTF1 | NM_001324 | -1.685 | -3.3255 |
| CSTF2 | NM_001325 | -0.7925 | -1.6185 |
| CSTF3 | NM_001326 | -3.1955 | -3.9795 |
| CTBP1 | NM_001328 | -0.1235 | -0.469 |
| CTBP2 | NM_001329 | -0.9365 | -1.7355 |
| CTF1 | NM_001330 | -0.1125 | -0.5205 |
| CTNND1 | NM_001331 | -0.546 | -1.48 |
| CTNND2 | NM_001332 | -0.277 | -0.817 |
| CTSL2 | NM_001333 | -0.622 | -1.111 |
| CTSO | NM_001334 | -0.297 | -1.018 |
| CTSW | NM_001335 | 0.082 | -0.368 |
| CTSZ | NM_001336 | -0.1835 | -0.16 |
| CX3CR1 | NM_001337 | -0.0255 | -0.0575 |
| CXADR | NM_001338 | -0.8815 | -1.0315 |
| CYLC2 | NM_001340 | -0.45 | -0.3105 |
| DAB2 | NM_001343 | 0.1555 | 0.2915 |
| DAD1 | NM_001344 | -0.2305 | -0.5285 |
| DGKA | NM_001345 | -0.0475 | -0.417 |
| DGKG | NM_001346 | 0.0795 | -0.2595 |
| DGKQ | NM_001347 | 0.368 | -0.759 |
| DAPK3 | NM_001348 | -0.5485 | -1.4875 |
| DARS | NM_001349 | -0.9145 | -0.5065 |
| DAXX | NM_001350 | -0.2725 | -0.882 |
| DAZL | NM_001351 | 0.037 | -0.12 |
| DBP | NM_001352 | -0.5965 | -1.5875 |
| AKR1C1 | NM_001353 | -0.133 | -0.3525 |

| | | | |
|---|---|---|---|
| AKR1C2 | NM_001354 | 0.161 | -0.448 |
| DDT | NM_001355 | 0.2315 | -0.3765 |
| DDX3X | NM_001356 | -1.372 | -2.2 |
| DHX9 | NM_001357 | -1.008 | -0.847 |
| DHX15 | NM_001358 | -1.204 | -1.7015 |
| DECR1 | NM_001359 | -0.5125 | -0.4435 |
| DHCR7 | NM_001360 | -0.636 | -1.5385 |
| DHODH | NM_001361 | -0.457 | -1.1225 |
| DIO3 | NM_001362 | 0.386 | -0.2285 |
| DKC1 | NM_001363 | -0.7235 | -0.959 |
| DLG2 | NM_001364 | -0.104 | -0.1645 |
| DLG4 | NM_001365 | -0.458 | -0.9215 |
| DNAH5 | NM_001369 | 0.1415 | -0.492 |
| DNAH6 | NM_001370 | 0.1265 | -0.404 |
| DNAH9 | NM_001372 | 0.3285 | -0.1595 |
| DNAH14 | NM_001373 | 0.163 | -0.345 |
| DNASE1L2 | NM_001374 | -0.3805 | -0.7655 |
| DNASE2 | NM_001375 | -0.083 | -0.6485 |
| DYNC1H1 | NM_001376 | -0.549 | -0.924 |
| DYNC2H1 | NM_001377 | 0.123 | -1.0755 |
| DYNC1I2 | NM_001378 | -0.8365 | -0.7655 |
| DNMT1 | NM_001379 | -0.1225 | -0.8115 |
| DOCK1 | NM_001380 | -1.303 | -0.9 |
| DOK1 | NM_001381 | -0.5415 | -2.156 |
| DPAGT1 | NM_001382 | -0.1295 | -0.848 |
| DPH1 | NM_001383 | -0.847 | -1.479 |
| DPH2 | NM_001384 | -0.7815 | -1.744 |
| DPYS | NM_001385 | 0.167 | 0.212 |
| DPYSL2 | NM_001386 | -0.1565 | -0.526 |
| DPYSL3 | NM_001387 | -0.0235 | -0.5435 |
| DRG2 | NM_001388 | -0.243 | -0.976 |
| DSCAM | NM_001389 | -0.022 | -1.071 |
| DTNA | NM_001390 | -0.5815 | -1.7065 |
| DTNA | NM_001391 | -0.5815 | -1.7065 |
| DTNA | NM_001392 | -0.5815 | -1.7065 |
| ECM2 | NM_001393 | 0.2135 | -0.0515 |
| DUSP4 | NM_001394 | -0.5185 | -1.986 |
| DUSP9 | NM_001395 | 0.0985 | -0.946 |
| DYRK1A | NM_001396 | 0.1795 | -0.404 |
| ECE1 | NM_001397 | -0.608 | -1.636 |
| EDA | NM_001399 | -0.0235 | 0.383 |
| S1PR1 | NM_001400 | -0.491 | -0.075 |
| LPAR1 | NM_001401 | -0.6865 | -2.2215 |

| | | | |
|---|---|---|---|
| EEF1A1 | NM_001402 | -0.2605 | -0.358 |
| EEF1G | NM_001404 | 0.134 | -0.317 |
| EFNA2 | NM_001405 | 0.0785 | -1.1135 |
| EFNB3 | NM_001406 | -0.1405 | -0.9845 |
| CELSR3 | NM_001407 | -0.338 | -1.447 |
| CELSR2 | NM_001408 | -0.266 | -0.8465 |
| MEGF6 | NM_001409 | 0.1725 | -0.4055 |
| MEGF8 | NM_001410 | -0.1855 | -0.7985 |
| EIF1AX | NM_001412 | -1.1405 | -1.1015 |
| EIF2B1 | NM_001414 | -0.4675 | -0.93 |
| EIF2S3 | NM_001415 | -0.6125 | -0.356 |
| EIF4A1 | NM_001416 | -0.018 | -0.2655 |
| EIF4B | NM_001417 | -0.903 | -0.7325 |
| EIF4G2 | NM_001418 | -0.732 | -0.4535 |
| ELAVL1 | NM_001419 | -0.267 | -1.588 |
| ELAVL3 | NM_001420 | -1.109 | -1.8905 |
| ELF4 | NM_001421 | -0.0105 | -1.7325 |
| ELF5 | NM_001422 | -0.3485 | -0.1935 |
| EMP1 | NM_001423 | 0.0635 | 0.173 |
| EMP2 | NM_001424 | -0.038 | -0.7745 |
| EMP3 | NM_001425 | 0.2095 | -0.043 |
| EN1 | NM_001426 | 0.2885 | -0.4225 |
| EN2 | NM_001427 | -0.1365 | -0.553 |
| ENO1 | NM_001428 | 0.1355 | -0.367 |
| EP300 | NM_001429 | -1.1545 | -2.566 |
| EPAS1 | NM_001430 | -0.0635 | -0.96 |
| EPB41L2 | NM_001431 | -0.624 | -0.5595 |
| EREG | NM_001432 | 0.1775 | -0.213 |
| ERN1 | NM_001433 | 0.045 | -0.276 |
| FBL | NM_001436 | 0.125 | -0.193 |
| ESR2 | NM_001437 | -0.0185 | -0.086 |
| ESRRG | NM_001438 | -1.2545 | -4.612 |
| EXTL2 | NM_001439 | -1.0695 | -1.201 |
| EXTL3 | NM_001440 | -0.6505 | -2.9975 |
| FAAH | NM_001441 | -0.0415 | -0.508 |
| FABP4 | NM_001442 | 0.0625 | 0.1975 |
| FABP1 | NM_001443 | 0.3495 | -0.2875 |
| FABP5 | NM_001444 | -0.3535 | -0.5185 |
| FABP7 | NM_001446 | -0.5165 | -0.452 |
| FAT2 | NM_001447 | -0.0835 | -0.5775 |
| GPC4 | NM_001448 | 0.219 | -0.2325 |
| FHL1 | NM_001449 | 0.1485 | -0.162 |
| FHL2 | NM_001450 | -0.5065 | -2.9275 |

| | | | |
|---|---|---|---|
| FOXF1 | NM_001451 | -0.1965 | -0.1905 |
| FOXF2 | NM_001452 | 0.1295 | -0.2025 |
| FOXC1 | NM_001453 | 0.81 | 0.8155 |
| FOXJ1 | NM_001454 | -0.1675 | -0.483 |
| FOXO3 | NM_001455 | -1.5695 | -2.4285 |
| FLNA | NM_001456 | 0.006 | -0.325 |
| FLNB | NM_001457 | -0.094 | -0.873 |
| FLNC | NM_001458 | -0.3105 | -0.923 |
| FLT3LG | NM_001459 | -0.495 | -0.258 |
| FMO2 | NM_001460 | -0.188 | -0.3305 |
| FMO5 | NM_001461 | -0.3265 | -0.0655 |
| FPR2 | NM_001462 | -0.0935 | -0.3865 |
| FRZB | NM_001463 | -0.21 | -1.8525 |
| ADAM2 | NM_001464 | 0.037 | -0.429 |
| FYB | NM_001465 | 0.1725 | -0.4775 |
| FZD2 | NM_001466 | -0.7345 | -0.9175 |
| SLC37A4 | NM_001467 | -0.327 | -1.19 |
| GAGE1 | NM_001468 | 0.0105 | -1.0725 |
| XRCC6 | NM_001469 | -0.136 | -0.212 |
| GABBR1 | NM_001470 | -0.616 | -1.272 |
| B4GALNT1 | NM_001478 | 0.2855 | -0.5015 |
| GALR1 | NM_001480 | 0.1365 | -0.6685 |
| GAS8 | NM_001481 | -0.722 | -1.1105 |
| GATM | NM_001482 | -0.5205 | -0.9495 |
| GBAS | NM_001483 | -1.23 | -1.315 |
| GBX2 | NM_001485 | -0.067 | 0.0625 |
| GCKR | NM_001486 | -0.089 | -0.51 |
| BLOC1S1 | NM_001487 | 0.306 | -0.1995 |
| TADA2A | NM_001488 | -1.5365 | -2.296 |
| GCNT1 | NM_001490 | -0.308 | -0.466 |
| GCNT2 | NM_001491 | -0.2005 | -2.219 |
| GDI1 | NM_001493 | 0.182 | -0.19 |
| GDI2 | NM_001494 | -0.8215 | -0.6675 |
| GFRA2 | NM_001495 | -0.694 | -2.2955 |
| GFRA3 | NM_001496 | 0.0225 | -0.11 |
| B4GALT1 | NM_001497 | 0.154 | -0.606 |
| GCLC | NM_001498 | -1.575 | -2.98 |
| GLE1 | NM_001499 | -0.827 | -1.7725 |
| GMDS | NM_001500 | -0.0135 | -0.8775 |
| GNRH2 | NM_001501 | 0.128 | -0.4425 |
| GP2 | NM_001502 | 0.1395 | -0.1095 |
| GPLD1 | NM_001503 | -0.2785 | -0.155 |
| GPER | NM_001505 | 0.005 | -0.5815 |

| | | | |
|---|---|---|---|
| GPR32 | NM_001506 | -0.636 | -0.3545 |
| MLNR | NM_001507 | 0.2745 | -0.195 |
| GPR39 | NM_001508 | -0.4175 | -0.3515 |
| GPX5 | NM_001509 | 0.37 | -0.2015 |
| GRID2 | NM_001510 | 0.2425 | -0.255 |
| CXCL1 | NM_001511 | 0.069 | -0.4495 |
| GSTA4 | NM_001512 | -0.6285 | -0.407 |
| GSTZ1 | NM_001513 | -0.004 | -0.4935 |
| GTF2B | NM_001514 | -1.2925 | -3.138 |
| GTF2H3 | NM_001516 | -0.858 | -1.3125 |
| GTF2H4 | NM_001517 | -0.2725 | -0.181 |
| GTF2I | NM_001518 | -0.1905 | -0.6505 |
| BRF1 | NM_001519 | -0.3265 | -1.5405 |
| GTF3C1 | NM_001520 | -0.369 | -1.541 |
| GTF3C2 | NM_001521 | -0.9595 | -2.184 |
| GUCY2F | NM_001522 | -0.0475 | 0.1035 |
| HAS1 | NM_001523 | 0.1525 | 0.17 |
| HCRT | NM_001524 | -0.1015 | 0.34 |
| HCRTR1 | NM_001525 | -0.0315 | -0.1765 |
| HCRTR2 | NM_001526 | -0.023 | -0.375 |
| HDAC2 | NM_001527 | -0.999 | -1.0355 |
| HGFAC | NM_001528 | -0.112 | -0.3725 |
| HIF1A | NM_001530 | -0.299 | -1.259 |
| MR1 | NM_001531 | -0.28 | -0.164 |
| SLC29A2 | NM_001532 | -0.1935 | -1.1475 |
| HNRNPL | NM_001533 | -0.005 | -0.5555 |
| PRMT2 | NM_001535 | -0.777 | -0.847 |
| PRMT1 | NM_001536 | -0.1365 | -0.383 |
| HSBP1 | NM_001537 | -0.2455 | -0.0845 |
| HSF4 | NM_001538 | -0.302 | -0.6445 |
| DNAJA1 | NM_001539 | -0.612 | -0.7595 |
| HSPB1 | NM_001540 | 0.1785 | -0.048 |
| HSPB2 | NM_001541 | 0.104 | -0.4675 |
| IGSF3 | NM_001542 | -0.3675 | -1.1225 |
| NDST1 | NM_001543 | 1.678 | 1.1745 |
| ICAM4 | NM_001544 | -0.2025 | -0.241 |
| ICT1 | NM_001545 | -0.7445 | -2.435 |
| ID4 | NM_001546 | 0.014 | 1.1935 |
| IFIT2 | NM_001547 | 0.024 | -0.364 |
| IFIT1 | NM_001548 | -0.226 | -0.554 |
| IFIT3 | NM_001549 | -0.1885 | -0.458 |
| IFRD1 | NM_001550 | -1.735 | -3.0135 |
| IGBP1 | NM_001551 | -0.935 | -1.0515 |

| | | | |
|---|---|---|---|
| IGFBP4 | NM_001552 | -0.4975 | -2.1005 |
| IGFBP7 | NM_001553 | -0.003 | -0.3225 |
| CYR61 | NM_001554 | 0.0575 | 0.524 |
| IGSF1 | NM_001555 | 0.3395 | -0.3795 |
| IKBKB | NM_001556 | -0.9915 | -2.568 |
| CXCR2 | NM_001557 | -0.132 | -0.1315 |
| IL10RA | NM_001558 | 0.042 | -0.6485 |
| IL12RB2 | NM_001559 | 0.263 | -0.015 |
| IL13RA1 | NM_001560 | -0.2145 | -0.404 |
| TNFRSF9 | NM_001561 | -0.143 | -0.2725 |
| IL18 | NM_001562 | 0.192 | -0.246 |
| IMPG1 | NM_001563 | -0.186 | 0.2755 |
| ING2 | NM_001564 | -1.1355 | -2.1765 |
| CXCL10 | NM_001565 | -0.446 | -0.2015 |
| INPP4A | NM_001566 | -0.6765 | -1.123 |
| INPPL1 | NM_001567 | -0.4525 | -0.849 |
| EIF3E | NM_001568 | -0.399 | -0.445 |
| IRAK2 | NM_001570 | -0.0425 | -0.3765 |
| IRF3 | NM_001571 | -0.355 | -1.256 |
| MPPED2 | NM_001584 | -1.0105 | -2.2935 |
| TEX28 | NM_001586 | -0.1235 | -0.4765 |
| OCRL | NM_001587 | -0.9435 | -0.8775 |
| PAX6 | NM_001604 | 0.165 | -0.474 |
| AARS | NM_001605 | -0.441 | -1.6995 |
| ABCA2 | NM_001606 | 0.2175 | -0.177 |
| ACAA1 | NM_001607 | -0.0955 | -1.153 |
| ACADL | NM_001608 | -0.168 | -0.6945 |
| ACADSB | NM_001609 | -1.393 | -1.6445 |
| ACP2 | NM_001610 | -0.4 | -1.225 |
| ACP5 | NM_001611 | -0.016 | -0.56 |
| ACRV1 | NM_001612 | -0.047 | -0.667 |
| ACTA2 | NM_001613 | -0.22 | -0.5805 |
| ACTG1 | NM_001614 | 0.067 | -0.427 |
| ACTG2 | NM_001615 | 0.1455 | 0.2195 |
| ACVR2A | NM_001616 | -0.9585 | -1.3285 |
| ADD2 | NM_001617 | 0.4305 | -0.786 |
| PARP1 | NM_001618 | -0.18 | -0.5615 |
| ADRBK1 | NM_001619 | -0.166 | -0.3185 |
| AHNAK | NM_001620 | -0.867 | -1.2525 |
| AHR | NM_001621 | 0.1665 | -0.1965 |
| AHSG | NM_001622 | 0.0275 | -0.299 |
| AIF1 | NM_001623 | -0.1545 | -0.3475 |
| AIM1 | NM_001624 | -0.264 | -0.0475 |

| | | | |
|---|---|---|---|
| AK2 | NM_001625 | -0.2575 | -1.04 |
| AKT2 | NM_001626 | -0.1 | -0.836 |
| ALCAM | NM_001627 | -0.614 | -1.0345 |
| AKR1B1 | NM_001628 | 0.3025 | -0.13 |
| ALOX5AP | NM_001629 | 0.145 | -0.292 |
| ALPI | NM_001631 | -0.159 | -0.188 |
| ALPP | NM_001632 | -0.1055 | -0.459 |
| AMBP | NM_001633 | -0.152 | -0.2585 |
| AMD1 | NM_001634 | -1.0395 | -2.7245 |
| AMPH | NM_001635 | -0.5445 | -0.85 |
| SLC25A6 | NM_001636 | 0.2365 | -0.3025 |
| AOAH | NM_001637 | -0.0155 | -0.3145 |
| APOF | NM_001638 | -0.052 | -0.1235 |
| APCS | NM_001639 | 0.1755 | -0.491 |
| APEH | NM_001640 | 0.136 | -0.685 |
| APEX1 | NM_001641 | -0.5705 | -1.4655 |
| APLP2 | NM_001642 | 0.0665 | -0.5475 |
| APOA2 | NM_001643 | -0.151 | -0.141 |
| APOBEC1 | NM_001644 | -0.005 | -0.1095 |
| APOC1 | NM_001645 | 0.45 | -0.221 |
| APOC4 | NM_001646 | 0.163 | -0.28 |
| APOD | NM_001647 | 0.004 | -0.478 |
| KLK3 | NM_001648 | -0.024 | -0.0285 |
| SHROOM2 | NM_001649 | -0.327 | -0.5655 |
| AQP4 | NM_001650 | -0.111 | -0.3035 |
| AQP5 | NM_001651 | 0.0065 | -0.1955 |
| AQP6 | NM_001652 | 0.0735 | -0.275 |
| ARAF | NM_001654 | -0.1105 | -0.7615 |
| ARCN1 | NM_001655 | -1.224 | -2.7005 |
| TRIM23 | NM_001656 | -0.8735 | -2.712 |
| AREG | NM_001657 | 0.4965 | -0.57 |
| ARF1 | NM_001658 | 0.118 | -0.4555 |
| ARF3 | NM_001659 | 0.1135 | -1.245 |
| ARF4 | NM_001660 | -0.542 | -1.1275 |
| ARL4D | NM_001661 | -0.3615 | -1.0995 |
| ARF5 | NM_001662 | 0.1955 | -0.4285 |
| ARF6 | NM_001663 | -0.78 | -2.69 |
| RHOA | NM_001664 | 0.042 | -0.43 |
| RHOG | NM_001665 | -0.4795 | -2.1235 |
| ARHGAP4 | NM_001666 | -0.0145 | -0.4515 |
| ARL2 | NM_001667 | -0.329 | -1.804 |
| ARNT | NM_001668 | -0.8185 | -2.4545 |
| ARSD | NM_001669 | -0.0115 | -0.832 |

| | | | |
|---|---|---|---|
| ARVCF | NM_001670 | -0.6015 | -1.353 |
| ASGR1 | NM_001671 | 0.0135 | -0.572 |
| ASIP | NM_001672 | -0.1785 | -0.1945 |
| ASNS | NM_001673 | -0.3555 | -0.499 |
| ATF3 | NM_001674 | -0.3345 | 1.001 |
| ATF4 | NM_001675 | -0.635 | -1.2975 |
| ATP12A | NM_001676 | 0.197 | 0.059 |
| ATP1B1 | NM_001677 | -0.602 | -0.7695 |
| ATP1B2 | NM_001678 | 0.3825 | -0.3485 |
| ATP1B3 | NM_001679 | -0.5235 | -0.973 |
| FXYD2 | NM_001680 | 0.1985 | -0.304 |
| ATP2A2 | NM_001681 | -0.248 | -1.0545 |
| ATP2B1 | NM_001682 | -0.169 | -1.4015 |
| ATP2B2 | NM_001683 | -1.334 | -2.4225 |
| ATP2B4 | NM_001684 | 0.034 | -0.262 |
| ATP5B | NM_001686 | 0.1445 | -0.2265 |
| ATP5D | NM_001687 | 0.2845 | -0.324 |
| ATP5F1 | NM_001688 | -0.507 | -0.8065 |
| ATP5G3 | NM_001689 | 0.0935 | -0.4055 |
| ATP6V1A | NM_001690 | -0.8505 | -0.6965 |
| ATP6V1B1 | NM_001692 | -0.4515 | -0.4075 |
| ATP6V1B2 | NM_001693 | -0.622 | -1.159 |
| ATP6V0C | NM_001694 | 0.266 | 0.008 |
| ATP6V1C1 | NM_001695 | -1.2525 | -2.7005 |
| ATP6V1E1 | NM_001696 | -1.1765 | -1.0575 |
| ATP5O | NM_001697 | -0.2575 | -0.293 |
| AUH | NM_001698 | -1.6875 | -2.5965 |
| AXL | NM_001699 | -0.0465 | 0.036 |
| AZU1 | NM_001700 | 0.214 | -0.134 |
| BAAT | NM_001701 | -0.0015 | -0.465 |
| BAI1 | NM_001702 | -0.348 | -0.3855 |
| BAI2 | NM_001703 | -0.177 | -0.7495 |
| BAI3 | NM_001704 | -1.912 | -4.0155 |
| BCL6 | NM_001706 | 0.303 | -0.525 |
| BCL7B | NM_001707 | -0.3765 | -1.8145 |
| OPN1SW | NM_001708 | -0.345 | -0.4945 |
| BDNF | NM_001709 | -0.605 | -1.907 |
| CFB | NM_001710 | 0.2415 | -0.288 |
| BGN | NM_001711 | -0.0345 | -0.558 |
| CEACAM1 | NM_001712 | 0.085 | -0.347 |
| BHMT | NM_001713 | 0.0735 | -0.0915 |
| BICD1 | NM_001714 | -0.8615 | -1.9545 |
| BLK | NM_001715 | -0.0295 | -0.431 |

| | | | |
|---|---|---|---|
| CXCR5 | NM_001716 | -0.002 | -0.348 |
| BNC1 | NM_001717 | -0.1505 | 0.2035 |
| BMP6 | NM_001718 | -0.017 | -0.1605 |
| BMP7 | NM_001719 | -0.3755 | -0.6695 |
| BMP8B | NM_001720 | 0.2415 | -0.709 |
| POLR3D | NM_001722 | -0.688 | -1.6105 |
| DST | NM_001723 | -0.3215 | -2.0285 |
| BPGM | NM_001724 | -1.2025 | -1.297 |
| BPI | NM_001725 | -0.0335 | -0.6595 |
| BRS3 | NM_001727 | -0.429 | 0.063 |
| BSG | NM_001728 | 0.389 | -0.097 |
| BTC | NM_001729 | -0.3345 | -0.3605 |
| KLF5 | NM_001730 | 0.1455 | 0.015 |
| BTG1 | NM_001731 | -0.1105 | -0.1475 |
| BTN1A1 | NM_001732 | -0.3365 | -0.574 |
| C1R | NM_001733 | -0.027 | -0.529 |
| C1S | NM_001734 | -0.1455 | -0.3475 |
| C5 | NM_001735 | -0.15 | -0.4245 |
| C5AR1 | NM_001736 | 0.2365 | 0.126 |
| C9 | NM_001737 | -0.0175 | -0.16 |
| CA1 | NM_001738 | 0.052 | -0.7995 |
| CA5A | NM_001739 | 0.0145 | -0.257 |
| CALB2 | NM_001740 | -0.1525 | -0.477 |
| CALCA | NM_001741 | 0.0585 | -0.5015 |
| CALCR | NM_001742 | 0.01 | 0.0765 |
| CALM2 | NM_001743 | -0.472 | -0.4125 |
| CAMK4 | NM_001744 | -0.6645 | -1.393 |
| CAMLG | NM_001745 | -1.0115 | -1.2075 |
| CANX | NM_001746 | -0.8695 | -0.3845 |
| CAPG | NM_001747 | 0.055 | -0.563 |
| CAPN2 | NM_001748 | -0.81 | -0.524 |
| CAPNS1 | NM_001749 | 0.2915 | -0.159 |
| CARS | NM_001751 | -0.4275 | -1.4015 |
| CAT | NM_001752 | -0.998 | -1.259 |
| CAV1 | NM_001753 | 0.087 | -0.168 |
| RUNX1 | NM_001754 | 0.0375 | -0.054 |
| CBFB | NM_001755 | -1.3565 | -2.781 |
| SERPINA6 | NM_001756 | -0.163 | -0.2825 |
| CBR1 | NM_001757 | -1.0745 | -2.953 |
| CCND2 | NM_001759 | 0.3135 | 0.418 |
| CCND3 | NM_001760 | -0.6295 | -1.572 |
| CCNF | NM_001761 | -1.081 | -3.2795 |
| CCT6A | NM_001762 | -1.4135 | -0.945 |

| | | | |
|---|---|---|---|
| CD1A | NM_001763 | 0.018 | -0.3 |
| CD1B | NM_001764 | 0.1755 | 0.1055 |
| CD1C | NM_001765 | -0.271 | -0.137 |
| CD1D | NM_001766 | -0.107 | -0.4355 |
| CD2 | NM_001767 | -0.1215 | -0.1375 |
| CD8A | NM_001768 | 0.0545 | -0.1035 |
| CD9 | NM_001769 | -0.5805 | -2.201 |
| CD19 | NM_001770 | -0.148 | -0.193 |
| CD22 | NM_001771 | 0.4995 | 0.1545 |
| CD33 | NM_001772 | 0.26 | -0.237 |
| CD34 | NM_001773 | -0.1735 | -0.332 |
| CD37 | NM_001774 | 0.5415 | 0.209 |
| CD38 | NM_001775 | -0.2025 | 0.1025 |
| ENTPD1 | NM_001776 | -0.2325 | -0.583 |
| CD47 | NM_001777 | -1.0665 | -1.2815 |
| CD48 | NM_001778 | 0.0945 | -0.3015 |
| CD58 | NM_001779 | -1.438 | -2.471 |
| CD63 | NM_001780 | 0.1715 | -0.2165 |
| CD69 | NM_001781 | 0.0085 | -0.007 |
| CD72 | NM_001782 | 0.5515 | -0.5475 |
| CD79A | NM_001783 | -0.0825 | -0.3615 |
| CD97 | NM_001784 | -0.193 | 0.051 |
| CDA | NM_001785 | 0.4245 | -0.173 |
| CDK1 | NM_001786 | -0.999 | -1.665 |
| 7-Sep | NM_001788 | -0.268 | -0.5055 |
| CDC25A | NM_001789 | -0.8005 | -3.334 |
| CDC25C | NM_001790 | -1.3315 | -2.71 |
| CDC42 | NM_001791 | -0.3145 | -0.3855 |
| CDH2 | NM_001792 | -0.5105 | -0.874 |
| CDH3 | NM_001793 | 0.024 | -0.6215 |
| CDH4 | NM_001794 | -0.473 | -0.5345 |
| CDH5 | NM_001795 | 0.044 | -0.679 |
| CDH8 | NM_001796 | 0.0445 | -0.5655 |
| CDH11 | NM_001797 | -1.549 | -3.119 |
| CDK2 | NM_001798 | -0.2675 | -1.1355 |
| CDK7 | NM_001799 | -1.2265 | -3.2165 |
| CDKN2D | NM_001800 | 0.674 | 0.2845 |
| CDO1 | NM_001801 | -0.294 | -0.806 |
| CDR2 | NM_001802 | -1.4425 | -2.851 |
| CD52 | NM_001803 | -0.129 | -0.237 |
| CDX1 | NM_001804 | -0.225 | -0.1095 |
| CEBPE | NM_001805 | 0.1745 | -0.461 |
| CEBPG | NM_001806 | -0.9955 | -1.152 |

| | | | |
|---|---|---|---|
| CEL | NM_001807 | -1.5195 | -0.82 |
| CENPA | NM_001809 | -1.5115 | -4.3695 |
| CENPB | NM_001810 | -0.4445 | -1.324 |
| CENPC1 | NM_001812 | -0.448 | -2.6685 |
| CENPE | NM_001813 | -0.233 | -1.807 |
| CTSC | NM_001814 | -0.4375 | -2.279 |
| CEACAM3 | NM_001815 | 0.148 | -0.014 |
| CEACAM8 | NM_001816 | 0.0585 | -0.3835 |
| CEACAM4 | NM_001817 | 0.0645 | -0.4525 |
| AKR1C4 | NM_001818 | -0.558 | 0.215 |
| CHGB | NM_001819 | -0.56 | -0.42 |
| CHML | NM_001821 | -0.0295 | -2.5655 |
| CHN1 | NM_001822 | -0.9215 | -1.0645 |
| CKM | NM_001824 | 0.135 | -0.0315 |
| CKMT2 | NM_001825 | -0.0045 | -0.5015 |
| CKS1B | NM_001826 | -0.0085 | -0.165 |
| CKS2 | NM_001827 | -0.6175 | -0.96 |
| CLC | NM_001828 | 0.196 | 0.039 |
| CLCN3 | NM_001829 | -1.6635 | -2.8755 |
| CLCN4 | NM_001830 | -1.034 | -1.8945 |
| CLPS | NM_001832 | -0.3115 | -0.3895 |
| CLTA | NM_001833 | -0.0495 | -0.5025 |
| CLTCL1 | NM_001835 | 0.2595 | -1.119 |
| CMA1 | NM_001836 | -0.3245 | 0.311 |
| CCR3 | NM_001837 | -0.0235 | -0.3235 |
| CCR7 | NM_001838 | -0.0715 | -0.216 |
| CNN3 | NM_001839 | -0.7835 | -0.5285 |
| CNR2 | NM_001841 | -0.0535 | -0.2885 |
| CNTN1 | NM_001843 | -0.5615 | -0.6425 |
| COL2A1 | NM_001844 | 0.0005 | 0.007 |
| COL4A1 | NM_001845 | -0.2945 | -1.0035 |
| COL4A2 | NM_001846 | 0.027 | -0.6135 |
| COL4A6 | NM_001847 | -0.0095 | -0.6015 |
| COL6A1 | NM_001848 | 0.0355 | -0.1465 |
| COL6A2 | NM_001849 | -0.0985 | -0.48 |
| COL8A1 | NM_001850 | -0.1245 | -0.139 |
| COL9A1 | NM_001851 | -0.2045 | -0.218 |
| COL9A2 | NM_001852 | -0.2185 | -0.4405 |
| COL9A3 | NM_001853 | 0.0455 | -0.5165 |
| COL11A1 | NM_001854 | 0.313 | -0.504 |
| COL15A1 | NM_001855 | 0.218 | 0.1895 |
| COL16A1 | NM_001856 | -0.163 | -0.7495 |
| COL19A1 | NM_001858 | -0.029 | -0.065 |

| | | | |
|---|---|---|---|
| SLC31A1 | NM_001859 | -0.69 | -1.265 |
| SLC31A2 | NM_001860 | -0.2265 | -1.0525 |
| COX4I1 | NM_001861 | 0.282 | -0.083 |
| COX5B | NM_001862 | 0.308 | -0.2395 |
| COX6B1 | NM_001863 | 0.314 | -0.209 |
| COX7A1 | NM_001864 | 0.4455 | -0.098 |
| COX7A2 | NM_001865 | 0.1805 | -0.1835 |
| COX7B | NM_001866 | 0.0365 | -0.0935 |
| COX7C | NM_001867 | 0.06 | -0.2205 |
| CPA1 | NM_001868 | 0.1765 | -0.29 |
| CPA2 | NM_001869 | -0.097 | 0.0035 |
| CPA3 | NM_001870 | 0.217 | -0.064 |
| CPB1 | NM_001871 | 0.0015 | -0.1895 |
| CPB2 | NM_001872 | -0.172 | 0.391 |
| CPE | NM_001873 | -0.6275 | -0.3355 |
| CPM | NM_001874 | 0.222 | -0.293 |
| CPS1 | NM_001875 | -0.208 | 0.025 |
| CPT1A | NM_001876 | -0.816 | -1.0275 |
| CR2 | NM_001877 | 0.1755 | 0.042 |
| CRABP2 | NM_001878 | -0.0495 | -0.416 |
| MASP1 | NM_001879 | 0.035 | -0.4165 |
| ATF2 | NM_001880 | -1.587 | -3.1055 |
| CREM | NM_001881 | -0.2665 | -1.346 |
| CRHBP | NM_001882 | -0.0605 | -0.662 |
| CRHR2 | NM_001883 | -0.442 | -0.621 |
| HAPLN1 | NM_001884 | 0.6405 | -0.193 |
| CRYAB | NM_001885 | 0.0865 | 0.0015 |
| CRYBA4 | NM_001886 | 0.131 | 0.303 |
| CRYBB1 | NM_001887 | 0.0205 | -0.262 |
| CRYM | NM_001888 | 0.1085 | -0.3055 |
| CSN1S1 | NM_001890 | 0.015 | -0.3585 |
| CSN2 | NM_001891 | -0.2775 | -0.283 |
| CSNK1A1 | NM_001892 | -1.33 | -2.16 |
| CSNK1D | NM_001893 | -1.008 | -2.954 |
| CSNK1E | NM_001894 | -0.36 | -0.916 |
| CSNK2A1 | NM_001895 | -0.677 | -2.1985 |
| CSNK2A2 | NM_001896 | -0.8955 | -1.6775 |
| CSPG4 | NM_001897 | -0.1365 | -0.2255 |
| CST1 | NM_001898 | -0.225 | -0.358 |
| CST4 | NM_001899 | 0.107 | -0.554 |
| CST5 | NM_001900 | 0.2 | 0.0285 |
| CTGF | NM_001901 | 0.1375 | 0.5745 |
| CTNNA1 | NM_001903 | -0.248 | -0.5355 |

| | | | |
|---|---|---|---|
| CTNNB1 | NM_001904 | -0.3905 | -0.266 |
| CTPS | NM_001905 | -0.2485 | -1.5395 |
| CTRL | NM_001907 | -0.1935 | -0.562 |
| CTSB | NM_001908 | 0.0525 | -0.4655 |
| CTSD | NM_001909 | 0.062 | -0.4735 |
| CTSE | NM_001910 | 0.02 | 0.024 |
| CTSG | NM_001911 | 0.169 | -0.0845 |
| CTSL1 | NM_001912 | -0.21 | -0.7485 |
| CUX1 | NM_001913 | 0.1975 | -0.1795 |
| CYB5A | NM_001914 | -0.0135 | -0.7865 |
| CYB561 | NM_001915 | -0.8475 | -0.8595 |
| CYC1 | NM_001916 | 0.134 | -0.4315 |
| DAO | NM_001917 | 0.057 | -0.0265 |
| DBT | NM_001918 | -1.291 | -1.8585 |
| ECI1 | NM_001919 | 0.0735 | -0.7085 |
| DCN | NM_001920 | 0.3285 | 0.0705 |
| DCTD | NM_001921 | -0.0205 | -0.7945 |
| DCT | NM_001922 | 0.051 | -0.506 |
| DDB1 | NM_001923 | 0.209 | -0.441 |
| GADD45A | NM_001924 | -0.0925 | 0.158 |
| DEFA4 | NM_001925 | -0.155 | -0.335 |
| DEFA6 | NM_001926 | 0.241 | -0.3195 |
| DES | NM_001927 | 0.033 | -0.5725 |
| CFD | NM_001928 | 0.2615 | -0.277 |
| DHPS | NM_001930 | -0.027 | -0.3955 |
| DLAT | NM_001931 | -0.8935 | -2.024 |
| MPP3 | NM_001932 | -0.767 | -1.4395 |
| DLST | NM_001933 | -0.164 | -1.708 |
| DLX4 | NM_001934 | 0.0375 | -0.259 |
| DPP4 | NM_001935 | -0.0675 | -0.2185 |
| DPP6 | NM_001936 | -0.537 | -0.9885 |
| DPT | NM_001937 | -0.352 | -0.71 |
| DR1 | NM_001938 | -0.713 | -0.91 |
| DRP2 | NM_001939 | 0.095 | -0.083 |
| ATN1 | NM_001940 | -0.3765 | -0.8135 |
| DSC3 | NM_001941 | -0.6245 | -0.519 |
| DSG1 | NM_001942 | 0.1285 | -0.108 |
| DSG2 | NM_001943 | -0.5875 | 0.3745 |
| DSG3 | NM_001944 | -0.092 | -0.5865 |
| HBEGF | NM_001945 | 0.004 | 1.937 |
| DUSP6 | NM_001946 | -0.338 | 0.281 |
| DUSP7 | NM_001947 | -0.564 | -1.0695 |
| DUT | NM_001948 | -0.175 | -0.6775 |

| | | | |
|---|---|---|---|
| E2F3 | NM_001949 | -1.7195 | -3.6495 |
| E2F4 | NM_001950 | -0.8195 | -2.173 |
| E2F5 | NM_001951 | -2.6775 | -4.7945 |
| TYMP | NM_001953 | -0.071 | -0.255 |
| DDR1 | NM_001954 | -0.0885 | -0.7975 |
| EDN1 | NM_001955 | 0.329 | -0.3645 |
| EDN2 | NM_001956 | -0.0235 | -0.72 |
| EDNRA | NM_001957 | -1.017 | -1.892 |
| EEF1A2 | NM_001958 | 0.251 | -0.247 |
| EEF1B2 | NM_001959 | -0.0455 | -0.2865 |
| EEF1D | NM_001960 | 0.3765 | -0.1945 |
| EEF2 | NM_001961 | 0.1325 | -0.359 |
| EFNA5 | NM_001962 | -0.998 | -2.2275 |
| EGF | NM_001963 | 0.5775 | 0.32 |
| EGR1 | NM_001964 | -0.4435 | 1.364 |
| EGR4 | NM_001965 | -0.303 | -0.3505 |
| EHHADH | NM_001966 | -0.193 | -0.6 |
| EIF4A2 | NM_001967 | -0.9805 | -0.318 |
| EIF4E | NM_001968 | -1.155 | -1.9695 |
| EIF5 | NM_001969 | -1.4435 | -2.6125 |
| EIF5A | NM_001970 | 0.2005 | -0.1815 |
| CELA1 | NM_001971 | 0.3075 | -0.2365 |
| ELANE | NM_001972 | 0.1205 | -0.135 |
| ELK4 | NM_001973 | 0.202 | -0.5775 |
| EMR1 | NM_001974 | 0.148 | -0.179 |
| ENO2 | NM_001975 | 0.1285 | -0.27 |
| ENO3 | NM_001976 | 0.087 | -0.3825 |
| ENPEP | NM_001977 | 0.068 | 0.349 |
| EPB49 | NM_001978 | 0.133 | -0.7355 |
| EPHX2 | NM_001979 | -0.034 | -0.426 |
| STX2 | NM_001980 | -1.258 | -1.53 |
| EPS15 | NM_001981 | -0.9405 | -1.0495 |
| ERBB3 | NM_001982 | 0.1765 | -0.3605 |
| ERCC1 | NM_001983 | -0.223 | -0.697 |
| ESD | NM_001984 | -0.165 | -0.4005 |
| ETFB | NM_001985 | 0.1225 | -0.4855 |
| ETV4 | NM_001986 | 0.369 | -0.18 |
| ETV6 | NM_001987 | -1.0035 | -1.8855 |
| EVPL | NM_001988 | -0.2215 | 0.285 |
| EVX1 | NM_001989 | -0.042 | -0.123 |
| EYA3 | NM_001990 | -1.6865 | -2.994 |
| EZH1 | NM_001991 | -1.0635 | -2.3405 |
| F2R | NM_001992 | -0.7415 | -1.412 |

| | | | |
|---|---|---|---|
| F3 | NM_001993 | -0.706 | -1.129 |
| F13B | NM_001994 | 0.0605 | -0.066 |
| ACSL1 | NM_001995 | -1.0115 | -1.697 |
| FBLN1 | NM_001996 | 0.1535 | -0.176 |
| FAU | NM_001997 | 0.198 | -0.266 |
| FBLN2 | NM_001998 | 0.09 | -0.6295 |
| FBN2 | NM_001999 | 0.082 | -0.28 |
| FCAR | NM_002000 | -0.5115 | -0.1385 |
| FCER1A | NM_002001 | 0.0475 | 0.3295 |
| FCER2 | NM_002002 | 0.1375 | -0.3115 |
| FCN1 | NM_002003 | 0.113 | -0.189 |
| FDPS | NM_002004 | -0.2515 | -0.6715 |
| FES | NM_002005 | -0.0495 | -0.585 |
| FGF2 | NM_002006 | -1.2305 | -2.226 |
| FGF4 | NM_002007 | -0.066 | 0.002 |
| FGF7 | NM_002009 | 0.156 | -0.3155 |
| FGF9 | NM_002010 | -0.2895 | -0.388 |
| FGFR4 | NM_002011 | -0.136 | -0.67 |
| FHIT | NM_002012 | 0.2265 | -0.349 |
| FKBP3 | NM_002013 | -0.691 | -0.3325 |
| FKBP4 | NM_002014 | -0.085 | -0.4615 |
| FOXO1 | NM_002015 | -0.4425 | -0.6415 |
| FLG | NM_002016 | 0.117 | -0.8585 |
| FLI1 | NM_002017 | 0.058 | 0.076 |
| FLII | NM_002018 | 0.2405 | -0.533 |
| FLT1 | NM_002019 | -0.074 | 0.037 |
| FLT4 | NM_002020 | 0.0755 | -0.3575 |
| FMO1 | NM_002021 | -0.1185 | -0.0115 |
| FMO4 | NM_002022 | -0.1785 | -0.259 |
| FMOD | NM_002023 | -0.183 | -0.317 |
| FMR1 | NM_002024 | -1.4225 | -1.6075 |
| AFF2 | NM_002025 | -0.2825 | -1.332 |
| FN1 | NM_002026 | -0.124 | -0.5965 |
| FNTA | NM_002027 | -1.82 | -2.6545 |
| FNTB | NM_002028 | -0.044 | -1.358 |
| FPR1 | NM_002029 | 0.088 | -0.5575 |
| FPR3 | NM_002030 | -0.4035 | -0.5965 |
| FRK | NM_002031 | 0.338 | -0.356 |
| FTH1 | NM_002032 | 0.15 | -0.1535 |
| FUT4 | NM_002033 | -1.0155 | -1.3935 |
| FUT5 | NM_002034 | -0.305 | -0.3985 |
| KDSR | NM_002035 | -0.8155 | -1.5315 |
| DARC | NM_002036 | -0.175 | -0.4375 |

| | | | |
|---|---|---|---|
| FYN | NM_002037 | -0.603 | -1.9005 |
| GAB1 | NM_002039 | -0.474 | -1.571 |
| GABPA | NM_002040 | -0.8585 | -1.919 |
| GABPB1 | NM_002041 | -2.2725 | -4.4245 |
| GABRR1 | NM_002042 | 0.5445 | -0.6195 |
| GABRR2 | NM_002043 | 0.215 | -0.324 |
| GALK2 | NM_002044 | -1.106 | -2.3835 |
| GAP43 | NM_002045 | 0.1145 | -0.4145 |
| GAPDH | NM_002046 | 0.1095 | -0.307 |
| GARS | NM_002047 | -0.512 | -1.563 |
| GAS1 | NM_002048 | -0.3645 | -0.146 |
| GATA1 | NM_002049 | -0.1075 | -0.7325 |
| GATA3 | NM_002051 | -0.7485 | -1.9475 |
| GATA4 | NM_002052 | 0.0505 | -0.547 |
| GBP1 | NM_002053 | 0.026 | 0.085 |
| GCG | NM_002054 | 0.1555 | -0.4185 |
| GFAP | NM_002055 | 0.0515 | -0.47 |
| GFPT1 | NM_002056 | -0.9885 | -2.1945 |
| GH2 | NM_002059 | 0.284 | 0.0655 |
| GJA4 | NM_002060 | 0.133 | 0.02 |
| GCLM | NM_002061 | -0.893 | -2.835 |
| GLP1R | NM_002062 | 0.0145 | -0.2095 |
| GLRA2 | NM_002063 | -0.0015 | -0.655 |
| GLRX | NM_002064 | -0.7235 | -0.853 |
| GLUL | NM_002065 | -0.3065 | -1.226 |
| GML | NM_002066 | -0.188 | -0.238 |
| GNA11 | NM_002067 | -0.513 | -1.5865 |
| GNA15 | NM_002068 | -0.0105 | -0.206 |
| GNAI1 | NM_002069 | -1.4015 | -2.8915 |
| GNAI2 | NM_002070 | 0.096 | -0.5715 |
| GNAL | NM_002071 | 0.2605 | -0.6505 |
| GNAQ | NM_002072 | -0.8825 | -1.679 |
| GNAZ | NM_002073 | -1.47 | -1.7895 |
| GNB1 | NM_002074 | 0.0555 | -0.643 |
| GNB3 | NM_002075 | -0.717 | -1.3885 |
| GNS | NM_002076 | -0.51 | -1.1805 |
| GOLGA1 | NM_002077 | -1.1975 | -2.214 |
| GOLGA4 | NM_002078 | 0.329 | -0.7345 |
| GOT1 | NM_002079 | 0.0725 | -0.7185 |
| GOT2 | NM_002080 | -0.052 | -0.8495 |
| GPC1 | NM_002081 | -0.1985 | -0.892 |
| GRK6 | NM_002082 | -0.6775 | -1.1955 |
| GPX2 | NM_002083 | -0.2225 | -0.2465 |

| | | | |
|---|---|---|---|
| GPX3 | NM_002084 | 0.4565 | -0.205 |
| GPX4 | NM_002085 | 0.345 | -0.2735 |
| GRB2 | NM_002086 | 0.131 | -0.862 |
| GRN | NM_002087 | 0.079 | -0.516 |
| GRIK5 | NM_002088 | -0.3005 | -0.7415 |
| CXCL2 | NM_002089 | -0.193 | -0.1395 |
| CXCL3 | NM_002090 | -0.176 | -0.1435 |
| GRP | NM_002091 | -0.3465 | -0.1805 |
| GRSF1 | NM_002092 | -1.0765 | -1.905 |
| GSK3B | NM_002093 | -1.4495 | -3.4265 |
| GSPT1 | NM_002094 | -0.6225 | -1.308 |
| GTF2E2 | NM_002095 | -1.191 | -2.2135 |
| GTF2F1 | NM_002096 | -0.283 | -0.915 |
| GTF3A | NM_002097 | -0.1865 | -0.3615 |
| GUCA1B | NM_002098 | -0.3395 | -0.9375 |
| GYPA | NM_002099 | 0.3675 | -0.005 |
| GYPB | NM_002100 | 0.0285 | -0.237 |
| GYPC | NM_002101 | 0.1955 | -0.386 |
| GYPE | NM_002102 | 0.3465 | -0.119 |
| GYS1 | NM_002103 | -0.6365 | -1.3365 |
| GZMK | NM_002104 | -0.04 | 0.1345 |
| H2AFX | NM_002105 | -0.5175 | -1.278 |
| H2AFZ | NM_002106 | -0.185 | -0.449 |
| H3F3A | NM_002107 | -0.1385 | -0.403 |
| HAL | NM_002108 | -0.1705 | -0.3185 |
| HARS | NM_002109 | -0.481 | -1.7155 |
| HCK | NM_002110 | -0.386 | 0.01 |
| HTT | NM_002111 | -0.0455 | -0.7275 |
| HDC | NM_002112 | -0.214 | 0.4475 |
| HIVEP1 | NM_002114 | -1.1185 | -2.9285 |
| HK3 | NM_002115 | 0.118 | -0.6265 |
| HLA-A | NM_002116 | 0.184 | -0.362 |
| HLA-C | NM_002117 | 0.372 | -0.3075 |
| HLA-DMB | NM_002118 | -0.057 | -0.3545 |
| HLA-DOA | NM_002119 | -0.05 | -0.178 |
| HLA-DOB | NM_002120 | 0.257 | -0.451 |
| HLA-DPB1 | NM_002121 | 0.0715 | -0.363 |
| HLA-DQA1 | NM_002122 | 0.148 | -0.026 |
| HLA-DQB1 | NM_002123 | 0.0105 | -0.2455 |
| HLA-DRB1 | NM_002124 | -0.23 | -0.436 |
| HLA-DRB5 | NM_002125 | 0.0155 | -0.27 |
| HLF | NM_002126 | -0.16 | -0.467 |
| HLA-G | NM_002127 | -0.17 | -0.5605 |

| | | | |
|---|---|---|---|
| HMGB1 | NM_002128 | -0.565 | -0.101 |
| HMGB2 | NM_002129 | -1.223 | -0.61 |
| HMGCS1 | NM_002130 | -1.2025 | -1.4315 |
| HMGA1 | NM_002131 | 0.274 | -0.169 |
| HMOX1 | NM_002133 | -0.055 | -0.1115 |
| HMOX2 | NM_002134 | -0.2595 | -1.9745 |
| NR4A1 | NM_002135 | -0.7825 | -1.903 |
| HNRNPA1 | NM_002136 | -0.242 | -0.3295 |
| HNRNPA2B1 | NM_002137 | -0.454 | -0.449 |
| HNRNPD | NM_002138 | -0.4805 | -1.078 |
| RBMX | NM_002139 | -0.7635 | -1.229 |
| HNRNPK | NM_002140 | -0.667 | -0.5185 |
| HOXA4 | NM_002141 | 0.0585 | -0.461 |
| HPCA | NM_002143 | -0.0525 | -0.5225 |
| HOXB1 | NM_002144 | 0.135 | -0.1145 |
| HOXB2 | NM_002145 | -0.077 | -0.199 |
| HOXB3 | NM_002146 | 0.4295 | -0.478 |
| HOXB5 | NM_002147 | 0.137 | -0.102 |
| HOXD10 | NM_002148 | -0.084 | -0.296 |
| HPCAL1 | NM_002149 | -0.144 | -1.042 |
| HPD | NM_002150 | 0.2355 | 0.1415 |
| HPN | NM_002151 | -0.4155 | -0.4165 |
| HRC | NM_002152 | -0.067 | -0.3885 |
| HSD17B2 | NM_002153 | 0.053 | -0.391 |
| HSPA4 | NM_002154 | -0.886 | -1.791 |
| HSPA6 | NM_002155 | -0.067 | -0.239 |
| HSPD1 | NM_002156 | -0.792 | -0.697 |
| HSPE1 | NM_002157 | -0.7495 | -0.7135 |
| FOXN2 | NM_002158 | -1.0025 | -3.0285 |
| HTN1 | NM_002159 | 0.044 | -0.498 |
| TNC | NM_002160 | -0.016 | -0.485 |
| IARS | NM_002161 | -0.9215 | -1.016 |
| ICAM3 | NM_002162 | -0.046 | -0.3805 |
| IRF8 | NM_002163 | 0.0605 | -0.2745 |
| IDO1 | NM_002164 | -0.2665 | -0.286 |
| ID1 | NM_002165 | 0.417 | 0.7295 |
| ID2 | NM_002166 | -0.096 | 1.0025 |
| ID3 | NM_002167 | -0.1165 | 0.2485 |
| IDH2 | NM_002168 | 0.1165 | -0.643 |
| IFNA5 | NM_002169 | 0.148 | -0.1695 |
| IFNA8 | NM_002170 | 0.0595 | -0.212 |
| IFNA10 | NM_002171 | 0.236 | -0.251 |
| IFNA14 | NM_002172 | 0.031 | -0.0995 |

| | | | |
|---|---|---|---|
| IFNA16 | NM_002173 | -0.326 | -0.061 |
| IFNA21 | NM_002175 | 0.5775 | -0.233 |
| IFNB1 | NM_002176 | 0.1535 | 0.1455 |
| IFNW1 | NM_002177 | -0.1285 | 0.0315 |
| IGFBP6 | NM_002178 | 0.04 | -0.317 |
| IGHMBP2 | NM_002180 | -0.5695 | -1.9815 |
| IHH | NM_002181 | -0.2835 | -0.709 |
| IL1RAP | NM_002182 | -0.273 | -0.634 |
| IL3RA | NM_002183 | 0.054 | -0.381 |
| IL6ST | NM_002184 | -0.292 | -1.308 |
| IL7R | NM_002185 | 0.145 | 0.0585 |
| IL9R | NM_002186 | -0.131 | -0.3465 |
| IL12B | NM_002187 | -0.095 | -0.5145 |
| IL13 | NM_002188 | -0.302 | -0.7055 |
| IL15RA | NM_002189 | -0.133 | -0.2235 |
| IL17A | NM_002190 | -0.199 | -0.1815 |
| INHA | NM_002191 | -0.0365 | -0.302 |
| INHBA | NM_002192 | 0.278 | -0.4595 |
| INHBB | NM_002193 | 0.1685 | -0.186 |
| INPP1 | NM_002194 | -0.6145 | -2.4895 |
| INSL4 | NM_002195 | 0.512 | 0.3065 |
| INSM1 | NM_002196 | 0.6555 | 0.2315 |
| ACO1 | NM_002197 | -0.0135 | -1.061 |
| IRF1 | NM_002198 | 0.0445 | 0.085 |
| IRF2 | NM_002199 | -1.362 | -3.0975 |
| ISG20 | NM_002201 | -0.12 | -0.8055 |
| ISL1 | NM_002202 | -0.8165 | -2.7295 |
| ITGA2 | NM_002203 | -0.324 | -0.4975 |
| ITGA3 | NM_002204 | -0.1555 | -0.415 |
| ITGA5 | NM_002205 | -0.009 | 0.4385 |
| ITGA7 | NM_002206 | 0.174 | -0.207 |
| ITGA9 | NM_002207 | -1.5695 | -1.32 |
| ITGAE | NM_002208 | -0.2345 | -0.542 |
| ITGAL | NM_002209 | -0.0375 | -0.0335 |
| ITGAV | NM_002210 | -1.023 | -1.5975 |
| ITGB1 | NM_002211 | -0.47 | -0.3105 |
| EIF6 | NM_002212 | -0.1485 | -1.276 |
| ITGB5 | NM_002213 | -0.356 | -0.6655 |
| ITGB8 | NM_002214 | 0.223 | 0.0485 |
| ITIH1 | NM_002215 | -0.223 | -0.309 |
| ITIH2 | NM_002216 | 0.113 | -0.097 |
| ITIH3 | NM_002217 | -0.1515 | -0.72 |
| ITPKA | NM_002220 | 0.304 | -0.454 |

| | | | |
|---|---|---|---|
| ITPKB | NM_002221 | -0.39 | -2.109 |
| ITPR1 | NM_002222 | -1.355 | -2.454 |
| ITPR2 | NM_002223 | -0.3835 | -1.2685 |
| ITPR3 | NM_002224 | 0.2125 | 0.146 |
| IVD | NM_002225 | -1.077 | -1.877 |
| JAG2 | NM_002226 | -0.2195 | -0.753 |
| JAK1 | NM_002227 | -0.743 | -0.853 |
| JUN | NM_002228 | 0.2045 | 0.553 |
| JUNB | NM_002229 | 0.7575 | 1.6895 |
| JUP | NM_002230 | -0.2395 | -1.001 |
| CD82 | NM_002231 | 0.1735 | -0.383 |
| KCNA3 | NM_002232 | -0.0785 | -0.346 |
| KCNA4 | NM_002233 | 0.3865 | -0.38 |
| KCNA5 | NM_002234 | -0.2505 | 0.2325 |
| KCNA6 | NM_002235 | -0.161 | -0.508 |
| KCNF1 | NM_002236 | -0.1605 | -0.684 |
| KCNG1 | NM_002237 | -1.2145 | -2.4805 |
| KCNH1 | NM_002238 | 0.204 | -0.494 |
| KCNJ3 | NM_002239 | 0.136 | -0.073 |
| KCNJ6 | NM_002240 | -0.6555 | -0.732 |
| KCNJ10 | NM_002241 | -0.0505 | -0.5315 |
| KCNJ13 | NM_002242 | -0.021 | -0.533 |
| KCNJ15 | NM_002243 | -0.0605 | 0.296 |
| KCNK1 | NM_002245 | 0.0375 | -1.527 |
| KCNK3 | NM_002246 | -1.2095 | -1.5315 |
| KCNMA1 | NM_002247 | 0.253 | -0.204 |
| KCNN1 | NM_002248 | -0.363 | -1.051 |
| KCNN3 | NM_002249 | -0.193 | 0.091 |
| KCNN4 | NM_002250 | -0.2225 | 0.066 |
| KCNS1 | NM_002251 | 0.4395 | -0.2485 |
| KCNS3 | NM_002252 | 0.1245 | -0.69 |
| KDR | NM_002253 | 0.226 | -0.62 |
| KIF3C | NM_002254 | -1.0015 | -1.668 |
| KIR2DL4 | NM_002255 | 0.013 | -0.415 |
| KISS1 | NM_002256 | -0.072 | -0.5115 |
| KLK1 | NM_002257 | -0.0605 | -0.555 |
| KLRB1 | NM_002258 | -0.037 | -0.0835 |
| KLRC2 | NM_002260 | 0.505 | -0.583 |
| KLRC3 | NM_002261 | 0.347 | -0.265 |
| KLRD1 | NM_002262 | -0.25 | -0.061 |
| KIFC1 | NM_002263 | -0.5885 | -2.1675 |
| KPNA1 | NM_002264 | -1.143 | -2.0435 |
| KPNB1 | NM_002265 | -0.7565 | -0.4265 |

| | | | |
|---|---|---|---|
| KPNA3 | NM_002267 | -1.039 | -3.227 |
| KPNA4 | NM_002268 | -1.325 | -1.857 |
| KPNA5 | NM_002269 | -0.604 | -1.46 |
| TNPO1 | NM_002270 | -0.5835 | -0.4805 |
| IPO5 | NM_002271 | -0.626 | -0.4485 |
| KRT4 | NM_002272 | -0.0835 | -0.5215 |
| KRT8 | NM_002273 | 0.0785 | -0.051 |
| KRT13 | NM_002274 | -0.427 | -0.0955 |
| KRT15 | NM_002275 | 0.077 | -0.2755 |
| KRT19 | NM_002276 | 0.0655 | -0.057 |
| KRT31 | NM_002277 | 0.0045 | -0.0545 |
| KRT32 | NM_002278 | -0.087 | -0.1995 |
| KRT33B | NM_002279 | 0.095 | -0.0695 |
| KRT35 | NM_002280 | 0.1825 | -0.1115 |
| KRT81 | NM_002281 | -0.072 | -0.551 |
| KRT83 | NM_002282 | 0.0075 | -0.6865 |
| KRT85 | NM_002283 | -0.1265 | 0.443 |
| KRT86 | NM_002284 | 0.1815 | 0.124 |
| AFF3 | NM_002285 | -1.1575 | -2.253 |
| LAG3 | NM_002286 | 0.0715 | 0.138 |
| LAIR1 | NM_002287 | -0.079 | -0.3735 |
| LAIR2 | NM_002288 | 0.5505 | 0.1355 |
| LALBA | NM_002289 | 0.222 | -0.413 |
| LAMA4 | NM_002290 | 0.188 | -0.2665 |
| LAMB1 | NM_002291 | -1.0325 | -1.7045 |
| LAMB2 | NM_002292 | -0.156 | -0.0775 |
| LAMC1 | NM_002293 | -0.798 | -1.1415 |
| LAMP2 | NM_002294 | -0.9845 | -0.8595 |
| LBR | NM_002296 | -1.042 | -1.4675 |
| LCN1 | NM_002297 | 0.0515 | -0.291 |
| LCP1 | NM_002298 | -0.9575 | -0.3725 |
| LCT | NM_002299 | -0.143 | -0.195 |
| LDHB | NM_002300 | -0.503 | -0.293 |
| LDHC | NM_002301 | -0.3495 | -0.3005 |
| LECT2 | NM_002302 | -0.261 | -0.2875 |
| LEPR | NM_002303 | 0.109 | 0.0965 |
| LFNG | NM_002304 | -0.3805 | -1.4655 |
| LGALS1 | NM_002305 | 0.588 | -0.368 |
| LGALS3 | NM_002306 | 0.069 | -0.728 |
| LGALS7 | NM_002307 | 0.0545 | 0.3975 |
| LGALS9 | NM_002308 | 0.329 | -0.164 |
| LIF | NM_002309 | -0.243 | 0.253 |
| LIFR | NM_002310 | -0.1865 | -0.709 |

| | | | |
|---|---|---|---|
| LIG3 | NM_002311 | -0.383 | -1.802 |
| LIG4 | NM_002312 | -1.0465 | -1.0575 |
| ABLIM1 | NM_002313 | -0.6685 | -1.1735 |
| LIMK1 | NM_002314 | -0.4925 | -0.764 |
| LMO1 | NM_002315 | -0.7815 | -1.4955 |
| LMX1B | NM_002316 | -0.136 | -0.3925 |
| LOX | NM_002317 | 0.059 | -0.4155 |
| LOXL2 | NM_002318 | 0.0865 | -0.541 |
| LRCH4 | NM_002319 | -0.301 | -1.181 |
| LRP1 | NM_002332 | -0.035 | -0.652 |
| LRP3 | NM_002333 | 0.049 | -0.731 |
| LRP4 | NM_002334 | -0.1895 | -0.727 |
| LRP5 | NM_002335 | -0.4925 | -1.6685 |
| LRP6 | NM_002336 | -0.71 | -2.22 |
| LRPAP1 | NM_002337 | -0.3415 | -1.0905 |
| LSAMP | NM_002338 | -0.1565 | -1.1835 |
| LSP1 | NM_002339 | 0.2225 | -0.3575 |
| LSS | NM_002340 | -0.3615 | -0.8805 |
| LTB | NM_002341 | -0.078 | -0.6505 |
| LTBR | NM_002342 | -0.774 | -0.2285 |
| LTF | NM_002343 | 0.4125 | 0.1915 |
| LTK | NM_002344 | -0.3495 | -0.2165 |
| LUM | NM_002345 | -0.356 | -0.2965 |
| LY6E | NM_002346 | 0.0875 | -0.805 |
| LY9 | NM_002348 | -0.1835 | -0.696 |
| LY75 | NM_002349 | -0.364 | 0.3725 |
| LYN | NM_002350 | -0.7165 | -1.669 |
| SH2D1A | NM_002351 | 0.25 | 0.3765 |
| TACSTD2 | NM_002353 | -0.34 | 0.077 |
| EPCAM | NM_002354 | -0.098 | -0.289 |
| M6PR | NM_002355 | -0.4675 | -1.9385 |
| MARCKS | NM_002356 | 0.099 | -0.531 |
| MXD1 | NM_002357 | -0.965 | -1.2205 |
| MAD2L1 | NM_002358 | -0.9925 | -1.01 |
| MAFG | NM_002359 | 0.033 | -0.1765 |
| MAFK | NM_002360 | -0.352 | -0.671 |
| MAGEA4 | NM_002362 | -0.2955 | -0.294 |
| MAGEB1 | NM_002363 | 0.0075 | -0.4555 |
| MAGEB2 | NM_002364 | 0.2575 | 0.0115 |
| MAGEB3 | NM_002365 | -0.077 | -0.199 |
| MAGEB4 | NM_002367 | -0.1795 | -0.6845 |
| MAGOH | NM_002370 | -1.0405 | -1.9735 |
| MAL | NM_002371 | -0.044 | -0.317 |

| | | | |
|---|---|---|---|
| MAN2A1 | NM_002372 | -0.794 | -2.275 |
| MAP1A | NM_002373 | -0.5425 | -1.5135 |
| MAP2 | NM_002374 | -1.005 | -1.082 |
| MAP4 | NM_002375 | -0.397 | -1.094 |
| MARK3 | NM_002376 | -1.1385 | -2.5945 |
| MAS1 | NM_002377 | -0.347 | -0.197 |
| MATN1 | NM_002379 | 0.0455 | -0.2855 |
| MATN2 | NM_002380 | 0.0845 | -0.6185 |
| MATN3 | NM_002381 | 0.134 | -0.2725 |
| MAX | NM_002382 | -0.588 | -2.2715 |
| MAZ | NM_002383 | 0.0975 | -0.1115 |
| MBD1 | NM_002384 | -0.503 | -2.472 |
| MBP | NM_002385 | 0.0715 | -0.0095 |
| MC1R | NM_002386 | -0.512 | -0.723 |
| MCC | NM_002387 | -1.1835 | -2.942 |
| MCM3 | NM_002388 | -0.4835 | -1.3595 |
| CD46 | NM_002389 | -1.213 | -1.466 |
| ADAM11 | NM_002390 | -0.1605 | -0.4635 |
| MDK | NM_002391 | 0.314 | -0.1 |
| MDM2 | NM_002392 | -1.401 | -2.95 |
| MDM4 | NM_002393 | -1.3115 | -1.2645 |
| SLC3A2 | NM_002394 | -0.5325 | -1.3455 |
| ME1 | NM_002395 | -0.6075 | -0.903 |
| ME2 | NM_002396 | -0.871 | -2.002 |
| MEF2C | NM_002397 | -1.515 | -0.257 |
| MEIS1 | NM_002398 | -1.502 | -3.5845 |
| MEIS2 | NM_002399 | -0.9845 | -2.763 |
| MAP3K3 | NM_002401 | -0.9385 | -3.338 |
| MEST | NM_002402 | -0.675 | -0.0765 |
| MFAP2 | NM_002403 | 0.051 | -0.4275 |
| MFAP4 | NM_002404 | -0.052 | -0.436 |
| MFNG | NM_002405 | -0.214 | -0.9 |
| MGAT1 | NM_002406 | -0.446 | -1.8305 |
| SCGB2A1 | NM_002407 | -0.039 | -0.1765 |
| MGAT2 | NM_002408 | -1.888 | -2.296 |
| MGAT3 | NM_002409 | -0.756 | -1.4135 |
| MGAT5 | NM_002410 | 0.063 | -0.148 |
| SCGB2A2 | NM_002411 | -0.3515 | 0.059 |
| MGMT | NM_002412 | 0.233 | -0.509 |
| MGST2 | NM_002413 | 0.0645 | -0.026 |
| CD99 | NM_002414 | -0.2785 | -0.45 |
| MIF | NM_002415 | 0.1485 | -0.3095 |
| CXCL9 | NM_002416 | 0.0475 | -0.2935 |

| | | | |
|---|---|---|---|
| MKI67 | NM_002417 | -0.799 | -2.2785 |
| MLN | NM_002418 | 0.1215 | -0.576 |
| MAP3K11 | NM_002419 | -0.5755 | -1.3885 |
| TRPM1 | NM_002420 | -0.016 | 0.0465 |
| MMP1 | NM_002421 | 0.2665 | -0.448 |
| MMP3 | NM_002422 | 0.084 | 0.2375 |
| MMP7 | NM_002423 | -0.2515 | -0.635 |
| MMP8 | NM_002424 | 0.275 | -0.34 |
| MMP10 | NM_002425 | 0.3255 | 0.1905 |
| MMP12 | NM_002426 | -0.1315 | -0.562 |
| MMP13 | NM_002427 | 0 | -0.408 |
| MMP15 | NM_002428 | -0.3685 | -0.937 |
| MMP19 | NM_002429 | 0.2215 | -0.5055 |
| MN1 | NM_002430 | 0.052 | -0.881 |
| MNAT1 | NM_002431 | -1.198 | -2.753 |
| MNDA | NM_002432 | 0.264 | -0.175 |
| MOG | NM_002433 | -0.501 | 0.329 |
| MPG | NM_002434 | -0.247 | -1.226 |
| MPI | NM_002435 | -0.63 | -1.3115 |
| MPP1 | NM_002436 | -0.1515 | -0.836 |
| MPV17 | NM_002437 | 0.007 | -0.4475 |
| MSH3 | NM_002439 | -0.652 | -1.5785 |
| MSH4 | NM_002440 | 0.1325 | -0.382 |
| MSH5 | NM_002441 | -0.695 | -1.045 |
| MSI1 | NM_002442 | -0.628 | -1.5615 |
| MSMB | NM_002443 | -0.277 | -0.6 |
| MSN | NM_002444 | 0.236 | -0.3815 |
| MSR1 | NM_002445 | -0.4005 | -0.451 |
| MAP3K10 | NM_002446 | -0.1845 | -0.6185 |
| MST1R | NM_002447 | -0.0595 | -0.449 |
| MSX1 | NM_002448 | 0.274 | 0.5035 |
| MSX2 | NM_002449 | 0.146 | -0.422 |
| MTAP | NM_002451 | -1.303 | -1.876 |
| NUDT1 | NM_002452 | 0.161 | -0.4465 |
| MTRR | NM_002454 | -1.2755 | -2.7305 |
| MTX1 | NM_002455 | -0.207 | -1.141 |
| MUC1 | NM_002456 | -0.1105 | -0.321 |
| MUC2 | NM_002457 | -0.0885 | 0.072 |
| MUC5B | NM_002458 | 0.116 | 0.0815 |
| IRF4 | NM_002460 | 0.1955 | -0.1095 |
| MVD | NM_002461 | -0.2255 | -0.6555 |
| MX2 | NM_002463 | 0.48 | 0.6475 |
| MYBPC1 | NM_002465 | 0.0155 | 0.076 |

| | | | |
|---|---|---|---|
| MYBL2 | NM_002466 | 0.0755 | -0.644 |
| MYC | NM_002467 | 0.2045 | 2.0635 |
| MYD88 | NM_002468 | -0.4585 | -1.2695 |
| MYF6 | NM_002469 | -0.2165 | -0.2345 |
| MYH3 | NM_002470 | -0.7275 | -1.1355 |
| MYH6 | NM_002471 | -0.079 | -0.564 |
| MYH8 | NM_002472 | 0.6345 | -0.526 |
| MYH9 | NM_002473 | 0.0245 | -1.0695 |
| MYH11 | NM_002474 | -0.05 | -0.205 |
| MYL6B | NM_002475 | 0.374 | -0.2435 |
| MYL5 | NM_002477 | 0.155 | -0.076 |
| MYOD1 | NM_002478 | -0.4065 | -0.2265 |
| MYOG | NM_002479 | -0.066 | -0.2275 |
| PPP1R12A | NM_002480 | -0.6635 | -2.7075 |
| PPP1R12B | NM_002481 | 0.8205 | 0.5085 |
| NASP | NM_002482 | -0.7075 | -1.004 |
| CEACAM6 | NM_002483 | -0.0965 | -0.3905 |
| NUBP1 | NM_002484 | -0.796 | -3.334 |
| NBN | NM_002485 | -1.2345 | -3.0115 |
| NCBP1 | NM_002486 | -1.1795 | -1.815 |
| NDN | NM_002487 | 0.1 | -0.3375 |
| NDUFA2 | NM_002488 | 0.2915 | -0.197 |
| NDUFA4 | NM_002489 | 0.184 | -0.052 |
| NDUFA6 | NM_002490 | -0.269 | -0.6775 |
| NDUFB3 | NM_002491 | -0.306 | -0.6325 |
| NDUFB5 | NM_002492 | -0.591 | -0.991 |
| NDUFB6 | NM_002493 | -0.3485 | -0.7025 |
| NDUFC1 | NM_002494 | -0.1845 | -1.0305 |
| NDUFS4 | NM_002495 | -0.7305 | -0.9795 |
| NDUFS8 | NM_002496 | 0.112 | -0.387 |
| NEK2 | NM_002497 | -1.1985 | -1.2175 |
| NEK3 | NM_002498 | -1.0365 | -1.524 |
| NEO1 | NM_002499 | -0.5275 | -1.356 |
| NEUROD1 | NM_002500 | -0.2985 | -0.792 |
| NFIX | NM_002501 | -0.0075 | -1.099 |
| NFKBIB | NM_002503 | -0.0265 | -0.657 |
| NFX1 | NM_002504 | -1.692 | -2.744 |
| NFYA | NM_002505 | -0.905 | -1.2805 |
| NGF | NM_002506 | 0.006 | -1.541 |
| NGFR | NM_002507 | -0.106 | -0.5015 |
| NID1 | NM_002508 | -0.0925 | -0.411 |
| NKX2-2 | NM_002509 | 0.2985 | -0.0445 |
| GPNMB | NM_002510 | -0.148 | -0.0895 |

| | | | |
|---|---|---|---|
| NMBR | NM_002511 | -0.212 | -0.4005 |
| NME3 | NM_002513 | -0.0775 | -0.5505 |
| NOV | NM_002514 | -0.1155 | -0.545 |
| NOVA1 | NM_002515 | -1.614 | -2.334 |
| NOVA2 | NM_002516 | -0.094 | -0.776 |
| NPAS1 | NM_002517 | -0.051 | -0.2625 |
| NPAS2 | NM_002518 | -0.1315 | -0.312 |
| NPAT | NM_002519 | -0.8655 | -2.9085 |
| NPM1 | NM_002520 | -0.3505 | -0.38 |
| NPPB | NM_002521 | -0.617 | -0.5645 |
| NPTX1 | NM_002522 | -0.6495 | -1.11 |
| NPTX2 | NM_002523 | -0.418 | -0.1545 |
| NRAS | NM_002524 | -1.2105 | -1.6585 |
| NRD1 | NM_002525 | -1.2875 | -0.8 |
| NT5E | NM_002526 | -0.5125 | -0.52 |
| NTF3 | NM_002527 | -0.5465 | -0.176 |
| NTHL1 | NM_002528 | -0.3225 | -1.3285 |
| NTRK3 | NM_002530 | -0.273 | -0.8245 |
| NTSR1 | NM_002531 | -0.3615 | -0.245 |
| NUP88 | NM_002532 | -1.321 | -1.902 |
| NVL | NM_002533 | -1.339 | -1.9675 |
| OAS1 | NM_002534 | -0.025 | -0.293 |
| OAS2 | NM_002535 | -0.0745 | -0.4055 |
| TBC1D25 | NM_002536 | -0.396 | -1.448 |
| OAZ2 | NM_002537 | -0.0225 | -0.777 |
| OCLN | NM_002538 | -0.7025 | -1.199 |
| ODC1 | NM_002539 | -0.093 | -0.43 |
| ODF2 | NM_002540 | -0.4545 | -0.8675 |
| OGDH | NM_002541 | -0.7535 | -1.7415 |
| OGG1 | NM_002542 | -0.6175 | -0.361 |
| OLR1 | NM_002543 | -0.316 | -0.3935 |
| OMG | NM_002544 | -0.4685 | 0.039 |
| OPCML | NM_002545 | -0.0225 | -0.0685 |
| TNFRSF11B | NM_002546 | 0.119 | 0.085 |
| OPHN1 | NM_002547 | -0.3245 | -0.175 |
| OR1D2 | NM_002548 | 0.1405 | -0.0345 |
| OR3A1 | NM_002550 | -0.1685 | 0.073 |
| OR3A2 | NM_002551 | 0.159 | -0.132 |
| ORC4 | NM_002552 | -0.643 | -2.021 |
| ORC5 | NM_002553 | -2.057 | -3.528 |
| SLC22A18 | NM_002555 | -0.084 | -0.4625 |
| OSBP | NM_002556 | -1.1435 | -2.125 |
| OVGP1 | NM_002557 | -0.1145 | -0.5485 |

| | | | |
|---|---|---|---|
| P2RX1 | NM_002558 | 0.094 | 0.1665 |
| P2RX3 | NM_002559 | -0.022 | -0.4025 |
| P2RX4 | NM_002560 | -0.3685 | -0.827 |
| P2RX5 | NM_002561 | -0.376 | -1.21 |
| P2RX7 | NM_002562 | -0.1055 | -0.2805 |
| P2RY1 | NM_002563 | 0.0385 | -0.04 |
| P2RY2 | NM_002564 | -0.097 | -0.3115 |
| P2RY4 | NM_002565 | -0.218 | -0.2715 |
| P2RY11 | NM_002566 | 0.005 | -0.276 |
| PEBP1 | NM_002567 | 0.359 | -0.098 |
| PABPC1 | NM_002568 | -0.102 | -0.081 |
| FURIN | NM_002569 | -0.1115 | -0.5845 |
| PCSK6 | NM_002570 | -0.1975 | -0.7225 |
| PAFAH1B2 | NM_002572 | -1.1985 | -1.8985 |
| PRDX1 | NM_002574 | -0.138 | 0.063 |
| PAK1 | NM_002576 | -0.294 | -1.475 |
| PAK2 | NM_002577 | -0.921 | -0.964 |
| PAK3 | NM_002578 | -0.866 | -1.7585 |
| PALM | NM_002579 | -0.1285 | -0.802 |
| PAPPA | NM_002581 | -0.124 | -0.715 |
| PARN | NM_002582 | -1.372 | -2.0105 |
| PAWR | NM_002583 | -0.6315 | -0.949 |
| PAX7 | NM_002584 | 0.2385 | -0.4755 |
| PBX1 | NM_002585 | -0.9275 | -0.778 |
| PBX2 | NM_002586 | -0.246 | -1.5235 |
| PCDH1 | NM_002587 | -0.3335 | -0.8565 |
| PCDH7 | NM_002589 | -0.437 | -0.099 |
| PCDH8 | NM_002590 | 0.301 | 0.1355 |
| PCK1 | NM_002591 | 0.1935 | 0.0665 |
| PCNA | NM_002592 | -0.695 | -0.909 |
| PCOLCE | NM_002593 | 0.1745 | -0.276 |
| PCSK2 | NM_002594 | -0.133 | -0.3435 |
| CDK17 | NM_002595 | -1.3335 | -2.9845 |
| CDK18 | NM_002596 | -0.501 | -0.8975 |
| PDC | NM_002597 | 0.058 | 0.2335 |
| PDCD2 | NM_002598 | -0.53 | -1.437 |
| PDE2A | NM_002599 | -0.4595 | -1.1105 |
| PDE4B | NM_002600 | -1.5795 | -3.566 |
| PDE6D | NM_002601 | -1.0375 | -2.6325 |
| PDE6G | NM_002602 | 0.1145 | -0.102 |
| PDE7A | NM_002603 | -1.3 | -2.545 |
| PDE8A | NM_002605 | -1.3115 | -2.6945 |
| PDGFA | NM_002607 | 0.163 | -0.0895 |

| | | | |
|---|---|---|---|
| PDGFB | NM_002608 | -0.2605 | -0.377 |
| PDGFRB | NM_002609 | -0.841 | -1.1055 |
| PDK1 | NM_002610 | -0.852 | -0.67 |
| PDK2 | NM_002611 | -0.2915 | -0.6255 |
| PDK4 | NM_002612 | -0.4985 | -0.842 |
| PDPK1 | NM_002613 | -1.505 | -2.511 |
| SERPINF1 | NM_002615 | 0.0185 | -0.455 |
| PER1 | NM_002616 | -0.593 | -1.1675 |
| PEX10 | NM_002617 | -0.5425 | -0.8795 |
| PEX13 | NM_002618 | -1.8775 | -3.228 |
| PF4 | NM_002619 | 0.105 | 0.0675 |
| PF4V1 | NM_002620 | -0.015 | -0.3695 |
| CFP | NM_002621 | 0.1695 | -0.2925 |
| PFDN1 | NM_002622 | -0.561 | -2.1835 |
| PFDN4 | NM_002623 | -0.576 | -0.6655 |
| PFDN5 | NM_002624 | 0.2505 | -0.0945 |
| PFKFB1 | NM_002625 | -0.0375 | -0.2475 |
| PFKL | NM_002626 | 0.2015 | -0.435 |
| PFKP | NM_002627 | -0.7765 | -0.843 |
| PFN2 | NM_002628 | -0.9655 | -1.668 |
| PGAM1 | NM_002629 | 0.0955 | -0.366 |
| PGC | NM_002630 | 0.1475 | -0.5335 |
| PGD | NM_002631 | 0.0965 | -0.5995 |
| PGF | NM_002632 | -0.8635 | -3.248 |
| PGM1 | NM_002633 | 0.017 | -0.4035 |
| PHB | NM_002634 | 0.1755 | -0.3305 |
| SLC25A3 | NM_002635 | 0.0645 | -0.4875 |
| PHF1 | NM_002636 | -0.1685 | -0.2715 |
| PHKA1 | NM_002637 | -0.603 | -0.5625 |
| PI3 | NM_002638 | 0.2225 | -0.1595 |
| SERPINB5 | NM_002639 | -0.099 | 0.2285 |
| SERPINB8 | NM_002640 | -0.041 | -0.4225 |
| PIGA | NM_002641 | -1.511 | -2.627 |
| PIGF | NM_002643 | -0.8915 | -0.8195 |
| PIGR | NM_002644 | -0.133 | -0.175 |
| PIK3C2A | NM_002645 | -0.524 | -2.6135 |
| PIK3C2B | NM_002646 | -0.6235 | -2.1695 |
| PIK3C3 | NM_002647 | -0.919 | -2.8745 |
| PIM1 | NM_002648 | -0.2235 | -0.804 |
| PIK3CG | NM_002649 | 0.08 | -0.515 |
| PI4KB | NM_002651 | -0.2755 | -1.6825 |
| PIP | NM_002652 | 0.2795 | -0.451 |
| PITX1 | NM_002653 | -0.2835 | -0.732 |

| | | | |
|---|---|---|---|
| PKM2 | NM_002654 | 0.2085 | -0.32 |
| PLAG1 | NM_002655 | 0.097 | -0.1135 |
| PLAGL1 | NM_002656 | -0.0175 | -0.963 |
| PLAGL2 | NM_002657 | -0.8175 | -2.903 |
| PLAU | NM_002658 | -0.1455 | 0.363 |
| PLAUR | NM_002659 | -0.253 | -1.3375 |
| PLCG1 | NM_002660 | 0.02 | -0.672 |
| PLCG2 | NM_002661 | -0.0175 | -0.3515 |
| PLD1 | NM_002662 | -0.092 | 0.1985 |
| PLD2 | NM_002663 | -0.2145 | -0.564 |
| PLEK | NM_002664 | -0.0315 | -0.0395 |
| PLGLB2 | NM_002665 | -0.349 | -1.5855 |
| PLIN1 | NM_002666 | -0.0525 | -0.5135 |
| PLN | NM_002667 | -0.1415 | 0.3595 |
| PLP2 | NM_002668 | 0.0025 | -0.716 |
| PLRG1 | NM_002669 | -1.224 | -2.0575 |
| PLS1 | NM_002670 | -1.5265 | -2.7355 |
| PLXNB1 | NM_002673 | -0.3655 | -0.81 |
| PMCH | NM_002674 | -0.258 | -0.983 |
| PML | NM_002675 | -0.6635 | -1.394 |
| PMM1 | NM_002676 | -0.388 | -1.377 |
| PMP2 | NM_002677 | 0.066 | -0.2695 |
| EXOSC10 | NM_002685 | -1.145 | -1.8775 |
| PNMT | NM_002686 | -0.09 | -0.572 |
| PNN | NM_002687 | -0.9545 | -2.0865 |
| POLA2 | NM_002689 | -1.1575 | -3.147 |
| POLB | NM_002690 | -0.5065 | -1.1065 |
| POLD1 | NM_002691 | 0.215 | -0.506 |
| POLE2 | NM_002692 | -1.604 | -3.319 |
| POLG | NM_002693 | -0.2095 | -1.5865 |
| POLR2E | NM_002695 | 0.1325 | -0.422 |
| POLR2G | NM_002696 | -0.0045 | -0.585 |
| POU2F1 | NM_002697 | -0.4685 | -1.654 |
| POU2F2 | NM_002698 | -0.15 | -0.759 |
| POU3F1 | NM_002699 | -0.4395 | -0.239 |
| POU4F3 | NM_002700 | -0.121 | -0.1825 |
| POU5F1 | NM_002701 | -0.0935 | 0.0605 |
| POU6F1 | NM_002702 | -0.1525 | -0.187 |
| PPAT | NM_002703 | -1.1855 | -2.408 |
| PPBP | NM_002704 | -0.025 | 0.275 |
| PPL | NM_002705 | 0.178 | -0.2385 |
| PPM1B | NM_002706 | -1.071 | -2.7885 |
| PPP1CA | NM_002708 | 0.144 | -0.4505 |

| | | | |
|---|---|---|---|
| PPP1CB | NM_002709 | -0.855 | -1.057 |
| PPP1CC | NM_002710 | -0.986 | -1.684 |
| PPP1R3A | NM_002711 | -0.1225 | -0.3015 |
| PPP1R7 | NM_002712 | -0.4255 | -0.7865 |
| PPP1R8 | NM_002713 | -1.4105 | -3.159 |
| PPP1R10 | NM_002714 | -2.553 | -4.4345 |
| PPP2CA | NM_002715 | -0.8165 | -1.3485 |
| PPP2R1B | NM_002716 | -0.8865 | -1.919 |
| PPP2R2A | NM_002717 | -1.072 | -2.6155 |
| PPP2R3A | NM_002718 | -0.5405 | -2.0855 |
| PPP2R5C | NM_002719 | -2.0235 | -3.1815 |
| PPP4C | NM_002720 | 0.1745 | -0.7335 |
| PPP6C | NM_002721 | -0.4145 | -1.4785 |
| PPY | NM_002722 | -0.332 | -0.4665 |
| PRB4 | NM_002723 | -0.156 | -0.28 |
| PRELP | NM_002725 | -0.4875 | -0.388 |
| PREP | NM_002726 | -0.6275 | -1.881 |
| SRGN | NM_002727 | 0.21 | 0.1935 |
| PRG2 | NM_002728 | -0.332 | -0.044 |
| HHEX | NM_002729 | -0.173 | 1.5725 |
| PRKACA | NM_002730 | 0.133 | -0.491 |
| PRKACB | NM_002731 | -1.8395 | -2.3 |
| PRKACG | NM_002732 | -0.251 | -0.4125 |
| PRKAG1 | NM_002733 | -0.4985 | -1.3715 |
| PRKAR1A | NM_002734 | -0.918 | -0.517 |
| PRKAR1B | NM_002735 | -0.343 | -1.096 |
| PRKAR2B | NM_002736 | -0.853 | -0.889 |
| PRKCA | NM_002737 | -0.796 | -1.618 |
| PRKCB | NM_002738 | -0.2515 | -0.2715 |
| PRKCG | NM_002739 | -0.035 | -0.0915 |
| PRKCI | NM_002740 | -1.3485 | -1.6105 |
| PKN1 | NM_002741 | 0.0515 | -0.439 |
| PRKD1 | NM_002742 | -0.1085 | -0.2035 |
| PRKCSH | NM_002743 | 0.0595 | -0.344 |
| PRKCZ | NM_002744 | -0.1325 | -0.866 |
| MAPK1 | NM_002745 | -1.176 | -2.003 |
| MAPK3 | NM_002746 | 0.1245 | -0.463 |
| MAPK4 | NM_002747 | -0.04 | -1.723 |
| MAPK6 | NM_002748 | -0.9475 | -2.4 |
| MAPK7 | NM_002749 | -0.8975 | -2.7945 |
| MAPK8 | NM_002750 | -0.5925 | -0.8075 |
| MAPK11 | NM_002751 | -0.024 | -0.439 |
| MAPK9 | NM_002752 | -0.5785 | -1.1935 |

| | | | |
|---|---|---|---|
| MAPK10 | NM_002753 | -0.778 | -0.715 |
| MAPK13 | NM_002754 | 0.171 | -0.442 |
| MAP2K1 | NM_002755 | -0.4335 | -2.013 |
| MAP2K3 | NM_002756 | -0.55 | -1.4165 |
| MAP2K5 | NM_002757 | -0.421 | -0.4285 |
| MAP2K6 | NM_002758 | -0.6285 | -1.3115 |
| EIF2AK2 | NM_002759 | -0.96 | -1.591 |
| PRM1 | NM_002761 | 0.034 | -0.2515 |
| PRM2 | NM_002762 | 0.2135 | 0.0615 |
| PROX1 | NM_002763 | -1.0545 | -2.0055 |
| PRPS1 | NM_002764 | -0.2775 | -0.446 |
| PRPS2 | NM_002765 | -1.4695 | -1.6375 |
| PRPSAP1 | NM_002766 | -0.551 | -1.9405 |
| PRPSAP2 | NM_002767 | -1.0105 | -1.3285 |
| CHMP1A | NM_002768 | -0.5105 | -1.38 |
| PRSS1 | NM_002769 | -0.216 | -0.836 |
| PRSS3 | NM_002771 | 0.118 | -0.569 |
| TMPRSS15 | NM_002772 | -0.0355 | -0.3505 |
| PRSS8 | NM_002773 | 0.116 | -0.597 |
| KLK6 | NM_002774 | -0.04 | -0.6725 |
| HTRA1 | NM_002775 | -0.28 | -0.5175 |
| KLK10 | NM_002776 | 0.0135 | -0.1875 |
| PRTN3 | NM_002777 | -0.343 | -0.1855 |
| PSAP | NM_002778 | 0.31 | -0.2785 |
| PSD | NM_002779 | -0.1065 | -0.354 |
| PSG4 | NM_002780 | -0.2255 | -0.084 |
| PSG5 | NM_002781 | 0.1185 | -0.0425 |
| PSG6 | NM_002782 | -0.107 | -0.493 |
| PSG7 | NM_002783 | 0.11 | -0.0205 |
| PSG9 | NM_002784 | -0.072 | -0.5515 |
| PSMA1 | NM_002786 | -0.522 | -0.8925 |
| PSMA2 | NM_002787 | -0.818 | -1.247 |
| PSMA3 | NM_002788 | -1.15 | -1.0895 |
| PSMA4 | NM_002789 | -0.9245 | -0.6685 |
| PSMA5 | NM_002790 | -0.9455 | -1.464 |
| PSMA6 | NM_002791 | -0.564 | -0.4505 |
| PSMA7 | NM_002792 | 0.158 | -0.3075 |
| PSMB1 | NM_002793 | -0.5195 | -0.6955 |
| PSMB2 | NM_002794 | -0.0845 | -1.1205 |
| PSMB3 | NM_002795 | 0.2375 | -0.2565 |
| PSMB4 | NM_002796 | 0.084 | -0.6965 |
| PSMB5 | NM_002797 | -0.148 | -1.1285 |
| PSMB6 | NM_002798 | 0.2155 | -0.3055 |

| | | | |
|---|---|---|---|
| PSMB7 | NM_002799 | 0.143 | -0.1075 |
| PSMB9 | NM_002800 | 0.0445 | -0.2125 |
| PSMB10 | NM_002801 | -0.0565 | -0.47 |
| PSMC1 | NM_002802 | -0.7325 | -0.6265 |
| PSMC2 | NM_002803 | -1.085 | -1.7615 |
| PSMC3 | NM_002804 | -0.0035 | -0.5505 |
| PSMC5 | NM_002805 | -0.3105 | -0.5585 |
| PSMC6 | NM_002806 | -1.3275 | -2.7345 |
| PSMD1 | NM_002807 | -0.6925 | -1.134 |
| PSMD2 | NM_002808 | 0.2575 | -0.1145 |
| PSMD3 | NM_002809 | -0.1515 | -1.065 |
| PSMD4 | NM_002810 | -0.019 | -0.7115 |
| PSMD7 | NM_002811 | -1.0785 | -1.981 |
| PSMD8 | NM_002812 | -0.0345 | -0.466 |
| PSMD9 | NM_002813 | -0.043 | -0.9355 |
| PSMD10 | NM_002814 | -0.738 | -0.97 |
| PSMD11 | NM_002815 | -0.2725 | -1.041 |
| PSMD12 | NM_002816 | -0.8505 | -2.519 |
| PSMD13 | NM_002817 | -0.0685 | -1.097 |
| PSME2 | NM_002818 | -0.0835 | -0.125 |
| PTBP1 | NM_002819 | -0.09 | -0.805 |
| PTHLH | NM_002820 | 0.03 | 0.2685 |
| PTK7 | NM_002821 | 0.131 | -0.5825 |
| TWF1 | NM_002822 | -0.72 | -1.393 |
| PTMA | NM_002823 | 0.0515 | -0.2865 |
| PTMS | NM_002824 | 0.1995 | -0.233 |
| PTN | NM_002825 | -0.2525 | -0.1795 |
| QSOX1 | NM_002826 | -0.385 | -1.049 |
| PTPN1 | NM_002827 | -0.582 | -1.548 |
| PTPN2 | NM_002828 | 0.016 | -1.1395 |
| PTPN3 | NM_002829 | -0.329 | -1.728 |
| PTPN4 | NM_002830 | -0.5505 | -1.935 |
| PTPN7 | NM_002832 | 0.149 | -0.398 |
| PTPN9 | NM_002833 | -1.005 | -2.4505 |
| PTPN11 | NM_002834 | -1.16 | -0.947 |
| PTPN12 | NM_002835 | -1.377 | -2.534 |
| PTPRA | NM_002836 | -0.569 | -1.255 |
| PTPRB | NM_002837 | 0.138 | -0.102 |
| PTPRC | NM_002838 | 0.147 | -0.4865 |
| PTPRD | NM_002839 | -0.843 | -2.8715 |
| PTPRF | NM_002840 | -0.0035 | -0.384 |
| PTPRG | NM_002841 | -0.5015 | -0.696 |
| PTPRH | NM_002842 | -0.338 | -0.687 |

| | | | |
|---|---|---|---|
| PTPRJ | NM_002843 | 0.667 | 0.2955 |
| PTPRK | NM_002844 | -1.262 | -1.9395 |
| PTPRM | NM_002845 | -0.134 | -0.431 |
| PTPRN | NM_002846 | 0.0625 | -0.4285 |
| PTPRN2 | NM_002847 | -0.063 | -0.502 |
| PTPRO | NM_002848 | 0.452 | -0.3665 |
| PTPRR | NM_002849 | -0.95 | -1.8585 |
| PTPRS | NM_002850 | -0.073 | -0.5195 |
| PTPRZ1 | NM_002851 | 0.0475 | -0.2985 |
| PTX3 | NM_002852 | -0.7805 | -2.587 |
| RAD1 | NM_002853 | -1.464 | -2.0355 |
| PVALB | NM_002854 | -0.053 | -0.242 |
| PVRL1 | NM_002855 | -0.899 | -2.0275 |
| PVRL2 | NM_002856 | -0.35 | -0.48 |
| PEX19 | NM_002857 | -0.245 | -0.8145 |
| ABCD3 | NM_002858 | -0.8785 | -1.197 |
| PXN | NM_002859 | -0.134 | -1.135 |
| ALDH18A1 | NM_002860 | -0.039 | -0.9145 |
| PCYT2 | NM_002861 | -0.4005 | -0.8435 |
| PYGB | NM_002862 | -0.254 | -0.6985 |
| PYGL | NM_002863 | -0.5495 | -0.5935 |
| PZP | NM_002864 | 0.186 | 0.093 |
| RAB2A | NM_002865 | -1.148 | -2.2375 |
| RAB3A | NM_002866 | 0.098 | -1.1285 |
| RAB3B | NM_002867 | -0.6425 | -0.8805 |
| RAB5B | NM_002868 | -0.341 | -1.307 |
| RAB6A | NM_002869 | -1.152 | -2.154 |
| RAB13 | NM_002870 | -0.066 | -0.443 |
| RABIF | NM_002871 | -0.624 | -2.639 |
| RAC2 | NM_002872 | -0.6495 | -0.69 |
| RAD17 | NM_002873 | -1.675 | -2.954 |
| RAD23B | NM_002874 | -0.688 | -1.143 |
| RAD51 | NM_002875 | -0.7715 | -2.366 |
| RAD51C | NM_002876 | -2.1425 | -3.151 |
| RAF1 | NM_002880 | -0.3805 | -1.5195 |
| RALB | NM_002881 | -0.2795 | -0.901 |
| RANBP1 | NM_002882 | -0.0395 | -0.4465 |
| RANGAP1 | NM_002883 | -0.9995 | -1.7455 |
| RAP1A | NM_002884 | -1.231 | -2.913 |
| RAP2B | NM_002886 | -0.698 | -2.1125 |
| RARS | NM_002887 | -1.27 | -3.3605 |
| RARRES1 | NM_002888 | -0.4085 | -0.595 |
| RARRES2 | NM_002889 | 0.1185 | -0.4125 |

| | | | |
|---|---|---|---|
| RASA1 | NM_002890 | -1.316 | -2.3375 |
| RASGRF1 | NM_002891 | -0.0115 | 0.137 |
| ARID4A | NM_002892 | -1.0025 | -2.4305 |
| RBBP7 | NM_002893 | -0.82 | -0.9305 |
| RBBP8 | NM_002894 | -0.8455 | -2.2825 |
| RBL1 | NM_002895 | -0.455 | -1.483 |
| RBM4 | NM_002896 | -0.0515 | -0.2005 |
| RBMS1 | NM_002897 | -0.7015 | -1.7475 |
| RBMS2 | NM_002898 | -0.2805 | -0.772 |
| RBP1 | NM_002899 | -0.0345 | -0.1625 |
| RBP3 | NM_002900 | -0.4205 | -0.314 |
| RCN1 | NM_002901 | -0.8635 | -0.64 |
| RCN2 | NM_002902 | -0.982 | -0.536 |
| RCVRN | NM_002903 | -0.2935 | -0.1575 |
| RDBP | NM_002904 | -0.2 | -0.6575 |
| RDH5 | NM_002905 | -0.2815 | -0.639 |
| RDX | NM_002906 | -0.338 | -1.2195 |
| RECQL | NM_002907 | -0.5055 | -2.111 |
| REL | NM_002908 | -0.4205 | -1.6465 |
| REG1A | NM_002909 | 0.26 | -0.287 |
| RENBP | NM_002910 | 0.0805 | 0.2075 |
| UPF1 | NM_002911 | -0.4355 | -1.7095 |
| REV3L | NM_002912 | -1.0355 | -3.2965 |
| RFC1 | NM_002913 | -0.627 | -2.491 |
| RFC2 | NM_002914 | -0.2625 | -1.3135 |
| RFC3 | NM_002915 | -0.7505 | -0.9725 |
| RFC4 | NM_002916 | -0.563 | -0.6235 |
| RFNG | NM_002917 | -0.594 | -1.4045 |
| RFX1 | NM_002918 | -1.23 | -2.2375 |
| RFX3 | NM_002919 | -0.2065 | 0.1065 |
| RGR | NM_002921 | -0.239 | -0.106 |
| RGS1 | NM_002922 | -0.269 | -0.036 |
| RGS2 | NM_002923 | -1.1165 | 0.233 |
| RGS7 | NM_002924 | -1.2365 | -2.3065 |
| RGS10 | NM_002925 | -0.199 | -0.911 |
| RGS12 | NM_002926 | 0.0255 | -0.857 |
| RGS13 | NM_002927 | -0.5995 | 0.3025 |
| RGS16 | NM_002928 | 0.429 | 2.631 |
| GRK1 | NM_002929 | -0.343 | -0.0965 |
| RIT2 | NM_002930 | -0.4165 | -0.093 |
| RING1 | NM_002931 | -0.131 | -0.6385 |
| RNASE2 | NM_002934 | -0.2855 | 0.033 |
| RNASE3 | NM_002935 | 0.2375 | -0.088 |

| | | | |
|---|---|---|---|
| RNASEH1 | NM_002936 | -0.6545 | -1.305 |
| RNASE4 | NM_002937 | -0.421 | -0.3605 |
| RNF4 | NM_002938 | -0.83 | -3.0035 |
| RNH1 | NM_002939 | -0.33 | -0.6735 |
| ABCE1 | NM_002940 | -0.9235 | -2.7145 |
| ROBO1 | NM_002941 | -0.903 | -1.004 |
| ROBO2 | NM_002942 | -1.674 | -1.748 |
| RORA | NM_002943 | -0.4935 | -1.1875 |
| ROS1 | NM_002944 | 0.145 | -0.0545 |
| RPA1 | NM_002945 | -0.7325 | -0.5125 |
| RPA2 | NM_002946 | -0.583 | -1.475 |
| RPA3 | NM_002947 | -0.1165 | -1.5125 |
| RPL15 | NM_002948 | 0.0585 | -0.3655 |
| MRPL12 | NM_002949 | -0.436 | -1.565 |
| RPN1 | NM_002950 | 0.103 | -0.3055 |
| RPN2 | NM_002951 | -0.1615 | -0.345 |
| RPS2 | NM_002952 | 0.091 | -0.3165 |
| RPS6KA1 | NM_002953 | -0.2155 | -1.1745 |
| RPS27A | NM_002954 | -0.0905 | -0.196 |
| CLIP1 | NM_002956 | -0.823 | -2.401 |
| RXRA | NM_002957 | -0.6885 | -1.402 |
| RYK | NM_002958 | -1.2015 | -1.8335 |
| SORT1 | NM_002959 | -1.048 | -1.288 |
| S100A3 | NM_002960 | 0.2965 | 0.0865 |
| S100A4 | NM_002961 | 0.4775 | -0.0265 |
| S100A5 | NM_002962 | -0.2555 | -0.618 |
| S100A7 | NM_002963 | -0.0775 | -0.2325 |
| S100A8 | NM_002964 | -0.22 | -0.356 |
| S100A9 | NM_002965 | 0.2725 | -0.548 |
| S100A10 | NM_002966 | -0.0565 | -0.847 |
| SAFB | NM_002967 | -0.7525 | -1.85 |
| SALL1 | NM_002968 | -0.299 | -0.479 |
| MAPK12 | NM_002969 | -0.2995 | -0.39 |
| SAT1 | NM_002970 | 0.8915 | 1.859 |
| SATB1 | NM_002971 | -1.235 | -2.3285 |
| SBF1 | NM_002972 | 0.1035 | -0.678 |
| ATXN2 | NM_002973 | -0.773 | -1.8045 |
| SERPINB4 | NM_002974 | 0.0045 | 0.1045 |
| CLEC11A | NM_002975 | 0.0385 | -0.5375 |
| SCN7A | NM_002976 | -0.126 | -0.045 |
| SCN9A | NM_002977 | -0.581 | -0.8735 |
| SCP2 | NM_002979 | -0.5505 | -0.4715 |
| SCTR | NM_002980 | 0.13 | -0.166 |

| | | | |
|---|---|---|---|
| CCL1 | NM_002981 | -0.084 | -0.492 |
| CCL2 | NM_002982 | 0.541 | 0.2015 |
| CCL3 | NM_002983 | 0.0075 | -0.463 |
| CCL4 | NM_002984 | 0.4585 | -0.47 |
| CCL5 | NM_002985 | -0.066 | -0.401 |
| CCL11 | NM_002986 | 0.214 | -0.367 |
| CCL17 | NM_002987 | -0.1005 | -0.2645 |
| CCL18 | NM_002988 | -0.4215 | -0.535 |
| CCL21 | NM_002989 | 0.021 | -0.3205 |
| CCL22 | NM_002990 | 0.287 | -0.6205 |
| CCL24 | NM_002991 | -0.024 | -0.4085 |
| CXCL6 | NM_002993 | -0.0535 | 0.0295 |
| CXCL5 | NM_002994 | 0.31 | -0.337 |
| XCL1 | NM_002995 | -0.536 | -0.149 |
| CX3CL1 | NM_002996 | -0.575 | -1.1055 |
| SDC1 | NM_002997 | -0.717 | -1.498 |
| SDC2 | NM_002998 | -0.506 | -1.4345 |
| SDC4 | NM_002999 | -0.741 | -0.7745 |
| SDHB | NM_003000 | -0.274 | -1.512 |
| SDHC | NM_003001 | 0.1615 | -0.5105 |
| SDHD | NM_003002 | -1.497 | -1.84 |
| SEC14L1 | NM_003003 | 0.111 | -0.986 |
| SECTM1 | NM_003004 | -0.0965 | -1.249 |
| SELP | NM_003005 | 0.122 | -0.2715 |
| SELPLG | NM_003006 | 0.181 | -0.117 |
| SEMG1 | NM_003007 | 0.7045 | 0.4565 |
| SEMG2 | NM_003008 | -0.322 | -0.309 |
| SEPW1 | NM_003009 | 0.34 | -0.3825 |
| MAP2K4 | NM_003010 | -1.281 | -3.69 |
| SET | NM_003011 | -0.3805 | -0.4045 |
| SFRP1 | NM_003012 | -0.8185 | -1.0895 |
| SFRP2 | NM_003013 | 0.029 | -0.0445 |
| SFRP4 | NM_003014 | 0.237 | -0.137 |
| SFRP5 | NM_003015 | -0.2015 | -0.543 |
| SRSF2 | NM_003016 | -0.693 | -0.8275 |
| SRSF3 | NM_003017 | -1.308 | -1.6985 |
| SFTPC | NM_003018 | 0.128 | -0.5065 |
| SFTPD | NM_003019 | 0.0015 | -0.075 |
| SCG5 | NM_003020 | -0.294 | -0.6975 |
| SGTA | NM_003021 | -0.4495 | -1.107 |
| SH3BGRL | NM_003022 | -0.152 | 0.0185 |
| SH3BP2 | NM_003023 | -0.299 | -1.011 |
| ITSN1 | NM_003024 | -0.605 | -1.252 |

| | | | |
|---|---|---|---|
| SH3GL1 | NM_003025 | -0.39 | -1.18 |
| SH3GL2 | NM_003026 | -0.089 | -0.048 |
| SH3GL3 | NM_003027 | -0.5055 | -0.8405 |
| SHB | NM_003028 | -0.687 | -1.9865 |
| SHC1 | NM_003029 | -0.1715 | -1.0885 |
| SHOX2 | NM_003030 | -1.196 | -0.9625 |
| SIAH1 | NM_003031 | -1.064 | -2.284 |
| ST6GAL1 | NM_003032 | -1.305 | -3.5355 |
| ST3GAL1 | NM_003033 | -0.807 | -1.611 |
| ST8SIA1 | NM_003034 | -0.6865 | -1.5515 |
| STIL | NM_003035 | -1.287 | -2.6155 |
| SKI | NM_003036 | -0.6385 | -2.62 |
| SLAMF1 | NM_003037 | -0.0335 | -0.236 |
| SLC1A4 | NM_003038 | -0.0625 | -0.888 |
| SLC2A5 | NM_003039 | -0.306 | -0.32 |
| SLC4A2 | NM_003040 | -0.0725 | -0.5895 |
| SLC5A2 | NM_003041 | 0.354 | -0.772 |
| SLC6A1 | NM_003042 | 0.1225 | 0.4315 |
| SLC6A6 | NM_003043 | -1.0605 | -1.7335 |
| SLC6A12 | NM_003044 | 0.0565 | -0.349 |
| SLC7A1 | NM_003045 | -0.787 | -2.4975 |
| SLC7A2 | NM_003046 | -1.211 | -2.2195 |
| SLC9A1 | NM_003047 | -0.3075 | -1.075 |
| SLC9A2 | NM_003048 | 0.0255 | 0.1485 |
| SLC10A1 | NM_003049 | -0.0085 | -0.0015 |
| SLC16A1 | NM_003051 | -1.225 | -1.0865 |
| SLC34A1 | NM_003052 | 0.189 | -0.0565 |
| SLC18A1 | NM_003053 | -0.188 | -0.27 |
| SLC18A2 | NM_003054 | -1.112 | -0.686 |
| SLC18A3 | NM_003055 | -0.211 | -1.302 |
| SLC22A1 | NM_003057 | -0.1825 | -0.231 |
| SLC22A2 | NM_003058 | -0.134 | -0.858 |
| SLC22A4 | NM_003059 | -0.289 | -0.8745 |
| SLC22A5 | NM_003060 | -0.1055 | -1.072 |
| SLIT1 | NM_003061 | -0.4825 | -0.848 |
| SLIT3 | NM_003062 | -0.7025 | -2.1355 |
| SLN | NM_003063 | -0.098 | -0.4705 |
| SLPI | NM_003064 | 0.024 | -0.512 |
| SNAI2 | NM_003068 | -0.729 | -1.3445 |
| SMARCA1 | NM_003069 | -0.4755 | -0.8945 |
| SMARCA2 | NM_003070 | 0.4585 | -0.9245 |
| HLTF | NM_003071 | -0.3175 | -2.3365 |
| SMARCA4 | NM_003072 | -0.2165 | -0.7055 |

| | | | |
|---|---|---|---|
| SMARCB1 | NM_003073 | -0.373 | -1.1505 |
| SMARCC1 | NM_003074 | -0.547 | -0.8535 |
| SMARCC2 | NM_003075 | 0.0015 | -0.5455 |
| SMARCD1 | NM_003076 | -0.1235 | -0.787 |
| SMARCD3 | NM_003078 | 0.0975 | -0.4005 |
| SMARCE1 | NM_003079 | -0.5935 | -0.468 |
| SMPD2 | NM_003080 | -0.256 | -0.7645 |
| SNAP25 | NM_003081 | -0.733 | -0.7435 |
| SNAPC1 | NM_003082 | -1.4215 | -3.8545 |
| SNAPC2 | NM_003083 | -0.015 | -0.3405 |
| SNCB | NM_003085 | 0.0735 | -0.398 |
| SNAPC4 | NM_003086 | -0.8115 | -1.651 |
| SNCG | NM_003087 | 0.288 | -0.2215 |
| FSCN1 | NM_003088 | 0.1195 | -0.1785 |
| SNRNP70 | NM_003089 | -0.127 | -0.7965 |
| SNRPA1 | NM_003090 | -0.887 | -1.3325 |
| SNRPB | NM_003091 | 0.208 | -0.577 |
| SNRPB2 | NM_003092 | -0.9335 | -0.602 |
| SNRPC | NM_003093 | 0.0895 | -0.37 |
| SNRPE | NM_003094 | -0.509 | -0.368 |
| SNRPF | NM_003095 | -0.0935 | -0.691 |
| SNRPG | NM_003096 | -0.2755 | -0.8865 |
| SNRPN | NM_003097 | -0.837 | -1.1145 |
| SNTA1 | NM_003098 | -0.044 | -0.7095 |
| SNX1 | NM_003099 | -0.4635 | -1.3525 |
| SNX2 | NM_003100 | -0.8735 | -2.267 |
| SOAT1 | NM_003101 | -0.61 | -1.001 |
| SOD3 | NM_003102 | -0.0445 | -0.537 |
| SORD | NM_003104 | -0.214 | -0.916 |
| SORL1 | NM_003105 | -0.468 | -0.6335 |
| SOX2 | NM_003106 | -0.309 | 0.8865 |
| SOX4 | NM_003107 | 0.8445 | 0.774 |
| SOX11 | NM_003108 | -1.852 | -2.0655 |
| SP1 | NM_003109 | -1.318 | -2.7675 |
| SP2 | NM_003110 | -0.703 | -2.376 |
| SP3 | NM_003111 | -0.703 | -2.25 |
| SP4 | NM_003112 | -1.7995 | -3.381 |
| SP100 | NM_003113 | 0.134 | -0.2535 |
| SPAG1 | NM_003114 | -0.917 | -0.9835 |
| UAP1 | NM_003115 | -1.2045 | -2.0475 |
| SPAG4 | NM_003116 | -0.232 | 0.1035 |
| SPAM1 | NM_003117 | -0.0085 | -0.13 |
| SPARC | NM_003118 | 0.1105 | -0.2985 |

| | | | |
|---|---|---|---|
| SPG7 | NM_003119 | -0.82 | -1.242 |
| SPIB | NM_003121 | -0.0165 | 0.049 |
| SPINK1 | NM_003122 | 0.0935 | 0.24 |
| SPN | NM_003123 | 0.1035 | -0.475 |
| SPR | NM_003124 | -0.483 | -1.0675 |
| SPRR1B | NM_003125 | -0.245 | -0.1175 |
| SPTA1 | NM_003126 | -0.3185 | -0.364 |
| SPTAN1 | NM_003127 | -0.1575 | -0.6805 |
| SPTBN1 | NM_003128 | -0.57 | -0.798 |
| SQLE | NM_003129 | -1.174 | -3.2555 |
| SRI | NM_003130 | -0.443 | -1.4995 |
| SRF | NM_003131 | -0.5605 | -1.9375 |
| SRM | NM_003132 | 0.135 | -0.338 |
| SRP9 | NM_003133 | -0.7285 | -0.369 |
| SRP14 | NM_003134 | -0.4445 | 0.038 |
| SRP19 | NM_003135 | -1.294 | -3.035 |
| SRP54 | NM_003136 | -1.039 | -1.845 |
| SRPK1 | NM_003137 | -1.339 | -1.697 |
| SRPR | NM_003139 | -0.098 | -0.921 |
| SRY | NM_003140 | -0.316 | -0.4795 |
| TRIM21 | NM_003141 | -0.1775 | -0.254 |
| SSB | NM_003142 | -0.4775 | -0.8855 |
| SSBP1 | NM_003143 | -0.568 | -1.092 |
| SSR1 | NM_003144 | -1.043 | -2.0215 |
| SSR2 | NM_003145 | -0.154 | -1.0095 |
| SSRP1 | NM_003146 | 0.051 | -0.2795 |
| SSX2 | NM_003147 | -0.0125 | -0.2755 |
| STAC | NM_003149 | -0.144 | -0.4565 |
| STAT3 | NM_003150 | 0.0245 | -0.7415 |
| STAT4 | NM_003151 | -0.3535 | -0.386 |
| STAT5A | NM_003152 | -0.269 | -1.0575 |
| STAT6 | NM_003153 | -0.142 | 0.0855 |
| STATH | NM_003154 | 0.0715 | 0.0645 |
| STC1 | NM_003155 | -0.85 | -1.242 |
| STIM1 | NM_003156 | 0.139 | -0.6545 |
| NEK4 | NM_003157 | -1.744 | -3.005 |
| CDKL5 | NM_003159 | 0.259 | -0.0665 |
| AURKC | NM_003160 | -0.348 | -0.578 |
| RPS6KB1 | NM_003161 | -0.9435 | -2.6725 |
| STRN | NM_003162 | -0.4775 | -0.405 |
| STX5 | NM_003164 | -0.5045 | -1.1935 |
| STXBP1 | NM_003165 | -1.2315 | -1.146 |
| SULT2A1 | NM_003167 | -0.015 | -0.204 |

| | | | |
|---|---|---|---|
| SUPT4H1 | NM_003168 | -0.347 | -1.288 |
| SUPT5H | NM_003169 | -0.0065 | -0.121 |
| SUPT6H | NM_003170 | -0.6585 | -1.5045 |
| SUPV3L1 | NM_003171 | -1.2755 | -2.2375 |
| SURF1 | NM_003172 | 0.1555 | -0.547 |
| SUV39H1 | NM_003173 | -0.9635 | -2.636 |
| SVIL | NM_003174 | -1.2575 | -2.2735 |
| SYCP1 | NM_003176 | 0.6615 | 0.202 |
| SYK | NM_003177 | -0.2105 | -0.4485 |
| SYN2 | NM_003178 | 0.1435 | -2.0985 |
| SYP | NM_003179 | -0.276 | -0.937 |
| SYT5 | NM_003180 | 0.106 | -0.003 |
| T | NM_003181 | 0.2475 | -0.0195 |
| TAC1 | NM_003182 | -0.0685 | 0.202 |
| ADAM17 | NM_003183 | -0.879 | -0.992 |
| TAF2 | NM_003184 | -1.5325 | -3.064 |
| TAF4 | NM_003185 | -1.187 | -3.488 |
| TAGLN | NM_003186 | 0.149 | -0.3335 |
| TAF9 | NM_003187 | -0.354 | 0.077 |
| MAP3K7 | NM_003188 | -1.443 | -2.156 |
| TAL1 | NM_003189 | 0.091 | -0.305 |
| TAPBP | NM_003190 | -0.0195 | -0.866 |
| TBCC | NM_003192 | -0.2765 | -0.4465 |
| TBCE | NM_003193 | -1.225 | -1.8775 |
| TBP | NM_003194 | -1.5125 | -3.825 |
| TCEA2 | NM_003195 | -0.1375 | -0.5745 |
| TCEA3 | NM_003196 | -0.2925 | -0.4475 |
| TCEB3 | NM_003198 | -0.191 | -0.754 |
| TCF4 | NM_003199 | -1.0505 | -2.2855 |
| TCF3 | NM_003200 | -0.076 | -1.1245 |
| TFAM | NM_003201 | -0.7265 | -0.403 |
| TCF7 | NM_003202 | -0.874 | -1.8875 |
| C2orf3 | NM_003203 | 0.155 | -1.125 |
| NFE2L1 | NM_003204 | -0.634 | -2.0955 |
| TCF12 | NM_003205 | -0.8565 | -1.8315 |
| TCF21 | NM_003206 | -0.2535 | -0.3065 |
| TDG | NM_003211 | -1.927 | -1.927 |
| TEAD3 | NM_003214 | -0.38 | -1.1875 |
| TEC | NM_003215 | -0.248 | -0.319 |
| TEF | NM_003216 | -0.8255 | -1.627 |
| TMBIM6 | NM_003217 | 0.1265 | -0.491 |
| TERF1 | NM_003218 | -0.858 | -1.243 |
| TFAP2A | NM_003220 | 0.0025 | -0.092 |

| | | | |
|---|---|---|---|
| TFAP2B | NM_003221 | -1.026 | -2.5585 |
| TFAP2C | NM_003222 | 0.1745 | -0.254 |
| TFAP4 | NM_003223 | -0.506 | -1.252 |
| ARFRP1 | NM_003224 | -0.7575 | -1.4965 |
| TFF1 | NM_003225 | 0.0865 | -0.044 |
| TFF3 | NM_003226 | -0.2685 | -0.437 |
| TFR2 | NM_003227 | -0.1785 | -0.6055 |
| TFRC | NM_003234 | -0.9985 | -0.5185 |
| TG | NM_003235 | 0.1255 | -0.392 |
| TGFA | NM_003236 | -0.284 | -0.276 |
| TGFB2 | NM_003238 | 0.1295 | -0.414 |
| TGFB3 | NM_003239 | -0.2115 | -1.1435 |
| LEFTY2 | NM_003240 | -0.145 | 0.346 |
| TGM4 | NM_003241 | 0.3225 | -0.1565 |
| TGFBR2 | NM_003242 | 0.172 | -0.011 |
| TGFBR3 | NM_003243 | -0.867 | -0.873 |
| TGIF1 | NM_003244 | -0.406 | -1.9425 |
| TGM3 | NM_003245 | 0.0855 | -0.0885 |
| THBS1 | NM_003246 | 0.3345 | 0.103 |
| THBS2 | NM_003247 | -1.1045 | -0.9375 |
| THBS4 | NM_003248 | -0.432 | -0.2525 |
| THOP1 | NM_003249 | -0.5325 | -1.204 |
| THRA | NM_003250 | -0.483 | -3.231 |
| THRSP | NM_003251 | 0.1055 | -0.2315 |
| TIAL1 | NM_003252 | -0.8975 | -1.4415 |
| TIAM1 | NM_003253 | -0.7035 | -1.8625 |
| TIMP1 | NM_003254 | 0.4155 | -0.224 |
| TIMP2 | NM_003255 | -0.2075 | -0.6005 |
| TIMP4 | NM_003256 | -0.186 | -0.047 |
| TJP1 | NM_003257 | -0.981 | -2.6965 |
| TK1 | NM_003258 | -0.7685 | -1.7805 |
| ICAM5 | NM_003259 | -0.128 | -0.511 |
| TLE2 | NM_003260 | 0.052 | -0.3565 |
| SEC62 | NM_003262 | -0.3935 | -0.925 |
| TLR1 | NM_003263 | 0.3975 | 0.3385 |
| TLR2 | NM_003264 | 0.022 | -0.324 |
| TLR3 | NM_003265 | 0.8335 | -0.0635 |
| TLR5 | NM_003268 | 0.0655 | 0.2595 |
| NR2E1 | NM_003269 | -0.0525 | -0.281 |
| TSPAN6 | NM_003270 | -0.7715 | -0.5445 |
| TSPAN4 | NM_003271 | 0.2605 | -0.4945 |
| GPR137B | NM_003272 | 0.124 | -1.419 |
| TM7SF2 | NM_003273 | 0.1545 | -0.5615 |

| | | | |
|---|---|---|---|
| TRAPPC10 | NM_003274 | 0.451 | -0.1865 |
| TMOD1 | NM_003275 | -0.4135 | -0.9585 |
| TMPO | NM_003276 | -0.235 | 0.663 |
| CLEC3B | NM_003278 | -0.0605 | -0.3485 |
| TNNC2 | NM_003279 | 0.4625 | -0.0525 |
| TNNC1 | NM_003280 | 0.068 | 0.0315 |
| TNNI1 | NM_003281 | -0.032 | 0.5045 |
| TNNI2 | NM_003282 | 0.3555 | -0.17 |
| TNNT1 | NM_003283 | 0.244 | -0.08 |
| TNP1 | NM_003284 | -0.0695 | -0.17 |
| TNR | NM_003285 | -0.1945 | -0.146 |
| TOP1 | NM_003286 | -0.9825 | -1.589 |
| TPD52L2 | NM_003288 | -0.269 | -0.5175 |
| TPM2 | NM_003289 | 0.087 | -0.087 |
| TPM4 | NM_003290 | -0.74 | -0.729 |
| TPP2 | NM_003291 | -0.7235 | -1.5545 |
| TPR | NM_003292 | -0.0675 | -2.144 |
| TPSAB1 | NM_003294 | 0.0225 | -0.288 |
| TPT1 | NM_003295 | -0.0485 | -0.317 |
| CRISP2 | NM_003296 | 0.2505 | 0.003 |
| NR2C1 | NM_003297 | -0.899 | -2.0125 |
| NR2C2 | NM_003298 | -1.5305 | -2.7175 |
| HSP90B1 | NM_003299 | -0.245 | -0.169 |
| TRAF3 | NM_003300 | -0.419 | -1.483 |
| TRHR | NM_003301 | 0.145 | -0.158 |
| TRIP6 | NM_003302 | -0.3085 | -0.5225 |
| TRPC1 | NM_003304 | -1.7495 | -2.4435 |
| TRPC3 | NM_003305 | -0.2595 | -0.5005 |
| TRPC4 | NM_003306 | 0.2865 | -0.5715 |
| TRPM2 | NM_003307 | -0.067 | -0.4225 |
| TSPYL1 | NM_003309 | -0.7395 | -1.5415 |
| TSSC1 | NM_003310 | -0.228 | -1.2775 |
| PHLDA2 | NM_003311 | 0.095 | 0.98 |
| TST | NM_003312 | -0.7905 | -0.977 |
| TSTA3 | NM_003313 | 0.217 | -0.214 |
| TTC1 | NM_003314 | -1.5255 | -2.739 |
| DNAJC7 | NM_003315 | -0.7385 | -1.0455 |
| TTC3 | NM_003316 | -0.1875 | -0.796 |
| NKX2-1 | NM_003317 | 0.0405 | -0.429 |
| TTK | NM_003318 | -0.6665 | -3.175 |
| TTN | NM_003319 | 0.34 | 0.126 |
| TUB | NM_003320 | -0.339 | -0.781 |
| TUFM | NM_003321 | 0.2635 | -0.3435 |

| | | | |
|---|---|---|---|
| TULP1 | NM_003322 | 0.1525 | -0.0005 |
| TULP2 | NM_003323 | 0.1925 | 0.0035 |
| TULP3 | NM_003324 | -1.0875 | -1.925 |
| HIRA | NM_003325 | -0.693 | -2.1255 |
| TNFSF4 | NM_003326 | -0.4075 | -0.917 |
| TNFRSF4 | NM_003327 | -0.029 | -0.36 |
| TXK | NM_003328 | -0.0975 | -0.169 |
| TXN | NM_003329 | -0.204 | -0.298 |
| TXNRD1 | NM_003330 | -1.0215 | -1.544 |
| TYK2 | NM_003331 | -0.0635 | -0.967 |
| TYROBP | NM_003332 | -0.0705 | -0.519 |
| UBA52 | NM_003333 | 0.2265 | -0.1225 |
| UBA1 | NM_003334 | 0.1535 | -0.356 |
| UBA7 | NM_003335 | -0.065 | -0.125 |
| UBE2A | NM_003336 | -0.7045 | -1.541 |
| UBE2B | NM_003337 | -1.0075 | -1.919 |
| UBE2D1 | NM_003338 | -1.0135 | -1.645 |
| UBE2D2 | NM_003339 | -0.733 | -1.5415 |
| UBE2D3 | NM_003340 | -0.7795 | -0.961 |
| UBE2E1 | NM_003341 | -0.9535 | -0.9985 |
| UBE2G1 | NM_003342 | -0.686 | -1.9795 |
| UBE2G2 | NM_003343 | -0.591 | -0.8965 |
| UBE2H | NM_003344 | -1.0205 | -1.6345 |
| UBE2I | NM_003345 | -0.544 | -0.675 |
| UBE2L3 | NM_003347 | -0.436 | -1.5275 |
| UBE2N | NM_003348 | -0.8345 | -0.679 |
| UBE2V2 | NM_003350 | -1.6095 | -1.769 |
| SUMO1 | NM_003352 | -0.9765 | -0.777 |
| UCN | NM_003353 | -0.4355 | -0.602 |
| UCP2 | NM_003355 | -0.227 | -0.9305 |
| UCP3 | NM_003356 | -0.3545 | -1.058 |
| SCGB1A1 | NM_003357 | 0.105 | -0.393 |
| UGCG | NM_003358 | -0.6435 | -1.2615 |
| UGDH | NM_003359 | -1.1075 | -1.923 |
| UGT8 | NM_003360 | -0.725 | -2.499 |
| UMOD | NM_003361 | 0.0715 | -0.3695 |
| UNG | NM_003362 | -0.5145 | -1.653 |
| USP4 | NM_003363 | -0.3965 | -1.4 |
| UPP1 | NM_003364 | -0.087 | -0.3655 |
| UQCRC1 | NM_003365 | 0.2495 | -0.3565 |
| UQCRC2 | NM_003366 | -0.711 | -0.2595 |
| USF2 | NM_003367 | -0.065 | -0.674 |
| USP1 | NM_003368 | -0.701 | -2.4895 |

| | | | |
|---|---|---|---|
| UVRAG | NM_003369 | -0.9275 | -1.302 |
| VASP | NM_003370 | -0.412 | -1.6655 |
| VAV2 | NM_003371 | -0.723 | -1.7655 |
| VBP1 | NM_003372 | -0.7005 | -0.274 |
| VCL | NM_003373 | -0.7635 | -1.1865 |
| VDAC1 | NM_003374 | -0.189 | -0.3715 |
| VDAC2 | NM_003375 | -0.0035 | -0.4675 |
| VEGFA | NM_003376 | -0.674 | -0.6745 |
| VEGFB | NM_003377 | -0.0235 | -0.382 |
| VGF | NM_003378 | 0.256 | 0.6615 |
| EZR | NM_003379 | -0.026 | -1.0215 |
| VIM | NM_003380 | -0.0435 | -0.4405 |
| VIP | NM_003381 | -0.2855 | -0.3655 |
| VIPR2 | NM_003382 | -0.3905 | -0.7015 |
| VLDLR | NM_003383 | -0.4475 | -2.851 |
| VRK1 | NM_003384 | -1.065 | -2.303 |
| VSNL1 | NM_003385 | -0.8375 | -1.3105 |
| ZAN | NM_003386 | -0.182 | -0.2715 |
| WIPF1 | NM_003387 | -1.691 | -2.723 |
| CORO2A | NM_003389 | -1.4295 | -1.8165 |
| WEE1 | NM_003390 | -1.576 | -2.758 |
| WNT2 | NM_003391 | -0.0895 | -0.0155 |
| WNT5A | NM_003392 | -0.5275 | -0.7545 |
| WNT8B | NM_003393 | -0.0375 | -0.5195 |
| WNT10B | NM_003394 | 0.1025 | -0.26 |
| WNT9A | NM_003395 | 0.0395 | -0.3305 |
| WNT9B | NM_003396 | -0.0285 | -0.324 |
| XPNPEP2 | NM_003399 | 0.022 | -0.2985 |
| XPO1 | NM_003400 | -1.1475 | -2.4275 |
| XRCC4 | NM_003401 | -1.0375 | -1.7165 |
| YY1 | NM_003403 | -0.8685 | -1.114 |
| YWHAB | NM_003404 | 0.0485 | -0.286 |
| YWHAH | NM_003405 | -0.9285 | -1.4355 |
| YWHAZ | NM_003406 | -0.584 | -0.5065 |
| ZFP36 | NM_003407 | 0.229 | 1.724 |
| ZFP37 | NM_003408 | -0.1725 | -2.3955 |
| ZFP161 | NM_003409 | -1.4405 | -2.7725 |
| ZFX | NM_003410 | -1.5795 | -2.8065 |
| ZFY | NM_003411 | 0.373 | -1.1505 |
| ZIC1 | NM_003412 | 0.113 | -0.13 |
| ZIC3 | NM_003413 | -0.9685 | -2.3815 |
| ZNF267 | NM_003414 | -0.6095 | -2.289 |
| ZNF268 | NM_003415 | -0.732 | -2.6285 |

| | | | |
|---|---|---|---|
| ZNF7 | NM_003416 | -1.8325 | -3.62 |
| ZNF264 | NM_003417 | -1.5605 | -3.056 |
| CNBP | NM_003418 | -0.682 | -0.669 |
| ZNF345 | NM_003419 | -1.543 | -1.7015 |
| ZNF35 | NM_003420 | -0.7205 | -2.361 |
| ZNF37A | NM_003421 | -0.73 | -1.6925 |
| MZF1 | NM_003422 | -0.6255 | -0.777 |
| ZNF43 | NM_003423 | -1.048 | -2.862 |
| ZNF45 | NM_003425 | -1.5725 | -3.3005 |
| ZNF74 | NM_003426 | -0.7685 | -2.8165 |
| ZNF76 | NM_003427 | -0.383 | -1.3285 |
| ZNF84 | NM_003428 | -1.1965 | -2.2425 |
| ZNF85 | NM_003429 | -0.827 | -1.546 |
| ZNF91 | NM_003430 | -0.477 | -1.7885 |
| ZNF124 | NM_003431 | -0.9225 | -1.566 |
| ZNF131 | NM_003432 | -1.378 | -2.3595 |
| ZNF132 | NM_003433 | -1.3305 | -1.5895 |
| ZNF133 | NM_003434 | -1.0865 | -2.068 |
| ZNF134 | NM_003435 | -1.067 | -2.1295 |
| ZNF135 | NM_003436 | -0.054 | -0.975 |
| ZNF136 | NM_003437 | -0.234 | -2.0415 |
| ZKSCAN1 | NM_003439 | -1.577 | -2.466 |
| ZNF140 | NM_003440 | -1.799 | -4.4555 |
| ZNF141 | NM_003441 | -0.3705 | -1.1765 |
| ZNF143 | NM_003442 | -1.0555 | -2.757 |
| ZBTB17 | NM_003443 | 0.044 | -0.912 |
| ZNF155 | NM_003445 | -1.396 | -2.096 |
| ZNF157 | NM_003446 | -0.9305 | -0.6535 |
| ZNF165 | NM_003447 | -0.2965 | -0.844 |
| TRIM26 | NM_003449 | -0.7365 | -1.5965 |
| ZNF174 | NM_003450 | -0.865 | -2.879 |
| ZNF177 | NM_003451 | -0.9255 | -2.1965 |
| ZNF189 | NM_003452 | -1.9665 | -3.5655 |
| ZMYM2 | NM_003453 | -0.9155 | -2.786 |
| ZNF200 | NM_003454 | -1.8995 | -3.8065 |
| ZNF202 | NM_003455 | -1.555 | -3.5815 |
| ZNF205 | NM_003456 | -0.871 | -1.846 |
| ZNF207 | NM_003457 | -0.662 | -1.4985 |
| BSN | NM_003458 | -0.829 | -1.9855 |
| SLC30A3 | NM_003459 | -0.169 | -0.7355 |
| ZP2 | NM_003460 | 0.218 | -0.3735 |
| ZYX | NM_003461 | -0.415 | -0.8865 |
| DNALI1 | NM_003462 | -0.288 | -0.048 |

| | | | |
|---|---|---|---|
| PTP4A1 | NM_003463 | -0.6835 | -0.7895 |
| CHIT1 | NM_003465 | 0.005 | -0.0545 |
| PAX8 | NM_003466 | 0.1665 | -0.088 |
| CXCR4 | NM_003467 | 0.077 | -0.3235 |
| FZD5 | NM_003468 | -0.095 | -0.5925 |
| SCG2 | NM_003469 | -0.994 | -2.497 |
| USP7 | NM_003470 | -0.9855 | -2.015 |
| KCNAB1 | NM_003471 | -1.2125 | -1.0695 |
| DEK | NM_003472 | -0.6915 | -1.241 |
| STAM | NM_003473 | -1.072 | -2.389 |
| ADAM12 | NM_003474 | -0.6285 | -0.705 |
| RASSF7 | NM_003475 | -0.4195 | -1.298 |
| CSRP3 | NM_003476 | 0.044 | 0.0145 |
| PDHX | NM_003477 | -0.9945 | -2.4695 |
| CUL5 | NM_003478 | -0.8325 | -2.387 |
| MFAP5 | NM_003480 | 0.1685 | -0.3595 |
| MLL2 | NM_003482 | -0.65 | -1.5935 |
| HMGA2 | NM_003483 | 0.246 | -0.136 |
| HMGA2 | NM_003484 | 0.246 | -0.136 |
| GPR68 | NM_003485 | -0.255 | -0.473 |
| SLC7A5 | NM_003486 | -0.581 | -1.131 |
| TAF15 | NM_003487 | -0.404 | -0.7105 |
| AKAP1 | NM_003488 | -0.1795 | -1.1685 |
| NRIP1 | NM_003489 | -0.154 | -3.257 |
| SYN3 | NM_003490 | -0.103 | -0.304 |
| NAA10 | NM_003491 | 0.3345 | -0.119 |
| TMEM187 | NM_003492 | -0.671 | -1.4855 |
| HIST3H3 | NM_003493 | 0.405 | -0.1425 |
| DYSF | NM_003494 | -0.1995 | -0.4835 |
| HIST1H4I | NM_003495 | 0.1825 | -0.326 |
| TRRAP | NM_003496 | -0.6835 | -1.599 |
| SNN | NM_003498 | -0.6315 | -2.2015 |
| ACOX2 | NM_003500 | -0.1395 | 0.5655 |
| ACOX3 | NM_003501 | -0.232 | -1.5875 |
| AXIN1 | NM_003502 | -0.544 | -1.683 |
| CDC7 | NM_003503 | -1.6505 | -2.831 |
| CDC45 | NM_003504 | -0.455 | -1.2005 |
| FZD1 | NM_003505 | -0.3385 | -0.729 |
| FZD6 | NM_003506 | -1.104 | -0.792 |
| FZD7 | NM_003507 | -0.2145 | -0.3375 |
| FZD9 | NM_003508 | -0.034 | -0.857 |
| HIST1H2AI | NM_003509 | 0.506 | 0.899 |
| HIST1H2AK | NM_003510 | 0.51 | 1 |

| | | | |
|---|---|---|---|
| HIST1H2AL | NM_003511 | 0.332 | 0.396 |
| HIST1H2AC | NM_003512 | -0.3645 | 0.0705 |
| HIST1H2AB | NM_003513 | 0.615 | 0.168 |
| HIST1H2AM | NM_003514 | 1.752 | 1.4955 |
| HIST2H2AA3 | NM_003516 | 0.778 | 0.407 |
| HIST2H2AC | NM_003517 | 0.7615 | 0.089 |
| HIST1H2BG | NM_003518 | 1.9035 | 1.5515 |
| HIST1H2BL | NM_003519 | 0.135 | 0.018 |
| HIST1H2BN | NM_003520 | 1.4955 | 1.676 |
| HIST1H2BM | NM_003521 | 0.4775 | 0.131 |
| HIST1H2BF | NM_003522 | 0.3485 | 0.9115 |
| HIST1H2BE | NM_003523 | 0.448 | 0.219 |
| HIST1H2BH | NM_003524 | 0.5585 | -0.039 |
| HIST1H2BI | NM_003525 | -0.37 | 0.2495 |
| HIST1H2BC | NM_003526 | 0.7925 | 0.4285 |
| HIST1H2BO | NM_003527 | -0.013 | 0.449 |
| HIST2H2BE | NM_003528 | 1.216 | 1.3175 |
| HIST1H3A | NM_003529 | 1.473 | 2.183 |
| HIST1H3D | NM_003530 | 0.949 | 0.826 |
| HIST1H3C | NM_003531 | -0.2715 | 0.7135 |
| HIST1H3E | NM_003532 | 0.109 | 0.1445 |
| HIST1H3I | NM_003533 | 0.4885 | 0.2105 |
| HIST1H3G | NM_003534 | 0.916 | 0.387 |
| HIST1H3J | NM_003535 | 0.8065 | 0.3585 |
| HIST1H3H | NM_003536 | -0.2105 | 0.0255 |
| HIST1H3B | NM_003537 | 0.799 | 0.3475 |
| HIST1H4A | NM_003538 | 0.2785 | 0.798 |
| HIST1H4D | NM_003539 | 0.9915 | 1.6175 |
| HIST1H4F | NM_003540 | 0.145 | 0.088 |
| HIST1H4K | NM_003541 | 0.3675 | 0.549 |
| HIST1H4C | NM_003542 | 0.241 | -0.258 |
| HIST1H4H | NM_003543 | 0.226 | 0.1305 |
| HIST1H4B | NM_003544 | 0.8055 | 1.14 |
| HIST1H4E | NM_003545 | 0.5485 | 1.0805 |
| HIST1H4L | NM_003546 | -0.1365 | 0.5175 |
| HIST1H4G | NM_003547 | -0.04 | -0.5125 |
| HYAL3 | NM_003549 | -0.227 | -0.372 |
| MAD1L1 | NM_003550 | -0.696 | -2.1325 |
| NME5 | NM_003551 | -0.035 | 0.041 |
| OR1E1 | NM_003553 | -0.385 | -0.8185 |
| OR1E2 | NM_003554 | -0.012 | 0.1145 |
| OR1G1 | NM_003555 | -0.205 | -0.3455 |
| PIP5K1A | NM_003557 | -0.1475 | -1.0495 |

| | | | |
|---|---|---|---|
| PIP5K1B | NM_003558 | 0.2135 | -0.2835 |
| PIP4K2B | NM_003559 | -0.884 | -1.2485 |
| PLA2G6 | NM_003560 | -0.287 | -0.6265 |
| PLA2G10 | NM_003561 | 0.0825 | -0.5835 |
| SLC25A11 | NM_003562 | -0.9435 | -1.366 |
| SPOP | NM_003563 | -0.491 | -2.0825 |
| TAGLN2 | NM_003564 | 0.4755 | -0.093 |
| ULK1 | NM_003565 | -0.656 | -0.9465 |
| EEA1 | NM_003566 | -0.433 | -2.114 |
| BCAR3 | NM_003567 | -1.1075 | -1.542 |
| ANXA9 | NM_003568 | 0.3045 | -0.292 |
| STX7 | NM_003569 | -0.7055 | -1.469 |
| BFSP2 | NM_003571 | 0.138 | -0.358 |
| LTBP4 | NM_003573 | -0.3845 | -0.861 |
| VAPA | NM_003574 | -0.824 | -1.753 |
| ZNF282 | NM_003575 | -0.5965 | -2.126 |
| STK24 | NM_003576 | -0.0975 | -1.0485 |
| UTF1 | NM_003577 | -0.0095 | -0.252 |
| SOAT2 | NM_003578 | -0.0225 | -0.326 |
| RAD54L | NM_003579 | -0.519 | -1.697 |
| NSMAF | NM_003580 | -1.0595 | -1.7115 |
| NCK2 | NM_003581 | -0.2055 | -2.32 |
| DYRK3 | NM_003582 | -1.1385 | -2.4665 |
| DYRK2 | NM_003583 | -1.1665 | -2.362 |
| DUSP11 | NM_003584 | -1.2345 | -3.7785 |
| DOC2B | NM_003585 | -0.727 | -0.9545 |
| DOC2A | NM_003586 | -0.4 | -0.852 |
| DHX16 | NM_003587 | -1.009 | -2.9925 |
| CUL4B | NM_003588 | -1.0075 | -0.7675 |
| CUL4A | NM_003589 | -1.5185 | -1.7745 |
| CUL3 | NM_003590 | -1.1015 | -2.595 |
| CUL2 | NM_003591 | -0.918 | -2.743 |
| CUL1 | NM_003592 | -1.713 | -3.63 |
| FOXN1 | NM_003593 | -0.128 | -0.3515 |
| TTF2 | NM_003594 | -1.239 | -0.964 |
| TPST1 | NM_003596 | -0.8785 | -2.4285 |
| KLF11 | NM_003597 | -0.6115 | -0.926 |
| TEAD2 | NM_003598 | -0.3115 | -1.2075 |
| SUPT3H | NM_003599 | -0.378 | -0.3605 |
| SMARCA5 | NM_003601 | -0.6385 | -1.0365 |
| FKBP6 | NM_003602 | -0.0565 | 0.116 |
| SORBS2 | NM_003603 | -0.5915 | -1.332 |
| IRS4 | NM_003604 | -0.09 | 0.06 |

| | | | |
|---|---|---|---|
| CDC42BPA | NM_003607 | 0.1535 | -1.323 |
| GPR65 | NM_003608 | -0.4065 | -0.3025 |
| HIRIP3 | NM_003609 | -0.6655 | -1.381 |
| OFD1 | NM_003611 | -1.2245 | -2.328 |
| SEMA7A | NM_003612 | 0.068 | -0.58 |
| CILP | NM_003613 | 0.0565 | -0.1005 |
| GALR3 | NM_003614 | 0.071 | -0.494 |
| SLC4A7 | NM_003615 | -1.2395 | -2.311 |
| RGS5 | NM_003617 | -0.1725 | -0.09 |
| MAP4K3 | NM_003618 | -1.1505 | -2.5855 |
| PRSS12 | NM_003619 | 0.102 | -0.5055 |
| PPM1D | NM_003620 | -1.5285 | -3.71 |
| PPFIBP2 | NM_003621 | -0.4265 | -1.261 |
| PPFIBP1 | NM_003622 | -0.7885 | -1.0285 |
| RANBP3 | NM_003624 | -0.5885 | -1.8495 |
| PPFIA2 | NM_003625 | -0.055 | -0.2715 |
| PPFIA1 | NM_003626 | -1.8235 | -2.609 |
| SLC43A1 | NM_003627 | -0.5665 | -1.581 |
| PKP4 | NM_003628 | -1.0155 | -2.478 |
| PIK3R3 | NM_003629 | -1.1455 | -3.346 |
| PEX3 | NM_003630 | -1.267 | -2.354 |
| PARG | NM_003631 | -1.5215 | -3.3545 |
| CNTNAP1 | NM_003632 | -0.7235 | -1.169 |
| ENC1 | NM_003633 | -0.37 | 0.212 |
| NIPSNAP1 | NM_003634 | -0.137 | -0.607 |
| NDST2 | NM_003635 | -0.6795 | -1.967 |
| KCNAB2 | NM_003636 | -0.5345 | -0.7975 |
| ITGA10 | NM_003637 | 0.03 | -0.5635 |
| ITGA8 | NM_003638 | -0.001 | -0.101 |
| IKBKG | NM_003639 | -0.315 | -0.9625 |
| IKBKAP | NM_003640 | -1.0295 | -1.101 |
| IFITM1 | NM_003641 | 0.112 | -0.265 |
| HAT1 | NM_003642 | -0.601 | -2.193 |
| GCM1 | NM_003643 | 0.125 | -0.222 |
| GAS7 | NM_003644 | -0.1265 | -0.5135 |
| SLC27A2 | NM_003645 | -0.097 | -1.474 |
| DGKZ | NM_003646 | -0.2985 | -0.85 |
| DGKE | NM_003647 | -0.4985 | -2.4025 |
| DDO | NM_003649 | -0.2045 | -0.4105 |
| CST7 | NM_003650 | 0.181 | -0.341 |
| CSDA | NM_003651 | 0.1425 | -0.201 |
| COPS3 | NM_003653 | -0.8075 | -1.259 |
| CHST1 | NM_003654 | -0.144 | -0.4775 |

| | | | |
|---|---|---|---|
| CBX4 | NM_003655 | 0.079 | 0.644 |
| CAMK1 | NM_003656 | 0.137 | -0.2465 |
| BCAS1 | NM_003657 | 0.0955 | -0.138 |
| BARX2 | NM_003658 | -0.4565 | -0.042 |
| AGPS | NM_003659 | -0.392 | -1.319 |
| PPFIA3 | NM_003660 | -0.7535 | -1.573 |
| APOL1 | NM_003661 | 0.0945 | -0.236 |
| PIR | NM_003662 | -0.2605 | -0.7305 |
| CGGBP1 | NM_003663 | -1.2975 | -1.9045 |
| AP3B1 | NM_003664 | -0.697 | -1.3905 |
| FCN3 | NM_003665 | 0.144 | 0.065 |
| BLZF1 | NM_003666 | -0.6045 | -2.4955 |
| LGR5 | NM_003667 | -1.454 | -2.6345 |
| MAPKAPK5 | NM_003668 | -1.1725 | -2.4585 |
| BHLHE40 | NM_003670 | 0.159 | 0.117 |
| CDC14B | NM_003671 | -0.3795 | -1.1515 |
| CDC14A | NM_003672 | -0.1155 | -0.0625 |
| TCAP | NM_003673 | -0.2385 | -0.3405 |
| PRPF18 | NM_003675 | -0.421 | -1.7355 |
| DEGS1 | NM_003676 | -0.7635 | -1.569 |
| DENR | NM_003677 | -1.2855 | -1.573 |
| KMO | NM_003679 | -0.427 | -0.148 |
| YARS | NM_003680 | -0.4785 | -1.0125 |
| PDXK | NM_003681 | -0.163 | -0.902 |
| MADD | NM_003682 | -0.419 | -1.6545 |
| RRP1 | NM_003683 | -0.622 | -2.261 |
| MKNK1 | NM_003684 | -1.232 | -1.5305 |
| KHSRP | NM_003685 | 0.262 | -0.6375 |
| PDLIM4 | NM_003687 | -0.476 | -1.322 |
| CASK | NM_003688 | -1.1375 | -1.4375 |
| AKR7A2 | NM_003689 | 0.345 | -0.4655 |
| PRKRA | NM_003690 | -0.8245 | -1.3795 |
| TMEFF1 | NM_003692 | -0.5895 | -0.4615 |
| SCARF1 | NM_003693 | -0.059 | -0.074 |
| LY6D | NM_003695 | 0.015 | -0.6525 |
| OR6A2 | NM_003696 | -0.0005 | -0.128 |
| OR5F1 | NM_003697 | 0.0275 | -0.0395 |
| OR2D2 | NM_003700 | 0.1845 | 0.293 |
| TNFSF11 | NM_003701 | 0.0845 | -0.34 |
| NOP14 | NM_003703 | -0.2055 | -1.1095 |
| FAM193A | NM_003704 | -0.728 | -2.5165 |
| SLC25A12 | NM_003705 | -1.378 | -2.9365 |
| PLA2G4C | NM_003706 | -0.087 | -0.472 |

| | | | |
|---|---|---|---|
| RUVBL1 | NM_003707 | -0.3995 | -1.083 |
| RDH16 | NM_003708 | -0.087 | -0.4595 |
| KLF7 | NM_003709 | -1.46 | -2.9435 |
| SPINT1 | NM_003710 | 0.127 | -0.468 |
| PPAP2A | NM_003711 | -0.741 | -3.284 |
| PPAP2C | NM_003712 | 0.0025 | -0.527 |
| PPAP2B | NM_003713 | -0.4115 | -1.593 |
| STC2 | NM_003714 | -0.9395 | -0.3 |
| USO1 | NM_003715 | -0.8035 | -2.009 |
| CADPS | NM_003716 | -0.6905 | -0.4185 |
| NPFF | NM_003717 | 0.168 | -0.8445 |
| CDK13 | NM_003718 | -1.294 | -3.574 |
| PDE8B | NM_003719 | 0.316 | -1.1125 |
| PSMG1 | NM_003720 | -1.1155 | -2.581 |
| RFXANK | NM_003721 | -0.056 | -1.1185 |
| TP63 | NM_003722 | -0.1155 | 0.2695 |
| JRK | NM_003724 | -0.2325 | -1.3575 |
| SKAP1 | NM_003726 | 0.0145 | -0.096 |
| UNC5C | NM_003728 | -0.4325 | -0.14 |
| RTCD1 | NM_003729 | -1.0415 | -2.6275 |
| RNASET2 | NM_003730 | -0.183 | -0.8805 |
| SSNA1 | NM_003731 | -0.3285 | -1.0835 |
| OASL | NM_003733 | 0.086 | -0.2105 |
| AOC3 | NM_003734 | 0.1675 | -0.253 |
| PCDHGA12 | NM_003735 | -0.3165 | -0.698 |
| PCDHGB4 | NM_003736 | 0.1015 | -0.5365 |
| DCHS1 | NM_003737 | -0.1715 | -1.071 |
| PTCH2 | NM_003738 | -0.2615 | -0.5365 |
| AKR1C3 | NM_003739 | -0.037 | -0.5335 |
| KCNK5 | NM_003740 | -0.7 | -1.381 |
| CHRD | NM_003741 | -0.1395 | -0.596 |
| ABCB11 | NM_003742 | 0.2135 | -0.704 |
| NCOA1 | NM_003743 | -0.707 | -2.847 |
| NUMB | NM_003744 | -0.986 | -2.4315 |
| SOCS1 | NM_003745 | -0.045 | -0.044 |
| DYNLL1 | NM_003746 | -0.274 | -0.403 |
| TNKS | NM_003747 | -0.953 | -1.575 |
| ALDH4A1 | NM_003748 | -0.1265 | -0.537 |
| IRS2 | NM_003749 | -0.7305 | -1.5765 |
| EIF3A | NM_003750 | -0.6315 | -2.0615 |
| EIF3B | NM_003751 | -0.047 | -0.533 |
| EIF3D | NM_003753 | -0.1905 | -0.703 |
| EIF3F | NM_003754 | 0.3055 | -0.3635 |

| | | | |
|---|---|---|---|
| EIF3G | NM_003755 | 0.2605 | -0.324 |
| EIF3H | NM_003756 | -0.2895 | -0.066 |
| EIF3I | NM_003757 | 0.1825 | 0.06 |
| EIF3J | NM_003758 | -1.083 | -1.59 |
| SLC4A4 | NM_003759 | 0.0425 | -0.0635 |
| EIF4G3 | NM_003760 | -0.836 | -1.756 |
| VAMP8 | NM_003761 | 0.2405 | -0.044 |
| VAMP4 | NM_003762 | -1.2335 | -1.6575 |
| STX16 | NM_003763 | -1.417 | -1.407 |
| STX11 | NM_003764 | -0.0975 | -0.272 |
| STX10 | NM_003765 | -0.147 | -0.8965 |
| BECN1 | NM_003766 | -0.878 | -1.458 |
| PEA15 | NM_003768 | -0.323 | -0.907 |
| SRSF9 | NM_003769 | -0.0305 | -0.5725 |
| KRT37 | NM_003770 | -0.281 | 0.118 |
| KRT36 | NM_003771 | -0.092 | 0.1765 |
| JRKL | NM_003772 | -1.2795 | -1.824 |
| GALNT4 | NM_003774 | -0.2205 | 0.9325 |
| S1PR4 | NM_003775 | -0.048 | -0.2115 |
| MRPL40 | NM_003776 | -0.6045 | -1.02 |
| DNAH11 | NM_003777 | -0.134 | -1.05 |
| B4GALT4 | NM_003778 | -1.009 | -1.594 |
| B4GALT3 | NM_003779 | -0.472 | -1.9975 |
| B4GALT2 | NM_003780 | -0.36 | -1.042 |
| B3GALNT1 | NM_003781 | 0.0155 | -0.871 |
| B3GALT4 | NM_003782 | 0.1605 | 0.575 |
| B3GALT2 | NM_003783 | 0.153 | -2.72 |
| SERPINB7 | NM_003784 | 0.6645 | 0.064 |
| PAGE1 | NM_003785 | -0.167 | 0.055 |
| ABCC3 | NM_003786 | -0.3115 | -0.1385 |
| NOL4 | NM_003787 | -1.322 | -2.234 |
| TRADD | NM_003789 | -0.429 | -0.976 |
| TNFRSF25 | NM_003790 | -0.183 | -0.233 |
| MBTPS1 | NM_003791 | -0.6275 | -1.6805 |
| EDF1 | NM_003792 | 0.064 | -0.4445 |
| CTSF | NM_003793 | -0.144 | -0.5255 |
| SNX4 | NM_003794 | -1.2555 | -1.491 |
| SNX3 | NM_003795 | -0.557 | -1.264 |
| C19orf2 | NM_003796 | -0.8885 | -1.442 |
| EED | NM_003797 | -1.2235 | -2.812 |
| CTNNAL1 | NM_003798 | -0.5065 | -1.34 |
| RNMT | NM_003799 | -1.034 | -2.2755 |
| RNGTT | NM_003800 | -1.207 | -2.5355 |

| | | | |
|---|---|---|---|
| GPAA1 | NM_003801 | 0.096 | -0.306 |
| MYH13 | NM_003802 | -0.1055 | -0.365 |
| MYOM1 | NM_003803 | 0.203 | -0.317 |
| RIPK1 | NM_003804 | -0.5125 | -1.184 |
| CRADD | NM_003805 | -0.222 | -1.903 |
| HRK | NM_003806 | -0.124 | -0.813 |
| TNFSF14 | NM_003807 | -0.1025 | -0.107 |
| TNFSF12 | NM_003809 | 0.052 | -0.1405 |
| TNFSF10 | NM_003810 | -0.391 | -0.174 |
| TNFSF9 | NM_003811 | -0.0145 | -0.105 |
| ADAM23 | NM_003812 | 0.0205 | -0.2105 |
| ADAM21 | NM_003813 | 0.115 | -0.1505 |
| ADAM20 | NM_003814 | 0.283 | -0.658 |
| ADAM15 | NM_003815 | 0.1465 | -0.561 |
| ADAM9 | NM_003816 | -0.5215 | -1.088 |
| ADAM7 | NM_003817 | -0.142 | -0.5405 |
| CDS2 | NM_003818 | -0.3765 | -1.129 |
| PABPC4 | NM_003819 | 0.0305 | -0.3275 |
| TNFRSF14 | NM_003820 | 0.6975 | -0.121 |
| RIPK2 | NM_003821 | -0.385 | -1.3635 |
| NR5A2 | NM_003822 | -0.1685 | -0.2445 |
| TNFRSF6B | NM_003823 | -0.1955 | -0.5275 |
| FADD | NM_003824 | -1.2385 | -3.3885 |
| SNAP23 | NM_003825 | -1.021 | -1.399 |
| NAPG | NM_003826 | -1.2745 | -1.7735 |
| NAPA | NM_003827 | 0.1705 | -0.4475 |
| MTMR1 | NM_003828 | -1.245 | -1.977 |
| MPDZ | NM_003829 | -0.697 | -1.823 |
| SIGLEC5 | NM_003830 | -0.016 | -0.487 |
| RIOK3 | NM_003831 | -0.8475 | -1.2165 |
| MATN4 | NM_003833 | 0.0645 | -0.1435 |
| RGS11 | NM_003834 | -0.156 | -0.396 |
| RGS9 | NM_003835 | -0.004 | -0.4435 |
| DLK1 | NM_003836 | -0.0135 | -0.2945 |
| FBP2 | NM_003837 | 0.21 | -0.622 |
| FPGT | NM_003838 | -1.2485 | -2.245 |
| TNFRSF11A | NM_003839 | -0.0855 | -0.433 |
| TNFRSF10D | NM_003840 | 0.025 | 0.1175 |
| TNFRSF10C | NM_003841 | 0.166 | -0.27 |
| TNFRSF10B | NM_003842 | -0.8155 | -2.116 |
| SCEL | NM_003843 | -0.0095 | -0.0995 |
| TNFRSF10A | NM_003844 | -0.5005 | -0.36 |
| DYRK4 | NM_003845 | -0.472 | -0.2245 |

| | | | |
|---|---|---|---|
| PEX11B | NM_003846 | -0.4645 | -1.3595 |
| PEX11A | NM_003847 | -0.958 | -1.7435 |
| SUCLG2 | NM_003848 | -0.7795 | -0.8435 |
| SUCLG1 | NM_003849 | -0.4675 | -1.0805 |
| SUCLA2 | NM_003850 | -1.382 | -2.6615 |
| CREG1 | NM_003851 | -0.737 | -0.5345 |
| TRIM24 | NM_003852 | -1.127 | -2.6415 |
| IL18RAP | NM_003853 | -0.052 | -0.785 |
| IL1RL2 | NM_003854 | -0.239 | -0.4695 |
| IL18R1 | NM_003855 | 0.3245 | -0.191 |
| IL1RL1 | NM_003856 | 0.0475 | 0.073 |
| GALR2 | NM_003857 | -0.1275 | -0.627 |
| DPM1 | NM_003859 | -1.2485 | -2.1075 |
| BANF1 | NM_003860 | 0.2465 | -0.493 |
| DCAF5 | NM_003861 | -1.075 | -2.8315 |
| FGF18 | NM_003862 | -0.086 | -0.5625 |
| DPM2 | NM_003863 | -0.3505 | -1.73 |
| SAP30 | NM_003864 | -1.0535 | -3.4115 |
| HESX1 | NM_003865 | -0.974 | -0.876 |
| INPP4B | NM_003866 | -0.044 | -0.407 |
| FGF17 | NM_003867 | -0.3705 | -0.5315 |
| FGF16 | NM_003868 | 0.242 | 0.3645 |
| CES2 | NM_003869 | -0.5975 | -1.474 |
| IQGAP1 | NM_003870 | -0.469 | -0.9475 |
| NRP2 | NM_003872 | -0.568 | -1.518 |
| NRP1 | NM_003873 | -2.015 | -3.5045 |
| CD84 | NM_003874 | -0.316 | -0.0675 |
| GMPS | NM_003875 | -0.869 | -1.684 |
| TMEM11 | NM_003876 | -0.536 | -3.2155 |
| SOCS2 | NM_003877 | -1.162 | -1.845 |
| GGH | NM_003878 | -0.8925 | -0.674 |
| CFLAR | NM_003879 | -0.587 | -0.769 |
| WISP3 | NM_003880 | -0.3395 | -0.111 |
| WISP2 | NM_003881 | -0.1635 | -0.479 |
| WISP1 | NM_003882 | 0.096 | -0.1165 |
| HDAC3 | NM_003883 | 0.0105 | -0.7845 |
| KAT2B | NM_003884 | -0.953 | -1.356 |
| CDK5R1 | NM_003885 | -0.3255 | 0.3705 |
| AKAP4 | NM_003886 | -0.4505 | -0.346 |
| ALDH1A2 | NM_003888 | -0.0655 | 0.0445 |
| NR1I2 | NM_003889 | -0.2945 | -0.269 |
| FCGBP | NM_003890 | -0.0545 | -0.3785 |
| PROZ | NM_003891 | 0.055 | -0.1295 |

| | | | |
|---|---|---|---|
| LDB1 | NM_003893 | -0.446 | -1.63 |
| SYNJ1 | NM_003895 | -0.59 | -1.34 |
| ST3GAL5 | NM_003896 | -0.6065 | -2.274 |
| IER3 | NM_003897 | 1.7475 | 3.0745 |
| SYNJ2 | NM_003898 | -0.825 | -1.2495 |
| ARHGEF7 | NM_003899 | -1.0645 | -2.5795 |
| SQSTM1 | NM_003900 | -0.709 | -1.4875 |
| SGPL1 | NM_003901 | -0.8245 | -1.3465 |
| FUBP1 | NM_003902 | -1.018 | -1.5135 |
| CDC16 | NM_003903 | -1.309 | -2.5375 |
| ZNF259 | NM_003904 | -0.04 | -1.1085 |
| NAE1 | NM_003905 | -1.5405 | -1.984 |
| MCM3AP | NM_003906 | -0.801 | -2.261 |
| EIF2B5 | NM_003907 | -0.5125 | -1.679 |
| EIF2S2 | NM_003908 | -1.1075 | -2.194 |
| CPNE3 | NM_003909 | -1.059 | -0.8105 |
| BUD31 | NM_003910 | -0.4125 | -1.2905 |
| PRPF4B | NM_003913 | -0.675 | -1.2585 |
| CCNA1 | NM_003914 | 0.1145 | -0.8025 |
| CPNE1 | NM_003915 | -0.2365 | -0.7265 |
| AP1S2 | NM_003916 | -0.7145 | -0.3905 |
| AP1G2 | NM_003917 | -0.401 | -0.684 |
| GYG2 | NM_003918 | -0.1645 | -0.707 |
| SGCE | NM_003919 | -1.5125 | -1.894 |
| TIMELESS | NM_003920 | -0.0535 | -0.491 |
| BCL10 | NM_003921 | -1.375 | -2.475 |
| HERC1 | NM_003922 | -0.8555 | -1.939 |
| FOXH1 | NM_003923 | 0.243 | 0.0195 |
| PHOX2B | NM_003924 | -0.664 | -1.571 |
| MBD4 | NM_003925 | -1.3175 | -2.1845 |
| MBD3 | NM_003926 | -0.917 | -1.9515 |
| MBD2 | NM_003927 | -0.906 | -2.2875 |
| RAB7L1 | NM_003929 | -1.158 | -0.9295 |
| SKAP2 | NM_003930 | -0.386 | 0.2525 |
| WASF1 | NM_003931 | -0.4905 | -1.144 |
| ST13 | NM_003932 | -0.6685 | -0.448 |
| BAIAP3 | NM_003933 | 0.01 | 0.3965 |
| FUBP3 | NM_003934 | -0.8115 | -1.0295 |
| TOP3B | NM_003935 | -1.074 | -1.8375 |
| CDK5R2 | NM_003936 | -1.0195 | -1.98 |
| KYNU | NM_003937 | -0.2195 | -0.235 |
| AP3D1 | NM_003938 | -0.206 | -1.224 |
| BTRC | NM_003939 | -0.64 | -1.588 |

| | | | |
|---|---|---|---|
| USP13 | NM_003940 | -0.9245 | -1.7 |
| WASL | NM_003941 | -1.6465 | -3.217 |
| RPS6KA4 | NM_003942 | -0.4875 | -1.7495 |
| STBD1 | NM_003943 | -0.6245 | -0.6315 |
| SELENBP1 | NM_003944 | 0.232 | -0.279 |
| ATP6V0E1 | NM_003945 | -0.2545 | -2.3375 |
| NOL3 | NM_003946 | -0.1915 | -0.91 |
| KALRN | NM_003947 | 0.643 | 0.008 |
| CDKL2 | NM_003948 | -0.5325 | -0.7785 |
| F2RL3 | NM_003950 | -0.051 | -0.2495 |
| SLC25A14 | NM_003951 | -0.4775 | -0.985 |
| RPS6KB2 | NM_003952 | 0.086 | -0.6265 |
| MPZL1 | NM_003953 | -0.7115 | -1.592 |
| MAP3K14 | NM_003954 | -0.7805 | -2.148 |
| SOCS3 | NM_003955 | 0.293 | -0.048 |
| CH25H | NM_003956 | 0.363 | 0.1955 |
| BRSK2 | NM_003957 | -1.0855 | -1.566 |
| RNF8 | NM_003958 | -1.169 | -2.8165 |
| HIP1R | NM_003959 | 0.014 | -0.8535 |
| NAT8 | NM_003960 | 0.1225 | 0.03 |
| RHBDL1 | NM_003961 | -0.106 | -0.209 |
| TM4SF5 | NM_003963 | 0.0755 | -0.0725 |
| CCRL2 | NM_003965 | 0.1265 | -0.2325 |
| SEMA5A | NM_003966 | -0.236 | -0.829 |
| TAAR5 | NM_003967 | -0.06 | -0.1785 |
| UBA3 | NM_003968 | -1.104 | -2.075 |
| UBE2M | NM_003969 | -0.234 | -0.64 |
| MYOM2 | NM_003970 | 0.036 | -0.347 |
| SPAG9 | NM_003971 | -1.214 | -1.697 |
| BTAF1 | NM_003972 | -1.1825 | -2.963 |
| RPL14 | NM_003973 | -0.0915 | -0.2095 |
| DOK2 | NM_003974 | -0.1725 | -0.7265 |
| SH2D2A | NM_003975 | 0.388 | -1.1415 |
| AIP | NM_003977 | 0.0265 | -0.926 |
| PSTPIP1 | NM_003978 | 0.0045 | -0.6485 |
| GPRC5A | NM_003979 | -0.018 | -0.915 |
| MAP7 | NM_003980 | -0.406 | -0.8655 |
| PRC1 | NM_003981 | -0.9005 | -0.9005 |
| SLC7A6 | NM_003983 | -0.708 | -0.9895 |
| SLC13A2 | NM_003984 | 0.1895 | -0.123 |
| TNK1 | NM_003985 | -0.0795 | -0.341 |
| BBOX1 | NM_003986 | -0.116 | 0.3135 |
| PAX2 | NM_003987 | 0.2985 | -0.22 |

| | | | |
|---|---|---|---|
| PAX2 | NM_003988 | 0.2985 | -0.22 |
| PAX2 | NM_003989 | 0.2985 | -0.22 |
| PAX2 | NM_003990 | 0.2985 | -0.22 |
| EDNRB | NM_003991 | -0.4215 | 0.038 |
| CLK3 | NM_003992 | -0.3935 | -2.1085 |
| CLK2 | NM_003993 | -0.789 | -1.966 |
| KITLG | NM_003994 | -0.1075 | 0.2045 |
| NPR2 | NM_003995 | -0.4015 | -1.6885 |
| GPX5 | NM_003996 | 0.37 | -0.2015 |
| NFKB1 | NM_003998 | -0.6035 | -2.378 |
| OSMR | NM_003999 | -0.006 | -0.5335 |
| CHI3L2 | NM_004000 | -0.1455 | -0.1445 |
| FCGR2B | NM_004001 | -0.2175 | -0.407 |
| GJB2 | NM_004004 | -0.0765 | -0.079 |
| DMD | NM_004006 | -0.1335 | -0.38 |
| DMD | NM_004007 | -0.1335 | -0.38 |
| DMD | NM_004009 | -0.1335 | -0.38 |
| DMD | NM_004010 | -0.1335 | -0.38 |
| DMD | NM_004011 | -0.1335 | -0.38 |
| DMD | NM_004012 | -0.1335 | -0.38 |
| DMD | NM_004013 | -0.1335 | -0.38 |
| DMD | NM_004014 | -0.1335 | -0.38 |
| DMD | NM_004015 | -0.1335 | -0.38 |
| DMD | NM_004016 | -0.1335 | -0.38 |
| DMD | NM_004017 | -0.1335 | -0.38 |
| DMD | NM_004018 | -0.1335 | -0.38 |
| DMD | NM_004019 | -0.2615 | -0.2815 |
| DMD | NM_004020 | -0.1335 | -0.38 |
| DMD | NM_004021 | -1.2955 | -1.304 |
| DMD | NM_004022 | -0.1335 | -0.38 |
| DMD | NM_004023 | -0.1335 | -0.38 |
| INPP4A | NM_004027 | -0.6765 | -1.123 |
| AQP4 | NM_004028 | -0.111 | -0.3035 |
| IRF7 | NM_004031 | 0.082 | -0.0055 |
| ANXA7 | NM_004034 | -0.5935 | -1.062 |
| ACOX1 | NM_004035 | -1.53 | -1.926 |
| ADCY3 | NM_004036 | -0.4885 | -1.286 |
| AMPD2 | NM_004037 | 0.0185 | -0.7785 |
| ANXA2 | NM_004039 | -0.2605 | -0.2505 |
| RHOB | NM_004040 | 0.0975 | 0.6355 |
| ARRB1 | NM_004041 | -0.0295 | -1.2355 |
| ARSF | NM_004042 | -0.0895 | -0.4285 |
| ASMT | NM_004043 | 0.14 | -0.476 |

| | | | |
|---|---|---|---|
| ATIC | NM_004044 | -0.585 | -1.232 |
| ATOX1 | NM_004045 | 0.284 | -0.3225 |
| ATP5A1 | NM_004046 | -0.505 | -0.4375 |
| ATP6V0B | NM_004047 | 0.3195 | -0.0155 |
| B2M | NM_004048 | -0.5975 | -0.242 |
| BCL2A1 | NM_004049 | 0.1515 | 0.0765 |
| BCL2L2 | NM_004050 | -1.614 | -2.4985 |
| BDH1 | NM_004051 | -0.852 | -2.027 |
| BNIP3 | NM_004052 | -0.4515 | -1.2255 |
| BYSL | NM_004053 | -0.6335 | -3.2955 |
| C3AR1 | NM_004054 | -0.1775 | -0.353 |
| CAPN5 | NM_004055 | 0.0795 | -0.8705 |
| CA8 | NM_004056 | -0.357 | -0.3715 |
| S100G | NM_004057 | 0.086 | 0.0155 |
| CAPS | NM_004058 | 0.0155 | -0.219 |
| CCNG1 | NM_004060 | -0.771 | -1.4995 |
| CDH12 | NM_004061 | -0.183 | -0.166 |
| CDH16 | NM_004062 | 0.394 | -0.267 |
| CDH17 | NM_004063 | -0.21 | -0.1885 |
| CDKN1B | NM_004064 | -1.648 | -0.5545 |
| CDR1 | NM_004065 | -0.153 | -0.3155 |
| CETN1 | NM_004066 | -0.1435 | -0.351 |
| CHN2 | NM_004067 | 0.2065 | 0.1045 |
| AP2M1 | NM_004068 | 0.2455 | -0.3765 |
| AP2S1 | NM_004069 | 0.2195 | -0.465 |
| CLCNKA | NM_004070 | -0.1665 | -0.2825 |
| CLK1 | NM_004071 | -0.908 | -1.031 |
| CMKLR1 | NM_004072 | 0.234 | 0.0755 |
| PLK3 | NM_004073 | -0.5785 | -1.4605 |
| COX8A | NM_004074 | 0.2255 | -0.2595 |
| CRY1 | NM_004075 | -0.7955 | -1.1845 |
| CRYBB3 | NM_004076 | 0.136 | -0.1615 |
| CS | NM_004077 | 0.003 | -0.6385 |
| CSRP1 | NM_004078 | 0.314 | -0.5765 |
| CTSS | NM_004079 | 0.427 | -0.496 |
| DGKB | NM_004080 | 0.0735 | -0.6315 |
| DAZ1 | NM_004081 | 0.3975 | -0.2135 |
| DCTN1 | NM_004082 | 0.0625 | -0.901 |
| DDIT3 | NM_004083 | -0.677 | -1.316 |
| TIMM8A | NM_004085 | -1.1315 | -1.295 |
| COCH | NM_004086 | -1.0935 | -1.3035 |
| DLG1 | NM_004087 | -1.5135 | -3.089 |
| DNTT | NM_004088 | 0.181 | 0.254 |

| | | | |
|---|---|---|---|
| TSC22D3 | NM_004089 | 0.017 | 0.148 |
| DUSP3 | NM_004090 | -0.839 | -1.513 |
| E2F2 | NM_004091 | -0.936 | -2.5105 |
| ECHS1 | NM_004092 | -0.0105 | -1.0445 |
| EFNB2 | NM_004093 | -0.602 | -2.8595 |
| EIF2S1 | NM_004094 | -1.226 | -1.901 |
| EIF4EBP1 | NM_004095 | 0.015 | -0.797 |
| EIF4EBP2 | NM_004096 | -0.1215 | -0.908 |
| EMX1 | NM_004097 | -0.0015 | -0.119 |
| EMX2 | NM_004098 | 0.047 | -0.258 |
| STOM | NM_004099 | -0.2575 | -0.409 |
| EYA4 | NM_004100 | -1.1145 | -2.9115 |
| F2RL2 | NM_004101 | -0.5275 | -0.2265 |
| FABP3 | NM_004102 | 0.0525 | -0.125 |
| PTK2B | NM_004103 | -0.212 | -0.2605 |
| FASN | NM_004104 | 0.137 | -0.7625 |
| FCER1G | NM_004106 | -0.302 | -0.1125 |
| FCGRT | NM_004107 | -0.3565 | -0.9975 |
| FCN2 | NM_004108 | -0.484 | -1.436 |
| FDX1 | NM_004109 | -0.4155 | -1.9045 |
| FDXR | NM_004110 | -0.0465 | -0.6465 |
| FEN1 | NM_004111 | -0.577 | -1.8245 |
| FGF11 | NM_004112 | -0.2685 | -0.6865 |
| FGF12 | NM_004113 | 0.0225 | -0.931 |
| FGF13 | NM_004114 | -0.387 | -0.415 |
| FGF14 | NM_004115 | -0.8225 | -1.604 |
| FKBP5 | NM_004117 | -0.789 | -1.4185 |
| FOXS1 | NM_004118 | 0.2635 | -0.0145 |
| FLT3 | NM_004119 | -0.2715 | -0.211 |
| GBP2 | NM_004120 | 0.3475 | -0.5735 |
| GGT5 | NM_004121 | -0.1495 | -0.3845 |
| GHSR | NM_004122 | -0.005 | -0.298 |
| GIP | NM_004123 | -0.133 | -0.3475 |
| GMFB | NM_004124 | -0.668 | -0.5275 |
| GNG11 | NM_004126 | -0.1765 | -0.174 |
| GTF2F2 | NM_004128 | -0.966 | -1.4655 |
| GYG1 | NM_004130 | -0.614 | -1.262 |
| GZMB | NM_004131 | 0.149 | -0.1205 |
| HABP2 | NM_004132 | 0.177 | 0.3795 |
| HNF4G | NM_004133 | -0.628 | -1.8075 |
| HSPA9 | NM_004134 | -0.972 | -0.615 |
| IREB2 | NM_004136 | -1.267 | -2.696 |
| KCNMB1 | NM_004137 | 0.2655 | -0.1015 |

| | | | |
|---|---|---|---|
| KRT33A | NM_004138 | 0.1705 | -0.8645 |
| LBP | NM_004139 | -0.1165 | -0.105 |
| LLGL1 | NM_004140 | 0.2425 | -0.3155 |
| MYO9B | NM_004145 | -0.645 | -0.4465 |
| NDUFB7 | NM_004146 | 0.2615 | -0.274 |
| DRG1 | NM_004147 | -0.73 | -1.3345 |
| NINJ1 | NM_004148 | -0.186 | -0.8865 |
| OAZ1 | NM_004152 | 0.163 | -0.2625 |
| ORC1 | NM_004153 | -0.8935 | -2.2575 |
| P2RY6 | NM_004154 | -0.941 | -0.208 |
| SERPINB9 | NM_004155 | -0.3535 | -0.7205 |
| PRKAR2A | NM_004157 | -0.4265 | -1.0245 |
| PSPN | NM_004158 | -0.2975 | 0.268 |
| PSMB8 | NM_004159 | -0.037 | 0.2895 |
| PYY | NM_004160 | 0.5315 | -0.3865 |
| RAB1A | NM_004161 | -0.9995 | -1.639 |
| RAB5A | NM_004162 | -1.8595 | -3.5125 |
| RAB27B | NM_004163 | -0.4305 | -0.2255 |
| RBP2 | NM_004164 | 0.0925 | 0.0925 |
| SDHA | NM_004168 | -0.1955 | -1.4735 |
| SHMT1 | NM_004169 | -0.9225 | -2.2875 |
| SLC1A1 | NM_004170 | 0.8835 | -2.2065 |
| SLC1A2 | NM_004171 | 0.0675 | -0.4915 |
| SLC1A3 | NM_004172 | -0.2025 | -0.363 |
| SLC7A4 | NM_004173 | 0.2525 | -0.2825 |
| SLC9A3 | NM_004174 | 0.121 | -0.0855 |
| SNRPD3 | NM_004175 | -0.441 | -0.9425 |
| SREBF1 | NM_004176 | -0.514 | -1.0535 |
| STX3 | NM_004177 | -0.4325 | -1.7105 |
| TARBP2 | NM_004178 | -0.102 | -0.257 |
| TPH1 | NM_004179 | -0.1425 | -0.114 |
| TANK | NM_004180 | -2.529 | -4.724 |
| UCHL1 | NM_004181 | 0.0505 | -0.385 |
| UXT | NM_004182 | 0.0345 | -0.3155 |
| BEST1 | NM_004183 | 0.126 | -0.161 |
| WARS | NM_004184 | -0.249 | -0.9745 |
| WNT2B | NM_004185 | -0.034 | -0.266 |
| SEMA3F | NM_004186 | -0.8385 | -1.156 |
| KDM5C | NM_004187 | 0.0145 | -0.985 |
| GFI1B | NM_004188 | -0.163 | -0.3775 |
| SOX14 | NM_004189 | 0.024 | -0.469 |
| LIPF | NM_004190 | 0.173 | 0.088 |
| ASMTL | NM_004192 | -0.1645 | -0.841 |

| | | | |
|---|---|---|---|
| GBF1 | NM_004193 | -0.2285 | -1.1165 |
| ADAM22 | NM_004194 | -0.5105 | -0.9525 |
| TNFRSF18 | NM_004195 | -0.2905 | -1.0095 |
| CDKL1 | NM_004196 | -0.674 | -0.425 |
| STK19 | NM_004197 | -0.158 | -0.452 |
| CHRNA6 | NM_004198 | -0.0865 | -0.4445 |
| P4HA2 | NM_004199 | 0.109 | -0.906 |
| SYT7 | NM_004200 | -0.215 | -0.5635 |
| TMSB4Y | NM_004202 | 0.1465 | -0.2835 |
| PKMYT1 | NM_004203 | -0.6415 | -1.432 |
| PIGQ | NM_004204 | -0.1355 | -0.5885 |
| USP2 | NM_004205 | 0.16 | 0.0275 |
| SEC22C | NM_004206 | -0.698 | -1.946 |
| SLC16A3 | NM_004207 | 0.023 | -0.6925 |
| AIFM1 | NM_004208 | -0.571 | -1.3845 |
| SYNGR3 | NM_004209 | -0.2795 | 0.024 |
| NEURL | NM_004210 | -0.2725 | -0.6875 |
| SLC6A5 | NM_004211 | -0.142 | -0.1335 |
| SLC28A2 | NM_004212 | -0.0115 | -0.0855 |
| SLC28A1 | NM_004213 | -0.2805 | -0.0575 |
| FIBP | NM_004214 | 0.1345 | -0.427 |
| EBAG9 | NM_004215 | -1.5475 | -3.593 |
| AURKB | NM_004217 | -0.9835 | -2.3765 |
| RAB11B | NM_004218 | 0.0895 | -0.4945 |
| PTTG1 | NM_004219 | -0.494 | -1.003 |
| ZNF213 | NM_004220 | -0.112 | -0.505 |
| IL32 | NM_004221 | 0.0195 | -0.0945 |
| GPR50 | NM_004224 | -0.0875 | -0.395 |
| MFHAS1 | NM_004225 | -0.755 | -3.492 |
| STK17B | NM_004226 | -1.904 | -2.99 |
| CYTH3 | NM_004227 | -0.5215 | -0.8905 |
| CYTH2 | NM_004228 | -0.751 | -2.6665 |
| MED14 | NM_004229 | -0.9575 | -1.45 |
| S1PR2 | NM_004230 | -0.4475 | -1.4785 |
| ATP6V1F | NM_004231 | 0.0025 | -0.6315 |
| SOCS6 | NM_004232 | -0.5425 | -1.2655 |
| CD83 | NM_004233 | -0.552 | -1.466 |
| ZNF235 | NM_004234 | -0.7315 | -1.966 |
| KLF4 | NM_004235 | 0.0275 | 0.168 |
| COPS2 | NM_004236 | -0.4095 | -1.847 |
| TRIP13 | NM_004237 | -0.694 | -0.936 |
| TRIP12 | NM_004238 | -0.996 | -1.906 |
| TRIP11 | NM_004239 | -0.2415 | -0.7915 |

| | | | |
|---|---|---|---|
| TRIP10 | NM_004240 | 0.077 | -0.3965 |
| JMJD1C | NM_004241 | -0.3455 | -2.3015 |
| HMGN3 | NM_004242 | -0.5085 | -0.584 |
| CD163 | NM_004244 | -0.5645 | -0.5245 |
| GLP2R | NM_004246 | 0.244 | 0.0025 |
| EFTUD2 | NM_004247 | -0.1155 | -0.8155 |
| PRLHR | NM_004248 | -0.6755 | -0.3665 |
| RAB28 | NM_004249 | -0.961 | -2.559 |
| RAB9A | NM_004251 | -1.328 | -3.5535 |
| SLC9A3R1 | NM_004252 | -0.385 | -1.5295 |
| SLC22A8 | NM_004254 | -0.0395 | -0.388 |
| COX5A | NM_004255 | 0.1525 | -0.23 |
| SLC22A13 | NM_004256 | -0.3615 | -0.3715 |
| TGFBRAP1 | NM_004257 | -0.046 | -1.2695 |
| CD101 | NM_004258 | -0.095 | -0.4555 |
| RECQL5 | NM_004259 | -0.791 | -1.203 |
| RECQL4 | NM_004260 | -0.3445 | -1.062 |
| TMPRSS11D | NM_004262 | 0.112 | 0.0405 |
| SEMA4F | NM_004263 | -0.3475 | -1.378 |
| MED21 | NM_004264 | -1.1765 | -2.628 |
| FADS2 | NM_004265 | -0.2015 | -0.928 |
| CHST2 | NM_004267 | -0.9805 | -2.2035 |
| MED17 | NM_004268 | -1.575 | -3.574 |
| MED27 | NM_004269 | -0.547 | -2.436 |
| LY86 | NM_004271 | -0.1755 | -0.1455 |
| HOMER1 | NM_004272 | -1.2045 | -2.6075 |
| CHST3 | NM_004273 | -0.6305 | -2.181 |
| AKAP6 | NM_004274 | -0.1395 | -0.347 |
| MED20 | NM_004275 | -1.374 | -3.0635 |
| SLC25A27 | NM_004277 | -0.4455 | -0.4175 |
| PIGL | NM_004278 | -0.461 | -0.2085 |
| PMPCB | NM_004279 | -0.9565 | -1.647 |
| EEF1E1 | NM_004280 | -1.621 | -2.9335 |
| BAG3 | NM_004281 | -0.302 | -1.7045 |
| BAG2 | NM_004282 | -1.2055 | -2.8385 |
| RAB3D | NM_004283 | -0.1865 | -0.4205 |
| CHD1L | NM_004284 | -0.383 | -0.639 |
| H6PD | NM_004285 | -0.238 | -0.3975 |
| GTPBP1 | NM_004286 | -0.2105 | -0.796 |
| GOSR2 | NM_004287 | -0.48 | -1.608 |
| CYTIP | NM_004288 | 0.121 | -0.3275 |
| NFE2L3 | NM_004289 | -0.458 | -1.222 |
| RNF14 | NM_004290 | -1.725 | -3.6045 |

| | | | |
|---|---|---|---|
| CARTPT | NM_004291 | -0.2175 | -0.4545 |
| RIN1 | NM_004292 | -0.148 | -0.483 |
| GDA | NM_004293 | -0.1385 | -0.5655 |
| MTRF1 | NM_004294 | -0.9855 | -1.1705 |
| TRAF4 | NM_004295 | -1.1105 | -2.6405 |
| RGS6 | NM_004296 | -0.041 | -0.518 |
| GNA14 | NM_004297 | -0.015 | -0.401 |
| NUP155 | NM_004298 | -1.0925 | -1.9745 |
| ABCB7 | NM_004299 | -0.9605 | -1.8415 |
| ACP1 | NM_004300 | -0.709 | -0.655 |
| ACTL6A | NM_004301 | -1.089 | -0.766 |
| ACVR1B | NM_004302 | -0.7235 | -2.381 |
| ALK | NM_004304 | -0.7875 | -1.381 |
| BIN1 | NM_004305 | 0.2095 | -0.4445 |
| ANXA13 | NM_004306 | 0.2945 | 0.6705 |
| APBB2 | NM_004307 | -0.1085 | -0.418 |
| ARHGAP1 | NM_004308 | -0.4715 | -1.4755 |
| ARHGDIA | NM_004309 | -0.172 | -1.133 |
| RHOH | NM_004310 | 0.0865 | -0.0565 |
| ARL3 | NM_004311 | -0.572 | -0.832 |
| ARR3 | NM_004312 | -0.2245 | -0.5285 |
| ARRB2 | NM_004313 | 0.152 | -0.7385 |
| ART1 | NM_004314 | -0.176 | -0.563 |
| ASAH1 | NM_004315 | -0.843 | -0.7025 |
| ASCL1 | NM_004316 | -0.604 | -0.9755 |
| ASNA1 | NM_004317 | 0.0785 | -1.3 |
| ASPH | NM_004318 | -1.0635 | -0.5415 |
| ASTN1 | NM_004319 | -0.1935 | -0.8735 |
| KIF1A | NM_004321 | -0.346 | -0.6845 |
| BAD | NM_004322 | -0.2855 | -0.9855 |
| BAG1 | NM_004323 | -0.2585 | -1.336 |
| BAX | NM_004324 | -0.281 | -1.03 |
| BCL9 | NM_004326 | -0.771 | -2.217 |
| BCR | NM_004327 | -0.376 | -1.5265 |
| BCS1L | NM_004328 | -0.31 | -1.3055 |
| BMPR1A | NM_004329 | -1.248 | -1.9705 |
| BNIP2 | NM_004330 | -1.4115 | -3.4495 |
| BNIP3L | NM_004331 | -0.582 | -0.7645 |
| BPHL | NM_004332 | -0.3265 | -1.2945 |
| BRAF | NM_004333 | -0.4585 | -0.6405 |
| BST1 | NM_004334 | -0.8235 | -0.708 |
| BST2 | NM_004335 | 0.1065 | 0.0475 |
| BUB1 | NM_004336 | -1.484 | -2.998 |

| | | | |
|---|---|---|---|
| C18orf1 | NM_004338 | -0.1455 | -2.338 |
| PTTG1IP | NM_004339 | -0.088 | -0.753 |
| CAD | NM_004341 | -0.571 | -1.406 |
| CALD1 | NM_004342 | -0.6825 | -0.8615 |
| CALR | NM_004343 | -0.0315 | -0.5455 |
| CETN2 | NM_004344 | -0.3725 | -0.2745 |
| CAMP | NM_004345 | 0.128 | 0.1425 |
| CASP3 | NM_004346 | -1.112 | -2.1085 |
| CASP5 | NM_004347 | -0.085 | -0.228 |
| RUNX2 | NM_004348 | 0.37 | -0.2655 |
| RUNX1T1 | NM_004349 | -1.1215 | -2.571 |
| RUNX3 | NM_004350 | -0.0435 | -0.659 |
| CBLN1 | NM_004352 | -0.5905 | -1.5355 |
| CCNG2 | NM_004354 | -1.051 | -1.5245 |
| CD74 | NM_004355 | -0.1105 | -0.6615 |
| CD81 | NM_004356 | 0.186 | -0.198 |
| CD151 | NM_004357 | -0.2565 | -1.2915 |
| CDC25B | NM_004358 | -0.3235 | -1.0145 |
| CDC34 | NM_004359 | -0.4165 | -1.9645 |
| CDH1 | NM_004360 | 0.144 | 0.1925 |
| CDH7 | NM_004361 | 0.048 | -1.912 |
| CLGN | NM_004362 | -0.653 | -1.1445 |
| CEACAM5 | NM_004363 | 0.0455 | -0.269 |
| CEBPA | NM_004364 | -0.374 | 0.009 |
| CETN3 | NM_004365 | -2.003 | -2.5255 |
| CLCN2 | NM_004366 | -0.2475 | -0.5305 |
| CCR6 | NM_004367 | 0.1095 | -0.029 |
| CNN2 | NM_004368 | -0.067 | -0.928 |
| COL6A3 | NM_004369 | -0.2995 | -0.3935 |
| COL12A1 | NM_004370 | 0.2205 | 0.624 |
| COPA | NM_004371 | -0.594 | -0.618 |
| COX6A1 | NM_004373 | 0.248 | -0.1775 |
| COX6C | NM_004374 | -0.003 | -0.0785 |
| COX11 | NM_004375 | -1.2915 | -2.066 |
| COX15 | NM_004376 | -0.9595 | -1.182 |
| CRABP1 | NM_004378 | 0.3535 | -0.1665 |
| CREB1 | NM_004379 | -2.091 | -3.7615 |
| CREBBP | NM_004380 | -0.55 | -2.652 |
| ATF6B | NM_004381 | -0.2215 | -0.791 |
| CSK | NM_004383 | -0.28 | -1.5995 |
| CSNK1G3 | NM_004384 | -0.839 | -2.3445 |
| VCAN | NM_004385 | -0.48 | -1.449 |
| NCAN | NM_004386 | -0.6425 | -0.7315 |

| | | | |
|---|---|---|---|
| NKX2-5 | NM_004387 | 0.061 | -0.2805 |
| CTBS | NM_004388 | -0.6175 | -1.2095 |
| CTNNA2 | NM_004389 | -0.583 | -0.399 |
| CTSH | NM_004390 | 0.004 | -0.5165 |
| CYP8B1 | NM_004391 | -0.2335 | -0.387 |
| DACH1 | NM_004392 | -1.0525 | -2.4995 |
| DAG1 | NM_004393 | -0.512 | -1.5425 |
| DAP | NM_004394 | -0.2215 | -0.7095 |
| DBN1 | NM_004395 | -0.361 | -1.3155 |
| DDX5 | NM_004396 | -0.954 | -0.7895 |
| DDX6 | NM_004397 | -1.041 | -1.294 |
| DDX10 | NM_004398 | -0.972 | -2.563 |
| DDX11 | NM_004399 | -0.204 | -1.65 |
| DFFA | NM_004401 | -0.599 | -1.1795 |
| DFFB | NM_004402 | 0.23 | -0.9445 |
| 2-Sep | NM_004404 | -0.889 | -0.366 |
| DLX2 | NM_004405 | -0.076 | -0.4425 |
| DMBT1 | NM_004406 | -0.2315 | -0.2675 |
| DMP1 | NM_004407 | -0.186 | -0.0255 |
| DNM1 | NM_004408 | -0.395 | -0.52 |
| DMPK | NM_004409 | -0.673 | -0.582 |
| DYNC1I1 | NM_004411 | -0.923 | 0.481 |
| TRDMT1 | NM_004412 | -1.028 | -1.8355 |
| DPEP1 | NM_004413 | -0.1255 | -0.213 |
| RCAN1 | NM_004414 | -0.58 | -0.9125 |
| DSP | NM_004415 | -0.057 | 0.0875 |
| DTX1 | NM_004416 | -1.0815 | -2.2195 |
| DUSP1 | NM_004417 | -0.077 | -0.0045 |
| DUSP2 | NM_004418 | -0.438 | -0.4615 |
| DUSP5 | NM_004419 | -0.28 | -1.5745 |
| DUSP8 | NM_004420 | -0.216 | -0.736 |
| DVL1 | NM_004421 | -0.441 | -1.3535 |
| DVL2 | NM_004422 | -0.336 | -1.827 |
| DVL3 | NM_004423 | -0.5305 | -0.976 |
| E4F1 | NM_004424 | -0.6125 | -2.2025 |
| ECM1 | NM_004425 | -0.205 | -0.624 |
| PHC1 | NM_004426 | -0.384 | -0.8015 |
| PHC2 | NM_004427 | -1.0245 | -2.472 |
| EFNA1 | NM_004428 | -0.516 | -0.4145 |
| EFNB1 | NM_004429 | -0.112 | -0.461 |
| EGR3 | NM_004430 | -0.1135 | -0.415 |
| EPHA2 | NM_004431 | -0.3645 | -0.293 |
| ELAVL2 | NM_004432 | -1.319 | -2.33 |

| | | | |
|---|---|---|---|
| ELF3 | NM_004433 | 0.209 | -0.283 |
| EML1 | NM_004434 | -0.534 | -1.474 |
| ENDOG | NM_004435 | -0.6575 | -1.681 |
| ENSA | NM_004436 | 0.1805 | -0.5175 |
| EPB41 | NM_004437 | -0.672 | -0.825 |
| EPHA4 | NM_004438 | -0.234 | -0.276 |
| EPHA5 | NM_004439 | -0.0215 | -2.903 |
| EPHA7 | NM_004440 | 0.459 | 1.0405 |
| EPHB1 | NM_004441 | 0.215 | -0.428 |
| EPHB2 | NM_004442 | -0.259 | -0.745 |
| EPHB3 | NM_004443 | -0.5475 | -0.769 |
| EPHB4 | NM_004444 | -0.0145 | -0.41 |
| EPHB6 | NM_004445 | -0.204 | -0.0515 |
| EPRS | NM_004446 | -0.0055 | -1.0505 |
| EPS8 | NM_004447 | -0.6465 | -1.8885 |
| ERBB2 | NM_004448 | 0.113 | -0.579 |
| ERG | NM_004449 | -0.1435 | -0.526 |
| ERH | NM_004450 | -0.4215 | -0.3455 |
| ESRRA | NM_004451 | -0.894 | -1.8835 |
| ESRRB | NM_004452 | 0.452 | -0.4045 |
| ETFDH | NM_004453 | -0.9205 | -1.684 |
| ETV5 | NM_004454 | -0.624 | -2.5055 |
| EXTL1 | NM_004455 | 0.2085 | 0.0115 |
| EZH2 | NM_004456 | -0.954 | -1.97 |
| ACSL3 | NM_004457 | -0.737 | -1.3815 |
| ACSL4 | NM_004458 | -0.283 | -1.4105 |
| BPTF | NM_004459 | -1.2375 | -2.0965 |
| FAP | NM_004460 | -0.453 | -0.179 |
| FARSA | NM_004461 | -1.161 | -2.889 |
| FDFT1 | NM_004462 | -0.3445 | -1.4045 |
| FGD1 | NM_004463 | -0.2505 | -1.147 |
| FGF5 | NM_004464 | 0.176 | -0.331 |
| FGF10 | NM_004465 | -0.206 | -1.2575 |
| GPC5 | NM_004466 | -0.1995 | -0.172 |
| FGL1 | NM_004467 | 0.0905 | -0.006 |
| FHL3 | NM_004468 | -0.588 | -1.0375 |
| FIGF | NM_004469 | -0.8455 | -2.0485 |
| FKBP2 | NM_004470 | 0.2615 | -0.4 |
| FOXD1 | NM_004472 | -0.231 | -1.13 |
| FOXE1 | NM_004473 | 0.1475 | 0.201 |
| FOXD2 | NM_004474 | 0.1955 | -0.182 |
| FLOT2 | NM_004475 | -0.086 | -0.7775 |
| FOLH1 | NM_004476 | -0.4335 | -0.138 |

| | | | |
|---|---|---|---|
| FRG1 | NM_004477 | -0.5555 | -1.4505 |
| FUT7 | NM_004479 | 0.154 | -0.521 |
| FUT8 | NM_004480 | -1.1135 | -2.375 |
| GALNT2 | NM_004481 | -0.099 | -0.9605 |
| GALNT3 | NM_004482 | -0.076 | -0.088 |
| GCSH | NM_004483 | -0.2965 | -0.065 |
| GPC3 | NM_004484 | -0.0175 | -0.2355 |
| GNG4 | NM_004485 | 0.1385 | -0.3365 |
| GOLGA2 | NM_004486 | -0.6235 | -1.151 |
| GOLGB1 | NM_004487 | -0.347 | -2.005 |
| GP5 | NM_004488 | -0.1485 | -0.317 |
| GPS2 | NM_004489 | -0.1885 | -0.8215 |
| GRB14 | NM_004490 | 0.248 | -0.6005 |
| ARHGAP35 | NM_004491 | -1.4245 | -2.648 |
| GTF2A2 | NM_004492 | -0.8735 | -0.9155 |
| HSD17B10 | NM_004493 | 0.353 | -0.217 |
| HDGF | NM_004494 | 0.248 | -0.3115 |
| NRG1 | NM_004495 | -0.1445 | -0.887 |
| FOXA1 | NM_004496 | -0.0195 | -0.381 |
| FOXA3 | NM_004497 | -0.0265 | 0.0885 |
| ONECUT1 | NM_004498 | -0.3585 | -0.492 |
| HNRNPAB | NM_004499 | -0.2745 | -1.0395 |
| HNRNPC | NM_004500 | -0.1685 | -0.3245 |
| HNRNPU | NM_004501 | -0.587 | -0.5255 |
| HOXB7 | NM_004502 | -0.23 | -0.277 |
| HOXC6 | NM_004503 | -0.3895 | -0.19 |
| AGFG1 | NM_004504 | -0.5755 | -1.4755 |
| USP6 | NM_004505 | -1.339 | -2.143 |
| HSF2 | NM_004506 | -1.4485 | -3.69 |
| HUS1 | NM_004507 | -0.5675 | -0.9855 |
| IDI1 | NM_004508 | -1.698 | -2.7515 |
| SP110 | NM_004509 | -0.213 | -0.626 |
| SP110 | NM_004510 | -0.213 | -0.626 |
| IL16 | NM_004513 | -0.113 | -0.7195 |
| FOXK2 | NM_004514 | -0.943 | -2.812 |
| ILF2 | NM_004515 | -0.177 | -0.4865 |
| ILF3 | NM_004516 | -0.0295 | -0.4595 |
| ILK | NM_004517 | 0.0575 | -0.811 |
| KCNQ2 | NM_004518 | -1.0055 | -3.149 |
| KCNQ3 | NM_004519 | -0.239 | -0.413 |
| KIF2A | NM_004520 | -0.9335 | -2.404 |
| KIF5B | NM_004521 | -0.168 | -1.7265 |
| KIF5C | NM_004522 | -0.707 | -0.7365 |

| | | | |
|---|---|---|---|
| KIF11 | NM_004523 | -1.1175 | -3.604 |
| LRP2 | NM_004525 | 0.2095 | -0.089 |
| MCM2 | NM_004526 | -0.0895 | -0.616 |
| MEOX1 | NM_004527 | 0.0855 | 0.1905 |
| MGST3 | NM_004528 | 0.248 | -0.259 |
| MLLT3 | NM_004529 | -0.7325 | -1.1185 |
| MMP2 | NM_004530 | 0.2795 | -0.31 |
| MOCS2 | NM_004531 | -0.753 | -1.7415 |
| MUC4 | NM_004532 | 0.0875 | -0.2915 |
| MYBPC2 | NM_004533 | -0.2945 | -0.2435 |
| MYT1 | NM_004535 | -1.4835 | -1.758 |
| NAIP | NM_004536 | -0.9625 | -0.931 |
| NAP1L1 | NM_004537 | -0.2335 | -0.25 |
| NAP1L3 | NM_004538 | -0.877 | -0.547 |
| NARS | NM_004539 | -0.825 | -0.568 |
| NCAM2 | NM_004540 | 0.0755 | -0.5525 |
| NDUFA1 | NM_004541 | 0.157 | -0.162 |
| NDUFA3 | NM_004542 | 0.35 | -0.031 |
| NEB | NM_004543 | -0.594 | -0.371 |
| NDUFA10 | NM_004544 | 0.1285 | -0.346 |
| NDUFB1 | NM_004545 | 0.171 | -0.3445 |
| NDUFB2 | NM_004546 | 0.3825 | -0.0005 |
| NDUFB4 | NM_004547 | -0.1255 | -0.642 |
| NDUFB10 | NM_004548 | 0.055 | -0.4285 |
| NDUFC2 | NM_004549 | 0.1625 | -0.2805 |
| NDUFS2 | NM_004550 | -0.3225 | -1.2365 |
| NDUFS3 | NM_004551 | 0.026 | -0.303 |
| NDUFS5 | NM_004552 | 0.116 | -0.216 |
| NDUFS6 | NM_004553 | 0.2355 | -0.1335 |
| NFATC4 | NM_004554 | -0.1935 | -0.7425 |
| NFATC3 | NM_004555 | -1.19 | -2.157 |
| NFKBIE | NM_004556 | -0.9005 | -2.2485 |
| NOTCH4 | NM_004557 | -0.0855 | -0.684 |
| NRTN | NM_004558 | 0.0055 | -0.5745 |
| YBX1 | NM_004559 | 0.0065 | -0.1795 |
| ROR2 | NM_004560 | -0.596 | -2.6745 |
| OVOL1 | NM_004561 | -0.026 | -0.176 |
| PARK2 | NM_004562 | -0.0205 | -0.377 |
| PCK2 | NM_004563 | -0.545 | -1.3295 |
| PET112 | NM_004564 | -0.629 | -1.45 |
| PEX14 | NM_004565 | -0.7775 | -2.0705 |
| PFKFB3 | NM_004566 | -0.3905 | -2.155 |
| PFKFB4 | NM_004567 | -1.2355 | -1.8135 |

| | | | |
|---|---|---|---|
| SERPINB6 | NM_004568 | -0.41 | -0.8315 |
| PIGH | NM_004569 | -0.6135 | -0.8675 |
| PIK3C2G | NM_004570 | -0.083 | -0.265 |
| PKNOX1 | NM_004571 | -0.797 | -2.1875 |
| PKP2 | NM_004572 | -1.0725 | -0.488 |
| PLCB2 | NM_004573 | -0.169 | -0.2255 |
| 4-Sep | NM_004574 | 0.015 | -0.14 |
| POU4F2 | NM_004575 | -0.4365 | 0.245 |
| PPP2R2B | NM_004576 | -1.281 | -1.527 |
| PSPH | NM_004577 | -0.4615 | -1.314 |
| RAB4A | NM_004578 | -0.3345 | -0.858 |
| MAP4K2 | NM_004579 | 0.307 | -0.249 |
| RAB27A | NM_004580 | -0.468 | -1.317 |
| RABGGTA | NM_004581 | -0.1685 | -0.7645 |
| RABGGTB | NM_004582 | -1.83 | -2.905 |
| RAB5C | NM_004583 | 0.0955 | -0.6495 |
| RAD9A | NM_004584 | -1.096 | -1.762 |
| RARRES3 | NM_004585 | -0.0425 | -0.1055 |
| RPS6KA3 | NM_004586 | -1.3435 | -2.3055 |
| RRBP1 | NM_004587 | -0.296 | -0.8495 |
| SCN2B | NM_004588 | -0.0885 | -0.592 |
| SCO1 | NM_004589 | -0.304 | -1.368 |
| CCL16 | NM_004590 | -0.3005 | 0.3445 |
| CCL20 | NM_004591 | -0.173 | -0.025 |
| SFSWAP | NM_004592 | -0.9875 | -2.2325 |
| TRA2B | NM_004593 | -0.809 | -1.2385 |
| SLC9A5 | NM_004594 | -0.4285 | -0.8825 |
| SMS | NM_004595 | -0.3315 | -0.1075 |
| SNRPA | NM_004596 | -0.137 | -0.686 |
| SNRPD2 | NM_004597 | 0.221 | -0.117 |
| SPOCK1 | NM_004598 | -0.4825 | -0.0965 |
| SREBF2 | NM_004599 | -0.0125 | -0.683 |
| TROVE2 | NM_004600 | -0.8535 | -1.218 |
| STAU1 | NM_004602 | -0.7415 | -1.708 |
| STX1A | NM_004603 | -0.2475 | -0.9095 |
| STX4 | NM_004604 | -0.95 | -2.176 |
| TAF1 | NM_004606 | -1.2275 | -1.719 |
| TBCA | NM_004607 | -0.305 | -0.498 |
| TBX6 | NM_004608 | 0.05 | -0.566 |
| TCF15 | NM_004609 | -0.407 | -1.413 |
| TCP10 | NM_004610 | -0.27 | -0.2865 |
| TGFBR1 | NM_004612 | -0.846 | -1.0815 |
| TGM2 | NM_004613 | -0.1535 | -0.4795 |

| | | | |
|---|---|---|---|
| TK2 | NM_004614 | -0.4315 | -1.081 |
| TSPAN7 | NM_004615 | 0.045 | -0.249 |
| TSPAN8 | NM_004616 | 0.026 | 0.009 |
| TM4SF4 | NM_004617 | -0.097 | -0.1975 |
| TOP3A | NM_004618 | -0.6995 | -1.6355 |
| TRAF5 | NM_004619 | -1.117 | -1.329 |
| TRAF6 | NM_004620 | -1.2975 | -2.657 |
| TRPC6 | NM_004621 | 0.459 | 0.163 |
| TSN | NM_004622 | -0.8505 | -0.748 |
| TTC4 | NM_004623 | -1.2445 | -4.385 |
| VIPR1 | NM_004624 | 0.117 | -0.0895 |
| WNT7A | NM_004625 | 0.024 | -0.1525 |
| WNT11 | NM_004626 | -0.3485 | -2.2995 |
| WRB | NM_004627 | -1.1845 | -1.306 |
| XPC | NM_004628 | -1.0635 | -2.6125 |
| FANCG | NM_004629 | -0.357 | -1.5635 |
| SF1 | NM_004630 | -0.1995 | -1.228 |
| LRP8 | NM_004631 | -1.334 | -2.2045 |
| DAP3 | NM_004632 | -0.873 | -1.247 |
| IL1R2 | NM_004633 | -0.0255 | -0.209 |
| BRPF1 | NM_004634 | -0.693 | -2.1945 |
| MAPKAPK3 | NM_004635 | -0.147 | -0.8605 |
| SEMA3B | NM_004636 | -0.146 | -0.431 |
| RAB7A | NM_004637 | -0.298 | -1.124 |
| BAG6 | NM_004639 | 0.246 | -0.368 |
| DDX39B | NM_004640 | -0.0445 | -0.3975 |
| MLLT10 | NM_004641 | -1.468 | -2.454 |
| CDK2AP1 | NM_004642 | -0.3225 | -0.2635 |
| PABPN1 | NM_004643 | -0.198 | -1.0915 |
| AP3B2 | NM_004644 | -0.1565 | -0.759 |
| COIL | NM_004645 | -1.6145 | -3.1645 |
| NPHS1 | NM_004646 | -0.16 | -0.0015 |
| DPF1 | NM_004647 | -0.418 | -1.191 |
| C21orf33 | NM_004649 | -0.362 | -0.843 |
| PNPLA4 | NM_004650 | -0.3905 | -0.8045 |
| USP11 | NM_004651 | 0.0095 | -0.368 |
| KDM5D | NM_004653 | 0.0725 | -1.4715 |
| USP9Y | NM_004654 | -0.063 | -1.5645 |
| AXIN2 | NM_004655 | -0.645 | -1.0605 |
| BAP1 | NM_004656 | -0.3135 | -0.8965 |
| SDPR | NM_004657 | -0.046 | -0.347 |
| RASAL1 | NM_004658 | -0.1145 | 0.1045 |
| DDX3Y | NM_004660 | -0.423 | -0.929 |

| | | | |
|---|---|---|---|
| CDC23 | NM_004661 | -1.606 | -2.9445 |
| DNAH9 | NM_004662 | 0.234 | -0.2265 |
| RAB11A | NM_004663 | -0.3755 | -0.5395 |
| LIN7A | NM_004664 | -0.536 | -1.2495 |
| VNN2 | NM_004665 | -0.0095 | 0.0735 |
| VNN1 | NM_004666 | 0.0945 | -0.071 |
| HERC2 | NM_004667 | -0.7205 | -1.328 |
| MGAM | NM_004668 | 0.169 | -0.801 |
| CLIC3 | NM_004669 | -0.384 | 0.0785 |
| PAPSS2 | NM_004670 | -0.3295 | -1.385 |
| PIAS2 | NM_004671 | -0.739 | -0.86 |
| MAP3K6 | NM_004672 | 0.0175 | -0.182 |
| ANGPTL1 | NM_004673 | -0.0355 | 0.0785 |
| ASH2L | NM_004674 | -0.946 | -1.8355 |
| DIRAS3 | NM_004675 | -0.6185 | -0.466 |
| BPY2 | NM_004678 | 0.28 | 0.4805 |
| EIF1AY | NM_004681 | -0.1605 | -0.313 |
| RGN | NM_004683 | 0.3585 | -0.3015 |
| SPARCL1 | NM_004684 | -0.022 | -0.3485 |
| MTMR6 | NM_004685 | -1.302 | -2.523 |
| MTMR7 | NM_004686 | -0.5095 | -0.655 |
| MTMR4 | NM_004687 | -0.9055 | -2.09 |
| NMI | NM_004688 | -0.0665 | -0.6695 |
| MTA1 | NM_004689 | -0.1085 | -0.587 |
| LATS1 | NM_004690 | 0.3205 | -0.901 |
| ATP6V0D1 | NM_004691 | -0.0445 | -0.8915 |
| KRT75 | NM_004693 | -0.0735 | -0.561 |
| SLC16A6 | NM_004694 | -1.038 | -1.1295 |
| SLC16A5 | NM_004695 | -0.014 | -0.553 |
| SLC16A4 | NM_004696 | 0.437 | -0.4225 |
| PRPF4 | NM_004697 | -1.297 | -3.679 |
| PRPF3 | NM_004698 | -1.5175 | -2.714 |
| FAM50A | NM_004699 | -0.0635 | -0.6285 |
| KCNQ4 | NM_004700 | -0.6435 | -1.578 |
| CCNB2 | NM_004701 | -0.6895 | -1.5125 |
| RABEP1 | NM_004703 | -0.654 | -2.245 |
| RRP9 | NM_004704 | -0.474 | -1.3175 |
| PRKRIR | NM_004705 | -0.827 | -1.02 |
| ARHGEF1 | NM_004706 | -0.14 | -0.908 |
| ATG12 | NM_004707 | -1.3315 | -1.6715 |
| PDCD5 | NM_004708 | -0.3495 | -0.9275 |
| CXorf1 | NM_004709 | -0.106 | -0.4195 |
| SYNGR2 | NM_004710 | -0.348 | -1.9425 |

| | | | |
|---|---|---|---|
| SYNGR1 | NM_004711 | -0.0035 | -0.574 |
| HGS | NM_004712 | -0.0295 | -0.5405 |
| DYRK1B | NM_004714 | -0.123 | -0.4535 |
| CTDP1 | NM_004715 | -0.689 | -2.1235 |
| PCSK7 | NM_004716 | -0.423 | -1.4075 |
| DGKI | NM_004717 | -0.4745 | -0.7855 |
| COX7A2L | NM_004718 | 0.271 | -0.4335 |
| SCAF11 | NM_004719 | -0.5605 | -2.1695 |
| LPAR2 | NM_004720 | -0.4685 | -1.146 |
| MAP3K13 | NM_004721 | -0.6495 | -1.591 |
| AP4M1 | NM_004722 | -0.687 | -1.8 |
| ARHGEF2 | NM_004723 | -0.355 | -2.111 |
| ZW10 | NM_004724 | -1.5285 | -2.461 |
| BUB3 | NM_004725 | -0.8095 | -0.875 |
| REPS2 | NM_004726 | 0.017 | -0.4125 |
| SLC24A1 | NM_004727 | -0.66 | -1.394 |
| DDX21 | NM_004728 | -1.028 | -1.26 |
| ZBED1 | NM_004729 | -0.0825 | -1.462 |
| ETF1 | NM_004730 | -1.871 | -2.959 |
| SLC16A7 | NM_004731 | -0.451 | -1.6525 |
| KCNAB3 | NM_004732 | -0.5235 | -0.9065 |
| SLC33A1 | NM_004733 | -1.543 | -2.122 |
| DCLK1 | NM_004734 | -1.1205 | -1.1045 |
| LRRFIP1 | NM_004735 | -1.425 | -2.078 |
| XPR1 | NM_004736 | -0.751 | -0.929 |
| LARGE | NM_004737 | -0.2875 | -1.2755 |
| VAPB | NM_004738 | -0.3075 | -1.7435 |
| MTA2 | NM_004739 | 0.045 | -0.4825 |
| TIAF1 | NM_004740 | -0.313 | -0.564 |
| NOLC1 | NM_004741 | -0.681 | -0.5135 |
| MAGI1 | NM_004742 | -1.3115 | -3.687 |
| LRAT | NM_004744 | 0.226 | 0.1025 |
| DLGAP2 | NM_004745 | 0.0575 | -0.44 |
| DLGAP1 | NM_004746 | -0.322 | -0.227 |
| DLG5 | NM_004747 | -0.3995 | -0.98 |
| CCPG1 | NM_004748 | -0.52 | -1.423 |
| CRLF1 | NM_004750 | 0.0215 | -0.4695 |
| GCNT3 | NM_004751 | 0.171 | -0.359 |
| GCM2 | NM_004752 | -0.419 | -0.198 |
| DHRS3 | NM_004753 | 0.3775 | 0.124 |
| RPS6KA5 | NM_004755 | -0.7935 | -1.922 |
| NUMBL | NM_004756 | -1.1525 | -2.865 |
| AIMP1 | NM_004757 | -0.4575 | -0.4495 |

| | | | |
|---|---|---|---|
| BZRAP1 | NM_004758 | 0.1195 | -0.5245 |
| MAPKAPK2 | NM_004759 | 0.19 | -1.023 |
| STK17A | NM_004760 | -0.6455 | -1.785 |
| RGL2 | NM_004761 | -0.0725 | -0.774 |
| CYTH1 | NM_004762 | -0.4665 | -1.849 |
| ITGB1BP1 | NM_004763 | -0.568 | -0.5855 |
| PIWIL1 | NM_004764 | -0.097 | -0.1975 |
| BCL7C | NM_004765 | 0.3365 | -0.3765 |
| COPB2 | NM_004766 | -0.8695 | -1.5 |
| GPR37L1 | NM_004767 | -0.0285 | 0.0285 |
| SRSF11 | NM_004768 | -1.044 | -1.3515 |
| ACCN3 | NM_004769 | -0.226 | -0.094 |
| KCNB2 | NM_004770 | -0.1895 | -1.3255 |
| MMP20 | NM_004771 | -0.101 | -0.0925 |
| C5orf13 | NM_004772 | -0.2675 | -0.438 |
| ZNHIT3 | NM_004773 | -0.869 | -2.0875 |
| MED1 | NM_004774 | -1.8645 | -2.349 |
| B4GALT6 | NM_004775 | -1.6055 | -2.6635 |
| B4GALT5 | NM_004776 | -0.6595 | -2.056 |
| GPR44 | NM_004778 | -0.276 | -0.7965 |
| CNOT8 | NM_004779 | -0.9575 | -1.51 |
| TCEAL1 | NM_004780 | -0.9445 | -1.035 |
| VAMP3 | NM_004781 | -0.7705 | -1.5855 |
| SNAP29 | NM_004782 | -2.628 | -1.695 |
| TAOK2 | NM_004783 | -0.31 | -0.8235 |
| NDST3 | NM_004784 | 0.1345 | -0.322 |
| SLC9A3R2 | NM_004785 | 0.049 | -0.5035 |
| TXNL1 | NM_004786 | -0.9355 | -1.635 |
| SLIT2 | NM_004787 | -0.932 | -1.7625 |
| UBE4A | NM_004788 | -1.402 | -1.59 |
| LHX2 | NM_004789 | -0.145 | -0.13 |
| SLC22A6 | NM_004790 | 0.128 | 0.005 |
| ITGBL1 | NM_004791 | -0.115 | -0.0005 |
| PPIG | NM_004792 | 0.1205 | -0.688 |
| LONP1 | NM_004793 | -0.5855 | -1.278 |
| RAB33A | NM_004794 | -0.9075 | -2.2365 |
| KL | NM_004795 | 0.1675 | 0.1955 |
| NRXN3 | NM_004796 | -0.261 | -0.418 |
| ADIPOQ | NM_004797 | -0.0225 | -0.371 |
| KIF3B | NM_004798 | -1.079 | -2.3875 |
| ZFYVE9 | NM_004799 | -0.7965 | -1.929 |
| TM9SF2 | NM_004800 | -0.926 | -1.016 |
| NRXN1 | NM_004801 | -0.3315 | -0.859 |

| | | | |
|---|---|---|---|
| OTOF | NM_004802 | -0.2305 | -0.381 |
| SLC22A14 | NM_004803 | -0.102 | -0.21 |
| CIAO1 | NM_004804 | -1.0795 | -1.1425 |
| POLR2D | NM_004805 | -0.718 | -2.5655 |
| HS6ST1 | NM_004807 | -0.7145 | -1.8835 |
| NMT2 | NM_004808 | -0.2785 | -0.797 |
| STOML1 | NM_004809 | -0.668 | -1.2375 |
| GRAP2 | NM_004810 | 0.0515 | -0.2475 |
| LPXN | NM_004811 | -0.78 | -1.0555 |
| PEX16 | NM_004813 | -0.234 | -1.1585 |
| SNRNP40 | NM_004814 | -0.853 | -1.7535 |
| ARHGAP29 | NM_004815 | -0.418 | -3.218 |
| FAM189A2 | NM_004816 | -0.269 | -0.269 |
| TJP2 | NM_004817 | -0.9095 | -1.5395 |
| DDX23 | NM_004818 | -0.6785 | -1.8335 |
| SYMPK | NM_004819 | -0.364 | -0.8225 |
| CYP7B1 | NM_004820 | -0.3365 | -0.506 |
| HAND1 | NM_004821 | -0.49 | -2.5965 |
| NTN1 | NM_004822 | 0.0135 | -0.152 |
| KCNK6 | NM_004823 | 0.039 | 0.136 |
| CDYL | NM_004824 | -1.0205 | -0.886 |
| CDY2A | NM_004825 | 0.519 | -0.682 |
| ECEL1 | NM_004826 | 0.159 | -0.098 |
| ABCG2 | NM_004827 | -0.3485 | -0.1175 |
| NCR2 | NM_004828 | 0.114 | -0.0945 |
| NCR1 | NM_004829 | -0.3495 | -0.467 |
| MED23 | NM_004830 | -1.215 | -2.344 |
| MED26 | NM_004831 | -0.341 | -2.5785 |
| GSTO1 | NM_004832 | 0.0025 | -0.721 |
| AIM2 | NM_004833 | 0.533 | -0.793 |
| MAP4K4 | NM_004834 | -0.341 | -0.495 |
| AGTR1 | NM_004835 | -0.793 | -0.37 |
| EIF2AK3 | NM_004836 | -1.2335 | -3.903 |
| HOMER3 | NM_004838 | -0.0835 | -0.867 |
| HOMER2 | NM_004839 | 0.214 | -0.203 |
| ARHGEF6 | NM_004840 | -0.4145 | -0.603 |
| RASAL2 | NM_004841 | -0.3805 | -1.5385 |
| AKAP7 | NM_004842 | 0.197 | -0.225 |
| IL27RA | NM_004843 | -0.3945 | -0.7205 |
| SH3BP5 | NM_004844 | -0.358 | -0.844 |
| PCYT1B | NM_004845 | -0.7035 | -1.3885 |
| EIF4E2 | NM_004846 | 0.0155 | -0.5975 |
| AIF1 | NM_004847 | -0.1545 | -0.3475 |

| | | | |
|---|---|---|---|
| C1orf38 | NM_004848 | -0.032 | -0.253 |
| ATG5 | NM_004849 | -1.274 | -2.794 |
| ROCK2 | NM_004850 | -0.2765 | -2.3295 |
| NAPSA | NM_004851 | 0.022 | -0.619 |
| ONECUT2 | NM_004852 | -0.88 | -1.708 |
| STX8 | NM_004853 | -0.3485 | -0.0785 |
| CHST10 | NM_004854 | -0.618 | -2.295 |
| PIGB | NM_004855 | -1.536 | -2.2865 |
| KIF23 | NM_004856 | -1.2105 | -2.492 |
| AKAP5 | NM_004857 | -1.0795 | -1.381 |
| SLC4A8 | NM_004858 | 1.0945 | 0.3425 |
| CLTC | NM_004859 | -0.6085 | -1.304 |
| FXR2 | NM_004860 | -0.445 | -1.1855 |
| GAL3ST1 | NM_004861 | 0.0255 | -0.488 |
| LITAF | NM_004862 | -0.104 | -1.0455 |
| SPTLC2 | NM_004863 | -1.3195 | -1.8845 |
| GDF15 | NM_004864 | 0.1855 | 0.262 |
| TBPL1 | NM_004865 | -1.3185 | -3.6165 |
| SCAMP1 | NM_004866 | -1.017 | -1.4425 |
| ITM2A | NM_004867 | 0.0705 | 0.0155 |
| VPS4B | NM_004869 | -1.622 | -3.71 |
| MPDU1 | NM_004870 | -0.0295 | -0.893 |
| GOSR1 | NM_004871 | -1.479 | -2.7135 |
| TMEM59 | NM_004872 | -0.874 | -0.78 |
| BAG5 | NM_004873 | -0.861 | -0.89 |
| BAG4 | NM_004874 | -1.3245 | -2.9755 |
| GMFG | NM_004877 | -0.03 | -0.1385 |
| PTGES | NM_004878 | -0.116 | -0.435 |
| EI24 | NM_004879 | 0.1685 | -0.3725 |
| TP53I3 | NM_004881 | -0.4655 | -1.214 |
| CIR1 | NM_004882 | -0.3705 | -1.783 |
| NRG2 | NM_004883 | -0.0085 | -0.3145 |
| IGDCC3 | NM_004884 | 0.2075 | -1.672 |
| NPFFR2 | NM_004885 | 0.664 | -0.5265 |
| APBA3 | NM_004886 | -0.534 | -1.354 |
| CXCL14 | NM_004887 | 0.0545 | -0.6735 |
| ATP6V1G1 | NM_004888 | -0.912 | -0.464 |
| ATP5J2 | NM_004889 | 0.2425 | -0.187 |
| SPAG7 | NM_004890 | -0.4155 | -1.1005 |
| MRPL33 | NM_004891 | -0.124 | -0.4135 |
| SEC22B | NM_004892 | -0.921 | -1.336 |
| H2AFY | NM_004893 | 0.04 | -0.4995 |
| C14orf2 | NM_004894 | 0.2335 | -0.119 |

| | | | |
|---|---|---|---|
| NLRP3 | NM_004895 | -0.0565 | 0.0615 |
| VPS26A | NM_004896 | -0.4455 | -1.8035 |
| MINPP1 | NM_004897 | -1.683 | -4.1155 |
| CLOCK | NM_004898 | -0.7565 | -2.0625 |
| APOBEC3B | NM_004900 | 0.2585 | -0.263 |
| ENTPD4 | NM_004901 | -1.1605 | -2.0365 |
| RBM39 | NM_004902 | -1.0695 | -2.1315 |
| CREB5 | NM_004904 | -0.219 | -1.026 |
| PRDX6 | NM_004905 | 0.2105 | -0.2605 |
| WTAP | NM_004906 | -1.4565 | -1.782 |
| IER2 | NM_004907 | 0.918 | 2.093 |
| PITPNM1 | NM_004910 | -0.0625 | -0.589 |
| PDIA4 | NM_004911 | -0.513 | -1.668 |
| KRIT1 | NM_004912 | -1.236 | -1.9075 |
| C16orf7 | NM_004913 | -0.508 | -0.5985 |
| RAB36 | NM_004914 | -1.2665 | -1.109 |
| ABCG1 | NM_004915 | -0.1035 | -0.8765 |
| KLK4 | NM_004917 | 0.088 | -0.247 |
| TCL1B | NM_004918 | -0.0765 | 0.089 |
| SEC24C | NM_004922 | -0.3155 | -1.7295 |
| MTL5 | NM_004923 | -0.734 | -0.2215 |
| ACTN4 | NM_004924 | -0.033 | -0.8325 |
| AQP3 | NM_004925 | 0.4725 | 1.5675 |
| ZFP36L1 | NM_004926 | -0.349 | -0.4295 |
| MRPL49 | NM_004927 | -0.973 | -2.6095 |
| C21orf2 | NM_004928 | -0.576 | -0.539 |
| CALB1 | NM_004929 | -0.325 | -0.8865 |
| CAPZB | NM_004930 | 0.034 | -0.666 |
| CD8B | NM_004931 | 0.0235 | 0.1305 |
| CDH6 | NM_004932 | -0.076 | -0.4285 |
| CDH15 | NM_004933 | 0.0925 | -0.695 |
| CDH18 | NM_004934 | -0.238 | -0.5555 |
| CDK5 | NM_004935 | -0.0365 | -0.6165 |
| CDKN2B | NM_004936 | 0.0735 | 0.1195 |
| CTNS | NM_004937 | -1.33 | -2.5025 |
| DAPK1 | NM_004938 | -1.4585 | -2.8495 |
| DDX1 | NM_004939 | -0.6745 | -0.5465 |
| DHX8 | NM_004941 | -0.726 | -2.066 |
| DEFB4A | NM_004942 | -0.021 | -0.1015 |
| DMWD | NM_004943 | -0.1405 | -0.516 |
| DNASE1L3 | NM_004944 | 0.6405 | 0.2015 |
| DNM2 | NM_004945 | -0.1715 | -0.9265 |
| DOCK2 | NM_004946 | 0.126 | -0.4395 |

| | | | |
|---|---|---|---|
| DOCK3 | NM_004947 | -0.8515 | -0.1985 |
| DSC1 | NM_004948 | -0.114 | -0.114 |
| DSC2 | NM_004949 | -0.9115 | -0.5975 |
| EPYC | NM_004950 | -0.1135 | -0.4425 |
| GPR183 | NM_004951 | -0.1505 | -0.365 |
| EFNA3 | NM_004952 | -0.454 | -0.408 |
| EIF4G1 | NM_004953 | -0.1565 | -0.948 |
| MARK2 | NM_004954 | -0.758 | -2.485 |
| SLC29A1 | NM_004955 | 0.1325 | -0.6855 |
| ETV1 | NM_004956 | -1.24 | -2.281 |
| FPGS | NM_004957 | -0.0305 | -0.629 |
| MTOR | NM_004958 | -0.882 | -1.42 |
| NR5A1 | NM_004959 | 0.1715 | -0.0985 |
| FUS | NM_004960 | 0.078 | -0.447 |
| GABRE | NM_004961 | -0.015 | -0.5085 |
| GDF10 | NM_004962 | -0.2975 | -0.53 |
| GUCY2C | NM_004963 | -0.053 | 0.1555 |
| HDAC1 | NM_004964 | -0.145 | -0.3245 |
| HMGN1 | NM_004965 | -0.544 | -0.471 |
| HNRNPF | NM_004966 | -0.7185 | -2.124 |
| IBSP | NM_004967 | -0.1395 | -0.025 |
| ICA1 | NM_004968 | -0.608 | -1.057 |
| IDE | NM_004969 | -0.728 | -1.5025 |
| IGFALS | NM_004970 | -0.024 | -0.2015 |
| JAK2 | NM_004972 | -0.4355 | -1.2425 |
| JARID2 | NM_004973 | -0.5345 | -1.5335 |
| KCNA2 | NM_004974 | -0.035 | 0.022 |
| KCNB1 | NM_004975 | -1.1115 | -1.5775 |
| KCNC1 | NM_004976 | -0.449 | -1.628 |
| KCNC3 | NM_004977 | -0.094 | -0.533 |
| KCNC4 | NM_004978 | -0.2685 | -0.906 |
| KCND1 | NM_004979 | -0.251 | -0.0555 |
| KCND3 | NM_004980 | -0.308 | -0.221 |
| KCNJ4 | NM_004981 | -0.291 | -0.247 |
| KCNJ8 | NM_004982 | -1.691 | -3.133 |
| KCNJ9 | NM_004983 | -0.0025 | -0.2605 |
| KIF5A | NM_004984 | -0.367 | -1.4055 |
| KRAS | NM_004985 | -1.1215 | -2.1625 |
| KTN1 | NM_004986 | 0.2755 | -1.327 |
| LIMS1 | NM_004987 | -0.657 | -0.715 |
| MAGEA1 | NM_004988 | -0.2285 | -0.4875 |
| MARS | NM_004990 | -0.323 | -1.6505 |
| MECOM | NM_004991 | -0.401 | -0.379 |

| | | | |
|---|---|---|---|
| MECP2 | NM_004992 | -0.787 | -1.64 |
| ATXN3 | NM_004993 | -1.768 | -2.82 |
| MMP9 | NM_004994 | 0.1165 | -0.1945 |
| MMP14 | NM_004995 | 0.013 | -0.4985 |
| ABCC1 | NM_004996 | -0.2555 | -1.302 |
| MYBPH | NM_004997 | 0.028 | -0.508 |
| MYO1E | NM_004998 | 0.0625 | -0.516 |
| MYO6 | NM_004999 | -0.4295 | -0.748 |
| NDUFA5 | NM_005000 | -0.5935 | -0.617 |
| NDUFA7 | NM_005001 | 0.1415 | -0.4215 |
| NDUFA9 | NM_005002 | -0.0995 | -0.7275 |
| NDUFAB1 | NM_005003 | -0.143 | -0.675 |
| NDUFB8 | NM_005004 | 0.255 | -0.287 |
| NDUFB9 | NM_005005 | 0.2485 | -0.173 |
| NDUFS1 | NM_005006 | -0.957 | -1.692 |
| NFKBIL1 | NM_005007 | -0.7705 | -1.652 |
| NHP2L1 | NM_005008 | 0.016 | -0.547 |
| NME4 | NM_005009 | 0.259 | -0.4015 |
| NRCAM | NM_005010 | -1.005 | -0.5705 |
| NRF1 | NM_005011 | -0.9375 | -3.32 |
| ROR1 | NM_005012 | -0.561 | -2.197 |
| NUCB2 | NM_005013 | -0.1515 | -0.772 |
| OMD | NM_005014 | -0.089 | -0.46 |
| OXA1L | NM_005015 | -0.183 | -1.211 |
| PCBP2 | NM_005016 | 0.152 | -0.2755 |
| PCYT1A | NM_005017 | -1.7185 | -2.634 |
| PDCD1 | NM_005018 | 0.0415 | -0.259 |
| PDE1A | NM_005019 | -0.0115 | -0.9385 |
| PDE1C | NM_005020 | -0.003 | -0.273 |
| ENPP3 | NM_005021 | 0.1025 | -0.378 |
| PFN1 | NM_005022 | 0.357 | -0.005 |
| PGGT1B | NM_005023 | -0.579 | -1.5535 |
| SERPINB10 | NM_005024 | -0.3855 | -0.1095 |
| SERPINI1 | NM_005025 | -0.413 | -0.1745 |
| PIK3CD | NM_005026 | 0.118 | -0.7065 |
| PIK3R2 | NM_005027 | 0.1355 | -0.529 |
| PIP4K2A | NM_005028 | -0.74 | -1.937 |
| PITX3 | NM_005029 | -0.003 | -0.105 |
| PLK1 | NM_005030 | -0.6705 | -1.2735 |
| FXYD1 | NM_005031 | 0.3475 | -0.088 |
| PLS3 | NM_005032 | -0.444 | -0.2105 |
| EXOSC9 | NM_005033 | -1.131 | -2.6995 |
| POLR2K | NM_005034 | -0.719 | -0.325 |

| | | | |
|---|---|---|---|
| POLRMT | NM_005035 | -0.4425 | -1.208 |
| PPARA | NM_005036 | 0.0355 | -0.1175 |
| PPARG | NM_005037 | -0.758 | -1.625 |
| PPID | NM_005038 | -1.023 | -1.678 |
| PRB1 | NM_005039 | 0.0585 | 0.193 |
| PRCP | NM_005040 | -0.1975 | -0.6945 |
| PRF1 | NM_005041 | 0.172 | -0.109 |
| PRH2 | NM_005042 | -0.1825 | -0.4995 |
| PRKX | NM_005044 | -0.8615 | -1.6965 |
| RELN | NM_005045 | -0.3295 | -0.663 |
| KLK7 | NM_005046 | 0.075 | -0.4105 |
| PSMD5 | NM_005047 | -1.715 | -1.8975 |
| PTH2R | NM_005048 | -0.7325 | -0.5415 |
| PWP2 | NM_005049 | -0.4145 | -2.368 |
| ABCD4 | NM_005050 | -0.44 | -1.802 |
| QARS | NM_005051 | 0.1775 | -0.392 |
| RAC3 | NM_005052 | 0.3105 | -0.3255 |
| RAD23A | NM_005053 | -0.107 | -1.275 |
| RAPSN | NM_005055 | -0.0925 | -0.2695 |
| RBBP5 | NM_005057 | -0.7855 | -1.6135 |
| RLN2 | NM_005059 | -0.324 | 0.1365 |
| RORC | NM_005060 | 0.021 | -0.4155 |
| RPL3L | NM_005061 | 0.094 | 0.181 |
| SCD | NM_005063 | -0.506 | -0.6665 |
| CCL23 | NM_005064 | -0.337 | -0.426 |
| SEL1L | NM_005065 | -1.5285 | -3.0275 |
| SFPQ | NM_005066 | -0.3265 | -0.989 |
| SIAH2 | NM_005067 | -0.8 | -3.0015 |
| SIM1 | NM_005068 | -0.26 | -0.613 |
| SIM2 | NM_005069 | 0.469 | -0.227 |
| SLC4A3 | NM_005070 | -0.503 | -0.686 |
| SLC1A6 | NM_005071 | 0.304 | -0.246 |
| SLC12A4 | NM_005072 | 0.111 | -0.6285 |
| SLC15A1 | NM_005073 | -0.094 | -0.3515 |
| SLC17A1 | NM_005074 | -0.0975 | 0.047 |
| CNTN2 | NM_005076 | -0.572 | -0.6895 |
| TLE1 | NM_005077 | -0.4115 | -1.9615 |
| TLE3 | NM_005078 | -0.616 | -2.0775 |
| TPD52 | NM_005079 | -0.9245 | -0.882 |
| XBP1 | NM_005080 | -1.467 | -3.776 |
| TRIM25 | NM_005082 | -0.2165 | -0.8705 |
| PLA2G7 | NM_005084 | -0.4225 | 0.139 |
| NUP214 | NM_005085 | -0.5715 | -2.165 |

| | | | |
|---|---|---|---|
| SSPN | NM_005086 | -1.1245 | -1.207 |
| FXR1 | NM_005087 | -0.8035 | -1.643 |
| AKAP17A | NM_005088 | -0.198 | -0.887 |
| ZRSR2 | NM_005089 | -0.599 | -0.9265 |
| PGLYRP1 | NM_005091 | -0.15 | -0.323 |
| TNFSF18 | NM_005092 | -0.2895 | -0.016 |
| CBFA2T2 | NM_005093 | -0.7565 | -2.345 |
| SLC27A4 | NM_005094 | -0.5315 | -1.6775 |
| ZMYM4 | NM_005095 | -0.83 | -2.0545 |
| ZMYM3 | NM_005096 | -0.354 | -0.9715 |
| LGI1 | NM_005097 | 0.241 | -0.551 |
| MSC | NM_005098 | -0.517 | -0.5815 |
| ADAMTS4 | NM_005099 | -1.6645 | -3.187 |
| AKAP12 | NM_005100 | -0.705 | -1.101 |
| ISG15 | NM_005101 | -0.1685 | 0.038 |
| FEZ2 | NM_005102 | -0.769 | -1.257 |
| FEZ1 | NM_005103 | -0.0505 | -0.803 |
| BRD2 | NM_005104 | 0.032 | -1.0205 |
| RBM8A | NM_005105 | -1.03 | -1.6265 |
| EXOG | NM_005107 | -1.0555 | -2.0335 |
| XYLB | NM_005108 | -0.2585 | -0.594 |
| OXSR1 | NM_005109 | -0.9635 | -1.502 |
| GFPT2 | NM_005110 | -0.001 | -1.343 |
| WDR1 | NM_005112 | -0.141 | -1.273 |
| GOLGA5 | NM_005113 | -1.6925 | -3.2085 |
| HS3ST1 | NM_005114 | 0.072 | -0.265 |
| MVP | NM_005115 | -0.238 | -0.476 |
| SLC23A2 | NM_005116 | -0.275 | -1.355 |
| FGF19 | NM_005117 | -0.4155 | -0.335 |
| TNFSF15 | NM_005118 | -0.2215 | -0.1185 |
| THRAP3 | NM_005119 | -0.9265 | -2.4185 |
| MED12 | NM_005120 | -0.2875 | -0.901 |
| MED13 | NM_005121 | -1.064 | -4.377 |
| NR1I3 | NM_005122 | 0.0515 | 0.029 |
| NR1H4 | NM_005123 | -0.443 | -0.311 |
| NUP153 | NM_005124 | -1.5905 | -3.3005 |
| CCS | NM_005125 | -0.1485 | -0.9985 |
| NR1D2 | NM_005126 | -0.8765 | -2.094 |
| CLEC2B | NM_005127 | 0.1125 | -0.2335 |
| DOPEY2 | NM_005128 | -0.45 | -2.085 |
| FGFBP1 | NM_005130 | 0.166 | -0.3595 |
| THOC1 | NM_005131 | -1.3375 | -2.5765 |
| RCE1 | NM_005133 | -0.5625 | -2.7175 |

| | | | |
|---|---|---|---|
| PPP4R1 | NM_005134 | -1.1755 | -1.829 |
| SLC12A6 | NM_005135 | -0.356 | -0.688 |
| DGCR2 | NM_005137 | -0.9895 | -2.0125 |
| SCO2 | NM_005138 | 0.0935 | 0.202 |
| ANXA3 | NM_005139 | -0.0865 | 0.2895 |
| CNGA2 | NM_005140 | 0.3695 | -0.686 |
| FGB | NM_005141 | -0.06 | -0.553 |
| GIF | NM_005142 | 0.221 | 0.009 |
| HP | NM_005143 | 0.243 | -0.0175 |
| HR | NM_005144 | -0.0375 | -0.594 |
| SART1 | NM_005146 | -0.4325 | -1.0245 |
| DNAJA3 | NM_005147 | -1.437 | -3.122 |
| UNC119 | NM_005148 | -0.0365 | -0.315 |
| TBX19 | NM_005149 | -0.409 | -0.3405 |
| USP14 | NM_005151 | -0.734 | -1.408 |
| USP10 | NM_005153 | -1.001 | -3.0005 |
| USP8 | NM_005154 | -0.683 | -2.3825 |
| ROD1 | NM_005156 | -1.0745 | -1.801 |
| ABL1 | NM_005157 | -0.564 | -2.848 |
| ABL2 | NM_005158 | -1.382 | -2.9035 |
| ACTC1 | NM_005159 | 0.1515 | -0.023 |
| ADRBK2 | NM_005160 | -1.4285 | -2.1045 |
| APLNR | NM_005161 | 0.1735 | -0.1355 |
| AKT1 | NM_005163 | -0.054 | -0.615 |
| ABCD2 | NM_005164 | -0.1145 | -1.442 |
| ALDOC | NM_005165 | 0.1355 | -0.373 |
| APLP1 | NM_005166 | 0.297 | -0.3075 |
| PPM1J | NM_005167 | -0.066 | -0.263 |
| RND3 | NM_005168 | -1.357 | -1.5605 |
| PHOX2A | NM_005169 | -1.0165 | -2.7745 |
| ASCL2 | NM_005170 | 0.2125 | -0.0005 |
| ATF1 | NM_005171 | -0.8455 | -3.3715 |
| ATOH1 | NM_005172 | 0.097 | -0.4525 |
| ATP2A3 | NM_005173 | -0.0895 | -0.635 |
| ATP5C1 | NM_005174 | -0.432 | -0.5025 |
| ATP5G2 | NM_005176 | 0.2015 | -0.3115 |
| ATP6V0A1 | NM_005177 | -0.52 | -1.1825 |
| BCL3 | NM_005178 | -0.528 | -0.2505 |
| BMI1 | NM_005180 | -0.7775 | -1.228 |
| CA3 | NM_005181 | 0.195 | 0.025 |
| CA7 | NM_005182 | -0.2745 | -0.108 |
| CACNA1F | NM_005183 | 0.0525 | -0.057 |
| CALM3 | NM_005184 | 0.1525 | -0.4135 |

| | | | |
|---|---|---|---|
| CALML3 | NM_005185 | -0.171 | -0.026 |
| CAPN1 | NM_005186 | 0.0425 | -0.4865 |
| CBFA2T3 | NM_005187 | -0.1045 | -1.9075 |
| CBL | NM_005188 | -1.095 | -1.875 |
| CBX2 | NM_005189 | -1.2955 | -2.7045 |
| CCNC | NM_005190 | -1.755 | -3.482 |
| CD80 | NM_005191 | -0.132 | -0.4435 |
| CDKN3 | NM_005192 | -1.4765 | -2.2735 |
| CDX4 | NM_005193 | -0.303 | -0.191 |
| CEBPB | NM_005194 | -0.6375 | -1.4465 |
| CEBPD | NM_005195 | -0.323 | -0.5065 |
| FOXN3 | NM_005197 | -1.2655 | -1.661 |
| CHRNG | NM_005199 | -0.2345 | -0.518 |
| CCR8 | NM_005201 | -0.25 | -0.414 |
| COL8A2 | NM_005202 | 0.272 | 0.1555 |
| MAP3K8 | NM_005204 | -0.464 | -0.2965 |
| COX6A2 | NM_005205 | -0.1185 | -0.265 |
| CRK | NM_005206 | -1.721 | -2.55 |
| CRKL | NM_005207 | -0.9505 | -1.9865 |
| CRYBA1 | NM_005208 | 0.4475 | 0.193 |
| CRYGB | NM_005210 | 0.1855 | -0.4395 |
| CSF1R | NM_005211 | -0.1965 | -0.381 |
| CSN3 | NM_005212 | -0.0015 | -0.5895 |
| CSTA | NM_005213 | -0.2715 | -0.1835 |
| CTLA4 | NM_005214 | 0.019 | -0.2365 |
| DCC | NM_005215 | 0.3625 | -0.0125 |
| DDOST | NM_005216 | 0.314 | -0.35 |
| DEFA3 | NM_005217 | -0.284 | -0.398 |
| DEFB1 | NM_005218 | 0.0905 | 0.15 |
| DIAPH1 | NM_005219 | -0.243 | -0.9065 |
| DLX3 | NM_005220 | -0.076 | -0.452 |
| DLX5 | NM_005221 | -0.151 | 0.2335 |
| DLX6 | NM_005222 | -0.7565 | -2.24 |
| DNASE1 | NM_005223 | -0.1855 | -0.334 |
| ARID3A | NM_005224 | -1.0255 | -2.043 |
| E2F1 | NM_005225 | -0.693 | -2.0355 |
| S1PR3 | NM_005226 | -0.733 | -2.0415 |
| EFNA4 | NM_005227 | -0.984 | -0.7625 |
| EGFR | NM_005228 | -0.7905 | -0.81 |
| ELK1 | NM_005229 | -0.324 | -0.85 |
| ELK3 | NM_005230 | -1.386 | -1.6395 |
| CTTN | NM_005231 | -0.321 | -0.5175 |
| EPHA1 | NM_005232 | -0.3165 | -0.297 |

| | | | |
|---|---|---|---|
| EPHA3 | NM_005233 | 0.213 | -0.1 |
| NR2F6 | NM_005234 | -0.4545 | -1.128 |
| ERBB4 | NM_005235 | -0.45 | -0.549 |
| ERCC4 | NM_005236 | -0.1625 | -0.6155 |
| ETS1 | NM_005238 | -1.0855 | -2.968 |
| ETS2 | NM_005239 | -0.7175 | -2.521 |
| ETV3 | NM_005240 | 0.282 | 0.26 |
| MECOM | NM_005241 | -0.401 | -0.379 |
| F2RL1 | NM_005242 | 0.0675 | -0.365 |
| EWSR1 | NM_005243 | -0.296 | -0.8785 |
| EYA2 | NM_005244 | -0.37 | -1.9865 |
| FAT1 | NM_005245 | -0.89 | -1.661 |
| FER | NM_005246 | 0.0195 | -1.178 |
| FGF3 | NM_005247 | 0.4035 | -0.3705 |
| FGR | NM_005248 | -0.091 | -0.2515 |
| FOXG1 | NM_005249 | 0.4035 | -0.038 |
| FOXL1 | NM_005250 | 0.0685 | -0.1365 |
| FOXC2 | NM_005251 | 0.1325 | -0.137 |
| FOS | NM_005252 | 0.7745 | 1.369 |
| FOSL2 | NM_005253 | -0.8665 | -1.4985 |
| GABPB1 | NM_005254 | -2.2725 | -4.4245 |
| GAK | NM_005255 | -0.252 | -1.174 |
| GAS2 | NM_005256 | -0.19 | -0.1115 |
| GATA6 | NM_005257 | -0.693 | -0.4495 |
| GCHFR | NM_005258 | 0.0105 | -0.537 |
| MSTN | NM_005259 | 0.0705 | 0.26 |
| GDF9 | NM_005260 | -0.167 | -0.496 |
| GEM | NM_005261 | -1.376 | -1.4245 |
| GFER | NM_005262 | -0.1735 | -0.6485 |
| GFI1 | NM_005263 | 0.225 | -0.352 |
| GFRA1 | NM_005264 | -0.702 | -0.89 |
| GJA5 | NM_005266 | -0.0645 | -0.0995 |
| GJA8 | NM_005267 | 0.0495 | -0.066 |
| GJB5 | NM_005268 | 0.131 | -0.148 |
| GLI1 | NM_005269 | 0.0825 | -0.227 |
| GLI2 | NM_005270 | -1.0185 | -3.1695 |
| GLUD1 | NM_005271 | -0.851 | -0.4615 |
| GNAT2 | NM_005272 | -0.065 | 0.003 |
| GNB2 | NM_005273 | 0.2125 | -0.437 |
| GNG5 | NM_005274 | 0.162 | -0.6135 |
| GNL1 | NM_005275 | -0.313 | -1.2715 |
| GPD1 | NM_005276 | -0.0325 | -0.306 |
| GPM6A | NM_005277 | -0.5865 | -0.293 |

| | | | |
|---|---|---|---|
| GPM6B | NM_005278 | -1.02 | -0.3295 |
| GPR1 | NM_005279 | -0.39 | -0.3715 |
| GPR3 | NM_005281 | -0.051 | -0.1285 |
| GPR4 | NM_005282 | -0.236 | -0.503 |
| GPR6 | NM_005284 | 0.0425 | -0.504 |
| NPBWR1 | NM_005285 | -0.0505 | -0.2485 |
| NPBWR2 | NM_005286 | -0.1135 | -0.5095 |
| GPR12 | NM_005288 | -0.0695 | -0.4535 |
| GPR15 | NM_005290 | 0.249 | -0.797 |
| GPR17 | NM_005291 | 0.1635 | -0.439 |
| GPR18 | NM_005292 | 0.473 | 0.5005 |
| GPR20 | NM_005293 | 0.403 | -0.0595 |
| GPR21 | NM_005294 | -0.351 | -0.702 |
| GPR22 | NM_005295 | -0.423 | -1.0255 |
| LPAR4 | NM_005296 | 0.034 | -0.2975 |
| MCHR1 | NM_005297 | -0.1755 | -0.712 |
| GPR25 | NM_005298 | -0.0555 | -0.478 |
| GPR31 | NM_005299 | 0.081 | -0.0395 |
| GPR34 | NM_005300 | 0.3895 | -0.223 |
| GPR35 | NM_005301 | 0.0475 | -0.621 |
| GPR37 | NM_005302 | -0.2055 | -0.334 |
| FFAR1 | NM_005303 | 0.1735 | -0.013 |
| FFAR3 | NM_005304 | -0.0015 | -0.3895 |
| FFAR2 | NM_005306 | -0.1655 | -0.5915 |
| GRK5 | NM_005308 | -0.821 | -2.159 |
| GPT | NM_005309 | 0.114 | -0.3425 |
| GRB7 | NM_005310 | 0.1555 | -0.111 |
| GRB10 | NM_005311 | 0.1195 | -0.5235 |
| RAPGEF1 | NM_005312 | -0.599 | -1.5355 |
| PDIA3 | NM_005313 | -0.092 | -0.3675 |
| GRPR | NM_005314 | -0.232 | -0.276 |
| GSC2 | NM_005315 | 0.3615 | -0.1505 |
| GTF2H1 | NM_005316 | -1.641 | -3.4835 |
| GZMM | NM_005317 | -0.114 | -0.426 |
| H1F0 | NM_005318 | -0.1485 | -0.3985 |
| HIST1H1C | NM_005319 | 0.412 | -0.6865 |
| HIST1H1D | NM_005320 | 0.3005 | -0.406 |
| HIST1H1E | NM_005321 | 0.04 | 0.079 |
| HIST1H1B | NM_005322 | 0.4445 | 0.099 |
| HIST1H1T | NM_005323 | -0.186 | 0.2555 |
| H3F3B | NM_005324 | -0.555 | -0.49 |
| HIST1H1A | NM_005325 | 0.3195 | 0.0525 |
| HAGH | NM_005326 | -0.167 | -0.805 |

| | | | |
|---|---|---|---|
| HADH | NM_005327 | -0.8285 | -1.3545 |
| HAS2 | NM_005328 | 0.283 | -0.1355 |
| HAS3 | NM_005329 | -0.5985 | -1.2145 |
| HBE1 | NM_005330 | 0.099 | 0.0485 |
| HBQ1 | NM_005331 | -0.028 | -0.284 |
| HBZ | NM_005332 | 0.278 | 0.605 |
| HCCS | NM_005333 | -0.662 | -2.824 |
| HCFC1 | NM_005334 | -0.043 | -0.754 |
| HCLS1 | NM_005335 | 0.08 | -0.0445 |
| HDLBP | NM_005336 | -0.0575 | -0.3395 |
| NCKAP1L | NM_005337 | -0.004 | -0.252 |
| HIP1 | NM_005338 | -0.733 | -1.1245 |
| UBE2K | NM_005339 | -0.7565 | -1.5355 |
| HINT1 | NM_005340 | 0.007 | -0.322 |
| ZBTB48 | NM_005341 | -0.9715 | -1.8045 |
| HMGB3 | NM_005342 | -0.3895 | -0.6785 |
| HRAS | NM_005343 | -0.062 | -0.789 |
| HSPA1A | NM_005345 | 0.066 | 0.2735 |
| HSPA1B | NM_005346 | 0.1775 | 0.337 |
| HSPA5 | NM_005347 | -1.0145 | -1.1375 |
| HSP90AA1 | NM_005348 | 0.0215 | -0.1525 |
| RBPJ | NM_005349 | -0.7465 | -1.343 |
| ITGAD | NM_005353 | 0.3975 | -0.3385 |
| JUND | NM_005354 | -0.521 | -1.2395 |
| LCK | NM_005356 | -0.47 | -0.194 |
| LIPE | NM_005357 | -0.5625 | -1.546 |
| LMO7 | NM_005358 | -0.236 | -0.5035 |
| SMAD4 | NM_005359 | -0.309 | -1.551 |
| MAF | NM_005360 | -0.238 | -0.441 |
| MAGEA3 | NM_005362 | -0.081 | -0.2675 |
| MAGEA8 | NM_005364 | 0.2605 | 0.359 |
| MAGEA12 | NM_005367 | -0.0605 | -0.1465 |
| MCF2 | NM_005369 | 0.004 | -0.1765 |
| RAB8A | NM_005370 | -0.803 | -2.458 |
| METTL1 | NM_005371 | -1.1365 | -2.773 |
| MOS | NM_005372 | 0.2395 | -0.271 |
| MPL | NM_005373 | -0.5215 | -0.266 |
| MPP2 | NM_005374 | -0.1515 | -1.363 |
| MYB | NM_005375 | -0.286 | -1.0575 |
| MYCL1 | NM_005376 | 0.082 | -0.437 |
| MYCN | NM_005378 | -0.2965 | -0.847 |
| MYO1A | NM_005379 | -0.2485 | -0.0395 |
| NBL1 | NM_005380 | -0.0625 | -0.5305 |

| | | | |
|---|---|---|---|
| NCL | NM_005381 | -0.61 | -0.629 |
| NEFM | NM_005382 | -0.124 | 0.7635 |
| NEU2 | NM_005383 | -0.1365 | -0.5085 |
| NFIL3 | NM_005384 | -0.489 | -0.8085 |
| NKTR | NM_005385 | 0.0535 | -1.5855 |
| NNAT | NM_005386 | -0.263 | -0.458 |
| NUP98 | NM_005387 | -0.305 | -1.525 |
| PDCL | NM_005388 | -0.8985 | -1.6355 |
| PCMT1 | NM_005389 | -0.662 | -0.839 |
| PDHA2 | NM_005390 | 0.242 | -0.3305 |
| PDK3 | NM_005391 | 0.3405 | -0.721 |
| PHF2 | NM_005392 | -0.495 | -1.5065 |
| PLXNB3 | NM_005393 | -0.03 | -0.178 |
| PNLIPRP2 | NM_005396 | -0.0395 | -0.41 |
| PODXL | NM_005397 | -0.897 | -1.0695 |
| PPP1R3C | NM_005398 | -0.711 | -0.716 |
| PRKAB2 | NM_005399 | -1.128 | -1.1215 |
| PRKCE | NM_005400 | -0.9165 | -2.4965 |
| PTPN14 | NM_005401 | 0.0205 | -0.2215 |
| RALA | NM_005402 | -1.4005 | -2.125 |
| ROCK1 | NM_005406 | -0.142 | -2.3415 |
| SALL2 | NM_005407 | -0.915 | -1.8735 |
| CCL13 | NM_005408 | -0.104 | -0.298 |
| CXCL11 | NM_005409 | -0.475 | -0.5915 |
| SEPP1 | NM_005410 | -0.4645 | -0.542 |
| SHMT2 | NM_005412 | 0.348 | -0.3655 |
| SIX3 | NM_005413 | -0.3585 | -1.2865 |
| SKIL | NM_005414 | -0.6595 | -0.461 |
| SLC20A1 | NM_005415 | -1.2115 | -2.294 |
| SPRR3 | NM_005416 | -0.154 | -0.544 |
| SRC | NM_005417 | -0.4795 | -1.772 |
| ST5 | NM_005418 | -0.314 | -1.584 |
| STAT2 | NM_005419 | -0.1785 | -1.0725 |
| SULT1E1 | NM_005420 | 0.045 | -0.064 |
| TAL2 | NM_005421 | -0.343 | -0.7815 |
| TECTA | NM_005422 | -0.039 | -0.4155 |
| TFF2 | NM_005423 | 0.243 | 0.0825 |
| TIE1 | NM_005424 | -0.0115 | -0.1325 |
| TNP2 | NM_005425 | -0.2395 | 0.2165 |
| TP53BP2 | NM_005426 | -0.9605 | -1.718 |
| TP73 | NM_005427 | -0.0585 | -0.73 |
| VAV1 | NM_005428 | -0.494 | -0.081 |
| VEGFC | NM_005429 | -0.1775 | -2.5355 |

| | | | |
|---|---|---|---|
| WNT1 | NM_005430 | 0.1155 | -0.153 |
| XRCC2 | NM_005431 | -1.05 | -1.5355 |
| YES1 | NM_005433 | -1.0595 | -1.9905 |
| MALL | NM_005434 | 0.0565 | -0.288 |
| ARHGEF5 | NM_005435 | 0.3975 | -0.119 |
| CCDC6 | NM_005436 | -1.377 | -2.666 |
| NCOA4 | NM_005437 | -1.1285 | -1.1545 |
| FOSL1 | NM_005438 | 0.142 | -0.4265 |
| MLF2 | NM_005439 | 0.0285 | -0.5575 |
| RND2 | NM_005440 | 0.1975 | -0.6125 |
| CHAF1B | NM_005441 | -0.481 | -1.928 |
| EOMES | NM_005442 | -0.274 | -0.795 |
| PAPSS1 | NM_005443 | -0.48 | -0.262 |
| RQCD1 | NM_005444 | -0.927 | -2.6825 |
| SMC3 | NM_005445 | 0.583 | -1.2625 |
| P2RX6 | NM_005446 | -0.1535 | -0.2975 |
| RASSF9 | NM_005447 | 0.296 | -0.028 |
| BMP15 | NM_005448 | -0.442 | 0.135 |
| FAIM3 | NM_005449 | -0.066 | -0.257 |
| NOG | NM_005450 | -0.1155 | 1.029 |
| PDLIM7 | NM_005451 | -0.1715 | -0.419 |
| WDR46 | NM_005452 | -0.341 | -0.8595 |
| ZBTB22 | NM_005453 | -0.975 | -1.1915 |
| CER1 | NM_005454 | -0.0045 | -0.043 |
| ZRANB2 | NM_005455 | -0.83 | -0.944 |
| MAPK8IP1 | NM_005456 | -0.164 | -0.954 |
| GABBR2 | NM_005458 | 0.125 | -0.52 |
| GUCA1C | NM_005459 | -0.11 | -0.408 |
| SNCAIP | NM_005460 | -0.0295 | -0.8295 |
| MAFB | NM_005461 | 1.2385 | 1.6835 |
| MAGEC1 | NM_005462 | 0.318 | 0.3595 |
| AKT3 | NM_005465 | -0.707 | -1.202 |
| MED6 | NM_005466 | -0.715 | -1.865 |
| NAALAD2 | NM_005467 | -0.867 | -1.1435 |
| NAALADL1 | NM_005468 | 0.011 | -0.137 |
| ACOT8 | NM_005469 | -0.233 | -0.9085 |
| ABI1 | NM_005470 | -0.7035 | -1.5425 |
| GNPDA1 | NM_005471 | -0.1275 | -0.715 |
| KCNE3 | NM_005472 | -0.4285 | -0.1405 |
| HDAC5 | NM_005474 | -0.104 | -0.696 |
| SH2B3 | NM_005475 | -0.4205 | -1.6025 |
| GNE | NM_005476 | -1.8615 | -3.847 |
| HCN4 | NM_005477 | -0.191 | -1.407 |

| | | | |
|---|---|---|---|
| INSL5 | NM_005478 | -0.4585 | -0.063 |
| FRAT1 | NM_005479 | -0.0975 | -0.8765 |
| TROAP | NM_005480 | -1.283 | -2.8485 |
| MED16 | NM_005481 | -0.2415 | -1.1945 |
| PIGK | NM_005482 | -0.599 | -1.0095 |
| CHAF1A | NM_005483 | -1.407 | -3.3325 |
| PARP2 | NM_005484 | -0.9395 | -1.605 |
| PARP3 | NM_005485 | -0.193 | -0.867 |
| TOM1L1 | NM_005486 | -0.2865 | -0.9565 |
| TOM1 | NM_005488 | -0.1625 | -0.9325 |
| SH2D3C | NM_005489 | -0.452 | -1.414 |
| SH2D3A | NM_005490 | 0.1125 | -0.6465 |
| MAMLD1 | NM_005491 | -0.061 | -0.4465 |
| CST8 | NM_005492 | -0.0765 | -0.0615 |
| RANBP9 | NM_005493 | -1.0325 | -1.938 |
| DNAJB6 | NM_005494 | -0.3425 | -0.126 |
| SLC17A4 | NM_005495 | 0.2135 | -0.3715 |
| SMC4 | NM_005496 | -0.278 | -1.8885 |
| GJC1 | NM_005497 | -1.5865 | -2.9855 |
| AP1M2 | NM_005498 | 0.2055 | -0.42 |
| UBA2 | NM_005499 | -0.929 | -0.508 |
| SAE1 | NM_005500 | -0.134 | -0.7595 |
| ITGA3 | NM_005501 | -0.1555 | -0.415 |
| ABCA1 | NM_005502 | 0.1555 | -0.2175 |
| APBA2 | NM_005503 | -0.2215 | -0.696 |
| BCAT1 | NM_005504 | -0.5705 | -0.6075 |
| SCARB1 | NM_005505 | -0.3435 | -0.969 |
| SCARB2 | NM_005506 | -0.879 | -0.612 |
| CFL1 | NM_005507 | 0.305 | -0.138 |
| CCR4 | NM_005508 | -0.116 | -0.147 |
| DMXL1 | NM_005509 | 0.1755 | -0.6355 |
| DOM3Z | NM_005510 | -0.206 | -1.258 |
| MLANA | NM_005511 | 0.256 | -0.0595 |
| GTF2E1 | NM_005513 | -1.852 | -4.0515 |
| HLA-B | NM_005514 | 0.0005 | -0.348 |
| MNX1 | NM_005515 | 0.6805 | 1.474 |
| HLA-E | NM_005516 | 0.0395 | -0.4635 |
| HMGN2 | NM_005517 | -0.298 | -0.3465 |
| HMGCS2 | NM_005518 | 0.0265 | -0.384 |
| HMX2 | NM_005519 | 0.0905 | -1.162 |
| HNRNPH1 | NM_005520 | -0.6565 | -0.388 |
| TLX1 | NM_005521 | -0.356 | -0.2525 |
| HOXA1 | NM_005522 | -0.003 | -0.1385 |

| | | | |
|---|---|---|---|
| HOXA11 | NM_005523 | 0.2105 | -0.3125 |
| HES1 | NM_005524 | 0.2375 | -0.5015 |
| HSD11B1 | NM_005525 | 0.0395 | -0.0545 |
| HSF1 | NM_005526 | -0.064 | -0.6145 |
| HSPA1L | NM_005527 | -0.8465 | -0.358 |
| DNAJC4 | NM_005528 | 0.1795 | -0.2185 |
| HSPG2 | NM_005529 | -0.1085 | -0.2895 |
| IDH3A | NM_005530 | -1.4195 | -2.7825 |
| IFI16 | NM_005531 | -0.048 | -1.737 |
| IFI27 | NM_005532 | -0.162 | -0.0745 |
| IFI35 | NM_005533 | -0.2425 | -0.218 |
| IFNGR2 | NM_005534 | -0.857 | -2.0975 |
| IL12RB1 | NM_005535 | -0.022 | -0.187 |
| IMPA1 | NM_005536 | -0.9455 | -1.2825 |
| ING1 | NM_005537 | -0.47 | -1.281 |
| INHBC | NM_005538 | -0.028 | -0.038 |
| INPP5A | NM_005539 | -0.305 | -1.7 |
| INPP5B | NM_005540 | -0.3165 | -0.774 |
| INPP5D | NM_005541 | 0.219 | -0.327 |
| INSIG1 | NM_005542 | -0.7045 | -2.0735 |
| INSL3 | NM_005543 | -0.151 | -0.1585 |
| IRS1 | NM_005544 | -0.2985 | -0.985 |
| ISLR | NM_005545 | 0.056 | -0.523 |
| ITK | NM_005546 | 0.3175 | -0.389 |
| IVL | NM_005547 | 0.1375 | -0.3885 |
| KARS | NM_005548 | -0.4815 | -0.587 |
| KCNA10 | NM_005549 | -0.4845 | -0.1355 |
| KIFC3 | NM_005550 | -0.1945 | -0.42 |
| KLK2 | NM_005551 | 0.1735 | -0.465 |
| KLC1 | NM_005552 | -0.539 | -0.653 |
| KRTAP5-9 | NM_005553 | -0.028 | -0.32 |
| KRT6A | NM_005554 | 0.206 | -0.353 |
| KRT6B | NM_005555 | -0.2085 | -0.2445 |
| KRT7 | NM_005556 | 0.333 | -0.255 |
| LAD1 | NM_005558 | 0.1345 | -0.26 |
| LAMA1 | NM_005559 | -0.561 | -1.779 |
| LAMA5 | NM_005560 | -0.1975 | -0.742 |
| LAMP1 | NM_005561 | 0.1945 | -0.631 |
| LAMC2 | NM_005562 | -0.1985 | -0.162 |
| STMN1 | NM_005563 | -0.072 | -0.3225 |
| LCN2 | NM_005564 | 0.076 | 0.324 |
| LCP2 | NM_005565 | -0.245 | 0.0095 |
| LDHA | NM_005566 | -0.2955 | -0.612 |

| | | | |
|---|---|---|---|
| LGALS3BP | NM_005567 | -0.368 | -1.1995 |
| LHX1 | NM_005568 | 0.1815 | -0.4255 |
| LIMK2 | NM_005569 | -0.436 | -1.716 |
| LMAN1 | NM_005570 | -0.5385 | -0.865 |
| LMNA | NM_005572 | 0.481 | 0.1685 |
| LMNB1 | NM_005573 | -0.7785 | -2.475 |
| LMO2 | NM_005574 | 0.0105 | -0.5865 |
| LNPEP | NM_005575 | 0.0985 | -0.2655 |
| LOXL1 | NM_005576 | -0.3785 | -0.8445 |
| LPA | NM_005577 | -0.2325 | -0.4265 |
| LPP | NM_005578 | -0.087 | -1.078 |
| BCAM | NM_005581 | 0.147 | -0.4 |
| CD180 | NM_005582 | -0.3325 | -0.4675 |
| LYL1 | NM_005583 | -0.197 | -0.399 |
| MAB21L1 | NM_005584 | -0.6575 | -0.333 |
| SMAD6 | NM_005585 | -1.5125 | -2.861 |
| MDFI | NM_005586 | -0.7865 | -1.5565 |
| MEF2A | NM_005587 | -1.6265 | -3.195 |
| MEP1A | NM_005588 | 0.085 | -0.024 |
| ALDH6A1 | NM_005589 | -0.2825 | -0.768 |
| MRE11A | NM_005590 | -0.7355 | -1.597 |
| MRE11A | NM_005591 | -0.7355 | -1.597 |
| MUSK | NM_005592 | 0.263 | -0.379 |
| MYF5 | NM_005593 | -8.88E-16 | -0.2655 |
| NACA | NM_005594 | -0.099 | -0.188 |
| NFIA | NM_005595 | -0.9245 | -2.006 |
| NFIB | NM_005596 | -0.775 | -1.879 |
| NFIC | NM_005597 | -0.9405 | -2.6905 |
| NHLH1 | NM_005598 | -0.149 | -1.673 |
| NHLH2 | NM_005599 | -0.489 | -0.589 |
| NIT1 | NM_005600 | -0.8 | -1.634 |
| NKG7 | NM_005601 | 0.124 | -0.1 |
| CLDN11 | NM_005602 | 0.3 | -0.482 |
| ATP8B1 | NM_005603 | -0.464 | -0.545 |
| POU3F2 | NM_005604 | -1.357 | -1.0985 |
| PPP3CC | NM_005605 | -0.8545 | -1.943 |
| LGMN | NM_005606 | 0.082 | -1.197 |
| PTK2 | NM_005607 | -1.0095 | -1.4185 |
| PTPRCAP | NM_005608 | -0.3285 | -0.266 |
| PYGM | NM_005609 | 0.061 | 0.2365 |
| RBBP4 | NM_005610 | -0.5985 | -0.552 |
| RBL2 | NM_005611 | -1.723 | -1.443 |
| REST | NM_005612 | -0.0355 | -0.42 |

| | | | |
|---|---|---|---|
| RGS4 | NM_005613 | -1.2155 | -2.4785 |
| RHEB | NM_005614 | -0.283 | -0.5215 |
| RNASE6 | NM_005615 | -0.2555 | -0.2715 |
| RPS14 | NM_005617 | 0.09 | -0.273 |
| DLL1 | NM_005618 | -0.1015 | -0.6395 |
| S100A11 | NM_005620 | 0.434 | -0.1495 |
| S100A12 | NM_005621 | -0.1725 | -0.1635 |
| ACSM3 | NM_005622 | -0.1075 | -0.6615 |
| CCL8 | NM_005623 | -0.242 | -0.4955 |
| CCL25 | NM_005624 | 0.214 | -0.3075 |
| SDCBP | NM_005625 | -0.3655 | -0.421 |
| SRSF4 | NM_005626 | -0.7975 | -1.978 |
| SGK1 | NM_005627 | 1.649 | 2.3965 |
| SLC1A5 | NM_005628 | -0.705 | -2.222 |
| SLC6A8 | NM_005629 | -0.3295 | -0.8945 |
| SLCO2A1 | NM_005630 | 0.143 | -0.4075 |
| SMO | NM_005631 | -0.404 | -1.1695 |
| SOLH | NM_005632 | -0.165 | -0.731 |
| SOS1 | NM_005633 | -1.3535 | -3.625 |
| SOX3 | NM_005634 | -0.1585 | -0.385 |
| SSX1 | NM_005635 | -0.5435 | -0.3045 |
| SSX4 | NM_005636 | 0.073 | -0.3915 |
| SS18 | NM_005637 | -1.255 | -2.1105 |
| VAMP7 | NM_005638 | -0.849 | -0.845 |
| SYT1 | NM_005639 | -0.7285 | -1.8615 |
| TAF4B | NM_005640 | -0.5875 | -1.7495 |
| TAF6 | NM_005641 | -0.177 | -0.7425 |
| TAF7 | NM_005642 | -1.2125 | -1.126 |
| TAF11 | NM_005643 | -0.8 | -1.251 |
| TAF12 | NM_005644 | -1.0015 | -1.56 |
| TAF13 | NM_005645 | 0.061 | -0.738 |
| TARBP1 | NM_005646 | -1.7655 | -2.482 |
| TBL1X | NM_005647 | -0.5815 | -1.7555 |
| TCEB1 | NM_005648 | -0.623 | -1.14 |
| ZNF354A | NM_005649 | -1.0565 | -1.963 |
| TCF20 | NM_005650 | -0.7495 | -1.9245 |
| TDO2 | NM_005651 | 0.1815 | -0.0965 |
| TERF2 | NM_005652 | -1.656 | -2.2995 |
| TFCP2 | NM_005653 | -1.3895 | -2.062 |
| NR2F1 | NM_005654 | -0.387 | -0.337 |
| KLF10 | NM_005655 | -0.2625 | 0.0135 |
| TMPRSS2 | NM_005656 | 0.0215 | -0.5965 |
| TP53BP1 | NM_005657 | -0.8905 | -1.7795 |

| | | | |
|---|---|---|---|
| TRAF1 | NM_005658 | -0.021 | -0.542 |
| UFD1L | NM_005659 | 0.0395 | -0.415 |
| SLC35A2 | NM_005660 | -0.6345 | -1.8085 |
| VDAC3 | NM_005662 | -0.731 | -1.1245 |
| WHSC2 | NM_005663 | -0.4985 | -2.3635 |
| MKRN3 | NM_005664 | -0.5735 | -0.265 |
| EVI5 | NM_005665 | -0.837 | -2.149 |
| CFHR2 | NM_005666 | 0.8145 | -0.0945 |
| RNF103 | NM_005667 | -0.868 | -2.174 |
| ST8SIA4 | NM_005668 | -0.4765 | -2.272 |
| REEP5 | NM_005669 | -0.127 | -0.7535 |
| EPM2A | NM_005670 | -0.5975 | -1.5745 |
| UBXN8 | NM_005671 | -1.224 | -1.5955 |
| PSCA | NM_005672 | 0.0785 | -0.3215 |
| ZNF239 | NM_005674 | -1.5085 | -1.6205 |
| DGCR6 | NM_005675 | 0.085 | -0.7455 |
| RBM10 | NM_005676 | -0.748 | -1.5345 |
| COLQ | NM_005677 | -0.6275 | -0.8835 |
| TAF1C | NM_005679 | -0.0525 | -0.5505 |
| TAF1B | NM_005680 | -1.7325 | -4.043 |
| TAF1A | NM_005681 | -1.289 | -3.186 |
| GPR56 | NM_005682 | -0.1105 | -0.4785 |
| GPR55 | NM_005683 | -0.065 | -0.3975 |
| GPR52 | NM_005684 | 0.1105 | -0.4645 |
| GTF2IRD1 | NM_005685 | -0.802 | -2.0405 |
| SOX13 | NM_005686 | 0.122 | -0.127 |
| FARSB | NM_005687 | -0.587 | -0.7985 |
| ABCC5 | NM_005688 | -1.0025 | -1.894 |
| ABCB6 | NM_005689 | -0.7635 | -1.413 |
| DNM1L | NM_005690 | -0.55 | -1.307 |
| ABCC9 | NM_005691 | -0.1095 | -0.2225 |
| ABCF2 | NM_005692 | -0.32 | -1.22 |
| NR1H3 | NM_005693 | -0.6225 | -0.551 |
| COX17 | NM_005694 | 0.044 | -1.0635 |
| SCAMP2 | NM_005697 | -0.3735 | -1.9225 |
| SCAMP3 | NM_005698 | -0.0765 | -0.7375 |
| IL18BP | NM_005699 | -0.38 | -0.2835 |
| DPP3 | NM_005700 | 0.06 | -0.621 |
| SNUPN | NM_005701 | -0.6075 | -1.559 |
| ERAL1 | NM_005702 | -0.3705 | -1.512 |
| PTPRU | NM_005704 | 0.1825 | -0.509 |
| TSSC4 | NM_005706 | -0.6775 | -1.9125 |
| PDCD7 | NM_005707 | -0.9255 | -2.6995 |

| | | | |
|---|---|---|---|
| GPC6 | NM_005708 | -0.985 | -2.1285 |
| USH1C | NM_005709 | 0.2785 | -0.076 |
| PQBP1 | NM_005710 | -0.1635 | -0.5965 |
| EDIL3 | NM_005711 | -0.2005 | -0.259 |
| COL4A3BP | NM_005713 | -0.813 | -2.6395 |
| KCNK7 | NM_005714 | 0.0455 | -0.1555 |
| UST | NM_005715 | 0.118 | -0.1305 |
| GIPC1 | NM_005716 | -0.391 | -1.469 |
| ARPC5 | NM_005717 | -0.9625 | -1.127 |
| ARPC4 | NM_005718 | -0.0805 | -0.7065 |
| ARPC3 | NM_005719 | -0.3095 | -0.652 |
| ACTR3 | NM_005721 | -1.106 | -1.373 |
| ACTR2 | NM_005722 | -0.813 | -0.665 |
| TSPAN5 | NM_005723 | -0.6555 | -2.367 |
| TSPAN3 | NM_005724 | -0.7005 | -0.5225 |
| TSPAN2 | NM_005725 | 0.028 | -0.5665 |
| TSFM | NM_005726 | 0.118 | -1.6475 |
| TSPAN1 | NM_005727 | -0.021 | -0.3295 |
| PPIF | NM_005729 | -0.334 | -1.157 |
| CTDSP2 | NM_005730 | -0.61 | -3.32 |
| ARPC2 | NM_005731 | -0.509 | -0.566 |
| RAD50 | NM_005732 | -0.0235 | -2.058 |
| KIF20A | NM_005733 | -1.3475 | -2.359 |
| HIPK3 | NM_005734 | -0.2365 | -0.0915 |
| ACTR1B | NM_005735 | -0.0795 | -0.653 |
| ACTR1A | NM_005736 | -0.0905 | -0.8495 |
| ARL4C | NM_005737 | -0.9555 | -1.7075 |
| ARL4A | NM_005738 | -0.918 | -2.4755 |
| RASGRP1 | NM_005739 | 0.0795 | 0.168 |
| DNAL4 | NM_005740 | -0.8895 | -1.4935 |
| ZNF263 | NM_005741 | -0.549 | -1.547 |
| PDIA6 | NM_005742 | -0.477 | -0.413 |
| ARIH1 | NM_005744 | -0.913 | -2.6755 |
| BCAP31 | NM_005745 | 0.233 | -0.6095 |
| NAMPT | NM_005746 | -1.004 | -2.251 |
| CELA3A | NM_005747 | 0.091 | -0.449 |
| YAF2 | NM_005748 | -1.6075 | -2.688 |
| TOB1 | NM_005749 | -0.7865 | 0.262 |
| C4orf6 | NM_005750 | 0.03 | -0.379 |
| AKAP9 | NM_005751 | -0.4385 | -2.9245 |
| CLEC3A | NM_005752 | -0.5445 | 0.238 |
| G3BP1 | NM_005754 | -0.9005 | -1.1585 |
| EBI3 | NM_005755 | 0.1735 | -0.2355 |

| | | | |
|---|---|---|---|
| GPR64 | NM_005756 | -0.2675 | -0.304 |
| ABI2 | NM_005759 | -1.112 | -1.5695 |
| CEBPZ | NM_005760 | -0.922 | -3.271 |
| PLXNC1 | NM_005761 | -0.5835 | -0.6505 |
| TRIM28 | NM_005762 | 0.268 | -0.2175 |
| AASS | NM_005763 | -0.0735 | -1.154 |
| PDZK1IP1 | NM_005764 | 0.1275 | 0.011 |
| ATP6AP2 | NM_005765 | -0.711 | -0.5795 |
| FARP1 | NM_005766 | -0.2055 | -0.598 |
| LPAR6 | NM_005767 | 0.064 | 0.3105 |
| LPCAT3 | NM_005768 | -0.049 | -1.131 |
| CHST4 | NM_005769 | -0.3985 | -0.394 |
| DHRS9 | NM_005771 | 0.026 | -0.099 |
| RCL1 | NM_005772 | 0.516 | -0.7955 |
| ZNF256 | NM_005773 | -0.5075 | -1.2865 |
| SORBS3 | NM_005775 | -0.028 | -0.3915 |
| CNIH | NM_005776 | -0.2005 | -1.5395 |
| RBM6 | NM_005777 | -1.173 | -1.2265 |
| RBM5 | NM_005778 | -1.1865 | -2.983 |
| LHFPL2 | NM_005779 | -1.1405 | -2.2465 |
| LHFP | NM_005780 | -0.854 | -1.284 |
| TNK2 | NM_005781 | -0.8325 | -1.0255 |
| THOC4 | NM_005782 | -0.0775 | -0.317 |
| TXNDC9 | NM_005783 | -0.931 | -2.6175 |
| RNF41 | NM_005785 | -0.776 | -2.4925 |
| TSHZ1 | NM_005786 | -1.16 | -2.7755 |
| ALG3 | NM_005787 | -0.2825 | -1.162 |
| PRMT3 | NM_005788 | -0.884 | -1.36 |
| PSME3 | NM_005789 | -0.3 | -1.795 |
| MPHOSPH10 | NM_005791 | -0.899 | -2.9485 |
| MPHOSPH6 | NM_005792 | -1.1805 | -2.7825 |
| NME6 | NM_005793 | -0.8325 | -1.835 |
| DHRS2 | NM_005794 | -0.331 | -0.5265 |
| CALCRL | NM_005795 | 0.013 | -0.558 |
| NUTF2 | NM_005796 | 0.115 | -0.3875 |
| MPZL2 | NM_005797 | -0.14 | -0.4805 |
| TRIM13 | NM_005798 | -1.2885 | -3.7325 |
| USPL1 | NM_005800 | -0.743 | -2.402 |
| EIF1 | NM_005801 | -0.124 | -0.426 |
| TOPORS | NM_005802 | -0.643 | -1.386 |
| FLOT1 | NM_005803 | -0.108 | -0.861 |
| DDX39A | NM_005804 | -0.351 | -0.7765 |
| PSMD14 | NM_005805 | -1.3535 | -2.39 |

| | | | |
|---|---|---|---|
| OLIG2 | NM_005806 | -0.1715 | -0.303 |
| PRG4 | NM_005807 | -0.2445 | -0.2815 |
| CTDSPL | NM_005808 | -0.992 | -2.586 |
| PRDX2 | NM_005809 | 0.1945 | -0.2865 |
| KLRG1 | NM_005810 | -0.897 | -1.0865 |
| GDF11 | NM_005811 | -0.137 | -0.4405 |
| PRKD3 | NM_005813 | -1.329 | -3.0095 |
| GPA33 | NM_005814 | -0.1305 | -0.1835 |
| ZNF443 | NM_005815 | -0.767 | -2.7265 |
| CD96 | NM_005816 | -0.1315 | -0.111 |
| PLIN3 | NM_005817 | -0.1495 | -0.7095 |
| STX6 | NM_005819 | -1.353 | -3.8325 |
| RCAN2 | NM_005822 | 0.1425 | -0.3635 |
| HNRNPR | NM_005826 | -0.601 | -0.736 |
| SLC35B1 | NM_005827 | -0.288 | -1.3545 |
| DCAF7 | NM_005828 | -0.6635 | -1.2595 |
| AP3S2 | NM_005829 | -0.6365 | -1.77 |
| MRPS31 | NM_005830 | -1.7465 | -3.508 |
| CALCOCO2 | NM_005831 | -0.8515 | -1.3175 |
| KCNMB2 | NM_005832 | 0.0645 | -0.083 |
| RABEPK | NM_005833 | -0.829 | -1.7325 |
| TIMM17B | NM_005834 | 0.301 | -0.764 |
| SLC17A2 | NM_005835 | 0.2255 | 0.297 |
| HRSP12 | NM_005836 | -1.8725 | -1.752 |
| POP7 | NM_005837 | -0.3105 | -0.7875 |
| GLYAT | NM_005838 | 0.0055 | -0.2145 |
| SRRM1 | NM_005839 | -0.821 | -2.213 |
| SPRY3 | NM_005840 | -0.0495 | -0.06 |
| SPRY1 | NM_005841 | -0.4925 | -0.8795 |
| SPRY2 | NM_005842 | -0.45 | -0.5505 |
| STAM2 | NM_005843 | -1.419 | -2.486 |
| ABCC4 | NM_005845 | -1.143 | -1.335 |
| SLC23A1 | NM_005847 | 0.0435 | -0.3485 |
| DENND4A | NM_005848 | -0.6575 | -1.5325 |
| IGSF6 | NM_005849 | 0.779 | 0.3725 |
| SF3B4 | NM_005850 | -0.295 | -1.287 |
| CDK2AP2 | NM_005851 | 0.0315 | -0.4025 |
| CHD3 | NM_005852 | 0.1465 | -0.3305 |
| IRX5 | NM_005853 | -0.263 | -0.4775 |
| RAMP2 | NM_005854 | 0.1705 | -0.803 |
| RAMP1 | NM_005855 | -0.3845 | -0.8725 |
| RAMP3 | NM_005856 | -0.314 | -0.1045 |
| ZMPSTE24 | NM_005857 | -1.366 | -2.7595 |

| | | | |
|---|---|---|---|
| AKAP8 | NM_005858 | -0.287 | -1.5655 |
| PURA | NM_005859 | -0.539 | -0.071 |
| FSTL3 | NM_005860 | -0.298 | -0.4615 |
| STUB1 | NM_005861 | -0.105 | -0.7615 |
| STAG1 | NM_005862 | -0.5915 | -1.882 |
| NET1 | NM_005863 | -1.431 | -2.1175 |
| EFS | NM_005864 | -0.34 | -1.292 |
| PRSS16 | NM_005865 | 0.0935 | -0.0415 |
| SIGMAR1 | NM_005866 | -0.399 | -0.938 |
| DSCR4 | NM_005867 | 0.095 | -0.226 |
| BET1 | NM_005868 | -1.6625 | -3.15 |
| CWC27 | NM_005869 | -0.465 | -1.1905 |
| SAP18 | NM_005870 | -0.391 | -1.2465 |
| SMNDC1 | NM_005871 | -1.222 | -2.9715 |
| BCAS2 | NM_005872 | -1.465 | -2.755 |
| RGS19 | NM_005873 | -1.0635 | -2.5945 |
| LILRB2 | NM_005874 | -0.0155 | -0.304 |
| EIF1B | NM_005875 | -1.063 | -2.437 |
| SPEG | NM_005876 | -0.5125 | -1.422 |
| SF3A1 | NM_005877 | 0.1675 | -1.004 |
| TRAIP | NM_005879 | -0.993 | -2.5575 |
| DNAJA2 | NM_005880 | -1.3385 | -2.539 |
| BCKDK | NM_005881 | -0.2665 | -0.9875 |
| MAEA | NM_005882 | -0.588 | -1.705 |
| APC2 | NM_005883 | -0.9555 | -1.764 |
| PAK4 | NM_005884 | -0.203 | -0.8765 |
| 6-Mar | NM_005885 | -0.998 | -0.977 |
| KATNB1 | NM_005886 | -0.4715 | -1.539 |
| SLC25A3 | NM_005888 | 0.0495 | -0.527 |
| ACAT2 | NM_005891 | -0.327 | -1.1745 |
| FMNL1 | NM_005892 | -0.268 | -0.168 |
| CCIN | NM_005893 | 0.1355 | -0.292 |
| CD5L | NM_005894 | -0.1705 | -0.2955 |
| GOLGA3 | NM_005895 | -0.584 | -1.377 |
| IDH1 | NM_005896 | -1.173 | -1.882 |
| IPP | NM_005897 | -1.2605 | -0.6055 |
| CAPRIN1 | NM_005898 | -1.311 | -1.7165 |
| NBR1 | NM_005899 | -1.3145 | -2.0335 |
| SMAD1 | NM_005900 | -1.533 | -3.241 |
| SMAD2 | NM_005901 | -1.341 | -2.2695 |
| SMAD3 | NM_005902 | -0.758 | -2.003 |
| SMAD5 | NM_005903 | -1.112 | -1.9855 |
| SMAD7 | NM_005904 | -0.1785 | -0.69 |

| | | | |
|---|---|---|---|
| SMAD9 | NM_005905 | -0.631 | -1.7815 |
| MAK | NM_005906 | 0.084 | -0.5075 |
| MAN1A1 | NM_005907 | -1.409 | -2.613 |
| MANBA | NM_005908 | -0.8265 | -1.358 |
| MAP1B | NM_005909 | -0.1645 | -0.6505 |
| MAPT | NM_005910 | 0.0845 | -0.5055 |
| MAT2A | NM_005911 | -1.235 | -1.2425 |
| MC4R | NM_005912 | -0.2255 | -0.15 |
| MC5R | NM_005913 | 0.2075 | -0.265 |
| MCM4 | NM_005914 | -0.47 | -0.8205 |
| MCM6 | NM_005915 | -1.1835 | -1.486 |
| MCM7 | NM_005916 | -0.135 | -0.7985 |
| MDH1 | NM_005917 | -0.556 | -0.6175 |
| MDH2 | NM_005918 | 0.2775 | -0.258 |
| EF2BNB-MEF: | NM_005919 | 0.013 | -0.144 |
| MEF2D | NM_005920 | -0.7105 | -3.7415 |
| MAP3K1 | NM_005921 | -1.4455 | -2.752 |
| MAP3K5 | NM_005923 | -1.6295 | -1.6845 |
| MEOX2 | NM_005924 | -0.345 | -0.2735 |
| MEP1B | NM_005925 | 0.106 | -0.1305 |
| MFAP1 | NM_005926 | -1.177 | -3.284 |
| MFAP3 | NM_005927 | -1.5585 | -2.1475 |
| MFGE8 | NM_005928 | 0.0805 | -0.476 |
| MFI2 | NM_005929 | -0.3375 | -0.3425 |
| CTAGE5 | NM_005930 | -0.6255 | -1.4575 |
| MICB | NM_005931 | 0.09 | -0.7215 |
| MIPEP | NM_005932 | -0.377 | -1.412 |
| MLL | NM_005933 | -0.375 | -1.726 |
| MLLT1 | NM_005934 | -0.9845 | -1.867 |
| AFF1 | NM_005935 | -0.826 | -1.6925 |
| MLLT6 | NM_005937 | -0.7115 | -1.118 |
| FOXO4 | NM_005938 | -0.1415 | -0.472 |
| MMP11 | NM_005940 | -0.118 | -0.5255 |
| MMP16 | NM_005941 | -0.2735 | -1.3145 |
| MOCS1 | NM_005943 | -0.2925 | -1.621 |
| CD200 | NM_005944 | -1.6905 | -2.4175 |
| MT1A | NM_005946 | 0.1595 | -0.257 |
| MT1B | NM_005947 | 0.095 | -0.181 |
| MT1F | NM_005949 | 0.0125 | -0.138 |
| MT1G | NM_005950 | 0.074 | -0.1455 |
| MT1H | NM_005951 | -0.181 | 0.219 |
| MT1X | NM_005952 | 0.1495 | 0.059 |
| MT2A | NM_005953 | 1.123 | 0.6835 |

| | | | |
|---|---|---|---|
| MT3 | NM_005954 | -0.7615 | -0.559 |
| MTF1 | NM_005955 | -0.825 | -1.7295 |
| MTHFD1 | NM_005956 | -0.717 | -0.718 |
| MTHFR | NM_005957 | -1.108 | -1.5175 |
| MTNR1A | NM_005958 | -0.129 | -0.644 |
| MTNR1B | NM_005959 | -0.0575 | -0.2855 |
| MUC6 | NM_005961 | 0.1575 | -0.2045 |
| MXI1 | NM_005962 | -0.881 | -1.754 |
| MYH1 | NM_005963 | 0.12 | 0.164 |
| MYH10 | NM_005964 | -0.479 | -0.8705 |
| NAB1 | NM_005966 | -1.2565 | -1.4975 |
| NAB2 | NM_005967 | -0.993 | -3.0715 |
| HNRNPM | NM_005968 | 0.2625 | -0.1005 |
| NAP1L4 | NM_005969 | -0.5855 | -1.169 |
| FXYD3 | NM_005971 | 0.4865 | -0.123 |
| PPYR1 | NM_005972 | 0.105 | -0.1605 |
| PRCC | NM_005973 | -0.816 | -2.8485 |
| PTK6 | NM_005975 | -0.1875 | 0.0835 |
| RNF6 | NM_005977 | -0.7325 | -1.723 |
| S100A2 | NM_005978 | -0.0275 | -0.2105 |
| S100A13 | NM_005979 | 0.248 | -0.5225 |
| S100P | NM_005980 | -0.003 | -0.533 |
| TSPAN31 | NM_005981 | -0.08 | -0.329 |
| SIX1 | NM_005982 | 0.1235 | 0.0685 |
| SKP2 | NM_005983 | -0.9155 | -1.594 |
| SLC25A1 | NM_005984 | 0.177 | -0.354 |
| SNAI1 | NM_005985 | -0.0995 | -0.7625 |
| SOX1 | NM_005986 | -0.154 | -0.176 |
| SPRR1A | NM_005987 | 0.0425 | -0.465 |
| SPRR2A | NM_005988 | 0.0825 | -0.5695 |
| AKR1D1 | NM_005989 | -0.1975 | -0.5885 |
| STK10 | NM_005990 | -0.3175 | -0.577 |
| TBX1 | NM_005992 | 0.062 | -0.1275 |
| TBCD | NM_005993 | 0.1525 | -0.482 |
| TBX2 | NM_005994 | -0.787 | -1.834 |
| TBX10 | NM_005995 | 0.1555 | -0.4865 |
| TBX3 | NM_005996 | -0.494 | -1.367 |
| VPS72 | NM_005997 | -0.642 | -2.9095 |
| CCT3 | NM_005998 | -0.228 | -0.5705 |
| TSNAX | NM_005999 | -1.4785 | -2.3385 |
| TUBA4A | NM_006000 | 0.188 | -0.409 |
| TUBA3C | NM_006001 | -0.0635 | -0.6455 |
| UCHL3 | NM_006002 | -0.7775 | -1.3385 |

| | | | |
|---|---|---|---|
| UQCRFS1 | NM_006003 | -0.279 | -2.1905 |
| UQCRH | NM_006004 | 0.0655 | -0.156 |
| WFS1 | NM_006005 | -0.1785 | -1.0815 |
| ZBTB16 | NM_006006 | -0.258 | -0.3035 |
| ZFAND5 | NM_006007 | -1.4895 | -2.4895 |
| TUBA1A | NM_006009 | 0.066 | -0.4505 |
| MANF | NM_006010 | -1.1855 | -1.6105 |
| ST8SIA2 | NM_006011 | -1.788 | -2.5145 |
| CLPP | NM_006012 | -0.3045 | -1.094 |
| RPL10 | NM_006013 | 0.0705 | -0.347 |
| LAGE3 | NM_006014 | -0.017 | -0.8775 |
| ARID1A | NM_006015 | -0.576 | -2.6255 |
| CD164 | NM_006016 | -0.8875 | -1.047 |
| PROM1 | NM_006017 | -0.308 | -0.5795 |
| HCAR3 | NM_006018 | -0.1595 | -0.1575 |
| TCIRG1 | NM_006019 | -0.0585 | -0.26 |
| ALKBH1 | NM_006020 | -1.303 | -2.918 |
| TSC22D1 | NM_006022 | -0.405 | 0.533 |
| CDC123 | NM_006023 | -0.9645 | -1.6955 |
| TAX1BP1 | NM_006024 | -0.0635 | -0.6615 |
| ENDOU | NM_006025 | 0.2245 | 0.3265 |
| H1FX | NM_006026 | 0.2165 | -0.142 |
| HTR3B | NM_006028 | 0.0435 | -0.318 |
| PNMA1 | NM_006029 | -0.158 | -0.11 |
| CACNA2D2 | NM_006030 | -0.3505 | -0.4495 |
| PCNT | NM_006031 | -0.7765 | -1.7365 |
| CPNE6 | NM_006032 | -0.087 | -0.549 |
| LIPG | NM_006033 | -0.3155 | 0.136 |
| TP53I11 | NM_006034 | -0.6055 | -1.23 |
| CDC42BPB | NM_006035 | -0.2935 | -0.783 |
| PREPL | NM_006036 | -1.0425 | -0.942 |
| HDAC4 | NM_006037 | -1.569 | -3.3455 |
| SPATA2 | NM_006038 | -0.232 | -1.0245 |
| MRC2 | NM_006039 | -0.4935 | -0.8825 |
| HS3ST4 | NM_006040 | -0.633 | -1.7435 |
| HS3ST3B1 | NM_006041 | -0.5825 | -0.051 |
| HS3ST2 | NM_006043 | -0.632 | -1.4945 |
| HDAC6 | NM_006044 | -0.234 | -0.742 |
| ATP9A | NM_006045 | -0.358 | -0.8225 |
| RBM12 | NM_006047 | -0.381 | -1.116 |
| UBE4B | NM_006048 | -0.6625 | -1.442 |
| SNAPC5 | NM_006049 | -0.9325 | -1.0445 |
| APBB3 | NM_006051 | -0.3785 | -0.84 |

| | | | |
|---|---|---|---|
| DSCR3 | NM_006052 | -1.2695 | -2.7335 |
| TCIRG1 | NM_006053 | -0.0585 | -0.26 |
| RTN3 | NM_006054 | -0.3165 | -0.408 |
| LANCL1 | NM_006055 | -1.0955 | -1.029 |
| NMUR1 | NM_006056 | -0.0205 | -0.363 |
| B3GALT5 | NM_006057 | 0.26 | -0.06 |
| TNIP1 | NM_006058 | -0.5655 | -1.6915 |
| LAMC3 | NM_006059 | -0.0255 | -0.4305 |
| IKZF1 | NM_006060 | -0.056 | -0.385 |
| CRISP3 | NM_006061 | 0.2405 | -0.536 |
| SMYD5 | NM_006062 | -0.3225 | -1.87 |
| KBTBD10 | NM_006063 | -0.147 | -0.425 |
| SIRPB1 | NM_006065 | 0.046 | -0.012 |
| AKR1A1 | NM_006066 | 0.2825 | -0.4265 |
| COX4NB | NM_006067 | -0.415 | -2.2255 |
| TLR6 | NM_006068 | -0.046 | -0.408 |
| TFG | NM_006070 | -0.5535 | -1.65 |
| PKDREJ | NM_006071 | 0.0315 | 0.0515 |
| CCL26 | NM_006072 | 0.0835 | -0.389 |
| TRDN | NM_006073 | -0.1255 | -0.53 |
| TRIM22 | NM_006074 | -0.1175 | -0.168 |
| AGFG2 | NM_006076 | -0.066 | -0.8285 |
| CACNG2 | NM_006078 | 0.0845 | -0.5965 |
| CITED2 | NM_006079 | 0.405 | 0.737 |
| SEMA3A | NM_006080 | -0.188 | -1.382 |
| TUBA1B | NM_006082 | 0.1425 | -0.2915 |
| IK | NM_006083 | -0.929 | -1.341 |
| IRF9 | NM_006084 | -0.6655 | -2.1745 |
| BPNT1 | NM_006085 | -0.8575 | -1.8975 |
| TUBB3 | NM_006086 | -0.147 | -1.0135 |
| SCML2 | NM_006089 | -0.4055 | -1.253 |
| CEPT1 | NM_006090 | -1.63 | -3.414 |
| CORO2B | NM_006091 | 0.1565 | -1.74 |
| NOD1 | NM_006092 | -0.591 | -1.041 |
| PRG3 | NM_006093 | 0.0145 | -0.0215 |
| DLC1 | NM_006094 | -1.246 | -2.5865 |
| ATP8A1 | NM_006095 | -0.172 | -0.3635 |
| NDRG1 | NM_006096 | 0.1135 | -0.508 |
| MYL9 | NM_006097 | 0.184 | -0.4845 |
| GNB2L1 | NM_006098 | 0.054 | -0.386 |
| PIAS3 | NM_006099 | -0.4985 | -1.2595 |
| ST3GAL6 | NM_006100 | -1.3555 | -2.6455 |
| NDC80 | NM_006101 | -2.0705 | -4.027 |

| | | | |
|---|---|---|---|
| WFDC2 | NM_006103 | 0.206 | -0.382 |
| RAPGEF3 | NM_006105 | -0.4965 | -0.2995 |
| YAP1 | NM_006106 | -0.309 | -1.111 |
| LUC7L3 | NM_006107 | -0.2535 | -0.5065 |
| SPON1 | NM_006108 | -0.5115 | -0.273 |
| PRMT5 | NM_006109 | -0.319 | -1.413 |
| CD2BP2 | NM_006110 | -0.53 | -1.514 |
| ACAA2 | NM_006111 | -0.32 | -0.7885 |
| PPIE | NM_006112 | -0.108 | -0.9975 |
| VAV3 | NM_006113 | -0.119 | -0.537 |
| TOMM40 | NM_006114 | -0.439 | -1.1905 |
| TAB1 | NM_006116 | -1.4505 | -1.5455 |
| HAX1 | NM_006118 | -0.456 | -1.0295 |
| HLA-DMA | NM_006120 | -0.006 | 0.2205 |
| KRT1 | NM_006121 | -0.166 | -0.4045 |
| MAN2A2 | NM_006122 | -0.4625 | -0.98 |
| IDS | NM_006123 | -0.7255 | -0.5545 |
| ARHGAP6 | NM_006125 | 0.0155 | -0.2675 |
| BMP1 | NM_006129 | 0.233 | -0.124 |
| DAGLA | NM_006133 | -0.6395 | -1.7225 |
| TMEM50B | NM_006134 | -0.801 | -1.2535 |
| CAPZA1 | NM_006135 | -0.936 | -1.002 |
| CAPZA2 | NM_006136 | -0.681 | -1.34 |
| CD7 | NM_006137 | -0.021 | -0.567 |
| MS4A3 | NM_006138 | -0.2525 | -0.3235 |
| CD28 | NM_006139 | -0.539 | -1.0495 |
| CSF2RA | NM_006140 | -0.4185 | -0.0625 |
| DYNC1LI2 | NM_006141 | -1.049 | -1.3375 |
| SFN | NM_006142 | 0.143 | -0.154 |
| GPR19 | NM_006143 | -1.7245 | -4.1165 |
| GZMA | NM_006144 | -0.17 | -0.094 |
| DNAJB1 | NM_006145 | -0.907 | -3.3045 |
| IRF6 | NM_006147 | 0.3485 | -0.2805 |
| LASP1 | NM_006148 | -0.1595 | -1.396 |
| LGALS4 | NM_006149 | 0.1105 | -0.327 |
| PRICKLE3 | NM_006150 | 0.105 | -0.386 |
| LPO | NM_006151 | 0.0965 | -0.295 |
| LRMP | NM_006152 | -0.0245 | -0.224 |
| NCK1 | NM_006153 | -1.093 | -2.476 |
| NEDD4 | NM_006154 | -0.8155 | -1.2625 |
| NEDD8 | NM_006156 | 0.1985 | -0.2105 |
| NELL1 | NM_006157 | -0.071 | -0.3835 |
| NEFL | NM_006158 | -0.653 | 0.0945 |

| | | | |
|---|---|---|---|
| NELL2 | NM_006159 | -0.61 | -0.706 |
| NEUROD2 | NM_006160 | -0.0785 | 0.108 |
| NEUROG1 | NM_006161 | 0.154 | 0.419 |
| NFATC1 | NM_006162 | 0.305 | 0.07 |
| NFE2L2 | NM_006164 | -1.476 | -3.3855 |
| NFRKB | NM_006165 | -0.6905 | -2.1255 |
| NFYB | NM_006166 | -1.117 | -1.516 |
| NKX3-1 | NM_006167 | -0.0145 | -0.225 |
| NKX6-1 | NM_006168 | -0.4545 | -0.3885 |
| NNMT | NM_006169 | -0.1275 | 0.0045 |
| NOP2 | NM_006170 | -0.476 | -1.5105 |
| NPPA | NM_006172 | -0.081 | -0.0125 |
| NPY5R | NM_006174 | -0.1535 | -0.1055 |
| NRL | NM_006177 | -0.266 | -0.3795 |
| NSF | NM_006178 | -1.0375 | -1.021 |
| NTF4 | NM_006179 | 0.0675 | -0.0565 |
| NTRK2 | NM_006180 | -0.042 | -0.1995 |
| NTN3 | NM_006181 | -0.0285 | -0.3585 |
| DDR2 | NM_006182 | -1.2105 | -1.327 |
| NTS | NM_006183 | -0.528 | -0.052 |
| NUCB1 | NM_006184 | -0.0195 | -0.898 |
| NUMA1 | NM_006185 | -0.3535 | -1.077 |
| NR4A2 | NM_006186 | 0.0325 | 0.2465 |
| OAS3 | NM_006187 | -0.1215 | -0.73 |
| OCM2 | NM_006188 | 0.143 | 0.166 |
| OMP | NM_006189 | 0.0495 | -0.226 |
| ORC2 | NM_006190 | -1.803 | -3.6435 |
| PA2G4 | NM_006191 | 0.075 | -0.528 |
| PAX1 | NM_006192 | 0.0785 | -1.186 |
| PAX4 | NM_006193 | -0.069 | 0.0745 |
| PAX9 | NM_006194 | -0.251 | -0.6205 |
| PBX3 | NM_006195 | -1.0965 | -1.7535 |
| PCBP1 | NM_006196 | -0.4885 | -1.851 |
| PCM1 | NM_006197 | -0.316 | -2.9755 |
| PCP4 | NM_006198 | -0.213 | 0.07 |
| PCSK5 | NM_006200 | 0.0695 | -0.9235 |
| CDK16 | NM_006201 | 0.0835 | -0.881 |
| PDE4A | NM_006202 | -0.085 | -0.6775 |
| PDE4D | NM_006203 | -1.413 | -2.114 |
| PDE6C | NM_006204 | 0.134 | 0.3755 |
| PDE6H | NM_006205 | -0.186 | 0.4305 |
| PDGFRA | NM_006206 | -0.1435 | -3.2065 |
| PDGFRL | NM_006207 | -0.4765 | -0.4275 |

| | | | |
|---|---|---|---|
| ENPP1 | NM_006208 | 0.128 | -0.6025 |
| ENPP2 | NM_006209 | -0.193 | -0.4575 |
| PEG3 | NM_006210 | -0.7405 | -0.577 |
| PFKFB2 | NM_006212 | -0.753 | -0.1705 |
| PHKG1 | NM_006213 | 0.5835 | -0.406 |
| PHYH | NM_006214 | -0.801 | -0.252 |
| SERPINA4 | NM_006215 | -0.216 | -0.576 |
| SERPINE2 | NM_006216 | -0.353 | -0.7755 |
| SERPINI2 | NM_006217 | -0.3535 | -0.0285 |
| PIK3CA | NM_006218 | -0.5365 | -2.0955 |
| PIK3CB | NM_006219 | -1.3475 | -1.7415 |
| PIN1 | NM_006221 | 0.2155 | -0.4325 |
| PIN4 | NM_006223 | -0.824 | -1.142 |
| PITPNA | NM_006224 | -0.43 | -0.8675 |
| PLCL1 | NM_006226 | -0.228 | -0.9385 |
| PNOC | NM_006228 | -0.046 | -0.536 |
| PNLIPRP1 | NM_006229 | -0.4905 | 0.056 |
| POLD2 | NM_006230 | 0.156 | -0.415 |
| POLE | NM_006231 | -0.174 | -1.044 |
| POLR2H | NM_006232 | -0.3895 | -1.824 |
| POLR2I | NM_006233 | 0.131 | -0.293 |
| POLR2J | NM_006234 | 0.1305 | -0.5345 |
| POU2AF1 | NM_006235 | 0.712 | 0.0275 |
| POU3F3 | NM_006236 | 0.211 | -0.316 |
| POU4F1 | NM_006237 | -0.655 | -0.3445 |
| PPARD | NM_006238 | -0.3695 | -1.267 |
| PPEF2 | NM_006239 | 0.367 | -0.3855 |
| PPEF1 | NM_006240 | 0.221 | -0.6725 |
| PPP1R2 | NM_006241 | -0.929 | -1.4895 |
| PPP1R3D | NM_006242 | -0.299 | -0.469 |
| PPP2R5A | NM_006243 | -0.7905 | -1.555 |
| PPP2R5B | NM_006244 | -0.401 | -1.065 |
| PPP2R5D | NM_006245 | -0.236 | -1.1885 |
| PPP2R5E | NM_006246 | -1.1795 | -1.7435 |
| PPP5C | NM_006247 | 0.043 | -0.7645 |
| PRB2 | NM_006248 | -0.28 | -0.6635 |
| PRB3 | NM_006249 | 0.116 | -0.1345 |
| PRH1 | NM_006250 | -0.339 | -0.748 |
| PRKAA1 | NM_006251 | -0.0165 | -0.431 |
| PRKAA2 | NM_006252 | -0.632 | -1.3325 |
| PRKAB1 | NM_006253 | -0.986 | -3.0215 |
| PRKCD | NM_006254 | -0.3105 | -0.8455 |
| PRKCH | NM_006255 | -0.249 | -1.1135 |

| | | | |
|---|---|---|---|
| PKN2 | NM_006256 | -0.8115 | -2.684 |
| PRKCQ | NM_006257 | 0.083 | -0.2895 |
| PRKG1 | NM_006258 | 0.092 | -0.6315 |
| PRKG2 | NM_006259 | 0.602 | -0.371 |
| DNAJC3 | NM_006260 | -0.613 | -1.546 |
| PROP1 | NM_006261 | 0.221 | -0.0635 |
| PRPH | NM_006262 | 0.425 | -0.041 |
| PSME1 | NM_006263 | 0.19 | -0.3635 |
| PTPN13 | NM_006264 | -0.161 | -1.5885 |
| RAD21 | NM_006265 | -0.8125 | -0.755 |
| RALGDS | NM_006266 | -0.5895 | -1.053 |
| RANBP2 | NM_006267 | -0.2 | -2.538 |
| DPF2 | NM_006268 | -0.826 | -3.0735 |
| RP1 | NM_006269 | -0.425 | -0.5195 |
| RRAS | NM_006270 | -0.0205 | -0.543 |
| S100A1 | NM_006271 | -0.159 | -0.29 |
| S100B | NM_006272 | -0.063 | -0.436 |
| CCL7 | NM_006273 | -0.256 | -0.01 |
| CCL19 | NM_006274 | 0.2055 | -0.3235 |
| SRSF6 | NM_006275 | -1.3235 | -1.363 |
| ITSN2 | NM_006277 | -0.6735 | -1.5785 |
| ST3GAL4 | NM_006278 | -0.2045 | -1.463 |
| ST3GAL3 | NM_006279 | -0.993 | -1.0135 |
| SSR4 | NM_006280 | 0.271 | -0.15 |
| STK3 | NM_006281 | -0.9895 | -2.584 |
| STK4 | NM_006282 | -1.2775 | -2.237 |
| TACC1 | NM_006283 | -1.963 | -3.2825 |
| TAF10 | NM_006284 | 0.2345 | -0.286 |
| TESK1 | NM_006285 | -0.7185 | -2.2625 |
| TFDP2 | NM_006286 | -0.717 | -1.247 |
| TFPI | NM_006287 | -0.269 | -0.344 |
| THY1 | NM_006288 | -0.6285 | -1.704 |
| TLN1 | NM_006289 | 0.1015 | -0.729 |
| TNFAIP3 | NM_006290 | -0.0915 | -0.164 |
| TNFAIP2 | NM_006291 | 0.123 | -0.3775 |
| TSG101 | NM_006292 | -1.215 | -2.7885 |
| TYRO3 | NM_006293 | -0.2135 | -0.9695 |
| UQCRB | NM_006294 | -0.2305 | -0.266 |
| VARS | NM_006295 | 0.164 | -0.322 |
| VRK2 | NM_006296 | -0.494 | -1.714 |
| XRCC1 | NM_006297 | -0.2095 | -0.981 |
| ZNF192 | NM_006298 | -1.4065 | -1.7525 |
| ZNF193 | NM_006299 | -1.0725 | -2.599 |

| | | | |
|---|---|---|---|
| ZNF230 | NM_006300 | -0.58 | -1.135 |
| MAP3K12 | NM_006301 | -1.0245 | -1.4335 |
| MOGS | NM_006302 | -0.787 | -2.074 |
| AIMP2 | NM_006303 | -0.6655 | -2.9645 |
| SHFM1 | NM_006304 | -0.2725 | -0.276 |
| ANP32A | NM_006305 | -0.402 | -1.267 |
| SMC1A | NM_006306 | -0.751 | -0.449 |
| SRPX | NM_006307 | -0.095 | -0.891 |
| HSPB3 | NM_006308 | 0.0835 | -0.3015 |
| LRRFIP2 | NM_006309 | -1.1075 | -2.1585 |
| NPEPPS | NM_006310 | -0.5375 | -1.465 |
| NCOR1 | NM_006311 | -0.6605 | -1.2695 |
| NCOR2 | NM_006312 | -0.413 | -1.3165 |
| USP15 | NM_006313 | 0.3825 | 0.199 |
| CNKSR1 | NM_006314 | -0.6385 | -0.237 |
| PCGF3 | NM_006315 | -0.984 | -1.6115 |
| BASP1 | NM_006317 | -0.583 | -1.389 |
| CDIPT | NM_006319 | -0.4905 | -1.2615 |
| PGRMC2 | NM_006320 | -1.4335 | -3.096 |
| ARIH2 | NM_006321 | -0.4625 | -1.415 |
| TUBGCP3 | NM_006322 | -1.77 | -2.9005 |
| SEC24B | NM_006323 | -1.504 | -3.3245 |
| CFDP1 | NM_006324 | -0.9015 | -0.525 |
| RAN | NM_006325 | -0.544 | -0.839 |
| TIMM23 | NM_006327 | -0.4815 | -2.1705 |
| RBM14 | NM_006328 | 0.1865 | -1.0305 |
| FBLN5 | NM_006329 | 0.027 | 0.0485 |
| LYPLA1 | NM_006330 | -0.8995 | -1.0535 |
| EMG1 | NM_006331 | -0.9155 | -2.537 |
| IFI30 | NM_006332 | 0.033 | -0.3495 |
| C1D | NM_006333 | -0.6195 | -1.269 |
| OLFM1 | NM_006334 | -0.525 | -0.7545 |
| TIMM17A | NM_006335 | -1.01 | -1.847 |
| ZER1 | NM_006336 | -0.873 | -1.5295 |
| HMG20B | NM_006339 | 0.3175 | -0.463 |
| BAIAP2 | NM_006340 | -0.5985 | -1.2375 |
| MAD2L2 | NM_006341 | -0.0055 | -1.1685 |
| TACC3 | NM_006342 | -0.718 | -1.885 |
| MERTK | NM_006343 | -0.351 | -0.251 |
| CLEC10A | NM_006344 | -0.03 | -0.2195 |
| SLC30A9 | NM_006345 | -0.846 | -0.8915 |
| PIBF1 | NM_006346 | -0.045 | -0.256 |
| PPIH | NM_006347 | 0.0795 | -0.3965 |

| | | | |
|---|---|---|---|
| COG5 | NM_006348 | -0.827 | -1.1225 |
| ZNHIT1 | NM_006349 | 0.186 | -0.4515 |
| FST | NM_006350 | -0.0155 | -0.531 |
| TIMM44 | NM_006351 | -0.439 | -0.902 |
| ZNF238 | NM_006352 | -1.1315 | 0.106 |
| HMGN4 | NM_006353 | -1.0795 | -1.44 |
| TADA3 | NM_006354 | -0.294 | -0.809 |
| TRIM38 | NM_006355 | 0.1335 | -0.405 |
| ATP5H | NM_006356 | -0.167 | -0.152 |
| UBE2E3 | NM_006357 | -0.3 | -0.266 |
| SLC25A17 | NM_006358 | -0.9985 | -2.1725 |
| SLC9A6 | NM_006359 | -1.075 | -2.2795 |
| EIF3M | NM_006360 | -0.9535 | -1.3625 |
| HOXB13 | NM_006361 | 0.2895 | -0.213 |
| NXF1 | NM_006362 | 0.33 | -0.0235 |
| SEC23B | NM_006363 | -0.6955 | -1.268 |
| SEC23A | NM_006364 | -1.1605 | -1.085 |
| C1orf61 | NM_006365 | -0.5 | -1.4615 |
| CAP2 | NM_006366 | -1.0535 | -0.787 |
| CAP1 | NM_006367 | -0.699 | -0.8875 |
| CREB3 | NM_006368 | -0.429 | -1.084 |
| LRRC41 | NM_006369 | -0.38 | -1.672 |
| VTI1B | NM_006370 | -0.1715 | -0.5935 |
| CRTAP | NM_006371 | 0.149 | -0.243 |
| SYNCRIP | NM_006372 | -0.617 | -0.922 |
| VAT1 | NM_006373 | 0.225 | -0.384 |
| STK25 | NM_006374 | 0.0705 | -0.4515 |
| ENOX2 | NM_006375 | -0.821 | -1.953 |
| UNC13B | NM_006377 | -0.1045 | -0.371 |
| SEMA4D | NM_006378 | 0.2185 | -0.7145 |
| SEMA3C | NM_006379 | -0.124 | -0.716 |
| APPBP2 | NM_006380 | -1.334 | -3.331 |
| CDRT1 | NM_006382 | -0.0655 | -0.2075 |
| CIB2 | NM_006383 | 0.1295 | -0.3445 |
| CIB1 | NM_006384 | 0.1375 | -0.529 |
| ZNF211 | NM_006385 | -0.8455 | -0.756 |
| DDX17 | NM_006386 | -1.2945 | -2.045 |
| CHERP | NM_006387 | -0.209 | -0.9565 |
| KAT5 | NM_006388 | -0.5595 | -2.078 |
| HYOU1 | NM_006389 | 0.1975 | -0.405 |
| IPO8 | NM_006390 | -1.4055 | -2.075 |
| IPO7 | NM_006391 | -0.792 | -1.223 |
| NOP56 | NM_006392 | -0.3265 | -0.5735 |

| | | | |
|---|---|---|---|
| NEBL | NM_006393 | 0.1015 | 0.1055 |
| ATG7 | NM_006395 | -0.2245 | -0.7755 |
| SSSCA1 | NM_006396 | -0.7105 | -1.8175 |
| RNASEH2A | NM_006397 | -0.3925 | -0.9605 |
| UBD | NM_006398 | 0.0405 | 0.0855 |
| BATF | NM_006399 | -0.8 | -0.3005 |
| DCTN2 | NM_006400 | 0.062 | -0.541 |
| ANP32B | NM_006401 | -0.061 | -0.457 |
| HBXIP | NM_006402 | -0.218 | -0.6125 |
| NEDD9 | NM_006403 | -0.997 | -3.3505 |
| PROCR | NM_006404 | -0.1675 | -0.36 |
| TM9SF1 | NM_006405 | -0.3175 | -1.8215 |
| PRDX4 | NM_006406 | 0.0395 | -0.266 |
| ARL6IP5 | NM_006407 | -0.749 | -0.627 |
| AGR2 | NM_006408 | -0.0095 | -0.2205 |
| ARPC1A | NM_006409 | 0.1 | -0.845 |
| HTATIP2 | NM_006410 | 0.04 | -0.531 |
| AGPAT1 | NM_006411 | 0.216 | -0.2985 |
| AGPAT2 | NM_006412 | -0.481 | -0.96 |
| RPP30 | NM_006413 | -0.88 | -2.174 |
| RPP38 | NM_006414 | -0.959 | -2.8755 |
| SPTLC1 | NM_006415 | -1.402 | -1.596 |
| SLC35A1 | NM_006416 | -1.678 | -1.95 |
| IFI44 | NM_006417 | -0.301 | -1.237 |
| OLFM4 | NM_006418 | 0.2415 | -0.327 |
| CXCL13 | NM_006419 | 0.0885 | -0.262 |
| ARFGEF2 | NM_006420 | -1.486 | -3.4545 |
| ARFGEF1 | NM_006421 | -0.5785 | -2.712 |
| AKAP3 | NM_006422 | 0.107 | -0.3475 |
| RABAC1 | NM_006423 | 0.149 | -0.3215 |
| SLC34A2 | NM_006424 | 0.213 | -0.1625 |
| SLU7 | NM_006425 | -0.7325 | -1.2925 |
| DPYSL4 | NM_006426 | -0.1635 | -0.735 |
| SIVA1 | NM_006427 | 0.4625 | -0.3755 |
| MRPL28 | NM_006428 | -0.048 | -1.093 |
| CCT4 | NM_006430 | -0.6325 | -0.839 |
| CCT2 | NM_006431 | -1.284 | -1.2655 |
| NPC2 | NM_006432 | -0.0045 | -0.2715 |
| GNLY | NM_006433 | 0.087 | -0.188 |
| SORBS1 | NM_006434 | -0.3905 | -0.84 |
| IFITM2 | NM_006435 | -0.125 | -0.9545 |
| PARP4 | NM_006437 | -1.0335 | -1.339 |
| COLEC10 | NM_006438 | 0.1325 | -0.03 |

| | | | |
|---|---|---|---|
| MAB21L2 | NM_006439 | -0.0145 | -1.5575 |
| TXNRD2 | NM_006440 | -0.135 | -0.9505 |
| MTHFS | NM_006441 | -0.7945 | -3.214 |
| DRAP1 | NM_006442 | 0.16 | -0.5365 |
| SMC2 | NM_006444 | 0.068 | -1.8885 |
| PRPF8 | NM_006445 | -0.0905 | -0.567 |
| SLCO1B1 | NM_006446 | 0.1995 | -0.351 |
| USP16 | NM_006447 | -0.374 | -1.7405 |
| CDC42EP3 | NM_006449 | -1.1525 | -3.371 |
| PAIP1 | NM_006451 | -1.4565 | -2.8425 |
| PAICS | NM_006452 | -0.696 | -0.5225 |
| TBL3 | NM_006453 | -0.215 | -1.257 |
| MXD4 | NM_006454 | -0.0835 | -0.5615 |
| LEPREL4 | NM_006455 | -0.839 | -1.0715 |
| ST6GALNAC2 | NM_006456 | -0.362 | -0.059 |
| PDLIM5 | NM_006457 | -0.872 | -2.2955 |
| TRIM3 | NM_006458 | -0.031 | -0.7325 |
| ERLIN1 | NM_006459 | -0.82 | -1.909 |
| HEXIM1 | NM_006460 | 0.339 | 1.553 |
| SPAG5 | NM_006461 | -1.1855 | -1.9985 |
| RBCK1 | NM_006462 | -0.3195 | -1.0605 |
| TGOLN2 | NM_006464 | -1.6925 | -1.8035 |
| ARID3B | NM_006465 | -0.5455 | -3.5595 |
| POLR3F | NM_006466 | -1.628 | -3.2615 |
| POLR3G | NM_006467 | -1.123 | -1.228 |
| POLR3C | NM_006468 | -1.574 | -2.319 |
| IVNS1ABP | NM_006469 | -1.6 | -4.0765 |
| TRIM16 | NM_006470 | -0.4575 | -0.897 |
| MYL12A | NM_006471 | -1.001 | -1.624 |
| TXNIP | NM_006472 | 0.274 | 0.89 |
| TAF6L | NM_006473 | -0.4175 | -1.2755 |
| PDPN | NM_006474 | 0.369 | -0.079 |
| POSTN | NM_006475 | 0.2475 | -0.192 |
| ATP5L | NM_006476 | 0.153 | -0.1985 |
| GAS2L1 | NM_006478 | -0.2565 | -0.419 |
| RAD51AP1 | NM_006479 | -0.7315 | -1.125 |
| RGS14 | NM_006480 | -0.302 | -0.7595 |
| DYRK2 | NM_006482 | -1.62 | -3.0275 |
| FBLN1 | NM_006485 | 0.1535 | -0.176 |
| FBLN1 | NM_006486 | 0.1535 | -0.176 |
| FBLN1 | NM_006487 | 0.1535 | -0.176 |
| KHK | NM_006488 | -0.3475 | -0.618 |
| NOVA1 | NM_006489 | -0.2255 | -0.648 |

| | | | |
|---|---|---|---|
| NOVA1 | NM_006491 | -0.2255 | -0.648 |
| ALX3 | NM_006492 | -0.3935 | -0.178 |
| CLN5 | NM_006493 | -1.6095 | -1.904 |
| ERF | NM_006494 | -0.9785 | -2.126 |
| EVI2B | NM_006495 | -0.0385 | -0.1285 |
| GNAI3 | NM_006496 | -0.904 | -1.1405 |
| HIC1 | NM_006497 | -0.1405 | -0.188 |
| LGALS2 | NM_006498 | 0.06 | 0.007 |
| LGALS8 | NM_006499 | -0.262 | -0.614 |
| MCAM | NM_006500 | 0.0775 | -0.5645 |
| POLH | NM_006502 | -0.821 | -1.7215 |
| PSMC4 | NM_006503 | 0.0055 | -0.679 |
| PTPRE | NM_006504 | -0.093 | -0.598 |
| PVR | NM_006505 | -0.592 | -1.9265 |
| RASA2 | NM_006506 | 1.14 | 0.446 |
| REG1B | NM_006507 | 0.1705 | 0.114 |
| RELB | NM_006509 | 0.012 | -0.5195 |
| TRIM27 | NM_006510 | -0.9955 | -3.048 |
| RSC1A1 | NM_006511 | -0.63 | -1.5055 |
| SAA4 | NM_006512 | 0.0565 | -0.194 |
| SARS | NM_006513 | -0.3145 | -1.0165 |
| SCN10A | NM_006514 | -0.0595 | -0.3565 |
| SETMAR | NM_006515 | -1.5105 | -3.311 |
| SLC2A1 | NM_006516 | -0.706 | -2.5 |
| SLC16A2 | NM_006517 | 0.212 | -0.2525 |
| DYNLT1 | NM_006519 | -0.7835 | -0.874 |
| DYNLT3 | NM_006520 | -0.442 | -0.705 |
| TFE3 | NM_006521 | -0.308 | -0.9885 |
| WNT6 | NM_006522 | 0.049 | -0.658 |
| ZNF138 | NM_006524 | -0.8825 | -1.2765 |
| ZNF217 | NM_006526 | -1.5545 | -3.9355 |
| SLBP | NM_006527 | -1.037 | -1.8665 |
| TFPI2 | NM_006528 | -0.296 | -0.444 |
| GLRA3 | NM_006529 | 0.2565 | -0.1075 |
| YEATS4 | NM_006530 | -1.6295 | -2.889 |
| IFT88 | NM_006531 | -1.2905 | -2.2555 |
| ELL | NM_006532 | -0.2185 | -1.1975 |
| MIA | NM_006533 | -0.1785 | -0.4975 |
| NCOA3 | NM_006534 | -1.302 | -1.833 |
| CLCA2 | NM_006536 | 0.047 | -0.128 |
| USP3 | NM_006537 | -0.3215 | -1.1965 |
| BCL2L11 | NM_006538 | -0.4175 | -1.9595 |
| CACNG3 | NM_006539 | 0.231 | -0.067 |

| | | | |
|---|---|---|---|
| NCOA2 | NM_006540 | -0.28 | -0.85 |
| GLRX3 | NM_006541 | -0.5545 | -1.143 |
| EXOC5 | NM_006544 | -0.5935 | -1.277 |
| NPRL2 | NM_006545 | -0.28 | -0.959 |
| IGF2BP1 | NM_006546 | -0.893 | -3.278 |
| IGF2BP3 | NM_006547 | -1.2115 | -3.5485 |
| IGF2BP2 | NM_006548 | 0.0105 | -2.1195 |
| CAMKK2 | NM_006549 | -0.5185 | -0.88 |
| SCGB1D2 | NM_006551 | -0.188 | -0.315 |
| SCGB1D1 | NM_006552 | 0.427 | -0.3165 |
| SLMO1 | NM_006553 | -0.7705 | -1.0715 |
| MTX2 | NM_006554 | -1.468 | -2.8175 |
| YKT6 | NM_006555 | -0.143 | -0.6165 |
| PMVK | NM_006556 | -0.173 | -1.2215 |
| DMRT2 | NM_006557 | 0.0885 | -0.145 |
| KHDRBS3 | NM_006558 | -0.2035 | -1.4785 |
| KHDRBS1 | NM_006559 | -0.2365 | -0.4335 |
| CELF1 | NM_006560 | -0.732 | -1.313 |
| CELF2 | NM_006561 | -0.901 | -0.5255 |
| LBX1 | NM_006562 | -0.026 | -0.275 |
| KLF1 | NM_006563 | -0.466 | -0.823 |
| CXCR6 | NM_006564 | -0.103 | -0.813 |
| CTCF | NM_006565 | -1.0065 | -3.079 |
| CD226 | NM_006566 | 0.252 | -0.1485 |
| FARS2 | NM_006567 | -0.235 | -1.7535 |
| CGRRF1 | NM_006568 | -0.9605 | -2.441 |
| CGREF1 | NM_006569 | -0.422 | -0.6225 |
| RRAGA | NM_006570 | -1.199 | -2.1645 |
| DCTN6 | NM_006571 | -0.7605 | -0.9015 |
| GNA13 | NM_006572 | -1.1875 | -1.2315 |
| TNFSF13B | NM_006573 | -0.0015 | -0.0165 |
| CSPG5 | NM_006574 | -0.7315 | -1.1085 |
| MAP4K5 | NM_006575 | -1.049 | -1.9605 |
| AVIL | NM_006576 | -0.211 | 0.3145 |
| B3GNT2 | NM_006577 | -0.7915 | -1.435 |
| GNB5 | NM_006578 | -0.3375 | -1.0535 |
| EBP | NM_006579 | -0.225 | -0.938 |
| CLDN16 | NM_006580 | -0.4035 | -0.676 |
| FUT9 | NM_006581 | -0.854 | -1.608 |
| GMEB1 | NM_006582 | -0.787 | -1.7065 |
| RRH | NM_006583 | -0.1435 | -0.5915 |
| CCT6B | NM_006584 | -0.016 | -0.853 |
| CCT8 | NM_006585 | -0.645 | -0.4 |

| | | | |
|---|---|---|---|
| CNPY3 | NM_006586 | -0.3245 | -1.141 |
| CORIN | NM_006587 | -0.176 | -0.437 |
| SULT1C4 | NM_006588 | -0.991 | -1.699 |
| FAM189B | NM_006589 | -0.457 | -2.0255 |
| USP39 | NM_006590 | -0.196 | -0.964 |
| POLD3 | NM_006591 | -1.9065 | -4.0395 |
| TBR1 | NM_006593 | -0.4435 | 0.1065 |
| AP4B1 | NM_006594 | -1.012 | -2.0555 |
| API5 | NM_006595 | -1.267 | -1.653 |
| HSPA8 | NM_006597 | -0.527 | -0.638 |
| SLC12A7 | NM_006598 | -0.4 | -1.0115 |
| NFAT5 | NM_006599 | -1.0085 | -2.852 |
| NUDC | NM_006600 | -0.327 | -0.973 |
| PTGES3 | NM_006601 | -0.7895 | -0.616 |
| TCFL5 | NM_006602 | -1.5915 | -1.4975 |
| STAG2 | NM_006603 | -0.396 | -2.1605 |
| RFPL2 | NM_006605 | 0.159 | -0.118 |
| RBBP9 | NM_006606 | -1.053 | -1.594 |
| PTTG2 | NM_006607 | -0.309 | -0.3055 |
| PHTF1 | NM_006608 | -1.151 | -1.886 |
| MAP3K2 | NM_006609 | -0.3315 | -1.4355 |
| MASP2 | NM_006610 | -0.162 | 0.418 |
| KIF1C | NM_006612 | -0.197 | -1.4495 |
| GRAP | NM_006613 | -0.069 | -0.175 |
| CHL1 | NM_006614 | 0.14 | 0.104 |
| NES | NM_006617 | -0.2075 | -0.4165 |
| KDM5B | NM_006618 | -1.3685 | -2.982 |
| HBS1L | NM_006620 | -0.5595 | -0.957 |
| AHCYL1 | NM_006621 | -0.5435 | -1.022 |
| PLK2 | NM_006622 | -0.1035 | 3.4305 |
| PHGDH | NM_006623 | 0.203 | -0.45 |
| ZMYND11 | NM_006624 | -1.3235 | -2.104 |
| SRSF10 | NM_006625 | -1.748 | -1.245 |
| ZBTB6 | NM_006626 | -1.128 | -2.7415 |
| POP4 | NM_006627 | -1.3675 | -2.16 |
| ARPP19 | NM_006628 | -1.145 | -0.612 |
| ZNF234 | NM_006630 | -1.3605 | -2.1535 |
| ZNF266 | NM_006631 | -1.6575 | -2.9525 |
| SLC17A3 | NM_006632 | -0.089 | -0.248 |
| IQGAP2 | NM_006633 | -0.0035 | -0.48 |
| VAMP5 | NM_006634 | -0.127 | -0.083 |
| ZNF460 | NM_006635 | -0.1475 | -0.154 |
| MTHFD2 | NM_006636 | -0.8095 | -0.9245 |

| | | | |
|---|---|---|---|
| OR5I1 | NM_006637 | -0.11 | 0.0355 |
| RPP40 | NM_006638 | -0.9265 | -1.4975 |
| CYSLTR1 | NM_006639 | 0.264 | 0.0705 |
| SDCCAG8 | NM_006642 | -1.089 | -2.146 |
| SDCCAG3 | NM_006643 | -0.705 | -1.3425 |
| HSPH1 | NM_006644 | -0.4705 | -0.803 |
| STARD10 | NM_006645 | -0.378 | -1.3195 |
| WASF3 | NM_006646 | -0.9835 | -0.7755 |
| NOXA1 | NM_006647 | -0.0375 | -0.362 |
| WNK2 | NM_006648 | -0.136 | -0.655 |
| UTP14A | NM_006649 | -1.25 | -1.8015 |
| CPLX2 | NM_006650 | -0.204 | -0.964 |
| CPLX1 | NM_006651 | -0.1215 | -0.5875 |
| SPINT3 | NM_006652 | 0.197 | -0.3745 |
| FRS3 | NM_006653 | -0.6195 | -1.4025 |
| FRS2 | NM_006654 | -1.1355 | -2.8635 |
| NEU3 | NM_006656 | -0.0645 | -0.8985 |
| FTCD | NM_006657 | 0.1725 | -0.434 |
| PPP1R17 | NM_006658 | 0.3755 | -0.3135 |
| TUBGCP2 | NM_006659 | 0.011 | -0.633 |
| CLPX | NM_006660 | -1.392 | -3.7785 |
| PDE10A | NM_006661 | -1.016 | -2.6285 |
| SRCAP | NM_006662 | -0.332 | -2.227 |
| PPP1R13L | NM_006663 | -0.085 | 0.076 |
| CCL27 | NM_006664 | -0.512 | -0.4915 |
| HPSE | NM_006665 | -0.8815 | -0.394 |
| RUVBL2 | NM_006666 | -0.0655 | -0.68 |
| PGRMC1 | NM_006667 | -0.6265 | -0.294 |
| CYP46A1 | NM_006668 | 0.0785 | -0.686 |
| LILRB1 | NM_006669 | -0.3465 | -0.2005 |
| TPBG | NM_006670 | 0.253 | -0.037 |
| SLC1A7 | NM_006671 | 0.1485 | -0.4135 |
| SLC22A7 | NM_006672 | -0.15 | -0.295 |
| TSPAN9 | NM_006675 | -0.0465 | -1.23 |
| USP20 | NM_006676 | -0.5065 | -1.404 |
| USP19 | NM_006677 | -0.294 | -1.7655 |
| CD300C | NM_006678 | 0.212 | -0.312 |
| ME3 | NM_006680 | -0.046 | -0.678 |
| NMU | NM_006681 | -0.202 | -0.55 |
| FGL2 | NM_006682 | 0.096 | -0.0625 |
| EDDM3A | NM_006683 | -0.041 | 0.01 |
| CFHR4 | NM_006684 | 0.1145 | -0.1675 |
| ACTL7B | NM_006686 | -0.102 | -0.3585 |

| | | | |
|---|---|---|---|
| ACTL7A | NM_006687 | 0.29 | -0.1855 |
| C1QL1 | NM_006688 | 0.1685 | -0.467 |
| MMP24 | NM_006690 | -0.236 | -1.1825 |
| LYVE1 | NM_006691 | -0.1915 | -0.4005 |
| CPSF4 | NM_006693 | -0.489 | -0.9835 |
| JTB | NM_006694 | -0.312 | -0.9615 |
| RUNDC3A | NM_006695 | -0.1315 | -0.6425 |
| BRD8 | NM_006696 | -1.3885 | -2.5435 |
| BLCAP | NM_006698 | -0.487 | -1.549 |
| MAN1A2 | NM_006699 | -0.6275 | -1.3505 |
| TRAFD1 | NM_006700 | -0.581 | -1.643 |
| TXNL4A | NM_006701 | 0.1885 | -0.588 |
| PNPLA6 | NM_006702 | -0.249 | -0.7665 |
| NUDT3 | NM_006703 | -0.121 | -1.129 |
| SUGT1 | NM_006704 | -0.8635 | -1.433 |
| GADD45G | NM_006705 | -0.079 | 0.167 |
| TCERG1 | NM_006706 | -1.516 | -2.5565 |
| GLO1 | NM_006708 | -1.333 | -1.192 |
| EHMT2 | NM_006709 | -0.0595 | -0.4805 |
| COPS8 | NM_006710 | -0.568 | -0.578 |
| RNPS1 | NM_006711 | -0.511 | -1.249 |
| FASTK | NM_006712 | -0.279 | -0.8605 |
| SUB1 | NM_006713 | -0.718 | -1.167 |
| SMPDL3A | NM_006714 | -1.059 | -2.323 |
| MAN2C1 | NM_006715 | -0.0575 | -0.5625 |
| DBF4 | NM_006716 | -1.3165 | -3.3825 |
| SPIN1 | NM_006717 | -1.104 | -1.1515 |
| PLAGL1 | NM_006718 | -0.0175 | -0.963 |
| ABLIM1 | NM_006720 | -0.6685 | -1.1735 |
| ADK | NM_006721 | -0.902 | -0.845 |
| MITF | NM_006722 | -0.596 | -0.749 |
| MAP3K4 | NM_006724 | -1.1485 | -3.0815 |
| CD6 | NM_006725 | -0.109 | -0.1515 |
| LRBA | NM_006726 | -0.5395 | -0.953 |
| CDH10 | NM_006727 | -1.245 | -0.796 |
| DIAPH2 | NM_006729 | -0.0385 | -0.6995 |
| DNASE1L1 | NM_006730 | -0.4735 | -1.1035 |
| FKTN | NM_006731 | -0.97 | -1.501 |
| FOSB | NM_006732 | 0.038 | -0.2555 |
| CENPI | NM_006733 | -1.0995 | -1.429 |
| HIVEP2 | NM_006734 | -0.818 | -2.1355 |
| HOXA2 | NM_006735 | -0.1965 | 0.031 |
| DNAJB2 | NM_006736 | -0.4895 | -0.996 |

| | | | |
|---|---|---|---|
| AKAP13 | NM_006738 | -0.518 | -1.411 |
| MCM5 | NM_006739 | -0.001 | -0.8365 |
| PPP1R1A | NM_006741 | -0.0655 | -0.263 |
| PSKH1 | NM_006742 | -0.452 | -1.5025 |
| RBM3 | NM_006743 | -0.343 | -0.4015 |
| RBP4 | NM_006744 | 0.025 | -0.2465 |
| SCML1 | NM_006746 | -0.428 | -0.3135 |
| SLA | NM_006748 | 0.084 | -0.2425 |
| SLC20A2 | NM_006749 | -1.1545 | -2.6615 |
| SNTB2 | NM_006750 | -0.8595 | -1.976 |
| SSFA2 | NM_006751 | -1.764 | -3.4835 |
| SURF6 | NM_006753 | -0.8 | -2.245 |
| SYPL1 | NM_006754 | -0.4565 | -2.174 |
| TALDO1 | NM_006755 | 0.316 | -0.253 |
| TCEA1 | NM_006756 | -0.983 | -0.92 |
| TNNT3 | NM_006757 | 0.164 | -0.231 |
| UGP2 | NM_006759 | -0.8735 | -1.8615 |
| UPK2 | NM_006760 | -0.415 | -0.425 |
| YWHAE | NM_006761 | -0.454 | -0.622 |
| LAPTM5 | NM_006762 | -0.216 | 0.0565 |
| BTG2 | NM_006763 | -0.7035 | -0.724 |
| IFRD2 | NM_006764 | -0.7105 | -2.8245 |
| TUSC3 | NM_006765 | -0.377 | -0.2655 |
| LZTR1 | NM_006767 | -0.599 | -1.0155 |
| BRAP | NM_006768 | -1.5425 | -2.7555 |
| LMO4 | NM_006769 | -0.6465 | -2.0265 |
| MARCO | NM_006770 | -0.1555 | -0.395 |
| KRT38 | NM_006771 | -0.225 | -0.31 |
| SYNGAP1 | NM_006772 | -0.466 | 0.042 |
| DDX18 | NM_006773 | -0.8955 | -1.5885 |
| INMT | NM_006774 | 0.2125 | -0.1835 |
| QKI | NM_006775 | -1.192 | -2.0845 |
| ZBTB33 | NM_006777 | -1.261 | -2.149 |
| TRIM10 | NM_006778 | -0.2855 | -0.1865 |
| CDC42EP2 | NM_006779 | -0.055 | -0.6085 |
| C6orf10 | NM_006781 | 0.047 | -0.016 |
| ZFPL1 | NM_006782 | -0.5045 | -1.523 |
| GJB6 | NM_006783 | -0.523 | -0.47 |
| WDR3 | NM_006784 | -1.4865 | -3.2945 |
| MALT1 | NM_006785 | -0.4465 | -1.1365 |
| UTS2 | NM_006786 | 0.4405 | -0.346 |
| RALBP1 | NM_006788 | -0.8495 | -1.4535 |
| APOBEC2 | NM_006789 | -0.289 | -0.5675 |

| | | | |
|---|---|---|---|
| MYOT | NM_006790 | 0.0575 | -0.1395 |
| MORF4L1 | NM_006791 | -0.484 | -0.5235 |
| PRDX3 | NM_006793 | -0.8075 | -0.491 |
| GPR75 | NM_006794 | 0.064 | -0.9995 |
| EHD1 | NM_006795 | -1.1205 | -1.715 |
| AFG3L2 | NM_006796 | -0.8925 | -2.458 |
| UGT2A1 | NM_006798 | -0.001 | 0.2775 |
| PRSS21 | NM_006799 | 0.03 | -0.2765 |
| MSL3 | NM_006800 | -0.7495 | -0.6285 |
| KDELR1 | NM_006801 | 0.079 | -0.513 |
| SF3A3 | NM_006802 | -0.83 | -1.0465 |
| AP3M2 | NM_006803 | -1.4335 | -1.794 |
| STARD3 | NM_006804 | -0.175 | -0.9895 |
| HNRNPA0 | NM_006805 | -0.6165 | -0.7835 |
| BTG3 | NM_006806 | -0.6795 | -0.853 |
| CBX1 | NM_006807 | -0.7795 | -0.358 |
| SEC61B | NM_006808 | -0.037 | -0.4155 |
| TOMM34 | NM_006809 | -0.0345 | -0.61 |
| PDIA5 | NM_006810 | -0.2645 | -1.0645 |
| SERINC3 | NM_006811 | -1.2785 | -2.0455 |
| OS9 | NM_006812 | -0.0835 | -0.562 |
| PNRC1 | NM_006813 | -0.573 | -2.432 |
| PSMF1 | NM_006814 | -0.269 | -1.13 |
| TMED2 | NM_006815 | -1.1405 | -0.5155 |
| LMAN2 | NM_006816 | 0.0925 | -0.4105 |
| ERP29 | NM_006817 | 0.269 | -0.6125 |
| MLLT11 | NM_006818 | -0.59 | -0.6385 |
| STIP1 | NM_006819 | -0.6135 | -1.285 |
| IFI44L | NM_006820 | -0.062 | 0.1185 |
| RAB40B | NM_006822 | -0.963 | -1.985 |
| PKIA | NM_006823 | -1.2465 | -1.942 |
| EBNA1BP2 | NM_006824 | -0.383 | -1.1075 |
| CKAP4 | NM_006825 | -0.4305 | -1.3295 |
| YWHAQ | NM_006826 | -0.794 | -0.433 |
| TMED10 | NM_006827 | -0.993 | -1.336 |
| ASCC3 | NM_006828 | -0.5965 | -2.297 |
| C10orf116 | NM_006829 | 0.2825 | -0.3625 |
| UQCR11 | NM_006830 | 0.337 | -0.1805 |
| FERMT2 | NM_006832 | -0.7075 | -1.6395 |
| COPS6 | NM_006833 | 0.127 | -0.3965 |
| RAB32 | NM_006834 | -0.043 | -1.82 |
| CCNI | NM_006835 | -0.0365 | -0.218 |
| GCN1L1 | NM_006836 | -0.2405 | -1.178 |

| | | | |
|---|---|---|---|
| COPS5 | NM_006837 | -1.28 | -2.129 |
| METAP2 | NM_006838 | -0.893 | -1.5175 |
| IMMT | NM_006839 | -0.7055 | -0.813 |
| LILRB5 | NM_006840 | 0.386 | -0.259 |
| SLC38A3 | NM_006841 | -0.099 | 0.4015 |
| SF3B2 | NM_006842 | -0.096 | -0.556 |
| SDS | NM_006843 | -0.128 | -0.267 |
| ILVBL | NM_006844 | -0.0985 | -0.7345 |
| KIF2C | NM_006845 | -1.121 | -2.3395 |
| SPINK5 | NM_006846 | -0.4955 | -0.411 |
| LILRB4 | NM_006847 | 0.0845 | -0.528 |
| CCDC85B | NM_006848 | -0.7505 | -1.612 |
| PDIA2 | NM_006849 | 0.182 | -0.132 |
| IL24 | NM_006850 | -0.124 | -0.286 |
| GLIPR1 | NM_006851 | -0.042 | -0.546 |
| TLK2 | NM_006852 | -1.519 | -3.608 |
| KDELR2 | NM_006854 | -0.2355 | -0.555 |
| KDELR3 | NM_006855 | -0.4985 | -0.2345 |
| ATF7 | NM_006856 | -0.6065 | -0.2775 |
| SNRNP27 | NM_006857 | -0.67 | -1.356 |
| TMED1 | NM_006858 | -0.367 | -1.8565 |
| LIAS | NM_006859 | -0.459 | -0.9645 |
| IFT27 | NM_006860 | -0.287 | -1.2615 |
| RAB35 | NM_006861 | -0.742 | -2.223 |
| TDRKH | NM_006862 | -1.0575 | -0.972 |
| LILRA1 | NM_006863 | 0.08 | -0.4565 |
| LILRB3 | NM_006864 | 0.0225 | -0.585 |
| LILRA3 | NM_006865 | 0.32 | 0.398 |
| LILRA2 | NM_006866 | -0.151 | -0.7585 |
| RBPMS | NM_006867 | -0.1975 | -1.048 |
| RAB31 | NM_006868 | -0.1595 | -0.2145 |
| ADAP1 | NM_006869 | -0.1475 | -0.5105 |
| DSTN | NM_006870 | -0.358 | -0.732 |
| RIPK3 | NM_006871 | -0.139 | -0.2535 |
| GTF2A1L | NM_006872 | -0.29 | -0.145 |
| STON1 | NM_006873 | -1.346 | -2.0295 |
| ELF2 | NM_006874 | -1.2945 | -3.008 |
| PIM2 | NM_006875 | -0.5185 | -0.8965 |
| B3GNT1 | NM_006876 | -0.374 | -1.588 |
| GMPR | NM_006877 | -0.2665 | -0.941 |
| SHOX | NM_006883 | -0.0285 | -0.2935 |
| SHOX2 | NM_006884 | -0.6445 | -0.635 |
| ZFHX3 | NM_006885 | -0.4935 | -1.687 |

| | | | |
|---|---|---|---|
| ATP5E | NM_006886 | 0.103 | -0.0615 |
| ZFP36L2 | NM_006887 | -0.479 | -0.195 |
| CALM1 | NM_006888 | -0.2355 | -0.3665 |
| CD86 | NM_006889 | -0.29 | -0.3095 |
| CEACAM7 | NM_006890 | -0.2125 | -0.4885 |
| CRYGD | NM_006891 | -0.2165 | 0.065 |
| DNMT3B | NM_006892 | -0.217 | -0.679 |
| EIF2D | NM_006893 | 0.163 | -0.626 |
| FMO3 | NM_006894 | 0.064 | 0.5715 |
| HNMT | NM_006895 | 0.0555 | -0.3355 |
| HOXA7 | NM_006896 | -0.021 | -0.4135 |
| HOXC9 | NM_006897 | -0.091 | -0.2005 |
| HOXD3 | NM_006898 | -1.0285 | -1.8875 |
| IDH3B | NM_006899 | 0.021 | -0.479 |
| MYO9A | NM_006901 | -0.309 | -2.0535 |
| PRRX1 | NM_006902 | -0.269 | -0.55 |
| PPA2 | NM_006903 | -0.8515 | -0.9805 |
| PRKDC | NM_006904 | -0.496 | -0.521 |
| PSG1 | NM_006905 | -0.1935 | -0.236 |
| PTPN5 | NM_006906 | -0.1585 | -0.496 |
| PYCR1 | NM_006907 | -0.375 | -0.734 |
| RAC1 | NM_006908 | 0.0325 | -0.319 |
| RASGRF2 | NM_006909 | -0.011 | -0.4455 |
| RBBP6 | NM_006910 | -0.2585 | -0.423 |
| RLN1 | NM_006911 | 0.2705 | -0.2145 |
| RIT1 | NM_006912 | -0.7385 | -1.548 |
| RNF5 | NM_006913 | -0.0505 | -0.6515 |
| RORB | NM_006914 | 0.098 | -1.2745 |
| RP2 | NM_006915 | -1.2575 | -2.701 |
| RPE | NM_006916 | -1.193 | -2.2785 |
| RXRG | NM_006917 | -0.1035 | -0.325 |
| SC5DL | NM_006918 | -1.3535 | -1.9715 |
| SERPINB3 | NM_006919 | 0.3185 | 0.1095 |
| SCN1A | NM_006920 | -0.315 | -0.3145 |
| SCN3A | NM_006922 | -0.919 | -2.154 |
| SDF2 | NM_006923 | -1.1545 | -2.9895 |
| SRSF1 | NM_006924 | -1.407 | -1.605 |
| SRSF5 | NM_006925 | -0.813 | -0.776 |
| ST3GAL2 | NM_006927 | 0.147 | -0.318 |
| SKIV2L | NM_006929 | -0.179 | -0.718 |
| SKP1 | NM_006930 | -0.2765 | -0.1745 |
| SMTN | NM_006932 | -0.301 | -0.5865 |
| SLC5A3 | NM_006933 | -1.623 | -2.4265 |

| | | | |
|---|---|---|---|
| SLC6A9 | NM_006934 | 0.164 | -0.447 |
| SUMO3 | NM_006936 | 0.2025 | -0.267 |
| SUMO2 | NM_006937 | -0.327 | -0.5265 |
| SNRPD1 | NM_006938 | -1.196 | -1.4785 |
| SOS2 | NM_006939 | -1.5015 | -3.9455 |
| SOX5 | NM_006940 | 0.17 | -0.8475 |
| SOX10 | NM_006941 | 0.262 | -0.375 |
| SOX15 | NM_006942 | 0.2185 | -0.5895 |
| SOX12 | NM_006943 | -0.287 | -0.876 |
| SPP2 | NM_006944 | -0.222 | 0.2025 |
| SPRR2D | NM_006945 | -0.0055 | -0.291 |
| SPTBN2 | NM_006946 | 0.063 | -0.4065 |
| SRP72 | NM_006947 | -0.3855 | -0.5025 |
| HSPA13 | NM_006948 | -1.647 | -3.4745 |
| STXBP2 | NM_006949 | 0.1405 | -0.353 |
| SYN1 | NM_006950 | 0.1985 | -0.533 |
| TAF5 | NM_006951 | -1.58 | -3.679 |
| UPK1B | NM_006952 | -0.0625 | -0.0785 |
| UPK3A | NM_006953 | 0.0245 | -0.167 |
| ZNF33A | NM_006954 | 0.4915 | 0.8205 |
| ZNF33B | NM_006955 | -0.355 | -2.2915 |
| ZNF12 | NM_006956 | -0.728 | -2.594 |
| ZNF16 | NM_006958 | -1.0275 | -2.6235 |
| ZNF17 | NM_006959 | -0.675 | -1.983 |
| ZNF19 | NM_006961 | -0.453 | -0.9465 |
| ZNF182 | NM_006962 | -0.6305 | -1.2225 |
| ZNF22 | NM_006963 | -1.6115 | -1.731 |
| ZNF24 | NM_006965 | -0.8435 | -1.685 |
| ZNF28 | NM_006969 | 0.1305 | -0.2885 |
| ZNF32 | NM_006973 | -1.233 | -1.8655 |
| ZNF33A | NM_006974 | 0.4915 | 0.8205 |
| ZBTB25 | NM_006977 | -0.7335 | -1.405 |
| RNF113A | NM_006978 | -0.521 | -0.632 |
| SLC39A7 | NM_006979 | -0.223 | -1.2435 |
| MTERF | NM_006980 | -1.6185 | -3.336 |
| NR4A3 | NM_006981 | -1.053 | -0.0425 |
| ALX1 | NM_006982 | -0.856 | -2.568 |
| MMP23B | NM_006983 | 0.0435 | -0.4545 |
| CLDN10 | NM_006984 | 0.126 | -1.3265 |
| NPIP | NM_006985 | -1.0545 | -1.124 |
| MAGED1 | NM_006986 | 0.138 | -0.3165 |
| RPH3AL | NM_006987 | -0.2765 | -0.5205 |
| ADAMTS1 | NM_006988 | -0.793 | -0.22 |

| | | | |
|---|---|---|---|
| RASA4 | NM_006989 | -1.054 | -1.5585 |
| WASF2 | NM_006990 | -0.288 | -1.2595 |
| ZNF197 | NM_006991 | -0.9105 | -1.193 |
| LRRC23 | NM_006992 | 0.153 | -0.44 |
| NPM3 | NM_006993 | 0.4265 | -0.296 |
| BTN3A3 | NM_006994 | -0.139 | -0.7845 |
| BTN2A2 | NM_006995 | -1.0235 | -1.553 |
| SLC19A2 | NM_006996 | -1.674 | -2.487 |
| TACC2 | NM_006997 | -0.5305 | -1.198 |
| SCGN | NM_006998 | -0.044 | -0.1585 |
| PAPD7 | NM_006999 | -0.617 | -1.4495 |
| UPK1A | NM_007000 | -0.088 | -0.3485 |
| SLC35D2 | NM_007001 | -0.46 | -0.641 |
| ADRM1 | NM_007002 | 0.225 | -0.0155 |
| PAGE4 | NM_007003 | -0.039 | -0.1115 |
| TLE4 | NM_007005 | 0.277 | 0.4805 |
| NUDT21 | NM_007006 | -0.8885 | -0.419 |
| CPSF6 | NM_007007 | 0.0285 | 0.213 |
| RTN4 | NM_007008 | -0.2735 | -0.847 |
| ZPBP | NM_007009 | 0.037 | -0.3605 |
| DDX52 | NM_007010 | -1.6325 | -3.8975 |
| ABHD2 | NM_007011 | -1.5785 | -2.1755 |
| WWP1 | NM_007013 | -0.974 | -2.2 |
| WWP2 | NM_007014 | -0.4925 | -1.787 |
| LECT1 | NM_007015 | -0.4455 | -0.411 |
| SOX30 | NM_007017 | -0.0275 | 0.2485 |
| CNTRL | NM_007018 | -0.686 | -2.305 |
| C10orf10 | NM_007021 | 0.6425 | 0.534 |
| CYB561D2 | NM_007022 | -0.799 | -1.893 |
| RAPGEF4 | NM_007023 | -0.226 | -0.721 |
| TMEM115 | NM_007024 | -1.034 | -2.753 |
| DUSP14 | NM_007026 | -0.2925 | -0.6915 |
| TOPBP1 | NM_007027 | -1.1505 | -2.634 |
| TRIM31 | NM_007028 | 0.2025 | 0.073 |
| STMN2 | NM_007029 | -0.738 | -0.3235 |
| TPPP | NM_007030 | -0.936 | -2.0325 |
| HSF2BP | NM_007031 | 0.008 | -0.477 |
| TRIOBP | NM_007032 | -0.5405 | -0.9545 |
| RER1 | NM_007033 | -0.531 | -1.35 |
| DNAJB4 | NM_007034 | -1.489 | -2.1115 |
| KERA | NM_007035 | -0.119 | 0.1965 |
| ESM1 | NM_007036 | -0.14 | -0.407 |
| ADAMTS8 | NM_007037 | -0.0375 | -2.0065 |

| | | | |
|---|---|---|---|
| ADAMTS5 | NM_007038 | -0.6905 | -3.115 |
| PTPN21 | NM_007039 | -1.1595 | -2.6665 |
| HNRNPUL1 | NM_007040 | -0.1985 | -1.2905 |
| ATE1 | NM_007041 | -0.5925 | -1.1715 |
| RPP14 | NM_007042 | -1.4495 | -3.006 |
| KRR1 | NM_007043 | -0.605 | -1.841 |
| KATNA1 | NM_007044 | -1.009 | -3.057 |
| FGFR1OP | NM_007045 | -1.1925 | -1.378 |
| EMILIN1 | NM_007046 | -0.706 | -1.7775 |
| BTN3A2 | NM_007047 | -0.5525 | -0.435 |
| BTN3A1 | NM_007048 | -0.839 | -0.7685 |
| BTN2A1 | NM_007049 | -0.878 | -2.342 |
| PTPRT | NM_007050 | -0.017 | -0.1395 |
| FAF1 | NM_007051 | -1.012 | -1.8665 |
| NOX1 | NM_007052 | -0.263 | -0.528 |
| CD160 | NM_007053 | -0.0615 | -0.4495 |
| KIF3A | NM_007054 | -0.989 | -1.713 |
| POLR3A | NM_007055 | -0.4195 | -1.3805 |
| CLASRP | NM_007056 | -0.34 | -0.918 |
| ZWINT | NM_007057 | -0.5895 | -1.3875 |
| CAPN11 | NM_007058 | 0.008 | 0.0425 |
| KPTN | NM_007059 | -0.0325 | -0.468 |
| PWP1 | NM_007062 | -1.161 | -2.04 |
| KALRN | NM_007064 | -0.1745 | -0.7935 |
| CDC37 | NM_007065 | 0.225 | -0.4825 |
| PKIG | NM_007066 | -0.017 | -0.979 |
| KAT7 | NM_007067 | -0.3425 | -2.567 |
| DMC1 | NM_007068 | 0.5815 | -0.294 |
| PLA2G16 | NM_007069 | -0.3085 | -0.512 |
| HHLA2 | NM_007072 | 0.1735 | 0.22 |
| BVES | NM_007073 | -0.112 | -1.426 |
| CORO1A | NM_007074 | -0.0025 | -0.735 |
| WDR45 | NM_007075 | 0.257 | -0.379 |
| FICD | NM_007076 | 0.211 | 0.3435 |
| AP4S1 | NM_007077 | -1.2495 | -1.2885 |
| LDB3 | NM_007078 | -0.218 | -0.4 |
| PTP4A3 | NM_007079 | -0.1995 | -0.8785 |
| LSM6 | NM_007080 | -1.1485 | -1.967 |
| RABL2A | NM_007082 | -0.6055 | -0.751 |
| NUDT6 | NM_007083 | -1.166 | -2.557 |
| SOX21 | NM_007084 | -0.5585 | -0.5245 |
| FSTL1 | NM_007085 | -0.8185 | -0.692 |
| WDHD1 | NM_007086 | -0.756 | -2.1825 |

| | | | |
|---|---|---|---|
| CLTA | NM_007096 | -0.0495 | -0.5025 |
| CLTB | NM_007097 | -0.035 | -0.4675 |
| CLTCL1 | NM_007098 | 0.2595 | -1.119 |
| ACP1 | NM_007099 | -0.5725 | -0.618 |
| ATP5I | NM_007100 | 0.3375 | -0.145 |
| SARDH | NM_007101 | -0.15 | -0.133 |
| GUCA2B | NM_007102 | 0.15 | -0.249 |
| NDUFV1 | NM_007103 | 0.164 | -0.372 |
| RPL10A | NM_007104 | -0.1615 | -0.198 |
| SLC22A18AS | NM_007105 | 0.022 | -0.343 |
| UBL3 | NM_007106 | -0.8485 | -2.3075 |
| SSR3 | NM_007107 | -0.401 | -0.8365 |
| TCEB2 | NM_007108 | 0.3355 | -0.124 |
| TCF19 | NM_007109 | -0.791 | -1.8185 |
| TEP1 | NM_007110 | 0.096 | -0.268 |
| TFDP1 | NM_007111 | -0.7685 | -1.203 |
| THBS3 | NM_007112 | 0.1735 | -0.5815 |
| TCHH | NM_007113 | 0.4895 | -0.0625 |
| TMF1 | NM_007114 | -0.649 | -1.8195 |
| TNFAIP6 | NM_007115 | -0.3275 | -0.24 |
| TRH | NM_007117 | 0.0265 | 0.0565 |
| TRIO | NM_007118 | -1.1365 | -2.223 |
| UGT1A4 | NM_007120 | 0.1265 | 0.2685 |
| NR1H2 | NM_007121 | -0.617 | -1.936 |
| USF1 | NM_007122 | -0.2735 | -1.0115 |
| USH2A | NM_007123 | 0.18 | -0.139 |
| UTRN | NM_007124 | -0.9425 | -2.0505 |
| UTY | NM_007125 | 0.003 | -0.875 |
| VCP | NM_007126 | 0.012 | -1.105 |
| VIL1 | NM_007127 | 0.052 | -0.519 |
| VPREB1 | NM_007128 | -0.0315 | -0.3875 |
| ZIC2 | NM_007129 | -0.1805 | -1.6255 |
| ZNF41 | NM_007130 | -0.5895 | -2.1015 |
| ZNF75D | NM_007131 | -1.1875 | -1.536 |
| ZNF135 | NM_007134 | -0.054 | -0.975 |
| ZNF79 | NM_007135 | -1.5785 | -2.2065 |
| ZNF80 | NM_007136 | 0.1765 | -0.28 |
| ZNF81 | NM_007137 | 0.3075 | -0.084 |
| ZNF90 | NM_007138 | -0.2485 | -0.6765 |
| PCGF2 | NM_007144 | -0.55 | -1.4375 |
| ZNF146 | NM_007145 | -1.3365 | -3.687 |
| VEZF1 | NM_007146 | -1.476 | -2.6445 |
| ZNF175 | NM_007147 | -1.911 | -3.63 |

| | | | |
|---|---|---|---|
| RNF112 | NM_007148 | -0.58 | -1.3595 |
| ZNF184 | NM_007149 | -0.414 | -2.633 |
| ZNF185 | NM_007150 | 0.0615 | -0.333 |
| ZNF195 | NM_007152 | -1.2155 | -2.4815 |
| ZNF208 | NM_007153 | 0.2935 | 0.622 |
| ZP3 | NM_007155 | -0.6395 | -0.795 |
| ZXDA | NM_007156 | -0.1795 | -0.396 |
| ZXDB | NM_007157 | -0.235 | -1.04 |
| CSDE1 | NM_007158 | -0.4585 | -0.404 |
| SLMAP | NM_007159 | -0.938 | -2.915 |
| OR2H2 | NM_007160 | 0.1395 | 0.0615 |
| LST1 | NM_007161 | -0.04 | -0.218 |
| TFEB | NM_007162 | -0.194 | -1.0625 |
| SLC14A2 | NM_007163 | 0.0035 | -0.3915 |
| SF3A2 | NM_007165 | -0.0335 | -0.8365 |
| PICALM | NM_007166 | -0.851 | -1.871 |
| ZMYM6 | NM_007167 | -0.285 | -0.776 |
| ABCA8 | NM_007168 | -0.2825 | 0.212 |
| TESK2 | NM_007170 | -0.2195 | -0.1545 |
| POMT1 | NM_007171 | -0.4295 | -0.8615 |
| NUP50 | NM_007172 | -1.5965 | -1.9415 |
| PRSS23 | NM_007173 | -0.3325 | -0.368 |
| CIT | NM_007174 | -0.8815 | -1.848 |
| ERLIN2 | NM_007175 | -1.1865 | -2.204 |
| C14orf1 | NM_007176 | -0.248 | -1.8115 |
| FAM107A | NM_007177 | 0.159 | -0.139 |
| STRAP | NM_007178 | -0.7245 | -0.6295 |
| INSL6 | NM_007179 | 0.1395 | -0.052 |
| TREH | NM_007180 | 0.3155 | 0.014 |
| MAP4K1 | NM_007181 | 0.19 | -0.238 |
| RASSF1 | NM_007182 | 0.833 | 0.5995 |
| PKP3 | NM_007183 | 0.1675 | -0.3385 |
| NISCH | NM_007184 | -0.792 | -1.5815 |
| CELF3 | NM_007185 | -0.2615 | -0.9645 |
| CEP250 | NM_007186 | -0.443 | -1.4315 |
| WBP4 | NM_007187 | -0.8475 | -1.305 |
| ABCB8 | NM_007188 | 0.0185 | -0.5295 |
| ABCF2 | NM_007189 | -0.2255 | -1.1585 |
| SEC23IP | NM_007190 | -1.5755 | -2.992 |
| WIF1 | NM_007191 | -0.206 | -0.387 |
| SUPT16H | NM_007192 | -0.806 | -0.924 |
| ANXA10 | NM_007193 | -0.1285 | -0.72 |
| CHEK2 | NM_007194 | -1.0785 | -2.1535 |

| | | | |
|---|---|---|---|
| POLI | NM_007195 | -1.032 | -1.9375 |
| FZD10 | NM_007197 | 0.1545 | -0.05 |
| PROSC | NM_007198 | -1.5535 | -2.689 |
| IRAK3 | NM_007199 | -0.0905 | -0.294 |
| AKAP13 | NM_007200 | -0.518 | -1.411 |
| AKAP10 | NM_007202 | -0.9295 | -2.835 |
| PALM2-AKAP2 | NM_007203 | -0.6225 | -2.367 |
| DDX20 | NM_007204 | -1.448 | -3.973 |
| DUSP10 | NM_007207 | -0.4735 | -1.9265 |
| MRPL3 | NM_007208 | -1.163 | -1.598 |
| RPL35 | NM_007209 | 0.1645 | -0.2675 |
| GALNT6 | NM_007210 | -0.572 | -0.794 |
| RASSF8 | NM_007211 | -0.5525 | -1.803 |
| RNF2 | NM_007212 | -0.746 | -1.406 |
| PRAF2 | NM_007213 | -0.0025 | -0.462 |
| SEC63 | NM_007214 | -0.4015 | -0.8895 |
| POLG2 | NM_007215 | -1.785 | -2.5965 |
| HPS5 | NM_007216 | -1.3785 | -3.276 |
| PDCD10 | NM_007217 | -0.9 | -2.46 |
| RNF139 | NM_007218 | -1.32 | -2.551 |
| RNF24 | NM_007219 | -0.8565 | -2.1095 |
| CA5B | NM_007220 | -0.583 | -0.8995 |
| PMF1 | NM_007221 | -0.1515 | -1.0065 |
| ZHX1 | NM_007222 | -0.5085 | -2.0115 |
| GPR176 | NM_007223 | -0.336 | -1.182 |
| NXPH4 | NM_007224 | -0.16 | -0.4275 |
| NXPH3 | NM_007225 | -0.0655 | 0.1975 |
| NXPH2 | NM_007226 | -0.58 | -2.255 |
| GPR45 | NM_007227 | 0.1675 | 0.0775 |
| PACSIN2 | NM_007229 | -0.074 | -1.3545 |
| SLC6A14 | NM_007231 | -0.1505 | -0.121 |
| HRH3 | NM_007232 | -1.019 | -1.617 |
| XPOT | NM_007235 | -1.558 | -1.7375 |
| CHP | NM_007236 | -0.0695 | -0.835 |
| SP140 | NM_007237 | 0.2105 | 0.0325 |
| PXMP4 | NM_007238 | -0.191 | -1.313 |
| DUSP12 | NM_007240 | -0.823 | -1.2265 |
| SNF8 | NM_007241 | 0.206 | -0.1485 |
| DDX19B | NM_007242 | -0.313 | -1.2175 |
| NRM | NM_007243 | -0.1655 | -0.688 |
| PRR4 | NM_007244 | -0.476 | -0.797 |
| ATXN2L | NM_007245 | -0.151 | -0.819 |
| KLHL2 | NM_007246 | -0.98 | -1.7285 |

| | | | |
|---|---|---|---|
| SYNRG | NM_007247 | -0.907 | -2.1575 |
| KLF12 | NM_007249 | -0.4835 | -1.8545 |
| KLF8 | NM_007250 | -0.096 | -0.2575 |
| POU6F2 | NM_007252 | 0.0745 | -0.1245 |
| CYP4F8 | NM_007253 | -0.119 | -0.668 |
| PNKP | NM_007254 | -0.101 | -0.777 |
| B4GALT7 | NM_007255 | -0.5645 | -1.213 |
| SLCO2B1 | NM_007256 | 0.073 | -0.4525 |
| PNMA2 | NM_007257 | -1.5155 | -1.4135 |
| VPS45 | NM_007259 | -1.3675 | -1.13 |
| LYPLA2 | NM_007260 | -0.2045 | -1.63 |
| CD300A | NM_007261 | -0.3155 | -0.136 |
| PARK7 | NM_007262 | -0.047 | -0.2085 |
| COPE | NM_007263 | -0.389 | -0.6735 |
| GPR182 | NM_007264 | -0.3275 | 0.147 |
| ECD | NM_007265 | -1.0745 | -2.271 |
| GPN1 | NM_007266 | -0.849 | -1.145 |
| TMC6 | NM_007267 | -0.0285 | -0.358 |
| VSIG4 | NM_007268 | -0.1545 | 0.0275 |
| STXBP3 | NM_007269 | -1.4555 | -3.048 |
| FKBP9 | NM_007270 | -0.222 | -0.568 |
| STK38 | NM_007271 | -1.539 | -2.9085 |
| CTRC | NM_007272 | -0.091 | -0.026 |
| PHB2 | NM_007273 | 0.101 | -0.413 |
| ACOT7 | NM_007274 | 0.1665 | -0.2405 |
| TUSC2 | NM_007275 | -0.076 | -0.7435 |
| CBX3 | NM_007276 | -0.842 | -0.6445 |
| EXOC3 | NM_007277 | -0.307 | -1.6225 |
| GABARAP | NM_007278 | 0.161 | -0.253 |
| U2AF2 | NM_007279 | -0.24 | -0.0765 |
| OIP5 | NM_007280 | -1.126 | -1.466 |
| SCRG1 | NM_007281 | -0.4265 | -0.7365 |
| RNF13 | NM_007282 | -1.259 | -1.978 |
| MGLL | NM_007283 | -0.246 | -0.8355 |
| TWF2 | NM_007284 | -0.184 | -0.8285 |
| GABARAPL2 | NM_007285 | -0.181 | -0.435 |
| SYNPO | NM_007286 | -0.167 | -0.616 |
| MME | NM_007287 | 0.2055 | -0.4565 |
| MME | NM_007288 | -0.403 | -0.471 |
| MME | NM_007289 | -0.403 | -0.471 |
| ACOX1 | NM_007292 | -1.148 | -2.459 |
| BRCA1 | NM_007294 | -1.1685 | -2.067 |
| BRCA1 | NM_007297 | -1.1685 | -2.067 |

| | | | |
|---|---|---|---|
| BRCA1 | NM_007298 | -1.1685 | -2.067 |
| BRCA1 | NM_007299 | -1.1685 | -2.067 |
| BRCA1 | NM_007300 | -1.1685 | -2.067 |
| SNCA | NM_007308 | -0.884 | -1.472 |
| DIAPH2 | NM_007309 | -0.293 | -0.32 |
| COMT | NM_007310 | -0.6255 | -0.8935 |
| TSPO | NM_007311 | 0.078 | -0.354 |
| HYAL1 | NM_007312 | -0.0895 | 0.034 |
| ABL1 | NM_007313 | -0.564 | -2.848 |
| ABL2 | NM_007314 | -1.382 | -2.9035 |
| STAT1 | NM_007315 | -1.011 | -1.3815 |
| KIF22 | NM_007317 | -0.1145 | -1.225 |
| PSEN1 | NM_007318 | -1.579 | -2.5275 |
| RANBP3 | NM_007320 | -0.5885 | -1.8495 |
| RANBP3 | NM_007322 | -0.5885 | -1.8495 |
| ZFYVE9 | NM_007323 | -0.7965 | -1.929 |
| ZFYVE9 | NM_007324 | -0.7965 | -1.929 |
| GRIA3 | NM_007325 | 0.146 | 0.326 |
| CYB5R3 | NM_007326 | 0.0845 | -0.5685 |
| GRIN1 | NM_007327 | 0.049 | -0.318 |
| DMBT1 | NM_007329 | -0.033 | 0.151 |
| WHSC1 | NM_007331 | -0.668 | -1.475 |
| TRPA1 | NM_007332 | -0.272 | -0.531 |
| KLRC3 | NM_007333 | 0.0675 | 0.07 |
| KLRD1 | NM_007334 | 0.099 | -0.2005 |
| DLEC1 | NM_007335 | -0.1505 | 0.151 |
| DLEC1 | NM_007337 | -0.3435 | -0.158 |
| SH3BGR | NM_007341 | -0.992 | -1.53 |
| NUPL2 | NM_007342 | -1.103 | -1.487 |
| PRSS3 | NM_007343 | 0.253 | -0.5905 |
| TTF1 | NM_007344 | -0.692 | -1.0905 |
| ZNF236 | NM_007345 | -0.58 | -1.931 |
| OGFR | NM_007346 | -0.5985 | -0.6425 |
| AP4E1 | NM_007347 | -1.1035 | -2.5285 |
| ATF6 | NM_007348 | -1.486 | -1.96 |
| PAXIP1 | NM_007349 | -1.6585 | -3.378 |
| PHLDA1 | NM_007350 | -0.328 | -1.1505 |
| MMRN1 | NM_007351 | 0.5315 | -0.278 |
| CELA3B | NM_007352 | 0.0115 | -0.105 |
| GNA12 | NM_007353 | -0.4205 | -1.5605 |
| C3orf27 | NM_007354 | 0.1925 | -0.3 |
| HSP90AB1 | NM_007355 | -0.29 | -0.391 |
| LAMB4 | NM_007356 | -0.172 | -0.158 |

| | | | |
|---|---|---|---|
| COG2 | NM_007357 | -1.412 | -3.549 |
| MTF2 | NM_007358 | -1.0445 | -2.919 |
| CASC3 | NM_007359 | -0.155 | -0.9835 |
| KLRK1 | NM_007360 | -0.1085 | -0.237 |
| NID2 | NM_007361 | -0.5605 | -0.192 |
| NCBP2 | NM_007362 | -0.7395 | -0.5605 |
| NONO | NM_007363 | -0.0775 | -0.3525 |
| TMED3 | NM_007364 | 0.2835 | -0.689 |
| PADI2 | NM_007365 | 0.013 | -0.2545 |
| PLA2R1 | NM_007366 | 0.0735 | 0.058 |
| RALY | NM_007367 | 0.271 | -0.4725 |
| RASA3 | NM_007368 | -0.736 | -1.6965 |
| RFC5 | NM_007370 | -0.6595 | -1.6615 |
| BRD3 | NM_007371 | -0.992 | -2.186 |
| DDX42 | NM_007372 | -1.3155 | -1.7265 |
| SHOC2 | NM_007373 | -1.3595 | -4.216 |
| SIX6 | NM_007374 | -0.0395 | -0.2395 |
| TARDBP | NM_007375 | -1.23 | -1.5945 |
| AGTR1 | NM_009585 | -1.3285 | -0.891 |
| SIM2 | NM_009586 | 0.469 | -0.227 |
| LGALS9 | NM_009587 | -0.0175 | -0.322 |
| LTB | NM_009588 | -0.078 | -0.6505 |
| AOC2 | NM_009590 | -0.641 | -0.4625 |
| DNM1L | NM_012062 | -0.623 | -0.965 |
| DNM1L | NM_012063 | -0.623 | -0.965 |
| MIP | NM_012064 | 0.361 | -0.2805 |
| AKR7A3 | NM_012067 | -0.432 | 0.2785 |
| ATF5 | NM_012068 | 0.2865 | -0.5355 |
| ATP1B4 | NM_012069 | 0.03 | -0.2615 |
| COMMD3 | NM_012071 | -0.6595 | -1.2495 |
| CD93 | NM_012072 | 0.4915 | 0.114 |
| CCT5 | NM_012073 | -0.6325 | -0.7235 |
| DPF3 | NM_012074 | -0.0985 | -1.7785 |
| DGAT1 | NM_012079 | 0.1805 | -0.5425 |
| HDHD1 | NM_012080 | -1.5915 | -2.954 |
| ELL2 | NM_012081 | -0.4505 | -1.4685 |
| ZFPM2 | NM_012082 | -1.3055 | -3.156 |
| FRAT2 | NM_012083 | -0.823 | -1.953 |
| GLUD2 | NM_012084 | 0.4945 | -0.5785 |
| GTF3C3 | NM_012086 | -1.034 | -2.012 |
| GTF3C5 | NM_012087 | -0.9325 | -2.521 |
| PGLS | NM_012088 | -0.1715 | -0.7675 |
| ABCB10 | NM_012089 | -1.165 | -2.141 |

| | | | |
|---|---|---|---|
| MACF1 | NM_012090 | -1.015 | -2.023 |
| ADAT1 | NM_012091 | -0.7945 | -1.5755 |
| ICOS | NM_012092 | -0.0245 | -0.152 |
| AK5 | NM_012093 | -0.6805 | -1.1025 |
| PRDX5 | NM_012094 | 0.362 | -0.204 |
| AP3M1 | NM_012095 | -1.249 | -1.334 |
| APPL1 | NM_012096 | -1.1 | -2.6785 |
| ARL5A | NM_012097 | -0.956 | -1.7965 |
| ANGPTL2 | NM_012098 | -1.1455 | -3.385 |
| CD3EAP | NM_012099 | -0.3435 | -1.8005 |
| DNPEP | NM_012100 | -0.038 | -0.897 |
| TRIM29 | NM_012101 | -0.247 | 0.0275 |
| RERE | NM_012102 | -0.1985 | -1.225 |
| BACE1 | NM_012104 | -0.8535 | -1.567 |
| BACE2 | NM_012105 | 0.0685 | -0.327 |
| ARL2BP | NM_012106 | -0.4975 | -0.6225 |
| STAP1 | NM_012108 | -0.381 | -0.026 |
| TMEM59L | NM_012109 | 0.2645 | -0.256 |
| CHIC2 | NM_012110 | -0.915 | -3.211 |
| AHSA1 | NM_012111 | -0.3395 | -1.276 |
| TPX2 | NM_012112 | -0.9975 | -1.9625 |
| CA14 | NM_012113 | -0.305 | -0.591 |
| CASP14 | NM_012114 | -0.1005 | -0.716 |
| CASP8AP2 | NM_012115 | -1.7405 | -3.8585 |
| CBLC | NM_012116 | 0.8065 | -0.121 |
| CBX5 | NM_012117 | -1.341 | -1.0305 |
| CCRN4L | NM_012118 | -0.0705 | -1.147 |
| CDK20 | NM_012119 | -0.936 | -1.6585 |
| CD2AP | NM_012120 | -0.7685 | -1.0085 |
| CDC42EP4 | NM_012121 | -0.8505 | -2.82 |
| MTO1 | NM_012123 | -1.785 | -3.24 |
| CHORDC1 | NM_012124 | -1.0505 | -2.5735 |
| CHRM5 | NM_012125 | -0.0605 | -0.327 |
| CIZ1 | NM_012127 | 0.1205 | -0.4265 |
| CLCA4 | NM_012128 | -0.418 | -0.0855 |
| CLDN12 | NM_012129 | -1.07 | -1.5145 |
| CLDN17 | NM_012131 | -0.035 | -0.2455 |
| COPG2 | NM_012133 | -0.709 | -0.7395 |
| LMOD1 | NM_012134 | 0.0355 | -0.418 |
| FAM50B | NM_012135 | -0.2065 | -1.444 |
| DDAH1 | NM_012137 | -0.714 | -0.511 |
| AATF | NM_012138 | -0.753 | -2.135 |
| SERGEF | NM_012139 | -0.171 | -1.2165 |

| | | | |
|---|---|---|---|
| SLC25A10 | NM_012140 | -0.376 | -1.0595 |
| INTS6 | NM_012141 | -2.319 | -3.86 |
| CCNDBP1 | NM_012142 | -0.7355 | -1.5015 |
| DNAI1 | NM_012144 | -0.0485 | -0.4625 |
| DTYMK | NM_012145 | -0.53 | -1.349 |
| F8A1 | NM_012151 | -1.3155 | -2.8815 |
| LPAR3 | NM_012152 | -0.3445 | -1.643 |
| EHF | NM_012153 | -0.081 | -0.349 |
| EIF2C2 | NM_012154 | 0.2285 | -0.883 |
| EML2 | NM_012155 | -0.154 | -0.4775 |
| EPB41L1 | NM_012156 | -0.072 | -0.674 |
| FBXL2 | NM_012157 | -0.983 | -1.5135 |
| FBXL3 | NM_012158 | -1.2295 | -3.291 |
| FBXL21 | NM_012159 | 0.111 | -0.771 |
| FBXL4 | NM_012160 | -1.626 | -1.7445 |
| FBXL5 | NM_012161 | -1.564 | -3.956 |
| LRRC29 | NM_012163 | 0.0185 | -0.2165 |
| FBXW2 | NM_012164 | -1.9015 | -2.251 |
| FBXO10 | NM_012166 | -1.117 | -2.852 |
| FBXO2 | NM_012168 | 0.036 | -0.2165 |
| FBXO22 | NM_012170 | -0.893 | -2.3845 |
| TSPAN17 | NM_012171 | -0.0675 | -0.8525 |
| FBXO24 | NM_012172 | 0.3045 | -0.2115 |
| FBXO25 | NM_012173 | -0.3795 | -1.0575 |
| FBXW8 | NM_012174 | -1.0655 | -1.2495 |
| FBXO3 | NM_012175 | -0.942 | -1.617 |
| FBXO4 | NM_012176 | -0.386 | -1.5855 |
| FBXO7 | NM_012179 | -0.3045 | -1.4695 |
| FBXO8 | NM_012180 | -1.9105 | -4.466 |
| FKBP8 | NM_012181 | 0.084 | -0.6465 |
| FOXB1 | NM_012182 | -0.235 | -0.1 |
| FOXD3 | NM_012183 | 0.0245 | -0.1 |
| FOXD4L1 | NM_012184 | -0.6305 | -0.615 |
| FOXE3 | NM_012186 | 0.0085 | -0.276 |
| FOXI1 | NM_012188 | 0.059 | -0.239 |
| ALDH1L1 | NM_012190 | -0.017 | -0.33 |
| NAT6 | NM_012191 | -0.6955 | -1.457 |
| FXC1 | NM_012192 | -1.3915 | -2.896 |
| FZD4 | NM_012193 | -0.3415 | -0.6095 |
| C11orf41 | NM_012194 | -0.7965 | -1.5365 |
| RABGAP1 | NM_012197 | -1.338 | -2.034 |
| GCA | NM_012198 | -0.4845 | -0.656 |
| EIF2C1 | NM_012199 | -0.1185 | -1.295 |

| | | | |
|---|---|---|---|
| B3GAT3 | NM_012200 | -0.5505 | -0.902 |
| GLG1 | NM_012201 | -0.6955 | -1.193 |
| GNG3 | NM_012202 | -0.0315 | -0.433 |
| GRHPR | NM_012203 | -0.289 | -0.7825 |
| GTF3C4 | NM_012204 | 0.0125 | -1.3135 |
| HAAO | NM_012205 | 0.1225 | -0.245 |
| HAVCR1 | NM_012206 | -0.21 | -0.3655 |
| HNRNPH3 | NM_012207 | -0.8765 | -0.506 |
| HARS2 | NM_012208 | -0.8905 | -1.991 |
| TRIM32 | NM_012210 | -0.808 | -3.729 |
| PTGR1 | NM_012212 | -0.405 | -0.8225 |
| MLYCD | NM_012213 | -0.944 | -2.168 |
| MGAT4A | NM_012214 | -0.1515 | -0.028 |
| MGEA5 | NM_012215 | -1.325 | -1.762 |
| MID2 | NM_012216 | -0.432 | -0.697 |
| TPSD1 | NM_012217 | -0.016 | -0.395 |
| ILF3 | NM_012218 | -0.0295 | -0.4595 |
| MRAS | NM_012219 | -1.0805 | -1.3675 |
| MYO1B | NM_012223 | -0.744 | -1.4695 |
| NEK1 | NM_012224 | -0.1285 | -0.993 |
| NUBP2 | NM_012225 | -0.071 | -1.1085 |
| P2RX2 | NM_012226 | 0.1135 | -0.3945 |
| MSRB2 | NM_012228 | -0.148 | -1.109 |
| NT5C2 | NM_012229 | -1.418 | -1.362 |
| POMZP3 | NM_012230 | -0.2365 | -0.72 |
| PRDM2 | NM_012231 | -0.841 | -0.9575 |
| PTRF | NM_012232 | 0.0365 | -0.552 |
| RAB3GAP1 | NM_012233 | -0.928 | -1.5545 |
| RYBP | NM_012234 | -1.025 | -2.231 |
| SCAP | NM_012235 | 0.1055 | -0.959 |
| SCMH1 | NM_012236 | -0.3695 | -0.8695 |
| SIRT2 | NM_012237 | 0.186 | -0.3835 |
| SIRT1 | NM_012238 | -1.5965 | -3.54 |
| SIRT3 | NM_012239 | -0.296 | -1.67 |
| SIRT4 | NM_012240 | -1.058 | -1.55 |
| SIRT5 | NM_012241 | -0.7815 | -1.5085 |
| DKK1 | NM_012242 | -1.3225 | -3.347 |
| SLC35A3 | NM_012243 | -0.673 | -1.8725 |
| SLC7A8 | NM_012244 | -0.096 | -0.0705 |
| SNW1 | NM_012245 | -1.2085 | -2.682 |
| SEPHS1 | NM_012247 | -0.965 | -0.7845 |
| SEPHS2 | NM_012248 | -0.7385 | -1.3195 |
| RHOQ | NM_012249 | -0.793 | -1.29 |

| | | | |
|---|---|---|---|
| RRAS2 | NM_012250 | -0.9475 | -2.199 |
| TFEC | NM_012252 | -0.0955 | -0.0655 |
| TKTL1 | NM_012253 | 0.0995 | -0.0545 |
| SLC27A5 | NM_012254 | -0.001 | -0.609 |
| XRN2 | NM_012255 | -0.944 | -0.6985 |
| ZNF212 | NM_012256 | -0.938 | -2.923 |
| HBP1 | NM_012257 | -1.091 | -3.0115 |
| HEY1 | NM_012258 | 0.1855 | 0.746 |
| HEY2 | NM_012259 | -0.1195 | -0.605 |
| HACL1 | NM_012260 | -1.261 | -1.8215 |
| C20orf103 | NM_012261 | -0.226 | -0.581 |
| HS2ST1 | NM_012262 | -0.987 | -2.4845 |
| TTLL1 | NM_012263 | -0.535 | -1.084 |
| TMEM184B | NM_012264 | -0.557 | -1.984 |
| RHBDD3 | NM_012265 | -0.483 | -0.788 |
| DNAJB5 | NM_012266 | -0.865 | -2.141 |
| HSPBP1 | NM_012267 | -0.3365 | -0.8485 |
| PLD3 | NM_012268 | -0.0105 | -0.2835 |
| HYAL4 | NM_012269 | 0.116 | 0.0055 |
| CXorf27 | NM_012274 | -0.0245 | 0.0145 |
| LILRA4 | NM_012276 | -0.1725 | -0.174 |
| ITGB1BP2 | NM_012278 | -0.344 | -0.7155 |
| ZNF346 | NM_012279 | -1.5315 | -2.9825 |
| FTSJ1 | NM_012280 | -0.4045 | -1.59 |
| KCND2 | NM_012281 | -0.209 | -1.258 |
| KCNE1L | NM_012282 | 0.0635 | -0.413 |
| KCNG2 | NM_012283 | 0.1435 | -0.2225 |
| KCNH3 | NM_012284 | -0.098 | -0.7185 |
| KCNH4 | NM_012285 | -0.707 | -0.7345 |
| MORF4L2 | NM_012286 | -1.2745 | -0.9635 |
| ACAP2 | NM_012287 | -1.3835 | -1.554 |
| TRAM2 | NM_012288 | -0.62 | -2.065 |
| TLK1 | NM_012290 | -1.191 | -2.977 |
| ESPL1 | NM_012291 | -0.848 | -2.3795 |
| HMHA1 | NM_012292 | 0.2195 | 0.2795 |
| PXDN | NM_012293 | -0.453 | -1.035 |
| RAPGEF5 | NM_012294 | -1.729 | -4.0765 |
| CABIN1 | NM_012295 | -0.1205 | -0.971 |
| GAB2 | NM_012296 | -0.118 | -1.1545 |
| G3BP2 | NM_012297 | -0.6225 | -1.066 |
| CAND2 | NM_012298 | -0.7965 | -1.762 |
| FBXW11 | NM_012300 | -1.208 | -2.5455 |
| MAGI2 | NM_012301 | -1.4115 | -1.67 |

| | | | |
|---|---|---|---|
| LPHN2 | NM_012302 | -1.27 | -3.394 |
| FBXL7 | NM_012304 | -0.2265 | -1.6285 |
| AP2A2 | NM_012305 | -0.423 | -1.131 |
| FAIM2 | NM_012306 | 0.0295 | -0.4895 |
| EPB41L3 | NM_012307 | -0.9175 | -1.344 |
| KDM2A | NM_012308 | -0.944 | -2.957 |
| SHANK2 | NM_012309 | -0.894 | -1.6275 |
| KIF4A | NM_012310 | -0.9985 | -1.3455 |
| KIN | NM_012311 | -1.4775 | -3.935 |
| KIR2DS4 | NM_012314 | 0.458 | -0.6095 |
| KLK9 | NM_012315 | -0.2005 | -0.3785 |
| KPNA6 | NM_012316 | -0.5615 | -1.868 |
| LDOC1 | NM_012317 | 0.013 | -0.8085 |
| LETM1 | NM_012318 | -0.889 | -1.8265 |
| SLC39A6 | NM_012319 | -0.9015 | -0.6785 |
| PLA2G15 | NM_012320 | -0.443 | -1.2355 |
| LSM4 | NM_012321 | 0.1635 | -0.4385 |
| LSM5 | NM_012322 | -0.4875 | -0.5105 |
| MAFF | NM_012323 | 0.0575 | -0.5275 |
| MAPK8IP2 | NM_012324 | -0.3865 | -0.7985 |
| MAPRE1 | NM_012325 | -0.5355 | -0.532 |
| MAPRE3 | NM_012326 | -0.252 | -1.2915 |
| PIGN | NM_012327 | -0.524 | -0.642 |
| DNAJB9 | NM_012328 | -1.602 | -3.877 |
| MMD | NM_012329 | -1.601 | -1.267 |
| MSRA | NM_012331 | 0.296 | -0.4945 |
| MYCBP | NM_012333 | -0.5395 | -0.764 |
| MYO10 | NM_012334 | 0.5455 | 0.0715 |
| MYO1F | NM_012335 | -0.3105 | 0.032 |
| NARF | NM_012336 | -0.3915 | -1.5865 |
| CCDC19 | NM_012337 | 0.2285 | 0.203 |
| TSPAN12 | NM_012338 | -1.746 | -1.578 |
| TSPAN15 | NM_012339 | -0.2215 | -0.7015 |
| NFATC2 | NM_012340 | -0.2475 | -0.6905 |
| GTPBP4 | NM_012341 | -1.395 | -2.118 |
| BAMBI | NM_012342 | -0.042 | -0.7055 |
| NNT | NM_012343 | -0.9795 | -0.7045 |
| NTSR2 | NM_012344 | 0.488 | -1.211 |
| NUFIP1 | NM_012345 | -0.5945 | -1.673 |
| NUP62 | NM_012346 | -0.971 | -2.442 |
| FBXO9 | NM_012347 | -1.2715 | -2.089 |
| OR10J1 | NM_012351 | 0.2095 | -0.1495 |
| OR1A2 | NM_012352 | 0.361 | 0.1115 |

| | | | |
|---|---|---|---|
| OR1C1 | NM_012353 | 0.1435 | -0.4725 |
| OR1F1 | NM_012360 | -0.737 | -0.1405 |
| OR1N1 | NM_012363 | 0.1675 | -0.5275 |
| OR1Q1 | NM_012364 | 0.231 | 0.2105 |
| OR2A5 | NM_012365 | 0.093 | -0.1655 |
| OR2B6 | NM_012367 | -0.082 | -0.006 |
| OR2C1 | NM_012368 | -0.024 | 0.119 |
| OR2F1 | NM_012369 | -0.46 | -0.086 |
| OR3A3 | NM_012373 | 0.0715 | -0.6235 |
| OR4D1 | NM_012374 | 0.311 | -0.0565 |
| OR52A1 | NM_012375 | -0.379 | -0.161 |
| OR7C2 | NM_012377 | -0.199 | 0.256 |
| OR8B8 | NM_012378 | -0.0615 | -0.181 |
| ORC3 | NM_012381 | -1.2905 | -1.8755 |
| TTC33 | NM_012382 | -0.829 | -2.172 |
| OSTF1 | NM_012383 | -0.4025 | -1.419 |
| GMEB2 | NM_012384 | -1.091 | -2.578 |
| NUPR1 | NM_012385 | 0.222 | -0.0375 |
| PADI4 | NM_012387 | -0.2315 | -0.2535 |
| PLDN | NM_012388 | -1.245 | -1.094 |
| SMR3A | NM_012390 | 0.005 | 0.1095 |
| SPDEF | NM_012391 | -0.046 | -0.3155 |
| PEF1 | NM_012392 | -0.675 | -1.609 |
| PFAS | NM_012393 | -0.006 | -0.6895 |
| PFDN2 | NM_012394 | -0.2285 | -0.274 |
| CDK14 | NM_012395 | -0.3655 | -1.4705 |
| PHLDA3 | NM_012396 | -0.613 | -1.7665 |
| SERPINB13 | NM_012397 | -0.3655 | 0.0105 |
| PIP5K1C | NM_012398 | -0.7975 | -1.3155 |
| PITPNB | NM_012399 | -0.952 | -2.169 |
| PLA2G2D | NM_012400 | 0.2305 | -0.4935 |
| PLXNB2 | NM_012401 | -0.4645 | -0.7825 |
| ARFIP2 | NM_012402 | -0.4445 | -1.55 |
| ANP32C | NM_012403 | -0.228 | -0.018 |
| ANP32D | NM_012404 | 0.2325 | 0.3245 |
| ICMT | NM_012405 | -0.8515 | -2.3935 |
| PRDM4 | NM_012406 | -1.272 | -3.3605 |
| PICK1 | NM_012407 | -0.526 | -0.9615 |
| ZMYND8 | NM_012408 | -1.439 | -3.8555 |
| PRND | NM_012409 | -0.421 | -0.496 |
| SEZ6L2 | NM_012410 | -0.099 | -0.6895 |
| PTPN22 | NM_012411 | 0.167 | -0.009 |
| H2AFV | NM_012412 | 0.292 | -0.25 |

| | | | |
|---|---|---|---|
| QPCT | NM_012413 | -0.842 | -0.5905 |
| RAB3GAP2 | NM_012414 | -0.6565 | -1.503 |
| RAD54B | NM_012415 | -1.051 | -2.1605 |
| RANBP6 | NM_012416 | -1.1245 | -2.386 |
| PITPNC1 | NM_012417 | -0.4415 | -1.5865 |
| RGS17 | NM_012419 | -1.1265 | -2.3335 |
| IFIT5 | NM_012420 | -1.2085 | -2.906 |
| RLF | NM_012421 | -0.5975 | -2.0945 |
| RPL13A | NM_012423 | 0.08 | -0.341 |
| RPS6KC1 | NM_012424 | -0.8095 | -2.0805 |
| RSU1 | NM_012425 | -0.4885 | -0.733 |
| SF3B3 | NM_012426 | -0.4305 | -1.507 |
| KLK5 | NM_012427 | -0.0635 | -0.251 |
| NPTN | NM_012428 | -0.4955 | -1.3235 |
| SEC14L2 | NM_012429 | -0.0205 | -0.3355 |
| SEC22A | NM_012430 | -1.492 | -2.6455 |
| SEMA3E | NM_012431 | -0.013 | 0.157 |
| SETDB1 | NM_012432 | -0.321 | -0.8105 |
| SF3B1 | NM_012433 | -0.6705 | -1.226 |
| SLC17A5 | NM_012434 | -2.226 | -3.008 |
| SHC2 | NM_012435 | -0.0595 | -0.623 |
| SNAPIN | NM_012437 | -0.9925 | -2.2865 |
| SPAG6 | NM_012443 | -0.4005 | 0.1875 |
| SPO11 | NM_012444 | 0.152 | 0.11 |
| SPON2 | NM_012445 | 0.1585 | -0.062 |
| SSBP2 | NM_012446 | -0.3695 | -1.35 |
| STAG3 | NM_012447 | -0.1505 | -0.2015 |
| STAT5B | NM_012448 | -1.0355 | -2.1405 |
| STEAP1 | NM_012449 | -0.3475 | -0.0745 |
| SLC13A4 | NM_012450 | 0.0775 | 0.105 |
| SYNGR4 | NM_012451 | 0.2955 | -0.313 |
| TNFRSF13B | NM_012452 | -0.2255 | -0.5305 |
| TBL2 | NM_012453 | -1.8235 | -3.0415 |
| TIAM2 | NM_012454 | -0.8985 | -2.3525 |
| PSD4 | NM_012455 | 0.2255 | -0.5835 |
| TIMM10 | NM_012456 | -0.4615 | -1.1945 |
| TIMM13 | NM_012458 | -0.0065 | -0.8765 |
| TIMM8B | NM_012459 | -0.3465 | -0.3595 |
| TIMM9 | NM_012460 | -1.4515 | -2.8205 |
| TINF2 | NM_012461 | -1.1365 | -2.681 |
| ATP6V0A2 | NM_012463 | -0.9235 | -2.66 |
| TLL1 | NM_012464 | -0.2835 | -0.729 |
| TLL2 | NM_012465 | -0.281 | -0.3305 |

| | | | |
|---|---|---|---|
| TSPAN16 | NM_012466 | 0.0555 | -0.219 |
| TPSG1 | NM_012467 | -0.031 | -0.54 |
| PRPF6 | NM_012469 | -0.067 | -1.1105 |
| TNPO3 | NM_012470 | -0.7495 | -1.4865 |
| TRPC5 | NM_012471 | 0.232 | -0.0175 |
| LRRC6 | NM_012472 | -0.2535 | -1.117 |
| TXN2 | NM_012473 | -0.022 | -0.582 |
| UCK2 | NM_012474 | 0.216 | -0.9745 |
| USP21 | NM_012475 | -0.396 | -1.078 |
| VAX2 | NM_012476 | -0.377 | -0.4055 |
| WBP2 | NM_012478 | -0.627 | -1.2385 |
| YWHAG | NM_012479 | -0.8 | -1.1295 |
| IKZF3 | NM_012481 | -0.1495 | 0.048 |
| ZNF281 | NM_012482 | -1.737 | -2.3295 |
| GNLY | NM_012483 | 0.087 | -0.188 |
| HMMR | NM_012484 | -1.0135 | -2.5545 |
| HMMR | NM_012485 | -1.0135 | -2.5545 |
| PSEN2 | NM_012486 | -0.1655 | -0.7825 |
| ACAN | NM_013227 | 0.0585 | -0.1605 |
| APAF1 | NM_013229 | -0.295 | -1.123 |
| CD24 | NM_013230 | -1.045 | -1.133 |
| FLRT2 | NM_013231 | -1.8135 | -2.5105 |
| PDCD6 | NM_013232 | -0.5125 | -1.0595 |
| STK39 | NM_013233 | -1.1595 | -1.718 |
| EIF3K | NM_013234 | 0.1655 | -0.2425 |
| DROSHA | NM_013235 | -0.919 | -1.884 |
| ATXN10 | NM_013236 | -0.5975 | -0.7965 |
| PRELID1 | NM_013237 | 0.3205 | -0.2945 |
| DNAJC15 | NM_013238 | 0.484 | -0.894 |
| PPP2R3B | NM_013239 | -0.455 | -0.9995 |
| N6AMT1 | NM_013240 | -0.884 | -2.009 |
| FHOD1 | NM_013241 | -0.197 | -0.842 |
| C16orf80 | NM_013242 | -0.6775 | -1.7165 |
| SCG3 | NM_013243 | -0.2465 | -0.4465 |
| MGAT4C | NM_013244 | -0.215 | -0.4195 |
| VPS4A | NM_013245 | -0.1595 | -1.232 |
| CLCF1 | NM_013246 | -0.2025 | -0.206 |
| HTRA2 | NM_013247 | -0.78 | -1.9055 |
| NXT1 | NM_013248 | -0.505 | -2.1685 |
| ZNF214 | NM_013249 | -0.285 | 0.403 |
| ZNF215 | NM_013250 | -0.2225 | -0.4485 |
| TAC3 | NM_013251 | 0.1885 | -0.1635 |
| CLEC5A | NM_013252 | -0.1915 | -0.143 |

| | | | |
|---|---|---|---|
| DKK3 | NM_013253 | -0.871 | -1.1315 |
| TBK1 | NM_013254 | -1.035 | -2.7795 |
| MKLN1 | NM_013255 | -0.9615 | -1.6665 |
| ZNF180 | NM_013256 | -1.261 | -2.953 |
| PYCARD | NM_013258 | 0.103 | -0.324 |
| TAGLN3 | NM_013259 | -1.188 | -1.2195 |
| SAP30BP | NM_013260 | -0.2965 | -2.2135 |
| PPARGC1A | NM_013261 | -0.849 | -2.517 |
| MYLIP | NM_013262 | -0.133 | -0.991 |
| BRD7 | NM_013263 | -0.96 | -1.752 |
| DDX25 | NM_013264 | -1.022 | -1.3105 |
| C11orf2 | NM_013265 | 0.0755 | -0.422 |
| CTNNA3 | NM_013266 | -0.1425 | -0.549 |
| GLS2 | NM_013267 | 0.062 | -0.388 |
| LGALS13 | NM_013268 | 0.2565 | -0.0225 |
| CLEC2D | NM_013269 | -0.242 | -0.3335 |
| PRSS50 | NM_013270 | 0.506 | 0.2735 |
| PCSK1N | NM_013271 | 0.1345 | -0.215 |
| SLCO3A1 | NM_013272 | -0.267 | -0.953 |
| POLL | NM_013274 | -0.2525 | -0.865 |
| ANKRD11 | NM_013275 | -0.8125 | -1.842 |
| SHPK | NM_013276 | -0.0315 | -1.75 |
| RACGAP1 | NM_013277 | -1.4795 | -1.8495 |
| IL17C | NM_013278 | -0.4315 | -0.1985 |
| C11orf9 | NM_013279 | -0.2265 | -0.6235 |
| FLRT1 | NM_013280 | -0.0605 | -1.09 |
| FLRT3 | NM_013281 | -1.3695 | -3.3835 |
| UHRF1 | NM_013282 | -0.367 | -1.564 |
| MAT2B | NM_013283 | -1.459 | -2.274 |
| POLM | NM_013284 | -0.822 | -1.4875 |
| GNL2 | NM_013285 | -0.563 | -1.768 |
| RBM15B | NM_013286 | -1.545 | -2.9655 |
| KIR3DL1 | NM_013289 | -0.114 | -0.434 |
| CPSF1 | NM_013291 | -0.0415 | -0.4835 |
| MYLPF | NM_013292 | -0.07 | -0.152 |
| TRA2A | NM_013293 | -0.82 | -0.96 |
| GPSM2 | NM_013296 | -1.7055 | -2.675 |
| SAC3D1 | NM_013299 | -0.666 | -2.926 |
| C12orf24 | NM_013300 | -1.5985 | -1.9725 |
| CCDC106 | NM_013301 | 0.0565 | -0.4715 |
| EEF2K | NM_013302 | -0.369 | -1.0495 |
| ZDHHC1 | NM_013304 | -0.0635 | -0.5515 |
| ST8SIA5 | NM_013305 | -0.355 | -0.0525 |

| | | | |
|---|---|---|---|
| SNX15 | NM_013306 | -0.7675 | -1.5305 |
| GPR171 | NM_013308 | 0.1625 | 0.0775 |
| SLC30A4 | NM_013309 | -0.1975 | -0.873 |
| C2orf27A | NM_013310 | -0.625 | -0.6825 |
| HOOK2 | NM_013312 | -0.286 | -1.1375 |
| YPEL1 | NM_013313 | -0.1925 | -0.435 |
| BLNK | NM_013314 | -0.223 | -0.5565 |
| CNOT4 | NM_013316 | -1.293 | -3.4325 |
| PRRC2B | NM_013318 | -0.1525 | -0.9095 |
| UBIAD1 | NM_013319 | -0.8025 | -2.2195 |
| HCFC2 | NM_013320 | -0.6045 | -1.458 |
| SNX8 | NM_013321 | 0.115 | -0.8465 |
| SNX10 | NM_013322 | 0.488 | -0.142 |
| SNX11 | NM_013323 | -0.691 | -2.7925 |
| CISH | NM_013324 | -0.316 | -1.774 |
| ATG4B | NM_013325 | -0.367 | -0.835 |
| C18orf8 | NM_013326 | -0.8215 | -2.477 |
| PARVB | NM_013327 | 0.0445 | 0.1985 |
| PYCR2 | NM_013328 | -0.9545 | -1.5655 |
| GCFC1 | NM_013329 | -1.326 | -2.046 |
| NME7 | NM_013330 | -0.944 | -1.408 |
| C7orf68 | NM_013332 | -0.896 | -2.7265 |
| EPN1 | NM_013333 | 0.202 | -0.2085 |
| GMPPB | NM_013334 | -0.4325 | -1.648 |
| GMPPA | NM_013335 | -0.1755 | -0.796 |
| SEC61A1 | NM_013336 | 0.002 | -1.0615 |
| TIMM22 | NM_013337 | -0.791 | -2.0115 |
| ALG5 | NM_013338 | -1.1745 | -1.4165 |
| ALG6 | NM_013339 | -1.674 | -2.386 |
| PCDHB1 | NM_013340 | -0.049 | -0.103 |
| OLA1 | NM_013341 | -0.701 | -0.7115 |
| TFPT | NM_013342 | -0.645 | -2.405 |
| GPR132 | NM_013345 | -0.052 | -0.4905 |
| SNX12 | NM_013346 | 0.044 | -0.9105 |
| RPA4 | NM_013347 | -0.227 | 0.075 |
| KCNJ14 | NM_013348 | -0.9265 | -0.844 |
| NENF | NM_013349 | 0.3225 | -0.2065 |
| TBX21 | NM_013351 | -0.0155 | 0 |
| DSE | NM_013352 | -0.966 | -2.887 |
| TMOD4 | NM_013353 | -0.4025 | -0.4555 |
| CNOT7 | NM_013354 | -1.0595 | -1.475 |
| PKN3 | NM_013355 | -0.4755 | -1.206 |
| SLC16A8 | NM_013356 | -0.281 | -0.516 |

| | | | |
|---|---|---|---|
| PURG | NM_013357 | -0.066 | -0.5995 |
| PADI1 | NM_013358 | 0.077 | -0.3545 |
| ZNF221 | NM_013359 | -0.25 | -0.3735 |
| ZNF223 | NM_013361 | -1.091 | -2.338 |
| ZNF225 | NM_013362 | -0.941 | -2.7275 |
| PCOLCE2 | NM_013363 | -0.8945 | -1.622 |
| PNMA3 | NM_013364 | -0.353 | -0.846 |
| GGA1 | NM_013365 | -0.682 | -1.404 |
| ANAPC2 | NM_013366 | -0.37 | -1.1375 |
| ANAPC4 | NM_013367 | -1.1085 | -2.312 |
| SERTAD3 | NM_013368 | -0.5115 | -1.3795 |
| DNMT3L | NM_013369 | 0.005 | -0.076 |
| IL19 | NM_013371 | 0.0405 | -0.3105 |
| GREM1 | NM_013372 | 0.0545 | 0.0425 |
| ZDHHC8 | NM_013373 | -0.2145 | -0.858 |
| PDCD6IP | NM_013374 | -0.661 | -1.2085 |
| ABT1 | NM_013375 | -0.644 | -2.722 |
| SERTAD1 | NM_013376 | 0.634 | 1.249 |
| PDZRN4 | NM_013377 | 0.2225 | 0.181 |
| VPREB3 | NM_013378 | 0.0635 | 0.2375 |
| DPP7 | NM_013379 | 0.4075 | -0.117 |
| ZFP112 | NM_013380 | -0.293 | -1.8005 |
| TRHDE | NM_013381 | -0.1665 | -1.0905 |
| POMT2 | NM_013382 | 0.3285 | -0.6625 |
| CYTH4 | NM_013385 | 0.102 | -0.3995 |
| SLC25A24 | NM_013386 | -1.356 | -2.1885 |
| UQCR10 | NM_013387 | 0.0625 | -0.4215 |
| PREB | NM_013388 | -1.042 | -2.386 |
| NPC1L1 | NM_013389 | -0.0785 | -0.547 |
| TMEM2 | NM_013390 | -0.1435 | -1.1785 |
| DMGDH | NM_013391 | -0.0015 | -0.3095 |
| NRBP1 | NM_013392 | -0.152 | -1.6335 |
| FTSJ2 | NM_013393 | -1.0905 | -2.2925 |
| USP25 | NM_013396 | -0.427 | -1.906 |
| PRICKLE4 | NM_013397 | -0.228 | -0.36 |
| ZNF224 | NM_013398 | -1.059 | -2.2545 |
| C16orf5 | NM_013399 | -0.7265 | -2.1975 |
| REPIN1 | NM_013400 | -0.289 | -1.1165 |
| RAB3IL1 | NM_013401 | -0.315 | -1.262 |
| FADS1 | NM_013402 | -0.4245 | -0.4135 |
| STRN4 | NM_013403 | 0.038 | -0.7445 |
| MSLN | NM_013404 | 0.2505 | -0.518 |
| DHPS | NM_013406 | -0.027 | -0.3955 |

| | | | |
|---|---|---|---|
| FST | NM_013409 | -0.04 | -0.2225 |
| AK4 | NM_013410 | -0.494 | -0.979 |
| AK2 | NM_013411 | -0.2575 | -1.04 |
| RABL2A | NM_013412 | -0.6055 | -0.751 |
| NCF4 | NM_013416 | 0.422 | 0.2005 |
| IARS | NM_013417 | -0.9215 | -1.016 |
| ARHGAP6 | NM_013423 | 0.0155 | -0.2675 |
| ARHGAP6 | NM_013427 | -0.659 | -1.8955 |
| KLRC4 | NM_013431 | -0.311 | -0.2 |
| TONSL | NM_013432 | -0.1045 | -0.617 |
| TNPO2 | NM_013433 | -0.709 | -1.032 |
| KCNIP3 | NM_013434 | 0.0455 | -0.2635 |
| RAX | NM_013435 | -0.3645 | -0.4795 |
| NCKAP1 | NM_013436 | -0.764 | -1.344 |
| LRP12 | NM_013437 | -1.1615 | -2.206 |
| UBQLN1 | NM_013438 | -1.104 | -1.012 |
| PILRA | NM_013439 | 0.275 | -0.069 |
| RCAN3 | NM_013441 | -0.1435 | 0.322 |
| STOML2 | NM_013442 | -0.1175 | -0.5375 |
| ST6GALNAC6 | NM_013443 | -0.72 | -1.955 |
| UBQLN2 | NM_013444 | -0.6465 | -1.5035 |
| GAD1 | NM_013445 | 0.033 | 0.0685 |
| MKRN1 | NM_013446 | -0.3455 | -0.87 |
| EMR2 | NM_013447 | -0.016 | -0.484 |
| BAZ1A | NM_013448 | -1.241 | -4.5405 |
| BAZ2A | NM_013449 | -0.2955 | -1.654 |
| BAZ2B | NM_013450 | -0.4135 | -1.952 |
| MYOF | NM_013451 | -0.237 | -0.1815 |
| VCX | NM_013452 | -0.03 | -0.16 |
| SPANXA1 | NM_013453 | 0.1685 | -0.472 |
| OR12D2 | NM_013936 | -0.1355 | -0.4835 |
| OR11A1 | NM_013937 | -0.297 | -0.307 |
| OR10H3 | NM_013938 | 0.1725 | -0.4325 |
| OR10H2 | NM_013939 | -0.146 | -0.1255 |
| OR10H1 | NM_013940 | -0.2105 | -0.3525 |
| OR10C1 | NM_013941 | 0.09 | 0.155 |
| PAX3 | NM_013942 | 0.1345 | 0.0185 |
| CLIC4 | NM_013943 | -1.0075 | -0.7305 |
| PAX7 | NM_013945 | 0.2385 | -0.4755 |
| PAX8 | NM_013952 | 0.1665 | -0.088 |
| PAX8 | NM_013953 | 0.1665 | -0.088 |
| NOX1 | NM_013955 | -0.263 | -0.528 |
| NRG1 | NM_013956 | -0.1445 | -0.887 |

| | | | |
|---|---|---|---|
| NRG1 | NM_013957 | -0.1445 | -0.887 |
| NRG1 | NM_013958 | -0.1445 | -0.887 |
| NRG1 | NM_013959 | -0.1445 | -0.887 |
| NRG1 | NM_013960 | -0.1445 | -0.887 |
| NRG1 | NM_013962 | -0.1445 | -0.887 |
| NRG1 | NM_013964 | -0.1445 | -0.887 |
| DDAH2 | NM_013974 | 0.0135 | -0.9945 |
| LIG3 | NM_013975 | -0.383 | -1.802 |
| GCDH | NM_013976 | 0.018 | -0.555 |
| BNIP1 | NM_013978 | -0.6045 | -1.8855 |
| BNIP1 | NM_013979 | -0.5905 | -2.2155 |
| BNIP1 | NM_013980 | -0.6045 | -1.8855 |
| NRG2 | NM_013981 | -0.0085 | -0.3145 |
| NRG2 | NM_013982 | 0.1515 | -0.164 |
| NRG2 | NM_013983 | -0.0085 | -0.3145 |
| EWSR1 | NM_013986 | -0.296 | -0.8785 |
| PARK2 | NM_013987 | -0.0205 | -0.377 |
| PARK2 | NM_013988 | -0.0205 | -0.377 |
| DIO2 | NM_013989 | 0.0605 | -0.2345 |
| PAX8 | NM_013992 | 0.1665 | -0.088 |
| DDR1 | NM_013993 | 0.1375 | -0.6015 |
| DDR1 | NM_013994 | -0.0885 | -0.7975 |
| LAMP2 | NM_013995 | -0.942 | -0.8055 |
| MEOX1 | NM_013999 | 0.0855 | 0.1905 |
| VCL | NM_014000 | -0.7635 | -1.1865 |
| GGA3 | NM_014001 | -0.9555 | -2.739 |
| IKBKE | NM_014002 | -0.1855 | -0.3585 |
| DHX38 | NM_014003 | -0.1045 | -0.4435 |
| PCDHGA8 | NM_014004 | -0.296 | 0.1455 |
| PCDHA9 | NM_014005 | -0.4555 | -0.474 |
| ZBTB43 | NM_014007 | -0.8695 | -1.6645 |
| CCDC22 | NM_014008 | 0.209 | -0.469 |
| FOXP3 | NM_014009 | -0.14 | -0.278 |
| ASTN2 | NM_014010 | 0.029 | -1.152 |
| SOCS5 | NM_014011 | -0.0255 | 0.079 |
| REM1 | NM_014012 | -0.0155 | -0.109 |
| SNRNP200 | NM_014014 | -0.144 | -0.407 |
| DEXI | NM_014015 | -0.7185 | -2.7235 |
| SACM1L | NM_014016 | -1.3545 | -2.715 |
| LAMTOR2 | NM_014017 | 0.0715 | -0.512 |
| MRPS28 | NM_014018 | -0.721 | -2.325 |
| TMEM176B | NM_014020 | -0.045 | -0.207 |
| SSX2IP | NM_014021 | -0.6025 | -1.314 |

| | | | |
|---|---|---|---|
| WDR37 | NM_014023 | -1.4875 | -3.2925 |
| DCPS | NM_014026 | -0.053 | -1.442 |
| OSTM1 | NM_014028 | -1.7625 | -2.7425 |
| GIT1 | NM_014030 | -0.018 | -0.6135 |
| METTL7A | NM_014033 | -0.6725 | -2.2175 |
| ASF1A | NM_014034 | -1.2555 | -4.691 |
| SNX24 | NM_014035 | -0.669 | -0.7945 |
| SLC6A16 | NM_014037 | 0.136 | -0.1595 |
| BZW2 | NM_014038 | -0.725 | -0.9785 |
| C11orf54 | NM_014039 | -1.763 | -1.635 |
| SPCS1 | NM_014041 | -0.1755 | -0.711 |
| C11orf51 | NM_014042 | 0.344 | -0.327 |
| CHMP2B | NM_014043 | -1.2205 | -2.054 |
| UNC50 | NM_014044 | -1.5225 | -3.374 |
| LRP10 | NM_014045 | -0.363 | -0.5205 |
| MRPS18B | NM_014046 | -0.2875 | -1.761 |
| C19orf53 | NM_014047 | 0.013 | -0.6 |
| MKL2 | NM_014048 | -1.5885 | -4.1605 |
| ACAD9 | NM_014049 | -0.585 | -1.2965 |
| MRPL42 | NM_014050 | -0.924 | -1.098 |
| TMEM14A | NM_014051 | -0.2655 | -0.608 |
| FLVCR1 | NM_014053 | -0.7305 | -1.899 |
| IFT81 | NM_014055 | -0.4495 | -0.6725 |
| OGN | NM_014057 | 0.433 | -0.2595 |
| TMPRSS11E | NM_014058 | -0.0725 | -0.427 |
| C13orf15 | NM_014059 | -0.192 | -0.1005 |
| MCTS1 | NM_014060 | -0.325 | -0.86 |
| MAGEH1 | NM_014061 | -0.676 | -0.6565 |
| NOB1 | NM_014062 | -0.4755 | -2.1775 |
| DBNL | NM_014063 | -0.088 | -0.6955 |
| METTL11A | NM_014064 | -0.3815 | -2.0545 |
| ASTE1 | NM_014065 | -1.744 | -3.119 |
| COMMD5 | NM_014066 | -0.6445 | -2.1015 |
| MACROD1 | NM_014067 | 0.2905 | -0.298 |
| PSORS1C1 | NM_014068 | -0.1445 | -0.8155 |
| PSORS1C2 | NM_014069 | -0.188 | 0.0995 |
| C6orf15 | NM_014070 | 0.1955 | 0.0595 |
| NCOA6 | NM_014071 | -1.147 | -2.5645 |
| FAM32A | NM_014077 | -0.6 | -1.1575 |
| MRPL13 | NM_014078 | -1.1695 | -2.2045 |
| KLF15 | NM_014079 | -1.216 | -2.7905 |
| DUOX2 | NM_014080 | 0.0595 | 0.029 |
| NUPL1 | NM_014089 | -2.027 | -3.172 |

| | | | |
|---|---|---|---|
| SLC43A3 | NM_014096 | -0.6805 | -1.804 |
| PRDX3 | NM_014098 | -0.863 | -0.5005 |
| ZNF770 | NM_014106 | -1.459 | -2.8455 |
| ATAD2 | NM_014109 | -0.801 | -3.089 |
| PPP1R8 | NM_014110 | -1.4105 | -3.159 |
| TRPS1 | NM_014112 | -0.3805 | -1.3555 |
| C16orf72 | NM_014117 | -1.4885 | -3.823 |
| SCN11A | NM_014139 | -0.533 | -0.352 |
| SMARCAL1 | NM_014140 | -1.7225 | -3.108 |
| CNTNAP2 | NM_014141 | -0.575 | -0.5395 |
| NUDT5 | NM_014142 | -0.6215 | -1.0695 |
| CD274 | NM_014143 | 0.229 | -0.127 |
| LAT2 | NM_014146 | 0.1465 | -0.4915 |
| WDR91 | NM_014149 | -0.3385 | -1.49 |
| ZC3H7A | NM_014153 | -1.288 | -1.5825 |
| ARMC8 | NM_014154 | -0.332 | -1.113 |
| ZBTB44 | NM_014155 | -1.329 | -2.552 |
| CCDC113 | NM_014157 | -1.13 | -1.0975 |
| SETD2 | NM_014159 | -0.676 | -3.114 |
| MKRN2 | NM_014160 | -0.838 | -1.4815 |
| MRPL18 | NM_014161 | -1.236 | -3.6825 |
| FXYD5 | NM_014164 | -0.0945 | 0.022 |
| NDUFAF4 | NM_014165 | -0.952 | -2.3235 |
| MED4 | NM_014166 | -0.8475 | -1.04 |
| CCDC59 | NM_014167 | -0.7755 | -1.846 |
| METTL5 | NM_014168 | -1.6095 | -2.4705 |
| CHMP4A | NM_014169 | -0.4365 | -1.008 |
| GTPBP8 | NM_014170 | -0.908 | -1.5095 |
| CRIPT | NM_014171 | -0.5835 | -1.0815 |
| BABAM1 | NM_014173 | 0.164 | -0.498 |
| THYN1 | NM_014174 | -1.0725 | -1.3245 |
| MRPL15 | NM_014175 | -1.4585 | -2.9235 |
| UBE2T | NM_014176 | -1.4495 | -2.891 |
| C18orf55 | NM_014177 | -1.2895 | -3.4495 |
| STXBP6 | NM_014178 | 0.177 | -0.4215 |
| MRPL22 | NM_014180 | -0.6005 | -2.3645 |
| LGALSL | NM_014181 | -1.5025 | -1.7715 |
| ORMDL2 | NM_014182 | -0.7365 | -1.1355 |
| DYNLRB1 | NM_014183 | -0.2025 | -0.553 |
| CNIH4 | NM_014184 | -0.4475 | -1.686 |
| COMMD9 | NM_014186 | 0.121 | -0.64 |
| TMEM208 | NM_014187 | -0.056 | -0.7525 |
| SSU72 | NM_014188 | -1.021 | -1.5415 |

| | | | |
|---|---|---|---|
| ADD1 | NM_014189 | 0.0445 | -0.8975 |
| ADD1 | NM_014190 | 0.0445 | -0.8975 |
| SCN8A | NM_014191 | -0.232 | -0.931 |
| AP2A1 | NM_014203 | -0.0675 | -0.3815 |
| ZNHIT2 | NM_014205 | -0.7095 | -1.4105 |
| C11orf10 | NM_014206 | 0.164 | -0.317 |
| CD5 | NM_014207 | 0.4445 | -0.3625 |
| DSPP | NM_014208 | 0.0915 | 0.225 |
| ETV2 | NM_014209 | -0.3855 | -0.486 |
| EVI2A | NM_014210 | 0.3345 | -0.6005 |
| GABRP | NM_014211 | -0.002 | -0.1575 |
| HOXC11 | NM_014212 | 0.047 | 0.6415 |
| HOXD9 | NM_014213 | -0.1715 | -0.834 |
| IMPA2 | NM_014214 | -0.5875 | 0.048 |
| INSRR | NM_014215 | 0.0315 | -0.336 |
| ITPK1 | NM_014216 | 0.0235 | -0.9185 |
| KCNK2 | NM_014217 | 0.1875 | -1.1915 |
| KIR2DL1 | NM_014218 | -0.102 | -0.2335 |
| TM4SF1 | NM_014220 | -0.776 | -0.2825 |
| NDUFA8 | NM_014222 | -0.4755 | -1.511 |
| NFYC | NM_014223 | -0.3825 | -1.432 |
| PPP2R1A | NM_014225 | 0.2255 | -0.3285 |
| MOK | NM_014226 | -0.249 | -0.6315 |
| SLC5A4 | NM_014227 | 0.0335 | -0.291 |
| SLC6A7 | NM_014228 | -0.113 | -0.702 |
| SLC6A11 | NM_014229 | -0.136 | -0.4065 |
| SRP68 | NM_014230 | -1.186 | -3.3995 |
| VAMP1 | NM_014231 | 0.081 | -0.882 |
| VAMP2 | NM_014232 | 0.174 | -0.1055 |
| UBTF | NM_014233 | -0.4465 | -1.622 |
| HSD17B8 | NM_014234 | 0.1525 | -0.2845 |
| UBL4A | NM_014235 | -0.2025 | -1.0805 |
| GNPAT | NM_014236 | -0.877 | -0.709 |
| ADAM18 | NM_014237 | 0.1245 | -0.213 |
| KSR1 | NM_014238 | -0.171 | -0.762 |
| EIF2B2 | NM_014239 | -0.513 | -1.778 |
| LIMD1 | NM_014240 | -0.575 | -2.1465 |
| PTPLA | NM_014241 | -0.3555 | -0.5605 |
| ADAMTS3 | NM_014243 | -0.843 | -1.108 |
| ADAMTS2 | NM_014244 | -0.244 | -0.3745 |
| RNF7 | NM_014245 | -0.175 | -0.8935 |
| CELSR1 | NM_014246 | 0.253 | -0.1595 |
| RAPGEF2 | NM_014247 | -1.465 | -3.9435 |

| | | | |
|---|---|---|---|
| RBX1 | NM_014248 | -0.325 | -0.704 |
| NR2E3 | NM_014249 | 0.3865 | 0.558 |
| SLC25A13 | NM_014251 | -0.4575 | -1.218 |
| SLC25A15 | NM_014252 | -0.5945 | -1.901 |
| ODZ1 | NM_014253 | -0.0075 | -0.5005 |
| TMEM5 | NM_014254 | -1.3905 | -2.6595 |
| CNPY2 | NM_014255 | -0.398 | -0.8235 |
| B3GNT3 | NM_014256 | 0.093 | -0.053 |
| CLEC4M | NM_014257 | -0.3085 | -0.2745 |
| SYCP2 | NM_014258 | -0.726 | 0.5285 |
| PFDN6 | NM_014260 | -0.5205 | -1.434 |
| LEPREL2 | NM_014262 | -0.178 | -1.014 |
| YME1L1 | NM_014263 | -1.1755 | -1.5985 |
| PLK4 | NM_014264 | -1.158 | -3.2425 |
| ADAM28 | NM_014265 | 0.275 | 0.224 |
| HCST | NM_014266 | -0.164 | -0.1505 |
| C11orf58 | NM_014267 | -1.074 | -1.5065 |
| MAPRE2 | NM_014268 | -0.7635 | -1.343 |
| ADAM29 | NM_014269 | 0.0215 | -0.2305 |
| SLC7A9 | NM_014270 | 0.0585 | -0.8785 |
| IL1RAPL1 | NM_014271 | 0.198 | -0.4605 |
| ADAMTS7 | NM_014272 | -0.094 | -0.5925 |
| MGAT4B | NM_014275 | -0.016 | -0.7525 |
| RBPJL | NM_014276 | -0.036 | -0.3015 |
| HSPA4L | NM_014278 | -0.5575 | -1.761 |
| OLFM1 | NM_014279 | -0.525 | -0.7545 |
| DNAJC8 | NM_014280 | -0.164 | -0.544 |
| PUF60 | NM_014281 | 0.0035 | -0.874 |
| HABP4 | NM_014282 | -0.5235 | -1.5375 |
| C1orf9 | NM_014283 | -0.9445 | -2.206 |
| NCDN | NM_014284 | -0.191 | -0.6295 |
| EXOSC2 | NM_014285 | -1.61 | -2.542 |
| NCS1 | NM_014286 | -0.2275 | -0.618 |
| NOMO1 | NM_014287 | -0.2055 | -0.9065 |
| ITGB3BP | NM_014288 | -1.011 | -0.904 |
| CAPN6 | NM_014289 | -0.334 | -0.618 |
| TDRD7 | NM_014290 | 0.554 | -0.322 |
| GCAT | NM_014291 | -0.0065 | -0.5005 |
| CBX6 | NM_014292 | -1.2185 | -1.8525 |
| NPTXR | NM_014293 | -0.233 | -0.601 |
| TRAM1 | NM_014294 | -1.55 | -3.5275 |
| CAPN7 | NM_014296 | -1.1175 | -2.3685 |
| ETHE1 | NM_014297 | -0.063 | -0.4025 |

| | | | |
|---|---|---|---|
| QPRT | NM_014298 | 0.0605 | -0.545 |
| BRD4 | NM_014299 | 0.0445 | -0.87 |
| SEC11A | NM_014300 | -0.5125 | -0.9775 |
| ISCU | NM_014301 | -0.3305 | -0.766 |
| SEC61G | NM_014302 | -0.0735 | -0.3165 |
| PES1 | NM_014303 | -0.1135 | -1.4035 |
| TGDS | NM_014305 | -1.2345 | -2.161 |
| C22orf28 | NM_014306 | -0.3675 | -1.3905 |
| PIK3R5 | NM_014308 | 0.015 | -0.1775 |
| RBFOX2 | NM_014309 | -0.681 | -1.2645 |
| RASD2 | NM_014310 | -0.329 | 0.002 |
| SMUG1 | NM_014311 | 0.0505 | -0.914 |
| VSIG2 | NM_014312 | 0.267 | -0.1145 |
| TMEM50A | NM_014313 | -0.9305 | -1.1715 |
| DDX58 | NM_014314 | 0.0115 | -0.3465 |
| KLHDC2 | NM_014315 | -1.437 | -1.779 |
| CARHSP1 | NM_014316 | 0.125 | -0.8365 |
| PDSS1 | NM_014317 | -1.552 | -2.4065 |
| LEMD3 | NM_014319 | -1.4425 | -3.7715 |
| HEBP2 | NM_014320 | 0.149 | -0.474 |
| ORC6 | NM_014321 | -1.173 | -0.7775 |
| OPN3 | NM_014322 | -1.9625 | -2.348 |
| PATZ1 | NM_014323 | -1.123 | -3.614 |
| AMACR | NM_014324 | -0.4105 | -1.661 |
| CORO1C | NM_014325 | -0.6235 | -0.9055 |
| DAPK2 | NM_014326 | -0.347 | 0.1585 |
| RUSC1 | NM_014328 | -0.2105 | -0.7045 |
| EDC4 | NM_014329 | -0.172 | -0.995 |
| PPP1R15A | NM_014330 | -0.2005 | -0.8335 |
| SLC7A11 | NM_014331 | 0.256 | -0.2805 |
| SMPX | NM_014332 | -0.2665 | 0.083 |
| CADM1 | NM_014333 | -0.7415 | -0.5395 |
| C9orf4 | NM_014334 | 0.345 | -0.352 |
| EID1 | NM_014335 | -1.194 | -0.9605 |
| AIPL1 | NM_014336 | -0.0765 | -0.6315 |
| PPIL2 | NM_014337 | -0.5575 | -1.012 |
| PISD | NM_014338 | -0.559 | -1.836 |
| IL17RA | NM_014339 | 0 | -1.053 |
| MTCH1 | NM_014341 | 0.4985 | -0.154 |
| MTCH2 | NM_014342 | -0.12 | -1.165 |
| CLDN15 | NM_014343 | 0.1215 | -1.521 |
| FJX1 | NM_014344 | -0.856 | -1.705 |
| ZNF318 | NM_014345 | -1.3025 | -2.363 |

| | | | |
|---|---|---|---|
| TBC1D22A | NM_014346 | -0.3605 | -1.1045 |
| ZNF324 | NM_014347 | -0.5945 | -1.1625 |
| TNFAIP8 | NM_014350 | -0.4285 | -2.3435 |
| SULT4A1 | NM_014351 | 0.298 | -0.0445 |
| POU2F3 | NM_014352 | 0.138 | 0.3435 |
| RAB26 | NM_014353 | -0.0415 | -0.341 |
| LCE2B | NM_014357 | 0.1325 | -0.0505 |
| CLEC4E | NM_014358 | -0.2445 | -0.0885 |
| OPTC | NM_014359 | 0.137 | -0.348 |
| NKX2-8 | NM_014360 | -0.063 | -0.435 |
| CNTN5 | NM_014361 | -0.29 | -0.2995 |
| HIBCH | NM_014362 | -0.5965 | -0.9265 |
| SACS | NM_014363 | -0.815 | -3.6785 |
| GAPDHS | NM_014364 | -0.0165 | 0.0855 |
| HSPB8 | NM_014365 | -0.1465 | -0.1015 |
| GNL3 | NM_014366 | -1.5065 | -2.8865 |
| FAM162A | NM_014367 | -0.3525 | -0.7915 |
| LHX6 | NM_014368 | -0.398 | -0.4705 |
| PTPN18 | NM_014369 | -0.1995 | -0.329 |
| SRPK3 | NM_014370 | -0.1165 | -0.258 |
| AKAP8L | NM_014371 | -0.089 | -0.7975 |
| RNF11 | NM_014372 | -0.949 | -0.8795 |
| GPR160 | NM_014373 | -0.6715 | -0.4085 |
| REPIN1 | NM_014374 | -0.3255 | -0.718 |
| FETUB | NM_014375 | 0.233 | -0.377 |
| CYFIP2 | NM_014376 | -0.0385 | -0.562 |
| DNAJC2 | NM_014377 | -1.032 | -2.6015 |
| KCNV1 | NM_014379 | -0.1085 | -0.261 |
| MLH3 | NM_014381 | -0.898 | -0.9805 |
| ATP2C1 | NM_014382 | -0.972 | -1.569 |
| ZBTB32 | NM_014383 | 0.0355 | -0.507 |
| ACAD8 | NM_014384 | -0.541 | -1.5475 |
| PKD2L2 | NM_014386 | 0.028 | -0.433 |
| DIEXF | NM_014388 | -2.0135 | -2.2635 |
| PELP1 | NM_014389 | 0.08 | -1.0135 |
| SND1 | NM_014390 | 0.2685 | -0.2675 |
| ANKRD1 | NM_014391 | -0.3415 | 0.403 |
| D4S234E | NM_014392 | -0.0425 | -0.873 |
| STAU2 | NM_014393 | -0.786 | -0.834 |
| GHITM | NM_014394 | -0.1595 | -0.6575 |
| DAPP1 | NM_014395 | 0.0645 | -0.1695 |
| VPS41 | NM_014396 | -0.8055 | -0.468 |
| NEK6 | NM_014397 | -0.1015 | -0.482 |

| | | | |
|---|---|---|---|
| LAMP3 | NM_014398 | 0.2655 | -0.009 |
| TSPAN13 | NM_014399 | -0.301 | -0.111 |
| LYPD3 | NM_014400 | -0.205 | -0.5425 |
| UQCRQ | NM_014402 | 0.213 | -0.193 |
| CACNG4 | NM_014405 | -0.8975 | -1.3985 |
| KCNMB3 | NM_014407 | -0.5575 | -0.341 |
| TRAPPC3 | NM_014408 | -0.7005 | -1.709 |
| TAF5L | NM_014409 | -0.5965 | -1.8665 |
| CLUL1 | NM_014410 | 0.1115 | -0.3915 |
| CACYBP | NM_014412 | -0.6125 | -1.4635 |
| EIF2AK1 | NM_014413 | -0.508 | -0.8085 |
| ZBTB11 | NM_014415 | -0.9305 | -2.4075 |
| BBC3 | NM_014417 | 0.1715 | -0.1555 |
| DKKL1 | NM_014419 | 0.1385 | -0.281 |
| DKK4 | NM_014420 | -0.2015 | -0.266 |
| DKK2 | NM_014421 | -0.098 | -0.035 |
| AFF4 | NM_014423 | -1.4205 | -3.2535 |
| HSPB7 | NM_014424 | -0.187 | -0.4295 |
| INVS | NM_014425 | -0.6355 | -1.393 |
| SNX5 | NM_014426 | 0.9905 | 1.8715 |
| CPNE7 | NM_014427 | 0.1025 | -0.276 |
| TJP3 | NM_014428 | -0.12 | -0.2375 |
| MORC1 | NM_014429 | 0.0095 | -0.113 |
| CIDEB | NM_014430 | 0.0985 | -0.21 |
| KIAA1274 | NM_014431 | -0.185 | -1.113 |
| IL20RA | NM_014432 | 0.194 | 0.1475 |
| RTDR1 | NM_014433 | -0.0935 | -0.097 |
| NDOR1 | NM_014434 | -0.3475 | -0.622 |
| NAAA | NM_014435 | -0.619 | -0.863 |
| SLC39A1 | NM_014437 | -0.1725 | -0.623 |
| IL36B | NM_014438 | 0.2815 | 0.006 |
| IL37 | NM_014439 | 0.0805 | 0.0145 |
| IL36A | NM_014440 | 0.211 | -0.0045 |
| SIGLEC9 | NM_014441 | -0.375 | -0.288 |
| SIGLEC8 | NM_014442 | -0.2685 | -0.217 |
| IL17B | NM_014443 | 0.085 | -0.225 |
| TUBGCP4 | NM_014444 | -0.7335 | -1.2255 |
| SERP1 | NM_014445 | -0.357 | -1.519 |
| ARFIP1 | NM_014447 | -1.3235 | -3.1465 |
| ARHGEF16 | NM_014448 | -0.1985 | -0.3645 |
| GPR162 | NM_014449 | -0.5585 | -0.5425 |
| SIT1 | NM_014450 | -0.407 | -0.227 |
| BBS9 | NM_014451 | -0.615 | -1.0105 |

| | | | |
|---|---|---|---|
| TNFRSF21 | NM_014452 | -0.4265 | -2.0815 |
| CHMP2A | NM_014453 | 0.0915 | -0.2545 |
| SESN1 | NM_014454 | -1.329 | -2.1515 |
| RNF115 | NM_014455 | -0.696 | -2.1705 |
| PDCD4 | NM_014456 | -1.3635 | -3.0195 |
| KLHL20 | NM_014458 | -0.262 | -2.073 |
| CSDC2 | NM_014460 | -0.0345 | -0.4655 |
| CNTN6 | NM_014461 | -0.142 | -0.0985 |
| LSM1 | NM_014462 | -1.1905 | -2.1465 |
| LSM3 | NM_014463 | -0.193 | -0.3125 |
| TINAG | NM_014464 | 0.3305 | -0.4175 |
| SULT1B1 | NM_014465 | -0.193 | -0.1125 |
| TEKT2 | NM_014466 | -0.37 | -0.126 |
| SRPX2 | NM_014467 | -0.152 | 0.0005 |
| VENTX | NM_014468 | 0.1245 | -0.377 |
| RBMXL2 | NM_014469 | 0.049 | -0.198 |
| RND1 | NM_014470 | 0.19 | -0.179 |
| SPINK4 | NM_014471 | -0.3065 | -0.238 |
| C10orf28 | NM_014472 | -0.662 | -1.397 |
| SMPDL3B | NM_014474 | -0.055 | -0.3335 |
| DHDH | NM_014475 | -0.29 | -0.2625 |
| PDLIM3 | NM_014476 | -0.334 | -1.205 |
| TP53TG5 | NM_014477 | 0.287 | -0.478 |
| CRCP | NM_014478 | -0.735 | -1.8625 |
| ADAMDEC1 | NM_014479 | 0.052 | 0.2435 |
| ZNF544 | NM_014480 | -1.405 | -2.096 |
| APEX2 | NM_014481 | -1.0215 | -2.3495 |
| BMP10 | NM_014482 | 0.1715 | -0.3375 |
| RBMS3 | NM_014483 | -0.315 | -0.709 |
| MOCS3 | NM_014484 | -1.773 | -3.2665 |
| HPGDS | NM_014485 | -0.093 | -0.3395 |
| ZNF330 | NM_014487 | -0.771 | -1.4825 |
| RAB30 | NM_014488 | 0.329 | -1.043 |
| PGAP2 | NM_014489 | -0.393 | -1.112 |
| FOXP2 | NM_014491 | 0.8085 | -0.3895 |
| TNRC6A | NM_014494 | -1.863 | -3.251 |
| ANGPTL3 | NM_014495 | -0.7245 | 0.1905 |
| RPS6KA6 | NM_014496 | -0.4795 | -0.3825 |
| ZNF638 | NM_014497 | -1.0405 | -2.5595 |
| GOLIM4 | NM_014498 | -0.4535 | -0.813 |
| HTATSF1 | NM_014500 | -0.78 | -0.327 |
| UBE2S | NM_014501 | -0.405 | -0.89 |
| PRPF19 | NM_014502 | -0.0645 | -0.917 |

| | | | |
|---|---|---|---|
| UTP20 | NM_014503 | -0.284 | -1.489 |
| KCNMB4 | NM_014505 | -0.3395 | -1.4915 |
| TOR1B | NM_014506 | -1.3335 | -2.5545 |
| MCAT | NM_014507 | -0.875 | -2.853 |
| APOBEC3C | NM_014508 | -0.293 | -0.311 |
| SERHL2 | NM_014509 | -0.118 | -0.2715 |
| PCLO | NM_014510 | -1.0235 | -1.31 |
| CNOT2 | NM_014515 | -1.895 | -3.0975 |
| CNOT3 | NM_014516 | -0.383 | -1.5195 |
| UBP1 | NM_014517 | -0.966 | -0.97 |
| ZNF229 | NM_014518 | -0.9355 | -1.9275 |
| ZNF232 | NM_014519 | -1.7925 | -3.382 |
| MYBBP1A | NM_014520 | -0.534 | -0.937 |
| SH3BP4 | NM_014521 | -0.862 | -2.759 |
| PCDH11X | NM_014522 | -0.172 | -0.1865 |
| TMOD3 | NM_014547 | -0.3335 | -0.922 |
| TMOD2 | NM_014548 | -0.94 | -1.242 |
| CARD10 | NM_014550 | -0.219 | -0.4715 |
| NCAPH2 | NM_014551 | 0.2225 | -0.4435 |
| TFCP2L1 | NM_014553 | -0.0605 | -0.2145 |
| SENP1 | NM_014554 | -2.024 | -2.985 |
| TRPM5 | NM_014555 | 0.1625 | -0.263 |
| OTX1 | NM_014562 | -0.393 | -0.2325 |
| OR1A1 | NM_014565 | 0.1815 | -0.2455 |
| OR1D5 | NM_014566 | -0.206 | -0.2445 |
| BCAR1 | NM_014567 | -0.474 | -1.64 |
| GALNT5 | NM_014568 | -0.2005 | -0.4775 |
| ZKSCAN5 | NM_014569 | -1.448 | -2.2495 |
| ARFGAP3 | NM_014570 | -1.302 | -1.741 |
| HEYL | NM_014571 | 0.046 | -0.564 |
| LATS2 | NM_014572 | -0.2995 | -2.2445 |
| TMEM97 | NM_014573 | -0.97 | -1.2605 |
| STRN3 | NM_014574 | -1.472 | -3.2245 |
| SCHIP1 | NM_014575 | -1.05 | -2.186 |
| A1CF | NM_014576 | 0.0105 | -0.246 |
| BRD1 | NM_014577 | -0.549 | -2.1435 |
| RHOD | NM_014578 | 0.058 | -0.3555 |
| SLC39A2 | NM_014579 | 0.258 | -0.3115 |
| SLC2A8 | NM_014580 | -0.508 | -1.7525 |
| OBP2B | NM_014581 | 0.2205 | -0.461 |
| LMCD1 | NM_014583 | -0.867 | -1.1285 |
| ERO1L | NM_014584 | -0.3855 | -1.1015 |
| SLC40A1 | NM_014585 | -0.132 | -1.0305 |

| | | | |
|---|---|---|---|
| HUNK | NM_014586 | -0.3975 | -1.8935 |
| SOX8 | NM_014587 | 0.2765 | 0.384 |
| VSX1 | NM_014588 | 0.1785 | -0.178 |
| PLA2G2E | NM_014589 | -0.048 | -0.209 |
| KCNIP2 | NM_014591 | 0.244 | -0.242 |
| KCNIP1 | NM_014592 | -0.239 | -0.458 |
| CXXC1 | NM_014593 | 0.0585 | -0.5545 |
| ZNF354C | NM_014594 | -0.547 | -1.2665 |
| NT5C | NM_014595 | -0.6195 | -2.1445 |
| ZNRD1 | NM_014596 | -0.816 | -2.6705 |
| DNTTIP2 | NM_014597 | -0.76 | -2.5595 |
| SOCS7 | NM_014598 | 0.7745 | -1.03 |
| MAGED2 | NM_014599 | 0.016 | -0.594 |
| EHD3 | NM_014600 | -0.7905 | -1.9745 |
| EHD2 | NM_014601 | -0.0965 | -0.323 |
| PIK3R4 | NM_014602 | -1.6135 | -3.466 |
| CDR2L | NM_014603 | -1.1225 | -2.6125 |
| TAX1BP3 | NM_014604 | -0.203 | -0.7535 |
| HERC3 | NM_014606 | 0.005 | -0.3145 |
| UBXN4 | NM_014607 | -0.9675 | -2.278 |
| CYFIP1 | NM_014608 | -0.1425 | -0.522 |
| MDN1 | NM_014611 | -0.942 | -1.6625 |
| FAM120A | NM_014612 | -0.8685 | -2.1755 |
| FAF2 | NM_014613 | -1.892 | -3.0575 |
| PSME4 | NM_014614 | -0.7895 | -1.4205 |
| KIAA0182 | NM_014615 | -0.799 | -1.3995 |
| ATP11B | NM_014616 | -0.6415 | -1.6645 |
| CRYGA | NM_014617 | -0.1605 | -0.4985 |
| DBC1 | NM_014618 | 0.408 | -1.4245 |
| GRIK4 | NM_014619 | 0.05 | 0.065 |
| HOXC4 | NM_014620 | -0.56 | -1.449 |
| HOXD4 | NM_014621 | -0.289 | -0.9055 |
| VWA5A | NM_014622 | 0.0015 | -0.2135 |
| MEA1 | NM_014623 | -0.2 | -0.631 |
| S100A6 | NM_014624 | -0.335 | -0.423 |
| NPHS2 | NM_014625 | 0.136 | 0.0595 |
| ARHGEF10 | NM_014629 | -1.825 | -2.6115 |
| ZNF592 | NM_014630 | -0.0065 | -1.2415 |
| SH3PXD2A | NM_014631 | -0.623 | -1.7965 |
| MICAL2 | NM_014632 | -0.4585 | -0.823 |
| CTR9 | NM_014633 | -0.721 | -1.4095 |
| PPM1F | NM_014634 | -1.068 | -1.977 |
| RALGPS1 | NM_014636 | -0.989 | -0.613 |

| | | | |
|---|---|---|---|
| MTFR1 | NM_014637 | -0.7485 | -1.9155 |
| PLCH2 | NM_014638 | 0.221 | -0.9035 |
| TTC37 | NM_014639 | 0.2645 | -1.3995 |
| TTLL4 | NM_014640 | -0.181 | -2.2325 |
| MDC1 | NM_014641 | -0.7245 | -1.425 |
| ZNF516 | NM_014643 | -0.899 | -2.1795 |
| PDE4DIP | NM_014644 | -0.2405 | -0.902 |
| LPIN2 | NM_014646 | -0.823 | -1.873 |
| KIAA0430 | NM_014647 | -1.4 | -2.7775 |
| DZIP3 | NM_014648 | -0.5905 | -1.1135 |
| SAFB2 | NM_014649 | -0.5665 | -1.535 |
| ZNF432 | NM_014650 | -0.147 | -1.0575 |
| IPO13 | NM_014652 | -0.516 | -1.31 |
| WSCD2 | NM_014653 | -0.1685 | -0.4975 |
| SDC3 | NM_014654 | -0.408 | -0.7105 |
| SLC25A44 | NM_014655 | -0.6345 | -1.4645 |
| KIAA0040 | NM_014656 | -0.591 | 0.2585 |
| TTI1 | NM_014657 | -1.3095 | -3.02 |
| PPIP5K1 | NM_014659 | -0.2065 | -0.879 |
| PHF14 | NM_014660 | -0.3955 | -1.4025 |
| FAM53B | NM_014661 | -0.5845 | -1.8395 |
| DEPDC5 | NM_014662 | -1.6815 | -3.1645 |
| KDM4A | NM_014663 | -0.338 | -1.1265 |
| LRRC14 | NM_014665 | -0.7045 | -1.4705 |
| CLINT1 | NM_014666 | -1.044 | -2.019 |
| VGLL4 | NM_014667 | -0.198 | -1.1195 |
| GREB1 | NM_014668 | -0.43 | -0.7655 |
| NUP93 | NM_014669 | 0.009 | -1.033 |
| BZW1 | NM_014670 | -0.8115 | -0.797 |
| UBE3C | NM_014671 | -2.0255 | -4.0425 |
| KIAA0391 | NM_014672 | -1.316 | -2.2195 |
| TTC35 | NM_014673 | -1.6205 | -3.462 |
| EDEM1 | NM_014674 | -1.063 | -2.321 |
| CROCC | NM_014675 | -0.588 | -0.946 |
| PUM1 | NM_014676 | -1.262 | -2.858 |
| RIMS2 | NM_014677 | -1.074 | -2.011 |
| PPP6R2 | NM_014678 | -0.1485 | -0.811 |
| CEP57 | NM_014679 | -1.329 | -2.3185 |
| KIAA0100 | NM_014680 | -0.0855 | -1.168 |
| DHX34 | NM_014681 | -0.5575 | -1.105 |
| ST18 | NM_014682 | -0.485 | -0.272 |
| ULK2 | NM_014683 | -1.286 | -1.7435 |
| HERPUD1 | NM_014685 | -1.1025 | -2.921 |

| | | | |
|---|---|---|---|
| KIAA0355 | NM_014686 | -1.221 | -3.255 |
| KIAA0226 | NM_014687 | -0.9985 | -1.733 |
| USP6NL | NM_014688 | -0.8545 | -1.573 |
| DOCK10 | NM_014689 | -0.9735 | -3.332 |
| FAM131B | NM_014690 | 0.1315 | -0.118 |
| AQR | NM_014691 | -1.0245 | -2.1925 |
| SEC14L5 | NM_014692 | 0.027 | -0.2095 |
| ECE2 | NM_014693 | 0.059 | -0.2655 |
| CCDC144A | NM_014695 | -0.0975 | -0.886 |
| GPRIN2 | NM_014696 | -0.277 | -0.2285 |
| NOS1AP | NM_014697 | 0.02 | -0.5025 |
| TMEM63A | NM_014698 | -0.0895 | -0.268 |
| ZNF646 | NM_014699 | -0.885 | -2.2375 |
| RAB11FIP3 | NM_014700 | -0.67 | -1.525 |
| SECISBP2L | NM_014701 | -0.7085 | -2.452 |
| KIAA0408 | NM_014702 | -0.699 | -1.95 |
| VPRBP | NM_014703 | -1.292 | -3.776 |
| CEP104 | NM_014704 | -0.551 | -0.699 |
| DOCK4 | NM_014705 | -0.7385 | -2.801 |
| SART3 | NM_014706 | -1.008 | -2.39 |
| HDAC9 | NM_014707 | -0.716 | -2.2515 |
| KNTC1 | NM_014708 | -1.0915 | -1.78 |
| USP34 | NM_014709 | -0.7325 | -2.535 |
| GPRASP1 | NM_014710 | -0.836 | -0.7705 |
| CCP110 | NM_014711 | -1.219 | -2.1635 |
| SETD1A | NM_014712 | -0.3395 | -1.412 |
| LAPTM4A | NM_014713 | -0.0325 | -0.655 |
| IFT140 | NM_014714 | -0.1095 | -0.4015 |
| ARHGAP32 | NM_014715 | -1.033 | -2.2805 |
| ACAP1 | NM_014716 | 0.1025 | -0.131 |
| ZNF536 | NM_014717 | -0.2575 | -0.9565 |
| CLSTN3 | NM_014718 | -0.26 | -0.864 |
| FAM115A | NM_014719 | -0.791 | -1.2205 |
| SLK | NM_014720 | -0.5825 | -2.6205 |
| PHACTR2 | NM_014721 | -0.047 | -1.1355 |
| FAM65B | NM_014722 | -0.0715 | -0.292 |
| SNPH | NM_014723 | -0.009 | -0.8305 |
| STARD8 | NM_014725 | 0.104 | -0.245 |
| TBKBP1 | NM_014726 | -0.4355 | -0.983 |
| MLL4 | NM_014727 | -0.666 | -1.3995 |
| FRMPD4 | NM_014728 | 0.356 | -0.167 |
| TOX | NM_014729 | -0.5665 | -1.3735 |
| MLEC | NM_014730 | -0.0145 | -0.5055 |

| | | | |
|---|---|---|---|
| ProSAPiP1 | NM_014731 | -0.058 | -0.7815 |
| KIAA0513 | NM_014732 | -1.1815 | -1.639 |
| ZFYVE16 | NM_014733 | -0.7555 | -2.052 |
| KIAA0247 | NM_014734 | -0.5405 | -1.3995 |
| PHF16 | NM_014735 | -1.287 | -2.5855 |
| KIAA0101 | NM_014736 | -0.8535 | -0.4175 |
| RASSF2 | NM_014737 | -1.276 | -1.8795 |
| KIAA0195 | NM_014738 | -0.2465 | -0.656 |
| BCLAF1 | NM_014739 | 0.361 | -0.795 |
| EIF4A3 | NM_014740 | -0.5485 | -2.0635 |
| ATG13 | NM_014741 | -0.4675 | -1.8515 |
| TM9SF4 | NM_014742 | -0.3935 | -1.629 |
| KIAA0232 | NM_014743 | -1.492 | -3.0915 |
| TBC1D5 | NM_014744 | -0.458 | -0.404 |
| RNF144A | NM_014746 | -1.12 | -1.482 |
| RIMS3 | NM_014747 | -0.2895 | -0.7645 |
| SNX17 | NM_014748 | -0.2805 | -1.193 |
| KIAA0586 | NM_014749 | -0.7505 | -1.783 |
| DLGAP5 | NM_014750 | -0.8455 | -2.7575 |
| MTSS1 | NM_014751 | -0.773 | -1.7765 |
| SPCS2 | NM_014752 | -0.3755 | -0.5025 |
| BMS1 | NM_014753 | -0.576 | -1.419 |
| PTDSS1 | NM_014754 | -0.303 | -1.17 |
| SERTAD2 | NM_014755 | -0.708 | -1.2545 |
| CKAP5 | NM_014756 | -0.7565 | -1.4895 |
| MAML1 | NM_014757 | -0.5575 | -2.436 |
| SNX19 | NM_014758 | -0.7035 | -1.282 |
| PHYHIP | NM_014759 | 0.324 | -0.5405 |
| TATDN2 | NM_014760 | -0.199 | -1.156 |
| IST1 | NM_014761 | -0.5145 | -2.2235 |
| DHCR24 | NM_014762 | -0.15 | -0.9415 |
| MRPL19 | NM_014763 | -0.8845 | -2.718 |
| DAZAP2 | NM_014764 | -0.3285 | -0.667 |
| TOMM20 | NM_014765 | -0.9025 | -0.5105 |
| SCRN1 | NM_014766 | -1.1465 | -0.6265 |
| SPOCK2 | NM_014767 | 0.0785 | -0.178 |
| AGAP2 | NM_014770 | -0.2175 | -0.982 |
| RNF40 | NM_014771 | -0.418 | -1.3325 |
| CTIF | NM_014772 | -0.5645 | -1.402 |
| KIAA0141 | NM_014773 | -0.4585 | -0.8325 |
| KIAA0494 | NM_014774 | -1.138 | -2.6775 |
| SFI1 | NM_014775 | -0.3805 | -0.5215 |
| GIT2 | NM_014776 | -0.6875 | -2.828 |

| | | | |
|---|---|---|---|
| URB2 | NM_014777 | -0.5975 | -3.435 |
| TSC22D2 | NM_014779 | -1.224 | -2.9655 |
| CUL7 | NM_014780 | -0.176 | -0.486 |
| RB1CC1 | NM_014781 | -0.525 | -2.3125 |
| ARHGAP11A | NM_014783 | -0.7855 | -2.753 |
| ARHGEF11 | NM_014784 | -0.2695 | -0.661 |
| ARHGEF17 | NM_014786 | -0.6715 | -1.2165 |
| DNAJC6 | NM_014787 | -1.95 | -2.751 |
| TRIM14 | NM_014788 | -0.4795 | -1.407 |
| ZNF623 | NM_014789 | -1.27 | -2.375 |
| JAKMIP2 | NM_014790 | -0.846 | -3.0195 |
| MELK | NM_014791 | -1.328 | -2.1915 |
| LCMT2 | NM_014793 | -1.417 | -2.2225 |
| ZEB2 | NM_014795 | -1.3735 | -2.6525 |
| ZBTB24 | NM_014797 | -1.7475 | -3.846 |
| PLEKHM1 | NM_014798 | -1.263 | -2.93 |
| HEPH | NM_014799 | 0.1965 | 0.139 |
| ELMO1 | NM_014800 | -0.1695 | -0.73 |
| PCNXL2 | NM_014801 | -0.0765 | -0.7435 |
| KIAA0528 | NM_014802 | -1.076 | -1.4055 |
| ZNF518A | NM_014803 | -0.41 | -2.4735 |
| KIAA0753 | NM_014804 | -1.465 | -2.4815 |
| EPM2AIP1 | NM_014805 | -1.101 | -0.3605 |
| RUSC2 | NM_014806 | -0.534 | -2.126 |
| C2CD2L | NM_014807 | -0.2215 | -0.776 |
| FARP2 | NM_014808 | -0.4715 | -0.602 |
| KIAA0319 | NM_014809 | -0.3735 | -0.3995 |
| CEP350 | NM_014810 | -0.427 | -3.517 |
| PPP1R26 | NM_014811 | 0.3945 | -1.8495 |
| CEP170 | NM_014812 | -0.5965 | -3.3665 |
| LRIG2 | NM_014813 | -0.5595 | -1.1865 |
| PSMD6 | NM_014814 | -1.2795 | -2.732 |
| MED24 | NM_014815 | -0.0585 | -0.808 |
| TRIL | NM_014817 | -0.505 | -1.1265 |
| TRIM66 | NM_014818 | -0.527 | -0.4485 |
| PJA2 | NM_014819 | -0.498 | -1.6785 |
| TOMM70A | NM_014820 | -1.6325 | -2.293 |
| SEC24D | NM_014822 | -1.2345 | -2.4405 |
| WNK1 | NM_014823 | -0.809 | -1.332 |
| FCHSD2 | NM_014824 | -1.104 | -2.522 |
| URB1 | NM_014825 | -0.4125 | -1.5045 |
| CDC42BPA | NM_014826 | 0.1535 | -1.323 |
| ZC3H11A | NM_014827 | -2.055 | -2.2495 |

| | | | |
|---|---|---|---|
| TOX4 | NM_014828 | -0.7335 | -2.342 |
| DDX46 | NM_014829 | -0.6165 | -1.4935 |
| ZBTB39 | NM_014830 | -0.865 | -1.8055 |
| TRANK1 | NM_014831 | -0.489 | -1.5515 |
| TBC1D4 | NM_014832 | -1.4435 | -2.8615 |
| OSBPL2 | NM_014835 | -0.7155 | -2.481 |
| RHOBTB1 | NM_014836 | -1.7895 | -2.6595 |
| ZBED4 | NM_014838 | -0.801 | -1.9315 |
| LPPR4 | NM_014839 | -1.2525 | -1.894 |
| NUAK1 | NM_014840 | -0.03 | -1.05 |
| SNAP91 | NM_014841 | -1.1855 | -1.6045 |
| TECPR2 | NM_014844 | 0.061 | -1.837 |
| FIG4 | NM_014845 | -0.99 | -1.1415 |
| KIAA0196 | NM_014846 | -1.2615 | -2.437 |
| UBAP2L | NM_014847 | -0.3455 | -1.631 |
| SV2B | NM_014848 | -0.1735 | -0.309 |
| SV2A | NM_014849 | -0.265 | -1.2045 |
| SRGAP3 | NM_014850 | -0.975 | -0.936 |
| KLHL21 | NM_014851 | -0.6585 | -1.4195 |
| SGSM2 | NM_014853 | -0.731 | -0.735 |
| KIAA0415 | NM_014855 | -0.4695 | -1.2615 |
| DENND4B | NM_014856 | -0.834 | -1.634 |
| RABGAP1L | NM_014857 | -0.5975 | -0.655 |
| TMCC2 | NM_014858 | -0.5715 | -0.8105 |
| ARHGAP44 | NM_014859 | -0.301 | -2.018 |
| SUPT7L | NM_014860 | -0.6625 | -2.0985 |
| ATP2C2 | NM_014861 | -0.007 | -0.44 |
| ARNT2 | NM_014862 | -0.5195 | -1.6585 |
| CHST15 | NM_014863 | -1.6085 | -2.4965 |
| FAM20B | NM_014864 | -1.0655 | -1.488 |
| NCAPD2 | NM_014865 | -0.6385 | -1.6285 |
| SEC16A | NM_014866 | -0.5365 | -1.795 |
| KBTBD11 | NM_014867 | -1.0635 | -1.8455 |
| RNF10 | NM_014868 | -0.011 | -1.1465 |
| IQSEC1 | NM_014869 | -0.685 | -1.8495 |
| ZBTB40 | NM_014870 | -1.173 | -2.494 |
| ZBTB5 | NM_014872 | -1.223 | -2.47 |
| LPGAT1 | NM_014873 | -1.0925 | -1.817 |
| MFN2 | NM_014874 | -0.57 | -1.308 |
| KIF14 | NM_014875 | -0.947 | -2.8155 |
| JOSD1 | NM_014876 | -1.2005 | -3.332 |
| HELZ | NM_014877 | -0.7665 | -2.4625 |
| KIAA0020 | NM_014878 | -0.4095 | -1.9485 |

| | | | |
|---|---|---|---|
| P2RY14 | NM_014879 | 0.01 | -0.1365 |
| CD302 | NM_014880 | -0.3835 | -1.126 |
| DCLRE1A | NM_014881 | -1.178 | -2.193 |
| ARHGAP25 | NM_014882 | 0.145 | -0.1345 |
| FAM13A | NM_014883 | -1.3375 | -2.468 |
| SUGP2 | NM_014884 | -0.8145 | -0.844 |
| ANAPC10 | NM_014885 | -1.4705 | -3.122 |
| NSA2 | NM_014886 | -0.6925 | -0.354 |
| N4BP2L2 | NM_014887 | -1.705 | -2.0525 |
| FAM3C | NM_014888 | -1.714 | -2.532 |
| PITRM1 | NM_014889 | -0.1205 | -0.8345 |
| FILIP1L | NM_014890 | -0.1355 | -0.2175 |
| PDAP1 | NM_014891 | -0.052 | -0.5155 |
| SCAF8 | NM_014892 | -0.589 | -2.206 |
| NLGN4Y | NM_014893 | 0.0205 | -0.5245 |
| KIAA1009 | NM_014895 | -0.7705 | -1.6455 |
| ZNF652 | NM_014897 | 0.075 | -0.6565 |
| ZFP30 | NM_014898 | -1.2435 | -2.1225 |
| RHOBTB3 | NM_014899 | -1.3275 | -0.751 |
| COBLL1 | NM_014900 | 0.1485 | -0.8565 |
| RNF44 | NM_014901 | -0.644 | -1.687 |
| DLGAP4 | NM_014902 | -0.685 | -1.695 |
| NAV3 | NM_014903 | -0.133 | -0.017 |
| RAB11FIP2 | NM_014904 | -1.3855 | -2.656 |
| GLS | NM_014905 | -0.9865 | -1.583 |
| PPM1E | NM_014906 | -1.1635 | -1.337 |
| FRMPD1 | NM_014907 | 0.296 | -1.788 |
| DOLK | NM_014908 | -1.4035 | -2.8155 |
| VASH1 | NM_014909 | -0.6295 | -1.4555 |
| ZNF507 | NM_014910 | -1.163 | -3.222 |
| AAK1 | NM_014911 | -0.072 | -0.4475 |
| CPEB3 | NM_014912 | -1.214 | -2.07 |
| ADNP2 | NM_014913 | -1.3495 | -2.6265 |
| AGAP1 | NM_014914 | -0.997 | -1.5385 |
| ANKRD26 | NM_014915 | -0.306 | -0.8785 |
| LMTK2 | NM_014916 | -0.229 | -0.291 |
| NTNG1 | NM_014917 | -0.0105 | -1.3755 |
| CHSY1 | NM_014918 | -1.3345 | -2.2935 |
| ICK | NM_014920 | -1.2345 | -3.351 |
| LPHN1 | NM_014921 | 0.0045 | -1.145 |
| NLRP1 | NM_014922 | -0.5045 | -0.3005 |
| FNDC3A | NM_014923 | -1.2875 | -2.941 |
| ATG14 | NM_014924 | -1.1335 | -2.283 |

| | | | |
|---|---|---|---|
| R3HDM2 | NM_014925 | -0.651 | -0.73 |
| SLITRK3 | NM_014926 | -0.42 | -1.373 |
| CNKSR2 | NM_014927 | -1.3135 | -2.7785 |
| FASTKD2 | NM_014929 | -1.006 | -2.7225 |
| ZNF510 | NM_014930 | -0.788 | -2.107 |
| PPP6R1 | NM_014931 | -0.487 | -1.4375 |
| NLGN1 | NM_014932 | 0.0285 | 0.073 |
| SEC31A | NM_014933 | -0.6105 | -0.4885 |
| DZIP1 | NM_014934 | -1.1165 | -1.613 |
| PLEKHA6 | NM_014935 | -1.1955 | -1.778 |
| ENPP4 | NM_014936 | -0.1725 | -0.089 |
| INPP5F | NM_014937 | -0.915 | -1.909 |
| MLXIP | NM_014938 | -0.439 | -1.636 |
| TRAPPC8 | NM_014939 | -1.523 | -4.2285 |
| MON1B | NM_014940 | -0.5515 | -1.5065 |
| MORC2 | NM_014941 | -1.5295 | -2.9035 |
| ANKRD6 | NM_014942 | -0.4595 | -0.767 |
| ZHX2 | NM_014943 | -0.9445 | -1.4355 |
| CLSTN1 | NM_014944 | -0.0065 | -0.791 |
| ABLIM3 | NM_014945 | -0.05 | -0.5535 |
| SPAST | NM_014946 | -1.5625 | -2.017 |
| FOXJ3 | NM_014947 | -0.9695 | -1.7435 |
| UBOX5 | NM_014948 | -0.415 | -1.1325 |
| KIAA0907 | NM_014949 | -1.453 | -1.931 |
| ZBTB1 | NM_014950 | -1.133 | -2.026 |
| ZNF365 | NM_014951 | 0.1755 | -1.899 |
| BAHD1 | NM_014952 | -0.4265 | -1.6735 |
| DIS3 | NM_014953 | -0.638 | -1.895 |
| RPH3A | NM_014954 | 0.0905 | -0.5395 |
| METTL13 | NM_014955 | -0.7815 | -3.855 |
| CEP164 | NM_014956 | -0.73 | -1.6655 |
| DENND3 | NM_014957 | -0.281 | -0.722 |
| CARD8 | NM_014959 | -0.39 | -1.377 |
| ARSG | NM_014960 | 0 | -0.5085 |
| RUFY3 | NM_014961 | -1.1765 | -1.148 |
| BTBD3 | NM_014962 | -0.7955 | -0.8695 |
| SBNO2 | NM_014963 | -0.3235 | -1.3475 |
| EPN2 | NM_014964 | -0.616 | -1.7135 |
| TRAK1 | NM_014965 | -0.093 | -0.963 |
| DHX30 | NM_014966 | -0.6665 | -2.5445 |
| FAN1 | NM_014967 | -1.914 | -2.345 |
| WDR47 | NM_014969 | -0.4135 | -2.223 |
| KIFAP3 | NM_014970 | -0.737 | -1.367 |

| | | | |
|---|---|---|---|
| EFR3B | NM_014971 | -0.376 | -0.7415 |
| TCF25 | NM_014972 | 0.0315 | -0.782 |
| DIP2C | NM_014974 | -1.0305 | -2.4955 |
| MAST1 | NM_014975 | -0.757 | -1.3795 |
| PDCD11 | NM_014976 | -0.7345 | -2.0365 |
| ACIN1 | NM_014977 | -0.501 | -1.3165 |
| SORCS3 | NM_014978 | -0.039 | -0.2595 |
| SV2C | NM_014979 | -0.8955 | -0.55 |
| STXBP5L | NM_014980 | -0.233 | -1.279 |
| MYH15 | NM_014981 | -0.3225 | 0.1795 |
| PCNX | NM_014982 | -1.3125 | -3.228 |
| HMGXB3 | NM_014983 | -0.4005 | -1.7455 |
| CEP152 | NM_014985 | -0.4725 | -2.5485 |
| IGSF9B | NM_014987 | -0.2695 | -0.8635 |
| LIMCH1 | NM_014988 | -0.95 | -1.2835 |
| RIMS1 | NM_014989 | -0.247 | -1.09 |
| RALGAPA1 | NM_014990 | 0.107 | -0.3085 |
| WDFY3 | NM_014991 | -0.897 | -2.84 |
| DAAM1 | NM_014992 | -0.643 | -1.631 |
| MAPKBP1 | NM_014994 | -0.2935 | -1.0835 |
| PLCH1 | NM_014996 | 0.017 | -0.4845 |
| KLHDC10 | NM_014997 | -1.764 | -2.3505 |
| RAB21 | NM_014999 | -0.8365 | -2.908 |
| STK38L | NM_015000 | -1.084 | -2.41 |
| SPEN | NM_015001 | -1.1015 | -2.9335 |
| FBXO21 | NM_015002 | -1.3675 | -2.4855 |
| EXOSC7 | NM_015004 | -0.559 | -1.8515 |
| KIAA0284 | NM_015005 | 0.0785 | -0.256 |
| PDZRN3 | NM_015009 | -0.769 | -2.265 |
| MYO16 | NM_015011 | 0.0445 | -0.321 |
| TMEM41B | NM_015012 | -1.2185 | -1.4945 |
| KDM1A | NM_015013 | -0.33 | -1.0275 |
| RBM34 | NM_015014 | -1.4265 | -2.3475 |
| KDM4B | NM_015015 | -0.401 | -1.5045 |
| MAST3 | NM_015016 | -0.386 | -1.3245 |
| USP33 | NM_015017 | -1.4695 | -0.885 |
| DOPEY1 | NM_015018 | -0.2565 | -1.544 |
| PHLPP2 | NM_015020 | -1.6835 | -2.42 |
| ZNF292 | NM_015021 | -0.4125 | -2.4725 |
| WDTC1 | NM_015023 | -0.368 | -0.4105 |
| XPO7 | NM_015024 | -1.212 | -1.8465 |
| MYT1L | NM_015025 | -1.4525 | -2.0865 |
| MON2 | NM_015026 | -1.094 | -1.997 |

| | | | |
|---|---|---|---|
| PDXDC1 | NM_015027 | -0.8735 | -1.764 |
| TNIK | NM_015028 | -0.5465 | -1.6965 |
| POP1 | NM_015029 | -0.8435 | -2.6675 |
| FRYL | NM_015030 | -0.4815 | -2.5465 |
| PDS5B | NM_015032 | -0.4405 | -2.851 |
| FNBP1 | NM_015033 | -0.59 | -1.0255 |
| ZHX3 | NM_015035 | -0.3955 | -1.3935 |
| ENDOD1 | NM_015036 | -1.077 | -1.64 |
| KIAA0913 | NM_015037 | -0.167 | -1.371 |
| KIAA0754 | NM_015038 | -1.306 | 0.1595 |
| NMNAT2 | NM_015039 | -0.627 | -0.483 |
| PIKFYVE | NM_015040 | -1.2035 | -3.069 |
| CLUAP1 | NM_015041 | -1.01 | -1.4345 |
| ZNF609 | NM_015042 | -1.0425 | -3.187 |
| TBC1D9B | NM_015043 | -0.2625 | -0.6745 |
| GGA2 | NM_015044 | -0.248 | -1.3285 |
| WAPAL | NM_015045 | -1.9735 | -4.932 |
| SETX | NM_015046 | -0.677 | -2.2945 |
| KIAA0090 | NM_015047 | -0.0545 | -0.927 |
| SETD1B | NM_015048 | -0.9195 | -2.458 |
| TRAK2 | NM_015049 | -2.036 | -3.553 |
| FTSJD2 | NM_015050 | -0.2625 | -1.229 |
| ERP44 | NM_015051 | -1.124 | -2.1605 |
| HECW1 | NM_015052 | 0.03 | -0.739 |
| PPFIA4 | NM_015053 | -0.289 | -0.442 |
| UHRF1BP1L | NM_015054 | -0.4855 | -2.4045 |
| SWAP70 | NM_015055 | -0.405 | -1.5705 |
| RRP1B | NM_015056 | -1.403 | -2.116 |
| MYCBP2 | NM_015057 | -0.3445 | -2.5055 |
| KIAA0564 | NM_015058 | -0.501 | -1.349 |
| TLN2 | NM_015059 | -0.3475 | -0.9575 |
| AVL9 | NM_015060 | -1.1365 | -2.1795 |
| KDM4C | NM_015061 | -0.956 | -1.4015 |
| PPRC1 | NM_015062 | -1 | -3.1835 |
| SLC8A2 | NM_015063 | 0.105 | -0.698 |
| EXPH5 | NM_015065 | -0.0765 | -0.238 |
| PEG10 | NM_015068 | -0.4375 | 0.29 |
| ZNF423 | NM_015069 | -0.261 | -1.1635 |
| ZC3H13 | NM_015070 | 0.1405 | 0.025 |
| ARHGAP26 | NM_015071 | -0.5285 | -0.7645 |
| TTLL5 | NM_015072 | -0.8105 | -1.428 |
| SIPA1L3 | NM_015073 | -1.005 | -2.4965 |
| KIF1B | NM_015074 | -1.2905 | -1.539 |

| | | | |
|---|---|---|---|
| CDK19 | NM_015076 | -1.678 | -1.615 |
| SARM1 | NM_015077 | -0.9175 | -1.2485 |
| MCF2L2 | NM_015078 | 0.072 | 0.032 |
| TBC1D2B | NM_015079 | -0.7375 | -1.994 |
| NRXN2 | NM_015080 | -0.217 | -1.237 |
| FSTL4 | NM_015082 | 0.1355 | -0.8605 |
| CDK12 | NM_015083 | -1.0345 | -2.657 |
| MRPS27 | NM_015084 | -0.2305 | -1.6055 |
| RAP1GAP2 | NM_015085 | -1.2035 | -2.1815 |
| DDN | NM_015086 | -0.191 | -0.553 |
| SPG20 | NM_015087 | -1.3075 | -2.834 |
| TNRC6B | NM_015088 | -2.032 | -3.354 |
| CUL9 | NM_015089 | -0.431 | -0.6945 |
| NFASC | NM_015090 | -0.854 | -0.85 |
| FAM179B | NM_015091 | -1.755 | -4.4535 |
| SMG1 | NM_015092 | -0.812 | -1.8015 |
| TAB2 | NM_015093 | -0.8295 | -2.8545 |
| HIC2 | NM_015094 | -0.694 | -2.23 |
| CLASP2 | NM_015097 | -0.98 | -1.246 |
| CAMTA2 | NM_015099 | -0.529 | -1.2335 |
| POGZ | NM_015100 | -1.1695 | -2.5125 |
| GLT25D2 | NM_015101 | -0.243 | -0.6735 |
| NPHP4 | NM_015102 | -0.4145 | -0.928 |
| PLXND1 | NM_015103 | -0.797 | -1 |
| ATG2A | NM_015104 | -0.4595 | -1.3775 |
| RAD54L2 | NM_015106 | -0.4335 | -1.258 |
| PHF8 | NM_015107 | -0.6995 | -1.366 |
| SMC5 | NM_015110 | -0.4955 | -2.586 |
| N4BP3 | NM_015111 | -0.0385 | -0.4235 |
| MAST2 | NM_015112 | -0.3625 | -0.8825 |
| ZZEF1 | NM_015113 | -0.804 | -1.6165 |
| ANKLE2 | NM_015114 | -1.8935 | -3.07 |
| DCUN1D4 | NM_015115 | -1.0695 | -1.9695 |
| LRCH1 | NM_015116 | -0.158 | -1.2395 |
| ZC3H3 | NM_015117 | -0.3145 | -0.9725 |
| ALMS1 | NM_015120 | -0.4735 | -2.2895 |
| FCHO1 | NM_015122 | -0.267 | -0.7095 |
| FRMD4B | NM_015123 | -0.9805 | -3.4195 |
| GRAMD4 | NM_015124 | -0.6755 | -1.2785 |
| CIC | NM_015125 | -0.673 | -1.453 |
| CLCC1 | NM_015127 | -0.8185 | -1.2665 |
| TBC1D9 | NM_015130 | -1.6145 | -4.392 |
| WDR43 | NM_015131 | -1.342 | -2.8235 |

| | | | |
|---|---|---|---|
| SNX13 | NM_015132 | -0.6855 | -2.605 |
| MAPK8IP3 | NM_015133 | -0.066 | -0.89 |
| MPRIP | NM_015134 | 0.003 | -0.6485 |
| NUP205 | NM_015135 | -1.0205 | -1.7495 |
| STAB1 | NM_015136 | 0.0555 | -0.2685 |
| EFR3A | NM_015137 | -0.8925 | -0.814 |
| RTF1 | NM_015138 | -1.7435 | -2.4795 |
| SLC35D1 | NM_015139 | -1.221 | -2.461 |
| TTLL12 | NM_015140 | -0.9365 | -1.869 |
| GPD1L | NM_015141 | -0.5205 | -0.683 |
| METAP1 | NM_015143 | -0.6035 | -1.4285 |
| ZCCHC14 | NM_015144 | -1.6345 | -2.9 |
| 8-Sep | NM_015146 | -0.346 | -0.7465 |
| CEP68 | NM_015147 | -2.614 | -3.0085 |
| PASK | NM_015148 | -0.514 | -1.2115 |
| RGL1 | NM_015149 | -1.044 | -2.785 |
| RFTN1 | NM_015150 | -0.13 | -1.081 |
| DIP2A | NM_015151 | -1.243 | -1.7675 |
| PHF3 | NM_015153 | -0.2325 | -2.5795 |
| MESDC2 | NM_015154 | -0.7845 | -2.5525 |
| LARP4B | NM_015155 | -1.0095 | -2.3315 |
| RCOR1 | NM_015156 | -1.0795 | -2.4025 |
| PHLDB1 | NM_015157 | -0.1505 | -1.0515 |
| KANK1 | NM_015158 | -0.154 | -0.148 |
| FAM168A | NM_015159 | -0.175 | -0.9865 |
| PMPCA | NM_015160 | -0.091 | -1.361 |
| ARL6IP1 | NM_015161 | -0.7815 | 0.0845 |
| ACSBG1 | NM_015162 | -0.2995 | -0.4895 |
| TRIM9 | NM_015163 | -1.443 | -1.2735 |
| PLEKHM2 | NM_015164 | -0.457 | -1.7755 |
| MLC1 | NM_015166 | 0.4835 | -0.244 |
| JMJD6 | NM_015167 | -1.127 | -2.3835 |
| ZC3H4 | NM_015168 | -1.851 | -4.5355 |
| RRS1 | NM_015169 | -0.865 | -2.964 |
| SULF1 | NM_015170 | 0.0655 | -0.37 |
| XPO6 | NM_015171 | -0.5515 | -2.2905 |
| TBC1D1 | NM_015173 | -0.3495 | -0.901 |
| ZFR2 | NM_015174 | -0.1325 | -0.0495 |
| NBEAL2 | NM_015175 | -0.439 | -0.834 |
| FBXO28 | NM_015176 | -2.225 | -4.7375 |
| DTX4 | NM_015177 | -0.587 | 0.075 |
| RHOBTB2 | NM_015178 | -0.5215 | -1.47 |
| RRP12 | NM_015179 | -0.0985 | -1.081 |

| | | | |
|---|---|---|---|
| SYNE2 | NM_015180 | -0.116 | -0.9965 |
| MAST4 | NM_015183 | -0.147 | -0.7595 |
| ARHGEF9 | NM_015185 | 0.1965 | -0.1395 |
| VPS13A | NM_015186 | -0.1695 | -1.3065 |
| SEL1L3 | NM_015187 | -1.269 | -1.029 |
| TBC1D12 | NM_015188 | -0.929 | -2.425 |
| EXOC6B | NM_015189 | -0.3205 | -0.8315 |
| DNAJC9 | NM_015190 | -0.947 | -1.527 |
| SIK2 | NM_015191 | -2.303 | -4.4525 |
| PLCB1 | NM_015192 | -0.4485 | -2.5255 |
| ARC | NM_015193 | 1.203 | 2.628 |
| MYO1D | NM_015194 | -0.1945 | -0.4475 |
| KIAA0922 | NM_015196 | -0.758 | -2.888 |
| PACS2 | NM_015197 | -0.003 | -0.6055 |
| COBL | NM_015198 | -0.266 | -0.321 |
| ANKRD28 | NM_015199 | -1.155 | -2.4335 |
| BOP1 | NM_015201 | 0.1 | -0.6295 |
| KIAA0556 | NM_015202 | -0.1105 | -0.6705 |
| RPRD2 | NM_015203 | -0.4845 | -1.8235 |
| THSD7A | NM_015204 | -0.875 | -0.848 |
| ATP11A | NM_015205 | -1.094 | -1.963 |
| KIAA1024 | NM_015206 | -0.83 | -1.4195 |
| OTUD3 | NM_015207 | -1.237 | -1.691 |
| ANKRD12 | NM_015208 | 0.2205 | -0.6865 |
| KAZN | NM_015209 | -0.9125 | -1.916 |
| DENND5A | NM_015213 | -0.9235 | -2.7135 |
| DDHD2 | NM_015214 | -1.3185 | -1.3035 |
| CAMTA1 | NM_015215 | 0.0475 | -0.016 |
| PPIP5K2 | NM_015216 | -0.6045 | -1.3765 |
| EXOC7 | NM_015219 | -0.1865 | -0.535 |
| DNMBP | NM_015221 | -0.7505 | -2.6055 |
| PRUNE2 | NM_015225 | -0.4485 | -0.297 |
| CLEC16A | NM_015226 | -0.6945 | -1.2335 |
| POFUT2 | NM_015227 | -0.552 | -1.331 |
| KIAA0664 | NM_015229 | -0.402 | -1.266 |
| ARAP2 | NM_015230 | -0.0385 | 0.3465 |
| NUP160 | NM_015231 | -1.7245 | -2.7885 |
| IQSEC3 | NM_015232 | 0.5215 | 0.376 |
| MTUS2 | NM_015233 | -1.016 | -0.2905 |
| GPR116 | NM_015234 | -0.0625 | -0.218 |
| CSTF2T | NM_015235 | -1.749 | -3.0615 |
| LPHN3 | NM_015236 | -0.73 | -1.543 |
| KIAA1107 | NM_015237 | -0.31 | -0.5015 |

| | | | |
|---|---|---|---|
| WWC1 | NM_015238 | -0.3225 | -1.103 |
| AGTPBP1 | NM_015239 | -0.5425 | -1.8365 |
| MICAL3 | NM_015241 | -0.6905 | -1.79 |
| ARAP1 | NM_015242 | -0.278 | -0.885 |
| VPS13B | NM_015243 | -0.1415 | -0.4325 |
| ANKS1A | NM_015245 | -1.0325 | -2.5815 |
| CYLD | NM_015247 | -0.745 | -1.417 |
| BICD2 | NM_015250 | -0.4905 | -0.841 |
| ATMIN | NM_015251 | -1.0005 | -1.5305 |
| EHBP1 | NM_015252 | -1.1145 | -2.771 |
| WSCD1 | NM_015253 | 0.4245 | -0.146 |
| KIF13B | NM_015254 | -0.5175 | -1.564 |
| UBR2 | NM_015255 | -0.696 | -1.729 |
| ACSL6 | NM_015256 | -0.249 | -0.309 |
| TMEM194A | NM_015257 | -1.4055 | -1.613 |
| FKBP15 | NM_015258 | -1.2095 | -2.4395 |
| ICOSLG | NM_015259 | 0.0875 | -0.4455 |
| SIN3B | NM_015260 | -1.273 | -1.738 |
| NCAPD3 | NM_015261 | -1.182 | -1.5185 |
| DMXL2 | NM_015263 | -0.469 | -1.567 |
| KIAA0930 | NM_015264 | -0.5615 | -1.029 |
| SATB2 | NM_015265 | -0.9025 | -2.576 |
| SLC9A8 | NM_015266 | -0.136 | -0.5775 |
| CUX2 | NM_015267 | -1.4345 | -1.9075 |
| DNAJC13 | NM_015268 | -1.0435 | -2.8435 |
| ZCCHC11 | NM_015269 | -1.3105 | -2.9285 |
| ADCY6 | NM_015270 | -0.357 | -1.1495 |
| TRIM2 | NM_015271 | -1.667 | -2.998 |
| RPGRIP1L | NM_015272 | -1.289 | -2.3945 |
| MAN2B2 | NM_015274 | -0.385 | -0.7855 |
| KIAA1033 | NM_015275 | -0.531 | -2.1185 |
| USP22 | NM_015276 | -0.3775 | -0.9605 |
| NEDD4L | NM_015277 | -0.9225 | -2.059 |
| SASH1 | NM_015278 | 0.0725 | -3.4805 |
| TBC1D30 | NM_015279 | -0.7505 | -2.431 |
| CLASP1 | NM_015282 | -0.888 | -1.189 |
| DPY19L1 | NM_015283 | -0.6065 | -0.4965 |
| SZT2 | NM_015284 | 0.1815 | -0.314 |
| WDR7 | NM_015285 | -0.486 | -2.009 |
| SYNM | NM_015286 | -1.7265 | -2.62 |
| PHF15 | NM_015288 | -0.8835 | -2.0615 |
| VPS39 | NM_015289 | -0.3865 | -0.947 |
| DNAJC16 | NM_015291 | -1.7295 | -2.57 |

| | | | |
|---|---|---|---|
| ESYT1 | NM_015292 | -0.0745 | -0.8905 |
| TRIM37 | NM_015294 | -1.0025 | -0.965 |
| SMCHD1 | NM_015295 | -0.513 | -2.197 |
| DOCK9 | NM_015296 | -0.676 | -1.6925 |
| KIAA1045 | NM_015297 | -0.3805 | -0.2605 |
| KHNYN | NM_015299 | -1.0475 | -2.448 |
| HAUS5 | NM_015302 | -0.87 | -0.978 |
| ANGEL1 | NM_015305 | -0.418 | -1.4915 |
| USP24 | NM_015306 | -1.19 | -2.714 |
| FAM189A1 | NM_015307 | 0.0935 | -0.322 |
| FNBP4 | NM_015308 | -1.911 | -3.199 |
| PSD3 | NM_015310 | -0.646 | -1.435 |
| OBSL1 | NM_015311 | -0.39 | -0.521 |
| KIAA1109 | NM_015312 | -0.712 | -2.036 |
| ARHGEF12 | NM_015313 | -1.0745 | -1.5105 |
| KIAA0895 | NM_015314 | -2.1685 | -3.382 |
| LARP1 | NM_015315 | 0.306 | -0.739 |
| PPP1R13B | NM_015316 | -0.302 | -1.634 |
| PUM2 | NM_015317 | -1.849 | -3.5845 |
| TENC1 | NM_015319 | -0.3235 | -0.4435 |
| ARHGEF4 | NM_015320 | -0.09 | -0.835 |
| CRTC1 | NM_015321 | -0.4185 | -1.0555 |
| FEM1B | NM_015322 | -0.631 | -0.9395 |
| RRP8 | NM_015324 | -0.6995 | -1.9825 |
| KIAA0947 | NM_015325 | -0.88 | -3.04 |
| SRGAP2 | NM_015326 | -0.804 | -2.229 |
| SMG5 | NM_015327 | -0.7585 | -2.02 |
| AHCYL2 | NM_015328 | -0.866 | -1.1225 |
| MAU2 | NM_015329 | -2.193 | -3.109 |
| SPECC1L | NM_015330 | -1.5055 | -2.427 |
| NCSTN | NM_015331 | -0.127 | -1.061 |
| NUDCD3 | NM_015332 | -0.6095 | -1.506 |
| MED13L | NM_015335 | -0.9805 | -1.8775 |
| ZDHHC17 | NM_015336 | -0.786 | -1.835 |
| ASXL1 | NM_015338 | -1.0175 | -3.4285 |
| ADNP | NM_015339 | -1.0315 | -2.5785 |
| LARS2 | NM_015340 | -0.399 | -1.7985 |
| NCAPH | NM_015341 | -0.904 | -2.8005 |
| PPWD1 | NM_015342 | -1.2885 | -2.711 |
| CTDNEP1 | NM_015343 | -0.337 | -1.6245 |
| LEPROTL1 | NM_015344 | -1.0725 | -1.772 |
| DAAM2 | NM_015345 | -0.8795 | -1.056 |
| ZFYVE26 | NM_015346 | -1.2285 | -2.356 |

| | | | |
|---|---|---|---|
| RIMBP2 | NM_015347 | -1.7295 | -2.1745 |
| TMEM131 | NM_015348 | -1.4005 | -2.777 |
| KIAA0240 | NM_015349 | -1.745 | -2.5365 |
| LRRC8B | NM_015350 | -1.1245 | -2.2545 |
| TTC9 | NM_015351 | -0.258 | -1.4425 |
| POFUT1 | NM_015352 | -0.758 | -1.8095 |
| KCTD2 | NM_015353 | -0.635 | -1.752 |
| NUP188 | NM_015354 | -0.2545 | -0.823 |
| SUZ12 | NM_015355 | -1.3235 | -3.1 |
| SCRIB | NM_015356 | -0.3305 | -0.553 |
| MORC3 | NM_015358 | -0.8155 | -2.715 |
| SLC39A14 | NM_015359 | -0.8675 | -1.971 |
| SKIV2L2 | NM_015360 | -0.6845 | -2.1745 |
| R3HDM1 | NM_015361 | -0.6035 | -1.114 |
| C17orf81 | NM_015362 | -0.8065 | -1.7845 |
| ZIM2 | NM_015363 | -0.1365 | -0.1895 |
| LY96 | NM_015364 | 0.0895 | -0.009 |
| AMMECR1 | NM_015365 | -1.402 | -2.528 |
| PRR5 | NM_015366 | -0.5815 | -0.556 |
| BCL2L13 | NM_015367 | -0.2655 | -0.649 |
| PANX1 | NM_015368 | 1.1855 | 1.2825 |
| C22orf31 | NM_015370 | 0.067 | -0.4015 |
| C22orf24 | NM_015372 | 0.278 | 0.031 |
| SUN2 | NM_015374 | -0.535 | -1.1555 |
| DSTYK | NM_015375 | -1.07 | -2.544 |
| RASGRP3 | NM_015376 | -0.1455 | 0.0325 |
| VPS13D | NM_015378 | -0.7465 | -1.393 |
| BRI3 | NM_015379 | -0.1085 | -0.4535 |
| SAMM50 | NM_015380 | -0.2855 | -1.0715 |
| FAM19A5 | NM_015381 | -0.477 | -1.6575 |
| HECTD1 | NM_015382 | -1.1345 | -3.346 |
| NIPBL | NM_015384 | -0.323 | -2.366 |
| SORBS1 | NM_015385 | -0.3905 | -0.84 |
| COG4 | NM_015386 | -0.2175 | -0.8875 |
| YIPF3 | NM_015388 | -0.129 | -0.896 |
| ANAPC13 | NM_015391 | -1.338 | -1.6595 |
| NPDC1 | NM_015392 | 0.1275 | -0.2705 |
| PARM1 | NM_015393 | -0.137 | -1.4555 |
| ZNF10 | NM_015394 | -0.511 | -2.012 |
| TECPR1 | NM_015395 | -0.317 | -1.1025 |
| ARMC8 | NM_015396 | -0.332 | -1.113 |
| DCAF12 | NM_015397 | -0.8095 | -3.228 |
| FAM149A | NM_015398 | 0.214 | -0.4905 |

| | | | |
|---|---|---|---|
| BRMS1 | NM_015399 | -0.3835 | -1.673 |
| HDAC7 | NM_015401 | -0.093 | -0.681 |
| DFNB31 | NM_015404 | -0.0395 | -0.501 |
| ABHD14A | NM_015407 | -0.0955 | -0.746 |
| EP400 | NM_015409 | -0.744 | -1.063 |
| SUMF2 | NM_015411 | 0.3035 | -0.3715 |
| C3orf17 | NM_015412 | -0.8805 | -1.5895 |
| RPL36 | NM_015414 | 0.531 | 0.5715 |
| BRP44 | NM_015415 | -0.0225 | -0.882 |
| LETMD1 | NM_015416 | -0.3545 | -0.817 |
| SPEF1 | NM_015417 | -0.0125 | 0.356 |
| MXRA5 | NM_015419 | 0.208 | -1.468 |
| DCAF13 | NM_015420 | -0.7755 | -1.3955 |
| TMEM186 | NM_015421 | -1.288 | -2.4095 |
| AASDHPPT | NM_015423 | -1.2225 | -2.0485 |
| CHRDL2 | NM_015424 | 0.0015 | -0.283 |
| POLR1A | NM_015425 | -0.325 | -2.073 |
| POC1A | NM_015426 | -0.5565 | -2.072 |
| ZNF473 | NM_015428 | -1.374 | -2.3005 |
| ABI3BP | NM_015429 | -0.117 | 0.211 |
| PAMR1 | NM_015430 | 0.061 | -0.1 |
| TRIM58 | NM_015431 | 0.1565 | -0.2075 |
| INTS7 | NM_015434 | -1.678 | -2.6125 |
| RNF19A | NM_015435 | -0.4845 | -2.1575 |
| RCHY1 | NM_015436 | -0.779 | -2.1975 |
| CCDC28A | NM_015439 | -1.464 | -2.788 |
| MTHFD1L | NM_015440 | -0.4735 | -1.38 |
| OLFML2B | NM_015441 | -0.2 | -0.561 |
| CNOT10 | NM_015442 | -1.713 | -2.327 |
| KIAA1267 | NM_015443 | -0.9175 | -3.09 |
| TMEM158 | NM_015444 | 0.2225 | 0.0505 |
| AHCTF1 | NM_015446 | -0.378 | -1.453 |
| CAMSAP1 | NM_015447 | -0.7675 | -1.934 |
| DPCD | NM_015448 | -0.167 | -0.6145 |
| C1orf43 | NM_015449 | -0.496 | -1.4995 |
| POT1 | NM_015450 | -1.327 | -2.0685 |
| THUMPD3 | NM_015453 | -1.6515 | -3.024 |
| LARP7 | NM_015454 | -0.5895 | -1.99 |
| CNOT6 | NM_015455 | -1.6095 | -2.861 |
| COBRA1 | NM_015456 | -0.555 | -1.189 |
| ZDHHC5 | NM_015457 | -1.0265 | -3.7085 |
| MTMR9 | NM_015458 | -1.181 | -3.2235 |
| ATL3 | NM_015459 | 0.4705 | -0.3825 |

| | | | |
|---|---|---|---|
| MYRIP | NM_015460 | -0.986 | -3.1765 |
| ZNF521 | NM_015461 | -0.1185 | -1.743 |
| NOL11 | NM_015462 | -1.3945 | -3.09 |
| CNRIP1 | NM_015463 | -0.647 | -0.9645 |
| SOSTDC1 | NM_015464 | -0.1095 | -0.659 |
| GEMIN5 | NM_015465 | -1.2445 | -3.014 |
| PTPN23 | NM_015466 | -0.1375 | -0.925 |
| NIPSNAP3A | NM_015469 | -0.673 | -0.9795 |
| RAB11FIP5 | NM_015470 | -0.1605 | -0.951 |
| NSL1 | NM_015471 | -1.1745 | -2.4495 |
| WWTR1 | NM_015472 | -0.4415 | -0.417 |
| HEATR5A | NM_015473 | 0.2135 | -0.778 |
| SAMHD1 | NM_015474 | -0.5855 | -0.67 |
| FAM98A | NM_015475 | -1.1085 | -2.0515 |
| C18orf10 | NM_015476 | 0.0675 | -0.323 |
| SIN3A | NM_015477 | -1.285 | -3.2455 |
| L3MBTL1 | NM_015478 | -0.9435 | -0.831 |
| PVRL3 | NM_015480 | -0.6825 | -1.524 |
| ZNF385A | NM_015481 | 0.0465 | -0.701 |
| SLC22A23 | NM_015482 | -0.1425 | -0.583 |
| KBTBD2 | NM_015483 | -1.0685 | -2.51 |
| SYF2 | NM_015484 | -0.878 | -1.5465 |
| RWDD3 | NM_015485 | -1.2525 | -2.6935 |
| PNKD | NM_015488 | 0.0495 | -0.402 |
| SEC31B | NM_015490 | -0.6225 | -0.402 |
| C15orf39 | NM_015492 | -0.458 | -3.1405 |
| KANK2 | NM_015493 | -0.577 | -1.394 |
| KIAA1429 | NM_015496 | -1.681 | -4.115 |
| TMEM87A | NM_015497 | -1.5265 | -2.5045 |
| C2CD2 | NM_015500 | -1.7015 | -2.6495 |
| SH2B1 | NM_015503 | -0.333 | -0.928 |
| MMACHC | NM_015506 | -1.4145 | -1.9115 |
| EGFL6 | NM_015507 | -0.1755 | 0.005 |
| TIPARP | NM_015508 | -1.237 | -2.5035 |
| NECAP1 | NM_015509 | -0.373 | -1.4275 |
| DHRS7B | NM_015510 | -0.9445 | -2.6 |
| C20orf4 | NM_015511 | -0.5515 | -2.088 |
| DNAH1 | NM_015512 | -0.1835 | -0.3725 |
| CRELD1 | NM_015513 | -0.554 | -0.848 |
| NGDN | NM_015514 | -0.2205 | -0.723 |
| KRT23 | NM_015515 | 0.227 | 0.3115 |
| TSKU | NM_015516 | -0.3875 | -1.915 |
| HINFP | NM_015517 | -0.8985 | -2.719 |

| | | | |
|---|---|---|---|
| MAGI1 | NM_015520 | -1.3115 | -3.687 |
| DYNC2LI1 | NM_015522 | 0.1015 | -1.1405 |
| REXO2 | NM_015523 | -0.7885 | -1.4545 |
| IBTK | NM_015525 | -1.493 | -3.4055 |
| CLIP3 | NM_015526 | 0.1355 | -0.3365 |
| TBC1D10B | NM_015527 | -0.3525 | -1.3825 |
| RNF167 | NM_015528 | -0.2925 | -1.1725 |
| MOXD1 | NM_015529 | -0.0615 | -0.1105 |
| GORASP2 | NM_015530 | -0.741 | -2.1635 |
| C2CD3 | NM_015531 | -0.8385 | -0.912 |
| GRINL1A | NM_015532 | -0.1435 | -0.441 |
| DAK | NM_015533 | 0.073 | -0.4035 |
| ZZZ3 | NM_015534 | -1.0895 | -3.37 |
| SPATS2L | NM_015535 | -0.8245 | -2.3115 |
| NELF | NM_015537 | -0.0835 | -0.7205 |
| RPAP1 | NM_015540 | -0.652 | -1.362 |
| LRIG1 | NM_015541 | 0.37 | -0.342 |
| UPF2 | NM_015542 | -0.8095 | -2.8555 |
| RAB1A | NM_015543 | -0.9995 | -1.639 |
| TMEM98 | NM_015544 | 0.109 | -0.293 |
| PTCD1 | NM_015545 | -0.8405 | -2.3115 |
| ACOT11 | NM_015547 | -0.1735 | -0.2135 |
| DST | NM_015548 | -0.3215 | -2.0285 |
| PLEKHG3 | NM_015549 | -0.4595 | -1.787 |
| OSBPL3 | NM_015550 | -0.492 | -1.0435 |
| SUSD5 | NM_015551 | -0.8245 | -2.2435 |
| IPCEF1 | NM_015553 | 0.046 | 0.296 |
| GLCE | NM_015554 | -1.41 | -2.6595 |
| ZNF451 | NM_015555 | -1.3515 | -3.1765 |
| SIPA1L1 | NM_015556 | -0.292 | -0.836 |
| CHD5 | NM_015557 | -0.1685 | -0.2915 |
| SETBP1 | NM_015559 | -0.9825 | -3.0275 |
| OPA1 | NM_015560 | -0.7875 | -1.6125 |
| UBXN7 | NM_015562 | -0.9375 | -2.307 |
| LRRTM2 | NM_015564 | -0.451 | 0.006 |
| LTN1 | NM_015565 | -0.682 | -2.9975 |
| FAM169A | NM_015566 | -0.9935 | -2.805 |
| SLITRK5 | NM_015567 | -0.802 | -3.378 |
| PPP1R16B | NM_015568 | -0.5915 | -1.6745 |
| DNM3 | NM_015569 | -0.598 | -1.1115 |
| AUTS2 | NM_015570 | -0.148 | -2.2845 |
| SENP6 | NM_015571 | -1.6285 | -2.9675 |
| GIGYF2 | NM_015575 | -1.4825 | -2.957 |

| | | | |
|---|---|---|---|
| ERC2 | NM_015576 | -0.952 | -1.5875 |
| RAI14 | NM_015577 | -0.8165 | -1.194 |
| LSM14A | NM_015578 | -1.424 | -1.9535 |
| POLDIP2 | NM_015584 | -0.4505 | -1.229 |
| C20orf26 | NM_015585 | 0.0205 | -0.517 |
| SAMD4A | NM_015589 | -0.2185 | -0.4035 |
| GPATCH4 | NM_015590 | -1.2045 | -1.895 |
| TBC1D29 | NM_015594 | 0.099 | -0.0445 |
| ARHGEF26 | NM_015595 | 0.133 | -0.135 |
| KLK13 | NM_015596 | 0.035 | -0.2925 |
| GPSM1 | NM_015597 | -0.2785 | -0.3115 |
| PGM3 | NM_015599 | -0.843 | -1.77 |
| ABHD12 | NM_015600 | -0.0435 | -0.8505 |
| HERC4 | NM_015601 | 0.3045 | 0.105 |
| TOR1AIP1 | NM_015602 | -1.3245 | -2.3565 |
| CCDC9 | NM_015603 | -0.2395 | -0.3535 |
| DCAF4 | NM_015604 | -0.637 | -1.72 |
| CHTOP | NM_015607 | -0.83 | -2.324 |
| C10orf137 | NM_015608 | -1.7665 | -3.765 |
| C1orf144 | NM_015609 | 0.1575 | -0.953 |
| WIPI2 | NM_015610 | -0.1835 | -1.579 |
| LRIT1 | NM_015613 | 0.145 | -0.1165 |
| PYGO1 | NM_015617 | -0.4325 | -1.603 |
| CCDC69 | NM_015621 | 0.496 | 0.067 |
| WSB1 | NM_015626 | -1.4575 | -2.001 |
| LDLRAP1 | NM_015627 | -0.047 | -0.491 |
| PRPF31 | NM_015629 | -0.164 | -1.0755 |
| EPC2 | NM_015630 | -0.854 | -3.1885 |
| TCTN3 | NM_015631 | -0.857 | -2.2265 |
| FGFR1OP2 | NM_015633 | -1.003 | -1.799 |
| KIAA1279 | NM_015634 | -1.442 | -2.655 |
| GAPVD1 | NM_015635 | -1.499 | -3.3445 |
| EIF2B4 | NM_015636 | -0.8805 | -2.379 |
| TRPC4AP | NM_015638 | -0.211 | -1.096 |
| SERBP1 | NM_015640 | -1.056 | -1.278 |
| TES | NM_015641 | -0.178 | -1.1135 |
| ZBTB20 | NM_015642 | -0.114 | -0.5685 |
| RAP1B | NM_015646 | -0.98 | -1.353 |
| IRF2BP1 | NM_015649 | -0.6465 | -2.003 |
| TRAF3IP1 | NM_015650 | -1.1135 | -2.1735 |
| PHF19 | NM_015651 | -0.987 | -2.7695 |
| C10orf12 | NM_015652 | -0.643 | -0.652 |
| RIBC2 | NM_015653 | -0.572 | -0.7565 |

| | | | |
|---|---|---|---|
| NAT9 | NM_015654 | -1.0795 | -1.649 |
| ZNF337 | NM_015655 | -0.7415 | -1.6635 |
| KIF26A | NM_015656 | -1.061 | -1.9805 |
| ABCA12 | NM_015657 | 0.0545 | -0.1445 |
| NOC2L | NM_015658 | 0.304 | 0.062 |
| RSL1D1 | NM_015659 | -1.0665 | -1.087 |
| GIMAP2 | NM_015660 | -0.0025 | 0.2785 |
| IFT172 | NM_015662 | -0.938 | -1.033 |
| AAAS | NM_015665 | -0.0175 | -1.005 |
| GTPBP5 | NM_015666 | -0.7215 | -0.922 |
| RGS22 | NM_015668 | -0.058 | -0.2115 |
| PCDHB5 | NM_015669 | -0.4985 | -0.942 |
| SENP3 | NM_015670 | -0.119 | -1.276 |
| GADD45B | NM_015675 | -0.34 | 0.207 |
| SH3YL1 | NM_015677 | -1.7685 | -1.7025 |
| NBEA | NM_015678 | -1.4285 | -1.5005 |
| TRUB2 | NM_015679 | -0.3915 | -1.5365 |
| B9D1 | NM_015681 | -0.1795 | -0.535 |
| ATP5S | NM_015684 | -0.2015 | -0.9055 |
| SDCBP2 | NM_015685 | 0.0295 | -0.399 |
| FAM155B | NM_015686 | -0.714 | -2.0825 |
| FILIP1 | NM_015687 | 0.0425 | -0.5565 |
| FAM184B | NM_015688 | -0.104 | -0.1495 |
| DENND2A | NM_015689 | -0.7015 | -1.1655 |
| STK36 | NM_015690 | -0.5865 | -0.712 |
| WWC3 | NM_015691 | -0.3805 | -0.922 |
| CPAMD8 | NM_015692 | -0.0185 | -0.4445 |
| INTU | NM_015693 | -0.7005 | -1.628 |
| ZNF777 | NM_015694 | -0.5535 | -2.0885 |
| BRPF3 | NM_015695 | -0.4845 | -2.6015 |
| GPX7 | NM_015696 | -0.2205 | -0.9095 |
| COQ2 | NM_015697 | -0.5605 | -2.1905 |
| GPKOW | NM_015698 | -0.7045 | -2.564 |
| NFU1 | NM_015700 | -0.9095 | -1.7615 |
| ERLEC1 | NM_015701 | -1.4385 | -3.3535 |
| MMADHC | NM_015702 | -1.2105 | -1.9235 |
| RRP7A | NM_015703 | -0.61 | -1.5235 |
| PPPDE2 | NM_015704 | -0.735 | -1.5555 |
| SGSM3 | NM_015705 | 0.12 | -0.406 |
| GLTSCR2 | NM_015710 | 0.237 | -0.271 |
| GLTSCR1 | NM_015711 | -0.3445 | -1.414 |
| RRM2B | NM_015713 | -1.2105 | -2.8885 |
| G0S2 | NM_015714 | -0.19 | 0.0355 |

| | | | |
|---|---|---|---|
| PLA2G3 | NM_015715 | -0.576 | -0.266 |
| CD207 | NM_015717 | -0.2125 | -0.105 |
| NOX3 | NM_015718 | 0.161 | -0.3065 |
| COL5A3 | NM_015719 | -0.139 | -0.36 |
| PODXL2 | NM_015720 | -0.028 | -0.6695 |
| GEMIN4 | NM_015721 | -1.0665 | -3.1165 |
| CALY | NM_015722 | 0.193 | -0.3305 |
| PNPLA8 | NM_015723 | -0.7235 | -2.712 |
| RDH8 | NM_015725 | -0.1175 | 0.13 |
| DCAF8 | NM_015726 | -0.3115 | -1.5235 |
| TACR1 | NM_015727 | -0.1275 | -0.196 |
| ACHE | NM_015831 | 0.2575 | -0.317 |
| MBD2 | NM_015832 | -0.906 | -2.2875 |
| ADARB1 | NM_015833 | 0.249 | -0.6755 |
| ADARB1 | NM_015834 | 0.249 | -0.6755 |
| WARS2 | NM_015836 | -0.4355 | -1.2705 |
| FCN2 | NM_015837 | -0.484 | -1.436 |
| ADAR | NM_015840 | -0.4295 | -1.1245 |
| ADAR | NM_015841 | -0.567 | -1.21 |
| LMO7 | NM_015842 | -0.236 | -0.5035 |
| MBD1 | NM_015844 | -0.503 | -2.472 |
| MBD1 | NM_015845 | -0.503 | -2.472 |
| MBD1 | NM_015846 | -0.503 | -2.472 |
| MBD1 | NM_015847 | -0.503 | -2.472 |
| KRT76 | NM_015848 | -0.063 | -0.078 |
| CELA2B | NM_015849 | 0.444 | -0.131 |
| FGFR1 | NM_015850 | -0.0765 | -1.193 |
| ZNF117 | NM_015852 | -0.7625 | -1.2595 |
| UBXN1 | NM_015853 | -0.035 | -0.825 |
| GTF2A1 | NM_015859 | -0.3875 | -0.2745 |
| FAM65B | NM_015864 | -0.0715 | -0.292 |
| SLC14A1 | NM_015865 | -0.054 | 0.3315 |
| PRDM2 | NM_015866 | -0.841 | -0.9575 |
| KIR2DL3 | NM_015868 | 0.3925 | -0.292 |
| PPARG | NM_015869 | -0.758 | -1.625 |
| ZNF593 | NM_015871 | -0.274 | -1.135 |
| ZBTB7B | NM_015872 | -0.083 | -0.545 |
| VILL | NM_015873 | -0.257 | -0.9115 |
| RBPJ | NM_015874 | -0.7465 | -1.343 |
| AZIN1 | NM_015878 | -0.724 | -0.8905 |
| ST8SIA3 | NM_015879 | -1.5725 | -3.0165 |
| DKK3 | NM_015881 | -0.871 | -1.1315 |
| MBTPS2 | NM_015884 | -1.7635 | -2.2295 |

| | | | |
|---|---|---|---|
| PCF11 | NM_015885 | -1.597 | -2.6925 |
| PI15 | NM_015886 | -0.413 | -0.65 |
| HOOK1 | NM_015888 | -0.5315 | -1.349 |
| MED15 | NM_015889 | -0.209 | -0.705 |
| CDC40 | NM_015891 | -1.095 | -2.273 |
| CHST15 | NM_015892 | -1.6085 | -2.4965 |
| PRLH | NM_015893 | 0.0925 | -0.442 |
| STMN3 | NM_015894 | -0.1715 | -0.513 |
| GMNN | NM_015895 | -1.24 | -2.416 |
| ZMYND10 | NM_015896 | -0.3985 | 0.087 |
| PIAS4 | NM_015897 | -0.796 | -1.7915 |
| ZBTB7A | NM_015898 | -0.8965 | -1.1545 |
| PLA1A | NM_015900 | 0.1475 | 0.02 |
| NUDT13 | NM_015901 | -0.05 | -0.6585 |
| UBR5 | NM_015902 | -0.771 | -1.7755 |
| EIF5B | NM_015904 | -0.1275 | -1.6275 |
| TRIM24 | NM_015905 | -1.125 | -2.8775 |
| TRIM33 | NM_015906 | -0.8205 | -1.962 |
| LAP3 | NM_015907 | -0.842 | -1.388 |
| SRRT | NM_015908 | 0.2585 | 0.405 |
| NBAS | NM_015909 | -0.789 | -0.753 |
| WDPCP | NM_015910 | -0.427 | -1.0115 |
| ZNF691 | NM_015911 | -1.513 | -2.6305 |
| FAM135B | NM_015912 | -0.862 | -1.589 |
| TXNDC12 | NM_015913 | -0.8575 | -0.7625 |
| TXNDC11 | NM_015914 | -0.4435 | -1.941 |
| ATL1 | NM_015915 | -1.3525 | -2.599 |
| CALHM2 | NM_015916 | -0.449 | -1.0875 |
| GSTK1 | NM_015917 | 0.1115 | -0.66 |
| POP5 | NM_015918 | -0.5905 | -1.5755 |
| ZNF226 | NM_015919 | -0.596 | -0.881 |
| RPS27L | NM_015920 | -0.8565 | -0.8335 |
| CUTA | NM_015921 | 0.1705 | -0.2855 |
| TEX264 | NM_015926 | -0.387 | -0.9905 |
| TGFB1I1 | NM_015927 | -0.545 | -1.7215 |
| LIPT1 | NM_015929 | -1.095 | -1.639 |
| C3orf32 | NM_015931 | -0.7205 | -0.604 |
| POMP | NM_015932 | -0.5875 | -0.615 |
| CCDC72 | NM_015933 | 0.0235 | -0.1705 |
| NOP58 | NM_015934 | -0.6055 | -1.42 |
| METTL13 | NM_015935 | -0.7815 | -3.855 |
| PIGT | NM_015937 | 0.1725 | -0.2625 |
| NMD3 | NM_015938 | -0.194 | -1.1235 |

| | | | |
|---|---|---|---|
| TRMT6 | NM_015939 | -0.9995 | -2.2165 |
| ATP6V1H | NM_015941 | -0.878 | -1.8475 |
| MTERFD1 | NM_015942 | -1.1525 | -3.4605 |
| AMDHD2 | NM_015944 | -0.0785 | -0.3915 |
| SLC35C2 | NM_015945 | -0.676 | -1.7595 |
| PELO | NM_015946 | -0.4895 | -1.308 |
| ASCC1 | NM_015947 | -1.1025 | -1.866 |
| SLC35B3 | NM_015948 | -2.031 | -3.208 |
| GET4 | NM_015949 | -0.741 | -1.5385 |
| MRPL2 | NM_015950 | -0.8355 | -2.1335 |
| NOSIP | NM_015953 | -0.0985 | -0.765 |
| DERA | NM_015954 | -1.3325 | -2.072 |
| MRPL4 | NM_015956 | -0.268 | -1.166 |
| APIP | NM_015957 | -1.311 | -2.7835 |
| DPH5 | NM_015958 | -1.2555 | -1.7365 |
| TMX2 | NM_015959 | -0.595 | -0.747 |
| CUTC | NM_015960 | -2.0555 | -2.4225 |
| FCF1 | NM_015962 | -1.3235 | -1.6055 |
| THAP4 | NM_015963 | -0.718 | -2.4365 |
| NDUFA13 | NM_015965 | 0.377 | -0.173 |
| ERGIC3 | NM_015966 | 0.076 | -0.279 |
| PTPN22 | NM_015967 | 0.167 | -0.009 |
| MRPS17 | NM_015969 | -1.1275 | -2.25 |
| MRPS7 | NM_015971 | -0.2895 | -1.455 |
| POLR1D | NM_015972 | -0.115 | -1.0565 |
| GAL | NM_015973 | 0.286 | -0.2235 |
| CRYL1 | NM_015974 | 0.226 | -0.376 |
| TAF9B | NM_015975 | -0.6755 | -0.5065 |
| SNX7 | NM_015976 | -0.8015 | -1.3235 |
| TNNI3K | NM_015978 | 0.846 | 0.1005 |
| MED23 | NM_015979 | -0.715 | -1.414 |
| HMP19 | NM_015980 | -0.1835 | -0.429 |
| CAMK2A | NM_015981 | -0.1675 | -0.2045 |
| YBX2 | NM_015982 | -0.0025 | -0.3555 |
| UBE2D4 | NM_015983 | -0.0765 | -0.7595 |
| UCHL5 | NM_015984 | -1.1565 | -2.158 |
| ANGPT4 | NM_015985 | -0.1415 | -0.039 |
| CRLF3 | NM_015986 | -1.5055 | -1.3715 |
| HEBP1 | NM_015987 | -0.6195 | -1.624 |
| CSAD | NM_015989 | -0.7915 | -1.114 |
| KLHL5 | NM_015990 | -0.891 | -1.6235 |
| C1QA | NM_015991 | 0.0535 | -0.1915 |
| PLLP | NM_015993 | -0.148 | -0.748 |

| | | | |
|---|---|---|---|
| ATP6V1D | NM_015994 | -0.7945 | -1.2415 |
| KLF13 | NM_015995 | -0.4265 | -1.202 |
| ADIPOR1 | NM_015999 | -0.1255 | -2.337 |
| UTP18 | NM_016001 | -1.587 | -2.6815 |
| SCCPDH | NM_016002 | -1.153 | -1.177 |
| WIPI2 | NM_016003 | -0.1835 | -1.579 |
| IFT52 | NM_016004 | -0.522 | -0.54 |
| ABHD5 | NM_016006 | -0.57 | -1.763 |
| SH3GLB1 | NM_016009 | -0.6525 | -2.0345 |
| FAM164A | NM_016010 | -0.948 | -1.4735 |
| MECR | NM_016011 | -0.2225 | -0.797 |
| NDUFAF1 | NM_016013 | -1.432 | -2.349 |
| FAM108B1 | NM_016014 | -0.917 | -2.666 |
| PHF20L1 | NM_016018 | 0.045 | -0.5465 |
| LUC7L2 | NM_016019 | -0.965 | -1.8725 |
| TFB1M | NM_016020 | -0.9295 | -2.494 |
| UBE2J1 | NM_016021 | -0.2895 | -1.03 |
| APH1A | NM_016022 | -0.174 | -0.71 |
| OTUD6B | NM_016023 | -0.6665 | -0.8285 |
| RBMX2 | NM_016024 | -0.4935 | -1.3555 |
| METTL9 | NM_016025 | -0.547 | -0.784 |
| RDH11 | NM_016026 | -1.332 | -1.7405 |
| LACTB2 | NM_016027 | -2.04 | -2.718 |
| SUV420H1 | NM_016028 | 0.255 | 0.4305 |
| DHRS7 | NM_016029 | -0.461 | -1.161 |
| TTC15 | NM_016030 | -1.2835 | -2.358 |
| ZDHHC9 | NM_016032 | -0.174 | -1.033 |
| FAM82B | NM_016033 | -0.5875 | -1.0765 |
| MRPS2 | NM_016034 | -0.33 | -2.014 |
| COQ4 | NM_016035 | -0.204 | -0.7545 |
| UTP11L | NM_016037 | -1.052 | -1.8445 |
| SBDS | NM_016038 | -0.5705 | -0.828 |
| C14orf166 | NM_016039 | -0.7105 | -0.7715 |
| TMED5 | NM_016040 | -0.9455 | -1.8555 |
| DERL2 | NM_016041 | -0.6575 | -2.632 |
| EXOSC3 | NM_016042 | -1.556 | -3.747 |
| FAHD2A | NM_016044 | -0.042 | -0.612 |
| SLMO2 | NM_016045 | -0.777 | -1.1035 |
| EXOSC1 | NM_016046 | -0.26 | -0.7285 |
| SF3B14 | NM_016047 | -0.245 | -0.1405 |
| ISOC1 | NM_016048 | -0.2575 | -0.6595 |
| FAM158A | NM_016049 | -0.318 | -1.1525 |
| MRPL11 | NM_016050 | 0.2155 | -0.4795 |

| | | | |
|---|---|---|---|
| RRP15 | NM_016052 | -1.4175 | -2.274 |
| CCDC53 | NM_016053 | -0.839 | -0.9195 |
| MRPL48 | NM_016055 | -0.7055 | -0.3035 |
| TMBIM4 | NM_016056 | -0.9755 | -1.5875 |
| COPZ1 | NM_016057 | 0.005 | -0.421 |
| TPRKB | NM_016058 | -1.5945 | -1.911 |
| PPIL1 | NM_016059 | -1.102 | -2.0315 |
| MED31 | NM_016060 | -0.96 | -2.8605 |
| YPEL5 | NM_016061 | -0.6555 | -1.0835 |
| FAM96B | NM_016062 | 0.134 | -0.403 |
| HDDC2 | NM_016063 | -0.783 | -0.784 |
| MRPS16 | NM_016065 | -0.2895 | -0.8055 |
| GLRX2 | NM_016066 | -0.795 | -2.886 |
| MRPS18C | NM_016067 | -1.178 | -1.88 |
| FIS1 | NM_016068 | 0.2875 | -0.382 |
| PAM16 | NM_016069 | -0.2555 | -0.7795 |
| MRPS23 | NM_016070 | -0.7465 | -2.153 |
| GOLT1B | NM_016072 | -1.171 | -1.9945 |
| HDGFRP3 | NM_016073 | -0.7165 | -0.8585 |
| BOLA1 | NM_016074 | -1.0635 | -3.3845 |
| VPS36 | NM_016075 | -1.223 | -1.1345 |
| PPPDE1 | NM_016076 | -1.6275 | -2.1775 |
| PTRH2 | NM_016077 | -1.429 | -3.28 |
| CHMP3 | NM_016079 | -0.6405 | -0.5095 |
| GLOD4 | NM_016080 | -0.6715 | -1.1675 |
| PALLD | NM_016081 | -1.083 | -2.532 |
| CDK5RAP1 | NM_016082 | -0.7295 | -2.4375 |
| CNR1 | NM_016083 | 0.0025 | -0.3535 |
| RASD1 | NM_016084 | -0.0755 | 0.5325 |
| C2orf28 | NM_016085 | -0.192 | -0.8825 |
| STYXL1 | NM_016086 | -0.1785 | -0.7405 |
| WNT16 | NM_016087 | 0.074 | -0.401 |
| ZNF589 | NM_016089 | -0.7055 | -1.0635 |
| RBM7 | NM_016090 | -1.0005 | -3.0385 |
| EIF3L | NM_016091 | -0.276 | -0.2385 |
| RPL26L1 | NM_016093 | -0.3525 | -1.1385 |
| COMMD2 | NM_016094 | -1.03 | -2.386 |
| GINS2 | NM_016095 | -0.3 | -0.9965 |
| ZNF706 | NM_016096 | -0.6945 | -2.1645 |
| IER3IP1 | NM_016097 | -1.077 | -1.5525 |
| BRP44L | NM_016098 | -0.8315 | -1.92 |
| GOLGA7 | NM_016099 | -1.293 | -1.831 |
| NAA20 | NM_016100 | -1.117 | -3.8895 |

| | | | |
|---|---|---|---|
| NIP7 | NM_016101 | -1.349 | -2.368 |
| SAR1B | NM_016103 | -0.672 | -1.69 |
| RWDD1 | NM_016104 | -0.667 | -0.922 |
| SCFD1 | NM_016106 | -1.037 | -2.3905 |
| ZFR | NM_016107 | -1.1835 | -1.7605 |
| AIG1 | NM_016108 | -0.35 | -0.474 |
| TELO2 | NM_016111 | -0.498 | -1.244 |
| PKD2L1 | NM_016112 | -0.2505 | 0.0685 |
| TRPV2 | NM_016113 | -0.121 | -0.106 |
| ASB4 | NM_016116 | -0.0995 | -0.185 |
| NUB1 | NM_016118 | -1.1455 | -1.6815 |
| RLIM | NM_016120 | 0.477 | -0.521 |
| KCTD3 | NM_016121 | -1.7125 | -2.798 |
| CCDC41 | NM_016122 | -0.5175 | -2.417 |
| IRAK4 | NM_016123 | -0.9745 | -1.3175 |
| RHD | NM_016124 | 0.1115 | -0.5965 |
| RNFT1 | NM_016125 | -1.2565 | -2.021 |
| HSPB11 | NM_016126 | -0.564 | -1.121 |
| TMEM66 | NM_016127 | -0.712 | -0.6445 |
| COPG | NM_016128 | -0.218 | -0.853 |
| COPS4 | NM_016129 | -1.011 | -1.678 |
| RAB10 | NM_016131 | -1.245 | -2.4225 |
| MYEF2 | NM_016132 | -0.9895 | -0.9035 |
| INSIG2 | NM_016133 | -1.109 | -2.0035 |
| PGCP | NM_016134 | -0.348 | -0.4185 |
| ETV7 | NM_016135 | -0.1025 | -0.315 |
| COQ7 | NM_016138 | -0.613 | -1.8345 |
| CHCHD2 | NM_016139 | 0.1755 | -0.1355 |
| TPPP3 | NM_016140 | -0.347 | -0.4695 |
| DYNC1LI1 | NM_016141 | -0.8975 | -2.449 |
| HSD17B12 | NM_016142 | -0.74 | -0.973 |
| NSFL1C | NM_016143 | 0.013 | -0.898 |
| COMMD10 | NM_016144 | -0.7095 | -0.483 |
| C19orf56 | NM_016145 | 0.302 | -0.2215 |
| TRAPPC4 | NM_016146 | -0.8625 | -2.468 |
| PPME1 | NM_016147 | -0.6595 | -1.6425 |
| SHANK1 | NM_016148 | -0.657 | -0.517 |
| ASB2 | NM_016150 | 0.2445 | -0.029 |
| TAOK2 | NM_016151 | -0.517 | -0.6235 |
| RARB | NM_016152 | -0.117 | -0.6575 |
| RAB4B | NM_016154 | -0.464 | -0.8935 |
| MMP17 | NM_016155 | -0.2435 | -1.2505 |
| MTMR2 | NM_016156 | -0.759 | -1.315 |

| | | | |
|---|---|---|---|
| TRO | NM_016157 | -0.4355 | -1.2345 |
| A4GNT | NM_016161 | 0.03 | -0.2775 |
| ING4 | NM_016162 | -0.7445 | -2.024 |
| PIAS1 | NM_016166 | -0.794 | -2.256 |
| NOL7 | NM_016167 | -1.392 | -2.623 |
| SUFU | NM_016169 | -1.1665 | -1.821 |
| TLX2 | NM_016170 | -0.962 | -2.1755 |
| UBAC1 | NM_016172 | -0.067 | -0.8195 |
| HEMK1 | NM_016173 | -0.33 | -0.9365 |
| CERCAM | NM_016174 | -0.042 | 0.0895 |
| C5orf45 | NM_016175 | -0.613 | -0.845 |
| SDF4 | NM_016176 | -0.4455 | -0.161 |
| OAZ3 | NM_016178 | -0.202 | -0.6265 |
| TRPC4 | NM_016179 | 0.2865 | -0.5715 |
| SLC45A2 | NM_016180 | -0.083 | -0.1445 |
| MRTO4 | NM_016183 | -0.3865 | -1.6725 |
| CLEC4A | NM_016184 | -0.5095 | -0.591 |
| HN1 | NM_016185 | -0.35 | -1.5975 |
| ACTL6B | NM_016188 | 0.0915 | -0.5915 |
| CRNN | NM_016190 | -0.173 | -0.977 |
| TMEFF2 | NM_016192 | -0.943 | -2.1875 |
| GNB5 | NM_016194 | -0.3375 | -1.0535 |
| KIF20B | NM_016195 | -0.703 | -2.0725 |
| RBM19 | NM_016196 | -0.588 | -1.405 |
| LSM7 | NM_016199 | 0.089 | -0.2085 |
| NAA38 | NM_016200 | -1.3295 | -2.818 |
| AMOTL2 | NM_016201 | 0.0475 | -2.316 |
| ZNF580 | NM_016202 | 0.3085 | 0.086 |
| PRKAG2 | NM_016203 | -0.789 | -1.3805 |
| GDF2 | NM_016204 | -0.079 | 0.134 |
| PDGFC | NM_016205 | -0.002 | -1.482 |
| VGLL3 | NM_016206 | 0.318 | -0.1695 |
| CPSF3 | NM_016207 | -1.0225 | -1.881 |
| VPS28 | NM_016208 | 0.002 | -0.607 |
| TRAPPC2L | NM_016209 | 0.2535 | -0.394 |
| C3orf18 | NM_016210 | -0.561 | -1.213 |
| SEC31A | NM_016211 | -0.6105 | -0.4885 |
| TRIP4 | NM_016213 | -1.509 | -2.336 |
| EGFL7 | NM_016215 | 0.154 | -0.4955 |
| DBR1 | NM_016216 | -1.4455 | -2.5195 |
| HECA | NM_016217 | -1.0115 | -1.6625 |
| POLK | NM_016218 | -0.131 | -1.4805 |
| MAN1B1 | NM_016219 | -0.5265 | -1.357 |

| | | | |
|---|---|---|---|
| ZNF107 | NM_016220 | -1.044 | -2.142 |
| DCTN4 | NM_016221 | -0.835 | -0.785 |
| DDX41 | NM_016222 | 0.0885 | -0.486 |
| PACSIN3 | NM_016223 | -0.467 | -1.169 |
| SNX9 | NM_016224 | -0.7555 | -1.9485 |
| VPS29 | NM_016226 | -1.014 | -1.315 |
| C1orf9 | NM_016227 | -0.4165 | -1.6055 |
| AADAT | NM_016228 | -1.2535 | -2.1135 |
| CYB5R2 | NM_016229 | 0.236 | -0.4295 |
| CYB5R4 | NM_016230 | -1.265 | -1.4665 |
| NLK | NM_016231 | -1.6305 | -3.2635 |
| IL1RL1 | NM_016232 | 0.124 | -0.475 |
| PADI3 | NM_016233 | -0.149 | 0.121 |
| ACSL5 | NM_016234 | 0.026 | -0.1355 |
| GPRC5B | NM_016235 | -0.1335 | -1.013 |
| ANAPC5 | NM_016237 | -0.1575 | -0.765 |
| ANAPC7 | NM_016238 | -1.09 | -1.641 |
| MYO15A | NM_016239 | 0.0055 | -0.177 |
| SCARA3 | NM_016240 | -0.3365 | -1.055 |
| EMCN | NM_016242 | 0.0025 | -0.5065 |
| CYB5R1 | NM_016243 | -0.361 | -0.767 |
| HSD17B11 | NM_016245 | -1.0325 | -0.79 |
| HSD17B14 | NM_016246 | 0.259 | -0.6235 |
| IMPG2 | NM_016247 | -0.2965 | 0.087 |
| AKAP11 | NM_016248 | -0.538 | -2.547 |
| MAGEC2 | NM_016249 | 0.1495 | 0.097 |
| BIRC6 | NM_016252 | -0.3515 | -1.855 |
| FAM8A1 | NM_016255 | -1.155 | -1.9095 |
| NAGPA | NM_016256 | -0.448 | -1.1895 |
| HPCAL4 | NM_016257 | -0.816 | -0.5655 |
| YTHDF2 | NM_016258 | -1.191 | -3.014 |
| IKZF2 | NM_016260 | -0.0565 | 0.004 |
| TUBD1 | NM_016261 | -2.037 | -2.406 |
| TUBE1 | NM_016262 | -2.1285 | -2.5065 |
| FZR1 | NM_016263 | -0.7205 | -1.2115 |
| ZNF44 | NM_016264 | -0.9355 | -1.6565 |
| ZNF12 | NM_016265 | -0.728 | -2.594 |
| VGLL1 | NM_016267 | -0.0585 | -0.042 |
| LEF1 | NM_016269 | -0.7345 | -0.6265 |
| KLF2 | NM_016270 | 0.2565 | 0.116 |
| RNF138 | NM_016271 | -1.3675 | -2.6415 |
| TOB2 | NM_016272 | -0.7945 | -1.352 |
| PLEKHO1 | NM_016274 | -1.025 | -3.048 |

| | | | |
|---|---|---|---|
| SELT | NM_016275 | -1.342 | -3.0485 |
| SGK2 | NM_016276 | -0.0075 | 0.1375 |
| RAB23 | NM_016277 | -1.551 | -3.139 |
| CDH9 | NM_016279 | 0.4845 | -0.1755 |
| TAOK3 | NM_016281 | -1.383 | -2.623 |
| AK3 | NM_016282 | -1.0535 | -0.948 |
| TAF9 | NM_016283 | -0.354 | 0.077 |
| CNOT1 | NM_016284 | -1.325 | -2.607 |
| DCXR | NM_016286 | 0.31 | -0.1695 |
| HP1BP3 | NM_016287 | -0.474 | -0.879 |
| CAB39 | NM_016289 | -1.1265 | -2.199 |
| UIMC1 | NM_016290 | -1.672 | -2.1365 |
| IP6K2 | NM_016291 | 0.385 | -0.656 |
| TRAP1 | NM_016292 | -0.0755 | -0.863 |
| BIN2 | NM_016293 | -0.1275 | 0.066 |
| PCYOX1 | NM_016297 | -1.4665 | -2.0555 |
| FBXO40 | NM_016298 | -0.0905 | 0.0185 |
| HSPA14 | NM_016299 | -1.5955 | -3.1675 |
| ARPP21 | NM_016300 | -0.0715 | -0.5665 |
| GPN3 | NM_016301 | -0.998 | -1.9035 |
| CRBN | NM_016302 | -1.2105 | -2.0545 |
| WBP5 | NM_016303 | -1.3325 | -2.078 |
| RSL24D1 | NM_016304 | -0.7125 | -0.3565 |
| SS18L2 | NM_016305 | -1.3945 | -2.672 |
| DNAJB11 | NM_016306 | -1.0075 | -1.237 |
| PRRX2 | NM_016307 | -0.824 | -1.796 |
| CMPK1 | NM_016308 | -1.017 | -1.233 |
| LCMT1 | NM_016309 | -0.4625 | -1.4425 |
| POLR3K | NM_016310 | -0.558 | -1.5535 |
| ATPIF1 | NM_016311 | -0.1785 | -0.465 |
| WBP11 | NM_016312 | -0.222 | -1.4175 |
| GULP1 | NM_016315 | -0.694 | -1.231 |
| REV1 | NM_016316 | -0.8115 | -2.483 |
| P2RX2 | NM_016318 | 0.1135 | -0.3945 |
| COPS7A | NM_016319 | -0.3125 | -1.784 |
| NUP98 | NM_016320 | -1.3115 | -2.5025 |
| RHCG | NM_016321 | -0.0545 | -0.288 |
| RAB14 | NM_016322 | -1.0815 | -3.0385 |
| HERC5 | NM_016323 | -0.157 | -0.124 |
| ZNF274 | NM_016324 | -1.0695 | -3.114 |
| ZNF274 | NM_016325 | -1.0695 | -3.114 |
| UPB1 | NM_016327 | 0.008 | -0.4605 |
| GTF2IRD1 | NM_016328 | -0.802 | -2.0405 |

| | | | |
|---|---|---|---|
| SFMBT1 | NM_016329 | -0.647 | -1.472 |
| ZNF639 | NM_016331 | -1.56 | -3.0995 |
| SEPX1 | NM_016332 | -0.537 | -1.68 |
| SRRM2 | NM_016333 | -0.4205 | -0.574 |
| PRODH | NM_016335 | 0.306 | -0.3295 |
| EVL | NM_016337 | 0.359 | -0.1265 |
| IPO11 | NM_016338 | -0.7115 | -1.8105 |
| RAPGEFL1 | NM_016339 | -1.0295 | -1.0775 |
| RAPGEF6 | NM_016340 | -0.88 | -3.2735 |
| PLCE1 | NM_016341 | -0.0555 | -0.5115 |
| CENPF | NM_016343 | -0.132 | -2.055 |
| NR2E3 | NM_016346 | -0.009 | -0.2485 |
| NAT8B | NM_016347 | -0.17 | -0.2465 |
| NIN | NM_016350 | -0.7115 | -2.9645 |
| ADAM22 | NM_016351 | -0.5105 | -0.9525 |
| CPA4 | NM_016352 | -0.167 | -0.2055 |
| ZDHHC2 | NM_016353 | -0.085 | -0.4225 |
| SLCO4A1 | NM_016354 | -0.8635 | -1.724 |
| DDX47 | NM_016355 | -0.792 | -1.683 |
| DCDC2 | NM_016356 | -0.3545 | 0.037 |
| LIMA1 | NM_016357 | -1.0115 | -2.3605 |
| IRX4 | NM_016358 | 0.0735 | -0.442 |
| NUSAP1 | NM_016359 | -1.4285 | -1.266 |
| TACO1 | NM_016360 | -0.824 | -3.5335 |
| ACP6 | NM_016361 | -0.385 | -1.2665 |
| GHRL | NM_016362 | -0.155 | 0.1135 |
| GP6 | NM_016363 | -0.1205 | -0.242 |
| DUSP13 | NM_016364 | -0.0005 | -0.2145 |
| CABP2 | NM_016366 | 0.007 | 0.0405 |
| ISYNA1 | NM_016368 | 0.1685 | -0.36 |
| CLDN18 | NM_016369 | 0.037 | -0.2065 |
| RAB9B | NM_016370 | -1.1195 | -1.786 |
| HSD17B7 | NM_016371 | -0.873 | -1.7105 |
| TPRA1 | NM_016372 | -0.693 | -2.315 |
| WWOX | NM_016373 | -0.3165 | -0.2245 |
| ARID4B | NM_016374 | -0.587 | -4 |
| ANKFY1 | NM_016376 | -1.2295 | -2.2465 |
| AKAP7 | NM_016377 | 0.197 | -0.225 |
| VCX2 | NM_016378 | -0.158 | -0.6995 |
| VCX3A | NM_016379 | -0.2505 | -0.348 |
| TREX1 | NM_016381 | -0.561 | -1.188 |
| CD244 | NM_016382 | 0.081 | -0.3635 |
| LUZP4 | NM_016383 | -0.07 | 0.225 |

| | | | |
|---|---|---|---|
| TRAT1 | NM_016388 | 0.586 | 0.039 |
| C9orf114 | NM_016390 | -0.4705 | -0.8915 |
| NOP16 | NM_016391 | -0.7105 | -2.1755 |
| PTPLAD1 | NM_016395 | -0.9475 | -0.7435 |
| CTDSPL2 | NM_016396 | -1.2675 | -2.714 |
| TRIAP1 | NM_016399 | -0.9735 | -0.224 |
| C15orf63 | NM_016400 | 1.27 | 1.924 |
| C11orf73 | NM_016401 | -0.6245 | -1.0285 |
| CWC15 | NM_016403 | -1.1205 | -1.7885 |
| TRMT112 | NM_016404 | 0.2335 | -0.1695 |
| UFC1 | NM_016406 | -0.046 | -0.3985 |
| C20orf43 | NM_016407 | -0.0745 | -0.6405 |
| CDK5RAP1 | NM_016408 | -0.6295 | -2.4945 |
| CHMP5 | NM_016410 | -1.061 | -1.2815 |
| GLRX5 | NM_016417 | 0.14 | -0.985 |
| NF2 | NM_016418 | -0.0255 | -0.984 |
| RNF141 | NM_016422 | -1.074 | -0.7735 |
| ZNF219 | NM_016423 | -0.5185 | -1.2055 |
| LUC7L3 | NM_016424 | -0.271 | -0.521 |
| GTSE1 | NM_016426 | -0.7395 | -1.737 |
| TCEB3B | NM_016427 | 0.1195 | -1.0235 |
| ABI3 | NM_016428 | -0.0475 | -0.1865 |
| COPZ2 | NM_016429 | -0.026 | -0.4925 |
| MAPK8IP2 | NM_016431 | -0.1255 | -0.5995 |
| GLTP | NM_016433 | -0.4915 | -2.6435 |
| RTEL1 | NM_016434 | -0.338 | -0.541 |
| PHF20 | NM_016436 | -1.504 | -2.1425 |
| TUBG2 | NM_016437 | -0.4995 | -1.1865 |
| HIGD1B | NM_016438 | 0.185 | -0.2085 |
| VRK3 | NM_016440 | -0.092 | -1.031 |
| CRIM1 | NM_016441 | -0.265 | -0.745 |
| ERAP1 | NM_016442 | -0.239 | -0.649 |
| PLEK2 | NM_016445 | -1.0295 | 0.1775 |
| TMEM8B | NM_016446 | 0.0825 | -0.454 |
| MPP6 | NM_016447 | -0.486 | -0.8065 |
| DTL | NM_016448 | -1.568 | -3.5195 |
| C22orf43 | NM_016449 | -0.002 | -0.1085 |
| COPB1 | NM_016451 | -0.7555 | -1.998 |
| CAPN9 | NM_016452 | -0.0985 | -0.249 |
| NCKIPSD | NM_016453 | -0.674 | -1.7515 |
| TMEM85 | NM_016454 | -0.583 | -0.914 |
| TMEM9 | NM_016456 | -0.317 | -0.8935 |
| PRKD2 | NM_016457 | -0.3955 | -1.4515 |

| | | | |
|---|---|---|---|
| TMEM14C | NM_016462 | 0.0865 | -0.6195 |
| CXXC5 | NM_016463 | -0.5495 | -1.5915 |
| TMEM138 | NM_016464 | -1.2975 | -2.9305 |
| ANKRD39 | NM_016466 | -0.3775 | -2.16 |
| ORMDL1 | NM_016467 | -0.831 | -1.5085 |
| COX16 | NM_016468 | -0.1235 | -0.2095 |
| C20orf111 | NM_016470 | -0.4025 | -1.4545 |
| C14orf129 | NM_016472 | -1.8365 | -4.142 |
| C3orf19 | NM_016474 | -1.1515 | -2.6465 |
| JKAMP | NM_016475 | -1.635 | -2.8215 |
| ANAPC11 | NM_016476 | 0.2395 | -0.369 |
| ZC3HC1 | NM_016478 | -1.284 | -3.1565 |
| SHISA5 | NM_016479 | -0.3035 | -0.8105 |
| PAIP2 | NM_016480 | -0.7715 | -0.365 |
| C9orf156 | NM_016481 | -0.642 | -2.3005 |
| PHF7 | NM_016483 | 0.1165 | -0.0795 |
| PDZD11 | NM_016484 | -0.0735 | -0.3735 |
| VTA1 | NM_016485 | -0.897 | -1.186 |
| TMEM69 | NM_016486 | -1.4095 | -1.9715 |
| PPHLN1 | NM_016488 | -1.1515 | -2.0805 |
| MRPL37 | NM_016491 | -0.006 | -0.7505 |
| RANGRF | NM_016492 | -0.052 | -0.961 |
| RNF181 | NM_016494 | -0.0745 | -0.6455 |
| TBC1D7 | NM_016495 | -0.4225 | -1.3195 |
| 2-Mar | NM_016496 | 0.235 | -0.757 |
| MRPL51 | NM_016497 | -0.175 | -0.486 |
| MTFP1 | NM_016498 | -0.796 | -2.1015 |
| TMEM216 | NM_016499 | -1.035 | -2.735 |
| CXorf26 | NM_016500 | -0.918 | -1.1365 |
| MRPL27 | NM_016504 | -0.5625 | -2.046 |
| ZCCHC17 | NM_016505 | -0.4575 | -1.3625 |
| KBTBD4 | NM_016506 | -0.8915 | -2.4865 |
| CDK12 | NM_016507 | -1.0345 | -2.657 |
| CDKL3 | NM_016508 | -0.8285 | -1.061 |
| CLEC1B | NM_016509 | -0.0585 | -0.406 |
| SCLY | NM_016510 | -0.9965 | -1.8125 |
| CLEC1A | NM_016511 | -0.178 | 0.1815 |
| ICK | NM_016513 | 0.2655 | -0.2335 |
| VPS54 | NM_016516 | -1.312 | -3.535 |
| PIPOX | NM_016518 | -0.131 | -0.233 |
| AMBN | NM_016519 | 0.0485 | -0.5775 |
| C9orf78 | NM_016520 | -0.8175 | -0.645 |
| TFDP3 | NM_016521 | 0.0645 | -0.3995 |

| | | | |
|---|---|---|---|
| NTM | NM_016522 | -0.022 | -0.5335 |
| KLRF1 | NM_016523 | -0.39 | 0.2 |
| SYT17 | NM_016524 | 0.0945 | -2.4625 |
| UBAP1 | NM_016525 | -0.7235 | -2.371 |
| BET1L | NM_016526 | -0.779 | -2.2775 |
| HAO2 | NM_016527 | -0.137 | 0.3645 |
| ATP8A2 | NM_016529 | 0.093 | -0.427 |
| RAB8B | NM_016530 | -0.888 | -1.3255 |
| KLF3 | NM_016531 | -2.346 | -3.6335 |
| INPP5K | NM_016532 | -0.279 | -1.1715 |
| NINJ2 | NM_016533 | -0.033 | -0.44 |
| ZNF581 | NM_016535 | -0.7375 | -1.6205 |
| ZNF571 | NM_016536 | -0.2705 | -1.6195 |
| SIRT7 | NM_016538 | -0.2875 | -0.8365 |
| SIRT6 | NM_016539 | 0.287 | -0.2465 |
| GPR83 | NM_016540 | -0.4845 | -1.249 |
| GNG13 | NM_016541 | -0.0525 | -0.3255 |
| MST4 | NM_016542 | -0.986 | -1.184 |
| SIGLEC7 | NM_016543 | 0.0655 | -0.826 |
| DNAJC27 | NM_016544 | -1.253 | -1.7505 |
| IER5 | NM_016545 | 0.321 | 1.5 |
| C1RL | NM_016546 | -0.571 | -0.374 |
| SDF4 | NM_016547 | -0.0095 | -0.6655 |
| GOLM1 | NM_016548 | -0.204 | -0.727 |
| TM7SF3 | NM_016551 | -0.082 | -1.0335 |
| ANKMY1 | NM_016552 | -0.3115 | -0.423 |
| NUP62 | NM_016553 | -0.971 | -2.442 |
| PSMC3IP | NM_016556 | -0.975 | -0.868 |
| CCRL1 | NM_016557 | -0.5115 | 0.691 |
| SCAND1 | NM_016558 | -0.161 | -0.7845 |
| PEX5L | NM_016559 | -0.8255 | -1.889 |
| BFAR | NM_016561 | -1.538 | -3.4235 |
| TLR7 | NM_016562 | -0.344 | -0.207 |
| RASL12 | NM_016563 | 0.075 | -0.2885 |
| CEND1 | NM_016564 | -0.791 | -1.2965 |
| CHCHD8 | NM_016565 | -0.948 | -2.5375 |
| BCCIP | NM_016567 | -0.8725 | -1.1475 |
| RXFP3 | NM_016568 | 0.0335 | -0.811 |
| TBX3 | NM_016569 | -0.494 | -1.367 |
| ERGIC2 | NM_016570 | -0.7355 | -2.3375 |
| LGSN | NM_016571 | -0.26 | -0.779 |
| GMIP | NM_016573 | -0.0525 | -0.567 |
| DRD2 | NM_016574 | -0.448 | -0.4105 |

| | | | |
|---|---|---|---|
| RAB6B | NM_016577 | -0.3575 | -1.456 |
| RSF1 | NM_016578 | -0.1595 | -2.1765 |
| CD320 | NM_016579 | -0.3775 | -1.7225 |
| PCDH12 | NM_016580 | 0.5365 | -0.0885 |
| ECSIT | NM_016581 | -0.377 | -0.8845 |
| SLC15A3 | NM_016582 | -0.1425 | -0.2285 |
| IL23A | NM_016584 | -0.186 | -0.3075 |
| MBIP | NM_016586 | -1.276 | -2.536 |
| CBX3 | NM_016587 | -1.08 | -0.9355 |
| NRN1 | NM_016588 | 0.2565 | -0.91 |
| GCNT4 | NM_016591 | -0.3485 | 0.2355 |
| GNAS | NM_016592 | 0.0835 | -0.2445 |
| CYP39A1 | NM_016593 | 0.051 | -0.5705 |
| ZDHHC3 | NM_016598 | -0.351 | -1.815 |
| MYOZ2 | NM_016599 | 0.0305 | -0.4805 |
| KCNK9 | NM_016601 | -0.0355 | -0.279 |
| CCR10 | NM_016602 | -0.46 | -0.6385 |
| FAM13B | NM_016603 | -0.7795 | -1.69 |
| KDM3B | NM_016604 | -0.718 | -1.3045 |
| FAM53C | NM_016605 | -0.864 | -2.147 |
| REEP2 | NM_016606 | -0.267 | -0.787 |
| ARMCX3 | NM_016607 | -1.275 | -1.2435 |
| ARMCX1 | NM_016608 | -1.175 | -1.581 |
| SLC22A17 | NM_016609 | -0.0205 | -0.3835 |
| SLC25A37 | NM_016612 | -1.3295 | -1.5 |
| FAM198B | NM_016613 | -0.33 | -0.2505 |
| TDP2 | NM_016614 | -1.9755 | -3.2415 |
| SLC6A13 | NM_016615 | 0.1085 | -0.356 |
| TXNDC3 | NM_016616 | 0.5005 | -0.0415 |
| UFM1 | NM_016617 | -0.8365 | -1.262 |
| KRCC1 | NM_016618 | -0.7075 | -3.187 |
| PLAC8 | NM_016619 | 0.002 | -0.258 |
| ZNF644 | NM_016620 | -0.9615 | -3.751 |
| PHF21A | NM_016621 | -1.179 | -2.4045 |
| MRPL35 | NM_016622 | -0.7435 | -1.996 |
| FAM49B | NM_016623 | -0.878 | -1.141 |
| RSRC1 | NM_016625 | -0.884 | -1.048 |
| MEX3C | NM_016626 | -1.2765 | -2.006 |
| AMZ2 | NM_016627 | -0.9675 | -0.9535 |
| WAC | NM_016628 | -1.6395 | -3.798 |
| SPG21 | NM_016630 | -0.355 | -1.2155 |
| GCFC1 | NM_016631 | -1.5635 | -2.301 |
| ARL17A | NM_016632 | -0.4185 | -0.8025 |

| | | | |
|---|---|---|---|
| AHSP | NM_016633 | -0.2695 | -0.4025 |
| ARL6IP4 | NM_016638 | 0.269 | -0.3275 |
| TNFRSF12A | NM_016639 | -0.478 | -0.658 |
| MRPS30 | NM_016640 | -0.7795 | -2.802 |
| GDE1 | NM_016641 | -1.1175 | -1.573 |
| SPTBN5 | NM_016642 | 0.12 | -0.339 |
| ZNF771 | NM_016643 | -0.296 | -1.0545 |
| PRR16 | NM_016644 | -0.0685 | -0.494 |
| C8orf55 | NM_016647 | -0.815 | -2.1845 |
| LARP7 | NM_016648 | -0.5895 | -1.99 |
| ESF1 | NM_016649 | -0.224 | -0.622 |
| DACT1 | NM_016651 | -0.099 | -1.0915 |
| CRNKL1 | NM_016652 | -0.7385 | -2.0585 |
| ZAK | NM_016653 | -0.608 | -1.459 |
| GABPB1 | NM_016654 | -2.2725 | -4.4245 |
| GABPB1 | NM_016655 | -2.2725 | -4.4245 |
| RRAGB | NM_016656 | -0.83 | -0.6685 |
| KDELR3 | NM_016657 | -0.081 | -0.5915 |
| RALY | NM_016732 | 0.271 | -0.4725 |
| LIMK2 | NM_016733 | -0.9735 | -2.8255 |
| PAX5 | NM_016734 | -0.364 | -0.733 |
| OAS1 | NM_016816 | -0.025 | -0.293 |
| OAS2 | NM_016817 | -0.0745 | -0.4055 |
| ABCG1 | NM_016818 | -0.1035 | -0.8765 |
| OGG1 | NM_016819 | -0.6175 | -0.361 |
| OGG1 | NM_016820 | -0.6175 | -0.361 |
| OGG1 | NM_016821 | -0.6175 | -0.361 |
| CRK | NM_016823 | -1.688 | -2.3115 |
| ADD3 | NM_016824 | -0.794 | -1.0915 |
| OGG1 | NM_016826 | -0.6175 | -0.361 |
| OGG1 | NM_016827 | -0.6175 | -0.361 |
| OGG1 | NM_016828 | -0.6175 | -0.361 |
| OGG1 | NM_016829 | -0.6175 | -0.361 |
| VAMP1 | NM_016830 | 0.081 | -0.882 |
| PER3 | NM_016831 | -1.5795 | -1.17 |
| MAPT | NM_016834 | 0.0845 | -0.5055 |
| MAPT | NM_016835 | 0.0845 | -0.5055 |
| RBMS1 | NM_016836 | -0.7015 | -1.7475 |
| MAPT | NM_016841 | 0.0845 | -0.5055 |
| SHC3 | NM_016848 | -0.835 | -1.3195 |
| CLIC5 | NM_016929 | -0.1505 | -0.399 |
| STX18 | NM_016930 | -0.7665 | -2.2225 |
| NOX4 | NM_016931 | 0.0195 | -0.5715 |

| | | | |
|---|---|---|---|
| SIX2 | NM_016932 | -0.3175 | -1.6345 |
| UBN1 | NM_016936 | -0.6695 | -2.3755 |
| POLA1 | NM_016937 | -0.6235 | -0.9845 |
| EFEMP2 | NM_016938 | 0.092 | -0.442 |
| RWDD2B | NM_016940 | -1.2385 | -1.1435 |
| DLL3 | NM_016941 | -0.354 | -0.929 |
| TAS2R3 | NM_016943 | 0.1105 | -0.209 |
| TAS2R4 | NM_016944 | -0.337 | -0.3865 |
| TAS2R16 | NM_016945 | -0.0825 | -0.199 |
| F11R | NM_016946 | -0.4095 | -0.9595 |
| PARD6A | NM_016948 | -0.847 | -1.216 |
| SPOCK3 | NM_016950 | -0.533 | -0.107 |
| CKLF | NM_016951 | -0.3285 | -1.091 |
| CDON | NM_016952 | -1.268 | -2.232 |
| PDE11A | NM_016953 | -0.2435 | -0.2575 |
| TBX22 | NM_016954 | 0.1475 | 0.218 |
| SEPSECS | NM_016955 | -0.8405 | -1.672 |
| HOXC10 | NM_017409 | 0.1655 | -0.5575 |
| HOXC13 | NM_017410 | -0.225 | -0.207 |
| SMN2 | NM_017411 | -1.132 | -1.4585 |
| FZD3 | NM_017412 | -1.285 | -1.291 |
| APLN | NM_017413 | -0.071 | -0.286 |
| USP18 | NM_017414 | -0.4535 | -1.2775 |
| KLHL3 | NM_017415 | -0.7575 | -0.794 |
| IL1RAPL2 | NM_017416 | -0.6875 | 0.085 |
| GALNT8 | NM_017417 | -0.045 | -0.055 |
| 1-Dec | NM_017418 | 0.122 | -0.1795 |
| ACCN5 | NM_017419 | 0.032 | -0.2275 |
| SIX4 | NM_017420 | -1.8175 | -2.078 |
| COQ3 | NM_017421 | -1.1545 | -2.4565 |
| CALML5 | NM_017422 | 0.198 | -0.4255 |
| GALNT7 | NM_017423 | -0.773 | -1.948 |
| CECR1 | NM_017424 | 0.073 | -0.672 |
| SPA17 | NM_017425 | -0.2455 | 0.186 |
| NUP54 | NM_017426 | -1.4095 | -2.4985 |
| BCMO1 | NM_017429 | 0.1425 | -0.363 |
| PRKAG3 | NM_017431 | -0.051 | -0.582 |
| PTOV1 | NM_017432 | 0.2295 | -0.378 |
| MYO3A | NM_017433 | 0.2395 | -0.466 |
| DUOX1 | NM_017434 | -0.212 | -0.465 |
| SLCO1C1 | NM_017435 | 0.281 | -0.1675 |
| A4GALT | NM_017436 | -0.277 | -0.3585 |
| CPSF2 | NM_017437 | -1.561 | -2.366 |

| | | | |
|---|---|---|---|
| SETD4 | NM_017438 | -1.0665 | -1.798 |
| PION | NM_017439 | -0.0585 | -0.7965 |
| MDM1 | NM_017440 | -1.474 | -2.426 |
| TLR9 | NM_017442 | -0.293 | -0.231 |
| POLE3 | NM_017443 | -0.0985 | 0.0545 |
| CHRAC1 | NM_017444 | -0.8595 | -3.015 |
| MRPL39 | NM_017446 | -1.1185 | -2.554 |
| C21orf91 | NM_017447 | -0.4885 | -1.794 |
| EPHB2 | NM_017449 | -0.259 | -0.745 |
| BAIAP2 | NM_017450 | -0.5985 | -1.2375 |
| BAIAP2 | NM_017451 | -0.5985 | -1.2375 |
| STAU1 | NM_017452 | -0.7415 | -1.708 |
| STAU1 | NM_017453 | -0.7415 | -1.708 |
| STAU1 | NM_017454 | -0.7415 | -1.708 |
| NPTN | NM_017455 | -0.6125 | -1.858 |
| CYTH1 | NM_017456 | -0.4665 | -1.849 |
| CYTH2 | NM_017457 | -0.9945 | -0.868 |
| MVP | NM_017458 | -0.238 | -0.476 |
| MFAP2 | NM_017459 | 0.051 | -0.4275 |
| CYP3A4 | NM_017460 | -0.127 | 0.2815 |
| UBQLN3 | NM_017481 | -0.2675 | -0.157 |
| ADD2 | NM_017482 | -0.0145 | -0.5655 |
| ADD2 | NM_017488 | -0.0145 | -0.5655 |
| TERF1 | NM_017489 | -0.858 | -1.243 |
| MARK2 | NM_017490 | -0.475 | -2.8425 |
| WDR1 | NM_017491 | -0.141 | -1.273 |
| OTUD4 | NM_017493 | 0.89 | 0.9025 |
| RBM38 | NM_017495 | -0.825 | -1.768 |
| SURF2 | NM_017503 | -0.7245 | -1.715 |
| OR2M4 | NM_017504 | 0.324 | -0.5 |
| OR7A5 | NM_017506 | -0.545 | -0.6665 |
| SOX6 | NM_017508 | -0.07 | -0.12 |
| KLK15 | NM_017509 | -0.233 | -0.2765 |
| TMED9 | NM_017510 | -0.06 | -0.974 |
| ENOSF1 | NM_017512 | -0.779 | -0.6855 |
| PLXNA3 | NM_017514 | -0.246 | -0.9355 |
| SLC35F2 | NM_017515 | -0.7125 | -1.317 |
| RAB39 | NM_017516 | -0.9345 | -1.474 |
| HAUS7 | NM_017518 | -0.3185 | -1.473 |
| ARID1B | NM_017519 | -0.899 | -2.266 |
| MPHOSPH8 | NM_017520 | -0.6275 | -1.5785 |
| FEV | NM_017521 | -0.1595 | -0.7475 |
| LRP8 | NM_017522 | -1.334 | -2.2045 |

| | | | |
|---|---|---|---|
| XAF1 | NM_017523 | -0.0415 | -0.2355 |
| CDC42BPG | NM_017525 | 0.105 | 0.061 |
| LEPROT | NM_017526 | -0.848 | -1.1735 |
| LY6K | NM_017527 | -0.0195 | -0.6825 |
| WBSCR22 | NM_017528 | 0.402 | -0.37 |
| ZNF821 | NM_017530 | -0.663 | -1.671 |
| MYH4 | NM_017533 | -0.3645 | -0.633 |
| MYH2 | NM_017534 | -0.2985 | 0.202 |
| DNAH3 | NM_017539 | 0.1145 | -0.14 |
| CRYGS | NM_017541 | -0.958 | -0.737 |
| POGK | NM_017542 | -1.217 | -2.7465 |
| NKRF | NM_017544 | -1.094 | -2.1835 |
| HAO1 | NM_017545 | 0.1905 | 0.0045 |
| C2orf29 | NM_017546 | -0.693 | -2.4015 |
| FOXRED1 | NM_017547 | -0.207 | -1.103 |
| CDV3 | NM_017548 | -0.9495 | -1.161 |
| EPDR1 | NM_017549 | -0.183 | -2.312 |
| MIER2 | NM_017550 | -0.335 | -1.3575 |
| GRID1 | NM_017551 | -0.1025 | -1.4235 |
| ATAD2B | NM_017552 | -0.969 | -2.9155 |
| INO80 | NM_017553 | -0.8715 | -2.5615 |
| PARP14 | NM_017554 | -0.15 | -0.2525 |
| FBLIM1 | NM_017556 | -0.0575 | -0.515 |
| HYDIN | NM_017558 | -0.422 | -0.2415 |
| FNDC8 | NM_017559 | -0.2295 | 0.2375 |
| ZNF853 | NM_017560 | -0.8475 | -2.3245 |
| FAM22F | NM_017561 | 0.0795 | -0.2215 |
| IL17RD | NM_017563 | -1.7165 | -1.4005 |
| STAB2 | NM_017564 | -0.269 | -0.096 |
| FAM20A | NM_017565 | 0.0275 | -0.2675 |
| KLHDC4 | NM_017566 | -0.0635 | -0.5435 |
| NAGK | NM_017567 | -0.727 | -2.2825 |
| FAM48A | NM_017569 | -1.5075 | -2.5815 |
| OPLAH | NM_017570 | -0.0215 | -0.276 |
| MKNK2 | NM_017572 | -0.1155 | 0.9995 |
| PCSK4 | NM_017573 | -0.0705 | -0.3685 |
| SMG6 | NM_017575 | -0.4545 | -1.22 |
| KIF27 | NM_017576 | -0.3825 | -0.78 |
| GRAMD1C | NM_017577 | 0.271 | -0.3075 |
| ROPN1 | NM_017578 | 0.215 | -0.38 |
| DMBT1 | NM_017579 | -0.033 | 0.151 |
| ZRANB1 | NM_017580 | -1.3255 | -2.14 |
| CHRNA9 | NM_017581 | 0.0665 | -0.2085 |

| | | | |
|---|---|---|---|
| UBE2Q1 | NM_017582 | -0.279 | -0.9005 |
| TRIM44 | NM_017583 | -0.7 | -1.263 |
| MIOX | NM_017584 | 0.0035 | -0.0835 |
| SLC2A6 | NM_017585 | -0.533 | -1.219 |
| C9orf7 | NM_017586 | -0.5155 | -1.604 |
| WDR5 | NM_017588 | -1.04 | -1.632 |
| BTG4 | NM_017589 | 0.105 | 0.159 |
| ZC3H7B | NM_017590 | 0.0305 | -0.7045 |
| MED29 | NM_017592 | -1.626 | -3.0445 |
| BMP2K | NM_017593 | -0.288 | -0.703 |
| DIRAS2 | NM_017594 | -0.168 | -0.284 |
| NKIRAS2 | NM_017595 | -0.569 | -1.22 |
| KIF21B | NM_017596 | 0.44 | -0.5925 |
| VEZT | NM_017599 | -1.132 | -1.656 |
| OTUD5 | NM_017602 | -0.415 | -1.06 |
| PPP1R12C | NM_017607 | -0.9125 | -1.423 |
| RNF111 | NM_017610 | -1.9405 | -5.361 |
| SLC43A3 | NM_017611 | -0.6805 | -1.804 |
| ZCCHC8 | NM_017612 | -1.7385 | -3.6565 |
| DONSON | NM_017613 | -0.8515 | -1.5725 |
| BHMT2 | NM_017614 | 0.2245 | -0.583 |
| NSMCE4A | NM_017615 | -1.3165 | -2.2585 |
| NOTCH1 | NM_017617 | 0.019 | -0.4585 |
| RNPC3 | NM_017619 | -0.5755 | -1.3995 |
| ILF3 | NM_017620 | -0.0295 | -0.4595 |
| ALKBH4 | NM_017621 | -0.4695 | -1.861 |
| C17orf59 | NM_017622 | -0.706 | -1.5405 |
| CNNM3 | NM_017623 | -0.597 | -1.5755 |
| ITLN1 | NM_017625 | -0.386 | -0.114 |
| DNAJB12 | NM_017626 | -0.7045 | -2.1445 |
| TET2 | NM_017628 | 1.5025 | 0.8195 |
| EIF2C4 | NM_017629 | -1.948 | -2.635 |
| DDX60 | NM_017631 | 0.036 | -0.322 |
| CDKN2AIP | NM_017632 | -1.4925 | -1.8295 |
| FAM46A | NM_017633 | -0.31 | 0.7385 |
| KCTD9 | NM_017634 | -1.3865 | -1.909 |
| SUV420H1 | NM_017635 | 0.188 | 0.562 |
| TRPM4 | NM_017636 | 0.269 | -0.616 |
| BNC2 | NM_017637 | -0.5445 | -1.4825 |
| MED18 | NM_017638 | -0.273 | -0.435 |
| DCHS2 | NM_017639 | -0.6575 | -0.402 |
| LRRC16A | NM_017640 | -0.638 | -0.971 |
| KIF21A | NM_017641 | -0.5345 | -2.2535 |

| | | | |
|---|---|---|---|
| MBTD1 | NM_017643 | -1.706 | -2.7105 |
| KLHL24 | NM_017644 | -1.8915 | -2.731 |
| HAUS6 | NM_017645 | -0.807 | -2.1335 |
| TRIT1 | NM_017646 | -1.436 | -2.4865 |
| FTSJ3 | NM_017647 | -1.025 | -2.146 |
| CNNM2 | NM_017649 | -0.628 | -1.258 |
| PPP1R9A | NM_017650 | -0.6255 | -1.3355 |
| AHI1 | NM_017651 | -0.9195 | -1.136 |
| ZNF586 | NM_017652 | -1.2695 | -1.7975 |
| DYM | NM_017653 | -0.3455 | -0.591 |
| SAMD9 | NM_017654 | 0.4335 | -0.3945 |
| GIPC2 | NM_017655 | 0.057 | -0.422 |
| ZNF562 | NM_017656 | -1.0175 | -2.2645 |
| AFTPH | NM_017657 | -1.2 | -2.3835 |
| KLHL28 | NM_017658 | -1.226 | -1.811 |
| QPCTL | NM_017659 | -0.0475 | -0.4115 |
| GATAD2A | NM_017660 | 0.02 | -0.809 |
| ZNF280D | NM_017661 | -0.5185 | -0.6775 |
| TRPM6 | NM_017662 | -0.1945 | -0.5305 |
| ANKRD10 | NM_017664 | -0.033 | -0.175 |
| ZCCHC10 | NM_017665 | -1.8285 | -3.509 |
| ZNF280C | NM_017666 | -2.018 | -3.3675 |
| CCDC132 | NM_017667 | -1.095 | -2.395 |
| NDE1 | NM_017668 | -1.4135 | -2.1175 |
| ERCC6L | NM_017669 | -0.813 | -2.6405 |
| OTUB1 | NM_017670 | 0.175 | -0.559 |
| FERMT1 | NM_017671 | -0.2075 | -0.384 |
| TRPM7 | NM_017672 | -0.675 | -1.6775 |
| CDHR2 | NM_017675 | 0.3 | 0.2735 |
| GIN1 | NM_017676 | -1.4565 | -2.411 |
| MTMR8 | NM_017677 | -0.079 | -0.2815 |
| FAM55D | NM_017678 | -0.0105 | 0.323 |
| BCAS3 | NM_017679 | -0.0395 | -0.6885 |
| ASPN | NM_017680 | -0.352 | 0.1105 |
| NUP62CL | NM_017681 | 0.019 | 0.2085 |
| BEST2 | NM_017682 | 0.194 | -0.339 |
| VPS13C | NM_017684 | 0.0215 | -0.285 |
| GDAP2 | NM_017686 | 0.181 | -0.4085 |
| BSPRY | NM_017688 | -0.325 | -0.4765 |
| LRRC49 | NM_017691 | -0.469 | -1.252 |
| BIVM | NM_017693 | -1.2535 | -1.413 |
| MFSD6 | NM_017694 | -0.9315 | -2.234 |
| MCM9 | NM_017696 | -1.5535 | -3.3235 |

| | | | |
|---|---|---|---|
| ESRP1 | NM_017697 | 0.333 | -0.0835 |
| SIDT1 | NM_017699 | 0.0445 | -0.22 |
| ARHGEF38 | NM_017700 | -0.2565 | -0.236 |
| DEF8 | NM_017702 | 0.5695 | -0.1755 |
| FBXL12 | NM_017703 | -0.9375 | -2.7875 |
| ANKRD49 | NM_017704 | -1.6495 | -2.8625 |
| PAQR5 | NM_017705 | -0.6285 | -0.4595 |
| WDR55 | NM_017706 | -0.461 | -1.4065 |
| ASAP3 | NM_017707 | -0.017 | -0.986 |
| FAM83E | NM_017708 | 0.3275 | 0.111 |
| FAM46C | NM_017709 | -0.3015 | -0.2725 |
| GDPD2 | NM_017711 | 0.2655 | -0.345 |
| PGPEP1 | NM_017712 | -0.2155 | 0.079 |
| TASP1 | NM_017714 | -1.3165 | -2.207 |
| ZNF3 | NM_017715 | -1.195 | -2.632 |
| MS4A12 | NM_017716 | -0.181 | -0.158 |
| SNRK | NM_017719 | -1.5295 | -3.637 |
| STAP2 | NM_017720 | -0.1785 | -0.4 |
| CC2D1A | NM_017721 | -0.9225 | -1.7055 |
| TRMT1 | NM_017722 | -0.1185 | -0.694 |
| C9orf167 | NM_017723 | -0.2125 | -0.605 |
| LRRFIP2 | NM_017724 | -1.1075 | -2.1585 |
| PPP1R14D | NM_017726 | 0.1725 | -0.173 |
| TMEM214 | NM_017727 | -0.4525 | -1.1425 |
| TMEM104 | NM_017728 | -1.0045 | -3.1205 |
| EPS8L1 | NM_017729 | -0.2715 | -0.3745 |
| QRICH1 | NM_017730 | -1.0735 | -3.469 |
| PIGG | NM_017733 | -0.8 | -1.669 |
| PALMD | NM_017734 | -0.905 | -0.7645 |
| TTC27 | NM_017735 | -1.045 | -2.3645 |
| THUMPD1 | NM_017736 | -1.081 | -2.6455 |
| FNBP1L | NM_017737 | -0.7195 | -1.4815 |
| CNTLN | NM_017738 | -0.5065 | -1.2525 |
| POMGNT1 | NM_017739 | -0.079 | -0.597 |
| ZDHHC7 | NM_017740 | -0.6285 | -1.835 |
| DCAF16 | NM_017741 | -1.1315 | -1.425 |
| ZCCHC2 | NM_017742 | -1.3025 | -2.9805 |
| DPP8 | NM_017743 | -1.3795 | -1.5685 |
| ST7L | NM_017744 | -1.44 | -1.7125 |
| BCOR | NM_017745 | -1.034 | -3.5405 |
| TEX10 | NM_017746 | -1.609 | -3.454 |
| ANKHD1 | NM_017747 | 0.2395 | -0.272 |
| CWC25 | NM_017748 | -0.944 | -1.342 |

| | | | |
|---|---|---|---|
| AMBRA1 | NM_017749 | -0.0575 | -1.7855 |
| RETSAT | NM_017750 | -0.1485 | -0.924 |
| SMPD4 | NM_017751 | -0.4145 | -1.509 |
| TBC1D8B | NM_017752 | -0.34 | -0.3285 |
| LPPR1 | NM_017753 | 0.462 | -1.675 |
| UHRF1BP1 | NM_017754 | -1.8705 | -2.372 |
| NSUN2 | NM_017755 | -0.6355 | -1.641 |
| ZNF407 | NM_017757 | -0.7205 | -2.612 |
| ALKBH5 | NM_017758 | -0.0105 | -0.8195 |
| INO80D | NM_017759 | -0.9105 | -2.1245 |
| NCAPG2 | NM_017760 | -0.87 | -1.4985 |
| PNRC2 | NM_017761 | -2.0615 | -2.846 |
| MTMR10 | NM_017762 | -1.0665 | -2.4395 |
| RNF43 | NM_017763 | -0.0065 | -0.4885 |
| PQLC2 | NM_017765 | -0.583 | -0.741 |
| CASZ1 | NM_017766 | 0.257 | -0.387 |
| SLC39A4 | NM_017767 | 0.193 | -0.492 |
| LRRC40 | NM_017768 | -0.8745 | -1.5705 |
| G2E3 | NM_017769 | -0.843 | -2.347 |
| ELOVL2 | NM_017770 | -1.6805 | -1.578 |
| PXK | NM_017771 | -0.779 | -1.2035 |
| TBC1D22B | NM_017772 | -0.776 | -2.645 |
| LAX1 | NM_017773 | 0.0505 | -0.408 |
| CDKAL1 | NM_017774 | -0.7935 | -1.0245 |
| TTC19 | NM_017775 | -0.76 | -0.735 |
| MKS1 | NM_017777 | -0.4345 | -1.096 |
| WHSC1L1 | NM_017778 | 0.074 | -0.524 |
| DEPDC1 | NM_017779 | -1.5305 | -4.08 |
| CHD7 | NM_017780 | -0.997 | -2.557 |
| CYP2W1 | NM_017781 | 0.1585 | -0.444 |
| FAM208B | NM_017782 | -0.286 | -0.414 |
| OSBPL10 | NM_017784 | 0.0155 | -1.174 |
| CCDC99 | NM_017785 | -1.3055 | -3.506 |
| SYBU | NM_017786 | -0.7135 | -2.029 |
| C10orf26 | NM_017787 | -1.1695 | -3.171 |
| SEMA4C | NM_017789 | -0.673 | -1.485 |
| RGS3 | NM_017790 | -0.617 | -1.4575 |
| FLVCR2 | NM_017791 | 0.026 | -0.3215 |
| RPP25 | NM_017793 | -0.765 | -2.4035 |
| KIAA1797 | NM_017794 | -0.743 | -0.9965 |
| BTBD2 | NM_017797 | -0.459 | -0.935 |
| YTHDF1 | NM_017798 | -0.6195 | -1.6455 |
| C14orf101 | NM_017799 | -1.118 | -1.0625 |

| | | | |
|---|---|---|---|
| CMTM6 | NM_017801 | -0.723 | -1.084 |
| HEATR2 | NM_017802 | -1.4445 | -1.696 |
| DUS2L | NM_017803 | -2.1105 | -3.2785 |
| RASIP1 | NM_017805 | 0.0315 | -1.0025 |
| LIME1 | NM_017806 | -0.4565 | -0.704 |
| OSGEP | NM_017807 | -0.4265 | -1.4515 |
| ZNF434 | NM_017810 | -1.121 | -2.78 |
| UBE2R2 | NM_017811 | -0.3765 | -1.738 |
| CHCHD3 | NM_017812 | -0.632 | -1.3735 |
| IMPAD1 | NM_017813 | -1.2895 | -1.423 |
| TMEM161A | NM_017814 | -0.489 | -0.959 |
| HAUS4 | NM_017815 | -0.0665 | -0.817 |
| LYAR | NM_017816 | -1.0245 | -2.262 |
| RAB20 | NM_017817 | -0.1725 | -0.9125 |
| RG9MTD1 | NM_017819 | -1.5505 | -3.484 |
| EXD3 | NM_017820 | 0.164 | -0.224 |
| RHBDL2 | NM_017821 | -0.137 | -0.223 |
| C12orf41 | NM_017822 | -1.0615 | -3.009 |
| DUSP23 | NM_017823 | 0.0825 | -0.3905 |
| ADPRHL2 | NM_017825 | -0.6305 | -1.9845 |
| SOHLH2 | NM_017826 | -0.276 | -0.948 |
| SARS2 | NM_017827 | -0.3875 | -0.2805 |
| COMMD4 | NM_017828 | 0.0005 | -0.8035 |
| OCIAD1 | NM_017830 | -1.113 | -1.1115 |
| RNF125 | NM_017831 | -0.459 | -0.8475 |
| DNAJC28 | NM_017833 | -0.789 | -0.0755 |
| SLC41A3 | NM_017836 | 0.056 | -0.745 |
| PIGV | NM_017837 | -0.641 | -0.985 |
| NHP2 | NM_017838 | 0.164 | -0.3505 |
| LPCAT2 | NM_017839 | -1.374 | -1.798 |
| MRPL16 | NM_017840 | -0.5905 | -2.284 |
| SDHAF2 | NM_017841 | -0.8305 | -3.378 |
| SLC48A1 | NM_017842 | -0.6075 | -1.832 |
| BCAS4 | NM_017843 | -0.196 | -0.902 |
| ANKMY1 | NM_017844 | -0.3115 | -0.423 |
| COMMD8 | NM_017845 | -1.486 | -3.686 |
| TRNAU1AP | NM_017846 | -0.6925 | -1.475 |
| C1orf27 | NM_017847 | -0.6855 | -1.2995 |
| FAM120C | NM_017848 | 0.152 | -0.2185 |
| TMEM127 | NM_017849 | -0.429 | -1.5705 |
| C1orf109 | NM_017850 | -0.921 | -1.3965 |
| PARP16 | NM_017851 | -0.7785 | -2.7795 |
| NLRP2 | NM_017852 | 0.1685 | -0.2315 |

| | | | |
|---|---|---|---|
| TXNL4B | NM_017853 | -0.6745 | -1.6195 |
| TMEM160 | NM_017854 | 0.1745 | -0.7015 |
| ODAM | NM_017855 | 0.0355 | -0.249 |
| GEMIN8 | NM_017856 | -1.0135 | -1.2385 |
| SSH3 | NM_017857 | 0.0215 | -0.265 |
| TIPIN | NM_017858 | -1.1305 | -2.798 |
| UCKL1 | NM_017859 | 0.0965 | -0.5275 |
| C1orf56 | NM_017860 | -0.603 | -1.2435 |
| PIGX | NM_017861 | -1.06 | -1.315 |
| CXorf48 | NM_017863 | -0.076 | -0.495 |
| INTS8 | NM_017864 | -1.724 | -2.5685 |
| ZNF692 | NM_017865 | -0.526 | -1.4495 |
| TMEM70 | NM_017866 | -1.2235 | -4.072 |
| C4orf27 | NM_017867 | -0.8815 | -2.404 |
| TTC12 | NM_017868 | -0.8495 | -0.633 |
| BANP | NM_017869 | -0.715 | -1.8985 |
| TMEM132A | NM_017870 | -0.1165 | -0.6175 |
| CPSF3L | NM_017871 | -0.0295 | -0.6455 |
| THG1L | NM_017872 | -0.7225 | -1.2645 |
| ASB6 | NM_017873 | -0.696 | -2.7405 |
| SLC25A38 | NM_017875 | -0.638 | -1.808 |
| C2orf18 | NM_017877 | -0.0995 | -0.322 |
| HRASLS2 | NM_017878 | -0.108 | 0.157 |
| ZNF416 | NM_017879 | -0.769 | -2.2635 |
| C2orf42 | NM_017880 | -1.485 | -3.369 |
| C9orf95 | NM_017881 | 0.449 | -0.321 |
| CLN6 | NM_017882 | 0.031 | -0.716 |
| WDR13 | NM_017883 | -0.0425 | -0.8285 |
| PINX1 | NM_017884 | -0.558 | -2.0155 |
| HCFC1R1 | NM_017885 | 0.062 | -0.7925 |
| ULK4 | NM_017886 | 0.667 | -0.2675 |
| C1orf123 | NM_017887 | -0.2845 | -1.07 |
| ACSM5 | NM_017888 | 0.0075 | -0.174 |
| VPS13B | NM_017890 | -0.4705 | -1.3945 |
| C1orf159 | NM_017891 | -0.3105 | -0.711 |
| PRPF40A | NM_017892 | -0.0445 | -1.1935 |
| SEMA4G | NM_017893 | -0.042 | -0.667 |
| ZSCAN2 | NM_017894 | -1.2225 | -3.0835 |
| DDX27 | NM_017895 | -0.8005 | -2.192 |
| C20orf11 | NM_017896 | -0.579 | -1.343 |
| MOSC2 | NM_017898 | -0.744 | -1.335 |
| TESC | NM_017899 | -0.026 | -0.9045 |
| AURKAIP1 | NM_017900 | -0.1445 | -0.823 |

| | | | |
|---|---|---|---|
| TPCN1 | NM_017901 | -0.141 | -0.5965 |
| HIF1AN | NM_017902 | -1.194 | -2.4975 |
| TTC22 | NM_017904 | 0.0495 | 0.0465 |
| TMCO3 | NM_017905 | -0.9755 | -2.348 |
| PAK1IP1 | NM_017906 | -0.9905 | -1.438 |
| ZNF446 | NM_017908 | -0.473 | -0.788 |
| RMND1 | NM_017909 | -0.809 | -1.4065 |
| TRMT61B | NM_017910 | -1.8605 | -2.909 |
| FAM118A | NM_017911 | -0.8535 | -1.463 |
| HERC6 | NM_017912 | -0.026 | -0.6095 |
| CDC37L1 | NM_017913 | -1.0685 | -2.8365 |
| C19orf24 | NM_017914 | -0.311 | -1.252 |
| C12orf48 | NM_017915 | -1.3945 | -1.898 |
| PIH1D1 | NM_017916 | 0.0295 | -0.5855 |
| PPP2R3C | NM_017917 | -1.6795 | -2.799 |
| CCDC109B | NM_017918 | -0.503 | -0.5185 |
| STX17 | NM_017919 | -0.2195 | -1.264 |
| URGCP | NM_017920 | -0.426 | -2.246 |
| NPLOC4 | NM_017921 | -0.8255 | -1.923 |
| PRPF39 | NM_017922 | -0.96 | -1.4515 |
| 1-Mar | NM_017923 | 0.0775 | -0.266 |
| C14orf119 | NM_017924 | -1.2515 | -3.491 |
| DENND4C | NM_017925 | -1.157 | -2.591 |
| C14orf118 | NM_017926 | 0.0155 | -1.1965 |
| PEX26 | NM_017929 | 0.079 | -0.819 |
| TTC38 | NM_017931 | -0.2115 | -0.8235 |
| PID1 | NM_017933 | -0.054 | -0.927 |
| PHIP | NM_017934 | -0.5855 | -4.1325 |
| BANK1 | NM_017935 | 0.4275 | -0.518 |
| FAM70A | NM_017938 | -0.3695 | -0.6185 |
| NBPF1 | NM_017940 | -2.086 | -1.448 |
| C17orf80 | NM_017941 | -1.7265 | -3.598 |
| FBXO34 | NM_017943 | -1.46 | -3.041 |
| USP47 | NM_017944 | -0.634 | -2.2135 |
| SLC35A5 | NM_017945 | -1.593 | -2.9135 |
| FKBP14 | NM_017946 | -1.046 | -1.2865 |
| MOCOS | NM_017947 | 0.1725 | -0.2005 |
| NOL8 | NM_017948 | -0.8 | -2.761 |
| CUEDC1 | NM_017949 | -0.431 | -1.245 |
| CCDC40 | NM_017950 | -0.124 | -0.883 |
| SMPD4 | NM_017951 | -0.353 | -1.6095 |
| PTCD3 | NM_017952 | -1.1215 | -2.1375 |
| ZNHIT6 | NM_017953 | -1.1375 | -1.976 |

| | | | |
|---|---|---|---|
| CADPS2 | NM_017954 | -0.0335 | 0.042 |
| CDCA4 | NM_017955 | -0.87 | -3.7575 |
| TRMT12 | NM_017956 | -1.393 | -2.7585 |
| EPN3 | NM_017957 | -0.213 | -0.508 |
| PLEKHB2 | NM_017958 | -0.458 | -0.6465 |
| SLC30A6 | NM_017964 | -1.377 | -2.531 |
| VPS37C | NM_017966 | -0.5585 | -2.575 |
| IWS1 | NM_017969 | -1.0735 | -3.5385 |
| C14orf102 | NM_017970 | -0.706 | -2.3235 |
| MRPL20 | NM_017971 | -0.6195 | -1.8155 |
| C14orf118 | NM_017972 | -0.358 | -1.2615 |
| ATG16L1 | NM_017974 | -0.4715 | -2.0815 |
| ZWILCH | NM_017975 | -0.8695 | -0.606 |
| ANKHD1 | NM_017978 | 0.2395 | -0.272 |
| LIMS2 | NM_017980 | -0.007 | -0.366 |
| SUSD4 | NM_017982 | 0.037 | -0.27 |
| WIPI1 | NM_017983 | -1.2465 | -2.088 |
| ZCWPW1 | NM_017984 | -0.1175 | -0.483 |
| GPR172B | NM_017986 | 0.0285 | -0.614 |
| RUFY2 | NM_017987 | -0.3865 | -0.694 |
| SCYL2 | NM_017988 | -0.667 | -1.9165 |
| PDPR | NM_017990 | -0.7355 | -1.865 |
| KIAA1310 | NM_017991 | -1.004 | -1.6945 |
| ENOX1 | NM_017993 | -1.231 | -2.061 |
| C7orf42 | NM_017994 | -0.722 | -1.217 |
| DET1 | NM_017996 | -1.097 | -3.0075 |
| C9orf40 | NM_017998 | -0.6555 | -2.4475 |
| RNF31 | NM_017999 | -0.678 | -1.382 |
| MREG | NM_018000 | -0.3315 | -1.1115 |
| OXR1 | NM_018002 | -1.0475 | -1.3835 |
| UACA | NM_018003 | -0.0525 | -0.8955 |
| TMEM45A | NM_018004 | -0.718 | -0.6765 |
| TRMU | NM_018006 | -0.845 | -1.513 |
| FEZF2 | NM_018008 | -0.0905 | 0.1295 |
| TAPBPL | NM_018009 | -0.2825 | -0.623 |
| IFT57 | NM_018010 | -0.855 | -1.4865 |
| ARGLU1 | NM_018011 | -0.141 | -0.25 |
| KIF26B | NM_018012 | -0.806 | -1.809 |
| SOBP | NM_018013 | -1.404 | -2.773 |
| BCL11A | NM_018014 | -1.1425 | -2.253 |
| CXorf57 | NM_018015 | 0.0195 | -0.1775 |
| C10orf118 | NM_018017 | 0.2275 | -0.6395 |
| SLC38A4 | NM_018018 | 0.3245 | 0.3095 |

| | | | |
|---|---|---|---|
| MED9 | NM_018019 | -1.41 | -3.5485 |
| TMEM51 | NM_018022 | -0.569 | -2.0175 |
| YEATS2 | NM_018023 | -1.5315 | -3.0775 |
| WDYHV1 | NM_018024 | -0.439 | -1.3925 |
| GPATCH1 | NM_018025 | -1.5755 | -2.7905 |
| PACS1 | NM_018026 | 0.097 | -0.54 |
| FRMD4A | NM_018027 | -1.591 | -2.327 |
| SAMD4B | NM_018028 | -0.134 | -0.8765 |
| EBLN2 | NM_018029 | -1.1485 | -2.6255 |
| OSBPL1A | NM_018030 | -0.4005 | -0.736 |
| WDR6 | NM_018031 | -0.309 | -1.3145 |
| LUC7L | NM_018032 | -0.538 | -1.901 |
| WDR70 | NM_018034 | -0.8875 | -1.633 |
| ATP5SL | NM_018035 | -0.2885 | -1.115 |
| ATG2B | NM_018036 | -1.411 | -2.5815 |
| KDM4D | NM_018039 | -0.4055 | -1.837 |
| GPATCH2 | NM_018040 | -1.0355 | -2.1135 |
| SLFN12 | NM_018042 | 0.0995 | -0.322 |
| ANO1 | NM_018043 | -0.1205 | -0.181 |
| NSUN5 | NM_018044 | -0.653 | -2.114 |
| BSDC1 | NM_018045 | -0.936 | -1.5705 |
| AGGF1 | NM_018046 | -1.56 | -3.0535 |
| RBM22 | NM_018047 | -0.568 | -1.604 |
| MAGOHB | NM_018048 | -1.113 | -1.469 |
| PLEKHJ1 | NM_018049 | -0.359 | -1.835 |
| MANSC1 | NM_018050 | -1.0855 | -3.067 |
| WDR60 | NM_018051 | -1.2965 | -1.568 |
| VAC14 | NM_018052 | -0.4595 | -1.0385 |
| XKR8 | NM_018053 | -0.49 | -2.3375 |
| ARHGAP17 | NM_018054 | -0.481 | -1.689 |
| NODAL | NM_018055 | 0.016 | -0.3145 |
| TMEM39B | NM_018056 | -1.145 | -2.326 |
| SLC6A15 | NM_018057 | -0.975 | -1.2145 |
| CRTAC1 | NM_018058 | 0.1395 | -0.2175 |
| RADIL | NM_018059 | -0.9915 | -1.5795 |
| IARS2 | NM_018060 | -1.072 | -1.58 |
| PRPF38B | NM_018061 | -0.391 | 0.091 |
| FANCL | NM_018062 | -0.9345 | -1.58 |
| HELLS | NM_018063 | -0.5545 | -1.639 |
| AKIRIN2 | NM_018064 | -0.999 | -3.227 |
| GPN2 | NM_018066 | -0.8175 | -2.01 |
| MAP7D1 | NM_018067 | -0.553 | -2.0645 |
| PIWIL2 | NM_018068 | 0.1355 | -0.133 |

| | | | |
|---|---|---|---|
| SSBP3 | NM_018070 | -0.017 | -0.824 |
| ARHGEF40 | NM_018071 | -1.0585 | -1.5465 |
| HEATR1 | NM_018072 | -0.8105 | -2.48 |
| TRIM68 | NM_018073 | -0.746 | -2.457 |
| CCDC94 | NM_018074 | -0.6325 | -2.09 |
| ANO10 | NM_018075 | -0.105 | -0.7545 |
| ARMC4 | NM_018076 | 0.081 | 0.035 |
| RBM28 | NM_018077 | -0.7105 | -1.5725 |
| LARP1B | NM_018078 | -1.286 | -2.196 |
| SRBD1 | NM_018079 | -0.6735 | -1.9625 |
| VPS13C | NM_018080 | 0.0215 | -0.285 |
| WRAP53 | NM_018081 | -0.6655 | -1.67 |
| POLR3B | NM_018082 | -1.205 | -3.336 |
| ZNF358 | NM_018083 | 0.23 | -0.4795 |
| CCDC88A | NM_018084 | -0.354 | -2.62 |
| IPO9 | NM_018085 | -0.274 | -0.8665 |
| FIGN | NM_018086 | -0.007 | -0.85 |
| TMEM48 | NM_018087 | -1.147 | -1.555 |
| ANKZF1 | NM_018089 | -0.7855 | -1.479 |
| NECAP2 | NM_018090 | -0.6085 | -1.8115 |
| ELP3 | NM_018091 | -1.0015 | -1.84 |
| NETO2 | NM_018092 | -1.36 | -2.516 |
| WDR74 | NM_018093 | -0.0605 | -0.191 |
| GSPT2 | NM_018094 | -1.1105 | -1.7465 |
| KBTBD4 | NM_018095 | -0.8705 | -2.603 |
| NLE1 | NM_018096 | -0.5005 | -1.1845 |
| HAUS2 | NM_018097 | -1.8135 | -2.163 |
| ECT2 | NM_018098 | -0.852 | -1.7355 |
| FAR2 | NM_018099 | -1.0515 | -1.304 |
| EFHC1 | NM_018100 | -1.005 | -0.877 |
| CDCA8 | NM_018101 | -0.807 | -2.6225 |
| ZNF334 | NM_018102 | -0.7345 | -1.6945 |
| LRRC8D | NM_018103 | -1.705 | -3.6305 |
| THAP1 | NM_018105 | -1.3595 | -2.034 |
| ZDHHC4 | NM_018106 | 0.0195 | -0.739 |
| RBM23 | NM_018107 | -1.154 | -2.968 |
| MTPAP | NM_018109 | -1.2685 | -2.842 |
| DOK4 | NM_018110 | -0.5615 | -1.3555 |
| C19orf73 | NM_018111 | -0.38 | -0.8855 |
| TMEM38B | NM_018112 | -0.9665 | -1.915 |
| LMBR1L | NM_018113 | -0.848 | -1.927 |
| DALRD3 | NM_018114 | 0.1585 | -0.5235 |
| SDAD1 | NM_018115 | -0.182 | -0.6125 |

| | | | |
|---|---|---|---|
| MSTO1 | NM_018116 | -0.721 | -1.379 |
| WDR11 | NM_018117 | -0.9285 | -1.168 |
| POLR3E | NM_018119 | -0.9695 | -2.09 |
| ARMC1 | NM_018120 | -1.7865 | -3.082 |
| FAM178A | NM_018121 | -1.6445 | -3.148 |
| DARS2 | NM_018122 | -0.864 | -1.4725 |
| RFWD3 | NM_018124 | -1.078 | -2.9665 |
| ARHGEF10L | NM_018125 | -0.5445 | -1.896 |
| TMEM33 | NM_018126 | -0.5475 | -0.943 |
| ELAC2 | NM_018127 | -0.345 | -1.142 |
| TSR1 | NM_018128 | -0.924 | -1.818 |
| PNPO | NM_018129 | -0.442 | -0.8095 |
| SHQ1 | NM_018130 | -1.389 | -2.894 |
| CEP55 | NM_018131 | -1.8455 | -3.662 |
| CENPQ | NM_018132 | -1.4365 | -3.916 |
| MSL2 | NM_018133 | -0.771 | -1.2225 |
| IQCC | NM_018134 | -1.032 | -0.9575 |
| MRPS18A | NM_018135 | -0.711 | -2.2555 |
| ASPM | NM_018136 | -0.921 | -3.1225 |
| PRMT6 | NM_018137 | -1.469 | -1.8155 |
| TBCCD1 | NM_018138 | -1.582 | -3.1835 |
| DNAAF2 | NM_018139 | -2.073 | -3.3515 |
| CEP72 | NM_018140 | -0.8765 | -2.672 |
| MRPS10 | NM_018141 | -1.5105 | -1.876 |
| INTS10 | NM_018142 | -1.299 | -1.6655 |
| KLHL11 | NM_018143 | 0.1855 | 0.3525 |
| SEC61A2 | NM_018144 | -0.119 | -0.3935 |
| FAM82A2 | NM_018145 | -0.409 | -1.5505 |
| RNMTL1 | NM_018146 | -0.932 | -2.9395 |
| SMG8 | NM_018149 | -1.148 | -1.939 |
| RNF220 | NM_018150 | -1.0915 | -2.06 |
| RIF1 | NM_018151 | -0.23 | -2.372 |
| ANTXR1 | NM_018153 | -1.115 | -1.0975 |
| ASF1B | NM_018154 | -1.6695 | -2.5615 |
| SLC25A36 | NM_018155 | -0.9555 | -1.1085 |
| VPS13D | NM_018156 | -0.7465 | -1.393 |
| RIC8B | NM_018157 | -1.424 | -2.937 |
| SLC4A1AP | NM_018158 | -1.5105 | -3.074 |
| NUDT11 | NM_018159 | -0.919 | -1.054 |
| NADSYN1 | NM_018161 | -0.1695 | -0.7175 |
| DNAJC17 | NM_018163 | -1.0445 | -2.186 |
| C12orf11 | NM_018164 | -0.939 | -3.658 |
| PBRM1 | NM_018165 | -1.239 | -1.3545 |

| | | | |
|---|---|---|---|
| FAM176B | NM_018166 | -0.067 | -0.5655 |
| BTBD7 | NM_018167 | -1.448 | -3.0015 |
| C14orf105 | NM_018168 | -0.165 | -0.231 |
| C12orf35 | NM_018169 | -0.498 | -2.687 |
| RPRD1A | NM_018170 | -1.0905 | -0.647 |
| APPL2 | NM_018171 | -1.2225 | -1.2925 |
| PLEKHG6 | NM_018173 | 0.168 | -0.7195 |
| MAP1S | NM_018174 | -0.5795 | -1.2685 |
| LGI2 | NM_018176 | -0.1595 | -0.7795 |
| N4BP2 | NM_018177 | -0.043 | -0.5355 |
| GOLPH3L | NM_018178 | -2.4145 | -4.086 |
| ATF7IP | NM_018179 | -1.2775 | -2.845 |
| DHX32 | NM_018180 | -1.574 | -3.206 |
| ZNF532 | NM_018181 | -1.119 | -1.586 |
| C17orf63 | NM_018182 | -0.8405 | -2.8985 |
| SBNO1 | NM_018183 | -0.5575 | -1.7525 |
| ARL8B | NM_018184 | -1.2735 | -1.6875 |
| C1orf112 | NM_018186 | -1.5045 | -2.6305 |
| ATAD3A | NM_018188 | -0.3505 | -1.0495 |
| DPPA4 | NM_018189 | 0.005 | -0.525 |
| BBS7 | NM_018190 | -1.0085 | -1.5105 |
| RCBTB1 | NM_018191 | -1.1605 | -0.605 |
| LEPREL1 | NM_018192 | 0.0385 | -0.356 |
| FANCI | NM_018193 | -0.91 | -1.1085 |
| HHAT | NM_018194 | -0.2445 | -0.559 |
| C11orf57 | NM_018195 | -1.6205 | -2.729 |
| TMLHE | NM_018196 | -0.5795 | -1.2935 |
| ZFP64 | NM_018197 | -1.161 | -3.3265 |
| DNAJC11 | NM_018198 | -1.062 | -1.703 |
| EXD2 | NM_018199 | -0.8335 | -2.6185 |
| HMG20A | NM_018200 | -1.373 | -2.115 |
| TBC1D13 | NM_018201 | -0.754 | -2.0145 |
| TMEM57 | NM_018202 | -1.192 | -2.399 |
| KLHDC8A | NM_018203 | -0.338 | -1.3065 |
| CKAP2 | NM_018204 | -1.05 | -3.19 |
| VPS35 | NM_018206 | -0.733 | -0.8235 |
| TRIM62 | NM_018207 | 0.15 | -0.2325 |
| ETNK2 | NM_018208 | 0.305 | -0.3475 |
| ARFGAP1 | NM_018209 | -0.413 | -1.0065 |
| CARKD | NM_018210 | -0.31 | -1.2745 |
| RAVER2 | NM_018211 | -1.3365 | -2.1135 |
| ENAH | NM_018212 | -0.8325 | -0.2935 |
| LRRC1 | NM_018214 | -0.955 | -2.176 |

| | | | |
|---|---|---|---|
| PNMAL1 | NM_018215 | -1.3745 | -1.5295 |
| PANK4 | NM_018216 | -0.672 | -2.221 |
| EDEM2 | NM_018217 | -0.251 | -1.233 |
| USP40 | NM_018218 | -1.3315 | -1.213 |
| CCDC87 | NM_018219 | -0.3145 | -0.0415 |
| MOB1A | NM_018221 | -0.2885 | -1.1215 |
| PARVA | NM_018222 | 0.0025 | -0.4105 |
| CHFR | NM_018223 | -0.195 | -1.4025 |
| C7orf44 | NM_018224 | 0.0995 | -0.2085 |
| SMU1 | NM_018225 | -1.0315 | -1.0785 |
| RNPEPL1 | NM_018226 | -0.302 | -1.191 |
| UBA6 | NM_018227 | -0.592 | -1.916 |
| VRTN | NM_018228 | -0.187 | -0.1895 |
| MUDENG | NM_018229 | -1.1615 | -2.559 |
| NUP133 | NM_018230 | -1.41 | -1.8515 |
| SLC38A7 | NM_018231 | -0.676 | -1.5975 |
| OGFOD1 | NM_018233 | -1.238 | -2.1695 |
| STEAP3 | NM_018234 | -0.179 | -0.492 |
| CNDP2 | NM_018235 | -0.3655 | -0.9445 |
| CCAR1 | NM_018237 | -0.646 | -1.1765 |
| AGK | NM_018238 | -0.7355 | -1.4185 |
| KIRREL | NM_018240 | 0.0175 | -0.1765 |
| TMEM184C | NM_018241 | -0.867 | -2.543 |
| SLC47A1 | NM_018242 | -0.153 | -0.466 |
| UQCC | NM_018244 | -1.13 | -1.8685 |
| OGDHL | NM_018245 | -0.4505 | -0.9615 |
| CCDC25 | NM_018246 | -0.7905 | -1.248 |
| TMEM30A | NM_018247 | -1.066 | -1.1005 |
| NEIL3 | NM_018248 | -1.287 | -2.706 |
| CDK5RAP2 | NM_018249 | -0.822 | -1.2085 |
| INTS9 | NM_018250 | -0.6895 | -2.5525 |
| TMEM206 | NM_018252 | -0.7105 | -1.981 |
| YY1AP1 | NM_018253 | -0.6905 | -2.053 |
| RCOR3 | NM_018254 | -1.8 | -2.731 |
| ELP2 | NM_018255 | -1.1775 | -1.7635 |
| WDR12 | NM_018256 | -1.2885 | -2.3595 |
| PCMTD2 | NM_018257 | -1.248 | -0.864 |
| TTC17 | NM_018259 | -1.315 | -2.396 |
| ZNF701 | NM_018260 | -0.749 | -1.0985 |
| EXOC1 | NM_018261 | -1.285 | -2.101 |
| IFT122 | NM_018262 | 0.1445 | -0.996 |
| ASXL2 | NM_018263 | -1.4485 | -3.2245 |
| TYW1 | NM_018264 | -1.5435 | -2.4345 |

| | | | |
|---|---|---|---|
| C1orf106 | NM_018265 | -0.053 | -0.225 |
| TMEM39A | NM_018266 | -1.155 | -2.6795 |
| WDR41 | NM_018268 | -0.959 | -1.3025 |
| ADI1 | NM_018269 | 0.1725 | -0.3955 |
| C20orf20 | NM_018270 | -0.87 | -2.0345 |
| THNSL2 | NM_018271 | -0.258 | -0.7225 |
| CASC1 | NM_018272 | -0.017 | -0.1445 |
| TMEM143 | NM_018273 | -0.278 | -0.692 |
| C7orf43 | NM_018275 | -0.511 | -1.153 |
| TMEM19 | NM_018279 | -0.932 | -2.4575 |
| C22orf26 | NM_018280 | 0.3165 | -0.4505 |
| ECHDC2 | NM_018281 | -0.0225 | -0.5385 |
| NUDT15 | NM_018283 | -0.671 | -0.6065 |
| GBP3 | NM_018284 | -0.258 | 0.4765 |
| IMP3 | NM_018285 | -0.8335 | -1.969 |
| TMEM100 | NM_018286 | -0.7665 | -0.6835 |
| ARHGAP12 | NM_018287 | -0.507 | -2.2685 |
| PHF10 | NM_018288 | -0.9525 | -0.2705 |
| VPS53 | NM_018289 | -0.7235 | -0.988 |
| PGM2 | NM_018290 | -0.891 | -1.254 |
| QRSL1 | NM_018292 | -1.525 | -2.378 |
| ZNF654 | NM_018293 | -0.529 | -2.277 |
| CWF19L1 | NM_018294 | -0.727 | -1.917 |
| TMEM140 | NM_018295 | 0.2015 | -0.3365 |
| LRRC36 | NM_018296 | -0.6505 | -0.383 |
| NGLY1 | NM_018297 | -1.0265 | -1.912 |
| MCOLN3 | NM_018298 | -0.3135 | -0.844 |
| UBE2W | NM_018299 | -1.268 | -3.192 |
| RBM41 | NM_018301 | -0.9315 | -1.578 |
| C4orf19 | NM_018302 | -0.2505 | -0.039 |
| EXOC2 | NM_018303 | -1.2235 | -1.454 |
| PRR11 | NM_018304 | -1.079 | -1.896 |
| TMEM40 | NM_018306 | -0.345 | -0.2255 |
| RHOT1 | NM_018307 | -1.606 | -2.0195 |
| TBC1D23 | NM_018309 | -0.943 | -2.234 |
| BRF2 | NM_018310 | -0.2205 | -0.8705 |
| PPP6R3 | NM_018312 | -0.764 | -1.7485 |
| PBRM1 | NM_018313 | -1.239 | -1.3545 |
| UEVLD | NM_018314 | -0.1565 | -1.233 |
| FBXW7 | NM_018315 | -0.993 | -1.8715 |
| KLHL26 | NM_018316 | -0.8005 | -1.929 |
| TBC1D19 | NM_018317 | -0.7405 | -1.4255 |
| CCDC91 | NM_018318 | -0.868 | -1.848 |

| | | | |
|---|---|---|---|
| TDP1 | NM_018319 | -1.678 | -3.915 |
| RNF121 | NM_018320 | -0.4415 | -2.1355 |
| BRIX1 | NM_018321 | -1.31 | -2.254 |
| C6orf64 | NM_018322 | -1.2685 | -2.739 |
| PI4K2B | NM_018323 | -0.9435 | -1.72 |
| OLAH | NM_018324 | -0.186 | -0.3725 |
| C9orf72 | NM_018325 | -1.604 | -2.907 |
| GIMAP4 | NM_018326 | 0.201 | -0.3845 |
| SPTLC3 | NM_018327 | -0.409 | 0.1885 |
| MBD5 | NM_018328 | -0.9965 | -1.4885 |
| KIAA1598 | NM_018330 | 0.117 | -0.1755 |
| DDX19A | NM_018332 | -0.553 | -1.094 |
| LRRN3 | NM_018334 | -0.572 | -4.192 |
| ZNF839 | NM_018335 | -0.9215 | -2.0345 |
| ZNF444 | NM_018337 | -0.504 | -0.9145 |
| WDR52 | NM_018338 | -0.3865 | -0.9105 |
| RFK | NM_018339 | -1.2405 | -1.275 |
| CPPED1 | NM_018340 | -0.7375 | -1.844 |
| C6orf70 | NM_018341 | -0.8385 | -1.41 |
| TMEM144 | NM_018342 | -0.132 | -0.3655 |
| RIOK2 | NM_018343 | -1.0185 | -2.7605 |
| SLC29A3 | NM_018344 | -0.593 | -1.936 |
| RSAD1 | NM_018346 | -0.5255 | -1.179 |
| C20orf29 | NM_018347 | -0.8105 | -1.6785 |
| FTSJD1 | NM_018348 | -2.1685 | -3.794 |
| MCTP2 | NM_018349 | 0.006 | -0.7005 |
| FGD6 | NM_018351 | -0.752 | -2.033 |
| C4orf43 | NM_018352 | -1.0145 | -1.8675 |
| MIS18BP1 | NM_018353 | -0.8485 | -3.38 |
| C20orf46 | NM_018354 | -0.1225 | -0.3065 |
| ZNF415 | NM_018355 | 0.2285 | -2.842 |
| C5orf22 | NM_018356 | -1.5715 | -2.1675 |
| LARP6 | NM_018357 | -0.8985 | -3.2275 |
| ABCF3 | NM_018358 | -0.3005 | -1.963 |
| UFSP2 | NM_018359 | -0.796 | -1.4925 |
| TXLNG | NM_018360 | -1.8695 | -3.742 |
| AGPAT5 | NM_018361 | -1.4605 | -2.169 |
| LIN7C | NM_018362 | -1.1305 | -1.691 |
| RNLS | NM_018363 | 0.1675 | -0.7285 |
| RSBN1 | NM_018364 | -1.931 | -3.6505 |
| MNS1 | NM_018365 | -0.203 | -1.0705 |
| CNO | NM_018366 | -0.821 | -2.2655 |
| ACER3 | NM_018367 | -1.049 | -1.3075 |

| | | | |
|---|---|---|---|
| LMBRD1 | NM_018368 | -1.342 | -2.1665 |
| DEPDC1B | NM_018369 | -1.2525 | -1.4635 |
| DRAM1 | NM_018370 | -0.524 | -1.231 |
| CSGALNACT1 | NM_018371 | 0.0685 | 0.0865 |
| LRIF1 | NM_018372 | -1.8025 | -4.474 |
| SYNJ2BP | NM_018373 | -1.253 | -2.264 |
| TMEM106B | NM_018374 | -0.9485 | -1.4925 |
| SLC39A9 | NM_018375 | -1.168 | -1.6965 |
| NIPSNAP3B | NM_018376 | -0.5155 | -1.384 |
| FBXL8 | NM_018378 | -0.1005 | -0.1805 |
| FAM63A | NM_018379 | -0.6285 | -0.816 |
| DDX28 | NM_018380 | -0.524 | -1.2535 |
| C19orf66 | NM_018381 | 0.118 | -0.034 |
| WDR33 | NM_018383 | -1.522 | -3.302 |
| GIMAP5 | NM_018384 | 0.1545 | -0.4095 |
| LSG1 | NM_018385 | -0.942 | -1.249 |
| PCID2 | NM_018386 | -1.3035 | -2.211 |
| STRBP | NM_018387 | -0.697 | -1.6225 |
| MBNL3 | NM_018388 | 0.1225 | -0.466 |
| SLC35C1 | NM_018389 | -0.5575 | -1.1465 |
| PLCXD1 | NM_018390 | -0.7865 | -1.604 |
| C4orf21 | NM_018392 | -1.1805 | -2.8575 |
| TCP11L1 | NM_018393 | -0.449 | -1.4305 |
| ABHD10 | NM_018394 | -1.082 | -2.923 |
| METTL2B | NM_018396 | -1.1085 | -1.88 |
| CHDH | NM_018397 | -0.426 | -0.8695 |
| CACNA2D3 | NM_018398 | -0.2785 | -0.41 |
| SCN3B | NM_018400 | -0.7525 | -1.0675 |
| STK32B | NM_018401 | 0.065 | -0.705 |
| IL26 | NM_018402 | -0.019 | -0.112 |
| DCP1A | NM_018403 | -1.7765 | -4.168 |
| ADAP2 | NM_018404 | 0.308 | -0.456 |
| C17orf79 | NM_018405 | 0.1955 | -0.293 |
| MUC4 | NM_018406 | 0.0875 | -0.2915 |
| LAPTM4B | NM_018407 | -0.102 | -0.211 |
| LRP2BP | NM_018409 | -0.018 | -0.3055 |
| HJURP | NM_018410 | -1.237 | -3.2325 |
| HR | NM_018411 | 0.0995 | -0.9505 |
| ST7 | NM_018412 | -0.7215 | -2.056 |
| CHST11 | NM_018413 | -0.643 | -1.778 |
| ST6GALNAC1 | NM_018414 | 0.2525 | 0.1505 |
| FOXJ2 | NM_018416 | -0.558 | -1.533 |
| ADCY10 | NM_018417 | -0.397 | -0.2165 |

| | | | |
|---|---|---|---|
| SPATA7 | NM_018418 | -1.2045 | -1.5325 |
| SOX18 | NM_018419 | -0.087 | -0.3565 |
| SLC22A15 | NM_018420 | 0.044 | -2.1285 |
| TBC1D2 | NM_018421 | -0.29 | -0.759 |
| STYK1 | NM_018423 | -0.022 | -0.2205 |
| EPB41L4B | NM_018424 | -0.372 | -1.219 |
| PI4K2A | NM_018425 | -0.557 | -2.649 |
| TMEM63B | NM_018426 | -0.31 | -0.7105 |
| RRN3 | NM_018427 | -1.442 | -1.797 |
| UTP6 | NM_018428 | -1.5115 | -2.192 |
| BDP1 | NM_018429 | -0.953 | -3.545 |
| TSNAXIP1 | NM_018430 | -0.1365 | -0.63 |
| DOK5 | NM_018431 | 0.1075 | -0.2045 |
| KDM3A | NM_018433 | -0.9915 | -2.8725 |
| RNF130 | NM_018434 | -0.73 | -1.2715 |
| ALLC | NM_018436 | -0.161 | -0.265 |
| HEMGN | NM_018437 | 0.092 | -0.3815 |
| FBXO6 | NM_018438 | -0.186 | -0.316 |
| IMPACT | NM_018439 | -1.061 | -1.251 |
| PAG1 | NM_018440 | -0.878 | -2.664 |
| PECR | NM_018441 | -0.6065 | -1.4935 |
| DCAF6 | NM_018442 | -1.115 | -1.723 |
| ZNF302 | NM_018443 | -1.7395 | -2.6875 |
| PDP1 | NM_018444 | -1.2095 | -1.779 |
| SELS | NM_018445 | -1.0055 | -2.704 |
| GLT8D1 | NM_018446 | -1.635 | -2.303 |
| TMEM111 | NM_018447 | -0.6755 | -0.905 |
| CAND1 | NM_018448 | -0.8545 | -2.107 |
| UBAP2 | NM_018449 | -0.585 | -2.1855 |
| CENPJ | NM_018451 | -1.406 | -3.3495 |
| C6orf35 | NM_018452 | -1.768 | -2.1805 |
| EAPP | NM_018453 | -1.42 | -2.5355 |
| NUSAP1 | NM_018454 | -1.4285 | -1.266 |
| CENPN | NM_018455 | -0.332 | -0.2905 |
| EAF2 | NM_018456 | -1.002 | -1.568 |
| PRR13 | NM_018457 | -0.577 | -1.289 |
| ARHGAP15 | NM_018460 | 0.576 | -0.3055 |
| PPP2R2D | NM_018461 | -1.0415 | -2.9265 |
| ITFG2 | NM_018463 | -0.4815 | -1.542 |
| CISD1 | NM_018464 | -0.4965 | -0.9145 |
| C9orf46 | NM_018465 | -1.381 | -2.405 |
| ALG13 | NM_018466 | -0.6645 | -1.719 |
| USE1 | NM_018467 | -0.0315 | -0.517 |

| | | | |
|---|---|---|---|
| TEX2 | NM_018469 | -1.152 | -3.25 |
| ZC3H15 | NM_018471 | -0.436 | -1.1065 |
| ACOT13 | NM_018473 | -1.188 | -1.1875 |
| PLK1S1 | NM_018474 | -1.326 | -1.6115 |
| TMEM165 | NM_018475 | -1.6555 | -2.5805 |
| BEX1 | NM_018476 | -0.211 | -0.396 |
| ACTR10 | NM_018477 | -1.432 | -1.291 |
| ECHDC1 | NM_018479 | -1.0715 | -1.5455 |
| TMEM126B | NM_018480 | -1.144 | -2.1265 |
| ASAP1 | NM_018482 | -0.7935 | -1.952 |
| SLC22A11 | NM_018484 | 0.164 | -0.3515 |
| GPR77 | NM_018485 | -0.2495 | -0.132 |
| HDAC8 | NM_018486 | 0.4635 | -0.028 |
| TMEM176A | NM_018487 | -0.029 | -0.278 |
| TBX4 | NM_018488 | -0.243 | -0.0635 |
| ASH1L | NM_018489 | -0.3325 | -1.5655 |
| LGR4 | NM_018490 | -0.5375 | -1.304 |
| PBK | NM_018492 | -0.766 | -1.155 |
| TMCO6 | NM_018502 | -0.1355 | -0.641 |
| LRRC59 | NM_018509 | -0.181 | -1.138 |
| MCM10 | NM_018518 | -1.526 | -2.724 |
| NAA16 | NM_018527 | -1.0985 | -2.0525 |
| GSDMB | NM_018530 | -0.6985 | -0.7295 |
| NRP2 | NM_018534 | -0.568 | -1.518 |
| ERMAP | NM_018538 | -0.1405 | -0.2345 |
| C17orf85 | NM_018553 | -0.443 | -1.603 |
| ZNF331 | NM_018555 | -1.416 | -1.737 |
| SIRPG | NM_018556 | -0.05 | -0.39 |
| LRP1B | NM_018557 | -0.5755 | -0.481 |
| GABRQ | NM_018558 | 0.2255 | -0.5555 |
| KIAA1704 | NM_018559 | -1.405 | -2.152 |
| USP49 | NM_018561 | -0.3645 | -1.2295 |
| SERINC2 | NM_018565 | -0.1555 | -0.2945 |
| YOD1 | NM_018566 | -1.5785 | -1.863 |
| AP1AR | NM_018569 | -1.1245 | -2.4445 |
| STRADB | NM_018571 | -0.715 | -1.189 |
| CAMK2N1 | NM_018584 | -0.554 | -0.932 |
| CSGALNACT2 | NM_018590 | -1.3285 | -2.333 |
| SLC16A10 | NM_018593 | -0.882 | -2.611 |
| DNAJA4 | NM_018602 | 0.237 | -0.0035 |
| ETNK1 | NM_018638 | -1.594 | -2.5455 |
| WSB2 | NM_018639 | -0.215 | -0.81 |
| LMO3 | NM_018640 | -1.304 | -1.7735 |

| | | | |
|---|---|---|---|
| CHST12 | NM_018641 | -1.021 | -2.4235 |
| TREM1 | NM_018643 | 0.0775 | -0.248 |
| B3GAT1 | NM_018644 | -0.05 | -0.3115 |
| HES6 | NM_018645 | 0.143 | 0.155 |
| TRPV6 | NM_018646 | -0.018 | -0.087 |
| TNFRSF19 | NM_018647 | -0.231 | -0.731 |
| NOP10 | NM_018648 | -0.3015 | -1.147 |
| H2AFY2 | NM_018649 | -0.3525 | -1.1195 |
| MARK1 | NM_018650 | -1.278 | -2.307 |
| ZNF167 | NM_018651 | -1.339 | -2.7965 |
| GPRC5D | NM_018654 | 0.0735 | 0.008 |
| LENEP | NM_018655 | -0.0145 | -0.2185 |
| SLC35E3 | NM_018656 | -0.783 | -2.2245 |
| MYNN | NM_018657 | -1.564 | -3.3895 |
| KCNJ16 | NM_018658 | -0.2105 | -0.1965 |
| CYTL1 | NM_018659 | 0.016 | -0.164 |
| ZNF395 | NM_018660 | -0.4665 | -1.3905 |
| DISC1 | NM_018662 | -0.2995 | -0.02 |
| PXMP2 | NM_018663 | 0.309 | -0.268 |
| BATF3 | NM_018664 | -0.4545 | -1.7275 |
| DDX43 | NM_018665 | 0.099 | -0.0715 |
| SAGE1 | NM_018666 | 0.2025 | -0.2695 |
| SMPD3 | NM_018667 | -0.439 | -1.4615 |
| VPS33B | NM_018668 | -0.626 | -2.394 |
| WDR4 | NM_018669 | -0.162 | -0.803 |
| MESP1 | NM_018670 | -0.3665 | -0.7185 |
| UNC45A | NM_018671 | 0.0765 | -0.5065 |
| ABCA5 | NM_018672 | -0.3855 | -0.867 |
| ACCN4 | NM_018674 | -0.0145 | -0.812 |
| THSD1 | NM_018676 | -1.0675 | -2.3195 |
| ACSS2 | NM_018677 | -0.1015 | -0.494 |
| TCP11 | NM_018679 | -0.107 | -0.197 |
| MLL5 | NM_018682 | -0.3845 | -1.8195 |
| RNF114 | NM_018683 | -0.731 | -1.8735 |
| ZC4H2 | NM_018684 | -0.4615 | -0.892 |
| ANLN | NM_018685 | -0.906 | -2.8325 |
| CMAS | NM_018686 | -1.186 | -2.031 |
| BIN3 | NM_018688 | -0.8205 | -2.4605 |
| KIAA1199 | NM_018689 | -0.2225 | -0.129 |
| FAM114A2 | NM_018691 | -1.154 | -1.1065 |
| ARL6IP4 | NM_018694 | 0.269 | -0.3275 |
| ERBB2IP | NM_018695 | -0.84 | -3.27 |
| ELAC1 | NM_018696 | -1.3 | -3.23 |

| | | | |
|---|---|---|---|
| LANCL2 | NM_018697 | -0.5905 | -1.7295 |
| NXT2 | NM_018698 | -0.5415 | -0.357 |
| PRDM5 | NM_018699 | -0.0715 | -1.446 |
| TRIM36 | NM_018700 | -0.206 | -0.9195 |
| ADARB2 | NM_018702 | 0.5175 | -0.2305 |
| RBBP6 | NM_018703 | -0.2585 | -0.423 |
| CTTNBP2NL | NM_018704 | -1.6785 | -4.1615 |
| DHTKD1 | NM_018706 | -1.209 | -1.5035 |
| FEM1A | NM_018708 | -1.377 | -2.7315 |
| TMEM55A | NM_018710 | -0.828 | -1.427 |
| SVOP | NM_018711 | -0.616 | -1.3535 |
| ELMOD1 | NM_018712 | -1.019 | -0.601 |
| SLC30A10 | NM_018713 | 0.508 | 0.0555 |
| COG1 | NM_018714 | -0.611 | -1.958 |
| RCC2 | NM_018715 | -0.6465 | -1.2235 |
| MAML3 | NM_018717 | -1.1595 | -2.968 |
| CEP41 | NM_018718 | -1.298 | -1.3655 |
| CDCA7L | NM_018719 | -0.0355 | -1.774 |
| RBFOX1 | NM_018723 | -0.464 | -0.8375 |
| IL20 | NM_018724 | -0.0145 | 0.0585 |
| IL17RB | NM_018725 | -0.855 | -1.4025 |
| TRPV1 | NM_018727 | -0.4195 | -0.5215 |
| MYO5C | NM_018728 | -0.0865 | -0.6805 |
| TAP2 | NM_018833 | -0.2035 | -0.459 |
| MATR3 | NM_018834 | -0.217 | -0.35 |
| RC3H2 | NM_018835 | -0.4275 | -1.5575 |
| AJAP1 | NM_018836 | -0.089 | -0.822 |
| SULF2 | NM_018837 | -0.5715 | -1.1015 |
| NDUFA12 | NM_018838 | -0.3095 | -0.408 |
| NSFL1C | NM_018839 | 0.013 | -0.898 |
| C20orf24 | NM_018840 | -0.352 | -1.906 |
| GNG12 | NM_018841 | -1.0355 | -1.8205 |
| BAIAP2L1 | NM_018842 | -0.1445 | -0.3935 |
| SLC25A40 | NM_018843 | -1.7565 | -2.9135 |
| BCAP29 | NM_018844 | -1.4795 | -1.146 |
| KLHL7 | NM_018846 | 0.4635 | -0.0885 |
| KLHL9 | NM_018847 | -1.5225 | -2.7475 |
| MKKS | NM_018848 | -0.442 | -1.5535 |
| ABCB4 | NM_018849 | -0.215 | -0.6335 |
| ABCB4 | NM_018850 | -0.2445 | -0.103 |
| RAC1 | NM_018890 | -0.314 | -0.5775 |
| LAMC2 | NM_018891 | -0.1985 | -0.162 |
| DNAH7 | NM_018897 | -0.164 | -0.4945 |

| | | | |
|---|---|---|---|
| PCDHAC1 | NM_018898 | -0.0465 | -0.4465 |
| PCDHAC2 | NM_018899 | 0.01 | -0.7045 |
| PCDHA11 | NM_018902 | -1.2995 | -2.3275 |
| PCDHA12 | NM_018903 | 0.36 | 0.1045 |
| PCDHA13 | NM_018904 | -0.07 | -0.3675 |
| PCDHA2 | NM_018905 | -0.1675 | -0.1005 |
| PCDHA4 | NM_018907 | 0.004 | -0.42 |
| PCDHA6 | NM_018909 | 0.0855 | 0.252 |
| PCDHA7 | NM_018910 | -0.052 | -0.2255 |
| PCDHA8 | NM_018911 | 0.164 | -0.6945 |
| PCDHGA1 | NM_018912 | 0.5405 | 0.26 |
| PCDHGA10 | NM_018913 | -0.646 | -0.1015 |
| PCDHGA11 | NM_018914 | -0.1945 | -0.312 |
| PCDHGA2 | NM_018915 | -0.068 | -0.32 |
| PCDHGA5 | NM_018918 | -0.4245 | -0.1265 |
| PCDHGA6 | NM_018919 | -0.0425 | -0.048 |
| PCDHGA7 | NM_018920 | 0.166 | -0.0725 |
| PCDHGA9 | NM_018921 | -0.0035 | -0.306 |
| PCDHGB1 | NM_018922 | -0.442 | -0.217 |
| PCDHGB2 | NM_018923 | 0.1095 | -0.1725 |
| PCDHGB3 | NM_018924 | 0.0135 | -0.127 |
| PCDHGB5 | NM_018925 | -0.159 | -0.313 |
| PCDHGB6 | NM_018926 | -0.1495 | 0.195 |
| PCDHGC5 | NM_018929 | -0.3175 | 0.125 |
| PCDHB10 | NM_018930 | -0.886 | -1.6815 |
| PCDHB11 | NM_018931 | -0.2555 | -0.299 |
| PCDHB12 | NM_018932 | -0.726 | -0.879 |
| PCDHB13 | NM_018933 | -0.553 | -1.1345 |
| PCDHB14 | NM_018934 | -0.592 | -1.569 |
| PCDHB15 | NM_018935 | -0.9515 | -0.779 |
| PCDHB2 | NM_018936 | -0.416 | -1.5975 |
| PCDHB3 | NM_018937 | -0.077 | -0.604 |
| PCDHB4 | NM_018938 | 0.036 | -0.3325 |
| PCDHB6 | NM_018939 | -0.509 | -0.3625 |
| PCDHB7 | NM_018940 | -0.3455 | -0.065 |
| CLN8 | NM_018941 | -0.14 | -2.0035 |
| HMX1 | NM_018942 | -0.1275 | -1.711 |
| TUBA8 | NM_018943 | -0.343 | -1.4635 |
| MIS18A | NM_018944 | -0.8515 | -1.6615 |
| PDE7B | NM_018945 | -0.3075 | 0.0255 |
| NANS | NM_018946 | -0.237 | -1.0045 |
| CYCS | NM_018947 | -0.645 | -0.86 |
| ERRFI1 | NM_018948 | -0.5615 | -1.5535 |

| | | | |
|---|---|---|---|
| UTS2R | NM_018949 | -0.0095 | -0.478 |
| HLA-F | NM_018950 | 0.083 | -0.4835 |
| HOXA10 | NM_018951 | -0.0585 | -0.107 |
| HOXB6 | NM_018952 | 0.1365 | -0.3415 |
| HOXC5 | NM_018953 | 0.0565 | -0.3965 |
| UBB | NM_018955 | -0.0885 | -0.785 |
| C9orf9 | NM_018956 | -0.375 | -1.0745 |
| SH3BP1 | NM_018957 | -0.1885 | -1.1145 |
| C15orf2 | NM_018958 | 0.495 | -0.382 |
| DAZAP1 | NM_018959 | -0.089 | -0.763 |
| GNMT | NM_018960 | 0.1285 | -0.298 |
| UBASH3A | NM_018961 | -0.044 | -0.2395 |
| DSCR6 | NM_018962 | -0.2325 | -0.4765 |
| BRWD1 | NM_018963 | -0.9065 | -1.164 |
| SLC37A1 | NM_018964 | -0.4065 | -1.0545 |
| TREM2 | NM_018965 | 0.283 | -0.1255 |
| SNTG1 | NM_018967 | -0.015 | -0.452 |
| SNTG2 | NM_018968 | 0.397 | 0.016 |
| GPR173 | NM_018969 | -0.296 | -0.325 |
| GPR85 | NM_018970 | -0.6355 | -1.333 |
| GPR27 | NM_018971 | 0.0445 | -0.6915 |
| GDAP1 | NM_018972 | -0.695 | -0.7505 |
| UNC93A | NM_018974 | -0.4735 | -0.6625 |
| TERF2IP | NM_018975 | -0.4765 | -0.343 |
| SLC38A2 | NM_018976 | -1.697 | -2.8895 |
| NLGN3 | NM_018977 | -0.1905 | -0.3365 |
| WNK1 | NM_018979 | -0.809 | -1.332 |
| TAS2R5 | NM_018980 | -0.4825 | -0.5615 |
| DNAJC10 | NM_018981 | -0.4685 | -1.1455 |
| YIPF1 | NM_018982 | -0.3485 | -1.373 |
| GAR1 | NM_018983 | -0.933 | -3.1825 |
| SSH1 | NM_018984 | -1.212 | -2.3545 |
| SH3TC1 | NM_018986 | -0.1085 | -0.542 |
| GFOD1 | NM_018988 | -0.571 | -1.5355 |
| RBM27 | NM_018989 | -1.052 | -2.8815 |
| SASH3 | NM_018990 | 0.2925 | 0.011 |
| KCTD5 | NM_018992 | -0.9095 | -2.5415 |
| RIN2 | NM_018993 | 0.1275 | -0.4685 |
| FBXO42 | NM_018994 | -1.1725 | -1.964 |
| MOV10L1 | NM_018995 | 0.0735 | -0.502 |
| TNRC6C | NM_018996 | -1.0965 | -3.3925 |
| MRPS21 | NM_018997 | -0.4155 | -0.668 |
| FBXW5 | NM_018998 | 0.1475 | -0.415 |

| | | | |
|---|---|---|---|
| FAM190B | NM_018999 | -0.3565 | -1.378 |
| FAM134B | NM_019000 | -0.3635 | -0.695 |
| XRN1 | NM_019001 | -1.3255 | -2.6375 |
| ETAA1 | NM_019002 | -0.9785 | -1.9465 |
| ANKIB1 | NM_019004 | -1.485 | -3.5495 |
| MIOS | NM_019005 | -1.976 | -4.419 |
| ZFAND6 | NM_019006 | -0.836 | -0.9915 |
| ARMCX6 | NM_019007 | -0.5045 | -1.0455 |
| SMCR7L | NM_019008 | -0.8895 | -2.347 |
| TOLLIP | NM_019009 | 0.0255 | -0.897 |
| KRT20 | NM_019010 | 0.231 | 0.2045 |
| PLEKHA5 | NM_019012 | -1.354 | -3.347 |
| FAM64A | NM_019013 | -0.969 | -2.2775 |
| POLR1B | NM_019014 | -1.5995 | -3.9215 |
| CHPF2 | NM_019015 | -0.5705 | -1.6135 |
| KRT24 | NM_019016 | -0.1025 | -0.276 |
| FAM105A | NM_019018 | -0.8235 | -1.111 |
| TBC1D16 | NM_019020 | 0.149 | -0.9595 |
| C11orf71 | NM_019021 | -1.033 | -0.4135 |
| TMX3 | NM_019022 | -1.1225 | -2.211 |
| PRMT7 | NM_019023 | -0.4405 | -1.181 |
| HEATR5B | NM_019024 | -1.671 | -2.4295 |
| TMCO1 | NM_019026 | -0.684 | -0.4425 |
| RBM47 | NM_019027 | 0.1795 | 0.1535 |
| DHX29 | NM_019030 | -0.948 | -2.54 |
| ADAMTSL4 | NM_019032 | 0.2995 | 0.027 |
| RHOF | NM_019034 | -0.116 | -0.734 |
| PCDH18 | NM_019035 | -1.3425 | -1.3805 |
| HMGCLL1 | NM_019036 | 0.2295 | -0.454 |
| EXOSC4 | NM_019037 | -1.1035 | -3.0015 |
| ELP4 | NM_019040 | -0.9145 | -2.4735 |
| MTRF1L | NM_019041 | -0.924 | -1.3715 |
| PUS7 | NM_019042 | -1.751 | -2.204 |
| APBB1IP | NM_019043 | 0.0155 | -0.4305 |
| CCDC93 | NM_019044 | -1.3775 | -2.2775 |
| WDR44 | NM_019045 | -0.9015 | -2.158 |
| ANKRD16 | NM_019046 | -0.5055 | -1.1495 |
| ASNSD1 | NM_019048 | -1.0975 | -1.7785 |
| USP53 | NM_019050 | -0.538 | -1.0255 |
| MRPL50 | NM_019051 | -1.2165 | -3.093 |
| CCHCR1 | NM_019052 | -0.339 | -0.8795 |
| EXOC6 | NM_019053 | -1.4685 | -3.121 |
| FAM35A | NM_019054 | -1.362 | -2.256 |

| | | | |
|---|---|---|---|
| ROBO4 | NM_019055 | 0.2495 | -0.3865 |
| NDUFB11 | NM_019056 | 0.327 | -0.1905 |
| DDIT4 | NM_019058 | 0.051 | 1.014 |
| CRCT1 | NM_019060 | 0.131 | -0.549 |
| RNF186 | NM_019062 | 0.1985 | -0.232 |
| EML4 | NM_019063 | -1.083 | -1.7695 |
| NECAB2 | NM_019065 | -0.128 | -0.1535 |
| MAGEL2 | NM_019066 | 0.181 | -0.2945 |
| GNL3L | NM_019067 | -0.0475 | -0.7615 |
| WDR5B | NM_019069 | -1.2135 | -2.018 |
| DDX49 | NM_019070 | 0.071 | -0.8115 |
| ING3 | NM_019071 | -0.45 | 0.1745 |
| SGTB | NM_019072 | -1.168 | -2.3285 |
| SPATA6 | NM_019073 | -0.1805 | -0.551 |
| DLL4 | NM_019074 | 0.1545 | 0.0035 |
| UGT1A10 | NM_019075 | 0.2595 | 0.603 |
| L1TD1 | NM_019079 | -0.078 | 0.206 |
| NDFIP2 | NM_019080 | -0.695 | -0.857 |
| DDX56 | NM_019082 | -0.8075 | -2.205 |
| CCDC76 | NM_019083 | -1.205 | -1.927 |
| CCNJ | NM_019084 | -0.6185 | -1.9185 |
| VSIG10 | NM_019086 | -1.062 | -1.4025 |
| ARL15 | NM_019087 | -0.7885 | -1.3295 |
| PAF1 | NM_019088 | -0.6985 | -2.69 |
| HES2 | NM_019089 | -0.0735 | -0.1535 |
| KIAA1383 | NM_019090 | -0.1035 | -0.2555 |
| PLEKHA3 | NM_019091 | -1.425 | -2.6375 |
| NUDT4 | NM_019094 | -0.358 | -1.3375 |
| CRLS1 | NM_019095 | -0.914 | -2.4385 |
| GTPBP2 | NM_019096 | -1.1875 | -2.809 |
| CNGB3 | NM_019098 | 0.719 | -0.4315 |
| C1orf183 | NM_019099 | -0.616 | -1.673 |
| DMAP1 | NM_019100 | 0.1275 | -0.9345 |
| APOM | NM_019101 | 0.2325 | -0.206 |
| HOXA5 | NM_019102 | -0.011 | 0.081 |
| ZMAT5 | NM_019103 | -0.175 | -0.932 |
| LIN37 | NM_019104 | -1.173 | -2.575 |
| 3-Sep | NM_019106 | -0.1345 | -0.647 |
| C19orf10 | NM_019107 | -0.126 | -1.2095 |
| SMG9 | NM_019108 | -0.259 | -1.238 |
| ALG1 | NM_019109 | -0.2705 | -0.8615 |
| ZKSCAN4 | NM_019110 | -1.3155 | -2.663 |
| HLA-DRA | NM_019111 | 0.165 | -0.4035 |

| | | | |
|---|---|---|---|
| ABCA7 | NM_019112 | 0.1755 | -0.315 |
| FGF21 | NM_019113 | 0.0065 | -0.081 |
| EPB41L4B | NM_019114 | -0.372 | -1.219 |
| UBFD1 | NM_019116 | -0.3635 | -1.1215 |
| KLHL4 | NM_019117 | -0.174 | -0.3925 |
| PCDHB9 | NM_019119 | 0.172 | -0.5465 |
| PCDHB8 | NM_019120 | 0.21 | -0.7775 |
| S100A4 | NM_019554 | 0.4775 | -0.0265 |
| ARHGEF3 | NM_019555 | -0.9255 | -2.32 |
| MOSPD1 | NM_019556 | -1.868 | -3.8325 |
| FAM54B | NM_019557 | -0.6165 | -1.5125 |
| HOXD8 | NM_019558 | -0.9985 | -1.8955 |
| YLPM1 | NM_019589 | -0.5305 | -1.942 |
| KIAA1217 | NM_019590 | 0.0705 | -0.1695 |
| ZNF26 | NM_019591 | -0.586 | -1.695 |
| RNF20 | NM_019592 | -0.332 | -0.949 |
| GPCPD1 | NM_019593 | -0.5215 | -1.2455 |
| LRRC8A | NM_019594 | -0.714 | -1.6635 |
| ITSN2 | NM_019595 | -0.6735 | -1.5785 |
| C21orf62 | NM_019596 | -0.086 | -0.0875 |
| HNRNPH2 | NM_019597 | -0.7175 | -0.763 |
| KLK12 | NM_019598 | 0.148 | -0.2485 |
| TAS2R1 | NM_019599 | -0.03 | -0.0535 |
| KIAA1370 | NM_019600 | -1.1275 | -2.0975 |
| SUSD2 | NM_019601 | -0.0785 | -0.7775 |
| BTNL2 | NM_019602 | 0.051 | -0.4675 |
| CRTAM | NM_019604 | -0.169 | -0.104 |
| SERTAD4 | NM_019605 | -1.4255 | -2.8895 |
| MEPCE | NM_019606 | -0.7145 | -3.606 |
| C8orf44 | NM_019607 | -0.7505 | -0.7225 |
| CPXM1 | NM_019609 | -0.136 | -0.7355 |
| RBMXL1 | NM_019610 | -0.711 | -2.3305 |
| IRGC | NM_019612 | 0.057 | -0.3395 |
| WDR45L | NM_019613 | -0.4005 | -0.869 |
| F7 | NM_019616 | -0.2585 | -0.101 |
| GKN1 | NM_019617 | -0.32 | -0.6675 |
| PARD3 | NM_019619 | -0.282 | -0.83 |
| ABCB9 | NM_019624 | 0.0745 | -0.486 |
| ABCB9 | NM_019625 | 0.0745 | -0.486 |
| LTB4R2 | NM_019839 | -0.164 | -0.52 |
| TRPV5 | NM_019841 | -0.0355 | -0.491 |
| KCNQ5 | NM_019842 | 0.5015 | -0.1945 |
| EIF4ENIF1 | NM_019843 | -1.6685 | -4.82 |

| | | | |
|---|---|---|---|
| SLCO1B3 | NM_019844 | -0.0765 | -0.6935 |
| RPRM | NM_019845 | 0.0795 | -0.131 |
| SLC7A10 | NM_019849 | 0.1325 | 0.108 |
| NGEF | NM_019850 | -0.193 | -0.644 |
| FGF20 | NM_019851 | 0.032 | -0.3765 |
| METTL3 | NM_019852 | -1.0975 | -2.1155 |
| PRMT8 | NM_019854 | -0.144 | -0.2625 |
| CABP5 | NM_019855 | -0.147 | 0.12 |
| MYOM1 | NM_019856 | 0.203 | -0.317 |
| CTPS2 | NM_019857 | -1.088 | -1.1125 |
| GPR162 | NM_019858 | -0.337 | -1.7695 |
| HTR7 | NM_019859 | -0.0655 | -0.482 |
| HTR7 | NM_019860 | -0.0655 | -0.482 |
| ABCC1 | NM_019862 | -0.4025 | -0.9845 |
| F8 | NM_019863 | 0.245 | -0.3385 |
| GSK3A | NM_019884 | -0.1015 | -0.85 |
| CYP26B1 | NM_019885 | 0.0625 | -0.391 |
| CHST7 | NM_019886 | -0.6325 | -1.1125 |
| MC3R | NM_019888 | 0.064 | -0.17 |
| ERO1LB | NM_019891 | -0.1785 | -1.4335 |
| INPP5E | NM_019892 | -0.898 | -1.825 |
| TMPRSS4 | NM_019894 | -0.072 | -0.4105 |
| CLDND1 | NM_019895 | -1.2655 | -1.8485 |
| POLE4 | NM_019896 | 0.408 | -0.485 |
| OR2S2 | NM_019897 | -0.0865 | -0.2985 |
| ABCC1 | NM_019898 | -0.4025 | -0.9845 |
| ABCC1 | NM_019899 | -0.4025 | -0.9845 |
| ABCC1 | NM_019900 | -0.4025 | -0.9845 |
| ADD3 | NM_019903 | -0.794 | -1.0915 |
| ACCN2 | NM_020039 | 0.0515 | -0.481 |
| SLC2A9 | NM_020041 | -0.1645 | 0.0555 |
| HLA-DQA2 | NM_020056 | -0.091 | -0.568 |
| OPN1LW | NM_020061 | -0.1575 | -0.265 |
| SLC2A4RG | NM_020062 | -0.427 | -0.599 |
| BARHL2 | NM_020063 | -0.214 | -0.2475 |
| BARHL1 | NM_020064 | 0.2905 | 0.146 |
| FMN2 | NM_020066 | -1.612 | -3.1685 |
| IGLL1 | NM_020070 | 0.155 | -0.287 |
| FSTL5 | NM_020116 | -0.3675 | -0.3765 |
| LARS | NM_020117 | -0.908 | -0.8925 |
| ZC3HAV1 | NM_020119 | -1.124 | -2.0005 |
| UGGT1 | NM_020120 | -0.5005 | -0.923 |
| UGGT2 | NM_020121 | 0.802 | 0.069 |

| | | | |
|---|---|---|---|
| KCMF1 | NM_020122 | -0.688 | -2.5205 |
| TM9SF3 | NM_020123 | -0.667 | -1.2165 |
| IFNK | NM_020124 | 0.2005 | 0.2215 |
| SLAMF8 | NM_020125 | 0.0095 | -0.2585 |
| SPHK2 | NM_020126 | 0.0025 | -0.1975 |
| TUFT1 | NM_020127 | -0.0645 | -1.368 |
| MDM1 | NM_020128 | -1.0145 | -0.964 |
| C8orf4 | NM_020130 | 0.086 | 1.099 |
| UBQLN4 | NM_020131 | 0.07 | -1.0425 |
| AGPAT3 | NM_020132 | -0.856 | -1.2755 |
| AGPAT4 | NM_020133 | -0.438 | -1.5195 |
| DPYSL5 | NM_020134 | -0.804 | -1.0385 |
| WRNIP1 | NM_020135 | -0.348 | -1.0675 |
| GRIPAP1 | NM_020137 | -0.3665 | -0.801 |
| BDH2 | NM_020139 | -0.678 | -1.002 |
| ANKS1B | NM_020140 | -0.3535 | -0.4295 |
| TMEM167B | NM_020141 | -1.45 | -2.2435 |
| NDUFA4L2 | NM_020142 | 0.246 | -0.2655 |
| PNO1 | NM_020143 | -1.2055 | -4.063 |
| PAPOLB | NM_020144 | -0.4425 | -1.417 |
| SH3GLB2 | NM_020145 | 0.126 | -0.7045 |
| THAP10 | NM_020147 | -1.4185 | -3.087 |
| SPIRE1 | NM_020148 | -1.6745 | -2.523 |
| SAR1A | NM_020150 | -0.3255 | -0.8665 |
| STARD7 | NM_020151 | -0.35 | -1.243 |
| C21orf7 | NM_020152 | 0.0645 | -0.141 |
| IFT46 | NM_020153 | -1.0425 | -2.1835 |
| C15orf24 | NM_020154 | -0.8585 | -2.4375 |
| GPR137 | NM_020155 | 0.158 | -0.7635 |
| C1GALT1 | NM_020156 | -1.4375 | -1.4795 |
| OTOR | NM_020157 | 0.0425 | -0.3135 |
| EXOSC5 | NM_020158 | -0.053 | -0.6715 |
| SMARCAD1 | NM_020159 | -1.1235 | -4.0625 |
| MEIS3 | NM_020160 | 0.037 | -0.27 |
| C2orf83 | NM_020161 | 0.1715 | 0.206 |
| DHX33 | NM_020162 | -1.0555 | -1.858 |
| SEMA3G | NM_020163 | -0.107 | -0.633 |
| ASPH | NM_020164 | -1.0635 | -0.5415 |
| RAD18 | NM_020165 | -0.9145 | -2.9375 |
| MCCC1 | NM_020166 | -0.615 | -1.0165 |
| NMUR2 | NM_020167 | -0.042 | -0.084 |
| PAK6 | NM_020168 | -0.0785 | -0.326 |
| LXN | NM_020169 | 0.1675 | 0.497 |

| | | | |
|---|---|---|---|
| NCLN | NM_020170 | -0.7445 | -1.38 |
| DUS3L | NM_020175 | -0.094 | -1.0905 |
| FEM1C | NM_020177 | -1.019 | -2.806 |
| CA10 | NM_020178 | -0.378 | -0.3395 |
| C11orf75 | NM_020179 | -0.705 | -3.3525 |
| CELF4 | NM_020180 | -1.4965 | -1.4715 |
| PMEPA1 | NM_020182 | -0.854 | -3.3055 |
| ARNTL2 | NM_020183 | -0.255 | -0.831 |
| CNNM4 | NM_020184 | -0.0365 | -0.921 |
| DUSP22 | NM_020185 | 0.153 | -0.655 |
| ACN9 | NM_020186 | -1.224 | -3.2985 |
| C3orf37 | NM_020187 | -0.6235 | -1.7985 |
| C16orf61 | NM_020188 | -0.305 | -1.1305 |
| ENY2 | NM_020189 | -0.776 | -0.7585 |
| OLFML3 | NM_020190 | -0.0785 | -0.652 |
| MRPS22 | NM_020191 | -1.3635 | -2.4405 |
| C7orf36 | NM_020192 | -2.2205 | -4.3815 |
| C11orf30 | NM_020193 | -1.065 | -3.0805 |
| MFF | NM_020194 | -0.5365 | -1.267 |
| SDR39U1 | NM_020195 | -0.126 | -1.1425 |
| XAB2 | NM_020196 | -0.03 | -1.0255 |
| SMYD2 | NM_020197 | -0.5725 | -0.9975 |
| CCDC47 | NM_020198 | -0.844 | -1.7435 |
| C5orf15 | NM_020199 | -1.0215 | -2.379 |
| PRTFDC1 | NM_020200 | -0.72 | -0.5865 |
| NT5M | NM_020201 | -0.236 | -1.3875 |
| NIT2 | NM_020202 | -0.4095 | -0.7295 |
| MEPE | NM_020203 | -0.2565 | -0.5515 |
| LHX9 | NM_020204 | -1.053 | -0.939 |
| OTUD7B | NM_020205 | -0.946 | -2.9075 |
| SLC6A20 | NM_020208 | 0.176 | -0.1915 |
| SHD | NM_020209 | -0.647 | -1.3245 |
| SEMA4B | NM_020210 | -0.9245 | -2.87 |
| RGMA | NM_020211 | -0.8385 | -2.211 |
| WDR93 | NM_020212 | -0.2105 | -0.212 |
| PARP6 | NM_020214 | -0.499 | -0.6655 |
| RNPEP | NM_020216 | 0.0125 | -0.7475 |
| ATXN7L3 | NM_020218 | -0.599 | -1.7575 |
| CEACAM19 | NM_020219 | -0.466 | -0.4635 |
| FAM20C | NM_020223 | -1.708 | -3.38 |
| STOX2 | NM_020225 | -1.253 | -2.135 |
| PRDM8 | NM_020226 | -0.4005 | -1.3745 |
| PRDM9 | NM_020227 | 0.0405 | -0.119 |

| | | | |
|---|---|---|---|
| PRDM10 | NM_020228 | -1.265 | -2.4405 |
| PRDM11 | NM_020229 | 1.021 | -0.5505 |
| PPAN | NM_020230 | -1.0615 | -3.0595 |
| PSMG2 | NM_020232 | -1.1905 | -2.5655 |
| C17orf48 | NM_020233 | -0.9025 | -1.343 |
| DTWD1 | NM_020234 | -1.87 | -2.782 |
| BBX | NM_020235 | -0.131 | -1.1405 |
| MRPL1 | NM_020236 | -1.058 | -2.3655 |
| INCENP | NM_020238 | -0.169 | -0.73 |
| CDC42SE1 | NM_020239 | -0.952 | -2.505 |
| CDC42SE2 | NM_020240 | -1.5315 | -1.9045 |
| KIF15 | NM_020242 | -0.3565 | -2.578 |
| TOMM22 | NM_020243 | -0.086 | -0.7645 |
| CHPT1 | NM_020244 | -0.2535 | -0.752 |
| TULP4 | NM_020245 | -1.237 | -2.0945 |
| SLC12A9 | NM_020246 | -0.156 | -0.873 |
| ADCK3 | NM_020247 | -0.257 | -1.218 |
| CTNNBIP1 | NM_020248 | 0.0815 | -0.328 |
| ARRB1 | NM_020251 | -0.0295 | -1.2355 |
| ABCC9 | NM_020297 | -0.394 | 0.1585 |
| AKR1B10 | NM_020299 | -0.0705 | -0.277 |
| MGST1 | NM_020300 | -0.374 | -0.7395 |
| CCNL1 | NM_020307 | -1.5515 | -2.248 |
| SLC17A7 | NM_020309 | -0.077 | -0.265 |
| MNT | NM_020310 | -0.615 | -1.2685 |
| CXCR7 | NM_020311 | -0.162 | -0.2895 |
| COQ9 | NM_020312 | -1.009 | -2.111 |
| CIAPIN1 | NM_020313 | -0.505 | -1.6505 |
| C16orf62 | NM_020314 | -0.263 | -0.526 |
| PDXP | NM_020315 | 0.144 | -0.169 |
| C1orf63 | NM_020317 | -0.229 | 0.0085 |
| PAPPA2 | NM_020318 | 0.1595 | -0.366 |
| ANKMY2 | NM_020319 | -1.0785 | -1.6615 |
| RARS2 | NM_020320 | -0.686 | -0.5635 |
| ACCN3 | NM_020321 | -0.226 | -0.094 |
| ACCN3 | NM_020322 | -0.226 | -0.094 |
| ACVR1B | NM_020327 | -0.8805 | -2.2285 |
| ACVR1B | NM_020328 | -0.8805 | -2.2285 |
| VANGL2 | NM_020335 | -0.079 | -0.7255 |
| RALGAPB | NM_020336 | -1.7185 | -3.1855 |
| ANKRD50 | NM_020337 | -1.129 | -3.358 |
| ZMIZ1 | NM_020338 | -0.905 | -2.1885 |
| KIAA1244 | NM_020340 | -0.0875 | -0.522 |

| | | | |
|---|---|---|---|
| PAK7 | NM_020341 | -0.8485 | -1.6705 |
| SLC39A10 | NM_020342 | -1.039 | -1.637 |
| RALGAPA2 | NM_020343 | -0.764 | -0.824 |
| SLC24A2 | NM_020344 | 0.0105 | -0.2615 |
| NKIRAS1 | NM_020345 | -1.574 | -2.7155 |
| SLC17A6 | NM_020346 | -0.1175 | -0.36 |
| LZTFL1 | NM_020347 | -0.625 | -2.001 |
| CNNM1 | NM_020348 | -0.59 | -1.9745 |
| ANKRD2 | NM_020349 | -0.0185 | 0.0385 |
| AGTRAP | NM_020350 | -0.3905 | -0.525 |
| COL8A1 | NM_020351 | -0.1245 | -0.139 |
| PLSCR4 | NM_020353 | -0.1515 | -0.1005 |
| ENTPD7 | NM_020354 | 0.0865 | -0.6075 |
| CASS4 | NM_020356 | 0.2985 | -0.203 |
| PCNP | NM_020357 | -0.7025 | -0.6755 |
| PLSCR3 | NM_020360 | -0.2225 | -1.151 |
| CPA6 | NM_020361 | -0.361 | -0.2605 |
| PITHD1 | NM_020362 | -0.4035 | -1.1475 |
| EIF2B3 | NM_020365 | -1.4105 | -1.2655 |
| RPGRIP1 | NM_020366 | -0.087 | 0.106 |
| PARP11 | NM_020367 | -0.8105 | -1.582 |
| UTP3 | NM_020368 | -1.5435 | -2.3495 |
| FSCN3 | NM_020369 | -0.2295 | -0.488 |
| GPR84 | NM_020370 | -0.112 | -0.06 |
| AVEN | NM_020371 | 0.0675 | -0.4965 |
| SLC22A17 | NM_020372 | -0.051 | -0.6365 |
| ANO2 | NM_020373 | 0.041 | -0.205 |
| C12orf4 | NM_020374 | -1.1715 | -2.3695 |
| C12orf5 | NM_020375 | -1.8345 | -0.5495 |
| PNPLA2 | NM_020376 | 0.019 | -0.6005 |
| CYSLTR2 | NM_020377 | 0.108 | -0.2645 |
| NAT14 | NM_020378 | -0.3 | -0.6115 |
| MAN1C1 | NM_020379 | -0.037 | -0.5095 |
| PDSS2 | NM_020381 | -1.1705 | -2.5975 |
| SETD8 | NM_020382 | -0.814 | -1.6995 |
| XPNPEP1 | NM_020383 | -0.0015 | -0.6505 |
| CLDN2 | NM_020384 | -0.2865 | -0.2975 |
| REXO4 | NM_020385 | -0.9715 | -2.175 |
| HRASLS | NM_020386 | -0.3575 | -0.987 |
| RAB25 | NM_020387 | 0.0085 | -0.055 |
| TRPC7 | NM_020389 | 0.0815 | -0.262 |
| EIF5A2 | NM_020390 | 0.009 | -0.587 |
| PGLYRP4 | NM_020393 | 0.0955 | -0.6295 |

| | | | |
|---|---|---|---|
| ZNF695 | NM_020394 | -0.0785 | -0.464 |
| INTS12 | NM_020395 | -1.333 | -3.1315 |
| BCL2L10 | NM_020396 | -0.228 | -0.1015 |
| CAMK1D | NM_020397 | -0.4705 | -1.2695 |
| SPINLW1 | NM_020398 | -0.1845 | -0.347 |
| LPAR5 | NM_020400 | -0.143 | -0.3575 |
| NUP107 | NM_020401 | -0.854 | -1.415 |
| CHRNA10 | NM_020402 | -0.3775 | -1.657 |
| PCDH9 | NM_020403 | -0.0605 | -0.012 |
| CD248 | NM_020404 | -1.2395 | -3.565 |
| PLXDC1 | NM_020405 | -0.172 | -0.522 |
| CD177 | NM_020406 | -0.0295 | 0.191 |
| RHBG | NM_020407 | -0.9955 | -0.9725 |
| LYRM4 | NM_020408 | -0.0965 | -0.564 |
| MRPL47 | NM_020409 | -1.05 | -1.61 |
| ATP13A1 | NM_020410 | -0.1415 | -0.638 |
| CHMP1B | NM_020412 | -1.1515 | -1.743 |
| DDX24 | NM_020414 | -0.5515 | -1.1685 |
| RETN | NM_020415 | 0.076 | -0.2075 |
| PCBP4 | NM_020418 | -0.2155 | -0.7735 |
| ADCK1 | NM_020421 | -0.324 | -1.6805 |
| TMEM159 | NM_020422 | -0.015 | -0.5195 |
| SCYL3 | NM_020423 | -2.2215 | -3.259 |
| LYRM1 | NM_020424 | -0.986 | -0.867 |
| C6orf162 | NM_020425 | -0.46 | -1.2885 |
| LYZL6 | NM_020426 | 0.22 | -0.018 |
| SLURP1 | NM_020427 | -0.106 | -0.7235 |
| SLC44A2 | NM_020428 | -0.108 | -0.9795 |
| SMURF1 | NM_020429 | -0.526 | -1.2175 |
| TMEM63C | NM_020431 | -0.1315 | -1.083 |
| PHTF2 | NM_020432 | -1.417 | -2.733 |
| JPH2 | NM_020433 | 0.1585 | -0.19 |
| GJC2 | NM_020435 | 0.006 | -0.426 |
| SALL4 | NM_020436 | -0.322 | -0.6655 |
| ASPHD2 | NM_020437 | -1.033 | -2.7005 |
| DOLPP1 | NM_020438 | -1.037 | -3.314 |
| CAMK1G | NM_020439 | -0.631 | -0.367 |
| PTGFRN | NM_020440 | -0.983 | -2.7895 |
| CORO1B | NM_020441 | 0.083 | -0.4095 |
| VARS2 | NM_020442 | 0.05 | -0.762 |
| NAV1 | NM_020443 | -1.2845 | -1.6955 |
| KIAA1191 | NM_020444 | -0.09 | -0.759 |
| C15orf17 | NM_020447 | -0.7315 | -1.237 |

| | | | |
|---|---|---|---|
| NIPAL3 | NM_020448 | -1.271 | -0.5925 |
| ATP8B2 | NM_020452 | -1.1575 | -2.6785 |
| ATP10D | NM_020453 | -2.2525 | -2.455 |
| GPR126 | NM_020455 | 0.149 | 0.0315 |
| SPRYD7 | NM_020456 | -1.1495 | -2.5115 |
| THAP11 | NM_020457 | -0.71 | -1.1425 |
| TTC7A | NM_020458 | -0.349 | -0.7985 |
| PAIP2B | NM_020459 | -0.317 | -0.5505 |
| TUBGCP6 | NM_020461 | -0.358 | -0.646 |
| SMEK2 | NM_020463 | -0.4415 | -0.5825 |
| NHSL1 | NM_020464 | 0.085 | -0.484 |
| NDRG4 | NM_020465 | -0.529 | -0.8035 |
| LYRM2 | NM_020466 | -1.381 | -2.254 |
| SNX14 | NM_020468 | -1.274 | -1.7475 |
| ABO | NM_020469 | 0.2365 | -0.143 |
| YIF1A | NM_020470 | -0.198 | -0.864 |
| PIGA | NM_020473 | -0.841 | -1.258 |
| GALNT1 | NM_020474 | -0.651 | -1.2715 |
| ANK1 | NM_020475 | -0.59 | -0.8445 |
| ANK1 | NM_020476 | -0.1485 | -0.816 |
| ANK1 | NM_020477 | -0.1485 | -0.816 |
| ANK1 | NM_020478 | -0.1485 | -0.816 |
| ANK1 | NM_020480 | -0.1485 | -0.816 |
| FHL5 | NM_020482 | 0.146 | -0.066 |
| PBXIP1 | NM_020524 | -0.552 | -0.587 |
| IL22 | NM_020525 | -0.1995 | -0.252 |
| EPHA8 | NM_020526 | -0.0655 | -0.1775 |
| PCBP3 | NM_020528 | -0.5335 | -0.738 |
| NFKBIA | NM_020529 | -0.092 | -0.021 |
| OSM | NM_020530 | 0.1755 | -0.4755 |
| C20orf3 | NM_020531 | -0.109 | -0.9215 |
| RTN4 | NM_020532 | -0.2735 | -0.847 |
| MCOLN1 | NM_020533 | -0.433 | -1.079 |
| CSRP2BP | NM_020536 | -2.323 | -4.1075 |
| ADCY2 | NM_020546 | 0.011 | -0.2165 |
| AMHR2 | NM_020547 | 0.3215 | -0.3315 |
| DBI | NM_020548 | -0.241 | -0.2425 |
| CHAT | NM_020549 | 0.0335 | -0.886 |
| RET | NM_020630 | 0.021 | -0.6075 |
| PLEKHG5 | NM_020631 | -0.055 | -0.5065 |
| ATP6V0A4 | NM_020632 | -0.1235 | -0.142 |
| VN1R1 | NM_020633 | -0.0235 | -0.832 |
| GDF3 | NM_020634 | 0.104 | -0.6795 |

| | | | |
|---|---|---|---|
| FGF22 | NM_020637 | -0.011 | -0.242 |
| FGF23 | NM_020638 | -0.267 | -0.8345 |
| RIPK4 | NM_020639 | 0.0265 | -0.147 |
| DCUN1D1 | NM_020640 | -0.486 | -1.646 |
| C9orf11 | NM_020641 | -0.0335 | -0.1285 |
| C11orf16 | NM_020643 | -0.1135 | 0.4095 |
| TMEM9B | NM_020644 | -0.8955 | -1.0385 |
| NRIP3 | NM_020645 | 0.083 | 0.411 |
| ASCL3 | NM_020646 | -0.209 | -0.006 |
| JPH1 | NM_020647 | -1.7475 | -4.0365 |
| TWSG1 | NM_020648 | -1.2485 | -2.6075 |
| CBX8 | NM_020649 | -0.215 | -0.7695 |
| RCN3 | NM_020650 | -0.0555 | -0.7585 |
| PELI1 | NM_020651 | -1.1085 | -1.3875 |
| ZNF286A | NM_020652 | -1.7755 | -3.352 |
| ZNF287 | NM_020653 | -1.028 | -1.895 |
| SENP7 | NM_020654 | -0.6515 | -1.524 |
| JPH3 | NM_020655 | -0.312 | -0.4035 |
| ZNF304 | NM_020657 | -1.6735 | -2.8115 |
| TTYH1 | NM_020659 | 0.0605 | -0.2985 |
| GJD2 | NM_020660 | 0.0265 | -0.157 |
| AICDA | NM_020661 | 0.316 | -0.2355 |
| MRS2 | NM_020662 | -0.7565 | -1.2055 |
| RHOJ | NM_020663 | 0.065 | -0.6695 |
| DECR2 | NM_020664 | -0.2685 | -0.714 |
| TMEM27 | NM_020665 | -0.9405 | -0.878 |
| CLK4 | NM_020666 | -0.097 | -0.776 |
| S100A14 | NM_020672 | 0.434 | -0.197 |
| RAB22A | NM_020673 | -1.374 | -1.3975 |
| SPC25 | NM_020675 | -1.5375 | -2.838 |
| ABHD6 | NM_020676 | -1.1405 | -2.797 |
| NMRAL1 | NM_020677 | -0.6405 | -0.3855 |
| LRTM1 | NM_020678 | 0.301 | 0.2335 |
| MIF4GD | NM_020679 | -0.5145 | -1.258 |
| SCYL1 | NM_020680 | -0.2065 | -1.1025 |
| AS3MT | NM_020682 | -1.72 | -2.193 |
| ADORA3 | NM_020683 | 0.1545 | -0.062 |
| C3orf14 | NM_020685 | -1.161 | -1.7005 |
| ABAT | NM_020686 | -0.9025 | -1.074 |
| SLC24A3 | NM_020689 | 0.235 | -0.6445 |
| NKHD1-EIF4EB | NM_020690 | -0.243 | -1.321 |
| GALNTL1 | NM_020692 | -0.3345 | -0.8705 |
| DSCAML1 | NM_020693 | -0.1175 | -1.367 |

| | | | |
|---|---|---|---|
| REXO1 | NM_020695 | -0.1905 | -0.782 |
| KIAA1143 | NM_020696 | -2.1495 | -2.0675 |
| KCNS2 | NM_020697 | 0.0505 | -0.9885 |
| TMCC3 | NM_020698 | -0.346 | -0.933 |
| GATAD2B | NM_020699 | -0.5505 | -1.7325 |
| PPM1H | NM_020700 | -0.071 | -1.241 |
| ISY1 | NM_020701 | -0.5875 | -1.8275 |
| KIAA1161 | NM_020702 | 0.096 | -0.3955 |
| AMIGO1 | NM_020703 | -0.825 | -0.276 |
| FAM40B | NM_020704 | -0.407 | -0.353 |
| TBC1D24 | NM_020705 | -0.425 | -1.5425 |
| HHATL | NM_020707 | -0.1485 | -0.263 |
| SLC12A5 | NM_020708 | 0.072 | -0.159 |
| PNMAL2 | NM_020709 | -0.5715 | -0.96 |
| LRRC47 | NM_020710 | -0.579 | -1.857 |
| ERMN | NM_020711 | -0.109 | -0.1205 |
| ZNF512B | NM_020713 | -0.8225 | -1.0855 |
| ZNF490 | NM_020714 | -1.147 | -2.1105 |
| PLEKHH1 | NM_020715 | -0.1135 | -0.817 |
| GRAMD1B | NM_020716 | -0.285 | -0.367 |
| SHROOM4 | NM_020717 | -0.006 | -0.3345 |
| USP31 | NM_020718 | -1.256 | -2.1585 |
| PRR12 | NM_020719 | -0.786 | -2.2805 |
| KIAA1210 | NM_020721 | -0.1255 | 0.2225 |
| KIAA1211 | NM_020722 | -1.2095 | -1.569 |
| RNF150 | NM_020724 | -0.634 | -1.3835 |
| ATXN7L1 | NM_020725 | 0.3515 | -0.3115 |
| NLN | NM_020726 | -1.205 | -3.0915 |
| ZNF295 | NM_020727 | -1.4495 | -3.176 |
| ESYT2 | NM_020728 | -1.1365 | -1.0835 |
| ODF2L | NM_020729 | -0.615 | -1.1145 |
| DLG3 | NM_020730 | -0.8345 | -2.8375 |
| AHRR | NM_020731 | 0.1635 | -0.353 |
| ARID1B | NM_020732 | -0.755 | -2.4065 |
| HEG1 | NM_020733 | -0.905 | -2.233 |
| RIMKLB | NM_020734 | -0.7645 | -2.109 |
| LRFN2 | NM_020737 | -1.051 | -1.5975 |
| KIDINS220 | NM_020738 | -0.376 | -0.8055 |
| CCPG1 | NM_020739 | -0.52 | -1.423 |
| ANKFY1 | NM_020740 | -1.2295 | -2.2465 |
| KIAA1257 | NM_020741 | 0.394 | -0.357 |
| NLGN4X | NM_020742 | -1.0145 | -1.803 |
| MTA3 | NM_020744 | -0.688 | -0.931 |

| | | | |
|---|---|---|---|
| AARS2 | NM_020745 | -0.9405 | -1.724 |
| MAVS | NM_020746 | -1.15 | -1.6835 |
| ZNF608 | NM_020747 | -1.123 | -2.2685 |
| INTS2 | NM_020748 | -1.8635 | -2.95 |
| MTUS1 | NM_020749 | -0.8945 | -2.8875 |
| XPO5 | NM_020750 | -0.763 | -0.9265 |
| COG6 | NM_020751 | -1.2695 | -2.292 |
| GPR158 | NM_020752 | -1.2115 | -2.027 |
| CASKIN2 | NM_020753 | -0.576 | -2.4365 |
| ARHGAP31 | NM_020754 | -0.1335 | 0.0635 |
| SERINC1 | NM_020755 | -0.827 | -0.393 |
| STARD9 | NM_020759 | -0.798 | -1.5505 |
| HECW2 | NM_020760 | -0.849 | -2.786 |
| RPTOR | NM_020761 | -0.82 | -3.1755 |
| SRGAP1 | NM_020762 | -0.8335 | -1.726 |
| CASKIN1 | NM_020764 | -0.467 | -0.6925 |
| UBR4 | NM_020765 | -0.43 | -1.1225 |
| PCDH19 | NM_020766 | -1.295 | -2.411 |
| KCTD16 | NM_020768 | 0.3175 | -0.733 |
| RGAG1 | NM_020769 | -0.165 | -0.621 |
| CGN | NM_020770 | -0.564 | -0.373 |
| HACE1 | NM_020771 | -1.0505 | -1.581 |
| NUFIP2 | NM_020772 | -1.386 | -2.9785 |
| TBC1D14 | NM_020773 | 0.0385 | -0.9445 |
| MIB1 | NM_020774 | -0.8955 | -1.8085 |
| KIAA1324 | NM_020775 | -0.0615 | -0.294 |
| KIAA1328 | NM_020776 | -0.3035 | -0.863 |
| SORCS2 | NM_020777 | 0.0235 | -0.2835 |
| ALPK3 | NM_020778 | -0.099 | 0.108 |
| WDR35 | NM_020779 | -1.0685 | -2.47 |
| PTCHD2 | NM_020780 | -0.7485 | -1.005 |
| ZNF398 | NM_020781 | -1.4925 | -3.016 |
| KLHDC5 | NM_020782 | -2.117 | -2.1425 |
| SYT4 | NM_020783 | -0.7055 | -1.115 |
| TXNDC16 | NM_020784 | -0.5665 | -0.7755 |
| PDP2 | NM_020786 | -1.225 | -1.7215 |
| ZNF624 | NM_020787 | -0.462 | -1.983 |
| IGSF9 | NM_020789 | -0.745 | -1.3015 |
| TAOK1 | NM_020791 | -0.197 | -1.4335 |
| NCEH1 | NM_020792 | -0.9735 | -1.7315 |
| LRRC7 | NM_020794 | -0.175 | -0.1145 |
| NLGN2 | NM_020795 | -0.4075 | -0.8605 |
| SEMA6A | NM_020796 | -1.148 | -2.741 |

| | | | |
|---|---|---|---|
| USP35 | NM_020798 | -0.4565 | -1.3285 |
| STAMBPL1 | NM_020799 | -1.2535 | -2.537 |
| IFT80 | NM_020800 | -1.272 | -1.6675 |
| ARRDC3 | NM_020801 | -0.823 | -0.2825 |
| KIAA1377 | NM_020802 | -0.6375 | -0.946 |
| KLHL8 | NM_020803 | -1.462 | -2.5635 |
| PACSIN1 | NM_020804 | -0.0815 | -0.4885 |
| KLHL14 | NM_020805 | 0.253 | -0.3395 |
| GPHN | NM_020806 | -0.996 | -2.378 |
| ZNF319 | NM_020807 | -0.794 | -1.9455 |
| SIPA1L2 | NM_020808 | -1.113 | -3.229 |
| ARHGAP20 | NM_020809 | 0.027 | -0.572 |
| TRMT5 | NM_020810 | -1.2485 | -2.53 |
| CARNS1 | NM_020811 | -0.1435 | -0.3585 |
| DOCK6 | NM_020812 | 0.2155 | -0.4805 |
| ZNF471 | NM_020813 | 0.0975 | -0.5585 |
| 4-Mar | NM_020814 | -0.651 | -1.4985 |
| PCDH10 | NM_020815 | 0.234 | -0.244 |
| KIF17 | NM_020816 | -0.4085 | -0.335 |
| KIAA1407 | NM_020817 | 0.234 | -0.5795 |
| FAM135A | NM_020819 | -1.5035 | -3.084 |
| PREX1 | NM_020820 | -0.3265 | -0.9485 |
| VPS13C | NM_020821 | -0.3665 | -1.589 |
| KCNT1 | NM_020822 | 0.024 | -0.5635 |
| TMEM181 | NM_020823 | -1.099 | -1.173 |
| ARHGAP21 | NM_020824 | -0.9835 | -2.2885 |
| CRAMP1L | NM_020825 | -0.9585 | -2.18 |
| SYT13 | NM_020826 | -0.6095 | -0.4875 |
| KIAA1430 | NM_020827 | -0.995 | -1.3895 |
| ZFP28 | NM_020828 | -0.52 | -1.6485 |
| KIAA1432 | NM_020829 | -1.6935 | -2.2255 |
| WDFY1 | NM_020830 | -0.9 | -0.703 |
| MKL1 | NM_020831 | -0.5725 | -2.311 |
| ZNF687 | NM_020832 | -0.596 | -2.2225 |
| HOMEZ | NM_020834 | -0.763 | -1.5365 |
| BEGAIN | NM_020836 | -1.0055 | -1.2105 |
| WDR48 | NM_020839 | -0.7465 | -1.287 |
| FNIP2 | NM_020840 | -0.9855 | -2.3215 |
| OSBPL8 | NM_020841 | -0.0895 | -0.896 |
| SCAPER | NM_020843 | -0.67 | -1.512 |
| KIAA1456 | NM_020844 | -0.493 | -0.683 |
| PITPNM2 | NM_020845 | -0.1955 | -1.149 |
| SLAIN2 | NM_020846 | -1.2375 | -2.9 |

| | | | |
|---|---|---|---|
| KIAA1462 | NM_020848 | -0.032 | -0.322 |
| RANBP10 | NM_020850 | -0.423 | -1.5965 |
| ISLR2 | NM_020851 | -0.113 | -0.8215 |
| KIAA1467 | NM_020853 | -0.026 | -0.5775 |
| KIAA1468 | NM_020854 | -1.252 | -3.1965 |
| TSHZ3 | NM_020856 | -0.7555 | -3.072 |
| VPS18 | NM_020857 | -0.6945 | -1.8085 |
| SEMA6D | NM_020858 | -0.2595 | -0.628 |
| SHROOM3 | NM_020859 | -0.643 | -2.322 |
| STIM2 | NM_020860 | -1.165 | -2.3935 |
| ZBTB2 | NM_020861 | -0.853 | -1.6065 |
| LRFN1 | NM_020862 | -0.26 | -0.7935 |
| ZFAT | NM_020863 | -0.234 | -1.554 |
| KIAA1486 | NM_020864 | 0.1155 | -0.539 |
| DHX36 | NM_020865 | -1.176 | -2.453 |
| KLHL1 | NM_020866 | -0.2295 | -0.1435 |
| DPP10 | NM_020868 | -0.0165 | -0.1135 |
| SH3RF1 | NM_020870 | -0.689 | -2.1765 |
| LRCH2 | NM_020871 | -0.531 | -1.3595 |
| CNTN3 | NM_020872 | -0.3355 | -0.2575 |
| LRRN1 | NM_020873 | 0.1635 | -1.5925 |
| DNAH2 | NM_020877 | -0.036 | -0.517 |
| CCDC146 | NM_020879 | -0.1925 | -0.2335 |
| ZNF530 | NM_020880 | -0.4645 | -1.201 |
| COL20A1 | NM_020882 | -0.1405 | -0.412 |
| ZSWIM5 | NM_020883 | -0.6575 | -1.373 |
| MYH7B | NM_020884 | 0.333 | -0.29 |
| USP28 | NM_020886 | -1.113 | -1.876 |
| KIAA1522 | NM_020888 | -0.495 | -1.898 |
| PHF12 | NM_020889 | 0.8125 | -0.898 |
| KIAA1524 | NM_020890 | -0.774 | -2.402 |
| DTX2 | NM_020892 | -0.5255 | -1.3885 |
| KIAA1530 | NM_020894 | -1.1715 | -1.1095 |
| GRAMD1A | NM_020895 | -0.774 | -1.7275 |
| OSBPL5 | NM_020896 | -0.3035 | -0.723 |
| HCN3 | NM_020897 | -0.9665 | -1.323 |
| CALCOCO1 | NM_020898 | -0.241 | -0.7335 |
| ZBTB4 | NM_020899 | -1.105 | -2.014 |
| PHRF1 | NM_020901 | -0.333 | -2.273 |
| CAMSAP3 | NM_020902 | 0.1145 | -0.596 |
| USP29 | NM_020903 | -0.1985 | 0.119 |
| PLEKHA4 | NM_020904 | -0.592 | -0.4405 |
| RDH14 | NM_020905 | -2.1305 | -3.907 |

| | | | |
|---|---|---|---|
| TLE3 | NM_020908 | -0.616 | -2.0775 |
| EPB41L5 | NM_020909 | -0.93 | -1.41 |
| KIAA1549 | NM_020910 | -0.8425 | -3.1455 |
| PLXNA4 | NM_020911 | -0.565 | -0.9625 |
| FLYWCH1 | NM_020912 | -0.2205 | -1.24 |
| RNF213 | NM_020914 | -0.864 | -1.354 |
| ZFP14 | NM_020917 | -0.8425 | -1.5765 |
| GPAM | NM_020918 | -1.475 | -1.881 |
| ALS2 | NM_020919 | -1.659 | -2.0115 |
| CHD8 | NM_020920 | -0.9515 | -2.3895 |
| NIN | NM_020921 | -0.7115 | -2.9645 |
| WNK3 | NM_020922 | -0.384 | -1.3415 |
| ZDBF2 | NM_020923 | -1.1205 | -1.802 |
| ZBTB26 | NM_020924 | -0.7365 | -1.5085 |
| CACHD1 | NM_020925 | -0.785 | -2.899 |
| VAT1L | NM_020927 | -0.918 | -0.202 |
| ZSWIM6 | NM_020928 | -1.41 | -4.114 |
| LRRC4C | NM_020929 | -0.6 | -0.372 |
| KIAA1586 | NM_020931 | -1.2805 | -2.125 |
| MAGEE1 | NM_020932 | -0.1285 | -1.379 |
| ZNF317 | NM_020933 | -1.1135 | -2.6105 |
| USP37 | NM_020935 | -1.12 | -3.152 |
| DDX55 | NM_020936 | -1.221 | -2.547 |
| FANCM | NM_020937 | -1.0325 | -3.2935 |
| CPNE5 | NM_020939 | 0.0965 | 0.005 |
| FAM160B1 | NM_020940 | -1.039 | -2.6055 |
| CWC22 | NM_020943 | -1.0135 | -2.1395 |
| GBA2 | NM_020944 | -0.7475 | -1.6875 |
| WDFY4 | NM_020945 | -0.0565 | -0.4315 |
| DENND1A | NM_020946 | 0.73 | 0.0485 |
| KIAA1609 | NM_020947 | -0.312 | -0.7215 |
| MIER1 | NM_020948 | -0.8485 | -3.324 |
| SLC7A14 | NM_020949 | -1.4255 | -1.8675 |
| KIAA1614 | NM_020950 | -0.0685 | -0.0975 |
| ZNF529 | NM_020951 | -1.4525 | -2.2325 |
| TRPM3 | NM_020952 | 0.08 | 0.028 |
| RNF213 | NM_020954 | -0.864 | -1.354 |
| PRX | NM_020956 | 0.0905 | -0.625 |
| PCDHB16 | NM_020957 | -0.6335 | -1.1025 |
| PPP4R4 | NM_020958 | 0.202 | -0.4155 |
| ANO8 | NM_020959 | -0.123 | -0.657 |
| GPR107 | NM_020960 | -0.1445 | -0.608 |
| METTL14 | NM_020961 | -1.009 | -2.2515 |

| | | | |
|---|---|---|---|
| IGDCC4 | NM_020962 | -0.6425 | -1.465 |
| MOV10 | NM_020963 | -0.0085 | -0.941 |
| EPG5 | NM_020964 | -0.4695 | -2.1425 |
| NCOA5 | NM_020967 | -1.335 | -3.6335 |
| SPTBN4 | NM_020971 | -0.0315 | -0.6905 |
| ZFYVE28 | NM_020972 | -0.178 | -0.5695 |
| GBA3 | NM_020973 | 0.3935 | 0.4805 |
| SCUBE2 | NM_020974 | -0.1885 | -0.153 |
| RET | NM_020975 | -0.8315 | -0.911 |
| ANK2 | NM_020977 | -0.69 | -1.0435 |
| AMY2B | NM_020978 | -0.3155 | -0.4145 |
| SH2B2 | NM_020979 | -0.6435 | -1.369 |
| AQP9 | NM_020980 | 0.072 | -0.5475 |
| B3GALT1 | NM_020981 | 0.3525 | 0.0975 |
| CLDN9 | NM_020982 | -0.1145 | -0.393 |
| ADCY6 | NM_020983 | 0.0525 | -0.1035 |
| CHAT | NM_020984 | 0.0335 | -0.886 |
| CHAT | NM_020985 | 0.0335 | -0.886 |
| CHAT | NM_020986 | 0.0335 | -0.886 |
| ANK3 | NM_020987 | -0.2625 | -1.591 |
| GNAO1 | NM_020988 | -0.3355 | -1.044 |
| CRYGC | NM_020989 | 0.067 | -0.128 |
| CKMT1B | NM_020990 | -0.9695 | 0.2105 |
| CSH2 | NM_020991 | -0.1275 | 0.154 |
| PDLIM1 | NM_020992 | 0.1055 | -0.55 |
| BCL7A | NM_020993 | -0.735 | -1.6145 |
| CTAG2 | NM_020994 | 0.2195 | -0.398 |
| HPR | NM_020995 | 0.171 | 0.0625 |
| FGF6 | NM_020996 | -0.1905 | -0.18 |
| LEFTY1 | NM_020997 | -0.035 | -0.447 |
| MST1 | NM_020998 | 0.887 | -0.172 |
| NEUROG3 | NM_020999 | 0.142 | 0.2455 |
| IFNA6 | NM_021002 | 0.1935 | 0.024 |
| PPM1A | NM_021003 | -0.2755 | -0.7395 |
| DHRS4 | NM_021004 | -0.1325 | -0.558 |
| NR2F2 | NM_021005 | -0.3325 | -0.513 |
| SCN2A | NM_021007 | 0.1875 | 0.159 |
| DEAF1 | NM_021008 | -0.716 | -0.989 |
| UBC | NM_021009 | -0.1175 | -0.41 |
| DEFA5 | NM_021010 | 0.068 | -0.3225 |
| KCNJ12 | NM_021012 | -0.8045 | -1.5625 |
| KRT34 | NM_021013 | -0.0785 | -0.5005 |
| SSX3 | NM_021014 | -0.2805 | 0.6655 |

| | | | |
|---|---|---|---|
| SSX5 | NM_021015 | 0.044 | 0.0155 |
| PSG3 | NM_021016 | 0.054 | 0.1315 |
| HIST1H3F | NM_021018 | 0.6475 | 0.2525 |
| MYL6 | NM_021019 | 0.3225 | -0.0445 |
| LZTS1 | NM_021020 | -0.6815 | -2.1475 |
| SNTB1 | NM_021021 | 0.208 | -0.09 |
| CFHR3 | NM_021023 | -0.5635 | -0.217 |
| TLX3 | NM_021025 | -0.581 | -1.729 |
| RFPL1 | NM_021026 | 0.035 | 0.6965 |
| RPL36A | NM_021029 | -0.0595 | -0.52 |
| ZNF14 | NM_021030 | -0.395 | -0.733 |
| FGF12 | NM_021032 | 0.0225 | -0.931 |
| RAP2A | NM_021033 | -1.509 | -2.7395 |
| IFITM3 | NM_021034 | -0.1295 | -0.1545 |
| ZNFX1 | NM_021035 | -0.441 | -1.7085 |
| MBNL1 | NM_021038 | -1.4015 | -3.2555 |
| DHH | NM_021044 | -0.328 | -0.2575 |
| ZNF248 | NM_021045 | -1.2725 | -2.2985 |
| KRTAP5-8 | NM_021046 | -0.4 | -0.429 |
| ZNF253 | NM_021047 | -0.6275 | -1.3055 |
| MAGEA10 | NM_021048 | 0.033 | -0.2885 |
| MAGEA5 | NM_021049 | -0.153 | 0.011 |
| HIST1H2AE | NM_021052 | 1.6995 | 2.2615 |
| IFNA7 | NM_021057 | -0.0745 | -0.2425 |
| HIST1H2BJ | NM_021058 | 0.1665 | -0.321 |
| ZNF250 | NM_021061 | -1.084 | -2.694 |
| HIST1H2BB | NM_021062 | 0.0075 | 1.1555 |
| HIST1H2BD | NM_021063 | -0.181 | -0.3375 |
| HIST1H2AG | NM_021064 | -0.407 | 0.1515 |
| HIST1H2AD | NM_021065 | 1.0725 | 0.78 |
| HIST1H2AJ | NM_021066 | 0.1635 | 0.143 |
| GINS1 | NM_021067 | -1.1785 | -2.5105 |
| IFNA4 | NM_021068 | 0.2715 | 0.084 |
| SORBS2 | NM_021069 | -0.5915 | -1.332 |
| LTBP3 | NM_021070 | -0.3685 | -0.559 |
| ART4 | NM_021071 | -0.0675 | -0.355 |
| HCN1 | NM_021072 | 0.1145 | -0.1975 |
| BMP5 | NM_021073 | -0.0925 | -0.4415 |
| NDUFV2 | NM_021074 | -1.007 | -2.2055 |
| NDUFV3 | NM_021075 | -0.3885 | -1.2635 |
| NEFH | NM_021076 | -0.2475 | -0.585 |
| NMB | NM_021077 | 0.3615 | 1.1675 |
| KAT2A | NM_021078 | -0.358 | -0.933 |

| | | | |
|---|---|---|---|
| NMT1 | NM_021079 | -0.365 | -1.917 |
| DAB1 | NM_021080 | -0.7615 | -0.948 |
| GHRH | NM_021081 | -0.177 | -0.5755 |
| SLC15A2 | NM_021082 | -0.1275 | -0.373 |
| XK | NM_021083 | 0.093 | -0.8665 |
| ZNF2 | NM_021088 | -0.4865 | -1.942 |
| ZNF8 | NM_021089 | -0.575 | -2.4145 |
| MTMR3 | NM_021090 | -0.32 | -1.7465 |
| SLCO1A2 | NM_021094 | 0.443 | 0.065 |
| SLC5A6 | NM_021095 | -0.409 | -1.764 |
| CACNA1I | NM_021096 | -0.0065 | -0.2005 |
| SLC8A1 | NM_021097 | -1.2225 | -2.382 |
| CACNA1H | NM_021098 | -0.174 | -0.6275 |
| NFS1 | NM_021100 | -0.4815 | -1.573 |
| CLDN1 | NM_021101 | -0.083 | -0.187 |
| SPINT2 | NM_021102 | -0.0695 | -0.1925 |
| TMSB10 | NM_021103 | 0.3905 | -0.048 |
| RPL41 | NM_021104 | 0.038 | -0.227 |
| PLSCR1 | NM_021105 | -1.475 | -2.8165 |
| RGS3 | NM_021106 | -0.617 | -1.4575 |
| MRPS12 | NM_021107 | -0.5375 | -2.297 |
| TMSB4X | NM_021109 | 0.123 | -0.404 |
| COL14A1 | NM_021110 | -0.1555 | -0.5225 |
| RECK | NM_021111 | -0.043 | -0.11 |
| SPINK2 | NM_021114 | -0.1535 | -0.024 |
| SEZ6L | NM_021115 | -0.135 | -0.801 |
| ADCY1 | NM_021116 | -1.421 | -1.3255 |
| CRY2 | NM_021117 | -0.2015 | -0.1015 |
| CYLC1 | NM_021118 | 0.242 | 0.0035 |
| DLG3 | NM_021120 | -0.6285 | -1.61 |
| MPST | NM_021126 | -0.618 | -1.257 |
| PMAIP1 | NM_021127 | -0.886 | -0.182 |
| POLR2L | NM_021128 | 0.3995 | -0.3655 |
| PPA1 | NM_021129 | -0.6955 | -0.5245 |
| PPIA | NM_021130 | -0.1055 | -0.363 |
| PPP2R4 | NM_021131 | -0.1025 | -0.9615 |
| PPP3CB | NM_021132 | -1.485 | -2.206 |
| RNASEL | NM_021133 | -1.2785 | -1.05 |
| MRPL23 | NM_021134 | 0.329 | -0.226 |
| RPS6KA2 | NM_021135 | -0.6265 | -1.588 |
| RTN1 | NM_021136 | -0.235 | -0.1905 |
| TNFAIP1 | NM_021137 | -1.2225 | -2.8155 |
| TRAF2 | NM_021138 | -0.748 | -1.604 |

| | | | |
|---|---|---|---|
| UGT2B4 | NM_021139 | 0.101 | -0.3365 |
| KDM6A | NM_021140 | -1.1475 | -2.449 |
| XRCC5 | NM_021141 | -0.8175 | -0.5485 |
| PSIP1 | NM_021144 | -0.356 | -0.6365 |
| DMTF1 | NM_021145 | -1.3265 | -2.0925 |
| ANGPTL7 | NM_021146 | -0.0905 | -0.047 |
| CCNO | NM_021147 | -0.177 | -0.2965 |
| ZNF273 | NM_021148 | -0.1605 | -0.6125 |
| COTL1 | NM_021149 | 0.2965 | -0.261 |
| GRIP1 | NM_021150 | -0.785 | -2.721 |
| CROT | NM_021151 | -0.014 | 0.263 |
| CDH19 | NM_021153 | 0.3085 | 0.0925 |
| PSAT1 | NM_021154 | -0.7885 | -1.5255 |
| CD209 | NM_021155 | -0.1225 | -0.3 |
| TMX4 | NM_021156 | -1.3735 | -1.0825 |
| TRIB3 | NM_021158 | -0.8015 | -2.098 |
| RAP1GDS1 | NM_021159 | -0.9505 | -0.795 |
| ABHD16A | NM_021160 | -0.1625 | -1.177 |
| KCNK10 | NM_021161 | -0.468 | -0.758 |
| RBAK | NM_021163 | -0.753 | -3.574 |
| FAM5B | NM_021165 | 0.4915 | 0.2455 |
| GATAD1 | NM_021167 | -1.6535 | -3.1375 |
| RAB40C | NM_021168 | -0.574 | -1.8625 |
| HES4 | NM_021170 | -0.0085 | -0.7365 |
| POLD4 | NM_021173 | -0.1995 | -0.2095 |
| KIAA1967 | NM_021174 | -0.085 | -0.7915 |
| HAMP | NM_021175 | 0.14 | -0.322 |
| G6PC2 | NM_021176 | -0.3455 | -0.6325 |
| LSM2 | NM_021177 | -0.562 | -1.677 |
| CCNB1IP1 | NM_021178 | -1.152 | -0.782 |
| C1orf114 | NM_021179 | -0.5035 | -0.4935 |
| SLAMF7 | NM_021181 | -0.1355 | -0.234 |
| HMHB1 | NM_021182 | 0.2655 | -0.108 |
| RAP2C | NM_021183 | -1.3415 | -2.1965 |
| C6orf47 | NM_021184 | -0.7485 | -1.7685 |
| CATSPERG | NM_021185 | 0.0835 | -0.1505 |
| ZP4 | NM_021186 | 0.095 | -0.26 |
| CYP4F11 | NM_021187 | 0.001 | -0.2155 |
| ZNF410 | NM_021188 | -1.314 | -2.961 |
| CADM3 | NM_021189 | -0.083 | -0.5645 |
| PTBP2 | NM_021190 | -1.261 | -1.1895 |
| NEUROD4 | NM_021191 | 0.5625 | -1.6055 |
| HOXD11 | NM_021192 | -0.342 | -0.268 |

| | | | |
|---|---|---|---|
| HOXD12 | NM_021193 | -0.039 | 0.0305 |
| SLC30A1 | NM_021194 | -1.1375 | 0.0125 |
| CLDN6 | NM_021195 | -0.0235 | -0.0985 |
| SLC4A5 | NM_021196 | 0.018 | -0.3365 |
| WFDC1 | NM_021197 | -0.0935 | -0.7255 |
| CTDSP1 | NM_021198 | -0.5355 | -1.7085 |
| SQRDL | NM_021199 | -0.438 | -0.195 |
| PLEKHB1 | NM_021200 | 0.339 | -0.3015 |
| MS4A7 | NM_021201 | -0.248 | 0.2435 |
| TP53INP2 | NM_021202 | 0.2075 | -0.535 |
| SRPRB | NM_021203 | -0.7995 | -1.8415 |
| ENOPH1 | NM_021204 | -1.202 | -1.9045 |
| RHOU | NM_021205 | -1.713 | -0.8815 |
| NLRC4 | NM_021209 | 0.028 | -0.119 |
| TRAPPC1 | NM_021210 | 0.2105 | -0.481 |
| ZBED5 | NM_021211 | -1.885 | -4.337 |
| PCTP | NM_021213 | -0.9595 | -3.381 |
| FAM108C1 | NM_021214 | -0.906 | -2.93 |
| RPRD1B | NM_021215 | -0.7205 | -1.595 |
| ZNF71 | NM_021216 | -0.368 | -2.672 |
| ZNF77 | NM_021217 | -0.951 | -2.239 |
| C9orf80 | NM_021218 | -1.5395 | -2.295 |
| JAM2 | NM_021219 | -0.7705 | -1.111 |
| OVOL2 | NM_021220 | -0.1395 | -0.475 |
| LY6G5B | NM_021221 | -0.1255 | -0.515 |
| PRUNE | NM_021222 | -1.248 | -3.3855 |
| MYL7 | NM_021223 | 0.0115 | 0.07 |
| ZNF462 | NM_021224 | -0.9935 | -2.511 |
| PROL1 | NM_021225 | -0.025 | -0.4275 |
| ARHGAP22 | NM_021226 | -0.8585 | -1.96 |
| OSTC | NM_021227 | -0.6245 | -0.7435 |
| SCAF1 | NM_021228 | -0.081 | -0.5145 |
| NTN4 | NM_021229 | -0.273 | -1.0165 |
| C19orf29 | NM_021231 | -0.7265 | -1.556 |
| PRODH2 | NM_021232 | -0.054 | -0.1565 |
| DNASE2B | NM_021233 | 0.074 | 0.0685 |
| EPS15L1 | NM_021235 | -0.2005 | -1.1355 |
| SELK | NM_021237 | -0.331 | -0.5505 |
| FAM60A | NM_021238 | -0.9635 | -0.9235 |
| RBM25 | NM_021239 | -0.3715 | -1.5295 |
| DMRT3 | NM_021240 | -0.2095 | -0.047 |
| WIZ | NM_021241 | -0.2995 | -0.872 |
| MID1IP1 | NM_021242 | -0.5775 | -1.1735 |

| | | | |
|---|---|---|---|
| C6orf115 | NM_021243 | -0.454 | -1.7175 |
| RRAGD | NM_021244 | -0.581 | -1.706 |
| MYOZ1 | NM_021245 | 0.102 | 0.005 |
| LY6G6D | NM_021246 | 0.0285 | -0.1185 |
| PRM3 | NM_021247 | 0.1915 | -0.473 |
| CDH22 | NM_021248 | -0.4185 | -0.7045 |
| SNX6 | NM_021249 | -1.3295 | -1.4905 |
| RAB18 | NM_021252 | -1.1215 | -1.897 |
| TRIM39 | NM_021253 | -0.9615 | -2.4325 |
| C21orf59 | NM_021254 | -0.7175 | -1.257 |
| PELI2 | NM_021255 | -1.0455 | -1.9225 |
| NGB | NM_021257 | 0.246 | -0.332 |
| IL22RA1 | NM_021258 | -0.178 | -0.1325 |
| TMEM8A | NM_021259 | 0.1635 | -0.5245 |
| ZFYVE1 | NM_021260 | -0.7865 | -2.3925 |
| CERS1 | NM_021267 | 0.0675 | -0.6075 |
| IFNA17 | NM_021268 | 0.423 | -0.3175 |
| ZNF708 | NM_021269 | -1.1085 | -2.944 |
| LAIR2 | NM_021270 | 0.5505 | 0.1355 |
| GRIN1 | NM_021569 | 0.2565 | 0.227 |
| BARX1 | NM_021570 | 0.51 | 0.0475 |
| CARD18 | NM_021571 | -0.139 | -0.224 |
| ENPP5 | NM_021572 | -0.538 | -1.029 |
| BCR | NM_021574 | -0.376 | -1.5265 |
| AP2S1 | NM_021575 | 0.2355 | -0.4605 |
| ADAMTS2 | NM_021599 | 0.4125 | -0.358 |
| CD79A | NM_021601 | -0.0825 | -0.3615 |
| CD79B | NM_021602 | 0.0665 | -0.1825 |
| FXYD2 | NM_021603 | 0.1985 | -0.304 |
| KCNN2 | NM_021614 | 0.2205 | -1.617 |
| CHST6 | NM_021615 | -0.28 | -0.2375 |
| PRDM12 | NM_021619 | 0.121 | 1.042 |
| PRDM13 | NM_021620 | 0.4275 | -0.2305 |
| PLEKHA1 | NM_021622 | -1.3095 | -1.491 |
| PLEKHA2 | NM_021623 | -0.8255 | -1.5435 |
| HRH4 | NM_021624 | 0.2355 | -0.062 |
| TRPV4 | NM_021625 | -0.186 | -0.2965 |
| SCPEP1 | NM_021626 | -0.3455 | -0.9195 |
| SENP2 | NM_021627 | -1.0345 | -1.848 |
| ALOXE3 | NM_021628 | 0.1215 | -0.1975 |
| GNB4 | NM_021629 | -1.1785 | -1.474 |
| PDLIM2 | NM_021630 | -0.171 | -0.878 |
| ZNF350 | NM_021632 | -0.0835 | -1.4355 |

| | | | |
|---|---|---|---|
| KLHL12 | NM_021633 | -1.668 | -2.703 |
| RXFP1 | NM_021634 | 0.007 | -0.1485 |
| PBOV1 | NM_021635 | 0.22 | -0.332 |
| LGR6 | NM_021636 | -0.0305 | -0.0485 |
| TMEM35 | NM_021637 | -0.1965 | -0.5975 |
| GPBP1L1 | NM_021639 | -1.5915 | -3.0115 |
| C12orf10 | NM_021640 | -0.2855 | -1.1105 |
| ADAM12 | NM_021641 | -0.698 | -0.496 |
| FCGR2A | NM_021642 | 0.0625 | -0.108 |
| TRIB2 | NM_021643 | -1.0095 | -1.803 |
| HNRNPH3 | NM_021644 | -0.7305 | -1.108 |
| UTP14C | NM_021645 | -1.509 | -2.878 |
| ZNF500 | NM_021646 | -0.5345 | -1.6055 |
| MFAP3L | NM_021647 | -1.2035 | -2.7615 |
| TSPYL4 | NM_021648 | -0.954 | -1.291 |
| NFYA | NM_021705 | -0.905 | -1.2805 |
| LAIR1 | NM_021706 | -0.079 | -0.3735 |
| SIVA1 | NM_021709 | -0.3035 | -1.184 |
| ADAM22 | NM_021721 | -0.5105 | -0.9525 |
| ADAM22 | NM_021722 | -0.5105 | -0.9525 |
| ADAM22 | NM_021723 | -0.5105 | -0.9525 |
| NR1D1 | NM_021724 | -0.107 | -0.5465 |
| FADS3 | NM_021727 | -0.221 | -0.9235 |
| OTX2 | NM_021728 | -0.079 | -0.323 |
| VPS11 | NM_021729 | -0.211 | -1.5425 |
| AVPI1 | NM_021732 | -0.458 | -1.7745 |
| TSKS | NM_021733 | 0.1755 | -0.158 |
| SLC25A19 | NM_021734 | -1.207 | -3.072 |
| CLCN6 | NM_021735 | 0.216 | -0.4645 |
| CLCN6 | NM_021736 | 0.216 | -0.4645 |
| CLCN6 | NM_021737 | 0.216 | -0.4645 |
| SVIL | NM_021738 | -1.2135 | -2.329 |
| ADAM28 | NM_021777 | 0.275 | 0.224 |
| EDA2R | NM_021783 | -1.048 | -1.686 |
| FOXA2 | NM_021784 | 0.1275 | -0.073 |
| RAI2 | NM_021785 | -0.295 | -0.8365 |
| ADAM30 | NM_021794 | 0.186 | 0.0405 |
| ELK4 | NM_021795 | -0.003 | -0.1535 |
| PLAC1 | NM_021796 | 0.019 | -0.176 |
| IL21R | NM_021798 | 0.2435 | -0.3815 |
| DNAJC12 | NM_021800 | -1.516 | -2.466 |
| MMP26 | NM_021801 | -0.0645 | -0.092 |
| IL21 | NM_021803 | -0.084 | -0.094 |

| | | | |
|---|---|---|---|
| ACE2 | NM_021804 | -0.4635 | -0.2225 |
| SIGIRR | NM_021805 | 0.337 | -0.445 |
| FAM3A | NM_021806 | -0.427 | -1.367 |
| EXOC4 | NM_021807 | -0.5215 | -0.564 |
| GALNT9 | NM_021808 | 0.0435 | -0.297 |
| TGIF2 | NM_021809 | -1.0185 | -3.2745 |
| CDH26 | NM_021810 | -0.076 | -0.0065 |
| BACH2 | NM_021813 | -0.841 | -3.054 |
| ELOVL5 | NM_021814 | -1.283 | -1.5085 |
| SLC5A7 | NM_021815 | 0.01 | -0.1465 |
| HAPLN2 | NM_021817 | -0.085 | -0.568 |
| SAV1 | NM_021818 | -1.0755 | -1.9595 |
| LMAN1L | NM_021819 | 0.242 | -0.4565 |
| MRPS35 | NM_021821 | -1.125 | -2.825 |
| APOBEC3G | NM_021822 | -0.1945 | -0.2605 |
| PPCDC | NM_021823 | -0.864 | -1.5835 |
| NIF3L1 | NM_021824 | -1.771 | -4.249 |
| CCDC90B | NM_021825 | -2.12 | -2.4905 |
| FASTKD5 | NM_021826 | -0.614 | -1.724 |
| CCDC81 | NM_021827 | -0.467 | -0.517 |
| HPSE2 | NM_021828 | -0.2705 | -0.563 |
| C10orf2 | NM_021830 | -1.032 | -2.465 |
| AGBL5 | NM_021831 | -0.251 | -0.614 |
| UCP1 | NM_021833 | 0.513 | -0.146 |
| FGG | NM_021870 | 0.258 | -0.323 |
| FGA | NM_021871 | -0.1405 | -0.1215 |
| CDC25B | NM_021872 | -0.3235 | -1.0145 |
| CDC25B | NM_021873 | -0.396 | -1.11 |
| FXYD1 | NM_021902 | 0.3475 | -0.088 |
| GABBR1 | NM_021903 | -0.497 | -1.0825 |
| GABBR1 | NM_021904 | -0.497 | -1.0825 |
| DTNB | NM_021907 | -0.2295 | -0.905 |
| ST7 | NM_021908 | -0.9475 | -2.294 |
| FXYD3 | NM_021910 | 0.4865 | -0.123 |
| GABRB2 | NM_021911 | -0.188 | -0.324 |
| GABRB3 | NM_021912 | -0.9845 | -1.441 |
| AXL | NM_021913 | -0.0465 | 0.036 |
| CFL2 | NM_021914 | -0.8905 | -2.2675 |
| ZNF69 | NM_021915 | -0.415 | -1.3455 |
| ZNF70 | NM_021916 | -0.2325 | -0.4335 |
| SCT | NM_021920 | 0.3165 | -0.1805 |
| FANCE | NM_021922 | -0.624 | -2.391 |
| FGFRL1 | NM_021923 | -0.1025 | -0.5225 |

| | | | |
|---|---|---|---|
| CDHR5 | NM_021924 | -0.0115 | -0.41 |
| C2orf43 | NM_021925 | -1.3905 | -1.395 |
| ALX4 | NM_021926 | -0.014 | -0.255 |
| GUF1 | NM_021927 | -0.972 | -1.554 |
| SPCS3 | NM_021928 | -1.37 | -1.4725 |
| RINT1 | NM_021930 | -1.173 | -2.5065 |
| DHX35 | NM_021931 | -0.867 | -1.835 |
| RIC8A | NM_021932 | -0.6535 | -1.918 |
| MIIP | NM_021933 | -0.253 | -1.0045 |
| C12orf44 | NM_021934 | -0.803 | -1.8865 |
| PROK2 | NM_021935 | -0.4945 | -0.454 |
| PAPPA2 | NM_021936 | 0.1595 | -0.366 |
| EEFSEC | NM_021937 | -0.465 | -1.384 |
| CELF5 | NM_021938 | 0.0545 | -1.067 |
| FKBP10 | NM_021939 | 0.061 | -0.519 |
| SMAP1 | NM_021940 | -0.7115 | -1.1265 |
| C4orf41 | NM_021942 | -1.5025 | -2.356 |
| ZFAND3 | NM_021943 | 0.0295 | -0.839 |
| C14orf93 | NM_021944 | -0.785 | -1.9555 |
| SLC22A23 | NM_021945 | 0.3215 | 0.0895 |
| BCORL1 | NM_021946 | -0.0445 | -1.166 |
| SRR | NM_021947 | -0.641 | -0.9855 |
| BCAN | NM_021948 | 0.014 | 0.1105 |
| ATP2B3 | NM_021949 | 0.0715 | -0.5615 |
| MS4A1 | NM_021950 | -0.2405 | -0.1505 |
| DMRT1 | NM_021951 | 0.143 | 0.025 |
| ELAVL4 | NM_021952 | -1.8505 | -0.916 |
| FOXM1 | NM_021953 | -0.3635 | -0.9255 |
| GJA3 | NM_021954 | -0.245 | 0.116 |
| GNGT1 | NM_021955 | -0.3105 | 0.267 |
| GRIK2 | NM_021956 | -0.8425 | -1.477 |
| GYS2 | NM_021957 | -0.0035 | -0.3495 |
| HLX | NM_021958 | -0.696 | -2.607 |
| PPP1R11 | NM_021959 | -0.389 | -1.929 |
| MCL1 | NM_021960 | -1.401 | -1.656 |
| TEAD1 | NM_021961 | -0.7415 | -1.6805 |
| ABR | NM_021962 | -0.7775 | -0.998 |
| NAP1L2 | NM_021963 | 0.439 | 0.139 |
| ZNF148 | NM_021964 | -0.5705 | -2.4965 |
| PGM5 | NM_021965 | 0.113 | -0.1765 |
| TCL1A | NM_021966 | -0.0035 | 0.0105 |
| HIST1H4J | NM_021968 | 0.448 | 0.242 |
| NR0B2 | NM_021969 | 0.319 | -0.4375 |

| | | | |
|---|---|---|---|
| GMPPB | NM_021971 | -0.4325 | -1.648 |
| SPHK1 | NM_021972 | -0.1685 | -0.8105 |
| HAND2 | NM_021973 | -1.0535 | -1.817 |
| POLR2F | NM_021974 | 0.1755 | -0.388 |
| RELA | NM_021975 | -0.7155 | -2.538 |
| RXRB | NM_021976 | -0.364 | -1.582 |
| SLC22A3 | NM_021977 | 0.0115 | 0.0125 |
| ST14 | NM_021978 | -0.211 | -0.43 |
| HSPA2 | NM_021979 | -0.3245 | -1.0375 |
| OPTN | NM_021980 | -0.034 | -0.4645 |
| SEC24A | NM_021982 | -1.497 | -3.46 |
| UBE2V1 | NM_021988 | -0.2425 | -1.1055 |
| TMSB15A | NM_021992 | 0.045 | -0.13 |
| ZNF277 | NM_021994 | -0.2965 | -1.7915 |
| GBGT1 | NM_021996 | -0.634 | -0.772 |
| ZNF711 | NM_021998 | -0.7785 | -2.873 |
| ITM2B | NM_021999 | -0.666 | -0.722 |
| NR1I2 | NM_022002 | -0.2945 | -0.269 |
| FXYD6 | NM_022003 | 0.187 | -0.2205 |
| FXYD7 | NM_022006 | 0.323 | 0.2515 |
| CUZD1 | NM_022034 | -0.6315 | -0.9935 |
| GPRC5C | NM_022036 | -0.875 | -1.927 |
| TIA1 | NM_022037 | -1.508 | -2.606 |
| FBXW4 | NM_022039 | -0.3175 | -1.128 |
| GAN | NM_022041 | -0.605 | -0.6635 |
| SLC26A1 | NM_022042 | 0.0175 | -0.086 |
| SDF2L1 | NM_022044 | -0.851 | -2.811 |
| MTBP | NM_022045 | -0.588 | -1.3865 |
| KLK14 | NM_022046 | -0.2335 | -0.278 |
| DEF6 | NM_022047 | -0.122 | 0.148 |
| CSNK1G1 | NM_022048 | -1.2695 | -2.26 |
| GPR88 | NM_022049 | 0.0505 | -0.2515 |
| EGLN1 | NM_022051 | -1.483 | -2.3515 |
| NXF3 | NM_022052 | 0.29 | 0.129 |
| NXF2 | NM_022053 | 0.134 | 0.02 |
| KCNK13 | NM_022054 | -0.061 | 0.02 |
| KCNK12 | NM_022055 | 0.131 | -0.903 |
| SLC4A10 | NM_022058 | 0.189 | 0.0315 |
| CXCL16 | NM_022059 | -0.004 | -0.244 |
| ABHD4 | NM_022060 | -0.3975 | -0.466 |
| MRPL17 | NM_022061 | -1.2495 | -3.995 |
| PKNOX2 | NM_022062 | -0.594 | 0.2425 |
| FAM204A | NM_022063 | -0.788 | -0.5045 |

| | | | |
|---|---|---|---|
| RNF123 | NM_022064 | -0.1325 | -0.471 |
| THADA | NM_022065 | -1.246 | -1.931 |
| UBE2O | NM_022066 | -0.538 | -2.146 |
| PIEZO2 | NM_022068 | -0.7465 | 0.11 |
| HEATR6 | NM_022070 | -0.8835 | -2.5625 |
| SH2D4A | NM_022071 | -0.1745 | -0.106 |
| NSUN3 | NM_022072 | -0.9725 | -2.01 |
| EGLN3 | NM_022073 | -0.221 | -1.6905 |
| FAM111A | NM_022074 | -0.195 | -0.1145 |
| DUSP21 | NM_022076 | -0.159 | 0.1065 |
| MANBAL | NM_022077 | -0.36 | -1.3845 |
| GPATCH3 | NM_022078 | -0.4095 | -1.1075 |
| HERC4 | NM_022079 | 0.3045 | 0.105 |
| NAPB | NM_022080 | -0.79 | -0.9205 |
| HPS4 | NM_022081 | -0.954 | -2.2725 |
| SLC17A9 | NM_022082 | -0.017 | -0.4985 |
| GALNT11 | NM_022087 | -0.2815 | -1.1375 |
| ZFP64 | NM_022088 | -0.438 | -1.059 |
| ATP13A2 | NM_022089 | -0.065 | -0.5495 |
| C5orf54 | NM_022090 | -1.3205 | -2.4765 |
| ASCC3 | NM_022091 | 0.6105 | 0.461 |
| CHTF18 | NM_022092 | -0.134 | -0.711 |
| TNN | NM_022093 | -0.206 | -0.4645 |
| CIDEC | NM_022094 | 0.0575 | -0.2065 |
| ZNF335 | NM_022095 | -0.7895 | -1.39 |
| ANKRD5 | NM_022096 | -0.7305 | -1.24 |
| CHP2 | NM_022097 | 0.1925 | 0.0685 |
| XPNPEP3 | NM_022098 | -1.33 | -2.076 |
| MRPS14 | NM_022100 | -0.373 | -1.458 |
| CXorf56 | NM_022101 | -0.398 | -1.031 |
| ZNF667 | NM_022103 | -1.3515 | -1.919 |
| PCIF1 | NM_022104 | -0.603 | -1.798 |
| DIDO1 | NM_022105 | 0.457 | -1.4845 |
| C20orf177 | NM_022106 | -1.3265 | -1.6035 |
| GPSM3 | NM_022107 | 0.152 | 0.1135 |
| FKBPL | NM_022110 | -0.6695 | -1.9005 |
| CLSPN | NM_022111 | -0.8865 | -1.5745 |
| TP53AIP1 | NM_022112 | 0.029 | -0.2835 |
| KIF13A | NM_022113 | -0.256 | -0.6685 |
| PRDM16 | NM_022114 | 0.2685 | -0.5095 |
| PRDM15 | NM_022115 | -1.0505 | -2.1465 |
| FIGNL1 | NM_022116 | -1.3355 | -2.4095 |
| TSPYL2 | NM_022117 | -0.143 | 0.14 |

| | | | |
|---|---|---|---|
| RBM26 | NM_022118 | -0.7885 | -2.0765 |
| PRSS22 | NM_022119 | -0.0715 | -0.499 |
| OXCT2 | NM_022120 | 0.175 | -0.3395 |
| PERP | NM_022121 | -0.1655 | -0.4095 |
| MMP27 | NM_022122 | 0.197 | 0.259 |
| NPAS3 | NM_022123 | 0.0345 | -0.345 |
| CDH23 | NM_022124 | -0.124 | -0.5565 |
| LHPP | NM_022126 | -0.0585 | -0.051 |
| SLC28A3 | NM_022127 | 0.148 | -0.155 |
| RBKS | NM_022128 | -0.6305 | -1.0165 |
| PBLD | NM_022129 | -0.898 | -0.767 |
| GOLPH3 | NM_022130 | -1.7665 | -4.0455 |
| CLSTN2 | NM_022131 | 0.2025 | -0.2435 |
| MCCC2 | NM_022132 | -0.16 | -0.8575 |
| SNX16 | NM_022133 | -1.217 | -2.706 |
| GAL3ST2 | NM_022134 | 0.006 | -0.433 |
| POPDC2 | NM_022135 | -0.293 | -1.7815 |
| SAMSN1 | NM_022136 | 0.3835 | 0.772 |
| SMOC1 | NM_022137 | -0.018 | -0.241 |
| SMOC2 | NM_022138 | -0.2995 | -0.7415 |
| GFRA4 | NM_022139 | -0.3155 | -0.15 |
| EPB41L4A | NM_022140 | -0.436 | -0.198 |
| PARVG | NM_022141 | 0.088 | -0.186 |
| ELSPBP1 | NM_022142 | 0.065 | -0.2995 |
| LRRC4 | NM_022143 | -0.3465 | -0.4575 |
| TNMD | NM_022144 | -0.2865 | -0.2515 |
| CENPK | NM_022145 | -1.3045 | -1.5795 |
| NPFFR1 | NM_022146 | 0.1515 | -0.1615 |
| RTP4 | NM_022147 | -0.1075 | -0.3295 |
| CRLF2 | NM_022148 | 0.159 | -0.3395 |
| MAGEF1 | NM_022149 | -0.8795 | -2.4365 |
| NPVF | NM_022150 | -0.552 | 0.0295 |
| MOAP1 | NM_022151 | -0.7305 | -0.768 |
| TMBIM1 | NM_022152 | 0.112 | -0.2465 |
| C10orf54 | NM_022153 | -0.491 | -0.149 |
| SLC39A8 | NM_022154 | -0.067 | -0.998 |
| DUS1L | NM_022156 | 0.118 | -0.849 |
| RRAGC | NM_022157 | -1.0815 | -2.097 |
| FN3K | NM_022158 | -0.1865 | -0.202 |
| ELTD1 | NM_022159 | -0.271 | -0.163 |
| DMRTA1 | NM_022160 | -0.432 | -1.1535 |
| BIRC7 | NM_022161 | 0.062 | -0.5875 |
| NOD2 | NM_022162 | 0.279 | -0.28 |

| | | | |
|---|---|---|---|
| MRPL46 | NM_022163 | -0.7945 | -3.4435 |
| TINAGL1 | NM_022164 | -0.337 | -0.2875 |
| LIN7B | NM_022165 | -0.49 | -1.429 |
| XYLT1 | NM_022166 | -1.196 | -2.4905 |
| XYLT2 | NM_022167 | -0.901 | -1.678 |
| IFIH1 | NM_022168 | -0.0295 | -0.341 |
| ABCG4 | NM_022169 | 0.021 | -0.364 |
| EIF4H | NM_022170 | -0.1795 | -1.0705 |
| TCTA | NM_022171 | -0.252 | -0.8175 |
| PC | NM_022172 | 0.171 | -0.143 |
| TIA1 | NM_022173 | -1.508 | -2.606 |
| HRH2 | NM_022304 | -0.311 | -0.232 |
| ICA1 | NM_022307 | -0.608 | -1.057 |
| ITGB1BP1 | NM_022334 | -0.568 | -0.5855 |
| EDAR | NM_022336 | -0.2035 | -0.319 |
| RAB38 | NM_022337 | -0.3955 | -0.8045 |
| C11orf24 | NM_022338 | -0.317 | -1.4935 |
| ZFYVE20 | NM_022340 | -1.236 | -2.6105 |
| KIF9 | NM_022342 | -0.163 | -0.91 |
| GLIPR2 | NM_022343 | -0.103 | -1.2705 |
| C17orf75 | NM_022344 | -1.3585 | -1.922 |
| NCAPG | NM_022346 | -0.8145 | -2.332 |
| TOR1AIP2 | NM_022347 | -0.5465 | -1.3555 |
| MS4A6A | NM_022349 | -0.1165 | 0.223 |
| ERAP2 | NM_022350 | -0.0005 | -0.449 |
| NECAB1 | NM_022351 | -0.23 | -0.807 |
| OSGEPL1 | NM_022353 | -2.1445 | -2.8285 |
| DPEP2 | NM_022355 | 0.1925 | 0.0045 |
| LEPRE1 | NM_022356 | -0.778 | -1.6275 |
| DPEP3 | NM_022357 | 0.098 | -0.4035 |
| KCNK15 | NM_022358 | 0.0115 | 0.0495 |
| PDE4DIP | NM_022359 | -0.9925 | -0.862 |
| EDDM3B | NM_022360 | 0.1745 | 0.0475 |
| POPDC3 | NM_022361 | -0.6305 | -1.7685 |
| MMS19 | NM_022362 | -0.0215 | -1.414 |
| LHX5 | NM_022363 | 0.0585 | -0.3415 |
| DNAJC1 | NM_022365 | -0.0765 | -0.753 |
| TFB2M | NM_022366 | -1.479 | -2.67 |
| SEMA4A | NM_022367 | -0.0655 | -0.5805 |
| PJA1 | NM_022368 | -0.633 | -1.645 |
| STRA6 | NM_022369 | -0.683 | -1.1345 |
| ROBO3 | NM_022370 | -0.1785 | -0.829 |
| TOR3A | NM_022371 | -0.3095 | -1.164 |

| | | | |
|---|---|---|---|
| MLST8 | NM_022372 | -0.377 | -0.9365 |
| HERPUD2 | NM_022373 | -1.8605 | -4.692 |
| ATL2 | NM_022374 | -1.648 | -2.5255 |
| OCLM | NM_022375 | 0.035 | -0.354 |
| ICAM4 | NM_022377 | -0.2025 | -0.241 |
| SLC6A20 | NM_022405 | 0.176 | -0.1915 |
| XRCC4 | NM_022406 | -1.0375 | -1.7165 |
| ABCG5 | NM_022436 | -0.117 | -0.0555 |
| ABCG8 | NM_022437 | -0.0535 | -0.2375 |
| MAL | NM_022438 | -0.044 | -0.317 |
| MAL | NM_022439 | -0.044 | -0.317 |
| MAL | NM_022440 | -0.044 | -0.317 |
| UBE2V1 | NM_022442 | -0.2425 | -1.1055 |
| MLF1 | NM_022443 | -0.534 | -0.676 |
| SLC13A1 | NM_022444 | -0.119 | -0.1515 |
| TPK1 | NM_022445 | -0.135 | -0.782 |
| RAB17 | NM_022449 | -0.463 | -0.871 |
| RHBDF1 | NM_022450 | 0.128 | -0.3425 |
| NOC3L | NM_022451 | -0.608 | -2.411 |
| RNF25 | NM_022453 | -0.4635 | -2.144 |
| SOX17 | NM_022454 | -0.0565 | -0.2385 |
| NSD1 | NM_022455 | 0.1895 | 1.22 |
| RAB3IP | NM_022456 | -0.321 | -0.6525 |
| RFWD2 | NM_022457 | -1.255 | -2.924 |
| LMBR1 | NM_022458 | -0.9255 | -0.944 |
| XPO4 | NM_022459 | -0.8215 | -2.1935 |
| HS1BP3 | NM_022460 | -1.003 | -1.4235 |
| AZI2 | NM_022461 | -0.797 | -1.6665 |
| HIF3A | NM_022462 | -0.243 | -0.193 |
| NXN | NM_022463 | -0.3995 | -1.138 |
| SIL1 | NM_022464 | -0.2665 | -1.1565 |
| IKZF4 | NM_022465 | -0.7695 | -1.593 |
| IKZF5 | NM_022466 | -1.024 | -2.1895 |
| CHST8 | NM_022467 | -0.963 | -2.104 |
| MMP25 | NM_022468 | -0.5045 | -0.1115 |
| GREM2 | NM_022469 | -0.2885 | -0.834 |
| ZMAT3 | NM_022470 | -0.9225 | -1.074 |
| ZFP106 | NM_022473 | -1.1305 | -2.206 |
| MPP5 | NM_022474 | -1.0365 | -2.5645 |
| HHIP | NM_022475 | 0.0655 | -0.3415 |
| AKTIP | NM_022476 | -1.0405 | -1.765 |
| NDRG3 | NM_022477 | -0.0405 | -0.4 |
| CDH24 | NM_022478 | 0.176 | -0.2255 |

| | | | |
|---|---|---|---|
| WBSCR17 | NM_022479 | -0.4055 | -1.4005 |
| KLHL25 | NM_022480 | -0.514 | -2.2525 |
| ARAP3 | NM_022481 | 0.051 | -1.0585 |
| GZF1 | NM_022482 | -1.622 | -2.505 |
| C5orf28 | NM_022483 | -1.533 | -1.9275 |
| TMEM168 | NM_022484 | -0.591 | -2.003 |
| MTMR14 | NM_022485 | -0.595 | -1.782 |
| SUSD1 | NM_022486 | -0.1215 | -0.9525 |
| DCLRE1C | NM_022487 | -1.3045 | -1.667 |
| ATG3 | NM_022488 | -1.2035 | -2.9645 |
| INF2 | NM_022489 | -0.3 | -0.759 |
| POLR1E | NM_022490 | -0.611 | -2.114 |
| SUDS3 | NM_022491 | -0.9325 | -1.747 |
| TTC31 | NM_022492 | -0.5155 | -1.621 |
| NARFL | NM_022493 | -0.2415 | -0.943 |
| ZDHHC6 | NM_022494 | -1.685 | -3.0015 |
| C14orf135 | NM_022495 | -1.5335 | -2.7225 |
| ACTR6 | NM_022496 | -1.5045 | -2.0355 |
| MRPS25 | NM_022497 | -0.299 | -1.0115 |
| FEZ1 | NM_022549 | -0.066 | -0.5695 |
| XRCC4 | NM_022550 | -1.0375 | -1.7165 |
| RPS18 | NM_022551 | 0.01 | -0.303 |
| DNMT3A | NM_022552 | -0.1735 | -1.304 |
| VPS52 | NM_022553 | -0.22 | -1.242 |
| GH2 | NM_022556 | 0.284 | 0.0655 |
| GH2 | NM_022557 | -0.152 | -0.172 |
| GH2 | NM_022558 | 0.284 | 0.0655 |
| GH1 | NM_022560 | 0.1075 | -0.174 |
| GH1 | NM_022562 | 0.317 | -0.477 |
| MESDC1 | NM_022566 | -0.5875 | -2.025 |
| NYX | NM_022567 | 0.143 | -0.2165 |
| ALDH8A1 | NM_022568 | -0.583 | -0.1985 |
| NDST4 | NM_022569 | 0.2625 | -0.157 |
| CLEC7A | NM_022570 | -0.082 | -0.414 |
| GPR135 | NM_022571 | -0.159 | -0.3685 |
| PNKD | NM_022572 | -0.237 | -0.377 |
| TSPY2 | NM_022573 | 0.441 | 0.04 |
| GIGYF1 | NM_022574 | -0.4715 | -0.8185 |
| VPS16 | NM_022575 | -0.0095 | -0.5665 |
| CSHL1 | NM_022579 | -0.2255 | -0.2515 |
| CSHL1 | NM_022580 | -0.2255 | -0.2515 |
| CSHL1 | NM_022581 | -0.058 | -0.3085 |
| CSH2 | NM_022644 | 0.0535 | -0.081 |

| | | | |
|---|---|---|---|
| CSH2 | NM_022645 | 0.174 | -0.4825 |
| TNS1 | NM_022648 | -0.0425 | -0.545 |
| RASA1 | NM_022650 | -0.9685 | -2.3215 |
| DUSP6 | NM_022652 | -0.338 | 0.281 |
| HOXC8 | NM_022658 | -0.507 | -0.858 |
| EBF2 | NM_022659 | 0.1185 | -0.0155 |
| SPANXC | NM_022661 | 0.0365 | -0.373 |
| ANAPC1 | NM_022662 | -1.4155 | -2.1175 |
| ECM1 | NM_022664 | -0.205 | -0.624 |
| PRRX1 | NM_022716 | 0.175 | -0.434 |
| SNRNP35 | NM_022717 | 0.0845 | -0.1885 |
| DGCR14 | NM_022719 | -0.3955 | -2.1105 |
| DGCR8 | NM_022720 | -0.62 | -1.337 |
| FANCF | NM_022725 | -1.3355 | -2.662 |
| ELOVL4 | NM_022726 | -0.809 | -3.2635 |
| TRMT2A | NM_022727 | -0.595 | -1.096 |
| NEUROD6 | NM_022728 | 0.1415 | -0.206 |
| COPS7B | NM_022730 | -0.093 | -1.1205 |
| NUCKS1 | NM_022731 | -0.8515 | -0.7005 |
| SMAP2 | NM_022733 | -0.39 | -1.859 |
| ACBD3 | NM_022735 | -1.6855 | -3.4545 |
| MFSD1 | NM_022736 | -0.865 | -0.943 |
| LPPR2 | NM_022737 | 0.149 | -0.261 |
| SMURF2 | NM_022739 | -1.538 | -3.1845 |
| HIPK2 | NM_022740 | -1.617 | -1.9625 |
| CCDC136 | NM_022742 | -1.0535 | -1.186 |
| SMYD3 | NM_022743 | -0.0065 | -0.4605 |
| C16orf58 | NM_022744 | -0.403 | -1.0615 |
| ATPAF1 | NM_022745 | -0.403 | -0.375 |
| MOSC1 | NM_022746 | -0.618 | -1.5415 |
| TNS3 | NM_022748 | -0.618 | -2.853 |
| FAM160B2 | NM_022749 | -0.485 | -1.3065 |
| PARP12 | NM_022750 | -0.1925 | -0.8045 |
| FAM59A | NM_022751 | -0.3745 | -1.9465 |
| ZNF574 | NM_022752 | -0.2695 | -1.464 |
| S100PBP | NM_022753 | -2.2345 | -2.5455 |
| SFXN1 | NM_022754 | -0.5545 | -1.2445 |
| IPPK | NM_022755 | -0.873 | -2.106 |
| MEAF6 | NM_022756 | -0.5355 | -0.935 |
| CCDC14 | NM_022757 | -1.3075 | -0.9335 |
| C6orf106 | NM_022758 | -0.2165 | -1.6465 |
| FAM113A | NM_022760 | -0.331 | -0.8955 |
| C11orf1 | NM_022761 | -0.4855 | -0.4845 |

| | | | |
|---|---|---|---|
| RMND5B | NM_022762 | -0.6545 | -1.547 |
| FNDC3B | NM_022763 | -1.212 | -3.2945 |
| MTHFSD | NM_022764 | 0.244 | -0.499 |
| MICAL1 | NM_022765 | -0.1685 | -0.8575 |
| CERK | NM_022766 | -0.3995 | -0.8415 |
| AEN | NM_022767 | -0.9265 | -2.967 |
| RBM15 | NM_022768 | -0.741 | -1.098 |
| CRTC3 | NM_022769 | -0.951 | -3.1605 |
| TBC1D15 | NM_022771 | -0.7245 | -2.5675 |
| EPS8L2 | NM_022772 | -0.0205 | -0.131 |
| LMF1 | NM_022773 | 0.068 | -0.534 |
| DEM1 | NM_022774 | -1.252 | -2.612 |
| OSBPL11 | NM_022776 | -1.4695 | -4.0345 |
| RABL5 | NM_022777 | -0.4325 | -0.978 |
| CEP85 | NM_022778 | -1.307 | -2.797 |
| DDX31 | NM_022779 | -1.222 | -2.847 |
| RMND5A | NM_022780 | -1.3265 | -1.8345 |
| RNF38 | NM_022781 | -0.8165 | -1 |
| MPHOSPH9 | NM_022782 | -0.991 | -2.474 |
| IQCH | NM_022784 | 0.229 | -0.1345 |
| EFCAB6 | NM_022785 | -0.25 | -0.2285 |
| ARV1 | NM_022786 | -1.5275 | -2.8795 |
| NMNAT1 | NM_022787 | -0.433 | -0.605 |
| P2RY12 | NM_022788 | 0.4405 | -0.2335 |
| IL25 | NM_022789 | -0.1475 | -0.263 |
| CTBP2 | NM_022802 | -0.9365 | -1.7355 |
| UCP3 | NM_022803 | -0.058 | -0.9395 |
| SNURF | NM_022804 | 0.104 | -0.313 |
| SNRPN | NM_022805 | 0.3275 | -0.2535 |
| SNRPN | NM_022806 | -0.837 | -1.1145 |
| SNRPN | NM_022807 | -0.837 | -1.1145 |
| SNRPN | NM_022808 | -0.837 | -1.1145 |
| CDC25C | NM_022809 | -1.3315 | -2.71 |
| SLC25A14 | NM_022810 | -0.4775 | -0.985 |
| PER2 | NM_022817 | -1.378 | -3.0765 |
| MAP1LC3B | NM_022818 | -0.215 | -0.3765 |
| PLA2G2F | NM_022819 | 0.0295 | 0.183 |
| CYP3A43 | NM_022820 | 0.0215 | -0.44 |
| ELOVL1 | NM_022821 | -0.863 | -1.419 |
| KLC2 | NM_022822 | -0.104 | -0.639 |
| FNDC4 | NM_022823 | 0.011 | -0.596 |
| 7-Mar | NM_022826 | -1.498 | -4.1035 |
| SPATA20 | NM_022827 | -0.541 | -0.3255 |

| | | | |
|---|---|---|---|
| YTHDC2 | NM_022828 | -1.3325 | -1.641 |
| SLC13A3 | NM_022829 | 0.38 | -0.2435 |
| TUT1 | NM_022830 | -0.509 | -1.5825 |
| AIDA | NM_022831 | -0.8095 | -1.07 |
| USP46 | NM_022832 | -0.482 | -1.186 |
| FAM129B | NM_022833 | -0.5435 | -1.9905 |
| VWA1 | NM_022834 | -0.1515 | -0.512 |
| PLEKHG2 | NM_022835 | -0.3425 | -0.5225 |
| DCLRE1B | NM_022836 | -1.3545 | -3.561 |
| ARMCX5 | NM_022838 | -1.5105 | -1.902 |
| MRPS11 | NM_022839 | -0.58 | -1.797 |
| METTL4 | NM_022840 | -1.5895 | -2.199 |
| RFX7 | NM_022841 | -1.021 | -3.295 |
| CDCP1 | NM_022842 | 0.152 | -0.2525 |
| PCDH20 | NM_022843 | -0.158 | -1.1885 |
| MYH11 | NM_022844 | -0.05 | -0.205 |
| CBFB | NM_022845 | -1.49 | -2.336 |
| IFI6 | NM_022873 | 0.224 | -0.6665 |
| SMN1 | NM_022874 | -1.2465 | -2.9415 |
| SMN2 | NM_022875 | -1.1185 | -1.331 |
| BCL11A | NM_022893 | -1.1425 | -2.253 |
| PAPOLG | NM_022894 | -1.329 | -4.017 |
| C12orf43 | NM_022895 | -1.163 | -2.618 |
| LPIN3 | NM_022896 | -0.1935 | -0.388 |
| RANBP17 | NM_022897 | -1.315 | -1.4495 |
| BCL11B | NM_022898 | -0.782 | -3.157 |
| ACTR8 | NM_022899 | -1.014 | -2.5905 |
| CASD1 | NM_022900 | -1.5065 | -2.5035 |
| LRRC19 | NM_022901 | 0.2045 | 0.059 |
| SLC30A5 | NM_022902 | -1.8635 | -2.8875 |
| CCDC71 | NM_022903 | -0.575 | -1.0895 |
| RASAL3 | NM_022904 | 0.1565 | -0.256 |
| TTC23 | NM_022905 | -0.5295 | -1.3105 |
| NT5DC2 | NM_022908 | 0.2165 | -0.149 |
| CENPH | NM_022909 | -0.488 | -0.213 |
| NDRG4 | NM_022910 | -0.529 | -0.8035 |
| SLC26A6 | NM_022911 | -0.1195 | -0.7715 |
| REEP1 | NM_022912 | -1.5215 | -1.2915 |
| GPBP1 | NM_022913 | -0.808 | -2.45 |
| ACD | NM_022914 | -0.4235 | -1.2055 |
| MRPL44 | NM_022915 | -0.5875 | -1.3895 |
| VPS33A | NM_022916 | -0.8685 | -2.551 |
| NOL6 | NM_022917 | -0.4965 | -1.276 |

| | | | |
|---|---|---|---|
| TMEM135 | NM_022918 | -0.9945 | -2.259 |
| FGFR4 | NM_022963 | -0.136 | -0.67 |
| FGFR3 | NM_022965 | 0.069 | 0.864 |
| FGFR2 | NM_022970 | -1.4165 | -3.3415 |
| ACSL4 | NM_022977 | -1.127 | -2.109 |
| SERF1B | NM_022978 | -0.557 | -2.199 |
| ARID4A | NM_023000 | -1.0025 | -2.4305 |
| ARID4A | NM_023001 | -1.0025 | -2.4305 |
| HAPLN4 | NM_023002 | 0.085 | -1.373 |
| TM6SF1 | NM_023003 | -0.2825 | -0.3745 |
| RTN4R | NM_023004 | -0.6095 | -1.9795 |
| JMJD4 | NM_023007 | -1.0045 | -1.808 |
| KRI1 | NM_023008 | -0.9375 | -2.073 |
| MARCKSL1 | NM_023009 | 0.06 | -0.343 |
| UPF3B | NM_023010 | -0.8705 | -1.3295 |
| UPF3A | NM_023011 | -1.2855 | -0.8695 |
| RSRC2 | NM_023012 | -1.086 | -2.288 |
| PRAMEF1 | NM_023013 | 0.136 | -0.412 |
| PRAMEF2 | NM_023014 | 0.0225 | -0.0275 |
| INTS3 | NM_023015 | -0.5055 | -1.3865 |
| ANKRD57 | NM_023016 | -0.6135 | -0.5495 |
| NADK | NM_023018 | 0.262 | -0.6915 |
| METTL1 | NM_023033 | -1.1365 | -2.773 |
| WHSC1L1 | NM_023034 | 0.074 | -0.524 |
| CACNA1A | NM_023035 | 0.246 | -0.356 |
| DNAI2 | NM_023036 | 0.384 | 0.2545 |
| FRY | NM_023037 | -1.5115 | -2.529 |
| ANKRA2 | NM_023039 | -2.04 | -2.665 |
| FOXL2 | NM_023067 | 0.2695 | -0.0835 |
| SIGLEC1 | NM_023068 | -0.0445 | -0.164 |
| ZNF643 | NM_023070 | -1.63 | -1.4125 |
| SPATS2 | NM_023071 | -1.1245 | -1.7995 |
| ZSWIM4 | NM_023072 | -0.773 | -1.6245 |
| C5orf42 | NM_023073 | -0.3555 | -1.526 |
| ZNF649 | NM_023074 | -0.22 | -1.248 |
| MPPE1 | NM_023075 | -1.091 | -1.281 |
| UNKL | NM_023076 | -1.258 | -1.8815 |
| SELRC1 | NM_023077 | -0.802 | -2.3235 |
| PYCRL | NM_023078 | -0.0015 | -0.5905 |
| UBE2Z | NM_023079 | -0.512 | -1.6895 |
| C8orf33 | NM_023080 | -0.0865 | -0.8765 |
| CAPN10 | NM_023083 | -0.345 | -1.3505 |
| CAPN10 | NM_023085 | -0.345 | -1.3505 |

| | | | |
|---|---|---|---|
| FGFR1 | NM_023105 | -0.0765 | -1.193 |
| FGFR1 | NM_023106 | -0.0765 | -1.193 |
| FGFR1 | NM_023110 | -0.0765 | -1.193 |
| OTUB2 | NM_023112 | -0.798 | -1.7985 |
| GPR87 | NM_023915 | 0.067 | 0.318 |
| TAS2R9 | NM_023917 | 0.1415 | -0.274 |
| TAS2R8 | NM_023918 | -0.1205 | -0.044 |
| TAS2R7 | NM_023919 | -0.1545 | -0.8525 |
| TAS2R13 | NM_023920 | 0.0035 | -0.038 |
| TAS2R10 | NM_023921 | -0.0185 | 0.1045 |
| TAS2R14 | NM_023922 | 0.7855 | -0.567 |
| PHACTR4 | NM_023923 | -1.2105 | -2.973 |
| BRD9 | NM_023924 | -0.3105 | -1.066 |
| CAPRIN2 | NM_023925 | -1.5535 | -2.0835 |
| ZSCAN18 | NM_023926 | -1.358 | -1.592 |
| GRAMD3 | NM_023927 | -0.033 | -0.2755 |
| AACS | NM_023928 | -0.3045 | -1.0795 |
| ZBTB10 | NM_023929 | -1.0225 | -1.6245 |
| KCTD14 | NM_023930 | -0.3215 | -0.261 |
| ZNF747 | NM_023931 | -0.8135 | -1.1595 |
| DLK2 | NM_023932 | -0.131 | -1.489 |
| FAM173A | NM_023933 | -0.1785 | -1.0785 |
| FUNDC2 | NM_023934 | 0.203 | -0.231 |
| DDRGK1 | NM_023935 | -0.2335 | -1.003 |
| MRPS34 | NM_023936 | -0.1955 | -1.0795 |
| MRPL34 | NM_023937 | -0.169 | -0.3405 |
| C1orf116 | NM_023938 | -0.096 | -0.247 |
| RASL11B | NM_023940 | -0.0875 | -0.971 |
| TMEM108 | NM_023943 | -0.148 | -2.2635 |
| CYP4F12 | NM_023944 | 0.081 | -0.3375 |
| MS4A5 | NM_023945 | 0.3105 | -0.132 |
| LYNX1 | NM_023946 | 0.0405 | -0.7195 |
| CHID1 | NM_023947 | 0.211 | -0.2895 |
| L1CAM | NM_024003 | 0.457 | -0.229 |
| VKORC1 | NM_024006 | 0.0205 | -0.64 |
| EBF1 | NM_024007 | -0.8785 | -1.3245 |
| GJB3 | NM_024009 | -0.3115 | -0.085 |
| MTRR | NM_024010 | -1.3955 | -3.014 |
| CDK11A | NM_024011 | -0.39 | -0.955 |
| HTR5A | NM_024012 | -0.2875 | -0.0285 |
| HOXA6 | NM_024014 | 0.0875 | -0.2845 |
| HOXB4 | NM_024015 | -0.2255 | -0.5255 |
| HOXB8 | NM_024016 | -0.0785 | -0.35 |

| | | | |
|---|---|---|---|
| HOXB9 | NM_024017 | -0.5685 | -0.408 |
| NEUROG2 | NM_024019 | -0.034 | 0.4495 |
| MS4A4A | NM_024021 | -0.252 | -0.338 |
| TMPRSS3 | NM_024022 | -0.3595 | -0.257 |
| DUSP26 | NM_024025 | -0.6355 | -1.5825 |
| MRP63 | NM_024026 | -0.984 | -1.9045 |
| PCYOX1L | NM_024028 | -0.998 | -2.027 |
| YIPF2 | NM_024029 | -0.16 | -0.642 |
| PRR14 | NM_024031 | -0.641 | -1.2585 |
| C17orf53 | NM_024032 | -0.793 | -2.348 |
| C7orf49 | NM_024033 | -1.075 | -2.6955 |
| GDAP1L1 | NM_024034 | 0.063 | -0.6595 |
| LRFN4 | NM_024036 | -0.061 | -1.239 |
| C1orf135 | NM_024037 | -0.53 | -2.154 |
| C19orf43 | NM_024038 | -0.0145 | -0.7525 |
| MIS12 | NM_024039 | -1.5915 | -2.7255 |
| CUEDC2 | NM_024040 | -0.037 | -0.532 |
| SCNM1 | NM_024041 | -0.742 | -1.738 |
| METRN | NM_024042 | 0.183 | -0.4395 |
| DBNDD1 | NM_024043 | -0.6185 | -1.1715 |
| SLX1B | NM_024044 | 0.181 | -0.7325 |
| DDX50 | NM_024045 | -1.3435 | -2.3475 |
| CAMKV | NM_024046 | -0.118 | -0.899 |
| NUDT9 | NM_024047 | -1.2055 | -1.7015 |
| DDA1 | NM_024050 | -0.082 | -0.7565 |
| GGCT | NM_024051 | -0.6585 | -1.0075 |
| C17orf39 | NM_024052 | -1.21 | -2.956 |
| SLC30A5 | NM_024055 | -0.1325 | -0.7165 |
| TMEM106C | NM_024056 | 0.009 | -0.2755 |
| NUP37 | NM_024057 | -1.8125 | -3.5525 |
| C20orf195 | NM_024059 | 0.11 | -0.5775 |
| AHNAK | NM_024060 | -0.867 | -1.2525 |
| ZNF655 | NM_024061 | 0.1805 | -0.147 |
| SPATA5L1 | NM_024063 | -1.133 | -2.7405 |
| PDCL3 | NM_024065 | -0.6595 | -1.2855 |
| ERI3 | NM_024066 | -0.12 | -0.977 |
| C7orf26 | NM_024067 | -0.746 | -2.299 |
| OBFC2B | NM_024068 | 0.0915 | -0.361 |
| C19orf50 | NM_024069 | -0.3865 | -1.2645 |
| PVRIG | NM_024070 | -0.2185 | 0.0905 |
| ZFYVE21 | NM_024071 | -0.025 | -0.635 |
| DDX54 | NM_024072 | 0.012 | -0.959 |
| TMEM38A | NM_024074 | -0.1345 | -0.556 |

| | | | |
|---|---|---|---|
| TSEN34 | NM_024075 | -0.4255 | -1.2195 |
| KCTD15 | NM_024076 | -0.9515 | -1.9305 |
| SECISBP2 | NM_024077 | -0.973 | -1.3355 |
| NOC4L | NM_024078 | -0.221 | -1.1045 |
| TRPM8 | NM_024080 | -0.222 | -0.1265 |
| PRRG4 | NM_024081 | -0.091 | -0.191 |
| PRRG3 | NM_024082 | -0.1105 | -0.311 |
| ASPSCR1 | NM_024083 | -0.345 | -1.0815 |
| ATG9A | NM_024085 | -0.346 | -1.619 |
| ASB9 | NM_024087 | 0.199 | -0.333 |
| KDELC1 | NM_024089 | -1.1165 | -1.78 |
| ELOVL6 | NM_024090 | -1.554 | -2.3615 |
| FASTKD3 | NM_024091 | -0.9645 | -1.674 |
| TMEM109 | NM_024092 | -0.62 | -1.8135 |
| C2orf49 | NM_024093 | -1.3265 | -2.0285 |
| DSCC1 | NM_024094 | -1.6065 | -3.401 |
| ASB8 | NM_024095 | -0.9825 | -3.0405 |
| DCTPP1 | NM_024096 | -0.29 | -0.949 |
| C1orf50 | NM_024097 | -0.5385 | -1.6855 |
| CCDC86 | NM_024098 | -0.569 | -2.1105 |
| C11orf48 | NM_024099 | -0.8095 | -1.5695 |
| WDR18 | NM_024100 | -0.0445 | -0.59 |
| MLPH | NM_024101 | 0.2095 | -0.456 |
| WDR77 | NM_024102 | -0.057 | -0.567 |
| SLC25A23 | NM_024103 | 0.233 | -0.0615 |
| C19orf42 | NM_024104 | -0.308 | -1.1095 |
| ALG12 | NM_024105 | -0.217 | -1.409 |
| ZNF426 | NM_024106 | -1.1635 | -2.067 |
| TRAPPC6A | NM_024108 | 0.2645 | -0.297 |
| CARD14 | NM_024110 | -0.2475 | -0.0885 |
| CHAC1 | NM_024111 | -0.766 | -0.7025 |
| C9orf16 | NM_024112 | 0.5165 | -0.126 |
| C11orf49 | NM_024113 | -0.31 | -0.7105 |
| TRIM48 | NM_024114 | -0.034 | -0.164 |
| TAF1D | NM_024116 | -0.52 | 0.076 |
| MAPKAP1 | NM_024117 | -0.387 | -1.259 |
| DHX58 | NM_024119 | -0.061 | 0.488 |
| C20orf7 | NM_024120 | -0.965 | -2.578 |
| TMEM185B | NM_024121 | -1.1335 | -2.4835 |
| APOO | NM_024122 | -0.64 | -0.5895 |
| TPSB2 | NM_024164 | -0.423 | -0.2305 |
| PHF1 | NM_024165 | -0.045 | -0.691 |
| UBL5 | NM_024292 | 0.2045 | -0.098 |

| | | | |
|---|---|---|---|
| FAM134A | NM_024293 | -0.76 | -1.195 |
| C6orf106 | NM_024294 | -0.2165 | -1.6465 |
| DERL1 | NM_024295 | -1.0645 | -1.3215 |
| CCDC28B | NM_024296 | 0.165 | -0.5575 |
| PHF23 | NM_024297 | -1.15 | -3.7635 |
| MBOAT7 | NM_024298 | -1.4555 | -1.2885 |
| PPDPF | NM_024299 | 0.173 | -0.2705 |
| CHCHD7 | NM_024300 | -0.5065 | -0.8085 |
| FKRP | NM_024301 | -1.1795 | -1.837 |
| MMP28 | NM_024302 | -0.2615 | -0.548 |
| ZSCAN5A | NM_024303 | -0.818 | -0.935 |
| FA2H | NM_024306 | -0.2135 | -0.538 |
| GDPD3 | NM_024307 | -0.2595 | -0.8625 |
| DHRS11 | NM_024308 | 0.132 | -0.8485 |
| TNIP2 | NM_024309 | -0.3345 | -1.457 |
| PLEKHF1 | NM_024310 | -0.1635 | -0.8735 |
| MFSD11 | NM_024311 | -0.679 | -1.3765 |
| GNPTAB | NM_024312 | -0.544 | -1.4135 |
| NOL12 | NM_024313 | -0.605 | -1.3265 |
| C7orf23 | NM_024315 | -1.1985 | -2.5605 |
| LENG1 | NM_024316 | -0.7105 | -2.0975 |
| LILRA6 | NM_024318 | -0.035 | 0.102 |
| C1orf35 | NM_024319 | -0.6475 | -1.9225 |
| PRR15L | NM_024320 | 0.468 | -0.3135 |
| RBM42 | NM_024321 | -0.5475 | -1.7445 |
| CENPO | NM_024322 | -0.4035 | -1.9545 |
| C19orf57 | NM_024323 | 0.0755 | -0.65 |
| CRELD2 | NM_024324 | -0.141 | -0.287 |
| ZNF343 | NM_024325 | -1.204 | -2.059 |
| FBXL15 | NM_024326 | -0.067 | -0.929 |
| ZNF576 | NM_024327 | -0.7 | -2.022 |
| THTPA | NM_024328 | -0.6035 | -0.637 |
| EFHD2 | NM_024329 | -0.4865 | -1.128 |
| SLC27A3 | NM_024330 | -0.309 | -0.909 |
| TTPAL | NM_024331 | -1.6715 | -3.6475 |
| BRCC3 | NM_024332 | -0.2775 | -1.067 |
| FSD1 | NM_024333 | -0.019 | -0.7495 |
| TMEM43 | NM_024334 | -1.2535 | -1.912 |
| IRX6 | NM_024335 | -0.143 | -0.1 |
| IRX3 | NM_024336 | -0.701 | -1.0075 |
| IRX1 | NM_024337 | 0.249 | 0.5075 |
| THOC6 | NM_024339 | -0.1825 | -1.684 |
| ZNF557 | NM_024341 | -1.963 | -3.434 |

| | | | |
|---|---|---|---|
| CAPN3 | NM_024344 | -0.009 | -0.367 |
| DCAF10 | NM_024345 | -1.4615 | -3.59 |
| DCTN3 | NM_024348 | 0.171 | -0.4205 |
| NDUFS7 | NM_024407 | 0.188 | -0.23 |
| NOTCH2 | NM_024408 | -0.0975 | -0.29 |
| NPPC | NM_024409 | -0.117 | 0.304 |
| ODF1 | NM_024410 | 0.08 | -0.1725 |
| PDYN | NM_024411 | -0.3215 | -0.2845 |
| DDX4 | NM_024415 | -0.1035 | -0.1605 |
| FDXR | NM_024417 | -0.0465 | -0.6465 |
| BZRAP1 | NM_024418 | 0.1195 | -0.5245 |
| PGS1 | NM_024419 | -1.0405 | -2.611 |
| PLA2G4A | NM_024420 | -0.1925 | 0.4515 |
| DSC2 | NM_024422 | -0.9115 | -0.5975 |
| DSC3 | NM_024423 | -0.6245 | -0.519 |
| WT1 | NM_024424 | 0.373 | -0.2905 |
| WT1 | NM_024426 | 0.373 | -0.2905 |
| PRKAG2 | NM_024429 | -0.789 | -1.3805 |
| PSTPIP2 | NM_024430 | -0.3565 | -1.2255 |
| ATP10A | NM_024490 | 0.318 | -0.148 |
| CEP70 | NM_024491 | 0.0215 | -0.2565 |
| ZKSCAN3 | NM_024493 | -0.6935 | -1.886 |
| WNT2B | NM_024494 | -0.034 | -0.266 |
| IRF2BPL | NM_024496 | 0.004 | 0.129 |
| HOXD1 | NM_024501 | -0.293 | 0.2155 |
| HIVEP3 | NM_024503 | 0.0365 | -0.5415 |
| PRDM14 | NM_024504 | -0.2215 | -0.2495 |
| NOX5 | NM_024505 | -0.2865 | -0.428 |
| GLB1L | NM_024506 | -0.041 | 0.161 |
| KREMEN2 | NM_024507 | -0.175 | -0.2055 |
| ZBED2 | NM_024508 | -0.155 | 0.1335 |
| LRFN3 | NM_024509 | -0.2755 | -1.522 |
| HAUS3 | NM_024511 | -0.9695 | -1.39 |
| LRRC2 | NM_024512 | 0.0445 | -1.5135 |
| FYCO1 | NM_024513 | 0.106 | -0.3215 |
| CYP2R1 | NM_024514 | -1.4195 | -2.7905 |
| WDR25 | NM_024515 | -0.0745 | -0.562 |
| C16orf53 | NM_024516 | -0.65 | -1.948 |
| ULBP3 | NM_024518 | -0.5785 | -0.728 |
| FAM65A | NM_024519 | -0.3885 | -1.086 |
| C2orf47 | NM_024520 | -0.96 | -3.0115 |
| NKAIN1 | NM_024522 | 0.1225 | -0.5825 |
| GCC1 | NM_024523 | -1.253 | -1.999 |

| | | | |
|---|---|---|---|
| ATP13A3 | NM_024524 | -1.131 | -1.935 |
| TTC13 | NM_024525 | -1.744 | -2.6015 |
| EPS8L3 | NM_024526 | 0.1025 | -0.1265 |
| ABHD8 | NM_024527 | -0.0665 | -0.5395 |
| NKAP | NM_024528 | -1.244 | -2.4 |
| CDC73 | NM_024529 | -1.012 | -2.281 |
| GPR172A | NM_024531 | -0.4355 | -1.02 |
| SPAG16 | NM_024532 | -0.8395 | -1.181 |
| CHST5 | NM_024533 | 0.4405 | -0.193 |
| CORO7 | NM_024535 | -0.336 | -0.551 |
| CHPF | NM_024536 | -0.043 | -0.6745 |
| CARS2 | NM_024537 | -0.1725 | -0.872 |
| RNF128 | NM_024539 | -0.0755 | -0.2425 |
| C10orf76 | NM_024541 | -0.897 | -2.267 |
| MUL1 | NM_024544 | -0.586 | -1.7175 |
| SAP130 | NM_024545 | -0.6475 | -2.1355 |
| RNF219 | NM_024546 | -1.025 | -3.7045 |
| CEP97 | NM_024548 | -0.7165 | -1.6625 |
| TCTN1 | NM_024549 | -0.269 | -0.5095 |
| ADIPOR2 | NM_024551 | -0.7505 | -2.8405 |
| CERS4 | NM_024552 | -0.1605 | -0.72 |
| CCDC132 | NM_024553 | -1.095 | -2.395 |
| PGBD5 | NM_024554 | -0.804 | -3.8775 |
| FBXL6 | NM_024555 | -0.133 | -0.58 |
| FAM118B | NM_024556 | -1.483 | -3.134 |
| RIC3 | NM_024557 | -0.736 | -1.2185 |
| METTL21D | NM_024558 | -1.9045 | -3.2035 |
| ACSS3 | NM_024560 | -0.401 | -0.1355 |
| NAA16 | NM_024561 | -1.0985 | -2.0525 |
| TMCO7 | NM_024562 | -0.5855 | -1.8195 |
| CCNJL | NM_024565 | -1.0205 | -3.353 |
| HMBOX1 | NM_024567 | -0.2745 | -1.3475 |
| MPZL1 | NM_024569 | -0.7115 | -1.592 |
| RNASEH2B | NM_024570 | -0.8505 | -1.7145 |
| SNRNP25 | NM_024571 | -0.044 | -0.4735 |
| GALNT14 | NM_024572 | -0.3725 | -1.7895 |
| C6orf211 | NM_024573 | -1.099 | -3.0505 |
| NDNF | NM_024574 | -0.516 | -1.743 |
| TNFAIP8L2 | NM_024575 | 0.2375 | -0.084 |
| OGFRL1 | NM_024576 | -0.536 | -0.436 |
| SH3TC2 | NM_024577 | 0.2125 | -0.2145 |
| OCEL1 | NM_024578 | -0.2745 | -0.7675 |
| C1orf54 | NM_024579 | -0.2615 | -0.1925 |

| | | | |
|---|---|---|---|
| EFTUD1 | NM_024580 | -0.902 | -2.5115 |
| FAM184A | NM_024581 | -0.605 | -2.9825 |
| FAT4 | NM_024582 | -0.386 | -2.182 |
| SCRN3 | NM_024583 | -1.4825 | -2.216 |
| ARMC7 | NM_024585 | -1.1415 | -1.973 |
| OSBPL9 | NM_024586 | -0.581 | -1.5355 |
| TMEM53 | NM_024587 | -0.1075 | -0.74 |
| ROGDI | NM_024589 | 0.044 | -0.561 |
| ARSJ | NM_024590 | -0.024 | -0.448 |
| CHMP6 | NM_024591 | -1.119 | -2.7295 |
| SRD5A3 | NM_024592 | -0.29 | -0.7245 |
| EFCAB1 | NM_024593 | 0.086 | -0.244 |
| PANK3 | NM_024594 | -0.3525 | -0.859 |
| AKIRIN1 | NM_024595 | -0.918 | -1.868 |
| MCPH1 | NM_024596 | -0.097 | -1.6855 |
| MAP7D3 | NM_024597 | -0.876 | -1.4865 |
| C16orf57 | NM_024598 | -0.0965 | -0.5025 |
| RHBDF2 | NM_024599 | -0.1065 | -0.425 |
| TMEM204 | NM_024600 | 0.1665 | -0.1375 |
| HECTD3 | NM_024602 | -0.157 | -1.069 |
| BEND5 | NM_024603 | -1.024 | -2.4895 |
| RPAP3 | NM_024604 | -0.424 | -2.4425 |
| ARHGAP10 | NM_024605 | -0.5775 | -2.3105 |
| PPP1R3B | NM_024607 | -0.7655 | -1.9515 |
| NEIL1 | NM_024608 | -0.3155 | -0.8255 |
| HSPBAP1 | NM_024610 | -1.3715 | -2.321 |
| NARG2 | NM_024611 | -0.979 | -2.575 |
| DHX40 | NM_024612 | -0.8125 | -0.81 |
| PLEKHF2 | NM_024613 | -1.113 | -1.369 |
| PARP8 | NM_024615 | -1.0815 | -1.577 |
| C3orf52 | NM_024616 | -0.2535 | -0.4715 |
| ZCCHC6 | NM_024617 | -0.4 | -2.221 |
| NLRX1 | NM_024618 | -0.67 | -1.5965 |
| FN3KRP | NM_024619 | -0.0985 | -1.3255 |
| ZNF329 | NM_024620 | -1.057 | -3.0745 |
| VEPH1 | NM_024621 | 0.139 | -0.02 |
| FASTKD1 | NM_024622 | -1.722 | -2.248 |
| OGFOD2 | NM_024623 | -0.154 | -1.04 |
| SMC6 | NM_024624 | -0.2945 | -2.4585 |
| ZC3HAV1 | NM_024625 | -1.124 | -2.0005 |
| VTCN1 | NM_024626 | 0.1005 | -0.237 |
| C22orf29 | NM_024627 | -1.04 | -1.675 |
| SLC12A8 | NM_024628 | 0.012 | -0.3145 |

| | | | |
|---|---|---|---|
| MLF1IP | NM_024629 | -0.515 | -1.39 |
| ZDHHC14 | NM_024630 | -0.303 | -1.449 |
| C11orf61 | NM_024631 | -0.699 | -2.6245 |
| SAP30L | NM_024632 | -0.9355 | -2.2765 |
| NAA35 | NM_024635 | -0.7375 | -2.189 |
| STEAP4 | NM_024636 | 0.0535 | -0.2615 |
| GAL3ST4 | NM_024637 | -0.5815 | -0.501 |
| QTRTD1 | NM_024638 | -1.854 | -2.867 |
| YRDC | NM_024640 | -0.6115 | -1.1655 |
| MANEA | NM_024641 | -0.2505 | -1.734 |
| GALNT12 | NM_024642 | -0.8075 | -1.0315 |
| FAM164C | NM_024643 | -0.749 | -0.7335 |
| C14orf169 | NM_024644 | -0.9915 | -2.2145 |
| ZMAT4 | NM_024645 | -1.0205 | -2.5145 |
| ZYG11B | NM_024646 | -1.3935 | -1.952 |
| C17orf101 | NM_024648 | 0.0565 | -0.7425 |
| BBS1 | NM_024649 | -0.365 | -0.8405 |
| C11orf80 | NM_024650 | 0.2015 | -0.3075 |
| LRRK1 | NM_024652 | -0.007 | -0.961 |
| PRKRIP1 | NM_024653 | -0.806 | -2.2815 |
| NOL9 | NM_024654 | 0.0575 | -1.293 |
| GLT25D1 | NM_024656 | -0.285 | -1.049 |
| MORC4 | NM_024657 | -0.482 | -0.7485 |
| IPO4 | NM_024658 | -0.0045 | -0.802 |
| GTDC1 | NM_024659 | -0.8465 | -1.3465 |
| IGFLR1 | NM_024660 | -0.198 | -0.6665 |
| CCDC51 | NM_024661 | -0.8725 | -2.238 |
| NAT10 | NM_024662 | -0.4635 | -2.4435 |
| NPEPL1 | NM_024663 | 0.113 | -0.2505 |
| PPCS | NM_024664 | -1.314 | -2.014 |
| TBL1XR1 | NM_024665 | -0.8815 | -1.618 |
| AAGAB | NM_024666 | -0.7395 | -0.9125 |
| VPS37B | NM_024667 | -0.6765 | -2.6645 |
| ANKHD1 | NM_024668 | 0.2675 | -0.5015 |
| ANKRD55 | NM_024669 | 0.4865 | -0.0975 |
| SUV39H2 | NM_024670 | -1.187 | -2.8455 |
| ZNF768 | NM_024671 | -0.224 | -0.789 |
| THAP9 | NM_024672 | -1.0365 | -2.716 |
| LIN28A | NM_024674 | -0.149 | -0.601 |
| PALB2 | NM_024675 | -1.8535 | -3.5655 |
| SH3D21 | NM_024676 | -0.4665 | -0.4465 |
| NSUN7 | NM_024677 | 0.4175 | -0.3635 |
| NARS2 | NM_024678 | -1.12 | -1.875 |

| | | | |
|---|---|---|---|
| E2F8 | NM_024680 | -1.4425 | -2.673 |
| KCTD17 | NM_024681 | -0.099 | -0.6515 |
| TBC1D17 | NM_024682 | -0.2975 | -0.8775 |
| C17orf42 | NM_024683 | -1.4835 | -3.186 |
| C11orf67 | NM_024684 | -0.096 | -0.672 |
| BBS10 | NM_024685 | -1.729 | -3.099 |
| TTLL7 | NM_024686 | -0.13 | -1.142 |
| ZBBX | NM_024687 | -0.29 | 0.098 |
| C10orf68 | NM_024688 | 0.298 | -0.0855 |
| CXorf36 | NM_024689 | 0.13 | -0.2245 |
| MUC16 | NM_024690 | -0.224 | -0.0125 |
| ZNF419 | NM_024691 | -1.2995 | -2.445 |
| CLIP4 | NM_024692 | -1.7015 | -2.757 |
| ECHDC3 | NM_024693 | -0.8245 | -2.69 |
| C6orf103 | NM_024694 | -0.3315 | -0.409 |
| ZNF385D | NM_024697 | -0.4385 | -0.566 |
| SLC25A22 | NM_024698 | -0.3325 | 0.2085 |
| ZFAND1 | NM_024699 | -1.4555 | -1.211 |
| SNIP1 | NM_024700 | -1.105 | -2.317 |
| ASB13 | NM_024701 | -0.681 | -1.707 |
| ZNF750 | NM_024702 | 0.303 | 1.1625 |
| KIF16B | NM_024704 | 0.0925 | -0.7335 |
| DHRS12 | NM_024705 | 0.052 | -0.6175 |
| ZNF668 | NM_024706 | -0.364 | -1.7575 |
| GEMIN7 | NM_024707 | -1.229 | -2.997 |
| ASB7 | NM_024708 | 0.0485 | -0.782 |
| C1orf115 | NM_024709 | -0.162 | -0.51 |
| GIMAP6 | NM_024711 | 0.054 | 0.0995 |
| ELMO3 | NM_024712 | -0.1835 | -1.209 |
| C15orf29 | NM_024713 | -0.98 | -1.9355 |
| TXNDC15 | NM_024715 | -0.1585 | -1.83 |
| MCTP1 | NM_024717 | -0.0905 | -0.036 |
| C9orf86 | NM_024718 | -0.3215 | -0.834 |
| GRTP1 | NM_024719 | -0.154 | -0.587 |
| ZFHX4 | NM_024721 | -0.974 | -3.626 |
| ACBD4 | NM_024722 | -0.031 | -0.6915 |
| CCDC82 | NM_024725 | -1.5 | -2.295 |
| IQCA1 | NM_024726 | -0.1445 | -0.0685 |
| LRRC31 | NM_024727 | 0.122 | -0.29 |
| C7orf10 | NM_024728 | -0.2055 | -0.7715 |
| MYH14 | NM_024729 | -0.1625 | 0.0935 |
| RERGL | NM_024730 | 0.095 | 0.0335 |
| KLHL36 | NM_024731 | 0.1455 | -0.5845 |

| | | | |
|---|---|---|---|
| ZNF665 | NM_024733 | -0.0275 | -0.728 |
| CLMN | NM_024734 | -0.32 | -0.7385 |
| FBXO31 | NM_024735 | -0.804 | -2.416 |
| GSDMD | NM_024736 | -0.0245 | -0.3735 |
| C12orf49 | NM_024738 | 0.497 | -0.3945 |
| ALG9 | NM_024740 | -0.122 | -0.8265 |
| ZNF408 | NM_024741 | -0.222 | -1.2965 |
| ARMC5 | NM_024742 | 0.003 | -0.978 |
| UGT2A3 | NM_024743 | -0.228 | -0.184 |
| ALS2CR8 | NM_024744 | -0.28 | -0.1025 |
| SHCBP1 | NM_024745 | -1.7 | -2.3605 |
| HHIPL2 | NM_024746 | -0.3565 | -0.3085 |
| HPS6 | NM_024747 | -1.245 | -3.0945 |
| VASH2 | NM_024749 | -1.685 | -2.4415 |
| GSTCD | NM_024751 | -1.968 | -2.4 |
| TTC21B | NM_024753 | -0.7355 | -1.9675 |
| PTCD2 | NM_024754 | -1.152 | -2.205 |
| SLTM | NM_024755 | -0.6445 | -1.596 |
| MMRN2 | NM_024756 | -0.0695 | -0.2265 |
| EHMT1 | NM_024757 | -0.5765 | -2.0305 |
| AGMAT | NM_024758 | 0.0545 | 0.096 |
| NIPAL2 | NM_024759 | -0.1045 | -0.7025 |
| TLE6 | NM_024760 | -0.071 | -0.606 |
| MOB3B | NM_024761 | -1.004 | -0.9665 |
| ZNF552 | NM_024762 | -1.0185 | -2.381 |
| WDR78 | NM_024763 | 0.079 | -0.3585 |
| CATSPERB | NM_024764 | -0.1785 | -0.265 |
| CAMKMT | NM_024766 | -0.5715 | -0.597 |
| DLC1 | NM_024767 | -1.246 | -2.5865 |
| CCDC48 | NM_024768 | -0.0075 | -1.157 |
| CLMP | NM_024769 | -0.9105 | -1.0705 |
| METTL8 | NM_024770 | -0.799 | -2.1005 |
| NAA40 | NM_024771 | -0.5215 | -1.5785 |
| ZMYM1 | NM_024772 | -0.8375 | -2.2895 |
| JMJD5 | NM_024773 | -0.358 | -1.3565 |
| GEMIN6 | NM_024775 | -1.6655 | -3.399 |
| LONRF3 | NM_024778 | 0.081 | -0.4965 |
| TMC5 | NM_024780 | 0.147 | 0.37 |
| CCDC102B | NM_024781 | 0.0095 | -0.082 |
| NHEJ1 | NM_024782 | -0.394 | -1.719 |
| AGBL2 | NM_024783 | -0.1885 | -0.5505 |
| ZBTB3 | NM_024784 | -0.836 | -2.606 |
| FAM124B | NM_024785 | 0.115 | -0.259 |

| | | | |
|---|---|---|---|
| ZDHHC11 | NM_024786 | -0.589 | -0.5895 |
| RNF122 | NM_024787 | -0.2415 | -0.92 |
| TMEM180 | NM_024789 | -0.4975 | -0.956 |
| CSPP1 | NM_024790 | -1.687 | -3.547 |
| PDZD3 | NM_024791 | -0.073 | -0.4995 |
| FAM57A | NM_024792 | -1.171 | -2.7655 |
| CLUAP1 | NM_024793 | -1.01 | -1.4345 |
| EPHX3 | NM_024794 | -0.3675 | -0.3435 |
| TM4SF20 | NM_024795 | 0.2685 | 0.106 |
| SNX22 | NM_024798 | -0.191 | -0.4375 |
| NEK11 | NM_024800 | -0.304 | -0.3635 |
| TUBAL3 | NM_024803 | -0.028 | -0.454 |
| ZNF669 | NM_024804 | -0.5315 | -1.281 |
| RBFA | NM_024805 | -0.1975 | -0.9555 |
| C11orf63 | NM_024806 | -0.461 | -1.26 |
| TREML2 | NM_024807 | 0.132 | -0.4435 |
| BORA | NM_024808 | -1.295 | -2.805 |
| TCTN2 | NM_024809 | -0.57 | -0.842 |
| CPSF7 | NM_024811 | -1.14 | -2.4285 |
| BAALC | NM_024812 | 0.117 | -0.706 |
| RPAP2 | NM_024813 | -1.218 | -2.2825 |
| CBLL1 | NM_024814 | -1.538 | -3.4125 |
| NUDT18 | NM_024815 | -0.1715 | -0.7735 |
| RABEP2 | NM_024816 | -0.3415 | -0.843 |
| THSD4 | NM_024817 | -0.008 | -0.1615 |
| UBA5 | NM_024818 | -0.5925 | -1.1045 |
| DCAKD | NM_024819 | -0.2265 | -0.7385 |
| DENND1A | NM_024820 | 0.73 | 0.0485 |
| CCDC134 | NM_024821 | -0.0175 | -0.31 |
| ZC3H14 | NM_024824 | -0.9785 | -2.06 |
| PODNL1 | NM_024825 | 0.1085 | -0.0735 |
| HDAC11 | NM_024827 | 0.1065 | -0.638 |
| C9orf82 | NM_024828 | -1.624 | -3.2525 |
| PLBD1 | NM_024829 | -0.4155 | -0.2155 |
| LPCAT1 | NM_024830 | -0.4765 | -1.1155 |
| TGS1 | NM_024831 | -0.837 | -3.884 |
| RIN3 | NM_024832 | -0.0825 | 0.0325 |
| ZNF671 | NM_024833 | -0.87 | -2.104 |
| GGNBP2 | NM_024835 | -1.492 | -2.301 |
| ZNF672 | NM_024836 | -0.931 | -3.2255 |
| ATP8B4 | NM_024837 | 0.091 | -0.1925 |
| THNSL1 | NM_024838 | -1.4215 | -3.4705 |
| ZNF613 | NM_024840 | -0.7815 | -2.612 |

| | | | |
|---|---|---|---|
| PRR5L | NM_024841 | 0.1745 | -0.223 |
| CYBRD1 | NM_024843 | -0.2815 | -0.442 |
| NUP85 | NM_024844 | -0.71 | -2.2875 |
| TMC7 | NM_024847 | -0.2535 | -0.4015 |
| MORN1 | NM_024848 | -0.292 | -0.242 |
| BTNL8 | NM_024850 | -0.0105 | 0.1015 |
| EIF2C3 | NM_024852 | -1.2195 | -1.746 |
| PYROXD1 | NM_024854 | -0.6295 | -1.92 |
| ACTR5 | NM_024855 | -0.59 | -2.09 |
| ATAD5 | NM_024857 | -0.937 | -2.0535 |
| MAGIX | NM_024859 | -0.2885 | -0.31 |
| SETD6 | NM_024860 | -0.143 | -0.7975 |
| C2orf54 | NM_024861 | 0.0125 | -0.4925 |
| TCEAL4 | NM_024863 | -0.824 | -1.113 |
| MRM1 | NM_024864 | -0.854 | -1.1165 |
| NANOG | NM_024865 | 0.087 | -0.337 |
| ADM2 | NM_024866 | 0.008 | -0.6315 |
| SPEF2 | NM_024867 | 0.008 | -0.2905 |
| GRRP1 | NM_024869 | -0.064 | 0.018 |
| PREX2 | NM_024870 | -0.197 | -0.3015 |
| MAP6D1 | NM_024871 | -0.242 | -1.1965 |
| DOK3 | NM_024872 | -0.0635 | -0.4125 |
| TNIP3 | NM_024873 | 0.3175 | -0.134 |
| KIAA0319L | NM_024874 | -0.136 | -0.8625 |
| SYNPO2L | NM_024875 | -0.0025 | 0.267 |
| ADCK4 | NM_024876 | -0.433 | -1.817 |
| CNTD2 | NM_024877 | 0.218 | -0.1265 |
| SLC35E1 | NM_024881 | -0.6185 | -1.6975 |
| L2HGDH | NM_024884 | -0.5335 | -1.3 |
| TAF7L | NM_024885 | -0.556 | -0.1925 |
| C10orf95 | NM_024886 | -0.095 | -0.1885 |
| DHDDS | NM_024887 | -0.572 | -1.322 |
| LPPR3 | NM_024888 | 0.2005 | -0.3615 |
| C10orf81 | NM_024889 | 0.1905 | -0.272 |
| SYNDIG1 | NM_024893 | -0.3345 | -1.2405 |
| NOL10 | NM_024894 | -1.2605 | -1.923 |
| PDZD7 | NM_024895 | -0.036 | -0.842 |
| ERMP1 | NM_024896 | -0.547 | -0.4235 |
| PAQR6 | NM_024897 | -0.157 | -0.2395 |
| DENND1C | NM_024898 | 0.16 | -0.412 |
| CEP76 | NM_024899 | -0.605 | -1.133 |
| PHF17 | NM_024900 | -1.568 | -2.9265 |
| DENND2D | NM_024901 | -0.46 | -0.0815 |

| | | | |
|---|---|---|---|
| DNAJC22 | NM_024902 | -0.0695 | -0.2795 |
| SCD5 | NM_024906 | 0.1855 | -0.498 |
| FBXO17 | NM_024907 | -0.078 | -0.654 |
| WDR76 | NM_024908 | -0.3815 | -2.414 |
| ATAT1 | NM_024909 | -0.7065 | -1.499 |
| WLS | NM_024911 | -0.092 | -0.74 |
| C7orf58 | NM_024913 | -0.0785 | -0.0015 |
| GRHL2 | NM_024915 | -0.121 | -0.3055 |
| TRMT2B | NM_024917 | 0.3785 | -0.6125 |
| DSN1 | NM_024918 | -1.969 | -2.419 |
| FRMD1 | NM_024919 | 0.17 | -0.1935 |
| POF1B | NM_024921 | 0.4555 | -0.6105 |
| CES3 | NM_024922 | -0.159 | -0.316 |
| NUP210 | NM_024923 | -0.416 | -1.4915 |
| TTC26 | NM_024926 | 0.077 | -0.6115 |
| PLEKHH3 | NM_024927 | -0.2425 | -0.835 |
| OBFC1 | NM_024928 | -0.7475 | -1.383 |
| ELOVL7 | NM_024930 | 0.237 | -1.5305 |
| ZCCHC4 | NM_024936 | -2.0625 | -3.1035 |
| ESRP2 | NM_024939 | -0.0305 | -0.441 |
| DOCK5 | NM_024940 | -1.014 | -2.254 |
| C5orf44 | NM_024941 | -1.5265 | -2.229 |
| C10orf88 | NM_024942 | -2.215 | -4.059 |
| TMEM156 | NM_024943 | 0.177 | -0.7755 |
| CHODL | NM_024944 | -0.281 | -1.2275 |
| RMI1 | NM_024945 | -1.381 | -2.978 |
| FAM192A | NM_024946 | -0.5095 | -1.0945 |
| PHC3 | NM_024947 | -1.3185 | -2.5795 |
| FAM188A | NM_024948 | -1.1 | -2.2415 |
| WWC2 | NM_024949 | -1.3455 | -2.032 |
| C14orf159 | NM_024952 | 0.069 | -0.2425 |
| NAA25 | NM_024953 | -0.9565 | -1.23 |
| UBTD1 | NM_024954 | -0.016 | -0.7905 |
| FOXRED2 | NM_024955 | -0.8835 | -1.66 |
| TMEM62 | NM_024956 | -0.2935 | -1.424 |
| NRSN2 | NM_024958 | -0.391 | -0.7165 |
| SLC24A6 | NM_024959 | -0.0985 | -0.876 |
| PANK2 | NM_024960 | 0.093 | -0.087 |
| FBXL18 | NM_024963 | -1.3185 | -2.0225 |
| SEMA6D | NM_024966 | -0.2595 | -0.628 |
| ZNF556 | NM_024967 | 0.4025 | -0.3945 |
| CSRNP3 | NM_024969 | -0.2085 | -0.7845 |
| TRPM3 | NM_024971 | 0.08 | 0.028 |

| | | | |
|---|---|---|---|
| MCF2L | NM_024979 | -0.714 | -1.049 |
| GPR157 | NM_024980 | 0.089 | -0.3545 |
| PGAP1 | NM_024989 | -0.681 | -1.2385 |
| SORBS1 | NM_024991 | -0.3905 | -0.84 |
| LRRTM4 | NM_024993 | 0.357 | -0.345 |
| GFM1 | NM_024996 | -0.4135 | -1.1175 |
| ATF7IP2 | NM_024997 | 0.041 | -2.9005 |
| DCAF17 | NM_025000 | -1.371 | -2.3295 |
| ADAMTS20 | NM_025003 | -0.333 | -0.536 |
| CCDC15 | NM_025004 | -0.446 | -1.529 |
| ADAMTSL4 | NM_025008 | -0.1785 | -0.505 |
| CEP135 | NM_025009 | -0.8545 | -2.7015 |
| KLHL18 | NM_025010 | -0.7055 | -3.103 |
| ARHGEF15 | NM_025014 | 0.3835 | 0.0975 |
| HSPA12A | NM_025015 | -1.383 | -1.5065 |
| ZNF606 | NM_025027 | -0.432 | -2.486 |
| MZT2B | NM_025029 | 0.177 | -0.2405 |
| C7orf69 | NM_025031 | -0.4595 | -0.2525 |
| ZNF614 | NM_025040 | -0.9695 | -2.586 |
| C3orf36 | NM_025041 | -0.0785 | -0.2935 |
| BAIAP2L2 | NM_025045 | 0.125 | -0.154 |
| ARL14 | NM_025047 | 0.17 | -0.683 |
| GPR110 | NM_025048 | 0.3135 | -0.1005 |
| PIF1 | NM_025049 | -1.0065 | -1.113 |
| YSK4 | NM_025052 | -0.136 | -0.2335 |
| VCPIP1 | NM_025054 | -1.1745 | -3.1775 |
| CCDC33 | NM_025055 | 0.1785 | -0.24 |
| C14orf45 | NM_025057 | 0.136 | -0.153 |
| TRIM46 | NM_025058 | -0.1665 | -0.4955 |
| C6orf97 | NM_025059 | -0.026 | 0.1555 |
| LRRC8E | NM_025061 | -0.1545 | -0.1545 |
| C1orf129 | NM_025063 | 0.465 | 0.2465 |
| RPF1 | NM_025065 | -1.1785 | -1.9685 |
| ZNF703 | NM_025069 | 0.2375 | -0.518 |
| PTGES2 | NM_025072 | 0.0425 | -0.892 |
| SIKE1 | NM_025073 | -1.0135 | -1.155 |
| FRAS1 | NM_025074 | -0.841 | -0.736 |
| THOC7 | NM_025075 | -0.8435 | -1.232 |
| UXS1 | NM_025076 | -0.6155 | -1.5585 |
| TOE1 | NM_025077 | -0.9365 | -1.7525 |
| PQLC1 | NM_025078 | -0.204 | -0.913 |
| ZC3H12A | NM_025079 | -0.163 | -0.302 |
| ASRGL1 | NM_025080 | -0.61 | -1.0175 |

| | | | |
|---|---|---|---|
| NYNRIN | NM_025081 | -0.4025 | -1.655 |
| CENPT | NM_025082 | -0.3895 | -1.168 |
| EDC3 | NM_025083 | -0.6485 | -2.21 |
| CWH43 | NM_025087 | 0.203 | -0.452 |
| USP36 | NM_025090 | -0.49 | -2.333 |
| ATHL1 | NM_025092 | -0.165 | -0.3915 |
| MOGAT2 | NM_025098 | -0.206 | -0.4525 |
| CTC1 | NM_025099 | -1.3515 | -1.003 |
| DBF4B | NM_025104 | -1.309 | -3.3195 |
| SPSB1 | NM_025106 | 0.5955 | -0.648 |
| MYCT1 | NM_025107 | -0.139 | -0.5035 |
| C16orf59 | NM_025108 | -0.655 | -1.717 |
| MYO19 | NM_025109 | -1.2365 | -2.001 |
| ZXDC | NM_025112 | -1.6455 | -2.1275 |
| KIAA0226L | NM_025113 | -0.2335 | -0.396 |
| CEP290 | NM_025114 | -0.4665 | -2.047 |
| TTI2 | NM_025115 | -0.5595 | -1.186 |
| TMEM134 | NM_025124 | 0.2025 | -0.49 |
| C10orf57 | NM_025125 | -0.3475 | -1.015 |
| RNF34 | NM_025126 | -1.419 | -3.199 |
| MUS81 | NM_025128 | -1.052 | -2.249 |
| FUZ | NM_025129 | 0.0225 | -0.4835 |
| HKDC1 | NM_025130 | 0.0765 | -0.0645 |
| WDR19 | NM_025132 | -1.391 | -1.576 |
| FBXO11 | NM_025133 | -1.3205 | -2.855 |
| CHD9 | NM_025134 | -0.889 | -3.344 |
| FHOD3 | NM_025135 | -0.929 | -2.5845 |
| OPA3 | NM_025136 | -0.6185 | -1.6835 |
| SPG11 | NM_025137 | -1.4955 | -1.5965 |
| ARMC9 | NM_025139 | -0.657 | -1.187 |
| CCDC92 | NM_025140 | -0.739 | -1.6315 |
| TM2D3 | NM_025141 | -0.646 | -1.265 |
| TAOK1 | NM_025142 | -0.206 | -1.1335 |
| ALPK1 | NM_025144 | -0.3575 | -0.2435 |
| NAA50 | NM_025146 | -0.88 | -1.524 |
| COQ10B | NM_025147 | -1.272 | -2.8995 |
| ACSF2 | NM_025149 | -0.5795 | -0.88 |
| TARS2 | NM_025150 | -0.3675 | -1.824 |
| RAB11FIP1 | NM_025151 | 0.2305 | -0.715 |
| NUBPL | NM_025152 | -0.9165 | -0.3035 |
| ATP10B | NM_025153 | -0.1255 | -0.539 |
| SUN1 | NM_025154 | -0.5265 | -1.293 |
| PAAF1 | NM_025155 | -1.0415 | -2.292 |

| | | | |
|---|---|---|---|
| PXN | NM_025157 | -0.252 | -0.7805 |
| RUFY1 | NM_025158 | -1.251 | -2.2055 |
| CXorf21 | NM_025159 | -0.625 | 0.021 |
| WDR26 | NM_025160 | -1.4135 | -3.47 |
| C17orf70 | NM_025161 | -0.6195 | -1.273 |
| PIGZ | NM_025163 | 0.058 | -0.233 |
| SIK3 | NM_025164 | -0.4645 | -2.073 |
| ELL3 | NM_025165 | -0.269 | -0.5465 |
| ZNF167 | NM_025169 | -0.9625 | -2.241 |
| PREX2 | NM_025170 | -0.197 | -0.3015 |
| NINL | NM_025176 | -1.227 | -2.283 |
| PLXNA2 | NM_025179 | -0.898 | -1.858 |
| CEP63 | NM_025180 | -0.6295 | -1.7645 |
| SLC35F5 | NM_025181 | -0.8675 | -2.158 |
| KIAA1539 | NM_025182 | -0.63 | -1.0725 |
| EFHC2 | NM_025184 | -0.209 | 0.133 |
| TANC2 | NM_025185 | -0.9915 | -2.972 |
| C16orf70 | NM_025187 | -0.5965 | -1.9875 |
| TRIM45 | NM_025188 | -0.9595 | -1.957 |
| ZNF430 | NM_025189 | -1.0255 | -1.9455 |
| ANKRD36B | NM_025190 | 0.1015 | -0.8405 |
| EDEM3 | NM_025191 | -1.958 | -3.6345 |
| HSD3B7 | NM_025193 | 0.085 | -0.3355 |
| ITPKC | NM_025194 | -0.6505 | -1.8075 |
| TRIB1 | NM_025195 | -0.6785 | -1.4725 |
| GRPEL1 | NM_025196 | -0.741 | -1.859 |
| MTERFD3 | NM_025198 | -1.8725 | -3.0075 |
| PLEKHO2 | NM_025201 | -0.493 | -0.994 |
| EFHD1 | NM_025202 | 0.0685 | -0.6375 |
| C2orf44 | NM_025203 | -1.976 | -4.4465 |
| TRABD | NM_025204 | -0.423 | -1.044 |
| MED28 | NM_025205 | -0.3895 | -0.8865 |
| FLAD1 | NM_025207 | -0.7995 | -1.8245 |
| PDGFD | NM_025208 | -0.256 | 0.0165 |
| EPC1 | NM_025209 | -1.1985 | -3.4285 |
| GKAP1 | NM_025211 | -1.581 | -3.605 |
| CXXC4 | NM_025212 | -0.753 | -0.9445 |
| SPTBN4 | NM_025213 | -0.5495 | -1.166 |
| CCDC68 | NM_025214 | -0.815 | -0.938 |
| PUS1 | NM_025215 | -1.048 | -2.034 |
| WNT10A | NM_025216 | -0.2 | -0.396 |
| ULBP2 | NM_025217 | -0.616 | -1.2345 |
| ULBP1 | NM_025218 | -0.777 | -0.7665 |

| | | | |
|---|---|---|---|
| DNAJC5 | NM_025219 | -0.9355 | -2.2865 |
| ADAM33 | NM_025220 | -0.0045 | -0.7295 |
| KCNIP4 | NM_025221 | -0.668 | 0.1505 |
| WDR82 | NM_025222 | -1.074 | -1.7655 |
| ZBTB46 | NM_025224 | -0.4415 | -1.4215 |
| PNPLA3 | NM_025225 | -0.909 | -1.3435 |
| BPIFB2 | NM_025227 | -0.24 | -0.4195 |
| TRAF3IP3 | NM_025228 | -0.117 | -0.469 |
| SEL1L2 | NM_025229 | 0.037 | -0.2575 |
| DCAF11 | NM_025230 | -0.0445 | -0.6475 |
| ZSCAN16 | NM_025231 | -1.5215 | -1.896 |
| REEP4 | NM_025232 | -0.3095 | -1.099 |
| COASY | NM_025233 | -0.0855 | -0.8935 |
| WDR61 | NM_025234 | -0.4475 | -1.006 |
| TNKS2 | NM_025235 | -1.639 | -3.998 |
| SOST | NM_025237 | 0.0295 | 0.0825 |
| BTBD1 | NM_025238 | -0.7935 | -0.87 |
| PDCD1LG2 | NM_025239 | 0.2145 | 0.3455 |
| CD276 | NM_025240 | -0.167 | -0.9 |
| UBXN6 | NM_025241 | 0.2825 | -0.2915 |
| SLC19A3 | NM_025243 | -0.399 | -1.108 |
| TSGA10 | NM_025244 | 0.211 | 0.173 |
| PBX4 | NM_025245 | -0.9845 | -1.5205 |
| TMEM22 | NM_025246 | -1.4185 | -3.657 |
| ACAD10 | NM_025247 | -0.4685 | -0.8675 |
| SRCIN1 | NM_025248 | -0.7455 | -1.112 |
| TTYH3 | NM_025250 | -0.3045 | -0.894 |
| ARHGAP39 | NM_025251 | -0.9135 | -2.1085 |
| SLC44A4 | NM_025257 | -0.0065 | -0.3865 |
| C6orf27 | NM_025258 | -0.161 | -0.459 |
| MSH5 | NM_025259 | -0.018 | -0.4245 |
| C6orf25 | NM_025260 | 0.1835 | -0.2545 |
| LY6G6C | NM_025261 | -0.0045 | -0.1785 |
| LY6G5C | NM_025262 | -0.3815 | 0.1635 |
| PRR3 | NM_025263 | -1.292 | -3.2065 |
| THUMPD2 | NM_025264 | -1.3675 | -2.615 |
| TSEN2 | NM_025265 | -1.9225 | -3.4725 |
| AARSD1 | NM_025267 | -0.415 | -0.5785 |
| TMEM121 | NM_025268 | -0.351 | -0.805 |
| PRR7 | NM_030567 | 0.023 | -0.5035 |
| KHDC1 | NM_030568 | -0.472 | -0.696 |
| ITIH5 | NM_030569 | -0.036 | -0.487 |
| NDFIP1 | NM_030571 | -0.319 | -0.514 |

| | | | |
|---|---|---|---|
| C12orf39 | NM_030572 | 0.1195 | 0.3415 |
| LIMD2 | NM_030576 | -0.134 | -1.199 |
| TMEM177 | NM_030577 | -1.05 | -3.27 |
| B9D2 | NM_030578 | -0.057 | -1.294 |
| CYB5B | NM_030579 | -0.829 | -1.8495 |
| ZNF34 | NM_030580 | -0.6625 | -1.1095 |
| WDR59 | NM_030581 | -1.2535 | -2.036 |
| COL18A1 | NM_030582 | -0.069 | -0.7035 |
| MATN2 | NM_030583 | 0.0845 | -0.6185 |
| B4GALT2 | NM_030587 | -0.36 | -1.042 |
| CYP2A7 | NM_030589 | -0.298 | -0.272 |
| MATN4 | NM_030590 | 0.0465 | -0.275 |
| SIRT2 | NM_030593 | 0.186 | -0.3835 |
| CPEB1 | NM_030594 | -0.2155 | -1.676 |
| ZFP2 | NM_030613 | -0.3355 | -1.5295 |
| KIF25 | NM_030615 | -0.2875 | -0.2625 |
| DICER1 | NM_030621 | 0.0765 | -1.67 |
| CYP2S1 | NM_030622 | -0.041 | -0.4445 |
| SPHKAP | NM_030623 | -0.1105 | -0.4535 |
| KLHL15 | NM_030624 | -1.43 | -2.2255 |
| TET1 | NM_030625 | -0.18 | -2.891 |
| LRRC27 | NM_030626 | -0.3155 | -1.0585 |
| CPEB4 | NM_030627 | -1.478 | -2.561 |
| INTS5 | NM_030628 | -0.697 | -1.3635 |
| CMIP | NM_030629 | -0.5915 | -1.692 |
| C17orf28 | NM_030630 | 0.027 | -0.299 |
| SLC25A21 | NM_030631 | -1.082 | 0.523 |
| ASXL3 | NM_030632 | -0.428 | -2.0715 |
| ZNF436 | NM_030634 | -1.4275 | -1.886 |
| EEPD1 | NM_030636 | -0.1485 | -1.6715 |
| DDHD1 | NM_030637 | -0.884 | -1.4545 |
| DUSP16 | NM_030640 | -1.983 | -2.6695 |
| APOL6 | NM_030641 | -0.072 | -0.383 |
| APOL5 | NM_030642 | 0.16 | -0.345 |
| APOL4 | NM_030643 | 0.0515 | -0.062 |
| SH3BP5L | NM_030645 | -0.3815 | -1.5585 |
| JHDM1D | NM_030647 | -0.927 | -1.683 |
| SETD7 | NM_030648 | -1.7335 | -2.1275 |
| ACAP3 | NM_030649 | -0.086 | -0.5705 |
| KIAA1715 | NM_030650 | -0.663 | -1.3315 |
| PRRT1 | NM_030651 | -0.0225 | -1.241 |
| DDX11 | NM_030653 | -0.526 | -1.5555 |
| LIM2 | NM_030657 | -0.0175 | -0.037 |

| | | | |
|---|---|---|---|
| ATXN3 | NM_030660 | -0.521 | -2.1695 |
| MAP2K2 | NM_030662 | -0.116 | -0.8235 |
| SMCP | NM_030663 | 0.019 | -0.295 |
| PTER | NM_030664 | 0.4395 | -0.1565 |
| RAI1 | NM_030665 | -0.907 | -2.7205 |
| SERPINB1 | NM_030666 | 0.0515 | -0.449 |
| PTPRO | NM_030667 | -1.2965 | -1.8335 |
| PTPRO | NM_030668 | 0.452 | -0.3665 |
| PTPRO | NM_030669 | 0.452 | -0.3665 |
| PTPRO | NM_030670 | 0.452 | -0.3665 |
| PTPRO | NM_030671 | 0.452 | -0.3665 |
| SLC38A1 | NM_030674 | -1.147 | -0.9795 |
| ZEB1 | NM_030751 | 0.001 | -1.4665 |
| TCP1 | NM_030752 | -0.597 | -0.7485 |
| WNT3 | NM_030753 | -1.035 | -1.463 |
| SAA2 | NM_030754 | -0.1435 | 0.1855 |
| TMX1 | NM_030755 | -1.36 | -2.073 |
| TCF7L2 | NM_030756 | -0.4665 | -1.848 |
| OSBP2 | NM_030758 | -0.0975 | -0.517 |
| NRBF2 | NM_030759 | -1.6105 | -3.606 |
| S1PR5 | NM_030760 | 0.3555 | -0.0875 |
| WNT4 | NM_030761 | 0.1075 | -0.2635 |
| BHLHE41 | NM_030762 | -0.0335 | -0.3855 |
| HMGN5 | NM_030763 | 0.1115 | 0.4135 |
| FCRL2 | NM_030764 | 0.0955 | -0.0255 |
| B3GNT4 | NM_030765 | -0.16 | -1.004 |
| BCL2L14 | NM_030766 | -0.4775 | -0.1325 |
| AKNA | NM_030767 | -0.2755 | -1.176 |
| ILKAP | NM_030768 | -0.5995 | -2.0485 |
| NPL | NM_030769 | -1.021 | -0.68 |
| TMPRSS5 | NM_030770 | 0.281 | -0.2085 |
| CCDC34 | NM_030771 | -0.9935 | -1.2645 |
| GJA9 | NM_030772 | 0.0145 | 0.026 |
| TUBB1 | NM_030773 | -0.1035 | -0.018 |
| OR51E2 | NM_030774 | -0.0025 | -0.1455 |
| WNT5B | NM_030775 | -0.249 | -2.1665 |
| ZBP1 | NM_030776 | -0.198 | 0.0605 |
| SLC2A10 | NM_030777 | 0.3325 | -0.3165 |
| KCNH6 | NM_030779 | 0.139 | -0.513 |
| SLC25A32 | NM_030780 | -1.6715 | -3.7155 |
| CLPTM1L | NM_030782 | 0.105 | -0.6725 |
| PTDSS2 | NM_030783 | -0.2585 | -1.0615 |
| GPR63 | NM_030784 | -0.7555 | -2.0085 |

| | | | |
|---|---|---|---|
| RSPH6A | NM_030785 | -0.024 | -0.3605 |
| SYNC | NM_030786 | -0.5325 | -0.3575 |
| CFHR5 | NM_030787 | 0.0995 | 0.113 |
| TM7SF4 | NM_030788 | 0.146 | -0.0695 |
| HM13 | NM_030789 | -0.503 | -1.167 |
| ITFG1 | NM_030790 | -0.949 | -0.8895 |
| SGPP1 | NM_030791 | -1.025 | -2.7955 |
| GDPD5 | NM_030792 | -0.598 | -0.8175 |
| FBXO38 | NM_030793 | -1.4205 | -3.1485 |
| STMN4 | NM_030795 | -0.2865 | -1.783 |
| VOPP1 | NM_030796 | -0.155 | -0.7635 |
| FAM49A | NM_030797 | -0.1565 | -0.467 |
| WBSCR16 | NM_030798 | -0.503 | -1.5295 |
| YIPF5 | NM_030799 | -1.051 | -1.5445 |
| FAM117A | NM_030802 | -0.862 | -3.421 |
| ATG16L1 | NM_030803 | -0.48 | -1.9875 |
| LMAN2L | NM_030805 | -1.639 | -3.117 |
| C1orf21 | NM_030806 | -0.5845 | -1.7825 |
| SLC2A11 | NM_030807 | -0.092 | -0.506 |
| NDEL1 | NM_030808 | -0.6695 | -2.2225 |
| CSRNP2 | NM_030809 | -1.49 | -3.156 |
| TXNDC5 | NM_030810 | -0.1275 | -0.4245 |
| MRPS26 | NM_030811 | -0.623 | -1.798 |
| ACTL8 | NM_030812 | 0.3015 | -0.645 |
| CLPB | NM_030813 | 0.3375 | -0.475 |
| PDRG1 | NM_030815 | -0.6745 | -2.32 |
| ANKRD13C | NM_030816 | -1.508 | -3.0565 |
| APOLD1 | NM_030817 | 0.0915 | -0.177 |
| CCDC130 | NM_030818 | -0.385 | -1.4785 |
| GFOD2 | NM_030819 | 0.353 | -0.8485 |
| COL21A1 | NM_030820 | -0.0265 | -0.0675 |
| PLA2G12A | NM_030821 | -0.953 | -0.9085 |
| ZNF442 | NM_030824 | -0.217 | -0.4385 |
| OR5V1 | NM_030876 | -0.0705 | -0.1545 |
| CTNNBL1 | NM_030877 | -0.2655 | -1.0685 |
| APOL2 | NM_030882 | -0.548 | -0.541 |
| OR2H1 | NM_030883 | -0.009 | -0.4775 |
| MAP4 | NM_030885 | -0.397 | -1.094 |
| LRRC3 | NM_030891 | -0.0815 | -0.421 |
| CD1E | NM_030893 | 0.0335 | -0.3035 |
| ZNF696 | NM_030895 | -0.7505 | -1.1455 |
| TBRG4 | NM_030900 | -0.44 | -2.4025 |
| OR7A17 | NM_030901 | 0.327 | -0.2 |

| | | | |
|---|---|---|---|
| OR2W1 | NM_030903 | -0.2385 | 0.1915 |
| OR2T1 | NM_030904 | 0.0435 | -0.2665 |
| OR2J2 | NM_030905 | 0.389 | -0.278 |
| STK33 | NM_030906 | -0.0785 | -0.4535 |
| RSG1 | NM_030907 | -0.1945 | -0.554 |
| OR2A4 | NM_030908 | 0.2265 | 0.4365 |
| CDADC1 | NM_030911 | -0.5785 | -1.921 |
| TRIM8 | NM_030912 | -0.8065 | -2.32 |
| SEMA6C | NM_030913 | 0.038 | -0.625 |
| URM1 | NM_030914 | -0.096 | -0.913 |
| LBH | NM_030915 | 0.1415 | -0.66 |
| PVRL4 | NM_030916 | 0.38 | -0.21 |
| FIP1L1 | NM_030917 | -1.338 | -2.24 |
| SNX27 | NM_030918 | -1.1755 | -2.4335 |
| FAM83D | NM_030919 | -0.747 | -3.0275 |
| ANP32E | NM_030920 | -0.689 | -0.6605 |
| NIPA2 | NM_030922 | -1.056 | -2.0195 |
| TMEM163 | NM_030923 | -0.1395 | -0.614 |
| ACSBG2 | NM_030924 | 0.28 | -0.0155 |
| CAB39L | NM_030925 | -0.5415 | -1.0815 |
| ITM2C | NM_030926 | -0.229 | -0.813 |
| TSPAN14 | NM_030927 | 0.418 | -1.1895 |
| CDT1 | NM_030928 | -0.4945 | -2.3235 |
| KAZALD1 | NM_030929 | 0.2295 | -0.105 |
| UNC93B1 | NM_030930 | -0.159 | -0.348 |
| DEFB126 | NM_030931 | 0.131 | 0.0215 |
| DIAPH3 | NM_030932 | -0.799 | -2.15 |
| SHCBP1L | NM_030933 | 0.3015 | 0.1705 |
| TRMT1L | NM_030934 | -2.3555 | -3.3645 |
| TSC22D4 | NM_030935 | -0.2335 | -0.6475 |
| RNF32 | NM_030936 | 0.085 | -0.3895 |
| CCNL2 | NM_030937 | -0.676 | -1.05 |
| VMP1 | NM_030938 | -1.0365 | -1.7405 |
| C6orf62 | NM_030939 | -1.3115 | -1.6935 |
| ISCA1 | NM_030940 | -1.3695 | -1.928 |
| LOC81691 | NM_030941 | -0.6175 | -1.064 |
| AMN | NM_030943 | -0.0545 | -0.2155 |
| C1QTNF3 | NM_030945 | -1.4 | -0.993 |
| OR14J1 | NM_030946 | 0.1785 | 0.1385 |
| PHACTR1 | NM_030948 | -0.2605 | -0.5105 |
| PPP1R14C | NM_030949 | -0.0755 | -0.033 |
| NUAK2 | NM_030952 | -0.2055 | 0.354 |
| TIGD6 | NM_030953 | -0.9915 | -1.5385 |

| | | | |
|---|---|---|---|
| RNF170 | NM_030954 | -0.4265 | -1.0105 |
| ADAMTS12 | NM_030955 | 0.255 | -0.1335 |
| TLR10 | NM_030956 | 0.3965 | 0.028 |
| ADAMTS10 | NM_030957 | -0.07 | -0.0865 |
| SLCO5A1 | NM_030958 | -0.1355 | -1.0045 |
| OR12D3 | NM_030959 | 0.0105 | -0.0225 |
| SPACA1 | NM_030960 | 0.0145 | -0.238 |
| TRIM56 | NM_030961 | -0.108 | 0.2135 |
| SBF2 | NM_030962 | -1.0395 | -1.6385 |
| RNF146 | NM_030963 | -2.1675 | -4.566 |
| SPRY4 | NM_030964 | -0.172 | -0.609 |
| ST6GALNAC5 | NM_030965 | -1.79 | -3.5825 |
| KRTAP1-3 | NM_030966 | -0.495 | -0.233 |
| KRTAP1-1 | NM_030967 | 0.0245 | -0.1745 |
| C1QTNF1 | NM_030968 | -0.0505 | -0.258 |
| TMEM14B | NM_030969 | -0.15 | -0.5515 |
| SFXN3 | NM_030971 | -0.1055 | -0.65 |
| ZNF611 | NM_030972 | -1.295 | -2.064 |
| MED25 | NM_030973 | -0.0765 | -0.5915 |
| SHARPIN | NM_030974 | -0.181 | -0.4985 |
| KRTAP9-9 | NM_030975 | 0.6285 | -0.326 |
| ARPC5L | NM_030978 | -0.116 | -1.4385 |
| PABPC3 | NM_030979 | -0.0955 | -0.1365 |
| ISG20L2 | NM_030980 | -0.981 | -2.887 |
| RAB1B | NM_030981 | 0.135 | -0.6675 |
| TBXAS1 | NM_030984 | 0.099 | -0.327 |
| HNRNPA1 | NM_031157 | -0.491 | -0.279 |
| CCR9 | NM_031200 | -0.162 | 0.0795 |
| HNRNPM | NM_031203 | 0.2625 | -0.1005 |
| LAS1L | NM_031206 | -0.227 | -0.8355 |
| HYI | NM_031207 | 0.2225 | -0.255 |
| FAHD1 | NM_031208 | -1.4295 | -2.738 |
| QTRT1 | NM_031209 | -0.0615 | -0.7025 |
| SLIRP | NM_031210 | -0.4425 | -0.6875 |
| SLC25A28 | NM_031212 | -0.7125 | -1.7085 |
| FAM108A1 | NM_031213 | 0.02 | -0.8525 |
| CABLES2 | NM_031215 | -0.8755 | -1.8535 |
| SEH1L | NM_031216 | -1.094 | -1.427 |
| KIF18A | NM_031217 | -0.942 | -3.036 |
| ZNF93 | NM_031218 | -0.3645 | -2.595 |
| HDHD3 | NM_031219 | -0.2875 | -1.722 |
| PITPNM3 | NM_031220 | -0.873 | -1.0645 |
| CYP19A1 | NM_031226 | -0.045 | -0.0015 |

| | | | |
|---|---|---|---|
| RBCK1 | NM_031229 | -0.3195 | -1.0605 |
| NECAB3 | NM_031232 | -0.271 | -0.841 |
| HNRNPA2B1 | NM_031243 | -0.454 | -0.449 |
| SIRT5 | NM_031244 | -0.262 | -0.7645 |
| PSG2 | NM_031246 | -0.1955 | -0.081 |
| HNRNPK | NM_031262 | -0.673 | -0.383 |
| HNRNPK | NM_031263 | -0.667 | -0.5185 |
| CDHR5 | NM_031264 | -0.0115 | -0.41 |
| HNRNPAB | NM_031266 | -0.4475 | -1.099 |
| CDK13 | NM_031267 | -1.294 | -3.574 |
| PDPK1 | NM_031268 | -1.16 | -1.9625 |
| TEX15 | NM_031271 | -0.18 | -1.72 |
| TEX13B | NM_031273 | -0.2515 | -0.147 |
| TEX13A | NM_031274 | 0.275 | 0.0825 |
| TEX12 | NM_031275 | 0.5105 | -0.268 |
| RNF17 | NM_031277 | 0.1105 | -0.457 |
| AGXT2L1 | NM_031279 | 0.017 | -0.49 |
| MRPS15 | NM_031280 | -0.106 | -0.899 |
| FCRL5 | NM_031281 | -0.185 | -0.4305 |
| FCRL4 | NM_031282 | 0.1205 | -0.1505 |
| TCF7L1 | NM_031283 | -0.128 | -0.777 |
| ADPGK | NM_031284 | -0.269 | -1.5775 |
| SH3BGRL3 | NM_031286 | 0.342 | -0.3285 |
| SF3B5 | NM_031287 | -0.2475 | -0.9515 |
| INO80B | NM_031288 | -0.548 | -1.259 |
| GSG1 | NM_031289 | -0.1 | 0.0575 |
| CCDC70 | NM_031290 | 0.2035 | 0.1565 |
| SLC25A31 | NM_031291 | -0.497 | 0.458 |
| PUS7L | NM_031292 | -0.454 | -0.847 |
| PMFBP1 | NM_031293 | -0.027 | -0.218 |
| LRRC48 | NM_031294 | 0.0285 | -0.669 |
| RAB33B | NM_031296 | -0.8605 | -1.368 |
| RNF208 | NM_031297 | -0.3695 | -0.7935 |
| CDCA3 | NM_031299 | -1.442 | -2.462 |
| MXD3 | NM_031300 | -0.6585 | -1.719 |
| APH1B | NM_031301 | -1.2 | -1.694 |
| GLT8D2 | NM_031302 | -0.036 | -0.0205 |
| KATNAL2 | NM_031303 | -0.4075 | -1.0395 |
| DOHH | NM_031304 | -0.2785 | -1.1675 |
| ARHGAP24 | NM_031305 | -0.2485 | -0.394 |
| PUS3 | NM_031307 | -0.718 | -2.1425 |
| EPPK1 | NM_031308 | 0.03 | -0.566 |
| SCRT1 | NM_031309 | -0.8215 | -0.7315 |

| | | | |
|---|---|---|---|
| PLVAP | NM_031310 | 0.214 | 0.1465 |
| CPVL | NM_031311 | -1.037 | -0.6645 |
| ALPPL2 | NM_031313 | -0.102 | -0.181 |
| HNRNPC | NM_031314 | -0.1685 | -0.3245 |
| COL4A3BP | NM_031361 | -0.8425 | -1.933 |
| HNRNPD | NM_031369 | -0.4805 | -1.078 |
| HNRNPD | NM_031370 | -0.4805 | -1.078 |
| ARID4B | NM_031371 | -0.3485 | -1.9785 |
| HNRPDL | NM_031372 | -1.353 | -0.9925 |
| HUWE1 | NM_031407 | -0.623 | -1.119 |
| CCR6 | NM_031409 | 0.1095 | -0.029 |
| PCDHA1 | NM_031410 | -0.0305 | -0.4335 |
| GABARAPL1 | NM_031412 | -0.367 | 0.0035 |
| CECR2 | NM_031413 | 0.0075 | -2.035 |
| STK31 | NM_031414 | -0.3935 | -0.1425 |
| GSDMC | NM_031415 | -0.1265 | -0.522 |
| MARK4 | NM_031417 | -0.1 | -0.61 |
| ANO3 | NM_031418 | -0.033 | -0.144 |
| NFKBIZ | NM_031419 | -0.336 | 0.606 |
| MRPL9 | NM_031420 | -0.4595 | -1.5225 |
| TTC25 | NM_031421 | -0.428 | -0.4585 |
| CHST9 | NM_031422 | -0.094 | 0.45 |
| AIF1L | NM_031426 | 0.253 | -0.097 |
| DNAL1 | NM_031427 | -0.7275 | -1.4465 |
| RTBDN | NM_031429 | -0.2505 | -0.4875 |
| RILP | NM_031430 | 0.024 | -0.4205 |
| COG3 | NM_031431 | -0.4955 | -1.4795 |
| UCK1 | NM_031432 | -1.2175 | -2.1285 |
| MFRP | NM_031433 | -0.091 | -0.2585 |
| TMUB1 | NM_031434 | -0.2665 | -0.8895 |
| THAP2 | NM_031435 | -1.3715 | -2.5965 |
| NUDT12 | NM_031438 | -0.9815 | -1.6645 |
| SOX7 | NM_031439 | -0.084 | -0.22 |
| RTP3 | NM_031440 | 0.025 | -0.076 |
| TMEM47 | NM_031442 | -0.121 | 0.2945 |
| CCM2 | NM_031443 | -1.528 | -2.997 |
| C22orf13 | NM_031444 | -0.87 | -2.349 |
| AMMECR1L | NM_031445 | -0.9475 | -3.702 |
| C18orf21 | NM_031446 | -1.18 | -2.666 |
| C19orf12 | NM_031448 | -0.531 | -1.53 |
| ZMIZ2 | NM_031449 | -0.078 | -0.729 |
| TEX101 | NM_031451 | 0.254 | -0.3075 |
| FAM103A1 | NM_031452 | -0.887 | -2.401 |

| | | | |
|---|---|---|---|
| FAM107B | NM_031453 | -1.2425 | -1.297 |
| SELO | NM_031454 | -0.9795 | -2.2025 |
| CCDC3 | NM_031455 | 0.1905 | -0.4545 |
| FBXW10 | NM_031456 | -0.202 | -0.3125 |
| MS4A8B | NM_031457 | -0.122 | -0.33 |
| PARP9 | NM_031458 | -0.0065 | -1.1455 |
| SESN2 | NM_031459 | -0.5545 | -2.188 |
| KCNK17 | NM_031460 | -0.6405 | -0.1825 |
| CRISPLD1 | NM_031461 | -1.1535 | -2.645 |
| CD99L2 | NM_031462 | -0.596 | -1.175 |
| HSDL1 | NM_031463 | -1.411 | -1.7925 |
| RPS6KL1 | NM_031464 | -0.7295 | -1.5455 |
| TRAPPC9 | NM_031466 | 0.242 | -0.2615 |
| SLC4A9 | NM_031467 | 0.118 | -0.678 |
| CALN1 | NM_031468 | -0.072 | -1.047 |
| SH3BGRL2 | NM_031469 | -1.0375 | -1.5715 |
| FERMT3 | NM_031471 | -0.01 | -0.329 |
| TRPT1 | NM_031472 | -0.1225 | -0.352 |
| IFT81 | NM_031473 | -0.4495 | -0.6725 |
| NRIP2 | NM_031474 | -0.0075 | -0.054 |
| ESPN | NM_031475 | -0.005 | -0.21 |
| CRISPLD2 | NM_031476 | -0.0635 | -0.6925 |
| YPEL3 | NM_031477 | 0.2165 | -0.294 |
| FAM57B | NM_031478 | -0.138 | -0.663 |
| INHBE | NM_031479 | -0.2215 | -1.0575 |
| RIOK1 | NM_031480 | -0.967 | -1.894 |
| SLC25A18 | NM_031481 | -0.0925 | -0.0875 |
| ATG10 | NM_031482 | -0.8715 | -1.924 |
| ITCH | NM_031483 | -0.7815 | -1.709 |
| GRWD1 | NM_031485 | -0.645 | -2.5765 |
| ZNF484 | NM_031486 | -0.14 | -1.2435 |
| L3MBTL2 | NM_031488 | -0.7755 | -2.15 |
| LONP2 | NM_031490 | -1.242 | -1.9455 |
| RBP5 | NM_031491 | 0.2 | 0.1675 |
| RBM4B | NM_031492 | -0.4965 | -2.0615 |
| PCDHA2 | NM_031495 | 0.051 | -0.2185 |
| PCDHA3 | NM_031497 | 0.038 | -0.3565 |
| GNGT2 | NM_031498 | -0.4605 | -0.2865 |
| PCDHA4 | NM_031500 | 0.004 | -0.42 |
| PCDHA5 | NM_031501 | -0.158 | -0.4805 |
| HNRNPU | NM_031844 | -0.587 | -0.5255 |
| MAP2 | NM_031845 | -1.2885 | -1.3145 |
| MAP2 | NM_031847 | -1.005 | -1.082 |

| | | | |
|---|---|---|---|
| PCDHA6 | NM_031848 | 0.0855 | 0.252 |
| PCDHA6 | NM_031849 | 0.0855 | 0.252 |
| AGTR1 | NM_031850 | -0.793 | -0.37 |
| PCDHA7 | NM_031852 | -0.052 | -0.2255 |
| KRTAP4-12 | NM_031854 | 0.014 | -0.045 |
| PCDHA8 | NM_031856 | 0.164 | -0.6945 |
| NBR1 | NM_031858 | -1.298 | -1.909 |
| PCDHA10 | NM_031859 | 0.3345 | -0.393 |
| PCDHA11 | NM_031861 | 0.2255 | -0.36 |
| NBR1 | NM_031862 | -1.2645 | -1.5055 |
| PCDHA12 | NM_031864 | 0.36 | 0.1045 |
| PCDHA13 | NM_031865 | -0.07 | -0.3675 |
| FZD8 | NM_031866 | -0.263 | -0.361 |
| PCDHAC1 | NM_031882 | -0.0465 | -0.4465 |
| PCDHAC2 | NM_031883 | 0.01 | -0.7045 |
| BBS2 | NM_031885 | -1.0525 | -1.1295 |
| KCNA7 | NM_031886 | 0.156 | -0.227 |
| ENAM | NM_031889 | -0.081 | -0.2805 |
| CECR6 | NM_031890 | -0.595 | -1.523 |
| CDH20 | NM_031891 | -0.0945 | -0.6385 |
| SH3KBP1 | NM_031892 | -0.134 | -0.166 |
| FTHL17 | NM_031894 | -0.6205 | -0.106 |
| CACNG8 | NM_031895 | -0.2315 | -0.739 |
| CACNG7 | NM_031896 | 0.0255 | -0.455 |
| CACNG6 | NM_031897 | 0.2045 | -0.323 |
| TEKT3 | NM_031898 | 0.0225 | -0.3035 |
| GORASP1 | NM_031899 | -0.6075 | -2.0225 |
| AGXT2 | NM_031900 | 0.556 | 0.0425 |
| MRPS21 | NM_031901 | -0.4155 | -0.668 |
| MRPS5 | NM_031902 | -0.279 | -1.5895 |
| MRPL32 | NM_031903 | -0.8025 | -2.1475 |
| FRMD8 | NM_031904 | -0.0835 | -0.39 |
| ARMC10 | NM_031905 | -0.149 | -0.8295 |
| USP26 | NM_031907 | 0.2535 | -0.3685 |
| C1QTNF2 | NM_031908 | -0.1065 | -0.504 |
| C1QTNF4 | NM_031909 | -0.0345 | -0.3105 |
| C1QTNF6 | NM_031910 | 0.3685 | -0.212 |
| C1QTNF7 | NM_031911 | 0.1575 | 0.0505 |
| SYT15 | NM_031912 | 0.115 | -0.499 |
| ESYT3 | NM_031913 | -0.132 | -0.2855 |
| SYT16 | NM_031914 | 0.0725 | -0.3665 |
| SETDB2 | NM_031915 | -0.928 | -2.114 |
| ROPN1L | NM_031916 | -0.1375 | -0.373 |

| | | | |
|---|---|---|---|
| ANGPTL6 | NM_031917 | 0.1815 | -0.267 |
| KLF16 | NM_031918 | -0.3265 | -0.91 |
| FSD1L | NM_031919 | 0.7015 | -0.1525 |
| ATAD3B | NM_031921 | -0.4465 | -1.0355 |
| REPS1 | NM_031922 | -0.734 | -2.017 |
| TAF3 | NM_031923 | -0.365 | -1.853 |
| RSPH3 | NM_031924 | -0.4305 | -0.8115 |
| TMEM120A | NM_031925 | 0.228 | -0.2265 |
| HMCN1 | NM_031935 | -0.358 | -0.343 |
| GPR61 | NM_031936 | -0.331 | -0.3365 |
| TBC1D10A | NM_031937 | -0.7015 | -1.9335 |
| BCO2 | NM_031938 | 0.077 | 0.07 |
| MRO | NM_031939 | 0.0155 | 0.159 |
| TM2D2 | NM_031940 | -0.8485 | -2.6585 |
| USHBP1 | NM_031941 | -0.1625 | -0.465 |
| CDCA7 | NM_031942 | -0.9375 | -0.757 |
| MIXL1 | NM_031944 | 0.1045 | 0.0305 |
| TSPAN10 | NM_031945 | 0.1105 | -0.4285 |
| AGAP3 | NM_031946 | -1.072 | -2.155 |
| SLC25A2 | NM_031947 | -0.1735 | -0.339 |
| PRSS27 | NM_031948 | 0.1575 | -0.533 |
| TTLL2 | NM_031949 | 0.352 | 0.3265 |
| FGFBP2 | NM_031950 | 0.1715 | -0.4925 |
| WDR87 | NM_031951 | 0.068 | -0.1875 |
| SPATA9 | NM_031952 | 0.251 | -0.2395 |
| SNX25 | NM_031953 | -0.7805 | -2.2065 |
| KCTD10 | NM_031954 | -0.656 | -2.3975 |
| SPATA16 | NM_031955 | -0.1685 | -0.645 |
| TTC29 | NM_031956 | -0.562 | -0.0485 |
| KRTAP1-5 | NM_031957 | 0.1765 | -0.1875 |
| KRTAP3-1 | NM_031958 | -0.197 | -0.5665 |
| KRTAP3-2 | NM_031959 | 0.0245 | -0.6115 |
| KRTAP4-8 | NM_031960 | 0.0365 | -0.073 |
| KRTAP9-2 | NM_031961 | -0.2855 | -0.2975 |
| KRTAP9-3 | NM_031962 | 0.27 | -0.8265 |
| KRTAP9-8 | NM_031963 | 0.001 | -0.0995 |
| KRTAP17-1 | NM_031964 | 0.168 | -0.559 |
| GSG2 | NM_031965 | -1.079 | -1.8385 |
| CCNB1 | NM_031966 | -1.749 | -3.218 |
| NARF | NM_031968 | -0.3915 | -1.5865 |
| PCBP2 | NM_031989 | 0.152 | -0.2755 |
| PTBP1 | NM_031990 | -0.09 | -0.805 |
| PTBP1 | NM_031991 | -0.09 | -0.805 |

| | | | |
|---|---|---|---|
| EIF4H | NM_031992 | -0.1795 | -1.0705 |
| PCDHGA1 | NM_031993 | 0.5405 | 0.26 |
| PCDHGA2 | NM_032009 | -0.068 | -0.32 |
| PCDHGA3 | NM_032011 | -0.359 | -0.0265 |
| C9orf5 | NM_032012 | -1.3455 | -2.079 |
| NDRG3 | NM_032013 | -0.0405 | -0.4 |
| MRPS24 | NM_032014 | 0.1445 | -0.4495 |
| RNF26 | NM_032015 | -1.0515 | -2.8035 |
| STARD3NL | NM_032016 | -0.797 | -1.931 |
| STK40 | NM_032017 | -1.011 | -1.9705 |
| C1orf124 | NM_032018 | -0.6125 | -1.391 |
| HDAC10 | NM_032019 | -0.0905 | -0.133 |
| FUCA2 | NM_032020 | 0.267 | -0.293 |
| TMEM133 | NM_032021 | -0.479 | -2.0065 |
| RASSF4 | NM_032023 | -0.166 | -0.654 |
| C10orf11 | NM_032024 | -0.3985 | 0.158 |
| EIF2A | NM_032025 | -1.033 | -1.9645 |
| TATDN1 | NM_032026 | -0.833 | -1.039 |
| TM2D1 | NM_032027 | -0.8915 | -1.309 |
| TSSK1B | NM_032028 | -0.029 | -0.325 |
| FKSG83 | NM_032030 | 0.244 | 0.3205 |
| SLC4A11 | NM_032034 | 0.1205 | -0.5975 |
| IFI27L2 | NM_032036 | 0.146 | -0.572 |
| TSSK6 | NM_032037 | 0.3195 | 0.294 |
| SPNS1 | NM_032038 | 0.117 | -0.5125 |
| ITFG3 | NM_032039 | -0.3785 | -1.2125 |
| CCDC8 | NM_032040 | 0.0405 | -0.304 |
| NCALD | NM_032041 | -1.0055 | -1.9685 |
| FAM172A | NM_032042 | -0.6135 | -0.6715 |
| BRIP1 | NM_032043 | -0.7595 | -2.2585 |
| REG4 | NM_032044 | -0.144 | -0.446 |
| KREMEN1 | NM_032045 | -0.393 | -1.4955 |
| B3GNT5 | NM_032047 | -1.0315 | -2.63 |
| EMILIN2 | NM_032048 | -0.863 | -1.9795 |
| AGTR1 | NM_032049 | -0.793 | -0.37 |
| PATZ1 | NM_032050 | -1.123 | -3.614 |
| PATZ1 | NM_032051 | -1.123 | -3.614 |
| PATZ1 | NM_032052 | -1.123 | -3.614 |
| PCDHGA4 | NM_032053 | -0.4285 | -0.062 |
| PCDHGA5 | NM_032054 | -0.4245 | -0.1265 |
| PCDHGA6 | NM_032086 | -0.0425 | -0.048 |
| PCDHGA7 | NM_032087 | 0.166 | -0.0725 |
| PCDHGA8 | NM_032088 | -0.478 | -0.455 |

| | | | |
|---|---|---|---|
| PCDHGA9 | NM_032089 | -0.0035 | -0.306 |
| PCDHGA10 | NM_032090 | -0.646 | -0.1015 |
| PCDHGA11 | NM_032091 | -0.1945 | -0.312 |
| PCDHGA11 | NM_032092 | -0.1945 | -0.312 |
| PCDHGA12 | NM_032094 | -0.3165 | -0.698 |
| PCDHGB1 | NM_032095 | -0.442 | -0.217 |
| PCDHGB2 | NM_032096 | 0.1095 | -0.1725 |
| PCDHGB3 | NM_032097 | 0.2075 | -0.5765 |
| PCDHGB4 | NM_032098 | 0.1015 | -0.5365 |
| PCDHGB5 | NM_032099 | -0.159 | -0.313 |
| PCDHGB6 | NM_032100 | -0.1495 | 0.195 |
| PCDHGB7 | NM_032101 | 0.086 | -0.497 |
| SRSF8 | NM_032102 | -1.52 | -1.554 |
| PPP1R12B | NM_032103 | 0.8205 | 0.5085 |
| PPP1R12B | NM_032104 | 0.8205 | 0.5085 |
| L3MBTL1 | NM_032107 | -0.9435 | -0.831 |
| SEMA6B | NM_032108 | 0.0655 | -0.139 |
| OTP | NM_032109 | -0.038 | 0.929 |
| DMRTA2 | NM_032110 | 0.267 | -0.1365 |
| MRPL14 | NM_032111 | -0.1975 | -1.7615 |
| MRPL43 | NM_032112 | -0.429 | -1.4575 |
| KATNAL1 | NM_032116 | -1.173 | -2.6555 |
| MND1 | NM_032117 | -1.9525 | -2.8125 |
| WDR54 | NM_032118 | 0.0275 | -0.5465 |
| GPR98 | NM_032119 | 0.077 | 0.0655 |
| C7orf64 | NM_032120 | -0.9225 | -1.7565 |
| MAGT1 | NM_032121 | -1.0965 | -1.418 |
| DTNBP1 | NM_032122 | -0.451 | -2.306 |
| KIRREL2 | NM_032123 | -0.4785 | -0.524 |
| HDHD2 | NM_032124 | -0.714 | -0.6585 |
| TMEM222 | NM_032125 | -0.3725 | -1.792 |
| C1orf49 | NM_032126 | -0.119 | -0.0755 |
| FAM160A2 | NM_032127 | -0.1085 | -1.0235 |
| SLC10A7 | NM_032128 | -0.6675 | -0.9285 |
| PLEKHN1 | NM_032129 | -0.0885 | -0.185 |
| FAM186B | NM_032130 | -0.119 | -0.397 |
| ARMC2 | NM_032131 | 0.1 | -0.051 |
| HORMAD1 | NM_032132 | 0.0315 | -0.6115 |
| MYCBPAP | NM_032133 | 0.2725 | 0.055 |
| QRICH2 | NM_032134 | -0.1865 | -0.2895 |
| FSCB | NM_032135 | -0.059 | -0.0015 |
| TKTL2 | NM_032136 | -0.392 | 0.1655 |
| C3orf20 | NM_032137 | -0.101 | -0.0275 |

| | | | |
|---|---|---|---|
| KBTBD7 | NM_032138 | -0.918 | -0.6675 |
| ANKRD27 | NM_032139 | -1.516 | -2.762 |
| C16orf48 | NM_032140 | -0.6215 | -1.47 |
| NSRP1 | NM_032141 | -1.4125 | -3.1965 |
| CEP192 | NM_032142 | -1.4515 | -2.237 |
| ZRANB3 | NM_032143 | -0.5775 | -1.7825 |
| FBXO30 | NM_032145 | -0.768 | -2.4725 |
| ARL6 | NM_032146 | -0.4705 | -2.1995 |
| USP44 | NM_032147 | -0.22 | -2.7055 |
| SLC41A2 | NM_032148 | -0.4185 | -0.7115 |
| C4orf17 | NM_032149 | -0.1675 | -0.3355 |
| PCBD2 | NM_032151 | -0.2995 | -0.5795 |
| PRAM1 | NM_032152 | 0.106 | -0.2815 |
| ZIC4 | NM_032153 | 0.116 | -0.0595 |
| CAPRIN2 | NM_032156 | -1.5535 | -2.0835 |
| DSEL | NM_032160 | -0.1665 | -0.38 |
| ZNF394 | NM_032164 | -0.922 | -2.117 |
| SNX29 | NM_032167 | -0.3415 | -0.9405 |
| WDR75 | NM_032168 | -1.1825 | -3.5545 |
| CEP78 | NM_032171 | -0.1155 | -1.0435 |
| USP42 | NM_032172 | -1.1685 | -3.38 |
| ZNRF3 | NM_032173 | -1.2 | -4.1155 |
| TOMM40L | NM_032174 | -1.3525 | -2.3385 |
| UTP15 | NM_032175 | -1.132 | -2.3205 |
| PHAX | NM_032177 | -0.9195 | -1.9665 |
| SLC7A6OS | NM_032178 | -0.216 | -0.5685 |
| FAM161A | NM_032180 | -2.0005 | -1.9825 |
| FAM176A | NM_032181 | 0.1435 | -0.388 |
| FAM175B | NM_032182 | -1.197 | -2.904 |
| SYDE2 | NM_032184 | -0.3445 | -0.712 |
| ZNF644 | NM_032186 | -0.7555 | -3.4265 |
| KAT8 | NM_032188 | -0.4675 | -1.37 |
| ATP11A | NM_032189 | -1.094 | -1.963 |
| PPP1R1B | NM_032192 | -0.207 | -0.123 |
| RNASEH2C | NM_032193 | 0.2275 | -0.2895 |
| RPF2 | NM_032194 | -1.028 | -2.5455 |
| SON | NM_032195 | -1.7915 | -2.9265 |
| ARID5B | NM_032199 | -0.737 | -1.2855 |
| ASCC2 | NM_032204 | -0.348 | -1.204 |
| PHF20L1 | NM_032205 | 0.045 | -0.5465 |
| NLRC5 | NM_032206 | -0.36 | -0.5935 |
| C19orf44 | NM_032207 | -0.6535 | -0.783 |
| ANTXR1 | NM_032208 | -1.115 | -1.0975 |

| | | | |
|---|---|---|---|
| LOXL4 | NM_032211 | -0.1755 | -0.5335 |
| ELMOD3 | NM_032213 | -0.546 | -1.328 |
| SLA2 | NM_032214 | 0.1045 | -0.285 |
| ANKRD17 | NM_032217 | -1.696 | -4.3965 |
| MFSD7 | NM_032219 | -0.067 | -0.387 |
| CHD6 | NM_032221 | -1.557 | -3.3025 |
| FAM188B | NM_032222 | -0.7035 | -0.739 |
| PCNXL3 | NM_032223 | -0.1075 | -1.194 |
| ZCCHC7 | NM_032226 | -1.216 | -2.3005 |
| TMEM164 | NM_032227 | 0.1245 | -0.43 |
| FAR1 | NM_032228 | -0.819 | -1.679 |
| SLITRK6 | NM_032229 | 0.0875 | -0.1625 |
| C12orf26 | NM_032230 | -0.9345 | -1.7175 |
| FAM96A | NM_032231 | -0.712 | -1.636 |
| SETD3 | NM_032233 | -1.2935 | -2.999 |
| USP48 | NM_032236 | -0.76 | -1.467 |
| SGK196 | NM_032237 | -0.5285 | -0.7815 |
| LARP1B | NM_032239 | -0.1655 | -0.745 |
| PLXNA1 | NM_032242 | -0.662 | -1.344 |
| TXNDC2 | NM_032243 | 0.188 | 0.1015 |
| MEX3B | NM_032246 | 0.2125 | 0.6555 |
| CCDC88B | NM_032251 | -0.057 | -0.1515 |
| TMEM117 | NM_032256 | -0.9695 | -1.484 |
| ZMYND12 | NM_032257 | 0.2445 | -0.3005 |
| WDR24 | NM_032259 | -0.1655 | -0.935 |
| RGPD5 | NM_032260 | 0.368 | 0.876 |
| C21orf56 | NM_032261 | -0.3795 | -0.951 |
| IQCG | NM_032263 | -0.6465 | -0.368 |
| NBPF3 | NM_032264 | -0.719 | -0.7575 |
| C2orf16 | NM_032266 | -0.128 | -0.5755 |
| ZNRF1 | NM_032268 | -0.016 | -1.327 |
| CCDC135 | NM_032269 | 0.1605 | -0.212 |
| LRRC8C | NM_032270 | -0.804 | -1.4625 |
| TRAF7 | NM_032271 | 0.1805 | -0.417 |
| MAF1 | NM_032272 | -0.109 | -0.9975 |
| TMEM126A | NM_032273 | -0.7475 | -1.0555 |
| RHBDD1 | NM_032276 | -1.1225 | -1.4295 |
| ATP13A4 | NM_032279 | 0.155 | -0.4255 |
| ELAVL3 | NM_032281 | -1.109 | -1.8905 |
| ZDHHC18 | NM_032283 | -0.2235 | -0.612 |
| MRI1 | NM_032285 | -0.7605 | -1.3515 |
| MED10 | NM_032286 | -0.828 | -2.4115 |
| LDOC1L | NM_032287 | -1.3945 | -3.2735 |

| | | | |
|---|---|---|---|
| FYTTD1 | NM_032288 | -0.7315 | -1.058 |
| PSD2 | NM_032289 | -0.953 | -0.8535 |
| ANKRD32 | NM_032290 | -0.7465 | -2.7685 |
| SGIP1 | NM_032291 | -1.238 | -1.317 |
| GON4L | NM_032292 | -1.1995 | -2.121 |
| GARNL3 | NM_032293 | 0.102 | -0.7955 |
| CAMKK1 | NM_032294 | -1.274 | -1.892 |
| SLC37A3 | NM_032295 | -1.665 | -1.8665 |
| FLYWCH1 | NM_032296 | -0.2205 | -1.24 |
| SYT3 | NM_032298 | -0.6255 | -0.2585 |
| DCUN1D5 | NM_032299 | -1.3285 | -3.452 |
| TCHP | NM_032300 | -1.0595 | -1.8355 |
| FBXW9 | NM_032301 | -0.128 | -0.7225 |
| PSMG3 | NM_032302 | -0.468 | -1.438 |
| HSDL2 | NM_032303 | -0.684 | -0.6045 |
| HAGHL | NM_032304 | 0.2435 | -0.2785 |
| POLR3GL | NM_032305 | -0.815 | -1.339 |
| ALKBH7 | NM_032306 | 0.1345 | -0.7375 |
| C9orf64 | NM_032307 | -1.988 | -3.6255 |
| CHCHD5 | NM_032309 | -0.187 | -0.81 |
| C9orf89 | NM_032310 | -0.017 | -0.971 |
| POLDIP3 | NM_032311 | -0.468 | -1.283 |
| YIPF4 | NM_032312 | -1.4405 | -3.006 |
| COQ5 | NM_032314 | -1.3375 | -1.5795 |
| SLC25A33 | NM_032315 | -0.511 | -2.124 |
| NICN1 | NM_032316 | -0.622 | -1.2425 |
| DNAJC30 | NM_032317 | 0.459 | 0.459 |
| PRADC1 | NM_032319 | -0.3525 | -1.4285 |
| BTBD10 | NM_032320 | -1.564 | -3.398 |
| C2orf88 | NM_032321 | 0.3055 | -0.476 |
| RNF135 | NM_032322 | -0.099 | -0.3995 |
| TMEM79 | NM_032323 | -0.886 | -1.435 |
| EIF1AD | NM_032325 | -0.888 | -3.158 |
| TMEM175 | NM_032326 | -0.236 | -0.702 |
| ZDHHC16 | NM_032327 | -0.4315 | -1.684 |
| EFCAB2 | NM_032328 | -0.875 | -1.0975 |
| ING5 | NM_032329 | -1.717 | -1.8055 |
| CAPNS2 | NM_032330 | -0.1535 | -0.1545 |
| ECE2 | NM_032331 | -0.19 | -1.4955 |
| C10orf58 | NM_032333 | -0.401 | -0.456 |
| UTP23 | NM_032334 | -1.6515 | -3.4465 |
| PHF6 | NM_032335 | -0.8655 | -2.384 |
| GINS4 | NM_032336 | -1.127 | -1.5235 |

| | | | |
|---|---|---|---|
| LLPH | NM_032338 | -0.805 | -1.7195 |
| MIEN1 | NM_032339 | -0.0885 | -0.497 |
| C6orf125 | NM_032340 | -0.2015 | -0.532 |
| DDI2 | NM_032341 | -0.046 | -0.593 |
| C9orf125 | NM_032342 | -0.682 | -2.392 |
| CHCHD6 | NM_032343 | 0.197 | -0.2625 |
| NUDT22 | NM_032344 | -0.336 | -1.1265 |
| WIBG | NM_032345 | -0.67 | -3.0195 |
| PDCD2L | NM_032346 | -0.7425 | -2.3095 |
| ZNF397 | NM_032347 | -1.106 | -1.745 |
| MXRA8 | NM_032348 | 0.0495 | -0.678 |
| NUDT16L1 | NM_032349 | -0.3915 | -0.98 |
| C7orf50 | NM_032350 | -0.5965 | -1.261 |
| MRPL45 | NM_032351 | -1.225 | -1.88 |
| BRMS1L | NM_032352 | -0.457 | -0.8755 |
| VPS25 | NM_032353 | -0.448 | -2.009 |
| TMEM107 | NM_032354 | -0.1865 | -0.5385 |
| MON1A | NM_032355 | -0.303 | -1.817 |
| LSMD1 | NM_032356 | 0.317 | -0.263 |
| CCDC115 | NM_032357 | -0.299 | -0.652 |
| CCDC77 | NM_032358 | -2.026 | -3.523 |
| C3orf26 | NM_032359 | -0.6975 | -0.7255 |
| ACBD6 | NM_032360 | -0.392 | -1.409 |
| THOC3 | NM_032361 | -0.517 | -0.564 |
| DNAJC14 | NM_032364 | -1.2475 | -2.991 |
| C16orf13 | NM_032366 | 0.23 | -0.2425 |
| ZBED3 | NM_032367 | -0.688 | -1.8995 |
| LZIC | NM_032368 | -1.2 | -1.3215 |
| HVCN1 | NM_032369 | 0.058 | -0.405 |
| ZNF414 | NM_032370 | -0.465 | -0.793 |
| DYDC2 | NM_032372 | 0.013 | -0.498 |
| PCGF5 | NM_032373 | 0.376 | 0.1555 |
| AKT1S1 | NM_032375 | -0.1365 | -0.59 |
| TMEM101 | NM_032376 | -0.367 | -1.693 |
| ELOF1 | NM_032377 | -0.175 | -1.117 |
| EEF1D | NM_032378 | 0.3765 | -0.1945 |
| SYTL2 | NM_032379 | -0.046 | -0.2065 |
| COG8 | NM_032382 | -0.4285 | -2.135 |
| HPS3 | NM_032383 | -1.5905 | -1.915 |
| C5orf4 | NM_032385 | -0.388 | -0.4285 |
| WNK4 | NM_032387 | -0.144 | 0.1815 |
| ARFGAP2 | NM_032389 | -0.3405 | -1.365 |
| MKI67IP | NM_032390 | -1.7765 | -3.2375 |

| | | | |
|---|---|---|---|
| PRAC | NM_032391 | -0.1845 | -0.1955 |
| PCDHGC3 | NM_032402 | 0.013 | 0.1875 |
| TMPRSS3 | NM_032405 | 0.071 | -0.348 |
| PCDHGC4 | NM_032406 | 0.2155 | -0.455 |
| PCDHGC5 | NM_032407 | -0.3175 | 0.125 |
| BAZ1B | NM_032408 | -1.5205 | -2.863 |
| PINK1 | NM_032409 | -0.119 | -0.583 |
| HOOK3 | NM_032410 | -1.651 | -2.746 |
| C2orf40 | NM_032411 | -0.446 | -0.214 |
| C5orf32 | NM_032412 | -0.052 | -1.198 |
| C15orf48 | NM_032413 | -0.329 | -0.363 |
| PROK1 | NM_032414 | -0.201 | -0.4185 |
| CARD11 | NM_032415 | 0.1785 | -0.27 |
| PCDH1 | NM_032420 | -0.473 | -0.6605 |
| CLIP2 | NM_032421 | -0.4475 | -1.0325 |
| ZNF528 | NM_032423 | -0.2585 | -0.7625 |
| KIAA1826 | NM_032424 | -1.3955 | -3.18 |
| HHIPL1 | NM_032425 | -0.0845 | -0.19 |
| MAML2 | NM_032427 | -0.924 | -1.7735 |
| LZTS2 | NM_032429 | -0.275 | -1.267 |
| BRSK1 | NM_032430 | -0.3585 | -0.356 |
| SYVN1 | NM_032431 | -0.0205 | -0.998 |
| ABLIM2 | NM_032432 | 0.0865 | 0.1585 |
| ZNF333 | NM_032433 | 0.2425 | -0.0895 |
| ZNF512 | NM_032434 | -0.7415 | -1.0105 |
| KIAA1804 | NM_032435 | -0.4925 | -3.092 |
| CHAMP1 | NM_032436 | -1.117 | -3.371 |
| EFCAB7 | NM_032437 | -0.881 | -2.2105 |
| L3MBTL3 | NM_032438 | -0.453 | -1.039 |
| PHYHIPL | NM_032439 | -0.879 | -2.4345 |
| LCOR | NM_032440 | -0.375 | -0.826 |
| NEURL4 | NM_032442 | -0.773 | -1.3315 |
| SLX4 | NM_032444 | -0.994 | -2.134 |
| MEGF11 | NM_032445 | -0.0065 | -0.2995 |
| MEGF10 | NM_032446 | 0.3845 | -0.2885 |
| FBN3 | NM_032447 | 0.1805 | 0.152 |
| FAM120B | NM_032448 | -1.36 | -2.4865 |
| CC2D1B | NM_032449 | -0.559 | -1.5645 |
| HEATR7A | NM_032450 | -0.095 | -0.565 |
| SPIRE2 | NM_032451 | -0.916 | -0.6 |
| JPH4 | NM_032452 | -0.662 | -1.1085 |
| ZNF527 | NM_032453 | -0.732 | -1.6045 |
| STK19 | NM_032454 | -0.8375 | -1.4225 |

| | | | |
|---|---|---|---|
| PCDH7 | NM_032456 | -0.437 | -0.099 |
| PCDH7 | NM_032457 | -0.437 | -0.099 |
| PHF6 | NM_032458 | -0.995 | -2.543 |
| EFS | NM_032459 | -1.2695 | -2.6785 |
| LAT2 | NM_032463 | 0.1465 | -0.4915 |
| LAT2 | NM_032464 | 0.1465 | -0.4915 |
| ASPH | NM_032466 | -1.3145 | -1.287 |
| ASPH | NM_032467 | -1.0635 | -0.5415 |
| ASPH | NM_032468 | -1.0635 | -0.5415 |
| PKIB | NM_032471 | -0.7525 | -1.06 |
| PPIL3 | NM_032472 | -1.016 | -2.1165 |
| MRPS6 | NM_032476 | -0.884 | -1.62 |
| MRPL41 | NM_032477 | 0.1215 | -0.3165 |
| MRPL38 | NM_032478 | -0.361 | -1.2915 |
| MRPL36 | NM_032479 | -1.0465 | -2.88 |
| DOT1L | NM_032482 | -0.5555 | -1.232 |
| PPAPDC1B | NM_032483 | -0.047 | -0.366 |
| MCM8 | NM_032485 | -0.878 | -1.377 |
| DCTN5 | NM_032486 | -0.685 | -1.5425 |
| ARPM1 | NM_032487 | -0.715 | -0.4725 |
| CNFN | NM_032488 | 0.1785 | -0.158 |
| ACRBP | NM_032489 | 0.0095 | -0.0755 |
| C14orf142 | NM_032490 | -1.133 | -1.752 |
| RFX4 | NM_032491 | 0.2 | -0.0625 |
| JAGN1 | NM_032492 | -1.3235 | -3.228 |
| AP1M1 | NM_032493 | -0.332 | -1.168 |
| ZC3H8 | NM_032494 | -1.202 | -2.394 |
| HOPX | NM_032495 | -0.0895 | -0.5305 |
| ARHGAP9 | NM_032496 | -0.1585 | -0.6555 |
| ZNF559 | NM_032497 | -1.098 | -3.5055 |
| C15orf41 | NM_032499 | -0.9165 | -1.5505 |
| ACSS1 | NM_032501 | 0 | -0.4265 |
| MCHR2 | NM_032503 | 0.2015 | -0.241 |
| UNC80 | NM_032504 | -1.145 | -1.494 |
| KBTBD8 | NM_032505 | -0.743 | -1.157 |
| KIAA1841 | NM_032506 | -0.3325 | 0.0965 |
| PGBD1 | NM_032507 | -1.4745 | -2.4205 |
| TMEM185A | NM_032508 | 0.696 | -0.1825 |
| MAK16 | NM_032509 | -1.2045 | -2.724 |
| PARD6G | NM_032510 | -0.723 | -2.9475 |
| C6orf168 | NM_032511 | -1.074 | -2.6035 |
| PDZD4 | NM_032512 | -0.079 | -0.561 |
| SLC30A2 | NM_032513 | -0.0905 | 0.188 |

| | | | |
|---|---|---|---|
| MAP1LC3A | NM_032514 | 0.0915 | -0.4175 |
| BOK | NM_032515 | -0.344 | -0.6505 |
| COL25A1 | NM_032518 | -0.2135 | -0.8615 |
| GNPTG | NM_032520 | 0.1235 | -0.553 |
| PARD6B | NM_032521 | -1.7285 | -2.8255 |
| ZBTB37 | NM_032522 | -0.079 | -0.437 |
| OSBPL6 | NM_032523 | -0.6335 | -1.7275 |
| KRTAP4-4 | NM_032524 | -0.397 | -0.132 |
| TUBB6 | NM_032525 | -0.163 | -1.2575 |
| NT5C1A | NM_032526 | -0.4385 | -0.052 |
| ST6GAL2 | NM_032528 | -1.1305 | -2.5 |
| ZNF594 | NM_032530 | -1.398 | -3.1845 |
| KIRREL3 | NM_032531 | -0.179 | -0.3365 |
| FNDC1 | NM_032532 | -0.0145 | 0.668 |
| KRBA1 | NM_032534 | -0.751 | -0.8875 |
| NTNG2 | NM_032536 | -0.0975 | -0.518 |
| TTBK1 | NM_032538 | -0.409 | -0.9775 |
| SLITRK2 | NM_032539 | -0.1435 | -0.487 |
| TRIM54 | NM_032546 | 0.1885 | 0.204 |
| SCOC | NM_032547 | -0.5325 | -1.0695 |
| ABTB1 | NM_032548 | 0.3145 | -0.5195 |
| IMMP2L | NM_032549 | -0.2665 | -0.753 |
| AFAP1L2 | NM_032550 | 0.0205 | -0.0525 |
| KISS1R | NM_032551 | -0.1315 | -0.4575 |
| DAB2IP | NM_032552 | -0.722 | -2.202 |
| GPR174 | NM_032553 | 0.371 | -0.059 |
| HCAR1 | NM_032554 | 0.0135 | -0.3925 |
| IL1F10 | NM_032556 | -0.0685 | 0.4065 |
| USP38 | NM_032557 | -1.6865 | -4.9395 |
| HIATL1 | NM_032558 | -0.4245 | -1.7555 |
| KIF2B | NM_032559 | -0.056 | 0.1735 |
| SMEK1 | NM_032560 | -1.6185 | -4.2415 |
| C22orf23 | NM_032561 | -0.267 | -0.3445 |
| PLA2G12B | NM_032562 | -0.079 | -0.345 |
| LCE3D | NM_032563 | 0.359 | -0.4775 |
| DGAT2 | NM_032564 | -0.4965 | -0.612 |
| EBPL | NM_032565 | -0.0875 | -0.3545 |
| SPINK7 | NM_032566 | -0.146 | -0.5745 |
| SPZ1 | NM_032567 | 0.432 | -0.045 |
| GLYR1 | NM_032569 | -0.531 | -0.87 |
| EMR3 | NM_032571 | 0.0345 | -0.1835 |
| RNASE7 | NM_032572 | -0.191 | -0.286 |
| DPY30 | NM_032574 | -1.0135 | -1.393 |

| | | | |
|---|---|---|---|
| GLIS2 | NM_032575 | -0.538 | -2.486 |
| CYorf15B | NM_032576 | 0.2495 | -0.895 |
| MYPN | NM_032578 | -0.141 | -0.5765 |
| RETNLB | NM_032579 | -0.0875 | -0.057 |
| HES7 | NM_032580 | -0.429 | -0.801 |
| FAM126A | NM_032581 | -1.011 | -2.037 |
| USP32 | NM_032582 | -1.4375 | -2.4295 |
| ABCC11 | NM_032583 | 0.1905 | -0.542 |
| ZNF347 | NM_032584 | -1.1355 | -2.425 |
| CARD6 | NM_032587 | 0.416 | -0.576 |
| TRIM63 | NM_032588 | -0.2965 | -0.653 |
| KDM2B | NM_032590 | -0.1165 | -0.655 |
| SLC9A7 | NM_032591 | 0.2005 | -0.027 |
| ACCS | NM_032592 | 0.1525 | -0.293 |
| HINT2 | NM_032593 | 0.14 | -0.612 |
| INSM2 | NM_032594 | -0.663 | -0.6125 |
| PPP1R9B | NM_032595 | -0.577 | -1.3035 |
| C9orf24 | NM_032596 | 0.0805 | -0.0955 |
| MS4A14 | NM_032597 | 0.1455 | -0.2355 |
| SPATA22 | NM_032598 | -0.29 | -0.129 |
| FAM71F1 | NM_032599 | -0.1375 | 0.1645 |
| CCDC54 | NM_032600 | -0.139 | -0.096 |
| MCEE | NM_032601 | -1.471 | -2.08 |
| GJA10 | NM_032602 | -0.046 | -0.05 |
| LOXL3 | NM_032603 | -0.4685 | -1.908 |
| ABHD1 | NM_032604 | -0.021 | -0.188 |
| CAPS2 | NM_032606 | -0.376 | 0.3325 |
| CREB3L3 | NM_032607 | -0.081 | -0.614 |
| MYO18B | NM_032608 | -0.1795 | -0.424 |
| COX4I2 | NM_032609 | 0.4165 | -0.2635 |
| GTPBP3 | NM_032620 | 0.1175 | -0.088 |
| BEX2 | NM_032621 | -0.2965 | -0.7945 |
| LNX1 | NM_032622 | -0.3485 | -0.7315 |
| C4orf49 | NM_032623 | -0.6935 | -0.917 |
| RBBP6 | NM_032626 | -0.2585 | -0.423 |
| SSBP4 | NM_032627 | 0.106 | -0.328 |
| CINP | NM_032630 | 0.2125 | -0.6645 |
| HDGFRP2 | NM_032631 | -0.176 | -1.313 |
| PAPOLA | NM_032632 | 0.144 | -0.3525 |
| TMEM147 | NM_032635 | 0.16 | -0.479 |
| SKP2 | NM_032637 | -0.9155 | -1.594 |
| GATA2 | NM_032638 | -0.829 | -1.959 |
| PLEKHA8 | NM_032639 | -0.9615 | -1.962 |

| | | | |
|---|---|---|---|
| SPSB2 | NM_032641 | -0.369 | -0.71 |
| WNT5B | NM_032642 | -0.2895 | -1.3225 |
| IRF5 | NM_032643 | -0.1315 | -0.227 |
| TTYH2 | NM_032646 | -0.5955 | -1.363 |
| CBX2 | NM_032647 | -0.4735 | -0.6685 |
| FAM167B | NM_032648 | -0.3325 | -0.3 |
| CNDP1 | NM_032649 | -0.1735 | -0.2625 |
| DHX37 | NM_032656 | -0.709 | -1.67 |
| USP30 | NM_032663 | -1.4215 | -3.1045 |
| FUT10 | NM_032664 | 0.3355 | -0.057 |
| BSCL2 | NM_032667 | -0.107 | -0.9375 |
| PCGF1 | NM_032673 | -0.1435 | -0.5195 |
| ZNF577 | NM_032679 | -0.4305 | -1.437 |
| EFCAB4B | NM_032680 | 0.1915 | -0.324 |
| SPRYD5 | NM_032681 | -0.2525 | -0.1605 |
| FOXP1 | NM_032682 | -1.1875 | -1.91 |
| MPV17L2 | NM_032683 | -0.156 | -1.379 |
| CYHR1 | NM_032687 | -0.7155 | -1.125 |
| ZNF607 | NM_032689 | -1.8145 | -3.423 |
| NAA11 | NM_032693 | -0.1725 | -0.257 |
| SUV420H2 | NM_032701 | -0.4245 | -1.2325 |
| TUBA1C | NM_032704 | -0.142 | -1.604 |
| PYROXD2 | NM_032709 | -0.321 | -0.727 |
| MAFG | NM_032711 | 0.033 | -0.1765 |
| INF2 | NM_032714 | -0.3 | -0.759 |
| AGPAT9 | NM_032717 | -0.58 | -0.305 |
| MFSD9 | NM_032718 | -0.1405 | -0.7025 |
| BUD13 | NM_032725 | -0.7175 | -3.378 |
| PLCD4 | NM_032726 | -0.355 | -0.4475 |
| INA | NM_032727 | -0.4405 | -0.386 |
| PPAPDC3 | NM_032728 | -1.1985 | -1.952 |
| RTN4IP1 | NM_032730 | -1.607 | -2.9405 |
| TXNDC17 | NM_032731 | -0.165 | -0.5195 |
| IL17RC | NM_032732 | -0.0415 | -0.5445 |
| BEST3 | NM_032735 | -0.005 | -0.125 |
| LMNB2 | NM_032737 | -0.4945 | -0.697 |
| FCRLA | NM_032738 | -0.2045 | -0.5575 |
| SFT2D3 | NM_032740 | -0.6835 | -0.8125 |
| AGPAT1 | NM_032741 | -0.031 | -0.4585 |
| C6orf105 | NM_032744 | -0.1875 | -0.553 |
| USMG5 | NM_032747 | 0.12 | 0.041 |
| ABHD14B | NM_032750 | 0.244 | -0.4485 |
| ZNF496 | NM_032752 | -0.5655 | -1.499 |

| | | | |
|---|---|---|---|
| RAX2 | NM_032753 | 0.0685 | -0.189 |
| HPDL | NM_032756 | -0.7215 | -1.5155 |
| PHF5A | NM_032758 | -0.8095 | -2.5555 |
| TRIM52 | NM_032765 | -1.366 | -1.7985 |
| ZNF503 | NM_032772 | 0.5985 | 0.791 |
| LRCH3 | NM_032773 | -0.992 | -1.771 |
| KLHL22 | NM_032775 | -1.12 | -2.0645 |
| JMJD1C | NM_032776 | -0.5435 | -2.673 |
| GPR124 | NM_032777 | -0.9375 | -1.827 |
| MINA | NM_032778 | -0.971 | -2.2405 |
| CCDC142 | NM_032779 | -0.7095 | -1.694 |
| TMEM25 | NM_032780 | -0.588 | -2.037 |
| PTPN5 | NM_032781 | -0.1585 | -0.496 |
| HAVCR2 | NM_032782 | -0.0695 | -0.148 |
| CBR4 | NM_032783 | -1.421 | -2.0685 |
| RSPO3 | NM_032784 | 0.1985 | 0.103 |
| AGBL4 | NM_032785 | -0.1645 | -0.6325 |
| ZC3H10 | NM_032786 | -0.993 | -1.621 |
| GPR128 | NM_032787 | -0.002 | -0.2305 |
| ZNF514 | NM_032788 | -1.3445 | -2.025 |
| PARP10 | NM_032789 | 0.0775 | -0.56 |
| ORAI1 | NM_032790 | -1.187 | -2.444 |
| ZBTB45 | NM_032792 | -0.836 | -2.254 |
| MFSD2A | NM_032793 | -0.4865 | -1.5115 |
| RPUSD4 | NM_032795 | -0.904 | -2.3975 |
| SYAP1 | NM_032796 | -0.6075 | -0.857 |
| AIFM2 | NM_032797 | -0.4 | -0.3975 |
| ZDHHC12 | NM_032799 | -0.2825 | -1.0875 |
| C1orf198 | NM_032800 | 0.0345 | -0.926 |
| JAM3 | NM_032801 | -0.324 | -1.404 |
| SPPL2A | NM_032802 | -0.474 | -0.982 |
| SLC7A3 | NM_032803 | 0.278 | 0.405 |
| ADO | NM_032804 | -1.082 | -2.0125 |
| ZSCAN10 | NM_032805 | -0.264 | -0.2205 |
| C3orf39 | NM_032806 | 0.0195 | -0.874 |
| FBXO18 | NM_032807 | -0.8625 | -2.041 |
| LINGO1 | NM_032808 | -0.3675 | -0.6305 |
| FAM73B | NM_032809 | -0.7005 | -1.7885 |
| ATAD1 | NM_032810 | -1.2355 | -1.843 |
| TBRG1 | NM_032811 | -0.443 | -0.445 |
| PLXDC2 | NM_032812 | 0.3025 | -0.4865 |
| TMTC4 | NM_032813 | -1.7565 | -2.2445 |
| RNFT2 | NM_032814 | -0.645 | -1.206 |

| | | | |
|---|---|---|---|
| NFATC2IP | NM_032815 | -0.815 | -1.6975 |
| ITIH5 | NM_032817 | -0.036 | -0.487 |
| C9orf100 | NM_032818 | -0.949 | -2.1295 |
| ZNF341 | NM_032819 | -0.731 | -1.769 |
| HYDIN | NM_032821 | 0.1905 | 0.063 |
| FAM136A | NM_032822 | -1.1395 | -1.5875 |
| C9orf3 | NM_032823 | -0.654 | -2.7235 |
| TMEM87B | NM_032824 | -0.131 | -0.6935 |
| ZNF382 | NM_032825 | -1.027 | -2.703 |
| SLC35B4 | NM_032826 | -1.8255 | -1.937 |
| ATOH8 | NM_032827 | -0.2845 | -2.2415 |
| ZNF587 | NM_032828 | -1.1555 | -1.527 |
| C12orf34 | NM_032829 | -0.676 | -2.7835 |
| CIRH1A | NM_032830 | -0.546 | -1.589 |
| ORAI2 | NM_032831 | -0.5505 | -1.0075 |
| LRP11 | NM_032832 | -1.486 | -2.282 |
| PPP1R15B | NM_032833 | -1.5115 | -2.177 |
| ALG10 | NM_032834 | -0.807 | -2.109 |
| FIZ1 | NM_032836 | -1.172 | -1.9105 |
| FAM104A | NM_032837 | -0.643 | -2.5405 |
| ZNF566 | NM_032838 | -1.2225 | -2.2965 |
| DIRC2 | NM_032839 | -0.6205 | -2.416 |
| SPRYD3 | NM_032840 | -0.3435 | -1.093 |
| TMEM209 | NM_032842 | -1.3985 | -2.191 |
| FIBCD1 | NM_032843 | -0.067 | -0.416 |
| MASTL | NM_032844 | -1.912 | -4.0075 |
| RAB2B | NM_032846 | -0.589 | -0.8845 |
| C8orf76 | NM_032847 | -1.5325 | -3.9615 |
| C12orf52 | NM_032848 | -0.6895 | -1.9635 |
| C13orf33 | NM_032849 | -0.015 | -0.127 |
| ATG4C | NM_032852 | -1.534 | -3.445 |
| MUM1 | NM_032853 | -0.873 | -1.7175 |
| CORO6 | NM_032854 | -0.2575 | -0.897 |
| HSH2D | NM_032855 | 0.0755 | -0.5215 |
| WDR73 | NM_032856 | -0.957 | -1.7515 |
| LACTB | NM_032857 | -1.5135 | -2.677 |
| MAEL | NM_032858 | 0.169 | -0.358 |
| ABHD13 | NM_032859 | -1.0665 | -1.2585 |
| LTV1 | NM_032860 | -1.184 | -3.3135 |
| SERAC1 | NM_032861 | -1.0955 | -1.4745 |
| TIGD5 | NM_032862 | -1.007 | -2.194 |
| PRPF38A | NM_032864 | -0.775 | -1.212 |
| TNS4 | NM_032865 | 0.14 | -0.282 |

| | | | |
|---|---|---|---|
| CGNL1 | NM_032866 | -0.4065 | -0.174 |
| MICALCL | NM_032867 | 0.085 | -0.213 |
| MPND | NM_032868 | 0.319 | -0.278 |
| NUDCD1 | NM_032869 | -1.199 | -2.8695 |
| PNISR | NM_032870 | -0.2355 | -1.0545 |
| RELT | NM_032871 | -0.2845 | -1.263 |
| SYTL1 | NM_032872 | -0.042 | -0.6325 |
| UBASH3B | NM_032873 | -0.243 | -0.437 |
| FBXL20 | NM_032875 | -0.9725 | -1.695 |
| JUB | NM_032876 | -0.8655 | -1.3835 |
| ALKBH6 | NM_032878 | -0.4385 | -1.0345 |
| IGSF21 | NM_032880 | -0.094 | -0.4625 |
| LSM10 | NM_032881 | -1.07 | -2.9 |
| PNMA6A | NM_032882 | 0.0155 | -0.6115 |
| TOX2 | NM_032883 | -0.1435 | -0.454 |
| ATG4D | NM_032885 | -0.292 | -1.4895 |
| COL27A1 | NM_032888 | -0.2525 | -0.0015 |
| MFSD5 | NM_032889 | -0.5305 | -1.9465 |
| DISP1 | NM_032890 | -1.412 | -2.7215 |
| FRMD5 | NM_032892 | 0.257 | -0.747 |
| CEP19 | NM_032898 | -1.6165 | -1.785 |
| FAM83A | NM_032899 | 0.082 | -0.2355 |
| ARHGAP19 | NM_032900 | -1.4255 | -1.9875 |
| C12orf62 | NM_032901 | -0.084 | -0.8965 |
| PPP1R16A | NM_032902 | -0.358 | -0.6595 |
| RBM17 | NM_032905 | -0.789 | -1.2275 |
| PIGY | NM_032906 | -0.9375 | -1.269 |
| RERG | NM_032918 | -1.9705 | -0.6555 |
| ZNF3 | NM_032924 | -1.195 | -2.632 |
| TMEM128 | NM_032927 | -0.597 | -0.922 |
| TMEM141 | NM_032928 | 0.1 | -0.457 |
| C11orf70 | NM_032930 | -1.025 | -1.08 |
| RAB11FIP4 | NM_032932 | -0.244 | -0.93 |
| C18orf45 | NM_032933 | -0.1355 | -1.2625 |
| MT4 | NM_032935 | 0.0585 | -0.4035 |
| TMEM60 | NM_032936 | -1.957 | -4.0565 |
| C9orf37 | NM_032937 | -0.7785 | -1.593 |
| POLR2C | NM_032940 | -0.1965 | -0.93 |
| RECQL | NM_032941 | -0.493 | -2.1555 |
| SYTL2 | NM_032943 | -0.046 | -0.2065 |
| STK31 | NM_032944 | -0.3935 | -0.1425 |
| NXF5 | NM_032946 | -0.106 | 0.186 |
| C5orf62 | NM_032947 | -0.3995 | -0.854 |

| | | | |
|---|---|---|---|
| PCDH8 | NM_032949 | 0.0205 | 0.261 |
| MLXIPL | NM_032951 | 0.136 | -0.012 |
| MLXIPL | NM_032952 | 0.136 | -0.012 |
| MLXIPL | NM_032953 | 0.136 | -0.012 |
| MLXIPL | NM_032954 | 0.136 | -0.012 |
| AIF1 | NM_032955 | -0.1545 | -0.3475 |
| RTEL1 | NM_032957 | -0.338 | -0.541 |
| MAPKAPK2 | NM_032960 | 0.2005 | -0.8085 |
| PCDH10 | NM_032961 | 0.041 | -1.4465 |
| CCL14 | NM_032962 | -0.0075 | -0.4705 |
| CCL15 | NM_032965 | 0.1725 | -0.5095 |
| CXCR5 | NM_032966 | -0.002 | -0.348 |
| SEC22C | NM_032970 | -0.698 | -1.946 |
| PCDH11Y | NM_032972 | -0.1165 | -0.178 |
| CASP10 | NM_032974 | 0.2235 | 0.0155 |
| DTNA | NM_032975 | -0.5815 | -1.7065 |
| CASP10 | NM_032976 | 0.2235 | 0.0155 |
| CASP10 | NM_032977 | 0.2235 | 0.0155 |
| DTNA | NM_032978 | -0.5815 | -1.7065 |
| DTNA | NM_032979 | -0.5815 | -1.7065 |
| DTNA | NM_032980 | -0.5815 | -1.7065 |
| DTNA | NM_032981 | -0.5815 | -1.7065 |
| CASP2 | NM_032982 | -0.539 | -2.0025 |
| CASP2 | NM_032983 | -0.7065 | -1.2805 |
| SEC23B | NM_032985 | -0.6955 | -1.268 |
| SEC23B | NM_032986 | -0.6955 | -1.268 |
| BAD | NM_032989 | -0.2855 | -0.9855 |
| CASP3 | NM_032991 | -1.314 | -2.373 |
| CASP6 | NM_032992 | -0.571 | -1.925 |
| GAR1 | NM_032993 | -0.933 | -3.1825 |
| ARHGEF4 | NM_032995 | -0.2845 | -1.205 |
| CASP9 | NM_032996 | -0.281 | -0.646 |
| ZWINT | NM_032997 | -0.5895 | -1.3875 |
| DEDD | NM_032998 | -1.01 | -2.161 |
| GTF2I | NM_032999 | -0.1905 | -0.6505 |
| GTF2I | NM_033000 | -0.1905 | -0.6505 |
| GTF2I | NM_033001 | -0.1905 | -0.6505 |
| NLRP1 | NM_033004 | -0.5045 | -0.3005 |
| NLRP1 | NM_033006 | -0.5045 | -0.3005 |
| NLRP1 | NM_033007 | -0.5045 | -0.3005 |
| PLAT | NM_033011 | -0.6655 | -1.295 |
| TNFSF11 | NM_033012 | 0.0845 | -0.34 |
| NR1I2 | NM_033013 | -0.2945 | -0.269 |

| | | | |
|---|---|---|---|
| OGN | NM_033014 | 0.254 | 0.329 |
| FASTK | NM_033015 | -0.279 | -0.8605 |
| PDGFB | NM_033016 | 0.062 | -0.184 |
| TRIM4 | NM_033017 | -1.2 | -3.2785 |
| CDK16 | NM_033018 | -0.188 | -1.0355 |
| TRIM33 | NM_033020 | -0.8205 | -1.962 |
| RPS24 | NM_033022 | -0.179 | -0.225 |
| PDGFA | NM_033023 | 0.163 | -0.0895 |
| SYDE1 | NM_033025 | -0.4335 | -1.0355 |
| PCLO | NM_033026 | -0.1595 | -0.499 |
| CSRNP1 | NM_033027 | -0.342 | -1.299 |
| BBS4 | NM_033028 | -0.8125 | -0.7225 |
| LMLN | NM_033029 | -0.6 | -0.8235 |
| BOLL | NM_033030 | -0.1165 | -0.074 |
| CCNB3 | NM_033031 | 0.126 | -0.4125 |
| KRT82 | NM_033033 | 0.0055 | -0.0895 |
| TRIM5 | NM_033034 | -0.0275 | 0.15 |
| TSLP | NM_033035 | -0.155 | -0.498 |
| GAL3ST3 | NM_033036 | -0.574 | -1.5005 |
| KRT84 | NM_033045 | -0.1705 | -0.002 |
| RTKN | NM_033046 | -0.986 | -1.5025 |
| CPXCR1 | NM_033048 | 0.1525 | -0.0345 |
| MUC13 | NM_033049 | 0.103 | 0 |
| SUCNR1 | NM_033050 | -0.2795 | -0.1885 |
| SLC46A2 | NM_033051 | -0.317 | -0.083 |
| MYO1G | NM_033054 | -0.073 | -0.3015 |
| HIAT1 | NM_033055 | -1.51 | -2.2465 |
| PCDH15 | NM_033056 | 0.3695 | 0.4005 |
| OR2B2 | NM_033057 | 0.153 | -0.4475 |
| TRIM55 | NM_033058 | -0.77 | 0.197 |
| KRTAP4-11 | NM_033059 | 0.0335 | -0.3275 |
| KRTAP4-1 | NM_033060 | 0.1485 | 0.184 |
| KRTAP4-7 | NM_033061 | 0.18 | -0.0035 |
| KRTAP4-2 | NM_033062 | 0.468 | -0.2975 |
| MAP6 | NM_033063 | -0.048 | -0.706 |
| ATCAY | NM_033064 | -1.0415 | -1.7345 |
| MPP4 | NM_033066 | 0.166 | 0.062 |
| DMRTB1 | NM_033067 | -0.0615 | -0.4445 |
| ACPT | NM_033068 | -0.0855 | -0.0065 |
| CECR5 | NM_033070 | -0.0635 | -0.9545 |
| SYNE1 | NM_033071 | -0.7175 | -1.7235 |
| DIDO1 | NM_033081 | 0.457 | -1.4845 |
| SARNP | NM_033082 | -0.892 | -0.6475 |

| | | | |
|---|---|---|---|
| EAF1 | NM_033083 | -0.6605 | -1.12 |
| FANCD2 | NM_033084 | -0.816 | -0.939 |
| FATE1 | NM_033085 | -0.1375 | -0.5955 |
| FGD3 | NM_033086 | 0.0665 | -0.804 |
| ALG2 | NM_033087 | -1.2275 | -2.6445 |
| FAM40A | NM_033088 | -0.3095 | -1.644 |
| ZCCHC3 | NM_033089 | -0.2495 | -0.3805 |
| GREB1 | NM_033090 | -0.43 | -0.7655 |
| TRIM4 | NM_033091 | -1.2 | -3.2785 |
| TRIM5 | NM_033092 | -0.0275 | 0.15 |
| TRIM5 | NM_033093 | -0.0275 | 0.15 |
| CDHR1 | NM_033100 | -0.241 | -0.3995 |
| SLC45A3 | NM_033102 | -0.233 | -0.3795 |
| RHPN2 | NM_033103 | -0.73 | -1.7825 |
| STON2 | NM_033104 | -0.1445 | -0.1745 |
| DNAJC5B | NM_033105 | -0.2585 | -0.041 |
| GALP | NM_033106 | -0.0485 | -0.0565 |
| GTPBP10 | NM_033107 | -1.3095 | -2.358 |
| HSFY1 | NM_033108 | 0.213 | 0.1055 |
| PNPT1 | NM_033109 | -1.1765 | -2.089 |
| A2LD1 | NM_033110 | -0.3365 | -0.382 |
| N4BP2L2 | NM_033111 | -0.8585 | -1.5195 |
| RRP36 | NM_033112 | -1.236 | -2.599 |
| ZNF628 | NM_033113 | 0.0155 | -1.086 |
| ZCRB1 | NM_033114 | -0.6115 | -0.706 |
| TBCK | NM_033115 | -0.9635 | -1.4765 |
| NEK9 | NM_033116 | -1.1855 | -1.142 |
| RBM18 | NM_033117 | -0.3785 | -1.0235 |
| MYLK2 | NM_033118 | -0.0185 | -0.571 |
| NKD1 | NM_033119 | 0.0525 | -0.564 |
| NKD2 | NM_033120 | -0.3895 | -0.373 |
| ANKRD13A | NM_033121 | -0.5685 | -1.214 |
| CABS1 | NM_033122 | -0.002 | -0.229 |
| PLCZ1 | NM_033123 | -0.183 | -0.7045 |
| CCDC65 | NM_033124 | -0.0725 | -0.3615 |
| SLC22A16 | NM_033125 | -0.0645 | -0.47 |
| PSKH2 | NM_033126 | -0.1585 | -0.3705 |
| SEC16B | NM_033127 | -0.0565 | -0.4745 |
| SCIN | NM_033128 | 0.2455 | -0.4325 |
| SCRT2 | NM_033129 | -1.081 | -1.866 |
| SIGLEC10 | NM_033130 | 0.156 | -0.129 |
| WNT3A | NM_033131 | 0.17 | -0.2845 |
| ZIC5 | NM_033132 | -0.357 | -1.235 |

| | | | |
|---|---|---|---|
| CNP | NM_033133 | -0.2045 | -1.0625 |
| PDGFD | NM_033135 | -0.256 | 0.0165 |
| FGF1 | NM_033136 | 0.2095 | -0.095 |
| FGF1 | NM_033137 | 0.2095 | -0.095 |
| CALD1 | NM_033138 | -0.6825 | -0.8615 |
| CALD1 | NM_033139 | -0.6825 | -0.8615 |
| CALD1 | NM_033140 | -0.403 | -0.6285 |
| MAP3K9 | NM_033141 | -0.212 | -0.791 |
| FGF5 | NM_033143 | 0.176 | -0.331 |
| DTNB | NM_033147 | -0.2295 | -0.905 |
| DTNB | NM_033148 | -0.2295 | -0.905 |
| COL2A1 | NM_033150 | 0.0005 | 0.007 |
| ABCC11 | NM_033151 | 0.1905 | -0.542 |
| CALD1 | NM_033157 | -0.6825 | -0.8615 |
| HYAL2 | NM_033158 | -0.542 | -1.406 |
| HYAL1 | NM_033159 | -0.0895 | 0.034 |
| ZNF658 | NM_033160 | -0.1415 | 0.1185 |
| SURF4 | NM_033161 | -0.2335 | -1.1565 |
| FGF8 | NM_033164 | -0.044 | -0.1865 |
| B3GALNT1 | NM_033167 | 0.0155 | -0.871 |
| B3GALNT1 | NM_033168 | 0.0155 | -0.871 |
| B3GALNT1 | NM_033169 | 0.0155 | -0.871 |
| B3GALT5 | NM_033170 | 0.26 | -0.06 |
| B3GALT5 | NM_033171 | 0.26 | -0.06 |
| B3GALT5 | NM_033172 | 0.26 | -0.06 |
| B3GALT5 | NM_033173 | 0.26 | -0.06 |
| NKX2-4 | NM_033176 | -0.046 | -0.449 |
| OR51B4 | NM_033179 | -0.5105 | -0.1855 |
| OR51B2 | NM_033180 | 0.0145 | -0.1345 |
| CNR1 | NM_033181 | 0.0025 | -0.3535 |
| FBXO44 | NM_033182 | -0.2985 | -0.644 |
| KRTAP2-4 | NM_033184 | 0.033 | -0.1165 |
| KRTAP3-3 | NM_033185 | -0.407 | -0.2205 |
| KRTAP4-3 | NM_033187 | -0.1665 | -0.7195 |
| KRTAP4-5 | NM_033188 | -0.0615 | -0.333 |
| KRTAP9-4 | NM_033191 | -0.1135 | -0.1355 |
| HSPB9 | NM_033194 | 0.566 | 0.6125 |
| LDHAL6B | NM_033195 | 0.032 | -0.1485 |
| ZNF682 | NM_033196 | -1.334 | -1.4795 |
| PIGS | NM_033198 | -0.222 | -0.8975 |
| UCN2 | NM_033199 | -0.2165 | 0.0005 |
| LMF2 | NM_033200 | -0.3035 | -0.963 |
| C16orf45 | NM_033201 | -0.59 | -1.1025 |

| | | | |
|---|---|---|---|
| ZNF101 | NM_033204 | -0.6765 | -1.3215 |
| OPALIN | NM_033207 | -0.561 | 0.612 |
| TIGD7 | NM_033208 | -0.798 | -1.709 |
| ZNF502 | NM_033210 | -0.9835 | -2.178 |
| C5orf30 | NM_033211 | -0.8445 | -1.5395 |
| CCDC102A | NM_033212 | -0.8125 | -2.238 |
| ZNF670 | NM_033213 | -0.7045 | -2.6505 |
| GK2 | NM_033214 | -0.0535 | -0.006 |
| PPP1R3F | NM_033215 | -0.6525 | -1.2665 |
| TRIM14 | NM_033219 | -0.6075 | -1.466 |
| TRIM14 | NM_033220 | -0.6075 | -1.466 |
| PSIP1 | NM_033222 | -0.356 | -0.6365 |
| GABRG3 | NM_033223 | -0.1305 | -0.304 |
| PURB | NM_033224 | -1.9065 | -2.102 |
| CSMD1 | NM_033225 | 0.968 | -0.2815 |
| ABCC12 | NM_033226 | 0.028 | -0.17 |
| TRIM23 | NM_033227 | -0.8735 | -2.712 |
| TRIM23 | NM_033228 | -0.8735 | -2.712 |
| TRIM15 | NM_033229 | -0.0795 | -0.4355 |
| PML | NM_033238 | -0.6635 | -1.394 |
| PML | NM_033239 | -0.6635 | -1.394 |
| PML | NM_033240 | -0.6635 | -1.394 |
| PML | NM_033244 | -0.6635 | -1.394 |
| PML | NM_033246 | -0.6635 | -1.394 |
| PML | NM_033247 | -0.6635 | -1.394 |
| PML | NM_033249 | -0.6635 | -1.394 |
| PML | NM_033250 | -0.6635 | -1.394 |
| RPL13 | NM_033251 | 0.0625 | -0.329 |
| NT5C1B | NM_033253 | -0.0925 | -0.479 |
| BOC | NM_033254 | -0.082 | -0.5275 |
| PPP1R14A | NM_033256 | -0.473 | -0.927 |
| DGCR6L | NM_033257 | 0.028 | -0.7495 |
| GNG8 | NM_033258 | -0.036 | -0.168 |
| CAMK2N2 | NM_033259 | -0.358 | -1.2615 |
| FOXQ1 | NM_033260 | 0.271 | 0.211 |
| IDI2 | NM_033261 | -0.1845 | -0.5565 |
| ERN2 | NM_033266 | 0.1385 | -0.468 |
| IRX2 | NM_033267 | 0.0735 | -0.2405 |
| BTBD6 | NM_033271 | -0.533 | -1.6295 |
| KCNH7 | NM_033272 | -0.1805 | -0.3045 |
| ZNF479 | NM_033273 | -0.32 | 0.1435 |
| ADAM19 | NM_033274 | -0.3945 | -1.2 |
| XRCC6BP1 | NM_033276 | -0.318 | -1.441 |

| | | | |
|---|---|---|---|
| LACRT | NM_033277 | -0.077 | 0.1735 |
| TRIM3 | NM_033278 | -0.428 | -1.2885 |
| SEC11C | NM_033280 | -0.318 | -0.4295 |
| MRPS36 | NM_033281 | -0.8425 | -1.6885 |
| OPN4 | NM_033282 | -0.226 | -0.2685 |
| TBL1Y | NM_033284 | 0.1395 | -0.4755 |
| TP53INP1 | NM_033285 | 0.155 | -0.151 |
| C15orf23 | NM_033286 | -2.1465 | -3.481 |
| ZNF160 | NM_033288 | 0.66 | 0.363 |
| MID1 | NM_033289 | -0.5825 | -1.5365 |
| MID1 | NM_033290 | -0.5825 | -1.5365 |
| CASP1 | NM_033292 | 0.024 | 0.318 |
| CASP1 | NM_033293 | 0.024 | 0.318 |
| CASP1 | NM_033294 | 0.024 | 0.318 |
| CASP1 | NM_033295 | 0.024 | 0.318 |
| MRFAP1 | NM_033296 | -1.087 | -1.314 |
| NLRP12 | NM_033297 | -0.3515 | -0.143 |
| LRP8 | NM_033300 | -1.334 | -2.2045 |
| RPL8 | NM_033301 | 0.2675 | -0.1125 |
| ADRA1A | NM_033302 | -0.2555 | -0.1675 |
| ADRA1A | NM_033303 | -0.2555 | -0.1675 |
| ADRA1A | NM_033304 | -0.2825 | 0.174 |
| VPS13A | NM_033305 | -0.1695 | -1.3065 |
| CASP4 | NM_033306 | -0.092 | -0.2085 |
| B3GNT9 | NM_033309 | -0.3855 | -0.7175 |
| KCNK4 | NM_033310 | -0.009 | -0.411 |
| CDC14A | NM_033312 | -0.1155 | -0.0625 |
| CDC14A | NM_033313 | -0.1155 | -0.0625 |
| RASL10B | NM_033315 | -0.514 | -1.1135 |
| MFI2 | NM_033316 | -0.5155 | -0.656 |
| DMKN | NM_033317 | 0.229 | -0.1485 |
| C22orf32 | NM_033318 | 0.146 | -0.4335 |
| CENPL | NM_033319 | -1.4525 | -3.2925 |
| SOX6 | NM_033326 | -0.325 | 0.2215 |
| CAPZA3 | NM_033328 | -0.002 | -0.3675 |
| CDC14B | NM_033331 | -0.1575 | -1.4865 |
| NR6A1 | NM_033334 | -0.1805 | -0.4015 |
| CAV3 | NM_033337 | 0.2515 | -0.27 |
| CASP7 | NM_033338 | -1.071 | -1.6805 |
| CASP7 | NM_033339 | -1.071 | -1.6805 |
| CASP7 | NM_033340 | -1.071 | -1.6805 |
| BIRC8 | NM_033341 | 0.168 | 0.1715 |
| TRIM7 | NM_033342 | 0.3155 | -0.264 |

| | | | |
|---|---|---|---|
| LHX4 | NM_033343 | 0.0115 | -0.3475 |
| RGS8 | NM_033345 | 0.169 | -0.3095 |
| KCNK7 | NM_033347 | 0.0455 | -0.1555 |
| KCNK7 | NM_033348 | 0.0455 | -0.1555 |
| CASP8 | NM_033355 | 0.105 | 0.061 |
| CASP8 | NM_033356 | 0.105 | 0.061 |
| CASP8 | NM_033358 | 0.105 | 0.061 |
| KRAS | NM_033360 | -1.389 | -2.6755 |
| MRPS12 | NM_033362 | -0.5375 | -2.297 |
| MRPS12 | NM_033363 | -0.5375 | -2.297 |
| C3orf15 | NM_033364 | -0.25 | -0.1905 |
| MYO1C | NM_033375 | -0.007 | -0.775 |
| CGB1 | NM_033377 | 0.3295 | -0.109 |
| CDK1 | NM_033379 | -1.2655 | -2.246 |
| COL4A5 | NM_033380 | 0.0465 | -0.197 |
| SEC14L2 | NM_033382 | -0.0205 | -0.3355 |
| MICALL1 | NM_033386 | -0.3085 | -1.231 |
| FAM78A | NM_033387 | -1.091 | -0.8015 |
| ATG16L2 | NM_033388 | 0.3115 | -0.236 |
| SSH2 | NM_033389 | -1.2645 | -2.771 |
| ZC3H12C | NM_033390 | -1.5525 | -3.2535 |
| FHDC1 | NM_033393 | -1.0935 | -1.5715 |
| TANC1 | NM_033394 | -1.0495 | -1.5425 |
| KIAA1731 | NM_033395 | -0.69 | -2.0385 |
| TNKS1BP1 | NM_033396 | -0.435 | -1.174 |
| ITPRIP | NM_033397 | -0.2855 | -1.6135 |
| ZFHX2 | NM_033400 | -1.3585 | -2.285 |
| CNTNAP4 | NM_033401 | -0.513 | -0.5555 |
| LRRCC1 | NM_033402 | -0.2185 | -0.5705 |
| DCLK3 | NM_033403 | -0.7475 | -1.778 |
| PRIC285 | NM_033405 | 0.0455 | -0.2105 |
| FBXO3 | NM_033406 | -0.7955 | -1.247 |
| DOCK7 | NM_033407 | -0.335 | -1.058 |
| C20orf54 | NM_033409 | 0.0705 | -0.3915 |
| ZNF764 | NM_033410 | -1.6745 | -2.7765 |
| RWDD2A | NM_033411 | -1.1485 | -2.0475 |
| MCART1 | NM_033412 | 0.099 | -1.0575 |
| LRRC46 | NM_033413 | -0.3635 | -0.569 |
| ZNF622 | NM_033414 | -0.564 | -1.834 |
| ARMC6 | NM_033415 | -0.691 | -1.6735 |
| IMP4 | NM_033416 | -0.3085 | -1.429 |
| HAUS8 | NM_033417 | -1.191 | -2.6035 |
| METTL18 | NM_033418 | -1.7 | -2.479 |

| | | | |
|---|---|---|---|
| PGAP3 | NM_033419 | 0.105 | -0.7745 |
| C19orf6 | NM_033420 | -0.226 | -0.768 |
| SNX21 | NM_033421 | -0.511 | -0.79 |
| GZMH | NM_033423 | 0.002 | -0.208 |
| DIXDC1 | NM_033425 | -2.3785 | -4.033 |
| KIAA1737 | NM_033426 | -1.365 | -2.21 |
| CTTNBP2 | NM_033427 | 0.2215 | 0.061 |
| C9orf123 | NM_033428 | -0.016 | -0.8685 |
| CALML4 | NM_033429 | -0.737 | -1.1135 |
| PDE5A | NM_033430 | -0.0945 | -0.303 |
| PDE5A | NM_033437 | -0.0945 | -0.303 |
| SLAMF9 | NM_033438 | 0.2935 | -0.6235 |
| IL33 | NM_033439 | -0.0065 | 0.086 |
| CELA2A | NM_033440 | -0.2325 | 0.0335 |
| HIST3H2A | NM_033445 | -0.066 | -0.656 |
| FAM125B | NM_033446 | -0.507 | -1.376 |
| KRT71 | NM_033448 | 0.3455 | -0.103 |
| FCHSD1 | NM_033449 | -0.235 | -0.8025 |
| ABCC10 | NM_033450 | -0.3455 | -1.8015 |
| TRIM47 | NM_033452 | -0.1355 | -0.6705 |
| ITPA | NM_033453 | -0.0915 | -1.1425 |
| KCNK7 | NM_033455 | 0.0455 | -0.1555 |
| MMEL1 | NM_033467 | -0.2005 | -0.3725 |
| ZNF257 | NM_033468 | -0.7085 | 0.0805 |
| FBXO9 | NM_033480 | -1.2715 | -2.089 |
| FBXO9 | NM_033481 | -1.2715 | -2.089 |
| POM121L2 | NM_033482 | -0.1765 | -0.1515 |
| FBXO4 | NM_033484 | -0.386 | -1.5855 |
| CDK11B | NM_033486 | -0.2975 | -0.3865 |
| CDK11B | NM_033487 | -0.2975 | -0.3865 |
| CDK11B | NM_033488 | -0.2975 | -0.3865 |
| CDK11B | NM_033489 | -0.4425 | -0.604 |
| CDK11B | NM_033492 | -0.2975 | -0.3865 |
| CDK11B | NM_033493 | -0.2975 | -0.3865 |
| KLHL13 | NM_033495 | -1.243 | -1.7845 |
| HK1 | NM_033496 | 0.0465 | -0.666 |
| HK1 | NM_033497 | 0.0465 | -0.666 |
| HK1 | NM_033498 | 0.0465 | -0.666 |
| HK1 | NM_033500 | 0.0205 | -0.67 |
| TRERF1 | NM_033502 | -1.8095 | -3.278 |
| BMF | NM_033503 | 0.206 | -0.8675 |
| TMEM54 | NM_033504 | 0.1125 | -0.565 |
| EPT1 | NM_033505 | -2.012 | -2.27 |

| | | | |
|---|---|---|---|
| FBXO24 | NM_033506 | 0.3045 | -0.2115 |
| DISP2 | NM_033510 | -1.205 | -2.464 |
| TSPYL5 | NM_033512 | -0.1435 | -2.236 |
| C19orf20 | NM_033513 | -0.862 | -1.167 |
| LIMS3 | NM_033514 | 0.57 | -0.126 |
| ARHGAP18 | NM_033515 | 0.0275 | -0.005 |
| TPD52L3 | NM_033516 | -0.2735 | -0.293 |
| SLC38A5 | NM_033518 | 0.2595 | -0.688 |
| C19orf33 | NM_033520 | -0.109 | -0.242 |
| CDK11A | NM_033529 | -0.39 | -0.955 |
| MFN1 | NM_033540 | -1.0845 | -2.2525 |
| SYS1 | NM_033542 | -0.6285 | -1.7545 |
| CEACAM21 | NM_033543 | 0.322 | -0.306 |
| RCCD1 | NM_033544 | -0.805 | -2.1675 |
| MYL12B | NM_033546 | -1.3565 | -0.325 |
| INTS4 | NM_033547 | -0.759 | -1.535 |
| TRIM41 | NM_033549 | -0.936 | -1.7015 |
| TP53RK | NM_033550 | -0.8835 | -3.5145 |
| GUCA2A | NM_033553 | -0.1775 | -0.2825 |
| HLA-DPA1 | NM_033554 | -0.0475 | 0.0035 |
| YIF1B | NM_033557 | 0.239 | -0.397 |
| FBXO21 | NM_033624 | -1.3675 | -2.4855 |
| RPL34 | NM_033625 | -0.0055 | -0.284 |
| CCDC120 | NM_033626 | -0.284 | -0.137 |
| SCAND1 | NM_033630 | -0.161 | -0.7845 |
| LUZP1 | NM_033631 | -0.5905 | -1.251 |
| FBXW7 | NM_033632 | -1.1985 | -2.537 |
| BTRC | NM_033637 | -1.272 | -2.54 |
| COL4A6 | NM_033641 | -0.0095 | -0.6015 |
| FGF13 | NM_033642 | -0.0635 | -0.424 |
| RPL36 | NM_033643 | 0.531 | 0.5715 |
| FBXW11 | NM_033644 | -1.208 | -2.5455 |
| FBXW11 | NM_033645 | -1.208 | -2.5455 |
| CDH7 | NM_033646 | -0.2735 | 0.1975 |
| HELB | NM_033647 | -0.046 | -0.9105 |
| CNTNAP3 | NM_033655 | 0.5635 | -0.0765 |
| BRWD1 | NM_033656 | -0.9065 | -1.164 |
| DAP3 | NM_033657 | -0.766 | -1.349 |
| WDR4 | NM_033661 | -0.162 | -0.803 |
| DRD3 | NM_033663 | 0.596 | 0.507 |
| ITGB1 | NM_033668 | -0.47 | -0.3105 |
| CCNDBP1 | NM_037370 | -0.7355 | -1.5015 |
| CDC42 | NM_044472 | -0.3145 | -0.3855 |

| | | | |
|---|---|---|---|
| CTDP1 | NM_048368 | -0.689 | -2.1235 |
| TRIM13 | NM_052811 | -1.2885 | -3.7325 |
| CARD9 | NM_052813 | -0.364 | -0.6375 |
| CARD9 | NM_052814 | -0.3555 | -0.0455 |
| MID2 | NM_052817 | -0.432 | -0.697 |
| N4BP2L1 | NM_052818 | -1.521 | -2.131 |
| CARD14 | NM_052819 | -0.0635 | -0.802 |
| CORO2A | NM_052820 | -1.0215 | -1.256 |
| CDK2 | NM_052827 | -0.3105 | -1.065 |
| TRIM10 | NM_052828 | -0.2855 | -0.1865 |
| C6orf192 | NM_052831 | -0.68 | -0.8195 |
| SLC26A7 | NM_052832 | 0.1165 | -0.1585 |
| WDR7 | NM_052834 | -0.2335 | -1.0545 |
| CDH23 | NM_052836 | 0.637 | 0.184 |
| 1-Sep | NM_052838 | -0.0375 | -0.733 |
| PANX2 | NM_052839 | 0.004 | -0.707 |
| CELF6 | NM_052840 | 0.277 | -0.693 |
| TSSK3 | NM_052841 | 0.069 | -0.4705 |
| OBSCN | NM_052843 | 0.063 | -0.0625 |
| WDR34 | NM_052844 | -0.311 | -0.933 |
| MMAB | NM_052845 | -0.2855 | -1.347 |
| EMILIN3 | NM_052846 | -0.74 | -1.669 |
| GNG7 | NM_052847 | -0.4525 | -0.562 |
| CCDC97 | NM_052848 | -1.1775 | -2.037 |
| C15orf57 | NM_052849 | -0.812 | -2.1395 |
| GADD45GIP1 | NM_052850 | -0.103 | -0.884 |
| STARD13 | NM_052851 | -1.199 | -0.698 |
| ZNF486 | NM_052852 | 0.077 | -1.1295 |
| ADCK2 | NM_052853 | -0.6935 | -1.798 |
| CREB3L1 | NM_052854 | -0.244 | -0.699 |
| ANKRD40 | NM_052855 | -1.4795 | -2.728 |
| ZNF830 | NM_052857 | -1.382 | -2.6195 |
| MARVELD3 | NM_052858 | 0.176 | 0.4905 |
| RFT1 | NM_052859 | -0.8205 | -2.1365 |
| ZNF300 | NM_052860 | -1.0795 | -1.933 |
| RCSD1 | NM_052862 | -0.3715 | -0.3465 |
| SCGB3A1 | NM_052863 | -0.0605 | -0.221 |
| TIFA | NM_052864 | -0.8785 | -2.774 |
| C20orf72 | NM_052865 | -1.8675 | -2.856 |
| ADAMTSL1 | NM_052866 | -0.2905 | -1.0215 |
| NALCN | NM_052867 | -0.2675 | -0.2805 |
| IGSF8 | NM_052868 | 0.093 | -0.45 |
| TTYH2 | NM_052869 | 0.1055 | -0.266 |

| | | | |
|---|---|---|---|
| SNX18 | NM_052870 | -0.994 | -1.925 |
| IL17F | NM_052872 | 0.296 | 0.084 |
| STX1B | NM_052874 | -0.36 | -0.752 |
| VPS26B | NM_052875 | -0.573 | -1.323 |
| NACC1 | NM_052876 | -0.497 | -1.383 |
| MED8 | NM_052877 | -0.7625 | -2.161 |
| LARP4 | NM_052879 | -0.0595 | -1.7255 |
| PIK3IP1 | NM_052880 | -0.592 | -0.71 |
| ZIM3 | NM_052882 | 0.364 | -0.229 |
| TXNRD3 | NM_052883 | -0.673 | -1.183 |
| SIGLEC11 | NM_052884 | 0.1865 | -0.342 |
| SLC2A13 | NM_052885 | -1.5 | -3.056 |
| MAL2 | NM_052886 | -0.05 | -0.0635 |
| CARD16 | NM_052889 | 0.0185 | -0.255 |
| PGLYRP2 | NM_052890 | -0.2865 | -0.3245 |
| PGLYRP3 | NM_052891 | 0.0895 | -0.0825 |
| PKD1L2 | NM_052892 | 0.2595 | 0.559 |
| BTBD9 | NM_052893 | -0.3395 | -1.3525 |
| CSMD2 | NM_052896 | -0.096 | -0.4985 |
| MBD6 | NM_052897 | -0.4995 | -1.0275 |
| XKR4 | NM_052898 | -0.1805 | -0.3655 |
| GPRIN1 | NM_052899 | -0.549 | -1.6435 |
| CSMD3 | NM_052900 | 0.0275 | 0.0215 |
| SLC25A25 | NM_052901 | -0.6575 | -2.329 |
| STK11IP | NM_052902 | -0.1135 | -0.959 |
| TUBGCP5 | NM_052903 | -0.9525 | -1.5795 |
| KLHL32 | NM_052904 | -0.315 | -0.4525 |
| FMNL2 | NM_052905 | -0.8185 | -1.082 |
| ELFN2 | NM_052906 | -0.24 | -0.261 |
| TMEM132B | NM_052907 | 0.1545 | 0.006 |
| PLEKHG4B | NM_052909 | -0.29 | -0.942 |
| SLITRK1 | NM_052910 | -0.322 | -0.712 |
| ESCO1 | NM_052911 | -0.6365 | -2.7335 |
| TMEM200A | NM_052913 | -0.3165 | -1.683 |
| RNF157 | NM_052916 | -0.096 | -0.758 |
| GALNT13 | NM_052917 | -0.5525 | -1.296 |
| SORCS1 | NM_052918 | -0.958 | -0.506 |
| KLHL29 | NM_052920 | 0.6365 | -0.4405 |
| SCAND3 | NM_052923 | -0.718 | -2.232 |
| RHPN1 | NM_052924 | 0.077 | -0.293 |
| LENG8 | NM_052925 | -0.2975 | -0.5475 |
| PNMA5 | NM_052926 | -0.1525 | -0.2185 |
| PWWP2A | NM_052927 | -1.501 | -1.749 |

| | | | |
|---|---|---|---|
| SMYD4 | NM_052928 | -0.852 | -2.448 |
| FHAD1 | NM_052929 | -0.1315 | -0.349 |
| SLAMF6 | NM_052931 | -0.1415 | -0.109 |
| TMEM123 | NM_052932 | -0.844 | -1.0475 |
| TSGA13 | NM_052933 | 0.0175 | -0.482 |
| SLC26A9 | NM_052934 | -0.165 | -0.2665 |
| NT5C3L | NM_052935 | 0.011 | -0.7255 |
| ATG4A | NM_052936 | -0.8935 | -1.365 |
| PCMTD1 | NM_052937 | -1.579 | -3.0285 |
| FCRL1 | NM_052938 | -0.391 | -0.3955 |
| FCRL3 | NM_052939 | -0.1335 | -0.106 |
| LRRC42 | NM_052940 | -0.656 | -2.8735 |
| GBP4 | NM_052941 | -0.0215 | -0.409 |
| GBP5 | NM_052942 | 0.0225 | -0.258 |
| FAM46B | NM_052943 | -0.28 | -1.062 |
| SLC5A11 | NM_052944 | 0.0125 | -0.034 |
| TNFRSF13C | NM_052945 | 0.0985 | 0.0115 |
| ALPK2 | NM_052947 | -0.2455 | -0.3905 |
| ARHGAP33 | NM_052948 | -0.0135 | -0.299 |
| WDFY2 | NM_052950 | -0.3725 | -1.815 |
| DNTTIP1 | NM_052951 | -0.8375 | -2.4265 |
| DIRC1 | NM_052952 | 0.2855 | -0.3665 |
| LRRC3B | NM_052953 | 0.0735 | -0.0105 |
| CYYR1 | NM_052954 | -0.2905 | -3.3035 |
| TGM7 | NM_052955 | 0.2025 | 0.283 |
| ACSM1 | NM_052956 | 0.2345 | -0.2775 |
| ACRC | NM_052957 | 0.0815 | -0.5085 |
| C8orf34 | NM_052958 | 0.157 | -0.2925 |
| PANX3 | NM_052959 | -0.018 | 0.0795 |
| RBP7 | NM_052960 | -0.3665 | -0.0445 |
| SLC26A8 | NM_052961 | -0.062 | -0.094 |
| IL22RA2 | NM_052962 | 0.1725 | 0.0575 |
| TOP1MT | NM_052963 | 0.0045 | -0.906 |
| CLNK | NM_052964 | 0.429 | 0.0755 |
| TSEN15 | NM_052965 | -1.131 | -1.5755 |
| FAM129A | NM_052966 | 0.0705 | -0.2995 |
| MAS1L | NM_052967 | -0.085 | -0.439 |
| APOA5 | NM_052968 | 0.137 | -0.409 |
| RPL39L | NM_052969 | -0.9805 | -1.548 |
| HSPA12B | NM_052970 | -0.0325 | -0.061 |
| LEAP2 | NM_052971 | -0.7965 | -1.5695 |
| LRG1 | NM_052972 | 0.1555 | -0.219 |
| TRIM9 | NM_052978 | 0.7335 | -1.314 |

| | | | |
|---|---|---|---|
| IFT122 | NM_052985 | -0.014 | -0.803 |
| CDK10 | NM_052987 | -0.3685 | -0.763 |
| CDK10 | NM_052988 | -0.3685 | -0.763 |
| IFT122 | NM_052989 | 0.1445 | -0.996 |
| IFT122 | NM_052990 | 0.1445 | -0.996 |
| CLRN1 | NM_052995 | -0.0205 | -0.34 |
| PRDM7 | NM_052996 | 0.1405 | -0.159 |
| ANKRD30A | NM_052997 | -0.3305 | -0.115 |
| ADC | NM_052998 | 0.325 | -0.4385 |
| CMTM1 | NM_052999 | -0.184 | -0.0705 |
| OSR2 | NM_053001 | -0.4925 | -0.084 |
| MED12L | NM_053002 | 0.785 | -0.0685 |
| SIGLEC12 | NM_053003 | 0.0855 | -0.1495 |
| GNB1L | NM_053004 | -0.432 | -0.9385 |
| MOB2 | NM_053005 | -0.2085 | -1.389 |
| TSSK2 | NM_053006 | -0.0395 | -0.259 |
| ENO3 | NM_053013 | 0.181 | -0.1135 |
| PALM2 | NM_053016 | 0.215 | -0.353 |
| ART5 | NM_053017 | -0.121 | -0.4685 |
| ZFP91 | NM_053023 | -1.0505 | -2.0155 |
| PFN2 | NM_053024 | -0.7565 | -1.517 |
| MYLK | NM_053025 | -0.219 | -0.624 |
| MYLK | NM_053026 | -0.219 | -0.624 |
| MYLK | NM_053027 | -0.219 | -0.624 |
| MYLK | NM_053028 | -0.219 | -0.624 |
| MYLK | NM_053031 | -0.219 | -0.624 |
| MYLK | NM_053032 | -0.219 | -0.624 |
| ANTXR1 | NM_053034 | -1.115 | -1.0975 |
| MRPS33 | NM_053035 | -0.726 | -0.692 |
| NPFFR2 | NM_053036 | 0.664 | -0.5265 |
| UGT2B28 | NM_053039 | -0.4495 | -0.511 |
| COMMD7 | NM_053041 | -0.0995 | -0.9 |
| ZNF518B | NM_053042 | -1.7265 | -1.628 |
| RBM33 | NM_053043 | -0.456 | -2.4275 |
| HTRA3 | NM_053044 | 0.077 | -0.334 |
| TMEM203 | NM_053045 | -0.8835 | -2.7685 |
| EGLN2 | NM_053046 | -0.5655 | -2.1135 |
| UCN3 | NM_053049 | 0.364 | 0.0995 |
| MRPL53 | NM_053050 | -0.067 | -0.757 |
| CNTROB | NM_053051 | -0.435 | -0.7295 |
| SNAP47 | NM_053052 | -0.746 | -2.6475 |
| TADA1 | NM_053053 | -1.728 | -3.4165 |
| CATSPER1 | NM_053054 | 0.2145 | -0.2365 |

| | | | |
|---|---|---|---|
| THEM4 | NM_053055 | -0.3255 | -0.7505 |
| CCND1 | NM_053056 | -1.604 | -1.0455 |
| GNG2 | NM_053064 | -0.68 | -1.0385 |
| UBQLN1 | NM_053067 | -0.8815 | -1.414 |
| GLMN | NM_053274 | -1.1765 | -2.2175 |
| RPLP0 | NM_053275 | 0.125 | -0.2855 |
| VIT | NM_053276 | 0.118 | 0.1785 |
| CLIC6 | NM_053277 | -0.0305 | -0.3655 |
| TAAR8 | NM_053278 | 0.0115 | -0.2565 |
| FAM167A | NM_053279 | -0.7935 | -1.5715 |
| ODF3 | NM_053280 | 0.015 | -0.325 |
| DACH2 | NM_053281 | -0.0405 | -0.161 |
| SH2D1B | NM_053282 | -0.186 | -0.482 |
| DCD | NM_053283 | 0.139 | -0.0265 |
| WFIKKN1 | NM_053284 | -0.185 | -0.1285 |
| TEKT1 | NM_053285 | -0.195 | -0.17 |
| ASS1 | NM_054012 | 0.267 | -0.304 |
| MGAT4B | NM_054013 | -0.016 | -0.7525 |
| FKBP1A | NM_054014 | 0.2695 | -0.1105 |
| SRSF10 | NM_054016 | -1.748 | -1.245 |
| CATSPER2 | NM_054020 | -0.5565 | -0.5735 |
| GPR101 | NM_054021 | 0.0465 | -0.272 |
| GOSR2 | NM_054022 | -0.48 | -1.608 |
| SCGB3A2 | NM_054023 | 0.067 | -0.0035 |
| MIA2 | NM_054024 | -0.033 | 0.1605 |
| B3GAT1 | NM_054025 | -0.134 | -0.807 |
| CNOT7 | NM_054026 | -1.0595 | -1.475 |
| ANKH | NM_054027 | -0.546 | -1.8735 |
| MRGPRX2 | NM_054030 | 0.1305 | 0.015 |
| MRGPRX3 | NM_054031 | 0.1865 | -0.1875 |
| MRGPRX4 | NM_054032 | -0.266 | -0.632 |
| FKBP1B | NM_054033 | -0.007 | -0.6205 |
| FN1 | NM_054034 | -0.124 | -0.5965 |
| UNC119 | NM_054035 | 0.0325 | -0.1905 |
| OR6C3 | NM_054104 | 0.144 | -0.4895 |
| OR6C2 | NM_054105 | 0.302 | -0.175 |
| OR5AC2 | NM_054106 | -0.193 | -0.0885 |
| OR1J2 | NM_054107 | -0.1985 | -0.5015 |
| HRASLS5 | NM_054108 | -0.2215 | -0.361 |
| GALNTL2 | NM_054110 | -0.2095 | -0.3685 |
| IP6K3 | NM_054111 | -0.288 | -0.8155 |
| DEFB118 | NM_054112 | 0.071 | -0.3355 |
| CIB3 | NM_054113 | -0.1415 | -0.654 |

| | | | |
|---|---|---|---|
| TAGAP | NM_054114 | -0.0275 | 0.19 |
| KRT3 | NM_057088 | 0.511 | 0.208 |
| ARTN | NM_057090 | -0.0085 | -0.564 |
| ARTN | NM_057091 | -0.0085 | -0.564 |
| CRYBA2 | NM_057093 | -0.0435 | -0.093 |
| CYP3A43 | NM_057095 | 0.0215 | -0.44 |
| CYP3A43 | NM_057096 | 0.0215 | -0.44 |
| CYP26A1 | NM_057157 | 0.175 | -0.312 |
| DUSP4 | NM_057158 | -0.5185 | -1.986 |
| LPAR1 | NM_057159 | -0.6115 | -2.395 |
| KLHDC3 | NM_057161 | -0.0095 | -0.8585 |
| KLHL4 | NM_057162 | -0.174 | -0.3925 |
| COL6A3 | NM_057164 | -0.2995 | -0.3935 |
| COL6A3 | NM_057165 | -0.2995 | -0.3935 |
| COL6A3 | NM_057166 | -0.2995 | -0.3935 |
| COL6A3 | NM_057167 | -0.3445 | -0.2445 |
| WNT16 | NM_057168 | 0.074 | -0.401 |
| GIT2 | NM_057169 | -1.5725 | -3.2765 |
| GIT2 | NM_057170 | -0.6875 | -2.828 |
| PEX16 | NM_057174 | -0.234 | -1.1585 |
| NAA15 | NM_057175 | -0.8895 | -2.563 |
| BSND | NM_057176 | 0.042 | -0.195 |
| PARD3B | NM_057177 | 0 | -0.155 |
| TWIST2 | NM_057179 | 0.012 | -0.4085 |
| VPS29 | NM_057180 | -1.014 | -1.315 |
| CCNE2 | NM_057749 | -1.216 | -3.0345 |
| PI4KA | NM_058004 | -0.052 | -0.5565 |
| TSR2 | NM_058163 | -0.1405 | -0.6325 |
| OLFM2 | NM_058164 | 0.092 | -0.4105 |
| MOGAT1 | NM_058165 | -0.2185 | 0.413 |
| TRIM6 | NM_058166 | -0.17 | -0.341 |
| UBE2J2 | NM_058167 | -0.808 | -2.5555 |
| LOH12CR1 | NM_058169 | -0.552 | -2.1625 |
| OLFM3 | NM_058170 | -0.5845 | -0.7695 |
| ANTXR2 | NM_058172 | -1.623 | -2.2605 |
| MUCL1 | NM_058173 | 0.1485 | -0.4225 |
| COL6A2 | NM_058174 | -0.0985 | -0.48 |
| COL6A2 | NM_058175 | -0.0985 | -0.48 |
| HDAC9 | NM_058176 | -0.862 | -2.3525 |
| PSAT1 | NM_058179 | -0.7885 | -1.5255 |
| C21orf58 | NM_058180 | -0.03 | -0.5475 |
| YBEY | NM_058181 | -0.132 | -0.689 |
| FAM165B | NM_058182 | -0.2055 | -0.678 |

| | | | |
|---|---|---|---|
| FAM3B | NM_058186 | 0.032 | -0.6835 |
| C21orf63 | NM_058187 | 0.109 | -0.7605 |
| RPUSD1 | NM_058192 | -0.6055 | -1.119 |
| CDKN2A | NM_058195 | 0.204 | 0.138 |
| CDKN2A | NM_058197 | 0.204 | 0.138 |
| SPAG11B | NM_058203 | 0.136 | -0.481 |
| RAD51C | NM_058216 | -1.921 | -3.0235 |
| EXOSC6 | NM_058219 | -0.667 | -1.826 |
| TECTB | NM_058222 | 0.1195 | -0.485 |
| FBXO32 | NM_058229 | -0.13 | -0.686 |
| ZNF354B | NM_058230 | -0.269 | -1.0465 |
| PPP4R4 | NM_058237 | 0.202 | -0.4155 |
| WNT7B | NM_058238 | -0.2575 | -0.3415 |
| CCNT2 | NM_058241 | -0.1945 | -1.091 |
| BRD4 | NM_058243 | 0.0445 | -0.87 |
| WNT8A | NM_058244 | -0.1005 | 0.0185 |
| DNAJB6 | NM_058246 | -0.3425 | -0.126 |
| CDKN1A | NM_078467 | -0.645 | 0.0405 |
| BCCIP | NM_078468 | -0.647 | -1.616 |
| BCCIP | NM_078469 | -0.8725 | -1.1475 |
| COX15 | NM_078470 | -0.9595 | -1.182 |
| MYO18A | NM_078471 | -0.157 | -0.478 |
| TM2D2 | NM_078473 | -0.8485 | -2.6585 |
| TM2D3 | NM_078474 | -0.646 | -1.265 |
| BTN2A1 | NM_078476 | -1.4835 | -3.688 |
| PUF60 | NM_078480 | 0.0035 | -0.874 |
| CD97 | NM_078481 | -0.222 | -0.47 |
| SLC36A1 | NM_078483 | -0.7685 | -1.6555 |
| COL9A1 | NM_078485 | -0.2045 | -0.218 |
| CDKN2B | NM_078487 | 0.0735 | 0.1195 |
| VNN2 | NM_078488 | -0.0095 | 0.0735 |
| CDKN2C | NM_078626 | -0.597 | -2.3365 |
| MSL3 | NM_078628 | -1.0535 | -0.7935 |
| MSL3 | NM_078629 | -1.0535 | -0.7935 |
| MYL1 | NM_079420 | 0.4225 | 0.2945 |
| MYL1 | NM_079422 | 0.0175 | 0.243 |
| MYL6 | NM_079423 | 0.3205 | -0.139 |
| SCAMP4 | NM_079834 | -0.5255 | -1.34 |
| BANP | NM_079837 | -0.907 | -2.7155 |
| ABCA10 | NM_080282 | -1.9295 | -2.8555 |
| ABCA9 | NM_080283 | -0.121 | -0.29 |
| ABCA6 | NM_080284 | -0.05 | -0.2665 |
| TUBA3D | NM_080386 | 0.191 | 0.003 |

| | | | |
|---|---|---|---|
| CLEC4D | NM_080387 | 0.2045 | 0.387 |
| S100A16 | NM_080388 | 0.069 | -0.633 |
| DEFB104A | NM_080389 | 0.223 | -0.1035 |
| TCEAL2 | NM_080390 | -0.378 | -0.2055 |
| PTP4A2 | NM_080391 | -0.817 | -0.334 |
| 4-Sep | NM_080415 | -0.102 | -0.242 |
| PTPN2 | NM_080422 | 0.016 | -1.1395 |
| PTPN2 | NM_080423 | -1.271 | -2.5055 |
| SP110 | NM_080424 | -0.213 | -0.626 |
| GNAS | NM_080425 | 0.0755 | -0.2415 |
| GNAS | NM_080426 | 0.0835 | -0.2445 |
| AQP10 | NM_080429 | 0.321 | -0.1925 |
| SELM | NM_080430 | 0.2215 | -0.4335 |
| ACTRT2 | NM_080431 | -0.082 | -0.1295 |
| GATA5 | NM_080473 | -0.0115 | -0.11 |
| SERPINB12 | NM_080474 | 0.0695 | -0.2555 |
| SERPINB11 | NM_080475 | -0.2075 | 0.0295 |
| PIGU | NM_080476 | -0.717 | -1.9575 |
| SDCBP2 | NM_080489 | 0.0295 | -0.399 |
| GAB2 | NM_080491 | -0.2075 | -0.3365 |
| COLQ | NM_080538 | 0.0405 | -0.239 |
| COLQ | NM_080539 | 0.0405 | -0.239 |
| SLC44A1 | NM_080546 | -0.7075 | -1.205 |
| PTPN6 | NM_080548 | -0.259 | -0.0345 |
| PTPN6 | NM_080549 | -0.171 | -0.8015 |
| SYNRG | NM_080550 | -0.907 | -2.1575 |
| SLC32A1 | NM_080552 | -0.3035 | -0.43 |
| C20orf96 | NM_080571 | 0.097 | -0.4415 |
| BPIFA2 | NM_080574 | 0.0175 | -0.013 |
| PTPN7 | NM_080588 | -0.427 | 0.095 |
| CAPS | NM_080590 | 0.1875 | -0.5075 |
| PTGS1 | NM_080591 | 0.3395 | -0.4795 |
| C2orf28 | NM_080592 | -0.192 | -0.8825 |
| HIST1H2BK | NM_080593 | 0.018 | -0.7395 |
| RNPS1 | NM_080594 | -0.511 | -1.249 |
| HIST1H2AH | NM_080596 | -0.2605 | -0.2065 |
| OSBPL1A | NM_080597 | -0.34 | -1.1105 |
| DDX39B | NM_080598 | -0.188 | -0.702 |
| UPF2 | NM_080599 | -0.8095 | -2.8555 |
| MAG | NM_080600 | -0.3265 | -0.549 |
| ZSWIM1 | NM_080603 | -0.852 | -2.206 |
| TJAP1 | NM_080604 | -1.0215 | -2.418 |
| B3GALT6 | NM_080605 | -1.277 | -2.2755 |

| | | | |
|---|---|---|---|
| BHLHE23 | NM_080606 | -0.137 | -0.261 |
| VSTM2L | NM_080607 | -0.8735 | -0.839 |
| C20orf165 | NM_080608 | 0.1745 | 0.004 |
| CST9L | NM_080610 | 0.3535 | -0.0655 |
| DUSP15 | NM_080611 | 0.001 | -0.4515 |
| WFDC3 | NM_080614 | -0.1955 | -0.0935 |
| GCNT7 | NM_080615 | -0.1415 | -0.2365 |
| C20orf112 | NM_080616 | -1.556 | -1.9075 |
| CBLN4 | NM_080617 | 0.293 | -0.431 |
| CTCFL | NM_080618 | 0.014 | -0.3755 |
| SAMD10 | NM_080621 | -0.4385 | -0.836 |
| ABHD16B | NM_080622 | -0.2305 | -0.0365 |
| C20orf160 | NM_080625 | 0.0065 | -2.2045 |
| BRI3BP | NM_080626 | -0.327 | -1.475 |
| C20orf118 | NM_080628 | 0.066 | -0.6345 |
| COL11A1 | NM_080629 | 0.313 | -0.504 |
| COL11A1 | NM_080630 | 0.313 | -0.504 |
| VPS41 | NM_080631 | -0.8055 | -0.468 |
| UPF3B | NM_080632 | -0.8245 | -1.455 |
| COL12A1 | NM_080645 | 0.2205 | 0.624 |
| TBX1 | NM_080646 | -0.2025 | -0.905 |
| TBX1 | NM_080647 | 0.039 | -0.928 |
| ATPBD4 | NM_080650 | -0.9095 | -1.098 |
| MED30 | NM_080651 | -0.551 | -1.3755 |
| TMEM41A | NM_080652 | -0.835 | -1.8565 |
| ATP6V1E2 | NM_080653 | -0.367 | -0.8165 |
| CCDC34 | NM_080654 | -0.9935 | -1.2645 |
| C9orf30 | NM_080655 | -0.4765 | -1.0335 |
| CDKN2AIPNL | NM_080656 | -0.056 | -0.663 |
| RSAD2 | NM_080657 | -0.1795 | -0.0865 |
| ACY3 | NM_080658 | 0.562 | -0.0925 |
| C11orf52 | NM_080659 | 0.1255 | -0.1415 |
| ZC3HAV1L | NM_080660 | -1.069 | -2.066 |
| GLYATL1 | NM_080661 | -0.0535 | -0.276 |
| PEX11G | NM_080662 | -0.1115 | -0.585 |
| ERI2 | NM_080663 | -1.7085 | -2.577 |
| C14orf126 | NM_080664 | -1.165 | -2.393 |
| WDR89 | NM_080666 | -1.551 | -3.1125 |
| CCDC104 | NM_080667 | -1.038 | -0.753 |
| CDCA5 | NM_080668 | -0.6505 | -1.4545 |
| SLC46A1 | NM_080669 | -1.1485 | -1.6215 |
| SLC35A4 | NM_080670 | -0.9525 | -2.541 |
| KCNE4 | NM_080671 | -0.037 | -0.3845 |

| | | | |
|---|---|---|---|
| PHACTR3 | NM_080672 | -0.852 | -1.6995 |
| SUN5 | NM_080675 | -0.0325 | -0.1345 |
| MACROD2 | NM_080676 | -0.339 | -0.2965 |
| DYNLL2 | NM_080677 | -0.311 | -0.9655 |
| UBE2F | NM_080678 | -0.13 | -0.855 |
| COL11A2 | NM_080679 | 0.2385 | -0.5075 |
| COL11A2 | NM_080680 | 0.2385 | -0.5075 |
| COL11A2 | NM_080681 | 0.2385 | -0.5075 |
| VCAM1 | NM_080682 | -0.458 | -0.438 |
| PTPN13 | NM_080683 | -0.161 | -1.5885 |
| PTPN13 | NM_080684 | -0.161 | -1.5885 |
| PTPN13 | NM_080685 | -0.161 | -1.5885 |
| PRRC2A | NM_080686 | -0.3485 | -1.387 |
| UPF3A | NM_080687 | -1.2855 | -0.8695 |
| TREX2 | NM_080701 | -0.0405 | -0.078 |
| TRPV1 | NM_080704 | -0.4195 | -0.5215 |
| TRPV1 | NM_080705 | -0.4195 | -0.5215 |
| TRPV1 | NM_080706 | -1.1845 | -1.4055 |
| TBX5 | NM_080717 | 0.241 | -0.285 |
| TBX5 | NM_080718 | 0.241 | -0.285 |
| C20orf123 | NM_080721 | 0.1945 | -0.584 |
| ADAMTS14 | NM_080722 | 0.224 | -0.433 |
| NRSN1 | NM_080723 | -1.6615 | -2.8055 |
| SRXN1 | NM_080725 | -0.859 | -1.859 |
| IFFO1 | NM_080730 | -0.913 | -1.502 |
| EGLN2 | NM_080732 | -0.5655 | -2.1135 |
| SYTL4 | NM_080737 | 0.148 | -1.633 |
| EDARADD | NM_080738 | 0.379 | -0.522 |
| C20orf141 | NM_080739 | 0.017 | -0.074 |
| ZNF280A | NM_080740 | -0.9725 | -0.1725 |
| NEU4 | NM_080741 | -0.454 | -0.5785 |
| B3GAT2 | NM_080742 | -0.322 | -1.0725 |
| SRSF12 | NM_080743 | -0.799 | -1.899 |
| SRCRB4D | NM_080744 | -0.6665 | -0.9625 |
| RPL10L | NM_080746 | 0.167 | 0.0215 |
| KRT72 | NM_080747 | -0.041 | -0.643 |
| ROMO1 | NM_080748 | 0.256 | -0.2275 |
| NEURL2 | NM_080749 | -0.1285 | -0.158 |
| DPH3P1 | NM_080750 | -0.2165 | -0.2875 |
| TMC2 | NM_080751 | 0.014 | -0.2485 |
| ZSWIM3 | NM_080752 | -0.921 | -2.3475 |
| WFDC10A | NM_080753 | 0.4255 | 0.219 |
| DACH1 | NM_080759 | -1.4675 | -4.4805 |

| | | | |
|---|---|---|---|
| DACH1 | NM_080760 | -1.0525 | -2.4995 |
| ZNF280B | NM_080764 | -1.5185 | -2.2375 |
| SIRPA | NM_080792 | -0.4105 | -0.587 |
| MRPL39 | NM_080794 | -1.0565 | -2.581 |
| DIDO1 | NM_080796 | 0.457 | -1.4845 |
| DIDO1 | NM_080797 | -0.7975 | -1.187 |
| COL13A1 | NM_080798 | 0.2635 | -0.061 |
| COL13A1 | NM_080800 | 0.2635 | -0.061 |
| COL13A1 | NM_080801 | 0.2635 | -0.061 |
| COL13A1 | NM_080802 | 0.2635 | -0.061 |
| COL13A1 | NM_080805 | 0.2635 | -0.061 |
| SIRPG | NM_080816 | -0.05 | -0.39 |
| GPR82 | NM_080817 | -0.1735 | -0.0645 |
| OXGR1 | NM_080818 | -0.776 | 0.029 |
| GPR78 | NM_080819 | 0.195 | -0.1675 |
| DTD1 | NM_080820 | -0.0585 | -0.6055 |
| C20orf108 | NM_080821 | -0.7205 | -0.7785 |
| OVCA2 | NM_080822 | -0.624 | -1.586 |
| SRMS | NM_080823 | 0.22 | -0.3075 |
| C20orf144 | NM_080825 | -0.0725 | 0.0145 |
| ISM1 | NM_080826 | 0.1015 | 0.2475 |
| FAM65C | NM_080829 | 0.0105 | -0.523 |
| CST11 | NM_080830 | -0.0905 | -0.2355 |
| DEFB129 | NM_080831 | -0.41 | -0.7615 |
| PABPC5 | NM_080832 | -0.1115 | -1.384 |
| C20orf151 | NM_080833 | -0.074 | -0.298 |
| C20orf152 | NM_080834 | -0.0985 | -0.4025 |
| STK35 | NM_080836 | -0.888 | -2.7575 |
| WISP1 | NM_080838 | 0.096 | -0.1165 |
| PTPRA | NM_080840 | -0.569 | -1.255 |
| PTPRA | NM_080841 | -0.4485 | -1.0645 |
| OR1K1 | NM_080859 | -0.101 | -0.226 |
| RSPH1 | NM_080860 | 0.0255 | -0.1085 |
| SPSB3 | NM_080861 | 0.056 | -0.6515 |
| SPSB4 | NM_080862 | -0.625 | -1.592 |
| ASB16 | NM_080863 | 0.1155 | -0.476 |
| RLN3 | NM_080864 | 0.0635 | 0.0075 |
| GPR62 | NM_080865 | 0.1245 | -0.3015 |
| SLC22A9 | NM_080866 | -0.198 | -0.288 |
| SOCS4 | NM_080867 | -1.5885 | -2.4395 |
| ASB17 | NM_080868 | -0.109 | -0.164 |
| WFDC12 | NM_080869 | -0.0935 | -0.1055 |
| DPCR1 | NM_080870 | 0.1115 | -0.3185 |

| | | | |
|---|---|---|---|
| ASB10 | NM_080871 | 0.2975 | -0.385 |
| UNC5D | NM_080872 | -0.0975 | -0.3035 |
| ASB11 | NM_080873 | 0.5025 | 0.3885 |
| ASB5 | NM_080874 | -0.287 | -0.995 |
| MIB2 | NM_080875 | -0.306 | -0.029 |
| DUSP19 | NM_080876 | -0.4655 | -1.05 |
| SLC34A3 | NM_080877 | -0.3805 | -0.416 |
| ITLN2 | NM_080878 | -0.361 | -0.3525 |
| RAB40A | NM_080879 | -0.3505 | -0.1095 |
| DBN1 | NM_080881 | -0.3835 | -1.279 |
| UNG | NM_080911 | -0.5145 | -1.653 |
| ASGR2 | NM_080912 | 0.198 | -0.1445 |
| ASGR2 | NM_080913 | 0.198 | -0.1445 |
| ASGR2 | NM_080914 | 0.198 | -0.1445 |
| DGUOK | NM_080916 | 0.024 | -0.574 |
| PTPRC | NM_080921 | 0.147 | -0.4865 |
| PTPRC | NM_080923 | 0.147 | -0.4865 |
| DCBLD2 | NM_080927 | -1.624 | -2.1235 |
| ASB15 | NM_080928 | -0.942 | 0.09 |
| WAC | NM_100486 | -1.6395 | -3.798 |
| DYRK1A | NM_101395 | 0.1795 | -0.404 |
| ATRIP | NM_130384 | -1.0355 | -2.278 |
| MRVI1 | NM_130385 | 0.1675 | -0.759 |
| COLEC12 | NM_130386 | -0.1485 | -2.072 |
| ASB14 | NM_130387 | 0.496 | 0.148 |
| ASB12 | NM_130388 | -0.4425 | -0.065 |
| TRIM34 | NM_130390 | -0.137 | -0.0815 |
| PTPRD | NM_130391 | -0.291 | -1.5925 |
| PTPRD | NM_130392 | -0.291 | -1.5925 |
| PTPRD | NM_130393 | -0.291 | -1.5925 |
| WRNIP1 | NM_130395 | -0.309 | -0.621 |
| EXO1 | NM_130398 | -1.8755 | -4.103 |
| DPP8 | NM_130434 | -1.3795 | -1.5685 |
| PTPRE | NM_130435 | -0.093 | -0.598 |
| DYRK1A | NM_130436 | 0.1795 | -0.404 |
| DYRK1A | NM_130438 | 0.1795 | -0.404 |
| MXI1 | NM_130439 | -0.881 | -1.754 |
| PTPRF | NM_130440 | -0.0035 | -0.384 |
| CLEC4C | NM_130441 | 0.143 | 0.2565 |
| DPP3 | NM_130443 | 0.06 | -0.621 |
| COL18A1 | NM_130445 | -0.069 | -0.7035 |
| KLHL6 | NM_130446 | 0.228 | -0.354 |
| TOR2A | NM_130459 | -0.475 | -1.4345 |

| | | | |
|---|---|---|---|
| ATP6V1G2 | NM_130463 | 0.0255 | -0.322 |
| TSPAN17 | NM_130465 | -0.0675 | -0.8525 |
| UBE3B | NM_130466 | -0.991 | -2.0465 |
| PAGE5 | NM_130467 | -0.1615 | -0.341 |
| CHST14 | NM_130468 | -0.9095 | -2.418 |
| JDP2 | NM_130469 | -0.9605 | -2.184 |
| MADD | NM_130470 | -0.668 | -1.477 |
| MADD | NM_130471 | -0.668 | -1.477 |
| MADD | NM_130472 | -0.668 | -1.477 |
| MADD | NM_130473 | -0.668 | -1.477 |
| MADD | NM_130474 | -0.668 | -1.477 |
| MADD | NM_130475 | -0.668 | -1.477 |
| MADD | NM_130476 | -0.668 | -1.477 |
| GIMAP1 | NM_130759 | 0.2835 | -0.016 |
| MADCAM1 | NM_130760 | 0.215 | -0.235 |
| MADCAM1 | NM_130762 | 0.215 | -0.235 |
| INPP5K | NM_130766 | -0.279 | -1.1715 |
| ACOT12 | NM_130767 | -0.185 | 0.0485 |
| ASZ1 | NM_130768 | 0.0405 | -0.659 |
| GPHA2 | NM_130769 | 0.198 | -0.308 |
| HTR3C | NM_130770 | -0.1115 | -0.2555 |
| S100Z | NM_130772 | -0.224 | 0.1675 |
| CNTNAP5 | NM_130773 | 0.113 | -0.2825 |
| XAGE5 | NM_130775 | -0.1245 | 0.151 |
| XAGE3 | NM_130776 | -0.253 | -0.0745 |
| XAGE2 | NM_130777 | 0.073 | -0.2835 |
| RAB24 | NM_130781 | -0.8445 | -1.505 |
| RGS18 | NM_130782 | -0.091 | -0.557 |
| TSPAN18 | NM_130783 | -0.361 | -1.5675 |
| SYCE1 | NM_130784 | -0.0115 | 0.16 |
| TPTE2 | NM_130785 | -0.1185 | -0.1605 |
| A1BG | NM_130786 | -0.0845 | -0.7775 |
| AP2A1 | NM_130787 | 0.0425 | -0.412 |
| WWOX | NM_130791 | -0.0675 | -0.5235 |
| CST11 | NM_130794 | -0.0905 | -0.2355 |
| RGS3 | NM_130795 | -0.617 | -1.4575 |
| DPP6 | NM_130797 | -0.537 | -0.9885 |
| SNAP23 | NM_130798 | -1.021 | -1.399 |
| MEN1 | NM_130803 | -0.81 | -1.9455 |
| RXFP2 | NM_130806 | -0.215 | -0.3625 |
| MOB3A | NM_130807 | -1.2995 | -2.418 |
| CPNE4 | NM_130808 | -0.41 | -0.429 |
| PRRC1 | NM_130809 | -1.5815 | -3.3235 |

| | | | |
|---|---|---|---|
| DYX1C1 | NM_130810 | -0.6375 | -1.257 |
| SNAP25 | NM_130811 | -0.827 | -0.8375 |
| LRRC15 | NM_130830 | -0.0675 | -0.0795 |
| OPA1 | NM_130831 | -0.7875 | -1.6125 |
| OPA1 | NM_130832 | -0.7875 | -1.6125 |
| OPA1 | NM_130833 | -0.7875 | -1.6125 |
| OPA1 | NM_130834 | -0.7875 | -1.6125 |
| OPA1 | NM_130835 | -0.7875 | -1.6125 |
| OPA1 | NM_130836 | -0.7875 | -1.6125 |
| OPA1 | NM_130837 | -0.7875 | -1.6125 |
| UBE3A | NM_130838 | -1.2775 | -2.2785 |
| UBE3A | NM_130839 | -1.2775 | -2.2785 |
| ATP6V0A4 | NM_130840 | -0.341 | 0.0465 |
| ATP6V0A4 | NM_130841 | -0.341 | 0.0465 |
| PTPRN2 | NM_130842 | -0.063 | -0.502 |
| PTPRN2 | NM_130843 | -0.317 | -0.706 |
| WWOX | NM_130844 | -0.132 | -0.3865 |
| PTPRR | NM_130846 | -0.95 | -1.8585 |
| AMOTL1 | NM_130847 | -1.1865 | -2.466 |
| C5orf20 | NM_130848 | -0.251 | -0.118 |
| SLC39A4 | NM_130849 | 0.193 | -0.492 |
| BPIFA1 | NM_130852 | -0.3445 | -0.513 |
| PTPRS | NM_130853 | -0.0245 | -0.5485 |
| PTPRS | NM_130854 | -0.0245 | -0.5485 |
| PTPRS | NM_130855 | -0.0245 | -0.5485 |
| WFDC8 | NM_130896 | -0.0565 | -0.186 |
| DYNLRB2 | NM_130897 | 0.089 | 0.092 |
| CREB3L4 | NM_130898 | -0.901 | -1.6625 |
| FAM71B | NM_130899 | 0.221 | -0.2085 |
| RAET1L | NM_130900 | -0.1115 | 0.1835 |
| OTUD7A | NM_130901 | -0.131 | -1.079 |
| COX7B2 | NM_130902 | 0.098 | -0.215 |
| PPIL3 | NM_130906 | -1.016 | -2.1165 |
| PTPRT | NM_133170 | -0.017 | -0.1395 |
| ELMO2 | NM_133171 | -0.351 | -1.7265 |
| APBB3 | NM_133172 | -0.3785 | -0.84 |
| APBB3 | NM_133173 | -0.3785 | -0.84 |
| APBB3 | NM_133174 | -0.3785 | -0.84 |
| XAGE3 | NM_133179 | -0.253 | -0.0745 |
| EPS8L1 | NM_133180 | 0.5065 | 0.097 |
| EPS8L3 | NM_133181 | 0.1025 | -0.1265 |
| LRPPRC | NM_133259 | -0.5245 | -1.0905 |
| GIPC3 | NM_133261 | -0.12 | -0.558 |

| | | | |
|---|---|---|---|
| ATP6V1G3 | NM_133262 | 0.0445 | -0.346 |
| PPARGC1B | NM_133263 | -0.3485 | -0.471 |
| WIPF2 | NM_133264 | -0.5495 | -2.158 |
| AMOT | NM_133265 | -0.921 | -0.796 |
| SHANK2 | NM_133266 | -0.894 | -1.6275 |
| GSX2 | NM_133267 | 0.159 | -0.356 |
| FCAR | NM_133269 | -0.5115 | -0.1385 |
| FCAR | NM_133271 | -0.5115 | -0.1385 |
| FCAR | NM_133272 | -0.5115 | -0.1385 |
| FCAR | NM_133273 | -0.5115 | -0.1385 |
| FCAR | NM_133274 | -0.5115 | -0.1385 |
| FCAR | NM_133277 | -0.5115 | -0.1385 |
| FCAR | NM_133278 | -0.007 | -0.321 |
| FCAR | NM_133279 | -0.5115 | -0.1385 |
| PHF10 | NM_133325 | -0.9525 | -0.2705 |
| ATP6V1G3 | NM_133326 | 0.0445 | -0.346 |
| DEDD2 | NM_133328 | -0.3965 | -2.0385 |
| KCNG3 | NM_133329 | -0.587 | -1.081 |
| WHSC1 | NM_133330 | -0.7515 | -1.5505 |
| WHSC1 | NM_133331 | -0.668 | -1.475 |
| WHSC1 | NM_133334 | -0.668 | -1.475 |
| WHSC1 | NM_133335 | -0.668 | -1.475 |
| MYOF | NM_133337 | -0.237 | -0.1815 |
| RAD17 | NM_133338 | -1.675 | -2.954 |
| RAD17 | NM_133339 | -1.675 | -2.954 |
| RAD17 | NM_133340 | -1.675 | -2.954 |
| RAD17 | NM_133341 | -1.675 | -2.954 |
| RAD17 | NM_133342 | -1.675 | -2.954 |
| RAD17 | NM_133343 | -1.675 | -2.954 |
| RAD17 | NM_133344 | -1.675 | -2.954 |
| PAQR8 | NM_133367 | -1.3455 | -2.1265 |
| RSPRY1 | NM_133368 | -1.411 | -2.793 |
| UNC5A | NM_133369 | -0.3375 | -1.4995 |
| YTHDC1 | NM_133370 | -1.3355 | -2.657 |
| MYOZ3 | NM_133371 | -0.077 | -0.1185 |
| FNIP1 | NM_133372 | -1.496 | -2.1105 |
| PLCD3 | NM_133373 | 0.184 | -0.658 |
| ZNF618 | NM_133374 | -0.524 | -1.6785 |
| DIS3L | NM_133375 | -1.497 | -1.8245 |
| ITGB1 | NM_133376 | -0.47 | -0.3105 |
| TTN | NM_133378 | 0.34 | 0.126 |
| TTN | NM_133379 | 0.34 | 0.126 |
| TTN | NM_133432 | 0.34 | 0.126 |

| | | | |
|---|---|---|---|
| NIPBL | NM_133433 | -0.323 | -2.366 |
| ASNS | NM_133436 | -0.1835 | -0.436 |
| TTN | NM_133437 | 0.34 | 0.126 |
| TADA2A | NM_133439 | -1.5365 | -2.296 |
| GPT2 | NM_133443 | -1.159 | -2.125 |
| ZNF526 | NM_133444 | -1.1325 | -1.8465 |
| GRIN3A | NM_133445 | 0.2375 | -1.437 |
| AGAP4 | NM_133446 | -0.52 | -1.8425 |
| AGAP11 | NM_133447 | -0.7755 | -0.6115 |
| TMEM132D | NM_133448 | -0.33 | 0.026 |
| ANKS3 | NM_133450 | -0.58 | -1.421 |
| RAVER1 | NM_133452 | -0.46 | -1.1075 |
| SGSM1 | NM_133454 | -0.293 | -0.5015 |
| EMID1 | NM_133455 | -0.126 | -0.4225 |
| SHROOM1 | NM_133456 | -0.0935 | -0.1175 |
| EMID2 | NM_133457 | -0.118 | -0.2115 |
| ZFP90 | NM_133458 | -1.46 | -2.5325 |
| CCBE1 | NM_133459 | -0.3985 | -0.563 |
| ZNF418 | NM_133460 | -0.2155 | -1.356 |
| TTC14 | NM_133462 | -0.826 | -1.667 |
| AMZ1 | NM_133463 | 0.1235 | -0.1615 |
| ZNF483 | NM_133464 | 0.0615 | 0.1535 |
| KIAA1958 | NM_133465 | -0.882 | -1.9385 |
| ZFP82 | NM_133466 | -0.684 | -2.8615 |
| CITED4 | NM_133467 | 0.1265 | -0.058 |
| BMPER | NM_133468 | -0.209 | -0.0965 |
| ZNF431 | NM_133473 | -1.032 | -2.465 |
| ZNF721 | NM_133474 | -0.4725 | -1.94 |
| ANKRD24 | NM_133475 | -0.2055 | -0.3035 |
| ZNF384 | NM_133476 | -0.8115 | -2.357 |
| SYNPO2 | NM_133477 | -1.1055 | -2.166 |
| SLC4A5 | NM_133478 | -0.479 | -0.381 |
| TADA3 | NM_133480 | -0.294 | -0.809 |
| TANK | NM_133484 | -1.848 | -3.981 |
| MBNL3 | NM_133486 | 0.1225 | -0.466 |
| RAD51 | NM_133487 | -0.7715 | -2.366 |
| SLC26A10 | NM_133489 | -0.0795 | -0.5015 |
| SAT2 | NM_133491 | -0.0925 | -1.054 |
| ACER1 | NM_133492 | 0.4995 | 0.2245 |
| CD109 | NM_133493 | -0.1945 | 0.07 |
| NEK7 | NM_133494 | -0.8315 | -1.928 |
| SLC30A7 | NM_133496 | -1.33 | -2.6405 |
| KCNV2 | NM_133497 | -0.007 | -0.587 |

| | | | |
|---|---|---|---|
| SPACA4 | NM_133498 | -0.069 | -0.6095 |
| ZNF274 | NM_133502 | -1.0695 | -3.114 |
| DCN | NM_133503 | 0.3285 | 0.0705 |
| DCN | NM_133504 | 0.3285 | 0.0705 |
| DCN | NM_133505 | 0.3285 | 0.0705 |
| DCN | NM_133506 | 0.3285 | 0.0705 |
| DCN | NM_133507 | 0.3285 | 0.0705 |
| RAD51B | NM_133509 | -0.0015 | 0.1 |
| RAD51B | NM_133510 | -0.4165 | -0.168 |
| SYN2 | NM_133625 | 0.4845 | -0.931 |
| ROBO1 | NM_133631 | -0.903 | -1.004 |
| SYN3 | NM_133633 | -0.103 | -0.304 |
| POFUT2 | NM_133635 | -0.552 | -1.331 |
| HELQ | NM_133636 | -2.1665 | -2.831 |
| DQX1 | NM_133637 | 0.0595 | -0.368 |
| ADAMTS19 | NM_133638 | -0.976 | -0.6685 |
| RHOV | NM_133639 | -0.167 | -0.9735 |
| MED22 | NM_133640 | -1.496 | -2.073 |
| LARGE | NM_133642 | -0.342 | -1.4415 |
| GTPBP3 | NM_133644 | -0.705 | -1.389 |
| MTO1 | NM_133645 | -1.785 | -3.24 |
| ZAK | NM_133646 | -1.2735 | -2.3555 |
| SLC12A6 | NM_133647 | -0.356 | -0.688 |
| TBL1Y | NM_134258 | 0.1395 | -0.4755 |
| TBL1Y | NM_134259 | 0.1395 | -0.4755 |
| RORA | NM_134260 | -0.291 | -0.7685 |
| RORA | NM_134261 | -0.291 | -0.7685 |
| RORA | NM_134262 | -0.291 | -0.7685 |
| WSB1 | NM_134265 | -1.201 | -2.064 |
| SLC26A7 | NM_134266 | 0.1165 | -0.1585 |
| CYGB | NM_134268 | -0.9075 | -1.645 |
| SMTN | NM_134269 | -0.25 | -0.4715 |
| SMTN | NM_134270 | -0.232 | -0.64 |
| TARBP2 | NM_134323 | -0.134 | -1.4165 |
| TARBP2 | NM_134324 | -0.102 | -0.257 |
| SLC26A9 | NM_134325 | -0.083 | -0.4265 |
| HPCAL1 | NM_134421 | -0.2015 | -0.865 |
| RAD52 | NM_134424 | -1.302 | -1.306 |
| SLC26A1 | NM_134425 | 0.0175 | -0.086 |
| RGS3 | NM_134427 | -0.617 | -1.4575 |
| RFX3 | NM_134428 | -0.2065 | 0.1065 |
| SLCO1A2 | NM_134431 | 0.443 | 0.065 |
| RFX2 | NM_134433 | -0.166 | -0.346 |

| | | | |
|---|---|---|---|
| RLN2 | NM_134441 | -0.324 | 0.1365 |
| CREB1 | NM_134442 | -2.091 | -3.7615 |
| NLRP4 | NM_134444 | 0.0805 | -0.1545 |
| CD99L2 | NM_134445 | -0.5455 | -0.894 |
| CD99L2 | NM_134446 | -0.487 | -1.161 |
| C19orf2 | NM_134447 | -0.934 | -1.4935 |
| IL1RAP | NM_134470 | 0.0075 | -0.354 |
| ATRX | NM_138270 | 0.3615 | -1.5905 |
| C6orf25 | NM_138272 | 0.1835 | -0.2545 |
| C6orf25 | NM_138273 | 0.1835 | -0.2545 |
| C6orf25 | NM_138274 | 0.1835 | -0.2545 |
| C6orf25 | NM_138275 | 0.1835 | -0.2545 |
| C6orf25 | NM_138277 | 0.1835 | -0.2545 |
| BNIPL | NM_138278 | 0.2465 | 0.062 |
| DLX4 | NM_138281 | 0.0375 | -0.259 |
| ATP6V1G2 | NM_138282 | 0.0255 | -0.322 |
| CSTL1 | NM_138283 | 0.0765 | 0.046 |
| IL17D | NM_138284 | -0.299 | -0.7705 |
| NUP35 | NM_138285 | -0.906 | -1.9225 |
| ZNF681 | NM_138286 | -0.979 | -1.924 |
| DTX3L | NM_138287 | -0.0365 | -1.6985 |
| SPTSSA | NM_138288 | -1.12 | -1.1345 |
| ACTRT1 | NM_138289 | -0.0645 | 0.0795 |
| RUNDC3B | NM_138290 | -0.7425 | -1.8 |
| PATE1 | NM_138294 | 0.001 | -0.022 |
| PKD1L1 | NM_138295 | 0.0495 | -0.227 |
| PTCRA | NM_138296 | -0.0825 | -0.39 |
| MUC4 | NM_138297 | 0.0875 | -0.2915 |
| PYGO2 | NM_138300 | -0.609 | -1.4135 |
| PANK1 | NM_138316 | -2.021 | -4.2235 |
| KCNK10 | NM_138317 | -0.468 | -0.758 |
| KCNK10 | NM_138318 | -0.468 | -0.758 |
| PCSK6 | NM_138319 | -0.3075 | -0.326 |
| PCSK6 | NM_138320 | -0.3075 | -0.326 |
| PCSK6 | NM_138321 | -0.002 | -0.092 |
| PCSK6 | NM_138322 | -0.3075 | -0.326 |
| PCSK6 | NM_138323 | -0.3075 | -0.326 |
| PCSK6 | NM_138324 | -0.3075 | -0.326 |
| PCSK6 | NM_138325 | 0.418 | -0.237 |
| ACMSD | NM_138326 | 0.092 | -0.6955 |
| TAAR1 | NM_138327 | -0.252 | 0.365 |
| RHBDL3 | NM_138328 | -1.224 | -1.8175 |
| NLRP6 | NM_138329 | 0.3165 | -0.003 |

| | | | |
|---|---|---|---|
| ZNF675 | NM_138330 | -0.2605 | -0.8175 |
| RNASE8 | NM_138331 | 0.2155 | -0.532 |
| FAM122A | NM_138333 | -1.2375 | -3.118 |
| JOSD2 | NM_138334 | -0.2835 | -0.098 |
| GNPDA2 | NM_138335 | -0.8815 | -2.5435 |
| MYEOV2 | NM_138336 | 0.2875 | -0.5235 |
| CLEC12A | NM_138337 | 0.1505 | 0.021 |
| POLR3H | NM_138338 | -0.629 | -2.1645 |
| ABHD3 | NM_138340 | -1.3695 | -0.9655 |
| TMEM116 | NM_138341 | -0.7285 | -1.6775 |
| GLB1L2 | NM_138342 | -0.317 | -1.024 |
| KLC4 | NM_138343 | -0.2375 | -0.203 |
| FAM181A | NM_138344 | 0.007 | -0.388 |
| VWA5B2 | NM_138345 | -0.0485 | -0.5525 |
| KIAA2013 | NM_138346 | -0.0625 | -1.0175 |
| ZNF551 | NM_138347 | -1.2415 | -1.3335 |
| FAM105B | NM_138348 | -0.5835 | -1.3925 |
| TP53I13 | NM_138349 | -0.5175 | -0.9815 |
| THAP3 | NM_138350 | -0.4755 | -0.9105 |
| SAMD1 | NM_138352 | -0.0815 | -0.7265 |
| DCAF15 | NM_138353 | -0.1755 | -0.814 |
| SCRN2 | NM_138355 | -0.3315 | -1.237 |
| SHF | NM_138356 | -0.3605 | -1.784 |
| MCU | NM_138357 | -0.6125 | -1.4175 |
| C19orf52 | NM_138358 | -0.63 | -2.016 |
| LRRC16B | NM_138360 | -0.2935 | -0.2845 |
| LRSAM1 | NM_138361 | -0.2465 | -0.847 |
| FAM104B | NM_138362 | -0.595 | -0.7705 |
| PRMT10 | NM_138364 | -1.1005 | -2.578 |
| ZNF251 | NM_138367 | -0.861 | -1.5585 |
| DKFZp761E198 | NM_138368 | -0.1355 | -0.7445 |
| BOD1 | NM_138369 | -0.4935 | -1.3225 |
| PKDCC | NM_138370 | -0.9615 | -2.527 |
| FAM113B | NM_138371 | -0.1315 | -0.113 |
| MYADM | NM_138373 | -0.7145 | -2.059 |
| ZNF845 | NM_138374 | 0.025 | -0.861 |
| CABLES1 | NM_138375 | -0.5665 | -1.7955 |
| TTC5 | NM_138376 | -0.519 | -1.334 |
| FDXACB1 | NM_138378 | -1.303 | -2.3985 |
| TIMD4 | NM_138379 | 0.191 | 0.006 |
| OXNAD1 | NM_138381 | -0.6915 | -2.24 |
| RIPPLY1 | NM_138382 | 0.2465 | -0.1455 |
| MTSS1L | NM_138383 | -0.34 | -0.782 |

| | | | |
|---|---|---|---|
| MTG1 | NM_138384 | -0.0235 | -0.9215 |
| TMEM129 | NM_138385 | -0.466 | -0.71 |
| NAF1 | NM_138386 | -0.407 | -1.656 |
| G6PC3 | NM_138387 | -0.036 | -0.7355 |
| FAM114A1 | NM_138389 | -0.733 | -1.206 |
| TMEM169 | NM_138390 | -1.5175 | -2.4055 |
| TMEM183A | NM_138391 | -0.505 | -0.5955 |
| SHKBP1 | NM_138392 | 0.2525 | -0.4025 |
| REEP6 | NM_138393 | -0.099 | -0.491 |
| HNRPLL | NM_138394 | -1.6025 | -2.7035 |
| MARS2 | NM_138395 | -1.215 | -2.7665 |
| TMEM44 | NM_138399 | -1.785 | -1.482 |
| NOM1 | NM_138400 | -2.4105 | -2.738 |
| FAM125A | NM_138401 | 0.1645 | -0.4405 |
| SP140L | NM_138402 | -0.2525 | -0.112 |
| MYL10 | NM_138403 | 0.101 | -0.008 |
| GTF3C6 | NM_138408 | 0.158 | -0.626 |
| MRAP2 | NM_138409 | -0.474 | -1.0515 |
| CMTM7 | NM_138410 | -0.04 | -1.0945 |
| FAM71E1 | NM_138411 | 0.0035 | -0.5 |
| RDH13 | NM_138412 | -0.7765 | -1.8235 |
| HOGA1 | NM_138413 | -0.4235 | -0.4065 |
| CCDC101 | NM_138414 | -0.6935 | -2.332 |
| PHF21B | NM_138415 | -0.3435 | -0.999 |
| KTI12 | NM_138417 | -0.5505 | -2.0525 |
| FAM195A | NM_138418 | -0.5765 | -0.8195 |
| FAM54A | NM_138419 | -1.8985 | -3.635 |
| AHNAK2 | NM_138420 | -0.376 | -0.8195 |
| SAAL1 | NM_138421 | -0.6835 | -1.282 |
| ADAT3 | NM_138422 | -0.058 | -0.684 |
| CASC4 | NM_138423 | -1.4015 | -1.1535 |
| KIF12 | NM_138424 | -0.117 | -0.6755 |
| C12orf57 | NM_138425 | 0.5205 | -0.0605 |
| GLCCI1 | NM_138426 | -1.257 | -1.876 |
| C1orf212 | NM_138428 | -0.8555 | -2.974 |
| ADPRHL1 | NM_138430 | -0.1145 | -0.4825 |
| MFSD3 | NM_138431 | -0.3725 | -0.715 |
| SDSL | NM_138432 | -0.318 | -0.229 |
| KLHDC7B | NM_138433 | 0.0645 | -0.387 |
| C7orf29 | NM_138434 | 0.4715 | -0.343 |
| FAM83F | NM_138435 | -0.322 | -0.0435 |
| FLYWCH2 | NM_138439 | -0.5165 | -1.2625 |
| VASN | NM_138440 | -0.0075 | -0.7615 |

| | | | |
|---|---|---|---|
| CCDC124 | NM_138442 | -0.045 | -0.666 |
| HAUS1 | NM_138443 | -1.499 | -2.111 |
| KCTD12 | NM_138444 | -1.591 | -1.3935 |
| GPR146 | NM_138445 | -0.8025 | -0.6395 |
| C7orf30 | NM_138446 | -1.114 | -5.142 |
| ZNF689 | NM_138447 | -0.5505 | -0.997 |
| ACYP2 | NM_138448 | 0.1435 | -0.256 |
| ARL11 | NM_138450 | -0.2505 | -0.1915 |
| IQCD | NM_138451 | 0.0435 | -0.5535 |
| RAB3C | NM_138453 | -0.4295 | -0.775 |
| NXNL1 | NM_138454 | 0.0645 | -0.5115 |
| CTHRC1 | NM_138455 | -0.2645 | -1.579 |
| BATF2 | NM_138456 | -0.2435 | -0.4935 |
| FOXP4 | NM_138457 | -0.104 | -0.826 |
| WDR92 | NM_138458 | -0.508 | -0.935 |
| NUS1 | NM_138459 | -1.4495 | -2.2635 |
| CMTM5 | NM_138460 | -0.0685 | -0.6575 |
| TM4SF19 | NM_138461 | 0.172 | 0.042 |
| ZMYND19 | NM_138462 | -0.7605 | -2.1415 |
| TLCD1 | NM_138463 | -0.528 | -0.706 |
| C5orf55 | NM_138464 | -0.006 | 0.0935 |
| TYW3 | NM_138467 | -0.172 | -0.754 |
| ICA1L | NM_138468 | -0.5505 | -0.896 |
| C11orf84 | NM_138471 | -0.96 | -1.915 |
| SP1 | NM_138473 | -1.824 | -3.087 |
| MDP1 | NM_138476 | -0.2505 | -0.926 |
| CDAN1 | NM_138477 | -0.882 | -1.8195 |
| CHADL | NM_138481 | -0.4135 | -0.161 |
| SGOL1 | NM_138484 | -0.38 | -0.871 |
| GTPBP8 | NM_138485 | -0.6805 | -1.567 |
| PRELID2 | NM_138492 | -0.6515 | -1.0585 |
| CCDC167 | NM_138493 | -0.1315 | -0.659 |
| ZNF655 | NM_138494 | -0.9175 | -1.7805 |
| ATXN7L1 | NM_138495 | 0.3515 | -0.3115 |
| CYHR1 | NM_138496 | -0.473 | -0.8475 |
| PWWP2B | NM_138499 | -0.8345 | -1.8365 |
| TECR | NM_138501 | 0.1395 | -0.2815 |
| TLR4 | NM_138554 | -0.1665 | 0.27 |
| KIF23 | NM_138555 | -1.291 | -2.407 |
| PPP1R8 | NM_138558 | -1.4105 | -3.159 |
| BCL11A | NM_138559 | -1.1425 | -2.253 |
| KLK15 | NM_138563 | 0.4315 | -0.0445 |
| KLK15 | NM_138564 | -0.233 | -0.2765 |

| | | | |
|---|---|---|---|
| CTTN | NM_138565 | -0.321 | -0.5175 |
| SYT8 | NM_138567 | -0.325 | -0.05 |
| EXOC3L2 | NM_138568 | -0.0775 | -0.373 |
| MLIP | NM_138569 | -0.042 | -0.6155 |
| SLC38A10 | NM_138570 | -0.3355 | -0.5775 |
| HINT3 | NM_138571 | -0.756 | -0.783 |
| TAF8 | NM_138572 | -0.346 | -0.7715 |
| NRG4 | NM_138573 | 0.2625 | -0.0285 |
| HDGFL1 | NM_138574 | -0.3775 | -0.3715 |
| PGAM5 | NM_138575 | -0.229 | -1.028 |
| BCL2L1 | NM_138578 | 0.423 | 0.4545 |
| H2AFY | NM_138609 | 0.04 | -0.4995 |
| H2AFY | NM_138610 | 0.04 | -0.4995 |
| HAS3 | NM_138612 | 0.121 | -0.2645 |
| RHCE | NM_138616 | -0.4685 | -0.9525 |
| RHCE | NM_138617 | -0.242 | -0.775 |
| RHCE | NM_138618 | 0.143 | 0.187 |
| GGA3 | NM_138619 | -0.9555 | -2.739 |
| DDX31 | NM_138620 | -1.564 | -3.056 |
| BCL2L11 | NM_138621 | -0.4725 | -1.6275 |
| BCL2L11 | NM_138622 | -0.4175 | -1.9595 |
| BCL2L11 | NM_138623 | -0.4175 | -1.9595 |
| BCL2L11 | NM_138624 | -0.4175 | -1.9595 |
| BCL2L11 | NM_138625 | -0.4175 | -1.9595 |
| BCL2L11 | NM_138626 | -0.4175 | -1.9595 |
| BCL2L11 | NM_138627 | -0.4175 | -1.9595 |
| TRIOBP | NM_138632 | 0.027 | -0.347 |
| AKAP7 | NM_138633 | -0.358 | -2.2945 |
| MSMB | NM_138634 | -0.277 | -0.6 |
| H2AFV | NM_138635 | 0.292 | -0.25 |
| TLR8 | NM_138636 | -0.2885 | -0.0265 |
| CFL2 | NM_138638 | -0.8905 | -2.2675 |
| BCL2L12 | NM_138639 | -0.1495 | -0.81 |
| PPP1R14B | NM_138689 | 0.205 | -0.155 |
| TMC1 | NM_138691 | -0.27 | -0.192 |
| KLF14 | NM_138693 | -0.5465 | -0.325 |
| PKHD1 | NM_138694 | 0.0635 | -0.0495 |
| TAS1R1 | NM_138697 | -0.378 | -0.5595 |
| TRIM40 | NM_138700 | 0.125 | -0.368 |
| C7orf11 | NM_138701 | -0.425 | -2.454 |
| MAGEC3 | NM_138702 | -0.1615 | -0.3695 |
| MAGEE2 | NM_138703 | 0.038 | 0.0365 |
| NDNL2 | NM_138704 | -0.639 | -0.5155 |

| | | | |
|---|---|---|---|
| CALML6 | NM_138705 | 0.004 | -0.0725 |
| B3GNT6 | NM_138706 | 0.0145 | -0.0115 |
| DAB2IP | NM_138709 | -0.106 | -0.4885 |
| PPARG | NM_138711 | -0.8965 | -1.2445 |
| PPARG | NM_138712 | -0.758 | -1.625 |
| NFAT5 | NM_138713 | -1.0085 | -2.852 |
| NFAT5 | NM_138714 | -1.0085 | -2.852 |
| MSR1 | NM_138715 | -0.4005 | -0.451 |
| MSR1 | NM_138716 | -0.4005 | -0.451 |
| PPT2 | NM_138717 | -0.031 | -0.4985 |
| HIST1H2BD | NM_138720 | -1.0135 | -1.633 |
| BCL2L14 | NM_138722 | -0.154 | -0.094 |
| BCL2L14 | NM_138723 | -0.4775 | -0.1325 |
| ST7L | NM_138727 | -1.44 | -1.7125 |
| ST7L | NM_138728 | -1.44 | -1.7125 |
| ST7L | NM_138729 | -1.44 | -1.7125 |
| MIPOL1 | NM_138731 | -0.0125 | -0.2125 |
| NRXN2 | NM_138732 | -0.5115 | -1.693 |
| PGK2 | NM_138733 | -0.4085 | -0.5025 |
| NRXN2 | NM_138734 | -0.217 | -1.237 |
| NRXN1 | NM_138735 | -0.3315 | -0.859 |
| GNAO1 | NM_138736 | -0.3355 | -1.044 |
| HEPH | NM_138737 | 0.1965 | 0.139 |
| C1orf43 | NM_138740 | -0.496 | -1.4995 |
| BAX | NM_138764 | -0.4365 | -0.9995 |
| PAM | NM_138766 | -0.7815 | -0.7685 |
| MYEOV | NM_138768 | -0.3115 | -0.18 |
| RHOT2 | NM_138769 | -0.2515 | -1.0445 |
| CCDC74A | NM_138770 | 0.1965 | -0.245 |
| CCDC126 | NM_138771 | -1.499 | -2.8695 |
| SLC25A46 | NM_138773 | -0.978 | -1.7495 |
| C19orf22 | NM_138774 | -0.198 | -1.052 |
| ALKBH8 | NM_138775 | -0.526 | -1.5595 |
| MRRF | NM_138777 | -0.8035 | -1.159 |
| WDR85 | NM_138778 | -0.8105 | -2.356 |
| C13orf27 | NM_138779 | -1.9595 | -3.6375 |
| SYTL5 | NM_138780 | -0.244 | -0.192 |
| FCHO2 | NM_138782 | -1.051 | -1.705 |
| ZNF653 | NM_138783 | -0.468 | -1.1555 |
| C6orf72 | NM_138785 | -0.82 | -1.3895 |
| TM4SF18 | NM_138786 | -0.097 | 0.415 |
| C11orf74 | NM_138787 | -0.744 | -1.004 |
| TMEM45B | NM_138788 | -0.127 | -0.33 |

| | | | |
|---|---|---|---|
| PIH1D2 | NM_138789 | -0.0735 | -0.122 |
| PLD4 | NM_138790 | 0.468 | -0.0495 |
| LEO1 | NM_138792 | -0.6665 | -2.275 |
| CANT1 | NM_138793 | -0.562 | -3.138 |
| LYPLAL1 | NM_138794 | -1.329 | -2.2485 |
| ARL8A | NM_138795 | -0.379 | -1.503 |
| SPATA17 | NM_138796 | -0.0545 | -0.2645 |
| ANKRD54 | NM_138797 | -0.627 | -1.8055 |
| MITD1 | NM_138798 | -1.555 | -2.5485 |
| MBOAT2 | NM_138799 | -1.061 | -1.4405 |
| GALM | NM_138801 | -0.4325 | -1.049 |
| ZFAND2B | NM_138802 | -0.2605 | -0.9885 |
| CCDC148 | NM_138803 | 0.388 | -0.006 |
| C2orf65 | NM_138804 | -0.065 | -0.169 |
| FAM3D | NM_138805 | -0.2345 | -0.399 |
| CD200R1 | NM_138806 | 0.072 | -0.447 |
| TAMM41 | NM_138807 | -0.835 | -2.319 |
| CMBL | NM_138809 | -0.481 | -0.758 |
| TAGAP | NM_138810 | -0.0275 | 0.19 |
| C7orf31 | NM_138811 | -0.9955 | -1.984 |
| DYDC1 | NM_138812 | -0.408 | -0.439 |
| ATP8B3 | NM_138813 | -0.31 | -0.096 |
| PNPLA5 | NM_138814 | -0.0895 | -0.7535 |
| DPPA2 | NM_138815 | 0.1665 | -0.045 |
| SLC7A13 | NM_138817 | 0.1265 | -0.0215 |
| FAM122C | NM_138819 | -0.3205 | -0.2315 |
| HIGD2A | NM_138820 | 0.217 | -0.359 |
| PAM | NM_138821 | -0.739 | -0.8865 |
| PAM | NM_138822 | -0.739 | -0.8865 |
| TAF1 | NM_138923 | -0.827 | -0.987 |
| GAMT | NM_138924 | 0.0595 | -0.2325 |
| SON | NM_138927 | -1.7915 | -2.9265 |
| DIABLO | NM_138929 | -0.606 | -2.1555 |
| A1CF | NM_138932 | 0.0105 | -0.246 |
| A1CF | NM_138933 | 0.0105 | -0.246 |
| REG3A | NM_138938 | -0.0245 | 0.0265 |
| CD200R1 | NM_138939 | 0.072 | -0.447 |
| CD200R1 | NM_138940 | 0.072 | -0.447 |
| MAPK1 | NM_138957 | -1.176 | -2.003 |
| VANGL1 | NM_138959 | -0.187 | -1.193 |
| ESAM | NM_138961 | 0.005 | -0.4745 |
| MSI2 | NM_138962 | -0.7105 | -1.588 |
| PROKR1 | NM_138964 | 0.113 | -0.1385 |

| | | | |
|---|---|---|---|
| NETO1 | NM_138966 | -0.866 | -1.0225 |
| SCAMP5 | NM_138967 | -0.65 | -1.723 |
| SDR16C5 | NM_138969 | 0.017 | -0.857 |
| NRXN3 | NM_138970 | -0.261 | -0.418 |
| BACE1 | NM_138971 | -0.613 | -1.0805 |
| BACE1 | NM_138972 | -0.613 | -1.0805 |
| BACE1 | NM_138973 | -0.613 | -1.0805 |
| MAPK10 | NM_138980 | -0.772 | -0.7575 |
| MAPK10 | NM_138981 | -0.778 | -0.715 |
| MAPK10 | NM_138982 | -0.778 | -0.715 |
| OLIG1 | NM_138983 | 0.6465 | 0.2335 |
| BACE2 | NM_138991 | -0.189 | -0.442 |
| BACE2 | NM_138992 | -0.189 | -0.442 |
| MYO3B | NM_138995 | -0.0665 | 0.1395 |
| NETO1 | NM_138999 | -0.866 | -1.0225 |
| HFE | NM_139003 | 0.087 | -0.1325 |
| HFE | NM_139004 | 0.087 | -0.1325 |
| HFE | NM_139006 | 0.087 | -0.1325 |
| HFE | NM_139007 | 0.087 | -0.1325 |
| HFE | NM_139008 | 0.087 | -0.1325 |
| HFE | NM_139009 | 0.087 | -0.1325 |
| HFE | NM_139010 | 0.087 | -0.1325 |
| HFE | NM_139011 | 0.087 | -0.1325 |
| MAPK14 | NM_139012 | -0.9775 | -2.637 |
| MAPK14 | NM_139013 | -0.9775 | -2.637 |
| MAPK14 | NM_139014 | -0.9775 | -2.637 |
| SPPL3 | NM_139015 | -0.297 | -1.4765 |
| IL31RA | NM_139017 | 0.0715 | -0.4265 |
| CD300LF | NM_139018 | 0.067 | -0.2625 |
| MAPK15 | NM_139021 | -0.1595 | 0.033 |
| TSPAN32 | NM_139022 | -0.0785 | -0.01 |
| ADAMTS13 | NM_139025 | -0.6415 | -0.368 |
| ADAMTS13 | NM_139026 | -0.1505 | -0.132 |
| ADAMTS13 | NM_139027 | -0.1505 | -0.132 |
| CD151 | NM_139029 | -0.2565 | -1.2915 |
| CD151 | NM_139030 | -0.116 | -1.335 |
| MAPK7 | NM_139032 | -0.8975 | -2.7945 |
| MAPK7 | NM_139033 | -0.8975 | -2.7945 |
| MAPK7 | NM_139034 | -0.8975 | -2.7945 |
| SMARCA1 | NM_139035 | -0.4905 | -0.899 |
| SMARCA2 | NM_139045 | 0.0205 | -1.3125 |
| MAPK8 | NM_139046 | -0.5925 | -0.8075 |
| MAPK8 | NM_139047 | -0.5925 | -0.8075 |

| | | | |
|---|---|---|---|
| HLTF | NM_139048 | -0.4 | -1.894 |
| MAPK8 | NM_139049 | -0.5925 | -0.8075 |
| EPS8L3 | NM_139053 | 0.1025 | -0.1265 |
| ADAMTS15 | NM_139055 | -0.0265 | -0.3345 |
| ADAMTS16 | NM_139056 | 0.054 | -0.0995 |
| ADAMTS17 | NM_139057 | -0.8015 | -2.1255 |
| ARX | NM_139058 | 0.1545 | -0.072 |
| CSNK1D | NM_139062 | -1.008 | -2.954 |
| SMARCC2 | NM_139067 | 0.0015 | -0.5455 |
| MAPK9 | NM_139068 | -0.5785 | -1.1935 |
| MAPK9 | NM_139069 | -0.5785 | -1.1935 |
| MAPK9 | NM_139070 | -0.5785 | -1.1935 |
| SMARCD1 | NM_139071 | -0.1235 | -0.787 |
| DNER | NM_139072 | 0.073 | -0.4525 |
| SPATA3 | NM_139073 | 0.0895 | -0.0755 |
| DEFB127 | NM_139074 | -0.0655 | 0.231 |
| TPCN2 | NM_139075 | -0.4455 | -0.877 |
| FAM175A | NM_139076 | -0.6905 | -1.208 |
| MAPKAPK5 | NM_139078 | -1.1725 | -2.4585 |
| YY1AP1 | NM_139118 | -0.6905 | -2.053 |
| YY1AP1 | NM_139119 | -0.6905 | -2.053 |
| YY1AP1 | NM_139121 | -0.6905 | -2.053 |
| MASP1 | NM_139125 | 0.167 | -0.0785 |
| PPIL4 | NM_139126 | -1.025 | -2.3735 |
| NUP98 | NM_139131 | -1.3115 | -2.5025 |
| NUP98 | NM_139132 | -1.4615 | -3.374 |
| ARID1A | NM_139135 | -0.6995 | -2.0615 |
| KCNC2 | NM_139136 | 0.1075 | -0.151 |
| KCNC2 | NM_139137 | 0.1075 | -0.151 |
| ADAMTS14 | NM_139155 | -0.2645 | -1.1575 |
| AMPD2 | NM_139156 | 0.231 | -1.033 |
| ST5 | NM_139157 | -0.314 | -1.584 |
| CDK15 | NM_139158 | -0.1875 | -0.1565 |
| DPP9 | NM_139159 | -0.374 | -1.3475 |
| DEPDC7 | NM_139160 | -0.928 | -2.211 |
| CRB3 | NM_139161 | 0.021 | -0.531 |
| ALS2CR12 | NM_139163 | -0.0675 | -0.913 |
| STARD4 | NM_139164 | -0.937 | -1.33 |
| RAET1E | NM_139165 | 0.1635 | -0.512 |
| ABRA | NM_139166 | -0.2515 | -0.5215 |
| SGCZ | NM_139167 | 0.579 | -0.375 |
| SREK1 | NM_139168 | -0.4285 | -1.9685 |
| TRUB1 | NM_139169 | -1.813 | -4.068 |

| | | | |
|---|---|---|---|
| C16orf71 | NM_139170 | 0.018 | 0.3145 |
| STARD6 | NM_139171 | 0.0305 | -0.023 |
| TMEM190 | NM_139172 | -0.1105 | -0.389 |
| ADAD2 | NM_139174 | 0.245 | -0.3945 |
| RNF133 | NM_139175 | 0.56 | 0.102 |
| SLC39A11 | NM_139177 | -0.8785 | -1.396 |
| ALKBH3 | NM_139178 | -0.584 | -1.57 |
| DAGLB | NM_139179 | -0.874 | -2.324 |
| BRD8 | NM_139199 | -1.2755 | -2.346 |
| GIT2 | NM_139201 | -0.6875 | -2.828 |
| MLC1 | NM_139202 | 0.4835 | -0.244 |
| NAP1L1 | NM_139207 | -0.2335 | -0.25 |
| MASP2 | NM_139208 | 0.3075 | 0.0555 |
| GRK7 | NM_139209 | 0.1985 | -0.3775 |
| HOPX | NM_139211 | -0.0895 | -0.5305 |
| HOPX | NM_139212 | -0.0895 | -0.5305 |
| TAF15 | NM_139215 | -0.404 | -0.7105 |
| NOL6 | NM_139235 | -0.4965 | -1.276 |
| NFKBID | NM_139239 | -0.226 | -0.771 |
| C1orf105 | NM_139240 | -0.28 | -0.0585 |
| FGD4 | NM_139241 | -0.5285 | -1.3445 |
| MTFMT | NM_139242 | -1.5675 | -3.3455 |
| ADAD1 | NM_139243 | 0.0575 | -0.476 |
| STXBP5 | NM_139244 | -0.8715 | -1.8025 |
| PPM1L | NM_139245 | 0.148 | -0.4985 |
| TSTD2 | NM_139246 | -1.377 | -2.9285 |
| ADCY4 | NM_139247 | 0.1855 | -0.08 |
| LIPH | NM_139248 | 0.1585 | -0.4535 |
| MS4A6E | NM_139249 | -0.1875 | -0.0635 |
| CTAG1A | NM_139250 | -0.009 | -0.366 |
| EHD4 | NM_139265 | -0.446 | -0.977 |
| STAT1 | NM_139266 | -1.011 | -1.3815 |
| CARS | NM_139273 | -0.4275 | -1.4015 |
| STAT3 | NM_139276 | 0.0245 | -0.7415 |
| KLK7 | NM_139277 | 0.075 | -0.4105 |
| LGI3 | NM_139278 | -0.0235 | -0.403 |
| MCFD2 | NM_139279 | -1.76 | -2.256 |
| ORMDL3 | NM_139280 | -0.3695 | -0.71 |
| WDR36 | NM_139281 | -1.343 | -3.2435 |
| RHOXF1 | NM_139282 | 0.497 | -0.419 |
| PPTC7 | NM_139283 | -1.4225 | -1.793 |
| LGI4 | NM_139284 | -0.078 | -0.423 |
| GAS2L2 | NM_139285 | -0.3525 | -0.235 |

| | | | |
|---|---|---|---|
| CDC26 | NM_139286 | -1.1085 | -1.238 |
| AKAP4 | NM_139289 | -0.4505 | -0.346 |
| YME1L1 | NM_139312 | -1.3785 | -2.0035 |
| ANGPTL4 | NM_139314 | -0.0155 | -0.2235 |
| TAF6 | NM_139315 | -0.139 | -0.7035 |
| AMPH | NM_139316 | -0.2975 | -0.748 |
| BIRC7 | NM_139317 | 0.062 | -0.5875 |
| KCNH5 | NM_139318 | 0.288 | -0.0545 |
| SLC17A8 | NM_139319 | -0.2775 | -0.166 |
| CHRFAM7A | NM_139320 | -0.143 | -0.791 |
| ATRN | NM_139321 | -0.9795 | -1.746 |
| ATRN | NM_139322 | 0.1085 | -0.3095 |
| YWHAB | NM_139323 | 0.0485 | -0.286 |
| BIN1 | NM_139343 | 0.2095 | -0.4445 |
| BIN1 | NM_139344 | 0.2095 | -0.4445 |
| BIN1 | NM_139345 | 0.2095 | -0.4445 |
| BIN1 | NM_139346 | 0.2095 | -0.4445 |
| BIN1 | NM_139347 | 0.2095 | -0.4445 |
| BIN1 | NM_139348 | 0.2095 | -0.4445 |
| BIN1 | NM_139349 | 0.2095 | -0.4445 |
| BIN1 | NM_139350 | 0.2095 | -0.4445 |
| BIN1 | NM_139351 | 0.2095 | -0.4445 |
| TAF1A | NM_139352 | -0.956 | -1.7995 |
| TAF1C | NM_139353 | -0.059 | -0.827 |
| MATK | NM_139354 | 0.3 | -0.2525 |
| RGS3 | NM_144488 | -0.617 | -1.4575 |
| RGS3 | NM_144489 | -0.617 | -1.4575 |
| CLDN14 | NM_144492 | 0.163 | -0.324 |
| PQBP1 | NM_144495 | -0.1635 | -0.5965 |
| AKAP12 | NM_144497 | -0.705 | -1.101 |
| OSBPL2 | NM_144498 | -0.7155 | -2.481 |
| GNAT1 | NM_144499 | -0.057 | -0.5365 |
| KLK8 | NM_144505 | 0.155 | -0.303 |
| KLK8 | NM_144506 | -0.0745 | -0.047 |
| KLK8 | NM_144507 | -0.2555 | 0.0025 |
| CASC5 | NM_144508 | -0.5275 | -2.5045 |
| RPIA | NM_144563 | -0.2245 | -1.1795 |
| SLC39A3 | NM_144564 | -0.72 | -1.6835 |
| DUOXA1 | NM_144565 | 0.149 | -0.5715 |
| ZNF700 | NM_144566 | -1.8205 | -3.4865 |
| ANGEL2 | NM_144567 | -1.554 | -2.25 |
| SPOCD1 | NM_144569 | 0.059 | -0.2195 |
| HN1L | NM_144570 | -0.387 | -1.077 |

| | | | |
|---|---|---|---|
| CNOT6L | NM_144571 | -1.0395 | -1.738 |
| TBC1D2B | NM_144572 | -0.5345 | -1.0485 |
| NEXN | NM_144573 | -0.2235 | 0.029 |
| WDR20 | NM_144574 | -1.131 | -2.623 |
| CAPN13 | NM_144575 | 0.046 | -0.327 |
| COQ10A | NM_144576 | -0.2365 | -1.337 |
| CCDC114 | NM_144577 | -0.1285 | -0.352 |
| MAPK1IP1L | NM_144578 | -0.352 | -1.2795 |
| SFXN5 | NM_144579 | -0.4835 | -1.069 |
| C1orf85 | NM_144580 | -0.364 | -0.282 |
| C14orf149 | NM_144581 | -0.7025 | -1.3315 |
| TEX261 | NM_144582 | -0.3715 | -1.4325 |
| ATP6V1C2 | NM_144583 | 0.134 | -0.2445 |
| HENMT1 | NM_144584 | -0.6495 | -0.431 |
| SLC22A12 | NM_144585 | 0.054 | -0.43 |
| LYPD1 | NM_144586 | -0.1915 | -1.6205 |
| BTBD16 | NM_144587 | 0.148 | -0.0175 |
| ZFYVE27 | NM_144588 | -0.2705 | -1.3395 |
| COMTD1 | NM_144589 | -0.076 | -0.701 |
| ANKRD22 | NM_144590 | -0.19 | -0.326 |
| C10orf32 | NM_144591 | -0.799 | -0.813 |
| RHEBL1 | NM_144593 | -0.849 | -1.2445 |
| GTSF1 | NM_144594 | 0.0075 | -0.2845 |
| SLAIN1 | NM_144595 | -1.978 | -4.0495 |
| TTC8 | NM_144596 | -1.1725 | -1.948 |
| C15orf40 | NM_144597 | -1.326 | -2.6275 |
| LRRC28 | NM_144598 | -0.176 | -1.307 |
| NIPA1 | NM_144599 | -0.871 | -1.961 |
| FOPNL | NM_144600 | -1.9875 | -2.4185 |
| CMTM3 | NM_144601 | -0.5575 | -2.2335 |
| C16orf78 | NM_144602 | 0.0785 | 0.2455 |
| ZC3H18 | NM_144604 | -0.733 | -2.664 |
| FLCN | NM_144606 | 0.3965 | -0.6245 |
| CYB5D1 | NM_144607 | -1.0435 | -1.5455 |
| HEXIM2 | NM_144608 | -1.611 | -2.949 |
| CCDC43 | NM_144609 | -0.964 | -1.675 |
| CYB5D2 | NM_144611 | -0.619 | -1.262 |
| LOXHD1 | NM_144612 | 0.357 | 0.1095 |
| COX6B2 | NM_144613 | -0.445 | -0.179 |
| MBD3L2 | NM_144614 | 0.35 | -0.316 |
| TMIGD2 | NM_144615 | -0.1935 | -0.163 |
| JSRP1 | NM_144616 | 0.037 | -0.531 |
| HSPB6 | NM_144617 | -0.026 | -0.3865 |

| | | | |
|---|---|---|---|
| GABPB2 | NM_144618 | -0.063 | -0.328 |
| LRRC39 | NM_144620 | -0.653 | -0.992 |
| DCST2 | NM_144622 | -0.228 | -0.4335 |
| UHMK1 | NM_144624 | -0.505 | -1.07 |
| WDR64 | NM_144625 | 0.1005 | -0.17 |
| TMEM125 | NM_144626 | 0.28 | -0.4215 |
| C1orf182 | NM_144627 | 0.176 | -0.456 |
| TBC1D20 | NM_144628 | -0.9225 | -2.524 |
| RFTN2 | NM_144629 | 0.322 | -0.111 |
| ZNF513 | NM_144631 | -0.522 | -1.2905 |
| TMEM182 | NM_144632 | -0.332 | -0.939 |
| KCNH8 | NM_144633 | -0.0885 | -2.1115 |
| LYZL4 | NM_144634 | -0.2305 | -0.3865 |
| FAM131A | NM_144635 | -0.9705 | -1.4145 |
| CHCHD4 | NM_144636 | -1.4695 | -3.395 |
| TMEM42 | NM_144638 | -0.702 | -1.8785 |
| UROC1 | NM_144639 | -0.0645 | -0.48 |
| PPM1M | NM_144641 | -0.252 | -0.72 |
| SYNPR | NM_144642 | -0.62 | 0.132 |
| SCLT1 | NM_144643 | -0.8875 | -1.838 |
| SPATA4 | NM_144644 | 0.0575 | 0.2275 |
| C4orf36 | NM_144645 | -0.1385 | -0.6 |
| IGJ | NM_144646 | -0.233 | -0.748 |
| CAPSL | NM_144647 | 0.09 | -0.561 |
| LRGUK | NM_144648 | -0.0675 | -0.1265 |
| TMEM71 | NM_144649 | -0.49 | -0.5615 |
| ADHFE1 | NM_144650 | -0.035 | -0.1985 |
| PXDNL | NM_144651 | -0.4295 | -0.154 |
| LETM2 | NM_144652 | -0.4165 | -0.652 |
| NACC2 | NM_144653 | 0.094 | -0.203 |
| HDX | NM_144657 | -0.3865 | -1.7905 |
| DOCK11 | NM_144658 | -0.191 | -1.272 |
| TCP10L | NM_144659 | 0.067 | -0.3745 |
| SAMD8 | NM_144660 | -0.2475 | -1.1105 |
| C10orf82 | NM_144661 | -0.1345 | -0.165 |
| PSMA8 | NM_144662 | 0.3565 | -0.195 |
| C11orf40 | NM_144663 | -0.6105 | -0.3245 |
| FAM76B | NM_144664 | -1.138 | -1.5115 |
| SESN3 | NM_144665 | 0.2175 | -0.2795 |
| DNHD1 | NM_144666 | -0.194 | -0.4025 |
| WDR66 | NM_144668 | 0.2105 | -0.496 |
| GLT1D1 | NM_144669 | -0.17 | -1.463 |
| A2ML1 | NM_144670 | -0.261 | -0.31 |

| | | | |
|---|---|---|---|
| FAM109A | NM_144671 | -0.196 | -1.0595 |
| CMTM2 | NM_144673 | 0.0175 | -0.446 |
| TEKT5 | NM_144674 | -0.149 | -0.201 |
| GSG1L | NM_144675 | -0.044 | -0.5835 |
| TMED6 | NM_144676 | -0.077 | -0.3085 |
| MGAT5B | NM_144677 | -0.0295 | -0.273 |
| C17orf56 | NM_144679 | -0.656 | -1.3695 |
| ZNF18 | NM_144680 | -0.7815 | -2.2335 |
| CCDC42 | NM_144681 | 0.136 | -0.0675 |
| SLFN13 | NM_144682 | -0.268 | -0.154 |
| DHRS13 | NM_144683 | -0.873 | -1.624 |
| ZNF480 | NM_144684 | -0.5605 | -1.2705 |
| HIPK4 | NM_144685 | -0.143 | -0.3125 |
| NLRP12 | NM_144687 | -0.3515 | -0.143 |
| CCDC155 | NM_144688 | -0.239 | -0.199 |
| ZNF420 | NM_144689 | -0.4855 | -2.0795 |
| ZNF582 | NM_144690 | -0.3375 | -1.628 |
| CAPN12 | NM_144691 | -0.04 | -0.533 |
| ZNF558 | NM_144693 | -0.992 | -2.0215 |
| ZNF570 | NM_144694 | -0.3695 | -1.1285 |
| BROX | NM_144695 | -0.697 | -0.7575 |
| AXDND1 | NM_144696 | 0.3705 | -0.1465 |
| C1orf51 | NM_144697 | -0.69 | -1.9195 |
| ANKRD35 | NM_144698 | -0.1025 | -0.283 |
| IL23R | NM_144701 | -0.451 | -0.073 |
| LSM14B | NM_144703 | -0.784 | -1.557 |
| AIFM3 | NM_144704 | -0.031 | -0.0575 |
| TEKT4 | NM_144705 | -0.007 | -0.1275 |
| C2orf15 | NM_144706 | -0.063 | 0.1975 |
| PROM2 | NM_144707 | 0.0475 | -0.427 |
| ANKAR | NM_144708 | 0.292 | -0.523 |
| PUS10 | NM_144709 | -0.3115 | -1.093 |
| KLHL23 | NM_144711 | -1.731 | -2.1635 |
| FAM82A1 | NM_144713 | -0.477 | -1.316 |
| EFHB | NM_144715 | -0.4445 | -0.6885 |
| CCDC12 | NM_144716 | -0.5255 | -2.6675 |
| IL20RB | NM_144717 | -0.3265 | -0.345 |
| SPICE1 | NM_144718 | -0.138 | -1.27 |
| CCDC13 | NM_144719 | -0.006 | 0.1225 |
| JAKMIP1 | NM_144720 | -1.466 | -2.789 |
| THAP6 | NM_144721 | -1.0365 | -2.0385 |
| SPEF2 | NM_144722 | 0.008 | -0.2905 |
| ZMAT2 | NM_144723 | -0.3625 | -0.4815 |

| | | | |
|---|---|---|---|
| TTC23L | NM_144725 | 0.0475 | -0.654 |
| RNF145 | NM_144726 | -1.6055 | -2.5945 |
| CRYGN | NM_144727 | 0.0495 | -0.2105 |
| DUSP10 | NM_144728 | -0.4735 | -1.9265 |
| DUSP10 | NM_144729 | -0.4735 | -1.9265 |
| HNRNPUL1 | NM_144732 | -0.1985 | -1.2905 |
| C2orf56 | NM_144736 | -1.2545 | -1.942 |
| MPZL2 | NM_144765 | -0.14 | -0.4805 |
| AKAP13 | NM_144767 | -0.518 | -1.411 |
| FOXI1 | NM_144769 | 0.1295 | -0.1705 |
| RBM11 | NM_144770 | 0.03 | -0.963 |
| APOA1BP | NM_144772 | 0.1085 | -0.4905 |
| PROKR2 | NM_144773 | -0.1945 | -0.462 |
| SMCR8 | NM_144775 | -1.6225 | -2.2475 |
| SCEL | NM_144777 | -0.0095 | -0.0995 |
| MBNL2 | NM_144778 | -0.506 | -1.9015 |
| FXYD5 | NM_144779 | -0.0945 | 0.022 |
| PDCD2 | NM_144781 | -0.431 | -1.326 |
| KLK11 | NM_144947 | -0.1455 | 0.559 |
| SOCS5 | NM_144949 | -0.0255 | 0.079 |
| PEBP4 | NM_144962 | 0.0345 | -0.3265 |
| FAM91A1 | NM_144963 | -1.2085 | -1.7495 |
| RG9MTD3 | NM_144964 | -0.5945 | -0.7925 |
| TTC16 | NM_144965 | -0.171 | 0.015 |
| FREM1 | NM_144966 | -0.323 | -0.3045 |
| ARHGAP36 | NM_144967 | -0.035 | -0.0345 |
| RIBC1 | NM_144968 | 0.1535 | -0.2195 |
| ZDHHC15 | NM_144969 | 0.114 | -0.7405 |
| CXorf38 | NM_144970 | -0.6255 | -0.9865 |
| LDHAL6A | NM_144972 | -0.3455 | 0.027 |
| DENND5B | NM_144973 | -1.789 | -3.244 |
| CCDC122 | NM_144974 | -0.01 | 0.136 |
| SLFN5 | NM_144975 | 0.1185 | -0.323 |
| ZNF564 | NM_144976 | -1.2495 | -2.601 |
| DENND1B | NM_144977 | 0.216 | -1.094 |
| CCDC138 | NM_144978 | -1.639 | -2.348 |
| RBM46 | NM_144979 | 0.517 | -0.572 |
| C6orf118 | NM_144980 | -0.031 | -0.032 |
| IMMP1L | NM_144981 | -0.8915 | -1.844 |
| ZFC3H1 | NM_144982 | -0.651 | -2.941 |
| VSTM4 | NM_144984 | -0.146 | -0.588 |
| CDH24 | NM_144985 | -0.056 | -0.4715 |
| U2AF1L4 | NM_144987 | -0.0975 | -0.698 |

| | | | |
|---|---|---|---|
| ALG14 | NM_144988 | -1.068 | -1.9355 |
| DNAH14 | NM_144989 | -0.407 | -0.5325 |
| SLFNL1 | NM_144990 | 0.2815 | -0.409 |
| TSPEAR | NM_144991 | -0.179 | -0.2745 |
| VWA3B | NM_144992 | -0.005 | -0.256 |
| TET3 | NM_144993 | -1.0995 | -3.093 |
| ANKRD23 | NM_144994 | -0.306 | -0.6545 |
| ARL13B | NM_144996 | -1.609 | -3.8075 |
| FLCN | NM_144997 | -0.4925 | -1.1205 |
| STRA13 | NM_144998 | 0.021 | -0.989 |
| LRRC45 | NM_144999 | -0.442 | -1.1465 |
| RANBP3L | NM_145000 | -0.434 | -0.025 |
| STK32A | NM_145001 | 0.074 | 0.4385 |
| TSNARE1 | NM_145003 | 0.4055 | 0.4 |
| ADAM32 | NM_145004 | 0.5155 | 0.0265 |
| C9orf72 | NM_145005 | -1.4165 | -2.6255 |
| SUSD3 | NM_145006 | -0.2915 | -0.5055 |
| NLRP11 | NM_145007 | -0.156 | -0.041 |
| YPEL4 | NM_145008 | -0.098 | -0.4965 |
| ENKUR | NM_145010 | 0.169 | -0.224 |
| ZNF25 | NM_145011 | -0.9495 | -1.221 |
| CCNY | NM_145012 | -0.0215 | -0.4895 |
| C11orf45 | NM_145013 | -0.1905 | -0.5275 |
| MRGPRF | NM_145015 | 0.1195 | 0.0555 |
| GLYATL2 | NM_145016 | -0.567 | 0.0145 |
| PPP1R32 | NM_145017 | -0.067 | -0.1395 |
| C11orf82 | NM_145018 | -0.941 | -2.6015 |
| FAM124A | NM_145019 | -1.583 | -2.2085 |
| CCDC11 | NM_145020 | 0.1505 | 0.0715 |
| 8-Mar | NM_145021 | -0.287 | -0.6155 |
| CCDC7 | NM_145023 | -0.0675 | -0.205 |
| CES5A | NM_145024 | -0.0455 | -0.294 |
| AKD1 | NM_145025 | -0.502 | -0.6475 |
| KIF6 | NM_145027 | -0.1745 | -0.383 |
| C6orf81 | NM_145028 | -0.118 | -0.3125 |
| C6orf136 | NM_145029 | -0.2985 | -1.448 |
| FBXL13 | NM_145032 | -0.0325 | 0.042 |
| TOR1AIP2 | NM_145034 | -0.5465 | -1.3555 |
| FAM55C | NM_145037 | -1.3355 | -1.7835 |
| CCDC164 | NM_145038 | -0.0075 | -0.091 |
| CENPBD1 | NM_145039 | -0.7785 | -1.3645 |
| PRKCDBP | NM_145040 | 0.483 | 0.277 |
| TMEM106A | NM_145041 | 0.383 | -0.333 |

| | | | |
|---|---|---|---|
| NEIL2 | NM_145043 | -0.38 | -1.2965 |
| ZNF501 | NM_145044 | -0.983 | -2.7205 |
| CCDC151 | NM_145045 | -0.0855 | -0.2115 |
| CALR3 | NM_145046 | -0.041 | 0.126 |
| OSCP1 | NM_145047 | -0.1875 | -0.6665 |
| PACRGL | NM_145048 | -1.421 | -2.629 |
| UBLCP1 | NM_145049 | -0.836 | -1.354 |
| RNF183 | NM_145051 | 0.124 | 0.142 |
| UPRT | NM_145052 | -0.9285 | -2.36 |
| UBQLNL | NM_145053 | 0.1825 | 0.0265 |
| WDR16 | NM_145054 | 0.4525 | -0.186 |
| C18orf25 | NM_145055 | -0.741 | -1.615 |
| DACT3 | NM_145056 | -0.2945 | -1.2155 |
| CDC42EP5 | NM_145057 | -0.113 | -0.331 |
| RILPL2 | NM_145058 | -0.574 | -1.3095 |
| FUK | NM_145059 | -0.2595 | -0.589 |
| SKA1 | NM_145060 | -0.7235 | -2.473 |
| SKA3 | NM_145061 | -2.0545 | -2.899 |
| ZUFSP | NM_145062 | -1.3145 | -2.447 |
| C6orf130 | NM_145063 | -1.054 | -1.4795 |
| STAC3 | NM_145064 | 0.096 | -0.033 |
| PELI3 | NM_145065 | 0.012 | -0.4545 |
| TRPV3 | NM_145068 | -0.084 | -0.0695 |
| CISH | NM_145071 | -0.316 | -1.774 |
| HTRA2 | NM_145074 | -0.78 | -1.9055 |
| NSMCE1 | NM_145080 | -0.1485 | -0.5495 |
| ZKSCAN5 | NM_145102 | -1.448 | -2.2495 |
| MAP2K3 | NM_145109 | -0.55 | -1.4165 |
| FAM200A | NM_145111 | -1.781 | -3.713 |
| MAX | NM_145112 | -0.588 | -2.2715 |
| MAX | NM_145113 | -0.4005 | -1.6465 |
| MAX | NM_145114 | -0.588 | -2.2715 |
| ZNF498 | NM_145115 | -0.816 | -3.2425 |
| MAX | NM_145116 | -0.588 | -2.2715 |
| NAV2 | NM_145117 | -0.2205 | -1.109 |
| PJA1 | NM_145119 | -0.633 | -1.645 |
| MAP2K5 | NM_145160 | -0.6025 | -1.96 |
| CHURC1 | NM_145165 | -1.208 | -0.718 |
| ZBTB47 | NM_145166 | -0.4495 | -1.1405 |
| PIGM | NM_145167 | -0.9865 | -1.174 |
| SDR42E1 | NM_145168 | -0.036 | -0.4655 |
| SFT2D1 | NM_145169 | -0.773 | -2.5155 |
| TTC18 | NM_145170 | 0.2405 | -0.229 |

| | | | |
|---|---|---|---|
| GPHB5 | NM_145171 | 0.264 | -0.6145 |
| WDR63 | NM_145172 | 0.594 | -0.1475 |
| DIRAS1 | NM_145173 | -0.6545 | -1.6635 |
| DNAJB7 | NM_145174 | 0.123 | 0.1605 |
| FAM84A | NM_145175 | -0.652 | -0.8315 |
| SLC2A12 | NM_145176 | -0.334 | -0.6865 |
| DHRSX | NM_145177 | -0.343 | -0.9585 |
| ATOH7 | NM_145178 | 0.611 | 1.459 |
| PYCARD | NM_145182 | 0.12 | -0.2045 |
| MAP2K7 | NM_145185 | -0.999 | -2.427 |
| ABCC11 | NM_145186 | 0.1905 | -0.542 |
| CABP4 | NM_145200 | -0.3525 | -0.6025 |
| NAPRT1 | NM_145201 | -0.4165 | -1.354 |
| PRAP1 | NM_145202 | -0.0205 | -0.1765 |
| CSNK1A1L | NM_145203 | -0.559 | -0.334 |
| SENP8 | NM_145204 | -0.575 | -1.7815 |
| HMGB4 | NM_145205 | -0.165 | 0.1325 |
| VTI1A | NM_145206 | -0.4145 | -0.902 |
| SPATA5 | NM_145207 | -1.617 | -3.0205 |
| MBD3L1 | NM_145208 | 0.0925 | -0.3215 |
| MRPL30 | NM_145212 | -0.6025 | -1.025 |
| TRIM11 | NM_145214 | -0.9685 | -1.8935 |
| ATP6V0E2 | NM_145230 | 0.172 | -0.544 |
| EFCAB11 | NM_145231 | -0.7665 | -1.5045 |
| CTU1 | NM_145232 | -0.509 | -0.992 |
| ZNF625 | NM_145233 | -0.2655 | -1.1325 |
| CHRDL1 | NM_145234 | 0.032 | 0.1005 |
| FANK1 | NM_145235 | 0.114 | -0.132 |
| B3GNT7 | NM_145236 | -0.6455 | -0.42 |
| ZSCAN20 | NM_145238 | -0.5165 | -0.423 |
| PRRT2 | NM_145239 | -0.394 | -0.48 |
| WDR31 | NM_145241 | -0.0035 | -0.5875 |
| OMA1 | NM_145243 | -2.1225 | -3.037 |
| DDIT4L | NM_145244 | -0.232 | -0.077 |
| EVI5L | NM_145245 | -0.2895 | -1.031 |
| FRA10AC1 | NM_145246 | -1.322 | -2.0735 |
| SPACA7 | NM_145248 | 0.252 | -0.601 |
| IFI27L1 | NM_145249 | 0.302 | -0.4565 |
| RNASE11 | NM_145250 | 0.4985 | -0.4855 |
| STYX | NM_145251 | -1.3385 | -2.7085 |
| ZG16B | NM_145252 | -0.133 | -0.576 |
| FAM100A | NM_145253 | -0.342 | -1.18 |
| TMEM170A | NM_145254 | -1.227 | -2.474 |

| | | | |
|---|---|---|---|
| MRPL10 | NM_145255 | -0.58 | -1.938 |
| LRRC25 | NM_145256 | -0.009 | -0.5585 |
| C1orf96 | NM_145257 | -1.2265 | -1.5715 |
| ACVR1C | NM_145259 | -0.0915 | -0.6225 |
| OSR1 | NM_145260 | -0.063 | -0.6095 |
| DNAJC19 | NM_145261 | -0.699 | -1.694 |
| GLYCTK | NM_145262 | 0.35 | -0.0325 |
| SPATA18 | NM_145263 | -0.7345 | -0.3205 |
| CCDC127 | NM_145265 | -0.853 | -2.796 |
| NUDCD2 | NM_145266 | -0.8805 | -2.1255 |
| C6orf57 | NM_145267 | -0.6295 | -1.7405 |
| C7orf45 | NM_145268 | -0.33 | 0.602 |
| FAM92A1 | NM_145269 | -0.759 | -1.338 |
| C16orf11 | NM_145270 | 0.017 | -0.5885 |
| ZNF688 | NM_145271 | -0.097 | -0.5025 |
| C17orf50 | NM_145272 | 0.3145 | -0.3595 |
| CD300LG | NM_145273 | -0.1235 | -0.406 |
| TMEM99 | NM_145274 | -0.0735 | -1.925 |
| ZNF563 | NM_145276 | -0.175 | -1.114 |
| HFE2 | NM_145277 | -0.084 | -0.3185 |
| C1orf150 | NM_145278 | -0.3155 | -0.24 |
| MOB3C | NM_145279 | -0.529 | -0.177 |
| SLC25A48 | NM_145282 | -0.278 | -0.3015 |
| NXNL2 | NM_145283 | 0.245 | -0.148 |
| FAM122B | NM_145284 | -1.2125 | -1.7475 |
| NKX2-3 | NM_145285 | -0.2685 | -0.4095 |
| STOML3 | NM_145286 | -0.122 | 0.043 |
| ZNF519 | NM_145287 | 0.0575 | -0.619 |
| ZNF296 | NM_145288 | 0.166 | -0.0655 |
| GPR125 | NM_145290 | -1.1845 | -1.9815 |
| ZBTB49 | NM_145291 | -0.2685 | -1.6775 |
| GALNTL5 | NM_145292 | 0.2175 | 0.0075 |
| WDR90 | NM_145294 | -0.368 | -0.5575 |
| ZNF627 | NM_145295 | -1.805 | -3.592 |
| CADM4 | NM_145296 | -0.2245 | -0.8465 |
| ZNF626 | NM_145297 | -0.4565 | -0.362 |
| APOBEC3F | NM_145298 | -0.457 | -0.5485 |
| FAM18B2 | NM_145301 | 0.1785 | -0.258 |
| C7orf33 | NM_145304 | 0.036 | 0.1045 |
| SLC25A43 | NM_145305 | -0.481 | -1.605 |
| C10orf35 | NM_145306 | -0.698 | -2.4045 |
| RTKN2 | NM_145307 | -0.1055 | -1.348 |
| LRTOMT | NM_145309 | -0.0875 | -0.4045 |

| | | | |
|---|---|---|---|
| ZNF485 | NM_145312 | -1.354 | -1.882 |
| RASGEF1A | NM_145313 | -0.6295 | 0.1065 |
| UCMA | NM_145314 | 0.4235 | -0.4515 |
| LACE1 | NM_145315 | -0.886 | -0.878 |
| TMEM217 | NM_145316 | -0.4475 | -0.3985 |
| OSBPL3 | NM_145320 | -0.492 | -1.0435 |
| OSBPL3 | NM_145321 | -0.492 | -1.0435 |
| OSBPL3 | NM_145322 | -0.492 | -1.0435 |
| ZNF493 | NM_145326 | -0.422 | -1.98 |
| MRPL33 | NM_145330 | -0.124 | -0.4135 |
| MAP3K7 | NM_145331 | -1.443 | -2.156 |
| MAP3K7 | NM_145332 | -1.443 | -2.156 |
| MAP3K7 | NM_145333 | -1.443 | -2.156 |
| PDCD4 | NM_145341 | -1.3635 | -3.0195 |
| APOL1 | NM_145343 | 0.0435 | -0.19 |
| UBXN11 | NM_145345 | 0.113 | -0.131 |
| SCARF1 | NM_145350 | -0.059 | -0.074 |
| SCARF1 | NM_145352 | -0.059 | -0.074 |
| APOL2 | NM_145637 | -0.548 | -0.541 |
| OSBPL5 | NM_145638 | -0.3035 | -0.723 |
| APOL3 | NM_145640 | 0.2035 | -0.1925 |
| APOL3 | NM_145641 | 0.2035 | -0.1925 |
| APOL3 | NM_145642 | 0.2035 | -0.1925 |
| MRPL35 | NM_145644 | -0.7435 | -1.996 |
| WDR67 | NM_145647 | -2.045 | -2.8845 |
| SLC15A4 | NM_145648 | -1.096 | -2.6585 |
| GCNT2 | NM_145649 | 0.0895 | -0.804 |
| MUC15 | NM_145650 | 0.2045 | -0.1605 |
| WFDC5 | NM_145652 | 0.402 | -0.367 |
| RDM1 | NM_145654 | -0.173 | -0.392 |
| GCNT2 | NM_145655 | 0.0895 | -0.804 |
| GSX1 | NM_145657 | -0.2105 | -0.292 |
| SPESP1 | NM_145658 | 0.1795 | 0.0915 |
| IL27 | NM_145659 | 0.1775 | -0.589 |
| APOL4 | NM_145660 | 0.0515 | -0.062 |
| DBF4B | NM_145663 | -1.309 | -3.3195 |
| SPANXB2 | NM_145664 | 0.3505 | -0.0585 |
| SPANXE | NM_145665 | -0.0265 | -0.2695 |
| BRF1 | NM_145685 | -0.009 | -0.1075 |
| MAP4K4 | NM_145686 | -0.7285 | -0.8655 |
| MAP4K4 | NM_145687 | -0.341 | -0.495 |
| APBB1 | NM_145689 | -0.006 | -1.067 |
| YWHAZ | NM_145690 | -0.584 | -0.5065 |

| | | | |
|---|---|---|---|
| ATPAF2 | NM_145691 | 0.309 | -0.5685 |
| LPIN1 | NM_145693 | -1.807 | -3.0105 |
| DGKB | NM_145695 | 0.0735 | -0.6315 |
| NUF2 | NM_145697 | -0.978 | -1.512 |
| ACBD5 | NM_145698 | -1.7095 | -2.396 |
| APOBEC3A | NM_145699 | -0.159 | -0.53 |
| CDCA4 | NM_145701 | -0.87 | -3.7575 |
| TIGD1 | NM_145702 | -1.237 | -1.744 |
| ATXN2L | NM_145714 | -0.151 | -0.819 |
| TIGD2 | NM_145715 | -1.45 | -2.4135 |
| SSBP3 | NM_145716 | -0.017 | -0.824 |
| TIGD3 | NM_145719 | -0.634 | -0.976 |
| TIGD4 | NM_145720 | -0.157 | -0.024 |
| TRAF3 | NM_145725 | -0.419 | -1.483 |
| TRAF3 | NM_145726 | -0.419 | -1.483 |
| SYNM | NM_145728 | -1.7265 | -2.62 |
| MRPL24 | NM_145729 | -0.1625 | -1.0565 |
| AP1B1 | NM_145730 | -0.03 | -0.202 |
| SYNGR1 | NM_145731 | -0.0035 | -0.574 |
| ARHGEF7 | NM_145735 | -0.5705 | -1.762 |
| SYNGR1 | NM_145738 | -0.0035 | -0.574 |
| OSBPL6 | NM_145739 | -0.6335 | -1.7275 |
| GSTA1 | NM_145740 | -0.0675 | -0.102 |
| PHLDB2 | NM_145753 | -0.3555 | -2.6365 |
| KIFC2 | NM_145754 | -0.408 | -0.406 |
| TTC21A | NM_145755 | -0.0765 | -0.3015 |
| ZNF396 | NM_145756 | -0.5655 | -0.492 |
| TRAF5 | NM_145759 | -0.905 | -1.337 |
| MGST1 | NM_145764 | -0.374 | -0.7395 |
| MGST1 | NM_145791 | -0.374 | -0.7395 |
| MGST1 | NM_145792 | -0.374 | -0.7395 |
| GFRA1 | NM_145793 | 0.0155 | -0.6755 |
| POGZ | NM_145796 | -1.1695 | -2.5125 |
| OSBPL7 | NM_145798 | -0.1495 | -0.3205 |
| 6-Sep | NM_145799 | -0.7035 | -0.663 |
| TRAF6 | NM_145803 | -1.2975 | -2.657 |
| ABTB2 | NM_145804 | 0.095 | -0.4965 |
| ISL2 | NM_145805 | 0.155 | -1.7105 |
| ZNF511 | NM_145806 | -0.2035 | -1.555 |
| NTN5 | NM_145807 | 0.039 | 0.118 |
| CDCA7 | NM_145810 | -0.9375 | -0.757 |
| CACNG6 | NM_145814 | 0.2045 | -0.323 |
| CACNG6 | NM_145815 | 0.2045 | -0.323 |

| | | | |
|---|---|---|---|
| CRYZL1 | NM_145858 | -1.4645 | -1.525 |
| PDCD10 | NM_145859 | -1.0075 | -2.663 |
| PDCD10 | NM_145860 | -1.0075 | -2.663 |
| EDARADD | NM_145861 | 0.379 | -0.522 |
| CHEK2 | NM_145862 | -1.0785 | -2.1535 |
| ANKS4B | NM_145865 | -0.2135 | -0.126 |
| FZD3 | NM_145866 | -1.285 | -1.291 |
| LTC4S | NM_145867 | 0.151 | -0.0605 |
| ANXA11 | NM_145868 | 0.1595 | -0.83 |
| ANXA11 | NM_145869 | 0.1595 | -0.83 |
| GSTZ1 | NM_145870 | -0.004 | -0.4935 |
| GSTZ1 | NM_145871 | -0.004 | -0.4935 |
| ASB4 | NM_145872 | -0.0995 | -0.185 |
| KLK10 | NM_145888 | 0.0135 | -0.1875 |
| RBFOX1 | NM_145891 | -0.464 | -0.8375 |
| RBFOX1 | NM_145892 | -0.464 | -0.8375 |
| RBFOX1 | NM_145893 | -0.464 | -0.8375 |
| KLK12 | NM_145894 | 0.039 | -0.4715 |
| KLK12 | NM_145895 | 0.148 | -0.2485 |
| PFDN5 | NM_145897 | 0.2505 | -0.0945 |
| CCL23 | NM_145898 | -0.337 | -0.426 |
| HMGA1 | NM_145899 | 0.274 | -0.169 |
| HMGA1 | NM_145901 | 0.274 | -0.169 |
| HMGA1 | NM_145902 | 0.274 | -0.169 |
| HMGA1 | NM_145903 | 0.274 | -0.169 |
| HMGA1 | NM_145905 | 0.274 | -0.169 |
| NEK11 | NM_145910 | 0.157 | -0.3875 |
| ZNF23 | NM_145911 | -0.8725 | -1.8865 |
| NFAM1 | NM_145912 | 0.198 | 0.297 |
| SLC5A8 | NM_145913 | -0.234 | -0.234 |
| ZSCAN21 | NM_145914 | -1.288 | -3.1675 |
| CTSL1 | NM_145918 | -0.21 | -0.7485 |
| MRPL4 | NM_146387 | -0.0745 | -0.487 |
| MRPL4 | NM_146388 | -0.087 | -1.3105 |
| EVC2 | NM_147127 | -0.6345 | -0.0795 |
| ZNRF2 | NM_147128 | -1.049 | -2.516 |
| ALS2CL | NM_147129 | 0.067 | -0.5855 |
| NFX1 | NM_147133 | -1.692 | -2.744 |
| NFX1 | NM_147134 | -1.692 | -2.744 |
| BVES | NM_147147 | -0.9615 | -1.9025 |
| GSTM4 | NM_147148 | 0.1655 | -0.5215 |
| ITSN2 | NM_147152 | -0.6735 | -1.5785 |
| SGMS1 | NM_147156 | -1.037 | -1.726 |

| | | | |
|---|---|---|---|
| SIGMAR1 | NM_147157 | -0.357 | -0.888 |
| ACOT11 | NM_147161 | -0.747 | -0.7725 |
| IL11RA | NM_147162 | 0.1245 | -0.261 |
| CNTFR | NM_147164 | -0.2485 | -0.715 |
| HS6ST2 | NM_147175 | -0.683 | -0.8165 |
| PPP3R2 | NM_147180 | -0.2505 | -0.2615 |
| KCNIP4 | NM_147181 | -0.668 | 0.1505 |
| KCNIP4 | NM_147182 | -0.668 | 0.1505 |
| KCNIP4 | NM_147183 | -0.668 | 0.1505 |
| TP53I3 | NM_147184 | -0.4655 | -1.214 |
| AKAP9 | NM_147185 | 0.394 | 0.152 |
| TNFRSF10B | NM_147187 | -0.3515 | -1.3295 |
| FBXO22 | NM_147188 | -0.893 | -2.3845 |
| FAM110B | NM_147189 | -0.4055 | -1.689 |
| CERS5 | NM_147190 | -0.346 | -1.1535 |
| MMP21 | NM_147191 | 0.2155 | -0.391 |
| DMBX1 | NM_147192 | -0.3425 | -0.472 |
| GLIS1 | NM_147193 | -0.3075 | -0.6575 |
| ANKRD18A | NM_147195 | 0.0275 | 0.17 |
| TMIE | NM_147196 | -0.1875 | -0.205 |
| WFDC11 | NM_147197 | 0.0565 | -0.0345 |
| WFDC9 | NM_147198 | -0.1135 | -0.365 |
| MRGPRX1 | NM_147199 | 0.1885 | -0.0495 |
| C9orf25 | NM_147202 | -0.7835 | -1.407 |
| FGL1 | NM_147203 | 0.0905 | -0.006 |
| TRPV4 | NM_147204 | 0.2725 | -0.2715 |
| NCOA1 | NM_147223 | -0.707 | -2.847 |
| NCOA1 | NM_147233 | -0.707 | -2.847 |
| TRAF3IP2 | NM_147686 | -0.354 | -0.4515 |
| SNX15 | NM_147777 | -0.698 | -1.5225 |
| CTSB | NM_147780 | 0.0525 | -0.4655 |
| CTSB | NM_147781 | 0.0525 | -0.4655 |
| CTSB | NM_147782 | 0.0525 | -0.4655 |
| CTSB | NM_147783 | 0.0525 | -0.4655 |
| CTSC | NM_148170 | -0.4375 | -2.279 |
| PEMT | NM_148172 | 0.1055 | -0.672 |
| AZIN1 | NM_148174 | -0.724 | -0.8905 |
| PPIL2 | NM_148175 | -0.5575 | -1.012 |
| PPIL2 | NM_148176 | -0.5575 | -1.012 |
| C9orf23 | NM_148179 | -0.982 | -2.2335 |
| ATXN2L | NM_148414 | -0.151 | -0.819 |
| ATXN2L | NM_148415 | -0.151 | -0.819 |
| ATXN2L | NM_148416 | -0.151 | -0.819 |

| | | | |
|---|---|---|---|
| CCL28 | NM_148672 | -0.0315 | 0.3345 |
| SMC1B | NM_148674 | -0.0575 | -0.1785 |
| MRPL10 | NM_148887 | -0.58 | -1.938 |
| SVIP | NM_148893 | -0.3945 | -1.0675 |
| BOD1L | NM_148894 | -0.263 | -2.0675 |
| NPB | NM_148896 | -0.0915 | -0.3705 |
| SDR9C7 | NM_148897 | -0.1725 | -0.029 |
| FOXP2 | NM_148898 | 0.8085 | -0.3895 |
| FOXP2 | NM_148899 | 0.8085 | -0.3895 |
| FOXP2 | NM_148900 | 0.8085 | -0.3895 |
| TNFRSF18 | NM_148901 | -0.2905 | -1.0095 |
| TNFRSF18 | NM_148902 | -0.2905 | -1.0095 |
| GREB1 | NM_148903 | -0.43 | -0.7655 |
| OSBPL9 | NM_148904 | -0.581 | -1.5355 |
| OSBPL9 | NM_148905 | -0.581 | -1.5355 |
| OSBPL9 | NM_148906 | -0.581 | -1.5355 |
| OSBPL9 | NM_148907 | -0.581 | -1.5355 |
| OSBPL9 | NM_148908 | -0.581 | -1.5355 |
| OSBPL9 | NM_148909 | -0.581 | -1.5355 |
| TIRAP | NM_148910 | -0.676 | -0.7195 |
| ABHD11 | NM_148912 | -0.2895 | -0.9 |
| ABHD11 | NM_148913 | -0.2895 | -0.9 |
| SHMT1 | NM_148918 | -0.5635 | -1.7005 |
| PSMB8 | NM_148919 | -0.5065 | -0.205 |
| PIGQ | NM_148920 | -0.1355 | -0.5885 |
| EPN2 | NM_148921 | -0.2555 | -0.8275 |
| CYB5A | NM_148923 | -0.0135 | -0.7865 |
| SNX1 | NM_148955 | -0.4635 | -1.3525 |
| NSUN5 | NM_148956 | -0.653 | -2.114 |
| TNFRSF19 | NM_148957 | -0.231 | -0.731 |
| HUS1B | NM_148959 | 0.536 | 0.1935 |
| CLDN19 | NM_148960 | 0.065 | -0.5455 |
| OTOS | NM_148961 | 0.08 | 0.0305 |
| OXER1 | NM_148962 | 0.2355 | -0.18 |
| GPRC6A | NM_148963 | -0.1735 | -0.109 |
| CTSE | NM_148964 | 0.02 | 0.024 |
| TNFRSF25 | NM_148965 | -1.163 | -0.7055 |
| TNFRSF25 | NM_148966 | -0.183 | -0.233 |
| TNFRSF25 | NM_148967 | -0.183 | -0.233 |
| TNFRSF25 | NM_148970 | -0.183 | -0.233 |
| MS4A4A | NM_148975 | -0.252 | -0.338 |
| PANK1 | NM_148977 | -2.021 | -4.2235 |
| PANK1 | NM_148978 | -2.021 | -4.2235 |

| | | | |
|---|---|---|---|
| TAGAP | NM_152133 | -0.0275 | 0.19 |
| GJD3 | NM_152219 | -0.155 | -0.224 |
| CSNK1E | NM_152221 | -0.36 | -0.916 |
| RELT | NM_152222 | -0.2845 | -1.263 |
| SNX5 | NM_152227 | 0.9905 | 1.8715 |
| TAS1R3 | NM_152228 | -0.0645 | -0.5645 |
| IPMK | NM_152230 | -0.1925 | -0.901 |
| FBXO34 | NM_152231 | -1.46 | -3.041 |
| TAS1R2 | NM_152232 | -0.175 | -0.3405 |
| SNX6 | NM_152233 | -1.3295 | -1.4905 |
| GAS2L1 | NM_152236 | -0.2565 | -0.419 |
| GAS2L1 | NM_152237 | -0.548 | -1.308 |
| SNX7 | NM_152238 | -0.8435 | -1.3125 |
| ZMAT3 | NM_152240 | -0.6285 | -1.103 |
| CDC42EP1 | NM_152243 | -0.013 | -0.6905 |
| SNX11 | NM_152244 | -0.6075 | -2.98 |
| C15orf42 | NM_152259 | -1.0715 | -3.6045 |
| RPUSD2 | NM_152260 | -0.9435 | -3.121 |
| C12orf23 | NM_152261 | -1.3005 | -2.026 |
| ZNF439 | NM_152262 | -0.23 | -1.6275 |
| TPM3 | NM_152263 | -0.2085 | -0.625 |
| BTF3L4 | NM_152265 | -1.462 | -2.232 |
| C19orf40 | NM_152266 | -0.491 | -0.909 |
| RNF185 | NM_152267 | -0.6015 | -2.192 |
| PARS2 | NM_152268 | -1.2825 | -2.747 |
| C12orf65 | NM_152269 | -1.857 | -3.858 |
| SLFN11 | NM_152270 | -0.0355 | 0.0725 |
| LONRF1 | NM_152271 | -1.414 | -4.709 |
| CHMP7 | NM_152272 | -0.922 | -3.2445 |
| TTC30A | NM_152275 | -0.8415 | -1.627 |
| UBTD2 | NM_152277 | -1.263 | -2.337 |
| TCEAL7 | NM_152278 | -1.8335 | -2.54 |
| ZNF585B | NM_152279 | -0.913 | -2.0645 |
| SYT11 | NM_152280 | -0.8915 | -1.344 |
| GORAB | NM_152281 | -2.007 | -4.367 |
| ACPL2 | NM_152282 | -1.316 | -2.3835 |
| ZFP62 | NM_152283 | -1.4165 | -2.934 |
| CHMP4C | NM_152284 | -0.0525 | -0.5355 |
| ARRDC1 | NM_152285 | 0.1555 | -0.632 |
| PNPLA7 | NM_152286 | -0.0645 | -0.3575 |
| ZNF276 | NM_152287 | -0.412 | -0.8895 |
| ORAI3 | NM_152288 | -0.085 | -0.46 |
| ZNF561 | NM_152289 | -2.1515 | -3.313 |

| | | | |
|---|---|---|---|
| C1orf158 | NM_152290 | 0.3575 | -0.1245 |
| RG9MTD2 | NM_152292 | -0.4635 | -0.608 |
| TADA2B | NM_152293 | -0.455 | -1.4205 |
| TARS | NM_152295 | 0.072 | -0.4705 |
| ATP1A3 | NM_152296 | -0.166 | -0.7015 |
| NASP | NM_152298 | -0.7075 | -1.004 |
| NCAPH2 | NM_152299 | 0.203 | -0.574 |
| RAB42 | NM_152304 | -0.221 | -0.54 |
| POGLUT1 | NM_152305 | -1.4555 | -1.7065 |
| TRMT61A | NM_152307 | -0.3365 | -1.29 |
| PIK3AP1 | NM_152309 | -0.804 | -0.7965 |
| ELOVL3 | NM_152310 | -0.084 | -0.1405 |
| CLRN3 | NM_152311 | -0.018 | -0.066 |
| GYLTL1B | NM_152312 | 0.1695 | -0.1645 |
| SLC36A4 | NM_152313 | -0.9055 | -1.6085 |
| FAM55A | NM_152315 | -0.109 | -0.8525 |
| C11orf46 | NM_152316 | -1.304 | -3.102 |
| DEPDC4 | NM_152317 | -0.357 | -1.3065 |
| C12orf45 | NM_152318 | -0.6695 | -1.6785 |
| C12orf54 | NM_152319 | -0.218 | -0.122 |
| ZNF641 | NM_152320 | -1.107 | -2.528 |
| ERP27 | NM_152321 | -0.059 | -0.3415 |
| SPIC | NM_152323 | -0.2295 | -0.232 |
| C13orf16 | NM_152324 | 0.0205 | -0.5055 |
| C13orf26 | NM_152325 | 0.303 | 0.2215 |
| ANKRD9 | NM_152326 | -0.2595 | -1.0805 |
| AK7 | NM_152327 | -0.5665 | -0.5295 |
| ADSSL1 | NM_152328 | 0.1895 | -0.508 |
| FRMD6 | NM_152330 | -1.082 | -4.15 |
| ACOT4 | NM_152331 | 0.409 | -0.3165 |
| TARSL2 | NM_152334 | -0.7635 | -1.449 |
| C15orf27 | NM_152335 | 0.2705 | -0.538 |
| AGBL1 | NM_152336 | 0.0475 | -0.409 |
| C16orf46 | NM_152337 | -0.007 | -0.036 |
| ZG16 | NM_152338 | 0.251 | -0.3775 |
| SPATA2L | NM_152339 | -0.2545 | -0.373 |
| PAQR4 | NM_152341 | -0.5535 | -1.447 |
| CDYL2 | NM_152342 | -0.161 | -0.176 |
| C17orf46 | NM_152343 | 0.0755 | -0.204 |
| LSM12 | NM_152344 | -0.3765 | -1.7505 |
| ANKRD13B | NM_152345 | -0.2345 | -0.976 |
| SLC43A2 | NM_152346 | -0.4045 | -1.097 |
| C17orf57 | NM_152347 | -0.3425 | -0.389 |

| | | | |
|---|---|---|---|
| WDR81 | NM_152348 | -0.9245 | -2.243 |
| KRT222 | NM_152349 | 0.1615 | -0.3085 |
| SLC5A10 | NM_152351 | 0.323 | -0.5975 |
| C18orf19 | NM_152352 | -1.4095 | -2.7925 |
| CLDND2 | NM_152353 | 0.042 | 0.165 |
| ZNF285 | NM_152354 | -1.1365 | -1.6495 |
| ZNF441 | NM_152355 | -0.3395 | -2.5195 |
| ZNF491 | NM_152356 | -0.326 | -0.624 |
| ZNF440 | NM_152357 | 0.0015 | -0.576 |
| IZUMO2 | NM_152358 | 0.5375 | 0.1675 |
| CPT1C | NM_152359 | -0.537 | -0.947 |
| ZNF573 | NM_152360 | -1.6785 | -1.9385 |
| EID2B | NM_152361 | -0.979 | -0.8505 |
| TNFAIP8L1 | NM_152362 | -0.725 | -1.731 |
| ANKLE1 | NM_152363 | 0.158 | -0.397 |
| C1orf172 | NM_152365 | 0.029 | -0.1895 |
| KLHDC9 | NM_152366 | -0.163 | -0.1045 |
| MAB21L3 | NM_152367 | -0.089 | 0.0875 |
| SLC44A3 | NM_152369 | 0.137 | -0.4 |
| C1orf93 | NM_152371 | -0.526 | -0.739 |
| MYOM3 | NM_152372 | 0.1585 | -0.3625 |
| ZNF684 | NM_152373 | -1.337 | -1.9685 |
| C1orf216 | NM_152374 | -0.0715 | -1.146 |
| KLHDC7A | NM_152375 | -0.0365 | 0.039 |
| UBXN10 | NM_152376 | 0.0565 | -0.3315 |
| C1orf87 | NM_152377 | 0.0185 | 0.053 |
| C1orf131 | NM_152379 | -1.5685 | -3.0415 |
| TBX15 | NM_152380 | 0.066 | -0.0695 |
| XIRP2 | NM_152381 | 0.0425 | -0.241 |
| DIS3L2 | NM_152383 | 0.181 | -0.711 |
| BBS5 | NM_152384 | -0.3635 | -1.17 |
| C2orf63 | NM_152385 | -1.5685 | -1.0725 |
| SGPP2 | NM_152386 | -0.301 | -0.6585 |
| KCTD18 | NM_152387 | -2.03 | -3.052 |
| TMEM237 | NM_152388 | -1.363 | -1.1065 |
| CCDC108 | NM_152389 | 0.0615 | 0.142 |
| TMEM178 | NM_152390 | -0.4395 | -1.705 |
| PQLC3 | NM_152391 | -0.541 | -0.488 |
| AHSA2 | NM_152392 | -1.08 | -0.493 |
| KBTBD5 | NM_152393 | -0.122 | -0.1545 |
| FAM194A | NM_152394 | -0.047 | 0.004 |
| NUDT16 | NM_152395 | -1.124 | -1.4325 |
| METTL6 | NM_152396 | -1.0155 | -2.2665 |

| | | | |
|---|---|---|---|
| IQCF1 | NM_152397 | 0.083 | -0.1725 |
| TMEM155 | NM_152399 | 0.5375 | -0.1175 |
| C4orf32 | NM_152400 | -0.617 | -0.477 |
| PDCL2 | NM_152401 | -0.1685 | -0.488 |
| TRAM1L1 | NM_152402 | -0.8505 | -1.7595 |
| EGFLAM | NM_152403 | -0.277 | -0.894 |
| UGT3A1 | NM_152404 | 0.2045 | -0.8015 |
| JMY | NM_152405 | -0.982 | -2.805 |
| AFAP1L1 | NM_152406 | -0.4675 | -0.085 |
| GRPEL2 | NM_152407 | -1.677 | -1.508 |
| C5orf24 | NM_152409 | -1.581 | -1.3705 |
| ZNF786 | NM_152411 | -0.9615 | -2.3335 |
| ZNF572 | NM_152412 | -0.85 | -1.873 |
| GOT1L1 | NM_152413 | 0.0885 | -0.05 |
| BHLHE22 | NM_152414 | -0.5235 | -1.375 |
| VPS37A | NM_152415 | -1.5415 | -3.6465 |
| C8orf38 | NM_152416 | -0.763 | -2.432 |
| TMEM68 | NM_152417 | -0.407 | -1.538 |
| DCAF4L2 | NM_152418 | -0.18 | 0.0015 |
| HGSNAT | NM_152419 | 0.0775 | -0.5515 |
| C9orf41 | NM_152420 | -1.9395 | -2.383 |
| FAM69B | NM_152421 | -0.586 | -1.4305 |
| PTPDC1 | NM_152422 | -0.464 | -1.953 |
| MUM1L1 | NM_152423 | 0.2975 | -0.1205 |
| FAM123B | NM_152424 | 0.0155 | -0.3845 |
| APOBEC3D | NM_152426 | -0.4085 | -0.47 |
| FGFBP3 | NM_152429 | -0.4145 | -0.619 |
| OR51E1 | NM_152430 | -0.4305 | -0.2625 |
| PIWIL4 | NM_152431 | -0.0915 | -0.802 |
| ARHGAP42 | NM_152432 | -0.288 | -1.189 |
| KBTBD3 | NM_152433 | -1.057 | -0.79 |
| CWF19L2 | NM_152434 | 0.0335 | -0.5185 |
| AMDHD1 | NM_152435 | 0.082 | 0.0025 |
| GLIPR1L2 | NM_152436 | 0.022 | -0.064 |
| ZNF664 | NM_152437 | -1.028 | -0.8335 |
| DDX11 | NM_152438 | -0.204 | -1.65 |
| BEST3 | NM_152439 | -0.005 | -0.125 |
| C12orf66 | NM_152440 | -1.12 | -2.444 |
| FBXL14 | NM_152441 | -0.2325 | -0.859 |
| RAD9B | NM_152442 | 0.022 | -0.309 |
| RDH12 | NM_152443 | -0.401 | -0.1585 |
| PTGR2 | NM_152444 | -1.0615 | -0.789 |
| FAM161B | NM_152445 | -0.51 | -0.45 |

| | | | |
|---|---|---|---|
| CEP128 | NM_152446 | -0.325 | -2.167 |
| LRFN5 | NM_152447 | -0.685 | -0.1525 |
| C15orf43 | NM_152448 | -0.292 | -0.655 |
| LYSMD4 | NM_152449 | -0.768 | -1.812 |
| FAM81A | NM_152450 | -0.159 | -0.4855 |
| GCOM1 | NM_152451 | -0.3855 | -0.511 |
| TMCO5A | NM_152453 | -0.332 | 0.445 |
| ZSCAN29 | NM_152455 | -1.5045 | -1.843 |
| IL34 | NM_152456 | 0.0565 | -0.3225 |
| ZNF597 | NM_152457 | -0.3295 | -1.005 |
| ZNF785 | NM_152458 | -0.0335 | -0.598 |
| C16orf89 | NM_152459 | -0.22 | -0.2345 |
| C17orf77 | NM_152460 | -0.2865 | -0.517 |
| SLC35G3 | NM_152462 | 0.066 | -0.22 |
| EME1 | NM_152463 | -0.725 | -1.8965 |
| TMEM199 | NM_152464 | -1.523 | -2.192 |
| PROCA1 | NM_152465 | -0.0265 | -0.3945 |
| KLHL10 | NM_152467 | 0.3285 | -0.265 |
| TMC8 | NM_152468 | 0.256 | -0.335 |
| RNF165 | NM_152470 | -0.889 | -1.658 |
| C19orf18 | NM_152474 | -0.173 | -0.3865 |
| ZNF417 | NM_152475 | -0.5205 | -0.876 |
| ZNF560 | NM_152476 | 0.1915 | -0.883 |
| ZNF565 | NM_152477 | -1.2075 | -1.6725 |
| ZNF583 | NM_152478 | -0.8585 | -1.5245 |
| TTC9B | NM_152479 | 0.4125 | 0.224 |
| FAM187B | NM_152481 | -0.008 | -0.192 |
| C19orf25 | NM_152482 | -0.579 | -1.75 |
| ZNF569 | NM_152484 | -0.004 | -1.5065 |
| C1orf74 | NM_152485 | -1.157 | -1.625 |
| SAMD11 | NM_152486 | -0.8185 | -1.642 |
| TMEM56 | NM_152487 | -0.4725 | -1.1695 |
| UBE2U | NM_152489 | -0.0225 | -0.1145 |
| B3GALNT2 | NM_152490 | -1.4605 | -0.9145 |
| PM20D1 | NM_152491 | -0.3535 | -0.355 |
| CCDC27 | NM_152492 | -0.0325 | -0.677 |
| ZNF362 | NM_152493 | -1.3885 | -2.5325 |
| DCST1 | NM_152494 | -0.0625 | -0.238 |
| CNIH3 | NM_152495 | -0.3185 | -0.47 |
| MANEAL | NM_152496 | -0.876 | -1.801 |
| WDR65 | NM_152498 | -0.0525 | -0.137 |
| CCDC24 | NM_152499 | -0.0825 | -0.2775 |
| PYHIN1 | NM_152501 | 0.134 | -0.378 |

| | | | |
|---|---|---|---|
| C20orf132 | NM_152503 | -0.328 | -0.737 |
| C20orf196 | NM_152504 | -0.3815 | -1.0405 |
| LCA5L | NM_152505 | 0.231 | -1.2325 |
| HORMAD2 | NM_152510 | 0.042 | -0.504 |
| DUSP18 | NM_152511 | -0.5715 | -0.9725 |
| ENTHD1 | NM_152512 | 0.359 | 0.03 |
| MEI1 | NM_152513 | 0.1375 | -0.5105 |
| CKAP2L | NM_152515 | -0.448 | -2.334 |
| COMMD1 | NM_152516 | -0.3655 | -1.0185 |
| TTC30B | NM_152517 | -1.337 | -1.894 |
| C2orf67 | NM_152519 | -0.4625 | -0.9165 |
| ZNF385B | NM_152520 | -0.432 | -1.3425 |
| ARL6IP6 | NM_152522 | -0.9535 | -1.4515 |
| CCNYL1 | NM_152523 | -0.841 | -1.267 |
| SGOL2 | NM_152524 | 0.0415 | -1.0965 |
| ALS2CR11 | NM_152525 | 0.262 | 0.2225 |
| PARD3B | NM_152526 | 0 | -0.155 |
| SLC16A14 | NM_152527 | -0.2115 | -3.3045 |
| GPR155 | NM_152529 | -0.756 | -0.979 |
| C3orf22 | NM_152533 | 0.196 | 0.2405 |
| FGD5 | NM_152536 | -0.087 | -0.8175 |
| IGSF11 | NM_152538 | 0.147 | -0.4015 |
| C3orf30 | NM_152539 | 0.351 | -0.0475 |
| SCFD2 | NM_152540 | -0.3685 | -1.2245 |
| PPM1K | NM_152542 | -0.872 | -1.374 |
| C4orf45 | NM_152543 | 0.3675 | 0.4285 |
| RASGEF1B | NM_152545 | 0.1985 | -0.5605 |
| SRFBP1 | NM_152546 | -0.455 | -2.33 |
| BTNL9 | NM_152547 | 0.1305 | -0.2195 |
| FAM81B | NM_152548 | -0.2575 | -0.114 |
| CCDC112 | NM_152549 | -0.5065 | -2.202 |
| SNRNP48 | NM_152551 | -1.0095 | -2.3815 |
| SAMD3 | NM_152552 | -0.158 | -0.0975 |
| RNF217 | NM_152553 | -0.4855 | -0.86 |
| C6orf195 | NM_152554 | 0.1505 | -0.0885 |
| C7orf60 | NM_152556 | -1.597 | -3.955 |
| ZNF746 | NM_152557 | -0.569 | -2.3685 |
| IQCE | NM_152558 | -0.1225 | -0.487 |
| WBSCR27 | NM_152559 | 0.032 | -1.226 |
| CDCA2 | NM_152562 | -0.956 | -2.426 |
| VPS13B | NM_152564 | -0.1415 | -0.4325 |
| ATP6V0D2 | NM_152565 | -0.17 | -0.114 |
| NKX6-3 | NM_152568 | -0.2905 | 0.026 |

| | | | |
|---|---|---|---|
| C9orf66 | NM_152569 | 0.3625 | -0.0215 |
| LINGO2 | NM_152570 | -0.286 | -1.613 |
| C9orf163 | NM_152571 | 0.016 | -0.4835 |
| AK8 | NM_152572 | 0.0275 | -0.171 |
| RASEF | NM_152573 | -0.205 | -0.1305 |
| TTC39B | NM_152574 | 0.2655 | -0.91 |
| ZNF645 | NM_152577 | -0.1975 | -0.283 |
| FMR1NB | NM_152578 | -0.06 | -0.409 |
| MOSPD2 | NM_152581 | -1.6895 | -2.4475 |
| HSFY1 | NM_152584 | 0.213 | 0.1055 |
| USP54 | NM_152586 | -1.5245 | -2.668 |
| C11orf65 | NM_152587 | -0.106 | -0.216 |
| TMTC2 | NM_152588 | -0.709 | -1.124 |
| C12orf50 | NM_152589 | 0.0145 | -0.0255 |
| IFLTD1 | NM_152590 | -0.0405 | -0.3735 |
| CCDC63 | NM_152591 | -0.1885 | -0.345 |
| C14orf49 | NM_152592 | -0.2395 | -0.1315 |
| SPRED1 | NM_152594 | -1.9955 | -4.2875 |
| PGBD4 | NM_152595 | -0.285 | 0.0635 |
| EXD1 | NM_152596 | -0.312 | -0.203 |
| FSIP1 | NM_152597 | -0.2435 | -0.711 |
| MFSD6L | NM_152599 | 0.1365 | -0.3595 |
| ZNF579 | NM_152600 | -0.655 | -1.6795 |
| ZNF709 | NM_152601 | -0.7855 | -2.388 |
| ZNF567 | NM_152603 | -0.8675 | -1.794 |
| ZNF383 | NM_152604 | -1.0485 | -1.4585 |
| ZNF781 | NM_152605 | -0.883 | -1.3405 |
| ZNF540 | NM_152606 | -0.205 | -1.361 |
| C1orf177 | NM_152607 | -0.029 | 0.0455 |
| C1orf55 | NM_152608 | -1.5985 | -3 |
| CNST | NM_152609 | -1.806 | -3.636 |
| C1orf65 | NM_152610 | -0.258 | -0.184 |
| LRRN4 | NM_152611 | -0.0835 | 0.184 |
| CCDC116 | NM_152612 | -0.059 | -0.216 |
| WBP2NL | NM_152613 | 0.3705 | 0.3685 |
| C2orf57 | NM_152614 | 0.0425 | -0.002 |
| PARP15 | NM_152615 | -0.1725 | -0.109 |
| TRIM42 | NM_152616 | 0.713 | 0.0295 |
| RNF168 | NM_152617 | -0.334 | -0.6825 |
| BBS12 | NM_152618 | -0.312 | -1.1115 |
| TRIM60 | NM_152620 | 0.185 | -0.0605 |
| SGMS2 | NM_152621 | 0.002 | 0.184 |
| MIER3 | NM_152622 | -1.142 | -2.859 |

| | | | |
|---|---|---|---|
| CDC20B | NM_152623 | -0.0955 | -0.157 |
| DCP2 | NM_152624 | -1.175 | -1.6745 |
| ZNF366 | NM_152625 | -0.253 | -0.162 |
| ZNF92 | NM_152626 | -1.184 | -3.9665 |
| SNX31 | NM_152628 | 0.3505 | -0.111 |
| GLIS3 | NM_152629 | 0.042 | -0.6525 |
| FAM46D | NM_152630 | 0.2295 | -0.07 |
| FAM47B | NM_152631 | 0.2165 | -0.911 |
| CXorf22 | NM_152632 | 0.2165 | -0.185 |
| TCEANC | NM_152634 | -0.5795 | -0.9015 |
| OIT3 | NM_152635 | 0.5475 | -0.49 |
| METTL15 | NM_152636 | -1.7935 | -2.5735 |
| METTL7B | NM_152637 | -0.3775 | -0.4125 |
| C12orf12 | NM_152638 | -0.1655 | -0.486 |
| DCP1B | NM_152640 | -1.8665 | -3.6205 |
| ARID2 | NM_152641 | -1.4525 | -3.829 |
| KNDC1 | NM_152643 | 0.309 | -0.1535 |
| FAM24B | NM_152644 | -0.15 | -0.791 |
| C15orf33 | NM_152647 | -0.5365 | -0.411 |
| MLKL | NM_152649 | 0.202 | 0.2045 |
| ZNF48 | NM_152652 | -0.438 | -0.9335 |
| UBE2E2 | NM_152653 | -0.329 | -1.3585 |
| DAND5 | NM_152654 | -0.042 | -0.379 |
| ZNF585A | NM_152655 | -0.5565 | -2.2025 |
| GGN | NM_152657 | 0.072 | -0.3695 |
| THAP8 | NM_152658 | -1.1855 | -2.052 |
| FAM76A | NM_152660 | -0.3435 | -0.4015 |
| RALGPS2 | NM_152663 | 0.247 | -0.895 |
| TCTEX1D1 | NM_152665 | -0.07 | 0.041 |
| PLD5 | NM_152666 | -0.466 | -1.648 |
| NANP | NM_152667 | -1.469 | -3.037 |
| C2orf51 | NM_152670 | -0.065 | -0.4735 |
| PIKFYVE | NM_152671 | 0.8725 | 0.09 |
| OSTalpha | NM_152672 | 0.1865 | -0.579 |
| MUC20 | NM_152673 | 0.0055 | -0.557 |
| FBXO15 | NM_152676 | -0.34 | -0.4035 |
| ZSCAN4 | NM_152677 | -0.2755 | 0.0815 |
| FAM116A | NM_152678 | -1.7015 | -3.6585 |
| SLC10A4 | NM_152679 | -0.75 | -3.6635 |
| TMEM154 | NM_152680 | -0.1695 | -0.2135 |
| RWDD4 | NM_152682 | -0.869 | -0.937 |
| CCDC111 | NM_152683 | -2.0495 | -2.5955 |
| SLC23A1 | NM_152685 | 0.0435 | -0.3485 |

| | | | |
|---|---|---|---|
| DNAJC18 | NM_152686 | -1.0095 | -1.323 |
| GAPT | NM_152687 | -0.147 | -0.373 |
| KHDRBS2 | NM_152688 | 0.165 | 0.0975 |
| C1GALT1C1 | NM_152692 | -1.4495 | -2.3095 |
| ZCCHC5 | NM_152694 | -0.0055 | -0.1135 |
| ZNF449 | NM_152695 | -0.7265 | -2.316 |
| HIPK1 | NM_152696 | 0.267 | -0.807 |
| SLC44A5 | NM_152697 | -0.405 | 0.1365 |
| FAM123C | NM_152698 | -0.124 | -0.984 |
| SENP5 | NM_152699 | -1.5085 | -2.4755 |
| ABCA13 | NM_152701 | 0.066 | 0.0095 |
| SAMD9L | NM_152703 | -0.075 | 0.116 |
| FAM123A | NM_152704 | -0.083 | -0.692 |
| POLR1D | NM_152705 | -0.681 | -1.811 |
| C7orf62 | NM_152706 | -0.1315 | -0.0385 |
| SLC25A16 | NM_152707 | -0.671 | -1.074 |
| STOX1 | NM_152709 | -0.5335 | -0.9015 |
| C10orf27 | NM_152710 | -0.0295 | -0.467 |
| STT3A | NM_152713 | -0.277 | -1.019 |
| TBCEL | NM_152715 | -1.1345 | -2.159 |
| PATL1 | NM_152716 | -0.8395 | -2.5995 |
| MS4A15 | NM_152717 | 0.428 | -0.3275 |
| VWCE | NM_152718 | -0.563 | -1.498 |
| SPERT | NM_152719 | -0.163 | -0.038 |
| NEK3 | NM_152720 | -0.764 | -1.376 |
| DOK6 | NM_152721 | -0.3845 | -1.055 |
| HEPACAM | NM_152722 | -0.48 | -0.155 |
| CCDC89 | NM_152723 | -0.268 | -0.5925 |
| RSU1 | NM_152724 | -0.4885 | -0.733 |
| SLC39A12 | NM_152725 | -0.3685 | -0.402 |
| EFHA1 | NM_152726 | -0.7 | -1.0625 |
| CPNE2 | NM_152727 | -0.1155 | -0.783 |
| NT5DC1 | NM_152729 | -1.189 | -1.8865 |
| C6orf170 | NM_152730 | -0.756 | -1.168 |
| BEND6 | NM_152731 | 0.0535 | -0.9965 |
| RSPH9 | NM_152732 | 0.3305 | -0.246 |
| BTBD9 | NM_152733 | -0.2065 | -0.124 |
| C6orf89 | NM_152734 | -0.0065 | -1.158 |
| ZBTB9 | NM_152735 | -0.901 | -1.524 |
| ZNF187 | NM_152736 | -1.4035 | -2.8105 |
| RNF182 | NM_152737 | -0.4445 | -0.8195 |
| HOXA9 | NM_152739 | -0.173 | -0.1545 |
| HIBADH | NM_152740 | -1.2455 | -1.686 |

| | | | |
|---|---|---|---|
| GPC2 | NM_152742 | -0.1415 | -1.029 |
| BRAT1 | NM_152743 | -0.472 | -1.13 |
| SDK1 | NM_152744 | -0.3905 | -0.688 |
| NXPH1 | NM_152745 | -0.6075 | -1.634 |
| KIAA1324L | NM_152748 | -1.0615 | -1.0995 |
| ATXN7L1 | NM_152749 | 0.3515 | -0.3115 |
| CDHR3 | NM_152750 | 0.089 | -0.3915 |
| BEND7 | NM_152751 | -0.119 | -0.3705 |
| SCUBE3 | NM_152753 | 0.033 | -0.3245 |
| SEMA3D | NM_152754 | -0.0565 | -0.007 |
| CNPY4 | NM_152755 | -0.7965 | -1.078 |
| RICTOR | NM_152756 | -0.3295 | -2.308 |
| YTHDF3 | NM_152758 | -1.151 | -3.347 |
| LRRC43 | NM_152759 | 0.1565 | 0.1715 |
| SNX32 | NM_152760 | -0.098 | -0.474 |
| CXorf58 | NM_152761 | 0.275 | -0.0525 |
| TSGA10IP | NM_152762 | -0.088 | -0.2625 |
| AKNAD1 | NM_152763 | 0.2575 | -0.1475 |
| C16orf73 | NM_152764 | -0.049 | -0.0535 |
| C8orf46 | NM_152765 | -0.279 | -0.3805 |
| C17orf61 | NM_152766 | 0.2665 | -0.3465 |
| C19orf26 | NM_152769 | -0.2405 | 0.291 |
| C4orf22 | NM_152770 | 0.5225 | 0.017 |
| TCP11L2 | NM_152772 | -0.0515 | -0.164 |
| TCTEX1D2 | NM_152773 | -0.226 | -0.595 |
| TMEM196 | NM_152774 | -0.136 | -1.125 |
| CCDC110 | NM_152775 | 0.6145 | -0.5245 |
| MFSD8 | NM_152778 | -1.933 | -3.15 |
| GLIPR1L1 | NM_152779 | -0.212 | -0.471 |
| MAP7D2 | NM_152780 | -0.6265 | -1.088 |
| C17orf66 | NM_152781 | 0.057 | -0.193 |
| SUN3 | NM_152782 | -0.274 | -0.2025 |
| D2HGDH | NM_152783 | -1.148 | -1.4655 |
| TMEM146 | NM_152784 | 0.044 | -0.272 |
| GCET2 | NM_152785 | 0.21 | 0.0825 |
| C9orf43 | NM_152786 | -0.3745 | -0.5695 |
| TAB3 | NM_152787 | -0.9615 | -1.843 |
| ANKS1B | NM_152788 | -0.3535 | -0.4295 |
| ZNF555 | NM_152791 | -0.8285 | -1.792 |
| ASPRV1 | NM_152792 | -0.2775 | -0.5655 |
| C7orf41 | NM_152793 | -1.031 | -0.8295 |
| HIF3A | NM_152794 | -0.243 | -0.193 |
| HIF3A | NM_152795 | -0.243 | -0.193 |

| | | | |
|---|---|---|---|
| HIF3A | NM_152796 | -0.243 | -0.193 |
| SNX3 | NM_152827 | -0.575 | -1.3135 |
| TES | NM_152829 | -0.323 | -1.45 |
| ACE | NM_152830 | 0.061 | -0.606 |
| FAM89B | NM_152832 | -0.0525 | -1.1165 |
| C9orf69 | NM_152833 | -1.366 | -2.2675 |
| TMEM18 | NM_152834 | -0.2895 | -1.302 |
| PDIK1L | NM_152835 | -1.7805 | -3.184 |
| SNX16 | NM_152836 | -1.142 | -2.185 |
| SNX16 | NM_152837 | -1.142 | -2.185 |
| RBM12 | NM_152838 | -0.381 | -1.116 |
| HPS4 | NM_152841 | -0.954 | -2.2725 |
| PIGO | NM_152850 | -0.908 | -1.5075 |
| MS4A6A | NM_152851 | -0.1165 | 0.223 |
| MS4A6A | NM_152852 | -0.1165 | 0.223 |
| CD40 | NM_152854 | 0.119 | -0.448 |
| IGLL1 | NM_152855 | 0.155 | -0.287 |
| RBM10 | NM_152856 | -0.748 | -1.5345 |
| WTAP | NM_152857 | -0.3035 | -0.7795 |
| WTAP | NM_152858 | -0.0215 | -0.59 |
| SP7 | NM_152860 | -0.002 | -0.1105 |
| ARPC2 | NM_152862 | -0.509 | -0.566 |
| NKAIN4 | NM_152864 | -0.071 | -0.5315 |
| MS4A1 | NM_152866 | -0.2405 | -0.1505 |
| KCNJ4 | NM_152868 | -0.291 | -0.247 |
| RGN | NM_152869 | 0.3585 | -0.3015 |
| FAS | NM_152871 | -0.4565 | -1.1795 |
| FAS | NM_152872 | -0.4565 | -1.1795 |
| DGKD | NM_152879 | -0.4465 | -1.269 |
| PTK7 | NM_152880 | 0.131 | -0.5825 |
| PTK7 | NM_152881 | 0.131 | -0.5825 |
| PTK7 | NM_152882 | 0.131 | -0.5825 |
| COL22A1 | NM_152888 | -0.0635 | -0.2565 |
| CHST13 | NM_152889 | -0.2505 | -0.5745 |
| COL24A1 | NM_152890 | 0.027 | 0.121 |
| PRSS33 | NM_152891 | -0.0555 | 0.226 |
| LRWD1 | NM_152892 | -0.2165 | -1.366 |
| UHRF2 | NM_152896 | -0.8125 | -2.4195 |
| SNX21 | NM_152897 | -0.511 | -0.79 |
| FERD3L | NM_152898 | -0.099 | -0.248 |
| MAGI3 | NM_152900 | -0.9535 | -2.801 |
| PYDC1 | NM_152901 | 0.304 | 0.106 |
| TIPRL | NM_152902 | -1.157 | -1.807 |

| | | | |
|---|---|---|---|
| KBTBD6 | NM_152903 | -1.4185 | -0.369 |
| SPECC1 | NM_152904 | -0.3345 | -0.4385 |
| NEDD1 | NM_152905 | -1.072 | -2.8115 |
| C22orf25 | NM_152906 | -0.3335 | -1.406 |
| SLC47A2 | NM_152908 | 0.3 | -0.191 |
| ZNF548 | NM_152909 | -1.348 | -2.2395 |
| DGKH | NM_152910 | -0.112 | -0.3965 |
| PAOX | NM_152911 | -0.336 | -0.675 |
| MTIF3 | NM_152912 | -1.0275 | -1.766 |
| TMEM130 | NM_152913 | -1.053 | -1.3895 |
| C17orf103 | NM_152914 | -0.586 | -1.251 |
| EMR2 | NM_152916 | -0.016 | -0.484 |
| EMR2 | NM_152917 | -0.016 | -0.484 |
| EMR2 | NM_152918 | -0.016 | -0.484 |
| EMR2 | NM_152919 | -0.016 | -0.484 |
| EMR2 | NM_152920 | -0.016 | -0.484 |
| EMR2 | NM_152921 | -0.016 | -0.484 |
| ABHD2 | NM_152924 | 0.1315 | -0.694 |
| CPNE1 | NM_152925 | -0.2265 | -0.82 |
| GLT8D1 | NM_152932 | -1.635 | -2.303 |
| TNFRSF8 | NM_152942 | 0.054 | -0.171 |
| ZNF268 | NM_152943 | -1.119 | -1.924 |
| RBM45 | NM_152945 | -1.06 | -1.572 |
| SPPL2B | NM_152988 | -0.3725 | -0.6545 |
| SOX5 | NM_152989 | 0.17 | -0.8475 |
| PXT1 | NM_152990 | 0.137 | -0.631 |
| EED | NM_152991 | -0.9755 | -2.1375 |
| POMZP3 | NM_152992 | -0.0595 | -0.4935 |
| PPP1R21 | NM_152994 | -1.53 | -1.514 |
| NFXL1 | NM_152995 | -0.402 | -2.1785 |
| ST6GALNAC3 | NM_152996 | -1.064 | -1.891 |
| C4orf7 | NM_152997 | 0.074 | 0.23 |
| EZH2 | NM_152998 | -1.045 | -1.709 |
| STEAP2 | NM_152999 | 0.2585 | -0.0985 |
| APCDD1 | NM_153000 | -0.515 | -0.873 |
| GPR156 | NM_153002 | -0.1445 | -0.1305 |
| RIOK1 | NM_153005 | -0.967 | -1.894 |
| NAGS | NM_153006 | -0.38 | -1.2595 |
| ODF4 | NM_153007 | 0.029 | -0.258 |
| TMEM74 | NM_153015 | -0.5105 | -0.6725 |
| ZFP3 | NM_153018 | -1.2685 | -2.4495 |
| RBM24 | NM_153020 | -0.1465 | -0.9645 |
| PLB1 | NM_153021 | -0.139 | -0.082 |

| | | | |
|---|---|---|---|
| C12orf59 | NM_153022 | -0.104 | -0.3075 |
| SPATA13 | NM_153023 | -0.197 | -0.9725 |
| C16orf55 | NM_153025 | 0.0765 | -0.5795 |
| PRICKLE1 | NM_153026 | -0.785 | -2.75 |
| C4orf39 | NM_153027 | -0.33 | 0.103 |
| ZNF75A | NM_153028 | -1.677 | -2.7425 |
| N4BP1 | NM_153029 | -1.69 | -3.878 |
| KCTD7 | NM_153033 | -0.747 | -1.0405 |
| ZNF488 | NM_153034 | -0.531 | -1.8765 |
| TCEANC2 | NM_153035 | -1.505 | -1.5455 |
| FAM26D | NM_153036 | 0.065 | -0.5095 |
| CCDC140 | NM_153038 | -0.1835 | 0.045 |
| C15orf32 | NM_153040 | 0.1295 | -0.1775 |
| KDM1B | NM_153042 | -1.049 | -0.7475 |
| C9orf91 | NM_153045 | -0.698 | -2.2295 |
| TDRD9 | NM_153046 | -0.044 | -0.195 |
| FYN | NM_153047 | -0.603 | -1.9005 |
| FYN | NM_153048 | -0.603 | -1.9005 |
| MTMR3 | NM_153050 | -0.32 | -1.7465 |
| MTMR3 | NM_153051 | -0.32 | -1.7465 |
| MINA | NM_153182 | -0.971 | -2.2405 |
| NUDT10 | NM_153183 | 0.1075 | -0.532 |
| CADM2 | NM_153184 | -0.113 | -0.487 |
| KANK1 | NM_153186 | -0.154 | -0.148 |
| TNPO1 | NM_153188 | -0.5835 | -0.4805 |
| SPAM1 | NM_153189 | 0.21 | 0.111 |
| EDF1 | NM_153200 | -0.1405 | -1.2265 |
| HSPA8 | NM_153201 | -0.63 | -0.691 |
| ADAM33 | NM_153202 | -0.0045 | -0.7295 |
| AMICA1 | NM_153206 | 0.1805 | 0.0645 |
| AEBP2 | NM_153207 | -1.5545 | -3.7785 |
| IQCK | NM_153208 | -0.7585 | -1.189 |
| KIF19 | NM_153209 | -0.014 | -0.5155 |
| USP43 | NM_153210 | 0.0745 | -0.0725 |
| TTC39C | NM_153211 | -1.234 | -2.3885 |
| GJB4 | NM_153212 | -0.178 | -0.177 |
| ARHGEF19 | NM_153213 | -0.203 | -0.7715 |
| FBLN7 | NM_153214 | -0.184 | -0.0455 |
| C3orf45 | NM_153215 | -0.0735 | -0.113 |
| POU5F2 | NM_153216 | 0.004 | -0.1585 |
| TMEM174 | NM_153217 | 0.216 | -0.1365 |
| ZNF524 | NM_153219 | 0.0365 | -0.731 |
| CILP2 | NM_153221 | -0.15 | -0.401 |

| | | | |
|---|---|---|---|
| CEP120 | NM_153223 | -1.6895 | -3.209 |
| C8orf84 | NM_153225 | -0.1545 | -0.1095 |
| ANKFN1 | NM_153228 | -0.0525 | 0.479 |
| TMEM92 | NM_153229 | 0.0865 | -0.5085 |
| FBXO39 | NM_153230 | -0.157 | 0.17 |
| EID2 | NM_153232 | -1.46 | -3.0435 |
| LIX1 | NM_153234 | -0.422 | -0.8745 |
| TXLNB | NM_153235 | -0.0465 | -0.073 |
| GIMAP7 | NM_153236 | 0.0255 | -0.769 |
| C9orf71 | NM_153237 | 0.177 | 0.141 |
| NPHP3 | NM_153240 | -0.8275 | -2.2675 |
| C10orf111 | NM_153244 | 0.165 | 0.069 |
| C6orf223 | NM_153246 | -0.2375 | -0.5085 |
| SLC29A4 | NM_153247 | -0.444 | -1.0065 |
| ZDHHC20 | NM_153251 | 0.0335 | -0.414 |
| BRWD3 | NM_153252 | -0.4075 | -0.606 |
| SIPA1 | NM_153253 | -0.5025 | -0.83 |
| TTLL10 | NM_153254 | -0.4335 | -0.3545 |
| MCM9 | NM_153255 | -1.5535 | -3.3235 |
| C10orf47 | NM_153256 | -0.215 | -1.2585 |
| ZNF461 | NM_153257 | -1.3995 | -2.181 |
| MCOLN2 | NM_153259 | -0.3485 | -0.2325 |
| LRRC57 | NM_153260 | -0.8175 | -1.2085 |
| TMEM188 | NM_153261 | -1.252 | -1.9425 |
| SYT14 | NM_153262 | -0.389 | -1.406 |
| ZNF549 | NM_153263 | -1.2245 | -3.2425 |
| COL6A5 | NM_153264 | -0.2925 | -0.197 |
| EML3 | NM_153265 | 0.104 | -0.825 |
| TMEM151A | NM_153266 | -0.388 | -0.8575 |
| MAMDC2 | NM_153267 | 0.011 | -0.648 |
| PLCXD2 | NM_153268 | -1.049 | -2.5005 |
| C20orf96 | NM_153269 | 0.097 | -0.4415 |
| KLHL34 | NM_153270 | 0.306 | -0.6235 |
| SNX33 | NM_153271 | -0.3175 | -0.9645 |
| IP6K1 | NM_153273 | -0.3725 | -1.5875 |
| BEST4 | NM_153274 | -0.0965 | -0.0955 |
| SLC22A6 | NM_153276 | 0.128 | 0.005 |
| SLC22A6 | NM_153277 | 0.128 | 0.005 |
| SLC22A6 | NM_153278 | 0.128 | 0.005 |
| UBA1 | NM_153280 | -0.54 | -0.1625 |
| HYAL1 | NM_153281 | -0.0895 | 0.034 |
| HYAL1 | NM_153282 | -0.0895 | 0.034 |
| HYAL1 | NM_153283 | -0.0895 | 0.034 |

| | | | |
|---|---|---|---|
| HYAL1 | NM_153285 | -0.0895 | 0.034 |
| DEFB119 | NM_153289 | 0.219 | -0.292 |
| SLC22A7 | NM_153320 | -0.15 | -0.295 |
| PMP22 | NM_153321 | -0.7095 | -1.6925 |
| PMP22 | NM_153322 | -0.7095 | -1.6925 |
| DEFB119 | NM_153323 | -0.0695 | -0.567 |
| DEFB123 | NM_153324 | -0.3445 | -0.052 |
| DEFB125 | NM_153325 | -0.038 | 0.04 |
| ALDH16A1 | NM_153329 | -0.082 | -0.603 |
| DNAJB8 | NM_153330 | -0.0485 | -0.3405 |
| KCTD6 | NM_153331 | -1.0575 | -2.174 |
| ERI1 | NM_153332 | -1.036 | -2.729 |
| TCEAL8 | NM_153333 | -1.8655 | -2.099 |
| SCARF2 | NM_153334 | -0.036 | -1.111 |
| STRADA | NM_153335 | -1.068 | -2.0825 |
| PSTK | NM_153336 | -0.4605 | -1.7805 |
| SNX20 | NM_153337 | 0.034 | -0.208 |
| GGT6 | NM_153338 | -0.2525 | -0.139 |
| PUSL1 | NM_153339 | 0.0105 | -1.1825 |
| ATXN7L2 | NM_153340 | -0.7575 | -1.4475 |
| ENPP6 | NM_153343 | -0.3165 | -0.247 |
| TMEM139 | NM_153345 | 0.2435 | -0.3775 |
| BEND2 | NM_153346 | -0.1025 | -0.1145 |
| TMEM86A | NM_153347 | 0.0555 | -1.0735 |
| FBXW8 | NM_153348 | -1.0655 | -1.2495 |
| FBXL16 | NM_153350 | -0.602 | -1.261 |
| LRRC34 | NM_153353 | -1.6075 | -1.4715 |
| TMEM161B | NM_153354 | -1.272 | -2.5815 |
| NKAIN2 | NM_153355 | -0.2975 | -0.4165 |
| TBC1D21 | NM_153356 | 0.4285 | -0.224 |
| SLC16A11 | NM_153357 | -0.1765 | -0.4205 |
| ZNF791 | NM_153358 | -1.4255 | -2.7285 |
| APCDD1L | NM_153360 | 0.1005 | -0.16 |
| NIM1 | NM_153361 | -0.2215 | -0.3525 |
| PRSS35 | NM_153362 | -0.1805 | -0.6685 |
| ZNF679 | NM_153363 | 0.0425 | -0.1305 |
| FAM71C | NM_153364 | -0.082 | -0.0985 |
| TAPT1 | NM_153365 | -1.174 | -2.364 |
| SVEP1 | NM_153366 | 0.1435 | -0.6835 |
| ZCCHC24 | NM_153367 | -0.4095 | -1.4265 |
| GJD4 | NM_153368 | 0.004 | -0.187 |
| KIAA1919 | NM_153369 | -0.8215 | -1.4165 |
| PI16 | NM_153370 | 0.0345 | -0.139 |

| | | | |
|---|---|---|---|
| LNX2 | NM_153371 | -1.3725 | -2.5695 |
| AGXT2L2 | NM_153373 | -0.352 | -0.3395 |
| LYSMD2 | NM_153374 | -0.236 | -0.595 |
| CCDC96 | NM_153376 | 0.0675 | -0.3535 |
| LRIG3 | NM_153377 | -0.4595 | -1.3465 |
| ZNF41 | NM_153380 | -0.3595 | -1.811 |
| ODF2 | NM_153432 | -0.4545 | -0.8675 |
| ODF2 | NM_153433 | -0.4545 | -0.8675 |
| ODF2 | NM_153435 | -0.4545 | -0.8675 |
| ODF2 | NM_153436 | -0.4545 | -0.8675 |
| ODF2 | NM_153437 | -0.4545 | -0.8675 |
| ODF2 | NM_153439 | -0.4545 | -0.8675 |
| ODF2 | NM_153440 | -0.4545 | -0.8675 |
| GPR26 | NM_153442 | -0.204 | -0.9985 |
| OR5P2 | NM_153444 | 0.165 | -0.101 |
| OR5P3 | NM_153445 | -0.19 | -0.119 |
| B4GALNT2 | NM_153446 | 0.1105 | -0.324 |
| NLRP5 | NM_153447 | -0.158 | -0.264 |
| ESX1 | NM_153448 | 0.168 | -0.2395 |
| SLC2A14 | NM_153449 | 0.1555 | -0.0745 |
| MED19 | NM_153450 | -0.4715 | -2.2235 |
| ORAOV1 | NM_153451 | -0.5265 | -1.352 |
| VGLL2 | NM_153453 | -0.441 | -0.2245 |
| HS6ST3 | NM_153456 | -0.6015 | -0.542 |
| IL17RC | NM_153460 | -0.0215 | -0.42 |
| IL17RC | NM_153461 | -0.0215 | -0.42 |
| ILF3 | NM_153464 | -0.0295 | -0.4595 |
| UXT | NM_153477 | 0.0345 | -0.3155 |
| CSAG1 | NM_153478 | 0.211 | -0.4865 |
| IL17RE | NM_153480 | 0.0985 | -0.376 |
| IL17RE | NM_153483 | 0.0985 | -0.376 |
| LDHD | NM_153486 | 0.0005 | 0.013 |
| MDGA1 | NM_153487 | -0.0705 | -0.4285 |
| KRT13 | NM_153490 | 0.2275 | -0.0255 |
| TAB1 | NM_153497 | -0.4285 | -0.8275 |
| CAMK1D | NM_153498 | -0.4705 | -1.2695 |
| CAMKK2 | NM_153499 | -0.5185 | -0.88 |
| CAMKK2 | NM_153500 | -0.5185 | -0.88 |
| COG7 | NM_153603 | -0.5715 | -1.745 |
| MYOCD | NM_153604 | 0.2005 | -0.0665 |
| FAM71A | NM_153606 | 0.037 | -0.046 |
| C5orf41 | NM_153607 | -0.2265 | -0.516 |
| ZNF114 | NM_153608 | -1.598 | -2.462 |

| | | | |
|---|---|---|---|
| TMPRSS6 | NM_153609 | 0.038 | -0.6105 |
| CMYA5 | NM_153610 | -0.0365 | 0.1055 |
| CYBASC3 | NM_153611 | -0.5905 | -1.257 |
| HS3ST5 | NM_153612 | -0.6975 | -2.06 |
| LPCAT4 | NM_153613 | -0.458 | -1.6885 |
| DNAJB13 | NM_153614 | 0.372 | -0.335 |
| RGL4 | NM_153615 | 0.1695 | -0.559 |
| SEMA6D | NM_153616 | -0.2595 | -0.628 |
| SEMA6D | NM_153617 | -0.2595 | -0.628 |
| SEMA6D | NM_153618 | -0.8435 | -2.907 |
| SEMA6D | NM_153619 | -0.2595 | -0.628 |
| HOXA1 | NM_153620 | 0.019 | -0.1235 |
| HOXA3 | NM_153631 | 0.2835 | 0.167 |
| CPNE8 | NM_153634 | -1.1645 | -0.5235 |
| CPNE9 | NM_153635 | 0.051 | -0.395 |
| PANK2 | NM_153638 | 0.093 | -0.087 |
| PANK2 | NM_153640 | -0.9945 | -2.538 |
| NUP50 | NM_153645 | -1.5965 | -1.9415 |
| SLC24A4 | NM_153646 | -0.0585 | -0.1895 |
| SLC24A4 | NM_153647 | -0.0585 | -0.1895 |
| SLC24A4 | NM_153648 | -0.0585 | -0.1895 |
| TPM3 | NM_153649 | -0.2085 | -0.625 |
| FOXA2 | NM_153675 | 0.1275 | -0.073 |
| USH1C | NM_153676 | 0.2785 | -0.076 |
| PIGP | NM_153681 | -1.094 | -1.2485 |
| PIGP | NM_153682 | -1.094 | -1.2485 |
| C12orf53 | NM_153685 | -0.122 | -0.6775 |
| LCORL | NM_153686 | -0.3705 | -1.0855 |
| IKBIP | NM_153687 | -1.6145 | -3.31 |
| ZFP1 | NM_153688 | -0.7545 | -2.4955 |
| C2orf69 | NM_153689 | -1.524 | -2.6145 |
| FAM43A | NM_153690 | -0.4395 | -0.436 |
| HTRA4 | NM_153692 | -0.162 | -0.163 |
| HOXC6 | NM_153693 | -0.3895 | -0.19 |
| SYCP3 | NM_153694 | 0.0495 | -0.38 |
| ZNF367 | NM_153695 | -1.856 | -3.5365 |
| ANKRD44 | NM_153697 | 0.0395 | -0.3575 |
| C9orf21 | NM_153698 | -0.3525 | -2.679 |
| GSTA5 | NM_153699 | 0.197 | -0.7345 |
| STRC | NM_153700 | -0.063 | -0.0775 |
| IL12RB1 | NM_153701 | -0.022 | -0.187 |
| ELMOD2 | NM_153702 | -0.93 | -1.378 |
| PODN | NM_153703 | 0.2825 | -0.503 |

| | | | |
|---|---|---|---|
| TMEM67 | NM_153704 | -1.1445 | -1.783 |
| KDELC2 | NM_153705 | -1.11 | -1.7085 |
| C5orf35 | NM_153706 | -0.549 | -0.6755 |
| FAM154A | NM_153707 | -0.074 | -0.4515 |
| RTP1 | NM_153708 | 0.034 | -0.7025 |
| C9orf96 | NM_153710 | -0.3655 | -0.3665 |
| FAM26E | NM_153711 | 0.0445 | -0.096 |
| TTL | NM_153712 | -1.2175 | -1.773 |
| LIX1L | NM_153713 | -0.0775 | -1.214 |
| C10orf67 | NM_153714 | 0.202 | -0.406 |
| EVC | NM_153717 | -0.0655 | -0.3575 |
| NUP62 | NM_153718 | -0.971 | -2.442 |
| NUP62 | NM_153719 | -1.2405 | -2.969 |
| DPM3 | NM_153741 | 0.2405 | -0.56 |
| CTH | NM_153742 | -0.6085 | -2.028 |
| PIGC | NM_153747 | -0.7405 | -2.225 |
| KCNC2 | NM_153748 | 0.1075 | -0.151 |
| FNDC5 | NM_153756 | -0.1825 | -0.6595 |
| NAP1L5 | NM_153757 | -1.256 | -0.455 |
| IL19 | NM_153758 | 0.147 | -0.0935 |
| DNMT3A | NM_153759 | -0.1735 | -1.304 |
| KCNJ1 | NM_153764 | -0.357 | -0.0905 |
| KCNJ1 | NM_153765 | -0.357 | -0.0905 |
| KCNJ1 | NM_153766 | -0.357 | -0.0905 |
| KCNJ1 | NM_153767 | -0.357 | -0.0905 |
| CABYR | NM_153770 | -0.825 | -1.8285 |
| TAF1L | NM_153809 | -0.3545 | 0.008 |
| C10orf46 | NM_153810 | -0.953 | -2.3395 |
| SLC38A6 | NM_153811 | -0.9925 | -1.0935 |
| PHF13 | NM_153812 | -0.602 | -1.6725 |
| ZFPM1 | NM_153813 | -1.1485 | -1.504 |
| RASGRF1 | NM_153815 | -0.0115 | 0.137 |
| SNX14 | NM_153816 | -1.274 | -1.7475 |
| PEX10 | NM_153818 | -0.687 | -0.8285 |
| GSG1 | NM_153823 | 0.109 | 0.452 |
| PYCR1 | NM_153824 | -0.2335 | -0.731 |
| CD46 | NM_153826 | -1.213 | -1.466 |
| MINK1 | NM_153827 | -0.1005 | -1.0415 |
| RTN4 | NM_153828 | -0.2735 | -0.847 |
| PTK2 | NM_153831 | -1.1985 | -1.656 |
| GPR161 | NM_153832 | -0.7185 | -1.698 |
| H1FOO | NM_153833 | -0.3535 | -0.282 |
| GPR112 | NM_153834 | 0.649 | -0.6845 |

| | | | |
|---|---|---|---|
| GPR113 | NM_153835 | -0.1235 | -0.5355 |
| CREG2 | NM_153836 | -0.013 | 0.1965 |
| GPR114 | NM_153837 | 0.108 | -0.2095 |
| GPR115 | NM_153838 | 0.268 | 0.2785 |
| GPR111 | NM_153839 | -0.0425 | -0.363 |
| GPR110 | NM_153840 | 0.3135 | -0.1005 |
| CSF3R | NM_156039 | 0.041 | 0.0825 |
| RGS20 | NM_170587 | -0.1465 | -2.1375 |
| CASC5 | NM_170589 | -0.5275 | -2.5045 |
| SH2D3C | NM_170600 | -0.452 | -1.414 |
| SIAE | NM_170601 | -0.0585 | -0.3865 |
| RASGRP4 | NM_170604 | -0.2685 | -0.173 |
| MLL3 | NM_170606 | -0.6775 | -2.1975 |
| MLX | NM_170607 | -0.974 | -2.578 |
| CRISP1 | NM_170609 | -0.225 | -0.141 |
| HIST1H2BA | NM_170610 | -0.031 | 0.0495 |
| CBLB | NM_170662 | -1.046 | -2.7255 |
| ATP2A2 | NM_170665 | -0.248 | -1.0545 |
| RASGRP3 | NM_170672 | -0.1455 | 0.0325 |
| MEIS2 | NM_170674 | -0.9845 | -2.763 |
| MEIS2 | NM_170675 | -0.9845 | -2.763 |
| MEIS2 | NM_170676 | -0.9845 | -2.763 |
| MEIS2 | NM_170677 | -1.661 | -2.875 |
| ITGB1BP3 | NM_170678 | 0.405 | -0.4685 |
| SKP1 | NM_170679 | -0.2765 | -0.1745 |
| GFM2 | NM_170681 | -1.157 | -1.939 |
| P2RX2 | NM_170682 | 0.1135 | -0.3945 |
| P2RX2 | NM_170683 | 0.1135 | -0.3945 |
| TAC4 | NM_170685 | -0.119 | -0.192 |
| ZNF398 | NM_170686 | -1.4475 | -2.925 |
| GFM2 | NM_170691 | -1.187 | -1.7915 |
| RASAL2 | NM_170692 | -0.3805 | -1.5385 |
| SGK2 | NM_170693 | -0.0075 | 0.1375 |
| TGIF1 | NM_170695 | -0.406 | -1.9425 |
| ALDH1A2 | NM_170696 | -0.0655 | 0.0445 |
| ALDH1A2 | NM_170697 | -0.0655 | 0.0445 |
| NMNAT2 | NM_170706 | -0.3125 | -0.6265 |
| LMNA | NM_170707 | 0.481 | 0.1685 |
| LMNA | NM_170708 | 0.481 | 0.1685 |
| WDR17 | NM_170710 | -1.221 | -0.6095 |
| DAZAP1 | NM_170711 | -0.089 | -0.763 |
| RASSF1 | NM_170712 | 0.833 | 0.5995 |
| RASSF1 | NM_170713 | 0.833 | 0.5995 |

| | | | |
|---|---|---|---|
| RASSF1 | NM_170714 | -1.0285 | -1.09 |
| PROSER1 | NM_170719 | -1.4335 | -2.4995 |
| KCNJ14 | NM_170720 | -0.9265 | -0.844 |
| MSI2 | NM_170721 | 0.472 | -0.3675 |
| NLRX1 | NM_170722 | -0.358 | -1.058 |
| PKHD1 | NM_170724 | 0.0635 | -0.0495 |
| PGBD2 | NM_170725 | -0.836 | -2.242 |
| ALDH4A1 | NM_170726 | -0.1265 | -0.537 |
| BDNF | NM_170731 | -0.605 | -1.907 |
| BDNF | NM_170732 | -0.605 | -1.907 |
| BDNF | NM_170733 | -0.605 | -1.907 |
| BDNF | NM_170734 | -0.605 | -1.907 |
| BDNF | NM_170735 | -0.605 | -1.907 |
| KCNJ15 | NM_170736 | 0.3025 | -0.278 |
| KCNJ15 | NM_170737 | -0.0605 | 0.296 |
| MRPL11 | NM_170738 | 0.2155 | -0.4795 |
| MRPL11 | NM_170739 | 0.2155 | -0.4795 |
| ALDH5A1 | NM_170740 | -0.6095 | -0.693 |
| KCNJ16 | NM_170741 | -0.2105 | -0.1965 |
| KCNJ16 | NM_170742 | -0.2105 | -0.1965 |
| IL28RA | NM_170743 | -0.0755 | -0.352 |
| UNC5B | NM_170744 | -0.3675 | -1.5 |
| HIST1H2AA | NM_170745 | -0.177 | -0.3745 |
| C11orf31 | NM_170746 | 0.1785 | -0.293 |
| PSMD10 | NM_170750 | -0.738 | -0.97 |
| PGBD3 | NM_170753 | -0.842 | -2.02 |
| TENC1 | NM_170754 | -0.3235 | -0.4435 |
| RNF39 | NM_170769 | -0.135 | -0.249 |
| ALDH8A1 | NM_170771 | -0.583 | -0.1985 |
| RASSF2 | NM_170774 | -1.276 | -1.8795 |
| KCNN2 | NM_170775 | 0.2205 | -1.617 |
| GPR97 | NM_170776 | -0.1745 | -0.4515 |
| CD200R1 | NM_170780 | 0.072 | -0.447 |
| KCNN3 | NM_170782 | -0.15 | -0.6095 |
| ZNRD1 | NM_170783 | -0.816 | -2.6705 |
| MKKS | NM_170784 | -0.442 | -1.5535 |
| CAMK2A | NM_171825 | -0.1005 | -0.2155 |
| CD8A | NM_171827 | 0.0545 | -0.1035 |
| KCNMB3 | NM_171828 | -0.5575 | -0.341 |
| KCNMB3 | NM_171829 | -0.5575 | -0.341 |
| KCNMB3 | NM_171830 | -0.4755 | -0.5265 |
| LACTB | NM_171846 | -1.109 | -1.7855 |
| TRIM35 | NM_171982 | -0.6625 | -3.1025 |

| | | | |
|---|---|---|---|
| USP2 | NM_171997 | 0.16 | 0.0275 |
| RAB39B | NM_171998 | -1.6135 | -2.915 |
| SALL3 | NM_171999 | -0.0225 | -0.135 |
| TEDDM1 | NM_172000 | 0.0885 | -0.164 |
| HSCB | NM_172002 | -0.721 | -1.409 |
| CLECL1 | NM_172004 | 0.6255 | -0.029 |
| WFDC13 | NM_172005 | -0.2725 | -0.0215 |
| WFDC10B | NM_172006 | 0.069 | -0.2815 |
| TNFSF14 | NM_172014 | -0.014 | -0.147 |
| TRIM39 | NM_172016 | -0.8185 | -1.834 |
| POM121 | NM_172020 | -0.8535 | -2.4815 |
| ABTB1 | NM_172027 | 0.3145 | -0.5195 |
| RDH10 | NM_172037 | -1.3045 | -1.7605 |
| KCNH2 | NM_172056 | 0.0625 | -0.671 |
| KCNH2 | NM_172057 | -0.4365 | -1.796 |
| EYA1 | NM_172058 | -1.4595 | -2.6105 |
| EYA1 | NM_172059 | -1.4595 | -2.6105 |
| EYA1 | NM_172060 | -1.833 | -3.9325 |
| PLEKHH2 | NM_172069 | 0.0435 | -1.1245 |
| UBR3 | NM_172070 | -0.376 | -1.757 |
| RC3H1 | NM_172071 | -1.4445 | -2.9735 |
| CAMK2B | NM_172078 | -0.105 | -0.256 |
| CAMK2B | NM_172079 | -0.105 | -0.256 |
| CAMK2B | NM_172080 | -0.105 | -0.256 |
| CAMK2B | NM_172081 | -0.105 | -0.256 |
| CAMK2B | NM_172082 | -0.105 | -0.256 |
| CAMK2B | NM_172083 | -0.105 | -0.256 |
| CAMK2B | NM_172084 | -0.105 | -0.256 |
| TNFSF13 | NM_172087 | -0.22 | -0.435 |
| CATSPER2 | NM_172095 | -0.5565 | -0.5735 |
| CATSPER2 | NM_172097 | -1.306 | -0.9335 |
| CD8B | NM_172101 | 0.0235 | 0.1305 |
| CD8B | NM_172102 | 0.0235 | 0.1305 |
| EYA4 | NM_172103 | -0.5495 | -1.353 |
| EYA4 | NM_172105 | -0.5495 | -1.353 |
| KCNQ2 | NM_172106 | -1.0055 | -3.149 |
| KCNQ2 | NM_172107 | -1.0055 | -3.149 |
| KCNQ2 | NM_172108 | -1.0055 | -3.149 |
| KCNQ2 | NM_172109 | 0.1585 | -0.3545 |
| EYA2 | NM_172110 | -0.37 | -1.9865 |
| CAMK2D | NM_172114 | -0.8975 | -1.093 |
| CAMK2D | NM_172115 | -0.8975 | -1.093 |
| CAMK2D | NM_172127 | -0.8975 | -1.093 |

| | | | |
|---|---|---|---|
| CAMK2D | NM_172128 | -0.8975 | -1.093 |
| CAMK2D | NM_172129 | -0.8975 | -1.093 |
| KCNAB2 | NM_172130 | -0.5345 | -0.7975 |
| WFDC10B | NM_172131 | 0.069 | -0.2815 |
| IL29 | NM_172140 | -0.351 | -0.181 |
| KCNAB1 | NM_172159 | -1.2125 | -1.0695 |
| KCNAB1 | NM_172160 | -1.2125 | -1.0695 |
| KCNQ4 | NM_172163 | -0.6435 | -1.578 |
| MSH5 | NM_172165 | -0.018 | -0.4245 |
| MSH5 | NM_172166 | -0.018 | -0.4245 |
| NOXO1 | NM_172168 | -0.083 | -0.334 |
| CAMK2G | NM_172169 | -0.476 | -0.1155 |
| CAMK2G | NM_172170 | -0.476 | -0.1155 |
| CAMK2G | NM_172171 | -0.302 | -0.896 |
| CAMK2G | NM_172173 | -0.476 | -0.1155 |
| IL15 | NM_172175 | 0.0515 | -0.575 |
| MRPL42 | NM_172177 | -0.924 | -1.098 |
| KLHDC1 | NM_172193 | -0.2155 | -1.0205 |
| OR4Q3 | NM_172194 | -0.168 | -0.496 |
| EIF2B4 | NM_172195 | -0.8805 | -2.379 |
| AGER | NM_172197 | -0.0285 | -0.2605 |
| KCND3 | NM_172198 | 0.1635 | 0.0225 |
| IL15RA | NM_172200 | -0.133 | -0.2235 |
| KCNE2 | NM_172201 | -0.225 | -0.3175 |
| CAMKK1 | NM_172207 | -0.0025 | -0.3055 |
| TAPBP | NM_172208 | -0.0195 | -0.866 |
| TAPBP | NM_172209 | -0.0195 | -0.866 |
| CSF1 | NM_172210 | -0.519 | -0.253 |
| CSF1 | NM_172212 | -0.0195 | -0.1285 |
| CD8B | NM_172213 | 0.0235 | 0.1305 |
| CAMKK2 | NM_172214 | -0.5185 | -0.88 |
| CAMKK2 | NM_172215 | -0.5185 | -0.88 |
| CAMKK2 | NM_172216 | -0.5185 | -0.88 |
| IL16 | NM_172217 | -0.113 | -0.7195 |
| SPAG1 | NM_172218 | -0.8815 | -0.9295 |
| CSF3 | NM_172219 | 0.093 | -0.3545 |
| CSF3 | NM_172220 | 0.093 | -0.3545 |
| CAMKK2 | NM_172226 | -0.5185 | -0.88 |
| KREMEN2 | NM_172229 | -0.038 | -0.6545 |
| SYVN1 | NM_172230 | -0.704 | -1.7565 |
| SUGP1 | NM_172231 | -0.665 | -1.796 |
| ABCA5 | NM_172232 | -0.3855 | -0.867 |
| POFUT1 | NM_172236 | -0.758 | -1.8095 |

| | | | |
|---|---|---|---|
| TFAP2D | NM_172238 | 0.3005 | -0.081 |
| REXO1L1 | NM_172239 | -0.166 | -0.159 |
| POC1B | NM_172240 | -1.0405 | -1.3155 |
| CTAGE1 | NM_172241 | -0.195 | -0.6995 |
| SPAG6 | NM_172242 | 0.123 | -0.0355 |
| SGCD | NM_172244 | 0.104 | -0.1815 |
| CSF2RA | NM_172245 | -0.4185 | -0.0625 |
| CSF2RA | NM_172246 | -0.4185 | -0.0625 |
| CSF2RA | NM_172247 | -0.4185 | -0.0625 |
| CSF2RA | NM_172249 | -0.4185 | -0.0625 |
| MMAA | NM_172250 | -0.841 | -1.2445 |
| MRPL54 | NM_172251 | 0.1245 | -0.4415 |
| SPAG8 | NM_172312 | -0.1515 | -0.1645 |
| CSF3R | NM_172313 | 0.177 | -0.543 |
| IL25 | NM_172314 | -0.1475 | -0.263 |
| MEIS2 | NM_172315 | -0.9845 | -2.763 |
| MEIS2 | NM_172316 | -0.9845 | -2.763 |
| PSENEN | NM_172341 | 0.061 | -0.6175 |
| KCNG3 | NM_172344 | -0.56 | -0.5145 |
| KCNG4 | NM_172347 | 0.069 | -0.0735 |
| NSD1 | NM_172349 | 0.1895 | 1.22 |
| CD46 | NM_172350 | -1.213 | -1.466 |
| CD46 | NM_172351 | -1.213 | -1.466 |
| CD46 | NM_172352 | -1.213 | -1.466 |
| CD46 | NM_172353 | -1.213 | -1.466 |
| CD46 | NM_172359 | -1.213 | -1.466 |
| CD46 | NM_172361 | -1.213 | -1.466 |
| KCNH1 | NM_172362 | -1.1505 | -1.795 |
| CACNA2D4 | NM_172364 | -0.176 | -0.4355 |
| PPP1R36 | NM_172365 | -0.2705 | -0.4005 |
| FBXO16 | NM_172366 | -1.134 | -1.5225 |
| TUSC5 | NM_172367 | -0.06 | -0.25 |
| C1QC | NM_172369 | -0.0585 | -0.569 |
| DAOA | NM_172370 | -0.1605 | -0.088 |
| ELF1 | NM_172373 | -0.088 | -1.1125 |
| IL4I1 | NM_172374 | -0.0955 | -0.233 |
| KCNH5 | NM_172375 | 0.288 | -0.0545 |
| KCNH5 | NM_172376 | 0.288 | -0.0545 |
| CTAG2 | NM_172377 | 0.2195 | -0.398 |
| NFATC1 | NM_172387 | 0.305 | 0.07 |
| NFATC1 | NM_172388 | 0.305 | 0.07 |
| NFATC1 | NM_172389 | 0.305 | 0.07 |
| NFATC1 | NM_172390 | 0.305 | 0.07 |

| | | | |
|---|---|---|---|
| AQP11 | NM_173039 | 0.039 | -0.332 |
| IL18BP | NM_173042 | -0.38 | -0.2835 |
| IL18BP | NM_173044 | -0.38 | -0.2835 |
| SCUBE1 | NM_173050 | -0.2945 | -0.341 |
| RELN | NM_173054 | -0.3295 | -0.663 |
| CAST | NM_173060 | -0.609 | -1.1605 |
| SLC35C2 | NM_173073 | -0.676 | -1.7595 |
| PIGF | NM_173074 | -0.884 | -0.907 |
| APBB2 | NM_173075 | -0.1085 | -0.418 |
| ABCA12 | NM_173076 | -0.052 | -0.2855 |
| CPO | NM_173077 | 0.0005 | -0.114 |
| SLITRK4 | NM_173078 | -0.1695 | -0.2385 |
| RUNDC1 | NM_173079 | -1.5135 | -3.4575 |
| SPRR4 | NM_173080 | 0.0945 | -0.2285 |
| ARMC3 | NM_173081 | -0.5805 | -0.2915 |
| SHPRH | NM_173082 | -0.2245 | -1.3585 |
| LIN9 | NM_173083 | -1.7325 | -4.067 |
| TRIM59 | NM_173084 | -0.8925 | -1.9285 |
| KRT6C | NM_173086 | -0.007 | -0.2735 |
| CAPN3 | NM_173087 | -0.009 | -0.367 |
| CAPN3 | NM_173088 | 0.3355 | -0.179 |
| CAPN3 | NM_173089 | -0.009 | -0.367 |
| CAPN3 | NM_173090 | -0.009 | -0.367 |
| NFATC2 | NM_173091 | -0.2475 | -0.6905 |
| KCNH6 | NM_173092 | 0.139 | -0.513 |
| SMG7 | NM_173156 | -0.58 | -1.4845 |
| NR4A1 | NM_173157 | -0.6525 | -1.619 |
| NPAS3 | NM_173159 | 0.0345 | -0.345 |
| FXYD4 | NM_173160 | -0.0385 | -0.202 |
| IL1F10 | NM_173161 | -0.0685 | 0.4065 |
| KCNH7 | NM_173162 | 0.031 | -0.6495 |
| NFATC3 | NM_173163 | -1.19 | -2.157 |
| NFATC3 | NM_173165 | -1.173 | -2.1255 |
| UNC45B | NM_173167 | 0.1015 | 0.0325 |
| IL36RN | NM_173170 | -0.109 | -0.2885 |
| PTK2B | NM_173174 | 0.259 | -0.493 |
| PTK2B | NM_173175 | -0.212 | -0.2605 |
| PTK2B | NM_173176 | -0.212 | -0.2605 |
| C1D | NM_173177 | -0.698 | -1.6015 |
| SLC35C2 | NM_173179 | -0.676 | -1.7595 |
| KCNIP2 | NM_173191 | 0.244 | -0.242 |
| KCNIP2 | NM_173192 | 0.244 | -0.242 |
| KCNIP2 | NM_173193 | 0.244 | -0.242 |

| | | | |
|---|---|---|---|
| KCNIP2 | NM_173194 | 0.244 | -0.242 |
| KCNIP2 | NM_173195 | 0.244 | -0.242 |
| KCNIP2 | NM_173197 | 0.244 | -0.242 |
| NR4A3 | NM_173199 | 0.4705 | -0.083 |
| NR4A3 | NM_173200 | -1.053 | -0.0425 |
| ATP2A1 | NM_173201 | 0.092 | -0.759 |
| PIAS2 | NM_173206 | -0.267 | -1.0175 |
| TGIF1 | NM_173207 | -0.406 | -1.9425 |
| TGIF1 | NM_173208 | -0.406 | -1.9425 |
| TGIF1 | NM_173209 | -0.406 | -1.9425 |
| TGIF1 | NM_173210 | -0.406 | -1.9425 |
| TGIF1 | NM_173211 | -0.406 | -1.9425 |
| NFAT5 | NM_173214 | -1.0085 | -2.852 |
| NFAT5 | NM_173215 | -1.0085 | -2.852 |
| ST6GAL1 | NM_173216 | -0.3555 | -1.579 |
| ST6GAL1 | NM_173217 | -0.3555 | -1.579 |
| ST3GAL1 | NM_173344 | -1.17 | -1.904 |
| FAM149B1 | NM_173348 | -0.6025 | -0.821 |
| OR6B3 | NM_173351 | 0.3985 | -0.077 |
| KRT78 | NM_173352 | 0.191 | -0.228 |
| TPH2 | NM_173353 | -0.002 | 0.1975 |
| SIK1 | NM_173354 | -0.302 | 0.378 |
| UPP2 | NM_173355 | 0.0085 | -0.036 |
| SSX7 | NM_173358 | 0.149 | -0.639 |
| EIF4E3 | NM_173359 | -0.939 | -0.9835 |
| RFESD | NM_173362 | -1.7005 | -2.2875 |
| PDE8A | NM_173454 | -0.804 | -2.083 |
| PAPLN | NM_173462 | -0.032 | -0.193 |
| CCDC149 | NM_173463 | -0.183 | -0.479 |
| L3MBTL4 | NM_173464 | 0.029 | -0.208 |
| COL23A1 | NM_173465 | 0.087 | -0.383 |
| MCAT | NM_173467 | -0.875 | -2.853 |
| MOB1B | NM_173468 | -1.412 | -2.409 |
| UBE2Q2 | NM_173469 | -0.8435 | -1.4045 |
| MMGT1 | NM_173470 | -0.7715 | -1.0585 |
| SLC25A26 | NM_173471 | -0.112 | -1.016 |
| C3orf24 | NM_173472 | -0.4145 | -0.0465 |
| ANAPC16 | NM_173473 | -0.5375 | -0.5575 |
| NTAN1 | NM_173474 | 0.0925 | -0.6495 |
| DCUN1D3 | NM_173475 | -0.584 | -1.415 |
| USH1G | NM_173477 | -0.111 | -0.2065 |
| CNTD1 | NM_173478 | -0.007 | -0.4605 |
| WDR88 | NM_173479 | 0.0915 | -0.352 |

| | | | |
|---|---|---|---|
| ZNF57 | NM_173480 | -0.9345 | -1.9025 |
| C19orf21 | NM_173481 | 0.255 | -0.571 |
| CCDC105 | NM_173482 | -0.0905 | -0.2085 |
| CYP4F22 | NM_173483 | 0.3035 | -0.3625 |
| KLF17 | NM_173484 | -0.0335 | 0.116 |
| TSHZ2 | NM_173485 | -0.311 | -0.505 |
| C4orf33 | NM_173487 | -0.813 | -1.0815 |
| SLCO6A1 | NM_173488 | -0.149 | -0.184 |
| HEATR7B2 | NM_173489 | 0.0925 | -0.303 |
| TMEM171 | NM_173490 | -0.0565 | -0.502 |
| LSM11 | NM_173491 | -0.1505 | -1.437 |
| PIP5KL1 | NM_173492 | -0.1435 | -0.3725 |
| PASD1 | NM_173493 | -0.5045 | -0.622 |
| CXorf41 | NM_173494 | -0.192 | -0.3135 |
| PTCHD1 | NM_173495 | 0.26 | -0.484 |
| MPP7 | NM_173496 | -0.0765 | -0.0475 |
| HECTD2 | NM_173497 | -0.1785 | 1.6195 |
| SPATA8 | NM_173499 | -0.0935 | -0.476 |
| TTBK2 | NM_173500 | -0.144 | -1.0095 |
| PRSS36 | NM_173502 | -0.3695 | 0.061 |
| EFCAB3 | NM_173503 | -0.874 | -0.955 |
| ANKRD29 | NM_173505 | -0.4295 | -0.217 |
| LYPD4 | NM_173506 | -0.106 | -0.2965 |
| SLC35F3 | NM_173508 | 0.2865 | -0.4195 |
| FAM163A | NM_173509 | -0.886 | -2.2625 |
| CCDC117 | NM_173510 | -1.8975 | -3.331 |
| FAM117B | NM_173511 | -1.0305 | -2.017 |
| SLC38A11 | NM_173512 | 0.096 | -0.3225 |
| SLC38A9 | NM_173514 | -2.0145 | -3.008 |
| CNKSR3 | NM_173515 | -0.794 | -2.92 |
| PNLDC1 | NM_173516 | 0.0235 | -0.579 |
| VKORC1L1 | NM_173517 | -1.8265 | -2.5735 |
| C8orf45 | NM_173518 | 0.395 | -0.1675 |
| CLVS1 | NM_173519 | -0.9575 | -1.8755 |
| C9orf84 | NM_173521 | 0.2575 | -0.4255 |
| MAGEB6 | NM_173523 | 0.082 | 0.5325 |
| C11orf42 | NM_173525 | 0.1375 | 0.0695 |
| FAM71D | NM_173526 | -0.086 | -0.8375 |
| REM2 | NM_173527 | 0.0885 | -0.126 |
| C15orf26 | NM_173528 | -0.3325 | -0.1335 |
| C18orf54 | NM_173529 | -0.9345 | -2.3105 |
| ZNF610 | NM_173530 | -0.783 | -2.4 |
| ZNF100 | NM_173531 | -1.052 | -1.887 |

| | | | |
|---|---|---|---|
| TDRD5 | NM_173533 | -0.375 | -0.2865 |
| CLEC4F | NM_173535 | -0.002 | -0.106 |
| GABRG1 | NM_173536 | 0.182 | -0.0345 |
| GTF2IRD2 | NM_173537 | -1.1395 | -1.0935 |
| CNBD1 | NM_173538 | -0.156 | -0.1075 |
| ZNF596 | NM_173539 | -0.0975 | -1.826 |
| FUT11 | NM_173540 | -0.346 | -0.2505 |
| C10orf91 | NM_173541 | 0.2745 | -0.1695 |
| PLBD2 | NM_173542 | 0.132 | -0.3055 |
| DZIP1L | NM_173543 | -0.59 | -1.1805 |
| FAM129C | NM_173544 | 0.0795 | -0.346 |
| APLF | NM_173545 | -1.164 | -0.9975 |
| KLHDC8B | NM_173546 | -1.132 | -2.266 |
| TRIM65 | NM_173547 | -1.1695 | -1.973 |
| ZNF584 | NM_173548 | -0.74 | -2.262 |
| C8orf47 | NM_173549 | 0.225 | -0.2335 |
| C9orf93 | NM_173550 | -0.9695 | -0.1165 |
| ANKS6 | NM_173551 | -0.31 | -1.0525 |
| C3orf58 | NM_173552 | -0.6545 | -0.446 |
| TRIML2 | NM_173553 | 0.1065 | -0.2865 |
| C10orf107 | NM_173554 | 0.032 | -0.213 |
| TYSND1 | NM_173555 | -0.482 | -1.451 |
| CCDC83 | NM_173556 | -0.1445 | 0.0895 |
| RNF152 | NM_173557 | 0.0195 | 0.035 |
| FGD2 | NM_173558 | 0.0445 | -0.206 |
| RFX6 | NM_173560 | -0.598 | -0.338 |
| UNC5CL | NM_173561 | -0.297 | -0.311 |
| KCTD20 | NM_173562 | -1.6555 | -0.939 |
| C6orf146 | NM_173563 | -0.155 | -0.254 |
| C7orf51 | NM_173564 | -0.335 | -1.1985 |
| RSPH10B | NM_173565 | -0.057 | -0.214 |
| PRR14L | NM_173566 | -1.382 | -2.8885 |
| EPHX4 | NM_173567 | -0.867 | -0.2535 |
| UMODL1 | NM_173568 | 0.2685 | -0.2605 |
| UBN2 | NM_173569 | -1.128 | -2.1135 |
| ZDHHC23 | NM_173570 | -0.01 | -0.977 |
| C10orf93 | NM_173572 | -0.3 | -0.1385 |
| C11orf35 | NM_173573 | 0.2 | -0.2105 |
| ZNF683 | NM_173574 | 0.188 | 0.039 |
| STK32C | NM_173575 | 0.0485 | -0.6085 |
| MKX | NM_173576 | 0.4215 | -0.2645 |
| PGM2L1 | NM_173582 | -0.564 | -0.9765 |
| EFCAB4A | NM_173584 | 0.1805 | -0.3795 |

| | | | |
|---|---|---|---|
| RCOR2 | NM_173587 | -0.354 | -1.2415 |
| IGSF22 | NM_173588 | 0.033 | 0.065 |
| DNHD1 | NM_173589 | -0.194 | -0.4025 |
| B4GALNT3 | NM_173593 | -0.0695 | -0.5745 |
| ANKRD52 | NM_173595 | -0.3205 | -1.0665 |
| KSR2 | NM_173598 | -0.1545 | -0.387 |
| GXYLT1 | NM_173601 | -0.238 | -1.7845 |
| DIP2B | NM_173602 | -1.198 | -2.1555 |
| KCNRG | NM_173605 | -0.711 | -0.512 |
| FAM177A1 | NM_173607 | -1.005 | -1.0975 |
| FAM98B | NM_173611 | -0.579 | -0.465 |
| VWA3A | NM_173615 | -0.135 | -0.2145 |
| INO80E | NM_173618 | -0.5945 | -1.1915 |
| HEXDC | NM_173620 | -0.425 | -0.722 |
| C17orf78 | NM_173625 | 0.231 | -0.0535 |
| SLC26A11 | NM_173626 | -0.6935 | -1.224 |
| ENDOV | NM_173627 | -0.125 | -0.823 |
| C18orf26 | NM_173629 | 0.2525 | -0.1375 |
| RTTN | NM_173630 | -1.3205 | -2.487 |
| ZNF547 | NM_173631 | -0.8075 | -1.978 |
| ZNF776 | NM_173632 | -1.005 | -2.581 |
| TMEM145 | NM_173633 | -0.2675 | -0.452 |
| C19orf75 | NM_173635 | -0.3975 | -0.43 |
| WDR62 | NM_173636 | -0.2145 | -0.33 |
| SLC25A41 | NM_173637 | -0.008 | -0.023 |
| EPHA10 | NM_173641 | -0.1765 | -0.426 |
| RIMKLA | NM_173642 | -0.208 | -1.2745 |
| C20orf197 | NM_173644 | -0.0185 | 0.2575 |
| RNF149 | NM_173647 | -1.1295 | -3.569 |
| CCDC141 | NM_173648 | -0.041 | -0.4445 |
| C2orf61 | NM_173649 | -0.086 | -0.235 |
| DNAJC5G | NM_173650 | 0.0215 | -0.55 |
| SLC9A9 | NM_173653 | -0.3445 | -0.3495 |
| C3orf64 | NM_173654 | -1.197 | -1.838 |
| EPHA6 | NM_173655 | 0.1685 | -0.066 |
| C3orf33 | NM_173657 | -0.4295 | -1.345 |
| ZNF660 | NM_173658 | -0.862 | -1.4265 |
| DOK7 | NM_173660 | -0.1075 | -0.3615 |
| RNF175 | NM_173662 | -0.05 | -0.3475 |
| FAM153A | NM_173663 | -0.1485 | 0.013 |
| ARL10 | NM_173664 | -0.3725 | -0.998 |
| DTWD2 | NM_173666 | -0.9175 | -1.021 |
| C5orf64 | NM_173667 | -0.1865 | 0.1035 |

| | | | |
|---|---|---|---|
| PPIL6 | NM_173672 | -0.089 | 0.242 |
| DCBLD1 | NM_173674 | -0.505 | -0.5045 |
| PNPLA1 | NM_173676 | -0.1375 | -0.1595 |
| FAM115C | NM_173678 | 0.0335 | -0.7205 |
| ZNF775 | NM_173680 | -1.038 | -2.287 |
| ATG9B | NM_173681 | -0.126 | -0.716 |
| XKR6 | NM_173683 | -0.313 | -0.7505 |
| NSMCE2 | NM_173685 | -0.85 | -1.3415 |
| C8orf58 | NM_173686 | -0.011 | -0.624 |
| C8orf31 | NM_173687 | -0.1575 | -0.7865 |
| NKAIN3 | NM_173688 | 0.332 | -0.428 |
| CRB2 | NM_173689 | -0.246 | 0.0855 |
| SCAI | NM_173690 | -1.277 | -1.986 |
| ATP11C | NM_173694 | -0.333 | -0.9285 |
| CXorf59 | NM_173695 | -0.039 | -0.3395 |
| FAM133A | NM_173698 | -0.052 | -0.2075 |
| MAGEB18 | NM_173699 | -0.029 | -0.127 |
| WARS | NM_173701 | -0.249 | -0.9745 |
| ELAC2 | NM_173717 | -0.345 | -1.142 |
| ARHGEF15 | NM_173728 | -0.086 | -0.205 |
| PDZD8 | NM_173791 | -0.5125 | -1.0125 |
| C22orf39 | NM_173793 | 0.1825 | -0.918 |
| FUNDC1 | NM_173794 | -1.2805 | -1.9015 |
| PAPD4 | NM_173797 | -1.2115 | -2.8295 |
| ZCCHC12 | NM_173798 | -0.2585 | 0.055 |
| TIGIT | NM_173799 | 0.045 | -0.2535 |
| AQPEP | NM_173800 | -0.3405 | -0.1695 |
| PLAC1L | NM_173801 | 0.0565 | -0.102 |
| MPV17L | NM_173803 | 0.1715 | -1.206 |
| TMEM86B | NM_173804 | -0.113 | -0.5565 |
| PDZD9 | NM_173806 | -0.151 | -0.2455 |
| C1orf101 | NM_173807 | -0.6315 | -0.537 |
| NEGR1 | NM_173808 | -0.7135 | -0.265 |
| BLOC1S2 | NM_173809 | -1.1145 | -0.969 |
| TTC9C | NM_173810 | -0.859 | -2.7455 |
| HARBI1 | NM_173811 | -0.5065 | -1.347 |
| DPY19L2 | NM_173812 | -0.921 | -1.772 |
| PRTG | NM_173814 | -1.333 | -2.1515 |
| CES4A | NM_173815 | -0.0045 | -0.428 |
| C2orf85 | NM_173821 | -0.2135 | -0.406 |
| FAM126B | NM_173822 | -1.4235 | -2.9145 |
| C3orf38 | NM_173824 | -1.477 | -2.617 |
| RABL3 | NM_173825 | -1.1525 | -1.545 |

| | | | |
|---|---|---|---|
| C3orf23 | NM_173826 | -1.564 | -3.481 |
| COX18 | NM_173827 | -1.0605 | -2.194 |
| RELL2 | NM_173828 | -0.307 | -0.804 |
| SREK1IP1 | NM_173829 | -0.4075 | -1.191 |
| CEP57L1 | NM_173830 | -1.0785 | -1.9845 |
| ZNF707 | NM_173831 | -0.0455 | -0.405 |
| ZFP41 | NM_173832 | -0.983 | -1.308 |
| SCARA5 | NM_173833 | 0.1055 | -0.33 |
| YIPF6 | NM_173834 | -1.451 | -1.335 |
| IL1RN | NM_173841 | -0.094 | -0.1175 |
| IL1RN | NM_173842 | -0.094 | -0.1175 |
| IL1RN | NM_173843 | -0.094 | -0.1175 |
| MALT1 | NM_173844 | -0.4465 | -1.1365 |
| TPPP2 | NM_173846 | 0.142 | -0.1625 |
| SPACA3 | NM_173847 | -0.0525 | 0.265 |
| RALYL | NM_173848 | 0.284 | -1.265 |
| GSC | NM_173849 | 0.2395 | -1.115 |
| SERPINA12 | NM_173850 | -0.0575 | -0.1595 |
| SLC30A8 | NM_173851 | -0.0215 | 0.104 |
| KRTCAP2 | NM_173852 | -0.0765 | -0.4795 |
| KRTCAP3 | NM_173853 | 0.1265 | -0.266 |
| SLC41A1 | NM_173854 | -0.669 | -2.082 |
| MORN3 | NM_173855 | 0.2945 | -0.237 |
| VN1R2 | NM_173856 | -0.2095 | 0.0455 |
| VN1R4 | NM_173857 | -0.161 | -0.362 |
| VN1R5 | NM_173858 | -0.0435 | -0.2075 |
| HOXC12 | NM_173860 | 0.221 | -0.1035 |
| CLCN3 | NM_173872 | -1.6635 | -2.8755 |
| IDH3B | NM_174855 | 0.021 | -0.479 |
| IDH3B | NM_174856 | 0.021 | -0.479 |
| AK5 | NM_174858 | -0.6805 | -1.1025 |
| IDH3G | NM_174869 | 0.187 | -0.2315 |
| PSMD12 | NM_174871 | -0.8505 | -2.519 |
| P2RX2 | NM_174872 | 0.1135 | -0.3945 |
| P2RX2 | NM_174873 | 0.1135 | -0.3945 |
| CLRN1 | NM_174878 | -0.0205 | -0.34 |
| CRB3 | NM_174881 | 0.0825 | -0.092 |
| TGIF1 | NM_174886 | -0.406 | -1.9425 |
| IFT20 | NM_174887 | -0.5875 | -0.9905 |
| NDUFAF2 | NM_174889 | -0.1465 | -1.096 |
| ANUBL1 | NM_174890 | -0.8995 | -2.39 |
| C14orf79 | NM_174891 | -0.248 | -0.547 |
| CD300LB | NM_174892 | 0.3705 | -0.258 |

| | | | |
|---|---|---|---|
| TRAPPC5 | NM_174894 | -0.4215 | -0.8795 |
| PCP2 | NM_174895 | 0.0765 | 0.1475 |
| C1orf162 | NM_174896 | -1.014 | -1.0885 |
| BPIFB6 | NM_174897 | 0.1535 | -0.096 |
| LYG1 | NM_174898 | 0.088 | -0.75 |
| FBXO36 | NM_174899 | -0.7875 | -1.415 |
| ZFP42 | NM_174900 | -0.603 | -0.3845 |
| FAM9C | NM_174901 | -0.2525 | -0.388 |
| LDLRAD3 | NM_174902 | 0.1865 | -0.547 |
| RNF151 | NM_174903 | 0.1155 | -0.503 |
| FAM98C | NM_174905 | 0.0045 | -0.5335 |
| PPP4R2 | NM_174907 | -1.485 | -3.5565 |
| CCDC50 | NM_174908 | -0.824 | -1.356 |
| TMEM167A | NM_174909 | -0.49 | -0.716 |
| TCTE3 | NM_174910 | -0.06 | -0.0115 |
| FAM84B | NM_174911 | -0.049 | -0.478 |
| FAAH2 | NM_174912 | 0.143 | -0.4335 |
| C14orf21 | NM_174913 | -0.346 | -1.064 |
| UGT3A2 | NM_174914 | -0.377 | -0.1205 |
| UBR1 | NM_174916 | -1.7675 | -3.656 |
| ACSF3 | NM_174917 | -0.204 | -0.9045 |
| C19orf59 | NM_174918 | -0.152 | -0.092 |
| ARHGAP27 | NM_174919 | -0.1505 | -0.415 |
| SAMD14 | NM_174920 | -0.193 | -1.0025 |
| C4orf34 | NM_174921 | -1.2655 | -1.052 |
| ADCK5 | NM_174922 | 0.0975 | -0.329 |
| CCDC107 | NM_174923 | -0.4295 | -0.869 |
| PDILT | NM_174924 | 0.0565 | -0.074 |
| TMEM136 | NM_174926 | -0.1555 | -0.9635 |
| SPATA19 | NM_174927 | 0.1815 | -0.2755 |
| N6AMT2 | NM_174928 | -1.043 | -1.466 |
| ZMIZ2 | NM_174929 | -0.078 | -0.729 |
| CCDC75 | NM_174931 | -1.525 | -2.4435 |
| BPIFC | NM_174932 | 0.2315 | -0.0335 |
| PHYHD1 | NM_174933 | 0.0375 | -0.1295 |
| SCN4B | NM_174934 | 0.382 | -0.2585 |
| PCSK9 | NM_174936 | -0.032 | -0.447 |
| TCERG1L | NM_174937 | 0.378 | 0.0165 |
| FRMD3 | NM_174938 | -0.405 | -1.2715 |
| TMEM80 | NM_174940 | -0.263 | -0.609 |
| CD163L1 | NM_174941 | -0.0125 | -0.805 |
| GAS2L3 | NM_174942 | -0.5565 | -1.166 |
| C14orf178 | NM_174943 | 0.063 | -0.36 |

| | | | |
|---|---|---|---|
| TSSK4 | NM_174944 | 0.044 | -0.053 |
| ZNF575 | NM_174945 | -0.127 | -0.3535 |
| FAM9A | NM_174951 | -0.1935 | -0.031 |
| C4orf37 | NM_174952 | -0.203 | -0.335 |
| ATP2A3 | NM_174953 | -0.0895 | -0.635 |
| ATP2A3 | NM_174954 | -0.0895 | -0.635 |
| ATP2A3 | NM_174955 | -0.0895 | -0.635 |
| ATP2A3 | NM_174956 | -0.0895 | -0.635 |
| ATP2A3 | NM_174957 | -0.0895 | -0.635 |
| ATP2A3 | NM_174958 | -0.0895 | -0.635 |
| SVOPL | NM_174959 | -0.282 | -0.3475 |
| ST3GAL3 | NM_174963 | -0.7235 | -0.9275 |
| ST3GAL3 | NM_174964 | -0.7235 | -0.9275 |
| ST3GAL3 | NM_174965 | -0.7235 | -0.9275 |
| ST3GAL3 | NM_174966 | -0.7235 | -0.9275 |
| ST3GAL3 | NM_174967 | -0.7235 | -0.9275 |
| ST3GAL3 | NM_174968 | -0.7235 | -0.9275 |
| ST3GAL3 | NM_174969 | -0.7235 | -0.9275 |
| ST3GAL3 | NM_174970 | -0.7235 | -0.9275 |
| ST3GAL3 | NM_174971 | -0.7235 | -0.9275 |
| SEC14L3 | NM_174975 | 0.086 | -0.1705 |
| ZDHHC22 | NM_174976 | 0.0445 | -0.713 |
| SEC14L4 | NM_174977 | 0.0305 | -0.5145 |
| C14orf39 | NM_174978 | -0.3225 | -0.814 |
| C19orf28 | NM_174983 | -0.02 | -0.6725 |
| CNTN1 | NM_175038 | -0.5615 | -0.6425 |
| ST6GALNAC4 | NM_175039 | -0.361 | -0.8175 |
| ST6GALNAC4 | NM_175040 | -0.361 | -0.8175 |
| ST8SIA4 | NM_175052 | -0.4765 | -2.272 |
| KRT74 | NM_175053 | -0.2695 | -0.3425 |
| HIST4H4 | NM_175054 | 0.2955 | 0.5995 |
| HIST3H2BB | NM_175055 | 0.865 | 0.2825 |
| ZPLD1 | NM_175056 | -0.0915 | -0.3955 |
| TAAR9 | NM_175057 | 0.1765 | -0.2405 |
| PLEKHA7 | NM_175058 | -0.603 | -0.2495 |
| CLEC14A | NM_175060 | -0.168 | -0.3465 |
| JAZF1 | NM_175061 | -1.631 | -2.78 |
| RASGEF1C | NM_175062 | 0.2795 | -0.2155 |
| C19orf63 | NM_175063 | 0.201 | -0.3595 |
| SPDYE1 | NM_175064 | -0.1045 | -1.307 |
| HIST2H2AB | NM_175065 | 0.3 | 0.1995 |
| DDX51 | NM_175066 | -0.2705 | -0.128 |
| TAAR6 | NM_175067 | 0.15 | -0.5585 |

| | | | |
|---|---|---|---|
| KRT73 | NM_175068 | -0.0705 | -0.4665 |
| APTX | NM_175069 | -0.647 | -1.9035 |
| APTX | NM_175073 | -0.647 | -1.9035 |
| C8orf42 | NM_175075 | -0.421 | -0.6815 |
| SLA2 | NM_175077 | 0.1045 | -0.285 |
| KRT77 | NM_175078 | -0.2635 | -0.2875 |
| P2RX5 | NM_175080 | -0.3315 | -1.107 |
| GART | NM_175085 | -0.557 | -1.4935 |
| CNTN5 | NM_175566 | -0.307 | -0.0135 |
| XG | NM_175569 | 0.1985 | -0.0995 |
| GIMAP8 | NM_175571 | 0.394 | -0.0065 |
| ADRM1 | NM_175573 | 0.2535 | -0.0435 |
| WFIKKN2 | NM_175575 | 0.033 | -0.824 |
| IFT88 | NM_175605 | -1.2905 | -2.2555 |
| CNTN4 | NM_175607 | 0.0425 | -1.246 |
| ARFGAP1 | NM_175609 | -0.413 | -1.0065 |
| TJP1 | NM_175610 | -0.4925 | -1.662 |
| GRIK1 | NM_175611 | 0.2995 | 0.483 |
| CNTN4 | NM_175613 | 0.195 | -0.4 |
| NDUFA11 | NM_175614 | 0.258 | -0.2365 |
| MT1E | NM_175617 | -0.0775 | -0.287 |
| ZAR1 | NM_175619 | 0.173 | 0.327 |
| RAB3IP | NM_175623 | -0.321 | -0.6525 |
| RAB3IP | NM_175624 | -0.321 | -0.6525 |
| RAB3IP | NM_175625 | -0.321 | -0.6525 |
| DNMT3A | NM_175629 | -0.4295 | -1.6295 |
| DNMT3A | NM_175630 | -0.1735 | -1.304 |
| RUNX1T1 | NM_175634 | -1.1215 | -2.571 |
| RUNX1T1 | NM_175635 | -1.1215 | -2.571 |
| RUNX1T1 | NM_175636 | -1.1215 | -2.571 |
| CBX7 | NM_175709 | -0.936 | -0.934 |
| CR1L | NM_175710 | -0.1335 | -0.4775 |
| SSX3 | NM_175711 | 0.5645 | 0.1315 |
| TPO | NM_175719 | 0.244 | -0.0135 |
| TPO | NM_175721 | 0.244 | -0.0135 |
| TPO | NM_175722 | 0.244 | -0.0135 |
| SSX5 | NM_175723 | 0.044 | 0.0155 |
| IL5RA | NM_175724 | 0.056 | -0.797 |
| IL5RA | NM_175725 | -0.036 | -0.4405 |
| IL5RA | NM_175726 | 0.056 | -0.797 |
| IL5RA | NM_175727 | 0.056 | -0.797 |
| IL5RA | NM_175728 | 0.056 | -0.797 |
| PTPMT1 | NM_175732 | -0.5465 | -1.355 |

| | | | |
|---|---|---|---|
| SYT9 | NM_175733 | -0.033 | -0.014 |
| C17orf74 | NM_175734 | -0.016 | -0.1025 |
| LYG2 | NM_175735 | 0.0705 | -0.6195 |
| FMNL3 | NM_175736 | -0.3395 | -1.0255 |
| KLB | NM_175737 | 0.0465 | -0.1915 |
| RAB37 | NM_175738 | -0.2305 | -0.536 |
| SERPINA9 | NM_175739 | 0.014 | 0.121 |
| C15orf55 | NM_175741 | 0.193 | -0.6695 |
| MAGEA2 | NM_175743 | -0.1855 | -0.8515 |
| RHOC | NM_175744 | 0.094 | -0.803 |
| OLIG3 | NM_175747 | 0.039 | 0.2355 |
| UBR7 | NM_175748 | -1.252 | -1.999 |
| IL6ST | NM_175767 | -0.292 | -1.308 |
| GRIK2 | NM_175768 | -0.8425 | -1.477 |
| TCF23 | NM_175769 | -0.066 | -0.1815 |
| KRT79 | NM_175834 | -0.007 | -0.651 |
| SMOX | NM_175839 | -0.241 | 0.003 |
| SMOX | NM_175840 | -0.132 | -0.287 |
| SMOX | NM_175841 | -0.132 | -0.287 |
| SMOX | NM_175842 | -0.132 | -0.287 |
| PTBP1 | NM_175847 | -0.1505 | -0.85 |
| DNMT3B | NM_175848 | -0.217 | -0.679 |
| DNMT3B | NM_175849 | -0.217 | -0.679 |
| DNMT3B | NM_175850 | -0.217 | -0.679 |
| TXLNA | NM_175852 | -0.667 | -1.5765 |
| PAN3 | NM_175854 | -1.0655 | -2.203 |
| CHSY3 | NM_175856 | -0.068 | -0.445 |
| KRTAP8-1 | NM_175857 | -0.0925 | -0.5605 |
| KRTAP11-1 | NM_175858 | -0.1515 | -0.05 |
| CTPS2 | NM_175859 | -1.088 | -1.1125 |
| TMTC1 | NM_175861 | -0.9805 | -1.379 |
| CD86 | NM_175862 | -0.29 | -0.3095 |
| UHMK1 | NM_175866 | -0.505 | -1.07 |
| C19orf39 | NM_175871 | -0.197 | -0.2745 |
| ZNF792 | NM_175872 | 0.0835 | -0.5185 |
| ANKRD43 | NM_175873 | -0.476 | -0.66 |
| C12orf60 | NM_175874 | -1.215 | -2.496 |
| SIX5 | NM_175875 | -0.482 | -0.375 |
| EXOC8 | NM_175876 | -1.2735 | -2.6535 |
| XKR3 | NM_175878 | -0.1765 | 0.0505 |
| ODF3L1 | NM_175881 | 0.1085 | -0.3995 |
| IMP5 | NM_175882 | -0.0035 | 0.0895 |
| OR7D2 | NM_175883 | -0.133 | -1.909 |

| | | | |
|---|---|---|---|
| C7orf74 | NM_175884 | 0.0065 | 1.2125 |
| FAM181B | NM_175885 | -0.044 | -0.5245 |
| PRPS1L1 | NM_175886 | -0.4555 | -0.512 |
| PRR15 | NM_175887 | -0.334 | 0.0015 |
| ZNF620 | NM_175888 | -1.7665 | -1.908 |
| C12orf61 | NM_175895 | 0.369 | 0.4 |
| C16orf54 | NM_175900 | 0.124 | -0.0785 |
| C17orf101 | NM_175902 | -0.5155 | -1.3315 |
| ZADH2 | NM_175907 | -0.783 | -2.3345 |
| ZNF493 | NM_175910 | -1.129 | -2.08 |
| OR2L13 | NM_175911 | -0.2815 | -0.53 |
| JPH2 | NM_175913 | 0.1585 | -0.19 |
| HNF4A | NM_175914 | -0.024 | 0.0685 |
| CRIPAK | NM_175918 | -0.826 | -1.1075 |
| LNPEP | NM_175920 | 0.0985 | -0.2655 |
| C5orf51 | NM_175921 | -1.8195 | -2.9305 |
| PRR18 | NM_175922 | -0.092 | 0.2815 |
| ILDR1 | NM_175924 | 0.4515 | 0.0865 |
| FGF14 | NM_175929 | -1.164 | -2.258 |
| CBFA2T3 | NM_175931 | 0.284 | -0.764 |
| PSMD13 | NM_175932 | -0.0685 | -1.097 |
| DUOX1 | NM_175940 | -0.212 | -0.465 |
| P2RY2 | NM_176071 | -0.097 | -0.3115 |
| P2RY2 | NM_176072 | -0.097 | -0.3115 |
| CDK5RAP3 | NM_176096 | -0.118 | -0.5915 |
| NHLRC4 | NM_176677 | -0.058 | -0.4895 |
| FAM151A | NM_176782 | -0.2245 | -0.429 |
| PSME1 | NM_176783 | 0.073 | -0.399 |
| PIGN | NM_176787 | -1.1315 | -0.6825 |
| GTSF1L | NM_176791 | -0.173 | 0.1785 |
| MRPL43 | NM_176792 | -0.429 | -1.4575 |
| MRPL43 | NM_176793 | -0.429 | -1.4575 |
| MRPL43 | NM_176794 | -0.429 | -1.4575 |
| HRAS | NM_176795 | -0.062 | -0.789 |
| P2RY6 | NM_176796 | -0.651 | -0.041 |
| P2RY6 | NM_176797 | -0.651 | -0.041 |
| P2RY6 | NM_176798 | -0.651 | -0.041 |
| ADD1 | NM_176801 | -0.126 | -0.66 |
| MRPS11 | NM_176805 | -0.58 | -1.797 |
| NLRP13 | NM_176810 | -0.158 | -0.205 |
| NLRP8 | NM_176811 | 0.106 | -0.0265 |
| CHMP4B | NM_176812 | -0.1875 | -1.0455 |
| AGR3 | NM_176813 | 0.1185 | -0.5505 |

| | | | |
|---|---|---|---|
| ZNF800 | NM_176814 | -0.1585 | -0.576 |
| DHFRL1 | NM_176815 | -0.342 | -0.103 |
| CCDC125 | NM_176816 | -0.9325 | -1.4255 |
| TAS2R38 | NM_176817 | 0.06 | -0.1435 |
| GATC | NM_176818 | -0.7385 | -1.2085 |
| CXorf36 | NM_176819 | 0.13 | -0.2245 |
| NLRP9 | NM_176820 | 0.199 | -0.1545 |
| NLRP10 | NM_176821 | -0.25 | 0.4535 |
| NLRP14 | NM_176822 | -0.141 | 0.2995 |
| S100A7A | NM_176823 | -0.3075 | -0.063 |
| BBS7 | NM_176824 | -1.0085 | -1.5105 |
| SULT1C2 | NM_176825 | 0.141 | -0.333 |
| PSME3 | NM_176863 | -0.3 | -1.795 |
| PPA2 | NM_176866 | -0.8515 | -0.9805 |
| PPA2 | NM_176867 | -0.8515 | -0.9805 |
| PPA2 | NM_176869 | -0.8515 | -0.9805 |
| MT1M | NM_176870 | -0.0535 | -0.2625 |
| PDLIM2 | NM_176871 | -0.0325 | -0.3215 |
| CCKBR | NM_176875 | -0.109 | -0.393 |
| P2RY12 | NM_176876 | 0.4405 | -0.2335 |
| INADL | NM_176877 | 0.1775 | -0.392 |
| NR2C2AP | NM_176880 | -0.649 | -2.8485 |
| TAS2R39 | NM_176881 | -0.156 | -0.494 |
| TAS2R40 | NM_176882 | -0.2785 | 0.1135 |
| TAS2R41 | NM_176883 | -0.462 | -0.308 |
| TAS2R43 | NM_176884 | 0.319 | -0.3085 |
| TAS2R31 | NM_176885 | 0.5075 | -0.7735 |
| TAS2R46 | NM_176887 | 0.084 | -0.001 |
| TAS2R19 | NM_176888 | -0.053 | -0.7305 |
| TAS2R20 | NM_176889 | -0.053 | -1.4645 |
| TAS2R50 | NM_176890 | 0.022 | -0.1685 |
| IFNE | NM_176891 | -0.0375 | -0.3815 |
| CD86 | NM_176892 | -0.29 | -0.3095 |
| P2RY13 | NM_176894 | 0.0585 | -0.1335 |
| PPAP2A | NM_176895 | -0.741 | -3.284 |
| LMX1A | NM_177398 | -0.0185 | -0.2275 |
| NKX6-2 | NM_177400 | 0.2155 | -0.0425 |
| MIDN | NM_177401 | -0.7325 | -0.305 |
| SYT2 | NM_177402 | -0.1195 | -0.265 |
| MAGEB1 | NM_177404 | 0.0075 | -0.4555 |
| CECR1 | NM_177405 | -0.2795 | -0.429 |
| MAGEB1 | NM_177415 | 0.0075 | -0.4555 |
| KLC3 | NM_177417 | 0.3905 | -0.3085 |

| | | | |
|---|---|---|---|
| EIF2C3 | NM_177422 | -1.2195 | -1.746 |
| PPFIA1 | NM_177423 | -1.3405 | -2.0065 |
| STX12 | NM_177424 | -1.745 | -1.7615 |
| MAGED2 | NM_177433 | 0.016 | -0.594 |
| FTSJ1 | NM_177434 | -0.4045 | -1.59 |
| PPARD | NM_177435 | -0.1275 | -0.4155 |
| TAS2R60 | NM_177437 | 0.07 | -0.1775 |
| DICER1 | NM_177438 | -0.733 | -2.665 |
| FTSJ1 | NM_177439 | -0.4045 | -1.59 |
| TMUB2 | NM_177441 | -0.7605 | -1.3775 |
| PPFIBP1 | NM_177444 | 0.068 | 0.023 |
| TRAPPC6B | NM_177452 | -1.033 | -1.6405 |
| FAM171B | NM_177454 | -0.4125 | -1.274 |
| BHLHA15 | NM_177455 | 0.1645 | 0.3375 |
| MAGEC3 | NM_177456 | -0.1615 | -0.3695 |
| LYNX1 | NM_177457 | 0.0405 | -0.7195 |
| LYNX1 | NM_177458 | 0.0405 | -0.7195 |
| LYNX1 | NM_177476 | -0.186 | -0.239 |
| LYNX1 | NM_177477 | 0.0405 | -0.7195 |
| FTMT | NM_177478 | -0.0125 | -0.651 |
| GPLD1 | NM_177483 | -0.188 | -0.6645 |
| MEST | NM_177524 | -0.705 | -0.2215 |
| MEST | NM_177525 | -0.705 | -0.2215 |
| PPAP2C | NM_177526 | 0.0965 | -0.5855 |
| SULT1A2 | NM_177528 | -1.3765 | -1.275 |
| PKHD1L1 | NM_177531 | -0.119 | 0.071 |
| RASSF6 | NM_177532 | -0.1675 | 0.0235 |
| NUDT14 | NM_177533 | 0.0605 | -0.322 |
| CYP20A1 | NM_177538 | -1.0145 | -1.8535 |
| TAS1R1 | NM_177540 | 0.0235 | -0.2275 |
| SNRPD2 | NM_177542 | 0.221 | -0.117 |
| PPAP2C | NM_177543 | 0.0025 | -0.527 |
| SLC13A5 | NM_177550 | 0.199 | -0.514 |
| GAS2 | NM_177553 | -0.19 | -0.1115 |
| TRO | NM_177556 | -0.4355 | -1.2345 |
| CSNK2A1 | NM_177559 | -0.677 | -2.1985 |
| CSNK2A1 | NM_177560 | -0.596 | -1.913 |
| ASAH1 | NM_177924 | -0.843 | -0.7025 |
| H2AFJ | NM_177925 | -0.0375 | 0.744 |
| GOLM1 | NM_177937 | 0.1595 | -0.661 |
| P4HTM | NM_177938 | -0.3255 | -0.924 |
| P4HTM | NM_177939 | -0.3255 | -0.924 |
| ARMCX3 | NM_177947 | -1.275 | -1.2435 |

| | | | |
|---|---|---|---|
| ARMCX3 | NM_177948 | -1.275 | -1.2435 |
| ARMCX2 | NM_177949 | -1.7295 | -2.8975 |
| PPM1A | NM_177951 | -0.2755 | -0.7395 |
| PPM1A | NM_177952 | -0.2755 | -0.7395 |
| SYT12 | NM_177963 | 0.1 | -0.4725 |
| LYPD6B | NM_177964 | -0.215 | -0.3975 |
| C8orf37 | NM_177965 | -0.617 | -0.3955 |
| PDE12 | NM_177966 | -1.269 | -2.657 |
| UBAC2 | NM_177967 | -0.317 | -1.719 |
| PPM1B | NM_177968 | -1.071 | -2.7885 |
| PPM1B | NM_177969 | -1.071 | -2.7885 |
| TUB | NM_177972 | 0.0995 | -0.711 |
| SULT2B1 | NM_177973 | 0.119 | -0.465 |
| CASC4 | NM_177974 | -1.4015 | -1.1535 |
| ARL6 | NM_177976 | -0.403 | -1.7965 |
| HAP1 | NM_177977 | -0.4085 | 0.046 |
| CDH26 | NM_177980 | -0.4745 | -0.0415 |
| PPM1G | NM_177983 | -0.3005 | -1.0765 |
| ARL5A | NM_177985 | -0.956 | -1.7965 |
| DSG4 | NM_177986 | 0.2905 | -0.5695 |
| TUBB8 | NM_177987 | -0.1185 | -0.211 |
| MRPL47 | NM_177988 | -1.05 | -1.61 |
| ACTL6A | NM_177989 | -1.089 | -0.766 |
| PAK7 | NM_177990 | -0.8485 | -1.6705 |
| PTPDC1 | NM_177995 | -1.2105 | -2.486 |
| EPB41L1 | NM_177996 | -0.202 | -0.718 |
| OTOP1 | NM_177998 | -1.254 | -0.237 |
| ASB6 | NM_177999 | -0.696 | -2.7405 |
| PPP2R4 | NM_178000 | -0.1025 | -0.9615 |
| PPP2R4 | NM_178001 | -0.255 | -1.167 |
| PPP2R4 | NM_178003 | -0.1025 | -0.9615 |
| STARD13 | NM_178006 | -1.199 | -0.698 |
| STARD13 | NM_178007 | -1.199 | -0.698 |
| DGKH | NM_178009 | -0.112 | -0.3965 |
| SOX5 | NM_178010 | 0.17 | -0.8475 |
| LRRTM3 | NM_178011 | 0.1755 | -0.257 |
| TUBB2B | NM_178012 | 0.186 | -0.6105 |
| PRIMA1 | NM_178013 | -1.1455 | -2.7435 |
| TUBB | NM_178014 | 0.038 | -0.6225 |
| CATSPER3 | NM_178019 | -0.0655 | -0.093 |
| GGT7 | NM_178026 | -0.3015 | -0.756 |
| TMEM132A | NM_178031 | 0.235 | -0.504 |
| CYP4X1 | NM_178033 | 0.793 | -0.19 |

| | | | |
|---|---|---|---|
| PLA2G4D | NM_178034 | -0.1705 | -0.417 |
| ERC1 | NM_178039 | -0.142 | -0.4435 |
| ERC1 | NM_178040 | -0.142 | -0.4435 |
| ACTL6A | NM_178042 | -1.089 | -0.766 |
| LARP1B | NM_178043 | -0.331 | -0.67 |
| DLX1 | NM_178120 | -0.274 | -0.749 |
| C1orf201 | NM_178122 | -0.447 | -0.256 |
| SESTD1 | NM_178123 | -0.127 | -0.883 |
| CXorf40A | NM_178124 | -0.523 | -2.002 |
| TRIM50 | NM_178125 | -0.057 | -0.297 |
| FAM134C | NM_178126 | -0.5375 | -2.8405 |
| ANGPTL5 | NM_178127 | 0.127 | 0.502 |
| FADS6 | NM_178128 | -0.0405 | -0.239 |
| P2RY8 | NM_178129 | -0.0755 | -0.403 |
| NME9 | NM_178130 | -0.1195 | 0.023 |
| CYP4Z1 | NM_178134 | -0.3065 | 0.688 |
| HSD17B13 | NM_178135 | 0.304 | -0.367 |
| POLDIP3 | NM_178136 | -0.1455 | -1.1875 |
| LHX3 | NM_178138 | 0.0565 | 0.226 |
| PDZD2 | NM_178140 | -0.9055 | -1.14 |
| SLC35B2 | NM_178148 | -0.815 | -2.009 |
| FBXO18 | NM_178150 | -0.8625 | -2.041 |
| DCX | NM_178151 | -0.96 | -0.4395 |
| DCX | NM_178152 | -0.96 | -0.4395 |
| DCX | NM_178153 | -0.96 | -0.4395 |
| FUT8 | NM_178155 | -1.1135 | -2.375 |
| FUT8 | NM_178156 | -1.0465 | -2.1975 |
| OTOP2 | NM_178160 | -0.0755 | -0.318 |
| PTF1A | NM_178161 | -0.0985 | 0.219 |
| ZNF598 | NM_178167 | -0.6355 | -1.096 |
| OR10A5 | NM_178168 | -0.3725 | -0.056 |
| RASSF3 | NM_178169 | -0.033 | -0.3865 |
| NEK8 | NM_178170 | -0.1045 | -0.249 |
| GSDMA | NM_178171 | 0.053 | -0.1535 |
| GPIHBP1 | NM_178172 | 0.08 | -0.608 |
| CCDC36 | NM_178173 | -0.2905 | -0.3135 |
| TREML1 | NM_178174 | -0.023 | -0.393 |
| LHFPL1 | NM_178175 | -0.0935 | -0.312 |
| MOGAT3 | NM_178176 | 0.038 | -0.097 |
| NMNAT3 | NM_178177 | -0.9115 | -1.598 |
| CDCP1 | NM_178181 | 0.152 | -0.2525 |
| ATPIF1 | NM_178190 | -0.366 | -0.641 |
| ATPIF1 | NM_178191 | -0.366 | -0.641 |

| | | | |
|---|---|---|---|
| IQGAP3 | NM_178229 | -0.278 | -0.927 |
| PPIAL4A | NM_178230 | 0.3735 | -0.7195 |
| ICA1L | NM_178231 | -0.5505 | -0.896 |
| HAPLN3 | NM_178232 | 0.136 | -0.079 |
| OTOP3 | NM_178233 | -0.279 | -0.2215 |
| TUSC3 | NM_178234 | -0.377 | -0.2655 |
| EXOC1 | NM_178237 | -1.285 | -2.101 |
| PILRB | NM_178238 | -1.2115 | -1.8945 |
| ATG4A | NM_178270 | -0.8935 | -1.365 |
| PILRA | NM_178272 | 0.275 | -0.069 |
| PILRA | NM_178273 | 0.275 | -0.069 |
| SERINC5 | NM_178276 | 0.0585 | -0.388 |
| SNAI3 | NM_178310 | 0.1185 | -0.477 |
| SPTBN1 | NM_178313 | -0.57 | -0.798 |
| RILPL1 | NM_178314 | -0.76 | -1.3935 |
| SPTLC1 | NM_178324 | -1.402 | -1.596 |
| ATG4B | NM_178326 | -0.367 | -0.835 |
| CCR3 | NM_178328 | -0.0235 | -0.3235 |
| CCR3 | NM_178329 | -0.0235 | -0.3235 |
| CCDC50 | NM_178335 | -0.824 | -1.356 |
| MRPL52 | NM_178336 | 0.071 | -0.4075 |
| C3orf35 | NM_178339 | 0.617 | -0.069 |
| C3orf35 | NM_178342 | -0.074 | -0.219 |
| LCE1A | NM_178348 | 0.0925 | -0.284 |
| LCE1B | NM_178349 | -0.2865 | -0.0775 |
| LCE1C | NM_178351 | 0.2635 | -0.636 |
| LCE1D | NM_178352 | 0.1225 | -0.546 |
| LCE1E | NM_178353 | -0.0235 | -0.548 |
| LCE1F | NM_178354 | 0.194 | -0.0955 |
| LCE4A | NM_178356 | 0.132 | -0.238 |
| PAQR7 | NM_178422 | -0.97 | -1.717 |
| HDAC9 | NM_178423 | -0.862 | -2.3525 |
| SOX30 | NM_178424 | -0.0275 | 0.2485 |
| HDAC9 | NM_178425 | -0.862 | -2.3525 |
| ARNT | NM_178427 | -0.8185 | -2.4545 |
| LCE2A | NM_178428 | -0.059 | -0.349 |
| LCE2C | NM_178429 | 0.4815 | -0.312 |
| LCE2D | NM_178430 | 0.1305 | -0.073 |
| LCE3A | NM_178431 | 0.1065 | -0.2765 |
| CDK20 | NM_178432 | -0.936 | -1.6585 |
| LCE3B | NM_178433 | 0.021 | -0.46 |
| LCE3C | NM_178434 | 0.2215 | 0.25 |
| LCE3E | NM_178435 | -0.468 | 0.007 |

| | | | |
|---|---|---|---|
| LCE5A | NM_178438 | -0.5675 | -0.891 |
| GMCL1 | NM_178439 | -1.223 | -3.123 |
| ZFYVE1 | NM_178441 | -0.7865 | -2.3925 |
| FERMT3 | NM_178443 | -0.01 | -0.329 |
| CCRL1 | NM_178445 | -0.5115 | 0.691 |
| C9orf140 | NM_178448 | -0.7905 | -1.765 |
| PTH2 | NM_178449 | -0.054 | -0.3235 |
| 3-Mar | NM_178450 | 0.2805 | -0.1365 |
| DNAAF1 | NM_178452 | -0.0545 | -0.4205 |
| DRAM2 | NM_178454 | -1.6105 | -1.453 |
| SPINT4 | NM_178455 | 0.3575 | -0.18 |
| C20orf85 | NM_178456 | -0.1305 | -0.2675 |
| ZNF831 | NM_178457 | 0.219 | 0.0375 |
| SIRPD | NM_178460 | -0.198 | -0.095 |
| C20orf166 | NM_178463 | -0.072 | -0.61 |
| BPIFA3 | NM_178466 | 0.176 | -0.055 |
| FAM83C | NM_178468 | 0.162 | -0.3905 |
| LCN8 | NM_178469 | 0.123 | 0.0865 |
| DCAF12L1 | NM_178470 | -0.0125 | -0.1885 |
| GPR119 | NM_178471 | -0.1235 | 0.109 |
| BANF2 | NM_178477 | 0.199 | -0.0455 |
| C20orf79 | NM_178483 | 0.0165 | -0.3025 |
| R3HDML | NM_178491 | -0.639 | -0.427 |
| NOTUM | NM_178493 | -0.4155 | -0.8105 |
| ITPRIPL1 | NM_178495 | -0.408 | -1.432 |
| MB21D2 | NM_178496 | -0.303 | -0.8615 |
| C4orf26 | NM_178497 | 0.299 | -0.2295 |
| SLC5A12 | NM_178498 | -0.048 | 0.041 |
| CCDC60 | NM_178499 | -0.154 | 0.0165 |
| PHOSPHO1 | NM_178500 | 0.03 | -0.443 |
| DTX3 | NM_178502 | -0.0915 | -0.073 |
| DNAH12 | NM_178504 | -0.0425 | -0.297 |
| TMEM26 | NM_178505 | 0.482 | 0.052 |
| OAF | NM_178507 | -0.9075 | -1.809 |
| C6orf1 | NM_178508 | -0.592 | -1.6475 |
| STXBP4 | NM_178509 | -0.3885 | -1.6885 |
| ANKK1 | NM_178510 | -0.069 | -0.036 |
| PRR24 | NM_178511 | 0.0855 | -0.69 |
| PIGW | NM_178517 | -0.4925 | -0.8155 |
| TMEM102 | NM_178518 | -0.27 | -0.392 |
| TMEM105 | NM_178520 | -0.0655 | -0.5635 |
| ZNF616 | NM_178523 | -0.4815 | -1.713 |
| ACTL9 | NM_178525 | 0.0785 | -0.3265 |

| | | | |
|---|---|---|---|
| SLC25A42 | NM_178526 | -0.359 | -0.812 |
| SLC9A11 | NM_178527 | -0.039 | -0.2285 |
| RNF180 | NM_178532 | -0.2875 | -0.358 |
| LCN12 | NM_178536 | -0.4405 | -0.923 |
| B4GALNT4 | NM_178537 | -0.245 | -0.782 |
| FAM19A2 | NM_178539 | -0.48 | -0.247 |
| C17orf65 | NM_178542 | -0.278 | -0.5785 |
| ENPP7 | NM_178543 | -0.054 | -0.4355 |
| ZNF546 | NM_178544 | -0.4505 | -1.118 |
| TMEM52 | NM_178545 | 0.042 | -0.3885 |
| ZBTB8OS | NM_178547 | -0.5835 | -0.719 |
| TFAP2E | NM_178548 | -0.0125 | -0.844 |
| ZNF678 | NM_178549 | -0.253 | -0.069 |
| C1orf110 | NM_178550 | -0.094 | -0.434 |
| C22orf33 | NM_178552 | 0.233 | -0.1295 |
| C2orf53 | NM_178553 | -0.0835 | -0.554 |
| KY | NM_178554 | 0.0405 | -0.2785 |
| TRIML1 | NM_178556 | 0.0525 | 0.246 |
| NAT8L | NM_178557 | -0.017 | -0.323 |
| ZNF680 | NM_178558 | -1.107 | -1.6155 |
| ABCB5 | NM_178559 | -0.0545 | -0.4635 |
| CTAGE6P | NM_178561 | 0.0665 | 0.248 |
| TSPAN33 | NM_178562 | -0.131 | -1.096 |
| AGBL3 | NM_178563 | -0.3555 | -0.383 |
| NRBP2 | NM_178564 | 0.058 | 0.127 |
| RSPO2 | NM_178565 | -0.2955 | -0.796 |
| ZDHHC21 | NM_178566 | -0.7765 | -0.7855 |
| RTN4RL1 | NM_178568 | 0.0415 | 0.021 |
| C5orf38 | NM_178569 | -0.004 | -0.436 |
| RTN4RL2 | NM_178570 | -0.2185 | -0.9115 |
| TBC1D26 | NM_178571 | -0.039 | -0.1245 |
| PSMF1 | NM_178578 | -0.269 | -1.13 |
| HM13 | NM_178580 | -0.445 | -1.202 |
| HM13 | NM_178581 | -0.4385 | -0.8515 |
| HM13 | NM_178582 | -0.4385 | -0.8515 |
| 10-Sep | NM_178584 | 0.0375 | -1.8735 |
| PPP2R5C | NM_178586 | -2.0235 | -3.1815 |
| PPP2R5C | NM_178587 | -1.081 | -2.396 |
| MTDH | NM_178812 | -0.539 | -1.4725 |
| AKAP14 | NM_178813 | 0.3755 | 0.045 |
| ARL5B | NM_178815 | -0.3295 | -1.134 |
| MRAP | NM_178817 | -0.2095 | -0.4715 |
| CMTM4 | NM_178818 | -0.635 | -0.685 |

| | | | |
|---|---|---|---|
| AGPAT6 | NM_178819 | -0.582 | -1.8105 |
| FBXO27 | NM_178820 | -0.214 | -0.369 |
| WDR69 | NM_178821 | -0.018 | -0.057 |
| IGSF10 | NM_178822 | -0.2155 | -0.419 |
| WDR49 | NM_178824 | 0.254 | 0.2185 |
| ANO4 | NM_178826 | -0.3795 | -1.4875 |
| IQUB | NM_178827 | -0.269 | -0.3795 |
| C9orf79 | NM_178828 | -0.0875 | 0.0535 |
| C7orf34 | NM_178829 | -0.247 | -0.2895 |
| C19orf47 | NM_178830 | -0.531 | -1.608 |
| GATS | NM_178831 | -0.1315 | -0.525 |
| MORN4 | NM_178832 | -0.883 | -1.3525 |
| SLC9B2 | NM_178833 | -0.5655 | -1.381 |
| LAYN | NM_178834 | -0.975 | -3.0185 |
| ZNF827 | NM_178835 | -1.6625 | -4.7035 |
| PLD6 | NM_178836 | -0.5505 | -1.6285 |
| C19orf51 | NM_178837 | 0.5315 | -0.192 |
| LRRTM1 | NM_178839 | -0.74 | -1.4445 |
| C1orf64 | NM_178840 | 0.2265 | -0.0185 |
| RNF166 | NM_178841 | -0.7085 | -0.3725 |
| CERS3 | NM_178842 | -0.327 | 0.0525 |
| NLRC3 | NM_178844 | 0.212 | -0.374 |
| HNF4A | NM_178849 | -0.024 | 0.0685 |
| HNF4A | NM_178850 | -0.024 | 0.0685 |
| RP1L1 | NM_178857 | 0.0105 | -0.1885 |
| SFXN2 | NM_178858 | -0.2695 | -1.427 |
| OSTBETA | NM_178859 | -0.0955 | -0.1745 |
| SEZ6 | NM_178860 | -0.4865 | -1.3495 |
| RNF113B | NM_178861 | 0.0135 | -0.135 |
| STT3B | NM_178862 | -1.007 | -2.0945 |
| KCTD13 | NM_178863 | -0.453 | -0.4875 |
| NPAS4 | NM_178864 | -0.08 | -0.2205 |
| SERINC2 | NM_178865 | 0.035 | -0.3205 |
| CMTM8 | NM_178868 | -0.4555 | -2.0775 |
| SNRNP35 | NM_180699 | -0.8075 | -2.5405 |
| PPP2R5D | NM_180976 | -0.5725 | -0.692 |
| PPP2R5D | NM_180977 | -0.5725 | -0.692 |
| MRPL52 | NM_180982 | 0.071 | -0.4075 |
| GPR180 | NM_180989 | -1.401 | -3.65 |
| ZACN | NM_180990 | -0.123 | -0.451 |
| SLCO4C1 | NM_180991 | 0.0635 | -0.5365 |
| PBRM1 | NM_181042 | -1.239 | -1.3545 |
| AXIN1 | NM_181050 | -0.544 | -1.683 |

| | | | |
|---|---|---|---|
| HIF1A | NM_181054 | -0.736 | -1.572 |
| GOLGA8A | NM_181077 | -1.2085 | -1.186 |
| IL21R | NM_181078 | 0.2435 | -0.3815 |
| IL21R | NM_181079 | 0.2435 | -0.3815 |
| SCYL3 | NM_181093 | -2.2215 | -3.259 |
| WDR17 | NM_181265 | -0.759 | -1.5425 |
| CMTM1 | NM_181268 | -0.184 | -0.0705 |
| CMTM1 | NM_181269 | -0.184 | -0.0705 |
| CMTM1 | NM_181270 | -0.184 | -0.0705 |
| CMTM1 | NM_181271 | -0.184 | -0.0705 |
| CMTM1 | NM_181272 | -0.184 | -0.0705 |
| CMTM1 | NM_181283 | -0.184 | -0.0705 |
| WDR20 | NM_181291 | -1.131 | -2.623 |
| CMTM1 | NM_181296 | -0.184 | -0.0705 |
| NLGN3 | NM_181303 | -0.1905 | -0.3365 |
| MRPL52 | NM_181304 | 0.071 | -0.4075 |
| MRPL52 | NM_181305 | 0.071 | -0.4075 |
| MRPL52 | NM_181306 | 0.071 | -0.4075 |
| MRPL52 | NM_181307 | 0.071 | -0.4075 |
| WDR20 | NM_181308 | -1.131 | -2.623 |
| IL22RA2 | NM_181309 | 0.1725 | 0.0575 |
| IL22RA2 | NM_181310 | 0.1725 | 0.0575 |
| NLGN4X | NM_181332 | -1.0885 | -2.285 |
| PRR5 | NM_181333 | -0.5815 | -0.556 |
| ARHGAP8 | NM_181335 | 0.261 | -0.4215 |
| LEMD2 | NM_181336 | -1.1375 | -2.1185 |
| KAAG1 | NM_181337 | 0.0145 | -0.402 |
| DCAF4 | NM_181340 | -0.5925 | -1.5045 |
| DCAF4 | NM_181341 | -0.5925 | -1.5045 |
| FKBP7 | NM_181342 | -1.426 | -1.331 |
| SMURF1 | NM_181349 | -0.018 | -0.859 |
| NCAM1 | NM_181351 | -0.7595 | -0.572 |
| ID1 | NM_181353 | 0.466 | 0.9325 |
| OXR1 | NM_181354 | -0.4415 | -0.855 |
| SUPT3H | NM_181356 | -0.378 | -0.3605 |
| DCAF11 | NM_181357 | -0.0445 | -0.6475 |
| HIPK1 | NM_181358 | 0.267 | -0.807 |
| IL6R | NM_181359 | -0.468 | -0.3015 |
| KCNMB2 | NM_181361 | 0.0645 | -0.083 |
| FXN | NM_181425 | 0.019 | -1.496 |
| CCDC39 | NM_181426 | -0.0265 | -0.2725 |
| GABPB1 | NM_181427 | -2.2725 | -4.4245 |
| TAS2R42 | NM_181429 | 0.0125 | -0.0765 |

| | | | |
|---|---|---|---|
| C1QTNF3 | NM_181435 | -1.4 | -0.993 |
| MRPL55 | NM_181441 | -0.57 | -1.937 |
| ADNP | NM_181442 | -1.5625 | -2.7285 |
| BTBD3 | NM_181443 | -0.7955 | -0.8695 |
| FSHR | NM_181446 | 0.354 | 0.2315 |
| CD300E | NM_181449 | -0.04 | -0.4155 |
| GCC2 | NM_181453 | -0.879 | -3.3185 |
| MRPL55 | NM_181454 | -0.57 | -1.937 |
| MRPL55 | NM_181455 | -0.57 | -1.937 |
| MRPL55 | NM_181456 | -0.57 | -1.937 |
| PAX3 | NM_181457 | 0.1345 | 0.0185 |
| PAX3 | NM_181458 | 0.1345 | 0.0185 |
| PAX3 | NM_181459 | 0.1345 | 0.0185 |
| PAX3 | NM_181460 | 0.1345 | 0.0185 |
| PAX3 | NM_181461 | 0.1345 | 0.0185 |
| MRPL55 | NM_181462 | -0.57 | -1.937 |
| MRPL55 | NM_181463 | -0.57 | -1.937 |
| MRPL55 | NM_181464 | -0.57 | -1.937 |
| MRPL55 | NM_181465 | -0.57 | -1.937 |
| EIF6 | NM_181466 | -0.1485 | -1.276 |
| EIF6 | NM_181468 | -0.15 | -1.103 |
| RFC2 | NM_181471 | -0.2625 | -1.3135 |
| C18orf1 | NM_181481 | -0.1455 | -2.338 |
| C18orf1 | NM_181482 | -0.1455 | -2.338 |
| C18orf1 | NM_181483 | -0.1455 | -2.338 |
| TBX5 | NM_181486 | 0.241 | -0.285 |
| ZNF445 | NM_181489 | -1.0225 | -3.209 |
| MED22 | NM_181491 | -1.496 | -2.073 |
| TCF20 | NM_181492 | -0.585 | -1.7695 |
| ITPA | NM_181493 | -0.0915 | -1.1425 |
| CUX1 | NM_181500 | 0.11 | -0.0355 |
| ITGA1 | NM_181501 | 0.211 | -1.1075 |
| EXOSC8 | NM_181503 | -1.2645 | -2.141 |
| PIK3R1 | NM_181504 | -0.308 | -1.1495 |
| LRRC70 | NM_181506 | -0.149 | 0.613 |
| HPS5 | NM_181507 | -1.3785 | -3.276 |
| HPS5 | NM_181508 | -1.3785 | -3.276 |
| WFDC8 | NM_181510 | -0.472 | -0.531 |
| MRPL21 | NM_181515 | -0.057 | -0.877 |
| SYT15 | NM_181519 | 0.115 | -0.499 |
| CMTM4 | NM_181521 | -0.641 | -1.0695 |
| PIK3R1 | NM_181523 | -0.308 | -1.1495 |
| PIK3R1 | NM_181524 | -0.308 | -1.1495 |

| | | | |
|---|---|---|---|
| MYL9 | NM_181526 | -0.3675 | -0.3385 |
| BTN2A2 | NM_181531 | -1.0235 | -1.553 |
| ERAS | NM_181532 | 0.222 | -0.319 |
| KRT25 | NM_181534 | -0.436 | -0.3275 |
| KRT28 | NM_181535 | -0.0795 | 0.0275 |
| PKD1L3 | NM_181536 | -0.2955 | 0.04 |
| KRT27 | NM_181537 | -0.224 | -0.4405 |
| GJC3 | NM_181538 | -0.0835 | -0.2775 |
| KRT26 | NM_181539 | -0.43 | -0.1345 |
| CUX1 | NM_181552 | -0.918 | -1.486 |
| CMTM3 | NM_181553 | -0.5575 | -2.2335 |
| RFC3 | NM_181558 | -0.911 | -1.1495 |
| CREM | NM_181571 | -1.5625 | -1.937 |
| RFC4 | NM_181573 | -1.67 | -3.476 |
| AUP1 | NM_181575 | -0.011 | -0.602 |
| RFC5 | NM_181578 | -0.6595 | -1.6615 |
| DUS4L | NM_181581 | -1.582 | -2.772 |
| UPP1 | NM_181597 | -0.087 | -0.3655 |
| ATL1 | NM_181598 | -1.3525 | -2.599 |
| KRTAP13-1 | NM_181599 | 0.076 | -0.2485 |
| KRTAP13-4 | NM_181600 | -0.156 | -0.6675 |
| KRTAP6-1 | NM_181602 | 0.4395 | 0.2435 |
| KRTAP6-2 | NM_181604 | -0.1185 | -0.3195 |
| KRTAP6-3 | NM_181605 | 0.34 | -0.1565 |
| KRTAP7-1 | NM_181606 | 0.2615 | -0.3765 |
| KRTAP19-1 | NM_181607 | 0.2735 | 1.157 |
| KRTAP19-2 | NM_181608 | -0.056 | -0.168 |
| KRTAP19-3 | NM_181609 | -0.072 | -0.218 |
| KRTAP19-4 | NM_181610 | 0.547 | -0.082 |
| KRTAP19-5 | NM_181611 | 0.377 | -0.4145 |
| KRTAP19-6 | NM_181612 | 0.0335 | 0.0785 |
| KRTAP19-7 | NM_181614 | -0.2415 | -0.21 |
| KRTAP20-1 | NM_181615 | 0.0785 | -0.277 |
| KRTAP20-2 | NM_181616 | -0.414 | -0.256 |
| KRTAP21-2 | NM_181617 | -0.145 | -0.2485 |
| KRTAP21-1 | NM_181619 | 0.388 | -0.3355 |
| KRTAP22-1 | NM_181620 | -0.1445 | -0.43 |
| KRTAP13-2 | NM_181621 | -0.504 | 0.0975 |
| KRTAP13-3 | NM_181622 | 0.1305 | -0.4285 |
| KRTAP15-1 | NM_181623 | -0.4525 | -0.034 |
| KRTAP23-1 | NM_181624 | 0.0295 | -0.3615 |
| CKLF | NM_181640 | -0.368 | -1.093 |
| SPINT1 | NM_181642 | -0.054 | -0.583 |

| | | | |
|---|---|---|---|
| C1orf88 | NM_181643 | 0.142 | 0.0595 |
| MFSD4 | NM_181644 | -0.0655 | -0.1375 |
| CCDC67 | NM_181645 | 0.0065 | 0.8195 |
| ZNF804B | NM_181646 | -0.114 | -0.2355 |
| PRDX5 | NM_181652 | 0.317 | -0.197 |
| CPLX4 | NM_181654 | 0.032 | -0.255 |
| C17orf58 | NM_181655 | -0.011 | 0.762 |
| C17orf58 | NM_181656 | -0.0225 | 1.0865 |
| LTB4R | NM_181657 | -0.785 | -0.6295 |
| NCOA3 | NM_181659 | -1.412 | -2.0115 |
| VPS13B | NM_181661 | -0.1415 | -0.4325 |
| ANKS1B | NM_181670 | -0.3535 | -0.4295 |
| PITPNC1 | NM_181671 | -0.4415 | -1.5865 |
| OGT | NM_181672 | -1.9915 | -1.1995 |
| OGT | NM_181673 | -1.9915 | -1.1995 |
| PPP2R2B | NM_181674 | -1.329 | -1.6335 |
| PPP2R2B | NM_181675 | -1.281 | -1.527 |
| PPP2R2B | NM_181676 | -1.281 | -1.527 |
| PPP2R2B | NM_181677 | -1.281 | -1.527 |
| PPP2R2B | NM_181678 | -1.281 | -1.527 |
| KRTAP12-2 | NM_181684 | -0.1495 | -0.55 |
| KRTAP12-1 | NM_181686 | -0.138 | -0.5455 |
| KRTAP10-10 | NM_181688 | 0.0235 | -0.5015 |
| NNAT | NM_181689 | -0.263 | -0.458 |
| AKT3 | NM_181690 | -0.707 | -1.202 |
| PRDX1 | NM_181696 | -0.138 | 0.063 |
| PRDX1 | NM_181697 | -0.138 | 0.063 |
| CCNY | NM_181698 | -0.0215 | -0.4895 |
| PPP2R1B | NM_181699 | -0.8865 | -1.919 |
| PPP2R1B | NM_181700 | -0.8865 | -1.919 |
| QSOX2 | NM_181701 | -0.6605 | -1.6905 |
| GJA5 | NM_181703 | -0.0645 | -0.0995 |
| LYRM7 | NM_181705 | -0.8015 | -1.0855 |
| DNAJC24 | NM_181706 | -1.187 | -2.6205 |
| C17orf64 | NM_181707 | -0.1805 | -0.4605 |
| BCDIN3D | NM_181708 | -1.5095 | -2.9405 |
| FAM101A | NM_181709 | -0.4085 | -2.3585 |
| ZNRF4 | NM_181710 | -0.301 | -0.357 |
| GRASP | NM_181711 | -0.74 | -0.738 |
| KANK4 | NM_181712 | -0.642 | -1.2855 |
| UBXN2A | NM_181713 | -1.3135 | -2.4565 |
| LCA5 | NM_181714 | -0.6 | -1.51 |
| CRTC2 | NM_181715 | -0.4855 | -0.362 |

| | | | |
|---|---|---|---|
| CENPV | NM_181716 | 0.0115 | -0.315 |
| ASPHD1 | NM_181718 | -0.3855 | -0.994 |
| TMCO4 | NM_181719 | -0.0445 | -0.401 |
| FOXR1 | NM_181721 | 0.1115 | -0.329 |
| EFHA2 | NM_181723 | -1.363 | -2.4305 |
| TMEM119 | NM_181724 | -0.621 | -1.085 |
| METTL2A | NM_181725 | -1.0755 | -2.9635 |
| ANKRD37 | NM_181726 | -0.8135 | -1.2705 |
| SPATA12 | NM_181727 | -0.224 | -0.2985 |
| COG5 | NM_181733 | -0.827 | -1.1225 |
| PRDX2 | NM_181738 | 0.212 | -0.236 |
| ORC4 | NM_181741 | -0.643 | -2.021 |
| ORC4 | NM_181742 | -0.643 | -2.021 |
| OPN5 | NM_181744 | -0.1195 | -0.059 |
| O3FAR1 | NM_181745 | 0.2145 | -0.185 |
| ORC5 | NM_181747 | -1.5 | -3.4645 |
| HSD11B1 | NM_181755 | 0.0395 | -0.0545 |
| ZNF233 | NM_181756 | -1.236 | -0.688 |
| UBE2A | NM_181762 | -0.7045 | -1.541 |
| APOBEC3H | NM_181773 | 0.034 | -0.5875 |
| SLC36A3 | NM_181774 | -0.198 | -0.1625 |
| PLXNA4 | NM_181775 | -0.048 | -0.433 |
| SLC36A2 | NM_181776 | 0.2735 | -0.031 |
| UBE2A | NM_181777 | -0.7045 | -1.541 |
| BTLA | NM_181780 | -0.0195 | 0.135 |
| NCOA7 | NM_181782 | -1.2485 | -2.957 |
| TMTC3 | NM_181783 | -0.694 | -2.8435 |
| SPRED2 | NM_181784 | -0.83 | -3.2795 |
| SLC46A3 | NM_181785 | -0.134 | -1.1445 |
| HKR1 | NM_181786 | -1.669 | -2.9065 |
| DPY19L4 | NM_181787 | -0.885 | -1.259 |
| H1FNT | NM_181788 | -0.3025 | -0.1405 |
| GLDN | NM_181789 | 0.0095 | -0.5335 |
| GPR142 | NM_181790 | -0.179 | -0.173 |
| GPR141 | NM_181791 | 0.0345 | 0.1165 |
| PKIB | NM_181794 | -0.7525 | -1.06 |
| PKIB | NM_181795 | -0.7525 | -1.06 |
| UBE2C | NM_181799 | -0.286 | -0.9185 |
| UBE2C | NM_181802 | 0.071 | -0.493 |
| UBE2C | NM_181803 | -0.0225 | -0.43 |
| PKIG | NM_181804 | -0.017 | -0.979 |
| PKIG | NM_181805 | -0.017 | -0.979 |
| AASDH | NM_181806 | -1.5945 | -2.7675 |

| | | | |
|---|---|---|---|
| DCDC1 | NM_181807 | -0.061 | -0.016 |
| POLN | NM_181808 | -0.135 | -0.287 |
| BMP8A | NM_181809 | -0.205 | -0.6085 |
| ABHD12B | NM_181814 | -0.282 | -0.1365 |
| NF2 | NM_181825 | -0.0255 | -0.984 |
| NF2 | NM_181828 | -0.0255 | -0.984 |
| NF2 | NM_181829 | -0.0255 | -0.984 |
| NF2 | NM_181830 | -0.0255 | -0.984 |
| NF2 | NM_181831 | -0.0255 | -0.984 |
| NF2 | NM_181832 | -0.6225 | -1.625 |
| NF2 | NM_181833 | -0.0255 | -0.984 |
| TMED7 | NM_181836 | -1.193 | -2.0785 |
| ORC3 | NM_181837 | -1.2905 | -1.8755 |
| UBE2D2 | NM_181838 | -0.733 | -1.5415 |
| PKIA | NM_181839 | -1.2255 | -1.749 |
| KCNK18 | NM_181840 | -0.001 | -0.43 |
| ZBTB12 | NM_181842 | -0.1305 | -0.483 |
| NUDT8 | NM_181843 | 0.027 | -0.4335 |
| BCL6B | NM_181844 | -0.016 | -0.4115 |
| ZNF283 | NM_181845 | -1.064 | -2.497 |
| ZSCAN22 | NM_181846 | -0.958 | -1.6475 |
| AMIGO2 | NM_181847 | 0.3685 | -1.7235 |
| APAF1 | NM_181861 | -0.295 | -1.123 |
| ACOT7 | NM_181864 | -0.0065 | -0.056 |
| ACOT7 | NM_181865 | -0.0065 | -0.056 |
| ACOT7 | NM_181866 | -0.0065 | -0.056 |
| APAF1 | NM_181868 | -0.295 | -1.123 |
| APAF1 | NM_181869 | -0.295 | -1.123 |
| PKLR | NM_181871 | 0.0545 | -0.254 |
| DMRT2 | NM_181872 | 0.0885 | -0.145 |
| MTMR11 | NM_181873 | -0.289 | -0.1505 |
| GRM7 | NM_181874 | 0.146 | -0.7625 |
| PPP2R2C | NM_181876 | 0.0455 | -0.444 |
| ZSCAN2 | NM_181877 | -1.2225 | -3.0835 |
| LILRA5 | NM_181879 | -0.368 | 0.141 |
| VCY1B | NM_181880 | 0.039 | -0.4535 |
| RXFP4 | NM_181885 | 0.1285 | -0.1535 |
| UBE2D3 | NM_181886 | -0.7795 | -0.961 |
| UBE2D3 | NM_181887 | -0.7795 | -0.961 |
| UBE2D3 | NM_181888 | -0.7795 | -0.961 |
| UBE2D3 | NM_181889 | -0.7795 | -0.961 |
| UBE2D3 | NM_181890 | -0.7795 | -0.961 |
| UBE2D3 | NM_181891 | -0.7795 | -0.961 |

| | | | |
|---|---|---|---|
| UBE2D3 | NM_181892 | -0.7795 | -0.961 |
| UBE2D3 | NM_181893 | -0.7795 | -0.961 |
| PPP2R3A | NM_181897 | -0.734 | -1.83 |
| STARD5 | NM_181900 | 0.1305 | -0.9885 |
| LILRA5 | NM_181985 | 0.0665 | 0.15 |
| LILRA5 | NM_181986 | -0.1945 | -0.364 |
| ENOX2 | NM_182314 | -0.821 | -1.953 |
| RPS6KA5 | NM_182398 | -0.7935 | -1.922 |
| PKM2 | NM_182470 | 0.2085 | -0.32 |
| PKM2 | NM_182471 | 0.2085 | -0.32 |
| EPHA5 | NM_182472 | 0.436 | -0.8175 |
| COQ6 | NM_182476 | -0.0835 | -1.088 |
| COQ6 | NM_182480 | -0.0835 | -1.088 |
| BAGE2 | NM_182482 | -0.3855 | -0.6895 |
| BAGE5 | NM_182484 | -0.396 | -0.7185 |
| CPEB2 | NM_182485 | -1.141 | -2.434 |
| C1QTNF6 | NM_182486 | 0.3685 | -0.212 |
| OLFML2A | NM_182487 | 0.1015 | -0.3855 |
| USP12 | NM_182488 | -1.2835 | -2.8445 |
| STRA8 | NM_182489 | -0.2955 | -0.193 |
| ZNF227 | NM_182490 | -0.927 | -2.697 |
| ZFAND2A | NM_182491 | -0.212 | -1.456 |
| LRP5L | NM_182492 | -0.1795 | -1.1655 |
| MYLK3 | NM_182493 | -0.046 | -0.4075 |
| FAM55B | NM_182495 | 0.2855 | -0.3365 |
| CCDC38 | NM_182496 | -0.077 | 0.0415 |
| KRT40 | NM_182497 | 0.0375 | -0.316 |
| ZNF428 | NM_182498 | 0.476 | -0.2585 |
| C2orf50 | NM_182500 | -0.0205 | -0.013 |
| MTERFD2 | NM_182501 | -0.592 | -1.698 |
| TMPRSS11B | NM_182502 | -0.0155 | -0.2435 |
| ADAT2 | NM_182503 | -1.2895 | -2.139 |
| WBSCR28 | NM_182504 | 0.177 | -0.1735 |
| C9orf85 | NM_182505 | -1.7995 | -2.1425 |
| MAGEB10 | NM_182506 | -0.573 | -0.2575 |
| KRT80 | NM_182507 | 0.1945 | 0.103 |
| C13orf30 | NM_182508 | 0.0225 | -0.092 |
| ISM2 | NM_182509 | -0.007 | -0.4875 |
| CBLN2 | NM_182511 | -0.0755 | -0.3895 |
| SPC24 | NM_182513 | -0.6215 | -1.5185 |
| ZNF714 | NM_182515 | -1.7035 | -2.4245 |
| C1orf210 | NM_182517 | 0.0595 | 0.5025 |
| BPIFB4 | NM_182519 | 0.281 | -0.4195 |

| | | | |
|---|---|---|---|
| C22orf15 | NM_182520 | -0.3875 | -0.2205 |
| ZSWIM2 | NM_182521 | -0.107 | -0.2145 |
| FAM19A4 | NM_182522 | -0.0795 | -1.037 |
| CMC1 | NM_182523 | -0.524 | -1.4405 |
| ZNF595 | NM_182524 | -0.94 | -2.665 |
| TMEM229B | NM_182526 | -0.188 | -0.331 |
| CABP7 | NM_182527 | 0.0165 | -2.1175 |
| C1QL2 | NM_182528 | -0.3845 | 0.258 |
| THAP5 | NM_182529 | -1.464 | -2.5725 |
| ZNF778 | NM_182531 | -0.847 | -1.2845 |
| TMEM61 | NM_182532 | 0.1445 | 0.044 |
| C1orf86 | NM_182533 | 0.1065 | -0.4025 |
| GKN2 | NM_182536 | -0.2305 | 0.3665 |
| HTR3D | NM_182537 | -0.136 | -0.3185 |
| SPNS3 | NM_182538 | -0.042 | -0.4125 |
| TCTE1 | NM_182539 | 0.125 | -0.3375 |
| DDX26B | NM_182540 | -0.9975 | -2.1325 |
| TMEM31 | NM_182541 | -0.2485 | -0.2225 |
| FAM194B | NM_182542 | 0.21 | 0.027 |
| NSUN6 | NM_182543 | 0.0175 | 0.5525 |
| VSTM2A | NM_182546 | -1.427 | -1.0845 |
| TMED4 | NM_182547 | -0.2215 | -0.6765 |
| LHFPL5 | NM_182548 | -0.019 | 0.0555 |
| LCLAT1 | NM_182551 | -1.444 | -2.4 |
| WDR27 | NM_182552 | -0.7935 | -0.7385 |
| CNIH2 | NM_182553 | 0.1485 | -0.4485 |
| C10orf53 | NM_182554 | -0.053 | -0.2365 |
| SLC25A45 | NM_182556 | -0.099 | -0.9425 |
| BCL9L | NM_182557 | -0.696 | -1.845 |
| TMPRSS12 | NM_182559 | 0.6655 | -0.1825 |
| C14orf177 | NM_182560 | 0.044 | 0.097 |
| FAM169B | NM_182562 | 0.1895 | -0.1625 |
| C16orf79 | NM_182563 | -0.352 | -0.7015 |
| FAM100B | NM_182565 | -0.508 | -2.0105 |
| VMO1 | NM_182566 | 0.0295 | -0.7035 |
| GDPD1 | NM_182569 | -2.4975 | -3.392 |
| ZSCAN1 | NM_182572 | 0.023 | -0.6595 |
| LYPD5 | NM_182573 | 0.2225 | -0.2255 |
| MAMSTR | NM_182574 | -0.139 | -0.5555 |
| IZUMO1 | NM_182575 | 0.0015 | -0.446 |
| ODF3L2 | NM_182577 | -0.1175 | -0.2405 |
| THEM5 | NM_182578 | -0.067 | 0.1895 |
| CYB561D1 | NM_182580 | 0.0715 | -0.298 |

| | | | |
|---|---|---|---|
| C1orf111 | NM_182581 | 0.006 | -0.045 |
| UNC80 | NM_182587 | -0.009 | -0.5335 |
| RGPD4 | NM_182588 | -0.65 | -2.405 |
| HTR3E | NM_182589 | -0.156 | -0.39 |
| YIPF7 | NM_182592 | -0.3655 | -0.4825 |
| ZNF454 | NM_182594 | -0.4255 | -0.699 |
| POM121L12 | NM_182595 | -0.2345 | -0.445 |
| C7orf53 | NM_182597 | 0.0505 | -0.9355 |
| C10orf81 | NM_182601 | 0.1905 | -0.272 |
| ANKRD42 | NM_182603 | -0.8885 | -1.5125 |
| TMPRSS11A | NM_182606 | -0.013 | 0.0625 |
| VSIG1 | NM_182607 | 0.1535 | -0.3755 |
| ANKRD33 | NM_182608 | 0.058 | -0.303 |
| ZNF677 | NM_182609 | -0.2965 | -1.089 |
| SAMD7 | NM_182610 | -0.0995 | -0.0725 |
| PDDC1 | NM_182612 | -0.056 | -0.7085 |
| FAM70B | NM_182614 | 0.114 | -0.3035 |
| C15orf38 | NM_182616 | -0.2865 | -0.7685 |
| ACSM2B | NM_182617 | -0.0065 | -0.023 |
| SKA2 | NM_182620 | -1.3445 | -0.85 |
| FAM131C | NM_182623 | -0.1275 | -0.3145 |
| GEN1 | NM_182625 | -0.458 | -2.017 |
| C2orf48 | NM_182626 | -0.392 | -0.33 |
| WDR53 | NM_182627 | -1.8485 | -3.1945 |
| CCDC37 | NM_182628 | -0.096 | -0.235 |
| SLC6A18 | NM_182632 | 0.212 | -0.125 |
| ZNF713 | NM_182633 | -0.878 | -1.1345 |
| HPS1 | NM_182639 | 0.005 | -0.6805 |
| MRPS9 | NM_182640 | -1.0855 | -2.878 |
| BPTF | NM_182641 | -1.2375 | -2.0965 |
| CTDSP1 | NM_182642 | -0.375 | -0.399 |
| DLC1 | NM_182643 | -1.246 | -2.5865 |
| EPHA3 | NM_182644 | 0.213 | -0.1 |
| VGLL2 | NM_182645 | -0.043 | -0.121 |
| CPEB2 | NM_182646 | 0.569 | -0.0055 |
| OPRL1 | NM_182647 | -1.03 | -1.4635 |
| BAZ1A | NM_182648 | -1.018 | -2.812 |
| PCNA | NM_182649 | -1.081 | -0.9455 |
| BPIFB3 | NM_182658 | -0.198 | -0.6705 |
| UTY | NM_182659 | 0.003 | -0.875 |
| UTY | NM_182660 | 0.003 | -0.875 |
| RASSF5 | NM_182663 | -0.189 | -0.6775 |
| RASSF5 | NM_182664 | -0.189 | -0.6775 |

| | | | |
|---|---|---|---|
| RASSF5 | NM_182665 | -0.861 | -0.2125 |
| UBE2E1 | NM_182666 | -0.9535 | -0.9985 |
| PLTP | NM_182676 | 0.1485 | -0.5285 |
| UBE2E3 | NM_182678 | -0.163 | -0.367 |
| GPATCH4 | NM_182679 | -1.387 | -2.406 |
| AMELX | NM_182680 | -0.0465 | -0.4 |
| AMELX | NM_182681 | -0.0465 | -0.4 |
| UPK3B | NM_182684 | 0.1935 | -0.125 |
| EFNA1 | NM_182685 | -0.516 | -0.4145 |
| PKMYT1 | NM_182687 | -0.537 | -1.4205 |
| UBE2G2 | NM_182688 | -0.591 | -0.8965 |
| SRPK2 | NM_182691 | -1.401 | -1.814 |
| SRPK2 | NM_182692 | -0.524 | -1.0745 |
| UBE2H | NM_182697 | -1.0205 | -1.6345 |
| DDX53 | NM_182699 | -0.085 | 0.0975 |
| SP8 | NM_182700 | 0.19 | 0.027 |
| GPX6 | NM_182701 | -0.0085 | -0.2335 |
| PRSS42 | NM_182702 | 0.0445 | -0.2715 |
| ANKDD1A | NM_182703 | -1.176 | -0.9235 |
| SELV | NM_182704 | 0.1675 | 0.1985 |
| FAM101B | NM_182705 | -0.8335 | -0.8615 |
| SCRIB | NM_182706 | -0.136 | -0.699 |
| PSG8 | NM_182707 | -0.1775 | -0.2295 |
| SYPL1 | NM_182715 | -0.4565 | -2.174 |
| CREM | NM_182717 | -1.5625 | -1.937 |
| CREM | NM_182718 | -1.5625 | -1.937 |
| CREM | NM_182719 | -1.5625 | -1.937 |
| CREM | NM_182720 | -1.5625 | -1.937 |
| CREM | NM_182721 | -1.5625 | -1.937 |
| CREM | NM_182722 | -1.5625 | -1.937 |
| CREM | NM_182723 | -1.5625 | -1.937 |
| CREM | NM_182724 | -1.5625 | -1.937 |
| CREM | NM_182725 | -1.5625 | -1.937 |
| SLC7A8 | NM_182728 | -0.096 | -0.0705 |
| TXNRD1 | NM_182729 | -1.177 | -1.714 |
| PLCB1 | NM_182734 | -0.1205 | -0.646 |
| NDUFB6 | NM_182739 | -0.3485 | -0.7025 |
| TXNRD1 | NM_182742 | -1.177 | -1.714 |
| TXNRD1 | NM_182743 | -1.177 | -1.714 |
| NBL1 | NM_182744 | -0.012 | -0.7585 |
| MCM4 | NM_182746 | -0.47 | -0.8205 |
| N6AMT1 | NM_182749 | -1.0305 | -2.752 |
| MCM10 | NM_182751 | -1.8195 | -3.7565 |

| | | | |
|---|---|---|---|
| TPRG1L | NM_182752 | -0.8145 | -1.273 |
| SPDYA | NM_182756 | -1.208 | -1.9585 |
| RNF144B | NM_182757 | -0.2435 | 0.212 |
| WDR72 | NM_182758 | -0.3445 | 0.142 |
| FAM19A3 | NM_182759 | -0.0445 | -0.258 |
| SUMF1 | NM_182760 | -0.7115 | -1.17 |
| FAM170A | NM_182761 | -0.2355 | -0.1995 |
| MACC1 | NM_182762 | 0.04 | -0.0065 |
| MCL1 | NM_182763 | 0.1675 | -0.209 |
| ELMO2 | NM_182764 | -0.2885 | -1.793 |
| HECTD2 | NM_182765 | -1.5305 | -1.5655 |
| SLC6A15 | NM_182767 | -0.975 | -1.2145 |
| CREM | NM_182769 | -1.5625 | -1.937 |
| CREM | NM_182770 | -1.5625 | -1.937 |
| CREM | NM_182771 | -1.5625 | -1.937 |
| CREM | NM_182772 | -1.5625 | -1.937 |
| ALS2CL | NM_182775 | 0.067 | -0.5855 |
| MCM7 | NM_182776 | -0.114 | -0.7625 |
| PAIP1 | NM_182789 | -1.355 | -2.3645 |
| CCDC33 | NM_182791 | 0.1785 | -0.24 |
| NPM2 | NM_182795 | -0.127 | -0.2395 |
| MAT2B | NM_182796 | -1.459 | -2.274 |
| PLCB4 | NM_182797 | -0.5485 | -1.781 |
| EGFLAM | NM_182798 | -0.277 | -0.894 |
| EGFLAM | NM_182801 | -0.277 | -0.894 |
| MCM8 | NM_182802 | -0.878 | -1.377 |
| ATF4 | NM_182810 | -0.635 | -1.2975 |
| PLCG1 | NM_182811 | 0.006 | -0.736 |
| SCARA3 | NM_182826 | -0.593 | -1.315 |
| GDF7 | NM_182828 | 0.1385 | -0.284 |
| MDGA2 | NM_182830 | -0.4705 | 0.1505 |
| PLAC4 | NM_182832 | 0.085 | 0.0645 |
| GDPD4 | NM_182833 | -0.194 | -0.386 |
| SCFD1 | NM_182835 | -0.807 | -2.047 |
| RABGGTA | NM_182836 | -0.2205 | -0.8645 |
| SLC35E2 | NM_182838 | -0.1305 | -0.6825 |
| ACCN4 | NM_182847 | 0.0265 | -0.755 |
| CLDN10 | NM_182848 | -0.342 | -0.061 |
| CCNB1IP1 | NM_182849 | -1.152 | -0.782 |
| CREM | NM_182850 | -1.5625 | -1.937 |
| CCNB1IP1 | NM_182851 | -0.5215 | -0.16 |
| CCNB1IP1 | NM_182852 | -1.152 | -0.782 |
| CREM | NM_182853 | -1.5625 | -1.937 |

| | | | |
|---|---|---|---|
| SNX20 | NM_182854 | -0.13 | -0.257 |
| VSX2 | NM_182894 | -0.0585 | -0.1265 |
| ARL13B | NM_182896 | -1.172 | -2.9525 |
| CREB5 | NM_182898 | -0.219 | -1.026 |
| CREB5 | NM_182899 | -0.219 | -1.026 |
| AKIP1 | NM_182901 | -0.5725 | -1.3675 |
| KIF9 | NM_182902 | -0.163 | -0.91 |
| P4HA3 | NM_182904 | -0.086 | -0.2785 |
| CLEC10A | NM_182906 | -0.03 | -0.2195 |
| PRDM1 | NM_182907 | 0 | -0.304 |
| DHRS2 | NM_182908 | -0.331 | -0.5265 |
| FILIP1L | NM_182909 | -0.1355 | -0.2175 |
| SYNE2 | NM_182910 | -0.116 | -0.9965 |
| TSGA10 | NM_182911 | 0.211 | 0.173 |
| SYNE2 | NM_182913 | -0.116 | -0.9965 |
| SYNE2 | NM_182914 | -0.1285 | -0.6255 |
| STEAP3 | NM_182915 | -0.614 | -0.933 |
| TRNT1 | NM_182916 | -1.2445 | -2.3095 |
| EIF4G1 | NM_182917 | -0.1565 | -0.948 |
| ERG | NM_182918 | -0.1435 | -0.526 |
| TICAM1 | NM_182919 | -0.409 | -2.1455 |
| ADAMTS9 | NM_182920 | -1.344 | -2.6965 |
| HEATR3 | NM_182922 | -0.7525 | -2.5665 |
| KLC1 | NM_182923 | -0.539 | -0.653 |
| MICALL2 | NM_182924 | -0.0485 | -0.089 |
| FLT4 | NM_182925 | 0.0755 | -0.3575 |
| KTN1 | NM_182926 | 0.2755 | -1.327 |
| MLL5 | NM_182931 | -0.3845 | -1.8195 |
| MOBP | NM_182935 | -0.3065 | -0.085 |
| PLOD2 | NM_182943 | -0.507 | -1.801 |
| NIN | NM_182944 | -0.7115 | -2.9645 |
| NIN | NM_182946 | -0.7115 | -2.9645 |
| ARHGEF25 | NM_182947 | -0.142 | -0.7815 |
| PRKACB | NM_182948 | -1.4275 | -1.8175 |
| PRELID2 | NM_182960 | -0.1895 | -0.604 |
| SYNE1 | NM_182961 | -0.778 | -1.851 |
| BIRC3 | NM_182962 | -0.24 | -0.0535 |
| NAV2 | NM_182964 | -0.733 | -3.3365 |
| SPHK1 | NM_182965 | -0.1685 | -0.8105 |
| NEDD9 | NM_182966 | -0.455 | -1.4445 |
| XRRA1 | NM_182969 | -1.166 | -2.0055 |
| RIMS4 | NM_182970 | -0.2505 | -0.408 |
| COX8C | NM_182971 | -0.629 | -0.779 |

| | | | |
|---|---|---|---|
| IRF2BP2 | NM_182972 | -0.8925 | -1.7395 |
| TMPRSS9 | NM_182973 | 0.0725 | -0.537 |
| GLT6D1 | NM_182974 | 0.355 | -0.2065 |
| ZNF326 | NM_182975 | -0.962 | -1.643 |
| ZNF326 | NM_182976 | -0.962 | -1.643 |
| NNT | NM_182977 | -0.662 | -0.6205 |
| GNAL | NM_182978 | 0.2605 | -0.6505 |
| OSGIN1 | NM_182981 | 0.0295 | -0.1365 |
| GRK4 | NM_182982 | -0.9455 | -1.2855 |
| HPN | NM_182983 | -0.136 | -0.594 |
| TRMT2A | NM_182984 | -0.595 | -1.096 |
| TRIM69 | NM_182985 | -0.0125 | -0.1875 |
| SHC1 | NM_183001 | -0.1715 | -1.0885 |
| SLC8A3 | NM_183002 | -1.663 | -2.5345 |
| EIF5 | NM_183004 | -1.4435 | -2.6125 |
| DLGAP4 | NM_183006 | -0.685 | -1.695 |
| UBXN11 | NM_183008 | 0.113 | -0.131 |
| KIAA1429 | NM_183009 | -0.584 | -2.4705 |
| CREM | NM_183011 | -1.5625 | -1.937 |
| CREM | NM_183012 | -1.5625 | -1.937 |
| CREM | NM_183013 | -1.5625 | -1.937 |
| DTNBP1 | NM_183040 | -0.446 | -1.0745 |
| RNF6 | NM_183043 | -0.3705 | -1.2255 |
| RNF6 | NM_183044 | -0.3705 | -1.2255 |
| ZMYND8 | NM_183047 | -1.439 | -3.8555 |
| ZMYND8 | NM_183048 | -1.439 | -3.8555 |
| BCKDHB | NM_183050 | -0.64 | -0.8255 |
| VPS28 | NM_183057 | 0.1485 | -0.6045 |
| LYZL2 | NM_183058 | -0.1805 | -0.2045 |
| RD3 | NM_183059 | -0.0705 | -0.56 |
| CREM | NM_183060 | -1.5625 | -1.937 |
| SLC9A10 | NM_183061 | -0.0025 | -0.065 |
| PRSS38 | NM_183062 | -0.177 | -0.3745 |
| TMEM107 | NM_183065 | -0.1865 | -0.5385 |
| CYP2U1 | NM_183075 | -1.3165 | -2.397 |
| RNF8 | NM_183078 | -1.169 | -2.8165 |
| PRNP | NM_183079 | -1.532 | -1.462 |
| RAB23 | NM_183227 | -1.1295 | -2.823 |
| IKZF3 | NM_183228 | -0.1495 | 0.048 |
| IKZF3 | NM_183229 | -0.1495 | 0.048 |
| IKZF3 | NM_183230 | -0.443 | -0.12 |
| IKZF3 | NM_183231 | -0.1495 | 0.048 |
| IKZF3 | NM_183232 | -0.1495 | 0.048 |

| | | | |
|---|---|---|---|
| SLC22A18 | NM_183233 | -0.084 | -0.4625 |
| RAB27A | NM_183234 | -0.468 | -1.317 |
| RAB27A | NM_183235 | -0.468 | -1.317 |
| RAB27A | NM_183236 | -0.468 | -1.317 |
| RNF7 | NM_183237 | -0.175 | -0.8935 |
| ZNF605 | NM_183238 | -0.958 | -1.7945 |
| GSTO2 | NM_183239 | -0.043 | -0.7895 |
| TMEM37 | NM_183240 | 0.1455 | -0.475 |
| C9orf142 | NM_183241 | -0.0815 | -0.6645 |
| BTBD8 | NM_183242 | 0.042 | -0.2555 |
| IMPDH1 | NM_183243 | -0.0955 | -1.069 |
| PHACTR3 | NM_183244 | -0.8465 | -1.5995 |
| INVS | NM_183245 | -0.6355 | -1.393 |
| PHACTR3 | NM_183246 | -0.8465 | -1.5995 |
| PAIP1 | NM_183323 | -1.355 | -2.3645 |
| RGS11 | NM_183337 | -0.156 | -0.396 |
| SEC13 | NM_183352 | 0.0055 | -1.0065 |
| RLIM | NM_183353 | -1.784 | -2.666 |
| ASNS | NM_183356 | -0.3555 | -0.499 |
| ADCY5 | NM_183357 | -0.1925 | -0.78 |
| DTNB | NM_183360 | -0.2295 | -0.905 |
| DTNB | NM_183361 | -0.2295 | -0.905 |
| C6orf145 | NM_183373 | -0.0865 | -1.108 |
| CYP26C1 | NM_183374 | 0.2245 | -0.1505 |
| PRSS48 | NM_183375 | -0.3305 | -0.257 |
| ARRDC4 | NM_183376 | -0.801 | -1.308 |
| ACCN1 | NM_183377 | -0.571 | -0.3355 |
| OVCH1 | NM_183378 | -0.3185 | -0.49 |
| RNF13 | NM_183381 | -1.259 | -1.978 |
| EML5 | NM_183387 | -1.0435 | -1.32 |
| CADPS | NM_183393 | -0.6905 | -0.4185 |
| CADPS | NM_183394 | -0.6905 | -0.4185 |
| NLRP3 | NM_183395 | -0.0565 | 0.0615 |
| PXMP4 | NM_183397 | -0.191 | -1.313 |
| RNF14 | NM_183398 | -1.5105 | -3.03 |
| RNF14 | NM_183399 | -1.5105 | -3.03 |
| RNF14 | NM_183400 | -1.5105 | -3.03 |
| RNF14 | NM_183401 | -1.5105 | -3.03 |
| RBL1 | NM_183404 | -0.599 | -1.504 |
| FBXO44 | NM_183412 | -0.2985 | -0.644 |
| FBXO44 | NM_183413 | -0.2985 | -0.644 |
| UBE3B | NM_183415 | -0.971 | -1.959 |
| KIF1B | NM_183416 | -1.0095 | -1.65 |

| | | | |
|---|---|---|---|
| RNF19A | NM_183419 | -1.2735 | -2.3865 |
| FBXO25 | NM_183420 | -0.3795 | -1.0575 |
| FBXO25 | NM_183421 | -0.3795 | -1.0575 |
| TSC22D1 | NM_183422 | -0.259 | 0.5975 |
| ALDOA | NM_184041 | 0.2905 | -0.155 |
| ALDOA | NM_184043 | 0.305 | -0.157 |
| TRIM55 | NM_184085 | -0.77 | 0.197 |
| TRIM55 | NM_184086 | -0.77 | 0.197 |
| TRIM55 | NM_184087 | -0.77 | 0.197 |
| NCKIPSD | NM_184231 | -0.373 | -0.437 |
| RBM39 | NM_184234 | -1.0695 | -2.1315 |
| TRIM54 | NM_187841 | -0.142 | -0.1635 |
| CREB3L2 | NM_194071 | -0.5945 | -1.7485 |
| HNRNPA3 | NM_194247 | -0.7865 | -0.4355 |
| OTOF | NM_194248 | -0.082 | 0.228 |
| DND1 | NM_194249 | 0.671 | 1.0065 |
| ZNF804A | NM_194250 | -0.7285 | -3.939 |
| GPR151 | NM_194251 | 0.005 | -0.406 |
| TTLL11 | NM_194252 | -0.663 | -2.154 |
| SLC19A1 | NM_194255 | -0.8135 | -1.8015 |
| UBE2I | NM_194259 | -0.544 | -0.675 |
| UBE2I | NM_194260 | -0.544 | -0.675 |
| UBE2I | NM_194261 | -0.544 | -0.675 |
| RNF34 | NM_194271 | -0.4835 | -1.1215 |
| RBPMS2 | NM_194272 | -0.128 | -0.901 |
| FRMD7 | NM_194277 | -0.15 | 0.0155 |
| C14orf43 | NM_194278 | -0.628 | -3.4255 |
| ISCA2 | NM_194279 | -1.5705 | -3.999 |
| TMEM219 | NM_194280 | 0.2185 | -0.4965 |
| LIN54 | NM_194282 | -0.743 | -2.6445 |
| DNAJC21 | NM_194283 | -1.107 | -2.3615 |
| CLDN23 | NM_194284 | -0.006 | -0.207 |
| SPTY2D1 | NM_194285 | -1.663 | -2.058 |
| SRRM4 | NM_194286 | -1.8135 | -2.6425 |
| C14orf166B | NM_194287 | 0.068 | -0.1285 |
| TMEM65 | NM_194291 | -0.922 | -1.2955 |
| SASS6 | NM_194292 | -1.028 | -2.6805 |
| XIRP1 | NM_194293 | -0.0875 | -0.3295 |
| IDO2 | NM_194294 | -0.058 | -0.599 |
| SPATA24 | NM_194296 | 0.0505 | -0.4045 |
| SLC16A9 | NM_194298 | -1.843 | -2.573 |
| CCDC129 | NM_194300 | 0.175 | -0.0535 |
| RALGAPA1 | NM_194301 | 0.107 | -0.3085 |

| | | | |
|---|---|---|---|
| CCDC108 | NM_194302 | 0.1525 | -0.128 |
| ESPNL | NM_194312 | -0.022 | -0.215 |
| KIF24 | NM_194313 | -1.0745 | -1.7295 |
| ZBTB41 | NM_194314 | -0.802 | -3.344 |
| UBE2J2 | NM_194315 | -0.808 | -2.5555 |
| LYPD6 | NM_194317 | -0.122 | -0.588 |
| B3GALTL | NM_194318 | -0.867 | -1.335 |
| ZNF169 | NM_194320 | -0.386 | -1.0635 |
| OTOF | NM_194322 | -0.2305 | -0.381 |
| OTOF | NM_194323 | -0.2305 | -0.381 |
| TMSB15B | NM_194324 | -0.176 | -0.2215 |
| ZNF30 | NM_194325 | -1.384 | -2.6945 |
| RPS19BP1 | NM_194326 | -0.554 | -1.664 |
| RNF38 | NM_194328 | -1.237 | -2.0815 |
| RNF38 | NM_194329 | -0.8165 | -1 |
| RNF38 | NM_194330 | -0.8165 | -1 |
| RNF38 | NM_194332 | -0.8165 | -1 |
| STX2 | NM_194356 | -1.258 | -1.53 |
| RNF41 | NM_194358 | -0.776 | -2.4925 |
| RNF41 | NM_194359 | -0.776 | -2.4925 |
| FGFR1OP | NM_194429 | -1.1925 | -1.378 |
| RNASE4 | NM_194431 | -0.332 | -0.1095 |
| VAPA | NM_194434 | -0.824 | -1.753 |
| VIP | NM_194435 | -0.2855 | -0.3655 |
| RNF212 | NM_194439 | -0.092 | -0.003 |
| BTN3A1 | NM_194441 | -0.9645 | -0.962 |
| LBR | NM_194442 | -0.771 | -1.3795 |
| PHLPP1 | NM_194449 | -0.537 | -2.153 |
| LIAS | NM_194451 | -0.459 | -0.9645 |
| KRIT1 | NM_194454 | -1.236 | -1.9075 |
| KRIT1 | NM_194455 | -1.236 | -1.9075 |
| KRIT1 | NM_194456 | -1.236 | -1.9075 |
| UBE2J2 | NM_194457 | -0.808 | -2.5555 |
| UBE2J2 | NM_194458 | -0.808 | -2.5555 |
| RNF126 | NM_194460 | -0.6815 | -1.725 |
| RNF135 | NM_197939 | -0.099 | -0.3995 |
| ADAMTS6 | NM_197941 | -0.0965 | -0.2195 |
| CLEC7A | NM_197947 | -0.082 | -0.414 |
| CLEC7A | NM_197948 | -0.082 | -0.414 |
| CLEC7A | NM_197949 | -0.082 | -0.414 |
| CLEC7A | NM_197950 | -0.082 | -0.414 |
| CLEC7A | NM_197954 | -0.082 | -0.414 |
| C15orf48 | NM_197955 | -0.329 | -0.363 |

| | | | |
|---|---|---|---|
| NAIF1 | NM_197956 | -0.8585 | -1.7065 |
| MAX | NM_197957 | -0.588 | -2.2715 |
| LARP6 | NM_197958 | 0.396 | -0.1955 |
| DPP8 | NM_197960 | -1.3795 | -1.5685 |
| DPP8 | NM_197961 | -1.3795 | -1.5685 |
| C7orf55 | NM_197964 | -0.394 | -1.126 |
| SLC10A6 | NM_197965 | 0.216 | 0.0345 |
| BID | NM_197966 | -0.028 | -1.151 |
| BID | NM_197967 | -0.028 | -1.151 |
| ZMYM2 | NM_197968 | -0.9155 | -2.786 |
| BOLL | NM_197970 | -0.1165 | -0.074 |
| NME7 | NM_197972 | -0.944 | -1.408 |
| BTN3A3 | NM_197974 | -0.139 | -0.7845 |
| BTNL3 | NM_197975 | 0.327 | -0.3925 |
| ZNF189 | NM_197977 | -1.65 | -3.72 |
| HEMGN | NM_197978 | 0.092 | -0.3815 |
| PHC2 | NM_198040 | -0.8175 | -1.99 |
| NUDT6 | NM_198041 | -1.166 | -2.557 |
| ZDHHC16 | NM_198043 | -0.4315 | -1.684 |
| ZDHHC16 | NM_198044 | -0.4315 | -1.684 |
| ZDHHC16 | NM_198045 | -0.4315 | -1.684 |
| ZDHHC16 | NM_198046 | -0.4315 | -1.684 |
| HIBCH | NM_198047 | -0.5965 | -0.9265 |
| CD247 | NM_198053 | -0.409 | -0.555 |
| MZF1 | NM_198055 | -0.8915 | -0.9335 |
| SCN5A | NM_198056 | 0.021 | -0.709 |
| TSC22D3 | NM_198057 | -0.063 | 0.017 |
| ZNF266 | NM_198058 | -1.6575 | -2.9525 |
| NRAP | NM_198060 | -0.291 | -0.437 |
| CES2 | NM_198061 | -0.5975 | -1.474 |
| GNPNAT1 | NM_198066 | -1.492 | -3.502 |
| OR2C3 | NM_198074 | -0.072 | -0.0785 |
| LRRC56 | NM_198075 | -0.1605 | -0.6555 |
| FAM36A | NM_198076 | -0.6465 | -1.142 |
| C1orf52 | NM_198077 | -1.532 | -2.2295 |
| MSRB3 | NM_198080 | -0.2365 | -0.701 |
| SCML4 | NM_198081 | 0.231 | 0.156 |
| CCDC57 | NM_198082 | -0.533 | -0.8895 |
| DHRS4L2 | NM_198083 | -0.148 | -0.4855 |
| RNF148 | NM_198085 | 0.0755 | 0.1295 |
| JUB | NM_198086 | -0.8655 | -1.3835 |
| ZNF200 | NM_198087 | -1.8995 | -3.8065 |
| ZNF200 | NM_198088 | -1.8995 | -3.8065 |

| | | | |
|---|---|---|---|
| ZNF155 | NM_198089 | -0.9545 | -1.674 |
| AQP1 | NM_198098 | -0.4675 | -0.332 |
| EBAG9 | NM_198120 | -1.5475 | -3.593 |
| CSMD3 | NM_198123 | 0.0275 | 0.0215 |
| CSMD3 | NM_198124 | 0.0275 | 0.0215 |
| TYROBP | NM_198125 | 0.058 | -0.44 |
| RNF138 | NM_198128 | -1.3675 | -2.6415 |
| CATSPER4 | NM_198137 | -0.309 | -0.0175 |
| GANC | NM_198141 | -0.195 | -0.401 |
| ABHD15 | NM_198147 | -0.624 | -0.7265 |
| CPXM2 | NM_198148 | -0.2165 | -0.3775 |
| SHISA4 | NM_198149 | -0.0965 | -0.5855 |
| ARSK | NM_198150 | -1.6745 | -2.8185 |
| HEPACAM2 | NM_198151 | 0.185 | 0.0755 |
| UTS2D | NM_198152 | 0.0985 | 0.102 |
| TREML4 | NM_198153 | 0.2135 | -0.3395 |
| TMEM95 | NM_198154 | -0.103 | -0.145 |
| C21orf33 | NM_198155 | -0.6465 | -1.044 |
| VHL | NM_198156 | -0.342 | -0.81 |
| MITF | NM_198158 | -0.596 | -0.749 |
| MITF | NM_198159 | -0.596 | -0.749 |
| GRHL3 | NM_198173 | 0.279 | -0.256 |
| MITF | NM_198177 | -0.596 | -0.749 |
| MITF | NM_198178 | -0.596 | -0.749 |
| QRFPR | NM_198179 | 0.3715 | -0.8925 |
| QRFP | NM_198180 | 0.045 | 0.0865 |
| GOLGA6L9 | NM_198181 | -0.643 | -0.6045 |
| GRHL1 | NM_198182 | -0.248 | -0.669 |
| UBE2L6 | NM_198183 | 0.0235 | -0.6875 |
| OSTN | NM_198184 | -0.2685 | 0.0675 |
| OVCH2 | NM_198185 | 0.0135 | 0.122 |
| ASTN2 | NM_198186 | 0.029 | -1.152 |
| ASTN2 | NM_198187 | 0.0335 | -0.978 |
| ASTN2 | NM_198188 | 0.029 | -1.152 |
| COPS8 | NM_198189 | -0.568 | -0.578 |
| STOM | NM_198194 | -0.2575 | -0.409 |
| UBA3 | NM_198195 | -1.104 | -2.075 |
| CD96 | NM_198196 | -0.1315 | -0.111 |
| MLX | NM_198204 | -0.974 | -2.578 |
| MLX | NM_198205 | -0.974 | -2.578 |
| CERS1 | NM_198207 | -0.445 | -0.7115 |
| CAV2 | NM_198212 | 0.498 | 0.241 |
| FAM13C | NM_198215 | -0.277 | -1.1595 |

| | | | |
|---|---|---|---|
| SNRPB | NM_198216 | 0.0995 | -0.723 |
| ING1 | NM_198217 | -0.47 | -1.281 |
| ING1 | NM_198218 | -0.47 | -1.281 |
| ING1 | NM_198219 | -0.6625 | -0.9285 |
| SNRPB2 | NM_198220 | -0.9335 | -0.602 |
| RGS12 | NM_198227 | 0.0255 | -0.857 |
| RGS12 | NM_198229 | 0.0255 | -0.857 |
| RNASE1 | NM_198235 | -0.0115 | -0.142 |
| ARHGEF11 | NM_198236 | -0.802 | -1.649 |
| WISP3 | NM_198239 | -0.3395 | -0.111 |
| CLIP1 | NM_198240 | -0.7815 | -1.605 |
| EIF4G1 | NM_198241 | -0.1565 | -0.948 |
| EIF4G1 | NM_198242 | -0.1565 | -0.948 |
| ASB7 | NM_198243 | -1.791 | -3.9145 |
| EIF4G1 | NM_198244 | -0.1565 | -0.948 |
| TERT | NM_198253 | -0.6095 | -0.8365 |
| E2F6 | NM_198256 | -1.5175 | -4.976 |
| ING3 | NM_198267 | -0.105 | 1.974 |
| HIPK1 | NM_198268 | -1.4105 | -1.615 |
| HIPK1 | NM_198269 | 0.267 | -0.807 |
| NHS | NM_198270 | -0.2435 | -0.3225 |
| LMOD3 | NM_198271 | -0.4285 | 0.245 |
| LYSMD3 | NM_198273 | -1.533 | -3.493 |
| SMYD1 | NM_198274 | 0.074 | -0.107 |
| MPZL3 | NM_198275 | -0.026 | -0.076 |
| TMEM17 | NM_198276 | -0.5805 | -1.2655 |
| SLC37A2 | NM_198277 | 0.045 | -0.496 |
| CXorf23 | NM_198279 | -0.8525 | -1.406 |
| GPRIN3 | NM_198281 | -0.321 | -0.3735 |
| TMEM173 | NM_198282 | -0.0815 | -0.373 |
| EYS | NM_198283 | 0.3525 | -0.5735 |
| WDR86 | NM_198285 | -0.316 | -0.4415 |
| TTC8 | NM_198309 | -1.174 | -2.0145 |
| TTC8 | NM_198310 | -1.1725 | -1.948 |
| VWA5A | NM_198315 | -0.06 | -0.016 |
| TENC1 | NM_198316 | -0.3235 | -0.4435 |
| KLHL17 | NM_198317 | -0.161 | -0.882 |
| PRMT1 | NM_198318 | -0.1365 | -0.383 |
| CPM | NM_198320 | 0.222 | -0.293 |
| GALNT10 | NM_198321 | -0.454 | -0.819 |
| UBA5 | NM_198329 | -0.6 | -1.1065 |
| STAT2 | NM_198332 | -0.1785 | -1.0725 |
| P2RY10 | NM_198333 | 0.181 | 0.33 |

| | | | |
|---|---|---|---|
| GANAB | NM_198334 | 0.0525 | -0.578 |
| GANAB | NM_198335 | 0.0525 | -0.578 |
| INSIG1 | NM_198336 | -1.733 | -4.4215 |
| INSIG1 | NM_198337 | -0.7045 | -2.0735 |
| KCTD8 | NM_198353 | 0.019 | -2.279 |
| ELF5 | NM_198381 | -0.3485 | -0.1935 |
| PDPN | NM_198389 | 0.369 | -0.079 |
| CMIP | NM_198390 | -0.2595 | -1.246 |
| FLRT3 | NM_198391 | -1.2565 | -3.191 |
| TCF21 | NM_198392 | -0.51 | 0.552 |
| TEX14 | NM_198393 | -0.487 | -0.4225 |
| G3BP1 | NM_198395 | -0.9005 | -1.1585 |
| CACNA1G | NM_198396 | -0.072 | -0.253 |
| CACNA1G | NM_198397 | 0.0745 | -0.3485 |
| ERGIC3 | NM_198398 | 0.1 | -0.3035 |
| ARPP21 | NM_198399 | -0.757 | 0.091 |
| NEDD4 | NM_198400 | -0.8155 | -1.2625 |
| ANKRD46 | NM_198401 | -1.677 | -2.2245 |
| PTPLB | NM_198402 | -1.3045 | -1.472 |
| MMD2 | NM_198403 | -0.222 | -0.371 |
| KCTD4 | NM_198404 | -0.3095 | -0.43 |
| PAQR6 | NM_198406 | -0.2915 | -0.0805 |
| GHSR | NM_198407 | -0.005 | -0.298 |
| CHMP2A | NM_198426 | 0.0915 | -0.2545 |
| BCAN | NM_198427 | -0.1545 | -0.0145 |
| BBS9 | NM_198428 | -0.615 | -1.0105 |
| AURKA | NM_198433 | -1.172 | -3.3945 |
| KBTBD3 | NM_198439 | -1.057 | -0.79 |
| DERL3 | NM_198440 | -0.0925 | -0.6215 |
| NRN1L | NM_198443 | -0.028 | -0.5015 |
| RINL | NM_198445 | 0.0075 | -0.277 |
| C1orf122 | NM_198446 | 0.1605 | -0.6715 |
| GOLT1A | NM_198447 | 0.3245 | -0.1365 |
| REG3G | NM_198448 | 0.1515 | -0.042 |
| EMB | NM_198449 | -1.132 | -2.023 |
| APOOL | NM_198450 | -0.6855 | -0.6505 |
| FOXR2 | NM_198451 | -0.099 | -0.076 |
| SSPO | NM_198455 | 0.08 | -0.565 |
| FAM120C | NM_198456 | 0.152 | -0.2185 |
| ZNF600 | NM_198457 | -0.792 | -0.9395 |
| ZNF497 | NM_198458 | -0.279 | -0.3395 |
| DENND2C | NM_198459 | -0.2635 | -0.1415 |
| GBP6 | NM_198460 | 0.612 | -0.139 |

| | | | |
|---|---|---|---|
| LONRF2 | NM_198461 | -1.109 | -1.1425 |
| C3orf67 | NM_198463 | 0.273 | -0.7875 |
| PRSS55 | NM_198464 | 0.351 | 0.246 |
| NRK | NM_198465 | -0.042 | -0.28 |
| RSBN1L | NM_198467 | -0.7115 | -1.7645 |
| MMS22L | NM_198468 | -1.8265 | -3.427 |
| MORN5 | NM_198469 | 0.026 | -0.079 |
| KANK3 | NM_198471 | 0.14 | -0.159 |
| C10orf125 | NM_198472 | -0.145 | -0.476 |
| OLFML1 | NM_198474 | -0.06 | -0.051 |
| FAM171A2 | NM_198475 | -0.0595 | -1.379 |
| C19orf54 | NM_198476 | -0.648 | -1.512 |
| CXCL17 | NM_198477 | 0.0075 | -0.179 |
| NKPD1 | NM_198478 | -0.017 | -0.2635 |
| TPRX1 | NM_198479 | -0.0815 | -0.2285 |
| ZNF615 | NM_198480 | -0.449 | -2.3075 |
| VSTM1 | NM_198481 | 0.0725 | -0.182 |
| SH2D6 | NM_198482 | 0.001 | -0.5335 |
| RUFY4 | NM_198483 | 0.1165 | -0.4695 |
| ZNF621 | NM_198484 | -1.192 | -1.952 |
| TPRG1 | NM_198485 | 0.011 | -0.31 |
| RPL7L1 | NM_198486 | -1.148 | -1.612 |
| FAM83H | NM_198488 | 0.0635 | 0.259 |
| CCDC84 | NM_198489 | -0.5405 | -1.361 |
| FAM92B | NM_198491 | 0.623 | -0.173 |
| CLEC4G | NM_198492 | 0.141 | -0.4265 |
| ANKRD45 | NM_198493 | 0.0595 | -0.3165 |
| ZNF642 | NM_198494 | -1.268 | -2.3565 |
| VWA2 | NM_198496 | -0.1045 | -0.13 |
| C11orf53 | NM_198498 | -0.2975 | -0.3215 |
| SMTNL2 | NM_198501 | -0.311 | -0.5095 |
| KCNT2 | NM_198503 | -1.162 | -2.6985 |
| PAQR9 | NM_198504 | -0.1175 | -0.8825 |
| ATP13A5 | NM_198505 | 0.1805 | -0.2795 |
| LRIT3 | NM_198506 | -0.513 | -0.7455 |
| FAM174A | NM_198507 | -0.7755 | -3.8195 |
| KLRG2 | NM_198508 | 0.0195 | -0.3425 |
| ITIH6 | NM_198510 | -0.2475 | -0.1515 |
| LANCL3 | NM_198511 | -0.3095 | -0.303 |
| DGAT2L6 | NM_198512 | -0.1205 | -0.2545 |
| PHF20L1 | NM_198513 | -0.019 | -0.505 |
| NHLRC2 | NM_198514 | -0.504 | -1.5375 |
| C10orf96 | NM_198515 | 0.3155 | 0.142 |

| | | | |
|---|---|---|---|
| GALNTL4 | NM_198516 | -0.091 | -1.376 |
| TBC1D10C | NM_198517 | 0.069 | -0.218 |
| TEX9 | NM_198524 | -0.42 | -0.155 |
| KIF7 | NM_198525 | -0.4295 | -1.5255 |
| ZNF710 | NM_198526 | -0.586 | -1.97 |
| HDDC3 | NM_198527 | -0.748 | -1.898 |
| EFCAB5 | NM_198529 | 0.022 | -0.161 |
| MXRA7 | NM_198530 | 0.173 | -0.265 |
| ATP9B | NM_198531 | -0.0965 | -0.6195 |
| C19orf35 | NM_198532 | -0.1905 | -0.5885 |
| HSD11B1L | NM_198533 | 0.229 | -0.244 |
| C19orf45 | NM_198534 | -0.211 | -0.2795 |
| ZNF699 | NM_198535 | -0.1595 | -0.312 |
| YJEFN3 | NM_198537 | -0.031 | -0.746 |
| SBSN | NM_198538 | -0.3535 | -0.274 |
| ZNF568 | NM_198539 | -0.113 | -0.859 |
| B3GNT8 | NM_198540 | -0.227 | -0.4365 |
| IGFL1 | NM_198541 | 0.256 | 0.1885 |
| ZNF773 | NM_198542 | -1.113 | -2.0395 |
| C1orf187 | NM_198545 | 0.199 | -0.725 |
| SPATA21 | NM_198546 | -0.4785 | -0.41 |
| FAM73A | NM_198549 | -0.324 | -1.1565 |
| MIA3 | NM_198551 | -0.8255 | -2.181 |
| FAM89A | NM_198552 | -1.0525 | -1.7095 |
| RBM43 | NM_198557 | -0.9005 | -1.8285 |
| C2orf62 | NM_198559 | -0.129 | -0.4765 |
| LHFPL4 | NM_198560 | -0.689 | -1.488 |
| C3orf62 | NM_198562 | -1.36 | -1.723 |
| TMEM110 | NM_198563 | -0.1525 | -1.5435 |
| DNAH12 | NM_198564 | -0.5515 | -0.1095 |
| LRRC33 | NM_198565 | -0.5025 | -0.876 |
| C5orf34 | NM_198566 | -1.535 | -3.0955 |
| C5orf25 | NM_198567 | -0.095 | -3.7265 |
| GJB7 | NM_198568 | -0.0925 | 0.429 |
| GPR126 | NM_198569 | 0.149 | 0.0315 |
| VWC2 | NM_198570 | 0.2455 | -0.0895 |
| C7orf52 | NM_198571 | 0.0475 | -0.406 |
| SPATC1 | NM_198572 | -0.0015 | -0.236 |
| ENHO | NM_198573 | -0.3355 | -0.434 |
| AGRN | NM_198576 | -0.23 | -0.321 |
| LRRK2 | NM_198578 | -0.267 | 0.196 |
| SLC27A1 | NM_198580 | -0.325 | -0.628 |
| ZC3H6 | NM_198581 | -1.0545 | -2.9915 |

| | | | |
|---|---|---|---|
| KLHL30 | NM_198582 | -0.65 | -0.133 |
| CA13 | NM_198584 | -0.221 | 0.219 |
| ENTPD8 | NM_198585 | 0.1655 | -0.5425 |
| NHLRC1 | NM_198586 | -0.273 | -0.434 |
| BSG | NM_198589 | 0.389 | -0.097 |
| BSG | NM_198591 | 0.389 | -0.097 |
| C1QTNF1 | NM_198593 | 0.2255 | 0.164 |
| C1QTNF1 | NM_198594 | 0.2255 | 0.164 |
| AFAP1 | NM_198595 | -0.6 | -1.938 |
| SULF2 | NM_198596 | -0.6765 | -1.0065 |
| SEC24C | NM_198597 | -0.3155 | -1.7295 |
| RAPGEF1 | NM_198679 | -0.599 | -1.5355 |
| PLEKHG5 | NM_198681 | -0.055 | -0.5065 |
| GYPE | NM_198682 | 0.3465 | -0.119 |
| RAB15 | NM_198686 | -0.1335 | -1.6055 |
| KRTAP10-4 | NM_198687 | 0.2925 | -0.5855 |
| KRTAP10-7 | NM_198689 | 0.2605 | -0.1765 |
| KRTAP10-9 | NM_198690 | 0.0275 | -0.3575 |
| KRTAP10-1 | NM_198691 | -0.228 | -0.049 |
| KRTAP10-11 | NM_198692 | -0.0315 | -0.741 |
| KRTAP10-2 | NM_198693 | -0.0805 | -0.3785 |
| KRTAP10-5 | NM_198694 | 0.1015 | -0.466 |
| KRTAP10-8 | NM_198695 | -0.133 | -0.4205 |
| KRTAP10-3 | NM_198696 | -0.0865 | -0.4655 |
| KRTAP12-3 | NM_198697 | -0.157 | -0.5895 |
| KRTAP12-4 | NM_198698 | 0.1925 | -0.5055 |
| KRTAP10-12 | NM_198699 | 0.115 | -0.3245 |
| CELF1 | NM_198700 | -0.732 | -1.313 |
| HSD11B1L | NM_198704 | 0.229 | -0.244 |
| HSD11B1L | NM_198705 | 0.229 | -0.244 |
| HSD11B1L | NM_198706 | 0.229 | -0.244 |
| HSD11B1L | NM_198707 | 0.229 | -0.244 |
| HSD11B1L | NM_198708 | 0.229 | -0.244 |
| ARSB | NM_198709 | 0.003 | -0.3315 |
| PTGER3 | NM_198714 | -0.2095 | -0.2735 |
| PTGER3 | NM_198715 | -0.2095 | -0.2735 |
| PTGER3 | NM_198716 | -0.2095 | -0.2735 |
| PTGER3 | NM_198717 | -0.2095 | -0.2735 |
| PTGER3 | NM_198718 | -0.2095 | -0.2735 |
| PTGER3 | NM_198719 | -0.2095 | -0.2735 |
| COL25A1 | NM_198721 | 0.119 | -0.206 |
| AMIGO3 | NM_198722 | -0.2095 | -0.8125 |
| TCEA2 | NM_198723 | -0.2305 | -0.6375 |

| | | | |
|---|---|---|---|
| CD47 | NM_198793 | -1.0665 | -1.2815 |
| MAP4K5 | NM_198794 | -0.781 | -1.6865 |
| TDRD1 | NM_198795 | -0.352 | -0.146 |
| ANKRD5 | NM_198798 | -1.118 | -0.9225 |
| GPR133 | NM_198827 | -0.047 | -0.0365 |
| MAST4 | NM_198828 | 0.335 | -0.1125 |
| ACLY | NM_198830 | -0.193 | -0.9385 |
| SERPINB8 | NM_198833 | -0.041 | -0.4225 |
| ACACA | NM_198834 | -0.9135 | -0.9015 |
| ACACA | NM_198836 | 1.4915 | 0.469 |
| ACACA | NM_198837 | 1.4915 | 0.469 |
| ACACA | NM_198838 | 1.4915 | 0.469 |
| ACACA | NM_198839 | 1.4915 | 0.469 |
| FAM120AOS | NM_198841 | -1.305 | -1.44 |
| SFTPB | NM_198843 | 0.1175 | -0.316 |
| ZPBP2 | NM_198844 | -0.068 | 0.0415 |
| SIGLEC6 | NM_198845 | 0.088 | 0.198 |
| SIGLEC6 | NM_198846 | 0.088 | 0.198 |
| FAM111A | NM_198847 | -0.195 | -0.1145 |
| SIAH3 | NM_198849 | -0.472 | -1.1485 |
| PHLDB3 | NM_198850 | -0.8535 | -0.87 |
| TRIM74 | NM_198853 | 0.051 | -0.073 |
| ZNF211 | NM_198855 | -0.964 | -0.463 |
| EFCAB6 | NM_198856 | -0.4925 | -0.1885 |
| PRICKLE2 | NM_198859 | -0.9695 | -0.715 |
| ALKBH6 | NM_198867 | -0.4385 | -1.0345 |
| TBC1D9B | NM_198868 | -0.2625 | -0.6745 |
| QRICH1 | NM_198880 | -1.117 | -3.5845 |
| TBC1D8B | NM_198881 | -0.34 | -0.3285 |
| SYNRG | NM_198882 | -0.907 | -2.1575 |
| NUP43 | NM_198887 | -1.127 | -1.55 |
| ANKRD17 | NM_198889 | -1.696 | -4.3965 |
| ATG16L1 | NM_198890 | -0.4715 | -2.0815 |
| BMP2K | NM_198892 | -0.2195 | -0.9395 |
| ZNF160 | NM_198893 | 0.66 | 0.363 |
| RAB6A | NM_198896 | -1.152 | -2.154 |
| FIBP | NM_198897 | 0.1345 | -0.427 |
| FMNL3 | NM_198900 | -0.3395 | -1.0255 |
| SRI | NM_198901 | -0.443 | -1.4995 |
| TSPAN3 | NM_198902 | -0.7005 | -0.5225 |
| GABRG2 | NM_198903 | 0.011 | -0.577 |
| GABRG2 | NM_198904 | 0.011 | -0.577 |
| UBE2CBP | NM_198920 | -1.1845 | -1.9915 |

| | | | |
|---|---|---|---|
| MRGPRD | NM_198923 | -0.1105 | -0.7275 |
| SEMA4B | NM_198925 | -0.503 | -0.8585 |
| C1orf183 | NM_198926 | -0.616 | -1.673 |
| PYHIN1 | NM_198928 | 0.134 | -0.378 |
| PYHIN1 | NM_198929 | 0.134 | -0.378 |
| PYHIN1 | NM_198930 | 0.134 | -0.378 |
| SS18L1 | NM_198935 | -2.12 | -2.0445 |
| SERINC3 | NM_198941 | -0.621 | -1.402 |
| OR7C1 | NM_198944 | 0.447 | 0.224 |
| LCN6 | NM_198946 | -0.0195 | -0.073 |
| FAM111B | NM_198947 | -0.364 | -0.31 |
| NUDT1 | NM_198948 | 0.161 | -0.4465 |
| NUDT1 | NM_198949 | 0.161 | -0.4465 |
| NUDT1 | NM_198950 | 0.161 | -0.4465 |
| TGM2 | NM_198951 | -0.1535 | -0.4795 |
| NUDT1 | NM_198952 | 0.161 | -0.4465 |
| NUDT1 | NM_198953 | 0.161 | -0.4465 |
| NUDT1 | NM_198954 | 0.161 | -0.4465 |
| MGAT5B | NM_198955 | -0.0295 | -0.273 |
| SP8 | NM_198956 | 0.19 | 0.027 |
| DHX57 | NM_198963 | -1.5035 | -3.1455 |
| PTHLH | NM_198964 | -0.169 | 0.58 |
| PTHLH | NM_198965 | 0.2015 | 0.0235 |
| PTHLH | NM_198966 | -0.169 | 0.58 |
| DZIP1 | NM_198968 | -1.1165 | -1.613 |
| AES | NM_198969 | 0.245 | -0.215 |
| AES | NM_198970 | 0.245 | -0.215 |
| HINFP | NM_198971 | -0.5445 | -2.3425 |
| MKNK1 | NM_198973 | -0.7305 | -0.8235 |
| TH1L | NM_198976 | -0.154 | -0.804 |
| ARHGEF1 | NM_198977 | -0.14 | -0.908 |
| LENG9 | NM_198988 | 0.2035 | -0.3925 |
| DLEU7 | NM_198989 | -0.124 | -0.591 |
| NAPEPLD | NM_198990 | -1.3605 | -2.712 |
| KCTD1 | NM_198991 | -1.2195 | -3.109 |
| SYT10 | NM_198992 | 0.2475 | -0.146 |
| STAC2 | NM_198993 | -0.0925 | -0.67 |
| TGM6 | NM_198994 | -0.3355 | -0.184 |
| C18orf34 | NM_198995 | -0.0855 | -0.242 |
| LIPI | NM_198996 | -0.1075 | -0.185 |
| SLC26A5 | NM_198999 | -0.0105 | -0.4415 |
| LHFPL3 | NM_199000 | 0.246 | -0.564 |
| C9orf169 | NM_199001 | 0.207 | -0.3105 |

| | | | |
|---|---|---|---|
| ARHGEF1 | NM_199002 | -0.14 | -0.908 |
| THAP1 | NM_199003 | -1.3595 | -2.034 |
| ARRB2 | NM_199004 | 0.152 | -0.7385 |
| SCN1B | NM_199037 | -0.114 | -0.5 |
| KLHL5 | NM_199039 | -0.8775 | -1.448 |
| NUDT4 | NM_199040 | -0.358 | -1.3375 |
| NSUN4 | NM_199044 | -0.693 | -2.063 |
| TEPP | NM_199046 | 0.06 | -0.122 |
| TBPL2 | NM_199047 | -0.001 | 0.008 |
| C2CD2 | NM_199050 | -1.7015 | -2.6495 |
| FAM5C | NM_199051 | 0.379 | -0.479 |
| C4orf41 | NM_199053 | -1.47 | -2.457 |
| MKNK2 | NM_199054 | -0.1155 | 0.9995 |
| NDUFAF3 | NM_199069 | -0.069 | -0.846 |
| NDUFAF3 | NM_199070 | -0.069 | -0.846 |
| MDFIC | NM_199072 | 0.2375 | -0.2935 |
| NDUFAF3 | NM_199073 | -0.069 | -0.846 |
| NDUFAF3 | NM_199074 | -0.069 | -0.846 |
| CNNM2 | NM_199077 | 0.1265 | -0.3605 |
| CNNM3 | NM_199078 | -0.597 | -1.5755 |
| VWA1 | NM_199121 | -0.1515 | -0.512 |
| SETD3 | NM_199123 | -1.2935 | -2.999 |
| C11orf63 | NM_199124 | -0.3885 | -0.5895 |
| ZNF585A | NM_199126 | -0.6415 | -2.359 |
| TMEM189 | NM_199129 | -0.8755 | -2.279 |
| VAX1 | NM_199131 | 0.106 | -0.047 |
| ZNF468 | NM_199132 | -0.6015 | -1.45 |
| FAM173B | NM_199133 | -0.585 | -1.2365 |
| C7orf46 | NM_199136 | -1.001 | -1.393 |
| FAM123A | NM_199138 | -0.083 | -0.692 |
| XAF1 | NM_199139 | -0.2265 | -0.051 |
| CARM1 | NM_199141 | -0.097 | -0.889 |
| UBE2V1 | NM_199144 | -0.345 | -1.404 |
| LHX6 | NM_199160 | -0.398 | -0.4705 |
| ADPRHL1 | NM_199162 | -0.116 | -0.5785 |
| ADSSL1 | NM_199165 | 0.1895 | -0.508 |
| ALAS1 | NM_199166 | -0.5745 | -2.806 |
| CLUL1 | NM_199167 | 0.1115 | -0.3915 |
| CXCL12 | NM_199168 | -0.2575 | -1.2595 |
| PMEPA1 | NM_199169 | -0.854 | -3.3055 |
| PMEPA1 | NM_199170 | -0.854 | -3.3055 |
| PMEPA1 | NM_199171 | -0.854 | -3.3055 |
| BGLAP | NM_199173 | 0.143 | -0.7135 |

| | | | |
|---|---|---|---|
| MRRF | NM_199177 | -0.8035 | -1.159 |
| KIRREL2 | NM_199179 | -0.4785 | -0.524 |
| KIRREL2 | NM_199180 | -0.7795 | -1.2475 |
| KIAA0889 | NM_199181 | -1.4965 | -1.86 |
| PRSS45 | NM_199183 | -0.178 | -0.073 |
| C6orf108 | NM_199184 | -0.035 | -0.5445 |
| NPM1 | NM_199185 | -0.3505 | -0.38 |
| BPGM | NM_199186 | -1.2225 | -1.197 |
| KRT18 | NM_199187 | 0.1865 | -0.2155 |
| LARP4 | NM_199188 | -1.5835 | -3.357 |
| MATR3 | NM_199189 | -0.217 | -0.35 |
| LARP4 | NM_199190 | -0.0595 | -1.7255 |
| BRE | NM_199191 | -0.0305 | -0.7825 |
| BRE | NM_199194 | -0.0755 | -1.067 |
| THEG | NM_199202 | -0.0515 | -0.291 |
| METAP1D | NM_199227 | -0.5165 | -0.4745 |
| RPE | NM_199229 | -1.193 | -2.2785 |
| GDNF | NM_199231 | -0.214 | -0.568 |
| COLEC11 | NM_199235 | 0.0725 | -0.781 |
| UNC13D | NM_199242 | -0.2205 | -0.071 |
| GPR150 | NM_199243 | 0.1625 | 0.046 |
| FOXD4L4 | NM_199244 | -0.2535 | -0.28 |
| VAMP1 | NM_199245 | 0.081 | -0.882 |
| CCNG1 | NM_199246 | -0.771 | -1.4995 |
| CACNB1 | NM_199247 | -0.268 | -1.1035 |
| CACNB1 | NM_199248 | -0.268 | -1.1035 |
| C19orf48 | NM_199250 | -0.879 | -2.0845 |
| TPTE2 | NM_199254 | -0.1185 | -0.1605 |
| TPTE | NM_199259 | -1.8605 | -0.212 |
| SP6 | NM_199262 | 0.0615 | -0.1325 |
| FAM179A | NM_199280 | -0.157 | -0.3935 |
| ARHGAP27 | NM_199282 | -0.7125 | -1.098 |
| PRR19 | NM_199285 | -0.409 | -0.5535 |
| DPPA3 | NM_199286 | -0.373 | -0.0855 |
| CCDC137 | NM_199287 | -1.1805 | -2.932 |
| NEK5 | NM_199289 | 0.1525 | -0.3165 |
| NACA2 | NM_199290 | -0.3795 | -0.1855 |
| TH | NM_199292 | 0.1285 | -0.3115 |
| TH | NM_199293 | 0.162 | -0.2025 |
| APITD1 | NM_199294 | -0.7235 | -0.719 |
| ISM2 | NM_199296 | -0.007 | -0.4875 |
| PHF17 | NM_199320 | -1.845 | -3.8595 |
| ZPBP2 | NM_199321 | 0.031 | -0.2125 |

| | | | |
|---|---|---|---|
| SPRY1 | NM_199327 | -0.548 | -1.166 |
| CLDN8 | NM_199328 | 0.308 | -0.463 |
| SLC43A3 | NM_199329 | -0.6805 | -1.804 |
| HOMER2 | NM_199330 | 0.3395 | -0.3195 |
| THRA | NM_199334 | -0.483 | -3.231 |
| FYB | NM_199335 | 0.1725 | -0.4775 |
| FAHD2B | NM_199336 | 0.1745 | -0.416 |
| TMEM179B | NM_199337 | -0.15 | -0.762 |
| SPEM1 | NM_199339 | -0.3085 | -0.5385 |
| LRRC37A3 | NM_199340 | -0.3925 | -0.4295 |
| CCDC23 | NM_199342 | -0.7005 | -0.9785 |
| SFT2D2 | NM_199344 | -0.0305 | -0.542 |
| PFN4 | NM_199346 | 0.22 | 0.516 |
| NEK10 | NM_199347 | 0.357 | -0.6695 |
| KCP | NM_199349 | 0.211 | -0.377 |
| C9orf50 | NM_199350 | 0.4885 | -0.529 |
| ILDR2 | NM_199351 | -0.1085 | -0.0405 |
| SLC22A25 | NM_199352 | -0.0965 | 0.135 |
| PRB1 | NM_199353 | 0.0585 | 0.193 |
| PRB1 | NM_199354 | 0.0585 | 0.193 |
| ADAMTS18 | NM_199355 | 0.019 | -0.0735 |
| ARHGAP11A | NM_199357 | -0.557 | -0.7665 |
| TPD52L2 | NM_199359 | -0.269 | -0.5175 |
| TPD52L2 | NM_199360 | -0.269 | -0.5175 |
| TPD52L2 | NM_199361 | -0.269 | -0.5175 |
| TPD52L2 | NM_199362 | -0.269 | -0.5175 |
| TPD52L2 | NM_199363 | -0.269 | -0.5175 |
| SPG7 | NM_199367 | 0.2705 | -0.432 |
| TRPC4AP | NM_199368 | -0.188 | -1.3235 |
| PRC1 | NM_199413 | -0.8885 | -0.7755 |
| PRC1 | NM_199414 | -0.8885 | -0.7755 |
| UBOX5 | NM_199415 | -0.5395 | -1.156 |
| PRCP | NM_199418 | -0.1975 | -0.6945 |
| POLQ | NM_199420 | -0.8255 | -2.184 |
| SOCS4 | NM_199421 | -0.0395 | -1.415 |
| WWP2 | NM_199423 | -0.5175 | -0.819 |
| WWP2 | NM_199424 | -0.5175 | -0.819 |
| VSX1 | NM_199425 | 0.1785 | -0.178 |
| ZFP64 | NM_199426 | -0.438 | -1.059 |
| ZFP64 | NM_199427 | -0.6155 | -1.587 |
| SPAST | NM_199436 | -1.0135 | -1.8355 |
| PRDM10 | NM_199437 | -0.1245 | -0.673 |
| PRDM10 | NM_199438 | -0.1245 | -0.673 |

| | | | |
|---|---|---|---|
| PRDM10 | NM_199439 | -0.1245 | -0.673 |
| HSPD1 | NM_199440 | -0.792 | -0.697 |
| ZNF334 | NM_199441 | -0.7345 | -1.6945 |
| COPE | NM_199442 | 0.096 | -0.324 |
| USP4 | NM_199443 | -0.3965 | -1.4 |
| COPE | NM_199444 | 0.2 | -0.3505 |
| ZNF365 | NM_199450 | -0.2075 | -0.5045 |
| ZNF365 | NM_199451 | -0.2075 | -0.5045 |
| ZNF365 | NM_199452 | -0.2075 | -0.5045 |
| HTR4 | NM_199453 | -0.427 | 0.4045 |
| PRDM16 | NM_199454 | 0.2685 | -0.5095 |
| TEPP | NM_199456 | 0.06 | -0.122 |
| C10orf71 | NM_199459 | 0.4625 | -0.345 |
| CACNA1C | NM_199460 | -0.2715 | -0.3655 |
| NANOS1 | NM_199461 | -0.627 | -1.055 |
| DSTYK | NM_199462 | -1.07 | -2.544 |
| KCNRG | NM_199464 | -0.5045 | -1.073 |
| PLP1 | NM_199478 | 0.0055 | -0.3675 |
| C20orf24 | NM_199483 | -0.1675 | -1.47 |
| UQCC | NM_199487 | -1.13 | -1.8685 |
| CCDC80 | NM_199511 | -0.315 | -0.463 |
| CCDC80 | NM_199512 | -0.315 | -0.463 |
| MAGED2 | NM_201222 | 0.016 | -0.594 |
| DDX47 | NM_201224 | -0.04 | 0.687 |
| CRB1 | NM_201253 | -0.674 | -0.994 |
| DNAJC12 | NM_201262 | -1.516 | -2.466 |
| WARS2 | NM_201263 | -0.4355 | -1.2705 |
| NRP2 | NM_201264 | 0.2155 | -0.6165 |
| UBL7 | NM_201265 | -0.032 | -0.6335 |
| NRP2 | NM_201266 | -1.5015 | -1.9995 |
| NRP2 | NM_201267 | -0.568 | -1.518 |
| ZNF644 | NM_201269 | -0.9615 | -3.751 |
| MPRIP | NM_201274 | 0.1685 | -0.438 |
| CNN2 | NM_201277 | -0.067 | -0.928 |
| MTMR2 | NM_201278 | -0.759 | -1.315 |
| NRP2 | NM_201279 | -0.568 | -1.518 |
| MUTED | NM_201280 | -1.377 | -2.427 |
| MTMR2 | NM_201281 | -0.759 | -1.315 |
| EGFR | NM_201282 | -0.735 | -0.8205 |
| EGFR | NM_201283 | -0.735 | -0.8205 |
| EGFR | NM_201284 | -0.735 | -0.8205 |
| USP51 | NM_201286 | 0.4785 | 0.173 |
| PRELP | NM_201348 | -0.4875 | -0.388 |

| | | | |
|---|---|---|---|
| PLEC | NM_201380 | -0.405 | -0.896 |
| GPX1 | NM_201397 | 0.216 | -0.2285 |
| FLAD1 | NM_201398 | -0.7995 | -1.8245 |
| FAM86A | NM_201400 | -0.2495 | -1.3925 |
| LUC7L | NM_201412 | -0.552 | -1.7705 |
| APP | NM_201413 | -0.3025 | -0.3875 |
| APP | NM_201414 | -0.3025 | -0.3875 |
| RTN3 | NM_201428 | -0.3165 | -0.408 |
| RTN3 | NM_201429 | -0.3165 | -0.408 |
| RTN3 | NM_201430 | -0.3165 | -0.408 |
| RASSF6 | NM_201431 | -0.1675 | 0.0235 |
| GAS7 | NM_201432 | -0.1265 | -0.5135 |
| GAS7 | NM_201433 | -0.1265 | -0.5135 |
| RAB5C | NM_201434 | 0.0955 | -0.6495 |
| CCDC62 | NM_201435 | 0.1065 | -0.452 |
| H2AFV | NM_201436 | 0.292 | -0.25 |
| TCEA1 | NM_201437 | -1.112 | -0.845 |
| PPHLN1 | NM_201438 | -1.1515 | -2.0805 |
| PPHLN1 | NM_201439 | -1.1515 | -2.0805 |
| PPHLN1 | NM_201440 | -1.1515 | -2.0805 |
| C1S | NM_201442 | -0.1455 | -0.3475 |
| TEAD4 | NM_201443 | -0.1755 | -1.7605 |
| DGKA | NM_201444 | -0.0475 | -0.417 |
| DGKA | NM_201445 | -0.0475 | -0.417 |
| EGFL7 | NM_201446 | 0.154 | -0.4955 |
| PPHLN1 | NM_201515 | -1.1515 | -2.0805 |
| H2AFV | NM_201516 | 0.292 | -0.25 |
| H2AFV | NM_201517 | 0.292 | -0.25 |
| SLC25A35 | NM_201520 | -0.2945 | -0.7625 |
| KLC4 | NM_201523 | 0.2645 | -0.3475 |
| GPR56 | NM_201524 | -0.1105 | -0.4785 |
| GPR56 | NM_201525 | -0.1105 | -0.4785 |
| ISLR | NM_201526 | 0.056 | -0.523 |
| DGKZ | NM_201532 | -0.2985 | -0.85 |
| DGKZ | NM_201533 | -0.2985 | -0.85 |
| NDRG2 | NM_201535 | 0.245 | -0.252 |
| NDRG2 | NM_201540 | 0.3755 | -0.2795 |
| MED8 | NM_201542 | -1.0935 | -2.279 |
| LGALS8 | NM_201543 | -0.262 | -0.614 |
| LGALS8 | NM_201544 | -0.262 | -0.614 |
| LGALS8 | NM_201545 | -0.1575 | -0.886 |
| CDCP2 | NM_201546 | 0.0375 | -0.396 |
| CERKL | NM_201548 | -0.0855 | -1.3575 |

| | | | |
|---|---|---|---|
| LRRC10 | NM_201550 | -0.292 | -0.112 |
| NELL1 | NM_201551 | -0.071 | -0.3835 |
| FGL1 | NM_201552 | 0.0905 | -0.006 |
| FGL1 | NM_201553 | 0.0905 | -0.006 |
| DGKA | NM_201554 | -0.0475 | -0.417 |
| FHL2 | NM_201555 | -0.5065 | -2.9275 |
| FHL2 | NM_201557 | -0.5065 | -2.9275 |
| FOXO3 | NM_201559 | -1.568 | -2.087 |
| SLC16A13 | NM_201566 | -0.0885 | -0.1715 |
| CDC25A | NM_201567 | -0.768 | -2.423 |
| SMG7 | NM_201568 | -0.2385 | -1.0785 |
| SMG7 | NM_201569 | -0.2385 | -1.0785 |
| CACNB2 | NM_201570 | 0.037 | -0.4365 |
| CACNB2 | NM_201571 | 0.037 | -0.4365 |
| CACNB2 | NM_201572 | 0.037 | -0.4365 |
| SLC4A3 | NM_201574 | -0.503 | -0.686 |
| SEZ6L2 | NM_201575 | -0.099 | -0.6895 |
| MAFA | NM_201589 | -0.073 | -0.0815 |
| CACNB2 | NM_201590 | 0.037 | -0.4365 |
| GPM6A | NM_201591 | -0.5865 | -0.293 |
| GPM6A | NM_201592 | -0.5865 | -0.293 |
| CACNB2 | NM_201593 | 0.037 | -0.4365 |
| CACNB2 | NM_201596 | 0.037 | -0.4365 |
| CACNB2 | NM_201597 | 0.037 | -0.4365 |
| FAM86A | NM_201598 | -0.2495 | -1.3925 |
| ZMYM3 | NM_201599 | -0.313 | -0.944 |
| IKBIP | NM_201612 | -1.6145 | -3.31 |
| IKBIP | NM_201613 | -1.6145 | -3.31 |
| CLEC12A | NM_201623 | -0.1225 | 0.2765 |
| USP33 | NM_201626 | -0.4475 | -0.817 |
| TRIM41 | NM_201627 | -0.936 | -1.7015 |
| TJP2 | NM_201629 | -0.8935 | -1.5035 |
| LRRN2 | NM_201630 | -0.843 | -1.412 |
| TGM5 | NM_201631 | -0.07 | -0.1115 |
| TCF7 | NM_201632 | -0.874 | -1.8875 |
| TCF7 | NM_201634 | -0.874 | -1.8875 |
| TBXA2R | NM_201636 | 0.088 | -0.4375 |
| GLYAT | NM_201648 | 0.0055 | -0.2145 |
| SLC6A9 | NM_201649 | -0.49 | -0.254 |
| LRRC23 | NM_201650 | 0.153 | -0.44 |
| SLC28A1 | NM_201651 | -0.2805 | -0.0575 |
| CHIA | NM_201653 | 0.11 | 0.1955 |
| SF1 | NM_201995 | -0.0435 | -0.7715 |

| | | | |
|---|---|---|---|
| SF1 | NM_201997 | -0.0435 | -0.7715 |
| SF1 | NM_201998 | -0.0435 | -0.7715 |
| ELF2 | NM_201999 | -1.2945 | -3.008 |
| ACSM3 | NM_202000 | 0.401 | -0.0685 |
| ERCC1 | NM_202001 | -0.1885 | -0.6235 |
| FOXM1 | NM_202002 | -0.183 | -0.867 |
| FOXM1 | NM_202003 | -0.183 | -0.867 |
| HFE2 | NM_202004 | -0.084 | -0.3185 |
| GIPC1 | NM_202467 | -0.391 | -1.469 |
| GIPC1 | NM_202468 | -0.41 | -1.537 |
| GIPC1 | NM_202469 | -0.391 | -1.469 |
| GIPC1 | NM_202470 | -0.4145 | -1.9685 |
| GIPC1 | NM_202494 | -0.391 | -1.469 |
| ENOSF1 | NM_202758 | -0.779 | -0.6855 |
| BMX | NM_203281 | 0.051 | -0.1895 |
| ZNF254 | NM_203282 | -0.24 | 0.185 |
| RBPJ | NM_203283 | -0.7465 | -1.343 |
| RBPJ | NM_203284 | -0.7465 | -1.343 |
| PVRL1 | NM_203285 | 0.7915 | 0.4105 |
| PVRL1 | NM_203286 | -0.171 | -0.828 |
| PSG11 | NM_203287 | -0.3235 | -0.3255 |
| POU5F1 | NM_203289 | -0.0935 | 0.0605 |
| POLR1C | NM_203290 | -0.683 | -1.894 |
| TRIM7 | NM_203293 | 0.3155 | -0.264 |
| TRIM7 | NM_203294 | 0.3155 | -0.264 |
| TRIM7 | NM_203295 | 0.3155 | -0.264 |
| TRIM7 | NM_203296 | 0.3155 | -0.264 |
| TRIM7 | NM_203297 | 0.3155 | -0.264 |
| CHCHD1 | NM_203298 | -0.329 | -1.27 |
| FBXO33 | NM_203301 | -1.1465 | -2.2705 |
| ZCCHC13 | NM_203303 | 0.0895 | -0.3105 |
| MEX3D | NM_203304 | -0.7905 | -2.4225 |
| FAM102A | NM_203305 | -0.216 | -0.969 |
| HEATR4 | NM_203309 | -0.427 | -0.3535 |
| BDH1 | NM_203314 | -0.852 | -2.027 |
| BDH1 | NM_203315 | -0.852 | -2.027 |
| MYO18A | NM_203318 | -0.157 | -0.478 |
| SLC23A2 | NM_203327 | -0.0075 | -0.5465 |
| CD59 | NM_203329 | -0.806 | -0.8585 |
| CD59 | NM_203330 | -0.806 | -0.8585 |
| CD59 | NM_203331 | -0.806 | -0.8585 |
| CLU | NM_203339 | -0.006 | -0.3765 |
| EPB41 | NM_203342 | -0.672 | -0.825 |

| | | | |
|---|---|---|---|
| EPB41 | NM_203343 | -0.672 | -0.825 |
| HDLBP | NM_203346 | -0.0575 | -0.3395 |
| LCN15 | NM_203347 | -0.0355 | -0.1085 |
| SHC4 | NM_203349 | -0.56 | -0.8725 |
| ZRANB2 | NM_203350 | -0.934 | -1.13 |
| MAP3K3 | NM_203351 | -0.9385 | -3.338 |
| PDLIM7 | NM_203352 | -0.1715 | -0.419 |
| CTAGE5 | NM_203354 | -0.6255 | -1.4575 |
| CTAGE5 | NM_203355 | -0.6255 | -1.4575 |
| CTAGE5 | NM_203356 | -0.6255 | -1.4575 |
| CAPRIN1 | NM_203364 | -1.311 | -1.7165 |
| RAPH1 | NM_203365 | -1.0855 | -2.13 |
| C3orf54 | NM_203370 | -0.39 | -0.658 |
| FIBIN | NM_203371 | 0.0615 | -0.466 |
| ACSL3 | NM_203372 | -0.737 | -1.3815 |
| FBXL22 | NM_203373 | 0.2855 | 0.5815 |
| ZNF784 | NM_203374 | -0.8665 | -2.1885 |
| TMEM81 | NM_203376 | -0.8615 | -1.283 |
| MB | NM_203377 | 0.0875 | -0.632 |
| MB | NM_203378 | 0.2025 | 0.154 |
| ACSL5 | NM_203379 | 0.386 | -0.1935 |
| ACSL5 | NM_203380 | 0.386 | -0.1935 |
| AMACR | NM_203382 | -0.4105 | -1.661 |
| RNH1 | NM_203383 | -0.33 | -0.6735 |
| RNH1 | NM_203384 | -0.33 | -0.6735 |
| RNH1 | NM_203385 | -0.33 | -0.6735 |
| RNH1 | NM_203386 | -0.33 | -0.6735 |
| RNH1 | NM_203387 | -0.33 | -0.6735 |
| RNH1 | NM_203388 | -0.33 | -0.6735 |
| RNH1 | NM_203389 | -0.33 | -0.6735 |
| RBM12B | NM_203390 | -1.652 | -2.5035 |
| GK | NM_203391 | -1.097 | -1.8855 |
| E2F7 | NM_203394 | -1.918 | -4.025 |
| IYD | NM_203395 | -0.101 | 0.0085 |
| MBLAC1 | NM_203397 | -0.057 | -0.66 |
| STMN1 | NM_203399 | -0.072 | -0.3225 |
| RPRML | NM_203400 | -0.13 | -0.855 |
| STMN1 | NM_203401 | -0.072 | -0.3225 |
| FITM1 | NM_203402 | 0.052 | -0.4675 |
| C9orf150 | NM_203403 | -0.4365 | -1.141 |
| AMPD2 | NM_203404 | 0.0185 | -0.7785 |
| KRTAP26-1 | NM_203405 | -0.3735 | -0.5495 |
| MBLAC2 | NM_203406 | -1.673 | -3.2825 |

| | | | |
|---|---|---|---|
| FAM47A | NM_203408 | -0.126 | 0.5735 |
| TMEM88 | NM_203411 | -0.2305 | -0.836 |
| UBL4B | NM_203412 | -0.315 | -0.4465 |
| C17orf81 | NM_203413 | -0.8065 | -1.7845 |
| C17orf81 | NM_203414 | -0.749 | -1.8625 |
| C17orf81 | NM_203415 | -0.8065 | -1.7845 |
| CD163 | NM_203416 | -0.5645 | -0.5245 |
| RCAN1 | NM_203417 | -0.58 | -0.9125 |
| RCAN1 | NM_203418 | -0.7185 | -1.035 |
| LRRN4CL | NM_203422 | -0.06 | -0.152 |
| IQCF2 | NM_203424 | 0.2165 | -0.107 |
| C17orf82 | NM_203425 | 0.142 | -0.324 |
| IER5L | NM_203434 | 1.151 | 2.294 |
| ASCL4 | NM_203436 | 0.181 | -0.0495 |
| AFTPH | NM_203437 | -1.2 | -2.3835 |
| ABCB9 | NM_203444 | 0.0745 | -0.486 |
| SYNJ1 | NM_203446 | -1.344 | -3.2155 |
| DOCK8 | NM_203447 | 0.0285 | 0.1265 |
| SERTM1 | NM_203451 | -0.509 | -0.6545 |
| PPAPDC2 | NM_203453 | -1.368 | -2.5135 |
| APOBEC4 | NM_203454 | 0.2995 | 0.0655 |
| PPIE | NM_203456 | -0.108 | -0.9975 |
| NOTCH2NL | NM_203458 | -0.2545 | -0.3035 |
| MRFAP1L1 | NM_203462 | -1.2935 | -1.9335 |
| AK4 | NM_203464 | -0.494 | -0.979 |
| ENTPD2 | NM_203468 | 0.242 | -0.3105 |
| LGALS14 | NM_203471 | 0.056 | -0.0055 |
| SELS | NM_203472 | -1.0055 | -2.704 |
| PORCN | NM_203475 | -0.255 | -0.7455 |
| DLL3 | NM_203486 | -0.354 | -0.929 |
| PCDH9 | NM_203487 | -0.0605 | -0.012 |
| ACYP1 | NM_203488 | -0.9395 | -0.955 |
| USP50 | NM_203494 | -0.0875 | -0.3325 |
| COMMD6 | NM_203495 | -0.07 | -0.1775 |
| COMMD6 | NM_203497 | -0.07 | -0.1775 |
| DDX42 | NM_203499 | -1.3155 | -1.7265 |
| KEAP1 | NM_203500 | -0.8335 | -3.2125 |
| CLEC4C | NM_203503 | 0.373 | -0.1555 |
| G3BP2 | NM_203504 | -0.6225 | -1.066 |
| G3BP2 | NM_203505 | -0.9885 | -1.231 |
| GRB2 | NM_203506 | -0.0425 | -1.049 |
| LYPD2 | NM_205545 | -0.2315 | -0.41 |
| FAM151B | NM_205548 | -0.3225 | -0.6765 |

| | | | |
|---|---|---|---|
| C19orf70 | NM_205767 | 0.071 | -0.649 |
| ZNF238 | NM_205768 | -1.1955 | 0.114 |
| IGSF1 | NM_205833 | 0.418 | -0.425 |
| LSR | NM_205834 | 0.2225 | -0.1815 |
| FBXO38 | NM_205836 | -1.4205 | -3.1485 |
| LST1 | NM_205837 | -0.04 | -0.218 |
| LST1 | NM_205838 | -0.04 | -0.218 |
| LST1 | NM_205839 | 0.051 | -0.149 |
| LST1 | NM_205840 | -0.04 | -0.218 |
| SPINK6 | NM_205841 | -0.2105 | -0.109 |
| NCKAP1 | NM_205842 | -0.4995 | -1.1715 |
| NFIC | NM_205843 | -0.9405 | -2.6905 |
| AKR1C2 | NM_205845 | 0.259 | -0.458 |
| PRELID2 | NM_205846 | -0.1895 | -0.604 |
| GMPPA | NM_205847 | -0.1755 | -0.796 |
| SYT6 | NM_205848 | -0.822 | -0.3285 |
| FAM9B | NM_205849 | -0.096 | -0.278 |
| SLC24A5 | NM_205850 | -0.858 | -0.7375 |
| CLEC12B | NM_205852 | -0.0965 | 0.042 |
| MUSTN1 | NM_205853 | 0.0495 | -0.461 |
| SFTA2 | NM_205854 | 0.2155 | -0.3715 |
| FAM180A | NM_205855 | -0.2675 | -0.1595 |
| OR2K2 | NM_205859 | -0.153 | -0.003 |
| NR5A2 | NM_205860 | -0.1685 | -0.2445 |
| DHDDS | NM_205861 | -0.5955 | -1.0075 |
| UGT1A6 | NM_205862 | -0.1565 | -0.129 |
| PARD3B | NM_205863 | 0.064 | -0.034 |
| CAGE1 | NM_205864 | 0.414 | 0.325 |
| ESRRG | NM_206594 | -1.3465 | -4.87 |
| ESRRG | NM_206595 | -1.2545 | -4.612 |
| CLYBL | NM_206808 | 0.3105 | -0.0905 |
| MOG | NM_206809 | -0.501 | 0.329 |
| MOG | NM_206810 | -0.501 | 0.329 |
| MOG | NM_206811 | -0.501 | 0.329 |
| MOG | NM_206812 | -0.501 | 0.329 |
| MOG | NM_206814 | -0.501 | 0.329 |
| OSCAR | NM_206818 | -0.083 | -0.2545 |
| MYBPC1 | NM_206819 | -0.102 | -0.558 |
| MYBPC1 | NM_206820 | 0.1965 | -0.4485 |
| MYBPC1 | NM_206821 | 0.0155 | 0.076 |
| GNL3 | NM_206825 | -1.288 | -2.3675 |
| GNL3 | NM_206826 | -1.288 | -2.3675 |
| RASL11A | NM_206827 | -0.0405 | -0.0885 |

| | | | |
|---|---|---|---|
| DPH3 | NM_206831 | -0.619 | -1.8035 |
| TMIGD1 | NM_206832 | 0.219 | -0.3905 |
| CTXN1 | NM_206833 | 0.096 | -0.547 |
| OSCP1 | NM_206837 | -0.2075 | -0.871 |
| MORF4L1 | NM_206839 | -0.5475 | -0.524 |
| NVL | NM_206840 | -1.339 | -1.9675 |
| RTN1 | NM_206852 | -0.235 | -0.1905 |
| QKI | NM_206853 | -1.192 | -2.0845 |
| QKI | NM_206854 | -1.192 | -2.0845 |
| QKI | NM_206855 | -1.192 | -2.0845 |
| TACC2 | NM_206860 | -0.5305 | -1.198 |
| TACC2 | NM_206861 | -0.5305 | -1.198 |
| TACC2 | NM_206862 | -1.188 | -4.026 |
| BACH1 | NM_206866 | -0.896 | -2.286 |
| PPP1CA | NM_206873 | 0.144 | -0.4505 |
| PPP1CB | NM_206876 | -0.855 | -1.057 |
| OR2V2 | NM_206880 | -0.2295 | -0.333 |
| SLC26A5 | NM_206883 | -0.0105 | -0.4415 |
| SLC26A5 | NM_206884 | -0.0905 | -0.276 |
| SLC26A5 | NM_206885 | -0.137 | -0.3365 |
| CCDC18 | NM_206886 | -0.0035 | -0.321 |
| DIP2A | NM_206889 | -1.243 | -1.7675 |
| DIP2A | NM_206890 | -1.243 | -1.7675 |
| DIP2A | NM_206891 | -1.243 | -1.7675 |
| MS4A10 | NM_206893 | 0.287 | -0.4875 |
| ZNF790 | NM_206894 | -1.034 | -2.2625 |
| C2orf82 | NM_206895 | 0.1055 | -0.061 |
| MRAP | NM_206898 | -0.2015 | -0.423 |
| OR10P1 | NM_206899 | 0.179 | 0.04 |
| RTN2 | NM_206900 | -0.527 | -1.3695 |
| PRKAA1 | NM_206907 | -0.0165 | -0.431 |
| PSD3 | NM_206909 | -0.646 | -1.435 |
| METTL21B | NM_206914 | -0.9435 | -0.77 |
| NGFRAP1 | NM_206917 | -0.2035 | -0.489 |
| DEGS2 | NM_206918 | 0.3175 | -0.4035 |
| ARL9 | NM_206919 | -0.017 | 0.034 |
| MAMDC4 | NM_206920 | -0.1615 | -0.5435 |
| C6orf204 | NM_206921 | -0.733 | -2.4905 |
| CRIP3 | NM_206922 | 0.0145 | -0.393 |
| YY2 | NM_206923 | -0.2895 | -0.409 |
| CA12 | NM_206925 | -0.858 | -1.8465 |
| SEPN1 | NM_206926 | 0.1265 | -0.265 |
| SYTL2 | NM_206927 | -0.046 | -0.2065 |

| | | | |
|---|---|---|---|
| SYTL2 | NM_206928 | -0.046 | -0.2065 |
| SYTL2 | NM_206929 | -0.046 | -0.2065 |
| SYTL2 | NM_206930 | -0.046 | -0.2065 |
| USH2A | NM_206933 | -0.1165 | 0.2205 |
| MS4A7 | NM_206938 | -0.248 | 0.2435 |
| MS4A7 | NM_206939 | -0.248 | 0.2435 |
| MS4A7 | NM_206940 | -0.248 | 0.2435 |
| LTBP1 | NM_206943 | -0.0975 | -0.986 |
| TRPM3 | NM_206944 | 0.08 | 0.028 |
| TRPM3 | NM_206945 | 0.08 | 0.028 |
| TRPM3 | NM_206946 | 0.08 | 0.028 |
| TRPM3 | NM_206947 | 0.08 | 0.028 |
| TRPM3 | NM_206948 | -0.424 | -0.351 |
| IFI27L1 | NM_206949 | 0.302 | -0.4565 |
| PRAME | NM_206953 | -0.5185 | -1.3285 |
| LTK | NM_206961 | 0.1185 | -0.382 |
| PRMT2 | NM_206962 | -0.777 | -0.847 |
| RARRES1 | NM_206963 | 0.5315 | -0.751 |
| FAM3B | NM_206964 | 0.032 | -0.6835 |
| FTCD | NM_206965 | -0.033 | -0.4415 |
| C5orf46 | NM_206966 | -0.104 | -0.0905 |
| C16orf74 | NM_206967 | -0.3755 | -0.135 |
| SPAG17 | NM_206996 | 0.03 | -0.4255 |
| GPR152 | NM_206997 | 0.095 | -0.291 |
| SCGB1D4 | NM_206998 | -0.1245 | 0.018 |
| CNOT1 | NM_206999 | -1.325 | -2.607 |
| BCL2L11 | NM_207002 | -0.4175 | -1.9595 |
| BCL2L11 | NM_207003 | -0.4175 | -1.9595 |
| FAM83A | NM_207006 | -0.0925 | -0.111 |
| FAM45A | NM_207009 | -1.67 | -3.228 |
| AP3M1 | NM_207012 | -1.249 | -1.334 |
| TCEB2 | NM_207013 | 0.359 | -0.11 |
| WDR78 | NM_207014 | 0.079 | -0.3585 |
| NAALADL2 | NM_207015 | 0.25 | 0.04 |
| EDN3 | NM_207032 | 0.004 | -0.3385 |
| EDN3 | NM_207033 | 0.004 | -0.3385 |
| EDN3 | NM_207034 | 0.004 | -0.3385 |
| TCF12 | NM_207036 | -0.9775 | -1.674 |
| TCF12 | NM_207037 | -0.9775 | -1.674 |
| TCF12 | NM_207038 | -0.9775 | -1.674 |
| TCF12 | NM_207040 | -0.9775 | -1.674 |
| ENSA | NM_207042 | 0.342 | -0.1555 |
| ENSA | NM_207043 | 0.1805 | -0.5175 |

| | | | |
|---|---|---|---|
| ENSA | NM_207044 | 0.1805 | -0.5175 |
| ENSA | NM_207045 | 0.1805 | -0.5175 |
| ENSA | NM_207046 | 0.18 | -0.417 |
| ENSA | NM_207047 | 0.1805 | -0.5175 |
| FBXW12 | NM_207102 | 0.1725 | -0.029 |
| C17orf87 | NM_207103 | 0.1115 | -0.273 |
| ASTN1 | NM_207108 | -0.645 | -0.389 |
| RNF216 | NM_207111 | -0.7985 | -2.7505 |
| SLC37A3 | NM_207113 | -1.439 | -2.024 |
| RNF216 | NM_207116 | -0.427 | -1.6315 |
| SLC25A47 | NM_207117 | -0.0455 | -0.472 |
| GTF2H5 | NM_207118 | -0.9525 | -0.9375 |
| LRRC20 | NM_207119 | -0.834 | -1.7645 |
| EXT2 | NM_207122 | -0.1755 | -1.341 |
| GAB1 | NM_207123 | -0.474 | -1.571 |
| PAOX | NM_207127 | -0.336 | -0.675 |
| PAOX | NM_207128 | -0.336 | -0.675 |
| LMOD2 | NM_207163 | 0.328 | 0.4805 |
| ENSA | NM_207168 | 0.1805 | -0.5175 |
| SYF2 | NM_207170 | -0.878 | -1.5465 |
| POGZ | NM_207171 | -1.1695 | -2.5125 |
| NPSR1 | NM_207172 | 0.022 | -0.369 |
| NPSR1 | NM_207173 | 0.022 | -0.369 |
| ABCG1 | NM_207174 | -0.1035 | -0.8765 |
| NPHP1 | NM_207181 | -0.3955 | -0.1105 |
| OR10A4 | NM_207186 | -0.104 | -0.4575 |
| BRDT | NM_207189 | -0.0665 | -0.0145 |
| ADAM15 | NM_207191 | 0.1465 | -0.561 |
| ADAM15 | NM_207194 | 0.1465 | -0.561 |
| ADAM15 | NM_207195 | 0.1465 | -0.561 |
| ADAM15 | NM_207196 | 0.1465 | -0.561 |
| ADAM15 | NM_207197 | 0.1465 | -0.561 |
| USF2 | NM_207291 | -0.0375 | -0.607 |
| MBNL1 | NM_207292 | -1.4015 | -3.2555 |
| MBNL1 | NM_207293 | -1.4015 | -3.2555 |
| MBNL1 | NM_207294 | -1.4015 | -3.2555 |
| MBNL1 | NM_207295 | -1.4015 | -3.2555 |
| MBNL1 | NM_207296 | -1.4015 | -3.2555 |
| MBNL1 | NM_207297 | -1.4015 | -3.2555 |
| LPPR1 | NM_207299 | 0.462 | -1.675 |
| ATRNL1 | NM_207303 | -0.103 | -1.443 |
| MBNL2 | NM_207304 | -1.218 | -2.9915 |
| FOXD4 | NM_207305 | -0.3905 | -0.8225 |

| | | | |
|---|---|---|---|
| C3orf25 | NM_207307 | 0.014 | -0.4785 |
| NUP210L | NM_207308 | 0.2135 | 0.277 |
| UAP1L1 | NM_207309 | -0.691 | -1.2155 |
| CCDC74B | NM_207310 | 0.18 | -0.2055 |
| CCDC64 | NM_207311 | -0.229 | -0.8515 |
| TUBA3E | NM_207312 | -0.0805 | -0.2775 |
| TMEM132E | NM_207313 | -0.027 | -0.3345 |
| CMPK2 | NM_207315 | 0.209 | 0.027 |
| TMEM207 | NM_207316 | 0.7 | -0.113 |
| ZNF474 | NM_207317 | -0.083 | -0.1415 |
| FAM199X | NM_207318 | -0.027 | -0.7705 |
| OTUD6A | NM_207320 | -0.346 | -0.425 |
| C10orf129 | NM_207321 | -0.025 | -0.146 |
| C2CD4A | NM_207322 | -1.323 | -0.0225 |
| DPY19L3 | NM_207325 | 1.985 | 2.916 |
| C22orf40 | NM_207327 | -0.852 | -2.386 |
| GPAT2 | NM_207328 | 0.018 | -0.2965 |
| NIPAL1 | NM_207330 | 0.27 | -0.576 |
| ERICH1 | NM_207332 | -0.369 | -1.3095 |
| ZNF320 | NM_207333 | -1.6725 | -2.168 |
| FAM43B | NM_207334 | -0.1175 | -1.1185 |
| KBTBD12 | NM_207335 | -0.172 | -0.044 |
| ZNF467 | NM_207336 | 0.0375 | -0.4585 |
| LCTL | NM_207338 | -0.6635 | -0.218 |
| PAGE2 | NM_207339 | 0.2355 | -0.412 |
| ZDHHC24 | NM_207340 | -0.076 | -0.6485 |
| ZP1 | NM_207341 | 0.2455 | -0.428 |
| RNF214 | NM_207343 | -1.114 | -2.7785 |
| SPRYD4 | NM_207344 | -0.593 | -1.474 |
| CLEC9A | NM_207345 | -0.3605 | 0.1375 |
| TSEN54 | NM_207346 | -0.5745 | -2.1875 |
| SLC25A34 | NM_207348 | -0.3865 | -0.335 |
| PRRT3 | NM_207351 | -0.5055 | -0.0455 |
| CYP4V2 | NM_207352 | 0.2295 | -1.0845 |
| ANKRD13D | NM_207354 | -0.456 | -1.279 |
| POTEB | NM_207355 | -0.2195 | -0.495 |
| C1orf174 | NM_207356 | -0.438 | -0.6515 |
| GADL1 | NM_207359 | 0.2895 | -0.05 |
| ZC3H12D | NM_207360 | -0.057 | -0.5275 |
| FREM2 | NM_207361 | -0.2645 | -0.226 |
| C2orf55 | NM_207362 | -0.086 | -0.38 |
| NCKAP5 | NM_207363 | -0.803 | -2.0515 |
| GPR148 | NM_207364 | -0.0815 | -0.214 |

| | | | |
|---|---|---|---|
| AADACL2 | NM_207365 | -0.4915 | -0.0015 |
| 14-Sep | NM_207366 | -0.676 | -0.0885 |
| FAM195B | NM_207368 | 0.155 | -0.4665 |
| GPR153 | NM_207370 | -0.329 | -0.675 |
| C10orf140 | NM_207371 | -1.5725 | -2.6225 |
| SH2D4B | NM_207372 | -0.2765 | -0.2505 |
| C10orf99 | NM_207373 | -0.303 | -0.11 |
| OR10W1 | NM_207374 | 0.215 | -0.093 |
| TOMM20L | NM_207377 | 0.054 | -0.138 |
| TMEM179 | NM_207379 | -0.597 | -0.1715 |
| C15orf52 | NM_207380 | 0.0595 | -0.406 |
| TNFAIP8L3 | NM_207381 | -0.072 | -0.1705 |
| SHISA6 | NM_207386 | 0.3635 | 0.0285 |
| CLEC17A | NM_207390 | 0.187 | -0.0875 |
| RGS9BP | NM_207391 | -0.054 | 0.0275 |
| KRTDAP | NM_207392 | 0.1795 | -0.053 |
| IGFL3 | NM_207393 | -0.0985 | -0.1215 |
| ZNF324B | NM_207395 | -0.652 | -1.3005 |
| RNF207 | NM_207396 | -0.5845 | -0.5685 |
| CD164L2 | NM_207397 | -0.011 | -0.4775 |
| GBP7 | NM_207398 | -0.1565 | -0.067 |
| C1orf229 | NM_207401 | 0.1095 | -0.3225 |
| ZNF662 | NM_207404 | -0.299 | -0.5375 |
| BEND4 | NM_207406 | -1.4905 | -2.1415 |
| TMPRSS11F | NM_207407 | -0.182 | 0.1645 |
| C5orf48 | NM_207408 | 0.0675 | 0.007 |
| C6orf126 | NM_207409 | 0.259 | -0.4485 |
| GFRAL | NM_207410 | -0.0085 | -0.126 |
| XKR5 | NM_207411 | -1.0255 | -2.223 |
| C8orf86 | NM_207412 | 0.1155 | -0.115 |
| FAM150A | NM_207413 | -0.218 | 0.178 |
| FLJ43860 | NM_207414 | 0.02 | -0.2065 |
| FAM75D3 | NM_207416 | 0.021 | -0.183 |
| C9orf171 | NM_207417 | -0.118 | 0.25 |
| FAM72D | NM_207418 | -1.913 | -3.148 |
| C1QTNF8 | NM_207419 | -0.0595 | -0.071 |
| SLC2A7 | NM_207420 | -0.0875 | -0.0685 |
| PADI6 | NM_207421 | -0.1045 | -0.3655 |
| FLJ44635 | NM_207422 | -0.194 | -0.4055 |
| FOXI2 | NM_207426 | -0.1415 | -0.3975 |
| C11orf88 | NM_207430 | 0.1165 | -0.021 |
| C12orf76 | NM_207435 | -0.7535 | -1.4425 |
| DNAH10 | NM_207437 | 0.007 | -0.1335 |

| | | | |
|---|---|---|---|
| C13orf35 | NM_207440 | -0.236 | -0.608 |
| C15orf53 | NM_207444 | -0.118 | -0.5275 |
| C15orf54 | NM_207445 | -0.2235 | -0.4885 |
| FAM174B | NM_207446 | -0.3085 | -1.168 |
| C17orf102 | NM_207454 | -0.0155 | -0.105 |
| TEX19 | NM_207459 | -1.582 | -1.393 |
| FAM177B | NM_207468 | -0.111 | 0.111 |
| DEFB132 | NM_207469 | 0.071 | -0.2855 |
| FAM190A | NM_207491 | -0.49 | 0.246 |
| C6orf138 | NM_207499 | -0.345 | -0.487 |
| SAMD12 | NM_207506 | 0.158 | -0.0775 |
| LCNL1 | NM_207510 | -0.152 | -0.2425 |
| C9orf139 | NM_207511 | 0.064 | -0.387 |
| DEF8 | NM_207514 | -0.915 | -1.911 |
| ADAMTSL3 | NM_207517 | -0.214 | -0.2025 |
| PRKACA | NM_207518 | 0.133 | -0.491 |
| ZAP70 | NM_207519 | 0.2835 | 0.2875 |
| RTN4 | NM_207520 | -0.2735 | -0.847 |
| RTN4 | NM_207521 | -0.2735 | -0.847 |
| MAP6 | NM_207577 | 0.1135 | -0.455 |
| PRKACB | NM_207578 | -1.4275 | -1.8175 |
| DUOXA2 | NM_207581 | -0.2335 | -0.3045 |
| ERVFRD-1 | NM_207582 | -0.1145 | -0.053 |
| IFNAR2 | NM_207584 | -0.982 | -3.069 |
| IFNAR2 | NM_207585 | -0.982 | -3.069 |
| ABCG1 | NM_207627 | -0.1035 | -0.8765 |
| ABCG1 | NM_207628 | -0.1035 | -0.8765 |
| ABCG1 | NM_207629 | -0.1035 | -0.8765 |
| C22orf36 | NM_207644 | -1.072 | -1.9245 |
| C11orf87 | NM_207645 | 0.1625 | 0.0775 |
| FSD1L | NM_207647 | -0.1835 | -2.04 |
| ZC3H14 | NM_207660 | -0.9785 | -2.06 |
| ZC3H14 | NM_207661 | -0.9785 | -2.06 |
| ZC3H14 | NM_207662 | -0.9785 | -2.06 |
| GRIPAP1 | NM_207672 | -0.3665 | -0.801 |
| ARL4A | NM_212460 | -0.918 | -2.4755 |
| CHKA | NM_212469 | -0.757 | -1.9725 |
| PRKAR1A | NM_212471 | -0.848 | -0.987 |
| PRKAR1A | NM_212472 | -0.848 | -0.987 |
| FN1 | NM_212474 | -0.124 | -0.5965 |
| FN1 | NM_212476 | -0.124 | -0.5965 |
| FN1 | NM_212478 | -0.124 | -0.5965 |
| ZMYND11 | NM_212479 | -1.3235 | -2.104 |

| | | | |
|---|---|---|---|
| ARID5A | NM_212481 | 0.056 | -0.588 |
| FN1 | NM_212482 | -0.124 | -0.5965 |
| GPS1 | NM_212492 | 0.158 | -0.4545 |
| CDK18 | NM_212502 | -0.3015 | -0.6405 |
| CDK18 | NM_212503 | -0.4835 | -0.749 |
| ABCA2 | NM_212533 | 0.2175 | -0.177 |
| PRKCB | NM_212535 | -0.2515 | -0.2715 |
| PRKCD | NM_212539 | -0.3105 | -0.8455 |
| B4GALT4 | NM_212543 | -1.009 | -1.594 |
| BLOC1S3 | NM_212550 | 0.1555 | -0.495 |
| LYSMD1 | NM_212551 | -0.795 | -2.516 |
| METTL10 | NM_212554 | -1.1925 | -1.195 |
| PATE2 | NM_212555 | 0.072 | -0.5935 |
| ASB18 | NM_212556 | 0.129 | 0.085 |
| AMTN | NM_212557 | -0.1915 | -0.146 |
| TMEM215 | NM_212558 | -0.0085 | -0.1135 |
| XKRX | NM_212559 | -0.293 | -0.2435 |
| DFFA | NM_213566 | -0.599 | -1.1795 |
| SLC39A3 | NM_213568 | -0.152 | -1.62 |
| NEBL | NM_213569 | 0.1015 | 0.1055 |
| RAPH1 | NM_213589 | -1.534 | -3.2 |
| TRIM13 | NM_213590 | -1.2885 | -3.7325 |
| DIO1 | NM_213593 | -0.1625 | -0.1355 |
| RFX4 | NM_213594 | 0.2 | -0.0625 |
| ISCU | NM_213595 | -0.1485 | -0.663 |
| FOXN4 | NM_213596 | -1.099 | -1.315 |
| KRBA2 | NM_213597 | -0.0365 | -0.2705 |
| ZNF543 | NM_213598 | -1.1775 | -2.238 |
| ANO5 | NM_213599 | -1.1555 | -2.011 |
| PLA2G4F | NM_213600 | 0.027 | -0.2105 |
| TMED8 | NM_213601 | -0.236 | -0.603 |
| SIGLEC15 | NM_213602 | 0.114 | -0.361 |
| ZNF789 | NM_213603 | -1.5275 | -1.364 |
| ADAMTSL5 | NM_213604 | -0.03 | -0.2785 |
| ZNF517 | NM_213605 | 0.07 | -0.889 |
| SLC16A12 | NM_213606 | 0.1765 | 0.2855 |
| CCDC103 | NM_213607 | -0.052 | -0.167 |
| C2orf66 | NM_213608 | -0.366 | -0.103 |
| FAM19A1 | NM_213609 | -0.0135 | -0.4425 |
| SLC25A3 | NM_213611 | 0.0495 | -0.527 |
| SLC26A1 | NM_213613 | -0.054 | -0.206 |
| ST5 | NM_213618 | -0.314 | -1.584 |
| ATP6V1H | NM_213619 | -0.878 | -1.8475 |

| | | | |
|---|---|---|---|
| ATP6V1H | NM_213620 | -0.878 | -1.8475 |
| HTR3A | NM_213621 | -0.0885 | -0.5375 |
| STAMBP | NM_213622 | -1.1125 | -1.6665 |
| C20orf132 | NM_213631 | -0.168 | -0.8015 |
| PSG4 | NM_213633 | -0.2255 | -0.084 |
| PDLIM7 | NM_213636 | -0.1715 | -0.419 |
| WARS | NM_213645 | -0.249 | -0.9745 |
| WARS | NM_213646 | -0.249 | -0.9745 |
| FGFR4 | NM_213647 | -0.136 | -0.67 |
| TCF7 | NM_213648 | -0.874 | -1.8875 |
| SFXN4 | NM_213649 | -0.54 | -1.5445 |
| SLC25A24 | NM_213651 | -0.829 | -1.949 |
| HFE2 | NM_213652 | -0.084 | -0.3185 |
| HFE2 | NM_213653 | -0.084 | -0.3185 |
| ARMC8 | NM_213654 | -0.332 | -1.113 |
| WNK1 | NM_213655 | -0.809 | -1.332 |
| KRT39 | NM_213656 | -0.2745 | -0.0045 |
| KLRC1 | NM_213658 | -0.282 | -0.5655 |
| STAT3 | NM_213662 | 0.0245 | -0.7415 |
| TPM2 | NM_213674 | 0.087 | -0.087 |
| CHCHD10 | NM_213720 | 0.0655 | -0.999 |
| RPLP1 | NM_213725 | 0.069 | -0.3185 |
| INCA1 | NM_213726 | -0.1405 | -0.3445 |
| C2orf27B | NM_214461 | -0.42 | -0.714 |
| DACT2 | NM_214462 | 0.0485 | -0.288 |
| C4orf40 | NM_214711 | -0.088 | -0.417 |
| SNORD15A | NR_000005 | -0.38 | -0.783 |
| SNORD15B | NR_000025 | 0.9775 | 0.246 |
| RAB9BP1 | NR_000039 | 0.078 | 0.1635 |
| CELP | NR_001275 | -0.167 | -0.0295 |
| CYP2B7P1 | NR_001278 | -0.023 | -0.3895 |
| PCDHB17 | NR_001280 | 0.2185 | -0.9215 |
| PCDHB18 | NR_001281 | -0.221 | -0.395 |
| TFAMP1 | NR_001288 | -0.156 | 0.017 |
| HLA-H | NR_001434 | -0.3625 | -0.316 |
| HLA-DPB2 | NR_001435 | 0.3055 | 0.6485 |
| LOC339240 | NR_001443 | -0.026 | -0.109 |
| ANXA2P3 | NR_001446 | -0.236 | -0.272 |
| TTTY6 | NR_001527 | -0.093 | -0.138 |
| TTTY13 | NR_001537 | 0.0195 | -0.1535 |
| TTTY5 | NR_001541 | 0.212 | 0.113 |
| TTTY10 | NR_001542 | 0.1285 | -0.1665 |
| TTTY14 | NR_001543 | -0.312 | -0.41 |

| | | | |
|---|---|---|---|
| TTTY11 | NR_001548 | -0.1795 | 0.1555 |
| TTTY12 | NR_001551 | 0.0335 | 0.183 |
| ANXA2P1 | NR_001562 | -0.1035 | 0.0525 |
| XIST | NR_001564 | -0.178 | -0.467 |
| ADAM3A | NR_001569 | -0.107 | -0.226 |
| TDH | NR_001578 | -0.8075 | -1.091 |
| OR6W1P | NR_002140 | 0.0885 | 0.0625 |
| TTTY6B | NR_002175 | -0.0575 | -0.4445 |
| PPY2 | NR_002181 | 0.462 | 0.249 |
| NACAP1 | NR_002182 | -0.11 | 0.435 |
| [illegible]KFZP586I142[illegible] | NR_002186 | -0.638 | -1.242 |
| PPP1R2P9 | NR_002191 | -0.092 | -0.4825 |
| PMS2CL | NR_002217 | -1.24 | -1.148 |
| INGX | NR_002226 | -0.309 | -0.2935 |
| DISC2 | NR_002227 | -0.123 | -0.615 |
| SNORA6 | NR_002325 | 1.3095 | 1.1335 |
| SNORA10 | NR_002327 | -0.1685 | -0.0725 |
| GNRHR2 | NR_002328 | 0.019 | -0.6825 |
| SNORA33 | NR_002436 | 0.2475 | 0.9175 |
| SNORD16 | NR_002440 | -0.247 | -0.086 |
| SNORA65 | NR_002449 | 4.085 | 3.705 |
| ABCA11P | NR_002451 | -0.454 | -1 |
| SNORA21 | NR_002576 | 1.3455 | 0.1895 |
| SNORA3 | NR_002580 | 0.698 | 1.482 |
| SNORA72 | NR_002581 | -0.3905 | 0.595 |
| SNORA52 | NR_002585 | -0.07 | -0.5495 |
| SNORA41 | NR_002590 | 1.1615 | 2.103 |
| DLEU1 | NR_002605 | -1.118 | -2.1965 |
| DLEU2 | NR_002612 | -0.6665 | -1.563 |
| NPY6R | NR_002713 | -0.1895 | 0.097 |
| REG1P | NR_002714 | 0.068 | -0.4235 |
| ATXN8OS | NR_002717 | 0.1175 | -0.3525 |
| ZNF204P | NR_002722 | -0.044 | -0.588 |
| MBL1P | NR_002724 | 0.0545 | -0.4625 |
| KCNQ1OT1 | NR_002728 | -0.6125 | -0.2505 |
| MST1P9 | NR_002729 | -0.017 | -0.293 |
| BRD7P3 | NR_002730 | -0.027 | 0.018 |
| PTTG3P | NR_002734 | -0.353 | -0.4615 |
| HTR7P1 | NR_002774 | -0.005 | -0.1115 |
| CYP4Z2P | NR_002788 | -0.143 | -0.4275 |
| NAPSB | NR_002798 | -0.282 | -0.4365 |
| RPL13P5 | NR_002803 | -0.4205 | -0.701 |
| SIGLECP3 | NR_002804 | -0.103 | -0.2345 |

| | | | |
|---|---|---|---|
| MALAT1 | NR_002819 | 0.2665 | 0.0905 |
| SIGLEC16 | NR_002825 | -0.1785 | -0.722 |
| SNORA71A | NR_002911 | -0.015 | -0.5965 |
| SNORA74A | NR_002915 | 2.047 | 2.428 |
| SNORD8 | NR_002916 | -0.1935 | -0.6965 |
| SNORA19 | NR_002917 | -0.1565 | 1.206 |
| SNORA5A | NR_002919 | 0.4655 | 0.4935 |
| SNORA13 | NR_002922 | 1.0765 | 0.7695 |
| BCORP1 | NR_002923 | 0.2355 | -0.1125 |
| GSTM2P1 | NR_002932 | 0.2965 | -0.266 |
| LOC493754 | NR_002933 | -0.597 | -1.3005 |
| LRRC37A4 | NR_002940 | -0.212 | -0.956 |
| KLKP1 | NR_002948 | 0.014 | -0.109 |
| NUDT16P1 | NR_002949 | -0.1275 | -0.527 |
| SNORA2A | NR_002950 | 1.3205 | 1.983 |
| SNORA12 | NR_002954 | 0.9185 | -1.5735 |
| SNORA14B | NR_002956 | 0.32 | 0.0925 |
| SNORA15 | NR_002957 | 0.5485 | 0.2555 |
| SNORA20 | NR_002960 | 0.444 | 1.8325 |
| SNORA23 | NR_002962 | -0.027 | -0.2015 |
| SNORA28 | NR_002964 | 0.671 | 1.508 |
| SNORA29 | NR_002965 | -0.4625 | 0.175 |
| SNORA30 | NR_002966 | 0.6735 | 0.5625 |
| SNORA34 | NR_002968 | -0.1895 | -0.087 |
| SNORA37 | NR_002970 | 0.455 | 1.2205 |
| SNORA42 | NR_002974 | 0.359 | 0.7245 |
| SNORA45 | NR_002977 | 0.5515 | 1.946 |
| SNORA46 | NR_002978 | -0.2705 | -0.0515 |
| SNORA49 | NR_002979 | -0.5665 | -1.128 |
| SNORA50 | NR_002980 | -0.2735 | 0.5515 |
| SNORA54 | NR_002982 | 0.204 | 0.9835 |
| SNORA56 | NR_002984 | -0.5145 | -1.4965 |
| SNORA58 | NR_002985 | -0.3985 | -0.33 |
| SNORA60 | NR_002986 | 0.483 | 0.5035 |
| SNORA74B | NR_002988 | 0.0365 | -0.4785 |
| SNORA35 | NR_002993 | 0.1355 | -0.2955 |
| SNORA76 | NR_002995 | 1.695 | 2.4285 |
| SNORA80 | NR_002996 | -0.5975 | 0.4265 |
| SCARNA20 | NR_002999 | 1.741 | -0.569 |
| SNORA47 | NR_003014 | 0.107 | -1.171 |
| SNORA53 | NR_003015 | 1.0705 | 1.2235 |
| SNORA26 | NR_003016 | 1.5375 | 1.351 |
| SNORA71C | NR_003017 | -0.1925 | -0.3285 |

| | | | |
|---|---|---|---|
| SNORA71D | NR_003018 | 0.143 | 0.2885 |
| SNORA77 | NR_003019 | -0.1975 | -0.5625 |
| SNORA78 | NR_003020 | -0.1725 | -0.6145 |
| SNORD9 | NR_003029 | -0.1425 | -0.134 |
| SNORD17 | NR_003045 | -0.283 | 0.774 |
| SNORD67 | NR_003056 | -0.1175 | -0.3525 |
| LILRP2 | NR_003061 | 0.042 | -0.5825 |
| PYY2 | NR_003064 | 0.1525 | -0.0745 |
| SNORD89 | NR_003070 | -0.5015 | -0.9755 |
| GSTTP1 | NR_003081 | -0.0155 | -0.372 |
| HSD17B7P2 | NR_003086 | -0.5475 | -0.0255 |
| ABCC13 | NR_003087 | -0.4795 | -0.4185 |
| ABCC13 | NR_003088 | -0.4795 | -0.4185 |
| OTUB1 | NR_003089 | 0.175 | -0.559 |
| MOBP | NR_003090 | -0.3065 | -0.085 |
| E2F6 | NR_003092 | -1.5175 | -4.976 |
| E2F6 | NR_003093 | -1.5175 | -4.976 |
| E2F6 | NR_003094 | -1.5175 | -4.976 |
| E2F6 | NR_003095 | -1.5175 | -4.976 |
| SNHG1 | NR_003098 | -1.0005 | -0.1375 |
| ZNF273 | NR_003099 | -0.1605 | -0.6125 |
| POT1 | NR_003102 | -1.327 | -2.0685 |
| POT1 | NR_003103 | -1.327 | -2.0685 |
| POT1 | NR_003104 | -1.327 | -2.0685 |
| ZWILCH | NR_003105 | -0.8695 | -0.606 |
| NBR2 | NR_003108 | -0.9545 | -1.186 |
| TRNAU1AP | NR_003109 | -0.6925 | -1.475 |
| ZNF542 | NR_003127 | -0.957 | -1.783 |
| HSP90B3P | NR_003130 | 0.252 | -0.1655 |
| HSP90AB2P | NR_003132 | 0.531 | 0.2115 |
| SNHG9 | NR_003142 | 1.071 | 1.2245 |
| LOC162632 | NR_003190 | -0.4775 | -0.851 |
| LGALS3 | NR_003225 | 0.069 | -0.728 |
| SNHG11 | NR_003239 | -0.2355 | -0.6045 |
| HNRPDL | NR_003249 | -1.353 | -0.9925 |
| ZNF74 | NR_003253 | -0.7685 | -2.8165 |
| TSIX | NR_003255 | -0.4375 | 0.033 |
| ABCD4 | NR_003256 | -0.44 | -1.802 |
| GNAS | NR_003259 | 0.0835 | -0.2445 |
| SDHAP3 | NR_003263 | -1.598 | -1.7745 |
| SDHAP2 | NR_003265 | 0.201 | -0.4405 |
| LOC220729 | NR_003266 | -0.315 | -0.1595 |
| PSPC1 | NR_003272 | -0.1795 | -1.0695 |

| | | | |
|---|---|---|---|
| CES1P1 | NR_003276 | 0.074 | -0.317 |
| EP400NL | NR_003290 | -0.2415 | -2.1235 |
| RRN3P1 | NR_003370 | -0.257 | -0.9795 |
| EARS2 | NR_003501 | -0.651 | -1.5685 |
| PPP4R1L | NR_003505 | -0.168 | 0.0295 |
| HSFY1 | NR_003510 | 0.213 | 0.1055 |
| CDKN2B-AS | NR_003529 | 0.0765 | -0.4035 |
| SLED1 | NR_003542 | 0.1475 | 0.6735 |
| C5orf44 | NR_003545 | -1.5265 | -2.229 |
| DPY19L2P2 | NR_003561 | -0.152 | -0.203 |
| MPP3 | NR_003562 | -0.767 | -1.4395 |
| ABCC6P1 | NR_003569 | -0.288 | -0.1255 |
| ANXA2P2 | NR_003573 | -0.399 | -0.507 |
| SNHG8 | NR_003584 | -0.327 | -0.153 |
| SCAND2 | NR_003654 | -0.2815 | 0.192 |
| MT1DP | NR_003658 | -0.233 | -0.35 |
| WASH3P | NR_003659 | -0.4645 | -1.2975 |
| DEFB109P1B | NR_003668 | -0.0665 | -0.22 |
| MT1IP | NR_003669 | 0.219 | -0.2465 |
| KGFLP2 | NR_003670 | 0.1235 | -0.22 |
| SNHG7 | NR_003672 | -0.7035 | -1.727 |
| NBLA00301 | NR_003679 | -0.7025 | -1.3895 |
| SNORD12B | NR_003695 | 1.057 | 0.8285 |
| PI4KAP2 | NR_003700 | 0.0515 | -0.5305 |
| SNORA84 | NR_003704 | 0.7925 | 1.8115 |
| SNORA70B | NR_003707 | 0 | -0.069 |
| SNORA70C | NR_003708 | -0.504 | -0.4775 |
| LOC728613 | NR_003713 | -0.092 | -0.65 |
| POM121L9P | NR_003714 | 0.156 | -0.3735 |
| CSN1S2AP | NR_003720 | 0.0785 | -0.113 |
| LOC389705 | NR_003920 | 0.0495 | -0.2675 |
| PMCHL1 | NR_003921 | -0.335 | 0.101 |
| PMCHL2 | NR_003922 | -0.1845 | 0.051 |
| GUCY1B2 | NR_003923 | -0.012 | 0.1815 |
| FKBP9L | NR_003949 | 0.0825 | 0.07 |
| SNORD94 | NR_004378 | -0.247 | -0.6185 |
| SCARNA10 | NR_004387 | 0.1305 | -0.141 |
| SCARNA14 | NR_004388 | 1.2615 | -1.2175 |
| SNORA16B | NR_004389 | -0.594 | 0.122 |
| SEC1 | NR_004401 | -0.0215 | -0.271 |
| SNORD97 | NR_004403 | 0.54 | 0.639 |
| HPVC1 | NR_004422 | 0.0015 | -0.1065 |
| AMN1 | NR_004854 | -1.468 | -2.112 |

| | | | |
|---|---|---|---|
| POFUT2 | NR_004858 | -0.552 | -1.331 |
| SCAND2 | NR_004859 | -0.2815 | 0.192 |
| SERPINA13 | NR_015340 | -0.552 | -0.611 |
| AAA1 | NR_015356 | -0.452 | -0.817 |
| LOC388796 | NR_015366 | 2.057 | 2.6115 |
| FLJ42709 | NR_015369 | -0.4155 | -0.187 |
| LOC401588 | NR_015378 | -0.161 | -0.0125 |
| UCA1 | NR_015379 | 0.0085 | -0.1655 |
| TP53TG1 | NR_015381 | 0.0395 | -0.5265 |
| .OC10012855 | NR_015398 | -0.2745 | -0.168 |
| LOC150568 | NR_015399 | -0.2325 | -0.45 |
| LOC149837 | NR_015406 | -0.537 | -1.388 |
| LOC154761 | NR_015421 | 0.065 | -0.035 |
| ANKRD36BP2 | NR_015424 | 0.087 | -0.2415 |
| C8ORFK29 | NR_015428 | -0.133 | -0.096 |
| LOC148413 | NR_015434 | -0.9325 | -1.725 |
| LOC652276 | NR_015441 | -0.8915 | -1.098 |
| TBRG1 | NR_016021 | -0.443 | -0.445 |
| FLJ42709 | NR_021490 | -0.4155 | -0.187 |
| FLJ42709 | NR_021491 | -0.4155 | -0.187 |
| ZNF137P | NR_023311 | -0.566 | -1.1135 |
| CIRBP | NR_023312 | -0.5905 | -2.0015 |
| CIRBP | NR_023313 | -0.5905 | -2.0015 |
| THTPA | NR_023314 | -0.4225 | -0.666 |
| ASZ1 | NR_023315 | 0.0405 | -0.659 |
| ACSF3 | NR_023316 | -0.204 | -0.9045 |
| ADPGK | NR_023318 | -0.269 | -1.5775 |
| ADPGK | NR_023319 | -0.269 | -1.5775 |
| KRTAP20-4 | NR_023342 | -0.0985 | -0.138 |
| CNKSR1 | NR_023345 | -0.6385 | -0.237 |
| CYB5D2 | NR_023346 | -0.619 | -1.262 |
| CYB5D2 | NR_023347 | -0.619 | -1.262 |
| GAS8 | NR_023348 | -0.722 | -1.1105 |
| TSEN15 | NR_023349 | -1.017 | -1.3245 |
| COPB2 | NR_023350 | -0.8695 | -1.5 |
| EXOSC7 | NR_023353 | 0.043 | 0.1435 |
| SCARNA9L | NR_023358 | 0.5865 | 0.5025 |
| CABLES1 | NR_023359 | -0.5665 | -1.7955 |
| CHRAC1 | NR_023360 | -0.8595 | -3.015 |
| AP3S2 | NR_023361 | -0.6365 | -1.77 |
| CCDC144C | NR_023380 | -0.576 | -0.913 |
| ZNF815 | NR_023382 | -0.5565 | 0.0115 |
| ZNF252 | NR_023392 | -1.5165 | -2.0155 |

| | | | |
|---|---|---|---|
| IPW | NR_023915 | -0.779 | -0.9055 |
| PIN1P1 | NR_023916 | 0.1775 | 0.0075 |
| WT1-AS | NR_023920 | -0.1715 | -0.3035 |
| DDX11L2 | NR_024004 | 0.2785 | -0.1595 |
| DDX11L2 | NR_024005 | 0.2785 | -0.1595 |
| SMCR5 | NR_024007 | 0.1325 | 0.229 |
| UGT2A3 | NR_024010 | -0.228 | -0.184 |
| FKSG29 | NR_024013 | -0.106 | -0.486 |
| TDRG1 | NR_024015 | -0.0795 | -0.2805 |
| C3orf55 | NR_024016 | -0.011 | -0.2205 |
| ZNF30 | NR_024018 | -1.384 | -2.6945 |
| FAM193B | NR_024019 | -0.862 | -1.8655 |
| NOL8 | NR_024020 | -0.8 | -2.761 |
| SETMAR | NR_024022 | -1.5105 | -3.311 |
| ACAA1 | NR_024024 | 0.064 | -0.877 |
| IL9R | NR_024033 | -0.131 | -0.3465 |
| AACSP1 | NR_024035 | -0.611 | -1.1165 |
| MPST | NR_024038 | -0.618 | -1.257 |
| CSGALNACT1 | NR_024040 | 0.0685 | 0.0865 |
| GUCY2E | NR_024042 | -0.2145 | -0.366 |
| LGALS9 | NR_024043 | 0.329 | -0.164 |
| WNT2 | NR_024047 | -0.0895 | -0.0155 |
| ZNF542 | NR_024055 | -0.957 | -1.783 |
| ZNF542 | NR_024056 | -0.957 | -1.783 |
| ZNF542 | NR_024057 | -0.957 | -1.783 |
| YWHAE | NR_024058 | -0.454 | -0.622 |
| WASH2P | NR_024077 | -0.672 | -1.9005 |
| ACP1 | NR_024080 | -0.5725 | -0.618 |
| C14orf165 | NR_024081 | 0.007 | -0.772 |
| SAP30L | NR_024084 | -0.9355 | -2.2765 |
| C4orf11 | NR_024087 | -0.3045 | -0.2465 |
| C4orf11 | NR_024088 | -0.3045 | -0.2465 |
| ADAM3A | NR_024106 | -0.107 | -0.226 |
| ADAM3A | NR_024107 | -0.107 | -0.226 |
| PFKL | NR_024108 | 0.193 | -0.366 |
| C1orf52 | NR_024113 | -1.532 | -2.2295 |
| ZNF620 | NR_024114 | -1.7665 | -1.908 |
| GIMAP6 | NR_024115 | 0.054 | 0.0995 |
| PBX3 | NR_024122 | -1.0965 | -1.7535 |
| PBX3 | NR_024123 | -1.0965 | -1.7535 |
| SNHG12 | NR_024127 | -0.911 | -2.034 |
| GHRL | NR_024132 | -0.155 | 0.1135 |
| GHRL | NR_024133 | -0.155 | 0.1135 |

| | | | |
|---|---|---|---|
| GHRL | NR_024134 | -0.155 | 0.1135 |
| GHRL | NR_024135 | -0.155 | 0.1135 |
| GHRL | NR_024136 | -0.155 | 0.1135 |
| GHRL | NR_024137 | -0.155 | 0.1135 |
| GHRL | NR_024138 | -0.155 | 0.1135 |
| RNF121 | NR_024147 | -0.4415 | -2.1355 |
| RNF121 | NR_024148 | -0.4415 | -2.1355 |
| ACBD5 | NR_024150 | -1.221 | -1.9905 |
| DGCR9 | NR_024159 | -0.261 | -0.505 |
| FKBP1AP1 | NR_024162 | 0.001 | -0.395 |
| CKS1B | NR_024163 | -0.0085 | -0.165 |
| TLR4 | NR_024168 | -0.1665 | 0.27 |
| TLR4 | NR_024169 | -0.1665 | 0.27 |
| MDM4 | NR_024171 | -1.3115 | -1.2645 |
| MKNK1 | NR_024174 | -0.7305 | -0.8235 |
| MKNK1 | NR_024176 | -0.7305 | -0.8235 |
| LOC284100 | NR_024178 | -0.278 | -0.23 |
| PDLIM5 | NR_024179 | 1.1995 | 2.0065 |
| ZNF577 | NR_024181 | -0.4305 | -1.437 |
| ATL2 | NR_024191 | -1.434 | -2.358 |
| HILS1 | NR_024192 | -0.106 | -0.307 |
| HILS1 | NR_024193 | -0.106 | -0.307 |
| TOM1 | NR_024194 | -0.1625 | -0.9325 |
| TOM1 | NR_024195 | -0.1625 | -0.9325 |
| CBLL1 | NR_024199 | -0.714 | -2.346 |
| ECD | NR_024203 | -1.0745 | -2.271 |
| KIAA1875 | NR_024207 | 0.4325 | -0.5905 |
| RNF185 | NR_024209 | -0.576 | -1.27 |
| RNF185 | NR_024210 | -0.576 | -1.27 |
| RNF185 | NR_024211 | -0.576 | -1.27 |
| RNF185 | NR_024212 | -0.576 | -1.27 |
| KBTBD4 | NR_024222 | -0.8915 | -2.4865 |
| CASP5 | NR_024239 | -0.085 | -0.228 |
| MUM1 | NR_024247 | -0.7325 | -1.595 |
| NECAP1 | NR_024260 | -0.556 | -1.704 |
| POGLUT1 | NR_024265 | -1.4555 | -1.7065 |
| SEC13 | NR_024272 | 0.0055 | -1.0065 |
| SEC13 | NR_024273 | 0.0055 | -1.0065 |
| C9orf53 | NR_024274 | -0.1735 | -0.3735 |
| LITAF | NR_024320 | -0.104 | -1.0455 |
| FLJ10661 | NR_024361 | -0.13 | -0.6275 |
| FLJ10661 | NR_024362 | -0.13 | -0.6275 |
| FLJ10661 | NR_024363 | -0.13 | -0.6275 |

| | | | |
|---|---|---|---|
| FLJ45340 | NR_024368 | -1.2345 | -1.519 |
| LOC642846 | NR_024374 | -0.1535 | -0.717 |
| FER1L4 | NR_024377 | 0.222 | -0.1795 |
| PLEKHM1P | NR_024386 | 0.164 | -0.011 |
| FLJ16779 | NR_024389 | 0.5915 | 0.461 |
| MGC23270 | NR_024396 | 0 | -0.545 |
| FLJ42393 | NR_024413 | -0.0385 | -0.2225 |
| OBFC2A | NR_024415 | -1.992 | -1.8 |
| PARVG | NR_024426 | 0.088 | -0.186 |
| PARVG | NR_024427 | 0.088 | -0.186 |
| PARVG | NR_024428 | 0.088 | -0.186 |
| .OC10013399[illegible] | NR_024434 | 0.546 | -0.4115 |
| .OC10013399[illegible] | NR_024435 | 0.546 | -0.4115 |
| LOC728989 | NR_024442 | -0.3495 | -0.1835 |
| .OC10013425[illegible] | NR_024452 | -0.184 | -0.106 |
| .OC10019094[illegible] | NR_024457 | -0.343 | -0.202 |
| LSM5 | NR_024466 | -0.4875 | -0.5105 |
| .OC10012788[illegible] | NR_024470 | 0.1255 | -0.6915 |
| .OC10013172[illegible] | NR_024479 | 0.115 | -0.2065 |
| LOC400657 | NR_024484 | -0.4535 | -1.2475 |
| LOC646813 | NR_024504 | 0.052 | 0.4045 |
| TTL | NR_024505 | -1.2175 | -1.773 |
| TTL | NR_024506 | -1.2175 | -1.773 |
| TTL | NR_024507 | -1.2175 | -1.773 |
| ADAMTS13 | NR_024514 | -0.1505 | -0.132 |
| APLP2 | NR_024515 | -0.3565 | -0.861 |
| APLP2 | NR_024516 | -0.3565 | -0.861 |
| DAXX | NR_024517 | -0.2725 | -0.882 |
| LMAN2L | NR_024518 | -1.639 | -3.117 |
| LMAN2L | NR_024519 | -1.639 | -3.117 |
| LMAN2L | NR_024520 | -1.639 | -3.117 |
| LMAN2L | NR_024521 | -1.639 | -3.117 |
| TMEM67 | NR_024522 | -1.1445 | -1.783 |
| ARL6IP6 | NR_024526 | -0.9535 | -1.4515 |
| SCN4B | NR_024527 | 0.382 | -0.2585 |
| ALG2 | NR_024532 | -1.2275 | -2.6445 |
| ALG3 | NR_024533 | -0.2825 | -1.162 |
| ALG3 | NR_024534 | -0.2825 | -1.162 |
| GSTM3 | NR_024537 | -0.0165 | -0.132 |
| GSTM4 | NR_024538 | 0.1655 | -0.5215 |
| POPDC3 | NR_024539 | -0.6305 | -1.7685 |
| WASH7P | NR_024540 | -0.4405 | -1.163 |
| SNHG7 | NR_024542 | -0.7035 | -1.727 |

| | | | |
|---|---|---|---|
| SNHG7 | NR_024543 | -0.7035 | -1.727 |
| TXNL1 | NR_024546 | -0.9355 | -1.635 |
| CRCP | NR_024548 | -0.735 | -1.8625 |
| DMTF1 | NR_024549 | -1.036 | -1.9595 |
| DMTF1 | NR_024550 | -1.036 | -1.9595 |
| CALHM2 | NR_024552 | -0.449 | -1.0875 |
| ZNF204P | NR_024553 | -0.044 | -0.588 |
| RHOBTB1 | NR_024554 | -0.933 | -1.701 |
| RHOBTB1 | NR_024555 | -0.933 | -1.701 |
| WAC | NR_024557 | -1.6395 | -3.798 |
| .OC10012867! | NR_024561 | 0.336 | 0.449 |
| .OC10012867! | NR_024562 | 0.336 | 0.449 |
| ZNF271 | NR_024565 | -0.3165 | -1.569 |
| ZNF271 | NR_024566 | -0.3165 | -1.569 |
| .OC10013055 | NR_024567 | -0.474 | -0.293 |
| FBXO31 | NR_024568 | -0.804 | -2.416 |
| .OC10013087 | NR_024569 | 0.069 | -0.3935 |
| SEC61A2 | NR_024576 | -0.119 | -0.3935 |
| SEC61A2 | NR_024577 | -0.119 | -0.3935 |
| ASB13 | NR_024581 | -0.681 | -1.707 |
| POM121L8P | NR_024583 | -0.068 | -0.4525 |
| .OC10012829 | NR_024585 | -0.1905 | -0.346 |
| DNM1P35 | NR_024595 | -0.035 | -0.4695 |
| C11orf73 | NR_024596 | -0.6245 | -1.0285 |
| C11orf73 | NR_024597 | -0.6245 | -1.0285 |
| C11orf73 | NR_024598 | -0.6245 | -1.0285 |
| CLCA4 | NR_024602 | -0.418 | -0.0855 |
| MPDU1 | NR_024603 | 0.003 | -0.951 |
| CLCA3P | NR_024604 | 0.424 | 0.754 |
| EFCAB1 | NR_024605 | -0.043 | -0.168 |
| LOC442459 | NR_024608 | -0.125 | -0.0715 |
| HINT1 | NR_024610 | 0.007 | -0.322 |
| HINT1 | NR_024611 | 0.007 | -0.322 |
| ATXN1L | NR_024612 | -0.9495 | -1.768 |
| C3orf51 | NR_024615 | -0.017 | -0.0565 |
| C11orf21 | NR_024621 | -0.0635 | -0.2615 |
| API5 | NR_024625 | -1.267 | -1.653 |
| C14orf162 | NR_024630 | -0.822 | -1.8645 |
| C21orf88 | NR_026542 | -0.082 | -0.2755 |
| C21orf88 | NR_026543 | -0.082 | -0.2755 |
| C21orf90 | NR_026547 | 0.1695 | -0.1175 |
| C21orf90 | NR_026548 | 0.1695 | -0.1175 |
| RHBG | NR_026549 | -0.9955 | -0.9725 |

| | | | |
|---|---|---|---|
| BRP44 | NR_026550 | -0.0225 | -0.882 |
| CALCOCO1 | NR_026554 | -0.1165 | -0.447 |
| GDEP | NR_026555 | -0.0475 | 0.096 |
| LRRC27 | NR_026559 | -0.3155 | -1.0585 |
| C20orf24 | NR_026562 | -0.1675 | -1.47 |
| ESPNP | NR_026567 | -0.044 | -0.2925 |
| MAPRE2 | NR_026570 | -0.7635 | -1.343 |
| NOX4 | NR_026571 | 0.0195 | -0.5715 |
| GK3P | NR_026575 | -0.416 | -0.3985 |
| SAA3P | NR_026576 | 0.109 | -0.0475 |
| TFDP1 | NR_026580 | -0.7685 | -1.203 |
| MLF2 | NR_026581 | 0.0285 | -0.5575 |
| ID2B | NR_026582 | -0.5385 | -0.335 |
| EFCAB2 | NR_026586 | 0.225 | -0.224 |
| EFCAB2 | NR_026587 | 0.225 | -0.224 |
| EFCAB2 | NR_026588 | 0.225 | -0.224 |
| CDYL | NR_026590 | -1.21 | -1.1755 |
| RAD1 | NR_026591 | -1.464 | -2.0355 |
| BMS1P4 | NR_026592 | -0.401 | -1.059 |
| TMX2 | NR_026593 | -0.195 | -0.7195 |
| NENF | NR_026598 | 0.3225 | -0.2065 |
| INTS2 | NR_026641 | -0.949 | -1.6555 |
| UBAC2 | NR_026644 | -0.317 | -1.719 |
| DET1 | NR_026645 | -1.097 | -3.0075 |
| BPHL | NR_026648 | -0.3265 | -1.2945 |
| BPHL | NR_026649 | -0.3265 | -1.2945 |
| BPHL | NR_026650 | -0.3265 | -1.2945 |
| CD58 | NR_026665 | -1.438 | -2.471 |
| RPS2P32 | NR_026676 | 0.6345 | 0.6035 |
| .OC10026816 | NR_026682 | 0.4385 | -0.4915 |
| .OC10026816 | NR_026683 | 0.4385 | -0.4915 |
| IL10RA | NR_026691 | 0.3595 | -0.2605 |
| CD209 | NR_026692 | 0.0665 | -0.056 |
| ZNF487P | NR_026693 | -0.346 | -0.923 |
| FGF1 | NR_026695 | 0.2095 | -0.095 |
| FGF1 | NR_026696 | 0.2095 | -0.095 |
| FAM134C | NR_026697 | -0.6525 | -1.5615 |
| KPNA1 | NR_026698 | -1.143 | -2.0435 |
| GLYCTK | NR_026699 | 0.35 | -0.0325 |
| GLYCTK | NR_026700 | 0.35 | -0.0325 |
| GLYCTK | NR_026701 | 0.35 | -0.0325 |
| GLYCTK | NR_026702 | 0.35 | -0.0325 |
| CLEC4M | NR_026707 | -0.3085 | -0.2745 |

| | | | |
|---|---|---|---|
| CLEC4M | NR_026708 | -0.3085 | -0.2745 |
| CLEC4M | NR_026709 | -0.3085 | -0.2745 |
| KIR3DX1 | NR_026716 | 0.035 | 0.0225 |
| STK19 | NR_026717 | -0.158 | -0.452 |
| DSCR9 | NR_026719 | 0.192 | -0.3195 |
| SARNP | NR_026722 | -0.557 | -0.6715 |
| SARNP | NR_026723 | -0.557 | -0.6715 |
| EDEM2 | NR_026728 | -0.12 | 0.143 |
| TPTE2P1 | NR_026730 | 0.1475 | -0.0835 |
| TMPRSS7 | NR_026734 | -0.056 | -0.2755 |
| C6orf52 | NR_026736 | 0.019 | -0.386 |
| C6orf52 | NR_026737 | 0.019 | -0.386 |
| EFCAB5 | NR_026738 | 0.022 | -0.161 |
| tAKR | NR_026743 | -0.0515 | 0.3345 |
| COG6 | NR_026745 | -1.2695 | -2.292 |
| ZNF619 | NR_026746 | -0.387 | -1.166 |
| LOC440905 | NR_026758 | -0.15 | -0.3335 |
| PPBPL2 | NR_026769 | 0.093 | -0.084 |
| C17orf88 | NR_026770 | -0.153 | -0.8765 |
| C6orf123 | NR_026773 | -0.14 | -0.262 |
| BPESC1 | NR_026783 | 0.4705 | -0.229 |
| LMLN | NR_026786 | -0.6 | -0.8235 |
| LMLN | NR_026787 | -0.6 | -0.8235 |
| SH3GL3 | NR_026799 | -0.5055 | -0.8405 |
| C15orf5 | NR_026813 | 0.587 | -0.5345 |
| PSORS1C3 | NR_026816 | -0.3465 | -0.188 |
| INTS9 | NR_026826 | -0.9445 | -2.658 |
| LOC84856 | NR_026827 | 0.481 | 0.1705 |
| FLJ32063 | NR_026830 | 0.2245 | -0.179 |
| DSCR8 | NR_026838 | -0.31 | -0.2955 |
| DSCR8 | NR_026839 | -0.31 | -0.2955 |
| DSCR8 | NR_026840 | -0.31 | -0.2955 |
| DSCR8 | NR_026841 | -0.31 | -0.2955 |
| DSCR8 | NR_026842 | -0.31 | -0.2955 |
| C18orf18 | NR_026849 | -0.3435 | -0.3185 |
| FAM75D5 | NR_026851 | 0.342 | -0.259 |
| PPP1R3E | NR_026862 | -0.7715 | -1.2315 |
| C7orf13 | NR_026865 | -0.5595 | -1.3955 |
| NEURL3 | NR_026875 | 0.0545 | 0.2895 |
| FAM185A | NR_026879 | -0.621 | -2.041 |
| PRG1 | NR_026881 | -0.0385 | 0.2885 |
| HSPC072 | NR_026883 | -0.0785 | -0.347 |
| HSPC072 | NR_026884 | -0.0785 | -0.347 |

| | | | |
|---|---|---|---|
| RPSAP9 | NR_026890 | 0.321 | 0.426 |
| POU6F1 | NR_026893 | -0.1525 | -0.187 |
| MUDENG | NR_026895 | -1.1615 | -2.559 |
| RPL10 | NR_026898 | 0.0705 | -0.347 |
| PABPC1P2 | NR_026904 | -0.021 | -0.2915 |
| STK16 | NR_026909 | -0.356 | -1.018 |
| ABHD11 | NR_026910 | -0.2895 | -0.9 |
| ABHD11 | NR_026912 | -0.2895 | -0.9 |
| ANKRD30BP2 | NR_026916 | 0.245 | 0.0085 |
| PTPN3 | NR_026918 | -0.329 | -1.728 |
| LOC202181 | NR_026921 | 0.0415 | -0.259 |
| C5orf27 | NR_026936 | 0.1025 | -0.314 |
| CASC2 | NR_026939 | 0.079 | -0.283 |
| CASC2 | NR_026940 | 0.079 | -0.283 |
| CASC2 | NR_026941 | 0.079 | -0.283 |
| LOC642852 | NR_026943 | -1.1975 | -1.4755 |
| LOC284440 | NR_026956 | -0.0225 | -0.2935 |
| LOC284749 | NR_026958 | 0.13 | -0.1385 |
| .OC10013069: | NR_026966 | -0.098 | -0.348 |
| LOC285456 | NR_026968 | 0.361 | -0.0865 |
| RPL23AP82 | NR_026981 | -0.701 | -1.7645 |
| RPL23AP82 | NR_026982 | -0.701 | -1.7645 |
| C22orf34 | NR_026997 | -0.044 | -0.056 |
| PGAP2 | NR_027014 | -0.393 | -1.112 |
| PGAP2 | NR_027015 | -0.393 | -1.112 |
| PGAP2 | NR_027016 | -0.393 | -1.112 |
| PGAP2 | NR_027017 | -0.393 | -1.112 |
| PGAP2 | NR_027018 | -0.393 | -1.112 |
| GUSBP1 | NR_027026 | -0.212 | -0.7985 |
| GUSBP1 | NR_027027 | -0.212 | -0.7985 |
| GUSBP1 | NR_027028 | -0.212 | -0.7985 |
| .OC10013077( | NR_027032 | -1.0455 | -1.7135 |
| LOC401093 | NR_027037 | -0.3785 | 0.2305 |
| LOC401093 | NR_027038 | -0.3785 | 0.2305 |
| LOC440040 | NR_027044 | 0.3705 | -0.334 |
| FLJ34503 | NR_027060 | 0.101 | -0.285 |
| .OC10013221! | NR_027069 | -0.0245 | 0.033 |
| PISRT1 | NR_027070 | -0.4515 | -0.4805 |
| AFMID | NR_027083 | -0.3455 | -1.381 |
| MGC45800 | NR_027107 | -0.8325 | -1.0085 |
| SHROOM4 | NR_027121 | -0.006 | -0.3345 |
| RBM18 | NR_027125 | -0.3785 | -1.0235 |
| RBM18 | NR_027126 | -0.3785 | -1.0235 |

| | | | |
|---|---|---|---|
| C21orf67 | NR_027128 | -0.06 | -0.5535 |
| C21orf67 | NR_027129 | -0.06 | -0.5535 |
| ZNF738 | NR_027130 | -1.034 | -0.9015 |
| ZSCAN18 | NR_027135 | -0.326 | -0.8385 |
| C1orf126 | NR_027136 | -0.853 | -0.063 |
| SPIN3 | NR_027139 | -0.8325 | -0.9125 |
| TNFRSF10B | NR_027140 | -0.3515 | -1.3295 |
| C10orf108 | NR_027151 | 0.628 | -0.528 |
| C10orf108 | NR_027152 | 0.628 | -0.528 |
| TUBBP5 | NR_027156 | -0.0855 | -0.6415 |
| LOC729678 | NR_027183 | -0.54 | -0.581 |
| LOC728743 | NR_027237 | -0.087 | -0.7375 |
| ZNF529 | NR_027239 | -1.4525 | -2.2325 |
| LOC388796 | NR_027241 | 2.057 | 2.6115 |
| GNN | NR_027249 | 0.0845 | -0.241 |
| SSX8 | NR_027250 | 0.1115 | -0.3635 |
| .OC10012972( | NR_027251 | 0.474 | 0.265 |
| LOC285696 | NR_027253 | -0.082 | -0.491 |
| LOC388387 | NR_027254 | -0.045 | -0.2295 |
| POLE3 | NR_027261 | -0.0985 | 0.0545 |
| GLG1 | NR_027264 | -0.6955 | -1.193 |
| GLG1 | NR_027265 | -0.6955 | -1.193 |
| SAE1 | NR_027280 | -0.134 | -0.7595 |
| LOC399815 | NR_027282 | -0.7085 | -0.366 |
| IFLTD1 | NR_027296 | -0.7185 | -0.891 |
| HOMER3 | NR_027297 | -0.1185 | -0.629 |
| XPC | NR_027299 | -1.0635 | -2.6125 |
| TRIB2 | NR_027303 | -0.996 | -1.6585 |
| LOC148145 | NR_027318 | 0.234 | -0.0475 |
| C7orf54 | NR_027330 | -0.791 | -0.809 |
| HPGD | NR_027332 | -0.3415 | -0.8265 |
| .OC10013001! | NR_027335 | 0.1075 | -0.0095 |
| .OC10013001! | NR_027336 | 0.1075 | -0.0095 |
| FKBP9L | NR_027339 | 0.0825 | 0.07 |
| FKBP9L | NR_027340 | 0.0825 | 0.07 |
| FKBP9L | NR_027342 | 0.0825 | 0.07 |
| TMPRSS3 | NR_027348 | -0.3595 | -0.257 |
| MIR17HG | NR_027349 | -0.8005 | 0.205 |
| MIR17HG | NR_027350 | -0.8005 | 0.205 |
| CD8A | NR_027353 | 0.0545 | -0.1035 |
| C12orf32 | NR_027363 | -1.5995 | -2.9275 |
| C12orf32 | NR_027364 | -1.5995 | -2.9275 |
| C12orf32 | NR_027365 | -1.5995 | -2.9275 |

| | | | |
|---|---|---|---|
| POP4 | NR_027368 | -1.3675 | -2.16 |
| RDH13 | NR_027381 | -0.7765 | -1.8235 |
| RDH13 | NR_027382 | -0.7765 | -1.8235 |
| AKAP17A | NR_027383 | 0.028 | -0.5075 |
| SLC13A2 | NR_027384 | 0.1895 | -0.123 |
| IGFALS | NR_027389 | -0.024 | -0.2015 |
| GRINL1A | NR_027390 | -0.1435 | -0.441 |
| SLC15A3 | NR_027391 | -0.1425 | -0.2285 |
| TJAP1 | NR_027394 | -1.0215 | -2.418 |
| GORAB | NR_027397 | -2.007 | -4.367 |
| GFOD2 | NR_027398 | 0.353 | -0.8485 |
| MTHFD2 | NR_027405 | -0.8095 | -0.9245 |
| GOLGA8A | NR_027409 | -1.2085 | -1.186 |
| NEK3 | NR_027415 | -0.764 | -1.376 |
| AVPR2 | NR_027419 | 0.0455 | -0.3515 |
| LOC389834 | NR_027420 | -0.427 | -0.963 |
| FAM66D | NR_027425 | -0.41 | -0.2565 |
| TATDN1 | NR_027427 | -0.833 | -1.039 |
| LOC219347 | NR_027428 | 0.0315 | -0.3485 |
| LOC219347 | NR_027429 | 0.0315 | -0.3485 |
| LOC219347 | NR_027430 | 0.0315 | -0.3485 |
| LOC219347 | NR_027431 | 0.0315 | -0.3485 |
| LOC219347 | NR_027432 | 0.0315 | -0.3485 |
| .OC10012971( | NR_027435 | -0.4615 | -0.348 |
| IP6K2 | NR_027437 | 0.385 | -0.656 |
| IP6K2 | NR_027438 | 0.385 | -0.656 |
| FLJ36777 | NR_027441 | 0.3465 | -0.2 |
| .OC10028884: | NR_027442 | -0.4485 | -0.38 |
| .OC10013395 | NR_027444 | 0.3295 | 0.201 |
| TBC1D15 | NR_027449 | -0.7245 | -2.5675 |
| LOC647979 | NR_027451 | -1.091 | -0.619 |
| SYT14 | NR_027458 | -0.4435 | -1.3615 |
| SYT14 | NR_027459 | -0.4435 | -1.3615 |
| SHBG | NR_027462 | -0.0225 | -0.5525 |
| SHBG | NR_027463 | -0.0225 | -0.5525 |
| AIFM3 | NR_027464 | -0.0965 | 0.04 |
| LOC440173 | NR_027471 | -0.076 | -0.0675 |
| KDM2A | NR_027473 | -0.944 | -2.957 |
| AGXT2L1 | NR_027474 | 0.017 | -0.49 |
| AGXT2L1 | NR_027475 | 0.017 | -0.49 |
| ABHD14B | NR_027476 | 0.21 | -0.294 |
| POTEG | NR_027480 | -0.008 | -0.311 |
| TPI1 | NR_027483 | 0.128 | -0.2305 |

| | | | |
|---|---|---|---|
| TBC1D3P2 | NR_027486 | -0.043 | -0.572 |
| MTHFSD | NR_027489 | 0.244 | -0.499 |
| MTHFSD | NR_027490 | 0.244 | -0.499 |
| CCNDBP1 | NR_027513 | -0.7355 | -1.5015 |
| CCNDBP1 | NR_027514 | -0.7355 | -1.5015 |
| FHL1 | NR_027621 | -0.136 | -0.286 |
| BZW2 | NR_027624 | -0.725 | -0.9785 |
| BRD9 | NR_027633 | -0.3105 | -1.066 |
| SPATA5L1 | NR_027635 | -1.133 | -2.7405 |
| ADAM9 | NR_027638 | -0.5215 | -1.088 |
| ADAM9 | NR_027639 | -0.5215 | -1.088 |
| LPO | NR_027647 | 0.0965 | -0.295 |
| C15orf40 | NR_027649 | 0.7175 | 0.2555 |
| C15orf40 | NR_027650 | 0.7175 | 0.2555 |
| KLF6 | NR_027653 | -0.1815 | 0.227 |
| SMAD6 | NR_027654 | -0.6655 | -1.897 |
| BACH1 | NR_027655 | -0.896 | -2.286 |
| ZNF138 | NR_027660 | -0.8825 | -1.2765 |
| ZNF138 | NR_027661 | -0.8825 | -1.2765 |
| SLC25A13 | NR_027662 | -0.4575 | -1.218 |
| RNF170 | NR_027668 | -0.4265 | -1.0105 |
| RNF170 | NR_027669 | -0.4265 | -1.0105 |
| UGGT1 | NR_027671 | -0.5005 | -0.923 |
| ADARB1 | NR_027672 | 0.249 | -0.6755 |
| ADARB1 | NR_027673 | 0.249 | -0.6755 |
| ADARB1 | NR_027674 | 0.249 | -0.6755 |
| BRCA1 | NR_027676 | -1.1685 | -2.067 |
| RPAIN | NR_027678 | -1.303 | -1.4555 |
| RPAIN | NR_027679 | -1.303 | -1.4555 |
| RPAIN | NR_027680 | -1.303 | -1.4555 |
| RPAIN | NR_027681 | -1.303 | -1.4555 |
| RPAIN | NR_027682 | -1.303 | -1.4555 |
| RPAIN | NR_027683 | -1.303 | -1.4555 |
| RPAIN | NR_027684 | -1.303 | -1.4555 |
| RPAIN | NR_027685 | -1.303 | -1.4555 |
| CERKL | NR_027689 | -0.0855 | -1.3575 |
| CERKL | NR_027690 | -0.0855 | -1.3575 |
| PANX2 | NR_027691 | 0.004 | -0.707 |
| PHF1 | NR_027692 | -0.1685 | -0.2715 |
| C1orf170 | NR_027693 | -0.4505 | -0.476 |
| DSCR10 | NR_027695 | -0.08 | -0.136 |
| NOP56 | NR_027700 | -0.3265 | -0.5735 |
| CDK10 | NR_027702 | -0.3685 | -0.763 |

| | | | |
|---|---|---|---|
| CDK10 | NR_027703 | -0.3685 | -0.763 |
| FLJ11235 | NR_027706 | 0.146 | -0.3545 |
| FLJ14107 | NR_027715 | -0.5565 | -1.1185 |
| H2AFJ | NR_027716 | -0.0375 | 0.744 |
| TTC31 | NR_027749 | -0.5155 | -1.621 |
| FAM20A | NR_027751 | 0.0275 | -0.2675 |
| ALDH1L2 | NR_027752 | -0.29 | -0.4185 |
| HHATL | NR_027753 | -0.1485 | -0.263 |
| CEACAM22P | NR_027754 | -0.118 | -0.2085 |
| CSF2RA | NR_027760 | -0.4185 | -0.0625 |
| RBM34 | NR_027762 | 0.392 | 1.2725 |
| GYS1 | NR_027763 | -0.446 | -0.8565 |
| NUDT16P1 | NR_027766 | -0.1275 | -0.527 |
| TNIK | NR_027767 | -0.5465 | -1.6965 |
| DPY19L2P2 | NR_027768 | -0.152 | -0.203 |
| HMGXB4 | NR_027780 | -1.501 | -2.162 |
| MT1DP | NR_027781 | -0.233 | -0.35 |
| SAT1 | NR_027783 | 0.8915 | 1.859 |
| SERHL | NR_027786 | -0.371 | -0.229 |
| ZNF767 | NR_027788 | -0.7595 | -1.5115 |
| ZNF767 | NR_027789 | -0.7595 | -1.5115 |
| C3orf17 | NR_027794 | -0.8805 | -1.5895 |
| C3orf17 | NR_027796 | -0.8805 | -1.5895 |
| PTGES2 | NR_027811 | 0.0425 | -0.892 |
| PTGES2 | NR_027812 | 0.0425 | -0.892 |
| SDK1 | NR_027816 | -0.468 | -0.79 |
| HLA-L | NR_027822 | 0.25 | -0.1165 |
| LIMS2 | NR_027823 | -0.007 | -0.366 |
| BCAP29 | NR_027830 | -1.4795 | -1.146 |
| APOL3 | NR_027833 | 0.2035 | -0.1925 |
| APOL3 | NR_027834 | 0.2035 | -0.1925 |
| APOL3 | NR_027835 | 0.2035 | -0.1925 |
| FYTTD1 | NR_027840 | -0.7315 | -1.058 |
| MTX2 | NR_027850 | -1.468 | -2.8175 |
| CLK1 | NR_027855 | -0.908 | -1.031 |
| CLK1 | NR_027856 | -0.908 | -1.031 |
| HAT1 | NR_027862 | -0.601 | -2.193 |
| PDCD6IP | NR_027867 | -0.661 | -1.2085 |
| PDCD6IP | NR_027868 | -0.661 | -1.2085 |
| GCFC1 | NR_027873 | -1.326 | -2.046 |
| ADAM9 | NR_027878 | -0.5215 | -1.088 |
| TMEM189 | NR_027889 | -0.8755 | -2.279 |
| C6orf124 | NR_027906 | 0.086 | -0.0975 |

| | | | |
|---|---|---|---|
| CCDC62 | NR_027918 | 0.1065 | -0.452 |
| COX11 | NR_027941 | -1.2915 | -2.066 |
| COX11 | NR_027942 | -1.2915 | -2.066 |
| ERC1 | NR_027946 | -0.142 | -0.4435 |
| ERC1 | NR_027948 | -0.142 | -0.4435 |
| ERC1 | NR_027949 | -0.142 | -0.4435 |
| TYW3 | NR_027962 | -0.172 | -0.754 |
| APLNR | NR_027991 | 0.1735 | -0.1355 |
| RDM1 | NR_027996 | -0.173 | -0.392 |
| RDM1 | NR_027997 | -0.173 | -0.392 |
| RDM1 | NR_027998 | -0.173 | -0.392 |
| RDM1 | NR_027999 | -0.173 | -0.392 |
| RDM1 | NR_028000 | -0.173 | -0.392 |
| CREBZF | NR_028024 | -1.9725 | -2.7395 |
| CREBZF | NR_028025 | -1.9725 | -2.7395 |
| CREBZF | NR_028026 | -1.9725 | -2.7395 |
| CREBZF | NR_028027 | -1.9725 | -2.7395 |
| ITPRIPL2 | NR_028028 | -1.2635 | -1.6325 |
| HCG9 | NR_028032 | 0.3 | 0.095 |
| FAS | NR_028033 | -0.4565 | -1.1795 |
| FAS | NR_028034 | -0.4565 | -1.1795 |
| FAS | NR_028035 | -0.4565 | -1.1795 |
| FAS | NR_028036 | -0.4565 | -1.1795 |
| GATS | NR_028038 | -0.0295 | -0.654 |
| GATS | NR_028039 | -0.0295 | -0.654 |
| GATS | NR_028040 | -0.0295 | -0.654 |
| GRM8 | NR_028041 | -1.312 | -1.6935 |
| KLRAP1 | NR_028045 | -0.138 | -1.095 |
| MTERFD2 | NR_028049 | -0.592 | -1.698 |
| MTERFD2 | NR_028050 | -0.592 | -1.698 |
| MTERFD2 | NR_028051 | -0.592 | -1.698 |
| NGRN | NR_028052 | 0.24 | -0.0685 |
| PLCXD1 | NR_028057 | -0.0525 | -1.1695 |
| GCC2 | NR_028063 | -0.3565 | -2.198 |
| RAB2B | NR_028074 | -0.589 | -0.8845 |
| SCARF1 | NR_028075 | -0.059 | -0.074 |
| SCARF1 | NR_028076 | -0.059 | -0.074 |
| ZSCAN12 | NR_028077 | 0.159 | -0.6535 |
| FAM172A | NR_028080 | -0.6135 | -0.6715 |
| UFSP2 | NR_028085 | -0.796 | -1.4925 |
| NXF5 | NR_028089 | -0.106 | 0.186 |
| LPAL2 | NR_028092 | -0.157 | -0.3945 |
| LPAL2 | NR_028093 | -0.157 | -0.3945 |

| | | | |
|---|---|---|---|
| DCAF11 | NR_028099 | -0.0445 | -0.6475 |
| DCAF11 | NR_028100 | -0.0445 | -0.6475 |
| THUMPD2 | NR_028102 | -1.3675 | -2.615 |
| DCAF8 | NR_028103 | -0.5305 | -1.7445 |
| DCAF8 | NR_028104 | -0.5305 | -1.7445 |
| DCAF8 | NR_028105 | -0.5305 | -1.7445 |
| DCAF8 | NR_028106 | -0.5305 | -1.7445 |
| .OC10013395[illegible] | NR_028119 | 0.3295 | 0.201 |
| CFL2 | NR_028130 | -0.8905 | -2.2675 |
| CFL2 | NR_028131 | -0.8905 | -2.2675 |
| EXOC7 | NR_028133 | -0.1865 | -0.535 |
| KCNMB3 | NR_028135 | -0.5575 | -0.341 |
| KALRN | NR_028136 | -0.1745 | -0.7935 |
| FLJ42289 | NR_028139 | -0.015 | -0.7505 |
| FLJ42289 | NR_028140 | -0.015 | -0.7505 |
| GABARAPL3 | NR_028287 | -0.1445 | 0.2485 |
| TCL6 | NR_028288 | 0.0765 | -0.298 |
| VNN3 | NR_028290 | -0.0745 | -0.197 |
| VNN3 | NR_028291 | -0.0745 | -0.197 |
| PTGER3 | NR_028292 | -0.2095 | -0.2735 |
| PTGER3 | NR_028293 | -0.2095 | -0.2735 |
| PTGER3 | NR_028294 | -0.2095 | -0.2735 |
| NLGN4Y | NR_028319 | 0.0205 | -0.5245 |
| SLC5A6 | NR_028323 | -0.328 | -1.5345 |
| FLJ45445 | NR_028324 | 0.1375 | -0.6145 |
| .OC10013206[illegible] | NR_028325 | -1.1635 | -1.1645 |
| C15orf37 | NR_028330 | -0.0285 | -0.1955 |
| TRAF3IP2 | NR_028338 | 0.058 | -0.0505 |
| LOC285501 | NR_028342 | 0.0255 | 0.345 |
| FLJ33360 | NR_028351 | 0.0535 | -0.0525 |
| PIGP | NR_028352 | -1.094 | -1.2485 |
| MRPL30 | NR_028356 | -0.6025 | -1.025 |
| GGNBP1 | NR_028361 | 0.0775 | -0.2865 |
| TIMM8B | NR_028383 | -0.3465 | -0.3595 |
| FUS | NR_028388 | 0.078 | -0.447 |
| CSRP2BP | NR_028402 | -2.323 | -4.1075 |
| FXYD5 | NR_028406 | -0.0945 | 0.022 |
| ARMCX4 | NR_028407 | -0.275 | -0.669 |
| APOC1P1 | NR_028412 | 0.1925 | -0.429 |
| APOC1P1 | NR_028413 | 0.1925 | -0.429 |
| APOC1P1 | NR_028414 | 0.1925 | -0.429 |
| AMT | NR_028435 | -0.056 | -0.721 |
| UBE2L3 | NR_028436 | -0.436 | -1.5275 |

| | | | |
|---|---|---|---|
| UBE2L3 | NR_028437 | -0.436 | -1.5275 |
| ATXN3 | NR_028453 | -0.521 | -2.1695 |
| ATXN3 | NR_028454 | -0.521 | -2.1695 |
| ATXN3 | NR_028455 | -0.521 | -2.1695 |
| ATXN3 | NR_028456 | -0.521 | -2.1695 |
| ATXN3 | NR_028457 | -0.521 | -2.1695 |
| ATXN3 | NR_028458 | -0.521 | -2.1695 |
| ATXN3 | NR_028459 | -0.521 | -2.1695 |
| ATXN3 | NR_028460 | -0.521 | -2.1695 |
| ATXN3 | NR_028461 | -0.521 | -2.1695 |
| ATXN3 | NR_028462 | -0.521 | -2.1695 |
| ATXN3 | NR_028463 | -0.521 | -2.1695 |
| ATXN3 | NR_028464 | -0.521 | -2.1695 |
| ATXN3 | NR_028465 | -0.521 | -2.1695 |
| ATXN3 | NR_028466 | -0.521 | -2.1695 |
| ATXN3 | NR_028467 | -0.521 | -2.1695 |
| ATXN3 | NR_028468 | -0.521 | -2.1695 |
| ATXN3 | NR_028469 | -0.521 | -2.1695 |
| ATXN3 | NR_028470 | -0.521 | -2.1695 |
| TMEM138 | NR_028473 | -1.2975 | -2.9305 |
| SLC25A26 | NR_028475 | -0.112 | -1.016 |
| RBMX | NR_028476 | -0.7635 | -1.229 |
| RBMX | NR_028477 | -0.7635 | -1.229 |
| DCAF17 | NR_028482 | -0.496 | -0.617 |
| TMEM41B | NR_028491 | -1.2185 | -1.4945 |
| LYRM2 | NR_028493 | -1.381 | -2.254 |
| LYRM2 | NR_028494 | -1.381 | -2.254 |
| LYRM2 | NR_028495 | -1.381 | -2.254 |
| LDHA | NR_028500 | -0.2955 | -0.612 |
| ZNF833P | NR_028594 | -0.851 | -2.317 |
| EDNRA | NR_028596 | -0.9585 | -0.9835 |
| .OC10028721 | NR_029193 | 0.223 | -0.4095 |
| C20orf7 | NR_029377 | -0.965 | -2.578 |
| .OC10028951 | NR_029378 | -0.0185 | 0.299 |
| FLJ41350 | NR_029380 | 0.0155 | -0.869 |
| TBXAS1 | NR_029394 | 0.099 | -0.327 |
| IGLL3P | NR_029395 | 0.29 | -0.17 |
| LOC643406 | NR_029405 | -0.5915 | 0.0535 |
| .OC10013309 | NR_029411 | 0.3225 | -0.0265 |
| MFNG | NR_029413 | -0.0455 | -0.46 |
| SHMT2 | NR_029415 | 0.348 | -0.3655 |
| SHMT2 | NR_029416 | 0.348 | -0.3655 |
| SHMT2 | NR_029417 | 0.348 | -0.3655 |

| | | | |
|---|---|---|---|
| WDR13 | NR_029427 | -0.0425 | -0.8285 |
| DNAJC7 | NR_029431 | -0.7385 | -1.0455 |
| LST1 | NR_029461 | -0.04 | -0.218 |
| LST1 | NR_029462 | -0.04 | -0.218 |
| COX7A2 | NR_029466 | 0.154 | -0.262 |
| SELL | NR_029467 | 0.016 | 0.0455 |
| SNRPC | NR_029472 | 0.0895 | -0.37 |
| MIR21 | NR_029493 | -1.6515 | -1.8285 |
| ANO1 | NR_030691 | -0.1205 | -0.181 |
| FAM104B | NR_030722 | -0.595 | -0.7705 |
| XPNPEP1 | NR_030724 | -0.154 | -0.6625 |
| RCC1 | NR_030725 | -0.3655 | -1.5915 |
| RCC1 | NR_030726 | -0.3655 | -1.5915 |
| TTC3P1 | NR_030737 | 0.246 | -0.469 |
| ZNF667 | NR_030740 | -1.3515 | -1.919 |
| TMED5 | NR_030761 | -0.9455 | -1.8555 |
| ATP5SL | NR_030765 | -0.2885 | -1.115 |
| ANKRD13D | NR_030767 | -0.456 | -1.279 |
| CCM2 | NR_030770 | -1.528 | -2.997 |
| UPRT | NR_030774 | -0.9285 | -2.36 |
| SHISA4 | NR_030775 | -0.0965 | -0.5855 |
| ATXN3 | NR_031765 | -0.521 | -2.1695 |
| ZIC4 | NR_033118 | 0.116 | -0.0595 |
| ZIC4 | NR_033119 | 0.116 | -0.0595 |
| PDZD3 | NR_033122 | -0.073 | -0.4995 |
| ALG13 | NR_033124 | -0.6645 | -1.719 |
| ALG13 | NR_033125 | -0.6645 | -1.719 |
| ALG13 | NR_033127 | -0.6645 | -1.719 |
| ALG13 | NR_033128 | -0.6645 | -1.719 |
| ALG13 | NR_033129 | -0.6645 | -1.719 |
| ALG13 | NR_033131 | -0.6645 | -1.719 |
| ALG13 | NR_033132 | -0.6645 | -1.719 |
| ALG13 | NR_033134 | -0.6645 | -1.719 |
| TMEM48 | NR_033142 | -1.147 | -1.555 |
| TMEM135 | NR_033149 | -0.9945 | -2.259 |
| PLA2G4F | NR_033151 | 0.027 | -0.2105 |
| TMEM150A | NR_033179 | -0.119 | -0.5455 |
| SYAP1 | NR_033181 | -0.6075 | -0.857 |
| MIB2 | NR_033183 | -0.306 | -0.029 |
| C1orf220 | NR_033186 | -0.1385 | -0.0155 |
| PIN4 | NR_033187 | -0.824 | -1.142 |
| PPP2R2D | NR_033191 | -1.0415 | -2.9265 |
| CCDC59 | NR_033192 | -0.7755 | -1.846 |

| | | | |
|---|---|---|---|
| ZFAND1 | NR_033193 | -1.4555 | -1.211 |
| ZFAND1 | NR_033194 | -1.4555 | -1.211 |
| ZFAND1 | NR_033195 | -1.4555 | -1.211 |
| ZFAND1 | NR_033196 | -1.4555 | -1.211 |
| LARP4 | NR_033200 | -0.0595 | -1.7255 |
| LOC404266 | NR_033201 | -0.1375 | -0.279 |
| LOC404266 | NR_033202 | -0.1375 | -0.279 |
| LOC404266 | NR_033203 | -0.1375 | -0.279 |
| LOC404266 | NR_033204 | -0.1375 | -0.279 |
| LOC404266 | NR_033205 | -0.1375 | -0.279 |
| SMPD4 | NR_033230 | -0.4145 | -1.509 |
| SMPD4 | NR_033231 | -0.4145 | -1.509 |
| SMPD4 | NR_033232 | -0.4145 | -1.509 |
| MOCS1 | NR_033233 | -0.2925 | -1.621 |
| STRBP | NR_033234 | -0.697 | -1.6225 |
| FAM3A | NR_033237 | -0.1035 | -0.785 |
| UBAP1 | NR_033243 | -0.7235 | -2.371 |
| GCSH | NR_033249 | -0.2965 | -0.065 |
| ZEB2 | NR_033258 | -1.3735 | -2.6525 |
| NUDT16 | NR_033268 | -1.109 | -1.566 |
| FUZ | NR_033269 | -0.147 | -0.6085 |
| GK5 | NR_033289 | -0.1945 | -0.088 |
| FAM114A1 | NR_033290 | -0.733 | -1.206 |
| CCDC163P | NR_033296 | -0.2785 | -0.6135 |
| CCDC163P | NR_033297 | -0.2785 | -0.6135 |
| CCDC163P | NR_033298 | -0.2785 | -0.6135 |
| DHX9 | NR_033302 | -1.008 | -0.847 |
| NSUN5P1 | NR_033322 | -0.8125 | -1.5445 |
| NSUN5P2 | NR_033323 | -0.5825 | -0.7965 |
| EFHC1 | NR_033327 | -1.005 | -0.877 |
| KLHL7 | NR_033328 | 0.4635 | -0.0885 |
| KLHL7 | NR_033329 | 0.4635 | -0.0885 |
| TMEM70 | NR_033334 | -1.2235 | -4.072 |
| C17orf70 | NR_033338 | -0.6195 | -1.273 |
| C4orf42 | NR_033339 | -0.8565 | -0.882 |
| SLC38A6 | NR_033344 | -0.9925 | -1.0935 |
| GGA3 | NR_033345 | -0.9555 | -2.739 |
| RAI2 | NR_033348 | -0.295 | -0.8365 |
| RAI2 | NR_033349 | -0.295 | -0.8365 |
| PRCD | NR_033357 | -0.012 | -0.3815 |
| TMCC1 | NR_033361 | -0.9735 | -1.5765 |
| ATG12 | NR_033362 | -1.3315 | -1.6715 |
| ATG12 | NR_033363 | -1.3315 | -1.6715 |

| | | | |
|---|---|---|---|
| BOD1P | NR_033372 | 0.5005 | 0.1975 |
| LOC401242 | NR_033379 | -0.323 | -0.6485 |
| PDHB | NR_033384 | -1.0295 | -2.649 |
| KLHDC9 | NR_033385 | -0.163 | -0.1045 |
| KLHDC9 | NR_033386 | -0.163 | -0.1045 |
| .OC10013221! | NR_033389 | -0.0245 | 0.033 |
| TYROBP | NR_033390 | 0.058 | -0.44 |
| TROVE2 | NR_033393 | -0.8535 | -1.218 |
| SLC22A20 | NR_033396 | 0.445 | -0.183 |
| PRMT1 | NR_033397 | -0.1365 | -0.383 |
| BMP1 | NR_033403 | 0.233 | -0.124 |
| BMP1 | NR_033404 | 0.233 | -0.124 |
| POLL | NR_033406 | -0.2525 | -0.865 |
| FBXW4P1 | NR_033408 | -0.203 | -0.684 |
| SLC5A1 | NR_033411 | -0.121 | -0.1215 |
| SLC29A3 | NR_033413 | -0.593 | -1.936 |
| SLC29A3 | NR_033414 | -0.593 | -1.936 |
| ZNF542 | NR_033418 | -0.957 | -1.783 |
| SLC11A2 | NR_033421 | -0.876 | -1.3705 |
| SLC11A2 | NR_033422 | -0.876 | -1.3705 |
| TSLP | NR_033425 | -0.155 | -0.498 |
| ACAD9 | NR_033426 | -0.585 | -1.2965 |
| ARL13B | NR_033427 | -1.172 | -2.9525 |
| ABTB1 | NR_033429 | 0.3145 | -0.5195 |
| UMPS | NR_033434 | -1.5695 | -5.0975 |
| PHF14 | NR_033435 | -0.3955 | -1.4025 |
| PHF14 | NR_033436 | -0.3955 | -1.4025 |
| UMPS | NR_033437 | -1.5695 | -5.0975 |
| PLSCR4 | NR_033438 | -0.1515 | -0.1005 |
| PLSCR4 | NR_033439 | -0.1515 | -0.1005 |
| PAM | NR_033440 | -0.739 | -0.8865 |
| HDAC2 | NR_033441 | -0.999 | -1.0355 |
| TCF7 | NR_033449 | -0.874 | -1.8875 |
| PAGE3 | NR_033460 | 0.074 | 0.0865 |
| NT5C3L | NR_033464 | 0.011 | -0.7255 |
| NT5C3L | NR_033465 | 0.011 | -0.7255 |
| PPP2R1A | NR_033500 | 0.2255 | -0.3285 |
| MASP1 | NR_033519 | 0.167 | -0.0785 |
| LMF1 | NR_033645 | 0.068 | -0.534 |
| LMF1 | NR_033646 | 0.068 | -0.534 |
| SF1 | NR_033649 | -0.0435 | -0.7715 |
| SF1 | NR_033650 | -0.0435 | -0.7715 |
| .OC10013289: | NR_033651 | -0.274 | 0.098 |

| | | | |
|---|---|---|---|
| .OC10013289: | NR_033652 | -0.274 | 0.098 |
| TAC3 | NR_033654 | 0.1885 | -0.1635 |
| AGA | NR_033655 | -0.795 | -0.6705 |
| STAT6 | NR_033659 | -0.142 | 0.0855 |
| SCAMP5 | NR_033660 | -0.65 | -1.723 |
| IGLL5 | NR_033661 | 0.017 | -0.0175 |
| CSF3 | NR_033662 | 0.093 | -0.3545 |
| EPOR | NR_033663 | -0.3795 | -0.8455 |
| AAA1 | NR_033664 | -0.452 | -0.817 |
| AAA1 | NR_033665 | -0.452 | -0.817 |
| ARMCX6 | NR_033669 | -0.5045 | -1.0455 |
| ARMCX6 | NR_033670 | -0.5045 | -1.0455 |
| NOX5 | NR_033671 | -0.027 | -0.1445 |
| NOX5 | NR_033672 | -0.027 | -0.1445 |
| DGCR2 | NR_033674 | -0.5775 | -1.2855 |
| VDAC2 | NR_033675 | -0.0035 | -0.4675 |
| NUP62CL | NR_033676 | 0.019 | 0.2085 |
| DDX49 | NR_033677 | 0.071 | -0.8115 |
| CARD8 | NR_033678 | -0.39 | -1.377 |
| CARD8 | NR_033679 | -0.39 | -1.377 |
| CARD8 | NR_033680 | -0.39 | -1.377 |
| PCYT2 | NR_033681 | -0.4005 | -0.8435 |
| PCYT2 | NR_033682 | -0.4005 | -0.8435 |
| PCYT2 | NR_033683 | -0.4005 | -0.8435 |
| PCYT2 | NR_033685 | -0.4005 | -0.8435 |
| PEF1 | NR_033686 | -0.675 | -1.609 |
| SLC25A1 | NR_033687 | 0.1415 | -0.5055 |
| PEF1 | NR_033688 | -0.675 | -1.609 |
| PDZD9 | NR_033694 | -0.151 | -0.2455 |
| NDUFA2 | NR_033697 | 0.2915 | -0.197 |
| FNTA | NR_033698 | -1.82 | -2.6545 |
| FMR1 | NR_033699 | -1.4225 | -1.6075 |
| FMR1 | NR_033700 | -1.4225 | -1.6075 |
| AIF1L | NR_033701 | 0.253 | -0.097 |
| MYNN | NR_033702 | -1.564 | -3.3895 |
| MYNN | NR_033703 | -1.564 | -3.3895 |
| VAMP4 | NR_033704 | -1.2335 | -1.6575 |
| VAMP7 | NR_033714 | -0.849 | -0.845 |
| VAMP7 | NR_033715 | -0.849 | -0.845 |
| SNX7 | NR_033716 | -0.8435 | -1.3125 |
| SCML2 | NR_033717 | -0.4055 | -1.253 |
| DNAJC19 | NR_033721 | -0.699 | -1.694 |
| DNAJC19 | NR_033722 | -0.699 | -1.694 |

| | | | |
|---|---|---|---|
| DNAJC19 | NR_033723 | -0.699 | -1.694 |
| DNAJC19 | NR_033724 | -0.699 | -1.694 |
| DNAJC19 | NR_033725 | -0.699 | -1.694 |
| ZNF630 | NR_033730 | -0.4205 | -1.2405 |
| BCORP1 | NR_033732 | 0.2355 | -0.1125 |
| DMKN | NR_033746 | 0.229 | -0.1485 |
| FLJ43826 | NR_033753 | -0.2865 | -0.2635 |
| ATP5L | NR_033759 | 0.153 | -0.1985 |
| ATP5S | NR_033761 | -0.2015 | -0.9055 |
| FOXP2 | NR_033766 | 0.8085 | -0.3895 |
| FOXP2 | NR_033767 | 0.8085 | -0.3895 |
| SPRNP1 | NR_033789 | -0.059 | -0.365 |
| TAF6 | NR_033792 | -0.073 | -0.646 |
| DSTNP2 | NR_033796 | -0.6245 | -1.3385 |
| CAST | NR_033798 | -0.609 | -1.1605 |
| OPN5 | NR_033806 | -0.1195 | -0.059 |
| CYP3A5 | NR_033807 | -0.131 | 0.34 |
| CYP3A5 | NR_033808 | -0.131 | 0.34 |
| CYP3A5 | NR_033809 | -0.131 | 0.34 |
| CYP3A5 | NR_033810 | -0.131 | 0.34 |
| CYP3A5 | NR_033811 | -0.131 | 0.34 |
| CYP3A5 | NR_033812 | -0.131 | 0.34 |
| DLST | NR_033814 | -0.164 | -1.708 |
| ALS2CL | NR_033815 | 0.067 | -0.5855 |
| IKBKB | NR_033818 | -0.9915 | -2.568 |
| IKBKB | NR_033819 | -0.9915 | -2.568 |
| ATAT1 | NR_033821 | -0.7065 | -1.499 |
| ATAT1 | NR_033822 | -0.7065 | -1.499 |
| ATAT1 | NR_033823 | -0.7065 | -1.499 |
| KLHL22 | NR_033825 | -0.638 | -2.23 |
| SNRPD2P2 | NR_033826 | -0.073 | -0.19 |
| UNQ6975 | NR_033831 | -0.327 | -0.541 |
| PIGA | NR_033835 | -0.841 | -1.258 |
| PIGA | NR_033836 | -0.841 | -1.258 |
| MGC39372 | NR_033851 | -0.2135 | -0.422 |
| .OC10028897 | NR_033857 | -0.0585 | 0.4225 |
| CYP4F24P | NR_033864 | 0.0395 | 0.051 |
| LOC338579 | NR_033891 | -0.3055 | 0.141 |
| LOC440419 | NR_033892 | -0.1025 | -0.4355 |
| KIAA1967 | NR_033902 | -0.03 | -0.797 |
| DHX35 | NR_033905 | -0.844 | -1.921 |
| ORC2 | NR_033915 | -1.703 | -2.959 |
| P2RX7 | NR_033948 | -0.1055 | -0.2805 |

| | | | |
|---|---|---|---|
| P2RX7 | NR_033949 | -0.1055 | -0.2805 |
| P2RX7 | NR_033950 | -0.1055 | -0.2805 |
| P2RX7 | NR_033951 | -0.1055 | -0.2805 |
| P2RX7 | NR_033952 | -0.1055 | -0.2805 |
| P2RX7 | NR_033953 | -0.1055 | -0.2805 |
| P2RX7 | NR_033954 | -0.1055 | -0.2805 |
| P2RX7 | NR_033955 | -0.1055 | -0.2805 |
| P2RX7 | NR_033956 | -0.1055 | -0.2805 |
| LOC729862 | NR_033961 | 0.0985 | -0.018 |
| TNFSF10 | NR_033994 | -0.391 | -0.174 |
| YAF2 | NR_034000 | -1.6075 | -2.688 |
| SRSF6 | NR_034009 | -0.6525 | -1.656 |
| SNHG8 | NR_034010 | -0.327 | -0.153 |
| SNHG8 | NR_034011 | -0.327 | -0.153 |
| .OC10013106( | NR_034014 | 0.034 | -0.442 |
| .OC10013106( | NR_034015 | 0.034 | -0.442 |
| SRSF10 | NR_034035 | -1.748 | -1.245 |
| SORD | NR_034039 | -0.214 | -0.916 |
| SRSF1 | NR_034041 | -1.407 | -1.605 |
| TNPO3 | NR_034053 | -0.7495 | -1.4865 |
| PSMD1 | NR_034059 | -0.6925 | -1.134 |
| ZNF85 | NR_034060 | -0.827 | -1.546 |
| CASP12 | NR_034061 | -0.2185 | -0.359 |
| CASP12 | NR_034063 | -0.2185 | -0.359 |
| CASP12 | NR_034064 | -0.2185 | -0.359 |
| CASP12 | NR_034065 | -0.2185 | -0.359 |
| CASP12 | NR_034066 | -0.2185 | -0.359 |
| CASP12 | NR_034067 | -0.2185 | -0.359 |
| CASP12 | NR_034068 | -0.2185 | -0.359 |
| SRPK1 | NR_034069 | -0.864 | -1.4045 |
| CASP12 | NR_034070 | -0.2185 | -0.359 |
| CASP12 | NR_034071 | -0.2185 | -0.359 |
| SARS | NR_034072 | -0.3145 | -1.0165 |
| LOC729852 | NR_034084 | -0.9115 | -1.2945 |
| C9orf29 | NR_034087 | 0.171 | -0.2165 |
| LOC401109 | NR_034088 | -0.1025 | -0.3 |
| .OC10012982 | NR_034093 | 0.0795 | -0.1805 |
| .OC10012982 | NR_034094 | 0.0795 | -0.1805 |
| .OC10012808 | NR_034097 | -0.113 | -0.7195 |
| EML2 | NR_034098 | -0.154 | -0.4775 |
| LOC388948 | NR_034099 | -0.3575 | 0.048 |
| .OC10028834( | NR_034101 | -0.3805 | 0.017 |
| CRNDE | NR_034105 | -0.2615 | -0.38 |

| | | | |
|---|---|---|---|
| CRNDE | NR_034106 | -0.2615 | -0.38 |
| LOC284751 | NR_034124 | 0.049 | -0.521 |
| C12orf33 | NR_034140 | 1.1125 | -0.0115 |
| SIRT2 | NR_034146 | 0.186 | -0.3835 |
| LOC200261 | NR_034149 | 0.1545 | -0.327 |
| LOC149950 | NR_034152 | -0.31 | -0.139 |
| C11orf92 | NR_034154 | 0.239 | 0.1595 |
| LGALS17A | NR_034156 | 0.2045 | -0.137 |
| EXD2 | NR_034164 | -0.9125 | -1.425 |
| EXD2 | NR_034165 | -0.9125 | -1.425 |
| VNN2 | NR_034173 | -0.0095 | 0.0735 |
| VNN2 | NR_034174 | -0.0095 | 0.0735 |
| LOC653712 | NR_034179 | 0.0695 | -0.3635 |
| SUV39H2 | NR_034181 | -1.187 | -2.8455 |
| SRGN | NR_036430 | 0.21 | 0.1935 |
| ZMAT1 | NR_036431 | 0.1925 | -0.226 |
| RSRC2 | NR_036434 | -1.086 | -2.288 |
| RSRC2 | NR_036435 | -1.086 | -2.288 |
| RSRC2 | NR_036436 | -1.086 | -2.288 |
| KCNC4 | NR_036437 | -0.0495 | -0.3955 |
| CDADC1 | NR_036439 | -0.5785 | -1.921 |
| POU5F1P3 | NR_036440 | -0.0455 | 0.161 |
| VRK2 | NR_036441 | -0.494 | -1.714 |
| LMF1 | NR_036442 | 0.068 | -0.534 |
| HERC2P9 | NR_036443 | -0.196 | -0.598 |
| PKD1P1 | NR_036447 | -1.5485 | -1.7565 |
| DTNBP1 | NR_036448 | -0.281 | -0.619 |
| ZNF826P | NR_036455 | -0.1905 | -1.38 |
| ADAT1 | NR_036460 | -0.537 | -1.0965 |
| HIST2H2BC | NR_036461 | -0.1435 | 0.643 |
| APITD1 | NR_036462 | -0.7235 | -0.719 |
| FBXL5 | NR_036464 | -1.564 | -3.956 |
| TMEM126B | NR_036465 | -1.144 | -2.1265 |
| IFFO1 | NR_036467 | -0.913 | -1.502 |
| CIDEA | NR_036468 | -0.2375 | -0.5555 |
| ENY2 | NR_036471 | -0.716 | -0.688 |
| ENY2 | NR_036472 | -0.716 | -0.688 |
| GPATCH8 | NR_036474 | -1.026 | -2.1815 |
| LOC440104 | NR_036476 | -0.1175 | -0.4945 |
| LOC440104 | NR_036477 | -0.1175 | -0.4945 |
| LOC440104 | NR_036478 | -0.1175 | -0.4945 |
| LOC440104 | NR_036479 | -0.1175 | -0.4945 |
| LOC730441 | NR_036483 | 0.0865 | -0.0735 |

| | | | |
|---|---|---|---|
| PEX19 | NR_036492 | -0.245 | -0.8145 |
| PEX19 | NR_036493 | -0.245 | -0.8145 |
| LOC646938 | NR_036495 | 0.126 | -0.0875 |
| LOC339803 | NR_036496 | -0.711 | -0.402 |
| LOC728752 | NR_036504 | 0.1635 | -0.3795 |
| .OC10019220 | NR_036506 | 0.301 | -0.236 |
| UROD | NR_036510 | 0.1625 | -0.6445 |
| C17orf62 | NR_036514 | -0.531 | -1.6595 |
| C17orf62 | NR_036516 | -0.531 | -1.6595 |
| C17orf62 | NR_036517 | -0.531 | -1.6595 |
| C17orf62 | NR_036518 | -0.531 | -1.6595 |
| C17orf62 | NR_036519 | -0.531 | -1.6595 |
| .OC10028704 | NR_036520 | 0.162 | -0.639 |
| .OC10012825 | NR_036521 | -0.483 | -1.0795 |
| .OC10012825 | NR_036522 | -0.483 | -1.0795 |
| MICA | NR_036523 | -0.0455 | -0.166 |
| MICA | NR_036524 | -0.0455 | -0.166 |
| .OC10049948 | NR_036526 | 0.3655 | 0.3925 |
| MATR3 | NR_036535 | -0.217 | -0.35 |
| GPR89A | NR_036541 | -0.846 | -1.2885 |
| CARS | NR_036542 | -0.4275 | -1.4015 |
| PPIE | NR_036543 | -0.108 | -0.9975 |
| PPIE | NR_036544 | -0.108 | -0.9975 |
| .OC10027170 | NR_036546 | 0.156 | -0.306 |
| .OC10012996 | NR_036549 | -0.0665 | -0.316 |
| C21orf59 | NR_036552 | -0.7175 | -1.257 |
| C12orf36 | NR_036555 | -0.166 | -0.4515 |
| ANKRD54 | NR_036556 | -0.627 | -1.8055 |
| CASP5 | NR_036562 | -0.085 | -0.228 |
| CCT3 | NR_036564 | -0.166 | -0.5775 |
| CCT3 | NR_036565 | -0.166 | -0.5775 |
| PILRB | NR_036569 | -1.0325 | -1.552 |
| PILRB | NR_036570 | -1.0325 | -1.552 |
| APTX | NR_036576 | -0.647 | -1.9035 |
| APTX | NR_036577 | -0.647 | -1.9035 |
| APTX | NR_036578 | -0.647 | -1.9035 |
| APTX | NR_036579 | -0.647 | -1.9035 |
| IRF4 | NR_036585 | -0.5695 | -0.3175 |
| ZNF28 | NR_036599 | 0.1305 | -0.2885 |
| ZNF28 | NR_036600 | 0.1305 | -0.2885 |
| GGPS1 | NR_036605 | -1.767 | -2.246 |
| SRSF2 | NR_036608 | -0.693 | -0.8275 |
| SRSF3 | NR_036610 | -1.308 | -1.6985 |

| | | | |
|---|---|---|---|
| DCLK2 | NR_036614 | -0.185 | -1.0295 |
| DBF4B | NR_036623 | -1.309 | -3.3195 |
| RBM5 | NR_036627 | -1.0975 | -2.9845 |
| VAPB | NR_036633 | -0.3075 | -1.7435 |
| TGFBR3 | NR_036634 | -0.535 | -0.8875 |
| C1orf93 | NR_036637 | 0.09 | -0.5055 |
| C1orf93 | NR_036638 | 0.09 | -0.5055 |
| YIPF1 | NR_036639 | -0.3485 | -1.373 |
| YIPF1 | NR_036640 | -0.3485 | -1.373 |
| PDGFC | NR_036641 | -0.1085 | -0.6905 |
| HSDL2 | NR_036651 | -0.789 | -0.6675 |
| OSBPL9 | NR_036662 | -0.581 | -1.5355 |
| MT1JP | NR_036677 | 0.241 | -0.2445 |
| BCL7B | NR_036682 | -0.3765 | -1.8145 |
| CES2 | NR_036684 | -0.5975 | -1.474 |
| HSP90AA4P | NR_036692 | 0.392 | -0.143 |
| CLEC2D | NR_036693 | 0.0845 | -0.721 |
| HSP90AA6P | NR_036751 | -0.1305 | -0.3505 |
| NOL4 | NR_036752 | -1.113 | -1.8675 |
| DNAJC25 | NR_037148 | -0.946 | -1.437 |
| TTLL3 | NR_037162 | -0.09 | -0.7 |
| POU2F1 | NR_037163 | -0.422 | -1.2635 |
| ASB13 | NR_037164 | -0.681 | -1.707 |
| PRPF38B | NR_037185 | -0.391 | 0.091 |
| SLC3A2 | NR_037193 | -0.5325 | -1.3455 |
| NFS1 | NR_037570 | -0.4815 | -1.573 |
| DCTN5 | NR_037573 | -0.685 | -1.5425 |
| MRPL10 | NR_037575 | -0.58 | -1.938 |
| TMEM222 | NR_037576 | -0.5635 | -1.9395 |
| TMEM222 | NR_037577 | -0.5635 | -1.9395 |
| TMEM222 | NR_037580 | -0.5635 | -1.9395 |
| AP3S2 | NR_037582 | -0.6365 | -1.77 |
| C21orf37 | NR_037585 | 0.1775 | 0.1795 |
| C21orf37 | NR_037586 | 0.1775 | 0.1795 |
| AK2 | NR_037591 | -0.2575 | -1.04 |
| AK2 | NR_037592 | -0.2575 | -1.04 |
| LOC442028 | NR_037597 | -0.0485 | -0.0325 |
| MRPS14 | NR_037606 | -0.3425 | -1.3925 |
| UCHL5 | NR_037607 | -0.881 | -1.905 |
| CMTM3 | NR_037613 | -0.5575 | -2.2335 |
| CNOT2 | NR_037615 | -1.895 | -3.0975 |
| CMTM3 | NR_037619 | -0.5575 | -2.2335 |
| FBXO22 | NR_037623 | -0.893 | -2.3845 |

| | | | |
|---|---|---|---|
| BRD2 | NR_037625 | 0.032 | -1.0205 |
| ZSWIM3 | NR_037628 | -0.822 | -2.049 |
| MYCBP | NR_037632 | -0.5395 | -0.764 |
| SLC28A3 | NR_037638 | 0.148 | -0.155 |
| RWDD3 | NR_037643 | -1.2525 | -2.6935 |
| TMX2 | NR_037645 | -0.195 | -0.7195 |
| FOXRED1 | NR_037647 | -0.207 | -1.103 |
| FOXRED1 | NR_037648 | -0.207 | -1.103 |
| CCNT2 | NR_037649 | -0.1945 | -1.091 |
| ST20 | NR_037652 | -0.3235 | -1.081 |
| ST20 | NR_037653 | -0.3235 | -1.081 |
| MTHFS | NR_037654 | -0.7945 | -3.214 |
| POC1B | NR_037659 | -0.9035 | -1.528 |
| POC1B | NR_037660 | -0.9035 | -1.528 |
| PTPN7 | NR_037663 | -0.427 | 0.095 |
| PTPN7 | NR_037664 | -0.427 | 0.095 |
| INTS7 | NR_037667 | -1.678 | -2.6125 |
| SCNN1D | NR_037668 | -0.3945 | -0.488 |
| SNX10 | NR_037670 | 0.488 | -0.142 |
| SERF2 | NR_037672 | 0.172 | -0.2695 |
| PLXNC1 | NR_037687 | -0.5835 | -0.6505 |
| ACTG1 | NR_037688 | 0.067 | -0.427 |
| SAFB | NR_037699 | -0.7525 | -1.85 |
| RNF7 | NR_037702 | -0.175 | -0.8935 |
| RNF7 | NR_037703 | -0.175 | -0.8935 |
| ZNF778 | NR_037705 | -0.847 | -1.2845 |
| FAM161A | NR_037710 | -2.0005 | -1.9825 |
| RNASEK | NR_037715 | 0.035 | -0.358 |
| RNASEK | NR_037716 | 0.035 | -0.358 |
| EFEMP2 | NR_037718 | 0.092 | -0.442 |
| SEC23IP | NR_037771 | -1.488 | -2.929 |
| STAM | NR_037774 | -1.0975 | -2.3155 |
| WBSCR22 | NR_037776 | 0.266 | -0.35 |
| ZNF625 | NR_037801 | -0.2655 | -1.1325 |
| IL17RC | NR_037807 | -0.0215 | -0.42 |
| IL15 | NR_037840 | 0.0515 | -0.575 |
| SNRNP27 | NR_037862 | -0.67 | -1.356 |
| ZHX1 | NR_037873 | -0.5085 | -2.0115 |
| ZHX1 | NR_037874 | -0.5085 | -2.0115 |
| .OC10028701( | NR_037885 | 0.035 | -0.496 |
| ZNF695 | NR_037892 | -0.0785 | -0.464 |
| PRLR | NR_037910 | -0.028 | 0.0215 |
| FAM24B | NR_037911 | 0.246 | -0.287 |

| | | | |
|---|---|---|---|
| RCHY1 | NR_037913 | -0.779 | -2.1975 |
| RCHY1 | NR_037914 | -0.779 | -2.1975 |
| XIAP | NR_037916 | -1.9625 | -2.4475 |
| HUS1 | NR_037917 | -0.5675 | -0.9855 |
| MAVS | NR_037921 | -1.15 | -1.6835 |
| SSR4 | NR_037927 | 0.271 | -0.15 |
| SCNM1 | NR_037937 | -0.1915 | -0.9175 |
| HOXA10 | NR_037939 | -0.0585 | -0.107 |
| STX16 | NR_037941 | -1.417 | -1.407 |
| STX16 | NR_037942 | -1.417 | -1.407 |
| STX16 | NR_037943 | -1.417 | -1.407 |
| WISP1 | NR_037944 | 0.096 | -0.1165 |
| NSUN2 | NR_037947 | -0.8135 | -1.825 |
| BSCL2 | NR_037948 | -0.107 | -0.9375 |
| BSCL2 | NR_037949 | -0.107 | -0.9375 |
| RHOU | NR_037962 | -0.617 | -0.894 |
| TTC19 | NR_037985 | -1.0405 | -0.905 |
| PLIN2 | NR_038064 | -0.946 | -1.7575 |
| TECR | NR_038103 | 0.2725 | -0.131 |
| TECR | NR_038104 | 0.2725 | -0.131 |
| MMAB | NR_038118 | -0.2855 | -1.347 |
| ELMO1 | NR_038120 | -0.1695 | -0.73 |
| ELMO1 | NR_038121 | -0.1695 | -0.73 |
| CAPZB | NR_038125 | 0.034 | -0.666 |
| CHN1 | NR_038133 | -0.9215 | -1.0645 |
| LOC641515 | NR_038135 | 0.416 | 0.3625 |
| UBR7 | NR_038150 | -0.472 | -1.452 |
| MRPL42 | NR_038159 | -0.924 | -1.098 |
| MRPL42 | NR_038160 | -0.924 | -1.098 |
| MRPL42 | NR_038161 | -0.924 | -1.098 |
| NSFL1C | NR_038164 | 0.013 | -0.898 |
| FUT8 | NR_038167 | -1.1135 | -2.375 |
| FUT8 | NR_038170 | -1.1135 | -2.375 |
| CAPRIN2 | NR_038177 | -1.5535 | -2.0835 |
| AGER | NR_038190 | -0.0285 | -0.2605 |
| DHPS | NR_038192 | -0.027 | -0.3955 |
| PBX4 | NR_038198 | 0.301 | -0.4265 |
| VEZT | NR_038241 | -1.132 | -1.656 |
| VEZT | NR_038242 | -1.132 | -1.656 |
| .OC10028863 | NR_038253 | -0.968 | -0.7705 |
| .OC10028863 | NR_038254 | -0.968 | -0.7705 |
| .OC10028863 | NR_038255 | -0.968 | -0.7705 |
| TRPM2 | NR_038257 | -0.067 | -0.4225 |

| | | | |
|---|---|---|---|
| HIVEP3 | NR_038260 | 0.0365 | -0.5415 |
| HIVEP3 | NR_038261 | 0.0365 | -0.5415 |
| PPFIA2 | NR_038265 | -0.055 | -0.2715 |
| SHARPIN | NR_038270 | -0.141 | -0.3865 |
| OR51B5 | NR_038321 | 0.2095 | -0.6415 |
| MAP3K13 | NR_038322 | -0.6495 | -1.591 |
| CLU | NR_038335 | -0.006 | -0.3765 |
| DCP2 | NR_038352 | -0.864 | -1.3175 |
| FAM153C | NR_038353 | 0.0485 | -0.3625 |
| WDR20 | NR_038354 | -1.131 | -2.623 |
| MGC45922 | NR_038359 | -0.134 | -0.3695 |
| ZNF345 | NR_038362 | -0.843 | -1.283 |
| FDXACB1 | NR_038364 | -0.681 | -0.9495 |
| NDRG3 | NR_038370 | -0.0405 | -0.4 |
| MGC39584 | NR_038377 | 0.0075 | -0.303 |
| LOC441242 | NR_038378 | 0.064 | -0.303 |
| ANAPC16 | NR_038391 | -0.5375 | -0.5575 |
| ANAPC16 | NR_038392 | -0.5375 | -0.5575 |
| MGC16075 | NR_038407 | -0.2505 | -0.818 |
| PIN1 | NR_038422 | 0.2155 | -0.4325 |
| NAPA | NR_038456 | 0.1705 | -0.4475 |
| NAPA | NR_038457 | 0.1705 | -0.4475 |
| GFOD1 | NR_038459 | -0.571 | -1.5355 |
| PIN1 | NR_038830 | 0.2155 | -0.4325 |
| .OC10012984 | NR_038910 | -0.184 | -0.246 |
| LOC284889 | NR_038911 | 0.198 | -0.284 |
| CAMTA1 | NR_038934 | 0.0475 | -0.016 |
| .OC10012826 | NR_038945 | 0.0915 | 0.309 |
| .OC10013298 | NR_038985 | 0 | -0.141 |
| TMEM139 | NR_040003 | 0.2435 | -0.3775 |
| CLPTM1 | NR_040006 | -0.1805 | -1.3915 |
| CLPTM1 | NR_040007 | -0.1805 | -1.3915 |
| CAPN1 | NR_040008 | 0.0425 | -0.4865 |
| IKBKB | NR_040009 | -0.9915 | -2.568 |
| TMEM50B | NR_040016 | -0.801 | -1.2535 |
| RNF41 | NR_040053 | -0.776 | -2.4925 |
| NDRG4 | NR_040072 | -0.529 | -0.8035 |
| .OC10013123 | NR_040073 | -0.393 | -0.5335 |
| .OC10028722 | NR_040074 | -0.1935 | -0.3385 |
| .OC10028722 | NR_040075 | -0.1935 | -0.3385 |
| FLJ33534 | NR_040080 | 0.0505 | 0.003 |
| ZNF195 | NR_040083 | -1.2155 | -2.4815 |
| .OC10013328 | NR_040084 | 0.191 | -0.2205 |

| | | | |
|---|---|---|---|
| LOC348120 | NR_040094 | -0.0245 | -0.348 |
| KLHDC3 | NR_040101 | -0.0095 | -0.8585 |
| CARKD | NR_040103 | -0.5245 | -1.336 |
| CARKD | NR_040104 | -0.5245 | -1.336 |
| LOC728175 | NR_040108 | 0.093 | -0.368 |
| NSF | NR_040116 | -1.0695 | -1.094 |
| MPPE1 | NR_040241 | -1.091 | -1.281 |
| MPPE1 | NR_040242 | -1.091 | -1.281 |
| MPPE1 | NR_040243 | -1.091 | -1.281 |
| .OC10028731 | NR_040245 | -0.016 | -0.0935 |
| ZNF410 | NR_040251 | -1.314 | -2.961 |
| ANKS3 | NR_040252 | -0.58 | -1.421 |
| LOC282980 | NR_040253 | 0.5055 | 0.1135 |
| NFKBIB | NR_040515 | -0.0265 | -0.657 |
| PDE4C | NR_040546 | -0.183 | -0.1575 |
| STAG3L1 | NR_040583 | -0.1405 | -0.6375 |
| STAG3L4 | NR_040585 | -0.195 | -0.874 |
| STAG3L4 | NR_040586 | -0.195 | -0.874 |
| HCP5 | NR_040662 | -0.298 | -0.395 |
| KCNQ1 | NR_040711 | 0.1635 | -0.2765 |
| CHRAC1 | NR_040712 | -0.8595 | -3.015 |
| RASSF3 | NR_040718 | -0.033 | -0.3865 |
| RBM39 | NR_040722 | -1.0695 | -2.1315 |
| RBM39 | NR_040723 | -1.0695 | -2.1315 |
| RBM39 | NR_040724 | -1.0695 | -2.1315 |
| C1orf50 | NR_040733 | -0.5385 | -1.6855 |
| ZIC4 | NR_040762 | 0.116 | -0.0595 |
| FLJ41484 | NR_044995 | -0.4425 | -0.307 |
| PSPC1 | NR_044998 | -0.1795 | -1.0695 |
| LMO3 | NR_045012 | -1.304 | -1.7735 |
| LMO3 | NR_045013 | -1.304 | -1.7735 |
| LMO3 | NR_045014 | -1.304 | -1.7735 |
| LETMD1 | NR_045017 | -0.3545 | -0.817 |
| LETMD1 | NR_045018 | -0.3545 | -0.817 |
| LETMD1 | NR_045019 | -0.3545 | -0.817 |
| LETMD1 | NR_045020 | -0.3545 | -0.817 |
| ATPAF1 | NR_045030 | -0.403 | -0.375 |
| APH1A | NR_045033 | -0.174 | -0.71 |
| APH1A | NR_045034 | -0.174 | -0.71 |
| APH1A | NR_045035 | -0.174 | -0.71 |
| SMUG1 | NR_045039 | 0.0505 | -0.914 |
| ZC4H2 | NR_045044 | -0.3345 | -0.7665 |
| UBTF | NR_045058 | -0.4465 | -1.622 |

| | | | |
|---|---|---|---|
| CHST9 | NR_045066 | -0.443 | -0.03 |
| CSRNP2 | NR_045072 | -0.295 | -1.872 |
| TSLP | NR_045089 | -0.155 | -0.498 |
| TPD52L2 | NR_045090 | -0.269 | -0.5175 |
| RNF146 | NR_045102 | -2.1675 | -4.566 |
| RNF146 | NR_045103 | -2.1675 | -4.566 |
| C15orf44 | NR_045104 | -1.312 | -3.044 |
| C15orf44 | NR_045105 | -1.312 | -3.044 |
| C15orf44 | NR_045106 | -1.312 | -3.044 |
| C15orf44 | NR_045107 | -1.312 | -3.044 |
| EGFL7 | NR_045110 | 0.154 | -0.4955 |
| EGFL7 | NR_045111 | 0.154 | -0.4955 |
| C5orf56 | NR_045116 | -0.0585 | -0.602 |
| ITGB3BP | NR_045147 | -0.905 | -1.0665 |
| CHEK1 | NR_045204 | -1.066 | -1.552 |
| CHEK1 | NR_045205 | -1.066 | -1.552 |
| DLST | NR_045209 | -0.164 | -1.708 |
| FCGR1B | NR_045213 | -0.1445 | -0.729 |
| TRIM41 | NR_045218 | -0.936 | -1.7015 |
| NRP1 | NR_045259 | -2.015 | -3.5045 |
| NACA | NR_045277 | -0.099 | -0.188 |
| PPFIA1 | NR_045286 | -1.176 | -1.92 |
| ASPSCR1 | NR_045351 | -0.623 | -1.1805 |
| RIC3 | NR_045405 | -0.736 | -1.2185 |
