## Supplementary material for "CDK12 loss in cancer cells affects DNA damage response genes through premature cleavage and polyadenylation"

**Supplementary Table 2: TT-seq expression changes with 2hours THZ531 treatment**

| gene_id | baseMean | log2FoldChange | lfcSE | stat | pvalue | padj | hgnc_symbol | gene_biotype | gene_gc(%) | PI | absPI | ensembl_gene_id | significant |
|---|---|---|---|---|---|---|---|---|---|---|---|---|---|
| ENSG00000261326 | 163.723856 | -6.465310374 | 0.404415 | -15.98682 | 1.58E-57 | 1.10E-53 | LINC01355 | lincRNA | 36 | -342.396062 | 342.396062 | ENSG00000261326 | down |
| ENSG00000186834 | 1220.130784 | 3.429670215 | 0.2061844 | 16.63399 | 3.95E-62 | 5.50E-58 | HEXIM1 | protein_coding | 48.22 | 196.3804416 | 196.3804416 | ENSG00000186834 | up |
| ENSG00000279838 | 913.827209 | -3.475681868 | 0.220685 | -15.74952 | 6.92E-56 | 3.21E-52 | | TEC | 41.03 | -178.9740616 | 178.9740616 | ENSG00000279838 | down |
| ENSG00000189367 | 377.1392595 | -4.221858291 | 0.3014854 | -14.00352 | 1.48E-44 | 4.13E-41 | KIAA0408 | protein_coding | 35.14 | -170.4955298 | 170.4955298 | ENSG00000189367 | down |
| ENSG00000280184 | 139.3176808 | -4.266808087 | 0.3260291 | -13.0872 | 3.90E-39 | 6.03E-36 | | TEC | 45.01 | -150.2759662 | 150.2759662 | ENSG00000280184 | down |
| ENSG00000239653 | 126.0461765 | -3.885859023 | 0.2838573 | -13.68948 | 1.17E-42 | 2.72E-39 | PSMD6-AS2 | antisense | 38.82 | -149.858025 | 149.858025 | ENSG00000239653 | down |
| ENSG00000170345 | 387.0461916 | 3.639377416 | 0.2592672 | 14.03717 | 9.23E-45 | 3.21E-41 | FOS | protein_coding | 53.92 | 147.3692995 | 147.3692995 | ENSG00000170345 | up |
| ENSG00000280347 | 127.4655583 | -3.637614254 | 0.27428 | -13.26241 | 3.82E-40 | 6.66E-37 | | TEC | 37.71 | -131.5973726 | 131.5973726 | ENSG00000280347 | down |
| ENSG00000279951 | 51.74314919 | -4.78462654 | 0.4227602 | -11.31759 | 1.07E-29 | 8.31E-27 | | TEC | 40.65 | -124.7859332 | 124.7859332 | ENSG00000279951 | down |
| ENSG00000213976 | 88.24768281 | -3.759165199 | 0.3057813 | -12.29364 | 9.80E-35 | 1.36E-31 | | unprocessed_pseudogene | 37.56 | -116.0269338 | 116.0269338 | ENSG00000213976 | down |
| ENSG00000250001 | 68.11348091 | -4.590094364 | 0.4195352 | -10.9409 | 7.35E-28 | 3.53E-25 | | sense_intronic | 46.18 | -112.2397 | 112.2397 | ENSG00000250001 | down |
| ENSG00000279145 | 421.4349563 | -2.998832719 | 0.2225805 | -13.47302 | 2.25E-41 | 4.48E-38 | | TEC | 38.91 | -112.0013926 | 112.0013926 | ENSG00000279145 | down |
| ENSG00000212126 | 43.39918852 | -4.725658913 | 0.4512493 | -10.47239 | 1.16E-25 | 4.35E-23 | TAS2R50 | protein_coding | 38.5 | -105.6713754 | 105.6713754 | ENSG00000212126 | down |
| ENSG00000121101 | 103.5112638 | 3.604913662 | 0.3022165 | 11.92825 | 8.43E-33 | 9.78E-30 | TEX14 | protein_coding | 45.25 | 104.5766403 | 104.5766403 | ENSG00000121101 | up |
| ENSG00000279880 | 124.1805623 | -3.58166688 | 0.2996829 | -11.95152 | 6.37E-33 | 8.07E-30 | | TEC | 40.21 | -104.202187 | 104.202187 | ENSG00000279880 | down |
| ENSG00000237260 | 35.25750145 | -5.242281446 | 0.5462912 | -9.59613 | 8.30E-22 | 1.93E-19 | | processed_pseudogene | 45.35 | -98.11101475 | 98.11101475 | ENSG00000237260 | down |
| ENSG00000257818 | 65.32770988 | -3.796907901 | 0.3380352 | -11.23229 | 2.83E-29 | 1.97E-26 | C1GALT1P1 | processed_pseudogene | 37.83 | -97.6011046 | 97.6011046 | ENSG00000257818 | down |
| ENSG00000270137 | 38.0445613 | -4.483052609 | 0.4419211 | -10.14446 | 3.51E-24 | 1.22E-21 | | sense_intronic | 36.55 | -93.75585228 | 93.75585228 | ENSG00000270137 | down |
| ENSG00000259326 | 76.20241285 | -3.724641007 | 0.3388373 | -10.99242 | 4.16E-28 | 2.07E-25 | | antisense | 36.31 | -91.94164995 | 91.94164995 | ENSG00000259326 | down |
| ENSG00000242539 | 54.73829865 | -3.621589563 | 0.3259791 | -11.10988 | 1.12E-28 | 6.80E-26 | | antisense | 38.14 | -91.14665332 | 91.14665332 | ENSG00000242539 | down |
| ENSG00000116652 | 105.5091216 | -3.399122071 | 0.2974219 | -11.42862 | 3.01E-30 | 2.46E-27 | DLEU2L | sense_intronic | 35.86 | -90.4451163 | 90.4451163 | ENSG00000116652 | down |
| ENSG00000274995 | 73.23910803 | -3.407313901 | 0.3017272 | -11.2927 | 1.43E-29 | 1.04E-26 | | sense_intronic | 47.1 | -88.52515036 | 88.52515036 | ENSG00000274995 | down |
| ENSG00000127364 | 66.05128477 | -3.474792599 | 0.3114363 | -11.15731 | 6.60E-29 | 4.17E-26 | TAS2R4 | protein_coding | 41.46 | -88.18831465 | 88.18831465 | ENSG00000127364 | down |
| ENSG00000249655 | 36.94557321 | -6.311151964 | 0.7629395 | -8.272152 | 1.32E-16 | 1.32E-14 | | antisense | 43.27 | -87.59974277 | 87.59974277 | ENSG00000249655 | down |
| ENSG00000277151 | 246.921163 | -3.05612776 | 0.2581367 | -11.83918 | 2.45E-32 | 2.43E-29 | | lincRNA | 34.33 | -87.44664425 | 87.44664425 | ENSG00000277151 | down |
| ENSG00000267088 | 37.57221587 | -4.538847895 | 0.4706622 | -9.643536 | 5.24E-22 | 1.30E-19 | | processed_pseudogene | 32.88 | -85.71847229 | 85.71847229 | ENSG00000267088 | down |
| ENSG00000127362 | 36.96032622 | -4.419191537 | 0.4548961 | -9.714726 | 2.61E-22 | 7.41E-20 | TAS2R3 | protein_coding | 45.69 | -84.53872431 | 84.53872431 | ENSG00000127362 | down |
| ENSG00000279598 | 265.7687229 | -2.89347097 | 0.2454536 | -11.78826 | 4.49E-32 | 4.16E-29 | | TEC | 39.95 | -82.11782153 | 82.11782153 | ENSG00000279598 | down |
| ENSG00000279764 | 57.66792 | -3.452893006 | 0.3209864 | -10.75713 | 5.49E-27 | 2.46E-24 | | TEC | 55.67 | -81.51744957 | 81.51744957 | ENSG00000279764 | down |
| ENSG00000216809 | 235.8830665 | -3.234417426 | 0.2923707 | -11.06273 | 1.90E-28 | 1.01E-25 | | processed_pseudogene | 44.93 | -80.85233499 | 80.85233499 | ENSG00000216809 | down |

| | | | | | | | | | | | | | |
|---|---|---|---|---|---|---|---|---|---|---|---|---|---|
| ENSG00000256019 | 68.18850293 | -3.668294644 | 0.353399 | -10.38004 | 3.06E-25 | 1.12E-22 | | processed_pseudogene | 39.68 | -80.52204023 | 80.52204023 | ENSG00000256019 | down |
| ENSG00000258441 | 1144.38401 | -2.898367008 | 0.2488702 | -11.6461 | 2.40E-31 | 2.09E-28 | LINC00641 | processed_transcript | 41.26 | -80.22622614 | 80.22622614 | ENSG00000258441 | down |
| ENSG00000254665 | 37.87312049 | -3.924865152 | 0.3941634 | -9.957457 | 2.34E-23 | 6.93E-21 | | antisense | 47.55 | -79.12215623 | 79.12215623 | ENSG00000254665 | down |
| ENSG00000234383 | 35.75790545 | -4.159677952 | 0.4310484 | -9.650142 | 4.91E-22 | 1.24E-19 | CTBP2P8 | processed_pseudogene | 55.86 | -78.64144823 | 78.64144823 | ENSG00000234383 | down |
| ENSG00000279900 | 35.89458419 | -4.092473432 | 0.4219507 | -9.698938 | 3.05E-22 | 8.16E-20 | | TEC | 40.27 | -78.11903444 | 78.11903444 | ENSG00000279900 | down |
| ENSG00000267405 | 54.71965876 | -3.725558356 | 0.3703288 | -10.06014 | 8.29E-24 | 2.68E-21 | | lincRNA | 47.65 | -76.63946396 | 76.63946396 | ENSG00000267405 | down |
| ENSG00000118515 | 273.9506882 | 2.931243157 | 0.2621015 | 11.18362 | 4.90E-29 | 3.25E-26 | SGK1 | protein_coding | 42.27 | 74.71112991 | 74.71112991 | ENSG00000118515 | up |
| ENSG00000119737 | 114.4328548 | -3.57591595 | 0.3530709 | -10.12804 | 4.15E-24 | 1.41E-21 | GPR75 | protein_coding | 44.92 | -74.56184913 | 74.56184913 | ENSG00000119737 | down |
| ENSG00000145632 | 644.0971346 | 2.56536216 | 0.2161315 | 11.86945 | 1.71E-32 | 1.83E-29 | PLK2 | protein_coding | 42.35 | 73.72404547 | 73.72404547 | ENSG00000145632 | up |
| ENSG00000207697 | 59.66853761 | -3.87893845 | 0.4014709 | -9.661817 | 4.38E-22 | 1.13E-19 | MIR573 | miRNA | 43.43 | -73.49487537 | 73.49487537 | ENSG00000207697 | down |
| ENSG00000218996 | 37.68813654 | -4.117588826 | 0.4395315 | -9.368131 | 7.38E-21 | 1.43E-18 | ARL4AP5 | processed_pseudogene | 39.73 | -73.47993397 | 73.47993397 | ENSG00000218996 | down |
| ENSG00000276136 | 33.4876425 | -4.045914035 | 0.4307438 | -9.392854 | 5.84E-21 | 1.15E-18 | | lincRNA | 42.69 | -72.5883266 | 72.5883266 | ENSG00000276136 | down |
| ENSG00000197019 | 197.5824863 | 2.863497173 | 0.2586711 | 11.07003 | 1.75E-28 | 9.76E-26 | SERTAD1 | protein_coding | 52.57 | 71.61701007 | 71.61701007 | ENSG00000197019 | up |
| ENSG00000234473 | 39.26156325 | -3.870315864 | 0.4075325 | -9.49695 | 2.16E-21 | 4.56E-19 | ZNF101P2 | processed_pseudogene | 44.02 | -70.98585382 | 70.98585382 | ENSG00000234473 | down |
| ENSG00000279425 | 100.5735269 | -3.04185058 | 0.2888747 | -10.53 | 6.28E-26 | 2.50E-23 | | TEC | 43.76 | -68.75232676 | 68.75232676 | ENSG00000279425 | down |
| ENSG00000237481 | 23.93055581 | -4.835028373 | 0.5783238 | -8.360417 | 6.25E-17 | 6.75E-15 | | antisense | 38.05 | -68.51718535 | 68.51718535 | ENSG00000237481 | down |
| ENSG00000229087 | 236.8497872 | -2.709762149 | 0.244506 | -11.0826 | 1.52E-28 | 8.84E-26 | | processed_pseudogene | 48.85 | -67.88915774 | 67.88915774 | ENSG00000229087 | down |
| ENSG00000254783 | 21.45059159 | -4.962407894 | 0.6087517 | -8.151776 | 3.59E-16 | 3.24E-14 | | processed_pseudogene | 48.11 | -66.93870139 | 66.93870139 | ENSG00000254783 | down |
| ENSG00000273338 | 182.330262 | 2.779014679 | 0.2560927 | 10.8516 | 1.96E-27 | 9.10E-25 | | antisense | 28.71 | 66.81083575 | 66.81083575 | ENSG00000273338 | up |
| ENSG00000280010 | 61.16568441 | -3.211229982 | 0.317853 | -10.10288 | 5.36E-24 | 1.78E-21 | | TEC | 38.67 | -66.6329781 | 66.6329781 | ENSG00000280010 | down |
| ENSG00000230832 | 29.25846492 | 4.200063831 | 0.4762597 | 8.818852 | 1.16E-18 | 1.75E-16 | | processed_pseudogene | 35.29 | 66.18018904 | 66.18018904 | ENSG00000230832 | up |
| ENSG00000267397 | 41.51962665 | -3.517444966 | 0.3664541 | -9.598596 | 8.10E-22 | 1.91E-19 | | lincRNA | 37.35 | -65.84099432 | 65.84099432 | ENSG00000267397 | down |
| ENSG00000187536 | 21.32062343 | -4.771070435 | 0.5792316 | -8.236896 | 1.77E-16 | 1.71E-14 | TPM3P7 | processed_pseudogene | 48.5 | -65.68477925 | 65.68477925 | ENSG00000187536 | down |
| ENSG00000280551 | 212.5011536 | -2.777184868 | 0.2603826 | -10.66579 | 1.47E-26 | 6.21E-24 | | TEC | 38.4 | -64.45019808 | 64.45019808 | ENSG00000280551 | down |
| ENSG00000254676 | 59.76502898 | -3.184663269 | 0.3193092 | -9.973604 | 1.99E-23 | 6.15E-21 | | sense_overlapping | 35.34 | -64.36488108 | 64.36488108 | ENSG00000254676 | down |
| ENSG00000279801 | 74.1523446 | -2.929510404 | 0.2827402 | -10.36114 | 3.72E-25 | 1.33E-22 | | TEC | 53.69 | -64.08657352 | 64.08657352 | ENSG00000279801 | down |
| ENSG00000250764 | 49.37962601 | -3.398639676 | 0.3539883 | -9.600995 | 7.92E-22 | 1.90E-19 | | antisense | 41.29 | -63.62627427 | 63.62627427 | ENSG00000250764 | down |
| ENSG00000260743 | 53.70396518 | -3.32089828 | 0.3419459 | -9.711765 | 2.69E-22 | 7.48E-20 | | lincRNA | 34.74 | -63.51570426 | 63.51570426 | ENSG00000260743 | down |
| ENSG00000237179 | 116.4250186 | 2.720574563 | 0.2554745 | 10.64911 | 1.76E-26 | 7.21E-24 | LINC01797 | lincRNA | 36.28 | 62.95985145 | 62.95985145 | ENSG00000237179 | up |
| ENSG00000232795 | 33.16677602 | -3.869302955 | 0.4333925 | -8.927942 | 4.34E-19 | 6.87E-17 | SCAND3P1 | processed_pseudogene | 36.22 | -62.54052145 | 62.54052145 | ENSG00000232795 | down |
| ENSG00000172183 | 20.58612156 | -4.218992871 | 0.4966829 | -8.494339 | 1.99E-17 | 2.47E-15 | ISG20 | protein_coding | 52.19 | -61.62469737 | 61.62469737 | ENSG00000172183 | down |
| ENSG00000263720 | 27.70127487 | -4.254032594 | 0.504498 | -8.432209 | 3.39E-17 | 3.94E-15 | | sense_intronic | 40.05 | -61.27929764 | 61.27929764 | ENSG00000263720 | down |

| | | | | | | | | | | | | | |
|---|---|---|---|---|---|---|---|---|---|---|---|---|---|
| ENSG00000201944 | 80.30486835 | -3.032731619 | 0.304191 | -9.969827 | 2.07E-23 | 6.25E-21 | | snoRNA | 35.61 | -61.2730692 | 61.2730692 | ENSG00000201944 | down |
| ENSG00000255330 | 317.1729459 | -2.997687625 | 0.2998 | -9.998959 | 1.54E-23 | 4.87E-21 | SOGA3 | protein_coding | 38.15 | -60.88956582 | 60.88956582 | ENSG00000255330 | down |
| ENSG00000259719 | 29.40513018 | -4.28159252 | 0.5134809 | -8.338367 | 7.53E-17 | 8.01E-15 | LINC02284 | lincRNA | 42.62 | -60.355938 | 60.355938 | ENSG00000259719 | down |
| ENSG00000142871 | 216.8153901 | 2.589117866 | 0.2422203 | 10.6891 | 1.14E-26 | 4.98E-24 | CYR61 | protein_coding | 50.25 | 60.33346223 | 60.33346223 | ENSG00000142871 | up |
| ENSG00000257752 | 24.04998864 | -5.608453549 | 0.7710933 | -7.273378 | 3.51E-13 | 1.84E-11 | | processed_pseudogene | 43.24 | -60.20502829 | 60.20502829 | ENSG00000257752 | down |
| ENSG00000281026 | 341.077209 | -2.662430712 | 0.2530016 | -10.52338 | 6.74E-26 | 2.61E-23 | N4BP2L2-IT2 | sense_intronic | 38.1 | -60.1278658 | 60.1278658 | ENSG00000281026 | down |
| ENSG00000242318 | 22.64576895 | -4.265749398 | 0.5158271 | -8.269726 | 1.34E-16 | 1.34E-14 | | processed_pseudogene | 40.72 | -59.18482731 | 59.18482731 | ENSG00000242318 | down |
| ENSG00000279616 | 46.1023127 | -3.106277136 | 0.3253652 | -9.547047 | 1.33E-21 | 3.00E-19 | | TEC | 42.03 | -57.53867313 | 57.53867313 | ENSG00000279616 | down |
| ENSG00000214756 | 578.3743645 | 2.299387221 | 0.2078921 | 11.06048 | 1.95E-28 | 1.01E-25 | METTL12 | protein_coding | 52.54 | 57.47892166 | 57.47892166 | ENSG00000214756 | up |
| ENSG00000251348 | 145.2563813 | -3.129533132 | 0.3303736 | -9.472709 | 2.73E-21 | 5.50E-19 | HSPD1P11 | processed_pseudogene | 40.36 | -57.14359555 | 57.14359555 | ENSG00000251348 | down |
| ENSG00000279070 | 81.7629378 | 2.915961831 | 0.2967965 | 9.824786 | 8.81E-23 | 2.55E-20 | | lincRNA | 44.39 | 57.13163968 | 57.13163968 | ENSG00000279070 | up |
| ENSG00000261542 | 105.0129269 | -3.091813762 | 0.3245515 | -9.526421 | 1.63E-21 | 3.54E-19 | | lincRNA | 32.38 | -57.04641599 | 57.04641599 | ENSG00000261542 | down |
| ENSG00000271862 | 80.45478942 | -2.978058615 | 0.3101313 | -9.602573 | 7.80E-22 | 1.90E-19 | | lincRNA | 40.34 | -55.75253405 | 55.75253405 | ENSG00000271862 | down |
| ENSG00000229065 | 19.17089843 | -5.147085985 | 0.7048789 | -7.302086 | 2.83E-13 | 1.51E-11 | | antisense | 47.48 | -55.69455619 | 55.69455619 | ENSG00000229065 | down |
| ENSG00000225401 | 16.18749697 | -5.143859114 | 0.7058699 | -7.287262 | 3.16E-13 | 1.67E-11 | TGIF2P1 | processed_pseudogene | 59.42 | -55.43921682 | 55.43921682 | ENSG00000225401 | down |
| ENSG00000255837 | 75.40858158 | -3.107271799 | 0.3335847 | -9.31479 | 1.22E-20 | 2.26E-18 | TAS2R20 | protein_coding | 34.76 | -54.83133955 | 54.83133955 | ENSG00000255837 | down |
| ENSG00000229591 | 24.77396511 | -4.059054877 | 0.4980698 | -8.14957 | 3.65E-16 | 3.27E-14 | | antisense | 43.97 | -54.73870826 | 54.73870826 | ENSG00000229591 | down |
| ENSG00000254463 | 40.62757213 | -3.488883868 | 0.3969286 | -8.789702 | 1.50E-18 | 2.22E-16 | | processed_pseudogene | 45.21 | -54.6143643 | 54.6143643 | ENSG00000254463 | down |
| ENSG00000223375 | 19.52546428 | -7.327484866 | 1.2095274 | -6.058139 | 1.38E-09 | 3.70E-08 | | processed_pseudogene | 44.91 | -54.45527151 | 54.45527151 | ENSG00000223375 | down |
| ENSG00000232623 | 17.41952447 | -5.426341532 | 0.7730223 | -7.019644 | 2.22E-12 | 9.80E-11 | | antisense | 48.46 | -54.31090993 | 54.31090993 | ENSG00000232623 | down |
| ENSG00000147437 | 47.01562507 | -3.044430789 | 0.3269205 | -9.31245 | 1.25E-20 | 2.26E-18 | GNRH1 | protein_coding | 39.78 | -53.72243856 | 53.72243856 | ENSG00000147437 | down |
| ENSG00000279698 | 21.15366529 | -5.413220149 | 0.7773796 | -6.963419 | 3.32E-12 | 1.41E-10 | | TEC | 41.33 | -53.3176758 | 53.3176758 | ENSG00000279698 | down |
| ENSG00000261229 | 26.86730636 | -3.649968244 | 0.4306208 | -8.476061 | 2.33E-17 | 2.84E-15 | | lincRNA | 40.63 | -53.09709648 | 53.09709648 | ENSG00000261229 | down |
| ENSG00000240770 | 22.26690468 | -4.881753755 | 0.666854 | -7.320574 | 2.47E-13 | 1.33E-11 | C21orf91-OT1 | lincRNA | 39.23 | -53.0876431 | 53.0876431 | ENSG00000240770 | down |
| ENSG00000266288 | 21.20041978 | -4.227224079 | 0.537442 | -7.865452 | 3.68E-15 | 2.77E-13 | | lincRNA | 37.89 | -53.08490787 | 53.08490787 | ENSG00000266288 | down |
| ENSG00000252960 | 20.35287188 | -4.808824324 | 0.6539058 | -7.354002 | 1.92E-13 | 1.06E-11 | RN7SKP258 | misc_RNA | 44.08 | -52.76989467 | 52.76989467 | ENSG00000252960 | down |
| ENSG00000278765 | 72.19942326 | -2.816339157 | 0.2946962 | -9.556753 | 1.22E-21 | 2.77E-19 | | lincRNA | 38.47 | -52.26276875 | 52.26276875 | ENSG00000278765 | down |
| ENSG00000260955 | 31.91738834 | -3.540255704 | 0.4150661 | -8.529378 | 1.47E-17 | 1.85E-15 | | lincRNA | 43.05 | -52.16165841 | 52.16165841 | ENSG00000260955 | down |
| ENSG00000227773 | 26.63112005 | -3.610978701 | 0.4281683 | -8.43355 | 3.35E-17 | 3.92E-15 | ASH1L-IT1 | sense_intronic | 43.24 | -52.02096968 | 52.02096968 | ENSG00000227773 | down |
| ENSG00000213090 | 75.81629656 | -2.929115339 | 0.3143564 | -9.317816 | 1.19E-20 | 2.23E-18 | | processed_pseudogene | 38.19 | -51.701313 | 51.701313 | ENSG00000213090 | down |
| ENSG00000251213 | 108.938352 | -2.819872631 | 0.297434 | -9.480666 | 2.53E-21 | 5.17E-19 | | lincRNA | 35.61 | -51.56484308 | 51.56484308 | ENSG00000251213 | down |
| ENSG00000214842 | 78.84556372 | 2.693326177 | 0.2776424 | 9.700703 | 2.99E-22 | 8.16E-20 | RAD51AP2 | protein_coding | 34.45 | 51.41146152 | 51.41146152 | ENSG00000214842 | up |

| | | | | | | | | | | | | | |
|---|---|---|---|---|---|---|---|---|---|---|---|---|---|
| ENSG00000268392 | 24.77313402 | -3.95851854 | 0.4955195 | -7.988624 | 1.36E-15 | 1.09E-13 | | lincRNA | 41.22 | -51.31950392 | 51.31950392 | ENSG00000268392 | down |
| ENSG00000260198 | 65.40522756 | -2.85664834 | 0.3077412 | -9.282633 | 1.65E-20 | 2.95E-18 | | sense_intronic | 49.45 | -50.07699394 | 50.07699394 | ENSG00000260198 | down |
| ENSG00000277039 | 16.32999292 | -6.520966993 | 1.0597165 | -6.153501 | 7.58E-10 | 2.14E-08 | | misc_RNA | 55.7 | -50.01828149 | 50.01828149 | ENSG00000277039 | down |
| ENSG00000223476 | 23.9823678 | -3.675326064 | 0.4497322 | -8.172255 | 3.03E-16 | 2.83E-14 | VN1R42P | processed_pseudogene | 38.37 | -49.79506101 | 49.79506101 | ENSG00000223476 | down |
| ENSG00000226977 | 9.542898072 | -7.152471193 | 1.2207799 | -5.858936 | 4.66E-09 | 1.13E-07 | HMGN1P24 | processed_pseudogene | 47.65 | -49.69859098 | 49.69859098 | ENSG00000226977 | down |
| ENSG00000279499 | 16.34760538 | -5.275977257 | 0.7749418 | -6.808224 | 9.88E-12 | 4.01E-10 | | TEC | 29.16 | -49.577072 | 49.577072 | ENSG00000279499 | down |
| ENSG00000257246 | 39.78428961 | -3.184602636 | 0.3647141 | -8.731778 | 2.51E-18 | 3.64E-16 | PHBP19 | processed_pseudogene | 51.59 | -49.16830582 | 49.16830582 | ENSG00000257246 | down |
| ENSG00000123870 | 52.17075463 | -3.003779678 | 0.335521 | -8.952582 | 3.47E-19 | 5.56E-17 | ZNF137P | transcribed_unprocessed_pseud | 46.4 | -48.82682664 | 48.82682664 | ENSG00000123870 | down |
| ENSG00000118491 | 36.7533909 | -3.382080347 | 0.4022166 | -8.408604 | 4.15E-17 | 4.58E-15 | ZC2HC1B | protein_coding | 39.82 | -48.49454954 | 48.49454954 | ENSG00000118491 | down |
| ENSG00000279589 | 104.0630579 | -2.956642174 | 0.3289232 | -8.988852 | 2.50E-19 | 4.14E-17 | | TEC | 36.46 | -48.43839396 | 48.43839396 | ENSG00000279589 | down |
| ENSG00000231333 | 29.2677336 | -3.318663572 | 0.3908866 | -8.490094 | 2.06E-17 | 2.54E-15 | RPL34P6 | processed_pseudogene | 42.61 | -48.43419587 | 48.43419587 | ENSG00000231333 | down |
| ENSG00000163352 | 20.04170213 | -3.912710828 | 0.5018756 | -7.796176 | 6.38E-15 | 4.60E-13 | LENEP | protein_coding | 54.66 | -48.27082771 | 48.27082771 | ENSG00000163352 | down |
| ENSG00000226555 | 60.62235126 | -2.828438253 | 0.3099633 | -9.125074 | 7.17E-20 | 1.22E-17 | AGKP1 | processed_pseudogene | 44.81 | -47.84200714 | 47.84200714 | ENSG00000226555 | down |
| ENSG00000258739 | 91.72350045 | -2.740620792 | 0.2962243 | -9.251845 | 2.21E-20 | 3.89E-18 | ENO1P2 | transcribed_processed_pseudog | 43.23 | -47.71472193 | 47.71472193 | ENSG00000258739 | down |
| ENSG00000259416 | 30.69894982 | -3.457468236 | 0.419398 | -8.243883 | 1.67E-16 | 1.62E-14 | | antisense | 55.74 | -47.67725153 | 47.67725153 | ENSG00000259416 | down |
| ENSG00000258900 | 44.37735231 | -2.975351945 | 0.3353202 | -8.873166 | 7.11E-19 | 1.10E-16 | HNRNPCP1 | processed_pseudogene | 45.95 | -47.48265824 | 47.48265824 | ENSG00000258900 | down |
| ENSG00000127366 | 32.21227528 | -3.309002936 | 0.3938235 | -8.402249 | 4.38E-17 | 4.80E-15 | TAS2R5 | protein_coding | 48.39 | -47.38025889 | 47.38025889 | ENSG00000127366 | down |
| ENSG00000261167 | 365.5399263 | -2.661334358 | 0.2845714 | -9.352078 | 8.59E-21 | 1.64E-18 | | sense_overlapping | 39.27 | -47.33285341 | 47.33285341 | ENSG00000261167 | down |
| ENSG00000261056 | 167.2258326 | -2.542391488 | 0.2666097 | -9.536007 | 1.48E-21 | 3.28E-19 | | transcribed_processed_pseudog | 45.32 | -46.9937226 | 46.9937226 | ENSG00000261056 | down |
| ENSG00000226761 | 98.39988224 | -2.799613264 | 0.3079517 | -9.091079 | 9.81E-20 | 1.64E-17 | TAS2R46 | protein_coding | 38.71 | -46.9882854 | 46.9882854 | ENSG00000226761 | down |
| ENSG00000266533 | 31.14585663 | -3.080398508 | 0.3560106 | -8.652547 | 5.04E-18 | 7.01E-16 | MIR3619 | miRNA | 65.06 | -46.68084681 | 46.68084681 | ENSG00000266533 | down |
| ENSG00000248559 | 14.49209925 | -5.627298782 | 0.8849104 | -6.359173 | 2.03E-10 | 6.46E-09 | | antisense | 44.13 | -46.0852689 | 46.0852689 | ENSG00000248559 | down |
| ENSG00000207737 | 6.554937549 | -6.985845574 | 1.2241457 | -5.706711 | 1.15E-08 | 2.54E-07 | MIR181B2 | miRNA | 46.07 | -46.0786003 | 46.0786003 | ENSG00000207737 | down |
| ENSG00000207864 | 51.11138552 | -2.97529854 | 0.3423784 | -8.690089 | 3.62E-18 | 5.15E-16 | MIR27B | miRNA | 47.42 | -45.48814316 | 45.48814316 | ENSG00000207864 | down |
| ENSG00000225595 | 16.33208989 | 4.568005958 | 0.6536793 | 6.988146 | 2.79E-12 | 1.21E-10 | LINC01623 | lincRNA | 44.93 | 45.30493625 | 45.30493625 | ENSG00000225595 | up |
| ENSG00000266258 | 36.70402408 | -2.955743374 | 0.3422931 | -8.635123 | 5.87E-18 | 7.93E-16 | LINC01909 | lincRNA | 39.71 | -44.63389857 | 44.63389857 | ENSG00000266258 | down |
| ENSG00000273786 | 16.8608013 | -3.797981139 | 0.5004969 | -7.588421 | 3.24E-14 | 2.03E-12 | | sense_intronic | 33.48 | -44.40716272 | 44.40716272 | ENSG00000273786 | down |
| ENSG00000258525 | 39.61956503 | -3.044172939 | 0.3591771 | -8.475409 | 2.34E-17 | 2.84E-15 | | antisense | 39.27 | -44.2844248 | 44.2844248 | ENSG00000258525 | down |
| ENSG00000270012 | 87.67422139 | -2.495876986 | 0.2679908 | -9.313292 | 1.24E-20 | 2.26E-18 | | lincRNA | 49.5 | -44.04258376 | 44.04258376 | ENSG00000270012 | down |
| ENSG00000233875 | 15.89622765 | -4.330543571 | 0.6140558 | -7.052362 | 1.76E-12 | 8.03E-11 | | antisense | 53.59 | -43.71807284 | 43.71807284 | ENSG00000233875 | down |
| ENSG00000279587 | 32.3253011 | -3.032397827 | 0.3598874 | -8.425963 | 3.58E-17 | 4.08E-15 | | TEC | 30.42 | -43.63314836 | 43.63314836 | ENSG00000279587 | down |
| ENSG00000249002 | 18.72325115 | -4.092371562 | 0.5667779 | -7.220415 | 5.18E-13 | 2.61E-11 | | processed_pseudogene | 41.62 | -43.30773039 | 43.30773039 | ENSG00000249002 | down |

| | | | | | | | | | | | | | |
|---|---|---|---|---|---|---|---|---|---|---|---|---|---|
| ENSG00000232882 | 25.40162108 | -3.623510866 | 0.4735271 | -7.652172 | 1.98E-14 | 1.28E-12 | PHKA1P1 | processed_pseudogene | 49.11 | -43.09236092 | 43.09236092 | ENSG00000232882 | down |
| ENSG00000233956 | 17.68298067 | -4.044507428 | 0.5598202 | -7.224655 | 5.02E-13 | 2.58E-11 | BTF3P6 | processed_pseudogene | 40.28 | -42.82408166 | 42.82408166 | ENSG00000233956 | down |
| ENSG00000225693 | 30.69492626 | -3.056800389 | 0.3685791 | -8.293472 | 1.10E-16 | 1.14E-14 | LAGE3P1 | processed_pseudogene | 62.19 | -42.6179724 | 42.6179724 | ENSG00000225693 | down |
| ENSG00000274825 | 25.32643709 | -3.229627694 | 0.4007255 | -8.059451 | 7.66E-16 | 6.55E-14 | | lincRNA | 46.08 | -42.57945053 | 42.57945053 | ENSG00000274825 | down |
| ENSG00000271924 | 26.01934418 | -3.261976997 | 0.4069004 | -8.016647 | 1.09E-15 | 8.85E-14 | RNA5SP108 | rRNA | 47.86 | -42.57907331 | 42.57907331 | ENSG00000271924 | down |
| ENSG00000232450 | 25.71539287 | -3.330632438 | 0.4216097 | -7.8998 | 2.79E-15 | 2.13E-13 | | transcribed_processed_pseudog | 54.14 | -42.20766722 | 42.20766722 | ENSG00000232450 | down |
| ENSG00000120738 | 379.4916135 | 2.474308283 | 0.27089 | 9.133996 | 6.60E-20 | 1.15E-17 | EGR1 | protein_coding | 56.56 | 41.91405689 | 41.91405689 | ENSG00000120738 | up |
| ENSG00000253180 | 15.58800622 | -4.167985067 | 0.5927965 | -7.031055 | 2.05E-12 | 9.18E-11 | | processed_pseudogene | 41.97 | -41.83540257 | 41.83540257 | ENSG00000253180 | down |
| ENSG00000162396 | 93.74958646 | -2.76014963 | 0.3193793 | -8.64223 | 5.51E-18 | 7.60E-16 | PARS2 | protein_coding | 49.77 | -41.73133413 | 41.73133413 | ENSG00000162396 | down |
| ENSG00000272650 | 15.21147444 | -4.696005871 | 0.709386 | -6.619818 | 3.60E-11 | 1.31E-09 | | lincRNA | 28.52 | -41.71208662 | 41.71208662 | ENSG00000272650 | down |
| ENSG00000237162 | 14.49375299 | -5.4138882 | 0.8792557 | -6.157354 | 7.40E-10 | 2.09E-08 | | processed_pseudogene | 42.78 | -41.574188 | 41.574188 | ENSG00000237162 | down |
| ENSG00000266289 | 20.47165566 | -3.458760691 | 0.449825 | -7.689125 | 1.48E-14 | 9.87E-13 | | lincRNA | 31.92 | -41.52493958 | 41.52493958 | ENSG00000266289 | down |
| ENSG00000231697 | 28.50422399 | -3.180091265 | 0.3972916 | -8.004425 | 1.20E-15 | 9.66E-14 | NANOGP5 | transcribed_processed_pseudog | 46.86 | -41.38900942 | 41.38900942 | ENSG00000231697 | down |
| ENSG00000224324 | 13.67143955 | -4.118757567 | 0.5865475 | -7.022036 | 2.19E-12 | 9.76E-11 | THAP5P1 | processed_pseudogene | 40.08 | -41.2314698 | 41.2314698 | ENSG00000224324 | down |
| ENSG00000258808 | 15.72676083 | 4.273521922 | 0.6199226 | 6.893638 | 5.44E-12 | 2.26E-10 | LINC02310 | lincRNA | 44.19 | 41.22173113 | 41.22173113 | ENSG00000258808 | up |
| ENSG00000271840 | 25.03017009 | -3.357666628 | 0.4317907 | -7.776145 | 7.48E-15 | 5.34E-13 | | lincRNA | 43.32 | -41.20724736 | 41.20724736 | ENSG00000271840 | down |
| ENSG00000166343 | 85.68544245 | -2.624602823 | 0.2986367 | -8.788615 | 1.51E-18 | 2.22E-16 | MSS51 | protein_coding | 42.14 | -41.08506333 | 41.08506333 | ENSG00000166343 | down |
| ENSG00000226769 | 33.63760509 | -3.263213116 | 0.4155205 | -7.853315 | 4.05E-15 | 3.00E-13 | GAPDHP54 | processed_pseudogene | 51.18 | -40.86448302 | 40.86448302 | ENSG00000226769 | down |
| ENSG00000236605 | 807.1930687 | 2.142284089 | 0.2215677 | 9.668757 | 4.09E-22 | 1.08E-19 | | lincRNA | 46.76 | 40.63587314 | 40.63587314 | ENSG00000236605 | up |
| ENSG00000278899 | 47.63300687 | 2.660661951 | 0.3067821 | 8.672806 | 4.22E-18 | 5.93E-16 | | antisense | 59.54 | 40.5139081 | 40.5139081 | ENSG00000278899 | up |
| ENSG00000232142 | 14.95910923 | -4.669242247 | 0.7149176 | -6.531161 | 6.53E-11 | 2.27E-09 | RPS25P9 | processed_pseudogene | 45.63 | -40.36444426 | 40.36444426 | ENSG00000232142 | down |
| ENSG00000279573 | 57.77103526 | -2.693262167 | 0.3131883 | -8.599497 | 8.01E-18 | 1.06E-15 | | TEC | 50.46 | -40.32892719 | 40.32892719 | ENSG00000279573 | down |
| ENSG00000279250 | 26.49498519 | -3.295875061 | 0.4247082 | -7.760329 | 8.47E-15 | 5.93E-13 | | TEC | 48.4 | -40.29928597 | 40.29928597 | ENSG00000279250 | down |
| ENSG00000225750 | 16.31977351 | -3.857173799 | 0.5379343 | -7.170344 | 7.48E-13 | 3.63E-11 | | lincRNA | 47.46 | -40.26962969 | 40.26962969 | ENSG00000225750 | down |
| ENSG00000222898 | 13.19560535 | -4.657071502 | 0.7130309 | -6.531374 | 6.52E-11 | 2.27E-09 | RN7SKP97 | misc_RNA | 49.85 | -40.25923118 | 40.25923118 | ENSG00000222898 | down |
| ENSG00000277840 | 38.85082009 | -2.827462975 | 0.338034 | -8.364434 | 6.04E-17 | 6.57E-15 | | sense_intronic | 38.84 | -40.10023583 | 40.10023583 | ENSG00000277840 | down |
| ENSG00000253463 | 15.99814114 | -6.055451196 | 1.059447 | -5.715672 | 1.09E-08 | 2.42E-07 | HMGB1P19 | processed_pseudogene | 43.15 | -40.06535021 | 40.06535021 | ENSG00000253463 | down |
| ENSG00000279265 | 13.58820167 | -4.65747002 | 0.7151875 | -6.512236 | 7.40E-11 | 2.55E-09 | | TEC | 39.48 | -40.02248708 | 40.02248708 | ENSG00000279265 | down |
| ENSG00000275106 | 34.8712451 | -2.853223605 | 0.3444753 | -8.282811 | 1.20E-16 | 1.22E-14 | | TEC | 40.38 | -39.69609837 | 39.69609837 | ENSG00000275106 | down |
| ENSG00000236514 | 39.48281366 | 2.753685677 | 0.3267602 | 8.427238 | 3.54E-17 | 4.07E-15 | | antisense | 49.23 | 39.62593957 | 39.62593957 | ENSG00000236514 | up |
| ENSG00000227740 | 27.00476322 | -3.32511715 | 0.4344133 | -7.654272 | 1.94E-14 | 1.27E-12 | | lincRNA | 38.64 | -39.55534755 | 39.55534755 | ENSG00000227740 | down |
| ENSG00000185338 | 36.46491527 | 2.859043161 | 0.350198 | 8.164076 | 3.24E-16 | 2.97E-14 | SOCS1 | protein_coding | 68.56 | 38.67309886 | 38.67309886 | ENSG00000185338 | up |

| | | | | | | | | | | | | | |
|---|---|---|---|---|---|---|---|---|---|---|---|---|---|
| ENSG00000238735 | 17.8034434 | -5.647681984 | 0.9728013 | -5.805586 | 6.41E-09 | 1.50E-07 | RNU7-113P | snRNA | 37.1 | -38.53366345 | 38.53366345 | ENSG00000238735 | down |
| ENSG00000267274 | 25.46712277 | -3.309727521 | 0.4425624 | -7.478556 | 7.51E-14 | 4.34E-12 | | sense_overlapping | 45.7 | -37.6064191 | 37.6064191 | ENSG00000267274 | down |
| ENSG00000264395 | 159.2401502 | 2.210744358 | 0.2421079 | 9.131235 | 6.77E-20 | 1.16E-17 | MIR3193 | miRNA | 56.36 | 37.43679354 | 37.43679354 | ENSG00000264395 | up |
| ENSG00000223825 | 14.2026464 | -3.974679339 | 0.5848707 | -6.795826 | 1.08E-11 | 4.30E-10 | DAZAP2P1 | processed_pseudogene | 54.87 | -37.22878322 | 37.22878322 | ENSG00000223825 | down |
| ENSG00000157450 | 687.410453 | -2.366841747 | 0.2688374 | -8.803991 | 1.32E-18 | 1.98E-16 | RNF111 | protein_coding | 39.52 | -37.16902461 | 37.16902461 | ENSG00000157450 | down |
| ENSG00000197182 | 1238.253786 | -2.043420577 | 0.2161918 | -9.451889 | 3.33E-21 | 6.62E-19 | MIRLET7BHG | lincRNA | 56.82 | -37.14779089 | 37.14779089 | ENSG00000197182 | down |
| ENSG00000231888 | 26.11946106 | -2.999325521 | 0.3842276 | -7.806117 | 5.90E-15 | 4.28E-13 | MTND5P15 | processed_pseudogene | 44.99 | -37.09831966 | 37.09831966 | ENSG00000231888 | down |
| ENSG00000159905 | 247.3797253 | -2.404287774 | 0.2762568 | -8.703091 | 3.23E-18 | 4.64E-16 | ZNF221 | protein_coding | 41.33 | -36.86706496 | 36.86706496 | ENSG00000159905 | down |
| ENSG00000270062 | 26.50707019 | -2.998766577 | 0.3870339 | -7.748073 | 9.33E-15 | 6.43E-13 | | sense_intronic | 32.06 | -36.56020624 | 36.56020624 | ENSG00000270062 | down |
| ENSG00000213613 | 58.44040818 | -2.624152564 | 0.3165958 | -8.288651 | 1.15E-16 | 1.18E-14 | RPL11P3 | processed_pseudogene | 46.27 | -36.55123552 | 36.55123552 | ENSG00000213613 | down |
| ENSG00000270878 | 40.37156413 | -2.700413406 | 0.3307802 | -8.163769 | 3.25E-16 | 2.97E-14 | | processed_pseudogene | 60.03 | -36.52737952 | 36.52737952 | ENSG00000270878 | down |
| ENSG00000258215 | 36.48672743 | -2.870879516 | 0.3629078 | -7.910769 | 2.56E-15 | 1.97E-13 | | processed_pseudogene | 51.99 | -36.47750311 | 36.47750311 | ENSG00000258215 | down |
| ENSG00000178662 | 817.3447094 | -2.224871218 | 0.2483274 | -8.959427 | 3.26E-19 | 5.35E-17 | CSRNP3 | protein_coding | 36.48 | -36.20305105 | 36.20305105 | ENSG00000178662 | down |
| ENSG00000279235 | 100.1339755 | -2.50893364 | 0.2982907 | -8.411036 | 4.06E-17 | 4.56E-15 | | TEC | 44.07 | -35.97993573 | 35.97993573 | ENSG00000279235 | down |
| ENSG00000245059 | 30.30232303 | 2.879301422 | 0.3670671 | 7.844074 | 4.36E-15 | 3.21E-13 | | lincRNA | 48.28 | 35.97135273 | 35.97135273 | ENSG00000245059 | up |
| ENSG00000198576 | 404.9206871 | 2.481125676 | 0.2933352 | 8.458329 | 2.71E-17 | 3.23E-15 | ARC | protein_coding | 66.75 | 35.95425665 | 35.95425665 | ENSG00000198576 | up |
| ENSG00000206712 | 16.47647731 | -3.834574445 | 0.5641871 | -6.796637 | 1.07E-11 | 4.30E-10 | RNU6-26P | snRNA | 44.86 | -35.91649252 | 35.91649252 | ENSG00000206712 | down |
| ENSG00000269939 | 63.74910513 | -2.427074592 | 0.2839538 | -8.547427 | 1.26E-17 | 1.61E-15 | PCF11-AS1 | lincRNA | 35.62 | -35.90564758 | 35.90564758 | ENSG00000269939 | down |
| ENSG00000229330 | 31.65607738 | -2.702570393 | 0.33422 | -8.086201 | 6.16E-16 | 5.29E-14 | | sense_intronic | 51.89 | -35.88073579 | 35.88073579 | ENSG00000229330 | down |
| ENSG00000239577 | 18.05261615 | -3.814535494 | 0.5603868 | -6.806968 | 9.97E-12 | 4.03E-10 | RN7SL388P | misc_RNA | 52.8 | -35.83462898 | 35.83462898 | ENSG00000239577 | down |
| ENSG00000172201 | 431.3673321 | 2.222060041 | 0.2492576 | 8.914714 | 4.89E-19 | 7.65E-17 | ID4 | protein_coding | 53.08 | 35.81134255 | 35.81134255 | ENSG00000172201 | up |
| ENSG00000260966 | 289.1681836 | -2.604790857 | 0.3165603 | -8.228421 | 1.90E-16 | 1.82E-14 | | sense_overlapping | 34.44 | -35.78871193 | 35.78871193 | ENSG00000260966 | down |
| ENSG00000279544 | 213.3077795 | -2.642557081 | 0.3240226 | -8.155471 | 3.48E-16 | 3.17E-14 | | TEC | 37.73 | -35.67346664 | 35.67346664 | ENSG00000279544 | down |
| ENSG00000279602 | 2286.880872 | 1.94970736 | 0.2055995 | 9.483036 | 2.47E-21 | 5.13E-19 | | TEC | 46.4 | 35.65949145 | 35.65949145 | ENSG00000279602 | up |
| ENSG00000255067 | 30.77278927 | -2.910741067 | 0.3751635 | -7.758593 | 8.59E-15 | 5.98E-13 | | sense_intronic | 47.1 | -35.57921005 | 35.57921005 | ENSG00000255067 | down |
| ENSG00000233484 | 6.59290916 | -6.42268241 | 1.2317688 | -5.214195 | 1.85E-07 | 3.06E-06 | | transcribed_processed_pseudog | 46.46 | -35.41541391 | 35.41541391 | ENSG00000233484 | down |
| ENSG00000231066 | 42.2399077 | -2.603673469 | 0.3184351 | -8.176466 | 2.92E-16 | 2.75E-14 | NPM1P9 | processed_pseudogene | 41.35 | -35.30766983 | 35.30766983 | ENSG00000231066 | down |
| ENSG00000254500 | 12.08379418 | -4.735279843 | 0.7817709 | -6.05712 | 1.39E-09 | 3.72E-08 | RANP3 | processed_pseudogene | 41.67 | -35.18186298 | 35.18186298 | ENSG00000254500 | down |
| ENSG00000273444 | 5.537584915 | -6.400171659 | 1.2312952 | -5.197918 | 2.02E-07 | 3.32E-06 | | unprocessed_pseudogene | 40.12 | -35.06840763 | 35.06840763 | ENSG00000273444 | down |
| ENSG00000259820 | 386.7621775 | -2.134256065 | 0.2383121 | -8.955719 | 3.38E-19 | 5.46E-17 | | lincRNA | 36.21 | -34.70823905 | 34.70823905 | ENSG00000259820 | down |
| ENSG00000279355 | 92.7742352 | -2.454743079 | 0.2941294 | -8.345794 | 7.07E-17 | 7.58E-15 | AGPAT4-IT1 | TEC | 51.33 | -34.66235659 | 34.66235659 | ENSG00000279355 | down |
| ENSG00000165507 | 161.6070079 | 2.182871166 | 0.2469648 | 8.838793 | 9.68E-19 | 1.48E-16 | C10orf10 | protein_coding | 57.22 | 34.55402609 | 34.55402609 | ENSG00000165507 | up |

| | | | | | | | | | | | | | |
|---|---|---|---|---|---|---|---|---|---|---|---|---|---|
| ENSG00000212195 | 1065.698238 | 1.881387261 | 0.1980799 | 9.498122 | 2.14E-21 | 4.56E-19 | | snoRNA | 43.37 | 34.50671361 | 34.50671361 | ENSG00000212195 | up |
| ENSG00000222627 | 8.288037561 | -6.319943832 | 1.2191076 | -5.184074 | 2.17E-07 | 3.56E-06 | RNU2-37P | snRNA | 45.2 | -34.43765438 | 34.43765438 | ENSG00000222627 | down |
| ENSG00000207597 | 42.54256422 | -2.797448885 | 0.3600579 | -7.769442 | 7.88E-15 | 5.57E-13 | MIR490 | miRNA | 51.56 | -34.28001543 | 34.28001543 | ENSG00000207597 | down |
| ENSG00000272576 | 62.5617702 | -2.418664729 | 0.2902177 | -8.333966 | 7.82E-17 | 8.25E-15 | | lincRNA | 35.08 | -34.06387754 | 34.06387754 | ENSG00000272576 | down |
| ENSG00000180626 | 956.720429 | -2.273113841 | 0.265837 | -8.550782 | 1.22E-17 | 1.58E-15 | ZNF594 | protein_coding | 43.95 | -33.64757598 | 33.64757598 | ENSG00000180626 | down |
| ENSG00000228956 | 505.7685127 | -2.26008666 | 0.2636317 | -8.572895 | 1.01E-17 | 1.31E-15 | SATB1-AS1 | processed_transcript | 37.1 | -33.63391813 | 33.63391813 | ENSG00000228956 | down |
| ENSG00000231241 | 11.78931792 | -4.697124219 | 0.7909011 | -5.938953 | 2.87E-09 | 7.21E-08 | RPS3AP3 | processed_pseudogene | 35.95 | -33.54744542 | 33.54744542 | ENSG00000231241 | down |
| ENSG00000201012 | 4.364326325 | -6.296434564 | 1.2338577 | -5.103047 | 3.34E-07 | 5.20E-06 | | misc_RNA | 36.79 | -33.27066604 | 33.27066604 | ENSG00000201012 | down |
| ENSG00000270607 | 97.52600933 | 2.241087708 | 0.2626303 | 8.533242 | 1.42E-17 | 1.80E-15 | | lincRNA | 47.57 | 33.0435994 | 33.0435994 | ENSG00000270607 | up |
| ENSG00000277423 | 11.88625664 | -4.350719432 | 0.7120082 | -6.11049 | 9.93E-10 | 2.74E-08 | | lincRNA | 37.98 | -32.89856246 | 32.89856246 | ENSG00000277423 | down |
| ENSG00000254409 | 17.20246828 | -3.277975142 | 0.4669296 | -7.020277 | 2.21E-12 | 9.79E-11 | | sense_overlapping | 48.33 | -32.81038142 | 32.81038142 | ENSG00000254409 | down |
| ENSG00000225030 | 26.31439437 | -2.894836865 | 0.3877332 | -7.466054 | 8.26E-14 | 4.73E-12 | | sense_intronic | 52.3 | -32.78318022 | 32.78318022 | ENSG00000225030 | down |
| ENSG00000184068 | 86.33992757 | 2.257836215 | 0.2666919 | 8.466085 | 2.54E-17 | 3.05E-15 | | antisense | 56.07 | 32.77535821 | 32.77535821 | ENSG00000184068 | up |
| ENSG00000272672 | 18.30845786 | -3.317555222 | 0.475703 | -6.974006 | 3.08E-12 | 1.32E-10 | | lincRNA | 44.06 | -32.77148155 | 32.77148155 | ENSG00000272672 | down |
| ENSG00000271991 | 187.1111886 | 2.166307508 | 0.2515976 | 8.610206 | 7.29E-18 | 9.76E-16 | | lincRNA | 46.06 | 32.51713503 | 32.51713503 | ENSG00000271991 | up |
| ENSG00000272386 | 13.65123845 | -3.535089844 | 0.5249449 | -6.734211 | 1.65E-11 | 6.39E-10 | | sense_intronic | 38.35 | -32.50283852 | 32.50283852 | ENSG00000272386 | down |
| ENSG00000259407 | 32.55434063 | -2.821750059 | 0.3747985 | -7.528713 | 5.12E-14 | 3.09E-12 | | antisense | 48.91 | -32.47903865 | 32.47903865 | ENSG00000259407 | down |
| ENSG00000232015 | 17.47507573 | -3.40387097 | 0.4963675 | -6.857562 | 7.00E-12 | 2.90E-10 | HSPE1P25 | processed_pseudogene | 44.71 | -32.46348442 | 32.46348442 | ENSG00000232015 | down |
| ENSG00000254180 | 14.32833697 | -3.258752434 | 0.4662972 | -6.988574 | 2.78E-12 | 1.21E-10 | | lincRNA | 38.39 | -32.31991654 | 32.31991654 | ENSG00000254180 | down |
| ENSG00000213139 | 54.93819411 | -2.32885189 | 0.2818023 | -8.264133 | 1.41E-16 | 1.39E-14 | CRYGS | protein_coding | 37.09 | -32.27125371 | 32.27125371 | ENSG00000213139 | down |
| ENSG00000225342 | 72.77870278 | 2.357711698 | 0.2877642 | 8.193206 | 2.54E-16 | 2.41E-14 | | antisense | 36.31 | 32.10764576 | 32.10764576 | ENSG00000225342 | up |
| ENSG00000261799 | 214.9497996 | -2.239231218 | 0.2662881 | -8.409056 | 4.13E-17 | 4.58E-15 | | sense_overlapping | 39.78 | -32.10760777 | 32.10760777 | ENSG00000261799 | down |
| ENSG00000279281 | 302.4588072 | -2.283458246 | 0.2748737 | -8.307299 | 9.79E-17 | 1.02E-14 | | TEC | 45.42 | -31.94401312 | 31.94401312 | ENSG00000279281 | down |
| ENSG00000265010 | 10.23896558 | -4.312247173 | 0.7130897 | -6.047271 | 1.47E-09 | 3.92E-08 | | sense_intronic | 44.73 | -31.9386972 | 31.9386972 | ENSG00000265010 | down |
| ENSG00000270883 | 21.38911669 | -3.032344327 | 0.4224011 | -7.178827 | 7.03E-13 | 3.43E-11 | | lincRNA | 41.47 | -31.7345039 | 31.7345039 | ENSG00000270883 | down |
| ENSG00000236153 | 37.34533871 | 2.64363388 | 0.343996 | 7.685072 | 1.53E-14 | 1.01E-12 | | antisense | 46.54 | 31.70787314 | 31.70787314 | ENSG00000236153 | up |
| ENSG00000253257 | 12.54024252 | -4.106096303 | 0.6667277 | -6.15858 | 7.34E-10 | 2.08E-08 | MTND4P7 | processed_pseudogene | 35.38 | -31.54160664 | 31.54160664 | ENSG00000253257 | down |
| ENSG00000230673 | 47.85241269 | -2.4524979 | 0.308444 | -7.951192 | 1.85E-15 | 1.44E-13 | PABPC1P3 | processed_pseudogene | 46.62 | -31.49046791 | 31.49046791 | ENSG00000230673 | down |
| ENSG00000275569 | 73.89955696 | -2.390056436 | 0.29667 | -8.05628 | 7.87E-16 | 6.68E-14 | | antisense | 32.5 | -31.48996471 | 31.48996471 | ENSG00000275569 | down |
| ENSG00000235313 | 20.26239874 | -3.044894347 | 0.4266973 | -7.135959 | 9.61E-13 | 4.63E-11 | HM13-IT1 | sense_intronic | 52.69 | -31.46707933 | 31.46707933 | ENSG00000235313 | down |
| ENSG00000253784 | 79.34962008 | 2.240221509 | 0.2702943 | 8.288083 | 1.15E-16 | 1.18E-14 | | lincRNA | 42.76 | 31.20354553 | 31.20354553 | ENSG00000253784 | up |
| ENSG00000207763 | 12.50454915 | -4.27678994 | 0.7143388 | -5.987061 | 2.14E-09 | 5.49E-08 | MIR617 | miRNA | 45.36 | -31.05175527 | 31.05175527 | ENSG00000207763 | down |

| | | | | | | | | | | | | | |
|---|---|---|---|---|---|---|---|---|---|---|---|---|---|
| ENSG00000261351 | 79.66267118 | -2.304504136 | 0.2840543 | -8.1129 | 4.94E-16 | 4.36E-14 | | antisense | 45.05 | -30.79041127 | 30.79041127 | ENSG00000261351 | down |
| ENSG00000171223 | 1388.699705 | 2.052274965 | 0.2389004 | 8.590506 | 8.66E-18 | 1.14E-15 | JUNB | protein_coding | 64.89 | 30.66944484 | 30.66944484 | ENSG00000171223 | up |
| ENSG00000240032 | 45.08850945 | -2.584731102 | 0.3392284 | -7.619442 | 2.55E-14 | 1.61E-12 | LNCSRLR | antisense | 34.2 | -30.48052673 | 30.48052673 | ENSG00000240032 | down |
| ENSG00000260949 | 132.5443358 | -2.326902685 | 0.2897835 | -8.029797 | 9.76E-16 | 8.09E-14 | | sense_overlapping | 46.98 | -30.4637505 | 30.4637505 | ENSG00000260949 | down |
| ENSG00000272777 | 74.59139209 | -2.328760241 | 0.2902354 | -8.023694 | 1.03E-15 | 8.40E-14 | | lincRNA | 35.4 | -30.44974866 | 30.44974866 | ENSG00000272777 | down |
| ENSG00000246863 | 135.4750363 | 2.097775074 | 0.2491896 | 8.418389 | 3.82E-17 | 4.32E-15 | | lincRNA | 41.23 | 30.13339114 | 30.13339114 | ENSG00000246863 | up |
| ENSG00000275029 | 35.22789614 | -2.749200251 | 0.374199 | -7.346894 | 2.03E-13 | 1.11E-11 | HMGB1P24 | transcribed_processed_pseudog | 44.34 | -30.11444656 | 30.11444656 | ENSG00000275029 | down |
| ENSG00000255115 | 11.57289001 | -3.735760398 | 0.5921475 | -6.308834 | 2.81E-10 | 8.74E-09 | | processed_pseudogene | 41.26 | -30.10505047 | 30.10505047 | ENSG00000255115 | down |
| ENSG00000252768 | 10.68150361 | -4.209533561 | 0.7102888 | -5.92651 | 3.09E-09 | 7.69E-08 | RNU6-856P | snRNA | 40.82 | -29.94597564 | 29.94597564 | ENSG00000252768 | down |
| ENSG00000229692 | 60.7324052 | -2.619796637 | 0.3497059 | -7.491427 | 6.81E-14 | 3.97E-12 | SOS1-IT1 | sense_intronic | 38.13 | -29.86918112 | 29.86918112 | ENSG00000229692 | down |
| ENSG00000270135 | 11.86728925 | -3.615951647 | 0.5668998 | -6.378467 | 1.79E-10 | 5.79E-09 | | antisense | 29.45 | -29.78531869 | 29.78531869 | ENSG00000270135 | down |
| ENSG00000266680 | 39.05016829 | 2.562506864 | 0.3387268 | 7.565113 | 3.88E-14 | 2.40E-12 | | antisense | 45.23 | 29.77671092 | 29.77671092 | ENSG00000266680 | up |
| ENSG00000274315 | 68.27442201 | -2.221634661 | 0.2735124 | -8.122611 | 4.56E-16 | 4.05E-14 | | antisense | 33.4 | -29.7542502 | 29.7542502 | ENSG00000274315 | down |
| ENSG00000233072 | 14.81811334 | -3.302950877 | 0.4970833 | -6.644662 | 3.04E-11 | 1.12E-09 | RPS15AP6 | processed_pseudogene | 41.3 | -29.56853317 | 29.56853317 | ENSG00000233072 | down |
| ENSG00000206786 | 4.618405356 | -6.050051622 | 1.237431 | -4.889203 | 1.01E-06 | 1.38E-05 | RNU6-701P | snRNA | 40.19 | -29.40533418 | 29.40533418 | ENSG00000206786 | down |
| ENSG00000259959 | 242.4497406 | -2.114580305 | 0.2556088 | -8.272722 | 1.31E-16 | 1.32E-14 | | lincRNA | 37.48 | -29.35069411 | 29.35069411 | ENSG00000259959 | down |
| ENSG00000237370 | 12.73484399 | 3.529886793 | 0.5516317 | 6.398992 | 1.56E-10 | 5.11E-09 | | antisense | 58.51 | 29.26778716 | 29.26778716 | ENSG00000237370 | up |
| ENSG00000184530 | 191.1303528 | 1.936864559 | 0.2242154 | 8.63841 | 5.70E-18 | 7.78E-16 | C6orf58 | protein_coding | 35.52 | 29.26406293 | 29.26406293 | ENSG00000184530 | up |
| ENSG00000271789 | 58.08649013 | -2.368067146 | 0.3038732 | -7.792944 | 6.55E-15 | 4.70E-13 | | lincRNA | 39.49 | -29.19366382 | 29.19366382 | ENSG00000271789 | down |
| ENSG00000224635 | 132.6635685 | -2.016943036 | 0.2386618 | -8.451053 | 2.89E-17 | 3.41E-15 | | lincRNA | 48.32 | -29.18056852 | 29.18056852 | ENSG00000224635 | down |
| ENSG00000266676 | 12.4865056 | -3.994858868 | 0.665351 | -6.004138 | 1.92E-09 | 4.98E-08 | MIR4297 | miRNA | 68.42 | -29.17426968 | 29.17426968 | ENSG00000266676 | down |
| ENSG00000273449 | 164.1207349 | -2.131669045 | 0.2595843 | -8.211857 | 2.18E-16 | 2.08E-14 | | lincRNA | 33.87 | -29.16671788 | 29.16671788 | ENSG00000273449 | down |
| ENSG00000145012 | 17.67406955 | -3.229737838 | 0.4856392 | -6.650489 | 2.92E-11 | 1.08E-09 | LPP | protein_coding | 39.17 | -28.9649222 | 28.9649222 | ENSG00000145012 | down |
| ENSG00000200651 | 12.87729214 | -3.436721639 | 0.533206 | -6.445392 | 1.15E-10 | 3.80E-09 | | misc_RNA | 42.86 | -28.93631366 | 28.93631366 | ENSG00000200651 | down |
| ENSG00000270574 | 158.1867893 | -2.237204947 | 0.2814517 | -7.948805 | 1.88E-15 | 1.46E-13 | | antisense | 40.39 | -28.71279413 | 28.71279413 | ENSG00000270574 | down |
| ENSG00000166603 | 37.38600675 | 2.469613561 | 0.3266942 | 7.559405 | 4.05E-14 | 2.47E-12 | MC4R | protein_coding | 43.7 | 28.66440011 | 28.66440011 | ENSG00000166603 | up |
| ENSG00000188394 | 34.71238511 | -2.589930029 | 0.3509351 | -7.380083 | 1.58E-13 | 8.88E-12 | GPR21 | protein_coding | 45.81 | -28.62270924 | 28.62270924 | ENSG00000188394 | down |
| ENSG00000187987 | 352.9475559 | -2.109145939 | 0.2583335 | -8.16443 | 3.23E-16 | 2.97E-14 | ZSCAN23 | protein_coding | 39.86 | -28.52954811 | 28.52954811 | ENSG00000187987 | down |
| ENSG00000260278 | 94.59994703 | 2.173422057 | 0.2705887 | 8.032197 | 9.57E-16 | 7.98E-14 | | antisense | 45.09 | 28.46721877 | 28.46721877 | ENSG00000260278 | up |
| ENSG00000207563 | 25.98121263 | -2.65014306 | 0.364874 | -7.263174 | 3.78E-13 | 1.97E-11 | MIR23B | miRNA | 50.52 | -28.37020435 | 28.37020435 | ENSG00000207563 | down |
| ENSG00000242999 | 11.04143097 | -3.407006628 | 0.5341305 | -6.378603 | 1.79E-10 | 5.79E-09 | RN7SL239P | misc_RNA | 53.67 | -28.06419668 | 28.06419668 | ENSG00000242999 | down |
| ENSG00000274308 | 32.27142146 | -2.618826792 | 0.3607743 | -7.258907 | 3.90E-13 | 2.01E-11 | | processed_pseudogene | 55.77 | -28.01178308 | 28.01178308 | ENSG00000274308 | down |

| | | | | | | | | | | | | | |
|---|---|---|---|---|---|---|---|---|---|---|---|---|---|
| ENSG00000143970 | 1083.881102 | -2.230560708 | 0.2837332 | -7.861473 | 3.80E-15 | 2.84E-13 | ASXL2 | protein_coding | 39.53 | -27.98551571 | 27.98551571 | ENSG00000143970 | down |
| ENSG00000235369 | 8.743060718 | -4.391633083 | 0.7837728 | -5.603196 | 2.10E-08 | 4.39E-07 | RPL36AP15 | processed_pseudogene | 44.74 | -27.91836574 | 27.91836574 | ENSG00000235369 | down |
| ENSG00000273243 | 27.89136538 | -2.749691805 | 0.3898688 | -7.052865 | 1.75E-12 | 8.03E-11 | | lincRNA | 43.97 | -27.75932128 | 27.75932128 | ENSG00000273243 | down |
| ENSG00000223787 | 15.09698743 | -3.450221635 | 0.5479534 | -6.29656 | 3.04E-10 | 9.35E-09 | | processed_pseudogene | 40.75 | -27.70193296 | 27.70193296 | ENSG00000223787 | down |
| ENSG00000263755 | 11.51244434 | -3.426126345 | 0.5437587 | -6.300821 | 2.96E-10 | 9.14E-09 | RN7SL498P | misc_RNA | 54.36 | -27.54278603 | 27.54278603 | ENSG00000263755 | down |
| ENSG00000257999 | 36.01901565 | 2.43374141 | 0.3263443 | 7.457587 | 8.81E-14 | 5.03E-12 | | antisense | 39.27 | 27.49781544 | 27.49781544 | ENSG00000257999 | up |
| ENSG00000235518 | 47.00196133 | -2.519522346 | 0.3440768 | -7.322558 | 2.43E-13 | 1.32E-11 | | lincRNA | 41.75 | -27.40833613 | 27.40833613 | ENSG00000235518 | down |
| ENSG00000283236 | 157.0694302 | -2.228086131 | 0.2875191 | -7.749349 | 9.24E-15 | 6.40E-13 | | transcribed_unprocessed_pseud | 38.76 | -27.16918402 | 27.16918402 | ENSG00000283236 | down |
| ENSG00000199630 | 17.19110508 | -2.995555904 | 0.4477978 | -6.689528 | 2.24E-11 | 8.52E-10 | | misc_RNA | 40.62 | -27.16882731 | 27.16882731 | ENSG00000199630 | down |
| ENSG00000270015 | 161.0249652 | -2.223224107 | 0.287318 | -7.737852 | 1.01E-14 | 6.90E-13 | | sense_intronic | 51.16 | -27.0368531 | 27.0368531 | ENSG00000270015 | down |
| ENSG00000241030 | 14.92492809 | -3.448146482 | 0.5541745 | -6.222131 | 4.90E-10 | 1.45E-08 | | processed_pseudogene | 54.23 | -27.03215011 | 27.03215011 | ENSG00000241030 | down |
| ENSG00000256274 | 170.4469715 | -2.358587609 | 0.3143358 | -7.503401 | 6.22E-14 | 3.68E-12 | | transcribed_processed_pseudog | 33.67 | -26.96729826 | 26.96729826 | ENSG00000256274 | down |
| ENSG00000230325 | 15.76393098 | -2.936308085 | 0.4363505 | -6.729243 | 1.71E-11 | 6.60E-10 | | antisense | 43.55 | -26.9574272 | 26.9574272 | ENSG00000230325 | down |
| ENSG00000268912 | 125.2606965 | -2.191528324 | 0.2818592 | -7.77526 | 7.53E-15 | 5.35E-13 | | lincRNA | 45.69 | -26.8939309 | 26.8939309 | ENSG00000268912 | down |
| ENSG00000264370 | 9.132718697 | -4.079247164 | 0.7174572 | -5.685701 | 1.30E-08 | 2.82E-07 | MIR3125 | miRNA | 57.69 | -26.71748146 | 26.71748146 | ENSG00000264370 | down |
| ENSG00000279233 | 92.0945554 | -2.064761993 | 0.2587967 | -7.978315 | 1.48E-15 | 1.17E-13 | | TEC | 53.04 | -26.6984237 | 26.6984237 | ENSG00000279233 | down |
| ENSG00000274115 | 73.88855007 | -2.238570384 | 0.2928033 | -7.645304 | 2.08E-14 | 1.33E-12 | MIR6081 | miRNA | 64.95 | -26.58462665 | 26.58462665 | ENSG00000274115 | down |
| ENSG00000252452 | 12.75138457 | -3.313692904 | 0.5266802 | -6.29166 | 3.14E-10 | 9.59E-09 | RNU6-107P | snRNA | 44.12 | -26.56978384 | 26.56978384 | ENSG00000252452 | down |
| ENSG00000280028 | 116.2159107 | -2.167556655 | 0.2790905 | -7.7665 | 8.07E-15 | 5.67E-13 | | TEC | 58.77 | -26.54420576 | 26.54420576 | ENSG00000280028 | down |
| ENSG00000242689 | 114.9822087 | -2.189016399 | 0.283542 | -7.720255 | 1.16E-14 | 7.88E-13 | CNTF | protein_coding | 41.59 | -26.4941014 | 26.4941014 | ENSG00000242689 | down |
| ENSG00000225527 | 32.29398957 | -2.518773158 | 0.3502314 | -7.191741 | 6.40E-13 | 3.17E-11 | | antisense | 37.37 | -26.44458499 | 26.44458499 | ENSG00000225527 | down |
| ENSG00000253057 | 16.6000114 | -3.198101821 | 0.5017568 | -6.373809 | 1.84E-10 | 5.94E-09 | RN7SKP57 | misc_RNA | 44.71 | -26.30762604 | 26.30762604 | ENSG00000253057 | down |
| ENSG00000199781 | 11.45966629 | -3.46978431 | 0.5676633 | -6.112399 | 9.81E-10 | 2.72E-08 | | misc_RNA | 36.36 | -26.25227929 | 26.25227929 | ENSG00000199781 | down |
| ENSG00000207759 | 8.916595342 | -4.697046508 | 0.8953803 | -5.245868 | 1.56E-07 | 2.61E-06 | MIR181A1 | miRNA | 41.82 | -26.2258638 | 26.2258638 | ENSG00000207759 | down |
| ENSG00000204103 | 162.2599208 | 1.942934195 | 0.2384197 | 8.14922 | 3.66E-16 | 3.27E-14 | MAFB | protein_coding | 56.44 | 26.20159404 | 26.20159404 | ENSG00000204103 | up |
| ENSG00000240531 | 17.91889137 | -3.077794581 | 0.4749514 | -6.48023 | 9.16E-11 | 3.09E-09 | | transcribed_processed_pseudog | 41.28 | -26.19327289 | 26.19327289 | ENSG00000240531 | down |
| ENSG00000258640 | 22.0889825 | -2.850657927 | 0.4238663 | -6.72537 | 1.75E-11 | 6.76E-10 | RPL21P5 | processed_pseudogene | 45.75 | -26.14159751 | 26.14159751 | ENSG00000258640 | down |
| ENSG00000251754 | 3.289310102 | -5.830537458 | 1.2431449 | -4.690151 | 2.73E-06 | 3.34E-05 | RNU6-999P | snRNA | 44.76 | -26.09666868 | 26.09666868 | ENSG00000251754 | down |
| ENSG00000180787 | 181.8969891 | -2.094181467 | 0.2673479 | -7.83317 | 4.76E-15 | 3.49E-13 | ZFP3 | protein_coding | 45.78 | -26.08862312 | 26.08862312 | ENSG00000180787 | down |
| ENSG00000227583 | 37.20641194 | -2.582631147 | 0.3669655 | -7.037803 | 1.95E-12 | 8.83E-11 | RPS3AP37 | processed_pseudogene | 39.6 | -25.96610217 | 25.96610217 | ENSG00000227583 | down |
| ENSG00000214773 | 102.1454725 | -2.047052546 | 0.2593629 | -7.892619 | 2.96E-15 | 2.24E-13 | | sense_intronic | 45.6 | -25.8950612 | 25.8950612 | ENSG00000214773 | down |
| ENSG00000131849 | 292.8114371 | -1.947387059 | 0.240784 | -8.087691 | 6.08E-16 | 5.26E-14 | ZNF132 | protein_coding | 48.36 | -25.85963438 | 25.85963438 | ENSG00000131849 | down |

| | | | | | | | | | | | | | |
|---|---|---|---|---|---|---|---|---|---|---|---|---|---|
| ENSG00000137331 | 82.37703562 | 2.150899132 | 0.2796223 | 7.692159 | 1.45E-14 | 9.68E-13 | IER3 | protein_coding | 58.55 | 25.84078374 | 25.84078374 | ENSG00000137331 | up |
| ENSG00000213842 | 37.93585712 | -2.468267671 | 0.3438301 | -7.178743 | 7.04E-13 | 3.43E-11 | SUGT1P2 | processed_pseudogene | 38.41 | -25.83125186 | 25.83125186 | ENSG00000213842 | down |
| ENSG00000180938 | 213.8207728 | -1.864556495 | 0.2259412 | -8.252399 | 1.55E-16 | 1.52E-14 | ZNF572 | protein_coding | 39.84 | -25.76357344 | 25.76357344 | ENSG00000180938 | down |
| ENSG00000220744 | 39.82749217 | -2.507010288 | 0.3524349 | -7.113399 | 1.13E-12 | 5.34E-11 | RPL5P18 | processed_pseudogene | 40.52 | -25.75243615 | 25.75243615 | ENSG00000220744 | down |
| ENSG00000264814 | 7.975240942 | -4.287646563 | 0.78785 | -5.442212 | 5.26E-08 | 1.01E-06 | MIR1273C | miRNA | 49.35 | -25.71141047 | 25.71141047 | ENSG00000264814 | down |
| ENSG00000240729 | 33.1550168 | -2.494543205 | 0.3503212 | -7.120731 | 1.07E-12 | 5.12E-11 | | processed_pseudogene | 42.23 | -25.67090289 | 25.67090289 | ENSG00000240729 | down |
| ENSG00000253476 | 17.02038868 | -2.93579741 | 0.4470259 | -6.567399 | 5.12E-11 | 1.83E-09 | | lincRNA | 40.57 | -25.6531332 | 25.6531332 | ENSG00000253476 | down |
| ENSG00000256064 | 25.38882772 | -2.629799349 | 0.3794865 | -6.929889 | 4.21E-12 | 1.77E-10 | | lincRNA | 37.64 | -25.64484733 | 25.64484733 | ENSG00000256064 | down |
| ENSG00000273320 | 51.81977802 | 2.200838698 | 0.2911096 | 7.560172 | 4.03E-14 | 2.47E-12 | | antisense | 47.11 | 25.54620775 | 25.54620775 | ENSG00000273320 | up |
| ENSG00000278743 | 87.97979585 | 2.063441506 | 0.2642522 | 7.808608 | 5.78E-15 | 4.22E-13 | | lincRNA | 60.14 | 25.53549781 | 25.53549781 | ENSG00000278743 | up |
| ENSG00000235613 | 53.12830399 | -2.21611301 | 0.2954681 | -7.500347 | 6.36E-14 | 3.75E-12 | NSRP1P1 | processed_pseudogene | 41.35 | -25.32226464 | 25.32226464 | ENSG00000235613 | down |
| ENSG00000135747 | 117.6832769 | -2.170979536 | 0.286768 | -7.570507 | 3.72E-14 | 2.31E-12 | ZNF670-ZNF695 | protein_coding | 42.2 | -25.26204296 | 25.26204296 | ENSG00000135747 | down |
| ENSG00000261072 | 126.9439348 | -2.274947354 | 0.3085363 | -7.373353 | 1.66E-13 | 9.27E-12 | | processed_pseudogene | 44.95 | -25.09968365 | 25.09968365 | ENSG00000261072 | down |
| ENSG00000229816 | 91.91605166 | -2.131056848 | 0.2805819 | -7.595132 | 3.07E-14 | 1.94E-12 | DDX50P1 | processed_pseudogene | 42.67 | -24.9607383 | 24.9607383 | ENSG00000229816 | down |
| ENSG00000276116 | 54.05623109 | 2.227223019 | 0.3002895 | 7.416918 | 1.20E-13 | 6.78E-12 | FUT8-AS1 | antisense | 56.67 | 24.8746673 | 24.8746673 | ENSG00000276116 | up |
| ENSG00000267632 | 117.9785127 | -1.955206924 | 0.2470851 | -7.913091 | 2.51E-15 | 1.94E-13 | | sense_intronic | 50.58 | -24.85407102 | 24.85407102 | ENSG00000267632 | down |
| ENSG00000260369 | 206.506143 | 1.89313107 | 0.2358371 | 8.027284 | 9.97E-16 | 8.21E-14 | | lincRNA | 61.81 | 24.77286722 | 24.77286722 | ENSG00000260369 | up |
| ENSG00000144802 | 130.1778436 | 1.907528163 | 0.2386243 | 7.993854 | 1.31E-15 | 1.05E-13 | NFKBIZ | protein_coding | 40.23 | 24.76021367 | 24.76021367 | ENSG00000144802 | up |
| ENSG00000260350 | 35.83306665 | -2.465934741 | 0.3506274 | -7.032922 | 2.02E-12 | 9.11E-11 | | antisense | 57.21 | -24.75880122 | 24.75880122 | ENSG00000260350 | down |
| ENSG00000261002 | 22.11119763 | 2.621742185 | 0.3843115 | 6.82192 | 8.98E-12 | 3.67E-10 | | antisense | 38.94 | 24.73769275 | 24.73769275 | ENSG00000261002 | up |
| ENSG00000268598 | 10.123599 | -3.402596122 | 0.5684854 | -5.985371 | 2.16E-09 | 5.52E-08 | VN1R80P | unprocessed_pseudogene | 40.83 | -24.69746064 | 24.69746064 | ENSG00000268598 | down |
| ENSG00000163263 | 10.24205247 | -3.524329843 | 0.5996171 | -5.877634 | 4.16E-09 | 1.01E-07 | C1orf189 | protein_coding | 46.57 | -24.64786298 | 24.64786298 | ENSG00000163263 | down |
| ENSG00000168209 | 1287.650492 | 2.150576196 | 0.2873313 | 7.484658 | 7.17E-14 | 4.16E-12 | DDIT4 | protein_coding | 58.72 | 24.47517912 | 24.47517912 | ENSG00000168209 | up |
| ENSG00000248476 | 35.63939964 | -2.320484718 | 0.3221418 | -7.203303 | 5.88E-13 | 2.94E-11 | BACH1-IT1 | sense_intronic | 43.74 | -24.43736521 | 24.43736521 | ENSG00000248476 | down |
| ENSG00000259548 | 15.90034505 | -2.798623381 | 0.4264728 | -6.562255 | 5.30E-11 | 1.89E-09 | | lincRNA | 46.83 | -24.41566276 | 24.41566276 | ENSG00000259548 | down |
| ENSG00000256658 | 66.99325873 | 2.052275452 | 0.2684292 | 7.6455 | 2.08E-14 | 1.33E-12 | | antisense | 51.61 | 24.37224091 | 24.37224091 | ENSG00000256658 | up |
| ENSG00000259469 | 26.01171204 | -2.473847292 | 0.355343 | -6.961857 | 3.36E-12 | 1.43E-10 | | lincRNA | 35.16 | -24.35759948 | 24.35759948 | ENSG00000259469 | down |
| ENSG00000276081 | 4.215759924 | -5.696951639 | 1.2472172 | -4.56773 | 4.93E-06 | 5.63E-05 | MIR7156 | miRNA | 55 | -24.20751416 | 24.20751416 | ENSG00000276081 | down |
| ENSG00000252277 | 11.55384355 | -4.217553776 | 0.7941255 | -5.310941 | 1.09E-07 | 1.93E-06 | SNORD116-30 | snoRNA | 35.29 | -24.10384844 | 24.10384844 | ENSG00000252277 | down |
| ENSG00000279696 | 327.7591307 | -1.844961631 | 0.2302671 | -8.012266 | 1.13E-15 | 9.12E-14 | | TEC | 37.34 | -24.05867725 | 24.05867725 | ENSG00000279696 | down |
| ENSG00000279348 | 750.5156812 | -1.969117762 | 0.2543795 | -7.740866 | 9.87E-15 | 6.77E-13 | | TEC | 40.08 | -23.96270683 | 23.96270683 | ENSG00000279348 | down |
| ENSG00000270750 | 48.22341135 | -2.23700585 | 0.3081283 | -7.259981 | 3.87E-13 | 2.00E-11 | | lincRNA | 34.73 | -23.93181416 | 23.93181416 | ENSG00000270750 | down |

| | | | | | | | | | | | | | |
|---|---|---|---|---|---|---|---|---|---|---|---|---|---|
| ENSG00000276672 | 581.548511 | -1.903352271 | 0.2422066 | -7.858384 | 3.89E-15 | 2.90E-13 | | sense_intronic | 39.6 | -23.86427623 | 23.86427623 | ENSG00000276672 | down |
| ENSG00000277806 | 35.93171898 | -2.34505125 | 0.3313241 | -7.077817 | 1.46E-12 | 6.82E-11 | | sense_intronic | 29.65 | -23.84042484 | 23.84042484 | ENSG00000277806 | down |
| ENSG00000199172 | 11.22100338 | -3.613181963 | 0.6330112 | -5.707927 | 1.14E-08 | 2.52E-07 | MIR331 | miRNA | 51.06 | -23.83985385 | 23.83985385 | ENSG00000199172 | down |
| ENSG00000272077 | 160.4844952 | -2.003781801 | 0.2618939 | -7.651121 | 1.99E-14 | 1.28E-12 | | lincRNA | 41.53 | -23.82768375 | 23.82768375 | ENSG00000272077 | down |
| ENSG00000257511 | 38.13316014 | -2.387902069 | 0.3408025 | -7.006703 | 2.44E-12 | 1.07E-10 | | processed_pseudogene | 39.4 | -23.80723771 | 23.80723771 | ENSG00000257511 | down |
| ENSG00000199962 | 4.187936619 | -6.147773108 | 1.4131444 | -4.350421 | 1.36E-05 | 0.000135503 | | misc_RNA | 39.45 | -23.77989419 | 23.77989419 | ENSG00000199962 | down |
| ENSG00000230498 | 40.45739099 | -2.423084948 | 0.3493521 | -6.935938 | 4.04E-12 | 1.70E-10 | | antisense | 34.89 | -23.6708788 | 23.6708788 | ENSG00000230498 | down |
| ENSG00000091844 | 239.2241295 | -2.176667189 | 0.2974098 | -7.318747 | 2.50E-13 | 1.34E-11 | RGS17 | protein_coding | 37.43 | -23.66643748 | 23.66643748 | ENSG00000091844 | down |
| ENSG00000229127 | 188.9909464 | -2.036508443 | 0.2693581 | -7.5606 | 4.01E-14 | 2.47E-12 | | antisense | 37.9 | -23.63874637 | 23.63874637 | ENSG00000229127 | down |
| ENSG00000251988 | 20.53476206 | -2.703493166 | 0.4124883 | -6.554108 | 5.60E-11 | 1.98E-09 | RNU4ATAC18P | snRNA | 47.62 | -23.5275887 | 23.5275887 | ENSG00000251988 | down |
| ENSG00000207417 | 7.308150537 | -3.920110974 | 0.7203845 | -5.441692 | 5.28E-08 | 1.01E-06 | | misc_RNA | 41.12 | -23.5048129 | 23.5048129 | ENSG00000207417 | down |
| ENSG00000222724 | 321.8993503 | 1.762450968 | 0.2176158 | 8.098911 | 5.55E-16 | 4.83E-14 | RNU2-63P | snRNA | 38.66 | 23.46960841 | 23.46960841 | ENSG00000222724 | up |
| ENSG00000104369 | 254.4214475 | -1.970983378 | 0.2574186 | -7.656725 | 1.91E-14 | 1.25E-12 | JPH1 | protein_coding | 39.71 | -23.45898553 | 23.45898553 | ENSG00000104369 | down |
| ENSG00000252326 | 8.338623481 | -3.923635956 | 0.7223478 | -5.431782 | 5.58E-08 | 1.06E-06 | SNORD116-25 | snoRNA | 43.48 | -23.44145104 | 23.44145104 | ENSG00000252326 | down |
| ENSG00000256973 | 54.40198399 | -2.272009292 | 0.3189358 | -7.123719 | 1.05E-12 | 5.04E-11 | | lincRNA | 42.08 | -23.39546103 | 23.39546103 | ENSG00000256973 | down |
| ENSG00000271815 | 23.9537415 | -2.335065321 | 0.3325763 | -7.021141 | 2.20E-12 | 9.78E-11 | | lincRNA | 48.24 | -23.37332923 | 23.37332923 | ENSG00000271815 | down |
| ENSG00000261334 | 61.57961085 | 2.056369016 | 0.2751499 | 7.473631 | 7.80E-14 | 4.49E-12 | | lincRNA | 59.15 | 23.33552009 | 23.33552009 | ENSG00000261334 | up |
| ENSG00000260583 | 30.44130422 | -2.341535478 | 0.3343805 | -7.002607 | 2.51E-12 | 1.10E-10 | | antisense | 44.17 | -23.31841963 | 23.31841963 | ENSG00000260583 | down |
| ENSG00000273199 | 19.00863062 | 2.653961768 | 0.4037639 | 6.573053 | 4.93E-11 | 1.76E-09 | | antisense | 67.35 | 23.23125262 | 23.23125262 | ENSG00000273199 | up |
| ENSG00000280176 | 13.65182454 | -2.867396322 | 0.4549414 | -6.302782 | 2.92E-10 | 9.05E-09 | | TEC | 47.22 | -23.06411994 | 23.06411994 | ENSG00000280176 | down |
| ENSG00000251374 | 9.770409358 | -3.569048448 | 0.6318976 | -5.648143 | 1.62E-08 | 3.45E-07 | | processed_pseudogene | 39.81 | -23.06255163 | 23.06255163 | ENSG00000251374 | down |
| ENSG00000272102 | 25.01695721 | -2.533031184 | 0.378204 | -6.697525 | 2.12E-11 | 8.09E-10 | | lincRNA | 39.93 | -23.03099934 | 23.03099934 | ENSG00000272102 | down |
| ENSG00000256229 | 639.0795192 | -2.068688327 | 0.2794266 | -7.403334 | 1.33E-13 | 7.49E-12 | ZNF486 | protein_coding | 40.9 | -23.01569088 | 23.01569088 | ENSG00000256229 | down |
| ENSG00000217331 | 62.42700091 | -2.237530835 | 0.314417 | -7.116444 | 1.11E-12 | 5.24E-11 | | processed_pseudogene | 30.92 | -23.00244769 | 23.00244769 | ENSG00000217331 | down |
| ENSG00000270100 | 29.76893402 | -2.485593809 | 0.3681031 | -6.75244 | 1.45E-11 | 5.72E-10 | | lincRNA | 40.06 | -22.97376394 | 22.97376394 | ENSG00000270100 | down |
| ENSG00000222222 | 49.31724827 | 2.181141807 | 0.303275 | 7.191961 | 6.39E-13 | 3.17E-11 | RNU2-17P | snRNA | 42.63 | 22.89979537 | 22.89979537 | ENSG00000222222 | up |
| ENSG00000244356 | 15.35257372 | -3.00937209 | 0.4929963 | -6.104249 | 1.03E-09 | 2.84E-08 | RN7SL398P | misc_RNA | 53.36 | -22.70986748 | 22.70986748 | ENSG00000244356 | down |
| ENSG00000206922 | 16.33843451 | -2.945898703 | 0.4778239 | -6.165239 | 7.04E-10 | 2.01E-08 | RNU6-80P | snRNA | 41.12 | -22.67534066 | 22.67534066 | ENSG00000206922 | down |
| ENSG00000146426 | 289.4563043 | -1.898192192 | 0.2476297 | -7.665447 | 1.78E-14 | 1.18E-12 | TIAM2 | protein_coding | 43.49 | -22.64470841 | 22.64470841 | ENSG00000146426 | down |
| ENSG00000276696 | 7.239094718 | -4.463810858 | 0.8970785 | -4.975942 | 6.49E-07 | 9.19E-06 | | misc_RNA | 35.38 | -22.4833668 | 22.4833668 | ENSG00000276696 | down |
| ENSG00000133740 | 434.7147176 | -2.100301353 | 0.2892099 | -7.262205 | 3.81E-13 | 1.98E-11 | E2F5 | protein_coding | 39.49 | -22.48093509 | 22.48093509 | ENSG00000133740 | down |
| ENSG00000203506 | 193.7705031 | 1.769605679 | 0.223901 | 7.903517 | 2.71E-15 | 2.07E-13 | RBMS3-AS2 | antisense | 36.04 | 22.44415887 | 22.44415887 | ENSG00000203506 | up |

| | | | | | | | | | | | | | |
|---|---|---|---|---|---|---|---|---|---|---|---|---|---|
| ENSG00000199122 | 5.361896316 | -5.50202836 | 1.2300368 | -4.47306 | 7.71E-06 | 8.34E-05 | MIR148B | miRNA | 43.43 | -22.44137141 | 22.44137141 | ENSG00000199122 | down |
| ENSG00000199664 | 3.134662649 | -5.558970649 | 1.2503704 | -4.445859 | 8.75E-06 | 9.30E-05 | RNU6-1266P | snRNA | 39.05 | -22.41002008 | 22.41002008 | ENSG00000199664 | down |
| ENSG00000186594 | 53.82664368 | 2.124306973 | 0.2954343 | 7.190456 | 6.46E-13 | 3.18E-11 | MIR22HG | lincRNA | 56.95 | 22.30094638 | 22.30094638 | ENSG00000186594 | up |
| ENSG00000207082 | 13.03597951 | -3.133888505 | 0.5287926 | -5.926498 | 3.09E-09 | 7.69E-08 | RNU6-171P | snRNA | 43.93 | -22.29400181 | 22.29400181 | ENSG00000207082 | down |
| ENSG00000278861 | 21.68597242 | -2.565031754 | 0.3921056 | -6.541686 | 6.08E-11 | 2.13E-09 | | TEC | 37.44 | -22.2412539 | 22.2412539 | ENSG00000278861 | down |
| ENSG00000259818 | 88.19070429 | 1.932390439 | 0.2568237 | 7.52419 | 5.30E-14 | 3.18E-12 | | antisense | 47.12 | 22.21685425 | 22.21685425 | ENSG00000259818 | up |
| ENSG00000206676 | 9.368629208 | -4.099702742 | 0.7957404 | -5.15206 | 2.58E-07 | 4.14E-06 | | misc_RNA | 38.83 | -22.06981895 | 22.06981895 | ENSG00000206676 | down |
| ENSG00000196267 | 148.4559479 | -2.098687592 | 0.2919289 | -7.189036 | 6.53E-13 | 3.20E-11 | ZNF836 | protein_coding | 43.18 | -22.0257314 | 22.0257314 | ENSG00000196267 | down |
| ENSG00000222445 | 11.79319266 | -3.197627584 | 0.5491217 | -5.823168 | 5.77E-09 | 1.36E-07 | RN7SKP56 | misc_RNA | 45.62 | -21.95370724 | 21.95370724 | ENSG00000222445 | down |
| ENSG00000259707 | 17.81023575 | -2.565512659 | 0.3949045 | -6.49654 | 8.22E-11 | 2.80E-09 | | processed_pseudogene | 52.2 | -21.94056718 | 21.94056718 | ENSG00000259707 | down |
| ENSG00000000457 | 418.1321252 | -2.07257964 | 0.2869669 | -7.222365 | 5.11E-13 | 2.60E-11 | SCYL3 | protein_coding | 40.14 | -21.93965095 | 21.93965095 | ENSG00000000457 | down |
| ENSG00000237887 | 43.62714216 | -2.159964051 | 0.3057621 | -7.064199 | 1.62E-12 | 7.45E-11 | RPL23AP32 | processed_pseudogene | 48.8 | -21.87607103 | 21.87607103 | ENSG00000237887 | down |
| ENSG00000276216 | 751.9746772 | 1.630308003 | 0.2012004 | 8.102908 | 5.37E-16 | 4.70E-14 | | lincRNA | 49.57 | 21.72875837 | 21.72875837 | ENSG00000276216 | up |
| ENSG00000203721 | 57.55954304 | -2.169365836 | 0.3104245 | -6.988384 | 2.78E-12 | 1.21E-10 | LINC00862 | lincRNA | 42.07 | -21.51551066 | 21.51551066 | ENSG00000203721 | down |
| ENSG00000249212 | 69.72479258 | -2.167374044 | 0.3101822 | -6.987421 | 2.80E-12 | 1.21E-10 | ATP1B1P1 | unprocessed_pseudogene | 43.17 | -21.49382576 | 21.49382576 | ENSG00000249212 | down |
| ENSG00000161551 | 332.3515189 | -2.128826515 | 0.3019506 | -7.050247 | 1.79E-12 | 8.13E-11 | ZNF577 | protein_coding | 41 | -21.48008815 | 21.48008815 | ENSG00000161551 | down |
| ENSG00000067798 | 46.12474458 | 2.167518142 | 0.3103751 | 6.983545 | 2.88E-12 | 1.24E-10 | NAV3 | protein_coding | 35.89 | 21.47217973 | 21.47217973 | ENSG00000067798 | up |
| ENSG00000252549 | 11.7651219 | -3.03674954 | 0.5145739 | -5.901484 | 3.60E-09 | 8.85E-08 | RNU6-759P | snRNA | 39.81 | -21.41895461 | 21.41895461 | ENSG00000252549 | down |
| ENSG00000272418 | 13.48693826 | -3.028731516 | 0.5126481 | -5.908013 | 3.46E-09 | 8.53E-08 | | sense_intronic | 43.51 | -21.40984723 | 21.40984723 | ENSG00000272418 | down |
| ENSG00000260855 | 153.5032236 | -2.076648228 | 0.2915803 | -7.122045 | 1.06E-12 | 5.09E-11 | | antisense | 38.91 | -21.37592862 | 21.37592862 | ENSG00000260855 | down |
| ENSG00000278492 | 15.53029432 | -2.666583197 | 0.4239616 | -6.28968 | 3.18E-10 | 9.69E-09 | | sense_intronic | 32.64 | -21.36890945 | 21.36890945 | ENSG00000278492 | down |
| ENSG00000179029 | 876.9072595 | 1.617508283 | 0.2009542 | 8.049141 | 8.34E-16 | 7.04E-14 | TMEM107 | protein_coding | 49.51 | 21.27460755 | 21.27460755 | ENSG00000179029 | up |
| ENSG00000261226 | 154.9277207 | 1.898113728 | 0.2555156 | 7.428563 | 1.10E-13 | 6.24E-12 | | sense_intronic | 61.68 | 21.26816371 | 21.26816371 | ENSG00000261226 | up |
| ENSG00000222522 | 8.307713218 | -3.598099461 | 0.6673226 | -5.391845 | 6.97E-08 | 1.29E-06 | RNU6-519P | snRNA | 39.81 | -21.19133815 | 21.19133815 | ENSG00000222522 | down |
| ENSG00000274349 | 141.1081314 | -1.922732209 | 0.2612556 | -7.359582 | 1.84E-13 | 1.02E-11 | ZNF658 | protein_coding | 38.33 | -21.1307786 | 21.1307786 | ENSG00000274349 | down |
| ENSG00000207004 | 24.95801567 | -2.493032354 | 0.3856208 | -6.464985 | 1.01E-10 | 3.39E-09 | RNU6-301P | snRNA | 44.86 | -21.11526664 | 21.11526664 | ENSG00000207004 | down |
| ENSG00000233012 | 114.4381669 | -1.930574132 | 0.2630683 | -7.33868 | 2.16E-13 | 1.18E-11 | HDAC1P2 | transcribed_processed_pseudog | 46.87 | -21.09911753 | 21.09911753 | ENSG00000233012 | down |
| ENSG00000207134 | 4.850406087 | -4.916676691 | 1.0752134 | -4.572745 | 4.81E-06 | 5.54E-05 | RNU6-106P | snRNA | 37.38 | -20.92640265 | 20.92640265 | ENSG00000207134 | down |
| ENSG00000167554 | 218.7348522 | -1.792518016 | 0.2366166 | -7.575621 | 3.57E-14 | 2.23E-12 | ZNF610 | protein_coding | 44.88 | -20.88534753 | 20.88534753 | ENSG00000167554 | down |
| ENSG00000243845 | 11.95626933 | -3.868878301 | 0.749938 | -5.158931 | 2.48E-07 | 4.00E-06 | RN7SL30P | misc_RNA | 55.56 | -20.88496005 | 20.88496005 | ENSG00000243845 | down |
| ENSG00000227630 | 22.67648464 | 2.446487168 | 0.3770792 | 6.487993 | 8.70E-11 | 2.95E-09 | LINC01132 | lincRNA | 50.62 | 20.86754493 | 20.86754493 | ENSG00000227630 | up |
| ENSG00000267334 | 106.5195072 | 1.89051791 | 0.256332 | 7.375272 | 1.64E-13 | 9.17E-12 | | lincRNA | 49.61 | 20.86677467 | 20.86677467 | ENSG00000267334 | up |

| | | | | | | | | | | | | | |
|---|---|---|---|---|---|---|---|---|---|---|---|---|---|
| ENSG00000208037 | 11.06465665 | 3.221779454 | 0.5702891 | 5.649379 | 1.61E-08 | 3.43E-07 | MIR320A | miRNA | 59.76 | 20.82647988 | 20.82647988 | ENSG00000208037 | up |
| ENSG00000237021 | 1189.086467 | 1.608687032 | 0.2016658 | 7.976995 | 1.50E-15 | 1.18E-13 | | lincRNA | 34.41 | 20.79762807 | 20.79762807 | ENSG00000237021 | up |
| ENSG00000231307 | 34.90793866 | -2.215918603 | 0.3260656 | -6.795929 | 1.08E-11 | 4.30E-10 | RPS3P2 | processed_pseudogene | 46.93 | -20.75537327 | 20.75537327 | ENSG00000231307 | down |
| ENSG00000167625 | 216.1325404 | -1.781826191 | 0.2366238 | -7.530208 | 5.07E-14 | 3.07E-12 | ZNF526 | protein_coding | 54.5 | -20.51475803 | 20.51475803 | ENSG00000167625 | down |
| ENSG00000264229 | 321.6100673 | 1.693796704 | 0.2194643 | 7.717868 | 1.18E-14 | 8.00E-13 | RNU4ATAC | snRNA | 45.67 | 20.49016431 | 20.49016431 | ENSG00000264229 | up |
| ENSG00000265452 | 16.86614771 | -2.666106024 | 0.433228 | -6.154048 | 7.55E-10 | 2.13E-08 | MIR3682 | miRNA | 29.76 | -20.45169092 | 20.45169092 | ENSG00000265452 | down |
| ENSG00000272788 | 18.35469517 | -2.501681422 | 0.3943402 | -6.343968 | 2.24E-10 | 7.07E-09 | | sense_intronic | 45.54 | -20.39023085 | 20.39023085 | ENSG00000272788 | down |
| ENSG00000118260 | 1854.780452 | -1.990851812 | 0.2806064 | -7.094819 | 1.30E-12 | 6.09E-11 | CREB1 | protein_coding | 37.47 | -20.33698932 | 20.33698932 | ENSG00000118260 | down |
| ENSG00000231010 | 71.70141699 | 2.007281302 | 0.2841235 | 7.064819 | 1.61E-12 | 7.44E-11 | | lincRNA | 46.96 | 20.33070896 | 20.33070896 | ENSG00000231010 | up |
| ENSG00000255647 | 132.7198766 | 1.779145552 | 0.237216 | 7.500108 | 6.38E-14 | 3.75E-12 | | processed_transcript | 46.57 | 20.32928568 | 20.32928568 | ENSG00000255647 | up |
| ENSG00000260686 | 90.5223772 | -1.918530945 | 0.2656339 | -7.222463 | 5.11E-13 | 2.60E-11 | | sense_overlapping | 31.22 | -20.30894178 | 20.30894178 | ENSG00000260686 | down |
| ENSG00000116741 | 1574.90265 | 1.54934238 | 0.192861 | 8.033468 | 9.48E-16 | 7.95E-14 | RGS2 | protein_coding | 40.83 | 20.29603163 | 20.29603163 | ENSG00000116741 | up |
| ENSG00000207975 | 15.61864885 | -2.730337535 | 0.4513847 | -6.048804 | 1.46E-09 | 3.89E-08 | MIR181B1 | miRNA | 44.55 | -20.23127375 | 20.23127375 | ENSG00000207975 | down |
| ENSG00000261762 | 206.3247296 | -1.996727968 | 0.2825738 | -7.066219 | 1.59E-12 | 7.39E-11 | | antisense | 39.46 | -20.22967241 | 20.22967241 | ENSG00000261762 | down |
| ENSG00000200550 | 9.668705999 | -3.291665112 | 0.5993369 | -5.492179 | 3.97E-08 | 7.79E-07 | RNU6-137P | snRNA | 37.96 | -20.10788986 | 20.10788986 | ENSG00000200550 | down |
| ENSG00000188483 | 561.8933519 | 1.749772942 | 0.2327114 | 7.519067 | 5.52E-14 | 3.28E-12 | IER5L | protein_coding | 64.99 | 20.09402928 | 20.09402928 | ENSG00000188483 | up |
| ENSG00000260452 | 39.10097658 | -2.142594537 | 0.3152874 | -6.795688 | 1.08E-11 | 4.30E-10 | TPRKBP2 | processed_pseudogene | 33.68 | -20.06858434 | 20.06858434 | ENSG00000260452 | down |
| ENSG00000239466 | 11.49019245 | -3.546239061 | 0.6713773 | -5.282036 | 1.28E-07 | 2.20E-06 | RN7SL552P | misc_RNA | 52.54 | -20.06402976 | 20.06402976 | ENSG00000239466 | down |
| ENSG00000252034 | 7.12095432 | -3.543736908 | 0.671498 | -5.27736 | 1.31E-07 | 2.24E-06 | RNY4P37 | misc_RNA | 40.62 | -20.01819477 | 20.01819477 | ENSG00000252034 | down |
| ENSG00000239494 | 10.12801674 | -2.978330711 | 0.5175012 | -5.755215 | 8.65E-09 | 1.96E-07 | RN7SL333P | misc_RNA | 54.88 | -19.97644167 | 19.97644167 | ENSG00000239494 | down |
| ENSG00000271843 | 34.1664078 | -2.205743425 | 0.330285 | -6.678303 | 2.42E-11 | 9.05E-10 | | lincRNA | 36.82 | -19.94761384 | 19.94761384 | ENSG00000271843 | down |
| ENSG00000130544 | 534.6384422 | -1.879379699 | 0.2602875 | -7.220399 | 5.18E-13 | 2.61E-11 | ZNF557 | protein_coding | 45.81 | -19.88863134 | 19.88863134 | ENSG00000130544 | down |
| ENSG00000250569 | 34.26070636 | -2.327816273 | 0.3596051 | -6.473257 | 9.59E-11 | 3.22E-09 | NTAN1P2 | processed_pseudogene | 48.2 | -19.76761518 | 19.76761518 | ENSG00000250569 | down |
| ENSG00000212124 | 183.2008244 | -2.001265431 | 0.2872485 | -6.967019 | 3.24E-12 | 1.39E-10 | TAS2R19 | protein_coding | 40.52 | -19.72841491 | 19.72841491 | ENSG00000212124 | down |
| ENSG00000273391 | 112.2931284 | 1.837684001 | 0.2527605 | 7.270456 | 3.58E-13 | 1.88E-11 | | antisense | 62.06 | 19.71270796 | 19.71270796 | ENSG00000273391 | up |
| ENSG00000270019 | 418.6187365 | 1.65746164 | 0.216604 | 7.652037 | 1.98E-14 | 1.28E-12 | | lincRNA | 44.12 | 19.71125183 | 19.71125183 | ENSG00000270019 | up |
| ENSG00000237631 | 52.78591847 | -2.121488603 | 0.314418 | -6.747351 | 1.51E-11 | 5.87E-10 | | processed_pseudogene | 44.82 | -19.5838892 | 19.5838892 | ENSG00000237631 | down |
| ENSG00000207245 | 9.271686303 | -4.272618788 | 0.9013027 | -4.740493 | 2.13E-06 | 2.69E-05 | SNORD116-29 | snoRNA | 40.96 | -19.52947244 | 19.52947244 | ENSG00000207245 | down |
| ENSG00000278784 | 35.93924237 | -2.155609011 | 0.3226966 | -6.679987 | 2.39E-11 | 8.97E-10 | | sense_intronic | 44.67 | -19.50245944 | 19.50245944 | ENSG00000278784 | down |
| ENSG00000271011 | 41.65508761 | -2.154215179 | 0.3227176 | -6.675233 | 2.47E-11 | 9.21E-10 | | antisense | 39.81 | -19.46454336 | 19.46454336 | ENSG00000271011 | down |
| ENSG00000197013 | 283.5781515 | -1.797086316 | 0.2463988 | -7.293405 | 3.02E-13 | 1.61E-11 | ZNF429 | protein_coding | 41.06 | -19.39819565 | 19.39819565 | ENSG00000197013 | down |
| ENSG00000273437 | 25.22193273 | -2.351528746 | 0.3692342 | -6.368665 | 1.91E-10 | 6.12E-09 | | lincRNA | 40 | -19.31416596 | 19.31416596 | ENSG00000273437 | down |

| | | | | | | | | | | | | | |
|---|---|---|---|---|---|---|---|---|---|---|---|---|---|
| ENSG00000151687 | 79.26154747 | -2.000447679 | 0.290123 | -6.895171 | 5.38E-12 | 2.24E-10 | ANKAR | protein_coding | 37.84 | -19.30277993 | 19.30277993 | ENSG00000151687 | down |
| ENSG00000273702 | 229.5423056 | 1.663347372 | 0.220668 | 7.537782 | 4.78E-14 | 2.91E-12 | | lincRNA | 49.57 | 19.18945185 | 19.18945185 | ENSG00000273702 | up |
| ENSG00000254905 | 24.86788954 | -2.267134406 | 0.3508407 | -6.462005 | 1.03E-10 | 3.45E-09 | | antisense | 50.41 | -19.1849457 | 19.1849457 | ENSG00000254905 | down |
| ENSG00000270361 | 20.49626535 | 2.460136137 | 0.3974975 | 6.18906 | 6.05E-10 | 1.75E-08 | | lincRNA | 45.8 | 19.08201014 | 19.08201014 | ENSG00000270361 | up |
| ENSG00000267264 | 17.62883114 | -2.582986962 | 0.4278464 | -6.037184 | 1.57E-09 | 4.16E-08 | | processed_pseudogene | 43.98 | -19.06503913 | 19.06503913 | ENSG00000267264 | down |
| ENSG00000263361 | 3.752405349 | -5.228437474 | 1.2418243 | -4.210288 | 2.55E-05 | 0.000232853 | MIR378H | miRNA | 51.81 | -18.99448976 | 18.99448976 | ENSG00000263361 | down |
| ENSG00000198799 | 1049.526109 | -1.85613856 | 0.2619032 | -7.087116 | 1.37E-12 | 6.40E-11 | LRIG2 | protein_coding | 41.19 | -18.92141284 | 18.92141284 | ENSG00000198799 | down |
| ENSG00000202361 | 10.53538194 | -3.002225318 | 0.5384198 | -5.575993 | 2.46E-08 | 5.04E-07 | | misc_RNA | 38.38 | -18.90689477 | 18.90689477 | ENSG00000202361 | down |
| ENSG00000199855 | 5.987223278 | -4.224863713 | 0.9010714 | -4.688711 | 2.75E-06 | 3.36E-05 | RNU6-490P | snRNA | 39.25 | -18.90021257 | 18.90021257 | ENSG00000199855 | down |
| ENSG00000221203 | 7.230674472 | -3.897356798 | 0.8006155 | -4.867951 | 1.13E-06 | 1.52E-05 | MIR1262 | miRNA | 35.48 | -18.78154382 | 18.78154382 | ENSG00000221203 | down |
| ENSG00000226216 | 72.03732239 | -2.076993527 | 0.3116217 | -6.665112 | 2.64E-11 | 9.82E-10 | RPS12P5 | processed_pseudogene | 44.74 | -18.70941859 | 18.70941859 | ENSG00000226216 | down |
| ENSG00000252236 | 16.88869603 | 2.553739867 | 0.4249763 | 6.009135 | 1.87E-09 | 4.84E-08 | | snoRNA | 47.52 | 18.68194981 | 18.68194981 | ENSG00000252236 | up |
| ENSG00000249685 | 12.83342639 | -2.758066349 | 0.477617 | -5.774641 | 7.71E-09 | 1.77E-07 | | lincRNA | 41.75 | -18.62132559 | 18.62132559 | ENSG00000249685 | down |
| ENSG00000207654 | 8.924134493 | -3.462792991 | 0.6725491 | -5.148758 | 2.62E-07 | 4.20E-06 | MIR128-1 | miRNA | 50 | -18.61987711 | 18.61987711 | ENSG00000207654 | down |
| ENSG00000111679 | 35.41108007 | 2.157422715 | 0.3310793 | 6.516332 | 7.20E-11 | 2.49E-09 | PTPN6 | protein_coding | 57.71 | 18.56236377 | 18.56236377 | ENSG00000111679 | up |
| ENSG00000268117 | 9.93698424 | -3.035417221 | 0.5528973 | -5.49002 | 4.02E-08 | 7.87E-07 | VN1R84P | unprocessed_pseudogene | 41.28 | -18.52828293 | 18.52828293 | ENSG00000268117 | down |
| ENSG00000138166 | 153.4619162 | 1.839293398 | 0.2612466 | 7.04045 | 1.92E-12 | 8.69E-11 | DUSP5 | protein_coding | 49.09 | 18.50506376 | 18.50506376 | ENSG00000138166 | up |
| ENSG00000102786 | 4400.680358 | -1.559742683 | 0.2042001 | -7.638303 | 2.20E-14 | 1.40E-12 | INTS6 | protein_coding | 36.13 | -18.4893239 | 18.4893239 | ENSG00000102786 | down |
| ENSG00000238731 | 3.880168107 | -5.171631164 | 1.2390371 | -4.173911 | 2.99E-05 | 0.000268259 | RNU7-90P | snRNA | 43.55 | -18.47019953 | 18.47019953 | ENSG00000238731 | down |
| ENSG00000256188 | 290.0231648 | -1.998680443 | 0.2961021 | -6.749972 | 1.48E-11 | 5.78E-10 | TAS2R30 | protein_coding | 35.98 | -18.46345976 | 18.46345976 | ENSG00000256188 | down |
| ENSG00000136826 | 78.8147324 | 1.980673611 | 0.2922143 | 6.778155 | 1.22E-11 | 4.81E-10 | KLF4 | protein_coding | 56.93 | 18.45481952 | 18.45481952 | ENSG00000136826 | up |
| ENSG00000236670 | 38.74585925 | -2.099150044 | 0.3186965 | -6.586674 | 4.50E-11 | 1.62E-09 | KRT18P5 | processed_pseudogene | 54.69 | -18.45120474 | 18.45120474 | ENSG00000236670 | down |
| ENSG00000277568 | 9.943199689 | -3.019892849 | 0.5501856 | -5.488862 | 4.05E-08 | 7.91E-07 | | misc_RNA | 57.05 | -18.42676341 | 18.42676341 | ENSG00000277568 | down |
| ENSG00000225183 | 6.559135144 | -3.849934821 | 0.7937234 | -4.850474 | 1.23E-06 | 1.65E-05 | | processed_pseudogene | 44.54 | -18.41590011 | 18.41590011 | ENSG00000225183 | down |
| ENSG00000100815 | 1367.316094 | -1.887242771 | 0.2725004 | -6.925651 | 4.34E-12 | 1.82E-10 | TRIP11 | protein_coding | 37.97 | -18.38163298 | 18.38163298 | ENSG00000100815 | down |
| ENSG00000141232 | 1879.744833 | 1.600089945 | 0.2127661 | 7.520419 | 5.46E-14 | 3.26E-12 | TOB1 | protein_coding | 50.17 | 18.3793085 | 18.3793085 | ENSG00000141232 | up |
| ENSG00000221788 | 27.60877959 | -2.343442445 | 0.3771425 | -6.213679 | 5.18E-10 | 1.52E-08 | MIR1252 | miRNA | 26.15 | -18.32333191 | 18.32333191 | ENSG00000221788 | down |
| ENSG00000257354 | 662.0976562 | -1.776805045 | 0.2496615 | -7.116857 | 1.10E-12 | 5.24E-11 | | lincRNA | 40.48 | -18.2660567 | 18.2660567 | ENSG00000257354 | down |
| ENSG00000143333 | 75.60933065 | 2.014238514 | 0.301339 | 6.684295 | 2.32E-11 | 8.76E-10 | RGS16 | protein_coding | 50.88 | 18.24443273 | 18.24443273 | ENSG00000143333 | up |
| ENSG00000221949 | 158.732989 | 1.728350114 | 0.2396672 | 7.211458 | 5.54E-13 | 2.78E-11 | LINC01465 | lincRNA | 58.2 | 18.24374457 | 18.24374457 | ENSG00000221949 | up |
| ENSG00000264448 | 25.70775771 | 2.309545696 | 0.3715823 | 6.215435 | 5.12E-10 | 1.51E-08 | | lincRNA | 52.48 | 18.0652773 | 18.0652773 | ENSG00000264448 | up |
| ENSG00000218682 | 14.1046701 | -2.497779236 | 0.4181646 | -5.973197 | 2.33E-09 | 5.88E-08 | | processed_pseudogene | 53.56 | -18.06073555 | 18.06073555 | ENSG00000218682 | down |

| | | | | | | | | | | | | | |
|---|---|---|---|---|---|---|---|---|---|---|---|---|---|
| ENSG00000250838 | 275.5619204 | 1.575451355 | 0.2102995 | 7.491465 | 6.81E-14 | 3.97E-12 | | antisense | 61.43 | 17.96224989 | 17.96224989 | ENSG00000250838 | up |
| ENSG00000260920 | 78.95614147 | -1.91908357 | 0.2825631 | -6.791698 | 1.11E-11 | 4.41E-10 | | sense_overlapping | 37.89 | -17.95439413 | 17.95439413 | ENSG00000260920 | down |
| ENSG00000224046 | 147.8362717 | 1.705636403 | 0.2369249 | 7.19906 | 6.06E-13 | 3.03E-11 | | antisense | 47.9 | 17.9419158 | 17.9419158 | ENSG00000224046 | up |
| ENSG00000213131 | 55.21168201 | -2.034995538 | 0.3085338 | -6.595697 | 4.23E-11 | 1.53E-09 | YWHAZP4 | processed_pseudogene | 46.71 | -17.93872799 | 17.93872799 | ENSG00000213131 | down |
| ENSG00000225215 | 42.70784915 | -2.03624564 | 0.3092411 | -6.584653 | 4.56E-11 | 1.64E-09 | SMARCE1P1 | processed_pseudogene | 47.22 | -17.88854478 | 17.88854478 | ENSG00000225215 | down |
| ENSG00000275709 | 160.7552433 | 1.655625403 | 0.2271349 | 7.289171 | 3.12E-13 | 1.65E-11 | | lincRNA | 40.69 | 17.85137064 | 17.85137064 | ENSG00000275709 | up |
| ENSG00000222267 | 6.150232059 | -3.832882842 | 0.8013542 | -4.783007 | 1.73E-06 | 2.23E-05 | RNU6-892P | snRNA | 34.34 | -17.83217442 | 17.83217442 | ENSG00000222267 | down |
| ENSG00000269973 | 110.1799393 | -1.919805371 | 0.2837236 | -6.766464 | 1.32E-11 | 5.21E-10 | | lincRNA | 42.62 | -17.82265528 | 17.82265528 | ENSG00000269973 | down |
| ENSG00000228825 | 12.40362922 | 2.746118979 | 0.4864474 | 5.645254 | 1.65E-08 | 3.51E-07 | LAMTOR3P1 | processed_pseudogene | 39.46 | 17.72671589 | 17.72671589 | ENSG00000228825 | up |
| ENSG00000276702 | 16.75788362 | -2.396447265 | 0.3971494 | -6.03412 | 1.60E-09 | 4.22E-08 | | antisense | 27.46 | -17.67239906 | 17.67239906 | ENSG00000276702 | down |
| ENSG00000249725 | 14.63785555 | 2.566330057 | 0.4400197 | 5.832308 | 5.47E-09 | 1.30E-07 | | lincRNA | 38.54 | 17.66908746 | 17.66908746 | ENSG00000249725 | up |
| ENSG00000254512 | 40.1736475 | -2.918950366 | 0.5344421 | -5.461678 | 4.72E-08 | 9.11E-07 | PHBP2 | processed_pseudogene | 53.1 | -17.63214497 | 17.63214497 | ENSG00000254512 | down |
| ENSG00000212306 | 5.336783863 | -4.654439178 | 1.0823361 | -4.300364 | 1.71E-05 | 0.000164869 | | misc_RNA | 48.42 | -17.60709262 | 17.60709262 | ENSG00000212306 | down |
| ENSG00000261291 | 39.13316134 | -2.115831231 | 0.3300884 | -6.409892 | 1.46E-10 | 4.78E-09 | | sense_intronic | 44.24 | -17.60457181 | 17.60457181 | ENSG00000261291 | down |
| ENSG00000276103 | 5.502145891 | -4.123271533 | 0.9022073 | -4.570204 | 4.87E-06 | 5.59E-05 | | snRNA | 44.44 | -17.53370388 | 17.53370388 | ENSG00000276103 | down |
| ENSG00000235990 | 19.83929274 | -2.385576361 | 0.3961026 | -6.022622 | 1.72E-09 | 4.49E-08 | RPL23AP20 | processed_pseudogene | 49.64 | -17.52832197 | 17.52832197 | ENSG00000235990 | down |
| ENSG00000266501 | 17.40116128 | -2.409985489 | 0.4027214 | -5.984251 | 2.17E-09 | 5.54E-08 | | processed_pseudogene | 47.49 | -17.48739316 | 17.48739316 | ENSG00000266501 | down |
| ENSG00000204388 | 269.6414577 | 1.607946086 | 0.219669 | 7.319858 | 2.48E-13 | 1.33E-11 | HSPA1B | protein_coding | 58.75 | 17.48594301 | 17.48594301 | ENSG00000204388 | up |
| ENSG00000267766 | 11.29944631 | -2.745346823 | 0.4899645 | -5.603155 | 2.10E-08 | 4.39E-07 | | sense_intronic | 45.41 | -17.45264125 | 17.45264125 | ENSG00000267766 | down |
| ENSG00000255374 | 64.71261034 | -2.053119607 | 0.3171784 | -6.473074 | 9.60E-11 | 3.22E-09 | TAS2R43 | protein_coding | 38.95 | -17.43491477 | 17.43491477 | ENSG00000255374 | down |
| ENSG00000224738 | 67.81315147 | 1.8531601 | 0.2720405 | 6.812075 | 9.62E-12 | 3.92E-10 | | antisense | 42.19 | 17.43288573 | 17.43288573 | ENSG00000224738 | up |
| ENSG00000197837 | 857.575929 | 1.587074823 | 0.2158828 | 7.351559 | 1.96E-13 | 1.08E-11 | HIST4H4 | protein_coding | 49.57 | 17.4059801 | 17.4059801 | ENSG00000197837 | up |
| ENSG00000162783 | 3803.088443 | 1.442430602 | 0.1871417 | 7.70769 | 1.28E-14 | 8.62E-13 | IER5 | protein_coding | 49.76 | 17.40240207 | 17.40240207 | ENSG00000162783 | up |
| ENSG00000249274 | 38.60441198 | -2.029809538 | 0.3128402 | -6.488328 | 8.68E-11 | 2.95E-09 | PDLIM1P4 | processed_pseudogene | 45.03 | -17.31345347 | 17.31345347 | ENSG00000249274 | down |
| ENSG00000126803 | 244.5911181 | 1.587537681 | 0.2168715 | 7.320177 | 2.48E-13 | 1.33E-11 | HSPA2 | protein_coding | 45.54 | 17.26400758 | 17.26400758 | ENSG00000126803 | up |
| ENSG00000264638 | 7.367610365 | -5.04400428 | 1.2372723 | -4.076713 | 4.57E-05 | 0.000389256 | MIR3152 | miRNA | 45.95 | -17.19887018 | 17.19887018 | ENSG00000264638 | down |
| ENSG00000253168 | 21.87785958 | -3.520117096 | 0.7190786 | -4.895316 | 9.81E-07 | 1.34E-05 | | antisense | 38.68 | -17.15348364 | 17.15348364 | ENSG00000253168 | down |
| ENSG00000284031 | 1.703142715 | -5.115339335 | 1.266561 | -4.038763 | 5.37E-05 | 0.000447986 | MIR4761 | miRNA | 62.2 | -17.12991732 | 17.12991732 | ENSG00000284031 | down |
| ENSG00000265972 | 3713.492097 | 1.710514284 | 0.2436339 | 7.020839 | 2.21E-12 | 9.78E-11 | TXNIP | protein_coding | 42.49 | 17.12175379 | 17.12175379 | ENSG00000265972 | up |
| ENSG00000243370 | 17.8457022 | -2.48089438 | 0.4255866 | -5.829352 | 5.56E-09 | 1.32E-07 | RN7SL775P | misc_RNA | 53.57 | -17.06362303 | 17.06362303 | ENSG00000243370 | down |
| ENSG00000259232 | 27.92079528 | -2.194451154 | 0.3541809 | -6.195848 | 5.80E-10 | 1.68E-08 | | processed_pseudogene | 53.87 | -17.06029687 | 17.06029687 | ENSG00000259232 | down |
| ENSG00000274270 | 89.69731857 | 1.73608304 | 0.2496258 | 6.954742 | 3.53E-12 | 1.49E-10 | | sense_intronic | 58.61 | 17.05776574 | 17.05776574 | ENSG00000274270 | up |

| | | | | | | | | | | | | | |
|---|---|---|---|---|---|---|---|---|---|---|---|---|---|
| ENSG00000264160 | 11.14939267 | -2.743748241 | 0.4952862 | -5.539723 | 3.03E-08 | 6.08E-07 | MIR4762 | miRNA | 44 | -17.05584014 | 17.05584014 | ENSG00000264160 | down |
| ENSG00000238523 | 1.876994582 | -5.105343317 | 1.2671001 | -4.029156 | 5.60E-05 | 0.000463087 | RNU7-107P | snRNA | 32.79 | -17.02293701 | 17.02293701 | ENSG00000238523 | down |
| ENSG00000202382 | 4.797181049 | -4.59178792 | 1.0802008 | -4.250865 | 2.13E-05 | 0.000199626 | | misc_RNA | 44.12 | -16.98861517 | 16.98861517 | ENSG00000202382 | down |
| ENSG00000221044 | 17.31638363 | -2.451536986 | 0.419044 | -5.850309 | 4.91E-09 | 1.18E-07 | | snoRNA | 50.49 | -16.98465264 | 16.98465264 | ENSG00000221044 | down |
| ENSG00000236711 | 115.8942202 | -1.86574692 | 0.2785001 | -6.699269 | 2.09E-11 | 8.01E-10 | SMAD9-IT1 | sense_intronic | 42.29 | -16.97131537 | 16.97131537 | ENSG00000236711 | down |
| ENSG00000266072 | 5.929869241 | -4.075173572 | 0.9022461 | -4.516699 | 6.28E-06 | 6.95E-05 | MIR5693 | miRNA | 46.58 | -16.94345069 | 16.94345069 | ENSG00000266072 | down |
| ENSG00000160888 | 986.5842813 | 1.622960425 | 0.2267967 | 7.156014 | 8.31E-13 | 4.02E-11 | IER2 | protein_coding | 62.33 | 16.87275989 | 16.87275989 | ENSG00000160888 | up |
| ENSG00000228412 | 63.49495821 | 1.894014462 | 0.2859026 | 6.624684 | 3.48E-11 | 1.27E-09 | | antisense | 38.79 | 16.84843765 | 16.84843765 | ENSG00000228412 | up |
| ENSG00000109787 | 629.8989405 | -1.704579103 | 0.2447172 | -6.965507 | 3.27E-12 | 1.40E-10 | KLF3 | protein_coding | 41.91 | -16.79801559 | 16.79801559 | ENSG00000109787 | down |
| ENSG00000010539 | 359.7873879 | -1.597852599 | 0.2222154 | -7.190558 | 6.45E-13 | 3.18E-11 | ZNF200 | protein_coding | 43.67 | -16.77423536 | 16.77423536 | ENSG00000010539 | down |
| ENSG00000257027 | 129.5984592 | -1.93120696 | 0.2949498 | -6.547579 | 5.85E-11 | 2.07E-09 | | sense_intronic | 41.43 | -16.7721 | 16.7721 | ENSG00000257027 | down |
| ENSG00000229521 | 54.99001706 | -1.990064391 | 0.3093416 | -6.433225 | 1.25E-10 | 4.11E-09 | MYCBP2-AS2 | sense_intronic | 42.35 | -16.68859993 | 16.68859993 | ENSG00000229521 | down |
| ENSG00000229623 | 34.83062724 | -2.024005325 | 0.3171604 | -6.381645 | 1.75E-10 | 5.70E-09 | METTL21AP1 | processed_pseudogene | 46.93 | -16.68627917 | 16.68627917 | ENSG00000229623 | down |
| ENSG00000238015 | 13.64113943 | -2.512018426 | 0.4389455 | -5.722848 | 1.05E-08 | 2.34E-07 | | processed_pseudogene | 42.92 | -16.65791385 | 16.65791385 | ENSG00000238015 | down |
| ENSG00000231128 | 54.8109462 | 1.828995027 | 0.2742055 | 6.670161 | 2.56E-11 | 9.51E-10 | | antisense | 38.96 | 16.50066134 | 16.50066134 | ENSG00000231128 | up |
| ENSG00000207162 | 7.60332166 | -3.758783416 | 0.8107402 | -4.636237 | 3.55E-06 | 4.23E-05 | RNU6-549P | snRNA | 39.25 | -16.44124464 | 16.44124464 | ENSG00000207162 | down |
| ENSG00000265281 | 20.97070154 | -2.250335397 | 0.3752836 | -5.996359 | 2.02E-09 | 5.19E-08 | MIR3935 | miRNA | 46.15 | -16.39271179 | 16.39271179 | ENSG00000265281 | down |
| ENSG00000260288 | 79.89203562 | 1.734351869 | 0.2541167 | 6.825021 | 8.79E-12 | 3.60E-10 | | processed_transcript | 49.1 | 16.37869071 | 16.37869071 | ENSG00000260288 | up |
| ENSG00000244314 | 22.89057273 | -2.233862688 | 0.3714765 | -6.01347 | 1.82E-09 | 4.73E-08 | RN7SL36P | misc_RNA | 54.71 | -16.36420424 | 16.36420424 | ENSG00000244314 | down |
| ENSG00000280440 | 13.55816902 | 2.581105534 | 0.4619763 | 5.587095 | 2.31E-08 | 4.78E-07 | | TEC | 43.35 | 16.31480802 | 16.31480802 | ENSG00000280440 | up |
| ENSG00000204611 | 667.1189458 | -1.716643741 | 0.2510819 | -6.836987 | 8.09E-12 | 3.33E-10 | ZNF616 | protein_coding | 42.23 | -16.26925843 | 16.26925843 | ENSG00000204611 | down |
| ENSG00000236577 | 6.195203456 | -3.71608537 | 0.8017515 | -4.634959 | 3.57E-06 | 4.24E-05 | SNRPGP14 | processed_pseudogene | 41.27 | -16.24726952 | 16.24726952 | ENSG00000236577 | down |
| ENSG00000270117 | 26.67301694 | 2.123658538 | 0.3468298 | 6.123058 | 9.18E-10 | 2.55E-08 | | lincRNA | 52.78 | 16.12553124 | 16.12553124 | ENSG00000270117 | up |
| ENSG00000274159 | 26.34759467 | -2.120099986 | 0.3464905 | -6.118782 | 9.43E-10 | 2.62E-08 | | sense_intronic | 38.52 | -16.07563503 | 16.07563503 | ENSG00000274159 | down |
| ENSG00000199331 | 16.64885213 | -2.33847657 | 0.4017142 | -5.821245 | 5.84E-09 | 1.38E-07 | | misc_RNA | 44.12 | -16.04513063 | 16.04513063 | ENSG00000199331 | down |
| ENSG00000206609 | 3.842543044 | -4.934714806 | 1.2419703 | -3.973295 | 7.09E-05 | 0.000566226 | SNORD116-11 | snoRNA | 40.22 | -16.0230683 | 16.0230683 | ENSG00000206609 | down |
| ENSG00000283615 | 4.231863347 | -4.508115078 | 1.0832386 | -4.161701 | 3.16E-05 | 0.000280848 | MIR924 | miRNA | 43.4 | -16.01069964 | 16.01069964 | ENSG00000283615 | down |
| ENSG00000197362 | 132.1357035 | -1.784646496 | 0.2682996 | -6.651693 | 2.90E-11 | 1.07E-09 | ZNF786 | protein_coding | 46.13 | -16.0093499 | 16.0093499 | ENSG00000197362 | down |
| ENSG00000236047 | 33.25954517 | -2.883183994 | 0.5522441 | -5.220851 | 1.78E-07 | 2.96E-06 | | processed_pseudogene | 50.08 | -15.93916964 | 15.93916964 | ENSG00000236047 | down |
| ENSG00000275413 | 23.81626704 | -2.415179339 | 0.423457 | -5.703482 | 1.17E-08 | 2.57E-07 | | antisense | 32.33 | -15.91795225 | 15.91795225 | ENSG00000275413 | down |
| ENSG00000225568 | 42.02157857 | -2.055196671 | 0.3323187 | -6.184414 | 6.23E-10 | 1.80E-08 | | processed_pseudogene | 42.25 | -15.91666063 | 15.91666063 | ENSG00000225568 | down |
| ENSG00000111215 | 140.1193014 | -2.06895657 | 0.3356881 | -6.163331 | 7.12E-10 | 2.03E-08 | PRR4 | protein_coding | 36.64 | -15.91629547 | 15.91629547 | ENSG00000111215 | down |

| | | | | | | | | | | | | | |
|---|---|---|---|---|---|---|---|---|---|---|---|---|---|
| ENSG00000270141 | 41.93188242 | 2.069879233 | 0.336022 | 6.159951 | 7.28E-10 | 2.07E-08 | TERC | lincRNA | 66.73 | 15.90604087 | 15.90604087 | ENSG00000270141 | up |
| ENSG00000253726 | 264.2580833 | -1.727650785 | 0.2564022 | -6.738051 | 1.61E-11 | 6.24E-10 | | sense_intronic | 38 | -15.90234555 | 15.90234555 | ENSG00000253726 | down |
| ENSG00000267127 | 36.84574443 | -2.031223502 | 0.3270531 | -6.210684 | 5.28E-10 | 1.54E-08 | | protein_coding | 46.26 | -15.86898864 | 15.86898864 | ENSG00000267127 | down |
| ENSG00000179774 | 48.03790908 | 1.874090372 | 0.290259 | 6.456613 | 1.07E-10 | 3.56E-09 | ATOH7 | protein_coding | 54.26 | 15.83382365 | 15.83382365 | ENSG00000179774 | up |
| ENSG00000163110 | 148.7330064 | 1.688263108 | 0.2487596 | 6.786726 | 1.15E-11 | 4.55E-10 | PDLIM5 | protein_coding | 36.89 | 15.7717268 | 15.7717268 | ENSG00000163110 | up |
| ENSG00000235651 | 93.07036764 | -1.928889665 | 0.3038291 | -6.3486 | 2.17E-10 | 6.89E-09 | | processed_pseudogene | 47.42 | -15.74303108 | 15.74303108 | ENSG00000235651 | down |
| ENSG00000277728 | 30.16370233 | -1.979186225 | 0.3164243 | -6.254849 | 3.98E-10 | 1.19E-08 | | antisense | 44.61 | -15.68296135 | 15.68296135 | ENSG00000277728 | down |
| ENSG00000219902 | 53.00422145 | -1.932475795 | 0.3056305 | -6.322916 | 2.57E-10 | 8.03E-09 | RPL35P3 | processed_pseudogene | 56.6 | -15.6438716 | 15.6438716 | ENSG00000219902 | down |
| ENSG00000240996 | 26.53510332 | -2.156969668 | 0.3609307 | -5.976133 | 2.28E-09 | 5.78E-08 | | lincRNA | 36.1 | -15.61160589 | 15.61160589 | ENSG00000240996 | down |
| ENSG00000272335 | 165.9493328 | -1.885680501 | 0.2954777 | -6.381802 | 1.75E-10 | 5.70E-09 | | lincRNA | 35.2 | -15.5459034 | 15.5459034 | ENSG00000272335 | down |
| ENSG00000255046 | 52.06608272 | 1.838286871 | 0.2848766 | 6.452923 | 1.10E-10 | 3.64E-09 | | lincRNA | 47.88 | 15.51377863 | 15.51377863 | ENSG00000255046 | up |
| ENSG00000232454 | 21.44284625 | -2.18447164 | 0.3690976 | -5.918412 | 3.25E-09 | 8.07E-08 | | processed_pseudogene | 37.61 | -15.49502283 | 15.49502283 | ENSG00000232454 | down |
| ENSG00000266415 | 10.93631583 | -2.89947573 | 0.5651387 | -5.130556 | 2.89E-07 | 4.55E-06 | MIR4636 | miRNA | 45 | -15.48897308 | 15.48897308 | ENSG00000266415 | down |
| ENSG00000224349 | 88.9887477 | 1.687180406 | 0.2509303 | 6.723701 | 1.77E-11 | 6.81E-10 | | antisense | 64.38 | 15.46570148 | 15.46570148 | ENSG00000224349 | up |
| ENSG00000253509 | 8.532193141 | -2.872685908 | 0.5580996 | -5.147264 | 2.64E-07 | 4.22E-06 | | sense_intronic | 48.86 | -15.44012028 | 15.44012028 | ENSG00000253509 | down |
| ENSG00000211520 | 3.496073065 | -4.873175351 | 1.2423354 | -3.922592 | 8.76E-05 | 0.000680621 | MIR216B | miRNA | 45.12 | -15.43380867 | 15.43380867 | ENSG00000211520 | down |
| ENSG00000277202 | 5.421913776 | -4.876133183 | 1.2452939 | -3.915648 | 9.02E-05 | 0.000696626 | MIR8063 | miRNA | 38.27 | -15.39395482 | 15.39395482 | ENSG00000277202 | down |
| ENSG00000279792 | 214.7221855 | -1.521100608 | 0.2155934 | -7.055414 | 1.72E-12 | 7.91E-11 | | TEC | 56.49 | -15.36609985 | 15.36609985 | ENSG00000279792 | down |
| ENSG00000273456 | 117.3771181 | 1.616371564 | 0.2366481 | 6.830276 | 8.48E-12 | 3.48E-10 | | lincRNA | 51.56 | 15.28815811 | 15.28815811 | ENSG00000273456 | up |
| ENSG00000222043 | 39.29218634 | 1.935713445 | 0.310173 | 6.240754 | 4.35E-10 | 1.29E-08 | | antisense | 68.45 | 15.26984728 | 15.26984728 | ENSG00000222043 | up |
| ENSG00000253327 | 53.53912832 | 1.790293416 | 0.275971 | 6.487251 | 8.74E-11 | 2.96E-09 | RAD21-AS1 | antisense | 42.45 | 15.26854332 | 15.26854332 | ENSG00000253327 | up |
| ENSG00000272428 | 11.99597423 | -2.645735634 | 0.4960294 | -5.333828 | 9.62E-08 | 1.72E-06 | | lincRNA | 32.23 | -15.25212313 | 15.25212313 | ENSG00000272428 | down |
| ENSG00000261578 | 175.7835 | -1.661080181 | 0.2472485 | -6.718261 | 1.84E-11 | 7.05E-10 | | sense_overlapping | 47.28 | -15.20150391 | 15.20150391 | ENSG00000261578 | down |
| ENSG00000226853 | 61.49374141 | 1.769280168 | 0.2717653 | 6.510325 | 7.50E-11 | 2.58E-09 | | lincRNA | 45.09 | 15.19586753 | 15.19586753 | ENSG00000226853 | up |
| ENSG00000270638 | 500.8852496 | 1.485298085 | 0.2095212 | 7.08901 | 1.35E-12 | 6.33E-11 | | lincRNA | 41.96 | 15.14773925 | 15.14773925 | ENSG00000270638 | up |
| ENSG00000222624 | 8.456494066 | -3.412406396 | 0.7311215 | -4.667359 | 3.05E-06 | 3.69E-05 | RNU2-15P | snRNA | 38.74 | -15.12558748 | 15.12558748 | ENSG00000222624 | down |
| ENSG00000264864 | 18.02574357 | 2.255247562 | 0.3919146 | 5.754436 | 8.69E-09 | 1.97E-07 | MIR3613 | miRNA | 35.63 | 15.12360989 | 15.12360989 | ENSG00000264864 | up |
| ENSG00000259946 | 88.09591248 | -2.855488535 | 0.5590307 | -5.107928 | 3.26E-07 | 5.07E-06 | | lincRNA | 32.1 | -15.1191606 | 15.1191606 | ENSG00000259946 | down |
| ENSG00000281195 | 389.80023 | -1.668483607 | 0.2496482 | -6.683339 | 2.34E-11 | 8.79E-10 | | processed_transcript | 36.8 | -15.10990878 | 15.10990878 | ENSG00000281195 | down |
| ENSG00000208032 | 9.119389876 | -2.789996104 | 0.5403075 | -5.163719 | 2.42E-07 | 3.92E-06 | MIR548A3 | miRNA | 36.08 | -15.08385861 | 15.08385861 | ENSG00000208032 | down |
| ENSG00000162772 | 495.5344289 | 1.76719433 | 0.2724888 | 6.485382 | 8.85E-11 | 2.99E-09 | ATF3 | protein_coding | 43.16 | 15.06389332 | 15.06389332 | ENSG00000162772 | up |
| ENSG00000249140 | 27.41192718 | -2.007397394 | 0.3300205 | -6.082644 | 1.18E-09 | 3.21E-08 | PRDX2P3 | processed_pseudogene | 54.61 | -15.04114658 | 15.04114658 | ENSG00000249140 | down |

| | | | | | | | | | | | | | |
|---|---|---|---|---|---|---|---|---|---|---|---|---|---|
| ENSG00000267793 | 38.83027479 | -2.051854628 | 0.3412497 | -6.012765 | 1.82E-09 | 4.74E-08 | | transcribed_processed_pseudog | 35.26 | -15.02869177 | 15.02869177 | ENSG00000267793 | down |
| ENSG00000198182 | 277.7975545 | -1.943780079 | 0.3146589 | -6.177419 | 6.52E-10 | 1.87E-08 | ZNF607 | protein_coding | 41.55 | -15.02161483 | 15.02161483 | ENSG00000198182 | down |
| ENSG00000251323 | 78.99360996 | -1.929120319 | 0.3113504 | -6.195978 | 5.79E-10 | 1.68E-08 | | lincRNA | 39.08 | -14.99753836 | 14.99753836 | ENSG00000251323 | down |
| ENSG00000230359 | 72.79679904 | 1.728135729 | 0.2640889 | 6.543765 | 6.00E-11 | 2.11E-09 | TPI1P2 | transcribed_processed_pseudog | 50.02 | 14.99257669 | 14.99257669 | ENSG00000230359 | up |
| ENSG00000272910 | 14.93176671 | -2.29371353 | 0.404054 | -5.676749 | 1.37E-08 | 2.96E-07 | | antisense | 33.73 | -14.97387786 | 14.97387786 | ENSG00000272910 | down |
| ENSG00000224660 | 224.6665527 | -1.682029239 | 0.2541968 | -6.617035 | 3.66E-11 | 1.33E-09 | SH3BP5-AS1 | antisense | 45.6 | -14.9287188 | 14.9287188 | ENSG00000224660 | down |
| ENSG00000223039 | 27.29348743 | -2.231945529 | 0.3886879 | -5.742256 | 9.34E-09 | 2.09E-07 | RN7SKP268 | misc_RNA | 52.85 | -14.90694886 | 14.90694886 | ENSG00000223039 | down |
| ENSG00000279786 | 58.52923906 | -1.805857458 | 0.2833097 | -6.374146 | 1.84E-10 | 5.94E-09 | | TEC | 43.34 | -14.8550063 | 14.8550063 | ENSG00000279786 | down |
| ENSG00000222160 | 4.290070214 | -3.917129008 | 0.9107306 | -4.301084 | 1.70E-05 | 0.000164554 | | misc_RNA | 45.19 | -14.82120985 | 14.82120985 | ENSG00000222160 | down |
| ENSG00000264536 | 3.48937158 | -4.809048468 | 1.2444168 | -3.8645 | 0.000111317 | 0.000839581 | MIR5706 | miRNA | 45 | -14.79233136 | 14.79233136 | ENSG00000264536 | down |
| ENSG00000233133 | 4.804095046 | -3.386097274 | 0.7312113 | -4.630805 | 3.64E-06 | 4.32E-05 | | unprocessed_pseudogene | 68.18 | -14.77750399 | 14.77750399 | ENSG00000233133 | down |
| ENSG00000199788 | 7.845721989 | -2.903453065 | 0.5806872 | -5.000029 | 5.73E-07 | 8.24E-06 | RNY3P2 | misc_RNA | 44.12 | -14.76194692 | 14.76194692 | ENSG00000199788 | down |
| ENSG00000249771 | 62.34749236 | 1.720047743 | 0.2645633 | 6.50146 | 7.95E-11 | 2.72E-09 | | lincRNA | 41.45 | 14.73264258 | 14.73264258 | ENSG00000249771 | up |
| ENSG00000201557 | 9.231046982 | -3.656744156 | 0.8235803 | -4.440058 | 8.99E-06 | 9.54E-05 | SNORD114-15 | snoRNA | 45.83 | -14.70232262 | 14.70232262 | ENSG00000201557 | down |
| ENSG00000257023 | 36.08890788 | 1.981995889 | 0.3279934 | 6.042792 | 1.51E-09 | 4.02E-08 | | antisense | 38.09 | 14.65739799 | 14.65739799 | ENSG00000257023 | up |
| ENSG00000243373 | 19.71683926 | -2.212896649 | 0.3871506 | -5.715854 | 1.09E-08 | 2.42E-07 | RN7SL173P | misc_RNA | 51.71 | -14.64143238 | 14.64143238 | ENSG00000243373 | down |
| ENSG00000273768 | 775.7121105 | 1.458430509 | 0.2074215 | 7.031241 | 2.05E-12 | 9.18E-11 | | snRNA | 54.27 | 14.63873466 | 14.63873466 | ENSG00000273768 | up |
| ENSG00000266618 | 16.69511564 | -2.363153286 | 0.4277347 | -5.524811 | 3.30E-08 | 6.56E-07 | MIR4742 | miRNA | 34.12 | -14.61152056 | 14.61152056 | ENSG00000266618 | down |
| ENSG00000164442 | 1338.99321 | 1.379603458 | 0.1909852 | 7.223615 | 5.06E-13 | 2.59E-11 | CITED2 | protein_coding | 50.32 | 14.60515034 | 14.60515034 | ENSG00000164442 | up |
| ENSG00000272367 | 53.49798998 | -1.940841418 | 0.3184258 | -6.095114 | 1.09E-09 | 2.99E-08 | | antisense | 32.76 | -14.603139 | 14.603139 | ENSG00000272367 | down |
| ENSG00000207305 | 3.665837469 | -4.366996723 | 1.0825505 | -4.033989 | 5.48E-05 | 0.000456098 | | misc_RNA | 38.89 | -14.58988077 | 14.58988077 | ENSG00000207305 | down |
| ENSG00000265201 | 7.795742131 | -2.887100905 | 0.579423 | -4.982717 | 6.27E-07 | 8.90E-06 | MIR4677 | miRNA | 46.25 | -14.58184091 | 14.58184091 | ENSG00000265201 | down |
| ENSG00000252186 | 75.02451185 | 1.808140045 | 0.2870865 | 6.298241 | 3.01E-10 | 9.27E-09 | RNU6-781P | snRNA | 48.51 | 14.52438716 | 14.52438716 | ENSG00000252186 | up |
| ENSG00000186638 | 334.815898 | -1.717057446 | 0.2657885 | -6.460239 | 1.05E-10 | 3.48E-09 | KIF24 | protein_coding | 42.81 | -14.52316246 | 14.52316246 | ENSG00000186638 | down |
| ENSG00000259788 | 38.12782407 | 1.884526859 | 0.3056267 | 6.166106 | 7.00E-10 | 2.00E-08 | | antisense | 47.6 | 14.50849696 | 14.50849696 | ENSG00000259788 | up |
| ENSG00000176018 | 196.558976 | -1.823193515 | 0.2909642 | -6.26604 | 3.70E-10 | 1.12E-08 | LYSMD3 | protein_coding | 35.92 | -14.49686874 | 14.49686874 | ENSG00000176018 | down |
| ENSG00000206965 | 4.240049802 | -4.356841888 | 1.0839778 | -4.019309 | 5.84E-05 | 0.000479453 | RNU6-5P | snRNA | 45.79 | -14.46146534 | 14.46146534 | ENSG00000206965 | down |
| ENSG00000261770 | 232.6028995 | -1.778970911 | 0.2809723 | -6.331481 | 2.43E-10 | 7.63E-09 | | lincRNA | 46.86 | -14.44059566 | 14.44059566 | ENSG00000261770 | down |
| ENSG00000264572 | 11.39483432 | -2.449909509 | 0.4544092 | -5.391417 | 6.99E-08 | 1.29E-06 | MIR4296 | miRNA | 56.82 | -14.4278486 | 14.4278486 | ENSG00000264572 | down |
| ENSG00000235672 | 17.28102883 | -2.221728723 | 0.3923748 | -5.662261 | 1.49E-08 | 3.21E-07 | | transcribed_processed_pseudog | 47.87 | -14.4268353 | 14.4268353 | ENSG00000235672 | down |
| ENSG00000264522 | 688.0559321 | -1.649879014 | 0.2514589 | -6.561228 | 5.34E-11 | 1.90E-09 | OTUD7B | protein_coding | 42.46 | -14.39071373 | 14.39071373 | ENSG00000264522 | down |
| ENSG00000229388 | 49.28696716 | 1.764125213 | 0.2784257 | 6.336073 | 2.36E-10 | 7.42E-09 | LINC01715 | lincRNA | 47.96 | 14.34116945 | 14.34116945 | ENSG00000229388 | up |

| | | | | | | | | | | | | | |
|---|---|---|---|---|---|---|---|---|---|---|---|---|---|
| ENSG00000222714 | 9.993772573 | -2.483276672 | 0.4652193 | -5.337862 | 9.40E-08 | 1.69E-06 | RN7SKP38 | misc_RNA | 49.51 | -14.33678724 | 14.33678724 | ENSG00000222714 | down |
| ENSG00000259183 | 10.09101093 | -2.512098247 | 0.4740061 | -5.299717 | 1.16E-07 | 2.02E-06 | MED28P6 | processed_pseudogene | 40.69 | -14.30494785 | 14.30494785 | ENSG00000259183 | down |
| ENSG00000253008 | 98.0053074 | -1.799304554 | 0.2872387 | -6.264144 | 3.75E-10 | 1.13E-08 | MIR2355 | miRNA | 41.38 | -14.30017209 | 14.30017209 | ENSG00000253008 | down |
| ENSG00000167074 | 175.611331 | -1.816515753 | 0.291668 | -6.228026 | 4.72E-10 | 1.40E-08 | TEF | protein_coding | 51.78 | -14.26877815 | 14.26877815 | ENSG00000167074 | down |
| ENSG00000230714 | 7.782278061 | 2.92037049 | 0.5967437 | 4.893844 | 9.89E-07 | 1.35E-05 | | lincRNA | 32.75 | 14.22267709 | 14.22267709 | ENSG00000230714 | up |
| ENSG00000275185 | 58.87907865 | -1.887390517 | 0.310384 | -6.080825 | 1.20E-09 | 3.24E-08 | | antisense | 37.27 | -14.13584468 | 14.13584468 | ENSG00000275185 | down |
| ENSG00000275004 | 183.2271472 | -1.636556588 | 0.2506866 | -6.528296 | 6.65E-11 | 2.30E-09 | ZNF280B | protein_coding | 41.55 | -14.13580077 | 14.13580077 | ENSG00000275004 | down |
| ENSG00000230815 | 22.51201428 | -2.312635489 | 0.4213664 | -5.48842 | 4.06E-08 | 7.92E-07 | | sense_intronic | 39.44 | -14.11012184 | 14.11012184 | ENSG00000230815 | down |
| ENSG00000274978 | 355.7648693 | 1.479655372 | 0.2160391 | 6.849015 | 7.44E-12 | 3.07E-10 | RNU11 | snRNA | 51.49 | 14.07530573 | 14.07530573 | ENSG00000274978 | up |
| ENSG00000208018 | 8.935318131 | -2.891087539 | 0.5916262 | -4.886679 | 1.03E-06 | 1.40E-05 | MIR645 | miRNA | 56.38 | -14.03728254 | 14.03728254 | ENSG00000208018 | down |
| ENSG00000240589 | 27.44188649 | -1.92834926 | 0.3221618 | -5.985654 | 2.16E-09 | 5.52E-08 | RN7SL258P | misc_RNA | 53.33 | -13.99676254 | 13.99676254 | ENSG00000240589 | down |
| ENSG00000247735 | 86.11180709 | 1.612636595 | 0.2466275 | 6.538754 | 6.20E-11 | 2.17E-09 | | antisense | 47.76 | 13.97111771 | 13.97111771 | ENSG00000247735 | up |
| ENSG00000152443 | 954.9756791 | -1.558467861 | 0.2348001 | -6.637423 | 3.19E-11 | 1.17E-09 | ZNF776 | protein_coding | 43.81 | -13.92018434 | 13.92018434 | ENSG00000152443 | down |
| ENSG00000256436 | 189.0177217 | -1.820709852 | 0.296818 | -6.134094 | 8.56E-10 | 2.39E-08 | TAS2R31 | protein_coding | 40.45 | -13.87683586 | 13.87683586 | ENSG00000256436 | down |
| ENSG00000279372 | 23.22669024 | 2.102723586 | 0.3684726 | 5.706594 | 1.15E-08 | 2.54E-07 | | TEC | 51.01 | 13.86955361 | 13.86955361 | ENSG00000279372 | up |
| ENSG00000221783 | 4.456582915 | -3.82507793 | 0.9097304 | -4.204628 | 2.62E-05 | 0.000237665 | MIR1183 | miRNA | 48.31 | -13.86221472 | 13.86221472 | ENSG00000221783 | down |
| ENSG00000198551 | 537.9379281 | -1.615748695 | 0.2483281 | -6.506509 | 7.69E-11 | 2.64E-09 | ZNF627 | protein_coding | 48.23 | -13.86113428 | 13.86113428 | ENSG00000198551 | down |
| ENSG00000272716 | 81.11627354 | 1.741790514 | 0.2780627 | 6.26402 | 3.75E-10 | 1.13E-08 | | lincRNA | 47.89 | 13.84307288 | 13.84307288 | ENSG00000272716 | up |
| ENSG00000211491 | 5.892627491 | -3.162002548 | 0.6817081 | -4.638353 | 3.51E-06 | 4.19E-05 | MIR320D1 | miRNA | 47.92 | -13.84257496 | 13.84257496 | ENSG00000211491 | down |
| ENSG00000266146 | 9.673692773 | -2.643078411 | 0.520397 | -5.078965 | 3.79E-07 | 5.83E-06 | MIR5190 | miRNA | 61.25 | -13.83563442 | 13.83563442 | ENSG00000266146 | down |
| ENSG00000177873 | 216.7730611 | -1.473236833 | 0.2166085 | -6.801382 | 1.04E-11 | 4.18E-10 | ZNF619 | protein_coding | 45.23 | -13.81695883 | 13.81695883 | ENSG00000177873 | down |
| ENSG00000200090 | 22.15801654 | -2.113693314 | 0.3727267 | -5.670893 | 1.42E-08 | 3.06E-07 | | misc_RNA | 44.44 | -13.7686904 | 13.7686904 | ENSG00000200090 | down |
| ENSG00000202119 | 8.541143209 | -2.85228904 | 0.5855488 | -4.871138 | 1.11E-06 | 1.50E-05 | RNU6-302P | snRNA | 45.79 | -13.76169829 | 13.76169829 | ENSG00000202119 | down |
| ENSG00000172086 | 265.514732 | -1.804616668 | 0.2941777 | -6.134444 | 8.55E-10 | 2.39E-08 | KRCC1 | protein_coding | 38.55 | -13.75432874 | 13.75432874 | ENSG00000172086 | down |
| ENSG00000228988 | 11.86992713 | 2.46019265 | 0.4687478 | 5.248435 | 1.53E-07 | 2.58E-06 | | antisense | 36.76 | 13.74874359 | 13.74874359 | ENSG00000228988 | up |
| ENSG00000280186 | 110.7990779 | 1.587186996 | 0.2430085 | 6.531405 | 6.52E-11 | 2.27E-09 | | TEC | 42.04 | 13.72083898 | 13.72083898 | ENSG00000280186 | up |
| ENSG00000272936 | 59.19137603 | -1.73434524 | 0.2776154 | -6.247294 | 4.18E-10 | 1.25E-08 | | antisense | 26.74 | -13.70964908 | 13.70964908 | ENSG00000272936 | down |
| ENSG00000269889 | 85.4181246 | 1.693753817 | 0.2680578 | 6.318615 | 2.64E-10 | 8.24E-09 | | antisense | 21.04 | 13.6925295 | 13.6925295 | ENSG00000269889 | up |
| ENSG00000208028 | 27.86256564 | 2.012507158 | 0.3476135 | 5.789497 | 7.06E-09 | 1.64E-07 | MIR616 | miRNA | 40.21 | 13.6561261 | 13.6561261 | ENSG00000208028 | up |
| ENSG00000200413 | 7.794823466 | -3.330618287 | 0.7430884 | -4.48213 | 7.39E-06 | 8.03E-05 | SNORD114-26 | snoRNA | 47.22 | -13.63941793 | 13.63941793 | ENSG00000200413 | down |
| ENSG00000244086 | 9.538213672 | 2.574806773 | 0.5040784 | 5.107949 | 3.26E-07 | 5.07E-06 | | processed_pseudogene | 44.54 | 13.63301468 | 13.63301468 | ENSG00000244086 | up |
| ENSG00000207943 | 3.693346272 | -4.286251039 | 1.0903773 | -3.93098 | 8.46E-05 | 0.000659883 | MIR634 | miRNA | 52.58 | -13.63256272 | 13.63256272 | ENSG00000207943 | down |

| | | | | | | | | | | | | | |
|---|---|---|---|---|---|---|---|---|---|---|---|---|---|
| ENSG00000207117 | 19.25557626 | -2.344481059 | 0.4375549 | -5.358141 | 8.41E-08 | 1.53E-06 | | misc_RNA | 42.72 | -13.63101574 | 13.63101574 | ENSG00000207117 | down |
| ENSG00000202408 | 3.882002695 | 4.636633447 | 1.2278642 | 3.776178 | 0.000159253 | 0.001152435 | RNU1-122P | snRNA | 48.72 | 13.62420743 | 13.62420743 | ENSG00000202408 | up |
| ENSG00000284059 | 5.251209733 | -3.513466309 | 0.8065639 | -4.356092 | 1.32E-05 | 0.000132816 | MIR5572 | miRNA | 59.12 | -13.62083444 | 13.62083444 | ENSG00000284059 | down |
| ENSG00000275441 | 28.70580172 | 1.924463272 | 0.3255433 | 5.911542 | 3.39E-09 | 8.40E-08 | | antisense | 53.24 | 13.61732395 | 13.61732395 | ENSG00000275441 | up |
| ENSG00000252985 | 11.98093851 | -2.420369212 | 0.4601639 | -5.259798 | 1.44E-07 | 2.44E-06 | | snoRNA | 43.59 | -13.58343032 | 13.58343032 | ENSG00000252985 | down |
| ENSG00000273682 | 29.10572934 | -2.105517002 | 0.3732369 | -5.641236 | 1.69E-08 | 3.58E-07 | | processed_pseudogene | 56.59 | -13.57294609 | 13.57294609 | ENSG00000273682 | down |
| ENSG00000200608 | 8.208468801 | -3.304627767 | 0.7367557 | -4.485378 | 7.28E-06 | 7.93E-05 | SNORD114-11 | snoRNA | 49.33 | -13.55073656 | 13.55073656 | ENSG00000200608 | down |
| ENSG00000147459 | 988.4931239 | -1.506324697 | 0.2262199 | -6.658674 | 2.76E-11 | 1.02E-09 | DOCK5 | protein_coding | 42.37 | -13.54194863 | 13.54194863 | ENSG00000147459 | down |
| ENSG00000233644 | 27.22623945 | -2.06236113 | 0.3624274 | -5.690412 | 1.27E-08 | 2.75E-07 | ARHGEF7-IT1 | sense_intronic | 43.09 | -13.53083442 | 13.53083442 | ENSG00000233644 | down |
| ENSG00000139725 | 579.3112233 | 1.392058324 | 0.20127 | 6.916373 | 4.63E-12 | 1.94E-10 | RHOF | protein_coding | 56.89 | 13.52078206 | 13.52078206 | ENSG00000139725 | up |
| ENSG00000178607 | 175.5212705 | 1.659776518 | 0.2622732 | 6.328427 | 2.48E-10 | 7.77E-09 | ERN1 | protein_coding | 45.1 | 13.46040859 | 13.46040859 | ENSG00000178607 | up |
| ENSG00000149308 | 887.6463472 | -1.683540818 | 0.2681337 | -6.278737 | 3.41E-10 | 1.04E-08 | NPAT | protein_coding | 40.27 | -13.4429058 | 13.4429058 | ENSG00000149308 | down |
| ENSG00000277440 | 33.89798715 | -1.852208046 | 0.3096894 | -5.980856 | 2.22E-09 | 5.65E-08 | | sense_intronic | 43.81 | -13.42473852 | 13.42473852 | ENSG00000277440 | down |
| ENSG00000196150 | 624.8660953 | -1.716472236 | 0.2762405 | -6.213689 | 5.18E-10 | 1.52E-08 | ZNF250 | protein_coding | 47.13 | -13.42106377 | 13.42106377 | ENSG00000196150 | down |
| ENSG00000275202 | 52.24874806 | 1.732274982 | 0.2801461 | 6.183471 | 6.27E-10 | 1.81E-08 | | antisense | 61.76 | 13.41282492 | 13.41282492 | ENSG00000275202 | up |
| ENSG00000216977 | 40.48633793 | -1.950694057 | 0.3348016 | -5.826418 | 5.66E-09 | 1.35E-07 | RPL21P65 | processed_pseudogene | 45.49 | -13.40345338 | 13.40345338 | ENSG00000216977 | down |
| ENSG00000272345 | 26.82769332 | -1.925402835 | 0.3285129 | -5.860966 | 4.60E-09 | 1.11E-07 | | antisense | 35.29 | -13.387335 | 13.387335 | ENSG00000272345 | down |
| ENSG00000262202 | 315.7603146 | 1.445493248 | 0.2141052 | 6.751321 | 1.47E-11 | 5.75E-10 | | lincRNA | 56.2 | 13.35728848 | 13.35728848 | ENSG00000262202 | up |
| ENSG00000253925 | 78.6025224 | 1.684816908 | 0.2693081 | 6.256095 | 3.95E-10 | 1.18E-08 | | antisense | 44.09 | 13.35466514 | 13.35466514 | ENSG00000253925 | up |
| ENSG00000268439 | 73.19050735 | -1.747621086 | 0.2847087 | -6.138278 | 8.34E-10 | 2.34E-08 | EMG1 | protein_coding | 45.52 | -13.33670544 | 13.33670544 | ENSG00000268439 | down |
| ENSG00000164011 | 107.0841308 | -1.683310761 | 0.2695072 | -6.245883 | 4.21E-10 | 1.25E-08 | ZNF691 | protein_coding | 47.25 | -13.30119565 | 13.30119565 | ENSG00000164011 | down |
| ENSG00000225742 | 31.56492615 | 1.927706802 | 0.3309801 | 5.824237 | 5.74E-09 | 1.36E-07 | LINC02036 | lincRNA | 44.61 | 13.23884969 | 13.23884969 | ENSG00000225742 | up |
| ENSG00000145868 | 1148.166217 | -1.76003516 | 0.288884 | -6.092533 | 1.11E-09 | 3.03E-08 | FBXO38 | protein_coding | 37.05 | -13.233399 | 13.233399 | ENSG00000145868 | down |
| ENSG00000224223 | 178.6204704 | -1.848657787 | 0.3118619 | -5.927808 | 3.07E-09 | 7.66E-08 | VSTM2A-OT1 | sense_overlapping | 35.79 | -13.15459803 | 13.15459803 | ENSG00000224223 | down |
| ENSG00000276137 | 3.735289018 | -3.770585758 | 0.914478 | -4.123211 | 3.74E-05 | 0.000325126 | MIR6133 | miRNA | 50.93 | -13.15160906 | 13.15160906 | ENSG00000276137 | down |
| ENSG00000241217 | 46.42509324 | -1.816300391 | 0.3038853 | -5.976928 | 2.27E-09 | 5.77E-08 | RN7SL809P | misc_RNA | 50.84 | -13.14833881 | 13.14833881 | ENSG00000241217 | down |
| ENSG00000275401 | 109.9728466 | -1.559797855 | 0.241862 | -6.449123 | 1.12E-10 | 3.72E-09 | | lincRNA | 50.69 | -13.14816135 | 13.14816135 | ENSG00000275401 | down |
| ENSG00000271646 | 49.60297391 | 1.742963277 | 0.2857177 | 6.100299 | 1.06E-09 | 2.90E-08 | | lincRNA | 50.34 | 13.13733784 | 13.13733784 | ENSG00000271646 | up |
| ENSG00000212447 | 3.996865564 | -4.226595069 | 1.0881158 | -3.884325 | 0.000102615 | 0.000781995 | SNORD90 | snoRNA | 33.33 | -13.13116877 | 13.13116877 | ENSG00000212447 | down |
| ENSG00000227036 | 53.80252306 | 1.815924728 | 0.3041905 | 5.969696 | 2.38E-09 | 6.00E-08 | LINC00511 | lincRNA | 47.33 | 13.11494539 | 13.11494539 | ENSG00000227036 | up |
| ENSG00000202187 | 3.450333279 | -4.220965918 | 1.0873537 | -3.88187 | 0.000103656 | 0.000788636 | RNA5SP355 | rRNA | 47.41 | -13.09817746 | 13.09817746 | ENSG00000202187 | down |
| ENSG00000271265 | 26.97047153 | -1.952260021 | 0.3393826 | -5.752388 | 8.80E-09 | 1.99E-07 | | lincRNA | 36.1 | -13.08288693 | 13.08288693 | ENSG00000271265 | down |

| | | | | | | | | | | | | | |
|---|---|---|---|---|---|---|---|---|---|---|---|---|---|
| ENSG00000141040 | 306.1634949 | -1.475351942 | 0.2231447 | -6.611637 | 3.80E-11 | 1.38E-09 | ZNF287 | protein_coding | 40.28 | -13.07266544 | 13.07266544 | ENSG00000141040 | down |
| ENSG00000208002 | 12.08057242 | -3.345167515 | 0.7652311 | -4.371447 | 1.23E-05 | 0.000124984 | MIR643 | miRNA | 38.14 | -13.05667699 | 13.05667699 | ENSG00000208002 | down |
| ENSG00000264210 | 1.58653943 | 4.627463636 | 1.2513383 | 3.698012 | 0.000217295 | 0.001508173 | MIR4716 | miRNA | 45.24 | 13.05661422 | 13.05661422 | ENSG00000264210 | up |
| ENSG00000222881 | 19.38442386 | -2.092252306 | 0.3774635 | -5.542927 | 2.97E-08 | 5.98E-07 | | misc_RNA | 38.32 | -13.01997896 | 13.01997896 | ENSG00000222881 | down |
| ENSG00000277324 | 39.97219232 | 1.828974173 | 0.3084615 | 5.929343 | 3.04E-09 | 7.62E-08 | | antisense | 51.28 | 13.01910344 | 13.01910344 | ENSG00000277324 | up |
| ENSG00000277476 | 107.1927263 | -1.625001218 | 0.2584311 | -6.287949 | 3.22E-10 | 9.78E-09 | | lincRNA | 59.74 | -13.01576776 | 13.01576776 | ENSG00000277476 | down |
| ENSG00000274213 | 21.33478837 | 2.152181893 | 0.3940958 | 5.461063 | 4.73E-08 | 9.12E-07 | | lincRNA | 63.51 | 12.99961772 | 12.99961772 | ENSG00000274213 | up |
| ENSG00000280088 | 127.6947645 | -1.762242768 | 0.2920024 | -6.035028 | 1.59E-09 | 4.21E-08 | | TEC | 39.54 | -12.99836164 | 12.99836164 | ENSG00000280088 | down |
| ENSG00000274799 | 10.630284 | -2.489073001 | 0.4911384 | -5.067967 | 4.02E-07 | 6.10E-06 | | misc_RNA | 55.32 | -12.97881408 | 12.97881408 | ENSG00000274799 | down |
| ENSG00000273175 | 166.5811009 | 1.577594892 | 0.2476139 | 6.371189 | 1.88E-10 | 6.03E-09 | | lincRNA | 45.05 | 12.9671919 | 12.9671919 | ENSG00000273175 | up |
| ENSG00000265565 | 38.39255273 | 1.914676625 | 0.3314111 | 5.777346 | 7.59E-09 | 1.75E-07 | MIR3143 | miRNA | 41.27 | 12.93909348 | 12.93909348 | ENSG00000265565 | up |
| ENSG00000261447 | 22.11056841 | 2.047347843 | 0.3674925 | 5.571128 | 2.53E-08 | 5.17E-07 | | lincRNA | 53.89 | 12.87119908 | 12.87119908 | ENSG00000261447 | up |
| ENSG00000268521 | 47.6421059 | -1.95619461 | 0.3436682 | -5.692102 | 1.25E-08 | 2.73E-07 | VN1R83P | unprocessed_pseudogene | 35.96 | -12.8401819 | 12.8401819 | ENSG00000268521 | down |
| ENSG00000206828 | 529.8120688 | 1.416714141 | 0.2118939 | 6.685959 | 2.29E-11 | 8.68E-10 | | snRNA | 45.12 | 12.83754025 | 12.83754025 | ENSG00000206828 | up |
| ENSG00000146267 | 1024.405821 | -1.571315417 | 0.2474794 | -6.349278 | 2.16E-10 | 6.88E-09 | FAXC | protein_coding | 41.71 | -12.8260661 | 12.8260661 | ENSG00000146267 | down |
| ENSG00000259291 | 122.083504 | -1.629622245 | 0.2614139 | -6.233878 | 4.55E-10 | 1.35E-08 | ZNF710-AS1 | antisense | 53.18 | -12.82566215 | 12.82566215 | ENSG00000259291 | down |
| ENSG00000206784 | 4.326781027 | -3.726928519 | 0.9100214 | -4.09543 | 4.21E-05 | 0.000362158 | | misc_RNA | 40.71 | -12.82474147 | 12.82474147 | ENSG00000206784 | down |
| ENSG00000266063 | 4.227674026 | -4.592544609 | 1.2500437 | -3.673907 | 0.000238869 | 0.001635092 | MIR4771-2 | miRNA | 44.59 | -12.79693132 | 12.79693132 | ENSG00000266063 | down |
| ENSG00000173218 | 392.0940419 | -1.471213632 | 0.2248305 | -6.543658 | 6.00E-11 | 2.11E-09 | VANGL1 | protein_coding | 45.32 | -12.76362895 | 12.76362895 | ENSG00000173218 | down |
| ENSG00000270964 | 82.88188402 | 1.611523863 | 0.2577853 | 6.251419 | 4.07E-10 | 1.22E-08 | | lincRNA | 46.77 | 12.75575426 | 12.75575426 | ENSG00000270964 | up |
| ENSG00000264885 | 200.6267165 | -1.537413143 | 0.2402079 | -6.400343 | 1.55E-10 | 5.08E-09 | | sense_intronic | 46.1 | -12.75168282 | 12.75168282 | ENSG00000264885 | down |
| ENSG00000238658 | 13.86915194 | -2.205026034 | 0.4126949 | -5.342993 | 9.14E-08 | 1.65E-06 | RNU6-625P | snRNA | 38.32 | -12.75127526 | 12.75127526 | ENSG00000238658 | down |
| ENSG00000254028 | 22.13261908 | 1.981729192 | 0.3523444 | 5.624409 | 1.86E-08 | 3.91E-07 | | lincRNA | 43.09 | 12.69748532 | 12.69748532 | ENSG00000254028 | up |
| ENSG00000275769 | 22.26598378 | -2.015657162 | 0.3622167 | -5.564783 | 2.62E-08 | 5.35E-07 | | sense_intronic | 34.05 | -12.64138303 | 12.64138303 | ENSG00000275769 | down |
| ENSG00000236095 | 25.81002273 | -2.031542287 | 0.3670245 | -5.535169 | 3.11E-08 | 6.22E-07 | | sense_intronic | 42.14 | -12.60818897 | 12.60818897 | ENSG00000236095 | down |
| ENSG00000260400 | 147.5878597 | -1.881785004 | 0.3276725 | -5.742884 | 9.31E-09 | 2.09E-07 | | sense_overlapping | 33.03 | -12.56997204 | 12.56997204 | ENSG00000260400 | down |
| ENSG00000238250 | 20.7589253 | -2.130565421 | 0.3953646 | -5.388863 | 7.09E-08 | 1.31E-06 | ST6GAL2-IT1 | sense_intronic | 39.64 | -12.53526194 | 12.53526194 | ENSG00000238250 | down |
| ENSG00000233220 | 52.48998556 | 1.700929934 | 0.2821096 | 6.029324 | 1.65E-09 | 4.33E-08 | LINC00167 | lincRNA | 54.19 | 12.52423557 | 12.52423557 | ENSG00000233220 | up |
| ENSG00000237373 | 23.35286525 | -1.983385861 | 0.3555308 | -5.578662 | 2.42E-08 | 4.97E-07 | BRWD1-IT1 | sense_overlapping | 37.38 | -12.50256372 | 12.50256372 | ENSG00000237373 | down |
| ENSG00000152782 | 440.0716984 | -1.687152104 | 0.2796331 | -6.03345 | 1.60E-09 | 4.23E-08 | PANK1 | protein_coding | 39.91 | -12.44010994 | 12.44010994 | ENSG00000152782 | down |
| ENSG00000268105 | 52.19339384 | -1.763180336 | 0.298716 | -5.902531 | 3.58E-09 | 8.81E-08 | | processed_pseudogene | 36.78 | -12.43966203 | 12.43966203 | ENSG00000268105 | down |
| ENSG00000185716 | 272.8967459 | -1.801210604 | 0.3083799 | -5.840881 | 5.19E-09 | 1.24E-07 | C16orf52 | protein_coding | 39.3 | -12.43881959 | 12.43881959 | ENSG00000185716 | down |

| | | | | | | | | | | | | | |
|---|---|---|---|---|---|---|---|---|---|---|---|---|---|
| ENSG00000267543 | 19.43560888 | -1.966976802 | 0.3519853 | -5.588235 | 2.29E-08 | 4.75E-07 | | sense_intronic | 46.92 | -12.43731926 | 12.43731926 | ENSG00000267543 | down |
| ENSG00000279089 | 376.274008 | -1.691602032 | 0.280796 | -6.024309 | 1.70E-09 | 4.46E-08 | | TEC | 37.78 | -12.43415399 | 12.43415399 | ENSG00000279089 | down |
| ENSG00000199460 | 6.062683748 | -2.945538697 | 0.6474007 | -4.549792 | 5.37E-06 | 6.09E-05 | RNU6-1216P | snRNA | 40.19 | -12.41622746 | 12.41622746 | ENSG00000199460 | down |
| ENSG00000257680 | 52.2654836 | -1.995956116 | 0.3605088 | -5.536498 | 3.09E-08 | 6.18E-07 | | processed_pseudogene | 41.86 | -12.39266203 | 12.39266203 | ENSG00000257680 | down |
| ENSG00000130921 | 399.6939754 | -1.61274076 | 0.2623508 | -6.147268 | 7.88E-10 | 2.22E-08 | C12orf65 | protein_coding | 47.41 | -12.34422108 | 12.34422108 | ENSG00000130921 | down |
| ENSG00000273306 | 76.36794574 | -1.782478927 | 0.3060352 | -5.824425 | 5.73E-09 | 1.36E-07 | | antisense | 44.41 | -12.2414729 | 12.2414729 | ENSG00000273306 | down |
| ENSG00000263235 | 41.98021627 | -1.748216981 | 0.2972708 | -5.880891 | 4.08E-09 | 9.97E-08 | | sense_overlapping | 50.84 | -12.23999006 | 12.23999006 | ENSG00000263235 | down |
| ENSG00000206778 | 5.811170335 | -3.047521158 | 0.6868748 | -4.436793 | 9.13E-06 | 9.68E-05 | RNU6-213P | snRNA | 41.35 | -12.23381012 | 12.23381012 | ENSG00000206778 | down |
| ENSG00000252112 | 9.683194746 | -2.352723852 | 0.4652503 | -5.0569 | 4.26E-07 | 6.41E-06 | | snoRNA | 43.75 | -12.21849129 | 12.21849129 | ENSG00000252112 | down |
| ENSG00000273156 | 64.63271804 | -1.799092046 | 0.3112391 | -5.780418 | 7.45E-09 | 1.72E-07 | | lincRNA | 30.83 | -12.17096159 | 12.17096159 | ENSG00000273156 | down |
| ENSG00000269961 | 30.19703628 | 1.844805751 | 0.3233942 | 5.704511 | 1.17E-08 | 2.55E-07 | | lincRNA | 48.32 | 12.16231701 | 12.16231701 | ENSG00000269961 | up |
| ENSG00000206727 | 4.464364842 | -4.518977781 | 1.2534564 | -3.605213 | 0.000311897 | 0.002064925 | SNORD116-9 | snoRNA | 45.26 | -12.13388803 | 12.13388803 | ENSG00000206727 | down |
| ENSG00000224152 | 125.9428897 | 1.551196993 | 0.2497173 | 6.211812 | 5.24E-10 | 1.53E-08 | | antisense | 51.32 | 12.12219145 | 12.12219145 | ENSG00000224152 | up |
| ENSG00000139800 | 1001.483158 | -1.378053428 | 0.2094822 | -6.578379 | 4.76E-11 | 1.71E-09 | ZIC5 | protein_coding | 51.78 | -12.08257071 | 12.08257071 | ENSG00000139800 | down |
| ENSG00000141639 | 250.2078667 | -1.553890845 | 0.2507748 | -6.19636 | 5.78E-10 | 1.68E-08 | MAPK4 | protein_coding | 45.75 | -12.08039608 | 12.08039608 | ENSG00000141639 | down |
| ENSG00000261758 | 86.59879589 | 1.602547213 | 0.2630145 | 6.092998 | 1.11E-09 | 3.03E-08 | | antisense | 43.73 | 12.04993424 | 12.04993424 | ENSG00000261758 | up |
| ENSG00000279948 | 119.3438404 | -1.585348327 | 0.2587892 | -6.126021 | 9.01E-10 | 2.51E-08 | | TEC | 46.34 | -12.04943084 | 12.04943084 | ENSG00000279948 | down |
| ENSG00000273783 | 172.1531682 | 1.494222388 | 0.2367735 | 6.310766 | 2.78E-10 | 8.65E-09 | | antisense | 46.82 | 12.04801105 | 12.04801105 | ENSG00000273783 | up |
| ENSG00000271576 | 696.8178268 | 1.323626649 | 0.1978823 | 6.688959 | 2.25E-11 | 8.53E-10 | | lincRNA | 43.79 | 12.00424379 | 12.00424379 | ENSG00000271576 | up |
| ENSG00000283078 | 106.6335789 | 1.614138922 | 0.2665027 | 6.056746 | 1.39E-09 | 3.72E-08 | | lincRNA | 51.24 | 11.9923399 | 11.9923399 | ENSG00000283078 | up |
| ENSG00000176840 | 10.05508018 | 2.38868143 | 0.4809293 | 4.966804 | 6.81E-07 | 9.57E-06 | MIR7-3HG | lincRNA | 51.53 | 11.9887361 | 11.9887361 | ENSG00000176840 | up |
| ENSG00000268713 | 52.84346253 | -1.797543121 | 0.314306 | -5.719087 | 1.07E-08 | 2.38E-07 | | lincRNA | 40.14 | -11.9052362 | 11.9052362 | ENSG00000268713 | down |
| ENSG00000242791 | 27.68916145 | -1.890879736 | 0.3391871 | -5.57474 | 2.48E-08 | 5.07E-07 | | lincRNA | 37.87 | -11.9033476 | 11.9033476 | ENSG00000242791 | down |
| ENSG00000109674 | 496.3885167 | -1.615454311 | 0.2682159 | -6.022962 | 1.71E-09 | 4.49E-08 | NEIL3 | protein_coding | 37.34 | -11.86991001 | 11.86991001 | ENSG00000109674 | down |
| ENSG00000272662 | 44.36147305 | 1.676124888 | 0.2836939 | 5.908216 | 3.46E-09 | 8.53E-08 | | lincRNA | 42.86 | 11.84838523 | 11.84838523 | ENSG00000272662 | up |
| ENSG00000115221 | 16.10962515 | -2.091736933 | 0.395932 | -5.283072 | 1.27E-07 | 2.19E-06 | ITGB6 | protein_coding | 40.66 | -11.83758924 | 11.83758924 | ENSG00000115221 | down |
| ENSG00000283736 | 26.46514747 | 1.890497349 | 0.3405502 | 5.551303 | 2.84E-08 | 5.74E-07 | MIR484 | miRNA | 59.49 | 11.79902462 | 11.79902462 | ENSG00000283736 | up |
| ENSG00000252153 | 7.998113151 | -2.721464007 | 0.5903938 | -4.609574 | 4.03E-06 | 4.74E-05 | MIR2278 | miRNA | 58.33 | -11.76801075 | 11.76801075 | ENSG00000252153 | down |
| ENSG00000248521 | 136.3565334 | -1.728767829 | 0.2983028 | -5.795346 | 6.82E-09 | 1.59E-07 | | processed_pseudogene | 55.01 | -11.75190992 | 11.75190992 | ENSG00000248521 | down |
| ENSG00000255571 | 758.1175585 | -1.557978223 | 0.2553096 | -6.10231 | 1.05E-09 | 2.87E-08 | MIR9-3HG | lincRNA | 53.82 | -11.75021927 | 11.75021927 | ENSG00000255571 | down |
| ENSG00000252317 | 5.231884894 | -2.999664006 | 0.68582 | -4.373836 | 1.22E-05 | 0.000123979 | | misc_RNA | 37.63 | -11.71864244 | 11.71864244 | ENSG00000252317 | down |
| ENSG00000277590 | 57.80642837 | 1.629276785 | 0.2738706 | 5.949074 | 2.70E-09 | 6.79E-08 | MIR7845 | miRNA | 72.73 | 11.67893474 | 11.67893474 | ENSG00000277590 | up |

| | | | | | | | | | | | | | |
|---|---|---|---|---|---|---|---|---|---|---|---|---|---|
| ENSG00000225423 | 69.19149252 | -1.677880956 | 0.286772 | -5.850924 | 4.89E-09 | 1.18E-07 | TNPO1P1 | processed_pseudogene | 41.12 | -11.62607081 | 11.62607081 | ENSG00000225423 | down |
| ENSG00000274220 | 56.85954464 | 1.738753345 | 0.3026589 | 5.744927 | 9.20E-09 | 2.07E-07 | | lincRNA | 46.42 | 11.62244627 | 11.62244627 | ENSG00000274220 | up |
| ENSG00000263766 | 33.7769026 | -1.764935458 | 0.3096321 | -5.700105 | 1.20E-08 | 2.61E-07 | | antisense | 43.3 | -11.61834361 | 11.61834361 | ENSG00000263766 | down |
| ENSG00000237605 | 17.04986574 | 2.030211866 | 0.3827991 | 5.303597 | 1.14E-07 | 1.98E-06 | | antisense | 56.26 | 11.57741545 | 11.57741545 | ENSG00000237605 | up |
| ENSG00000256294 | 223.2486174 | -1.529502397 | 0.2504569 | -6.106848 | 1.02E-09 | 2.80E-08 | ZNF225 | protein_coding | 41.4 | -11.5517035 | 11.5517035 | ENSG00000256294 | down |
| ENSG00000258001 | 19.27753674 | 2.011101704 | 0.3778551 | 5.322415 | 1.02E-07 | 1.82E-06 | | lincRNA | 43.48 | 11.54319116 | 11.54319116 | ENSG00000258001 | up |
| ENSG00000196152 | 172.201909 | -1.438556392 | 0.2285935 | -6.293077 | 3.11E-10 | 9.52E-09 | ZNF79 | protein_coding | 47.15 | -11.53893864 | 11.53893864 | ENSG00000196152 | down |
| ENSG00000177614 | 358.9189377 | -1.646631436 | 0.2798865 | -5.883212 | 4.02E-09 | 9.85E-08 | PGBD5 | protein_coding | 48.34 | -11.53752136 | 11.53752136 | ENSG00000177614 | down |
| ENSG00000207174 | 6.747883458 | -2.986310574 | 0.6878856 | -4.341289 | 1.42E-05 | 0.000140569 | SNORD116-15 | snoRNA | 41.3 | -11.50360115 | 11.50360115 | ENSG00000207174 | down |
| ENSG00000270742 | 96.40405095 | -1.687460173 | 0.2909186 | -5.800455 | 6.61E-09 | 1.55E-07 | | sense_intronic | 41.28 | -11.49219585 | 11.49219585 | ENSG00000270742 | down |
| ENSG00000207093 | 4.538609191 | -4.048136763 | 1.0920942 | -3.706765 | 0.000209923 | 0.001463578 | SNORD116-8 | snoRNA | 44.21 | -11.47478481 | 11.47478481 | ENSG00000207093 | down |
| ENSG00000232906 | 82.79603994 | -1.792185719 | 0.3187999 | -5.621663 | 1.89E-08 | 3.97E-07 | | processed_pseudogene | 33.95 | -11.47182344 | 11.47182344 | ENSG00000232906 | down |
| ENSG00000222958 | 3.368031767 | -4.047916984 | 1.0932771 | -3.702554 | 0.00021344 | 0.001482898 | MIR1913 | miRNA | 63.75 | -11.45110662 | 11.45110662 | ENSG00000222958 | down |
| ENSG00000208012 | 2.763822447 | -4.43401957 | 1.2557278 | -3.531036 | 0.000413936 | 0.002643683 | MIRLET7F2 | miRNA | 42.17 | -11.42997418 | 11.42997418 | ENSG00000208012 | down |
| ENSG00000270277 | 163.6683147 | -1.538722647 | 0.2541217 | -6.055062 | 1.40E-09 | 3.75E-08 | | antisense | 39.77 | -11.42632215 | 11.42632215 | ENSG00000270277 | down |
| ENSG00000283676 | 2.731289528 | 4.380183888 | 1.233917 | 3.549821 | 0.000385494 | 0.002481383 | MIR5087 | miRNA | 42.11 | 11.41172044 | 11.41172044 | ENSG00000283676 | up |
| ENSG00000199204 | 8.17752673 | -2.858749098 | 0.6471948 | -4.417139 | 1.00E-05 | 0.000104529 | | misc_RNA | 40.2 | -11.380004 | 11.380004 | ENSG00000199204 | down |
| ENSG00000241525 | 35.98027945 | 1.837689958 | 0.33246 | 5.527552 | 3.25E-08 | 6.47E-07 | | antisense | 50.98 | 11.37387005 | 11.37387005 | ENSG00000241525 | up |
| ENSG00000265745 | 19.92520694 | -1.961053542 | 0.3666876 | -5.348022 | 8.89E-08 | 1.61E-06 | RN7SL375P | misc_RNA | 52.68 | -11.36075745 | 11.36075745 | ENSG00000265745 | down |
| ENSG00000163684 | 532.0847647 | -1.586813102 | 0.2676845 | -5.927924 | 3.07E-09 | 7.66E-08 | RPP14 | protein_coding | 44.14 | -11.2913751 | 11.2913751 | ENSG00000163684 | down |
| ENSG00000218730 | 18.77108254 | -2.060728897 | 0.3965169 | -5.197077 | 2.02E-07 | 3.33E-06 | | processed_pseudogene | 44.33 | -11.28834154 | 11.28834154 | ENSG00000218730 | down |
| ENSG00000238300 | 10.41952262 | -2.397637649 | 0.4983976 | -4.810692 | 1.50E-06 | 1.96E-05 | SNORD121B | snoRNA | 40.74 | -11.28508561 | 11.28508561 | ENSG00000238300 | down |
| ENSG00000125846 | 323.3663284 | -1.673589534 | 0.2900265 | -5.770472 | 7.90E-09 | 1.81E-07 | ZNF133 | protein_coding | 40.41 | -11.28379261 | 11.28379261 | ENSG00000125846 | down |
| ENSG00000165655 | 311.0763285 | 1.375174319 | 0.2162188 | 6.360106 | 2.02E-10 | 6.44E-09 | ZNF503 | protein_coding | 61.2 | 11.26437067 | 11.26437067 | ENSG00000165655 | up |
| ENSG00000118596 | 41.11444633 | -1.744767283 | 0.3092009 | -5.642828 | 1.67E-08 | 3.55E-07 | SLC16A7 | protein_coding | 35.78 | -11.25327389 | 11.25327389 | ENSG00000118596 | down |
| ENSG00000232082 | 44.61334581 | -1.893262069 | 0.3498508 | -5.411626 | 6.25E-08 | 1.17E-06 | RPS6KA2-IT1 | sense_intronic | 53.76 | -11.22802671 | 11.22802671 | ENSG00000232082 | down |
| ENSG00000251774 | 11.14482311 | -2.382495814 | 0.4948977 | -4.814118 | 1.48E-06 | 1.94E-05 | RNU6-377P | snRNA | 40.2 | -11.22669558 | 11.22669558 | ENSG00000251774 | down |
| ENSG00000207309 | 19.9718555 | -2.168517435 | 0.4308685 | -5.032898 | 4.83E-07 | 7.09E-06 | RNU4-70P | snRNA | 41.01 | -11.16687764 | 11.16687764 | ENSG00000207309 | down |
| ENSG00000261428 | 278.1890744 | -1.731683095 | 0.3071893 | -5.637186 | 1.73E-08 | 3.65E-07 | | antisense | 40.23 | -11.14767583 | 11.14767583 | ENSG00000261428 | down |
| ENSG00000117616 | 528.5979559 | 1.396346772 | 0.2226751 | 6.27078 | 3.59E-10 | 1.09E-08 | RSRP1 | protein_coding | 46.4 | 11.11999799 | 11.11999799 | ENSG00000117616 | up |
| ENSG00000184557 | 109.2446673 | 1.57924714 | 0.267899 | 5.894935 | 3.75E-09 | 9.19E-08 | SOCS3 | protein_coding | 62.24 | 11.11281648 | 11.11281648 | ENSG00000184557 | up |
| ENSG00000224142 | 24.93481463 | -1.956870806 | 0.3699962 | -5.288895 | 1.23E-07 | 2.13E-06 | AMMECR1-IT1 | sense_intronic | 43.78 | -11.10033798 | 11.10033798 | ENSG00000224142 | down |

| | | | | | | | | | | | | | |
|---|---|---|---|---|---|---|---|---|---|---|---|---|---|
| ENSG00000105072 | 91.81539351 | -1.729952448 | 0.3073954 | -5.627776 | 1.83E-08 | 3.85E-07 | C19orf44 | protein_coding | 51.63 | -11.09780494 | 11.09780494 | ENSG00000105072 | down |
| ENSG00000278099 | 202.4581274 | 1.436846659 | 0.2326752 | 6.175332 | 6.60E-10 | 1.89E-08 | | snRNA | 51.52 | 11.09704954 | 11.09704954 | ENSG00000278099 | up |
| ENSG00000215930 | 10.006492 | -2.383511212 | 0.4982295 | -4.783963 | 1.72E-06 | 2.22E-05 | MIR942 | miRNA | 44.19 | -11.09305698 | 11.09305698 | ENSG00000215930 | down |
| ENSG00000275765 | 107.4923891 | 1.478011074 | 0.2430571 | 6.080921 | 1.19E-09 | 3.24E-08 | | lincRNA | 61.96 | 11.06974672 | 11.06974672 | ENSG00000275765 | up |
| ENSG00000259562 | 67.55977007 | -1.590930194 | 0.2713647 | -5.862702 | 4.55E-09 | 1.10E-07 | | transcribed_processed_pseudog | 41.58 | -11.06776968 | 11.06776968 | ENSG00000259562 | down |
| ENSG00000206621 | 10.15851403 | -2.344951966 | 0.487155 | -4.813564 | 1.48E-06 | 1.94E-05 | SNORD116-14 | snoRNA | 41.3 | -11.04791745 | 11.04791745 | ENSG00000206621 | down |
| ENSG00000272991 | 27.24706604 | 1.871648807 | 0.3467146 | 5.398241 | 6.73E-08 | 1.26E-06 | | antisense | 34.57 | 11.04461274 | 11.04461274 | ENSG00000272991 | up |
| ENSG00000141582 | 631.5809595 | 1.510153759 | 0.2517444 | 5.998759 | 1.99E-09 | 5.13E-08 | CBX4 | protein_coding | 62.53 | 11.00928762 | 11.00928762 | ENSG00000141582 | up |
| ENSG00000275908 | 2.762640222 | -3.99266022 | 1.0964356 | -3.641491 | 0.000271064 | 0.001824963 | | snRNA | 38.95 | -10.93488226 | 10.93488226 | ENSG00000275908 | down |
| ENSG00000225218 | 12.44309996 | -2.215497447 | 0.4501597 | -4.921581 | 8.58E-07 | 1.18E-05 | | lincRNA | 51.1 | -10.91734567 | 10.91734567 | ENSG00000225218 | down |
| ENSG00000207205 | 332.4423336 | 1.335603829 | 0.2104348 | 6.346877 | 2.20E-10 | 6.95E-09 | RNVU1-15 | snRNA | 52.73 | 10.89563038 | 10.89563038 | ENSG00000207205 | up |
| ENSG00000203872 | 204.453752 | -1.614616371 | 0.2798698 | -5.76917 | 7.97E-09 | 1.82E-07 | C6orf163 | protein_coding | 40.23 | -10.88191551 | 10.88191551 | ENSG00000203872 | down |
| ENSG00000251748 | 15.80064763 | -2.116050387 | 0.4211709 | -5.024209 | 5.06E-07 | 7.35E-06 | RNU4ATAC11P | snRNA | 49.21 | -10.86343128 | 10.86343128 | ENSG00000251748 | down |
| ENSG00000199914 | 6.387236942 | -3.977519364 | 1.093745 | -3.636606 | 0.000276254 | 0.001856969 | SNORD114-16 | snoRNA | 47.14 | -10.86338191 | 10.86338191 | ENSG00000199914 | down |
| ENSG00000222266 | 4.174381406 | -3.268980682 | 0.8134358 | -4.018732 | 5.85E-05 | 0.000479532 | RNU6-757P | snRNA | 38 | -10.85034307 | 10.85034307 | ENSG00000222266 | down |
| ENSG00000153767 | 251.0904748 | -1.64030018 | 0.2875134 | -5.705126 | 1.16E-08 | 2.55E-07 | GTF2E1 | protein_coding | 38.04 | -10.81446833 | 10.81446833 | ENSG00000153767 | down |
| ENSG00000139675 | 240.5322028 | -1.728974453 | 0.3112955 | -5.554125 | 2.79E-08 | 5.65E-07 | HNRNPA1L2 | protein_coding | 41.98 | -10.80196015 | 10.80196015 | ENSG00000139675 | down |
| ENSG00000279059 | 203.1273883 | -1.452723468 | 0.2397468 | -6.059407 | 1.37E-09 | 3.68E-08 | | TEC | 39.83 | -10.79987991 | 10.79987991 | ENSG00000279059 | down |
| ENSG00000231466 | 72.63340035 | -1.668842885 | 0.2953491 | -5.650407 | 1.60E-08 | 3.42E-07 | | processed_pseudogene | 41.79 | -10.7904678 | 10.7904678 | ENSG00000231466 | down |
| ENSG00000206717 | 5.091238304 | -3.268300065 | 0.8161581 | -4.004494 | 6.22E-05 | 0.000504266 | | misc_RNA | 44.9 | -10.77669753 | 10.77669753 | ENSG00000206717 | down |
| ENSG00000225526 | 11.69167501 | 2.230943314 | 0.4579363 | 4.871733 | 1.11E-06 | 1.49E-05 | MKRN2OS | protein_coding | 43.71 | 10.76581358 | 10.76581358 | ENSG00000225526 | up |
| ENSG00000260475 | 38.70197563 | 1.700835404 | 0.3046766 | 5.582428 | 2.37E-08 | 4.88E-07 | | lincRNA | 59.61 | 10.73528545 | 10.73528545 | ENSG00000260475 | up |
| ENSG00000279031 | 24.30495061 | -1.869631776 | 0.3512333 | -5.323048 | 1.02E-07 | 1.82E-06 | | TEC | 47.62 | -10.73297741 | 10.73297741 | ENSG00000279031 | down |
| ENSG00000102554 | 29.39268092 | 1.793842983 | 0.3302585 | 5.431633 | 5.58E-08 | 1.06E-06 | KLF5 | protein_coding | 38.99 | 10.71717227 | 10.71717227 | ENSG00000102554 | up |
| ENSG00000200769 | 14.65197395 | -2.074571542 | 0.4118807 | -5.036827 | 4.73E-07 | 6.98E-06 | | misc_RNA | 42.39 | -10.69667915 | 10.69667915 | ENSG00000200769 | down |
| ENSG00000272735 | 37.04162967 | 1.693780881 | 0.3033649 | 5.583312 | 2.36E-08 | 4.86E-07 | | lincRNA | 53.38 | 10.69340969 | 10.69340969 | ENSG00000272735 | up |
| ENSG00000238311 | 6.472953478 | -2.633957296 | 0.5912885 | -4.454606 | 8.40E-06 | 8.99E-05 | SNORD13E | snoRNA | 42.31 | -10.65792151 | 10.65792151 | ENSG00000238311 | down |
| ENSG00000200997 | 91.77367591 | 1.553478682 | 0.2670467 | 5.817255 | 5.98E-09 | 1.41E-07 | RNU1-85P | snRNA | 41.46 | 10.64401565 | 10.64401565 | ENSG00000200997 | up |
| ENSG00000142166 | 460.5662765 | -1.534240866 | 0.2621321 | -5.85293 | 4.83E-09 | 1.17E-07 | IFNAR1 | protein_coding | 41.3 | -10.63767076 | 10.63767076 | ENSG00000142166 | down |
| ENSG00000272148 | 25.51995539 | -1.827933126 | 0.3410711 | -5.359391 | 8.35E-08 | 1.53E-06 | | antisense | 42.5 | -10.63116728 | 10.63116728 | ENSG00000272148 | down |
| ENSG00000200522 | 6.215984248 | -2.572842282 | 0.57172 | -4.500179 | 6.79E-06 | 7.46E-05 | RNU6-957P | snRNA | 37.37 | -10.6186291 | 10.6186291 | ENSG00000200522 | down |
| ENSG00000224687 | 72.9672438 | 1.56535768 | 0.2704939 | 5.787034 | 7.16E-09 | 1.66E-07 | RASAL2-AS1 | lincRNA | 45.58 | 10.61310367 | 10.61310367 | ENSG00000224687 | up |

| | | | | | | | | | | | | | |
|---|---|---|---|---|---|---|---|---|---|---|---|---|---|
| ENSG00000236073 | 6.960767559 | -2.902337171 | 0.6871462 | -4.223755 | 2.40E-05 | 0.000221391 | | processed_pseudogene | 41.74 | -10.60757835 | 10.60757835 | ENSG00000236073 | down |
| ENSG00000206737 | 61.31507959 | 1.586517742 | 0.2764179 | 5.739563 | 9.49E-09 | 2.12E-07 | RNVU1-18 | snRNA | 54.27 | 10.58634831 | 10.58634831 | ENSG00000206737 | up |
| ENSG00000200406 | 6.633851254 | -3.252880976 | 0.8179019 | -3.977104 | 6.98E-05 | 0.000558519 | SNORD114-23 | snoRNA | 45.83 | -10.58149924 | 10.58149924 | ENSG00000200406 | down |
| ENSG00000276449 | 123.5000229 | -1.596687657 | 0.2793872 | -5.714964 | 1.10E-08 | 2.42E-07 | | antisense | 41.89 | -10.56255785 | 10.56255785 | ENSG00000276449 | down |
| ENSG00000207263 | 5.767920507 | -2.789680982 | 0.6488907 | -4.299154 | 1.71E-05 | 0.000165542 | SNORD116-16 | snoRNA | 43.48 | -10.54803787 | 10.54803787 | ENSG00000207263 | down |
| ENSG00000238964 | 6.709943422 | -2.701700288 | 0.6180347 | -4.371438 | 1.23E-05 | 0.000124984 | RNU7-125P | snRNA | 35.48 | -10.5451305 | 10.5451305 | ENSG00000238964 | down |
| ENSG00000270022 | 465.1173141 | 1.314063859 | 0.2087918 | 6.293658 | 3.10E-10 | 9.51E-09 | | lincRNA | 49.25 | 10.54123866 | 10.54123866 | ENSG00000270022 | up |
| ENSG00000264292 | 5.855943179 | -2.698064138 | 0.6169023 | -4.373568 | 1.22E-05 | 0.000124041 | MIR2467 | miRNA | 60.49 | -10.53981334 | 10.53981334 | ENSG00000264292 | down |
| ENSG00000185278 | 1224.484266 | -1.552176698 | 0.2683097 | -5.785018 | 7.25E-09 | 1.68E-07 | ZBTB37 | protein_coding | 40.41 | -10.5167708 | 10.5167708 | ENSG00000185278 | down |
| ENSG00000234911 | 126.8650628 | 1.448098904 | 0.2418937 | 5.986509 | 2.14E-09 | 5.50E-08 | TEX21P | unitary_pseudogene | 42.85 | 10.51299188 | 10.51299188 | ENSG00000234911 | up |
| ENSG00000273680 | 101.0827305 | -1.473071305 | 0.2481678 | -5.935788 | 2.92E-09 | 7.34E-08 | | antisense | 36.71 | -10.50966552 | 10.50966552 | ENSG00000273680 | down |
| ENSG00000272477 | 569.1527998 | -1.430467773 | 0.2377329 | -6.017123 | 1.78E-09 | 4.64E-08 | | sense_intronic | 31.69 | -10.49060364 | 10.49060364 | ENSG00000272477 | down |
| ENSG00000187772 | 2063.793102 | -1.54103806 | 0.2659591 | -5.794267 | 6.86E-09 | 1.60E-07 | LIN28B | protein_coding | 36.8 | -10.47257002 | 10.47257002 | ENSG00000187772 | down |
| ENSG00000201302 | 109.2180505 | 1.432988686 | 0.2387312 | 6.002519 | 1.94E-09 | 5.02E-08 | SNORA65 | snoRNA | 54.62 | 10.46000032 | 10.46000032 | ENSG00000201302 | up |
| ENSG00000237738 | 9.793655988 | -2.28328861 | 0.4811678 | -4.745306 | 2.08E-06 | 2.63E-05 | RNF216-IT1 | sense_intronic | 48.47 | -10.45656131 | 10.45656131 | ENSG00000237738 | down |
| ENSG00000256667 | 210.5519904 | -1.641573622 | 0.2932435 | -5.597987 | 2.17E-08 | 4.52E-07 | KLRA1P | transcribed_unprocessed_pseud | 33.5 | -10.41557983 | 10.41557983 | ENSG00000256667 | down |
| ENSG00000199568 | 740.6958151 | 1.266905321 | 0.1989813 | 6.366957 | 1.93E-10 | 6.17E-09 | RNU5A-1 | snRNA | 41.38 | 10.40080652 | 10.40080652 | ENSG00000199568 | up |
| ENSG00000145723 | 112.1274533 | -1.499665273 | 0.2563526 | -5.85001 | 4.92E-09 | 1.18E-07 | GIN1 | protein_coding | 36.01 | -10.3898806 | 10.3898806 | ENSG00000145723 | down |
| ENSG00000196511 | 43.19101845 | 1.60590298 | 0.2842175 | 5.65026 | 1.60E-08 | 3.42E-07 | TPK1 | protein_coding | 37.29 | 10.38350857 | 10.38350857 | ENSG00000196511 | up |
| ENSG00000212429 | 10.36406456 | -2.244517708 | 0.4708175 | -4.767277 | 1.87E-06 | 2.39E-05 | RNU11-6P | snRNA | 43.61 | -10.37343598 | 10.37343598 | ENSG00000212429 | down |
| ENSG00000207336 | 7.383754322 | -2.543875542 | 0.5688066 | -4.472303 | 7.74E-06 | 8.36E-05 | RNU6-658P | snRNA | 38.68 | -10.3727684 | 10.3727684 | ENSG00000207336 | down |
| ENSG00000272440 | 13.14366964 | -2.010360195 | 0.3995181 | -5.031962 | 4.85E-07 | 7.10E-06 | | lincRNA | 26.98 | -10.3507998 | 10.3507998 | ENSG00000272440 | down |
| ENSG00000200999 | 6.543109092 | -2.806597163 | 0.6617605 | -4.241107 | 2.22E-05 | 0.000206701 | | snoRNA | 42.5 | -10.34135051 | 10.34135051 | ENSG00000200999 | down |
| ENSG00000258655 | 119.4484232 | 1.410027339 | 0.2343489 | 6.016786 | 1.78E-09 | 4.64E-08 | ARHGAP5-AS1 | antisense | 46 | 10.34057238 | 10.34057238 | ENSG00000258655 | up |
| ENSG00000161298 | 474.9475256 | -1.538895933 | 0.2675402 | -5.752017 | 8.82E-09 | 1.99E-07 | ZNF382 | protein_coding | 41.92 | -10.31238373 | 10.31238373 | ENSG00000161298 | down |
| ENSG00000200683 | 2.828865435 | -3.907555147 | 1.0941752 | -3.571234 | 0.000355304 | 0.002310553 | RNU6-379P | snRNA | 38.46 | -10.30142522 | 10.30142522 | ENSG00000200683 | down |
| ENSG00000260273 | 39.00474511 | 1.72287534 | 0.3172275 | 5.43104 | 5.60E-08 | 1.06E-06 | | antisense | 47.71 | 10.29171471 | 10.29171471 | ENSG00000260273 | up |
| ENSG00000207739 | 60.02679604 | -1.526940557 | 0.2652435 | -5.756751 | 8.57E-09 | 1.95E-07 | MIR218-2 | miRNA | 55.45 | -10.24653662 | 10.24653662 | ENSG00000207739 | down |
| ENSG00000259826 | 75.11984999 | 1.518188272 | 0.2629912 | 5.772772 | 7.80E-09 | 1.79E-07 | | antisense | 47.18 | 10.24394944 | 10.24394944 | ENSG00000259826 | up |
| ENSG00000222112 | 26.11760102 | -1.905659024 | 0.3702387 | -5.147109 | 2.65E-07 | 4.22E-06 | RN7SKP16 | misc_RNA | 54.18 | -10.24254147 | 10.24254147 | ENSG00000222112 | down |
| ENSG00000227172 | 20.98624164 | 1.902513977 | 0.369661 | 5.146645 | 2.65E-07 | 4.22E-06 | | antisense | 40.78 | 10.22454254 | 10.22454254 | ENSG00000227172 | up |
| ENSG00000155542 | 314.1161521 | 1.367136397 | 0.2250572 | 6.074617 | 1.24E-09 | 3.36E-08 | SETD9 | protein_coding | 38.6 | 10.21751028 | 10.21751028 | ENSG00000155542 | up |

| | | | | | | | | | | | | | |
|---|---|---|---|---|---|---|---|---|---|---|---|---|---|
| ENSG00000244232 | 13.59817985 | -2.126426386 | 0.437489 | -4.860525 | 1.17E-06 | 1.57E-05 | RN7SL698P | misc_RNA | 52.98 | -10.21624547 | 10.21624547 | ENSG00000244232 | down |
| ENSG00000251726 | 6.558859522 | -2.59460507 | 0.590961 | -4.390484 | 1.13E-05 | 0.000116384 | RNU7-41P | snRNA | 44.26 | -10.20745687 | 10.20745687 | ENSG00000251726 | down |
| ENSG00000208883 | 4.309899014 | -3.907812935 | 1.0997861 | -3.553248 | 0.000380505 | 0.00245381 | SNORD96B | snoRNA | 50.63 | -10.20001325 | 10.20001325 | ENSG00000208883 | down |
| ENSG00000138756 | 328.160297 | -1.516985249 | 0.2633082 | -5.761253 | 8.35E-09 | 1.90E-07 | BMP2K | protein_coding | 35.68 | -10.19622574 | 10.19622574 | ENSG00000138756 | down |
| ENSG00000171016 | 1435.296779 | -1.56642043 | 0.2763359 | -5.668537 | 1.44E-08 | 3.10E-07 | PYGO1 | protein_coding | 39.31 | -10.1954264 | 10.1954264 | ENSG00000171016 | down |
| ENSG00000272906 | 34.46798685 | 1.688295814 | 0.3094296 | 5.456155 | 4.87E-08 | 9.36E-07 | | lincRNA | 54.1 | 10.17852228 | 10.17852228 | ENSG00000272906 | up |
| ENSG00000276030 | 12.83015079 | -2.735779322 | 0.6421216 | -4.260532 | 2.04E-05 | 0.000192507 | | lincRNA | 32.28 | -10.16493576 | 10.16493576 | ENSG00000276030 | down |
| ENSG00000227962 | 21.8388481 | -1.950708848 | 0.3855336 | -5.059764 | 4.20E-07 | 6.33E-06 | | processed_pseudogene | 40.43 | -10.14158307 | 10.14158307 | ENSG00000227962 | down |
| ENSG00000265093 | 20.40540444 | -1.894767645 | 0.3690223 | -5.134562 | 2.83E-07 | 4.46E-06 | RN7SL246P | misc_RNA | 50 | -10.1384187 | 10.1384187 | ENSG00000265093 | down |
| ENSG00000215218 | 152.7403041 | -1.577273459 | 0.2801588 | -5.629927 | 1.80E-08 | 3.80E-07 | UBE2QL1 | protein_coding | 47.32 | -10.1258584 | 10.1258584 | ENSG00000215218 | down |
| ENSG00000258650 | 7.801902021 | -2.378659673 | 0.5202241 | -4.572375 | 4.82E-06 | 5.54E-05 | HSBP1P1 | processed_pseudogene | 46.39 | -10.12395451 | 10.12395451 | ENSG00000258650 | down |
| ENSG00000271797 | 23.15096618 | 1.812596746 | 0.3456444 | 5.244108 | 1.57E-07 | 2.63E-06 | | antisense | 53.6 | 10.11403116 | 10.11403116 | ENSG00000271797 | up |
| ENSG00000175595 | 196.6483366 | -1.537114509 | 0.2701248 | -5.690386 | 1.27E-08 | 2.75E-07 | ERCC4 | protein_coding | 39.57 | -10.08477207 | 10.08477207 | ENSG00000175595 | down |
| ENSG00000240207 | 158.1723508 | -1.597182242 | 0.2860495 | -5.583588 | 2.36E-08 | 4.86E-07 | | antisense | 43.38 | -10.08362375 | 10.08362375 | ENSG00000240207 | down |
| ENSG00000276326 | 3.63048874 | -3.465561295 | 0.923704 | -3.751809 | 0.000175563 | 0.001255451 | MIR2909 | miRNA | 52.17 | -10.05428729 | 10.05428729 | ENSG00000276326 | down |
| ENSG00000201558 | 254.3454157 | 1.312869029 | 0.2136461 | 6.145065 | 7.99E-10 | 2.24E-08 | RNVU1-6 | snRNA | 50.61 | 10.04218016 | 10.04218016 | ENSG00000201558 | up |
| ENSG00000256588 | 31.70906132 | -1.710281398 | 0.3178399 | -5.380953 | 7.41E-08 | 1.36E-06 | | sense_intronic | 41.14 | -10.03376732 | 10.03376732 | ENSG00000256588 | down |
| ENSG00000185379 | 439.8785428 | -1.476855744 | 0.2549481 | -5.792769 | 6.92E-09 | 1.61E-07 | RAD51D | protein_coding | 47.17 | -10.03175168 | 10.03175168 | ENSG00000185379 | down |
| ENSG00000247363 | 31.82545171 | 1.701077816 | 0.3153738 | 5.393846 | 6.90E-08 | 1.28E-06 | | lincRNA | 42.44 | 10.02393589 | 10.02393589 | ENSG00000247363 | up |
| ENSG00000243951 | 18.96431323 | -1.916687992 | 0.377626 | -5.075625 | 3.86E-07 | 5.90E-06 | RN7SL308P | misc_RNA | 52.53 | -10.02317188 | 10.02317188 | ENSG00000243951 | down |
| ENSG00000235688 | 14.47778155 | 2.072240767 | 0.425615 | 4.868816 | 1.12E-06 | 1.51E-05 | | antisense | 57.25 | 9.988423219 | 9.988423219 | ENSG00000235688 | up |
| ENSG00000228701 | 63.08968688 | 1.50701465 | 0.2634779 | 5.719701 | 1.07E-08 | 2.38E-07 | TNKS2-AS1 | antisense | 38.44 | 9.982368215 | 9.982368215 | ENSG00000228701 | up |
| ENSG00000252503 | 4.719326979 | -2.993596245 | 0.7430711 | -4.028681 | 5.61E-05 | 0.000463549 | RNU6-531P | snRNA | 46.15 | -9.980362447 | 9.980362447 | ENSG00000252503 | down |
| ENSG00000222343 | 36.83462695 | -1.925911497 | 0.3813837 | -5.0498 | 4.42E-07 | 6.64E-06 | RN7SKP139 | misc_RNA | 48.81 | -9.972609074 | 9.972609074 | ENSG00000222343 | down |
| ENSG00000268362 | 281.81184 | -1.428390204 | 0.2432234 | -5.87275 | 4.29E-09 | 1.04E-07 | | lincRNA | 45.45 | -9.972430303 | 9.972430303 | ENSG00000268362 | down |
| ENSG00000215952 | 13.75371236 | -1.993942023 | 0.4026552 | -4.951983 | 7.35E-07 | 1.02E-05 | MIR921 | miRNA | 55.36 | -9.950191803 | 9.950191803 | ENSG00000215952 | down |
| ENSG00000153814 | 132.9411446 | -1.633192686 | 0.2978527 | -5.483223 | 4.18E-08 | 8.13E-07 | JAZF1 | protein_coding | 41.44 | -9.945761361 | 9.945761361 | ENSG00000153814 | down |
| ENSG00000047056 | 328.3336158 | -1.43858849 | 0.24617 | -5.843881 | 5.10E-09 | 1.22E-07 | WDR37 | protein_coding | 45.5 | -9.94479913 | 9.94479913 | ENSG00000047056 | down |
| ENSG00000206763 | 22.38822785 | -1.926021604 | 0.382044 | -5.04136 | 4.62E-07 | 6.87E-06 | RNU6-10P | snRNA | 44.86 | -9.944328339 | 9.944328339 | ENSG00000206763 | down |
| ENSG00000232519 | 41.74289946 | 1.598703929 | 0.2889331 | 5.533129 | 3.15E-08 | 6.28E-07 | | antisense | 64.13 | 9.914819944 | 9.914819944 | ENSG00000232519 | up |
| ENSG00000110768 | 814.6017764 | -1.588839537 | 0.2863055 | -5.549455 | 2.87E-08 | 5.78E-07 | GTF2H1 | protein_coding | 40.71 | -9.911019779 | 9.911019779 | ENSG00000110768 | down |
| ENSG00000249493 | 21.4142032 | -1.827150257 | 0.3533073 | -5.171561 | 2.32E-07 | 3.78E-06 | ANKRD20A18P | unprocessed_pseudogene | 35.73 | -9.907667726 | 9.907667726 | ENSG00000249493 | down |

| | | | | | | | | | | | | | |
|---|---|---|---|---|---|---|---|---|---|---|---|---|---|
| ENSG00000267144 | 19.50567187 | 1.919605865 | 0.3811031 | 5.036972 | 4.73E-07 | 6.98E-06 | | lincRNA | 39.12 | 9.89766206 | 9.89766206 | ENSG00000267144 | up |
| ENSG00000281468 | 35.19887801 | -1.830428077 | 0.3545048 | -5.163339 | 2.43E-07 | 3.92E-06 | | sense_intronic | 30.71 | -9.896041887 | 9.896041887 | ENSG00000281468 | down |
| ENSG00000207993 | 3.41326656 | -3.876858562 | 1.1038557 | -3.512106 | 0.00044457 | 0.002813522 | MIR134 | miRNA | 60.27 | -9.888895998 | 9.888895998 | ENSG00000207993 | down |
| ENSG00000275097 | 75.36764893 | -1.558624901 | 0.2791044 | -5.58438 | 2.35E-08 | 4.84E-07 | | 3prime_overlapping_ncRNA | 31.22 | -9.842278029 | 9.842278029 | ENSG00000275097 | down |
| ENSG00000201318 | 4.214479533 | -3.446119739 | 0.9262129 | -3.720656 | 0.000198706 | 0.00139515 | SNORD114-30 | snoRNA | 45.83 | -9.839978607 | 9.839978607 | ENSG00000201318 | down |
| ENSG00000240399 | 51.54913587 | -1.590873603 | 0.2882151 | -5.519745 | 3.39E-08 | 6.74E-07 | | transcribed_processed_pseudog | 43.65 | -9.817528614 | 9.817528614 | ENSG00000240399 | down |
| ENSG00000280040 | 23.04828173 | 1.848312587 | 0.3613322 | 5.115272 | 3.13E-07 | 4.90E-06 | | TEC | 34.38 | 9.814908863 | 9.814908863 | ENSG00000280040 | up |
| ENSG00000151065 | 444.318317 | -1.585321525 | 0.2867827 | -5.527955 | 3.24E-08 | 6.46E-07 | DCP1B | protein_coding | 38.72 | -9.812501014 | 9.812501014 | ENSG00000151065 | down |
| ENSG00000252908 | 40.13200544 | -1.613384321 | 0.2945481 | -5.47749 | 4.31E-08 | 8.38E-07 | RNU6-1003P | snRNA | 37.14 | -9.804382294 | 9.804382294 | ENSG00000252908 | down |
| ENSG00000259891 | 77.3786405 | -1.442986145 | 0.2491686 | -5.791203 | 6.99E-09 | 1.62E-07 | | antisense | 48.41 | -9.796890286 | 9.796890286 | ENSG00000259891 | down |
| ENSG00000136099 | 1511.683219 | 1.217485469 | 0.1931461 | 6.303443 | 2.91E-10 | 9.03E-09 | PCDH8 | protein_coding | 56.58 | 9.794016439 | 9.794016439 | ENSG00000136099 | up |
| ENSG00000262172 | 25.69953224 | -1.820207664 | 0.3534287 | -5.15014 | 2.60E-07 | 4.18E-06 | | sense_intronic | 62.58 | -9.791491879 | 9.791491879 | ENSG00000262172 | down |
| ENSG00000261526 | 211.3403962 | 1.433319317 | 0.2467754 | 5.808193 | 6.32E-09 | 1.48E-07 | | lincRNA | 55.35 | 9.788057624 | 9.788057624 | ENSG00000261526 | up |
| ENSG00000227331 | 68.02050128 | -1.584720495 | 0.2874642 | -5.512758 | 3.53E-08 | 6.97E-07 | | processed_pseudogene | 50.75 | -9.757102022 | 9.757102022 | ENSG00000227331 | down |
| ENSG00000225026 | 10.70456885 | -2.746618648 | 0.6602527 | -4.159951 | 3.18E-05 | 0.000282648 | | processed_pseudogene | 40.81 | -9.747074302 | 9.747074302 | ENSG00000225026 | down |
| ENSG00000256442 | 27.68758675 | 1.700792763 | 0.3203925 | 5.308466 | 1.11E-07 | 1.95E-06 | | antisense | 50.9 | 9.713019395 | 9.713019395 | ENSG00000256442 | up |
| ENSG00000266126 | 21.41007312 | -1.924375986 | 0.386357 | -4.980823 | 6.33E-07 | 8.98E-06 | | sense_intronic | 39.89 | -9.712088595 | 9.712088595 | ENSG00000266126 | down |
| ENSG00000226276 | 16.07132921 | -1.95636029 | 0.3964956 | -4.934129 | 8.05E-07 | 1.11E-05 | | lincRNA | 37.65 | -9.692420186 | 9.692420186 | ENSG00000226276 | down |
| ENSG00000226766 | 59.878209 | -1.64296248 | 0.3046819 | -5.392386 | 6.95E-08 | 1.29E-06 | FABP7P1 | processed_pseudogene | 40.41 | -9.677581158 | 9.677581158 | ENSG00000226766 | down |
| ENSG00000272192 | 41.06789022 | 1.69295648 | 0.3189538 | 5.307843 | 1.11E-07 | 1.95E-06 | | lincRNA | 39.66 | 9.667610982 | 9.667610982 | ENSG00000272192 | up |
| ENSG00000223225 | 6.387060532 | -2.485134774 | 0.5697977 | -4.361434 | 1.29E-05 | 0.000129894 | RNU6-1054P | snRNA | 40.38 | -9.658254277 | 9.658254277 | ENSG00000223225 | down |
| ENSG00000241400 | 17.76735947 | -1.849229562 | 0.3646431 | -5.071341 | 3.95E-07 | 6.00E-06 | | processed_pseudogene | 45.3 | -9.655826411 | 9.655826411 | ENSG00000241400 | down |
| ENSG00000183309 | 522.3797236 | -1.509522146 | 0.2687654 | -5.616505 | 1.95E-08 | 4.09E-07 | ZNF623 | protein_coding | 47.94 | -9.643909393 | 9.643909393 | ENSG00000183309 | down |
| ENSG00000197016 | 462.0325534 | -1.424754785 | 0.2464183 | -5.781856 | 7.39E-09 | 1.71E-07 | ZNF470 | protein_coding | 40.11 | -9.642812866 | 9.642812866 | ENSG00000197016 | down |
| ENSG00000198920 | 391.4370937 | -1.519898284 | 0.2722772 | -5.582172 | 2.38E-08 | 4.88E-07 | KIAA0753 | protein_coding | 41.58 | -9.593251583 | 9.593251583 | ENSG00000198920 | down |
| ENSG00000207197 | 4.817993932 | -3.409450978 | 0.9235768 | -3.691573 | 0.000222872 | 0.001541502 | SNORD116-12 | snoRNA | 35.87 | -9.587567089 | 9.587567089 | ENSG00000207197 | down |
| ENSG00000198464 | 1008.886071 | -1.528801296 | 0.2748878 | -5.561547 | 2.67E-08 | 5.44E-07 | ZNF480 | protein_coding | 42.41 | -9.576676031 | 9.576676031 | ENSG00000198464 | down |
| ENSG00000273063 | 20.56297065 | 1.855003328 | 0.3685131 | 5.033751 | 4.81E-07 | 7.08E-06 | | antisense | 55.92 | 9.553335945 | 9.553335945 | ENSG00000273063 | up |
| ENSG00000221639 | 21.1214891 | -1.882215525 | 0.3769019 | -4.993913 | 5.92E-07 | 8.48E-06 | | snoRNA | 52.8 | -9.546308463 | 9.546308463 | ENSG00000221639 | down |
| ENSG00000221763 | 35.21072197 | 1.598315827 | 0.2945265 | 5.426731 | 5.74E-08 | 1.08E-06 | MIR1289-1 | miRNA | 41.67 | 9.533724817 | 9.533724817 | ENSG00000221763 | up |
| ENSG00000255893 | 25.78609155 | -1.851258527 | 0.3681157 | -5.029013 | 4.93E-07 | 7.20E-06 | | antisense | 39.22 | -9.520945914 | 9.520945914 | ENSG00000255893 | down |
| ENSG00000181218 | 766.9934705 | 1.231677519 | 0.1993571 | 6.178248 | 6.48E-10 | 1.86E-08 | HIST3H2A | protein_coding | 57.65 | 9.520159433 | 9.520159433 | ENSG00000181218 | up |

| | | | | | | | | | | | | | |
|---|---|---|---|---|---|---|---|---|---|---|---|---|---|
| ENSG00000253513 | 14.75131265 | 1.916143375 | 0.3878143 | 4.940879 | 7.78E-07 | 1.08E-05 | | antisense | 49.42 | 9.519488158 | 9.519488158 | ENSG00000253513 | up |
| ENSG00000206756 | 8.284586056 | -2.466199999 | 0.5680593 | -4.341448 | 1.42E-05 | 0.000140568 | | misc_RNA | 41.18 | -9.500086116 | 9.500086116 | ENSG00000206756 | down |
| ENSG00000259151 | 74.60765583 | -1.520387248 | 0.2739482 | -5.549908 | 2.86E-08 | 5.78E-07 | CAP2P1 | transcribed_processed_pseudog | 48.92 | -9.484775759 | 9.484775759 | ENSG00000259151 | down |
| ENSG00000274210 | 6.903121358 | 2.469523645 | 0.5700075 | 4.332441 | 1.47E-05 | 0.000145614 | | snRNA | 51.52 | 9.475058991 | 9.475058991 | ENSG00000274210 | up |
| ENSG00000136280 | 172.2237188 | -1.599613335 | 0.2961733 | -5.400937 | 6.63E-08 | 1.24E-06 | CCM2 | protein_coding | 46.33 | -9.448838529 | 9.448838529 | ENSG00000136280 | down |
| ENSG00000256304 | 171.696724 | -1.552921248 | 0.2833979 | -5.47965 | 4.26E-08 | 8.29E-07 | CCDC150P1 | processed_pseudogene | 44.83 | -9.444244178 | 9.444244178 | ENSG00000256304 | down |
| ENSG00000235092 | 48.58450729 | 1.538187073 | 0.2796533 | 5.500337 | 3.79E-08 | 7.45E-07 | ID2-AS1 | antisense | 48.44 | 9.425371125 | 9.425371125 | ENSG00000235092 | up |
| ENSG00000162997 | 53.4811968 | 1.508609319 | 0.2718695 | 5.549021 | 2.87E-08 | 5.79E-07 | PRORSD1P | transcribed_unitary_pseudogene | 44.56 | 9.409873357 | 9.409873357 | ENSG00000162997 | up |
| ENSG00000198039 | 634.4332496 | -1.596716026 | 0.29599 | -5.394493 | 6.87E-08 | 1.28E-06 | ZNF273 | protein_coding | 44.96 | -9.409612407 | 9.409612407 | ENSG00000198039 | down |
| ENSG00000199237 | 6.327909962 | -2.703978006 | 0.6569691 | -4.115837 | 3.86E-05 | 0.000334482 | RNU6-834P | snRNA | 40 | -9.398018639 | 9.398018639 | ENSG00000199237 | down |
| ENSG00000228950 | 51.69442505 | 1.639301968 | 0.3087849 | 5.308881 | 1.10E-07 | 1.94E-06 | | lincRNA | 46.07 | 9.362572838 | 9.362572838 | ENSG00000228950 | up |
| ENSG00000107929 | 869.5080819 | -1.411810164 | 0.2469373 | -5.717282 | 1.08E-08 | 2.40E-07 | LARP4B | protein_coding | 42.7 | -9.344971388 | 9.344971388 | ENSG00000107929 | down |
| ENSG00000181619 | 187.0888456 | -1.387135454 | 0.2405271 | -5.767064 | 8.07E-09 | 1.84E-07 | GPR135 | protein_coding | 40.34 | -9.343226653 | 9.343226653 | ENSG00000181619 | down |
| ENSG00000283193 | 25.51761214 | -1.836948676 | 0.3673164 | -5.000998 | 5.70E-07 | 8.20E-06 | MIR1243 | miRNA | 30.11 | -9.342732642 | 9.342732642 | ENSG00000283193 | down |
| ENSG00000269887 | 37.23153928 | -1.739084534 | 0.3379924 | -5.145336 | 2.67E-07 | 4.25E-06 | | sense_intronic | 33.15 | -9.341832181 | 9.341832181 | ENSG00000269887 | down |
| ENSG00000251787 | 2.377733032 | -4.161206951 | 1.2668868 | -3.284593 | 0.001021299 | 0.005742951 | RNU7-47P | snRNA | 45.16 | -9.324702474 | 9.324702474 | ENSG00000251787 | down |
| ENSG00000208004 | 3.089536342 | -4.158732038 | 1.2671149 | -3.282048 | 0.001030561 | 0.005777126 | MIR323B | miRNA | 48.78 | -9.308440643 | 9.308440643 | ENSG00000208004 | down |
| ENSG00000271936 | 60.54147865 | -1.600617259 | 0.2987228 | -5.358203 | 8.41E-08 | 1.53E-06 | | antisense | 53.25 | -9.306127244 | 9.306127244 | ENSG00000271936 | down |
| ENSG00000279584 | 16.57588221 | 1.929395926 | 0.3964956 | 4.866122 | 1.14E-06 | 1.53E-05 | | TEC | 56.12 | 9.292523805 | 9.292523805 | ENSG00000279584 | up |
| ENSG00000200612 | 7.384098445 | -2.937511523 | 0.7505275 | -3.913929 | 9.08E-05 | 0.000700441 | SNORD114-25 | snoRNA | 47.22 | -9.266757976 | 9.266757976 | ENSG00000200612 | down |
| ENSG00000207698 | 5.05901423 | -2.786410577 | 0.6931917 | -4.019682 | 5.83E-05 | 0.000478976 | MIR32 | miRNA | 38.57 | -9.250008295 | 9.250008295 | ENSG00000207698 | down |
| ENSG00000207978 | 2.634238102 | -3.778220496 | 1.0990591 | -3.437686 | 0.000586707 | 0.003565568 | MIR154 | miRNA | 40.48 | -9.248597573 | 9.248597573 | ENSG00000207978 | down |
| ENSG00000251595 | 179.5944219 | -1.635688203 | 0.3099391 | -5.27745 | 1.31E-07 | 2.24E-06 | ABCA11P | transcribed_processed_pseudog | 39.69 | -9.239829562 | 9.239829562 | ENSG00000251595 | down |
| ENSG00000256400 | 8.765552199 | -2.446589362 | 0.5698743 | -4.293209 | 1.76E-05 | 0.000168985 | | processed_pseudogene | 36.46 | -9.228908331 | 9.228908331 | ENSG00000256400 | down |
| ENSG00000185798 | 222.5951072 | -1.445308157 | 0.2575406 | -5.611962 | 2.00E-08 | 4.19E-07 | WDR53 | protein_coding | 43.07 | -9.218117799 | 9.218117799 | ENSG00000185798 | down |
| ENSG00000271870 | 101.5836661 | -1.609556052 | 0.3028264 | -5.315111 | 1.07E-07 | 1.89E-06 | | antisense | 33.81 | -9.213929402 | 9.213929402 | ENSG00000271870 | down |
| ENSG00000261202 | 50.49612924 | 1.58723767 | 0.2971747 | 5.341092 | 9.24E-08 | 1.66E-06 | | antisense | 46.62 | 9.173267396 | 9.173267396 | ENSG00000261202 | up |
| ENSG00000269867 | 38.8236883 | -1.538283217 | 0.2839143 | -5.418125 | 6.02E-08 | 1.13E-06 | | sense_intronic | 46.52 | -9.146178803 | 9.146178803 | ENSG00000269867 | down |
| ENSG00000202191 | 11.33025233 | -2.321649699 | 0.5291616 | -4.387412 | 1.15E-05 | 0.000117173 | SNORD113-1 | snoRNA | 33.33 | -9.126802427 | 9.126802427 | ENSG00000202191 | down |
| ENSG00000143878 | 4358.802345 | 1.22213614 | 0.2013363 | 6.070124 | 1.28E-09 | 3.45E-08 | RHOB | protein_coding | 57.34 | 9.120005181 | 9.120005181 | ENSG00000143878 | up |
| ENSG00000272008 | 310.1900656 | 1.286177745 | 0.2175989 | 5.910773 | 3.41E-09 | 8.42E-08 | | antisense | 45.53 | 9.099260068 | 9.099260068 | ENSG00000272008 | up |
| ENSG00000264857 | 6.557378963 | 2.623088176 | 0.6380714 | 4.110963 | 3.94E-05 | 0.000340101 | MIR5696 | miRNA | 41.18 | 9.097899034 | 9.097899034 | ENSG00000264857 | up |

| | | | | | | | | | | | | | |
|---|---|---|---|---|---|---|---|---|---|---|---|---|---|
| ENSG00000272945 | 24.08698315 | -1.779026792 | 0.3553476 | -5.00644 | 5.54E-07 | 8.02E-06 | | sense_intronic | 49.8 | -9.065964581 | 9.065964581 | ENSG00000272945 | down |
| ENSG00000223486 | 72.778828 | 1.456347968 | 0.2628443 | 5.540726 | 3.01E-08 | 6.05E-07 | | lincRNA | 56.45 | 9.055739749 | 9.055739749 | ENSG00000223486 | up |
| ENSG00000222020 | 65.82223084 | 1.470184752 | 0.2666748 | 5.513025 | 3.53E-08 | 6.97E-07 | | antisense | 64.05 | 9.051970216 | 9.051970216 | ENSG00000222020 | up |
| ENSG00000198948 | 71.12988735 | -1.536543082 | 0.2849264 | -5.392772 | 6.94E-08 | 1.29E-06 | MFAP3L | protein_coding | 42.91 | -9.051282434 | 9.051282434 | ENSG00000198948 | down |
| ENSG00000200367 | 10.25138222 | -2.447656299 | 0.5762484 | -4.247572 | 2.16E-05 | 0.00020193 | SNORD113-8 | snoRNA | 39.19 | -9.043596255 | 9.043596255 | ENSG00000200367 | down |
| ENSG00000273165 | 27.85164078 | -1.7351473 | 0.3427808 | -5.061973 | 4.15E-07 | 6.27E-06 | | lincRNA | 34.15 | -9.02799439 | 9.02799439 | ENSG00000273165 | down |
| ENSG00000275175 | 11.11897489 | 2.114173834 | 0.4618152 | 4.577965 | 4.70E-06 | 5.42E-05 | | lincRNA | 36.71 | 9.018977547 | 9.018977547 | ENSG00000275175 | up |
| ENSG00000136848 | 625.5375938 | -1.309106758 | 0.2244497 | -5.832517 | 5.46E-09 | 1.30E-07 | DAB2IP | protein_coding | 53.14 | -9.013151578 | 9.013151578 | ENSG00000136848 | down |
| ENSG00000251609 | 44.51253022 | -1.747544174 | 0.3468598 | -5.038186 | 4.70E-07 | 6.96E-06 | SETP12 | processed_pseudogene | 40.75 | -9.012660785 | 9.012660785 | ENSG00000251609 | down |
| ENSG00000252305 | 10.05049173 | -2.06362193 | 0.4455395 | -4.631737 | 3.63E-06 | 4.31E-05 | | snoRNA | 53.89 | -9.009265909 | 9.009265909 | ENSG00000252305 | down |
| ENSG00000203327 | 29.50764765 | -1.746091236 | 0.346711 | -5.036157 | 4.75E-07 | 7.00E-06 | | antisense | 37.18 | -9.001157198 | 9.001157198 | ENSG00000203327 | down |
| ENSG00000157741 | 987.9845461 | -1.386937482 | 0.2454052 | -5.651622 | 1.59E-08 | 3.40E-07 | UBN2 | protein_coding | 39.02 | -8.97063385 | 8.97063385 | ENSG00000157741 | down |
| ENSG00000268357 | 65.90084735 | -1.669259713 | 0.325017 | -5.135915 | 2.81E-07 | 4.43E-06 | VN1R81P | unprocessed_pseudogene | 39.25 | -8.9361791 | 8.9361791 | ENSG00000268357 | down |
| ENSG00000148926 | 208.6266295 | 1.300110873 | 0.2231839 | 5.825289 | 5.70E-09 | 1.35E-07 | ADM | protein_coding | 60.01 | 8.930364966 | 8.930364966 | ENSG00000148926 | up |
| ENSG00000244357 | 23.03394806 | -1.735067535 | 0.3449876 | -5.029361 | 4.92E-07 | 7.19E-06 | RN7SL145P | misc_RNA | 50.68 | -8.923960132 | 8.923960132 | ENSG00000244357 | down |
| ENSG00000224707 | 73.62384179 | -1.562299584 | 0.2944184 | -5.306392 | 1.12E-07 | 1.96E-06 | E2F3-IT1 | sense_intronic | 39.27 | -8.917806018 | 8.917806018 | ENSG00000224707 | down |
| ENSG00000076650 | 281.092388 | -1.489453205 | 0.2742744 | -5.430522 | 5.62E-08 | 1.06E-06 | GPATCH1 | protein_coding | 45.71 | -8.8963548 | 8.8963548 | ENSG00000076650 | down |
| ENSG00000108423 | 313.5328276 | -1.519398522 | 0.2828052 | -5.372597 | 7.76E-08 | 1.42E-06 | TUBD1 | protein_coding | 45.29 | -8.884164022 | 8.884164022 | ENSG00000108423 | down |
| ENSG00000101445 | 304.9790089 | -1.451211116 | 0.2640252 | -5.496488 | 3.87E-08 | 7.61E-07 | PPP1R16B | protein_coding | 47.52 | -8.87955483 | 8.87955483 | ENSG00000101445 | down |
| ENSG00000165115 | 174.3994857 | -1.440380921 | 0.2612131 | -5.514199 | 3.50E-08 | 6.93E-07 | KIF27 | protein_coding | 38.37 | -8.871754849 | 8.871754849 | ENSG00000165115 | down |
| ENSG00000265100 | 63.56335272 | 1.435526734 | 0.2601488 | 5.518098 | 3.43E-08 | 6.80E-07 | | antisense | 57.86 | 8.853904258 | 8.853904258 | ENSG00000265100 | up |
| ENSG00000188315 | 186.9401875 | -1.396801434 | 0.2497262 | -5.593332 | 2.23E-08 | 4.62E-07 | C3orf62 | protein_coding | 48.07 | -8.848970701 | 8.848970701 | ENSG00000188315 | down |
| ENSG00000227470 | 47.89386074 | -1.71006159 | 0.3387835 | -5.047653 | 4.47E-07 | 6.69E-06 | | processed_pseudogene | 43.79 | -8.848966017 | 8.848966017 | ENSG00000227470 | down |
| ENSG00000204842 | 570.5494882 | -1.317741088 | 0.2293149 | -5.746427 | 9.11E-09 | 2.05E-07 | ATXN2 | protein_coding | 41.99 | -8.81239787 | 8.81239787 | ENSG00000204842 | down |
| ENSG00000232536 | 21.85057229 | 1.795636464 | 0.3657953 | 4.908857 | 9.16E-07 | 1.26E-05 | | sense_intronic | 45.61 | 8.800815155 | 8.800815155 | ENSG00000232536 | up |
| ENSG00000201500 | 8.414397757 | -2.649327528 | 0.6590023 | -4.02021 | 5.81E-05 | 0.000478187 | | snoRNA | 39.74 | -8.796832715 | 8.796832715 | ENSG00000201500 | down |
| ENSG00000083097 | 1165.166107 | -1.56336986 | 0.2970071 | -5.263746 | 1.41E-07 | 2.40E-06 | DOPEY1 | protein_coding | 35.96 | -8.786766559 | 8.786766559 | ENSG00000083097 | down |
| ENSG00000166153 | 53.92521409 | -1.516263098 | 0.2838147 | -5.342441 | 9.17E-08 | 1.65E-06 | DEPDC4 | protein_coding | 39.88 | -8.767126561 | 8.767126561 | ENSG00000166153 | down |
| ENSG00000200563 | 4.34912851 | -2.741101016 | 0.6957547 | -3.939752 | 8.16E-05 | 0.000639079 | RNU6-640P | snRNA | 36.19 | -8.756298574 | 8.756298574 | ENSG00000200563 | down |
| ENSG00000244184 | 40.94845402 | 1.527448109 | 0.2872175 | 5.318089 | 1.05E-07 | 1.86E-06 | | antisense | 50.54 | 8.753908696 | 8.753908696 | ENSG00000244184 | up |
| ENSG00000144227 | 54.48544145 | -1.551220726 | 0.2943473 | -5.270036 | 1.36E-07 | 2.32E-06 | NXPH2 | protein_coding | 38.25 | -8.739086261 | 8.739086261 | ENSG00000144227 | down |
| ENSG00000274421 | 88.17296412 | -1.439890907 | 0.2633547 | -5.467496 | 4.56E-08 | 8.83E-07 | | sense_intronic | 39.25 | -8.717412494 | 8.717412494 | ENSG00000274421 | down |

| | | | | | | | | | | | | | |
|---|---|---|---|---|---|---|---|---|---|---|---|---|---|
| ENSG00000212125 | 64.62320392 | -1.540495212 | 0.2922222 | -5.271657 | 1.35E-07 | 2.31E-06 | TAS2R15P | processed_pseudogene | 38.47 | -8.683755294 | 8.683755294 | ENSG00000212125 | down |
| ENSG00000259642 | 138.60147 | 1.288745126 | 0.2234421 | 5.767691 | 8.04E-09 | 1.83E-07 | ST20-AS1 | antisense | 56.17 | 8.681670688 | 8.681670688 | ENSG00000259642 | up |
| ENSG00000264112 | 1737.862429 | -1.45213336 | 0.2674466 | -5.429619 | 5.65E-08 | 1.07E-06 | | lincRNA | 40.9 | -8.671112817 | 8.671112817 | ENSG00000264112 | down |
| ENSG00000265558 | 28.62197764 | -1.724158213 | 0.3475987 | -4.960197 | 7.04E-07 | 9.85E-06 | MIR3918 | miRNA | 62.37 | -8.631798458 | 8.631798458 | ENSG00000265558 | down |
| ENSG00000272663 | 39.4041655 | 1.516719877 | 0.2864113 | 5.2956 | 1.19E-07 | 2.06E-06 | | lincRNA | 47.03 | 8.622820987 | 8.622820987 | ENSG00000272663 | up |
| ENSG00000275673 | 38.81046122 | -1.564094025 | 0.3003236 | -5.208029 | 1.91E-07 | 3.15E-06 | | sense_intronic | 52.38 | -8.604686348 | 8.604686348 | ENSG00000275673 | down |
| ENSG00000207626 | 6.037390272 | -2.616846545 | 0.6545795 | -3.997752 | 6.39E-05 | 0.000517034 | MIR562 | miRNA | 37.89 | -8.600216664 | 8.600216664 | ENSG00000207626 | down |
| ENSG00000221808 | 5.159661188 | -3.03146818 | 0.8181951 | -3.705068 | 0.000211334 | 0.001470468 | MIR1256 | miRNA | 42.02 | -8.586768179 | 8.586768179 | ENSG00000221808 | down |
| ENSG00000243854 | 16.19856258 | -1.855460544 | 0.3891161 | -4.768398 | 1.86E-06 | 2.38E-05 | RN7SL67P | misc_RNA | 53.38 | -8.579079679 | 8.579079679 | ENSG00000243854 | down |
| ENSG00000258101 | 41.06412607 | 1.530447443 | 0.2914841 | 5.250535 | 1.52E-07 | 2.55E-06 | | antisense | 46.26 | 8.559650223 | 8.559650223 | ENSG00000258101 | up |
| ENSG00000201710 | 7.70872613 | -2.743098865 | 0.7046807 | -3.892684 | 9.91E-05 | 0.000758016 | | snoRNA | 36 | -8.55935138 | 8.55935138 | ENSG00000201710 | down |
| ENSG00000258284 | 7.590022604 | -2.336853937 | 0.5540691 | -4.217622 | 2.47E-05 | 0.00022615 | POLR2KP1 | processed_pseudogene | 41.07 | -8.519241646 | 8.519241646 | ENSG00000258284 | down |
| ENSG00000269906 | 73.25574351 | 1.465957823 | 0.2741024 | 5.348213 | 8.88E-08 | 1.61E-06 | | sense_intronic | 35.18 | 8.492573458 | 8.492573458 | ENSG00000269906 | up |
| ENSG00000207761 | 3.847068845 | -4.041259802 | 1.2724303 | -3.176017 | 0.001493124 | 0.007925568 | MIR329-1 | miRNA | 42.5 | -8.490564053 | 8.490564053 | ENSG00000207761 | down |
| ENSG00000265967 | 17.71905219 | -2.504246653 | 0.6164114 | -4.062622 | 4.85E-05 | 0.000410953 | | antisense | 42.96 | -8.479899864 | 8.479899864 | ENSG00000265967 | down |
| ENSG00000180221 | 29.36580613 | -1.747396214 | 0.3578639 | -4.882851 | 1.05E-06 | 1.42E-05 | TPT1P10 | processed_pseudogene | 42.14 | -8.47217945 | 8.47217945 | ENSG00000180221 | down |
| ENSG00000196214 | 749.5081318 | -1.462975236 | 0.273683 | -5.345511 | 9.02E-08 | 1.63E-06 | ZNF766 | protein_coding | 44.77 | -8.467643543 | 8.467643543 | ENSG00000196214 | down |
| ENSG00000131127 | 1442.044204 | -1.439701544 | 0.2674102 | -5.383869 | 7.29E-08 | 1.34E-06 | ZNF141 | protein_coding | 40.31 | -8.455653596 | 8.455653596 | ENSG00000131127 | down |
| ENSG00000266498 | 29.21825965 | 1.599496286 | 0.3135668 | 5.100974 | 3.38E-07 | 5.25E-06 | | lincRNA | 48.69 | 8.444980274 | 8.444980274 | ENSG00000266498 | up |
| ENSG00000200795 | 695.0087459 | 1.16506118 | 0.1948326 | 5.979805 | 2.23E-09 | 5.68E-08 | RNU4-1 | snRNA | 48.23 | 8.441980532 | 8.441980532 | ENSG00000200795 | up |
| ENSG00000122035 | 109.8548967 | 1.304357713 | 0.2307159 | 5.653524 | 1.57E-08 | 3.37E-07 | RASL11A | protein_coding | 53.83 | 8.441911032 | 8.441911032 | ENSG00000122035 | up |
| ENSG00000183475 | 392.6668489 | -1.424353511 | 0.2632902 | -5.409824 | 6.31E-08 | 1.18E-06 | ASB7 | protein_coding | 40.07 | -8.441763781 | 8.441763781 | ENSG00000183475 | down |
| ENSG00000275127 | 6.335523304 | -2.513546934 | 0.6216571 | -4.043301 | 5.27E-05 | 0.000440454 | SNORD116-22 | snoRNA | 43.48 | -8.435714459 | 8.435714459 | ENSG00000275127 | down |
| ENSG00000275810 | 11.0334814 | -2.162106891 | 0.4948791 | -4.36896 | 1.25E-05 | 0.000125952 | MIR8082 | miRNA | 50.62 | -8.431772044 | 8.431772044 | ENSG00000275810 | down |
| ENSG00000114315 | 53.4117627 | 1.617884836 | 0.3194594 | 5.064445 | 4.10E-07 | 6.20E-06 | HES1 | protein_coding | 53.32 | 8.425465088 | 8.425465088 | ENSG00000114315 | up |
| ENSG00000252212 | 14.53659731 | -1.894501843 | 0.4055465 | -4.671479 | 2.99E-06 | 3.63E-05 | RNU2-58P | snRNA | 43.72 | -8.411064448 | 8.411064448 | ENSG00000252212 | down |
| ENSG00000122741 | 626.9195707 | -1.50426295 | 0.2866556 | -5.247631 | 1.54E-07 | 2.59E-06 | DCAF10 | protein_coding | 38.56 | -8.404485248 | 8.404485248 | ENSG00000122741 | down |
| ENSG00000261242 | 46.99436572 | 1.49820258 | 0.2849582 | 5.257622 | 1.46E-07 | 2.47E-06 | | antisense | 47.72 | 8.401993077 | 8.401993077 | ENSG00000261242 | up |
| ENSG00000125398 | 118.0716963 | 1.382320195 | 0.2523836 | 5.477061 | 4.32E-08 | 8.39E-07 | SOX9 | protein_coding | 53.34 | 8.399609653 | 8.399609653 | ENSG00000125398 | up |
| ENSG00000070269 | 57.08576775 | -1.65566075 | 0.3315664 | -4.993452 | 5.93E-07 | 8.48E-06 | TMEM260 | protein_coding | 39.13 | -8.397015701 | 8.397015701 | ENSG00000070269 | down |
| ENSG00000207279 | 11.84130554 | -2.131210453 | 0.4856307 | -4.388541 | 1.14E-05 | 0.000116986 | SNORD116-24 | snoRNA | 40.22 | -8.379634319 | 8.379634319 | ENSG00000207279 | down |
| ENSG00000135315 | 721.8788286 | -1.471936946 | 0.2779707 | -5.295295 | 1.19E-07 | 2.07E-06 | CEP162 | protein_coding | 37 | -8.367953246 | 8.367953246 | ENSG00000135315 | down |

| | | | | | | | | | | | | | |
|---|---|---|---|---|---|---|---|---|---|---|---|---|---|
| ENSG00000207009 | 14.33119144 | 1.93277705 | 0.4190169 | 4.612647 | 3.98E-06 | 4.68E-05 | | misc_RNA | 46.36 | 8.367894268 | 8.367894268 | ENSG00000207009 | up |
| ENSG00000260060 | 94.04480796 | -1.39700783 | 0.2569009 | -5.437925 | 5.39E-08 | 1.03E-06 | | antisense | 50.06 | -8.364401929 | 8.364401929 | ENSG00000260060 | down |
| ENSG00000135913 | 1868.225903 | -1.317825281 | 0.2354244 | -5.597659 | 2.17E-08 | 4.52E-07 | USP37 | protein_coding | 39.62 | -8.361208827 | 8.361208827 | ENSG00000135913 | down |
| ENSG00000255100 | 13.07045516 | 1.940250251 | 0.4221712 | 4.595885 | 4.31E-06 | 5.02E-05 | | antisense | 50.41 | 8.341605183 | 8.341605183 | ENSG00000255100 | up |
| ENSG00000196482 | 1933.26341 | -1.380256943 | 0.2527465 | -5.461033 | 4.73E-08 | 9.12E-07 | ESRRG | protein_coding | 38.06 | -8.337033535 | 8.337033535 | ENSG00000196482 | down |
| ENSG00000201003 | 9.614582815 | -2.058960796 | 0.462368 | -4.453078 | 8.46E-06 | 9.04E-05 | | snoRNA | 47.79 | -8.325603529 | 8.325603529 | ENSG00000201003 | down |
| ENSG00000271387 | 78.7802561 | 1.389420327 | 0.2555298 | 5.437409 | 5.41E-08 | 1.03E-06 | | lincRNA | 56.72 | 8.318505005 | 8.318505005 | ENSG00000271387 | up |
| ENSG00000012174 | 319.8195245 | -1.450318906 | 0.2726363 | -5.31961 | 1.04E-07 | 1.85E-06 | MBTPS2 | protein_coding | 38.49 | -8.315536976 | 8.315536976 | ENSG00000012174 | down |
| ENSG00000254884 | 71.0363869 | 1.459288357 | 0.2753277 | 5.300188 | 1.16E-07 | 2.02E-06 | PRR13P2 | processed_pseudogene | 56.65 | 8.31064407 | 8.31064407 | ENSG00000254884 | up |
| ENSG00000104722 | 710.6929588 | 1.192552379 | 0.2033976 | 5.863158 | 4.54E-09 | 1.10E-07 | NEFM | protein_coding | 49.83 | 8.296856173 | 8.296856173 | ENSG00000104722 | up |
| ENSG00000279865 | 96.65490587 | 1.404843625 | 0.2602784 | 5.397465 | 6.76E-08 | 1.26E-06 | | TEC | 38.59 | 8.288172505 | 8.288172505 | ENSG00000279865 | up |
| ENSG00000276524 | 63.2619842 | 1.434839546 | 0.2687576 | 5.338787 | 9.36E-08 | 1.68E-06 | | lincRNA | 58.04 | 8.285382046 | 8.285382046 | ENSG00000276524 | up |
| ENSG00000214432 | 19.09595889 | 1.715521188 | 0.3519983 | 4.873663 | 1.10E-06 | 1.48E-05 | | antisense | 47.04 | 8.285109464 | 8.285109464 | ENSG00000214432 | up |
| ENSG00000251972 | 16.80561998 | -2.021880688 | 0.4510453 | -4.482656 | 7.37E-06 | 8.02E-05 | RNU6-123P | snRNA | 33.33 | -8.281407611 | 8.281407611 | ENSG00000251972 | down |
| ENSG00000136870 | 738.7198794 | -1.449378513 | 0.2730984 | -5.307166 | 1.11E-07 | 1.95E-06 | ZNF189 | protein_coding | 39.9 | -8.275119039 | 8.275119039 | ENSG00000136870 | down |
| ENSG00000164074 | 279.6089216 | -1.466998338 | 0.2782699 | -5.271854 | 1.35E-07 | 2.31E-06 | ABHD18 | protein_coding | 39.82 | -8.269454187 | 8.269454187 | ENSG00000164074 | down |
| ENSG00000196456 | 115.7495285 | -1.533072375 | 0.2974626 | -5.153832 | 2.55E-07 | 4.10E-06 | ZNF775 | protein_coding | 52.5 | -8.258469868 | 8.258469868 | ENSG00000196456 | down |
| ENSG00000266910 | 42.41545877 | -1.595549791 | 0.3162282 | -5.045565 | 4.52E-07 | 6.75E-06 | | processed_pseudogene | 42.23 | -8.250324204 | 8.250324204 | ENSG00000266910 | down |
| ENSG00000278475 | 67.07081616 | -1.594098672 | 0.3159765 | -5.044991 | 4.54E-07 | 6.76E-06 | | sense_intronic | 33.05 | -8.241483704 | 8.241483704 | ENSG00000278475 | down |
| ENSG00000230734 | 29.42650953 | -1.629940669 | 0.3271102 | -4.982848 | 6.27E-07 | 8.90E-06 | RPL10P3 | processed_pseudogene | 50.85 | -8.232318962 | 8.232318962 | ENSG00000230734 | down |
| ENSG00000212283 | 17.70821592 | -1.816294069 | 0.3848366 | -4.71965 | 2.36E-06 | 2.96E-05 | SNORD89 | snoRNA | 47.37 | -8.226703206 | 8.226703206 | ENSG00000212283 | down |
| ENSG00000207063 | 8.539040427 | -2.14299918 | 0.4949483 | -4.329743 | 1.49E-05 | 0.000146992 | SNORD116-1 | snoRNA | 42.11 | -8.213484139 | 8.213484139 | ENSG00000207063 | down |
| ENSG00000155016 | 311.7335121 | -1.385388296 | 0.2563074 | -5.405182 | 6.47E-08 | 1.21E-06 | CYP2U1 | protein_coding | 40.45 | -8.19604879 | 8.19604879 | ENSG00000155016 | down |
| ENSG00000261135 | 30.89038478 | 1.585222633 | 0.3142483 | 5.04449 | 4.55E-07 | 6.77E-06 | | lincRNA | 41.26 | 8.194527317 | 8.194527317 | ENSG00000261135 | up |
| ENSG00000231125 | 137.4214663 | -1.511750127 | 0.2924827 | -5.168682 | 2.36E-07 | 3.83E-06 | | sense_intronic | 40.43 | -8.18807521 | 8.18807521 | ENSG00000231125 | down |
| ENSG00000205181 | 329.3015303 | -1.456292062 | 0.2768227 | -5.260739 | 1.43E-07 | 2.43E-06 | LINC00654 | lincRNA | 50.26 | -8.175369624 | 8.175369624 | ENSG00000205181 | down |
| ENSG00000275710 | 13.06019625 | -2.781184426 | 0.7366005 | -3.775703 | 0.000159557 | 0.001154034 | | antisense | 25.73 | -8.170512362 | 8.170512362 | ENSG00000275710 | down |
| ENSG00000212535 | 15.83705881 | -1.922685928 | 0.4209523 | -4.567467 | 4.94E-06 | 5.63E-05 | RNU6-808P | snRNA | 32.41 | -8.169886243 | 8.169886243 | ENSG00000212535 | down |
| ENSG00000208025 | 8.810996213 | -2.28619155 | 0.5476737 | -4.174368 | 2.99E-05 | 0.000267894 | MIR591 | miRNA | 41.05 | -8.166362211 | 8.166362211 | ENSG00000208025 | down |
| ENSG00000274344 | 10.45695235 | -2.041414567 | 0.4614832 | -4.423595 | 9.71E-06 | 0.000101772 | | misc_RNA | 37.5 | -8.150085364 | 8.150085364 | ENSG00000274344 | down |
| ENSG00000207290 | 4.195205285 | -2.687084744 | 0.7001696 | -3.837763 | 0.00012416 | 0.000927407 | | misc_RNA | 40.35 | -8.149202024 | 8.149202024 | ENSG00000207290 | down |
| ENSG00000151151 | 433.7273088 | -1.358652684 | 0.2498764 | -5.437299 | 5.41E-08 | 1.03E-06 | IPMK | protein_coding | 38.62 | -8.134298122 | 8.134298122 | ENSG00000151151 | down |

| | | | | | | | | | | | | | |
|---|---|---|---|---|---|---|---|---|---|---|---|---|---|
| ENSG00000166292 | 62.82774099 | 1.414782104 | 0.265676 | 5.325216 | 1.01E-07 | 1.80E-06 | TMEM100 | protein_coding | 40.18 | 8.128369722 | 8.128369722 | ENSG00000166292 | up |
| ENSG00000107890 | 794.0272668 | -1.477439532 | 0.2837412 | -5.206997 | 1.92E-07 | 3.17E-06 | ANKRD26 | protein_coding | 38.15 | -8.125161622 | 8.125161622 | ENSG00000107890 | down |
| ENSG00000260519 | 19.61692635 | 1.718418427 | 0.3563367 | 4.822457 | 1.42E-06 | 1.87E-05 | | antisense | 54.97 | 8.123597548 | 8.123597548 | ENSG00000260519 | up |
| ENSG00000199015 | 3.036550621 | -3.621733261 | 1.1032711 | -3.282723 | 0.001028097 | 0.005767204 | MIR377 | miRNA | 39.13 | -8.109186476 | 8.109186476 | ENSG00000199015 | down |
| ENSG00000266649 | 393.7753442 | -1.334374394 | 0.2437078 | -5.475305 | 4.37E-08 | 8.46E-07 | MIR3126 | miRNA | 40.54 | -8.103326817 | 8.103326817 | ENSG00000266649 | down |
| ENSG00000179134 | 132.0335964 | -1.487761266 | 0.2870935 | -5.182149 | 2.19E-07 | 3.59E-06 | SAMD4B | protein_coding | 49.82 | -8.100966276 | 8.100966276 | ENSG00000179134 | down |
| ENSG00000267692 | 7.924906906 | -2.207540035 | 0.5217158 | -4.231307 | 2.32E-05 | 0.000214653 | | processed_pseudogene | 43.31 | -8.097835774 | 8.097835774 | ENSG00000267692 | down |
| ENSG00000272205 | 36.76517595 | 1.542604247 | 0.303308 | 5.085934 | 3.66E-07 | 5.64E-06 | | lincRNA | 58.21 | 8.096645234 | 8.096645234 | ENSG00000272205 | up |
| ENSG00000141905 | 16.44307117 | -1.802588074 | 0.3838559 | -4.696002 | 2.65E-06 | 3.26E-05 | NFIC | protein_coding | 55.9 | -8.087771795 | 8.087771795 | ENSG00000141905 | down |
| ENSG00000088876 | 349.2962554 | -1.395421986 | 0.260893 | -5.348637 | 8.86E-08 | 1.61E-06 | ZNF343 | protein_coding | 43.04 | -8.084425602 | 8.084425602 | ENSG00000088876 | down |
| ENSG00000254936 | 61.55101864 | -1.494731129 | 0.2894022 | -5.164892 | 2.41E-07 | 3.90E-06 | | sense_intronic | 43.59 | -8.084252991 | 8.084252991 | ENSG00000254936 | down |
| ENSG00000237440 | 828.3590942 | -1.40302648 | 0.2631936 | -5.330777 | 9.78E-08 | 1.75E-06 | ZNF737 | protein_coding | 39.46 | -8.078700682 | 8.078700682 | ENSG00000237440 | down |
| ENSG00000275939 | 11.85582134 | -2.129151804 | 0.4945266 | -4.305435 | 1.67E-05 | 0.000161925 | | misc_RNA | 57.23 | -8.070947469 | 8.070947469 | ENSG00000275939 | down |
| ENSG00000105085 | 66.23880241 | 1.414479954 | 0.2666981 | 5.303676 | 1.13E-07 | 1.98E-06 | MED26 | protein_coding | 52.52 | 8.066164102 | 8.066164102 | ENSG00000105085 | up |
| ENSG00000243173 | 28.92566296 | -1.549198906 | 0.3061321 | -5.060557 | 4.18E-07 | 6.31E-06 | RN7SL861P | misc_RNA | 52.28 | -8.056235057 | 8.056235057 | ENSG00000243173 | down |
| ENSG00000140557 | 403.8328673 | -1.418114536 | 0.2685017 | -5.281585 | 1.28E-07 | 2.20E-06 | ST8SIA2 | protein_coding | 48 | -8.022700216 | 8.022700216 | ENSG00000140557 | down |
| ENSG00000229447 | 17.99720492 | -1.749651422 | 0.3685778 | -4.747034 | 2.06E-06 | 2.62E-05 | | processed_pseudogene | 39.13 | -8.017795467 | 8.017795467 | ENSG00000229447 | down |
| ENSG00000122778 | 846.0390148 | -1.275725953 | 0.2294406 | -5.560159 | 2.70E-08 | 5.48E-07 | KIAA1549 | protein_coding | 44.25 | -7.987769763 | 7.987769763 | ENSG00000122778 | down |
| ENSG00000249471 | 113.1539627 | -1.420265538 | 0.269772 | -5.264688 | 1.40E-07 | 2.39E-06 | ZNF324B | protein_coding | 58.99 | -7.984871958 | 7.984871958 | ENSG00000249471 | down |
| ENSG00000181827 | 2186.749013 | -1.338351774 | 0.2467576 | -5.423751 | 5.84E-08 | 1.10E-06 | RFX7 | protein_coding | 37.13 | -7.974167632 | 7.974167632 | ENSG00000181827 | down |
| ENSG00000234160 | 99.28931829 | -1.473920358 | 0.2854572 | -5.163367 | 2.43E-07 | 3.92E-06 | | antisense | 48.87 | -7.968615534 | 7.968615534 | ENSG00000234160 | down |
| ENSG00000207605 | 4.620630294 | 2.591930109 | 0.6717047 | 3.858735 | 0.000113976 | 0.000857311 | MIR191 | miRNA | 59.78 | 7.949091336 | 7.949091336 | ENSG00000207605 | up |
| ENSG00000197020 | 976.5589983 | -1.321696803 | 0.2426525 | -5.446871 | 5.13E-08 | 9.84E-07 | ZNF100 | protein_coding | 38.56 | -7.939163357 | 7.939163357 | ENSG00000197020 | down |
| ENSG00000265435 | 6.574596788 | -2.28475396 | 0.5552143 | -4.115085 | 3.87E-05 | 0.000335119 | MIR3121 | miRNA | 35.06 | -7.939065218 | 7.939065218 | ENSG00000265435 | down |
| ENSG00000242889 | 56.87653691 | -1.516246905 | 0.2986518 | -5.076972 | 3.83E-07 | 5.86E-06 | RN7SL449P | misc_RNA | 55.94 | -7.933029563 | 7.933029563 | ENSG00000242889 | down |
| ENSG00000201231 | 5.256932929 | -2.980402121 | 0.8313469 | -3.585028 | 0.000337042 | 0.002204149 | RNU4-50P | snRNA | 41.13 | -7.918210059 | 7.918210059 | ENSG00000201231 | down |
| ENSG00000266900 | 158.2326708 | -1.470880464 | 0.2857214 | -5.147954 | 2.63E-07 | 4.21E-06 | | sense_intronic | 34.37 | -7.907106035 | 7.907106035 | ENSG00000266900 | down |
| ENSG00000200257 | 5.188204336 | -2.399299033 | 0.5993862 | -4.002927 | 6.26E-05 | 0.000507028 | RNU6-97P | snRNA | 41.35 | -7.905613585 | 7.905613585 | ENSG00000200257 | down |
| ENSG00000276292 | 14.06749282 | -1.883909219 | 0.4151484 | -4.537918 | 5.68E-06 | 6.39E-05 | | antisense | 35.68 | -7.902141094 | 7.902141094 | ENSG00000276292 | down |
| ENSG00000260951 | 146.3367605 | -1.511128649 | 0.2977815 | -5.074623 | 3.88E-07 | 5.92E-06 | | lincRNA | 40.38 | -7.899645084 | 7.899645084 | ENSG00000260951 | down |
| ENSG00000265917 | 25.08322408 | -1.763424632 | 0.3758375 | -4.691987 | 2.71E-06 | 3.32E-05 | MIR3685 | miRNA | 45.16 | -7.899039562 | 7.899039562 | ENSG00000265917 | down |
| ENSG00000264099 | 37.53421613 | -1.655496668 | 0.3420466 | -4.839974 | 1.30E-06 | 1.73E-05 | MIR4803 | miRNA | 35.14 | -7.884030625 | 7.884030625 | ENSG00000264099 | down |

| | | | | | | | | | | | | | |
|---|---|---|---|---|---|---|---|---|---|---|---|---|---|
| ENSG00000269001 | 148.7121749 | -1.450393287 | 0.2802058 | -5.176171 | 2.26E-07 | 3.69E-06 | | transcribed_unprocessed_pseud | 44.81 | -7.879527891 | 7.879527891 | ENSG00000269001 | down |
| ENSG00000230376 | 8.927320566 | 2.131606536 | 0.5017634 | 4.248231 | 2.15E-05 | 0.000201473 | MEMO1P4 | processed_pseudogene | 44.04 | 7.877957159 | 7.877957159 | ENSG00000230376 | up |
| ENSG00000206638 | 2.617606554 | -3.204967257 | 0.9304102 | -3.444682 | 0.000571731 | 0.003486734 | RNU6-672P | snRNA | 39.25 | -7.876467162 | 7.876467162 | ENSG00000206638 | down |
| ENSG00000270231 | 193.3701737 | -1.479322097 | 0.2889321 | -5.119964 | 3.06E-07 | 4.79E-06 | NBPF8 | unprocessed_pseudogene | 44.63 | -7.869305487 | 7.869305487 | ENSG00000270231 | down |
| ENSG00000245888 | 96.77448813 | 1.371352028 | 0.2578206 | 5.319017 | 1.04E-07 | 1.85E-06 | | antisense | 44.09 | 7.861592013 | 7.861592013 | ENSG00000245888 | up |
| ENSG00000275745 | 7.732027488 | 2.39611473 | 0.5998854 | 3.994287 | 6.49E-05 | 0.00052374 | | misc_RNA | 41.44 | 7.861375698 | 7.861375698 | ENSG00000275745 | up |
| ENSG00000207493 | 15.29446125 | -1.828076896 | 0.3980107 | -4.593035 | 4.37E-06 | 5.08E-05 | SNORA46 | snoRNA | 43.7 | -7.849659931 | 7.849659931 | ENSG00000207493 | down |
| ENSG00000237200 | 9.216407962 | -2.123651056 | 0.5001897 | -4.245692 | 2.18E-05 | 0.000203259 | ZBTB40-IT1 | sense_intronic | 44.07 | -7.840415364 | 7.840415364 | ENSG00000237200 | down |
| ENSG00000140948 | 481.8831812 | -1.349463632 | 0.252056 | -5.353825 | 8.61E-08 | 1.57E-06 | ZCCHC14 | protein_coding | 48.02 | -7.833439678 | 7.833439678 | ENSG00000140948 | down |
| ENSG00000165801 | 228.8645673 | -1.407517242 | 0.2688661 | -5.235013 | 1.65E-07 | 2.75E-06 | ARHGEF40 | protein_coding | 53.84 | -7.826554455 | 7.826554455 | ENSG00000165801 | down |
| ENSG00000126749 | 1176.192769 | -1.393070655 | 0.2650314 | -5.256247 | 1.47E-07 | 2.48E-06 | EMG1 | protein_coding | 48.3 | -7.808622234 | 7.808622234 | ENSG00000126749 | down |
| ENSG00000254331 | 11.4073613 | -1.985229412 | 0.4523282 | -4.388913 | 1.14E-05 | 0.000116934 | CKS1BP7 | processed_pseudogene | 45.96 | -7.80603889 | 7.80603889 | ENSG00000254331 | down |
| ENSG00000238812 | 3.931102269 | -2.774429552 | 0.7513486 | -3.6926 | 0.000221973 | 0.001536815 | RNU7-127P | snRNA | 46.77 | -7.805520579 | 7.805520579 | ENSG00000238812 | down |
| ENSG00000272843 | 41.09284179 | 1.491100933 | 0.293846 | 5.074431 | 3.89E-07 | 5.92E-06 | | antisense | 51.27 | 7.79494728 | 7.79494728 | ENSG00000272843 | up |
| ENSG00000278770 | 9.802025239 | -1.961291156 | 0.4446639 | -4.410727 | 1.03E-05 | 0.000107205 | | misc_RNA | 53.02 | -7.78590337 | 7.78590337 | ENSG00000278770 | down |
| ENSG00000233924 | 50.23675393 | -1.548138656 | 0.3116382 | -4.967744 | 6.77E-07 | 9.55E-06 | RPSAP13 | processed_pseudogene | 49.6 | -7.771969383 | 7.771969383 | ENSG00000233924 | down |
| ENSG00000103494 | 382.8558829 | -1.449268317 | 0.2819832 | -5.139556 | 2.75E-07 | 4.36E-06 | RPGRIP1L | protein_coding | 37.27 | -7.769242054 | 7.769242054 | ENSG00000103494 | down |
| ENSG00000125449 | 125.3808262 | -1.356610854 | 0.2555717 | -5.308142 | 1.11E-07 | 1.95E-06 | ARMC7 | protein_coding | 51.99 | -7.747136496 | 7.747136496 | ENSG00000125449 | down |
| ENSG00000236756 | 16.87666576 | 1.748419911 | 0.3747046 | 4.666129 | 3.07E-06 | 3.71E-05 | DNAJC9-AS1 | antisense | 41.21 | 7.746036522 | 7.746036522 | ENSG00000236756 | up |
| ENSG00000196081 | 1068.421426 | -1.456765492 | 0.284631 | -5.118084 | 3.09E-07 | 4.83E-06 | ZNF724 | protein_coding | 40.39 | -7.743723272 | 7.743723272 | ENSG00000196081 | down |
| ENSG00000271270 | 212.8570164 | -1.479892795 | 0.2914914 | -5.076969 | 3.84E-07 | 5.86E-06 | TMCC1-AS1 | antisense | 44.04 | -7.74282424 | 7.74282424 | ENSG00000271270 | down |
| ENSG00000124216 | 1107.161698 | 1.269824043 | 0.2314431 | 5.48655 | 4.10E-08 | 7.99E-07 | SNAI1 | protein_coding | 55.16 | 7.742536319 | 7.742536319 | ENSG00000124216 | up |
| ENSG00000219395 | 64.99492898 | -1.634800887 | 0.3387357 | -4.826184 | 1.39E-06 | 1.84E-05 | HSPA8P15 | processed_pseudogene | 47.04 | -7.739658492 | 7.739658492 | ENSG00000219395 | down |
| ENSG00000253633 | 59.38751021 | 1.44325404 | 0.2807639 | 5.140454 | 2.74E-07 | 4.34E-06 | | lincRNA | 43.71 | 7.739284028 | 7.739284028 | ENSG00000253633 | up |
| ENSG00000167528 | 54.07167329 | -1.506147697 | 0.2997801 | -5.024175 | 5.06E-07 | 7.35E-06 | ZNF641 | protein_coding | 46.15 | -7.732297919 | 7.732297919 | ENSG00000167528 | down |
| ENSG00000198743 | 2080.33468 | -1.430026478 | 0.277084 | -5.160985 | 2.46E-07 | 3.97E-06 | SLC5A3 | protein_coding | 39.03 | -7.724217575 | 7.724217575 | ENSG00000198743 | down |
| ENSG00000268996 | 64.45143605 | 1.358907788 | 0.2568494 | 5.290679 | 1.22E-07 | 2.11E-06 | MAN1B1-AS1 | antisense | 60.58 | 7.712686856 | 7.712686856 | ENSG00000268996 | up |
| ENSG00000197302 | 467.3881823 | -1.399528124 | 0.2685603 | -5.211225 | 1.88E-07 | 3.10E-06 | ZNF720 | protein_coding | 38.78 | -7.709091407 | 7.709091407 | ENSG00000197302 | down |
| ENSG00000075151 | 181.0625411 | 1.250761973 | 0.2267971 | 5.514893 | 3.49E-08 | 6.91E-07 | EIF4G3 | protein_coding | 39.36 | 7.705202872 | 7.705202872 | ENSG00000075151 | up |
| ENSG00000251369 | 351.5848546 | -1.394898111 | 0.2673217 | -5.218051 | 1.81E-07 | 3.00E-06 | ZNF550 | protein_coding | 44.48 | -7.703009523 | 7.703009523 | ENSG00000251369 | down |
| ENSG00000228293 | 30.44471473 | 1.55375031 | 0.3148045 | 4.935603 | 7.99E-07 | 1.10E-05 | | antisense | 45.57 | 7.702193516 | 7.702193516 | ENSG00000228293 | up |
| ENSG00000154319 | 348.833369 | -1.448326973 | 0.2830556 | -5.116758 | 3.11E-07 | 4.86E-06 | FAM167A | protein_coding | 52.59 | -7.695151125 | 7.695151125 | ENSG00000154319 | down |

| | | | | | | | | | | | | | |
|---|---|---|---|---|---|---|---|---|---|---|---|---|---|
| ENSG00000166833 | 1727.013264 | -1.212616466 | 0.21665 | -5.597122 | 2.18E-08 | 4.53E-07 | NAV2 | protein_coding | 44.51 | -7.692845323 | 7.692845323 | ENSG00000166833 | down |
| ENSG00000137992 | 768.9775155 | -1.395905285 | 0.2678148 | -5.212203 | 1.87E-07 | 3.09E-06 | DBT | protein_coding | 39.15 | -7.691612587 | 7.691612587 | ENSG00000137992 | down |
| ENSG00000236723 | 75.0036555 | -1.367177673 | 0.2596063 | -5.266351 | 1.39E-07 | 2.37E-06 | | antisense | 47.26 | -7.691055923 | 7.691055923 | ENSG00000236723 | down |
| ENSG00000112715 | 984.0231281 | 1.280457842 | 0.2351309 | 5.445724 | 5.16E-08 | 9.89E-07 | VEGFA | protein_coding | 56.4 | 7.688632788 | 7.688632788 | ENSG00000112715 | up |
| ENSG00000047932 | 1919.261972 | -1.476078111 | 0.2915437 | -5.062974 | 4.13E-07 | 6.24E-06 | GOPC | protein_coding | 36.66 | -7.682725697 | 7.682725697 | ENSG00000047932 | down |
| ENSG00000276365 | 3.711823768 | -2.761153525 | 0.7522099 | -3.670722 | 0.000241866 | 0.001654793 | MIR145 | miRNA | 47.73 | -7.679475911 | 7.679475911 | ENSG00000276365 | down |
| ENSG00000200832 | 6.56418448 | -2.375639088 | 0.5994591 | -3.962971 | 7.40E-05 | 0.000586195 | SNORD114-4 | snoRNA | 42.67 | -7.677964957 | 7.677964957 | ENSG00000200832 | down |
| ENSG00000168517 | 130.9608474 | -1.302013236 | 0.2416793 | -5.387359 | 7.15E-08 | 1.32E-06 | HEXIM2 | protein_coding | 53.43 | -7.657210171 | 7.657210171 | ENSG00000168517 | down |
| ENSG00000273329 | 430.0515187 | -1.163924769 | 0.204291 | -5.697387 | 1.22E-08 | 2.65E-07 | | lincRNA | 46.42 | -7.654700471 | 7.654700471 | ENSG00000273329 | down |
| ENSG00000090447 | 456.4902426 | -1.300938481 | 0.2414576 | -5.387854 | 7.13E-08 | 1.31E-06 | TFAP4 | protein_coding | 56.39 | -7.65169795 | 7.65169795 | ENSG00000090447 | down |
| ENSG00000104728 | 120.271576 | -1.500481858 | 0.2996883 | -5.006808 | 5.53E-07 | 8.01E-06 | ARHGEF10 | protein_coding | 50 | -7.647057903 | 7.647057903 | ENSG00000104728 | down |
| ENSG00000199347 | 388.7089256 | 1.158411597 | 0.2030564 | 5.704875 | 1.16E-08 | 2.55E-07 | RNU5E-1 | snRNA | 40 | 7.637385939 | 7.637385939 | ENSG00000199347 | up |
| ENSG00000266993 | 15.18445199 | -1.870716912 | 0.4181392 | -4.47391 | 7.68E-06 | 8.32E-05 | | antisense | 42.62 | -7.632776275 | 7.632776275 | ENSG00000266993 | down |
| ENSG00000164031 | 570.202118 | -1.497256212 | 0.299121 | -5.005521 | 5.57E-07 | 8.05E-06 | DNAJB14 | protein_coding | 34.8 | -7.627625445 | 7.627625445 | ENSG00000164031 | down |
| ENSG00000202523 | 11.94160076 | -2.019408281 | 0.4702055 | -4.294735 | 1.75E-05 | 0.000168058 | | misc_RNA | 34.31 | -7.622339663 | 7.622339663 | ENSG00000202523 | down |
| ENSG00000272599 | 139.6788602 | -1.436330837 | 0.2809067 | -5.113195 | 3.17E-07 | 4.94E-06 | | antisense | 40.62 | -7.621039729 | 7.621039729 | ENSG00000272599 | down |
| ENSG00000202189 | 6.11506757 | -2.382701266 | 0.6048115 | -3.939576 | 8.16E-05 | 0.000639186 | SNORA30B | snoRNA | 53.49 | -7.611235681 | 7.611235681 | ENSG00000202189 | down |
| ENSG00000222164 | 52.88413104 | -1.498418179 | 0.2998518 | -4.997195 | 5.82E-07 | 8.35E-06 | RN7SKP266 | misc_RNA | 43.55 | -7.609474672 | 7.609474672 | ENSG00000222164 | down |
| ENSG00000231188 | 34.41658277 | 1.493722928 | 0.29857 | 5.002924 | 5.65E-07 | 8.13E-06 | | antisense | 48.9 | 7.602896869 | 7.602896869 | ENSG00000231188 | up |
| ENSG00000258099 | 57.58180381 | 1.397420273 | 0.2698406 | 5.178689 | 2.23E-07 | 3.65E-06 | ATXN2-AS | antisense | 61.79 | 7.599219546 | 7.599219546 | ENSG00000258099 | up |
| ENSG00000225808 | 61.14599445 | -2.186317883 | 0.5315226 | -4.113311 | 3.90E-05 | 0.000337286 | DNAJC19P5 | processed_pseudogene | 41.14 | -7.590899358 | 7.590899358 | ENSG00000225808 | down |
| ENSG00000275318 | 43.550788 | -1.479969913 | 0.2947982 | -5.020281 | 5.16E-07 | 7.48E-06 | | unprocessed_pseudogene | 35.56 | -7.586212703 | 7.586212703 | ENSG00000275318 | down |
| ENSG00000224165 | 67.83987247 | 1.401843848 | 0.2713929 | 5.165366 | 2.40E-07 | 3.90E-06 | DNAJC27-AS1 | antisense | 45.18 | 7.582703583 | 7.582703583 | ENSG00000224165 | up |
| ENSG00000259408 | 65.62120168 | -1.472012411 | 0.2924592 | -5.033223 | 4.82E-07 | 7.08E-06 | | antisense | 42.23 | -7.580495709 | 7.580495709 | ENSG00000259408 | down |
| ENSG00000200855 | 9.930363223 | -2.054344866 | 0.4844622 | -4.240465 | 2.23E-05 | 0.000206904 | | misc_RNA | 44.12 | -7.568682503 | 7.568682503 | ENSG00000200855 | down |
| ENSG00000221164 | 37.60310149 | -1.5157089 | 0.3060541 | -4.952422 | 7.33E-07 | 1.02E-05 | SNORA11F | snoRNA | 50.78 | -7.563874379 | 7.563874379 | ENSG00000221164 | down |
| ENSG00000212017 | 15.5978652 | -1.847702239 | 0.4123473 | -4.480937 | 7.43E-06 | 8.07E-05 | MIR548U | miRNA | 33.33 | -7.562777967 | 7.562777967 | ENSG00000212017 | down |
| ENSG00000163026 | 730.6143188 | -1.449330815 | 0.2859905 | -5.067758 | 4.03E-07 | 6.11E-06 | WDCP | protein_coding | 43.29 | -7.557264713 | 7.557264713 | ENSG00000163026 | down |
| ENSG00000185875 | 375.1794177 | -1.489683285 | 0.2982878 | -4.994114 | 5.91E-07 | 8.48E-06 | THNSL1 | protein_coding | 37.6 | -7.555453283 | 7.555453283 | ENSG00000185875 | down |
| ENSG00000273723 | 333.0080822 | 1.168167894 | 0.2071142 | 5.64021 | 1.70E-08 | 3.59E-07 | | lincRNA | 38.77 | 7.528195525 | 7.528195525 | ENSG00000273723 | up |
| ENSG00000224977 | 22.137511 | 1.728962302 | 0.3738987 | 4.624147 | 3.76E-06 | 4.44E-05 | | lincRNA | 54.51 | 7.525182112 | 7.525182112 | ENSG00000224977 | up |
| ENSG00000139318 | 294.0251401 | 1.199994129 | 0.2160481 | 5.554291 | 2.79E-08 | 5.65E-07 | DUSP6 | protein_coding | 46.57 | 7.497096754 | 7.497096754 | ENSG00000139318 | up |

| | | | | | | | | | | | | | |
|---|---|---|---|---|---|---|---|---|---|---|---|---|---|
| ENSG00000283200 | 8.604780258 | -2.117012558 | 0.5101355 | -4.149902 | 3.33E-05 | 0.000293289 | MIR1206 | miRNA | 42.37 | -7.478780599 | 7.478780599 | ENSG00000283200 | down |
| ENSG00000258634 | 261.2694541 | -1.38003436 | 0.267499 | -5.159026 | 2.48E-07 | 4.00E-06 | | antisense | 53.13 | -7.449694778 | 7.449694778 | ENSG00000258634 | down |
| ENSG00000205683 | 81.52746946 | -1.442802 | 0.2866559 | -5.03322 | 4.82E-07 | 7.08E-06 | DPF3 | protein_coding | 45.75 | -7.4300694 | 7.4300694 | ENSG00000205683 | down |
| ENSG00000207034 | 16.8395774 | -1.757415287 | 0.385851 | -4.554647 | 5.25E-06 | 5.97E-05 | | misc_RNA | 41.03 | -7.422887364 | 7.422887364 | ENSG00000207034 | down |
| ENSG00000270640 | 57.86285406 | 1.353814453 | 0.2605094 | 5.196798 | 2.03E-07 | 3.33E-06 | | sense_intronic | 39.86 | 7.415790535 | 7.415790535 | ENSG00000270640 | up |
| ENSG00000269846 | 37.53044521 | 1.488233458 | 0.3007835 | 4.947857 | 7.50E-07 | 1.04E-05 | | antisense | 55.52 | 7.415474857 | 7.415474857 | ENSG00000269846 | up |
| ENSG00000272055 | 5.597382938 | -2.499655717 | 0.6588087 | -3.794206 | 0.000148117 | 0.001083105 | RNU6-6P | snRNA | 45.79 | -7.412302954 | 7.412302954 | ENSG00000272055 | down |
| ENSG00000261839 | 78.4881662 | 1.304118716 | 0.2463341 | 5.294105 | 1.20E-07 | 2.08E-06 | | lincRNA | 43.07 | 7.41092478 | 7.41092478 | ENSG00000261839 | up |
| ENSG00000199325 | 4.539565226 | 2.549376046 | 0.6788668 | 3.75534 | 0.000173106 | 0.001239792 | RNU4-39P | snRNA | 45.67 | 7.410146929 | 7.410146929 | ENSG00000199325 | up |
| ENSG00000212207 | 5.788144074 | -2.512028951 | 0.6644231 | -3.780767 | 0.000156346 | 0.001134343 | RNU6-1321P | snRNA | 33.64 | -7.398567274 | 7.398567274 | ENSG00000212207 | down |
| ENSG00000260404 | 187.769258 | -1.302418135 | 0.2462047 | -5.289982 | 1.22E-07 | 2.12E-06 | | transcribed_unprocessed_pseud | 46.61 | -7.39061768 | 7.39061768 | ENSG00000260404 | down |
| ENSG00000278333 | 4.820747689 | -2.727098815 | 0.7527929 | -3.622641 | 0.00029161 | 0.001950088 | | misc_RNA | 55.29 | -7.390289693 | 7.390289693 | ENSG00000278333 | down |
| ENSG00000269837 | 313.5013521 | -1.409813997 | 0.2773901 | -5.082423 | 3.73E-07 | 5.73E-06 | IPO5P1 | transcribed_processed_pseudog | 46.62 | -7.389700543 | 7.389700543 | ENSG00000269837 | down |
| ENSG00000180953 | 119.2775243 | -1.604118749 | 0.3372524 | -4.756434 | 1.97E-06 | 2.50E-05 | ST20 | protein_coding | 41.13 | -7.381380032 | 7.381380032 | ENSG00000180953 | down |
| ENSG00000171488 | 308.7818175 | -1.314920681 | 0.250224 | -5.254975 | 1.48E-07 | 2.50E-06 | LRRC8C | protein_coding | 39.69 | -7.367308642 | 7.367308642 | ENSG00000171488 | down |
| ENSG00000261575 | 39.52380446 | 1.454791106 | 0.2915894 | 4.989176 | 6.06E-07 | 8.64E-06 | | transcribed_unprocessed_pseud | 52.83 | 7.366221521 | 7.366221521 | ENSG00000261575 | up |
| ENSG00000197937 | 477.3217069 | -1.466140404 | 0.295215 | -4.966348 | 6.82E-07 | 9.58E-06 | ZNF347 | protein_coding | 44.35 | -7.358236682 | 7.358236682 | ENSG00000197937 | down |
| ENSG00000062725 | 1897.352029 | -1.450827535 | 0.2905443 | -4.993481 | 5.93E-07 | 8.48E-06 | APPBP2 | protein_coding | 39.69 | -7.358162953 | 7.358162953 | ENSG00000062725 | down |
| ENSG00000276730 | 4.282701559 | -2.724584709 | 0.7542447 | -3.612335 | 0.000303452 | 0.002015727 | | misc_RNA | 46.75 | -7.344303919 | 7.344303919 | ENSG00000276730 | down |
| ENSG00000256464 | 31.11101793 | -1.636976246 | 0.348724 | -4.694189 | 2.68E-06 | 3.29E-05 | YWHABP2 | processed_pseudogene | 43.2 | -7.339032097 | 7.339032097 | ENSG00000256464 | down |
| ENSG00000129757 | 148.6811463 | 1.366013317 | 0.2658286 | 5.138699 | 2.77E-07 | 4.37E-06 | CDKN1C | protein_coding | 67.44 | 7.320898756 | 7.320898756 | ENSG00000129757 | up |
| ENSG00000264676 | 8.612650148 | -1.97398716 | 0.4640854 | -4.253499 | 2.10E-05 | 0.000197985 | RN7SL204P | misc_RNA | 58.4 | -7.310402061 | 7.310402061 | ENSG00000264676 | down |
| ENSG00000138696 | 304.5048197 | -1.298913797 | 0.2469632 | -5.259544 | 1.44E-07 | 2.44E-06 | BMPR1B | protein_coding | 36.7 | -7.289582078 | 7.289582078 | ENSG00000138696 | down |
| ENSG00000198346 | 290.9859922 | -1.458962514 | 0.294609 | -4.9522 | 7.34E-07 | 1.02E-05 | ZNF813 | protein_coding | 43.99 | -7.28060152 | 7.28060152 | ENSG00000198346 | down |
| ENSG00000138246 | 1890.5949 | -1.381889067 | 0.2711981 | -5.095498 | 3.48E-07 | 5.39E-06 | DNAJC13 | protein_coding | 36.8 | -7.280261943 | 7.280261943 | ENSG00000138246 | down |
| ENSG00000265333 | 7.054508569 | -2.190651408 | 0.5451125 | -4.018715 | 5.85E-05 | 0.000479532 | MIR3137 | miRNA | 54.67 | -7.27117155 | 7.27117155 | ENSG00000265333 | down |
| ENSG00000083814 | 142.1243869 | -1.347496207 | 0.2616676 | -5.149648 | 2.61E-07 | 4.18E-06 | ZNF671 | protein_coding | 49.44 | -7.247760791 | 7.247760791 | ENSG00000083814 | down |
| ENSG00000238386 | 7.377222539 | -2.529471701 | 0.6786501 | -3.72721 | 0.000193611 | 0.001364582 | RNU7-48P | snRNA | 33.33 | -7.246937122 | 7.246937122 | ENSG00000238386 | down |
| ENSG00000197933 | 213.7703456 | -1.435861574 | 0.2883872 | -4.978936 | 6.39E-07 | 9.06E-06 | ZNF823 | protein_coding | 44.81 | -7.241170018 | 7.241170018 | ENSG00000197933 | down |
| ENSG00000120129 | 608.7002946 | 1.108397883 | 0.195198 | 5.678325 | 1.36E-08 | 2.94E-07 | DUSP1 | protein_coding | 52.99 | 7.239555184 | 7.239555184 | ENSG00000120129 | up |
| ENSG00000129292 | 3709.879826 | -1.388734021 | 0.2741226 | -5.066106 | 4.06E-07 | 6.15E-06 | PHF20L1 | protein_coding | 34.77 | -7.236719432 | 7.236719432 | ENSG00000129292 | down |
| ENSG00000236860 | 13.80313929 | -1.857999715 | 0.4256789 | -4.364792 | 1.27E-05 | 0.0001281 | RPL39P29 | processed_pseudogene | 40.13 | -7.232174893 | 7.232174893 | ENSG00000236860 | down |

| | | | | | | | | | | | | | |
|---|---|---|---|---|---|---|---|---|---|---|---|---|---|
| ENSG00000207136 | 6.837913174 | -2.185122115 | 0.54458 | -4.01249 | 6.01E-05 | 0.000490445 | RNU6-769P | snRNA | 36.54 | -7.231464762 | 7.231464762 | ENSG00000207136 | down |
| ENSG00000213865 | 121.8631972 | -1.439669698 | 0.2898287 | -4.967313 | 6.79E-07 | 9.56E-06 | C8orf44 | protein_coding | 40.76 | -7.226676017 | 7.226676017 | ENSG00000213865 | down |
| ENSG00000220515 | 59.39514629 | -1.557453425 | 0.3262614 | -4.773637 | 1.81E-06 | 2.32E-05 | PGAM1P10 | processed_pseudogene | 53.09 | -7.217535109 | 7.217535109 | ENSG00000220515 | down |
| ENSG00000184205 | 334.996755 | 1.246341836 | 0.2331628 | 5.345371 | 9.02E-08 | 1.63E-06 | TSPYL2 | protein_coding | 52.46 | 7.213777881 | 7.213777881 | ENSG00000184205 | up |
| ENSG00000232503 | 28.99601595 | -1.689861935 | 0.3690901 | -4.578453 | 4.68E-06 | 5.41E-05 | CYCSP8 | processed_pseudogene | 46.47 | -7.209985716 | 7.209985716 | ENSG00000232503 | down |
| ENSG00000152193 | 1021.765443 | -1.35212195 | 0.2639429 | -5.122782 | 3.01E-07 | 4.73E-06 | RNF219 | protein_coding | 38.52 | -7.200114091 | 7.200114091 | ENSG00000152193 | down |
| ENSG00000163393 | 327.3272167 | -1.394675943 | 0.2768089 | -5.038408 | 4.69E-07 | 6.96E-06 | SLC22A15 | protein_coding | 39.5 | -7.192860948 | 7.192860948 | ENSG00000163393 | down |
| ENSG00000259378 | 75.27112398 | -1.488771321 | 0.3058851 | -4.867093 | 1.13E-06 | 1.52E-05 | DCAF13P3 | processed_pseudogene | 39.43 | -7.172277287 | 7.172277287 | ENSG00000259378 | down |
| ENSG00000275291 | 132.2251171 | 1.237621915 | 0.2313612 | 5.349306 | 8.83E-08 | 1.60E-06 | | snRNA | 46.91 | 7.171492097 | 7.171492097 | ENSG00000275291 | up |
| ENSG00000143322 | 1617.252663 | -1.298479167 | 0.249031 | -5.214127 | 1.85E-07 | 3.06E-06 | ABL2 | protein_coding | 41.31 | -7.159964361 | 7.159964361 | ENSG00000143322 | down |
| ENSG00000280129 | 172.2148999 | 1.214433543 | 0.2251388 | 5.394154 | 6.88E-08 | 1.28E-06 | | TEC | 42.77 | 7.156490585 | 7.156490585 | ENSG00000280129 | up |
| ENSG00000160229 | 356.7281479 | -1.389513902 | 0.2763942 | -5.02729 | 4.97E-07 | 7.25E-06 | ZNF66 | protein_coding | 38.46 | -7.14142338 | 7.14142338 | ENSG00000160229 | down |
| ENSG00000275611 | 14.90597055 | -1.889743978 | 0.4399618 | -4.295246 | 1.74E-05 | 0.000167787 | | sense_intronic | 36.59 | -7.13423905 | 7.13423905 | ENSG00000275611 | down |
| ENSG00000105750 | 266.0113842 | -1.423890347 | 0.2870109 | -4.961103 | 7.01E-07 | 9.83E-06 | ZNF85 | protein_coding | 37.87 | -7.130180084 | 7.130180084 | ENSG00000105750 | down |
| ENSG00000262074 | 492.0616034 | 1.158486939 | 0.21056 | 5.501933 | 3.76E-08 | 7.40E-07 | SNORD3B-2 | snoRNA | 56.13 | 7.102571737 | 7.102571737 | ENSG00000262074 | up |
| ENSG00000213096 | 1011.672129 | -1.423214099 | 0.2875057 | -4.950212 | 7.41E-07 | 1.03E-05 | ZNF254 | protein_coding | 41.43 | -7.097129879 | 7.097129879 | ENSG00000213096 | down |
| ENSG00000278916 | 157.4185339 | 1.241764568 | 0.2338599 | 5.309866 | 1.10E-07 | 1.94E-06 | CEP83-AS1 | lincRNA | 44.48 | 7.094342319 | 7.094342319 | ENSG00000278916 | up |
| ENSG00000125347 | 144.0357105 | 1.251881318 | 0.2370434 | 5.281232 | 1.28E-07 | 2.20E-06 | IRF1 | protein_coding | 53.85 | 7.081892869 | 7.081892869 | ENSG00000125347 | up |
| ENSG00000164683 | 110.0871498 | 1.310220918 | 0.2539004 | 5.160374 | 2.46E-07 | 3.98E-06 | HEY1 | protein_coding | 46.45 | 7.075895901 | 7.075895901 | ENSG00000164683 | up |
| ENSG00000173275 | 173.6005322 | -1.364488482 | 0.2703174 | -5.047727 | 4.47E-07 | 6.69E-06 | ZNF449 | protein_coding | 37.9 | -7.060746984 | 7.060746984 | ENSG00000173275 | down |
| ENSG00000185658 | 4493.74781 | -1.358824065 | 0.2686702 | -5.057591 | 4.25E-07 | 6.39E-06 | BRWD1 | protein_coding | 38.65 | -7.058335694 | 7.058335694 | ENSG00000185658 | down |
| ENSG00000216031 | 3.440342558 | -3.46329255 | 1.1106886 | -3.118149 | 0.001819907 | 0.009398579 | MIR298 | miRNA | 53.41 | -7.01987854 | 7.01987854 | ENSG00000216031 | down |
| ENSG00000283162 | 81.95281344 | -1.315640066 | 0.2565986 | -5.127229 | 2.94E-07 | 4.63E-06 | | transcribed_unprocessed_pseud | 41.54 | -7.018687003 | 7.018687003 | ENSG00000283162 | down |
| ENSG00000228079 | 14.86302936 | -2.282065643 | 0.5909998 | -3.861365 | 0.000112755 | 0.000848592 | | lincRNA | 39.55 | -7.008911217 | 7.008911217 | ENSG00000228079 | down |
| ENSG00000223023 | 6.794645593 | -2.151092585 | 0.5409139 | -3.976774 | 6.99E-05 | 0.000558971 | | misc_RNA | 37.96 | -6.996666327 | 6.996666327 | ENSG00000223023 | down |
| ENSG00000207642 | 28.69827787 | -1.582835832 | 0.3397743 | -4.658492 | 3.19E-06 | 3.83E-05 | MIR571 | miRNA | 50 | -6.990506237 | 6.990506237 | ENSG00000207642 | down |
| ENSG00000252148 | 6.585625836 | -2.307572617 | 0.6020917 | -3.832593 | 0.000126799 | 0.000944079 | RNU6-1206P | snRNA | 30.1 | -6.980387628 | 6.980387628 | ENSG00000252148 | down |
| ENSG00000277453 | 17.2911027 | 1.624852978 | 0.353768 | 4.59299 | 4.37E-06 | 5.08E-05 | | lincRNA | 49.15 | 6.977027798 | 6.977027798 | ENSG00000277453 | up |
| ENSG00000240291 | 102.2803766 | -1.483605704 | 0.3087524 | -4.805163 | 1.55E-06 | 2.01E-05 | | antisense | 39.11 | -6.968763774 | 6.968763774 | ENSG00000240291 | down |
| ENSG00000272854 | 13.92327753 | 1.815158998 | 0.4188588 | 4.333582 | 1.47E-05 | 0.000145067 | | antisense | 49.1 | 6.96736378 | 6.96736378 | ENSG00000272854 | up |
| ENSG00000179943 | 143.9022202 | -1.308525988 | 0.2555666 | -5.120098 | 3.05E-07 | 4.79E-06 | FIZ1 | protein_coding | 57.1 | -6.96074963 | 6.96074963 | ENSG00000179943 | down |
| ENSG00000228606 | 11.04542084 | 1.837667391 | 0.4272709 | 4.300942 | 1.70E-05 | 0.000164554 | | antisense | 58.74 | 6.953167482 | 6.953167482 | ENSG00000228606 | up |

| | | | | | | | | | | | | | |
|---|---|---|---|---|---|---|---|---|---|---|---|---|---|
| ENSG00000156097 | 22.562853 | 1.633191405 | 0.3573802 | 4.569898 | 4.88E-06 | 5.60E-05 | GPR61 | protein_coding | 49.33 | 6.94449469 | 6.94449469 | ENSG00000156097 | up |
| ENSG00000118762 | 977.6014356 | -1.363311183 | 0.2724393 | -5.004091 | 5.61E-07 | 8.09E-06 | PKD2 | protein_coding | 40.51 | -6.942086268 | 6.942086268 | ENSG00000118762 | down |
| ENSG00000272829 | 43.72845595 | 1.472758486 | 0.3060181 | 4.812652 | 1.49E-06 | 1.95E-05 | | lincRNA | 52.91 | 6.936380705 | 6.936380705 | ENSG00000272829 | up |
| ENSG00000279319 | 157.4212109 | -1.319538846 | 0.2598815 | -5.077463 | 3.83E-07 | 5.86E-06 | | TEC | 34.93 | -6.903849655 | 6.903849655 | ENSG00000279319 | down |
| ENSG00000234284 | 202.618195 | -1.255010396 | 0.241034 | -5.206778 | 1.92E-07 | 3.17E-06 | ZNF879 | protein_coding | 43.57 | -6.901915159 | 6.901915159 | ENSG00000234284 | down |
| ENSG00000200394 | 46.30862609 | -1.370489309 | 0.2755871 | -4.972981 | 6.59E-07 | 9.31E-06 | SNORA38B | snoRNA | 48.09 | -6.895004896 | 6.895004896 | ENSG00000200394 | down |
| ENSG00000273373 | 742.6362229 | -1.428886646 | 0.2934966 | -4.868495 | 1.12E-06 | 1.51E-05 | | antisense | 34.98 | -6.886981858 | 6.886981858 | ENSG00000273373 | down |
| ENSG00000188613 | 377.5815589 | 1.264672114 | 0.2440723 | 5.181547 | 2.20E-07 | 3.60E-06 | NANOS1 | protein_coding | 50.86 | 6.885102221 | 6.885102221 | ENSG00000188613 | up |
| ENSG00000213062 | 42.57075873 | 1.40622433 | 0.2870197 | 4.899399 | 9.61E-07 | 1.31E-05 | | lincRNA | 46.98 | 6.863999969 | 6.863999969 | ENSG00000213062 | up |
| ENSG00000253764 | 26.9576369 | 1.496712682 | 0.3155403 | 4.743333 | 2.10E-06 | 2.65E-05 | | lincRNA | 65.14 | 6.849767156 | 6.849767156 | ENSG00000253764 | up |
| ENSG00000212354 | 3.800172385 | -2.68148355 | 0.7634205 | -3.51246 | 0.00044398 | 0.002811063 | RNU6-1242P | snRNA | 33.01 | -6.840811388 | 6.840811388 | ENSG00000212354 | down |
| ENSG00000258365 | 52.63595292 | -1.453151134 | 0.302196 | -4.808638 | 1.52E-06 | 1.98E-05 | | antisense | 45.53 | -6.834910721 | 6.834910721 | ENSG00000258365 | down |
| ENSG00000198429 | 284.2080675 | -1.376862156 | 0.2788953 | -4.936842 | 7.94E-07 | 1.10E-05 | ZNF69 | protein_coding | 43.23 | -6.828532211 | 6.828532211 | ENSG00000198429 | down |
| ENSG00000221287 | 3.503790813 | -2.664682322 | 0.7575187 | -3.517645 | 0.000435394 | 0.002762984 | MIR1289-2 | miRNA | 53.15 | -6.8179138 | 6.8179138 | ENSG00000221287 | down |
| ENSG00000272800 | 50.50450281 | -1.614059645 | 0.3545278 | -4.552703 | 5.30E-06 | 6.02E-05 | | lincRNA | 30.68 | -6.812053682 | 6.812053682 | ENSG00000272800 | down |
| ENSG00000272807 | 119.1181076 | -1.397379387 | 0.2860495 | -4.885096 | 1.03E-06 | 1.40E-05 | | antisense | 31.62 | -6.781459196 | 6.781459196 | ENSG00000272807 | down |
| ENSG00000198538 | 434.3479771 | -1.467453212 | 0.3081522 | -4.762106 | 1.92E-06 | 2.44E-05 | ZNF28 | protein_coding | 45.1 | -6.768672029 | 6.768672029 | ENSG00000198538 | down |
| ENSG00000178882 | 164.9260949 | -1.317145845 | 0.2621367 | -5.024652 | 5.04E-07 | 7.35E-06 | RFLNA | protein_coding | 47.18 | -6.762221815 | 6.762221815 | ENSG00000178882 | down |
| ENSG00000051341 | 1739.756998 | -1.259026678 | 0.2447239 | -5.144683 | 2.68E-07 | 4.26E-06 | POLQ | protein_coding | 39.06 | -6.761829697 | 6.761829697 | ENSG00000051341 | down |
| ENSG00000206987 | 5.608191708 | 2.588851172 | 0.7287615 | 3.552398 | 0.000381737 | 0.002460612 | | snoRNA | 44.36 | 6.754201222 | 6.754201222 | ENSG00000206987 | up |
| ENSG00000198435 | 127.9194962 | 1.283862731 | 0.2524012 | 5.086596 | 3.65E-07 | 5.63E-06 | NRARP | protein_coding | 67.29 | 6.739919724 | 6.739919724 | ENSG00000198435 | up |
| ENSG00000200942 | 4.562808758 | -2.657656534 | 0.7594874 | -3.499277 | 0.000466522 | 0.002927167 | RNU6-501P | snRNA | 39 | -6.733312293 | 6.733312293 | ENSG00000200942 | down |
| ENSG00000207375 | 3.996597458 | -3.047041021 | 0.9352082 | -3.258142 | 0.001121443 | 0.006183705 | SNORD116-23 | snoRNA | 44.57 | -6.730155704 | 6.730155704 | ENSG00000207375 | down |
| ENSG00000264802 | 7.10995154 | -2.086608314 | 0.5271911 | -3.957974 | 7.56E-05 | 0.000597285 | MIR5191 | miRNA | 56.67 | -6.726846468 | 6.726846468 | ENSG00000264802 | down |
| ENSG00000272426 | 33.06695795 | 1.518250272 | 0.3254938 | 4.664452 | 3.09E-06 | 3.73E-05 | | lincRNA | 40.87 | 6.723224318 | 6.723224318 | ENSG00000272426 | up |
| ENSG00000250600 | 4.344747319 | 2.467934032 | 0.6798123 | 3.630317 | 0.000283074 | 0.001897562 | ROPN1L-AS1 | antisense | 57.06 | 6.717232693 | 6.717232693 | ENSG00000250600 | up |
| ENSG00000277918 | 232.9457866 | 1.178611084 | 0.2227609 | 5.290923 | 1.22E-07 | 2.11E-06 | | snRNA | 53.66 | 6.689432458 | 6.689432458 | ENSG00000277918 | up |
| ENSG00000257433 | 35.54263276 | 1.419260521 | 0.2949772 | 4.811425 | 1.50E-06 | 1.96E-05 | | antisense | 50.18 | 6.681789015 | 6.681789015 | ENSG00000257433 | up |
| ENSG00000265195 | 24.24199473 | -1.548291451 | 0.3364175 | -4.602292 | 4.18E-06 | 4.88E-05 | MIR4312 | miRNA | 59.21 | -6.675466037 | 6.675466037 | ENSG00000265195 | down |
| ENSG00000213183 | 34.75742238 | -1.544738462 | 0.3352592 | -4.607594 | 4.07E-06 | 4.77E-05 | RPS7P8 | processed_pseudogene | 46.13 | -6.674987047 | 6.674987047 | ENSG00000213183 | down |
| ENSG00000172403 | 2526.613088 | -1.319923636 | 0.2647584 | -4.985388 | 6.18E-07 | 8.80E-06 | SYNPO2 | protein_coding | 38.36 | -6.672681106 | 6.672681106 | ENSG00000172403 | down |
| ENSG00000144711 | 459.6278641 | -1.320299388 | 0.2648962 | -4.984215 | 6.22E-07 | 8.85E-06 | IQSEC1 | protein_coding | 54.36 | -6.671687368 | 6.671687368 | ENSG00000144711 | down |

| | | | | | | | | | | | | | |
|---|---|---|---|---|---|---|---|---|---|---|---|---|---|
| ENSG00000199471 | 9.73328357 | -1.98053221 | 0.4888856 | -4.051116 | 5.10E-05 | 0.000428828 | | misc_RNA | 43.56 | -6.669871923 | 6.669871923 | ENSG00000199471 | down |
| ENSG00000259931 | 11.17530213 | -1.874226627 | 0.4501069 | -4.163959 | 3.13E-05 | 0.000278617 | | processed_pseudogene | 39.11 | -6.662860565 | 6.662860565 | ENSG00000259931 | down |
| ENSG00000200291 | 6.031476833 | -2.081420434 | 0.5285131 | -3.938257 | 8.21E-05 | 0.000641988 | | misc_RNA | 45.1 | -6.644878716 | 6.644878716 | ENSG00000200291 | down |
| ENSG00000207248 | 12.55144994 | -1.720657803 | 0.3959849 | -4.345261 | 1.39E-05 | 0.000138345 | RNU6-1005P | snRNA | 43.93 | -6.640081989 | 6.640081989 | ENSG00000207248 | down |
| ENSG00000276517 | 360.9312108 | -1.366038449 | 0.2795501 | -4.88656 | 1.03E-06 | 1.40E-05 | | sense_intronic | 38.95 | -6.632613993 | 6.632613993 | ENSG00000276517 | down |
| ENSG00000146278 | 1221.925216 | 1.148351936 | 0.2152255 | 5.335575 | 9.52E-08 | 1.70E-06 | PNRC1 | protein_coding | 48.53 | 6.624172837 | 6.624172837 | ENSG00000146278 | up |
| ENSG00000119547 | 1176.154061 | -1.323082154 | 0.266732 | -4.960342 | 7.04E-07 | 9.85E-06 | ONECUT2 | protein_coding | 44.08 | -6.623857607 | 6.623857607 | ENSG00000119547 | down |
| ENSG00000130066 | 196.2982952 | 1.192356133 | 0.2280867 | 5.227645 | 1.72E-07 | 2.86E-06 | SAT1 | protein_coding | 43.75 | 6.61011913 | 6.61011913 | ENSG00000130066 | up |
| ENSG00000252282 | 3.692990154 | -3.03886089 | 0.9397958 | -3.233533 | 0.001222692 | 0.006681138 | RNU6-1089P | snRNA | 43.27 | -6.609976937 | 6.609976937 | ENSG00000252282 | down |
| ENSG00000197989 | 636.6971921 | 1.110302634 | 0.2048353 | 5.420466 | 5.94E-08 | 1.12E-06 | SNHG12 | antisense | 51.03 | 6.607194384 | 6.607194384 | ENSG00000197989 | up |
| ENSG00000272159 | 22.82391887 | -1.513108722 | 0.3268678 | -4.629115 | 3.67E-06 | 4.34E-05 | | antisense | 42.59 | -6.600893745 | 6.600893745 | ENSG00000272159 | down |
| ENSG00000116514 | 123.3173202 | 1.229063495 | 0.2389295 | 5.144042 | 2.69E-07 | 4.27E-06 | RNF19B | protein_coding | 44.34 | 6.599692848 | 6.599692848 | ENSG00000116514 | up |
| ENSG00000234779 | 18.27812568 | 1.634374119 | 0.3675915 | 4.44617 | 8.74E-06 | 9.30E-05 | | antisense | 61.04 | 6.589178571 | 6.589178571 | ENSG00000234779 | up |
| ENSG00000214188 | 103.5332938 | 1.401581702 | 0.2916164 | 4.806251 | 1.54E-06 | 2.00E-05 | ST7-OT4 | sense_intronic | 38.94 | 6.586224542 | 6.586224542 | ENSG00000214188 | up |
| ENSG00000200169 | 182.0134115 | 1.226585186 | 0.2385031 | 5.142849 | 2.71E-07 | 4.29E-06 | RNU5D-1 | snRNA | 37.93 | 6.58360813 | 6.58360813 | ENSG00000200169 | up |
| ENSG00000243094 | 23.38298255 | -1.652866573 | 0.3743137 | -4.415726 | 1.01E-05 | 0.000104993 | | processed_pseudogene | 46.04 | -6.576494334 | 6.576494334 | ENSG00000243094 | down |
| ENSG00000260804 | 861.1427107 | -1.248089383 | 0.2449086 | -5.096143 | 3.47E-07 | 5.38E-06 | LINC01963 | lincRNA | 44.66 | -6.576404279 | 6.576404279 | ENSG00000260804 | down |
| ENSG00000184635 | 983.0639064 | -1.282289276 | 0.2553875 | -5.020956 | 5.14E-07 | 7.46E-06 | ZNF93 | protein_coding | 40 | -6.574292981 | 6.574292981 | ENSG00000184635 | down |
| ENSG00000156239 | 446.0418754 | -1.389235425 | 0.2882273 | -4.81993 | 1.44E-06 | 1.89E-05 | N6AMT1 | protein_coding | 38.62 | -6.562154116 | 6.562154116 | ENSG00000156239 | down |
| ENSG00000262904 | 79.65042251 | 1.300096823 | 0.2614996 | 4.971698 | 6.64E-07 | 9.36E-06 | TMPOP2 | processed_pseudogene | 43.63 | 6.53769116 | 6.53769116 | ENSG00000262904 | up |
| ENSG00000122435 | 1148.373361 | -1.34754495 | 0.2760869 | -4.880872 | 1.06E-06 | 1.43E-05 | TRMT13 | protein_coding | 35.8 | -6.528212699 | 6.528212699 | ENSG00000122435 | down |
| ENSG00000271474 | 366.5004323 | 1.121645838 | 0.2093844 | 5.356874 | 8.47E-08 | 1.54E-06 | | antisense | 44.58 | 6.518567875 | 6.518567875 | ENSG00000271474 | up |
| ENSG00000125968 | 4353.291705 | 1.317719117 | 0.2673549 | 4.928726 | 8.28E-07 | 1.14E-05 | ID1 | protein_coding | 58.56 | 6.513127836 | 6.513127836 | ENSG00000125968 | up |
| ENSG00000102189 | 1440.629017 | -1.332992131 | 0.2719555 | -4.901509 | 9.51E-07 | 1.30E-05 | EEA1 | protein_coding | 36.99 | -6.512189382 | 6.512189382 | ENSG00000102189 | down |
| ENSG00000151116 | 180.6521739 | -1.428149736 | 0.3016157 | -4.734999 | 2.19E-06 | 2.76E-05 | UEVLD | protein_coding | 40.83 | -6.512161175 | 6.512161175 | ENSG00000151116 | down |
| ENSG00000212615 | 41.37913406 | 1.443531602 | 0.3067322 | 4.706162 | 2.52E-06 | 3.13E-05 | | snoRNA | 33.33 | 6.502970976 | 6.502970976 | ENSG00000212615 | up |
| ENSG00000228808 | 18.7946577 | -1.687550324 | 0.3888274 | -4.340101 | 1.42E-05 | 0.000141197 | HMGB3P4 | processed_pseudogene | 51.56 | -6.497361667 | 6.497361667 | ENSG00000228808 | down |
| ENSG00000277688 | 19.30888252 | 1.680442554 | 0.3865361 | 4.34744 | 1.38E-05 | 0.000137076 | | antisense | 39.22 | 6.491614413 | 6.491614413 | ENSG00000277688 | up |
| ENSG00000283515 | 58.54914813 | -1.402585817 | 0.2941065 | -4.768973 | 1.85E-06 | 2.37E-05 | | protein_coding | 47.26 | -6.486303141 | 6.486303141 | ENSG00000283515 | down |
| ENSG00000132466 | 3022.775382 | -1.287793258 | 0.2588379 | -4.975288 | 6.52E-07 | 9.21E-06 | ANKRD17 | protein_coding | 36.17 | -6.485049372 | 6.485049372 | ENSG00000132466 | down |
| ENSG00000227986 | 58.82462478 | -1.441279325 | 0.3065023 | -4.702344 | 2.57E-06 | 3.17E-05 | TRIM60P18 | unprocessed_pseudogene | 40.73 | -6.483335752 | 6.483335752 | ENSG00000227986 | down |
| ENSG00000206645 | 4.879367334 | -2.502885141 | 0.7071202 | -3.539547 | 0.000400815 | 0.00257524 | | misc_RNA | 39.22 | -6.480425889 | 6.480425889 | ENSG00000206645 | down |

| | | | | | | | | | | | | | |
|---|---|---|---|---|---|---|---|---|---|---|---|---|---|
| ENSG00000202293 | 3.536594366 | -3.379633085 | 1.1166887 | -3.026477 | 0.002474214 | 0.012104172 | SNORD114-22 | snoRNA | 45.83 | -6.478975952 | 6.478975952 | ENSG00000202293 | down |
| ENSG00000162994 | 75.8217272 | -1.411080077 | 0.297051 | -4.750296 | 2.03E-06 | 2.58E-05 | CLHC1 | protein_coding | 39.9 | -6.475614582 | 6.475614582 | ENSG00000162994 | down |
| ENSG00000274964 | 187.6171249 | -1.438904898 | 0.3061857 | -4.699451 | 2.61E-06 | 3.21E-05 | | sense_intronic | 34.71 | -6.464906877 | 6.464906877 | ENSG00000274964 | down |
| ENSG00000178171 | 2618.344439 | -1.232842265 | 0.2425929 | -5.081938 | 3.74E-07 | 5.74E-06 | AMER3 | protein_coding | 55.57 | -6.461307829 | 6.461307829 | ENSG00000178171 | down |
| ENSG00000206585 | 99.60074956 | 1.295772853 | 0.2618736 | 4.948085 | 7.49E-07 | 1.04E-05 | RNVU1-7 | snRNA | 54.88 | 6.456593064 | 6.456593064 | ENSG00000206585 | up |
| ENSG00000206870 | 7.392673512 | -2.085689246 | 0.5382462 | -3.874972 | 0.000106637 | 0.000807782 | RNU6-398P | snRNA | 42.2 | -6.450422783 | 6.450422783 | ENSG00000206870 | down |
| ENSG00000203865 | 75.83330211 | -1.308625206 | 0.2659504 | -4.920561 | 8.63E-07 | 1.18E-05 | ATP1A1-AS1 | processed_transcript | 45.35 | -6.446695515 | 6.446695515 | ENSG00000203865 | down |
| ENSG00000124225 | 324.0416 | -1.245638429 | 0.2467578 | -5.048021 | 4.46E-07 | 6.69E-06 | PMEPA1 | protein_coding | 54.59 | -6.445739844 | 6.445739844 | ENSG00000124225 | down |
| ENSG00000278356 | 33.84736249 | -1.529135697 | 0.3360975 | -4.549679 | 5.37E-06 | 6.09E-05 | | sense_intronic | 44.42 | -6.445712853 | 6.445712853 | ENSG00000278356 | down |
| ENSG00000165449 | 393.131663 | -1.251206603 | 0.2486141 | -5.032726 | 4.84E-07 | 7.09E-06 | SLC16A9 | protein_coding | 40.06 | -6.443144434 | 6.443144434 | ENSG00000165449 | down |
| ENSG00000201036 | 10.19015993 | -2.229597181 | 0.595695 | -3.74285 | 0.000181945 | 0.001295763 | | snoRNA | 38.2 | -6.437904943 | 6.437904943 | ENSG00000201036 | down |
| ENSG00000170734 | 487.7179533 | -1.224402114 | 0.2408074 | -5.084569 | 3.68E-07 | 5.67E-06 | POLH | protein_coding | 43.39 | -6.423267186 | 6.423267186 | ENSG00000170734 | down |
| ENSG00000273271 | 31.44045003 | -2.313967294 | 0.6310583 | -3.666804 | 0.000245601 | 0.001677869 | | antisense | 35.77 | -6.421818839 | 6.421818839 | ENSG00000273271 | down |
| ENSG00000206846 | 16.8466547 | -1.567816126 | 0.3501223 | -4.47791 | 7.54E-06 | 8.17E-05 | | misc_RNA | 43.62 | -6.40872735 | 6.40872735 | ENSG00000206846 | down |
| ENSG00000162231 | 995.4898208 | 1.148211487 | 0.2190582 | 5.241581 | 1.59E-07 | 2.66E-06 | NXF1 | protein_coding | 50.07 | 6.400625081 | 6.400625081 | ENSG00000162231 | up |
| ENSG00000251745 | 17.93116881 | -1.527905206 | 0.3372424 | -4.530585 | 5.88E-06 | 6.57E-05 | RNU7-124P | snRNA | 37.1 | -6.390836926 | 6.390836926 | ENSG00000251745 | down |
| ENSG00000173889 | 2056.388147 | -1.238514056 | 0.2457789 | -5.039139 | 4.68E-07 | 6.94E-06 | PHC3 | protein_coding | 37.75 | -6.388958569 | 6.388958569 | ENSG00000173889 | down |
| ENSG00000266402 | 244.3066544 | 1.131957622 | 0.2146606 | 5.273245 | 1.34E-07 | 2.29E-06 | SNHG25 | lincRNA | 63.71 | 6.383879415 | 6.383879415 | ENSG00000266402 | up |
| ENSG00000207650 | 10.92383937 | -1.844797472 | 0.4496136 | -4.103073 | 4.08E-05 | 0.000351044 | MIR570 | miRNA | 37.11 | -6.373107554 | 6.373107554 | ENSG00000207650 | down |
| ENSG00000239039 | 207.5229095 | 1.165074372 | 0.2243678 | 5.192698 | 2.07E-07 | 3.40E-06 | SNORD13 | snoRNA | 49.04 | 6.371373843 | 6.371373843 | ENSG00000239039 | up |
| ENSG00000212499 | 3.405630285 | -2.609978087 | 0.7604668 | -3.432074 | 0.000598985 | 0.00362279 | RNA5SP300 | rRNA | 54.21 | -6.37084389 | 6.37084389 | ENSG00000212499 | down |
| ENSG00000275857 | 16.21661812 | 1.644248579 | 0.3776299 | 4.354127 | 1.34E-05 | 0.000133627 | | antisense | 50.27 | 6.369993142 | 6.369993142 | ENSG00000275857 | up |
| ENSG00000260310 | 21.91120491 | 1.632969267 | 0.3738047 | 4.368509 | 1.25E-05 | 0.000126121 | | sense_intronic | 56.74 | 6.367294387 | 6.367294387 | ENSG00000260310 | up |
| ENSG00000277610 | 23.90309153 | 1.593345004 | 0.3601499 | 4.424116 | 9.68E-06 | 0.000101603 | RNVU1-4 | snRNA | 43.9 | 6.362374483 | 6.362374483 | ENSG00000277610 | up |
| ENSG00000105793 | 660.6668095 | -1.389531159 | 0.2928991 | -4.74406 | 2.09E-06 | 2.64E-05 | GTPBP10 | protein_coding | 38.02 | -6.36086682 | 6.36086682 | ENSG00000105793 | down |
| ENSG00000226609 | 38.15428305 | 1.413762961 | 0.3007606 | 4.700625 | 2.59E-06 | 3.20E-05 | | antisense | 43.09 | 6.354931499 | 6.354931499 | ENSG00000226609 | up |
| ENSG00000168040 | 190.2930497 | -1.214498567 | 0.2393659 | -5.073817 | 3.90E-07 | 5.93E-06 | FADD | protein_coding | 55.01 | -6.347841851 | 6.347841851 | ENSG00000168040 | down |
| ENSG00000118412 | 2693.775664 | -1.272921995 | 0.2573891 | -4.945516 | 7.59E-07 | 1.05E-05 | CASP8AP2 | protein_coding | 42.57 | -6.336541498 | 6.336541498 | ENSG00000118412 | down |
| ENSG00000135837 | 3652.058706 | -1.284297971 | 0.2609982 | -4.920716 | 8.62E-07 | 1.18E-05 | CEP350 | protein_coding | 36.47 | -6.326851976 | 6.326851976 | ENSG00000135837 | down |
| ENSG00000200170 | 5.210873389 | -2.139205422 | 0.5654935 | -3.7829 | 0.000155012 | 0.001126424 | | misc_RNA | 47.52 | -6.307015322 | 6.307015322 | ENSG00000200170 | down |
| ENSG00000078687 | 334.9080467 | -1.222839309 | 0.2427587 | -5.037262 | 4.72E-07 | 6.98E-06 | TNRC6C | protein_coding | 42.22 | -6.305151458 | 6.305151458 | ENSG00000078687 | down |
| ENSG00000087074 | 511.7705659 | 1.197430515 | 0.2352793 | 5.089401 | 3.59E-07 | 5.55E-06 | PPP1R15A | protein_coding | 56.68 | 6.29295067 | 6.29295067 | ENSG00000087074 | up |

| | | | | | | | | | | | | | |
|---|---|---|---|---|---|---|---|---|---|---|---|---|---|
| ENSG00000270558 | 32.53811672 | -1.46035717 | 0.3177293 | -4.59623 | 4.30E-06 | 5.02E-05 | | processed_pseudogene | 36.11 | -6.27894897 | 6.27894897 | ENSG00000270558 | down |
| ENSG00000237082 | 26.6881807 | -1.496783315 | 0.3300608 | -4.534872 | 5.76E-06 | 6.45E-05 | COX5BP6 | processed_pseudogene | 52.25 | -6.272083694 | 6.272083694 | ENSG00000237082 | down |
| ENSG00000274191 | 28.50137774 | -1.575788389 | 0.3568811 | -4.415444 | 1.01E-05 | 0.000105051 | | lincRNA | 43.93 | -6.269433175 | 6.269433175 | ENSG00000274191 | down |
| ENSG00000154734 | 930.1910578 | 1.058919508 | 0.1959785 | 5.403243 | 6.54E-08 | 1.22E-06 | ADAMTS1 | protein_coding | 45.12 | 6.260281884 | 6.260281884 | ENSG00000154734 | up |
| ENSG00000019549 | 808.7269944 | 1.211525413 | 0.2402708 | 5.042333 | 4.60E-07 | 6.84E-06 | SNAI2 | protein_coding | 41.13 | 6.25739428 | 6.25739428 | ENSG00000019549 | up |
| ENSG00000222365 | 419.3767898 | 1.105168189 | 0.2092082 | 5.282625 | 1.27E-07 | 2.19E-06 | SNORD12B | snoRNA | 41.76 | 6.253807787 | 6.253807787 | ENSG00000222365 | up |
| ENSG00000278576 | 118.9102565 | -1.268206373 | 0.2576219 | -4.922742 | 8.53E-07 | 1.18E-05 | | sense_intronic | 43.2 | -6.25208827 | 6.25208827 | ENSG00000278576 | down |
| ENSG00000204256 | 7056.854316 | 1.080492863 | 0.2024197 | 5.337883 | 9.40E-08 | 1.69E-06 | BRD2 | protein_coding | 51.46 | 6.238046877 | 6.238046877 | ENSG00000204256 | up |
| ENSG00000101138 | 890.2041345 | -1.224776906 | 0.2447245 | -5.004718 | 5.59E-07 | 8.07E-06 | CSTF1 | protein_coding | 41.14 | -6.237839619 | 6.237839619 | ENSG00000101138 | down |
| ENSG00000272247 | 27.94747583 | 1.442891616 | 0.3132048 | 4.606863 | 4.09E-06 | 4.79E-05 | | antisense | 51.84 | 6.233219852 | 6.233219852 | ENSG00000272247 | up |
| ENSG00000219150 | 8.95150461 | -2.845129342 | 0.8768278 | -3.244798 | 0.001175338 | 0.006447686 | | processed_pseudogene | 47.6 | -6.232529239 | 6.232529239 | ENSG00000219150 | down |
| ENSG00000104047 | 661.6667057 | -1.295802938 | 0.267155 | -4.850378 | 1.23E-06 | 1.65E-05 | DTWD1 | protein_coding | 36.31 | -6.198384798 | 6.198384798 | ENSG00000104047 | down |
| ENSG00000226419 | 815.4852608 | -1.113185574 | 0.2125962 | -5.23615 | 1.64E-07 | 2.74E-06 | SLC16A1-AS1 | antisense | 43.76 | -6.192308827 | 6.192308827 | ENSG00000226419 | down |
| ENSG00000234134 | 53.82323804 | 1.31039287 | 0.2724805 | 4.809125 | 1.52E-06 | 1.98E-05 | | sense_intronic | 57.31 | 6.164298193 | 6.164298193 | ENSG00000234134 | up |
| ENSG00000077782 | 192.5598832 | 1.234567396 | 0.2492637 | 4.952856 | 7.31E-07 | 1.02E-05 | FGFR1 | protein_coding | 50.55 | 6.161546004 | 6.161546004 | ENSG00000077782 | up |
| ENSG00000261335 | 185.7248208 | 1.176946046 | 0.2318042 | 5.077329 | 3.83E-07 | 5.86E-06 | | antisense | 46.77 | 6.157801705 | 6.157801705 | ENSG00000261335 | up |
| ENSG00000198440 | 340.7707938 | -1.295903078 | 0.2680676 | -4.83424 | 1.34E-06 | 1.77E-05 | ZNF583 | protein_coding | 38.95 | -6.156908977 | 6.156908977 | ENSG00000198440 | down |
| ENSG00000264249 | 8.46749762 | 1.898525473 | 0.4783575 | 3.968843 | 7.22E-05 | 0.000574931 | MIR3912 | miRNA | 31.43 | 6.151951584 | 6.151951584 | ENSG00000264249 | up |
| ENSG00000183850 | 219.6892576 | -1.299504406 | 0.2693227 | -4.825084 | 1.40E-06 | 1.85E-05 | ZNF730 | protein_coding | 41.52 | -6.149679668 | 6.149679668 | ENSG00000183850 | down |
| ENSG00000278514 | 20.04021216 | 1.587763119 | 0.3649791 | 4.350285 | 1.36E-05 | 0.000135503 | | lincRNA | 37.7 | 6.141547241 | 6.141547241 | ENSG00000278514 | up |
| ENSG00000221214 | 12.95026679 | -1.724700323 | 0.4138904 | -4.167046 | 3.09E-05 | 0.000275577 | MIR548E | miRNA | 35.23 | -6.139512877 | 6.139512877 | ENSG00000221214 | down |
| ENSG00000252759 | 123.4354147 | 1.196565069 | 0.2385744 | 5.015479 | 5.29E-07 | 7.66E-06 | | misc_RNA | 39.58 | 6.121055571 | 6.121055571 | ENSG00000252759 | up |
| ENSG00000185453 | 15.29428558 | -1.680107209 | 0.3985043 | -4.216033 | 2.49E-05 | 0.000227599 | C19orf68 | protein_coding | 50.04 | -6.120344019 | 6.120344019 | ENSG00000185453 | down |
| ENSG00000266490 | 72.45399866 | -1.468464448 | 0.3247281 | -4.522135 | 6.12E-06 | 6.80E-05 | | lincRNA | 37.01 | -6.119580141 | 6.119580141 | ENSG00000266490 | down |
| ENSG00000252906 | 25.87341241 | -1.525291911 | 0.3440561 | -4.433265 | 9.28E-06 | 9.82E-05 | SCARNA3 | scaRNA | 46.85 | -6.113214508 | 6.113214508 | ENSG00000252906 | down |
| ENSG00000185730 | 155.6790847 | -1.18699804 | 0.2358657 | -5.032516 | 4.84E-07 | 7.09E-06 | ZNF696 | protein_coding | 58.48 | -6.112478923 | 6.112478923 | ENSG00000185730 | down |
| ENSG00000236255 | 12.24604461 | -1.696524176 | 0.4048113 | -4.190901 | 2.78E-05 | 0.0002512 | | lincRNA | 48.08 | -6.107453168 | 6.107453168 | ENSG00000236255 | down |
| ENSG00000266579 | 42.82215251 | -1.46419533 | 0.323693 | -4.523408 | 6.09E-06 | 6.77E-05 | | lincRNA | 34.79 | -6.105106282 | 6.105106282 | ENSG00000266579 | down |
| ENSG00000031691 | 598.0822541 | -1.321453733 | 0.2772779 | -4.765811 | 1.88E-06 | 2.40E-05 | CENPQ | protein_coding | 37.99 | -6.104210007 | 6.104210007 | ENSG00000031691 | down |
| ENSG00000230251 | 55.08895654 | -1.496507982 | 0.3348536 | -4.469142 | 7.85E-06 | 8.47E-05 | PHBP4 | processed_pseudogene | 50.66 | -6.093982798 | 6.093982798 | ENSG00000230251 | down |
| ENSG00000270419 | 90.64388684 | 1.200365567 | 0.2405637 | 4.989803 | 6.04E-07 | 8.62E-06 | CAHM | lincRNA | 60.94 | 6.079113724 | 6.079113724 | ENSG00000270419 | up |
| ENSG00000199787 | 6.163090542 | -2.012801048 | 0.5256419 | -3.829225 | 0.000128547 | 0.000956071 | SNORA80C | snoRNA | 54.81 | -6.077672364 | 6.077672364 | ENSG00000199787 | down |

| | | | | | | | | | | | | | |
|---|---|---|---|---|---|---|---|---|---|---|---|---|---|
| ENSG00000175387 | 3669.023971 | -1.238281223 | 0.2520899 | -4.912061 | 9.01E-07 | 1.24E-05 | SMAD2 | protein_coding | 37.52 | -6.077351605 | 6.077351605 | ENSG00000175387 | down |
| ENSG00000230695 | 25.83421414 | 1.467771977 | 0.3257264 | 4.50615 | 6.60E-06 | 7.27E-05 | | antisense | 51.89 | 6.073923498 | 6.073923498 | ENSG00000230695 | up |
| ENSG00000201240 | 6.207298105 | -2.349055237 | 0.6649682 | -3.532583 | 0.000411521 | 0.002634304 | SNORD114-9 | snoRNA | 47.22 | -6.058998445 | 6.058998445 | ENSG00000201240 | down |
| ENSG00000201899 | 6.148629222 | -2.349430304 | 0.6652192 | -3.531814 | 0.00041272 | 0.002639547 | SNORD114-24 | snoRNA | 44.44 | -6.057936802 | 6.057936802 | ENSG00000201899 | down |
| ENSG00000265986 | 9.6637305 | -1.831179153 | 0.4568535 | -4.008242 | 6.12E-05 | 0.00049691 | MIR4735 | miRNA | 37.68 | -6.049706567 | 6.049706567 | ENSG00000265986 | down |
| ENSG00000272644 | 29.29711015 | -1.418842061 | 0.310048 | -4.576202 | 4.73E-06 | 5.46E-05 | | lincRNA | 44.96 | -6.048041387 | 6.048041387 | ENSG00000272644 | down |
| ENSG00000141026 | 436.4170777 | -1.084541825 | 0.2069384 | -5.240893 | 1.60E-07 | 2.67E-06 | MED9 | protein_coding | 47.9 | -6.04451151 | 6.04451151 | ENSG00000141026 | down |
| ENSG00000149474 | 522.7062951 | -1.248974236 | 0.2561123 | -4.876667 | 1.08E-06 | 1.46E-05 | KAT14 | protein_coding | 42.25 | -6.039648919 | 6.039648919 | ENSG00000149474 | down |
| ENSG00000204149 | 259.793381 | -1.340201669 | 0.2849169 | -4.703834 | 2.55E-06 | 3.16E-05 | AGAP6 | protein_coding | 41.1 | -6.031870196 | 6.031870196 | ENSG00000204149 | down |
| ENSG00000207450 | 4.616657708 | -2.206710795 | 0.6067791 | -3.636762 | 0.000276087 | 0.001856969 | | misc_RNA | 35.64 | -6.026957997 | 6.026957997 | ENSG00000207450 | down |
| ENSG00000235381 | 221.8503137 | -1.274221953 | 0.2645776 | -4.816061 | 1.46E-06 | 1.92E-05 | | antisense | 41.83 | -6.009196344 | 6.009196344 | ENSG00000235381 | down |
| ENSG00000204920 | 93.44860395 | -1.436595965 | 0.3170757 | -4.530767 | 5.88E-06 | 6.57E-05 | ZNF155 | protein_coding | 40.41 | -6.008913711 | 6.008913711 | ENSG00000204920 | down |
| ENSG00000186105 | 158.544576 | -1.349950877 | 0.2888955 | -4.6728 | 2.97E-06 | 3.61E-05 | LRRC70 | protein_coding | 32.72 | -5.996670927 | 5.996670927 | ENSG00000186105 | down |
| ENSG00000181450 | 1802.019313 | -1.327575207 | 0.2817995 | -4.711062 | 2.46E-06 | 3.06E-05 | ZNF678 | protein_coding | 40.32 | -5.992914826 | 5.992914826 | ENSG00000181450 | down |
| ENSG00000278231 | 15.76568563 | 1.670249153 | 0.3993024 | 4.182918 | 2.88E-05 | 0.000258845 | | lincRNA | 47.64 | 5.991117658 | 5.991117658 | ENSG00000278231 | up |
| ENSG00000204713 | 2017.161272 | -1.243915399 | 0.2556193 | -4.866282 | 1.14E-06 | 1.53E-05 | TRIM27 | protein_coding | 44.66 | -5.991053107 | 5.991053107 | ENSG00000204713 | down |
| ENSG00000197863 | 257.0519278 | -1.221479614 | 0.2493927 | -4.897817 | 9.69E-07 | 1.32E-05 | ZNF790 | protein_coding | 42.57 | -5.958482125 | 5.958482125 | ENSG00000197863 | down |
| ENSG00000222145 | 12.54413362 | -2.232267161 | 0.6211966 | -3.593495 | 0.000326272 | 0.002151909 | | snoRNA | 47.62 | -5.953849541 | 5.953849541 | ENSG00000222145 | down |
| ENSG00000273402 | 63.11101551 | -1.288206688 | 0.2704177 | -4.763767 | 1.90E-06 | 2.42E-05 | | antisense | 27.52 | -5.945473459 | 5.945473459 | ENSG00000273402 | down |
| ENSG00000242876 | 9.110902168 | -1.814652951 | 0.454829 | -3.989748 | 6.61E-05 | 0.00053263 | RN7SL812P | misc_RNA | 52.69 | -5.94040165 | 5.94040165 | ENSG00000242876 | down |
| ENSG00000207223 | 12.31285451 | -2.380159685 | 0.6856999 | -3.471139 | 0.000518255 | 0.003201274 | | misc_RNA | 39.22 | -5.93773003 | 5.93773003 | ENSG00000207223 | down |
| ENSG00000199591 | 4.751306593 | -2.256069501 | 0.6328894 | -3.564714 | 0.000364254 | 0.002361036 | | misc_RNA | 42.99 | -5.926463028 | 5.926463028 | ENSG00000199591 | down |
| ENSG00000266276 | 2.732573885 | -2.708071969 | 0.8350759 | -3.242905 | 0.001183177 | 0.006485577 | MIR4743 | miRNA | 53.62 | -5.925400545 | 5.925400545 | ENSG00000266276 | down |
| ENSG00000254324 | 6.821169822 | -2.034510353 | 0.5416176 | -3.756359 | 0.000172403 | 0.001236562 | MIR151A | miRNA | 52.22 | -5.915916549 | 5.915916549 | ENSG00000254324 | down |
| ENSG00000225156 | 120.1869976 | -1.355321929 | 0.2927519 | -4.629593 | 3.66E-06 | 4.34E-05 | | lincRNA | 50.97 | -5.912912092 | 5.912912092 | ENSG00000225156 | down |
| ENSG00000142544 | 87.85770641 | -1.33182686 | 0.2852446 | -4.66907 | 3.03E-06 | 3.67E-05 | CTU1 | protein_coding | 54.31 | -5.907170162 | 5.907170162 | ENSG00000142544 | down |
| ENSG00000231473 | 52.93800943 | 1.293651502 | 0.2732657 | 4.734043 | 2.20E-06 | 2.76E-05 | LINC00441 | lincRNA | 39.88 | 5.897015297 | 5.897015297 | ENSG00000231473 | up |
| ENSG00000277342 | 379.9301733 | 1.119199068 | 0.2196508 | 5.095356 | 3.48E-07 | 5.39E-06 | | sense_intronic | 30.36 | 5.896321619 | 5.896321619 | ENSG00000277342 | up |
| ENSG00000108599 | 628.7021355 | -1.265544775 | 0.2644674 | -4.785257 | 1.71E-06 | 2.21E-05 | AKAP10 | protein_coding | 41.26 | -5.892981839 | 5.892981839 | ENSG00000108599 | down |
| ENSG00000201136 | 5.842873256 | -2.040425502 | 0.546156 | -3.735975 | 0.000186989 | 0.001325275 | RNU6-353P | snRNA | 38.89 | -5.871720288 | 5.871720288 | ENSG00000201136 | down |
| ENSG00000260339 | 116.6066645 | 1.177878634 | 0.2379914 | 4.94925 | 7.45E-07 | 1.04E-05 | HEXA-AS1 | antisense | 41.18 | 5.871699147 | 5.871699147 | ENSG00000260339 | up |
| ENSG00000202142 | 4.062152412 | -3.268744875 | 1.1209008 | -2.916177 | 0.003543496 | 0.016222411 | SNORD114-18 | snoRNA | 37.5 | -5.850676152 | 5.850676152 | ENSG00000202142 | down |

| | | | | | | | | | | | | | |
|---|---|---|---|---|---|---|---|---|---|---|---|---|---|
| ENSG00000211514 | 43.22247741 | -1.353590471 | 0.2938543 | -4.606332 | 4.10E-06 | 4.80E-05 | MIR454 | miRNA | 39.13 | -5.846435605 | 5.846435605 | ENSG00000211514 | down |
| ENSG00000119866 | 1102.246356 | -1.128609816 | 0.2236648 | -5.045988 | 4.51E-07 | 6.74E-06 | BCL11A | protein_coding | 44.56 | -5.836412428 | 5.836412428 | ENSG00000119866 | down |
| ENSG00000243107 | 107.256653 | -1.467039014 | 0.3323672 | -4.41391 | 1.02E-05 | 0.000105719 | | sense_overlapping | 38.79 | -5.832721898 | 5.832721898 | ENSG00000243107 | down |
| ENSG00000256006 | 54.50803502 | -1.263980646 | 0.2654023 | -4.762508 | 1.91E-06 | 2.44E-05 | | sense_intronic | 43.46 | -5.830740784 | 5.830740784 | ENSG00000256006 | down |
| ENSG00000026652 | 960.4576244 | -1.32694207 | 0.2857047 | -4.644454 | 3.41E-06 | 4.08E-05 | AGPAT4 | protein_coding | 45.36 | -5.825099998 | 5.825099998 | ENSG00000026652 | down |
| ENSG00000173295 | 162.8089999 | -1.301926115 | 0.2777061 | -4.688144 | 2.76E-06 | 3.37E-05 | FAM86B3P | transcribed_unprocessed_pseud | 52.58 | -5.823182506 | 5.823182506 | ENSG00000173295 | down |
| ENSG00000207234 | 17.81746482 | -1.541897918 | 0.3589426 | -4.295667 | 1.74E-05 | 0.000167701 | RNU6-125P | snRNA | 41.12 | -5.821380408 | 5.821380408 | ENSG00000207234 | down |
| ENSG00000228886 | 13.3998864 | -1.62531372 | 0.3891884 | -4.176162 | 2.96E-05 | 0.000266306 | | lincRNA | 44.26 | -5.809878187 | 5.809878187 | ENSG00000228886 | down |
| ENSG00000196247 | 1482.964225 | -1.267526785 | 0.2670342 | -4.746683 | 2.07E-06 | 2.62E-05 | ZNF107 | protein_coding | 39.91 | -5.808000125 | 5.808000125 | ENSG00000196247 | down |
| ENSG00000243314 | 19.46823607 | -1.643359825 | 0.3963206 | -4.146542 | 3.38E-05 | 0.000297248 | | processed_pseudogene | 42.78 | -5.795933572 | 5.795933572 | ENSG00000243314 | down |
| ENSG00000197779 | 336.8412892 | -1.339521861 | 0.2905482 | -4.610325 | 4.02E-06 | 4.73E-05 | ZNF81 | protein_coding | 39.93 | -5.793901512 | 5.793901512 | ENSG00000197779 | down |
| ENSG00000168675 | 69.03122175 | -1.279932775 | 0.2714566 | -4.715055 | 2.42E-06 | 3.01E-05 | LDLRAD4 | protein_coding | 46.44 | -5.786756532 | 5.786756532 | ENSG00000168675 | down |
| ENSG00000272263 | 93.45710071 | -1.229071086 | 0.2554423 | -4.811541 | 1.50E-06 | 1.96E-05 | | antisense | 43.14 | -5.786389152 | 5.786389152 | ENSG00000272263 | down |
| ENSG00000231074 | 1667.604863 | -1.271460714 | 0.2690801 | -4.725213 | 2.30E-06 | 2.88E-05 | HCG18 | antisense | 42.72 | -5.773054472 | 5.773054472 | ENSG00000231074 | down |
| ENSG00000199942 | 5.234042596 | -2.54103207 | 0.7679453 | -3.308871 | 0.000936729 | 0.0053495 | SNORD114-19 | snoRNA | 40 | -5.772429126 | 5.772429126 | ENSG00000199942 | down |
| ENSG00000282988 | 150.0545347 | 1.294847951 | 0.2767309 | 4.679088 | 2.88E-06 | 3.51E-05 | HIST1H3D | protein_coding | 47.53 | 5.767651678 | 5.767651678 | ENSG00000282988 | up |
| ENSG00000259732 | 52.42167097 | -1.273019794 | 0.2699852 | -4.715146 | 2.42E-06 | 3.01E-05 | | sense_overlapping | 42.36 | -5.755501972 | 5.755501972 | ENSG00000259732 | down |
| ENSG00000223734 | 26.16251465 | 1.41382166 | 0.3164799 | 4.467335 | 7.92E-06 | 8.53E-05 | | lincRNA | 44.27 | 5.752562913 | 5.752562913 | ENSG00000223734 | up |
| ENSG00000239272 | 20.55712325 | -1.522032584 | 0.3543514 | -4.295263 | 1.74E-05 | 0.000167787 | RPL21P10 | processed_pseudogene | 47.08 | -5.746039899 | 5.746039899 | ENSG00000239272 | down |
| ENSG00000135048 | 52.01082972 | 1.284327877 | 0.2739138 | 4.688803 | 2.75E-06 | 3.36E-05 | TMEM2 | protein_coding | 40.46 | 5.745527319 | 5.745527319 | ENSG00000135048 | up |
| ENSG00000235088 | 37.93137543 | -1.41846012 | 0.3182763 | -4.456695 | 8.32E-06 | 8.91E-05 | | processed_pseudogene | 47.74 | -5.744642711 | 5.744642711 | ENSG00000235088 | down |
| ENSG00000200871 | 3.785250762 | -2.399747696 | 0.7076004 | -3.391388 | 0.000695395 | 0.004125257 | RNU6-810P | snRNA | 38.32 | -5.722315956 | 5.722315956 | ENSG00000200871 | down |
| ENSG00000199289 | 2.69031148 | 3.186222227 | 1.0921444 | 2.9174 | 0.003529628 | 0.016176104 | RNU6-502P | snRNA | 44.44 | 5.706925653 | 5.706925653 | ENSG00000199289 | up |
| ENSG00000239197 | 19.62545993 | -1.555616082 | 0.367736 | -4.230252 | 2.33E-05 | 0.00021552 | | snoRNA | 41.96 | -5.703685707 | 5.703685707 | ENSG00000239197 | down |
| ENSG00000168807 | 442.4182509 | -1.237621442 | 0.2600203 | -4.759712 | 1.94E-06 | 2.47E-05 | SNTB2 | protein_coding | 41.92 | -5.703171626 | 5.703171626 | ENSG00000168807 | down |
| ENSG00000278928 | 390.4078141 | -1.355697682 | 0.2983081 | -4.544622 | 5.50E-06 | 6.22E-05 | | TEC | 41.49 | -5.701921773 | 5.701921773 | ENSG00000278928 | down |
| ENSG00000166068 | 783.2142856 | -1.209681263 | 0.2514878 | -4.8101 | 1.51E-06 | 1.97E-05 | SPRED1 | protein_coding | 35.39 | -5.692605286 | 5.692605286 | ENSG00000166068 | down |
| ENSG00000082556 | 218.2060132 | -1.322617771 | 0.2880185 | -4.592128 | 4.39E-06 | 5.09E-05 | OPRK1 | protein_coding | 42.12 | -5.677826496 | 5.677826496 | ENSG00000082556 | down |
| ENSG00000265688 | 74.54118998 | -1.332928012 | 0.2917104 | -4.569354 | 4.89E-06 | 5.61E-05 | MAFG-AS1 | antisense | 65.54 | -5.666718481 | 5.666718481 | ENSG00000265688 | down |
| ENSG00000282527 | 11.70531697 | 1.773190289 | 0.4502698 | 3.938062 | 8.21E-05 | 0.000642149 | | antisense | 36.18 | 5.660668839 | 5.660668839 | ENSG00000282527 | up |
| ENSG00000121075 | 99.33882929 | 1.213019065 | 0.2533887 | 4.787187 | 1.69E-06 | 2.19E-05 | TBX4 | protein_coding | 52.81 | 5.652970715 | 5.652970715 | ENSG00000121075 | up |
| ENSG00000231961 | 28.26671282 | -1.507482838 | 0.3522656 | -4.279392 | 1.87E-05 | 0.000178714 | | processed_pseudogene | 44.88 | -5.649805776 | 5.649805776 | ENSG00000231961 | down |

| | | | | | | | | | | | | | |
|---|---|---|---|---|---|---|---|---|---|---|---|---|---|
| ENSG00000221036 | 2.988053631 | -2.89308324 | 0.946953 | -3.05515 | 0.002249482 | 0.011237722 | MIR1193 | miRNA | 55.13 | -5.639549994 | 5.639549994 | ENSG00000221036 | down |
| ENSG00000209042 | 270.5460129 | 1.071370374 | 0.2105152 | 5.089278 | 3.59E-07 | 5.55E-06 | SNORD12C | snoRNA | 41.57 | 5.63045691 | 5.63045691 | ENSG00000209042 | up |
| ENSG00000146648 | 455.9786122 | -1.175494769 | 0.2421967 | -4.853471 | 1.21E-06 | 1.62E-05 | EGFR | protein_coding | 44.57 | -5.62988093 | 5.62988093 | ENSG00000146648 | down |
| ENSG00000251733 | 19.19702847 | -1.579829054 | 0.3792223 | -4.165971 | 3.10E-05 | 0.000276525 | SCARNA8 | scaRNA | 47.33 | -5.621451518 | 5.621451518 | ENSG00000251733 | down |
| ENSG00000169955 | 251.4317499 | -1.083114088 | 0.2143041 | -5.054099 | 4.32E-07 | 6.49E-06 | ZNF747 | protein_coding | 57.48 | -5.618583435 | 5.618583435 | ENSG00000169955 | down |
| ENSG00000283563 | 566.6439955 | -1.272974791 | 0.2733859 | -4.656329 | 3.22E-06 | 3.87E-05 | | protein_coding | 37.06 | -5.616693341 | 5.616693341 | ENSG00000283563 | down |
| ENSG00000134852 | 822.3995935 | -1.19478189 | 0.248573 | -4.806563 | 1.54E-06 | 2.00E-05 | CLOCK | protein_coding | 37.44 | -5.614768801 | 5.614768801 | ENSG00000134852 | down |
| ENSG00000100234 | 1749.022104 | -1.177622398 | 0.2433968 | -4.838283 | 1.31E-06 | 1.74E-05 | TIMP3 | protein_coding | 47.18 | -5.604860188 | 5.604860188 | ENSG00000100234 | down |
| ENSG00000221406 | 13.8822119 | -2.201007779 | 0.6280862 | -3.504308 | 0.000457795 | 0.002885414 | MIR320B2 | miRNA | 36.96 | -5.590101591 | 5.590101591 | ENSG00000221406 | down |
| ENSG00000216179 | 3.331122473 | -2.873719222 | 0.9430007 | -3.04742 | 0.00230815 | 0.011444577 | MIR541 | miRNA | 53.57 | -5.579039232 | 5.579039232 | ENSG00000216179 | down |
| ENSG00000222767 | 3.644834891 | -2.87567289 | 0.9439676 | -3.046368 | 0.002316239 | 0.011480596 | | misc_RNA | 44.21 | -5.57890771 | 5.57890771 | ENSG00000222767 | down |
| ENSG00000239183 | 33.92260615 | 1.392108738 | 0.3140449 | 4.432833 | 9.30E-06 | 9.83E-05 | SNORA84 | snoRNA | 55.64 | 5.578677309 | 5.578677309 | ENSG00000239183 | up |
| ENSG00000254678 | 10.23412912 | 1.870206652 | 0.4927159 | 3.79571 | 0.000147222 | 0.001077123 | | processed_pseudogene | 58.8 | 5.550276908 | 5.550276908 | ENSG00000254678 | up |
| ENSG00000152749 | 710.6059751 | -1.326737901 | 0.2928468 | -4.530484 | 5.88E-06 | 6.57E-05 | GPR180 | protein_coding | 36.49 | -5.549405511 | 5.549405511 | ENSG00000152749 | down |
| ENSG00000187479 | 454.1287506 | 1.295377663 | 0.2828888 | 4.579105 | 4.67E-06 | 5.40E-05 | C11orf96 | protein_coding | 53.45 | 5.528159689 | 5.528159689 | ENSG00000187479 | up |
| ENSG00000253854 | 35.6791495 | -1.36247499 | 0.3054387 | -4.460715 | 8.17E-06 | 8.78E-05 | | antisense | 40.97 | -5.527180089 | 5.527180089 | ENSG00000253854 | down |
| ENSG00000204335 | 69.78437858 | 1.339445104 | 0.2979766 | 4.495135 | 6.95E-06 | 7.63E-05 | SP5 | protein_coding | 66.42 | 5.515276057 | 5.515276057 | ENSG00000204335 | up |
| ENSG00000135457 | 709.5828077 | -1.220513399 | 0.2589626 | -4.713088 | 2.44E-06 | 3.04E-05 | TFCP2 | protein_coding | 41.94 | -5.513942386 | 5.513942386 | ENSG00000135457 | down |
| ENSG00000256525 | 714.1143627 | -1.226961107 | 0.2611276 | -4.698703 | 2.62E-06 | 3.22E-05 | POLG2 | protein_coding | 41.34 | -5.511175571 | 5.511175571 | ENSG00000256525 | down |
| ENSG00000256037 | 23.98718993 | -1.703570786 | 0.4298846 | -3.962856 | 7.41E-05 | 0.000586195 | MRPL40P1 | processed_pseudogene | 50.99 | -5.505868659 | 5.505868659 | ENSG00000256037 | down |
| ENSG00000239419 | 112.0479406 | 1.149729934 | 0.2373057 | 4.844932 | 1.27E-06 | 1.69E-05 | RN7SL535P | misc_RNA | 53.72 | 5.486430413 | 5.486430413 | ENSG00000239419 | up |
| ENSG00000162086 | 1243.547908 | -1.161332853 | 0.2409386 | -4.820036 | 1.44E-06 | 1.89E-05 | ZNF75A | protein_coding | 45.02 | -5.485639815 | 5.485639815 | ENSG00000162086 | down |
| ENSG00000182903 | 1076.114261 | -1.214611125 | 0.2578355 | -4.710799 | 2.47E-06 | 3.06E-05 | ZNF721 | protein_coding | 41.11 | -5.482763717 | 5.482763717 | ENSG00000182903 | down |
| ENSG00000101463 | 239.9842134 | -1.236413968 | 0.265064 | -4.664587 | 3.09E-06 | 3.73E-05 | SYNDIG1 | protein_coding | 46.16 | -5.47517666 | 5.47517666 | ENSG00000101463 | down |
| ENSG00000233895 | 44.39234923 | -1.302269397 | 0.2871342 | -4.535403 | 5.75E-06 | 6.44E-05 | | lincRNA | 44.35 | -5.457966948 | 5.457966948 | ENSG00000233895 | down |
| ENSG00000138685 | 586.6045553 | -1.248970334 | 0.2697941 | -4.629347 | 3.67E-06 | 4.34E-05 | FGF2 | protein_coding | 38.74 | -5.448744788 | 5.448744788 | ENSG00000138685 | down |
| ENSG00000234345 | 41.00879599 | -1.432507314 | 0.3322043 | -4.312127 | 1.62E-05 | 0.000157539 | | processed_pseudogene | 44.18 | -5.447270779 | 5.447270779 | ENSG00000234345 | down |
| ENSG00000171295 | 396.4226146 | -1.231471782 | 0.2642791 | -4.659739 | 3.17E-06 | 3.81E-05 | ZNF440 | protein_coding | 44.1 | -5.441504094 | 5.441504094 | ENSG00000171295 | down |
| ENSG00000235159 | 194.090048 | 1.138518613 | 0.2351347 | 4.841984 | 1.29E-06 | 1.71E-05 | | lincRNA | 41.72 | 5.426538085 | 5.426538085 | ENSG00000235159 | up |
| ENSG00000234215 | 19.13718223 | 1.479028743 | 0.3500622 | 4.225045 | 2.39E-05 | 0.000220272 | | lincRNA | 44.2 | 5.40886887 | 5.40886887 | ENSG00000234215 | up |
| ENSG00000151490 | 549.7631922 | -1.186860243 | 0.2509524 | -4.729423 | 2.25E-06 | 2.82E-05 | PTPRO | protein_coding | 37.5 | -5.39914801 | 5.39914801 | ENSG00000151490 | down |
| ENSG00000267376 | 9.620440597 | -1.704798474 | 0.434851 | -3.92042 | 8.84E-05 | 0.000685638 | | processed_pseudogene | 47.06 | -5.39382091 | 5.39382091 | ENSG00000267376 | down |

| | | | | | | | | | | | | | |
|---|---|---|---|---|---|---|---|---|---|---|---|---|---|
| ENSG00000201689 | 4.324645098 | -2.627265189 | 0.8371563 | -3.138321 | 0.001699187 | 0.008870558 | SNORD114-29 | snoRNA | 42.86 | -5.391277095 | 5.391277095 | ENSG00000201689 | down |
| ENSG00000168661 | 109.0787114 | -1.33191592 | 0.2989913 | -4.454698 | 8.40E-06 | 8.99E-05 | ZNF30 | protein_coding | 42.7 | -5.389402233 | 5.389402233 | ENSG00000168661 | down |
| ENSG00000252636 | 25.48402814 | -1.417972457 | 0.3292251 | -4.306999 | 1.65E-05 | 0.000160896 | RNU6-826P | snRNA | 47.12 | -5.379013006 | 5.379013006 | ENSG00000252636 | down |
| ENSG00000249492 | 72.49271158 | 1.179286294 | 0.2491118 | 4.733964 | 2.20E-06 | 2.76E-05 | | processed_transcript | 36.71 | 5.375689899 | 5.375689899 | ENSG00000249492 | up |
| ENSG00000276115 | 228.3065644 | -1.266737347 | 0.2776703 | -4.562019 | 5.07E-06 | 5.78E-05 | | sense_intronic | 36.19 | -5.368790638 | 5.368790638 | ENSG00000276115 | down |
| ENSG00000130956 | 248.0710952 | -1.18684281 | 0.2517758 | -4.713888 | 2.43E-06 | 3.03E-05 | HABP4 | protein_coding | 43.78 | -5.363392228 | 5.363392228 | ENSG00000130956 | down |
| ENSG00000176732 | 84.57574645 | 1.17230777 | 0.2473786 | 4.738921 | 2.15E-06 | 2.70E-05 | PFN4 | protein_coding | 44.06 | 5.354947242 | 5.354947242 | ENSG00000176732 | up |
| ENSG00000162490 | 260.5677053 | -1.215480167 | 0.2615331 | -4.647519 | 3.36E-06 | 4.02E-05 | DRAXIN | protein_coding | 53.39 | -5.342729986 | 5.342729986 | ENSG00000162490 | down |
| ENSG00000146414 | 4471.130365 | -1.269927672 | 0.2794058 | -4.545101 | 5.49E-06 | 6.22E-05 | SHPRH | protein_coding | 35.43 | -5.341988808 | 5.341988808 | ENSG00000146414 | down |
| ENSG00000262251 | 196.1508059 | 1.099061877 | 0.2249258 | 4.886331 | 1.03E-06 | 1.40E-05 | | sense_intronic | 49.82 | 5.336256769 | 5.336256769 | ENSG00000262251 | up |
| ENSG00000187626 | 182.7437131 | -1.154809929 | 0.2422669 | -4.766684 | 1.87E-06 | 2.39E-05 | ZKSCAN4 | protein_coding | 44.69 | -5.336142213 | 5.336142213 | ENSG00000187626 | down |
| ENSG00000251445 | 4.384105133 | 2.40588114 | 0.736826 | 3.265196 | 0.001093884 | 0.006058136 | | antisense | 53.73 | 5.33542874 | 5.33542874 | ENSG00000251445 | up |
| ENSG00000127152 | 708.1998918 | -1.147919788 | 0.2401164 | -4.780681 | 1.75E-06 | 2.25E-05 | BCL11B | protein_coding | 50.77 | -5.335294153 | 5.335294153 | ENSG00000127152 | down |
| ENSG00000234707 | 28.93764331 | 1.360794026 | 0.3106109 | 4.381025 | 1.18E-05 | 0.000120309 | | antisense | 42.96 | 5.333909053 | 5.333909053 | ENSG00000234707 | up |
| ENSG00000125848 | 573.1138353 | -1.063670353 | 0.2144931 | -4.958997 | 7.09E-07 | 9.91E-06 | FLRT3 | protein_coding | 35.11 | -5.322751826 | 5.322751826 | ENSG00000125848 | down |
| ENSG00000260708 | 167.3140681 | 1.117536028 | 0.2309171 | 4.839555 | 1.30E-06 | 1.73E-05 | | antisense | 51.64 | 5.321523872 | 5.321523872 | ENSG00000260708 | up |
| ENSG00000180488 | 700.4646213 | -1.292741698 | 0.2878043 | -4.491738 | 7.06E-06 | 7.73E-05 | MIGA1 | protein_coding | 39.53 | -5.315337185 | 5.315337185 | ENSG00000180488 | down |
| ENSG00000245975 | 41.21997584 | 1.308597048 | 0.2932095 | 4.463011 | 8.08E-06 | 8.70E-05 | | lincRNA | 45.68 | 5.313825481 | 5.313825481 | ENSG00000245975 | up |
| ENSG00000259415 | 38.11007237 | 1.350867671 | 0.3078958 | 4.387418 | 1.15E-05 | 0.000117173 | | sense_intronic | 45.47 | 5.31049208 | 5.31049208 | ENSG00000259415 | up |
| ENSG00000107290 | 2825.757393 | -1.293878112 | 0.2885203 | -4.48453 | 7.31E-06 | 7.95E-05 | SETX | protein_coding | 41 | -5.304085216 | 5.304085216 | ENSG00000107290 | down |
| ENSG00000241634 | 145.6322786 | -1.338341728 | 0.3038117 | -4.405169 | 1.06E-05 | 0.000109582 | | processed_pseudogene | 50.49 | -5.300183739 | 5.300183739 | ENSG00000241634 | down |
| ENSG00000185112 | 1181.358443 | 1.045317156 | 0.2094197 | 4.991493 | 5.99E-07 | 8.56E-06 | FAM43A | protein_coding | 60.99 | 5.297397258 | 5.297397258 | ENSG00000185112 | up |
| ENSG00000221614 | 2.916551157 | -3.167541765 | 1.1265744 | -2.811658 | 0.004928689 | 0.021311223 | MIR1185-2 | miRNA | 38.37 | -5.294202765 | 5.294202765 | ENSG00000221614 | down |
| ENSG00000261061 | 76.47042861 | -1.309556258 | 0.2942294 | -4.450801 | 8.56E-06 | 9.13E-05 | | sense_intronic | 43.44 | -5.289784315 | 5.289784315 | ENSG00000261061 | down |
| ENSG00000201221 | 8.423535498 | -1.78905413 | 0.4723079 | -3.787898 | 0.000151927 | 0.001106321 | RNU4-40P | snRNA | 37.06 | -5.288656965 | 5.288656965 | ENSG00000201221 | down |
| ENSG00000138735 | 1124.091088 | -1.241632953 | 0.2715811 | -4.571867 | 4.83E-06 | 5.55E-05 | PDE5A | protein_coding | 36.45 | -5.283726665 | 5.283726665 | ENSG00000138735 | down |
| ENSG00000221176 | 8.134279512 | -1.782925639 | 0.4702203 | -3.791681 | 0.000149631 | 0.001092454 | MIR1207 | miRNA | 62.07 | -5.280307086 | 5.280307086 | ENSG00000221176 | down |
| ENSG00000165891 | 1102.003667 | -1.116229558 | 0.2314663 | -4.822429 | 1.42E-06 | 1.87E-05 | E2F7 | protein_coding | 40.18 | -5.276828717 | 5.276828717 | ENSG00000165891 | down |
| ENSG00000206656 | 5.914184675 | -2.688711253 | 0.8773058 | -3.064737 | 0.002178618 | 0.010934714 | SNORD116-17 | snoRNA | 42.39 | -5.273080556 | 5.273080556 | ENSG00000206656 | down |
| ENSG00000175906 | 784.9441331 | 1.135606809 | 0.2377671 | 4.776131 | 1.79E-06 | 2.30E-05 | ARL4D | protein_coding | 55.99 | 5.26827572 | 5.26827572 | ENSG00000175906 | up |
| ENSG00000272155 | 25.36213811 | -2.35057302 | 0.7155557 | -3.284962 | 0.001019963 | 0.005737753 | | antisense | 39.55 | -5.268240911 | 5.268240911 | ENSG00000272155 | down |
| ENSG00000178177 | 772.2797968 | -1.225840321 | 0.2669037 | -4.592818 | 4.37E-06 | 5.08E-05 | LCORL | protein_coding | 36.02 | -5.263689765 | 5.263689765 | ENSG00000178177 | down |

| | | | | | | | | | | | | | |
|---|---|---|---|---|---|---|---|---|---|---|---|---|---|
| ENSG00000207385 | 7.863008964 | -1.853681237 | 0.4996971 | -3.70961 | 0.000207579 | 0.001450138 | RNU6-310P | snRNA | 38.32 | -5.261842492 | 5.261842492 | ENSG00000207385 | down |
| ENSG00000234946 | 39.74674935 | 1.311080045 | 0.2956272 | 4.434911 | 9.21E-06 | 9.75E-05 | SDHCP3 | processed_pseudogene | 49.01 | 5.258589851 | 5.258589851 | ENSG00000234946 | up |
| ENSG00000180357 | 1562.371819 | -1.175115349 | 0.2506796 | -4.687719 | 2.76E-06 | 3.37E-05 | ZNF609 | protein_coding | 42.67 | -5.25537926 | 5.25537926 | ENSG00000180357 | down |
| ENSG00000151612 | 247.2524736 | -1.16600729 | 0.2477802 | -4.705813 | 2.53E-06 | 3.13E-05 | ZNF827 | protein_coding | 41.53 | -5.252332725 | 5.252332725 | ENSG00000151612 | down |
| ENSG00000172244 | 330.5348911 | -1.19342244 | 0.2566359 | -4.650255 | 3.32E-06 | 3.98E-05 | C5orf34 | protein_coding | 37.92 | -5.251440319 | 5.251440319 | ENSG00000172244 | down |
| ENSG00000143756 | 1030.313528 | -1.293794009 | 0.2901142 | -4.459603 | 8.21E-06 | 8.81E-05 | FBXO28 | protein_coding | 40.97 | -5.246078048 | 5.246078048 | ENSG00000143756 | down |
| ENSG00000274292 | 176.7236334 | 1.137770316 | 0.2390066 | 4.760414 | 1.93E-06 | 2.46E-05 | | lincRNA | 50.9 | 5.244308477 | 5.244308477 | ENSG00000274292 | up |
| ENSG00000273061 | 51.43868023 | 1.245260596 | 0.2738874 | 4.546616 | 5.45E-06 | 6.18E-05 | CDC37L1-AS1 | lincRNA | 41.23 | 5.241675728 | 5.241675728 | ENSG00000273061 | up |
| ENSG00000261474 | 24.83300961 | -1.396019501 | 0.3259869 | -4.28244 | 1.85E-05 | 0.000176738 | | lincRNA | 48.37 | -5.238800729 | 5.238800729 | ENSG00000261474 | down |
| ENSG00000226754 | 51.36102362 | 1.298841271 | 0.2920729 | 4.446977 | 8.71E-06 | 9.27E-05 | | antisense | 44.9 | 5.238122827 | 5.238122827 | ENSG00000226754 | up |
| ENSG00000274428 | 394.5015754 | 1.155030381 | 0.2448097 | 4.718074 | 2.38E-06 | 2.98E-05 | | snRNA | 45.62 | 5.228147877 | 5.228147877 | ENSG00000274428 | up |
| ENSG00000147813 | 69.37525593 | -1.297547567 | 0.2920965 | -4.442188 | 8.90E-06 | 9.46E-05 | NAPRT | protein_coding | 67.89 | -5.221644972 | 5.221644972 | ENSG00000147813 | down |
| ENSG00000239884 | 15.91244638 | -1.550428339 | 0.3829438 | -4.04871 | 5.15E-05 | 0.000431955 | RN7SL608P | misc_RNA | 55.2 | -5.216512234 | 5.216512234 | ENSG00000239884 | down |
| ENSG00000249085 | 32.69756309 | 1.346159212 | 0.3094575 | 4.350061 | 1.36E-05 | 0.000135545 | | antisense | 38.46 | 5.206832986 | 5.206832986 | ENSG00000249085 | up |
| ENSG00000270395 | 4.089938332 | -2.336775205 | 0.7141162 | -3.272262 | 0.001066906 | 0.005937065 | MREGP1 | processed_pseudogene | 51.76 | -5.202662174 | 5.202662174 | ENSG00000270395 | down |
| ENSG00000196437 | 560.6267905 | -1.182176118 | 0.2542243 | -4.65013 | 3.32E-06 | 3.98E-05 | ZNF569 | protein_coding | 39.27 | -5.201952906 | 5.201952906 | ENSG00000196437 | down |
| ENSG00000178386 | 151.9602337 | -1.26426738 | 0.2815119 | -4.49099 | 7.09E-06 | 7.75E-05 | ZNF223 | protein_coding | 43.75 | -5.196763228 | 5.196763228 | ENSG00000178386 | down |
| ENSG00000121297 | 633.2918926 | -1.31593706 | 0.2991751 | -4.398551 | 1.09E-05 | 0.000112641 | TSHZ3 | protein_coding | 46.88 | -5.195718764 | 5.195718764 | ENSG00000121297 | down |
| ENSG00000141179 | 328.5861517 | -1.248462064 | 0.276372 | -4.517324 | 6.26E-06 | 6.95E-05 | PCTP | protein_coding | 40.32 | -5.191299855 | 5.191299855 | ENSG00000141179 | down |
| ENSG00000279133 | 514.9976268 | -1.153785278 | 0.2453536 | -4.70254 | 2.57E-06 | 3.17E-05 | | TEC | 40.51 | -5.190131129 | 5.190131129 | ENSG00000279133 | down |
| ENSG00000201690 | 3.702575248 | -2.242405569 | 0.6714559 | -3.339617 | 0.000838939 | 0.004850727 | RNY1P2 | misc_RNA | 42.74 | -5.189359563 | 5.189359563 | ENSG00000201690 | down |
| ENSG00000274954 | 19.31714965 | 1.553453759 | 0.3851981 | 4.032869 | 5.51E-05 | 0.00045773 | | sense_intronic | 26.62 | 5.187588385 | 5.187588385 | ENSG00000274954 | up |
| ENSG00000163625 | 2255.487943 | -1.18495781 | 0.255489 | -4.637999 | 3.52E-06 | 4.19E-05 | WDFY3 | protein_coding | 36.88 | -5.187052584 | 5.187052584 | ENSG00000163625 | down |
| ENSG00000173334 | 831.1106232 | 1.005468155 | 0.1995859 | 5.037772 | 4.71E-07 | 6.97E-06 | TRIB1 | protein_coding | 51.37 | 5.185050351 | 5.185050351 | ENSG00000173334 | up |
| ENSG00000188234 | 88.31168348 | -1.403941241 | 0.3308136 | -4.243905 | 2.20E-05 | 0.000204574 | AGAP4 | protein_coding | 41.11 | -5.179350245 | 5.179350245 | ENSG00000188234 | down |
| ENSG00000234816 | 167.2875682 | 1.168249691 | 0.2502902 | 4.66758 | 3.05E-06 | 3.69E-05 | HIST1H2APS5 | unprocessed_pseudogene | 54.47 | 5.178405095 | 5.178405095 | ENSG00000234816 | up |
| ENSG00000240823 | 10.66451274 | -1.718176774 | 0.4492107 | -3.824879 | 0.000130836 | 0.000968954 | RN7SL23P | misc_RNA | 57.09 | -5.178063737 | 5.178063737 | ENSG00000240823 | down |
| ENSG00000120647 | 987.4088075 | -1.262757952 | 0.281835 | -4.480487 | 7.45E-06 | 8.08E-05 | CCDC77 | protein_coding | 43.8 | -5.167831146 | 5.167831146 | ENSG00000120647 | down |
| ENSG00000256771 | 1144.335392 | -1.208467526 | 0.2636486 | -4.583629 | 4.57E-06 | 5.29E-05 | ZNF253 | protein_coding | 38.27 | -5.167774064 | 5.167774064 | ENSG00000256771 | down |
| ENSG00000169379 | 285.1125804 | -1.230209519 | 0.2712245 | -4.535761 | 5.74E-06 | 6.43E-05 | ARL13B | protein_coding | 36.2 | -5.156431194 | 5.156431194 | ENSG00000169379 | down |
| ENSG00000259802 | 193.743049 | 1.080734017 | 0.2231154 | 4.843834 | 1.27E-06 | 1.70E-05 | | antisense | 54.16 | 5.155041373 | 5.155041373 | ENSG00000259802 | up |
| ENSG00000136169 | 335.4550958 | -1.234834657 | 0.272829 | -4.526039 | 6.01E-06 | 6.69E-05 | SETDB2 | protein_coding | 37.41 | -5.155006727 | 5.155006727 | ENSG00000136169 | down |

| | | | | | | | | | | | | | |
|---|---|---|---|---|---|---|---|---|---|---|---|---|---|
| ENSG00000116205 | 538.8102733 | -1.110315898 | 0.2324295 | -4.777001 | 1.78E-06 | 2.29E-05 | TCEANC2 | protein_coding | 42.31 | -5.152587255 | 5.152587255 | ENSG00000116205 | down |
| ENSG00000273181 | 16.30962333 | 1.529417776 | 0.3774836 | 4.051614 | 5.09E-05 | 0.000428175 | | lincRNA | 33.06 | 5.151658548 | 5.151658548 | ENSG00000273181 | up |
| ENSG00000163848 | 1602.249964 | -1.210071652 | 0.2646398 | -4.572523 | 4.82E-06 | 5.54E-05 | ZNF148 | protein_coding | 36.86 | -5.150257726 | 5.150257726 | ENSG00000163848 | down |
| ENSG00000274898 | 19.99560814 | 1.454907608 | 0.3503252 | 4.15302 | 3.28E-05 | 0.000289503 | | lincRNA | 43.78 | 5.147968104 | 5.147968104 | ENSG00000274898 | up |
| ENSG00000263002 | 490.4865568 | -1.148065276 | 0.2446268 | -4.69313 | 2.69E-06 | 3.30E-05 | ZNF234 | protein_coding | 41.72 | -5.144961158 | 5.144961158 | ENSG00000263002 | down |
| ENSG00000064393 | 897.2576349 | -1.178576847 | 0.2545321 | -4.630367 | 3.65E-06 | 4.33E-05 | HIPK2 | protein_coding | 45.62 | -5.142862555 | 5.142862555 | ENSG00000064393 | down |
| ENSG00000249459 | 468.8198721 | -1.086984385 | 0.2255168 | -4.81997 | 1.44E-06 | 1.89E-05 | ZNF286B | protein_coding | 42.4 | -5.134449444 | 5.134449444 | ENSG00000249459 | down |
| ENSG00000274565 | 33.71762371 | 1.321665708 | 0.3032561 | 4.358249 | 1.31E-05 | 0.000131703 | | antisense | 59.01 | 5.128596656 | 5.128596656 | ENSG00000274565 | up |
| ENSG00000241157 | 28.21520843 | -1.426943627 | 0.3411251 | -4.183051 | 2.88E-05 | 0.000258845 | | processed_pseudogene | 46.39 | -5.118390357 | 5.118390357 | ENSG00000241157 | down |
| ENSG00000273117 | 82.4526964 | 1.163847248 | 0.2503634 | 4.648631 | 3.34E-06 | 4.00E-05 | | lincRNA | 56.34 | 5.118062918 | 5.118062918 | ENSG00000273117 | up |
| ENSG00000167395 | 640.7171561 | -1.019405598 | 0.2052636 | -4.966325 | 6.82E-07 | 9.58E-06 | ZNF646 | protein_coding | 58.57 | -5.116172805 | 5.116172805 | ENSG00000167395 | down |
| ENSG00000106701 | 1510.377397 | -1.203084156 | 0.2633072 | -4.569127 | 4.90E-06 | 5.61E-05 | FSD1L | protein_coding | 37.37 | -5.114571945 | 5.114571945 | ENSG00000106701 | down |
| ENSG00000130349 | 318.923648 | -1.230075787 | 0.272345 | -4.516608 | 6.28E-06 | 6.95E-05 | C6orf203 | protein_coding | 47.04 | -5.114316749 | 5.114316749 | ENSG00000130349 | down |
| ENSG00000167562 | 326.5934315 | -1.126222899 | 0.2390187 | -4.711861 | 2.45E-06 | 3.05E-05 | ZNF701 | protein_coding | 45.33 | -5.085454433 | 5.085454433 | ENSG00000167562 | down |
| ENSG00000256628 | 107.5596049 | -1.123759672 | 0.2382295 | -4.71713 | 2.39E-06 | 2.99E-05 | ZBTB11-AS1 | antisense | 52.17 | -5.084777405 | 5.084777405 | ENSG00000256628 | down |
| ENSG00000185633 | 27.66128101 | 1.355657886 | 0.3168221 | 4.278924 | 1.88E-05 | 0.000178968 | NDUFA4L2 | protein_coding | 62.05 | 5.079953939 | 5.079953939 | ENSG00000185633 | up |
| ENSG00000226964 | 14.80938717 | 1.566841675 | 0.3947166 | 3.969535 | 7.20E-05 | 0.000573919 | RHEBP2 | processed_pseudogene | 40.67 | 5.078368186 | 5.078368186 | ENSG00000226964 | up |
| ENSG00000144182 | 83.40790842 | 1.139813218 | 0.2437346 | 4.676452 | 2.92E-06 | 3.56E-05 | LIPT1 | protein_coding | 41.3 | 5.071151507 | 5.071151507 | ENSG00000144182 | up |
| ENSG00000231908 | 20.92661591 | 1.45615845 | 0.3536801 | 4.117162 | 3.84E-05 | 0.000332942 | IDH1-AS1 | antisense | 52.09 | 5.063982479 | 5.063982479 | ENSG00000231908 | up |
| ENSG00000135637 | 276.214891 | -1.090578095 | 0.2282502 | -4.777995 | 1.77E-06 | 2.28E-05 | CCDC142 | protein_coding | 52.12 | -5.062892606 | 5.062892606 | ENSG00000135637 | down |
| ENSG00000266270 | 2.124265498 | -4.892039079 | 2.2585616 | -2.165998 | 0.030311366 | 0.092407523 | MIR5096 | miRNA | 54.29 | -5.059800176 | 5.059800176 | ENSG00000266270 | down |
| ENSG00000265112 | 167.4964183 | 1.080862992 | 0.2254614 | 4.794004 | 1.63E-06 | 2.12E-05 | MIR3153 | miRNA | 40.24 | 5.051730243 | 5.051730243 | ENSG00000265112 | up |
| ENSG00000184347 | 4846.850281 | -1.082049608 | 0.2259721 | -4.788421 | 1.68E-06 | 2.18E-05 | SLIT3 | protein_coding | 45.01 | -5.045073612 | 5.045073612 | ENSG00000184347 | down |
| ENSG00000135437 | 45.59140228 | -1.299336741 | 0.2981798 | -4.357561 | 1.32E-05 | 0.000132022 | RDH5 | protein_coding | 52.34 | -5.040586077 | 5.040586077 | ENSG00000135437 | down |
| ENSG00000271749 | 13.89113106 | -1.608434854 | 0.4126306 | -3.898001 | 9.70E-05 | 0.000742481 | | unprocessed_pseudogene | 43.53 | -5.033299142 | 5.033299142 | ENSG00000271749 | down |
| ENSG00000199646 | 6.694123743 | -2.31344231 | 0.7155949 | -3.232894 | 0.001225431 | 0.006693478 | RNU6-1272P | snRNA | 43.93 | -5.030229089 | 5.030229089 | ENSG00000199646 | down |
| ENSG00000231842 | 23.6744247 | -1.466035209 | 0.3586753 | -4.08736 | 4.36E-05 | 0.00037383 | | lincRNA | 35.46 | -5.024579739 | 5.024579739 | ENSG00000231842 | down |
| ENSG00000070159 | 86.14746311 | -1.328464565 | 0.3089679 | -4.299684 | 1.71E-05 | 0.000165261 | PTPN3 | protein_coding | 45.34 | -5.024026467 | 5.024026467 | ENSG00000070159 | down |
| ENSG00000219529 | 54.13251423 | 1.287045355 | 0.2944584 | 4.37089 | 1.24E-05 | 0.000125025 | | processed_pseudogene | 46.09 | 5.023340537 | 5.023340537 | ENSG00000219529 | up |
| ENSG00000283409 | 7.928780298 | -2.000615586 | 0.5746021 | -3.481741 | 0.000498166 | 0.00310195 | MIR3662 | miRNA | 37.89 | -5.018274613 | 5.018274613 | ENSG00000283409 | down |
| ENSG00000109066 | 299.020557 | -1.14745883 | 0.2478451 | -4.629741 | 3.66E-06 | 4.34E-05 | TMEM104 | protein_coding | 54.36 | -5.006060215 | 5.006060215 | ENSG00000109066 | down |
| ENSG00000143473 | 210.9584119 | -1.23251829 | 0.276265 | -4.461363 | 8.14E-06 | 8.76E-05 | KCNH1 | protein_coding | 41.06 | -5.001187294 | 5.001187294 | ENSG00000143473 | down |

| | | | | | | | | | | | | | |
|---|---|---|---|---|---|---|---|---|---|---|---|---|---|
| ENSG00000264895 | 770.0445929 | -1.12781671 | 0.2417143 | -4.665908 | 3.07E-06 | 3.71E-05 | | sense_intronic | 36.41 | -4.996472811 | 4.996472811 | ENSG00000264895 | down |
| ENSG00000253878 | 265.2774401 | 1.055696993 | 0.2190115 | 4.820282 | 1.43E-06 | 1.89E-05 | | sense_intronic | 46.4 | 4.986661183 | 4.986661183 | ENSG00000253878 | up |
| ENSG00000130222 | 805.6212826 | 1.119820747 | 0.2395832 | 4.674037 | 2.95E-06 | 3.59E-05 | GADD45G | protein_coding | 66.23 | 4.976905925 | 4.976905925 | ENSG00000130222 | up |
| ENSG00000015532 | 283.7771361 | -1.182866665 | 0.2605015 | -4.540729 | 5.61E-06 | 6.31E-05 | XYLT2 | protein_coding | 56.54 | -4.967604868 | 4.967604868 | ENSG00000015532 | down |
| ENSG00000154114 | 367.2134245 | -1.297417893 | 0.2998238 | -4.327267 | 1.51E-05 | 0.000148549 | TBCEL | protein_coding | 36.59 | -4.966685517 | 4.966685517 | ENSG00000154114 | down |
| ENSG00000187098 | 20.7507372 | 1.461467942 | 0.3591219 | 4.06956 | 4.71E-05 | 0.000400367 | MITF | protein_coding | 38.57 | 4.965397529 | 4.965397529 | ENSG00000187098 | up |
| ENSG00000077514 | 962.3944335 | -1.148039473 | 0.2491618 | -4.607607 | 4.07E-06 | 4.77E-05 | POLD3 | protein_coding | 41.72 | -4.960806505 | 4.960806505 | ENSG00000077514 | down |
| ENSG00000198093 | 755.1592619 | -1.199927717 | 0.2666392 | -4.500193 | 6.79E-06 | 7.46E-05 | ZNF649 | protein_coding | 43.38 | -4.95233908 | 4.95233908 | ENSG00000198093 | down |
| ENSG00000196757 | 582.7365759 | -1.250852565 | 0.2840806 | -4.40316 | 1.07E-05 | 0.000110519 | ZNF700 | protein_coding | 44.6 | -4.949075787 | 4.949075787 | ENSG00000196757 | down |
| ENSG00000273064 | 98.10764475 | 1.114623943 | 0.238708 | 4.669403 | 3.02E-06 | 3.67E-05 | | antisense | 46.27 | 4.944152011 | 4.944152011 | ENSG00000273064 | up |
| ENSG00000279421 | 16.24117858 | -1.494160529 | 0.3724847 | -4.011334 | 6.04E-05 | 0.000491919 | | TEC | 51.64 | -4.942841436 | 4.942841436 | ENSG00000279421 | down |
| ENSG00000214425 | 571.3438711 | -1.183186541 | 0.2613915 | -4.526492 | 6.00E-06 | 6.68E-05 | LRRC37A4P | transcribed_unprocessed_pseud | 42.83 | -4.940084976 | 4.940084976 | ENSG00000214425 | down |
| ENSG00000271141 | 95.22505615 | -1.30104335 | 0.3019402 | -4.308944 | 1.64E-05 | 0.000159711 | | lincRNA | 38.31 | -4.939625766 | 4.939625766 | ENSG00000271141 | down |
| ENSG00000279453 | 133.4003543 | -1.335434406 | 0.3143136 | -4.248732 | 2.15E-05 | 0.000201157 | | TEC | 38.44 | -4.936385907 | 4.936385907 | ENSG00000279453 | down |
| ENSG00000202314 | 58.25849828 | -1.222349541 | 0.2747099 | -4.449601 | 8.60E-06 | 9.17E-05 | SNORD6 | snoRNA | 45.21 | -4.935310105 | 4.935310105 | ENSG00000202314 | down |
| ENSG00000201217 | 22.23226912 | -1.475988294 | 0.3658117 | -4.034831 | 5.46E-05 | 0.000454737 | | misc_RNA | 41.59 | -4.93310625 | 4.93310625 | ENSG00000201217 | down |
| ENSG00000152439 | 219.1255481 | -1.174458201 | 0.2586345 | -4.540995 | 5.60E-06 | 6.31E-05 | ZNF773 | protein_coding | 43.76 | -4.932524206 | 4.932524206 | ENSG00000152439 | down |
| ENSG00000234183 | 104.8860742 | 1.11375298 | 0.2387703 | 4.664537 | 3.09E-06 | 3.73E-05 | LINC01952 | antisense | 49.87 | 4.93200051 | 4.93200051 | ENSG00000234183 | up |
| ENSG00000240328 | 43.37407538 | -1.412898281 | 0.3426193 | -4.123814 | 3.73E-05 | 0.000324682 | | processed_pseudogene | 44.15 | -4.92895532 | 4.92895532 | ENSG00000240328 | down |
| ENSG00000138744 | 67.50406484 | -1.194874786 | 0.265626 | -4.498336 | 6.85E-06 | 7.52E-05 | NAAA | protein_coding | 44.41 | -4.927394877 | 4.927394877 | ENSG00000138744 | down |
| ENSG00000172748 | 239.6961776 | -1.232501467 | 0.2784309 | -4.426597 | 9.57E-06 | 0.000100746 | ZNF596 | protein_coding | 40.95 | -4.926027328 | 4.926027328 | ENSG00000172748 | down |
| ENSG00000107282 | 644.3402746 | -1.133574696 | 0.2454477 | -4.618396 | 3.87E-06 | 4.56E-05 | APBA1 | protein_coding | 41.44 | -4.920570663 | 4.920570663 | ENSG00000107282 | down |
| ENSG00000165495 | 29.29919214 | -1.384423819 | 0.3325512 | -4.163039 | 3.14E-05 | 0.000279564 | PKNOX2 | protein_coding | 49.03 | -4.91957685 | 4.91957685 | ENSG00000165495 | down |
| ENSG00000212127 | 110.0721521 | -1.443705886 | 0.3545103 | -4.072395 | 4.65E-05 | 0.000395766 | TAS2R14 | protein_coding | 36.14 | -4.912298601 | 4.912298601 | ENSG00000212127 | down |
| ENSG00000048991 | 804.6654213 | -1.170843648 | 0.2581102 | -4.536217 | 5.73E-06 | 6.43E-05 | R3HDM1 | protein_coding | 37.98 | -4.908286994 | 4.908286994 | ENSG00000048991 | down |
| ENSG00000231729 | 12.6316712 | -1.565076013 | 0.4011171 | -3.901793 | 9.55E-05 | 0.000733667 | ARHGEF9-IT1 | sense_intronic | 43.53 | -4.905732104 | 4.905732104 | ENSG00000231729 | down |
| ENSG00000272195 | 79.22152903 | 1.15163375 | 0.2520847 | 4.56844 | 4.91E-06 | 5.62E-05 | | antisense | 62.79 | 4.894617647 | 4.894617647 | ENSG00000272195 | up |
| ENSG00000257004 | 16.35537493 | 1.507947166 | 0.3801083 | 3.967152 | 7.27E-05 | 0.000578033 | | lincRNA | 41.37 | 4.882804793 | 4.882804793 | ENSG00000257004 | up |
| ENSG00000138411 | 598.9189928 | -1.11433298 | 0.2403851 | -4.635615 | 3.56E-06 | 4.23E-05 | HECW2 | protein_coding | 39.16 | -4.873147466 | 4.873147466 | ENSG00000138411 | down |
| ENSG00000152804 | 246.4787855 | 1.158686837 | 0.2550097 | 4.543697 | 5.53E-06 | 6.25E-05 | HHEX | protein_coding | 46.38 | 4.871513594 | 4.871513594 | ENSG00000152804 | up |
| ENSG00000247595 | 43.32306154 | -1.305327538 | 0.3059136 | -4.266981 | 1.98E-05 | 0.000187281 | SPTY2D1-AS1 | protein_coding | 43.98 | -4.86561651 | 4.86561651 | ENSG00000247595 | down |
| ENSG00000274380 | 7.745330075 | -1.929396282 | 0.553009 | -3.488906 | 0.000485002 | 0.003025397 | MIR6801 | miRNA | 59.49 | -4.860569255 | 4.860569255 | ENSG00000274380 | down |

| | | | | | | | | | | | | | |
|---|---|---|---|---|---|---|---|---|---|---|---|---|---|
| ENSG00000283929 | 24.77012993 | -1.438006683 | 0.3544632 | -4.056858 | 4.97E-05 | 0.000419947 | MIR5010 | miRNA | 54.17 | -4.855868162 | 4.855868162 | ENSG00000283929 | down |
| ENSG00000133858 | 395.7816266 | -1.254542071 | 0.2883418 | -4.350885 | 1.36E-05 | 0.000135424 | ZFC3H1 | protein_coding | 36.14 | -4.852952304 | 4.852952304 | ENSG00000133858 | down |
| ENSG00000261423 | 99.89168112 | 1.157537319 | 0.255172 | 4.536303 | 5.72E-06 | 6.43E-05 | TMEM202-AS1 | lincRNA | 40.16 | 4.852505609 | 4.852505609 | ENSG00000261423 | up |
| ENSG00000270829 | 14.24278108 | 1.572430563 | 0.4063339 | 3.869799 | 0.000108925 | 0.000823198 | | lincRNA | 47.19 | 4.850155202 | 4.850155202 | ENSG00000270829 | up |
| ENSG00000224020 | 165.8831823 | 1.036262941 | 0.2163284 | 4.790231 | 1.67E-06 | 2.16E-05 | MIR181A2HG | antisense | 38.12 | 4.835232132 | 4.835232132 | ENSG00000224020 | up |
| ENSG00000116406 | 1334.54016 | -1.2605368 | 0.2910892 | -4.330414 | 1.49E-05 | 0.000146753 | EDEM3 | protein_coding | 34.39 | -4.832159421 | 4.832159421 | ENSG00000116406 | down |
| ENSG00000164603 | 349.4850745 | -1.191820547 | 0.2674026 | -4.457027 | 8.31E-06 | 8.91E-05 | BMT2 | protein_coding | 36.97 | -4.827176388 | 4.827176388 | ENSG00000164603 | down |
| ENSG00000200888 | 44.29979263 | -1.280328299 | 0.2987519 | -4.285591 | 1.82E-05 | 0.00017452 | | misc_RNA | 43.14 | -4.811670923 | 4.811670923 | ENSG00000200888 | down |
| ENSG00000252859 | 6.264454388 | -2.013676091 | 0.5932443 | -3.394346 | 0.000687928 | 0.004082704 | RNU6-375P | snRNA | 33 | -4.810777056 | 4.810777056 | ENSG00000252859 | down |
| ENSG00000159110 | 167.9413632 | -1.220764273 | 0.277918 | -4.392533 | 1.12E-05 | 0.000115462 | IFNAR2 | protein_coding | 44.16 | -4.806831617 | 4.806831617 | ENSG00000159110 | down |
| ENSG00000273066 | 336.1729307 | 1.024652682 | 0.2134057 | 4.80143 | 1.58E-06 | 2.04E-05 | | processed_transcript | 60.91 | 4.805094684 | 4.805094684 | ENSG00000273066 | up |
| ENSG00000118263 | 3447.907984 | -1.120990311 | 0.2443288 | -4.588039 | 4.47E-06 | 5.19E-05 | KLF7 | protein_coding | 41.5 | -4.803540375 | 4.803540375 | ENSG00000118263 | down |
| ENSG00000101624 | 530.786147 | 1.065078908 | 0.2263128 | 4.706224 | 2.52E-06 | 3.13E-05 | CEP76 | protein_coding | 40.84 | 4.798078005 | 4.798078005 | ENSG00000101624 | up |
| ENSG00000270194 | 104.161058 | 1.144102139 | 0.2525241 | 4.530666 | 5.88E-06 | 6.57E-05 | | antisense | 53.41 | 4.785486804 | 4.785486804 | ENSG00000270194 | up |
| ENSG00000271907 | 3.581090075 | -2.11394444 | 0.6401259 | -3.302389 | 0.000958652 | 0.005454561 | SNORA35B | snoRNA | 43.41 | -4.78436407 | 4.78436407 | ENSG00000271907 | down |
| ENSG00000272092 | 53.08493913 | 1.198432616 | 0.2714 | 4.415744 | 1.01E-05 | 0.000104993 | | lincRNA | 65.03 | 4.768373589 | 4.768373589 | ENSG00000272092 | up |
| ENSG00000165511 | 88.21264907 | 1.148865455 | 0.2547102 | 4.510481 | 6.47E-06 | 7.14E-05 | C10orf25 | antisense | 44.41 | 4.763432882 | 4.763432882 | ENSG00000165511 | up |
| ENSG00000070778 | 462.2418581 | -1.133505409 | 0.2495667 | -4.541893 | 5.58E-06 | 6.29E-05 | PTPN21 | protein_coding | 41.28 | -4.762227056 | 4.762227056 | ENSG00000070778 | down |
| ENSG00000278238 | 29.82165998 | 1.296009763 | 0.3061227 | 4.233629 | 2.30E-05 | 0.000212731 | | lincRNA | 45.06 | 4.759167883 | 4.759167883 | ENSG00000278238 | up |
| ENSG00000173258 | 443.4507849 | -1.126635079 | 0.2474332 | -4.553289 | 5.28E-06 | 6.01E-05 | ZNF483 | protein_coding | 42.33 | -4.755868311 | 4.755868311 | ENSG00000173258 | down |
| ENSG00000271818 | 40.11589602 | 1.239951314 | 0.2863793 | 4.329752 | 1.49E-05 | 0.000146992 | RN7SKP4 | misc_RNA | 52.8 | 4.75236787 | 4.75236787 | ENSG00000271818 | up |
| ENSG00000244405 | 458.9768025 | -1.193361435 | 0.2701742 | -4.417007 | 1.00E-05 | 0.000104529 | ETV5 | protein_coding | 42.69 | -4.750489615 | 4.750489615 | ENSG00000244405 | down |
| ENSG00000234676 | 17.82793569 | 1.439445958 | 0.3594777 | 4.00427 | 6.22E-05 | 0.000504449 | IFT74-AS1 | antisense | 39.53 | 4.746116448 | 4.746116448 | ENSG00000234676 | up |
| ENSG00000207014 | 4.280060039 | -2.785876275 | 0.980937 | -2.840016 | 0.004511133 | 0.019807161 | SNORD116-3 | snoRNA | 45.26 | -4.74484252 | 4.74484252 | ENSG00000207014 | down |
| ENSG00000177842 | 437.090827 | -1.082527564 | 0.2333747 | -4.638582 | 3.51E-06 | 4.19E-05 | ZNF620 | protein_coding | 43.94 | -4.739193763 | 4.739193763 | ENSG00000177842 | down |
| ENSG00000268403 | 537.331819 | 1.000831371 | 0.2076687 | 4.819366 | 1.44E-06 | 1.89E-05 | | antisense | 49.35 | 4.726680348 | 4.726680348 | ENSG00000268403 | up |
| ENSG00000129810 | 524.9466182 | -1.130560159 | 0.2496152 | -4.529213 | 5.92E-06 | 6.60E-05 | SGO1 | protein_coding | 37.06 | -4.726283811 | 4.726283811 | ENSG00000129810 | down |
| ENSG00000110921 | 183.6284649 | -1.274167542 | 0.2993341 | -4.256673 | 2.07E-05 | 0.000195452 | MVK | protein_coding | 52.45 | -4.725835338 | 4.725835338 | ENSG00000110921 | down |
| ENSG00000213540 | 40.49925622 | -1.260282013 | 0.294939 | -4.273026 | 1.93E-05 | 0.000183268 | | processed_pseudogene | 53.75 | -4.709563876 | 4.709563876 | ENSG00000213540 | down |
| ENSG00000264482 | 5.086829756 | -2.10112985 | 0.640336 | -3.281293 | 0.001033325 | 0.005783274 | MIR4705 | miRNA | 38.03 | -4.701963524 | 4.701963524 | ENSG00000264482 | down |
| ENSG00000222675 | 3.429056874 | 2.454422782 | 0.8113898 | 3.024961 | 0.00248665 | 0.012147941 | RNA5SP146 | rRNA | 55.28 | 4.701440276 | 4.701440276 | ENSG00000222675 | up |
| ENSG00000168411 | 1740.594429 | -1.174113 | 0.2651718 | -4.427745 | 9.52E-06 | 0.000100287 | RFWD3 | protein_coding | 43.53 | -4.694989493 | 4.694989493 | ENSG00000168411 | down |

| | | | | | | | | | | | | | |
|---|---|---|---|---|---|---|---|---|---|---|---|---|---|
| ENSG00000155729 | 231.2848033 | -1.193671019 | 0.272021 | -4.388158 | 1.14E-05 | 0.00011703 | KCTD18 | protein_coding | 40.26 | -4.693160376 | 4.693160376 | ENSG00000155729 | down |
| ENSG00000265817 | 328.5438573 | -1.133767889 | 0.2516876 | -4.504664 | 6.65E-06 | 7.32E-05 | FSBP | protein_coding | 35.6 | -4.688899735 | 4.688899735 | ENSG00000265817 | down |
| ENSG00000120370 | 323.1976971 | -1.188553356 | 0.2706723 | -4.391116 | 1.13E-05 | 0.000116132 | GORAB | protein_coding | 36.49 | -4.677016407 | 4.677016407 | ENSG00000120370 | down |
| ENSG00000241772 | 51.95688626 | -1.339413986 | 0.3248294 | -4.123439 | 3.73E-05 | 0.000325008 | | 3prime_overlapping_ncRNA | 32.74 | -4.672018763 | 4.672018763 | ENSG00000241772 | down |
| ENSG00000259804 | 93.99076764 | 1.07862528 | 0.2338918 | 4.611643 | 3.99E-06 | 4.70E-05 | | antisense | 59.84 | 4.668005914 | 4.668005914 | ENSG00000259804 | up |
| ENSG00000137936 | 47.91364714 | -1.256034749 | 0.2949669 | -4.258222 | 2.06E-05 | 0.000194374 | BCAR3 | protein_coding | 45.08 | -4.661598936 | 4.661598936 | ENSG00000137936 | down |
| ENSG00000139618 | 2045.63482 | -1.19268499 | 0.2728623 | -4.371014 | 1.24E-05 | 0.000125025 | BRCA2 | protein_coding | 38.11 | -4.655051848 | 4.655051848 | ENSG00000139618 | down |
| ENSG00000105771 | 178.3127525 | -1.191741373 | 0.2727426 | -4.369472 | 1.25E-05 | 0.000125748 | SMG9 | protein_coding | 50.25 | -4.648384096 | 4.648384096 | ENSG00000105771 | down |
| ENSG00000173875 | 759.890837 | -1.128020504 | 0.2510407 | -4.493376 | 7.01E-06 | 7.68E-05 | ZNF791 | protein_coding | 45.22 | -4.641440676 | 4.641440676 | ENSG00000173875 | down |
| ENSG00000111879 | 733.8636337 | -1.183367637 | 0.2701336 | -4.380675 | 1.18E-05 | 0.000120358 | FAM184A | protein_coding | 39.55 | -4.638237133 | 4.638237133 | ENSG00000111879 | down |
| ENSG00000253540 | 89.36167845 | -1.10389887 | 0.2431501 | -4.539989 | 5.63E-06 | 6.33E-05 | FAM86HP | transcribed_unprocessed_pseud | 53.04 | -4.63467419 | 4.63467419 | ENSG00000253540 | down |
| ENSG00000276934 | 31.44408646 | -1.319905294 | 0.3190505 | -4.136979 | 3.52E-05 | 0.000308929 | | sense_intronic | 38.72 | -4.633054456 | 4.633054456 | ENSG00000276934 | down |
| ENSG00000266897 | 16.76926725 | -1.487845798 | 0.3828844 | -3.885887 | 0.000101957 | 0.000777407 | | lincRNA | 51.19 | -4.626235932 | 4.626235932 | ENSG00000266897 | down |
| ENSG00000252874 | 21.70943273 | -1.425974024 | 0.3591782 | -3.970102 | 7.18E-05 | 0.000572884 | | misc_RNA | 38.89 | -4.622912161 | 4.622912161 | ENSG00000252874 | down |
| ENSG00000170088 | 534.7843713 | -1.174641573 | 0.2676443 | -4.388816 | 1.14E-05 | 0.000116934 | TMEM192 | protein_coding | 41.86 | -4.618759798 | 4.618759798 | ENSG00000170088 | down |
| ENSG00000178665 | 271.1650628 | -1.209489354 | 0.2799041 | -4.321084 | 1.55E-05 | 0.000152236 | ZNF713 | protein_coding | 43 | -4.617205774 | 4.617205774 | ENSG00000178665 | down |
| ENSG00000259659 | 69.54783952 | -1.313031533 | 0.317727 | -4.132578 | 3.59E-05 | 0.000313523 | | sense_intronic | 46.69 | -4.600508631 | 4.600508631 | ENSG00000259659 | down |
| ENSG00000255153 | 23.20147353 | 1.399327818 | 0.3499684 | 3.99844 | 6.38E-05 | 0.000515833 | TOLLIP-AS1 | antisense | 60.38 | 4.600277613 | 4.600277613 | ENSG00000255153 | up |
| ENSG00000261970 | 44.41329497 | -1.389281465 | 0.3461841 | -4.013129 | 5.99E-05 | 0.000489585 | | processed_pseudogene | 37.06 | -4.59876044 | 4.59876044 | ENSG00000261970 | down |
| ENSG00000270894 | 51.6827068 | -1.203228799 | 0.2785041 | -4.320327 | 1.56E-05 | 0.000152651 | | sense_overlapping | 35.62 | -4.591881477 | 4.591881477 | ENSG00000270894 | down |
| ENSG00000263968 | 53.51775404 | -1.191778667 | 0.274485 | -4.341871 | 1.41E-05 | 0.000140397 | RN7SL381P | misc_RNA | 56.71 | -4.591496677 | 4.591496677 | ENSG00000263968 | down |
| ENSG00000181072 | 1530.50763 | -1.021143273 | 0.2173445 | -4.69827 | 2.62E-06 | 3.23E-05 | CHRM2 | protein_coding | 37.16 | -4.586150406 | 4.586150406 | ENSG00000181072 | down |
| ENSG00000166845 | 475.2856503 | -1.108241778 | 0.2459447 | -4.506061 | 6.60E-06 | 7.27E-05 | C18orf54 | protein_coding | 34.77 | -4.586118202 | 4.586118202 | ENSG00000166845 | down |
| ENSG00000261455 | 237.3570989 | 1.093296848 | 0.2409982 | 4.536536 | 5.72E-06 | 6.43E-05 | LINC01003 | lincRNA | 60.83 | 4.583203494 | 4.583203494 | ENSG00000261455 | up |
| ENSG00000145014 | 98.50127036 | -1.227833184 | 0.2875483 | -4.270007 | 1.95E-05 | 0.000185259 | TMEM44 | protein_coding | 50.87 | -4.582544276 | 4.582544276 | ENSG00000145014 | down |
| ENSG00000198496 | 190.4606597 | -1.101912038 | 0.2439467 | -4.517019 | 6.27E-06 | 6.95E-05 | NBR2 | transcribed_unprocessed_pseud | 47.53 | -4.581614113 | 4.581614113 | ENSG00000198496 | down |
| ENSG00000263627 | 20.36920636 | 1.415349946 | 0.3569007 | 3.965668 | 7.32E-05 | 0.000580976 | PPP4R1-AS1 | lincRNA | 42.71 | 4.579848305 | 4.579848305 | ENSG00000263627 | up |
| ENSG00000270130 | 27.89278569 | -1.366027568 | 0.3381866 | -4.039272 | 5.36E-05 | 0.000447283 | | sense_intronic | 42.05 | -4.575397096 | 4.575397096 | ENSG00000270130 | down |
| ENSG00000267213 | 134.1178793 | 1.040431025 | 0.2239297 | 4.64624 | 3.38E-06 | 4.04E-05 | | antisense | 44.48 | 4.570875359 | 4.570875359 | ENSG00000267213 | up |
| ENSG00000225484 | 238.2907499 | -1.161938528 | 0.2646872 | -4.389855 | 1.13E-05 | 0.000116635 | NUTM2B-AS1 | antisense | 39.68 | -4.570104159 | 4.570104159 | ENSG00000225484 | down |
| ENSG00000260274 | 60.57277952 | 1.144500367 | 0.2587645 | 4.422941 | 9.74E-06 | 0.000102004 | | lincRNA | 54.55 | 4.568140111 | 4.568140111 | ENSG00000260274 | up |
| ENSG00000198521 | 2129.169509 | -1.17281966 | 0.2686746 | -4.365205 | 1.27E-05 | 0.00012795 | ZNF43 | protein_coding | 41.78 | -4.565739729 | 4.565739729 | ENSG00000198521 | down |

| | | | | | | | | | | | | | |
|---|---|---|---|---|---|---|---|---|---|---|---|---|---|
| ENSG00000230513 | 113.5999332 | 1.073317596 | 0.2349852 | 4.567597 | 4.93E-06 | 5.63E-05 | THAP7-AS1 | antisense | 50.16 | 4.560746264 | 4.560746264 | ENSG00000230513 | up |
| ENSG00000274248 | 20.12533895 | 1.437651799 | 0.3663154 | 3.924629 | 8.69E-05 | 0.000676022 | | lincRNA | 63.23 | 4.557412704 | 4.557412704 | ENSG00000274248 | up |
| ENSG00000114491 | 1594.64204 | -1.175415203 | 0.2699193 | -4.354692 | 1.33E-05 | 0.000133475 | UMPS | protein_coding | 42.81 | -4.554264821 | 4.554264821 | ENSG00000114491 | down |
| ENSG00000241462 | 21.70775454 | -1.502616131 | 0.3919053 | -3.834131 | 0.000126009 | 0.0009392 | | processed_pseudogene | 43.35 | -4.548782587 | 4.548782587 | ENSG00000241462 | down |
| ENSG00000272602 | 613.1662536 | -1.216277065 | 0.2845368 | -4.274586 | 1.91E-05 | 0.000182239 | ZNF595 | protein_coding | 38.31 | -4.548097631 | 4.548097631 | ENSG00000272602 | down |
| ENSG00000272533 | 100.6271449 | 1.096263992 | 0.2432659 | 4.506444 | 6.59E-06 | 7.27E-05 | SNORA28 | snoRNA | 41.27 | 4.536656916 | 4.536656916 | ENSG00000272533 | up |
| ENSG00000167106 | 77.52982672 | 1.137868273 | 0.2574657 | 4.419494 | 9.89E-06 | 0.000103489 | FAM102A | protein_coding | 55.6 | 4.534526747 | 4.534526747 | ENSG00000167106 | up |
| ENSG00000233560 | 135.4716886 | -1.288900681 | 0.3113514 | -4.139698 | 3.48E-05 | 0.000305483 | KRT8P39 | processed_pseudogene | 55.13 | -4.530501776 | 4.530501776 | ENSG00000233560 | down |
| ENSG00000268375 | 21.23804743 | -1.337215565 | 0.32913 | -4.06288 | 4.85E-05 | 0.000410749 | | antisense | 49.56 | -4.528377544 | 4.528377544 | ENSG00000268375 | down |
| ENSG00000145685 | 259.5191545 | -1.140392819 | 0.2587576 | -4.407186 | 1.05E-05 | 0.000108809 | LHFPL2 | protein_coding | 43.31 | -4.519756704 | 4.519756704 | ENSG00000145685 | down |
| ENSG00000086102 | 1610.546778 | -1.141861405 | 0.2593663 | -4.402504 | 1.07E-05 | 0.000110772 | NFX1 | protein_coding | 42.57 | -4.516714014 | 4.516714014 | ENSG00000086102 | down |
| ENSG00000281469 | 24.89694497 | -1.353258849 | 0.3358452 | -4.029413 | 5.59E-05 | 0.000462856 | | TEC | 29.71 | -4.512514903 | 4.512514903 | ENSG00000281469 | down |
| ENSG00000261071 | 94.51263768 | 1.106153242 | 0.2473829 | 4.471422 | 7.77E-06 | 8.39E-05 | | antisense | 54.16 | 4.508782786 | 4.508782786 | ENSG00000261071 | up |
| ENSG00000250568 | 126.8646404 | 1.066423715 | 0.2340748 | 4.555909 | 5.22E-06 | 5.94E-05 | | processed_pseudogene | 46.98 | 4.506711778 | 4.506711778 | ENSG00000250568 | up |
| ENSG00000272267 | 37.94971163 | 1.222316834 | 0.2881218 | 4.242362 | 2.21E-05 | 0.00020571 | | antisense | 55.56 | 4.506369288 | 4.506369288 | ENSG00000272267 | up |
| ENSG00000133639 | 3984.676329 | 0.991329728 | 0.2099697 | 4.7213 | 2.34E-06 | 2.93E-05 | BTG1 | protein_coding | 42.46 | 4.493223075 | 4.493223075 | ENSG00000133639 | n.s. |
| ENSG00000265083 | 2.701850503 | -2.685684917 | 0.9554185 | -2.811004 | 0.004938722 | 0.021334729 | MIR3691 | miRNA | 57.78 | -4.487545502 | 4.487545502 | ENSG00000265083 | down |
| ENSG00000111254 | 14.13207506 | 1.485481735 | 0.3881112 | 3.827464 | 0.00012947 | 0.000961906 | AKAP3 | protein_coding | 38.99 | 4.481501285 | 4.481501285 | ENSG00000111254 | up |
| ENSG00000171161 | 329.6759162 | -1.185111793 | 0.2757391 | -4.297947 | 1.72E-05 | 0.000166101 | ZNF672 | protein_coding | 52.37 | -4.479282532 | 4.479282532 | ENSG00000171161 | down |
| ENSG00000275632 | 70.67398898 | -1.232004201 | 0.2926804 | -4.209384 | 2.56E-05 | 0.000233479 | | lincRNA | 40.65 | -4.474333036 | 4.474333036 | ENSG00000275632 | down |
| ENSG00000197841 | 486.6092333 | -1.126482133 | 0.2556698 | -4.406004 | 1.05E-05 | 0.000109242 | ZNF181 | protein_coding | 39.74 | -4.462684142 | 4.462684142 | ENSG00000197841 | down |
| ENSG00000149476 | 198.4129185 | 1.029151711 | 0.2230364 | 4.614278 | 3.94E-06 | 4.65E-05 | TKFC | protein_coding | 54.64 | 4.458809048 | 4.458809048 | ENSG00000149476 | up |
| ENSG00000266887 | 1.839162996 | -3.648621137 | 1.5377759 | -2.372661 | 0.017660456 | 0.060205966 | MIR4535 | miRNA | 55.93 | -4.452633003 | 4.452633003 | ENSG00000266887 | down |
| ENSG00000167380 | 869.7865898 | -1.040518114 | 0.2270155 | -4.583467 | 4.57E-06 | 5.29E-05 | ZNF226 | protein_coding | 40.39 | -4.449571382 | 4.449571382 | ENSG00000167380 | down |
| ENSG00000149639 | 786.3132004 | -1.097090583 | 0.2465006 | -4.450661 | 8.56E-06 | 9.13E-05 | SOGA1 | protein_coding | 51.57 | -4.431556509 | 4.431556509 | ENSG00000149639 | down |
| ENSG00000196922 | 723.8175744 | -1.20192317 | 0.2834066 | -4.240985 | 2.23E-05 | 0.000206701 | ZNF252P | transcribed_unitary_pseudogene | 41.65 | -4.428675746 | 4.428675746 | ENSG00000196922 | down |
| ENSG00000115884 | 373.0587236 | -1.125285166 | 0.2565044 | -4.387001 | 1.15E-05 | 0.000117309 | SDC1 | protein_coding | 57.56 | -4.423124327 | 4.423124327 | ENSG00000115884 | down |
| ENSG00000207637 | 5.2411198 | -1.917894385 | 0.5753384 | -3.333507 | 0.000857587 | 0.004946222 | MIR595 | miRNA | 53.12 | -4.422139758 | 4.422139758 | ENSG00000207637 | down |
| ENSG00000147124 | 253.6077657 | -1.201977009 | 0.2836692 | -4.237249 | 2.26E-05 | 0.00020961 | ZNF41 | protein_coding | 43.28 | -4.421578783 | 4.421578783 | ENSG00000147124 | down |
| ENSG00000233060 | 34.62778311 | 1.296170675 | 0.3179475 | 4.076682 | 4.57E-05 | 0.000389256 | | lincRNA | 46.88 | 4.419637637 | 4.419637637 | ENSG00000233060 | up |
| ENSG00000188399 | 56.43259778 | -1.294775341 | 0.3175087 | -4.077921 | 4.54E-05 | 0.000388262 | ANKRD36P1 | unprocessed_pseudogene | 34.53 | -4.416316719 | 4.416316719 | ENSG00000188399 | down |
| ENSG00000155090 | 1022.52029 | 1.126264778 | 0.2571019 | 4.380616 | 1.18E-05 | 0.000120358 | KLF10 | protein_coding | 48.47 | 4.414421143 | 4.414421143 | ENSG00000155090 | up |

| | | | | | | | | | | | | | |
|---|---|---|---|---|---|---|---|---|---|---|---|---|---|
| ENSG00000119965 | 351.1456043 | -1.193058224 | 0.2806475 | -4.251091 | 2.13E-05 | 0.000199626 | C10orf88 | protein_coding | 36.24 | -4.414055569 | 4.414055569 | ENSG00000119965 | down |
| ENSG00000133624 | 383.6507417 | -1.125628871 | 0.2568738 | -4.38203 | 1.18E-05 | 0.000119842 | ZNF767P | transcribed_unprocessed_pseud | 46.36 | -4.414028987 | 4.414028987 | ENSG00000133624 | down |
| ENSG00000259663 | 640.1431404 | -1.04635222 | 0.2299801 | -4.549751 | 5.37E-06 | 6.09E-05 | | antisense | 41.79 | -4.410652348 | 4.410652348 | ENSG00000259663 | down |
| ENSG00000177606 | 2056.877255 | 0.928382314 | 0.1920788 | 4.83334 | 1.34E-06 | 1.78E-05 | JUN | protein_coding | 56.67 | 4.409356735 | 4.409356735 | ENSG00000177606 | n.s. |
| ENSG00000140471 | 869.254997 | -1.179211408 | 0.2759094 | -4.273908 | 1.92E-05 | 0.000182669 | LINS1 | protein_coding | 40.18 | -4.408286471 | 4.408286471 | ENSG00000140471 | down |
| ENSG00000186104 | 213.7605193 | -1.165479622 | 0.2711377 | -4.298478 | 1.72E-05 | 0.000165817 | CYP2R1 | protein_coding | 39.86 | -4.405944225 | 4.405944225 | ENSG00000186104 | down |
| ENSG00000173041 | 866.1025073 | -1.15302833 | 0.2667629 | -4.322297 | 1.54E-05 | 0.000151508 | ZNF680 | protein_coding | 38.98 | -4.40406756 | 4.40406756 | ENSG00000173041 | down |
| ENSG00000243319 | 153.6068414 | -1.153884674 | 0.2672092 | -4.318283 | 1.57E-05 | 0.000153747 | FGF14-IT1 | sense_intronic | 37.7 | -4.399983852 | 4.399983852 | ENSG00000243319 | down |
| ENSG00000240463 | 24.91125492 | -1.421067248 | 0.3668633 | -3.87356 | 0.000107257 | 0.000812037 | RPS19P3 | processed_pseudogene | 56.24 | -4.391700576 | 4.391700576 | ENSG00000240463 | down |
| ENSG00000115902 | 619.2489541 | -1.108384404 | 0.2515372 | -4.406443 | 1.05E-05 | 0.000109102 | SLC1A4 | protein_coding | 45.52 | -4.391605295 | 4.391605295 | ENSG00000115902 | down |
| ENSG00000169951 | 185.6024553 | -1.01894976 | 0.2215138 | -4.599938 | 4.23E-06 | 4.93E-05 | ZNF764 | protein_coding | 58.06 | -4.388577196 | 4.388577196 | ENSG00000169951 | down |
| ENSG00000235786 | 98.90819339 | -1.186094858 | 0.2792788 | -4.246992 | 2.17E-05 | 0.000202318 | ZNRF3-IT1 | sense_intronic | 45.22 | -4.381394346 | 4.381394346 | ENSG00000235786 | down |
| ENSG00000198853 | 329.0719405 | -1.051613275 | 0.2326636 | -4.519888 | 6.19E-06 | 6.87E-05 | RUSC2 | protein_coding | 44.22 | -4.377937777 | 4.377937777 | ENSG00000198853 | down |
| ENSG00000111145 | 379.2618587 | -1.111508853 | 0.2530221 | -4.392931 | 1.12E-05 | 0.000115337 | ELK3 | protein_coding | 45.11 | -4.377158001 | 4.377158001 | ENSG00000111145 | down |
| ENSG00000170915 | 251.6811108 | -1.065685121 | 0.2374017 | -4.488953 | 7.16E-06 | 7.82E-05 | PAQR8 | protein_coding | 42.53 | -4.37679192 | 4.37679192 | ENSG00000170915 | down |
| ENSG00000199899 | 3.942939184 | -1.992640479 | 0.6134897 | -3.248042 | 0.001162021 | 0.006378041 | | misc_RNA | 37.37 | -4.37446897 | 4.37446897 | ENSG00000199899 | down |
| ENSG00000251072 | 36.61296061 | 1.215761578 | 0.290236 | 4.188872 | 2.80E-05 | 0.000253292 | | lincRNA | 44.31 | 4.372339125 | 4.372339125 | ENSG00000251072 | up |
| ENSG00000264402 | 10.83388252 | 1.608407102 | 0.4433964 | 3.62747 | 0.000286212 | 0.001916756 | MIR548AL | miRNA | 32.99 | 4.370738924 | 4.370738924 | ENSG00000264402 | up |
| ENSG00000266236 | 39.46701914 | -1.23499099 | 0.2973361 | -4.153518 | 3.27E-05 | 0.000289213 | NARF-IT1 | sense_intronic | 49.03 | -4.37036484 | 4.37036484 | ENSG00000266236 | down |
| ENSG00000172216 | 236.4983419 | 1.153356033 | 0.2679908 | 4.303716 | 1.68E-05 | 0.000162959 | CEBPB | protein_coding | 66.16 | 4.368820985 | 4.368820985 | ENSG00000172216 | up |
| ENSG00000222790 | 5.007734727 | -2.047536981 | 0.63974 | -3.200577 | 0.001371528 | 0.007353664 | RNU4-14P | snRNA | 37.7 | -4.368412359 | 4.368412359 | ENSG00000222790 | down |
| ENSG00000256944 | 10.00626546 | 1.636025664 | 0.4551032 | 3.594846 | 0.000324584 | 0.002141791 | | antisense | 48.7 | 4.366917061 | 4.366917061 | ENSG00000256944 | up |
| ENSG00000252494 | 24.83355031 | -1.353076141 | 0.3416929 | -3.959919 | 7.50E-05 | 0.000592777 | RNU6-126P | snRNA | 36.89 | -4.366523802 | 4.366523802 | ENSG00000252494 | down |
| ENSG00000265185 | 645.7145974 | 0.953116441 | 0.2008621 | 4.745129 | 2.08E-06 | 2.63E-05 | SNORD3B-1 | snoRNA | 55.28 | 4.364897392 | 4.364897392 | ENSG00000265185 | n.s. |
| ENSG00000272969 | 32.6493241 | 1.291525316 | 0.3182725 | 4.057923 | 4.95E-05 | 0.000418545 | | antisense | 50.77 | 4.36310594 | 4.36310594 | ENSG00000272969 | up |
| ENSG00000261560 | 95.70440032 | -1.201628508 | 0.2857298 | -4.205471 | 2.61E-05 | 0.000236935 | | sense_intronic | 43.21 | -4.35634808 | 4.35634808 | ENSG00000261560 | down |
| ENSG00000206859 | 6.982532874 | -1.756842407 | 0.5077617 | -3.459974 | 0.000540227 | 0.00331932 | RNU6-767P | snRNA | 33.64 | -4.355126117 | 4.355126117 | ENSG00000206859 | down |
| ENSG00000270993 | 3.838560243 | 2.385055159 | 0.8092601 | 2.947205 | 0.00320661 | 0.014936643 | | processed_pseudogene | 60 | 4.354507296 | 4.354507296 | ENSG00000270993 | up |
| ENSG00000103647 | 254.1324569 | -1.08728336 | 0.2454362 | -4.430004 | 9.42E-06 | 9.94E-05 | CORO2B | protein_coding | 49.34 | -4.35183547 | 4.35183547 | ENSG00000103647 | down |
| ENSG00000233203 | 25.46294051 | 1.292965362 | 0.3193691 | 4.048499 | 5.15E-05 | 0.000432084 | | antisense | 47.36 | 4.350093381 | 4.350093381 | ENSG00000233203 | up |
| ENSG00000202270 | 4.770020785 | -2.120954958 | 0.6766872 | -3.134321 | 0.001722522 | 0.008968837 | SNORD114-12 | snoRNA | 44 | -4.342154501 | 4.342154501 | ENSG00000202270 | down |
| ENSG00000206622 | 16.61317749 | 1.47719772 | 0.3914613 | 3.773547 | 0.000160943 | 0.001163451 | SNORA69 | snoRNA | 36.36 | 4.334469983 | 4.334469983 | ENSG00000206622 | up |

| | | | | | | | | | | | | | |
|---|---|---|---|---|---|---|---|---|---|---|---|---|---|
| ENSG00000276213 | 8.108283601 | 1.707702796 | 0.4886905 | 3.494446 | 0.000475046 | 0.002972616 | | misc_RNA | 66.31 | 4.315127917 | 4.315127917 | ENSG00000276213 | up |
| ENSG00000200879 | 116.0934091 | 1.05045705 | 0.2339801 | 4.489515 | 7.14E-06 | 7.80E-05 | SNORD14E | snoRNA | 44.71 | 4.315095754 | 4.315095754 | ENSG00000200879 | up |
| ENSG00000233230 | 60.62241349 | 1.150964653 | 0.2690377 | 4.27808 | 1.89E-05 | 0.000179525 | | antisense | 56.26 | 4.311370072 | 4.311370072 | ENSG00000233230 | up |
| ENSG00000265800 | 117.8722984 | -1.09531327 | 0.2495658 | -4.388875 | 1.14E-05 | 0.000116934 | | sense_intronic | 43.68 | -4.306836243 | 4.306836243 | ENSG00000265800 | down |
| ENSG00000168813 | 1232.386784 | -1.090151975 | 0.2477996 | -4.399328 | 1.09E-05 | 0.000112322 | ZNF507 | protein_coding | 39.82 | -4.305595406 | 4.305595406 | ENSG00000168813 | down |
| ENSG00000246379 | 154.5230821 | 1.00223356 | 0.2183162 | 4.590742 | 4.42E-06 | 5.12E-05 | | lincRNA | 38.46 | 4.299930707 | 4.299930707 | ENSG00000246379 | up |
| ENSG00000173320 | 857.9552171 | -1.089545623 | 0.2478177 | -4.396561 | 1.10E-05 | 0.000113594 | STOX2 | protein_coding | 42.71 | -4.297871693 | 4.297871693 | ENSG00000173320 | down |
| ENSG00000200156 | 569.478352 | 0.935458316 | 0.1968635 | 4.751811 | 2.02E-06 | 2.56E-05 | RNU5B-1 | snRNA | 39.66 | 4.295604236 | 4.295604236 | ENSG00000200156 | n.s. |
| ENSG00000060709 | 700.5522407 | -1.107112606 | 0.2542143 | -4.355037 | 1.33E-05 | 0.000133361 | RIMBP2 | protein_coding | 50.38 | -4.290031084 | 4.290031084 | ENSG00000060709 | down |
| ENSG00000280904 | 5.868511481 | 1.859712929 | 0.5579141 | 3.333332 | 0.000858124 | 0.004947271 | | snoRNA | 43.75 | 4.287817951 | 4.287817951 | ENSG00000280904 | up |
| ENSG00000258789 | 26.48445014 | 1.332192189 | 0.3370052 | 3.953031 | 7.72E-05 | 0.000607693 | | antisense | 54.07 | 4.284751113 | 4.284751113 | ENSG00000258789 | up |
| ENSG00000213600 | 15.42105421 | 1.428083405 | 0.3742948 | 3.815397 | 0.000135964 | 0.001002609 | | sense_intronic | 57.49 | 4.282634489 | 4.282634489 | ENSG00000213600 | up |
| ENSG00000242540 | 112.8752958 | -1.145892772 | 0.2682212 | -4.272193 | 1.94E-05 | 0.000183704 | | 3prime_overlapping_ncRNA | 46.07 | -4.280919289 | 4.280919289 | ENSG00000242540 | down |
| ENSG00000276744 | 98.58931294 | 1.037109754 | 0.2307692 | 4.494142 | 6.99E-06 | 7.66E-05 | | sense_intronic | 57.72 | 4.268639246 | 4.268639246 | ENSG00000276744 | up |
| ENSG00000242182 | 15.40928706 | -1.441047034 | 0.3801462 | -3.790771 | 0.000150181 | 0.001095166 | RN7SL745P | misc_RNA | 50.84 | -4.266248802 | 4.266248802 | ENSG00000242182 | down |
| ENSG00000143457 | 275.2541018 | -1.174961827 | 0.2792009 | -4.208303 | 2.57E-05 | 0.000234139 | GOLPH3L | protein_coding | 40.09 | -4.265728813 | 4.265728813 | ENSG00000143457 | down |
| ENSG00000229419 | 325.709752 | -1.129397789 | 0.2630032 | -4.294237 | 1.75E-05 | 0.00016832 | RALGAPA1P1 | processed_pseudogene | 40.13 | -4.262194145 | 4.262194145 | ENSG00000229419 | down |
| ENSG00000121931 | 556.2855292 | -1.186738253 | 0.2835692 | -4.185004 | 2.85E-05 | 0.00025731 | LRIF1 | protein_coding | 36.58 | -4.259847573 | 4.259847573 | ENSG00000121931 | down |
| ENSG00000102780 | 1266.655024 | -1.183121674 | 0.2824636 | -4.18858 | 2.81E-05 | 0.000253454 | DGKH | protein_coding | 38.24 | -4.254625578 | 4.254625578 | ENSG00000102780 | down |
| ENSG00000146247 | 7157.283858 | -1.139455742 | 0.2669488 | -4.268443 | 1.97E-05 | 0.000186311 | PHIP | protein_coding | 34.29 | -4.249898226 | 4.249898226 | ENSG00000146247 | down |
| ENSG00000280365 | 54.26001935 | -1.172308217 | 0.2788761 | -4.203689 | 2.63E-05 | 0.000238343 | | TEC | 46.56 | -4.247035989 | 4.247035989 | ENSG00000280365 | down |
| ENSG00000180257 | 291.9074617 | -1.209658614 | 0.2925572 | -4.134776 | 3.55E-05 | 0.000311515 | ZNF816 | protein_coding | 43.93 | -4.241693178 | 4.241693178 | ENSG00000180257 | down |
| ENSG00000165244 | 966.9143564 | -1.129580882 | 0.2637775 | -4.282325 | 1.85E-05 | 0.000176738 | ZNF367 | protein_coding | 42.86 | -4.238944473 | 4.238944473 | ENSG00000165244 | down |
| ENSG00000130749 | 547.3334701 | -1.007880264 | 0.2218756 | -4.542546 | 5.56E-06 | 6.28E-05 | ZC3H4 | protein_coding | 51.95 | -4.235436359 | 4.235436359 | ENSG00000130749 | down |
| ENSG00000279702 | 19.49518785 | -1.918799524 | 0.5892255 | -3.256477 | 0.00112804 | 0.006217615 | | TEC | 32.92 | -4.233593533 | 4.233593533 | ENSG00000279702 | down |
| ENSG00000213793 | 178.1436271 | -1.161160335 | 0.2752986 | -4.217821 | 2.47E-05 | 0.000226099 | ZNF888 | protein_coding | 45.68 | -4.233243093 | 4.233243093 | ENSG00000213793 | down |
| ENSG00000204172 | 108.603303 | -1.075747133 | 0.2451787 | -4.387604 | 1.15E-05 | 0.000117173 | AGAP9 | protein_coding | 40.48 | -4.228946146 | 4.228946146 | ENSG00000204172 | down |
| ENSG00000231856 | 190.9373072 | 1.016994545 | 0.225178 | 4.516402 | 6.29E-06 | 6.96E-05 | | antisense | 47.93 | 4.228306072 | 4.228306072 | ENSG00000231856 | up |
| ENSG00000222259 | 22.19865898 | 1.297295724 | 0.3258929 | 3.980742 | 6.87E-05 | 0.000551624 | RN7SKP114 | misc_RNA | 54.12 | 4.227052271 | 4.227052271 | ENSG00000222259 | up |
| ENSG00000271155 | 81.72671537 | 1.074158558 | 0.2447723 | 4.388399 | 1.14E-05 | 0.000116986 | | antisense | 62.05 | 4.223447713 | 4.223447713 | ENSG00000271155 | up |
| ENSG00000144426 | 484.3617034 | -1.229836919 | 0.3006073 | -4.091174 | 4.29E-05 | 0.000368414 | NBEAL1 | protein_coding | 37.7 | -4.222846775 | 4.222846775 | ENSG00000144426 | down |
| ENSG00000185085 | 202.1292687 | -1.070602669 | 0.2435563 | -4.395709 | 1.10E-05 | 0.000113956 | INTS5 | protein_coding | 54.61 | -4.221668294 | 4.221668294 | ENSG00000185085 | down |

| | | | | | | | | | | | | | |
|---|---|---|---|---|---|---|---|---|---|---|---|---|---|
| ENSG00000182022 | 260.6091833 | -1.099563971 | 0.2536937 | -4.334218 | 1.46E-05 | 0.000144854 | CHST15 | protein_coding | 50.24 | -4.221303519 | 4.221303519 | ENSG00000182022 | down |
| ENSG00000284428 | 580.2907886 | -0.976108331 | 0.2118413 | -4.607734 | 4.07E-06 | 4.77E-05 | | processed_transcript | 45.13 | -4.217872881 | 4.217872881 | ENSG00000284428 | n.s. |
| ENSG00000274093 | 32.46154203 | -1.270651185 | 0.3162716 | -4.017595 | 5.88E-05 | 0.000481533 | | sense_intronic | 31.79 | -4.215225539 | 4.215225539 | ENSG00000274093 | down |
| ENSG00000254996 | 720.6002779 | -1.113306219 | 0.2588228 | -4.301422 | 1.70E-05 | 0.000164426 | ANKHD1-EIF4EBP3 | protein_coding | 38.97 | -4.212782658 | 4.212782658 | ENSG00000254996 | down |
| ENSG00000272746 | 22.83034264 | -1.413201791 | 0.3716648 | -3.802356 | 0.000143327 | 0.001051389 | | sense_intronic | 38.95 | -4.208849088 | 4.208849088 | ENSG00000272746 | down |
| ENSG00000278876 | 46.02168807 | -1.298193328 | 0.3274041 | -3.96511 | 7.34E-05 | 0.000581344 | | TEC | 45.85 | -4.200391349 | 4.200391349 | ENSG00000278876 | down |
| ENSG00000105778 | 987.7351786 | -1.173389212 | 0.280832 | -4.178261 | 2.94E-05 | 0.000264031 | AVL9 | protein_coding | 40.32 | -4.198792348 | 4.198792348 | ENSG00000105778 | down |
| ENSG00000217512 | 124.1463092 | -1.145976679 | 0.2710467 | -4.227968 | 2.36E-05 | 0.000217575 | | processed_pseudogene | 36.91 | -4.197014911 | 4.197014911 | ENSG00000217512 | down |
| ENSG00000135870 | 1668.736945 | -1.072984663 | 0.245441 | -4.37166 | 1.23E-05 | 0.000124984 | RC3H1 | protein_coding | 38.58 | -4.188015727 | 4.188015727 | ENSG00000135870 | down |
| ENSG00000135482 | 517.7138644 | -0.951003669 | 0.2044914 | -4.650581 | 3.31E-06 | 3.98E-05 | ZC3H10 | protein_coding | 48.69 | -4.184911255 | 4.184911255 | ENSG00000135482 | n.s. |
| ENSG00000157259 | 662.0326102 | -1.110195931 | 0.2586952 | -4.291521 | 1.77E-05 | 0.000170158 | GATAD1 | protein_coding | 42.23 | -4.184493579 | 4.184493579 | ENSG00000157259 | down |
| ENSG00000137496 | 145.4472563 | -1.044368593 | 0.2357531 | -4.429924 | 9.43E-06 | 9.94E-05 | IL18BP | protein_coding | 57.03 | -4.180069756 | 4.180069756 | ENSG00000137496 | down |
| ENSG00000274124 | 10.66550783 | -1.518048912 | 0.4157525 | -3.651328 | 0.000260887 | 0.001769548 | | antisense | 43.27 | -4.177879712 | 4.177879712 | ENSG00000274124 | down |
| ENSG00000213799 | 432.3341169 | -1.143740376 | 0.2710353 | -4.219894 | 2.44E-05 | 0.000224474 | ZNF845 | protein_coding | 44.17 | -4.173319532 | 4.173319532 | ENSG00000213799 | down |
| ENSG00000166398 | 410.9842474 | -1.086508062 | 0.2507324 | -4.333338 | 1.47E-05 | 0.000145125 | KIAA0355 | protein_coding | 44.95 | -4.170298664 | 4.170298664 | ENSG00000166398 | down |
| ENSG00000199030 | 65.24885217 | 1.125128403 | 0.2646216 | 4.251838 | 2.12E-05 | 0.000199325 | MIRLET7C | miRNA | 47.62 | 4.16346915 | 4.16346915 | ENSG00000199030 | up |
| ENSG00000056586 | 1601.144862 | -1.124752514 | 0.2646258 | -4.25035 | 2.13E-05 | 0.000199844 | RC3H2 | protein_coding | 40.5 | -4.160807925 | 4.160807925 | ENSG00000056586 | down |
| ENSG00000171649 | 1062.120847 | -1.113341324 | 0.260594 | -4.272322 | 1.93E-05 | 0.000183704 | ZIK1 | protein_coding | 47.45 | -4.159310946 | 4.159310946 | ENSG00000171649 | down |
| ENSG00000188419 | 389.0645073 | -1.142437442 | 0.271443 | -4.208756 | 2.57E-05 | 0.000233823 | CHM | protein_coding | 36.96 | -4.148318294 | 4.148318294 | ENSG00000188419 | down |
| ENSG00000162695 | 1265.405593 | -1.106794156 | 0.2586201 | -4.279613 | 1.87E-05 | 0.00017866 | SLC30A7 | protein_coding | 35.85 | -4.148235495 | 4.148235495 | ENSG00000162695 | down |
| ENSG00000164066 | 751.7434462 | -1.164160894 | 0.2793329 | -4.167646 | 3.08E-05 | 0.000275029 | INTU | protein_coding | 37.11 | -4.145135812 | 4.145135812 | ENSG00000164066 | down |
| ENSG00000257702 | 119.2228052 | -1.118598375 | 0.2631208 | -4.251273 | 2.13E-05 | 0.000199626 | LBX2-AS1 | antisense | 60.7 | -4.138570345 | 4.138570345 | ENSG00000257702 | down |
| ENSG00000221886 | 294.6308663 | -0.959302292 | 0.2084307 | -4.602499 | 4.17E-06 | 4.88E-05 | ZBED8 | protein_coding | 40 | -4.136102315 | 4.136102315 | ENSG00000221886 | n.s. |
| ENSG00000150510 | 299.0931994 | -1.014865549 | 0.2270136 | -4.470505 | 7.80E-06 | 8.42E-05 | FAM124A | protein_coding | 44.27 | -4.135137868 | 4.135137868 | ENSG00000150510 | down |
| ENSG00000222276 | 7.46616611 | -1.633861106 | 0.4675262 | -3.494695 | 0.000474605 | 0.002971187 | RNU2-33P | snRNA | 38.02 | -4.128881331 | 4.128881331 | ENSG00000222276 | down |
| ENSG00000189180 | 1724.835463 | -1.13668106 | 0.2700642 | -4.208929 | 2.57E-05 | 0.000233797 | ZNF33A | protein_coding | 39.26 | -4.127472809 | 4.127472809 | ENSG00000189180 | down |
| ENSG00000221520 | 11.08234544 | -1.549125051 | 0.4314411 | -3.590583 | 0.00032994 | 0.002168909 | MIR1285-1 | miRNA | 47.62 | -4.126495231 | 4.126495231 | ENSG00000221520 | down |
| ENSG00000263883 | 77.44664849 | 1.17205449 | 0.2829024 | 4.142964 | 3.43E-05 | 0.000301735 | EEF1DP7 | transcribed_processed_pseudog | 55.75 | 4.126070149 | 4.126070149 | ENSG00000263883 | up |
| ENSG00000252427 | 8.359530311 | -1.599119399 | 0.4526161 | -3.533059 | 0.000410781 | 0.002630775 | | snoRNA | 45.37 | -4.125594173 | 4.125594173 | ENSG00000252427 | down |
| ENSG00000186566 | 2192.664773 | -1.117660359 | 0.2633081 | -4.244687 | 2.19E-05 | 0.000203998 | GPATCH8 | protein_coding | 40.95 | -4.124585645 | 4.124585645 | ENSG00000186566 | down |
| ENSG00000115993 | 975.0594164 | -1.134764745 | 0.2694912 | -4.210767 | 2.55E-05 | 0.000232511 | TRAK2 | protein_coding | 37.73 | -4.123231472 | 4.123231472 | ENSG00000115993 | down |
| ENSG00000241549 | 91.59084693 | -1.186367499 | 0.2884721 | -4.112591 | 3.91E-05 | 0.000337981 | GUSBP2 | transcribed_unprocessed_pseud | 39.56 | -4.118009054 | 4.118009054 | ENSG00000241549 | down |

| | | | | | | | | | | | | | |
|---|---|---|---|---|---|---|---|---|---|---|---|---|---|
| ENSG00000206992 | 28.39801499 | 1.280171867 | 0.3238609 | 3.952844 | 7.72E-05 | 0.000607825 | RNU6-574P | snRNA | 42.99 | 4.117316646 | 4.117316646 | ENSG00000206992 | up |
| ENSG00000261529 | 126.1184167 | -1.08766287 | 0.2530154 | -4.298801 | 1.72E-05 | 0.000165691 | | sense_intronic | 37.99 | -4.112127347 | 4.112127347 | ENSG00000261529 | down |
| ENSG00000164506 | 514.4498 | -1.108288812 | 0.2603246 | -4.257334 | 2.07E-05 | 0.000195016 | STXBP5 | protein_coding | 34.37 | -4.111674266 | 4.111674266 | ENSG00000164506 | down |
| ENSG00000200571 | 8.82391111 | -1.664758211 | 0.4822491 | -3.452071 | 0.000556301 | 0.003406059 | RNU6-1284P | snRNA | 38 | -4.1082034 | 4.1082034 | ENSG00000200571 | down |
| ENSG00000201801 | 91.90372796 | 1.088533185 | 0.2537564 | 4.289677 | 1.79E-05 | 0.000171458 | RNU5E-4P | snRNA | 33.33 | 4.099244378 | 4.099244378 | ENSG00000201801 | up |
| ENSG00000265713 | 52.17684182 | -1.122421469 | 0.2660532 | -4.218786 | 2.46E-05 | 0.000225431 | | transcribed_processed_pseudog | 42.83 | -4.093456862 | 4.093456862 | ENSG00000265713 | down |
| ENSG00000253730 | 25.93725161 | 1.291356995 | 0.3293396 | 3.921049 | 8.82E-05 | 0.000684231 | | antisense | 42.46 | 4.086883029 | 4.086883029 | ENSG00000253730 | up |
| ENSG00000207996 | 92.86271736 | 1.072066374 | 0.2484171 | 4.315589 | 1.59E-05 | 0.000155382 | MIR301A | miRNA | 40.7 | 4.083071922 | 4.083071922 | ENSG00000207996 | up |
| ENSG00000128607 | 772.7787054 | -1.181471447 | 0.2879276 | -4.103363 | 4.07E-05 | 0.000350821 | KLHDC10 | protein_coding | 40.14 | -4.081883018 | 4.081883018 | ENSG00000128607 | down |
| ENSG00000273888 | 278.7856059 | 0.958268225 | 0.2094779 | 4.574554 | 4.77E-06 | 5.50E-05 | FRMD6-AS1 | antisense | 52.67 | 4.081841342 | 4.081841342 | ENSG00000273888 | n.s. |
| ENSG00000154222 | 101.4788656 | -1.211726391 | 0.2991622 | -4.0504 | 5.11E-05 | 0.000429364 | CC2D1B | protein_coding | 51.22 | -4.080094196 | 4.080094196 | ENSG00000154222 | down |
| ENSG00000132846 | 272.4608766 | -1.106840922 | 0.2610128 | -4.240563 | 2.23E-05 | 0.000206904 | ZBED3 | protein_coding | 43.7 | -4.077858426 | 4.077858426 | ENSG00000132846 | down |
| ENSG00000166025 | 1698.233364 | -1.017530721 | 0.2296475 | -4.430838 | 9.39E-06 | 9.92E-05 | AMOTL1 | protein_coding | 41.79 | -4.073854313 | 4.073854313 | ENSG00000166025 | down |
| ENSG00000236778 | 199.6440594 | 0.996056319 | 0.2224439 | 4.477786 | 7.54E-06 | 8.17E-05 | INTS6-AS1 | antisense | 40.22 | 4.071557415 | 4.071557415 | ENSG00000236778 | n.s. |
| ENSG00000122483 | 780.9494813 | -1.118665322 | 0.2654585 | -4.214088 | 2.51E-05 | 0.000229418 | CCDC18 | protein_coding | 36.98 | -4.071239153 | 4.071239153 | ENSG00000122483 | down |
| ENSG00000235319 | 27.66853057 | 1.323673824 | 0.3425737 | 3.86391 | 0.000111586 | 0.000841158 | | lincRNA | 48.96 | 4.07045939 | 4.07045939 | ENSG00000235319 | up |
| ENSG00000231327 | 52.48862558 | 1.113771764 | 0.263792 | 4.222159 | 2.42E-05 | 0.000222523 | LINC01816 | lincRNA | 57.49 | 4.068190367 | 4.068190367 | ENSG00000231327 | up |
| ENSG00000167232 | 1891.026939 | -1.096957846 | 0.2580512 | -4.250931 | 2.13E-05 | 0.000199626 | ZNF91 | protein_coding | 40.56 | -4.058505101 | 4.058505101 | ENSG00000167232 | down |
| ENSG00000280165 | 173.0616163 | 1.014381991 | 0.2291659 | 4.426409 | 9.58E-06 | 0.000100758 | PCDH20 | protein_coding | 44.3 | 4.054202831 | 4.054202831 | ENSG00000280165 | up |
| ENSG00000279659 | 80.3397868 | -1.155516976 | 0.2795948 | -4.132827 | 3.58E-05 | 0.000313459 | | TEC | 36.03 | -4.048721895 | 4.048721895 | ENSG00000279659 | down |
| ENSG00000175414 | 597.6163364 | -1.146361479 | 0.2763768 | -4.14782 | 3.36E-05 | 0.000295781 | ARL10 | protein_coding | 50.75 | -4.045544559 | 4.045544559 | ENSG00000175414 | down |
| ENSG00000246308 | 83.64524035 | 1.106482751 | 0.2620772 | 4.221972 | 2.42E-05 | 0.00022256 | | antisense | 39.66 | 4.041485866 | 4.041485866 | ENSG00000246308 | up |
| ENSG00000137764 | 190.2216544 | -1.057629934 | 0.2448795 | -4.318981 | 1.57E-05 | 0.000153369 | MAP2K5 | protein_coding | 39.55 | -4.034077155 | 4.034077155 | ENSG00000137764 | down |
| ENSG00000271040 | 18.08224076 | 1.335562729 | 0.3490898 | 3.825843 | 0.000130326 | 0.000965685 | | lincRNA | 52.72 | 4.026941195 | 4.026941195 | ENSG00000271040 | up |
| ENSG00000144026 | 897.289836 | -1.048550979 | 0.2420897 | -4.331249 | 1.48E-05 | 0.000146301 | ZNF514 | protein_coding | 46.72 | -4.020934392 | 4.020934392 | ENSG00000144026 | down |
| ENSG00000136010 | 1217.673264 | -1.061062708 | 0.2465872 | -4.302993 | 1.69E-05 | 0.000163378 | ALDH1L2 | protein_coding | 39.68 | -4.018037703 | 4.018037703 | ENSG00000136010 | down |
| ENSG00000272764 | 207.8889598 | 1.005410777 | 0.2272108 | 4.425012 | 9.64E-06 | 0.000101259 | | lincRNA | 46.83 | 4.016181993 | 4.016181993 | ENSG00000272764 | up |
| ENSG00000108510 | 5489.946741 | -1.110436893 | 0.264559 | -4.197312 | 2.70E-05 | 0.000244673 | MED13 | protein_coding | 38.73 | -4.010247178 | 4.010247178 | ENSG00000108510 | down |
| ENSG00000278987 | 122.8893666 | -1.075352165 | 0.2520003 | -4.267266 | 1.98E-05 | 0.000187169 | | TEC | 45 | -4.008661399 | 4.008661399 | ENSG00000278987 | down |
| ENSG00000178404 | 274.904188 | -0.99695373 | 0.2245359 | -4.440065 | 8.99E-06 | 9.54E-05 | CEP295NL | protein_coding | 53.83 | -4.008356819 | 4.008356819 | ENSG00000178404 | n.s. |
| ENSG00000167632 | 246.9613772 | -1.04726003 | 0.2421567 | -4.32472 | 1.53E-05 | 0.000149957 | TRAPPC9 | protein_coding | 47.51 | -4.004755741 | 4.004755741 | ENSG00000167632 | down |
| ENSG00000237406 | 68.14396945 | -1.138557884 | 0.2749535 | -4.140911 | 3.46E-05 | 0.000304256 | NDUFA9P1 | processed_pseudogene | 47.05 | -4.004035415 | 4.004035415 | ENSG00000237406 | down |

| | | | | | | | | | | | | | |
|---|---|---|---|---|---|---|---|---|---|---|---|---|---|
| ENSG00000201377 | 5.112008484 | -1.942031434 | 0.6175773 | -3.144596 | 0.001663162 | 0.008725019 | RNY4P23 | misc_RNA | 39.18 | -3.999096384 | 3.999096384 | ENSG00000201377 | down |
| ENSG00000200953 | 15.64820642 | -1.449542191 | 0.3965086 | -3.655765 | 0.000256416 | 0.001744908 | | misc_RNA | 38.18 | -3.998166931 | 3.998166931 | ENSG00000200953 | down |
| ENSG00000166135 | 826.5349971 | -1.04069102 | 0.2401449 | -4.333597 | 1.47E-05 | 0.000145067 | HIF1AN | protein_coding | 45.24 | -3.994621369 | 3.994621369 | ENSG00000166135 | down |
| ENSG00000207022 | 6.031848635 | -1.816298489 | 0.5587582 | -3.250598 | 0.001151625 | 0.006330074 | | snoRNA | 54.84 | -3.993297918 | 3.993297918 | ENSG00000207022 | down |
| ENSG00000261220 | 99.89220612 | 1.081557997 | 0.2547449 | 4.245651 | 2.18E-05 | 0.000203259 | | lincRNA | 46.54 | 3.993058988 | 3.993058988 | ENSG00000261220 | up |
| ENSG00000124164 | 1343.574727 | -1.126971975 | 0.2711501 | -4.156266 | 3.23E-05 | 0.000286512 | VAPB | protein_coding | 42.86 | -3.992700542 | 3.992700542 | ENSG00000124164 | down |
| ENSG00000201209 | 9.21364952 | -1.596973328 | 0.4596872 | -3.474043 | 0.000512678 | 0.003176687 | | snoRNA | 30.88 | -3.989280307 | 3.989280307 | ENSG00000201209 | down |
| ENSG00000129317 | 1705.456913 | -1.173128467 | 0.2883196 | -4.068847 | 4.72E-05 | 0.00040086 | PUS7L | protein_coding | 35.9 | -3.985126191 | 3.985126191 | ENSG00000129317 | down |
| ENSG00000199202 | 8.631196656 | -1.662933568 | 0.4894245 | -3.397732 | 0.000679468 | 0.004039385 | RNA5SP289 | rRNA | 47.45 | -3.980538788 | 3.980538788 | ENSG00000199202 | down |
| ENSG00000178440 | 36.08814503 | -1.755566362 | 0.5313841 | -3.303762 | 0.000953969 | 0.005432356 | LINC00843 | lincRNA | 43.38 | -3.976378423 | 3.976378423 | ENSG00000178440 | down |
| ENSG00000214770 | 230.995201 | 0.985292673 | 0.2215019 | 4.448236 | 8.66E-06 | 9.22E-05 | | antisense | 41.85 | 3.975787904 | 3.975787904 | ENSG00000214770 | n.s. |
| ENSG00000264966 | 3.538603779 | -1.934497074 | 0.6160008 | -3.140413 | 0.001687097 | 0.008820673 | MIR5094 | miRNA | 38.82 | -3.9744209 | 3.9744209 | ENSG00000264966 | down |
| ENSG00000197008 | 629.8401979 | -1.190355795 | 0.2952761 | -4.031332 | 5.55E-05 | 0.000460187 | ZNF138 | protein_coding | 39.99 | -3.972295946 | 3.972295946 | ENSG00000197008 | down |
| ENSG00000275591 | 231.0896629 | -1.188788805 | 0.2947516 | -4.033189 | 5.50E-05 | 0.000457381 | XKR5 | protein_coding | 49.65 | -3.970224176 | 3.970224176 | ENSG00000275591 | down |
| ENSG00000272081 | 15.07104181 | 1.455013311 | 0.4003775 | 3.634104 | 0.000278948 | 0.00187171 | | lincRNA | 54.66 | 3.968929069 | 3.968929069 | ENSG00000272081 | up |
| ENSG00000121542 | 317.3921543 | -1.236430845 | 0.3133313 | -3.946082 | 7.94E-05 | 0.000623829 | SEC22A | protein_coding | 37.72 | -3.962679825 | 3.962679825 | ENSG00000121542 | down |
| ENSG00000200632 | 3.089426752 | -2.573382422 | 0.9573907 | -2.687912 | 0.007190026 | 0.028991233 | SNORD113-7 | snoRNA | 41.56 | -3.957175885 | 3.957175885 | ENSG00000200632 | down |
| ENSG00000240857 | 451.6042193 | -1.098787646 | 0.2621561 | -4.191349 | 2.77E-05 | 0.000250868 | RDH14 | protein_coding | 41.88 | -3.95624426 | 3.95624426 | ENSG00000240857 | down |
| ENSG00000235919 | 172.8641288 | 1.059460912 | 0.2481256 | 4.269858 | 1.96E-05 | 0.000185259 | ASH1L-AS1 | antisense | 62.06 | 3.954141817 | 3.954141817 | ENSG00000235919 | up |
| ENSG00000265144 | 2.198486068 | -2.57480615 | 0.958781 | -2.6855 | 0.007242141 | 0.029150716 | MIR548AM | miRNA | 37.84 | -3.953230595 | 3.953230595 | ENSG00000265144 | down |
| ENSG00000166963 | 1657.109315 | -0.962971186 | 0.2146402 | -4.486443 | 7.24E-06 | 7.90E-05 | MAP1A | protein_coding | 52.57 | -3.950345121 | 3.950345121 | ENSG00000166963 | n.s. |
| ENSG00000162601 | 1967.893223 | -1.140709748 | 0.277841 | -4.10562 | 4.03E-05 | 0.000347628 | MYSM1 | protein_coding | 36.33 | -3.945584251 | 3.945584251 | ENSG00000162601 | down |
| ENSG00000283414 | 4.25016827 | -2.574310763 | 0.9596571 | -2.682532 | 0.007306716 | 0.029334317 | | snRNA | 35.56 | -3.945450489 | 3.945450489 | ENSG00000283414 | down |
| ENSG00000269053 | 12.09425106 | 1.483223783 | 0.4134689 | 3.587268 | 0.000334161 | 0.002189191 | | antisense | 47.45 | 3.94495442 | 3.94495442 | ENSG00000269053 | up |
| ENSG00000267681 | 13.17858978 | -1.482193436 | 0.4132818 | -3.586399 | 0.000335276 | 0.002194661 | | transcribed_processed_pseudog | 47 | -3.940607703 | 3.940607703 | ENSG00000267681 | down |
| ENSG00000095951 | 955.7391444 | -1.029739303 | 0.2380593 | -4.325558 | 1.52E-05 | 0.0001496 | HIVEP1 | protein_coding | 39.37 | -3.93882312 | 3.93882312 | ENSG00000095951 | down |
| ENSG00000252965 | 3.903037895 | -1.986499243 | 0.6445334 | -3.082073 | 0.002055642 | 0.010434087 | | misc_RNA | 49.06 | -3.936338753 | 3.936338753 | ENSG00000252965 | down |
| ENSG00000012048 | 3778.134549 | -1.106187863 | 0.2656465 | -4.164135 | 3.13E-05 | 0.00027858 | BRCA1 | protein_coding | 44.09 | -3.932553019 | 3.932553019 | ENSG00000012048 | down |
| ENSG00000167840 | 350.7287946 | -1.030025015 | 0.2384501 | -4.319666 | 1.56E-05 | 0.000153002 | ZNF232 | protein_coding | 50.38 | -3.92985855 | 3.92985855 | ENSG00000167840 | down |
| ENSG00000278129 | 856.6944057 | -1.122915461 | 0.2718913 | -4.130016 | 3.63E-05 | 0.000316592 | ZNF8 | protein_coding | 48.3 | -3.929642322 | 3.929642322 | ENSG00000278129 | down |
| ENSG00000283210 | 9.096824384 | -1.64650995 | 0.4855902 | -3.390739 | 0.000697044 | 0.004132849 | MIR1322 | miRNA | 33.8 | -3.92486881 | 3.92486881 | ENSG00000283210 | down |
| ENSG00000130818 | 1005.579678 | -1.0081126 | 0.2309887 | -4.364337 | 1.28E-05 | 0.000128274 | ZNF426 | protein_coding | 43.51 | -3.923436163 | 3.923436163 | ENSG00000130818 | down |

| | | | | | | | | | | | | | |
|---|---|---|---|---|---|---|---|---|---|---|---|---|---|
| ENSG00000207001 | 8.459848145 | -1.677182777 | 0.4994811 | -3.357851 | 0.00078551 | 0.004585602 | SNORD116-2 | snoRNA | 42.11 | -3.922265733 | 3.922265733 | ENSG00000207001 | down |
| ENSG00000183530 | 1455.947464 | -1.101359258 | 0.2643485 | -4.166315 | 3.10E-05 | 0.000276284 | PRR14L | protein_coding | 45.25 | -3.919345184 | 3.919345184 | ENSG00000183530 | down |
| ENSG00000226824 | 38.18227995 | -1.206062301 | 0.3034357 | -3.974688 | 7.05E-05 | 0.00056357 | | sense_intronic | 50.03 | -3.918559285 | 3.918559285 | ENSG00000226824 | down |
| ENSG00000204599 | 235.4330498 | -1.077726427 | 0.2560085 | -4.209729 | 2.56E-05 | 0.000233276 | TRIM39 | protein_coding | 44.36 | -3.914441841 | 3.914441841 | ENSG00000204599 | down |
| ENSG00000103248 | 67.70740249 | -1.082325375 | 0.2577303 | -4.199449 | 2.68E-05 | 0.000242691 | MTHFSD | protein_coding | 49.14 | -3.912548015 | 3.912548015 | ENSG00000103248 | down |
| ENSG00000176024 | 186.4159963 | -1.146855523 | 0.2812552 | -4.077634 | 4.55E-05 | 0.000388379 | ZNF613 | protein_coding | 41.55 | -3.911631283 | 3.911631283 | ENSG00000176024 | down |
| ENSG00000239151 | 1.984665061 | -2.566721206 | 0.9599165 | -2.673901 | 0.007497468 | 0.029953297 | RNU7-195P | snRNA | 38.71 | -3.910541861 | 3.910541861 | ENSG00000239151 | down |
| ENSG00000257180 | 44.55670613 | -1.187736173 | 0.2968448 | -4.001203 | 6.30E-05 | 0.000510143 | | sense_intronic | 41.44 | -3.91039384 | 3.91039384 | ENSG00000257180 | down |
| ENSG00000125462 | 164.5919463 | -1.046651019 | 0.2451175 | -4.269998 | 1.95E-05 | 0.000185259 | C1orf61 | protein_coding | 51.53 | -3.906332473 | 3.906332473 | ENSG00000125462 | down |
| ENSG00000139083 | 303.4575706 | -1.025925318 | 0.2378862 | -4.312674 | 1.61E-05 | 0.00015726 | ETV6 | protein_coding | 43.19 | -3.901985925 | 3.901985925 | ENSG00000139083 | down |
| ENSG00000179862 | 250.2027036 | 1.051932467 | 0.2471338 | 4.256531 | 2.08E-05 | 0.000195452 | CITED4 | protein_coding | 73.48 | 3.901574527 | 3.901574527 | ENSG00000179862 | up |
| ENSG00000234006 | 38.14424275 | 1.197117768 | 0.3007657 | 3.980233 | 6.88E-05 | 0.00055217 | DDX39B-AS1 | antisense | 55.21 | 3.900122617 | 3.900122617 | ENSG00000234006 | up |
| ENSG00000279901 | 112.9029678 | 1.018700264 | 0.2355207 | 4.32531 | 1.52E-05 | 0.000149662 | | TEC | 47.26 | 3.896413947 | 3.896413947 | ENSG00000279901 | up |
| ENSG00000252568 | 6.457466071 | -1.924970046 | 0.6181098 | -3.114285 | 0.001843914 | 0.009501412 | RNU7-28P | snRNA | 40.32 | -3.892697238 | 3.892697238 | ENSG00000252568 | down |
| ENSG00000196865 | 1529.553874 | -1.099732473 | 0.2647792 | -4.153395 | 3.28E-05 | 0.000289213 | NHLRC2 | protein_coding | 35.52 | -3.891714332 | 3.891714332 | ENSG00000196865 | down |
| ENSG00000201839 | 9.635495718 | -1.654958505 | 0.4916446 | -3.366169 | 0.000762201 | 0.004470147 | SNORD114-3 | snoRNA | 48 | -3.888619907 | 3.888619907 | ENSG00000201839 | down |
| ENSG00000239486 | 14.59080309 | 1.388652358 | 0.37712 | 3.682256 | 0.000231179 | 0.001590269 | | unprocessed_pseudogene | 43.29 | 3.886184365 | 3.886184365 | ENSG00000239486 | up |
| ENSG00000207692 | 3.656989677 | -2.233373007 | 0.7783944 | -2.869205 | 0.004115051 | 0.01835753 | MIR592 | miRNA | 45.36 | -3.877550373 | 3.877550373 | ENSG00000207692 | down |
| ENSG00000204576 | 582.6084135 | -1.000688381 | 0.2298073 | -4.354468 | 1.33E-05 | 0.000133515 | PRR3 | protein_coding | 44.95 | -3.877135929 | 3.877135929 | ENSG00000204576 | down |
| ENSG00000225193 | 30.97618014 | 1.22767046 | 0.3139687 | 3.910169 | 9.22E-05 | 0.000710649 | | transcribed_processed_pseudog | 47.47 | 3.865130033 | 3.865130033 | ENSG00000225193 | up |
| ENSG00000266173 | 625.0196052 | -1.049163103 | 0.247491 | -4.239197 | 2.24E-05 | 0.000207937 | STRADA | protein_coding | 48.88 | -3.863090134 | 3.863090134 | ENSG00000266173 | down |
| ENSG00000197299 | 2049.702648 | -1.076790075 | 0.2573781 | -4.183689 | 2.87E-05 | 0.000258469 | BLM | protein_coding | 40.73 | -3.863083538 | 3.863083538 | ENSG00000197299 | down |
| ENSG00000239344 | 38.62714474 | -1.276133432 | 0.3329255 | -3.83309 | 0.000126544 | 0.000942681 | | processed_pseudogene | 43.26 | -3.86111443 | 3.86111443 | ENSG00000239344 | down |
| ENSG00000204977 | 1456.337329 | -1.027126867 | 0.2399889 | -4.279893 | 1.87E-05 | 0.000178558 | TRIM13 | protein_coding | 41.26 | -3.84989926 | 3.84989926 | ENSG00000204977 | down |
| ENSG00000272474 | 4.551946671 | -2.225930533 | 0.7777685 | -2.861945 | 0.0042105 | 0.018687534 | SNORD113-5 | snoRNA | 41.03 | -3.847405183 | 3.847405183 | ENSG00000272474 | down |
| ENSG00000115970 | 1830.147524 | -1.061223411 | 0.2524274 | -4.204074 | 2.62E-05 | 0.000238092 | THADA | protein_coding | 40.47 | -3.845082305 | 3.845082305 | ENSG00000115970 | down |
| ENSG00000265520 | 11.24788967 | -1.464379447 | 0.4109428 | -3.563463 | 0.000365995 | 0.002371217 | MIR548V | miRNA | 37.5 | -3.844037967 | 3.844037967 | ENSG00000265520 | down |
| ENSG00000226756 | 32.74056306 | 1.199973408 | 0.304206 | 3.944608 | 7.99E-05 | 0.000627324 | | lincRNA | 40.49 | 3.842924562 | 3.842924562 | ENSG00000226756 | up |
| ENSG00000125945 | 426.8666505 | -1.010476591 | 0.2347139 | -4.305141 | 1.67E-05 | 0.000162027 | ZNF436 | protein_coding | 46 | -3.830123895 | 3.830123895 | ENSG00000125945 | down |
| ENSG00000198185 | 442.7567953 | -1.073260403 | 0.2572776 | -4.171604 | 3.02E-05 | 0.000270642 | ZNF334 | protein_coding | 40.56 | -3.828969684 | 3.828969684 | ENSG00000198185 | down |
| ENSG00000267481 | 97.49376479 | -1.172674725 | 0.2943128 | -3.984451 | 6.76E-05 | 0.000544022 | | sense_intronic | 40.74 | -3.828060067 | 3.828060067 | ENSG00000267481 | down |
| ENSG00000229273 | 57.59245147 | 1.102264841 | 0.2681655 | 4.11039 | 3.95E-05 | 0.000340735 | | processed_pseudogene | 62.66 | 3.822195129 | 3.822195129 | ENSG00000229273 | up |

| | | | | | | | | | | | | | |
|---|---|---|---|---|---|---|---|---|---|---|---|---|---|
| ENSG00000118518 | 727.2403258 | -1.177553526 | 0.2964599 | -3.97205 | 7.13E-05 | 0.000568543 | RNF146 | protein_coding | 36.49 | -3.821440091 | 3.821440091 | ENSG00000118518 | down |
| ENSG00000143493 | 2158.004608 | -1.119997357 | 0.2746541 | -4.077846 | 4.55E-05 | 0.000388262 | INTS7 | protein_coding | 38.12 | -3.82017088 | 3.82017088 | ENSG00000143493 | down |
| ENSG00000102935 | 355.1603702 | -0.986311237 | 0.2267206 | -4.350338 | 1.36E-05 | 0.000135503 | ZNF423 | protein_coding | 50.74 | -3.815101247 | 3.815101247 | ENSG00000102935 | n.s. |
| ENSG00000131797 | 308.9443918 | -1.046345226 | 0.2480575 | -4.218155 | 2.46E-05 | 0.000225913 | CLUHP3 | transcribed_unprocessed_pseud | 55.17 | -3.815035729 | 3.815035729 | ENSG00000131797 | down |
| ENSG00000205765 | 662.7236666 | -1.088773613 | 0.2635831 | -4.130665 | 3.62E-05 | 0.000315944 | C5orf51 | protein_coding | 36.01 | -3.811131454 | 3.811131454 | ENSG00000205765 | down |
| ENSG00000164815 | 609.201078 | -1.131698587 | 0.2794028 | -4.05042 | 5.11E-05 | 0.000429364 | ORC5 | protein_coding | 36.03 | -3.810626616 | 3.810626616 | ENSG00000164815 | down |
| ENSG00000200215 | 5.68691321 | -2.118400174 | 0.7243748 | -2.924453 | 0.003450622 | 0.015892493 | SNORD113-6 | snoRNA | 40 | -3.810595117 | 3.810595117 | ENSG00000200215 | down |
| ENSG00000273017 | 18.52064003 | -1.355875493 | 0.3676071 | -3.688383 | 0.000225684 | 0.001556315 | | sense_intronic | 27.88 | -3.807166047 | 3.807166047 | ENSG00000273017 | down |
| ENSG00000112218 | 410.6984069 | -1.072656193 | 0.2578529 | -4.159954 | 3.18E-05 | 0.000282648 | GPR63 | protein_coding | 36.26 | -3.806593106 | 3.806593106 | ENSG00000112218 | down |
| ENSG00000223044 | 12.46616477 | -1.830934055 | 0.5796524 | -3.158676 | 0.001584877 | 0.008358423 | RNU6-130P | snRNA | 39.81 | -3.804453262 | 3.804453262 | ENSG00000223044 | down |
| ENSG00000165494 | 3336.203844 | -0.935691275 | 0.2098924 | -4.457957 | 8.27E-06 | 8.88E-05 | PCF11 | protein_coding | 38.25 | -3.791236359 | 3.791236359 | ENSG00000165494 | n.s. |
| ENSG00000199047 | 2.459153682 | 2.474954006 | 0.9232283 | 2.68076 | 0.007345508 | 0.029473058 | MIR378A | miRNA | 53.03 | 3.788102242 | 3.788102242 | ENSG00000199047 | up |
| ENSG00000135083 | 207.6269193 | -1.036855637 | 0.2455517 | -4.222556 | 2.42E-05 | 0.000222278 | CCNJL | protein_coding | 46.79 | -3.78774154 | 3.78774154 | ENSG00000135083 | down |
| ENSG00000261777 | 55.42096659 | 1.102744633 | 0.2697864 | 4.087473 | 4.36E-05 | 0.00037383 | | processed_transcript | 46.95 | 3.779464711 | 3.779464711 | ENSG00000261777 | up |
| ENSG00000242699 | 50.29209961 | -1.217898096 | 0.3138006 | -3.881121 | 0.000103976 | 0.000790639 | RN7SL516P | misc_RNA | 49.16 | -3.777946657 | 3.777946657 | ENSG00000242699 | down |
| ENSG00000226498 | 50.29127688 | -1.160214764 | 0.2916497 | -3.97811 | 6.95E-05 | 0.000556801 | RPSAP21 | transcribed_processed_pseudog | 51.75 | -3.775687245 | 3.775687245 | ENSG00000226498 | down |
| ENSG00000215158 | 778.6243201 | -1.085556656 | 0.263787 | -4.115277 | 3.87E-05 | 0.000335048 | | transcribed_unprocessed_pseud | 38.71 | -3.772192558 | 3.772192558 | ENSG00000215158 | down |
| ENSG00000254531 | 47.88427733 | 1.122074877 | 0.2774234 | 4.044629 | 5.24E-05 | 0.000439018 | | lincRNA | 62.32 | 3.767386002 | 3.767386002 | ENSG00000254531 | up |
| ENSG00000279088 | 373.7903445 | 0.942839666 | 0.2131855 | 4.422626 | 9.75E-06 | 0.000102076 | | TEC | 42.48 | 3.762944444 | 3.762944444 | ENSG00000279088 | n.s. |
| ENSG00000158691 | 1108.902743 | -0.941745426 | 0.2128194 | -4.425092 | 9.64E-06 | 0.000101259 | ZSCAN12 | protein_coding | 41.13 | -3.761866405 | 3.761866405 | ENSG00000158691 | n.s. |
| ENSG00000138738 | 130.1956682 | -1.152877269 | 0.2894905 | -3.982435 | 6.82E-05 | 0.000548024 | PRDM5 | protein_coding | 37.44 | -3.759763793 | 3.759763793 | ENSG00000138738 | down |
| ENSG00000228084 | 38.06585389 | 1.165452067 | 0.294613 | 3.955874 | 7.63E-05 | 0.000601532 | | antisense | 43.18 | 3.753619775 | 3.753619775 | ENSG00000228084 | up |
| ENSG00000176095 | 674.297131 | -1.098656744 | 0.269636 | -4.074592 | 4.61E-05 | 0.000392287 | IP6K1 | protein_coding | 44.81 | -3.742459815 | 3.742459815 | ENSG00000176095 | down |
| ENSG00000272688 | 35.58033346 | 1.177694839 | 0.2997484 | 3.928945 | 8.53E-05 | 0.000665119 | | lincRNA | 46.94 | 3.74165515 | 3.74165515 | ENSG00000272688 | up |
| ENSG00000160445 | 200.3470464 | -1.119099752 | 0.2773241 | -4.03535 | 5.45E-05 | 0.000454005 | ZER1 | protein_coding | 51.42 | -3.741082292 | 3.741082292 | ENSG00000160445 | down |
| ENSG00000196670 | 1073.980509 | -1.118329414 | 0.2774835 | -4.030255 | 5.57E-05 | 0.000461474 | ZFP62 | protein_coding | 42.93 | -3.730581933 | 3.730581933 | ENSG00000196670 | down |
| ENSG00000278434 | 103.2934306 | -1.064612399 | 0.2576027 | -4.132769 | 3.58E-05 | 0.000313459 | | sense_intronic | 52.78 | -3.730208743 | 3.730208743 | ENSG00000278434 | down |
| ENSG00000114853 | 186.7656017 | -1.020362531 | 0.2416235 | -4.222945 | 2.41E-05 | 0.000222041 | ZBTB47 | protein_coding | 62.11 | -3.727962012 | 3.727962012 | ENSG00000114853 | down |
| ENSG00000152464 | 276.5722318 | -1.126205889 | 0.2806787 | -4.012439 | 6.01E-05 | 0.000490445 | RPP38 | protein_coding | 43.41 | -3.727076918 | 3.727076918 | ENSG00000152464 | down |
| ENSG00000280269 | 66.55155914 | -1.168245117 | 0.2969276 | -3.934444 | 8.34E-05 | 0.000651532 | | TEC | 38.04 | -3.722103806 | 3.722103806 | ENSG00000280269 | down |
| ENSG00000251821 | 28.03022797 | -1.286796695 | 0.3437443 | -3.743471 | 0.000181495 | 0.001293225 | RNU6-583P | snRNA | 40.22 | -3.716688318 | 3.716688318 | ENSG00000251821 | down |
| ENSG00000263412 | 448.6999123 | 0.928956207 | 0.2097697 | 4.428458 | 9.49E-06 | 0.000100032 | | processed_transcript | 49.4 | 3.715697258 | 3.715697258 | ENSG00000263412 | n.s. |

| | | | | | | | | | | | | | |
|---|---|---|---|---|---|---|---|---|---|---|---|---|---|
| ENSG00000148339 | 444.4895377 | -1.083197981 | 0.2651181 | -4.085718 | 4.39E-05 | 0.000376021 | SLC25A25 | protein_coding | 49.9 | -3.709723216 | 3.709723216 | ENSG00000148339 | down |
| ENSG00000240775 | 32.52566481 | -1.236109917 | 0.3238569 | -3.81684 | 0.000135172 | 0.000997765 | | processed_pseudogene | 47.13 | -3.709530668 | 3.709530668 | ENSG00000240775 | down |
| ENSG00000160551 | 3276.970852 | -1.018168334 | 0.2416621 | -4.213191 | 2.52E-05 | 0.000230181 | TAOK1 | protein_coding | 41.03 | -3.704026343 | 3.704026343 | ENSG00000160551 | down |
| ENSG00000157426 | 561.518396 | -1.171320067 | 0.2991014 | -3.916131 | 9.00E-05 | 0.00069562 | AASDH | protein_coding | 38.98 | -3.69859313 | 3.69859313 | ENSG00000157426 | down |
| ENSG00000215012 | 454.4097439 | -0.973740612 | 0.225992 | -4.30874 | 1.64E-05 | 0.000159747 | C22orf29 | protein_coding | 55.11 | -3.696871496 | 3.696871496 | ENSG00000215012 | n.s. |
| ENSG00000231663 | 70.25327347 | 1.080641555 | 0.264644 | 4.083379 | 4.44E-05 | 0.000379595 | | antisense | 48.54 | 3.696528744 | 3.696528744 | ENSG00000231663 | up |
| ENSG00000083817 | 279.640906 | -1.040822815 | 0.2503085 | -4.15816 | 3.21E-05 | 0.000284492 | ZNF416 | protein_coding | 48.61 | -3.690684777 | 3.690684777 | ENSG00000083817 | down |
| ENSG00000224016 | 14.16285995 | -2.091918681 | 0.7231744 | -2.892689 | 0.00381959 | 0.017243888 | | unprocessed_pseudogene | 39.96 | -3.688815757 | 3.688815757 | ENSG00000224016 | down |
| ENSG00000239627 | 90.17264036 | -1.267074455 | 0.3372199 | -3.757413 | 0.000171679 | 0.001234015 | RPL12P20 | processed_pseudogene | 46.35 | -3.685513792 | 3.685513792 | ENSG00000239627 | down |
| ENSG00000135976 | 1766.861095 | -1.094768373 | 0.2707461 | -4.043525 | 5.27E-05 | 0.000440298 | ANKRD36 | protein_coding | 35.92 | -3.674320116 | 3.674320116 | ENSG00000135976 | down |
| ENSG00000123200 | 2513.884978 | -1.055745206 | 0.2563576 | -4.118252 | 3.82E-05 | 0.000331578 | ZC3H13 | protein_coding | 34.9 | -3.673375021 | 3.673375021 | ENSG00000123200 | down |
| ENSG00000277782 | 57.93485144 | 1.102657785 | 0.2739576 | 4.024921 | 5.70E-05 | 0.000469825 | | lincRNA | 47.56 | 3.669716051 | 3.669716051 | ENSG00000277782 | up |
| ENSG00000224728 | 79.3577426 | -1.129443369 | 0.2841962 | -3.974167 | 7.06E-05 | 0.00056448 | IMPDH1P8 | processed_pseudogene | 57.11 | -3.668829136 | 3.668829136 | ENSG00000224728 | down |
| ENSG00000177352 | 258.2528293 | -0.938846101 | 0.2147904 | -4.370987 | 1.24E-05 | 0.000125025 | CCDC71 | protein_coding | 56.27 | -3.66431817 | 3.66431817 | ENSG00000177352 | n.s. |
| ENSG00000257167 | 678.9380506 | 0.899217483 | 0.2013439 | 4.466077 | 7.97E-06 | 8.58E-05 | TMPO-AS1 | antisense | 49.57 | 3.656746555 | 3.656746555 | ENSG00000257167 | n.s. |
| ENSG00000278963 | 1508.090498 | 0.880144329 | 0.195043 | 4.512567 | 6.40E-06 | 7.08E-05 | | TEC | 40.35 | 3.652716595 | 3.652716595 | ENSG00000278963 | n.s. |
| ENSG00000165102 | 118.248467 | -1.046171415 | 0.2536824 | -4.123941 | 3.72E-05 | 0.000324682 | HGSNAT | protein_coding | 44.1 | -3.649613159 | 3.649613159 | ENSG00000165102 | down |
| ENSG00000106003 | 343.264052 | 1.016820651 | 0.2430044 | 4.184371 | 2.86E-05 | 0.000257862 | LFNG | protein_coding | 62.83 | 3.648976309 | 3.648976309 | ENSG00000106003 | up |
| ENSG00000221821 | 36.09830105 | 1.137880069 | 0.2882764 | 3.947184 | 7.91E-05 | 0.000621316 | C6orf226 | protein_coding | 59.6 | 3.64882614 | 3.64882614 | ENSG00000221821 | up |
| ENSG00000024526 | 2754.423902 | -1.20259827 | 0.3133885 | -3.837403 | 0.000124342 | 0.000927768 | DEPDC1 | protein_coding | 34.94 | -3.646952019 | 3.646952019 | ENSG00000024526 | down |
| ENSG00000130766 | 205.767379 | 1.102199816 | 0.2747727 | 4.011315 | 6.04E-05 | 0.000491919 | SESN2 | protein_coding | 50.06 | 3.646193844 | 3.646193844 | ENSG00000130766 | up |
| ENSG00000180263 | 576.9938813 | 0.956739938 | 0.2216994 | 4.315483 | 1.59E-05 | 0.000155382 | FGD6 | protein_coding | 41.01 | 3.643839668 | 3.643839668 | ENSG00000180263 | n.s. |
| ENSG00000277948 | 28.5863444 | -1.307018941 | 0.3557414 | -3.674071 | 0.000238717 | 0.00163485 | | misc_RNA | 40.64 | -3.642036998 | 3.642036998 | ENSG00000277948 | down |
| ENSG00000124120 | 375.5742338 | -1.092422909 | 0.271287 | -4.026816 | 5.65E-05 | 0.000466608 | TTPAL | protein_coding | 46.55 | -3.638913366 | 3.638913366 | ENSG00000124120 | down |
| ENSG00000200241 | 19.88687882 | -1.291136306 | 0.3494993 | -3.694245 | 0.000220541 | 0.001528417 | | misc_RNA | 43.24 | -3.635527533 | 3.635527533 | ENSG00000200241 | down |
| ENSG00000254144 | 27.85740627 | 1.277870753 | 0.3440979 | 3.713684 | 0.000204264 | 0.001429128 | | antisense | 54.3 | 3.635451409 | 3.635451409 | ENSG00000254144 | up |
| ENSG00000075336 | 730.6413461 | -1.012928922 | 0.2421293 | -4.183422 | 2.87E-05 | 0.000258606 | TIMM21 | protein_coding | 42.31 | -3.633742147 | 3.633742147 | ENSG00000075336 | down |
| ENSG00000265355 | 12.09690776 | 1.496279593 | 0.4371829 | 3.422549 | 0.00062037 | 0.003732676 | MIR3136 | miRNA | 38.46 | 3.63293648 | 3.63293648 | ENSG00000265355 | up |
| ENSG00000139132 | 2030.973038 | -1.016004544 | 0.2433589 | -4.174922 | 2.98E-05 | 0.000267553 | FGD4 | protein_coding | 40.83 | -3.629767928 | 3.629767928 | ENSG00000139132 | down |
| ENSG00000241395 | 21.30639698 | -1.257744606 | 0.336155 | -3.741562 | 0.00018288 | 0.00130043 | RN7SL344P | misc_RNA | 55.07 | -3.629741592 | 3.629741592 | ENSG00000241395 | down |
| ENSG00000186260 | 267.2430269 | -1.067406278 | 0.2623182 | -4.069128 | 4.72E-05 | 0.000400702 | MKL2 | protein_coding | 38.53 | -3.626170157 | 3.626170157 | ENSG00000186260 | down |
| ENSG00000196705 | 2082.385141 | -1.079383765 | 0.2669214 | -4.043826 | 5.26E-05 | 0.000440179 | ZNF431 | protein_coding | 39.12 | -3.622811789 | 3.622811789 | ENSG00000196705 | down |

| | | | | | | | | | | | | | |
|---|---|---|---|---|---|---|---|---|---|---|---|---|---|
| ENSG00000207105 | 6.856407754 | -1.683617557 | 0.5238742 | -3.213782 | 0.00130999 | 0.007072119 | | misc_RNA | 39.64 | -3.620536145 | 3.620536145 | ENSG00000207105 | down |
| ENSG00000106771 | 1472.172887 | -1.018654326 | 0.244949 | -4.158639 | 3.20E-05 | 0.000284095 | TMEM245 | protein_coding | 39.52 | -3.612695102 | 3.612695102 | ENSG00000106771 | down |
| ENSG00000207582 | 4.550918924 | -1.832920987 | 0.5968538 | -3.070971 | 0.002133637 | 0.010749525 | MIR30B | miRNA | 43.18 | -3.608307931 | 3.608307931 | ENSG00000207582 | down |
| ENSG00000151240 | 592.9320855 | -1.024986845 | 0.2475516 | -4.140498 | 3.47E-05 | 0.000304612 | DIP2C | protein_coding | 48.43 | -3.604112414 | 3.604112414 | ENSG00000151240 | down |
| ENSG00000207042 | 7.154693562 | -1.537038205 | 0.4571999 | -3.361851 | 0.000774218 | 0.004529176 | RNU6-196P | snRNA | 39.25 | -3.60278798 | 3.60278798 | ENSG00000207042 | down |
| ENSG00000176049 | 1061.609995 | -1.033621811 | 0.2508758 | -4.120054 | 3.79E-05 | 0.000329202 | JAKMIP2 | protein_coding | 38.29 | -3.599627174 | 3.599627174 | ENSG00000176049 | down |
| ENSG00000163521 | 17.22662204 | 1.327625615 | 0.366557 | 3.62188 | 0.00029247 | 0.001953781 | GLB1L | protein_coding | 49.58 | 3.596702481 | 3.596702481 | ENSG00000163521 | up |
| ENSG00000212370 | 4.770392265 | -1.752169669 | 0.5589094 | -3.13498 | 0.001718661 | 0.008953037 | RNU6-482P | snRNA | 42.99 | -3.588495389 | 3.588495389 | ENSG00000212370 | down |
| ENSG00000104490 | 71.11030915 | 1.040903809 | 0.2539634 | 4.098637 | 4.16E-05 | 0.000357618 | NCALD | protein_coding | 40.58 | 3.58755922 | 3.58755922 | ENSG00000104490 | up |
| ENSG00000128596 | 209.0148526 | -1.11220071 | 0.2810451 | -3.957375 | 7.58E-05 | 0.000598445 | CCDC136 | protein_coding | 48.94 | -3.584596121 | 3.584596121 | ENSG00000128596 | down |
| ENSG00000183814 | 990.5542797 | -1.1058837 | 0.2787107 | -3.967855 | 7.25E-05 | 0.000576659 | LIN9 | protein_coding | 39.7 | -3.582046961 | 3.582046961 | ENSG00000183814 | down |
| ENSG00000227374 | 16.06528341 | -1.909324085 | 0.638978 | -2.98809 | 0.002807266 | 0.013321595 | | lincRNA | 60.47 | -3.580829949 | 3.580829949 | ENSG00000227374 | down |
| ENSG00000212385 | 9.050304033 | -2.21901721 | 0.8046732 | -2.757662 | 0.005821627 | 0.024458213 | RNU6-817P | snRNA | 44.23 | -3.576113277 | 3.576113277 | ENSG00000212385 | down |
| ENSG00000215630 | 25.88938018 | -1.218226556 | 0.3229356 | -3.772351 | 0.000161717 | 0.001167832 | GUSBP9 | unprocessed_pseudogene | 50.96 | -3.57259532 | 3.57259532 | ENSG00000215630 | down |
| ENSG00000270039 | 23.57664146 | -1.273566776 | 0.3457296 | -3.683707 | 0.000229866 | 0.001582334 | | lincRNA | 37.89 | -3.566880731 | 3.566880731 | ENSG00000270039 | down |
| ENSG00000146587 | 1096.445126 | -1.075999844 | 0.2680992 | -4.01344 | 5.98E-05 | 0.000489229 | RBAK | protein_coding | 40.26 | -3.562084723 | 3.562084723 | ENSG00000146587 | down |
| ENSG00000237280 | 56.30260971 | -1.093253614 | 0.2746706 | -3.980236 | 6.88E-05 | 0.00055217 | | sense_intronic | 42.61 | -3.561740759 | 3.561740759 | ENSG00000237280 | down |
| ENSG00000256361 | 38.70645706 | -1.297349526 | 0.3558858 | -3.64541 | 0.000266966 | 0.001799115 | | processed_pseudogene | 37.64 | -3.56114802 | 3.56114802 | ENSG00000256361 | down |
| ENSG00000088888 | 775.0345723 | -1.023244569 | 0.2486131 | -4.115811 | 3.86E-05 | 0.000334482 | MAVS | protein_coding | 52.52 | -3.556416328 | 3.556416328 | ENSG00000088888 | down |
| ENSG00000103528 | 528.0852861 | -1.022970635 | 0.2486299 | -4.114431 | 3.88E-05 | 0.000335861 | SYT17 | protein_coding | 45.9 | -3.553636168 | 3.553636168 | ENSG00000103528 | down |
| ENSG00000152219 | 1208.173435 | -1.14617732 | 0.2955089 | -3.878656 | 0.000105035 | 0.000797482 | ARL14EP | protein_coding | 36.48 | -3.551177144 | 3.551177144 | ENSG00000152219 | down |
| ENSG00000173404 | 1847.085553 | 0.907471501 | 0.2073832 | 4.37582 | 1.21E-05 | 0.000122946 | INSM1 | protein_coding | 63.13 | 3.548472717 | 3.548472717 | ENSG00000173404 | n.s. |
| ENSG00000205054 | 17.55295347 | 1.286313246 | 0.3518409 | 3.655951 | 0.00025623 | 0.001744908 | LINC01121 | processed_transcript | 44.5 | 3.547944387 | 3.547944387 | ENSG00000205054 | up |
| ENSG00000251768 | 8.077195464 | -1.591572011 | 0.4863415 | -3.27254 | 0.001065859 | 0.005935981 | RNA5SP217 | rRNA | 49.45 | -3.54364701 | 3.54364701 | ENSG00000251768 | down |
| ENSG00000226801 | 38.15904852 | -1.178306544 | 0.3087209 | -3.816737 | 0.000135228 | 0.000997765 | OSTCP8 | processed_pseudogene | 44.2 | -3.53606439 | 3.53606439 | ENSG00000226801 | down |
| ENSG00000206824 | 22.54944568 | -1.233043113 | 0.3307167 | -3.728397 | 0.000192702 | 0.001362374 | | misc_RNA | 44.12 | -3.533536196 | 3.533536196 | ENSG00000206824 | down |
| ENSG00000119669 | 405.5868137 | 0.990360912 | 0.2374548 | 4.170735 | 3.04E-05 | 0.000271502 | IRF2BPL | protein_coding | 62.81 | 3.531851385 | 3.531851385 | ENSG00000119669 | n.s. |
| ENSG00000265213 | 32.49313733 | 1.19213767 | 0.3144086 | 3.791683 | 0.00014963 | 0.001092454 | MIR3684 | miRNA | 36.49 | 3.530631255 | 3.530631255 | ENSG00000265213 | up |
| ENSG00000140265 | 1281.025659 | -1.025164107 | 0.2502312 | -4.096867 | 4.19E-05 | 0.00036014 | ZSCAN29 | protein_coding | 44.13 | -3.530181965 | 3.530181965 | ENSG00000140265 | down |
| ENSG00000143614 | 752.9829863 | -1.008514463 | 0.2441979 | -4.129906 | 3.63E-05 | 0.000316592 | GATAD2B | protein_coding | 45.03 | -3.529296061 | 3.529296061 | ENSG00000143614 | down |
| ENSG00000275902 | 36.59816164 | -1.209173007 | 0.3213491 | -3.762802 | 0.00016802 | 0.001209589 | | antisense | 54.49 | -3.527595595 | 3.527595595 | ENSG00000275902 | down |
| ENSG00000267270 | 72.71775863 | -1.011008304 | 0.2452941 | -4.121616 | 3.76E-05 | 0.00032718 | PARD6G-AS1 | antisense | 48.33 | -3.523579186 | 3.523579186 | ENSG00000267270 | down |

| | | | | | | | | | | | | | |
|---|---|---|---|---|---|---|---|---|---|---|---|---|---|
| ENSG00000200176 | 10.1701232 | -1.443549976 | 0.4207869 | -3.430597 | 0.000602255 | 0.003635057 | RNU1-19P | snRNA | 52.73 | -3.521523985 | 3.521523985 | ENSG00000200176 | down |
| ENSG00000261609 | 750.2297873 | -1.059277642 | 0.2635753 | -4.01888 | 5.85E-05 | 0.000479532 | GAN | protein_coding | 41.53 | -3.515935681 | 3.515935681 | ENSG00000261609 | down |
| ENSG00000230551 | 1143.937173 | -0.993040223 | 0.2390546 | -4.15403 | 3.27E-05 | 0.000288776 | | processed_transcript | 37.36 | -3.514805296 | 3.514805296 | ENSG00000230551 | n.s. |
| ENSG00000271938 | 6.040458484 | 1.796350418 | 0.5869416 | 3.060527 | 0.002209482 | 0.011065689 | | lincRNA | 39.73 | 3.513700115 | 3.513700115 | ENSG00000271938 | up |
| ENSG00000139190 | 200.0247607 | 0.95191291 | 0.2243384 | 4.243202 | 2.20E-05 | 0.000205079 | VAMP1 | protein_coding | 48.87 | 3.510730506 | 3.510730506 | ENSG00000139190 | n.s. |
| ENSG00000081386 | 1030.199769 | -1.078811474 | 0.2712098 | -3.977775 | 6.96E-05 | 0.000557265 | ZNF510 | protein_coding | 38.98 | -3.510385569 | 3.510385569 | ENSG00000081386 | down |
| ENSG00000234233 | 423.1730308 | 0.911650018 | 0.2100578 | 4.339996 | 1.42E-05 | 0.000141197 | KCNH1-IT1 | sense_intronic | 47.01 | 3.510010809 | 3.510010809 | ENSG00000234233 | n.s. |
| ENSG00000175105 | 743.5032716 | -1.066115097 | 0.266762 | -3.996502 | 6.43E-05 | 0.000519468 | ZNF654 | protein_coding | 35.51 | -3.50159203 | 3.50159203 | ENSG00000175105 | down |
| ENSG00000064042 | 4094.00339 | -1.035392794 | 0.2551179 | -4.058487 | 4.94E-05 | 0.000417788 | LIMCH1 | protein_coding | 40.5 | -3.498637591 | 3.498637591 | ENSG00000064042 | down |
| ENSG00000280432 | 100.0900429 | -1.767755508 | 0.5739824 | -3.079808 | 0.002071342 | 0.01050229 | | TEC | 41.42 | -3.497886089 | 3.497886089 | ENSG00000280432 | down |
| ENSG00000197951 | 581.5101137 | -1.079793913 | 0.272307 | -3.965354 | 7.33E-05 | 0.000581188 | ZNF71 | protein_coding | 46.22 | -3.493871067 | 3.493871067 | ENSG00000197951 | down |
| ENSG00000129493 | 19.65903418 | 1.248321082 | 0.3390998 | 3.68128 | 0.000232066 | 0.001594797 | HEATR5A | protein_coding | 39.35 | 3.491921731 | 3.491921731 | ENSG00000129493 | up |
| ENSG00000249020 | 42.93443949 | -1.177446833 | 0.3104112 | -3.793184 | 0.000148728 | 0.001087 | SNORA58 | snoRNA | 48.91 | -3.489682352 | 3.489682352 | ENSG00000249020 | down |
| ENSG00000152495 | 2226.790623 | -1.035315572 | 0.2554763 | -4.052491 | 5.07E-05 | 0.00042683 | CAMK4 | protein_coding | 37.31 | -3.488749825 | 3.488749825 | ENSG00000152495 | down |
| ENSG00000272986 | 22.12956994 | 1.220389716 | 0.3280826 | 3.719764 | 0.000199409 | 0.001399382 | | antisense | 41.48 | 3.483069822 | 3.483069822 | ENSG00000272986 | up |
| ENSG00000276791 | 265.9264723 | 0.966598139 | 0.2305698 | 4.192215 | 2.76E-05 | 0.000250075 | | lincRNA | 58.86 | 3.481619117 | 3.481619117 | ENSG00000276791 | n.s. |
| ENSG00000242493 | 30.54621804 | 1.176309967 | 0.3105672 | 3.787618 | 0.000152099 | 0.001106988 | RN7SL37P | misc_RNA | 50.49 | 3.477004159 | 3.477004159 | ENSG00000242493 | up |
| ENSG00000185650 | 134.3457553 | 0.961670963 | 0.2290767 | 4.198031 | 2.69E-05 | 0.000244057 | ZFP36L1 | protein_coding | 53.26 | 3.474045152 | 3.474045152 | ENSG00000185650 | n.s. |
| ENSG00000163155 | 196.5233093 | -1.021580425 | 0.2510592 | -4.069081 | 4.72E-05 | 0.000400702 | LYSMD1 | protein_coding | 46.07 | -3.470491533 | 3.470491533 | ENSG00000163155 | down |
| ENSG00000251805 | 4.151919333 | -1.774428684 | 0.5798502 | -3.06015 | 0.002212261 | 0.011072704 | | scaRNA | 47.66 | -3.470332353 | 3.470332353 | ENSG00000251805 | down |
| ENSG00000272146 | 59.87033291 | 1.046557028 | 0.2606001 | 4.01595 | 5.92E-05 | 0.000484335 | ARF4-AS1 | antisense | 43.98 | 3.469184317 | 3.469184317 | ENSG00000272146 | up |
| ENSG00000272711 | 229.9739131 | -1.048331183 | 0.2615086 | -4.008783 | 6.10E-05 | 0.000496476 | | lincRNA | 51.84 | -3.463793263 | 3.463793263 | ENSG00000272711 | down |
| ENSG00000205763 | 154.2705728 | -1.083291033 | 0.2748791 | -3.940972 | 8.12E-05 | 0.000636193 | RP9P | transcribed_unprocessed_pseud | 43.81 | -3.462643356 | 3.462643356 | ENSG00000205763 | down |
| ENSG00000279035 | 66.2874215 | 1.041197164 | 0.2588015 | 4.02315 | 5.74E-05 | 0.000473092 | | TEC | 60.02 | 3.462036837 | 3.462036837 | ENSG00000279035 | up |
| ENSG00000202444 | 12.21962913 | 1.408966779 | 0.4094836 | 3.440838 | 0.000579915 | 0.003532002 | RNU5E-6P | snRNA | 37.39 | 3.454757007 | 3.454757007 | ENSG00000202444 | up |
| ENSG00000233718 | 1544.367815 | 0.841230148 | 0.187554 | 4.485268 | 7.28E-06 | 7.93E-05 | MYCNOS | protein_coding | 59.15 | 3.449492325 | 3.449492325 | ENSG00000233718 | n.s. |
| ENSG00000170832 | 1632.997768 | -1.05527071 | 0.2647475 | -3.985952 | 6.72E-05 | 0.000540905 | USP32 | protein_coding | 40.19 | -3.447441422 | 3.447441422 | ENSG00000170832 | down |
| ENSG00000156398 | 465.1423389 | -0.973147603 | 0.2342034 | -4.155138 | 3.25E-05 | 0.000287746 | SFXN2 | protein_coding | 48.44 | -3.445907155 | 3.445907155 | ENSG00000156398 | n.s. |
| ENSG00000200731 | 12.83908947 | -1.425152213 | 0.4175948 | -3.412763 | 0.000643078 | 0.003849342 | RNU1-124P | snRNA | 48.28 | -3.441191774 | 3.441191774 | ENSG00000200731 | down |
| ENSG00000275457 | 110.0740452 | 1.050522206 | 0.2632558 | 3.9905 | 6.59E-05 | 0.000531381 | | antisense | 63.04 | 3.440033331 | 3.440033331 | ENSG00000275457 | up |
| ENSG00000169116 | 752.7340931 | -1.049238219 | 0.2629379 | -3.990442 | 6.60E-05 | 0.000531381 | PARM1 | protein_coding | 40.47 | -3.435828795 | 3.435828795 | ENSG00000169116 | down |
| ENSG00000197062 | 315.720259 | -1.039870556 | 0.2594274 | -4.00833 | 6.11E-05 | 0.00049691 | ZSCAN26 | protein_coding | 41.65 | -3.43544307 | 3.43544307 | ENSG00000197062 | down |

| | | | | | | | | | | | | | |
|---|---|---|---|---|---|---|---|---|---|---|---|---|---|
| ENSG00000166004 | 1665.634795 | -1.029403162 | 0.2553802 | -4.030865 | 5.56E-05 | 0.000460827 | CEP295 | protein_coding | 38.58 | -3.434564471 | 3.434564471 | ENSG00000166004 | down |
| ENSG00000181274 | 202.5175918 | 0.958329695 | 0.2295508 | 4.174805 | 2.98E-05 | 0.000267553 | FRAT2 | protein_coding | 60.23 | 3.423719325 | 3.423719325 | ENSG00000181274 | n.s. |
| ENSG00000221184 | 41.47933779 | -1.181499505 | 0.315187 | -3.748567 | 0.000177848 | 0.001271138 | MIR1254-1 | miRNA | 50.52 | -3.421395066 | 3.421395066 | ENSG00000221184 | down |
| ENSG00000228505 | 1171.496624 | -1.104085124 | 0.2846588 | -3.878626 | 0.000105048 | 0.000797482 | | lincRNA | 40.25 | -3.420763775 | 3.420763775 | ENSG00000228505 | down |
| ENSG00000138376 | 1039.049323 | -1.03777362 | 0.2592789 | -4.002538 | 6.27E-05 | 0.000507566 | BARD1 | protein_coding | 36.39 | -3.418953213 | 3.418953213 | ENSG00000138376 | down |
| ENSG00000277138 | 2.278812297 | -2.459109227 | 0.9666169 | -2.544037 | 0.010957942 | 0.040935719 | MIR6509 | miRNA | 37.65 | -3.412991743 | 3.412991743 | ENSG00000277138 | down |
| ENSG00000261783 | 23.53152884 | -1.260020665 | 0.3479902 | -3.620851 | 0.000293636 | 0.001959516 | | sense_intronic | 40.26 | -3.41194834 | 3.41194834 | ENSG00000261783 | down |
| ENSG00000259287 | 35.55545367 | -1.150857022 | 0.3036021 | -3.790675 | 0.000150238 | 0.001095166 | | antisense | 43.53 | -3.407135419 | 3.407135419 | ENSG00000259287 | down |
| ENSG00000280239 | 123.110397 | -1.020518952 | 0.2532023 | -4.030449 | 5.57E-05 | 0.000461368 | | TEC | 54.27 | -3.40440304 | 3.40440304 | ENSG00000280239 | down |
| ENSG00000207719 | 8.175494264 | -1.646996546 | 0.52309 | -3.148591 | 0.001640594 | 0.008619619 | MIR623 | miRNA | 52.04 | -3.40024403 | 3.40024403 | ENSG00000207719 | down |
| ENSG00000176476 | 196.8177272 | -1.118698413 | 0.2912077 | -3.841583 | 0.000122244 | 0.000914069 | SGF29 | protein_coding | 48.31 | -3.399747935 | 3.399747935 | ENSG00000176476 | down |
| ENSG00000260526 | 108.753224 | 1.003712527 | 0.2470957 | 4.06204 | 4.86E-05 | 0.000411729 | | antisense | 35.9 | 3.397957295 | 3.397957295 | ENSG00000260526 | up |
| ENSG00000128965 | 239.9764687 | 1.173277837 | 0.3130411 | 3.747999 | 0.000178251 | 0.001273364 | CHAC1 | protein_coding | 57.9 | 3.396694888 | 3.396694888 | ENSG00000128965 | up |
| ENSG00000186666 | 144.4485332 | -0.958709803 | 0.2307819 | -4.154181 | 3.26E-05 | 0.000288769 | BCDIN3D | protein_coding | 45.14 | -3.393304549 | 3.393304549 | ENSG00000186666 | n.s. |
| ENSG00000241745 | 41.81172694 | 1.068488251 | 0.2720388 | 3.927706 | 8.58E-05 | 0.00066818 | RN7SL788P | misc_RNA | 55.06 | 3.392564083 | 3.392564083 | ENSG00000241745 | up |
| ENSG00000207779 | 39.96101573 | 1.106577121 | 0.2869255 | 3.85667 | 0.000114942 | 0.000864114 | MIR15B | miRNA | 36.73 | 3.389920535 | 3.389920535 | ENSG00000207779 | up |
| ENSG00000212445 | 9.786544918 | -1.507163066 | 0.4580593 | -3.290323 | 0.001000725 | 0.005654664 | | snoRNA | 46.67 | -3.387489432 | 3.387489432 | ENSG00000212445 | down |
| ENSG00000261431 | 34.03872475 | 1.166573216 | 0.3108667 | 3.752648 | 0.000174976 | 0.001252543 | | antisense | 55.08 | 3.385637239 | 3.385637239 | ENSG00000261431 | up |
| ENSG00000272686 | 532.1743589 | 0.922004222 | 0.2178134 | 4.233 | 2.31E-05 | 0.000213185 | | antisense | 41.64 | 3.384902514 | 3.384902514 | ENSG00000272686 | n.s. |
| ENSG00000278864 | 185.610656 | -1.003316708 | 0.2474324 | -4.054913 | 5.02E-05 | 0.0004232 | | TEC | 53.36 | -3.384642602 | 3.384642602 | ENSG00000278864 | down |
| ENSG00000162650 | 716.9425297 | 0.887400653 | 0.2055375 | 4.317463 | 1.58E-05 | 0.000154211 | ATXN7L2 | protein_coding | 56.87 | 3.382668493 | 3.382668493 | ENSG00000162650 | n.s. |
| ENSG00000273727 | 7.845248744 | 1.518285956 | 0.4642423 | 3.27046 | 0.001073727 | 0.005970246 | | snRNA | 45.68 | 3.376679713 | 3.376679713 | ENSG00000273727 | up |
| ENSG00000266852 | 39.26893254 | 1.142251692 | 0.3021932 | 3.779872 | 0.000156909 | 0.001137834 | MIR4482 | miRNA | 48.57 | 3.362698651 | 3.362698651 | ENSG00000266852 | up |
| ENSG00000202358 | 10.15405941 | -1.436828788 | 0.427803 | -3.358622 | 0.00078332 | 0.004578576 | RNU6-652P | snRNA | 36.45 | -3.361129925 | 3.361129925 | ENSG00000202358 | down |
| ENSG00000136709 | 3913.271892 | -1.093099826 | 0.2829939 | -3.862627 | 0.000112174 | 0.000845132 | WDR33 | protein_coding | 41.82 | -3.359178419 | 3.359178419 | ENSG00000136709 | down |
| ENSG00000243403 | 14.72890173 | -1.754854001 | 0.5803214 | -3.023935 | 0.002495104 | 0.012184962 | | processed_pseudogene | 52.59 | -3.359099071 | 3.359099071 | ENSG00000243403 | down |
| ENSG00000202566 | 11.99782531 | -1.388878691 | 0.4072403 | -3.410465 | 0.000648523 | 0.003874191 | MIR421 | miRNA | 38.82 | -3.349724037 | 3.349724037 | ENSG00000202566 | down |
| ENSG00000270522 | 73.91678277 | 1.003505384 | 0.2489055 | 4.031672 | 5.54E-05 | 0.000459794 | | lincRNA | 47.71 | 3.34913527 | 3.34913527 | ENSG00000270522 | up |
| ENSG00000272768 | 77.4114459 | 1.046406381 | 0.2653962 | 3.942808 | 8.05E-05 | 0.000631696 | | antisense | 63.71 | 3.347968605 | 3.347968605 | ENSG00000272768 | up |
| ENSG00000116717 | 2155.841475 | 0.853194716 | 0.1946472 | 4.383287 | 1.17E-05 | 0.00011924 | GADD45A | protein_coding | 52.9 | 3.347576335 | 3.347576335 | ENSG00000116717 | n.s. |
| ENSG00000008196 | 288.7889944 | -0.991256999 | 0.2445422 | -4.053521 | 5.05E-05 | 0.000425469 | TFAP2B | protein_coding | 44.14 | -3.341658467 | 3.341658467 | ENSG00000008196 | n.s. |
| ENSG00000160207 | 31.85918621 | 1.171632476 | 0.315228 | 3.716778 | 0.00020178 | 0.001413879 | HSF2BP | protein_coding | 43.01 | 3.338669435 | 3.338669435 | ENSG00000160207 | up |

| | | | | | | | | | | | | | |
|---|---|---|---|---|---|---|---|---|---|---|---|---|---|
| ENSG00000136205 | 535.6009406 | -0.995145092 | 0.246223 | -4.041641 | 5.31E-05 | 0.000443317 | TNS3 | protein_coding | 48.46 | -3.337005547 | 3.337005547 | ENSG00000136205 | n.s. |
| ENSG00000187607 | 1788.740644 | -0.978222816 | 0.2399307 | -4.077105 | 4.56E-05 | 0.000389024 | ZNF286A | protein_coding | 39.97 | -3.33576326 | 3.33576326 | ENSG00000187607 | n.s. |
| ENSG00000279873 | 332.2152776 | 0.900615263 | 0.2118723 | 4.250745 | 2.13E-05 | 0.000199626 | | lincRNA | 56.62 | 3.332080311 | 3.332080311 | ENSG00000279873 | n.s. |
| ENSG00000232956 | 3537.540691 | 0.839119566 | 0.1903138 | 4.409136 | 1.04E-05 | 0.000107915 | SNHG15 | lincRNA | 50.27 | 3.328718311 | 3.328718311 | ENSG00000232956 | n.s. |
| ENSG00000224418 | 67.04865838 | 1.116255335 | 0.2935402 | 3.802734 | 0.000143108 | 0.001050337 | STK24-AS1 | antisense | 55.01 | 3.324957702 | 3.324957702 | ENSG00000224418 | up |
| ENSG00000155097 | 785.6181777 | -1.113846308 | 0.2926722 | -3.805781 | 0.000141357 | 0.001038585 | ATP6V1C1 | protein_coding | 38.39 | -3.323225055 | 3.323225055 | ENSG00000155097 | down |
| ENSG00000251934 | 15.212436 | -1.296360771 | 0.3682015 | -3.520792 | 0.000430261 | 0.002736646 | RNU6-1143P | snRNA | 34.62 | -3.322289338 | 3.322289338 | ENSG00000251934 | down |
| ENSG00000197566 | 64.70253139 | -1.026575347 | 0.2588891 | -3.96531 | 7.33E-05 | 0.000581188 | ZNF624 | protein_coding | 37.68 | -3.321672646 | 3.321672646 | ENSG00000197566 | down |
| ENSG00000282978 | 36.44860303 | -1.243230249 | 0.3457998 | -3.595231 | 0.000324104 | 0.00213964 | | processed_pseudogene | 43.09 | -3.319001251 | 3.319001251 | ENSG00000282978 | down |
| ENSG00000135297 | 988.5012955 | -0.945687441 | 0.2286351 | -4.136231 | 3.53E-05 | 0.000309742 | MTO1 | protein_coding | 42.61 | -3.318417143 | 3.318417143 | ENSG00000135297 | n.s. |
| ENSG00000238259 | 24.34386526 | -1.205753285 | 0.3303335 | -3.650109 | 0.000262129 | 0.001773381 | | processed_pseudogene | 46.67 | -3.317266045 | 3.317266045 | ENSG00000238259 | down |
| ENSG00000198984 | 8.829279863 | 1.471770066 | 0.4466985 | 3.294773 | 0.000985012 | 0.005579462 | MIR345 | miRNA | 60.2 | 3.316498021 | 3.316498021 | ENSG00000198984 | up |
| ENSG00000111249 | 470.319453 | -0.964590682 | 0.2356473 | -4.093366 | 4.25E-05 | 0.000365172 | CUX2 | protein_coding | 49.28 | -3.315783286 | 3.315783286 | ENSG00000111249 | n.s. |
| ENSG00000223013 | 3.603497027 | -1.776944496 | 0.5965201 | -2.978851 | 0.002893314 | 0.013650835 | RNA5SP344 | rRNA | 47.12 | -3.313718593 | 3.313718593 | ENSG00000223013 | down |
| ENSG00000254293 | 257.7541589 | 0.88236334 | 0.2059883 | 4.283561 | 1.84E-05 | 0.000176 | | lincRNA | 47.45 | 3.312822461 | 3.312822461 | ENSG00000254293 | n.s. |
| ENSG00000252751 | 4.395731507 | -1.747576148 | 0.5815008 | -3.005286 | 0.002653312 | 0.012747435 | RNU6-143P | snRNA | 41.94 | -3.310917927 | 3.310917927 | ENSG00000252751 | down |
| ENSG00000155744 | 975.8444972 | -1.043389662 | 0.2658795 | -3.924296 | 8.70E-05 | 0.000676201 | FAM126B | protein_coding | 39.16 | -3.30746608 | 3.30746608 | ENSG00000155744 | down |
| ENSG00000011523 | 1726.897895 | -1.090710989 | 0.2843398 | -3.835943 | 0.000125084 | 0.000932801 | CEP68 | protein_coding | 47.81 | -3.305084415 | 3.305084415 | ENSG00000011523 | down |
| ENSG00000181007 | 655.1910002 | -1.051700258 | 0.2696521 | -3.900212 | 9.61E-05 | 0.000737661 | ZFP82 | protein_coding | 40.74 | -3.294075799 | 3.294075799 | ENSG00000181007 | down |
| ENSG00000270083 | 134.3643524 | 0.965698262 | 0.2368984 | 4.076423 | 4.57E-05 | 0.00038945 | | sense_intronic | 45.11 | 3.292594801 | 3.292594801 | ENSG00000270083 | n.s. |
| ENSG00000162852 | 907.3854993 | -1.063648629 | 0.2743876 | -3.876446 | 0.000105993 | 0.000803782 | CNST | protein_coding | 41.32 | -3.291845305 | 3.291845305 | ENSG00000162852 | down |
| ENSG00000185920 | 3381.778598 | -0.93797079 | 0.2268867 | -4.134094 | 3.56E-05 | 0.000312244 | PTCH1 | protein_coding | 47.49 | -3.288061653 | 3.288061653 | ENSG00000185920 | n.s. |
| ENSG00000184588 | 1557.962327 | -1.048281031 | 0.2689301 | -3.897969 | 9.70E-05 | 0.000742481 | PDE4B | protein_coding | 37.45 | -3.280401441 | 3.280401441 | ENSG00000184588 | down |
| ENSG00000101680 | 395.8463621 | -0.935479498 | 0.2263101 | -4.133619 | 3.57E-05 | 0.000312695 | LAMA1 | protein_coding | 43.14 | -3.278742776 | 3.278742776 | ENSG00000101680 | n.s. |
| ENSG00000164252 | 1001.5895 | -1.046676757 | 0.2684675 | -3.898709 | 9.67E-05 | 0.000741438 | AGGF1 | protein_coding | 39.02 | -3.276020057 | 3.276020057 | ENSG00000164252 | down |
| ENSG00000181896 | 283.2225418 | -1.061901441 | 0.2744141 | -3.869704 | 0.000108967 | 0.000823198 | ZNF101 | protein_coding | 46.28 | -3.275430356 | 3.275430356 | ENSG00000181896 | down |
| ENSG00000119684 | 254.5460722 | -1.056997554 | 0.2726081 | -3.877351 | 0.0001056 | 0.000801235 | MLH3 | protein_coding | 41.37 | -3.272718154 | 3.272718154 | ENSG00000119684 | down |
| ENSG00000183741 | 1183.991103 | -0.879507492 | 0.2063182 | -4.26287 | 2.02E-05 | 0.000190632 | CBX6 | protein_coding | 58.47 | -3.271596131 | 3.271596131 | ENSG00000183741 | n.s. |
| ENSG00000229659 | 18.94072835 | -1.730040255 | 0.5764417 | -3.00124 | 0.002688822 | 0.012886909 | RPL26P6 | processed_pseudogene | 42.01 | -3.269518722 | 3.269518722 | ENSG00000229659 | down |
| ENSG00000206633 | 205.6271336 | 0.920110001 | 0.2210712 | 4.162052 | 3.15E-05 | 0.000280595 | SNORA80B | snoRNA | 58.82 | 3.268157016 | 3.268157016 | ENSG00000206633 | n.s. |
| ENSG00000221539 | 27.57989832 | 1.172541154 | 0.3191382 | 3.674086 | 0.000238703 | 0.00163485 | SNORD99 | snoRNA | 47.3 | 3.267311688 | 3.267311688 | ENSG00000221539 | up |
| ENSG00000260296 | 48.65462147 | -1.586692952 | 0.5053622 | -3.139714 | 0.001691127 | 0.008838426 | | sense_overlapping | 32.58 | -3.258472396 | 3.258472396 | ENSG00000260296 | down |

| | | | | | | | | | | | | | |
|---|---|---|---|---|---|---|---|---|---|---|---|---|---|
| ENSG00000236780 | 6973.332257 | -1.089597599 | 0.2860432 | -3.809207 | 0.000139413 | 0.001025926 | LINC01829 | lincRNA | 36.25 | -3.256680953 | 3.256680953 | ENSG00000236780 | down |
| ENSG00000165312 | 102.7655007 | 1.021477881 | 0.2597405 | 3.932687 | 8.40E-05 | 0.000655948 | OTUD1 | protein_coding | 49.91 | 3.251497226 | 3.251497226 | ENSG00000165312 | up |
| ENSG00000212246 | 14.60761595 | 1.331430209 | 0.3880003 | 3.431519 | 0.000600211 | 0.003627059 | RNU6-360P | snRNA | 42.31 | 3.249282654 | 3.249282654 | ENSG00000212246 | up |
| ENSG00000261407 | 13.1287907 | 1.326737177 | 0.3859751 | 3.437365 | 0.000587404 | 0.003566694 | | antisense | 52.27 | 3.247499868 | 3.247499868 | ENSG00000261407 | up |
| ENSG00000171067 | 279.24756 | -1.101402047 | 0.2911288 | -3.783213 | 0.000154817 | 0.001126184 | C11orf24 | protein_coding | 52.05 | -3.247363671 | 3.247363671 | ENSG00000171067 | down |
| ENSG00000135835 | 791.4080423 | -0.977740645 | 0.2432824 | -4.018954 | 5.85E-05 | 0.000479532 | KIAA1614 | protein_coding | 54.17 | -3.245299518 | 3.245299518 | ENSG00000135835 | n.s. |
| ENSG00000115738 | 4943.691125 | 1.059168583 | 0.2746712 | 3.856133 | 0.000115195 | 0.000865548 | ID2 | protein_coding | 49.26 | 3.243925161 | 3.243925161 | ENSG00000115738 | up |
| ENSG00000099725 | 528.5817315 | -0.969324567 | 0.240111 | -4.036985 | 5.41E-05 | 0.000451124 | PRKY | transcribed_unprocessed_pseud | 44.3 | -3.243073282 | 3.243073282 | ENSG00000099725 | n.s. |
| ENSG00000185219 | 1356.042316 | -0.963192282 | 0.2378981 | -4.048759 | 5.15E-05 | 0.000431955 | ZNF445 | protein_coding | 44.45 | -3.240720125 | 3.240720125 | ENSG00000185219 | n.s. |
| ENSG00000200183 | 8.482830919 | -1.517667781 | 0.4740094 | -3.201767 | 0.001365873 | 0.007333995 | RNU6-238P | snRNA | 41.51 | -3.239703803 | 3.239703803 | ENSG00000200183 | down |
| ENSG00000184939 | 701.9551786 | -1.094327496 | 0.2887224 | -3.790241 | 0.000150501 | 0.00109651 | ZFP90 | protein_coding | 44.64 | -3.239195541 | 3.239195541 | ENSG00000184939 | down |
| ENSG00000140006 | 583.092479 | -1.046755202 | 0.2701126 | -3.875255 | 0.000106513 | 0.000807285 | WDR89 | protein_coding | 41.2 | -3.237585749 | 3.237585749 | ENSG00000140006 | down |
| ENSG00000157653 | 93.1927318 | 0.970574529 | 0.2409525 | 4.028074 | 5.62E-05 | 0.000464395 | C9orf43 | protein_coding | 43.34 | 3.235033772 | 3.235033772 | ENSG00000157653 | n.s. |
| ENSG00000232821 | 362.2045024 | 0.879517603 | 0.2076724 | 4.23512 | 2.28E-05 | 0.000211465 | | antisense | 39.99 | 3.232018091 | 3.232018091 | ENSG00000232821 | n.s. |
| ENSG00000171467 | 606.073375 | -0.978884022 | 0.2443672 | -4.005792 | 6.18E-05 | 0.000501797 | ZNF318 | protein_coding | 42.92 | -3.229800233 | 3.229800233 | ENSG00000171467 | n.s. |
| ENSG00000177599 | 204.8267345 | -0.962209939 | 0.2379512 | -4.043729 | 5.26E-05 | 0.000440179 | ZNF491 | protein_coding | 44.18 | -3.229533021 | 3.229533021 | ENSG00000177599 | n.s. |
| ENSG00000037637 | 397.3277456 | -0.969506638 | 0.2408244 | -4.025783 | 5.68E-05 | 0.000468384 | FBXO42 | protein_coding | 43.85 | -3.227873088 | 3.227873088 | ENSG00000037637 | n.s. |
| ENSG00000264720 | 6.756495262 | -1.63163936 | 0.5299177 | -3.079043 | 0.002076667 | 0.010525461 | MIR3117 | miRNA | 46.15 | -3.226989236 | 3.226989236 | ENSG00000264720 | down |
| ENSG00000158636 | 263.1893295 | -1.028414395 | 0.2635676 | -3.9019 | 9.54E-05 | 0.000733667 | EMSY | protein_coding | 37.68 | -3.223565801 | 3.223565801 | ENSG00000158636 | down |
| ENSG00000207573 | 10.12587388 | 1.482288218 | 0.4588039 | 3.230767 | 0.001234587 | 0.006722391 | MIR550A2 | miRNA | 53.61 | 3.220235905 | 3.220235905 | ENSG00000207573 | up |
| ENSG00000266962 | 256.9966657 | -0.936654664 | 0.2290187 | -4.089861 | 4.32E-05 | 0.000370277 | | antisense | 63.34 | -3.214105355 | 3.214105355 | ENSG00000266962 | n.s. |
| ENSG00000279354 | 55.83531955 | 1.05904484 | 0.2759781 | 3.837424 | 0.000124332 | 0.000927768 | | TEC | 41.63 | 3.211617559 | 3.211617559 | ENSG00000279354 | up |
| ENSG00000252233 | 25.6326888 | -1.185965327 | 0.3275447 | -3.620774 | 0.000293723 | 0.001959516 | RN7SKP253 | misc_RNA | 42.07 | -3.21141751 | 3.21141751 | ENSG00000252233 | down |
| ENSG00000183963 | 26.80495062 | 1.14599232 | 0.3110069 | 3.684781 | 0.000228899 | 0.001576926 | SMTN | protein_coding | 54.79 | 3.211286746 | 3.211286746 | ENSG00000183963 | up |
| ENSG00000250461 | 106.7287907 | -1.141699187 | 0.3093596 | -3.690525 | 0.000223792 | 0.001546328 | | processed_pseudogene | 37.77 | -3.208971918 | 3.208971918 | ENSG00000250461 | down |
| ENSG00000153234 | 143.9301687 | 0.935645357 | 0.2289712 | 4.086302 | 4.38E-05 | 0.000375308 | NR4A2 | protein_coding | 45.15 | 3.20515824 | 3.20515824 | ENSG00000153234 | n.s. |
| ENSG00000136935 | 197.4976433 | -1.012855013 | 0.2584619 | -3.918779 | 8.90E-05 | 0.000689554 | GOLGA1 | protein_coding | 44.12 | -3.202071879 | 3.202071879 | ENSG00000136935 | down |
| ENSG00000161960 | 1312.507569 | 0.826897206 | 0.1899747 | 4.352669 | 1.34E-05 | 0.000134422 | EIF4A1 | protein_coding | 52.92 | 3.201355871 | 3.201355871 | ENSG00000161960 | n.s. |
| ENSG00000143507 | 436.0326118 | 0.980519043 | 0.2461203 | 3.983901 | 6.78E-05 | 0.000544967 | DUSP10 | protein_coding | 41.35 | 3.200051127 | 3.200051127 | ENSG00000143507 | n.s. |
| ENSG00000152969 | 208.201803 | -1.071523881 | 0.2814266 | -3.807472 | 0.000140395 | 0.001032602 | JAKMIP1 | protein_coding | 49.33 | -3.199642296 | 3.199642296 | ENSG00000152969 | down |
| ENSG00000004139 | 958.6821803 | -0.994353074 | 0.2514966 | -3.953743 | 7.69E-05 | 0.000606229 | SARM1 | protein_coding | 51.55 | -3.199194894 | 3.199194894 | ENSG00000004139 | n.s. |
| ENSG00000102870 | 953.6613052 | -0.985021328 | 0.2481683 | -3.969166 | 7.21E-05 | 0.00057448 | ZNF629 | protein_coding | 57.75 | -3.192183486 | 3.192183486 | ENSG00000102870 | n.s. |

| | | | | | | | | | | | | | |
|---|---|---|---|---|---|---|---|---|---|---|---|---|---|
| ENSG00000118200 | 3465.604283 | -1.027677982 | 0.2648493 | -3.880237 | 0.000104355 | 0.000793084 | CAMSAP2 | protein_coding | 37.69 | -3.186501626 | 3.186501626 | ENSG00000118200 | down |
| ENSG00000226253 | 31.80466549 | -1.157801114 | 0.3171135 | -3.651062 | 0.000261158 | 0.001770256 | MRPL35P3 | processed_pseudogene | 45.6 | -3.186226781 | 3.186226781 | ENSG00000226253 | down |
| ENSG00000167196 | 2024.987722 | -1.044163352 | 0.2713337 | -3.848262 | 0.000118959 | 0.000891902 | FBXO22 | protein_coding | 39.46 | -3.184366914 | 3.184366914 | ENSG00000167196 | down |
| ENSG00000242970 | 16.11401442 | -1.302116319 | 0.3790721 | -3.435009 | 0.000592534 | 0.003593872 | | processed_pseudogene | 51.88 | -3.182941889 | 3.182941889 | ENSG00000242970 | down |
| ENSG00000071242 | 950.2332201 | -0.959208119 | 0.2387627 | -4.017412 | 5.88E-05 | 0.000481622 | RPS6KA2 | protein_coding | 46.48 | -3.181975107 | 3.181975107 | ENSG00000071242 | n.s. |
| ENSG00000166123 | 829.9534214 | -0.983175546 | 0.2481681 | -3.961733 | 7.44E-05 | 0.000588625 | GPT2 | protein_coding | 51.96 | -3.175815482 | 3.175815482 | ENSG00000166123 | n.s. |
| ENSG00000150995 | 1536.896752 | -0.945043735 | 0.2338703 | -4.040887 | 5.32E-05 | 0.000444479 | ITPR1 | protein_coding | 42.9 | -3.167927529 | 3.167927529 | ENSG00000150995 | n.s. |
| ENSG00000282386 | 30.80578967 | 1.110598693 | 0.2986862 | 3.718279 | 0.000200584 | 0.00140692 | | antisense | 45.63 | 3.167128422 | 3.167128422 | ENSG00000282386 | up |
| ENSG00000081692 | 652.9176883 | -0.938744644 | 0.2317341 | -4.050957 | 5.10E-05 | 0.000428861 | JMJD4 | protein_coding | 61.26 | -3.161395104 | 3.161395104 | ENSG00000081692 | n.s. |
| ENSG00000247416 | 50.34717953 | 1.108467279 | 0.2981785 | 3.717463 | 0.000201234 | 0.001410764 | | antisense | 53.01 | 3.159736538 | 3.159736538 | ENSG00000247416 | up |
| ENSG00000275854 | 67.22735069 | -1.148403653 | 0.314596 | -3.650407 | 0.000261825 | 0.001773047 | | sense_intronic | 29.27 | -3.159579689 | 3.159579689 | ENSG00000275854 | down |
| ENSG00000161904 | 631.6929547 | -1.038141734 | 0.2701493 | -3.842844 | 0.000121617 | 0.00091036 | LEMD2 | protein_coding | 53.18 | -3.156767544 | 3.156767544 | ENSG00000161904 | down |
| ENSG00000177511 | 1315.623529 | -1.016668574 | 0.261901 | -3.881882 | 0.000103651 | 0.000788636 | ST8SIA3 | protein_coding | 41.88 | -3.154847886 | 3.154847886 | ENSG00000177511 | down |
| ENSG00000204120 | 2620.324721 | -0.978736773 | 0.2473418 | -3.957022 | 7.59E-05 | 0.00059899 | GIGYF2 | protein_coding | 39.44 | -3.154058194 | 3.154058194 | ENSG00000204120 | n.s. |
| ENSG00000170011 | 767.0564733 | -0.979909978 | 0.2477986 | -3.954462 | 7.67E-05 | 0.000604752 | MYRIP | protein_coding | 40.16 | -3.153764411 | 3.153764411 | ENSG00000170011 | n.s. |
| ENSG00000196236 | 508.5030723 | -0.970358174 | 0.2442342 | -3.973065 | 7.10E-05 | 0.00056645 | XPNPEP3 | protein_coding | 41.48 | -3.15059655 | 3.15059655 | ENSG00000196236 | n.s. |
| ENSG00000272516 | 44.83034955 | -1.09004279 | 0.2912591 | -3.742519 | 0.000182185 | 0.001296808 | | lincRNA | 33.22 | -3.147088883 | 3.147088883 | ENSG00000272516 | down |
| ENSG00000143995 | 21763.09322 | -0.88703976 | 0.2133318 | -4.158028 | 3.21E-05 | 0.000284492 | MEIS1 | protein_coding | 39.2 | -3.14538084 | 3.14538084 | ENSG00000143995 | n.s. |
| ENSG00000124224 | 71.04003857 | 1.009206689 | 0.2595258 | 3.888657 | 0.000100801 | 0.000770278 | PPP4R1L | transcribed_unprocessed_pseud | 41.96 | 3.142015932 | 3.142015932 | ENSG00000124224 | up |
| ENSG00000149646 | 84.54228436 | 1.025725972 | 0.2662161 | 3.852982 | 0.000116688 | 0.000875821 | CNBD2 | protein_coding | 45.62 | 3.136244114 | 3.136244114 | ENSG00000149646 | up |
| ENSG00000212443 | 58.16780023 | 1.009504877 | 0.2599313 | 3.883738 | 0.000102863 | 0.000783458 | SNORA53 | snoRNA | 39.36 | 3.135506392 | 3.135506392 | ENSG00000212443 | up |
| ENSG00000225213 | 27.84137474 | -1.140461769 | 0.3127716 | -3.646309 | 0.000266034 | 0.00179544 | | sense_intronic | 39.83 | -3.13151306 | 3.13151306 | ENSG00000225213 | down |
| ENSG00000080839 | 944.1926789 | -1.043672933 | 0.2736512 | -3.81388 | 0.000136802 | 0.001007772 | RBL1 | protein_coding | 43.07 | -3.127509522 | 3.127509522 | ENSG00000080839 | down |
| ENSG00000270108 | 45.20602353 | -1.111736352 | 0.3011548 | -3.691578 | 0.000222867 | 0.001541502 | | sense_intronic | 35.33 | -3.126264883 | 3.126264883 | ENSG00000270108 | down |
| ENSG00000207701 | 6.75624785 | -1.60553819 | 0.5261886 | -3.05126 | 0.00227883 | 0.011347696 | MIR597 | miRNA | 44.33 | -3.122920007 | 3.122920007 | ENSG00000207701 | down |
| ENSG00000066777 | 1388.33013 | -1.022361728 | 0.2655577 | -3.849867 | 0.000118182 | 0.000886555 | ARFGEF1 | protein_coding | 38.36 | -3.120548763 | 3.120548763 | ENSG00000066777 | down |
| ENSG00000100731 | 1482.245862 | -0.971053029 | 0.2458365 | -3.949995 | 7.82E-05 | 0.00061441 | PCNX1 | protein_coding | 36.87 | -3.118577405 | 3.118577405 | ENSG00000100731 | n.s. |
| ENSG00000223006 | 22.44256122 | -1.247788913 | 0.3596947 | -3.469022 | 0.000522357 | 0.003223749 | RN7SKP137 | misc_RNA | 52.12 | -3.109039259 | 3.109039259 | ENSG00000223006 | down |
| ENSG00000198604 | 4097.35924 | -1.067350698 | 0.284141 | -3.756412 | 0.000172367 | 0.001236562 | BAZ1A | protein_coding | 39.75 | -3.103625228 | 3.103625228 | ENSG00000198604 | down |
| ENSG00000266709 | 122.8628064 | 0.975199296 | 0.248048 | 3.931494 | 8.44E-05 | 0.000658842 | | lincRNA | 52.3 | 3.102322196 | 3.102322196 | ENSG00000266709 | n.s. |
| ENSG00000263276 | 235.4265943 | -0.996404742 | 0.2562649 | -3.888182 | 0.000100998 | 0.000771361 | | sense_overlapping | 41.07 | -3.101550935 | 3.101550935 | ENSG00000263276 | n.s. |
| ENSG00000111817 | 1057.345033 | -1.075120261 | 0.287408 | -3.740746 | 0.000183475 | 0.001303991 | DSE | protein_coding | 38.2 | -3.101426832 | 3.101426832 | ENSG00000111817 | down |

| | | | | | | | | | | | | | |
|---|---|---|---|---|---|---|---|---|---|---|---|---|---|
| ENSG00000236617 | 23.5193924 | 1.165732028 | 0.3249522 | 3.587395 | 0.000333998 | 0.002189191 | | antisense | 52.43 | 3.100516436 | 3.100516436 | ENSG00000236617 | up |
| ENSG00000239396 | 20.45608316 | -1.204085561 | 0.3412465 | -3.528492 | 0.000417935 | 0.002666777 | RN7SL414P | misc_RNA | 53.06 | -3.099332186 | 3.099332186 | ENSG00000239396 | down |
| ENSG00000163946 | 814.9859829 | -1.054072407 | 0.2793714 | -3.773015 | 0.000161286 | 0.00116533 | FAM208A | protein_coding | 38.7 | -3.09217538 | 3.09217538 | ENSG00000163946 | down |
| ENSG00000188848 | 1078.83988 | -1.055833807 | 0.2801158 | -3.769276 | 0.000163722 | 0.001181089 | BEND4 | protein_coding | 42.3 | -3.091182979 | 3.091182979 | ENSG00000188848 | down |
| ENSG00000227627 | 243.0139366 | -0.923921522 | 0.2293382 | -4.028642 | 5.61E-05 | 0.000463549 | | antisense | 42.78 | -3.080265644 | 3.080265644 | ENSG00000227627 | n.s. |
| ENSG00000138829 | 743.0239957 | -0.982643505 | 0.2518215 | -3.902143 | 9.53E-05 | 0.000733417 | FBN2 | protein_coding | 37.12 | -3.080242269 | 3.080242269 | ENSG00000138829 | n.s. |
| ENSG00000162971 | 1004.662012 | -1.019850548 | 0.2666677 | -3.824425 | 0.000131078 | 0.000970225 | TYW5 | protein_coding | 37.41 | -3.072939989 | 3.072939989 | ENSG00000162971 | down |
| ENSG00000165417 | 1368.44387 | -1.024543732 | 0.268537 | -3.81528 | 0.000136029 | 0.001002609 | GTF2A1 | protein_coding | 37.75 | -3.072472032 | 3.072472032 | ENSG00000165417 | down |
| ENSG00000102710 | 1529.135069 | -1.012088808 | 0.2636455 | -3.838824 | 0.000123625 | 0.000923902 | SUPT20H | protein_coding | 35.69 | -3.071056055 | 3.071056055 | ENSG00000102710 | down |
| ENSG00000237984 | 202.9897275 | 0.910996627 | 0.2247595 | 4.053207 | 5.05E-05 | 0.000425783 | PTENP1 | transcribed_processed_pseudog | 41.99 | 3.070798289 | 3.070798289 | ENSG00000237984 | n.s. |
| ENSG00000268201 | 27.54485675 | -1.139256296 | 0.3154572 | -3.611445 | 0.000304496 | 0.002021696 | | sense_intronic | 48.14 | -3.069480243 | 3.069480243 | ENSG00000268201 | down |
| ENSG00000232671 | 23.94553662 | 1.146795651 | 0.3185693 | 3.599831 | 0.000318424 | 0.002106138 | | antisense | 54.93 | 3.069413648 | 3.069413648 | ENSG00000232671 | up |
| ENSG00000257803 | 111.1026099 | 0.993885599 | 0.2566766 | 3.872132 | 0.000107888 | 0.000816369 | | processed_pseudogene | 41.61 | 3.069231285 | 3.069231285 | ENSG00000257803 | n.s. |
| ENSG00000276334 | 194.6451109 | -1.055257821 | 0.280927 | -3.756341 | 0.000172416 | 0.001236562 | | sense_intronic | 32.88 | -3.068461754 | 3.068461754 | ENSG00000276334 | down |
| ENSG00000248161 | 21.74243587 | 1.207648029 | 0.3449959 | 3.50047 | 0.000464438 | 0.002920166 | | lincRNA | 38.25 | 3.060895659 | 3.060895659 | ENSG00000248161 | up |
| ENSG00000274364 | 14.13141822 | -1.329501308 | 0.399386 | -3.328863 | 0.000872013 | 0.005006616 | | sense_intronic | 41.76 | -3.058458896 | 3.058458896 | ENSG00000274364 | down |
| ENSG00000120539 | 1498.721619 | -1.067377815 | 0.2863372 | -3.727695 | 0.000193239 | 0.001364335 | MASTL | protein_coding | 41.49 | -3.058121837 | 3.058121837 | ENSG00000120539 | down |
| ENSG00000260576 | 17.41506433 | -1.239147086 | 0.3590775 | -3.450918 | 0.000558683 | 0.003417639 | EIF5A2P1 | processed_pseudogene | 40.87 | -3.056076076 | 3.056076076 | ENSG00000260576 | down |
| ENSG00000272707 | 7.406907251 | -1.455401325 | 0.4584757 | -3.174435 | 0.001501286 | 0.007962819 | | lincRNA | 41.25 | -3.054790122 | 3.054790122 | ENSG00000272707 | down |
| ENSG00000227124 | 516.1928372 | -0.941332033 | 0.2372131 | -3.968297 | 7.24E-05 | 0.00057592 | ZNF717 | protein_coding | 41.48 | -3.049575065 | 3.049575065 | ENSG00000227124 | n.s. |
| ENSG00000179406 | 47.47976881 | 1.033527559 | 0.2735513 | 3.778186 | 0.000157975 | 0.001143777 | LINC00174 | processed_transcript | 44.08 | 3.040285154 | 3.040285154 | ENSG00000179406 | up |
| ENSG00000085231 | 407.462889 | -1.124119093 | 0.31082 | -3.616624 | 0.00029847 | 0.001986423 | AK6 | protein_coding | 41.64 | -3.037289184 | 3.037289184 | ENSG00000085231 | down |
| ENSG00000162687 | 1331.951393 | -1.070804825 | 0.2888737 | -3.706827 | 0.000209873 | 0.001463578 | KCNT2 | protein_coding | 33.86 | -3.035286519 | 3.035286519 | ENSG00000162687 | down |
| ENSG00000159882 | 211.3671161 | -0.969054769 | 0.2485685 | -3.898542 | 9.68E-05 | 0.00074154 | ZNF230 | protein_coding | 40.86 | -3.033010887 | 3.033010887 | ENSG00000159882 | n.s. |
| ENSG00000007541 | 7.16921802 | 1.561748429 | 0.512375 | 3.048058 | 0.002303258 | 0.011432535 | PIGQ | protein_coding | 64.36 | 3.032692818 | 3.032692818 | ENSG00000007541 | up |
| ENSG00000109920 | 2381.686052 | -0.997670126 | 0.2596681 | -3.842097 | 0.000121987 | 0.000912644 | FNBP4 | protein_coding | 43.42 | -3.032616473 | 3.032616473 | ENSG00000109920 | n.s. |
| ENSG00000261131 | 17.22649458 | -1.214105985 | 0.3496053 | -3.472791 | 0.000515077 | 0.00318588 | | sense_intronic | 43.42 | -3.031344076 | 3.031344076 | ENSG00000261131 | down |
| ENSG00000235423 | 134.0881302 | -0.958194338 | 0.2445262 | -3.918576 | 8.91E-05 | 0.000689752 | | antisense | 46.01 | -3.029146415 | 3.029146415 | ENSG00000235423 | n.s. |
| ENSG00000162222 | 197.612833 | -0.954030887 | 0.2430369 | -3.925456 | 8.66E-05 | 0.000674078 | TTC9C | protein_coding | 47.12 | -3.025508645 | 3.025508645 | ENSG00000162222 | n.s. |
| ENSG00000198707 | 2152.785829 | -1.068687432 | 0.2885879 | -3.703161 | 0.00021293 | 0.001480089 | CEP290 | protein_coding | 33.9 | -3.024077712 | 3.024077712 | ENSG00000198707 | down |
| ENSG00000162063 | 477.3487265 | -0.958451258 | 0.2448537 | -3.914383 | 9.06E-05 | 0.000699901 | CCNF | protein_coding | 52.76 | -3.023878392 | 3.023878392 | ENSG00000162063 | n.s. |
| ENSG00000270313 | 7.220586 | 1.517226373 | 0.4909731 | 3.090243 | 0.001999926 | 0.010188427 | COX6CP16 | processed_pseudogene | 49.07 | 3.022152414 | 3.022152414 | ENSG00000270313 | up |

| | | | | | | | | | | | | | |
|---|---|---|---|---|---|---|---|---|---|---|---|---|---|
| ENSG00000008853 | 202.4666387 | -0.920503526 | 0.2303914 | -3.99539 | 6.46E-05 | 0.000521611 | RHOBTB2 | protein_coding | 53.51 | -3.021693661 | 3.021693661 | ENSG00000008853 | n.s. |
| ENSG00000272086 | 79.85842696 | 1.015051839 | 0.2670513 | 3.800962 | 0.000144135 | 0.001056764 | | lincRNA | 52.49 | 3.020816649 | 3.020816649 | ENSG00000272086 | up |
| ENSG00000082269 | 391.8559757 | -0.963492951 | 0.2470262 | -3.900367 | 9.60E-05 | 0.000737596 | FAM135A | protein_coding | 36.09 | -3.017834698 | 3.017834698 | ENSG00000082269 | n.s. |
| ENSG00000168769 | 885.9728157 | -0.915518686 | 0.229236 | -3.993783 | 6.50E-05 | 0.000524552 | TET2 | protein_coding | 36.29 | -3.003094524 | 3.003094524 | ENSG00000168769 | n.s. |
| ENSG00000273210 | 29.51067267 | 1.140982118 | 0.3200483 | 3.565031 | 0.000363813 | 0.002360377 | | antisense | 47.15 | 2.997381296 | 2.997381296 | ENSG00000273210 | up |
| ENSG00000166669 | 415.3082216 | 0.901948415 | 0.2242878 | 4.021388 | 5.79E-05 | 0.000476121 | ATF7IP2 | protein_coding | 41.4 | 2.99652734 | 2.99652734 | ENSG00000166669 | n.s. |
| ENSG00000244710 | 31.0925912 | -1.160103623 | 0.3281437 | -3.535353 | 0.000407231 | 0.002614054 | RN7SL47P | misc_RNA | 54.85 | -2.996182799 | 2.996182799 | ENSG00000244710 | down |
| ENSG00000250938 | 27.94698941 | 1.121206971 | 0.312091 | 3.592564 | 0.00032744 | 0.002157573 | | antisense | 36.15 | 2.98917656 | 2.98917656 | ENSG00000250938 | up |
| ENSG00000103423 | 876.3915161 | -1.085970579 | 0.2974211 | -3.65129 | 0.000260927 | 0.001769548 | DNAJA3 | protein_coding | 49.02 | -2.988740622 | 2.988740622 | ENSG00000103423 | down |
| ENSG00000176155 | 102.3779014 | -0.943423554 | 0.2408874 | -3.916451 | 8.99E-05 | 0.000695083 | CCDC57 | protein_coding | 51.82 | -2.979296886 | 2.979296886 | ENSG00000176155 | n.s. |
| ENSG00000265727 | 36.58038318 | -1.084151003 | 0.2971944 | -3.647953 | 0.000264338 | 0.001786594 | RN7SL648P | misc_RNA | 53.16 | -2.9792189 | 2.9792189 | ENSG00000265727 | down |
| ENSG00000253816 | 47.28264586 | -1.071529392 | 0.2920212 | -3.669355 | 0.000243163 | 0.00166285 | | unprocessed_pseudogene | 40.49 | -2.977937614 | 2.977937614 | ENSG00000253816 | down |
| ENSG00000199753 | 97.84251611 | 0.940539535 | 0.2398497 | 3.92137 | 8.80E-05 | 0.000683702 | SNORD104 | snoRNA | 51.43 | 2.976932705 | 2.976932705 | ENSG00000199753 | n.s. |
| ENSG00000219085 | 44.27442247 | -1.074744896 | 0.2935988 | -3.66059 | 0.000251635 | 0.001716569 | NPM1P37 | processed_pseudogene | 39.24 | -2.972033804 | 2.972033804 | ENSG00000219085 | down |
| ENSG00000276900 | 69.36232332 | 0.985290909 | 0.2574878 | 3.826554 | 0.00012995 | 0.000964244 | | antisense | 46.74 | 2.971453174 | 2.971453174 | ENSG00000276900 | n.s. |
| ENSG00000208308 | 19.2627502 | -1.213108178 | 0.3527744 | -3.438765 | 0.000584375 | 0.003554759 | SNORA40B | snoRNA | 32.03 | -2.971132171 | 2.971132171 | ENSG00000208308 | down |
| ENSG00000276770 | 3.471647074 | -2.181356718 | 0.8673019 | -2.515107 | 0.011899643 | 0.043865164 | MIR6780B | miRNA | 59.49 | -2.962021182 | 2.962021182 | ENSG00000276770 | down |
| ENSG00000196584 | 1659.897086 | -1.045683697 | 0.2822672 | -3.704588 | 0.000211734 | 0.001472517 | XRCC2 | protein_coding | 42.57 | -2.961313291 | 2.961313291 | ENSG00000196584 | down |
| ENSG00000156509 | 272.3555229 | -0.958041631 | 0.2477751 | -3.866578 | 0.000110373 | 0.000832915 | FBXO43 | protein_coding | 38.91 | -2.950192909 | 2.950192909 | ENSG00000156509 | n.s. |
| ENSG00000272143 | 258.5339776 | 0.84956055 | 0.2065776 | 4.11255 | 3.91E-05 | 0.000337981 | FGF14-AS2 | lincRNA | 51.49 | 2.948915948 | 2.948915948 | ENSG00000272143 | n.s. |
| ENSG00000227354 | 247.4529973 | 0.887325857 | 0.2206527 | 4.021369 | 5.79E-05 | 0.000476121 | RBM26-AS1 | antisense | 38.6 | 2.947947072 | 2.947947072 | ENSG00000227354 | n.s. |
| ENSG00000151136 | 875.697865 | -0.889831518 | 0.2217879 | -4.012083 | 6.02E-05 | 0.000490897 | BTBD11 | protein_coding | 46.12 | -2.944461341 | 2.944461341 | ENSG00000151136 | n.s. |
| ENSG00000151491 | 122.0755354 | 0.974706806 | 0.2545399 | 3.829289 | 0.000128514 | 0.000956071 | EPS8 | protein_coding | 37.25 | 2.943136692 | 2.943136692 | ENSG00000151491 | n.s. |
| ENSG00000230149 | 39.32890006 | -1.078418002 | 0.2966849 | -3.634894 | 0.000278095 | 0.001866883 | | antisense | 57.97 | -2.942876439 | 2.942876439 | ENSG00000230149 | down |
| ENSG00000239407 | 72.36305663 | -1.136877604 | 0.3212832 | -3.538553 | 0.000402326 | 0.002583757 | | lincRNA | 38.6 | -2.941953045 | 2.941953045 | ENSG00000239407 | down |
| ENSG00000272948 | 64.30181808 | 1.020180866 | 0.2729575 | 3.737508 | 0.000185853 | 0.001318517 | | antisense | 57.28 | 2.938033325 | 2.938033325 | ENSG00000272948 | up |
| ENSG00000127603 | 52.05217035 | -1.045404355 | 0.2833469 | -3.689486 | 0.000224708 | 0.001550351 | MACF1 | protein_coding | 42.57 | -2.937136744 | 2.937136744 | ENSG00000127603 | down |
| ENSG00000174669 | 309.8463851 | -0.925310342 | 0.2357887 | -3.92432 | 8.70E-05 | 0.000676201 | SLC29A2 | protein_coding | 58.27 | -2.933163591 | 2.933163591 | ENSG00000174669 | n.s. |
| ENSG00000207931 | 4.343466497 | -2.246847547 | 0.9126676 | -2.461847 | 0.01382238 | 0.049510934 | MIR577 | miRNA | 39.58 | -2.932807589 | 2.932807589 | ENSG00000207931 | down |
| ENSG00000260018 | 22.63190398 | 1.155121031 | 0.330035 | 3.499995 | 0.000465266 | 0.002920606 | | antisense | 65.64 | 2.927685464 | 2.927685464 | ENSG00000260018 | up |
| ENSG00000252128 | 18.92004068 | 1.210917391 | 0.3549465 | 3.411549 | 0.00064595 | 0.003864873 | | snoRNA | 38.89 | 2.921779971 | 2.921779971 | ENSG00000252128 | up |
| ENSG00000264931 | 6.445776623 | -1.476127935 | 0.4795036 | -3.07845 | 0.002080805 | 0.010538756 | MIR3138 | miRNA | 67.07 | -2.918615845 | 2.918615845 | ENSG00000264931 | down |

| | | | | | | | | | | | | | |
|---|---|---|---|---|---|---|---|---|---|---|---|---|---|
| ENSG00000139746 | 2808.913999 | -1.017303 | 0.2728304 | -3.728702 | 0.000192469 | 0.001362003 | RBM26 | protein_coding | 34.26 | -2.915409389 | 2.915409389 | ENSG00000139746 | down |
| ENSG00000275662 | 4.400464521 | -2.036301037 | 0.7874883 | -2.585818 | 0.009714833 | 0.037098087 | SNORD112 | snoRNA | 37.66 | -2.913230988 | 2.913230988 | ENSG00000275662 | down |
| ENSG00000178193 | 47.857534 | 1.107808471 | 0.3106351 | 3.56627 | 0.000362099 | 0.002351446 | | antisense | 39.54 | 2.912057287 | 2.912057287 | ENSG00000178193 | up |
| ENSG00000270115 | 29.83243598 | -1.110326044 | 0.3125928 | -3.551989 | 0.000382331 | 0.002463302 | | lincRNA | 48.19 | -2.896265967 | 2.896265967 | ENSG00000270115 | down |
| ENSG00000199798 | 5.641885542 | -1.943841847 | 0.7357048 | -2.642149 | 0.008238178 | 0.032373738 | SNORD114-5 | snoRNA | 44.29 | -2.895949485 | 2.895949485 | ENSG00000199798 | down |
| ENSG00000151012 | 402.6646246 | -1.026627559 | 0.2777209 | -3.696616 | 0.000218493 | 0.001514978 | SLC7A11 | protein_coding | 35.52 | -2.894672552 | 2.894672552 | ENSG00000151012 | down |
| ENSG00000236184 | 30.62084832 | -1.130827347 | 0.3213971 | -3.518474 | 0.000434036 | 0.002755624 | TCEA1P4 | processed_pseudogene | 41.59 | -2.894669207 | 2.894669207 | ENSG00000236184 | down |
| ENSG00000169188 | 241.6005152 | -0.971927241 | 0.2557855 | -3.799774 | 0.000144828 | 0.001060167 | APEX2 | protein_coding | 49.09 | -2.891119594 | 2.891119594 | ENSG00000169188 | n.s. |
| ENSG00000216099 | 2.331990284 | -2.155648038 | 0.8629081 | -2.49812 | 0.012485392 | 0.045601815 | MIR889 | miRNA | 35.44 | -2.89076254 | 2.89076254 | ENSG00000216099 | down |
| ENSG00000251839 | 4.804720005 | -2.09002077 | 0.8231897 | -2.53893 | 0.01111922 | 0.041419122 | RNU7-12P | snRNA | 41.94 | -2.890078865 | 2.890078865 | ENSG00000251839 | down |
| ENSG00000222872 | 4.433130951 | -1.748520673 | 0.6255073 | -2.795364 | 0.005184128 | 0.02224988 | RNU4-78P | snRNA | 42.06 | -2.889731721 | 2.889731721 | ENSG00000222872 | down |
| ENSG00000259554 | 72.14959465 | -1.721725527 | 0.6113889 | -2.816089 | 0.004861221 | 0.021058738 | | sense_intronic | 39.21 | -2.886589322 | 2.886589322 | ENSG00000259554 | down |
| ENSG00000279347 | 80.75664842 | -1.125178915 | 0.3194915 | -3.52178 | 0.00042866 | 0.00272771 | | TEC | 44.12 | -2.885185763 | 2.885185763 | ENSG00000279347 | down |
| ENSG00000177200 | 6241.500967 | -0.979038275 | 0.2590277 | -3.779667 | 0.000157038 | 0.001138181 | CHD9 | protein_coding | 38.92 | -2.882081865 | 2.882081865 | ENSG00000177200 | n.s. |
| ENSG00000074603 | 1931.73854 | -0.910808603 | 0.2324629 | -3.918081 | 8.93E-05 | 0.000690784 | DPP8 | protein_coding | 40.97 | -2.878754297 | 2.878754297 | ENSG00000074603 | n.s. |
| ENSG00000164053 | 383.6094877 | -0.88768616 | 0.2237764 | -3.966844 | 7.28E-05 | 0.000578448 | ATRIP | protein_coding | 47.47 | -2.874093106 | 2.874093106 | ENSG00000164053 | n.s. |
| ENSG00000196505 | 884.7874335 | -0.966009649 | 0.2542148 | -3.799974 | 0.000144711 | 0.001060167 | GDAP2 | protein_coding | 36.55 | -2.873516974 | 2.873516974 | ENSG00000196505 | n.s. |
| ENSG00000163510 | 1189.88179 | -0.980814698 | 0.2602637 | -3.768543 | 0.000164203 | 0.001183949 | CWC22 | protein_coding | 35.03 | -2.870518187 | 2.870518187 | ENSG00000163510 | n.s. |
| ENSG00000142528 | 421.2984809 | -0.993795768 | 0.2656843 | -3.740514 | 0.000183644 | 0.001304529 | ZNF473 | protein_coding | 48.48 | -2.86664986 | 2.86664986 | ENSG00000142528 | n.s. |
| ENSG00000235499 | 116.2990853 | 0.984949164 | 0.2621549 | 3.757127 | 0.000171875 | 0.001234787 | | lincRNA | 69.31 | 2.864633854 | 2.864633854 | ENSG00000235499 | n.s. |
| ENSG00000260260 | 361.325808 | 0.866730271 | 0.216211 | 4.008724 | 6.10E-05 | 0.000496476 | SNHG19 | lincRNA | 54.09 | 2.863765308 | 2.863765308 | ENSG00000260260 | n.s. |
| ENSG00000201569 | 7.304026315 | -1.596889792 | 0.5469592 | -2.919577 | 0.003505065 | 0.016111264 | SNORD114-17 | snoRNA | 42.67 | -2.863016431 | 2.863016431 | ENSG00000201569 | down |
| ENSG00000165660 | 531.6279616 | -1.058498365 | 0.2926516 | -3.616923 | 0.000298126 | 0.001985081 | FAM175B | protein_coding | 42.62 | -2.860297292 | 2.860297292 | ENSG00000165660 | down |
| ENSG00000090905 | 1980.242622 | -0.931069779 | 0.2411064 | -3.861656 | 0.000112621 | 0.000848039 | TNRC6A | protein_coding | 42.59 | -2.859859026 | 2.859859026 | ENSG00000090905 | n.s. |
| ENSG00000154642 | 1275.22306 | -0.939444731 | 0.2443419 | -3.844796 | 0.000120653 | 0.000903629 | C21orf91 | protein_coding | 37.16 | -2.859679045 | 2.859679045 | ENSG00000154642 | n.s. |
| ENSG00000048649 | 2211.464676 | -0.992011776 | 0.2653141 | -3.739008 | 0.000184748 | 0.001311699 | RSF1 | protein_coding | 39.04 | -2.859142368 | 2.859142368 | ENSG00000048649 | n.s. |
| ENSG00000213693 | 109.0437464 | -1.035050343 | 0.2831215 | -3.655853 | 0.000256329 | 0.001744908 | SEC14L1P1 | processed_pseudogene | 50.02 | -2.854904174 | 2.854904174 | ENSG00000213693 | down |
| ENSG00000187801 | 167.1932187 | -1.060180584 | 0.2936694 | -3.610117 | 0.000306059 | 0.002029174 | ZFP69B | protein_coding | 41.8 | -2.854727717 | 2.854727717 | ENSG00000187801 | down |
| ENSG00000197563 | 366.8903916 | -1.045844674 | 0.2877057 | -3.63512 | 0.000277851 | 0.001866145 | PIGN | protein_coding | 38.03 | -2.854167237 | 2.854167237 | ENSG00000197563 | down |
| ENSG00000214067 | 45.26202842 | -1.058374736 | 0.2932782 | -3.608773 | 0.000307648 | 0.002038736 | | processed_pseudogene | 43.08 | -2.847704363 | 2.847704363 | ENSG00000214067 | down |
| ENSG00000268093 | 20.08864938 | 1.172395033 | 0.3425192 | 3.422859 | 0.000619662 | 0.003731642 | | antisense | 50.5 | 2.846692412 | 2.846692412 | ENSG00000268093 | up |
| ENSG00000236636 | 32.92030405 | -1.203971345 | 0.356848 | -3.373905 | 0.000741098 | 0.004361079 | | processed_pseudogene | 40.74 | -2.841861293 | 2.841861293 | ENSG00000236636 | down |

| | | | | | | | | | | | | | |
|---|---|---|---|---|---|---|---|---|---|---|---|---|---|
| ENSG00000180855 | 530.1418692 | -1.007690771 | 0.272573 | -3.696958 | 0.000218198 | 0.001513691 | ZNF443 | protein_coding | 41.62 | -2.841650595 | 2.841650595 | ENSG00000180855 | down |
| ENSG00000251756 | 11.89445254 | -1.469143083 | 0.4835093 | -3.0385 | 0.002377589 | 0.011717936 | RNA5SP385 | rRNA | 48.18 | -2.837134008 | 2.837134008 | ENSG00000251756 | down |
| ENSG00000101384 | 153.0443612 | 0.918840099 | 0.2374288 | 3.86996 | 0.000108853 | 0.000823198 | JAG1 | protein_coding | 47.52 | 2.834158273 | 2.834158273 | ENSG00000101384 | n.s. |
| ENSG00000113163 | 563.4961927 | -0.949948488 | 0.2500006 | -3.799784 | 0.000144822 | 0.001060167 | COL4A3BP | protein_coding | 37.13 | -2.825741034 | 2.825741034 | ENSG00000113163 | n.s. |
| ENSG00000143952 | 1047.002055 | -1.016868705 | 0.2772612 | -3.667548 | 0.000244888 | 0.001673819 | VPS54 | protein_coding | 36.99 | -2.823123943 | 2.823123943 | ENSG00000143952 | down |
| ENSG00000242125 | 3746.829312 | 0.790070762 | 0.1893232 | 4.173132 | 3.00E-05 | 0.000269006 | SNHG3 | sense_intronic | 46.41 | 2.820740705 | 2.820740705 | ENSG00000242125 | n.s. |
| ENSG00000259740 | 38.01889206 | -1.105961186 | 0.3149942 | -3.511053 | 0.000446336 | 0.002822134 | | sense_intronic | 39.95 | -2.819562196 | 2.819562196 | ENSG00000259740 | down |
| ENSG00000252255 | 12.448976 | -1.31676127 | 0.4111293 | -3.202791 | 0.001361025 | 0.007313608 | RNU2-35P | snRNA | 35.21 | -2.812428721 | 2.812428721 | ENSG00000252255 | down |
| ENSG00000091136 | 5684.58254 | -0.953694388 | 0.2520946 | -3.783081 | 0.000154899 | 0.001126191 | LAMB1 | protein_coding | 42.42 | -2.811861073 | 2.811861073 | ENSG00000091136 | n.s. |
| ENSG00000255568 | 105.7340002 | 0.917799063 | 0.2380924 | 3.854802 | 0.000115823 | 0.000869799 | BRWD1-AS2 | antisense | 59.24 | 2.808998596 | 2.808998596 | ENSG00000255568 | n.s. |
| ENSG00000182253 | 669.4622148 | -0.892426901 | 0.2283195 | -3.908675 | 9.28E-05 | 0.000714373 | SYNM | protein_coding | 49.91 | -2.807642169 | 2.807642169 | ENSG00000182253 | n.s. |
| ENSG00000177666 | 164.3143355 | 0.889693668 | 0.2273625 | 3.913106 | 9.11E-05 | 0.000702445 | PNPLA2 | protein_coding | 64.55 | 2.80554889 | 2.80554889 | ENSG00000177666 | n.s. |
| ENSG00000002016 | 254.7032365 | -0.988126005 | 0.2664688 | -3.708224 | 0.000208718 | 0.001456632 | RAD52 | protein_coding | 45.17 | -2.802967735 | 2.802967735 | ENSG00000002016 | n.s. |
| ENSG00000265291 | 3.393086517 | 1.948118839 | 0.7511663 | 2.593459 | 0.009501588 | 0.036423626 | MIR4710 | miRNA | 64.29 | 2.802596522 | 2.802596522 | ENSG00000265291 | up |
| ENSG00000158079 | 594.9782706 | -1.066139833 | 0.2990811 | -3.564718 | 0.000364248 | 0.002361036 | PTPDC1 | protein_coding | 39.39 | -2.800639918 | 2.800639918 | ENSG00000158079 | down |
| ENSG00000228629 | 16.07163444 | -1.213182428 | 0.36374 | -3.335301 | 0.000852071 | 0.004916446 | | antisense | 43.73 | -2.80044812 | 2.80044812 | ENSG00000228629 | down |
| ENSG00000146833 | 372.6352223 | -0.930642972 | 0.2435299 | -3.821473 | 0.000132657 | 0.000980871 | TRIM4 | protein_coding | 43.25 | -2.799735071 | 2.799735071 | ENSG00000146833 | n.s. |
| ENSG00000010818 | 31.00926225 | -1.132960566 | 0.3280556 | -3.453563 | 0.000553233 | 0.003390257 | HIVEP2 | protein_coding | 38.72 | -2.798149061 | 2.798149061 | ENSG00000010818 | down |
| ENSG00000047621 | 838.1215814 | -1.030146574 | 0.2840731 | -3.626343 | 0.000287464 | 0.00192421 | C12orf4 | protein_coding | 36.55 | -2.797618046 | 2.797618046 | ENSG00000047621 | down |
| ENSG00000203875 | 4425.002676 | 0.824601262 | 0.2028975 | 4.064128 | 4.82E-05 | 0.000408807 | SNHG5 | processed_transcript | 40.9 | 2.794146015 | 2.794146015 | ENSG00000203875 | n.s. |
| ENSG00000178966 | 447.8488918 | -0.97408002 | 0.261348 | -3.727138 | 0.000193666 | 0.001364582 | RMI1 | protein_coding | 38.66 | -2.790739527 | 2.790739527 | ENSG00000178966 | n.s. |
| ENSG00000118454 | 1199.914736 | -0.964651208 | 0.2575453 | -3.74556 | 0.000179992 | 0.001283826 | ANKRD13C | protein_coding | 37.55 | -2.789282966 | 2.789282966 | ENSG00000118454 | n.s. |
| ENSG00000234884 | 39.20791924 | 1.013367873 | 0.2775343 | 3.651325 | 0.000260891 | 0.001769548 | | antisense | 45.01 | 2.788927974 | 2.788927974 | ENSG00000234884 | up |
| ENSG00000137872 | 1053.307265 | -0.926607978 | 0.2424183 | -3.822352 | 0.000132185 | 0.0009779 | SEMA6D | protein_coding | 38.82 | -2.788817349 | 2.788817349 | ENSG00000137872 | n.s. |
| ENSG00000123908 | 1578.320777 | -0.896097334 | 0.2305591 | -3.886627 | 0.000101647 | 0.000775467 | AGO2 | protein_coding | 51.18 | -2.787253867 | 2.787253867 | ENSG00000123908 | n.s. |
| ENSG00000253797 | 806.3283176 | -1.051911158 | 0.2938335 | -3.579957 | 0.000343651 | 0.002241054 | UTP14C | protein_coding | 41.96 | -2.787088859 | 2.787088859 | ENSG00000253797 | down |
| ENSG00000251722 | 11.88500095 | -1.479602305 | 0.4942623 | -2.993557 | 0.002757462 | 0.013155947 | RNU5E-3P | snRNA | 32.35 | -2.782951262 | 2.782951262 | ENSG00000251722 | down |
| ENSG00000117597 | 1845.691863 | -0.925523983 | 0.242313 | -3.81954 | 0.000133701 | 0.000988068 | DIEXF | protein_coding | 41.44 | -2.781396974 | 2.781396974 | ENSG00000117597 | n.s. |
| ENSG00000111678 | 3138.721865 | 0.971732788 | 0.2608883 | 3.724709 | 0.000195541 | 0.001375703 | C12orf57 | protein_coding | 52.58 | 2.780589417 | 2.780589417 | ENSG00000111678 | n.s. |
| ENSG00000186448 | 961.2784502 | -0.960905964 | 0.2564582 | -3.746833 | 0.000179081 | 0.001278639 | ZNF197 | protein_coding | 40.2 | -2.780143278 | 2.780143278 | ENSG00000186448 | n.s. |
| ENSG00000241764 | 26.80138017 | 1.122385219 | 0.324577 | 3.457993 | 0.000544215 | 0.003339403 | | antisense | 56.77 | 2.779397538 | 2.779397538 | ENSG00000241764 | up |
| ENSG00000232184 | 76.50030241 | -1.136330523 | 0.3307053 | -3.436082 | 0.000590192 | 0.00358206 | | sense_intronic | 35.88 | -2.779313557 | 2.779313557 | ENSG00000232184 | down |

| | | | | | | | | | | | | | |
|---|---|---|---|---|---|---|---|---|---|---|---|---|---|
| ENSG00000103707 | 375.5319514 | -1.011130367 | 0.2772459 | -3.647053 | 0.000265265 | 0.001791121 | MTFMT | protein_coding | 40 | -2.777448801 | 2.777448801 | ENSG00000103707 | down |
| ENSG00000151458 | 1534.208447 | -0.920383061 | 0.240546 | -3.826225 | 0.000130124 | 0.000964702 | ANKRD50 | protein_coding | 35.15 | -2.775513615 | 2.775513615 | ENSG00000151458 | n.s. |
| ENSG00000227987 | 186.7802213 | -0.929103381 | 0.2440724 | -3.806672 | 0.000140849 | 0.001035399 | | sense_intronic | 51.01 | -2.773273305 | 2.773273305 | ENSG00000227987 | n.s. |
| ENSG00000151881 | 174.3148944 | -1.074858168 | 0.3043031 | -3.532196 | 0.000412124 | 0.00263695 | TMEM267 | protein_coding | 38.29 | -2.771949616 | 2.771949616 | ENSG00000151881 | down |
| ENSG00000158161 | 279.9109849 | -0.950781832 | 0.2527668 | -3.761499 | 0.000168898 | 0.001215283 | EYA3 | protein_coding | 39.52 | -2.771835887 | 2.771835887 | ENSG00000158161 | n.s. |
| ENSG00000068654 | 2544.210368 | -0.918728681 | 0.2400538 | -3.827178 | 0.000129621 | 0.000962511 | POLR1A | protein_coding | 46.69 | -2.77143159 | 2.77143159 | ENSG00000068654 | n.s. |
| ENSG00000279207 | 213.0389244 | -0.967671829 | 0.2597964 | -3.724732 | 0.000195523 | 0.001375703 | | TEC | 36.67 | -2.768969083 | 2.768969083 | ENSG00000279207 | n.s. |
| ENSG00000090612 | 2562.928464 | -1.04047312 | 0.2900538 | -3.587173 | 0.000334283 | 0.002189191 | ZNF268 | protein_coding | 41.98 | -2.76736328 | 2.76736328 | ENSG00000090612 | down |
| ENSG00000180096 | 17.41596073 | 1.17693266 | 0.3495051 | 3.367426 | 0.000758733 | 0.004453559 | 1-Sep | protein_coding | 52.75 | 2.767313264 | 2.767313264 | ENSG00000180096 | up |
| ENSG00000180998 | 247.8714395 | -1.039209822 | 0.289498 | -3.589696 | 0.000331064 | 0.002175276 | GPR137C | protein_coding | 37.31 | -2.766881269 | 2.766881269 | ENSG00000180998 | down |
| ENSG00000134255 | 1215.127227 | -0.987401651 | 0.2682105 | -3.681443 | 0.000231918 | 0.001594564 | CEPT1 | protein_coding | 36.54 | -2.762116074 | 2.762116074 | ENSG00000134255 | n.s. |
| ENSG00000266097 | 14.36412922 | 1.270506839 | 0.3930968 | 3.232046 | 0.001229074 | 0.006708116 | MIR5192 | miRNA | 55.43 | 2.761318865 | 2.761318865 | ENSG00000266097 | up |
| ENSG00000078549 | 1486.378497 | -0.859758589 | 0.2176494 | -3.950201 | 7.81E-05 | 0.00061423 | ADCYAP1R1 | protein_coding | 51.89 | -2.761259728 | 2.761259728 | ENSG00000078549 | n.s. |
| ENSG00000111596 | 3474.749996 | -1.002556447 | 0.2745507 | -3.651626 | 0.000260585 | 0.001769548 | CNOT2 | protein_coding | 35.15 | -2.759173442 | 2.759173442 | ENSG00000111596 | down |
| ENSG00000265421 | 3.453025256 | -1.774133133 | 0.6562661 | -2.703375 | 0.00686393 | 0.027910774 | MIR4459 | miRNA | 65.15 | -2.757407595 | 2.757407595 | ENSG00000265421 | down |
| ENSG00000155816 | 420.7399435 | -0.893611818 | 0.2309663 | -3.869014 | 0.000109277 | 0.000825085 | FMN2 | protein_coding | 39.28 | -2.755453018 | 2.755453018 | ENSG00000155816 | n.s. |
| ENSG00000178295 | 2889.189353 | -0.964030186 | 0.2590145 | -3.721916 | 0.000197717 | 0.001389606 | GEN1 | protein_coding | 35.87 | -2.754338666 | 2.754338666 | ENSG00000178295 | n.s. |
| ENSG00000198987 | 31.37420118 | 1.104967731 | 0.3186109 | 3.468079 | 0.000524193 | 0.003232214 | MIR16-2 | miRNA | 37.04 | 2.751922072 | 2.751922072 | ENSG00000198987 | up |
| ENSG00000268895 | 133.2292255 | -0.984535873 | 0.2675583 | -3.679705 | 0.000233503 | 0.001603882 | A1BG-AS1 | antisense | 55.91 | -2.751607968 | 2.751607968 | ENSG00000268895 | n.s. |
| ENSG00000161692 | 405.843727 | -0.959696935 | 0.2573854 | -3.728638 | 0.000192517 | 0.001362003 | DBF4B | protein_coding | 47.89 | -2.750320656 | 2.750320656 | ENSG00000161692 | n.s. |
| ENSG00000141378 | 2033.549753 | -0.954489668 | 0.2554908 | -3.735906 | 0.000187041 | 0.001325275 | PTRH2 | protein_coding | 39.24 | -2.746729219 | 2.746729219 | ENSG00000141378 | n.s. |
| ENSG00000263816 | 4.680538557 | 1.657539281 | 0.5922068 | 2.79892 | 0.005127386 | 0.022033518 | MIR548AW | miRNA | 41.54 | 2.746403609 | 2.746403609 | ENSG00000263816 | up |
| ENSG00000170949 | 667.3923323 | -1.034111579 | 0.288529 | -3.584081 | 0.000338266 | 0.002210082 | ZNF160 | protein_coding | 45.26 | -2.746178088 | 2.746178088 | ENSG00000170949 | down |
| ENSG00000122557 | 281.183118 | -0.951755104 | 0.2546537 | -3.737448 | 0.000185897 | 0.001318517 | HERPUD2 | protein_coding | 38.32 | -2.740973004 | 2.740973004 | ENSG00000122557 | n.s. |
| ENSG00000178878 | 350.8038511 | 0.828610997 | 0.2065831 | 4.011029 | 6.05E-05 | 0.000492228 | APOLD1 | protein_coding | 42.82 | 2.740907104 | 2.740907104 | ENSG00000178878 | n.s. |
| ENSG00000255062 | 55.29208883 | -0.996245098 | 0.2729249 | -3.650254 | 0.000261981 | 0.001773243 | | antisense | 45.33 | -2.740901251 | 2.740901251 | ENSG00000255062 | n.s. |
| ENSG00000199977 | 28.14683019 | -1.090102246 | 0.3129307 | -3.483526 | 0.000494856 | 0.003082718 | | snoRNA | 46.83 | -2.737318967 | 2.737318967 | ENSG00000199977 | down |
| ENSG00000157985 | 615.052072 | -0.897549264 | 0.233246 | -3.848081 | 0.000119047 | 0.000892083 | AGAP1 | protein_coding | 47.33 | -2.737161524 | 2.737161524 | ENSG00000157985 | n.s. |
| ENSG00000244748 | 31.35950156 | -1.09691647 | 0.3159965 | -3.471293 | 0.000517959 | 0.003200861 | RN7SL153P | misc_RNA | 49.66 | -2.73651392 | 2.73651392 | ENSG00000244748 | down |
| ENSG00000273107 | 27.46128708 | 1.076233276 | 0.3069449 | 3.506275 | 0.000454426 | 0.002866881 | | lincRNA | 49.08 | 2.736420353 | 2.736420353 | ENSG00000273107 | up |
| ENSG00000256044 | 6.276829541 | 1.46026769 | 0.4898026 | 2.981339 | 0.002869908 | 0.0135726 | | processed_pseudogene | 50 | 2.726811795 | 2.726811795 | ENSG00000256044 | up |
| ENSG00000274929 | 71.16166853 | 0.974442192 | 0.2649318 | 3.678086 | 0.00023499 | 0.00161171 | | sense_intronic | 49.57 | 2.721337493 | 2.721337493 | ENSG00000274929 | n.s. |

| | | | | | | | | | | | | | |
|---|---|---|---|---|---|---|---|---|---|---|---|---|---|
| ENSG00000272990 | 29.2432246 | -1.594591998 | 0.5609505 | -2.842661 | 0.004473868 | 0.019680778 | | antisense | 38.19 | -2.720306556 | 2.720306556 | ENSG00000272990 | down |
| ENSG00000268584 | 82.1653388 | 0.904678774 | 0.2369056 | 3.818731 | 0.00013414 | 0.000990787 | | antisense | 49.31 | 2.717672857 | 2.717672857 | ENSG00000268584 | n.s. |
| ENSG00000117625 | 1324.017689 | -0.985788546 | 0.2697839 | -3.653994 | 0.000258193 | 0.001755281 | RCOR3 | protein_coding | 37.38 | -2.716491391 | 2.716491391 | ENSG00000117625 | n.s. |
| ENSG00000060749 | 2690.660251 | -0.953092421 | 0.2565654 | -3.714813 | 0.000203354 | 0.001423502 | QSER1 | protein_coding | 38.54 | -2.713112919 | 2.713112919 | ENSG00000060749 | n.s. |
| ENSG00000137868 | 67.54104639 | -0.964792821 | 0.2613628 | -3.691394 | 0.000223029 | 0.001541822 | STRA6 | protein_coding | 55.21 | -2.712964485 | 2.712964485 | ENSG00000137868 | n.s. |
| ENSG00000207380 | 17.23264547 | -1.217930611 | 0.3724365 | -3.27017 | 0.001074828 | 0.005971601 | | misc_RNA | 38.05 | -2.708567138 | 2.708567138 | ENSG00000207380 | down |
| ENSG00000267280 | 346.6840497 | -0.897419577 | 0.2344341 | -3.828025 | 0.000129176 | 0.000960231 | TBX2-AS1 | antisense | 62.75 | -2.708075024 | 2.708075024 | ENSG00000267280 | n.s. |
| ENSG00000146966 | 158.6064654 | -1.041894474 | 0.2938664 | -3.545469 | 0.000391915 | 0.002520382 | DENND2A | protein_coding | 47.56 | -2.70739779 | 2.70739779 | ENSG00000146966 | down |
| ENSG00000166037 | 1172.558679 | -1.018878583 | 0.2844395 | -3.582058 | 0.000340898 | 0.002226233 | CEP57 | protein_coding | 37.69 | -2.70250344 | 2.70250344 | ENSG00000166037 | down |
| ENSG00000137275 | 218.2996659 | -1.026129123 | 0.2878104 | -3.565296 | 0.000363446 | 0.002359095 | RIPK1 | protein_coding | 45.29 | -2.695902482 | 2.695902482 | ENSG00000137275 | down |
| ENSG00000130021 | 160.486338 | -1.093093091 | 0.3169137 | -3.449182 | 0.000562287 | 0.003435157 | PUDP | protein_coding | 43.76 | -2.693439737 | 2.693439737 | ENSG00000130021 | down |
| ENSG00000207368 | 9.122326897 | -1.36862266 | 0.4458582 | -3.069637 | 0.002143194 | 0.010780236 | | misc_RNA | 43.24 | -2.692589542 | 2.692589542 | ENSG00000207368 | down |
| ENSG00000254491 | 61.26770687 | -0.981739347 | 0.2694306 | -3.643755 | 0.000268689 | 0.001809848 | | processed_pseudogene | 48.99 | -2.692280741 | 2.692280741 | ENSG00000254491 | n.s. |
| ENSG00000066455 | 542.9322257 | -1.012945372 | 0.2826006 | -3.584371 | 0.000337891 | 0.002208665 | GOLGA5 | protein_coding | 39.61 | -2.690251368 | 2.690251368 | ENSG00000066455 | down |
| ENSG00000131080 | 118.1414531 | -0.971623806 | 0.2654066 | -3.660888 | 0.000251343 | 0.001715415 | EDA2R | protein_coding | 39.73 | -2.687152786 | 2.687152786 | ENSG00000131080 | n.s. |
| ENSG00000221909 | 276.206488 | -0.858403968 | 0.2201627 | -3.898953 | 9.66E-05 | 0.000741098 | FAM200A | protein_coding | 44.02 | -2.686910995 | 2.686910995 | ENSG00000221909 | n.s. |
| ENSG00000149548 | 194.2840151 | -0.974708687 | 0.267009 | -3.650471 | 0.00026176 | 0.001773047 | CCDC15 | protein_coding | 37.57 | -2.68169625 | 2.68169625 | ENSG00000149548 | n.s. |
| ENSG00000081665 | 1047.742539 | -0.928091073 | 0.2479977 | -3.742338 | 0.000182316 | 0.00129708 | ZNF506 | protein_coding | 40.49 | -2.679429754 | 2.679429754 | ENSG00000081665 | n.s. |
| ENSG00000204138 | 297.374142 | -0.931059091 | 0.2492285 | -3.735765 | 0.000187145 | 0.001325343 | PHACTR4 | protein_coding | 42.87 | -2.679282547 | 2.679282547 | ENSG00000204138 | n.s. |
| ENSG00000129473 | 338.7427228 | -0.909064597 | 0.2403649 | -3.782019 | 0.000155562 | 0.001129242 | BCL2L2 | protein_coding | 51.4 | -2.679206851 | 2.679206851 | ENSG00000129473 | n.s. |
| ENSG00000267758 | 59.62472734 | -1.055403327 | 0.3014849 | -3.500684 | 0.000464066 | 0.002919654 | | antisense | 38.82 | -2.675097746 | 2.675097746 | ENSG00000267758 | down |
| ENSG00000252367 | 68.60532508 | 0.942428597 | 0.2541059 | 3.708803 | 0.000208241 | 0.001454034 | | misc_RNA | 37.36 | 2.674070892 | 2.674070892 | ENSG00000252367 | n.s. |
| ENSG00000202297 | 19.15213087 | -1.247243195 | 0.3888139 | -3.207815 | 0.001337474 | 0.007200945 | RNY3P12 | misc_RNA | 39.22 | -2.672356366 | 2.672356366 | ENSG00000202297 | down |
| ENSG00000238778 | 10.55989177 | -1.309780362 | 0.4188839 | -3.126834 | 0.001767 | 0.009162732 | RNU7-80P | snRNA | 40.98 | -2.669299674 | 2.669299674 | ENSG00000238778 | down |
| ENSG00000206630 | 45.75281291 | 0.990245831 | 0.2742967 | 3.610127 | 0.000306047 | 0.002029174 | SNORD60 | snoRNA | 42.17 | 2.666415764 | 2.666415764 | ENSG00000206630 | n.s. |
| ENSG00000170871 | 1265.914882 | -0.913986245 | 0.2431816 | -3.758451 | 0.000170969 | 0.001229543 | KIAA0232 | protein_coding | 42.27 | -2.659934281 | 2.659934281 | ENSG00000170871 | n.s. |
| ENSG00000260806 | 104.536043 | 0.901792596 | 0.2384201 | 3.782369 | 0.000155343 | 0.001128243 | | antisense | 52.38 | 2.658121633 | 2.658121633 | ENSG00000260806 | n.s. |
| ENSG00000227811 | 271.6792404 | -0.881043246 | 0.2302495 | -3.826473 | 0.000129993 | 0.000964244 | FAM212B-AS1 | antisense | 49.47 | -2.657061712 | 2.657061712 | ENSG00000227811 | n.s. |
| ENSG00000207023 | 48.31816413 | 0.992967857 | 0.2759152 | 3.598816 | 0.000319669 | 0.00211337 | RNU6-975P | snRNA | 38.1 | 2.656213292 | 2.656213292 | ENSG00000207023 | n.s. |
| ENSG00000118655 | 509.536606 | -0.913314886 | 0.2431595 | -3.756032 | 0.000172629 | 0.001237011 | DCLRE1B | protein_coding | 45.65 | -2.655578632 | 2.655578632 | ENSG00000118655 | n.s. |
| ENSG00000163312 | 300.6262904 | -0.912521706 | 0.2429341 | -3.756251 | 0.000172478 | 0.001236562 | HELQ | protein_coding | 38.46 | -2.653415971 | 2.653415971 | ENSG00000163312 | n.s. |
| ENSG00000011258 | 2550.357364 | -0.965966165 | 0.264952 | -3.645816 | 0.000266545 | 0.001797331 | MBTD1 | protein_coding | 39.5 | -2.651936294 | 2.651936294 | ENSG00000011258 | n.s. |

| | | | | | | | | | | | | | |
|---|---|---|---|---|---|---|---|---|---|---|---|---|---|
| ENSG00000206418 | 491.3820905 | -0.99547094 | 0.2773531 | -3.589183 | 0.000331716 | 0.002177676 | RAB12 | protein_coding | 42.23 | -2.649950261 | 2.649950261 | ENSG00000206418 | n.s. |
| ENSG00000240445 | 161.1899404 | -0.943017888 | 0.2555696 | -3.689868 | 0.000224371 | 0.001548791 | FOXO3B | processed_pseudogene | 60.32 | -2.649886909 | 2.649886909 | ENSG00000240445 | n.s. |
| ENSG00000278236 | 43.58513314 | -1.046221355 | 0.29906 | -3.498366 | 0.000468118 | 0.00293586 | | misc_RNA | 52.33 | -2.649309386 | 2.649309386 | ENSG00000278236 | down |
| ENSG00000213699 | 478.1745687 | -0.884990932 | 0.2322965 | -3.809747 | 0.000139109 | 0.001024228 | SLC35F6 | protein_coding | 52.66 | -2.645771892 | 2.645771892 | ENSG00000213699 | n.s. |
| ENSG00000141219 | 1208.863479 | -1.024705114 | 0.2900262 | -3.533147 | 0.000410644 | 0.002630775 | C17orf80 | protein_coding | 44.12 | -2.643653407 | 2.643653407 | ENSG00000141219 | down |
| ENSG00000040199 | 388.8423726 | -0.925035295 | 0.2485983 | -3.721004 | 0.000198432 | 0.00139393 | PHLPP2 | protein_coding | 41.82 | -2.641677946 | 2.641677946 | ENSG00000040199 | n.s. |
| ENSG00000273080 | 85.61839858 | 0.931788072 | 0.251404 | 3.706338 | 0.000210278 | 0.001465313 | | antisense | 51.74 | 2.640752161 | 2.640752161 | ENSG00000273080 | n.s. |
| ENSG00000275485 | 30.10820693 | -1.085177654 | 0.3167346 | -3.426142 | 0.000612221 | 0.003690023 | | sense_intronic | 43.4 | -2.640205721 | 2.640205721 | ENSG00000275485 | down |
| ENSG00000178403 | 1439.424785 | 0.781708621 | 0.1926661 | 4.057324 | 4.96E-05 | 0.000419364 | NEUROG2 | protein_coding | 56.57 | 2.640149371 | 2.640149371 | ENSG00000178403 | n.s. |
| ENSG00000038427 | 2136.372383 | -0.944715727 | 0.2567893 | -3.678953 | 0.000234193 | 0.001607036 | VCAN | protein_coding | 37.22 | -2.639511593 | 2.639511593 | ENSG00000038427 | n.s. |
| ENSG00000100503 | 2545.585372 | -0.901716136 | 0.2393159 | -3.767891 | 0.000164633 | 0.00118643 | NIN | protein_coding | 41.15 | -2.638203023 | 2.638203023 | ENSG00000100503 | n.s. |
| ENSG00000132604 | 271.2504711 | -0.935607125 | 0.2533173 | -3.69342 | 0.000221258 | 0.001532624 | TERF2 | protein_coding | 44.29 | -2.633326619 | 2.633326619 | ENSG00000132604 | n.s. |
| ENSG00000065526 | 1070.184574 | -0.918449239 | 0.2464204 | -3.727165 | 0.000193646 | 0.001364582 | SPEN | protein_coding | 42.99 | -2.631357324 | 2.631357324 | ENSG00000065526 | n.s. |
| ENSG00000197312 | 471.1741973 | -0.898281033 | 0.2383039 | -3.769477 | 0.00016359 | 0.001180749 | DDI2 | protein_coding | 42.4 | -2.630025349 | 2.630025349 | ENSG00000197312 | n.s. |
| ENSG00000108175 | 1705.786765 | -0.89970535 | 0.2388985 | -3.766057 | 0.000165846 | 0.001194555 | ZMIZ1 | protein_coding | 55.32 | -2.629653257 | 2.629653257 | ENSG00000108175 | n.s. |
| ENSG00000249709 | 353.9659988 | -0.969526597 | 0.2676482 | -3.622392 | 0.000291891 | 0.001951035 | ZNF564 | protein_coding | 44.24 | -2.627160168 | 2.627160168 | ENSG00000249709 | n.s. |
| ENSG00000200087 | 398.4771717 | 0.831418748 | 0.2124098 | 3.91422 | 9.07E-05 | 0.000699986 | SNORA73B | snoRNA | 54.9 | 2.623052071 | 2.623052071 | ENSG00000200087 | n.s. |
| ENSG00000199762 | 7.92927454 | 1.444237087 | 0.4915275 | 2.938263 | 0.00330057 | 0.015307741 | | misc_RNA | 38.46 | 2.621418714 | 2.621418714 | ENSG00000199762 | up |
| ENSG00000121621 | 1107.614409 | -1.018385359 | 0.2888308 | -3.52589 | 0.000422063 | 0.002691883 | KIF18A | protein_coding | 34.02 | -2.617193139 | 2.617193139 | ENSG00000121621 | down |
| ENSG00000173064 | 30.60884739 | -1.079072828 | 0.3155537 | -3.419617 | 0.000627093 | 0.003764993 | HECTD4 | protein_coding | 43.29 | -2.615927066 | 2.615927066 | ENSG00000173064 | down |
| ENSG00000029153 | 602.7281222 | -0.972450017 | 0.2695308 | -3.607936 | 0.000308643 | 0.002044354 | ARNTL2 | protein_coding | 39.26 | -2.615349861 | 2.615349861 | ENSG00000029153 | n.s. |
| ENSG00000122644 | 660.4770703 | 0.826996836 | 0.2110209 | 3.919029 | 8.89E-05 | 0.000689223 | ARL4A | protein_coding | 42.68 | 2.614666535 | 2.614666535 | ENSG00000122644 | n.s. |
| ENSG00000184208 | 363.7605602 | -0.903393263 | 0.2412043 | -3.745345 | 0.000180146 | 0.001284267 | C22orf46 | protein_coding | 51.21 | -2.612021446 | 2.612021446 | ENSG00000184208 | n.s. |
| ENSG00000197256 | 220.3837796 | -0.966828098 | 0.2674036 | -3.615614 | 0.000299637 | 0.001993235 | KANK2 | protein_coding | 52.98 | -2.610862781 | 2.610862781 | ENSG00000197256 | n.s. |
| ENSG00000156787 | 527.7309469 | -1.000064759 | 0.2816514 | -3.550718 | 0.000384182 | 0.002474082 | TBC1D31 | protein_coding | 39.82 | -2.606754686 | 2.606754686 | ENSG00000156787 | down |
| ENSG00000201649 | 4.06304601 | -1.698384253 | 0.6326267 | -2.684655 | 0.007260481 | 0.029199215 | RNY4P34 | misc_RNA | 40 | -2.606389434 | 2.606389434 | ENSG00000201649 | down |
| ENSG00000145348 | 820.549177 | -1.026635379 | 0.2932366 | -3.501047 | 0.000463433 | 0.002918309 | TBCK | protein_coding | 35.98 | -2.602385926 | 2.602385926 | ENSG00000145348 | down |
| ENSG00000127980 | 718.1996788 | -1.078168532 | 0.3160568 | -3.411313 | 0.000646509 | 0.003866558 | PEX1 | protein_coding | 37.11 | -2.601270782 | 2.601270782 | ENSG00000127980 | down |
| ENSG00000087258 | 377.7387864 | -0.835966512 | 0.2150507 | -3.8873 | 0.000101365 | 0.000773744 | GNAO1 | protein_coding | 49.48 | -2.601028489 | 2.601028489 | ENSG00000087258 | n.s. |
| ENSG00000121361 | 332.9048625 | -0.823473179 | 0.2106803 | -3.908639 | 9.28E-05 | 0.000714373 | KCNJ8 | protein_coding | 40.62 | -2.590708574 | 2.590708574 | ENSG00000121361 | n.s. |
| ENSG00000157184 | 366.1571205 | -1.024819704 | 0.2933697 | -3.49327 | 0.000477144 | 0.002981721 | CPT2 | protein_coding | 44.68 | -2.588215981 | 2.588215981 | ENSG00000157184 | down |
| ENSG00000159433 | 1446.897322 | -0.905493733 | 0.243268 | -3.722207 | 0.000197489 | 0.001388709 | STARD9 | protein_coding | 43.49 | -2.587347527 | 2.587347527 | ENSG00000159433 | n.s. |

| | | | | | | | | | | | | | |
|---|---|---|---|---|---|---|---|---|---|---|---|---|---|
| ENSG00000267682 | 43.95426373 | -1.006832231 | 0.2858023 | -3.522827 | 0.000426969 | 0.002718196 | | sense_intronic | 51.38 | -2.583248886 | 2.583248886 | ENSG00000267682 | down |
| ENSG00000163872 | 2220.913358 | -0.932220497 | 0.254471 | -3.663367 | 0.000248922 | 0.001699725 | YEATS2 | protein_coding | 42.58 | -2.58189774 | 2.58189774 | ENSG00000163872 | n.s. |
| ENSG00000274054 | 8.478865002 | 1.349551149 | 0.4464341 | 3.022957 | 0.002503174 | 0.012215806 | MIR4727 | miRNA | 50.91 | 2.581796433 | 2.581796433 | ENSG00000274054 | up |
| ENSG00000149970 | 404.203802 | -0.984422825 | 0.2764871 | -3.560466 | 0.000370198 | 0.002395104 | CNKSR2 | protein_coding | 34.96 | -2.579852878 | 2.579852878 | ENSG00000149970 | n.s. |
| ENSG00000130684 | 677.1020079 | -0.965693588 | 0.2689492 | -3.590617 | 0.000329897 | 0.002168909 | ZNF337 | protein_coding | 42.68 | -2.572374634 | 2.572374634 | ENSG00000130684 | n.s. |
| ENSG00000122484 | 1491.762049 | -0.973058698 | 0.2721226 | -3.575809 | 0.000349146 | 0.002273696 | RPAP2 | protein_coding | 38.47 | -2.572054623 | 2.572054623 | ENSG00000122484 | n.s. |
| ENSG00000230177 | 75.25751665 | -0.994186825 | 0.2813917 | -3.533107 | 0.000410706 | 0.002630775 | | antisense | 41.18 | -2.564918776 | 2.564918776 | ENSG00000230177 | n.s. |
| ENSG00000075461 | 251.8407088 | -0.927158263 | 0.2536516 | -3.655243 | 0.000256938 | 0.001747605 | CACNG4 | protein_coding | 52.37 | -2.556691401 | 2.556691401 | ENSG00000075461 | n.s. |
| ENSG00000252122 | 5.923904346 | -1.512591847 | 0.5349561 | -2.827506 | 0.00469121 | 0.020455908 | | snoRNA | 56.3 | -2.555041761 | 2.555041761 | ENSG00000252122 | down |
| ENSG00000267526 | 151.1888422 | -1.045642726 | 0.3046654 | -3.432102 | 0.000598923 | 0.00362279 | | processed_pseudogene | 38.22 | -2.552368776 | 2.552368776 | ENSG00000267526 | down |
| ENSG00000075420 | 86.80705668 | -1.029519019 | 0.297646 | -3.458871 | 0.000542444 | 0.003331475 | FNDC3B | protein_coding | 39.76 | -2.550492749 | 2.550492749 | ENSG00000075420 | down |
| ENSG00000207588 | 6.915881317 | -1.4619049 | 0.5081506 | -2.876913 | 0.004015868 | 0.018024802 | MIR593 | miRNA | 61 | -2.549751474 | 2.549751474 | ENSG00000207588 | down |
| ENSG00000151835 | 6293.614711 | -0.985969658 | 0.279056 | -3.533232 | 0.000410512 | 0.002630775 | SACS | protein_coding | 40.65 | -2.543719172 | 2.543719172 | ENSG00000151835 | n.s. |
| ENSG00000199179 | 86.61306031 | 0.893883715 | 0.2408046 | 3.712071 | 0.00020557 | 0.001436823 | MIRLET7I | miRNA | 53.57 | 2.540950952 | 2.540950952 | ENSG00000199179 | n.s. |
| ENSG00000068024 | 56.59484143 | -1.020986746 | 0.294818 | -3.463109 | 0.000533972 | 0.003285238 | HDAC4 | protein_coding | 51.33 | -2.53555237 | 2.53555237 | ENSG00000068024 | down |
| ENSG00000182568 | 7736.578032 | -0.85629416 | 0.2258863 | -3.790819 | 0.000150151 | 0.001095166 | SATB1 | protein_coding | 36.43 | -2.535076127 | 2.535076127 | ENSG00000182568 | n.s. |
| ENSG00000207185 | 5.154095453 | -1.560513666 | 0.5633845 | -2.769891 | 0.005607506 | 0.023744923 | RNU6-1157P | snRNA | 39.62 | -2.534944027 | 2.534944027 | ENSG00000207185 | down |
| ENSG00000128564 | 165.7651266 | 0.968875501 | 0.2723636 | 3.557287 | 0.000374705 | 0.002420892 | VGF | protein_coding | 68.27 | 2.534602162 | 2.534602162 | ENSG00000128564 | n.s. |
| ENSG00000140526 | 773.0152775 | -0.890146266 | 0.239621 | -3.714809 | 0.000203357 | 0.001423502 | ABHD2 | protein_coding | 44.01 | -2.533927748 | 2.533927748 | ENSG00000140526 | n.s. |
| ENSG00000164916 | 581.9868991 | -0.86074276 | 0.2277615 | -3.779141 | 0.00015737 | 0.001139994 | FOXK1 | protein_coding | 52.09 | -2.533249677 | 2.533249677 | ENSG00000164916 | n.s. |
| ENSG00000251013 | 78.72776995 | -0.954908528 | 0.2667027 | -3.580423 | 0.000343038 | 0.002238109 | GAPDHP62 | processed_pseudogene | 50.81 | -2.530621004 | 2.530621004 | ENSG00000251013 | n.s. |
| ENSG00000101003 | 779.5520687 | -0.959553348 | 0.2686654 | -3.571555 | 0.000354868 | 0.0023088 | GINS1 | protein_coding | 41.38 | -2.529971549 | 2.529971549 | ENSG00000101003 | n.s. |
| ENSG00000165480 | 638.6773819 | -0.960152905 | 0.2689954 | -3.569403 | 0.000357796 | 0.002324589 | SKA3 | protein_coding | 39.84 | -2.528710358 | 2.528710358 | ENSG00000165480 | n.s. |
| ENSG00000117971 | 598.7488636 | -0.918764415 | 0.25162 | -3.651397 | 0.000260818 | 0.001769548 | CHRNB4 | protein_coding | 46.53 | -2.528566226 | 2.528566226 | ENSG00000117971 | n.s. |
| ENSG00000163576 | 23.73677072 | 1.089687849 | 0.3259903 | 3.342701 | 0.000829674 | 0.004801142 | EFHB | protein_coding | 40.39 | 2.526610692 | 2.526610692 | ENSG00000163576 | up |
| ENSG00000081014 | 949.6763445 | -0.950351013 | 0.2650643 | -3.58536 | 0.000336614 | 0.002202383 | AP4E1 | protein_coding | 37.92 | -2.525184514 | 2.525184514 | ENSG00000081014 | n.s. |
| ENSG00000155850 | 600.0088705 | -0.977604998 | 0.2766282 | -3.534003 | 0.000409316 | 0.002626227 | SLC26A2 | protein_coding | 41.19 | -2.522873617 | 2.522873617 | ENSG00000155850 | n.s. |
| ENSG00000202242 | 2.420370431 | 1.992689495 | 0.8244448 | 2.417008 | 0.015648684 | 0.054441934 | RNU6-1276P | snRNA | 33.65 | 2.518891944 | 2.518891944 | ENSG00000202242 | up |
| ENSG00000257870 | 135.5668914 | 0.867808988 | 0.2312667 | 3.752417 | 0.000175138 | 0.001253055 | | lincRNA | 42.38 | 2.518407704 | 2.518407704 | ENSG00000257870 | n.s. |
| ENSG00000160993 | 163.7217594 | -1.058816159 | 0.3126546 | -3.386536 | 0.00070781 | 0.004184642 | ALKBH4 | protein_coding | 54.42 | -2.518226616 | 2.518226616 | ENSG00000160993 | down |
| ENSG00000113593 | 983.1663226 | -0.992893866 | 0.2836518 | -3.500396 | 0.000464567 | 0.002920166 | PPWD1 | protein_coding | 36.46 | -2.516581364 | 2.516581364 | ENSG00000113593 | n.s. |
| ENSG00000199032 | 4.159485136 | 1.657025137 | 0.62101 | 2.668275 | 0.00762419 | 0.030372417 | MIR425 | miRNA | 57.47 | 2.514569855 | 2.514569855 | ENSG00000199032 | up |

| | | | | | | | | | | | | | |
|---|---|---|---|---|---|---|---|---|---|---|---|---|---|
| ENSG00000186017 | 596.8880926 | -0.948755725 | 0.2650374 | -3.579705 | 0.000343982 | 0.002242164 | ZNF566 | protein_coding | 40.96 | -2.513569569 | 2.513569569 | ENSG00000186017 | n.s. |
| ENSG00000196141 | 79.90903847 | -1.048877115 | 0.308648 | -3.398296 | 0.000678071 | 0.004033046 | SPATS2L | protein_coding | 40.19 | -2.51139653 | 2.51139653 | ENSG00000196141 | down |
| ENSG00000278022 | 34.97079952 | 1.017451903 | 0.2947731 | 3.451644 | 0.000557182 | 0.003409954 | | sense_intronic | 45.1 | 2.510309726 | 2.510309726 | ENSG00000278022 | up |
| ENSG00000184349 | 712.5072028 | -0.87514887 | 0.2347467 | -3.728056 | 0.000192962 | 0.001363071 | EFNA5 | protein_coding | 38.27 | -2.507722908 | 2.507722908 | ENSG00000184349 | n.s. |
| ENSG00000280378 | 53.84316798 | -0.978807085 | 0.2787055 | -3.511976 | 0.000444788 | 0.002813625 | | TEC | 47.81 | -2.49667636 | 2.49667636 | ENSG00000280378 | n.s. |
| ENSG00000155592 | 977.7282094 | -0.881172579 | 0.2378143 | -3.705297 | 0.000211143 | 0.001469873 | ZKSCAN2 | protein_coding | 43.2 | -2.496115296 | 2.496115296 | ENSG00000155592 | n.s. |
| ENSG00000274023 | 53.07377328 | -0.992039605 | 0.2846484 | -3.48514 | 0.000491879 | 0.003065545 | | sense_overlapping | 41.8 | -2.493483902 | 2.493483902 | ENSG00000274023 | n.s. |
| ENSG00000162669 | 427.9974521 | -1.022896868 | 0.2984278 | -3.427619 | 0.0006089 | 0.003671594 | HFM1 | protein_coding | 37.16 | -2.490902553 | 2.490902553 | ENSG00000162669 | down |
| ENSG00000237298 | 1421.920646 | -0.90573421 | 0.2481982 | -3.649238 | 0.000263019 | 0.001778541 | TTN-AS1 | antisense | 37.97 | -2.490711109 | 2.490711109 | ENSG00000237298 | n.s. |
| ENSG00000172379 | 1216.29374 | -0.912852845 | 0.2512391 | -3.633402 | 0.000279708 | 0.001875904 | ARNT2 | protein_coding | 45.48 | -2.489157384 | 2.489157384 | ENSG00000172379 | n.s. |
| ENSG00000154240 | 284.5734153 | -0.934947738 | 0.2604919 | -3.589162 | 0.000331743 | 0.002177676 | CEP112 | protein_coding | 37.96 | -2.488837093 | 2.488837093 | ENSG00000154240 | n.s. |
| ENSG00000165030 | 314.8683148 | 0.933636939 | 0.2599055 | 3.592217 | 0.000327876 | 0.002159423 | NFIL3 | protein_coding | 41.93 | 2.48876079 | 2.48876079 | ENSG00000165030 | n.s. |
| ENSG00000087448 | 2118.328359 | -0.910747841 | 0.2504452 | -3.636515 | 0.000276351 | 0.001856969 | KLHL42 | protein_coding | 43.03 | -2.487430159 | 2.487430159 | ENSG00000087448 | n.s. |
| ENSG00000261420 | 109.8207514 | 0.877893922 | 0.2368979 | 3.705791 | 0.000210732 | 0.001467745 | | antisense | 54.52 | 2.487380215 | 2.487380215 | ENSG00000261420 | n.s. |
| ENSG00000241839 | 143.5895812 | 0.934189638 | 0.2603421 | 3.588315 | 0.000332822 | 0.002182702 | PLEKHO2 | protein_coding | 53.96 | 2.485883722 | 2.485883722 | ENSG00000241839 | n.s. |
| ENSG00000266573 | 7.929917414 | 1.414901239 | 0.4901209 | 2.886842 | 0.003891302 | 0.017544881 | | lincRNA | 41.83 | 2.484353725 | 2.484353725 | ENSG00000266573 | up |
| ENSG00000100441 | 743.0506096 | -0.918455445 | 0.2538039 | -3.61876 | 0.000296018 | 0.001972903 | KHNYN | protein_coding | 52.2 | -2.484324865 | 2.484324865 | ENSG00000100441 | n.s. |
| ENSG00000124766 | 4640.24432 | 0.777824844 | 0.197461 | 3.939131 | 8.18E-05 | 0.000640014 | SOX4 | protein_coding | 50.73 | 2.484224962 | 2.484224962 | ENSG00000124766 | n.s. |
| ENSG00000219433 | 43.59675889 | -0.991006187 | 0.2849589 | -3.477716 | 0.000505706 | 0.003139494 | BTBD10P2 | processed_pseudogene | 44.41 | -2.480627501 | 2.480627501 | ENSG00000219433 | n.s. |
| ENSG00000110925 | 655.1889757 | -0.882663603 | 0.2392083 | -3.689937 | 0.00022431 | 0.001548791 | CSRNP2 | protein_coding | 46.64 | -2.480290942 | 2.480290942 | ENSG00000110925 | n.s. |
| ENSG00000268129 | 56.22267363 | 0.947202043 | 0.26614 | 3.559037 | 0.000372217 | 0.002407047 | | lincRNA | 48.66 | 2.480263121 | 2.480263121 | ENSG00000268129 | n.s. |
| ENSG00000223658 | 90.08568028 | 0.989775538 | 0.2845235 | 3.478712 | 0.000503829 | 0.00313441 | C1GALT1C1L | protein_coding | 38.99 | 2.478243736 | 2.478243736 | ENSG00000223658 | n.s. |
| ENSG00000219608 | 13.4867872 | -1.213054133 | 0.3877159 | -3.128719 | 0.001755703 | 0.009121138 | HIGD1AP16 | processed_pseudogene | 39.92 | -2.474570973 | 2.474570973 | ENSG00000219608 | down |
| ENSG00000224505 | 115.2516165 | -0.883100847 | 0.2397348 | -3.683657 | 0.000229912 | 0.001582334 | | antisense | 46.18 | -2.47330211 | 2.47330211 | ENSG00000224505 | n.s. |
| ENSG00000201047 | 3.477276868 | -1.850184554 | 0.742212 | -2.492798 | 0.012674097 | 0.046121655 | | misc_RNA | 38.24 | -2.472022548 | 2.472022548 | ENSG00000201047 | down |
| ENSG00000268043 | 403.3966977 | -0.928763866 | 0.2588021 | -3.588703 | 0.000332328 | 0.002180489 | NBPF12 | protein_coding | 43.89 | -2.471854919 | 2.471854919 | ENSG00000268043 | n.s. |
| ENSG00000201317 | 9.2979953 | -1.298646638 | 0.4310777 | -3.012558 | 0.002590556 | 0.012545499 | RNU4-59P | snRNA | 37.07 | -2.469392225 | 2.469392225 | ENSG00000201317 | down |
| ENSG00000186272 | 309.914774 | -0.912688138 | 0.2522185 | -3.61864 | 0.000296155 | 0.001972903 | ZNF17 | protein_coding | 47.68 | -2.468724911 | 2.468724911 | ENSG00000186272 | n.s. |
| ENSG00000201574 | 7.829737938 | -2.039962803 | 0.8638479 | -2.361484 | 0.018201971 | 0.061735943 | RNU1-93P | snRNA | 44.97 | -2.467257313 | 2.467257313 | ENSG00000201574 | down |
| ENSG00000255081 | 19.77173908 | -1.135118365 | 0.3512712 | -3.231458 | 0.001231602 | 0.006713341 | | sense_intronic | 45.8 | -2.466681779 | 2.466681779 | ENSG00000255081 | down |
| ENSG00000279765 | 593.8243726 | -0.827903244 | 0.2177038 | -3.802888 | 0.000143019 | 0.001050238 | | protein_coding | 38.36 | -2.4660855 | 2.4660855 | ENSG00000279765 | n.s. |
| ENSG00000278318 | 414.7179443 | -0.987083432 | 0.2841605 | -3.473682 | 0.000513369 | 0.003179551 | ZNF229 | protein_coding | 41.83 | -2.465373363 | 2.465373363 | ENSG00000278318 | n.s. |

| | | | | | | | | | | | | | |
|---|---|---|---|---|---|---|---|---|---|---|---|---|---|
| ENSG00000116497 | 55.188256 | 0.957931154 | 0.2717191 | 3.525446 | 0.00042277 | 0.002695161 | S100PBP | protein_coding | 41.44 | 2.461323006 | 2.461323006 | ENSG00000116497 | n.s. |
| ENSG00000100629 | 1194.762327 | -0.922104767 | 0.2566657 | -3.59263 | 0.000327358 | 0.002157573 | CEP128 | protein_coding | 36.62 | -2.458363198 | 2.458363198 | ENSG00000100629 | n.s. |
| ENSG00000079335 | 188.5379789 | -0.907233648 | 0.2504939 | -3.62178 | 0.000292583 | 0.001953781 | CDC14A | protein_coding | 38.39 | -2.457808494 | 2.457808494 | ENSG00000079335 | n.s. |
| ENSG00000204439 | 290.063006 | -0.863784301 | 0.2326085 | -3.713468 | 0.000204438 | 0.001429631 | C6orf47 | protein_coding | 57.01 | -2.457272758 | 2.457272758 | ENSG00000204439 | n.s. |
| ENSG00000266144 | 3.401677986 | 1.865462289 | 0.7543199 | 2.473039 | 0.013396966 | 0.048173025 | MIR4654 | miRNA | 56.58 | 2.457179624 | 2.457179624 | ENSG00000266144 | up |
| ENSG00000170242 | 3086.113694 | -0.938744152 | 0.2640129 | -3.555676 | 0.000377009 | 0.002433517 | USP47 | protein_coding | 35.32 | -2.45365714 | 2.45365714 | ENSG00000170242 | n.s. |
| ENSG00000111266 | 498.5831932 | -0.973913291 | 0.2793037 | -3.486933 | 0.000488594 | 0.003046436 | DUSP16 | protein_coding | 40.28 | -2.450568319 | 2.450568319 | ENSG00000111266 | n.s. |
| ENSG00000198646 | 713.3679047 | -0.876573128 | 0.23823 | -3.679524 | 0.00023367 | 0.001604232 | NCOA6 | protein_coding | 43.48 | -2.449787752 | 2.449787752 | ENSG00000198646 | n.s. |
| ENSG00000143367 | 226.0565223 | 0.88674051 | 0.2423534 | 3.658873 | 0.000253327 | 0.001727261 | TUFT1 | protein_coding | 45.93 | 2.449746551 | 2.449746551 | ENSG00000143367 | n.s. |
| ENSG00000279161 | 99.15676666 | 1.056367955 | 0.3165225 | 3.337419 | 0.000845605 | 0.00488521 | | TEC | 56.04 | 2.441389682 | 2.441389682 | ENSG00000279161 | up |
| ENSG00000106692 | 113.6426909 | -0.994623009 | 0.2890132 | -3.441444 | 0.000578618 | 0.003527185 | FKTN | protein_coding | 37.44 | -2.439384245 | 2.439384245 | ENSG00000106692 | n.s. |
| ENSG00000264421 | 12.67819034 | 1.293082852 | 0.4313002 | 2.998104 | 0.002716648 | 0.012988971 | | antisense | 38.18 | 2.439304129 | 2.439304129 | ENSG00000264421 | up |
| ENSG00000279384 | 37.2472595 | -1.036961668 | 0.3078749 | -3.368127 | 0.000756808 | 0.00444601 | | TEC | 38.1 | -2.438964482 | 2.438964482 | ENSG00000279384 | down |
| ENSG00000168811 | 64.49115331 | 0.962197058 | 0.2748509 | 3.500796 | 0.000463871 | 0.002919654 | IL12A | protein_coding | 43.49 | 2.438850736 | 2.438850736 | ENSG00000168811 | n.s. |
| ENSG00000275523 | 3.028753159 | 1.768651126 | 0.6983728 | 2.532531 | 0.011324223 | 0.04208947 | MIR6508 | miRNA | 48.33 | 2.433357168 | 2.433357168 | ENSG00000275523 | up |
| ENSG00000261120 | 21.8599665 | 1.103521603 | 0.3390797 | 3.254461 | 0.001136079 | 0.0062545 | | antisense | 59.54 | 2.43194906 | 2.43194906 | ENSG00000261120 | up |
| ENSG00000119888 | 55.4530826 | 0.958363245 | 0.2738183 | 3.499997 | 0.000465264 | 0.002920606 | EPCAM | protein_coding | 44.35 | 2.428997538 | 2.428997538 | ENSG00000119888 | n.s. |
| ENSG00000199879 | 5.898558384 | 1.509729177 | 0.5479374 | 2.755295 | 0.005863925 | 0.02461325 | RNU1-120P | snRNA | 42.68 | 2.428899146 | 2.428899146 | ENSG00000199879 | up |
| ENSG00000207762 | 2.533197171 | -2.462625504 | 1.1677314 | -2.108897 | 0.034953449 | 0.103236503 | MIR329-2 | miRNA | 41.67 | -2.428559309 | 2.428559309 | ENSG00000207762 | n.s. |
| ENSG00000227210 | 15.38367059 | 1.195695413 | 0.3830944 | 3.121151 | 0.001801458 | 0.00932405 | | antisense | 39.4 | 2.427734476 | 2.427734476 | ENSG00000227210 | up |
| ENSG00000157193 | 2421.55969 | -0.984498425 | 0.2853324 | -3.450357 | 0.000559847 | 0.003423251 | LRP8 | protein_coding | 51.78 | -2.427341128 | 2.427341128 | ENSG00000157193 | n.s. |
| ENSG00000122863 | 264.9943528 | -0.86909675 | 0.2363236 | -3.677571 | 0.000235466 | 0.001614176 | CHST3 | protein_coding | 51.39 | -2.426560861 | 2.426560861 | ENSG00000122863 | n.s. |
| ENSG00000147874 | 2577.096993 | -0.977043593 | 0.2822129 | -3.46208 | 0.000536017 | 0.003296364 | HAUS6 | protein_coding | 39.7 | -2.424987818 | 2.424987818 | ENSG00000147874 | n.s. |
| ENSG00000108306 | 456.3080451 | -0.89408728 | 0.2469508 | -3.620508 | 0.000294025 | 0.001960592 | FBXL20 | protein_coding | 43.28 | -2.420842117 | 2.420842117 | ENSG00000108306 | n.s. |
| ENSG00000270487 | 3.659162121 | 1.695635908 | 0.6571881 | 2.580138 | 0.009876091 | 0.037528608 | | antisense | 42.64 | 2.417362213 | 2.417362213 | ENSG00000270487 | up |
| ENSG00000184857 | 107.7051886 | -0.947258331 | 0.2696372 | -3.513085 | 0.000442936 | 0.002805729 | TMEM186 | protein_coding | 47.7 | -2.417360001 | 2.417360001 | ENSG00000184857 | n.s. |
| ENSG00000110436 | 612.8689783 | -1.002456766 | 0.2942915 | -3.40634 | 0.000658402 | 0.0039242 | SLC1A2 | protein_coding | 42.2 | -2.412160504 | 2.412160504 | ENSG00000110436 | down |
| ENSG00000238835 | 35.4293506 | -1.063302673 | 0.3215919 | -3.306372 | 0.000945124 | 0.005390805 | SCARNA18 | snoRNA | 40.3 | -2.411938742 | 2.411938742 | ENSG00000238835 | down |
| ENSG00000101331 | 127.4849849 | -0.9875187 | 0.2878401 | -3.430789 | 0.000601829 | 0.003635057 | CCM2L | protein_coding | 50.16 | -2.409040799 | 2.409040799 | ENSG00000101331 | n.s. |
| ENSG00000178502 | 463.6995021 | 0.857217437 | 0.2325976 | 3.685409 | 0.000228335 | 0.001573819 | KLHL11 | protein_coding | 47.67 | 2.402819298 | 2.402819298 | ENSG00000178502 | n.s. |
| ENSG00000259767 | 53.43750819 | -1.049398711 | 0.3162011 | -3.31877 | 0.00090415 | 0.005184711 | | sense_intronic | 42.08 | -2.398165127 | 2.398165127 | ENSG00000259767 | down |
| ENSG00000088035 | 466.6515324 | -1.00907945 | 0.2981363 | -3.384624 | 0.000712758 | 0.004210321 | ALG6 | protein_coding | 37.93 | -2.39725467 | 2.39725467 | ENSG00000088035 | down |

| | | | | | | | | | | | | | |
|---|---|---|---|---|---|---|---|---|---|---|---|---|---|
| ENSG00000163755 | 659.1752152 | -0.945421244 | 0.2701191 | -3.500017 | 0.000465229 | 0.002920606 | HPS3 | protein_coding | 38.69 | -2.396195687 | 2.396195687 | ENSG00000163755 | n.s. |
| ENSG00000275092 | 42.56007769 | 0.977167236 | 0.2841639 | 3.438745 | 0.000584417 | 0.003554759 | | sense_intronic | 46.68 | 2.393268024 | 2.393268024 | ENSG00000275092 | n.s. |
| ENSG00000231249 | 24.03261644 | 1.061610543 | 0.3221769 | 3.295117 | 0.000983808 | 0.00557718 | ITPR1-AS1 | antisense | 47.6 | 2.392429919 | 2.392429919 | ENSG00000231249 | up |
| ENSG00000272469 | 16.66702189 | -1.179327549 | 0.3780612 | -3.119409 | 0.00181214 | 0.009361941 | RN7SL760P | misc_RNA | 54.44 | -2.392424075 | 2.392424075 | ENSG00000272469 | down |
| ENSG00000204859 | 109.5120453 | -0.954191997 | 0.2743143 | -3.478463 | 0.000504298 | 0.003134528 | ZBTB48 | protein_coding | 57.78 | -2.389132459 | 2.389132459 | ENSG00000204859 | n.s. |
| ENSG00000119596 | 1800.43984 | -0.86336259 | 0.2359929 | -3.658426 | 0.000253769 | 0.001729429 | YLPM1 | protein_coding | 39.67 | -2.384691519 | 2.384691519 | ENSG00000119596 | n.s. |
| ENSG00000233010 | 36.11341727 | -1.051896417 | 0.3184798 | -3.302867 | 0.000957019 | 0.005447495 | RPEP4 | processed_pseudogene | 47.34 | -2.381286399 | 2.381286399 | ENSG00000233010 | down |
| ENSG00000165272 | 64.09179362 | 1.023981348 | 0.3061773 | 3.344407 | 0.000824587 | 0.004775678 | AQP3 | protein_coding | 55.65 | 2.376624832 | 2.376624832 | ENSG00000165272 | up |
| ENSG00000252577 | 15.51188144 | -1.223575596 | 0.4014765 | -3.047689 | 0.002306081 | 0.011438392 | SCARNA20 | scaRNA | 61.07 | -2.375737255 | 2.375737255 | ENSG00000252577 | down |
| ENSG00000272183 | 63.38943688 | -0.934156763 | 0.2665907 | -3.504086 | 0.000458177 | 0.002886518 | | antisense | 57.94 | -2.37240861 | 2.37240861 | ENSG00000272183 | n.s. |
| ENSG00000267680 | 306.8681126 | -0.907888344 | 0.2554498 | -3.554077 | 0.000379308 | 0.00244722 | ZNF224 | protein_coding | 39.97 | -2.370793325 | 2.370793325 | ENSG00000267680 | n.s. |
| ENSG00000200816 | 31.46610182 | 1.055635672 | 0.3211673 | 3.286872 | 0.001013069 | 0.005712825 | SNORA38 | snoRNA | 44.7 | 2.36794817 | 2.36794817 | ENSG00000200816 | up |
| ENSG00000232545 | 38.97368949 | -1.027536098 | 0.3086112 | -3.329549 | 0.000869867 | 0.004996352 | | sense_intronic | 55.52 | -2.364717113 | 2.364717113 | ENSG00000232545 | down |
| ENSG00000207956 | 9.706669922 | -1.286823057 | 0.4352669 | -2.9564 | 0.003112533 | 0.014571551 | MIR579 | miRNA | 32.65 | -2.363243113 | 2.363243113 | ENSG00000207956 | down |
| ENSG00000269893 | 2149.949091 | 0.756607252 | 0.1943113 | 3.89379 | 9.87E-05 | 0.00075498 | SNHG8 | lincRNA | 45.83 | 2.362176494 | 2.362176494 | ENSG00000269893 | n.s. |
| ENSG00000239774 | 24.59379569 | 1.070384912 | 0.3287186 | 3.256235 | 0.001129003 | 0.006218 | | lincRNA | 38.78 | 2.36164295 | 2.36164295 | ENSG00000239774 | up |
| ENSG00000102908 | 787.8293955 | -0.934380348 | 0.2674199 | -3.494057 | 0.00047574 | 0.002974281 | NFAT5 | protein_coding | 38.64 | -2.360822195 | 2.360822195 | ENSG00000102908 | n.s. |
| ENSG00000091732 | 458.6321744 | -0.928359238 | 0.2647716 | -3.506265 | 0.000454443 | 0.002866881 | ZC3HC1 | protein_coding | 44.46 | -2.36043725 | 2.36043725 | ENSG00000091732 | n.s. |
| ENSG00000127720 | 171.8158123 | -1.055026621 | 0.3218953 | -3.277546 | 0.001047139 | 0.005848098 | METTL25 | protein_coding | 35.3 | -2.355858962 | 2.355858962 | ENSG00000127720 | down |
| ENSG00000091436 | 900.7494376 | -0.960206978 | 0.2790397 | -3.441112 | 0.000579328 | 0.003529973 | MAP3K20 | protein_coding | 38.91 | -2.354647019 | 2.354647019 | ENSG00000091436 | n.s. |
| ENSG00000283101 | 48.69666353 | 0.962999544 | 0.2806152 | 3.431744 | 0.000599714 | 0.003625624 | BMS1P21 | transcribed_unprocessed_pseud | 36.5 | 2.350313299 | 2.350313299 | ENSG00000283101 | n.s. |
| ENSG00000263439 | 12.451267 | -1.206144196 | 0.3949023 | -3.054285 | 0.002255977 | 0.011262087 | MIR4753 | miRNA | 38.55 | -2.350028571 | 2.350028571 | ENSG00000263439 | down |
| ENSG00000246090 | 64.25555253 | 0.919818814 | 0.2617816 | 3.513688 | 0.000441932 | 0.002800647 | | antisense | 38.9 | 2.348059825 | 2.348059825 | ENSG00000246090 | n.s. |
| ENSG00000270820 | 54.33159353 | 0.915755948 | 0.2602246 | 3.519099 | 0.000433015 | 0.002751652 | | antisense | 44.62 | 2.344707455 | 2.344707455 | ENSG00000270820 | n.s. |
| ENSG00000071205 | 525.129962 | -0.915552349 | 0.2601945 | -3.518723 | 0.000433629 | 0.002754297 | ARHGAP10 | protein_coding | 40.94 | -2.343804171 | 2.343804171 | ENSG00000071205 | n.s. |
| ENSG00000278177 | 45.62069886 | 0.957538915 | 0.2785615 | 3.437442 | 0.000587237 | 0.003566694 | | antisense | 52.69 | 2.34380068 | 2.34380068 | ENSG00000278177 | n.s. |
| ENSG00000235090 | 22.71927399 | 1.101759689 | 0.3447807 | 3.195538 | 0.001395706 | 0.007465391 | RPL7L1P3 | processed_pseudogene | 46.3 | 2.343384956 | 2.343384956 | ENSG00000235090 | up |
| ENSG00000206652 | 157.7347653 | 0.914034998 | 0.259704 | 3.519526 | 0.000432319 | 0.002748481 | RNU1-1 | snRNA | 54.27 | 2.340758872 | 2.340758872 | ENSG00000206652 | n.s. |
| ENSG00000273162 | 102.6711051 | 0.864270108 | 0.2390846 | 3.614913 | 0.000300448 | 0.001997679 | | lincRNA | 51.28 | 2.333075029 | 2.333075029 | ENSG00000273162 | n.s. |
| ENSG00000281881 | 16.96572868 | 1.128138422 | 0.3580136 | 3.151105 | 0.001626541 | 0.00855224 | | lincRNA | 47.65 | 2.332900153 | 2.332900153 | ENSG00000281881 | up |
| ENSG00000279197 | 90.06404146 | 1.048222798 | 0.3205272 | 3.270308 | 0.001074303 | 0.005971066 | | TEC | 36.38 | 2.331193117 | 2.331193117 | ENSG00000279197 | up |
| ENSG00000075292 | 6917.93531 | -0.943245284 | 0.2731438 | -3.453292 | 0.000553788 | 0.003392166 | ZNF638 | protein_coding | 38.16 | -2.329365785 | 2.329365785 | ENSG00000075292 | n.s. |

| | | | | | | | | | | | | | |
|---|---|---|---|---|---|---|---|---|---|---|---|---|---|
| ENSG00000228203 | 264.7535291 | -0.896714263 | 0.2529884 | -3.544487 | 0.000393377 | 0.002528622 | RNF144A-AS1 | processed_transcript | 51.22 | -2.328871064 | 2.328871064 | ENSG00000228203 | n.s. |
| ENSG00000261186 | 41.22418757 | 0.95951046 | 0.2804096 | 3.421817 | 0.000622042 | 0.003741116 | LINC01238 | antisense | 48.42 | 2.32873074 | 2.32873074 | ENSG00000261186 | n.s. |
| ENSG00000264049 | 8.239763216 | -1.312095319 | 0.4523645 | -2.900527 | 0.003725359 | 0.016867697 | MIR4737 | miRNA | 62.96 | -2.326271794 | 2.326271794 | ENSG00000264049 | down |
| ENSG00000158987 | 1112.541239 | -0.960746506 | 0.2811551 | -3.417141 | 0.000632825 | 0.003795979 | RAPGEF6 | protein_coding | 37.88 | -2.325656224 | 2.325656224 | ENSG00000158987 | n.s. |
| ENSG00000260318 | 47.25194359 | -0.99927481 | 0.2984711 | -3.347978 | 0.000814034 | 0.004733776 | COX6CP1 | processed_pseudogene | 45.58 | -2.323106341 | 2.323106341 | ENSG00000260318 | n.s. |
| ENSG00000174473 | 117.1298809 | 0.877194957 | 0.2451621 | 3.57802 | 0.000346206 | 0.002255606 | GALNTL6 | protein_coding | 36.53 | 2.32170408 | 2.32170408 | ENSG00000174473 | n.s. |
| ENSG00000125319 | 316.3874342 | -0.928008189 | 0.2670263 | -3.475343 | 0.0005102 | 0.003165559 | C17orf53 | protein_coding | 50.23 | -2.319602443 | 2.319602443 | ENSG00000125319 | n.s. |
| ENSG00000200428 | 10.0626578 | -1.765846004 | 0.7151184 | -2.469306 | 0.013537548 | 0.048615756 | | misc_RNA | 34.19 | -2.318949506 | 2.318949506 | ENSG00000200428 | down |
| ENSG00000257151 | 379.4714583 | -1.033193512 | 0.314805 | -3.282011 | 0.001030696 | 0.005777126 | PWAR6 | lincRNA | 34.45 | -2.312584796 | 2.312584796 | ENSG00000257151 | down |
| ENSG00000201950 | 8.312420053 | -1.469483868 | 0.5402119 | -2.720199 | 0.006524269 | 0.026740474 | SNORD113-9 | snoRNA | 41.67 | -2.311249629 | 2.311249629 | ENSG00000201950 | down |
| ENSG00000139767 | 683.1049278 | -0.816220839 | 0.220457 | -3.702404 | 0.000213566 | 0.001483032 | SRRM4 | protein_coding | 44.14 | -2.308965792 | 2.308965792 | ENSG00000139767 | n.s. |
| ENSG00000138363 | 1004.858662 | -0.861142151 | 0.2390357 | -3.602567 | 0.00031509 | 0.002085072 | ATIC | protein_coding | 41.16 | -2.308617771 | 2.308617771 | ENSG00000138363 | n.s. |
| ENSG00000248794 | 20.83291289 | -1.123211902 | 0.3576343 | -3.140671 | 0.00168561 | 0.008816208 | | processed_pseudogene | 44.44 | -2.307883887 | 2.307883887 | ENSG00000248794 | down |
| ENSG00000202124 | 5.011646621 | -1.526193734 | 0.5731884 | -2.662639 | 0.007753054 | 0.030788867 | RNY4P3 | misc_RNA | 43.16 | -2.307004049 | 2.307004049 | ENSG00000202124 | down |
| ENSG00000201662 | 27.94130174 | -1.126420799 | 0.3593108 | -3.134948 | 0.001718844 | 0.008953037 | RNU6-60P | snRNA | 41.75 | -2.306943165 | 2.306943165 | ENSG00000201662 | down |
| ENSG00000251865 | 8.646519392 | -1.340872317 | 0.4698631 | -2.853751 | 0.004320635 | 0.019091148 | RNU6-518P | snRNA | 36.46 | -2.305184723 | 2.305184723 | ENSG00000251865 | down |
| ENSG00000132549 | 1102.948808 | -0.881832792 | 0.2479909 | -3.555908 | 0.000376676 | 0.002432497 | VPS13B | protein_coding | 36.65 | -2.305064753 | 2.305064753 | ENSG00000132549 | n.s. |
| ENSG00000164327 | 1179.947804 | -0.952217149 | 0.278659 | -3.417141 | 0.000632826 | 0.003795979 | RICTOR | protein_coding | 35.53 | -2.305009413 | 2.305009413 | ENSG00000164327 | n.s. |
| ENSG00000276712 | 73.47483005 | 0.905236332 | 0.2579828 | 3.508901 | 0.000449962 | 0.002843766 | MIR7111 | miRNA | 58.33 | 2.304827789 | 2.304827789 | ENSG00000276712 | n.s. |
| ENSG00000168152 | 402.1578927 | -1.0351599 | 0.3163308 | -3.272396 | 0.0010664 | 0.005936618 | THAP9 | protein_coding | 37.02 | -2.304743023 | 2.304743023 | ENSG00000168152 | down |
| ENSG00000197162 | 432.0629661 | -0.838649049 | 0.2299442 | -3.647185 | 0.000265129 | 0.001791069 | ZNF785 | protein_coding | 49.75 | -2.303674672 | 2.303674672 | ENSG00000197162 | n.s. |
| ENSG00000207632 | 4.38754281 | -1.5868794 | 0.6093352 | -2.60428 | 0.009206759 | 0.035449588 | MIR588 | miRNA | 44.58 | -2.30159213 | 2.30159213 | ENSG00000207632 | down |
| ENSG00000236358 | 15.13489971 | -1.164690511 | 0.3788565 | -3.074226 | 0.002110497 | 0.010661992 | | antisense | 40.86 | -2.296957954 | 2.296957954 | ENSG00000236358 | down |
| ENSG00000265094 | 11.40381882 | -1.283223829 | 0.440074 | -2.915928 | 0.003546327 | 0.016225932 | | sense_intronic | 38.9 | -2.29670164 | 2.29670164 | ENSG00000265094 | down |
| ENSG00000258738 | 160.6453547 | 0.84047519 | 0.2310936 | 3.636947 | 0.000275889 | 0.001856547 | | antisense | 52.29 | 2.295584793 | 2.295584793 | ENSG00000258738 | n.s. |
| ENSG00000134698 | 286.2760187 | -0.860646454 | 0.2396594 | -3.591124 | 0.000329255 | 0.002167479 | AGO4 | protein_coding | 41.53 | -2.292800992 | 2.292800992 | ENSG00000134698 | n.s. |
| ENSG00000272654 | 86.34255872 | 0.960798248 | 0.2833669 | 3.390651 | 0.000697268 | 0.004132849 | | lincRNA | 44.22 | 2.290303241 | 2.290303241 | ENSG00000272654 | n.s. |
| ENSG00000228015 | 6.130730798 | 1.412441451 | 0.5104177 | 2.767227 | 0.005653541 | 0.023918035 | | processed_pseudogene | 65.38 | 2.2899553 | 2.2899553 | ENSG00000228015 | up |
| ENSG00000065802 | 829.1664532 | -0.920489874 | 0.2656094 | -3.465577 | 0.000529094 | 0.003258106 | ASB1 | protein_coding | 47.31 | -2.289290308 | 2.289290308 | ENSG00000065802 | n.s. |
| ENSG00000200150 | 11.7653288 | -1.807870193 | 0.7472232 | -2.419451 | 0.015543938 | 0.054172277 | | snoRNA | 35.62 | -2.289166652 | 2.289166652 | ENSG00000200150 | down |
| ENSG00000089876 | 327.2623434 | -1.054573252 | 0.3265866 | -3.229077 | 0.001241903 | 0.006756941 | DHX32 | protein_coding | 41.45 | -2.288687469 | 2.288687469 | ENSG00000089876 | down |
| ENSG00000207326 | 8.402546913 | -1.276395851 | 0.4376775 | -2.916293 | 0.003542175 | 0.016222411 | | misc_RNA | 39.47 | -2.284601293 | 2.284601293 | ENSG00000207326 | down |

| | | | | | | | | | | | | | |
|---|---|---|---|---|---|---|---|---|---|---|---|---|---|
| ENSG00000237765 | 1127.02011 | -0.913979956 | 0.2630438 | -3.47463 | 0.000511559 | 0.003171162 | FAM200B | protein_coding | 40.86 | -2.283836233 | 2.283836233 | ENSG00000237765 | n.s. |
| ENSG00000271533 | 258.9677708 | -1.050331095 | 0.3251153 | -3.230642 | 0.001235127 | 0.006722701 | | sense_intronic | 32.02 | -2.281798285 | 2.281798285 | ENSG00000271533 | down |
| ENSG00000144791 | 502.6820231 | -0.795498603 | 0.2134131 | -3.727505 | 0.000193385 | 0.001364582 | LIMD1 | protein_coding | 46.55 | -2.279103716 | 2.279103716 | ENSG00000144791 | n.s. |
| ENSG00000152217 | 1477.942224 | -0.86781032 | 0.2437016 | -3.560954 | 0.000369509 | 0.002391761 | SETBP1 | protein_coding | 42.22 | -2.274775811 | 2.274775811 | ENSG00000152217 | n.s. |
| ENSG00000276645 | 16.45277874 | -1.173374063 | 0.3855208 | -3.043607 | 0.002337599 | 0.011561775 | | misc_RNA | 49.86 | -2.27279679 | 2.27279679 | ENSG00000276645 | down |
| ENSG00000258890 | 3677.680442 | -0.898497627 | 0.2569874 | -3.496271 | 0.000471809 | 0.002955015 | CEP95 | protein_coding | 40.77 | -2.272696087 | 2.272696087 | ENSG00000258890 | n.s. |
| ENSG00000141371 | 9.939708042 | 1.255778877 | 0.4280487 | 2.933729 | 0.003349164 | 0.015502129 | C17orf64 | protein_coding | 45.51 | 2.272468322 | 2.272468322 | ENSG00000141371 | up |
| ENSG00000238886 | 11.34433354 | -1.199023954 | 0.399181 | -3.00371 | 0.002667094 | 0.012804808 | SNORD121A | snoRNA | 39.02 | -2.269305026 | 2.269305026 | ENSG00000238886 | down |
| ENSG00000109079 | 269.8592974 | -0.976126309 | 0.2915646 | -3.34789 | 0.000814293 | 0.004733776 | TNFAIP1 | protein_coding | 52.61 | -2.269290886 | 2.269290886 | ENSG00000109079 | n.s. |
| ENSG00000125898 | 161.5572531 | 0.83462512 | 0.2300444 | 3.628104 | 0.00028551 | 0.001912972 | FAM110A | protein_coding | 51.88 | 2.268754268 | 2.268754268 | ENSG00000125898 | n.s. |
| ENSG00000259775 | 45.20150236 | 0.966010889 | 0.2871072 | 3.364635 | 0.000766451 | 0.004491286 | | antisense | 35.16 | 2.267835426 | 2.267835426 | ENSG00000259775 | n.s. |
| ENSG00000080298 | 111.9591974 | -0.968067554 | 0.2883509 | -3.357255 | 0.000787203 | 0.004591636 | RFX3 | protein_coding | 35 | -2.263373486 | 2.263373486 | ENSG00000080298 | n.s. |
| ENSG00000224967 | 9.542304898 | -1.186668106 | 0.3931987 | -3.017986 | 0.00254461 | 0.012378968 | | processed_pseudogene | 46.41 | -2.263350557 | 2.263350557 | ENSG00000224967 | down |
| ENSG00000275770 | 4.016825642 | -1.584750257 | 0.6136624 | -2.582447 | 0.009810255 | 0.037331711 | MIR6505 | miRNA | 50.7 | -2.262899914 | 2.262899914 | ENSG00000275770 | down |
| ENSG00000268798 | 164.6682426 | 0.84929307 | 0.2365258 | 3.5907 | 0.000329791 | 0.002168909 | | lincRNA | 57 | 2.262311748 | 2.262311748 | ENSG00000268798 | n.s. |
| ENSG00000118922 | 882.5357905 | -0.985715038 | 0.2964733 | -3.324802 | 0.000884814 | 0.005075924 | KLF12 | protein_coding | 37.64 | -2.261708233 | 2.261708233 | ENSG00000118922 | n.s. |
| ENSG00000135749 | 751.5846964 | -0.929938702 | 0.2714913 | -3.425298 | 0.000614126 | 0.003699902 | PCNX2 | protein_coding | 40.87 | -2.261434052 | 2.261434052 | ENSG00000135749 | n.s. |
| ENSG00000252414 | 5.068560869 | -1.490503562 | 0.5588741 | -2.666976 | 0.007653722 | 0.030472626 | RNU6-100P | snRNA | 40.57 | -2.259737725 | 2.259737725 | ENSG00000252414 | down |
| ENSG00000275494 | 22.4779105 | 1.080155292 | 0.3409996 | 3.167614 | 0.001536952 | 0.008130312 | | sense_intronic | 64.04 | 2.257408729 | 2.257408729 | ENSG00000275494 | up |
| ENSG00000162337 | 406.1017991 | -0.83643562 | 0.2316717 | -3.610434 | 0.000305685 | 0.002028621 | LRP5 | protein_coding | 54.62 | -2.252352985 | 2.252352985 | ENSG00000162337 | n.s. |
| ENSG00000112874 | 434.1510709 | -1.016409014 | 0.311532 | -3.262615 | 0.001103894 | 0.006099268 | NUDT12 | protein_coding | 34.05 | -2.251063694 | 2.251063694 | ENSG00000112874 | down |
| ENSG00000113761 | 265.5607361 | -0.921493302 | 0.2684471 | -3.432681 | 0.000597644 | 0.003618924 | ZNF346 | protein_coding | 46.38 | -2.249752655 | 2.249752655 | ENSG00000113761 | n.s. |
| ENSG00000171634 | 6232.458085 | -0.900662644 | 0.2593037 | -3.473389 | 0.00051393 | 0.003180218 | BPTF | protein_coding | 40.86 | -2.249443721 | 2.249443721 | ENSG00000171634 | n.s. |
| ENSG00000212333 | 2.452781064 | -1.720561159 | 0.6986603 | -2.462658 | 0.013791148 | 0.049424492 | RNA5SP213 | rRNA | 46.23 | -2.247152294 | 2.247152294 | ENSG00000212333 | down |
| ENSG00000173705 | 698.1254454 | -0.91729669 | 0.2667842 | -3.438348 | 0.000585276 | 0.003558424 | SUSD5 | protein_coding | 41.61 | -2.246223178 | 2.246223178 | ENSG00000173705 | n.s. |
| ENSG00000198795 | 729.4442451 | -0.900004941 | 0.2592548 | -3.471508 | 0.000517545 | 0.003199722 | ZNF521 | protein_coding | 38.07 | -2.245411359 | 2.245411359 | ENSG00000198795 | n.s. |
| ENSG00000265442 | 4.013811217 | -1.612999598 | 0.6330783 | -2.547867 | 0.010838364 | 0.040560995 | MIR3941 | miRNA | 40.78 | -2.245120274 | 2.245120274 | ENSG00000265442 | down |
| ENSG00000269825 | 301.9558279 | -0.905434649 | 0.261674 | -3.460163 | 0.000539849 | 0.003318461 | | sense_intronic | 41.71 | -2.2446298 | 2.2446298 | ENSG00000269825 | n.s. |
| ENSG00000233654 | 68.08396685 | 0.898087096 | 0.2585623 | 3.473387 | 0.000513933 | 0.003180218 | | antisense | 39.04 | 2.243011178 | 2.243011178 | ENSG00000233654 | n.s. |
| ENSG00000257298 | 213.2950657 | -0.860803261 | 0.2425757 | -3.548596 | 0.000387291 | 0.002491796 | | sense_intronic | 43.93 | -2.241090482 | 2.241090482 | ENSG00000257298 | n.s. |
| ENSG00000101849 | 85.74760796 | -0.921952083 | 0.2694483 | -3.421628 | 0.000622473 | 0.003742095 | TBL1X | protein_coding | 45.23 | -2.237471825 | 2.237471825 | ENSG00000101849 | n.s. |
| ENSG00000280128 | 441.7883562 | -0.963698062 | 0.2881333 | -3.344626 | 0.000823937 | 0.004773896 | | TEC | 41.69 | -2.236865633 | 2.236865633 | ENSG00000280128 | n.s. |

| | | | | | | | | | | | | | |
|---|---|---|---|---|---|---|---|---|---|---|---|---|---|
| ENSG00000279520 | 39.87101138 | 0.958177191 | 0.2865246 | 3.344136 | 0.000825393 | 0.004778354 | | TEC | 54.69 | 2.223662537 | 2.223662537 | ENSG00000279520 | n.s. |
| ENSG00000269082 | 57.56849657 | -1.016217927 | 0.3135361 | -3.241151 | 0.001190479 | 0.00651791 | | processed_transcript | 44.76 | -2.22134229 | 2.22134229 | ENSG00000269082 | down |
| ENSG00000272562 | 119.4786552 | 0.837769162 | 0.2339622 | 3.580789 | 0.000342559 | 0.002236025 | | antisense | 41.12 | 2.220526657 | 2.220526657 | ENSG00000272562 | n.s. |
| ENSG00000163328 | 62.09548936 | 0.931805172 | 0.2749242 | 3.389316 | 0.000700671 | 0.004149489 | GPR155 | protein_coding | 39.18 | 2.219564911 | 2.219564911 | ENSG00000163328 | n.s. |
| ENSG00000278351 | 23.1858013 | -1.071136515 | 0.3396595 | -3.153559 | 0.001612924 | 0.008483845 | | sense_intronic | 29.92 | -2.218759943 | 2.218759943 | ENSG00000278351 | down |
| ENSG00000147526 | 147.7427856 | 0.841873539 | 0.2357845 | 3.570522 | 0.000356271 | 0.002315762 | TACC1 | protein_coding | 42.76 | 2.218594409 | 2.218594409 | ENSG00000147526 | n.s. |
| ENSG00000163884 | 166.2271623 | -0.99069346 | 0.3019276 | -3.281229 | 0.001033559 | 0.005783274 | KLF15 | protein_coding | 57.83 | -2.216999826 | 2.216999826 | ENSG00000163884 | n.s. |
| ENSG00000101191 | 2338.240575 | -0.87925429 | 0.251927 | -3.490115 | 0.000482813 | 0.003013092 | DIDO1 | protein_coding | 49.19 | -2.216589188 | 2.216589188 | ENSG00000101191 | n.s. |
| ENSG00000262333 | 64.0464911 | -0.961057976 | 0.2886174 | -3.329869 | 0.00086887 | 0.004992685 | HNRNPA1P16 | processed_pseudogene | 46.28 | -2.212034298 | 2.212034298 | ENSG00000262333 | n.s. |
| ENSG00000200556 | 20.503545 | 1.180071155 | 0.3951411 | 2.986455 | 0.002822321 | 0.013383917 | RNU6-103P | snRNA | 45.79 | 2.210765103 | 2.210765103 | ENSG00000200556 | up |
| ENSG00000133302 | 931.5458931 | -0.917462572 | 0.2691363 | -3.408914 | 0.00065222 | 0.003894025 | SLF1 | protein_coding | 36.06 | -2.210719004 | 2.210719004 | ENSG00000133302 | n.s. |
| ENSG00000047346 | 289.7503269 | -0.910553398 | 0.2662003 | -3.420558 | 0.000624928 | 0.003754209 | FAM214A | protein_coding | 37.21 | -2.208530512 | 2.208530512 | ENSG00000047346 | n.s. |
| ENSG00000283327 | 4.136472878 | 1.646225726 | 0.6591266 | 2.497587 | 0.012504194 | 0.045634572 | MIR6841 | miRNA | 45.83 | 2.207104737 | 2.207104737 | ENSG00000283327 | up |
| ENSG00000242971 | 16.93061255 | -1.146703722 | 0.3781137 | -3.032696 | 0.0024238 | 0.011899354 | RN7SL233P | misc_RNA | 52.71 | -2.206804483 | 2.206804483 | ENSG00000242971 | down |
| ENSG00000248540 | 200.706071 | -0.880959095 | 0.2532196 | -3.479032 | 0.000503228 | 0.003132069 | | lincRNA | 56.18 | -2.206070162 | 2.206070162 | ENSG00000248540 | n.s. |
| ENSG00000241975 | 43.38370828 | -1.070354824 | 0.3403308 | -3.145042 | 0.001660628 | 0.00871501 | ELOCP19 | processed_pseudogene | 42.14 | -2.204644195 | 2.204644195 | ENSG00000241975 | down |
| ENSG00000273759 | 155.1241154 | 0.85478176 | 0.2420592 | 3.531293 | 0.000413534 | 0.002643539 | | lincRNA | 47.71 | 2.203468569 | 2.203468569 | ENSG00000273759 | n.s. |
| ENSG00000166169 | 183.4929351 | -0.881387743 | 0.2536231 | -3.475187 | 0.000510497 | 0.003165994 | POLL | protein_coding | 52.96 | -2.203019777 | 2.203019777 | ENSG00000166169 | n.s. |
| ENSG00000154229 | 1497.39039 | -0.831352265 | 0.2325176 | -3.575438 | 0.000349642 | 0.00227586 | PRKCA | protein_coding | 43.87 | -2.197143104 | 2.197143104 | ENSG00000154229 | n.s. |
| ENSG00000119397 | 1696.224049 | -0.922713177 | 0.2723496 | -3.387973 | 0.000704111 | 0.004166313 | CNTRL | protein_coding | 38.88 | -2.196286317 | 2.196286317 | ENSG00000119397 | n.s. |
| ENSG00000271855 | 91.75151326 | 0.854896037 | 0.2426184 | 3.523624 | 0.000425687 | 0.002711273 | | lincRNA | 60.09 | 2.194370034 | 2.194370034 | ENSG00000271855 | n.s. |
| ENSG00000229618 | 40.13079156 | -1.011593226 | 0.3133869 | -3.227937 | 0.001246864 | 0.006773348 | | lincRNA | 35.74 | -2.194344602 | 2.194344602 | ENSG00000229618 | down |
| ENSG00000168140 | 74.11133308 | 0.943977368 | 0.2820132 | 3.34728 | 0.000816086 | 0.004742224 | VASN | protein_coding | 63.46 | 2.193820324 | 2.193820324 | ENSG00000168140 | n.s. |
| ENSG00000151474 | 515.5196158 | -0.867507778 | 0.2483826 | -3.492627 | 0.000478293 | 0.002987562 | FRMD4A | protein_coding | 43.78 | -2.190182167 | 2.190182167 | ENSG00000151474 | n.s. |
| ENSG00000267030 | 31.80939465 | -0.988264567 | 0.3029172 | -3.262491 | 0.001104378 | 0.006099268 | | antisense | 63.89 | -2.188731559 | 2.188731559 | ENSG00000267030 | n.s. |
| ENSG00000221771 | 8.527647357 | -1.83256217 | 0.7811968 | -2.345839 | 0.018984282 | 0.063956665 | MIR1205 | miRNA | 52.38 | -2.188288491 | 2.188288491 | ENSG00000221771 | down |
| ENSG00000204131 | 602.1966538 | -0.848461843 | 0.2402813 | -3.53112 | 0.000413805 | 0.002643683 | NHSL2 | protein_coding | 45.33 | -2.187157004 | 2.187157004 | ENSG00000204131 | n.s. |
| ENSG00000006025 | 86.48848942 | 0.910447117 | 0.2674991 | 3.403552 | 0.000665158 | 0.003962768 | OSBPL7 | protein_coding | 57.86 | 2.186895214 | 2.186895214 | ENSG00000006025 | n.s. |
| ENSG00000253596 | 25.45351789 | 1.03587673 | 0.3257696 | 3.179783 | 0.001473853 | 0.00784421 | | antisense | 36.36 | 2.180987503 | 2.180987503 | ENSG00000253596 | up |
| ENSG00000134982 | 1875.700006 | -0.891371762 | 0.2595193 | -3.434703 | 0.000593203 | 0.003595634 | APC | protein_coding | 37.63 | -2.178712761 | 2.178712761 | ENSG00000134982 | n.s. |
| ENSG00000249853 | 209.3288413 | -0.874969192 | 0.2523962 | -3.46665 | 0.000526987 | 0.003248004 | HS3ST5 | protein_coding | 37.56 | -2.177258822 | 2.177258822 | ENSG00000249853 | n.s. |
| ENSG00000266643 | 6.387918962 | -1.712868448 | 0.7075515 | -2.420839 | 0.01548472 | 0.054006451 | MIR3677 | miRNA | 71.67 | -2.171153865 | 2.171153865 | ENSG00000266643 | down |

| | | | | | | | | | | | | | |
|---|---|---|---|---|---|---|---|---|---|---|---|---|---|
| ENSG00000068784 | 1082.916008 | -0.879911507 | 0.2550994 | -3.449288 | 0.000562067 | 0.003435157 | SRBD1 | protein_coding | 36.55 | -2.168148931 | 2.168148931 | ENSG00000068784 | n.s. |
| ENSG00000101004 | 1035.740008 | -0.848755907 | 0.2414848 | -3.514738 | 0.000440188 | 0.002790864 | NINL | protein_coding | 49.07 | -2.167944443 | 2.167944443 | ENSG00000101004 | n.s. |
| ENSG00000149295 | 55.11028925 | 0.938231657 | 0.2811961 | 3.336575 | 0.000848177 | 0.004896006 | DRD2 | protein_coding | 48.35 | 2.167463417 | 2.167463417 | ENSG00000149295 | n.s. |
| ENSG00000224553 | 15.67729616 | -1.159295198 | 0.3886719 | -2.982709 | 0.002857094 | 0.013516591 | | processed_pseudogene | 50.86 | -2.166876726 | 2.166876726 | ENSG00000224553 | down |
| ENSG00000122783 | 731.7139325 | -0.824695552 | 0.2314689 | -3.562879 | 0.00036681 | 0.002375392 | C7orf49 | protein_coding | 40.03 | -2.164219415 | 2.164219415 | ENSG00000122783 | n.s. |
| ENSG00000196182 | 162.3544218 | -0.884457586 | 0.2576385 | -3.432941 | 0.000597073 | 0.003617512 | STK40 | protein_coding | 52.28 | -2.159482831 | 2.159482831 | ENSG00000196182 | n.s. |
| ENSG00000260565 | 620.0397983 | -0.807211973 | 0.2244921 | -3.595725 | 0.000323489 | 0.002136596 | ERVK13-1 | processed_transcript | 47.26 | -2.155480048 | 2.155480048 | ENSG00000260565 | n.s. |
| ENSG00000136897 | 1493.796911 | -0.967184489 | 0.2956156 | -3.271764 | 0.001068789 | 0.005945167 | MRPL50 | protein_coding | 35.67 | -2.152794026 | 2.152794026 | ENSG00000136897 | n.s. |
| ENSG00000175054 | 1839.436295 | -0.959717753 | 0.2924949 | -3.281144 | 0.00103387 | 0.005783274 | ATR | protein_coding | 37.14 | -2.147681576 | 2.147681576 | ENSG00000175054 | n.s. |
| ENSG00000271931 | 15.69108766 | 1.119746983 | 0.3699132 | 3.027053 | 0.002469507 | 0.012085392 | | antisense | 55.95 | 2.147382759 | 2.147382759 | ENSG00000271931 | up |
| ENSG00000169087 | 257.4499626 | -0.947671131 | 0.2870658 | -3.301233 | 0.00096261 | 0.005474842 | HSPBAP1 | protein_coding | 39.97 | -2.143279974 | 2.143279974 | ENSG00000169087 | n.s. |
| ENSG00000251493 | 622.8314763 | 0.859185836 | 0.2476175 | 3.46981 | 0.000520827 | 0.003215729 | FOXD1 | protein_coding | 53.9 | 2.141710241 | 2.141710241 | ENSG00000251493 | n.s. |
| ENSG00000166233 | 3130.345151 | -0.882962129 | 0.2581372 | -3.420515 | 0.000625027 | 0.003754209 | ARIH1 | protein_coding | 38.73 | -2.141608398 | 2.141608398 | ENSG00000166233 | n.s. |
| ENSG00000137802 | 149.610295 | 0.800406959 | 0.2225409 | 3.596674 | 0.000322312 | 0.002129829 | MAPKBP1 | protein_coding | 49.09 | 2.138411509 | 2.138411509 | ENSG00000137802 | n.s. |
| ENSG00000245281 | 42.97901474 | 0.915094908 | 0.2729403 | 3.352729 | 0.000800189 | 0.004657622 | | antisense | 40.7 | 2.133851018 | 2.133851018 | ENSG00000245281 | n.s. |
| ENSG00000198482 | 358.741952 | -0.908473218 | 0.2699708 | -3.36508 | 0.000765216 | 0.004485937 | ZNF808 | protein_coding | 44.95 | -2.133228484 | 2.133228484 | ENSG00000198482 | n.s. |
| ENSG00000076053 | 96.38650661 | 0.892431171 | 0.2628517 | 3.395189 | 0.000685814 | 0.00407189 | RBM7 | protein_coding | 38.27 | 2.133092493 | 2.133092493 | ENSG00000076053 | n.s. |
| ENSG00000186185 | 1316.812667 | -0.82696901 | 0.23422 | -3.530736 | 0.000414406 | 0.002645469 | KIF18B | protein_coding | 49.25 | -2.131510388 | 2.131510388 | ENSG00000186185 | n.s. |
| ENSG00000118058 | 2114.263371 | -0.845349635 | 0.2421953 | -3.490364 | 0.000482363 | 0.003011633 | KMT2A | protein_coding | 41.25 | -2.1312938 | 2.1312938 | ENSG00000118058 | n.s. |
| ENSG00000197056 | 277.0861214 | -1.03599686 | 0.3297529 | -3.141737 | 0.001679488 | 0.008787491 | ZMYM1 | protein_coding | 41.67 | -2.130149536 | 2.130149536 | ENSG00000197056 | down |
| ENSG00000264678 | 14.96897846 | -1.106274889 | 0.3647368 | -3.033077 | 0.002420736 | 0.011896894 | MIR3140 | miRNA | 36.67 | -2.129099482 | 2.129099482 | ENSG00000264678 | down |
| ENSG00000233170 | 44.44386111 | 0.923876478 | 0.2773014 | 3.331669 | 0.00086327 | 0.004966654 | | processed_pseudogene | 49.33 | 2.128552351 | 2.128552351 | ENSG00000233170 | n.s. |
| ENSG00000162929 | 366.5183698 | -0.897120707 | 0.2653047 | -3.381473 | 0.000720982 | 0.004255291 | KIAA1841 | protein_coding | 39.67 | -2.127136624 | 2.127136624 | ENSG00000162929 | n.s. |
| ENSG00000164187 | 485.5511208 | -1.011320371 | 0.3182502 | -3.177752 | 0.001484216 | 0.007887304 | LMBRD2 | protein_coding | 35.54 | -2.126878959 | 2.126878959 | ENSG00000164187 | down |
| ENSG00000177463 | 1444.239206 | -0.854899347 | 0.2467756 | -3.464278 | 0.000531657 | 0.00327244 | NR2C2 | protein_coding | 42.09 | -2.124534615 | 2.124534615 | ENSG00000177463 | n.s. |
| ENSG00000011454 | 975.3444451 | -0.899453552 | 0.2665877 | -3.37395 | 0.000740979 | 0.004361079 | RABGAP1 | protein_coding | 38.49 | -2.123075641 | 2.123075641 | ENSG00000011454 | n.s. |
| ENSG00000183960 | 491.6446777 | -0.890577156 | 0.2627588 | -3.389333 | 0.000700628 | 0.004149489 | KCNH8 | protein_coding | 36.03 | -2.121359557 | 2.121359557 | ENSG00000183960 | n.s. |
| ENSG00000180660 | 6881.774731 | 0.731830806 | 0.1953308 | 3.746622 | 0.000179232 | 0.001279059 | MAB21L1 | protein_coding | 46.24 | 2.117266584 | 2.117266584 | ENSG00000180660 | n.s. |
| ENSG00000171492 | 688.1673198 | -0.894490648 | 0.2649059 | -3.376635 | 0.000733783 | 0.004323515 | LRRC8D | protein_coding | 40.31 | -2.114721719 | 2.114721719 | ENSG00000171492 | n.s. |
| ENSG00000200480 | 5.380005325 | -1.934722749 | 0.8663837 | -2.233101 | 0.025542257 | 0.080732088 | SNORD114-28 | snoRNA | 47.22 | -2.114562604 | 2.114562604 | ENSG00000200480 | down |
| ENSG00000265008 | 19.16722007 | 1.073933752 | 0.349723 | 3.070812 | 0.002134772 | 0.010749525 | | antisense | 50.91 | 2.114157514 | 2.114157514 | ENSG00000265008 | up |
| ENSG00000167384 | 651.8817128 | -0.94280559 | 0.2870636 | -3.28431 | 0.001022326 | 0.005744084 | ZNF180 | protein_coding | 44.11 | -2.112619142 | 2.112619142 | ENSG00000167384 | n.s. |

| | | | | | | | | | | | | | |
|---|---|---|---|---|---|---|---|---|---|---|---|---|---|
| ENSG00000250853 | 70.71751346 | 0.911286991 | 0.2726839 | 3.341917 | 0.000832019 | 0.004812714 | RNF138P1 | processed_pseudogene | 37.73 | 2.112007768 | 2.112007768 | ENSG00000250853 | n.s. |
| ENSG00000164051 | 244.8193075 | -0.984036526 | 0.3065407 | -3.210133 | 0.001326734 | 0.007151422 | CCDC51 | protein_coding | 51.92 | -2.111356236 | 2.111356236 | ENSG00000164051 | n.s. |
| ENSG00000278520 | 8.536431378 | -1.264713285 | 0.4502482 | -2.808925 | 0.004970729 | 0.021446377 | MIR7851 | miRNA | 59.38 | -2.110358841 | 2.110358841 | ENSG00000278520 | down |
| ENSG00000257531 | 120.8225432 | 0.857398993 | 0.2487569 | 3.446735 | 0.000567406 | 0.003463389 | | transcribed_processed_pseudog | 51.61 | 2.109629159 | 2.109629159 | ENSG00000257531 | n.s. |
| ENSG00000214796 | 103.6644666 | -0.942083713 | 0.2872366 | -3.279818 | 0.001038742 | 0.005805864 | | transcribed_unprocessed_pseud | 51.87 | -2.106624602 | 2.106624602 | ENSG00000214796 | n.s. |
| ENSG00000023608 | 300.8686876 | -0.947241415 | 0.2897216 | -3.269488 | 0.001077422 | 0.005983625 | SNAPC1 | protein_coding | 39.61 | -2.105751434 | 2.105751434 | ENSG00000023608 | n.s. |
| ENSG00000159388 | 493.4198846 | 0.775661711 | 0.2140082 | 3.624449 | 0.000289579 | 0.001937435 | BTG2 | protein_coding | 53.33 | 2.104194 | 2.104194 | ENSG00000159388 | n.s. |
| ENSG00000104142 | 364.2276278 | -0.831445261 | 0.2378068 | -3.496306 | 0.000471748 | 0.002955015 | VPS18 | protein_coding | 53.64 | -2.103091132 | 2.103091132 | ENSG00000104142 | n.s. |
| ENSG00000276644 | 1645.171833 | -0.83576062 | 0.2402611 | -3.478552 | 0.000504131 | 0.003134528 | DACH1 | protein_coding | 33.93 | -2.092600683 | 2.092600683 | ENSG00000276644 | n.s. |
| ENSG00000260727 | 174.7152133 | 0.828126834 | 0.2370081 | 3.494087 | 0.000475686 | 0.002974281 | SLC7A5P1 | unprocessed_pseudogene | 69.27 | 2.092360154 | 2.092360154 | ENSG00000260727 | n.s. |
| ENSG00000005700 | 1823.552722 | -0.990645638 | 0.311049 | -3.184854 | 0.001448271 | 0.007725778 | IBTK | protein_coding | 36.07 | -2.092300843 | 2.092300843 | ENSG00000005700 | n.s. |
| ENSG00000143633 | 814.3344106 | -0.834955756 | 0.240058 | -3.478142 | 0.000504903 | 0.003136889 | C1orf131 | protein_coding | 43.5 | -2.090312475 | 2.090312475 | ENSG00000143633 | n.s. |
| ENSG00000180592 | 695.7630636 | 0.729147138 | 0.1955702 | 3.728313 | 0.000192766 | 0.001362374 | SKIDA1 | protein_coding | 45.4 | 2.089519643 | 2.089519643 | ENSG00000180592 | n.s. |
| ENSG00000106246 | 350.2010399 | -0.928774916 | 0.2822472 | -3.290643 | 0.000999586 | 0.005650519 | PTCD1 | protein_coding | 53.4 | -2.087803942 | 2.087803942 | ENSG00000106246 | n.s. |
| ENSG00000107731 | 2201.251111 | -0.821411547 | 0.2343241 | -3.505451 | 0.000455834 | 0.002874357 | UNC5B | protein_coding | 54.37 | -2.087583953 | 2.087583953 | ENSG00000107731 | n.s. |
| ENSG00000028839 | 1031.295735 | -0.932135578 | 0.284132 | -3.280643 | 0.001035709 | 0.005791235 | TBPL1 | protein_coding | 37.6 | -2.085400561 | 2.085400561 | ENSG00000028839 | n.s. |
| ENSG00000266405 | 78.71390504 | 0.897511997 | 0.2681834 | 3.346635 | 0.000817989 | 0.004749318 | | transcribed_processed_pseudog | 40.82 | 2.085251341 | 2.085251341 | ENSG00000266405 | n.s. |
| ENSG00000066739 | 1194.605815 | -0.868450985 | 0.2552201 | -3.402752 | 0.000667107 | 0.003972681 | ATG2B | protein_coding | 37.31 | -2.085078131 | 2.085078131 | ENSG00000066739 | n.s. |
| ENSG00000207031 | 25.34328014 | 1.021108756 | 0.3263074 | 3.129285 | 0.001752325 | 0.009107879 | SNORD59A | snoRNA | 37.33 | 2.083656924 | 2.083656924 | ENSG00000207031 | up |
| ENSG00000200502 | 19.81112568 | -1.07183823 | 0.3515204 | -3.049149 | 0.002294903 | 0.01140326 | | misc_RNA | 46.53 | -2.082550573 | 2.082550573 | ENSG00000200502 | down |
| ENSG00000275764 | 194.7280949 | -0.912290437 | 0.2756081 | -3.3101 | 0.000932626 | 0.005328254 | | lincRNA | 32.72 | -2.074014858 | 2.074014858 | ENSG00000275764 | n.s. |
| ENSG00000246067 | 587.2128034 | -0.724181098 | 0.1943484 | -3.7262 | 0.000194388 | 0.001368977 | RAB30-AS1 | lincRNA | 40.04 | -2.073767787 | 2.073767787 | ENSG00000246067 | n.s. |
| ENSG00000112167 | 262.4174304 | -0.791650121 | 0.2224464 | -3.558835 | 0.000372504 | 0.002407784 | SAYSD1 | protein_coding | 41.97 | -2.072842775 | 2.072842775 | ENSG00000112167 | n.s. |
| ENSG00000155970 | 83.66287896 | -0.958593221 | 0.2974878 | -3.222294 | 0.001271687 | 0.006894743 | MICU3 | protein_coding | 35.97 | -2.07198192 | 2.07198192 | ENSG00000155970 | n.s. |
| ENSG00000271798 | 9.637857567 | 1.242668906 | 0.4426064 | 2.807616 | 0.004990968 | 0.02152036 | SNORA51 | snoRNA | 50.76 | 2.071716041 | 2.071716041 | ENSG00000271798 | up |
| ENSG00000272137 | 30.16223099 | -1.056991703 | 0.3450237 | -3.063534 | 0.002187394 | 0.010966903 | | lincRNA | 29.94 | -2.071614959 | 2.071614959 | ENSG00000272137 | down |
| ENSG00000277182 | 60.39637653 | 0.909986142 | 0.274848 | 3.310871 | 0.000930061 | 0.005315779 | | antisense | 56.2 | 2.069702598 | 2.069702598 | ENSG00000277182 | n.s. |
| ENSG00000276058 | 19.41403778 | -1.109460776 | 0.3724032 | -2.979192 | 0.002890092 | 0.013644878 | STMN1P1 | processed_pseudogene | 43.86 | -2.06917802 | 2.06917802 | ENSG00000276058 | down |
| ENSG00000116128 | 635.2698423 | -0.809833637 | 0.2303824 | -3.515171 | 0.000439471 | 0.00278759 | BCL9 | protein_coding | 42.09 | -2.068939563 | 2.068939563 | ENSG00000116128 | n.s. |
| ENSG00000265321 | 7.613228911 | -1.363945238 | 0.5116617 | -2.665717 | 0.007682439 | 0.030543291 | MIR4263 | miRNA | 45.78 | -2.066491844 | 2.066491844 | ENSG00000265321 | down |
| ENSG00000151131 | 2292.364747 | -0.856246166 | 0.2510674 | -3.410424 | 0.00064862 | 0.003874191 | C12orf45 | protein_coding | 39.89 | -2.065110784 | 2.065110784 | ENSG00000151131 | n.s. |
| ENSG00000187257 | 1083.254116 | -0.863875601 | 0.2544256 | -3.395396 | 0.000685294 | 0.004070542 | RSBN1L | protein_coding | 39.26 | -2.0649631 | 2.0649631 | ENSG00000187257 | n.s. |

| | | | | | | | | | | | | | |
|---|---|---|---|---|---|---|---|---|---|---|---|---|---|
| ENSG00000265527 | 20.97766771 | -1.160031578 | 0.3990845 | -2.906732 | 0.003652264 | 0.016606948 | MIR5690 | miRNA | 30.14 | -2.064520005 | 2.064520005 | ENSG00000265527 | down |
| ENSG00000170471 | 1658.994948 | -0.861760167 | 0.2535872 | -3.398279 | 0.000678113 | 0.004033046 | RALGAPB | protein_coding | 39.37 | -2.063369923 | 2.063369923 | ENSG00000170471 | n.s. |
| ENSG00000240718 | 16.91123968 | -1.091804618 | 0.3638754 | -3.000491 | 0.00269545 | 0.012905349 | RN7SL851P | misc_RNA | 54.79 | -2.062670308 | 2.062670308 | ENSG00000240718 | down |
| ENSG00000212102 | 25.95376951 | 0.990481814 | 0.3132938 | 3.161511 | 0.001569527 | 0.008286891 | MIR301B | miRNA | 52.56 | 2.061795251 | 2.061795251 | ENSG00000212102 | n.s. |
| ENSG00000207194 | 2.5651467 | -2.34471205 | 1.1850554 | -1.978567 | 0.047864734 | 0.132147668 | RNU6-1026P | snRNA | 37.96 | -2.060862377 | 2.060862377 | ENSG00000207194 | n.s. |
| ENSG00000164663 | 519.3595374 | -0.814575053 | 0.2329787 | -3.49635 | 0.000471669 | 0.002955015 | USP49 | protein_coding | 41.51 | -2.06041895 | 2.06041895 | ENSG00000164663 | n.s. |
| ENSG00000143376 | 726.0236758 | -0.821746305 | 0.2362915 | -3.47768 | 0.000505774 | 0.003139494 | SNX27 | protein_coding | 40.41 | -2.056946274 | 2.056946274 | ENSG00000143376 | n.s. |
| ENSG00000160352 | 1735.284649 | -0.885366111 | 0.2646573 | -3.34533 | 0.000821846 | 0.004767735 | ZNF714 | protein_coding | 38.78 | -2.055543807 | 2.055543807 | ENSG00000160352 | n.s. |
| ENSG00000147592 | 184.9131825 | -0.964043669 | 0.3013238 | -3.199362 | 0.001377323 | 0.007378401 | LACTB2 | protein_coding | 38.38 | -2.055377504 | 2.055377504 | ENSG00000147592 | n.s. |
| ENSG00000082805 | 859.1674902 | -0.824655476 | 0.2377755 | -3.468211 | 0.000523936 | 0.003232062 | ERC1 | protein_coding | 39.35 | -2.053821193 | 2.053821193 | ENSG00000082805 | n.s. |
| ENSG00000280798 | 210.5262191 | -0.910960029 | 0.2765332 | -3.294216 | 0.000986966 | 0.005585984 | LINC00294 | lincRNA | 45.1 | -2.052302098 | 2.052302098 | ENSG00000280798 | n.s. |
| ENSG00000023041 | 1005.315954 | -0.865426618 | 0.2560022 | -3.380544 | 0.000723425 | 0.004267899 | ZDHHC6 | protein_coding | 38.61 | -2.05087578 | 2.05087578 | ENSG00000023041 | n.s. |
| ENSG00000278396 | 28.37594467 | 1.028273451 | 0.3327268 | 3.090444 | 0.001998576 | 0.010185275 | | sense_intronic | 37.36 | 2.048348715 | 2.048348715 | ENSG00000278396 | up |
| ENSG00000265942 | 54.21577738 | 0.916870306 | 0.2798889 | 3.275837 | 0.001053493 | 0.005878871 | RN7SL577P | misc_RNA | 52.15 | 2.045268184 | 2.045268184 | ENSG00000265942 | n.s. |
| ENSG00000184992 | 949.0985949 | -0.913795199 | 0.2784766 | -3.281407 | 0.001032904 | 0.005783274 | BRI3BP | protein_coding | 48.32 | -2.044914879 | 2.044914879 | ENSG00000184992 | n.s. |
| ENSG00000200823 | 11.20064633 | -1.278279505 | 0.4656072 | -2.745403 | 0.006043659 | 0.025155714 | SNORD114-2 | snoRNA | 42.31 | -2.044433387 | 2.044433387 | ENSG00000200823 | down |
| ENSG00000065559 | 893.8837277 | -0.903701397 | 0.2737998 | -3.300592 | 0.00096481 | 0.00548288 | MAP2K4 | protein_coding | 37.09 | -2.04326097 | 2.04326097 | ENSG00000065559 | n.s. |
| ENSG00000165195 | 248.1441876 | -0.911604714 | 0.2775062 | -3.284989 | 0.001019864 | 0.005737753 | PIGA | protein_coding | 39.56 | -2.043141484 | 2.043141484 | ENSG00000165195 | n.s. |
| ENSG00000201121 | 8.272153315 | -1.285891831 | 0.4703555 | -2.733872 | 0.006259436 | 0.025829915 | RNY1P12 | misc_RNA | 43.81 | -2.041838096 | 2.041838096 | ENSG00000201121 | down |
| ENSG00000181744 | 1672.359707 | 0.83692765 | 0.2439618 | 3.430568 | 0.000602318 | 0.003635057 | C3orf58 | protein_coding | 37.13 | 2.041675622 | 2.041675622 | ENSG00000181744 | n.s. |
| ENSG00000170340 | 640.0353085 | 0.906638057 | 0.2753156 | 3.293086 | 0.000990942 | 0.005606211 | B3GNT2 | protein_coding | 45.4 | 2.041141989 | 2.041141989 | ENSG00000170340 | n.s. |
| ENSG00000062650 | 1893.403437 | -0.897280316 | 0.2709912 | -3.311105 | 0.000929283 | 0.005313513 | WAPL | protein_coding | 39.87 | -2.04097021 | 2.04097021 | ENSG00000062650 | n.s. |
| ENSG00000232065 | 20.91340197 | 1.038680055 | 0.3387939 | 3.065817 | 0.002170762 | 0.010899838 | LINC01063 | antisense | 50.92 | 2.038492672 | 2.038492672 | ENSG00000232065 | up |
| ENSG00000244026 | 153.3930464 | -0.865880933 | 0.2571246 | -3.367554 | 0.000758381 | 0.004453372 | FAM86DP | transcribed_unprocessed_pseud | 52.79 | -2.035955417 | 2.035955417 | ENSG00000244026 | n.s. |
| ENSG00000156017 | 407.3083797 | -0.848387697 | 0.2494629 | -3.400857 | 0.000671749 | 0.003998616 | CARNMT1 | protein_coding | 37.59 | -2.034510279 | 2.034510279 | ENSG00000156017 | n.s. |
| ENSG00000141682 | 2778.816926 | 0.817512607 | 0.2358545 | 3.466173 | 0.000527923 | 0.003252332 | PMAIP1 | protein_coding | 39.83 | 2.033812068 | 2.033812068 | ENSG00000141682 | n.s. |
| ENSG00000199730 | 23.69923507 | -1.033093153 | 0.3364148 | -3.070891 | 0.002134212 | 0.010749525 | RN7SKP95 | misc_RNA | 50.5 | -2.033758271 | 2.033758271 | ENSG00000199730 | down |
| ENSG00000207653 | 13.71805865 | -1.557455387 | 0.6328001 | -2.461212 | 0.013846834 | 0.049555743 | MIR558 | miRNA | 42.55 | -2.032332898 | 2.032332898 | ENSG00000207653 | down |
| ENSG00000270722 | 281.5241338 | 0.752850089 | 0.2083755 | 3.612949 | 0.000302734 | 0.00201192 | | snRNA | 53.66 | 2.029976987 | 2.029976987 | ENSG00000270722 | n.s. |
| ENSG00000099910 | 390.3905995 | -0.937599646 | 0.2908046 | -3.224157 | 0.00126344 | 0.006852696 | KLHL22 | protein_coding | 49.11 | -2.029095534 | 2.029095534 | ENSG00000099910 | n.s. |
| ENSG00000078699 | 640.5405221 | -0.86523754 | 0.2574196 | -3.361195 | 0.000776061 | 0.004538051 | CBFA2T2 | protein_coding | 42.36 | -2.02736455 | 2.02736455 | ENSG00000078699 | n.s. |
| ENSG00000088854 | 437.8787899 | -0.869383534 | 0.2596412 | -3.348404 | 0.000812786 | 0.004728965 | C20orf194 | protein_coding | 43.79 | -2.02152007 | 2.02152007 | ENSG00000088854 | n.s. |

| | | | | | | | | | | | | | |
|---|---|---|---|---|---|---|---|---|---|---|---|---|---|
| ENSG00000188313 | 587.9551268 | -0.836592564 | 0.2450521 | -3.413937 | 0.000640313 | 0.003836091 | PLSCR1 | protein_coding | 36.31 | -2.021300557 | 2.021300557 | ENSG00000188313 | n.s. |
| ENSG00000099810 | 1449.789133 | -0.879189068 | 0.264224 | -3.327438 | 0.000876484 | 0.00503021 | MTAP | protein_coding | 40.09 | -2.020740333 | 2.020740333 | ENSG00000099810 | n.s. |
| ENSG00000197024 | 642.2392104 | -0.848584166 | 0.2505091 | -3.387438 | 0.000705486 | 0.004172679 | ZNF398 | protein_coding | 45.92 | -2.019278203 | 2.019278203 | ENSG00000197024 | n.s. |
| ENSG00000218690 | 28.12269983 | 1.013050357 | 0.3277269 | 3.091142 | 0.001993882 | 0.010165075 | HIST1H2APS4 | transcribed_unprocessed_pseud | 51.4 | 2.018897343 | 2.018897343 | ENSG00000218690 | up |
| ENSG00000278642 | 7.041160228 | 1.265249192 | 0.4616705 | 2.740589 | 0.006132914 | 0.025443553 | | lincRNA | 50.84 | 2.017341457 | 2.017341457 | ENSG00000278642 | up |
| ENSG00000042088 | 667.7927398 | -0.898714058 | 0.2737429 | -3.283059 | 0.001026871 | 0.005762647 | TDP1 | protein_coding | 42.14 | -2.012560442 | 2.012560442 | ENSG00000042088 | n.s. |
| ENSG00000274884 | 5.595755641 | 1.427786411 | 0.5567788 | 2.564369 | 0.01033635 | 0.039021965 | | misc_RNA | 68.97 | 2.011309675 | 2.011309675 | ENSG00000274884 | up |
| ENSG00000167524 | 142.1519434 | -0.889151217 | 0.2693813 | -3.300716 | 0.000964384 | 0.005482696 | | protein_coding | 45.98 | -2.010376115 | 2.010376115 | ENSG00000167524 | n.s. |
| ENSG00000260830 | 24.39870358 | 1.003086758 | 0.3235836 | 3.099931 | 0.001935656 | 0.00991543 | | antisense | 54.69 | 2.009873349 | 2.009873349 | ENSG00000260830 | up |
| ENSG00000272892 | 57.74783587 | 0.856197291 | 0.2545952 | 3.362975 | 0.000771074 | 0.004516474 | | lincRNA | 50.33 | 2.00795428 | 2.00795428 | ENSG00000272892 | n.s. |
| ENSG00000136925 | 833.1761774 | -0.923383432 | 0.2857654 | -3.231263 | 0.001232443 | 0.006713341 | TSTD2 | protein_coding | 42.44 | -2.006568793 | 2.006568793 | ENSG00000136925 | n.s. |
| ENSG00000112425 | 82.72900826 | 0.87931578 | 0.2651503 | 3.316291 | 0.000912207 | 0.005226607 | EPM2A | protein_coding | 37.55 | 2.006405305 | 2.006405305 | ENSG00000112425 | n.s. |
| ENSG00000202538 | 610.1381262 | 0.730759323 | 0.2004393 | 3.645789 | 0.000266573 | 0.001797331 | RNU4-2 | snRNA | 45.39 | 2.006206057 | 2.006206057 | ENSG00000202538 | n.s. |
| ENSG00000142556 | 749.738418 | -0.857799138 | 0.2554383 | -3.358146 | 0.000784671 | 0.004584548 | ZNF614 | protein_coding | 41.38 | -2.006137824 | 2.006137824 | ENSG00000142556 | n.s. |
| ENSG00000197343 | 2038.934959 | -0.859873741 | 0.2563944 | -3.353715 | 0.000797345 | 0.004644951 | ZNF655 | protein_coding | 42.73 | -2.006101679 | 2.006101679 | ENSG00000197343 | n.s. |
| ENSG00000177888 | 1008.469892 | -0.869485992 | 0.2609051 | -3.332576 | 0.000860461 | 0.004954588 | ZBTB41 | protein_coding | 33.4 | -2.004158647 | 2.004158647 | ENSG00000177888 | n.s. |
| ENSG00000155966 | 74.77549877 | -0.923778109 | 0.2861684 | -3.228092 | 0.001246188 | 0.006773348 | AFF2 | protein_coding | 38.84 | -2.003856348 | 2.003856348 | ENSG00000155966 | n.s. |
| ENSG00000186532 | 364.0576925 | -0.841424848 | 0.2484989 | -3.38603 | 0.000709116 | 0.004190589 | SMYD4 | protein_coding | 46.16 | -2.000676845 | 2.000676845 | ENSG00000186532 | n.s. |
| ENSG00000200164 | 10.39025173 | 1.178024731 | 0.4156142 | 2.834419 | 0.004590908 | 0.020097878 | | misc_RNA | 41.58 | 1.998931014 | 1.998931014 | ENSG00000200164 | up |
| ENSG00000179743 | 115.4740513 | 0.818744567 | 0.2385203 | 3.432599 | 0.000597826 | 0.003618924 | | antisense | 50.48 | 1.998899783 | 1.998899783 | ENSG00000179743 | n.s. |
| ENSG00000207863 | 14.9445389 | -1.14824967 | 0.3995617 | -2.873773 | 0.004056002 | 0.018163947 | MIR125B2 | miRNA | 49.44 | -1.998861288 | 1.998861288 | ENSG00000207863 | down |
| ENSG00000145715 | 823.0280188 | -0.845391743 | 0.2503878 | -3.37633 | 0.000734599 | 0.004326487 | RASA1 | protein_coding | 35.77 | -1.998391621 | 1.998391621 | ENSG00000145715 | n.s. |
| ENSG00000171813 | 188.1657519 | -0.90758395 | 0.279333 | -3.249111 | 0.001157664 | 0.006358249 | PWWP2B | protein_coding | 62.25 | -1.993655575 | 1.993655575 | ENSG00000171813 | n.s. |
| ENSG00000238741 | 1413.288382 | -0.782983718 | 0.2231995 | -3.508 | 0.00045149 | 0.002852127 | SCARNA7 | scaRNA | 37.27 | -1.992561295 | 1.992561295 | ENSG00000238741 | n.s. |
| ENSG00000155666 | 37.98767131 | 0.94520879 | 0.2970718 | 3.181752 | 0.001463871 | 0.007800028 | KDM8 | protein_coding | 51.92 | 1.992409239 | 1.992409239 | ENSG00000155666 | n.s. |
| ENSG00000157657 | 667.8937343 | -0.8150015 | 0.2372671 | -3.434954 | 0.000592655 | 0.003593872 | ZNF618 | protein_coding | 45.61 | -1.992220184 | 1.992220184 | ENSG00000157657 | n.s. |
| ENSG00000254615 | 149.7370558 | 0.844859175 | 0.2506548 | 3.370609 | 0.000750023 | 0.004409868 | | lincRNA | 57.15 | 1.990128661 | 1.990128661 | ENSG00000254615 | n.s. |
| ENSG00000119537 | 109.4530269 | 0.807012038 | 0.2339996 | 3.448775 | 0.000563136 | 0.003438833 | KDSR | protein_coding | 42.29 | 1.988145865 | 1.988145865 | ENSG00000119537 | n.s. |
| ENSG00000096968 | 230.3559883 | -0.886225076 | 0.2696422 | -3.28667 | 0.001013795 | 0.005714602 | JAK2 | protein_coding | 37.67 | -1.987815292 | 1.987815292 | ENSG00000096968 | n.s. |
| ENSG00000159082 | 217.3889226 | -0.836978583 | 0.2473577 | -3.383677 | 0.00071522 | 0.004223073 | SYNJ1 | protein_coding | 38.08 | -1.987298044 | 1.987298044 | ENSG00000159082 | n.s. |
| ENSG00000232466 | 40.35137717 | -0.992822018 | 0.3205851 | -3.096906 | 0.001955517 | 0.009998774 | | processed_pseudogene | 48.66 | -1.985696895 | 1.985696895 | ENSG00000232466 | n.s. |
| ENSG00000167257 | 301.8014493 | -0.872974098 | 0.2638894 | -3.308106 | 0.000939292 | 0.005361939 | RNF214 | protein_coding | 42.99 | -1.982243219 | 1.982243219 | ENSG00000167257 | n.s. |

| | | | | | | | | | | | | | |
|---|---|---|---|---|---|---|---|---|---|---|---|---|---|
| ENSG00000197852 | 1215.087738 | -0.873128994 | 0.2642104 | -3.304674 | 0.000950871 | 0.005416928 | FAM212B | protein_coding | 43.96 | -1.978725909 | 1.978725909 | ENSG00000197852 | n.s. |
| ENSG00000225793 | 83.00784076 | -0.812168674 | 0.2369037 | -3.428265 | 0.000607452 | 0.003664455 | | lincRNA | 37.06 | -1.978435304 | 1.978435304 | ENSG00000225793 | n.s. |
| ENSG00000165934 | 1832.367326 | -0.892505781 | 0.2738341 | -3.259294 | 0.001116898 | 0.006163521 | CPSF2 | protein_coding | 39.6 | -1.972590475 | 1.972590475 | ENSG00000165934 | n.s. |
| ENSG00000224078 | 2199.35192 | -0.866846735 | 0.2617529 | -3.311698 | 0.000927314 | 0.005304437 | SNHG14 | processed_transcript | 42.05 | -1.972389031 | 1.972389031 | ENSG00000224078 | n.s. |
| ENSG00000074047 | 59.27588341 | -0.914728431 | 0.2843919 | -3.216437 | 0.001297931 | 0.007012455 | GLI2 | protein_coding | 52.42 | -1.970443856 | 1.970443856 | ENSG00000074047 | n.s. |
| ENSG00000110851 | 1125.413324 | -0.896526846 | 0.2758812 | -3.249685 | 0.001155329 | 0.006347928 | PRDM4 | protein_coding | 41.21 | -1.969999408 | 1.969999408 | ENSG00000110851 | n.s. |
| ENSG00000238959 | 11.95917901 | -1.148763299 | 0.4033057 | -2.848369 | 0.0043944 | 0.01938632 | RNU7-19P | snRNA | 42.42 | -1.967262451 | 1.967262451 | ENSG00000238959 | down |
| ENSG00000265778 | 8.329712317 | 1.204911688 | 0.4337459 | 2.777921 | 0.005470799 | 0.023300683 | | antisense | 55.01 | 1.967176592 | 1.967176592 | ENSG00000265778 | up |
| ENSG00000148300 | 434.6447415 | -0.886098555 | 0.2713083 | -3.26602 | 0.001090704 | 0.006050002 | REXO4 | protein_coding | 51.72 | -1.96558324 | 1.96558324 | ENSG00000148300 | n.s. |
| ENSG00000181222 | 969.3573915 | 0.82505179 | 0.243457 | 3.388902 | 0.000701732 | 0.004154003 | POLR2A | protein_coding | 49.69 | 1.964888201 | 1.964888201 | ENSG00000181222 | n.s. |
| ENSG00000278716 | 172.5057079 | -0.901076804 | 0.2785049 | -3.235407 | 0.001214694 | 0.006645258 | | lincRNA | 57.9 | -1.962084061 | 1.962084061 | ENSG00000278716 | n.s. |
| ENSG00000276823 | 7.214556463 | 1.269457179 | 0.4710827 | 2.694765 | 0.007043833 | 0.028525678 | | misc_RNA | 47.4 | 1.961011778 | 1.961011778 | ENSG00000276823 | up |
| ENSG00000196876 | 344.8979479 | -0.796616853 | 0.2311308 | -3.446606 | 0.000567677 | 0.003463525 | SCN8A | protein_coding | 41.03 | -1.960061184 | 1.960061184 | ENSG00000196876 | n.s. |
| ENSG00000198265 | 3802.958657 | -0.811497826 | 0.2377536 | -3.413188 | 0.000642077 | 0.003845006 | HELZ | protein_coding | 38.17 | -1.959850804 | 1.959850804 | ENSG00000198265 | n.s. |
| ENSG00000158966 | 1079.667223 | -0.813907103 | 0.2388623 | -3.407432 | 0.000655772 | 0.0039102 | CACHD1 | protein_coding | 39.92 | -1.959726391 | 1.959726391 | ENSG00000158966 | n.s. |
| ENSG00000151967 | 1064.905721 | -0.826860905 | 0.2446285 | -3.380068 | 0.000724679 | 0.00427349 | SCHIP1 | protein_coding | 39.91 | -1.959013202 | 1.959013202 | ENSG00000151967 | n.s. |
| ENSG00000163611 | 468.9627849 | -0.880270269 | 0.269564 | -3.265534 | 0.00109258 | 0.006053317 | SPICE1 | protein_coding | 38.02 | -1.952445266 | 1.952445266 | ENSG00000163611 | n.s. |
| ENSG00000196867 | 429.7811996 | -0.875367953 | 0.2672807 | -3.275089 | 0.001056287 | 0.0058921 | ZFP28 | protein_coding | 40.78 | -1.951834103 | 1.951834103 | ENSG00000196867 | n.s. |
| ENSG00000101407 | 923.1424932 | -0.872175241 | 0.2658124 | -3.281168 | 0.001033782 | 0.005783274 | TTI1 | protein_coding | 43.33 | -1.951776646 | 1.951776646 | ENSG00000101407 | n.s. |
| ENSG00000006652 | 3180.322995 | 0.803510744 | 0.2347675 | 3.42258 | 0.000620297 | 0.003732676 | IFRD1 | protein_coding | 39.45 | 1.950907777 | 1.950907777 | ENSG00000006652 | n.s. |
| ENSG00000146243 | 1329.454447 | -0.875829947 | 0.268142 | -3.266292 | 0.001089658 | 0.006046756 | IRAK1BP1 | protein_coding | 37.25 | -1.943009082 | 1.943009082 | ENSG00000146243 | n.s. |
| ENSG00000176842 | 173.2105296 | 0.805664809 | 0.2363498 | 3.408781 | 0.000652539 | 0.003894254 | IRX5 | protein_coding | 59.38 | 1.941310339 | 1.941310339 | ENSG00000176842 | n.s. |
| ENSG00000163873 | 488.1455215 | -0.835423536 | 0.2497683 | -3.344794 | 0.000823437 | 0.004772984 | GRIK3 | protein_coding | 49.57 | -1.939193448 | 1.939193448 | ENSG00000163873 | n.s. |
| ENSG00000155363 | 63.07681339 | -0.863689424 | 0.2626944 | -3.287811 | 0.001009698 | 0.005696118 | MOV10 | protein_coding | 48.61 | -1.938482695 | 1.938482695 | ENSG00000155363 | n.s. |
| ENSG00000100842 | 354.4225934 | -0.781346363 | 0.225965 | -3.45782 | 0.000544564 | 0.003340073 | EFS | protein_coding | 54.28 | -1.934804267 | 1.934804267 | ENSG00000100842 | n.s. |
| ENSG00000242931 | 86.36591539 | -0.827732564 | 0.2467223 | -3.354916 | 0.000793892 | 0.004628709 | RPL7P49 | processed_pseudogene | 43.64 | -1.932374915 | 1.932374915 | ENSG00000242931 | n.s. |
| ENSG00000277382 | 10.8933227 | 1.22828468 | 0.4515138 | 2.72037 | 0.00652089 | 0.026740474 | | antisense | 58.74 | 1.931884093 | 1.931884093 | ENSG00000277382 | up |
| ENSG00000207609 | 10.30955769 | -1.191474489 | 0.4306687 | -2.766568 | 0.005664972 | 0.023951837 | MIR491 | miRNA | 51.19 | -1.930976448 | 1.930976448 | ENSG00000207609 | down |
| ENSG00000228939 | 73.78780396 | -1.015786065 | 0.3374877 | -3.009846 | 0.0026138 | 0.012622941 | AKT3-IT1 | sense_intronic | 43.31 | -1.92881464 | 1.92881464 | ENSG00000228939 | down |
| ENSG00000274695 | 26.90630397 | 1.019850744 | 0.3398044 | 3.001287 | 0.002688408 | 0.012886909 | | lincRNA | 38.31 | 1.927366193 | 1.927366193 | ENSG00000274695 | up |
| ENSG00000264961 | 3.582729744 | -2.383873602 | 1.2631563 | -1.887236 | 0.059128633 | 0.15543495 | MIR4730 | miRNA | 65.79 | -1.927245764 | 1.927245764 | ENSG00000264961 | n.s. |
| ENSG00000119401 | 362.5090584 | -0.85184743 | 0.2581481 | -3.29984 | 0.000967398 | 0.005495343 | TRIM32 | protein_coding | 43.3 | -1.92517962 | 1.92517962 | ENSG00000119401 | n.s. |

| | | | | | | | | | | | | | |
|---|---|---|---|---|---|---|---|---|---|---|---|---|---|
| ENSG00000225591 | 18.63240525 | -1.049106023 | 0.3551774 | -2.953753 | 0.003139356 | 0.014667537 | | processed_pseudogene | 47.45 | -1.923685723 | 1.923685723 | ENSG00000225591 | down |
| ENSG00000264470 | 5.669703357 | -1.277922128 | 0.4812564 | -2.655387 | 0.00792174 | 0.031333632 | MIR4794 | miRNA | 38.96 | -1.921981151 | 1.921981151 | ENSG00000264470 | down |
| ENSG00000189164 | 431.7514182 | -0.843697548 | 0.2546242 | -3.313502 | 0.000921356 | 0.005272519 | ZNF527 | protein_coding | 40.73 | -1.921927665 | 1.921927665 | ENSG00000189164 | n.s. |
| ENSG00000225830 | 338.1014981 | -0.841352171 | 0.2535921 | -3.317738 | 0.000907494 | 0.005201745 | ERCC6 | protein_coding | 40.54 | -1.921522925 | 1.921522925 | ENSG00000225830 | n.s. |
| ENSG00000112182 | 157.3895098 | -0.783562314 | 0.2277546 | -3.44038 | 0.000580898 | 0.003536439 | BACH2 | protein_coding | 40.5 | -1.920851139 | 1.920851139 | ENSG00000112182 | n.s. |
| ENSG00000273882 | 14.38449797 | -1.141193743 | 0.4043945 | -2.821981 | 0.0047728 | 0.020740229 | MIR8066 | miRNA | 39.74 | -1.920841847 | 1.920841847 | ENSG00000273882 | down |
| ENSG00000260947 | 191.4950326 | -0.945815483 | 0.3031634 | -3.119821 | 0.001809612 | 0.009352349 | | lincRNA | 38.24 | -1.919134621 | 1.919134621 | ENSG00000260947 | n.s. |
| ENSG00000186615 | 116.8816319 | -0.878546774 | 0.2714843 | -3.236087 | 0.001211806 | 0.006632065 | KTN1-AS1 | antisense | 40.78 | -1.913783457 | 1.913783457 | ENSG00000186615 | n.s. |
| ENSG00000117569 | 4037.083054 | -0.887521062 | 0.275877 | -3.21709 | 0.001294979 | 0.00700466 | PTBP2 | protein_coding | 35.67 | -1.912264367 | 1.912264367 | ENSG00000117569 | n.s. |
| ENSG00000085274 | 1413.02488 | -0.812614934 | 0.2416669 | -3.362541 | 0.000772285 | 0.004521669 | MYNN | protein_coding | 37.29 | -1.905339203 | 1.905339203 | ENSG00000085274 | n.s. |
| ENSG00000107651 | 1096.758709 | -0.863490061 | 0.2651729 | -3.256329 | 0.001128629 | 0.006218 | SEC23IP | protein_coding | 40.11 | -1.905160649 | 1.905160649 | ENSG00000107651 | n.s. |
| ENSG00000054598 | 225.2434428 | 0.804490595 | 0.2380975 | 3.378829 | 0.000727952 | 0.004290975 | FOXC1 | protein_coding | 56.32 | 1.904586462 | 1.904586462 | ENSG00000054598 | n.s. |
| ENSG00000252021 | 3.027931472 | -1.528314793 | 0.6372394 | -2.398337 | 0.016469713 | 0.056771132 | | misc_RNA | 49.53 | -1.904085291 | 1.904085291 | ENSG00000252021 | down |
| ENSG00000138398 | 3197.567144 | -0.859402259 | 0.2633324 | -3.263564 | 0.001100203 | 0.006088285 | PPIG | protein_coding | 37.96 | -1.904010019 | 1.904010019 | ENSG00000138398 | n.s. |
| ENSG00000181638 | 174.2402429 | -0.768472006 | 0.2221939 | -3.458565 | 0.000543062 | 0.003333796 | ZFP41 | protein_coding | 57.2 | -1.903552009 | 1.903552009 | ENSG00000181638 | n.s. |
| ENSG00000164970 | 322.6038719 | -0.860369122 | 0.2639443 | -3.259662 | 0.001115452 | 0.006157987 | FAM219A | protein_coding | 47.16 | -1.901898614 | 1.901898614 | ENSG00000164970 | n.s. |
| ENSG00000205643 | 83.64813733 | -0.923883298 | 0.2939509 | -3.142985 | 0.001672343 | 0.008757021 | CDPF1 | protein_coding | 57.88 | -1.901022557 | 1.901022557 | ENSG00000205643 | n.s. |
| ENSG00000199886 | 9.672513992 | -1.159412792 | 0.4165079 | -2.783652 | 0.005375075 | 0.022949147 | RNU6-249P | snRNA | 43.14 | -1.900548234 | 1.900548234 | ENSG00000199886 | down |
| ENSG00000281910 | 25.28392305 | -0.959510684 | 0.3114013 | -3.081267 | 0.002061218 | 0.010454766 | SNORA50A | snoRNA | 43.38 | -1.900489078 | 1.900489078 | ENSG00000281910 | n.s. |
| ENSG00000252941 | 3.745908065 | -1.457410498 | 0.592775 | -2.458623 | 0.013947082 | 0.049776304 | RNA5SP340 | rRNA | 41.12 | -1.898972879 | 1.898972879 | ENSG00000252941 | down |
| ENSG00000221634 | 11.33211012 | -1.184145358 | 0.4307485 | -2.749041 | 0.005976984 | 0.024968684 | MIR1276 | miRNA | 56.63 | -1.897716496 | 1.897716496 | ENSG00000221634 | down |
| ENSG00000230778 | 58.21862714 | -0.839555739 | 0.2545737 | -3.297889 | 0.000974146 | 0.005526109 | ANKRD63 | protein_coding | 73.84 | -1.895364703 | 1.895364703 | ENSG00000230778 | n.s. |
| ENSG00000213860 | 32.89212167 | 1.066152192 | 0.3672385 | 2.90316 | 0.00369418 | 0.016759228 | RPL21P75 | processed_pseudogene | 43.06 | 1.893215483 | 1.893215483 | ENSG00000213860 | up |
| ENSG00000269399 | 130.9157927 | 0.790273899 | 0.2326485 | 3.396858 | 0.000681644 | 0.004050589 | | lincRNA | 47.66 | 1.890715975 | 1.890715975 | ENSG00000269399 | n.s. |
| ENSG00000269189 | 49.23240449 | -0.885169458 | 0.2765008 | -3.201327 | 0.001367964 | 0.007339561 | | sense_intronic | 40.82 | -1.889243642 | 1.889243642 | ENSG00000269189 | n.s. |
| ENSG00000115289 | 2320.137174 | 0.780338385 | 0.2283671 | 3.417035 | 0.000633072 | 0.003795979 | PCGF1 | protein_coding | 56.33 | 1.888946575 | 1.888946575 | ENSG00000115289 | n.s. |
| ENSG00000100485 | 978.4977938 | -0.818795056 | 0.2458922 | -3.329894 | 0.00086879 | 0.004992685 | SOS2 | protein_coding | 38.27 | -1.884592596 | 1.884592596 | ENSG00000100485 | n.s. |
| ENSG00000278175 | 186.7956707 | 0.762209436 | 0.2206172 | 3.454896 | 0.000550505 | 0.003375023 | GLIDR | lincRNA | 40.28 | 1.883970839 | 1.883970839 | ENSG00000278175 | n.s. |
| ENSG00000272525 | 95.18492711 | 0.812582257 | 0.243495 | 3.337162 | 0.000846386 | 0.004887697 | | lincRNA | 48.25 | 1.877792894 | 1.877792894 | ENSG00000272525 | n.s. |
| ENSG00000108312 | 5261.092078 | -0.807652869 | 0.2414048 | -3.345638 | 0.000820936 | 0.004764437 | UBTF | protein_coding | 59.52 | -1.87536057 | 1.87536057 | ENSG00000108312 | n.s. |
| ENSG00000255142 | 13.30302608 | -1.069306183 | 0.3706635 | -2.884844 | 0.003916079 | 0.01763946 | | lincRNA | 59.44 | -1.875044134 | 1.875044134 | ENSG00000255142 | down |
| ENSG00000159217 | 4913.172881 | -0.801359332 | 0.2386636 | -3.357693 | 0.000785957 | 0.004586289 | IGF2BP1 | protein_coding | 47.35 | -1.8740097 | 1.8740097 | ENSG00000159217 | n.s. |

| | | | | | | | | | | | | | |
|---|---|---|---|---|---|---|---|---|---|---|---|---|---|
| ENSG00000197782 | 735.1618818 | -0.826195683 | 0.249903 | -3.306065 | 0.00094616 | 0.005394505 | ZNF780A | protein_coding | 38.52 | -1.873851765 | 1.873851765 | ENSG00000197782 | n.s. |
| ENSG00000120149 | 216.8335627 | 0.893645176 | 0.2817262 | 3.172034 | 0.001513752 | 0.008022827 | MSX2 | protein_coding | 47.07 | 1.872787693 | 1.872787693 | ENSG00000120149 | n.s. |
| ENSG00000100095 | 1169.456497 | -0.811581709 | 0.2435102 | -3.332845 | 0.000859628 | 0.004953892 | SEZ6L | protein_coding | 44.56 | -1.870739259 | 1.870739259 | ENSG00000100095 | n.s. |
| ENSG00000276174 | 76.23800046 | -0.87592956 | 0.2735549 | -3.202025 | 0.001364654 | 0.007330275 | | sense_intronic | 31.9 | -1.870004259 | 1.870004259 | ENSG00000276174 | n.s. |
| ENSG00000066735 | 2757.426942 | -0.793861117 | 0.2355641 | -3.370042 | 0.000751566 | 0.004417078 | KIF26A | protein_coding | 63.77 | -1.869435735 | 1.869435735 | ENSG00000066735 | n.s. |
| ENSG00000234709 | 28.42507676 | -1.01050311 | 0.3407081 | -2.965891 | 0.003018072 | 0.014181782 | UPF3AP3 | transcribed_processed_pseudog | 52.28 | -1.867681779 | 1.867681779 | ENSG00000234709 | down |
| ENSG00000199357 | 11.09581751 | 1.175450195 | 0.4297796 | 2.735007 | 0.006237898 | 0.02577159 | | misc_RNA | 39.18 | 1.867624375 | 1.867624375 | ENSG00000199357 | up |
| ENSG00000197050 | 543.2811853 | -0.826792171 | 0.2507091 | -3.297815 | 0.000974402 | 0.005526109 | ZNF420 | protein_coding | 41.84 | -1.866549918 | 1.866549918 | ENSG00000197050 | n.s. |
| ENSG00000279329 | 82.35578191 | 0.84109785 | 0.2575545 | 3.265708 | 0.001091908 | 0.006052006 | | TEC | 49.92 | 1.865639676 | 1.865639676 | ENSG00000279329 | n.s. |
| ENSG00000225900 | 30.2874423 | -0.946814928 | 0.3084276 | -3.069813 | 0.002141929 | 0.010777768 | HSPE1P13 | processed_pseudogene | 38.38 | -1.862831082 | 1.862831082 | ENSG00000225900 | n.s. |
| ENSG00000120693 | 3816.436763 | -0.856626311 | 0.2651029 | -3.231297 | 0.001232297 | 0.006713341 | SMAD9 | protein_coding | 41.07 | -1.861501476 | 1.861501476 | ENSG00000120693 | n.s. |
| ENSG00000051108 | 1624.744293 | 0.731666175 | 0.2086424 | 3.506795 | 0.000453539 | 0.002863773 | HERPUD1 | protein_coding | 43.16 | 1.860671986 | 1.860671986 | ENSG00000051108 | n.s. |
| ENSG00000253395 | 13.14380595 | 1.131644886 | 0.4060292 | 2.787102 | 0.005318175 | 0.022769049 | | lincRNA | 46.39 | 1.858902257 | 1.858902257 | ENSG00000253395 | up |
| ENSG00000232995 | 324.1361382 | -0.768825192 | 0.2251859 | -3.41418 | 0.000639742 | 0.003834322 | | antisense | 37.85 | -1.857721099 | 1.857721099 | ENSG00000232995 | n.s. |
| ENSG00000257303 | 35.7741425 | 0.900229759 | 0.286025 | 3.147382 | 0.001647397 | 0.008652095 | | antisense | 46.83 | 1.857064814 | 1.857064814 | ENSG00000257303 | n.s. |
| ENSG00000207268 | 11.78796804 | -1.107765738 | 0.3940635 | -2.811135 | 0.004936702 | 0.021332618 | SNORA70C | snoRNA | 50.37 | -1.851027625 | 1.851027625 | ENSG00000207268 | down |
| ENSG00000243960 | 43.8572297 | -0.92419083 | 0.2982717 | -3.098487 | 0.001945117 | 0.009956569 | | sense_overlapping | 41.95 | -1.850128669 | 1.850128669 | ENSG00000243960 | n.s. |
| ENSG00000111490 | 867.1360667 | -0.802387502 | 0.2407674 | -3.332625 | 0.000860307 | 0.004954588 | TBC1D30 | protein_coding | 41.2 | -1.849497134 | 1.849497134 | ENSG00000111490 | n.s. |
| ENSG00000198964 | 114.4413053 | -0.885430397 | 0.2796089 | -3.166674 | 0.00154193 | 0.008153548 | SGMS1 | protein_coding | 39.37 | -1.849357171 | 1.849357171 | ENSG00000198964 | n.s. |
| ENSG00000227161 | 25.51489794 | -1.022216522 | 0.3486108 | -2.932257 | 0.003365081 | 0.015570629 | | sense_overlapping | 38.3 | -1.847854502 | 1.847854502 | ENSG00000227161 | down |
| ENSG00000105829 | 165.6825323 | -0.908302438 | 0.2910652 | -3.120615 | 0.00180474 | 0.009337568 | BET1 | protein_coding | 35.67 | -1.84364163 | 1.84364163 | ENSG00000105829 | n.s. |
| ENSG00000260038 | 24.94799941 | 0.98422497 | 0.3299066 | 2.983344 | 0.002851175 | 0.013493169 | | sense_intronic | 40.86 | 1.840388531 | 1.840388531 | ENSG00000260038 | n.s. |
| ENSG00000276853 | 180.7864383 | 0.857344133 | 0.2672304 | 3.208258 | 0.001335418 | 0.007192658 | | sense_intronic | 32.32 | 1.837383311 | 1.837383311 | ENSG00000276853 | n.s. |
| ENSG00000112234 | 934.8781971 | -0.872546979 | 0.274402 | -3.179812 | 0.001473705 | 0.00784421 | FBXL4 | protein_coding | 36.71 | -1.837104745 | 1.837104745 | ENSG00000112234 | n.s. |
| ENSG00000160318 | 14.12428792 | 1.078875405 | 0.380102 | 2.838384 | 0.004534259 | 0.019883617 | CLDND2 | protein_coding | 66.05 | 1.835711477 | 1.835711477 | ENSG00000160318 | up |
| ENSG00000106683 | 205.4241975 | -0.823495302 | 0.2515233 | -3.274031 | 0.001060249 | 0.005907099 | LIMK1 | protein_coding | 54.59 | -1.83526284 | 1.83526284 | ENSG00000106683 | n.s. |
| ENSG00000229891 | 102.0746613 | 0.87602198 | 0.2762679 | 3.170915 | 0.001519596 | 0.008050736 | LINC01315 | lincRNA | 55.65 | 1.834534043 | 1.834534043 | ENSG00000229891 | n.s. |
| ENSG00000249605 | 28.2446612 | -0.973456077 | 0.3252489 | -2.992957 | 0.002762888 | 0.013168232 | | sense_intronic | 37.3 | -1.8305574 | 1.8305574 | ENSG00000249605 | n.s. |
| ENSG00000114988 | 344.3772673 | -0.947633318 | 0.3118965 | -3.038294 | 0.002379216 | 0.011721807 | LMAN2L | protein_coding | 44.97 | -1.829885075 | 1.829885075 | ENSG00000114988 | n.s. |
| ENSG00000073008 | 201.2138573 | -0.848596574 | 0.2636877 | -3.218188 | 0.001290031 | 0.006980608 | PVR | protein_coding | 50.36 | -1.82966481 | 1.82966481 | ENSG00000073008 | n.s. |
| ENSG00000145362 | 5668.014981 | -0.802203002 | 0.2420649 | -3.314 | 0.000919715 | 0.005265291 | ANK2 | protein_coding | 37.08 | -1.827881797 | 1.827881797 | ENSG00000145362 | n.s. |
| ENSG00000049618 | 4906.878231 | -0.811110206 | 0.2462113 | -3.294366 | 0.000986441 | 0.005585285 | ARID1B | protein_coding | 40.87 | -1.827394533 | 1.827394533 | ENSG00000049618 | n.s. |

| | | | | | | | | | | | | | |
|---|---|---|---|---|---|---|---|---|---|---|---|---|---|
| ENSG00000143387 | 75.63171283 | -0.901055951 | 0.2889865 | -3.117986 | 0.001820915 | 0.009400297 | CTSK | protein_coding | 43 | -1.826312842 | 1.826312842 | ENSG00000143387 | n.s. |
| ENSG00000080845 | 204.1282489 | -0.890030413 | 0.283634 | -3.137954 | 0.001701317 | 0.008878352 | DLGAP4 | protein_coding | 51.09 | -1.826046593 | 1.826046593 | ENSG00000080845 | n.s. |
| ENSG00000152926 | 968.1266368 | -0.870298155 | 0.2742604 | -3.173254 | 0.001507404 | 0.007992224 | ZNF117 | protein_coding | 39.14 | -1.825304465 | 1.825304465 | ENSG00000152926 | n.s. |
| ENSG00000022976 | 223.2690279 | -0.892524814 | 0.2849215 | -3.132528 | 0.001733077 | 0.009020424 | ZNF839 | protein_coding | 49.04 | -1.825010697 | 1.825010697 | ENSG00000022976 | n.s. |
| ENSG00000267160 | 175.8178738 | -0.791366816 | 0.2374806 | -3.332342 | 0.000861182 | 0.004956694 | | antisense | 29.78 | -1.823948493 | 1.823948493 | ENSG00000267160 | n.s. |
| ENSG00000113300 | 1607.603971 | -0.850473677 | 0.2650106 | -3.209206 | 0.00133102 | 0.007171744 | CNOT6 | protein_coding | 42.07 | -1.823734686 | 1.823734686 | ENSG00000113300 | n.s. |
| ENSG00000206775 | 10.99317871 | 1.168902143 | 0.4315856 | 2.70839 | 0.006761051 | 0.027543286 | SNORD37 | snoRNA | 43.94 | 1.823468937 | 1.823468937 | ENSG00000206775 | up |
| ENSG00000134444 | 1493.55527 | -0.811238225 | 0.2466563 | -3.288942 | 0.001005648 | 0.005677872 | KIAA1468 | protein_coding | 34.63 | -1.821890491 | 1.821890491 | ENSG00000134444 | n.s. |
| ENSG00000167281 | 403.5642442 | -0.82165529 | 0.2518467 | -3.262522 | 0.001104257 | 0.006099268 | RBFOX3 | protein_coding | 55.28 | -1.819738282 | 1.819738282 | ENSG00000167281 | n.s. |
| ENSG00000254985 | 52.16524776 | -0.906538171 | 0.2924598 | -3.099701 | 0.001937158 | 0.009919473 | RSF1-IT2 | sense_intronic | 39 | -1.816259554 | 1.816259554 | ENSG00000254985 | n.s. |
| ENSG00000248240 | 27.63555971 | 0.93218437 | 0.3052989 | 3.05335 | 0.002263018 | 0.01129319 | | antisense | 45.83 | 1.815133904 | 1.815133904 | ENSG00000248240 | n.s. |
| ENSG00000266441 | 40.45514199 | 0.897065533 | 0.2879769 | 3.115061 | 0.001839069 | 0.009479954 | | antisense | 59.4 | 1.81493739 | 1.81493739 | ENSG00000266441 | n.s. |
| ENSG00000079387 | 979.7555994 | -0.77560673 | 0.2309737 | -3.357987 | 0.000785124 | 0.004585267 | SENP1 | protein_coding | 38.62 | -1.813861356 | 1.813861356 | ENSG00000079387 | n.s. |
| ENSG00000268987 | 27.0976162 | 0.940953179 | 0.3101379 | 3.033983 | 0.002413482 | 0.011865434 | | sense_intronic | 47.59 | 1.812008956 | 1.812008956 | ENSG00000268987 | n.s. |
| ENSG00000264405 | 2.607395897 | 1.788597437 | 0.8356402 | 2.140392 | 0.032323124 | 0.097199754 | MIR3913-1 | miRNA | 33.33 | 1.81065949 | 1.81065949 | ENSG00000264405 | up |
| ENSG00000196912 | 586.9211033 | -0.881677391 | 0.2808913 | -3.138856 | 0.001696086 | 0.008857689 | ANKRD36B | protein_coding | 35.8 | -1.809801161 | 1.809801161 | ENSG00000196912 | n.s. |
| ENSG00000263540 | 3.374080339 | -1.559228373 | 0.675221 | -2.309212 | 0.020931821 | 0.069155748 | MIR5582 | miRNA | 32.35 | -1.808972658 | 1.808972658 | ENSG00000263540 | down |
| ENSG00000122481 | 424.0241363 | -0.941705786 | 0.3108665 | -3.029294 | 0.002451263 | 0.012012853 | RWDD3 | protein_coding | 40.77 | -1.808408327 | 1.808408327 | ENSG00000122481 | n.s. |
| ENSG00000264575 | 167.833312 | 0.771021138 | 0.229307 | 3.362397 | 0.000772688 | 0.004522127 | LINC00526 | lincRNA | 54.56 | 1.8077803 | 1.8077803 | ENSG00000264575 | n.s. |
| ENSG00000198863 | 490.841854 | -0.780994849 | 0.2339213 | -3.338707 | 0.000841693 | 0.004864627 | RUNDC1 | protein_coding | 47.47 | -1.806402383 | 1.806402383 | ENSG00000198863 | n.s. |
| ENSG00000189007 | 390.8435673 | -0.784644751 | 0.2356272 | -3.330026 | 0.000868378 | 0.004992685 | ADAT2 | protein_coding | 38.36 | -1.805990007 | 1.805990007 | ENSG00000189007 | n.s. |
| ENSG00000236977 | 72.61116521 | 0.894705994 | 0.2877584 | 3.109226 | 0.001875779 | 0.00964063 | ANKRD44-IT1 | sense_intronic | 40.66 | 1.80363298 | 1.80363298 | ENSG00000236977 | n.s. |
| ENSG00000136643 | 702.4987153 | -0.83224382 | 0.2579664 | -3.226172 | 0.00125458 | 0.006807293 | RPS6KC1 | protein_coding | 37.51 | -1.803493633 | 1.803493633 | ENSG00000136643 | n.s. |
| ENSG00000279203 | 37.92952578 | -0.932319014 | 0.3067381 | -3.039463 | 0.002370006 | 0.011689621 | | TEC | 39.46 | -1.80142641 | 1.80142641 | ENSG00000279203 | n.s. |
| ENSG00000163029 | 2324.788112 | -0.876298403 | 0.2793344 | -3.137095 | 0.00170631 | 0.008901072 | SMC6 | protein_coding | 36.86 | -1.796900437 | 1.796900437 | ENSG00000163029 | n.s. |
| ENSG00000202479 | 12.93603673 | 1.099596598 | 0.3956537 | 2.779189 | 0.005449477 | 0.023224079 | | snoRNA | 40 | 1.796808468 | 1.796808468 | ENSG00000202479 | up |
| ENSG00000278840 | 44.56527361 | -0.938405249 | 0.3105849 | -3.021413 | 0.002515979 | 0.01227026 | | sense_intronic | 29.41 | -1.793429655 | 1.793429655 | ENSG00000278840 | n.s. |
| ENSG00000169410 | 622.495303 | -0.846957372 | 0.2656391 | -3.188376 | 0.001430741 | 0.007641047 | PTPN9 | protein_coding | 45.12 | -1.792879262 | 1.792879262 | ENSG00000169410 | n.s. |
| ENSG00000173214 | 674.004072 | -0.808170704 | 0.2475758 | -3.264336 | 0.00109721 | 0.006074137 | MFSD4B | protein_coding | 43.27 | -1.791322812 | 1.791322812 | ENSG00000173214 | n.s. |
| ENSG00000236654 | 25.65192395 | -0.968811534 | 0.3266047 | -2.966313 | 0.003013939 | 0.014167141 | | processed_pseudogene | 46.86 | -1.791059109 | 1.791059109 | ENSG00000236654 | n.s. |
| ENSG00000214140 | 103.6433292 | -0.975010089 | 0.3298593 | -2.955836 | 0.00311823 | 0.014588411 | PRCD | protein_coding | 55 | -1.790110718 | 1.790110718 | ENSG00000214140 | n.s. |
| ENSG00000264911 | 28.42020118 | -1.055849426 | 0.37276 | -2.832518 | 0.004618301 | 0.020201258 | | processed_pseudogene | 39.06 | -1.789265231 | 1.789265231 | ENSG00000264911 | down |

| | | | | | | | | | | | | | |
|---|---|---|---|---|---|---|---|---|---|---|---|---|---|
| ENSG00000198974 | 6.03031577 | -1.339492003 | 0.5374324 | -2.492392 | 0.012688603 | 0.046162377 | MIR30E | miRNA | 51.09 | -1.789175317 | 1.789175317 | ENSG00000198974 | down |
| ENSG00000185298 | 882.4903365 | -0.823888624 | 0.2552855 | -3.227322 | 0.001249546 | 0.006785269 | CCDC137 | protein_coding | 57.88 | -1.786547224 | 1.786547224 | ENSG00000185298 | n.s. |
| ENSG00000146373 | 120.5178918 | -0.828442831 | 0.2575328 | -3.216844 | 0.001296091 | 0.007005234 | RNF217 | protein_coding | 36.66 | -1.784944175 | 1.784944175 | ENSG00000146373 | n.s. |
| ENSG00000117481 | 201.0115073 | -0.805625981 | 0.2468805 | -3.263222 | 0.001101531 | 0.006093213 | NSUN4 | protein_coding | 47.35 | -1.784585353 | 1.784585353 | ENSG00000117481 | n.s. |
| ENSG00000277383 | 26.45762692 | 0.926351832 | 0.3052856 | 3.034378 | 0.00241032 | 0.011854077 | | antisense | 65.46 | 1.784276145 | 1.784276145 | ENSG00000277383 | n.s. |
| ENSG00000124459 | 571.7934829 | -0.763795839 | 0.2277572 | -3.353553 | 0.000797811 | 0.004645725 | ZNF45 | protein_coding | 41.36 | -1.781894841 | 1.781894841 | ENSG00000124459 | n.s. |
| ENSG00000075303 | 550.0579553 | -0.892897472 | 0.2887069 | -3.092747 | 0.001983131 | 0.010119648 | SLC25A40 | protein_coding | 35.47 | -1.781182763 | 1.781182763 | ENSG00000075303 | n.s. |
| ENSG00000146757 | 1600.214236 | -0.871080322 | 0.2781323 | -3.131892 | 0.00173684 | 0.00903663 | ZNF92 | protein_coding | 39.16 | -1.780482513 | 1.780482513 | ENSG00000146757 | n.s. |
| ENSG00000179886 | 268.037026 | -0.739106015 | 0.2168865 | -3.4078 | 0.000654888 | 0.0039066 | TIGD5 | protein_coding | 64.53 | -1.779915907 | 1.779915907 | ENSG00000179886 | n.s. |
| ENSG00000146802 | 806.514966 | -0.897415055 | 0.2911136 | -3.082697 | 0.002051338 | 0.010416038 | TMEM168 | protein_coding | 35.46 | -1.778943575 | 1.778943575 | ENSG00000146802 | n.s. |
| ENSG00000249690 | 152.5578907 | 0.765909401 | 0.2289655 | 3.345086 | 0.00082257 | 0.004769947 | | antisense | 45.58 | 1.778048325 | 1.778048325 | ENSG00000249690 | n.s. |
| ENSG00000073614 | 4008.110861 | -0.793218112 | 0.2414479 | -3.285256 | 0.001018897 | 0.005736392 | KDM5A | protein_coding | 39.46 | -1.7778883 | 1.7778883 | ENSG00000073614 | n.s. |
| ENSG00000202222 | 27.5184574 | 0.922710331 | 0.3040726 | 3.034506 | 0.002409298 | 0.011853236 | | misc_RNA | 41.59 | 1.777290547 | 1.777290547 | ENSG00000202222 | n.s. |
| ENSG00000148143 | 2841.308576 | -0.752403162 | 0.2230992 | -3.372505 | 0.000744877 | 0.004381463 | ZNF462 | protein_coding | 40.08 | -1.774453225 | 1.774453225 | ENSG00000148143 | n.s. |
| ENSG00000174306 | 37.84106603 | -0.925283248 | 0.3057561 | -3.026214 | 0.002476372 | 0.012110476 | ZHX3 | protein_coding | 41.13 | -1.773618814 | 1.773618814 | ENSG00000174306 | n.s. |
| ENSG00000284154 | 5.651086087 | -1.392209442 | 0.57363 | -2.427016 | 0.015223567 | 0.05325571 | MIR3605 | miRNA | 55 | -1.773165028 | 1.773165028 | ENSG00000284154 | down |
| ENSG00000227726 | 20.11487158 | -1.011426548 | 0.3506501 | -2.884433 | 0.003921197 | 0.017656802 | | sense_intronic | 48.1 | -1.773119705 | 1.773119705 | ENSG00000227726 | down |
| ENSG00000183691 | 61.23457822 | 0.860408259 | 0.2735271 | 3.145605 | 0.001657436 | 0.008701541 | NOG | protein_coding | 59.46 | 1.77278846 | 1.77278846 | ENSG00000183691 | n.s. |
| ENSG00000205268 | 1716.360086 | -0.813836374 | 0.2516018 | -3.23462 | 0.001218046 | 0.006660978 | PDE7A | protein_coding | 37.7 | -1.771283959 | 1.771283959 | ENSG00000205268 | n.s. |
| ENSG00000272248 | 9.973831593 | 1.091288288 | 0.3942709 | 2.767864 | 0.005642497 | 0.023878568 | | lincRNA | 47.76 | 1.770060569 | 1.770060569 | ENSG00000272248 | up |
| ENSG00000203684 | 22.69264647 | -0.963507884 | 0.3258453 | -2.956949 | 0.003106996 | 0.014556087 | IBA57-AS1 | lincRNA | 52.98 | -1.769920957 | 1.769920957 | ENSG00000203684 | n.s. |
| ENSG00000244968 | 113.3440631 | 0.788847928 | 0.2402002 | 3.284127 | 0.001022989 | 0.005745492 | LIFR-AS1 | antisense | 41.55 | 1.767550098 | 1.767550098 | ENSG00000244968 | n.s. |
| ENSG00000123178 | 231.9595467 | -0.898433491 | 0.2927365 | -3.069086 | 0.00214715 | 0.010796235 | SPRYD7 | protein_coding | 40.33 | -1.766974014 | 1.766974014 | ENSG00000123178 | n.s. |
| ENSG00000270673 | 38.33686347 | 0.89734022 | 0.2922425 | 3.070533 | 0.00213677 | 0.010755692 | YTHDF3-AS1 | lincRNA | 55.29 | 1.766290086 | 1.766290086 | ENSG00000270673 | n.s. |
| ENSG00000198909 | 823.9552604 | -0.787824457 | 0.2398677 | -3.284412 | 0.001021955 | 0.005744084 | MAP3K3 | protein_coding | 45.21 | -1.765340644 | 1.765340644 | ENSG00000198909 | n.s. |
| ENSG00000283785 | 7.702943268 | -1.201568086 | 0.4583147 | -2.62171 | 0.008748986 | 0.034006737 | MIR15A | miRNA | 42.17 | -1.764424672 | 1.764424672 | ENSG00000283785 | down |
| ENSG00000111802 | 552.7124859 | -0.830352737 | 0.2601164 | -3.192235 | 0.001411765 | 0.00754549 | TDP2 | protein_coding | 39.02 | -1.762268013 | 1.762268013 | ENSG00000111802 | n.s. |
| ENSG00000145147 | 62.09922227 | 0.905171475 | 0.2968321 | 3.04944 | 0.002292687 | 0.011400388 | SLIT2 | protein_coding | 36.01 | 1.758820936 | 1.758820936 | ENSG00000145147 | n.s. |
| ENSG00000100592 | 314.2353571 | -0.784865884 | 0.2389981 | -3.283984 | 0.001023509 | 0.005746092 | DAAM1 | protein_coding | 39.75 | -1.758592002 | 1.758592002 | ENSG00000100592 | n.s. |
| ENSG00000249855 | 90.71445901 | -0.892439601 | 0.2906027 | -3.070995 | 0.002133464 | 0.010749525 | EEF1A1P19 | processed_pseudogene | 47.68 | -1.756866179 | 1.756866179 | ENSG00000249855 | n.s. |
| ENSG00000229955 | 33.63795215 | -0.964347574 | 0.3277097 | -2.942688 | 0.003253761 | 0.015115819 | | sense_intronic | 36.7 | -1.755660652 | 1.755660652 | ENSG00000229955 | n.s. |
| ENSG00000225447 | 15.72103444 | -1.047488916 | 0.3720772 | -2.815246 | 0.004873994 | 0.021107501 | RPS15AP10 | processed_pseudogene | 43.19 | -1.755133856 | 1.755133856 | ENSG00000225447 | down |

| | | | | | | | | | | | | | |
|---|---|---|---|---|---|---|---|---|---|---|---|---|---|
| ENSG00000196712 | 2041.08768 | -0.833046851 | 0.262028 | -3.179228 | 0.001476678 | 0.007856242 | NF1 | protein_coding | 37.62 | -1.753384638 | 1.753384638 | ENSG00000196712 | n.s. |
| ENSG00000144712 | 258.4277967 | -0.81610767 | 0.2541325 | -3.211347 | 0.001321145 | 0.00712681 | CAND2 | protein_coding | 50.87 | -1.752268712 | 1.752268712 | ENSG00000144712 | n.s. |
| ENSG00000066933 | 110.061532 | -0.901078114 | 0.2953836 | -3.050536 | 0.002284334 | 0.011371036 | MYO9A | protein_coding | 37.93 | -1.751876054 | 1.751876054 | ENSG00000066933 | n.s. |
| ENSG00000197275 | 235.0340773 | -0.830242655 | 0.2608056 | -3.183377 | 0.001455678 | 0.007762317 | RAD54B | protein_coding | 36.72 | -1.751819166 | 1.751819166 | ENSG00000197275 | n.s. |
| ENSG00000271659 | 46.71162169 | -0.870128562 | 0.2801048 | -3.106439 | 0.001893552 | 0.009717628 | | antisense | 41.18 | -1.751081307 | 1.751081307 | ENSG00000271659 | n.s. |
| ENSG00000182771 | 544.5008517 | -0.784959092 | 0.2397287 | -3.274364 | 0.001059002 | 0.005902515 | GRID1 | protein_coding | 45.04 | -1.749644692 | 1.749644692 | ENSG00000182771 | n.s. |
| ENSG00000167985 | 311.5245489 | -0.934803413 | 0.3131995 | -2.98469 | 0.002838658 | 0.013449151 | SDHAF2 | protein_coding | 45.09 | -1.749302432 | 1.749302432 | ENSG00000167985 | n.s. |
| ENSG00000135698 | 435.7781078 | -0.915920487 | 0.3033038 | -3.019812 | 0.002529319 | 0.012323031 | MPHOSPH6 | protein_coding | 40.61 | -1.748750916 | 1.748750916 | ENSG00000135698 | n.s. |
| ENSG00000064195 | 81.9121673 | 0.810319171 | 0.251674 | 3.219717 | 0.001283172 | 0.006948892 | DLX3 | protein_coding | 57.41 | 1.748737169 | 1.748737169 | ENSG00000064195 | n.s. |
| ENSG00000090776 | 169.2298439 | 0.789387152 | 0.2419293 | 3.262884 | 0.001102848 | 0.006098075 | EFNB1 | protein_coding | 56.43 | 1.748340374 | 1.748340374 | ENSG00000090776 | n.s. |
| ENSG00000198010 | 46.27496241 | -0.876047735 | 0.2832815 | -3.092499 | 0.001984786 | 0.010122409 | DLGAP2 | protein_coding | 47.91 | -1.747466544 | 1.747466544 | ENSG00000198010 | n.s. |
| ENSG00000200063 | 9.089552789 | -1.166089653 | 0.4399791 | -2.650329 | 0.008041334 | 0.03173455 | | snoRNA | 59.69 | -1.747347635 | 1.747347635 | ENSG00000200063 | down |
| ENSG00000265458 | 89.41532749 | 0.782960077 | 0.2389597 | 3.276536 | 0.001050888 | 0.005866684 | | antisense | 48.51 | 1.747259437 | 1.747259437 | ENSG00000265458 | n.s. |
| ENSG00000128594 | 188.1166234 | 0.770139772 | 0.2329054 | 3.306664 | 0.00094414 | 0.005387404 | LRRC4 | protein_coding | 56.07 | 1.747154856 | 1.747154856 | ENSG00000128594 | n.s. |
| ENSG00000199530 | 7.956461391 | -1.496201736 | 0.6458197 | -2.316748 | 0.020517449 | 0.068080183 | | misc_RNA | 38.89 | -1.74603643 | 1.74603643 | ENSG00000199530 | down |
| ENSG00000117318 | 1055.045334 | 0.778541 | 0.2369136 | 3.286181 | 0.001015558 | 0.005719908 | ID3 | protein_coding | 57.27 | 1.745964606 | 1.745964606 | ENSG00000117318 | n.s. |
| ENSG00000276859 | 10.26277385 | 1.14758417 | 0.429468 | 2.672107 | 0.007537671 | 0.030105275 | | misc_RNA | 65.05 | 1.745885677 | 1.745885677 | ENSG00000276859 | up |
| ENSG00000221650 | 11.17149176 | -1.098108927 | 0.401367 | -2.735923 | 0.006220566 | 0.025722883 | MIR1267 | miRNA | 44.87 | -1.745642192 | 1.745642192 | ENSG00000221650 | down |
| ENSG00000136478 | 799.3351961 | -0.841699174 | 0.2668506 | -3.154196 | 0.001609411 | 0.008468565 | TEX2 | protein_coding | 43.58 | -1.744160756 | 1.744160756 | ENSG00000136478 | n.s. |
| ENSG00000275191 | 17.22621033 | -1.318720899 | 0.5319771 | -2.478905 | 0.013178631 | 0.047596908 | | sense_intronic | 35.29 | -1.743904552 | 1.743904552 | ENSG00000275191 | down |
| ENSG00000112541 | 642.6137856 | -0.788633194 | 0.2419832 | -3.259041 | 0.001117894 | 0.006166575 | PDE10A | protein_coding | 40.93 | -1.742844643 | 1.742844643 | ENSG00000112541 | n.s. |
| ENSG00000142751 | 206.3968455 | -0.776100851 | 0.236052 | -3.287838 | 0.001009599 | 0.005696118 | GPN2 | protein_coding | 48.74 | -1.741897061 | 1.741897061 | ENSG00000142751 | n.s. |
| ENSG00000070495 | 503.0946774 | -0.805847272 | 0.2501097 | -3.221976 | 0.001273099 | 0.006897024 | JMJD6 | protein_coding | 48.83 | -1.741708549 | 1.741708549 | ENSG00000070495 | n.s. |
| ENSG00000076770 | 1082.368188 | -0.838480713 | 0.2655592 | -3.157415 | 0.001591746 | 0.008385381 | MBNL3 | protein_coding | 36.96 | -1.741086087 | 1.741086087 | ENSG00000076770 | n.s. |
| ENSG00000173812 | 5438.356227 | 0.676543209 | 0.1919696 | 3.524221 | 0.00042473 | 0.002706415 | EIF1 | protein_coding | 49.15 | 1.737096178 | 1.737096178 | ENSG00000173812 | n.s. |
| ENSG00000172469 | 762.4115 | -0.92136244 | 0.3076011 | -2.995315 | 0.002741612 | 0.013085865 | MANEA | protein_coding | 33.8 | -1.735107512 | 1.735107512 | ENSG00000172469 | n.s. |
| ENSG00000071189 | 1398.602462 | -0.850029819 | 0.2718118 | -3.127273 | 0.001764358 | 0.009155853 | SNX13 | protein_coding | 35.09 | -1.732616788 | 1.732616788 | ENSG00000071189 | n.s. |
| ENSG00000118231 | 120.8894412 | 0.900636398 | 0.2972205 | 3.030196 | 0.00244395 | 0.011981378 | CRYGD | protein_coding | 37.37 | 1.730566743 | 1.730566743 | ENSG00000118231 | n.s. |
| ENSG00000272583 | 30.52857063 | 0.91549375 | 0.305053 | 3.001098 | 0.002690082 | 0.012888512 | | lincRNA | 38.79 | 1.730097529 | 1.730097529 | ENSG00000272583 | n.s. |
| ENSG00000220323 | 25.89840104 | 0.933399928 | 0.3142966 | 2.969806 | 0.00297988 | 0.014035476 | | processed_pseudogene | 63.29 | 1.729378047 | 1.729378047 | ENSG00000220323 | n.s. |
| ENSG00000170681 | 322.4470862 | -0.79026058 | 0.2437175 | -3.242528 | 0.001184745 | 0.006489064 | CAVIN4 | protein_coding | 40.73 | -1.728946266 | 1.728946266 | ENSG00000170681 | n.s. |
| ENSG00000167981 | 193.804017 | -0.785174603 | 0.2415103 | -3.251102 | 0.001149588 | 0.006323869 | ZNF597 | protein_coding | 42.53 | -1.72661243 | 1.72661243 | ENSG00000167981 | n.s. |

| | | | | | | | | | | | | | |
|---|---|---|---|---|---|---|---|---|---|---|---|---|---|
| ENSG00000213197 | 46.66882513 | -0.891490077 | 0.2933071 | -3.039443 | 0.002370164 | 0.011689621 | | processed_pseudogene | 52.88 | -1.722536756 | 1.722536756 | ENSG00000213197 | n.s. |
| ENSG00000273008 | 118.8043795 | 0.76589325 | 0.2327003 | 3.291329 | 0.000997152 | 0.005639052 | | lincRNA | 54.4 | 1.722336082 | 1.722336082 | ENSG00000273008 | n.s. |
| ENSG00000270696 | 262.9270881 | -0.806615873 | 0.2519446 | -3.201561 | 0.001366853 | 0.007336426 | | antisense | 39.27 | -1.721734072 | 1.721734072 | ENSG00000270696 | n.s. |
| ENSG00000167695 | 176.9833354 | -0.827940084 | 0.2620671 | -3.159268 | 0.001581662 | 0.008344631 | FAM57A | protein_coding | 49.91 | -1.720950341 | 1.720950341 | ENSG00000167695 | n.s. |
| ENSG00000282100 | 131.176648 | -0.885693011 | 0.2905196 | -3.048651 | 0.002298713 | 0.011414043 | | processed_pseudogene | 45.94 | -1.720512267 | 1.720512267 | ENSG00000282100 | n.s. |
| ENSG00000040531 | 148.0246344 | -0.903955895 | 0.2999118 | -3.014072 | 0.002577665 | 0.012504817 | CTNS | protein_coding | 54.27 | -1.720158167 | 1.720158167 | ENSG00000040531 | n.s. |
| ENSG00000119321 | 590.8494683 | -0.769549315 | 0.2347132 | -3.278679 | 0.001042942 | 0.005826998 | FKBP15 | protein_coding | 41.19 | -1.719600383 | 1.719600383 | ENSG00000119321 | n.s. |
| ENSG00000174206 | 224.6600232 | -0.735987249 | 0.2194532 | -3.353732 | 0.000797295 | 0.004644951 | C12orf66 | protein_coding | 39.91 | -1.717072154 | 1.717072154 | ENSG00000174206 | n.s. |
| ENSG00000153404 | 485.34681 | -0.810677051 | 0.2542717 | -3.188231 | 0.001431459 | 0.00764195 | PLEKHG4B | protein_coding | 49.1 | -1.716037789 | 1.716037789 | ENSG00000153404 | n.s. |
| ENSG00000119820 | 2116.170277 | -0.909342015 | 0.3033976 | -2.997195 | 0.00272476 | 0.013014713 | YIPF4 | protein_coding | 41.2 | -1.714623853 | 1.714623853 | ENSG00000119820 | n.s. |
| ENSG00000178397 | 186.9509334 | -0.859395246 | 0.2778796 | -3.092689 | 0.001983518 | 0.010119648 | FAM220A | protein_coding | 48.26 | -1.714351363 | 1.714351363 | ENSG00000178397 | n.s. |
| ENSG00000143373 | 685.8750146 | -0.796863948 | 0.247911 | -3.214315 | 0.001307561 | 0.007061741 | ZNF687 | protein_coding | 57.99 | -1.714124641 | 1.714124641 | ENSG00000143373 | n.s. |
| ENSG00000113360 | 1268.482144 | -0.792944137 | 0.2460978 | -3.22207 | 0.001272682 | 0.006897024 | DROSHA | protein_coding | 39.75 | -1.713820511 | 1.713820511 | ENSG00000113360 | n.s. |
| ENSG00000100601 | 317.325535 | -0.815524796 | 0.2568166 | -3.175514 | 0.001495712 | 0.007936277 | ALKBH1 | protein_coding | 42.35 | -1.712914555 | 1.712914555 | ENSG00000100601 | n.s. |
| ENSG00000169609 | 782.1797604 | -0.782816114 | 0.2414115 | -3.242663 | 0.001184183 | 0.00648854 | C15orf40 | protein_coding | 44.6 | -1.712686607 | 1.712686607 | ENSG00000169609 | n.s. |
| ENSG00000212248 | 27.92985668 | 0.993936425 | 0.3479835 | 2.856274 | 0.004286445 | 0.018964147 | RNU6-750P | snRNA | 43.93 | 1.711624817 | 1.711624817 | ENSG00000212248 | n.s. |
| ENSG00000180425 | 239.7859074 | 0.750840988 | 0.2265455 | 3.314305 | 0.000918713 | 0.005261718 | C11orf71 | protein_coding | 39.34 | 1.711070811 | 1.711070811 | ENSG00000180425 | n.s. |
| ENSG00000114279 | 232.2412597 | -0.867030417 | 0.2819469 | -3.075155 | 0.002103933 | 0.010636548 | FGF12 | protein_coding | 37.5 | -1.710823819 | 1.710823819 | ENSG00000114279 | n.s. |
| ENSG00000204498 | 155.2735761 | -0.7887418 | 0.2444089 | -3.22714 | 0.001250342 | 0.006786942 | NFKBIL1 | protein_coding | 48.1 | -1.710249266 | 1.710249266 | ENSG00000204498 | n.s. |
| ENSG00000100105 | 1004.6029 | -0.757492117 | 0.229691 | -3.297874 | 0.000974199 | 0.005526109 | PATZ1 | protein_coding | 52.8 | -1.710099465 | 1.710099465 | ENSG00000100105 | n.s. |
| ENSG00000184677 | 692.4182879 | -0.80380583 | 0.2518597 | -3.191483 | 0.001415445 | 0.007562259 | ZBTB40 | protein_coding | 42.7 | -1.705152266 | 1.705152266 | ENSG00000184677 | n.s. |
| ENSG00000153339 | 185.7760983 | -0.860520173 | 0.2792156 | -3.08192 | 0.002056699 | 0.010435649 | TRAPPC8 | protein_coding | 38.41 | -1.705103978 | 1.705103978 | ENSG00000153339 | n.s. |
| ENSG00000272853 | 69.92446638 | 0.801853918 | 0.250976 | 3.194943 | 0.001398587 | 0.007477926 | | lincRNA | 56.86 | 1.704916889 | 1.704916889 | ENSG00000272853 | n.s. |
| ENSG00000235560 | 185.8903639 | 0.725465503 | 0.2155011 | 3.366412 | 0.000761528 | 0.004468082 | | antisense | 49.9 | 1.704756082 | 1.704756082 | ENSG00000235560 | n.s. |
| ENSG00000136059 | 234.5944145 | -0.84912341 | 0.273723 | -3.102126 | 0.00192136 | 0.009849445 | VILL | protein_coding | 55.98 | -1.703841032 | 1.703841032 | ENSG00000136059 | n.s. |
| ENSG00000064313 | 1184.562719 | -0.819496515 | 0.2595474 | -3.157406 | 0.001591795 | 0.008385381 | TAF2 | protein_coding | 37.5 | -1.701665833 | 1.701665833 | ENSG00000064313 | n.s. |
| ENSG00000243302 | 71.44434729 | -0.854224647 | 0.2764892 | -3.089541 | 0.002004662 | 0.010205085 | | processed_pseudogene | 58.96 | -1.700917905 | 1.700917905 | ENSG00000243302 | n.s. |
| ENSG00000109089 | 252.3407059 | -0.798739409 | 0.2498931 | -3.196324 | 0.001391908 | 0.007447939 | CDR2L | protein_coding | 53.8 | -1.699688605 | 1.699688605 | ENSG00000109089 | n.s. |
| ENSG00000264901 | 4.430056201 | -1.349170092 | 0.5598295 | -2.409966 | 0.015953998 | 0.055300861 | MIR3616 | miRNA | 54.35 | -1.696268526 | 1.696268526 | ENSG00000264901 | down |
| ENSG00000227540 | 130.2338253 | 0.781364473 | 0.242059 | 3.227992 | 0.001246627 | 0.006773348 | | antisense | 42.27 | 1.694933172 | 1.694933172 | ENSG00000227540 | n.s. |
| ENSG00000165118 | 374.522892 | -0.845268185 | 0.2726047 | -3.100711 | 0.001930568 | 0.009893004 | C9orf64 | protein_coding | 44.72 | -1.694485308 | 1.694485308 | ENSG00000165118 | n.s. |
| ENSG00000227666 | 25.8872329 | -0.997747405 | 0.3518898 | -2.835397 | 0.004576873 | 0.020045234 | CYCSP24 | processed_pseudogene | 43.67 | -1.69416398 | 1.69416398 | ENSG00000227666 | n.s. |

| | | | | | | | | | | | | | |
|---|---|---|---|---|---|---|---|---|---|---|---|---|---|
| ENSG00000225513 | 56.94318441 | 0.837465109 | 0.2690738 | 3.1124 | 0.001855731 | 0.00954817 | | processed_pseudogene | 54.22 | 1.691746387 | 1.691746387 | ENSG00000225513 | n.s. |
| ENSG00000129007 | 52.05925609 | -0.889117863 | 0.2952412 | -3.011496 | 0.002599634 | 0.012575946 | CALML4 | protein_coding | 50.69 | -1.689732341 | 1.689732341 | ENSG00000129007 | n.s. |
| ENSG00000277558 | 32.90760852 | -0.95044693 | 0.3272109 | -2.904692 | 0.003676144 | 0.016693721 | | sense_intronic | 50.6 | -1.68936889 | 1.68936889 | ENSG00000277558 | n.s. |
| ENSG00000168802 | 734.9864715 | -0.889564754 | 0.2957872 | -3.007448 | 0.002634508 | 0.012674586 | CHTF8 | protein_coding | 48.17 | -1.687563247 | 1.687563247 | ENSG00000168802 | n.s. |
| ENSG00000154945 | 1475.562889 | -0.819819682 | 0.2608625 | -3.142728 | 0.001673815 | 0.0087611 | ANKRD40 | protein_coding | 44.45 | -1.686730942 | 1.686730942 | ENSG00000154945 | n.s. |
| ENSG00000278281 | 2.186165299 | -1.570594744 | 0.7102838 | -2.211221 | 0.027020507 | 0.084381375 | MIR7856 | miRNA | 42.86 | -1.686431452 | 1.686431452 | ENSG00000278281 | down |
| ENSG00000147548 | 2849.501654 | -0.80113075 | 0.2521939 | -3.176647 | 0.001489885 | 0.00791139 | NSD3 | protein_coding | 38.79 | -1.68377432 | 1.68377432 | ENSG00000147548 | n.s. |
| ENSG00000122970 | 1159.472082 | -0.833631177 | 0.2679305 | -3.111371 | 0.001862211 | 0.00957443 | IFT81 | protein_coding | 39.9 | -1.683007193 | 1.683007193 | ENSG00000122970 | n.s. |
| ENSG00000237436 | 31.43253307 | 0.900919771 | 0.3022862 | 2.980354 | 0.002879155 | 0.013611708 | | antisense | 59.77 | 1.681195003 | 1.681195003 | ENSG00000237436 | n.s. |
| ENSG00000035403 | 859.5011222 | -0.852295214 | 0.2773574 | -3.072913 | 0.002119803 | 0.010701239 | VCL | protein_coding | 41.45 | -1.679503918 | 1.679503918 | ENSG00000035403 | n.s. |
| ENSG00000143458 | 412.7730366 | -0.835936786 | 0.2693491 | -3.103544 | 0.00191218 | 0.009809612 | GABPB2 | protein_coding | 44.85 | -1.678852098 | 1.678852098 | ENSG00000143458 | n.s. |
| ENSG00000169118 | 553.5626927 | -0.783512059 | 0.2442597 | -3.207701 | 0.001338005 | 0.007201021 | CSNK1G1 | protein_coding | 40.28 | -1.678757554 | 1.678757554 | ENSG00000169118 | n.s. |
| ENSG00000228043 | 83.82640779 | 0.787360642 | 0.2461669 | 3.198483 | 0.001381525 | 0.007398066 | | lincRNA | 38.27 | 1.677772476 | 1.677772476 | ENSG00000228043 | n.s. |
| ENSG00000155858 | 389.1508319 | -0.833224583 | 0.2681685 | -3.107093 | 0.00188937 | 0.009699741 | LSM11 | protein_coding | 43.31 | -1.67748095 | 1.67748095 | ENSG00000155858 | n.s. |
| ENSG00000148296 | 801.3379859 | -0.784401896 | 0.2447972 | -3.204293 | 0.001353946 | 0.007281187 | SURF6 | protein_coding | 53.39 | -1.676892681 | 1.676892681 | ENSG00000148296 | n.s. |
| ENSG00000215973 | 53.42463177 | 0.871076075 | 0.2872181 | 3.032804 | 0.00242293 | 0.011899354 | MIR933 | miRNA | 66.23 | 1.676365524 | 1.676365524 | ENSG00000215973 | n.s. |
| ENSG00000254027 | 25.87957294 | -0.940449945 | 0.3232578 | -2.909288 | 0.003622533 | 0.016514909 | | antisense | 64.09 | -1.675998241 | 1.675998241 | ENSG00000254027 | n.s. |
| ENSG00000208024 | 79.89763593 | 0.799826409 | 0.2522917 | 3.170245 | 0.001523103 | 0.008066248 | MIR199A2 | miRNA | 53.64 | 1.674299348 | 1.674299348 | ENSG00000208024 | n.s. |
| ENSG00000240233 | 10.98899001 | -1.059549579 | 0.3887027 | -2.725861 | 0.006413403 | 0.026355906 | RN7SL587P | misc_RNA | 55.87 | -1.673158103 | 1.673158103 | ENSG00000240233 | down |
| ENSG00000146463 | 1096.035994 | -0.876097731 | 0.2901896 | -3.019053 | 0.00253566 | 0.012344054 | ZMYM4 | protein_coding | 38.64 | -1.672069489 | 1.672069489 | ENSG00000146463 | n.s. |
| ENSG00000153363 | 272.0443355 | -0.834885868 | 0.2695698 | -3.097105 | 0.00195421 | 0.009995761 | LINC00467 | lincRNA | 39.05 | -1.669925484 | 1.669925484 | ENSG00000153363 | n.s. |
| ENSG00000129355 | 82.64182718 | 0.913651613 | 0.3099389 | 2.947844 | 0.003199981 | 0.014910754 | CDKN2D | protein_coding | 58.55 | 1.668785027 | 1.668785027 | ENSG00000129355 | n.s. |
| ENSG00000243508 | 52.76248626 | -0.834397464 | 0.2695542 | -3.095472 | 0.001964997 | 0.010039871 | | processed_pseudogene | 41.37 | -1.667352986 | 1.667352986 | ENSG00000243508 | n.s. |
| ENSG00000065060 | 989.3243937 | -0.811791453 | 0.2585831 | -3.139383 | 0.001693041 | 0.008845105 | UHRF1BP1 | protein_coding | 42.65 | -1.666848953 | 1.666848953 | ENSG00000065060 | n.s. |
| ENSG00000152683 | 893.5062848 | -0.874252 | 0.2897594 | -3.017165 | 0.002551509 | 0.012403862 | SLC30A6 | protein_coding | 39.74 | -1.66671166 | 1.66671166 | ENSG00000152683 | n.s. |
| ENSG00000170604 | 260.6527856 | -0.773040698 | 0.2403103 | -3.216844 | 0.001296092 | 0.007005234 | IRF2BP1 | protein_coding | 68.49 | -1.665575995 | 1.665575995 | ENSG00000170604 | n.s. |
| ENSG00000232587 | 68.85631703 | -0.847113889 | 0.2761079 | -3.068053 | 0.002154582 | 0.010829693 | EEF1A1P3 | processed_pseudogene | 45.49 | -1.664903965 | 1.664903965 | ENSG00000232587 | n.s. |
| ENSG00000228446 | 102.9009945 | -0.889787272 | 0.2981703 | -2.984158 | 0.002843603 | 0.013466491 | | processed_pseudogene | 41.88 | -1.664565584 | 1.664565584 | ENSG00000228446 | n.s. |
| ENSG00000136158 | 297.3784567 | 0.738678536 | 0.2241898 | 3.294881 | 0.000984634 | 0.005579462 | SPRY2 | protein_coding | 47.95 | 1.664543911 | 1.664543911 | ENSG00000136158 | n.s. |
| ENSG00000167378 | 810.4922674 | -0.779916853 | 0.2437285 | -3.199941 | 0.001374555 | 0.007366411 | IRGQ | protein_coding | 52.71 | -1.663363027 | 1.663363027 | ENSG00000167378 | n.s. |
| ENSG00000124160 | 777.5628121 | -0.82778817 | 0.2668 | -3.102654 | 0.001917938 | 0.009835528 | NCOA5 | protein_coding | 41.78 | -1.661538353 | 1.661538353 | ENSG00000124160 | n.s. |
| ENSG00000278013 | 19.82423 | 0.953375511 | 0.3319701 | 2.871871 | 0.004080489 | 0.01822868 | | lincRNA | 45.68 | 1.658153376 | 1.658153376 | ENSG00000278013 | n.s. |

| | | | | | | | | | | | | | |
|---|---|---|---|---|---|---|---|---|---|---|---|---|---|
| ENSG00000267904 | 50.18374866 | -0.914315155 | 0.3114407 | -2.93576 | 0.003327316 | 0.015411253 | | sense_intronic | 47.01 | -1.656887226 | 1.656887226 | ENSG00000267904 | n.s. |
| ENSG00000100906 | 953.019559 | 0.766972893 | 0.2382411 | 3.219314 | 0.001284975 | 0.006955953 | NFKBIA | protein_coding | 56.53 | 1.65485396 | 1.65485396 | ENSG00000100906 | n.s. |
| ENSG00000104880 | 324.7552896 | -0.846419981 | 0.2767138 | -3.058828 | 0.002222048 | 0.011112632 | ARHGEF18 | protein_coding | 51.44 | -1.654059594 | 1.654059594 | ENSG00000104880 | n.s. |
| ENSG00000196653 | 509.2041795 | -0.876948109 | 0.2925611 | -2.997488 | 0.00272215 | 0.013010811 | ZNF502 | protein_coding | 40.62 | -1.653657227 | 1.653657227 | ENSG00000196653 | n.s. |
| ENSG00000263727 | 26.54270083 | -0.922467271 | 0.3160314 | -2.91891 | 0.003512571 | 0.016129789 | | sense_intronic | 47.33 | -1.653403872 | 1.653403872 | ENSG00000263727 | n.s. |
| ENSG00000276926 | 15.63808125 | -1.012906281 | 0.3646845 | -2.777487 | 0.005478109 | 0.02331935 | MIR6797 | miRNA | 65.28 | -1.653350266 | 1.653350266 | ENSG00000276926 | down |
| ENSG00000275713 | 1844.203383 | 0.711276293 | 0.2125071 | 3.347071 | 0.000816703 | 0.004743827 | HIST1H2BH | protein_coding | 41.55 | 1.652914435 | 1.652914435 | ENSG00000275713 | n.s. |
| ENSG00000232034 | 21.04572631 | 0.961574595 | 0.3371048 | 2.85245 | 0.004338363 | 0.019157317 | | sense_intronic | 43.86 | 1.651663321 | 1.651663321 | ENSG00000232034 | n.s. |
| ENSG00000100294 | 273.8484117 | -0.851864526 | 0.2799492 | -3.042926 | 0.002342903 | 0.011583892 | MCAT | protein_coding | 52.52 | -1.649333679 | 1.649333679 | ENSG00000100294 | n.s. |
| ENSG00000200834 | 17.90499034 | -1.010265475 | 0.3637826 | -2.777114 | 0.005484401 | 0.023322943 | | misc_RNA | 41.58 | -1.648972121 | 1.648972121 | ENSG00000200834 | down |
| ENSG00000236536 | 26.26851773 | -0.950361456 | 0.3315981 | -2.866004 | 0.004156893 | 0.018508611 | | antisense | 37.7 | -1.646621125 | 1.646621125 | ENSG00000236536 | n.s. |
| ENSG00000203326 | 112.1601884 | -0.860991187 | 0.2851171 | -3.019781 | 0.002529571 | 0.012323031 | ZNF525 | protein_coding | 43.64 | -1.643875367 | 1.643875367 | ENSG00000203326 | n.s. |
| ENSG00000278700 | 13.25986181 | -1.007861414 | 0.3631647 | -2.775219 | 0.005516456 | 0.023430634 | | misc_RNA | 54.82 | -1.643031769 | 1.643031769 | ENSG00000278700 | down |
| ENSG00000267194 | 153.155824 | -0.824959736 | 0.2670072 | -3.089653 | 0.002003904 | 0.010204956 | | lincRNA | 36.86 | -1.642650627 | 1.642650627 | ENSG00000267194 | n.s. |
| ENSG00000140534 | 2096.262959 | -0.724173044 | 0.2190809 | -3.305506 | 0.000948052 | 0.005403079 | TICRR | protein_coding | 42.67 | -1.641960048 | 1.641960048 | ENSG00000140534 | n.s. |
| ENSG00000183496 | 2475.169985 | 0.680574616 | 0.1995296 | 3.410895 | 0.0006475 | 0.003870821 | MEX3B | protein_coding | 55.84 | 1.641680003 | 1.641680003 | ENSG00000183496 | n.s. |
| ENSG00000189144 | 98.8487495 | -0.837752922 | 0.2737879 | -3.059861 | 0.002214398 | 0.011078356 | ZNF573 | protein_coding | 41.38 | -1.63824652 | 1.63824652 | ENSG00000189144 | n.s. |
| ENSG00000108379 | 230.6507701 | -0.784709932 | 0.2478858 | -3.165611 | 0.001547575 | 0.008179071 | WNT3 | protein_coding | 53.82 | -1.637921929 | 1.637921929 | ENSG00000108379 | n.s. |
| ENSG00000213967 | 162.7531612 | -0.766054393 | 0.2390681 | -3.204336 | 0.001353744 | 0.007281187 | ZNF726 | protein_coding | 37.37 | -1.637669428 | 1.637669428 | ENSG00000213967 | n.s. |
| ENSG00000164144 | 589.4737437 | -0.890607185 | 0.3012907 | -2.955973 | 0.003116845 | 0.014586835 | ARFIP1 | protein_coding | 37.33 | -1.635189468 | 1.635189468 | ENSG00000164144 | n.s. |
| ENSG00000234689 | 11.49053251 | -1.167338593 | 0.4566242 | -2.556453 | 0.010574522 | 0.039799656 | LINC00444 | lincRNA | 41.89 | -1.634414903 | 1.634414903 | ENSG00000234689 | down |
| ENSG00000163933 | 480.6419372 | -0.840010145 | 0.275388 | -3.050278 | 0.002286293 | 0.011372656 | RFT1 | protein_coding | 45.77 | -1.633095722 | 1.633095722 | ENSG00000163933 | n.s. |
| ENSG00000183808 | 2589.542114 | -0.783670731 | 0.2478206 | -3.16225 | 0.00156555 | 0.008269027 | RBM12B | protein_coding | 39 | -1.632030009 | 1.632030009 | ENSG00000183808 | n.s. |
| ENSG00000273797 | 22.64196722 | 0.96452128 | 0.3408702 | 2.829585 | 0.004660841 | 0.020348978 | | sense_intronic | 36.05 | 1.631446663 | 1.631446663 | ENSG00000273797 | n.s. |
| ENSG00000145725 | 2572.605101 | -0.852568099 | 0.2819979 | -3.023314 | 0.002500228 | 0.012205705 | PPIP5K2 | protein_coding | 34.29 | -1.631335467 | 1.631335467 | ENSG00000145725 | n.s. |
| ENSG00000122966 | 1627.052991 | -0.764184359 | 0.2387665 | -3.200551 | 0.001371649 | 0.007353664 | CIT | protein_coding | 43.74 | -1.630384422 | 1.630384422 | ENSG00000122966 | n.s. |
| ENSG00000239763 | 23.70522782 | -0.943889293 | 0.329929 | -2.860886 | 0.004224591 | 0.018744098 | | processed_pseudogene | 50.13 | -1.630224668 | 1.630224668 | ENSG00000239763 | n.s. |
| ENSG00000215883 | 293.0060271 | -0.823261671 | 0.2674657 | -3.078008 | 0.002083894 | 0.010550567 | CYB5RL | protein_coding | 48.99 | -1.627361257 | 1.627361257 | ENSG00000215883 | n.s. |
| ENSG00000232693 | 88.91235633 | -0.910894658 | 0.312734 | -2.912682 | 0.003583392 | 0.016379372 | | sense_intronic | 43.37 | -1.626587107 | 1.626587107 | ENSG00000232693 | n.s. |
| ENSG00000236404 | 194.5454177 | -0.783776086 | 0.2483321 | -3.156161 | 0.001598608 | 0.008414904 | VLDLR-AS1 | antisense | 39.93 | -1.62629685 | 1.62629685 | ENSG00000236404 | n.s. |
| ENSG00000121690 | 386.0210072 | 0.796946628 | 0.2546761 | 3.129256 | 0.001752496 | 0.009107879 | DEPDC7 | protein_coding | 38.52 | 1.62623555 | 1.62623555 | ENSG00000121690 | n.s. |
| ENSG00000116580 | 1596.740902 | -0.773167121 | 0.2433213 | -3.177556 | 0.001485219 | 0.007889625 | GON4L | protein_coding | 44.03 | -1.625926874 | 1.625926874 | ENSG00000116580 | n.s. |

| | | | | | | | | | | | | | |
|---|---|---|---|---|---|---|---|---|---|---|---|---|---|
| ENSG00000189308 | 1001.598315 | -0.772640967 | 0.24306 | -3.178808 | 0.001478822 | 0.007864643 | LIN54 | protein_coding | 39.03 | -1.625884617 | 1.625884617 | ENSG00000189308 | n.s. |
| ENSG00000207815 | 13.27646486 | -1.320466597 | 0.5540378 | -2.383351 | 0.017155824 | 0.058731382 | MIR563 | miRNA | 44.3 | -1.625665748 | 1.625665748 | ENSG00000207815 | down |
| ENSG00000148444 | 735.3238344 | 0.747885903 | 0.2314174 | 3.231761 | 0.001230297 | 0.006712158 | COMMD3 | protein_coding | 41.08 | 1.625259133 | 1.625259133 | ENSG00000148444 | n.s. |
| ENSG00000115998 | 236.4955054 | -0.724345149 | 0.2204142 | -3.286291 | 0.001015161 | 0.005719908 | C2orf42 | protein_coding | 43.21 | -1.624424394 | 1.624424394 | ENSG00000115998 | n.s. |
| ENSG00000184708 | 504.4270254 | -0.755877275 | 0.2352831 | -3.212628 | 0.001315264 | 0.00709784 | EIF4ENIF1 | protein_coding | 44.22 | -1.624284866 | 1.624284866 | ENSG00000184708 | n.s. |
| ENSG00000109756 | 1070.312633 | -0.788469977 | 0.2507734 | -3.144154 | 0.001665679 | 0.008734934 | RAPGEF2 | protein_coding | 36.57 | -1.623254992 | 1.623254992 | ENSG00000109756 | n.s. |
| ENSG00000231165 | 23.12685973 | -0.994011258 | 0.3579282 | -2.777124 | 0.005484219 | 0.023322943 | TRBV26OR9-2 | TR_V_pseudogene | 44.37 | -1.622441718 | 1.622441718 | ENSG00000231165 | n.s. |
| ENSG00000265444 | 2.335305819 | 1.862921074 | 0.9468815 | 1.967428 | 0.049133897 | 0.134772436 | MIR4733 | miRNA | 52.63 | 1.621484492 | 1.621484492 | ENSG00000265444 | n.s. |
| ENSG00000197321 | 38.81381396 | -0.923893899 | 0.3201408 | -2.885898 | 0.003902984 | 0.017591855 | SVIL | protein_coding | 43.61 | -1.621145872 | 1.621145872 | ENSG00000197321 | n.s. |
| ENSG00000188636 | 1073.78716 | -0.745408468 | 0.2305954 | -3.232538 | 0.001226957 | 0.006699187 | RTL6 | protein_coding | 56.28 | -1.620501534 | 1.620501534 | ENSG00000188636 | n.s. |
| ENSG00000160062 | 281.4578509 | -0.852534465 | 0.2831023 | -3.011401 | 0.002600456 | 0.012575946 | ZBTB8A | protein_coding | 43.45 | -1.620207082 | 1.620207082 | ENSG00000160062 | n.s. |
| ENSG00000279146 | 92.30480986 | -0.822628489 | 0.2678723 | -3.070973 | 0.002133627 | 0.010749525 | | TEC | 55.12 | -1.619435275 | 1.619435275 | ENSG00000279146 | n.s. |
| ENSG00000209482 | 67.60942901 | 0.786747805 | 0.2503181 | 3.142992 | 0.001672303 | 0.008757021 | SNORD83A | snoRNA | 54.74 | 1.618846588 | 1.618846588 | ENSG00000209482 | n.s. |
| ENSG00000221264 | 3.418453288 | 1.542445407 | 0.7066947 | 2.182619 | 0.029063857 | 0.089229565 | MIR1284 | miRNA | 30 | 1.618782917 | 1.618782917 | ENSG00000221264 | up |
| ENSG00000198252 | 268.4775699 | -0.853563037 | 0.2838877 | -3.006692 | 0.002641068 | 0.012699308 | STYX | protein_coding | 38.27 | -1.61854325 | 1.61854325 | ENSG00000198252 | n.s. |
| ENSG00000196792 | 915.1175312 | -0.81028608 | 0.2618464 | -3.094509 | 0.00197139 | 0.010068841 | STRN3 | protein_coding | 38.51 | -1.618157915 | 1.618157915 | ENSG00000196792 | n.s. |
| ENSG00000001629 | 1023.31198 | -0.848747058 | 0.2813446 | -3.016752 | 0.002554985 | 0.012416426 | ANKIB1 | protein_coding | 36.77 | -1.617714779 | 1.617714779 | ENSG00000001629 | n.s. |
| ENSG00000263593 | 54.12557684 | -0.965943853 | 0.3432549 | -2.814072 | 0.004891834 | 0.021171588 | MIR4804 | miRNA | 45.21 | -1.617228177 | 1.617228177 | ENSG00000263593 | n.s. |
| ENSG00000196937 | 587.8152667 | -0.818202282 | 0.2659636 | -3.07637 | 0.002095378 | 0.010601 | FAM3C | protein_coding | 35.92 | -1.615665725 | 1.615665725 | ENSG00000196937 | n.s. |
| ENSG00000221869 | 145.0123366 | 0.791627771 | 0.2531515 | 3.127091 | 0.001765451 | 0.009158113 | CEBPD | protein_coding | 70.02 | 1.613490985 | 1.613490985 | ENSG00000221869 | n.s. |
| ENSG00000174953 | 3220.040354 | -0.830240099 | 0.2723196 | -3.048771 | 0.002297795 | 0.011413555 | DHX36 | protein_coding | 35.49 | -1.612807038 | 1.612807038 | ENSG00000174953 | n.s. |
| ENSG00000138336 | 1196.821811 | -0.789856464 | 0.2523593 | -3.129889 | 0.001748726 | 0.009095076 | TET1 | protein_coding | 42.09 | -1.612250017 | 1.612250017 | ENSG00000138336 | n.s. |
| ENSG00000182168 | 1219.011411 | -0.77211534 | 0.2439124 | -3.165544 | 0.001547932 | 0.008179071 | UNC5C | protein_coding | 37.41 | -1.611633288 | 1.611633288 | ENSG00000182168 | n.s. |
| ENSG00000174516 | 120.1633732 | -0.837269615 | 0.2762687 | -3.030635 | 0.002440402 | 0.011968197 | PELI3 | protein_coding | 56.57 | -1.609208139 | 1.609208139 | ENSG00000174516 | n.s. |
| ENSG00000119986 | 83.09005066 | 0.835707782 | 0.2755572 | 3.032793 | 0.00242302 | 0.011899354 | AVPI1 | protein_coding | 49.03 | 1.608300072 | 1.608300072 | ENSG00000119986 | n.s. |
| ENSG00000280000 | 68.75940721 | -0.874723804 | 0.2960554 | -2.954595 | 0.003130795 | 0.01463736 | | TEC | 38.91 | -1.604713403 | 1.604713403 | ENSG00000280000 | n.s. |
| ENSG00000179151 | 676.3464083 | -0.753475967 | 0.2356927 | -3.196857 | 0.001389338 | 0.007437042 | EDC3 | protein_coding | 43.7 | -1.603848758 | 1.603848758 | ENSG00000179151 | n.s. |
| ENSG00000000460 | 861.6232583 | -0.72924096 | 0.2242986 | -3.251206 | 0.001149166 | 0.006323869 | C1orf112 | protein_coding | 39.22 | -1.603613389 | 1.603613389 | ENSG00000000460 | n.s. |
| ENSG00000138286 | 237.3763471 | -0.843601061 | 0.2800832 | -3.011966 | 0.002595613 | 0.012565619 | FAM149B1 | protein_coding | 39.89 | -1.603530483 | 1.603530483 | ENSG00000138286 | n.s. |
| ENSG00000119906 | 548.7382367 | -0.720763079 | 0.2206926 | -3.265914 | 0.001091112 | 0.006050002 | SLF2 | protein_coding | 39.04 | -1.598828731 | 1.598828731 | ENSG00000119906 | n.s. |
| ENSG00000185722 | 946.9706155 | -0.784137746 | 0.2508189 | -3.12631 | 0.001770147 | 0.009175638 | ANKFY1 | protein_coding | 44.91 | -1.597573802 | 1.597573802 | ENSG00000185722 | n.s. |
| ENSG00000139977 | 533.3837102 | -0.830827412 | 0.274136 | -3.030712 | 0.002439778 | 0.011968197 | NAA30 | protein_coding | 37.28 | -1.596826411 | 1.596826411 | ENSG00000139977 | n.s. |

| | | | | | | | | | | | | | |
|---|---|---|---|---|---|---|---|---|---|---|---|---|---|
| ENSG00000263648 | 27.7656966 | -0.894059158 | 0.3070092 | -2.912158 | 0.003589413 | 0.016391004 | | processed_pseudogene | 43.95 | -1.596248244 | 1.596248244 | ENSG00000263648 | n.s. |
| ENSG00000103199 | 249.4561034 | -0.786196823 | 0.2519624 | -3.120294 | 0.001806706 | 0.009342117 | ZNF500 | protein_coding | 54.32 | -1.595629449 | 1.595629449 | ENSG00000103199 | n.s. |
| ENSG00000127418 | 118.38036 | -0.80385264 | 0.2605985 | -3.084641 | 0.002037982 | 0.010355776 | FGFRL1 | protein_coding | 66.74 | -1.595500655 | 1.595500655 | ENSG00000127418 | n.s. |
| ENSG00000180447 | 233.9304824 | 0.708823369 | 0.2154691 | 3.289675 | 0.001003033 | 0.005665406 | GAS1 | protein_coding | 57.73 | 1.592562366 | 1.592562366 | ENSG00000180447 | n.s. |
| ENSG00000162600 | 212.6315931 | -0.903333262 | 0.3124465 | -2.891161 | 0.003838211 | 0.017316724 | OMA1 | protein_coding | 39.98 | -1.591252484 | 1.591252484 | ENSG00000162600 | n.s. |
| ENSG00000107186 | 2079.536123 | -0.834527394 | 0.2767624 | -3.01532 | 0.00256708 | 0.012462155 | MPDZ | protein_coding | 35.81 | -1.589279689 | 1.589279689 | ENSG00000107186 | n.s. |
| ENSG00000140259 | 1072.241903 | -0.837425532 | 0.2784482 | -3.007473 | 0.002634295 | 0.012674586 | MFAP1 | protein_coding | 43.98 | -1.588651691 | 1.588651691 | ENSG00000140259 | n.s. |
| ENSG00000238227 | 294.9751087 | -0.722402217 | 0.2222143 | -3.250926 | 0.0011503 | 0.006325287 | TMEM250 | protein_coding | 65.35 | -1.588504527 | 1.588504527 | ENSG00000238227 | n.s. |
| ENSG00000238390 | 39.74777215 | -0.848812384 | 0.2844977 | -2.983548 | 0.002849276 | 0.013488768 | | snoRNA | 41.24 | -1.58730267 | 1.58730267 | ENSG00000238390 | n.s. |
| ENSG00000206675 | 11.93899556 | -1.079463612 | 0.4114661 | -2.623457 | 0.008704251 | 0.033851754 | RNU6-32P | snRNA | 44.86 | -1.587263607 | 1.587263607 | ENSG00000206675 | down |
| ENSG00000239291 | 113.0271509 | -0.798991344 | 0.2589058 | -3.086031 | 0.002028474 | 0.01031876 | | processed_pseudogene | 58.28 | -1.587094417 | 1.587094417 | ENSG00000239291 | n.s. |
| ENSG00000142065 | 397.4112574 | -0.833442729 | 0.2764289 | -3.015035 | 0.002569498 | 0.012469545 | ZFP14 | protein_coding | 42.69 | -1.58699946 | 1.58699946 | ENSG00000142065 | n.s. |
| ENSG00000170776 | 107.4410211 | 0.762618981 | 0.2413331 | 3.160026 | 0.001577549 | 0.008326084 | AKAP13 | protein_coding | 39.92 | 1.585911341 | 1.585911341 | ENSG00000170776 | n.s. |
| ENSG00000201044 | 3.642522015 | -1.846390289 | 0.9458918 | -1.95201 | 0.050937023 | 0.138541936 | RNU6-268P | snRNA | 42.99 | -1.584976042 | 1.584976042 | ENSG00000201044 | n.s. |
| ENSG00000283749 | 2.008539756 | 1.709989018 | 0.8402395 | 2.035121 | 0.041838701 | 0.118470661 | MIR4781 | miRNA | 53.95 | 1.584115339 | 1.584115339 | ENSG00000283749 | n.s. |
| ENSG00000108799 | 785.8230514 | -0.815881989 | 0.2679131 | -3.045324 | 0.002324301 | 0.011508267 | EZH1 | protein_coding | 45.44 | -1.581987071 | 1.581987071 | ENSG00000108799 | n.s. |
| ENSG00000189079 | 3389.282669 | -0.799627895 | 0.2602562 | -3.072464 | 0.002122994 | 0.010709581 | ARID2 | protein_coding | 36.1 | -1.575448875 | 1.575448875 | ENSG00000189079 | n.s. |
| ENSG00000151883 | 193.592587 | 0.748741584 | 0.2355666 | 3.178471 | 0.001480539 | 0.007870769 | PARP8 | protein_coding | 35.7 | 1.57533943 | 1.57533943 | ENSG00000151883 | n.s. |
| ENSG00000224255 | 25.49061388 | -0.924207402 | 0.3252637 | -2.84141 | 0.004491457 | 0.019745677 | PDCL3P6 | processed_pseudogene | 44.76 | -1.575337369 | 1.575337369 | ENSG00000224255 | n.s. |
| ENSG00000214029 | 1603.50971 | -0.809812537 | 0.2653954 | -3.051344 | 0.002278195 | 0.011347696 | ZNF891 | protein_coding | 39.65 | -1.57516015 | 1.57516015 | ENSG00000214029 | n.s. |
| ENSG00000166167 | 424.3458714 | -0.819681654 | 0.2706521 | -3.028544 | 0.002457355 | 0.012034382 | BTRC | protein_coding | 38.96 | -1.57344139 | 1.57344139 | ENSG00000166167 | n.s. |
| ENSG00000215126 | 43.98844273 | -0.932924847 | 0.3302612 | -2.82481 | 0.004730873 | 0.020590169 | CBWD6 | protein_coding | 36.14 | -1.573228577 | 1.573228577 | ENSG00000215126 | n.s. |
| ENSG00000103429 | 990.7576339 | -0.872439868 | 0.2980069 | -2.927583 | 0.00341608 | 0.015750894 | BFAR | protein_coding | 46.07 | -1.572742809 | 1.572742809 | ENSG00000103429 | n.s. |
| ENSG00000252291 | 4.726798548 | -1.374565679 | 0.600263 | -2.289939 | 0.022024844 | 0.071916487 | | snoRNA | 44.34 | -1.571364355 | 1.571364355 | ENSG00000252291 | down |
| ENSG00000256087 | 446.6311391 | -0.827437666 | 0.2748965 | -3.009997 | 0.002612501 | 0.012621047 | ZNF432 | protein_coding | 41.15 | -1.571225201 | 1.571225201 | ENSG00000256087 | n.s. |
| ENSG00000260903 | 1065.677936 | -0.778248342 | 0.250121 | -3.111487 | 0.001861474 | 0.00957418 | XKR7 | protein_coding | 51.87 | -1.571204336 | 1.571204336 | ENSG00000260903 | n.s. |
| ENSG00000035115 | 490.4009503 | -0.812053626 | 0.2669977 | -3.041425 | 0.00235461 | 0.01163764 | SH3YL1 | protein_coding | 40.78 | -1.570621412 | 1.570621412 | ENSG00000035115 | n.s. |
| ENSG00000253276 | 375.9170136 | 0.721316078 | 0.2230447 | 3.233953 | 0.001220895 | 0.006673936 | CCDC71L | protein_coding | 43.76 | 1.56930822 | 1.56930822 | ENSG00000253276 | n.s. |
| ENSG00000130227 | 1616.595609 | -0.796061348 | 0.2590503 | -3.072999 | 0.002119193 | 0.010701239 | XPO7 | protein_coding | 39.92 | -1.568691378 | 1.568691378 | ENSG00000130227 | n.s. |
| ENSG00000242474 | 159.866167 | 0.722828863 | 0.2238781 | 3.228671 | 0.001243669 | 0.006763909 | | lincRNA | 48.7 | 1.568395706 | 1.568395706 | ENSG00000242474 | n.s. |
| ENSG00000107758 | 625.0603521 | -0.883505231 | 0.3044603 | -2.901873 | 0.003709387 | 0.016806312 | PPP3CB | protein_coding | 38.91 | -1.567804406 | 1.567804406 | ENSG00000107758 | n.s. |
| ENSG00000198105 | 858.8942081 | -0.803866883 | 0.2631587 | -3.054685 | 0.002252972 | 0.011251123 | ZNF248 | protein_coding | 39.14 | -1.566579103 | 1.566579103 | ENSG00000198105 | n.s. |

| | | | | | | | | | | | | | |
|---|---|---|---|---|---|---|---|---|---|---|---|---|---|
| ENSG00000235194 | 142.9966908 | -0.731969636 | 0.2284786 | -3.203668 | 0.001356889 | 0.007294195 | PPP1R3E | protein_coding | 51.79 | -1.564235671 | 1.564235671 | ENSG00000235194 | n.s. |
| ENSG00000113838 | 274.3659692 | -0.827313989 | 0.275597 | -3.001898 | 0.002683024 | 0.01286798 | TBCCD1 | protein_coding | 40.74 | -1.564028501 | 1.564028501 | ENSG00000113838 | n.s. |
| ENSG00000180917 | 380.9427419 | -0.776011551 | 0.2496498 | -3.108401 | 0.001881029 | 0.00966048 | CMTR2 | protein_coding | 39.62 | -1.563664292 | 1.563664292 | ENSG00000180917 | n.s. |
| ENSG00000134247 | 1260.98291 | -0.772304794 | 0.2478653 | -3.115824 | 0.001834317 | 0.009465975 | PTGFRN | protein_coding | 44.19 | -1.563017203 | 1.563017203 | ENSG00000134247 | n.s. |
| ENSG00000268670 | 91.43508166 | 0.73958649 | 0.2321965 | 3.185175 | 0.001446665 | 0.007720167 | | sense_intronic | 63.52 | 1.562282778 | 1.562282778 | ENSG00000268670 | n.s. |
| ENSG00000171303 | 277.3276036 | -0.844592199 | 0.2847111 | -2.966488 | 0.003012219 | 0.014163839 | KCNK3 | protein_coding | 53.76 | -1.561498127 | 1.561498127 | ENSG00000171303 | n.s. |
| ENSG00000126653 | 1011.041427 | -0.718230004 | 0.2222591 | -3.231498 | 0.001231431 | 0.006713341 | NSRP1 | protein_coding | 38.57 | -1.560757819 | 1.560757819 | ENSG00000126653 | n.s. |
| ENSG00000170456 | 1035.834348 | -0.803360261 | 0.2637407 | -3.046023 | 0.002318898 | 0.011489684 | DENND5B | protein_coding | 40.95 | -1.558271416 | 1.558271416 | ENSG00000170456 | n.s. |
| ENSG00000111058 | 112.3650978 | 0.750418726 | 0.2377139 | 3.156814 | 0.00159503 | 0.008399244 | ACSS3 | protein_coding | 35.63 | 1.557689023 | 1.557689023 | ENSG00000111058 | n.s. |
| ENSG00000039650 | 83.27513184 | 0.825228935 | 0.2752093 | 2.998551 | 0.002712671 | 0.01297476 | PNKP | protein_coding | 59.53 | 1.557125034 | 1.557125034 | ENSG00000039650 | n.s. |
| ENSG00000162139 | 553.5393731 | -0.79386749 | 0.2591493 | -3.06336 | 0.002188666 | 0.01096933 | NEU3 | protein_coding | 43.57 | -1.555837303 | 1.555837303 | ENSG00000162139 | n.s. |
| ENSG00000135778 | 1366.971363 | -0.829104972 | 0.2774196 | -2.988631 | 0.002802301 | 0.013309037 | NTPCR | protein_coding | 42.72 | -1.555279362 | 1.555279362 | ENSG00000135778 | n.s. |
| ENSG00000165259 | 338.7465712 | -0.817604047 | 0.2716019 | -3.010303 | 0.002609868 | 0.012612701 | HDX | protein_coding | 34.93 | -1.552786978 | 1.552786978 | ENSG00000165259 | n.s. |
| ENSG00000255135 | 353.5519111 | 0.687144212 | 0.2082538 | 3.299551 | 0.000968395 | 0.005498762 | | lincRNA | 43.63 | 1.552763905 | 1.552763905 | ENSG00000255135 | n.s. |
| ENSG00000269921 | 75.26809897 | 0.782390128 | 0.2536939 | 3.083993 | 0.002042424 | 0.010374559 | | lincRNA | 42.72 | 1.552285767 | 1.552285767 | ENSG00000269921 | n.s. |
| ENSG00000083223 | 819.1449517 | -0.786646241 | 0.2558273 | -3.074911 | 0.00210565 | 0.010641368 | ZCCHC6 | protein_coding | 37.99 | -1.552055032 | 1.552055032 | ENSG00000083223 | n.s. |
| ENSG00000076706 | 185.2697722 | 0.780525634 | 0.2528275 | 3.087186 | 0.00202061 | 0.010282511 | MCAM | protein_coding | 59.59 | 1.551607543 | 1.551607543 | ENSG00000076706 | n.s. |
| ENSG00000139044 | 242.4055703 | -0.757376991 | 0.2415468 | -3.135529 | 0.001715442 | 0.008942008 | B4GALNT3 | protein_coding | 49.06 | -1.551535938 | 1.551535938 | ENSG00000139044 | n.s. |
| ENSG00000278482 | 9.295557189 | -1.139231022 | 0.4522462 | -2.519051 | 0.011767176 | 0.043468929 | | misc_RNA | 41.74 | -1.551428799 | 1.551428799 | ENSG00000278482 | down |
| ENSG00000107789 | 353.9175037 | -0.831871904 | 0.2792427 | -2.979029 | 0.002891637 | 0.013647544 | MINPP1 | protein_coding | 38.05 | -1.551395759 | 1.551395759 | ENSG00000107789 | n.s. |
| ENSG00000206897 | 23.65852643 | -0.956825024 | 0.3459579 | -2.765727 | 0.00567961 | 0.023997016 | SNORA9B | snoRNA | 50.38 | -1.549906077 | 1.549906077 | ENSG00000206897 | n.s. |
| ENSG00000120688 | 511.3868111 | -0.792084344 | 0.2588408 | -3.060121 | 0.002212473 | 0.011072704 | WBP4 | protein_coding | 38.92 | -1.549116034 | 1.549116034 | ENSG00000120688 | n.s. |
| ENSG00000174501 | 1646.96615 | -0.815193155 | 0.270953 | -3.008614 | 0.00262442 | 0.012652287 | ANKRD36C | protein_coding | 35.73 | -1.547098814 | 1.547098814 | ENSG00000174501 | n.s. |
| ENSG00000145246 | 360.0007876 | -0.84218495 | 0.2848235 | -2.956866 | 0.003107835 | 0.014556087 | ATP10D | protein_coding | 37.6 | -1.547056146 | 1.547056146 | ENSG00000145246 | n.s. |
| ENSG00000018869 | 189.9687678 | -0.795291103 | 0.2606146 | -3.051598 | 0.002276264 | 0.011346781 | ZNF582 | protein_coding | 40.84 | -1.546942451 | 1.546942451 | ENSG00000018869 | n.s. |
| ENSG00000115977 | 393.7825329 | -0.784599118 | 0.2553584 | -3.07254 | 0.002122451 | 0.010709581 | AAK1 | protein_coding | 41.95 | -1.545838765 | 1.545838765 | ENSG00000115977 | n.s. |
| ENSG00000177082 | 1133.917064 | -0.772321185 | 0.2493356 | -3.097516 | 0.001951498 | 0.009985557 | WDR73 | protein_coding | 48.66 | -1.545127152 | 1.545127152 | ENSG00000177082 | n.s. |
| ENSG00000135476 | 1137.093616 | -0.719213489 | 0.2239957 | -3.210836 | 0.001323493 | 0.007136716 | ESPL1 | protein_coding | 51.69 | -1.543792907 | 1.543792907 | ENSG00000135476 | n.s. |
| ENSG00000175727 | 565.089158 | -0.749868854 | 0.2385305 | -3.143702 | 0.001668253 | 0.008745138 | MLXIP | protein_coding | 50.38 | -1.543405054 | 1.543405054 | ENSG00000175727 | n.s. |
| ENSG00000245904 | 28.90221206 | 0.86008302 | 0.2946138 | 2.919358 | 0.003507535 | 0.016117295 | | antisense | 39.81 | 1.541877571 | 1.541877571 | ENSG00000245904 | n.s. |
| ENSG00000165521 | 6123.032039 | -0.81494604 | 0.2714644 | -3.002036 | 0.002681802 | 0.012866552 | EML5 | protein_coding | 37.41 | -1.540686315 | 1.540686315 | ENSG00000165521 | n.s. |
| ENSG00000171853 | 725.2682281 | -0.806120879 | 0.2668646 | -3.020711 | 0.002521818 | 0.012293863 | TRAPPC12 | protein_coding | 47.39 | -1.539942101 | 1.539942101 | ENSG00000171853 | n.s. |

| | | | | | | | | | | | | | |
|---|---|---|---|---|---|---|---|---|---|---|---|---|---|
| ENSG00000101346 | 634.5513807 | -0.79102668 | 0.2591447 | -3.052451 | 0.002269805 | 0.01132041 | POFUT1 | protein_coding | 47.65 | -1.539446958 | 1.539446958 | ENSG00000101346 | n.s. |
| ENSG00000173681 | 284.8162739 | -0.792008865 | 0.2597349 | -3.049297 | 0.002293772 | 0.01140171 | CXorf23 | protein_coding | 39.56 | -1.538896993 | 1.538896993 | ENSG00000173681 | n.s. |
| ENSG00000207333 | 9.344559007 | -1.169446547 | 0.4733829 | -2.470403 | 0.013496098 | 0.048479404 | RNU6-680P | snRNA | 40.95 | -1.537170508 | 1.537170508 | ENSG00000207333 | down |
| ENSG00000221439 | 43.29636215 | -0.914252744 | 0.3242651 | -2.81946 | 0.004810445 | 0.020890776 | RNU4ATAC16P | snRNA | 43.65 | -1.535986145 | 1.535986145 | ENSG00000221439 | n.s. |
| ENSG00000095787 | 3555.839966 | -0.789765287 | 0.2589117 | -3.050327 | 0.002285921 | 0.011372656 | WAC | protein_coding | 37.79 | -1.53541278 | 1.53541278 | ENSG00000095787 | n.s. |
| ENSG00000113595 | 619.5034937 | -0.794052308 | 0.2612923 | -3.038942 | 0.002374104 | 0.011704904 | TRIM23 | protein_coding | 37.19 | -1.533816958 | 1.533816958 | ENSG00000113595 | n.s. |
| ENSG00000145545 | 601.229308 | -0.81542782 | 0.2725214 | -2.99216 | 0.002770107 | 0.013185711 | SRD5A1 | protein_coding | 44.23 | -1.532919863 | 1.532919863 | ENSG00000145545 | n.s. |
| ENSG00000129173 | 368.3020847 | -0.697945284 | 0.2148845 | -3.248002 | 0.001162184 | 0.006378041 | E2F8 | protein_coding | 41.92 | -1.532208152 | 1.532208152 | ENSG00000129173 | n.s. |
| ENSG00000183798 | 152.8614393 | -0.829347614 | 0.2797646 | -2.964448 | 0.003032269 | 0.014243685 | EMILIN3 | protein_coding | 62.66 | -1.531288887 | 1.531288887 | ENSG00000183798 | n.s. |
| ENSG00000113742 | 68.19825377 | 0.81590853 | 0.2730354 | 2.988288 | 0.002805448 | 0.013317509 | CPEB4 | protein_coding | 37.31 | 1.530299271 | 1.530299271 | ENSG00000113742 | n.s. |
| ENSG00000170085 | 111.1785919 | -0.812568979 | 0.2712775 | -2.995342 | 0.002741374 | 0.013085865 | SIMC1 | protein_coding | 42.19 | -1.53022793 | 1.53022793 | ENSG00000170085 | n.s. |
| ENSG00000228638 | 46.44208035 | -0.937919841 | 0.3378932 | -2.775787 | 0.005506819 | 0.023396839 | FCF1P2 | processed_pseudogene | 40.48 | -1.529599834 | 1.529599834 | ENSG00000228638 | n.s. |
| ENSG00000140479 | 43.53725037 | -0.896524527 | 0.3155837 | -2.840846 | 0.004499409 | 0.019768152 | PCSK6 | protein_coding | 48.11 | -1.527708221 | 1.527708221 | ENSG00000140479 | n.s. |
| ENSG00000276656 | 3.917633929 | -1.39582844 | 0.6248571 | -2.233836 | 0.02549386 | 0.08061572 | MIR6083 | miRNA | 50 | -1.526450433 | 1.526450433 | ENSG00000276656 | down |
| ENSG00000160293 | 490.1683112 | -0.755191854 | 0.2425764 | -3.113212 | 0.001850629 | 0.009532483 | VAV2 | protein_coding | 56.31 | -1.52608714 | 1.52608714 | ENSG00000160293 | n.s. |
| ENSG00000107984 | 1498.682711 | 0.682838518 | 0.2085043 | 3.274937 | 0.001056854 | 0.005892904 | DKK1 | protein_coding | 43.26 | 1.522504967 | 1.522504967 | ENSG00000107984 | n.s. |
| ENSG00000127314 | 2241.149251 | -0.838868291 | 0.285581 | -2.93741 | 0.003309666 | 0.015344816 | RAP1B | protein_coding | 38.96 | -1.521739215 | 1.521739215 | ENSG00000127314 | n.s. |
| ENSG00000266570 | 24.68685605 | -0.927860703 | 0.333239 | -2.78437 | 0.005363177 | 0.022912402 | MIR5579 | miRNA | 46.55 | -1.521626022 | 1.521626022 | ENSG00000266570 | n.s. |
| ENSG00000199545 | 29.4161571 | 0.957570751 | 0.3500912 | 2.735204 | 0.006234164 | 0.025763807 | RNA5SP195 | rRNA | 50.43 | 1.521570317 | 1.521570317 | ENSG00000199545 | n.s. |
| ENSG00000139651 | 365.0287547 | -0.742179723 | 0.2366213 | -3.136572 | 0.001709354 | 0.008913607 | ZNF740 | protein_coding | 50.62 | -1.521428746 | 1.521428746 | ENSG00000139651 | n.s. |
| ENSG00000236337 | 25.81450188 | -0.896718495 | 0.3166973 | -2.831469 | 0.004633475 | 0.020261265 | FMR1-IT1 | sense_intronic | 45.88 | -1.518443417 | 1.518443417 | ENSG00000236337 | n.s. |
| ENSG00000112584 | 1138.960114 | -0.819899257 | 0.2761419 | -2.969123 | 0.00298651 | 0.014061949 | FAM120B | protein_coding | 44.49 | -1.518416111 | 1.518416111 | ENSG00000112584 | n.s. |
| ENSG00000137337 | 620.3207307 | -0.753500889 | 0.2423928 | -3.108594 | 0.001879796 | 0.00965771 | MDC1 | protein_coding | 48.84 | -1.518399114 | 1.518399114 | ENSG00000137337 | n.s. |
| ENSG00000148925 | 379.7022927 | -0.841781001 | 0.2874399 | -2.928546 | 0.003405517 | 0.015715947 | BTBD10 | protein_coding | 36.22 | -1.518286253 | 1.518286253 | ENSG00000148925 | n.s. |
| ENSG00000108001 | 4194.071909 | -0.760621467 | 0.2459106 | -3.093082 | 0.001980895 | 0.010113679 | EBF3 | protein_coding | 48.8 | -1.517508911 | 1.517508911 | ENSG00000108001 | n.s. |
| ENSG00000072840 | 318.9701798 | -0.74965338 | 0.240648 | -3.115145 | 0.001838547 | 0.009479954 | EVC | protein_coding | 47.4 | -1.516694041 | 1.516694041 | ENSG00000072840 | n.s. |
| ENSG00000100575 | 393.593884 | -0.849710354 | 0.2918682 | -2.911281 | 0.003599497 | 0.016420643 | TIMM9 | protein_coding | 40.88 | -1.516401457 | 1.516401457 | ENSG00000100575 | n.s. |
| ENSG00000204311 | 102.3937704 | 0.763405527 | 0.2474218 | 3.085442 | 0.002032501 | 0.010335468 | DFNB59 | protein_coding | 39.52 | 1.515871353 | 1.515871353 | ENSG00000204311 | n.s. |
| ENSG00000150961 | 50.54231253 | -0.864045741 | 0.2997122 | -2.882918 | 0.003940095 | 0.017730426 | SEC24D | protein_coding | 38.92 | -1.513186743 | 1.513186743 | ENSG00000150961 | n.s. |
| ENSG00000104133 | 1566.705697 | -0.828397259 | 0.2813055 | -2.944832 | 0.003231303 | 0.015016498 | SPG11 | protein_coding | 41.28 | -1.510525524 | 1.510525524 | ENSG00000104133 | n.s. |
| ENSG00000224003 | 218.4176701 | -0.812929897 | 0.2733766 | -2.973663 | 0.002942683 | 0.013864966 | YES1P1 | processed_pseudogene | 42.44 | -1.510489748 | 1.510489748 | ENSG00000224003 | n.s. |
| ENSG00000148541 | 563.2756741 | -0.746624673 | 0.2398504 | -3.112877 | 0.001852735 | 0.009539803 | FAM13C | protein_coding | 40.06 | -1.508525733 | 1.508525733 | ENSG00000148541 | n.s. |

| | | | | | | | | | | | | | |
|---|---|---|---|---|---|---|---|---|---|---|---|---|---|
| ENSG00000267672 | 55.78860116 | 0.801756292 | 0.2679049 | 2.99269 | 0.002765302 | 0.013168232 | | sense_intronic | 41.34 | 1.507680674 | 1.507680674 | ENSG00000267672 | n.s. |
| ENSG00000138686 | 474.8561913 | -0.83595478 | 0.2855479 | -2.927546 | 0.003416483 | 0.015750894 | BBS7 | protein_coding | 35.63 | -1.50697133 | 1.50697133 | ENSG00000138686 | n.s. |
| ENSG00000143643 | 434.7599562 | -0.850686509 | 0.2934382 | -2.899031 | 0.003743176 | 0.016937355 | TTC13 | protein_coding | 39.73 | -1.506697157 | 1.506697157 | ENSG00000143643 | n.s. |
| ENSG00000081087 | 419.4301576 | -0.849975281 | 0.2930865 | -2.900083 | 0.00373064 | 0.016886119 | OSTM1 | protein_coding | 42.48 | -1.506555819 | 1.506555819 | ENSG00000081087 | n.s. |
| ENSG00000207217 | 27.73629902 | -0.92135441 | 0.331465 | -2.779643 | 0.005441872 | 0.023198769 | SNORA80D | snoRNA | 58.52 | -1.505986088 | 1.505986088 | ENSG00000207217 | n.s. |
| ENSG00000104343 | 687.7735308 | -0.799515052 | 0.2671034 | -2.99328 | 0.002759968 | 0.013159943 | UBE2W | protein_coding | 39.29 | -1.503684727 | 1.503684727 | ENSG00000104343 | n.s. |
| ENSG00000134461 | 162.3746264 | -0.737345016 | 0.2357586 | -3.127543 | 0.001762741 | 0.009150872 | ANKRD16 | protein_coding | 46.86 | -1.50310548 | 1.50310548 | ENSG00000134461 | n.s. |
| ENSG00000058056 | 1268.862469 | -0.78215609 | 0.2582063 | -3.029191 | 0.002452096 | 0.012012853 | USP13 | protein_coding | 42.73 | -1.502016455 | 1.502016455 | ENSG00000058056 | n.s. |
| ENSG00000177189 | 859.2034696 | -0.803231407 | 0.269718 | -2.978042 | 0.002900966 | 0.013682299 | RPS6KA3 | protein_coding | 37.57 | -1.497095574 | 1.497095574 | ENSG00000177189 | n.s. |
| ENSG00000006756 | 219.9228005 | -0.857526453 | 0.2980626 | -2.877001 | 0.004014745 | 0.018024802 | ARSD | protein_coding | 48.88 | -1.49563719 | 1.49563719 | ENSG00000006756 | n.s. |
| ENSG00000165821 | 216.0340316 | -0.714526441 | 0.2255658 | -3.167708 | 0.00153646 | 0.008130312 | SALL2 | protein_coding | 48.58 | -1.493283639 | 1.493283639 | ENSG00000165821 | n.s. |
| ENSG00000124370 | 222.6212982 | -0.771860923 | 0.2538107 | -3.041089 | 0.002357243 | 0.011646521 | MCEE | protein_coding | 41.39 | -1.492627565 | 1.492627565 | ENSG00000124370 | n.s. |
| ENSG00000165476 | 788.0064856 | -0.823890991 | 0.2807654 | -2.934446 | 0.003341439 | 0.015471517 | REEP3 | protein_coding | 38.34 | -1.491627533 | 1.491627533 | ENSG00000165476 | n.s. |
| ENSG00000102804 | 2940.74945 | 0.680927552 | 0.2098646 | 3.244604 | 0.001176141 | 0.006449554 | TSC22D1 | protein_coding | 38.08 | 1.491551582 | 1.491551582 | ENSG00000102804 | n.s. |
| ENSG00000239908 | 19.08821556 | -0.96120379 | 0.3559175 | -2.700637 | 0.006920693 | 0.028100556 | RN7SL75P | misc_RNA | 50.89 | -1.49110111 | 1.49110111 | ENSG00000239908 | n.s. |
| ENSG00000084774 | 1624.445435 | -0.724409448 | 0.2304853 | -3.142974 | 0.001672407 | 0.008757021 | CAD | protein_coding | 51.17 | -1.490576467 | 1.490576467 | ENSG00000084774 | n.s. |
| ENSG00000143157 | 2820.969738 | -0.784390165 | 0.2609193 | -3.006256 | 0.002644863 | 0.012711226 | POGK | protein_coding | 47.02 | -1.487056733 | 1.487056733 | ENSG00000143157 | n.s. |
| ENSG00000184786 | 590.5853854 | 0.6698741 | 0.205054 | 3.266818 | 0.001087634 | 0.00603793 | TCTE3 | protein_coding | 38.54 | 1.486525617 | 1.486525617 | ENSG00000184786 | n.s. |
| ENSG00000112144 | 1067.6786 | -0.791745768 | 0.2648422 | -2.9895 | 0.002794345 | 0.013278385 | ICK | protein_coding | 40.86 | -1.485991813 | 1.485991813 | ENSG00000112144 | n.s. |
| ENSG00000233937 | 399.1769616 | -0.703568748 | 0.221153 | -3.181367 | 0.00146582 | 0.007807425 | | processed_transcript | 48.01 | -1.48276564 | 1.48276564 | ENSG00000233937 | n.s. |
| ENSG00000177302 | 1513.374187 | -0.754858366 | 0.2462191 | -3.0658 | 0.002170886 | 0.010899838 | TOP3A | protein_coding | 48.23 | -1.48146991 | 1.48146991 | ENSG00000177302 | n.s. |
| ENSG00000238829 | 2.754504529 | -1.458198226 | 0.6799287 | -2.144634 | 0.031982119 | 0.096464091 | RNU7-45P | snRNA | 39.68 | -1.480996172 | 1.480996172 | ENSG00000238829 | down |
| ENSG00000203819 | 71.02146141 | 0.785957766 | 0.2627545 | 2.991224 | 0.002778614 | 0.01320814 | | processed_pseudogene | 60 | 1.476939094 | 1.476939094 | ENSG00000203819 | n.s. |
| ENSG00000166024 | 605.3448057 | -0.774891003 | 0.2569444 | -3.015792 | 0.002563086 | 0.012447104 | R3HCC1L | protein_coding | 37.83 | -1.47611443 | 1.47611443 | ENSG00000166024 | n.s. |
| ENSG00000259196 | 23.98114676 | -0.91308005 | 0.3307575 | -2.760573 | 0.005770008 | 0.024270641 | HMBOX1-IT1 | sense_intronic | 38.15 | -1.474550092 | 1.474550092 | ENSG00000259196 | n.s. |
| ENSG00000182223 | 208.1276161 | -0.725284205 | 0.2323723 | -3.121216 | 0.001801059 | 0.00932405 | ZAR1 | protein_coding | 53.24 | -1.47261372 | 1.47261372 | ENSG00000182223 | n.s. |
| ENSG00000118707 | 474.7039361 | -0.791730592 | 0.2661412 | -2.974852 | 0.002931303 | 0.013816022 | TGIF2 | protein_coding | 51.95 | -1.47231566 | 1.47231566 | ENSG00000118707 | n.s. |
| ENSG00000173894 | 3381.225322 | -0.745284287 | 0.2422245 | -3.076832 | 0.00209213 | 0.010588412 | CBX2 | protein_coding | 58.13 | -1.472062536 | 1.472062536 | ENSG00000173894 | n.s. |
| ENSG00000279672 | 65.04435913 | -0.788597157 | 0.2646853 | -2.979376 | 0.002888358 | 0.01364132 | | TEC | 62.07 | -1.470846934 | 1.470846934 | ENSG00000279672 | n.s. |
| ENSG00000281128 | 121.3017326 | 0.72654286 | 0.233204 | 3.115482 | 0.001836443 | 0.009473433 | | antisense | 36.98 | 1.470154111 | 1.470154111 | ENSG00000281128 | n.s. |
| ENSG00000227034 | 20.70539597 | -1.022348988 | 0.3945269 | -2.591329 | 0.00956061 | 0.036609563 | | processed_pseudogene | 43.97 | -1.468507663 | 1.468507663 | ENSG00000227034 | down |
| ENSG00000136152 | 923.6574023 | -0.763217272 | 0.2517413 | -3.031753 | 0.002431382 | 0.011932371 | COG3 | protein_coding | 39.15 | -1.467875361 | 1.467875361 | ENSG00000136152 | n.s. |

| | | | | | | | | | | | | | |
|---|---|---|---|---|---|---|---|---|---|---|---|---|---|
| ENSG00000169604 | 24650.14562 | -0.755540206 | 0.2479013 | -3.047746 | 0.00230565 | 0.011438392 | ANTXR1 | protein_coding | 41.88 | -1.46698334 | 1.46698334 | ENSG00000169604 | n.s. |
| ENSG00000272886 | 885.0064266 | -0.769903644 | 0.2552804 | -3.015914 | 0.002562057 | 0.012446449 | DCP1A | protein_coding | 41.19 | -1.466631449 | 1.466631449 | ENSG00000272886 | n.s. |
| ENSG00000148120 | 4157.85343 | -0.725609114 | 0.2331194 | -3.112607 | 0.001854428 | 0.009544989 | C9orf3 | protein_coding | 42.89 | -1.465893363 | 1.465893363 | ENSG00000148120 | n.s. |
| ENSG00000157570 | 669.9952183 | -0.745965485 | 0.2432607 | -3.066527 | 0.002165611 | 0.010881198 | TSPAN18 | protein_coding | 50.12 | -1.464571423 | 1.464571423 | ENSG00000157570 | n.s. |
| ENSG00000129673 | 68.0927712 | 0.782044128 | 0.2619862 | 2.985058 | 0.002835242 | 0.013440609 | AANAT | protein_coding | 53.01 | 1.463658975 | 1.463658975 | ENSG00000129673 | n.s. |
| ENSG00000151693 | 809.4779486 | -0.77087812 | 0.256176 | -3.009174 | 0.002619594 | 0.012633396 | ASAP2 | protein_coding | 46.56 | -1.463496617 | 1.463496617 | ENSG00000151693 | n.s. |
| ENSG00000222011 | 133.7794296 | -0.856278791 | 0.3012022 | -2.84287 | 0.004470927 | 0.019680778 | FAM185A | protein_coding | 37.7 | -1.460775429 | 1.460775429 | ENSG00000222011 | n.s. |
| ENSG00000100281 | 1052.621295 | -0.744851727 | 0.243106 | -3.063897 | 0.002184743 | 0.010957554 | HMGXB4 | protein_coding | 41.49 | -1.460122689 | 1.460122689 | ENSG00000100281 | n.s. |
| ENSG00000228218 | 52.74242379 | -0.854732956 | 0.3004726 | -2.844628 | 0.004446329 | 0.019591687 | ATF4P3 | processed_pseudogene | 52.08 | -1.459822486 | 1.459822486 | ENSG00000228218 | n.s. |
| ENSG00000225746 | 292.8316784 | -0.874046473 | 0.311293 | -2.807794 | 0.004988219 | 0.021515172 | MEG8 | lincRNA | 38.95 | -1.457258508 | 1.457258508 | ENSG00000225746 | n.s. |
| ENSG00000254708 | 30.34510749 | 0.905289837 | 0.3286252 | 2.754779 | 0.005873176 | 0.024637612 | | processed_pseudogene | 39.73 | 1.456069425 | 1.456069425 | ENSG00000254708 | n.s. |
| ENSG00000127080 | 260.5877843 | -0.775903556 | 0.2596222 | -2.988587 | 0.002802707 | 0.013309037 | IPPK | protein_coding | 47.93 | -1.455481307 | 1.455481307 | ENSG00000127080 | n.s. |
| ENSG00000151461 | 1259.287839 | -0.770906275 | 0.2570937 | -2.998542 | 0.002712743 | 0.01297476 | UPF2 | protein_coding | 40.23 | -1.454623571 | 1.454623571 | ENSG00000151461 | n.s. |
| ENSG00000119630 | 2207.669293 | -0.712827024 | 0.2278415 | -3.128609 | 0.00175636 | 0.009121147 | PGF | protein_coding | 56.15 | -1.45413189 | 1.45413189 | ENSG00000119630 | n.s. |
| ENSG00000057019 | 1520.046395 | -0.751573896 | 0.2471752 | -3.040652 | 0.002360665 | 0.011655157 | DCBLD2 | protein_coding | 38.58 | -1.453154505 | 1.453154505 | ENSG00000057019 | n.s. |
| ENSG00000149292 | 67.759325 | -0.765244358 | 0.254303 | -3.009183 | 0.002619513 | 0.012633396 | TTC12 | protein_coding | 43.07 | -1.452801034 | 1.452801034 | ENSG00000149292 | n.s. |
| ENSG00000201785 | 35.17525722 | 0.860696713 | 0.3046848 | 2.824876 | 0.004729889 | 0.020590169 | SNORD117 | snoRNA | 44.74 | 1.451427378 | 1.451427378 | ENSG00000201785 | n.s. |
| ENSG00000240225 | 949.0379527 | -0.748241549 | 0.2456718 | -3.045696 | 0.002321428 | 0.011498127 | ZNF542P | transcribed_unprocessed_pseud | 40.44 | -1.451119373 | 1.451119373 | ENSG00000240225 | n.s. |
| ENSG00000124831 | 1400.383881 | -0.766115185 | 0.2550678 | -3.003575 | 0.002668276 | 0.012806067 | LRRFIP1 | protein_coding | 45.52 | -1.449937517 | 1.449937517 | ENSG00000124831 | n.s. |
| ENSG00000105523 | 35.19801583 | 0.83616314 | 0.2916571 | 2.866939 | 0.004144623 | 0.018459881 | FAM83E | protein_coding | 56.4 | 1.449715498 | 1.449715498 | ENSG00000105523 | n.s. |
| ENSG00000161149 | 102.7752875 | -0.764409648 | 0.25424 | -3.006646 | 0.002641471 | 0.012699308 | TUBA3FP | processed_transcript | 54.66 | -1.449488816 | 1.449488816 | ENSG00000161149 | n.s. |
| ENSG00000111052 | 1111.919005 | -0.857628563 | 0.3033555 | -2.82714 | 0.004696578 | 0.020472904 | LIN7A | protein_coding | 36.73 | -1.448380739 | 1.448380739 | ENSG00000111052 | n.s. |
| ENSG00000112877 | 222.5862257 | -0.790128498 | 0.2675597 | -2.953092 | 0.003146078 | 0.014694009 | CEP72 | protein_coding | 53.81 | -1.448194676 | 1.448194676 | ENSG00000112877 | n.s. |
| ENSG00000144057 | 662.2927659 | -0.762577579 | 0.2533822 | -3.009595 | 0.002615965 | 0.01262464 | ST6GAL2 | protein_coding | 41.34 | -1.447967813 | 1.447967813 | ENSG00000144057 | n.s. |
| ENSG00000073849 | 650.8710063 | -0.804969381 | 0.2752679 | -2.924313 | 0.00345218 | 0.01589441 | ST6GAL1 | protein_coding | 44.38 | -1.447943172 | 1.447943172 | ENSG00000073849 | n.s. |
| ENSG00000145860 | 1860.487285 | -0.769017408 | 0.256967 | -2.99267 | 0.00276549 | 0.013168232 | RNF145 | protein_coding | 36.97 | -1.446116102 | 1.446116102 | ENSG00000145860 | n.s. |
| ENSG00000228878 | 30.38843924 | 0.900760162 | 0.3273626 | 2.751567 | 0.005931093 | 0.024828205 | SEPT7-AS1 | lincRNA | 39.03 | 1.445769319 | 1.445769319 | ENSG00000228878 | n.s. |
| ENSG00000265135 | 6.720377649 | -1.319235877 | 0.5900577 | -2.235774 | 0.025366563 | 0.080338673 | MIR5687 | miRNA | 36.36 | -1.444662683 | 1.444662683 | ENSG00000265135 | down |
| ENSG00000100038 | 516.7056689 | -0.791681726 | 0.2687808 | -2.945455 | 0.003224798 | 0.01499628 | TOP3B | protein_coding | 54.98 | -1.444040499 | 1.444040499 | ENSG00000100038 | n.s. |
| ENSG00000166881 | 1622.432037 | -0.826726569 | 0.287181 | -2.878764 | 0.003992364 | 0.01794895 | NEMP1 | protein_coding | 44.16 | -1.443432308 | 1.443432308 | ENSG00000166881 | n.s. |
| ENSG00000201847 | 5.811283644 | -1.237251281 | 0.5341646 | -2.316236 | 0.020545383 | 0.068156629 | SNORD31B | snoRNA | 44.93 | -1.443243594 | 1.443243594 | ENSG00000201847 | down |
| ENSG00000204217 | 1327.934549 | -0.806028764 | 0.276404 | -2.916125 | 0.003544083 | 0.016222411 | BMPR2 | protein_coding | 41.04 | -1.442698481 | 1.442698481 | ENSG00000204217 | n.s. |

| | | | | | | | | | | | | | |
|---|---|---|---|---|---|---|---|---|---|---|---|---|---|
| ENSG00000004897 | 3993.349543 | -0.792381708 | 0.2693077 | -2.942291 | 0.003257937 | 0.015130174 | CDC27 | protein_coding | 38.66 | -1.442258384 | 1.442258384 | ENSG00000004897 | n.s. |
| ENSG00000164543 | 257.1210937 | 0.800169645 | 0.2733763 | 2.926989 | 0.003422606 | 0.015773898 | STK17A | protein_coding | 39.76 | 1.441954479 | 1.441954479 | ENSG00000164543 | n.s. |
| ENSG00000162065 | 390.3564487 | -0.784844954 | 0.2654118 | -2.957084 | 0.003105634 | 0.014556087 | TBC1D24 | protein_coding | 55.83 | -1.441725132 | 1.441725132 | ENSG00000162065 | n.s. |
| ENSG00000206699 | 4.752244315 | -1.649426485 | 0.8365153 | -1.971783 | 0.048634419 | 0.133689439 | | misc_RNA | 37.25 | -1.441438585 | 1.441438585 | ENSG00000206699 | n.s. |
| ENSG00000135093 | 350.1377906 | -0.799050459 | 0.2728307 | -2.928741 | 0.003403378 | 0.015711284 | USP30 | protein_coding | 42.28 | -1.44131789 | 1.44131789 | ENSG00000135093 | n.s. |
| ENSG00000107338 | 274.0699702 | -0.720453362 | 0.232652 | -3.096699 | 0.001956884 | 0.010002093 | SHB | protein_coding | 50.89 | -1.440841258 | 1.440841258 | ENSG00000107338 | n.s. |
| ENSG00000197603 | 1195.09052 | -0.765683019 | 0.2559238 | -2.99184 | 0.002773015 | 0.013190534 | C5orf42 | protein_coding | 38.94 | -1.43928319 | 1.43928319 | ENSG00000197603 | n.s. |
| ENSG00000187013 | 191.4082786 | 0.688207931 | 0.2172145 | 3.168333 | 0.001533159 | 0.008116415 | C17orf82 | lincRNA | 70.85 | 1.438792121 | 1.438792121 | ENSG00000187013 | n.s. |
| ENSG00000237457 | 25.72294507 | 0.88468451 | 0.3192905 | 2.770782 | 0.005592179 | 0.023701647 | LINC01351 | lincRNA | 37.76 | 1.437808263 | 1.437808263 | ENSG00000237457 | n.s. |
| ENSG00000280037 | 26.38780978 | -0.861051558 | 0.3064457 | -2.809801 | 0.00495721 | 0.021401314 | | TEC | 45.03 | -1.437576862 | 1.437576862 | ENSG00000280037 | n.s. |
| ENSG00000196378 | 87.98568416 | -0.828281989 | 0.2887624 | -2.868386 | 0.004125718 | 0.018399224 | ZNF34 | protein_coding | 49.73 | -1.437235321 | 1.437235321 | ENSG00000196378 | n.s. |
| ENSG00000254685 | 333.5365623 | -0.778401435 | 0.2626249 | -2.963929 | 0.003037387 | 0.014262913 | FPGT | protein_coding | 34.35 | -1.436766957 | 1.436766957 | ENSG00000254685 | n.s. |
| ENSG00000123213 | 1487.812906 | -0.819050295 | 0.283881 | -2.885189 | 0.003911785 | 0.017625821 | NLN | protein_coding | 40.37 | -1.436491898 | 1.436491898 | ENSG00000123213 | n.s. |
| ENSG00000258813 | 82.88749197 | 0.760204096 | 0.2533349 | 3.000788 | 0.002692823 | 0.012897204 | | antisense | 53.45 | 1.43640902 | 1.43640902 | ENSG00000258813 | n.s. |
| ENSG00000213801 | 91.34299479 | -0.904303177 | 0.3307097 | -2.734432 | 0.006248805 | 0.025808992 | ZNF321P | transcribed_processed_pseudog | 46.09 | -1.436240493 | 1.436240493 | ENSG00000213801 | n.s. |
| ENSG00000271886 | 13.43136795 | -0.995401537 | 0.3834204 | -2.59611 | 0.009428593 | 0.036203612 | MIR98 | miRNA | 42.86 | -1.434620567 | 1.434620567 | ENSG00000271886 | n.s. |
| ENSG00000122257 | 2569.496831 | -0.766519505 | 0.2568201 | -2.984656 | 0.002838975 | 0.013449151 | RBBP6 | protein_coding | 37.61 | -1.434391889 | 1.434391889 | ENSG00000122257 | n.s. |
| ENSG00000092148 | 3774.365157 | -0.775193654 | 0.2612139 | -2.967658 | 0.003000779 | 0.014119585 | HECTD1 | protein_coding | 39.35 | -1.434246294 | 1.434246294 | ENSG00000092148 | n.s. |
| ENSG00000227910 | 39.70786335 | 0.81345927 | 0.281267 | 2.892125 | 0.003826456 | 0.017269286 | | transcribed_processed_pseudog | 51.45 | 1.433905502 | 1.433905502 | ENSG00000227910 | n.s. |
| ENSG00000134046 | 630.8448632 | -0.743666807 | 0.2449777 | -3.035651 | 0.002400172 | 0.011812511 | MBD2 | protein_coding | 39.24 | -1.43353509 | 1.43353509 | ENSG00000134046 | n.s. |
| ENSG00000137478 | 224.5562823 | -0.75529263 | 0.2511053 | -3.007873 | 0.002630834 | 0.012670045 | FCHSD2 | protein_coding | 38.87 | -1.43295768 | 1.43295768 | ENSG00000137478 | n.s. |
| ENSG00000167635 | 4868.431452 | -0.804453925 | 0.2767532 | -2.906755 | 0.003651987 | 0.016606948 | ZNF146 | protein_coding | 42.84 | -1.431694838 | 1.431694838 | ENSG00000167635 | n.s. |
| ENSG00000130779 | 1402.88547 | -0.780878246 | 0.2644598 | -2.95273 | 0.003149778 | 0.014706356 | CLIP1 | protein_coding | 43.88 | -1.430955422 | 1.430955422 | ENSG00000130779 | n.s. |
| ENSG00000176994 | 808.7906093 | -0.678437879 | 0.2131833 | -3.182416 | 0.001460521 | 0.007785159 | SMCR8 | protein_coding | 52.68 | -1.430644026 | 1.430644026 | ENSG00000176994 | n.s. |
| ENSG00000183570 | 77.86465858 | -0.751438316 | 0.2493267 | -3.01387 | 0.002579382 | 0.012507298 | PCBP3 | protein_coding | 46.92 | -1.429864269 | 1.429864269 | ENSG00000183570 | n.s. |
| ENSG00000140718 | 111.9823325 | -0.780569273 | 0.2646758 | -2.949152 | 0.003186469 | 0.014862717 | FTO | protein_coding | 40.2 | -1.426803972 | 1.426803972 | ENSG00000140718 | n.s. |
| ENSG00000198169 | 278.3493517 | -0.79836919 | 0.2741381 | -2.912289 | 0.003587908 | 0.016391004 | ZNF251 | protein_coding | 47.51 | -1.425403906 | 1.425403906 | ENSG00000198169 | n.s. |
| ENSG00000058729 | 1070.360658 | -0.781291204 | 0.2652157 | -2.945871 | 0.003220466 | 0.014987546 | RIOK2 | protein_coding | 36.71 | -1.425285691 | 1.425285691 | ENSG00000058729 | n.s. |
| ENSG00000063015 | 320.1340544 | -0.736986787 | 0.2423672 | -3.040785 | 0.002359619 | 0.011654124 | SEZ6 | protein_coding | 54.21 | -1.424978969 | 1.424978969 | ENSG00000063015 | n.s. |
| ENSG00000141084 | 233.8257648 | -0.701454633 | 0.224807 | -3.120252 | 0.001806961 | 0.009342117 | RANBP10 | protein_coding | 49.73 | -1.42364054 | 1.42364054 | ENSG00000141084 | n.s. |
| ENSG00000117114 | 5150.317033 | -0.784197059 | 0.2670251 | -2.936791 | 0.003316272 | 0.015365212 | ADGRL2 | protein_coding | 36.78 | -1.422111133 | 1.422111133 | ENSG00000117114 | n.s. |
| ENSG00000201264 | 168.0417401 | 0.698247829 | 0.2234011 | 3.125535 | 0.001774822 | 0.009196444 | SNORD73B | snoRNA | 27.78 | 1.421897947 | 1.421897947 | ENSG00000201264 | n.s. |

| | | | | | | | | | | | | | |
|---|---|---|---|---|---|---|---|---|---|---|---|---|---|
| ENSG00000152503 | 215.3515524 | 0.71734883 | 0.2329866 | 3.078928 | 0.002077469 | 0.010525692 | TRIM36 | protein_coding | 40.18 | 1.418736167 | 1.418736167 | ENSG00000152503 | n.s. |
| ENSG00000144749 | 31.30531851 | 0.834927175 | 0.2944252 | 2.835787 | 0.004571295 | 0.020027103 | LRIG1 | protein_coding | 46.07 | 1.418025175 | 1.418025175 | ENSG00000144749 | n.s. |
| ENSG00000144283 | 1060.296502 | -0.785625991 | 0.2682824 | -2.928355 | 0.003407607 | 0.015720382 | PKP4 | protein_coding | 38.59 | -1.41690546 | 1.41690546 | ENSG00000144283 | n.s. |
| ENSG00000197927 | 81.11521988 | -0.860294626 | 0.3082837 | -2.790594 | 0.005261139 | 0.0225526 | C2orf27A | transcribed_unprocessed_pseud | 40.76 | -1.416736101 | 1.416736101 | ENSG00000197927 | n.s. |
| ENSG00000275714 | 1484.034393 | 0.717782942 | 0.233407 | 3.075241 | 0.002103323 | 0.010636548 | HIST1H3A | protein_coding | 60.58 | 1.416328805 | 1.416328805 | ENSG00000275714 | n.s. |
| ENSG00000265145 | 58.98760586 | -0.774324887 | 0.2626587 | -2.948027 | 0.003198089 | 0.014909606 | SNORD53 | snoRNA | 46.15 | -1.414330599 | 1.414330599 | ENSG00000265145 | n.s. |
| ENSG00000275964 | 145.3698112 | 0.684354652 | 0.2172464 | 3.150131 | 0.001631972 | 0.008577555 | | lincRNA | 65.99 | 1.414312303 | 1.414312303 | ENSG00000275964 | n.s. |
| ENSG00000111725 | 431.7376159 | -0.732685236 | 0.24122 | -3.037415 | 0.002386168 | 0.011747743 | PRKAB1 | protein_coding | 47.82 | -1.414115876 | 1.414115876 | ENSG00000111725 | n.s. |
| ENSG00000162869 | 84.49314228 | -0.826808106 | 0.2908495 | -2.842735 | 0.004472825 | 0.019680778 | PPP1R21 | protein_coding | 39.73 | -1.410499686 | 1.410499686 | ENSG00000162869 | n.s. |
| ENSG00000272301 | 54.44817306 | -0.843703128 | 0.3000966 | -2.811438 | 0.004932058 | 0.021319168 | | lincRNA | 33.68 | -1.410021763 | 1.410021763 | ENSG00000272301 | n.s. |
| ENSG00000198876 | 616.5907288 | -0.771264321 | 0.2616254 | -2.947972 | 0.003198664 | 0.014909606 | DCAF12 | protein_coding | 45.99 | -1.408740371 | 1.408740371 | ENSG00000198876 | n.s. |
| ENSG00000020256 | 146.0614305 | -0.779534726 | 0.2659244 | -2.931415 | 0.003374219 | 0.015602542 | ZFP64 | protein_coding | 44.05 | -1.408466957 | 1.408466957 | ENSG00000020256 | n.s. |
| ENSG00000207357 | 25.05588726 | 0.867351248 | 0.3133204 | 2.768257 | 0.005635701 | 0.02385706 | RNU6-2 | snRNA | 45.79 | 1.407175989 | 1.407175989 | ENSG00000207357 | n.s. |
| ENSG00000133059 | 452.2418455 | -0.723041093 | 0.2368753 | -3.052413 | 0.002270099 | 0.01132041 | DSTYK | protein_coding | 43.55 | -1.407137634 | 1.407137634 | ENSG00000133059 | n.s. |
| ENSG00000080371 | 1522.713126 | -0.773128504 | 0.2629305 | -2.940429 | 0.003277582 | 0.015206188 | RAB21 | protein_coding | 37.14 | -1.405531881 | 1.405531881 | ENSG00000080371 | n.s. |
| ENSG00000140386 | 771.849753 | -0.806757195 | 0.2806894 | -2.874199 | 0.004050542 | 0.018151172 | SCAPER | protein_coding | 37.7 | -1.404641187 | 1.404641187 | ENSG00000140386 | n.s. |
| ENSG00000177105 | 67.10044309 | -0.806231172 | 0.2804828 | -2.874441 | 0.004047437 | 0.018143101 | RHOG | protein_coding | 51.22 | -1.403881059 | 1.403881059 | ENSG00000177105 | n.s. |
| ENSG00000224985 | 21.87671878 | 0.89002061 | 0.3271222 | 2.720758 | 0.006513234 | 0.026718843 | | antisense | 46.94 | 1.400164722 | 1.400164722 | ENSG00000224985 | n.s. |
| ENSG00000136381 | 3172.516403 | -0.788831116 | 0.2718266 | -2.901964 | 0.00370831 | 0.016806312 | IREB2 | protein_coding | 39.42 | -1.399802576 | 1.399802576 | ENSG00000136381 | n.s. |
| ENSG00000103111 | 345.6300535 | -0.804453449 | 0.2800197 | -2.872846 | 0.004067919 | 0.018199754 | MON1B | protein_coding | 49.31 | -1.399696304 | 1.399696304 | ENSG00000103111 | n.s. |
| ENSG00000266038 | 10.66339491 | -1.045168787 | 0.4188926 | -2.495076 | 0.01259303 | 0.045898629 | MIR4659A | miRNA | 38.27 | -1.398645172 | 1.398645172 | ENSG00000266038 | down |
| ENSG00000107854 | 1374.362999 | -0.744066654 | 0.2487257 | -2.991515 | 0.002775966 | 0.01320006 | TNKS2 | protein_coding | 37.63 | -1.398416801 | 1.398416801 | ENSG00000107854 | n.s. |
| ENSG00000207352 | 7.870607528 | -1.405765102 | 0.6636033 | -2.118382 | 0.034142745 | 0.101314883 | RNU6-540P | snRNA | 40.59 | -1.397789848 | 1.397789848 | ENSG00000207352 | n.s. |
| ENSG00000187824 | 29.46955566 | -0.927676468 | 0.3490672 | -2.657587 | 0.007870224 | 0.031191895 | TMEM220 | protein_coding | 40.31 | -1.397042019 | 1.397042019 | ENSG00000187824 | n.s. |
| ENSG00000213468 | 240.5440999 | -0.719277978 | 0.2362406 | -3.044684 | 0.00232925 | 0.011528672 | FIRRE | lincRNA | 40.91 | -1.394119478 | 1.394119478 | ENSG00000213468 | n.s. |
| ENSG00000138688 | 2534.551889 | -0.784638578 | 0.2702943 | -2.902905 | 0.003697189 | 0.016767414 | KIAA1109 | protein_coding | 34.76 | -1.393152406 | 1.393152406 | ENSG00000138688 | n.s. |
| ENSG00000149716 | 415.2107616 | -0.710080055 | 0.2317372 | -3.064161 | 0.002182815 | 0.010951832 | ORAOV1 | protein_coding | 53.52 | -1.392121347 | 1.392121347 | ENSG00000149716 | n.s. |
| ENSG00000168710 | 2152.34697 | -0.775233037 | 0.265392 | -2.921087 | 0.003488124 | 0.016038688 | AHCYL1 | protein_coding | 40.97 | -1.391412407 | 1.391412407 | ENSG00000168710 | n.s. |
| ENSG00000196387 | 1488.532224 | -0.713687767 | 0.2335793 | -3.055441 | 0.002247301 | 0.011230859 | ZNF140 | protein_coding | 41.08 | -1.391396411 | 1.391396411 | ENSG00000196387 | n.s. |
| ENSG00000151422 | 308.8081062 | -0.780369191 | 0.268226 | -2.909372 | 0.003621552 | 0.016514909 | FER | protein_coding | 35.73 | -1.390714516 | 1.390714516 | ENSG00000151422 | n.s. |
| ENSG00000091972 | 447.7864606 | -0.769268865 | 0.2623603 | -2.932108 | 0.003366697 | 0.01557293 | CD200 | protein_coding | 38.61 | -1.39055323 | 1.39055323 | ENSG00000091972 | n.s. |
| ENSG00000225697 | 165.9377318 | 0.712712713 | 0.2332018 | 3.056206 | 0.002241572 | 0.011206251 | SLC26A6 | protein_coding | 59.37 | 1.390174404 | 1.390174404 | ENSG00000225697 | n.s. |

| | | | | | | | | | | | | | |
|---|---|---|---|---|---|---|---|---|---|---|---|---|---|
| ENSG00000198626 | 2614.842421 | -0.780093875 | 0.2682389 | -2.908205 | 0.003635096 | 0.016550502 | RYR2 | protein_coding | 38.53 | -1.389494494 | 1.389494494 | ENSG00000198626 | n.s. |
| ENSG00000125633 | 621.3628562 | -0.724960072 | 0.2395872 | -3.025871 | 0.002479179 | 0.012119949 | CCDC93 | protein_coding | 42.78 | -1.389385411 | 1.389385411 | ENSG00000125633 | n.s. |
| ENSG00000220771 | 9.928333116 | -1.053903795 | 0.4260687 | -2.473554 | 0.013377663 | 0.048128477 | BOLA2P3 | processed_pseudogene | 60.94 | -1.388621404 | 1.388621404 | ENSG00000220771 | down |
| ENSG00000198131 | 998.4061366 | -0.731151159 | 0.2429299 | -3.009721 | 0.002614876 | 0.01262376 | ZNF544 | protein_coding | 47.34 | -1.388318061 | 1.388318061 | ENSG00000198131 | n.s. |
| ENSG00000269971 | 11.07324206 | 1.036404426 | 0.4157208 | 2.49303 | 0.012665804 | 0.046103527 | | sense_intronic | 41.85 | 1.384911839 | 1.384911839 | ENSG00000269971 | up |
| ENSG00000272579 | 54.70078 | -0.938946147 | 0.3575056 | -2.626381 | 0.008629808 | 0.03359978 | | antisense | 29.38 | -1.383690729 | 1.383690729 | ENSG00000272579 | n.s. |
| ENSG00000089775 | 542.985078 | -0.735409721 | 0.2456708 | -2.993477 | 0.002758185 | 0.013155947 | ZBTB25 | protein_coding | 42.25 | -1.383215885 | 1.383215885 | ENSG00000089775 | n.s. |
| ENSG00000280406 | 114.8417765 | 0.728688277 | 0.2422096 | 3.008503 | 0.002625381 | 0.012652536 | | TEC | 30.49 | 1.382920954 | 1.382920954 | ENSG00000280406 | n.s. |
| ENSG00000247828 | 267.2749033 | -0.792651673 | 0.2754692 | -2.87746 | 0.004008908 | 0.018005168 | TMEM161B-AS1 | antisense | 37.29 | -1.382862344 | 1.382862344 | ENSG00000247828 | n.s. |
| ENSG00000089335 | 1652.307833 | -0.757911327 | 0.2572608 | -2.946082 | 0.003218273 | 0.014985956 | ZNF302 | protein_coding | 39.83 | -1.382669409 | 1.382669409 | ENSG00000089335 | n.s. |
| ENSG00000244018 | 20.94376279 | -0.88369396 | 0.326067 | -2.710161 | 0.006725059 | 0.027418155 | RPL35P6 | processed_pseudogene | 48.39 | -1.380296185 | 1.380296185 | ENSG00000244018 | n.s. |
| ENSG00000072518 | 71.08239198 | -0.75654825 | 0.2568681 | -2.94528 | 0.00322663 | 0.014999791 | MARK2 | protein_coding | 49.65 | -1.379879536 | 1.379879536 | ENSG00000072518 | n.s. |
| ENSG00000118007 | 1578.384119 | -0.793589435 | 0.2763795 | -2.871376 | 0.004086889 | 0.018249442 | STAG1 | protein_coding | 38.94 | -1.379853955 | 1.379853955 | ENSG00000118007 | n.s. |
| ENSG00000137094 | 230.7256417 | -0.888375837 | 0.3288538 | -2.701431 | 0.006904177 | 0.028050931 | DNAJB5 | protein_coding | 55.15 | -1.378806136 | 1.378806136 | ENSG00000137094 | n.s. |
| ENSG00000144191 | 1671.91767 | -0.770789393 | 0.2644539 | -2.914645 | 0.00356093 | 0.016282044 | CNGA3 | protein_coding | 47.34 | -1.378395793 | 1.378395793 | ENSG00000144191 | n.s. |
| ENSG00000166435 | 30.97216469 | -0.86028462 | 0.3130322 | -2.74823 | 0.00599179 | 0.025007101 | XRRA1 | protein_coding | 41.82 | -1.378121459 | 1.378121459 | ENSG00000166435 | n.s. |
| ENSG00000274934 | 54.37315922 | 0.776634962 | 0.2677171 | 2.900954 | 0.00372028 | 0.016850181 | | misc_RNA | 42.51 | 1.377280893 | 1.377280893 | ENSG00000274934 | n.s. |
| ENSG00000188428 | 271.1240635 | -0.856326706 | 0.3110442 | -2.753071 | 0.005903916 | 0.024744198 | BLOC1S5 | protein_coding | 40 | -1.375711647 | 1.375711647 | ENSG00000188428 | n.s. |
| ENSG00000132535 | 94.13999875 | 0.719719029 | 0.2382072 | 3.021399 | 0.002516095 | 0.01227026 | DLG4 | protein_coding | 55.08 | 1.375488309 | 1.375488309 | ENSG00000132535 | n.s. |
| ENSG00000145882 | 304.5647845 | -0.812451817 | 0.2869984 | -2.830859 | 0.004642324 | 0.020293587 | PCYOX1L | protein_coding | 50.16 | -1.375189405 | 1.375189405 | ENSG00000145882 | n.s. |
| ENSG00000001631 | 955.78959 | -0.745592821 | 0.2518762 | -2.960156 | 0.00307483 | 0.014419287 | KRIT1 | protein_coding | 36.48 | -1.372678293 | 1.372678293 | ENSG00000001631 | n.s. |
| ENSG00000258311 | 134.1870489 | -0.779791728 | 0.2698775 | -2.889428 | 0.00385943 | 0.017406818 | | protein_coding | 53.8 | -1.371872472 | 1.371872472 | ENSG00000258311 | n.s. |
| ENSG00000163104 | 2943.818865 | -0.789714277 | 0.275219 | -2.869403 | 0.00411247 | 0.018351898 | SMARCAD1 | protein_coding | 36.02 | -1.371195913 | 1.371195913 | ENSG00000163104 | n.s. |
| ENSG00000160392 | 153.0546283 | -0.76431769 | 0.2618868 | -2.918504 | 0.003517154 | 0.016145512 | C19orf47 | protein_coding | 50.5 | -1.369617701 | 1.369617701 | ENSG00000160392 | n.s. |
| ENSG00000138623 | 106.2915142 | 0.742976409 | 0.2508403 | 2.96195 | 0.00305697 | 0.014345196 | SEMA7A | protein_coding | 57.93 | 1.369523586 | 1.369523586 | ENSG00000138623 | n.s. |
| ENSG00000243609 | 55.78518824 | -0.79321525 | 0.2774324 | -2.85913 | 0.004248044 | 0.018842154 | | processed_pseudogene | 55.79 | -1.368192755 | 1.368192755 | ENSG00000243609 | n.s. |
| ENSG00000118482 | 4465.041644 | -0.752986868 | 0.2563906 | -2.936874 | 0.003315393 | 0.015365212 | PHF3 | protein_coding | 36.07 | -1.365512655 | 1.365512655 | ENSG00000118482 | n.s. |
| ENSG00000212384 | 8.352661296 | -1.132967982 | 0.4814225 | -2.353375 | 0.018603837 | 0.062884491 | SNORD113-2 | snoRNA | 31.94 | -1.361211688 | 1.361211688 | ENSG00000212384 | down |
| ENSG00000160336 | 391.082352 | -0.782894404 | 0.2727937 | -2.869913 | 0.004105841 | 0.01832819 | ZNF761 | protein_coding | 44.23 | -1.35979396 | 1.35979396 | ENSG00000160336 | n.s. |
| ENSG00000106780 | 566.3623016 | -0.741335188 | 0.2509431 | -2.954197 | 0.003134841 | 0.014651354 | MEGF9 | protein_coding | 38.18 | -1.359699344 | 1.359699344 | ENSG00000106780 | n.s. |
| ENSG00000040633 | 471.8219443 | -0.804600238 | 0.284496 | -2.82816 | 0.004681645 | 0.020426997 | PHF23 | protein_coding | 59.51 | -1.359609843 | 1.359609843 | ENSG00000040633 | n.s. |
| ENSG00000196659 | 294.3274155 | 0.654740846 | 0.2073548 | 3.157588 | 0.001590804 | 0.008385381 | TTC30B | protein_coding | 43.32 | 1.359554443 | 1.359554443 | ENSG00000196659 | n.s. |

| | | | | | | | | | | | | | |
|---|---|---|---|---|---|---|---|---|---|---|---|---|---|
| ENSG00000065413 | 9.278558984 | -1.354627138 | 0.6360368 | -2.129794 | 0.033188654 | 0.099181291 | ANKRD44 | protein_coding | 40.24 | -1.359463485 | 1.359463485 | ENSG00000065413 | down |
| ENSG00000146555 | 301.4633398 | -0.714916725 | 0.2373043 | -3.012658 | 0.002589706 | 0.012545499 | SDK1 | protein_coding | 43.12 | -1.359422765 | 1.359422765 | ENSG00000146555 | n.s. |
| ENSG00000164114 | 615.2195487 | -0.798922424 | 0.2814959 | -2.838132 | 0.004537847 | 0.019893088 | MAP9 | protein_coding | 33.75 | -1.359204956 | 1.359204956 | ENSG00000164114 | n.s. |
| ENSG00000147862 | 9121.783917 | -0.722905962 | 0.2416123 | -2.992009 | 0.002771485 | 0.013187761 | NFIB | protein_coding | 38.29 | -1.358939534 | 1.358939534 | ENSG00000147862 | n.s. |
| ENSG00000196455 | 1152.512515 | -0.756162292 | 0.2587587 | -2.922268 | 0.00347492 | 0.015983257 | PIK3R4 | protein_coding | 36.67 | -1.358320584 | 1.358320584 | ENSG00000196455 | n.s. |
| ENSG00000155287 | 225.5921123 | -0.790467121 | 0.2772645 | -2.85095 | 0.004358881 | 0.019241818 | SLC25A28 | protein_coding | 45.02 | -1.356247043 | 1.356247043 | ENSG00000155287 | n.s. |
| ENSG00000113812 | 841.8859614 | -0.759611564 | 0.2608419 | -2.912153 | 0.003589469 | 0.016391004 | ACTR8 | protein_coding | 42.11 | -1.356206258 | 1.356206258 | ENSG00000113812 | n.s. |
| ENSG00000005436 | 1353.279788 | -0.779186616 | 0.2711964 | -2.873145 | 0.004064071 | 0.018188383 | GCFC2 | protein_coding | 36.39 | -1.355945162 | 1.355945162 | ENSG00000005436 | n.s. |
| ENSG00000270988 | 54.317867 | 0.795117672 | 0.2799583 | 2.840129 | 0.004509527 | 0.019806356 | | lincRNA | 46.28 | 1.354240785 | 1.354240785 | ENSG00000270988 | n.s. |
| ENSG00000141646 | 3116.696664 | -0.762879137 | 0.2628318 | -2.902538 | 0.003701525 | 0.016781612 | SMAD4 | protein_coding | 37.94 | -1.354237358 | 1.354237358 | ENSG00000141646 | n.s. |
| ENSG00000197044 | 488.8337634 | -0.885268812 | 0.3305206 | -2.678408 | 0.007397315 | 0.029621171 | ZNF441 | protein_coding | 41.78 | -1.353042882 | 1.353042882 | ENSG00000197044 | n.s. |
| ENSG00000120451 | 991.8372271 | -0.716964846 | 0.2390879 | -2.99875 | 0.002710901 | 0.01297476 | SNX19 | protein_coding | 42.46 | -1.352841451 | 1.352841451 | ENSG00000120451 | n.s. |
| ENSG00000115107 | 195.8312935 | -0.827242996 | 0.2975363 | -2.780309 | 0.005430718 | 0.023165405 | STEAP3 | protein_coding | 51.67 | -1.352674738 | 1.352674738 | ENSG00000115107 | n.s. |
| ENSG00000009335 | 1615.463538 | -0.765205275 | 0.2642864 | -2.895364 | 0.003787194 | 0.017114283 | UBE3C | protein_coding | 42.12 | -1.35184324 | 1.35184324 | ENSG00000009335 | n.s. |
| ENSG00000277478 | 4.348549582 | 1.272710146 | 0.5799987 | 2.194333 | 0.028211491 | 0.087150785 | MIR6165 | miRNA | 66.67 | 1.348727465 | 1.348727465 | ENSG00000277478 | up |
| ENSG00000144357 | 2267.470312 | -0.77720542 | 0.2709775 | -2.868154 | 0.00412874 | 0.018406803 | UBR3 | protein_coding | 36.88 | -1.348468225 | 1.348468225 | ENSG00000144357 | n.s. |
| ENSG00000103319 | 527.4892789 | -0.728507858 | 0.245429 | -2.968303 | 0.002994487 | 0.014094738 | EEF2K | protein_coding | 47.84 | -1.348426507 | 1.348426507 | ENSG00000103319 | n.s. |
| ENSG00000253696 | 63.59255861 | 0.757527241 | 0.2606261 | 2.906567 | 0.003654188 | 0.016610273 | KBTBD11-OT1 | lincRNA | 54.8 | 1.348113081 | 1.348113081 | ENSG00000253696 | n.s. |
| ENSG00000259623 | 548.0957453 | 0.746373459 | 0.2547206 | 2.930165 | 0.003387821 | 0.015651449 | | sense_overlapping | 34.51 | 1.34753656 | 1.34753656 | ENSG00000259623 | n.s. |
| ENSG00000255423 | 276.7060673 | -0.819014029 | 0.2936414 | -2.789164 | 0.00528443 | 0.022631536 | EBLN2 | protein_coding | 41.75 | -1.347512295 | 1.347512295 | ENSG00000255423 | n.s. |
| ENSG00000004777 | 56.01080248 | 0.790659294 | 0.2782987 | 2.841046 | 0.004496583 | 0.019761969 | ARHGAP33 | protein_coding | 60.93 | 1.347417674 | 1.347417674 | ENSG00000004777 | n.s. |
| ENSG00000153956 | 7151.990028 | -0.778027666 | 0.2716578 | -2.863999 | 0.004183293 | 0.018608305 | CACNA2D1 | protein_coding | 34.98 | -1.346215975 | 1.346215975 | ENSG00000153956 | n.s. |
| ENSG00000070718 | 1204.698627 | -0.76986403 | 0.267364 | -2.87946 | 0.003983564 | 0.017920247 | AP3M2 | protein_coding | 41.4 | -1.344687633 | 1.344687633 | ENSG00000070718 | n.s. |
| ENSG00000212420 | 12.86527322 | -0.982319298 | 0.3890102 | -2.525176 | 0.011564039 | 0.042809389 | RNU6-1111P | snRNA | 39.25 | -1.344265621 | 1.344265621 | ENSG00000212420 | n.s. |
| ENSG00000181027 | 261.1412609 | -0.774719291 | 0.270126 | -2.867993 | 0.004130852 | 0.018410326 | FKRP | protein_coding | 51.13 | -1.344090338 | 1.344090338 | ENSG00000181027 | n.s. |
| ENSG00000198040 | 4843.551575 | -0.735249757 | 0.249589 | -2.945842 | 0.003220767 | 0.014987546 | ZNF84 | protein_coding | 39.16 | -1.341293686 | 1.341293686 | ENSG00000198040 | n.s. |
| ENSG00000162409 | 859.5919608 | -0.801104101 | 0.2847555 | -2.813305 | 0.004903505 | 0.021215508 | PRKAA2 | protein_coding | 36.59 | -1.340524793 | 1.340524793 | ENSG00000162409 | n.s. |
| ENSG00000164116 | 464.5338389 | -0.748197316 | 0.2564225 | -2.91783 | 0.003524764 | 0.016161494 | GUCY1A3 | protein_coding | 36.88 | -1.340409333 | 1.340409333 | ENSG00000164116 | n.s. |
| ENSG00000135045 | 353.8782933 | -0.754313716 | 0.2597634 | -2.90385 | 0.003686053 | 0.016727808 | C9orf40 | protein_coding | 42.71 | -1.340084304 | 1.340084304 | ENSG00000135045 | n.s. |
| ENSG00000135956 | 305.5780138 | -0.693819146 | 0.2284042 | -3.037681 | 0.002384061 | 0.011741521 | TMEM127 | protein_coding | 51.14 | -1.33926219 | 1.33926219 | ENSG00000135956 | n.s. |
| ENSG00000262633 | 67.96405848 | -0.813824924 | 0.2917634 | -2.789332 | 0.005281686 | 0.022626744 | | protein_coding | 45.82 | -1.339049578 | 1.339049578 | ENSG00000262633 | n.s. |
| ENSG00000279583 | 34.54417764 | 0.791478078 | 0.2796902 | 2.829838 | 0.004657153 | 0.020339253 | | TEC | 63.49 | 1.338915768 | 1.338915768 | ENSG00000279583 | n.s. |

| | | | | | | | | | | | | | |
|---|---|---|---|---|---|---|---|---|---|---|---|---|---|
| ENSG00000160716 | 876.1300119 | -0.692033887 | 0.2276227 | -3.040268 | 0.002363679 | 0.011665902 | CHRNB2 | protein_coding | 54.69 | -1.337758037 | 1.337758037 | ENSG00000160716 | n.s. |
| ENSG00000198841 | 551.9777941 | 0.683402187 | 0.2232443 | 3.06123 | 0.002204299 | 0.011043707 | KTI12 | protein_coding | 55.37 | 1.337339581 | 1.337339581 | ENSG00000198841 | n.s. |
| ENSG00000232640 | 16.72496069 | 0.967727629 | 0.3812913 | 2.538027 | 0.011147948 | 0.041511868 | | antisense | 47.25 | 1.337232862 | 1.337232862 | ENSG00000232640 | n.s. |
| ENSG00000122870 | 260.0175634 | -0.773203748 | 0.2701291 | -2.862349 | 0.004205134 | 0.018675622 | BICC1 | protein_coding | 37.96 | -1.336656595 | 1.336656595 | ENSG00000122870 | n.s. |
| ENSG00000175449 | 126.1309418 | 0.704334573 | 0.2341899 | 3.007536 | 0.002633751 | 0.012674586 | RFESD | protein_coding | 37.57 | 1.336169328 | 1.336169328 | ENSG00000175449 | n.s. |
| ENSG00000279561 | 244.2617163 | 0.704305808 | 0.2344651 | 3.003883 | 0.002665574 | 0.012801928 | | TEC | 42.28 | 1.333056939 | 1.333056939 | ENSG00000279561 | n.s. |
| ENSG00000250420 | 562.5147245 | -0.746279437 | 0.2562777 | -2.911995 | 0.003591284 | 0.016393919 | AACSP1 | transcribed_unprocessed_pseud | 48.81 | -1.332345525 | 1.332345525 | ENSG00000250420 | n.s. |
| ENSG00000090470 | 680.3877226 | -0.740813585 | 0.2533728 | -2.923808 | 0.003457774 | 0.015909647 | PDCD7 | protein_coding | 45.72 | -1.332234293 | 1.332234293 | ENSG00000090470 | n.s. |
| ENSG00000112365 | 1207.406904 | -0.700915817 | 0.2327653 | -3.011255 | 0.002601702 | 0.012577603 | ZBTB24 | protein_coding | 43.86 | -1.332021905 | 1.332021905 | ENSG00000112365 | n.s. |
| ENSG00000205269 | 729.35137 | -0.843634119 | 0.3096222 | -2.724721 | 0.006435586 | 0.026423669 | TMEM170B | protein_coding | 36.55 | -1.33126045 | 1.33126045 | ENSG00000205269 | n.s. |
| ENSG00000230119 | 11.51863643 | -0.995569028 | 0.3993123 | -2.493209 | 0.012659437 | 0.046093441 | | processed_pseudogene | 39.58 | -1.330439495 | 1.330439495 | ENSG00000230119 | n.s. |
| ENSG00000163960 | 1314.946463 | -0.709321073 | 0.2373999 | -2.987874 | 0.002809254 | 0.013326489 | UBXN7 | protein_coding | 42.28 | -1.33017865 | 1.33017865 | ENSG00000163960 | n.s. |
| ENSG00000116141 | 1265.227207 | -0.747661304 | 0.2572754 | -2.906074 | 0.003659952 | 0.016631041 | MARK1 | protein_coding | 37.1 | -1.330149705 | 1.330149705 | ENSG00000116141 | n.s. |
| ENSG00000163528 | 250.757037 | -0.830656572 | 0.302415 | -2.746744 | 0.006019016 | 0.025083139 | CHCHD4 | protein_coding | 48.18 | -1.329563963 | 1.329563963 | ENSG00000163528 | n.s. |
| ENSG00000151247 | 2129.908327 | -0.763498494 | 0.2656713 | -2.873846 | 0.004055065 | 0.018163947 | EIF4E | protein_coding | 39.19 | -1.329090374 | 1.329090374 | ENSG00000151247 | n.s. |
| ENSG00000167291 | 2258.996076 | -0.719823362 | 0.2429615 | -2.962705 | 0.003049487 | 0.014314903 | TBC1D16 | protein_coding | 52.48 | -1.327506591 | 1.327506591 | ENSG00000167291 | n.s. |
| ENSG00000103091 | 22.77016583 | -0.874243201 | 0.3275793 | -2.668799 | 0.007612308 | 0.030359831 | WDR59 | protein_coding | 44.73 | -1.326839475 | 1.326839475 | ENSG00000103091 | n.s. |
| ENSG00000113194 | 1430.326464 | -0.801553861 | 0.2865237 | -2.797513 | 0.005149763 | 0.022116025 | FAF2 | protein_coding | 42.96 | -1.326806434 | 1.326806434 | ENSG00000113194 | n.s. |
| ENSG00000155849 | 39.98878768 | -0.787929452 | 0.2791491 | -2.822612 | 0.004763424 | 0.020718887 | ELMO1 | protein_coding | 41.67 | -1.326584487 | 1.326584487 | ENSG00000155849 | n.s. |
| ENSG00000207231 | 5.71361409 | -1.174635223 | 0.5171006 | -2.27158 | 0.023111919 | 0.074436096 | | misc_RNA | 43.86 | -1.325242739 | 1.325242739 | ENSG00000207231 | down |
| ENSG00000243667 | 409.4055913 | -0.768776804 | 0.2693019 | -2.854703 | 0.00430771 | 0.019046125 | WDR92 | protein_coding | 39.84 | -1.322444761 | 1.322444761 | ENSG00000243667 | n.s. |
| ENSG00000266262 | 4.767531331 | -1.148481329 | 0.499676 | -2.298452 | 0.021536061 | 0.070679478 | MIR4428 | miRNA | 58.9 | -1.321565121 | 1.321565121 | ENSG00000266262 | down |
| ENSG00000134318 | 2473.18661 | -0.73159719 | 0.249618 | -2.930867 | 0.003380177 | 0.015624903 | ROCK2 | protein_coding | 36.75 | -1.321398171 | 1.321398171 | ENSG00000134318 | n.s. |
| ENSG00000159640 | 77.26606874 | 0.759319269 | 0.2643783 | 2.872093 | 0.004077623 | 0.01822868 | ACE | protein_coding | 58.35 | 1.320642072 | 1.320642072 | ENSG00000159640 | n.s. |
| ENSG00000185760 | 1394.554362 | -0.747098966 | 0.2579806 | -2.895951 | 0.003780119 | 0.017087858 | KCNQ5 | protein_coding | 37.35 | -1.320357254 | 1.320357254 | ENSG00000185760 | n.s. |
| ENSG00000156384 | 288.1460811 | 0.681293048 | 0.2237935 | 3.044293 | 0.002332279 | 0.011539558 | SFR1 | protein_coding | 37.96 | 1.320217047 | 1.320217047 | ENSG00000156384 | n.s. |
| ENSG00000130522 | 670.5290981 | 0.693784059 | 0.2302025 | 3.013799 | 0.00257999 | 0.012507298 | JUND | protein_coding | 69.67 | 1.320157644 | 1.320157644 | ENSG00000130522 | n.s. |
| ENSG00000168297 | 369.2662694 | -0.739306517 | 0.2539218 | -2.911552 | 0.003596382 | 0.01641181 | PXK | protein_coding | 43.61 | -1.319546439 | 1.319546439 | ENSG00000168297 | n.s. |
| ENSG00000081138 | 816.4417963 | -0.79425397 | 0.2835596 | -2.801012 | 0.005094262 | 0.021904699 | CDH7 | protein_coding | 35.27 | -1.31803485 | 1.31803485 | ENSG00000081138 | n.s. |
| ENSG00000123636 | 2319.480195 | -0.784424755 | 0.2782404 | -2.819234 | 0.004813845 | 0.020899022 | BAZ2B | protein_coding | 36.16 | -1.317734788 | 1.317734788 | ENSG00000123636 | n.s. |
| ENSG00000138433 | 743.1325616 | -0.763807721 | 0.2671772 | -2.858806 | 0.004252387 | 0.018855408 | CIR1 | protein_coding | 37.69 | -1.317235347 | 1.317235347 | ENSG00000138433 | n.s. |
| ENSG00000275377 | 6.03180386 | -1.140382655 | 0.4951244 | -2.303224 | 0.021266217 | 0.069980983 | MIR1299 | miRNA | 50.6 | -1.31716473 | 1.31716473 | ENSG00000275377 | down |

| | | | | | | | | | | | | | |
|---|---|---|---|---|---|---|---|---|---|---|---|---|---|
| ENSG00000278223 | 4.633995764 | -1.205523763 | 0.5404266 | -2.230689 | 0.025701712 | 0.081088815 | MIR6783 | miRNA | 59.38 | -1.315273498 | 1.315273498 | ENSG00000278223 | down |
| ENSG00000257135 | 60.43327192 | 0.753510692 | 0.2622573 | 2.873174 | 0.004063708 | 0.018188383 | | lincRNA | 54.18 | 1.311263767 | 1.311263767 | ENSG00000257135 | n.s. |
| ENSG00000162624 | 55.33422519 | 0.788113038 | 0.2811233 | 2.803443 | 0.005056017 | 0.021746963 | LHX8 | protein_coding | 39.78 | 1.310317823 | 1.310317823 | ENSG00000162624 | n.s. |
| ENSG00000243926 | 25.2288065 | 0.880526455 | 0.3337357 | 2.638394 | 0.008329962 | 0.032669877 | TIPARP-AS1 | antisense | 43.54 | 1.308332434 | 1.308332434 | ENSG00000243926 | n.s. |
| ENSG00000143340 | 585.0950814 | -0.76523851 | 0.2690157 | -2.844587 | 0.004446904 | 0.019591687 | FAM163A | protein_coding | 47.32 | -1.306972402 | 1.306972402 | ENSG00000143340 | n.s. |
| ENSG00000110200 | 589.8348596 | 0.658282456 | 0.2133863 | 3.084934 | 0.002035976 | 0.010349357 | ANAPC15 | protein_coding | 52.32 | 1.306747701 | 1.306747701 | ENSG00000110200 | n.s. |
| ENSG00000062370 | 615.392397 | -0.766401796 | 0.2697043 | -2.841638 | 0.004488248 | 0.019737801 | ZNF112 | protein_coding | 38.31 | -1.306486091 | 1.306486091 | ENSG00000062370 | n.s. |
| ENSG00000264812 | 19.50525732 | 0.893603616 | 0.3421218 | 2.611946 | 0.009002859 | 0.034789566 | | sense_intronic | 50.59 | 1.30336644 | 1.30336644 | ENSG00000264812 | n.s. |
| ENSG00000160213 | 267.0982638 | 0.709422888 | 0.2403448 | 2.951688 | 0.003160423 | 0.014751115 | CSTB | protein_coding | 54.21 | 1.299077564 | 1.299077564 | ENSG00000160213 | n.s. |
| ENSG00000143079 | 727.8382448 | -0.710624484 | 0.2409539 | -2.949213 | 0.003185843 | 0.014862717 | CTTNBP2NL | protein_coding | 39.63 | -1.298951766 | 1.298951766 | ENSG00000143079 | n.s. |
| ENSG00000253320 | 142.336905 | 0.680677712 | 0.2255108 | 3.018381 | 0.00254129 | 0.012367137 | AZIN1-AS1 | antisense | 44.04 | 1.298549843 | 1.298549843 | ENSG00000253320 | n.s. |
| ENSG00000251550 | 122.9300426 | -0.728637319 | 0.2504643 | -2.909147 | 0.003624165 | 0.016516941 | | processed_pseudogene | 38.31 | -1.29848298 | 1.29848298 | ENSG00000251550 | n.s. |
| ENSG00000044459 | 1173.498941 | -0.796681206 | 0.2871299 | -2.774637 | 0.005526335 | 0.023465434 | CNTLN | protein_coding | 36.43 | -1.29824892 | 1.29824892 | ENSG00000044459 | n.s. |
| ENSG00000207971 | 34.19433116 | 0.813424785 | 0.2966672 | 2.741876 | 0.006108938 | 0.025359196 | MIR125B1 | miRNA | 53.41 | 1.298115749 | 1.298115749 | ENSG00000207971 | n.s. |
| ENSG00000272215 | 8.452778436 | -1.067615237 | 0.4508927 | -2.367781 | 0.017895118 | 0.060851002 | | snRNA | 46.03 | -1.297934294 | 1.297934294 | ENSG00000272215 | down |
| ENSG00000143515 | 789.0237118 | -0.749173837 | 0.2614219 | -2.865766 | 0.004160018 | 0.018516603 | ATP8B2 | protein_coding | 52.37 | -1.297897733 | 1.297897733 | ENSG00000143515 | n.s. |
| ENSG00000133316 | 1948.678458 | 0.638888791 | 0.2046682 | 3.121583 | 0.001798812 | 0.009317286 | WDR74 | protein_coding | 51.65 | 1.29739822 | 1.29739822 | ENSG00000133316 | n.s. |
| ENSG00000221230 | 3.56122469 | -1.316006683 | 0.6243889 | -2.107672 | 0.035059405 | 0.103395911 | MIR548L | miRNA | 39.53 | -1.296920159 | 1.296920159 | ENSG00000221230 | n.s. |
| ENSG00000108587 | 975.341746 | -0.694269792 | 0.2330159 | -2.979495 | 0.002887242 | 0.013640674 | GOSR1 | protein_coding | 39.72 | -1.294927146 | 1.294927146 | ENSG00000108587 | n.s. |
| ENSG00000223960 | 425.5805575 | -0.702025314 | 0.2370263 | -2.961804 | 0.003058427 | 0.014347198 | | antisense | 39.56 | -1.29399617 | 1.29399617 | ENSG00000223960 | n.s. |
| ENSG00000168172 | 984.9059843 | -0.750654506 | 0.2627392 | -2.857033 | 0.004276218 | 0.018936954 | HOOK3 | protein_coding | 40.46 | -1.293144925 | 1.293144925 | ENSG00000168172 | n.s. |
| ENSG00000134343 | 675.1586005 | -0.813115676 | 0.2973762 | -2.7343 | 0.006251309 | 0.025811676 | ANO3 | protein_coding | 36.36 | -1.291377136 | 1.291377136 | ENSG00000134343 | n.s. |
| ENSG00000177182 | 92.22447048 | -0.74216081 | 0.2583472 | -2.872726 | 0.004069468 | 0.018200837 | CLVS1 | protein_coding | 39.85 | -1.291292003 | 1.291292003 | ENSG00000177182 | n.s. |
| ENSG00000237732 | 243.4290328 | 0.663762817 | 0.2175202 | 3.051499 | 0.002277017 | 0.011346781 | | transcribed_unprocessed_pseud | 39.28 | 1.291103189 | 1.291103189 | ENSG00000237732 | n.s. |
| ENSG00000135336 | 1366.93426 | -0.81382621 | 0.2978598 | -2.732246 | 0.006290423 | 0.025919373 | ORC3 | protein_coding | 38.95 | -1.29103397 | 1.29103397 | ENSG00000135336 | n.s. |
| ENSG00000270095 | 28.69049074 | 0.835598604 | 0.3104459 | 2.691607 | 0.007110859 | 0.028730263 | | sense_intronic | 35.27 | 1.288209279 | 1.288209279 | ENSG00000270095 | n.s. |
| ENSG00000114993 | 214.8318933 | -0.743737633 | 0.2598025 | -2.862704 | 0.004200425 | 0.018660664 | RTKN | protein_coding | 53.87 | -1.285976583 | 1.285976583 | ENSG00000114993 | n.s. |
| ENSG00000136448 | 2838.259058 | -0.744039204 | 0.2603157 | -2.858219 | 0.004260263 | 0.018878307 | NMT1 | protein_coding | 48.75 | -1.282751079 | 1.282751079 | ENSG00000136448 | n.s. |
| ENSG00000162769 | 1326.494849 | -0.739842093 | 0.2580578 | -2.866963 | 0.004144314 | 0.018459881 | FLVCR1 | protein_coding | 40.27 | -1.282716849 | 1.282716849 | ENSG00000162769 | n.s. |
| ENSG00000251648 | 33.83209485 | -0.845318109 | 0.3170405 | -2.666278 | 0.007669624 | 0.030509765 | | processed_pseudogene | 51.61 | -1.281131276 | 1.281131276 | ENSG00000251648 | n.s. |
| ENSG00000273344 | 292.9872344 | 0.66854021 | 0.2209741 | 3.025423 | 0.002482858 | 0.012133673 | PAXIP1-AS1 | lincRNA | 48.32 | 1.280928207 | 1.280928207 | ENSG00000273344 | n.s. |
| ENSG00000109572 | 1016.282967 | -0.752044024 | 0.2648176 | -2.839857 | 0.004513377 | 0.019810767 | CLCN3 | protein_coding | 37.1 | -1.280805214 | 1.280805214 | ENSG00000109572 | n.s. |

| | | | | | | | | | | | | | |
|---|---|---|---|---|---|---|---|---|---|---|---|---|---|
| ENSG00000162923 | 2287.193351 | -0.713863983 | 0.2445429 | -2.919176 | 0.003509575 | 0.016121348 | WDR26 | protein_coding | 38.39 | -1.279671608 | 1.279671608 | ENSG00000162923 | n.s. |
| ENSG00000252782 | 50.01767224 | 0.800770346 | 0.2920259 | 2.742121 | 0.006104392 | 0.025359196 | RNU6-341P | snRNA | 43.75 | 1.277920979 | 1.277920979 | ENSG00000252782 | n.s. |
| ENSG00000070882 | 283.9849836 | -0.713282136 | 0.2444595 | -2.917793 | 0.003525181 | 0.016161494 | OSBPL3 | protein_coding | 40.72 | -1.277858141 | 1.277858141 | ENSG00000070882 | n.s. |
| ENSG00000176407 | 1195.804824 | -0.755671713 | 0.2671594 | -2.828542 | 0.004676052 | 0.020408989 | KCMF1 | protein_coding | 41.72 | -1.277220084 | 1.277220084 | ENSG00000176407 | n.s. |
| ENSG00000140382 | 1524.677797 | -0.757466086 | 0.2683766 | -2.8224 | 0.004766566 | 0.020726079 | HMG20A | protein_coding | 38.22 | -1.275181171 | 1.275181171 | ENSG00000140382 | n.s. |
| ENSG00000281508 | 268.57521 | -0.670120625 | 0.2223585 | -3.013695 | 0.002580871 | 0.012507298 | CDR1 | antisense | 44.31 | -1.275129997 | 1.275129997 | ENSG00000281508 | n.s. |
| ENSG00000237004 | 29.0967644 | 0.830714711 | 0.3094504 | 2.684485 | 0.007264172 | 0.029205623 | ZNRF2P1 | processed_pseudogene | 75.62 | 1.274759569 | 1.274759569 | ENSG00000237004 | n.s. |
| ENSG00000119938 | 202.4874875 | 0.699407049 | 0.2377293 | 2.942032 | 0.003260664 | 0.015137788 | PPP1R3C | protein_coding | 40.41 | 1.272877178 | 1.272877178 | ENSG00000119938 | n.s. |
| ENSG00000246985 | 114.6499862 | -0.78986404 | 0.2865185 | -2.756765 | 0.005837635 | 0.024518068 | SOCS2-AS1 | processed_transcript | 42.92 | -1.272086896 | 1.272086896 | ENSG00000246985 | n.s. |
| ENSG00000140992 | 472.8394394 | -0.757388322 | 0.268856 | -2.817078 | 0.004846281 | 0.021007088 | PDPK1 | protein_coding | 52.51 | -1.270620511 | 1.270620511 | ENSG00000140992 | n.s. |
| ENSG00000082482 | 253.8434942 | -0.720106432 | 0.2489178 | -2.892949 | 0.00381643 | 0.017235209 | KCNK2 | protein_coding | 36.37 | -1.269967788 | 1.269967788 | ENSG00000082482 | n.s. |
| ENSG00000134077 | 1033.278359 | -0.754274024 | 0.2672664 | -2.82218 | 0.004769837 | 0.020733824 | THUMPD3 | protein_coding | 40.09 | -1.269684996 | 1.269684996 | ENSG00000134077 | n.s. |
| ENSG00000144229 | 4338.220358 | -0.7598373 | 0.2704338 | -2.809698 | 0.004958806 | 0.021401568 | THSD7B | protein_coding | 37.67 | -1.268589713 | 1.268589713 | ENSG00000144229 | n.s. |
| ENSG00000151093 | 213.6117046 | -0.665139641 | 0.2204156 | -3.017662 | 0.002547328 | 0.012387864 | OXSM | protein_coding | 42 | -1.268423675 | 1.268423675 | ENSG00000151093 | n.s. |
| ENSG00000175104 | 533.0909597 | -0.776844851 | 0.2797297 | -2.777126 | 0.005484183 | 0.023322943 | TRAF6 | protein_coding | 38.03 | -1.267979094 | 1.267979094 | ENSG00000175104 | n.s. |
| ENSG00000172113 | 252.3118601 | -0.707649654 | 0.2425087 | -2.918038 | 0.003522414 | 0.016161494 | NME6 | protein_coding | 47.93 | -1.267767446 | 1.267767446 | ENSG00000172113 | n.s. |
| ENSG00000174529 | 73.18624548 | -0.776455923 | 0.2796204 | -2.776821 | 0.005489336 | 0.023336803 | TMEM81 | protein_coding | 47.52 | -1.267143955 | 1.267143955 | ENSG00000174529 | n.s. |
| ENSG00000125703 | 437.7996211 | -0.795304364 | 0.2903083 | -2.739517 | 0.006152959 | 0.025511511 | ATG4C | protein_coding | 35.44 | -1.267129666 | 1.267129666 | ENSG00000125703 | n.s. |
| ENSG00000196361 | 927.4140046 | -0.707409927 | 0.2426278 | -2.915617 | 0.003549856 | 0.016236744 | ELAVL3 | protein_coding | 53.1 | -1.265910804 | 1.265910804 | ENSG00000196361 | n.s. |
| ENSG00000132953 | 1863.07604 | -0.725461446 | 0.2521981 | -2.876554 | 0.00402044 | 0.018039505 | XPO4 | protein_coding | 38.5 | -1.26504181 | 1.26504181 | ENSG00000132953 | n.s. |
| ENSG00000079156 | 73.60840771 | -0.727153351 | 0.253186 | -2.872012 | 0.00407867 | 0.01822868 | OSBPL6 | protein_coding | 39.34 | -1.264697667 | 1.264697667 | ENSG00000079156 | n.s. |
| ENSG00000207137 | 14.27870777 | -1.000365581 | 0.4138481 | -2.417229 | 0.015639186 | 0.05442249 | SNORD116-13 | snoRNA | 42.39 | -1.264683765 | 1.264683765 | ENSG00000207137 | down |
| ENSG00000197885 | 342.0901985 | -0.724253542 | 0.251586 | -2.878752 | 0.003992523 | 0.01794895 | NKIRAS1 | protein_coding | 42.91 | -1.264518406 | 1.264518406 | ENSG00000197885 | n.s. |
| ENSG00000276653 | 8.389379312 | -1.009570723 | 0.4197683 | -2.405067 | 0.016169519 | 0.055918582 | RN7SL856P | misc_RNA | 52.68 | -1.264430647 | 1.264430647 | ENSG00000276653 | down |
| ENSG00000238123 | 20.92633023 | -0.861728938 | 0.328859 | -2.62036 | 0.008783698 | 0.034103576 | MID1IP1-AS1 | antisense | 46.07 | -1.264328767 | 1.264328767 | ENSG00000238123 | n.s. |
| ENSG00000199575 | 14.64552844 | -1.385838405 | 0.6866401 | -2.018289 | 0.04356115 | 0.122426703 | SNORD114-1 | snoRNA | 48.61 | -1.264056257 | 1.264056257 | ENSG00000199575 | n.s. |
| ENSG00000168175 | 2155.136932 | -0.742134222 | 0.2613442 | -2.839681 | 0.004515869 | 0.019815456 | MAPK1IP1L | protein_coding | 40.17 | -1.263851562 | 1.263851562 | ENSG00000168175 | n.s. |
| ENSG00000139990 | 624.6755865 | -0.715923806 | 0.2474325 | -2.89341 | 0.00381083 | 0.017215506 | DCAF5 | protein_coding | 40.67 | -1.262947018 | 1.262947018 | ENSG00000139990 | n.s. |
| ENSG00000188878 | 62.4721391 | 0.774825446 | 0.2793735 | 2.773439 | 0.005546718 | 0.023537627 | FBF1 | protein_coding | 53.45 | 1.261599707 | 1.261599707 | ENSG00000188878 | n.s. |
| ENSG00000221083 | 3.32078375 | -1.307255797 | 0.6273089 | -2.083911 | 0.03716829 | 0.10839843 | | snoRNA | 55.2 | -1.261471787 | 1.261471787 | ENSG00000221083 | n.s. |
| ENSG00000222932 | 3.304635556 | 1.293197131 | 0.6172368 | 2.095139 | 0.036158611 | 0.105986599 | RNU6-172P | snRNA | 39.78 | 1.260542667 | 1.260542667 | ENSG00000222932 | n.s. |
| ENSG00000175066 | 761.8385582 | -0.750228493 | 0.2662657 | -2.817593 | 0.004838507 | 0.02098646 | GK5 | protein_coding | 38.79 | -1.258929038 | 1.258929038 | ENSG00000175066 | n.s. |

| | | | | | | | | | | | | | |
|---|---|---|---|---|---|---|---|---|---|---|---|---|---|
| ENSG00000150977 | 157.7524655 | 0.674939211 | 0.2265167 | 2.979644 | 0.002885831 | 0.013638639 | RILPL2 | protein_coding | 49.36 | 1.258916158 | 1.258916158 | ENSG00000150977 | n.s. |
| ENSG00000029639 | 388.1247543 | -0.705216092 | 0.2424721 | -2.908442 | 0.003632343 | 0.016543379 | TFB1M | protein_coding | 38.35 | -1.25625487 | 1.25625487 | ENSG00000029639 | n.s. |
| ENSG00000204681 | 21.99738659 | -0.941358942 | 0.3779149 | -2.490928 | 0.012740977 | 0.046304521 | GABBR1 | protein_coding | 45.01 | -1.256127349 | 1.256127349 | ENSG00000204681 | n.s. |
| ENSG00000274750 | 763.1373767 | 0.660302223 | 0.219234 | 3.011861 | 0.002596517 | 0.012565626 | HIST1H3E | protein_coding | 42.63 | 1.255112934 | 1.255112934 | ENSG00000274750 | n.s. |
| ENSG00000269958 | 107.3627681 | -0.74632439 | 0.2646291 | -2.820266 | 0.004798383 | 0.020844895 | | sense_intronic | 49.08 | -1.254571521 | 1.254571521 | ENSG00000269958 | n.s. |
| ENSG00000143355 | 773.6970215 | -0.704637666 | 0.24238 | -2.907161 | 0.003647254 | 0.016600431 | LHX9 | protein_coding | 43.13 | -1.254170938 | 1.254170938 | ENSG00000143355 | n.s. |
| ENSG00000178031 | 740.4397552 | -0.727308262 | 0.2544371 | -2.8585 | 0.004256495 | 0.018867615 | ADAMTSL1 | protein_coding | 38.59 | -1.254085266 | 1.254085266 | ENSG00000178031 | n.s. |
| ENSG00000174871 | 116.0855527 | -0.746143129 | 0.2647544 | -2.818246 | 0.004828678 | 0.020950354 | CNIH2 | protein_coding | 61.39 | -1.252631521 | 1.252631521 | ENSG00000174871 | n.s. |
| ENSG00000111540 | 528.7449153 | -0.772937387 | 0.2794991 | -2.765437 | 0.005684652 | 0.024004879 | RAB5B | protein_coding | 46.01 | -1.251927063 | 1.251927063 | ENSG00000111540 | n.s. |
| ENSG00000206734 | 8.007411792 | -1.006630742 | 0.4201574 | -2.395842 | 0.016582247 | 0.057104779 | | misc_RNA | 43.14 | -1.251571732 | 1.251571732 | ENSG00000206734 | down |
| ENSG00000264349 | 41.4422426 | 0.773549128 | 0.2799581 | 2.763089 | 0.005725721 | 0.02412809 | MIR4258 | miRNA | 68.13 | 1.25119797 | 1.25119797 | ENSG00000264349 | n.s. |
| ENSG00000214401 | 45.73692031 | 0.777354612 | 0.2821408 | 2.755201 | 0.005865601 | 0.02461325 | KANSL1-AS1 | antisense | 45.58 | 1.25063222 | 1.25063222 | ENSG00000214401 | n.s. |
| ENSG00000149256 | 435.0787177 | -0.651898337 | 0.2153171 | -3.02762 | 0.002464882 | 0.012066999 | TENM4 | protein_coding | 44.45 | -1.250602236 | 1.250602236 | ENSG00000149256 | n.s. |
| ENSG00000153207 | 4024.949269 | -0.73819647 | 0.2607841 | -2.830681 | 0.004644908 | 0.020298509 | AHCTF1 | protein_coding | 38.75 | -1.249424003 | 1.249424003 | ENSG00000153207 | n.s. |
| ENSG00000118420 | 361.8758556 | -0.725684221 | 0.254382 | -2.852734 | 0.004334485 | 0.019146267 | UBE3D | protein_coding | 38.49 | -1.246664453 | 1.246664453 | ENSG00000118420 | n.s. |
| ENSG00000121486 | 392.0228553 | -0.720749747 | 0.2517326 | -2.863156 | 0.004194435 | 0.018646237 | TRMT1L | protein_coding | 35.55 | -1.246470953 | 1.246470953 | ENSG00000121486 | n.s. |
| ENSG00000116473 | 840.2461297 | -0.753371695 | 0.2694211 | -2.796261 | 0.005169765 | 0.02219508 | RAP1A | protein_coding | 39.31 | -1.245883388 | 1.245883388 | ENSG00000116473 | n.s. |
| ENSG00000212526 | 4.617296533 | -1.166922074 | 0.5295148 | -2.203757 | 0.027541427 | 0.08563182 | RNU6-466P | snRNA | 39.81 | -1.245531578 | 1.245531578 | ENSG00000212526 | down |
| ENSG00000258913 | 44.6988611 | -0.765925491 | 0.276357 | -2.771507 | 0.005579745 | 0.023656149 | | lincRNA | 53.02 | -1.245437703 | 1.245437703 | ENSG00000258913 | n.s. |
| ENSG00000157106 | 3415.712098 | -0.703756429 | 0.2430117 | -2.895977 | 0.003779801 | 0.017087858 | SMG1 | protein_coding | 39.95 | -1.243757452 | 1.243757452 | ENSG00000157106 | n.s. |
| ENSG00000175470 | 462.0556617 | -0.776767349 | 0.2828166 | -2.746541 | 0.00602273 | 0.025091105 | PPP2R2D | protein_coding | 48.11 | -1.243200768 | 1.243200768 | ENSG00000175470 | n.s. |
| ENSG00000173065 | 42.77231818 | -0.776231187 | 0.2824953 | -2.747766 | 0.006000276 | 0.025027512 | FAM222B | protein_coding | 45.72 | -1.243198141 | 1.243198141 | ENSG00000173065 | n.s. |
| ENSG00000137145 | 850.558847 | -0.714420926 | 0.2487679 | -2.871837 | 0.004080931 | 0.01822868 | DENND4C | protein_coding | 38.75 | -1.242552863 | 1.242552863 | ENSG00000137145 | n.s. |
| ENSG00000266668 | 12.15645054 | -0.993916743 | 0.413742 | -2.402262 | 0.01629404 | 0.056279315 | MIR5692C2 | miRNA | 32.47 | -1.242049246 | 1.242049246 | ENSG00000266668 | n.s. |
| ENSG00000269946 | 44.4095957 | 0.798683711 | 0.2954157 | 2.703593 | 0.006859429 | 0.02790062 | | lincRNA | 40.89 | 1.241462891 | 1.241462891 | ENSG00000269946 | n.s. |
| ENSG00000080345 | 8855.735412 | -0.709545136 | 0.2464947 | -2.878541 | 0.003995196 | 0.017955168 | RIF1 | protein_coding | 38.04 | -1.238731375 | 1.238731375 | ENSG00000080345 | n.s. |
| ENSG00000252658 | 13.05464844 | -0.971816923 | 0.4004488 | -2.426819 | 0.015231844 | 0.053271279 | RNU6-786P | snRNA | 36.67 | -1.23761553 | 1.23761553 | ENSG00000252658 | n.s. |
| ENSG00000125741 | 276.2776427 | -0.73758154 | 0.2619015 | -2.816255 | 0.004858704 | 0.021054382 | OPA3 | protein_coding | 49.78 | -1.236671615 | 1.236671615 | ENSG00000125741 | n.s. |
| ENSG00000152147 | 432.6875427 | -0.717952392 | 0.251358 | -2.856294 | 0.004286177 | 0.018964147 | GEMIN6 | protein_coding | 45.01 | -1.236361905 | 1.236361905 | ENSG00000152147 | n.s. |
| ENSG00000172915 | 2914.333939 | -0.770476977 | 0.2802443 | -2.749305 | 0.005972181 | 0.024962678 | NBEA | protein_coding | 35.01 | -1.234850245 | 1.234850245 | ENSG00000172915 | n.s. |
| ENSG00000196295 | 162.8654843 | -0.776737554 | 0.2839002 | -2.735953 | 0.00621999 | 0.025722883 | | processed_transcript | 42.17 | -1.234764432 | 1.234764432 | ENSG00000196295 | n.s. |
| ENSG00000228170 | 27.2156676 | 0.801815115 | 0.298156 | 2.689247 | 0.007161335 | 0.028915303 | | antisense | 48.47 | 1.233891032 | 1.233891032 | ENSG00000228170 | n.s. |

| | | | | | | | | | | | | | |
|---|---|---|---|---|---|---|---|---|---|---|---|---|---|
| ENSG00000116127 | 4445.29303 | -0.688499088 | 0.2359664 | -2.917785 | 0.003525279 | 0.016161494 | ALMS1 | protein_coding | 38.28 | -1.233458852 | 1.233458852 | ENSG00000116127 | n.s. |
| ENSG00000271971 | 37.64256296 | -0.864538509 | 0.3349896 | -2.580792 | 0.009857387 | 0.037477991 | | antisense | 31.39 | -1.233025313 | 1.233025313 | ENSG00000271971 | n.s. |
| ENSG00000141446 | 706.8746447 | -0.769387736 | 0.2798762 | -2.749029 | 0.005977205 | 0.024968684 | ESCO1 | protein_coding | 41.11 | -1.23302413 | 1.23302413 | ENSG00000141446 | n.s. |
| ENSG00000273893 | 55.83311252 | -0.764699749 | 0.2772344 | -2.758315 | 0.005810014 | 0.024416789 | | sense_intronic | 46 | -1.232934166 | 1.232934166 | ENSG00000273893 | n.s. |
| ENSG00000235652 | 63.97792353 | 0.711985207 | 0.2486719 | 2.863152 | 0.004194499 | 0.018646237 | | antisense | 36.99 | 1.23131348 | 1.23131348 | ENSG00000235652 | n.s. |
| ENSG00000154721 | 210.0606511 | -0.728527829 | 0.2576167 | -2.827952 | 0.004684679 | 0.020433832 | JAM2 | protein_coding | 39.65 | -1.230957185 | 1.230957185 | ENSG00000154721 | n.s. |
| ENSG00000242294 | 239.5692951 | 0.669733088 | 0.226634 | 2.955131 | 0.003125363 | 0.014616873 | STAG3L5P | transcribed_unprocessed_pseud | 47.29 | 1.229057683 | 1.229057683 | ENSG00000242294 | n.s. |
| ENSG00000200882 | 40.58458007 | -0.856991126 | 0.3312274 | -2.58732 | 0.009672585 | 0.036957027 | RNU6-681P | snRNA | 35.58 | -1.227470935 | 1.227470935 | ENSG00000200882 | n.s. |
| ENSG00000238966 | 48.86618911 | 0.786425488 | 0.2903697 | 2.708359 | 0.006761686 | 0.027543286 | | snoRNA | 36.11 | 1.226811378 | 1.226811378 | ENSG00000238966 | n.s. |
| ENSG00000007968 | 445.6350977 | -0.671665105 | 0.2280135 | -2.945725 | 0.003221983 | 0.014988198 | E2F2 | protein_coding | 51.66 | -1.225285461 | 1.225285461 | ENSG00000007968 | n.s. |
| ENSG00000207606 | 5.901691805 | 1.113909711 | 0.4974041 | 2.239446 | 0.025126902 | 0.079745122 | MIR554 | miRNA | 48.96 | 1.223402437 | 1.223402437 | ENSG00000207606 | up |
| ENSG00000131389 | 459.8065052 | -0.701765493 | 0.244062 | -2.875357 | 0.004035706 | 0.018096338 | SLC6A6 | protein_coding | 52.21 | -1.222762715 | 1.222762715 | ENSG00000131389 | n.s. |
| ENSG00000252615 | 9.21907582 | -1.062070101 | 0.4629405 | -2.294183 | 0.021780002 | 0.071351287 | | misc_RNA | 41 | -1.217767657 | 1.217767657 | ENSG00000252615 | down |
| ENSG00000279098 | 18.57479361 | -0.916422441 | 0.3687307 | -2.485344 | 0.01294264 | 0.046914963 | | TEC | 27.38 | -1.217640072 | 1.217640072 | ENSG00000279098 | n.s. |
| ENSG00000261765 | 34.30084399 | 0.783329602 | 0.2898558 | 2.70248 | 0.006882427 | 0.027977818 | | sense_intronic | 49.38 | 1.216656696 | 1.216656696 | ENSG00000261765 | n.s. |
| ENSG00000153391 | 269.8322517 | 0.668422576 | 0.2272954 | 2.940766 | 0.003274016 | 0.015194706 | INO80C | protein_coding | 42.29 | 1.215397917 | 1.215397917 | ENSG00000153391 | n.s. |
| ENSG00000264235 | 53.95869787 | 0.769747818 | 0.2824415 | 2.725335 | 0.006423622 | 0.026382327 | | antisense | 40.41 | 1.215190792 | 1.215190792 | ENSG00000264235 | n.s. |
| ENSG00000149115 | 844.964842 | -0.674725043 | 0.2306058 | -2.925881 | 0.003434822 | 0.015824958 | TNKS1BP1 | protein_coding | 55.55 | -1.214948665 | 1.214948665 | ENSG00000149115 | n.s. |
| ENSG00000270157 | 59.61673619 | -0.85128344 | 0.3300589 | -2.579187 | 0.009903327 | 0.037601315 | | sense_intronic | 42.92 | -1.212906058 | 1.212906058 | ENSG00000270157 | n.s. |
| ENSG00000170234 | 944.9101216 | -0.709999844 | 0.2496036 | -2.844509 | 0.004447987 | 0.019591687 | PWWP2A | protein_coding | 42.31 | -1.212628728 | 1.212628728 | ENSG00000170234 | n.s. |
| ENSG00000274322 | 17.76348787 | 0.898317949 | 0.358458 | 2.506062 | 0.012208413 | 0.044778117 | | protein_coding | 45.09 | 1.211771778 | 1.211771778 | ENSG00000274322 | n.s. |
| ENSG00000113356 | 197.6059987 | -0.773258758 | 0.2848821 | -2.714312 | 0.006641366 | 0.027132554 | POLR3G | protein_coding | 37.84 | -1.211317058 | 1.211317058 | ENSG00000113356 | n.s. |
| ENSG00000196417 | 406.1973652 | -0.73257439 | 0.262089 | -2.795136 | 0.005187784 | 0.022251855 | ZNF765 | protein_coding | 44.73 | -1.210677183 | 1.210677183 | ENSG00000196417 | n.s. |
| ENSG00000256981 | 296.1947558 | -0.718592854 | 0.2545298 | -2.823217 | 0.004754431 | 0.020686233 | | processed_pseudogene | 39.01 | -1.2103393 | 1.2103393 | ENSG00000256981 | n.s. |
| ENSG00000140470 | 215.1435427 | -0.675312938 | 0.231581 | -2.916098 | 0.003544392 | 0.016222411 | ADAMTS17 | protein_coding | 46.62 | -1.208732235 | 1.208732235 | ENSG00000140470 | n.s. |
| ENSG00000177051 | 144.6448971 | -0.725911435 | 0.2588503 | -2.804368 | 0.005041537 | 0.021698089 | FBXO46 | protein_coding | 51.45 | -1.207610665 | 1.207610665 | ENSG00000177051 | n.s. |
| ENSG00000163297 | 2513.365177 | -0.725608197 | 0.2587329 | -2.804468 | 0.005039965 | 0.021698032 | ANTXR2 | protein_coding | 35.81 | -1.207107028 | 1.207107028 | ENSG00000163297 | n.s. |
| ENSG00000182963 | 3083.434558 | -0.751726857 | 0.2732168 | -2.751393 | 0.005934244 | 0.024833927 | GJC1 | protein_coding | 44.78 | -1.206487474 | 1.206487474 | ENSG00000182963 | n.s. |
| ENSG00000266922 | 19.302089 | -0.864478054 | 0.3390425 | -2.549763 | 0.010779616 | 0.040377882 | | sense_intronic | 41.07 | -1.20495832 | 1.20495832 | ENSG00000266922 | n.s. |
| ENSG00000268543 | 21.20291532 | -0.861168357 | 0.3371378 | -2.554352 | 0.010638579 | 0.039946315 | | TEC | 47.14 | -1.204363989 | 1.204363989 | ENSG00000268543 | n.s. |
| ENSG00000076321 | 810.6981094 | -0.730572323 | 0.2618378 | -2.790171 | 0.005268019 | 0.022575139 | KLHL20 | protein_coding | 38.33 | -1.20279204 | 1.20279204 | ENSG00000076321 | n.s. |
| ENSG00000259682 | 26.90814338 | -0.85982454 | 0.3365768 | -2.554617 | 0.010630484 | 0.039926687 | | sense_intronic | 45.14 | -1.202668161 | 1.202668161 | ENSG00000259682 | n.s. |

| | | | | | | | | | | | | | |
|---|---|---|---|---|---|---|---|---|---|---|---|---|---|
| ENSG00000258325 | 46.23908773 | 0.759432056 | 0.2783012 | 2.728813 | 0.006356273 | 0.026167471 | ITFG2-AS1 | antisense | 47.69 | 1.201602448 | 1.201602448 | ENSG00000258325 | n.s. |
| ENSG00000188227 | 726.190083 | -0.741859202 | 0.2682693 | -2.765352 | 0.005686137 | 0.024004879 | ZNF793 | protein_coding | 42.72 | -1.201589711 | 1.201589711 | ENSG00000188227 | n.s. |
| ENSG00000172831 | 378.8207475 | -0.711518391 | 0.2516993 | -2.826859 | 0.004700706 | 0.020478076 | CES2 | protein_coding | 55.49 | -1.201548828 | 1.201548828 | ENSG00000172831 | n.s. |
| ENSG00000244044 | 39.61913588 | -0.828133 | 0.3179549 | -2.604561 | 0.009199206 | 0.035430302 | RN7SL735P | misc_RNA | 55.82 | -1.201310555 | 1.201310555 | ENSG00000244044 | n.s. |
| ENSG00000266640 | 9.397714978 | 0.99583809 | 0.4223147 | 2.358047 | 0.018371356 | 0.06218925 | MIR4754 | miRNA | 69.66 | 1.201264234 | 1.201264234 | ENSG00000266640 | n.s. |
| ENSG00000132205 | 260.477355 | -0.718574128 | 0.2557185 | -2.81002 | 0.004953845 | 0.021393417 | EMILIN2 | protein_coding | 47.46 | -1.199817477 | 1.199817477 | ENSG00000132205 | n.s. |
| ENSG00000052126 | 975.3424012 | -0.706479173 | 0.2492554 | -2.834358 | 0.00459178 | 0.020097878 | PLEKHA5 | protein_coding | 38.09 | -1.198789035 | 1.198789035 | ENSG00000052126 | n.s. |
| ENSG00000164080 | 1080.954982 | -0.652464316 | 0.2206634 | -2.956831 | 0.003108184 | 0.014556087 | RAD54L2 | protein_coding | 44.38 | -1.198547812 | 1.198547812 | ENSG00000164080 | n.s. |
| ENSG00000095739 | 1233.45729 | 0.663387329 | 0.2264018 | 2.930133 | 0.003388168 | 0.015651449 | BAMBI | protein_coding | 47.11 | 1.19770963 | 1.19770963 | ENSG00000095739 | n.s. |
| ENSG00000144824 | 2159.290458 | -0.764545655 | 0.2817704 | -2.713364 | 0.006660385 | 0.027186319 | PHLDB2 | protein_coding | 38.78 | -1.197010592 | 1.197010592 | ENSG00000144824 | n.s. |
| ENSG00000103671 | 295.4934999 | -0.788782237 | 0.2956682 | -2.667796 | 0.00763507 | 0.030407061 | TRIP4 | protein_coding | 42.11 | -1.196602811 | 1.196602811 | ENSG00000103671 | n.s. |
| ENSG00000269038 | 21.36955967 | -0.873405449 | 0.3457595 | -2.526049 | 0.011535342 | 0.042748621 | | lincRNA | 48.67 | -1.195760094 | 1.195760094 | ENSG00000269038 | n.s. |
| ENSG00000090615 | 1453.636966 | -0.694980295 | 0.2434297 | -2.854953 | 0.004304322 | 0.019037191 | GOLGA3 | protein_coding | 53.56 | -1.195642113 | 1.195642113 | ENSG00000090615 | n.s. |
| ENSG00000199279 | 13.66197616 | -0.963438919 | 0.4026806 | -2.392564 | 0.01673113 | 0.057546325 | RNU6-661P | snRNA | 39.25 | -1.194647311 | 1.194647311 | ENSG00000199279 | n.s. |
| ENSG00000101187 | 112.7155697 | -0.77648083 | 0.288785 | -2.688785 | 0.007171263 | 0.028940723 | SLCO4A1 | protein_coding | 58.22 | -1.19460848 | 1.19460848 | ENSG00000101187 | n.s. |
| ENSG00000222831 | 2.304952857 | -1.462544376 | 0.7705112 | -1.898148 | 0.057676563 | 0.152522466 | MIR1537 | miRNA | 40.98 | -1.19441053 | 1.19441053 | ENSG00000222831 | n.s. |
| ENSG00000273173 | 69.38387545 | 0.76162039 | 0.2804593 | 2.715619 | 0.006615209 | 0.027033623 | SNURF | protein_coding | 40.75 | 1.19429369 | 1.19429369 | ENSG00000273173 | n.s. |
| ENSG00000132330 | 127.493915 | -0.748879236 | 0.2732767 | -2.74037 | 0.006136999 | 0.025452916 | SCLY | protein_coding | 46.25 | -1.193910046 | 1.193910046 | ENSG00000132330 | n.s. |
| ENSG00000078124 | 496.72777 | -0.74207571 | 0.2694831 | -2.753701 | 0.005892563 | 0.024704053 | ACER3 | protein_coding | 38.74 | -1.192687674 | 1.192687674 | ENSG00000078124 | n.s. |
| ENSG00000101181 | 226.761839 | -0.764044733 | 0.2820466 | -2.70893 | 0.00675005 | 0.027511985 | MTG2 | protein_coding | 49.97 | -1.192275055 | 1.192275055 | ENSG00000101181 | n.s. |
| ENSG00000182010 | 727.7593445 | -0.780910978 | 0.2917693 | -2.676467 | 0.007440285 | 0.029759001 | RTKN2 | protein_coding | 35.12 | -1.191968184 | 1.191968184 | ENSG00000182010 | n.s. |
| ENSG00000078142 | 5802.834352 | -0.715833654 | 0.2551572 | -2.805462 | 0.005024458 | 0.021637961 | PIK3C3 | protein_coding | 35.33 | -1.191708181 | 1.191708181 | ENSG00000078142 | n.s. |
| ENSG00000273983 | 3307.320531 | 0.682544835 | 0.2371271 | 2.878393 | 0.003997073 | 0.01795781 | HIST1H3G | protein_coding | 42.68 | 1.191550342 | 1.191550342 | ENSG00000273983 | n.s. |
| ENSG00000164684 | 2415.98056 | -0.725424527 | 0.260297 | -2.786911 | 0.005321309 | 0.022774813 | ZNF704 | protein_coding | 40.26 | -1.191542559 | 1.191542559 | ENSG00000164684 | n.s. |
| ENSG00000165832 | 953.4007672 | -0.749725871 | 0.274095 | -2.735277 | 0.006232774 | 0.025763807 | TRUB1 | protein_coding | 36.89 | -1.191306888 | 1.191306888 | ENSG00000165832 | n.s. |
| ENSG00000055917 | 4159.020927 | -0.736145736 | 0.2665952 | -2.761287 | 0.00575741 | 0.024246953 | PUM2 | protein_coding | 37.42 | -1.18912774 | 1.18912774 | ENSG00000055917 | n.s. |
| ENSG00000228623 | 193.552713 | -0.789272952 | 0.2970172 | -2.657331 | 0.007876204 | 0.031205632 | ZNF883 | protein_coding | 38.73 | -1.18846118 | 1.18846118 | ENSG00000228623 | n.s. |
| ENSG00000168228 | 334.5421496 | -0.738753242 | 0.2681812 | -2.75468 | 0.005874959 | 0.024637669 | ZCCHC4 | protein_coding | 38.93 | -1.188211002 | 1.188211002 | ENSG00000168228 | n.s. |
| ENSG00000274471 | 113.9192062 | 0.740927199 | 0.2695652 | 2.748602 | 0.005985009 | 0.02498629 | | unprocessed_pseudogene | 45.2 | 1.187186329 | 1.187186329 | ENSG00000274471 | n.s. |
| ENSG00000154920 | 703.8318017 | -0.666136117 | 0.2289937 | -2.908971 | 0.003626201 | 0.016520809 | EME1 | protein_coding | 47.88 | -1.187033702 | 1.187033702 | ENSG00000154920 | n.s. |
| ENSG00000275054 | 7.626967378 | -1.60868585 | 0.8997849 | -1.787856 | 0.073799251 | 0.183051554 | | misc_RNA | 32.43 | -1.186287702 | 1.186287702 | ENSG00000275054 | n.s. |
| ENSG00000271895 | 33.1343645 | -0.816198065 | 0.3133499 | -2.60475 | 0.009194133 | 0.035420563 | | antisense | 42.99 | -1.18409489 | 1.18409489 | ENSG00000271895 | n.s. |

| | | | | | | | | | | | | | |
|---|---|---|---|---|---|---|---|---|---|---|---|---|---|
| ENSG00000177283 | 738.798999 | 0.623234092 | 0.207174 | 3.008264 | 0.002627449 | 0.012658121 | FZD8 | protein_coding | 64.57 | 1.182668175 | 1.182668175 | ENSG00000177283 | n.s. |
| ENSG00000265871 | 14.42695938 | 0.921127993 | 0.3777614 | 2.438385 | 0.014753034 | 0.052014814 | MIR3174 | miRNA | 45.98 | 1.182611316 | 1.182611316 | ENSG00000265871 | n.s. |
| ENSG00000104427 | 492.1827566 | -0.764955157 | 0.2838876 | -2.69457 | 0.00704795 | 0.028534054 | ZC2HC1A | protein_coding | 34.79 | -1.181577673 | 1.181577673 | ENSG00000104427 | n.s. |
| ENSG00000272518 | 62.36920681 | -0.824109154 | 0.3186896 | -2.58593 | 0.009711662 | 0.03709615 | | sense_intronic | 40.02 | -1.179029198 | 1.179029198 | ENSG00000272518 | n.s. |
| ENSG00000139734 | 1085.768624 | -0.722855367 | 0.2604132 | -2.775801 | 0.005506582 | 0.023396839 | DIAPH3 | protein_coding | 35.91 | -1.178863482 | 1.178863482 | ENSG00000139734 | n.s. |
| ENSG00000090432 | 160.1587922 | -0.76014164 | 0.2816035 | -2.699333 | 0.006947866 | 0.028186228 | MUL1 | protein_coding | 47.52 | -1.178191451 | 1.178191451 | ENSG00000090432 | n.s. |
| ENSG00000265140 | 3.862594228 | 1.28640892 | 0.6364174 | 2.021329 | 0.04324571 | 0.121687556 | MIR4301 | miRNA | 56.06 | 1.176747489 | 1.176747489 | ENSG00000265140 | n.s. |
| ENSG00000143398 | 1543.792298 | -0.747784096 | 0.2748087 | -2.721108 | 0.006506358 | 0.026698504 | PIP5K1A | protein_coding | 45.21 | -1.176648052 | 1.176648052 | ENSG00000143398 | n.s. |
| ENSG00000143416 | 174.5241965 | 0.791384426 | 0.2998866 | 2.638946 | 0.008316426 | 0.032644376 | SELENBP1 | protein_coding | 54.96 | 1.176148913 | 1.176148913 | ENSG00000143416 | n.s. |
| ENSG00000151692 | 1190.953854 | -0.736771823 | 0.2687046 | -2.74194 | 0.006107756 | 0.025359196 | RNF144A | protein_coding | 44.89 | -1.175788006 | 1.175788006 | ENSG00000151692 | n.s. |
| ENSG00000167555 | 1568.978141 | -0.728456434 | 0.2640305 | -2.758985 | 0.005798112 | 0.024374128 | ZNF528 | protein_coding | 43.66 | -1.175051889 | 1.175051889 | ENSG00000167555 | n.s. |
| ENSG00000138592 | 1964.321429 | -0.700152546 | 0.2485168 | -2.817325 | 0.004842549 | 0.020997448 | USP8 | protein_coding | 39.77 | -1.174739387 | 1.174739387 | ENSG00000138592 | n.s. |
| ENSG00000112210 | 650.3252478 | -0.714956558 | 0.2566753 | -2.785452 | 0.005345319 | 0.022857149 | RAB23 | protein_coding | 37.66 | -1.173227936 | 1.173227936 | ENSG00000112210 | n.s. |
| ENSG00000263004 | 37.87234816 | 0.79156795 | 0.300508 | 2.634099 | 0.008436081 | 0.032983868 | | lincRNA | 45.5 | 1.172864977 | 1.172864977 | ENSG00000263004 | n.s. |
| ENSG00000253739 | 15.93546245 | -0.916970367 | 0.3769812 | -2.432403 | 0.014998992 | 0.052698493 | RNU6-323P | snRNA | 42.06 | -1.172073176 | 1.172073176 | ENSG00000253739 | n.s. |
| ENSG00000200974 | 8.422266257 | 1.0506034 | 0.4653301 | 2.257759 | 0.023960673 | 0.076673054 | RNU4-87P | snRNA | 35.37 | 1.171798108 | 1.171798108 | ENSG00000200974 | up |
| ENSG00000091428 | 206.8503071 | -0.730300742 | 0.2655115 | -2.750542 | 0.005949668 | 0.024883516 | RAPGEF4 | protein_coding | 40.56 | -1.171466847 | 1.171466847 | ENSG00000091428 | n.s. |
| ENSG00000104299 | 311.1136734 | -0.737899677 | 0.2699557 | -2.73341 | 0.006268217 | 0.025858484 | INTS9 | protein_coding | 42.01 | -1.171339694 | 1.171339694 | ENSG00000104299 | n.s. |
| ENSG00000087586 | 739.6025922 | -0.755981774 | 0.2803052 | -2.696996 | 0.00699682 | 0.028360037 | AURKA | protein_coding | 42.93 | -1.169725462 | 1.169725462 | ENSG00000087586 | n.s. |
| ENSG00000083535 | 1178.511453 | -0.764792178 | 0.2854719 | -2.679046 | 0.007383231 | 0.02957328 | PIBF1 | protein_coding | 36.99 | -1.169444109 | 1.169444109 | ENSG00000083535 | n.s. |
| ENSG00000122482 | 5205.166548 | -0.723375997 | 0.2618866 | -2.762172 | 0.005741818 | 0.024188603 | ZNF644 | protein_coding | 36.39 | -1.169257158 | 1.169257158 | ENSG00000122482 | n.s. |
| ENSG00000134463 | 111.1610341 | -0.770123972 | 0.2885823 | -2.668646 | 0.00761577 | 0.030360932 | ECHDC3 | protein_coding | 47.51 | -1.168805524 | 1.168805524 | ENSG00000134463 | n.s. |
| ENSG00000173567 | 25.9847433 | 0.803930329 | 0.3083052 | 2.60758 | 0.009118489 | 0.035168068 | ADGRF3 | protein_coding | 45.62 | 1.168795306 | 1.168795306 | ENSG00000173567 | n.s. |
| ENSG00000280123 | 339.395639 | -0.714637029 | 0.2570719 | -2.779911 | 0.005437372 | 0.023186688 | | TEC | 30.55 | -1.168260943 | 1.168260943 | ENSG00000280123 | n.s. |
| ENSG00000256223 | 717.9667836 | -0.739512829 | 0.271346 | -2.72535 | 0.006423344 | 0.026382327 | ZNF10 | protein_coding | 40.03 | -1.167459211 | 1.167459211 | ENSG00000256223 | n.s. |
| ENSG00000255883 | 22.74221458 | -0.899368195 | 0.3665689 | -2.453477 | 0.014148269 | 0.050354385 | FUNDC2P1 | processed_pseudogene | 51.05 | -1.167346373 | 1.167346373 | ENSG00000255883 | n.s. |
| ENSG00000198355 | 585.8629938 | 0.738296929 | 0.2707367 | 2.726993 | 0.006391444 | 0.026281178 | PIM3 | protein_coding | 69.51 | 1.166771371 | 1.166771371 | ENSG00000198355 | n.s. |
| ENSG00000228175 | 60.88870741 | 0.733428484 | 0.2680342 | 2.736325 | 0.00621297 | 0.025714381 | GEMIN8P4 | processed_pseudogene | 53.77 | 1.166022151 | 1.166022151 | ENSG00000228175 | n.s. |
| ENSG00000166529 | 283.2618737 | -0.769341514 | 0.2885982 | -2.665787 | 0.007680828 | 0.030543291 | ZSCAN21 | protein_coding | 47.87 | -1.165617152 | 1.165617152 | ENSG00000166529 | n.s. |
| ENSG00000152422 | 182.1133855 | -0.767130158 | 0.2872647 | -2.670465 | 0.00757463 | 0.030244213 | XRCC4 | protein_coding | 35.2 | -1.165545121 | 1.165545121 | ENSG00000152422 | n.s. |
| ENSG00000222095 | 4.15956088 | -1.306697946 | 0.655193 | -1.994371 | 0.046111512 | 0.128248218 | | snoRNA | 47.3 | -1.165507483 | 1.165507483 | ENSG00000222095 | n.s. |
| ENSG00000119844 | 795.9401861 | -0.721509941 | 0.2613106 | -2.76112 | 0.005760358 | 0.024251456 | AFTPH | protein_coding | 37.29 | -1.165427726 | 1.165427726 | ENSG00000119844 | n.s. |

| | | | | | | | | | | | | | |
|---|---|---|---|---|---|---|---|---|---|---|---|---|---|
| ENSG00000005483 | 1648.446298 | -0.67934328 | 0.2383821 | -2.849809 | 0.004374553 | 0.01930488 | KMT2E | protein_coding | 37.46 | -1.164620535 | 1.164620535 | ENSG00000005483 | n.s. |
| ENSG00000260083 | 205.3433256 | 0.664655973 | 0.2304881 | 2.883689 | 0.00393047 | 0.017692834 | MIR762HG | antisense | 52.7 | 1.16461193 | 1.16461193 | ENSG00000260083 | n.s. |
| ENSG00000101574 | 244.136636 | -1.065157618 | 0.4770649 | -2.232731 | 0.025566692 | 0.080790979 | METTL4 | protein_coding | 36.78 | -1.16383076 | 1.16383076 | ENSG00000101574 | down |
| ENSG00000152518 | 826.6749025 | 0.618889426 | 0.2067954 | 2.992762 | 0.002764649 | 0.013168232 | ZFP36L2 | protein_coding | 55.96 | 1.163804557 | 1.163804557 | ENSG00000152518 | n.s. |
| ENSG00000270956 | 52.0253851 | -0.773836747 | 0.2915535 | -2.654185 | 0.007950029 | 0.031409832 | | antisense | 36.15 | -1.163025859 | 1.163025859 | ENSG00000270956 | n.s. |
| ENSG00000206770 | 4.989510292 | 1.164255389 | 0.5483072 | 2.123363 | 0.033723422 | 0.100327181 | | misc_RNA | 37.27 | 1.162603768 | 1.162603768 | ENSG00000206770 | n.s. |
| ENSG00000092439 | 3435.524357 | -0.727880006 | 0.2653241 | -2.743362 | 0.006081362 | 0.025282413 | TRPM7 | protein_coding | 38.89 | -1.162556468 | 1.162556468 | ENSG00000092439 | n.s. |
| ENSG00000201942 | 6.398427436 | -1.218729611 | 0.5889891 | -2.069189 | 0.038528363 | 0.111315709 | RNA5SP363 | rRNA | 41.74 | -1.161989879 | 1.161989879 | ENSG00000201942 | n.s. |
| ENSG00000153317 | 953.1659725 | -0.674212347 | 0.2359793 | -2.857082 | 0.004275549 | 0.018936954 | ASAP1 | protein_coding | 42.15 | -1.161458791 | 1.161458791 | ENSG00000153317 | n.s. |
| ENSG00000152578 | 144.4711417 | 0.674393305 | 0.2361093 | 2.856276 | 0.00428643 | 0.018964147 | GRIA4 | protein_coding | 35.32 | 1.161350251 | 1.161350251 | ENSG00000152578 | n.s. |
| ENSG00000169814 | 80.37495471 | 0.706927551 | 0.2536797 | 2.786694 | 0.005324881 | 0.022776751 | BTD | protein_coding | 41.69 | 1.16113432 | 1.16113432 | ENSG00000169814 | n.s. |
| ENSG00000179912 | 111.7170864 | -0.752239194 | 0.2793038 | -2.693265 | 0.007075605 | 0.028621046 | R3HDM2 | protein_coding | 43.16 | -1.16094165 | 1.16094165 | ENSG00000179912 | n.s. |
| ENSG00000272870 | 75.96885972 | 0.726191533 | 0.2645728 | 2.74477 | 0.006055334 | 0.025196775 | | antisense | 38.2 | 1.16092976 | 1.16092976 | ENSG00000272870 | n.s. |
| ENSG00000138190 | 587.1493442 | -0.788515386 | 0.3003991 | -2.624892 | 0.008667636 | 0.033718774 | EXOC6 | protein_coding | 37.36 | -1.160795752 | 1.160795752 | ENSG00000138190 | n.s. |
| ENSG00000159267 | 385.6938141 | -0.725229785 | 0.2641058 | -2.745982 | 0.006033001 | 0.025118862 | HLCS | protein_coding | 44.35 | -1.16036769 | 1.16036769 | ENSG00000159267 | n.s. |
| ENSG00000176102 | 1639.675573 | -0.767573735 | 0.2883888 | -2.661593 | 0.007777177 | 0.030867057 | CSTF3 | protein_coding | 37.22 | -1.159423792 | 1.159423792 | ENSG00000176102 | n.s. |
| ENSG00000269293 | 150.3167601 | -0.728829073 | 0.266277 | -2.737109 | 0.006198178 | 0.02566079 | ZSCAN16-AS1 | antisense | 40.71 | -1.159370253 | 1.159370253 | ENSG00000269293 | n.s. |
| ENSG00000103343 | 148.2736038 | -0.721298874 | 0.262043 | -2.752598 | 0.005912449 | 0.024772505 | ZNF174 | protein_coding | 46.45 | -1.158427686 | 1.158427686 | ENSG00000103343 | n.s. |
| ENSG00000115368 | 2807.576643 | -0.749941553 | 0.2784152 | -2.693609 | 0.007068309 | 0.028599846 | WDR75 | protein_coding | 37.03 | -1.15763701 | 1.15763701 | ENSG00000115368 | n.s. |
| ENSG00000254485 | 35.76371536 | 0.790999834 | 0.3022808 | 2.616772 | 0.008876559 | 0.03441613 | | antisense | 43.09 | 1.157420985 | 1.157420985 | ENSG00000254485 | n.s. |
| ENSG00000148297 | 518.5073953 | -0.752836243 | 0.2801207 | -2.687542 | 0.007198002 | 0.029014992 | MED22 | protein_coding | 56.94 | -1.157393531 | 1.157393531 | ENSG00000148297 | n.s. |
| ENSG00000107485 | 1476.987101 | -0.606178721 | 0.2007798 | -3.019123 | 0.002535078 | 0.012344054 | GATA3 | protein_coding | 48.73 | -1.156917668 | 1.156917668 | ENSG00000107485 | n.s. |
| ENSG00000183150 | 618.8086094 | -0.649828923 | 0.2235396 | -2.906997 | 0.003649166 | 0.016603703 | GPR19 | protein_coding | 41.27 | -1.156562289 | 1.156562289 | ENSG00000183150 | n.s. |
| ENSG00000114648 | 411.796836 | -0.727740163 | 0.2665293 | -2.730432 | 0.006325145 | 0.026047024 | KLHL18 | protein_coding | 44.87 | -1.152916454 | 1.152916454 | ENSG00000114648 | n.s. |
| ENSG00000117153 | 1066.889737 | -0.73496067 | 0.2705619 | -2.716423 | 0.006599152 | 0.026999706 | KLHL12 | protein_coding | 41.92 | -1.152889416 | 1.152889416 | ENSG00000117153 | n.s. |
| ENSG00000107614 | 46.20229165 | 0.753436624 | 0.2811301 | 2.680029 | 0.007361589 | 0.029529069 | TRDMT1 | protein_coding | 36.72 | 1.152569857 | 1.152569857 | ENSG00000107614 | n.s. |
| ENSG00000260000 | 199.8333804 | 0.647312816 | 0.2228424 | 2.904801 | 0.003674872 | 0.016693392 | | antisense | 42.32 | 1.150569664 | 1.150569664 | ENSG00000260000 | n.s. |
| ENSG00000083812 | 275.046243 | -0.677231301 | 0.2387742 | -2.836284 | 0.004564187 | 0.020002257 | ZNF324 | protein_coding | 57.75 | -1.150562471 | 1.150562471 | ENSG00000083812 | n.s. |
| ENSG00000130348 | 826.9495275 | -0.765402919 | 0.2884465 | -2.653535 | 0.007965349 | 0.031461432 | QRSL1 | protein_coding | 39.53 | -1.149804734 | 1.149804734 | ENSG00000130348 | n.s. |
| ENSG00000146707 | 141.9315652 | 0.680470126 | 0.2407014 | 2.82703 | 0.004698197 | 0.02047355 | POMZP3 | protein_coding | 49.24 | 1.149182605 | 1.149182605 | ENSG00000146707 | n.s. |
| ENSG00000198597 | 541.2494803 | -0.715105312 | 0.2598311 | -2.752193 | 0.005919757 | 0.024795661 | ZNF536 | protein_coding | 46.3 | -1.14819047 | 1.14819047 | ENSG00000198597 | n.s. |
| ENSG00000104412 | 668.1343355 | -0.777286458 | 0.2956375 | -2.629188 | 0.008558912 | 0.033361067 | EMC2 | protein_coding | 34.69 | -1.147865603 | 1.147865603 | ENSG00000104412 | n.s. |

| | | | | | | | | | | | | | |
|---|---|---|---|---|---|---|---|---|---|---|---|---|---|
| ENSG00000088812 | 745.3898126 | -0.710500479 | 0.2573318 | -2.761029 | 0.005761963 | 0.024251456 | ATRN | protein_coding | 40.34 | -1.147644559 | 1.147644559 | ENSG00000088812 | n.s. |
| ENSG00000167548 | 235.0711448 | -0.671699368 | 0.2360921 | -2.845074 | 0.004440115 | 0.019575591 | KMT2D | protein_coding | 54.13 | -1.147454028 | 1.147454028 | ENSG00000167548 | n.s. |
| ENSG00000177628 | 177.1403912 | 0.637139937 | 0.2178882 | 2.92416 | 0.003453875 | 0.015896958 | GBA | protein_coding | 51.52 | 1.146014671 | 1.146014671 | ENSG00000177628 | n.s. |
| ENSG00000108433 | 1523.351454 | -0.714033349 | 0.2595539 | -2.751003 | 0.005941315 | 0.024856048 | GOSR2 | protein_coding | 47.31 | -1.145715009 | 1.145715009 | ENSG00000108433 | n.s. |
| ENSG00000136536 | 3810.873489 | -0.741184077 | 0.2750859 | -2.694373 | 0.007052113 | 0.028542606 | 7-Mar | protein_coding | 38.09 | -1.144763533 | 1.144763533 | ENSG00000136536 | n.s. |
| ENSG00000237125 | 12160.9605 | -0.65622927 | 0.2281669 | -2.876094 | 0.004026304 | 0.018059997 | HAND2-AS1 | antisense | 41 | -1.143992896 | 1.143992896 | ENSG00000237125 | n.s. |
| ENSG00000124198 | 1160.778996 | -0.717852299 | 0.2622432 | -2.737354 | 0.006193569 | 0.025649336 | ARFGEF2 | protein_coding | 43.19 | -1.142048362 | 1.142048362 | ENSG00000124198 | n.s. |
| ENSG00000226686 | 131.0089192 | -0.663934986 | 0.2326489 | -2.853806 | 0.004319885 | 0.019091148 | LINC01535 | lincRNA | 44.46 | -1.141415754 | 1.141415754 | ENSG00000226686 | n.s. |
| ENSG00000157764 | 979.4258585 | -0.737985058 | 0.2737088 | -2.696241 | 0.007012684 | 0.028416065 | BRAF | protein_coding | 37.97 | -1.141246711 | 1.141246711 | ENSG00000157764 | n.s. |
| ENSG00000171928 | 655.7427107 | -0.712522479 | 0.2593407 | -2.747438 | 0.006006295 | 0.025045119 | TVP23B | protein_coding | 41.13 | -1.140945792 | 1.140945792 | ENSG00000171928 | n.s. |
| ENSG00000277369 | 112.6246689 | 0.658609056 | 0.2299381 | 2.864289 | 0.004179471 | 0.018597243 | | lincRNA | 58.79 | 1.139756843 | 1.139756843 | ENSG00000277369 | n.s. |
| ENSG00000119541 | 965.4511084 | -0.768892743 | 0.2918783 | -2.634292 | 0.008431294 | 0.032979202 | VPS4B | protein_coding | 39.19 | -1.139314403 | 1.139314403 | ENSG00000119541 | n.s. |
| ENSG00000065491 | 257.8197801 | -0.683729493 | 0.2436501 | -2.806194 | 0.005013051 | 0.021595517 | TBC1D22B | protein_coding | 43.34 | -1.13884473 | 1.13884473 | ENSG00000065491 | n.s. |
| ENSG00000200785 | 11.94972321 | -0.982608044 | 0.4258709 | -2.307291 | 0.021038607 | 0.069346716 | SNORD8 | snoRNA | 50 | -1.138817276 | 1.138817276 | ENSG00000200785 | n.s. |
| ENSG00000139350 | 1443.874503 | -0.694346936 | 0.2493654 | -2.784456 | 0.005361767 | 0.022912402 | NEDD1 | protein_coding | 35.02 | -1.138679936 | 1.138679936 | ENSG00000139350 | n.s. |
| ENSG00000189319 | 711.7328577 | -0.658674617 | 0.2301439 | -2.862013 | 0.004209603 | 0.018687534 | FAM53B | protein_coding | 52.68 | -1.138484826 | 1.138484826 | ENSG00000189319 | n.s. |
| ENSG00000116285 | 247.8747907 | 0.742100546 | 0.2765758 | 2.683172 | 0.007292741 | 0.029286653 | ERRFI1 | protein_coding | 42.8 | 1.137885216 | 1.137885216 | ENSG00000116285 | n.s. |
| ENSG00000214050 | 18.12610441 | -0.899382061 | 0.3720354 | -2.417463 | 0.015629104 | 0.05442249 | FBXO16 | protein_coding | 46.01 | -1.137018218 | 1.137018218 | ENSG00000214050 | n.s. |
| ENSG00000106261 | 1805.477025 | -0.693347094 | 0.2490487 | -2.783982 | 0.005369604 | 0.02293282 | ZKSCAN1 | protein_coding | 44.83 | -1.13677205 | 1.13677205 | ENSG00000106261 | n.s. |
| ENSG00000253106 | 205.0493085 | -0.714340984 | 0.260935 | -2.73762 | 0.006188549 | 0.025636171 | | antisense | 40.87 | -1.136621403 | 1.136621403 | ENSG00000253106 | n.s. |
| ENSG00000160007 | 1500.020972 | -0.6819526 | 0.2430032 | -2.806353 | 0.005010582 | 0.021591562 | ARHGAP35 | protein_coding | 47.96 | -1.135939316 | 1.135939316 | ENSG00000160007 | n.s. |
| ENSG00000187079 | 3235.506642 | -0.697705335 | 0.2516602 | -2.77241 | 0.005564285 | 0.023597787 | TEAD1 | protein_coding | 42.75 | -1.13525639 | 1.13525639 | ENSG00000187079 | n.s. |
| ENSG00000196628 | 16621.75386 | -0.68416467 | 0.244306 | -2.800441 | 0.005103276 | 0.021936683 | TCF4 | protein_coding | 37.78 | -1.134912221 | 1.134912221 | ENSG00000196628 | n.s. |
| ENSG00000186001 | 918.1695954 | -0.710858279 | 0.2591844 | -2.742674 | 0.006094112 | 0.025327857 | LRCH3 | protein_coding | 40.91 | -1.134815257 | 1.134815257 | ENSG00000186001 | n.s. |
| ENSG00000119283 | 442.3125808 | -0.664422235 | 0.2337161 | -2.84286 | 0.004471073 | 0.019680778 | TRIM67 | protein_coding | 48.93 | -1.133476252 | 1.133476252 | ENSG00000119283 | n.s. |
| ENSG00000143469 | 1974.025825 | -0.753317316 | 0.2836184 | -2.656095 | 0.007905131 | 0.03129461 | SYT14 | protein_coding | 36.49 | -1.133388842 | 1.133388842 | ENSG00000143469 | n.s. |
| ENSG00000089048 | 2581.460598 | -0.723950073 | 0.26684 | -2.713049 | 0.006666727 | 0.027196261 | ESF1 | protein_coding | 37.97 | -1.133337175 | 1.133337175 | ENSG00000089048 | n.s. |
| ENSG00000125351 | 611.1576952 | -0.73074164 | 0.2706176 | -2.700273 | 0.00692825 | 0.028123039 | UPF3B | protein_coding | 42.87 | -1.133334791 | 1.133334791 | ENSG00000125351 | n.s. |
| ENSG00000135540 | 93.54781152 | 0.691257313 | 0.2483558 | 2.783334 | 0.005380335 | 0.022964564 | NHSL1 | protein_coding | 40.97 | 1.132930508 | 1.132930508 | ENSG00000135540 | n.s. |
| ENSG00000061337 | 629.6423984 | -0.716252169 | 0.2625236 | -2.728335 | 0.006365501 | 0.026197715 | LZTS1 | protein_coding | 50.43 | -1.13292226 | 1.13292226 | ENSG00000061337 | n.s. |
| ENSG00000107669 | 741.9556896 | -0.735829594 | 0.2737002 | -2.688451 | 0.007178431 | 0.028952866 | ATE1 | protein_coding | 39.1 | -1.131932878 | 1.131932878 | ENSG00000107669 | n.s. |
| ENSG00000171617 | 3378.061509 | 0.673412928 | 0.2389047 | 2.818751 | 0.004821094 | 0.020923967 | ENC1 | protein_coding | 45.65 | 1.130900017 | 1.130900017 | ENSG00000171617 | n.s. |

| | | | | | | | | | | | | | |
|---|---|---|---|---|---|---|---|---|---|---|---|---|---|
| ENSG00000237491 | 64.68418805 | 0.707128224 | 0.2577239 | 2.743744 | 0.006074293 | 0.025261127 | | lincRNA | 40.42 | 1.12967077 | 1.12967077 | ENSG00000237491 | n.s. |
| ENSG00000163659 | 927.5609979 | 0.674135369 | 0.239647 | 2.813035 | 0.004907631 | 0.021226762 | TIPARP | protein_coding | 38.27 | 1.127906847 | 1.127906847 | ENSG00000163659 | n.s. |
| ENSG00000118965 | 1594.517057 | -0.755698241 | 0.2858507 | -2.643681 | 0.008200985 | 0.032245783 | WDR35 | protein_coding | 36.9 | -1.127144386 | 1.127144386 | ENSG00000118965 | n.s. |
| ENSG00000128578 | 220.4557011 | -0.759682272 | 0.2881757 | -2.636177 | 0.008384598 | 0.032865641 | STRIP2 | protein_coding | 43.81 | -1.126804725 | 1.126804725 | ENSG00000128578 | n.s. |
| ENSG00000145734 | 3713.837185 | -0.68683119 | 0.2466331 | -2.784829 | 0.005355593 | 0.02289405 | BDP1 | protein_coding | 39.84 | -1.126593664 | 1.126593664 | ENSG00000145734 | n.s. |
| ENSG00000208022 | 43.7364498 | 0.746622363 | 0.280735 | 2.659528 | 0.007825032 | 0.031048141 | MIR618 | miRNA | 45.92 | 1.125879946 | 1.125879946 | ENSG00000208022 | n.s. |
| ENSG00000270381 | 18.11527167 | -0.890753737 | 0.3688472 | -2.414967 | 0.015736633 | 0.054679595 | | processed_pseudogene | 42.48 | -1.124286839 | 1.124286839 | ENSG00000270381 | n.s. |
| ENSG00000116604 | 131.0318692 | 0.671244903 | 0.2385182 | 2.814229 | 0.004889434 | 0.021167785 | MEF2D | protein_coding | 54.23 | 1.123881848 | 1.123881848 | ENSG00000116604 | n.s. |
| ENSG00000283204 | 11.55916083 | 1.011686072 | 0.4490808 | 2.252793 | 0.024272199 | 0.077474053 | MIR4434 | miRNA | 33.96 | 1.123825122 | 1.123825122 | ENSG00000283204 | up |
| ENSG00000083828 | 522.5491236 | -0.697990582 | 0.2532531 | -2.756099 | 0.005849528 | 0.02456061 | ZNF586 | protein_coding | 46.13 | -1.123597918 | 1.123597918 | ENSG00000083828 | n.s. |
| ENSG00000202058 | 30.31164243 | -0.795886741 | 0.310148 | -2.566151 | 0.010283392 | 0.038832564 | RN7SKP80 | misc_RNA | 56.89 | -1.122840144 | 1.122840144 | ENSG00000202058 | n.s. |
| ENSG00000203804 | 404.0738657 | 0.607229114 | 0.2046957 | 2.966497 | 0.00301213 | 0.014163839 | ADAMTSL4-AS1 | processed_transcript | 48.94 | 1.122656739 | 1.122656739 | ENSG00000203804 | n.s. |
| ENSG00000168280 | 6674.786753 | -0.692424744 | 0.2503654 | -2.765657 | 0.005680828 | 0.023997016 | KIF5C | protein_coding | 43.05 | -1.121619201 | 1.121619201 | ENSG00000168280 | n.s. |
| ENSG00000078246 | 528.2761218 | -0.734782633 | 0.2746211 | -2.675623 | 0.00745904 | 0.029816884 | TULP3 | protein_coding | 45 | -1.120938641 | 1.120938641 | ENSG00000078246 | n.s. |
| ENSG00000235354 | 29.93788589 | 0.779518228 | 0.300713 | 2.592233 | 0.00953551 | 0.036533546 | RPS29P16 | processed_pseudogene | 50.29 | 1.120407919 | 1.120407919 | ENSG00000235354 | n.s. |
| ENSG00000112139 | 7.358694784 | -1.03934138 | 0.4691443 | -2.215398 | 0.026732759 | 0.083696919 | MDGA1 | protein_coding | 52.18 | -1.119672625 | 1.119672625 | ENSG00000112139 | down |
| ENSG00000234608 | 773.2430402 | 0.675550373 | 0.2413972 | 2.798501 | 0.005134037 | 0.022055293 | MAPKAPK5-AS1 | lincRNA | 53.55 | 1.119040526 | 1.119040526 | ENSG00000234608 | n.s. |
| ENSG00000162241 | 16.88584382 | 0.870570089 | 0.3567169 | 2.440507 | 0.014666664 | 0.051840005 | SLC25A45 | protein_coding | 59.38 | 1.118974177 | 1.118974177 | ENSG00000162241 | n.s. |
| ENSG00000234072 | 374.1932214 | 0.601185101 | 0.2020083 | 2.976041 | 0.002919957 | 0.013767205 | | antisense | 49.35 | 1.118898186 | 1.118898186 | ENSG00000234072 | n.s. |
| ENSG00000174371 | 3310.194655 | -0.654118402 | 0.2298147 | -2.846287 | 0.004423235 | 0.019507348 | EXO1 | protein_coding | 39.58 | -1.118412796 | 1.118412796 | ENSG00000174371 | n.s. |
| ENSG00000182841 | 539.4686296 | -0.629343399 | 0.2167112 | -2.904066 | 0.00368351 | 0.016721717 | RRP7BP | transcribed_unprocessed_pseud | 54.88 | -1.118166458 | 1.118166458 | ENSG00000182841 | n.s. |
| ENSG00000167785 | 447.206776 | -0.745461975 | 0.2814099 | -2.649025 | 0.008072428 | 0.031830194 | ZNF558 | protein_coding | 44.39 | -1.116076382 | 1.116076382 | ENSG00000167785 | n.s. |
| ENSG00000260007 | 146.2359748 | -0.689268033 | 0.2493469 | -2.764294 | 0.005704614 | 0.024053718 | | protein_coding | 50.05 | -1.115799348 | 1.115799348 | ENSG00000260007 | n.s. |
| ENSG00000220804 | 61.57148253 | -0.767423939 | 0.2943045 | -2.607585 | 0.009118356 | 0.035168068 | LINC01881 | transcribed_unprocessed_pseud | 39.11 | -1.115720437 | 1.115720437 | ENSG00000220804 | n.s. |
| ENSG00000056277 | 812.114403 | -0.699688422 | 0.2552895 | -2.740765 | 0.006129638 | 0.025437542 | ZNF280C | protein_coding | 38.03 | -1.115670582 | 1.115670582 | ENSG00000056277 | n.s. |
| ENSG00000265806 | 16.07416302 | -1.157928134 | 0.5563207 | -2.081404 | 0.037396935 | 0.108791793 | MIR4292 | miRNA | 70.15 | -1.115552441 | 1.115552441 | ENSG00000265806 | n.s. |
| ENSG00000230184 | 18.70004361 | -0.892165745 | 0.371494 | -2.401561 | 0.016325266 | 0.056372874 | SMYD3-IT1 | sense_intronic | 46.52 | -1.114252406 | 1.114252406 | ENSG00000230184 | n.s. |
| ENSG00000176783 | 503.6567857 | -0.723688461 | 0.2691117 | -2.689175 | 0.007162887 | 0.028915303 | RUFY1 | protein_coding | 47.71 | -1.113664092 | 1.113664092 | ENSG00000176783 | n.s. |
| ENSG00000204406 | 420.6614406 | -0.695102392 | 0.2529459 | -2.748028 | 0.005995495 | 0.025015065 | MBD5 | protein_coding | 36.28 | -1.113413873 | 1.113413873 | ENSG00000204406 | n.s. |
| ENSG00000036549 | 3045.133318 | -0.713763368 | 0.2636864 | -2.706864 | 0.006792202 | 0.027651413 | ZZZ3 | protein_coding | 37.17 | -1.112245083 | 1.112245083 | ENSG00000036549 | n.s. |
| ENSG00000260160 | 40.01809356 | -0.763075166 | 0.2922697 | -2.610859 | 0.009031502 | 0.034880889 | | sense_overlapping | 42.96 | -1.112115249 | 1.112115249 | ENSG00000260160 | n.s. |
| ENSG00000204381 | 1114.433057 | -0.717320972 | 0.2658085 | -2.698638 | 0.006962376 | 0.028236867 | LAYN | protein_coding | 44.3 | -1.111261812 | 1.111261812 | ENSG00000204381 | n.s. |

| | | | | | | | | | | | | | |
|---|---|---|---|---|---|---|---|---|---|---|---|---|---|
| ENSG00000102606 | 1373.509386 | -0.712531673 | 0.2633113 | -2.706043 | 0.006809019 | 0.027711772 | ARHGEF7 | protein_coding | 44.55 | -1.109651008 | 1.109651008 | ENSG00000102606 | n.s. |
| ENSG00000144792 | 1200.966026 | -0.654166071 | 0.2308687 | -2.833498 | 0.004604158 | 0.020145725 | ZNF660 | protein_coding | 45.57 | -1.109346011 | 1.109346011 | ENSG00000144792 | n.s. |
| ENSG00000013561 | 683.4934468 | -0.7435551 | 0.2812517 | -2.643735 | 0.008199672 | 0.032245783 | RNF14 | protein_coding | 42.04 | -1.109032562 | 1.109032562 | ENSG00000013561 | n.s. |
| ENSG00000252916 | 12.32096502 | -0.942641517 | 0.4052765 | -2.325922 | 0.020022724 | 0.066797184 | RNU6-762P | snRNA | 40.2 | -1.107831755 | 1.107831755 | ENSG00000252916 | n.s. |
| ENSG00000239827 | 122.2797387 | 0.724754505 | 0.2706245 | 2.678081 | 0.007404524 | 0.029641514 | SUGT1P3 | transcribed_unprocessed_pseud | 40.63 | 1.107497081 | 1.107497081 | ENSG00000239827 | n.s. |
| ENSG00000160688 | 1745.884447 | -0.685439321 | 0.2483712 | -2.759738 | 0.00578478 | 0.024325427 | FLAD1 | protein_coding | 51.25 | -1.106257611 | 1.106257611 | ENSG00000160688 | n.s. |
| ENSG00000115170 | 243.2129872 | -0.675828765 | 0.2430298 | -2.780847 | 0.005421732 | 0.023134164 | ACVR1 | protein_coding | 40.31 | -1.105484328 | 1.105484328 | ENSG00000115170 | n.s. |
| ENSG00000153107 | 1879.513955 | -0.722771646 | 0.269748 | -2.679433 | 0.007374705 | 0.029564075 | ANAPC1 | protein_coding | 39.07 | -1.10528821 | 1.10528821 | ENSG00000153107 | n.s. |
| ENSG00000207638 | 69.0260393 | 0.729453267 | 0.273743 | 2.664737 | 0.007704849 | 0.030623642 | MIR99A | miRNA | 53.09 | 1.104350788 | 1.104350788 | ENSG00000207638 | n.s. |
| ENSG00000113272 | 382.9616198 | -0.750738129 | 0.2862471 | -2.622693 | 0.008723789 | 0.033918267 | THG1L | protein_coding | 43.11 | -1.103259491 | 1.103259491 | ENSG00000113272 | n.s. |
| ENSG00000133961 | 80.86514166 | -0.736266456 | 0.2779498 | -2.648919 | 0.008074975 | 0.031831222 | NUMB | protein_coding | 40.56 | -1.10229889 | 1.10229889 | ENSG00000133961 | n.s. |
| ENSG00000202169 | 7.013171894 | -1.51502797 | 0.8551508 | -1.77165 | 0.076452676 | 0.188098711 | | misc_RNA | 36.46 | -1.099325779 | 1.099325779 | ENSG00000202169 | n.s. |
| ENSG00000280227 | 81.98855268 | -0.741506647 | 0.2814876 | -2.634243 | 0.008432519 | 0.032979202 | | TEC | 28.6 | -1.098734785 | 1.098734785 | ENSG00000280227 | n.s. |
| ENSG00000230715 | 40.49725545 | -0.758439431 | 0.2914968 | -2.601879 | 0.009271454 | 0.035678955 | | processed_pseudogene | 65.26 | -1.09790772 | 1.09790772 | ENSG00000230715 | n.s. |
| ENSG00000109861 | 2239.271595 | -0.741482693 | 0.2817698 | -2.63152 | 0.008500391 | 0.033157995 | CTSC | protein_coding | 38.6 | -1.0969582 | 1.0969582 | ENSG00000109861 | n.s. |
| ENSG00000116095 | 1003.459881 | -0.714540634 | 0.2662716 | -2.683503 | 0.007285535 | 0.029266156 | PLEKHA3 | protein_coding | 37.26 | -1.095844033 | 1.095844033 | ENSG00000116095 | n.s. |
| ENSG00000201616 | 14.42744711 | -0.918678184 | 0.3920277 | -2.343401 | 0.019108814 | 0.064263771 | RNU1-91P | snRNA | 46.01 | -1.095095441 | 1.095095441 | ENSG00000201616 | n.s. |
| ENSG00000152133 | 899.2455983 | -0.755028051 | 0.2900173 | -2.603389 | 0.009230704 | 0.035531959 | GPATCH11 | protein_coding | 38.7 | -1.094323195 | 1.094323195 | ENSG00000152133 | n.s. |
| ENSG00000272894 | 69.98321206 | 0.711907456 | 0.2649338 | 2.687114 | 0.007207233 | 0.029043791 | | lincRNA | 41.47 | 1.09416381 | 1.09416381 | ENSG00000272894 | n.s. |
| ENSG00000144485 | 2316.263884 | 0.696538868 | 0.2562812 | 2.717869 | 0.006570379 | 0.026905703 | HES6 | protein_coding | 70.08 | 1.093674442 | 1.093674442 | ENSG00000144485 | n.s. |
| ENSG00000207554 | 2.861227553 | 1.350968503 | 0.7154287 | 1.888334 | 0.058981094 | 0.155147132 | MIR647 | miRNA | 66.67 | 1.093279702 | 1.093279702 | ENSG00000207554 | n.s. |
| ENSG00000251718 | 9.439855557 | -0.965271106 | 0.4247317 | -2.272661 | 0.023046623 | 0.074315952 | RNU2-13P | snRNA | 38.05 | -1.08971188 | 1.08971188 | ENSG00000251718 | n.s. |
| ENSG00000135968 | 1630.059281 | -0.717535571 | 0.26878 | -2.669602 | 0.007594119 | 0.030309868 | GCC2 | protein_coding | 38.51 | -1.089517455 | 1.089517455 | ENSG00000135968 | n.s. |
| ENSG00000115840 | 699.1770785 | -0.743060229 | 0.2836758 | -2.6194 | 0.008808468 | 0.034180684 | SLC25A12 | protein_coding | 40.22 | -1.08948921 | 1.08948921 | ENSG00000115840 | n.s. |
| ENSG00000206878 | 8.830956158 | -0.975328501 | 0.4316591 | -2.259488 | 0.023853062 | 0.076398937 | | snoRNA | 47.2 | -1.089356776 | 1.089356776 | ENSG00000206878 | n.s. |
| ENSG00000152767 | 1574.829587 | -0.645635835 | 0.2285256 | -2.825223 | 0.004724771 | 0.020576472 | FARP1 | protein_coding | 44.52 | -1.088948175 | 1.088948175 | ENSG00000152767 | n.s. |
| ENSG00000274211 | 902.0053505 | -0.653699102 | 0.2329254 | -2.806474 | 0.0050087 | 0.021590133 | SOCS7 | protein_coding | 44.99 | -1.088895803 | 1.088895803 | ENSG00000274211 | n.s. |
| ENSG00000065328 | 2118.993795 | -0.702425995 | 0.2606129 | -2.695285 | 0.007032854 | 0.028489505 | MCM10 | protein_coding | 43.59 | -1.085469492 | 1.085469492 | ENSG00000065328 | n.s. |
| ENSG00000212389 | 20.30823004 | 0.822878576 | 0.3324686 | 2.475056 | 0.013321515 | 0.047988474 | RNU6-1275P | snRNA | 39.25 | 1.085264157 | 1.085264157 | ENSG00000212389 | n.s. |
| ENSG00000148730 | 1582.067915 | -0.675392931 | 0.2456444 | -2.749474 | 0.005969105 | 0.024957311 | EIF4EBP2 | protein_coding | 43.66 | -1.082521285 | 1.082521285 | ENSG00000148730 | n.s. |
| ENSG00000071537 | 969.356934 | -0.701182742 | 0.2603685 | -2.69304 | 0.007080387 | 0.028632072 | SEL1L | protein_coding | 38.48 | -1.082028197 | 1.082028197 | ENSG00000071537 | n.s. |
| ENSG00000103546 | 882.1608777 | -0.705235275 | 0.2627931 | -2.683614 | 0.007283117 | 0.029266156 | SLC6A2 | protein_coding | 48.47 | -1.081573016 | 1.081573016 | ENSG00000103546 | n.s. |

| | | | | | | | | | | | | | |
|---|---|---|---|---|---|---|---|---|---|---|---|---|---|
| ENSG00000112763 | 634.5351226 | -0.662102174 | 0.2383839 | -2.777462 | 0.005478532 | 0.02331935 | BTN2A1 | protein_coding | 45.31 | -1.080738491 | 1.080738491 | ENSG00000112763 | n.s. |
| ENSG00000061987 | 724.3410117 | -0.727147611 | 0.2755722 | -2.638683 | 0.008322877 | 0.032660491 | MON2 | protein_coding | 36.21 | -1.080524836 | 1.080524836 | ENSG00000061987 | n.s. |
| ENSG00000174469 | 14.05621198 | 0.871425705 | 0.364302 | 2.392042 | 0.016754937 | 0.057599748 | CNTNAP2 | protein_coding | 37.52 | 1.080201364 | 1.080201364 | ENSG00000174469 | n.s. |
| ENSG00000234167 | 14.03456235 | -0.884661996 | 0.3728694 | -2.372579 | 0.017664395 | 0.060205966 | ELOBP4 | processed_pseudogene | 58.45 | -1.079606529 | 1.079606529 | ENSG00000234167 | n.s. |
| ENSG00000157895 | 421.4880818 | -0.75079008 | 0.2897739 | -2.590951 | 0.009571113 | 0.036619566 | C12orf43 | protein_coding | 49.65 | -1.078349882 | 1.078349882 | ENSG00000157895 | n.s. |
| ENSG00000119778 | 1433.888275 | -0.681936908 | 0.249986 | -2.7279 | 0.006373891 | 0.026224493 | ATAD2B | protein_coding | 37.98 | -1.078341992 | 1.078341992 | ENSG00000119778 | n.s. |
| ENSG00000131711 | 12065.35358 | -0.666845158 | 0.2415527 | -2.760661 | 0.005768452 | 0.024270641 | MAP1B | protein_coding | 42.29 | -1.076900749 | 1.076900749 | ENSG00000131711 | n.s. |
| ENSG00000255717 | 14102.47064 | 0.570914765 | 0.1904835 | 2.997187 | 0.002724836 | 0.013014713 | SNHG1 | processed_transcript | 41.79 | 1.076497135 | 1.076497135 | ENSG00000255717 | n.s. |
| ENSG00000180035 | 395.1135291 | -0.661089296 | 0.2384242 | -2.772744 | 0.005558578 | 0.023580768 | ZNF48 | protein_coding | 53.07 | -1.075884522 | 1.075884522 | ENSG00000180035 | n.s. |
| ENSG00000258986 | 355.837882 | -0.672151116 | 0.244756 | -2.746209 | 0.006028834 | 0.02510902 | TMEM179 | protein_coding | 53.82 | -1.075556205 | 1.075556205 | ENSG00000258986 | n.s. |
| ENSG00000107779 | 1375.246025 | -0.74544462 | 0.2870095 | -2.597282 | 0.009396482 | 0.036091206 | BMPR1A | protein_coding | 39.9 | -1.075377368 | 1.075377368 | ENSG00000107779 | n.s. |
| ENSG00000199875 | 6.450871593 | -1.578670007 | 0.9258911 | -1.705028 | 0.088189235 | 0.208642093 | | misc_RNA | 37.17 | -1.074439759 | 1.074439759 | ENSG00000199875 | n.s. |
| ENSG00000223599 | 158.3428309 | -0.725797038 | 0.2757788 | -2.631808 | 0.008493176 | 0.033141952 | | processed_pseudogene | 49.2 | -1.073905201 | 1.073905201 | ENSG00000223599 | n.s. |
| ENSG00000120832 | 765.2670085 | -0.675040513 | 0.2467053 | -2.736222 | 0.006214907 | 0.025714754 | MTERF2 | protein_coding | 41.69 | -1.073191306 | 1.073191306 | ENSG00000120832 | n.s. |
| ENSG00000103047 | 418.4759824 | -0.71775249 | 0.2711422 | -2.647144 | 0.008117473 | 0.031980639 | TANGO6 | protein_coding | 43.08 | -1.07312098 | 1.07312098 | ENSG00000103047 | n.s. |
| ENSG00000236088 | 194.760808 | -0.739104367 | 0.2837149 | -2.605095 | 0.009184874 | 0.035394685 | COX10-AS1 | processed_transcript | 40.99 | -1.072486227 | 1.072486227 | ENSG00000236088 | n.s. |
| ENSG00000234912 | 513.7342767 | 0.656550319 | 0.2363175 | 2.778255 | 0.005465177 | 0.023283863 | SNHG20 | processed_transcript | 49.6 | 1.072110535 | 1.072110535 | ENSG00000234912 | n.s. |
| ENSG00000132182 | 758.5573765 | -0.644784187 | 0.2299915 | -2.803514 | 0.005054911 | 0.021746963 | NUP210 | protein_coding | 52.6 | -1.072019078 | 1.072019078 | ENSG00000132182 | n.s. |
| ENSG00000254860 | 40.93641571 | 0.731117037 | 0.27928 | 2.617864 | 0.00884821 | 0.034315773 | TMEM9B-AS1 | antisense | 43.8 | 1.070725443 | 1.070725443 | ENSG00000254860 | n.s. |
| ENSG00000202392 | 15.56792649 | -0.880720941 | 0.3719819 | -2.367645 | 0.017901724 | 0.060851002 | RN7SKP292 | misc_RNA | 53.63 | -1.070720867 | 1.070720867 | ENSG00000202392 | n.s. |
| ENSG00000134954 | 248.4486224 | -0.682099663 | 0.2510869 | -2.716587 | 0.006595876 | 0.026994234 | ETS1 | protein_coding | 42.5 | -1.070029522 | 1.070029522 | ENSG00000134954 | n.s. |
| ENSG00000245317 | 63.76299866 | 0.732736456 | 0.2803834 | 2.613338 | 0.008966263 | 0.034686656 | | lincRNA | 46.28 | 1.069676204 | 1.069676204 | ENSG00000245317 | n.s. |
| ENSG00000203965 | 412.1696355 | -0.772526567 | 0.3040954 | -2.540409 | 0.011072295 | 0.041284834 | EFCAB7 | protein_coding | 35.15 | -1.069338584 | 1.069338584 | ENSG00000203965 | n.s. |
| ENSG00000133958 | 1082.045808 | -0.673330585 | 0.2462705 | -2.734109 | 0.006254924 | 0.025818948 | UNC79 | protein_coding | 39.32 | -1.069290356 | 1.069290356 | ENSG00000133958 | n.s. |
| ENSG00000115020 | 1952.512424 | -0.680837231 | 0.2505221 | -2.717674 | 0.006574264 | 0.026913695 | PIKFYVE | protein_coding | 37.08 | -1.068932612 | 1.068932612 | ENSG00000115020 | n.s. |
| ENSG00000263731 | 639.6698056 | -0.669186874 | 0.2440552 | -2.741949 | 0.006107587 | 0.025359196 | | lincRNA | 43.7 | -1.067931584 | 1.067931584 | ENSG00000263731 | n.s. |
| ENSG00000183137 | 316.2055213 | -0.762567287 | 0.2983262 | -2.556153 | 0.010583662 | 0.039804517 | CEP57L1 | protein_coding | 36.59 | -1.06764578 | 1.06764578 | ENSG00000183137 | n.s. |
| ENSG00000106125 | 222.7166955 | -0.691215994 | 0.2566962 | -2.692739 | 0.007086768 | 0.028644403 | MINDY4 | protein_coding | 48.22 | -1.066518789 | 1.066518789 | ENSG00000106125 | n.s. |
| ENSG00000109171 | 289.993299 | -0.67197644 | 0.2459249 | -2.732445 | 0.006286608 | 0.025911323 | SLAIN2 | protein_coding | 37.94 | -1.066097621 | 1.066097621 | ENSG00000109171 | n.s. |
| ENSG00000078403 | 1467.460742 | -0.679550563 | 0.2502199 | -2.715813 | 0.006611316 | 0.027025647 | MLLT10 | protein_coding | 39.77 | -1.065687433 | 1.065687433 | ENSG00000078403 | n.s. |
| ENSG00000167912 | 66.78405469 | 0.693952227 | 0.2583692 | 2.685894 | 0.007233613 | 0.029133234 | | antisense | 54.4 | 1.065640884 | 1.065640884 | ENSG00000167912 | n.s. |
| ENSG00000274904 | 19.22879095 | 0.892224583 | 0.3803757 | 2.345641 | 0.018994413 | 0.063971748 | | antisense | 61.54 | 1.065326661 | 1.065326661 | ENSG00000274904 | n.s. |

| | | | | | | | | | | | | | |
|---|---|---|---|---|---|---|---|---|---|---|---|---|---|
| ENSG00000101928 | 194.7554432 | -0.831443401 | 0.3415726 | -2.434163 | 0.014926281 | 0.052519235 | MOSPD1 | protein_coding | 40.84 | -1.063982831 | 1.063982831 | ENSG00000101928 | n.s. |
| ENSG00000188042 | 492.9504836 | 0.647707378 | 0.232418 | 2.786821 | 0.005322793 | 0.022774813 | ARL4C | protein_coding | 50.16 | 1.063888629 | 1.063888629 | ENSG00000188042 | n.s. |
| ENSG00000161813 | 2320.922061 | -0.685539133 | 0.253834 | -2.700738 | 0.006918584 | 0.028100189 | LARP4 | protein_coding | 41.45 | -1.063470518 | 1.063470518 | ENSG00000161813 | n.s. |
| ENSG00000167157 | 270.1953616 | -0.817795381 | 0.3329597 | -2.456139 | 0.014043872 | 0.050085253 | PRRX2 | protein_coding | 54.69 | -1.063371259 | 1.063371259 | ENSG00000167157 | n.s. |
| ENSG00000279656 | 36.95890314 | -0.726187185 | 0.2775533 | -2.616388 | 0.00888654 | 0.034445239 | | TEC | 46.87 | -1.062318026 | 1.062318026 | ENSG00000279656 | n.s. |
| ENSG00000065609 | 32.19086281 | -0.813118855 | 0.3301319 | -2.463012 | 0.013777536 | 0.049411814 | SNAP91 | protein_coding | 36.62 | -1.062070807 | 1.062070807 | ENSG00000065609 | n.s. |
| ENSG00000152954 | 210.5607637 | -0.742161647 | 0.287086 | -2.585154 | 0.009733547 | 0.037150891 | NRSN1 | protein_coding | 40.3 | -1.061313989 | 1.061313989 | ENSG00000152954 | n.s. |
| ENSG00000179454 | 633.4009822 | -0.651257427 | 0.2347509 | -2.774249 | 0.005532937 | 0.023486306 | KLHL28 | protein_coding | 37.29 | -1.061019021 | 1.061019021 | ENSG00000179454 | n.s. |
| ENSG00000269940 | 87.83831948 | -0.689250592 | 0.2562954 | -2.689282 | 0.007160583 | 0.028915303 | | sense_intronic | 51.23 | -1.060668611 | 1.060668611 | ENSG00000269940 | n.s. |
| ENSG00000182504 | 1295.388122 | -0.677343412 | 0.2496199 | -2.713499 | 0.006657669 | 0.027183203 | CEP97 | protein_coding | 40.61 | -1.060516157 | 1.060516157 | ENSG00000182504 | n.s. |
| ENSG00000152582 | 49.13079116 | -0.784328166 | 0.3126276 | -2.508825 | 0.012113331 | 0.044511456 | SPEF2 | protein_coding | 37.81 | -1.060041631 | 1.060041631 | ENSG00000152582 | n.s. |
| ENSG00000271780 | 40.11201243 | 0.74439471 | 0.2887516 | 2.577976 | 0.009938099 | 0.037723053 | | lincRNA | 58.94 | 1.059566345 | 1.059566345 | ENSG00000271780 | n.s. |
| ENSG00000140650 | 900.1496858 | -0.654557355 | 0.2368156 | -2.763995 | 0.005709835 | 0.024068433 | PMM2 | protein_coding | 46.16 | -1.059435239 | 1.059435239 | ENSG00000140650 | n.s. |
| ENSG00000238832 | 7.441029852 | -1.384070851 | 0.7570209 | -1.828313 | 0.067502655 | 0.171785681 | | snoRNA | 32.41 | -1.058832248 | 1.058832248 | ENSG00000238832 | n.s. |
| ENSG00000112419 | 1038.014872 | -0.732638243 | 0.2819486 | -2.598481 | 0.009363714 | 0.035994199 | PHACTR2 | protein_coding | 40.02 | -1.057759276 | 1.057759276 | ENSG00000112419 | n.s. |
| ENSG00000105011 | 331.3869151 | -0.762397459 | 0.2997426 | -2.543508 | 0.01097457 | 0.040986839 | ASF1B | protein_coding | 51.11 | -1.057716366 | 1.057716366 | ENSG00000105011 | n.s. |
| ENSG00000160999 | 87.00953978 | -0.70440338 | 0.265535 | -2.652771 | 0.007983405 | 0.031523809 | SH2B2 | protein_coding | 56.99 | -1.057563984 | 1.057563984 | ENSG00000160999 | n.s. |
| ENSG00000198369 | 542.8321074 | -0.675199713 | 0.2487834 | -2.714007 | 0.006647484 | 0.027149582 | SPRED2 | protein_coding | 43.76 | -1.057522677 | 1.057522677 | ENSG00000198369 | n.s. |
| ENSG00000196652 | 606.7166581 | -0.695910046 | 0.260685 | -2.669544 | 0.007595427 | 0.030309868 | ZKSCAN5 | protein_coding | 46.16 | -1.056680914 | 1.056680914 | ENSG00000196652 | n.s. |
| ENSG00000066279 | 8671.591698 | -0.69557839 | 0.2605917 | -2.669227 | 0.007602616 | 0.030329862 | ASPM | protein_coding | 34.66 | -1.055978124 | 1.055978124 | ENSG00000066279 | n.s. |
| ENSG00000276641 | 3.473183062 | -1.22945771 | 0.6305592 | -1.949789 | 0.051201215 | 0.139070331 | MIR6769A | miRNA | 64.38 | -1.053356966 | 1.053356966 | ENSG00000276641 | n.s. |
| ENSG00000198814 | 211.9065468 | -0.769057301 | 0.3044818 | -2.525791 | 0.011543814 | 0.042757256 | GK | protein_coding | 38.63 | -1.05283189 | 1.05283189 | ENSG00000198814 | n.s. |
| ENSG00000061918 | 169.8047554 | -0.793037951 | 0.3194524 | -2.482491 | 0.013046722 | 0.047230763 | GUCY1B3 | protein_coding | 36.52 | -1.051389916 | 1.051389916 | ENSG00000061918 | n.s. |
| ENSG00000065833 | 217.3464286 | -0.707666586 | 0.2682537 | -2.63805 | 0.008338434 | 0.032693894 | ME1 | protein_coding | 36.91 | -1.051262311 | 1.051262311 | ENSG00000065833 | n.s. |
| ENSG00000174132 | 113.170488 | -0.842472369 | 0.3509939 | -2.400248 | 0.016383965 | 0.05654394 | FAM174A | protein_coding | 35.6 | -1.051080266 | 1.051080266 | ENSG00000174132 | n.s. |
| ENSG00000207756 | 5.051770299 | -1.15946074 | 0.576575 | -2.010945 | 0.044331225 | 0.124189869 | MIR580 | miRNA | 29.9 | -1.05037152 | 1.05037152 | ENSG00000207756 | n.s. |
| ENSG00000150756 | 282.3171803 | -0.740365883 | 0.2878223 | -2.572302 | 0.010102472 | 0.038242719 | FAM173B | protein_coding | 42.65 | -1.049432537 | 1.049432537 | ENSG00000150756 | n.s. |
| ENSG00000128805 | 43.62878898 | -0.770145624 | 0.3057514 | -2.518863 | 0.011773462 | 0.043480614 | ARHGAP22 | protein_coding | 47.75 | -1.048710634 | 1.048710634 | ENSG00000128805 | n.s. |
| ENSG00000278526 | 12.4121277 | -0.897801499 | 0.3873846 | -2.317597 | 0.02047121 | 0.067943692 | | misc_RNA | 59.12 | -1.048498249 | 1.048498249 | ENSG00000278526 | n.s. |
| ENSG00000141568 | 2375.443156 | -0.664536236 | 0.2441584 | -2.721742 | 0.006493882 | 0.026655163 | FOXK2 | protein_coding | 49.72 | -1.04612534 | 1.04612534 | ENSG00000141568 | n.s. |
| ENSG00000177410 | 3139.46919 | 0.590634316 | 0.2037657 | 2.898595 | 0.00374839 | 0.016955436 | ZFAS1 | antisense | 45.74 | 1.045830894 | 1.045830894 | ENSG00000177410 | n.s. |
| ENSG00000163806 | 134.6766296 | -0.683196502 | 0.2549397 | -2.679835 | 0.007365838 | 0.029537606 | SPDYA | protein_coding | 39.58 | -1.045034246 | 1.045034246 | ENSG00000163806 | n.s. |

| | | | | | | | | | | | | | |
|---|---|---|---|---|---|---|---|---|---|---|---|---|---|
| ENSG00000172785 | 323.3375999 | -0.742128634 | 0.2894841 | -2.563625 | 0.010358536 | 0.039084524 | CBWD1 | protein_coding | 36.26 | -1.044913534 | 1.044913534 | ENSG00000172785 | n.s. |
| ENSG00000197763 | 236.3525325 | -0.703051516 | 0.2664679 | -2.638409 | 0.008329598 | 0.032669877 | TXNRD3 | protein_coding | 39.52 | -1.044630851 | 1.044630851 | ENSG00000197763 | n.s. |
| ENSG00000167264 | 137.5627315 | -0.633044441 | 0.2267242 | -2.792134 | 0.005236165 | 0.022452456 | DUS2 | protein_coding | 48.49 | -1.043723257 | 1.043723257 | ENSG00000167264 | n.s. |
| ENSG00000241127 | 864.1385503 | -0.672184154 | 0.248949 | -2.700087 | 0.006932125 | 0.028130567 | YAE1D1 | protein_coding | 40.07 | -1.042437642 | 1.042437642 | ENSG00000241127 | n.s. |
| ENSG00000243260 | 34.64868034 | -0.780377365 | 0.3132326 | -2.491367 | 0.012725251 | 0.046271525 | RN7SL558P | misc_RNA | 49.5 | -1.041558888 | 1.041558888 | ENSG00000243260 | n.s. |
| ENSG00000215939 | 11.05432878 | -0.981236398 | 0.446777 | -2.196255 | 0.028073686 | 0.08687929 | MIR873 | miRNA | 46.75 | -1.041173984 | 1.041173984 | ENSG00000215939 | n.s. |
| ENSG00000232044 | 131.2589991 | 0.64237535 | 0.2322445 | 2.765944 | 0.005675823 | 0.023990434 | LINC01105 | processed_transcript | 43.52 | 1.040623591 | 1.040623591 | ENSG00000232044 | n.s. |
| ENSG00000252947 | 45.05469926 | 0.748967728 | 0.2945001 | 2.543184 | 0.010984752 | 0.041013865 | SCARNA1 | scaRNA | 40.36 | 1.038870146 | 1.038870146 | ENSG00000252947 | n.s. |
| ENSG00000232546 | 13.02167823 | -0.881002308 | 0.3781728 | -2.329629 | 0.01982576 | 0.066258774 | | unprocessed_pseudogene | 41.56 | -1.038487287 | 1.038487287 | ENSG00000232546 | n.s. |
| ENSG00000243398 | 17.32961174 | -0.841206119 | 0.3525832 | -2.385837 | 0.017040307 | 0.058407728 | RN7SL141P | misc_RNA | 54.51 | -1.037652718 | 1.037652718 | ENSG00000243398 | n.s. |
| ENSG00000139546 | 827.2522181 | 0.653396392 | 0.2391 | 2.732733 | 0.006281124 | 0.025896386 | TARBP2 | protein_coding | 57.56 | 1.036783805 | 1.036783805 | ENSG00000139546 | n.s. |
| ENSG00000118873 | 2852.281301 | -0.693425318 | 0.2619838 | -2.646825 | 0.008125137 | 0.031992725 | RAB3GAP2 | protein_coding | 37.8 | -1.036635321 | 1.036635321 | ENSG00000118873 | n.s. |
| ENSG00000202400 | 40.36540002 | 0.755531865 | 0.2990212 | 2.526683 | 0.011514537 | 0.042694249 | SNORD82 | snoRNA | 37.14 | 1.034799577 | 1.034799577 | ENSG00000202400 | n.s. |
| ENSG00000207155 | 5.95703207 | -1.08694596 | 0.526337 | -2.065114 | 0.038912191 | 0.112145401 | RNY1P14 | misc_RNA | 47.22 | -1.032836194 | 1.032836194 | ENSG00000207155 | n.s. |
| ENSG00000119946 | 330.8837396 | -0.656466364 | 0.2413265 | -2.720241 | 0.006523434 | 0.026740474 | CNNM1 | protein_coding | 44.48 | -1.03251058 | 1.03251058 | ENSG00000119946 | n.s. |
| ENSG00000250366 | 119.7641172 | -0.745478916 | 0.2934587 | -2.54032 | 0.011075117 | 0.041284834 | TUNAR | protein_coding | 47.34 | -1.031898968 | 1.031898968 | ENSG00000250366 | n.s. |
| ENSG00000181192 | 1158.589095 | -0.721932839 | 0.2794846 | -2.583086 | 0.009792096 | 0.037280653 | DHTKD1 | protein_coding | 46.21 | -1.031292963 | 1.031292963 | ENSG00000181192 | n.s. |
| ENSG00000238365 | 8.243648086 | -1.238205227 | 0.6452687 | -1.918899 | 0.054997166 | 0.146945877 | RNU7-57P | snRNA | 40 | -1.031230055 | 1.031230055 | ENSG00000238365 | n.s. |
| ENSG00000174521 | 149.0841299 | 0.758541639 | 0.3014311 | 2.516468 | 0.011853773 | 0.043719227 | TTC9B | protein_coding | 63.92 | 1.031106531 | 1.031106531 | ENSG00000174521 | n.s. |
| ENSG00000260257 | 143.7124774 | -0.671470844 | 0.2503386 | -2.68225 | 0.007312868 | 0.029350553 | | lincRNA | 40.4 | -1.028950989 | 1.028950989 | ENSG00000260257 | n.s. |
| ENSG00000200354 | 101.1622064 | 0.635548192 | 0.2299134 | 2.764294 | 0.005704617 | 0.024053718 | SNORA71D | snoRNA | 53.73 | 1.028836713 | 1.028836713 | ENSG00000200354 | n.s. |
| ENSG00000134504 | 166.5200568 | -0.646508553 | 0.23615 | -2.737703 | 0.006186999 | 0.025636171 | KCTD1 | protein_coding | 42.61 | -1.028690046 | 1.028690046 | ENSG00000134504 | n.s. |
| ENSG00000266240 | 9.855867809 | 0.928861645 | 0.4129614 | 2.24927 | 0.024495321 | 0.078078837 | MIR5091 | miRNA | 63.44 | 1.02868331 | 1.02868331 | ENSG00000266240 | n.s. |
| ENSG00000271983 | 21.43393927 | 0.846541866 | 0.3578733 | 2.365479 | 0.018006759 | 0.061148317 | | lincRNA | 43.09 | 1.02737632 | 1.02737632 | ENSG00000271983 | n.s. |
| ENSG00000238998 | 21.6270846 | -0.846259961 | 0.3577227 | -2.365687 | 0.017996657 | 0.061129596 | RNU7-187P | snRNA | 29.41 | -1.027146737 | 1.027146737 | ENSG00000238998 | n.s. |
| ENSG00000175087 | 240.811222 | -0.711753938 | 0.274039 | -2.597273 | 0.009396726 | 0.036091206 | PDIK1L | protein_coding | 43.1 | -1.026775238 | 1.026775238 | ENSG00000175087 | n.s. |
| ENSG00000230749 | 48.12788505 | 0.702175391 | 0.2684768 | 2.615405 | 0.008912179 | 0.03451579 | MEIS1-AS2 | antisense | 50.78 | 1.026567914 | 1.026567914 | ENSG00000230749 | n.s. |
| ENSG00000132688 | 3572.082048 | -0.629839768 | 0.2273357 | -2.770527 | 0.005596565 | 0.023713019 | NES | protein_coding | 59 | -1.023497903 | 1.023497903 | ENSG00000132688 | n.s. |
| ENSG00000174989 | 796.7715396 | -0.65246131 | 0.2402021 | -2.716302 | 0.006601577 | 0.027001689 | FBXW8 | protein_coding | 43.26 | -1.023456725 | 1.023456725 | ENSG00000174989 | n.s. |
| ENSG00000126461 | 294.9995067 | -0.647058983 | 0.2373318 | -2.726389 | 0.006403142 | 0.026321506 | SCAF1 | protein_coding | 58.2 | -1.022152132 | 1.022152132 | ENSG00000126461 | n.s. |
| ENSG00000011376 | 867.092549 | -0.664970791 | 0.2475079 | -2.686665 | 0.007216935 | 0.029074475 | LARS2 | protein_coding | 42.15 | -1.021719719 | 1.021719719 | ENSG00000011376 | n.s. |
| ENSG00000102763 | 885.4804136 | -0.692543132 | 0.2635476 | -2.627772 | 0.00859461 | 0.033481464 | VWA8 | protein_coding | 37.16 | -1.021636552 | 1.021636552 | ENSG00000102763 | n.s. |

| | | | | | | | | | | | | | |
|---|---|---|---|---|---|---|---|---|---|---|---|---|---|
| ENSG00000170161 | 191.6356532 | 0.649422857 | 0.2388051 | 2.719468 | 0.006538698 | 0.026791728 | | lincRNA | 48.19 | 1.020892261 | 1.020892261 | ENSG00000170161 | n.s. |
| ENSG00000040275 | 1460.822039 | -0.686851128 | 0.2602642 | -2.639054 | 0.008313778 | 0.032643184 | SPDL1 | protein_coding | 37.28 | -1.020803288 | 1.020803288 | ENSG00000040275 | n.s. |
| ENSG00000213066 | 419.3565456 | -0.717247466 | 0.2782788 | -2.577442 | 0.009953472 | 0.037760814 | FGFR1OP | protein_coding | 41.03 | -1.020613487 | 1.020613487 | ENSG00000213066 | n.s. |
| ENSG00000126003 | 320.4930822 | -0.648742895 | 0.2385914 | -2.719054 | 0.006546892 | 0.026817409 | PLAGL2 | protein_coding | 48.56 | -1.019553423 | 1.019553423 | ENSG00000126003 | n.s. |
| ENSG00000259926 | 23.91056314 | -0.798215481 | 0.3284027 | -2.430599 | 0.015073869 | 0.052847235 | | sense_intronic | 40.21 | -1.019303391 | 1.019303391 | ENSG00000259926 | n.s. |
| ENSG00000203724 | 103.9163592 | 0.755642934 | 0.3015456 | 2.505899 | 0.012214042 | 0.044786965 | C1orf53 | protein_coding | 40.35 | 1.019247738 | 1.019247738 | ENSG00000203724 | n.s. |
| ENSG00000174197 | 2546.583534 | -0.663756309 | 0.2471775 | -2.685343 | 0.007245544 | 0.029155987 | MGA | protein_coding | 40.41 | -1.019046649 | 1.019046649 | ENSG00000174197 | n.s. |
| ENSG00000137834 | 4497.286551 | -0.564228034 | 0.1925837 | -2.929781 | 0.003392007 | 0.015663987 | SMAD6 | protein_coding | 50.83 | -1.018486724 | 1.018486724 | ENSG00000137834 | n.s. |
| ENSG00000072071 | 778.4070462 | -0.61597659 | 0.2203737 | -2.795145 | 0.00518764 | 0.022251855 | ADGRL1 | protein_coding | 56.42 | -1.017983719 | 1.017983719 | ENSG00000072071 | n.s. |
| ENSG00000075407 | 1569.93711 | -0.724331804 | 0.2829002 | -2.560379 | 0.010455811 | 0.039399662 | ZNF37A | protein_coding | 39.11 | -1.017329452 | 1.017329452 | ENSG00000075407 | n.s. |
| ENSG00000180098 | 151.9249034 | -0.71486903 | 0.2773774 | -2.577243 | 0.009959197 | 0.03777224 | TRNAU1AP | protein_coding | 47.61 | -1.01713514 | 1.01713514 | ENSG00000180098 | n.s. |
| ENSG00000139323 | 390.0481626 | -0.770653308 | 0.3112839 | -2.475725 | 0.013296585 | 0.047918373 | POC1B | protein_coding | 39.79 | -1.016875452 | 1.016875452 | ENSG00000139323 | n.s. |
| ENSG00000210191 | 20.07020371 | 0.883851306 | 0.3846426 | 2.297851 | 0.021570267 | 0.070747426 | MT-TL2 | Mt_tRNA | 35.21 | 1.016684748 | 1.016684748 | ENSG00000210191 | n.s. |
| ENSG00000120658 | 1142.501253 | -0.685454225 | 0.2600987 | -2.635362 | 0.008404758 | 0.03293539 | ENOX1 | protein_coding | 39.04 | -1.016074287 | 1.016074287 | ENSG00000120658 | n.s. |
| ENSG00000254662 | 13.16949374 | 0.875237257 | 0.3791155 | 2.308629 | 0.020964159 | 0.069232445 | | antisense | 50.38 | 1.015004183 | 1.015004183 | ENSG00000254662 | n.s. |
| ENSG00000106404 | 123.7709869 | -0.725426527 | 0.2839207 | -2.555033 | 0.010617785 | 0.039900519 | CLDN15 | protein_coding | 59.59 | -1.014887274 | 1.014887274 | ENSG00000106404 | n.s. |
| ENSG00000139324 | 1119.395654 | -0.724841067 | 0.283602 | -2.555839 | 0.010593212 | 0.039829677 | TMTC3 | protein_coding | 34.6 | -1.014627609 | 1.014627609 | ENSG00000139324 | n.s. |
| ENSG00000154822 | 884.4751783 | -0.650583168 | 0.2403353 | -2.706982 | 0.006789803 | 0.027649731 | PLCL2 | protein_coding | 39.07 | -1.01380966 | 1.01380966 | ENSG00000154822 | n.s. |
| ENSG00000116750 | 2263.970409 | -0.714857572 | 0.2780283 | -2.571169 | 0.010135594 | 0.038347249 | UCHL5 | protein_coding | 35 | -1.012428319 | 1.012428319 | ENSG00000116750 | n.s. |
| ENSG00000188542 | 330.9474263 | 0.584917275 | 0.204305 | 2.862961 | 0.004197022 | 0.018651498 | DUSP28 | protein_coding | 53.35 | 1.011489424 | 1.011489424 | ENSG00000188542 | n.s. |
| ENSG00000267796 | 67.65545752 | -0.781010257 | 0.3188979 | -2.449092 | 0.014321691 | 0.050867577 | LIN37 | protein_coding | 54.65 | -1.010282805 | 1.010282805 | ENSG00000267796 | n.s. |
| ENSG00000105173 | 106.2321751 | 0.645833883 | 0.2381965 | 2.711349 | 0.006701003 | 0.027328078 | CCNE1 | protein_coding | 47.46 | 1.009690822 | 1.009690822 | ENSG00000105173 | n.s. |
| ENSG00000259519 | 7.747250395 | 0.966293327 | 0.4437672 | 2.177478 | 0.029444903 | 0.090180682 | | antisense | 50.21 | 1.009666836 | 1.009666836 | ENSG00000259519 | n.s. |
| ENSG00000096060 | 1879.24523 | -0.638227903 | 0.2339833 | -2.727664 | 0.00637845 | 0.026235497 | FKBP5 | protein_coding | 41.97 | -1.009108968 | 1.009108968 | ENSG00000096060 | n.s. |
| ENSG00000152292 | 27.43825714 | 0.746171411 | 0.2973151 | 2.509699 | 0.012083415 | 0.044424975 | SH2D6 | protein_coding | 54.62 | 1.009101927 | 1.009101927 | ENSG00000152292 | n.s. |
| ENSG00000239912 | 11.73569747 | -0.910337679 | 0.4050464 | -2.24749 | 0.024608725 | 0.078348144 | RPL39P36 | processed_pseudogene | 37.82 | -1.006807335 | 1.006807335 | ENSG00000239912 | n.s. |
| ENSG00000106484 | 217.1110186 | 0.750131582 | 0.3003009 | 2.497933 | 0.012491966 | 0.0456019 | MEST | protein_coding | 44.94 | 1.00593925 | 1.00593925 | ENSG00000106484 | n.s. |
| ENSG00000160094 | 57.86464219 | -0.731380869 | 0.2889936 | -2.530785 | 0.011380749 | 0.042265717 | ZNF362 | protein_coding | 50.56 | -1.004925917 | 1.004925917 | ENSG00000160094 | n.s. |
| ENSG00000137193 | 383.5271616 | 0.618215865 | 0.2231825 | 2.770001 | 0.005605604 | 0.02374409 | PIM1 | protein_coding | 53.6 | 1.004257345 | 1.004257345 | ENSG00000137193 | n.s. |
| ENSG00000256299 | 40.94020299 | -0.733524512 | 0.2905079 | -2.524972 | 0.011570737 | 0.042822799 | | lincRNA | 43.4 | -1.003699894 | 1.003699894 | ENSG00000256299 | n.s. |
| ENSG00000268205 | 762.736292 | -0.716432608 | 0.2802027 | -2.556837 | 0.010562877 | 0.039769317 | | lincRNA | 36.36 | -1.003329386 | 1.003329386 | ENSG00000268205 | n.s. |
| ENSG00000102595 | 1276.921667 | -0.706928423 | 0.2747552 | -2.572939 | 0.010083905 | 0.038191613 | UGGT2 | protein_coding | 37.01 | -1.002447127 | 1.002447127 | ENSG00000102595 | n.s. |

| | | | | | | | | | | | | | |
|---|---|---|---|---|---|---|---|---|---|---|---|---|---|
| ENSG00000107957 | 1105.521866 | -0.664765558 | 0.249993 | -2.659136 | 0.007834123 | 0.031075355 | SH3PXD2A | protein_coding | 51.58 | -1.002189857 | 1.002189857 | ENSG00000107957 | n.s. |
| ENSG00000163041 | 2609.777271 | 0.638681978 | 0.2351517 | 2.716042 | 0.006606756 | 0.027014938 | H3F3A | protein_coding | 44.56 | 1.00170628 | 1.00170628 | ENSG00000163041 | n.s. |
| ENSG00000258450 | 54.69882107 | -0.701169095 | 0.2714442 | -2.583105 | 0.00979154 | 0.037280653 | | sense_intronic | 50.17 | -1.001631613 | 1.001631613 | ENSG00000258450 | n.s. |
| ENSG00000144935 | 369.1446443 | -0.699005121 | 0.2704036 | -2.585044 | 0.009736666 | 0.037150891 | TRPC1 | protein_coding | 35.58 | -0.99959883 | 0.99959883 | ENSG00000144935 | n.s. |
| ENSG00000131378 | 207.0153673 | -0.659213091 | 0.2471929 | -2.666797 | 0.007657798 | 0.030480136 | RFTN1 | protein_coding | 43.5 | -0.999355907 | 0.999355907 | ENSG00000131378 | n.s. |
| ENSG00000182827 | 1329.340946 | -0.654164107 | 0.2443103 | -2.677596 | 0.007415267 | 0.029675989 | ACBD3 | protein_coding | 41.45 | -0.99929769 | 0.99929769 | ENSG00000182827 | n.s. |
| ENSG00000135905 | 734.4550347 | -0.699125322 | 0.2705194 | -2.584381 | 0.009755396 | 0.037212158 | DOCK10 | protein_coding | 39.37 | -0.999270409 | 0.999270409 | ENSG00000135905 | n.s. |
| ENSG00000277634 | 17.05058505 | -0.864359173 | 0.3751498 | -2.304037 | 0.02122056 | 0.069863765 | | misc_RNA | 38.71 | -0.998981398 | 0.998981398 | ENSG00000277634 | n.s. |
| ENSG00000174485 | 902.3491865 | -0.719910801 | 0.2830927 | -2.543021 | 0.010989849 | 0.041020695 | DENND4A | protein_coding | 37.31 | -0.998514113 | 0.998514113 | ENSG00000174485 | n.s. |
| ENSG00000184675 | 485.5909213 | -0.666500162 | 0.2516271 | -2.648761 | 0.008078736 | 0.031837033 | AMER1 | protein_coding | 50.95 | -0.997795674 | 0.997795674 | ENSG00000184675 | n.s. |
| ENSG00000109458 | 54.47258355 | -0.747410012 | 0.3000298 | -2.491119 | 0.012734129 | 0.046291718 | GAB1 | protein_coding | 38.61 | -0.997416198 | 0.997416198 | ENSG00000109458 | n.s. |
| ENSG00000059588 | 1686.114379 | -0.686555603 | 0.2634735 | -2.605786 | 0.009166369 | 0.035333157 | TARBP1 | protein_coding | 40.27 | -0.996753482 | 0.996753482 | ENSG00000059588 | n.s. |
| ENSG00000188229 | 3272.680503 | 0.714322498 | 0.2800135 | 2.551029 | 0.01074055 | 0.040264049 | TUBB4B | protein_coding | 63.25 | 0.996538856 | 0.996538856 | ENSG00000188229 | n.s. |
| ENSG00000254635 | 519.9528855 | -0.672799752 | 0.2554974 | -2.633294 | 0.008456115 | 0.033052916 | WAC-AS1 | antisense | 43.31 | -0.996275296 | 0.996275296 | ENSG00000254635 | n.s. |
| ENSG00000283122 | 201.2251336 | 0.62357443 | 0.227272 | 2.743737 | 0.006074427 | 0.025261127 | HYMAI | non_coding | 40.45 | 0.996189635 | 0.996189635 | ENSG00000283122 | n.s. |
| ENSG00000175137 | 614.0475628 | -0.714751612 | 0.2804886 | -2.548238 | 0.010826855 | 0.040533019 | SH3BP5L | protein_coding | 52.31 | -0.995070798 | 0.995070798 | ENSG00000175137 | n.s. |
| ENSG00000224348 | 11.88629119 | -0.889118164 | 0.3931159 | -2.26172 | 0.023714684 | 0.076008183 | | processed_pseudogene | 59.41 | -0.995047392 | 0.995047392 | ENSG00000224348 | n.s. |
| ENSG00000143363 | 322.3539127 | -0.744455612 | 0.2985943 | -2.493201 | 0.012659723 | 0.046093441 | PRUNE1 | protein_coding | 46.61 | -0.994861351 | 0.994861351 | ENSG00000143363 | n.s. |
| ENSG00000238197 | 122.0139427 | 0.642289002 | 0.2380365 | 2.69828 | 0.006969884 | 0.028259082 | PAXBP1-AS1 | antisense | 39.6 | 0.994804148 | 0.994804148 | ENSG00000238197 | n.s. |
| ENSG00000101333 | 73.69367219 | -0.701178619 | 0.272529 | -2.572859 | 0.010086229 | 0.038191613 | PLCB4 | protein_coding | 39.14 | -0.994293721 | 0.994293721 | ENSG00000101333 | n.s. |
| ENSG00000188215 | 269.8357165 | -0.634731105 | 0.233945 | -2.713164 | 0.006664419 | 0.027194815 | DCUN1D3 | protein_coding | 43.13 | -0.993680363 | 0.993680363 | ENSG00000188215 | n.s. |
| ENSG00000235109 | 63.08874422 | -0.738410191 | 0.2950778 | -2.502425 | 0.012334559 | 0.04513379 | ZSCAN31 | protein_coding | 41.29 | -0.993529578 | 0.993529578 | ENSG00000235109 | n.s. |
| ENSG00000144674 | 2311.29977 | -0.680188544 | 0.2604117 | -2.611974 | 0.009002121 | 0.034789566 | GOLGA4 | protein_coding | 37.82 | -0.992089675 | 0.992089675 | ENSG00000144674 | n.s. |
| ENSG00000278540 | 3147.426294 | -0.673441216 | 0.2564758 | -2.62575 | 0.008645841 | 0.033652793 | ACACA | protein_coding | 41.27 | -0.991964695 | 0.991964695 | ENSG00000278540 | n.s. |
| ENSG00000147912 | 278.6602631 | -0.646566511 | 0.2409351 | -2.683571 | 0.00728404 | 0.029266156 | FBXO10 | protein_coding | 46.45 | -0.991596585 | 0.991596585 | ENSG00000147912 | n.s. |
| ENSG00000135541 | 1185.438886 | -0.693262017 | 0.2681772 | -2.585089 | 0.009735381 | 0.037150891 | AHI1 | protein_coding | 36.31 | -0.991386015 | 0.991386015 | ENSG00000135541 | n.s. |
| ENSG00000270048 | 20.0908919 | -0.881413794 | 0.3886721 | -2.267757 | 0.023344046 | 0.075044828 | | sense_intronic | 44.87 | -0.99130779 | 0.99130779 | ENSG00000270048 | n.s. |
| ENSG00000227345 | 1027.239558 | -0.699546963 | 0.2719981 | -2.571882 | 0.010114748 | 0.038278782 | PARG | protein_coding | 38.39 | -0.99128735 | 0.99128735 | ENSG00000227345 | n.s. |
| ENSG00000105948 | 316.0885463 | -0.70535593 | 0.2754903 | -2.560366 | 0.010456206 | 0.039399662 | TTC26 | protein_coding | 38.13 | -0.990677695 | 0.990677695 | ENSG00000105948 | n.s. |
| ENSG00000035687 | 1412.071816 | -0.69017796 | 0.2665298 | -2.589496 | 0.009611654 | 0.036744389 | ADSS | protein_coding | 38.3 | -0.990273526 | 0.990273526 | ENSG00000035687 | n.s. |
| ENSG00000100150 | 128.5705556 | 0.641380059 | 0.238192 | 2.692702 | 0.007087551 | 0.028644403 | DEPDC5 | protein_coding | 45.23 | 0.989623922 | 0.989623922 | ENSG00000100150 | n.s. |
| ENSG00000222679 | 33.32014569 | 0.768982944 | 0.3149156 | 2.44187 | 0.014611419 | 0.051685669 | RNU6-1267P | snRNA | 41.12 | 0.989396418 | 0.989396418 | ENSG00000222679 | n.s. |

| | | | | | | | | | | | | | | |
|---|---|---|---|---|---|---|---|---|---|---|---|---|---|---|
| ENSG00000234444 | 922.5886954 | -0.667437349 | 0.25329 | -2.635072 | 0.008411945 | 0.032935745 | ZNF736 | | protein_coding | 38.55 | -0.989364081 | 0.989364081 | ENSG00000234444 | n.s. |
| ENSG00000189184 | 333.497353 | 0.583467618 | 0.2061579 | 2.830197 | 0.00465193 | 0.020322818 | PCDH18 | | protein_coding | 36.59 | 0.987236586 | 0.987236586 | ENSG00000189184 | n.s. |
| ENSG00000013441 | 3926.619666 | 0.614749635 | 0.2233915 | 2.751894 | 0.005925172 | 0.02481088 | CLK1 | | protein_coding | 41.85 | 0.986893142 | 0.986893142 | ENSG00000013441 | n.s. |
| ENSG00000137073 | 1044.517699 | -0.645247511 | 0.2408231 | -2.679343 | 0.007376686 | 0.029564075 | UBAP2 | | protein_coding | 43.26 | -0.986735535 | 0.986735535 | ENSG00000137073 | n.s. |
| ENSG00000257093 | 912.9604135 | -0.691515856 | 0.2679578 | -2.580689 | 0.009860328 | 0.037478938 | KIAA1147 | | protein_coding | 43.51 | -0.986248711 | 0.986248711 | ENSG00000257093 | n.s. |
| ENSG00000174365 | 290.7054285 | 0.609073019 | 0.2203299 | 2.764368 | 0.00570332 | 0.024053718 | SNHG11 | | processed_transcript | 53.2 | 0.985978233 | 0.985978233 | ENSG00000174365 | n.s. |
| ENSG00000213337 | 166.4480269 | -0.729958157 | 0.2912027 | -2.506701 | 0.012186372 | 0.044720837 | ANKRD39 | | protein_coding | 51.61 | -0.985071287 | 0.985071287 | ENSG00000213337 | n.s. |
| ENSG00000067369 | 3052.148593 | -0.680255642 | 0.2616638 | -2.599731 | 0.009329674 | 0.035873255 | TP53BP1 | | protein_coding | 41.72 | -0.983125331 | 0.983125331 | ENSG00000067369 | n.s. |
| ENSG00000162733 | 4501.200878 | -0.688164664 | 0.2664112 | -2.583092 | 0.009791911 | 0.037280653 | DDR2 | | protein_coding | 40.47 | -0.983054569 | 0.983054569 | ENSG00000162733 | n.s. |
| ENSG00000175029 | 1618.324543 | -0.708128484 | 0.2783152 | -2.54434 | 0.010948436 | 0.040911186 | CTBP2 | | protein_coding | 50.11 | -0.982994169 | 0.982994169 | ENSG00000175029 | n.s. |
| ENSG00000112249 | 4033.952219 | -0.700246265 | 0.2736059 | -2.559325 | 0.010487573 | 0.039507165 | ASCC3 | | protein_coding | 34.11 | -0.982672485 | 0.982672485 | ENSG00000112249 | n.s. |
| ENSG00000206715 | 6.061491856 | -1.008207628 | 0.4812974 | -2.094771 | 0.036191387 | 0.106015712 | RNU6-444P | | snRNA | 41.67 | -0.982629166 | 0.982629166 | ENSG00000206715 | n.s. |
| ENSG00000227542 | 101.048949 | -0.649454208 | 0.2438629 | -2.663194 | 0.007740273 | 0.03074688 | | | lincRNA | 42.43 | -0.982103969 | 0.982103969 | ENSG00000227542 | n.s. |
| ENSG00000205060 | 1298.220457 | -0.687649716 | 0.2662806 | -2.582425 | 0.009810869 | 0.037331711 | SLC35B4 | | protein_coding | 40.49 | -0.981910226 | 0.981910226 | ENSG00000205060 | n.s. |
| ENSG00000264529 | 49.93731615 | -0.756604798 | 0.3085068 | -2.452474 | 0.014187777 | 0.050469193 | | | transcribed_processed_pseudog | 39.88 | -0.981296479 | 0.981296479 | ENSG00000264529 | n.s. |
| ENSG00000199266 | 13.62962224 | 0.885277601 | 0.3935509 | 2.249461 | 0.024483154 | 0.078057924 | SNORA60 | | snoRNA | 50.75 | 0.980518422 | 0.980518422 | ENSG00000199266 | n.s. |
| ENSG00000197959 | 128.5615074 | -0.661395905 | 0.250988 | -2.635169 | 0.008409539 | 0.032935745 | DNM3 | | protein_coding | 37.28 | -0.980408652 | 0.980408652 | ENSG00000197959 | n.s. |
| ENSG00000271452 | 32.15388273 | -0.776070919 | 0.3216933 | -2.412456 | 0.015845445 | 0.054989064 | | | sense_intronic | 34.76 | -0.977634906 | 0.977634906 | ENSG00000271452 | n.s. |
| ENSG00000105229 | 235.6081397 | -0.704567247 | 0.2771946 | -2.541778 | 0.01102902 | 0.041146047 | PIAS4 | | protein_coding | 59.38 | -0.976299025 | 0.976299025 | ENSG00000105229 | n.s. |
| ENSG00000172432 | 701.2705355 | -0.684100493 | 0.2650343 | -2.581177 | 0.009846411 | 0.037456717 | GTPBP2 | | protein_coding | 46.56 | -0.975849027 | 0.975849027 | ENSG00000172432 | n.s. |
| ENSG00000148606 | 948.3885854 | -0.663980746 | 0.2532839 | -2.621488 | 0.008754691 | 0.034019415 | POLR3A | | protein_coding | 44.7 | -0.974905107 | 0.974905107 | ENSG00000148606 | n.s. |
| ENSG00000116574 | 1168.079906 | -0.699902524 | 0.2746106 | -2.548709 | 0.010812255 | 0.040489249 | RHOU | | protein_coding | 46.54 | -0.974726448 | 0.974726448 | ENSG00000116574 | n.s. |
| ENSG00000260219 | 27.33104768 | 0.766304935 | 0.3161583 | 2.423802 | 0.015358997 | 0.053635144 | | | lincRNA | 58.47 | 0.973629154 | 0.973629154 | ENSG00000260219 | n.s. |
| ENSG00000124126 | 264.6525606 | -0.642158979 | 0.2408416 | -2.666313 | 0.007668828 | 0.030509765 | PREX1 | | protein_coding | 50.46 | -0.973231194 | 0.973231194 | ENSG00000124126 | n.s. |
| ENSG00000236287 | 2581.66195 | -0.632141104 | 0.2351188 | -2.688603 | 0.007175164 | 0.028948077 | ZBED5 | | protein_coding | 40.77 | -0.972473413 | 0.972473413 | ENSG00000236287 | n.s. |
| ENSG00000152270 | 804.9351979 | -0.67352753 | 0.2592517 | -2.597967 | 0.009377745 | 0.036038187 | PDE3B | | protein_coding | 36.96 | -0.972059899 | 0.972059899 | ENSG00000152270 | n.s. |
| ENSG00000100167 | 647.2071924 | -0.662996188 | 0.2531349 | -2.619141 | 0.008815141 | 0.034197048 | | 3-Sep | protein_coding | 51.53 | -0.97195996 | 0.97195996 | ENSG00000100167 | n.s. |
| ENSG00000072422 | 634.9114419 | -0.704351871 | 0.2778641 | -2.534879 | 0.011248609 | 0.041843676 | RHOBTB1 | | protein_coding | 38.94 | -0.970857607 | 0.970857607 | ENSG00000072422 | n.s. |
| ENSG00000198818 | 389.5123112 | -0.693318027 | 0.2712329 | -2.556172 | 0.010583069 | 0.039804517 | SFT2D1 | | protein_coding | 43.24 | -0.970692132 | 0.970692132 | ENSG00000198818 | n.s. |
| ENSG00000169372 | 187.3212041 | 0.678621763 | 0.2626297 | 2.583949 | 0.009767634 | 0.037238437 | CRADD | | protein_coding | 40.55 | 0.969756309 | 0.969756309 | ENSG00000169372 | n.s. |
| ENSG00000215966 | 4.50842649 | -1.563805129 | 0.97004 | -1.612104 | 0.106939354 | 0.240147843 | MIR876 | | miRNA | 37.04 | -0.968810601 | 0.968810601 | ENSG00000215966 | n.s. |
| ENSG00000158615 | 4514.037375 | 0.603880262 | 0.219689 | 2.748796 | 0.005981456 | 0.024978949 | PPP1R15B | | protein_coding | 40.44 | 0.967673338 | 0.967673338 | ENSG00000158615 | n.s. |

| | | | | | | | | | | | | | |
|---|---|---|---|---|---|---|---|---|---|---|---|---|---|
| ENSG00000101337 | 433.1061322 | -0.652483005 | 0.2476094 | -2.63513 | 0.008410513 | 0.032935745 | TM9SF4 | protein_coding | 47.45 | -0.96719677 | 0.96719677 | ENSG00000101337 | n.s. |
| ENSG00000251869 | 19.29105065 | -0.836909871 | 0.3634206 | -2.302868 | 0.021286246 | 0.070025225 | SCARNA23 | scaRNA | 34.62 | -0.966417895 | 0.966417895 | ENSG00000251869 | n.s. |
| ENSG00000165275 | 19.74610042 | -1.136601623 | 0.5860766 | -1.93934 | 0.052459982 | 0.141825306 | TRMT10B | protein_coding | 42.56 | -0.964118088 | 0.964118088 | ENSG00000165275 | n.s. |
| ENSG00000229276 | 37.16479261 | -0.719408817 | 0.2881428 | -2.49671 | 0.012535158 | 0.045735589 | REV3L-IT1 | sense_intronic | 37.95 | -0.963824884 | 0.963824884 | ENSG00000229276 | n.s. |
| ENSG00000224870 | 148.9951506 | -0.676210703 | 0.2621204 | -2.579771 | 0.009886581 | 0.037555079 | | processed_transcript | 59.37 | -0.963824291 | 0.963824291 | ENSG00000224870 | n.s. |
| ENSG00000128731 | 3080.021262 | -0.642469734 | 0.2422843 | -2.651719 | 0.008008323 | 0.031613234 | HERC2 | protein_coding | 43.74 | -0.963788812 | 0.963788812 | ENSG00000128731 | n.s. |
| ENSG00000223111 | 20.96147375 | 0.852716819 | 0.3749989 | 2.273918 | 0.022970888 | 0.074132153 | | snoRNA | 52.24 | 0.963564364 | 0.963564364 | ENSG00000223111 | n.s. |
| ENSG00000271966 | 52.51285058 | 0.692226223 | 0.2716946 | 2.54781 | 0.010840154 | 0.040560995 | | lincRNA | 42.32 | 0.96350373 | 0.96350373 | ENSG00000271966 | n.s. |
| ENSG00000273018 | 138.8809702 | -0.749983996 | 0.30752 | -2.438814 | 0.014735553 | 0.052005857 | | processed_transcript | 39.96 | -0.962940258 | 0.962940258 | ENSG00000273018 | n.s. |
| ENSG00000121749 | 1351.006893 | -0.66991908 | 0.2585399 | -2.591163 | 0.009565213 | 0.036617118 | TBC1D15 | protein_coding | 37.01 | -0.962215389 | 0.962215389 | ENSG00000121749 | n.s. |
| ENSG00000206903 | 41.05689241 | 0.727073562 | 0.2932641 | 2.479245 | 0.013166093 | 0.047563962 | SNORA24B | snoRNA | 43.51 | 0.961716182 | 0.961716182 | ENSG00000206903 | n.s. |
| ENSG00000197372 | 1058.079853 | -0.650179014 | 0.2471347 | -2.630869 | 0.008516693 | 0.0332058 | ZNF675 | protein_coding | 41.44 | -0.961475659 | 0.961475659 | ENSG00000197372 | n.s. |
| ENSG00000125863 | 617.1108953 | -0.737212899 | 0.2997214 | -2.45966 | 0.013906852 | 0.049708856 | MKKS | protein_coding | 38.18 | -0.96100584 | 0.96100584 | ENSG00000125863 | n.s. |
| ENSG00000150433 | 327.9327909 | -0.72719093 | 0.2935679 | -2.477079 | 0.013246252 | 0.047791544 | TMEM218 | protein_coding | 41.01 | -0.960363929 | 0.960363929 | ENSG00000150433 | n.s. |
| ENSG00000178694 | 402.9020798 | -0.638093327 | 0.2402152 | -2.65634 | 0.007899384 | 0.031280753 | NSUN3 | protein_coding | 37.97 | -0.96015358 | 0.96015358 | ENSG00000178694 | n.s. |
| ENSG00000272472 | 69.64068013 | -0.788293753 | 0.3326169 | -2.369975 | 0.01778927 | 0.060587086 | | sense_intronic | 31.94 | -0.95984219 | 0.95984219 | ENSG00000272472 | n.s. |
| ENSG00000152061 | 606.1126865 | -0.640701977 | 0.2418566 | -2.649099 | 0.008070678 | 0.031830194 | RABGAP1L | protein_coding | 37.91 | -0.959233828 | 0.959233828 | ENSG00000152061 | n.s. |
| ENSG00000198466 | 576.0501779 | -0.658965092 | 0.2525858 | -2.608876 | 0.009084015 | 0.035064246 | ZNF587 | protein_coding | 46.53 | -0.958883492 | 0.958883492 | ENSG00000198466 | n.s. |
| ENSG00000088179 | 1896.431192 | -0.6871727 | 0.2693463 | -2.551261 | 0.010733395 | 0.040258907 | PTPN4 | protein_coding | 37.04 | -0.958700758 | 0.958700758 | ENSG00000088179 | n.s. |
| ENSG00000212241 | 3.521642093 | -1.672546396 | 1.0877988 | -1.537551 | 0.124158383 | 0.267304341 | | misc_RNA | 40.43 | -0.95835903 | 0.95835903 | ENSG00000212241 | n.s. |
| ENSG00000249421 | 34.19530622 | 0.756058349 | 0.3124265 | 2.419956 | 0.015522398 | 0.054110754 | ADAMTS19-AS1 | antisense | 63.75 | 0.957711523 | 0.957711523 | ENSG00000249421 | n.s. |
| ENSG00000122687 | 668.077646 | -0.676627873 | 0.2633261 | -2.569543 | 0.010183267 | 0.038496234 | MRM2 | protein_coding | 49.95 | -0.957145445 | 0.957145445 | ENSG00000122687 | n.s. |
| ENSG00000172007 | 354.6407314 | 0.675355116 | 0.2626885 | 2.570936 | 0.010142421 | 0.038362654 | RAB33B | protein_coding | 41.25 | 0.956364543 | 0.956364543 | ENSG00000172007 | n.s. |
| ENSG00000024048 | 954.2385533 | -0.649292409 | 0.2472924 | -2.625606 | 0.008649492 | 0.033657597 | UBR2 | protein_coding | 39.1 | -0.956353761 | 0.956353761 | ENSG00000024048 | n.s. |
| ENSG00000067646 | 424.0206604 | -0.744501462 | 0.3052894 | -2.438674 | 0.014741243 | 0.052008141 | ZFY | protein_coding | 37.27 | -0.955886771 | 0.955886771 | ENSG00000067646 | n.s. |
| ENSG00000275667 | 3.893764575 | -1.163503256 | 0.6111332 | -1.903845 | 0.05693031 | 0.150979181 | MIR6839 | miRNA | 49.56 | -0.955332668 | 0.955332668 | ENSG00000275667 | n.s. |
| ENSG00000176268 | 12.28787392 | -0.901285683 | 0.4106938 | -2.194544 | 0.028196326 | 0.087123267 | CYCSP34 | processed_pseudogene | 46.71 | -0.955241914 | 0.955241914 | ENSG00000176268 | n.s. |
| ENSG00000075188 | 851.5130078 | -0.692627529 | 0.2732606 | -2.534678 | 0.011255086 | 0.041843676 | NUP37 | protein_coding | 36.52 | -0.954697123 | 0.954697123 | ENSG00000075188 | n.s. |
| ENSG00000086712 | 950.5985476 | -0.679355045 | 0.2653118 | -2.560591 | 0.010449433 | 0.039395465 | TXLNG | protein_coding | 42.93 | -0.954190693 | 0.954190693 | ENSG00000086712 | n.s. |
| ENSG00000141449 | 204.2528297 | -0.666990515 | 0.258165 | -2.583583 | 0.009778007 | 0.037261168 | GREB1L | protein_coding | 40.54 | -0.952958388 | 0.952958388 | ENSG00000141449 | n.s. |
| ENSG00000198363 | 1964.478923 | -0.707529741 | 0.2825829 | -2.503795 | 0.012286912 | 0.044994918 | ASPH | protein_coding | 38.17 | -0.952926902 | 0.952926902 | ENSG00000198363 | n.s. |
| ENSG00000082397 | 2098.764534 | -0.645240883 | 0.2454513 | -2.628794 | 0.008568829 | 0.033390375 | EPB41L3 | protein_coding | 39.83 | -0.952619897 | 0.952619897 | ENSG00000082397 | n.s. |

| | | | | | | | | | | | | | |
|---|---|---|---|---|---|---|---|---|---|---|---|---|---|
| ENSG00000135473 | 135.5560431 | 0.632068445 | 0.2378208 | 2.657751 | 0.007866391 | 0.031191895 | PAN2 | protein_coding | 47.67 | 0.951868681 | 0.951868681 | ENSG00000135473 | n.s. |
| ENSG00000153827 | 3580.508622 | -0.668523613 | 0.2594146 | -2.577047 | 0.009964842 | 0.037783358 | TRIP12 | protein_coding | 38.19 | -0.951108174 | 0.951108174 | ENSG00000153827 | n.s. |
| ENSG00000148773 | 8696.546591 | -0.619632064 | 0.2308005 | -2.684709 | 0.007259306 | 0.029199215 | MKI67 | protein_coding | 43.51 | -0.950905227 | 0.950905227 | ENSG00000148773 | n.s. |
| ENSG00000198740 | 1232.795828 | -0.637806309 | 0.2413172 | -2.64302 | 0.008217013 | 0.03229968 | ZNF652 | protein_coding | 41.52 | -0.950842783 | 0.950842783 | ENSG00000198740 | n.s. |
| ENSG00000187391 | 9.421688979 | -0.937493693 | 0.4375788 | -2.142457 | 0.032156724 | 0.096803907 | MAGI2 | protein_coding | 36.83 | -0.950719026 | 0.950719026 | ENSG00000187391 | n.s. |
| ENSG00000138459 | 638.7634791 | -0.745305336 | 0.3069241 | -2.428305 | 0.015169586 | 0.053093551 | SLC35A5 | protein_coding | 38.43 | -0.950233171 | 0.950233171 | ENSG00000138459 | n.s. |
| ENSG00000189362 | 417.1244411 | -0.650399305 | 0.2488415 | -2.613709 | 0.008956534 | 0.034658651 | NEMP2 | protein_coding | 40.26 | -0.949705482 | 0.949705482 | ENSG00000189362 | n.s. |
| ENSG00000132952 | 896.1439496 | -0.669960218 | 0.2606159 | -2.57068 | 0.010149893 | 0.03838049 | USPL1 | protein_coding | 38.94 | -0.948589628 | 0.948589628 | ENSG00000132952 | n.s. |
| ENSG00000162775 | 4068.175108 | 0.556851803 | 0.1961175 | 2.839379 | 0.004520143 | 0.019827961 | RBM15 | protein_coding | 44.01 | 0.948163781 | 0.948163781 | ENSG00000162775 | n.s. |
| ENSG00000171105 | 422.5486121 | -0.645512952 | 0.2462632 | -2.621232 | 0.008761267 | 0.034035469 | INSR | protein_coding | 47.45 | -0.947657076 | 0.947657076 | ENSG00000171105 | n.s. |
| ENSG00000200972 | 195.3838449 | 0.629303736 | 0.2369784 | 2.655532 | 0.007918332 | 0.031329053 | RNU5A-8P | snRNA | 39.47 | 0.946506006 | 0.946506006 | ENSG00000200972 | n.s. |
| ENSG00000187240 | 694.5717337 | -0.700571483 | 0.2793504 | -2.507859 | 0.012146503 | 0.044598039 | DYNC2H1 | protein_coding | 35.1 | -0.94625086 | 0.94625086 | ENSG00000187240 | n.s. |
| ENSG00000134815 | 326.136783 | -0.644524729 | 0.2459529 | -2.620521 | 0.008779549 | 0.034096977 | DHX34 | protein_coding | 54.23 | -0.945700902 | 0.945700902 | ENSG00000134815 | n.s. |
| ENSG00000110060 | 299.0386462 | -0.689402398 | 0.272735 | -2.527737 | 0.011480027 | 0.04257763 | PUS3 | protein_coding | 40.24 | -0.945045575 | 0.945045575 | ENSG00000110060 | n.s. |
| ENSG00000214944 | 81.15456027 | -0.638488541 | 0.2426372 | -2.631454 | 0.00850205 | 0.033157995 | ARHGEF28 | protein_coding | 39.36 | -0.944587443 | 0.944587443 | ENSG00000214944 | n.s. |
| ENSG00000175155 | 365.8247466 | 0.582735903 | 0.2106172 | 2.766801 | 0.005660924 | 0.023941993 | YPEL2 | protein_coding | 46.62 | 0.944521487 | 0.944521487 | ENSG00000175155 | n.s. |
| ENSG00000206660 | 3.946915793 | 1.239186153 | 0.6793229 | 1.824149 | 0.06812955 | 0.17293851 | | misc_RNA | 50 | 0.944394035 | 0.944394035 | ENSG00000206660 | n.s. |
| ENSG00000218823 | 183.9296843 | -0.670576139 | 0.261639 | -2.562982 | 0.01037773 | 0.039146339 | PAPOLB | protein_coding | 40.47 | -0.943707734 | 0.943707734 | ENSG00000218823 | n.s. |
| ENSG00000252685 | 8.234134623 | -1.122202948 | 0.581313 | -1.930463 | 0.053549557 | 0.144238825 | RNU6-928P | snRNA | 40.38 | -0.943680462 | 0.943680462 | ENSG00000252685 | n.s. |
| ENSG00000187815 | 129.1998275 | -0.724217251 | 0.2945612 | -2.458631 | 0.013946805 | 0.049776304 | ZFP69 | protein_coding | 41.16 | -0.943638679 | 0.943638679 | ENSG00000187815 | n.s. |
| ENSG00000159885 | 203.4444943 | -0.710379659 | 0.2859795 | -2.484023 | 0.012990733 | 0.047052542 | ZNF222 | protein_coding | 42.04 | -0.942969975 | 0.942969975 | ENSG00000159885 | n.s. |
| ENSG00000205133 | 100.3546014 | 0.659872484 | 0.2554132 | 2.583549 | 0.009778949 | 0.037261168 | TRIQK | protein_coding | 34.5 | 0.942788547 | 0.942788547 | ENSG00000205133 | n.s. |
| ENSG00000175274 | 1361.892774 | -0.614012152 | 0.2286254 | -2.685669 | 0.007238476 | 0.029144392 | TP53I11 | protein_coding | 54.85 | -0.942781891 | 0.942781891 | ENSG00000175274 | n.s. |
| ENSG00000162849 | 1418.083758 | -0.62624502 | 0.2359449 | -2.6542 | 0.007949674 | 0.031409832 | KIF26B | protein_coding | 44.8 | -0.941205177 | 0.941205177 | ENSG00000162849 | n.s. |
| ENSG00000023902 | 1952.646319 | -0.629778696 | 0.2379973 | -2.646159 | 0.008141159 | 0.032037691 | PLEKHO1 | protein_coding | 51.63 | -0.941102733 | 0.941102733 | ENSG00000023902 | n.s. |
| ENSG00000074755 | 1078.269268 | -0.653277072 | 0.2517828 | -2.594605 | 0.009469959 | 0.036312375 | ZZEF1 | protein_coding | 44.49 | -0.940683281 | 0.940683281 | ENSG00000074755 | n.s. |
| ENSG00000168944 | 1934.117913 | -0.654930725 | 0.2528044 | -2.590662 | 0.009579163 | 0.036640297 | CEP120 | protein_coding | 36.19 | -0.940507383 | 0.940507383 | ENSG00000168944 | n.s. |
| ENSG00000079313 | 325.837789 | -0.616884505 | 0.2306519 | -2.674526 | 0.0074835 | 0.029906077 | REXO1 | protein_coding | 60.94 | -0.940280379 | 0.940280379 | ENSG00000079313 | n.s. |
| ENSG00000121671 | 148.2879647 | 0.62523929 | 0.2354934 | 2.655019 | 0.007930403 | 0.03135899 | CRY2 | protein_coding | 49.13 | 0.940133517 | 0.940133517 | ENSG00000121671 | n.s. |
| ENSG00000237036 | 203.676854 | -0.643331244 | 0.2461154 | -2.613941 | 0.008950447 | 0.034644724 | ZEB1-AS1 | antisense | 38.57 | -0.939497077 | 0.939497077 | ENSG00000237036 | n.s. |
| ENSG00000187764 | 47.00759133 | -0.694499656 | 0.2766872 | -2.510053 | 0.012071293 | 0.044403862 | SEMA4D | protein_coding | 51.34 | -0.939365825 | 0.939365825 | ENSG00000187764 | n.s. |
| ENSG00000283154 | 723.223335 | 0.586764512 | 0.2136068 | 2.746937 | 0.006015471 | 0.02507587 | IQCJ-SCHIP1 | protein_coding | 39 | 0.939259765 | 0.939259765 | ENSG00000283154 | n.s. |

| | | | | | | | | | | | | | |
|---|---|---|---|---|---|---|---|---|---|---|---|---|---|
| ENSG00000242073 | 29.74828665 | 0.730823082 | 0.299699 | 2.438523 | 0.014747407 | 0.052008141 | | transcribed_unprocessed_pseud | 41.4 | 0.938324707 | 0.938324707 | ENSG00000242073 | n.s. |
| ENSG00000167193 | 961.448625 | -0.731253148 | 0.3000478 | -2.437122 | 0.014804691 | 0.052170515 | CRK | protein_coding | 49.13 | -0.937886913 | 0.937886913 | ENSG00000167193 | n.s. |
| ENSG00000115464 | 4021.141728 | -0.670146858 | 0.2622574 | -2.555302 | 0.01060956 | 0.039880374 | USP34 | protein_coding | 38.76 | -0.937696808 | 0.937696808 | ENSG00000115464 | n.s. |
| ENSG00000075340 | 1782.140408 | -0.622462004 | 0.2342207 | -2.657587 | 0.007870223 | 0.031191895 | ADD2 | protein_coding | 44.81 | -0.937401782 | 0.937401782 | ENSG00000075340 | n.s. |
| ENSG00000183647 | 435.4379799 | -0.625469468 | 0.236049 | -2.649744 | 0.00805527 | 0.031780538 | ZNF530 | protein_coding | 46.12 | -0.936852409 | 0.936852409 | ENSG00000183647 | n.s. |
| ENSG00000115459 | 323.5424608 | -0.626570585 | 0.2367172 | -2.646917 | 0.008122939 | 0.031992725 | ELMOD3 | protein_coding | 46.14 | -0.936690921 | 0.936690921 | ENSG00000115459 | n.s. |
| ENSG00000115421 | 1131.415961 | -0.642367622 | 0.2459782 | -2.611482 | 0.009015066 | 0.034827071 | PAPOLG | protein_coding | 39.6 | -0.93662534 | 0.93662534 | ENSG00000115421 | n.s. |
| ENSG00000128694 | 507.3396303 | -0.632596197 | 0.2402561 | -2.633008 | 0.008463226 | 0.033071429 | OSGEPL1 | protein_coding | 39.12 | -0.936588424 | 0.936588424 | ENSG00000128694 | n.s. |
| ENSG00000137310 | 383.0133437 | -0.648039639 | 0.2492624 | -2.599829 | 0.009327031 | 0.035872998 | TCF19 | protein_coding | 51.1 | -0.936567833 | 0.936567833 | ENSG00000137310 | n.s. |
| ENSG00000175395 | 305.3798591 | -0.695902082 | 0.2782226 | -2.501242 | 0.012375869 | 0.045261159 | ZNF25 | protein_coding | 40.15 | -0.935483308 | 0.935483308 | ENSG00000175395 | n.s. |
| ENSG00000251868 | 10.05624101 | -0.969036383 | 0.4650305 | -2.083812 | 0.037177238 | 0.108401818 | RNU7-71P | snRNA | 43.33 | -0.935084677 | 0.935084677 | ENSG00000251868 | n.s. |
| ENSG00000273355 | 19.23172624 | 0.796113833 | 0.3425742 | 2.323917 | 0.020129967 | 0.06706972 | | lincRNA | 38.92 | 0.93421849 | 0.93421849 | ENSG00000273355 | n.s. |
| ENSG00000167302 | 523.1133554 | 0.639944496 | 0.2449063 | 2.613018 | 0.008974652 | 0.034709466 | TEPSIN | protein_coding | 62.93 | 0.934032309 | 0.934032309 | ENSG00000167302 | n.s. |
| ENSG00000206848 | 3.977706583 | -1.129631682 | 0.5912657 | -1.910531 | 0.056064823 | 0.149166927 | RNU6-890P | snRNA | 36.45 | -0.933445675 | 0.933445675 | ENSG00000206848 | n.s. |
| ENSG00000087116 | 431.6091183 | -0.637906043 | 0.243857 | -2.615902 | 0.008899202 | 0.034475123 | ADAMTS2 | protein_coding | 53.87 | -0.932933876 | 0.932933876 | ENSG00000087116 | n.s. |
| ENSG00000207561 | 6.422562878 | -1.075791381 | 0.5481153 | -1.96271 | 0.049679865 | 0.13578578 | MIR635 | miRNA | 43.88 | -0.932867878 | 0.932867878 | ENSG00000207561 | n.s. |
| ENSG00000229422 | 56.81439761 | 0.72333635 | 0.2958615 | 2.444848 | 0.014491325 | 0.051378333 | | processed_pseudogene | 69.6 | 0.93253968 | 0.93253968 | ENSG00000229422 | n.s. |
| ENSG00000146263 | 4131.121372 | -0.653377351 | 0.2531447 | -2.581043 | 0.009850218 | 0.037460964 | MMS22L | protein_coding | 35.58 | -0.931991205 | 0.931991205 | ENSG00000146263 | n.s. |
| ENSG00000121741 | 5124.915292 | -0.699544943 | 0.2811601 | -2.488066 | 0.012843993 | 0.04661807 | ZMYM2 | protein_coding | 37.62 | -0.931406112 | 0.931406112 | ENSG00000121741 | n.s. |
| ENSG00000134574 | 589.3052041 | 0.574707221 | 0.2078335 | 2.765229 | 0.00568828 | 0.024006646 | DDB2 | protein_coding | 47.15 | 0.930835184 | 0.930835184 | ENSG00000134574 | n.s. |
| ENSG00000252690 | 431.6255255 | -0.612456078 | 0.2295087 | -2.668552 | 0.007617888 | 0.030360932 | SCARNA15 | processed_transcript | 38.13 | -0.929515342 | 0.929515342 | ENSG00000252690 | n.s. |
| ENSG00000235127 | 429.78773 | -0.646686425 | 0.2495887 | -2.591008 | 0.009569516 | 0.036619566 | | lincRNA | 41.51 | -0.928827177 | 0.928827177 | ENSG00000235127 | n.s. |
| ENSG00000183853 | 287.4706997 | -0.622024937 | 0.2352096 | -2.644556 | 0.008179808 | 0.032180692 | KIRREL | protein_coding | 49.16 | -0.928312894 | 0.928312894 | ENSG00000183853 | n.s. |
| ENSG00000104218 | 736.9190694 | -0.659830141 | 0.2575461 | -2.561988 | 0.010407486 | 0.03924795 | CSPP1 | protein_coding | 38.01 | -0.92784194 | 0.92784194 | ENSG00000104218 | n.s. |
| ENSG00000227704 | 126.0789771 | 0.616899487 | 0.2323719 | 2.654794 | 0.007935678 | 0.031370937 | | antisense | 40 | 0.927491418 | 0.927491418 | ENSG00000227704 | n.s. |
| ENSG00000101361 | 6262.954185 | 0.622107065 | 0.2354797 | 2.641872 | 0.008244933 | 0.03239114 | NOP56 | protein_coding | 54.33 | 0.926674363 | 0.926674363 | ENSG00000101361 | n.s. |
| ENSG00000065029 | 196.9572866 | -0.710430828 | 0.2887932 | -2.459998 | 0.013893774 | 0.0496772 | ZNF76 | protein_coding | 48.23 | -0.926290182 | 0.926290182 | ENSG00000065029 | n.s. |
| ENSG00000251862 | 8.703528407 | -0.98535413 | 0.4803084 | -2.051503 | 0.040217994 | 0.115146028 | MIR1343 | miRNA | 79.76 | -0.925002211 | 0.925002211 | ENSG00000251862 | n.s. |
| ENSG00000251804 | 3.954143748 | -1.654862218 | 1.0920014 | -1.51544 | 0.129661108 | 0.276289203 | RNU6-1294P | snRNA | 37.04 | -0.924465754 | 0.924465754 | ENSG00000251804 | n.s. |
| ENSG00000175782 | 639.4652936 | -0.683990963 | 0.2726239 | -2.508917 | 0.012110177 | 0.044511456 | SLC35E3 | protein_coding | 43.41 | -0.924433072 | 0.924433072 | ENSG00000175782 | n.s. |
| ENSG00000151572 | 104.0041921 | -0.669601895 | 0.2641609 | -2.534826 | 0.011250318 | 0.041843676 | ANO4 | protein_coding | 39.06 | -0.922959277 | 0.922959277 | ENSG00000151572 | n.s. |
| ENSG00000188803 | 66.56579443 | -0.663724212 | 0.260672 | -2.546204 | 0.010890142 | 0.040726148 | SHISA6 | protein_coding | 43.68 | -0.922660726 | 0.922660726 | ENSG00000188803 | n.s. |

| | | | | | | | | | | | | | |
|---|---|---|---|---|---|---|---|---|---|---|---|---|---|
| ENSG00000164841 | 113.4755435 | 0.622644977 | 0.2363521 | 2.634396 | 0.008428712 | 0.032979202 | TMEM74 | protein_coding | 37.41 | 0.922610335 | 0.922610335 | ENSG00000164841 | n.s. |
| ENSG00000273363 | 25.98465167 | 0.763905324 | 0.3237447 | 2.359592 | 0.018295068 | 0.061991297 | | sense_intronic | 53.32 | 0.922544992 | 0.922544992 | ENSG00000273363 | n.s. |
| ENSG00000153721 | 1234.147019 | -0.675738863 | 0.2679824 | -2.521579 | 0.011682948 | 0.043192162 | CNKSR3 | protein_coding | 42.73 | -0.922109914 | 0.922109914 | ENSG00000153721 | n.s. |
| ENSG00000165061 | 321.0090313 | -0.702793226 | 0.2847405 | -2.468188 | 0.013579882 | 0.048755208 | ZMAT4 | protein_coding | 40.92 | -0.922049945 | 0.922049945 | ENSG00000165061 | n.s. |
| ENSG00000253352 | 4283.977887 | -0.639798819 | 0.2465078 | -2.595451 | 0.0094467 | 0.036243155 | TUG1 | antisense | 39.53 | -0.921805509 | 0.921805509 | ENSG00000253352 | n.s. |
| ENSG00000189042 | 544.9391652 | -0.688837504 | 0.2761552 | -2.494385 | 0.012617555 | 0.045971594 | ZNF567 | protein_coding | 40.54 | -0.921327354 | 0.921327354 | ENSG00000189042 | n.s. |
| ENSG00000108557 | 948.6977099 | -0.60222213 | 0.2249851 | -2.67672 | 0.007434681 | 0.029745133 | RAI1 | protein_coding | 56.45 | -0.919342722 | 0.919342722 | ENSG00000108557 | n.s. |
| ENSG00000094916 | 8814.107976 | -0.629450567 | 0.240786 | -2.614149 | 0.008945002 | 0.034633276 | CBX5 | protein_coding | 42.96 | -0.919316595 | 0.919316595 | ENSG00000094916 | n.s. |
| ENSG00000063601 | 474.1044201 | -0.69450574 | 0.2800586 | -2.479858 | 0.013143456 | 0.047506837 | MTMR1 | protein_coding | 42.48 | -0.919000474 | 0.919000474 | ENSG00000063601 | n.s. |
| ENSG00000132842 | 919.3616668 | -0.689253357 | 0.2768389 | -2.489727 | 0.012784128 | 0.046437102 | AP3B1 | protein_coding | 36.53 | -0.918867696 | 0.918867696 | ENSG00000132842 | n.s. |
| ENSG00000115942 | 1357.47487 | -0.684086843 | 0.2736204 | -2.500131 | 0.012414726 | 0.045379423 | ORC2 | protein_coding | 39.05 | -0.918825105 | 0.918825105 | ENSG00000115942 | n.s. |
| ENSG00000153922 | 4487.072488 | -0.639321191 | 0.2470953 | -2.587347 | 0.009671825 | 0.036957027 | CHD1 | protein_coding | 35.16 | -0.91570164 | 0.91570164 | ENSG00000153922 | n.s. |
| ENSG00000176473 | 108.8926085 | -0.67768402 | 0.2701943 | -2.508136 | 0.012136995 | 0.044574883 | WDR25 | protein_coding | 46.07 | -0.915489973 | 0.915489973 | ENSG00000176473 | n.s. |
| ENSG00000083799 | 688.3691208 | -0.665908615 | 0.2630384 | -2.531602 | 0.011354267 | 0.042189875 | CYLD | protein_coding | 36.42 | -0.915485679 | 0.915485679 | ENSG00000083799 | n.s. |
| ENSG00000212597 | 5.02314133 | -1.204143927 | 0.661974 | -1.81902 | 0.068908403 | 0.174058725 | RNU6-876P | snRNA | 36.54 | -0.914311544 | 0.914311544 | ENSG00000212597 | n.s. |
| ENSG00000080986 | 1347.953695 | -0.672652228 | 0.2673169 | -2.51631 | 0.011859071 | 0.043727183 | NDC80 | protein_coding | 36.36 | -0.914301531 | 0.914301531 | ENSG00000080986 | n.s. |
| ENSG00000131149 | 3205.992443 | -0.617306726 | 0.2345385 | -2.632006 | 0.008488248 | 0.033132007 | GSE1 | protein_coding | 56.58 | -0.913461017 | 0.913461017 | ENSG00000131149 | n.s. |
| ENSG00000179818 | 1188.707242 | 0.575634621 | 0.2106963 | 2.732058 | 0.006294004 | 0.025926455 | PCBP1-AS1 | processed_transcript | 45.84 | 0.913104367 | 0.913104367 | ENSG00000179818 | n.s. |
| ENSG00000137955 | 11571.3115 | 0.57344637 | 0.2094066 | 2.738435 | 0.006173246 | 0.025588002 | RABGGTB | protein_coding | 37.86 | 0.912905764 | 0.912905764 | ENSG00000137955 | n.s. |
| ENSG00000103024 | 169.1853333 | 0.718163947 | 0.2963522 | 2.423346 | 0.015378271 | 0.053675523 | NME3 | protein_coding | 69.2 | 0.912228878 | 0.912228878 | ENSG00000103024 | n.s. |
| ENSG00000243353 | 9.362582917 | -0.894226208 | 0.4160614 | -2.149265 | 0.031613392 | 0.095477929 | RPS29P19 | processed_pseudogene | 47.95 | -0.91219748 | 0.91219748 | ENSG00000243353 | n.s. |
| ENSG00000102390 | 384.6166338 | 0.627276406 | 0.2405396 | 2.607788 | 0.009112925 | 0.03516609 | PBDC1 | protein_coding | 42.82 | 0.911982061 | 0.911982061 | ENSG00000102390 | n.s. |
| ENSG00000235908 | 46.69638982 | -0.731389161 | 0.3048964 | -2.398812 | 0.016448362 | 0.056713855 | RHOA-IT1 | sense_intronic | 47.91 | -0.911538233 | 0.911538233 | ENSG00000235908 | n.s. |
| ENSG00000124508 | 238.5535561 | -0.674997488 | 0.2692013 | -2.507408 | 0.012162024 | 0.044643252 | BTN2A2 | protein_coding | 45.78 | -0.911411427 | 0.911411427 | ENSG00000124508 | n.s. |
| ENSG00000125107 | 3723.630491 | -0.629792977 | 0.2421259 | -2.601097 | 0.009292605 | 0.035750466 | CNOT1 | protein_coding | 41.44 | -0.911133019 | 0.911133019 | ENSG00000125107 | n.s. |
| ENSG00000174373 | 84.97364571 | 0.639113931 | 0.2477468 | 2.579706 | 0.009888452 | 0.037555079 | RALGAPA1 | protein_coding | 36.27 | 0.9109491 | 0.9109491 | ENSG00000174373 | n.s. |
| ENSG00000103148 | 103.8874306 | -0.658765272 | 0.2596013 | -2.537604 | 0.011161435 | 0.041550979 | NPRL3 | protein_coding | 51.3 | -0.910030693 | 0.910030693 | ENSG00000103148 | n.s. |
| ENSG00000239899 | 32.03579838 | -0.702388569 | 0.2868437 | -2.448681 | 0.014338046 | 0.050899698 | RN7SL674P | misc_RNA | 54.18 | -0.908388456 | 0.908388456 | ENSG00000239899 | n.s. |
| ENSG00000169016 | 1025.254869 | -0.688237747 | 0.2784667 | -2.471526 | 0.01345378 | 0.048352345 | E2F6 | protein_coding | 41.96 | -0.905433507 | 0.905433507 | ENSG00000169016 | n.s. |
| ENSG00000163655 | 3037.664347 | -0.673222585 | 0.2692103 | -2.500732 | 0.012393706 | 0.045314489 | GMPS | protein_coding | 38.88 | -0.90465154 | 0.90465154 | ENSG00000163655 | n.s. |
| ENSG00000139517 | 200.1822434 | -0.683383545 | 0.2755505 | -2.480066 | 0.013135799 | 0.047491489 | LNX2 | protein_coding | 37.54 | -0.904378997 | 0.904378997 | ENSG00000139517 | n.s. |
| ENSG00000115904 | 250.8401727 | -0.669156728 | 0.266738 | -2.508666 | 0.012118791 | 0.044519769 | SOS1 | protein_coding | 37.8 | -0.904329916 | 0.904329916 | ENSG00000115904 | n.s. |

| | | | | | | | | | | | | | |
|---|---|---|---|---|---|---|---|---|---|---|---|---|---|
| ENSG00000170364 | 973.8583813 | -0.640318715 | 0.2494101 | -2.567333 | 0.010248408 | 0.038710956 | SETMAR | protein_coding | 40.2 | -0.904236387 | 0.904236387 | ENSG00000170364 | n.s. |
| ENSG00000200597 | 13.99165026 | -1.101615093 | 0.5791415 | -1.902152 | 0.057151323 | 0.151421097 | RNU1-87P | snRNA | 45.29 | -0.903119049 | 0.903119049 | ENSG00000200597 | n.s. |
| ENSG00000174446 | 342.2587644 | -0.609432055 | 0.2313351 | -2.634413 | 0.008428296 | 0.032979202 | SNAPC5 | protein_coding | 44.54 | -0.903031956 | 0.903031956 | ENSG00000174446 | n.s. |
| ENSG00000091127 | 1464.872475 | -0.681819119 | 0.2751107 | -2.478345 | 0.013199348 | 0.047659366 | PUS7 | protein_coding | 43 | -0.901263788 | 0.901263788 | ENSG00000091127 | n.s. |
| ENSG00000198000 | 1601.062213 | -0.646222392 | 0.2533947 | -2.55026 | 0.010764251 | 0.040342037 | NOL8 | protein_coding | 38.87 | -0.900990516 | 0.900990516 | ENSG00000198000 | n.s. |
| ENSG00000100722 | 1753.490476 | -0.648817373 | 0.2549774 | -2.544607 | 0.010940065 | 0.040890881 | ZC3H14 | protein_coding | 40.2 | -0.90080087 | 0.90080087 | ENSG00000100722 | n.s. |
| ENSG00000136492 | 3828.512005 | -0.634039845 | 0.2462069 | -2.575231 | 0.010017305 | 0.037961605 | BRIP1 | protein_coding | 35.92 | -0.900752154 | 0.900752154 | ENSG00000136492 | n.s. |
| ENSG00000118193 | 2310.564048 | -0.664844775 | 0.2647476 | -2.51124 | 0.012030794 | 0.044278283 | KIF14 | protein_coding | 37.83 | -0.900072989 | 0.900072989 | ENSG00000118193 | n.s. |
| ENSG00000198824 | 625.4275709 | -0.646652965 | 0.2539287 | -2.546593 | 0.010878033 | 0.040691792 | CHAMP1 | protein_coding | 42.93 | -0.899166544 | 0.899166544 | ENSG00000198824 | n.s. |
| ENSG00000227946 | 99.78440316 | 0.660640498 | 0.2624389 | 2.517312 | 0.01182542 | 0.043637774 | | lincRNA | 63.47 | 0.898561857 | 0.898561857 | ENSG00000227946 | n.s. |
| ENSG00000276575 | 3.23940813 | -1.198975816 | 0.6643356 | -1.804774 | 0.071110019 | 0.178076974 | MIR6895 | miRNA | 53.85 | -0.898503165 | 0.898503165 | ENSG00000276575 | n.s. |
| ENSG00000090861 | 1025.388948 | -0.674499258 | 0.2710684 | -2.488299 | 0.012835584 | 0.046599697 | AARS | protein_coding | 49.47 | -0.898174618 | 0.898174618 | ENSG00000090861 | n.s. |
| ENSG00000155868 | 396.5189626 | -0.625473195 | 0.2414926 | -2.59003 | 0.009596747 | 0.036697474 | MED7 | protein_coding | 41.05 | -0.897781599 | 0.897781599 | ENSG00000155868 | n.s. |
| ENSG00000138180 | 1098.306068 | -0.702584028 | 0.2890357 | -2.430787 | 0.015066089 | 0.05283757 | CEP55 | protein_coding | 40.68 | -0.897239968 | 0.897239968 | ENSG00000138180 | n.s. |
| ENSG00000102760 | 62.74132807 | 0.731347851 | 0.3074984 | 2.37838 | 0.017388915 | 0.059441657 | RGCC | protein_coding | 45.66 | 0.896565979 | 0.896565979 | ENSG00000102760 | n.s. |
| ENSG00000133026 | 3578.107352 | -0.621591601 | 0.2394888 | -2.595493 | 0.00944553 | 0.036243155 | MYH10 | protein_coding | 42 | -0.895573023 | 0.895573023 | ENSG00000133026 | n.s. |
| ENSG00000272578 | 275.3153998 | -0.651964321 | 0.2577204 | -2.529735 | 0.011414865 | 0.042369811 | | transcribed_unprocessed_pseud | 49.78 | -0.89511016 | 0.89511016 | ENSG00000272578 | n.s. |
| ENSG00000096070 | 167.3471317 | -0.621606085 | 0.2395643 | -2.594736 | 0.009466371 | 0.036308617 | BRPF3 | protein_coding | 47.04 | -0.895106726 | 0.895106726 | ENSG00000096070 | n.s. |
| ENSG00000264201 | 1.259778855 | -1.47145971 | 0.9229762 | -1.594255 | 0.110878843 | 0.246450533 | MIR4701 | miRNA | 58.73 | -0.89504515 | 0.89504515 | ENSG00000264201 | n.s. |
| ENSG00000101751 | 545.5841915 | -0.601614514 | 0.2279597 | -2.639127 | 0.008311981 | 0.032643184 | POLI | protein_coding | 36.47 | -0.894123994 | 0.894123994 | ENSG00000101751 | n.s. |
| ENSG00000165055 | 605.2921758 | -0.651861518 | 0.2578111 | -2.528447 | 0.011456853 | 0.042503001 | METTL2B | protein_coding | 45.29 | -0.894080488 | 0.894080488 | ENSG00000165055 | n.s. |
| ENSG00000174483 | 76.35521358 | -0.693987289 | 0.2841384 | -2.442427 | 0.014588876 | 0.051619041 | BBS1 | protein_coding | 49.91 | -0.893293548 | 0.893293548 | ENSG00000174483 | n.s. |
| ENSG00000147576 | 27.75154886 | 0.791600557 | 0.3484547 | 2.271746 | 0.02310184 | 0.074420853 | ADHFE1 | protein_coding | 43.01 | 0.893167151 | 0.893167151 | ENSG00000147576 | n.s. |
| ENSG00000165006 | 351.7579719 | -0.693561286 | 0.2838985 | -2.44299 | 0.014566129 | 0.051575249 | UBAP1 | protein_coding | 42.8 | -0.893000847 | 0.893000847 | ENSG00000165006 | n.s. |
| ENSG00000143951 | 188.4420789 | -0.684867976 | 0.2785107 | -2.459037 | 0.013931035 | 0.049759314 | WDPCP | protein_coding | 37.72 | -0.892469002 | 0.892469002 | ENSG00000143951 | n.s. |
| ENSG00000113597 | 383.7398141 | -0.782946373 | 0.3427976 | -2.28399 | 0.022372097 | 0.072709315 | TRAPPC13 | protein_coding | 36.8 | -0.891313937 | 0.891313937 | ENSG00000113597 | n.s. |
| ENSG00000170919 | 37.36152774 | -0.701998466 | 0.2897783 | -2.422536 | 0.015412588 | 0.053768345 | TPT1-AS1 | processed_transcript | 48.88 | -0.891168333 | 0.891168333 | ENSG00000170919 | n.s. |
| ENSG00000186063 | 414.22057 | -0.672259009 | 0.2708765 | -2.481792 | 0.013072357 | 0.047311261 | AIDA | protein_coding | 38.46 | -0.890767034 | 0.890767034 | ENSG00000186063 | n.s. |
| ENSG00000130147 | 992.1298653 | -0.630525746 | 0.2455845 | -2.567449 | 0.01024499 | 0.038708549 | SH3BP4 | protein_coding | 50.77 | -0.890424115 | 0.890424115 | ENSG00000130147 | n.s. |
| ENSG00000114030 | 1685.065159 | -0.68185556 | 0.2769934 | -2.461632 | 0.013830668 | 0.049518973 | KPNA1 | protein_coding | 38.27 | -0.889977219 | 0.889977219 | ENSG00000114030 | n.s. |
| ENSG00000185670 | 111.0887569 | -0.634913074 | 0.2482669 | -2.557381 | 0.010546372 | 0.039717916 | ZBTB3 | protein_coding | 51.23 | -0.889521815 | 0.889521815 | ENSG00000185670 | n.s. |
| ENSG00000067141 | 163.3835022 | -0.655946309 | 0.261058 | -2.512646 | 0.011982949 | 0.044125522 | NEO1 | protein_coding | 38.73 | -0.889010688 | 0.889010688 | ENSG00000067141 | n.s. |

| | | | | | | | | | | | | | |
|---|---|---|---|---|---|---|---|---|---|---|---|---|---|
| ENSG00000108256 | 4044.670366 | -0.642714795 | 0.2530979 | -2.539392 | 0.011104526 | 0.041383383 | NUFIP2 | protein_coding | 41.64 | -0.8889864 | 0.8889864 | ENSG00000108256 | n.s. |
| ENSG00000136463 | 606.0840144 | -0.689597221 | 0.2821558 | -2.44403 | 0.014524207 | 0.051481808 | TACO1 | protein_coding | 49.33 | -0.888439968 | 0.888439968 | ENSG00000136463 | n.s. |
| ENSG00000138380 | 218.4663179 | -0.617259149 | 0.2380291 | -2.593209 | 0.009508501 | 0.036440092 | CARF | protein_coding | 38.49 | -0.887878233 | 0.887878233 | ENSG00000138380 | n.s. |
| ENSG00000206562 | 425.0368531 | -0.624953069 | 0.2427548 | -2.574421 | 0.010040798 | 0.03804028 | METTL6 | protein_coding | 42.52 | -0.887281055 | 0.887281055 | ENSG00000206562 | n.s. |
| ENSG00000013364 | 31.99043955 | -0.762035233 | 0.3294105 | -2.31333 | 0.020704515 | 0.068570162 | MVP | protein_coding | 52.54 | -0.886906004 | 0.886906004 | ENSG00000013364 | n.s. |
| ENSG00000164040 | 872.4823261 | -0.669658778 | 0.2699976 | -2.48024 | 0.013129391 | 0.047480652 | PGRMC2 | protein_coding | 38.64 | -0.886282227 | 0.886282227 | ENSG00000164040 | n.s. |
| ENSG00000092445 | 449.9250065 | -0.652429263 | 0.2594313 | -2.514844 | 0.011908512 | 0.043886238 | TYRO3 | protein_coding | 53.19 | -0.885784714 | 0.885784714 | ENSG00000092445 | n.s. |
| ENSG00000178691 | 1429.446648 | -0.646528982 | 0.2559091 | -2.526401 | 0.011523796 | 0.042717201 | SUZ12 | protein_coding | 41 | -0.885354985 | 0.885354985 | ENSG00000178691 | n.s. |
| ENSG00000175309 | 165.4881761 | -0.632820994 | 0.2477023 | -2.554765 | 0.010625961 | 0.039920469 | PHYKPL | protein_coding | 51.43 | -0.885192767 | 0.885192767 | ENSG00000175309 | n.s. |
| ENSG00000128652 | 237.08809 | 0.591356051 | 0.2234747 | 2.646188 | 0.008140446 | 0.032037691 | HOXD3 | protein_coding | 51.11 | 0.883686284 | 0.883686284 | ENSG00000128652 | n.s. |
| ENSG00000269609 | 208.0780602 | -0.666464974 | 0.2686787 | -2.480528 | 0.013118797 | 0.047454666 | RPARP-AS1 | processed_transcript | 51.6 | -0.882213738 | 0.882213738 | ENSG00000269609 | n.s. |
| ENSG00000144681 | 1064.5114 | -0.658863241 | 0.2639958 | -2.495734 | 0.012569678 | 0.045825511 | STAC | protein_coding | 39.86 | -0.882147174 | 0.882147174 | ENSG00000144681 | n.s. |
| ENSG00000265690 | 108.7838944 | 0.646071276 | 0.2562281 | 2.521469 | 0.011686585 | 0.043194141 | | protein_coding | 67.54 | 0.881612815 | 0.881612815 | ENSG00000265690 | n.s. |
| ENSG00000267575 | 401.4726982 | -0.58546713 | 0.2204435 | -2.65586 | 0.007910647 | 0.031307543 | | lincRNA | 43.58 | -0.880748069 | 0.880748069 | ENSG00000267575 | n.s. |
| ENSG00000172175 | 655.2858748 | -0.654964355 | 0.2618091 | -2.501687 | 0.012360314 | 0.045216146 | MALT1 | protein_coding | 38.97 | -0.880734798 | 0.880734798 | ENSG00000172175 | n.s. |
| ENSG00000241357 | 24.07694626 | -0.776046766 | 0.3404609 | -2.279401 | 0.022643252 | 0.073333799 | | antisense | 66.67 | -0.880577018 | 0.880577018 | ENSG00000241357 | n.s. |
| ENSG00000087502 | 2542.110446 | -0.662009597 | 0.2664038 | -2.484986 | 0.012955668 | 0.046949964 | ERGIC2 | protein_coding | 34.04 | -0.879390202 | 0.879390202 | ENSG00000087502 | n.s. |
| ENSG00000255270 | 104.5843043 | -0.687770259 | 0.2827595 | -2.432351 | 0.015001167 | 0.052698493 | NAV2-IT1 | sense_intronic | 44.27 | -0.879109185 | 0.879109185 | ENSG00000255270 | n.s. |
| ENSG00000154027 | 105.0800019 | -0.697819226 | 0.2892507 | -2.412507 | 0.015843243 | 0.054989064 | AK5 | protein_coding | 39.61 | -0.879059396 | 0.879059396 | ENSG00000154027 | n.s. |
| ENSG00000185615 | 83.67579919 | 0.643488458 | 0.2550904 | 2.52259 | 0.011649402 | 0.043091026 | PDIA2 | protein_coding | 64.25 | 0.878756312 | 0.878756312 | ENSG00000185615 | n.s. |
| ENSG00000267041 | 1628.871276 | -0.660593898 | 0.2660253 | -2.4832 | 0.013020803 | 0.047149189 | ZNF850 | protein_coding | 41.15 | -0.876294829 | 0.876294829 | ENSG00000267041 | n.s. |
| ENSG00000126773 | 1424.145488 | -0.68861936 | 0.2838949 | -2.425613 | 0.01528254 | 0.053432908 | PCNX4 | protein_coding | 37.99 | -0.876055483 | 0.876055483 | ENSG00000126773 | n.s. |
| ENSG00000186350 | 381.0324501 | -0.593642327 | 0.2259864 | -2.626894 | 0.008616813 | 0.033558568 | RXRA | protein_coding | 62.16 | -0.875145485 | 0.875145485 | ENSG00000186350 | n.s. |
| ENSG00000100867 | 173.5221759 | 0.576490403 | 0.2160355 | 2.668498 | 0.007619126 | 0.030360932 | DHRS2 | protein_coding | 49.5 | 0.87493078 | 0.87493078 | ENSG00000100867 | n.s. |
| ENSG00000186446 | 506.0434766 | -0.56191608 | 0.2077367 | -2.704944 | 0.006831586 | 0.02779549 | ZNF501 | protein_coding | 39.42 | -0.87435584 | 0.87435584 | ENSG00000186446 | n.s. |
| ENSG00000100350 | 476.0730233 | -0.578712481 | 0.217429 | -2.661616 | 0.007776662 | 0.030867057 | FOXRED2 | protein_coding | 52.7 | -0.874147967 | 0.874147967 | ENSG00000100350 | n.s. |
| ENSG00000090989 | 949.771387 | -0.682134147 | 0.2801578 | -2.434821 | 0.014899163 | 0.052450326 | EXOC1 | protein_coding | 34.99 | -0.873303474 | 0.873303474 | ENSG00000090989 | n.s. |
| ENSG00000212190 | 5.872305858 | -1.025225515 | 0.5276123 | -1.943142 | 0.051999008 | 0.140825168 | RNU6-298P | snRNA | 34.38 | -0.872794701 | 0.872794701 | ENSG00000212190 | n.s. |
| ENSG00000247240 | 253.6727278 | 0.579493804 | 0.2180805 | 2.657247 | 0.007878173 | 0.031205632 | UBL7-AS1 | antisense | 43.8 | 0.872582658 | 0.872582658 | ENSG00000247240 | n.s. |
| ENSG00000172731 | 449.2183183 | -0.656517865 | 0.2641143 | -2.485734 | 0.01292845 | 0.04687573 | LRRC20 | protein_coding | 49.41 | -0.87254635 | 0.87254635 | ENSG00000172731 | n.s. |
| ENSG00000215712 | 486.9255989 | -0.661677365 | 0.267454 | -2.473985 | 0.013361522 | 0.048082831 | TMEM242 | protein_coding | 43.34 | -0.872097366 | 0.872097366 | ENSG00000215712 | n.s. |
| ENSG00000189043 | 4192.667844 | 0.680285431 | 0.2792391 | 2.436211 | 0.014842018 | 0.052275594 | NDUFA4 | protein_coding | 37.82 | 0.871922534 | 0.871922534 | ENSG00000189043 | n.s. |

| | | | | | | | | | | | | | |
|---|---|---|---|---|---|---|---|---|---|---|---|---|---|
| ENSG00000256594 | 170.2365378 | -0.595337087 | 0.2275743 | -2.616012 | 0.008896339 | 0.034473623 | | transcribed_unprocessed_pseud | 39.19 | -0.87068827 | 0.87068827 | ENSG00000256594 | n.s. |
| ENSG00000166526 | 1631.254602 | -0.622088661 | 0.2436303 | -2.553412 | 0.010667316 | 0.040043419 | ZNF3 | protein_coding | 48.95 | -0.869349525 | 0.869349525 | ENSG00000166526 | n.s. |
| ENSG00000163389 | 663.2952736 | -0.644358537 | 0.2572211 | -2.505077 | 0.012242476 | 0.044879408 | POGLUT1 | protein_coding | 41.18 | -0.868564944 | 0.868564944 | ENSG00000163389 | n.s. |
| ENSG00000111912 | 87.43031371 | -0.690423372 | 0.2865258 | -2.409638 | 0.015968355 | 0.055319086 | NCOA7 | protein_coding | 39.05 | -0.867948484 | 0.867948484 | ENSG00000111912 | n.s. |
| ENSG00000206149 | 514.3512051 | -0.609740077 | 0.2365657 | -2.577466 | 0.009952779 | 0.037760814 | HERC2P9 | transcribed_unprocessed_pseud | 42.75 | -0.867634919 | 0.867634919 | ENSG00000206149 | n.s. |
| ENSG00000106299 | 947.5055562 | -0.635004759 | 0.2516926 | -2.522938 | 0.011637894 | 0.0430599 | WASL | protein_coding | 35.43 | -0.867370133 | 0.867370133 | ENSG00000106299 | n.s. |
| ENSG00000143624 | 838.7731123 | -0.633446239 | 0.2508289 | -2.525411 | 0.011556297 | 0.042792108 | INTS3 | protein_coding | 48.32 | -0.866957531 | 0.866957531 | ENSG00000143624 | n.s. |
| ENSG00000237731 | 48.24383018 | -0.690466617 | 0.2867228 | -2.408133 | 0.016034337 | 0.055528587 | RNGTTP1 | processed_pseudogene | 41.35 | -0.866869363 | 0.866869363 | ENSG00000237731 | n.s. |
| ENSG00000173480 | 625.4936796 | -0.649137546 | 0.2605177 | -2.491722 | 0.012712565 | 0.046237473 | ZNF417 | protein_coding | 45.26 | -0.866602459 | 0.866602459 | ENSG00000173480 | n.s. |
| ENSG00000054654 | 5880.40443 | -0.655854527 | 0.2648229 | -2.476578 | 0.013264854 | 0.047846259 | SYNE2 | protein_coding | 40.76 | -0.865827671 | 0.865827671 | ENSG00000054654 | n.s. |
| ENSG00000272606 | 82.70924767 | 0.661787911 | 0.2685707 | 2.464111 | 0.013735375 | 0.049275347 | | antisense | 46.45 | 0.865201864 | 0.865201864 | ENSG00000272606 | n.s. |
| ENSG00000210184 | 3.227035635 | -1.151529596 | 0.637163 | -1.807276 | 0.070719192 | 0.177404599 | MT-TS2 | Mt_tRNA | 45.76 | -0.864839175 | 0.864839175 | ENSG00000210184 | n.s. |
| ENSG00000241472 | 152.6644858 | 0.605074831 | 0.2341442 | 2.584198 | 0.009760585 | 0.037221755 | PTPRG-AS1 | processed_transcript | 37.73 | 0.864774859 | 0.864774859 | ENSG00000241472 | n.s. |
| ENSG00000147256 | 131.3970914 | 0.664458033 | 0.2704648 | 2.456726 | 0.014020955 | 0.050016337 | ARHGAP36 | protein_coding | 46.18 | 0.86438556 | 0.86438556 | ENSG00000147256 | n.s. |
| ENSG00000200089 | 4.521759268 | -1.498710449 | 0.9712326 | -1.543102 | 0.122806143 | 0.265024713 | SNORD114-31 | snoRNA | 50.67 | -0.86432674 | 0.86432674 | ENSG00000200089 | n.s. |
| ENSG00000272168 | 1003.340946 | 0.556792526 | 0.2061112 | 2.701418 | 0.006904442 | 0.028050931 | CASC15 | lincRNA | 40.31 | 0.864171356 | 0.864171356 | ENSG00000272168 | n.s. |
| ENSG00000136527 | 25464.59639 | 0.571234825 | 0.2144721 | 2.663446 | 0.00773448 | 0.030732639 | TRA2B | protein_coding | 38.67 | 0.863935633 | 0.863935633 | ENSG00000136527 | n.s. |
| ENSG00000253043 | 8.404502435 | 0.919886684 | 0.4482974 | 2.051956 | 0.040173897 | 0.115043432 | RNU7-181P | snRNA | 36.49 | 0.863900699 | 0.863900699 | ENSG00000253043 | n.s. |
| ENSG00000265342 | 16.53984697 | 0.81420404 | 0.3708493 | 2.195512 | 0.028126911 | 0.087005328 | | antisense | 48.16 | 0.863426055 | 0.863426055 | ENSG00000265342 | n.s. |
| ENSG00000169045 | 25957.53593 | 0.582935397 | 0.221464 | 2.63219 | 0.008483629 | 0.033123267 | HNRNPH1 | protein_coding | 45.97 | 0.86266676 | 0.86266676 | ENSG00000169045 | n.s. |
| ENSG00000159556 | 154.6685339 | 0.644908012 | 0.25855 | 2.494326 | 0.012619653 | 0.045971594 | ISL2 | protein_coding | 59.5 | 0.8625712 | 0.8625712 | ENSG00000159556 | n.s. |
| ENSG00000240106 | 21.15571238 | 0.75755929 | 0.3317281 | 2.283676 | 0.0223906 | 0.072735495 | RN7SL146P | misc_RNA | 56.9 | 0.862294592 | 0.862294592 | ENSG00000240106 | n.s. |
| ENSG00000151148 | 644.0972816 | -0.621686857 | 0.2444758 | -2.542938 | 0.010992474 | 0.041020695 | UBE3B | protein_coding | 46.32 | -0.8622778 | 0.8622778 | ENSG00000151148 | n.s. |
| ENSG00000166133 | 396.2746641 | -0.598370566 | 0.2305321 | -2.595607 | 0.009442407 | 0.036243155 | RPUSD2 | protein_coding | 48.69 | -0.862116759 | 0.862116759 | ENSG00000166133 | n.s. |
| ENSG00000068796 | 1776.933582 | -0.651576715 | 0.2633139 | -2.474525 | 0.013341353 | 0.048047505 | KIF2A | protein_coding | 38.86 | -0.85899258 | 0.85899258 | ENSG00000068796 | n.s. |
| ENSG00000138621 | 99.66917538 | -0.742876516 | 0.3224513 | -2.303841 | 0.021231554 | 0.069883434 | PPCDC | protein_coding | 47.94 | -0.858487238 | 0.858487238 | ENSG00000138621 | n.s. |
| ENSG00000198270 | 329.2088755 | 0.562301371 | 0.2101069 | 2.676263 | 0.007444825 | 0.02976861 | TMEM116 | protein_coding | 40.27 | 0.858207659 | 0.858207659 | ENSG00000198270 | n.s. |
| ENSG00000197497 | 101.8094029 | -0.643255193 | 0.2586492 | -2.48698 | 0.012883278 | 0.046748469 | ZNF665 | protein_coding | 45.26 | -0.855679031 | 0.855679031 | ENSG00000197497 | n.s. |
| ENSG00000228415 | 14.7727698 | -0.825516629 | 0.3807388 | -2.168197 | 0.030143716 | 0.091997142 | PTMAP1 | processed_pseudogene | 50.46 | -0.855421519 | 0.855421519 | ENSG00000228415 | n.s. |
| ENSG00000136682 | 426.7823681 | -0.645539581 | 0.2601461 | -2.48145 | 0.013084901 | 0.047344354 | CBWD2 | protein_coding | 36.27 | -0.855166812 | 0.855166812 | ENSG00000136682 | n.s. |
| ENSG00000006831 | 1851.245931 | -0.630745564 | 0.2511006 | -2.511924 | 0.012007505 | 0.044204254 | ADIPOR2 | protein_coding | 39.4 | -0.854367534 | 0.854367534 | ENSG00000006831 | n.s. |
| ENSG00000103126 | 385.1544624 | -0.62124278 | 0.2454273 | -2.53127 | 0.011365018 | 0.042218554 | AXIN1 | protein_coding | 54.36 | -0.853896115 | 0.853896115 | ENSG00000103126 | n.s. |

| | | | | | | | | | | | | | |
|---|---|---|---|---|---|---|---|---|---|---|---|---|---|
| ENSG00000276722 | 9.750063237 | 0.9101813 | 0.4439991 | 2.049962 | 0.04036815 | 0.115504675 | | misc_RNA | 56.14 | 0.853204348 | 0.853204348 | ENSG00000276722 | n.s. |
| ENSG00000169925 | 722.1800198 | -0.609846983 | 0.2389325 | -2.552382 | 0.010698904 | 0.040151168 | BRD3 | protein_coding | 57.61 | -0.851530448 | 0.851530448 | ENSG00000169925 | n.s. |
| ENSG00000252779 | 10.17900767 | -0.866853864 | 0.4120713 | -2.10365 | 0.035408952 | 0.104228084 | RNU6-182P | snRNA | 33.33 | -0.851263711 | 0.851263711 | ENSG00000252779 | n.s. |
| ENSG00000244300 | 153.7278328 | -0.669055687 | 0.2758381 | -2.425538 | 0.015285734 | 0.053432908 | GATA2-AS1 | antisense | 55.65 | -0.85116675 | 0.85116675 | ENSG00000244300 | n.s. |
| ENSG00000206838 | 10.16183976 | -0.884267942 | 0.42538 | -2.078772 | 0.037638319 | 0.109222827 | SNORA5A | snoRNA | 52.24 | -0.850388618 | 0.850388618 | ENSG00000206838 | n.s. |
| ENSG00000140396 | 1044.277739 | -0.626405779 | 0.249196 | -2.513707 | 0.011946974 | 0.044016334 | NCOA2 | protein_coding | 38.24 | -0.849648113 | 0.849648113 | ENSG00000140396 | n.s. |
| ENSG00000125841 | 279.8166324 | -0.626281999 | 0.2494595 | -2.510555 | 0.012054142 | 0.044352491 | NRSN2 | protein_coding | 51.94 | -0.84741089 | 0.84741089 | ENSG00000125841 | n.s. |
| ENSG00000159596 | 179.2821984 | -0.616896927 | 0.2438681 | -2.529633 | 0.011418174 | 0.042370798 | TMEM69 | protein_coding | 44.47 | -0.84695837 | 0.84695837 | ENSG00000159596 | n.s. |
| ENSG00000175536 | 88.49218903 | 0.608631893 | 0.2390887 | 2.545632 | 0.01090801 | 0.040782016 | LIPT2 | protein_coding | 55.44 | 0.845713072 | 0.845713072 | ENSG00000175536 | n.s. |
| ENSG00000110619 | 1257.041916 | -0.674077423 | 0.2801732 | -2.405931 | 0.016131315 | 0.055800322 | CARS | protein_coding | 52.02 | -0.844863902 | 0.844863902 | ENSG00000110619 | n.s. |
| ENSG00000107833 | 449.3751215 | 0.762449053 | 0.3389899 | 2.249179 | 0.024501086 | 0.078079336 | NPM3 | protein_coding | 57.44 | 0.844384794 | 0.844384794 | ENSG00000107833 | n.s. |
| ENSG00000167964 | 193.5098743 | 0.674801033 | 0.2811457 | 2.400183 | 0.016386899 | 0.05654394 | RAB26 | protein_coding | 58.26 | 0.841891171 | 0.841891171 | ENSG00000167964 | n.s. |
| ENSG00000185591 | 2105.305673 | -0.627709119 | 0.2512914 | -2.497934 | 0.012491957 | 0.0456019 | SP1 | protein_coding | 45.65 | -0.841768638 | 0.841768638 | ENSG00000185591 | n.s. |
| ENSG00000265210 | 3.526453033 | -1.07568438 | 0.5812262 | -1.850716 | 0.064210489 | 0.165188959 | MIR4486 | miRNA | 57.14 | -0.841205607 | 0.841205607 | ENSG00000265210 | n.s. |
| ENSG00000109654 | 884.9638021 | -0.686077193 | 0.2885232 | -2.377893 | 0.017411879 | 0.059505546 | TRIM2 | protein_coding | 42.05 | -0.840748189 | 0.840748189 | ENSG00000109654 | n.s. |
| ENSG00000147894 | 1147.104552 | -0.691529358 | 0.2920621 | -2.367748 | 0.017896737 | 0.060851002 | C9orf72 | protein_coding | 35.84 | -0.840714554 | 0.840714554 | ENSG00000147894 | n.s. |
| ENSG00000231113 | 70.94919921 | -0.706709727 | 0.3019545 | -2.340451 | 0.019260458 | 0.06467507 | | antisense | 48.07 | -0.840463795 | 0.840463795 | ENSG00000231113 | n.s. |
| ENSG00000201772 | 28.77428074 | -0.749692589 | 0.3311442 | -2.263946 | 0.023577454 | 0.075698439 | SNORA5C | snoRNA | 59.12 | -0.840340228 | 0.840340228 | ENSG00000201772 | n.s. |
| ENSG00000089057 | 554.8838123 | -0.63064136 | 0.2533994 | -2.488725 | 0.012820217 | 0.046556046 | SLC23A2 | protein_coding | 44.93 | -0.840029371 | 0.840029371 | ENSG00000089057 | n.s. |
| ENSG00000196911 | 1377.492918 | -0.657909395 | 0.2708278 | -2.429254 | 0.015129941 | 0.052981432 | KPNA5 | protein_coding | 35.56 | -0.839411009 | 0.839411009 | ENSG00000196911 | n.s. |
| ENSG00000214654 | 65.65956099 | 0.62226457 | 0.2484418 | 2.504669 | 0.012256602 | 0.044907543 | | protein_coding | 41.27 | 0.838613953 | 0.838613953 | ENSG00000214654 | n.s. |
| ENSG00000170222 | 214.5425417 | -0.657262238 | 0.270602 | -2.428889 | 0.015145183 | 0.053021469 | ADPRM | protein_coding | 40.57 | -0.838369694 | 0.838369694 | ENSG00000170222 | n.s. |
| ENSG00000171517 | 342.8236187 | -0.705241731 | 0.3014753 | -2.339302 | 0.019319798 | 0.064832379 | LPAR3 | protein_coding | 41.6 | -0.837973894 | 0.837973894 | ENSG00000171517 | n.s. |
| ENSG00000157107 | 429.7120253 | -0.70358587 | 0.3003951 | -2.342202 | 0.019170354 | 0.064424051 | FCHO2 | protein_coding | 37.04 | -0.837936978 | 0.837936978 | ENSG00000157107 | n.s. |
| ENSG00000196741 | 120.253956 | 0.593325799 | 0.2310911 | 2.567497 | 0.010243563 | 0.038708549 | LINC01560 | lincRNA | 39.98 | 0.837890606 | 0.837890606 | ENSG00000196741 | n.s. |
| ENSG00000108861 | 390.6194645 | -0.666228846 | 0.2764591 | -2.409864 | 0.015958462 | 0.055300861 | DUSP3 | protein_coding | 50.91 | -0.837628279 | 0.837628279 | ENSG00000108861 | n.s. |
| ENSG00000279366 | 36.50783113 | -0.730598368 | 0.3186016 | -2.293141 | 0.021839882 | 0.071513801 | | TEC | 38.59 | -0.836980898 | 0.836980898 | ENSG00000279366 | n.s. |
| ENSG00000164209 | 1740.331252 | -0.704183123 | 0.3010161 | -2.339354 | 0.019317131 | 0.064832379 | SLC25A46 | protein_coding | 35.23 | -0.836716047 | 0.836716047 | ENSG00000164209 | n.s. |
| ENSG00000188629 | 170.4317169 | -0.658208433 | 0.2715053 | -2.424293 | 0.015338216 | 0.053576012 | ZNF177 | protein_coding | 38.88 | -0.836602415 | 0.836602415 | ENSG00000188629 | n.s. |
| ENSG00000080503 | 125.8282901 | -0.653574463 | 0.2685857 | -2.433392 | 0.014958078 | 0.052605258 | SMARCA2 | protein_coding | 40.37 | -0.835902683 | 0.835902683 | ENSG00000080503 | n.s. |
| ENSG00000179909 | 399.0970746 | -0.638538159 | 0.2590532 | -2.464892 | 0.013705442 | 0.049180636 | ZNF154 | protein_coding | 46.08 | -0.835339364 | 0.835339364 | ENSG00000179909 | n.s. |
| ENSG00000075539 | 2523.803226 | -0.646535893 | 0.2642739 | -2.446461 | 0.014426622 | 0.051175138 | FRYL | protein_coding | 36.94 | -0.834639678 | 0.834639678 | ENSG00000075539 | n.s. |

| | | | | | | | | | | | | | |
|---|---|---|---|---|---|---|---|---|---|---|---|---|---|
| ENSG00000162599 | 2009.1133 | -0.59824665 | 0.2345035 | -2.551121 | 0.010737715 | 0.040264049 | NFIA | protein_coding | 40.05 | -0.834603465 | 0.834603465 | ENSG00000162599 | n.s. |
| ENSG00000143013 | 1851.071073 | -0.651351283 | 0.2674393 | -2.435511 | 0.014870773 | 0.052363625 | LMO4 | protein_coding | 42.55 | -0.834361644 | 0.834361644 | ENSG00000143013 | n.s. |
| ENSG00000169914 | 630.1867449 | -0.595815918 | 0.2330699 | -2.556383 | 0.010576652 | 0.039799656 | OTUD3 | protein_coding | 41.28 | -0.834214188 | 0.834214188 | ENSG00000169914 | n.s. |
| ENSG00000188486 | 1003.377561 | 0.571786688 | 0.2189021 | 2.612065 | 0.008999708 | 0.034789566 | H2AFX | protein_coding | 66.42 | 0.833980041 | 0.833980041 | ENSG00000188486 | n.s. |
| ENSG00000257267 | 1185.114855 | -0.643788773 | 0.2627625 | -2.450078 | 0.014282512 | 0.0507449 | ZNF271P | transcribed_unitary_pseudogene | 38.12 | -0.83345384 | 0.83345384 | ENSG00000257267 | n.s. |
| ENSG00000196693 | 1302.616062 | -0.637974003 | 0.25903 | -2.462934 | 0.013780512 | 0.049411814 | ZNF33B | protein_coding | 40.2 | -0.833301996 | 0.833301996 | ENSG00000196693 | n.s. |
| ENSG00000177683 | 890.5395017 | -0.612103802 | 0.243083 | -2.518085 | 0.011799469 | 0.043553554 | THAP5 | protein_coding | 39.2 | -0.833058827 | 0.833058827 | ENSG00000177683 | n.s. |
| ENSG00000114026 | 175.3289976 | 0.591039105 | 0.2305027 | 2.564131 | 0.010343448 | 0.039038175 | OGG1 | protein_coding | 49.74 | 0.832484779 | 0.832484779 | ENSG00000114026 | n.s. |
| ENSG00000170113 | 421.5653101 | -0.662077916 | 0.2747383 | -2.409849 | 0.015959122 | 0.055300861 | NIPA1 | protein_coding | 46.41 | -0.832409447 | 0.832409447 | ENSG00000170113 | n.s. |
| ENSG00000116641 | 2329.749723 | -0.6591811 | 0.272946 | -2.415061 | 0.01573258 | 0.05467916 | DOCK7 | protein_coding | 36.06 | -0.83200399 | 0.83200399 | ENSG00000116641 | n.s. |
| ENSG00000156531 | 1564.535137 | -0.647476085 | 0.2653649 | -2.439946 | 0.014689463 | 0.051882648 | PHF6 | protein_coding | 37.47 | -0.831992441 | 0.831992441 | ENSG00000156531 | n.s. |
| ENSG00000225791 | 263.6883639 | 0.562130456 | 0.2135591 | 2.632201 | 0.008483365 | 0.033123267 | TRAM2-AS1 | lincRNA | 44.52 | 0.831878217 | 0.831878217 | ENSG00000225791 | n.s. |
| ENSG00000138764 | 536.8935906 | 0.647369203 | 0.2653655 | 2.439538 | 0.014706058 | 0.05191492 | CCNG2 | protein_coding | 43.19 | 0.831680272 | 0.831680272 | ENSG00000138764 | n.s. |
| ENSG00000104517 | 4472.163885 | -0.626375957 | 0.2521592 | -2.484049 | 0.012989789 | 0.047052542 | UBR5 | protein_coding | 38.38 | -0.83146204 | 0.83146204 | ENSG00000104517 | n.s. |
| ENSG00000198554 | 1821.946285 | -0.666397898 | 0.2777704 | -2.399096 | 0.016435595 | 0.056683873 | WDHD1 | protein_coding | 39.43 | -0.830691958 | 0.830691958 | ENSG00000198554 | n.s. |
| ENSG00000119509 | 642.6116129 | -0.650281938 | 0.26748 | -2.431143 | 0.015051288 | 0.052812267 | INVS | protein_coding | 38.92 | -0.830582482 | 0.830582482 | ENSG00000119509 | n.s. |
| ENSG00000228889 | 163.8910125 | 0.584200938 | 0.2268073 | 2.57576 | 0.010002019 | 0.037913995 | UBAC2-AS1 | lincRNA | 44.85 | 0.830266635 | 0.830266635 | ENSG00000228889 | n.s. |
| ENSG00000048028 | 101.5296 | -0.673825716 | 0.2825969 | -2.384406 | 0.017106728 | 0.058620964 | USP28 | protein_coding | 41.45 | -0.8301176 | 0.8301176 | ENSG00000048028 | n.s. |
| ENSG00000199075 | 2.772314178 | -1.137444346 | 0.6401922 | -1.776723 | 0.07561378 | 0.186562229 | MIR26A1 | miRNA | 55.84 | -0.829397435 | 0.829397435 | ENSG00000199075 | n.s. |
| ENSG00000130584 | 263.886111 | -0.652798352 | 0.2693684 | -2.42344 | 0.01537428 | 0.05367505 | ZBTB46 | protein_coding | 53.89 | -0.829202446 | 0.829202446 | ENSG00000130584 | n.s. |
| ENSG00000113638 | 307.4011981 | -0.731346442 | 0.3211217 | -2.277474 | 0.022757923 | 0.073619556 | TTC33 | protein_coding | 38.63 | -0.828620496 | 0.828620496 | ENSG00000113638 | n.s. |
| ENSG00000265401 | 28.01342361 | 0.792261233 | 0.3637819 | 2.177847 | 0.029417428 | 0.090116357 | | antisense | 54.44 | 0.828068566 | 0.828068566 | ENSG00000265401 | n.s. |
| ENSG00000114982 | 1682.412765 | -0.650821706 | 0.2683419 | -2.425345 | 0.015293827 | 0.053447779 | KANSL3 | protein_coding | 44.51 | -0.827890988 | 0.827890988 | ENSG00000114982 | n.s. |
| ENSG00000164329 | 1034.64484 | -0.634260236 | 0.2577182 | -2.461061 | 0.013852686 | 0.049555743 | PAPD4 | protein_coding | 36.28 | -0.827649995 | 0.827649995 | ENSG00000164329 | n.s. |
| ENSG00000121067 | 1773.503757 | -0.655203816 | 0.2712165 | -2.415796 | 0.015700856 | 0.05459616 | SPOP | protein_coding | 40.44 | -0.827416213 | 0.827416213 | ENSG00000121067 | n.s. |
| ENSG00000132434 | 765.178452 | -0.661949218 | 0.275643 | -2.401473 | 0.016329225 | 0.056372874 | LANCL2 | protein_coding | 42.91 | -0.826728119 | 0.826728119 | ENSG00000132434 | n.s. |
| ENSG00000046651 | 171.443684 | 0.567853369 | 0.2176251 | 2.60932 | 0.009072243 | 0.035028518 | OFD1 | protein_coding | 39.8 | 0.826555007 | 0.826555007 | ENSG00000046651 | n.s. |
| ENSG00000113369 | 4404.683404 | 0.520353666 | 0.1904102 | 2.732803 | 0.006279787 | 0.025896386 | ARRDC3 | protein_coding | 35.81 | 0.825676818 | 0.825676818 | ENSG00000113369 | n.s. |
| ENSG00000264916 | 19.71435059 | -0.755414558 | 0.3384834 | -2.231763 | 0.025630661 | 0.080969578 | RN7SL230P | misc_RNA | 55.12 | -0.824669548 | 0.824669548 | ENSG00000264916 | n.s. |
| ENSG00000181264 | 315.0602753 | -0.661108889 | 0.2756618 | -2.398261 | 0.016473129 | 0.056771132 | TMEM136 | protein_coding | 43.45 | -0.823657348 | 0.823657348 | ENSG00000181264 | n.s. |
| ENSG00000173928 | 54.11920878 | 0.643304741 | 0.2642209 | 2.434723 | 0.014903197 | 0.052451266 | SWSAP1 | protein_coding | 53.77 | 0.823587049 | 0.823587049 | ENSG00000173928 | n.s. |
| ENSG00000082497 | 600.1740932 | -0.624279157 | 0.2522985 | -2.474368 | 0.01334723 | 0.048056239 | SERTAD4 | protein_coding | 43.15 | -0.822956135 | 0.822956135 | ENSG00000082497 | n.s. |

| | | | | | | | | | | | | | |
|---|---|---|---|---|---|---|---|---|---|---|---|---|---|
| ENSG00000122779 | 2965.636637 | -0.618432798 | 0.2487425 | -2.486237 | 0.012910201 | 0.046821757 | TRIM24 | protein_coding | 39.19 | -0.822238745 | 0.822238745 | ENSG00000122779 | n.s. |
| ENSG00000183783 | 110.7026814 | 0.639654072 | 0.2621297 | 2.44022 | 0.014678313 | 0.051856421 | KCTD8 | protein_coding | 35.64 | 0.822081787 | 0.822081787 | ENSG00000183783 | n.s. |
| ENSG00000226360 | 84.52726618 | 0.751442154 | 0.3363233 | 2.234285 | 0.025464338 | 0.080555929 | | processed_pseudogene | 51.69 | 0.822004443 | 0.822004443 | ENSG00000226360 | n.s. |
| ENSG00000150764 | 673.1837889 | -0.702862939 | 0.3030238 | -2.319497 | 0.020368079 | 0.067697484 | DIXDC1 | protein_coding | 43.3 | -0.821947229 | 0.821947229 | ENSG00000150764 | n.s. |
| ENSG00000123066 | 2639.573392 | -0.59435766 | 0.2341002 | -2.538903 | 0.011120066 | 0.041419122 | MED13L | protein_coding | 37.29 | -0.821877244 | 0.821877244 | ENSG00000123066 | n.s. |
| ENSG00000236609 | 179.9901558 | -0.655093296 | 0.2721478 | -2.407123 | 0.016078733 | 0.055646084 | ZNF853 | protein_coding | 57.53 | -0.821857377 | 0.821857377 | ENSG00000236609 | n.s. |
| ENSG00000070444 | 388.9202948 | 0.571779726 | 0.2206261 | 2.591623 | 0.009552444 | 0.036588355 | MNT | protein_coding | 61.67 | 0.821451411 | 0.821451411 | ENSG00000070444 | n.s. |
| ENSG00000230397 | 65.76461863 | -0.687589875 | 0.2930578 | -2.34626 | 0.018962878 | 0.063927396 | SPTLC1P1 | transcribed_processed_pseudog | 44.83 | -0.821197518 | 0.821197518 | ENSG00000230397 | n.s. |
| ENSG00000014138 | 1116.558148 | -0.622926115 | 0.2517892 | -2.473999 | 0.013361012 | 0.048082831 | POLA2 | protein_coding | 47.9 | -0.82102283 | 0.82102283 | ENSG00000014138 | n.s. |
| ENSG00000147439 | 115.1727728 | -0.678570599 | 0.2873912 | -2.361139 | 0.01821889 | 0.061778278 | BIN3 | protein_coding | 49.55 | -0.820503277 | 0.820503277 | ENSG00000147439 | n.s. |
| ENSG00000170946 | 330.851491 | -0.648798634 | 0.2683987 | -2.417294 | 0.015636369 | 0.05442249 | DNAJC24 | protein_coding | 36.59 | -0.820225241 | 0.820225241 | ENSG00000170946 | n.s. |
| ENSG00000238531 | 4.668553462 | 1.05752283 | 0.5741911 | 1.841761 | 0.065510082 | 0.167849994 | SNORD105B | snoRNA | 55.7 | 0.819663388 | 0.819663388 | ENSG00000238531 | n.s. |
| ENSG00000168116 | 2093.083818 | -0.636033148 | 0.2602696 | -2.443747 | 0.014535597 | 0.051496345 | KIAA1586 | protein_coding | 37.98 | -0.81935291 | 0.81935291 | ENSG00000168116 | n.s. |
| ENSG00000210174 | 3.800498299 | -1.132283305 | 0.6401703 | -1.768722 | 0.076940292 | 0.188997828 | MT-TR | Mt_tRNA | 23.08 | -0.81925587 | 0.81925587 | ENSG00000210174 | n.s. |
| ENSG00000214875 | 27.75421912 | 0.70614416 | 0.3057833 | 2.309296 | 0.02092714 | 0.069155748 | MED28P1 | processed_pseudogene | 47.99 | 0.819248483 | 0.819248483 | ENSG00000214875 | n.s. |
| ENSG00000115760 | 6485.97271 | -0.634140478 | 0.2592385 | -2.446166 | 0.014438438 | 0.051203863 | BIRC6 | protein_coding | 37.56 | -0.818483383 | 0.818483383 | ENSG00000115760 | n.s. |
| ENSG00000101882 | 317.0479541 | -0.68101005 | 0.2893299 | -2.35375 | 0.018585125 | 0.06285176 | NKAP | protein_coding | 41.82 | -0.818357897 | 0.818357897 | ENSG00000101882 | n.s. |
| ENSG00000118620 | 888.18969 | -0.630504418 | 0.257012 | -2.45321 | 0.01415875 | 0.050378809 | ZNF430 | protein_coding | 39.05 | -0.818238435 | 0.818238435 | ENSG00000118620 | n.s. |
| ENSG00000212027 | 9.656538551 | -0.907167456 | 0.4521929 | -2.006152 | 0.044840081 | 0.125363143 | MIR374B | miRNA | 41.67 | -0.818110942 | 0.818110942 | ENSG00000212027 | n.s. |
| ENSG00000141699 | 826.2521596 | -0.606327685 | 0.2420646 | -2.504817 | 0.01225146 | 0.04490052 | RETREG3 | protein_coding | 46.4 | -0.817177308 | 0.817177308 | ENSG00000141699 | n.s. |
| ENSG00000065883 | 1296.728442 | -0.606712494 | 0.2423811 | -2.503134 | 0.012309888 | 0.045055354 | CDK13 | protein_coding | 38.87 | -0.816788874 | 0.816788874 | ENSG00000065883 | n.s. |
| ENSG00000233016 | 1581.319992 | 0.551873498 | 0.2096511 | 2.632342 | 0.008479848 | 0.033123267 | SNHG7 | antisense | 58.94 | 0.816699286 | 0.816699286 | ENSG00000233016 | n.s. |
| ENSG00000155621 | 191.3946533 | -0.657774582 | 0.2747771 | -2.393848 | 0.016672671 | 0.057373604 | C9orf85 | protein_coding | 38.63 | -0.816487609 | 0.816487609 | ENSG00000155621 | n.s. |
| ENSG00000273284 | 40.44267188 | -0.721532689 | 0.3170245 | -2.275952 | 0.022848861 | 0.073862247 | | sense_intronic | 32.4 | -0.816470146 | 0.816470146 | ENSG00000273284 | n.s. |
| ENSG00000258568 | 84.67341813 | 0.59783348 | 0.2370111 | 2.522385 | 0.01165619 | 0.043104682 | RHOQP1 | processed_pseudogene | 48.88 | 0.816327 | 0.816327 | ENSG00000258568 | n.s. |
| ENSG00000076685 | 767.7763081 | -0.694679922 | 0.2987218 | -2.325508 | 0.0200448 | 0.066846407 | NT5C2 | protein_coding | 40.09 | -0.816194677 | 0.816194677 | ENSG00000076685 | n.s. |
| ENSG00000219470 | 61.74370863 | -0.634720563 | 0.2601736 | -2.439604 | 0.014703356 | 0.05191492 | | processed_pseudogene | 60.77 | -0.815430466 | 0.815430466 | ENSG00000219470 | n.s. |
| ENSG00000234684 | 66.37180894 | -0.645789057 | 0.2674367 | -2.414736 | 0.015746633 | 0.054700691 | SDCBP2-AS1 | antisense | 43.98 | -0.814990434 | 0.814990434 | ENSG00000234684 | n.s. |
| ENSG00000119772 | 2331.441021 | -0.606863954 | 0.2427568 | -2.499885 | 0.012423365 | 0.045399084 | DNMT3A | protein_coding | 53.43 | -0.814989716 | 0.814989716 | ENSG00000119772 | n.s. |
| ENSG00000261115 | 495.4300467 | -0.599055528 | 0.2380393 | -2.516625 | 0.011848496 | 0.043711344 | TMEM178B | protein_coding | 43.27 | -0.814359585 | 0.814359585 | ENSG00000261115 | n.s. |
| ENSG00000140987 | 417.6915992 | -0.659782693 | 0.276454 | -2.386591 | 0.017005401 | 0.058345539 | ZSCAN32 | protein_coding | 45.48 | -0.814166791 | 0.814166791 | ENSG00000140987 | n.s. |
| ENSG00000224790 | 89.14436033 | -0.636825031 | 0.2618213 | -2.432289 | 0.015003722 | 0.052698493 | | lincRNA | 47.75 | -0.813990903 | 0.813990903 | ENSG00000224790 | n.s. |

| | | | | | | | | | | | | | |
|---|---|---|---|---|---|---|---|---|---|---|---|---|---|
| ENSG00000186812 | 1878.720377 | -0.532267532 | 0.1986703 | -2.679149 | 0.007380944 | 0.029572629 | ZNF397 | protein_coding | 41.49 | -0.813895641 | 0.813895641 | ENSG00000186812 | n.s. |
| ENSG00000233250 | 10.66869364 | 0.822640432 | 0.3895556 | 2.111741 | 0.034708674 | 0.102647111 | | antisense | 45.34 | 0.813306153 | 0.813306153 | ENSG00000233250 | n.s. |
| ENSG00000173451 | 194.5157957 | -0.559616148 | 0.2146779 | -2.606772 | 0.009140031 | 0.035241389 | THAP2 | protein_coding | 37.49 | -0.813092594 | 0.813092594 | ENSG00000173451 | n.s. |
| ENSG00000116675 | 991.9792123 | -0.656511477 | 0.2747444 | -2.389535 | 0.016869711 | 0.057941564 | DNAJC6 | protein_coding | 40.01 | -0.812111122 | 0.812111122 | ENSG00000116675 | n.s. |
| ENSG00000166225 | 1013.381864 | -0.61538994 | 0.248575 | -2.475671 | 0.013298613 | 0.047918373 | FRS2 | protein_coding | 37.91 | -0.812005758 | 0.812005758 | ENSG00000166225 | n.s. |
| ENSG00000136861 | 2056.788514 | -0.616077932 | 0.2490975 | -2.47324 | 0.013389402 | 0.048158267 | CDK5RAP2 | protein_coding | 43.21 | -0.811577419 | 0.811577419 | ENSG00000136861 | n.s. |
| ENSG00000143379 | 1633.63898 | -0.622083656 | 0.2529634 | -2.459185 | 0.013925295 | 0.049751572 | SETDB1 | protein_coding | 44.16 | -0.810695186 | 0.810695186 | ENSG00000143379 | n.s. |
| ENSG00000129003 | 2634.975795 | -0.633420213 | 0.2604728 | -2.431809 | 0.015023616 | 0.052728463 | VPS13C | protein_coding | 34.88 | -0.809482458 | 0.809482458 | ENSG00000129003 | n.s. |
| ENSG00000177646 | 701.2994223 | -0.63259114 | 0.2602189 | -2.430996 | 0.015057362 | 0.052820269 | ACAD9 | protein_coding | 49.2 | -0.807945021 | 0.807945021 | ENSG00000177646 | n.s. |
| ENSG00000169136 | 440.4962966 | 0.618794169 | 0.2513204 | 2.462173 | 0.013809817 | 0.049478663 | ATF5 | protein_coding | 55.56 | 0.807886553 | 0.807886553 | ENSG00000169136 | n.s. |
| ENSG00000170260 | 241.1088488 | -0.660047029 | 0.277957 | -2.374637 | 0.017566198 | 0.05994465 | ZNF212 | protein_coding | 45.75 | -0.806742197 | 0.806742197 | ENSG00000170260 | n.s. |
| ENSG00000116539 | 3641.32401 | -0.594887799 | 0.2367271 | -2.512969 | 0.011971982 | 0.044096799 | ASH1L | protein_coding | 41.38 | -0.806425695 | 0.806425695 | ENSG00000116539 | n.s. |
| ENSG00000213742 | 74.44563739 | 0.62621589 | 0.2563366 | 2.442944 | 0.014568007 | 0.051575249 | ZNF337-AS1 | antisense | 41.18 | 0.8062897 | 0.8062897 | ENSG00000213742 | n.s. |
| ENSG00000168488 | 808.8570073 | 0.6004701 | 0.2403392 | 2.498427 | 0.012474578 | 0.045574272 | ATXN2L | protein_coding | 52.64 | 0.805398693 | 0.805398693 | ENSG00000168488 | n.s. |
| ENSG00000163584 | 5324.442356 | 0.703498243 | 0.3069474 | 2.291918 | 0.021910401 | 0.071677283 | RPL22L1 | protein_coding | 40.36 | 0.805237078 | 0.805237078 | ENSG00000163584 | n.s. |
| ENSG00000124786 | 356.7759205 | -0.638639235 | 0.264613 | -2.413484 | 0.015800829 | 0.05486158 | SLC35B3 | protein_coding | 34.49 | -0.80515272 | 0.80515272 | ENSG00000124786 | n.s. |
| ENSG00000148123 | 194.8695288 | -0.662464846 | 0.2798501 | -2.367214 | 0.017922586 | 0.060907044 | PLPPR1 | protein_coding | 38.59 | -0.805115042 | 0.805115042 | ENSG00000148123 | n.s. |
| ENSG00000227836 | 19.75982006 | -0.747749881 | 0.3377932 | -2.213632 | 0.026854094 | 0.083993063 | | processed_pseudogene | 45.37 | -0.804396857 | 0.804396857 | ENSG00000227836 | n.s. |
| ENSG00000198918 | 5588.447586 | 0.69428075 | 0.3009794 | 2.306738 | 0.021069426 | 0.069415431 | RPL39 | protein_coding | 46.11 | 0.80435478 | 0.80435478 | ENSG00000198918 | n.s. |
| ENSG00000239249 | 14.30263802 | -0.823855581 | 0.3928873 | -2.096926 | 0.036000152 | 0.105633322 | RN7SL757P | misc_RNA | 54.49 | -0.804247046 | 0.804247046 | ENSG00000239249 | n.s. |
| ENSG00000254791 | 108.6384567 | 0.635775321 | 0.2629222 | 2.418112 | 0.015601255 | 0.054358427 | FAR1-IT1 | sense_intronic | 44.96 | 0.804086103 | 0.804086103 | ENSG00000254791 | n.s. |
| ENSG00000103381 | 226.27343 | -0.696226422 | 0.3023382 | -2.302807 | 0.021289721 | 0.070025225 | CPPED1 | protein_coding | 44.42 | -0.803964318 | 0.803964318 | ENSG00000103381 | n.s. |
| ENSG00000135655 | 3518.510516 | -0.621288806 | 0.2538719 | -2.447253 | 0.014394962 | 0.051088721 | USP15 | protein_coding | 34.93 | -0.8025032 | 0.8025032 | ENSG00000135655 | n.s. |
| ENSG00000217767 | 20.39148483 | -0.761326431 | 0.347984 | -2.18782 | 0.028682705 | 0.088323649 | NDUFAB1P1 | processed_pseudogene | 47.39 | -0.802379435 | 0.802379435 | ENSG00000217767 | n.s. |
| ENSG00000276573 | 71.32621694 | 0.628525479 | 0.2586859 | 2.429686 | 0.015111906 | 0.05294491 | | sense_intronic | 39.01 | 0.802108998 | 0.802108998 | ENSG00000276573 | n.s. |
| ENSG00000206932 | 2.713335749 | 1.211958301 | 0.7232719 | 1.675661 | 0.093804654 | 0.217942059 | RNU6-4P | snRNA | 45.79 | 0.801903057 | 0.801903057 | ENSG00000206932 | n.s. |
| ENSG00000125249 | 1163.115293 | -0.683346001 | 0.2940632 | -2.323807 | 0.020135849 | 0.06706972 | RAP2A | protein_coding | 36.06 | -0.801888427 | 0.801888427 | ENSG00000125249 | n.s. |
| ENSG00000281156 | 5.126392915 | 0.963467633 | 0.5027532 | 1.916383 | 0.055316375 | 0.147513864 | MIR3651 | miRNA | 53.33 | 0.800802657 | 0.800802657 | ENSG00000281156 | n.s. |
| ENSG00000119771 | 446.57777 | -0.616938315 | 0.2514163 | -2.453852 | 0.014133531 | 0.050348555 | KLHL29 | protein_coding | 50.5 | -0.800793947 | 0.800793947 | ENSG00000119771 | n.s. |
| ENSG00000048828 | 1260.047598 | -0.593226071 | 0.2369402 | -2.503695 | 0.012290392 | 0.044995826 | FAM120A | protein_coding | 42.4 | -0.798973347 | 0.798973347 | ENSG00000048828 | n.s. |
| ENSG00000212135 | 18.74305249 | -0.738419458 | 0.3325138 | -2.220718 | 0.026370054 | 0.082878163 | SNORD67 | snoRNA | 51.35 | -0.798644863 | 0.798644863 | ENSG00000212135 | n.s. |
| ENSG00000180376 | 996.1036499 | -0.582162638 | 0.2304861 | -2.525804 | 0.011543391 | 0.042757256 | CCDC66 | protein_coding | 39.8 | -0.796974933 | 0.796974933 | ENSG00000180376 | n.s. |

| | | | | | | | | | | | | | |
|---|---|---|---|---|---|---|---|---|---|---|---|---|---|
| ENSG00000168303 | 843.6720358 | -0.618956817 | 0.2534043 | -2.442567 | 0.014583242 | 0.051612221 | MPLKIP | protein_coding | 40.64 | -0.796750583 | 0.796750583 | ENSG00000168303 | n.s. |
| ENSG00000198885 | 622.7041486 | -0.584784682 | 0.2321364 | -2.519143 | 0.011764102 | 0.043468929 | ITPRIPL1 | protein_coding | 50.7 | -0.796372096 | 0.796372096 | ENSG00000198885 | n.s. |
| ENSG00000141013 | 254.1045983 | -0.629554938 | 0.2603913 | -2.417726 | 0.015617812 | 0.054402501 | GAS8 | protein_coding | 52.63 | -0.795997388 | 0.795997388 | ENSG00000141013 | n.s. |
| ENSG00000178425 | 798.0701877 | -0.662297835 | 0.2813505 | -2.353996 | 0.018572832 | 0.062825447 | NT5DC1 | protein_coding | 36.29 | -0.795992199 | 0.795992199 | ENSG00000178425 | n.s. |
| ENSG00000138018 | 2701.911077 | -0.641448489 | 0.2682283 | -2.391427 | 0.016783019 | 0.057669188 | SELENOI | protein_coding | 42.63 | -0.794790704 | 0.794790704 | ENSG00000138018 | n.s. |
| ENSG00000103653 | 279.9153562 | -0.614159146 | 0.2507985 | -2.448815 | 0.014332692 | 0.050893669 | CSK | protein_coding | 56.64 | -0.79431428 | 0.79431428 | ENSG00000103653 | n.s. |
| ENSG00000154767 | 941.7619607 | -0.670892622 | 0.287348 | -2.334774 | 0.019555229 | 0.065511901 | XPC | protein_coding | 45.2 | -0.794122043 | 0.794122043 | ENSG00000154767 | n.s. |
| ENSG00000100796 | 2359.039637 | -0.600000339 | 0.2419712 | -2.479635 | 0.013151678 | 0.047524217 | PPP4R3A | protein_coding | 40.03 | -0.793851466 | 0.793851466 | ENSG00000100796 | n.s. |
| ENSG00000007047 | 272.1360165 | -0.616772253 | 0.2526749 | -2.440971 | 0.014647815 | 0.051788097 | MARK4 | protein_coding | 53.57 | -0.793027289 | 0.793027289 | ENSG00000007047 | n.s. |
| ENSG00000164741 | 3709.792533 | -0.581857365 | 0.2310157 | -2.518691 | 0.011779182 | 0.043490201 | DLC1 | protein_coding | 38.54 | -0.792261985 | 0.792261985 | ENSG00000164741 | n.s. |
| ENSG00000251728 | 4.848104477 | -1.028135504 | 0.5601282 | -1.835536 | 0.066426301 | 0.16960451 | | misc_RNA | 36.46 | -0.792242769 | 0.792242769 | ENSG00000251728 | n.s. |
| ENSG00000102781 | 881.290896 | -0.612948105 | 0.2505449 | -2.44646 | 0.014426662 | 0.051175138 | KATNAL1 | protein_coding | 38.41 | -0.791279826 | 0.791279826 | ENSG00000102781 | n.s. |
| ENSG00000101493 | 801.1069506 | -0.622168473 | 0.2565839 | -2.424815 | 0.015316199 | 0.053512532 | ZNF516 | protein_coding | 47.61 | -0.791114901 | 0.791114901 | ENSG00000101493 | n.s. |
| ENSG00000185000 | 209.8551712 | 0.63480481 | 0.2647338 | 2.397898 | 0.01648944 | 0.056799225 | DGAT1 | protein_coding | 63.02 | 0.790749428 | 0.790749428 | ENSG00000185000 | n.s. |
| ENSG00000112039 | 280.5120077 | -0.662226652 | 0.2823627 | -2.345305 | 0.01901152 | 0.06401388 | FANCE | protein_coding | 50.42 | -0.790517072 | 0.790517072 | ENSG00000112039 | n.s. |
| ENSG00000126391 | 175.1333642 | -0.598118984 | 0.2414127 | -2.477578 | 0.013227737 | 0.047737113 | FRMD8 | protein_coding | 56.27 | -0.790201219 | 0.790201219 | ENSG00000126391 | n.s. |
| ENSG00000099875 | 309.2376986 | 0.566786531 | 0.2222894 | 2.549769 | 0.010779433 | 0.040377882 | MKNK2 | protein_coding | 64.23 | 0.790019068 | 0.790019068 | ENSG00000099875 | n.s. |
| ENSG00000116830 | 2895.494203 | -0.639027297 | 0.267735 | -2.38679 | 0.016996183 | 0.058328287 | TTF2 | protein_coding | 41.16 | -0.788636867 | 0.788636867 | ENSG00000116830 | n.s. |
| ENSG00000206698 | 18.46618606 | -0.796848121 | 0.3772685 | -2.112151 | 0.034673502 | 0.10258376 | RNU1-73P | snRNA | 45.12 | -0.78802015 | 0.78802015 | ENSG00000206698 | n.s. |
| ENSG00000133422 | 954.6936274 | -0.604536022 | 0.2458783 | -2.45868 | 0.013944887 | 0.049776304 | MORC2 | protein_coding | 45.11 | -0.787696748 | 0.787696748 | ENSG00000133422 | n.s. |
| ENSG00000186665 | 205.2627624 | 0.626144026 | 0.2598163 | 2.409949 | 0.015954771 | 0.055300861 | C17orf58 | protein_coding | 52.41 | 0.787230914 | 0.787230914 | ENSG00000186665 | n.s. |
| ENSG00000252438 | 6.83249241 | -0.905849978 | 0.4608342 | -1.965674 | 0.049336242 | 0.135191597 | | snoRNA | 39.73 | -0.787229198 | 0.787229198 | ENSG00000252438 | n.s. |
| ENSG00000074054 | 2553.04379 | -0.589451689 | 0.2366525 | -2.49079 | 0.012745959 | 0.046310541 | CLASP1 | protein_coding | 41.05 | -0.786517262 | 0.786517262 | ENSG00000074054 | n.s. |
| ENSG00000163481 | 321.031759 | -0.676932982 | 0.2929681 | -2.310603 | 0.020854794 | 0.068969429 | RNF25 | protein_coding | 49.77 | -0.786151628 | 0.786151628 | ENSG00000163481 | n.s. |
| ENSG00000237976 | 63.16260356 | 0.658325494 | 0.280914 | 2.343512 | 0.019103124 | 0.064260157 | | antisense | 51.56 | 0.784762314 | 0.784762314 | ENSG00000237976 | n.s. |
| ENSG00000197238 | 106.3893423 | 0.600734592 | 0.2440932 | 2.461087 | 0.013851692 | 0.049555743 | HIST1H4J | protein_coding | 62.2 | 0.783902181 | 0.783902181 | ENSG00000197238 | n.s. |
| ENSG00000156802 | 3791.982577 | -0.64119534 | 0.2701022 | -2.373899 | 0.017601364 | 0.060020523 | ATAD2 | protein_coding | 40.58 | -0.78334849 | 0.78334849 | ENSG00000156802 | n.s. |
| ENSG00000271347 | 75.04064435 | -0.686267258 | 0.3000796 | -2.286951 | 0.022198682 | 0.072297602 | | sense_intronic | 26.49 | -0.782945917 | 0.782945917 | ENSG00000271347 | n.s. |
| ENSG00000151849 | 1699.895224 | -0.655454337 | 0.2795708 | -2.344502 | 0.019052505 | 0.064120867 | CENPJ | protein_coding | 41.37 | -0.781957427 | 0.781957427 | ENSG00000151849 | n.s. |
| ENSG00000128915 | 3091.87632 | -0.624927323 | 0.2599839 | -2.403715 | 0.016229398 | 0.05608387 | ICE2 | protein_coding | 36.91 | -0.781885334 | 0.781885334 | ENSG00000128915 | n.s. |
| ENSG00000153982 | 286.3722586 | -0.685230266 | 0.2996383 | -2.286858 | 0.022204091 | 0.072298307 | GDPD1 | protein_coding | 40.89 | -0.781759937 | 0.781759937 | ENSG00000153982 | n.s. |
| ENSG00000112079 | 698.4172676 | -0.660844311 | 0.2831698 | -2.333739 | 0.019609411 | 0.065646028 | STK38 | protein_coding | 41.52 | -0.781641063 | 0.781641063 | ENSG00000112079 | n.s. |

| | | | | | | | | | | | | | |
|---|---|---|---|---|---|---|---|---|---|---|---|---|---|
| ENSG00000066135 | 569.3452526 | -0.599216932 | 0.2435668 | -2.460175 | 0.013886943 | 0.049665531 | KDM4A | protein_coding | 46.75 | -0.781345872 | 0.781345872 | ENSG00000066135 | n.s. |
| ENSG00000100523 | 1459.231307 | -0.620641985 | 0.2574473 | -2.410754 | 0.015919592 | 0.055218855 | DDHD1 | protein_coding | 38.26 | -0.780713374 | 0.780713374 | ENSG00000100523 | n.s. |
| ENSG00000186868 | 65.10640728 | -0.681204126 | 0.297155 | -2.29242 | 0.021881429 | 0.071616158 | MAPT | protein_coding | 49.7 | -0.779971218 | 0.779971218 | ENSG00000186868 | n.s. |
| ENSG00000154359 | 592.7579959 | -0.636907033 | 0.2679694 | -2.37679 | 0.017464008 | 0.059625155 | LONRF1 | protein_coding | 38.66 | -0.779937548 | 0.779937548 | ENSG00000154359 | n.s. |
| ENSG00000265820 | 2.38216763 | -1.175156045 | 0.7002522 | -1.67819 | 0.093310074 | 0.217068699 | MIR3177 | miRNA | 63.41 | -0.779601806 | 0.779601806 | ENSG00000265820 | n.s. |
| ENSG00000166130 | 718.4178058 | -0.680679076 | 0.2969094 | -2.292548 | 0.021874037 | 0.071608799 | IKBIP | protein_coding | 40.95 | -0.779400418 | 0.779400418 | ENSG00000166130 | n.s. |
| ENSG00000138658 | 1744.47719 | -0.650756824 | 0.2769814 | -2.34946 | 0.018800644 | 0.063488084 | ZGRF1 | protein_coding | 37.99 | -0.779156208 | 0.779156208 | ENSG00000138658 | n.s. |
| ENSG00000165506 | 646.6150201 | -0.604669755 | 0.2474357 | -2.443745 | 0.014535706 | 0.051496345 | DNAAF2 | protein_coding | 46.14 | -0.778949847 | 0.778949847 | ENSG00000165506 | n.s. |
| ENSG00000277767 | 70.61611826 | -0.614793608 | 0.2540468 | -2.420001 | 0.015520464 | 0.054110754 | | lincRNA | 38.32 | -0.778769156 | 0.778769156 | ENSG00000277767 | n.s. |
| ENSG00000127452 | 118.232764 | 0.591516338 | 0.2392464 | 2.472414 | 0.013420385 | 0.048244777 | FBXL12 | protein_coding | 50.56 | 0.778760655 | 0.778760655 | ENSG00000127452 | n.s. |
| ENSG00000138802 | 1008.270271 | -0.609975444 | 0.2510819 | -2.429389 | 0.015124313 | 0.052975047 | SEC24B | protein_coding | 37.89 | -0.778285146 | 0.778285146 | ENSG00000138802 | n.s. |
| ENSG00000189060 | 1988.165858 | 0.626092231 | 0.2613803 | 2.39533 | 0.016605393 | 0.057170348 | H1F0 | protein_coding | 53.2 | 0.778125685 | 0.778125685 | ENSG00000189060 | n.s. |
| ENSG00000133863 | 742.0104741 | -0.652782702 | 0.2786178 | -2.342933 | 0.019132825 | 0.064313453 | TEX15 | protein_coding | 35.93 | -0.77791995 | 0.77791995 | ENSG00000133863 | n.s. |
| ENSG00000265282 | 34.34852433 | 0.707300578 | 0.3156161 | 2.241016 | 0.025025059 | 0.079476253 | | lincRNA | 48.05 | 0.777862734 | 0.777862734 | ENSG00000265282 | n.s. |
| ENSG00000164088 | 129.5729709 | 0.649739308 | 0.2766008 | 2.349014 | 0.018823177 | 0.063533357 | PPM1M | protein_coding | 58.15 | 0.77773678 | 0.77773678 | ENSG00000164088 | n.s. |
| ENSG00000099250 | 505.73689 | -0.598845268 | 0.2440545 | -2.453736 | 0.014138061 | 0.050348555 | NRP1 | protein_coding | 41.34 | -0.777308937 | 0.777308937 | ENSG00000099250 | n.s. |
| ENSG00000078725 | 138.5118289 | -0.593316247 | 0.2405464 | -2.466536 | 0.0136427 | 0.048968115 | BRINP1 | protein_coding | 40.76 | -0.777295675 | 0.777295675 | ENSG00000078725 | n.s. |
| ENSG00000249055 | 30.12682679 | -0.705740683 | 0.314777 | -2.242034 | 0.024959173 | 0.07932129 | TBCAP3 | transcribed_processed_pseudog | 40.71 | -0.776745418 | 0.776745418 | ENSG00000249055 | n.s. |
| ENSG00000283982 | 296.1041781 | -0.555796844 | 0.2178305 | -2.551511 | 0.010725698 | 0.040240878 | | lincRNA | 39.73 | -0.775521429 | 0.775521429 | ENSG00000283982 | n.s. |
| ENSG00000183354 | 593.8784415 | -0.642594375 | 0.2724683 | -2.358419 | 0.018352977 | 0.062157262 | KIAA2026 | protein_coding | 37.58 | -0.775295333 | 0.775295333 | ENSG00000183354 | n.s. |
| ENSG00000144366 | 2750.723359 | -0.629084771 | 0.2638815 | -2.383967 | 0.017127144 | 0.058662047 | GULP1 | protein_coding | 35.36 | -0.774807723 | 0.774807723 | ENSG00000144366 | n.s. |
| ENSG00000109436 | 828.658648 | -0.621468704 | 0.2591655 | -2.39796 | 0.016486646 | 0.056799225 | TBC1D9 | protein_coding | 41.09 | -0.774137207 | 0.774137207 | ENSG00000109436 | n.s. |
| ENSG00000182621 | 751.8876959 | -0.629267888 | 0.2641533 | -2.382208 | 0.017209197 | 0.058899617 | PLCB1 | protein_coding | 38.19 | -0.773928727 | 0.773928727 | ENSG00000182621 | n.s. |
| ENSG00000240370 | 65.21120811 | -0.639724208 | 0.2710639 | -2.360049 | 0.018272502 | 0.061944983 | RPL13P5 | transcribed_processed_pseudog | 48.71 | -0.772782918 | 0.772782918 | ENSG00000240370 | n.s. |
| ENSG00000218336 | 206.0070979 | -0.633523673 | 0.2670959 | -2.371896 | 0.017697075 | 0.060287834 | TENM3 | protein_coding | 38.92 | -0.772753374 | 0.772753374 | ENSG00000218336 | n.s. |
| ENSG00000162437 | 1147.590354 | -0.642815415 | 0.2731398 | -2.35343 | 0.018601089 | 0.062884491 | RAVER2 | protein_coding | 37.2 | -0.772314726 | 0.772314726 | ENSG00000162437 | n.s. |
| ENSG00000109790 | 615.5415346 | -0.658035929 | 0.2831702 | -2.323818 | 0.020135264 | 0.06706972 | KLHL5 | protein_coding | 36.93 | -0.772187728 | 0.772187728 | ENSG00000109790 | n.s. |
| ENSG00000182158 | 403.6874877 | -0.598338346 | 0.2449278 | -2.442917 | 0.014569091 | 0.051575249 | CREB3L2 | protein_coding | 43.66 | -0.770395726 | 0.770395726 | ENSG00000182158 | n.s. |
| ENSG00000213024 | 1359.062092 | -0.602100005 | 0.247554 | -2.432197 | 0.015007538 | 0.0526986 | NUP62 | protein_coding | 54.29 | -0.769604782 | 0.769604782 | ENSG00000213024 | n.s. |
| ENSG00000117533 | 486.5144381 | -0.663603294 | 0.287536 | -2.307897 | 0.021004879 | 0.069316591 | VAMP4 | protein_coding | 36.68 | -0.769224255 | 0.769224255 | ENSG00000117533 | n.s. |
| ENSG00000166263 | 913.7719243 | -0.637947061 | 0.2706374 | -2.357202 | 0.018413203 | 0.062315757 | STXBP4 | protein_coding | 36.22 | -0.768982743 | 0.768982743 | ENSG00000166263 | n.s. |
| ENSG00000171574 | 160.5818259 | -0.62077224 | 0.2598008 | -2.389416 | 0.016875174 | 0.057941564 | ZNF584 | protein_coding | 51.81 | -0.767901337 | 0.767901337 | ENSG00000171574 | n.s. |

| | | | | | | | | | | | | | |
|---|---|---|---|---|---|---|---|---|---|---|---|---|---|
| ENSG00000254330 | 68.26164916 | -0.716996685 | 0.3247677 | -2.207722 | 0.027263679 | 0.084996013 | | sense_intronic | 36.98 | -0.767617686 | 0.767617686 | ENSG00000254330 | n.s. |
| ENSG00000238113 | 41.47120353 | 0.652509618 | 0.2803933 | 2.327123 | 0.019958741 | 0.066623243 | LINC01410 | lincRNA | 40.99 | 0.767595503 | 0.767595503 | ENSG00000238113 | n.s. |
| ENSG00000266017 | 15.46812488 | -0.758682172 | 0.3547082 | -2.138891 | 0.032444509 | 0.097480556 | MIR4477B | miRNA | 41.98 | -0.76708989 | 0.76708989 | ENSG00000266017 | n.s. |
| ENSG00000238363 | 142.4502304 | 0.572501502 | 0.2293197 | 2.496521 | 0.012541811 | 0.045747875 | SNORA13 | snoRNA | 40.6 | 0.766939658 | 0.766939658 | ENSG00000238363 | n.s. |
| ENSG00000226642 | 185.1782085 | -0.661349947 | 0.2865792 | -2.307739 | 0.021013675 | 0.069316591 | ACTG1P12 | processed_pseudogene | 56.43 | -0.766612259 | 0.766612259 | ENSG00000226642 | n.s. |
| ENSG00000143353 | 603.7101396 | -0.595194971 | 0.2437823 | -2.441502 | 0.014626299 | 0.051725161 | LYPLAL1 | protein_coding | 33.96 | -0.765598187 | 0.765598187 | ENSG00000143353 | n.s. |
| ENSG00000122512 | 434.7599137 | -0.654216192 | 0.2819624 | -2.320225 | 0.020328725 | 0.067615106 | PMS2 | protein_coding | 45.05 | -0.765404336 | 0.765404336 | ENSG00000122512 | n.s. |
| ENSG00000232931 | 184.5923467 | -0.643481657 | 0.2749282 | -2.340545 | 0.01925564 | 0.06467507 | LINC00342 | lincRNA | 39.66 | -0.765268985 | 0.765268985 | ENSG00000232931 | n.s. |
| ENSG00000082213 | 706.552895 | -0.596735589 | 0.2449111 | -2.436539 | 0.014828564 | 0.052241422 | C5orf22 | protein_coding | 39.88 | -0.765006081 | 0.765006081 | ENSG00000082213 | n.s. |
| ENSG00000133114 | 834.9576993 | -0.660031743 | 0.2860443 | -2.307446 | 0.021029965 | 0.069346716 | GPALPP1 | protein_coding | 38.86 | -0.764959696 | 0.764959696 | ENSG00000133114 | n.s. |
| ENSG00000205791 | 55.94898184 | 0.623733942 | 0.2623667 | 2.377337 | 0.017438159 | 0.059566117 | LOH12CR2 | lincRNA | 45.19 | 0.764074521 | 0.764074521 | ENSG00000205791 | n.s. |
| ENSG00000130449 | 1657.14168 | -0.623670323 | 0.262358 | -2.377173 | 0.017445927 | 0.059578032 | ZSWIM6 | protein_coding | 38.17 | -0.763942415 | 0.763942415 | ENSG00000130449 | n.s. |
| ENSG00000128045 | 4563.07644 | 0.520718343 | 0.1987738 | 2.619653 | 0.008801931 | 0.034164842 | RASL11B | protein_coding | 47.73 | 0.763592097 | 0.763592097 | ENSG00000128045 | n.s. |
| ENSG00000084710 | 310.4417935 | -0.618683956 | 0.2592834 | -2.38613 | 0.017026734 | 0.058389957 | EFR3B | protein_coding | 48.38 | -0.763246792 | 0.763246792 | ENSG00000084710 | n.s. |
| ENSG00000108506 | 1424.132227 | -0.587857245 | 0.239517 | -2.454345 | 0.014114159 | 0.050323033 | INTS2 | protein_coding | 39.68 | -0.763175789 | 0.763175789 | ENSG00000108506 | n.s. |
| ENSG00000003393 | 1258.944059 | -0.609604438 | 0.2535896 | -2.403901 | 0.016221148 | 0.056083198 | ALS2 | protein_coding | 38.06 | -0.762717096 | 0.762717096 | ENSG00000003393 | n.s. |
| ENSG00000043093 | 1305.871879 | -0.61641941 | 0.25797 | -2.3895 | 0.016871321 | 0.057941564 | DCUN1D1 | protein_coding | 39.56 | -0.762516843 | 0.762516843 | ENSG00000043093 | n.s. |
| ENSG00000106049 | 248.7824949 | -0.660580971 | 0.287031 | -2.301427 | 0.021367505 | 0.07021472 | HIBADH | protein_coding | 36.88 | -0.762027599 | 0.762027599 | ENSG00000106049 | n.s. |
| ENSG00000107554 | 437.580268 | -0.563475019 | 0.2248283 | -2.506246 | 0.01220208 | 0.044766684 | DNMBP | protein_coding | 43.7 | -0.760153199 | 0.760153199 | ENSG00000107554 | n.s. |
| ENSG00000138640 | 1369.079482 | -0.644393781 | 0.2765661 | -2.329981 | 0.019807141 | 0.066212444 | FAM13A | protein_coding | 38.27 | -0.759779177 | 0.759779177 | ENSG00000138640 | n.s. |
| ENSG00000147536 | 632.704546 | -0.586096627 | 0.2390774 | -2.451494 | 0.014226471 | 0.05059391 | GINS4 | protein_coding | 46.76 | -0.75952365 | 0.75952365 | ENSG00000147536 | n.s. |
| ENSG00000213949 | 2288.070559 | -0.643580226 | 0.2761452 | -2.330586 | 0.019775185 | 0.066137377 | ITGA1 | protein_coding | 36.8 | -0.759137007 | 0.759137007 | ENSG00000213949 | n.s. |
| ENSG00000225975 | 51.52817738 | 0.644224314 | 0.2766162 | 2.328947 | 0.019861891 | 0.066336347 | LINC01534 | lincRNA | 47.16 | 0.7590563 | 0.7590563 | ENSG00000225975 | n.s. |
| ENSG00000145740 | 1003.258798 | -0.67974658 | 0.3009265 | -2.258846 | 0.023892971 | 0.076473986 | SLC30A5 | protein_coding | 39.35 | -0.758927727 | 0.758927727 | ENSG00000145740 | n.s. |
| ENSG00000198331 | 213.2047063 | -0.672398924 | 0.2960231 | -2.27144 | 0.023120327 | 0.074445956 | HYLS1 | protein_coding | 41.34 | -0.758572821 | 0.758572821 | ENSG00000198331 | n.s. |
| ENSG00000156103 | 1667.910643 | -0.655164422 | 0.2841938 | -2.305344 | 0.021147305 | 0.069639055 | MMP16 | protein_coding | 35.15 | -0.758121631 | 0.758121631 | ENSG00000156103 | n.s. |
| ENSG00000117151 | 335.9883637 | -0.633359511 | 0.2696653 | -2.348687 | 0.018839723 | 0.063573792 | CTBS | protein_coding | 36.59 | -0.757955188 | 0.757955188 | ENSG00000117151 | n.s. |
| ENSG00000135951 | 31.94447377 | 0.662745604 | 0.2895157 | 2.289152 | 0.022070502 | 0.072031786 | TSGA10 | protein_coding | 37.63 | 0.757170825 | 0.757170825 | ENSG00000135951 | n.s. |
| ENSG00000252622 | 35.15164326 | 0.663988856 | 0.2904405 | 2.286144 | 0.022245826 | 0.072417263 | RNU6-881P | snRNA | 40.2 | 0.757052137 | 0.757052137 | ENSG00000252622 | n.s. |
| ENSG00000138614 | 1099.495286 | -0.658478953 | 0.2866666 | -2.29702 | 0.021617638 | 0.070852724 | INTS14 | protein_coding | 40.67 | -0.757016014 | 0.757016014 | ENSG00000138614 | n.s. |
| ENSG00000124151 | 392.9636816 | -0.66142322 | 0.288715 | -2.290921 | 0.021967979 | 0.071831886 | NCOA3 | protein_coding | 41.15 | -0.756458315 | 0.756458315 | ENSG00000124151 | n.s. |
| ENSG00000207988 | 15.43785059 | -0.785312606 | 0.3773897 | -2.080906 | 0.037442468 | 0.10885602 | MIR576 | miRNA | 36.73 | -0.756371913 | 0.756371913 | ENSG00000207988 | n.s. |

| | | | | | | | | | | | | | |
|---|---|---|---|---|---|---|---|---|---|---|---|---|---|
| ENSG00000166012 | 4553.450095 | -0.558661689 | 0.2225865 | -2.509863 | 0.01207779 | 0.044416025 | TAF1D | protein_coding | 41.79 | -0.755567763 | 0.755567763 | ENSG00000166012 | n.s. |
| ENSG00000147421 | 488.6268381 | -0.600231581 | 0.248978 | -2.410782 | 0.015918378 | 0.055218855 | HMBOX1 | protein_coding | 37.72 | -0.755038869 | 0.755038869 | ENSG00000147421 | n.s. |
| ENSG00000153253 | 531.0724189 | -0.648293193 | 0.2802059 | -2.313631 | 0.020687953 | 0.068531614 | SCN3A | protein_coding | 35.07 | -0.754683967 | 0.754683967 | ENSG00000153253 | n.s. |
| ENSG00000154743 | 596.3455349 | -0.57846431 | 0.2351878 | -2.459585 | 0.013909768 | 0.049708856 | TSEN2 | protein_coding | 43.16 | -0.75406654 | 0.75406654 | ENSG00000154743 | n.s. |
| ENSG00000145375 | 1189.405991 | -0.581867777 | 0.2374116 | -2.450881 | 0.01425069 | 0.050654162 | SPATA5 | protein_coding | 37.39 | -0.753742712 | 0.753742712 | ENSG00000145375 | n.s. |
| ENSG00000025293 | 1378.158513 | -0.628460434 | 0.2672772 | -2.351344 | 0.018705755 | 0.063213648 | PHF20 | protein_coding | 44.08 | -0.753642929 | 0.753642929 | ENSG00000025293 | n.s. |
| ENSG00000144840 | 493.0227587 | -0.643200788 | 0.2770945 | -2.321232 | 0.020274312 | 0.067450239 | RABL3 | protein_coding | 37.93 | -0.75319874 | 0.75319874 | ENSG00000144840 | n.s. |
| ENSG00000271851 | 60.34832014 | 0.646938008 | 0.2796115 | 2.313703 | 0.020684032 | 0.068531614 | | sense_intronic | 40.23 | 0.753106384 | 0.753106384 | ENSG00000271851 | n.s. |
| ENSG00000267340 | 17.67745355 | -0.748365457 | 0.3509301 | -2.13252 | 0.032964109 | 0.098637285 | | processed_pseudogene | 50.51 | -0.752824886 | 0.752824886 | ENSG00000267340 | n.s. |
| ENSG00000047644 | 286.7990459 | -0.603293171 | 0.251398 | -2.399753 | 0.016406127 | 0.056596261 | WWC3 | protein_coding | 46.21 | -0.752434636 | 0.752434636 | ENSG00000047644 | n.s. |
| ENSG00000199674 | 9.135617618 | -0.860671858 | 0.4365656 | -1.971461 | 0.048671209 | 0.133764162 | RNU6-862P | snRNA | 40.95 | -0.751934771 | 0.751934771 | ENSG00000199674 | n.s. |
| ENSG00000170881 | 910.2220932 | 0.636003221 | 0.2726357 | 2.332795 | 0.019658908 | 0.065795906 | RNF139 | protein_coding | 39.17 | 0.751629324 | 0.751629324 | ENSG00000170881 | n.s. |
| ENSG00000196345 | 773.2837858 | -0.560767953 | 0.2246906 | -2.495734 | 0.012569676 | 0.045825511 | ZKSCAN7 | protein_coding | 45.82 | -0.75080811 | 0.75080811 | ENSG00000196345 | n.s. |
| ENSG00000199065 | 4.706631532 | -1.249843134 | 0.7905649 | -1.58095 | 0.113889584 | 0.251036202 | MIR101-2 | miRNA | 40.51 | -0.750235394 | 0.750235394 | ENSG00000199065 | n.s. |
| ENSG00000018189 | 1653.536465 | -0.60806022 | 0.2548109 | -2.386319 | 0.017017967 | 0.058374268 | RUFY3 | protein_coding | 39.91 | -0.750211659 | 0.750211659 | ENSG00000018189 | n.s. |
| ENSG00000164366 | 546.864271 | -0.617563523 | 0.2610509 | -2.365683 | 0.017996855 | 0.061129596 | CCDC127 | protein_coding | 51.76 | -0.749566784 | 0.749566784 | ENSG00000164366 | n.s. |
| ENSG00000227051 | 5108.551535 | -0.61557681 | 0.259803 | -2.369399 | 0.01781704 | 0.060666827 | C14orf132 | protein_coding | 47.76 | -0.74918697 | 0.74918697 | ENSG00000227051 | n.s. |
| ENSG00000112941 | 1038.592804 | -0.556241269 | 0.2221494 | -2.503906 | 0.012283056 | 0.04499263 | PAPD7 | protein_coding | 47.97 | -0.749178346 | 0.749178346 | ENSG00000112941 | n.s. |
| ENSG00000257261 | 16.30346518 | -0.748736371 | 0.3522695 | -2.125464 | 0.033547878 | 0.099944762 | | lincRNA | 38.48 | -0.748916041 | 0.748916041 | ENSG00000257261 | n.s. |
| ENSG00000096872 | 409.3166328 | -0.656208397 | 0.2868931 | -2.287292 | 0.022178761 | 0.072249623 | IFT74 | protein_coding | 37.56 | -0.748841669 | 0.748841669 | ENSG00000096872 | n.s. |
| ENSG00000198932 | 402.9133587 | -0.589163369 | 0.2431524 | -2.423021 | 0.015392023 | 0.05371006 | GPRASP1 | protein_coding | 48.65 | -0.748204698 | 0.748204698 | ENSG00000198932 | n.s. |
| ENSG00000089486 | 398.1331313 | -0.566477793 | 0.2287737 | -2.476149 | 0.01328083 | 0.047891478 | CDIP1 | protein_coding | 51.97 | -0.747604401 | 0.747604401 | ENSG00000089486 | n.s. |
| ENSG00000274015 | 55.33406443 | 0.640138821 | 0.2762084 | 2.317593 | 0.020471435 | 0.067943692 | | lincRNA | 43.32 | 0.747586669 | 0.747586669 | ENSG00000274015 | n.s. |
| ENSG00000199390 | 10.6968358 | -1.104076663 | 0.6496136 | -1.69959 | 0.089208112 | 0.210337771 | SNORD114-7 | snoRNA | 40.26 | -0.747551244 | 0.747551244 | ENSG00000199390 | n.s. |
| ENSG00000100154 | 1551.467552 | -0.542248228 | 0.2138845 | -2.535239 | 0.011237074 | 0.04182138 | TTC28 | protein_coding | 39.91 | -0.747544299 | 0.747544299 | ENSG00000100154 | n.s. |
| ENSG00000264559 | 4.815665243 | -1.197609307 | 0.7399448 | -1.618512 | 0.105552373 | 0.237684892 | MIR3162 | miRNA | 48.78 | -0.747306319 | 0.747306319 | ENSG00000264559 | n.s. |
| ENSG00000205559 | 41.25464947 | 0.654309592 | 0.2859667 | 2.288062 | 0.022133903 | 0.072120367 | CHKB-AS1 | antisense | 63.15 | 0.74718365 | 0.74718365 | ENSG00000205559 | n.s. |
| ENSG00000207766 | 4.286617317 | -0.963738116 | 0.5236457 | -1.840439 | 0.065703792 | 0.168176688 | MIR626 | miRNA | 31.91 | -0.746159014 | 0.746159014 | ENSG00000207766 | n.s. |
| ENSG00000120925 | 314.1912635 | -0.702275685 | 0.3195727 | -2.197546 | 0.027981495 | 0.086671047 | RNF170 | protein_coding | 42.83 | -0.745905233 | 0.745905233 | ENSG00000120925 | n.s. |
| ENSG00000178951 | 252.4711284 | -0.58343223 | 0.2398518 | -2.43247 | 0.014996242 | 0.052698493 | ZBTB7A | protein_coding | 62.12 | -0.745744129 | 0.745744129 | ENSG00000178951 | n.s. |
| ENSG00000088826 | 117.3224874 | 0.623491695 | 0.2655791 | 2.347669 | 0.018891321 | 0.063717022 | SMOX | protein_coding | 49.87 | 0.745536784 | 0.745536784 | ENSG00000088826 | n.s. |
| ENSG00000086758 | 3850.476165 | -0.579906051 | 0.2376249 | -2.440426 | 0.014669943 | 0.051840005 | HUWE1 | protein_coding | 41.2 | -0.745373525 | 0.745373525 | ENSG00000086758 | n.s. |

| | | | | | | | | | | | | | |
|---|---|---|---|---|---|---|---|---|---|---|---|---|---|
| ENSG00000064933 | 1573.190994 | -0.629853515 | 0.2702821 | -2.330356 | 0.019787344 | 0.06616215 | PMS1 | protein_coding | 36.81 | -0.742843185 | 0.742843185 | ENSG00000064933 | n.s. |
| ENSG00000266124 | 9.985841506 | 0.802364501 | 0.3944952 | 2.033902 | 0.041961479 | 0.118745868 | MIR5587 | miRNA | 64.15 | 0.742493259 | 0.742493259 | ENSG00000266124 | n.s. |
| ENSG00000172986 | 159.2578396 | -0.61173769 | 0.2586662 | -2.36497 | 0.018031535 | 0.061217524 | GXYLT2 | protein_coding | 43.26 | -0.74211382 | 0.74211382 | ENSG00000172986 | n.s. |
| ENSG00000136715 | 518.5545881 | -0.590888835 | 0.2453779 | -2.408077 | 0.016036806 | 0.055528587 | SAP130 | protein_coding | 43.39 | -0.741851112 | 0.741851112 | ENSG00000136715 | n.s. |
| ENSG00000271737 | 52.88921195 | 0.648344814 | 0.2832513 | 2.288938 | 0.022082929 | 0.072038572 | | lincRNA | 62.26 | 0.740691747 | 0.740691747 | ENSG00000271737 | n.s. |
| ENSG00000159579 | 1599.904438 | -0.611720616 | 0.2589861 | -2.361983 | 0.018177493 | 0.061676605 | RSPRY1 | protein_coding | 41.63 | -0.74010826 | 0.74010826 | ENSG00000159579 | n.s. |
| ENSG00000110888 | 221.4596215 | -0.640714933 | 0.278373 | -2.301642 | 0.021355383 | 0.070191454 | CAPRIN2 | protein_coding | 37.58 | -0.739202915 | 0.739202915 | ENSG00000110888 | n.s. |
| ENSG00000273841 | 1352.961591 | 0.5980409 | 0.2504403 | 2.387958 | 0.016942274 | 0.058157613 | TAF9 | protein_coding | 44.77 | 0.738815796 | 0.738815796 | ENSG00000273841 | n.s. |
| ENSG00000143494 | 609.17526 | -0.607606435 | 0.2566228 | -2.367702 | 0.017898936 | 0.060851002 | VASH2 | protein_coding | 48.23 | -0.738686749 | 0.738686749 | ENSG00000143494 | n.s. |
| ENSG00000145916 | 174.6880351 | -0.628110592 | 0.2699646 | -2.326641 | 0.0199844 | 0.066692906 | RMND5B | protein_coding | 49.48 | -0.738608031 | 0.738608031 | ENSG00000145916 | n.s. |
| ENSG00000251666 | 31.7864335 | -0.745792722 | 0.3529502 | -2.113025 | 0.034598621 | 0.102405741 | ZNF346-IT1 | sense_intronic | 44.4 | -0.738092932 | 0.738092932 | ENSG00000251666 | n.s. |
| ENSG00000136141 | 2582.210398 | -0.577145017 | 0.2373175 | -2.431953 | 0.015017657 | 0.052720835 | LRCH1 | protein_coding | 40.02 | -0.737601558 | 0.737601558 | ENSG00000136141 | n.s. |
| ENSG00000101353 | 86.88191448 | -0.618117337 | 0.2636275 | -2.344662 | 0.019044356 | 0.064108938 | MROH8 | protein_coding | 45.33 | -0.737464307 | 0.737464307 | ENSG00000101353 | n.s. |
| ENSG00000135341 | 2109.612577 | -0.624072245 | 0.2676059 | -2.332057 | 0.019697693 | 0.065894037 | MAP3K7 | protein_coding | 35.81 | -0.737125357 | 0.737125357 | ENSG00000135341 | n.s. |
| ENSG00000075568 | 2990.608278 | -0.576632509 | 0.2370393 | -2.432645 | 0.014988991 | 0.052698493 | TMEM131 | protein_coding | 39.29 | -0.737052713 | 0.737052713 | ENSG00000075568 | n.s. |
| ENSG00000138346 | 1340.639534 | -0.601559548 | 0.2530295 | -2.377428 | 0.017433826 | 0.059565928 | DNA2 | protein_coding | 42.26 | -0.736911703 | 0.736911703 | ENSG00000138346 | n.s. |
| ENSG00000168522 | 1834.000324 | -0.632532662 | 0.2733671 | -2.313858 | 0.020675521 | 0.068523037 | FNTA | protein_coding | 42.58 | -0.736371374 | 0.736371374 | ENSG00000168522 | n.s. |
| ENSG00000128872 | 3520.356432 | -0.627750085 | 0.2702647 | -2.322723 | 0.020194044 | 0.067247471 | TMOD2 | protein_coding | 41.52 | -0.735926519 | 0.735926519 | ENSG00000128872 | n.s. |
| ENSG00000115145 | 497.3434219 | -0.613537968 | 0.2609801 | -2.350899 | 0.018728112 | 0.063273841 | STAM2 | protein_coding | 39.09 | -0.735494469 | 0.735494469 | ENSG00000115145 | n.s. |
| ENSG00000171792 | 928.654427 | -0.64296923 | 0.2809609 | -2.288465 | 0.02211047 | 0.072077752 | RHNO1 | protein_coding | 47.54 | -0.734398662 | 0.734398662 | ENSG00000171792 | n.s. |
| ENSG00000135945 | 2031.244196 | -0.60380976 | 0.2549343 | -2.368492 | 0.017860771 | 0.060785996 | REV1 | protein_coding | 39.33 | -0.734351296 | 0.734351296 | ENSG00000135945 | n.s. |
| ENSG00000141314 | 251.3251028 | -0.63230537 | 0.2736479 | -2.310653 | 0.020852038 | 0.068969429 | RHBDL3 | protein_coding | 51.16 | -0.734323647 | 0.734323647 | ENSG00000141314 | n.s. |
| ENSG00000152763 | 32.71347107 | -0.674547875 | 0.3028484 | -2.227345 | 0.025924243 | 0.081661367 | WDR78 | protein_coding | 36.44 | -0.733896862 | 0.733896862 | ENSG00000152763 | n.s. |
| ENSG00000138073 | 503.9047959 | -0.69073942 | 0.3143425 | -2.19741 | 0.027991166 | 0.086681719 | PREB | protein_coding | 57.47 | -0.73361533 | 0.73361533 | ENSG00000138073 | n.s. |
| ENSG00000213703 | 56.3124632 | -0.633592109 | 0.2747577 | -2.306003 | 0.021110441 | 0.069534108 | | processed_pseudogene | 50.73 | -0.733574283 | 0.733574283 | ENSG00000213703 | n.s. |
| ENSG00000176490 | 205.0633161 | -0.644514181 | 0.2822477 | -2.283506 | 0.02240059 | 0.07275097 | DIRAS1 | protein_coding | 61.18 | -0.73356104 | 0.73356104 | ENSG00000176490 | n.s. |
| ENSG00000173230 | 3006.412879 | -0.595257058 | 0.2497337 | -2.383568 | 0.017145735 | 0.05871128 | GOLGB1 | protein_coding | 38.07 | -0.732927187 | 0.732927187 | ENSG00000173230 | n.s. |
| ENSG00000230910 | 298.1036514 | 0.555219726 | 0.224241 | 2.475996 | 0.013286502 | 0.047899525 | | antisense | 37.68 | 0.732706146 | 0.732706146 | ENSG00000230910 | n.s. |
| ENSG00000243911 | 9.665854304 | -0.819601306 | 0.4105099 | -1.996544 | 0.045874701 | 0.127666093 | RN7SL430P | misc_RNA | 52.89 | -0.732661632 | 0.732661632 | ENSG00000243911 | n.s. |
| ENSG00000139154 | 1455.98528 | -0.605469189 | 0.256321 | -2.362152 | 0.018169208 | 0.061669889 | AEBP2 | protein_coding | 40.33 | -0.732573419 | 0.732573419 | ENSG00000139154 | n.s. |
| ENSG00000181852 | 586.1872963 | -0.672558644 | 0.3017827 | -2.228619 | 0.025839284 | 0.081452317 | RNF41 | protein_coding | 47.86 | -0.732481304 | 0.732481304 | ENSG00000181852 | n.s. |
| ENSG00000139926 | 2641.773559 | -0.680139139 | 0.3072792 | -2.213424 | 0.026868408 | 0.084008273 | FRMD6 | protein_coding | 39.42 | -0.731610671 | 0.731610671 | ENSG00000139926 | n.s. |

| | | | | | | | | | | | | | |
|---|---|---|---|---|---|---|---|---|---|---|---|---|---|
| ENSG00000104812 | 200.5575119 | -0.635913892 | 0.2767712 | -2.297616 | 0.021583661 | 0.070774684 | GYS1 | protein_coding | 52.54 | -0.731378597 | 0.731378597 | ENSG00000104812 | n.s. |
| ENSG00000151657 | 613.3653389 | -0.616519948 | 0.263926 | -2.335958 | 0.019493429 | 0.065336307 | KIN | protein_coding | 39.94 | -0.730480836 | 0.730480836 | ENSG00000151657 | n.s. |
| ENSG00000186684 | 207.7067909 | -0.61110407 | 0.2604961 | -2.345924 | 0.018979959 | 0.063956665 | CYP27C1 | protein_coding | 45.93 | -0.729728042 | 0.729728042 | ENSG00000186684 | n.s. |
| ENSG00000176595 | 560.0978879 | -0.603091897 | 0.2553385 | -2.361931 | 0.018180046 | 0.061676605 | KBTBD11 | protein_coding | 50.18 | -0.72966855 | 0.72966855 | ENSG00000176595 | n.s. |
| ENSG00000136859 | 359.3471121 | -0.632287338 | 0.2746878 | -2.30184 | 0.021344206 | 0.070171284 | ANGPTL2 | protein_coding | 49.92 | -0.729558788 | 0.729558788 | ENSG00000136859 | n.s. |
| ENSG00000259581 | 126.5532795 | -0.561525961 | 0.2288667 | -2.453507 | 0.014147097 | 0.050354385 | TYRO3P | processed_pseudogene | 57.26 | -0.728839754 | 0.728839754 | ENSG00000259581 | n.s. |
| ENSG00000168826 | 106.0673646 | -0.627668397 | 0.2717049 | -2.31011 | 0.020882044 | 0.069026755 | ZBTB49 | protein_coding | 43.97 | -0.72871205 | 0.72871205 | ENSG00000168826 | n.s. |
| ENSG00000172456 | 52.76125532 | -0.644167273 | 0.2832168 | -2.274467 | 0.022937945 | 0.07404707 | FGGY | protein_coding | 40.3 | -0.728226029 | 0.728226029 | ENSG00000172456 | n.s. |
| ENSG00000163686 | 162.4522376 | -0.679839197 | 0.3079144 | -2.207884 | 0.027252356 | 0.084982861 | ABHD6 | protein_coding | 45.11 | -0.727882516 | 0.727882516 | ENSG00000163686 | n.s. |
| ENSG00000272523 | 39.13214189 | 0.637614863 | 0.2791844 | 2.283848 | 0.022380457 | 0.072719511 | LINC01023 | lincRNA | 63.53 | 0.725828272 | 0.725828272 | ENSG00000272523 | n.s. |
| ENSG00000104312 | 255.1736565 | 0.591139038 | 0.2483906 | 2.379876 | 0.017318445 | 0.059215302 | RIPK2 | protein_coding | 36.01 | 0.725662212 | 0.725662212 | ENSG00000104312 | n.s. |
| ENSG00000182544 | 225.8779079 | 0.574940924 | 0.2381108 | 2.414594 | 0.015752757 | 0.054708316 | MFSD5 | protein_coding | 57.75 | 0.725544775 | 0.725544775 | ENSG00000182544 | n.s. |
| ENSG00000213185 | 51.15727993 | -0.727631389 | 0.3432917 | -2.119571 | 0.034042201 | 0.101038067 | FAM24B | protein_coding | 41.03 | -0.724367942 | 0.724367942 | ENSG00000213185 | n.s. |
| ENSG00000164576 | 401.1738976 | -0.648564256 | 0.2871128 | -2.258918 | 0.023888477 | 0.076473986 | SAP30L | protein_coding | 45.76 | -0.72411309 | 0.72411309 | ENSG00000164576 | n.s. |
| ENSG00000123395 | 432.2219136 | 0.617349292 | 0.2658648 | 2.322042 | 0.020230679 | 0.067321163 | ATG101 | protein_coding | 51.45 | 0.723439772 | 0.723439772 | ENSG00000123395 | n.s. |
| ENSG00000102100 | 105.7936839 | -0.665543524 | 0.299014 | -2.225794 | 0.026028008 | 0.081951144 | SLC35A2 | protein_coding | 52.04 | -0.723076419 | 0.723076419 | ENSG00000102100 | n.s. |
| ENSG00000016864 | 466.2762008 | -0.649489044 | 0.2879955 | -2.255206 | 0.02412041 | 0.07708757 | GLT8D1 | protein_coding | 44.49 | -0.722891467 | 0.722891467 | ENSG00000016864 | n.s. |
| ENSG00000144161 | 1375.157092 | -0.640619871 | 0.2820951 | -2.270936 | 0.02315087 | 0.074527068 | ZC3H8 | protein_coding | 38.5 | -0.722418037 | 0.722418037 | ENSG00000144161 | n.s. |
| ENSG00000274066 | 3.474433073 | 1.090631774 | 0.6505376 | 1.676509 | 0.093638627 | 0.217723882 | MIR6514 | miRNA | 48.57 | 0.722100678 | 0.722100678 | ENSG00000274066 | n.s. |
| ENSG00000156299 | 1940.379393 | -0.577679616 | 0.2404493 | -2.4025 | 0.016283422 | 0.056256595 | TIAM1 | protein_coding | 43.86 | -0.721999326 | 0.721999326 | ENSG00000156299 | n.s. |
| ENSG00000199593 | 13.5926777 | -1.14544033 | 0.703307 | -1.628649 | 0.103387325 | 0.234287391 | SNORD114-14 | snoRNA | 46.67 | -0.72191501 | 0.72191501 | ENSG00000199593 | n.s. |
| ENSG00000116191 | 553.6803488 | -0.666380113 | 0.2999057 | -2.221966 | 0.026285626 | 0.082668798 | RALGPS2 | protein_coding | 36.82 | -0.721462011 | 0.721462011 | ENSG00000116191 | n.s. |
| ENSG00000278291 | 159.4824554 | 0.55679062 | 0.2272565 | 2.450053 | 0.014283507 | 0.0507449 | | antisense | 45.9 | 0.72082537 | 0.72082537 | ENSG00000278291 | n.s. |
| ENSG00000234685 | 43.6267181 | 0.656517837 | 0.2933573 | 2.237946 | 0.025224587 | 0.079982219 | NUS1P2 | processed_pseudogene | 54.41 | 0.720204367 | 0.720204367 | ENSG00000234685 | n.s. |
| ENSG00000141642 | 332.1135962 | 0.524306393 | 0.2072478 | 2.529853 | 0.01141103 | 0.042366872 | ELAC1 | protein_coding | 41.78 | 0.719858835 | 0.719858835 | ENSG00000141642 | n.s. |
| ENSG00000279342 | 394.2628025 | -0.600900797 | 0.2557885 | -2.349209 | 0.018813333 | 0.063515527 | | sense_intronic | 39.09 | -0.719350419 | 0.719350419 | ENSG00000279342 | n.s. |
| ENSG00000120868 | 235.4408383 | -0.604748375 | 0.2583964 | -2.34039 | 0.01926363 | 0.06467507 | APAF1 | protein_coding | 39.48 | -0.719204922 | 0.719204922 | ENSG00000120868 | n.s. |
| ENSG00000127993 | 602.4768818 | -0.554842785 | 0.2263408 | -2.45136 | 0.014231746 | 0.050599744 | RBM48 | protein_coding | 39.51 | -0.718993954 | 0.718993954 | ENSG00000127993 | n.s. |
| ENSG00000151718 | 481.3980225 | -0.595373423 | 0.252408 | -2.358774 | 0.018335407 | 0.062112864 | WWC2 | protein_coding | 39.31 | -0.718507627 | 0.718507627 | ENSG00000151718 | n.s. |
| ENSG00000084070 | 161.0841634 | -0.612885898 | 0.2638987 | -2.322429 | 0.02020985 | 0.067284012 | SMAP2 | protein_coding | 41.95 | -0.718356282 | 0.718356282 | ENSG00000084070 | n.s. |
| ENSG00000235034 | 26.16678655 | 0.710525298 | 0.3327763 | 2.135144 | 0.032749249 | 0.098099782 | C19orf81 | protein_coding | 50.87 | 0.716445364 | 0.716445364 | ENSG00000235034 | n.s. |
| ENSG00000249850 | 220.1806485 | 0.586866257 | 0.2473744 | 2.372381 | 0.017673846 | 0.060223435 | KRT18P31 | processed_pseudogene | 55.88 | 0.71611444 | 0.71611444 | ENSG00000249850 | n.s. |

| | | | | | | | | | | | | | |
|---|---|---|---|---|---|---|---|---|---|---|---|---|---|
| ENSG00000202031 | 5.665234558 | 0.928662367 | 0.5059309 | 1.835552 | 0.066423964 | 0.16960451 | SNORD38A | snoRNA | 47.14 | 0.71559249 | 0.71559249 | ENSG00000202031 | n.s. |
| ENSG00000103540 | 1434.426383 | -0.616814718 | 0.2672828 | -2.307723 | 0.021014532 | 0.069316591 | CCP110 | protein_coding | 37.85 | -0.714988678 | 0.714988678 | ENSG00000103540 | n.s. |
| ENSG00000148835 | 454.1715294 | -0.633092405 | 0.278545 | -2.272855 | 0.02303493 | 0.074308465 | TAF5 | protein_coding | 43.36 | -0.714737083 | 0.714737083 | ENSG00000148835 | n.s. |
| ENSG00000159459 | 1830.32707 | -0.613254047 | 0.2649662 | -2.314462 | 0.020642421 | 0.068433296 | UBR1 | protein_coding | 39.81 | -0.714276945 | 0.714276945 | ENSG00000159459 | n.s. |
| ENSG00000160818 | 1644.399068 | -0.539750576 | 0.2178396 | -2.477743 | 0.013221628 | 0.047727439 | GPATCH4 | protein_coding | 48.64 | -0.713135664 | 0.713135664 | ENSG00000160818 | n.s. |
| ENSG00000200755 | 4.466845972 | -0.937309025 | 0.5145224 | -1.821707 | 0.06849948 | 0.173466398 | RNA5SP68 | rRNA | 50.42 | -0.713090308 | 0.713090308 | ENSG00000200755 | n.s. |
| ENSG00000129680 | 530.4001641 | -0.647842855 | 0.2890438 | -2.241331 | 0.02500462 | 0.079429461 | MAP7D3 | protein_coding | 40.66 | -0.712639052 | 0.712639052 | ENSG00000129680 | n.s. |
| ENSG00000048342 | 183.6903304 | 0.573780629 | 0.2396331 | 2.394413 | 0.016646999 | 0.057299422 | CC2D2A | protein_coding | 40.21 | 0.712549335 | 0.712549335 | ENSG00000048342 | n.s. |
| ENSG00000013293 | 1128.646495 | -0.649963287 | 0.2907321 | -2.235609 | 0.025377396 | 0.080338673 | SLC7A14 | protein_coding | 42.15 | -0.71175877 | 0.71175877 | ENSG00000013293 | n.s. |
| ENSG00000234222 | 53.31341833 | 0.604063829 | 0.259375 | 2.328921 | 0.019863264 | 0.066336347 | LIX1L-AS1 | antisense | 44.35 | 0.711737271 | 0.711737271 | ENSG00000234222 | n.s. |
| ENSG00000163877 | 221.5838029 | -0.622165726 | 0.2716401 | -2.290404 | 0.02199789 | 0.071895921 | SNIP1 | protein_coding | 43.23 | -0.711319431 | 0.711319431 | ENSG00000163877 | n.s. |
| ENSG00000008869 | 906.5262445 | -0.623799582 | 0.272887 | -2.285927 | 0.022258553 | 0.072441756 | HEATR5B | protein_coding | 38.59 | -0.711138406 | 0.711138406 | ENSG00000008869 | n.s. |
| ENSG00000131381 | 1019.303336 | -0.608195876 | 0.262317 | -2.318553 | 0.020419271 | 0.067835246 | RBSN | protein_coding | 44.4 | -0.710704005 | 0.710704005 | ENSG00000131381 | n.s. |
| ENSG00000275453 | 7.870379037 | -0.872602271 | 0.4605432 | -1.894724 | 0.058128955 | 0.153398302 | MIR548O2 | miRNA | 41.43 | -0.710454836 | 0.710454836 | ENSG00000275453 | n.s. |
| ENSG00000097046 | 2890.062924 | -0.634482034 | 0.2804366 | -2.262479 | 0.023667796 | 0.075910324 | CDC7 | protein_coding | 36.61 | -0.710428997 | 0.710428997 | ENSG00000097046 | n.s. |
| ENSG00000066044 | 865.0584157 | -0.625037741 | 0.2739994 | -2.281164 | 0.022538727 | 0.073086879 | ELAVL1 | protein_coding | 52.21 | -0.710143244 | 0.710143244 | ENSG00000066044 | n.s. |
| ENSG00000181788 | 533.0796808 | -0.597560561 | 0.2555224 | -2.338584 | 0.019356976 | 0.064941487 | SIAH2 | protein_coding | 44.98 | -0.709589882 | 0.709589882 | ENSG00000181788 | n.s. |
| ENSG00000257475 | 149.4693951 | -0.595097446 | 0.2539935 | -2.342963 | 0.019131273 | 0.064313453 | | sense_intronic | 44.99 | -0.709176537 | 0.709176537 | ENSG00000257475 | n.s. |
| ENSG00000077684 | 1511.799416 | -0.572198666 | 0.2392717 | -2.391418 | 0.016783419 | 0.057669188 | JADE1 | protein_coding | 44.21 | -0.708986284 | 0.708986284 | ENSG00000077684 | n.s. |
| ENSG00000283734 | 7.789649061 | 0.901342535 | 0.4854404 | 1.856752 | 0.063346389 | 0.163631126 | MIR4785 | miRNA | 80.82 | 0.708576085 | 0.708576085 | ENSG00000283734 | n.s. |
| ENSG00000259396 | 42.32334042 | -0.65739442 | 0.2967196 | -2.215541 | 0.026722941 | 0.083684999 | | sense_intronic | 45.55 | -0.708245446 | 0.708245446 | ENSG00000259396 | n.s. |
| ENSG00000244642 | 15.77965816 | -0.732299432 | 0.3510544 | -2.086 | 0.036978637 | 0.107989627 | RN7SL396P | misc_RNA | 56.27 | -0.707853783 | 0.707853783 | ENSG00000244642 | n.s. |
| ENSG00000252546 | 3.752786815 | 1.216987421 | 0.7842827 | 1.55172 | 0.120729166 | 0.262314635 | RNA5SP466 | rRNA | 43.52 | 0.707285681 | 0.707285681 | ENSG00000252546 | n.s. |
| ENSG00000198865 | 141.2608788 | -0.675672725 | 0.3100744 | -2.179066 | 0.029326733 | 0.089917662 | CCDC152 | protein_coding | 35.74 | -0.706858395 | 0.706858395 | ENSG00000198865 | n.s. |
| ENSG00000137601 | 778.9233991 | -0.609457414 | 0.2640138 | -2.30843 | 0.020975232 | 0.069252587 | NEK1 | protein_coding | 37.07 | -0.706704878 | 0.706704878 | ENSG00000137601 | n.s. |
| ENSG00000185088 | 2372.549615 | 0.512334881 | 0.2021233 | 2.534764 | 0.011252298 | 0.041843676 | RPS27L | protein_coding | 39.85 | 0.706187115 | 0.706187115 | ENSG00000185088 | n.s. |
| ENSG00000140443 | 142.016771 | -0.558582517 | 0.2310787 | -2.417282 | 0.015636889 | 0.05442249 | IGF1R | protein_coding | 44.52 | -0.706172078 | 0.706172078 | ENSG00000140443 | n.s. |
| ENSG00000107130 | 2006.873258 | -0.56858169 | 0.237537 | -2.393655 | 0.016681438 | 0.057389588 | NCS1 | protein_coding | 54.7 | -0.705704771 | 0.705704771 | ENSG00000107130 | n.s. |
| ENSG00000124302 | 523.2260403 | -0.619953002 | 0.2718003 | -2.280913 | 0.022553582 | 0.073111415 | CHST8 | protein_coding | 50.64 | -0.704275793 | 0.704275793 | ENSG00000124302 | n.s. |
| ENSG00000142039 | 162.5975616 | -0.628253778 | 0.2775089 | -2.263905 | 0.023579985 | 0.075698439 | CCDC97 | protein_coding | 53.51 | -0.704217877 | 0.704217877 | ENSG00000142039 | n.s. |
| ENSG00000235272 | 15.73979541 | -0.775739739 | 0.3857246 | -2.011124 | 0.044312408 | 0.124162137 | FAM103A2P | processed_pseudogene | 47.74 | -0.702828598 | 0.702828598 | ENSG00000235272 | n.s. |
| ENSG00000005889 | 216.7714069 | 0.528426402 | 0.2124988 | 2.486727 | 0.01289243 | 0.04676949 | ZFX | protein_coding | 40.77 | 0.702826859 | 0.702826859 | ENSG00000005889 | n.s. |

| | | | | | | | | | | | | | |
|---|---|---|---|---|---|---|---|---|---|---|---|---|---|
| ENSG00000277586 | 774.8380127 | 0.561565565 | 0.2336175 | 2.403783 | 0.016226424 | 0.05608387 | NEFL | protein_coding | 44.93 | 0.702609509 | 0.702609509 | ENSG00000277586 | n.s. |
| ENSG00000068097 | 1372.310369 | -0.593293844 | 0.2541222 | -2.33468 | 0.019560159 | 0.065512651 | HEATR6 | protein_coding | 40.4 | -0.70226699 | 0.70226699 | ENSG00000068097 | n.s. |
| ENSG00000207933 | 7.645949066 | 0.86361787 | 0.4565827 | 1.891482 | 0.058560036 | 0.154214372 | MIR9-1 | miRNA | 40.45 | 0.701149888 | 0.701149888 | ENSG00000207933 | n.s. |
| ENSG00000244754 | 3298.075335 | -0.618134494 | 0.2713116 | -2.278319 | 0.022707571 | 0.073525001 | N4BP2L2 | protein_coding | 39.02 | -0.700695602 | 0.700695602 | ENSG00000244754 | n.s. |
| ENSG00000145390 | 202.5770673 | 0.596160542 | 0.2564089 | 2.325039 | 0.020069888 | 0.066914046 | USP53 | protein_coding | 36.03 | 0.70018025 | 0.70018025 | ENSG00000145390 | n.s. |
| ENSG00000204623 | 210.4555034 | -0.53589972 | 0.217602 | -2.462752 | 0.013787542 | 0.04942429 | ZNRD1ASP | transcribed_unitary_pseudogene | 41.59 | -0.699916953 | 0.699916953 | ENSG00000204623 | n.s. |
| ENSG00000264585 | 41.05065163 | 0.622323349 | 0.2744823 | 2.267262 | 0.023374236 | 0.075124535 | MIR4449 | miRNA | 86.36 | 0.699627236 | 0.699627236 | ENSG00000264585 | n.s. |
| ENSG00000278948 | 81.06409327 | -0.603498205 | 0.2615562 | -2.307336 | 0.021036071 | 0.069346716 | | TEC | 52.24 | -0.69943879 | 0.69943879 | ENSG00000278948 | n.s. |
| ENSG00000186230 | 439.104287 | -0.593475682 | 0.2547914 | -2.329261 | 0.019845219 | 0.066307891 | ZNF749 | protein_coding | 48.04 | -0.699372389 | 0.699372389 | ENSG00000186230 | n.s. |
| ENSG00000100490 | 30.46844124 | 0.643266933 | 0.2890356 | 2.225563 | 0.026043487 | 0.081981342 | CDKL1 | protein_coding | 43.04 | 0.698771207 | 0.698771207 | ENSG00000100490 | n.s. |
| ENSG00000170385 | 903.835327 | 0.557092464 | 0.2315276 | 2.40616 | 0.016121185 | 0.055779139 | SLC30A1 | protein_coding | 38.51 | 0.698331113 | 0.698331113 | ENSG00000170385 | n.s. |
| ENSG00000126814 | 977.2189 | -0.577810278 | 0.244865 | -2.359709 | 0.018289261 | 0.061986705 | TRMT5 | protein_coding | 37.97 | -0.69782231 | 0.69782231 | ENSG00000126814 | n.s. |
| ENSG00000108784 | 602.6201944 | -0.537412381 | 0.219021 | -2.453702 | 0.014139399 | 0.050348555 | NAGLU | protein_coding | 57.01 | -0.697568251 | 0.697568251 | ENSG00000108784 | n.s. |
| ENSG00000114541 | 755.76409 | -0.6158978 | 0.2705362 | -2.276582 | 0.022811194 | 0.073774739 | FRMD4B | protein_coding | 41.42 | -0.697253168 | 0.697253168 | ENSG00000114541 | n.s. |
| ENSG00000220842 | 31.49426021 | 0.941581465 | 0.5254025 | 1.792115 | 0.073114613 | 0.181846153 | RPL21P16 | processed_pseudogene | 43.27 | 0.697048882 | 0.697048882 | ENSG00000220842 | n.s. |
| ENSG00000078018 | 3974.381255 | -0.596416766 | 0.2572566 | -2.318373 | 0.020429061 | 0.067851578 | MAP2 | protein_coding | 35.9 | -0.696877235 | 0.696877235 | ENSG00000078018 | n.s. |
| ENSG00000165392 | 1380.728451 | -0.613176978 | 0.2688041 | -2.28113 | 0.022540764 | 0.073086879 | WRN | protein_coding | 35.97 | -0.696667512 | 0.696667512 | ENSG00000165392 | n.s. |
| ENSG00000114999 | 2643.365359 | -0.588761128 | 0.2522656 | -2.333894 | 0.019601272 | 0.065634562 | TTL | protein_coding | 42.38 | -0.696426349 | 0.696426349 | ENSG00000114999 | n.s. |
| ENSG00000196367 | 1774.81398 | -0.545318937 | 0.2243826 | -2.430308 | 0.015085981 | 0.052867382 | TRRAP | protein_coding | 46.21 | -0.696269869 | 0.696269869 | ENSG00000196367 | n.s. |
| ENSG00000152348 | 192.3846405 | 0.536204848 | 0.2185083 | 2.453933 | 0.014130323 | 0.050348555 | ATG10 | protein_coding | 38.03 | 0.696000857 | 0.696000857 | ENSG00000152348 | n.s. |
| ENSG00000179833 | 648.9264754 | 0.5681673 | 0.2390619 | 2.376654 | 0.017470479 | 0.059632626 | SERTAD2 | protein_coding | 40.14 | 0.695729987 | 0.695729987 | ENSG00000179833 | n.s. |
| ENSG00000178381 | 233.7876615 | 0.59311953 | 0.255424 | 2.322098 | 0.020227656 | 0.067321163 | ZFAND2A | protein_coding | 54.85 | 0.695046165 | 0.695046165 | ENSG00000178381 | n.s. |
| ENSG00000183891 | 816.2504214 | 0.54172013 | 0.2222764 | 2.437146 | 0.014803706 | 0.052170515 | TTC32 | protein_coding | 39.39 | 0.694796626 | 0.694796626 | ENSG00000183891 | n.s. |
| ENSG00000276007 | 49.32111389 | -0.642391463 | 0.2894056 | -2.219692 | 0.026439663 | 0.083037633 | | sense_intronic | 37.34 | -0.694248539 | 0.694248539 | ENSG00000276007 | n.s. |
| ENSG00000280968 | 4.623633847 | -0.940168946 | 0.5254056 | -1.789415 | 0.073547929 | 0.18263025 | MIR3653 | miRNA | 56.36 | -0.694246403 | 0.694246403 | ENSG00000280968 | n.s. |
| ENSG00000165185 | 936.1842583 | -0.609062948 | 0.2665044 | -2.285377 | 0.022290731 | 0.072512582 | KIAA1958 | protein_coding | 38.95 | -0.694079991 | 0.694079991 | ENSG00000165185 | n.s. |
| ENSG00000199377 | 29.50172179 | 0.665431262 | 0.3058706 | 2.175532 | 0.029590278 | 0.090564313 | RNU5F-1 | snRNA | 35.04 | 0.694073356 | 0.694073356 | ENSG00000199377 | n.s. |
| ENSG00000185252 | 516.4403115 | -0.579711548 | 0.2469002 | -2.347959 | 0.018876606 | 0.063682818 | ZNF74 | protein_coding | 50.81 | -0.693322099 | 0.693322099 | ENSG00000185252 | n.s. |
| ENSG00000148153 | 564.9389838 | -0.632764496 | 0.2830292 | -2.235687 | 0.025372304 | 0.080338673 | INIP | protein_coding | 40.14 | -0.692924798 | 0.692924798 | ENSG00000148153 | n.s. |
| ENSG00000170846 | 507.4958065 | 0.500016518 | 0.1967315 | 2.541619 | 0.011034038 | 0.041153741 | | lincRNA | 53.75 | 0.692818227 | 0.692818227 | ENSG00000170846 | n.s. |
| ENSG00000163214 | 1479.995115 | -0.61251281 | 0.2692416 | -2.274956 | 0.022908588 | 0.073986609 | DHX57 | protein_coding | 41.54 | -0.692658199 | 0.692658199 | ENSG00000163214 | n.s. |
| ENSG00000092421 | 155.3370749 | -0.623997845 | 0.2771375 | -2.251582 | 0.024348675 | 0.077700345 | SEMA6A | protein_coding | 40.58 | -0.692373692 | 0.692373692 | ENSG00000092421 | n.s. |

| | | | | | | | | | | | | | |
|---|---|---|---|---|---|---|---|---|---|---|---|---|---|
| ENSG00000101294 | 1087.173439 | -0.561161075 | 0.2352032 | -2.385856 | 0.017039404 | 0.058407728 | HM13 | protein_coding | 49.84 | -0.692208844 | 0.692208844 | ENSG00000101294 | n.s. |
| ENSG00000163638 | 2211.680824 | -0.584233678 | 0.2501689 | -2.335357 | 0.019524803 | 0.065425713 | ADAMTS9 | protein_coding | 40.09 | -0.69187963 | 0.69187963 | ENSG00000163638 | n.s. |
| ENSG00000158195 | 309.6492009 | -0.538413789 | 0.2207949 | -2.438524 | 0.014747363 | 0.052008141 | WASF2 | protein_coding | 44 | -0.6912849 | 0.6912849 | ENSG00000158195 | n.s. |
| ENSG00000186310 | 733.979158 | 0.604832849 | 0.2641845 | 2.289433 | 0.022054185 | 0.071995409 | NAP1L3 | protein_coding | 42.29 | 0.691139592 | 0.691139592 | ENSG00000186310 | n.s. |
| ENSG00000199865 | 4.371710927 | 0.987466397 | 0.5698765 | 1.732772 | 0.083136116 | 0.199741872 | RNU6-495P | snRNA | 44.86 | 0.690763241 | 0.690763241 | ENSG00000199865 | n.s. |
| ENSG00000087299 | 368.822019 | -0.584630848 | 0.2506576 | -2.332389 | 0.019680255 | 0.065851524 | L2HGDH | protein_coding | 40.78 | -0.690702858 | 0.690702858 | ENSG00000087299 | n.s. |
| ENSG00000100580 | 870.6798146 | -0.586124168 | 0.2517175 | -2.328499 | 0.0198856 | 0.066395015 | TMED8 | protein_coding | 41.16 | -0.690374868 | 0.690374868 | ENSG00000100580 | n.s. |
| ENSG00000126870 | 299.5689529 | -0.607019279 | 0.2660625 | -2.281491 | 0.022519415 | 0.073051681 | WDR60 | protein_coding | 46.03 | -0.689798368 | 0.689798368 | ENSG00000126870 | n.s. |
| ENSG00000126062 | 368.5361419 | -0.545680395 | 0.2258199 | -2.41644 | 0.01567309 | 0.054513223 | TMEM115 | protein_coding | 57.53 | -0.689466166 | 0.689466166 | ENSG00000126062 | n.s. |
| ENSG00000204315 | 64.08201762 | -0.602606651 | 0.2631254 | -2.290188 | 0.022010401 | 0.071903054 | FKBPL | protein_coding | 54.57 | -0.688931658 | 0.688931658 | ENSG00000204315 | n.s. |
| ENSG00000212464 | 96.45769506 | 0.916143973 | 0.5070987 | 1.806638 | 0.070818691 | 0.177531264 | SNORA12 | snoRNA | 42.18 | 0.687772327 | 0.687772327 | ENSG00000212464 | n.s. |
| ENSG00000198805 | 1297.018026 | 0.576359314 | 0.2458451 | 2.3444 | 0.019057724 | 0.064122933 | PNP | protein_coding | 47.11 | 0.68758895 | 0.68758895 | ENSG00000198805 | n.s. |
| ENSG00000107821 | 67.14767589 | -0.592080272 | 0.2562661 | -2.310412 | 0.020865334 | 0.0689879 | KAZALD1 | protein_coding | 58.78 | -0.68753964 | 0.68753964 | ENSG00000107821 | n.s. |
| ENSG00000144868 | 688.3029336 | -0.562692575 | 0.2370172 | -2.374058 | 0.017593764 | 0.060015421 | TMEM108 | protein_coding | 40.21 | -0.687462419 | 0.687462419 | ENSG00000144868 | n.s. |
| ENSG00000188994 | 4376.874806 | -0.59735694 | 0.2598963 | -2.298443 | 0.021536583 | 0.070679478 | ZNF292 | protein_coding | 37.67 | -0.687382612 | 0.687382612 | ENSG00000188994 | n.s. |
| ENSG00000157851 | 818.9526516 | -0.587462003 | 0.2532241 | -2.31993 | 0.020344691 | 0.067635897 | DPYSL5 | protein_coding | 46.42 | -0.687226414 | 0.687226414 | ENSG00000157851 | n.s. |
| ENSG00000204116 | 326.3522721 | -0.617465417 | 0.2740403 | -2.253192 | 0.02424707 | 0.077411591 | CHIC1 | protein_coding | 37.63 | -0.68612387 | 0.68612387 | ENSG00000204116 | n.s. |
| ENSG00000266758 | 12.22030489 | 0.766114308 | 0.3834927 | 1.997728 | 0.045746111 | 0.127435595 | MIR3680-2 | miRNA | 43.68 | 0.685449557 | 0.685449557 | ENSG00000266758 | n.s. |
| ENSG00000165121 | 465.6393918 | -0.626239209 | 0.2801959 | -2.235005 | 0.025417008 | 0.080445783 | | transcribed_unprocessed_pseud | 37.75 | -0.685416759 | 0.685416759 | ENSG00000165121 | n.s. |
| ENSG00000135211 | 216.0822848 | -0.682296234 | 0.3202328 | -2.130626 | 0.033120004 | 0.099018641 | TMEM60 | protein_coding | 42.55 | -0.685218537 | 0.685218537 | ENSG00000135211 | n.s. |
| ENSG00000229992 | 28.94681683 | -0.692158516 | 0.3278036 | -2.111504 | 0.034729057 | 0.102682663 | HMGB3P9 | processed_pseudogene | 44.08 | -0.684200683 | 0.684200683 | ENSG00000229992 | n.s. |
| ENSG00000103994 | 1428.50906 | -0.602420616 | 0.2642431 | -2.279797 | 0.022619753 | 0.073274738 | ZNF106 | protein_coding | 41.35 | -0.683774944 | 0.683774944 | ENSG00000103994 | n.s. |
| ENSG00000126883 | 425.4345033 | 0.523868655 | 0.212816 | 2.461604 | 0.013831738 | 0.049518973 | NUP214 | protein_coding | 42.09 | 0.683768229 | 0.683768229 | ENSG00000126883 | n.s. |
| ENSG00000167702 | 146.5948801 | 0.622613125 | 0.2782369 | 2.237709 | 0.025240039 | 0.080012991 | KIFC2 | protein_coding | 63.6 | 0.682906666 | 0.682906666 | ENSG00000167702 | n.s. |
| ENSG00000201586 | 10.83659403 | -0.80966249 | 0.4188787 | -1.932928 | 0.053245042 | 0.143585265 | RNU6-593P | snRNA | 36.61 | -0.682456519 | 0.682456519 | ENSG00000201586 | n.s. |
| ENSG00000166507 | 217.6183423 | -0.565472978 | 0.2397966 | -2.358136 | 0.018366974 | 0.06218925 | NDST2 | protein_coding | 53.04 | -0.682121392 | 0.682121392 | ENSG00000166507 | n.s. |
| ENSG00000198951 | 202.7956985 | -0.601613958 | 0.2641224 | -2.277785 | 0.022739395 | 0.073576713 | NAGA | protein_coding | 53.29 | -0.681784803 | 0.681784803 | ENSG00000198951 | n.s. |
| ENSG00000228265 | 43.99837406 | 0.63571 | 0.2876803 | 2.209779 | 0.027120475 | 0.084644332 | RALY-AS1 | lincRNA | 49.5 | 0.68173675 | 0.68173675 | ENSG00000228265 | n.s. |
| ENSG00000146350 | 1038.655182 | -0.616253115 | 0.2742605 | -2.246963 | 0.024642388 | 0.078386101 | TBC1D32 | protein_coding | 35.75 | -0.681428624 | 0.681428624 | ENSG00000146350 | n.s. |
| ENSG00000173588 | 1224.279186 | -0.596342385 | 0.2605292 | -2.288966 | 0.022081333 | 0.072038572 | CEP83 | protein_coding | 37.69 | -0.681282357 | 0.681282357 | ENSG00000173588 | n.s. |
| ENSG00000196177 | 398.9216686 | -0.661840067 | 0.3063805 | -2.16019 | 0.030757954 | 0.093462025 | ACADSB | protein_coding | 40.72 | -0.681274879 | 0.681274879 | ENSG00000196177 | n.s. |
| ENSG00000039523 | 38.4922349 | 0.633543277 | 0.2863783 | 2.21226 | 0.026948721 | 0.084221559 | RIPOR1 | protein_coding | 57.05 | 0.680790858 | 0.680790858 | ENSG00000039523 | n.s. |

| | | | | | | | | | | | | | |
|---|---|---|---|---|---|---|---|---|---|---|---|---|---|
| ENSG00000261514 | 223.9267013 | 0.646195932 | 0.2954862 | 2.18689 | 0.028750522 | 0.088462655 | LINC01976 | lincRNA | 63.99 | 0.680599422 | 0.680599422 | ENSG00000261514 | n.s. |
| ENSG00000115966 | 1991.198611 | -0.59042917 | 0.2566579 | -2.300452 | 0.021422612 | 0.070379194 | ATF2 | protein_coding | 36.05 | -0.680502513 | 0.680502513 | ENSG00000115966 | n.s. |
| ENSG00000176624 | 4551.952633 | 0.517031223 | 0.2092571 | 2.470794 | 0.013481339 | 0.048438885 | MEX3C | protein_coding | 41.02 | 0.679795683 | 0.679795683 | ENSG00000176624 | n.s. |
| ENSG00000131626 | 1081.333275 | -0.589817413 | 0.2565161 | -2.299339 | 0.021485716 | 0.070553212 | PPFIA1 | protein_coding | 45 | -0.679164849 | 0.679164849 | ENSG00000131626 | n.s. |
| ENSG00000222511 | 6.139199828 | -0.891823323 | 0.4892734 | -1.822751 | 0.068341181 | 0.173254601 | | misc_RNA | 33.98 | -0.678958672 | 0.678958672 | ENSG00000222511 | n.s. |
| ENSG00000132475 | 37912.76415 | 0.563716533 | 0.2393426 | 2.35527 | 0.018509236 | 0.062625541 | H3F3B | protein_coding | 47.38 | 0.678291079 | 0.678291079 | ENSG00000132475 | n.s. |
| ENSG00000240771 | 758.6441182 | 0.583739901 | 0.2527957 | 2.309137 | 0.020935968 | 0.069155748 | ARHGEF25 | protein_coding | 56.4 | 0.677238522 | 0.677238522 | ENSG00000240771 | n.s. |
| ENSG00000073969 | 1019.095997 | -0.633648366 | 0.2874277 | -2.204548 | 0.027485803 | 0.085497059 | NSF | protein_coding | 38.62 | -0.676767393 | 0.676767393 | ENSG00000073969 | n.s. |
| ENSG00000112357 | 253.4890339 | -0.615631328 | 0.2749296 | -2.239232 | 0.025140795 | 0.079771031 | PEX7 | protein_coding | 38.37 | -0.676058496 | 0.676058496 | ENSG00000112357 | n.s. |
| ENSG00000277977 | 12.77363265 | -0.768154407 | 0.3880261 | -1.979646 | 0.047743298 | 0.131838543 | | unprocessed_pseudogene | 45.36 | -0.675943312 | 0.675943312 | ENSG00000277977 | n.s. |
| ENSG00000178773 | 100.3285079 | 0.59892018 | 0.2636008 | 2.272073 | 0.023082094 | 0.07438007 | CPNE7 | protein_coding | 60.32 | 0.675907426 | 0.675907426 | ENSG00000178773 | n.s. |
| ENSG00000266383 | 9.492976019 | 0.821886762 | 0.4314474 | 1.904953 | 0.056786231 | 0.150683189 | MIR5002 | miRNA | 48.45 | 0.675537658 | 0.675537658 | ENSG00000266383 | n.s. |
| ENSG00000172954 | 689.2007633 | -0.585011247 | 0.2540724 | -2.302538 | 0.021304877 | 0.070058527 | LCLAT1 | protein_coding | 37.54 | -0.6754183 | 0.6754183 | ENSG00000172954 | n.s. |
| ENSG00000105880 | 70.95182715 | 0.586915281 | 0.2553589 | 2.298394 | 0.021539397 | 0.070679478 | DLX5 | protein_coding | 55.55 | 0.675367326 | 0.675367326 | ENSG00000105880 | n.s. |
| ENSG00000277681 | 13.13237793 | -0.729283011 | 0.358477 | -2.034393 | 0.041912033 | 0.118630053 | MIR6739 | miRNA | 38.67 | -0.675174059 | 0.675174059 | ENSG00000277681 | n.s. |
| ENSG00000170396 | 693.1625526 | -0.549133522 | 0.2305557 | -2.381783 | 0.01722905 | 0.058953077 | ZNF804A | protein_coding | 34.04 | -0.67515611 | 0.67515611 | ENSG00000170396 | n.s. |
| ENSG00000198420 | 1135.125593 | -0.597156904 | 0.2626192 | -2.273851 | 0.022974951 | 0.074132153 | TCAF1 | protein_coding | 39.07 | -0.674783351 | 0.674783351 | ENSG00000198420 | n.s. |
| ENSG00000118257 | 3188.045368 | -0.581880359 | 0.2522457 | -2.3068 | 0.021065978 | 0.069415431 | NRP2 | protein_coding | 45.88 | -0.674133985 | 0.674133985 | ENSG00000118257 | n.s. |
| ENSG00000269044 | 92.46432403 | -0.638949212 | 0.291927 | -2.188729 | 0.028616529 | 0.088167278 | | sense_intronic | 46.78 | -0.673894984 | 0.673894984 | ENSG00000269044 | n.s. |
| ENSG00000165322 | 874.836974 | -0.604609519 | 0.2679133 | -2.256736 | 0.024024596 | 0.076824633 | ARHGAP12 | protein_coding | 36.94 | -0.67383701 | 0.67383701 | ENSG00000165322 | n.s. |
| ENSG00000265420 | 17.4691184 | 0.725127706 | 0.3559487 | 2.037169 | 0.041633117 | 0.118008528 | MIR4779 | miRNA | 36.14 | 0.672981313 | 0.672981313 | ENSG00000265420 | n.s. |
| ENSG00000249502 | 18.74084823 | 0.705807945 | 0.3411258 | 2.069055 | 0.038540968 | 0.111329023 | | antisense | 43.81 | 0.672911366 | 0.672911366 | ENSG00000249502 | n.s. |
| ENSG00000086289 | 270.7213785 | -0.636266488 | 0.2903473 | -2.191398 | 0.028422979 | 0.087687379 | EPDR1 | protein_coding | 38.37 | -0.672573707 | 0.672573707 | ENSG00000086289 | n.s. |
| ENSG00000163738 | 548.7698732 | 0.528000623 | 0.2175504 | 2.427027 | 0.015223117 | 0.05325571 | MTHFD2L | protein_coding | 36.15 | 0.672479449 | 0.672479449 | ENSG00000163738 | n.s. |
| ENSG00000117298 | 245.7464762 | -0.552405064 | 0.2331556 | -2.369255 | 0.017823973 | 0.060675594 | ECE1 | protein_coding | 52.19 | -0.672269209 | 0.672269209 | ENSG00000117298 | n.s. |
| ENSG00000135334 | 1035.167999 | 0.53550169 | 0.222397 | 2.407864 | 0.016046155 | 0.055547144 | AKIRIN2 | protein_coding | 40.89 | 0.672235764 | 0.672235764 | ENSG00000135334 | n.s. |
| ENSG00000034693 | 529.8140295 | -0.623121988 | 0.2810262 | -2.217309 | 0.026601982 | 0.083418781 | PEX3 | protein_coding | 35.11 | -0.67218422 | 0.67218422 | ENSG00000034693 | n.s. |
| ENSG00000160972 | 27.43924861 | 0.714925963 | 0.3482639 | 2.052828 | 0.040089218 | 0.114871821 | PPP1R16A | protein_coding | 57.4 | 0.671877765 | 0.671877765 | ENSG00000160972 | n.s. |
| ENSG00000241956 | 47.84377234 | 0.642130584 | 0.2946569 | 2.179248 | 0.029313213 | 0.089896006 | | antisense | 35.56 | 0.671835289 | 0.671835289 | ENSG00000241956 | n.s. |
| ENSG00000070367 | 2696.140615 | -0.622652835 | 0.2808774 | -2.216814 | 0.02663582 | 0.083487284 | EXOC5 | protein_coding | 35.97 | -0.671456155 | 0.671456155 | ENSG00000070367 | n.s. |
| ENSG00000037749 | 901.71952 | -0.58941572 | 0.2579953 | -2.284598 | 0.022336393 | 0.072610225 | MFAP3 | protein_coding | 43.54 | -0.671345812 | 0.671345812 | ENSG00000037749 | n.s. |
| ENSG00000008283 | 76.53463508 | 0.593397819 | 0.260774 | 2.275525 | 0.022874444 | 0.073927782 | CYB561 | protein_coding | 57.41 | 0.671247059 | 0.671247059 | ENSG00000008283 | n.s. |

| | | | | | | | | | | | | | |
|---|---|---|---|---|---|---|---|---|---|---|---|---|---|
| ENSG00000169071 | 380.6719756 | -0.600617682 | 0.2657472 | -2.260109 | 0.023814496 | 0.076292964 | ROR2 | protein_coding | 44.96 | -0.671199576 | 0.671199576 | ENSG00000169071 | n.s. |
| ENSG00000081377 | 351.5189499 | -0.583412742 | 0.2539688 | -2.297183 | 0.021608339 | 0.070838923 | CDC14B | protein_coding | 42.84 | -0.670765992 | 0.670765992 | ENSG00000081377 | n.s. |
| ENSG00000135972 | 735.5620596 | -0.613382162 | 0.2746192 | -2.233574 | 0.025511148 | 0.080652069 | MRPS9 | protein_coding | 36.63 | -0.670662539 | 0.670662539 | ENSG00000135972 | n.s. |
| ENSG00000093217 | 406.9989627 | -0.598647229 | 0.2645499 | -2.26289 | 0.023642499 | 0.075864141 | XYLB | protein_coding | 44.83 | -0.670463018 | 0.670463018 | ENSG00000093217 | n.s. |
| ENSG00000158411 | 358.3280551 | -0.65407984 | 0.3035222 | -2.154966 | 0.031164519 | 0.094532375 | MITD1 | protein_coding | 39.09 | -0.670052098 | 0.670052098 | ENSG00000158411 | n.s. |
| ENSG00000284415 | 4.187823838 | -1.055906922 | 0.6458165 | -1.634995 | 0.102050048 | 0.232239755 | MIR1287 | miRNA | 51.11 | -0.66951197 | 0.66951197 | ENSG00000284415 | n.s. |
| ENSG00000270072 | 28.65133724 | -0.678324168 | 0.3215255 | -2.109705 | 0.034883738 | 0.103052467 | | sense_intronic | 43.27 | -0.669466344 | 0.669466344 | ENSG00000270072 | n.s. |
| ENSG00000243702 | 21.19259101 | 0.730803618 | 0.3614087 | 2.022097 | 0.043166298 | 0.12153779 | RN7SL638P | misc_RNA | 51.72 | 0.668896269 | 0.668896269 | ENSG00000243702 | n.s. |
| ENSG00000174606 | 2023.621859 | -0.580748117 | 0.2525594 | -2.299451 | 0.021479321 | 0.070548854 | ANGEL2 | protein_coding | 38.68 | -0.668737286 | 0.668737286 | ENSG00000174606 | n.s. |
| ENSG00000113108 | 127.8252091 | 0.572167583 | 0.2467557 | 2.318762 | 0.02040797 | 0.067813881 | APBB3 | protein_coding | 53.75 | 0.668681611 | 0.668681611 | ENSG00000113108 | n.s. |
| ENSG00000277130 | 117.4270605 | -0.625473107 | 0.2835802 | -2.20563 | 0.027409915 | 0.085353922 | | sense_intronic | 33.28 | -0.668490969 | 0.668490969 | ENSG00000277130 | n.s. |
| ENSG00000067842 | 322.6124693 | -0.57466574 | 0.2486547 | -2.3111 | 0.02082734 | 0.068911373 | ATP2B3 | protein_coding | 58.21 | -0.667594417 | 0.667594417 | ENSG00000067842 | n.s. |
| ENSG00000119723 | 48.09713171 | -0.635042733 | 0.2906598 | -2.184831 | 0.028901216 | 0.08890668 | COQ6 | protein_coding | 45.54 | -0.667471577 | 0.667471577 | ENSG00000119723 | n.s. |
| ENSG00000160199 | 392.7281259 | -0.595389146 | 0.2630431 | -2.263466 | 0.023606965 | 0.075767584 | PKNOX1 | protein_coding | 46.99 | -0.667143398 | 0.667143398 | ENSG00000160199 | n.s. |
| ENSG00000119402 | 1455.449551 | -0.573007553 | 0.2477327 | -2.313007 | 0.020722236 | 0.068610475 | FBXW2 | protein_coding | 41.5 | -0.666757077 | 0.666757077 | ENSG00000119402 | n.s. |
| ENSG00000048471 | 148.0752453 | -0.582258657 | 0.2540073 | -2.292291 | 0.02188886 | 0.071623641 | SNX29 | protein_coding | 47.13 | -0.666653326 | 0.666653326 | ENSG00000048471 | n.s. |
| ENSG00000264070 | 32.51062785 | 0.645574278 | 0.2983961 | 2.163481 | 0.030504198 | 0.092865728 | | processed_pseudogene | 65.34 | 0.66632596 | 0.66632596 | ENSG00000264070 | n.s. |
| ENSG00000243005 | 114.7989496 | -0.585618099 | 0.2565375 | -2.282778 | 0.022443452 | 0.072873178 | RN7SL16P | misc_RNA | 53.16 | -0.666100936 | 0.666100936 | ENSG00000243005 | n.s. |
| ENSG00000124356 | 1128.341836 | -0.585984161 | 0.2568017 | -2.281855 | 0.022497929 | 0.07299899 | STAMBP | protein_coding | 42.1 | -0.666078323 | 0.666078323 | ENSG00000124356 | n.s. |
| ENSG00000115042 | 412.6565436 | -0.591687965 | 0.2608827 | -2.268023 | 0.023327793 | 0.075019383 | FAHD2A | protein_coding | 49.55 | -0.665546308 | 0.665546308 | ENSG00000115042 | n.s. |
| ENSG00000196458 | 1409.710406 | -0.615112405 | 0.2769422 | -2.221086 | 0.026345168 | 0.082837348 | ZNF605 | protein_coding | 46.54 | -0.665412493 | 0.665412493 | ENSG00000196458 | n.s. |
| ENSG00000185220 | 196.6086162 | -0.664800152 | 0.3127326 | -2.125778 | 0.033521754 | 0.099919372 | PGBD2 | protein_coding | 46.25 | -0.665033035 | 0.665033035 | ENSG00000185220 | n.s. |
| ENSG00000227372 | 300.226863 | -0.583273484 | 0.2552922 | -2.284729 | 0.022328722 | 0.072602242 | TP73-AS1 | transcribed_unitary_pseudogene | 54.46 | -0.664377642 | 0.664377642 | ENSG00000227372 | n.s. |
| ENSG00000187231 | 1570.429222 | -0.578796832 | 0.2522198 | -2.294811 | 0.021743961 | 0.07124998 | SESTD1 | protein_coding | 36.9 | -0.664004555 | 0.664004555 | ENSG00000187231 | n.s. |
| ENSG00000211459 | 632.1566697 | -0.593052212 | 0.2621592 | -2.262184 | 0.023686054 | 0.075951389 | MT-RNR1 | Mt_rRNA | 45.49 | -0.663900768 | 0.663900768 | ENSG00000211459 | n.s. |
| ENSG00000104356 | 1071.635721 | -0.605266277 | 0.2706232 | -2.236565 | 0.025314784 | 0.080213414 | POP1 | protein_coding | 41.8 | -0.663222337 | 0.663222337 | ENSG00000104356 | n.s. |
| ENSG00000154832 | 551.3814043 | 0.591698308 | 0.2616067 | 2.261786 | 0.023710621 | 0.076008183 | CXXC1 | protein_coding | 59.29 | 0.662193038 | 0.662193038 | ENSG00000154832 | n.s. |
| ENSG00000114127 | 1773.622706 | -0.618109339 | 0.279931 | -2.208078 | 0.027238853 | 0.084975701 | XRN1 | protein_coding | 36.98 | -0.661812911 | 0.661812911 | ENSG00000114127 | n.s. |
| ENSG00000180011 | 871.9708797 | -0.562386062 | 0.2417955 | -2.325875 | 0.020025242 | 0.066797184 | ZADH2 | protein_coding | 46.75 | -0.660939632 | 0.660939632 | ENSG00000180011 | n.s. |
| ENSG00000110514 | 123.0852907 | 0.565800728 | 0.2441139 | 2.317774 | 0.020461614 | 0.067943489 | MADD | protein_coding | 47.1 | 0.660771598 | 0.660771598 | ENSG00000110514 | n.s. |
| ENSG00000279192 | 628.7773536 | -0.587070018 | 0.2588521 | -2.267975 | 0.023330743 | 0.075019383 | PWAR5 | TEC | 35.82 | -0.66035192 | 0.66035192 | ENSG00000279192 | n.s. |
| ENSG00000157212 | 935.6584693 | -0.536908817 | 0.2254852 | -2.381127 | 0.017259766 | 0.059043667 | PAXIP1 | protein_coding | 40.81 | -0.659767879 | 0.659767879 | ENSG00000157212 | n.s. |

| | | | | | | | | | | | | | |
|---|---|---|---|---|---|---|---|---|---|---|---|---|---|
| ENSG00000234741 | 19172.00629 | 0.506189538 | 0.2058898 | 2.458546 | 0.013950092 | 0.049776304 | GAS5 | processed_transcript | 46.16 | 0.659553507 | 0.659553507 | ENSG00000234741 | n.s. |
| ENSG00000140950 | 20.40285159 | -0.94354276 | 0.5449987 | -1.731275 | 0.083402668 | 0.200244068 | TLDC1 | protein_coding | 49.64 | -0.659008327 | 0.659008327 | ENSG00000140950 | n.s. |
| ENSG00000103657 | 3883.263655 | -0.566615826 | 0.2450992 | -2.311782 | 0.020789721 | 0.068803253 | HERC1 | protein_coding | 38.81 | -0.658629154 | 0.658629154 | ENSG00000103657 | n.s. |
| ENSG00000205937 | 3252.689784 | 0.534498111 | 0.2241784 | 2.384253 | 0.017113836 | 0.058630891 | RNPS1 | protein_coding | 49.19 | 0.658434063 | 0.658434063 | ENSG00000205937 | n.s. |
| ENSG00000089022 | 4181.552092 | -0.589811451 | 0.2613215 | -2.257034 | 0.024005945 | 0.076782627 | MAPKAPK5 | protein_coding | 43.78 | -0.657484665 | 0.657484665 | ENSG00000089022 | n.s. |
| ENSG00000229797 | 38.64984904 | 0.614077099 | 0.2784231 | 2.205554 | 0.027415265 | 0.085353922 | | antisense | 45.12 | 0.656311184 | 0.656311184 | ENSG00000229797 | n.s. |
| ENSG00000188859 | 398.2361852 | -0.579850496 | 0.2549084 | -2.27474 | 0.022921519 | 0.074011203 | FAM78B | protein_coding | 45.08 | -0.655638427 | 0.655638427 | ENSG00000188859 | n.s. |
| ENSG00000267152 | 61.31357745 | -0.655209978 | 0.3081873 | -2.126012 | 0.033502221 | 0.099882532 | | lincRNA | 35.52 | -0.655544434 | 0.655544434 | ENSG00000267152 | n.s. |
| ENSG00000239911 | 140.7862771 | 0.608249568 | 0.2744998 | 2.215847 | 0.026701977 | 0.083675809 | PRKAG2-AS1 | antisense | 60.13 | 0.655328134 | 0.655328134 | ENSG00000239911 | n.s. |
| ENSG00000089916 | 2394.99743 | -0.573244594 | 0.2504532 | -2.288829 | 0.022089292 | 0.072042449 | GPATCH2L | protein_coding | 39.76 | -0.654881234 | 0.654881234 | ENSG00000089916 | n.s. |
| ENSG00000173933 | 1976.551199 | 0.518743737 | 0.2147717 | 2.415327 | 0.015721106 | 0.054652922 | RBM4 | protein_coding | 45.16 | 0.654855147 | 0.654855147 | ENSG00000173933 | n.s. |
| ENSG00000273204 | 21.75505206 | 0.70148216 | 0.3431126 | 2.044467 | 0.04090748 | 0.116640353 | | antisense | 54.08 | 0.654588903 | 0.654588903 | ENSG00000273204 | n.s. |
| ENSG00000131055 | 28.24382604 | 0.703502921 | 0.3450259 | 2.038986 | 0.041451445 | 0.117672915 | COX4I2 | protein_coding | 52.35 | 0.653781788 | 0.653781788 | ENSG00000131055 | n.s. |
| ENSG00000133789 | 551.1467037 | -0.588011295 | 0.2608938 | -2.253834 | 0.024206644 | 0.077317988 | SWAP70 | protein_coding | 39.97 | -0.653703598 | 0.653703598 | ENSG00000133789 | n.s. |
| ENSG00000112592 | 699.4949241 | -0.556828461 | 0.2395826 | -2.324161 | 0.020116884 | 0.067054675 | TBP | protein_coding | 39.46 | -0.6534777 | 0.6534777 | ENSG00000112592 | n.s. |
| ENSG00000100077 | 3125.284403 | -0.57082631 | 0.2490713 | -2.291819 | 0.021916105 | 0.0716791 | GRK3 | protein_coding | 40.55 | -0.653372051 | 0.653372051 | ENSG00000100077 | n.s. |
| ENSG00000157514 | 410.5121865 | 0.613260116 | 0.2787909 | 2.199714 | 0.027827212 | 0.08625073 | TSC22D3 | protein_coding | 48.75 | 0.652654242 | 0.652654242 | ENSG00000157514 | n.s. |
| ENSG00000023330 | 464.5214494 | -0.621067669 | 0.2844666 | -2.183271 | 0.029015867 | 0.089121536 | ALAS1 | protein_coding | 45.47 | -0.652131821 | 0.652131821 | ENSG00000023330 | n.s. |
| ENSG00000138606 | 1810.031156 | -0.536241714 | 0.2264502 | -2.368034 | 0.017882906 | 0.060846456 | SHF | protein_coding | 50.88 | -0.651943753 | 0.651943753 | ENSG00000138606 | n.s. |
| ENSG00000112159 | 6977.692261 | -0.559240977 | 0.2416311 | -2.314441 | 0.020643528 | 0.068433296 | MDN1 | protein_coding | 41.54 | -0.651366165 | 0.651366165 | ENSG00000112159 | n.s. |
| ENSG00000117600 | 1091.078029 | -0.631819606 | 0.2923418 | -2.161236 | 0.030677092 | 0.093277388 | PLPPR4 | protein_coding | 36.26 | -0.650915485 | 0.650915485 | ENSG00000117600 | n.s. |
| ENSG00000256603 | 25.1801522 | 0.684278821 | 0.3312743 | 2.065596 | 0.038866629 | 0.112083694 | | antisense | 44.96 | 0.65037798 | 0.65037798 | ENSG00000256603 | n.s. |
| ENSG00000165304 | 864.090554 | -0.60259379 | 0.2717372 | -2.217561 | 0.026584797 | 0.083383673 | MELK | protein_coding | 41.98 | -0.650149877 | 0.650149877 | ENSG00000165304 | n.s. |
| ENSG00000258297 | 214.823398 | -0.662509583 | 0.315085 | -2.102638 | 0.035497461 | 0.104422386 | | antisense | 39.85 | -0.650058635 | 0.650058635 | ENSG00000258297 | n.s. |
| ENSG00000162980 | 1779.753133 | -0.623323363 | 0.2865733 | -2.175092 | 0.029623208 | 0.090607188 | ARL5A | protein_coding | 37.35 | -0.650024883 | 0.650024883 | ENSG00000162980 | n.s. |
| ENSG00000138468 | 24.63623813 | -0.716297868 | 0.3559512 | -2.012349 | 0.044183199 | 0.123874884 | SENP7 | protein_coding | 37.56 | -0.649694155 | 0.649694155 | ENSG00000138468 | n.s. |
| ENSG00000105613 | 371.5452669 | -0.587916907 | 0.2618835 | -2.244956 | 0.024770949 | 0.078759061 | MAST1 | protein_coding | 52.4 | -0.648883581 | 0.648883581 | ENSG00000105613 | n.s. |
| ENSG00000196199 | 939.2849074 | -0.590390135 | 0.2635757 | -2.239926 | 0.025095718 | 0.079664316 | MPHOSPH8 | protein_coding | 43.3 | -0.648682996 | 0.648682996 | ENSG00000196199 | n.s. |
| ENSG00000136842 | 409.2555157 | -0.634476977 | 0.2947951 | -2.152264 | 0.031376557 | 0.095007489 | TMOD1 | protein_coding | 46.76 | -0.648589115 | 0.648589115 | ENSG00000136842 | n.s. |
| ENSG00000186871 | 578.0392886 | -0.595367802 | 0.2671632 | -2.228479 | 0.025848561 | 0.081452317 | ERCC6L | protein_coding | 43.97 | -0.648413024 | 0.648413024 | ENSG00000186871 | n.s. |
| ENSG00000255284 | 37.22224349 | -0.633481765 | 0.2942068 | -2.153185 | 0.031304105 | 0.094893764 | | lincRNA | 59.11 | -0.647901283 | 0.647901283 | ENSG00000255284 | n.s. |
| ENSG00000272661 | 20.52845906 | -0.751940928 | 0.3846025 | -1.955112 | 0.050569846 | 0.137597024 | | antisense | 51.39 | -0.647715118 | 0.647715118 | ENSG00000272661 | n.s. |

| | | | | | | | | | | | | | |
|---|---|---|---|---|---|---|---|---|---|---|---|---|---|
| ENSG00000104976 | 78.09649358 | 0.632936771 | 0.2940737 | 2.152306 | 0.031373227 | 0.095007489 | SNAPC2 | protein_coding | 64.16 | 0.647014652 | 0.647014652 | ENSG00000104976 | n.s. |
| ENSG00000235173 | 154.3835701 | -0.571085624 | 0.251028 | -2.274988 | 0.022906647 | 0.073986609 | HGH1 | protein_coding | 64.59 | -0.645810395 | 0.645810395 | ENSG00000235173 | n.s. |
| ENSG00000259746 | 22.94010407 | -0.724989964 | 0.3639966 | -1.99175 | 0.046398548 | 0.128789271 | HSPE1P3 | processed_pseudogene | 41.42 | -0.645328295 | 0.645328295 | ENSG00000259746 | n.s. |
| ENSG00000143630 | 187.236902 | -0.589393475 | 0.2637607 | -2.234577 | 0.025445157 | 0.080516572 | HCN3 | protein_coding | 53.79 | -0.644864079 | 0.644864079 | ENSG00000143630 | n.s. |
| ENSG00000142731 | 1933.412518 | -0.582452138 | 0.2590404 | -2.248499 | 0.024544375 | 0.078199389 | PLK4 | protein_coding | 36.34 | -0.644656068 | 0.644656068 | ENSG00000142731 | n.s. |
| ENSG00000070413 | 528.8909471 | -0.606963051 | 0.2762305 | -2.197306 | 0.02799859 | 0.086685427 | DGCR2 | protein_coding | 48.62 | -0.644627479 | 0.644627479 | ENSG00000070413 | n.s. |
| ENSG00000252585 | 20.59565239 | -0.703494163 | 0.3477681 | -2.022883 | 0.043085197 | 0.121358528 | | misc_RNA | 43.36 | -0.644351198 | 0.644351198 | ENSG00000252585 | n.s. |
| ENSG00000255857 | 133.9825555 | -0.646013532 | 0.3047131 | -2.120071 | 0.034000048 | 0.100955997 | PXN-AS1 | antisense | 49.38 | -0.643344126 | 0.643344126 | ENSG00000255857 | n.s. |
| ENSG00000157152 | 2238.716224 | -0.562398052 | 0.2455058 | -2.290773 | 0.021976546 | 0.071843027 | SYN2 | protein_coding | 40.96 | -0.64316706 | 0.64316706 | ENSG00000157152 | n.s. |
| ENSG00000166685 | 670.5244239 | -0.549732403 | 0.2369562 | -2.319974 | 0.020342262 | 0.067635897 | COG1 | protein_coding | 49.15 | -0.64308947 | 0.64308947 | ENSG00000166685 | n.s. |
| ENSG00000170265 | 434.5218996 | -0.569649693 | 0.2506556 | -2.272639 | 0.023047927 | 0.074315952 | ZNF282 | protein_coding | 51.85 | -0.643087765 | 0.643087765 | ENSG00000170265 | n.s. |
| ENSG00000130717 | 404.5389777 | -0.581388579 | 0.2587278 | -2.247105 | 0.024633299 | 0.078379576 | UCK1 | protein_coding | 57.27 | -0.642897798 | 0.642897798 | ENSG00000130717 | n.s. |
| ENSG00000165527 | 1554.328859 | 0.495267327 | 0.2018259 | 2.453933 | 0.014130333 | 0.050348555 | ARF6 | protein_coding | 43.17 | 0.642863423 | 0.642863423 | ENSG00000165527 | n.s. |
| ENSG00000275966 | 22.01165421 | 0.690477928 | 0.3384067 | 2.040379 | 0.041312592 | 0.117362827 | | lincRNA | 72.3 | 0.642468609 | 0.642468609 | ENSG00000275966 | n.s. |
| ENSG00000173113 | 1196.975529 | 0.683526339 | 0.3329822 | 2.052741 | 0.040097658 | 0.114872365 | TRMT112 | protein_coding | 56.49 | 0.642367416 | 0.642367416 | ENSG00000173113 | n.s. |
| ENSG00000202184 | 3.733229065 | -1.03994289 | 0.6461763 | -1.609379 | 0.10753339 | 0.241209504 | RNU6-1283P | snRNA | 35.24 | -0.642274536 | 0.642274536 | ENSG00000202184 | n.s. |
| ENSG00000137075 | 887.2533361 | -0.579162032 | 0.2573107 | -2.250828 | 0.024396446 | 0.077825605 | RNF38 | protein_coding | 41.31 | -0.642219743 | 0.642219743 | ENSG00000137075 | n.s. |
| ENSG00000185515 | 513.2478637 | -0.62383739 | 0.2888373 | -2.159822 | 0.030786417 | 0.093507698 | BRCC3 | protein_coding | 37.76 | -0.642023894 | 0.642023894 | ENSG00000185515 | n.s. |
| ENSG00000176903 | 1088.368317 | 0.566725101 | 0.2489814 | 2.276174 | 0.022835588 | 0.073836481 | PNMA1 | protein_coding | 53.75 | 0.641379237 | 0.641379237 | ENSG00000176903 | n.s. |
| ENSG00000256571 | 23.45787711 | -0.72014589 | 0.3618323 | -1.990275 | 0.046560634 | 0.129161927 | | sense_intronic | 40.99 | -0.640112825 | 0.640112825 | ENSG00000256571 | n.s. |
| ENSG00000070018 | 3805.064234 | -0.576781763 | 0.256268 | -2.250698 | 0.024404697 | 0.077825605 | LRP6 | protein_coding | 38.53 | -0.639580316 | 0.639580316 | ENSG00000070018 | n.s. |
| ENSG00000143067 | 393.210121 | -0.559043949 | 0.2441017 | -2.290209 | 0.022009201 | 0.071903054 | ZNF697 | protein_coding | 44.33 | -0.639128483 | 0.639128483 | ENSG00000143067 | n.s. |
| ENSG00000069275 | 11688.08786 | -0.598722679 | 0.271806 | -2.202757 | 0.027611848 | 0.085775952 | NUCKS1 | protein_coding | 39.57 | -0.638618239 | 0.638618239 | ENSG00000069275 | n.s. |
| ENSG00000207233 | 45.73726669 | 0.623881519 | 0.2897886 | 2.152885 | 0.031327699 | 0.094923951 | SNORA37 | snoRNA | 54.26 | 0.637996334 | 0.637996334 | ENSG00000207233 | n.s. |
| ENSG00000188211 | 831.9167472 | -0.584452868 | 0.261971 | -2.230983 | 0.025682278 | 0.081064239 | NCR3LG1 | protein_coding | 44.5 | -0.637737841 | 0.637737841 | ENSG00000188211 | n.s. |
| ENSG00000109686 | 217.4406601 | -0.592387546 | 0.2675051 | -2.214491 | 0.026795042 | 0.083854208 | SH3D19 | protein_coding | 40.79 | -0.637690467 | 0.637690467 | ENSG00000109686 | n.s. |
| ENSG00000166165 | 2866.112528 | 0.619363581 | 0.2866848 | 2.160434 | 0.030739084 | 0.093425073 | CKB | protein_coding | 70.4 | 0.637657446 | 0.637657446 | ENSG00000166165 | n.s. |
| ENSG00000229320 | 114.8300383 | -0.560654063 | 0.2456614 | -2.282223 | 0.022476173 | 0.072947837 | KRT8P12 | transcribed_processed_pseudog | 50.42 | -0.637456707 | 0.637456707 | ENSG00000229320 | n.s. |
| ENSG00000135116 | 467.6460399 | -0.57569094 | 0.2560759 | -2.248126 | 0.024568157 | 0.078239352 | HRK | protein_coding | 49 | -0.637045063 | 0.637045063 | ENSG00000135116 | n.s. |
| ENSG00000239306 | 2039.615496 | 0.497888351 | 0.2046071 | 2.433387 | 0.014958286 | 0.052605258 | RBM14 | protein_coding | 50.54 | 0.636784684 | 0.636784684 | ENSG00000239306 | n.s. |
| ENSG00000175643 | 332.6760763 | -0.615533561 | 0.2841627 | -2.16613 | 0.030301223 | 0.09239683 | RMI2 | protein_coding | 50.87 | -0.636672787 | 0.636672787 | ENSG00000175643 | n.s. |
| ENSG00000186814 | 569.4916427 | -0.612366898 | 0.2819767 | -2.171693 | 0.029878821 | 0.091328832 | ZSCAN30 | protein_coding | 42.35 | -0.636489313 | 0.636489313 | ENSG00000186814 | n.s. |

| | | | | | | | | | | | | | |
|---|---|---|---|---|---|---|---|---|---|---|---|---|---|
| ENSG00000064309 | 635.4834216 | -0.596747243 | 0.2710165 | -2.201886 | 0.027673392 | 0.085888685 | CDON | protein_coding | 39.95 | -0.636170781 | 0.636170781 | ENSG00000064309 | n.s. |
| ENSG00000252311 | 7.205673497 | -1.096611188 | 0.7085444 | -1.547696 | 0.121695544 | 0.263714718 | RNU1-103P | snRNA | 48.8 | -0.634790527 | 0.634790527 | ENSG00000252311 | n.s. |
| ENSG00000207092 | 5.147825009 | -1.301993664 | 0.9339464 | -1.394078 | 0.163294186 | 0.325640425 | | misc_RNA | 40.2 | -0.63441163 | 0.63441163 | ENSG00000207092 | n.s. |
| ENSG00000110092 | 10369.08561 | -0.55854141 | 0.2449509 | -2.280218 | 0.022594752 | 0.073210783 | CCND1 | protein_coding | 58.58 | -0.634181852 | 0.634181852 | ENSG00000110092 | n.s. |
| ENSG00000198648 | 871.6992576 | -0.611225469 | 0.2817565 | -2.169339 | 0.030056936 | 0.091792657 | STK39 | protein_coding | 40.16 | -0.633958204 | 0.633958204 | ENSG00000198648 | n.s. |
| ENSG00000157404 | 585.6776838 | -0.593354019 | 0.2691244 | -2.204758 | 0.027471095 | 0.085470403 | KIT | protein_coding | 40.87 | -0.633811416 | 0.633811416 | ENSG00000157404 | n.s. |
| ENSG00000264566 | 4.096126805 | -0.952003738 | 0.5667087 | -1.679882 | 0.092980285 | 0.216636219 | MIR23C | miRNA | 45 | -0.632386507 | 0.632386507 | ENSG00000264566 | n.s. |
| ENSG00000136878 | 229.4305459 | -0.60024426 | 0.2744636 | -2.186972 | 0.02874458 | 0.088462655 | USP20 | protein_coding | 52.14 | -0.632201281 | 0.632201281 | ENSG00000136878 | n.s. |
| ENSG00000200403 | 5.010836078 | -0.935808814 | 0.5515715 | -1.696623 | 0.089767979 | 0.211157217 | RNU6-1099P | snRNA | 40.19 | -0.632039723 | 0.632039723 | ENSG00000200403 | n.s. |
| ENSG00000071246 | 329.998149 | -0.55442892 | 0.2425655 | -2.285687 | 0.022272577 | 0.072470459 | VASH1 | protein_coding | 54.83 | -0.631959701 | 0.631959701 | ENSG00000071246 | n.s. |
| ENSG00000164654 | 1570.468954 | -0.567340209 | 0.2514842 | -2.255968 | 0.024072656 | 0.076960642 | MIOS | protein_coding | 36.45 | -0.63186456 | 0.63186456 | ENSG00000164654 | n.s. |
| ENSG00000211575 | 4.718051987 | 1.304211048 | 0.9397798 | 1.387784 | 0.165202955 | 0.328152674 | MIR760 | miRNA | 63.75 | 0.631139096 | 0.631139096 | ENSG00000211575 | n.s. |
| ENSG00000207342 | 18.63688525 | -0.712943463 | 0.3591867 | -1.984883 | 0.047157526 | 0.130453853 | | misc_RNA | 37.37 | -0.630629214 | 0.630629214 | ENSG00000207342 | n.s. |
| ENSG00000089123 | 257.3659325 | -0.568684317 | 0.2526697 | -2.250702 | 0.024404387 | 0.077825605 | TASP1 | protein_coding | 37.32 | -0.630601241 | 0.630601241 | ENSG00000089123 | n.s. |
| ENSG00000243725 | 977.363639 | -0.577926045 | 0.2591114 | -2.230416 | 0.025719856 | 0.0811093 | TTC4 | protein_coding | 42.06 | -0.630476483 | 0.630476483 | ENSG00000243725 | n.s. |
| ENSG00000231992 | 13.14754252 | 0.75027095 | 0.3888032 | 1.929693 | 0.05364484 | 0.144439587 | | sense_intronic | 42.34 | 0.630463004 | 0.630463004 | ENSG00000231992 | n.s. |
| ENSG00000100207 | 1223.292917 | -0.525756704 | 0.2237568 | -2.349679 | 0.018789588 | 0.063466142 | TCF20 | protein_coding | 49.75 | -0.62957152 | 0.62957152 | ENSG00000100207 | n.s. |
| ENSG00000160877 | 421.2863215 | -0.539685046 | 0.2330448 | -2.3158 | 0.020569215 | 0.068219433 | NACC1 | protein_coding | 55.78 | -0.629322359 | 0.629322359 | ENSG00000160877 | n.s. |
| ENSG00000054148 | 530.8601796 | 0.675511669 | 0.3309093 | 2.04138 | 0.041213056 | 0.117175695 | PHPT1 | protein_coding | 66.23 | 0.62901111 | 0.62901111 | ENSG00000054148 | n.s. |
| ENSG00000252641 | 8.286399363 | -0.876692064 | 0.4984849 | -1.758713 | 0.078626207 | 0.191886536 | RNU6-678P | snRNA | 37.61 | -0.628549195 | 0.628549195 | ENSG00000252641 | n.s. |
| ENSG00000168348 | 123.2669374 | 0.525594511 | 0.2239775 | 2.34664 | 0.018943541 | 0.063877674 | INSM2 | protein_coding | 58.21 | 0.627901958 | 0.627901958 | ENSG00000168348 | n.s. |
| ENSG00000177951 | 234.5372759 | -0.548721605 | 0.2396158 | -2.290006 | 0.02202095 | 0.071916487 | BET1L | protein_coding | 48.36 | -0.627282918 | 0.627282918 | ENSG00000177951 | n.s. |
| ENSG00000264358 | 49.85998903 | -0.665513016 | 0.323681 | -2.056077 | 0.039775113 | 0.114167392 | MIR3122 | miRNA | 50.68 | -0.627218011 | 0.627218011 | ENSG00000264358 | n.s. |
| ENSG00000196597 | 422.0110116 | -0.558603288 | 0.2465725 | -2.265473 | 0.023483696 | 0.075441509 | ZNF782 | protein_coding | 38.77 | -0.626970539 | 0.626970539 | ENSG00000196597 | n.s. |
| ENSG00000184402 | 798.7165414 | -0.606839743 | 0.2803804 | -2.164345 | 0.030437918 | 0.092691849 | SS18L1 | protein_coding | 55.02 | -0.626840242 | 0.626840242 | ENSG00000184402 | n.s. |
| ENSG00000136051 | 2727.135569 | -0.596692065 | 0.2732615 | -2.183594 | 0.028992123 | 0.089068255 | WASHC4 | protein_coding | 35.27 | -0.626691984 | 0.626691984 | ENSG00000136051 | n.s. |
| ENSG00000182150 | 3123.782722 | -0.567231359 | 0.2525737 | -2.245805 | 0.024716506 | 0.078603908 | ERCC6L2 | protein_coding | 37.1 | -0.62653872 | 0.62653872 | ENSG00000182150 | n.s. |
| ENSG00000127191 | 217.4607391 | -0.580827925 | 0.2619775 | -2.217091 | 0.026616874 | 0.083446686 | TRAF2 | protein_coding | 55.12 | -0.626475718 | 0.626475718 | ENSG00000127191 | n.s. |
| ENSG00000064999 | 366.2244061 | -0.559485759 | 0.247279 | -2.262569 | 0.023662265 | 0.075910073 | ANKS1A | protein_coding | 44.93 | -0.626456537 | 0.626456537 | ENSG00000064999 | n.s. |
| ENSG00000116044 | 2953.863656 | -0.582948 | 0.2635509 | -2.211899 | 0.02697362 | 0.084261546 | NFE2L2 | protein_coding | 40.63 | -0.62630218 | 0.62630218 | ENSG00000116044 | n.s. |
| ENSG00000155957 | 157.325674 | 0.570350534 | 0.2547909 | 2.238504 | 0.0251882 | 0.079885035 | TMBIM4 | protein_coding | 38.11 | 0.625979428 | 0.625979428 | ENSG00000155957 | n.s. |
| ENSG00000173409 | 356.1447965 | -0.668861037 | 0.3266906 | -2.047384 | 0.040620394 | 0.116035648 | ARV1 | protein_coding | 41.75 | -0.625658345 | 0.625658345 | ENSG00000173409 | n.s. |

| | | | | | | | | | | | | | |
|---|---|---|---|---|---|---|---|---|---|---|---|---|---|
| ENSG00000163807 | 1092.674812 | -0.633598307 | 0.3003044 | -2.109853 | 0.034870983 | 0.103036649 | KIAA1143 | protein_coding | 41.53 | -0.62536677 | 0.62536677 | ENSG00000163807 | n.s. |
| ENSG00000135643 | 161.5015199 | -0.58843472 | 0.2677175 | -2.197969 | 0.027951302 | 0.086596791 | KCNMB4 | protein_coding | 39.69 | -0.625210832 | 0.625210832 | ENSG00000135643 | n.s. |
| ENSG00000011143 | 419.9137877 | -0.595841617 | 0.2731273 | -2.181553 | 0.029142548 | 0.089431715 | MKS1 | protein_coding | 47.46 | -0.624744964 | 0.624744964 | ENSG00000011143 | n.s. |
| ENSG00000139192 | 109.5734058 | -0.549468283 | 0.2407615 | -2.28221 | 0.022476925 | 0.072947837 | TAPBPL | protein_coding | 50.43 | -0.624738613 | 0.624738613 | ENSG00000139192 | n.s. |
| ENSG00000146856 | 395.276433 | 0.508065491 | 0.2134565 | 2.380183 | 0.017304047 | 0.059180606 | AGBL3 | protein_coding | 36.82 | 0.623813273 | 0.623813273 | ENSG00000146856 | n.s. |
| ENSG00000101945 | 232.8527188 | -0.584563425 | 0.2653087 | -2.203333 | 0.027571264 | 0.085705449 | SUV39H1 | protein_coding | 54.35 | -0.623724246 | 0.623724246 | ENSG00000101945 | n.s. |
| ENSG00000228649 | 501.6131762 | -0.488239314 | 0.2008771 | -2.430538 | 0.015076436 | 0.052847235 | SNHG26 | processed_transcript | 43.38 | -0.62347073 | 0.62347073 | ENSG00000228649 | n.s. |
| ENSG00000279259 | 55.04949887 | 0.588412948 | 0.2682435 | 2.193578 | 0.028265761 | 0.087279704 | | TEC | 57.68 | 0.623180348 | 0.623180348 | ENSG00000279259 | n.s. |
| ENSG00000115935 | 151.6132838 | -0.576759863 | 0.2599684 | -2.218577 | 0.026515526 | 0.083203893 | WIPF1 | protein_coding | 41.82 | -0.622817802 | 0.622817802 | ENSG00000115935 | n.s. |
| ENSG00000171827 | 427.9104271 | -0.592597891 | 0.2714475 | -2.183103 | 0.029028214 | 0.089139795 | ZNF570 | protein_coding | 41.65 | -0.622185338 | 0.622185338 | ENSG00000171827 | n.s. |
| ENSG00000126005 | 90.74001002 | 0.596432366 | 0.2741842 | 2.175298 | 0.029607796 | 0.090579949 | MMP24-AS1 | protein_coding | 47.16 | 0.622059829 | 0.622059829 | ENSG00000126005 | n.s. |
| ENSG00000198142 | 612.900432 | 0.520693387 | 0.2219666 | 2.345819 | 0.018985341 | 0.063956665 | SOWAHC | protein_coding | 47.58 | 0.621767361 | 0.621767361 | ENSG00000198142 | n.s. |
| ENSG00000085511 | 1291.544666 | -0.55199188 | 0.2432514 | -2.269224 | 0.023254695 | 0.074826696 | MAP3K4 | protein_coding | 38.79 | -0.621511632 | 0.621511632 | ENSG00000085511 | n.s. |
| ENSG00000107829 | 202.4368143 | -0.580450216 | 0.2631351 | -2.205902 | 0.027390847 | 0.085316055 | FBXW4 | protein_coding | 45.8 | -0.620483422 | 0.620483422 | ENSG00000107829 | n.s. |
| ENSG00000204604 | 335.2622626 | -0.574033498 | 0.2586259 | -2.219551 | 0.02644923 | 0.083037633 | ZNF468 | protein_coding | 43.76 | -0.620372374 | 0.620372374 | ENSG00000204604 | n.s. |
| ENSG00000178202 | 438.3909883 | -0.613055508 | 0.2865071 | -2.139757 | 0.032374449 | 0.097312057 | KDELC2 | protein_coding | 41.21 | -0.620310007 | 0.620310007 | ENSG00000178202 | n.s. |
| ENSG00000144036 | 784.0115157 | -0.576420628 | 0.2604017 | -2.213583 | 0.02685751 | 0.083993063 | EXOC6B | protein_coding | 37.12 | -0.620088286 | 0.620088286 | ENSG00000144036 | n.s. |
| ENSG00000136827 | 544.9518816 | -0.644899257 | 0.310291 | -2.078369 | 0.037675353 | 0.109304842 | TOR1A | protein_coding | 52.06 | -0.619980737 | 0.619980737 | ENSG00000136827 | n.s. |
| ENSG00000186318 | 300.5333182 | -0.597205934 | 0.2752057 | -2.170034 | 0.030004237 | 0.09165182 | BACE1 | protein_coding | 49.51 | -0.619815499 | 0.619815499 | ENSG00000186318 | n.s. |
| ENSG00000174839 | 1087.008714 | -0.634323019 | 0.3024399 | -2.097353 | 0.035962371 | 0.10556696 | DENND6A | protein_coding | 39.72 | -0.619398656 | 0.619398656 | ENSG00000174839 | n.s. |
| ENSG00000274737 | 50.70064799 | 0.596190241 | 0.2747047 | 2.170295 | 0.029984475 | 0.091611553 | | sense_intronic | 56.98 | 0.618875133 | 0.618875133 | ENSG00000274737 | n.s. |
| ENSG00000111731 | 2028.159985 | -0.606433506 | 0.2820684 | -2.149952 | 0.031559013 | 0.095396469 | C2CD5 | protein_coding | 34.95 | -0.618845806 | 0.618845806 | ENSG00000111731 | n.s. |
| ENSG00000182541 | 405.4371023 | -0.57314079 | 0.2583891 | -2.218131 | 0.026545904 | 0.083280446 | LIMK2 | protein_coding | 47.35 | -0.618680814 | 0.618680814 | ENSG00000182541 | n.s. |
| ENSG00000120798 | 916.6991046 | -0.570135581 | 0.2563911 | -2.223695 | 0.026168969 | 0.082339109 | NR2C1 | protein_coding | 39.56 | -0.618251511 | 0.618251511 | ENSG00000120798 | n.s. |
| ENSG00000283146 | 3.623767888 | -1.00030707 | 0.6213849 | -1.609803 | 0.107440955 | 0.241040994 | MIR548AU | miRNA | 35.19 | -0.618098832 | 0.618098832 | ENSG00000283146 | n.s. |
| ENSG00000224817 | 11.58953719 | -0.776453627 | 0.4152695 | -1.869758 | 0.061517387 | 0.159975081 | | lincRNA | 39.76 | -0.618016448 | 0.618016448 | ENSG00000224817 | n.s. |
| ENSG00000235904 | 54.46180872 | 0.57919922 | 0.2630009 | 2.202271 | 0.027646203 | 0.08584257 | RBMS3-AS3 | antisense | 37.6 | 0.617598557 | 0.617598557 | ENSG00000235904 | n.s. |
| ENSG00000101290 | 625.9737394 | -0.585418389 | 0.2674322 | -2.189035 | 0.028594325 | 0.088118369 | CDS2 | protein_coding | 44.84 | -0.617577499 | 0.617577499 | ENSG00000101290 | n.s. |
| ENSG00000162688 | 1685.147031 | -0.597572657 | 0.2762158 | -2.163427 | 0.030508356 | 0.092865728 | AGL | protein_coding | 36.47 | -0.616781349 | 0.616781349 | ENSG00000162688 | n.s. |
| ENSG00000278619 | 92.08641226 | -0.545077991 | 0.2395831 | -2.27511 | 0.022899331 | 0.073986609 | MRM1 | protein_coding | 52.69 | -0.616399744 | 0.616399744 | ENSG00000278619 | n.s. |
| ENSG00000244265 | 20.65960475 | 0.664556805 | 0.3263552 | 2.036299 | 0.041720334 | 0.118207613 | SIAH2-AS1 | antisense | 56.64 | 0.616279779 | 0.616279779 | ENSG00000244265 | n.s. |
| ENSG00000117408 | 187.1784498 | -0.590792786 | 0.2715723 | -2.175453 | 0.029596181 | 0.090564313 | IPO13 | protein_coding | 51.66 | -0.616222224 | 0.616222224 | ENSG00000117408 | n.s. |

| | | | | | | | | | | | | | |
|---|---|---|---|---|---|---|---|---|---|---|---|---|---|
| ENSG00000231584 | 100.2648564 | -0.622836765 | 0.2948154 | -2.112633 | 0.03463218 | 0.102483282 | FAHD2CP | transcribed_unprocessed_pseud | 49.42 | -0.616201671 | 0.616201671 | ENSG00000231584 | n.s. |
| ENSG00000170445 | 1543.574889 | -0.585985151 | 0.2682417 | -2.184542 | 0.028922472 | 0.088908852 | HARS | protein_coding | 46.01 | -0.615902622 | 0.615902622 | ENSG00000170445 | n.s. |
| ENSG00000148690 | 339.3253725 | -0.576945154 | 0.2619165 | -2.202783 | 0.027610058 | 0.085775952 | FRA10AC1 | protein_coding | 34.82 | -0.61538958 | 0.61538958 | ENSG00000148690 | n.s. |
| ENSG00000163808 | 2565.970678 | -0.577737555 | 0.2625746 | -2.20028 | 0.02778705 | 0.086145423 | KIF15 | protein_coding | 41.48 | -0.615156339 | 0.615156339 | ENSG00000163808 | n.s. |
| ENSG00000185986 | 234.5051029 | -0.591895172 | 0.2725852 | -2.171414 | 0.02989992 | 0.091373262 | SDHAP3 | transcribed_unprocessed_pseud | 47.9 | -0.61508614 | 0.61508614 | ENSG00000185986 | n.s. |
| ENSG00000200397 | 41.79838394 | 0.59512722 | 0.2749319 | 2.164635 | 0.030415649 | 0.092644296 | | misc_RNA | 48.04 | 0.614874324 | 0.614874324 | ENSG00000200397 | n.s. |
| ENSG00000174720 | 1794.456216 | -0.599609696 | 0.2782167 | -2.155189 | 0.031147019 | 0.09449988 | LARP7 | protein_coding | 37.92 | -0.614341353 | 0.614341353 | ENSG00000174720 | n.s. |
| ENSG00000233797 | 30.37187995 | -0.638635878 | 0.3072193 | -2.078762 | 0.037639239 | 0.109222827 | UFL1-AS1 | antisense | 37.42 | -0.614167557 | 0.614167557 | ENSG00000233797 | n.s. |
| ENSG00000103253 | 69.27967213 | 0.635735548 | 0.3048998 | 2.085064 | 0.037063535 | 0.108180565 | HAGHL | protein_coding | 65.72 | 0.614025659 | 0.614025659 | ENSG00000103253 | n.s. |
| ENSG00000153714 | 71.34724551 | 0.571780932 | 0.2586422 | 2.210703 | 0.027056429 | 0.084463377 | LURAP1L | protein_coding | 36.85 | 0.613710519 | 0.613710519 | ENSG00000153714 | n.s. |
| ENSG00000234665 | 84.74571885 | -0.60246632 | 0.2807662 | -2.145794 | 0.031889425 | 0.096228103 | | lincRNA | 38.27 | -0.612526347 | 0.612526347 | ENSG00000234665 | n.s. |
| ENSG00000229325 | 82.4935823 | -0.616721655 | 0.2914893 | -2.115761 | 0.034365139 | 0.101875597 | ACAP2-IT1 | sense_intronic | 33.33 | -0.611744608 | 0.611744608 | ENSG00000229325 | n.s. |
| ENSG00000257923 | 282.6386369 | -0.536725585 | 0.2348811 | -2.285095 | 0.022307281 | 0.072549469 | CUX1 | protein_coding | 48.61 | -0.611526754 | 0.611526754 | ENSG00000257923 | n.s. |
| ENSG00000200636 | 4.48136199 | -1.362420019 | 1.0286616 | -1.324459 | 0.185350673 | 0.355753712 | SNORD114-27 | snoRNA | 45.71 | -0.611522988 | 0.611522988 | ENSG00000200636 | n.s. |
| ENSG00000111203 | 596.4412149 | -0.572671517 | 0.2600292 | -2.202335 | 0.027641636 | 0.08584257 | ITFG2 | protein_coding | 50.2 | -0.610638084 | 0.610638084 | ENSG00000111203 | n.s. |
| ENSG00000182263 | 656.7556304 | -0.54314601 | 0.2396485 | -2.266428 | 0.023425209 | 0.075270986 | FIGN | protein_coding | 36.3 | -0.610155236 | 0.610155236 | ENSG00000182263 | n.s. |
| ENSG00000196715 | 808.9504984 | -0.618551171 | 0.2933195 | -2.108796 | 0.034962157 | 0.103237219 | VKORC1L1 | protein_coding | 41.16 | -0.609992713 | 0.609992713 | ENSG00000196715 | n.s. |
| ENSG00000259318 | 51.32492201 | 0.608260002 | 0.2857099 | 2.128943 | 0.033258987 | 0.099348825 | | processed_pseudogene | 45.92 | 0.609985797 | 0.609985797 | ENSG00000259318 | n.s. |
| ENSG00000196781 | 816.3227735 | 0.542106943 | 0.2389757 | 2.268461 | 0.023301125 | 0.074958772 | TLE1 | protein_coding | 43.43 | 0.609966556 | 0.609966556 | ENSG00000196781 | n.s. |
| ENSG00000066583 | 370.092532 | 0.607094593 | 0.2850936 | 2.129457 | 0.033216454 | 0.099243065 | ISOC1 | protein_coding | 38.93 | 0.609097902 | 0.609097902 | ENSG00000066583 | n.s. |
| ENSG00000111269 | 667.7213941 | -0.633936684 | 0.3051198 | -2.077665 | 0.037740223 | 0.109470234 | CREBL2 | protein_coding | 40.66 | -0.609025478 | 0.609025478 | ENSG00000111269 | n.s. |
| ENSG00000251739 | 6.848214611 | -0.841156595 | 0.4754726 | -1.769096 | 0.076877886 | 0.188944538 | RNU6-1053P | snRNA | 46.53 | -0.60871614 | 0.60871614 | ENSG00000251739 | n.s. |
| ENSG00000259797 | 29.32613019 | -0.674213678 | 0.3358549 | -2.007455 | 0.044701208 | 0.125075344 | | sense_intronic | 48.39 | -0.608699184 | 0.608699184 | ENSG00000259797 | n.s. |
| ENSG00000166166 | 304.0636067 | -0.556574368 | 0.2493949 | -2.231699 | 0.025634842 | 0.080969578 | TRMT61A | protein_coding | 64.22 | -0.607600063 | 0.607600063 | ENSG00000166166 | n.s. |
| ENSG00000035862 | 2305.102121 | -0.534701733 | 0.2344801 | -2.280372 | 0.022585645 | 0.07319831 | TIMP2 | protein_coding | 52.6 | -0.607153254 | 0.607153254 | ENSG00000035862 | n.s. |
| ENSG00000141551 | 902.4949668 | -0.580124425 | 0.2662454 | -2.178909 | 0.029338465 | 0.089933827 | CSNK1D | protein_coding | 55.31 | -0.606854774 | 0.606854774 | ENSG00000141551 | n.s. |
| ENSG00000243701 | 165.3319305 | -0.597372257 | 0.2786513 | -2.143798 | 0.032049041 | 0.096584155 | DUBR | lincRNA | 38.56 | -0.606389062 | 0.606389062 | ENSG00000243701 | n.s. |
| ENSG00000078747 | 1398.117269 | -0.568934265 | 0.2585736 | -2.20028 | 0.027787033 | 0.086145423 | ITCH | protein_coding | 40.53 | -0.60578288 | 0.60578288 | ENSG00000078747 | n.s. |
| ENSG00000111224 | 59.99493325 | -0.607545395 | 0.2864265 | -2.121122 | 0.033911554 | 0.100757696 | PARP11 | protein_coding | 39.15 | -0.605553722 | 0.605553722 | ENSG00000111224 | n.s. |
| ENSG00000179218 | 3770.90551 | 0.571286371 | 0.2605906 | 2.192275 | 0.028359639 | 0.087511358 | CALR | protein_coding | 53.01 | 0.604384176 | 0.604384176 | ENSG00000179218 | n.s. |
| ENSG00000206937 | 9.608562395 | -0.794751663 | 0.4366682 | -1.820036 | 0.068753586 | 0.173856915 | SNORA70B | snoRNA | 49.63 | -0.603858696 | 0.603858696 | ENSG00000206937 | n.s. |
| ENSG00000125247 | 1152.290237 | -0.620515077 | 0.2965389 | -2.092525 | 0.036391572 | 0.106512478 | TMTC4 | protein_coding | 44.69 | -0.60351266 | 0.60351266 | ENSG00000125247 | n.s. |

| | | | | | | | | | | | | | |
|---|---|---|---|---|---|---|---|---|---|---|---|---|---|
| ENSG00000279638 | 8.241201156 | -0.77387769 | 0.4190457 | -1.846762 | 0.064781621 | 0.166381573 | | TEC | 32.61 | -0.60276929 | 0.60276929 | ENSG00000279638 | n.s. |
| ENSG00000173898 | 2805.917276 | -0.519991635 | 0.2253064 | -2.307931 | 0.021003 | 0.069316591 | SPTBN2 | protein_coding | 55.04 | -0.602754961 | 0.602754961 | ENSG00000173898 | n.s. |
| ENSG00000134146 | 161.3457972 | -0.626114121 | 0.3011274 | -2.079234 | 0.037595892 | 0.10918807 | DPH6 | protein_coding | 36.45 | -0.602212095 | 0.602212095 | ENSG00000134146 | n.s. |
| ENSG00000234624 | 20.20074864 | -0.65456634 | 0.3228609 | -2.027394 | 0.042622133 | 0.120346371 | | processed_pseudogene | 54.98 | -0.601917606 | 0.601917606 | ENSG00000234624 | n.s. |
| ENSG00000133703 | 1290.688553 | -0.589623162 | 0.2742165 | -2.15021 | 0.031538595 | 0.095355454 | KRAS | protein_coding | 36.37 | -0.601801512 | 0.601801512 | ENSG00000133703 | n.s. |
| ENSG00000120137 | 8890.412385 | -0.569933314 | 0.2602688 | -2.189788 | 0.028539636 | 0.088008273 | PANK3 | protein_coding | 37.86 | -0.60155122 | 0.60155122 | ENSG00000120137 | n.s. |
| ENSG00000203761 | 193.4867323 | 0.526342737 | 0.2300098 | 2.288349 | 0.022117213 | 0.072082856 | MSTO2P | unprocessed_pseudogene | 52.8 | 0.601171838 | 0.601171838 | ENSG00000203761 | n.s. |
| ENSG00000198846 | 298.7534435 | -0.561199932 | 0.2541816 | -2.20787 | 0.027253356 | 0.084982861 | TOX | protein_coding | 37.57 | -0.600859175 | 0.600859175 | ENSG00000198846 | n.s. |
| ENSG00000166783 | 800.8585012 | -0.564889588 | 0.2569079 | -2.198802 | 0.027892001 | 0.086432302 | MARF1 | protein_coding | 44.18 | -0.60066061 | 0.60066061 | ENSG00000166783 | n.s. |
| ENSG00000119004 | 521.7407193 | -0.571113533 | 0.2614304 | -2.184572 | 0.028920247 | 0.088908852 | CYP20A1 | protein_coding | 42.38 | -0.600271734 | 0.600271734 | ENSG00000119004 | n.s. |
| ENSG00000180008 | 2648.836639 | -0.592270919 | 0.2765431 | -2.141695 | 0.032218039 | 0.096942783 | SOCS4 | protein_coding | 37.83 | -0.600257406 | 0.600257406 | ENSG00000180008 | n.s. |
| ENSG00000201263 | 8.291572112 | -0.843018528 | 0.4813244 | -1.751456 | 0.079867415 | 0.194111575 | SNORD114-6 | snoRNA | 40.28 | -0.600185833 | 0.600185833 | ENSG00000201263 | n.s. |
| ENSG00000158545 | 572.3022963 | -0.547430056 | 0.2447416 | -2.236767 | 0.025301546 | 0.080189715 | ZC3H18 | protein_coding | 51.75 | -0.599918378 | 0.599918378 | ENSG00000158545 | n.s. |
| ENSG00000123643 | 37.34105147 | -0.658906302 | 0.3270938 | -2.014426 | 0.043964805 | 0.123411689 | SLC36A1 | protein_coding | 42.72 | -0.598711063 | 0.598711063 | ENSG00000123643 | n.s. |
| ENSG00000228485 | 72.42812334 | 0.57880471 | 0.2672797 | 2.165539 | 0.030346412 | 0.092494111 | GRK5-IT1 | sense_intronic | 39.16 | 0.598418039 | 0.598418039 | ENSG00000228485 | n.s. |
| ENSG00000278847 | 30.26015945 | 0.643495974 | 0.3157443 | 2.038029 | 0.041547022 | 0.11784686 | | lincRNA | 32.79 | 0.597603118 | 0.597603118 | ENSG00000278847 | n.s. |
| ENSG00000153006 | 773.9118138 | -0.602316804 | 0.2846859 | -2.115724 | 0.034368288 | 0.101875597 | SREK1IP1 | protein_coding | 36.21 | -0.597456007 | 0.597456007 | ENSG00000153006 | n.s. |
| ENSG00000183878 | 224.0262246 | 0.539957703 | 0.2402573 | 2.247414 | 0.024613573 | 0.078348144 | UTY | protein_coding | 37.47 | 0.597177716 | 0.597177716 | ENSG00000183878 | n.s. |
| ENSG00000048052 | 1948.249353 | -0.526648404 | 0.231168 | -2.278207 | 0.022714271 | 0.073529596 | HDAC9 | protein_coding | 36.13 | -0.596975885 | 0.596975885 | ENSG00000048052 | n.s. |
| ENSG00000075711 | 1938.689372 | -0.576624538 | 0.2660778 | -2.167127 | 0.030225135 | 0.092225413 | DLG1 | protein_coding | 36.59 | -0.596892541 | 0.596892541 | ENSG00000075711 | n.s. |
| ENSG00000188582 | 104.8104702 | 0.552041166 | 0.248743 | 2.219323 | 0.026464754 | 0.083063293 | PAQR9 | protein_coding | 41.7 | 0.596530635 | 0.596530635 | ENSG00000188582 | n.s. |
| ENSG00000186472 | 175.3709729 | -0.570082341 | 0.261714 | -2.178265 | 0.029386316 | 0.090060681 | PCLO | protein_coding | 33.81 | -0.596001006 | 0.596001006 | ENSG00000186472 | n.s. |
| ENSG00000175061 | 9416.061098 | 0.604180085 | 0.2865251 | 2.108646 | 0.034975125 | 0.103237219 | LRRC75A-AS1 | processed_transcript | 52.34 | 0.595820469 | 0.595820469 | ENSG00000175061 | n.s. |
| ENSG00000099968 | 607.851542 | -0.619217071 | 0.2977528 | -2.079634 | 0.037559073 | 0.109126663 | BCL2L13 | protein_coding | 42.8 | -0.595729624 | 0.595729624 | ENSG00000099968 | n.s. |
| ENSG00000135164 | 3376.078672 | -0.55791159 | 0.2531473 | -2.203901 | 0.027531309 | 0.085619479 | DMTF1 | protein_coding | 35.84 | -0.595530128 | 0.595530128 | ENSG00000135164 | n.s. |
| ENSG00000091009 | 687.4122731 | -0.560841109 | 0.2554645 | -2.195378 | 0.028136529 | 0.087015748 | RBM27 | protein_coding | 38.7 | -0.594717112 | 0.594717112 | ENSG00000091009 | n.s. |
| ENSG00000038532 | 428.9989234 | -0.558483857 | 0.2537926 | -2.200552 | 0.027767764 | 0.086143176 | CLEC16A | protein_coding | 48.06 | -0.594661949 | 0.594661949 | ENSG00000038532 | n.s. |
| ENSG00000009954 | 5229.758357 | -0.549555474 | 0.2474417 | -2.220949 | 0.026354418 | 0.082847721 | BAZ1B | protein_coding | 43.19 | -0.594464836 | 0.594464836 | ENSG00000009954 | n.s. |
| ENSG00000145198 | 236.0454294 | -0.557116177 | 0.2529186 | -2.202749 | 0.027612427 | 0.085775952 | VWA5B2 | protein_coding | 58.09 | -0.59423931 | 0.59423931 | ENSG00000145198 | n.s. |
| ENSG00000128266 | 217.8033479 | -0.566181085 | 0.2594609 | -2.182144 | 0.029098925 | 0.089317533 | GNAZ | protein_coding | 57.44 | -0.593959786 | 0.593959786 | ENSG00000128266 | n.s. |
| ENSG00000173166 | 537.4411303 | -0.581950551 | 0.2707432 | -2.149456 | 0.031598296 | 0.095453043 | RAPH1 | protein_coding | 39.16 | -0.5937119 | 0.5937119 | ENSG00000173166 | n.s. |
| ENSG00000162402 | 4365.273633 | -0.534147946 | 0.2370466 | -2.253346 | 0.024237364 | 0.077398352 | USP24 | protein_coding | 37.78 | -0.593581674 | 0.593581674 | ENSG00000162402 | n.s. |

| | | | | | | | | | | | | | |
|---|---|---|---|---|---|---|---|---|---|---|---|---|---|
| ENSG00000263069 | 19.33562997 | 0.666701071 | 0.3346739 | 1.992092 | 0.046361006 | 0.128736396 | | antisense | 48.14 | 0.593563065 | 0.593563065 | ENSG00000263069 | n.s. |
| ENSG00000140941 | 502.4308405 | 0.559941908 | 0.2551461 | 2.194593 | 0.028192797 | 0.087123267 | MAP1LC3B | protein_coding | 47.47 | 0.593463304 | 0.593463304 | ENSG00000140941 | n.s. |
| ENSG00000222345 | 66.87049804 | 0.58049333 | 0.2697868 | 2.151675 | 0.031422989 | 0.095047204 | | snoRNA | 35.53 | 0.593299392 | 0.593299392 | ENSG00000222345 | n.s. |
| ENSG00000149532 | 1523.289127 | -0.559654125 | 0.2551455 | -2.19347 | 0.028273501 | 0.087284246 | CPSF7 | protein_coding | 44.18 | -0.592709611 | 0.592709611 | ENSG00000149532 | n.s. |
| ENSG00000162407 | 603.4972684 | 0.583239343 | 0.2719738 | 2.144469 | 0.031995344 | 0.096464091 | PLPP3 | protein_coding | 41.42 | 0.592357897 | 0.592357897 | ENSG00000162407 | n.s. |
| ENSG00000199867 | 4.192266027 | 1.108120829 | 0.7513039 | 1.47493 | 0.140231285 | 0.292760715 | | misc_RNA | 39.22 | 0.59116828 | 0.59116828 | ENSG00000199867 | n.s. |
| ENSG00000272617 | 434.7508113 | -0.558340973 | 0.2546587 | -2.192507 | 0.028342946 | 0.087479238 | COG8 | protein_coding | 51.52 | -0.590777798 | 0.590777798 | ENSG00000272617 | n.s. |
| ENSG00000171451 | 446.8904472 | 0.482742439 | 0.2033438 | 2.374021 | 0.017595558 | 0.060015421 | DSEL | protein_coding | 41.51 | 0.589784368 | 0.589784368 | ENSG00000171451 | n.s. |
| ENSG00000248593 | 38.20942762 | -0.632697598 | 0.3097857 | -2.042372 | 0.041114674 | 0.116967636 | DSTNP2 | transcribed_unprocessed_pseud | 46.24 | -0.589632585 | 0.589632585 | ENSG00000248593 | n.s. |
| ENSG00000117791 | 11.27725095 | -0.744641714 | 0.3996843 | -1.863075 | 0.062451762 | 0.161890873 | 2-Mar | protein_coding | 44.02 | -0.588846013 | 0.588846013 | ENSG00000117791 | n.s. |
| ENSG00000109762 | 436.9293958 | -0.562706411 | 0.2581877 | -2.179447 | 0.029298487 | 0.08987064 | SNX25 | protein_coding | 41.36 | -0.588805953 | 0.588805953 | ENSG00000109762 | n.s. |
| ENSG00000138080 | 1086.171579 | -0.541404738 | 0.2431409 | -2.226712 | 0.025966532 | 0.081776075 | EMILIN1 | protein_coding | 61.63 | -0.588709294 | 0.588709294 | ENSG00000138080 | n.s. |
| ENSG00000150401 | 720.0525693 | -0.554892769 | 0.2526834 | -2.196 | 0.028091951 | 0.086916495 | DCUN1D2 | protein_coding | 44.89 | -0.588684515 | 0.588684515 | ENSG00000150401 | n.s. |
| ENSG00000163781 | 3768.37544 | -0.579490852 | 0.2702084 | -2.144607 | 0.031984256 | 0.096464091 | TOPBP1 | protein_coding | 37.82 | -0.5885508 | 0.5885508 | ENSG00000163781 | n.s. |
| ENSG00000154040 | 83.74536019 | 0.546587308 | 0.2469585 | 2.213276 | 0.026878619 | 0.084021331 | CABYR | protein_coding | 43.36 | 0.587915018 | 0.587915018 | ENSG00000154040 | n.s. |
| ENSG00000196656 | 12.63238529 | 0.744842751 | 0.4002605 | 1.860895 | 0.062759038 | 0.162525408 | | transcribed_processed_pseudog | 35.46 | 0.587739575 | 0.587739575 | ENSG00000196656 | n.s. |
| ENSG00000178385 | 730.4363488 | -0.543065436 | 0.2446758 | -2.219531 | 0.026450614 | 0.083037633 | PLEKHM3 | protein_coding | 41.34 | -0.586904414 | 0.586904414 | ENSG00000178385 | n.s. |
| ENSG00000170889 | 6783.651551 | 0.620349215 | 0.3012527 | 2.059232 | 0.039472024 | 0.113476976 | RPS9 | protein_coding | 51.79 | 0.586287234 | 0.586287234 | ENSG00000170889 | n.s. |
| ENSG00000256663 | 59.62836418 | -0.578328617 | 0.2700659 | -2.141435 | 0.032238975 | 0.096967648 | | processed_pseudogene | 62.26 | -0.586062685 | 0.586062685 | ENSG00000256663 | n.s. |
| ENSG00000115361 | 173.456662 | -0.590911002 | 0.2793743 | -2.115123 | 0.034419488 | 0.101978428 | ACADL | protein_coding | 36.42 | -0.585883347 | 0.585883347 | ENSG00000115361 | n.s. |
| ENSG00000176371 | 772.6611608 | -0.536927728 | 0.2406892 | -2.230792 | 0.025694889 | 0.081085661 | ZSCAN2 | protein_coding | 48.03 | -0.585818185 | 0.585818185 | ENSG00000176371 | n.s. |
| ENSG00000135912 | 577.360267 | -0.533291028 | 0.2382096 | -2.238747 | 0.025172363 | 0.079852997 | TTLL4 | protein_coding | 44.96 | -0.585398242 | 0.585398242 | ENSG00000135912 | n.s. |
| ENSG00000105327 | 77.10631585 | 0.575004007 | 0.2678476 | 2.146758 | 0.031812558 | 0.096016961 | BBC3 | protein_coding | 59.72 | 0.585154003 | 0.585154003 | ENSG00000105327 | n.s. |
| ENSG00000212371 | 28.5553833 | -0.604815639 | 0.2897993 | -2.087016 | 0.036886726 | 0.10784836 | | snoRNA | 41.83 | -0.584969493 | 0.584969493 | ENSG00000212371 | n.s. |
| ENSG00000055130 | 1595.789301 | -0.61036688 | 0.2942676 | -2.07419 | 0.038061672 | 0.110173109 | CUL1 | protein_coding | 41.22 | -0.584685325 | 0.584685325 | ENSG00000055130 | n.s. |
| ENSG00000091656 | 1042.6477 | -0.53600677 | 0.2405273 | -2.228465 | 0.025849502 | 0.081452317 | ZFHX4 | protein_coding | 36.38 | -0.583763128 | 0.583763128 | ENSG00000091656 | n.s. |
| ENSG00000211997 | 22.37419477 | -0.670631501 | 0.3409617 | -1.966882 | 0.049196793 | 0.13488912 | MIR708 | miRNA | 48.86 | -0.583464883 | 0.583464883 | ENSG00000211997 | n.s. |
| ENSG00000270061 | 78.64172942 | 0.569361785 | 0.2641682 | 2.1553 | 0.031138366 | 0.094494219 | | sense_intronic | 45.63 | 0.583365102 | 0.583365102 | ENSG00000270061 | n.s. |
| ENSG00000183576 | 918.7834963 | -0.590870045 | 0.2800518 | -2.10986 | 0.03487045 | 0.103036649 | SETD3 | protein_coding | 42.67 | -0.583193623 | 0.583193623 | ENSG00000183576 | n.s. |
| ENSG00000034152 | 41.98345585 | 0.58290998 | 0.2742493 | 2.125474 | 0.033547042 | 0.099944762 | MAP2K3 | protein_coding | 56.97 | 0.583049858 | 0.583049858 | ENSG00000034152 | n.s. |
| ENSG00000182405 | 286.2695305 | 0.491290377 | 0.2103003 | 2.336138 | 0.019484065 | 0.065320643 | PGBD4 | protein_coding | 42.18 | 0.582154311 | 0.582154311 | ENSG00000182405 | n.s. |
| ENSG00000207997 | 15.92831801 | -0.691697093 | 0.3582218 | -1.930918 | 0.05349314 | 0.144114743 | MIR644A | miRNA | 32.98 | -0.581918945 | 0.581918945 | ENSG00000207997 | n.s. |

| | | | | | | | | | | | | | |
|---|---|---|---|---|---|---|---|---|---|---|---|---|---|
| ENSG00000159792 | 204.8519087 | -0.57363346 | 0.2676869 | -2.142927 | 0.032118985 | 0.096732128 | PSKH1 | protein_coding | 52.29 | -0.581910565 | 0.581910565 | ENSG00000159792 | n.s. |
| ENSG00000111785 | 129.4068821 | -0.575779043 | 0.269301 | -2.13805 | 0.032512676 | 0.097664292 | RIC8B | protein_coding | 38.48 | -0.58168895 | 0.58168895 | ENSG00000111785 | n.s. |
| ENSG00000172572 | 392.7331015 | -0.581124995 | 0.2734169 | -2.125417 | 0.033551807 | 0.099944762 | PDE3A | protein_coding | 35.95 | -0.581264444 | 0.581264444 | ENSG00000172572 | n.s. |
| ENSG00000163162 | 928.6317016 | -0.578134931 | 0.2712427 | -2.13143 | 0.033053706 | 0.098841655 | RNF149 | protein_coding | 41.19 | -0.58106029 | 0.58106029 | ENSG00000163162 | n.s. |
| ENSG00000196440 | 43.10553341 | -0.583365719 | 0.2751885 | -2.119877 | 0.034016418 | 0.100983068 | ARMCX4 | protein_coding | 41.06 | -0.580887255 | 0.580887255 | ENSG00000196440 | n.s. |
| ENSG00000127334 | 7715.710628 | -0.539119362 | 0.2433207 | -2.215674 | 0.026713815 | 0.083682409 | DYRK2 | protein_coding | 43.53 | -0.580828778 | 0.580828778 | ENSG00000127334 | n.s. |
| ENSG00000165113 | 471.7378084 | -0.588968184 | 0.2793371 | -2.10845 | 0.034992125 | 0.10326311 | GKAP1 | protein_coding | 38.75 | -0.580754906 | 0.580754906 | ENSG00000165113 | n.s. |
| ENSG00000141569 | 778.0535435 | -0.60491973 | 0.29133 | -2.076408 | 0.037856272 | 0.109761116 | TRIM65 | protein_coding | 54.93 | -0.580451626 | 0.580451626 | ENSG00000141569 | n.s. |
| ENSG00000198837 | 564.7003341 | -0.536020575 | 0.2414564 | -2.219948 | 0.026422306 | 0.083004911 | DENND4B | protein_coding | 56.48 | -0.57938261 | 0.57938261 | ENSG00000198837 | n.s. |
| ENSG00000100211 | 154.4622499 | -0.588393012 | 0.2793247 | -2.106484 | 0.035162361 | 0.103602259 | CBY1 | protein_coding | 47.17 | -0.579349867 | 0.579349867 | ENSG00000100211 | n.s. |
| ENSG00000164418 | 482.8700401 | -0.548565707 | 0.2505752 | -2.189226 | 0.028580431 | 0.088114558 | GRIK2 | protein_coding | 33.94 | -0.578710673 | 0.578710673 | ENSG00000164418 | n.s. |
| ENSG00000121406 | 347.5926121 | -0.571743825 | 0.2671552 | -2.140119 | 0.032345151 | 0.097244986 | ZNF549 | protein_coding | 43.95 | -0.578680668 | 0.578680668 | ENSG00000121406 | n.s. |
| ENSG00000143344 | 199.7719228 | -0.510174192 | 0.2239623 | -2.277947 | 0.022729742 | 0.073562574 | RGL1 | protein_coding | 39.98 | -0.578202392 | 0.578202392 | ENSG00000143344 | n.s. |
| ENSG00000281649 | 723.345915 | -0.558464395 | 0.2577088 | -2.167037 | 0.03023204 | 0.09222627 | EBLN3P | lincRNA | 38.61 | -0.578091826 | 0.578091826 | ENSG00000281649 | n.s. |
| ENSG00000234465 | 160.2265775 | 0.517644188 | 0.2291559 | 2.258917 | 0.023888532 | 0.076473986 | PINLYP | protein_coding | 51.61 | 0.577942631 | 0.577942631 | ENSG00000234465 | n.s. |
| ENSG00000221066 | 25.25743497 | -0.660816474 | 0.3349202 | -1.973056 | 0.04848914 | 0.133493301 | SNORD111 | snoRNA | 37.23 | -0.577910788 | 0.577910788 | ENSG00000221066 | n.s. |
| ENSG00000252161 | 3.445859428 | -0.978541162 | 0.6250887 | -1.565444 | 0.117478889 | 0.256936628 | RNA5SP318 | rRNA | 50 | -0.577509531 | 0.577509531 | ENSG00000252161 | n.s. |
| ENSG00000280120 | 279.2868987 | -0.590990064 | 0.2816974 | -2.097961 | 0.035908634 | 0.105431445 | | TEC | 39.37 | -0.577414928 | 0.577414928 | ENSG00000280120 | n.s. |
| ENSG00000088986 | 2499.281496 | 0.646576024 | 0.3243012 | 1.993751 | 0.046179229 | 0.128282801 | DYNLL1 | protein_coding | 47.04 | 0.576636908 | 0.576636908 | ENSG00000088986 | n.s. |
| ENSG00000119392 | 488.0842305 | -0.575436842 | 0.2704151 | -2.127976 | 0.033339056 | 0.09954448 | GLE1 | protein_coding | 45.27 | -0.57657783 | 0.57657783 | ENSG00000119392 | n.s. |
| ENSG00000246465 | 227.3688288 | -0.538334257 | 0.243928 | -2.206939 | 0.027318299 | 0.085109124 | | lincRNA | 49.16 | -0.576030511 | 0.576030511 | ENSG00000246465 | n.s. |
| ENSG00000198911 | 1454.353928 | -0.547158617 | 0.2504858 | -2.18439 | 0.028933602 | 0.088908852 | SREBF2 | protein_coding | 48.21 | -0.575093799 | 0.575093799 | ENSG00000198911 | n.s. |
| ENSG00000196470 | 1237.304569 | 0.574193227 | 0.2704035 | 2.123468 | 0.033714629 | 0.100322456 | SIAH1 | protein_coding | 45.2 | 0.573390415 | 0.573390415 | ENSG00000196470 | n.s. |
| ENSG00000145780 | 616.2756066 | -0.550481355 | 0.2532541 | -2.173632 | 0.029732754 | 0.090922278 | FEM1C | protein_coding | 35.62 | -0.57323258 | 0.57323258 | ENSG00000145780 | n.s. |
| ENSG00000117748 | 578.0007636 | -0.640362655 | 0.3204907 | -1.998069 | 0.045709138 | 0.127383573 | RPA2 | protein_coding | 44.15 | -0.573051943 | 0.573051943 | ENSG00000117748 | n.s. |
| ENSG00000114904 | 798.3008283 | -0.55572232 | 0.2570175 | -2.162197 | 0.030603019 | 0.093072485 | NEK4 | protein_coding | 42.79 | -0.573049015 | 0.573049015 | ENSG00000114904 | n.s. |
| ENSG00000188641 | 818.1063593 | -0.582516732 | 0.2767597 | -2.104775 | 0.035310927 | 0.104005509 | DPYD | protein_coding | 35.09 | -0.572581127 | 0.572581127 | ENSG00000188641 | n.s. |
| ENSG00000130338 | 2476.110965 | -0.531368125 | 0.2398259 | -2.215641 | 0.026716103 | 0.083682409 | TULP4 | protein_coding | 43.15 | -0.572477859 | 0.572477859 | ENSG00000130338 | n.s. |
| ENSG00000163449 | 402.4440864 | -0.60415973 | 0.2930107 | -2.061903 | 0.039216969 | 0.112883576 | TMEM169 | protein_coding | 43.95 | -0.572362345 | 0.572362345 | ENSG00000163449 | n.s. |
| ENSG00000010165 | 1155.533347 | -0.539235066 | 0.2454821 | -2.196637 | 0.02804635 | 0.086813992 | METTL13 | protein_coding | 43.61 | -0.572349639 | 0.572349639 | ENSG00000010165 | n.s. |
| ENSG00000230732 | 137.9458785 | -0.535418255 | 0.2428291 | -2.204918 | 0.027459861 | 0.085454547 | | sense_intronic | 39.19 | -0.571968488 | 0.571968488 | ENSG00000230732 | n.s. |
| ENSG00000139352 | 1477.051738 | 0.481245966 | 0.2059048 | 2.337226 | 0.019427459 | 0.065146558 | ASCL1 | protein_coding | 51.29 | 0.570809941 | 0.570809941 | ENSG00000139352 | n.s. |

| | | | | | | | | | | | | | |
|---|---|---|---|---|---|---|---|---|---|---|---|---|---|
| ENSG00000139116 | 15678.41325 | -0.573037294 | 0.2705217 | -2.118268 | 0.034152411 | 0.101321973 | KIF21A | protein_coding | 36.22 | -0.569768896 | 0.569768896 | ENSG00000139116 | n.s. |
| ENSG00000121274 | 703.0713678 | -0.552475614 | 0.2554932 | -2.162389 | 0.030588213 | 0.093047781 | PAPD5 | protein_coding | 41.51 | -0.569764775 | 0.569764775 | ENSG00000121274 | n.s. |
| ENSG00000073060 | 1517.080666 | -0.552399321 | 0.2554252 | -2.162666 | 0.030566886 | 0.093023553 | SCARB1 | protein_coding | 51.45 | -0.569748568 | 0.569748568 | ENSG00000073060 | n.s. |
| ENSG00000198793 | 1182.632272 | -0.537313658 | 0.2447848 | -2.195045 | 0.028160386 | 0.087070189 | MTOR | protein_coding | 44.54 | -0.569622603 | 0.569622603 | ENSG00000198793 | n.s. |
| ENSG00000140332 | 2417.415024 | 0.498482714 | 0.2178233 | 2.288472 | 0.022110022 | 0.072077752 | TLE3 | protein_coding | 52.15 | 0.569366341 | 0.569366341 | ENSG00000140332 | n.s. |
| ENSG00000137221 | 271.967209 | -0.569203906 | 0.2678679 | -2.124943 | 0.033591389 | 0.100041265 | TJAP1 | protein_coding | 51.18 | -0.569101918 | 0.569101918 | ENSG00000137221 | n.s. |
| ENSG00000186767 | 389.3036322 | 0.538296524 | 0.2459006 | 2.189081 | 0.028590921 | 0.088118369 | SPIN4 | protein_coding | 37.43 | 0.567867063 | 0.567867063 | ENSG00000186767 | n.s. |
| ENSG00000100345 | 1422.964329 | -0.47855752 | 0.2047443 | -2.337342 | 0.019421418 | 0.06514199 | MYH9 | protein_coding | 52.37 | -0.56763573 | 0.56763573 | ENSG00000100345 | n.s. |
| ENSG00000221680 | 20.02539234 | 0.681526367 | 0.3551376 | 1.919049 | 0.05497816 | 0.146945877 | MIR1278 | miRNA | 37.04 | 0.567604189 | 0.567604189 | ENSG00000221680 | n.s. |
| ENSG00000165490 | 753.1455699 | -0.575531225 | 0.273013 | -2.108072 | 0.035024741 | 0.103315565 | DDIAS | protein_coding | 39.78 | -0.567378392 | 0.567378392 | ENSG00000165490 | n.s. |
| ENSG00000145833 | 2959.363349 | -0.554307185 | 0.2573456 | -2.153941 | 0.031244827 | 0.094734696 | DDX46 | protein_coding | 42.1 | -0.567328377 | 0.567328377 | ENSG00000145833 | n.s. |
| ENSG00000167173 | 289.9127729 | 0.494707604 | 0.2156773 | 2.29374 | 0.021805425 | 0.07141777 | C15orf39 | protein_coding | 58.35 | 0.567030747 | 0.567030747 | ENSG00000167173 | n.s. |
| ENSG00000066322 | 98.16207235 | -0.562133094 | 0.2632414 | -2.135428 | 0.032726029 | 0.098072429 | ELOVL1 | protein_coding | 58.74 | -0.566884844 | 0.566884844 | ENSG00000066322 | n.s. |
| ENSG00000198146 | 2074.620785 | -0.539312637 | 0.2468943 | -2.184387 | 0.028933851 | 0.088908852 | ZNF770 | protein_coding | 37.91 | -0.566847242 | 0.566847242 | ENSG00000198146 | n.s. |
| ENSG00000087263 | 984.5838394 | -0.588301918 | 0.2826144 | -2.081642 | 0.037375177 | 0.108780649 | OGFOD1 | protein_coding | 40.39 | -0.566798516 | 0.566798516 | ENSG00000087263 | n.s. |
| ENSG00000077232 | 3518.146632 | -0.569982184 | 0.2691414 | -2.117779 | 0.034193759 | 0.101423031 | DNAJC10 | protein_coding | 36.39 | -0.56648444 | 0.56648444 | ENSG00000077232 | n.s. |
| ENSG00000110713 | 2448.255271 | -0.565458076 | 0.2657963 | -2.127412 | 0.033385905 | 0.09962108 | NUP98 | protein_coding | 41.69 | -0.566390376 | 0.566390376 | ENSG00000110713 | n.s. |
| ENSG00000168096 | 63.2860415 | -0.567927538 | 0.2677866 | -2.120821 | 0.033936854 | 0.100811353 | ANKS3 | protein_coding | 49.79 | -0.565934429 | 0.565934429 | ENSG00000168096 | n.s. |
| ENSG00000174628 | 29.29411883 | -0.638583613 | 0.3214307 | -1.986691 | 0.046956601 | 0.129975498 | IQCK | protein_coding | 43.85 | -0.565873532 | 0.565873532 | ENSG00000174628 | n.s. |
| ENSG00000174928 | 121.6711314 | -0.595506851 | 0.2884158 | -2.064751 | 0.038946531 | 0.112221141 | C3orf33 | protein_coding | 40.75 | -0.565687035 | 0.565687035 | ENSG00000174928 | n.s. |
| ENSG00000136720 | 414.5192077 | -0.552710606 | 0.256825 | -2.152091 | 0.03139022 | 0.095007489 | HS6ST1 | protein_coding | 53.08 | -0.565004084 | 0.565004084 | ENSG00000136720 | n.s. |
| ENSG00000167619 | 137.8079575 | 0.552298335 | 0.2565551 | 2.152747 | 0.031338535 | 0.094936124 | TMEM145 | protein_coding | 57.14 | 0.564762883 | 0.564762883 | ENSG00000167619 | n.s. |
| ENSG00000104419 | 142.9498416 | 0.559684317 | 0.2620322 | 2.135937 | 0.032684554 | 0.098032538 | NDRG1 | protein_coding | 50.32 | 0.564514254 | 0.564514254 | ENSG00000104419 | n.s. |
| ENSG00000172888 | 901.0940702 | -0.565080002 | 0.2662026 | -2.122744 | 0.033775327 | 0.100460132 | ZNF621 | protein_coding | 41.97 | -0.563953378 | 0.563953378 | ENSG00000172888 | n.s. |
| ENSG00000256747 | 23.6586331 | 0.644903911 | 0.326943 | 1.972527 | 0.048549463 | 0.133561385 | | antisense | 73.54 | 0.563851798 | 0.563851798 | ENSG00000256747 | n.s. |
| ENSG00000184083 | 117.830834 | -0.544945205 | 0.2517298 | -2.164803 | 0.030402811 | 0.092643111 | FAM120C | protein_coding | 42.69 | -0.56303023 | 0.56303023 | ENSG00000184083 | n.s. |
| ENSG00000202082 | 7.488066313 | -1.133555264 | 0.8039107 | -1.410051 | 0.158524578 | 0.318674227 | RNU6-290P | snRNA | 33.66 | -0.562983689 | 0.562983689 | ENSG00000202082 | n.s. |
| ENSG00000177679 | 367.2321175 | -0.562388874 | 0.2646883 | -2.124721 | 0.033609883 | 0.100074936 | SRRM3 | protein_coding | 52.93 | -0.562205916 | 0.562205916 | ENSG00000177679 | n.s. |
| ENSG00000147475 | 624.7442029 | -0.572832787 | 0.2724021 | -2.102894 | 0.035475034 | 0.104378464 | ERLIN2 | protein_coding | 44.02 | -0.562171852 | 0.562171852 | ENSG00000147475 | n.s. |
| ENSG00000164099 | 2162.725936 | -0.550398115 | 0.2560631 | -2.149463 | 0.03159773 | 0.095453043 | PRSS12 | protein_coding | 36.51 | -0.561521782 | 0.561521782 | ENSG00000164099 | n.s. |
| ENSG00000271869 | 17.32499598 | 0.683599018 | 0.3589991 | 1.90418 | 0.056886721 | 0.150892326 | | lincRNA | 54.09 | 0.561462327 | 0.561462327 | ENSG00000271869 | n.s. |
| ENSG00000185189 | 150.6833227 | 0.533638222 | 0.2442955 | 2.184396 | 0.028933151 | 0.088908852 | NRBP2 | protein_coding | 61.1 | 0.560883121 | 0.560883121 | ENSG00000185189 | n.s. |

| | | | | | | | | | | | | | |
|---|---|---|---|---|---|---|---|---|---|---|---|---|---|
| ENSG00000108187 | 230.6271976 | -0.604674884 | 0.2969646 | -2.036185 | 0.041731792 | 0.118216022 | PBLD | protein_coding | 43.3 | -0.560729327 | 0.560729327 | ENSG00000108187 | n.s. |
| ENSG00000197329 | 1018.210836 | 0.55580251 | 0.2602743 | 2.135449 | 0.032724328 | 0.098072429 | PELI1 | protein_coding | 36.9 | 0.560500747 | 0.560500747 | ENSG00000197329 | n.s. |
| ENSG00000267500 | 118.0281418 | -0.568677342 | 0.2698303 | -2.107537 | 0.035071061 | 0.103408383 | ZNF887P | transcribed_unprocessed_pseud | 43.51 | -0.560399818 | 0.560399818 | ENSG00000267500 | n.s. |
| ENSG00000173221 | 267.1987535 | 0.561934732 | 0.2648395 | 2.121793 | 0.033855098 | 0.100632905 | GLRX | protein_coding | 41.1 | 0.560395025 | 0.560395025 | ENSG00000173221 | n.s. |
| ENSG00000154370 | 823.0270862 | -0.503432302 | 0.223236 | -2.255158 | 0.024123432 | 0.07708757 | TRIM11 | protein_coding | 56.97 | -0.560328028 | 0.560328028 | ENSG00000154370 | n.s. |
| ENSG00000204387 | 1741.839554 | 0.520272708 | 0.2349922 | 2.214 | 0.026828797 | 0.083940975 | C6orf48 | protein_coding | 48.85 | 0.559826956 | 0.559826956 | ENSG00000204387 | n.s. |
| ENSG00000204084 | 255.7279961 | -0.528339102 | 0.2408145 | -2.193967 | 0.028237762 | 0.087212591 | INPP5B | protein_coding | 46.64 | -0.559733441 | 0.559733441 | ENSG00000204084 | n.s. |
| ENSG00000187122 | 318.4476565 | -0.560702419 | 0.2642094 | -2.122189 | 0.03382184 | 0.100555516 | SLIT1 | protein_coding | 50.45 | -0.559353426 | 0.559353426 | ENSG00000187122 | n.s. |
| ENSG00000267698 | 38.5901992 | -0.596698386 | 0.2911476 | -2.04947 | 0.040416133 | 0.115618208 | | antisense | 51.74 | -0.559090734 | 0.559090734 | ENSG00000267698 | n.s. |
| ENSG00000132394 | 156.2929078 | -0.542775217 | 0.2512312 | -2.160461 | 0.030737004 | 0.093425073 | EEFSEC | protein_coding | 47.87 | -0.558806926 | 0.558806926 | ENSG00000132394 | n.s. |
| ENSG00000136153 | 278.0617067 | -0.596316915 | 0.2910154 | -2.049091 | 0.04045325 | 0.1156531 | LMO7 | protein_coding | 38.67 | -0.558655162 | 0.558655162 | ENSG00000136153 | n.s. |
| ENSG00000137818 | 13134.4157 | 0.599426413 | 0.2934605 | 2.042613 | 0.041090718 | 0.116967636 | RPLP1 | protein_coding | 47.38 | 0.558626027 | 0.558626027 | ENSG00000137818 | n.s. |
| ENSG00000206704 | 3.017465882 | -1.046642874 | 0.7095964 | -1.474983 | 0.140217032 | 0.292760715 | | misc_RNA | 45.54 | -0.558370578 | 0.558370578 | ENSG00000206704 | n.s. |
| ENSG00000232046 | 22795.15616 | 0.450417574 | 0.1882963 | 2.392068 | 0.016753741 | 0.057599748 | LINC01798 | lincRNA | 39.07 | 0.558328352 | 0.558328352 | ENSG00000232046 | n.s. |
| ENSG00000165282 | 340.2621979 | -0.504706354 | 0.2246051 | -2.247083 | 0.024634707 | 0.078379576 | PIGO | protein_coding | 53.46 | -0.558102817 | 0.558102817 | ENSG00000165282 | n.s. |
| ENSG00000033800 | 2031.48924 | -0.521188002 | 0.2361032 | -2.207458 | 0.027282073 | 0.085015287 | PIAS1 | protein_coding | 38.09 | -0.557933306 | 0.557933306 | ENSG00000033800 | n.s. |
| ENSG00000156599 | 842.4285142 | -0.522720521 | 0.2373924 | -2.201926 | 0.027670508 | 0.085888685 | ZDHHC5 | protein_coding | 45.38 | -0.557253554 | 0.557253554 | ENSG00000156599 | n.s. |
| ENSG00000139687 | 1150.343151 | -0.580999722 | 0.2799154 | -2.075626 | 0.037928544 | 0.109921336 | RB1 | protein_coding | 36.91 | -0.557131093 | 0.557131093 | ENSG00000139687 | n.s. |
| ENSG00000029725 | 2046.455885 | -0.566045491 | 0.269117 | -2.103344 | 0.035435738 | 0.104284883 | RABEP1 | protein_coding | 41.37 | -0.555731373 | 0.555731373 | ENSG00000029725 | n.s. |
| ENSG00000151445 | 347.5548179 | -0.566790649 | 0.2699097 | -2.099927 | 0.035735279 | 0.104966728 | VIPAS39 | protein_coding | 41.62 | -0.554858766 | 0.554858766 | ENSG00000151445 | n.s. |
| ENSG00000229770 | 20.1067468 | -0.692056241 | 0.3686007 | -1.877523 | 0.060446449 | 0.157868301 | | antisense | 44.21 | -0.554824993 | 0.554824993 | ENSG00000229770 | n.s. |
| ENSG00000170464 | 453.1132487 | -0.597247672 | 0.2932056 | -2.036959 | 0.041654181 | 0.118044203 | DNAJC18 | protein_coding | 44.33 | -0.554219164 | 0.554219164 | ENSG00000170464 | n.s. |
| ENSG00000163053 | 290.4328045 | -0.596616279 | 0.2927602 | -2.037901 | 0.041559812 | 0.117848731 | SLC16A14 | protein_coding | 42.86 | -0.554062677 | 0.554062677 | ENSG00000163053 | n.s. |
| ENSG00000261008 | 38.63563288 | -0.605792716 | 0.2997679 | -2.020872 | 0.043292971 | 0.12177132 | LINC01572 | lincRNA | 37.41 | -0.553970171 | 0.553970171 | ENSG00000261008 | n.s. |
| ENSG00000010803 | 42.90679479 | -0.611959807 | 0.304497 | -2.00974 | 0.04445874 | 0.124446931 | SCMH1 | protein_coding | 39.5 | -0.553833299 | 0.553833299 | ENSG00000010803 | n.s. |
| ENSG00000197081 | 1221.330598 | -0.506247116 | 0.2265887 | -2.234211 | 0.025469166 | 0.080555929 | IGF2R | protein_coding | 45.55 | -0.55378498 | 0.55378498 | ENSG00000197081 | n.s. |
| ENSG00000280828 | 95.45752822 | 0.582173891 | 0.2817362 | 2.06638 | 0.038792645 | 0.111893516 | | unprocessed_pseudogene | 41.25 | 0.553760941 | 0.553760941 | ENSG00000280828 | n.s. |
| ENSG00000102384 | 419.8608514 | -0.567483701 | 0.2707153 | -2.096238 | 0.036061108 | 0.105745326 | CENPI | protein_coding | 40.62 | -0.553715876 | 0.553715876 | ENSG00000102384 | n.s. |
| ENSG00000151729 | 751.7372887 | 0.515506287 | 0.2331374 | 2.21117 | 0.0270241 | 0.084381375 | SLC25A4 | protein_coding | 46.96 | 0.55352663 | 0.55352663 | ENSG00000151729 | n.s. |
| ENSG00000212694 | 387.7381485 | 0.481078911 | 0.2093289 | 2.298197 | 0.021550596 | 0.070699564 | LINC01089 | lincRNA | 58.94 | 0.553521342 | 0.553521342 | ENSG00000212694 | n.s. |
| ENSG00000131023 | 1606.43346 | -0.535724643 | 0.2474791 | -2.164727 | 0.030408606 | 0.092643111 | LATS1 | protein_coding | 41.22 | -0.553503666 | 0.553503666 | ENSG00000131023 | n.s. |
| ENSG00000105879 | 809.5919696 | -0.541424456 | 0.251579 | -2.152105 | 0.031389107 | 0.095007489 | CBLL1 | protein_coding | 36.56 | -0.553466906 | 0.553466906 | ENSG00000105879 | n.s. |

| | | | | | | | | | | | | | |
|---|---|---|---|---|---|---|---|---|---|---|---|---|---|
| ENSG00000186153 | 123.126885 | -0.595531818 | 0.292116 | -2.038683 | 0.041481691 | 0.117699121 | WWOX | protein_coding | 42.64 | -0.553384113 | 0.553384113 | ENSG00000186153 | n.s. |
| ENSG00000242086 | 742.7367263 | -0.51915323 | 0.2358495 | -2.201205 | 0.027721498 | 0.086018813 | LINC00969 | lincRNA | 48.33 | -0.553109254 | 0.553109254 | ENSG00000242086 | n.s. |
| ENSG00000161021 | 933.8078486 | -0.531573672 | 0.2446515 | -2.172779 | 0.029796948 | 0.091098573 | MAML1 | protein_coding | 49.38 | -0.553096253 | 0.553096253 | ENSG00000161021 | n.s. |
| ENSG00000117020 | 4040.618693 | -0.548687768 | 0.2570336 | -2.134693 | 0.032786087 | 0.098189005 | AKT3 | protein_coding | 37.34 | -0.55304278 | 0.55304278 | ENSG00000117020 | n.s. |
| ENSG00000179456 | 7305.983959 | 0.474972374 | 0.2053554 | 2.312929 | 0.020726543 | 0.068610475 | ZBTB18 | protein_coding | 44.35 | 0.552682403 | 0.552682403 | ENSG00000179456 | n.s. |
| ENSG00000123104 | 1961.573333 | -0.571363144 | 0.2737863 | -2.086894 | 0.036897682 | 0.107853384 | ITPR2 | protein_coding | 38.25 | -0.552603134 | 0.552603134 | ENSG00000123104 | n.s. |
| ENSG00000174718 | 2299.573002 | -0.543590141 | 0.2534803 | -2.144507 | 0.031992303 | 0.096464091 | KIAA1551 | protein_coding | 41.61 | -0.552088806 | 0.552088806 | ENSG00000174718 | n.s. |
| ENSG00000136560 | 1159.589698 | -0.51513951 | 0.2332666 | -2.208372 | 0.027218319 | 0.084930672 | TANK | protein_coding | 35.76 | -0.551681158 | 0.551681158 | ENSG00000136560 | n.s. |
| ENSG00000161048 | 163.5475022 | -0.500621932 | 0.2231945 | -2.242985 | 0.024897809 | 0.079144337 | NAPEPLD | protein_coding | 40.35 | -0.551475185 | 0.551475185 | ENSG00000161048 | n.s. |
| ENSG00000256325 | 82.78934841 | -0.556166922 | 0.2629638 | -2.114994 | 0.03443046 | 0.101978428 | | antisense | 37.61 | -0.55143488 | 0.55143488 | ENSG00000256325 | n.s. |
| ENSG00000119522 | 223.2205803 | -0.558082587 | 0.2643915 | -2.110819 | 0.034787862 | 0.102834692 | DENND1A | protein_coding | 42.52 | -0.551307659 | 0.551307659 | ENSG00000119522 | n.s. |
| ENSG00000092847 | 690.5578703 | -0.50100076 | 0.2235096 | -2.241518 | 0.02499255 | 0.079409235 | AGO1 | protein_coding | 44.84 | -0.551165459 | 0.551165459 | ENSG00000092847 | n.s. |
| ENSG00000159208 | 183.6306262 | 0.570189573 | 0.2733454 | 2.085967 | 0.03698158 | 0.107989627 | CIART | protein_coding | 47.53 | 0.551155482 | 0.551155482 | ENSG00000159208 | n.s. |
| ENSG00000162377 | 2079.970864 | -0.575900994 | 0.2778417 | -2.072767 | 0.038193983 | 0.110487187 | COA7 | protein_coding | 46.05 | -0.550957619 | 0.550957619 | ENSG00000162377 | n.s. |
| ENSG00000198453 | 214.2233098 | -0.536762128 | 0.2488625 | -2.156862 | 0.031016385 | 0.094165096 | ZNF568 | protein_coding | 40.05 | -0.550777012 | 0.550777012 | ENSG00000198453 | n.s. |
| ENSG00000206535 | 323.0266032 | 0.4975438 | 0.2212935 | 2.248343 | 0.024554328 | 0.078213202 | LNP1 | protein_coding | 39.31 | 0.550641643 | 0.550641643 | ENSG00000206535 | n.s. |
| ENSG00000125629 | 182.9859632 | -0.565518266 | 0.2701562 | -2.093301 | 0.036322253 | 0.106331944 | INSIG2 | protein_coding | 37.96 | -0.550439426 | 0.550439426 | ENSG00000125629 | n.s. |
| ENSG00000239883 | 502.5048651 | -0.555596213 | 0.262906 | -2.113289 | 0.034576054 | 0.102360705 | PARGP1 | transcribed_unprocessed_pseud | 38.95 | -0.549966208 | 0.549966208 | ENSG00000239883 | n.s. |
| ENSG00000183155 | 284.8365781 | -0.596899952 | 0.2940951 | -2.029615 | 0.042395651 | 0.11985272 | RABIF | protein_coding | 45.67 | -0.549955028 | 0.549955028 | ENSG00000183155 | n.s. |
| ENSG00000143942 | 279.6260144 | -0.5764126 | 0.2785048 | -2.069668 | 0.038483417 | 0.111208928 | CHAC2 | protein_coding | 39.11 | -0.549817123 | 0.549817123 | ENSG00000143942 | n.s. |
| ENSG00000163006 | 1498.533541 | -0.570029298 | 0.2736433 | -2.083111 | 0.037241131 | 0.108565382 | CCDC138 | protein_coding | 37.69 | -0.549684174 | 0.549684174 | ENSG00000163006 | n.s. |
| ENSG00000103404 | 888.5893732 | -0.504463559 | 0.2263765 | -2.228427 | 0.025852028 | 0.081452317 | USP31 | protein_coding | 41.37 | -0.549409526 | 0.549409526 | ENSG00000103404 | n.s. |
| ENSG00000155903 | 624.1657153 | -0.544765456 | 0.2551199 | -2.135331 | 0.032733974 | 0.098075128 | RASA2 | protein_coding | 36.35 | -0.549363884 | 0.549363884 | ENSG00000155903 | n.s. |
| ENSG00000148396 | 938.5400407 | -0.523979757 | 0.2402167 | -2.18128 | 0.029162749 | 0.089473986 | SEC16A | protein_coding | 52.76 | -0.549289664 | 0.549289664 | ENSG00000148396 | n.s. |
| ENSG00000116903 | 989.2772443 | 0.525255024 | 0.2411712 | 2.177934 | 0.029410953 | 0.09011635 | EXOC8 | protein_coding | 43.25 | 0.548994664 | 0.548994664 | ENSG00000116903 | n.s. |
| ENSG00000168538 | 1272.100302 | -0.570394944 | 0.2743555 | -2.079036 | 0.037614022 | 0.109195169 | TRAPPC11 | protein_coding | 38.51 | -0.548603904 | 0.548603904 | ENSG00000168538 | n.s. |
| ENSG00000252198 | 6.292249872 | -0.814436793 | 0.4808216 | -1.693844 | 0.090294909 | 0.212181606 | | misc_RNA | 53.02 | -0.548353996 | 0.548353996 | ENSG00000252198 | n.s. |
| ENSG00000138182 | 2810.932187 | -0.537202985 | 0.2500229 | -2.148615 | 0.0316649 | 0.09559202 | KIF20B | protein_coding | 35.15 | -0.547720539 | 0.547720539 | ENSG00000138182 | n.s. |
| ENSG00000131323 | 273.7891983 | -0.514300054 | 0.2337253 | -2.200446 | 0.027775232 | 0.086145423 | TRAF3 | protein_coding | 45.83 | -0.547610132 | 0.547610132 | ENSG00000131323 | n.s. |
| ENSG00000251807 | 33.77092785 | 0.626058046 | 0.3174141 | 1.97237 | 0.048567392 | 0.133574428 | RNU6-202P | snRNA | 44.79 | 0.547347948 | 0.547347948 | ENSG00000251807 | n.s. |
| ENSG00000134909 | 201.13843 | -0.519182443 | 0.2373085 | -2.187795 | 0.02868453 | 0.088323649 | ARHGAP32 | protein_coding | 36.86 | -0.547178317 | 0.547178317 | ENSG00000134909 | n.s. |
| ENSG00000197622 | 1376.740914 | -0.576755333 | 0.2795731 | -2.062986 | 0.039113964 | 0.112610363 | CDC42SE1 | protein_coding | 46.96 | -0.547007241 | 0.547007241 | ENSG00000197622 | n.s. |

| | | | | | | | | | | | | | |
|---|---|---|---|---|---|---|---|---|---|---|---|---|---|
| ENSG00000104897 | 729.7251104 | 0.503590309 | 0.226375 | 2.224585 | 0.026109141 | 0.082169431 | SF3A2 | protein_coding | 58.67 | 0.546541385 | 0.546541385 | ENSG00000104897 | n.s. |
| ENSG00000167202 | 583.7199188 | -0.545407956 | 0.2563136 | -2.127893 | 0.033345935 | 0.09954448 | TBC1D2B | protein_coding | 47.99 | -0.546489401 | 0.546489401 | ENSG00000167202 | n.s. |
| ENSG00000278991 | 706.4077965 | -0.505334624 | 0.2275919 | -2.220354 | 0.026394757 | 0.08293708 | | TEC | 37.19 | -0.546393699 | 0.546393699 | ENSG00000278991 | n.s. |
| ENSG00000274070 | 96.24326938 | -0.541044266 | 0.253224 | -2.136623 | 0.032628659 | 0.097928179 | CASTOR2 | protein_coding | 52.51 | -0.545963615 | 0.545963615 | ENSG00000274070 | n.s. |
| ENSG00000261801 | 126.8806179 | -0.55550261 | 0.2640379 | -2.103874 | 0.035389422 | 0.104192624 | LOXL1-AS1 | antisense | 50.5 | -0.545594123 | 0.545594123 | ENSG00000261801 | n.s. |
| ENSG00000220205 | 232.7741577 | 0.499721861 | 0.2239546 | 2.231353 | 0.025657742 | 0.081005156 | VAMP2 | protein_coding | 58.14 | 0.545440085 | 0.545440085 | ENSG00000220205 | n.s. |
| ENSG00000120458 | 311.3008279 | -0.565614043 | 0.2717895 | -2.081074 | 0.037427146 | 0.108834201 | MSANTD2 | protein_coding | 40.09 | -0.544819021 | 0.544819021 | ENSG00000120458 | n.s. |
| ENSG00000204764 | 889.9685481 | -0.563139 | 0.2698544 | -2.086825 | 0.036903937 | 0.107853384 | RANBP17 | protein_coding | 37.4 | -0.54464902 | 0.54464902 | ENSG00000204764 | n.s. |
| ENSG00000154122 | 242.9037261 | 0.482576923 | 0.2123812 | 2.27222 | 0.023073223 | 0.07438007 | ANKH | protein_coding | 43.86 | 0.544609009 | 0.544609009 | ENSG00000154122 | n.s. |
| ENSG00000150776 | 643.6502306 | -0.565434115 | 0.2718013 | -2.080321 | 0.037496066 | 0.108979102 | C11orf57 | protein_coding | 40.81 | -0.544318981 | 0.544318981 | ENSG00000150776 | n.s. |
| ENSG00000146063 | 606.4382221 | -0.529465679 | 0.2454199 | -2.157387 | 0.030975497 | 0.09406147 | TRIM41 | protein_coding | 52.09 | -0.543543238 | 0.543543238 | ENSG00000146063 | n.s. |
| ENSG00000133884 | 468.938622 | -0.558022218 | 0.266541 | -2.09357 | 0.036298285 | 0.106284127 | DPF2 | protein_coding | 48.63 | -0.543252256 | 0.543252256 | ENSG00000133884 | n.s. |
| ENSG00000176871 | 849.2860973 | -0.569841422 | 0.2754418 | -2.068827 | 0.038562357 | 0.111344607 | WSB2 | protein_coding | 46.42 | -0.543247386 | 0.543247386 | ENSG00000176871 | n.s. |
| ENSG00000159899 | 274.774903 | -0.531370778 | 0.2469243 | -2.151958 | 0.031400648 | 0.095007489 | NPR2 | protein_coding | 48.3 | -0.543189612 | 0.543189612 | ENSG00000159899 | n.s. |
| ENSG00000156052 | 1206.199505 | -0.569786409 | 0.2756679 | -2.06693 | 0.038740708 | 0.111790028 | GNAQ | protein_coding | 39.13 | -0.542207001 | 0.542207001 | ENSG00000156052 | n.s. |
| ENSG00000165240 | 451.9771938 | -0.581484215 | 0.2846164 | -2.043045 | 0.041047943 | 0.116921135 | ATP7A | protein_coding | 38.61 | -0.542005494 | 0.542005494 | ENSG00000165240 | n.s. |
| ENSG00000132823 | 503.3618179 | 0.569165953 | 0.2753548 | 2.067028 | 0.038731557 | 0.111790028 | OSER1 | protein_coding | 40.61 | 0.541616577 | 0.541616577 | ENSG00000132823 | n.s. |
| ENSG00000211456 | 1614.640418 | -0.529875243 | 0.2463265 | -2.151109 | 0.031467598 | 0.095161461 | SACM1L | protein_coding | 37.99 | -0.541288172 | 0.541288172 | ENSG00000211456 | n.s. |
| ENSG00000165288 | 548.0439945 | -0.545917334 | 0.2581203 | -2.114973 | 0.034432277 | 0.101978428 | BRWD3 | protein_coding | 36.15 | -0.541272499 | 0.541272499 | ENSG00000165288 | n.s. |
| ENSG00000163918 | 1924.646869 | -0.574251389 | 0.2792667 | -2.056283 | 0.039755256 | 0.114149809 | RFC4 | protein_coding | 40.5 | -0.541246181 | 0.541246181 | ENSG00000163918 | n.s. |
| ENSG00000170891 | 72.1790853 | 0.618619613 | 0.3135943 | 1.972675 | 0.048532581 | 0.133561385 | CYTL1 | protein_coding | 43.07 | 0.540870935 | 0.540870935 | ENSG00000170891 | n.s. |
| ENSG00000185869 | 671.9601021 | -0.569575598 | 0.2760606 | -2.063227 | 0.039091105 | 0.112567828 | ZNF829 | protein_coding | 39.33 | -0.540291274 | 0.540291274 | ENSG00000185869 | n.s. |
| ENSG00000104714 | 296.7677233 | -0.595389237 | 0.2957915 | -2.012868 | 0.044128541 | 0.123771489 | ERICH1 | protein_coding | 51.17 | -0.54024392 | 0.54024392 | ENSG00000104714 | n.s. |
| ENSG00000068615 | 382.3449906 | -0.568756302 | 0.2755412 | -2.064143 | 0.039004199 | 0.112364051 | REEP1 | protein_coding | 46.05 | -0.539961655 | 0.539961655 | ENSG00000068615 | n.s. |
| ENSG00000180801 | 53.30905372 | 0.587745255 | 0.2900824 | 2.026132 | 0.042751281 | 0.120660197 | ARSJ | protein_coding | 36.4 | 0.539806383 | 0.539806383 | ENSG00000180801 | n.s. |
| ENSG00000260329 | 280.0167008 | 0.478994851 | 0.2110039 | 2.270076 | 0.023202959 | 0.074677485 | | antisense | 45.75 | 0.53973634 | 0.53973634 | ENSG00000260329 | n.s. |
| ENSG00000006704 | 553.2219941 | -0.531602696 | 0.2479836 | -2.143701 | 0.032056882 | 0.096586881 | GTF2IRD1 | protein_coding | 52.72 | -0.539620254 | 0.539620254 | ENSG00000006704 | n.s. |
| ENSG00000116353 | 128.2582609 | -0.58002722 | 0.2843327 | -2.039959 | 0.041354372 | 0.117433597 | MECR | protein_coding | 48.21 | -0.539545749 | 0.539545749 | ENSG00000116353 | n.s. |
| ENSG00000170633 | 999.8675671 | -0.562487797 | 0.2710414 | -2.075284 | 0.037960248 | 0.109925235 | RNF34 | protein_coding | 43.68 | -0.539371009 | 0.539371009 | ENSG00000170633 | n.s. |
| ENSG00000136630 | 297.0641556 | 0.48369978 | 0.2142894 | 2.257227 | 0.023993888 | 0.076761695 | HLX | protein_coding | 54.42 | 0.539255333 | 0.539255333 | ENSG00000136630 | n.s. |
| ENSG00000011405 | 2889.379752 | -0.540805873 | 0.2549551 | -2.121181 | 0.033906585 | 0.100757696 | PIK3C2A | protein_coding | 39.84 | -0.539032988 | 0.539032988 | ENSG00000011405 | n.s. |
| ENSG00000170017 | 347.5441425 | -0.60718686 | 0.3053993 | -1.988174 | 0.046792461 | 0.12957268 | ALCAM | protein_coding | 35.29 | -0.538870177 | 0.538870177 | ENSG00000170017 | n.s. |

| | | | | | | | | | | | | | |
|---|---|---|---|---|---|---|---|---|---|---|---|---|---|
| ENSG00000221883 | 55.38766396 | 0.552449375 | 0.263701 | 2.094984 | 0.036172381 | 0.10599844 | ARIH2OS | protein_coding | 61.45 | 0.538472696 | 0.538472696 | ENSG00000221883 | n.s. |
| ENSG00000106211 | 270.6143648 | 0.579749597 | 0.2845387 | 2.037507 | 0.041599228 | 0.117936479 | HSPB1 | protein_coding | 63.41 | 0.538211606 | 0.538211606 | ENSG00000106211 | n.s. |
| ENSG00000205730 | 812.1825786 | -0.502543351 | 0.2276426 | -2.207598 | 0.027272294 | 0.085003839 | ITPRIPL2 | protein_coding | 49.31 | -0.538003543 | 0.538003543 | ENSG00000205730 | n.s. |
| ENSG00000267909 | 217.1137065 | -0.476570934 | 0.2097542 | -2.272044 | 0.023083839 | 0.07438007 | CCDC177 | protein_coding | 62.35 | -0.53783099 | 0.53783099 | ENSG00000267909 | n.s. |
| ENSG00000089280 | 3950.281685 | 0.507448404 | 0.2312284 | 2.194576 | 0.028193992 | 0.087123267 | FUS | protein_coding | 46.87 | 0.537827232 | 0.537827232 | ENSG00000089280 | n.s. |
| ENSG00000226314 | 94.92708788 | 0.519270925 | 0.2396439 | 2.166844 | 0.030246769 | 0.092250988 | ZNF192P1 | transcribed_unprocessed_pseud | 42.26 | 0.537460454 | 0.537460454 | ENSG00000226314 | n.s. |
| ENSG00000005810 | 8968.627124 | -0.548027278 | 0.2607941 | -2.101379 | 0.035607688 | 0.10465819 | MYCBP2 | protein_coding | 35.72 | -0.537191015 | 0.537191015 | ENSG00000005810 | n.s. |
| ENSG00000175764 | 109.2471992 | -0.548417155 | 0.2610868 | -2.100517 | 0.035683423 | 0.104858654 | TTLL11 | protein_coding | 44.23 | -0.537117414 | 0.537117414 | ENSG00000175764 | n.s. |
| ENSG00000173846 | 67.71515979 | 0.551245983 | 0.2632252 | 2.094199 | 0.036242253 | 0.106142384 | PLK3 | protein_coding | 59.69 | 0.536974865 | 0.536974865 | ENSG00000173846 | n.s. |
| ENSG00000182095 | 784.7465898 | -0.491738126 | 0.220464 | -2.230469 | 0.025716351 | 0.0811093 | TNRC18 | protein_coding | 54.68 | -0.536451554 | 0.536451554 | ENSG00000182095 | n.s. |
| ENSG00000256269 | 147.435369 | -0.58291188 | 0.2874266 | -2.028037 | 0.042556441 | 0.12018526 | HMBS | protein_coding | 50.91 | -0.536365664 | 0.536365664 | ENSG00000256269 | n.s. |
| ENSG00000272669 | 35.66758028 | -0.621278075 | 0.3174565 | -1.95705 | 0.050341643 | 0.137029658 | | antisense | 56.82 | -0.53627818 | 0.53627818 | ENSG00000272669 | n.s. |
| ENSG00000169519 | 136.2569088 | -0.568751526 | 0.2765409 | -2.056663 | 0.039718655 | 0.114091776 | METTL15 | protein_coding | 36.8 | -0.536188033 | 0.536188033 | ENSG00000169519 | n.s. |
| ENSG00000261173 | 37.07925187 | 0.601078134 | 0.3014592 | 1.993895 | 0.046163478 | 0.128282801 | | antisense | 43.26 | 0.536060453 | 0.536060453 | ENSG00000261173 | n.s. |
| ENSG00000141252 | 80.38366976 | 0.53140498 | 0.2487697 | 2.136132 | 0.03266862 | 0.09800586 | VPS53 | protein_coding | 45.86 | 0.536053687 | 0.536053687 | ENSG00000141252 | n.s. |
| ENSG00000224114 | 222.5119597 | 0.656266529 | 0.3461976 | 1.895641 | 0.058007497 | 0.153193928 | | processed_pseudogene | 54.27 | 0.534698699 | 0.534698699 | ENSG00000224114 | n.s. |
| ENSG00000185947 | 721.7094242 | -0.559983065 | 0.2704384 | -2.070649 | 0.038391608 | 0.110989692 | ZNF267 | protein_coding | 38.82 | -0.534625551 | 0.534625551 | ENSG00000185947 | n.s. |
| ENSG00000105722 | 251.2083768 | -0.542455533 | 0.2575156 | -2.106496 | 0.035161332 | 0.103602259 | ERF | protein_coding | 64.09 | -0.534118411 | 0.534118411 | ENSG00000105722 | n.s. |
| ENSG00000123485 | 1538.344576 | -0.526178223 | 0.2454799 | -2.143468 | 0.032075537 | 0.096622178 | HJURP | protein_coding | 48.25 | -0.534030474 | 0.534030474 | ENSG00000123485 | n.s. |
| ENSG00000171791 | 4064.631601 | 0.48888229 | 0.2190794 | 2.23153 | 0.025646012 | 0.080986489 | BCL2 | protein_coding | 42.74 | 0.533657761 | 0.533657761 | ENSG00000171791 | n.s. |
| ENSG00000236296 | 142.7168015 | 0.511355634 | 0.2350288 | 2.175715 | 0.02957658 | 0.090544135 | GUSBP5 | transcribed_processed_pseudog | 39.63 | 0.53341535 | 0.53341535 | ENSG00000236296 | n.s. |
| ENSG00000148218 | 241.1987086 | -0.608677213 | 0.3081022 | -1.975569 | 0.048203563 | 0.13295131 | ALAD | protein_coding | 51.45 | -0.533388333 | 0.533388333 | ENSG00000148218 | n.s. |
| ENSG00000252192 | 26.23469809 | 0.621293245 | 0.3184731 | 1.95085 | 0.051074902 | 0.138854282 | | snoRNA | 46.51 | 0.532722129 | 0.532722129 | ENSG00000252192 | n.s. |
| ENSG00000117707 | 866.8398092 | -0.529754835 | 0.2484709 | -2.13206 | 0.033001949 | 0.098729293 | PROX1 | protein_coding | 41.17 | -0.532697079 | 0.532697079 | ENSG00000117707 | n.s. |
| ENSG00000078053 | 259.3889308 | -0.594930211 | 0.2976882 | -1.998501 | 0.045662368 | 0.127278709 | AMPH | protein_coding | 39.15 | -0.532607844 | 0.532607844 | ENSG00000078053 | n.s. |
| ENSG00000117899 | 1101.507129 | -0.563266673 | 0.2735006 | -2.059471 | 0.03944913 | 0.11343458 | MESDC2 | protein_coding | 45.39 | -0.532430376 | 0.532430376 | ENSG00000117899 | n.s. |
| ENSG00000185477 | 2474.557062 | -0.558176404 | 0.2696925 | -2.069677 | 0.038482597 | 0.111208928 | GPRIN3 | protein_coding | 39.05 | -0.532422338 | 0.532422338 | ENSG00000185477 | n.s. |
| ENSG00000173349 | 1453.228791 | 0.447132283 | 0.1909665 | 2.341418 | 0.019210658 | 0.064543916 | SFT2D3 | protein_coding | 48.54 | 0.532152111 | 0.532152111 | ENSG00000173349 | n.s. |
| ENSG00000162714 | 1050.879552 | -0.542660618 | 0.2581819 | -2.101854 | 0.035566053 | 0.104602062 | ZNF496 | protein_coding | 48.34 | -0.532056897 | 0.532056897 | ENSG00000162714 | n.s. |
| ENSG00000122386 | 138.8149035 | -0.578371895 | 0.2853536 | -2.02686 | 0.04267672 | 0.12047607 | ZNF205 | protein_coding | 58.51 | -0.531581151 | 0.531581151 | ENSG00000122386 | n.s. |
| ENSG00000265491 | 1362.299532 | -0.531105801 | 0.2497727 | -2.126356 | 0.03347359 | 0.099818545 | RNF115 | protein_coding | 41.91 | -0.531524719 | 0.531524719 | ENSG00000265491 | n.s. |
| ENSG00000259467 | 45.59871255 | 0.563602882 | 0.2740748 | 2.056383 | 0.039745581 | 0.114145569 | NDUFAF4P1 | processed_pseudogene | 42 | 0.531218791 | 0.531218791 | ENSG00000259467 | n.s. |

| | | | | | | | | | | | | | |
|---|---|---|---|---|---|---|---|---|---|---|---|---|---|
| ENSG00000138443 | 2065.139326 | -0.541171995 | 0.257504 | -2.101606 | 0.035587816 | 0.10463667 | ABI2 | protein_coding | 39.51 | -0.530519615 | 0.530519615 | ENSG00000138443 | n.s. |
| ENSG00000162374 | 4826.200004 | -0.547124669 | 0.2618492 | -2.089465 | 0.036665913 | 0.107270331 | ELAVL4 | protein_coding | 40.1 | -0.530448516 | 0.530448516 | ENSG00000162374 | n.s. |
| ENSG00000088756 | 254.2502822 | -0.557115684 | 0.2695703 | -2.066681 | 0.038764252 | 0.111834788 | ARHGAP28 | protein_coding | 39.1 | -0.530052721 | 0.530052721 | ENSG00000088756 | n.s. |
| ENSG00000277581 | 109.0187599 | 0.503106488 | 0.2300159 | 2.187269 | 0.028722916 | 0.088416793 | | sense_intronic | 57.47 | 0.530005197 | 0.530005197 | ENSG00000277581 | n.s. |
| ENSG00000197323 | 4096.023084 | -0.565542555 | 0.275987 | -2.049164 | 0.040446088 | 0.1156531 | TRIM33 | protein_coding | 38.2 | -0.529824426 | 0.529824426 | ENSG00000197323 | n.s. |
| ENSG00000145908 | 2346.299794 | -0.56678243 | 0.2770312 | -2.045915 | 0.040764711 | 0.116304728 | ZNF300 | protein_coding | 36.58 | -0.529602993 | 0.529602993 | ENSG00000145908 | n.s. |
| ENSG00000105136 | 520.0233424 | -0.510821513 | 0.2355601 | -2.16854 | 0.030117641 | 0.091957876 | ZNF419 | protein_coding | 46.6 | -0.52942107 | 0.52942107 | ENSG00000105136 | n.s. |
| ENSG00000169738 | 1058.868881 | 0.584388789 | 0.2906574 | 2.010576 | 0.044370248 | 0.124224205 | DCXR | protein_coding | 61.99 | 0.529335728 | 0.529335728 | ENSG00000169738 | n.s. |
| ENSG00000116266 | 641.2644179 | -0.631779674 | 0.328119 | -1.925459 | 0.054171937 | 0.145352837 | STXBP3 | protein_coding | 36.03 | -0.529163799 | 0.529163799 | ENSG00000116266 | n.s. |
| ENSG00000159921 | 863.678133 | -0.567493247 | 0.2780783 | -2.040768 | 0.041273875 | 0.117324654 | GNE | protein_coding | 43.82 | -0.528115297 | 0.528115297 | ENSG00000159921 | n.s. |
| ENSG00000254726 | 2367.486844 | -0.524164447 | 0.2457624 | -2.13281 | 0.032940346 | 0.098608564 | MEX3A | protein_coding | 53.44 | -0.52735418 | 0.52735418 | ENSG00000254726 | n.s. |
| ENSG00000124201 | 445.5740512 | -0.520195056 | 0.2428975 | -2.141624 | 0.032223745 | 0.096942783 | ZNFX1 | protein_coding | 45.85 | -0.527209635 | 0.527209635 | ENSG00000124201 | n.s. |
| ENSG00000247626 | 682.785347 | 0.48124199 | 0.2152149 | 2.2361 | 0.025345249 | 0.080291674 | MARS2 | protein_coding | 48.39 | 0.527118542 | 0.527118542 | ENSG00000247626 | n.s. |
| ENSG00000108106 | 3411.961207 | 0.590265218 | 0.2958025 | 1.995471 | 0.045991578 | 0.127965775 | UBE2S | protein_coding | 54.62 | 0.527051454 | 0.527051454 | ENSG00000108106 | n.s. |
| ENSG00000130748 | 220.5957894 | 0.630353427 | 0.3277166 | 1.923471 | 0.054420877 | 0.145855454 | TMEM160 | protein_coding | 61.6 | 0.527024208 | 0.527024208 | ENSG00000130748 | n.s. |
| ENSG00000164611 | 2305.15542 | 0.57024622 | 0.2807105 | 2.031439 | 0.04221051 | 0.119353561 | PTTG1 | protein_coding | 39.4 | 0.526431134 | 0.526431134 | ENSG00000164611 | n.s. |
| ENSG00000176749 | 709.0822449 | 0.469282169 | 0.2072159 | 2.264701 | 0.023531031 | 0.075576134 | CDK5R1 | protein_coding | 58.04 | 0.526354073 | 0.526354073 | ENSG00000176749 | n.s. |
| ENSG00000063587 | 325.3816098 | -0.523871278 | 0.2458955 | -2.130463 | 0.033133433 | 0.099037523 | ZNF275 | protein_coding | 49.08 | -0.526071657 | 0.526071657 | ENSG00000063587 | n.s. |
| ENSG00000141576 | 1247.410715 | -0.513450931 | 0.2384846 | -2.152973 | 0.031320767 | 0.094923603 | RNF157 | protein_coding | 44.99 | -0.525068162 | 0.525068162 | ENSG00000141576 | n.s. |
| ENSG00000232040 | 346.2362569 | -0.532781894 | 0.2528328 | -2.10725 | 0.035095947 | 0.10345985 | ZBED9 | protein_coding | 39.71 | -0.524911722 | 0.524911722 | ENSG00000232040 | n.s. |
| ENSG00000103723 | 1045.889282 | -0.55050844 | 0.2660857 | -2.068914 | 0.038554185 | 0.111344102 | AP3B2 | protein_coding | 48.94 | -0.524817743 | 0.524817743 | ENSG00000103723 | n.s. |
| ENSG00000263553 | 23.63958663 | 0.672582205 | 0.3639181 | 1.848169 | 0.06457791 | 0.165919586 | | lincRNA | 42.41 | 0.524682956 | 0.524682956 | ENSG00000263553 | n.s. |
| ENSG00000170153 | 1696.325955 | -0.521020594 | 0.2442494 | -2.13315 | 0.03291242 | 0.098546156 | RNF150 | protein_coding | 38.46 | -0.524334448 | 0.524334448 | ENSG00000170153 | n.s. |
| ENSG00000165525 | 2385.961603 | -0.513612215 | 0.2389244 | -2.149685 | 0.031580103 | 0.095439498 | NEMF | protein_coding | 40.35 | -0.524024087 | 0.524024087 | ENSG00000165525 | n.s. |
| ENSG00000151914 | 12031.34742 | -0.531359908 | 0.2519921 | -2.108637 | 0.034975936 | 0.103237219 | DST | protein_coding | 38.07 | -0.524007855 | 0.524007855 | ENSG00000151914 | n.s. |
| ENSG00000198382 | 146.7563878 | -0.548251112 | 0.2647507 | -2.07082 | 0.038375597 | 0.110966447 | UVRAG | protein_coding | 38.73 | -0.523474723 | 0.523474723 | ENSG00000198382 | n.s. |
| ENSG00000161267 | 570.2633349 | -0.52355503 | 0.2464297 | -2.124561 | 0.033623259 | 0.100093357 | BDH1 | protein_coding | 48.52 | -0.523342858 | 0.523342858 | ENSG00000161267 | n.s. |
| ENSG00000255020 | 32.39786757 | 0.579986284 | 0.2890687 | 2.006396 | 0.044814033 | 0.125340654 | | antisense | 53.9 | 0.523094295 | 0.523094295 | ENSG00000255020 | n.s. |
| ENSG00000277583 | 7.082331992 | 0.804261024 | 0.4850099 | 1.658236 | 0.097269804 | 0.223849171 | | misc_RNA | 52.25 | 0.522805462 | 0.522805462 | ENSG00000277583 | n.s. |
| ENSG00000270181 | 782.8806744 | -0.551585319 | 0.2675647 | -2.061502 | 0.03925515 | 0.112946777 | BIVM-ERCC5 | protein_coding | 40.01 | -0.522420883 | 0.522420883 | ENSG00000270181 | n.s. |
| ENSG00000168763 | 218.4389134 | -0.551722013 | 0.2680553 | -2.058239 | 0.039567156 | 0.113726984 | CNNM3 | protein_coding | 53.5 | -0.520900878 | 0.520900878 | ENSG00000168763 | n.s. |
| ENSG00000136122 | 690.1850977 | -0.4845447 | 0.2190391 | -2.212138 | 0.026957148 | 0.084228988 | BORA | protein_coding | 36.79 | -0.520661894 | 0.520661894 | ENSG00000136122 | n.s. |

| | | | | | | | | | | | | | |
|---|---|---|---|---|---|---|---|---|---|---|---|---|---|
| ENSG00000278834 | 83.87564579 | 0.520117595 | 0.2445898 | 2.126489 | 0.033462563 | 0.099807041 | | antisense | 50.53 | 0.520553879 | 0.520553879 | ENSG00000278834 | n.s. |
| ENSG00000242615 | 72.76349735 | -0.566745301 | 0.2798026 | -2.025518 | 0.042814153 | 0.120766096 | | transcribed_processed_pseudog | 38.76 | -0.520303342 | 0.520303342 | ENSG00000242615 | n.s. |
| ENSG00000171940 | 1097.781111 | -0.479949776 | 0.2159314 | -2.222696 | 0.026236309 | 0.082532338 | ZNF217 | protein_coding | 44.04 | -0.519965999 | 0.519965999 | ENSG00000171940 | n.s. |
| ENSG00000150477 | 145.6886142 | -0.535847297 | 0.2564942 | -2.089121 | 0.036696868 | 0.107315792 | KIAA1328 | protein_coding | 37.13 | -0.51941627 | 0.51941627 | ENSG00000150477 | n.s. |
| ENSG00000247728 | 40.55658249 | 0.567478413 | 0.2806532 | 2.021991 | 0.043177253 | 0.121544055 | | antisense | 41.21 | 0.519393856 | 0.519393856 | ENSG00000247728 | n.s. |
| ENSG00000148516 | 4819.632855 | -0.539480939 | 0.2593317 | -2.080273 | 0.037500458 | 0.108979102 | ZEB1 | protein_coding | 36.04 | -0.51933498 | 0.51933498 | ENSG00000148516 | n.s. |
| ENSG00000166452 | 193.8953249 | -0.584970479 | 0.2941713 | -1.988537 | 0.046752325 | 0.129538831 | AKIP1 | protein_coding | 46.42 | -0.519219812 | 0.519219812 | ENSG00000166452 | n.s. |
| ENSG00000175198 | 267.0217318 | -0.578905514 | 0.2896679 | -1.998514 | 0.04566093 | 0.127278709 | PCCA | protein_coding | 39.84 | -0.518261826 | 0.518261826 | ENSG00000175198 | n.s. |
| ENSG00000137267 | 779.0120665 | 0.56617126 | 0.2801928 | 2.020649 | 0.043316128 | 0.121811846 | TUBB2A | protein_coding | 56.17 | 0.517656313 | 0.517656313 | ENSG00000137267 | n.s. |
| ENSG00000136021 | 1529.160277 | -0.552252533 | 0.2695733 | -2.048617 | 0.040499547 | 0.115737931 | SCYL2 | protein_coding | 37.97 | -0.51719791 | 0.51719791 | ENSG00000136021 | n.s. |
| ENSG00000203497 | 44.32837605 | 0.578017108 | 0.2893785 | 1.997443 | 0.045777092 | 0.127496391 | PDCD4-AS1 | antisense | 55.62 | 0.517037523 | 0.517037523 | ENSG00000203497 | n.s. |
| ENSG00000207460 | 44.29735182 | -0.606688656 | 0.3121019 | -1.94388 | 0.051909902 | 0.140665933 | SNORD116-19 | snoRNA | 42.39 | -0.516784112 | 0.516784112 | ENSG00000207460 | n.s. |
| ENSG00000134419 | 18413.09697 | 0.615326033 | 0.3190805 | 1.928435 | 0.053801042 | 0.144664332 | RPS15A | protein_coding | 46.41 | 0.516651448 | 0.516651448 | ENSG00000134419 | n.s. |
| ENSG00000130695 | 648.3451835 | -0.552494809 | 0.270028 | -2.046065 | 0.040750002 | 0.116304728 | CEP85 | protein_coding | 44.77 | -0.516252603 | 0.516252603 | ENSG00000130695 | n.s. |
| ENSG00000229204 | 15.93103911 | 0.676918061 | 0.3711423 | 1.823877 | 0.068170638 | 0.172948214 | PTGES3P3 | processed_pseudogene | 37.76 | 0.515868369 | 0.515868369 | ENSG00000229204 | n.s. |
| ENSG00000049540 | 121.5072058 | -0.547097444 | 0.2660916 | -2.056049 | 0.039777779 | 0.114167392 | ELN | protein_coding | 54.95 | -0.515616318 | 0.515616318 | ENSG00000049540 | n.s. |
| ENSG00000149311 | 2471.436494 | -0.537370756 | 0.2589066 | -2.075539 | 0.037936605 | 0.109921336 | ATM | protein_coding | 37.52 | -0.515294492 | 0.515294492 | ENSG00000149311 | n.s. |
| ENSG00000166813 | 553.0734141 | -0.543772293 | 0.2637715 | -2.061528 | 0.039252675 | 0.112946777 | KIF7 | protein_coding | 49.35 | -0.515020961 | 0.515020961 | ENSG00000166813 | n.s. |
| ENSG00000138767 | 680.2515815 | -0.493544823 | 0.2268258 | -2.175876 | 0.029564502 | 0.090527065 | CNOT6L | protein_coding | 36.25 | -0.514876601 | 0.514876601 | ENSG00000138767 | n.s. |
| ENSG00000205078 | 65.29956574 | 0.570947378 | 0.284878 | 2.004182 | 0.045050551 | 0.12582524 | SYCE1L | protein_coding | 46.1 | 0.513985233 | 0.513985233 | ENSG00000205078 | n.s. |
| ENSG00000140525 | 4134.541495 | -0.557734568 | 0.2748237 | -2.029427 | 0.042414843 | 0.119858302 | FANCI | protein_coding | 40.11 | -0.513858636 | 0.513858636 | ENSG00000140525 | n.s. |
| ENSG00000136111 | 458.5957224 | -0.548855728 | 0.2679938 | -2.048017 | 0.040558344 | 0.115882172 | TBC1D4 | protein_coding | 38.08 | -0.513719836 | 0.513719836 | ENSG00000136111 | n.s. |
| ENSG00000120526 | 1627.309645 | -0.580369068 | 0.2923894 | -1.984918 | 0.047153583 | 0.130453853 | NUDCD1 | protein_coding | 36.6 | -0.513361449 | 0.513361449 | ENSG00000120526 | n.s. |
| ENSG00000118495 | 321.169623 | -0.563467134 | 0.2794379 | -2.016431 | 0.043754965 | 0.122872203 | PLAGL1 | protein_coding | 40.32 | -0.513062946 | 0.513062946 | ENSG00000118495 | n.s. |
| ENSG00000055332 | 1491.482265 | -0.548295457 | 0.2679826 | -2.046011 | 0.04075525 | 0.116304728 | EIF2AK2 | protein_coding | 41.14 | -0.512328717 | 0.512328717 | ENSG00000055332 | n.s. |
| ENSG00000076003 | 3556.998206 | -0.551936341 | 0.2711458 | -2.03557 | 0.04179352 | 0.118366797 | MCM6 | protein_coding | 40.28 | -0.511518101 | 0.511518101 | ENSG00000076003 | n.s. |
| ENSG00000120324 | 197.081432 | -0.520270609 | 0.2471975 | -2.104676 | 0.035319552 | 0.10400891 | PCDHB10 | protein_coding | 48.48 | -0.511389307 | 0.511389307 | ENSG00000120324 | n.s. |
| ENSG00000157240 | 291.572155 | 0.474860517 | 0.2144083 | 2.214748 | 0.026777338 | 0.083817645 | FZD1 | protein_coding | 47.74 | 0.511265484 | 0.511265484 | ENSG00000157240 | n.s. |
| ENSG00000139218 | 4455.615646 | -0.546749594 | 0.2672107 | -2.046137 | 0.040742915 | 0.116304728 | SCAF11 | protein_coding | 36.52 | -0.510884259 | 0.510884259 | ENSG00000139218 | n.s. |
| ENSG00000264247 | 719.6860299 | -0.464733302 | 0.2074252 | -2.240486 | 0.02505939 | 0.079567133 | LINC00909 | lincRNA | 43.19 | -0.510865652 | 0.510865652 | ENSG00000264247 | n.s. |
| ENSG00000167996 | 2508.460768 | 0.506029259 | 0.2367699 | 2.13722 | 0.03258014 | 0.097828226 | FTH1 | protein_coding | 50.42 | 0.510854664 | 0.510854664 | ENSG00000167996 | n.s. |
| ENSG00000171469 | 939.2150212 | -0.483540038 | 0.2207156 | -2.190783 | 0.028467497 | 0.087805264 | ZNF561 | protein_coding | 44.62 | -0.510850111 | 0.510850111 | ENSG00000171469 | n.s. |

| | | | | | | | | | | | | | |
|---|---|---|---|---|---|---|---|---|---|---|---|---|---|
| ENSG00000283183 | 229.3682749 | -0.522081933 | 0.2486944 | -2.099291 | 0.035791237 | 0.10510892 | | lincRNA | 45.49 | -0.510784301 | 0.510784301 | ENSG00000283183 | n.s. |
| ENSG00000106868 | 501.6895454 | -0.52338709 | 0.2497418 | -2.095713 | 0.036107654 | 0.105859523 | SUSD1 | protein_coding | 42.78 | -0.510443763 | 0.510443763 | ENSG00000106868 | n.s. |
| ENSG00000112137 | 88.09596997 | -0.542033106 | 0.2637636 | -2.054996 | 0.039879427 | 0.114388395 | PHACTR1 | protein_coding | 41.2 | -0.510388148 | 0.510388148 | ENSG00000112137 | n.s. |
| ENSG00000099954 | 1427.343799 | -0.499243053 | 0.231998 | -2.151928 | 0.031403036 | 0.095007489 | CECR2 | protein_coding | 44.03 | -0.510347297 | 0.510347297 | ENSG00000099954 | n.s. |
| ENSG00000250539 | 117.1355498 | -0.528745905 | 0.2537158 | -2.084008 | 0.037159395 | 0.108395193 | KRT8P33 | processed_pseudogene | 58.61 | -0.510234499 | 0.510234499 | ENSG00000250539 | n.s. |
| ENSG00000154655 | 155.6310726 | -0.574302383 | 0.2887904 | -1.988648 | 0.046740105 | 0.12953075 | L3MBTL4 | protein_coding | 39.96 | -0.50976637 | 0.50976637 | ENSG00000154655 | n.s. |
| ENSG00000224885 | 63.89031696 | 0.551462961 | 0.2712879 | 2.032759 | 0.04207684 | 0.119023943 | TSSC1-IT1 | sense_intronic | 47.97 | 0.509753428 | 0.509753428 | ENSG00000224885 | n.s. |
| ENSG00000267108 | 7.480421426 | -0.762058973 | 0.4523406 | -1.684702 | 0.092046144 | 0.215170997 | | lincRNA | 52.44 | -0.508458144 | 0.508458144 | ENSG00000267108 | n.s. |
| ENSG00000258469 | 44.20724824 | 0.570630145 | 0.2864804 | 1.991864 | 0.046385943 | 0.128779956 | CHMP4BP1 | processed_pseudogene | 53.57 | 0.507947409 | 0.507947409 | ENSG00000258469 | n.s. |
| ENSG00000201990 | 5.264649322 | -0.841305688 | 0.5300971 | -1.587078 | 0.112494876 | 0.249086483 | | misc_RNA | 44.66 | -0.507854043 | 0.507854043 | ENSG00000201990 | n.s. |
| ENSG00000111726 | 797.9663985 | -0.57990495 | 0.2938301 | -1.973607 | 0.048426485 | 0.133406808 | CMAS | protein_coding | 37.51 | -0.507313612 | 0.507313612 | ENSG00000111726 | n.s. |
| ENSG00000169131 | 512.3816738 | -0.573960851 | 0.2894513 | -1.982928 | 0.047375528 | 0.130952843 | ZNF354A | protein_coding | 46.36 | -0.506741465 | 0.506741465 | ENSG00000169131 | n.s. |
| ENSG00000206754 | 197.4335442 | 0.474044396 | 0.2149878 | 2.204983 | 0.027455269 | 0.085454547 | SNORD101 | snoRNA | 36.99 | 0.506404954 | 0.506404954 | ENSG00000206754 | n.s. |
| ENSG00000100304 | 730.278417 | -0.54361178 | 0.2662315 | -2.041876 | 0.041163796 | 0.117083458 | TTLL12 | protein_coding | 58.31 | -0.506376798 | 0.506376798 | ENSG00000100304 | n.s. |
| ENSG00000168876 | 402.2455073 | -0.538229001 | 0.2620277 | -2.054092 | 0.039966805 | 0.114615413 | ANKRD49 | protein_coding | 37.68 | -0.506342688 | 0.506342688 | ENSG00000168876 | n.s. |
| ENSG00000138463 | 376.5224131 | -0.539760712 | 0.2632398 | -2.050453 | 0.040320288 | 0.115415165 | DIRC2 | protein_coding | 38.14 | -0.506153657 | 0.506153657 | ENSG00000138463 | n.s. |
| ENSG00000152904 | 845.1960159 | -0.555947457 | 0.2757415 | -2.01619 | 0.04378006 | 0.122917883 | GGPS1 | protein_coding | 43.08 | -0.506126186 | 0.506126186 | ENSG00000152904 | n.s. |
| ENSG00000219626 | 120.3535808 | 0.490515019 | 0.2268318 | 2.162461 | 0.030582641 | 0.093047781 | FAM228B | protein_coding | 43.18 | 0.505865187 | 0.505865187 | ENSG00000219626 | n.s. |
| ENSG00000083093 | 925.4213728 | -0.493243751 | 0.2287495 | -2.156262 | 0.031063226 | 0.094286743 | PALB2 | protein_coding | 43.53 | -0.505845816 | 0.505845816 | ENSG00000083093 | n.s. |
| ENSG00000280280 | 22.36908967 | -0.677062526 | 0.3758665 | -1.801338 | 0.071649637 | 0.179098366 | | TEC | 41.07 | -0.505703672 | 0.505703672 | ENSG00000280280 | n.s. |
| ENSG00000135821 | 8552.047004 | 0.502574298 | 0.2356672 | 2.132559 | 0.032960882 | 0.098637285 | GLUL | protein_coding | 47.77 | 0.505569084 | 0.505569084 | ENSG00000135821 | n.s. |
| ENSG00000153558 | 170.3535934 | -0.537798 | 0.2620408 | -2.052344 | 0.040136204 | 0.114959138 | FBXL2 | protein_coding | 38.73 | -0.505237827 | 0.505237827 | ENSG00000153558 | n.s. |
| ENSG00000067191 | 19.91188708 | -0.641344085 | 0.3450524 | -1.858686 | 0.063071678 | 0.163072791 | CACNB1 | protein_coding | 54.39 | -0.505134464 | 0.505134464 | ENSG00000067191 | n.s. |
| ENSG00000134202 | 1094.489423 | 0.535392494 | 0.2602814 | 2.056976 | 0.039688559 | 0.114028851 | GSTM3 | protein_coding | 47.16 | 0.504867226 | 0.504867226 | ENSG00000134202 | n.s. |
| ENSG00000103222 | 1006.088919 | -0.503104741 | 0.2363283 | -2.128839 | 0.03326762 | 0.099353299 | ABCC1 | protein_coding | 48.48 | -0.504522343 | 0.504522343 | ENSG00000103222 | n.s. |
| ENSG00000265462 | 31.04785139 | 0.60101771 | 0.3117502 | 1.927883 | 0.053869741 | 0.144793128 | MIR3680-1 | miRNA | 43.68 | 0.504405344 | 0.504405344 | ENSG00000265462 | n.s. |
| ENSG00000198298 | 207.413624 | -0.501746909 | 0.2353819 | -2.131629 | 0.033037324 | 0.09881389 | ZNF485 | protein_coding | 40.92 | -0.504346962 | 0.504346962 | ENSG00000198298 | n.s. |
| ENSG00000118246 | 2107.444395 | -0.54138395 | 0.2653195 | -2.040498 | 0.041300733 | 0.117362827 | FASTKD2 | protein_coding | 37.72 | -0.50374122 | 0.50374122 | ENSG00000118246 | n.s. |
| ENSG00000166189 | 212.5071137 | -0.52102093 | 0.2497197 | -2.086423 | 0.036940293 | 0.107936977 | HPS6 | protein_coding | 64.21 | -0.503738535 | 0.503738535 | ENSG00000166189 | n.s. |
| ENSG00000175575 | 484.9975801 | -0.505457685 | 0.2383693 | -2.120481 | 0.03396547 | 0.100874838 | PAAF1 | protein_coding | 42.99 | -0.503545614 | 0.503545614 | ENSG00000175575 | n.s. |
| ENSG00000129521 | 493.5156697 | 0.498368639 | 0.2331871 | 2.137205 | 0.032581303 | 0.097828226 | EGLN3 | protein_coding | 41.68 | 0.503120994 | 0.503120994 | ENSG00000129521 | n.s. |
| ENSG00000135245 | 894.2981297 | 0.452202052 | 0.2005899 | 2.254361 | 0.024173487 | 0.077229797 | HILPDA | protein_coding | 51.52 | 0.502945953 | 0.502945953 | ENSG00000135245 | n.s. |

| | | | | | | | | | | | | | |
|---|---|---|---|---|---|---|---|---|---|---|---|---|---|
| ENSG00000132716 | 2020.634305 | -0.531175656 | 0.2577598 | -2.060739 | 0.039327971 | 0.113109552 | DCAF8 | protein_coding | 42.25 | -0.50275814 | 0.50275814 | ENSG00000132716 | n.s. |
| ENSG00000276740 | 35.36895761 | -0.606450971 | 0.3169037 | -1.913676 | 0.055661612 | 0.148292503 | | sense_intronic | 44.14 | -0.502675569 | 0.502675569 | ENSG00000276740 | n.s. |
| ENSG00000252050 | 7.112341548 | -0.791559587 | 0.4844299 | -1.634002 | 0.102258346 | 0.232485786 | | snoRNA | 40 | -0.501535 | 0.501535 | ENSG00000252050 | n.s. |
| ENSG00000117174 | 1235.924849 | -0.52046387 | 0.2500287 | -2.081617 | 0.037377478 | 0.108780649 | ZNHIT6 | protein_coding | 36.46 | -0.50144006 | 0.50144006 | ENSG00000117174 | n.s. |
| ENSG00000008277 | 1961.17465 | -0.533188546 | 0.2596892 | -2.05318 | 0.040055138 | 0.114797794 | ADAM22 | protein_coding | 37.29 | -0.501232667 | 0.501232667 | ENSG00000008277 | n.s. |
| ENSG00000167615 | 1087.610348 | 0.475352979 | 0.2172773 | 2.187771 | 0.028686313 | 0.088323649 | LENG8 | protein_coding | 58.96 | 0.500985438 | 0.500985438 | ENSG00000167615 | n.s. |
| ENSG00000172167 | 1018.672624 | -0.523177045 | 0.252326 | -2.073417 | 0.038133497 | 0.110358072 | MTBP | protein_coding | 35.31 | -0.500782921 | 0.500782921 | ENSG00000172167 | n.s. |
| ENSG00000275055 | 64.88661556 | 0.545045561 | 0.2690183 | 2.026054 | 0.042759277 | 0.120660197 | | antisense | 45.72 | 0.500589448 | 0.500589448 | ENSG00000275055 | n.s. |
| ENSG00000223551 | 18.05029439 | 0.624548275 | 0.3327535 | 1.876909 | 0.060530504 | 0.157980801 | TMSB4XP4 | processed_pseudogene | 43.04 | 0.500510295 | 0.500510295 | ENSG00000223551 | n.s. |
| ENSG00000081019 | 1543.292142 | -0.518093196 | 0.2484816 | -2.085036 | 0.037066028 | 0.108180565 | RSBN1 | protein_coding | 37.22 | -0.500400704 | 0.500400704 | ENSG00000081019 | n.s. |
| ENSG00000103356 | 846.4496681 | -0.528395923 | 0.2563837 | -2.060957 | 0.039307105 | 0.113072897 | EARS2 | protein_coding | 47.93 | -0.500201498 | 0.500201498 | ENSG00000103356 | n.s. |
| ENSG00000158710 | 120.2243443 | 0.599942052 | 0.3127462 | 1.918303 | 0.055072595 | 0.14703274 | TAGLN2 | protein_coding | 56.56 | 0.499503323 | 0.499503323 | ENSG00000158710 | n.s. |
| ENSG00000128928 | 778.4733098 | -0.495035049 | 0.2318036 | -2.135579 | 0.032713723 | 0.098072429 | IVD | protein_coding | 50.9 | -0.499219614 | 0.499219614 | ENSG00000128928 | n.s. |
| ENSG00000272953 | 57.43866308 | 0.537498471 | 0.2637392 | 2.037992 | 0.041550689 | 0.11784686 | | lincRNA | 54.91 | 0.499165147 | 0.499165147 | ENSG00000272953 | n.s. |
| ENSG00000135775 | 1986.574859 | -0.566568939 | 0.2860308 | -1.980797 | 0.047614022 | 0.131533735 | COG2 | protein_coding | 39.42 | -0.499126185 | 0.499126185 | ENSG00000135775 | n.s. |
| ENSG00000166224 | 851.8369585 | -0.520206922 | 0.2506558 | -2.075383 | 0.037951006 | 0.109921336 | SGPL1 | protein_coding | 41.07 | -0.498835783 | 0.498835783 | ENSG00000166224 | n.s. |
| ENSG00000149636 | 746.0355195 | -0.541567816 | 0.2669506 | -2.028719 | 0.042486905 | 0.120013224 | DSN1 | protein_coding | 44.84 | -0.498659883 | 0.498659883 | ENSG00000149636 | n.s. |
| ENSG00000185753 | 167.8320527 | -0.546428161 | 0.2707026 | -2.018555 | 0.043533443 | 0.122373536 | CXorf38 | protein_coding | 45.19 | -0.498513236 | 0.498513236 | ENSG00000185753 | n.s. |
| ENSG00000197958 | 17002.35077 | 0.594722622 | 0.3087274 | 1.926368 | 0.054058417 | 0.145160141 | RPL12 | protein_coding | 51.23 | 0.498468323 | 0.498468323 | ENSG00000197958 | n.s. |
| ENSG00000006607 | 659.8753674 | -0.51029171 | 0.2433894 | -2.096606 | 0.036028446 | 0.105672424 | FARP2 | protein_coding | 45.6 | -0.498064269 | 0.498064269 | ENSG00000006607 | n.s. |
| ENSG00000083642 | 2199.116502 | -0.517697458 | 0.2490047 | -2.079067 | 0.037611166 | 0.109195169 | PDS5B | protein_coding | 36.62 | -0.497919642 | 0.497919642 | ENSG00000083642 | n.s. |
| ENSG00000180287 | 45.38398097 | -0.58462635 | 0.3008619 | -1.943172 | 0.051995407 | 0.140825168 | PLD5 | protein_coding | 40.38 | -0.497703942 | 0.497703942 | ENSG00000180287 | n.s. |
| ENSG00000026559 | 2636.772813 | -0.568865863 | 0.2884487 | -1.972156 | 0.048591804 | 0.133598675 | KCNG1 | protein_coding | 52.21 | -0.497301313 | 0.497301313 | ENSG00000026559 | n.s. |
| ENSG00000158417 | 3020.051396 | -0.497459216 | 0.2342202 | -2.123896 | 0.033678864 | 0.100237458 | EIF5B | protein_coding | 38.48 | -0.496946812 | 0.496946812 | ENSG00000158417 | n.s. |
| ENSG00000227097 | 299.7583114 | 0.588261856 | 0.3040397 | 1.934819 | 0.053012494 | 0.143068997 | RPS28P7 | processed_pseudogene | 61.9 | 0.496760353 | 0.496760353 | ENSG00000227097 | n.s. |
| ENSG00000172466 | 4672.610623 | -0.544798111 | 0.2700727 | -2.017227 | 0.043671798 | 0.122688145 | ZNF24 | protein_coding | 39.65 | -0.496418623 | 0.496418623 | ENSG00000172466 | n.s. |
| ENSG00000181541 | 7426.554704 | 0.490944434 | 0.2295168 | 2.139035 | 0.032432856 | 0.097466577 | MAB21L2 | protein_coding | 52.09 | 0.496415659 | 0.496415659 | ENSG00000181541 | n.s. |
| ENSG00000172493 | 1282.802598 | -0.497103118 | 0.2342205 | -2.122372 | 0.03380647 | 0.100531285 | AFF1 | protein_coding | 41.19 | -0.495959167 | 0.495959167 | ENSG00000172493 | n.s. |
| ENSG00000074657 | 2679.184195 | -0.517145305 | 0.2492038 | -2.07519 | 0.037968911 | 0.10992746 | ZNF532 | protein_coding | 43.34 | -0.49588743 | 0.49588743 | ENSG00000074657 | n.s. |
| ENSG00000115233 | 3603.850797 | -0.567604518 | 0.2879876 | -1.970934 | 0.048731417 | 0.133903201 | PSMD14 | protein_coding | 35.42 | -0.495637397 | 0.495637397 | ENSG00000115233 | n.s. |
| ENSG00000171033 | 1193.760246 | -0.606602342 | 0.3194955 | -1.898625 | 0.057613748 | 0.152385298 | PKIA | protein_coding | 34.75 | -0.495628648 | 0.495628648 | ENSG00000171033 | n.s. |
| ENSG00000198900 | 2033.222684 | -0.488177261 | 0.2278712 | -2.142338 | 0.032166283 | 0.096811749 | TOP1 | protein_coding | 39.86 | -0.495046858 | 0.495046858 | ENSG00000198900 | n.s. |

| | | | | | | | | | | | | | |
|---|---|---|---|---|---|---|---|---|---|---|---|---|---|
| ENSG00000167889 | 271.4707099 | -0.512097678 | 0.2454856 | -2.08606 | 0.03697319 | 0.107989627 | MGAT5B | protein_coding | 54.13 | -0.495002813 | 0.495002813 | ENSG00000167889 | n.s. |
| ENSG00000148950 | 314.9472116 | -0.589970452 | 0.3062241 | -1.926597 | 0.054029828 | 0.145138792 | IMMP1L | protein_coding | 34.91 | -0.494522966 | 0.494522966 | ENSG00000148950 | n.s. |
| ENSG00000244270 | 61.26771249 | -0.567882013 | 0.2887064 | -1.966988 | 0.049184578 | 0.13488219 | | processed_pseudogene | 46.9 | -0.494083128 | 0.494083128 | ENSG00000244270 | n.s. |
| ENSG00000127914 | 4384.094801 | -0.548730975 | 0.2738826 | -2.003526 | 0.045120842 | 0.125971021 | AKAP9 | protein_coding | 36.72 | -0.493709364 | 0.493709364 | ENSG00000127914 | n.s. |
| ENSG00000215030 | 240.7614329 | 0.61891445 | 0.3307238 | 1.871394 | 0.061290551 | 0.159534182 | | processed_pseudogene | 61.32 | 0.493365333 | 0.493365333 | ENSG00000215030 | n.s. |
| ENSG00000012232 | 582.2644811 | -0.530025754 | 0.2597862 | -2.040239 | 0.041326574 | 0.117378599 | EXTL3 | protein_coding | 44.36 | -0.493141832 | 0.493141832 | ENSG00000012232 | n.s. |
| ENSG00000052841 | 1814.008218 | -0.542877529 | 0.2695454 | -2.014049 | 0.044004415 | 0.123497977 | TTC17 | protein_coding | 37.89 | -0.493117459 | 0.493117459 | ENSG00000052841 | n.s. |
| ENSG00000168038 | 170.1143694 | -0.53293935 | 0.2620745 | -2.033542 | 0.041997825 | 0.118824573 | ULK4 | protein_coding | 39.39 | -0.493018855 | 0.493018855 | ENSG00000168038 | n.s. |
| ENSG00000271335 | 120.2904266 | -0.51163461 | 0.2457772 | -2.081701 | 0.037369783 | 0.108780649 | | antisense | 42.38 | -0.492933523 | 0.492933523 | ENSG00000271335 | n.s. |
| ENSG00000248909 | 37.98754086 | 0.568775911 | 0.2899888 | 1.961372 | 0.049835665 | 0.136078046 | HMGB1P21 | processed_pseudogene | 42.69 | 0.492680482 | 0.492680482 | ENSG00000248909 | n.s. |
| ENSG00000275180 | 942.9819955 | 0.440016807 | 0.1945962 | 2.261179 | 0.02374819 | 0.076098056 | | lincRNA | 36.25 | 0.492214434 | 0.492214434 | ENSG00000275180 | n.s. |
| ENSG00000233461 | 42.22899939 | 0.627973348 | 0.338979 | 1.852544 | 0.063947805 | 0.164756715 | | antisense | 41.53 | 0.491801646 | 0.491801646 | ENSG00000233461 | n.s. |
| ENSG00000173153 | 245.9435521 | -0.56137651 | 0.2842752 | -1.974764 | 0.048294916 | 0.133124157 | ESRRA | protein_coding | 57.05 | -0.491621623 | 0.491621623 | ENSG00000173153 | n.s. |
| ENSG00000167904 | 65.72214864 | -0.599836297 | 0.3155246 | -1.901076 | 0.057292012 | 0.151729543 | TMEM68 | protein_coding | 38.91 | -0.491223849 | 0.491223849 | ENSG00000167904 | n.s. |
| ENSG00000114480 | 205.1781935 | -0.549286872 | 0.2750627 | -1.996952 | 0.045830425 | 0.127593883 | GBE1 | protein_coding | 35.29 | -0.491155922 | 0.491155922 | ENSG00000114480 | n.s. |
| ENSG00000163412 | 701.3880253 | -0.553355013 | 0.2782964 | -1.988366 | 0.046771259 | 0.129551083 | EIF4E3 | protein_coding | 41.79 | -0.491135196 | 0.491135196 | ENSG00000163412 | n.s. |
| ENSG00000103642 | 209.5521747 | 0.5466812 | 0.2731163 | 2.001642 | 0.045323224 | 0.126510676 | LACTB | protein_coding | 40 | 0.490850193 | 0.490850193 | ENSG00000103642 | n.s. |
| ENSG00000163507 | 2555.620998 | -0.526222905 | 0.2575277 | -2.043364 | 0.041016393 | 0.116879092 | KIAA1524 | protein_coding | 34.15 | -0.490578234 | 0.490578234 | ENSG00000163507 | n.s. |
| ENSG00000067955 | 901.64844 | -0.525982472 | 0.2575143 | -2.042537 | 0.041098305 | 0.116967636 | CBFB | protein_coding | 40.45 | -0.4901811 | 0.4901811 | ENSG00000067955 | n.s. |
| ENSG00000152454 | 674.8549135 | 0.481509122 | 0.2245563 | 2.144269 | 0.032011357 | 0.096491474 | ZNF256 | protein_coding | 48.53 | 0.488977838 | 0.488977838 | ENSG00000152454 | n.s. |
| ENSG00000182973 | 962.4363318 | -0.583257833 | 0.3028757 | -1.925734 | 0.054137649 | 0.145316847 | CNOT10 | protein_coding | 41.97 | -0.488585774 | 0.488585774 | ENSG00000182973 | n.s. |
| ENSG00000207139 | 4.205899525 | -0.83232276 | 0.5334161 | -1.560363 | 0.118674145 | 0.25890005 | | misc_RNA | 38.61 | -0.488463482 | 0.488463482 | ENSG00000207139 | n.s. |
| ENSG00000187742 | 2792.676453 | -0.50675505 | 0.2433635 | -2.082297 | 0.037315362 | 0.108713494 | SECISBP2 | protein_coding | 42.36 | -0.488368227 | 0.488368227 | ENSG00000187742 | n.s. |
| ENSG00000122884 | 337.6954238 | -0.56251526 | 0.2864106 | -1.964017 | 0.049528159 | 0.13553486 | P4HA1 | protein_coding | 38.79 | -0.488234552 | 0.488234552 | ENSG00000122884 | n.s. |
| ENSG00000228439 | 42.15317472 | -0.603970595 | 0.3205464 | -1.884191 | 0.059539168 | 0.156201965 | TSTD3 | protein_coding | 40.48 | -0.486989648 | 0.486989648 | ENSG00000228439 | n.s. |
| ENSG00000206751 | 8.601388583 | -0.732850198 | 0.4361404 | -1.680308 | 0.092897414 | 0.216579153 | | misc_RNA | 42.59 | -0.486893469 | 0.486893469 | ENSG00000206751 | n.s. |
| ENSG00000252473 | 42.58367994 | -0.592611563 | 0.3111976 | -1.904294 | 0.056871949 | 0.150881887 | | snoRNA | 45.19 | -0.486749148 | 0.486749148 | ENSG00000252473 | n.s. |
| ENSG00000128590 | 600.5067362 | 0.474648721 | 0.2203395 | 2.15417 | 0.031226821 | 0.094700726 | DNAJB9 | protein_coding | 38.78 | 0.485872594 | 0.485872594 | ENSG00000128590 | n.s. |
| ENSG00000138593 | 1061.718473 | -0.497695981 | 0.237382 | -2.096604 | 0.036028657 | 0.105672424 | SECISBP2L | protein_coding | 36.79 | -0.485770354 | 0.485770354 | ENSG00000138593 | n.s. |
| ENSG00000276529 | 16.42319861 | -0.691325651 | 0.3979224 | -1.737338 | 0.08232758 | 0.198518688 | | lincRNA | 55.76 | -0.485447907 | 0.485447907 | ENSG00000276529 | n.s. |
| ENSG00000036257 | 2486.370379 | -0.529326538 | 0.2617332 | -2.02239 | 0.043136108 | 0.121477353 | CUL3 | protein_coding | 37.23 | -0.484600923 | 0.484600923 | ENSG00000036257 | n.s. |
| ENSG00000065243 | 2148.193377 | -0.572009774 | 0.2951759 | -1.937861 | 0.052640228 | 0.142174567 | PKN2 | protein_coding | 36.25 | -0.484594146 | 0.484594146 | ENSG00000065243 | n.s. |

| | | | | | | | | | | | | | |
|---|---|---|---|---|---|---|---|---|---|---|---|---|---|
| ENSG00000090581 | 324.8369634 | 0.545049409 | 0.2738554 | 1.990282 | 0.046559893 | 0.129161927 | GNPTG | protein_coding | 56.48 | 0.484475607 | 0.484475607 | ENSG00000090581 | n.s. |
| ENSG00000254694 | 50.17325525 | 0.568445771 | 0.2923397 | 1.94447 | 0.051838764 | 0.140516697 | | antisense | 35.49 | 0.484470453 | 0.484470453 | ENSG00000254694 | n.s. |
| ENSG00000164330 | 570.3562246 | -0.521264812 | 0.2556671 | -2.038842 | 0.04146583 | 0.1176781 | EBF1 | protein_coding | 39.97 | -0.484413659 | 0.484413659 | ENSG00000164330 | n.s. |
| ENSG00000128585 | 2301.906466 | -0.534085769 | 0.2654631 | -2.011902 | 0.044230255 | 0.123956892 | MKLN1 | protein_coding | 39.28 | -0.484271227 | 0.484271227 | ENSG00000128585 | n.s. |
| ENSG00000252485 | 2.528897948 | -0.93880417 | 0.6501481 | -1.443985 | 0.148743189 | 0.304910397 | | misc_RNA | 48.42 | -0.484261258 | 0.484261258 | ENSG00000252485 | n.s. |
| ENSG00000164823 | 659.9713058 | -0.548615223 | 0.276724 | -1.982535 | 0.047419339 | 0.131039604 | OSGIN2 | protein_coding | 38.2 | -0.484206385 | 0.484206385 | ENSG00000164823 | n.s. |
| ENSG00000100479 | 699.5269775 | -0.556535617 | 0.2830269 | -1.96637 | 0.049255882 | 0.13499796 | POLE2 | protein_coding | 41.33 | -0.484003884 | 0.484003884 | ENSG00000100479 | n.s. |
| ENSG00000164323 | 1672.421657 | -0.564145814 | 0.2892035 | -1.950688 | 0.051094181 | 0.138860879 | CFAP97 | protein_coding | 41.18 | -0.483709955 | 0.483709955 | ENSG00000164323 | n.s. |
| ENSG00000278818 | 22.16952641 | 0.606415655 | 0.3239279 | 1.87207 | 0.061196971 | 0.15937999 | | misc_RNA | 51.51 | 0.483656633 | 0.483656633 | ENSG00000278818 | n.s. |
| ENSG00000207442 | 20.82140924 | -0.631957521 | 0.3456833 | -1.82814 | 0.067528538 | 0.171820146 | SNORD116-6 | snoRNA | 41.67 | -0.483400686 | 0.483400686 | ENSG00000207442 | n.s. |
| ENSG00000145332 | 885.2393262 | -0.518690441 | 0.2539616 | -2.042397 | 0.04111218 | 0.116967636 | KLHL8 | protein_coding | 38.23 | -0.483385406 | 0.483385406 | ENSG00000145332 | n.s. |
| ENSG00000137285 | 2874.248125 | 0.543773714 | 0.2732797 | 1.989806 | 0.046612263 | 0.129227906 | TUBB2B | protein_coding | 56.75 | 0.483221082 | 0.483221082 | ENSG00000137285 | n.s. |
| ENSG00000148719 | 580.5430618 | -0.513019281 | 0.2496148 | -2.055244 | 0.039855471 | 0.114343235 | DNAJB12 | protein_coding | 53.03 | -0.483156185 | 0.483156185 | ENSG00000148719 | n.s. |
| ENSG00000207181 | 234.0069555 | 0.476206645 | 0.2222637 | 2.14253 | 0.032150849 | 0.096803907 | SNORA14B | snoRNA | 46.67 | 0.482924548 | 0.482924548 | ENSG00000207181 | n.s. |
| ENSG00000229989 | 1007.605058 | -0.499560053 | 0.2395984 | -2.084989 | 0.037070278 | 0.108180565 | MIR181A1HG | lincRNA | 35.03 | -0.482500454 | 0.482500454 | ENSG00000229989 | n.s. |
| ENSG00000151466 | 940.2077539 | -0.532607616 | 0.2648886 | -2.010685 | 0.044358732 | 0.124224205 | SCLT1 | protein_coding | 36.17 | -0.482432663 | 0.482432663 | ENSG00000151466 | n.s. |
| ENSG00000044574 | 1699.157606 | 0.494788025 | 0.2361844 | 2.094923 | 0.036177878 | 0.10599844 | HSPA5 | protein_coding | 43.98 | 0.482270148 | 0.482270148 | ENSG00000044574 | n.s. |
| ENSG00000239263 | 55.84454895 | -0.545067645 | 0.2746983 | -1.984241 | 0.047228976 | 0.130588341 | RBM43P1 | processed_pseudogene | 37.28 | -0.481891904 | 0.481891904 | ENSG00000239263 | n.s. |
| ENSG00000207622 | 4.23902392 | -1.163346347 | 0.9247502 | -1.258011 | 0.208387631 | 0.385411927 | MIR619 | miRNA | 56.57 | -0.481712464 | 0.481712464 | ENSG00000207622 | n.s. |
| ENSG00000240731 | 15.27064397 | 0.676697793 | 0.3864147 | 1.751222 | 0.079907699 | 0.194163158 | | sense_intronic | 47.97 | 0.481695936 | 0.481695936 | ENSG00000240731 | n.s. |
| ENSG00000198894 | 488.3945094 | -0.502267891 | 0.2420091 | -2.075409 | 0.037948639 | 0.109921336 | CIPC | protein_coding | 43.02 | -0.481633722 | 0.481633722 | ENSG00000198894 | n.s. |
| ENSG00000137571 | 543.1901697 | -0.514886183 | 0.2514988 | -2.047271 | 0.040631509 | 0.11604359 | SLCO5A1 | protein_coding | 40.04 | -0.481613643 | 0.481613643 | ENSG00000137571 | n.s. |
| ENSG00000168672 | 363.6462727 | 0.464525163 | 0.2142246 | 2.168402 | 0.030128099 | 0.09196964 | FAM84B | protein_coding | 50.4 | 0.481413214 | 0.481413214 | ENSG00000168672 | n.s. |
| ENSG00000253710 | 954.5982875 | -0.477143968 | 0.2233536 | -2.136272 | 0.032657248 | 0.097992857 | ALG11 | protein_coding | 41.48 | -0.481345496 | 0.481345496 | ENSG00000253710 | n.s. |
| ENSG00000171757 | 405.6383272 | -0.561703709 | 0.2880809 | -1.949812 | 0.051198512 | 0.139070331 | LRRC34 | protein_coding | 38.99 | -0.481248366 | 0.481248366 | ENSG00000171757 | n.s. |
| ENSG00000241868 | 16.49496993 | -0.635548915 | 0.3500375 | -1.815659 | 0.069422648 | 0.174979676 | RN7SL434P | misc_RNA | 54.61 | -0.481118405 | 0.481118405 | ENSG00000241868 | n.s. |
| ENSG00000275101 | 5.46384538 | -0.816752949 | 0.5225502 | -1.563013 | 0.118049452 | 0.257820004 | MIR6766 | miRNA | 54.17 | -0.480808894 | 0.480808894 | ENSG00000275101 | n.s. |
| ENSG00000162817 | 127.5890875 | 0.565336804 | 0.2911084 | 1.942015 | 0.052135295 | 0.14111192 | C1orf115 | protein_coding | 48.31 | 0.480782939 | 0.480782939 | ENSG00000162817 | n.s. |
| ENSG00000072756 | 1683.688917 | -0.476614518 | 0.2231919 | -2.135448 | 0.032724462 | 0.098072429 | TRNT1 | protein_coding | 38.89 | -0.480643373 | 0.480643373 | ENSG00000072756 | n.s. |
| ENSG00000143156 | 704.1846332 | -0.565070335 | 0.2912259 | -1.940316 | 0.052341291 | 0.141531909 | NME7 | protein_coding | 36.61 | -0.47982701 | 0.47982701 | ENSG00000143156 | n.s. |
| ENSG00000265725 | 46.47836303 | 0.593985689 | 0.3149701 | 1.885848 | 0.059315472 | 0.155732476 | MIR3144 | miRNA | 35.44 | 0.479715204 | 0.479715204 | ENSG00000265725 | n.s. |
| ENSG00000089157 | 37956.05813 | 0.577681569 | 0.3017753 | 1.914277 | 0.055584717 | 0.148144336 | RPLP0 | protein_coding | 48.92 | 0.479079961 | 0.479079961 | ENSG00000089157 | n.s. |

| | | | | | | | | | | | | | |
|---|---|---|---|---|---|---|---|---|---|---|---|---|---|
| ENSG00000135365 | 244.0465779 | -0.500589651 | 0.2414963 | -2.072867 | 0.038184646 | 0.11048313 | PHF21A | protein_coding | 38.79 | -0.478916136 | 0.478916136 | ENSG00000135365 | n.s. |
| ENSG00000054267 | 3087.07058 | -0.492822323 | 0.2356342 | -2.091472 | 0.036485772 | 0.106765744 | ARID4B | protein_coding | 39.95 | -0.478810433 | 0.478810433 | ENSG00000054267 | n.s. |
| ENSG00000197147 | 1570.23927 | -0.545316945 | 0.2758631 | -1.976766 | 0.048068045 | 0.13265637 | LRRC8B | protein_coding | 38.63 | -0.478391227 | 0.478391227 | ENSG00000197147 | n.s. |
| ENSG00000100162 | 608.0677668 | 0.464323062 | 0.2149672 | 2.159972 | 0.030774855 | 0.093492976 | CENPM | protein_coding | 53.29 | 0.477891053 | 0.477891053 | ENSG00000100162 | n.s. |
| ENSG00000109111 | 1521.520917 | -0.508142339 | 0.2474644 | -2.053396 | 0.040034211 | 0.114761441 | SUPT6H | protein_coding | 48.02 | -0.477757463 | 0.477757463 | ENSG00000109111 | n.s. |
| ENSG00000131495 | 812.9488075 | 0.593167256 | 0.3151137 | 1.882391 | 0.059782959 | 0.156634952 | NDUFA2 | protein_coding | 50.9 | 0.477565673 | 0.477565673 | ENSG00000131495 | n.s. |
| ENSG00000123975 | 1791.101 | 0.538484139 | 0.2709961 | 1.987055 | 0.046916271 | 0.129889688 | CKS2 | protein_coding | 38.43 | 0.477325978 | 0.477325978 | ENSG00000123975 | n.s. |
| ENSG00000139163 | 2975.790193 | -0.55196437 | 0.2817547 | -1.959025 | 0.050109897 | 0.136532309 | ETNK1 | protein_coding | 34.4 | -0.477319229 | 0.477319229 | ENSG00000139163 | n.s. |
| ENSG00000198121 | 634.0186813 | -0.505348738 | 0.2455422 | -2.058093 | 0.039581211 | 0.1137439 | LPAR1 | protein_coding | 38.26 | -0.477085535 | 0.477085535 | ENSG00000198121 | n.s. |
| ENSG00000175073 | 831.1420428 | -0.502060873 | 0.2431067 | -2.065187 | 0.038905315 | 0.112145401 | VCPIP1 | protein_coding | 39.75 | -0.477067546 | 0.477067546 | ENSG00000175073 | n.s. |
| ENSG00000235183 | 177.0303986 | 0.481898854 | 0.228201 | 2.111729 | 0.03470966 | 0.102647111 | | processed_pseudogene | 46.79 | 0.476430878 | 0.476430878 | ENSG00000235183 | n.s. |
| ENSG00000186487 | 1157.900707 | -0.480653526 | 0.2273072 | -2.114555 | 0.034467921 | 0.102062285 | MYT1L | protein_coding | 44.5 | -0.476392393 | 0.476392393 | ENSG00000186487 | n.s. |
| ENSG00000109062 | 184.1828843 | 0.511708818 | 0.2507867 | 2.040414 | 0.041309086 | 0.117362827 | SLC9A3R1 | protein_coding | 56.2 | 0.476129416 | 0.476129416 | ENSG00000109062 | n.s. |
| ENSG00000228801 | 87.25609782 | 0.545757405 | 0.2770733 | 1.969722 | 0.048870247 | 0.134231692 | | antisense | 40.39 | 0.475979557 | 0.475979557 | ENSG00000228801 | n.s. |
| ENSG00000085224 | 4947.311159 | -0.533356184 | 0.2675316 | -1.993619 | 0.046193705 | 0.128297416 | ATRX | protein_coding | 37.73 | -0.475637492 | 0.475637492 | ENSG00000085224 | n.s. |
| ENSG00000222297 | 6.111091155 | -0.919349722 | 0.6353675 | -1.446957 | 0.147908833 | 0.303871135 | RNU6-1156P | snRNA | 37 | -0.475589312 | 0.475589312 | ENSG00000222297 | n.s. |
| ENSG00000259972 | 119.0605729 | -0.537822224 | 0.2710099 | -1.984511 | 0.047198862 | 0.130542264 | | processed_transcript | 47.73 | -0.475568689 | 0.475568689 | ENSG00000259972 | n.s. |
| ENSG00000105656 | 143.5555255 | 0.487106004 | 0.2322879 | 2.096993 | 0.035994199 | 0.105633322 | ELL | protein_coding | 55.52 | 0.475512424 | 0.475512424 | ENSG00000105656 | n.s. |
| ENSG00000070770 | 990.2442375 | -0.493358517 | 0.236955 | -2.082077 | 0.037335475 | 0.10872659 | CSNK2A2 | protein_coding | 41.31 | -0.475431958 | 0.475431958 | ENSG00000070770 | n.s. |
| ENSG00000161217 | 659.1502001 | -0.524643128 | 0.2609375 | -2.010608 | 0.044366876 | 0.124224205 | PCYT1A | protein_coding | 46.69 | -0.47521848 | 0.47521848 | ENSG00000161217 | n.s. |
| ENSG00000151208 | 879.301867 | -0.473935861 | 0.2228437 | -2.126763 | 0.033439745 | 0.099760352 | DLG5 | protein_coding | 50 | -0.474429714 | 0.474429714 | ENSG00000151208 | n.s. |
| ENSG00000119912 | 966.6670926 | -0.546184043 | 0.2779399 | -1.965116 | 0.049400825 | 0.13534193 | IDE | protein_coding | 40.71 | -0.474397782 | 0.474397782 | ENSG00000119912 | n.s. |
| ENSG00000115808 | 801.2341883 | -0.504322737 | 0.2455792 | -2.053605 | 0.040013899 | 0.114726836 | STRN | protein_coding | 37.76 | -0.474232313 | 0.474232313 | ENSG00000115808 | n.s. |
| ENSG00000149294 | 2454.230749 | -0.516919148 | 0.2553694 | -2.024202 | 0.042949375 | 0.121089127 | NCAM1 | protein_coding | 41.71 | -0.473960506 | 0.473960506 | ENSG00000149294 | n.s. |
| ENSG00000116984 | 3123.175461 | -0.502736695 | 0.2445328 | -2.055907 | 0.039791458 | 0.114183115 | MTR | protein_coding | 40.73 | -0.473778113 | 0.473778113 | ENSG00000116984 | n.s. |
| ENSG00000152620 | 413.8922565 | -0.569248164 | 0.2968789 | -1.917443 | 0.055181736 | 0.147239422 | NADK2 | protein_coding | 35.98 | -0.473600751 | 0.473600751 | ENSG00000152620 | n.s. |
| ENSG00000136485 | 4325.358039 | -0.53273246 | 0.2676862 | -1.990138 | 0.046575721 | 0.129178041 | DCAF7 | protein_coding | 43.96 | -0.473498634 | 0.473498634 | ENSG00000136485 | n.s. |
| ENSG00000107371 | 1190.533113 | -0.463973262 | 0.2159126 | -2.148894 | 0.031642828 | 0.095546105 | EXOSC3 | protein_coding | 43.49 | -0.473153907 | 0.473153907 | ENSG00000107371 | n.s. |
| ENSG00000206755 | 9.651684496 | -0.709555009 | 0.4213933 | -1.683831 | 0.092214371 | 0.215463177 | SNORA30 | snoRNA | 51.94 | -0.473008485 | 0.473008485 | ENSG00000206755 | n.s. |
| ENSG00000136960 | 507.3448586 | 0.567001953 | 0.2956136 | 1.918051 | 0.055104572 | 0.147089908 | ENPP2 | protein_coding | 39.28 | 0.471982134 | 0.471982134 | ENSG00000136960 | n.s. |
| ENSG00000198690 | 1271.721646 | -0.489865907 | 0.2355845 | -2.079364 | 0.037583921 | 0.109176076 | FAN1 | protein_coding | 42.55 | -0.471188554 | 0.471188554 | ENSG00000198690 | n.s. |
| ENSG00000260625 | 41.0770239 | 0.561701588 | 0.2916957 | 1.925642 | 0.054149078 | 0.14531951 | | antisense | 48.85 | 0.470523982 | 0.470523982 | ENSG00000260625 | n.s. |

| | | | | | | | | | | | | | |
|---|---|---|---|---|---|---|---|---|---|---|---|---|---|
| ENSG00000136895 | 1154.827643 | -0.480475089 | 0.2287977 | -2.099999 | 0.035728896 | 0.104966728 | GARNL3 | protein_coding | 43.3 | -0.470360291 | 0.470360291 | ENSG00000136895 | n.s. |
| ENSG00000154548 | 459.5819672 | -0.559145276 | 0.2896579 | -1.930364 | 0.053561748 | 0.144243757 | SRSF12 | protein_coding | 40.77 | -0.470186926 | 0.470186926 | ENSG00000154548 | n.s. |
| ENSG00000107872 | 255.1609114 | 0.504854029 | 0.2472841 | 2.041595 | 0.041191731 | 0.117138984 | FBXL15 | protein_coding | 61.93 | 0.470169824 | 0.470169824 | ENSG00000107872 | n.s. |
| ENSG00000119929 | 365.9186615 | -0.603759401 | 0.3270078 | -1.846315 | 0.064846472 | 0.166487455 | CUTC | protein_coding | 40.02 | -0.470098231 | 0.470098231 | ENSG00000119929 | n.s. |
| ENSG00000222629 | 8.627300246 | -0.732933001 | 0.4456427 | -1.644665 | 0.100038896 | 0.228521994 | RNU2-42P | snRNA | 32.74 | -0.469862822 | 0.469862822 | ENSG00000222629 | n.s. |
| ENSG00000271147 | 170.2437678 | -0.560929363 | 0.2913835 | -1.925055 | 0.054222425 | 0.145460274 | ARMCX5-GPRASP2 | processed_transcript | 39.81 | -0.46964125 | 0.46964125 | ENSG00000271147 | n.s. |
| ENSG00000177335 | 81.99467491 | -0.536666046 | 0.2720644 | -1.97257 | 0.048544609 | 0.133561385 | C8orf31 | processed_transcript | 51.12 | -0.469217366 | 0.469217366 | ENSG00000177335 | n.s. |
| ENSG00000187514 | 25294.64372 | 0.582066078 | 0.3090013 | 1.883701 | 0.059605459 | 0.156312349 | PTMA | protein_coding | 61.71 | 0.469149164 | 0.469149164 | ENSG00000187514 | n.s. |
| ENSG00000099940 | 382.9599971 | -0.52282266 | 0.2613078 | -2.000792 | 0.045414795 | 0.126740868 | SNAP29 | protein_coding | 48.86 | -0.469015489 | 0.469015489 | ENSG00000099940 | n.s. |
| ENSG00000284138 | 23.99743748 | 0.625489406 | 0.3464592 | 1.805377 | 0.071015745 | 0.177897124 | | processed_pseudogene | 58.18 | 0.469011393 | 0.469011393 | ENSG00000284138 | n.s. |
| ENSG00000207780 | 11.86098485 | 0.678882176 | 0.3948519 | 1.719334 | 0.085553603 | 0.203861512 | MIR648 | miRNA | 63.83 | 0.468879995 | 0.468879995 | ENSG00000207780 | n.s. |
| ENSG00000173653 | 63.27411478 | 0.5169689 | 0.2569474 | 2.011964 | 0.04422371 | 0.123956892 | RCE1 | protein_coding | 60.51 | 0.46875086 | 0.46875086 | ENSG00000173653 | n.s. |
| ENSG00000227500 | 51.63759824 | 0.523688214 | 0.2622553 | 1.996864 | 0.045839951 | 0.12759489 | SCAMP4 | protein_coding | 60.16 | 0.468264572 | 0.468264572 | ENSG00000227500 | n.s. |
| ENSG00000063177 | 10134.62085 | 0.591331579 | 0.3171951 | 1.864252 | 0.062286302 | 0.161552196 | RPL18 | protein_coding | 56.83 | 0.468149604 | 0.468149604 | ENSG00000063177 | n.s. |
| ENSG00000180423 | 74.94113404 | -0.501961379 | 0.2456676 | -2.043255 | 0.041027248 | 0.116886102 | HARBI1 | protein_coding | 46.15 | -0.467947033 | 0.467947033 | ENSG00000180423 | n.s. |
| ENSG00000081760 | 709.3420854 | -0.5016184 | 0.2454689 | -2.043511 | 0.041001837 | 0.116861531 | AACS | protein_coding | 51.22 | -0.467673094 | 0.467673094 | ENSG00000081760 | n.s. |
| ENSG00000267655 | 38.15674102 | 0.574230913 | 0.3030331 | 1.894945 | 0.05809975 | 0.153350298 | | sense_intronic | 40.34 | 0.467605061 | 0.467605061 | ENSG00000267655 | n.s. |
| ENSG00000149927 | 45.63311916 | 0.608095873 | 0.331938 | 1.831956 | 0.06695797 | 0.170572803 | DOC2A | protein_coding | 55.16 | 0.467072489 | 0.467072489 | ENSG00000149927 | n.s. |
| ENSG00000113790 | 30.87844955 | -0.566070202 | 0.2966423 | -1.908258 | 0.056357818 | 0.149803342 | EHHADH | protein_coding | 39.78 | -0.466712709 | 0.466712709 | ENSG00000113790 | n.s. |
| ENSG00000242498 | 448.0869808 | -0.491721458 | 0.2383185 | -2.063296 | 0.039084543 | 0.112567828 | ARPIN | protein_coding | 49.81 | -0.466439949 | 0.466439949 | ENSG00000242498 | n.s. |
| ENSG00000154874 | 991.9963995 | -0.532043132 | 0.2693317 | -1.97542 | 0.04822051 | 0.13297171 | CCDC144B | transcribed_unprocessed_pseud | 38.34 | -0.46619787 | 0.46619787 | ENSG00000154874 | n.s. |
| ENSG00000203666 | 204.9014818 | -0.559763418 | 0.2917369 | -1.918727 | 0.055018951 | 0.146945877 | EFCAB2 | protein_coding | 42.37 | -0.466194818 | 0.466194818 | ENSG00000203666 | n.s. |
| ENSG00000066422 | 953.9228235 | -0.516881639 | 0.2577188 | -2.005604 | 0.044898566 | 0.125451081 | ZBTB11 | protein_coding | 40.45 | -0.465982027 | 0.465982027 | ENSG00000066422 | n.s. |
| ENSG00000100889 | 205.4114161 | -0.553075081 | 0.2862247 | -1.932311 | 0.053321184 | 0.143734917 | PCK2 | protein_coding | 49.2 | -0.465931309 | 0.465931309 | ENSG00000100889 | n.s. |
| ENSG00000263884 | 68.00307903 | 0.519430103 | 0.2597228 | 1.999941 | 0.045506676 | 0.126946393 | | lincRNA | 47.4 | 0.465606566 | 0.465606566 | ENSG00000263884 | n.s. |
| ENSG00000174599 | 293.1782014 | 0.469089661 | 0.2217376 | 2.115517 | 0.034385878 | 0.101906042 | TRAM1L1 | protein_coding | 40.63 | 0.465243157 | 0.465243157 | ENSG00000174599 | n.s. |
| ENSG00000157693 | 260.4337218 | -0.484581495 | 0.2334669 | -2.075589 | 0.037931946 | 0.109921336 | TMEM268 | protein_coding | 47.67 | -0.464673919 | 0.464673919 | ENSG00000157693 | n.s. |
| ENSG00000141027 | 2889.207667 | -0.488960231 | 0.2367599 | -2.065216 | 0.038902568 | 0.112145401 | NCOR1 | protein_coding | 39.72 | -0.464619072 | 0.464619072 | ENSG00000141027 | n.s. |
| ENSG00000241990 | 357.8668577 | 0.458879486 | 0.2147554 | 2.136754 | 0.032618001 | 0.097917298 | PRR34-AS1 | antisense | 55.48 | 0.46307391 | 0.46307391 | ENSG00000241990 | n.s. |
| ENSG00000258366 | 66.39058343 | 0.553265436 | 0.2875355 | 1.924164 | 0.054334051 | 0.145731649 | RTEL1 | protein_coding | 59.63 | 0.462776729 | 0.462776729 | ENSG00000258366 | n.s. |
| ENSG00000167523 | 161.8962463 | 0.476615021 | 0.2281324 | 2.089203 | 0.036689418 | 0.107315792 | SPATA33 | protein_coding | 52.08 | 0.462000271 | 0.462000271 | ENSG00000167523 | n.s. |
| ENSG00000197956 | 451.8100859 | 0.617380429 | 0.3423992 | 1.803101 | 0.071372314 | 0.178533367 | S100A6 | protein_coding | 58.51 | 0.461973801 | 0.461973801 | ENSG00000197956 | n.s. |

| | | | | | | | | | | | | |
|---|---|---|---|---|---|---|---|---|---|---|---|---|
| ENSG00000276584 | 2.829087739 | 0.923832222 | 0.6529259 | 1.414911 | 0.157094516 | 0.316368891 | MIR6737 | miRNA | 60 | 0.461737098 | 0.461737098 | ENSG00000276584 | n.s. |
| ENSG00000279799 | 59.53695564 | 0.530338951 | 0.2695264 | 1.967669 | 0.049106082 | 0.134746548 | | TEC | 63.46 | 0.461650697 | 0.461650697 | ENSG00000279799 | n.s. |
| ENSG00000168916 | 1362.032955 | 0.482442829 | 0.2328506 | 2.071898 | 0.038274914 | 0.110698303 | ZNF608 | protein_coding | 40.42 | 0.461147338 | 0.461147338 | ENSG00000168916 | n.s. |
| ENSG00000196376 | 210.6184026 | -0.491807016 | 0.2398845 | -2.050183 | 0.040346615 | 0.115466787 | SLC35F1 | protein_coding | 38.46 | -0.461090184 | 0.461090184 | ENSG00000196376 | n.s. |
| ENSG00000112559 | 139.2820808 | -0.619921162 | 0.3452355 | -1.795647 | 0.072550645 | 0.180960702 | MDFI | protein_coding | 58.13 | -0.460239221 | 0.460239221 | ENSG00000112559 | n.s. |
| ENSG00000198354 | 94.38713978 | 0.503112345 | 0.2491048 | 2.019682 | 0.0434164 | 0.122069172 | DCAF12L2 | protein_coding | 54.87 | 0.459539797 | 0.459539797 | ENSG00000198354 | n.s. |
| ENSG00000161920 | 112.11944 | -0.560082347 | 0.2943579 | -1.902726 | 0.057076322 | 0.15127996 | MED11 | protein_coding | 52.08 | -0.459389958 | 0.459389958 | ENSG00000161920 | n.s. |
| ENSG00000103064 | 97.02993431 | 0.501013142 | 0.2475923 | 2.023541 | 0.043017437 | 0.121216711 | SLC7A6 | protein_coding | 47.81 | 0.459147232 | 0.459147232 | ENSG00000103064 | n.s. |
| ENSG00000112701 | 4231.720363 | -0.512208698 | 0.256123 | -1.999854 | 0.045515985 | 0.126946928 | SENP6 | protein_coding | 37.44 | -0.459132508 | 0.459132508 | ENSG00000112701 | n.s. |
| ENSG00000179331 | 361.5054062 | -0.529023184 | 0.2694639 | -1.963244 | 0.049617875 | 0.135669615 | RAB39A | protein_coding | 42.88 | -0.458936821 | 0.458936821 | ENSG00000179331 | n.s. |
| ENSG00000231365 | 368.8976574 | -0.457909072 | 0.2152201 | -2.127631 | 0.033367654 | 0.099587963 | | antisense | 40.62 | -0.458730172 | 0.458730172 | ENSG00000231365 | n.s. |
| ENSG00000141367 | 8818.766884 | -0.518483391 | 0.2611927 | -1.985061 | 0.047137684 | 0.130450801 | CLTC | protein_coding | 38.54 | -0.458626169 | 0.458626169 | ENSG00000141367 | n.s. |
| ENSG00000129116 | 388.3899186 | -0.521681995 | 0.2636814 | -1.978456 | 0.047877318 | 0.132156205 | PALLD | protein_coding | 39.87 | -0.458512795 | 0.458512795 | ENSG00000129116 | n.s. |
| ENSG00000128944 | 1595.977379 | -0.522540006 | 0.26438 | -1.976473 | 0.048101184 | 0.13272152 | KNSTRN | protein_coding | 44.95 | -0.458298234 | 0.458298234 | ENSG00000128944 | n.s. |
| ENSG00000032742 | 397.2696247 | -0.51614321 | 0.259509 | -1.988922 | 0.0467098 | 0.129472536 | IFT88 | protein_coding | 38.74 | -0.458243476 | 0.458243476 | ENSG00000032742 | n.s. |
| ENSG00000175161 | 36.68166505 | 0.555923441 | 0.2914487 | 1.907449 | 0.056462524 | 0.150024375 | CADM2 | protein_coding | 34 | 0.457990952 | 0.457990952 | ENSG00000175161 | n.s. |
| ENSG00000258466 | 23.22148548 | -0.622818875 | 0.3489056 | -1.785064 | 0.074250907 | 0.184047602 | | protein_coding | 43.56 | -0.457815368 | 0.457815368 | ENSG00000258466 | n.s. |
| ENSG00000100263 | 483.2564359 | -0.502909896 | 0.2497738 | -2.013462 | 0.044066098 | 0.123621255 | RHBDD3 | protein_coding | 56.01 | -0.456595341 | 0.456595341 | ENSG00000100263 | n.s. |
| ENSG00000175895 | 137.992214 | 0.589358452 | 0.3201914 | 1.840644 | 0.065673723 | 0.168176429 | PLEKHF2 | protein_coding | 37.16 | 0.456301866 | 0.456301866 | ENSG00000175895 | n.s. |
| ENSG00000210164 | 7.299292563 | 0.745786623 | 0.4661611 | 1.599848 | 0.109632413 | 0.244480742 | MT-TG | Mt_tRNA | 26.47 | 0.456238953 | 0.456238953 | ENSG00000210164 | n.s. |
| ENSG00000274667 | 174.7558347 | 0.473398136 | 0.227367 | 2.082088 | 0.037334443 | 0.10872659 | | sense_intronic | 28.97 | 0.456196852 | 0.456196852 | ENSG00000274667 | n.s. |
| ENSG00000177119 | 3021.092738 | -0.536482633 | 0.2762815 | -1.941797 | 0.052161648 | 0.141155806 | ANO6 | protein_coding | 39.44 | -0.456171855 | 0.456171855 | ENSG00000177119 | n.s. |
| ENSG00000276533 | 24.08069109 | 0.600048688 | 0.3303269 | 1.81653 | 0.069289164 | 0.174795643 | | sense_intronic | 50.8 | 0.454518518 | 0.454518518 | ENSG00000276533 | n.s. |
| ENSG00000163660 | 10578.80389 | 0.457913429 | 0.216394 | 2.11611 | 0.034335478 | 0.101821696 | CCNL1 | protein_coding | 37.3 | 0.454323238 | 0.454323238 | ENSG00000163660 | n.s. |
| ENSG00000162734 | 380.5999124 | -0.536474171 | 0.2769534 | -1.937056 | 0.052738547 | 0.142384874 | PEA15 | protein_coding | 51.85 | -0.454144776 | 0.454144776 | ENSG00000162734 | n.s. |
| ENSG00000248283 | 67.38392471 | -0.541142755 | 0.2807408 | -1.927553 | 0.053910763 | 0.14487542 | CCNL2P1 | processed_pseudogene | 48.61 | -0.454021635 | 0.454021635 | ENSG00000248283 | n.s. |
| ENSG00000163635 | 238.197344 | -0.471226705 | 0.2264138 | -2.081263 | 0.037409825 | 0.108806559 | ATXN7 | protein_coding | 40.77 | -0.453953853 | 0.453953853 | ENSG00000163635 | n.s. |
| ENSG00000159842 | 457.7219063 | -0.529018879 | 0.27108 | -1.951523 | 0.050994845 | 0.138672114 | ABR | protein_coding | 55.2 | -0.453903946 | 0.453903946 | ENSG00000159842 | n.s. |
| ENSG00000258559 | 48.63245352 | -0.580280724 | 0.3136209 | -1.850262 | 0.064275855 | 0.165296034 | | sense_overlapping | 52.46 | -0.453627241 | 0.453627241 | ENSG00000258559 | n.s. |
| ENSG00000185808 | 160.4757955 | 0.523975155 | 0.2672881 | 1.960338 | 0.049956243 | 0.136246967 | PIGP | protein_coding | 42.2 | 0.453591227 | 0.453591227 | ENSG00000185808 | n.s. |
| ENSG00000137806 | 235.0431971 | -0.544556544 | 0.2838077 | -1.918752 | 0.055015752 | 0.146945877 | NDUFAF1 | protein_coding | 46.09 | -0.453529886 | 0.453529886 | ENSG00000137806 | n.s. |
| ENSG00000131016 | 9784.254643 | -0.492148925 | 0.2425032 | -2.029453 | 0.042412135 | 0.119858302 | AKAP12 | protein_coding | 43.87 | -0.453432493 | 0.453432493 | ENSG00000131016 | n.s. |

| | | | | | | | | | | | | | |
|---|---|---|---|---|---|---|---|---|---|---|---|---|---|
| ENSG00000156983 | 273.6583244 | 0.460281555 | 0.2185121 | 2.106435 | 0.035166579 | 0.103602259 | BRPF1 | protein_coding | 51.62 | 0.453207384 | 0.453207384 | ENSG00000156983 | n.s. |
| ENSG00000005156 | 2107.862833 | -0.493163729 | 0.2436602 | -2.023982 | 0.042972021 | 0.121113249 | LIG3 | protein_coding | 47.23 | -0.452136622 | 0.452136622 | ENSG00000005156 | n.s. |
| ENSG00000182986 | 754.4524686 | -0.548543603 | 0.2876229 | -1.907163 | 0.056499541 | 0.150094087 | ZNF320 | protein_coding | 43.93 | -0.451800488 | 0.451800488 | ENSG00000182986 | n.s. |
| ENSG00000199691 | 21.94148167 | -0.607441784 | 0.3380386 | -1.79696 | 0.072342017 | 0.180569721 | RN7SKP173 | misc_RNA | 48.25 | -0.451544932 | 0.451544932 | ENSG00000199691 | n.s. |
| ENSG00000139629 | 224.4954579 | -0.533416376 | 0.2753929 | -1.936929 | 0.052754069 | 0.142399166 | GALNT6 | protein_coding | 49.89 | -0.451532991 | 0.451532991 | ENSG00000139629 | n.s. |
| ENSG00000265828 | 40.59648257 | -0.557827835 | 0.2956623 | -1.886706 | 0.059199918 | 0.155491903 | MIR3939 | miRNA | 54.72 | -0.4508879 | 0.4508879 | ENSG00000265828 | n.s. |
| ENSG00000197728 | 4511.200451 | 0.543934066 | 0.2842891 | 1.913313 | 0.05570794 | 0.148359153 | RPS26 | protein_coding | 46.62 | 0.450750359 | 0.450750359 | ENSG00000197728 | n.s. |
| ENSG00000198315 | 1417.060232 | -0.501279242 | 0.2505605 | -2.000631 | 0.045432131 | 0.126763839 | ZKSCAN8 | protein_coding | 38.6 | -0.449649795 | 0.449649795 | ENSG00000198315 | n.s. |
| ENSG00000207965 | 6.54792424 | -0.768115424 | 0.4929769 | -1.558117 | 0.119205608 | 0.259855906 | MIR629 | miRNA | 59.79 | -0.449552923 | 0.449552923 | ENSG00000207965 | n.s. |
| ENSG00000104904 | 4851.544334 | 0.571940912 | 0.3081014 | 1.85634 | 0.063405087 | 0.163722001 | OAZ1 | protein_coding | 64.06 | 0.449484334 | 0.449484334 | ENSG00000104904 | n.s. |
| ENSG00000183762 | 1786.550973 | -0.525325658 | 0.2703006 | -1.943487 | 0.05195733 | 0.140767056 | KREMEN1 | protein_coding | 45.49 | -0.447314257 | 0.447314257 | ENSG00000183762 | n.s. |
| ENSG00000271833 | 98.25260125 | -0.514207576 | 0.2617075 | -1.964818 | 0.049435301 | 0.13540061 | | sense_intronic | 45.7 | -0.446527255 | 0.446527255 | ENSG00000271833 | n.s. |
| ENSG00000234740 | 96.50369977 | 0.494608422 | 0.2464612 | 2.006841 | 0.044766562 | 0.12523304 | | sense_intronic | 43.07 | 0.44627582 | 0.44627582 | ENSG00000234740 | n.s. |
| ENSG00000122912 | 520.4569123 | -0.52513583 | 0.2705795 | -1.940782 | 0.052284747 | 0.141406474 | SLC25A16 | protein_coding | 43.34 | -0.446119013 | 0.446119013 | ENSG00000122912 | n.s. |
| ENSG00000228506 | 257.4688403 | -0.462503951 | 0.2220696 | -2.082698 | 0.037278753 | 0.10865231 | | antisense | 42.95 | -0.445835788 | 0.445835788 | ENSG00000228506 | n.s. |
| ENSG00000128654 | 1052.582833 | -0.541330896 | 0.2838663 | -1.906992 | 0.056521564 | 0.150123947 | MTX2 | protein_coding | 35.02 | -0.445813074 | 0.445813074 | ENSG00000128654 | n.s. |
| ENSG00000123607 | 1860.737636 | -0.527965146 | 0.2728991 | -1.934653 | 0.053032866 | 0.14309624 | TTC21B | protein_coding | 36.17 | -0.445798869 | 0.445798869 | ENSG00000123607 | n.s. |
| ENSG00000165246 | 238.4048884 | -0.510663881 | 0.259395 | -1.968673 | 0.048990701 | 0.134456442 | NLGN4Y | protein_coding | 38.62 | -0.445001895 | 0.445001895 | ENSG00000165246 | n.s. |
| ENSG00000206948 | 17.10609115 | 0.638221446 | 0.3689951 | 1.72962 | 0.08369816 | 0.200849618 | SNORA36A | snoRNA | 40.15 | 0.444922672 | 0.444922672 | ENSG00000206948 | n.s. |
| ENSG00000139998 | 174.91462 | -0.51547972 | 0.2635164 | -1.956158 | 0.050446535 | 0.137288332 | RAB15 | protein_coding | 52.23 | -0.444532375 | 0.444532375 | ENSG00000139998 | n.s. |
| ENSG00000143384 | 9260.438954 | 0.460381256 | 0.2210774 | 2.082444 | 0.037301926 | 0.108697093 | MCL1 | protein_coding | 44.74 | 0.443707199 | 0.443707199 | ENSG00000143384 | n.s. |
| ENSG00000202078 | 17.67475088 | -0.608770034 | 0.3428736 | -1.775494 | 0.075816354 | 0.186896441 | | misc_RNA | 40.71 | -0.443427464 | 0.443427464 | ENSG00000202078 | n.s. |
| ENSG00000208005 | 3.932598557 | 0.862526931 | 0.5986579 | 1.440768 | 0.149650324 | 0.306183903 | MIR503 | miRNA | 59.15 | 0.443354062 | 0.443354062 | ENSG00000208005 | n.s. |
| ENSG00000188517 | 136.5380872 | 0.511605577 | 0.260795 | 1.961715 | 0.049795656 | 0.135995471 | COL25A1 | protein_coding | 36.92 | 0.443293726 | 0.443293726 | ENSG00000188517 | n.s. |
| ENSG00000213619 | 1243.664051 | 0.547702749 | 0.290229 | 1.88714 | 0.059141517 | 0.15543495 | NDUFS3 | protein_coding | 48.67 | 0.442791011 | 0.442791011 | ENSG00000213619 | n.s. |
| ENSG00000163644 | 550.914336 | -0.53811574 | 0.2823735 | -1.905688 | 0.05669074 | 0.150458477 | PPM1K | protein_coding | 39.9 | -0.442645042 | 0.442645042 | ENSG00000163644 | n.s. |
| ENSG00000273604 | 289.773873 | 0.472145162 | 0.230405 | 2.049196 | 0.040442912 | 0.1156531 | EPOP | protein_coding | 63.89 | 0.442325759 | 0.442325759 | ENSG00000273604 | n.s. |
| ENSG00000164077 | 170.0511082 | -0.531926594 | 0.2775662 | -1.916396 | 0.055314751 | 0.147513864 | MON1A | protein_coding | 51.16 | -0.442119917 | 0.442119917 | ENSG00000164077 | n.s. |
| ENSG00000223224 | 8.879320576 | -0.937372613 | 0.686757 | -1.364926 | 0.172276236 | 0.337926463 | SNORD71 | snoRNA | 48.84 | -0.441669162 | 0.441669162 | ENSG00000223224 | n.s. |
| ENSG00000206573 | 2654.0481 | -0.464058989 | 0.2245144 | -2.066945 | 0.038739319 | 0.111790028 | THUMPD3-AS1 | antisense | 41.7 | -0.441597113 | 0.441597113 | ENSG00000206573 | n.s. |
| ENSG00000258796 | 32.9658879 | -0.61125745 | 0.3459209 | -1.767044 | 0.077220847 | 0.189553219 | | processed_pseudogene | 37.17 | -0.441492207 | 0.441492207 | ENSG00000258796 | n.s. |
| ENSG00000178904 | 1965.699257 | -0.529767734 | 0.2760369 | -1.919192 | 0.054960037 | 0.146929455 | DPY19L3 | protein_coding | 40.45 | -0.441238848 | 0.441238848 | ENSG00000178904 | n.s. |

| | | | | | | | | | | | | | |
|---|---|---|---|---|---|---|---|---|---|---|---|---|---|
| ENSG00000236269 | 199.8799983 | 0.47233074 | 0.2308995 | 2.045611 | 0.040794611 | 0.11636619 | ENO1-IT1 | sense_intronic | 48.92 | 0.441238711 | 0.441238711 | ENSG00000236269 | n.s. |
| ENSG00000077235 | 1878.123992 | -0.477684179 | 0.2350907 | -2.031915 | 0.042162287 | 0.119241423 | GTF3C1 | protein_coding | 49.42 | -0.441176139 | 0.441176139 | ENSG00000077235 | n.s. |
| ENSG00000207750 | 36.04718396 | -0.594847871 | 0.3315277 | -1.794263 | 0.072771174 | 0.181380782 | MIR553 | miRNA | 32.35 | -0.441025404 | 0.441025404 | ENSG00000207750 | n.s. |
| ENSG00000116922 | 316.9679915 | -0.554206299 | 0.296493 | -1.869205 | 0.061594239 | 0.160145022 | C1orf109 | protein_coding | 48.39 | -0.44086366 | 0.44086366 | ENSG00000116922 | n.s. |
| ENSG00000201343 | 56.08963002 | -0.54237624 | 0.2866793 | -1.891927 | 0.058500716 | 0.154116456 | | misc_RNA | 40.91 | -0.440491398 | 0.440491398 | ENSG00000201343 | n.s. |
| ENSG00000077044 | 1130.450663 | -0.491235115 | 0.2459617 | -1.997202 | 0.045803285 | 0.127543827 | DGKD | protein_coding | 47.76 | -0.439331485 | 0.439331485 | ENSG00000077044 | n.s. |
| ENSG00000049167 | 445.2944317 | -0.55401249 | 0.2968912 | -1.866045 | 0.062035005 | 0.161110498 | ERCC8 | protein_coding | 37.16 | -0.439263296 | 0.439263296 | ENSG00000049167 | n.s. |
| ENSG00000125875 | 506.5933439 | -0.502087025 | 0.2543895 | -1.973694 | 0.048416544 | 0.133406766 | TBC1D20 | protein_coding | 46.39 | -0.439236848 | 0.439236848 | ENSG00000125875 | n.s. |
| ENSG00000123352 | 741.7791747 | -0.508739352 | 0.259923 | -1.95727 | 0.050315779 | 0.136986036 | SPATS2 | protein_coding | 41.68 | -0.43920674 | 0.43920674 | ENSG00000123352 | n.s. |
| ENSG00000264796 | 13.23040325 | -0.654283561 | 0.3871154 | -1.690151 | 0.090999059 | 0.213438182 | MIR5009 | miRNA | 34 | -0.43884623 | 0.43884623 | ENSG00000264796 | n.s. |
| ENSG00000169439 | 894.3769169 | 0.556212891 | 0.298961 | 1.860487 | 0.062816707 | 0.162624956 | SDC2 | protein_coding | 41.4 | 0.438747852 | 0.438747852 | ENSG00000169439 | n.s. |
| ENSG00000241547 | 46.45871163 | 0.524561762 | 0.2728058 | 1.922839 | 0.054500237 | 0.146008621 | ACTG1P20 | processed_pseudogene | 55.56 | 0.438335086 | 0.438335086 | ENSG00000241547 | n.s. |
| ENSG00000189339 | 407.954075 | -0.506819566 | 0.2587027 | -1.959081 | 0.050103278 | 0.136532309 | SLC35E2B | protein_coding | 53.52 | -0.438279602 | 0.438279602 | ENSG00000189339 | n.s. |
| ENSG00000170571 | 426.3391533 | -0.533485688 | 0.2802695 | -1.903474 | 0.056978688 | 0.151078703 | EMB | protein_coding | 36.85 | -0.437883321 | 0.437883321 | ENSG00000170571 | n.s. |
| ENSG00000136231 | 2322.940586 | -0.50521944 | 0.2575138 | -1.961912 | 0.049772738 | 0.135959552 | IGF2BP3 | protein_coding | 41.63 | -0.437818253 | 0.437818253 | ENSG00000136231 | n.s. |
| ENSG00000023318 | 732.2315136 | -0.51362754 | 0.2641545 | -1.944421 | 0.051844737 | 0.140516697 | ERP44 | protein_coding | 37.86 | -0.437750406 | 0.437750406 | ENSG00000023318 | n.s. |
| ENSG00000175662 | 28.73455666 | -0.55855074 | 0.3014641 | -1.852794 | 0.063911928 | 0.164694757 | TOM1L2 | protein_coding | 48.26 | -0.437524092 | 0.437524092 | ENSG00000175662 | n.s. |
| ENSG00000159884 | 123.7400485 | 0.531489755 | 0.2788211 | 1.906204 | 0.056623782 | 0.150309422 | CCDC107 | protein_coding | 52.82 | 0.437423401 | 0.437423401 | ENSG00000159884 | n.s. |
| ENSG00000147454 | 897.3057538 | -0.498630264 | 0.2523376 | -1.976044 | 0.048149756 | 0.132829216 | SLC25A37 | protein_coding | 48.45 | -0.437152339 | 0.437152339 | ENSG00000147454 | n.s. |
| ENSG00000197977 | 487.9789504 | -0.545126538 | 0.2903666 | -1.877373 | 0.060466951 | 0.157892228 | ELOVL2 | protein_coding | 39.64 | -0.436994827 | 0.436994827 | ENSG00000197977 | n.s. |
| ENSG00000156675 | 436.2912172 | 0.476790381 | 0.2356404 | 2.023381 | 0.043033886 | 0.121238525 | RAB11FIP1 | protein_coding | 45.87 | 0.436911327 | 0.436911327 | ENSG00000156675 | n.s. |
| ENSG00000141564 | 1464.134773 | -0.453233771 | 0.2177474 | -2.081466 | 0.037391272 | 0.108791793 | RPTOR | protein_coding | 49.12 | -0.436647167 | 0.436647167 | ENSG00000141564 | n.s. |
| ENSG00000038382 | 2964.404155 | -0.473845157 | 0.2335306 | -2.02905 | 0.042453231 | 0.11994244 | TRIO | protein_coding | 43.35 | -0.436424241 | 0.436424241 | ENSG00000038382 | n.s. |
| ENSG00000166199 | 493.6900248 | -0.483849919 | 0.2411718 | -2.006246 | 0.044830036 | 0.125360231 | ALKBH3 | protein_coding | 40.79 | -0.436355316 | 0.436355316 | ENSG00000166199 | n.s. |
| ENSG00000129696 | 330.1426746 | -0.500681725 | 0.254314 | -1.968754 | 0.048981329 | 0.134456442 | TTI2 | protein_coding | 42.98 | -0.43630326 | 0.43630326 | ENSG00000129696 | n.s. |
| ENSG00000279310 | 9.417952466 | 0.691227755 | 0.4239838 | 1.630317 | 0.103034612 | 0.233830627 | | TEC | 37.93 | 0.436232872 | 0.436232872 | ENSG00000279310 | n.s. |
| ENSG00000258682 | 52.15593381 | -0.551059159 | 0.2956175 | -1.864095 | 0.062308298 | 0.161579147 | | sense_overlapping | 37.85 | -0.436226525 | 0.436226525 | ENSG00000258682 | n.s. |
| ENSG00000109103 | 159.6448127 | 0.523148729 | 0.2724179 | 1.920391 | 0.054808576 | 0.146665349 | UNC119 | protein_coding | 57.56 | 0.436134699 | 0.436134699 | ENSG00000109103 | n.s. |
| ENSG00000110108 | 233.5844271 | -0.52641047 | 0.2751866 | -1.912922 | 0.055758013 | 0.148464107 | TMEM109 | protein_coding | 51.1 | -0.43606713 | 0.43606713 | ENSG00000110108 | n.s. |
| ENSG00000119782 | 81.24635272 | 0.580691436 | 0.321458 | 1.80643 | 0.07085119 | 0.177548797 | FKBP1B | protein_coding | 48.23 | 0.435914764 | 0.435914764 | ENSG00000119782 | n.s. |
| ENSG00000070950 | 1165.438522 | -0.501962471 | 0.2554825 | -1.964763 | 0.049441694 | 0.13540061 | RAD18 | protein_coding | 40.44 | -0.435893859 | 0.435893859 | ENSG00000070950 | n.s. |
| ENSG00000189241 | 1750.919813 | 0.482912401 | 0.2407267 | 2.006061 | 0.044849782 | 0.125365091 | TSPYL1 | protein_coding | 48.02 | 0.435501694 | 0.435501694 | ENSG00000189241 | n.s. |

| | | | | | | | | | | | | | |
|---|---|---|---|---|---|---|---|---|---|---|---|---|---|
| ENSG00000140743 | 298.3046635 | -0.523129843 | 0.2728221 | -1.917476 | 0.055177456 | 0.147239422 | CDR2 | protein_coding | 43.33 | -0.43523142 | 0.43523142 | ENSG00000140743 | n.s. |
| ENSG00000053254 | 1952.973415 | -0.502529003 | 0.2562522 | -1.961072 | 0.04987067 | 0.136126277 | FOXN3 | protein_coding | 44.31 | -0.435219279 | 0.435219279 | ENSG00000053254 | n.s. |
| ENSG00000102531 | 2590.256915 | -0.507432537 | 0.2601652 | -1.950425 | 0.051125542 | 0.138918992 | FNDC3A | protein_coding | 37.08 | -0.434990644 | 0.434990644 | ENSG00000102531 | n.s. |
| ENSG00000198780 | 908.8758782 | -0.507936536 | 0.2606272 | -1.9489 | 0.051307322 | 0.139249871 | FAM169A | protein_coding | 38.67 | -0.434897903 | 0.434897903 | ENSG00000198780 | n.s. |
| ENSG00000097033 | 2078.764888 | -0.535641653 | 0.2831762 | -1.891549 | 0.058551129 | 0.154214372 | SH3GLB1 | protein_coding | 36.11 | -0.434874148 | 0.434874148 | ENSG00000097033 | n.s. |
| ENSG00000212907 | 788.4732233 | -0.465277069 | 0.2274788 | -2.045365 | 0.040818912 | 0.116411657 | MT-ND4L | protein_coding | 43.1 | -0.434570423 | 0.434570423 | ENSG00000212907 | n.s. |
| ENSG00000185115 | 506.4844681 | 0.517097328 | 0.2679945 | 1.929507 | 0.053667958 | 0.144473894 | NSMCE3 | protein_coding | 41.84 | 0.434470668 | 0.434470668 | ENSG00000185115 | n.s. |
| ENSG00000131873 | 1299.41174 | -0.501412248 | 0.2556924 | -1.960998 | 0.049879293 | 0.136126277 | CHSY1 | protein_coding | 43.36 | -0.434252105 | 0.434252105 | ENSG00000131873 | n.s. |
| ENSG00000281780 | 17.52278143 | 0.607314624 | 0.3458381 | 1.756066 | 0.079077128 | 0.192851786 | | snoRNA | 35.96 | 0.434094121 | 0.434094121 | ENSG00000281780 | n.s. |
| ENSG00000109618 | 414.4414418 | -0.498667386 | 0.2534627 | -1.967419 | 0.049134877 | 0.134772436 | SEPSECS | protein_coding | 36.27 | -0.434039554 | 0.434039554 | ENSG00000109618 | n.s. |
| ENSG00000173598 | 1558.951814 | 0.472420093 | 0.2333919 | 2.02415 | 0.042954765 | 0.121089127 | NUDT4 | protein_coding | 41.97 | 0.433159551 | 0.433159551 | ENSG00000173598 | n.s. |
| ENSG00000145425 | 18896.36963 | 0.581229057 | 0.3230233 | 1.799341 | 0.07196479 | 0.1797893 | RPS3A | protein_coding | 41.56 | 0.43315291 | 0.43315291 | ENSG00000145425 | n.s. |
| ENSG00000158793 | 319.5862902 | -0.510893023 | 0.2635332 | -1.938629 | 0.052546589 | 0.1420043 | NIT1 | protein_coding | 49.13 | -0.433083251 | 0.433083251 | ENSG00000158793 | n.s. |
| ENSG00000144747 | 866.8018121 | -0.500751357 | 0.2556677 | -1.958602 | 0.050159404 | 0.136640458 | TMF1 | protein_coding | 36.59 | -0.432859833 | 0.432859833 | ENSG00000144747 | n.s. |
| ENSG00000222078 | 18.52515781 | -0.600344903 | 0.3402163 | -1.764598 | 0.077631378 | 0.190225567 | RN7SKP110 | misc_RNA | 50.88 | -0.432687251 | 0.432687251 | ENSG00000222078 | n.s. |
| ENSG00000127947 | 910.0926379 | -0.503363905 | 0.2577347 | -1.953031 | 0.050815937 | 0.138239604 | PTPN12 | protein_coding | 38.16 | -0.432574591 | 0.432574591 | ENSG00000127947 | n.s. |
| ENSG00000116977 | 37.61225605 | 0.539742646 | 0.287618 | 1.876595 | 0.060573584 | 0.158022486 | LGALS8 | protein_coding | 43.48 | 0.432485586 | 0.432485586 | ENSG00000116977 | n.s. |
| ENSG00000170903 | 1022.721076 | -0.5223612 | 0.273255 | -1.911625 | 0.055924256 | 0.148878282 | MSANTD4 | protein_coding | 38.77 | -0.432080809 | 0.432080809 | ENSG00000170903 | n.s. |
| ENSG00000180318 | 295.8139688 | -0.499978006 | 0.2553569 | -1.957958 | 0.050234999 | 0.13681773 | ALX1 | protein_coding | 31.86 | -0.43190981 | 0.43190981 | ENSG00000180318 | n.s. |
| ENSG00000112029 | 2698.219573 | -0.498891136 | 0.2544906 | -1.960352 | 0.04995463 | 0.136246967 | FBXO5 | protein_coding | 38.53 | -0.431876665 | 0.431876665 | ENSG00000112029 | n.s. |
| ENSG00000155760 | 299.0170313 | 0.447005217 | 0.2144617 | 2.084312 | 0.037131788 | 0.108337359 | FZD7 | protein_coding | 52.76 | 0.43145916 | 0.43145916 | ENSG00000155760 | n.s. |
| ENSG00000178700 | 476.2914496 | 0.436742511 | 0.2071526 | 2.108313 | 0.035003901 | 0.103275972 | DHFR2 | protein_coding | 35.09 | 0.430628426 | 0.430628426 | ENSG00000178700 | n.s. |
| ENSG00000125485 | 662.151202 | -0.480328317 | 0.2402047 | -1.999663 | 0.045536703 | 0.126979273 | DDX31 | protein_coding | 44.73 | -0.430502498 | 0.430502498 | ENSG00000125485 | n.s. |
| ENSG00000181798 | 62.227259 | 0.509801063 | 0.2635591 | 1.934295 | 0.053076837 | 0.143187135 | LINC00471 | lincRNA | 42.78 | 0.430321038 | 0.430321038 | ENSG00000181798 | n.s. |
| ENSG00000107185 | 443.8520616 | -0.475693408 | 0.2367586 | -2.009192 | 0.044516804 | 0.124584416 | RGP1 | protein_coding | 52.62 | -0.430281946 | 0.430281946 | ENSG00000107185 | n.s. |
| ENSG00000168077 | 278.6249473 | -0.502835069 | 0.2581096 | -1.948145 | 0.051397561 | 0.139413255 | SCARA3 | protein_coding | 50.25 | -0.430273924 | 0.430273924 | ENSG00000168077 | n.s. |
| ENSG00000162433 | 497.4347545 | -0.512302134 | 0.2656185 | -1.928714 | 0.0537664 | 0.144639071 | AK4 | protein_coding | 42.62 | -0.430187466 | 0.430187466 | ENSG00000162433 | n.s. |
| ENSG00000104375 | 67.64131859 | -0.606957338 | 0.3475209 | -1.746535 | 0.080718042 | 0.195563621 | STK3 | protein_coding | 39.36 | -0.430157906 | 0.430157906 | ENSG00000104375 | n.s. |
| ENSG00000105705 | 491.7912577 | -0.51965675 | 0.2718931 | -1.911254 | 0.055971977 | 0.148976836 | SUGP1 | protein_coding | 51.37 | -0.429694424 | 0.429694424 | ENSG00000105705 | n.s. |
| ENSG00000119574 | 109.1859926 | -0.467864554 | 0.2310167 | -2.025241 | 0.042842581 | 0.120821806 | ZBTB45 | protein_coding | 53.64 | -0.429431669 | 0.429431669 | ENSG00000119574 | n.s. |
| ENSG00000253347 | 18.37735238 | -0.885017793 | 0.6367619 | -1.389872 | 0.164567648 | 0.32718483 | | antisense | 38.81 | -0.429416688 | 0.429416688 | ENSG00000253347 | n.s. |
| ENSG00000136866 | 464.1345487 | -0.535675825 | 0.2854092 | -1.876869 | 0.060535984 | 0.157980801 | ZFP37 | protein_coding | 39.08 | -0.429288296 | 0.429288296 | ENSG00000136866 | n.s. |

| | | | | | | | | | | | | | |
|---|---|---|---|---|---|---|---|---|---|---|---|---|---|
| ENSG00000068028 | 317.8620448 | -0.460273099 | 0.2253361 | -2.042607 | 0.04109132 | 0.116967636 | RASSF1 | protein_coding | 54.57 | -0.428944283 | 0.428944283 | ENSG00000068028 | n.s. |
| ENSG00000259015 | 13.93708708 | -0.667825098 | 0.4058111 | -1.645655 | 0.099834751 | 0.228205424 | | processed_pseudogene | 51.75 | -0.428526027 | 0.428526027 | ENSG00000259015 | n.s. |
| ENSG00000277118 | 18.24661256 | -0.604137378 | 0.3459117 | -1.746507 | 0.080722842 | 0.195563621 | | misc_RNA | 54.55 | -0.428159367 | 0.428159367 | ENSG00000277118 | n.s. |
| ENSG00000136014 | 39.73089649 | 0.551731857 | 0.2994986 | 1.842185 | 0.065448063 | 0.167752832 | USP44 | protein_coding | 41.64 | 0.427774338 | 0.427774338 | ENSG00000136014 | n.s. |
| ENSG00000265625 | 35.59199677 | -0.536119699 | 0.2864362 | -1.87169 | 0.061249561 | 0.159487122 | | sense_intronic | 39.73 | -0.427434499 | 0.427434499 | ENSG00000265625 | n.s. |
| ENSG00000072415 | 482.472747 | -0.501872908 | 0.2583206 | -1.94283 | 0.052036733 | 0.140872532 | MPP5 | protein_coding | 37.63 | -0.427181009 | 0.427181009 | ENSG00000072415 | n.s. |
| ENSG00000184661 | 1017.677711 | -0.469003254 | 0.2325814 | -2.016513 | 0.043746409 | 0.122872203 | CDCA2 | protein_coding | 39.74 | -0.427049204 | 0.427049204 | ENSG00000184661 | n.s. |
| ENSG00000100209 | 130.5065235 | -0.548312963 | 0.2969772 | -1.846313 | 0.064846762 | 0.166487455 | HSCB | protein_coding | 46.7 | -0.426926609 | 0.426926609 | ENSG00000100209 | n.s. |
| ENSG00000213442 | 322.0060036 | 0.598793246 | 0.3416139 | 1.752836 | 0.079630143 | 0.193753682 | RPL18AP3 | processed_pseudogene | 60.23 | 0.426789907 | 0.426789907 | ENSG00000213442 | n.s. |
| ENSG00000157168 | 70.11373501 | -0.484271842 | 0.2445319 | -1.980404 | 0.047658169 | 0.131629576 | NRG1 | protein_coding | 37.9 | -0.426472311 | 0.426472311 | ENSG00000157168 | n.s. |
| ENSG00000207982 | 8.385281324 | -0.871971759 | 0.6239094 | -1.397594 | 0.16223517 | 0.324353859 | MIR548B | miRNA | 37.11 | -0.426377563 | 0.426377563 | ENSG00000207982 | n.s. |
| ENSG00000135932 | 1344.750694 | -0.517338835 | 0.2711815 | -1.907722 | 0.056427184 | 0.149959092 | CAB39 | protein_coding | 40.78 | -0.426301271 | 0.426301271 | ENSG00000135932 | n.s. |
| ENSG00000207133 | 23.0602406 | -0.607245512 | 0.3496576 | -1.736686 | 0.082442604 | 0.198624049 | SNORD116-7 | snoRNA | 44.21 | -0.42626702 | 0.42626702 | ENSG00000207133 | n.s. |
| ENSG00000114374 | 869.7333898 | -0.529433267 | 0.2814261 | -1.881251 | 0.05993776 | 0.156863428 | USP9Y | protein_coding | 36.81 | -0.425917574 | 0.425917574 | ENSG00000114374 | n.s. |
| ENSG00000144559 | 66.72124218 | -0.501241206 | 0.2582484 | -1.940927 | 0.052267155 | 0.14138636 | TAMM41 | protein_coding | 42.11 | -0.425850763 | 0.425850763 | ENSG00000144559 | n.s. |
| ENSG00000156475 | 231.1731882 | -0.510836261 | 0.2660362 | -1.920176 | 0.054835679 | 0.146709676 | PPP2R2B | protein_coding | 39.61 | -0.425803091 | 0.425803091 | ENSG00000156475 | n.s. |
| ENSG00000187605 | 1669.143801 | -0.459471888 | 0.2258514 | -2.034399 | 0.041911348 | 0.118630053 | TET3 | protein_coding | 46.75 | -0.425381497 | 0.425381497 | ENSG00000187605 | n.s. |
| ENSG00000137338 | 427.6678111 | -0.510698429 | 0.2660762 | -1.919369 | 0.054937686 | 0.146926125 | PGBD1 | protein_coding | 39.87 | -0.42536122 | 0.42536122 | ENSG00000137338 | n.s. |
| ENSG00000247556 | 3702.603328 | -0.51046462 | 0.2659248 | -1.919583 | 0.054910656 | 0.146882049 | OIP5-AS1 | processed_transcript | 41.46 | -0.425232995 | 0.425232995 | ENSG00000247556 | n.s. |
| ENSG00000147202 | 573.1802612 | -0.515318877 | 0.2699962 | -1.908615 | 0.056311731 | 0.149709419 | DIAPH2 | protein_coding | 37 | -0.425009694 | 0.425009694 | ENSG00000147202 | n.s. |
| ENSG00000104899 | 567.9986273 | 0.487459906 | 0.2475211 | 1.969367 | 0.048910996 | 0.13431712 | AMH | protein_coding | 71.68 | 0.425000999 | 0.425000999 | ENSG00000104899 | n.s. |
| ENSG00000142856 | 1145.749344 | -0.56172285 | 0.3096502 | -1.814056 | 0.069669068 | 0.175343897 | ITGB3BP | protein_coding | 36.68 | -0.424723897 | 0.424723897 | ENSG00000142856 | n.s. |
| ENSG00000196466 | 282.9518781 | -0.490045766 | 0.2498998 | -1.960969 | 0.049882659 | 0.136126277 | ZNF799 | protein_coding | 41.86 | -0.424408072 | 0.424408072 | ENSG00000196466 | n.s. |
| ENSG00000118961 | 1305.27157 | -0.555808378 | 0.3043822 | -1.826021 | 0.067847043 | 0.172409997 | LDAH | protein_coding | 38.21 | -0.424324989 | 0.424324989 | ENSG00000118961 | n.s. |
| ENSG00000154803 | 213.924962 | -0.507406185 | 0.2639225 | -1.922557 | 0.054535662 | 0.14604732 | FLCN | protein_coding | 53.29 | -0.423941119 | 0.423941119 | ENSG00000154803 | n.s. |
| ENSG00000156521 | 285.7106281 | -0.453491225 | 0.221651 | -2.045969 | 0.040759373 | 0.116304728 | TYSND1 | protein_coding | 53.24 | -0.423743393 | 0.423743393 | ENSG00000156521 | n.s. |
| ENSG00000085788 | 570.5233357 | 0.488133399 | 0.2485458 | 1.963957 | 0.049535027 | 0.13553486 | DDHD2 | protein_coding | 42.25 | 0.42367489 | 0.42367489 | ENSG00000085788 | n.s. |
| ENSG00000122299 | 1308.262542 | -0.494367127 | 0.2535904 | -1.949471 | 0.051239176 | 0.139130011 | ZC3H7A | protein_coding | 42.77 | -0.423464589 | 0.423464589 | ENSG00000122299 | n.s. |
| ENSG00000114933 | 1203.093709 | -0.487870063 | 0.2484165 | -1.96392 | 0.049539388 | 0.13553486 | INO80D | protein_coding | 40.81 | -0.423446328 | 0.423446328 | ENSG00000114933 | n.s. |
| ENSG00000247934 | 133.7562153 | 0.462733854 | 0.2288947 | 2.021602 | 0.043217502 | 0.121632765 | | antisense | 37.35 | 0.423378072 | 0.423378072 | ENSG00000247934 | n.s. |
| ENSG00000124177 | 3272.766603 | -0.46671585 | 0.2319038 | -2.012541 | 0.044162947 | 0.123843044 | CHD6 | protein_coding | 41.1 | -0.423371193 | 0.423371193 | ENSG00000124177 | n.s. |
| ENSG00000215193 | 1227.019791 | -0.508288055 | 0.2648388 | -1.919236 | 0.054954509 | 0.146929455 | PEX26 | protein_coding | 48.63 | -0.423348614 | 0.423348614 | ENSG00000215193 | n.s. |

| | | | | | | | | | | | | | |
|---|---|---|---|---|---|---|---|---|---|---|---|---|---|
| ENSG00000003987 | 193.7947142 | -0.508454087 | 0.2650533 | -1.918309 | 0.055071835 | 0.14703274 | MTMR7 | protein_coding | 40.36 | -0.423331728 | 0.423331728 | ENSG00000003987 | n.s. |
| ENSG00000067533 | 3152.227475 | -0.52409926 | 0.2779289 | -1.885731 | 0.059331183 | 0.155744356 | RRP15 | protein_coding | 37.64 | -0.423256107 | 0.423256107 | ENSG00000067533 | n.s. |
| ENSG00000166619 | 721.5601444 | -0.483877853 | 0.245245 | -1.973039 | 0.048491124 | 0.133493301 | BLCAP | protein_coding | 47.98 | -0.423170793 | 0.423170793 | ENSG00000166619 | n.s. |
| ENSG00000168564 | 1695.406045 | 0.43151768 | 0.2053332 | 2.101548 | 0.035592851 | 0.10463667 | CDKN2AIP | protein_coding | 43.85 | 0.423023726 | 0.423023726 | ENSG00000168564 | n.s. |
| ENSG00000252627 | 7.296491503 | -0.729185452 | 0.4704308 | -1.550038 | 0.121132446 | 0.263002074 | RNU6-122P | snRNA | 34.58 | -0.422957333 | 0.422957333 | ENSG00000252627 | n.s. |
| ENSG00000128791 | 515.9462827 | -0.523827779 | 0.2778376 | -1.885374 | 0.059379325 | 0.155841345 | TWSG1 | protein_coding | 38.45 | -0.422895235 | 0.422895235 | ENSG00000128791 | n.s. |
| ENSG00000235445 | 16.10616278 | 0.603491242 | 0.3479716 | 1.734312 | 0.082862768 | 0.199291471 | | sense_intronic | 53.98 | 0.422752427 | 0.422752427 | ENSG00000235445 | n.s. |
| ENSG00000078902 | 26.9400309 | 0.55633513 | 0.3057574 | 1.819531 | 0.068830412 | 0.173956403 | TOLLIP | protein_coding | 57.49 | 0.422569678 | 0.422569678 | ENSG00000078902 | n.s. |
| ENSG00000164180 | 1048.839969 | -0.493348835 | 0.2531259 | -1.949025 | 0.051292384 | 0.139236472 | TMEM161B | protein_coding | 35.47 | -0.422428455 | 0.422428455 | ENSG00000164180 | n.s. |
| ENSG00000188559 | 511.8919662 | -0.492926172 | 0.2530926 | -1.947612 | 0.051461438 | 0.139559331 | RALGAPA2 | protein_coding | 40.42 | -0.421570732 | 0.421570732 | ENSG00000188559 | n.s. |
| ENSG00000188322 | 280.4787903 | -0.452022211 | 0.2212963 | -2.042611 | 0.041090976 | 0.116967636 | SBK1 | protein_coding | 55.29 | -0.421254997 | 0.421254997 | ENSG00000188322 | n.s. |
| ENSG00000198399 | 207.328285 | 0.493161737 | 0.2534193 | 1.946031 | 0.051651046 | 0.140046254 | ITSN2 | protein_coding | 39.9 | 0.421026231 | 0.421026231 | ENSG00000198399 | n.s. |
| ENSG00000165355 | 215.1256892 | 0.507936894 | 0.2654177 | 1.913726 | 0.055655152 | 0.148292503 | FBXO33 | protein_coding | 38.98 | 0.421019142 | 0.421019142 | ENSG00000165355 | n.s. |
| ENSG00000106638 | 178.9688066 | -0.502241609 | 0.2606995 | -1.926515 | 0.054040041 | 0.145138792 | TBL2 | protein_coding | 52.83 | -0.420987202 | 0.420987202 | ENSG00000106638 | n.s. |
| ENSG00000134248 | 1692.604508 | 0.514125889 | 0.270478 | 1.900804 | 0.057327625 | 0.151772678 | LAMTOR5 | protein_coding | 42.55 | 0.420969569 | 0.420969569 | ENSG00000134248 | n.s. |
| ENSG00000101216 | 365.8828578 | -0.485076523 | 0.2470223 | -1.963695 | 0.049565435 | 0.13557948 | GMEB2 | protein_coding | 53.42 | -0.420952335 | 0.420952335 | ENSG00000101216 | n.s. |
| ENSG00000210194 | 61.70286619 | 0.555926399 | 0.3061981 | 1.815577 | 0.069435228 | 0.174979676 | MT-TE | Mt_tRNA | 39.13 | 0.420843174 | 0.420843174 | ENSG00000210194 | n.s. |
| ENSG00000266297 | 10.72982774 | -0.653904271 | 0.3968037 | -1.647929 | 0.099367274 | 0.227472962 | MIR744 | miRNA | 54.08 | -0.420506362 | 0.420506362 | ENSG00000266297 | n.s. |
| ENSG00000179981 | 299.8041802 | -0.493936447 | 0.2542108 | -1.943019 | 0.052013861 | 0.140837998 | TSHZ1 | protein_coding | 44.58 | -0.420478297 | 0.420478297 | ENSG00000179981 | n.s. |
| ENSG00000184897 | 2783.995966 | 0.485425647 | 0.2475778 | 1.960699 | 0.049914125 | 0.136185451 | H1FX | protein_coding | 68.61 | 0.420315158 | 0.420315158 | ENSG00000184897 | n.s. |
| ENSG00000177054 | 688.9488004 | -0.522084256 | 0.2773821 | -1.882184 | 0.059811006 | 0.15664948 | ZDHHC13 | protein_coding | 37.99 | -0.420314917 | 0.420314917 | ENSG00000177054 | n.s. |
| ENSG00000262380 | 16.49197817 | -0.580506234 | 0.3280395 | -1.769623 | 0.076789926 | 0.188795009 | | sense_intronic | 48.01 | -0.420292023 | 0.420292023 | ENSG00000262380 | n.s. |
| ENSG00000223546 | 116.7936503 | -0.514510914 | 0.271084 | -1.897976 | 0.057699278 | 0.152553561 | LINC00630 | lincRNA | 39.11 | -0.420138113 | 0.420138113 | ENSG00000223546 | n.s. |
| ENSG00000140043 | 136.3343407 | -0.554329866 | 0.3051712 | -1.816455 | 0.069300578 | 0.174795643 | PTGR2 | protein_coding | 42.1 | -0.419887909 | 0.419887909 | ENSG00000140043 | n.s. |
| ENSG00000009413 | 3440.158064 | -0.475727321 | 0.239966 | -1.982478 | 0.047425739 | 0.131039604 | REV3L | protein_coding | 36.9 | -0.419875711 | 0.419875711 | ENSG00000009413 | n.s. |
| ENSG00000156976 | 19331.30022 | 0.448226864 | 0.2187765 | 2.048789 | 0.040482719 | 0.115713589 | EIF4A2 | protein_coding | 41.94 | 0.41981629 | 0.41981629 | ENSG00000156976 | n.s. |
| ENSG00000187187 | 32.41505387 | -0.57168532 | 0.3205501 | -1.783451 | 0.074512918 | 0.184532793 | ZNF546 | protein_coding | 39.84 | -0.419574975 | 0.419574975 | ENSG00000187187 | n.s. |
| ENSG00000197122 | 298.9307383 | -0.468861023 | 0.2346731 | -1.997933 | 0.045723975 | 0.12739942 | SRC | protein_coding | 54.1 | -0.419552104 | 0.419552104 | ENSG00000197122 | n.s. |
| ENSG00000079215 | 89.06545169 | 0.482771071 | 0.2457677 | 1.964339 | 0.049490828 | 0.135508516 | SLC1A3 | protein_coding | 40.36 | 0.41906142 | 0.41906142 | ENSG00000079215 | n.s. |
| ENSG00000154710 | 359.9175809 | -0.498817803 | 0.258624 | -1.928737 | 0.053763467 | 0.144639071 | RABGEF1 | protein_coding | 44.49 | -0.41886448 | 0.41886448 | ENSG00000154710 | n.s. |
| ENSG00000252652 | 42.52149366 | -0.589736052 | 0.3373317 | -1.748238 | 0.08042288 | 0.195074523 | | misc_RNA | 39.58 | -0.418594322 | 0.418594322 | ENSG00000252652 | n.s. |
| ENSG00000089351 | 330.7406804 | -0.470980086 | 0.2366811 | -1.989935 | 0.046598053 | 0.129214239 | GRAMD1A | protein_coding | 55.68 | -0.418555117 | 0.418555117 | ENSG00000089351 | n.s. |

| | | | | | | | | | | | | | |
|---|---|---|---|---|---|---|---|---|---|---|---|---|---|
| ENSG00000125375 | 954.1515454 | -0.512597984 | 0.2700966 | -1.897832 | 0.057718204 | 0.152574627 | ATP5S | protein_coding | 40.64 | -0.418545319 | 0.418545319 | ENSG00000125375 | n.s. |
| ENSG00000063322 | 430.7805663 | -0.514098119 | 0.2713593 | -1.894529 | 0.058154815 | 0.153437462 | MED29 | protein_coding | 52.36 | -0.418511132 | 0.418511132 | ENSG00000063322 | n.s. |
| ENSG00000165675 | 562.2140953 | -0.547540244 | 0.2997095 | -1.826903 | 0.067714327 | 0.172197062 | ENOX2 | protein_coding | 38.87 | -0.418306654 | 0.418306654 | ENSG00000165675 | n.s. |
| ENSG00000112242 | 1226.226752 | -0.457414756 | 0.226336 | -2.020955 | 0.043284458 | 0.12177132 | E2F3 | protein_coding | 42.72 | -0.418285205 | 0.418285205 | ENSG00000112242 | n.s. |
| ENSG00000005175 | 1463.207121 | -0.506890694 | 0.265509 | -1.909128 | 0.056245553 | 0.149590607 | RPAP3 | protein_coding | 36.57 | -0.418233319 | 0.418233319 | ENSG00000005175 | n.s. |
| ENSG00000179941 | 592.187799 | -0.468580815 | 0.2349376 | -1.99449 | 0.046098485 | 0.128237604 | BBS10 | protein_coding | 37.34 | -0.417966873 | 0.417966873 | ENSG00000179941 | n.s. |
| ENSG00000144045 | 8.656253623 | 0.694473531 | 0.4386353 | 1.58326 | 0.113362287 | 0.250411411 | DQX1 | protein_coding | 49.43 | 0.4176188 | 0.4176188 | ENSG00000144045 | n.s. |
| ENSG00000230124 | 2195.462753 | -0.491492261 | 0.2532165 | -1.940996 | 0.052258736 | 0.14138636 | ACBD6 | protein_coding | 37.67 | -0.417568133 | 0.417568133 | ENSG00000230124 | n.s. |
| ENSG00000137814 | 824.1759675 | -0.532775994 | 0.2874106 | -1.85371 | 0.063780596 | 0.164539047 | HAUS2 | protein_coding | 42.11 | -0.417553075 | 0.417553075 | ENSG00000137814 | n.s. |
| ENSG00000207389 | 40.52027057 | 0.533719504 | 0.288291 | 1.851322 | 0.064123199 | 0.16506572 | RNU1-4 | snRNA | 54.27 | 0.417551777 | 0.417551777 | ENSG00000207389 | n.s. |
| ENSG00000160087 | 71.27824226 | -0.506350437 | 0.2656253 | -1.906258 | 0.056616694 | 0.150309422 | UBE2J2 | protein_coding | 56.33 | -0.416733395 | 0.416733395 | ENSG00000160087 | n.s. |
| ENSG00000174917 | 255.3414203 | 0.59105407 | 0.339721 | 1.739822 | 0.08189027 | 0.197709547 | C19orf70 | protein_coding | 61.19 | 0.416085728 | 0.416085728 | ENSG00000174917 | n.s. |
| ENSG00000102910 | 1408.298481 | -0.495544429 | 0.2569565 | -1.928515 | 0.053791079 | 0.144664332 | LONP2 | protein_coding | 39.56 | -0.416078197 | 0.416078197 | ENSG00000102910 | n.s. |
| ENSG00000214517 | 560.880809 | -0.514872528 | 0.2729926 | -1.886031 | 0.059290787 | 0.155697026 | PPME1 | protein_coding | 38.65 | -0.415872674 | 0.415872674 | ENSG00000214517 | n.s. |
| ENSG00000159307 | 72.47561233 | -0.528409734 | 0.2844111 | -1.857908 | 0.063182033 | 0.163327784 | SCUBE1 | protein_coding | 55.48 | -0.41582672 | 0.41582672 | ENSG00000159307 | n.s. |
| ENSG00000120254 | 1362.000733 | -0.495489265 | 0.2572255 | -1.926284 | 0.054068965 | 0.145160471 | MTHFD1L | protein_coding | 43.51 | -0.415295135 | 0.415295135 | ENSG00000120254 | n.s. |
| ENSG00000183864 | 317.256119 | -0.467777383 | 0.2352616 | -1.988329 | 0.046775357 | 0.129551083 | TOB2 | protein_coding | 51.46 | -0.415180004 | 0.415180004 | ENSG00000183864 | n.s. |
| ENSG00000101040 | 68.52713892 | -0.496535908 | 0.258147 | -1.923462 | 0.054422113 | 0.145855454 | ZMYND8 | protein_coding | 46.13 | -0.415142414 | 0.415142414 | ENSG00000101040 | n.s. |
| ENSG00000082516 | 1597.666049 | -0.475090189 | 0.2411488 | -1.970112 | 0.048825527 | 0.134135322 | GEMIN5 | protein_coding | 41.33 | -0.414495688 | 0.414495688 | ENSG00000082516 | n.s. |
| ENSG00000228906 | 95.93158243 | -0.493500769 | 0.2558393 | -1.928949 | 0.053737249 | 0.144632461 | | lincRNA | 45.9 | -0.414409485 | 0.414409485 | ENSG00000228906 | n.s. |
| ENSG00000230269 | 188.6752924 | -0.523012701 | 0.2804092 | -1.865177 | 0.06215666 | 0.161315948 | | lincRNA | 45.16 | -0.414394833 | 0.414394833 | ENSG00000230269 | n.s. |
| ENSG00000101935 | 669.2914726 | -0.538389257 | 0.2935651 | -1.833969 | 0.066658624 | 0.170041778 | AMMECR1 | protein_coding | 38.78 | -0.414260578 | 0.414260578 | ENSG00000101935 | n.s. |
| ENSG00000273951 | 52.58062866 | -0.520327 | 0.278465 | -1.868554 | 0.06168484 | 0.16035064 | | sense_intronic | 33.74 | -0.413623179 | 0.413623179 | ENSG00000273951 | n.s. |
| ENSG00000006377 | 118.0098194 | 0.47256276 | 0.2394307 | 1.973694 | 0.048416605 | 0.133406766 | DLX6 | protein_coding | 49.51 | 0.413408367 | 0.413408367 | ENSG00000006377 | n.s. |
| ENSG00000088038 | 209.4694431 | -0.494855624 | 0.2574669 | -1.922016 | 0.054603683 | 0.146173252 | CNOT3 | protein_coding | 57.91 | -0.413269812 | 0.413269812 | ENSG00000088038 | n.s. |
| ENSG00000007255 | 130.5875491 | 0.564258194 | 0.3168743 | 1.7807 | 0.074961394 | 0.185248047 | TRAPPC6A | protein_coding | 56.86 | 0.41317601 | 0.41317601 | ENSG00000007255 | n.s. |
| ENSG00000112977 | 314.3214057 | -0.527275457 | 0.2846603 | -1.852297 | 0.063983176 | 0.164817347 | DAP | protein_coding | 46.31 | -0.412855144 | 0.412855144 | ENSG00000112977 | n.s. |
| ENSG00000108590 | 277.254532 | -0.48147603 | 0.2468105 | -1.950792 | 0.051081781 | 0.138854282 | MED31 | protein_coding | 40.22 | -0.412837155 | 0.412837155 | ENSG00000108590 | n.s. |
| ENSG00000166250 | 561.2018419 | -0.520153552 | 0.2786139 | -1.866933 | 0.061910907 | 0.160818202 | CLMP | protein_coding | 45.02 | -0.412827563 | 0.412827563 | ENSG00000166250 | n.s. |
| ENSG00000103932 | 166.7855004 | -0.462889144 | 0.2321656 | -1.993789 | 0.04617517 | 0.128282801 | RPAP1 | protein_coding | 50.42 | -0.412819149 | 0.412819149 | ENSG00000103932 | n.s. |
| ENSG00000144655 | 408.9887833 | 0.509327564 | 0.2695954 | 1.88923 | 0.058861062 | 0.1549192 | CSRNP1 | protein_coding | 55.85 | 0.412501722 | 0.412501722 | ENSG00000144655 | n.s. |
| ENSG00000197124 | 282.916715 | -0.501758908 | 0.2636223 | -1.903325 | 0.05699815 | 0.151101532 | ZNF682 | protein_coding | 41.4 | -0.411809157 | 0.411809157 | ENSG00000197124 | n.s. |

| | | | | | | | | | | | | | |
|---|---|---|---|---|---|---|---|---|---|---|---|---|---|
| ENSG00000053900 | 752.5075933 | -0.516529291 | 0.2761797 | -1.870265 | 0.061447015 | 0.159851793 | ANAPC4 | protein_coding | 37.97 | -0.411303229 | 0.411303229 | ENSG00000053900 | n.s. |
| ENSG00000155561 | 5021.157908 | -0.498379077 | 0.2610841 | -1.908884 | 0.056277111 | 0.149645955 | NUP205 | protein_coding | 38.99 | -0.411130353 | 0.411130353 | ENSG00000155561 | n.s. |
| ENSG00000130150 | 110.9164775 | -0.485574992 | 0.2506749 | -1.937071 | 0.052736716 | 0.142384874 | MOSPD2 | protein_coding | 36.08 | -0.411056781 | 0.411056781 | ENSG00000130150 | n.s. |
| ENSG00000154814 | 559.024902 | -0.474111514 | 0.2416452 | -1.962015 | 0.049760751 | 0.13595348 | OXNAD1 | protein_coding | 42.08 | -0.410869624 | 0.410869624 | ENSG00000154814 | n.s. |
| ENSG00000104093 | 3205.120019 | -0.507088234 | 0.2684545 | -1.888917 | 0.058902993 | 0.154970971 | DMXL2 | protein_coding | 36.95 | -0.410614518 | 0.410614518 | ENSG00000104093 | n.s. |
| ENSG00000108733 | 158.9777532 | 0.447096341 | 0.2207211 | 2.025617 | 0.04280406 | 0.120762094 | PEX12 | protein_coding | 42.42 | 0.410465454 | 0.410465454 | ENSG00000108733 | n.s. |
| ENSG00000125834 | 449.9707676 | -0.449560507 | 0.222829 | -2.017514 | 0.043641918 | 0.122628946 | STK35 | protein_coding | 48.13 | -0.409732596 | 0.409732596 | ENSG00000125834 | n.s. |
| ENSG00000160194 | 601.6750016 | -0.518350509 | 0.2783462 | -1.862251 | 0.062567667 | 0.162130956 | NDUFV3 | protein_coding | 46.83 | -0.40956639 | 0.40956639 | ENSG00000160194 | n.s. |
| ENSG00000164961 | 924.9460096 | -0.509428251 | 0.2710317 | -1.879589 | 0.060164084 | 0.157219321 | WASHC5 | protein_coding | 40.15 | -0.409322586 | 0.409322586 | ENSG00000164961 | n.s. |
| ENSG00000239043 | 10.38472554 | -0.65941611 | 0.4086873 | -1.613498 | 0.106636368 | 0.23958337 | SNORD127 | snoRNA | 36.05 | -0.409196269 | 0.409196269 | ENSG00000239043 | n.s. |
| ENSG00000243446 | 8.78025693 | -0.672018524 | 0.4213115 | -1.595063 | 0.110698061 | 0.246171926 | RN7SL284P | misc_RNA | 53.82 | -0.409098989 | 0.409098989 | ENSG00000243446 | n.s. |
| ENSG00000262119 | 76.38321101 | 0.545966741 | 0.3025974 | 1.804268 | 0.071189316 | 0.178203677 | | sense_intronic | 31.17 | 0.408974589 | 0.408974589 | ENSG00000262119 | n.s. |
| ENSG00000154511 | 494.4150107 | -0.567395504 | 0.3215662 | -1.764475 | 0.077652027 | 0.190242685 | FAM69A | protein_coding | 39.43 | -0.40891742 | 0.40891742 | ENSG00000154511 | n.s. |
| ENSG00000078618 | 2788.064669 | -0.510096988 | 0.2717357 | -1.877181 | 0.06049333 | 0.157931489 | NRDC | protein_coding | 39.35 | -0.408858688 | 0.408858688 | ENSG00000078618 | n.s. |
| ENSG00000120963 | 1198.536474 | 0.470754594 | 0.2398146 | 1.962994 | 0.049646867 | 0.135722233 | ZNF706 | protein_coding | 40.04 | 0.408308528 | 0.408308528 | ENSG00000120963 | n.s. |
| ENSG00000262663 | 29.38053784 | -0.542995148 | 0.3002667 | -1.808376 | 0.07054801 | 0.17704397 | | lincRNA | 52.98 | -0.408288293 | 0.408288293 | ENSG00000262663 | n.s. |
| ENSG00000129911 | 232.2520913 | 0.469995755 | 0.239293 | 1.964102 | 0.049518294 | 0.13553486 | KLF16 | protein_coding | 65.43 | 0.407932341 | 0.407932341 | ENSG00000129911 | n.s. |
| ENSG00000265390 | 4.699455786 | 0.778870522 | 0.5340414 | 1.458446 | 0.144717673 | 0.299435899 | MIR4999 | miRNA | 28.57 | 0.407891481 | 0.407891481 | ENSG00000265390 | n.s. |
| ENSG00000213762 | 1424.405368 | -0.506546258 | 0.2691143 | -1.882272 | 0.059799142 | 0.156647876 | ZNF134 | protein_coding | 46.33 | -0.407807976 | 0.407807976 | ENSG00000213762 | n.s. |
| ENSG00000106392 | 199.1117794 | -0.519683761 | 0.2801926 | -1.854737 | 0.063633747 | 0.164251512 | C1GALT1 | protein_coding | 37.33 | -0.407687038 | 0.407687038 | ENSG00000106392 | n.s. |
| ENSG00000198755 | 12121.97963 | 0.546088126 | 0.3033673 | 1.800089 | 0.071846627 | 0.179526308 | RPL10A | protein_coding | 57.02 | 0.407311789 | 0.407311789 | ENSG00000198755 | n.s. |
| ENSG00000271918 | 175.2700028 | 0.452113669 | 0.2255383 | 2.004598 | 0.045006033 | 0.125726123 | | antisense | 52.67 | 0.407162031 | 0.407162031 | ENSG00000271918 | n.s. |
| ENSG00000042445 | 217.6280673 | -0.465435664 | 0.2359842 | -1.972317 | 0.048573391 | 0.133574428 | RETSAT | protein_coding | 51.7 | -0.406919546 | 0.406919546 | ENSG00000042445 | n.s. |
| ENSG00000164190 | 2428.906708 | -0.470080074 | 0.2398326 | -1.960034 | 0.049991783 | 0.136317195 | NIPBL | protein_coding | 35.04 | -0.406830499 | 0.406830499 | ENSG00000164190 | n.s. |
| ENSG00000199477 | 31.18763627 | 0.543092765 | 0.3010031 | 1.804276 | 0.071188016 | 0.178203677 | SNORA31 | snoRNA | 43.08 | 0.406821742 | 0.406821742 | ENSG00000199477 | n.s. |
| ENSG00000003096 | 1087.740371 | -0.531004874 | 0.2902667 | -1.829369 | 0.067344336 | 0.171476808 | KLHL13 | protein_coding | 36.56 | -0.40664067 | 0.40664067 | ENSG00000003096 | n.s. |
| ENSG00000095564 | 3365.913801 | -0.499283654 | 0.2635581 | -1.894397 | 0.058172366 | 0.153454689 | BTAF1 | protein_coding | 37.37 | -0.406426799 | 0.406426799 | ENSG00000095564 | n.s. |
| ENSG00000264653 | 11.57022335 | 0.648435645 | 0.3999694 | 1.621213 | 0.104971929 | 0.23681355 | MIR5194 | miRNA | 45 | 0.405657094 | 0.405657094 | ENSG00000264653 | n.s. |
| ENSG00000213965 | 281.0088664 | -0.473857096 | 0.2432064 | -1.948374 | 0.051370169 | 0.139366107 | NUDT19 | protein_coding | 46.13 | -0.405547199 | 0.405547199 | ENSG00000213965 | n.s. |
| ENSG00000283160 | 17.39206896 | -0.585049795 | 0.3396413 | -1.722552 | 0.084969531 | 0.20297005 | MIR4521 | miRNA | 41.67 | -0.405186791 | 0.405186791 | ENSG00000283160 | n.s. |
| ENSG00000093183 | 1376.783354 | -0.503295966 | 0.2674838 | -1.881594 | 0.059891179 | 0.156770989 | SEC22C | protein_coding | 43.29 | -0.405019528 | 0.405019528 | ENSG00000093183 | n.s. |
| ENSG00000238961 | 35.42775246 | -0.534210039 | 0.2939119 | -1.817585 | 0.069127516 | 0.174548859 | SNORA47 | snoRNA | 60.87 | -0.404975542 | 0.404975542 | ENSG00000238961 | n.s. |

| | | | | | | | | | | | | | |
|---|---|---|---|---|---|---|---|---|---|---|---|---|---|
| ENSG00000070371 | 314.2376795 | -0.520726266 | 0.2821959 | -1.845265 | 0.064999009 | 0.166847566 | CLTCL1 | protein_coding | 46.41 | -0.404958467 | 0.404958467 | ENSG00000070371 | n.s. |
| ENSG00000167526 | 15130.91085 | 0.569824684 | 0.3256821 | 1.749635 | 0.080181368 | 0.194692219 | RPL13 | protein_coding | 55.95 | 0.404946713 | 0.404946713 | ENSG00000167526 | n.s. |
| ENSG00000100650 | 7513.27707 | 0.467834027 | 0.2386984 | 1.959938 | 0.05000304 | 0.136321191 | SRSF5 | protein_coding | 43.68 | 0.404880702 | 0.404880702 | ENSG00000100650 | n.s. |
| ENSG00000266245 | 12.8801316 | -0.609324725 | 0.362842 | -1.679311 | 0.093091358 | 0.216743751 | MIR4644 | miRNA | 59.52 | -0.404624164 | 0.404624164 | ENSG00000266245 | n.s. |
| ENSG00000108788 | 1108.999512 | -0.499176216 | 0.2642479 | -1.889045 | 0.058885815 | 0.154955057 | MLX | protein_coding | 52.95 | -0.404230027 | 0.404230027 | ENSG00000108788 | n.s. |
| ENSG00000267422 | 12.4865975 | 0.617425723 | 0.3712161 | 1.663252 | 0.096262094 | 0.222043926 | | transcribed_processed_pseudog | 43.57 | 0.403525436 | 0.403525436 | ENSG00000267422 | n.s. |
| ENSG00000173542 | 960.9746205 | -0.503875777 | 0.26863 | -1.875724 | 0.060693188 | 0.158304845 | MOB1B | protein_coding | 40.16 | -0.403355479 | 0.403355479 | ENSG00000173542 | n.s. |
| ENSG00000155906 | 1036.452622 | -0.509041382 | 0.2730393 | -1.864352 | 0.062272237 | 0.161545809 | RMND1 | protein_coding | 42.3 | -0.403010254 | 0.403010254 | ENSG00000155906 | n.s. |
| ENSG00000134324 | 1759.24342 | -0.482522915 | 0.2510296 | -1.922175 | 0.054583724 | 0.146147921 | LPIN1 | protein_coding | 46.76 | -0.40300669 | 0.40300669 | ENSG00000134324 | n.s. |
| ENSG00000147364 | 688.9283719 | -0.496997001 | 0.2629695 | -1.889941 | 0.058765815 | 0.154715492 | FBXO25 | protein_coding | 41.96 | -0.402799269 | 0.402799269 | ENSG00000147364 | n.s. |
| ENSG00000183474 | 187.9416807 | -0.506647835 | 0.2712204 | -1.86803 | 0.061757887 | 0.160480602 | GTF2H2C | protein_coding | 38.56 | -0.402570948 | 0.402570948 | ENSG00000183474 | n.s. |
| ENSG00000162378 | 1847.177346 | -0.502854356 | 0.2681576 | -1.875219 | 0.060762567 | 0.15845612 | ZYG11B | protein_coding | 41.29 | -0.402329236 | 0.402329236 | ENSG00000162378 | n.s. |
| ENSG00000234329 | 38.15428123 | 0.514060234 | 0.277791 | 1.850529 | 0.064237365 | 0.165227569 | | processed_pseudogene | 50.13 | 0.401952686 | 0.401952686 | ENSG00000234329 | n.s. |
| ENSG00000181894 | 334.5662267 | -0.502893818 | 0.2684091 | -1.87361 | 0.060984265 | 0.158915202 | ZNF329 | protein_coding | 46.41 | -0.40172896 | 0.40172896 | ENSG00000181894 | n.s. |
| ENSG00000150787 | 548.0615827 | 0.513294059 | 0.2772104 | 1.851641 | 0.064077406 | 0.164999024 | PTS | protein_coding | 41.9 | 0.401662161 | 0.401662161 | ENSG00000150787 | n.s. |
| ENSG00000103174 | 76.51712675 | 0.510642235 | 0.2749498 | 1.85722 | 0.063279878 | 0.163489654 | NAGPA | protein_coding | 56.14 | 0.401625084 | 0.401625084 | ENSG00000103174 | n.s. |
| ENSG00000030066 | 2271.672082 | -0.518274646 | 0.2814659 | -1.841341 | 0.065571662 | 0.167945962 | NUP160 | protein_coding | 41.86 | -0.401574969 | 0.401574969 | ENSG00000030066 | n.s. |
| ENSG00000172209 | 481.9752519 | 0.442082187 | 0.219555 | 2.013537 | 0.044058168 | 0.123621255 | GPR22 | protein_coding | 30.79 | 0.401369447 | 0.401369447 | ENSG00000172209 | n.s. |
| ENSG00000158258 | 1683.346352 | -0.476721824 | 0.2468648 | -1.931105 | 0.053470071 | 0.144080471 | CLSTN2 | protein_coding | 42.71 | -0.401111302 | 0.401111302 | ENSG00000158258 | n.s. |
| ENSG00000116120 | 1689.571957 | -0.513204253 | 0.2774359 | -1.849812 | 0.064340646 | 0.165401554 | FARSB | protein_coding | 38.05 | -0.401048809 | 0.401048809 | ENSG00000116120 | n.s. |
| ENSG00000125931 | 147.8034869 | 0.485238395 | 0.2539928 | 1.910442 | 0.056076346 | 0.14916908 | CITED1 | protein_coding | 56.77 | 0.400962768 | 0.400962768 | ENSG00000125931 | n.s. |
| ENSG00000150867 | 415.8741108 | -0.475954188 | 0.2463765 | -1.931817 | 0.05338215 | 0.143871404 | PIP4K2A | protein_coding | 41.89 | -0.400765573 | 0.400765573 | ENSG00000150867 | n.s. |
| ENSG00000159167 | 1343.518971 | -0.429599303 | 0.2101776 | -2.043982 | 0.040955314 | 0.116752833 | STC1 | protein_coding | 43.81 | -0.400701263 | 0.400701263 | ENSG00000159167 | n.s. |
| ENSG00000006451 | 511.694888 | -0.510564331 | 0.2753751 | -1.854069 | 0.063729305 | 0.164437196 | RALA | protein_coding | 39.69 | -0.400282403 | 0.400282403 | ENSG00000006451 | n.s. |
| ENSG00000151948 | 192.9850753 | -0.518280507 | 0.282004 | -1.837848 | 0.066084765 | 0.168920041 | GLT1D1 | protein_coding | 45.85 | -0.40027779 | 0.40027779 | ENSG00000151948 | n.s. |
| ENSG00000253948 | 20.7482865 | -0.581769246 | 0.3392154 | -1.715044 | 0.086337233 | 0.205176131 | | lincRNA | 38.08 | -0.400183452 | 0.400183452 | ENSG00000253948 | n.s. |
| ENSG00000058272 | 3215.477761 | -0.494407039 | 0.2618518 | -1.888118 | 0.059010169 | 0.155164982 | PPP1R12A | protein_coding | 36.07 | -0.400077283 | 0.400077283 | ENSG00000058272 | n.s. |
| ENSG00000132664 | 693.7472871 | -0.491449378 | 0.2594646 | -1.894091 | 0.058212982 | 0.153532741 | POLR3F | protein_coding | 40.58 | -0.39994101 | 0.39994101 | ENSG00000132664 | n.s. |
| ENSG00000272407 | 15.95736281 | -0.601374756 | 0.35802 | -1.679724 | 0.093011056 | 0.216654815 | | transcribed_processed_pseudog | 44.18 | -0.399452149 | 0.399452149 | ENSG00000272407 | n.s. |
| ENSG00000248121 | 52.0425476 | 0.526712867 | 0.2899178 | 1.816766 | 0.069252917 | 0.17473928 | SMURF2P1 | transcribed_unprocessed_pseud | 37.46 | 0.39904265 | 0.39904265 | ENSG00000248121 | n.s. |
| ENSG00000153443 | 254.7085385 | 0.470477354 | 0.242659 | 1.938841 | 0.052520661 | 0.141961786 | UBALD1 | protein_coding | 64.7 | 0.398884131 | 0.398884131 | ENSG00000153443 | n.s. |
| ENSG00000100320 | 4651.678676 | -0.490759741 | 0.2593259 | -1.892444 | 0.058431805 | 0.153964046 | RBFOX2 | protein_coding | 39.31 | -0.398781886 | 0.398781886 | ENSG00000100320 | n.s. |

| | | | | | | | | | | | | | |
|---|---|---|---|---|---|---|---|---|---|---|---|---|---|
| ENSG00000040933 | 1228.695925 | -0.503293475 | 0.2698408 | -1.865149 | 0.062160458 | 0.161315948 | INPP4A | protein_coding | 47.33 | -0.398770842 | 0.398770842 | ENSG00000040933 | n.s. |
| ENSG00000169554 | 355.8123644 | 0.442193153 | 0.2204763 | 2.005627 | 0.044896046 | 0.125451081 | ZEB2 | protein_coding | 38.76 | 0.398648444 | 0.398648444 | ENSG00000169554 | n.s. |
| ENSG00000171466 | 1632.386424 | -0.492558823 | 0.2608727 | -1.888119 | 0.059009955 | 0.155164982 | ZNF562 | protein_coding | 43.08 | -0.398581695 | 0.398581695 | ENSG00000171466 | n.s. |
| ENSG00000034510 | 15652.37094 | 0.577180076 | 0.3358014 | 1.718814 | 0.085648321 | 0.204017378 | TMSB10 | protein_coding | 63.37 | 0.39844636 | 0.39844636 | ENSG00000034510 | n.s. |
| ENSG00000283871 | 9.333522936 | 0.647868923 | 0.4037383 | 1.604675 | 0.108565275 | 0.2429761 | MIR130B | miRNA | 57.32 | 0.398074276 | 0.398074276 | ENSG00000283871 | n.s. |
| ENSG00000160953 | 629.8500729 | -0.505259261 | 0.2717799 | -1.859075 | 0.063016468 | 0.163051159 | MUM1 | protein_coding | 52.99 | -0.397980649 | 0.397980649 | ENSG00000160953 | n.s. |
| ENSG00000103274 | 370.51378 | -0.525787584 | 0.2896929 | -1.814982 | 0.069526609 | 0.1751129 | NUBP1 | protein_coding | 50.62 | -0.397853926 | 0.397853926 | ENSG00000103274 | n.s. |
| ENSG00000125652 | 144.0345014 | 0.563687086 | 0.3235086 | 1.742417 | 0.081435431 | 0.196878886 | ALKBH7 | protein_coding | 64.35 | 0.397850828 | 0.397850828 | ENSG00000125652 | n.s. |
| ENSG00000153936 | 1773.365016 | -0.522539545 | 0.286673 | -1.822772 | 0.068337959 | 0.173254601 | HS2ST1 | protein_coding | 36.82 | -0.397817311 | 0.397817311 | ENSG00000153936 | n.s. |
| ENSG00000139405 | 351.6640098 | -0.478835122 | 0.2499623 | -1.91563 | 0.055412269 | 0.147713005 | RITA1 | protein_coding | 51.56 | -0.397711482 | 0.397711482 | ENSG00000139405 | n.s. |
| ENSG00000185344 | 1317.030453 | -0.498895592 | 0.2666543 | -1.870945 | 0.061352687 | 0.159636228 | ATP6V0A2 | protein_coding | 44.41 | -0.397554203 | 0.397554203 | ENSG00000185344 | n.s. |
| ENSG00000115758 | 10821.20618 | 0.524741165 | 0.2889503 | 1.816026 | 0.069366387 | 0.174921447 | ODC1 | protein_coding | 46.16 | 0.397311415 | 0.397311415 | ENSG00000115758 | n.s. |
| ENSG00000108439 | 392.542061 | -0.456348915 | 0.2320282 | -1.966782 | 0.049208363 | 0.134894277 | PNPO | protein_coding | 50.57 | -0.397026511 | 0.397026511 | ENSG00000108439 | n.s. |
| ENSG00000143702 | 6466.654943 | -0.491933559 | 0.2614193 | -1.88178 | 0.059865911 | 0.156734314 | CEP170 | protein_coding | 36.28 | -0.395925796 | 0.395925796 | ENSG00000143702 | n.s. |
| ENSG00000171681 | 3117.619038 | -0.485792471 | 0.2565678 | -1.893427 | 0.058301079 | 0.153706859 | ATF7IP | protein_coding | 36.1 | -0.395098297 | 0.395098297 | ENSG00000171681 | n.s. |
| ENSG00000089818 | 1114.669569 | 0.495096672 | 0.2644289 | 1.872325 | 0.061161713 | 0.159317966 | NECAP1 | protein_coding | 41.14 | 0.394956065 | 0.394956065 | ENSG00000089818 | n.s. |
| ENSG00000020922 | 1010.132942 | -0.491480296 | 0.2615142 | -1.879364 | 0.060194806 | 0.157240579 | MRE11 | protein_coding | 37.31 | -0.394872651 | 0.394872651 | ENSG00000020922 | n.s. |
| ENSG00000231871 | 54.92375273 | 0.500917024 | 0.2693516 | 1.859715 | 0.062925919 | 0.162847133 | IPO9-AS1 | antisense | 46.27 | 0.394832758 | 0.394832758 | ENSG00000231871 | n.s. |
| ENSG00000123415 | 402.440734 | -0.489935585 | 0.2601506 | -1.883277 | 0.059662874 | 0.156379177 | SMUG1 | protein_coding | 50.76 | -0.39480042 | 0.39480042 | ENSG00000123415 | n.s. |
| ENSG00000198794 | 1023.420117 | -0.475929446 | 0.2487351 | -1.913399 | 0.055696972 | 0.148358321 | SCAMP5 | protein_coding | 48.31 | -0.394397064 | 0.394397064 | ENSG00000198794 | n.s. |
| ENSG00000117523 | 7166.761735 | -0.482477831 | 0.2542783 | -1.89744 | 0.057769907 | 0.152662612 | PRRC2C | protein_coding | 38.3 | -0.393830881 | 0.393830881 | ENSG00000117523 | n.s. |
| ENSG00000011422 | 55.62184972 | -0.499254544 | 0.2685781 | -1.85888 | 0.063044151 | 0.163072791 | PLAUR | protein_coding | 49.94 | -0.393222113 | 0.393222113 | ENSG00000011422 | n.s. |
| ENSG00000145555 | 527.164936 | -0.44078752 | 0.2210751 | -1.993836 | 0.046170006 | 0.128282801 | MYO10 | protein_coding | 44.1 | -0.393108224 | 0.393108224 | ENSG00000145555 | n.s. |
| ENSG00000224043 | 31.08077342 | -0.602908934 | 0.3631444 | -1.660246 | 0.096865011 | 0.223151627 | CCNT2-AS1 | antisense | 41.05 | -0.392734844 | 0.392734844 | ENSG00000224043 | n.s. |
| ENSG00000073417 | 92.96602542 | -0.488428868 | 0.2597511 | -1.880373 | 0.060057271 | 0.157087616 | PDE8A | protein_coding | 41.8 | -0.392627478 | 0.392627478 | ENSG00000073417 | n.s. |
| ENSG00000198908 | 302.3894699 | -0.498138045 | 0.2680005 | -1.858721 | 0.063066755 | 0.163072791 | BHLHB9 | protein_coding | 42.71 | -0.392342738 | 0.392342738 | ENSG00000198908 | n.s. |
| ENSG00000137502 | 1270.038861 | -0.500773089 | 0.2702403 | -1.853066 | 0.06387291 | 0.164655161 | RAB30 | protein_coding | 41.21 | -0.392317982 | 0.392317982 | ENSG00000137502 | n.s. |
| ENSG00000185596 | 118.396416 | -0.48062311 | 0.2533563 | -1.897024 | 0.057824738 | 0.152740244 | WASH3P | transcribed_unprocessed_pseud | 54.04 | -0.392210816 | 0.392210816 | ENSG00000185596 | n.s. |
| ENSG00000259366 | 42.66027834 | -0.544854435 | 0.3090892 | -1.762774 | 0.07793863 | 0.190676426 | | sense_intronic | 40.61 | -0.39213337 | 0.39213337 | ENSG00000259366 | n.s. |
| ENSG00000135108 | 1919.951644 | -0.474227943 | 0.2481576 | -1.910995 | 0.056005197 | 0.149036766 | FBXO21 | protein_coding | 44.38 | -0.392047362 | 0.392047362 | ENSG00000135108 | n.s. |
| ENSG00000164244 | 1324.912378 | -0.51318952 | 0.2810288 | -1.82611 | 0.067833642 | 0.172407412 | PRRC1 | protein_coding | 36.29 | -0.391791495 | 0.391791495 | ENSG00000164244 | n.s. |
| ENSG00000225945 | 15.70121039 | 0.592338529 | 0.3535995 | 1.675168 | 0.093901269 | 0.218079627 | | antisense | 43.87 | 0.391763762 | 0.391763762 | ENSG00000225945 | n.s. |

| | | | | | | | | | | | | | |
|---|---|---|---|---|---|---|---|---|---|---|---|---|---|
| ENSG00000171103 | 723.3077018 | -0.496368109 | 0.2667439 | -1.860841 | 0.062766582 | 0.162525408 | TRMT61B | protein_coding | 41.37 | -0.39167352 | 0.39167352 | ENSG00000171103 | n.s. |
| ENSG00000144566 | 753.2762364 | -0.499861978 | 0.2697645 | -1.852957 | 0.063888518 | 0.164664908 | RAB5A | protein_coding | 39.8 | -0.391591346 | 0.391591346 | ENSG00000144566 | n.s. |
| ENSG00000180530 | 3750.797568 | -0.503683313 | 0.2730402 | -1.844722 | 0.065077948 | 0.166988623 | NRIP1 | protein_coding | 36.59 | -0.391519647 | 0.391519647 | ENSG00000180530 | n.s. |
| ENSG00000125966 | 487.5511934 | -0.50483587 | 0.2740133 | -1.842377 | 0.06542001 | 0.167711803 | MMP24 | protein_coding | 49.16 | -0.391468095 | 0.391468095 | ENSG00000125966 | n.s. |
| ENSG00000168434 | 187.8086916 | -0.495523781 | 0.2661462 | -1.861848 | 0.062624526 | 0.162248097 | COG7 | protein_coding | 45.97 | -0.391374785 | 0.391374785 | ENSG00000168434 | n.s. |
| ENSG00000140299 | 2150.930266 | -0.506657781 | 0.2757101 | -1.837647 | 0.066114461 | 0.168932214 | BNIP2 | protein_coding | 38.55 | -0.391285484 | 0.391285484 | ENSG00000140299 | n.s. |
| ENSG00000115750 | 1013.870787 | -0.491924179 | 0.2633106 | -1.868228 | 0.06173035 | 0.160438989 | TAF1B | protein_coding | 39.53 | -0.390927267 | 0.390927267 | ENSG00000115750 | n.s. |
| ENSG00000180901 | 781.0756763 | -0.46802161 | 0.2436565 | -1.920825 | 0.054753738 | 0.146546768 | KCTD2 | protein_coding | 49.14 | -0.390341141 | 0.390341141 | ENSG00000180901 | n.s. |
| ENSG00000020181 | 518.1766638 | -0.482761514 | 0.2558441 | -1.886936 | 0.059168925 | 0.155464983 | ADGRA2 | protein_coding | 55.25 | -0.390248678 | 0.390248678 | ENSG00000020181 | n.s. |
| ENSG00000199961 | 52.56924112 | 0.496314664 | 0.2673558 | 1.856383 | 0.063399044 | 0.163722001 | SNORD1B | snoRNA | 40.48 | 0.390050198 | 0.390050198 | ENSG00000199961 | n.s. |
| ENSG00000092208 | 315.0973813 | -0.545785893 | 0.3108813 | -1.755609 | 0.079155225 | 0.19300844 | GEMIN2 | protein_coding | 39.65 | -0.389922377 | 0.389922377 | ENSG00000092208 | n.s. |
| ENSG00000237953 | 25.54917394 | 0.567816927 | 0.3314817 | 1.712966 | 0.086718748 | 0.205862768 | | processed_pseudogene | 40.45 | 0.38976214 | 0.38976214 | ENSG00000237953 | n.s. |
| ENSG00000163125 | 1204.315971 | -0.436795903 | 0.2190617 | -1.99394 | 0.046158621 | 0.128282801 | RPRD2 | protein_coding | 41.76 | -0.389548374 | 0.389548374 | ENSG00000163125 | n.s. |
| ENSG00000144504 | 145.25623 | -0.444560206 | 0.2250574 | -1.975319 | 0.048231965 | 0.132976961 | ANKMY1 | protein_coding | 49.38 | -0.389534142 | 0.389534142 | ENSG00000144504 | n.s. |
| ENSG00000269918 | 177.9955016 | -0.542440808 | 0.3081601 | -1.760257 | 0.078364314 | 0.191486703 | | sense_intronic | 41.16 | -0.389397306 | 0.389397306 | ENSG00000269918 | n.s. |
| ENSG00000105640 | 8395.446259 | 0.50983477 | 0.2791802 | 1.826185 | 0.067822366 | 0.172407412 | RPL18A | protein_coding | 58.32 | 0.389230331 | 0.389230331 | ENSG00000105640 | n.s. |
| ENSG00000007923 | 597.8575108 | -0.508422094 | 0.2780034 | -1.828834 | 0.06742451 | 0.171618181 | DNAJC11 | protein_coding | 46.6 | -0.389164932 | 0.389164932 | ENSG00000007923 | n.s. |
| ENSG00000123689 | 27.9557749 | 0.574398289 | 0.3379511 | 1.699649 | 0.089196977 | 0.210337771 | G0S2 | protein_coding | 57.59 | 0.388915164 | 0.388915164 | ENSG00000123689 | n.s. |
| ENSG00000204366 | 115.6852784 | -0.503537375 | 0.2739827 | -1.837844 | 0.066085432 | 0.168920041 | ZBTB12 | protein_coding | 64.33 | -0.388891392 | 0.388891392 | ENSG00000204366 | n.s. |
| ENSG00000237753 | 66.89190256 | 0.478498751 | 0.2528343 | 1.892539 | 0.058419253 | 0.153960109 | | lincRNA | 48.77 | 0.388824157 | 0.388824157 | ENSG00000237753 | n.s. |
| ENSG00000023228 | 3431.834651 | -0.510646191 | 0.2802727 | -1.821962 | 0.068460798 | 0.173463092 | NDUFS1 | protein_coding | 40.81 | -0.388496005 | 0.388496005 | ENSG00000023228 | n.s. |
| ENSG00000149972 | 22.38354383 | 0.601089309 | 0.3638224 | 1.65215 | 0.09850392 | 0.225793559 | CNTN5 | protein_coding | 35.04 | 0.388477079 | 0.388477079 | ENSG00000149972 | n.s. |
| ENSG00000273343 | 37.42671488 | 0.513114774 | 0.2826191 | 1.81557 | 0.069436379 | 0.174979676 | | sense_intronic | 45.32 | 0.388434243 | 0.388434243 | ENSG00000273343 | n.s. |
| ENSG00000015171 | 1549.794963 | -0.514512529 | 0.2840089 | -1.811607 | 0.070046897 | 0.176072024 | ZMYND11 | protein_coding | 37.8 | -0.388101762 | 0.388101762 | ENSG00000015171 | n.s. |
| ENSG00000093167 | 427.0474628 | 0.457714614 | 0.2361202 | 1.938481 | 0.052564532 | 0.142025224 | LRRFIP2 | protein_coding | 38.6 | 0.387974705 | 0.387974705 | ENSG00000093167 | n.s. |
| ENSG00000279513 | 62.48357353 | -0.552370917 | 0.3180158 | -1.736929 | 0.082399668 | 0.198581339 | | TEC | 33.18 | -0.387798391 | 0.387798391 | ENSG00000279513 | n.s. |
| ENSG00000168803 | 516.0395622 | -0.509323352 | 0.279459 | -1.822533 | 0.068374125 | 0.173306561 | ADAL | protein_coding | 39.85 | -0.387689294 | 0.387689294 | ENSG00000168803 | n.s. |
| ENSG00000130827 | 381.6640777 | 0.447335015 | 0.2279889 | 1.962092 | 0.049751821 | 0.13595348 | PLXNA3 | protein_coding | 60.44 | 0.38766485 | 0.38766485 | ENSG00000130827 | n.s. |
| ENSG00000115548 | 2212.673419 | 0.473353846 | 0.249001 | 1.901012 | 0.057300433 | 0.151729543 | KDM3A | protein_coding | 40.15 | 0.387643595 | 0.387643595 | ENSG00000115548 | n.s. |
| ENSG00000130254 | 1023.597423 | 0.444701155 | 0.2258393 | 1.969105 | 0.048941069 | 0.134373202 | SAFB2 | protein_coding | 49.34 | 0.387640364 | 0.387640364 | ENSG00000130254 | n.s. |
| ENSG00000146592 | 133.8404234 | 0.511427795 | 0.2814188 | 1.817319 | 0.069168234 | 0.174620003 | CREB5 | protein_coding | 40.7 | 0.387614199 | 0.387614199 | ENSG00000146592 | n.s. |
| ENSG00000214223 | 70.38727997 | -0.531739223 | 0.2993635 | -1.776233 | 0.075694571 | 0.18668755 | HNRNPA1P10 | processed_pseudogene | 47.51 | -0.387576555 | 0.387576555 | ENSG00000214223 | n.s. |

| | | | | | | | | | | | | | |
|---|---|---|---|---|---|---|---|---|---|---|---|---|---|
| ENSG00000219438 | 1146.578828 | -0.529133504 | 0.2972493 | -1.7801 | 0.075059582 | 0.185424869 | FAM19A5 | protein_coding | 54.37 | -0.38723684 | 0.38723684 | ENSG00000219438 | n.s. |
| ENSG00000134874 | 1497.698319 | -0.472655933 | 0.2486218 | -1.901104 | 0.057288395 | 0.151729543 | DZIP1 | protein_coding | 41.42 | -0.387072053 | 0.387072053 | ENSG00000134874 | n.s. |
| ENSG00000168246 | 881.9583618 | -0.521225182 | 0.2902317 | -1.795894 | 0.072511441 | 0.180927742 | UBTD2 | protein_coding | 41.41 | -0.387007007 | 0.387007007 | ENSG00000168246 | n.s. |
| ENSG00000021762 | 103.6906077 | 0.462644954 | 0.2405538 | 1.92325 | 0.054448709 | 0.145898649 | OSBPL5 | protein_coding | 57 | 0.386747462 | 0.386747462 | ENSG00000021762 | n.s. |
| ENSG00000183049 | 433.1339308 | -0.474222906 | 0.250238 | -1.895088 | 0.058080799 | 0.153329346 | CAMK1D | protein_coding | 45.61 | -0.386195145 | 0.386195145 | ENSG00000183049 | n.s. |
| ENSG00000162623 | 1948.273068 | -0.461548739 | 0.2400501 | -1.922718 | 0.054515425 | 0.146021211 | TYW3 | protein_coding | 40.26 | -0.385662766 | 0.385662766 | ENSG00000162623 | n.s. |
| ENSG00000110700 | 10176.88015 | 0.537178066 | 0.3052014 | 1.760077 | 0.078394692 | 0.191522951 | RPS13 | protein_coding | 46.54 | 0.385575229 | 0.385575229 | ENSG00000110700 | n.s. |
| ENSG00000237651 | 333.5538573 | -0.534462049 | 0.302772 | -1.76523 | 0.077525178 | 0.190032228 | C2orf74 | protein_coding | 44.26 | -0.38543946 | 0.38543946 | ENSG00000237651 | n.s. |
| ENSG00000168264 | 4556.080014 | 0.401328349 | 0.1932004 | 2.077264 | 0.037777173 | 0.109554588 | IRF2BP2 | protein_coding | 48.01 | 0.385423478 | 0.385423478 | ENSG00000168264 | n.s. |
| ENSG00000173638 | 264.9776034 | -0.470733161 | 0.2477168 | -1.900288 | 0.057395394 | 0.151923207 | SLC19A1 | protein_coding | 59.34 | -0.385236665 | 0.385236665 | ENSG00000173638 | n.s. |
| ENSG00000154781 | 644.3451633 | -0.496370904 | 0.2693371 | -1.842936 | 0.065338405 | 0.167595175 | CCDC174 | protein_coding | 42.28 | -0.385054015 | 0.385054015 | ENSG00000154781 | n.s. |
| ENSG00000181026 | 2967.790544 | -0.413924602 | 0.2030092 | -2.038945 | 0.041455552 | 0.117672915 | AEN | protein_coding | 51.46 | -0.384669854 | 0.384669854 | ENSG00000181026 | n.s. |
| ENSG00000274561 | 83.64638954 | -0.511020516 | 0.2824413 | -1.809298 | 0.070404683 | 0.17681176 | | lincRNA | 33.41 | -0.384537242 | 0.384537242 | ENSG00000274561 | n.s. |
| ENSG00000226221 | 169.6476673 | 0.565583384 | 0.3319239 | 1.703955 | 0.088389541 | 0.209044942 | | processed_pseudogene | 43.84 | 0.38446115 | 0.38446115 | ENSG00000226221 | n.s. |
| ENSG00000139880 | 864.5678288 | 0.457573135 | 0.2372471 | 1.928678 | 0.053770871 | 0.144639071 | CDH24 | protein_coding | 57.26 | 0.384230739 | 0.384230739 | ENSG00000139880 | n.s. |
| ENSG00000005801 | 3170.155546 | -0.489709947 | 0.264063 | -1.854519 | 0.063664918 | 0.16430151 | ZNF195 | protein_coding | 41.43 | -0.384108132 | 0.384108132 | ENSG00000005801 | n.s. |
| ENSG00000144713 | 18368.86871 | 0.537901333 | 0.3066173 | 1.754309 | 0.079377659 | 0.193305557 | RPL32 | protein_coding | 47.4 | 0.383930121 | 0.383930121 | ENSG00000144713 | n.s. |
| ENSG00000186174 | 28.55924052 | 0.52339834 | 0.2937278 | 1.781916 | 0.074762923 | 0.184926792 | BCL9L | protein_coding | 59.25 | 0.383651069 | 0.383651069 | ENSG00000186174 | n.s. |
| ENSG00000262248 | 21.25813882 | -0.567786622 | 0.3346093 | -1.696865 | 0.089722279 | 0.211085381 | | processed_pseudogene | 49.54 | -0.383563621 | 0.383563621 | ENSG00000262248 | n.s. |
| ENSG00000134452 | 719.1426279 | -0.508873403 | 0.281036 | -1.810705 | 0.070186502 | 0.176391095 | FBXO18 | protein_coding | 45.82 | -0.383447988 | 0.383447988 | ENSG00000134452 | n.s. |
| ENSG00000243678 | 924.6270516 | 0.438393536 | 0.2221968 | 1.972997 | 0.04849595 | 0.133493301 | NME2 | protein_coding | 52.1 | 0.383392914 | 0.383392914 | ENSG00000243678 | n.s. |
| ENSG00000155329 | 465.4126505 | -0.519053633 | 0.289971 | -1.790019 | 0.073450826 | 0.182584512 | ZCCHC10 | protein_coding | 42.73 | -0.383339827 | 0.383339827 | ENSG00000155329 | n.s. |
| ENSG00000121005 | 383.8197172 | -0.504388982 | 0.2771146 | -1.820146 | 0.068736765 | 0.173845953 | CRISPLD1 | protein_coding | 35.22 | -0.383252611 | 0.383252611 | ENSG00000121005 | n.s. |
| ENSG00000186687 | 630.3363233 | -0.51483472 | 0.2864162 | -1.797506 | 0.072255402 | 0.180385864 | LYRM7 | protein_coding | 38.11 | -0.382932777 | 0.382932777 | ENSG00000186687 | n.s. |
| ENSG00000152457 | 330.4632108 | -0.475095428 | 0.2522232 | -1.883631 | 0.059614923 | 0.156312349 | DCLRE1C | protein_coding | 41.86 | -0.382930103 | 0.382930103 | ENSG00000152457 | n.s. |
| ENSG00000173575 | 6487.180563 | -0.433035787 | 0.2182427 | -1.984193 | 0.04723428 | 0.130588341 | CHD2 | protein_coding | 39.49 | -0.382845031 | 0.382845031 | ENSG00000173575 | n.s. |
| ENSG00000263652 | 2.8134631 | -0.913815692 | 0.7203982 | -1.268487 | 0.204624099 | 0.381594071 | MIR548AX | miRNA | 39.73 | -0.382339007 | 0.382339007 | ENSG00000263652 | n.s. |
| ENSG00000163618 | 209.571095 | -0.496278585 | 0.2705137 | -1.834578 | 0.066568197 | 0.169861118 | CADPS | protein_coding | 40 | -0.382087871 | 0.382087871 | ENSG00000163618 | n.s. |
| ENSG00000274051 | 19.65009399 | -0.578536941 | 0.3459177 | -1.67247 | 0.094431746 | 0.218952071 | | misc_RNA | 60.22 | -0.381632438 | 0.381632438 | ENSG00000274051 | n.s. |
| ENSG00000208009 | 10.34452887 | -0.631267403 | 0.3977734 | -1.587003 | 0.112512023 | 0.249086483 | MIR130A | miRNA | 50.56 | -0.381064466 | 0.381064466 | ENSG00000208009 | n.s. |
| ENSG00000124279 | 451.94326 | -0.47362804 | 0.2518579 | -1.880537 | 0.060034911 | 0.157058638 | FASTKD3 | protein_coding | 37.73 | -0.380767661 | 0.380767661 | ENSG00000124279 | n.s. |
| ENSG00000276131 | 25.59190226 | 0.533023566 | 0.303667 | 1.75529 | 0.079209798 | 0.193073882 | | antisense | 56.64 | 0.380726192 | 0.380726192 | ENSG00000276131 | n.s. |

| | | | | | | | | | | | | | |
|---|---|---|---|---|---|---|---|---|---|---|---|---|---|
| ENSG00000245857 | 49.72751999 | -0.505858975 | 0.2795635 | -1.80946 | 0.070379601 | 0.176780658 | | lincRNA | 46.12 | -0.380691888 | 0.380691888 | ENSG00000245857 | n.s. |
| ENSG00000121058 | 1874.305818 | -0.500640055 | 0.2749369 | -1.820927 | 0.068617907 | 0.173671535 | COIL | protein_coding | 43 | -0.380622294 | 0.380622294 | ENSG00000121058 | n.s. |
| ENSG00000129933 | 723.825207 | -0.481820246 | 0.2590476 | -1.859968 | 0.062890098 | 0.162784686 | MAU2 | protein_coding | 54.38 | -0.379860554 | 0.379860554 | ENSG00000129933 | n.s. |
| ENSG00000099889 | 99.23507661 | -0.512635478 | 0.2859195 | -1.792936 | 0.072983086 | 0.18174629 | ARVCF | protein_coding | 58.03 | -0.37962423 | 0.37962423 | ENSG00000099889 | n.s. |
| ENSG00000094804 | 2725.058038 | -0.455668599 | 0.2374676 | -1.918866 | 0.055001294 | 0.146945877 | CDC6 | protein_coding | 43.75 | -0.379500219 | 0.379500219 | ENSG00000094804 | n.s. |
| ENSG00000164897 | 148.9700653 | 0.479058793 | 0.256911 | 1.864687 | 0.062225229 | 0.161453944 | TMUB1 | protein_coding | 67.61 | 0.379391254 | 0.379391254 | ENSG00000164897 | n.s. |
| ENSG00000163281 | 1280.816383 | -0.474409826 | 0.2531164 | -1.874275 | 0.060892489 | 0.158705751 | GNPDA2 | protein_coding | 33.84 | -0.379246695 | 0.379246695 | ENSG00000163281 | n.s. |
| ENSG00000167716 | 384.106922 | -0.421236159 | 0.2102251 | -2.003738 | 0.04509811 | 0.125932809 | WDR81 | protein_coding | 57.66 | -0.379054038 | 0.379054038 | ENSG00000167716 | n.s. |
| ENSG00000156853 | 411.2352024 | -0.432988969 | 0.2193988 | -1.973525 | 0.048435784 | 0.133406808 | ZNF689 | protein_coding | 48.18 | -0.378788279 | 0.378788279 | ENSG00000156853 | n.s. |
| ENSG00000265681 | 2102.722684 | 0.436526225 | 0.2223231 | 1.963476 | 0.049590841 | 0.135622328 | RPL17 | protein_coding | 42.91 | 0.378760188 | 0.378760188 | ENSG00000265681 | n.s. |
| ENSG00000204104 | 330.8437742 | -0.467087535 | 0.2471505 | -1.889891 | 0.058772552 | 0.154715492 | TRAF3IP1 | protein_coding | 44.49 | -0.378558658 | 0.378558658 | ENSG00000204104 | n.s. |
| ENSG00000173327 | 391.7175838 | -0.437418764 | 0.2231951 | -1.959805 | 0.050018615 | 0.136336957 | MAP3K11 | protein_coding | 57.13 | -0.378536251 | 0.378536251 | ENSG00000173327 | n.s. |
| ENSG00000104290 | 1580.081492 | -0.495304269 | 0.2715618 | -1.82391 | 0.068165714 | 0.172948214 | FZD3 | protein_coding | 36.28 | -0.377463419 | 0.377463419 | ENSG00000104290 | n.s. |
| ENSG00000007168 | 2859.888861 | -0.495205644 | 0.2714938 | -1.824004 | 0.068151537 | 0.172948214 | PAFAH1B1 | protein_coding | 42.4 | -0.377388258 | 0.377388258 | ENSG00000007168 | n.s. |
| ENSG00000088387 | 174.9697413 | -0.509574416 | 0.2844311 | -1.791557 | 0.073204016 | 0.182035991 | DOCK9 | protein_coding | 41.78 | -0.377004931 | 0.377004931 | ENSG00000088387 | n.s. |
| ENSG00000104313 | 2157.294342 | -0.450777878 | 0.2343268 | -1.923714 | 0.054390407 | 0.145826619 | EYA1 | protein_coding | 37.53 | -0.37692387 | 0.37692387 | ENSG00000104313 | n.s. |
| ENSG00000051825 | 2852.799277 | -0.473305522 | 0.2530826 | -1.870162 | 0.061461285 | 0.159859046 | MPHOSPH9 | protein_coding | 42.99 | -0.376875571 | 0.376875571 | ENSG00000051825 | n.s. |
| ENSG00000272831 | 123.53789 | 0.483483568 | 0.261701 | 1.847466 | 0.064679659 | 0.16615035 | | antisense | 49.55 | 0.376874832 | 0.376874832 | ENSG00000272831 | n.s. |
| ENSG00000225573 | 46.37261953 | 0.743283236 | 0.5207 | 1.427469 | 0.153444689 | 0.311386031 | RPL35P5 | processed_pseudogene | 56.49 | 0.376622266 | 0.376622266 | ENSG00000225573 | n.s. |
| ENSG00000252157 | 14.5514227 | 0.624301195 | 0.393743 | 1.585555 | 0.112840312 | 0.249550887 | RNU6-479P | snRNA | 36.19 | 0.376354283 | 0.376354283 | ENSG00000252157 | n.s. |
| ENSG00000205542 | 5684.749144 | 0.532878826 | 0.305699 | 1.743149 | 0.081307632 | 0.196672369 | TMSB4X | protein_coding | 50.47 | 0.376349215 | 0.376349215 | ENSG00000205542 | n.s. |
| ENSG00000259744 | 31.15959578 | 0.560371408 | 0.331527 | 1.690274 | 0.090975546 | 0.213420813 | | sense_intronic | 37.84 | 0.375876536 | 0.375876536 | ENSG00000259744 | n.s. |
| ENSG00000173264 | 18.49280029 | -0.603382911 | 0.3735953 | -1.615071 | 0.106295316 | 0.239048568 | GPR137 | protein_coding | 56.27 | -0.375010838 | 0.375010838 | ENSG00000173264 | n.s. |
| ENSG00000143252 | 1431.669855 | -0.506480918 | 0.2825737 | -1.792385 | 0.073071236 | 0.181814996 | SDHC | protein_coding | 42.28 | -0.374983429 | 0.374983429 | ENSG00000143252 | n.s. |
| ENSG00000279500 | 215.1899413 | 0.444782267 | 0.2301067 | 1.932939 | 0.053243675 | 0.143585265 | | TEC | 46.58 | 0.374902581 | 0.374902581 | ENSG00000279500 | n.s. |
| ENSG00000033178 | 1284.454094 | -0.493063224 | 0.2709728 | -1.819604 | 0.068819357 | 0.173956403 | UBA6 | protein_coding | 35.52 | -0.374510895 | 0.374510895 | ENSG00000033178 | n.s. |
| ENSG00000225224 | 76.8682692 | -0.457201486 | 0.2406101 | -1.900175 | 0.057410094 | 0.151933234 | RPS27AP12 | processed_pseudogene | 44.09 | -0.374149564 | 0.374149564 | ENSG00000225224 | n.s. |
| ENSG00000279845 | 106.9023776 | -0.506504337 | 0.2830274 | -1.789594 | 0.073519132 | 0.18263025 | | TEC | 28.76 | -0.374016623 | 0.374016623 | ENSG00000279845 | n.s. |
| ENSG00000176148 | 471.085464 | -0.443821701 | 0.2296578 | -1.932535 | 0.053293498 | 0.143688104 | TCP11L1 | protein_coding | 42.1 | -0.373954927 | 0.373954927 | ENSG00000176148 | n.s. |
| ENSG00000159658 | 331.2834895 | -0.45886815 | 0.242103 | -1.895343 | 0.058047 | 0.15326918 | EFCAB14 | protein_coding | 40.17 | -0.373768833 | 0.373768833 | ENSG00000159658 | n.s. |
| ENSG00000188290 | 105.7839939 | 0.54004049 | 0.313814 | 1.720893 | 0.085270185 | 0.20336019 | HES4 | protein_coding | 74.32 | 0.373564402 | 0.373564402 | ENSG00000188290 | n.s. |
| ENSG00000078043 | 1760.416896 | -0.464052484 | 0.2465785 | -1.881966 | 0.059840608 | 0.156697532 | PIAS2 | protein_coding | 38.59 | -0.373533405 | 0.373533405 | ENSG00000078043 | n.s. |

| | | | | | | | | | | | | | |
|---|---|---|---|---|---|---|---|---|---|---|---|---|---|
| ENSG00000165775 | 858.9525312 | 0.530103197 | 0.3046118 | 1.740258 | 0.081813697 | 0.197587528 | FUNDC2 | protein_coding | 39.84 | 0.373320126 | 0.373320126 | ENSG00000165775 | n.s. |
| ENSG00000059728 | 364.2003463 | 0.419598851 | 0.2107656 | 1.990832 | 0.04649939 | 0.129043454 | MXD1 | protein_coding | 40.74 | 0.373134162 | 0.373134162 | ENSG00000059728 | n.s. |
| ENSG00000264947 | 4.589186448 | 0.790729412 | 0.5793783 | 1.364789 | 0.172319237 | 0.337930822 | MIR3181 | miRNA | 83.56 | 0.372569713 | 0.372569713 | ENSG00000264947 | n.s. |
| ENSG00000135148 | 576.542399 | -0.524244849 | 0.2996459 | -1.749548 | 0.080196405 | 0.194694777 | TRAFD1 | protein_coding | 44.88 | -0.372552348 | 0.372552348 | ENSG00000135148 | n.s. |
| ENSG00000118496 | 2031.248865 | -0.474259523 | 0.2555451 | -1.855874 | 0.063471559 | 0.163863252 | FBXO30 | protein_coding | 35.2 | -0.372539595 | 0.372539595 | ENSG00000118496 | n.s. |
| ENSG00000183579 | 1314.051081 | -0.483437547 | 0.2634364 | -1.835121 | 0.066487797 | 0.169730398 | ZNRF3 | protein_coding | 45.29 | -0.372363115 | 0.372363115 | ENSG00000183579 | n.s. |
| ENSG00000127804 | 951.5034509 | -0.489910588 | 0.2692801 | -1.819335 | 0.068860384 | 0.173991351 | METTL16 | protein_coding | 44.67 | -0.372073541 | 0.372073541 | ENSG00000127804 | n.s. |
| ENSG00000100246 | 103.7315377 | -0.551199785 | 0.3250131 | -1.695931 | 0.089898955 | 0.211358182 | DNAL4 | protein_coding | 51.3 | -0.372049346 | 0.372049346 | ENSG00000100246 | n.s. |
| ENSG00000197696 | 46.21663646 | 0.503900563 | 0.2816439 | 1.789141 | 0.073592126 | 0.182700722 | NMB | protein_coding | 56.33 | 0.372009497 | 0.372009497 | ENSG00000197696 | n.s. |
| ENSG00000176399 | 273.0420795 | 0.444536096 | 0.2310409 | 1.924058 | 0.054347263 | 0.14573901 | DMRTA1 | protein_coding | 37.55 | 0.371820742 | 0.371820742 | ENSG00000176399 | n.s. |
| ENSG00000232648 | 22.91253352 | -0.540485985 | 0.315148 | -1.715023 | 0.086341087 | 0.205176131 | | antisense | 40.1 | -0.371785805 | 0.371785805 | ENSG00000232648 | n.s. |
| ENSG00000203993 | 226.3042484 | -0.461003064 | 0.2447401 | -1.883644 | 0.059613195 | 0.156312349 | ARRDC1-AS1 | antisense | 56.78 | -0.371571563 | 0.371571563 | ENSG00000203993 | n.s. |
| ENSG00000100401 | 1338.357177 | -0.474113236 | 0.2557894 | -1.85353 | 0.063806434 | 0.16457521 | RANGAP1 | protein_coding | 53.26 | -0.371532001 | 0.371532001 | ENSG00000100401 | n.s. |
| ENSG00000267419 | 46.57069219 | -0.492037585 | 0.2715986 | -1.811635 | 0.070042626 | 0.176072024 | | transcribed_unprocessed_pseud | 41.25 | -0.371148695 | 0.371148695 | ENSG00000267419 | n.s. |
| ENSG00000207518 | 4.414636302 | -0.776508354 | 0.5639793 | -1.376838 | 0.168562205 | 0.332750674 | RNU6-59P | snRNA | 39.25 | -0.371078632 | 0.371078632 | ENSG00000207518 | n.s. |
| ENSG00000149273 | 18455.50501 | 0.510043883 | 0.2875464 | 1.773779 | 0.076099654 | 0.187462046 | RPS3 | protein_coding | 48.52 | 0.370846097 | 0.370846097 | ENSG00000149273 | n.s. |
| ENSG00000173114 | 963.2966024 | -0.48625707 | 0.2664497 | -1.824949 | 0.06800873 | 0.172757808 | LRRN3 | protein_coding | 34.94 | -0.370801317 | 0.370801317 | ENSG00000173114 | n.s. |
| ENSG00000106397 | 139.7137763 | -0.469057457 | 0.2520277 | -1.861135 | 0.062725185 | 0.162478651 | PLOD3 | protein_coding | 58.5 | -0.370181878 | 0.370181878 | ENSG00000106397 | n.s. |
| ENSG00000113569 | 3127.657599 | -0.472466168 | 0.2551394 | -1.851796 | 0.064055142 | 0.164972205 | NUP155 | protein_coding | 41.76 | -0.36974693 | 0.36974693 | ENSG00000113569 | n.s. |
| ENSG00000207012 | 4.922800574 | -0.888726608 | 0.7033112 | -1.263632 | 0.206362079 | 0.383807004 | RNU6-72P | snRNA | 39.24 | -0.369609935 | 0.369609935 | ENSG00000207012 | n.s. |
| ENSG00000260196 | 152.5493575 | -0.506257967 | 0.2848079 | -1.777542 | 0.075479162 | 0.186263093 | | antisense | 48.42 | -0.369504117 | 0.369504117 | ENSG00000260196 | n.s. |
| ENSG00000132646 | 7651.463538 | 0.494066548 | 0.2741473 | 1.802194 | 0.071514937 | 0.178857996 | PCNA | protein_coding | 42.54 | 0.369310614 | 0.369310614 | ENSG00000132646 | n.s. |
| ENSG00000257169 | 67.56022473 | 0.510438923 | 0.2886329 | 1.768471 | 0.076982184 | 0.189034031 | | processed_pseudogene | 41.64 | 0.369282146 | 0.369282146 | ENSG00000257169 | n.s. |
| ENSG00000232228 | 9.901074906 | -0.658474874 | 0.4336903 | -1.518307 | 0.128937143 | 0.27498287 | | processed_pseudogene | 46.86 | -0.369203148 | 0.369203148 | ENSG00000232228 | n.s. |
| ENSG00000198668 | 7099.748723 | 0.453796874 | 0.2396347 | 1.893702 | 0.058264537 | 0.153639611 | CALM1 | protein_coding | 47.09 | 0.369162305 | 0.369162305 | ENSG00000198668 | n.s. |
| ENSG00000184903 | 234.1541813 | 0.439848126 | 0.2282231 | 1.927273 | 0.053945649 | 0.144941195 | IMMP2L | protein_coding | 36.11 | 0.368948198 | 0.368948198 | ENSG00000184903 | n.s. |
| ENSG00000105321 | 84.84294847 | 0.48576637 | 0.2670107 | 1.819277 | 0.068869233 | 0.173991351 | CCDC9 | protein_coding | 56.02 | 0.368926122 | 0.368926122 | ENSG00000105321 | n.s. |
| ENSG00000153487 | 449.980028 | 0.443980128 | 0.2317176 | 1.91604 | 0.055360064 | 0.147602101 | ING1 | protein_coding | 49.86 | 0.368906401 | 0.368906401 | ENSG00000153487 | n.s. |
| ENSG00000116260 | 447.4055945 | -0.48681468 | 0.2679301 | -1.816947 | 0.069225301 | 0.174732391 | QSOX1 | protein_coding | 51.87 | -0.368823742 | 0.368823742 | ENSG00000116260 | n.s. |
| ENSG00000186815 | 50.07171079 | -0.508192082 | 0.2868232 | -1.771796 | 0.0764285 | 0.18807819 | TPCN1 | protein_coding | 50.56 | -0.368775461 | 0.368775461 | ENSG00000186815 | n.s. |
| ENSG00000250486 | 280.0769298 | 0.413435386 | 0.2073245 | 1.994146 | 0.046136082 | 0.128282801 | FAM218A | protein_coding | 39.77 | 0.368714728 | 0.368714728 | ENSG00000250486 | n.s. |
| ENSG00000100395 | 587.648716 | -0.504995316 | 0.2840677 | -1.777729 | 0.075448369 | 0.186230919 | L3MBTL2 | protein_coding | 52.42 | -0.368620429 | 0.368620429 | ENSG00000100395 | n.s. |

| | | | | | | | | | | | | | |
|---|---|---|---|---|---|---|---|---|---|---|---|---|---|
| ENSG00000210049 | 6.892244594 | -0.704188023 | 0.4833734 | -1.45682 | 0.14516608 | 0.300051563 | MT-TF | Mt_tRNA | 40.85 | -0.368152391 | 0.368152391 | ENSG00000210049 | n.s. |
| ENSG00000163888 | 109.1594746 | -0.486342932 | 0.2679605 | -1.81498 | 0.069526978 | 0.1751129 | CAMK2N2 | protein_coding | 68.28 | -0.368006874 | 0.368006874 | ENSG00000163888 | n.s. |
| ENSG00000119927 | 1082.030093 | -0.483124678 | 0.2649603 | -1.823385 | 0.068245109 | 0.173105602 | GPAM | protein_coding | 39.93 | -0.367990694 | 0.367990694 | ENSG00000119927 | n.s. |
| ENSG00000142541 | 24774.6162 | 0.551773137 | 0.3276251 | 1.68416 | 0.092150678 | 0.215379198 | RPL13A | protein_coding | 53.4 | 0.367920255 | 0.367920255 | ENSG00000142541 | n.s. |
| ENSG00000147419 | 958.1666582 | -0.502711168 | 0.2825398 | -1.779258 | 0.075197506 | 0.185732636 | CCDC25 | protein_coding | 40.07 | -0.367538055 | 0.367538055 | ENSG00000147419 | n.s. |
| ENSG00000092020 | 716.0265942 | -0.53849472 | 0.3153912 | -1.707387 | 0.087750213 | 0.207720748 | PPP2R3C | protein_coding | 42.2 | -0.367533482 | 0.367533482 | ENSG00000092020 | n.s. |
| ENSG00000213380 | 283.4244786 | -0.437702417 | 0.2270341 | -1.927915 | 0.053865719 | 0.144793128 | COG8 | protein_coding | 52.29 | -0.36734265 | 0.36734265 | ENSG00000213380 | n.s. |
| ENSG00000258978 | 14.29228653 | -0.623378717 | 0.3986119 | -1.563874 | 0.117847187 | 0.257539851 | HIF1AP1 | processed_pseudogene | 36 | -0.367267037 | 0.367267037 | ENSG00000258978 | n.s. |
| ENSG00000163069 | 1023.797166 | -0.507050667 | 0.2864563 | -1.77008 | 0.076713776 | 0.188674422 | SGCB | protein_coding | 38.68 | -0.367250192 | 0.367250192 | ENSG00000163069 | n.s. |
| ENSG00000111196 | 2043.431534 | -0.482916907 | 0.265187 | -1.821043 | 0.06860033 | 0.173658618 | MAGOHB | protein_coding | 36.42 | -0.367163492 | 0.367163492 | ENSG00000111196 | n.s. |
| ENSG00000120616 | 3276.378906 | -0.456067044 | 0.2425179 | -1.88055 | 0.060033195 | 0.157058638 | EPC1 | protein_coding | 38.9 | -0.366649706 | 0.366649706 | ENSG00000120616 | n.s. |
| ENSG00000011021 | 49.86862816 | -0.513731979 | 0.2929675 | -1.753546 | 0.079508302 | 0.193530435 | CLCN6 | protein_coding | 47.79 | -0.366419708 | 0.366419708 | ENSG00000011021 | n.s. |
| ENSG00000074964 | 189.1521472 | -0.477742679 | 0.2609678 | -1.830657 | 0.067151706 | 0.171017598 | ARHGEF10L | protein_coding | 54.36 | -0.366409141 | 0.366409141 | ENSG00000074964 | n.s. |
| ENSG00000176055 | 324.8298615 | -0.483908015 | 0.2666455 | -1.814799 | 0.069554822 | 0.175151346 | MBLAC2 | protein_coding | 36.86 | -0.366118282 | 0.366118282 | ENSG00000176055 | n.s. |
| ENSG00000140280 | 328.5119329 | 0.530125884 | 0.3084578 | 1.718633 | 0.085681179 | 0.204060735 | LYSMD2 | protein_coding | 41.49 | 0.365914384 | 0.365914384 | ENSG00000140280 | n.s. |
| ENSG00000114391 | 13658.40081 | 0.525202375 | 0.3039604 | 1.727865 | 0.084012502 | 0.201257066 | RPL24 | protein_coding | 42.52 | 0.365671557 | 0.365671557 | ENSG00000114391 | n.s. |
| ENSG00000120699 | 2134.069322 | 0.449586207 | 0.2374583 | 1.893327 | 0.05831436 | 0.153712767 | EXOSC8 | protein_coding | 40.36 | 0.365643993 | 0.365643993 | ENSG00000120699 | n.s. |
| ENSG00000124523 | 64.24040772 | -0.52637065 | 0.305034 | -1.725613 | 0.084417052 | 0.202017637 | SIRT5 | protein_coding | 42.7 | -0.365622693 | 0.365622693 | ENSG00000124523 | n.s. |
| ENSG00000148908 | 232.5451587 | 0.561795871 | 0.3386521 | 1.658918 | 0.097132412 | 0.223569941 | RGS10 | protein_coding | 49.42 | 0.365496859 | 0.365496859 | ENSG00000148908 | n.s. |
| ENSG00000111371 | 12086.45041 | -0.475819916 | 0.2597195 | -1.832053 | 0.06694349 | 0.170572803 | SLC38A1 | protein_coding | 39.96 | -0.36547262 | 0.36547262 | ENSG00000111371 | n.s. |
| ENSG00000136942 | 13339.74795 | 0.550334293 | 0.3277154 | 1.679305 | 0.093092554 | 0.216743751 | RPL35 | protein_coding | 56.22 | 0.365451366 | 0.365451366 | ENSG00000136942 | n.s. |
| ENSG00000107816 | 899.5195544 | -0.46087507 | 0.2470177 | -1.865757 | 0.062075356 | 0.161185227 | LZTS2 | protein_coding | 60.49 | -0.365324038 | 0.365324038 | ENSG00000107816 | n.s. |
| ENSG00000109680 | 347.3621174 | -0.487522328 | 0.2702535 | -1.803945 | 0.071239966 | 0.178298409 | TBC1D19 | protein_coding | 36.57 | -0.36508233 | 0.36508233 | ENSG00000109680 | n.s. |
| ENSG00000117598 | 1022.065032 | -0.516831691 | 0.2965512 | -1.742808 | 0.081367226 | 0.196762759 | PLPPR5 | protein_coding | 37.21 | -0.364912682 | 0.364912682 | ENSG00000117598 | n.s. |
| ENSG00000077454 | 156.740909 | 0.480099901 | 0.2637382 | 1.820365 | 0.068703409 | 0.173824743 | LRCH4 | protein_coding | 60.36 | 0.364822348 | 0.364822348 | ENSG00000077454 | n.s. |
| ENSG00000207771 | 14.11050985 | 0.615491715 | 0.3921059 | 1.569708 | 0.116483087 | 0.255481099 | MIR550A1 | miRNA | 50.52 | 0.364765763 | 0.364765763 | ENSG00000207771 | n.s. |
| ENSG00000132478 | 2158.762999 | -0.503727039 | 0.2847529 | -1.768997 | 0.07689434 | 0.188951622 | UNK | protein_coding | 50.87 | -0.364521758 | 0.364521758 | ENSG00000132478 | n.s. |
| ENSG00000261366 | 284.8641472 | 0.425812665 | 0.2185296 | 1.948535 | 0.051350959 | 0.139341142 | MANEA-AS1 | antisense | 32.72 | 0.364461845 | 0.364461845 | ENSG00000261366 | n.s. |
| ENSG00000128881 | 1192.114249 | -0.502403632 | 0.2836258 | -1.771361 | 0.076500752 | 0.188183741 | TTBK2 | protein_coding | 39.87 | -0.364452589 | 0.364452589 | ENSG00000128881 | n.s. |
| ENSG00000122406 | 30018.64984 | 0.523105998 | 0.3027099 | 1.728077 | 0.083974386 | 0.201200375 | RPL5 | protein_coding | 40.34 | 0.364275959 | 0.364275959 | ENSG00000122406 | n.s. |
| ENSG00000146530 | 254.2366395 | -0.48598442 | 0.2692801 | -1.804754 | 0.07111312 | 0.178076974 | VWDE | protein_coding | 33.94 | -0.36419295 | 0.36419295 | ENSG00000146530 | n.s. |
| ENSG00000226944 | 19.69322854 | -0.548197056 | 0.3263345 | -1.679862 | 0.09298413 | 0.216636219 | | antisense | 58.13 | -0.364150273 | 0.364150273 | ENSG00000226944 | n.s. |

| | | | | | | | | | | | | | |
|---|---|---|---|---|---|---|---|---|---|---|---|---|---|
| ENSG00000119725 | 294.0587719 | 0.453215756 | 0.2411413 | 1.879461 | 0.060181537 | 0.157235418 | ZNF410 | protein_coding | 41.75 | 0.364136028 | 0.364136028 | ENSG00000119725 | n.s. |
| ENSG00000164331 | 275.1352368 | -0.495551754 | 0.2776855 | -1.784579 | 0.074329571 | 0.184177009 | ANKRA2 | protein_coding | 37.28 | -0.364113879 | 0.364113879 | ENSG00000164331 | n.s. |
| ENSG00000173226 | 1171.488375 | -0.480653927 | 0.2648093 | -1.815095 | 0.069509338 | 0.1751129 | IQCB1 | protein_coding | 36.52 | -0.363702107 | 0.363702107 | ENSG00000173226 | n.s. |
| ENSG00000122873 | 716.4337378 | -0.528401141 | 0.3079825 | -1.715686 | 0.086219575 | 0.205027352 | CISD1 | protein_coding | 36.84 | -0.363639429 | 0.363639429 | ENSG00000122873 | n.s. |
| ENSG00000112312 | 2594.098071 | 0.454858647 | 0.2428001 | 1.873387 | 0.06101493 | 0.158965358 | GMNN | protein_coding | 36.82 | 0.363294468 | 0.363294468 | ENSG00000112312 | n.s. |
| ENSG00000165819 | 795.2731714 | -0.48023235 | 0.2648141 | -1.81347 | 0.069759363 | 0.17553942 | METTL3 | protein_coding | 42.49 | -0.362875733 | 0.362875733 | ENSG00000165819 | n.s. |
| ENSG00000273084 | 32.03335161 | 0.5344178 | 0.3139194 | 1.702404 | 0.088679617 | 0.209457066 | | lincRNA | 59.24 | 0.362818908 | 0.362818908 | ENSG00000273084 | n.s. |
| ENSG00000092964 | 3107.270174 | -0.427900591 | 0.2208083 | -1.937882 | 0.052637583 | 0.142174567 | DPYSL2 | protein_coding | 44.97 | -0.362508004 | 0.362508004 | ENSG00000092964 | n.s. |
| ENSG00000197905 | 157.9396285 | -0.462825414 | 0.2500669 | -1.850806 | 0.064197429 | 0.16518588 | TEAD4 | protein_coding | 51.6 | -0.361942006 | 0.361942006 | ENSG00000197905 | n.s. |
| ENSG00000078177 | 1845.696753 | -0.480545745 | 0.2655402 | -1.809691 | 0.070343681 | 0.176722316 | N4BP2 | protein_coding | 40.52 | -0.361710917 | 0.361710917 | ENSG00000078177 | n.s. |
| ENSG00000200742 | 18.01882903 | -0.573897825 | 0.3523677 | -1.62869 | 0.103378628 | 0.234287391 | | misc_RNA | 42.57 | -0.361699727 | 0.361699727 | ENSG00000200742 | n.s. |
| ENSG00000171365 | 515.2941153 | -0.493147227 | 0.2767867 | -1.781687 | 0.074800338 | 0.184970862 | CLCN5 | protein_coding | 39.57 | -0.361425965 | 0.361425965 | ENSG00000171365 | n.s. |
| ENSG00000128923 | 767.0123985 | -0.475248199 | 0.2610919 | -1.820234 | 0.068723443 | 0.17384384 | MINDY2 | protein_coding | 39.89 | -0.361112921 | 0.361112921 | ENSG00000128923 | n.s. |
| ENSG00000170629 | 257.2369248 | -0.454343142 | 0.2432491 | -1.86781 | 0.061788544 | 0.160530303 | DPY19L2P2 | transcribed_unprocessed_pseud | 36.36 | -0.360949717 | 0.360949717 | ENSG00000170629 | n.s. |
| ENSG00000115216 | 816.1555677 | 0.468726119 | 0.2555022 | 1.834528 | 0.066575603 | 0.169861118 | NRBP1 | protein_coding | 48.61 | 0.360875062 | 0.360875062 | ENSG00000115216 | n.s. |
| ENSG00000227598 | 55.04893878 | -0.47265083 | 0.2588928 | -1.825662 | 0.06790118 | 0.172516083 | | antisense | 43.79 | -0.360713128 | 0.360713128 | ENSG00000227598 | n.s. |
| ENSG00000162928 | 914.7861839 | -0.488886782 | 0.2734585 | -1.787792 | 0.073809626 | 0.183051554 | PEX13 | protein_coding | 39.79 | -0.360518107 | 0.360518107 | ENSG00000162928 | n.s. |
| ENSG00000136100 | 1939.140063 | -0.48546232 | 0.2703311 | -1.795806 | 0.072525326 | 0.180929961 | VPS36 | protein_coding | 39.35 | -0.36045068 | 0.36045068 | ENSG00000136100 | n.s. |
| ENSG00000177485 | 1334.734845 | -0.447841945 | 0.2382077 | -1.880048 | 0.060101474 | 0.157144192 | ZBTB33 | protein_coding | 39.37 | -0.359931317 | 0.359931317 | ENSG00000177485 | n.s. |
| ENSG00000189369 | 132.3396743 | 0.488183359 | 0.2731016 | 1.787552 | 0.073848316 | 0.183114889 | GSPT2 | protein_coding | 44.83 | 0.359926041 | 0.359926041 | ENSG00000189369 | n.s. |
| ENSG00000264706 | 26.53934041 | -0.523978264 | 0.3059428 | -1.712667 | 0.086773782 | 0.205923192 | RN7SL217P | misc_RNA | 54.55 | -0.359603526 | 0.359603526 | ENSG00000264706 | n.s. |
| ENSG00000183340 | 209.8079832 | -0.44439366 | 0.2355409 | -1.886694 | 0.059201507 | 0.155491903 | JRKL | protein_coding | 38.8 | -0.359199939 | 0.359199939 | ENSG00000183340 | n.s. |
| ENSG00000215421 | 1125.427536 | -0.460264849 | 0.248987 | -1.84855 | 0.06452286 | 0.165808745 | ZNF407 | protein_coding | 37.97 | -0.359187268 | 0.359187268 | ENSG00000215421 | n.s. |
| ENSG00000221500 | 143.1366616 | 0.436164537 | 0.2287966 | 1.906342 | 0.056605883 | 0.150309422 | SNORD100 | snoRNA | 50 | 0.358969431 | 0.358969431 | ENSG00000221500 | n.s. |
| ENSG00000172878 | 262.854984 | -0.496077574 | 0.2804867 | -1.768631 | 0.076955461 | 0.189001743 | METAP1D | protein_coding | 40.05 | -0.358929086 | 0.358929086 | ENSG00000172878 | n.s. |
| ENSG00000122335 | 330.7868032 | -0.508257714 | 0.2916373 | -1.742773 | 0.081373265 | 0.196762759 | SERAC1 | protein_coding | 39.81 | -0.358858965 | 0.358858965 | ENSG00000122335 | n.s. |
| ENSG00000259439 | 36.73173745 | -0.518012117 | 0.3007891 | -1.722177 | 0.085037452 | 0.203083437 | LINC01833 | lincRNA | 52.5 | -0.358632995 | 0.358632995 | ENSG00000259439 | n.s. |
| ENSG00000177706 | 377.9620526 | -0.457666614 | 0.2469401 | -1.853351 | 0.063832117 | 0.164610959 | FAM20C | protein_coding | 59.85 | -0.358600674 | 0.358600674 | ENSG00000177706 | n.s. |
| ENSG00000145391 | 1643.239367 | -0.513117485 | 0.2963764 | -1.731303 | 0.083397656 | 0.200244068 | SETD7 | protein_coding | 42.84 | -0.358381952 | 0.358381952 | ENSG00000145391 | n.s. |
| ENSG00000106100 | 48.24414787 | 0.497393387 | 0.2819377 | 1.764196 | 0.077698924 | 0.190324089 | NOD1 | protein_coding | 46.77 | 0.358375039 | 0.358375039 | ENSG00000106100 | n.s. |
| ENSG00000226085 | 43.93924352 | 0.515272717 | 0.2984161 | 1.726692 | 0.084222988 | 0.201633657 | UQCRFS1P1 | processed_pseudogene | 53.9 | 0.358339698 | 0.358339698 | ENSG00000226085 | n.s. |
| ENSG00000139174 | 225.2281202 | 0.418258506 | 0.2145534 | 1.949438 | 0.051243173 | 0.139130011 | PRICKLE1 | protein_coding | 41.36 | 0.358271528 | 0.358271528 | ENSG00000139174 | n.s. |

| | | | | | | | | | | | | | |
|---|---|---|---|---|---|---|---|---|---|---|---|---|---|
| ENSG00000162244 | 10745.73161 | 0.530037614 | 0.312323 | 1.697082 | 0.089681222 | 0.211085381 | RPL29 | protein_coding | 53.82 | 0.358062587 | 0.358062587 | ENSG00000162244 | n.s. |
| ENSG00000084628 | 220.4079998 | -0.463507308 | 0.2521702 | -1.838073 | 0.066051601 | 0.168920041 | NKAIN1 | protein_coding | 50.46 | -0.357975418 | 0.357975418 | ENSG00000084628 | n.s. |
| ENSG00000169139 | 1980.467993 | -0.537512227 | 0.3196031 | -1.681812 | 0.092605316 | 0.216151519 | UBE2V2 | protein_coding | 42.29 | -0.357575552 | 0.357575552 | ENSG00000169139 | n.s. |
| ENSG00000021776 | 2460.90545 | -0.460471809 | 0.2497652 | -1.843619 | 0.065238712 | 0.167370294 | AQR | protein_coding | 37.54 | -0.357474221 | 0.357474221 | ENSG00000021776 | n.s. |
| ENSG00000228659 | 22.12849524 | -0.592478241 | 0.3736276 | -1.585746 | 0.112797049 | 0.249550887 | LINC01201 | antisense | 41.35 | -0.357170106 | 0.357170106 | ENSG00000228659 | n.s. |
| ENSG00000136868 | 628.7563111 | -0.492594366 | 0.2784407 | -1.769118 | 0.076874266 | 0.188944538 | SLC31A1 | protein_coding | 41.28 | -0.356473626 | 0.356473626 | ENSG00000136868 | n.s. |
| ENSG00000167543 | 150.9027238 | 0.514860404 | 0.2990065 | 1.721904 | 0.085086949 | 0.203166796 | TP53I13 | protein_coding | 63.48 | 0.356359222 | 0.356359222 | ENSG00000167543 | n.s. |
| ENSG00000137871 | 1685.895085 | -0.488634717 | 0.2751057 | -1.776171 | 0.075704798 | 0.18668755 | ZNF280D | protein_coding | 37.52 | -0.356158342 | 0.356158342 | ENSG00000137871 | n.s. |
| ENSG00000145087 | 382.7924875 | -0.482138989 | 0.2694199 | -1.789545 | 0.073527072 | 0.18263025 | STXBP5L | protein_coding | 36.35 | -0.356024585 | 0.356024585 | ENSG00000145087 | n.s. |
| ENSG00000276390 | 64.2735444 | 0.532178655 | 0.3154423 | 1.687087 | 0.091586587 | 0.214528944 | | sense_intronic | 32.88 | 0.355768937 | 0.355768937 | ENSG00000276390 | n.s. |
| ENSG00000250510 | 123.2463105 | 0.542260687 | 0.3254409 | 1.666234 | 0.095666794 | 0.220963632 | GPR162 | protein_coding | 61.64 | 0.355549054 | 0.355549054 | ENSG00000250510 | n.s. |
| ENSG00000083720 | 1049.410209 | -0.481708533 | 0.2693585 | -1.788355 | 0.073718823 | 0.182924107 | OXCT1 | protein_coding | 37.48 | -0.355370381 | 0.355370381 | ENSG00000083720 | n.s. |
| ENSG00000263465 | 725.0644718 | 0.449112215 | 0.2410557 | 1.863106 | 0.062447392 | 0.161890873 | SRSF8 | protein_coding | 43.87 | 0.355147895 | 0.355147895 | ENSG00000263465 | n.s. |
| ENSG00000177990 | 175.9216004 | -0.449120781 | 0.2411278 | -1.862584 | 0.062520845 | 0.162039784 | DPY19L2 | protein_coding | 36.31 | -0.35497534 | 0.35497534 | ENSG00000177990 | n.s. |
| ENSG00000147251 | 527.3255877 | -0.490084329 | 0.2768886 | -1.76997 | 0.076732201 | 0.188686406 | DOCK11 | protein_coding | 39.68 | -0.354948176 | 0.354948176 | ENSG00000147251 | n.s. |
| ENSG00000128268 | 285.4883503 | -0.450802225 | 0.2426676 | -1.857694 | 0.063212475 | 0.163376144 | MGAT3 | protein_coding | 56.74 | -0.354696311 | 0.354696311 | ENSG00000128268 | n.s. |
| ENSG00000264384 | 32.49351659 | 0.509478871 | 0.2949598 | 1.727282 | 0.084116953 | 0.201437965 | RN7SL431P | misc_RNA | 51.8 | 0.354525292 | 0.354525292 | ENSG00000264384 | n.s. |
| ENSG00000229994 | 69.07297496 | -0.522071514 | 0.3068198 | -1.701557 | 0.088838374 | 0.209643505 | RPL5P4 | processed_pseudogene | 42.78 | -0.354235226 | 0.354235226 | ENSG00000229994 | n.s. |
| ENSG00000130382 | 530.662422 | -0.440710362 | 0.2344356 | -1.879878 | 0.060124677 | 0.157160745 | MLLT1 | protein_coding | 55.57 | -0.354179492 | 0.354179492 | ENSG00000130382 | n.s. |
| ENSG00000176542 | 633.4491164 | -0.482558599 | 0.2706659 | -1.782857 | 0.074609537 | 0.184706363 | USF3 | protein_coding | 40.61 | -0.353965488 | 0.353965488 | ENSG00000176542 | n.s. |
| ENSG00000117410 | 529.6187626 | 0.5300083 | 0.3143996 | 1.685779 | 0.091838368 | 0.21486567 | ATP6V0B | protein_coding | 57.08 | 0.353957016 | 0.353957016 | ENSG00000117410 | n.s. |
| ENSG00000272758 | 40.72704182 | -0.524224907 | 0.3090548 | -1.69622 | 0.089844251 | 0.21130093 | | antisense | 39.37 | -0.353903497 | 0.353903497 | ENSG00000272758 | n.s. |
| ENSG00000013523 | 423.8713453 | -0.432508546 | 0.2276982 | -1.899482 | 0.057501181 | 0.152116462 | ANGEL1 | protein_coding | 44.52 | -0.353715775 | 0.353715775 | ENSG00000013523 | n.s. |
| ENSG00000274712 | 147.6389144 | 0.458822088 | 0.2500414 | 1.834984 | 0.06650803 | 0.169750925 | | lincRNA | 51.97 | 0.353379198 | 0.353379198 | ENSG00000274712 | n.s. |
| ENSG00000166747 | 992.5989733 | -0.466167515 | 0.2565921 | -1.816765 | 0.069253131 | 0.17473928 | AP1G1 | protein_coding | 40.73 | -0.353172919 | 0.353172919 | ENSG00000166747 | n.s. |
| ENSG00000153094 | 283.2747375 | 0.424092761 | 0.2212036 | 1.917206 | 0.055211774 | 0.147291345 | BCL2L11 | protein_coding | 44.42 | 0.352770017 | 0.352770017 | ENSG00000153094 | n.s. |
| ENSG00000252213 | 64.63864284 | 0.453739972 | 0.2459636 | 1.844745 | 0.06507471 | 0.166988623 | SNORA74D | snoRNA | 53.19 | 0.352698032 | 0.352698032 | ENSG00000252213 | n.s. |
| ENSG00000100578 | 893.443856 | -0.442124479 | 0.2362944 | -1.871075 | 0.061334737 | 0.159619353 | KIAA0586 | protein_coding | 37.04 | -0.352335387 | 0.352335387 | ENSG00000100578 | n.s. |
| ENSG00000077279 | 935.1811837 | -0.46210561 | 0.2532868 | -1.824436 | 0.068086199 | 0.172859982 | DCX | protein_coding | 39.48 | -0.352265665 | 0.352265665 | ENSG00000077279 | n.s. |
| ENSG00000124596 | 824.2984784 | -0.458388809 | 0.2502032 | -1.832066 | 0.066941527 | 0.170572803 | OARD1 | protein_coding | 44.19 | -0.352083958 | 0.352083958 | ENSG00000124596 | n.s. |
| ENSG00000182606 | 550.6853267 | -0.460809213 | 0.2522438 | -1.82684 | 0.067723772 | 0.172197062 | TRAK1 | protein_coding | 44.94 | -0.352046379 | 0.352046379 | ENSG00000182606 | n.s. |
| ENSG00000106443 | 5528.464855 | -0.460501609 | 0.2520776 | -1.826825 | 0.067726145 | 0.172197062 | PHF14 | protein_coding | 34.81 | -0.351811378 | 0.351811378 | ENSG00000106443 | n.s. |

| | | | | | | | | | | | | | |
|---|---|---|---|---|---|---|---|---|---|---|---|---|---|
| ENSG00000101247 | 1079.266703 | -0.468075637 | 0.2589108 | -1.807865 | 0.070627551 | 0.17721164 | NDUFAF5 | protein_coding | 39.88 | -0.351762471 | 0.351762471 | ENSG00000101247 | n.s. |
| ENSG00000054282 | 575.0472854 | -0.457654139 | 0.2498235 | -1.83191 | 0.066964802 | 0.170572803 | SDCCAG8 | protein_coding | 39.81 | -0.351519666 | 0.351519666 | ENSG00000054282 | n.s. |
| ENSG00000263080 | 20.62274379 | 0.540832323 | 0.3261836 | 1.658061 | 0.097305184 | 0.223893584 | | antisense | 46.49 | 0.351518486 | 0.351518486 | ENSG00000263080 | n.s. |
| ENSG00000130706 | 961.4721863 | 0.461410408 | 0.2532809 | 1.821734 | 0.06849534 | 0.173466398 | ADRM1 | protein_coding | 64.09 | 0.351033951 | 0.351033951 | ENSG00000130706 | n.s. |
| ENSG00000198695 | 724.7055356 | 0.46631172 | 0.2577785 | 1.808963 | 0.070456767 | 0.176878755 | MT-ND6 | protein_coding | 42.67 | 0.350817652 | 0.350817652 | ENSG00000198695 | n.s. |
| ENSG00000277846 | 68.30826541 | 0.47248553 | 0.2631523 | 1.795483 | 0.072576747 | 0.180993382 | SNORD30 | snoRNA | 41.43 | 0.350743635 | 0.350743635 | ENSG00000277846 | n.s. |
| ENSG00000201376 | 35.42813736 | 0.507838536 | 0.2955531 | 1.718265 | 0.085748321 | 0.204185714 | | snoRNA | 45.93 | 0.350395715 | 0.350395715 | ENSG00000201376 | n.s. |
| ENSG00000229117 | 9615.553114 | 0.511147129 | 0.2986317 | 1.711631 | 0.086964738 | 0.206306023 | RPL41 | protein_coding | 46.1 | 0.35038527 | 0.35038527 | ENSG00000229117 | n.s. |
| ENSG00000080819 | 324.7135059 | -0.49177524 | 0.2806801 | -1.752085 | 0.079759247 | 0.193971702 | CPOX | protein_coding | 38.07 | -0.350272631 | 0.350272631 | ENSG00000080819 | n.s. |
| ENSG00000211643 | 55.32629283 | -0.490299531 | 0.2792973 | -1.755475 | 0.079178012 | 0.193030198 | IGLV5-52 | IG_V_gene | 53 | -0.350257602 | 0.350257602 | ENSG00000211643 | n.s. |
| ENSG00000130193 | 162.2170895 | -0.50247247 | 0.2907179 | -1.728385 | 0.083919261 | 0.201137522 | THEM6 | protein_coding | 52.84 | -0.349975544 | 0.349975544 | ENSG00000130193 | n.s. |
| ENSG00000087008 | 148.5093799 | -0.470248297 | 0.2615993 | -1.79759 | 0.072242043 | 0.180384858 | ACOX3 | protein_coding | 51.93 | -0.349770646 | 0.349770646 | ENSG00000087008 | n.s. |
| ENSG00000151332 | 512.3756109 | 0.458817596 | 0.2514831 | 1.824447 | 0.068084517 | 0.172859982 | MBIP | protein_coding | 35.27 | 0.349759193 | 0.349759193 | ENSG00000151332 | n.s. |
| ENSG00000166938 | 1022.044291 | -0.464682251 | 0.2568939 | -1.808849 | 0.070474506 | 0.176891391 | DIS3L | protein_coding | 42.92 | -0.349577345 | 0.349577345 | ENSG00000166938 | n.s. |
| ENSG00000092010 | 285.8122246 | 0.546801054 | 0.3330183 | 1.641955 | 0.100599315 | 0.229576178 | PSME1 | protein_coding | 57.4 | 0.349445887 | 0.349445887 | ENSG00000092010 | n.s. |
| ENSG00000118181 | 7860.673001 | 0.553609119 | 0.3395897 | 1.63023 | 0.103052979 | 0.233834197 | RPS25 | protein_coding | 47.62 | 0.349378272 | 0.349378272 | ENSG00000118181 | n.s. |
| ENSG00000213079 | 3150.679069 | -0.465661062 | 0.2578557 | -1.805898 | 0.070934341 | 0.177725181 | SCAF8 | protein_coding | 37.66 | -0.349362694 | 0.349362694 | ENSG00000213079 | n.s. |
| ENSG00000131368 | 1990.358416 | -0.471224357 | 0.2626956 | -1.793804 | 0.072844516 | 0.181498604 | MRPS25 | protein_coding | 47.97 | -0.349236958 | 0.349236958 | ENSG00000131368 | n.s. |
| ENSG00000120071 | 7771.911874 | -0.431846033 | 0.2288406 | -1.887104 | 0.059146331 | 0.15543495 | KANSL1 | protein_coding | 40.92 | -0.349126496 | 0.349126496 | ENSG00000120071 | n.s. |
| ENSG00000180957 | 1095.799496 | -0.450798287 | 0.2449463 | -1.840396 | 0.065710061 | 0.168176688 | PITPNB | protein_coding | 39.71 | -0.349023453 | 0.349023453 | ENSG00000180957 | n.s. |
| ENSG00000120800 | 3560.573035 | -0.454116126 | 0.2478268 | -1.832393 | 0.066892841 | 0.170514285 | UTP20 | protein_coding | 39.31 | -0.348869823 | 0.348869823 | ENSG00000120800 | n.s. |
| ENSG00000109881 | 447.1343928 | -0.483051784 | 0.2735846 | -1.765639 | 0.077456359 | 0.189936579 | CCDC34 | protein_coding | 37.62 | -0.348469397 | 0.348469397 | ENSG00000109881 | n.s. |
| ENSG00000197345 | 796.6097012 | 0.523604965 | 0.3113152 | 1.681913 | 0.092585803 | 0.216142209 | MRPL21 | protein_coding | 50.69 | 0.348333656 | 0.348333656 | ENSG00000197345 | n.s. |
| ENSG00000183513 | 930.9821931 | -0.487808379 | 0.2780268 | -1.754537 | 0.079338453 | 0.193285962 | COA5 | protein_coding | 42.89 | -0.348197468 | 0.348197468 | ENSG00000183513 | n.s. |
| ENSG00000213741 | 7744.334735 | 0.528845121 | 0.3166952 | 1.669887 | 0.094941764 | 0.219689908 | RPS29 | protein_coding | 43.43 | 0.348080514 | 0.348080514 | ENSG00000213741 | n.s. |
| ENSG00000197798 | 229.4023239 | -0.505092023 | 0.294261 | -1.716477 | 0.086074864 | 0.204753174 | FAM118B | protein_coding | 41.9 | -0.347891908 | 0.347891908 | ENSG00000197798 | n.s. |
| ENSG00000231728 | 57.8714773 | -0.463160379 | 0.2563083 | -1.807044 | 0.070755422 | 0.177404599 | | antisense | 52.99 | -0.347849713 | 0.347849713 | ENSG00000231728 | n.s. |
| ENSG00000149262 | 563.5844147 | -0.474051678 | 0.2660773 | -1.781631 | 0.074809375 | 0.184970862 | INTS4 | protein_coding | 40.73 | -0.347430901 | 0.347430901 | ENSG00000149262 | n.s. |
| ENSG00000151689 | 89.29405327 | -0.482029722 | 0.2732572 | -1.764015 | 0.077729476 | 0.190365436 | INPP1 | protein_coding | 40.14 | -0.347259949 | 0.347259949 | ENSG00000151689 | n.s. |
| ENSG00000183943 | 840.4992462 | -0.460189372 | 0.2539766 | -1.811936 | 0.069996088 | 0.176055903 | PRKX | protein_coding | 45.74 | -0.34714358 | 0.34714358 | ENSG00000183943 | n.s. |
| ENSG00000181826 | 304.0953852 | -0.447771479 | 0.243087 | -1.842022 | 0.065471958 | 0.16778319 | RELL1 | protein_coding | 42.83 | -0.347135534 | 0.347135534 | ENSG00000181826 | n.s. |
| ENSG00000156928 | 550.2147067 | -0.470037426 | 0.2627263 | -1.789076 | 0.073602553 | 0.182700722 | MALSU1 | protein_coding | 41.44 | -0.347009706 | 0.347009706 | ENSG00000156928 | n.s. |

| | | | | | | | | | | | | | |
|---|---|---|---|---|---|---|---|---|---|---|---|---|---|
| ENSG00000157538 | 562.055649 | -0.48568383 | 0.2768583 | -1.754269 | 0.0793845 | 0.193305557 | DSCR3 | protein_coding | 44.22 | -0.346659583 | 0.346659583 | ENSG00000157538 | n.s. |
| ENSG00000196323 | 1033.276543 | -0.472501956 | 0.265167 | -1.781903 | 0.074764987 | 0.184926792 | ZBTB44 | protein_coding | 39.13 | -0.346344012 | 0.346344012 | ENSG00000196323 | n.s. |
| ENSG00000124659 | 989.5241143 | 0.454879607 | 0.2495362 | 1.8229 | 0.068318501 | 0.173254601 | TBCC | protein_coding | 54.33 | 0.34630677 | 0.34630677 | ENSG00000124659 | n.s. |
| ENSG00000224905 | 80.08207098 | -0.486243826 | 0.2777593 | -1.750594 | 0.080015812 | 0.194391929 | | antisense | 42.94 | -0.345875819 | 0.345875819 | ENSG00000224905 | n.s. |
| ENSG00000165152 | 714.9173628 | -0.493371965 | 0.2844041 | -1.734757 | 0.082783889 | 0.199170569 | TMEM246 | protein_coding | 39.12 | -0.345742658 | 0.345742658 | ENSG00000165152 | n.s. |
| ENSG00000163681 | 843.2186616 | -0.43210012 | 0.2304519 | -1.875013 | 0.060790997 | 0.15850057 | SLMAP | protein_coding | 38.95 | -0.345666775 | 0.345666775 | ENSG00000163681 | n.s. |
| ENSG00000152700 | 728.2495366 | -0.490965864 | 0.2821953 | -1.739809 | 0.081892621 | 0.197709547 | SAR1B | protein_coding | 43.44 | -0.345626398 | 0.345626398 | ENSG00000152700 | n.s. |
| ENSG00000120029 | 388.3581736 | -0.472067854 | 0.2651487 | -1.780389 | 0.075012259 | 0.18534085 | C10orf76 | protein_coding | 42.35 | -0.345567288 | 0.345567288 | ENSG00000120029 | n.s. |
| ENSG00000171160 | 151.2935495 | -0.524969639 | 0.3143223 | -1.670164 | 0.094887013 | 0.219689908 | MORN4 | protein_coding | 46.51 | -0.345529711 | 0.345529711 | ENSG00000171160 | n.s. |
| ENSG00000182899 | 14432.88528 | 0.534198572 | 0.3232358 | 1.652659 | 0.098400252 | 0.225637619 | RPL35A | protein_coding | 44.91 | 0.345406649 | 0.345406649 | ENSG00000182899 | n.s. |
| ENSG00000054611 | 193.6785401 | -0.446575161 | 0.2428644 | -1.838784 | 0.065946969 | 0.168720993 | TBC1D22A | protein_coding | 49.42 | -0.345127073 | 0.345127073 | ENSG00000054611 | n.s. |
| ENSG00000114948 | 576.3179321 | -0.463117514 | 0.2574159 | -1.799102 | 0.072002568 | 0.17985141 | ADAM23 | protein_coding | 38.27 | -0.345062447 | 0.345062447 | ENSG00000114948 | n.s. |
| ENSG00000135535 | 2673.948208 | 0.459607037 | 0.2544141 | 1.806531 | 0.070835426 | 0.177541249 | CD164 | protein_coding | 38.11 | 0.345027338 | 0.345027338 | ENSG00000135535 | n.s. |
| ENSG00000112983 | 1439.469927 | -0.478052248 | 0.2707553 | -1.765624 | 0.077458873 | 0.189936579 | BRD8 | protein_coding | 43.02 | -0.344862776 | 0.344862776 | ENSG00000112983 | n.s. |
| ENSG00000054793 | 1073.619401 | -0.44080141 | 0.2382849 | -1.849892 | 0.064329051 | 0.165401554 | ATP9A | protein_coding | 46.69 | -0.344468853 | 0.344468853 | ENSG00000054793 | n.s. |
| ENSG00000166710 | 3861.23864 | 0.515065024 | 0.3053531 | 1.686785 | 0.091644663 | 0.214628873 | B2M | protein_coding | 42.63 | 0.344224062 | 0.344224062 | ENSG00000166710 | n.s. |
| ENSG00000134262 | 781.3072734 | -0.449490061 | 0.2458906 | -1.828008 | 0.067548353 | 0.171839159 | AP4B1 | protein_coding | 43.25 | -0.343804997 | 0.343804997 | ENSG00000134262 | n.s. |
| ENSG00000076043 | 423.0305925 | 0.48248823 | 0.275341 | 1.75233 | 0.079717164 | 0.193903228 | REXO2 | protein_coding | 39.61 | 0.343731833 | 0.343731833 | ENSG00000076043 | n.s. |
| ENSG00000138083 | 312.8698587 | -0.428795411 | 0.2284722 | -1.876795 | 0.060546258 | 0.157980801 | SIX3 | protein_coding | 57.2 | -0.343634793 | 0.343634793 | ENSG00000138083 | n.s. |
| ENSG00000245694 | 1236.574661 | 0.479027566 | 0.2723048 | 1.759159 | 0.07855044 | 0.191768854 | CRNDE | lincRNA | 39.73 | 0.343569074 | 0.343569074 | ENSG00000245694 | n.s. |
| ENSG00000152684 | 1361.679928 | 0.42621984 | 0.2262875 | 1.883533 | 0.05962822 | 0.156317777 | PELO | protein_coding | 38.85 | 0.343529623 | 0.343529623 | ENSG00000152684 | n.s. |
| ENSG00000053770 | 751.7395125 | -0.468550446 | 0.2630297 | -1.781359 | 0.074853799 | 0.18501499 | AP5M1 | protein_coding | 35.17 | -0.343350526 | 0.343350526 | ENSG00000053770 | n.s. |
| ENSG00000261596 | 94.14913715 | 0.486935666 | 0.2796267 | 1.741377 | 0.081617432 | 0.197250391 | | sense_intronic | 39.37 | 0.343280942 | 0.343280942 | ENSG00000261596 | n.s. |
| ENSG00000266308 | 11.59311294 | -0.61443561 | 0.4054373 | -1.515488 | 0.129648749 | 0.276289203 | RN7SL510P | misc_RNA | 57.41 | -0.343245905 | 0.343245905 | ENSG00000266308 | n.s. |
| ENSG00000101639 | 2166.471262 | -0.475240282 | 0.2690254 | -1.766526 | 0.077307682 | 0.189666054 | CEP192 | protein_coding | 40.44 | -0.343128422 | 0.343128422 | ENSG00000101639 | n.s. |
| ENSG00000174842 | 726.2888864 | -0.446591376 | 0.2436995 | -1.83255 | 0.066869576 | 0.170486195 | GLMN | protein_coding | 38 | -0.343120969 | 0.343120969 | ENSG00000174842 | n.s. |
| ENSG00000272447 | 141.4446234 | 0.451709245 | 0.2482389 | 1.819655 | 0.068811526 | 0.173956403 | | lincRNA | 63.73 | 0.343100084 | 0.343100084 | ENSG00000272447 | n.s. |
| ENSG00000112200 | 5075.276815 | -0.446489456 | 0.2436335 | -1.832628 | 0.066857936 | 0.170486195 | ZNF451 | protein_coding | 36.73 | -0.343042662 | 0.343042662 | ENSG00000112200 | n.s. |
| ENSG00000085760 | 1258.608195 | -0.476612646 | 0.2703679 | -1.76283 | 0.07792919 | 0.190676426 | MTIF2 | protein_coding | 39.6 | -0.34301955 | 0.34301955 | ENSG00000085760 | n.s. |
| ENSG00000237594 | 138.7959818 | 0.426667214 | 0.2269704 | 1.879836 | 0.06013038 | 0.157160745 | | antisense | 58.57 | 0.342893633 | 0.342893633 | ENSG00000237594 | n.s. |
| ENSG00000165671 | 2710.590738 | -0.424948879 | 0.2254596 | -1.884812 | 0.05945528 | 0.156011281 | NSD1 | protein_coding | 42.34 | -0.342867452 | 0.342867452 | ENSG00000165671 | n.s. |
| ENSG00000213390 | 485.2015031 | -0.504888821 | 0.2964994 | -1.702832 | 0.088599428 | 0.209398389 | ARHGAP19 | protein_coding | 42.77 | -0.342832972 | 0.342832972 | ENSG00000213390 | n.s. |

| | | | | | | | | | | | | | |
|---|---|---|---|---|---|---|---|---|---|---|---|---|---|
| ENSG00000204147 | 294.700764 | -0.471480631 | 0.2657775 | -1.773967 | 0.076068552 | 0.187451762 | ASAH2B | protein_coding | 38.24 | -0.342818505 | 0.342818505 | ENSG00000204147 | n.s. |
| ENSG00000101152 | 717.1837425 | -0.462729435 | 0.258192 | -1.792191 | 0.073102349 | 0.181846153 | DNAJC5 | protein_coding | 51.37 | -0.342556696 | 0.342556696 | ENSG00000101152 | n.s. |
| ENSG00000113273 | 307.8001483 | -0.442968095 | 0.2408532 | -1.839162 | 0.065891292 | 0.168609531 | ARSB | protein_coding | 41.01 | -0.342466554 | 0.342466554 | ENSG00000113273 | n.s. |
| ENSG00000120256 | 689.070257 | -0.485637626 | 0.2788496 | -1.741576 | 0.081582708 | 0.197200702 | LRP11 | protein_coding | 43.94 | -0.342418983 | 0.342418983 | ENSG00000120256 | n.s. |
| ENSG00000149806 | 4315.543921 | 0.537932043 | 0.3284204 | 1.637937 | 0.101434787 | 0.231103999 | FAU | protein_coding | 55.53 | 0.342228352 | 0.342228352 | ENSG00000149806 | n.s. |
| ENSG00000240808 | 25.73393512 | -0.522427736 | 0.3137372 | -1.665176 | 0.095877554 | 0.221340273 | | processed_pseudogene | 43.1 | -0.342158591 | 0.342158591 | ENSG00000240808 | n.s. |
| ENSG00000161533 | 1058.613645 | -0.470990507 | 0.2656858 | -1.772735 | 0.076272668 | 0.187788569 | ACOX1 | protein_coding | 44.33 | -0.342094934 | 0.342094934 | ENSG00000161533 | n.s. |
| ENSG00000198081 | 625.1641658 | -0.434161727 | 0.233583 | -1.858704 | 0.063069092 | 0.163072791 | ZBTB14 | protein_coding | 49 | -0.341953806 | 0.341953806 | ENSG00000198081 | n.s. |
| ENSG00000113615 | 681.3095771 | -0.444291344 | 0.2422802 | -1.833791 | 0.066684978 | 0.170077843 | SEC24A | protein_coding | 42.26 | -0.34181655 | 0.34181655 | ENSG00000113615 | n.s. |
| ENSG00000187741 | 458.4742542 | 0.434487382 | 0.2339107 | 1.857492 | 0.063241128 | 0.163419863 | FANCA | protein_coding | 50.45 | 0.341809119 | 0.341809119 | ENSG00000187741 | n.s. |
| ENSG00000153132 | 759.5117515 | -0.511719539 | 0.3035656 | -1.685697 | 0.091854195 | 0.214866591 | CLGN | protein_coding | 36.07 | -0.341742226 | 0.341742226 | ENSG00000153132 | n.s. |
| ENSG00000179988 | 338.0647749 | 0.472935816 | 0.2678833 | 1.765454 | 0.077487421 | 0.189973123 | PSTK | protein_coding | 40.38 | 0.341132311 | 0.341132311 | ENSG00000179988 | n.s. |
| ENSG00000178229 | 421.5444064 | -0.440303433 | 0.2392438 | -1.840397 | 0.065710017 | 0.168176688 | ZNF543 | protein_coding | 47.36 | -0.340897978 | 0.340897978 | ENSG00000178229 | n.s. |
| ENSG00000201247 | 13.05811498 | -0.65501653 | 0.4508235 | -1.452933 | 0.146242219 | 0.301783076 | SNORD114-13 | snoRNA | 45.95 | -0.340808454 | 0.340808454 | ENSG00000201247 | n.s. |
| ENSG00000196950 | 3092.598959 | -0.466746039 | 0.2624988 | -1.778088 | 0.075389362 | 0.186140465 | SLC39A10 | protein_coding | 38.48 | -0.340798917 | 0.340798917 | ENSG00000196950 | n.s. |
| ENSG00000226989 | 23.94861282 | 0.544989911 | 0.3364649 | 1.619752 | 0.105285458 | 0.237198936 | | processed_pseudogene | 50.63 | 0.340557254 | 0.340557254 | ENSG00000226989 | n.s. |
| ENSG00000156171 | 745.3812878 | -0.49234368 | 0.2861267 | -1.720719 | 0.085301767 | 0.203400667 | DRAM2 | protein_coding | 37.94 | -0.340528338 | 0.340528338 | ENSG00000156171 | n.s. |
| ENSG00000130294 | 4625.844882 | -0.4406114 | 0.2397283 | -1.837961 | 0.066068114 | 0.168920041 | KIF1A | protein_coding | 58.61 | -0.340292477 | 0.340292477 | ENSG00000130294 | n.s. |
| ENSG00000135127 | 127.8627652 | 0.448207175 | 0.2465645 | 1.817809 | 0.069093359 | 0.174494256 | BICDL1 | protein_coding | 43.81 | 0.339839136 | 0.339839136 | ENSG00000135127 | n.s. |
| ENSG00000183495 | 1299.594283 | -0.439493306 | 0.2391757 | -1.837533 | 0.066131187 | 0.168943949 | EP400 | protein_coding | 47.47 | -0.339401939 | 0.339401939 | ENSG00000183495 | n.s. |
| ENSG00000116793 | 772.4114323 | -0.454093032 | 0.2519963 | -1.801983 | 0.071548083 | 0.178908757 | PHTF1 | protein_coding | 39.1 | -0.339374784 | 0.339374784 | ENSG00000116793 | n.s. |
| ENSG00000138050 | 1636.279048 | -0.475774791 | 0.2714107 | -1.75297 | 0.079607186 | 0.193737258 | THUMPD2 | protein_coding | 36.6 | -0.339126015 | 0.339126015 | ENSG00000138050 | n.s. |
| ENSG00000188739 | 1506.497376 | -0.470718578 | 0.2669042 | -1.763624 | 0.077795302 | 0.190426159 | RBM34 | protein_coding | 44.48 | -0.339046067 | 0.339046067 | ENSG00000188739 | n.s. |
| ENSG00000177700 | 531.8089826 | 0.515658329 | 0.30907 | 1.668419 | 0.095232509 | 0.220289453 | POLR2L | protein_coding | 61.14 | 0.338790768 | 0.338790768 | ENSG00000177700 | n.s. |
| ENSG00000187961 | 255.8715162 | 0.447303282 | 0.246317 | 1.815966 | 0.069375582 | 0.174921447 | KLHL17 | protein_coding | 67.87 | 0.338678784 | 0.338678784 | ENSG00000187961 | n.s. |
| ENSG00000170264 | 459.6972116 | -0.472745287 | 0.2689286 | -1.757884 | 0.078767278 | 0.192163449 | FAM161A | protein_coding | 41.32 | -0.33864126 | 0.33864126 | ENSG00000170264 | n.s. |
| ENSG00000109971 | 27688.60715 | 0.473613571 | 0.2696889 | 1.756147 | 0.079063229 | 0.19285167 | HSPA8 | protein_coding | 49.25 | 0.338527896 | 0.338527896 | ENSG00000109971 | n.s. |
| ENSG00000133627 | 327.7711645 | -0.496313709 | 0.2909729 | -1.705704 | 0.088063137 | 0.208379172 | ACTR3B | protein_coding | 43.36 | -0.338061944 | 0.338061944 | ENSG00000133627 | n.s. |
| ENSG00000230630 | 1655.845426 | -0.391248869 | 0.1998329 | -1.95788 | 0.050244132 | 0.13681773 | DNM3OS | antisense | 33.12 | -0.337983317 | 0.337983317 | ENSG00000230630 | n.s. |
| ENSG00000090857 | 874.9009635 | -0.444075473 | 0.2437207 | -1.822067 | 0.068444817 | 0.173454167 | PDPR | protein_coding | 47.32 | -0.337859398 | 0.337859398 | ENSG00000090857 | n.s. |
| ENSG00000140497 | 273.6092824 | -0.470931488 | 0.267694 | -1.759216 | 0.078540891 | 0.191768854 | SCAMP2 | protein_coding | 51.05 | -0.337762389 | 0.337762389 | ENSG00000140497 | n.s. |
| ENSG00000173706 | 1161.646008 | -0.451749734 | 0.2507484 | -1.801606 | 0.071607473 | 0.179025111 | HEG1 | protein_coding | 44.98 | -0.337495923 | 0.337495923 | ENSG00000173706 | n.s. |

| | | | | | | | | | | | | | |
|---|---|---|---|---|---|---|---|---|---|---|---|---|---|
| ENSG00000277563 | 5.08460329 | -0.933347172 | 0.8096168 | -1.152826 | 0.248981859 | 0.435062051 | | misc_RNA | 43.18 | -0.337357213 | 0.337357213 | ENSG00000277563 | n.s. |
| ENSG00000220848 | 32.57720697 | -0.502765996 | 0.2974815 | -1.690075 | 0.09101361 | 0.213438182 | RPS18P9 | processed_pseudogene | 51.74 | -0.337219174 | 0.337219174 | ENSG00000220848 | n.s. |
| ENSG00000259706 | 87.63531244 | -0.478253351 | 0.2746769 | -1.741148 | 0.081657557 | 0.197278876 | HSP90B2P | processed_pseudogene | 40.31 | -0.337130073 | 0.337130073 | ENSG00000259706 | n.s. |
| ENSG00000167670 | 1897.762716 | -0.44025094 | 0.2406835 | -1.82917 | 0.06737419 | 0.171521456 | CHAF1A | protein_coding | 51.15 | -0.337092021 | 0.337092021 | ENSG00000167670 | n.s. |
| ENSG00000136636 | 3483.392599 | -0.463202324 | 0.2611573 | -1.773653 | 0.076120629 | 0.187480545 | KCTD3 | protein_coding | 36.04 | -0.336768375 | 0.336768375 | ENSG00000136636 | n.s. |
| ENSG00000162408 | 552.1233802 | -0.442381365 | 0.2428309 | -1.821767 | 0.068490335 | 0.173466398 | NOL9 | protein_coding | 47.78 | -0.336556947 | 0.336556947 | ENSG00000162408 | n.s. |
| ENSG00000110844 | 76.34575876 | -0.467995774 | 0.2657357 | -1.761133 | 0.078215963 | 0.191254102 | PRPF40B | protein_coding | 47.42 | -0.336203129 | 0.336203129 | ENSG00000110844 | n.s. |
| ENSG00000118985 | 48.84283553 | 0.546681743 | 0.3405751 | 1.605172 | 0.108455879 | 0.242770291 | ELL2 | protein_coding | 38.17 | 0.336102374 | 0.336102374 | ENSG00000118985 | n.s. |
| ENSG00000241015 | 302.1536718 | -0.468166928 | 0.2659478 | -1.760372 | 0.078344835 | 0.191486703 | TPM3P9 | transcribed_processed_pseudog | 45.51 | -0.336078957 | 0.336078957 | ENSG00000241015 | n.s. |
| ENSG00000107341 | 1102.866834 | -0.466572575 | 0.2646095 | -1.763249 | 0.077858446 | 0.190547222 | UBE2R2 | protein_coding | 42.11 | -0.335931031 | 0.335931031 | ENSG00000107341 | n.s. |
| ENSG00000165782 | 325.5941711 | -0.433385628 | 0.2358298 | -1.837705 | 0.066105867 | 0.168932214 | TMEM55B | protein_coding | 51.18 | -0.334698315 | 0.334698315 | ENSG00000165782 | n.s. |
| ENSG00000112902 | 2814.471489 | -0.462690146 | 0.2616713 | -1.768211 | 0.077025572 | 0.189107219 | SEMA5A | protein_coding | 40.18 | -0.334660031 | 0.334660031 | ENSG00000112902 | n.s. |
| ENSG00000117133 | 1809.005363 | 0.472803851 | 0.2709366 | 1.745072 | 0.08097235 | 0.196065743 | RPF1 | protein_coding | 37.23 | 0.334555191 | 0.334555191 | ENSG00000117133 | n.s. |
| ENSG00000211455 | 1466.440184 | -0.453006675 | 0.2531056 | -1.789794 | 0.073487114 | 0.18260951 | STK38L | protein_coding | 37.77 | -0.334534833 | 0.334534833 | ENSG00000211455 | n.s. |
| ENSG00000261934 | 100.0594394 | -0.455021972 | 0.2550454 | -1.784082 | 0.074410296 | 0.184311431 | PCDHGA9 | protein_coding | 45.5 | -0.334189854 | 0.334189854 | ENSG00000261934 | n.s. |
| ENSG00000162813 | 504.3278064 | -0.529221038 | 0.3246584 | -1.630086 | 0.103083386 | 0.233865078 | BPNT1 | protein_coding | 40.42 | -0.333956799 | 0.333956799 | ENSG00000162813 | n.s. |
| ENSG00000279306 | 28.75907425 | 0.505845323 | 0.3023414 | 1.673093 | 0.094308952 | 0.218734557 | | TEC | 58.41 | 0.333899696 | 0.333899696 | ENSG00000279306 | n.s. |
| ENSG00000182774 | 28459.52964 | 0.526991946 | 0.322628 | 1.633435 | 0.102377524 | 0.232618826 | RPS17 | protein_coding | 49.9 | 0.333773058 | 0.333773058 | ENSG00000182774 | n.s. |
| ENSG00000233927 | 2113.174825 | 0.504648236 | 0.3013294 | 1.67474 | 0.093985325 | 0.218202048 | RPS28 | protein_coding | 57.12 | 0.333643729 | 0.333643729 | ENSG00000233927 | n.s. |
| ENSG00000111684 | 117.5278074 | -0.488477447 | 0.2859541 | -1.708237 | 0.087592314 | 0.207511959 | LPCAT3 | protein_coding | 44.19 | -0.333609028 | 0.333609028 | ENSG00000111684 | n.s. |
| ENSG00000127526 | 651.1277823 | -0.45129101 | 0.2521963 | -1.789443 | 0.073543448 | 0.18263025 | SLC35E1 | protein_coding | 50.12 | -0.333245596 | 0.333245596 | ENSG00000127526 | n.s. |
| ENSG00000203791 | 629.536384 | -0.490646915 | 0.2880939 | -1.70308 | 0.088553117 | 0.209325135 | EEF1AKMT2 | protein_coding | 40.87 | -0.333236895 | 0.333236895 | ENSG00000203791 | n.s. |
| ENSG00000177600 | 5744.426943 | 0.499070643 | 0.2961292 | 1.685314 | 0.091928019 | 0.214967049 | RPLP2 | protein_coding | 57.05 | 0.333193588 | 0.333193588 | ENSG00000177600 | n.s. |
| ENSG00000100335 | 1034.223028 | -0.441697735 | 0.2437945 | -1.811762 | 0.070022925 | 0.176072024 | MIEF1 | protein_coding | 49.85 | -0.333176861 | 0.333176861 | ENSG00000100335 | n.s. |
| ENSG00000148634 | 557.5010327 | 0.461603892 | 0.2614287 | 1.765697 | 0.077446707 | 0.189936579 | HERC4 | protein_coding | 38 | 0.332997073 | 0.332997073 | ENSG00000148634 | n.s. |
| ENSG00000255150 | 247.5214054 | -0.456043073 | 0.2565351 | -1.777702 | 0.075452748 | 0.186230919 | EID3 | protein_coding | 44.72 | -0.332887827 | 0.332887827 | ENSG00000255150 | n.s. |
| ENSG00000153976 | 155.1050797 | 0.447002345 | 0.2485641 | 1.798338 | 0.07212341 | 0.180120939 | HS3ST3A1 | protein_coding | 40.94 | 0.332764547 | 0.332764547 | ENSG00000153976 | n.s. |
| ENSG00000119041 | 2837.440983 | -0.489867412 | 0.2877049 | -1.702673 | 0.088629226 | 0.209398389 | GTF3C3 | protein_coding | 36.72 | -0.332633034 | 0.332633034 | ENSG00000119041 | n.s. |
| ENSG00000075218 | 677.211815 | -0.465790961 | 0.2654292 | -1.754859 | 0.079283361 | 0.19318555 | GTSE1 | protein_coding | 50.13 | -0.332586559 | 0.332586559 | ENSG00000075218 | n.s. |
| ENSG00000160049 | 929.4327535 | -0.487188204 | 0.2853103 | -1.707573 | 0.087715555 | 0.207720748 | DFFA | protein_coding | 48.52 | -0.332515752 | 0.332515752 | ENSG00000160049 | n.s. |
| ENSG00000152359 | 111.2179537 | -0.482242454 | 0.280759 | -1.717639 | 0.085862527 | 0.204422703 | POC5 | protein_coding | 37.27 | -0.332492125 | 0.332492125 | ENSG00000152359 | n.s. |
| ENSG00000185187 | 85.06321031 | 0.513499613 | 0.3106014 | 1.653243 | 0.098281444 | 0.225543522 | SIGIRR | protein_coding | 65.22 | 0.332115961 | 0.332115961 | ENSG00000185187 | n.s. |

| | | | | | | | | | | | | | |
|---|---|---|---|---|---|---|---|---|---|---|---|---|---|
| ENSG00000112511 | 238.8499736 | 0.44783159 | 0.2495842 | 1.794311 | 0.07276358 | 0.181380782 | PHF1 | protein_coding | 57.49 | 0.332026251 | 0.332026251 | ENSG00000112511 | n.s. |
| ENSG00000164167 | 944.1831969 | -0.466205251 | 0.2661861 | -1.751426 | 0.079872528 | 0.194111575 | LSM6 | protein_coding | 39.71 | -0.33191416 | 0.33191416 | ENSG00000164167 | n.s. |
| ENSG00000164941 | 881.8747881 | -0.488645896 | 0.2869069 | -1.703151 | 0.088539746 | 0.209325135 | INTS8 | protein_coding | 40.21 | -0.331877846 | 0.331877846 | ENSG00000164941 | n.s. |
| ENSG00000258815 | 79.35200699 | -0.510052382 | 0.3074439 | -1.659009 | 0.097113907 | 0.223569941 | | lincRNA | 35.13 | -0.331833239 | 0.331833239 | ENSG00000258815 | n.s. |
| ENSG00000083844 | 245.3512666 | 0.404490114 | 0.2126004 | 1.902584 | 0.057094807 | 0.151300153 | ZNF264 | protein_coding | 44.47 | 0.331746868 | 0.331746868 | ENSG00000083844 | n.s. |
| ENSG00000145808 | 761.8078474 | -0.472381469 | 0.2721355 | -1.735832 | 0.082593577 | 0.198906024 | ADAMTS19 | protein_coding | 34.94 | -0.331305718 | 0.331305718 | ENSG00000145808 | n.s. |
| ENSG00000156011 | 2676.299999 | -0.451828806 | 0.2536189 | -1.781526 | 0.074826516 | 0.184980394 | PSD3 | protein_coding | 39.93 | -0.33113372 | 0.33113372 | ENSG00000156011 | n.s. |
| ENSG00000110696 | 4018.520301 | -0.508644808 | 0.3065675 | -1.659161 | 0.09708336 | 0.223567916 | C11orf58 | protein_coding | 38.23 | -0.330919491 | 0.330919491 | ENSG00000110696 | n.s. |
| ENSG00000100888 | 1921.548313 | -0.423462385 | 0.2289363 | -1.849696 | 0.064357435 | 0.165414172 | CHD8 | protein_coding | 42.52 | -0.330905061 | 0.330905061 | ENSG00000100888 | n.s. |
| ENSG00000141076 | 192.4134398 | 0.437244216 | 0.2410073 | 1.814236 | 0.069641329 | 0.175305771 | UTP4 | protein_coding | 43.98 | 0.330645732 | 0.330645732 | ENSG00000141076 | n.s. |
| ENSG00000141428 | 426.4701413 | -0.492116337 | 0.2908614 | -1.691927 | 0.090659846 | 0.212884438 | C18orf21 | protein_coding | 42.7 | -0.330631355 | 0.330631355 | ENSG00000141428 | n.s. |
| ENSG00000136603 | 2635.608918 | -0.466351553 | 0.2669409 | -1.747022 | 0.080633529 | 0.195449272 | SKIL | protein_coding | 39.63 | -0.330627369 | 0.330627369 | ENSG00000136603 | n.s. |
| ENSG00000054118 | 1442.235017 | -0.427586425 | 0.2328216 | -1.836541 | 0.066277643 | 0.169287034 | THRAP3 | protein_coding | 43.72 | -0.329830036 | 0.329830036 | ENSG00000054118 | n.s. |
| ENSG00000171453 | 671.3161289 | 0.448765164 | 0.2514309 | 1.784845 | 0.07428642 | 0.184102852 | POLR1C | protein_coding | 50.11 | 0.329815238 | 0.329815238 | ENSG00000171453 | n.s. |
| ENSG00000207005 | 191.0696858 | 0.421393001 | 0.2276311 | 1.85121 | 0.064139335 | 0.165066905 | RNU1-2 | snRNA | 54.27 | 0.329672597 | 0.329672597 | ENSG00000207005 | n.s. |
| ENSG00000275576 | 34.10285948 | -0.56325324 | 0.3615961 | -1.557686 | 0.119307693 | 0.25995634 | | sense_intronic | 44.66 | -0.329559237 | 0.329559237 | ENSG00000275576 | n.s. |
| ENSG00000157045 | 314.9143186 | 0.485677432 | 0.2854258 | 1.701589 | 0.088832365 | 0.209643505 | NTAN1 | protein_coding | 47.6 | 0.329541166 | 0.329541166 | ENSG00000157045 | n.s. |
| ENSG00000261270 | 77.87385604 | 0.453757704 | 0.2560616 | 1.772065 | 0.076383785 | 0.188028896 | | sense_intronic | 47.7 | 0.329326181 | 0.329326181 | ENSG00000261270 | n.s. |
| ENSG00000138092 | 1435.625684 | -0.436451635 | 0.2408814 | -1.811894 | 0.070002548 | 0.176055903 | CENPO | protein_coding | 43.93 | -0.329237033 | 0.329237033 | ENSG00000138092 | n.s. |
| ENSG00000144535 | 200.0046445 | -0.462229059 | 0.2639011 | -1.751524 | 0.079855674 | 0.194111575 | DIS3L2 | protein_coding | 42.48 | -0.329083316 | 0.329083316 | ENSG00000144535 | n.s. |
| ENSG00000104327 | 149.1491156 | -0.472386985 | 0.273306 | -1.728418 | 0.083913373 | 0.201137522 | CALB1 | protein_coding | 36.25 | -0.329020796 | 0.329020796 | ENSG00000104327 | n.s. |
| ENSG00000073536 | 732.1554321 | -0.407994381 | 0.216662 | -1.883091 | 0.059688012 | 0.156415619 | NLE1 | protein_coding | 52.13 | -0.328729185 | 0.328729185 | ENSG00000073536 | n.s. |
| ENSG00000027847 | 132.6314056 | -0.489133263 | 0.2890506 | -1.692207 | 0.090606582 | 0.212806249 | B4GALT7 | protein_coding | 57.92 | -0.328705194 | 0.328705194 | ENSG00000027847 | n.s. |
| ENSG00000228350 | 9.504818653 | -0.643179118 | 0.4483285 | -1.434616 | 0.15139663 | 0.308397261 | CELSR3-AS1 | antisense | 48.95 | -0.32859345 | 0.32859345 | ENSG00000228350 | n.s. |
| ENSG00000121578 | 339.9041461 | -0.509242569 | 0.3086516 | -1.649894 | 0.098964561 | 0.226625588 | B4GALT4 | protein_coding | 41.78 | -0.32830413 | 0.32830413 | ENSG00000121578 | n.s. |
| ENSG00000142937 | 12578.25806 | 0.51640973 | 0.3154568 | 1.637022 | 0.101625905 | 0.231425822 | RPS8 | protein_coding | 55.11 | 0.328223923 | 0.328223923 | ENSG00000142937 | n.s. |
| ENSG00000232346 | 98.69908398 | 0.560668671 | 0.3598864 | 1.557905 | 0.119255856 | 0.259884063 | | processed_pseudogene | 50.61 | 0.328114716 | 0.328114716 | ENSG00000232346 | n.s. |
| ENSG00000187164 | 277.4703705 | -0.455597165 | 0.2584735 | -1.762646 | 0.077960264 | 0.190695846 | SHTN1 | protein_coding | 39.09 | -0.327874494 | 0.327874494 | ENSG00000187164 | n.s. |
| ENSG00000130177 | 872.7283067 | -0.470308411 | 0.2719713 | -1.729258 | 0.083762969 | 0.200970501 | CDC16 | protein_coding | 41.32 | -0.327742734 | 0.327742734 | ENSG00000130177 | n.s. |
| ENSG00000184990 | 964.4151302 | 0.505638109 | 0.3054854 | 1.655196 | 0.097884808 | 0.224818572 | SIVA1 | protein_coding | 52.32 | 0.327738348 | 0.327738348 | ENSG00000184990 | n.s. |
| ENSG00000204386 | 222.8699187 | 0.456807238 | 0.2596167 | 1.759545 | 0.078485056 | 0.191676448 | NEU1 | protein_coding | 52.86 | 0.327727786 | 0.327727786 | ENSG00000204386 | n.s. |
| ENSG00000245556 | 142.6155496 | -0.421920322 | 0.2289864 | -1.842557 | 0.065393785 | 0.167706331 | SCAMP1-AS1 | lincRNA | 42.36 | -0.327178343 | 0.327178343 | ENSG00000245556 | n.s. |

| | | | | | | | | | | | | | |
|---|---|---|---|---|---|---|---|---|---|---|---|---|---|
| ENSG00000272006 | 9.26892647 | -0.769876805 | 0.6011437 | -1.280687 | 0.200303715 | 0.375903576 | | processed_pseudogene | 34.82 | -0.32713878 | 0.32713878 | ENSG00000272006 | n.s. |
| ENSG00000105409 | 916.8302471 | -0.445329691 | 0.2495941 | -1.784216 | 0.074388653 | 0.184290607 | ATP1A3 | protein_coding | 56.13 | -0.327093233 | 0.327093233 | ENSG00000105409 | n.s. |
| ENSG00000040608 | 129.3257444 | -0.434597362 | 0.2402131 | -1.809216 | 0.070417494 | 0.176812041 | RTN4R | protein_coding | 60.45 | -0.32702937 | 0.32702937 | ENSG00000040608 | n.s. |
| ENSG00000240877 | 13.79824999 | 0.563436892 | 0.3635062 | 1.550006 | 0.121140006 | 0.263002074 | RN7SL521P | misc_RNA | 67.34 | 0.326816401 | 0.326816401 | ENSG00000240877 | n.s. |
| ENSG00000139737 | 2640.071383 | -0.443117605 | 0.247822 | -1.788048 | 0.07376825 | 0.183014139 | SLAIN1 | protein_coding | 36.78 | -0.326806038 | 0.326806038 | ENSG00000139737 | n.s. |
| ENSG00000168887 | 830.5978345 | -0.431899268 | 0.2380161 | -1.81458 | 0.06958847 | 0.175204389 | C2orf68 | protein_coding | 51.15 | -0.326712369 | 0.326712369 | ENSG00000168887 | n.s. |
| ENSG00000106829 | 2452.43385 | 0.409832706 | 0.2189846 | 1.871514 | 0.061273875 | 0.159520598 | TLE4 | protein_coding | 39.28 | 0.32671176 | 0.32671176 | ENSG00000106829 | n.s. |
| ENSG00000115232 | 1791.367006 | -0.445642105 | 0.2500638 | -1.782114 | 0.074730671 | 0.184907612 | ITGA4 | protein_coding | 36.12 | -0.32667581 | 0.32667581 | ENSG00000115232 | n.s. |
| ENSG00000157911 | 130.7967854 | -0.49970578 | 0.3004785 | -1.663034 | 0.096305717 | 0.222107751 | PEX10 | protein_coding | 61.52 | -0.326525889 | 0.326525889 | ENSG00000157911 | n.s. |
| ENSG00000115239 | 251.2232769 | 0.408062044 | 0.2176901 | 1.874509 | 0.06086036 | 0.158651711 | ASB3 | protein_coding | 39.31 | 0.326268145 | 0.326268145 | ENSG00000115239 | n.s. |
| ENSG00000132199 | 76.84664543 | -0.483376359 | 0.2851488 | -1.695172 | 0.090042765 | 0.211624817 | ENOSF1 | protein_coding | 46.1 | -0.326005163 | 0.326005163 | ENSG00000132199 | n.s. |
| ENSG00000184924 | 523.1264085 | 0.486571121 | 0.2881021 | 1.688884 | 0.091241585 | 0.213864745 | PTRHD1 | protein_coding | 45.88 | 0.325934921 | 0.325934921 | ENSG00000184924 | n.s. |
| ENSG00000139496 | 3726.204105 | -0.451269631 | 0.2554552 | -1.766532 | 0.077306716 | 0.189666054 | NUP58 | protein_coding | 36.36 | -0.325821363 | 0.325821363 | ENSG00000139496 | n.s. |
| ENSG00000086200 | 1321.807341 | -0.470447911 | 0.2731161 | -1.72252 | 0.084975395 | 0.20297005 | IPO11 | protein_coding | 38.3 | -0.325817188 | 0.325817188 | ENSG00000086200 | n.s. |
| ENSG00000130643 | 65.90469261 | 0.550906251 | 0.3517614 | 1.566136 | 0.117316702 | 0.25667536 | CALY | protein_coding | 60.02 | 0.325373947 | 0.325373947 | ENSG00000130643 | n.s. |
| ENSG00000116337 | 88.97368179 | 0.481332788 | 0.2836523 | 1.696911 | 0.089713464 | 0.211085381 | AMPD2 | protein_coding | 56.74 | 0.325160439 | 0.325160439 | ENSG00000116337 | n.s. |
| ENSG00000076604 | 886.5886493 | 0.483588267 | 0.2857417 | 1.692397 | 0.090570366 | 0.212757079 | TRAF4 | protein_coding | 59.64 | 0.325027402 | 0.325027402 | ENSG00000076604 | n.s. |
| ENSG00000232838 | 139.6056043 | -0.452578218 | 0.257068 | -1.760539 | 0.078316492 | 0.19146629 | PET117 | protein_coding | 45.08 | -0.324909378 | 0.324909378 | ENSG00000232838 | n.s. |
| ENSG00000140995 | 177.3751676 | 0.46412996 | 0.267791 | 1.73318 | 0.083063713 | 0.19960236 | DEF8 | protein_coding | 57.08 | 0.324814079 | 0.324814079 | ENSG00000140995 | n.s. |
| ENSG00000252080 | 4.600443805 | -0.976260437 | 0.8970309 | -1.088324 | 0.276451987 | 0.465129063 | RNU7-99P | snRNA | 35.48 | -0.324534863 | 0.324534863 | ENSG00000252080 | n.s. |
| ENSG00000167637 | 169.635586 | -0.475251443 | 0.2783356 | -1.707476 | 0.087733568 | 0.207720748 | ZNF283 | protein_coding | 38.88 | -0.324368673 | 0.324368673 | ENSG00000167637 | n.s. |
| ENSG00000146476 | 652.6729779 | -0.463775654 | 0.2676833 | -1.732553 | 0.083175084 | 0.199801017 | ARMT1 | protein_coding | 39.28 | -0.324365761 | 0.324365761 | ENSG00000146476 | n.s. |
| ENSG00000134851 | 961.1187083 | -0.456365394 | 0.2612552 | -1.746819 | 0.080668805 | 0.195500743 | TMEM165 | protein_coding | 38.82 | -0.323495334 | 0.323495334 | ENSG00000134851 | n.s. |
| ENSG00000135090 | 317.8005044 | -0.433387492 | 0.2407115 | -1.800444 | 0.071790586 | 0.179418476 | TAOK3 | protein_coding | 39.71 | -0.32336464 | 0.32336464 | ENSG00000135090 | n.s. |
| ENSG00000229119 | 61.61170159 | 0.530568462 | 0.3324436 | 1.595965 | 0.110496497 | 0.245836167 | | processed_pseudogene | 55.25 | 0.323304134 | 0.323304134 | ENSG00000229119 | n.s. |
| ENSG00000142599 | 173.7157979 | -0.423989045 | 0.2323811 | -1.824542 | 0.068070163 | 0.172859982 | RERE | protein_coding | 41.9 | -0.323209196 | 0.323209196 | ENSG00000142599 | n.s. |
| ENSG00000172667 | 617.5334427 | -0.436086536 | 0.2432807 | -1.792524 | 0.073049072 | 0.181814996 | ZMAT3 | protein_coding | 40.73 | -0.32286552 | 0.32286552 | ENSG00000172667 | n.s. |
| ENSG00000210107 | 83.9832112 | 0.480573589 | 0.2840951 | 1.691594 | 0.090723386 | 0.212936901 | MT-TQ | Mt_tRNA | 48.61 | 0.322824862 | 0.322824862 | ENSG00000210107 | n.s. |
| ENSG00000205978 | 467.8468006 | -0.438373037 | 0.2454017 | -1.786349 | 0.074042826 | 0.183564507 | NYNRIN | protein_coding | 52.95 | -0.322735178 | 0.322735178 | ENSG00000205978 | n.s. |
| ENSG00000229054 | 22.28322494 | -0.521899071 | 0.3241014 | -1.610296 | 0.107333293 | 0.240838267 | RPS29P14 | processed_pseudogene | 52.63 | -0.322676891 | 0.322676891 | ENSG00000229054 | n.s. |
| ENSG00000112685 | 642.6872588 | -0.474890428 | 0.2789084 | -1.702675 | 0.088628889 | 0.209398389 | EXOC2 | protein_coding | 40.51 | -0.322463263 | 0.322463263 | ENSG00000112685 | n.s. |
| ENSG00000106785 | 189.0035902 | -0.471226084 | 0.275578 | -1.709955 | 0.087274117 | 0.206898949 | TRIM14 | protein_coding | 46.67 | -0.322432544 | 0.322432544 | ENSG00000106785 | n.s. |

| | | | | | | | | | | | | | |
|---|---|---|---|---|---|---|---|---|---|---|---|---|---|
| ENSG00000140450 | 278.6306143 | 0.435340354 | 0.2428872 | 1.792356 | 0.073075969 | 0.181814996 | ARRDC4 | protein_coding | 39.4 | 0.322313068 | 0.322313068 | ENSG00000140450 | n.s. |
| ENSG00000147687 | 565.8441898 | -0.476701748 | 0.2808466 | -1.697374 | 0.089625903 | 0.211036943 | TATDN1 | protein_coding | 40.76 | -0.32207949 | 0.32207949 | ENSG00000147687 | n.s. |
| ENSG00000171503 | 288.6934926 | -0.4783638 | 0.2824354 | -1.693711 | 0.090320264 | 0.212205373 | ETFDH | protein_coding | 38.19 | -0.322055378 | 0.322055378 | ENSG00000171503 | n.s. |
| ENSG00000116350 | 1494.35304 | -0.453032729 | 0.258967 | -1.749384 | 0.080224716 | 0.194729553 | SRSF4 | protein_coding | 43 | -0.32191062 | 0.32191062 | ENSG00000116350 | n.s. |
| ENSG00000092978 | 1301.200055 | -0.442702264 | 0.2495808 | -1.773783 | 0.076099007 | 0.187462046 | GPATCH2 | protein_coding | 36.36 | -0.321882905 | 0.321882905 | ENSG00000092978 | n.s. |
| ENSG00000130638 | 1980.889845 | -0.475548642 | 0.2799506 | -1.698688 | 0.089377996 | 0.210524418 | ATXN10 | protein_coding | 40.57 | -0.321802589 | 0.321802589 | ENSG00000130638 | n.s. |
| ENSG00000103351 | 456.8789344 | -0.466351617 | 0.271565 | -1.717274 | 0.085929035 | 0.204511104 | CLUAP1 | protein_coding | 45.59 | -0.321448292 | 0.321448292 | ENSG00000103351 | n.s. |
| ENSG00000071127 | 1059.599929 | -0.452784561 | 0.2589694 | -1.74841 | 0.080393116 | 0.195036304 | WDR1 | protein_coding | 54.98 | -0.321424761 | 0.321424761 | ENSG00000071127 | n.s. |
| ENSG00000249859 | 449.989996 | -0.410810126 | 0.2219041 | -1.851296 | 0.064127018 | 0.16506572 | PVT1 | lincRNA | 46.6 | -0.321394472 | 0.321394472 | ENSG00000249859 | n.s. |
| ENSG00000197417 | 306.9350826 | -0.464130655 | 0.2696521 | -1.72122 | 0.08521085 | 0.203308355 | SHPK | protein_coding | 50.5 | -0.321106368 | 0.321106368 | ENSG00000197417 | n.s. |
| ENSG00000243176 | 31.85164911 | 0.506468921 | 0.309926 | 1.634161 | 0.102225087 | 0.232459359 | | processed_transcript | 39.16 | 0.320925533 | 0.320925533 | ENSG00000243176 | n.s. |
| ENSG00000274475 | 23.75905691 | 0.526598545 | 0.3301449 | 1.595053 | 0.110700455 | 0.246171926 | RN7SL650P | misc_RNA | 54.85 | 0.320572908 | 0.320572908 | ENSG00000274475 | n.s. |
| ENSG00000060237 | 4328.8074 | -0.415008548 | 0.2257716 | -1.838179 | 0.06603603 | 0.168917811 | WNK1 | protein_coding | 38.72 | -0.320521292 | 0.320521292 | ENSG00000060237 | n.s. |
| ENSG00000167977 | 490.3778721 | -0.473141997 | 0.2784643 | -1.699112 | 0.089298124 | 0.2104623 | KCTD5 | protein_coding | 59.23 | -0.320234657 | 0.320234657 | ENSG00000167977 | n.s. |
| ENSG00000280033 | 41.80333672 | 0.461407398 | 0.2675854 | 1.724337 | 0.084647028 | 0.20238474 | | TEC | 61.64 | 0.320134713 | 0.320134713 | ENSG00000280033 | n.s. |
| ENSG00000119979 | 331.5435514 | -0.46243997 | 0.2688321 | -1.720181 | 0.08539947 | 0.203563914 | FAM45A | protein_coding | 45.15 | -0.319684384 | 0.319684384 | ENSG00000119979 | n.s. |
| ENSG00000228109 | 117.0127089 | 0.462345978 | 0.2687644 | 1.720265 | 0.085384281 | 0.203562558 | MELTF-AS1 | antisense | 61.61 | 0.319620745 | 0.319620745 | ENSG00000228109 | n.s. |
| ENSG00000155961 | 387.2560978 | -0.471717765 | 0.2774535 | -1.700169 | 0.089099201 | 0.210187762 | RAB39B | protein_coding | 41.08 | -0.319538111 | 0.319538111 | ENSG00000155961 | n.s. |
| ENSG00000203667 | 1081.638788 | 0.432679617 | 0.2417372 | 1.789876 | 0.07347381 | 0.182609043 | COX20 | protein_coding | 39.53 | 0.319524257 | 0.319524257 | ENSG00000203667 | n.s. |
| ENSG00000180370 | 1349.089467 | -0.460265532 | 0.2668607 | -1.724741 | 0.084574211 | 0.202317933 | PAK2 | protein_coding | 41.7 | -0.319408456 | 0.319408456 | ENSG00000180370 | n.s. |
| ENSG00000175093 | 519.6857647 | -0.451625565 | 0.2589339 | -1.744173 | 0.08112889 | 0.196367794 | SPSB4 | protein_coding | 47.04 | -0.319267542 | 0.319267542 | ENSG00000175093 | n.s. |
| ENSG00000203760 | 784.0142356 | 0.486232784 | 0.291579 | 1.667585 | 0.095398133 | 0.220562639 | CENPW | protein_coding | 35.46 | 0.319196288 | 0.319196288 | ENSG00000203760 | n.s. |
| ENSG00000164008 | 93.94879392 | -0.494840031 | 0.2998492 | -1.650296 | 0.098882348 | 0.226549109 | C1orf50 | protein_coding | 43.06 | -0.319091479 | 0.319091479 | ENSG00000164008 | n.s. |
| ENSG00000214389 | 248.8256943 | 0.551757735 | 0.3566942 | 1.546865 | 0.121895876 | 0.264047592 | RPS3AP26 | processed_pseudogene | 43.11 | 0.319091314 | 0.319091314 | ENSG00000214389 | n.s. |
| ENSG00000104852 | 3454.072819 | 0.437765712 | 0.2464494 | 1.77629 | 0.075685062 | 0.18668755 | SNRNP70 | protein_coding | 53.38 | 0.319080705 | 0.319080705 | ENSG00000104852 | n.s. |
| ENSG00000270704 | 50.4661245 | -0.513416231 | 0.3180104 | -1.614463 | 0.106426933 | 0.239228639 | | sense_intronic | 37.74 | -0.318927402 | 0.318927402 | ENSG00000270704 | n.s. |
| ENSG00000117475 | 839.5885118 | -0.458558538 | 0.2656375 | -1.726257 | 0.084301209 | 0.201775095 | BLZF1 | protein_coding | 35.78 | -0.318758915 | 0.318758915 | ENSG00000117475 | n.s. |
| ENSG00000162144 | 364.9202102 | 0.389738943 | 0.2051676 | 1.899612 | 0.05748401 | 0.152099938 | CYB561A3 | protein_coding | 52.14 | 0.318756164 | 0.318756164 | ENSG00000162144 | n.s. |
| ENSG00000128000 | 130.9647573 | -0.444776142 | 0.2529569 | -1.758308 | 0.078695083 | 0.192020967 | ZNF780B | protein_coding | 39.65 | -0.318749422 | 0.318749422 | ENSG00000128000 | n.s. |
| ENSG00000196510 | 1922.821233 | -0.485816364 | 0.2914265 | -1.667029 | 0.095508738 | 0.220745046 | ANAPC7 | protein_coding | 45.87 | -0.318748505 | 0.318748505 | ENSG00000196510 | n.s. |
| ENSG00000197149 | 345.4157171 | 0.517994695 | 0.3228807 | 1.604291 | 0.108649864 | 0.243048209 | | processed_pseudogene | 60.02 | 0.318208065 | 0.318208065 | ENSG00000197149 | n.s. |
| ENSG00000160298 | 405.3242509 | 0.389749713 | 0.2054111 | 1.897413 | 0.057773418 | 0.152662612 | C21orf58 | protein_coding | 52.19 | 0.31813995 | 0.31813995 | ENSG00000160298 | n.s. |

| | | | | | | | | | | | | | |
|---|---|---|---|---|---|---|---|---|---|---|---|---|---|
| ENSG00000186575 | 879.2380574 | -0.395488747 | 0.2103416 | -1.880221 | 0.060077954 | 0.157112201 | NF2 | protein_coding | 45.79 | -0.317889934 | 0.317889934 | ENSG00000186575 | n.s. |
| ENSG00000184305 | 618.5792535 | -0.484836086 | 0.2910356 | -1.665899 | 0.095733463 | 0.221080942 | CCSER1 | protein_coding | 35.16 | -0.317785179 | 0.317785179 | ENSG00000184305 | n.s. |
| ENSG00000117713 | 962.8435708 | -0.437559416 | 0.2469601 | -1.771782 | 0.076430826 | 0.18807819 | ARID1A | protein_coding | 46.19 | -0.31752005 | 0.31752005 | ENSG00000117713 | n.s. |
| ENSG00000247572 | 82.8211848 | -0.450477228 | 0.2588218 | -1.740492 | 0.081772668 | 0.197522701 | CKMT2-AS1 | antisense | 41.97 | -0.317308494 | 0.317308494 | ENSG00000247572 | n.s. |
| ENSG00000251682 | 211.1616931 | -0.463777512 | 0.2712747 | -1.709623 | 0.08733555 | 0.206974103 | | processed_pseudogene | 44.33 | -0.317262771 | 0.317262771 | ENSG00000251682 | n.s. |
| ENSG00000251730 | 3.485430112 | -0.762934793 | 0.6040659 | -1.262999 | 0.206589406 | 0.383846385 | | snoRNA | 37.78 | -0.317260746 | 0.317260746 | ENSG00000251730 | n.s. |
| ENSG00000178498 | 1166.322819 | 0.443814167 | 0.2528714 | 1.755098 | 0.079242515 | 0.193119821 | DTX3 | protein_coding | 55.6 | 0.316960161 | 0.316960161 | ENSG00000178498 | n.s. |
| ENSG00000109323 | 159.8888376 | -0.468280506 | 0.2756232 | -1.698988 | 0.089321391 | 0.2104623 | MANBA | protein_coding | 39.25 | -0.316944276 | 0.316944276 | ENSG00000109323 | n.s. |
| ENSG00000112062 | 1376.067793 | -0.437822847 | 0.2475165 | -1.768863 | 0.076916708 | 0.188973236 | MAPK14 | protein_coding | 39.88 | -0.316808481 | 0.316808481 | ENSG00000112062 | n.s. |
| ENSG00000278829 | 181.4668754 | 0.427052486 | 0.2379323 | 1.794849 | 0.072677795 | 0.181212919 | | sense_intronic | 36.65 | 0.316792165 | 0.316792165 | ENSG00000278829 | n.s. |
| ENSG00000175455 | 6162.236222 | -0.478253995 | 0.2854311 | -1.67555 | 0.093826381 | 0.217942059 | CCDC14 | protein_coding | 36.3 | -0.316441036 | 0.316441036 | ENSG00000175455 | n.s. |
| ENSG00000147457 | 666.055187 | -0.487921765 | 0.2947334 | -1.655468 | 0.097829577 | 0.224728792 | CHMP7 | protein_coding | 46.73 | -0.316339823 | 0.316339823 | ENSG00000147457 | n.s. |
| ENSG00000107625 | 1493.861386 | -0.489098757 | 0.2959204 | -1.652805 | 0.098370501 | 0.225636324 | DDX50 | protein_coding | 40.32 | -0.316246846 | 0.316246846 | ENSG00000107625 | n.s. |
| ENSG00000170473 | 395.3857441 | -0.512035293 | 0.3182415 | -1.608952 | 0.107626882 | 0.24130259 | PYM1 | protein_coding | 45.5 | -0.316150056 | 0.316150056 | ENSG00000170473 | n.s. |
| ENSG00000139291 | 618.7089281 | -0.45598998 | 0.265107 | -1.720022 | 0.085428408 | 0.203598037 | TMEM19 | protein_coding | 39.6 | -0.315192319 | 0.315192319 | ENSG00000139291 | n.s. |
| ENSG00000165568 | 43.92715407 | -0.54144077 | 0.3486693 | -1.552878 | 0.120452353 | 0.261794898 | AKR1E2 | protein_coding | 42.59 | -0.315139548 | 0.315139548 | ENSG00000165568 | n.s. |
| ENSG00000130255 | 8479.653865 | 0.514161393 | 0.3209538 | 1.60198 | 0.109160105 | 0.24383702 | RPL36 | protein_coding | 58.81 | 0.315129702 | 0.315129702 | ENSG00000130255 | n.s. |
| ENSG00000277452 | 15.50061787 | -0.566780794 | 0.3752699 | -1.510328 | 0.130959625 | 0.278289203 | RN7SL473P | misc_RNA | 52.84 | -0.314848796 | 0.314848796 | ENSG00000277452 | n.s. |
| ENSG00000198218 | 1150.974511 | -0.436960566 | 0.2477532 | -1.763693 | 0.077783688 | 0.190426159 | QRICH1 | protein_coding | 46.08 | -0.314731069 | 0.314731069 | ENSG00000198218 | n.s. |
| ENSG00000279691 | 68.7034456 | 0.442955799 | 0.2533412 | 1.748456 | 0.080385148 | 0.195036304 | | TEC | 58.13 | 0.314447474 | 0.314447474 | ENSG00000279691 | n.s. |
| ENSG00000073670 | 293.1030252 | 0.448989507 | 0.2589544 | 1.733855 | 0.082943747 | 0.199382905 | ADAM11 | protein_coding | 59.28 | 0.314432776 | 0.314432776 | ENSG00000073670 | n.s. |
| ENSG00000203705 | 608.1862532 | -0.470112247 | 0.2786457 | -1.687133 | 0.09157787 | 0.214528944 | TATDN3 | protein_coding | 40.66 | -0.314276668 | 0.314276668 | ENSG00000203705 | n.s. |
| ENSG00000186432 | 2129.456556 | -0.447859267 | 0.2580742 | -1.73539 | 0.082671799 | 0.198969655 | KPNA4 | protein_coding | 37.2 | -0.314044809 | 0.314044809 | ENSG00000186432 | n.s. |
| ENSG00000124614 | 2578.71363 | 0.498027652 | 0.3057239 | 1.629011 | 0.103310687 | 0.23422809 | RPS10 | protein_coding | 47.56 | 0.31393722 | 0.31393722 | ENSG00000124614 | n.s. |
| ENSG00000166949 | 243.2041881 | -0.436222335 | 0.2475271 | -1.762321 | 0.07801504 | 0.190796312 | SMAD3 | protein_coding | 48.34 | -0.313831445 | 0.313831445 | ENSG00000166949 | n.s. |
| ENSG00000188452 | 765.1623831 | -0.47345546 | 0.2824405 | -1.676301 | 0.093679163 | 0.217781764 | CERKL | protein_coding | 37.01 | -0.31341733 | 0.31341733 | ENSG00000188452 | n.s. |
| ENSG00000013810 | 861.7309932 | -0.4474639 | 0.2580546 | -1.733989 | 0.082919967 | 0.199382905 | TACC3 | protein_coding | 56.07 | -0.313364375 | 0.313364375 | ENSG00000013810 | n.s. |
| ENSG00000248206 | 42.50203524 | -0.474777564 | 0.2839077 | -1.672295 | 0.094466069 | 0.218989523 | | sense_intronic | 48.09 | -0.313152201 | 0.313152201 | ENSG00000248206 | n.s. |
| ENSG00000125354 | 805.6565506 | -0.44635904 | 0.2571799 | -1.735591 | 0.082636252 | 0.198918488 | 6-Sep | protein_coding | 47.21 | -0.313042687 | 0.313042687 | ENSG00000125354 | n.s. |
| ENSG00000014824 | 2779.964584 | -0.481015416 | 0.2899572 | -1.658918 | 0.097132223 | 0.223569941 | SLC30A9 | protein_coding | 36.98 | -0.312942178 | 0.312942178 | ENSG00000014824 | n.s. |
| ENSG00000177688 | 159.0061226 | -0.434039887 | 0.2459587 | -1.764686 | 0.0776166 | 0.190222835 | SUMO4 | protein_coding | 39.87 | -0.312828759 | 0.312828759 | ENSG00000177688 | n.s. |
| ENSG00000139437 | 725.5753213 | -0.422510859 | 0.2357052 | -1.792539 | 0.073046629 | 0.181814996 | TCHP | protein_coding | 46.02 | -0.312814492 | 0.312814492 | ENSG00000139437 | n.s. |

| | | | | | | | | | | | | | |
|---|---|---|---|---|---|---|---|---|---|---|---|---|---|
| ENSG00000100325 | 183.3363998 | -0.451817215 | 0.2624254 | -1.721698 | 0.085124298 | 0.203186286 | ASCC2 | protein_coding | 48.94 | -0.312705228 | 0.312705228 | ENSG00000100325 | n.s. |
| ENSG00000278012 | 254.8972849 | 0.429116702 | 0.2416783 | 1.77557 | 0.075803852 | 0.186896441 | | sense_intronic | 39.78 | 0.312568163 | 0.312568163 | ENSG00000278012 | n.s. |
| ENSG00000153395 | 673.7513189 | -0.468976375 | 0.2785625 | -1.683559 | 0.092266914 | 0.215542324 | LPCAT1 | protein_coding | 56.14 | -0.312557484 | 0.312557484 | ENSG00000153395 | n.s. |
| ENSG00000137876 | 6646.842342 | 0.496757149 | 0.3054773 | 1.626167 | 0.103914097 | 0.235188253 | RSL24D1 | protein_coding | 36.26 | 0.312253782 | 0.312253782 | ENSG00000137876 | n.s. |
| ENSG00000152520 | 674.1212655 | -0.454375257 | 0.2651075 | -1.713928 | 0.086541904 | 0.205583164 | PAN3 | protein_coding | 37.49 | -0.312161461 | 0.312161461 | ENSG00000152520 | n.s. |
| ENSG00000005302 | 88.0438135 | 0.464640598 | 0.274668 | 1.691645 | 0.090713737 | 0.212936901 | MSL3 | protein_coding | 42.85 | 0.31212189 | 0.31212189 | ENSG00000005302 | n.s. |
| ENSG00000074621 | 36.97450704 | 0.477284 | 0.2868484 | 1.66389 | 0.096134506 | 0.221859892 | SLC24A1 | protein_coding | 44.73 | 0.312106128 | 0.312106128 | ENSG00000074621 | n.s. |
| ENSG00000213221 | 142.5677557 | 0.460693537 | 0.2710143 | 1.699886 | 0.089152327 | 0.210277461 | DNLZ | protein_coding | 64.59 | 0.311985015 | 0.311985015 | ENSG00000213221 | n.s. |
| ENSG00000063176 | 219.064719 | -0.449132419 | 0.2604273 | -1.724598 | 0.084599882 | 0.202317933 | SPHK2 | protein_coding | 56.78 | -0.311682458 | 0.311682458 | ENSG00000063176 | n.s. |
| ENSG00000207047 | 13.17665501 | 0.608514101 | 0.4234502 | 1.437038 | 0.150707145 | 0.307532695 | SNORD51 | snoRNA | 38.75 | 0.311625369 | 0.311625369 | ENSG00000207047 | n.s. |
| ENSG00000137822 | 752.4560116 | -0.431344439 | 0.244156 | -1.766676 | 0.077282526 | 0.189666054 | TUBGCP4 | protein_coding | 42.65 | -0.311435167 | 0.311435167 | ENSG00000137822 | n.s. |
| ENSG00000207963 | 5.849510799 | -0.738397199 | 0.5795833 | -1.274014 | 0.202658448 | 0.378818842 | MIR569 | miRNA | 32.29 | -0.311284946 | 0.311284946 | ENSG00000207963 | n.s. |
| ENSG00000263895 | 16.96367314 | -0.729610272 | 0.5678828 | -1.28479 | 0.198865623 | 0.374454623 | | sense_intronic | 37.13 | -0.311252328 | 0.311252328 | ENSG00000263895 | n.s. |
| ENSG00000111653 | 386.9034941 | -0.524813359 | 0.3342838 | -1.569963 | 0.116423646 | 0.25539096 | ING4 | protein_coding | 47.73 | -0.311106462 | 0.311106462 | ENSG00000111653 | n.s. |
| ENSG00000258399 | 229.6885303 | -0.472000903 | 0.2824872 | -1.670875 | 0.094746349 | 0.219497105 | | lincRNA | 46.71 | -0.310846203 | 0.310846203 | ENSG00000258399 | n.s. |
| ENSG00000153885 | 1142.446012 | -0.443142499 | 0.2552684 | -1.735987 | 0.08256619 | 0.198887381 | KCTD15 | protein_coding | 56 | -0.310816945 | 0.310816945 | ENSG00000153885 | n.s. |
| ENSG00000167525 | 63.81134259 | 0.45002618 | 0.2618095 | 1.718907 | 0.085631268 | 0.204011661 | PROCA1 | protein_coding | 55.73 | 0.310673327 | 0.310673327 | ENSG00000167525 | n.s. |
| ENSG00000135316 | 11986.81082 | -0.445864969 | 0.257911 | -1.728755 | 0.083852905 | 0.201116968 | SYNCRIP | protein_coding | 40.05 | -0.310567818 | 0.310567818 | ENSG00000135316 | n.s. |
| ENSG00000147471 | 367.3575252 | -0.452452191 | 0.2641591 | -1.712802 | 0.086748944 | 0.205899343 | PLPBP | protein_coding | 43.59 | -0.310538317 | 0.310538317 | ENSG00000147471 | n.s. |
| ENSG00000272253 | 4.978031849 | -0.925780475 | 0.8463339 | -1.093871 | 0.274011355 | 0.461991017 | RNA5SP386 | rRNA | 53.27 | -0.310475729 | 0.310475729 | ENSG00000272253 | n.s. |
| ENSG00000207507 | 4.034994732 | 0.711806945 | 0.5468644 | 1.301615 | 0.193047983 | 0.366286055 | RNU6-9 | snRNA | 45.79 | 0.310475679 | 0.310475679 | ENSG00000207507 | n.s. |
| ENSG00000145920 | 3031.288111 | -0.445587991 | 0.2577329 | -1.728875 | 0.083831465 | 0.201100188 | CPLX2 | protein_coding | 50.92 | -0.310391035 | 0.310391035 | ENSG00000145920 | n.s. |
| ENSG00000141510 | 367.0420449 | -0.437350914 | 0.2502036 | -1.74798 | 0.080467465 | 0.195148669 | TP53 | protein_coding | 48.86 | -0.310359256 | 0.310359256 | ENSG00000141510 | n.s. |
| ENSG00000184752 | 1385.645389 | 0.495739383 | 0.3058221 | 1.621006 | 0.10501637 | 0.236860994 | NDUFA12 | protein_coding | 43.14 | 0.310088183 | 0.310088183 | ENSG00000184752 | n.s. |
| ENSG00000179104 | 335.6919822 | -0.424255546 | 0.2385734 | -1.778302 | 0.075354315 | 0.186086933 | TMTC2 | protein_coding | 37.42 | -0.309827089 | 0.309827089 | ENSG00000179104 | n.s. |
| ENSG00000158164 | 1359.293693 | 0.522125107 | 0.3324637 | 1.570473 | 0.116305191 | 0.25518176 | TMSB15A | protein_coding | 43.49 | 0.309698701 | 0.309698701 | ENSG00000158164 | n.s. |
| ENSG00000101888 | 137.7650327 | 0.491817252 | 0.3021605 | 1.627669 | 0.103595118 | 0.234672392 | NXT2 | protein_coding | 37.4 | 0.309617649 | 0.309617649 | ENSG00000101888 | n.s. |
| ENSG00000273139 | 12.93190709 | 0.565763766 | 0.3778876 | 1.497175 | 0.134347804 | 0.284015255 | | lincRNA | 56.13 | 0.309279477 | 0.309279477 | ENSG00000273139 | n.s. |
| ENSG00000235449 | 33.42427134 | 0.503728986 | 0.3143565 | 1.602413 | 0.109064315 | 0.243670433 | | processed_pseudogene | 47.37 | 0.308885186 | 0.308885186 | ENSG00000235449 | n.s. |
| ENSG00000156671 | 652.0142241 | -0.464807336 | 0.2765813 | -1.680545 | 0.092851372 | 0.216508064 | SAMD8 | protein_coding | 43.24 | -0.308876524 | 0.308876524 | ENSG00000156671 | n.s. |
| ENSG00000157869 | 365.2061862 | -0.492574675 | 0.3036465 | -1.622198 | 0.10476097 | 0.236476489 | RAB28 | protein_coding | 36.22 | -0.308456188 | 0.308456188 | ENSG00000157869 | n.s. |
| ENSG00000251216 | 34.21501668 | 0.527396556 | 0.3387334 | 1.556967 | 0.1194784 | 0.260171362 | | lincRNA | 36.87 | 0.308390131 | 0.308390131 | ENSG00000251216 | n.s. |

| | | | | | | | | | | | | | |
|---|---|---|---|---|---|---|---|---|---|---|---|---|---|
| ENSG00000280383 | 1356.657052 | 0.377636673 | 0.1990475 | 1.897219 | 0.057799012 | 0.15270126 | | sense_overlapping | 41.87 | 0.308210957 | 0.308210957 | ENSG00000280383 | n.s. |
| ENSG00000182272 | 1342.650512 | 0.447616661 | 0.2608484 | 1.716003 | 0.086161534 | 0.204924332 | B4GALNT4 | protein_coding | 65.98 | 0.308142207 | 0.308142207 | ENSG00000182272 | n.s. |
| ENSG00000164305 | 750.0601948 | -0.474157312 | 0.2860987 | -1.657321 | 0.097454679 | 0.224052426 | CASP3 | protein_coding | 40.48 | -0.308036463 | 0.308036463 | ENSG00000164305 | n.s. |
| ENSG00000118217 | 1293.273779 | -0.448434274 | 0.2617178 | -1.713427 | 0.086634095 | 0.205719674 | ATF6 | protein_coding | 37.21 | -0.307950656 | 0.307950656 | ENSG00000118217 | n.s. |
| ENSG00000115163 | 512.2806384 | -0.491122056 | 0.3025125 | -1.623477 | 0.10448753 | 0.236088912 | CENPA | protein_coding | 48.97 | -0.307896404 | 0.307896404 | ENSG00000115163 | n.s. |
| ENSG00000225178 | 817.7083284 | 0.526071384 | 0.3378713 | 1.557017 | 0.11946648 | 0.260171362 | RPSAP58 | processed_pseudogene | 52.36 | 0.307615249 | 0.307615249 | ENSG00000225178 | n.s. |
| ENSG00000142892 | 651.3618826 | -0.491747106 | 0.3034957 | -1.620277 | 0.105172794 | 0.237060194 | PIGK | protein_coding | 35.84 | -0.307411457 | 0.307411457 | ENSG00000142892 | n.s. |
| ENSG00000089234 | 1153.107528 | -0.446426165 | 0.2602964 | -1.715069 | 0.086332627 | 0.205176131 | BRAP | protein_coding | 44.41 | -0.307084579 | 0.307084579 | ENSG00000089234 | n.s. |
| ENSG00000157343 | 36.82788006 | 0.489764662 | 0.3019811 | 1.621839 | 0.104837856 | 0.236573334 | ARMC12 | protein_coding | 46.92 | 0.306609433 | 0.306609433 | ENSG00000157343 | n.s. |
| ENSG00000197283 | 37.86494952 | 0.480610017 | 0.2932555 | 1.638878 | 0.101238671 | 0.230732693 | SYNGAP1 | protein_coding | 53.86 | 0.30609613 | 0.30609613 | ENSG00000197283 | n.s. |
| ENSG00000127957 | 227.396693 | 0.452581671 | 0.2666548 | 1.697257 | 0.089648084 | 0.211053478 | PMS2P3 | transcribed_unprocessed_pseud | 42.92 | 0.305767565 | 0.305767565 | ENSG00000127957 | n.s. |
| ENSG00000174106 | 1066.219575 | -0.44983009 | 0.2640265 | -1.703731 | 0.08843138 | 0.209108374 | LEMD3 | protein_coding | 35.54 | -0.305717383 | 0.305717383 | ENSG00000174106 | n.s. |
| ENSG00000060339 | 3787.938641 | -0.458897871 | 0.2726801 | -1.682917 | 0.092391207 | 0.215796473 | CCAR1 | protein_coding | 41.75 | -0.305605634 | 0.305605634 | ENSG00000060339 | n.s. |
| ENSG00000164938 | 513.7976044 | -0.45757018 | 0.271428 | -1.685788 | 0.091836579 | 0.21486567 | TP53INP1 | protein_coding | 42.53 | -0.305580452 | 0.305580452 | ENSG00000164938 | n.s. |
| ENSG00000138814 | 1474.288202 | -0.438578603 | 0.2537184 | -1.728604 | 0.083880057 | 0.201129989 | PPP3CA | protein_coding | 35.79 | -0.305480159 | 0.305480159 | ENSG00000138814 | n.s. |
| ENSG00000262180 | 60.08481943 | -0.499885848 | 0.3128454 | -1.597869 | 0.110072213 | 0.245155324 | OCLM | protein_coding | 35.52 | -0.305209639 | 0.305209639 | ENSG00000262180 | n.s. |
| ENSG00000181873 | 584.1517857 | 0.435032221 | 0.2506262 | 1.735781 | 0.082602502 | 0.198906024 | IBA57 | protein_coding | 54.59 | 0.305110746 | 0.305110746 | ENSG00000181873 | n.s. |
| ENSG00000126215 | 318.2280504 | -0.431317334 | 0.2473527 | -1.743734 | 0.08120551 | 0.196493625 | XRCC3 | protein_coding | 59.79 | -0.304791056 | 0.304791056 | ENSG00000126215 | n.s. |
| ENSG00000031823 | 646.4424204 | -0.443329894 | 0.2586542 | -1.713987 | 0.086531109 | 0.205583164 | RANBP3 | protein_coding | 52.74 | -0.304573158 | 0.304573158 | ENSG00000031823 | n.s. |
| ENSG00000198718 | 556.010468 | 0.392744347 | 0.2131699 | 1.842401 | 0.065416539 | 0.167711803 | TOGARAM1 | protein_coding | 37.85 | 0.304548251 | 0.304548251 | ENSG00000198718 | n.s. |
| ENSG00000133424 | 181.3171544 | -0.412515763 | 0.2306207 | -1.78872 | 0.073659926 | 0.182810544 | LARGE1 | protein_coding | 43.98 | -0.304436121 | 0.304436121 | ENSG00000133424 | n.s. |
| ENSG00000268750 | 89.90478028 | 0.422615814 | 0.2396009 | 1.763832 | 0.077760253 | 0.190407316 | | protein_coding | 47.11 | 0.304417088 | 0.304417088 | ENSG00000268750 | n.s. |
| ENSG00000144857 | 62.97426472 | 0.448931832 | 0.263928 | 1.700963 | 0.088949904 | 0.209871127 | BOC | protein_coding | 47.04 | 0.304397019 | 0.304397019 | ENSG00000144857 | n.s. |
| ENSG00000107863 | 1573.110882 | -0.433312262 | 0.2493865 | -1.737513 | 0.082296687 | 0.198478569 | ARHGAP21 | protein_coding | 38.96 | -0.3043093 | 0.3043093 | ENSG00000107863 | n.s. |
| ENSG00000237729 | 11.75956526 | 0.62462693 | 0.4481422 | 1.393814 | 0.163373785 | 0.325741545 | | processed_pseudogene | 47.62 | 0.304272547 | 0.304272547 | ENSG00000237729 | n.s. |
| ENSG00000172661 | 556.7590544 | -0.453656418 | 0.2686487 | -1.688661 | 0.091284493 | 0.213929305 | WASHC2C | protein_coding | 42.53 | -0.303827186 | 0.303827186 | ENSG00000172661 | n.s. |
| ENSG00000221491 | 17.40221468 | -0.536877434 | 0.3516421 | -1.526772 | 0.126817651 | 0.271708712 | SNORA2C | snoRNA | 50.36 | -0.303817026 | 0.303817026 | ENSG00000221491 | n.s. |
| ENSG00000099995 | 1049.954031 | -0.427224651 | 0.2441573 | -1.749793 | 0.080154119 | 0.194692219 | SF3A1 | protein_coding | 49.04 | -0.303607799 | 0.303607799 | ENSG00000099995 | n.s. |
| ENSG00000170142 | 1345.595499 | 0.491391873 | 0.3053575 | 1.609234 | 0.107565091 | 0.241230784 | UBE2E1 | protein_coding | 41.45 | 0.303467539 | 0.303467539 | ENSG00000170142 | n.s. |
| ENSG00000265432 | 6.101273079 | -0.659631432 | 0.4905357 | -1.344717 | 0.178716783 | 0.34674941 | MIR4308 | miRNA | 40.74 | -0.303420082 | 0.303420082 | ENSG00000265432 | n.s. |
| ENSG00000272172 | 12.2600896 | 0.56764442 | 0.38437 | 1.476818 | 0.139724547 | 0.292099831 | | antisense | 66.37 | 0.303388172 | 0.303388172 | ENSG00000272172 | n.s. |
| ENSG00000170634 | 98.97994508 | 0.441617529 | 0.2577473 | 1.713374 | 0.086643695 | 0.205719674 | ACYP2 | protein_coding | 40.31 | 0.303269432 | 0.303269432 | ENSG00000170634 | n.s. |

| | | | | | | | | | | | | | |
|---|---|---|---|---|---|---|---|---|---|---|---|---|---|
| ENSG00000107036 | 1147.386575 | -0.431581772 | 0.2483077 | -1.738093 | 0.082194487 | 0.198300786 | RIC1 | protein_coding | 37.47 | -0.303261965 | 0.303261965 | ENSG00000107036 | n.s. |
| ENSG00000011198 | 385.2579243 | 0.431936955 | 0.2486874 | 1.736867 | 0.082410614 | 0.198581339 | ABHD5 | protein_coding | 40.26 | 0.303246335 | 0.303246335 | ENSG00000011198 | n.s. |
| ENSG00000280198 | 63.30862597 | 0.472170639 | 0.2870414 | 1.644957 | 0.099978747 | 0.22849706 | | TEC | 51.71 | 0.302717749 | 0.302717749 | ENSG00000280198 | n.s. |
| ENSG00000171126 | 158.7260784 | 0.439204739 | 0.2557994 | 1.716989 | 0.08598116 | 0.204565224 | KCNG3 | protein_coding | 42.42 | 0.302685937 | 0.302685937 | ENSG00000171126 | n.s. |
| ENSG00000197150 | 28.06668802 | -0.49402631 | 0.3084729 | -1.601523 | 0.109261175 | 0.243984498 | ABCB8 | protein_coding | 57.04 | -0.302659175 | 0.302659175 | ENSG00000197150 | n.s. |
| ENSG00000056736 | 154.9953641 | -0.455868778 | 0.27163 | -1.678271 | 0.093294156 | 0.217067936 | IL17RB | protein_coding | 45.37 | -0.302425318 | 0.302425318 | ENSG00000056736 | n.s. |
| ENSG00000222432 | 11.3361148 | -0.590100419 | 0.4105973 | -1.437176 | 0.15066808 | 0.307498046 | | misc_RNA | 40.71 | -0.302224438 | 0.302224438 | ENSG00000222432 | n.s. |
| ENSG00000198719 | 133.313532 | 0.428626701 | 0.246164 | 1.741224 | 0.081644331 | 0.197278876 | DLL1 | protein_coding | 53.53 | 0.302147284 | 0.302147284 | ENSG00000198719 | n.s. |
| ENSG00000269894 | 21.47135027 | 0.521786119 | 0.3370615 | 1.548044 | 0.121611694 | 0.263615073 | | sense_intronic | 41.41 | 0.302129692 | 0.302129692 | ENSG00000269894 | n.s. |
| ENSG00000170315 | 11400.00796 | 0.509251457 | 0.3242706 | 1.570452 | 0.116309951 | 0.25518176 | UBB | protein_coding | 51.44 | 0.302062691 | 0.302062691 | ENSG00000170315 | n.s. |
| ENSG00000156970 | 2290.666206 | -0.457016139 | 0.2729765 | -1.674196 | 0.094092058 | 0.218377016 | BUB1B | protein_coding | 38.74 | -0.301993099 | 0.301993099 | ENSG00000156970 | n.s. |
| ENSG00000148468 | 1302.950964 | -0.45847 | 0.2744113 | -1.67074 | 0.094772995 | 0.219518285 | FAM171A1 | protein_coding | 44.98 | -0.301915928 | 0.301915928 | ENSG00000148468 | n.s. |
| ENSG00000180398 | 1771.996194 | -0.45361728 | 0.2696936 | -1.681973 | 0.092574155 | 0.216142209 | MCFD2 | protein_coding | 45.38 | -0.30177362 | 0.30177362 | ENSG00000180398 | n.s. |
| ENSG00000146352 | 5112.247442 | -0.425327145 | 0.243385 | -1.747549 | 0.080542204 | 0.195261903 | CLVS2 | protein_coding | 36.42 | -0.301719626 | 0.301719626 | ENSG00000146352 | n.s. |
| ENSG00000100014 | 2027.561565 | -0.432533316 | 0.2499906 | -1.730198 | 0.083594856 | 0.200670894 | SPECC1L | protein_coding | 42.29 | -0.301698741 | 0.301698741 | ENSG00000100014 | n.s. |
| ENSG00000133612 | 860.2973726 | -0.446002569 | 0.2628239 | -1.696964 | 0.089703535 | 0.211085381 | AGAP3 | protein_coding | 57.73 | -0.301293397 | 0.301293397 | ENSG00000133612 | n.s. |
| ENSG00000161970 | 17021.12194 | 0.515483156 | 0.3312083 | 1.556372 | 0.119619758 | 0.260384131 | RPL26 | protein_coding | 43.78 | 0.301240875 | 0.301240875 | ENSG00000161970 | n.s. |
| ENSG00000179431 | 824.9530653 | 0.384370395 | 0.2074096 | 1.853195 | 0.063854432 | 0.16463801 | FJX1 | protein_coding | 63.8 | 0.301142632 | 0.301142632 | ENSG00000179431 | n.s. |
| ENSG00000147123 | 1692.171691 | 0.515963763 | 0.3318697 | 1.554718 | 0.120013244 | 0.261003499 | NDUFB11 | protein_coding | 52.3 | 0.300989355 | 0.300989355 | ENSG00000147123 | n.s. |
| ENSG00000182134 | 251.3526678 | -0.484898128 | 0.300479 | -1.61375 | 0.106581564 | 0.239537549 | TDRKH | protein_coding | 43.72 | -0.300940579 | 0.300940579 | ENSG00000182134 | n.s. |
| ENSG00000140836 | 836.6327688 | -0.42783883 | 0.2461062 | -1.738432 | 0.082134797 | 0.198225477 | ZFHX3 | protein_coding | 46.45 | -0.300702469 | 0.300702469 | ENSG00000140836 | n.s. |
| ENSG00000104497 | 167.8624649 | -0.454236782 | 0.2710364 | -1.675925 | 0.093752792 | 0.217880173 | SNX16 | protein_coding | 35.17 | -0.300605857 | 0.300605857 | ENSG00000104497 | n.s. |
| ENSG00000187951 | 966.3989023 | -0.41863526 | 0.2378277 | -1.760246 | 0.078366102 | 0.191486703 | ARHGAP11B | protein_coding | 40.92 | -0.300522085 | 0.300522085 | ENSG00000187951 | n.s. |
| ENSG00000100316 | 18670.09564 | 0.460870534 | 0.2774188 | 1.661281 | 0.096657099 | 0.222807416 | RPL3 | protein_coding | 53.41 | 0.300520014 | 0.300520014 | ENSG00000100316 | n.s. |
| ENSG00000123444 | 466.3820187 | -0.438042889 | 0.2558329 | -1.712222 | 0.0868557 | 0.206082466 | KBTBD4 | protein_coding | 48.95 | -0.300479444 | 0.300479444 | ENSG00000123444 | n.s. |
| ENSG00000224616 | 57.51546289 | 0.446784566 | 0.2641969 | 1.691105 | 0.090816808 | 0.213120245 | RTCA-AS1 | antisense | 42.8 | 0.299960119 | 0.299960119 | ENSG00000224616 | n.s. |
| ENSG00000104177 | 4631.537659 | -0.455656383 | 0.2728447 | -1.670021 | 0.094915163 | 0.219689908 | MYEF2 | protein_coding | 33.92 | -0.299908426 | 0.299908426 | ENSG00000104177 | n.s. |
| ENSG00000284231 | 3.175152082 | 0.745218181 | 0.6039863 | 1.233833 | 0.21726514 | 0.3960438 | MIR424 | miRNA | 51.02 | 0.299769067 | 0.299769067 | ENSG00000284231 | n.s. |
| ENSG00000230454 | 169.394049 | 0.400298709 | 0.2219837 | 1.80328 | 0.071344245 | 0.178501317 | | lincRNA | 58.51 | 0.299566972 | 0.299566972 | ENSG00000230454 | n.s. |
| ENSG00000184144 | 1149.694971 | -0.43071034 | 0.2494462 | -1.726666 | 0.084227634 | 0.201633657 | CNTN2 | protein_coding | 54.18 | -0.299531895 | 0.299531895 | ENSG00000184144 | n.s. |
| ENSG00000262265 | 86.89279011 | -0.432744357 | 0.2513832 | -1.721453 | 0.085168695 | 0.203257412 | | sense_intronic | 43.33 | -0.299439019 | 0.299439019 | ENSG00000262265 | n.s. |
| ENSG00000187676 | 193.9482982 | -0.457320556 | 0.2747868 | -1.664274 | 0.09605763 | 0.221719228 | B3GLCT | protein_coding | 41.08 | -0.299177572 | 0.299177572 | ENSG00000187676 | n.s. |

| | | | | | | | | | | | | | |
|---|---|---|---|---|---|---|---|---|---|---|---|---|---|
| ENSG00000137776 | 4843.94355 | -0.437406171 | 0.2558944 | -1.709323 | 0.087391159 | 0.207070644 | SLTM | protein_coding | 38.73 | -0.299133973 | 0.299133973 | ENSG00000137776 | n.s. |
| ENSG00000122126 | 646.3080545 | -0.431547508 | 0.2507286 | -1.721174 | 0.085219246 | 0.203308355 | OCRL | protein_coding | 39.83 | -0.298563888 | 0.298563888 | ENSG00000122126 | n.s. |
| ENSG00000162878 | 599.9870714 | -0.441052989 | 0.2596392 | -1.698715 | 0.08937294 | 0.210524418 | PKDCC | protein_coding | 56.22 | -0.298459466 | 0.298459466 | ENSG00000162878 | n.s. |
| ENSG00000137054 | 1569.820228 | -0.420480716 | 0.2405684 | -1.747863 | 0.080487683 | 0.195163708 | POLR1E | protein_coding | 46.29 | -0.298373511 | 0.298373511 | ENSG00000137054 | n.s. |
| ENSG00000065154 | 1071.158902 | 0.447017652 | 0.2653216 | 1.684814 | 0.092024407 | 0.215156309 | OAT | protein_coding | 42.49 | 0.298270703 | 0.298270703 | ENSG00000065154 | n.s. |
| ENSG00000134278 | 939.7978538 | -0.446515182 | 0.2648102 | -1.68617 | 0.091762979 | 0.214797572 | SPIRE1 | protein_coding | 40.66 | -0.298259028 | 0.298259028 | ENSG00000134278 | n.s. |
| ENSG00000188895 | 1198.64337 | -0.439173438 | 0.2579977 | -1.702238 | 0.088710783 | 0.209457066 | MSL1 | protein_coding | 45.42 | -0.298157036 | 0.298157036 | ENSG00000188895 | n.s. |
| ENSG00000105583 | 609.8912749 | 0.487246062 | 0.3046316 | 1.59946 | 0.10971848 | 0.244549375 | WDR83OS | protein_coding | 57.12 | 0.298015986 | 0.298015986 | ENSG00000105583 | n.s. |
| ENSG00000169057 | 193.2633787 | 0.417528856 | 0.2380103 | 1.754247 | 0.079388147 | 0.193305557 | MECP2 | protein_coding | 46.41 | 0.298013584 | 0.298013584 | ENSG00000169057 | n.s. |
| ENSG00000168724 | 1372.284241 | -0.438623785 | 0.2576774 | -1.70222 | 0.088714059 | 0.209457066 | DNAJC21 | protein_coding | 39.24 | -0.297783874 | 0.297783874 | ENSG00000168724 | n.s. |
| ENSG00000115875 | 11696.31752 | 0.394586227 | 0.2177733 | 1.811913 | 0.069999677 | 0.176055903 | SRSF7 | protein_coding | 41.45 | 0.297655887 | 0.297655887 | ENSG00000115875 | n.s. |
| ENSG00000145819 | 66.73168021 | 0.439729114 | 0.2588037 | 1.699084 | 0.089303416 | 0.2104623 | ARHGAP26 | protein_coding | 42.17 | 0.29761996 | 0.29761996 | ENSG00000145819 | n.s. |
| ENSG00000164032 | 10126.29475 | 0.484445827 | 0.3019514 | 1.604383 | 0.108629611 | 0.243041953 | H2AFZ | protein_coding | 49.59 | 0.297604157 | 0.297604157 | ENSG00000164032 | n.s. |
| ENSG00000159592 | 167.2629757 | 0.426662284 | 0.2466283 | 1.729981 | 0.083633672 | 0.200729463 | GPBP1L1 | protein_coding | 41.19 | 0.297549532 | 0.297549532 | ENSG00000159592 | n.s. |
| ENSG00000183763 | 384.3577088 | -0.434756394 | 0.2542476 | -1.709972 | 0.087270966 | 0.206898949 | TRAIP | protein_coding | 49.84 | -0.297478461 | 0.297478461 | ENSG00000183763 | n.s. |
| ENSG00000099194 | 6690.525169 | -0.43461229 | 0.2541395 | -1.710133 | 0.087241281 | 0.20689156 | SCD | protein_coding | 47.04 | -0.297386599 | 0.297386599 | ENSG00000099194 | n.s. |
| ENSG00000198792 | 189.8835907 | -0.4506996 | 0.2696182 | -1.671621 | 0.094599014 | 0.219224713 | TMEM184B | protein_coding | 53.51 | -0.297060834 | 0.297060834 | ENSG00000198792 | n.s. |
| ENSG00000151366 | 1690.369775 | 0.502193372 | 0.3203957 | 1.567416 | 0.117017479 | 0.256252639 | NDUFC2 | protein_coding | 44.63 | 0.296962837 | 0.296962837 | ENSG00000151366 | n.s. |
| ENSG00000006125 | 1374.013537 | -0.439956138 | 0.2594497 | -1.695728 | 0.089937412 | 0.211412898 | AP2B1 | protein_coding | 39.93 | -0.296912548 | 0.296912548 | ENSG00000006125 | n.s. |
| ENSG00000177173 | 48.25540459 | -0.473488653 | 0.2915562 | -1.624005 | 0.10437476 | 0.236025626 | NAP1L4P1 | processed_pseudogene | 42.35 | -0.296896724 | 0.296896724 | ENSG00000177173 | n.s. |
| ENSG00000166206 | 3131.916183 | -0.441608823 | 0.2610179 | -1.691872 | 0.090670453 | 0.212884438 | GABRB3 | protein_coding | 41.34 | -0.296697575 | 0.296697575 | ENSG00000166206 | n.s. |
| ENSG00000122026 | 9652.413904 | 0.497573287 | 0.3159233 | 1.574981 | 0.115260741 | 0.253398909 | RPL21 | protein_coding | 41.95 | 0.296650834 | 0.296650834 | ENSG00000122026 | n.s. |
| ENSG00000142875 | 855.947482 | -0.4716778 | 0.2900539 | -1.626173 | 0.103912837 | 0.235188253 | PRKACB | protein_coding | 34.7 | -0.296489295 | 0.296489295 | ENSG00000142875 | n.s. |
| ENSG00000212588 | 299.2198436 | 0.394600254 | 0.2183562 | 1.80714 | 0.070740437 | 0.177404599 | SNORA26 | snoRNA | 45.9 | 0.296358651 | 0.296358651 | ENSG00000212588 | n.s. |
| ENSG00000060069 | 136.7662736 | 0.407858812 | 0.230034 | 1.773037 | 0.076222507 | 0.187698261 | CTDP1 | protein_coding | 53.52 | 0.29632564 | 0.29632564 | ENSG00000060069 | n.s. |
| ENSG00000046647 | 173.7626268 | -0.42488258 | 0.245801 | -1.728564 | 0.083887226 | 0.201129989 | GEMIN8 | protein_coding | 40.79 | -0.295940562 | 0.295940562 | ENSG00000046647 | n.s. |
| ENSG00000148303 | 20635.51812 | 0.492739913 | 0.3117002 | 1.580814 | 0.113920635 | 0.251036202 | RPL7A | protein_coding | 50.42 | 0.295773856 | 0.295773856 | ENSG00000148303 | n.s. |
| ENSG00000284062 | 2.996847157 | 0.782437203 | 0.6590398 | 1.187238 | 0.235133718 | 0.418854499 | MIR4469 | miRNA | 72.15 | 0.295711825 | 0.295711825 | ENSG00000284062 | n.s. |
| ENSG00000087157 | 1008.912236 | -0.424696469 | 0.2458538 | -1.727435 | 0.084089602 | 0.20140711 | PGS1 | protein_coding | 52.04 | -0.295556976 | 0.295556976 | ENSG00000087157 | n.s. |
| ENSG00000252695 | 3.837128818 | 0.740606672 | 0.6037352 | 1.226708 | 0.219932476 | 0.399303693 | MIR2276 | miRNA | 57.3 | 0.295277416 | 0.295277416 | ENSG00000252695 | n.s. |
| ENSG00000184203 | 844.1458787 | 0.453822851 | 0.2737315 | 1.657913 | 0.09733511 | 0.223918187 | PPP1R2 | protein_coding | 39 | 0.294944295 | 0.294944295 | ENSG00000184203 | n.s. |
| ENSG00000197136 | 358.815386 | -0.392696966 | 0.2173101 | -1.807081 | 0.070749702 | 0.177404599 | PCNX3 | protein_coding | 60.5 | -0.294929215 | 0.294929215 | ENSG00000197136 | n.s. |

| | | | | | | | | | | | | | |
|---|---|---|---|---|---|---|---|---|---|---|---|---|---|
| ENSG00000202470 | 20.32499699 | -0.54385172 | 0.3652399 | -1.489026 | 0.13648053 | 0.287251639 | | misc_RNA | 40.78 | -0.294624863 | 0.294624863 | ENSG00000202470 | n.s. |
| ENSG00000119977 | 549.9626065 | -0.443866251 | 0.2644178 | -1.678655 | 0.09321926 | 0.216929929 | TCTN3 | protein_coding | 39.54 | -0.294585388 | 0.294585388 | ENSG00000119977 | n.s. |
| ENSG00000143476 | 3997.852661 | -0.440968937 | 0.2616192 | -1.685537 | 0.091884936 | 0.214902395 | DTL | protein_coding | 38.44 | -0.294460864 | 0.294460864 | ENSG00000143476 | n.s. |
| ENSG00000170631 | 343.7607605 | -0.414668897 | 0.2371615 | -1.748466 | 0.08038329 | 0.195036304 | ZNF16 | protein_coding | 47.4 | -0.294367039 | 0.294367039 | ENSG00000170631 | n.s. |
| ENSG00000070669 | 399.0521838 | 0.440559142 | 0.2612289 | 1.686487 | 0.091701997 | 0.214727028 | ASNS | protein_coding | 38.42 | 0.294343416 | 0.294343416 | ENSG00000070669 | n.s. |
| ENSG00000110900 | 951.9084498 | -0.43878643 | 0.259588 | -1.690319 | 0.090967033 | 0.213420813 | TSPAN11 | protein_coding | 51.34 | -0.294321803 | 0.294321803 | ENSG00000110900 | n.s. |
| ENSG00000265981 | 62.8125516 | 0.465787844 | 0.2855238 | 1.631345 | 0.102817564 | 0.233452209 | MIR544B | miRNA | 39.74 | 0.294285697 | 0.294285697 | ENSG00000265981 | n.s. |
| ENSG00000158882 | 325.7850851 | -0.451261638 | 0.2717458 | -1.660602 | 0.09679342 | 0.223047795 | TOMM40L | protein_coding | 52.77 | -0.294043015 | 0.294043015 | ENSG00000158882 | n.s. |
| ENSG00000134775 | 355.562575 | -0.4694765 | 0.2893973 | -1.622256 | 0.104748606 | 0.236476489 | FHOD3 | protein_coding | 43.06 | -0.293991833 | 0.293991833 | ENSG00000134775 | n.s. |
| ENSG00000168546 | 2768.337883 | -0.446631409 | 0.26746 | -1.6699 | 0.094939227 | 0.219689908 | GFRA2 | protein_coding | 52.15 | -0.29396828 | 0.29396828 | ENSG00000168546 | n.s. |
| ENSG00000202141 | 53.55549213 | 0.479848912 | 0.2996734 | 1.60124 | 0.109323832 | 0.244013714 | | misc_RNA | 42.48 | 0.293948613 | 0.293948613 | ENSG00000202141 | n.s. |
| ENSG00000198830 | 3199.506741 | 0.396403013 | 0.220948 | 1.794101 | 0.072797052 | 0.181412807 | HMGN2 | protein_coding | 48.34 | 0.293866262 | 0.293866262 | ENSG00000198830 | n.s. |
| ENSG00000226709 | 18.47832863 | -0.526162005 | 0.3473331 | -1.514863 | 0.129807104 | 0.27643076 | FGF12-AS3 | antisense | 34.08 | -0.293816036 | 0.293816036 | ENSG00000226709 | n.s. |
| ENSG00000153879 | 3375.340414 | 0.430446908 | 0.2520708 | 1.707643 | 0.087702612 | 0.207720748 | CEBPG | protein_coding | 44.81 | 0.293788676 | 0.293788676 | ENSG00000153879 | n.s. |
| ENSG00000100084 | 866.8535727 | -0.418853264 | 0.241318 | -1.73569 | 0.082618617 | 0.198910427 | HIRA | protein_coding | 46.11 | -0.293759575 | 0.293759575 | ENSG00000100084 | n.s. |
| ENSG00000167797 | 111.3792203 | 0.418976097 | 0.2415348 | 1.734641 | 0.082804433 | 0.199185578 | CDK2AP2 | protein_coding | 62.33 | 0.293594195 | 0.293594195 | ENSG00000167797 | n.s. |
| ENSG00000267216 | 511.2690097 | 0.370252139 | 0.1984708 | 1.865524 | 0.062107951 | 0.161239791 | | protein_coding | 47.15 | 0.293435103 | 0.293435103 | ENSG00000267216 | n.s. |
| ENSG00000182446 | 2442.060194 | -0.422613872 | 0.2450961 | -1.724278 | 0.084657669 | 0.20238474 | NPLOC4 | protein_coding | 50.06 | -0.293218902 | 0.293218902 | ENSG00000182446 | n.s. |
| ENSG00000073584 | 5634.22289 | -0.441109829 | 0.2626183 | -1.679661 | 0.093023234 | 0.216654815 | SMARCE1 | protein_coding | 37.72 | -0.292999111 | 0.292999111 | ENSG00000073584 | n.s. |
| ENSG00000100221 | 622.4846286 | -0.442427538 | 0.2638695 | -1.676691 | 0.093602979 | 0.217677346 | JOSD1 | protein_coding | 47.81 | -0.292969661 | 0.292969661 | ENSG00000100221 | n.s. |
| ENSG00000276564 | 137.7296521 | -0.440461098 | 0.2619774 | -1.681295 | 0.092705714 | 0.216313332 | | lincRNA | 32.83 | -0.292869946 | 0.292869946 | ENSG00000276564 | n.s. |
| ENSG00000157181 | 1568.781085 | -0.442492345 | 0.2640855 | -1.675565 | 0.093823504 | 0.217942059 | C1orf27 | protein_coding | 35.17 | -0.292779021 | 0.292779021 | ENSG00000157181 | n.s. |
| ENSG00000266555 | 6.138488251 | -0.656132689 | 0.497053 | -1.320046 | 0.186819765 | 0.357981225 | MIR3145 | miRNA | 31.71 | -0.292726873 | 0.292726873 | ENSG00000266555 | n.s. |
| ENSG00000149084 | 1669.274924 | -0.46836032 | 0.2891289 | -1.619901 | 0.105253493 | 0.237165299 | HSD17B12 | protein_coding | 38.16 | -0.292701246 | 0.292701246 | ENSG00000149084 | n.s. |
| ENSG00000258017 | 34.64201085 | 0.493060933 | 0.3138111 | 1.571203 | 0.116135507 | 0.254917798 | | antisense | 46.25 | 0.292680892 | 0.292680892 | ENSG00000258017 | n.s. |
| ENSG00000171604 | 375.8456619 | -0.451120492 | 0.2724933 | -1.655528 | 0.097817346 | 0.224728792 | CXXC5 | protein_coding | 58.2 | -0.292480039 | 0.292480039 | ENSG00000171604 | n.s. |
| ENSG00000234797 | 472.7711464 | 0.529903178 | 0.3520714 | 1.505101 | 0.132298048 | 0.280747708 | RPS3AP6 | processed_pseudogene | 42.26 | 0.292338988 | 0.292338988 | ENSG00000234797 | n.s. |
| ENSG00000163682 | 17572.98781 | 0.493300383 | 0.3143678 | 1.569182 | 0.11660547 | 0.255648723 | RPL9 | protein_coding | 41.12 | 0.292209626 | 0.292209626 | ENSG00000163682 | n.s. |
| ENSG00000160209 | 15.96388765 | 0.536068755 | 0.3587491 | 1.494272 | 0.135104555 | 0.285069603 | PDXK | protein_coding | 57.26 | 0.292183787 | 0.292183787 | ENSG00000160209 | n.s. |
| ENSG00000187498 | 732.731639 | -0.423769985 | 0.2467289 | -1.717553 | 0.08587822 | 0.204425108 | COL4A1 | protein_coding | 44.9 | -0.292174897 | 0.292174897 | ENSG00000187498 | n.s. |
| ENSG00000188760 | 243.946499 | -0.467410919 | 0.2885195 | -1.620033 | 0.105225283 | 0.237140113 | TMEM198 | protein_coding | 56.18 | -0.292129477 | 0.292129477 | ENSG00000188760 | n.s. |
| ENSG00000080802 | 841.7081304 | -0.420405647 | 0.2436654 | -1.72534 | 0.084466222 | 0.202100569 | CNOT4 | protein_coding | 37.19 | -0.29194333 | 0.29194333 | ENSG00000080802 | n.s. |

| | | | | | | | | | | | | | |
|---|---|---|---|---|---|---|---|---|---|---|---|---|---|
| ENSG00000171863 | 24354.37981 | 0.493628568 | 0.3149327 | 1.56741 | 0.11701891 | 0.256252639 | RPS7 | protein_coding | 44.76 | 0.291898197 | 0.291898197 | ENSG00000171863 | n.s. |
| ENSG00000272721 | 23.15009225 | 0.485352365 | 0.3065928 | 1.583052 | 0.113409548 | 0.250476069 | | antisense | 53.28 | 0.291810228 | 0.291810228 | ENSG00000272721 | n.s. |
| ENSG00000179335 | 1059.990584 | 0.393887197 | 0.2196715 | 1.793073 | 0.072961174 | 0.181724227 | CLK3 | protein_coding | 54.25 | 0.291707804 | 0.291707804 | ENSG00000179335 | n.s. |
| ENSG00000170291 | 475.4514112 | -0.487547415 | 0.3090945 | -1.577341 | 0.114717071 | 0.252522653 | ELP5 | protein_coding | 51.3 | -0.291406922 | 0.291406922 | ENSG00000170291 | n.s. |
| ENSG00000273802 | 2208.785392 | 0.383371575 | 0.2105708 | 1.82063 | 0.068663111 | 0.173754362 | HIST1H2BG | protein_coding | 48.76 | 0.291387042 | 0.291387042 | ENSG00000273802 | n.s. |
| ENSG00000171320 | 1309.375726 | -0.396866265 | 0.2225769 | -1.783052 | 0.07457777 | 0.184660553 | ESCO2 | protein_coding | 37.71 | -0.291151359 | 0.291151359 | ENSG00000171320 | n.s. |
| ENSG00000164587 | 15317.94367 | 0.475782755 | 0.29747 | 1.599431 | 0.109724912 | 0.244549375 | RPS14 | protein_coding | 48.97 | 0.291004645 | 0.291004645 | ENSG00000164587 | n.s. |
| ENSG00000138587 | 179.7516082 | -0.469675667 | 0.2913171 | -1.612249 | 0.106907852 | 0.240115828 | MNS1 | protein_coding | 35.19 | -0.291001279 | 0.291001279 | ENSG00000138587 | n.s. |
| ENSG00000184575 | 6094.204224 | -0.460358926 | 0.2822262 | -1.63117 | 0.102854418 | 0.23349781 | XPOT | protein_coding | 39.76 | -0.290816646 | 0.290816646 | ENSG00000184575 | n.s. |
| ENSG00000221990 | 110.8212438 | 0.417321566 | 0.2414688 | 1.728263 | 0.08394114 | 0.201155333 | EXOC3-AS1 | antisense | 55.32 | 0.290651303 | 0.290651303 | ENSG00000221990 | n.s. |
| ENSG00000104228 | 370.2892464 | -0.418774362 | 0.2428249 | -1.724594 | 0.084600661 | 0.202317933 | TRIM35 | protein_coding | 45.69 | -0.29061501 | 0.29061501 | ENSG00000104228 | n.s. |
| ENSG00000280789 | 547.5213698 | -0.464633281 | 0.2866789 | -1.620744 | 0.105072481 | 0.236910793 | PAGR1 | protein_coding | 54.75 | -0.290588694 | 0.290588694 | ENSG00000280789 | n.s. |
| ENSG00000272631 | 747.5072871 | 0.382078196 | 0.2098098 | 1.821069 | 0.06859636 | 0.173658618 | | antisense | 44.98 | 0.290495451 | 0.290495451 | ENSG00000272631 | n.s. |
| ENSG00000177551 | 3391.462428 | 0.374746165 | 0.203518 | 1.841342 | 0.065571467 | 0.167945962 | NHLH2 | protein_coding | 46.33 | 0.290364733 | 0.290364733 | ENSG00000177551 | n.s. |
| ENSG00000176124 | 150.6318503 | 0.436574427 | 0.2596284 | 1.681536 | 0.092658882 | 0.216240297 | DLEU1 | protein_coding | 39.72 | 0.29034966 | 0.29034966 | ENSG00000176124 | n.s. |
| ENSG00000183258 | 973.0848676 | 0.425244905 | 0.2490832 | 1.70724 | 0.087777383 | 0.207738315 | DDX41 | protein_coding | 59.55 | 0.290222587 | 0.290222587 | ENSG00000183258 | n.s. |
| ENSG00000125977 | 2855.103488 | -0.466125628 | 0.2883329 | -1.616623 | 0.10595963 | 0.238447701 | EIF2S2 | protein_coding | 42.27 | -0.290213013 | 0.290213013 | ENSG00000125977 | n.s. |
| ENSG00000161547 | 18423.38164 | 0.391651134 | 0.2183858 | 1.793391 | 0.072910476 | 0.181630445 | SRSF2 | protein_coding | 49.23 | 0.290139606 | 0.290139606 | ENSG00000161547 | n.s. |
| ENSG00000138794 | 79.92621135 | -0.487650099 | 0.3100275 | -1.572925 | 0.115736147 | 0.254283475 | CASP6 | protein_coding | 44.61 | -0.289996669 | 0.289996669 | ENSG00000138794 | n.s. |
| ENSG00000256053 | 135.2988212 | -0.501182126 | 0.3240075 | -1.546823 | 0.121906049 | 0.264047592 | APOPT1 | protein_coding | 48.97 | -0.289842539 | 0.289842539 | ENSG00000256053 | n.s. |
| ENSG00000004700 | 1888.928427 | -0.44011316 | 0.2634545 | -1.670547 | 0.094811198 | 0.219570245 | RECQL | protein_coding | 37.57 | -0.289782174 | 0.289782174 | ENSG00000004700 | n.s. |
| ENSG00000166595 | 628.3174128 | 0.513535792 | 0.3374198 | 1.521949 | 0.128021832 | 0.273507437 | FAM96B | protein_coding | 55.72 | 0.289136499 | 0.289136499 | ENSG00000166595 | n.s. |
| ENSG00000240972 | 4845.322083 | 0.510073274 | 0.3338558 | 1.527825 | 0.126555959 | 0.271249787 | MIF | protein_coding | 69.2 | 0.289023122 | 0.289023122 | ENSG00000240972 | n.s. |
| ENSG00000255198 | 73.91435722 | 0.473188359 | 0.2961287 | 1.597915 | 0.11006194 | 0.245155324 | SNHG9 | lincRNA | 66.42 | 0.288909256 | 0.288909256 | ENSG00000255198 | n.s. |
| ENSG00000170577 | 325.9604572 | 0.402483347 | 0.2287302 | 1.759642 | 0.078468454 | 0.191669523 | SIX2 | protein_coding | 59.16 | 0.288760445 | 0.288760445 | ENSG00000170577 | n.s. |
| ENSG00000186312 | 150.9466479 | -0.478573864 | 0.3018395 | -1.585524 | 0.112847259 | 0.249550887 | CA5BP1 | transcribed_unprocessed_pseud | 43 | -0.28850389 | 0.28850389 | ENSG00000186312 | n.s. |
| ENSG00000008952 | 3208.582255 | -0.445760107 | 0.2697533 | -1.652473 | 0.098438152 | 0.22567996 | SEC62 | protein_coding | 36.6 | -0.288187031 | 0.288187031 | ENSG00000008952 | n.s. |
| ENSG00000169905 | 3667.797298 | -0.40705494 | 0.2333872 | -1.744118 | 0.081138442 | 0.196367794 | TOR1AIP2 | protein_coding | 38.85 | -0.287759242 | 0.287759242 | ENSG00000169905 | n.s. |
| ENSG00000176444 | 1509.713968 | -0.425012408 | 0.250152 | -1.699017 | 0.089316067 | 0.2104623 | CLK2 | protein_coding | 50.41 | -0.287659315 | 0.287659315 | ENSG00000176444 | n.s. |
| ENSG00000171861 | 406.0488826 | -0.406407023 | 0.232851 | -1.745352 | 0.080923579 | 0.195981734 | MRM3 | protein_coding | 48.63 | -0.287648553 | 0.287648553 | ENSG00000171861 | n.s. |
| ENSG00000112851 | 1553.531311 | -0.444884321 | 0.2694097 | -1.65133 | 0.098671207 | 0.226102571 | ERBIN | protein_coding | 37.27 | -0.287259359 | 0.287259359 | ENSG00000112851 | n.s. |
| ENSG00000167377 | 370.08715 | -0.414627596 | 0.2406522 | -1.722933 | 0.084900602 | 0.202861005 | ZNF23 | protein_coding | 44.44 | -0.28725459 | 0.28725459 | ENSG00000167377 | n.s. |

| | | | | | | | | | | | | | |
|---|---|---|---|---|---|---|---|---|---|---|---|---|---|
| ENSG00000164951 | 1141.693136 | 0.41739129 | 0.2433002 | 1.71554 | 0.086246274 | 0.205055818 | PDP1 | protein_coding | 42.9 | 0.287218593 | 0.287218593 | ENSG00000164951 | n.s. |
| ENSG00000177192 | 575.7462727 | -0.386625266 | 0.2152342 | -1.7963 | 0.072446815 | 0.180798889 | PUS1 | protein_coding | 51.99 | -0.287186888 | 0.287186888 | ENSG00000177192 | n.s. |
| ENSG00000154001 | 1171.117792 | -0.4352597 | 0.2602516 | -1.672457 | 0.094434187 | 0.218952071 | PPP2R5E | protein_coding | 40.18 | -0.287119471 | 0.287119471 | ENSG00000154001 | n.s. |
| ENSG00000140993 | 621.9361224 | -0.433691307 | 0.2587706 | -1.675968 | 0.093744375 | 0.217880173 | TIGD7 | protein_coding | 44.07 | -0.287009226 | 0.287009226 | ENSG00000140993 | n.s. |
| ENSG00000100461 | 960.6063207 | -0.452902302 | 0.2771489 | -1.634148 | 0.102227841 | 0.232459359 | RBM23 | protein_coding | 45.96 | -0.286982886 | 0.286982886 | ENSG00000100461 | n.s. |
| ENSG00000252756 | 7.259923204 | -0.597630805 | 0.4328064 | -1.380827 | 0.167332094 | 0.331026533 | RNU6-577P | snRNA | 40.82 | -0.286944778 | 0.286944778 | ENSG00000252756 | n.s. |
| ENSG00000144815 | 905.1222024 | -0.428652715 | 0.2539262 | -1.6881 | 0.09139209 | 0.214145418 | NXPE3 | protein_coding | 39.47 | -0.286893496 | 0.286893496 | ENSG00000144815 | n.s. |
| ENSG00000196814 | 692.4453414 | -0.418500193 | 0.2445582 | -1.71125 | 0.087035024 | 0.206437587 | MVB12B | protein_coding | 47.23 | -0.28676103 | 0.28676103 | ENSG00000196814 | n.s. |
| ENSG00000138698 | 1095.017016 | -0.469766769 | 0.2940859 | -1.59738 | 0.110181094 | 0.245252019 | RAP1GDS1 | protein_coding | 36.01 | -0.286739721 | 0.286739721 | ENSG00000138698 | n.s. |
| ENSG00000132639 | 1146.110801 | -0.458282688 | 0.2828059 | -1.620485 | 0.1051281 | 0.236997819 | SNAP25 | protein_coding | 40.89 | -0.28654384 | 0.28654384 | ENSG00000132639 | n.s. |
| ENSG00000128973 | 1323.139729 | -0.414103567 | 0.2406173 | -1.721005 | 0.085249931 | 0.203346717 | CLN6 | protein_coding | 47.26 | -0.286461457 | 0.286461457 | ENSG00000128973 | n.s. |
| ENSG00000185008 | 684.4411523 | -0.445793064 | 0.2707398 | -1.646574 | 0.099645626 | 0.227847932 | ROBO2 | protein_coding | 35.65 | -0.286357363 | 0.286357363 | ENSG00000185008 | n.s. |
| ENSG00000279075 | 183.8111774 | 0.398974527 | 0.2268214 | 1.758981 | 0.078580705 | 0.191809109 | | TEC | 38.6 | 0.28611691 | 0.28611691 | ENSG00000279075 | n.s. |
| ENSG00000061676 | 5119.673104 | -0.44516505 | 0.2702929 | -1.646973 | 0.099563586 | 0.227735141 | NCKAP1 | protein_coding | 35.82 | -0.286049684 | 0.286049684 | ENSG00000061676 | n.s. |
| ENSG00000143776 | 3176.29811 | -0.432779242 | 0.2585372 | -1.673954 | 0.094139728 | 0.218451239 | CDC42BPA | protein_coding | 36.66 | -0.285913654 | 0.285913654 | ENSG00000143776 | n.s. |
| ENSG00000103363 | 2134.873196 | 0.494426055 | 0.3198181 | 1.54596 | 0.122114192 | 0.264210707 | ELOB | protein_coding | 55.52 | 0.285802777 | 0.285802777 | ENSG00000103363 | n.s. |
| ENSG00000158234 | 596.3287598 | -0.466532402 | 0.2913275 | -1.601402 | 0.109287977 | 0.244005213 | FAIM | protein_coding | 40.46 | -0.285798167 | 0.285798167 | ENSG00000158234 | n.s. |
| ENSG00000212402 | 18.74046612 | -0.523847814 | 0.3505048 | -1.494553 | 0.135031243 | 0.285027288 | SNORA74B | snoRNA | 54.73 | -0.285556546 | 0.285556546 | ENSG00000212402 | n.s. |
| ENSG00000122376 | 1041.441848 | -0.475649168 | 0.3008047 | -1.581256 | 0.113819596 | 0.251036202 | FAM35A | protein_coding | 40.46 | -0.285514903 | 0.285514903 | ENSG00000122376 | n.s. |
| ENSG00000185009 | 1326.186547 | -0.445272208 | 0.2708271 | -1.64412 | 0.100151409 | 0.228703964 | AP3M1 | protein_coding | 41.25 | -0.285297618 | 0.285297618 | ENSG00000185009 | n.s. |
| ENSG00000215041 | 404.2311577 | -0.437120756 | 0.2631935 | -1.660834 | 0.096746712 | 0.222977068 | NEURL4 | protein_coding | 57.23 | -0.284888994 | 0.284888994 | ENSG00000215041 | n.s. |
| ENSG00000105372 | 14656.2889 | 0.492447636 | 0.3184548 | 1.546366 | 0.122016231 | 0.264156622 | RPS19 | protein_coding | 53.52 | 0.284702937 | 0.284702937 | ENSG00000105372 | n.s. |
| ENSG00000182512 | 939.8501434 | 0.474279636 | 0.2999369 | 1.581265 | 0.113817551 | 0.251036202 | GLRX5 | protein_coding | 45.13 | 0.284692823 | 0.284692823 | ENSG00000182512 | n.s. |
| ENSG00000124541 | 765.7469254 | -0.452490826 | 0.2781204 | -1.62696 | 0.103745488 | 0.23490786 | RRP36 | protein_coding | 48.91 | -0.284663088 | 0.284663088 | ENSG00000124541 | n.s. |
| ENSG00000213853 | 548.9622196 | -0.405095231 | 0.2330935 | -1.737908 | 0.08222696 | 0.198344762 | EMP2 | protein_coding | 48.31 | -0.284611509 | 0.284611509 | ENSG00000213853 | n.s. |
| ENSG00000218510 | 169.6527835 | -0.404503676 | 0.2325646 | -1.739317 | 0.081978958 | 0.197883674 | LINC00339 | lincRNA | 47.63 | -0.284604756 | 0.284604756 | ENSG00000218510 | n.s. |
| ENSG00000203877 | 775.4771368 | 0.411175628 | 0.2388172 | 1.721717 | 0.085120743 | 0.203186286 | RIPPLY2 | protein_coding | 42.89 | 0.284576958 | 0.284576958 | ENSG00000203877 | n.s. |
| ENSG00000007392 | 2803.443533 | 0.410302903 | 0.238033 | 1.723723 | 0.084757839 | 0.20255465 | LUC7L | protein_coding | 47.33 | 0.284527741 | 0.284527741 | ENSG00000007392 | n.s. |
| ENSG00000180828 | 257.917686 | 0.387893756 | 0.2176515 | 1.782178 | 0.074720208 | 0.184907612 | BHLHE22 | protein_coding | 53.59 | 0.28434366 | 0.28434366 | ENSG00000180828 | n.s. |
| ENSG00000167491 | 1108.07039 | -0.38749343 | 0.2174038 | -1.782368 | 0.074689312 | 0.184870984 | GATAD2A | protein_coding | 50.87 | -0.284083541 | 0.284083541 | ENSG00000167491 | n.s. |
| ENSG00000135069 | 1842.163126 | -0.467113359 | 0.2930965 | -1.593718 | 0.110999125 | 0.246560437 | PSAT1 | protein_coding | 45.59 | -0.284040707 | 0.284040707 | ENSG00000135069 | n.s. |
| ENSG00000120709 | 1034.739171 | 0.398055755 | 0.2269645 | 1.753824 | 0.079460731 | 0.193448464 | FAM53C | protein_coding | 48.45 | 0.283986794 | 0.283986794 | ENSG00000120709 | n.s. |

| | | | | | | | | | | | | | |
|---|---|---|---|---|---|---|---|---|---|---|---|---|---|
| ENSG00000145022 | 161.219981 | -0.438963375 | 0.2656129 | -1.652643 | 0.098403478 | 0.225637619 | TCTA | protein_coding | 53.23 | -0.283828667 | 0.283828667 | ENSG00000145022 | n.s. |
| ENSG00000164169 | 382.7221492 | -0.420470563 | 0.2479168 | -1.696015 | 0.089883036 | 0.211356446 | PRMT9 | protein_coding | 39.69 | -0.283811114 | 0.283811114 | ENSG00000164169 | n.s. |
| ENSG00000172840 | 370.3579926 | -0.429351276 | 0.2565424 | -1.673607 | 0.094207828 | 0.218536418 | PDP2 | protein_coding | 48.25 | -0.28357629 | 0.28357629 | ENSG00000172840 | n.s. |
| ENSG00000007341 | 856.7845803 | -0.383008249 | 0.2136805 | -1.792434 | 0.073063411 | 0.181814996 | ST7L | protein_coding | 39.05 | -0.283567933 | 0.283567933 | ENSG00000007341 | n.s. |
| ENSG00000274925 | 222.9676395 | -0.376031776 | 0.2077653 | -1.809887 | 0.070313236 | 0.17667771 | | lincRNA | 48.73 | -0.283083579 | 0.283083579 | ENSG00000274925 | n.s. |
| ENSG00000267222 | 19.45961381 | -0.535671818 | 0.3653418 | -1.466221 | 0.142588097 | 0.296218154 | | sense_intronic | 38.89 | -0.283042736 | 0.283042736 | ENSG00000267222 | n.s. |
| ENSG00000120333 | 982.6109202 | -0.479401518 | 0.306292 | -1.565178 | 0.117541183 | 0.257032494 | MRPS14 | protein_coding | 40.16 | -0.282852634 | 0.282852634 | ENSG00000120333 | n.s. |
| ENSG00000150991 | 5471.998157 | 0.4615608 | 0.2881599 | 1.601752 | 0.109210358 | 0.243910139 | UBC | protein_coding | 50.7 | 0.282830679 | 0.282830679 | ENSG00000150991 | n.s. |
| ENSG00000252118 | 5.931496863 | 0.700984216 | 0.5669968 | 1.236311 | 0.216343083 | 0.395346468 | RNU6ATAC39P | snRNA | 37.58 | 0.282512156 | 0.282512156 | ENSG00000252118 | n.s. |
| ENSG00000185608 | 377.8034758 | 0.439871101 | 0.267194 | 1.646261 | 0.09970995 | 0.227957575 | MRPL40 | protein_coding | 51.63 | 0.282461456 | 0.282461456 | ENSG00000185608 | n.s. |
| ENSG00000085982 | 643.6496049 | -0.446601425 | 0.2736172 | -1.632212 | 0.102634772 | 0.233102736 | USP40 | protein_coding | 38.86 | -0.282454246 | 0.282454246 | ENSG00000085982 | n.s. |
| ENSG00000033100 | 622.7798759 | -0.402247252 | 0.2315845 | -1.736935 | 0.082398598 | 0.198581339 | CHPF2 | protein_coding | 56.82 | -0.282402336 | 0.282402336 | ENSG00000033100 | n.s. |
| ENSG00000197472 | 724.5656022 | -0.444101152 | 0.2712258 | -1.637385 | 0.101549957 | 0.231290701 | ZNF695 | protein_coding | 42.51 | -0.282378089 | 0.282378089 | ENSG00000197472 | n.s. |
| ENSG00000109501 | 130.9126437 | -0.435510614 | 0.2630507 | -1.655614 | 0.097799928 | 0.224728792 | WFS1 | protein_coding | 57.04 | -0.28235951 | 0.28235951 | ENSG00000109501 | n.s. |
| ENSG00000185787 | 6690.447443 | 0.425440183 | 0.2534365 | 1.678686 | 0.093213332 | 0.216929929 | MORF4L1 | protein_coding | 45.4 | 0.282356365 | 0.282356365 | ENSG00000185787 | n.s. |
| ENSG00000147324 | 650.9937923 | -0.377177648 | 0.2091638 | -1.803265 | 0.07134668 | 0.178501317 | MFHAS1 | protein_coding | 44.16 | -0.282264128 | 0.282264128 | ENSG00000147324 | n.s. |
| ENSG00000177613 | 1834.24332 | -0.381804877 | 0.2132388 | -1.790504 | 0.073372934 | 0.182423457 | CSTF2T | protein_coding | 43.36 | -0.282123 | 0.282123 | ENSG00000177613 | n.s. |
| ENSG00000166823 | 93.33546118 | 0.463864088 | 0.2910329 | 1.593855 | 0.110968596 | 0.246560437 | MESP1 | protein_coding | 60.98 | 0.282064902 | 0.282064902 | ENSG00000166823 | n.s. |
| ENSG00000115159 | 1279.304344 | -0.439830921 | 0.2675341 | -1.644018 | 0.100172436 | 0.228714469 | GPD2 | protein_coding | 37.9 | -0.281802468 | 0.281802468 | ENSG00000115159 | n.s. |
| ENSG00000186187 | 249.4444562 | 0.401095066 | 0.2313279 | 1.733881 | 0.08293923 | 0.199382905 | ZNRF1 | protein_coding | 45.85 | 0.28089172 | 0.28089172 | ENSG00000186187 | n.s. |
| ENSG00000092203 | 625.0667744 | -0.457554904 | 0.2854602 | -1.602868 | 0.108963862 | 0.24355496 | TOX4 | protein_coding | 43.81 | -0.28066556 | 0.28066556 | ENSG00000092203 | n.s. |
| ENSG00000178429 | 68.95757958 | 0.658277186 | 0.5127777 | 1.283748 | 0.199230239 | 0.374798354 | RPS3AP5 | processed_pseudogene | 42.39 | 0.280559267 | 0.280559267 | ENSG00000178429 | n.s. |
| ENSG00000168234 | 96.169492 | 0.445853418 | 0.2740195 | 1.627087 | 0.103718701 | 0.234885405 | TTC39C | protein_coding | 41.64 | 0.280505988 | 0.280505988 | ENSG00000168234 | n.s. |
| ENSG00000171914 | 42.81207845 | -0.456386703 | 0.2844318 | -1.604556 | 0.108591615 | 0.24299599 | TLN2 | protein_coding | 43.6 | -0.280404398 | 0.280404398 | ENSG00000171914 | n.s. |
| ENSG00000095485 | 613.3560366 | -0.460879152 | 0.2890648 | -1.59438 | 0.110850921 | 0.246427809 | CWF19L1 | protein_coding | 42.57 | -0.280357528 | 0.280357528 | ENSG00000095485 | n.s. |
| ENSG00000207445 | 126.3153357 | 0.398690885 | 0.2293537 | 1.738323 | 0.082153911 | 0.198237245 | SNORD15B | snoRNA | 47.95 | 0.280205833 | 0.280205833 | ENSG00000207445 | n.s. |
| ENSG00000163577 | 405.1907404 | -0.438535923 | 0.2672208 | -1.6411 | 0.100776747 | 0.229868061 | EIF5A2 | protein_coding | 38.29 | -0.280014557 | 0.280014557 | ENSG00000163577 | n.s. |
| ENSG00000147535 | 110.916917 | 0.43245296 | 0.2614991 | 1.653745 | 0.098179246 | 0.225346133 | PLPP5 | protein_coding | 45.35 | 0.279861885 | 0.279861885 | ENSG00000147535 | n.s. |
| ENSG00000178980 | 405.9176846 | 0.491235037 | 0.3205856 | 1.532305 | 0.125447114 | 0.269370784 | SELENOW | protein_coding | 54.44 | 0.279831798 | 0.279831798 | ENSG00000178980 | n.s. |
| ENSG00000115295 | 646.6573184 | -0.445576502 | 0.2741507 | -1.625298 | 0.104099157 | 0.235517154 | CLIP4 | protein_coding | 38.44 | -0.279811998 | 0.279811998 | ENSG00000115295 | n.s. |
| ENSG00000174796 | 181.9361327 | 0.411634455 | 0.2418222 | 1.702219 | 0.088714237 | 0.209457066 | THAP6 | protein_coding | 40.52 | 0.279460683 | 0.279460683 | ENSG00000174796 | n.s. |
| ENSG00000236144 | 292.542642 | -0.38377008 | 0.2161964 | -1.775099 | 0.075881533 | 0.187024002 | TMEM147-AS1 | antisense | 56.8 | -0.279424013 | 0.279424013 | ENSG00000236144 | n.s. |

| | | | | | | | | | | | | | |
|---|---|---|---|---|---|---|---|---|---|---|---|---|---|
| ENSG00000169155 | 430.6755924 | -0.451574974 | 0.2803703 | -1.610638 | 0.107258741 | 0.240709776 | ZBTB43 | protein_coding | 41.82 | -0.279301954 | 0.279301954 | ENSG00000169155 | n.s. |
| ENSG00000267430 | 89.44193455 | -0.42960537 | 0.2591364 | -1.657835 | 0.097350828 | 0.223918187 | | processed_pseudogene | 48.83 | -0.279205097 | 0.279205097 | ENSG00000267430 | n.s. |
| ENSG00000151893 | 1211.368921 | -0.419961517 | 0.2498387 | -1.68093 | 0.092776418 | 0.216406993 | CACUL1 | protein_coding | 39.56 | -0.279160477 | 0.279160477 | ENSG00000151893 | n.s. |
| ENSG00000141543 | 3679.497582 | 0.424526824 | 0.2543713 | 1.668926 | 0.095132091 | 0.220093735 | EIF4A3 | protein_coding | 46.23 | 0.279080674 | 0.279080674 | ENSG00000141543 | n.s. |
| ENSG00000136810 | 4205.338213 | 0.486762001 | 0.3166664 | 1.537145 | 0.124257903 | 0.267394558 | TXN | protein_coding | 42.28 | 0.278840364 | 0.278840364 | ENSG00000136810 | n.s. |
| ENSG00000133103 | 192.2442231 | 0.447899629 | 0.2771395 | 1.616152 | 0.106061451 | 0.238638264 | COG6 | protein_coding | 35.35 | 0.278709987 | 0.278709987 | ENSG00000133103 | n.s. |
| ENSG00000255112 | 1186.260804 | 0.427332298 | 0.2573952 | 1.660218 | 0.096870533 | 0.223151627 | CHMP1B | protein_coding | 41.52 | 0.278364234 | 0.278364234 | ENSG00000255112 | n.s. |
| ENSG00000103342 | 2874.888832 | -0.429898137 | 0.2599254 | -1.653929 | 0.09814197 | 0.225297717 | GSPT1 | protein_coding | 42.29 | -0.278248649 | 0.278248649 | ENSG00000103342 | n.s. |
| ENSG00000108298 | 20222.45707 | 0.484333966 | 0.3145747 | 1.539647 | 0.123646418 | 0.26640809 | RPL19 | protein_coding | 49.65 | 0.278226901 | 0.278226901 | ENSG00000108298 | n.s. |
| ENSG00000200455 | 12.24196334 | 0.561909591 | 0.3991428 | 1.407791 | 0.159193039 | 0.319879446 | RNU6-1112P | snRNA | 41.41 | 0.278152926 | 0.278152926 | ENSG00000200455 | n.s. |
| ENSG00000166479 | 1889.134248 | -0.435615413 | 0.2654807 | -1.640855 | 0.100827435 | 0.229908349 | TMX3 | protein_coding | 36.18 | -0.278116595 | 0.278116595 | ENSG00000166479 | n.s. |
| ENSG00000198862 | 1733.246756 | -0.427890321 | 0.258206 | -1.657167 | 0.0974858 | 0.224086972 | LTN1 | protein_coding | 36.74 | -0.277950446 | 0.277950446 | ENSG00000198862 | n.s. |
| ENSG00000040341 | 535.9423484 | -0.455151401 | 0.2848592 | -1.597812 | 0.11008483 | 0.245155324 | STAU2 | protein_coding | 38.25 | -0.277896635 | 0.277896635 | ENSG00000040341 | n.s. |
| ENSG00000177839 | 265.3448753 | -0.403094829 | 0.2347671 | -1.716999 | 0.085979388 | 0.204565224 | PCDHB9 | protein_coding | 46.34 | -0.277800135 | 0.277800135 | ENSG00000177839 | n.s. |
| ENSG00000175866 | 415.5843992 | 0.389662569 | 0.2223126 | 1.752768 | 0.079641767 | 0.193753682 | BAIAP2 | protein_coding | 60.69 | 0.277732009 | 0.277732009 | ENSG00000175866 | n.s. |
| ENSG00000116679 | 13744.98883 | -0.424870944 | 0.2554024 | -1.663536 | 0.096205263 | 0.22198639 | IVNS1ABP | protein_coding | 37.35 | -0.277726941 | 0.277726941 | ENSG00000116679 | n.s. |
| ENSG00000176974 | 410.8938036 | -0.43343919 | 0.2636928 | -1.643728 | 0.100232441 | 0.228813948 | SHMT1 | protein_coding | 49.9 | -0.277625388 | 0.277625388 | ENSG00000176974 | n.s. |
| ENSG00000136932 | 272.9806587 | -0.465450407 | 0.2955073 | -1.575089 | 0.115235851 | 0.253384201 | TRMO | protein_coding | 41.69 | -0.27751106 | 0.27751106 | ENSG00000136932 | n.s. |
| ENSG00000198160 | 1914.370012 | -0.422885616 | 0.2537576 | -1.666495 | 0.095614937 | 0.220880499 | MIER1 | protein_coding | 36.15 | -0.277346413 | 0.277346413 | ENSG00000198160 | n.s. |
| ENSG00000101126 | 2848.115473 | -0.419328665 | 0.2503401 | -1.675036 | 0.093927139 | 0.218103329 | ADNP | protein_coding | 41.52 | -0.277317857 | 0.277317857 | ENSG00000101126 | n.s. |
| ENSG00000110080 | 67.01157137 | -0.435159075 | 0.2654754 | -1.639169 | 0.101178004 | 0.23063218 | ST3GAL4 | protein_coding | 52.01 | -0.277231185 | 0.277231185 | ENSG00000110080 | n.s. |
| ENSG00000243749 | 41.19605301 | -0.474931904 | 0.3054433 | -1.554894 | 0.119971379 | 0.260953212 | TMEM35B | protein_coding | 51.77 | -0.277093013 | 0.277093013 | ENSG00000243749 | n.s. |
| ENSG00000135503 | 972.8089903 | -0.419329182 | 0.2505409 | -1.673696 | 0.094190403 | 0.218532409 | ACVR1B | protein_coding | 44.84 | -0.276960277 | 0.276960277 | ENSG00000135503 | n.s. |
| ENSG00000225783 | 1618.223219 | -0.407651735 | 0.2395314 | -1.701872 | 0.088779397 | 0.209575371 | MIAT | lincRNA | 49.99 | -0.276656827 | 0.276656827 | ENSG00000225783 | n.s. |
| ENSG00000207475 | 47.28015805 | -0.458178228 | 0.2886492 | -1.587318 | 0.112440555 | 0.249007452 | SNORA80E | snoRNA | 54.74 | -0.276642358 | 0.276642358 | ENSG00000207475 | n.s. |
| ENSG00000235843 | 41.10705687 | -0.456228061 | 0.2867376 | -1.5911 | 0.111587178 | 0.247629628 | | antisense | 42.48 | -0.276564262 | 0.276564262 | ENSG00000235843 | n.s. |
| ENSG00000133773 | 3289.114824 | 0.388977449 | 0.2223101 | 1.749706 | 0.080169013 | 0.194692219 | CCDC59 | protein_coding | 35.35 | 0.276427371 | 0.276427371 | ENSG00000133773 | n.s. |
| ENSG00000064419 | 1631.966466 | -0.414595819 | 0.2462242 | -1.683814 | 0.092217559 | 0.215463177 | TNPO3 | protein_coding | 39.8 | -0.276380742 | 0.276380742 | ENSG00000064419 | n.s. |
| ENSG00000224856 | 29.21490662 | 0.472577918 | 0.3037139 | 1.555997 | 0.119708857 | 0.260463575 | | processed_pseudogene | 45.34 | 0.276105068 | 0.276105068 | ENSG00000224856 | n.s. |
| ENSG00000101546 | 1079.710123 | 0.39337296 | 0.2267239 | 1.735031 | 0.082735316 | 0.19908811 | RBFA | protein_coding | 47.61 | 0.275736616 | 0.275736616 | ENSG00000101546 | n.s. |
| ENSG00000272371 | 73.32907913 | 0.414771476 | 0.246887 | 1.680006 | 0.092956216 | 0.216636219 | | lincRNA | 43.42 | 0.275519806 | 0.275519806 | ENSG00000272371 | n.s. |
| ENSG00000034677 | 616.7393161 | -0.43350937 | 0.264918 | -1.636391 | 0.101757798 | 0.231688279 | RNF19A | protein_coding | 38.61 | -0.27532004 | 0.27532004 | ENSG00000034677 | n.s. |

| | | | | | | | | | | | | | | |
|---|---|---|---|---|---|---|---|---|---|---|---|---|---|---|
| ENSG00000118473 | 162.7366017 | -0.440699673 | 0.2721438 | -1.619363 | 0.105369183 | 0.237349157 | SGIP1 | | protein_coding | 37.83 | -0.275266439 | 0.275266439 | ENSG00000118473 | n.s. |
| ENSG00000147654 | 405.8726425 | -0.439950602 | 0.2713762 | -1.621183 | 0.104978326 | 0.23681355 | EBAG9 | | protein_coding | 34.99 | -0.275230216 | 0.275230216 | ENSG00000147654 | n.s. |
| ENSG00000234936 | 29.52820062 | -0.488670267 | 0.3210171 | -1.522256 | 0.127944857 | 0.273426898 | | | antisense | 49.76 | -0.275198943 | 0.275198943 | ENSG00000234936 | n.s. |
| ENSG00000085382 | 151.81188 | 0.404106409 | 0.2371551 | 1.703975 | 0.088385739 | 0.209044942 | HACE1 | | protein_coding | 37.39 | 0.274695508 | 0.274695508 | ENSG00000085382 | n.s. |
| ENSG00000180822 | 778.8300387 | -0.418285073 | 0.2508714 | -1.667328 | 0.095449126 | 0.220643896 | PSMG4 | | protein_coding | 49.83 | -0.274523874 | 0.274523874 | ENSG00000180822 | n.s. |
| ENSG00000197744 | 189.9928387 | 0.514986022 | 0.34943 | 1.473789 | 0.140538473 | 0.293053342 | PTMAP2 | | processed_pseudogene | 51.99 | 0.27451501 | 0.27451501 | ENSG00000197744 | n.s. |
| ENSG00000168137 | 2657.240769 | -0.412578394 | 0.2454626 | -1.68082 | 0.092797979 | 0.216419808 | SETD5 | | protein_coding | 39.13 | -0.274242093 | 0.274242093 | ENSG00000168137 | n.s. |
| ENSG00000135002 | 672.8635325 | 0.454460024 | 0.2867725 | 1.584741 | 0.113025302 | 0.249746274 | RFK | | protein_coding | 40.38 | 0.273812611 | 0.273812611 | ENSG00000135002 | n.s. |
| ENSG00000003989 | 660.362001 | -0.442391245 | 0.2745707 | -1.611211 | 0.107133748 | 0.240506799 | SLC7A2 | | protein_coding | 40.13 | -0.273783838 | 0.273783838 | ENSG00000003989 | n.s. |
| ENSG00000115839 | 1962.857968 | -0.428640383 | 0.2611758 | -1.641195 | 0.100756927 | 0.229860509 | RAB3GAP1 | | protein_coding | 37.96 | -0.273702159 | 0.273702159 | ENSG00000115839 | n.s. |
| ENSG00000198060 | 943.6044306 | -0.449977185 | 0.2823388 | -1.593749 | 0.110992303 | 0.246560437 | | 5-Mar | protein_coding | 40.44 | -0.273620601 | 0.273620601 | ENSG00000198060 | n.s. |
| ENSG00000179950 | 2130.831713 | 0.443929593 | 0.2762524 | 1.606971 | 0.10806074 | 0.2420804 | PUF60 | | protein_coding | 62.04 | 0.273478552 | 0.273478552 | ENSG00000179950 | n.s. |
| ENSG00000171241 | 695.3820459 | -0.389379546 | 0.2241678 | -1.737 | 0.082387145 | 0.198581339 | SHCBP1 | | protein_coding | 39.54 | -0.273368414 | 0.273368414 | ENSG00000171241 | n.s. |
| ENSG00000123094 | 562.0130396 | -0.423957855 | 0.2569347 | -1.650061 | 0.098930492 | 0.226584839 | RASSF8 | | protein_coding | 38.09 | -0.273354947 | 0.273354947 | ENSG00000123094 | n.s. |
| ENSG00000064651 | 955.5603227 | -0.394010644 | 0.2285473 | -1.723978 | 0.084711751 | 0.202479263 | SLC12A2 | | protein_coding | 35.24 | -0.273293446 | 0.273293446 | ENSG00000064651 | n.s. |
| ENSG00000126790 | 250.1036625 | -0.383774548 | 0.2190738 | -1.751805 | 0.079807338 | 0.194054763 | L3HYPDH | | protein_coding | 42.53 | -0.273276528 | 0.273276528 | ENSG00000126790 | n.s. |
| ENSG00000155545 | 1239.225683 | -0.447269389 | 0.2799822 | -1.597492 | 0.110156014 | 0.245235399 | MIER3 | | protein_coding | 38.91 | -0.273020768 | 0.273020768 | ENSG00000155545 | n.s. |
| ENSG00000108021 | 3615.114764 | -0.435326329 | 0.2680339 | -1.624147 | 0.104344512 | 0.235995556 | FAM208B | | protein_coding | 38.83 | -0.272991476 | 0.272991476 | ENSG00000108021 | n.s. |
| ENSG00000254911 | 449.3130068 | -0.397199836 | 0.2317601 | -1.713841 | 0.086558008 | 0.205586343 | SCARNA9 | | antisense | 40.23 | -0.272878566 | 0.272878566 | ENSG00000254911 | n.s. |
| ENSG00000198886 | 3640.122206 | -0.385699883 | 0.221257 | -1.743221 | 0.081294949 | 0.196672369 | MT-ND4 | | protein_coding | 44.27 | -0.272403108 | 0.272403108 | ENSG00000198886 | n.s. |
| ENSG00000100003 | 8.460540895 | 0.56396751 | 0.4069855 | 1.385719 | 0.165832776 | 0.328901672 | SEC14L2 | | protein_coding | 50.43 | 0.27235904 | 0.27235904 | ENSG00000100003 | n.s. |
| ENSG00000012963 | 984.9564233 | -0.449856575 | 0.2832636 | -1.58812 | 0.112259214 | 0.248788962 | UBR7 | | protein_coding | 43.29 | -0.271789348 | 0.271789348 | ENSG00000012963 | n.s. |
| ENSG00000136295 | 572.3679355 | -0.404674718 | 0.2394872 | -1.689755 | 0.091074838 | 0.213509844 | TTYH3 | | protein_coding | 62.29 | -0.271367621 | 0.271367621 | ENSG00000136295 | n.s. |
| ENSG00000277368 | 16.48260702 | 0.50735327 | 0.3434052 | 1.477419 | 0.13956345 | 0.291900875 | | | antisense | 61.85 | 0.27131457 | 0.27131457 | ENSG00000277368 | n.s. |
| ENSG00000205220 | 130.614504 | 0.494479596 | 0.3296266 | 1.500121 | 0.133583185 | 0.282914312 | PSMB10 | | protein_coding | 61.37 | 0.271145455 | 0.271145455 | ENSG00000205220 | n.s. |
| ENSG00000073711 | 551.6926535 | -0.411722559 | 0.2465266 | -1.670094 | 0.094900806 | 0.219689908 | PPP2R3A | | protein_coding | 39.07 | -0.270991627 | 0.270991627 | ENSG00000073711 | n.s. |
| ENSG00000180182 | 966.3016288 | -0.407875733 | 0.2427538 | -1.680203 | 0.092917779 | 0.216590362 | MED14 | | protein_coding | 38.86 | -0.270976677 | 0.270976677 | ENSG00000180182 | n.s. |
| ENSG00000119787 | 1986.93496 | -0.442415808 | 0.2764132 | -1.60056 | 0.109474449 | 0.244186599 | ATL2 | | protein_coding | 39.46 | -0.270881543 | 0.270881543 | ENSG00000119787 | n.s. |
| ENSG00000133678 | 142.8342602 | -0.471627161 | 0.3064884 | -1.538809 | 0.123850915 | 0.266726677 | TMEM254 | | protein_coding | 42.93 | -0.270682649 | 0.270682649 | ENSG00000133678 | n.s. |
| ENSG00000101464 | 266.9515464 | -0.450689218 | 0.2850542 | -1.581065 | 0.113863165 | 0.251036202 | PIGU | | protein_coding | 44.54 | -0.270532352 | 0.270532352 | ENSG00000101464 | n.s. |
| ENSG00000068366 | 801.6098107 | -0.466403776 | 0.301115 | -1.548923 | 0.121400336 | 0.263320904 | ACSL4 | | protein_coding | 39.16 | -0.270287827 | 0.270287827 | ENSG00000068366 | n.s. |
| ENSG00000228409 | 611.4433764 | -0.409955163 | 0.2452041 | -1.671893 | 0.094545373 | 0.219136879 | CCT6P1 | | transcribed_unprocessed_pseud | 40.94 | -0.270277096 | 0.270277096 | ENSG00000228409 | n.s. |

| | | | | | | | | | | | | | |
|---|---|---|---|---|---|---|---|---|---|---|---|---|---|
| ENSG00000165138 | 594.5110223 | -0.431070625 | 0.2654642 | -1.623837 | 0.10441061 | 0.236068353 | ANKS6 | protein_coding | 46.95 | -0.270265 | 0.270265 | ENSG00000165138 | n.s. |
| ENSG00000276293 | 920.4907421 | -0.440217167 | 0.2746097 | -1.603065 | 0.108920294 | 0.243496669 | PIP4K2B | protein_coding | 49.36 | -0.270076303 | 0.270076303 | ENSG00000276293 | n.s. |
| ENSG00000125691 | 18597.74865 | 0.469318654 | 0.304546 | 1.541043 | 0.123306231 | 0.26583968 | RPL23 | protein_coding | 44.44 | 0.270036658 | 0.270036658 | ENSG00000125691 | n.s. |
| ENSG00000231500 | 18399.84122 | 0.465612614 | 0.300429 | 1.549826 | 0.121183383 | 0.26305523 | RPS18 | protein_coding | 46.95 | 0.27003346 | 0.27003346 | ENSG00000231500 | n.s. |
| ENSG00000168028 | 16926.00239 | 0.457025395 | 0.2918174 | 1.566135 | 0.117317023 | 0.25667536 | RPSA | protein_coding | 45.15 | 0.269926428 | 0.269926428 | ENSG00000168028 | n.s. |
| ENSG00000146826 | 130.4900773 | -0.411939458 | 0.2473813 | -1.665201 | 0.0958727 | 0.221340273 | C7orf43 | protein_coding | 60.47 | -0.269795447 | 0.269795447 | ENSG00000146826 | n.s. |
| ENSG00000155304 | 1627.383801 | -0.474064514 | 0.3096331 | -1.531052 | 0.125756445 | 0.269951739 | HSPA13 | protein_coding | 36.7 | -0.269607064 | 0.269607064 | ENSG00000155304 | n.s. |
| ENSG00000214435 | 427.7788158 | -0.448979768 | 0.2839645 | -1.581112 | 0.113852398 | 0.251036202 | AS3MT | protein_coding | 43.69 | -0.269506232 | 0.269506232 | ENSG00000214435 | n.s. |
| ENSG00000174607 | 365.9944434 | -0.444095076 | 0.2790951 | -1.591196 | 0.111565454 | 0.247620886 | UGT8 | protein_coding | 36.21 | -0.269216087 | 0.269216087 | ENSG00000174607 | n.s. |
| ENSG00000283196 | 46.92665357 | -0.468336154 | 0.3040892 | -1.540128 | 0.123529256 | 0.266196848 | | lincRNA | 46.72 | -0.269198259 | 0.269198259 | ENSG00000283196 | n.s. |
| ENSG00000140598 | 710.1922558 | -0.429321988 | 0.264402 | -1.623747 | 0.104429731 | 0.236070013 | EFL1 | protein_coding | 38.04 | -0.26916736 | 0.26916736 | ENSG00000140598 | n.s. |
| ENSG00000135119 | 242.1245684 | -0.384208616 | 0.221666 | -1.733277 | 0.08304641 | 0.199595231 | RNFT2 | protein_coding | 47.15 | -0.268888339 | 0.268888339 | ENSG00000135119 | n.s. |
| ENSG00000115282 | 483.9315166 | -0.424169478 | 0.259615 | -1.63384 | 0.102292355 | 0.232525137 | TTC31 | protein_coding | 50.85 | -0.268724128 | 0.268724128 | ENSG00000115282 | n.s. |
| ENSG00000095002 | 3046.96117 | -0.440195479 | 0.2757328 | -1.596456 | 0.11038692 | 0.245631627 | MSH2 | protein_coding | 44.66 | -0.268394101 | 0.268394101 | ENSG00000095002 | n.s. |
| ENSG00000121104 | 362.0448877 | -0.410771374 | 0.2470942 | -1.662408 | 0.096431014 | 0.222357625 | FAM117A | protein_coding | 46.7 | -0.268212336 | 0.268212336 | ENSG00000121104 | n.s. |
| ENSG00000175854 | 82.22001878 | -0.628730002 | 0.48948 | -1.284486 | 0.198972091 | 0.374464508 | SWI5 | protein_coding | 52.59 | -0.268209516 | 0.268209516 | ENSG00000175854 | n.s. |
| ENSG00000178971 | 312.7113724 | -0.427459449 | 0.2632986 | -1.623478 | 0.104487295 | 0.236088912 | CTC1 | protein_coding | 48.84 | -0.267984762 | 0.267984762 | ENSG00000178971 | n.s. |
| ENSG00000011243 | 1072.633993 | 0.392631873 | 0.2299642 | 1.707361 | 0.087755041 | 0.207720748 | AKAP8L | protein_coding | 48.86 | 0.267979154 | 0.267979154 | ENSG00000011243 | n.s. |
| ENSG00000115286 | 484.0975348 | 0.475060026 | 0.3119586 | 1.52283 | 0.127801163 | 0.273207684 | NDUFS7 | protein_coding | 59.57 | 0.267699693 | 0.267699693 | ENSG00000115286 | n.s. |
| ENSG00000214535 | 92.15450646 | 0.521907088 | 0.3628634 | 1.438302 | 0.150348484 | 0.307025805 | RPS15AP1 | processed_pseudogene | 46.29 | 0.267647065 | 0.267647065 | ENSG00000214535 | n.s. |
| ENSG00000132824 | 667.7415253 | -0.436410139 | 0.2723265 | -1.602525 | 0.109039473 | 0.243670433 | SERINC3 | protein_coding | 42.56 | -0.26760546 | 0.26760546 | ENSG00000132824 | n.s. |
| ENSG00000171208 | 2301.945341 | -0.429600899 | 0.2656682 | -1.617058 | 0.105865699 | 0.238313358 | NETO2 | protein_coding | 38.7 | -0.267577613 | 0.267577613 | ENSG00000171208 | n.s. |
| ENSG00000184445 | 5011.034034 | -0.447638203 | 0.2839727 | -1.576343 | 0.114946882 | 0.252829583 | KNTC1 | protein_coding | 41.24 | -0.26731705 | 0.26731705 | ENSG00000184445 | n.s. |
| ENSG00000171858 | 5397.606483 | 0.490321166 | 0.3281407 | 1.494241 | 0.135112714 | 0.285069603 | RPS21 | protein_coding | 59.5 | 0.267249105 | 0.267249105 | ENSG00000171858 | n.s. |
| ENSG00000150712 | 530.4899252 | -0.401941998 | 0.2391191 | -1.680928 | 0.09277694 | 0.216406993 | MTMR12 | protein_coding | 44.09 | -0.267182386 | 0.267182386 | ENSG00000150712 | n.s. |
| ENSG00000225479 | 48.93570927 | -0.492747338 | 0.3308903 | -1.489156 | 0.136446236 | 0.287251639 | PLCB1-IT1 | sense_intronic | 37.7 | -0.266939703 | 0.266939703 | ENSG00000225479 | n.s. |
| ENSG00000108591 | 367.2899378 | -0.461566362 | 0.2984594 | -1.546496 | 0.121984765 | 0.264135907 | DRG2 | protein_coding | 53.55 | -0.266865004 | 0.266865004 | ENSG00000108591 | n.s. |
| ENSG00000180616 | 12533.45517 | -0.428539275 | 0.2650697 | -1.616704 | 0.105942212 | 0.238447044 | SSTR2 | protein_coding | 47.17 | -0.266812005 | 0.266812005 | ENSG00000180616 | n.s. |
| ENSG00000165197 | 176.3805762 | 0.473858598 | 0.3114074 | 1.521668 | 0.12809238 | 0.273574199 | VEGFD | protein_coding | 39.3 | 0.266746787 | 0.266746787 | ENSG00000165197 | n.s. |
| ENSG00000173157 | 791.4103616 | -0.432098632 | 0.268859 | -1.607157 | 0.108019907 | 0.242027866 | ADAMTS20 | protein_coding | 35.33 | -0.26623093 | 0.26623093 | ENSG00000173157 | n.s. |
| ENSG00000089682 | 442.1547417 | -0.415928471 | 0.2531383 | -1.643088 | 0.100364796 | 0.229078533 | RBM41 | protein_coding | 36.02 | -0.266200713 | 0.266200713 | ENSG00000089682 | n.s. |
| ENSG00000214160 | 280.9376673 | -0.413919632 | 0.2512482 | -1.647453 | 0.099465007 | 0.227584435 | ALG3 | protein_coding | 51.41 | -0.266091316 | 0.266091316 | ENSG00000214160 | n.s. |

| | | | | | | | | | | | | | |
|---|---|---|---|---|---|---|---|---|---|---|---|---|---|
| ENSG00000201516 | 5.514058506 | -0.625533526 | 0.488312 | -1.281012 | 0.200189473 | 0.375790487 | | snoRNA | 46.34 | -0.265885664 | 0.265885664 | ENSG00000201516 | n.s. |
| ENSG00000276026 | 10.83958404 | 0.575450942 | 0.4267945 | 1.348309 | 0.177558957 | 0.345321045 | | antisense | 44.87 | 0.265729983 | 0.265729983 | ENSG00000276026 | n.s. |
| ENSG00000230071 | 78.38850248 | 0.419509479 | 0.2568312 | 1.633406 | 0.102383694 | 0.232618826 | RPL4P6 | processed_pseudogene | 46.63 | 0.265698485 | 0.265698485 | ENSG00000230071 | n.s. |
| ENSG00000253054 | 5.741792714 | 0.62715632 | 0.4906183 | 1.278298 | 0.201144467 | 0.377074783 | RNU7-77P | snRNA | 46.67 | 0.265646177 | 0.265646177 | ENSG00000253054 | n.s. |
| ENSG00000132359 | 913.9404442 | -0.418333142 | 0.2557357 | -1.635802 | 0.101880953 | 0.231930758 | RAP1GAP2 | protein_coding | 51.49 | -0.265491637 | 0.265491637 | ENSG00000132359 | n.s. |
| ENSG00000253190 | 49.36737785 | -0.438170318 | 0.275501 | -1.590449 | 0.111733659 | 0.247915177 | | lincRNA | 53.21 | -0.265398397 | 0.265398397 | ENSG00000253190 | n.s. |
| ENSG00000134802 | 373.5738666 | -0.404063602 | 0.2422381 | -1.668043 | 0.095307117 | 0.220388803 | SLC43A3 | protein_coding | 48.17 | -0.265393205 | 0.265393205 | ENSG00000134802 | n.s. |
| ENSG00000084073 | 682.6297625 | -0.44197945 | 0.2795544 | -1.581014 | 0.113874804 | 0.251036202 | ZMPSTE24 | protein_coding | 40.78 | -0.265304195 | 0.265304195 | ENSG00000084073 | n.s. |
| ENSG00000185837 | 25.2030273 | -0.477440862 | 0.3160199 | -1.510794 | 0.130841057 | 0.278249587 | HDHD5-AS1 | antisense | 49.27 | -0.265249658 | 0.265249658 | ENSG00000185837 | n.s. |
| ENSG00000108107 | 2407.449879 | 0.406233845 | 0.2443757 | 1.662333 | 0.096446003 | 0.222357625 | RPL28 | protein_coding | 52.88 | 0.265249565 | 0.265249565 | ENSG00000108107 | n.s. |
| ENSG00000278472 | 19.04426842 | -0.520335411 | 0.3631439 | -1.432863 | 0.151897006 | 0.309235563 | | sense_intronic | 35.67 | -0.265220457 | 0.265220457 | ENSG00000278472 | n.s. |
| ENSG00000260589 | 279.3425719 | 0.374767829 | 0.2148759 | 1.744113 | 0.081139404 | 0.196367794 | STAM-AS1 | antisense | 38.61 | 0.264934523 | 0.264934523 | ENSG00000260589 | n.s. |
| ENSG00000198887 | 2246.016925 | -0.410077798 | 0.2482984 | -1.651553 | 0.098625801 | 0.226035724 | SMC5 | protein_coding | 36.37 | -0.26483764 | 0.26483764 | ENSG00000198887 | n.s. |
| ENSG00000161016 | 11693.9741 | 0.479888229 | 0.318936 | 1.504654 | 0.132413168 | 0.280820798 | RPL8 | protein_coding | 60.89 | 0.264692301 | 0.264692301 | ENSG00000161016 | n.s. |
| ENSG00000207425 | 4.8773579 | -0.802464218 | 0.7431272 | -1.079848 | 0.280209993 | 0.467936452 | | misc_RNA | 46 | -0.26466323 | 0.26466323 | ENSG00000207425 | n.s. |
| ENSG00000111875 | 1520.757331 | -0.458787539 | 0.29732 | -1.543077 | 0.122812187 | 0.265024713 | ASF1A | protein_coding | 39.13 | -0.264589026 | 0.264589026 | ENSG00000111875 | n.s. |
| ENSG00000231416 | 58.97857675 | 0.418199961 | 0.2564351 | 1.630822 | 0.102927961 | 0.233626671 | | processed_pseudogene | 44.38 | 0.264083899 | 0.264083899 | ENSG00000231416 | n.s. |
| ENSG00000100288 | 248.2227558 | 0.441496979 | 0.279928 | 1.577181 | 0.114753982 | 0.252524055 | CHKB | protein_coding | 50.66 | 0.263881528 | 0.263881528 | ENSG00000100288 | n.s. |
| ENSG00000102738 | 557.4445083 | -0.427067533 | 0.2653949 | -1.609178 | 0.107577529 | 0.241230784 | MRPS31 | protein_coding | 39.19 | -0.263742931 | 0.263742931 | ENSG00000102738 | n.s. |
| ENSG00000175110 | 1137.675823 | 0.431075735 | 0.2694789 | 1.599664 | 0.10967315 | 0.244512294 | MRPS22 | protein_coding | 42.84 | 0.263688726 | 0.263688726 | ENSG00000175110 | n.s. |
| ENSG00000083290 | 705.9676278 | -0.41663683 | 0.2552158 | -1.632489 | 0.102576589 | 0.233019066 | ULK2 | protein_coding | 41.45 | -0.263568018 | 0.263568018 | ENSG00000083290 | n.s. |
| ENSG00000021574 | 1108.361186 | -0.454105469 | 0.2929176 | -1.550284 | 0.121073423 | 0.262974652 | SPAST | protein_coding | 39.98 | -0.263420275 | 0.263420275 | ENSG00000021574 | n.s. |
| ENSG00000089195 | 826.934681 | -0.432199252 | 0.2708282 | -1.595843 | 0.110523846 | 0.245857727 | TRMT6 | protein_coding | 42.56 | -0.263345979 | 0.263345979 | ENSG00000089195 | n.s. |
| ENSG00000175206 | 29.99061956 | 0.524553934 | 0.3695779 | 1.419333 | 0.155802063 | 0.314748871 | NPPA | protein_coding | 54.04 | 0.263344864 | 0.263344864 | ENSG00000175206 | n.s. |
| ENSG00000100418 | 492.0578207 | -0.460352627 | 0.2997082 | -1.536003 | 0.124537706 | 0.267913856 | DESI1 | protein_coding | 46.95 | -0.263323924 | 0.263323924 | ENSG00000100418 | n.s. |
| ENSG00000198791 | 3252.935027 | -0.425527722 | 0.2645499 | -1.608497 | 0.10772632 | 0.24144777 | CNOT7 | protein_coding | 36.74 | -0.262625838 | 0.262625838 | ENSG00000198791 | n.s. |
| ENSG00000102743 | 349.2723959 | -0.436915033 | 0.2760556 | -1.582707 | 0.113488369 | 0.250570657 | SLC25A15 | protein_coding | 46.36 | -0.262616427 | 0.262616427 | ENSG00000102743 | n.s. |
| ENSG00000174886 | 1008.274797 | 0.484474777 | 0.3253778 | 1.488961 | 0.136497654 | 0.287251639 | NDUFA11 | protein_coding | 59.05 | 0.262458147 | 0.262458147 | ENSG00000174886 | n.s. |
| ENSG00000130713 | 1686.864288 | -0.39283225 | 0.2329996 | -1.685978 | 0.091800007 | 0.214848125 | EXOSC2 | protein_coding | 47.74 | -0.262360259 | 0.262360259 | ENSG00000130713 | n.s. |
| ENSG00000198246 | 107.5378325 | -0.413323029 | 0.2527643 | -1.635211 | 0.102004721 | 0.23217455 | SLC29A3 | protein_coding | 51.86 | -0.262123425 | 0.262123425 | ENSG00000198246 | n.s. |
| ENSG00000108561 | 3786.926042 | 0.485168968 | 0.3265749 | 1.485628 | 0.137377473 | 0.288492693 | C1QBP | protein_coding | 46.67 | 0.261925833 | 0.261925833 | ENSG00000108561 | n.s. |
| ENSG00000144278 | 1568.53897 | -0.441949723 | 0.2816504 | -1.569143 | 0.116614597 | 0.255648723 | GALNT13 | protein_coding | 35.16 | -0.261791735 | 0.261791735 | ENSG00000144278 | n.s. |

| | | | | | | | | | | | | | |
|---|---|---|---|---|---|---|---|---|---|---|---|---|---|
| ENSG00000014919 | 692.8268789 | -0.434253925 | 0.2738757 | -1.585587 | 0.112832935 | 0.249550887 | COX15 | protein_coding | 42.42 | -0.26178602 | 0.26178602 | ENSG00000014919 | n.s. |
| ENSG00000243836 | 39.02481753 | 0.43610936 | 0.2758864 | 1.580757 | 0.113933618 | 0.251036202 | WDR86-AS1 | processed_transcript | 60.18 | 0.261780594 | 0.261780594 | ENSG00000243836 | n.s. |
| ENSG00000279957 | 69.73196604 | 0.458278623 | 0.2986068 | 1.534723 | 0.12485194 | 0.268465414 | | TEC | 35.72 | 0.261728263 | 0.261728263 | ENSG00000279957 | n.s. |
| ENSG00000135333 | 814.4330917 | -0.403297374 | 0.2435367 | -1.656003 | 0.097721278 | 0.224591178 | EPHA7 | protein_coding | 33.25 | -0.261581623 | 0.261581623 | ENSG00000135333 | n.s. |
| ENSG00000280376 | 39.72555771 | 0.453745638 | 0.2941486 | 1.542573 | 0.122934413 | 0.26520233 | | TEC | 43.92 | 0.26154927 | 0.26154927 | ENSG00000280376 | n.s. |
| ENSG00000115137 | 137.7679168 | 0.394974858 | 0.2355575 | 1.676766 | 0.093588159 | 0.217677346 | DNAJC27 | protein_coding | 40.71 | 0.261547124 | 0.261547124 | ENSG00000115137 | n.s. |
| ENSG00000115306 | 7194.541425 | -0.392632105 | 0.2333546 | -1.682556 | 0.092461088 | 0.215923471 | SPTBN1 | protein_coding | 42.78 | -0.26137525 | 0.26137525 | ENSG00000115306 | n.s. |
| ENSG00000120784 | 999.2312946 | -0.423722993 | 0.263535 | -1.607843 | 0.107869559 | 0.241729901 | ZFP30 | protein_coding | 41.09 | -0.2612971 | 0.2612971 | ENSG00000120784 | n.s. |
| ENSG00000135823 | 1575.61065 | -0.410066065 | 0.2502282 | -1.638768 | 0.101261556 | 0.230747077 | STX6 | protein_coding | 40.71 | -0.261156229 | 0.261156229 | ENSG00000135823 | n.s. |
| ENSG00000072134 | 662.0846836 | -0.441625392 | 0.2817332 | -1.56753 | 0.116990794 | 0.256252639 | EPN2 | protein_coding | 48.72 | -0.261147073 | 0.261147073 | ENSG00000072134 | n.s. |
| ENSG00000149150 | 117.8447842 | -0.432093117 | 0.2721202 | -1.587876 | 0.112314364 | 0.24882603 | SLC43A1 | protein_coding | 50.69 | -0.261029262 | 0.261029262 | ENSG00000149150 | n.s. |
| ENSG00000123349 | 3338.735232 | 0.489359537 | 0.3318451 | 1.474663 | 0.140303293 | 0.292826075 | PFDN5 | protein_coding | 47.28 | 0.261019608 | 0.261019608 | ENSG00000123349 | n.s. |
| ENSG00000112276 | 1718.646694 | -0.409407904 | 0.2497352 | -1.639368 | 0.101136697 | 0.23057577 | BVES | protein_coding | 40.62 | -0.260869121 | 0.260869121 | ENSG00000112276 | n.s. |
| ENSG00000180071 | 12.01338928 | -0.702381041 | 0.5986133 | -1.173347 | 0.240656752 | 0.425427116 | ANKRD18A | protein_coding | 36.89 | -0.260706166 | 0.260706166 | ENSG00000180071 | n.s. |
| ENSG00000273711 | 14.3978897 | 0.522643836 | 0.369858 | 1.413093 | 0.157628267 | 0.317101338 | | antisense | 32.32 | 0.260695752 | 0.260695752 | ENSG00000273711 | n.s. |
| ENSG00000167114 | 205.7490233 | -0.412898222 | 0.2533112 | -1.630004 | 0.103100753 | 0.23386637 | SLC27A4 | protein_coding | 54.67 | -0.260552084 | 0.260552084 | ENSG00000167114 | n.s. |
| ENSG00000182858 | 208.1099758 | -0.430426084 | 0.2708131 | -1.589384 | 0.11197372 | 0.248329102 | ALG12 | protein_coding | 55.69 | -0.260395893 | 0.260395893 | ENSG00000182858 | n.s. |
| ENSG00000174748 | 16116.94916 | 0.450536081 | 0.2914897 | 1.545633 | 0.122193233 | 0.264299579 | RPL15 | protein_coding | 46.46 | 0.260366391 | 0.260366391 | ENSG00000174748 | n.s. |
| ENSG00000008988 | 11975.95831 | 0.488298079 | 0.3313161 | 1.473813 | 0.140531817 | 0.293053342 | RPS20 | protein_coding | 41.97 | 0.260288913 | 0.260288913 | ENSG00000008988 | n.s. |
| ENSG00000236489 | 17.63133254 | -0.503317026 | 0.3480439 | -1.446131 | 0.148140505 | 0.304130383 | | processed_pseudogene | 45.15 | -0.2601848 | 0.2601848 | ENSG00000236489 | n.s. |
| ENSG00000005007 | 108.677007 | 0.413126337 | 0.2540535 | 1.626139 | 0.103919998 | 0.235188253 | UPF1 | protein_coding | 55.16 | 0.25968476 | 0.25968476 | ENSG00000005007 | n.s. |
| ENSG00000247271 | 94.47021393 | 0.404258636 | 0.245505 | 1.646641 | 0.099631832 | 0.227847932 | ZBED5-AS1 | antisense | 40.5 | 0.259677519 | 0.259677519 | ENSG00000247271 | n.s. |
| ENSG00000100324 | 265.4219599 | -0.449619957 | 0.2912446 | -1.543788 | 0.122639614 | 0.264812553 | TAB1 | protein_coding | 51.38 | -0.259458336 | 0.259458336 | ENSG00000100324 | n.s. |
| ENSG00000155093 | 163.6613254 | -0.431437746 | 0.2725744 | -1.582826 | 0.113461169 | 0.250550335 | PTPRN2 | protein_coding | 51.39 | -0.259339392 | 0.259339392 | ENSG00000155093 | n.s. |
| ENSG00000185950 | 241.801078 | 0.401437285 | 0.2432836 | 1.650079 | 0.098926731 | 0.226584839 | IRS2 | protein_coding | 46.75 | 0.258834378 | 0.258834378 | ENSG00000185950 | n.s. |
| ENSG00000179841 | 354.8597369 | 0.365266665 | 0.20919 | 1.7461 | 0.080793613 | 0.195701021 | AKAP5 | protein_coding | 37.55 | 0.258757429 | 0.258757429 | ENSG00000179841 | n.s. |
| ENSG00000202337 | 53.1134663 | 0.497996151 | 0.343123 | 1.451363 | 0.146678761 | 0.302414984 | RNU6-8 | snRNA | 46.73 | 0.258657555 | 0.258657555 | ENSG00000202337 | n.s. |
| ENSG00000131019 | 65.8829021 | 0.421511193 | 0.2629098 | 1.603254 | 0.108878587 | 0.243446014 | ULBP3 | protein_coding | 54.66 | 0.258638144 | 0.258638144 | ENSG00000131019 | n.s. |
| ENSG00000115297 | 5176.768473 | -0.404905022 | 0.2466908 | -1.641347 | 0.100725494 | 0.229826459 | TLX2 | protein_coding | 64.99 | -0.258572336 | 0.258572336 | ENSG00000115297 | n.s. |
| ENSG00000199053 | 3.559982344 | -0.957735957 | 1.0128533 | -0.945582 | 0.344361781 | 0.537144194 | MIR324 | miRNA | 61.45 | -0.258501664 | 0.258501664 | ENSG00000199053 | n.s. |
| ENSG00000044446 | 22.27604114 | -0.498160668 | 0.3434841 | -1.450317 | 0.146970223 | 0.302845422 | PHKA2 | protein_coding | 45.75 | -0.258435287 | 0.258435287 | ENSG00000044446 | n.s. |
| ENSG00000130477 | 993.3109509 | -0.407653011 | 0.2493819 | -1.634654 | 0.102121677 | 0.232326815 | UNC13A | protein_coding | 52.17 | -0.258411514 | 0.258411514 | ENSG00000130477 | n.s. |

| | | | | | | | | | | | | | |
|---|---|---|---|---|---|---|---|---|---|---|---|---|---|
| ENSG00000146535 | 394.1152916 | -0.435157111 | 0.2769302 | -1.57136 | 0.116099047 | 0.254879696 | GNA12 | protein_coding | 45.87 | -0.258337442 | 0.258337442 | ENSG00000146535 | n.s. |
| ENSG00000147403 | 13101.08745 | 0.447287879 | 0.2897085 | 1.543924 | 0.122606793 | 0.264782747 | RPL10 | protein_coding | 54.28 | 0.258134449 | 0.258134449 | ENSG00000147403 | n.s. |
| ENSG00000130935 | 5037.230595 | -0.42124459 | 0.2630893 | -1.601147 | 0.109344363 | 0.244013714 | NOL11 | protein_coding | 40.57 | -0.25804844 | 0.25804844 | ENSG00000130935 | n.s. |
| ENSG00000146834 | 572.2137485 | -0.377125317 | 0.2205575 | -1.709873 | 0.087289353 | 0.20689984 | MEPCE | protein_coding | 58.8 | -0.258044168 | 0.258044168 | ENSG00000146834 | n.s. |
| ENSG00000106367 | 339.1759797 | 0.470080203 | 0.313212 | 1.500837 | 0.133397729 | 0.282599396 | AP1S1 | protein_coding | 54.82 | 0.25799354 | 0.25799354 | ENSG00000106367 | n.s. |
| ENSG00000105193 | 3226.539866 | 0.437613664 | 0.2797804 | 1.564133 | 0.117786375 | 0.257487784 | RPS16 | protein_coding | 50.95 | 0.25786095 | 0.25786095 | ENSG00000105193 | n.s. |
| ENSG00000116747 | 4005.582359 | -0.446114273 | 0.2885944 | -1.545817 | 0.122148674 | 0.264244249 | TROVE2 | protein_coding | 35.99 | -0.257851576 | 0.257851576 | ENSG00000116747 | n.s. |
| ENSG00000143774 | 1593.111871 | 0.457982012 | 0.3010244 | 1.521412 | 0.128156546 | 0.273627295 | GUK1 | protein_coding | 61.76 | 0.257770862 | 0.257770862 | ENSG00000143774 | n.s. |
| ENSG00000279283 | 74.74812067 | 0.436415684 | 0.278662 | 1.566111 | 0.117322559 | 0.25667536 | | TEC | 57.82 | 0.257754007 | 0.257754007 | ENSG00000279283 | n.s. |
| ENSG00000204196 | 279.2320091 | 0.496931391 | 0.3428256 | 1.449517 | 0.147193321 | 0.303161628 | RPL12P16 | processed_pseudogene | 51.81 | 0.257572345 | 0.257572345 | ENSG00000204196 | n.s. |
| ENSG00000188566 | 248.7946657 | -0.410868924 | 0.2531778 | -1.622847 | 0.10462201 | 0.236316067 | NDOR1 | protein_coding | 59.32 | -0.257412154 | 0.257412154 | ENSG00000188566 | n.s. |
| ENSG00000277146 | 14.26195738 | -0.537747156 | 0.3903369 | -1.377649 | 0.168311697 | 0.332357277 | | processed_pseudogene | 43.24 | -0.257255448 | 0.257255448 | ENSG00000277146 | n.s. |
| ENSG00000006062 | 369.3665672 | -0.380408883 | 0.2240505 | -1.697871 | 0.089532146 | 0.210851839 | MAP3K14 | protein_coding | 49.97 | -0.257165005 | 0.257165005 | ENSG00000006062 | n.s. |
| ENSG00000147996 | 205.8178219 | -0.415338678 | 0.2577961 | -1.611113 | 0.107155037 | 0.240515812 | CBWD5 | protein_coding | 36.05 | -0.257034986 | 0.257034986 | ENSG00000147996 | n.s. |
| ENSG00000170540 | 6499.840272 | 0.447761358 | 0.2909798 | 1.538806 | 0.123851754 | 0.266726677 | ARL6IP1 | protein_coding | 40.91 | 0.256985264 | 0.256985264 | ENSG00000170540 | n.s. |
| ENSG00000119711 | 227.9599668 | -0.410742231 | 0.2536935 | -1.619049 | 0.105436641 | 0.237462691 | ALDH6A1 | protein_coding | 43.41 | -0.256469345 | 0.256469345 | ENSG00000119711 | n.s. |
| ENSG00000101412 | 1895.22755 | -0.440634515 | 0.283832 | -1.552448 | 0.120555039 | 0.261977182 | E2F1 | protein_coding | 55.58 | -0.256333195 | 0.256333195 | ENSG00000101412 | n.s. |
| ENSG00000033867 | 1005.253564 | -0.41653892 | 0.2593923 | -1.605826 | 0.108312189 | 0.242565645 | SLC4A7 | protein_coding | 36.85 | -0.256242543 | 0.256242543 | ENSG00000033867 | n.s. |
| ENSG00000153201 | 10812.61912 | -0.39838646 | 0.2418062 | -1.647544 | 0.099446239 | 0.227578893 | RANBP2 | protein_coding | 39.01 | -0.256109915 | 0.256109915 | ENSG00000153201 | n.s. |
| ENSG00000147118 | 204.5313043 | -0.422074675 | 0.2651251 | -1.591983 | 0.111388524 | 0.247307037 | ZNF182 | protein_coding | 40.39 | -0.256099518 | 0.256099518 | ENSG00000147118 | n.s. |
| ENSG00000179091 | 1091.687433 | 0.439811871 | 0.2831676 | 1.553186 | 0.120378779 | 0.261675839 | CYC1 | protein_coding | 63.31 | 0.25607447 | 0.25607447 | ENSG00000179091 | n.s. |
| ENSG00000265623 | 6.228939257 | 0.620130825 | 0.4937928 | 1.255852 | 0.209169534 | 0.386447375 | MIR3139 | miRNA | 46.05 | 0.256057994 | 0.256057994 | ENSG00000265623 | n.s. |
| ENSG00000007376 | 202.2166007 | -0.443955974 | 0.2877211 | -1.543008 | 0.122828865 | 0.265024713 | RPUSD1 | protein_coding | 67.17 | -0.25603546 | 0.25603546 | ENSG00000007376 | n.s. |
| ENSG00000164294 | 207.1554881 | -0.400549353 | 0.24411 | -1.640856 | 0.100827362 | 0.229908349 | GPX8 | protein_coding | 39.03 | -0.255728835 | 0.255728835 | ENSG00000164294 | n.s. |
| ENSG00000121879 | 1113.63288 | -0.3974588 | 0.2412186 | -1.647712 | 0.099411833 | 0.227537556 | PIK3CA | protein_coding | 35.78 | -0.255544907 | 0.255544907 | ENSG00000121879 | n.s. |
| ENSG00000136270 | 1363.428146 | -0.381714678 | 0.2263652 | -1.686278 | 0.091742201 | 0.214785046 | TBRG4 | protein_coding | 53.25 | -0.254983863 | 0.254983863 | ENSG00000136270 | n.s. |
| ENSG00000247315 | 973.1770682 | 0.380075563 | 0.2249036 | 1.689949 | 0.091037767 | 0.213458879 | ZCCHC3 | protein_coding | 55.16 | 0.254911274 | 0.254911274 | ENSG00000247315 | n.s. |
| ENSG00000164219 | 1592.558422 | -0.40693968 | 0.2507873 | -1.622649 | 0.104664472 | 0.236373633 | PGGT1B | protein_coding | 35.39 | -0.25490741 | 0.25490741 | ENSG00000164219 | n.s. |
| ENSG00000150990 | 756.0052997 | -0.402291402 | 0.2462414 | -1.633728 | 0.10231598 | 0.232540873 | DHX37 | protein_coding | 54.02 | -0.254851885 | 0.254851885 | ENSG00000150990 | n.s. |
| ENSG00000277170 | 113.265027 | 0.388290438 | 0.2329721 | 1.666682 | 0.095577638 | 0.220830974 | | sense_intronic | 45.3 | 0.254695235 | 0.254695235 | ENSG00000277170 | n.s. |
| ENSG00000197555 | 23.41658248 | 0.472879038 | 0.3188734 | 1.482968 | 0.138082945 | 0.289450292 | SIPA1L1 | protein_coding | 39.56 | 0.254610373 | 0.254610373 | ENSG00000197555 | n.s. |
| ENSG00000205544 | 187.7230765 | 0.485128714 | 0.3322351 | 1.460197 | 0.144235867 | 0.298794224 | TMEM256 | protein_coding | 57.95 | 0.254512011 | 0.254512011 | ENSG00000205544 | n.s. |

| | | | | | | | | | | | | | |
|---|---|---|---|---|---|---|---|---|---|---|---|---|---|
| ENSG00000129235 | 583.0725474 | 0.440406251 | 0.285038 | 1.545079 | 0.122327078 | 0.264465823 | TXNDC17 | protein_coding | 42.73 | 0.254392058 | 0.254392058 | ENSG00000129235 | n.s. |
| ENSG00000139514 | 1631.932419 | -0.396739049 | 0.241213 | -1.644766 | 0.10001798 | 0.228521994 | SLC7A1 | protein_coding | 47.44 | -0.254338294 | 0.254338294 | ENSG00000139514 | n.s. |
| ENSG00000167283 | 1656.076569 | 0.475803568 | 0.3224135 | 1.475756 | 0.140009439 | 0.29251972 | ATP5L | protein_coding | 42.83 | 0.254005284 | 0.254005284 | ENSG00000167283 | n.s. |
| ENSG00000152082 | 2208.248701 | 0.472983422 | 0.319448 | 1.480627 | 0.138705907 | 0.290406368 | MZT2B | protein_coding | 57.61 | 0.253989196 | 0.253989196 | ENSG00000152082 | n.s. |
| ENSG00000006576 | 1081.691095 | -0.447648431 | 0.2928738 | -1.528469 | 0.126396251 | 0.271032652 | PHTF2 | protein_coding | 37.65 | -0.253806985 | 0.253806985 | ENSG00000006576 | n.s. |
| ENSG00000279555 | 513.6193007 | -0.398936863 | 0.2435926 | -1.637721 | 0.101479831 | 0.231168797 | | TEC | 49.86 | -0.253752085 | 0.253752085 | ENSG00000279555 | n.s. |
| ENSG00000116918 | 1753.916375 | -0.430847792 | 0.2757166 | -1.562647 | 0.118135639 | 0.257927318 | TSNAX | protein_coding | 37.66 | -0.253555071 | 0.253555071 | ENSG00000116918 | n.s. |
| ENSG00000142534 | 25853.15953 | 0.463118035 | 0.3091267 | 1.49815 | 0.134094385 | 0.283651796 | RPS11 | protein_coding | 53.05 | 0.253424886 | 0.253424886 | ENSG00000142534 | n.s. |
| ENSG00000145979 | 434.9534566 | 0.389801468 | 0.2351623 | 1.657585 | 0.097401283 | 0.223966649 | TBC1D7 | protein_coding | 44.56 | 0.253299483 | 0.253299483 | ENSG00000145979 | n.s. |
| ENSG00000090061 | 220.4183651 | 0.402944083 | 0.2478662 | 1.625651 | 0.104023861 | 0.23538505 | CCNK | protein_coding | 50.14 | 0.253137985 | 0.253137985 | ENSG00000090061 | n.s. |
| ENSG00000161010 | 321.9557981 | -0.425466239 | 0.2704272 | -1.573312 | 0.115646693 | 0.254207279 | MRNIP | protein_coding | 49.4 | -0.253072433 | 0.253072433 | ENSG00000161010 | n.s. |
| ENSG00000156990 | 320.1222786 | -0.446851385 | 0.2927484 | -1.526401 | 0.12691003 | 0.271752671 | RPUSD3 | protein_coding | 52.21 | -0.252840212 | 0.252840212 | ENSG00000156990 | n.s. |
| ENSG00000143748 | 1613.163199 | -0.414424419 | 0.259556 | -1.596667 | 0.110340046 | 0.245566571 | NVL | protein_coding | 41.59 | -0.252728756 | 0.252728756 | ENSG00000143748 | n.s. |
| ENSG00000185090 | 490.0718264 | -0.42146084 | 0.2668012 | -1.579681 | 0.114179917 | 0.251459504 | MANEAL | protein_coding | 54.73 | -0.252679237 | 0.252679237 | ENSG00000185090 | n.s. |
| ENSG00000156508 | 123363.1782 | 0.453934366 | 0.3005019 | 1.510588 | 0.130893575 | 0.278289203 | EEF1A1 | protein_coding | 47.14 | 0.252162194 | 0.252162194 | ENSG00000156508 | n.s. |
| ENSG00000114416 | 5407.699842 | -0.387191241 | 0.2332956 | -1.65966 | 0.096982953 | 0.223373641 | FXR1 | protein_coding | 38.94 | -0.252049138 | 0.252049138 | ENSG00000114416 | n.s. |
| ENSG00000214087 | 1296.750179 | -0.406239414 | 0.2518199 | -1.613214 | 0.106698022 | 0.239683214 | ARL16 | protein_coding | 56.96 | -0.25201565 | 0.25201565 | ENSG00000214087 | n.s. |
| ENSG00000264701 | 15.68763813 | 0.497071368 | 0.3480019 | 1.428358 | 0.153188773 | 0.311194596 | | antisense | 51.52 | 0.251999254 | 0.251999254 | ENSG00000264701 | n.s. |
| ENSG00000146085 | 934.5416836 | -0.441651075 | 0.2880286 | -1.533358 | 0.125187576 | 0.268979418 | MUT | protein_coding | 34.4 | -0.251865195 | 0.251865195 | ENSG00000146085 | n.s. |
| ENSG00000155380 | 7640.976616 | 0.401907075 | 0.2477424 | 1.622278 | 0.104743772 | 0.236476489 | SLC16A1 | protein_coding | 40.76 | 0.251679046 | 0.251679046 | ENSG00000155380 | n.s. |
| ENSG00000173207 | 1787.132161 | 0.445756939 | 0.2924143 | 1.524402 | 0.1274083 | 0.272573104 | CKS1B | protein_coding | 44.4 | 0.25163737 | 0.25163737 | ENSG00000173207 | n.s. |
| ENSG00000152128 | 90.58927683 | -0.445170259 | 0.2918844 | -1.52516 | 0.127219276 | 0.272270001 | TMEM163 | protein_coding | 43.72 | -0.251521288 | 0.251521288 | ENSG00000152128 | n.s. |
| ENSG00000198563 | 2428.428344 | 0.359724648 | 0.2076923 | 1.732008 | 0.083272131 | 0.199999635 | DDX39B | protein_coding | 47.24 | 0.251437024 | 0.251437024 | ENSG00000198563 | n.s. |
| ENSG00000269938 | 104.4404609 | 0.447508721 | 0.2944574 | 1.519774 | 0.128567767 | 0.274379065 | | sense_intronic | 34.11 | 0.251342838 | 0.251342838 | ENSG00000269938 | n.s. |
| ENSG00000183337 | 1919.969159 | -0.391739319 | 0.2381776 | -1.644736 | 0.100024164 | 0.228521994 | BCOR | protein_coding | 53.23 | -0.251133107 | 0.251133107 | ENSG00000183337 | n.s. |
| ENSG00000134871 | 1479.18111 | -0.384098762 | 0.2309187 | -1.663351 | 0.09624221 | 0.222034845 | COL4A2 | protein_coding | 47.83 | -0.251038833 | 0.251038833 | ENSG00000134871 | n.s. |
| ENSG00000118046 | 366.6827265 | -0.416343776 | 0.262637 | -1.585244 | 0.112910908 | 0.249572722 | STK11 | protein_coding | 57.49 | -0.250973229 | 0.250973229 | ENSG00000118046 | n.s. |
| ENSG00000179387 | 2362.224476 | -0.427520712 | 0.2740454 | -1.560036 | 0.118751384 | 0.259027968 | ELMOD2 | protein_coding | 34.81 | -0.250806454 | 0.250806454 | ENSG00000179387 | n.s. |
| ENSG00000163939 | 3745.50647 | -0.411717934 | 0.2582868 | -1.594034 | 0.11092842 | 0.246521371 | PBRM1 | protein_coding | 40.81 | -0.250384379 | 0.250384379 | ENSG00000163939 | n.s. |
| ENSG00000265932 | 28.39433856 | 0.461323418 | 0.3095149 | 1.490473 | 0.136100037 | 0.286761625 | MIR3146 | miRNA | 37.97 | 0.250258251 | 0.250258251 | ENSG00000265932 | n.s. |
| ENSG00000111860 | 3071.779476 | -0.399635636 | 0.2463818 | -1.622018 | 0.104799589 | 0.236525316 | CEP85L | protein_coding | 38.17 | -0.250220811 | 0.250220811 | ENSG00000111860 | n.s. |
| ENSG00000086827 | 510.0414695 | -0.394663652 | 0.2414285 | -1.634702 | 0.102111548 | 0.232326815 | ZW10 | protein_coding | 39.15 | -0.25017755 | 0.25017755 | ENSG00000086827 | n.s. |

| | | | | | | | | | | | | | |
|---|---|---|---|---|---|---|---|---|---|---|---|---|---|
| ENSG00000144218 | 437.3634035 | -0.400929193 | 0.2478354 | -1.617724 | 0.10572214 | 0.23802868 | AFF3 | protein_coding | 41.63 | -0.249927516 | 0.249927516 | ENSG00000144218 | n.s. |
| ENSG00000076984 | 312.2190436 | -0.407760715 | 0.2546603 | -1.601195 | 0.109333779 | 0.244013714 | MAP2K7 | protein_coding | 63.14 | -0.249788409 | 0.249788409 | ENSG00000076984 | n.s. |
| ENSG00000153815 | 175.6063693 | -0.412708756 | 0.2596163 | -1.589688 | 0.111905222 | 0.248216728 | CMIP | protein_coding | 51.33 | -0.249758526 | 0.249758526 | ENSG00000153815 | n.s. |
| ENSG00000109929 | 491.4313774 | -0.441284759 | 0.2891145 | -1.526332 | 0.126927215 | 0.271752671 | SC5D | protein_coding | 38.78 | -0.249690469 | 0.249690469 | ENSG00000109929 | n.s. |
| ENSG00000181038 | 634.0564287 | -0.449360216 | 0.2975144 | -1.510381 | 0.13094614 | 0.278289203 | METTL23 | protein_coding | 47.85 | -0.249621237 | 0.249621237 | ENSG00000181038 | n.s. |
| ENSG00000198680 | 233.6000261 | 0.397808726 | 0.2448626 | 1.62462 | 0.104243522 | 0.235805451 | TUSC1 | protein_coding | 52.44 | 0.249603557 | 0.249603557 | ENSG00000198680 | n.s. |
| ENSG00000153250 | 2288.664679 | -0.393229754 | 0.2406345 | -1.634137 | 0.102230026 | 0.232459359 | RBMS1 | protein_coding | 39.2 | -0.2491712 | 0.2491712 | ENSG00000153250 | n.s. |
| ENSG00000156697 | 831.4887748 | -0.406539793 | 0.2539535 | -1.600843 | 0.109411609 | 0.244085535 | UTP14A | protein_coding | 45.09 | -0.248988531 | 0.248988531 | ENSG00000156697 | n.s. |
| ENSG00000180806 | 63.60834471 | 0.467202591 | 0.3170738 | 1.473482 | 0.140621101 | 0.293181729 | HOXC9 | protein_coding | 56.89 | 0.24895502 | 0.24895502 | ENSG00000180806 | n.s. |
| ENSG00000276023 | 260.9472111 | 0.471812425 | 0.3221448 | 1.464597 | 0.143030774 | 0.296960553 | DUSP14 | protein_coding | 46.28 | 0.248787275 | 0.248787275 | ENSG00000276023 | n.s. |
| ENSG00000072501 | 3374.108008 | -0.405306723 | 0.2528037 | -1.603247 | 0.10888015 | 0.243446014 | SMC1A | protein_coding | 45.54 | -0.248695124 | 0.248695124 | ENSG00000072501 | n.s. |
| ENSG00000269858 | 668.4146275 | -0.401761688 | 0.2493305 | -1.611362 | 0.107100836 | 0.240471687 | EGLN2 | protein_coding | 59.14 | -0.248664792 | 0.248664792 | ENSG00000269858 | n.s. |
| ENSG00000148429 | 860.7813494 | -0.400038734 | 0.2477043 | -1.614985 | 0.106314 | 0.239051973 | USP6NL | protein_coding | 38.22 | -0.24862714 | 0.24862714 | ENSG00000148429 | n.s. |
| ENSG00000164414 | 380.5832678 | -0.460416083 | 0.3100525 | -1.484962 | 0.137553963 | 0.288708717 | SLC35A1 | protein_coding | 39.25 | -0.248412941 | 0.248412941 | ENSG00000164414 | n.s. |
| ENSG00000130811 | 1377.937606 | 0.431504925 | 0.2799246 | 1.541504 | 0.123194102 | 0.265680217 | EIF3G | protein_coding | 59.24 | 0.248391833 | 0.248391833 | ENSG00000130811 | n.s. |
| ENSG00000197859 | 138.0209084 | -0.413457605 | 0.2613691 | -1.581891 | 0.113674355 | 0.250901719 | ADAMTSL2 | protein_coding | 59.43 | -0.248279788 | 0.248279788 | ENSG00000197859 | n.s. |
| ENSG00000279819 | 38.40611486 | -0.439446986 | 0.2883137 | -1.524197 | 0.127459432 | 0.2726406 | | TEC | 42.5 | -0.248028041 | 0.248028041 | ENSG00000279819 | n.s. |
| ENSG00000206503 | 559.7403082 | 0.427331823 | 0.2757495 | 1.54971 | 0.121211093 | 0.263074365 | HLA-A | protein_coding | 55.18 | 0.247818898 | 0.247818898 | ENSG00000206503 | n.s. |
| ENSG00000165684 | 345.7077732 | -0.372854782 | 0.2220484 | -1.67916 | 0.093120815 | 0.2167733 | SNAPC4 | protein_coding | 58.99 | -0.247573417 | 0.247573417 | ENSG00000165684 | n.s. |
| ENSG00000251485 | 58.3206145 | -0.445511748 | 0.2951977 | -1.509198 | 0.131248268 | 0.278774925 | | processed_pseudogene | 49.11 | -0.24714599 | 0.24714599 | ENSG00000251485 | n.s. |
| ENSG00000070614 | 1038.615727 | -0.383307312 | 0.2323679 | -1.649571 | 0.099030728 | 0.226739817 | NDST1 | protein_coding | 52.04 | -0.24703091 | 0.24703091 | ENSG00000070614 | n.s. |
| ENSG00000100393 | 1664.345501 | -0.392266001 | 0.2410075 | -1.627609 | 0.103607785 | 0.234672392 | EP300 | protein_coding | 42.33 | -0.246946354 | 0.246946354 | ENSG00000100393 | n.s. |
| ENSG00000159753 | 333.4214094 | 0.391202114 | 0.2401945 | 1.628689 | 0.103378926 | 0.234287391 | CARMIL2 | protein_coding | 62.9 | 0.246555556 | 0.246555556 | ENSG00000159753 | n.s. |
| ENSG00000165637 | 2281.122943 | 0.426298968 | 0.2755964 | 1.546823 | 0.121905816 | 0.264047592 | VDAC2 | protein_coding | 44.04 | 0.246536276 | 0.246536276 | ENSG00000165637 | n.s. |
| ENSG00000244627 | 89.0833403 | -0.423080969 | 0.2723685 | -1.553341 | 0.120341798 | 0.261636298 | | transcribed_unprocessed_pseud | 43.54 | -0.246360896 | 0.246360896 | ENSG00000244627 | n.s. |
| ENSG00000274340 | 90.51309451 | 0.39081672 | 0.2399052 | 1.629046 | 0.103303204 | 0.23422809 | | lincRNA | 37.51 | 0.246355628 | 0.246355628 | ENSG00000274340 | n.s. |
| ENSG00000137845 | 3569.423141 | -0.400587071 | 0.2495407 | -1.605298 | 0.108428318 | 0.242747624 | ADAM10 | protein_coding | 37.38 | -0.246298962 | 0.246298962 | ENSG00000137845 | n.s. |
| ENSG00000105810 | 12557.67286 | -0.418412656 | 0.2677927 | -1.56245 | 0.11818205 | 0.257988192 | CDK6 | protein_coding | 38.2 | -0.246194078 | 0.246194078 | ENSG00000105810 | n.s. |
| ENSG00000113558 | 5333.094207 | 0.419358784 | 0.2686862 | 1.560775 | 0.118576839 | 0.258809425 | SKP1 | protein_coding | 41.26 | 0.246171956 | 0.246171956 | ENSG00000113558 | n.s. |
| ENSG00000260852 | 158.2475833 | -0.40527233 | 0.2544535 | -1.592717 | 0.111223799 | 0.247020091 | FBXL19-AS1 | antisense | 52.62 | -0.246108804 | 0.246108804 | ENSG00000260852 | n.s. |
| ENSG00000198018 | 290.7497282 | -0.401102156 | 0.250314 | -1.602396 | 0.109068027 | 0.243670433 | ENTPD7 | protein_coding | 40.08 | -0.245954705 | 0.245954705 | ENSG00000198018 | n.s. |
| ENSG00000241859 | 203.6150149 | -0.404498442 | 0.2540104 | -1.592449 | 0.11128394 | 0.247114243 | ANOS2P | transcribed_unprocessed_pseud | 40.21 | -0.245571903 | 0.245571903 | ENSG00000241859 | n.s. |

| | | | | | | | | | | | | | |
|---|---|---|---|---|---|---|---|---|---|---|---|---|---|
| ENSG00000119943 | 58.78502204 | 0.41824891 | 0.2680129 | 1.560555 | 0.118628758 | 0.258882162 | PYROXD2 | protein_coding | 50.39 | 0.245469396 | 0.245469396 | ENSG00000119943 | n.s. |
| ENSG00000072121 | 413.651769 | -0.379428243 | 0.2295302 | -1.653065 | 0.098317705 | 0.225589553 | ZFYVE26 | protein_coding | 43 | -0.245369042 | 0.245369042 | ENSG00000072121 | n.s. |
| ENSG00000152558 | 2179.452418 | 0.414523862 | 0.2642908 | 1.568439 | 0.11677882 | 0.255928146 | TMEM123 | protein_coding | 40.62 | 0.245349191 | 0.245349191 | ENSG00000152558 | n.s. |
| ENSG00000112306 | 16677.84093 | 0.471985075 | 0.3252595 | 1.451103 | 0.146751145 | 0.302519423 | RPS12 | protein_coding | 42.96 | 0.24507671 | 0.24507671 | ENSG00000112306 | n.s. |
| ENSG00000264712 | 7.920418226 | 0.590909983 | 0.469294 | 1.259147 | 0.20797736 | 0.385062338 | MIR4504 | miRNA | 38.04 | 0.244913844 | 0.244913844 | ENSG00000264712 | n.s. |
| ENSG00000272667 | 137.4672042 | 0.3727548 | 0.2234636 | 1.668078 | 0.095300226 | 0.220388803 | | lincRNA | 51.9 | 0.244829257 | 0.244829257 | ENSG00000272667 | n.s. |
| ENSG00000068489 | 1491.450931 | -0.418302423 | 0.2686009 | -1.557338 | 0.119390262 | 0.260095544 | PRR11 | protein_coding | 44.47 | -0.24465132 | 0.24465132 | ENSG00000068489 | n.s. |
| ENSG00000171988 | 4270.300254 | -0.411527086 | 0.2616963 | -1.572537 | 0.115826054 | 0.254440857 | JMJD1C | protein_coding | 38.11 | -0.244617111 | 0.244617111 | ENSG00000171988 | n.s. |
| ENSG00000127125 | 136.8525844 | -0.41178475 | 0.2619714 | -1.571869 | 0.115980956 | 0.254660597 | PPCS | protein_coding | 42.3 | -0.244615891 | 0.244615891 | ENSG00000127125 | n.s. |
| ENSG00000117691 | 873.0269883 | 0.460292981 | 0.3129524 | 1.470808 | 0.141342978 | 0.294290157 | NENF | protein_coding | 47.72 | 0.2445188 | 0.2445188 | ENSG00000117691 | n.s. |
| ENSG00000010295 | 246.1435647 | -0.400094173 | 0.2504205 | -1.597689 | 0.110112138 | 0.245176923 | IFFO1 | protein_coding | 54.32 | -0.244265658 | 0.244265658 | ENSG00000010295 | n.s. |
| ENSG00000262050 | 22.5864566 | 0.477119097 | 0.3321365 | 1.436515 | 0.150855813 | 0.307700774 | | antisense | 57.9 | 0.244223626 | 0.244223626 | ENSG00000262050 | n.s. |
| ENSG00000088367 | 184.3581181 | -0.426061718 | 0.2771749 | -1.537158 | 0.124254556 | 0.267394558 | EPB41L1 | protein_coding | 49.98 | -0.244068363 | 0.244068363 | ENSG00000088367 | n.s. |
| ENSG00000133313 | 1008.280146 | -0.417366339 | 0.2681474 | -1.556481 | 0.119593689 | 0.260375751 | CNDP2 | protein_coding | 51.18 | -0.243908665 | 0.243908665 | ENSG00000133313 | n.s. |
| ENSG00000078140 | 1670.764559 | -0.428578582 | 0.2798338 | -1.531547 | 0.125634305 | 0.269731137 | UBE2K | protein_coding | 42.02 | -0.24389075 | 0.24389075 | ENSG00000078140 | n.s. |
| ENSG00000174444 | 48680.03595 | 0.44415795 | 0.2959527 | 1.500773 | 0.133414194 | 0.282599396 | RPL4 | protein_coding | 42.06 | 0.243766662 | 0.243766662 | ENSG00000174444 | n.s. |
| ENSG00000105426 | 1302.48784 | -0.385115095 | 0.235955 | -1.632155 | 0.102646906 | 0.233102736 | PTPRS | protein_coding | 52.47 | -0.243567054 | 0.243567054 | ENSG00000105426 | n.s. |
| ENSG00000082641 | 2436.631191 | -0.406432461 | 0.2573659 | -1.579201 | 0.114289974 | 0.251622282 | NFE2L1 | protein_coding | 54.35 | -0.243555019 | 0.243555019 | ENSG00000082641 | n.s. |
| ENSG00000148343 | 43.42228758 | 0.432171104 | 0.2837631 | 1.523 | 0.127758822 | 0.273207684 | MIGA2 | protein_coding | 53.65 | 0.243531482 | 0.243531482 | ENSG00000148343 | n.s. |
| ENSG00000213782 | 2124.858238 | 0.417786855 | 0.2688666 | 1.553881 | 0.120212691 | 0.261396424 | DDX47 | protein_coding | 40.84 | 0.24344455 | 0.24344455 | ENSG00000213782 | n.s. |
| ENSG00000166439 | 1271.798965 | -0.409050536 | 0.2600085 | -1.57322 | 0.115667871 | 0.254213697 | RNF169 | protein_coding | 39.49 | -0.24330369 | 0.24330369 | ENSG00000166439 | n.s. |
| ENSG00000119705 | 1738.912029 | 0.454957771 | 0.3081004 | 1.476654 | 0.139768294 | 0.292147419 | SLIRP | protein_coding | 41.29 | 0.243128498 | 0.243128498 | ENSG00000119705 | n.s. |
| ENSG00000106993 | 534.4616891 | -0.402623356 | 0.2536353 | -1.587411 | 0.112419633 | 0.249000725 | CDC37L1 | protein_coding | 36.8 | -0.243103736 | 0.243103736 | ENSG00000106993 | n.s. |
| ENSG00000099290 | 535.3435168 | -0.405527002 | 0.2567412 | -1.579517 | 0.114217559 | 0.251502621 | WASHC2A | protein_coding | 42.59 | -0.243096196 | 0.243096196 | ENSG00000099290 | n.s. |
| ENSG00000207816 | 25.63991962 | 0.469153938 | 0.323874 | 1.448569 | 0.147457905 | 0.303392407 | MIR124-2 | miRNA | 49.54 | 0.243019531 | 0.243019531 | ENSG00000207816 | n.s. |
| ENSG00000138778 | 3861.281671 | -0.412323919 | 0.2636367 | -1.563985 | 0.117820976 | 0.25752299 | CENPE | protein_coding | 34.67 | -0.242934654 | 0.242934654 | ENSG00000138778 | n.s. |
| ENSG00000175691 | 168.0424663 | -0.440306114 | 0.2926033 | -1.504788 | 0.132378472 | 0.280816379 | ZNF77 | protein_coding | 49.74 | -0.242862974 | 0.242862974 | ENSG00000175691 | n.s. |
| ENSG00000225470 | 981.4574426 | -0.409630844 | 0.2611273 | -1.568702 | 0.11671749 | 0.255834007 | JPX | lincRNA | 38.97 | -0.242518553 | 0.242518553 | ENSG00000225470 | n.s. |
| ENSG00000213935 | 26.63028635 | -0.450710994 | 0.3041158 | -1.482037 | 0.138330383 | 0.289794452 | | processed_pseudogene | 43.16 | -0.24244192 | 0.24244192 | ENSG00000213935 | n.s. |
| ENSG00000167770 | 901.5520764 | 0.404028348 | 0.2556692 | 1.580278 | 0.1140433 | 0.251198365 | OTUB1 | protein_coding | 59.12 | 0.242410217 | 0.242410217 | ENSG00000167770 | n.s. |
| ENSG00000204514 | 1295.641305 | -0.386274632 | 0.2379966 | -1.623026 | 0.104583936 | 0.236268398 | ZNF814 | protein_coding | 46.12 | -0.242037507 | 0.242037507 | ENSG00000204514 | n.s. |
| ENSG00000014164 | 173.6199542 | -0.376512443 | 0.2285745 | -1.64722 | 0.099512814 | 0.22765641 | ZC3H3 | protein_coding | 58.38 | -0.241992121 | 0.241992121 | ENSG00000014164 | n.s. |

| | | | | | | | | | | | | | |
|---|---|---|---|---|---|---|---|---|---|---|---|---|---|
| ENSG00000135537 | 56.83054278 | -0.41694108 | 0.2691573 | -1.549061 | 0.121367075 | 0.263289776 | AFG1L | protein_coding | 39.15 | -0.241644876 | 0.241644876 | ENSG00000135537 | n.s. |
| ENSG00000070601 | 304.421929 | -0.383017773 | 0.2350238 | -1.629698 | 0.103165417 | 0.233974931 | FRMPD1 | protein_coding | 44.15 | -0.24161936 | 0.24161936 | ENSG00000070601 | n.s. |
| ENSG00000143164 | 798.122319 | -0.427026495 | 0.2799315 | -1.525468 | 0.12714249 | 0.272171723 | DCAF6 | protein_coding | 35.91 | -0.241337012 | 0.241337012 | ENSG00000143164 | n.s. |
| ENSG00000101752 | 3383.376044 | -0.419375385 | 0.272134 | -1.541062 | 0.123301731 | 0.26583968 | MIB1 | protein_coding | 38.64 | -0.241300291 | 0.241300291 | ENSG00000101752 | n.s. |
| ENSG00000235655 | 286.331151 | 0.465055751 | 0.3208321 | 1.44953 | 0.147189642 | 0.303161628 | H3F3AP4 | processed_pseudogene | 51.58 | 0.24105038 | 0.24105038 | ENSG00000235655 | n.s. |
| ENSG00000110422 | 832.5349644 | -0.399112947 | 0.2516693 | -1.585863 | 0.11277045 | 0.249550887 | HIPK3 | protein_coding | 38.83 | -0.240601601 | 0.240601601 | ENSG00000110422 | n.s. |
| ENSG00000265584 | 3.115053594 | -0.711196261 | 0.6458973 | -1.101098 | 0.270853983 | 0.459062653 | MIR3978 | miRNA | 48.51 | -0.240475396 | 0.240475396 | ENSG00000265584 | n.s. |
| ENSG00000198467 | 156.9214548 | 0.407973187 | 0.2607806 | 1.564431 | 0.117716446 | 0.257375326 | TPM2 | protein_coding | 56.38 | 0.240472897 | 0.240472897 | ENSG00000198467 | n.s. |
| ENSG00000226950 | 4899.258447 | 0.37524923 | 0.2282225 | 1.644226 | 0.10012961 | 0.228691693 | DANCR | processed_transcript | 44.16 | 0.240440798 | 0.240440798 | ENSG00000226950 | n.s. |
| ENSG00000182141 | 882.9776574 | -0.415864604 | 0.2689539 | -1.54623 | 0.122049083 | 0.264156622 | ZNF708 | protein_coding | 39.17 | -0.240427337 | 0.240427337 | ENSG00000182141 | n.s. |
| ENSG00000114942 | 12877.27673 | 0.458115904 | 0.3138396 | 1.459714 | 0.144368785 | 0.29887322 | EEF1B2 | protein_coding | 45.25 | 0.240287746 | 0.240287746 | ENSG00000114942 | n.s. |
| ENSG00000196976 | 173.8517707 | 0.469650518 | 0.3270684 | 1.43594 | 0.151019463 | 0.307899251 | LAGE3 | protein_coding | 63.33 | 0.240269151 | 0.240269151 | ENSG00000196976 | n.s. |
| ENSG00000143578 | 902.9228376 | 0.398568556 | 0.2514046 | 1.585367 | 0.112882967 | 0.249572722 | CREB3L4 | protein_coding | 52.02 | 0.240258275 | 0.240258275 | ENSG00000143578 | n.s. |
| ENSG00000172262 | 1509.848236 | -0.396613509 | 0.2495963 | -1.58902 | 0.112055844 | 0.24843209 | ZNF131 | protein_coding | 44.17 | -0.2398688 | 0.2398688 | ENSG00000172262 | n.s. |
| ENSG00000156876 | 978.3160847 | -0.408587691 | 0.2618456 | -1.560414 | 0.118662001 | 0.25890005 | SASS6 | protein_coding | 36.48 | -0.239786987 | 0.239786987 | ENSG00000156876 | n.s. |
| ENSG00000272325 | 1832.638092 | -0.401776195 | 0.2549688 | -1.575785 | 0.115075254 | 0.253071041 | NUDT3 | protein_coding | 42.12 | -0.239762977 | 0.239762977 | ENSG00000272325 | n.s. |
| ENSG00000259363 | 52.23600746 | -0.445204361 | 0.3002073 | -1.48299 | 0.13807712 | 0.289450292 | | lincRNA | 36.24 | -0.239709607 | 0.239709607 | ENSG00000259363 | n.s. |
| ENSG00000126858 | 926.9804695 | -0.456447538 | 0.3124063 | -1.46107 | 0.143996221 | 0.298519861 | RHOT1 | protein_coding | 42.18 | -0.239647176 | 0.239647176 | ENSG00000126858 | n.s. |
| ENSG00000166503 | 4647.607858 | -0.421396782 | 0.2753079 | -1.530638 | 0.125858899 | 0.270088385 | | protein_coding | 39.61 | -0.239561581 | 0.239561581 | ENSG00000166503 | n.s. |
| ENSG00000229587 | 45.31013105 | -0.491175005 | 0.3521163 | -1.394923 | 0.16303921 | 0.325455995 | | sense_intronic | 38.86 | -0.239451609 | 0.239451609 | ENSG00000229587 | n.s. |
| ENSG00000149923 | 654.369001 | 0.446351466 | 0.3018285 | 1.478825 | 0.139187208 | 0.291238879 | PPP4C | protein_coding | 54.46 | 0.239133088 | 0.239133088 | ENSG00000149923 | n.s. |
| ENSG00000162959 | 486.2399168 | -0.42630499 | 0.2807469 | -1.518467 | 0.128896656 | 0.274954517 | MEMO1 | protein_coding | 38.06 | -0.239045894 | 0.239045894 | ENSG00000162959 | n.s. |
| ENSG00000103061 | 632.0726289 | -0.404492391 | 0.2581931 | -1.566628 | 0.117201747 | 0.256563235 | SLC7A6OS | protein_coding | 47.19 | -0.238976361 | 0.238976361 | ENSG00000103061 | n.s. |
| ENSG00000236552 | 659.6026225 | 0.479003454 | 0.3389583 | 1.413163 | 0.157607685 | 0.317101338 | RPL13AP5 | processed_pseudogene | 57.35 | 0.238927845 | 0.238927845 | ENSG00000236552 | n.s. |
| ENSG00000184271 | 46.40803197 | -0.455029973 | 0.3115065 | -1.46074 | 0.144086847 | 0.29861881 | POU6F1 | protein_coding | 50.08 | -0.238837425 | 0.238837425 | ENSG00000184271 | n.s. |
| ENSG00000070540 | 230.255981 | -0.448573026 | 0.3046325 | -1.472505 | 0.14088444 | 0.293554932 | WIPI1 | protein_coding | 46.58 | -0.238780185 | 0.238780185 | ENSG00000070540 | n.s. |
| ENSG00000156650 | 1268.263769 | -0.392189454 | 0.2459264 | -1.594743 | 0.110769645 | 0.246286452 | KAT6B | protein_coding | 40.6 | -0.23867061 | 0.23867061 | ENSG00000156650 | n.s. |
| ENSG00000128709 | 105.3766734 | 0.405341623 | 0.2593654 | 1.562821 | 0.118094743 | 0.257878468 | HOXD9 | protein_coding | 60.19 | 0.238577966 | 0.238577966 | ENSG00000128709 | n.s. |
| ENSG00000101901 | 489.9225117 | -0.435838332 | 0.2909617 | -1.497924 | 0.134153094 | 0.28370371 | ALG13 | protein_coding | 37.51 | -0.238462399 | 0.238462399 | ENSG00000101901 | n.s. |
| ENSG00000185946 | 1201.325137 | -0.404329815 | 0.2585681 | -1.563727 | 0.117881809 | 0.257575084 | RNPC3 | protein_coding | 35.03 | -0.238189138 | 0.238189138 | ENSG00000185946 | n.s. |
| ENSG00000079432 | 421.5749431 | -0.378042703 | 0.2322274 | -1.627899 | 0.10354637 | 0.234609619 | CIC | protein_coding | 61.52 | -0.23803617 | 0.23803617 | ENSG00000079432 | n.s. |
| ENSG00000005022 | 2068.789324 | 0.44305394 | 0.299262 | 1.480488 | 0.138742954 | 0.290440257 | SLC25A5 | protein_coding | 53.25 | 0.237894795 | 0.237894795 | ENSG00000005022 | n.s. |

| | | | | | | | | | | | | | |
|---|---|---|---|---|---|---|---|---|---|---|---|---|---|
| ENSG00000004961 | 272.5817008 | -0.405317355 | 0.2598886 | -1.559581 | 0.118858961 | 0.259222011 | HCCS | protein_coding | 42.99 | -0.237648965 | 0.237648965 | ENSG00000004961 | n.s. |
| ENSG00000119599 | 333.4754966 | 0.397357443 | 0.2519414 | 1.577182 | 0.114753632 | 0.252524055 | DCAF4 | protein_coding | 48.94 | 0.237499449 | 0.237499449 | ENSG00000119599 | n.s. |
| ENSG00000207221 | 6.463068924 | -0.752793361 | 0.7178425 | -1.048689 | 0.294321383 | 0.483674844 | SNORA70E | snoRNA | 49.63 | -0.237466031 | 0.237466031 | ENSG00000207221 | n.s. |
| ENSG00000149932 | 231.5781273 | 0.464340362 | 0.3235933 | 1.434951 | 0.151301164 | 0.308247895 | TMEM219 | protein_coding | 47.42 | 0.237324304 | 0.237324304 | ENSG00000149932 | n.s. |
| ENSG00000183617 | 417.1261482 | 0.483630027 | 0.3455025 | 1.399787 | 0.161577 | 0.323316552 | MRPL54 | protein_coding | 56.43 | 0.237158653 | 0.237158653 | ENSG00000183617 | n.s. |
| ENSG00000028116 | 344.7157664 | -0.392121715 | 0.2467839 | -1.588928 | 0.112076708 | 0.248438784 | VRK2 | protein_coding | 35.24 | -0.237147608 | 0.237147608 | ENSG00000028116 | n.s. |
| ENSG00000171714 | 531.1370773 | -0.441337494 | 0.2981279 | -1.480363 | 0.138776401 | 0.290466602 | ANO5 | protein_coding | 34.49 | -0.236955776 | 0.236955776 | ENSG00000171714 | n.s. |
| ENSG00000198700 | 6431.544572 | -0.404571353 | 0.2596777 | -1.557975 | 0.119239195 | 0.259884063 | IPO9 | protein_coding | 41.91 | -0.236763389 | 0.236763389 | ENSG00000198700 | n.s. |
| ENSG00000210151 | 33.29225414 | 0.442555362 | 0.2997312 | 1.476508 | 0.139807666 | 0.29218585 | MT-TS1 | Mt_tRNA | 49.28 | 0.236475395 | 0.236475395 | ENSG00000210151 | n.s. |
| ENSG00000036054 | 745.8433906 | -0.422879431 | 0.2790968 | -1.515171 | 0.129729082 | 0.276391432 | TBC1D23 | protein_coding | 36.46 | -0.236167769 | 0.236167769 | ENSG00000036054 | n.s. |
| ENSG00000111880 | 1547.858433 | -0.417377385 | 0.2736814 | -1.525048 | 0.127247066 | 0.272270001 | RNGTT | protein_coding | 36.64 | -0.235818309 | 0.235818309 | ENSG00000111880 | n.s. |
| ENSG00000171316 | 4528.394873 | -0.379059647 | 0.2346053 | -1.615734 | 0.106151942 | 0.238798478 | CHD7 | protein_coding | 39.98 | -0.23576314 | 0.23576314 | ENSG00000171316 | n.s. |
| ENSG00000275066 | 77.38315924 | -0.440913255 | 0.298486 | -1.477165 | 0.139631311 | 0.291948752 | SYNRG | protein_coding | 40.78 | -0.235753397 | 0.235753397 | ENSG00000275066 | n.s. |
| ENSG00000143621 | 10992.64051 | 0.426682382 | 0.2832859 | 1.50619 | 0.132018367 | 0.280279258 | ILF2 | protein_coding | 44.37 | 0.235703206 | 0.235703206 | ENSG00000143621 | n.s. |
| ENSG00000131263 | 3807.878094 | -0.399758326 | 0.2556697 | -1.563573 | 0.117917819 | 0.257613337 | RLIM | protein_coding | 38.47 | -0.235470309 | 0.235470309 | ENSG00000131263 | n.s. |
| ENSG00000267470 | 22.77134583 | -0.476048499 | 0.3384494 | -1.406557 | 0.159558696 | 0.320264977 | ZNF571-AS1 | antisense | 41.03 | -0.235401485 | 0.235401485 | ENSG00000267470 | n.s. |
| ENSG00000200922 | 19.18019743 | 0.482955444 | 0.3464382 | 1.394059 | 0.16329968 | 0.325640425 | | misc_RNA | 42.71 | 0.235325685 | 0.235325685 | ENSG00000200922 | n.s. |
| ENSG00000188167 | 199.5972551 | 0.362988189 | 0.2193399 | 1.654912 | 0.097942396 | 0.224913735 | TMPPE | protein_coding | 49.95 | 0.235210547 | 0.235210547 | ENSG00000188167 | n.s. |
| ENSG00000201208 | 31.34003901 | 0.470358421 | 0.3323622 | 1.415198 | 0.157010388 | 0.316269622 | | misc_RNA | 42.27 | 0.235152175 | 0.235152175 | ENSG00000201208 | n.s. |
| ENSG00000198046 | 976.5739402 | -0.377983828 | 0.2339495 | -1.615664 | 0.10616696 | 0.238798478 | ZNF667 | protein_coding | 43.97 | -0.235094014 | 0.235094014 | ENSG00000198046 | n.s. |
| ENSG00000169660 | 519.9887101 | 0.40225649 | 0.2584685 | 1.556308 | 0.119634941 | 0.260384131 | HEXDC | protein_coding | 55.13 | 0.235072855 | 0.235072855 | ENSG00000169660 | n.s. |
| ENSG00000154917 | 345.4185289 | -0.415350863 | 0.2720719 | -1.526621 | 0.126855175 | 0.271723785 | RAB6B | protein_coding | 49.59 | -0.235035567 | 0.235035567 | ENSG00000154917 | n.s. |
| ENSG00000228981 | 62.92685057 | 0.482189089 | 0.3457863 | 1.394471 | 0.1631753 | 0.325532268 | | processed_pseudogene | 59.55 | 0.235021834 | 0.235021834 | ENSG00000228981 | n.s. |
| ENSG00000147383 | 359.2595392 | -0.434948599 | 0.2928802 | -1.485073 | 0.137524373 | 0.288708717 | NSDHL | protein_coding | 45.97 | -0.234672212 | 0.234672212 | ENSG00000147383 | n.s. |
| ENSG00000130939 | 298.6322841 | -0.410916362 | 0.2678228 | -1.534285 | 0.124959622 | 0.268655469 | UBE4B | protein_coding | 44.12 | -0.234552832 | 0.234552832 | ENSG00000130939 | n.s. |
| ENSG00000106153 | 4861.198881 | 0.452735702 | 0.3126283 | 1.44816 | 0.14757244 | 0.303538349 | CHCHD2 | protein_coding | 45.29 | 0.234420403 | 0.234420403 | ENSG00000106153 | n.s. |
| ENSG00000136731 | 2893.070646 | -0.394149851 | 0.2505822 | -1.572936 | 0.115733588 | 0.254283475 | UGGT1 | protein_coding | 40.84 | -0.234393768 | 0.234393768 | ENSG00000136731 | n.s. |
| ENSG00000057663 | 1089.827822 | -0.433763014 | 0.2918461 | -1.486273 | 0.137206863 | 0.288221357 | ATG5 | protein_coding | 39.61 | -0.23435081 | 0.23435081 | ENSG00000057663 | n.s. |
| ENSG00000072182 | 554.7059754 | 0.373696438 | 0.2301547 | 1.623675 | 0.104445254 | 0.236070013 | ASIC4 | protein_coding | 56.96 | 0.234292411 | 0.234292411 | ENSG00000072182 | n.s. |
| ENSG00000109917 | 775.6693776 | 0.3879864 | 0.2445723 | 1.586388 | 0.112651425 | 0.249355451 | ZPR1 | protein_coding | 44.53 | 0.234026076 | 0.234026076 | ENSG00000109917 | n.s. |
| ENSG00000163743 | 684.9655353 | -0.421894589 | 0.2798127 | -1.507775 | 0.131612104 | 0.279462456 | RCHY1 | protein_coding | 34.99 | -0.233593131 | 0.233593131 | ENSG00000163743 | n.s. |
| ENSG00000172586 | 745.5987431 | 0.426388917 | 0.2845838 | 1.498289 | 0.13405809 | 0.283618111 | CHCHD1 | protein_coding | 45.52 | 0.233348174 | 0.233348174 | ENSG00000172586 | n.s. |

| | | | | | | | | | | | | | |
|---|---|---|---|---|---|---|---|---|---|---|---|---|---|
| ENSG00000260838 | 324.7803155 | -0.427269641 | 0.2856487 | -1.495787 | 0.134709105 | 0.284570848 | | lincRNA | 33.98 | -0.233207868 | 0.233207868 | ENSG00000260838 | n.s. |
| ENSG00000133030 | 3328.563211 | -0.379963456 | 0.236966 | -1.603451 | 0.108835082 | 0.243423429 | MPRIP | protein_coding | 51.29 | -0.233159873 | 0.233159873 | ENSG00000133030 | n.s. |
| ENSG00000111348 | 90.00097178 | 0.393789236 | 0.2511028 | 1.568239 | 0.116825298 | 0.25598971 | ARHGDIB | protein_coding | 41.12 | 0.233035606 | 0.233035606 | ENSG00000111348 | n.s. |
| ENSG00000156482 | 12647.95376 | 0.455406874 | 0.3171831 | 1.435785 | 0.151063423 | 0.307943783 | RPL30 | protein_coding | 42.56 | 0.232953622 | 0.232953622 | ENSG00000156482 | n.s. |
| ENSG00000196586 | 479.6745445 | -0.413202817 | 0.27134 | -1.522823 | 0.127803034 | 0.273207684 | MYO6 | protein_coding | 37.66 | -0.232842717 | 0.232842717 | ENSG00000196586 | n.s. |
| ENSG00000119950 | 717.9014128 | -0.413264006 | 0.2714146 | -1.52263 | 0.12785131 | 0.273268926 | MXI1 | protein_coding | 40.13 | -0.23283697 | 0.23283697 | ENSG00000119950 | n.s. |
| ENSG00000240342 | 3269.446973 | 0.449622711 | 0.310464 | 1.448228 | 0.147553313 | 0.303538349 | | processed_pseudogene | 61.21 | 0.232808539 | 0.232808539 | ENSG00000240342 | n.s. |
| ENSG00000205726 | 1042.044283 | -0.380950992 | 0.2384129 | -1.597862 | 0.110073645 | 0.245155324 | ITSN1 | protein_coding | 42.47 | -0.232592932 | 0.232592932 | ENSG00000205726 | n.s. |
| ENSG00000083845 | 9066.75654 | 0.438858197 | 0.2988763 | 1.468361 | 0.142006254 | 0.295273756 | RPS5 | protein_coding | 51.15 | 0.232496168 | 0.232496168 | ENSG00000083845 | n.s. |
| ENSG00000213930 | 151.1445457 | 0.402909619 | 0.2610193 | 1.543601 | 0.122685074 | 0.264869636 | GALT | protein_coding | 50.16 | 0.232465887 | 0.232465887 | ENSG00000213930 | n.s. |
| ENSG00000105258 | 534.0350636 | 0.451543654 | 0.3132395 | 1.441529 | 0.149435393 | 0.305879004 | POLR2I | protein_coding | 60.72 | 0.232296783 | 0.232296783 | ENSG00000105258 | n.s. |
| ENSG00000123106 | 1152.198289 | -0.428184748 | 0.2874101 | -1.489804 | 0.136275733 | 0.287002471 | CCDC91 | protein_coding | 35.52 | -0.232125102 | 0.232125102 | ENSG00000123106 | n.s. |
| ENSG00000205659 | 406.0641383 | -0.406683804 | 0.2651031 | -1.534059 | 0.125015187 | 0.268733433 | LIN52 | protein_coding | 40.2 | -0.232085622 | 0.232085622 | ENSG00000205659 | n.s. |
| ENSG00000154380 | 7160.896031 | -0.405861546 | 0.2644371 | -1.534813 | 0.124829754 | 0.268459169 | ENAH | protein_coding | 38.55 | -0.23179636 | 0.23179636 | ENSG00000154380 | n.s. |
| ENSG00000132661 | 448.5102465 | 0.430287504 | 0.2900632 | 1.483427 | 0.137961012 | 0.289281803 | NXT1 | protein_coding | 48.94 | 0.231786793 | 0.231786793 | ENSG00000132661 | n.s. |
| ENSG00000185800 | 128.9160513 | 0.372909523 | 0.2309609 | 1.614601 | 0.106397221 | 0.239200469 | DMWD | protein_coding | 57.55 | 0.231665556 | 0.231665556 | ENSG00000185800 | n.s. |
| ENSG00000196449 | 591.5784139 | 0.401308651 | 0.2598317 | 1.544495 | 0.122468575 | 0.264689533 | YRDC | protein_coding | 49.48 | 0.231660729 | 0.231660729 | ENSG00000196449 | n.s. |
| ENSG00000108094 | 785.8006413 | -0.405511055 | 0.2644762 | -1.533261 | 0.125211574 | 0.268989469 | CUL2 | protein_coding | 39.72 | -0.23124865 | 0.23124865 | ENSG00000108094 | n.s. |
| ENSG00000137692 | 658.8711713 | -0.416194172 | 0.2755614 | -1.51035 | 0.130954064 | 0.278289203 | DCUN1D5 | protein_coding | 36.06 | -0.231197379 | 0.231197379 | ENSG00000137692 | n.s. |
| ENSG00000144362 | 397.0741309 | -0.388938892 | 0.2473817 | -1.572222 | 0.115899063 | 0.254561075 | PHOSPHO2 | protein_coding | 37.24 | -0.231110602 | 0.231110602 | ENSG00000144362 | n.s. |
| ENSG00000134014 | 1075.500319 | -0.40010212 | 0.2590856 | -1.544286 | 0.122519181 | 0.264728452 | ELP3 | protein_coding | 40.16 | -0.230938695 | 0.230938695 | ENSG00000134014 | n.s. |
| ENSG00000123416 | 20787.39307 | 0.426793256 | 0.2869961 | 1.487105 | 0.136987087 | 0.28806392 | TUBA1B | protein_coding | 53.12 | 0.230686506 | 0.230686506 | ENSG00000123416 | n.s. |
| ENSG00000265110 | 2.737684496 | -0.698410991 | 0.6459355 | -1.081239 | 0.279590617 | 0.467427974 | MIR4731 | miRNA | 64.29 | -0.230674884 | 0.230674884 | ENSG00000265110 | n.s. |
| ENSG00000201176 | 5.55961105 | -0.609085522 | 0.5125269 | -1.188397 | 0.234676972 | 0.418254926 | RNU6-853P | snRNA | 39.25 | -0.230574766 | 0.230574766 | ENSG00000201176 | n.s. |
| ENSG00000168818 | 408.8475876 | -0.404092098 | 0.2635267 | -1.533401 | 0.125177111 | 0.268979418 | STX18 | protein_coding | 41.43 | -0.230446026 | 0.230446026 | ENSG00000168818 | n.s. |
| ENSG00000234127 | 375.4286283 | -0.413491493 | 0.2733471 | -1.512698 | 0.130356467 | 0.277346133 | TRIM26 | protein_coding | 46.9 | -0.230305618 | 0.230305618 | ENSG00000234127 | n.s. |
| ENSG00000277791 | 1246.283383 | 0.460163204 | 0.3249565 | 1.416076 | 0.156753185 | 0.315888638 | PSMB3 | protein_coding | 49.05 | 0.230296036 | 0.230296036 | ENSG00000277791 | n.s. |
| ENSG00000259637 | 40.76537893 | -0.4109766 | 0.2708457 | -1.517383 | 0.129170008 | 0.27536886 | DNM1P38 | unprocessed_pseudogene | 58.18 | -0.2301819 | 0.2301819 | ENSG00000259637 | n.s. |
| ENSG00000144647 | 216.445467 | 0.424546206 | 0.2851539 | 1.488832 | 0.136531639 | 0.287279736 | POMGNT2 | protein_coding | 47.68 | 0.22997456 | 0.22997456 | ENSG00000144647 | n.s. |
| ENSG00000135940 | 2055.964724 | 0.45569511 | 0.3200974 | 1.423614 | 0.154558215 | 0.312960156 | COX5B | protein_coding | 50.3 | 0.229903173 | 0.229903173 | ENSG00000135940 | n.s. |
| ENSG00000249784 | 225.2636126 | 0.602172592 | 0.5039658 | 1.194868 | 0.232138634 | 0.415272543 | SCARNA22 | scaRNA | 56 | 0.229829276 | 0.229829276 | ENSG00000249784 | n.s. |
| ENSG00000088448 | 2916.286444 | 0.381184898 | 0.2404753 | 1.585131 | 0.112936539 | 0.249589752 | ANKRD10 | protein_coding | 40 | 0.22976806 | 0.22976806 | ENSG00000088448 | n.s. |

| | | | | | | | | | | | | | |
|---|---|---|---|---|---|---|---|---|---|---|---|---|---|
| ENSG00000163867 | 46.59279688 | 0.409411033 | 0.2694969 | 1.519168 | 0.128720237 | 0.274662355 | ZMYM6 | protein_coding | 38.8 | 0.229761824 | 0.229761824 | ENSG00000163867 | n.s. |
| ENSG00000126457 | 3425.066422 | 0.435670272 | 0.2974474 | 1.464697 | 0.143003589 | 0.296948392 | PRMT1 | protein_coding | 58.23 | 0.229737222 | 0.229737222 | ENSG00000126457 | n.s. |
| ENSG00000173950 | 761.1495595 | -0.420893747 | 0.2814289 | -1.49556 | 0.134768455 | 0.284602032 | XXYLT1 | protein_coding | 48.89 | -0.229707815 | 0.229707815 | ENSG00000173950 | n.s. |
| ENSG00000010404 | 357.3691506 | -0.408823126 | 0.2689505 | -1.520068 | 0.128493796 | 0.274282156 | IDS | protein_coding | 45.34 | -0.229677831 | 0.229677831 | ENSG00000010404 | n.s. |
| ENSG00000168906 | 16848.62045 | 0.353393484 | 0.2131742 | 1.657769 | 0.097364124 | 0.223918187 | MAT2A | protein_coding | 44.46 | 0.229674182 | 0.229674182 | ENSG00000168906 | n.s. |
| ENSG00000162631 | 165.1835708 | -0.382606972 | 0.2420314 | -1.580815 | 0.113920265 | 0.251036202 | NTNG1 | protein_coding | 37.35 | -0.229665055 | 0.229665055 | ENSG00000162631 | n.s. |
| ENSG00000152455 | 1662.519718 | -0.408726525 | 0.2688932 | -1.520033 | 0.128502658 | 0.274282156 | SUV39H2 | protein_coding | 39.96 | -0.229623561 | 0.229623561 | ENSG00000152455 | n.s. |
| ENSG00000109390 | 512.1753961 | 0.444219398 | 0.3072972 | 1.445569 | 0.148298005 | 0.304356298 | NDUFC1 | protein_coding | 42.39 | 0.229491607 | 0.229491607 | ENSG00000109390 | n.s. |
| ENSG00000174547 | 1183.807351 | 0.456938478 | 0.3220299 | 1.418932 | 0.15591887 | 0.314890981 | MRPL11 | protein_coding | 48.04 | 0.229309904 | 0.229309904 | ENSG00000174547 | n.s. |
| ENSG00000267260 | 203.2287833 | -0.396848454 | 0.2568151 | -1.545269 | 0.122281105 | 0.26440749 | | lincRNA | 45.82 | -0.229269768 | 0.229269768 | ENSG00000267260 | n.s. |
| ENSG00000234851 | 270.6343908 | 0.449838371 | 0.3140064 | 1.432577 | 0.15197874 | 0.309266296 | RPL23AP42 | processed_pseudogene | 49.26 | 0.229267956 | 0.229267956 | ENSG00000234851 | n.s. |
| ENSG00000240668 | 57.69406207 | 0.423142581 | 0.2842367 | 1.488698 | 0.13656692 | 0.287310552 | KRT8P36 | processed_pseudogene | 56.53 | 0.229194512 | 0.229194512 | ENSG00000240668 | n.s. |
| ENSG00000169750 | 614.2283928 | 0.429565853 | 0.2913252 | 1.474523 | 0.14034074 | 0.292860329 | RAC3 | protein_coding | 68.77 | 0.229104417 | 0.229104417 | ENSG00000169750 | n.s. |
| ENSG00000176340 | 1981.020143 | 0.47061313 | 0.33844 | 1.390537 | 0.164366002 | 0.326976534 | COX8A | protein_coding | 57 | 0.228474869 | 0.228474869 | ENSG00000176340 | n.s. |
| ENSG00000071794 | 3321.159888 | -0.416921526 | 0.2780432 | -1.499485 | 0.133747935 | 0.283177084 | HLTF | protein_coding | 35.95 | -0.228448771 | 0.228448771 | ENSG00000071794 | n.s. |
| ENSG00000135953 | 374.9438536 | -0.396401611 | 0.2570496 | -1.542121 | 0.123044219 | 0.265398089 | MFSD9 | protein_coding | 41.84 | -0.228367843 | 0.228367843 | ENSG00000135953 | n.s. |
| ENSG00000133878 | 630.7501424 | -0.439155119 | 0.3024957 | -1.451773 | 0.146564648 | 0.302269232 | DUSP26 | protein_coding | 52.92 | -0.22818766 | 0.22818766 | ENSG00000133878 | n.s. |
| ENSG00000136877 | 531.192081 | 0.406293931 | 0.2675374 | 1.518643 | 0.128852328 | 0.274902078 | FPGS | protein_coding | 53.5 | 0.227858566 | 0.227858566 | ENSG00000136877 | n.s. |
| ENSG00000107263 | 553.6640339 | -0.376959362 | 0.2374041 | -1.587838 | 0.11232289 | 0.24882603 | RAPGEF1 | protein_coding | 48.49 | -0.22772273 | 0.22772273 | ENSG00000107263 | n.s. |
| ENSG00000201501 | 3.668107199 | -0.682160972 | 0.6255526 | -1.090493 | 0.275495863 | 0.463924638 | | misc_RNA | 42.16 | -0.22753654 | 0.22753654 | ENSG00000201501 | n.s. |
| ENSG00000173660 | 1861.901656 | 0.449595803 | 0.3152979 | 1.42594 | 0.1538857 | 0.312006786 | UQCRH | protein_coding | 46.3 | 0.227421725 | 0.227421725 | ENSG00000173660 | n.s. |
| ENSG00000109685 | 4247.296381 | -0.395358943 | 0.2566253 | -1.540608 | 0.123412247 | 0.265985868 | NSD2 | protein_coding | 47.14 | -0.227387311 | 0.227387311 | ENSG00000109685 | n.s. |
| ENSG00000144306 | 647.336234 | -0.418321702 | 0.2803746 | -1.49201 | 0.135696582 | 0.286128049 | SCRN3 | protein_coding | 37.9 | -0.227332564 | 0.227332564 | ENSG00000144306 | n.s. |
| ENSG00000198242 | 6899.567814 | 0.414853627 | 0.2766948 | 1.499318 | 0.133791091 | 0.283182329 | RPL23A | protein_coding | 49.99 | 0.227312345 | 0.227312345 | ENSG00000198242 | n.s. |
| ENSG00000137100 | 402.3701878 | 0.444257186 | 0.3094362 | 1.435699 | 0.151088016 | 0.307948829 | DCTN3 | protein_coding | 49.85 | 0.227247077 | 0.227247077 | ENSG00000137100 | n.s. |
| ENSG00000155636 | 386.3077735 | -0.403153554 | 0.2647168 | -1.522962 | 0.127768267 | 0.273207684 | RBM45 | protein_coding | 36.95 | -0.227179886 | 0.227179886 | ENSG00000155636 | n.s. |
| ENSG00000198663 | 656.5783 | -0.41579086 | 0.277805 | -1.4967 | 0.134471214 | 0.284232991 | C6orf89 | protein_coding | 42.84 | -0.227157156 | 0.227157156 | ENSG00000198663 | n.s. |
| ENSG00000112992 | 1747.366715 | -0.412140644 | 0.2740349 | -1.503972 | 0.132588691 | 0.281107407 | NNT | protein_coding | 37.88 | -0.22714213 | 0.22714213 | ENSG00000112992 | n.s. |
| ENSG00000185104 | 1622.305346 | -0.398308533 | 0.2597954 | -1.533163 | 0.125235795 | 0.268999995 | FAF1 | protein_coding | 38.27 | -0.227134536 | 0.227134536 | ENSG00000185104 | n.s. |
| ENSG00000110871 | 616.6176308 | -0.442996901 | 0.3080842 | -1.437909 | 0.150459957 | 0.307205922 | COQ5 | protein_coding | 45.87 | -0.227067106 | 0.227067106 | ENSG00000110871 | n.s. |
| ENSG00000180891 | 631.7640974 | -0.376611776 | 0.2375147 | -1.585636 | 0.112821937 | 0.249550887 | CUEDC1 | protein_coding | 51.38 | -0.227036975 | 0.227036975 | ENSG00000180891 | n.s. |
| ENSG00000062582 | 688.4178531 | 0.466386949 | 0.3349785 | 1.392289 | 0.163834837 | 0.326053807 | MRPS24 | protein_coding | 50.93 | 0.22699553 | 0.22699553 | ENSG00000062582 | n.s. |

| | | | | | | | | | | | | | |
|---|---|---|---|---|---|---|---|---|---|---|---|---|---|
| ENSG00000147669 | 1553.203903 | 0.424460765 | 0.2873163 | 1.477329 | 0.139587447 | 0.291900875 | POLR2K | protein_coding | 37.38 | 0.226986593 | 0.226986593 | ENSG00000147669 | n.s. |
| ENSG00000157837 | 441.2306046 | -0.391165734 | 0.2522687 | -1.550592 | 0.120999579 | 0.262861155 | SPPL3 | protein_coding | 40.88 | -0.226983144 | 0.226983144 | ENSG00000157837 | n.s. |
| ENSG00000053438 | 2821.335436 | 0.419034424 | 0.2815352 | 1.488391 | 0.136647884 | 0.287437452 | NNAT | protein_coding | 54.05 | 0.226888974 | 0.226888974 | ENSG00000053438 | n.s. |
| ENSG00000196526 | 2582.287468 | -0.379789399 | 0.240931 | -1.576341 | 0.1149473 | 0.252829583 | AFAP1 | protein_coding | 46.39 | -0.226799637 | 0.226799637 | ENSG00000196526 | n.s. |
| ENSG00000116761 | 485.9984116 | 0.39573487 | 0.2574175 | 1.537327 | 0.124213205 | 0.267381022 | CTH | protein_coding | 39.51 | 0.226704406 | 0.226704406 | ENSG00000116761 | n.s. |
| ENSG00000006459 | 533.0877759 | -0.397456474 | 0.259363 | -1.532433 | 0.125415571 | 0.269344593 | KDM7A | protein_coding | 39.18 | -0.22642767 | 0.22642767 | ENSG00000006459 | n.s. |
| ENSG00000123552 | 744.3404169 | -0.390708673 | 0.2523275 | -1.548419 | 0.121521543 | 0.263501704 | USP45 | protein_coding | 37.98 | -0.226304938 | 0.226304938 | ENSG00000123552 | n.s. |
| ENSG00000109265 | 1126.261633 | -0.390132787 | 0.2517954 | -1.549404 | 0.121284663 | 0.263193011 | KIAA1211 | protein_coding | 41.29 | -0.226169983 | 0.226169983 | ENSG00000109265 | n.s. |
| ENSG00000115084 | 742.928414 | -0.410198244 | 0.2728322 | -1.503482 | 0.132714793 | 0.281331921 | SLC35F5 | protein_coding | 36.74 | -0.225929394 | 0.225929394 | ENSG00000115084 | n.s. |
| ENSG00000123810 | 64.97520808 | 0.41970588 | 0.2830973 | 1.48255 | 0.138194026 | 0.289639534 | B9D2 | protein_coding | 52.34 | 0.225861429 | 0.225861429 | ENSG00000123810 | n.s. |
| ENSG00000115128 | 2484.402597 | 0.459676873 | 0.3281051 | 1.401005 | 0.161212615 | 0.322772972 | SF3B6 | protein_coding | 43.19 | 0.225748617 | 0.225748617 | ENSG00000115128 | n.s. |
| ENSG00000081177 | 45.83828707 | -0.467252147 | 0.3371752 | -1.385785 | 0.165812698 | 0.328901672 | EXD2 | protein_coding | 41.37 | -0.225651911 | 0.225651911 | ENSG00000081177 | n.s. |
| ENSG00000125817 | 860.3540122 | -0.389719405 | 0.2518028 | -1.547717 | 0.12169055 | 0.263714718 | CENPB | protein_coding | 66.2 | -0.225595169 | 0.225595169 | ENSG00000125817 | n.s. |
| ENSG00000204152 | 566.9178786 | 0.397620049 | 0.2600395 | 1.529076 | 0.126245702 | 0.270793238 | TIMM23B | protein_coding | 40.65 | 0.22559458 | 0.22559458 | ENSG00000204152 | n.s. |
| ENSG00000101109 | 927.10735 | -0.38950824 | 0.251611 | -1.548057 | 0.121608576 | 0.263615073 | STK4 | protein_coding | 38.46 | -0.225536863 | 0.225536863 | ENSG00000101109 | n.s. |
| ENSG00000109466 | 194.1826793 | 0.377373239 | 0.2393151 | 1.576889 | 0.114821177 | 0.252631991 | KLHL2 | protein_coding | 38.15 | 0.225484909 | 0.225484909 | ENSG00000109466 | n.s. |
| ENSG00000197756 | 35729.27615 | 0.42296039 | 0.2869487 | 1.473993 | 0.140483482 | 0.293026444 | RPL37A | protein_coding | 44.46 | 0.225477301 | 0.225477301 | ENSG00000197756 | n.s. |
| ENSG00000099385 | 260.5715348 | 0.438301606 | 0.3042435 | 1.440628 | 0.149689912 | 0.306219901 | BCL7C | protein_coding | 48.81 | 0.225272381 | 0.225272381 | ENSG00000099385 | n.s. |
| ENSG00000149480 | 1710.664279 | 0.392171875 | 0.2548181 | 1.539027 | 0.123797772 | 0.266692925 | MTA2 | protein_coding | 52.38 | 0.225102148 | 0.225102148 | ENSG00000149480 | n.s. |
| ENSG00000100138 | 2026.118796 | 0.428400389 | 0.293113 | 1.461554 | 0.143863576 | 0.298289288 | SNU13 | protein_coding | 46.79 | 0.225065432 | 0.225065432 | ENSG00000100138 | n.s. |
| ENSG00000120910 | 163.739949 | -0.431984491 | 0.2973881 | -1.452595 | 0.146336149 | 0.301932158 | PPP3CC | protein_coding | 39.2 | -0.224671085 | 0.224671085 | ENSG00000120910 | n.s. |
| ENSG00000126267 | 1716.103908 | 0.464093816 | 0.3352967 | 1.384129 | 0.166319053 | 0.329534801 | COX6B1 | protein_coding | 49.06 | 0.223739032 | 0.223739032 | ENSG00000126267 | n.s. |
| ENSG00000196787 | 1879.403166 | 0.345003686 | 0.2084992 | 1.6547 | 0.097985335 | 0.224975233 | HIST1H2AG | protein_coding | 46.8 | 0.223515913 | 0.223515913 | ENSG00000196787 | n.s. |
| ENSG00000038358 | 895.2953796 | 0.381528748 | 0.2448538 | 1.55819 | 0.119188252 | 0.259855906 | EDC4 | protein_coding | 57.32 | 0.223296341 | 0.223296341 | ENSG00000038358 | n.s. |
| ENSG00000101665 | 877.607336 | 0.362933972 | 0.2260698 | 1.605407 | 0.108404353 | 0.242733003 | SMAD7 | protein_coding | 52.06 | 0.223157636 | 0.223157636 | ENSG00000101665 | n.s. |
| ENSG00000115486 | 117.2552216 | 0.390177752 | 0.2540936 | 1.535567 | 0.124644628 | 0.268102449 | GGCX | protein_coding | 46.19 | 0.223064316 | 0.223064316 | ENSG00000115486 | n.s. |
| ENSG00000116161 | 4340.594439 | 0.405196081 | 0.2696833 | 1.502489 | 0.132970911 | 0.281746157 | CACYBP | protein_coding | 40.31 | 0.222915382 | 0.222915382 | ENSG00000116161 | n.s. |
| ENSG00000125991 | 1194.435134 | 0.426939127 | 0.2932796 | 1.455741 | 0.145464235 | 0.300547824 | ERGIC3 | protein_coding | 50.94 | 0.222899117 | 0.222899117 | ENSG00000125991 | n.s. |
| ENSG00000198625 | 3842.719218 | -0.395866865 | 0.2601947 | -1.521425 | 0.128153095 | 0.273627295 | MDM4 | protein_coding | 42.52 | -0.222809937 | 0.222809937 | ENSG00000198625 | n.s. |
| ENSG00000091592 | 132.7963577 | 0.433267996 | 0.3006546 | 1.441082 | 0.149561423 | 0.306046986 | NLRP1 | protein_coding | 46.74 | 0.222791556 | 0.222791556 | ENSG00000091592 | n.s. |
| ENSG00000280138 | 893.0384929 | 0.356026617 | 0.2196599 | 1.620809 | 0.105058652 | 0.236910793 | | TEC | 44.97 | 0.222664441 | 0.222664441 | ENSG00000280138 | n.s. |
| ENSG00000205758 | 473.7314494 | -0.426869521 | 0.2934012 | -1.4549 | 0.14569688 | 0.300880697 | CRYZL1 | protein_coding | 41.59 | -0.222657564 | 0.222657564 | ENSG00000205758 | n.s. |

| | | | | | | | | | | | | | |
|---|---|---|---|---|---|---|---|---|---|---|---|---|---|
| ENSG00000253437 | 4.878856457 | -0.595335869 | 0.5049061 | -1.179102 | 0.23835749 | 0.423027576 | RNU6-988P | snRNA | 35.51 | -0.222436127 | 0.222436127 | ENSG00000253437 | n.s. |
| ENSG00000235885 | 6831.927513 | -0.339035439 | 0.2033961 | -1.666873 | 0.095539706 | 0.220779972 | LINC01828 | lincRNA | 36.94 | -0.22242092 | 0.22242092 | ENSG00000235885 | n.s. |
| ENSG00000148187 | 513.7644026 | -0.392281467 | 0.2566882 | -1.528241 | 0.126452751 | 0.27111205 | MRRF | protein_coding | 41.15 | -0.222365212 | 0.222365212 | ENSG00000148187 | n.s. |
| ENSG00000167685 | 37.32013452 | -0.451081583 | 0.321311 | -1.403879 | 0.160355016 | 0.321564581 | ZNF444 | protein_coding | 54.02 | -0.222262237 | 0.222262237 | ENSG00000167685 | n.s. |
| ENSG00000172172 | 888.7009754 | -0.453971686 | 0.324672 | -1.398247 | 0.16203892 | 0.324008025 | MRPL13 | protein_coding | 35.86 | -0.222193823 | 0.222193823 | ENSG00000172172 | n.s. |
| ENSG00000141644 | 798.8982589 | -0.411864793 | 0.2774239 | -1.484604 | 0.137648667 | 0.288844369 | MBD1 | protein_coding | 49.32 | -0.222133549 | 0.222133549 | ENSG00000141644 | n.s. |
| ENSG00000257218 | 1105.701265 | -0.431341693 | 0.2991098 | -1.442085 | 0.14927844 | 0.305692561 | GATC | protein_coding | 46.68 | -0.222018095 | 0.222018095 | ENSG00000257218 | n.s. |
| ENSG00000188647 | 3588.621617 | -0.404989878 | 0.2702136 | -1.498777 | 0.133931547 | 0.283393453 | PTAR1 | protein_coding | 36.03 | -0.221776582 | 0.221776582 | ENSG00000188647 | n.s. |
| ENSG00000108474 | 202.1404645 | 0.362005305 | 0.2261056 | 1.601045 | 0.109366965 | 0.244025041 | PIGL | protein_coding | 44.78 | 0.221751999 | 0.221751999 | ENSG00000108474 | n.s. |
| ENSG00000207168 | 11.34490456 | -0.536202061 | 0.4265961 | -1.256931 | 0.20877849 | 0.385878522 | SNORA15 | snoRNA | 36.84 | -0.221746037 | 0.221746037 | ENSG00000207168 | n.s. |
| ENSG00000258982 | 22.9410921 | 0.462777062 | 0.3356469 | 1.378762 | 0.167968259 | 0.331907759 | | antisense | 48.69 | 0.221662158 | 0.221662158 | ENSG00000258982 | n.s. |
| ENSG00000182551 | 1352.039377 | 0.41115528 | 0.2770208 | 1.484203 | 0.137754947 | 0.289007271 | ADI1 | protein_coding | 44.71 | 0.221650206 | 0.221650206 | ENSG00000182551 | n.s. |
| ENSG00000133812 | 1097.819697 | -0.407537233 | 0.2730336 | -1.492626 | 0.135535005 | 0.28587394 | SBF2 | protein_coding | 38.31 | -0.221629112 | 0.221629112 | ENSG00000133812 | n.s. |
| ENSG00000066084 | 2140.142156 | -0.389772845 | 0.2546777 | -1.530455 | 0.125904086 | 0.270143719 | DIP2B | protein_coding | 41.84 | -0.221548883 | 0.221548883 | ENSG00000066084 | n.s. |
| ENSG00000139289 | 1605.553433 | 0.335193238 | 0.2002003 | 1.674289 | 0.094073785 | 0.218371009 | PHLDA1 | protein_coding | 45.13 | 0.221497374 | 0.221497374 | ENSG00000139289 | n.s. |
| ENSG00000121057 | 2486.853351 | 0.373672888 | 0.2381056 | 1.569358 | 0.116564608 | 0.255619632 | AKAP1 | protein_coding | 49.96 | 0.221365984 | 0.221365984 | ENSG00000121057 | n.s. |
| ENSG00000272288 | 17.17413058 | 0.651652598 | 0.5898375 | 1.1048 | 0.269246166 | 0.457439216 | | antisense | 62.48 | 0.221344627 | 0.221344627 | ENSG00000272288 | n.s. |
| ENSG00000154265 | 80.64891419 | 0.390651336 | 0.2557507 | 1.527469 | 0.126644384 | 0.271397531 | ABCA5 | protein_coding | 34.61 | 0.221262615 | 0.221262615 | ENSG00000154265 | n.s. |
| ENSG00000011451 | 470.0854377 | -0.368551008 | 0.2330582 | -1.581369 | 0.113793709 | 0.251036202 | WIZ | protein_coding | 58.16 | -0.221227771 | 0.221227771 | ENSG00000011451 | n.s. |
| ENSG00000139625 | 77.19959903 | -0.410448527 | 0.2766881 | -1.483434 | 0.137959106 | 0.289281803 | MAP3K12 | protein_coding | 52.61 | -0.221099955 | 0.221099955 | ENSG00000139625 | n.s. |
| ENSG00000122882 | 891.9168337 | -0.388659809 | 0.2539028 | -1.530743 | 0.125832991 | 0.270074415 | ECD | protein_coding | 39.47 | -0.220959536 | 0.220959536 | ENSG00000122882 | n.s. |
| ENSG00000198034 | 19575.82894 | 0.430632901 | 0.2992532 | 1.439025 | 0.150143394 | 0.306832009 | RPS4X | protein_coding | 46.44 | 0.220957456 | 0.220957456 | ENSG00000198034 | n.s. |
| ENSG00000073169 | 216.7374314 | -0.394466659 | 0.2599597 | -1.517415 | 0.129161988 | 0.27536886 | SELENOO | protein_coding | 55.78 | -0.220934927 | 0.220934927 | ENSG00000073169 | n.s. |
| ENSG00000200175 | 2.676832476 | -0.84960425 | 0.9217866 | -0.921693 | 0.356688746 | 0.549599094 | RNU6-1309P | snRNA | 37.96 | -0.220858016 | 0.220858016 | ENSG00000200175 | n.s. |
| ENSG00000163607 | 569.2250071 | 0.382359417 | 0.2474841 | 1.544986 | 0.122349711 | 0.264473689 | GTPBP8 | protein_coding | 37.14 | 0.220857501 | 0.220857501 | ENSG00000163607 | n.s. |
| ENSG00000136830 | 400.4156431 | -0.411059913 | 0.2775382 | -1.481093 | 0.138581763 | 0.290236756 | FAM129B | protein_coding | 54.31 | -0.220840947 | 0.220840947 | ENSG00000136830 | n.s. |
| ENSG00000238317 | 122.7271694 | 0.381840882 | 0.2469841 | 1.546014 | 0.12210122 | 0.264210707 | SNORD11 | snoRNA | 36.9 | 0.220722964 | 0.220722964 | ENSG00000238317 | n.s. |
| ENSG00000243147 | 1485.527359 | 0.433114966 | 0.3023997 | 1.43226 | 0.152069395 | 0.309360343 | MRPL33 | protein_coding | 39.12 | 0.22068739 | 0.22068739 | ENSG00000243147 | n.s. |
| ENSG00000201610 | 4.270781393 | -0.604081196 | 0.5208562 | -1.159785 | 0.246136331 | 0.432041872 | RNA5SP84 | rRNA | 57.94 | -0.220171987 | 0.220171987 | ENSG00000201610 | n.s. |
| ENSG00000121579 | 3308.987032 | -0.41448001 | 0.2818929 | -1.470345 | 0.141468223 | 0.29446286 | NAA50 | protein_coding | 37.41 | -0.220076233 | 0.220076233 | ENSG00000121579 | n.s. |
| ENSG00000106086 | 1193.16318 | -0.372480226 | 0.2377685 | -1.566567 | 0.117216027 | 0.256563235 | PLEKHA8 | protein_coding | 39.62 | -0.220063395 | 0.220063395 | ENSG00000106086 | n.s. |
| ENSG00000118564 | 1113.629907 | -0.407213788 | 0.2739465 | -1.486472 | 0.137154328 | 0.288197963 | FBXL5 | protein_coding | 37.87 | -0.220021312 | 0.220021312 | ENSG00000118564 | n.s. |

| | | | | | | | | | | | | | |
|---|---|---|---|---|---|---|---|---|---|---|---|---|---|
| ENSG00000221063 | 23.05116917 | -0.463806531 | 0.3383328 | -1.370859 | 0.170418911 | 0.335443596 | MIR1296 | miRNA | 57.61 | -0.220020772 | 0.220020772 | ENSG00000221063 | n.s. |
| ENSG00000135900 | 809.8004861 | 0.380759983 | 0.2463697 | 1.545482 | 0.122229652 | 0.264337285 | MRPL44 | protein_coding | 37.45 | 0.220018948 | 0.220018948 | ENSG00000135900 | n.s. |
| ENSG00000250220 | 61.48281416 | -0.41829601 | 0.2859914 | -1.462618 | 0.143572047 | 0.297862257 | | antisense | 39.01 | -0.220017228 | 0.220017228 | ENSG00000250220 | n.s. |
| ENSG00000100104 | 405.2736752 | -0.370239937 | 0.2355308 | -1.571938 | 0.115964875 | 0.254660597 | SRRD | protein_coding | 45.15 | -0.219936683 | 0.219936683 | ENSG00000100104 | n.s. |
| ENSG00000130810 | 642.7557109 | -0.366097869 | 0.2313917 | -1.582156 | 0.113613851 | 0.250807935 | PPAN | protein_coding | 58.7 | -0.21989988 | 0.21989988 | ENSG00000130810 | n.s. |
| ENSG00000104936 | 19.89032432 | -0.485946584 | 0.3650533 | -1.331166 | 0.183134366 | 0.353008414 | DMPK | protein_coding | 59.04 | -0.219752307 | 0.219752307 | ENSG00000104936 | n.s. |
| ENSG00000277872 | 2.23843749 | 0.721687027 | 0.7058831 | 1.022389 | 0.306596896 | 0.496308404 | MIR6817 | miRNA | 54.55 | 0.219572102 | 0.219572102 | ENSG00000277872 | n.s. |
| ENSG00000140988 | 22817.53474 | 0.437827243 | 0.3088228 | 1.41773 | 0.156269752 | 0.315233811 | RPS2 | protein_coding | 57.46 | 0.219512222 | 0.219512222 | ENSG00000140988 | n.s. |
| ENSG00000178096 | 275.0008644 | 0.372689208 | 0.2384415 | 1.563021 | 0.118047595 | 0.257820004 | BOLA1 | protein_coding | 45.46 | 0.219395946 | 0.219395946 | ENSG00000178096 | n.s. |
| ENSG00000151806 | 3234.073308 | -0.379862407 | 0.245983 | -1.544263 | 0.12252461 | 0.264728452 | GUF1 | protein_coding | 33.75 | -0.219256346 | 0.219256346 | ENSG00000151806 | n.s. |
| ENSG00000138835 | 210.193952 | 0.378001867 | 0.2440129 | 1.549106 | 0.121356284 | 0.263289776 | RGS3 | protein_coding | 48.79 | 0.219077032 | 0.219077032 | ENSG00000138835 | n.s. |
| ENSG00000198832 | 98.15596653 | 0.430293784 | 0.3006937 | 1.431004 | 0.15242912 | 0.309865768 | SELENOM | protein_coding | 49.48 | 0.218944837 | 0.218944837 | ENSG00000198832 | n.s. |
| ENSG00000139620 | 2470.259862 | 0.364683315 | 0.2306623 | 1.581027 | 0.113871869 | 0.251036202 | KANSL2 | protein_coding | 41.97 | 0.218906136 | 0.218906136 | ENSG00000139620 | n.s. |
| ENSG00000144893 | 378.4460914 | -0.358035184 | 0.2240317 | -1.598145 | 0.110010689 | 0.245147059 | MED12L | protein_coding | 38.81 | -0.218606728 | 0.218606728 | ENSG00000144893 | n.s. |
| ENSG00000164168 | 921.4753111 | -0.402749896 | 0.2701736 | -1.490708 | 0.136038224 | 0.286674769 | TMEM184C | protein_coding | 38.71 | -0.218536334 | 0.218536334 | ENSG00000164168 | n.s. |
| ENSG00000007866 | 200.9467512 | -0.383868521 | 0.2508382 | -1.530343 | 0.125931796 | 0.270161541 | TEAD3 | protein_coding | 56.75 | -0.218181837 | 0.218181837 | ENSG00000007866 | n.s. |
| ENSG00000099977 | 338.2408214 | 0.44809921 | 0.3218422 | 1.392295 | 0.163833219 | 0.326053807 | DDT | protein_coding | 54.22 | 0.218094691 | 0.218094691 | ENSG00000099977 | n.s. |
| ENSG00000134686 | 614.2623563 | -0.42177126 | 0.2916734 | -1.44604 | 0.148166084 | 0.304130383 | PHC2 | protein_coding | 45.03 | -0.218030516 | 0.218030516 | ENSG00000134686 | n.s. |
| ENSG00000142207 | 1410.086623 | -0.351126011 | 0.2175995 | -1.613634 | 0.106606787 | 0.239555568 | URB1 | protein_coding | 46.41 | -0.217906601 | 0.217906601 | ENSG00000142207 | n.s. |
| ENSG00000221983 | 6474.56464 | 0.427047144 | 0.2980581 | 1.432765 | 0.151925105 | 0.309247548 | UBA52 | protein_coding | 53.24 | 0.217663254 | 0.217663254 | ENSG00000221983 | n.s. |
| ENSG00000109475 | 11303.78287 | 0.428960722 | 0.3005149 | 1.427419 | 0.153459085 | 0.311386031 | RPL34 | protein_coding | 41.51 | 0.217354773 | 0.217354773 | ENSG00000109475 | n.s. |
| ENSG00000176208 | 2025.932717 | -0.370629721 | 0.2376929 | -1.55928 | 0.11893011 | 0.259336558 | ATAD5 | protein_coding | 41.02 | -0.217239517 | 0.217239517 | ENSG00000176208 | n.s. |
| ENSG00000131732 | 736.8901045 | -0.374960116 | 0.2421337 | -1.548567 | 0.121485954 | 0.263465566 | ZCCHC9 | protein_coding | 38.15 | -0.217205448 | 0.217205448 | ENSG00000131732 | n.s. |
| ENSG00000106537 | 485.7180896 | 0.442112359 | 0.315464 | 1.401467 | 0.161074503 | 0.322728494 | TSPAN13 | protein_coding | 39.23 | 0.217149095 | 0.217149095 | ENSG00000106537 | n.s. |
| ENSG00000198995 | 5.313779729 | -0.608421408 | 0.5330241 | -1.141452 | 0.253681921 | 0.43995753 | MIR340 | miRNA | 38.95 | -0.21695653 | 0.21695653 | ENSG00000198995 | n.s. |
| ENSG00000072041 | 3530.712598 | -0.410428101 | 0.279916 | -1.466255 | 0.142578966 | 0.296218154 | SLC6A15 | protein_coding | 34.54 | -0.216865418 | 0.216865418 | ENSG00000072041 | n.s. |
| ENSG00000092853 | 902.7065989 | -0.383758891 | 0.2516293 | -1.525096 | 0.127235084 | 0.272270001 | CLSPN | protein_coding | 42.79 | -0.216823853 | 0.216823853 | ENSG00000092853 | n.s. |
| ENSG00000138399 | 848.7640151 | -0.412599999 | 0.2825537 | -1.460253 | 0.144220446 | 0.298794224 | FASTKD1 | protein_coding | 37.97 | -0.216461431 | 0.216461431 | ENSG00000138399 | n.s. |
| ENSG00000199035 | 21.0384759 | -0.457831796 | 0.3347138 | -1.367831 | 0.171365047 | 0.336708375 | MIR103A1 | miRNA | 48.72 | -0.216438188 | 0.216438188 | ENSG00000199035 | n.s. |
| ENSG00000122678 | 57.59219479 | -0.394802741 | 0.2633133 | -1.499365 | 0.133779008 | 0.283182329 | POLM | protein_coding | 56.76 | -0.216325786 | 0.216325786 | ENSG00000122678 | n.s. |
| ENSG00000120519 | 125.8489086 | 0.393087599 | 0.2615851 | 1.502714 | 0.132912871 | 0.281666044 | SLC10A7 | protein_coding | 36.95 | 0.216302547 | 0.216302547 | ENSG00000120519 | n.s. |
| ENSG00000168476 | 140.3066087 | 0.374693375 | 0.2426511 | 1.544165 | 0.122548307 | 0.264738569 | REEP4 | protein_coding | 62.11 | 0.216266565 | 0.216266565 | ENSG00000168476 | n.s. |

| | | | | | | | | | | | | | |
|---|---|---|---|---|---|---|---|---|---|---|---|---|---|
| ENSG00000237854 | 221.6731205 | 0.360011513 | 0.2277903 | 1.580452 | 0.114003459 | 0.251150342 | LINC00674 | transcribed_unprocessed_pseud | 40.03 | 0.21603075 | 0.21603075 | ENSG00000237854 | n.s. |
| ENSG00000167130 | 373.8373501 | -0.407445113 | 0.2774467 | -1.468553 | 0.141954177 | 0.29520956 | DOLPP1 | protein_coding | 57.76 | -0.215892772 | 0.215892772 | ENSG00000167130 | n.s. |
| ENSG00000198585 | 1053.147311 | -0.384767291 | 0.2534269 | -1.518258 | 0.128949447 | 0.27498287 | NUDT16 | protein_coding | 48.83 | -0.21573685 | 0.21573685 | ENSG00000198585 | n.s. |
| ENSG00000108961 | 195.3512366 | 0.41780729 | 0.2891728 | 1.444836 | 0.148503977 | 0.304689194 | RANGRF | protein_coding | 59.59 | 0.215648286 | 0.215648286 | ENSG00000108961 | n.s. |
| ENSG00000177946 | 114.2633703 | 0.357606102 | 0.2255806 | 1.58527 | 0.112905 | 0.249572722 | CENPBD1 | protein_coding | 52.11 | 0.21556599 | 0.21556599 | ENSG00000177946 | n.s. |
| ENSG00000197746 | 2620.817082 | 0.42095646 | 0.2930695 | 1.436371 | 0.150896838 | 0.307739369 | PSAP | protein_coding | 47.72 | 0.215452647 | 0.215452647 | ENSG00000197746 | n.s. |
| ENSG00000159202 | 1205.237485 | -0.400836534 | 0.2706509 | -1.481009 | 0.13860405 | 0.290236756 | UBE2Z | protein_coding | 47.44 | -0.215348462 | 0.215348462 | ENSG00000159202 | n.s. |
| ENSG00000110955 | 13389.08669 | 0.416261253 | 0.2876348 | 1.447187 | 0.147844632 | 0.303871135 | ATP5B | protein_coding | 45.43 | 0.215336338 | 0.215336338 | ENSG00000110955 | n.s. |
| ENSG00000274012 | 22225.70825 | -0.326768528 | 0.1955683 | -1.670866 | 0.094748086 | 0.219497105 | RN7SL2 | misc_RNA | 62.67 | -0.215200343 | 0.215200343 | ENSG00000274012 | n.s. |
| ENSG00000136875 | 1066.795057 | -0.399133099 | 0.2689437 | -1.484077 | 0.13778857 | 0.289007271 | PRPF4 | protein_coding | 42.73 | -0.215169154 | 0.215169154 | ENSG00000136875 | n.s. |
| ENSG00000223086 | 7.904165353 | 0.597372107 | 0.5195274 | 1.149838 | 0.250210795 | 0.436497294 | RNA5SP155 | rRNA | 51.16 | 0.215064977 | 0.215064977 | ENSG00000223086 | n.s. |
| ENSG00000137074 | 89.2243248 | -0.372676997 | 0.241529 | -1.542991 | 0.122833044 | 0.265024713 | APTX | protein_coding | 42.68 | -0.214927903 | 0.214927903 | ENSG00000137074 | n.s. |
| ENSG00000200419 | 24.40897756 | 0.443022435 | 0.3188361 | 1.389499 | 0.164681035 | 0.327316781 | | misc_RNA | 43.14 | 0.214879937 | 0.214879937 | ENSG00000200419 | n.s. |
| ENSG00000163710 | 153.1624304 | 0.354130813 | 0.2224715 | 1.591803 | 0.11142893 | 0.247357302 | PCOLCE2 | protein_coding | 40.92 | 0.214842405 | 0.214842405 | ENSG00000163710 | n.s. |
| ENSG00000177954 | 22332.38036 | 0.439810872 | 0.3149276 | 1.396546 | 0.162550212 | 0.324750553 | RPS27 | protein_coding | 45.91 | 0.214825665 | 0.214825665 | ENSG00000177954 | n.s. |
| ENSG00000250571 | 77.21148874 | -0.408000947 | 0.2787966 | -1.463436 | 0.143348176 | 0.297530806 | GLI4 | protein_coding | 64.03 | -0.214799467 | 0.214799467 | ENSG00000250571 | n.s. |
| ENSG00000166140 | 147.8325106 | -0.393891304 | 0.2635287 | -1.494681 | 0.134997824 | 0.284999954 | ZFYVE19 | protein_coding | 53.53 | -0.214731896 | 0.214731896 | ENSG00000166140 | n.s. |
| ENSG00000198015 | 4030.543967 | -0.415370322 | 0.2872233 | -1.446158 | 0.148132886 | 0.304130383 | MRPL42 | protein_coding | 39.33 | -0.214721614 | 0.214721614 | ENSG00000198015 | n.s. |
| ENSG00000127252 | 148.2485506 | -0.406592339 | 0.2774125 | -1.46566 | 0.142740979 | 0.296491519 | HRASLS | protein_coding | 37.84 | -0.214675764 | 0.214675764 | ENSG00000127252 | n.s. |
| ENSG00000049130 | 462.5942965 | -0.379351212 | 0.2485329 | -1.526362 | 0.126919638 | 0.271752671 | KITLG | protein_coding | 36.84 | -0.214646846 | 0.214646846 | ENSG00000049130 | n.s. |
| ENSG00000223496 | 1100.349785 | 0.347545981 | 0.216023 | 1.608838 | 0.107651776 | 0.241319543 | EXOSC6 | protein_coding | 45.72 | 0.214577497 | 0.214577497 | ENSG00000223496 | n.s. |
| ENSG00000105202 | 1441.860723 | 0.410970791 | 0.2823151 | 1.455717 | 0.145470932 | 0.300547824 | FBL | protein_coding | 48.35 | 0.214562266 | 0.214562266 | ENSG00000105202 | n.s. |
| ENSG00000215251 | 815.7427364 | 0.399742634 | 0.2701877 | 1.4795 | 0.139006845 | 0.290905202 | FASTKD5 | protein_coding | 44.53 | 0.214361393 | 0.214361393 | ENSG00000215251 | n.s. |
| ENSG00000134222 | 525.2886229 | -0.388323006 | 0.2580665 | -1.50474 | 0.132390915 | 0.280816379 | PSRC1 | protein_coding | 53.65 | -0.214190258 | 0.214190258 | ENSG00000134222 | n.s. |
| ENSG00000276083 | 6.049949637 | 0.608077387 | 0.5378644 | 1.13054 | 0.258248656 | 0.444724309 | MIR7976 | miRNA | 33.33 | 0.213987985 | 0.213987985 | ENSG00000276083 | n.s. |
| ENSG00000151135 | 820.2156301 | 0.40650663 | 0.2778156 | 1.463225 | 0.143405872 | 0.297580668 | TMEM263 | protein_coding | 38.27 | 0.213983174 | 0.213983174 | ENSG00000151135 | n.s. |
| ENSG00000189091 | 3662.581771 | -0.368694008 | 0.2378339 | -1.550216 | 0.1210896 | 0.262974652 | SF3B3 | protein_coding | 43.72 | -0.213874273 | 0.213874273 | ENSG00000189091 | n.s. |
| ENSG00000099953 | 115.0930644 | 0.419946688 | 0.2933545 | 1.431533 | 0.152277435 | 0.309647834 | MMP11 | protein_coding | 57.83 | 0.213808277 | 0.213808277 | ENSG00000099953 | n.s. |
| ENSG00000244346 | 46.68412422 | -0.416883615 | 0.2899384 | -1.437835 | 0.150480815 | 0.307205922 | | processed_pseudogene | 39.53 | -0.213682208 | 0.213682208 | ENSG00000244346 | n.s. |
| ENSG00000205581 | 7565.531505 | 0.39522177 | 0.2657608 | 1.487133 | 0.136979646 | 0.28806392 | HMGN1 | protein_coding | 42.26 | 0.213621767 | 0.213621767 | ENSG00000205581 | n.s. |
| ENSG00000170854 | 2195.447747 | -0.389927581 | 0.2601295 | -1.498975 | 0.133880167 | 0.283327795 | RIOX2 | protein_coding | 41.54 | -0.213567555 | 0.213567555 | ENSG00000170854 | n.s. |
| ENSG00000136108 | 3182.933594 | -0.392180244 | 0.2626296 | -1.493282 | 0.135363267 | 0.285554965 | CKAP2 | protein_coding | 38.86 | -0.21346774 | 0.21346774 | ENSG00000136108 | n.s. |

| | | | | | | | | | | | | | |
|---|---|---|---|---|---|---|---|---|---|---|---|---|---|
| ENSG00000073464 | 281.6772461 | -0.416927076 | 0.2904353 | -1.435525 | 0.151137662 | 0.308004928 | CLCN4 | protein_coding | 45.24 | -0.213234158 | 0.213234158 | ENSG00000073464 | n.s. |
| ENSG00000153187 | 38646.0836 | -0.360344132 | 0.2298633 | -1.567645 | 0.116963994 | 0.256252639 | HNRNPU | protein_coding | 38.67 | -0.213082891 | 0.213082891 | ENSG00000153187 | n.s. |
| ENSG00000173273 | 1570.793355 | -0.378199112 | 0.248494 | -1.521965 | 0.128017927 | 0.273507437 | TNKS | protein_coding | 37.08 | -0.212937771 | 0.212937771 | ENSG00000173273 | n.s. |
| ENSG00000100097 | 69.0894711 | 0.474424307 | 0.358336 | 1.323965 | 0.185514711 | 0.356019479 | LGALS1 | protein_coding | 56.78 | 0.212791751 | 0.212791751 | ENSG00000100097 | n.s. |
| ENSG00000165175 | 832.8314409 | 0.360259087 | 0.2301226 | 1.56551 | 0.11746345 | 0.256936628 | MID1IP1 | protein_coding | 53.8 | 0.212615539 | 0.212615539 | ENSG00000165175 | n.s. |
| ENSG00000100744 | 420.443514 | -0.400525124 | 0.2724638 | -1.470012 | 0.141558435 | 0.29456256 | GSKIP | protein_coding | 38.94 | -0.212607729 | 0.212607729 | ENSG00000100744 | n.s. |
| ENSG00000108666 | 834.3240362 | -0.416141025 | 0.2900916 | -1.434516 | 0.151425057 | 0.308410044 | C17orf75 | protein_coding | 44.91 | -0.212594584 | 0.212594584 | ENSG00000108666 | n.s. |
| ENSG00000120889 | 1378.070335 | -0.393217587 | 0.2645236 | -1.486512 | 0.137143665 | 0.288197963 | TNFRSF10B | protein_coding | 43.23 | -0.212459038 | 0.212459038 | ENSG00000120889 | n.s. |
| ENSG00000181201 | 125.4412254 | 0.388130503 | 0.2592222 | 1.497289 | 0.134318151 | 0.283995691 | HIST3H2BA | unitary_pseudogene | 61.07 | 0.212186385 | 0.212186385 | ENSG00000181201 | n.s. |
| ENSG00000099795 | 813.3326796 | 0.442931195 | 0.321292 | 1.378594 | 0.168020011 | 0.331962908 | NDUFB7 | protein_coding | 57.48 | 0.212124373 | 0.212124373 | ENSG00000099795 | n.s. |
| ENSG00000137177 | 108.7768206 | -0.38995133 | 0.2613508 | -1.492061 | 0.135683177 | 0.286128049 | KIF13A | protein_coding | 40.89 | -0.211914982 | 0.211914982 | ENSG00000137177 | n.s. |
| ENSG00000103121 | 559.11332 | 0.403274748 | 0.2758983 | 1.461679 | 0.143829135 | 0.298262294 | CMC2 | protein_coding | 40.64 | 0.211881216 | 0.211881216 | ENSG00000103121 | n.s. |
| ENSG00000246705 | 92.70054036 | 0.420919795 | 0.2961099 | 1.421499 | 0.155171868 | 0.313837583 | H2AFJ | protein_coding | 45.61 | 0.211846849 | 0.211846849 | ENSG00000246705 | n.s. |
| ENSG00000103152 | 206.7106426 | 0.456625138 | 0.337581 | 1.352639 | 0.176171009 | 0.343625568 | MPG | protein_coding | 61.91 | 0.211835036 | 0.211835036 | ENSG00000103152 | n.s. |
| ENSG00000247092 | 620.3560856 | 0.327596385 | 0.1981972 | 1.652881 | 0.098355027 | 0.225636324 | SNHG10 | antisense | 47.24 | 0.211820869 | 0.211820869 | ENSG00000247092 | n.s. |
| ENSG00000232412 | 13.92158958 | 0.482565342 | 0.3690589 | 1.307557 | 0.191023744 | 0.36397816 | | lincRNA | 57.66 | 0.211809836 | 0.211809836 | ENSG00000232412 | n.s. |
| ENSG00000136631 | 1495.236362 | -0.403330359 | 0.2761841 | -1.460368 | 0.144189024 | 0.298786097 | VPS45 | protein_coding | 39.17 | -0.211603084 | 0.211603084 | ENSG00000136631 | n.s. |
| ENSG00000143303 | 92.45494665 | -0.402497809 | 0.27529 | -1.462086 | 0.143717517 | 0.298075225 | RRNAD1 | protein_coding | 51 | -0.211582681 | 0.211582681 | ENSG00000143303 | n.s. |
| ENSG00000107021 | 225.6422505 | -0.370906478 | 0.2418573 | -1.533575 | 0.125134146 | 0.268947626 | TBC1D13 | protein_coding | 52.07 | -0.211539939 | 0.211539939 | ENSG00000107021 | n.s. |
| ENSG00000179021 | 942.6666507 | 0.37873461 | 0.2499862 | 1.515022 | 0.129766889 | 0.276429681 | C3orf38 | protein_coding | 38.06 | 0.211491211 | 0.211491211 | ENSG00000179021 | n.s. |
| ENSG00000100916 | 637.1140764 | 0.396521634 | 0.268937 | 1.474404 | 0.140372968 | 0.292883685 | BRMS1L | protein_coding | 38.94 | 0.211466899 | 0.211466899 | ENSG00000100916 | n.s. |
| ENSG00000172071 | 727.4025082 | -0.349254862 | 0.2196625 | -1.589961 | 0.111843525 | 0.248119407 | EIF2AK3 | protein_coding | 38.57 | -0.211417682 | 0.211417682 | ENSG00000172071 | n.s. |
| ENSG00000257103 | 1916.385811 | -0.403839686 | 0.2769673 | -1.458077 | 0.144819216 | 0.299564426 | LSM14A | protein_coding | 39.83 | -0.211414018 | 0.211414018 | ENSG00000257103 | n.s. |
| ENSG00000103544 | 648.2614866 | -0.40121778 | 0.2742073 | -1.463192 | 0.143414945 | 0.297580668 | C16orf62 | protein_coding | 45.1 | -0.211199148 | 0.211199148 | ENSG00000103544 | n.s. |
| ENSG00000123219 | 1125.78901 | -0.391035717 | 0.2632616 | -1.48535 | 0.137451069 | 0.288603715 | CENPK | protein_coding | 37.27 | -0.211041227 | 0.211041227 | ENSG00000123219 | n.s. |
| ENSG00000125257 | 907.0823811 | -0.386659151 | 0.2585698 | -1.495376 | 0.134816306 | 0.284659907 | ABCC4 | protein_coding | 42.33 | -0.210989728 | 0.210989728 | ENSG00000125257 | n.s. |
| ENSG00000142507 | 1313.726285 | 0.416201294 | 0.2914344 | 1.428113 | 0.153259237 | 0.311280578 | PSMB6 | protein_coding | 52.71 | 0.210950784 | 0.210950784 | ENSG00000142507 | n.s. |
| ENSG00000168569 | 246.1655876 | -0.408466994 | 0.2826676 | -1.445044 | 0.148445637 | 0.304614385 | TMEM223 | protein_coding | 52.1 | -0.210870918 | 0.210870918 | ENSG00000168569 | n.s. |
| ENSG00000166847 | 2164.528635 | -0.39545517 | 0.2684886 | -1.472894 | 0.140779731 | 0.293424579 | DCTN5 | protein_coding | 42.94 | -0.210581266 | 0.210581266 | ENSG00000166847 | n.s. |
| ENSG00000141756 | 778.9608021 | 0.415043371 | 0.2906678 | 1.427896 | 0.153321746 | 0.311318615 | FKBP10 | protein_coding | 56.18 | 0.210341868 | 0.210341868 | ENSG00000141756 | n.s. |
| ENSG00000261824 | 674.4221759 | -0.379165628 | 0.2512289 | -1.509244 | 0.131236467 | 0.278774925 | LINC00662 | lincRNA | 42.21 | -0.210340726 | 0.210340726 | ENSG00000261824 | n.s. |
| ENSG00000185973 | 190.6256529 | -0.403148636 | 0.277067 | -1.455058 | 0.145653123 | 0.300834954 | TMLHE | protein_coding | 37.38 | -0.210311236 | 0.210311236 | ENSG00000185973 | n.s. |

| | | | | | | | | | | | | | |
|---|---|---|---|---|---|---|---|---|---|---|---|---|---|
| ENSG00000213137 | 23.81987178 | -0.466743175 | 0.3516032 | -1.327471 | 0.184352817 | 0.354522688 | ARF1P2 | processed_pseudogene | 55.2 | -0.210200574 | 0.210200574 | ENSG00000213137 | n.s. |
| ENSG00000147416 | 2474.713022 | 0.393952291 | 0.2671122 | 1.474857 | 0.140250949 | 0.292760715 | ATP6V1B2 | protein_coding | 39.51 | 0.210168505 | 0.210168505 | ENSG00000147416 | n.s. |
| ENSG00000276603 | 116.6359434 | 0.363332274 | 0.2349808 | 1.546221 | 0.12205125 | 0.264156622 | | sense_intronic | 43.68 | 0.210056375 | 0.210056375 | ENSG00000276603 | n.s. |
| ENSG00000231587 | 10.21536659 | -0.501686636 | 0.3954066 | -1.268787 | 0.204517197 | 0.381545992 | SNORD62B | snoRNA | 46.51 | -0.209932331 | 0.209932331 | ENSG00000231587 | n.s. |
| ENSG00000108515 | 54.60893186 | 0.450348315 | 0.3320393 | 1.35631 | 0.175000415 | 0.341921244 | ENO3 | protein_coding | 58.22 | 0.209895602 | 0.209895602 | ENSG00000108515 | n.s. |
| ENSG00000110107 | 1520.758341 | 0.358917379 | 0.2305252 | 1.556955 | 0.119481124 | 0.260171362 | PRPF19 | protein_coding | 46.52 | 0.209873531 | 0.209873531 | ENSG00000110107 | n.s. |
| ENSG00000162441 | 205.0122961 | -0.414142332 | 0.2901397 | -1.427389 | 0.153467711 | 0.311386031 | LZIC | protein_coding | 43.89 | -0.209846282 | 0.209846282 | ENSG00000162441 | n.s. |
| ENSG00000105856 | 481.4429595 | -0.408399877 | 0.2835596 | -1.440261 | 0.149793498 | 0.306341784 | HBP1 | protein_coding | 37.95 | -0.20983331 | 0.20983331 | ENSG00000105856 | n.s. |
| ENSG00000104763 | 236.2735056 | 0.417317129 | 0.2938026 | 1.4204 | 0.155491385 | 0.314346821 | ASAH1 | protein_coding | 40.73 | 0.209739803 | 0.209739803 | ENSG00000104763 | n.s. |
| ENSG00000130347 | 350.3086457 | -0.389715127 | 0.2629467 | -1.482107 | 0.138311916 | 0.289794452 | RTN4IP1 | protein_coding | 41.03 | -0.209631638 | 0.209631638 | ENSG00000130347 | n.s. |
| ENSG00000183779 | 181.9958574 | 0.354220521 | 0.2260105 | 1.567274 | 0.117050545 | 0.256281607 | ZNF703 | protein_coding | 65.89 | 0.209444416 | 0.209444416 | ENSG00000183779 | n.s. |
| ENSG00000171222 | 439.1735433 | 0.377028782 | 0.2498159 | 1.509226 | 0.13124098 | 0.278774925 | SCAND1 | protein_coding | 47.71 | 0.209155319 | 0.209155319 | ENSG00000171222 | n.s. |
| ENSG00000130005 | 237.4477068 | 0.426399478 | 0.3045675 | 1.400016 | 0.161508391 | 0.323225719 | GAMT | protein_coding | 64.25 | 0.209146424 | 0.209146424 | ENSG00000130005 | n.s. |
| ENSG00000115425 | 249.354672 | -0.391670166 | 0.2653846 | -1.475859 | 0.139981852 | 0.292505977 | PECR | protein_coding | 42.56 | -0.209099093 | 0.209099093 | ENSG00000115425 | n.s. |
| ENSG00000138942 | 351.3084585 | -0.418584681 | 0.2960636 | -1.413834 | 0.157410793 | 0.316801167 | RNF185 | protein_coding | 44.98 | -0.208963009 | 0.208963009 | ENSG00000138942 | n.s. |
| ENSG00000207951 | 69.22862485 | 0.394788517 | 0.269057 | 1.467304 | 0.142293306 | 0.295782278 | MIR561 | miRNA | 45.36 | 0.20885412 | 0.20885412 | ENSG00000207951 | n.s. |
| ENSG00000154518 | 3651.319794 | 0.395831893 | 0.2701989 | 1.464965 | 0.142930544 | 0.296840984 | ATP5G3 | protein_coding | 40.96 | 0.208791878 | 0.208791878 | ENSG00000154518 | n.s. |
| ENSG00000108518 | 3039.731601 | 0.401908203 | 0.277369 | 1.449002 | 0.147337077 | 0.303196555 | PFN1 | protein_coding | 57.98 | 0.20829927 | 0.20829927 | ENSG00000108518 | n.s. |
| ENSG00000132432 | 1335.294918 | 0.399907498 | 0.2750703 | 1.453838 | 0.145991325 | 0.30139935 | SEC61G | protein_coding | 39.24 | 0.208294895 | 0.208294895 | ENSG00000132432 | n.s. |
| ENSG00000197119 | 902.2724167 | -0.396819597 | 0.2717885 | -1.460031 | 0.144281649 | 0.298808427 | SLC25A29 | protein_coding | 57.04 | -0.208174401 | 0.208174401 | ENSG00000197119 | n.s. |
| ENSG00000013375 | 1851.222398 | -0.398998158 | 0.2743081 | -1.454562 | 0.14579059 | 0.301029569 | PGM3 | protein_coding | 38.66 | -0.208033986 | 0.208033986 | ENSG00000013375 | n.s. |
| ENSG00000205138 | 252.1605712 | -0.381095513 | 0.2547455 | -1.495985 | 0.134657531 | 0.284540416 | SDHAF1 | protein_coding | 63.38 | -0.208023287 | 0.208023287 | ENSG00000205138 | n.s. |
| ENSG00000188130 | 381.0233631 | 0.414733206 | 0.2925283 | 1.417754 | 0.156262588 | 0.315233811 | MAPK12 | protein_coding | 64.68 | 0.207933629 | 0.207933629 | ENSG00000188130 | n.s. |
| ENSG00000075415 | 9326.223731 | 0.385647909 | 0.2598473 | 1.484133 | 0.137773673 | 0.289007271 | SLC25A3 | protein_coding | 42.39 | 0.207899406 | 0.207899406 | ENSG00000075415 | n.s. |
| ENSG00000178105 | 1384.755573 | -0.384386998 | 0.2585037 | -1.486969 | 0.137023028 | 0.28806548 | DDX10 | protein_coding | 37.41 | -0.207764548 | 0.207764548 | ENSG00000178105 | n.s. |
| ENSG00000227008 | 131.6454813 | 0.431927165 | 0.312523 | 1.382065 | 0.16695166 | 0.330485957 | | processed_pseudogene | 42.31 | 0.207690877 | 0.207690877 | ENSG00000227008 | n.s. |
| ENSG00000171135 | 575.4467798 | -0.418284055 | 0.2968781 | -1.408942 | 0.158852348 | 0.319287028 | JAGN1 | protein_coding | 47.58 | -0.207393067 | 0.207393067 | ENSG00000171135 | n.s. |
| ENSG00000118689 | 963.0583302 | -0.37505741 | 0.2490227 | -1.506117 | 0.132037036 | 0.280279258 | FOXO3 | protein_coding | 41.71 | -0.207185104 | 0.207185104 | ENSG00000118689 | n.s. |
| ENSG00000198455 | 296.2872592 | 0.338633127 | 0.2116713 | 1.599807 | 0.109641439 | 0.244480742 | ZXDB | protein_coding | 46.1 | 0.207160625 | 0.207160625 | ENSG00000198455 | n.s. |
| ENSG00000145337 | 1132.816922 | 0.406947577 | 0.2843584 | 1.431108 | 0.152399206 | 0.309850197 | PYURF | protein_coding | 44.72 | 0.207074552 | 0.207074552 | ENSG00000145337 | n.s. |
| ENSG00000169019 | 287.2572855 | -0.432159952 | 0.3133441 | -1.379187 | 0.167837255 | 0.331790161 | COMMD8 | protein_coding | 36.41 | -0.207063606 | 0.207063606 | ENSG00000169019 | n.s. |
| ENSG00000084207 | 3949.789646 | 0.438838399 | 0.3214047 | 1.365376 | 0.172134823 | 0.337744242 | GSTP1 | protein_coding | 63.05 | 0.206873708 | 0.206873708 | ENSG00000084207 | n.s. |

| | | | | | | | | | | | | | |
|---|---|---|---|---|---|---|---|---|---|---|---|---|---|
| ENSG00000238793 | 3.233417365 | -0.926460931 | 1.1152971 | -0.830685 | 0.406151415 | 0.598079974 | SNORD124 | snoRNA | 49.04 | -0.206823823 | 0.206823823 | ENSG00000238793 | n.s. |
| ENSG00000170185 | 1253.726135 | -0.382709423 | 0.2574925 | -1.486293 | 0.137201527 | 0.288221357 | USP38 | protein_coding | 35.49 | -0.20676789 | 0.20676789 | ENSG00000170185 | n.s. |
| ENSG00000264546 | 149.641194 | -0.384179128 | 0.2593935 | -1.481067 | 0.138588691 | 0.290236756 | | sense_intronic | 42.53 | -0.20639931 | 0.20639931 | ENSG00000264546 | n.s. |
| ENSG00000131002 | 2532.991803 | -0.411066642 | 0.2900645 | -1.417156 | 0.156437312 | 0.315480401 | TXLNGY | transcribed_unprocessed_pseud | 36.42 | -0.20595574 | 0.20595574 | ENSG00000131002 | n.s. |
| ENSG00000118960 | 120.7243374 | -0.427053271 | 0.308552 | -1.384056 | 0.166341347 | 0.329534801 | HS1BP3 | protein_coding | 52.46 | -0.205881833 | 0.205881833 | ENSG00000118960 | n.s. |
| ENSG00000184009 | 32781.87995 | 0.39958047 | 0.2769337 | 1.442874 | 0.149055898 | 0.305371582 | ACTG1 | protein_coding | 61.09 | 0.205852423 | 0.205852423 | ENSG00000184009 | n.s. |
| ENSG00000104129 | 186.6419022 | -0.399873813 | 0.2772656 | -1.442205 | 0.149244608 | 0.305668237 | DNAJC17 | protein_coding | 50.19 | -0.205834921 | 0.205834921 | ENSG00000104129 | n.s. |
| ENSG00000278571 | 59.84346842 | -0.400443131 | 0.2780095 | -1.440394 | 0.149756084 | 0.306310263 | MIR7161 | miRNA | 52.38 | -0.205763078 | 0.205763078 | ENSG00000278571 | n.s. |
| ENSG00000143207 | 1519.05125 | -0.379441145 | 0.2546923 | -1.489802 | 0.136276186 | 0.287002471 | RFWD2 | protein_coding | 36.32 | -0.205700495 | 0.205700495 | ENSG00000143207 | n.s. |
| ENSG00000196428 | 2390.663804 | -0.352523387 | 0.2267052 | -1.554986 | 0.11994935 | 0.260946062 | TSC22D2 | protein_coding | 36.52 | -0.205679506 | 0.205679506 | ENSG00000196428 | n.s. |
| ENSG00000162729 | 111.729418 | 0.396739599 | 0.2737797 | 1.44912 | 0.147304138 | 0.303196555 | IGSF8 | protein_coding | 58.9 | 0.205620508 | 0.205620508 | ENSG00000162729 | n.s. |
| ENSG00000175334 | 727.7067997 | 0.430989317 | 0.3138544 | 1.373214 | 0.169685737 | 0.334426301 | BANF1 | protein_coding | 58.14 | 0.205021435 | 0.205021435 | ENSG00000175334 | n.s. |
| ENSG00000141971 | 169.8009907 | 0.443354526 | 0.3285446 | 1.34945 | 0.177192496 | 0.344940515 | MVB12A | protein_coding | 50.72 | 0.204943198 | 0.204943198 | ENSG00000141971 | n.s. |
| ENSG00000106415 | 1144.286441 | -0.390674055 | 0.2676416 | -1.459691 | 0.14437498 | 0.29887322 | GLCCI1 | protein_coding | 36.7 | -0.204913619 | 0.204913619 | ENSG00000106415 | n.s. |
| ENSG00000089639 | 161.5771797 | -0.386841676 | 0.2635868 | -1.467606 | 0.142211148 | 0.295655638 | GMIP | protein_coding | 56.14 | -0.204721967 | 0.204721967 | ENSG00000089639 | n.s. |
| ENSG00000131469 | 23358.6861 | 0.439257422 | 0.3240644 | 1.355463 | 0.17527 | 0.342171114 | RPL27 | protein_coding | 49.84 | 0.204587069 | 0.204587069 | ENSG00000131469 | n.s. |
| ENSG00000257509 | 45.11477872 | -0.414261759 | 0.2947362 | -1.405534 | 0.159862578 | 0.320747467 | | sense_intronic | 46.25 | -0.204577687 | 0.204577687 | ENSG00000257509 | n.s. |
| ENSG00000010810 | 4178.599105 | -0.383617017 | 0.2602474 | -1.474047 | 0.140468825 | 0.293026444 | FYN | protein_coding | 42.42 | -0.204503617 | 0.204503617 | ENSG00000010810 | n.s. |
| ENSG00000256073 | 64.93837789 | 0.4086869 | 0.2886716 | 1.41575 | 0.156848661 | 0.316035298 | URB1-AS1 | lincRNA | 54.63 | 0.20445151 | 0.20445151 | ENSG00000256073 | n.s. |
| ENSG00000264940 | 51.37795589 | -0.392327992 | 0.2699274 | -1.453458 | 0.146096735 | 0.301572253 | SNORD3C | snoRNA | 54.91 | -0.204249335 | 0.204249335 | ENSG00000264940 | n.s. |
| ENSG00000126226 | 1229.561898 | -0.406560891 | 0.2863163 | -1.419971 | 0.155616047 | 0.314507508 | PCID2 | protein_coding | 43.4 | -0.204243581 | 0.204243581 | ENSG00000126226 | n.s. |
| ENSG00000166886 | 553.7550999 | -0.39383755 | 0.2717993 | -1.449001 | 0.147337214 | 0.303196555 | NAB2 | protein_coding | 60.22 | -0.204116446 | 0.204116446 | ENSG00000166886 | n.s. |
| ENSG00000272163 | 172.7734891 | 0.343395304 | 0.2185662 | 1.571128 | 0.116153021 | 0.254917798 | | antisense | 56.17 | 0.203839398 | 0.203839398 | ENSG00000272163 | n.s. |
| ENSG00000153944 | 8456.89457 | -0.3644928 | 0.2405196 | -1.515439 | 0.129661255 | 0.276289203 | MSI2 | protein_coding | 47.5 | -0.203618831 | 0.203618831 | ENSG00000153944 | n.s. |
| ENSG00000010278 | 334.0952399 | -0.460248377 | 0.3505357 | -1.312986 | 0.189187705 | 0.361077507 | CD9 | protein_coding | 51.04 | -0.203613682 | 0.203613682 | ENSG00000010278 | n.s. |
| ENSG00000143811 | 1051.673543 | -0.358963556 | 0.2348164 | -1.528699 | 0.126339122 | 0.270951879 | PYCR2 | protein_coding | 54.97 | -0.203571044 | 0.203571044 | ENSG00000143811 | n.s. |
| ENSG00000159377 | 3002.927526 | 0.419212561 | 0.3016706 | 1.389637 | 0.164639138 | 0.327280229 | PSMB4 | protein_coding | 49.27 | 0.203351725 | 0.203351725 | ENSG00000159377 | n.s. |
| ENSG00000147852 | 2095.310782 | -0.364914331 | 0.2411849 | -1.513007 | 0.130277963 | 0.27722147 | VLDLR | protein_coding | 41.97 | -0.203320463 | 0.203320463 | ENSG00000147852 | n.s. |
| ENSG00000231485 | 24.66181426 | 0.434723263 | 0.3198656 | 1.359081 | 0.174120885 | 0.340680872 | | lincRNA | 73.9 | 0.203299312 | 0.203299312 | ENSG00000231485 | n.s. |
| ENSG00000125356 | 1001.542527 | 0.450045149 | 0.3383793 | 1.330002 | 0.183517548 | 0.35340454 | NDUFA1 | protein_coding | 43.57 | 0.203297939 | 0.203297939 | ENSG00000125356 | n.s. |
| ENSG00000128829 | 870.9990528 | -0.405971964 | 0.28662 | -1.416412 | 0.156654932 | 0.315782049 | EIF2AK4 | protein_coding | 40.17 | -0.203234665 | 0.203234665 | ENSG00000128829 | n.s. |
| ENSG00000164944 | 1773.663833 | -0.388237755 | 0.2664545 | -1.457051 | 0.145102403 | 0.300009023 | KIAA1429 | protein_coding | 38.74 | -0.202996199 | 0.202996199 | ENSG00000164944 | n.s. |

| | | | | | | | | | | | | | |
|---|---|---|---|---|---|---|---|---|---|---|---|---|---|
| ENSG00000187531 | 328.3233082 | 0.353910654 | 0.2300732 | 1.538253 | 0.123986883 | 0.266976395 | SIRT7 | protein_coding | 57.63 | 0.202977364 | 0.202977364 | ENSG00000187531 | n.s. |
| ENSG00000176978 | 1089.76827 | 0.40100239 | 0.2811417 | 1.426335 | 0.153771565 | 0.311820783 | DPP7 | protein_coding | 68.7 | 0.202945281 | 0.202945281 | ENSG00000176978 | n.s. |
| ENSG00000206914 | 13.92823314 | -0.470720155 | 0.3643228 | -1.292042 | 0.196342771 | 0.370618275 | | misc_RNA | 41.18 | -0.202914829 | 0.202914829 | ENSG00000206914 | n.s. |
| ENSG00000270659 | 32.9516411 | 0.416020659 | 0.2986832 | 1.392849 | 0.163665417 | 0.325949592 | | lincRNA | 30 | 0.202539474 | 0.202539474 | ENSG00000270659 | n.s. |
| ENSG00000017483 | 475.3618814 | 0.418760946 | 0.3020294 | 1.386491 | 0.165597173 | 0.32857481 | SLC38A5 | protein_coding | 54.89 | 0.202414692 | 0.202414692 | ENSG00000017483 | n.s. |
| ENSG00000184432 | 2717.919829 | -0.389443698 | 0.2682188 | -1.451963 | 0.146511985 | 0.302205388 | COPB2 | protein_coding | 37.63 | -0.202393032 | 0.202393032 | ENSG00000184432 | n.s. |
| ENSG00000204228 | 98.36018338 | 0.381100319 | 0.2591735 | 1.470445 | 0.141441276 | 0.294450791 | HSD17B8 | protein_coding | 56.62 | 0.20235942 | 0.20235942 | ENSG00000204228 | n.s. |
| ENSG00000120159 | 658.5129241 | -0.393189848 | 0.2727231 | -1.441718 | 0.149381876 | 0.305833568 | CAAP1 | protein_coding | 37.23 | -0.202302016 | 0.202302016 | ENSG00000120159 | n.s. |
| ENSG00000066382 | 50.45928118 | -0.40893519 | 0.2905814 | -1.4073 | 0.159338456 | 0.320125443 | MPPED2 | protein_coding | 40.01 | -0.202291981 | 0.202291981 | ENSG00000066382 | n.s. |
| ENSG00000106070 | 156.8724025 | -0.399879187 | 0.2806036 | -1.425068 | 0.15413755 | 0.312290033 | GRB10 | protein_coding | 45.52 | -0.202115687 | 0.202115687 | ENSG00000106070 | n.s. |
| ENSG00000234287 | 125.0745556 | 0.456941537 | 0.3481319 | 1.312553 | 0.189333634 | 0.361256981 | | processed_pseudogene | 49.41 | 0.202052123 | 0.202052123 | ENSG00000234287 | n.s. |
| ENSG00000125755 | 147.3214574 | -0.390259643 | 0.2695176 | -1.447993 | 0.147618982 | 0.303589229 | SYMPK | protein_coding | 51.56 | -0.20204274 | 0.20204274 | ENSG00000125755 | n.s. |
| ENSG00000065135 | 4733.577427 | -0.377781693 | 0.255713 | -1.477366 | 0.139577497 | 0.291900875 | GNAI3 | protein_coding | 39.95 | -0.202024277 | 0.202024277 | ENSG00000065135 | n.s. |
| ENSG00000272689 | 12.61895111 | -0.466063739 | 0.3594672 | -1.29654 | 0.194789422 | 0.36866264 | | sense_intronic | 35.95 | -0.201978449 | 0.201978449 | ENSG00000272689 | n.s. |
| ENSG00000100749 | 1856.356295 | -0.417676144 | 0.3011626 | -1.386879 | 0.165478595 | 0.328468611 | VRK1 | protein_coding | 38.99 | -0.201948974 | 0.201948974 | ENSG00000100749 | n.s. |
| ENSG00000161677 | 49.50617829 | 0.433866568 | 0.3202082 | 1.354951 | 0.175433011 | 0.342430088 | JOSD2 | protein_coding | 61.76 | 0.201933685 | 0.201933685 | ENSG00000161677 | n.s. |
| ENSG00000274494 | 3.655744083 | -0.628022699 | 0.5921711 | -1.060543 | 0.288897825 | 0.477838278 | MIR6832 | miRNA | 52.78 | -0.201418852 | 0.201418852 | ENSG00000274494 | n.s. |
| ENSG00000103194 | 1244.845398 | -0.379887649 | 0.258608 | -1.468971 | 0.141840713 | 0.295017664 | USP10 | protein_coding | 43.62 | -0.20139819 | 0.20139819 | ENSG00000103194 | n.s. |
| ENSG00000135776 | 577.6972745 | -0.399856075 | 0.2811455 | -1.422239 | 0.154956907 | 0.313585031 | ABCB10 | protein_coding | 43.62 | -0.201385384 | 0.201385384 | ENSG00000135776 | n.s. |
| ENSG00000215386 | 1214.402009 | -0.361029727 | 0.2384938 | -1.513791 | 0.130078895 | 0.276840179 | MIR99AHG | lincRNA | 35.47 | -0.201371868 | 0.201371868 | ENSG00000215386 | n.s. |
| ENSG00000164107 | 9541.918371 | 0.333154826 | 0.2097859 | 1.58807 | 0.112270419 | 0.248788962 | HAND2 | protein_coding | 55.84 | 0.201281782 | 0.201281782 | ENSG00000164107 | n.s. |
| ENSG00000167566 | 432.2959114 | -0.351304926 | 0.2286009 | -1.536761 | 0.124351888 | 0.267555453 | NCKAP5L | protein_coding | 52.89 | -0.20115235 | 0.20115235 | ENSG00000167566 | n.s. |
| ENSG00000165233 | 135.9250565 | 0.425318223 | 0.310984 | 1.367653 | 0.171420727 | 0.336770254 | CARD19 | protein_coding | 49.96 | 0.201033581 | 0.201033581 | ENSG00000165233 | n.s. |
| ENSG00000156162 | 923.3276725 | -0.41664642 | 0.3008924 | -1.384702 | 0.166143502 | 0.329376192 | DPY19L4 | protein_coding | 40.06 | -0.200951817 | 0.200951817 | ENSG00000156162 | n.s. |
| ENSG00000081791 | 952.3148512 | -0.36669406 | 0.2445383 | -1.499536 | 0.133734607 | 0.283177084 | KIAA0141 | protein_coding | 48.21 | -0.200927038 | 0.200927038 | ENSG00000081791 | n.s. |
| ENSG00000226210 | 232.7735488 | -0.38796103 | 0.2679711 | -1.447772 | 0.147680796 | 0.303671499 | | unprocessed_pseudogene | 54.31 | -0.200807064 | 0.200807064 | ENSG00000226210 | n.s. |
| ENSG00000137154 | 30073.52075 | 0.418520701 | 0.303362 | 1.379608 | 0.167707345 | 0.331674626 | RPS6 | protein_coding | 40.73 | 0.200591846 | 0.200591846 | ENSG00000137154 | n.s. |
| ENSG00000207652 | 45.88930209 | -0.434132088 | 0.3220184 | -1.348159 | 0.177607172 | 0.345366573 | MIR621 | miRNA | 55.21 | -0.200447337 | 0.200447337 | ENSG00000207652 | n.s. |
| ENSG00000172809 | 18595.22061 | 0.411134233 | 0.2950811 | 1.393292 | 0.16353147 | 0.325869279 | RPL38 | protein_coding | 48.98 | 0.20020452 | 0.20020452 | ENSG00000172809 | n.s. |
| ENSG00000229536 | 14.78753806 | 0.466865824 | 0.3624215 | 1.288185 | 0.197681721 | 0.372741413 | | lincRNA | 43.98 | 0.200095122 | 0.200095122 | ENSG00000229536 | n.s. |
| ENSG00000141741 | 217.3862868 | 0.418127478 | 0.3035104 | 1.377638 | 0.168315101 | 0.332357277 | MIEN1 | protein_coding | 57.02 | 0.200030015 | 0.200030015 | ENSG00000141741 | n.s. |
| ENSG00000183718 | 794.7240658 | 0.342168419 | 0.2199024 | 1.556002 | 0.119707704 | 0.260463575 | TRIM52 | protein_coding | 43.45 | 0.199912926 | 0.199912926 | ENSG00000183718 | n.s. |

| | | | | | | | | | | | | | |
|---|---|---|---|---|---|---|---|---|---|---|---|---|---|
| ENSG00000181220 | 413.6609294 | -0.360332335 | 0.2388733 | -1.508467 | 0.131435163 | 0.279129313 | ZNF746 | protein_coding | 46.99 | -0.199694217 | 0.199694217 | ENSG00000181220 | n.s. |
| ENSG00000077238 | 43.22491155 | -0.395410409 | 0.2776604 | -1.424079 | 0.154423604 | 0.312778562 | IL4R | protein_coding | 49.92 | -0.199588553 | 0.199588553 | ENSG00000077238 | n.s. |
| ENSG00000157064 | 1127.140273 | -0.357562505 | 0.2361723 | -1.51399 | 0.130028387 | 0.276774993 | NMNAT2 | protein_coding | 44.41 | -0.199474522 | 0.199474522 | ENSG00000157064 | n.s. |
| ENSG00000109132 | 8177.958108 | 0.32374262 | 0.2014727 | 1.606881 | 0.108080436 | 0.242085571 | PHOX2B | protein_coding | 52.1 | 0.199435521 | 0.199435521 | ENSG00000109132 | n.s. |
| ENSG00000263750 | 7.078636053 | -0.715848666 | 0.7415197 | -0.96538 | 0.334354288 | 0.526727173 | MIR548AV | miRNA | 38.71 | -0.199302488 | 0.199302488 | ENSG00000263750 | n.s. |
| ENSG00000140416 | 1867.515531 | -0.380664252 | 0.2610755 | -1.458062 | 0.144823515 | 0.299564426 | TPM1 | protein_coding | 45.06 | -0.199281452 | 0.199281452 | ENSG00000140416 | n.s. |
| ENSG00000212567 | 19.19454343 | 0.465122192 | 0.3612335 | 1.287594 | 0.197887282 | 0.373073026 | | snoRNA | 54.69 | 0.199168183 | 0.199168183 | ENSG00000212567 | n.s. |
| ENSG00000145041 | 1112.994222 | -0.388873233 | 0.270645 | -1.436839 | 0.150763776 | 0.307558102 | DCAF1 | protein_coding | 43.34 | -0.199131416 | 0.199131416 | ENSG00000145041 | n.s. |
| ENSG00000210082 | 5352.061086 | -0.387917983 | 0.2695545 | -1.439108 | 0.150120012 | 0.306829262 | MT-RNR2 | Mt_rRNA | 42.81 | -0.199041968 | 0.199041968 | ENSG00000210082 | n.s. |
| ENSG00000180611 | 229.524748 | 0.379967824 | 0.260474 | 1.458755 | 0.144632529 | 0.299348699 | MB21D2 | protein_coding | 42.73 | 0.199035743 | 0.199035743 | ENSG00000180611 | n.s. |
| ENSG00000238058 | 31.75963618 | -0.431282347 | 0.3200187 | -1.347679 | 0.17776179 | 0.345618964 | | antisense | 51.51 | -0.198994726 | 0.198994726 | ENSG00000238058 | n.s. |
| ENSG00000113318 | 962.0567749 | -0.351530533 | 0.2302509 | -1.526728 | 0.126828623 | 0.271708712 | MSH3 | protein_coding | 38.03 | -0.198929876 | 0.198929876 | ENSG00000113318 | n.s. |
| ENSG00000119616 | 649.4141023 | -0.384534222 | 0.2658728 | -1.446309 | 0.148090506 | 0.304130383 | FCF1 | protein_coding | 40.97 | -0.198781194 | 0.198781194 | ENSG00000119616 | n.s. |
| ENSG00000084733 | 1832.158075 | -0.407737384 | 0.2923565 | -1.394658 | 0.163118929 | 0.325513093 | RAB10 | protein_coding | 40.62 | -0.19874406 | 0.19874406 | ENSG00000084733 | n.s. |
| ENSG00000145901 | 58.61619194 | 0.407614868 | 0.2922163 | 1.394908 | 0.163043566 | 0.325455995 | TNIP1 | protein_coding | 50.01 | 0.198715397 | 0.198715397 | ENSG00000145901 | n.s. |
| ENSG00000160145 | 147.2539121 | -0.355432364 | 0.2346288 | -1.514871 | 0.129805006 | 0.27643076 | KALRN | protein_coding | 42.68 | -0.198478278 | 0.198478278 | ENSG00000160145 | n.s. |
| ENSG00000258315 | 295.5238323 | 0.417886598 | 0.3046252 | 1.371806 | 0.170123893 | 0.335026161 | C17orf49 | protein_coding | 58.07 | 0.198463237 | 0.198463237 | ENSG00000258315 | n.s. |
| ENSG00000167220 | 663.1793819 | -0.395597274 | 0.2788711 | -1.418567 | 0.156025355 | 0.31501465 | HDHD2 | protein_coding | 38.68 | -0.198458987 | 0.198458987 | ENSG00000167220 | n.s. |
| ENSG00000165406 | 106.8321231 | 0.378596586 | 0.2596709 | 1.457986 | 0.144844338 | 0.299564426 | 8-Mar | protein_coding | 40.34 | 0.198199008 | 0.198199008 | ENSG00000165406 | n.s. |
| ENSG00000230525 | 117.6642219 | -0.383986157 | 0.2657813 | -1.444745 | 0.148529557 | 0.304696219 | LINC01799 | lincRNA | 38.12 | -0.198187902 | 0.198187902 | ENSG00000230525 | n.s. |
| ENSG00000214655 | 45.56750246 | 0.402883298 | 0.2872986 | 1.402316 | 0.160821016 | 0.322452622 | ZSWIM8 | protein_coding | 54.46 | 0.198030874 | 0.198030874 | ENSG00000214655 | n.s. |
| ENSG00000181704 | 820.4725925 | -0.40673701 | 0.2921167 | -1.392379 | 0.163807799 | 0.326053807 | YIPF6 | protein_coding | 41.22 | -0.197963265 | 0.197963265 | ENSG00000181704 | n.s. |
| ENSG00000250170 | 31.76623737 | 0.411151788 | 0.2975446 | 1.381816 | 0.167028277 | 0.330542007 | RASA2-IT1 | sense_intronic | 40.44 | 0.197670819 | 0.197670819 | ENSG00000250170 | n.s. |
| ENSG00000164889 | 461.7771792 | 0.406816172 | 0.2925626 | 1.390527 | 0.164368941 | 0.326976534 | SLC4A2 | protein_coding | 59.83 | 0.197502504 | 0.197502504 | ENSG00000164889 | n.s. |
| ENSG00000147905 | 514.0922512 | -0.395920258 | 0.2802824 | -1.412576 | 0.157780385 | 0.3173615 | ZCCHC7 | protein_coding | 36.62 | -0.197344774 | 0.197344774 | ENSG00000147905 | n.s. |
| ENSG00000140264 | 4658.90658 | 0.422058164 | 0.3105896 | 1.358893 | 0.174180428 | 0.340749488 | SERF2 | protein_coding | 49.15 | 0.197339536 | 0.197339536 | ENSG00000140264 | n.s. |
| ENSG00000112294 | 234.647726 | -0.384128243 | 0.2667396 | -1.440087 | 0.149842781 | 0.306397568 | ALDH5A1 | protein_coding | 43.11 | -0.19733232 | 0.19733232 | ENSG00000112294 | n.s. |
| ENSG00000053501 | 155.2022466 | 0.353702053 | 0.2336207 | 1.514002 | 0.130025455 | 0.276774993 | USE1 | protein_coding | 55.26 | 0.197320879 | 0.197320879 | ENSG00000053501 | n.s. |
| ENSG00000270060 | 64.12153325 | 0.38458335 | 0.2673728 | 1.438379 | 0.150326652 | 0.307025805 | | sense_intronic | 48.71 | 0.197224003 | 0.197224003 | ENSG00000270060 | n.s. |
| ENSG00000100600 | 306.6647461 | 0.387230842 | 0.2704462 | 1.431822 | 0.152194669 | 0.309524741 | LGMN | protein_coding | 47.28 | 0.197218461 | 0.197218461 | ENSG00000100600 | n.s. |
| ENSG00000200534 | 255.8318062 | 0.32743306 | 0.2067076 | 1.58404 | 0.113184682 | 0.250058764 | SNORA33 | snoRNA | 40.77 | 0.197100924 | 0.197100924 | ENSG00000200534 | n.s. |
| ENSG00000268364 | 73.16633937 | 0.36523489 | 0.2459616 | 1.484927 | 0.137563286 | 0.288708717 | SMC5-AS1 | antisense | 39.57 | 0.19705887 | 0.19705887 | ENSG00000268364 | n.s. |

| | | | | | | | | | | | | | |
|---|---|---|---|---|---|---|---|---|---|---|---|---|---|
| ENSG00000221303 | 26.70834512 | -0.622157515 | 0.5916985 | -1.051477 | 0.293039435 | 0.482382645 | SNORA79 | snoRNA | 34.29 | -0.196980249 | 0.196980249 | ENSG00000221303 | n.s. |
| ENSG00000157833 | 386.0586359 | -0.359810686 | 0.2402104 | -1.497898 | 0.13415968 | 0.28370371 | GAREM2 | protein_coding | 58.42 | -0.196865014 | 0.196865014 | ENSG00000157833 | n.s. |
| ENSG00000272047 | 583.0202782 | -0.416684185 | 0.3050841 | -1.365801 | 0.172001449 | 0.337577696 | GTF2H5 | protein_coding | 40.89 | -0.196519202 | 0.196519202 | ENSG00000272047 | n.s. |
| ENSG00000167371 | 126.4631869 | 0.383817344 | 0.2673978 | 1.435379 | 0.151179023 | 0.308044129 | PRRT2 | protein_coding | 61.52 | 0.196279225 | 0.196279225 | ENSG00000167371 | n.s. |
| ENSG00000127586 | 597.7117025 | 0.369410269 | 0.2511421 | 1.470922 | 0.141312335 | 0.294270363 | CHTF18 | protein_coding | 66.52 | 0.19625049 | 0.19625049 | ENSG00000127586 | n.s. |
| ENSG00000059915 | 268.1617919 | 0.352720186 | 0.2333315 | 1.51167 | 0.130617917 | 0.277859932 | PSD | protein_coding | 59.67 | 0.196173823 | 0.196173823 | ENSG00000059915 | n.s. |
| ENSG00000143479 | 202.8200077 | -0.359675985 | 0.2406875 | -1.494369 | 0.135079086 | 0.285069603 | DYRK3 | protein_coding | 44.46 | -0.196041068 | 0.196041068 | ENSG00000143479 | n.s. |
| ENSG00000137038 | 1370.276758 | 0.381555248 | 0.2650048 | 1.439805 | 0.149922499 | 0.306515558 | TMEM261 | protein_coding | 38.15 | 0.195946735 | 0.195946735 | ENSG00000137038 | n.s. |
| ENSG00000112290 | 2792.687695 | -0.373156916 | 0.2556451 | -1.459668 | 0.144381332 | 0.29887322 | WASF1 | protein_coding | 36.65 | -0.195725653 | 0.195725653 | ENSG00000112290 | n.s. |
| ENSG00000279600 | 48.50047776 | -0.420067962 | 0.3099114 | -1.355445 | 0.175275758 | 0.342171114 | | TEC | 41.15 | -0.19564945 | 0.19564945 | ENSG00000279600 | n.s. |
| ENSG00000165898 | 431.7883739 | -0.37371827 | 0.2563518 | -1.457833 | 0.14488647 | 0.299607059 | ISCA2 | protein_coding | 45.59 | -0.195622066 | 0.195622066 | ENSG00000165898 | n.s. |
| ENSG00000188352 | 860.7995278 | -0.377059641 | 0.2602228 | -1.448988 | 0.147340939 | 0.303196555 | FOCAD | protein_coding | 36.92 | -0.195420863 | 0.195420863 | ENSG00000188352 | n.s. |
| ENSG00000134900 | 3011.515339 | -0.378277838 | 0.2618874 | -1.444429 | 0.148618289 | 0.30478393 | TPP2 | protein_coding | 37.69 | -0.195194366 | 0.195194366 | ENSG00000134900 | n.s. |
| ENSG00000119632 | 166.9680506 | 0.445332292 | 0.3409315 | 1.306222 | 0.191477067 | 0.364610572 | IFI27L2 | protein_coding | 54.59 | 0.195131581 | 0.195131581 | ENSG00000119632 | n.s. |
| ENSG00000240869 | 15.86465609 | -0.474239804 | 0.3785302 | -1.252845 | 0.210261987 | 0.387899346 | RN7SL128P | misc_RNA | 51.8 | -0.195045798 | 0.195045798 | ENSG00000240869 | n.s. |
| ENSG00000124193 | 7753.407179 | 0.35730717 | 0.2388779 | 1.495773 | 0.13471281 | 0.284570848 | SRSF6 | protein_coding | 41.69 | 0.195021681 | 0.195021681 | ENSG00000124193 | n.s. |
| ENSG00000264881 | 2.690765743 | -0.873571698 | 1.0515734 | -0.830728 | 0.406127174 | 0.598079974 | MIR1273D | miRNA | 54.65 | -0.195016792 | 0.195016792 | ENSG00000264881 | n.s. |
| ENSG00000267080 | 308.5712566 | -0.338845382 | 0.2199092 | -1.540842 | 0.123355192 | 0.265904063 | ASB16-AS1 | antisense | 56.18 | -0.194929286 | 0.194929286 | ENSG00000267080 | n.s. |
| ENSG00000263266 | 477.6556757 | 0.449833643 | 0.3470717 | 1.296083 | 0.194947019 | 0.368733506 | RPS7P1 | processed_pseudogene | 47.52 | 0.194907248 | 0.194907248 | ENSG00000263266 | n.s. |
| ENSG00000176953 | 618.7496577 | -0.379544297 | 0.263627 | -1.439702 | 0.149951692 | 0.306530231 | NFATC2IP | protein_coding | 49.47 | -0.194906126 | 0.194906126 | ENSG00000176953 | n.s. |
| ENSG00000123091 | 1463.562727 | 0.406434352 | 0.2945243 | 1.379969 | 0.167596183 | 0.331501869 | RNF11 | protein_coding | 41.25 | 0.19489097 | 0.19489097 | ENSG00000123091 | n.s. |
| ENSG00000196290 | 1041.08806 | -0.378262241 | 0.2621892 | -1.442707 | 0.149102932 | 0.305422998 | NIF3L1 | protein_coding | 41.02 | -0.194842224 | 0.194842224 | ENSG00000196290 | n.s. |
| ENSG00000070214 | 1023.796974 | -0.382150313 | 0.2667465 | -1.432635 | 0.151962307 | 0.309266296 | SLC44A1 | protein_coding | 38.86 | -0.194769559 | 0.194769559 | ENSG00000070214 | n.s. |
| ENSG00000121210 | 1240.398766 | -0.353078194 | 0.234638 | -1.504778 | 0.13238105 | 0.280816379 | KIAA0922 | protein_coding | 41.78 | -0.19475001 | 0.19475001 | ENSG00000121210 | n.s. |
| ENSG00000188807 | 215.3965232 | 0.367324001 | 0.2500358 | 1.469085 | 0.141809618 | 0.294997059 | TMEM201 | protein_coding | 59.34 | 0.194748689 | 0.194748689 | ENSG00000188807 | n.s. |
| ENSG00000025039 | 803.8710521 | -0.378116442 | 0.2623373 | -1.441337 | 0.149489522 | 0.305944821 | RRAGD | protein_coding | 42.25 | -0.194486799 | 0.194486799 | ENSG00000025039 | n.s. |
| ENSG00000272337 | 9.772336366 | -0.488908443 | 0.399393 | -1.224129 | 0.220903681 | 0.400295742 | RNU6-90P | snRNA | 40.57 | -0.194399301 | 0.194399301 | ENSG00000272337 | n.s. |
| ENSG00000131876 | 1699.628871 | 0.348267952 | 0.2299933 | 1.514253 | 0.129961789 | 0.276717844 | SNRPA1 | protein_coding | 42.28 | 0.194320574 | 0.194320574 | ENSG00000131876 | n.s. |
| ENSG00000156515 | 1375.944314 | -0.364152786 | 0.2468986 | -1.474908 | 0.140237169 | 0.292760715 | HK1 | protein_coding | 47.64 | -0.194270851 | 0.194270851 | ENSG00000156515 | n.s. |
| ENSG00000135622 | 340.6953597 | -0.389316432 | 0.2758398 | -1.411386 | 0.158130836 | 0.317928611 | SEMA4F | protein_coding | 44.97 | -0.19375126 | 0.19375126 | ENSG00000135622 | n.s. |
| ENSG00000065000 | 1081.089922 | -0.368914094 | 0.2525402 | -1.460813 | 0.144066667 | 0.29861881 | AP3D1 | protein_coding | 56.41 | -0.193636677 | 0.193636677 | ENSG00000065000 | n.s. |
| ENSG00000185641 | 137.9016884 | 0.435393274 | 0.3305994 | 1.316981 | 0.187844868 | 0.359253311 | | processed_pseudogene | 47.72 | 0.19357551 | 0.19357551 | ENSG00000185641 | n.s. |

| | | | | | | | | | | | | | |
|---|---|---|---|---|---|---|---|---|---|---|---|---|---|
| ENSG00000143971 | 9221.096998 | 0.31983049 | 0.201252 | 1.589204 | 0.112014282 | 0.248379494 | ETAA1 | protein_coding | 33.26 | 0.19346043 | 0.19346043 | ENSG00000143971 | n.s. |
| ENSG00000104332 | 890.7190213 | -0.370421827 | 0.2543959 | -1.456084 | 0.145369375 | 0.300427165 | SFRP1 | protein_coding | 48.62 | -0.1934568 | 0.1934568 | ENSG00000104332 | n.s. |
| ENSG00000102901 | 574.9285916 | 0.369624962 | 0.253696 | 1.45696 | 0.145127346 | 0.300016041 | CENPT | protein_coding | 53.77 | 0.193260453 | 0.193260453 | ENSG00000102901 | n.s. |
| ENSG00000196605 | 18.37252213 | -0.456846815 | 0.3578881 | -1.276507 | 0.201776263 | 0.377649 | ZNF846 | protein_coding | 45.78 | -0.193205845 | 0.193205845 | ENSG00000196605 | n.s. |
| ENSG00000272567 | 15.12664159 | -0.465467475 | 0.3696396 | -1.259247 | 0.207941255 | 0.385046694 | | lincRNA | 29.8 | -0.192930032 | 0.192930032 | ENSG00000272567 | n.s. |
| ENSG00000140691 | 185.4758931 | 0.361575041 | 0.2451115 | 1.475145 | 0.140173546 | 0.292760715 | ARMC5 | protein_coding | 58.18 | 0.192895657 | 0.192895657 | ENSG00000140691 | n.s. |
| ENSG00000119718 | 891.2286605 | -0.366550865 | 0.2506416 | -1.462451 | 0.143617796 | 0.29791278 | EIF2B2 | protein_coding | 44.08 | -0.192773083 | 0.192773083 | ENSG00000119718 | n.s. |
| ENSG00000173473 | 5159.916003 | -0.364115212 | 0.2482702 | -1.466609 | 0.142482487 | 0.296131314 | SMARCC1 | protein_coding | 41.6 | -0.192440594 | 0.192440594 | ENSG00000173473 | n.s. |
| ENSG00000202343 | 7.218090015 | -0.547842613 | 0.4854014 | -1.128638 | 0.259050455 | 0.445553983 | | snoRNA | 41.48 | -0.19234736 | 0.19234736 | ENSG00000202343 | n.s. |
| ENSG00000141504 | 343.7930201 | 0.412131686 | 0.303631 | 1.357344 | 0.174672033 | 0.341471317 | SAT2 | protein_coding | 56.73 | 0.192319508 | 0.192319508 | ENSG00000141504 | n.s. |
| ENSG00000067208 | 289.4890069 | -0.362201773 | 0.2463838 | -1.470071 | 0.141542472 | 0.29456256 | EVI5 | protein_coding | 38.19 | -0.192264834 | 0.192264834 | ENSG00000067208 | n.s. |
| ENSG00000111775 | 3019.369695 | 0.431251516 | 0.3269082 | 1.319182 | 0.187108168 | 0.358253346 | COX6A1 | protein_coding | 48.25 | 0.192256129 | 0.192256129 | ENSG00000111775 | n.s. |
| ENSG00000136098 | 319.9557495 | -0.371304007 | 0.2564888 | -1.447642 | 0.147717168 | 0.303701436 | NEK3 | protein_coding | 39.71 | -0.19216956 | 0.19216956 | ENSG00000136098 | n.s. |
| ENSG00000005339 | 403.5718939 | -0.351691519 | 0.2349983 | -1.49657 | 0.134505092 | 0.284261444 | CREBBP | protein_coding | 45.01 | -0.192122761 | 0.192122761 | ENSG00000005339 | n.s. |
| ENSG00000065615 | 1218.958923 | -0.378971351 | 0.2658197 | -1.425671 | 0.153963409 | 0.31211889 | CYB5R4 | protein_coding | 37.46 | -0.191638212 | 0.191638212 | ENSG00000065615 | n.s. |
| ENSG00000173818 | 68.70472764 | -0.405480821 | 0.2966566 | -1.366836 | 0.171676832 | 0.337130682 | ENDOV | protein_coding | 56.47 | -0.191468741 | 0.191468741 | ENSG00000173818 | n.s. |
| ENSG00000143126 | 579.0814797 | -0.337903807 | 0.2211628 | -1.527851 | 0.126549533 | 0.271249787 | CELSR2 | protein_coding | 59.75 | -0.191466635 | 0.191466635 | ENSG00000143126 | n.s. |
| ENSG00000254093 | 551.3525646 | -0.389326451 | 0.2776589 | -1.402175 | 0.160862884 | 0.322457766 | PINX1 | protein_coding | 43.17 | -0.191364525 | 0.191364525 | ENSG00000254093 | n.s. |
| ENSG00000165650 | 1475.485498 | -0.358868424 | 0.2438328 | -1.471781 | 0.141080095 | 0.293874651 | PDZD8 | protein_coding | 37.51 | -0.19085982 | 0.19085982 | ENSG00000165650 | n.s. |
| ENSG00000151239 | 1855.638748 | 0.376206414 | 0.2635615 | 1.427395 | 0.153466079 | 0.311386031 | TWF1 | protein_coding | 34.16 | 0.190624119 | 0.190624119 | ENSG00000151239 | n.s. |
| ENSG00000135838 | 108.8495104 | -0.377786968 | 0.2655245 | -1.422795 | 0.154795626 | 0.313304186 | NPL | protein_coding | 41.95 | -0.190417402 | 0.190417402 | ENSG00000135838 | n.s. |
| ENSG00000167930 | 281.4756899 | -0.377837596 | 0.2657362 | -1.421852 | 0.15506914 | 0.313720958 | FAM234A | protein_coding | 52.35 | -0.190224781 | 0.190224781 | ENSG00000167930 | n.s. |
| ENSG00000113460 | 2827.725973 | 0.371067431 | 0.258231 | 1.436959 | 0.150729545 | 0.307533331 | BRIX1 | protein_coding | 38.42 | 0.190026529 | 0.190026529 | ENSG00000113460 | n.s. |
| ENSG00000177971 | 1753.380063 | 0.368312986 | 0.2550552 | 1.444052 | 0.148724323 | 0.304910397 | IMP3 | protein_coding | 52.2 | 0.189986065 | 0.189986065 | ENSG00000177971 | n.s. |
| ENSG00000204560 | 662.50828 | -0.366543461 | 0.2529074 | -1.449319 | 0.147248563 | 0.303196555 | DHX16 | protein_coding | 49.99 | -0.189970582 | 0.189970582 | ENSG00000204560 | n.s. |
| ENSG00000107281 | 570.4774784 | 0.365742218 | 0.2519934 | 1.451396 | 0.146669624 | 0.302414984 | NPDC1 | protein_coding | 65.86 | 0.189965299 | 0.189965299 | ENSG00000107281 | n.s. |
| ENSG00000146282 | 1409.949147 | -0.379936117 | 0.2684474 | -1.41531 | 0.156977736 | 0.316249606 | RARS2 | protein_coding | 38.27 | -0.189956663 | 0.189956663 | ENSG00000146282 | n.s. |
| ENSG00000269968 | 77.06277493 | 0.378856732 | 0.267219 | 1.417776 | 0.15625605 | 0.315233811 | | antisense | 56.5 | 0.189946342 | 0.189946342 | ENSG00000269968 | n.s. |
| ENSG00000134779 | 1276.60145 | -0.38396253 | 0.2729301 | -1.406816 | 0.159481827 | 0.320264977 | TPGS2 | protein_coding | 42.82 | -0.189865843 | 0.189865843 | ENSG00000134779 | n.s. |
| ENSG00000269343 | 820.3314486 | -0.376272811 | 0.2643019 | -1.423648 | 0.154548452 | 0.312960156 | ZNF587B | protein_coding | 47.01 | -0.189833753 | 0.189833753 | ENSG00000269343 | n.s. |
| ENSG00000247400 | 135.8020356 | -0.350328536 | 0.2352621 | -1.489099 | 0.136461345 | 0.287251639 | DNAJC3-AS1 | lincRNA | 38.92 | -0.1897861 | 0.1897861 | ENSG00000247400 | n.s. |
| ENSG00000140367 | 1564.911387 | 0.383655968 | 0.2727691 | 1.406523 | 0.159568925 | 0.320264977 | UBE2Q2 | protein_coding | 38.58 | 0.189714252 | 0.189714252 | ENSG00000140367 | n.s. |

| | | | | | | | | | | | | | |
|---|---|---|---|---|---|---|---|---|---|---|---|---|---|
| ENSG00000163935 | 208.1367198 | -0.368631339 | 0.2556948 | -1.441685 | 0.14939123 | 0.305833568 | SFMBT1 | protein_coding | 40.71 | -0.189666298 | 0.189666298 | ENSG00000163935 | n.s. |
| ENSG00000130707 | 4611.759567 | -0.40522152 | 0.2981005 | -1.359345 | 0.174037134 | 0.340612738 | ASS1 | protein_coding | 55.65 | -0.189537955 | 0.189537955 | ENSG00000130707 | n.s. |
| ENSG00000187672 | 104.6701186 | -0.389418327 | 0.2796581 | -1.39248 | 0.163777026 | 0.326053807 | ERC2 | protein_coding | 40.68 | -0.189534076 | 0.189534076 | ENSG00000187672 | n.s. |
| ENSG00000173402 | 28.95524008 | 0.415873386 | 0.3110006 | 1.337211 | 0.181153816 | 0.350160291 | DAG1 | protein_coding | 47.51 | 0.18952727 | 0.18952727 | ENSG00000173402 | n.s. |
| ENSG00000196646 | 476.0877909 | -0.354157245 | 0.2396429 | -1.477854 | 0.139446878 | 0.291738375 | ZNF136 | protein_coding | 44.4 | -0.189476407 | 0.189476407 | ENSG00000196646 | n.s. |
| ENSG00000144451 | 525.9348382 | -0.390762374 | 0.2814145 | -1.388565 | 0.164964942 | 0.327787483 | SPAG16 | protein_coding | 35.89 | -0.189288277 | 0.189288277 | ENSG00000144451 | n.s. |
| ENSG00000169764 | 682.6772357 | -0.385508877 | 0.2752367 | -1.400645 | 0.161320301 | 0.322922584 | UGP2 | protein_coding | 38.65 | -0.189246917 | 0.189246917 | ENSG00000169764 | n.s. |
| ENSG00000140983 | 866.6796162 | 0.368016203 | 0.2558363 | 1.438483 | 0.150296956 | 0.307010639 | RHOT2 | protein_coding | 66.37 | 0.188735849 | 0.188735849 | ENSG00000140983 | n.s. |
| ENSG00000143919 | 143.6289248 | 0.371532824 | 0.2601966 | 1.427893 | 0.153322685 | 0.311318615 | CAMKMT | protein_coding | 37.93 | 0.188290944 | 0.188290944 | ENSG00000143919 | n.s. |
| ENSG00000149485 | 4696.471635 | -0.364960839 | 0.2527727 | -1.44383 | 0.148786812 | 0.304954922 | FADS1 | protein_coding | 50.81 | -0.188233791 | 0.188233791 | ENSG00000149485 | n.s. |
| ENSG00000121775 | 130.3543361 | -0.374870265 | 0.2640893 | -1.419483 | 0.155758286 | 0.314748871 | TMEM39B | protein_coding | 48.85 | -0.188198301 | 0.188198301 | ENSG00000121775 | n.s. |
| ENSG00000230735 | 16.52452976 | -0.443986656 | 0.3474602 | -1.277806 | 0.201317885 | 0.37727551 | | sense_intronic | 38.58 | -0.187957926 | 0.187957926 | ENSG00000230735 | n.s. |
| ENSG00000130540 | 375.8250189 | 0.352842718 | 0.2396007 | 1.472628 | 0.14085137 | 0.293529955 | SULT4A1 | protein_coding | 56.02 | 0.187834964 | 0.187834964 | ENSG00000130540 | n.s. |
| ENSG00000080644 | 4898.925858 | -0.359840636 | 0.2472284 | -1.455499 | 0.145531235 | 0.300627803 | CHRNA3 | protein_coding | 47.02 | -0.187826325 | 0.187826325 | ENSG00000080644 | n.s. |
| ENSG00000102984 | 80.63413225 | 0.388117565 | 0.2796937 | 1.387652 | 0.165243008 | 0.328152674 | ZNF821 | protein_coding | 45.9 | 0.187819425 | 0.187819425 | ENSG00000102984 | n.s. |
| ENSG00000143337 | 1230.619886 | 0.360601506 | 0.2481188 | 1.453342 | 0.146128713 | 0.301593547 | TOR1AIP1 | protein_coding | 38.76 | 0.187721194 | 0.187721194 | ENSG00000143337 | n.s. |
| ENSG00000156831 | 208.8853363 | -0.408068916 | 0.3034249 | -1.344876 | 0.178665187 | 0.346728182 | NSMCE2 | protein_coding | 39.89 | -0.187716132 | 0.187716132 | ENSG00000156831 | n.s. |
| ENSG00000114735 | 202.1258723 | -0.365856052 | 0.2542634 | -1.438886 | 0.150182795 | 0.306867486 | HEMK1 | protein_coding | 53.84 | -0.187702127 | 0.187702127 | ENSG00000114735 | n.s. |
| ENSG00000170160 | 542.0537694 | -0.409287343 | 0.3050378 | -1.341759 | 0.179674032 | 0.3480247 | CCDC144A | protein_coding | 38.37 | -0.187613198 | 0.187613198 | ENSG00000170160 | n.s. |
| ENSG00000139645 | 451.8950713 | -0.337726606 | 0.2236078 | -1.510353 | 0.130953437 | 0.278289203 | ANKRD52 | protein_coding | 53.52 | -0.18760836 | 0.18760836 | ENSG00000139645 | n.s. |
| ENSG00000186010 | 440.4285747 | 0.434987809 | 0.336598 | 1.292307 | 0.196250977 | 0.370522683 | NDUFA13 | protein_coding | 53.15 | 0.187560305 | 0.187560305 | ENSG00000186010 | n.s. |
| ENSG00000237892 | 610.6931881 | -0.363251665 | 0.251443 | -1.444668 | 0.14855117 | 0.304696219 | KLF7-IT1 | sense_intronic | 42.55 | -0.187486148 | 0.187486148 | ENSG00000237892 | n.s. |
| ENSG00000165792 | 1028.8237 | 0.357743313 | 0.2452535 | 1.458668 | 0.144656645 | 0.299354112 | METTL17 | protein_coding | 45.05 | 0.187391232 | 0.187391232 | ENSG00000165792 | n.s. |
| ENSG00000110315 | 633.6043869 | -0.389590141 | 0.281898 | -1.382025 | 0.166963888 | 0.330485957 | RNF141 | protein_coding | 36.1 | -0.187333247 | 0.187333247 | ENSG00000110315 | n.s. |
| ENSG00000117262 | 212.1817412 | -0.373106914 | 0.2628851 | -1.419277 | 0.155818142 | 0.314748871 | GPR89A | protein_coding | 38.91 | -0.187313035 | 0.187313035 | ENSG00000117262 | n.s. |
| ENSG00000186603 | 270.9338618 | 0.356946555 | 0.2444812 | 1.460016 | 0.144285646 | 0.298808427 | HPDL | protein_coding | 62.8 | 0.187256718 | 0.187256718 | ENSG00000186603 | n.s. |
| ENSG00000100034 | 331.3107393 | -0.362207241 | 0.2504802 | -1.446051 | 0.148162853 | 0.304130383 | PPM1F | protein_coding | 55.32 | -0.18723948 | 0.18723948 | ENSG00000100034 | n.s. |
| ENSG00000071655 | 800.5978465 | -0.352404862 | 0.239578 | -1.47094 | 0.141307317 | 0.294270363 | MBD3 | protein_coding | 58.04 | -0.187216308 | 0.187216308 | ENSG00000071655 | n.s. |
| ENSG00000109113 | 930.4120782 | 0.369916352 | 0.2593393 | 1.42638 | 0.153758669 | 0.311820783 | RAB34 | protein_coding | 58.35 | 0.187212795 | 0.187212795 | ENSG00000109113 | n.s. |
| ENSG00000264007 | 30.29159616 | 0.404384214 | 0.2993952 | 1.350671 | 0.176800987 | 0.344424255 | | sense_intronic | 53.29 | 0.18719199 | 0.18719199 | ENSG00000264007 | n.s. |
| ENSG00000065057 | 309.1555289 | 0.428668631 | 0.329392 | 1.301394 | 0.193123751 | 0.366379887 | NTHL1 | protein_coding | 61.44 | 0.186928834 | 0.186928834 | ENSG00000065057 | n.s. |
| ENSG00000164466 | 1476.806962 | -0.383807717 | 0.2754869 | -1.393198 | 0.163560055 | 0.325879599 | SFXN1 | protein_coding | 42.43 | -0.186892415 | 0.186892415 | ENSG00000164466 | n.s. |

| | | | | | | | | | | | | | |
|---|---|---|---|---|---|---|---|---|---|---|---|---|---|
| ENSG00000251141 | 37.97967944 | -0.428588839 | 0.3294873 | -1.300775 | 0.193335447 | 0.366681574 | | antisense | 36.75 | -0.186740835 | 0.186740835 | ENSG00000251141 | n.s. |
| ENSG00000133111 | 330.4955453 | -0.360904708 | 0.2493553 | -1.447351 | 0.147798569 | 0.303823931 | RFXAP | protein_coding | 38.45 | -0.186724164 | 0.186724164 | ENSG00000133111 | n.s. |
| ENSG00000164904 | 1988.784811 | -0.34394531 | 0.2309272 | -1.48941 | 0.136379391 | 0.287176386 | ALDH7A1 | protein_coding | 43.05 | -0.186367204 | 0.186367204 | ENSG00000164904 | n.s. |
| ENSG00000213553 | 843.8020751 | 0.422851336 | 0.3227425 | 1.310182 | 0.190134309 | 0.362635614 | RPLP0P6 | processed_pseudogene | 53.56 | 0.186278507 | 0.186278507 | ENSG00000213553 | n.s. |
| ENSG00000183971 | 181.8550962 | 0.389225185 | 0.2827424 | 1.376607 | 0.168633826 | 0.332750674 | NPW | protein_coding | 55.12 | 0.186003342 | 0.186003342 | ENSG00000183971 | n.s. |
| ENSG00000264357 | 4.280913313 | 0.595509292 | 0.5718512 | 1.041371 | 0.297703334 | 0.48717954 | MIR4648 | miRNA | 63.89 | 0.185984078 | 0.185984078 | ENSG00000264357 | n.s. |
| ENSG00000153933 | 1969.174328 | -0.367191981 | 0.2573107 | -1.427037 | 0.153569166 | 0.311546481 | DGKE | protein_coding | 38.37 | -0.185974348 | 0.185974348 | ENSG00000153933 | n.s. |
| ENSG00000060982 | 11131.86307 | -0.384974465 | 0.2777959 | -1.385818 | 0.165802574 | 0.328901672 | BCAT1 | protein_coding | 40.06 | -0.185917227 | 0.185917227 | ENSG00000060982 | n.s. |
| ENSG00000198786 | 1757.291999 | -0.339690806 | 0.2271223 | -1.495629 | 0.134750292 | 0.284602032 | MT-ND5 | protein_coding | 44.92 | -0.185390335 | 0.185390335 | ENSG00000198786 | n.s. |
| ENSG00000221916 | 27.65209039 | 0.426699896 | 0.3286682 | 1.298269 | 0.194194814 | 0.367944857 | C19orf73 | protein_coding | 63.04 | 0.185280461 | 0.185280461 | ENSG00000221916 | n.s. |
| ENSG00000185418 | 762.0691752 | -0.377128754 | 0.2691299 | -1.401289 | 0.161127683 | 0.322767464 | TARSL2 | protein_coding | 41.13 | -0.185211797 | 0.185211797 | ENSG00000185418 | n.s. |
| ENSG00000252498 | 94.17564842 | 0.406401099 | 0.3041193 | 1.336321 | 0.18144434 | 0.350478572 | RNU6-1016P | snRNA | 36.17 | 0.185050079 | 0.185050079 | ENSG00000252498 | n.s. |
| ENSG00000166548 | 22.34548543 | 0.428303891 | 0.3311214 | 1.293495 | 0.195839961 | 0.370020326 | TK2 | protein_coding | 49.54 | 0.184930663 | 0.184930663 | ENSG00000166548 | n.s. |
| ENSG00000115526 | 1103.832503 | 0.340400448 | 0.2282592 | 1.491289 | 0.1358857 | 0.286396701 | CHST10 | protein_coding | 48.91 | 0.184848333 | 0.184848333 | ENSG00000115526 | n.s. |
| ENSG00000198912 | 210.723687 | 0.363052045 | 0.2537204 | 1.430914 | 0.152454901 | 0.309872933 | C1orf174 | protein_coding | 49.32 | 0.184726819 | 0.184726819 | ENSG00000198912 | n.s. |
| ENSG00000231699 | 22.4032315 | -0.421943426 | 0.3235036 | -1.304293 | 0.192133754 | 0.36524826 | | processed_pseudogene | 40.49 | -0.184563052 | 0.184563052 | ENSG00000231699 | n.s. |
| ENSG00000225264 | 45.1778291 | 0.378985779 | 0.2721699 | 1.39246 | 0.163783043 | 0.326053807 | ZNRF2P2 | transcribed_processed_pseudog | 40.97 | 0.184456443 | 0.184456443 | ENSG00000225264 | n.s. |
| ENSG00000255864 | 39.11005697 | -0.393861766 | 0.2897884 | -1.359136 | 0.174103629 | 0.340680872 | | lincRNA | 37.59 | -0.184190341 | 0.184190341 | ENSG00000255864 | n.s. |
| ENSG00000105429 | 491.5559434 | -0.345046963 | 0.2338751 | -1.475347 | 0.140119187 | 0.292705092 | MEGF8 | protein_coding | 54.75 | -0.184106613 | 0.184106613 | ENSG00000105429 | n.s. |
| ENSG00000133247 | 241.6936598 | -0.385191535 | 0.2798053 | -1.376641 | 0.168623152 | 0.332750674 | KMT5C | protein_coding | 64.88 | -0.184075737 | 0.184075737 | ENSG00000133247 | n.s. |
| ENSG00000026036 | 376.8774509 | 0.361818869 | 0.25289 | 1.430736 | 0.152505835 | 0.309931213 | RTEL1-TNFRSF6B | protein_coding | 59.83 | 0.184069808 | 0.184069808 | ENSG00000026036 | n.s. |
| ENSG00000102221 | 357.3956454 | -0.380424483 | 0.2742957 | -1.386914 | 0.16546803 | 0.328468611 | JADE3 | protein_coding | 40.39 | -0.183937568 | 0.183937568 | ENSG00000102221 | n.s. |
| ENSG00000225930 | 270.8121051 | -0.36563763 | 0.2576272 | -1.419251 | 0.155825897 | 0.314748871 | LINC02249 | lincRNA | 40.51 | -0.183563187 | 0.183563187 | ENSG00000225930 | n.s. |
| ENSG00000169359 | 700.8316645 | -0.339375394 | 0.2283685 | -1.486087 | 0.137256135 | 0.288281366 | SLC33A1 | protein_coding | 39.65 | -0.183324964 | 0.183324964 | ENSG00000169359 | n.s. |
| ENSG00000269246 | 9.416285109 | 0.500188807 | 0.4295592 | 1.164423 | 0.244252472 | 0.430053217 | | lincRNA | 56.07 | 0.183308093 | 0.183308093 | ENSG00000269246 | n.s. |
| ENSG00000182372 | 333.2263137 | -0.375113285 | 0.2686625 | -1.396225 | 0.162646817 | 0.324896933 | CLN8 | protein_coding | 48.06 | -0.183150707 | 0.183150707 | ENSG00000182372 | n.s. |
| ENSG00000242622 | 411.1258699 | 0.316638901 | 0.2047177 | 1.54671 | 0.121933117 | 0.264065139 | | lincRNA | 41.62 | 0.183108771 | 0.183108771 | ENSG00000242622 | n.s. |
| ENSG00000259935 | 23.7768601 | -0.417577827 | 0.3195373 | -1.30682 | 0.191273697 | 0.364359514 | | sense_intronic | 30.82 | -0.183095302 | 0.183095302 | ENSG00000259935 | n.s. |
| ENSG00000138434 | 6400.808524 | -0.35366997 | 0.2444029 | -1.447078 | 0.147875106 | 0.303871135 | SSFA2 | protein_coding | 35.71 | -0.182957207 | 0.182957207 | ENSG00000138434 | n.s. |
| ENSG00000131013 | 1145.097557 | -0.374143479 | 0.2677602 | -1.397308 | 0.162320978 | 0.324478819 | PPIL4 | protein_coding | 38.44 | -0.182886439 | 0.182886439 | ENSG00000131013 | n.s. |
| ENSG00000110048 | 637.5631216 | -0.376940829 | 0.271375 | -1.389004 | 0.164831675 | 0.327569428 | OSBP | protein_coding | 42.8 | -0.182701952 | 0.182701952 | ENSG00000110048 | n.s. |
| ENSG00000076242 | 1611.302104 | -0.365040254 | 0.2580523 | -1.414598 | 0.157186364 | 0.316440969 | MLH1 | protein_coding | 41.89 | -0.182413277 | 0.182413277 | ENSG00000076242 | n.s. |

| | | | | | | | | | | | | | |
|---|---|---|---|---|---|---|---|---|---|---|---|---|---|
| ENSG00000166002 | 45.16062295 | -0.434356195 | 0.3416945 | -1.271183 | 0.203663592 | 0.380312258 | SMCO4 | protein_coding | 45.2 | -0.182368652 | 0.182368652 | ENSG00000166002 | n.s. |
| ENSG00000096080 | 573.7554975 | -0.384678185 | 0.2808017 | -1.369928 | 0.170709239 | 0.335751481 | MRPS18A | protein_coding | 49.03 | -0.18233056 | 0.18233056 | ENSG00000096080 | n.s. |
| ENSG00000204650 | 983.197971 | -0.373970512 | 0.2681155 | -1.394811 | 0.163072701 | 0.32546749 | LINC02210 | transcribed_unitary_pseudogene | 46.27 | -0.182307777 | 0.182307777 | ENSG00000204650 | n.s. |
| ENSG00000226525 | 80.64006102 | 0.421084802 | 0.3251221 | 1.295159 | 0.19526545 | 0.369185343 | RPS7P10 | processed_pseudogene | 45.77 | 0.182226785 | 0.182226785 | ENSG00000226525 | n.s. |
| ENSG00000201512 | 100.27276 | 0.351403747 | 0.2428462 | 1.447022 | 0.147890813 | 0.303871135 | SNORA71C | snoRNA | 57.46 | 0.181784866 | 0.181784866 | ENSG00000201512 | n.s. |
| ENSG00000018510 | 3358.978893 | -0.371816418 | 0.2662103 | -1.396702 | 0.162503187 | 0.324703196 | AGPS | protein_coding | 36.51 | -0.181637317 | 0.181637317 | ENSG00000018510 | n.s. |
| ENSG00000100596 | 793.4814491 | -0.358673484 | 0.2513749 | -1.426847 | 0.153624048 | 0.311612415 | SPTLC2 | protein_coding | 41.35 | -0.181626961 | 0.181626961 | ENSG00000100596 | n.s. |
| ENSG00000168300 | 283.7079709 | -0.370881241 | 0.2651929 | -1.398534 | 0.161952889 | 0.323929044 | PCMTD1 | protein_coding | 36.74 | -0.181564952 | 0.181564952 | ENSG00000168300 | n.s. |
| ENSG00000217930 | 432.4142916 | 0.414863397 | 0.3180084 | 1.304567 | 0.192040181 | 0.365120229 | PAM16 | protein_coding | 56.77 | 0.181529331 | 0.181529331 | ENSG00000217930 | n.s. |
| ENSG00000072736 | 228.8296778 | 0.334348381 | 0.2243379 | 1.490379 | 0.136124699 | 0.286770191 | NFATC3 | protein_coding | 41.53 | 0.181372627 | 0.181372627 | ENSG00000072736 | n.s. |
| ENSG00000100372 | 504.5542989 | -0.401032318 | 0.3013949 | -1.330588 | 0.183324751 | 0.353100739 | SLC25A17 | protein_coding | 41.77 | -0.181307262 | 0.181307262 | ENSG00000100372 | n.s. |
| ENSG00000004864 | 982.7790837 | -0.393391602 | 0.2920938 | -1.346799 | 0.178044965 | 0.346082207 | SLC25A13 | protein_coding | 39.76 | -0.181283005 | 0.181283005 | ENSG00000004864 | n.s. |
| ENSG00000178796 | 47.970447 | 0.435490539 | 0.3444453 | 1.264324 | 0.206113664 | 0.38349867 | RIIAD1 | protein_coding | 51.01 | 0.1812669 | 0.1812669 | ENSG00000178796 | n.s. |
| ENSG00000174177 | 236.6446232 | 0.375514431 | 0.2711209 | 1.385044 | 0.166038998 | 0.329216886 | CTU2 | protein_coding | 60.96 | 0.181192433 | 0.181192433 | ENSG00000174177 | n.s. |
| ENSG00000100416 | 277.8318785 | -0.358283164 | 0.2513367 | -1.425511 | 0.154009586 | 0.312132229 | TRMU | protein_coding | 52.44 | -0.181169961 | 0.181169961 | ENSG00000100416 | n.s. |
| ENSG00000136943 | 244.6944226 | 0.359683508 | 0.2529182 | 1.422134 | 0.154987382 | 0.313601121 | CTSV | protein_coding | 44.69 | 0.181144669 | 0.181144669 | ENSG00000136943 | n.s. |
| ENSG00000066427 | 156.5315351 | 0.366278395 | 0.2603384 | 1.406932 | 0.159447515 | 0.320264977 | ATXN3 | protein_coding | 42.78 | 0.181121206 | 0.181121206 | ENSG00000066427 | n.s. |
| ENSG00000167470 | 407.4415865 | 0.325882467 | 0.2156959 | 1.510842 | 0.130828683 | 0.278249587 | MIDN | protein_coding | 65.97 | 0.181049046 | 0.181049046 | ENSG00000167470 | n.s. |
| ENSG00000139793 | 1995.954612 | -0.373961304 | 0.2694573 | -1.387831 | 0.165188455 | 0.328152674 | MBNL2 | protein_coding | 40.24 | -0.18096887 | 0.18096887 | ENSG00000139793 | n.s. |
| ENSG00000174080 | 348.8761989 | 0.395556945 | 0.2951914 | 1.340002 | 0.180244775 | 0.34888753 | CTSF | protein_coding | 58.67 | 0.180893947 | 0.180893947 | ENSG00000174080 | n.s. |
| ENSG00000198589 | 6181.289819 | -0.368018316 | 0.2626529 | -1.401158 | 0.161166718 | 0.322772972 | LRBA | protein_coding | 36.23 | -0.18073484 | 0.18073484 | ENSG00000198589 | n.s. |
| ENSG00000064961 | 367.3700374 | 0.385113806 | 0.2827885 | 1.361844 | 0.173247148 | 0.339448359 | HMG20B | protein_coding | 65.34 | 0.180705521 | 0.180705521 | ENSG00000064961 | n.s. |
| ENSG00000125686 | 1359.242027 | -0.35885193 | 0.2524085 | -1.421711 | 0.155110235 | 0.313758507 | MED1 | protein_coding | 45.01 | -0.180647673 | 0.180647673 | ENSG00000125686 | n.s. |
| ENSG00000107897 | 816.6332468 | -0.379027427 | 0.2759148 | -1.373712 | 0.169531144 | 0.334189737 | ACBD5 | protein_coding | 40.95 | -0.180419667 | 0.180419667 | ENSG00000107897 | n.s. |
| ENSG00000155189 | 1883.883092 | -0.379743914 | 0.2767468 | -1.372171 | 0.170010172 | 0.334896947 | AGPAT5 | protein_coding | 38.61 | -0.180412084 | 0.180412084 | ENSG00000155189 | n.s. |
| ENSG00000184619 | 166.1972503 | -0.326828413 | 0.2171301 | -1.505219 | 0.132267715 | 0.280726128 | KRBA2 | protein_coding | 48.93 | -0.180316845 | 0.180316845 | ENSG00000184619 | n.s. |
| ENSG00000172053 | 1264.933105 | 0.376180097 | 0.2727059 | 1.379435 | 0.167760641 | 0.331697506 | QARS | protein_coding | 53.42 | 0.180287243 | 0.180287243 | ENSG00000172053 | n.s. |
| ENSG00000132740 | 102.9653063 | 0.364660208 | 0.259383 | 1.405876 | 0.159761046 | 0.320604358 | IGHMBP2 | protein_coding | 50.65 | 0.180153293 | 0.180153293 | ENSG00000132740 | n.s. |
| ENSG00000131845 | 439.4486761 | -0.353273299 | 0.2466314 | -1.432394 | 0.152031191 | 0.309327819 | ZNF304 | protein_coding | 47.4 | -0.180021368 | 0.180021368 | ENSG00000131845 | n.s. |
| ENSG00000136149 | 26.7058748 | 0.581855255 | 0.562313 | 1.034753 | 0.300784104 | 0.490501652 | RPL13AP25 | processed_pseudogene | 57.19 | 0.180002465 | 0.180002465 | ENSG00000136149 | n.s. |
| ENSG00000133704 | 1287.843428 | -0.366935744 | 0.2624192 | -1.398281 | 0.16202872 | 0.324008025 | IPO8 | protein_coding | 35.93 | -0.179594584 | 0.179594584 | ENSG00000133704 | n.s. |
| ENSG00000144524 | 152.4850347 | 0.360236758 | 0.255047 | 1.412433 | 0.157822572 | 0.317400502 | COPS7B | protein_coding | 44.9 | 0.179539259 | 0.179539259 | ENSG00000144524 | n.s. |

| | | | | | | | | | | | | | |
|---|---|---|---|---|---|---|---|---|---|---|---|---|---|
| ENSG00000272478 | 19.57105153 | -0.423582899 | 0.3312708 | -1.278661 | 0.201016567 | 0.376936521 | | antisense | 45.5 | -0.179485537 | 0.179485537 | ENSG00000272478 | n.s. |
| ENSG00000155111 | 1201.495644 | -0.356817567 | 0.2511045 | -1.420993 | 0.155318919 | 0.314089369 | CDK19 | protein_coding | 39.87 | -0.17946024 | 0.17946024 | ENSG00000155111 | n.s. |
| ENSG00000189343 | 285.9148726 | 0.418697892 | 0.3250175 | 1.288232 | 0.197665299 | 0.372741413 | | processed_pseudogene | 60.82 | 0.179450715 | 0.179450715 | ENSG00000189343 | n.s. |
| ENSG00000276234 | 206.488169 | -0.35794599 | 0.2525579 | -1.417283 | 0.156400294 | 0.31545144 | TADA2A | protein_coding | 42.79 | -0.179355097 | 0.179355097 | ENSG00000276234 | n.s. |
| ENSG00000255690 | 176.7755107 | 0.340354757 | 0.2325054 | 1.463857 | 0.143232945 | 0.297335961 | TRIL | protein_coding | 56.8 | 0.17928274 | 0.17928274 | ENSG00000255690 | n.s. |
| ENSG00000241360 | 1061.679524 | 0.387774619 | 0.2873722 | 1.349381 | 0.177214645 | 0.344940515 | PDXP | protein_coding | 55.3 | 0.179251064 | 0.179251064 | ENSG00000241360 | n.s. |
| ENSG00000242265 | 3993.711059 | 0.35956274 | 0.2546734 | 1.411858 | 0.157991694 | 0.317694736 | PEG10 | protein_coding | 43.22 | 0.179058642 | 0.179058642 | ENSG00000242265 | n.s. |
| ENSG00000135372 | 1125.569655 | -0.356681478 | 0.2514349 | -1.418584 | 0.156020434 | 0.31501465 | NAT10 | protein_coding | 45.79 | -0.178936129 | 0.178936129 | ENSG00000135372 | n.s. |
| ENSG00000163795 | 183.3891718 | -0.366565346 | 0.2626575 | -1.395602 | 0.162834379 | 0.325178293 | ZNF513 | protein_coding | 57.87 | -0.178839334 | 0.178839334 | ENSG00000163795 | n.s. |
| ENSG00000185046 | 17.30157354 | 0.442612351 | 0.3574755 | 1.238161 | 0.215656186 | 0.394661024 | ANKS1B | protein_coding | 37.84 | 0.178716139 | 0.178716139 | ENSG00000185046 | n.s. |
| ENSG00000272391 | 154.295476 | 0.36431818 | 0.2603211 | 1.399495 | 0.161664519 | 0.323445192 | POM121C | protein_coding | 45.74 | 0.178588517 | 0.178588517 | ENSG00000272391 | n.s. |
| ENSG00000126561 | 359.7896982 | -0.352928202 | 0.2475833 | -1.425493 | 0.154014825 | 0.312132229 | STAT5A | protein_coding | 54.63 | -0.178462164 | 0.178462164 | ENSG00000126561 | n.s. |
| ENSG00000105866 | 654.1735129 | -0.346138308 | 0.2397892 | -1.443511 | 0.148876659 | 0.305094161 | SP4 | protein_coding | 38.35 | -0.178457179 | 0.178457179 | ENSG00000105866 | n.s. |
| ENSG00000205213 | 641.9581281 | 0.356434643 | 0.2516294 | 1.416506 | 0.156627307 | 0.315772081 | LGR4 | protein_coding | 39.08 | 0.178440546 | 0.178440546 | ENSG00000205213 | n.s. |
| ENSG00000164405 | 2356.909932 | 0.401572163 | 0.3050696 | 1.316329 | 0.188063487 | 0.359523263 | UQCRQ | protein_coding | 51.29 | 0.178407671 | 0.178407671 | ENSG00000164405 | n.s. |
| ENSG00000225195 | 18.10576218 | -0.562408041 | 0.534133 | -1.052936 | 0.292370179 | 0.481793112 | | sense_intronic | 45.19 | -0.178361756 | 0.178361756 | ENSG00000225195 | n.s. |
| ENSG00000011566 | 1715.021182 | -0.353821809 | 0.2486433 | -1.423009 | 0.154733428 | 0.313269378 | MAP4K3 | protein_coding | 36.9 | -0.178355225 | 0.178355225 | ENSG00000011566 | n.s. |
| ENSG00000121851 | 182.6204347 | -0.377280496 | 0.2759724 | -1.367095 | 0.171595563 | 0.337025323 | POLR3GL | protein_coding | 44.9 | -0.178203714 | 0.178203714 | ENSG00000121851 | n.s. |
| ENSG00000102699 | 1199.131178 | -0.3784746 | 0.2774394 | -1.36417 | 0.172513964 | 0.338249812 | PARP4 | protein_coding | 42.88 | -0.178171624 | 0.178171624 | ENSG00000102699 | n.s. |
| ENSG00000173545 | 753.4685168 | 0.345240115 | 0.2390263 | 1.44436 | 0.148637713 | 0.30478393 | ZNF622 | protein_coding | 40.67 | 0.178146639 | 0.178146639 | ENSG00000173545 | n.s. |
| ENSG00000197579 | 1355.344154 | 0.360010007 | 0.2558943 | 1.40687 | 0.159465948 | 0.320264977 | TOPORS | protein_coding | 42.02 | 0.178021547 | 0.178021547 | ENSG00000197579 | n.s. |
| ENSG00000188976 | 744.65553 | 0.377395953 | 0.2765282 | 1.364765 | 0.172327001 | 0.337930822 | NOC2L | protein_coding | 59.55 | 0.17781848 | 0.17781848 | ENSG00000188976 | n.s. |
| ENSG00000007944 | 2143.250018 | 0.354298776 | 0.2497783 | 1.418453 | 0.156058607 | 0.315036101 | MYLIP | protein_coding | 40.19 | 0.177730323 | 0.177730323 | ENSG00000007944 | n.s. |
| ENSG00000135040 | 801.458296 | -0.361482497 | 0.2579721 | -1.401247 | 0.161140318 | 0.322767464 | NAA35 | protein_coding | 37.88 | -0.177527759 | 0.177527759 | ENSG00000135040 | n.s. |
| ENSG00000234202 | 7.782648966 | 0.500187242 | 0.4399745 | 1.136855 | 0.255598829 | 0.441696301 | | processed_pseudogene | 50.64 | 0.177504567 | 0.177504567 | ENSG00000234202 | n.s. |
| ENSG00000198917 | 576.3600511 | -0.353193827 | 0.2487238 | -1.420024 | 0.155600593 | 0.314507508 | SPOUT1 | protein_coding | 57.61 | -0.177433624 | 0.177433624 | ENSG00000198917 | n.s. |
| ENSG00000160741 | 188.8051779 | 0.366875125 | 0.2645783 | 1.386641 | 0.165551206 | 0.328530422 | CRTC2 | protein_coding | 53.2 | 0.177356391 | 0.177356391 | ENSG00000160741 | n.s. |
| ENSG00000223749 | 210.3032725 | -0.359078211 | 0.2554908 | -1.405445 | 0.15988913 | 0.320747467 | MIR503HG | lincRNA | 53.09 | -0.177326022 | 0.177326022 | ENSG00000223749 | n.s. |
| ENSG00000215417 | 8461.24257 | 0.30720834 | 0.1989722 | 1.543976 | 0.122594191 | 0.264782747 | MIR17HG | processed_transcript | 44.48 | 0.177293102 | 0.177293102 | ENSG00000215417 | n.s. |
| ENSG00000233597 | 23.16180583 | -0.425215901 | 0.3361039 | -1.265132 | 0.205823871 | 0.383010661 | | processed_pseudogene | 52.26 | -0.177225369 | 0.177225369 | ENSG00000233597 | n.s. |
| ENSG00000161036 | 198.7494595 | 0.373152917 | 0.272009 | 1.37184 | 0.170113128 | 0.335026161 | LRWD1 | protein_coding | 61.06 | 0.177218261 | 0.177218261 | ENSG00000161036 | n.s. |
| ENSG00000242071 | 266.7724067 | 0.426410835 | 0.3379206 | 1.261867 | 0.206996705 | 0.38424486 | RPL7AP6 | processed_pseudogene | 53.06 | 0.177127621 | 0.177127621 | ENSG00000242071 | n.s. |

| | | | | | | | | | | | | | |
|---|---|---|---|---|---|---|---|---|---|---|---|---|---|
| ENSG00000263072 | 355.1352143 | -0.33281384 | 0.2260534 | -1.47228 | 0.140945379 | 0.293637963 | ZNF213-AS1 | antisense | 53.29 | -0.177119465 | 0.177119465 | ENSG00000263072 | n.s. |
| ENSG00000115541 | 3630.734378 | 0.379504636 | 0.2796621 | 1.357011 | 0.174777656 | 0.341629834 | HSPE1 | protein_coding | 44.77 | 0.17701774 | 0.17701774 | ENSG00000115541 | n.s. |
| ENSG00000234420 | 1725.25924 | -0.372808615 | 0.2719764 | -1.370739 | 0.170456327 | 0.335443596 | ZNF37BP | transcribed_processed_pseudog | 39.07 | -0.176853135 | 0.176853135 | ENSG00000234420 | n.s. |
| ENSG00000140105 | 617.6984497 | -0.373527786 | 0.2729629 | -1.36842 | 0.171180696 | 0.336441112 | WARS | protein_coding | 46.89 | -0.176712612 | 0.176712612 | ENSG00000140105 | n.s. |
| ENSG00000060688 | 685.3327103 | -0.369758417 | 0.2685704 | -1.376765 | 0.168584815 | 0.332750674 | SNRNP40 | protein_coding | 42.01 | -0.176700543 | 0.176700543 | ENSG00000060688 | n.s. |
| ENSG00000109220 | 207.9960523 | -0.369743638 | 0.2686324 | -1.376393 | 0.168700065 | 0.332788798 | CHIC2 | protein_coding | 36.75 | -0.176675084 | 0.176675084 | ENSG00000109220 | n.s. |
| ENSG00000079102 | 2943.287847 | -0.349480108 | 0.245229 | -1.425117 | 0.154123304 | 0.312290033 | RUNX1T1 | protein_coding | 38.11 | -0.176641882 | 0.176641882 | ENSG00000079102 | n.s. |
| ENSG00000278771 | 266.1483605 | -0.320260728 | 0.2128948 | -1.504314 | 0.132500466 | 0.280963142 | RN7SL3 | misc_RNA | 62.21 | -0.17657596 | 0.17657596 | ENSG00000278771 | n.s. |
| ENSG00000221914 | 1976.939438 | -0.349305948 | 0.245166 | -1.424773 | 0.154222778 | 0.312417247 | PPP2R2A | protein_coding | 37.61 | -0.17649207 | 0.17649207 | ENSG00000221914 | n.s. |
| ENSG00000144231 | 1694.809377 | -0.358949654 | 0.2560322 | -1.401971 | 0.16092396 | 0.322519692 | POLR2D | protein_coding | 45.35 | -0.176403569 | 0.176403569 | ENSG00000144231 | n.s. |
| ENSG00000106803 | 1152.938858 | 0.415204633 | 0.3242831 | 1.280377 | 0.200412507 | 0.376006379 | SEC61B | protein_coding | 42.24 | 0.176380916 | 0.176380916 | ENSG00000106803 | n.s. |
| ENSG00000252062 | 8.328129216 | -0.482014441 | 0.4143483 | -1.163308 | 0.24470476 | 0.43061481 | RNU6-469P | snRNA | 42.57 | -0.176374404 | 0.176374404 | ENSG00000252062 | n.s. |
| ENSG00000181090 | 239.6792576 | -0.365063849 | 0.2635344 | -1.385261 | 0.16597274 | 0.329132383 | EHMT1 | protein_coding | 49.25 | -0.176190541 | 0.176190541 | ENSG00000181090 | n.s. |
| ENSG00000257815 | 160.6101542 | 0.343416689 | 0.2386843 | 1.43879 | 0.150209934 | 0.306877903 | | antisense | 38.83 | 0.176184569 | 0.176184569 | ENSG00000257815 | n.s. |
| ENSG00000146909 | 1148.295947 | -0.325386777 | 0.218829 | -1.486945 | 0.137029209 | 0.28806548 | NOM1 | protein_coding | 47.02 | -0.175874411 | 0.175874411 | ENSG00000146909 | n.s. |
| ENSG00000131174 | 2162.642685 | 0.42026984 | 0.3312705 | 1.268661 | 0.204562121 | 0.381545992 | COX7B | protein_coding | 40.81 | 0.17586322 | 0.17586322 | ENSG00000131174 | n.s. |
| ENSG00000197912 | 680.0538286 | -0.348747584 | 0.2451122 | -1.422808 | 0.154791908 | 0.313304186 | SPG7 | protein_coding | 52.29 | -0.175780571 | 0.175780571 | ENSG00000197912 | n.s. |
| ENSG00000206644 | 5.213785089 | -0.547332066 | 0.5162061 | -1.060298 | 0.289009275 | 0.477838278 | RNU6-1279P | snRNA | 49.51 | -0.175539827 | 0.175539827 | ENSG00000206644 | n.s. |
| ENSG00000124209 | 817.1448133 | -0.373954348 | 0.274718 | -1.36123 | 0.173441008 | 0.339589249 | RAB22A | protein_coding | 41.46 | -0.175401816 | 0.175401816 | ENSG00000124209 | n.s. |
| ENSG00000198075 | 185.6381893 | -0.322733743 | 0.2163368 | -1.491811 | 0.135748585 | 0.286194359 | SULT1C4 | protein_coding | 40.61 | -0.175353806 | 0.175353806 | ENSG00000198075 | n.s. |
| ENSG00000274997 | 806.390322 | 0.324480893 | 0.2182531 | 1.486718 | 0.137089292 | 0.288148282 | HIST1H2AH | protein_coding | 61.76 | 0.175344272 | 0.175344272 | ENSG00000274997 | n.s. |
| ENSG00000143324 | 1519.36476 | -0.35696635 | 0.2547074 | -1.401476 | 0.161071765 | 0.322728494 | XPR1 | protein_coding | 36.95 | -0.175328553 | 0.175328553 | ENSG00000143324 | n.s. |
| ENSG00000198836 | 4060.458389 | -0.364244702 | 0.2636057 | -1.381779 | 0.167039687 | 0.330542007 | OPA1 | protein_coding | 36.98 | -0.175119142 | 0.175119142 | ENSG00000198836 | n.s. |
| ENSG00000267034 | 294.7808719 | 0.336880433 | 0.2319811 | 1.452189 | 0.146449075 | 0.302120383 | | lincRNA | 41.22 | 0.175117177 | 0.175117177 | ENSG00000267034 | n.s. |
| ENSG00000119760 | 1717.978124 | -0.363139064 | 0.2623135 | -1.38437 | 0.166245046 | 0.329484665 | SUPT7L | protein_coding | 41.29 | -0.175092872 | 0.175092872 | ENSG00000119760 | n.s. |
| ENSG00000241370 | 220.1971655 | 0.400437369 | 0.3072224 | 1.303412 | 0.192434167 | 0.365569778 | RPP21 | protein_coding | 53.76 | 0.175003029 | 0.175003029 | ENSG00000241370 | n.s. |
| ENSG00000234456 | 2312.534064 | -0.375391817 | 0.2768486 | -1.355946 | 0.175116321 | 0.342003722 | MAGI2-AS3 | processed_transcript | 38.5 | -0.174921013 | 0.174921013 | ENSG00000234456 | n.s. |
| ENSG00000250988 | 79.49831542 | 0.354280533 | 0.2522108 | 1.4047 | 0.160110407 | 0.321120311 | SNHG21 | antisense | 40.87 | 0.174778003 | 0.174778003 | ENSG00000250988 | n.s. |
| ENSG00000101868 | 2080.016826 | -0.367380153 | 0.2675742 | -1.373003 | 0.169751495 | 0.334434706 | POLA1 | protein_coding | 38.8 | -0.174758572 | 0.174758572 | ENSG00000101868 | n.s. |
| ENSG00000197128 | 114.0868072 | -0.34939949 | 0.2469504 | -1.414857 | 0.157110475 | 0.316368891 | ZNF772 | protein_coding | 42.84 | -0.174632041 | 0.174632041 | ENSG00000197128 | n.s. |
| ENSG00000242372 | 993.8491469 | 0.411959873 | 0.3222531 | 1.278374 | 0.201117724 | 0.377074783 | EIF6 | protein_coding | 52.05 | 0.174494878 | 0.174494878 | ENSG00000242372 | n.s. |
| ENSG00000186468 | 17696.47259 | 0.405058198 | 0.3137088 | 1.291192 | 0.196637277 | 0.371093526 | RPS23 | protein_coding | 44.77 | 0.174384287 | 0.174384287 | ENSG00000186468 | n.s. |

| | | | | | | | | | | | | | |
|---|---|---|---|---|---|---|---|---|---|---|---|---|---|
| ENSG00000067900 | 1959.733169 | -0.338478454 | 0.2345751 | -1.442943 | 0.149036658 | 0.305371582 | ROCK1 | protein_coding | 36.92 | -0.174374413 | 0.174374413 | ENSG00000067900 | n.s. |
| ENSG00000245534 | 56.27095414 | -0.360290353 | 0.2596885 | -1.387394 | 0.165321512 | 0.328261752 | RORA-AS1 | antisense | 40.18 | -0.174301165 | 0.174301165 | ENSG00000245534 | n.s. |
| ENSG00000101460 | 64.50860602 | 0.415668468 | 0.327642 | 1.268667 | 0.204560007 | 0.381545992 | MAP1LC3A | protein_coding | 52.56 | 0.173937761 | 0.173937761 | ENSG00000101460 | n.s. |
| ENSG00000241468 | 2188.83651 | 0.420719883 | 0.334958 | 1.256038 | 0.209102252 | 0.386374341 | ATP5J2 | protein_coding | 53.67 | 0.173753829 | 0.173753829 | ENSG00000241468 | n.s. |
| ENSG00000126581 | 1348.365762 | -0.362495355 | 0.262789 | -1.379416 | 0.167766562 | 0.331697506 | BECN1 | protein_coding | 44.97 | -0.173728724 | 0.173728724 | ENSG00000126581 | n.s. |
| ENSG00000189136 | 181.0939793 | 0.340230768 | 0.2373102 | 1.433696 | 0.151659008 | 0.308841358 | UBE2Q2P1 | transcribed_unprocessed_pseud | 39.14 | 0.173607698 | 0.173607698 | ENSG00000189136 | n.s. |
| ENSG00000131931 | 363.140065 | 0.334799137 | 0.2309374 | 1.44974 | 0.147131043 | 0.30312304 | THAP1 | protein_coding | 41.03 | 0.173553529 | 0.173553529 | ENSG00000131931 | n.s. |
| ENSG00000113522 | 215.1820562 | -0.338670176 | 0.2355838 | -1.437578 | 0.150553805 | 0.30730987 | RAD50 | protein_coding | 38.81 | -0.173542554 | 0.173542554 | ENSG00000113522 | n.s. |
| ENSG00000134323 | 48939.41536 | 0.31523913 | 0.2097706 | 1.50278 | 0.132895704 | 0.281666044 | MYCN | protein_coding | 53.76 | 0.17346522 | 0.17346522 | ENSG00000134323 | n.s. |
| ENSG00000196821 | 584.6807713 | -0.356232385 | 0.2557326 | -1.392988 | 0.163623466 | 0.325912664 | C6orf106 | protein_coding | 41.83 | -0.173449108 | 0.173449108 | ENSG00000196821 | n.s. |
| ENSG00000047578 | 46.58538996 | -0.398862458 | 0.3069275 | -1.299533 | 0.193761047 | 0.367438717 | KIAA0556 | protein_coding | 47.84 | -0.173431413 | 0.173431413 | ENSG00000047578 | n.s. |
| ENSG00000126777 | 5218.033323 | -0.359508839 | 0.2597674 | -1.383964 | 0.16636943 | 0.329543544 | KTN1 | protein_coding | 37.04 | -0.173314607 | 0.173314607 | ENSG00000126777 | n.s. |
| ENSG00000169612 | 409.830947 | 0.376602115 | 0.2801755 | 1.344165 | 0.178895075 | 0.346998625 | FAM103A1 | protein_coding | 41.1 | 0.173113541 | 0.173113541 | ENSG00000169612 | n.s. |
| ENSG00000080854 | 68.81996313 | -0.386325947 | 0.2919688 | -1.323176 | 0.185777 | 0.3564737 | IGSF9B | protein_coding | 56.66 | -0.173063402 | 0.173063402 | ENSG00000080854 | n.s. |
| ENSG00000125885 | 2791.654905 | -0.34505834 | 0.2433051 | -1.418213 | 0.156128718 | 0.315131942 | MCM8 | protein_coding | 39.4 | -0.173049371 | 0.173049371 | ENSG00000125885 | n.s. |
| ENSG00000188321 | 578.3858147 | -0.392390043 | 0.2994113 | -1.310538 | 0.190013759 | 0.362455346 | ZNF559 | protein_coding | 40.31 | -0.172944142 | 0.172944142 | ENSG00000188321 | n.s. |
| ENSG00000164932 | 72.97588133 | 0.372811075 | 0.2756689 | 1.352387 | 0.176251494 | 0.343656347 | CTHRC1 | protein_coding | 39.94 | 0.172937973 | 0.172937973 | ENSG00000164932 | n.s. |
| ENSG00000174021 | 675.5339486 | 0.368359109 | 0.270531 | 1.361615 | 0.173319344 | 0.339501233 | GNG5 | protein_coding | 40.66 | 0.17281886 | 0.17281886 | ENSG00000174021 | n.s. |
| ENSG00000148408 | 805.0770748 | -0.343708754 | 0.2419333 | -1.420676 | 0.155410996 | 0.31422993 | CACNA1B | protein_coding | 49.94 | -0.172800418 | 0.172800418 | ENSG00000148408 | n.s. |
| ENSG00000226137 | 194.0733236 | -0.354100219 | 0.2537608 | -1.39541 | 0.162892353 | 0.325247417 | BAIAP2-AS1 | lincRNA | 59.08 | -0.172725181 | 0.172725181 | ENSG00000226137 | n.s. |
| ENSG00000106399 | 1045.326804 | 0.352401936 | 0.2518363 | 1.399329 | 0.161714294 | 0.323498293 | RPA3 | protein_coding | 37.19 | 0.172722059 | 0.172722059 | ENSG00000106399 | n.s. |
| ENSG00000138604 | 1771.125533 | -0.364009119 | 0.2655528 | -1.37076 | 0.170449748 | 0.335443596 | GLCE | protein_coding | 36.64 | -0.172678826 | 0.172678826 | ENSG00000138604 | n.s. |
| ENSG00000141349 | 686.5494827 | 0.306691058 | 0.2015344 | 1.52178 | 0.128064098 | 0.27355576 | G6PC3 | protein_coding | 53.16 | 0.172653 | 0.172653 | ENSG00000141349 | n.s. |
| ENSG00000133393 | 978.1192778 | -0.413723128 | 0.3269396 | -1.265442 | 0.205712925 | 0.383009756 | FOPNL | protein_coding | 41.7 | -0.17243573 | 0.17243573 | ENSG00000133393 | n.s. |
| ENSG00000159873 | 780.075138 | -0.353857424 | 0.2539584 | -1.393368 | 0.163508593 | 0.325869279 | CCDC117 | protein_coding | 44.33 | -0.172313201 | 0.172313201 | ENSG00000159873 | n.s. |
| ENSG00000134824 | 2414.949395 | -0.365495877 | 0.2676846 | -1.365398 | 0.172128091 | 0.337744242 | FADS2 | protein_coding | 51.23 | -0.172299159 | 0.172299159 | ENSG00000134824 | n.s. |
| ENSG00000006015 | 327.7047876 | 0.411601332 | 0.3243231 | 1.269109 | 0.204402226 | 0.381435758 | C19orf60 | protein_coding | 60.24 | 0.172287508 | 0.172287508 | ENSG00000006015 | n.s. |
| ENSG00000128482 | 339.9292404 | -0.369060953 | 0.2718858 | -1.357412 | 0.174650406 | 0.341471317 | RNF112 | protein_coding | 60.03 | -0.172220733 | 0.172220733 | ENSG00000128482 | n.s. |
| ENSG00000273568 | 62.24601903 | -0.385036399 | 0.2913499 | -1.32156 | 0.186314649 | 0.357259173 | | sense_intronic | 53.66 | -0.172117666 | 0.172117666 | ENSG00000273568 | n.s. |
| ENSG00000132471 | 314.1050212 | -0.371701743 | 0.2756333 | -1.348537 | 0.177485721 | 0.345275073 | WBP2 | protein_coding | 55.06 | -0.171664791 | 0.171664791 | ENSG00000132471 | n.s. |
| ENSG00000143947 | 18638.53327 | 0.404958414 | 0.3171748 | 1.276767 | 0.201684454 | 0.377649 | RPS27A | protein_coding | 45.81 | 0.171261635 | 0.171261635 | ENSG00000143947 | n.s. |
| ENSG00000109255 | 132.8083313 | 0.406244279 | 0.3188801 | 1.273972 | 0.202673386 | 0.378818842 | NMU | protein_coding | 38.09 | 0.171259762 | 0.171259762 | ENSG00000109255 | n.s. |

| | | | | | | | | | | | | | |
|---|---|---|---|---|---|---|---|---|---|---|---|---|---|
| ENSG00000101098 | 265.9676464 | -0.342616199 | 0.2421662 | -1.414798 | 0.157127837 | 0.316368891 | RIMS4 | protein_coding | 49.54 | -0.17124171 | 0.17124171 | ENSG00000101098 | n.s. |
| ENSG00000254172 | 4.044169025 | -0.566054934 | 0.5563476 | -1.017448 | 0.30894019 | 0.498767888 | RNU5A-3P | snRNA | 36.84 | -0.171006053 | 0.171006053 | ENSG00000254172 | n.s. |
| ENSG00000108091 | 837.1407379 | -0.330546899 | 0.2285383 | -1.446352 | 0.148078473 | 0.304130383 | CCDC6 | protein_coding | 40.92 | -0.170872977 | 0.170872977 | ENSG00000108091 | n.s. |
| ENSG00000120437 | 2506.57235 | -0.374795204 | 0.280254 | -1.337341 | 0.1811113 | 0.350126719 | ACAT2 | protein_coding | 44.57 | -0.170822189 | 0.170822189 | ENSG00000120437 | n.s. |
| ENSG00000150760 | 1367.87374 | -0.350264016 | 0.2511799 | -1.394475 | 0.16317433 | 0.325532268 | DOCK1 | protein_coding | 45.16 | -0.170720768 | 0.170720768 | ENSG00000150760 | n.s. |
| ENSG00000052795 | 1001.254128 | -0.329232947 | 0.2272058 | -1.449052 | 0.147323071 | 0.303196555 | FNIP2 | protein_coding | 40.56 | -0.170633448 | 0.170633448 | ENSG00000052795 | n.s. |
| ENSG00000229676 | 510.6048828 | -0.360125023 | 0.2630378 | -1.3691 | 0.170968001 | 0.33616544 | ZNF492 | protein_coding | 39.03 | -0.170500089 | 0.170500089 | ENSG00000229676 | n.s. |
| ENSG00000239900 | 507.8310846 | -0.347310882 | 0.2479729 | -1.4006 | 0.161333728 | 0.322922584 | ADSL | protein_coding | 44.97 | -0.170495461 | 0.170495461 | ENSG00000239900 | n.s. |
| ENSG00000025796 | 2698.092726 | -0.349634512 | 0.2507903 | -1.394131 | 0.163278107 | 0.325640425 | SEC63 | protein_coding | 39.82 | -0.170363502 | 0.170363502 | ENSG00000025796 | n.s. |
| ENSG00000148498 | 1297.173284 | -0.357781196 | 0.2604375 | -1.37377 | 0.169513078 | 0.334189737 | PARD3 | protein_coding | 40.01 | -0.170306315 | 0.170306315 | ENSG00000148498 | n.s. |
| ENSG00000131374 | 1314.255552 | -0.34305514 | 0.2435412 | -1.408613 | 0.158949764 | 0.319436715 | TBC1D5 | protein_coding | 37.62 | -0.170023331 | 0.170023331 | ENSG00000131374 | n.s. |
| ENSG00000278462 | 71.7731517 | -0.361752033 | 0.2654639 | -1.362717 | 0.172971834 | 0.339052068 | | sense_intronic | 32.33 | -0.169927086 | 0.169927086 | ENSG00000278462 | n.s. |
| ENSG00000039319 | 1186.272981 | -0.358098681 | 0.2611702 | -1.371132 | 0.170333959 | 0.335344982 | ZFYVE16 | protein_coding | 37.65 | -0.169920756 | 0.169920756 | ENSG00000039319 | n.s. |
| ENSG00000215114 | 615.5595249 | -0.355390929 | 0.2581441 | -1.376715 | 0.168600307 | 0.332750674 | UBXN2B | protein_coding | 38.03 | -0.169834593 | 0.169834593 | ENSG00000215114 | n.s. |
| ENSG00000065882 | 631.5735734 | -0.364407343 | 0.2687318 | -1.356026 | 0.175090903 | 0.342003722 | TBC1D1 | protein_coding | 42.84 | -0.169802586 | 0.169802586 | ENSG00000065882 | n.s. |
| ENSG00000283149 | 26.86858723 | 0.438929098 | 0.3644929 | 1.204218 | 0.228505188 | 0.410407344 | | protein_coding | 41.04 | 0.169771137 | 0.169771137 | ENSG00000283149 | n.s. |
| ENSG00000251050 | 13.26263293 | -0.585686155 | 0.5919119 | -0.989482 | 0.322427371 | 0.51309452 | | processed_pseudogene | 47.44 | -0.169733384 | 0.169733384 | ENSG00000251050 | n.s. |
| ENSG00000118579 | 2068.12407 | -0.362688644 | 0.2667802 | -1.359504 | 0.173987093 | 0.340601248 | MED28 | protein_coding | 44.87 | -0.169648979 | 0.169648979 | ENSG00000118579 | n.s. |
| ENSG00000146676 | 1803.302625 | -0.338241616 | 0.2385219 | -1.418073 | 0.156169334 | 0.315168234 | PURB | protein_coding | 40.58 | -0.169613816 | 0.169613816 | ENSG00000146676 | n.s. |
| ENSG00000111450 | 1103.054603 | -0.366058842 | 0.2709216 | -1.351162 | 0.176643679 | 0.34426231 | STX2 | protein_coding | 45.87 | -0.169525701 | 0.169525701 | ENSG00000111450 | n.s. |
| ENSG00000151779 | 49123.49551 | -0.349398483 | 0.2519421 | -1.386821 | 0.165496467 | 0.328468611 | NBAS | protein_coding | 38.12 | -0.168936307 | 0.168936307 | ENSG00000151779 | n.s. |
| ENSG00000075975 | 966.7105253 | 0.361183091 | 0.2658132 | 1.358785 | 0.174214639 | 0.340768534 | MKRN2 | protein_coding | 45.18 | 0.168867728 | 0.168867728 | ENSG00000075975 | n.s. |
| ENSG00000106615 | 1541.270899 | 0.380870185 | 0.2897435 | 1.314508 | 0.188675215 | 0.360395805 | RHEB | protein_coding | 42 | 0.168809388 | 0.168809388 | ENSG00000106615 | n.s. |
| ENSG00000153048 | 2612.692593 | 0.358858027 | 0.2634028 | 1.362393 | 0.173073986 | 0.339156805 | CARHSP1 | protein_coding | 55.97 | 0.168519537 | 0.168519537 | ENSG00000153048 | n.s. |
| ENSG00000156467 | 2796.008845 | 0.395095671 | 0.3077016 | 1.284022 | 0.199134172 | 0.374668254 | UQCRB | protein_coding | 38.18 | 0.168450265 | 0.168450265 | ENSG00000156467 | n.s. |
| ENSG00000162736 | 620.2450719 | -0.362274234 | 0.2675934 | -1.353823 | 0.175792768 | 0.342988047 | NCSTN | protein_coding | 46.85 | -0.16835645 | 0.16835645 | ENSG00000162736 | n.s. |
| ENSG00000056097 | 3100.534794 | -0.365693689 | 0.2719248 | -1.344834 | 0.178678871 | 0.346728182 | ZFR | protein_coding | 38.33 | -0.168223067 | 0.168223067 | ENSG00000056097 | n.s. |
| ENSG00000124789 | 3435.594219 | -0.335774188 | 0.2369663 | -1.41697 | 0.156491694 | 0.315544368 | NUP153 | protein_coding | 40.18 | -0.168202576 | 0.168202576 | ENSG00000124789 | n.s. |
| ENSG00000196562 | 2372.158169 | -0.363299118 | 0.2690308 | -1.3504 | 0.176887771 | 0.344545108 | SULF2 | protein_coding | 50.48 | -0.168118088 | 0.168118088 | ENSG00000196562 | n.s. |
| ENSG00000144120 | 374.0961464 | -0.345982391 | 0.2488747 | -1.390187 | 0.164472065 | 0.327119714 | TMEM177 | protein_coding | 52.47 | -0.16790293 | 0.16790293 | ENSG00000144120 | n.s. |
| ENSG00000233822 | 606.3572301 | 0.309121121 | 0.2072776 | 1.491339 | 0.135872497 | 0.286396701 | HIST1H2BN | protein_coding | 40.18 | 0.167862657 | 0.167862657 | ENSG00000233822 | n.s. |
| ENSG00000255310 | 187.9891132 | -0.372819167 | 0.2809223 | -1.327125 | 0.184467184 | 0.354640458 | | sense_intronic | 42.08 | -0.167847559 | 0.167847559 | ENSG00000255310 | n.s. |

| | | | | | | | | | | | | | |
|---|---|---|---|---|---|---|---|---|---|---|---|---|---|
| ENSG00000172869 | 556.9112222 | -0.348562337 | 0.2520239 | -1.383053 | 0.166648639 | 0.330002703 | DMXL1 | protein_coding | 37.58 | -0.167826666 | 0.167826666 | ENSG00000172869 | n.s. |
| ENSG00000104872 | 572.2537995 | 0.387204108 | 0.2989191 | 1.295347 | 0.195200419 | 0.369112514 | PIH1D1 | protein_coding | 55.81 | 0.167597903 | 0.167597903 | ENSG00000104872 | n.s. |
| ENSG00000243679 | 64.74553655 | -0.375797179 | 0.2847989 | -1.319518 | 0.186996046 | 0.358220411 | | processed_pseudogene | 67.24 | -0.167549049 | 0.167549049 | ENSG00000243679 | n.s. |
| ENSG00000115414 | 89.26406204 | 0.361320565 | 0.2676329 | 1.35006 | 0.176996666 | 0.344708991 | FN1 | protein_coding | 39.44 | 0.167127884 | 0.167127884 | ENSG00000115414 | n.s. |
| ENSG00000169618 | 201.0436403 | -0.369979604 | 0.2782983 | -1.329435 | 0.183704368 | 0.353568693 | PROKR1 | protein_coding | 46.47 | -0.167055484 | 0.167055484 | ENSG00000169618 | n.s. |
| ENSG00000124608 | 914.9536388 | -0.327507332 | 0.2284896 | -1.433358 | 0.151755593 | 0.308992853 | AARS2 | protein_coding | 54.66 | -0.167045628 | 0.167045628 | ENSG00000124608 | n.s. |
| ENSG00000164830 | 567.9010557 | -0.385001163 | 0.2970432 | -1.296111 | 0.194937084 | 0.368733506 | OXR1 | protein_coding | 36.58 | -0.166816152 | 0.166816152 | ENSG00000164830 | n.s. |
| ENSG00000237172 | 116.1169599 | -0.333763173 | 0.2359721 | -1.414418 | 0.157239287 | 0.316501749 | B3GNT9 | protein_coding | 62.77 | -0.166756054 | 0.166756054 | ENSG00000237172 | n.s. |
| ENSG00000158092 | 445.0559937 | -0.365945353 | 0.2740053 | -1.335541 | 0.181699339 | 0.350793455 | NCK1 | protein_coding | 39.43 | -0.166486296 | 0.166486296 | ENSG00000158092 | n.s. |
| ENSG00000261188 | 309.3318113 | 0.309386563 | 0.2087241 | 1.482275 | 0.138267023 | 0.28974891 | | antisense | 42.72 | 0.166443223 | 0.166443223 | ENSG00000261188 | n.s. |
| ENSG00000086015 | 282.6483182 | -0.332386798 | 0.2347147 | -1.416131 | 0.156737179 | 0.315888638 | MAST2 | protein_coding | 41.32 | -0.166348289 | 0.166348289 | ENSG00000086015 | n.s. |
| ENSG00000249637 | 30.01931987 | 0.420547512 | 0.3448321 | 1.219572 | 0.222627237 | 0.402319608 | | antisense | 44.25 | 0.166296598 | 0.166296598 | ENSG00000249637 | n.s. |
| ENSG00000120733 | 1777.896412 | -0.336078775 | 0.2389131 | -1.406699 | 0.159516715 | 0.320264977 | KDM3B | protein_coding | 43.18 | -0.166187779 | 0.166187779 | ENSG00000120733 | n.s. |
| ENSG00000112149 | 159.4530606 | 0.366941687 | 0.2755472 | 1.331684 | 0.182964122 | 0.352777935 | CD83 | protein_coding | 45.18 | 0.166040595 | 0.166040595 | ENSG00000112149 | n.s. |
| ENSG00000109736 | 369.4297225 | 0.340397753 | 0.244344 | 1.393109 | 0.163586897 | 0.325886445 | MFSD10 | protein_coding | 67.55 | 0.165751138 | 0.165751138 | ENSG00000109736 | n.s. |
| ENSG00000130312 | 446.690594 | 0.404559124 | 0.3237717 | 1.24952 | 0.211475064 | 0.389312087 | MRPL34 | protein_coding | 54.25 | 0.165748728 | 0.165748728 | ENSG00000130312 | n.s. |
| ENSG00000071051 | 1224.965542 | -0.359858428 | 0.2672941 | -1.346302 | 0.17820529 | 0.346287822 | NCK2 | protein_coding | 44.54 | -0.165737398 | 0.165737398 | ENSG00000071051 | n.s. |
| ENSG00000207508 | 6.958669052 | -0.523592299 | 0.498145 | -1.051084 | 0.293219953 | 0.482508144 | RNU6-1237P | snRNA | 38.32 | -0.165714529 | 0.165714529 | ENSG00000207508 | n.s. |
| ENSG00000183011 | 536.9129511 | 0.395796464 | 0.3118267 | 1.269283 | 0.204340061 | 0.381408866 | NAA38 | protein_coding | 44.28 | 0.165684047 | 0.165684047 | ENSG00000183011 | n.s. |
| ENSG00000172943 | 389.487904 | -0.335447249 | 0.238684 | -1.405403 | 0.15990147 | 0.320747467 | PHF8 | protein_coding | 43.07 | -0.165656184 | 0.165656184 | ENSG00000172943 | n.s. |
| ENSG00000156875 | 229.5673252 | 0.34992032 | 0.2556248 | 1.368883 | 0.171035992 | 0.336236689 | MFSD14A | protein_coding | 39.5 | 0.165636498 | 0.165636498 | ENSG00000156875 | n.s. |
| ENSG00000131242 | 52.87360607 | 0.3604056 | 0.2683338 | 1.343124 | 0.179231814 | 0.347397896 | RAB11FIP4 | protein_coding | 51.38 | 0.165488454 | 0.165488454 | ENSG00000131242 | n.s. |
| ENSG00000100099 | 225.0248962 | 0.313075287 | 0.2135104 | 1.466323 | 0.142560188 | 0.296218154 | HPS4 | protein_coding | 46.81 | 0.165425327 | 0.165425327 | ENSG00000100099 | n.s. |
| ENSG00000115268 | 6226.372564 | 0.365821344 | 0.2748362 | 1.331052 | 0.183171791 | 0.353031678 | RPS15 | protein_coding | 62.7 | 0.165419408 | 0.165419408 | ENSG00000115268 | n.s. |
| ENSG00000178343 | 116.6805026 | 0.355556407 | 0.262495 | 1.354527 | 0.175568422 | 0.342646361 | SHISA3 | protein_coding | 50.29 | 0.165388441 | 0.165388441 | ENSG00000178343 | n.s. |
| ENSG00000112578 | 658.6022278 | -0.342928135 | 0.2477357 | -1.38425 | 0.166281972 | 0.329510945 | BYSL | protein_coding | 48.76 | -0.165335995 | 0.165335995 | ENSG00000112578 | n.s. |
| ENSG00000169020 | 1428.303292 | 0.410828426 | 0.3328687 | 1.234206 | 0.217126219 | 0.39599795 | ATP5I | protein_coding | 57.33 | 0.165279177 | 0.165279177 | ENSG00000169020 | n.s. |
| ENSG00000125818 | 437.3961437 | -0.359346104 | 0.2672663 | -1.344525 | 0.17877875 | 0.34682131 | PSMF1 | protein_coding | 45.44 | -0.165261198 | 0.165261198 | ENSG00000125818 | n.s. |
| ENSG00000212534 | 102.1630044 | 0.338084005 | 0.2420345 | 1.396842 | 0.162460957 | 0.32466541 | SNORD70 | snoRNA | 37.5 | 0.165175667 | 0.165175667 | ENSG00000212534 | n.s. |
| ENSG00000123575 | 439.1967147 | -0.367902652 | 0.2777321 | -1.324667 | 0.185281485 | 0.355718991 | FAM199X | protein_coding | 36.81 | -0.165148906 | 0.165148906 | ENSG00000123575 | n.s. |
| ENSG00000093144 | 2147.470057 | -0.341269458 | 0.2459268 | -1.387687 | 0.165232305 | 0.328152674 | ECHDC1 | protein_coding | 36 | -0.165148499 | 0.165148499 | ENSG00000093144 | n.s. |
| ENSG00000146094 | 26.13924754 | -0.398740595 | 0.3169163 | -1.258189 | 0.208323448 | 0.385411927 | DOK3 | protein_coding | 58.13 | -0.165108452 | 0.165108452 | ENSG00000146094 | n.s. |

| | | | | | | | | | | | | | |
|---|---|---|---|---|---|---|---|---|---|---|---|---|---|
| ENSG00000163166 | 2439.261667 | -0.347620512 | 0.2534423 | -1.371596 | 0.1701891 | 0.335107176 | IWS1 | protein_coding | 41.3 | -0.165055876 | 0.165055876 | ENSG00000163166 | n.s. |
| ENSG00000162642 | 836.3891782 | -0.359539562 | 0.2677869 | -1.342633 | 0.179390742 | 0.347572682 | C1orf52 | protein_coding | 39.66 | -0.165012251 | 0.165012251 | ENSG00000162642 | n.s. |
| ENSG00000108061 | 813.8593832 | -0.367337971 | 0.2771644 | -1.325343 | 0.185057258 | 0.355533627 | SHOC2 | protein_coding | 35.08 | -0.164978578 | 0.164978578 | ENSG00000108061 | n.s. |
| ENSG00000136279 | 723.1388468 | -0.367040642 | 0.2770277 | -1.324924 | 0.185196401 | 0.355604674 | DBNL | protein_coding | 53.2 | -0.164813191 | 0.164813191 | ENSG00000136279 | n.s. |
| ENSG00000125743 | 3779.844749 | 0.401385956 | 0.3209084 | 1.25078 | 0.21101462 | 0.38874967 | SNRPD2 | protein_coding | 52.25 | 0.164700686 | 0.164700686 | ENSG00000125743 | n.s. |
| ENSG00000166260 | 2640.431369 | -0.364147702 | 0.2735422 | -1.331231 | 0.183113175 | 0.353008414 | COX11 | protein_coding | 41.57 | -0.164673032 | 0.164673032 | ENSG00000166260 | n.s. |
| ENSG00000114767 | 763.3018075 | 0.376684609 | 0.289126 | 1.302839 | 0.192629689 | 0.365641875 | RRP9 | protein_coding | 55.15 | 0.164590107 | 0.164590107 | ENSG00000114767 | n.s. |
| ENSG00000104447 | 83.07315417 | 0.355366962 | 0.2629933 | 1.35124 | 0.176618675 | 0.344261768 | TRPS1 | protein_coding | 35.81 | 0.164574422 | 0.164574422 | ENSG00000104447 | n.s. |
| ENSG00000151327 | 423.6421265 | -0.380563288 | 0.2939929 | -1.294464 | 0.195505063 | 0.36948785 | FAM177A1 | protein_coding | 41.39 | -0.164555504 | 0.164555504 | ENSG00000151327 | n.s. |
| ENSG00000169398 | 1950.656787 | -0.356244071 | 0.2641931 | -1.348423 | 0.177522335 | 0.345298055 | PTK2 | protein_coding | 40.07 | -0.164515603 | 0.164515603 | ENSG00000169398 | n.s. |
| ENSG00000225138 | 199.0088177 | 0.365197778 | 0.2750715 | 1.327647 | 0.184294844 | 0.354460162 | SLC9A3-AS1 | processed_transcript | 65.93 | 0.164496973 | 0.164496973 | ENSG00000225138 | n.s. |
| ENSG00000201644 | 8.985774791 | 0.495967276 | 0.4570632 | 1.085118 | 0.277869588 | 0.465952991 | | misc_RNA | 36.17 | 0.164491463 | 0.164491463 | ENSG00000201644 | n.s. |
| ENSG00000197498 | 1343.450238 | -0.360358507 | 0.2692877 | -1.338191 | 0.180834082 | 0.349736364 | RPF2 | protein_coding | 42.58 | -0.164416887 | 0.164416887 | ENSG00000197498 | n.s. |
| ENSG00000167468 | 1203.431843 | 0.405819725 | 0.3271121 | 1.240614 | 0.214748462 | 0.393638487 | GPX4 | protein_coding | 69.47 | 0.164317399 | 0.164317399 | ENSG00000167468 | n.s. |
| ENSG00000227383 | 33.13304208 | 0.395799715 | 0.3140613 | 1.260263 | 0.207574588 | 0.384662305 | | processed_pseudogene | 42.81 | 0.164225364 | 0.164225364 | ENSG00000227383 | n.s. |
| ENSG00000212309 | 11.40136941 | 0.592577867 | 0.6163902 | 0.961368 | 0.336367145 | 0.528463074 | | snoRNA | 36.47 | 0.16413539 | 0.16413539 | ENSG00000212309 | n.s. |
| ENSG00000174684 | 265.3226555 | 0.378507312 | 0.2918788 | 1.296796 | 0.194701354 | 0.368619384 | B4GAT1 | protein_coding | 57.6 | 0.164053332 | 0.164053332 | ENSG00000174684 | n.s. |
| ENSG00000124222 | 513.9250769 | -0.344275644 | 0.2504916 | -1.3744 | 0.169317498 | 0.333863125 | STX16 | protein_coding | 41.01 | -0.164023778 | 0.164023778 | ENSG00000124222 | n.s. |
| ENSG00000031003 | 1092.948436 | -0.343067179 | 0.2492514 | -1.37639 | 0.168700951 | 0.332788798 | FAM13B | protein_coding | 40.05 | -0.163928237 | 0.163928237 | ENSG00000031003 | n.s. |
| ENSG00000275835 | 928.4274617 | -0.355922909 | 0.2643584 | -1.346365 | 0.17818474 | 0.346287822 | TUBGCP5 | protein_coding | 43.36 | -0.163924845 | 0.163924845 | ENSG00000275835 | n.s. |
| ENSG00000137815 | 1308.930789 | -0.348441684 | 0.2555746 | -1.363366 | 0.172767249 | 0.338698734 | RTF1 | protein_coding | 45.65 | -0.16383255 | 0.16383255 | ENSG00000137815 | n.s. |
| ENSG00000115109 | 620.9683409 | 0.343334674 | 0.2497771 | 1.374564 | 0.169266617 | 0.333810072 | EPB41L5 | protein_coding | 38.5 | 0.163599166 | 0.163599166 | ENSG00000115109 | n.s. |
| ENSG00000077348 | 478.1647042 | 0.377954148 | 0.2919691 | 1.2945 | 0.195492632 | 0.36948785 | EXOSC5 | protein_coding | 52.53 | 0.163427312 | 0.163427312 | ENSG00000077348 | n.s. |
| ENSG00000169967 | 2581.441025 | -0.3458995 | 0.2530199 | -1.367084 | 0.171598974 | 0.337025323 | MAP3K2 | protein_coding | 38.69 | -0.163381293 | 0.163381293 | ENSG00000169967 | n.s. |
| ENSG00000105186 | 498.9186954 | -0.344331229 | 0.2512456 | -1.370497 | 0.17053192 | 0.33554493 | ANKRD27 | protein_coding | 48.98 | -0.163298851 | 0.163298851 | ENSG00000105186 | n.s. |
| ENSG00000113657 | 820.5002591 | -0.346104339 | 0.2534045 | -1.365818 | 0.171996188 | 0.337577696 | DPYSL3 | protein_coding | 41.43 | -0.163231893 | 0.163231893 | ENSG00000113657 | n.s. |
| ENSG00000218175 | 110.7659782 | 0.514072517 | 0.4878229 | 1.05381 | 0.291970055 | 0.481408223 | | processed_pseudogene | 53.91 | 0.163211082 | 0.163211082 | ENSG00000218175 | n.s. |
| ENSG00000113119 | 112.1315057 | -0.348877784 | 0.2566332 | -1.359441 | 0.1740068 | 0.340601248 | TMCO6 | protein_coding | 51.35 | -0.163188897 | 0.163188897 | ENSG00000113119 | n.s. |
| ENSG00000099999 | 106.3408035 | -0.371010198 | 0.2835784 | -1.308316 | 0.190766116 | 0.363691173 | RNF215 | protein_coding | 53.32 | -0.162972626 | 0.162972626 | ENSG00000099999 | n.s. |
| ENSG00000171703 | 140.2636461 | 0.38475328 | 0.301135 | 1.277677 | 0.201363386 | 0.37727551 | TCEA2 | protein_coding | 61.58 | 0.162881987 | 0.162881987 | ENSG00000171703 | n.s. |
| ENSG00000120662 | 195.3984155 | -0.339027456 | 0.2454778 | -1.381092 | 0.167250579 | 0.330912295 | MTRF1 | protein_coding | 39.67 | -0.162830513 | 0.162830513 | ENSG00000120662 | n.s. |
| ENSG00000153975 | 815.0980103 | -0.350907928 | 0.2594382 | -1.352568 | 0.176193559 | 0.343625568 | ZUFSP | protein_coding | 36.54 | -0.162791286 | 0.162791286 | ENSG00000153975 | n.s. |

| | | | | | | | | | | | | | |
|---|---|---|---|---|---|---|---|---|---|---|---|---|---|
| ENSG00000095139 | 1097.322689 | -0.363866393 | 0.2752833 | -1.321789 | 0.186238411 | 0.357162175 | ARCN1 | protein_coding | 42.33 | -0.162697232 | 0.162697232 | ENSG00000095139 | n.s. |
| ENSG00000174010 | 1157.654535 | 0.364828332 | 0.2765067 | 1.31942 | 0.187028889 | 0.358234037 | KLHL15 | protein_coding | 41.85 | 0.162652566 | 0.162652566 | ENSG00000174010 | n.s. |
| ENSG00000274591 | 67.78608075 | -0.371420845 | 0.2845241 | -1.305411 | 0.191753019 | 0.364873211 | | sense_intronic | 35.05 | -0.162629596 | 0.162629596 | ENSG00000274591 | n.s. |
| ENSG00000233098 | 114.0213988 | -0.403921644 | 0.3270302 | -1.23512 | 0.216785766 | 0.395791794 | CCDC144NL-AS1 | antisense | 43.23 | -0.162591878 | 0.162591878 | ENSG00000233098 | n.s. |
| ENSG00000253284 | 377.1865243 | -0.358133774 | 0.2685126 | -1.333769 | 0.182279519 | 0.351652729 | | sense_intronic | 38.08 | -0.162551909 | 0.162551909 | ENSG00000253284 | n.s. |
| ENSG00000128159 | 565.3194494 | 0.339243976 | 0.2460386 | 1.378824 | 0.167948887 | 0.331907759 | TUBGCP6 | protein_coding | 56.93 | 0.16249196 | 0.16249196 | ENSG00000128159 | n.s. |
| ENSG00000078070 | 571.1048391 | -0.357919771 | 0.2683444 | -1.333808 | 0.182266938 | 0.351652729 | MCCC1 | protein_coding | 41.62 | -0.162454776 | 0.162454776 | ENSG00000078070 | n.s. |
| ENSG00000074935 | 2028.431277 | -0.337661215 | 0.2442491 | -1.382446 | 0.166834724 | 0.330324213 | TUBE1 | protein_coding | 34.64 | -0.162435166 | 0.162435166 | ENSG00000074935 | n.s. |
| ENSG00000117016 | 1799.845113 | -0.347471967 | 0.2558018 | -1.358364 | 0.17434824 | 0.340934066 | RIMS3 | protein_coding | 51.39 | -0.162383936 | 0.162383936 | ENSG00000117016 | n.s. |
| ENSG00000109118 | 402.7417077 | -0.320213022 | 0.2241857 | -1.428338 | 0.153194553 | 0.311194596 | PHF12 | protein_coding | 46.18 | -0.16233774 | 0.16233774 | ENSG00000109118 | n.s. |
| ENSG00000199135 | 17.92253874 | -0.444243905 | 0.3824398 | -1.161605 | 0.245396095 | 0.431231836 | MIR101-1 | miRNA | 45.33 | -0.162277494 | 0.162277494 | ENSG00000199135 | n.s. |
| ENSG00000272455 | 83.09600314 | -0.345679237 | 0.2538765 | -1.361604 | 0.173322902 | 0.339501233 | | lincRNA | 49.44 | -0.162178402 | 0.162178402 | ENSG00000272455 | n.s. |
| ENSG00000166532 | 1787.033676 | -0.359438115 | 0.2705092 | -1.328746 | 0.183931714 | 0.353880085 | RIMKLB | protein_coding | 41.05 | -0.162158305 | 0.162158305 | ENSG00000166532 | n.s. |
| ENSG00000165661 | 776.3428903 | -0.358653539 | 0.2695463 | -1.330582 | 0.183326551 | 0.353100739 | QSOX2 | protein_coding | 54.55 | -0.162147757 | 0.162147757 | ENSG00000165661 | n.s. |
| ENSG00000231622 | 14.9359981 | 0.450220368 | 0.391512 | 1.149953 | 0.25016327 | 0.436469073 | RPS29P7 | processed_pseudogene | 49.71 | 0.162100278 | 0.162100278 | ENSG00000231622 | n.s. |
| ENSG00000124641 | 312.5736808 | -0.375795742 | 0.290804 | -1.292265 | 0.196265516 | 0.370522683 | MED20 | protein_coding | 46.57 | -0.162037562 | 0.162037562 | ENSG00000124641 | n.s. |
| ENSG00000164070 | 534.2708656 | -0.343152281 | 0.2510738 | -1.366739 | 0.171707209 | 0.337135305 | HSPA4L | protein_coding | 35.4 | -0.162035054 | 0.162035054 | ENSG00000164070 | n.s. |
| ENSG00000264268 | 5.454097444 | 0.542052895 | 0.5365915 | 1.010178 | 0.31241004 | 0.50255744 | MIR4767 | miRNA | 82.05 | 0.161973153 | 0.161973153 | ENSG00000264268 | n.s. |
| ENSG00000114739 | 2959.212307 | -0.339517739 | 0.24688 | -1.375234 | 0.169058879 | 0.333447623 | ACVR2B | protein_coding | 47.64 | -0.161940582 | 0.161940582 | ENSG00000114739 | n.s. |
| ENSG00000169981 | 940.7062831 | -0.344952975 | 0.2533698 | -1.36146 | 0.173368246 | 0.339542282 | ZNF35 | protein_coding | 42.6 | -0.161819557 | 0.161819557 | ENSG00000169981 | n.s. |
| ENSG00000145982 | 272.7596967 | -0.369543777 | 0.2831224 | -1.305244 | 0.191809871 | 0.364881656 | FARS2 | protein_coding | 41.44 | -0.161803992 | 0.161803992 | ENSG00000145982 | n.s. |
| ENSG00000228532 | 31.59861767 | 0.590607254 | 0.6183635 | 0.955113 | 0.339520345 | 0.532215916 | | processed_pseudogene | 45.49 | 0.161774498 | 0.161774498 | ENSG00000228532 | n.s. |
| ENSG00000110583 | 717.9068954 | -0.357155858 | 0.268107 | -1.332139 | 0.182814394 | 0.352538062 | NAA40 | protein_coding | 50.29 | -0.161718025 | 0.161718025 | ENSG00000110583 | n.s. |
| ENSG00000072958 | 429.1129027 | -0.36993605 | 0.2838126 | -1.303452 | 0.192420584 | 0.365569778 | AP1M1 | protein_coding | 51.59 | -0.161673047 | 0.161673047 | ENSG00000072958 | n.s. |
| ENSG00000200959 | 21.83317026 | 0.406534698 | 0.3321453 | 1.223966 | 0.220964866 | 0.400324506 | SNORA74A | snoRNA | 48.48 | 0.161633246 | 0.161633246 | ENSG00000200959 | n.s. |
| ENSG00000256092 | 67.77068092 | 0.351202736 | 0.2610425 | 1.345385 | 0.178500817 | 0.346571869 | | lincRNA | 59.97 | 0.161625849 | 0.161625849 | ENSG00000256092 | n.s. |
| ENSG00000103018 | 1497.987054 | -0.382150652 | 0.2993323 | -1.276677 | 0.20171628 | 0.377649 | CYB5B | protein_coding | 39.78 | -0.161615967 | 0.161615967 | ENSG00000103018 | n.s. |
| ENSG00000152990 | 2833.24219 | -0.354816746 | 0.2654584 | -1.336619 | 0.181346987 | 0.350388906 | ADGRA3 | protein_coding | 39.05 | -0.161601164 | 0.161601164 | ENSG00000152990 | n.s. |
| ENSG00000148158 | 798.165359 | -0.341041831 | 0.2491485 | -1.36883 | 0.171052537 | 0.336236689 | SNX30 | protein_coding | 44.26 | -0.161433822 | 0.161433822 | ENSG00000148158 | n.s. |
| ENSG00000108055 | 6987.15587 | -0.344789855 | 0.2538154 | -1.358428 | 0.174327993 | 0.340934066 | SMC3 | protein_coding | 36.86 | -0.161130506 | 0.161130506 | ENSG00000108055 | n.s. |
| ENSG00000188186 | 339.9000661 | 0.389228495 | 0.3094897 | 1.257646 | 0.208519856 | 0.385502849 | LAMTOR4 | protein_coding | 53.24 | 0.161129857 | 0.161129857 | ENSG00000188186 | n.s. |
| ENSG00000181751 | 1085.240654 | 0.361394101 | 0.2739581 | 1.319158 | 0.187116162 | 0.358253346 | C5orf30 | protein_coding | 37.54 | 0.161113012 | 0.161113012 | ENSG00000181751 | n.s. |

| | | | | | | | | | | | | | |
|---|---|---|---|---|---|---|---|---|---|---|---|---|---|
| ENSG00000127995 | 1759.58936 | -0.375699134 | 0.2915862 | -1.288467 | 0.19758347 | 0.372657093 | CASD1 | protein_coding | 34.66 | -0.16105869 | 0.16105869 | ENSG00000127995 | n.s. |
| ENSG00000100379 | 64.90161679 | 0.368416514 | 0.2826355 | 1.303504 | 0.192402762 | 0.365569778 | KCTD17 | protein_coding | 60.2 | 0.161008964 | 0.161008964 | ENSG00000100379 | n.s. |
| ENSG00000141552 | 1089.174328 | 0.400215737 | 0.3243434 | 1.233926 | 0.217230522 | 0.396032547 | ANAPC11 | protein_coding | 55.3 | 0.160994433 | 0.160994433 | ENSG00000141552 | n.s. |
| ENSG00000153707 | 114.0005764 | -0.343217743 | 0.2521282 | -1.361283 | 0.173424387 | 0.339589249 | PTPRD | protein_coding | 35.53 | -0.160984933 | 0.160984933 | ENSG00000153707 | n.s. |
| ENSG00000278259 | 241.7534989 | 0.302154144 | 0.2051036 | 1.473178 | 0.14070292 | 0.293308392 | MYO19 | protein_coding | 47.58 | 0.160950113 | 0.160950113 | ENSG00000278259 | n.s. |
| ENSG00000106346 | 453.4506143 | -0.303232342 | 0.2063159 | -1.469748 | 0.141630054 | 0.294667551 | USP42 | protein_coding | 45.58 | -0.160915606 | 0.160915606 | ENSG00000106346 | n.s. |
| ENSG00000085662 | 1749.615154 | 0.388625002 | 0.3090687 | 1.257407 | 0.208606476 | 0.385611785 | AKR1B1 | protein_coding | 50.04 | 0.160832341 | 0.160832341 | ENSG00000085662 | n.s. |
| ENSG00000252338 | 10.93022526 | -0.486426432 | 0.4495689 | -1.081984 | 0.27925959 | 0.467267308 | RNU6-503P | snRNA | 37.14 | -0.160732125 | 0.160732125 | ENSG00000252338 | n.s. |
| ENSG00000174740 | 113.452889 | 0.329052128 | 0.2357093 | 1.396008 | 0.162712084 | 0.324980684 | PABPC5 | protein_coding | 41.28 | 0.160624312 | 0.160624312 | ENSG00000174740 | n.s. |
| ENSG00000197601 | 2767.052678 | -0.357555916 | 0.2698308 | -1.325111 | 0.185134174 | 0.355604674 | FAR1 | protein_coding | 36.03 | -0.160554241 | 0.160554241 | ENSG00000197601 | n.s. |
| ENSG00000184900 | 1399.833244 | 0.398884765 | 0.3230599 | 1.234708 | 0.216939112 | 0.395916012 | SUMO3 | protein_coding | 54.29 | 0.160510006 | 0.160510006 | ENSG00000184900 | n.s. |
| ENSG00000198276 | 450.3200971 | 0.390446979 | 0.3117849 | 1.252296 | 0.210462007 | 0.388085178 | UCKL1 | protein_coding | 62.54 | 0.160502189 | 0.160502189 | ENSG00000198276 | n.s. |
| ENSG00000231767 | 90.933923 | 0.412955047 | 0.3421097 | 1.207084 | 0.227399842 | 0.408792574 | | processed_pseudogene | 42.92 | 0.160431798 | 0.160431798 | ENSG00000231767 | n.s. |
| ENSG00000173581 | 85.54136716 | 0.369444247 | 0.2845783 | 1.298217 | 0.19421294 | 0.367944857 | CCDC106 | protein_coding | 61.17 | 0.16041907 | 0.16041907 | ENSG00000173581 | n.s. |
| ENSG00000267637 | 13.45183885 | -0.442557995 | 0.3834416 | -1.154173 | 0.248429211 | 0.434517146 | | sense_intronic | 21.08 | -0.160202933 | 0.160202933 | ENSG00000267637 | n.s. |
| ENSG00000160097 | 732.0135936 | 0.372054992 | 0.2881965 | 1.290977 | 0.196711663 | 0.371125889 | FNDC5 | protein_coding | 54.01 | 0.160161767 | 0.160161767 | ENSG00000160097 | n.s. |
| ENSG00000142168 | 3698.377644 | 0.384748204 | 0.3045086 | 1.263505 | 0.206407673 | 0.383840528 | SOD1 | protein_coding | 44.51 | 0.159997222 | 0.159997222 | ENSG00000142168 | n.s. |
| ENSG00000144580 | 2314.338124 | -0.343105266 | 0.2528939 | -1.356716 | 0.174871331 | 0.341764956 | CNOT9 | protein_coding | 41.17 | -0.15998054 | 0.15998054 | ENSG00000144580 | n.s. |
| ENSG00000067048 | 3868.332996 | 0.328106061 | 0.2354561 | 1.393491 | 0.163471234 | 0.325842518 | DDX3Y | protein_coding | 37.7 | 0.159785108 | 0.159785108 | ENSG00000067048 | n.s. |
| ENSG00000230409 | 51.10064196 | 0.385843154 | 0.3066989 | 1.258052 | 0.208372898 | 0.385411927 | TCEA1P2 | processed_pseudogene | 41.72 | 0.159767946 | 0.159767946 | ENSG00000230409 | n.s. |
| ENSG00000176387 | 81.75776207 | 0.355792803 | 0.2685078 | 1.325074 | 0.185146483 | 0.355604674 | HSD11B2 | protein_coding | 59.84 | 0.159762546 | 0.159762546 | ENSG00000176387 | n.s. |
| ENSG00000270140 | 39.69048539 | -0.384830073 | 0.3052464 | -1.260719 | 0.207409987 | 0.384523202 | | sense_intronic | 40.27 | -0.159734285 | 0.159734285 | ENSG00000270140 | n.s. |
| ENSG00000150627 | 695.2388679 | -0.360008323 | 0.2737516 | -1.315091 | 0.188479175 | 0.360169579 | WDR17 | protein_coding | 34.28 | -0.159661161 | 0.159661161 | ENSG00000150627 | n.s. |
| ENSG00000164087 | 257.9824688 | -0.357653068 | 0.2711468 | -1.319039 | 0.187156177 | 0.35828069 | POC1A | protein_coding | 51.67 | -0.159433368 | 0.159433368 | ENSG00000164087 | n.s. |
| ENSG00000125944 | 5117.897437 | -0.349998425 | 0.2618539 | -1.336617 | 0.181347587 | 0.350388906 | HNRNPR | protein_coding | 38.85 | -0.159406661 | 0.159406661 | ENSG00000125944 | n.s. |
| ENSG00000117450 | 3259.074477 | 0.383223263 | 0.3033646 | 1.263243 | 0.206501724 | 0.383841982 | PRDX1 | protein_coding | 47.46 | 0.159362446 | 0.159362446 | ENSG00000117450 | n.s. |
| ENSG00000197982 | 165.553928 | 0.381281866 | 0.3008922 | 1.267171 | 0.205094212 | 0.382112501 | C1orf122 | protein_coding | 63.25 | 0.159302901 | 0.159302901 | ENSG00000197982 | n.s. |
| ENSG00000215612 | 705.8394524 | 0.348117367 | 0.2597008 | 1.340455 | 0.180097374 | 0.348747669 | HMX1 | protein_coding | 59.36 | 0.159259757 | 0.159259757 | ENSG00000215612 | n.s. |
| ENSG00000235912 | 11.37125358 | 0.620496676 | 0.6796935 | 0.912907 | 0.36129168 | 0.553871842 | | processed_pseudogene | 53.65 | 0.159213685 | 0.159213685 | ENSG00000235912 | n.s. |
| ENSG00000279528 | 25.05825189 | -0.420336861 | 0.3536934 | -1.188422 | 0.234667371 | 0.418254926 | | TEC | 42.12 | -0.159122274 | 0.159122274 | ENSG00000279528 | n.s. |
| ENSG00000213339 | 580.3774007 | 0.355868931 | 0.2693629 | 1.321151 | 0.186451139 | 0.357471662 | QTRT1 | protein_coding | 52.81 | 0.158987427 | 0.158987427 | ENSG00000213339 | n.s. |
| ENSG00000204628 | 29735.13885 | 0.374086147 | 0.292096 | 1.280696 | 0.200300517 | 0.375903576 | RACK1 | protein_coding | 52.35 | 0.158958011 | 0.158958011 | ENSG00000204628 | n.s. |

| | | | | | | | | | | | | | |
|---|---|---|---|---|---|---|---|---|---|---|---|---|---|
| ENSG00000269190 | 188.2981556 | 0.405892261 | 0.3346664 | 1.212827 | 0.225196129 | 0.406035444 | FBXO17 | protein_coding | 49.36 | 0.158880865 | 0.158880865 | ENSG00000269190 | n.s. |
| ENSG00000273148 | 104.4830425 | 0.327315356 | 0.2354754 | 1.390019 | 0.164523078 | 0.327142932 | | lincRNA | 41.13 | 0.158833854 | 0.158833854 | ENSG00000273148 | n.s. |
| ENSG00000152256 | 922.8507513 | -0.34494503 | 0.256336 | -1.345676 | 0.17840719 | 0.34653506 | PDK1 | protein_coding | 39.52 | -0.158761923 | 0.158761923 | ENSG00000152256 | n.s. |
| ENSG00000160959 | 466.820538 | -0.311598388 | 0.2176013 | -1.43197 | 0.152152508 | 0.309484203 | LRRC14 | protein_coding | 63.14 | -0.158716227 | 0.158716227 | ENSG00000160959 | n.s. |
| ENSG00000186283 | 321.5788184 | -0.354835876 | 0.2684236 | -1.321925 | 0.186193107 | 0.357124485 | TOR3A | protein_coding | 52.61 | -0.158675639 | 0.158675639 | ENSG00000186283 | n.s. |
| ENSG00000277072 | 173.5646384 | 0.326803017 | 0.2350873 | 1.390135 | 0.164487906 | 0.327119714 | STAG3L2 | transcribed_unprocessed_pseud | 47.41 | 0.158595309 | 0.158595309 | ENSG00000277072 | n.s. |
| ENSG00000250950 | 86.10411881 | 0.336508824 | 0.246519 | 1.365042 | 0.172239816 | 0.33790263 | | lincRNA | 44.72 | 0.158565794 | 0.158565794 | ENSG00000250950 | n.s. |
| ENSG00000168824 | 452.4589734 | -0.36604614 | 0.2825527 | -1.295497 | 0.1951489 | 0.369065218 | | protein_coding | 50.8 | -0.158460233 | 0.158460233 | ENSG00000168824 | n.s. |
| ENSG00000067715 | 9115.341527 | -0.343644021 | 0.2551589 | -1.346784 | 0.178049763 | 0.346082207 | SYT1 | protein_coding | 35.61 | -0.158358288 | 0.158358288 | ENSG00000067715 | n.s. |
| ENSG00000137288 | 1057.469035 | -0.379877198 | 0.3007834 | -1.262959 | 0.206603807 | 0.383846385 | UQCC2 | protein_coding | 49.22 | -0.157969101 | 0.157969101 | ENSG00000137288 | n.s. |
| ENSG00000166860 | 1554.716709 | 0.308628907 | 0.2148909 | 1.436212 | 0.15094203 | 0.307786451 | ZBTB39 | protein_coding | 49.81 | 0.157940998 | 0.157940998 | ENSG00000166860 | n.s. |
| ENSG00000134882 | 472.1528557 | -0.380329656 | 0.3015498 | -1.26125 | 0.207218847 | 0.384476015 | UBAC2 | protein_coding | 41.75 | -0.157886535 | 0.157886535 | ENSG00000134882 | n.s. |
| ENSG00000135679 | 4536.524349 | -0.342751646 | 0.2547588 | -1.345396 | 0.178497226 | 0.346571869 | MDM2 | protein_coding | 39.81 | -0.157736601 | 0.157736601 | ENSG00000135679 | n.s. |
| ENSG00000213923 | 1567.552732 | 0.322742042 | 0.2310431 | 1.396891 | 0.162446297 | 0.32466541 | CSNK1E | protein_coding | 55.79 | 0.157680136 | 0.157680136 | ENSG00000213923 | n.s. |
| ENSG00000120942 | 162.0195159 | -0.345275674 | 0.2580246 | -1.33815 | 0.180847431 | 0.349736364 | UBIAD1 | protein_coding | 48.21 | -0.157535206 | 0.157535206 | ENSG00000120942 | n.s. |
| ENSG00000176225 | 1053.05061 | -0.329151891 | 0.2388565 | -1.378032 | 0.168193362 | 0.33221112 | RTTN | protein_coding | 36.79 | -0.157527433 | 0.157527433 | ENSG00000176225 | n.s. |
| ENSG00000233844 | 324.4674557 | -0.355315106 | 0.2702882 | -1.314579 | 0.188651567 | 0.360395805 | KCNQ5-IT1 | sense_intronic | 36.35 | -0.157482858 | 0.157482858 | ENSG00000233844 | n.s. |
| ENSG00000166326 | 2697.623825 | -0.348134227 | 0.2616452 | -1.330558 | 0.183334428 | 0.353100739 | TRIM44 | protein_coding | 39.96 | -0.157391962 | 0.157391962 | ENSG00000166326 | n.s. |
| ENSG00000254004 | 1783.993456 | -0.355677673 | 0.2708341 | -1.313268 | 0.189092722 | 0.361044702 | ZNF260 | protein_coding | 40.16 | -0.157365682 | 0.157365682 | ENSG00000254004 | n.s. |
| ENSG00000106133 | 180.8645411 | 0.353956834 | 0.268873 | 1.316446 | 0.188024376 | 0.359497857 | NSUN5P2 | transcribed_unprocessed_pseud | 52.77 | 0.157264329 | 0.157264329 | ENSG00000106133 | n.s. |
| ENSG00000167515 | 445.8836726 | 0.368415636 | 0.2866646 | 1.28518 | 0.198729317 | 0.374260555 | TRAPPC2L | protein_coding | 58.39 | 0.157249351 | 0.157249351 | ENSG00000167515 | n.s. |
| ENSG00000004866 | 574.0668547 | -0.332489786 | 0.2430269 | -1.36812 | 0.17127467 | 0.336578296 | ST7 | protein_coding | 39.15 | -0.157239039 | 0.157239039 | ENSG00000004866 | n.s. |
| ENSG00000180354 | 982.8250842 | -0.344566419 | 0.2575015 | -1.338114 | 0.180859141 | 0.349736364 | MTURN | protein_coding | 46.82 | -0.157211602 | 0.157211602 | ENSG00000180354 | n.s. |
| ENSG00000171763 | 900.7680203 | -0.332898529 | 0.243583 | -1.366674 | 0.171727615 | 0.337135305 | SPATA5L1 | protein_coding | 41.47 | -0.157193276 | 0.157193276 | ENSG00000171763 | n.s. |
| ENSG00000245614 | 45.61521417 | 0.368193709 | 0.2866298 | 1.284562 | 0.198945491 | 0.374464508 | DDX11-AS1 | antisense | 44.82 | 0.157067511 | 0.157067511 | ENSG00000245614 | n.s. |
| ENSG00000156502 | 922.3023325 | -0.351699105 | 0.2662885 | -1.320745 | 0.186586521 | 0.357632728 | SUPV3L1 | protein_coding | 41.8 | -0.157055717 | 0.157055717 | ENSG00000156502 | n.s. |
| ENSG00000221716 | 5.481836491 | -0.531702048 | 0.5304658 | -1.002331 | 0.316183985 | 0.506702304 | SNORA11 | snoRNA | 53.91 | -0.156983499 | 0.156983499 | ENSG00000221716 | n.s. |
| ENSG00000109381 | 367.958774 | -0.338618578 | 0.2505084 | -1.351726 | 0.176463105 | 0.344006694 | ELF2 | protein_coding | 39.88 | -0.15692706 | 0.15692706 | ENSG00000109381 | n.s. |
| ENSG00000210117 | 151.6910088 | -0.363804232 | 0.2815203 | -1.292284 | 0.196258669 | 0.370522683 | MT-TW | Mt_tRNA | 35.29 | -0.156867001 | 0.156867001 | ENSG00000210117 | n.s. |
| ENSG00000200869 | 3.930955833 | 0.658743615 | 0.758368 | 0.868633 | 0.385047781 | 0.577945263 | RNU6-457P | snRNA | 40.19 | 0.15685561 | 0.15685561 | ENSG00000200869 | n.s. |
| ENSG00000181991 | 1041.690633 | -0.340931184 | 0.2536811 | -1.343936 | 0.178969018 | 0.347045353 | MRPS11 | protein_coding | 45.97 | -0.156696666 | 0.156696666 | ENSG00000181991 | n.s. |
| ENSG00000111615 | 4392.400712 | -0.343030587 | 0.2562886 | -1.338454 | 0.180748414 | 0.349667941 | KRR1 | protein_coding | 35.47 | -0.156540013 | 0.156540013 | ENSG00000111615 | n.s. |

| | | | | | | | | | | | | | |
|---|---|---|---|---|---|---|---|---|---|---|---|---|---|
| ENSG00000241343 | 9610.612669 | 0.358024499 | 0.2747566 | 1.30306 | 0.192554114 | 0.365622391 | RPL36A | protein_coding | 41.83 | 0.156444968 | 0.156444968 | ENSG00000241343 | n.s. |
| ENSG00000243224 | 85.46034207 | 0.341436542 | 0.2545729 | 1.341213 | 0.179851236 | 0.348319482 | | antisense | 54.35 | 0.156385528 | 0.156385528 | ENSG00000243224 | n.s. |
| ENSG00000082014 | 114.651733 | 0.395037457 | 0.3236295 | 1.220647 | 0.222219657 | 0.401970156 | SMARCD3 | protein_coding | 56.14 | 0.156358271 | 0.156358271 | ENSG00000082014 | n.s. |
| ENSG00000006282 | 328.1324877 | 0.360059505 | 0.2773333 | 1.298292 | 0.194187163 | 0.367944857 | SPATA20 | protein_coding | 57.05 | 0.156344053 | 0.156344053 | ENSG00000006282 | n.s. |
| ENSG00000121410 | 151.4216294 | 0.389085446 | 0.3157038 | 1.232438 | 0.217785364 | 0.396682343 | A1BG | protein_coding | 55.83 | 0.156240035 | 0.156240035 | ENSG00000121410 | n.s. |
| ENSG00000250021 | 237.1031528 | -0.374750747 | 0.2961068 | -1.265593 | 0.205658825 | 0.383009756 | C15orf38-AP3S2 | protein_coding | 45.41 | -0.156192425 | 0.156192425 | ENSG00000250021 | n.s. |
| ENSG00000105255 | 478.1281575 | 0.374299256 | 0.2956523 | 1.266012 | 0.205508861 | 0.382782591 | FSD1 | protein_coding | 54.71 | 0.156100689 | 0.156100689 | ENSG00000105255 | n.s. |
| ENSG00000150403 | 567.4287532 | -0.351470291 | 0.2670652 | -1.316047 | 0.188158318 | 0.35965517 | TMCO3 | protein_coding | 48.38 | -0.156092769 | 0.156092769 | ENSG00000150403 | n.s. |
| ENSG00000175183 | 688.4745618 | 0.378320789 | 0.3014311 | 1.255082 | 0.209448965 | 0.386894622 | CSRP2 | protein_coding | 42.39 | 0.156022258 | 0.156022258 | ENSG00000175183 | n.s. |
| ENSG00000133641 | 1424.932158 | -0.347539835 | 0.2623825 | -1.324554 | 0.185319068 | 0.35574209 | C12orf29 | protein_coding | 36.21 | -0.15599838 | 0.15599838 | ENSG00000133641 | n.s. |
| ENSG00000120549 | 84.24678554 | -0.349588254 | 0.2648291 | -1.320052 | 0.186817536 | 0.357981225 | KIAA1217 | protein_coding | 40.56 | -0.155965216 | 0.155965216 | ENSG00000120549 | n.s. |
| ENSG00000211450 | 720.7091738 | 0.350087993 | 0.2655297 | 1.318451 | 0.187352581 | 0.358508794 | SELENOH | protein_coding | 51.67 | 0.155964266 | 0.155964266 | ENSG00000211450 | n.s. |
| ENSG00000135441 | 289.8416515 | 0.383224925 | 0.3085166 | 1.242153 | 0.214180003 | 0.392889089 | BLOC1S1 | protein_coding | 55.05 | 0.155485861 | 0.155485861 | ENSG00000135441 | n.s. |
| ENSG00000101473 | 178.9342015 | -0.349170747 | 0.2649519 | -1.317864 | 0.187549023 | 0.358835377 | ACOT8 | protein_coding | 49.06 | -0.155417557 | 0.155417557 | ENSG00000101473 | n.s. |
| ENSG00000169371 | 240.0254202 | -0.368326348 | 0.2889295 | -1.274797 | 0.202381246 | 0.378577735 | SNUPN | protein_coding | 47.36 | -0.155376603 | 0.155376603 | ENSG00000169371 | n.s. |
| ENSG00000185043 | 151.5163715 | 0.37946988 | 0.3040087 | 1.248221 | 0.211950311 | 0.389671752 | CIB1 | protein_coding | 56.35 | 0.15531743 | 0.15531743 | ENSG00000185043 | n.s. |
| ENSG00000207547 | 73.79534934 | -0.36869142 | 0.2895459 | -1.273344 | 0.202896168 | 0.379082575 | MIR25 | miRNA | 63.1 | -0.155317225 | 0.155317225 | ENSG00000207547 | n.s. |
| ENSG00000232638 | 69.12841797 | 0.341184113 | 0.2554513 | 1.335613 | 0.181675763 | 0.350793455 | | lincRNA | 57.97 | 0.155221207 | 0.155221207 | ENSG00000232638 | n.s. |
| ENSG00000200913 | 47.77338147 | 0.353399647 | 0.2702112 | 1.307865 | 0.190919265 | 0.363833689 | SNORD46 | snoRNA | 48.08 | 0.155176757 | 0.155176757 | ENSG00000200913 | n.s. |
| ENSG00000283575 | 4.37169037 | 0.526651813 | 0.5262103 | 1.000839 | 0.316904629 | 0.507444026 | | snRNA | 42.71 | 0.155157868 | 0.155157868 | ENSG00000283575 | n.s. |
| ENSG00000169375 | 1636.001051 | -0.333609422 | 0.2464791 | -1.3535 | 0.175895954 | 0.343141287 | SIN3A | protein_coding | 44.13 | -0.154970592 | 0.154970592 | ENSG00000169375 | n.s. |
| ENSG00000172115 | 6844.759905 | 0.337772383 | 0.2516406 | 1.342281 | 0.179504983 | 0.347745635 | CYCS | protein_coding | 43.01 | 0.154949133 | 0.154949133 | ENSG00000172115 | n.s. |
| ENSG00000239218 | 66.72055375 | 0.363486986 | 0.2832807 | 1.283133 | 0.199445312 | 0.374899025 | RPS20P22 | transcribed_processed_pseudog | 47.12 | 0.154876603 | 0.154876603 | ENSG00000239218 | n.s. |
| ENSG00000161265 | 112.4409312 | 0.354913062 | 0.2727847 | 1.301074 | 0.193233102 | 0.366537395 | U2AF1L4 | protein_coding | 55.84 | 0.154700113 | 0.154700113 | ENSG00000161265 | n.s. |
| ENSG00000239569 | 70.64418136 | 0.33874241 | 0.25303 | 1.338744 | 0.180654026 | 0.349533909 | KMT2E-AS1 | antisense | 64.44 | 0.15463953 | 0.15463953 | ENSG00000239569 | n.s. |
| ENSG00000204347 | 121.0012463 | -0.375607043 | 0.2996914 | -1.253313 | 0.210091901 | 0.387688475 | BTBD17 | protein_coding | 58.45 | -0.154568725 | 0.154568725 | ENSG00000204347 | n.s. |
| ENSG00000178950 | 600.9922435 | -0.333887153 | 0.2473308 | -1.349962 | 0.177028154 | 0.344722097 | GAK | protein_coding | 56.5 | -0.154433118 | 0.154433118 | ENSG00000178950 | n.s. |
| ENSG00000132612 | 441.7586297 | -0.355956895 | 0.2745253 | -1.296627 | 0.194759641 | 0.36866264 | VPS4A | protein_coding | 54.49 | -0.15426135 | 0.15426135 | ENSG00000132612 | n.s. |
| ENSG00000143514 | 3233.698513 | 0.33593976 | 0.2501434 | 1.342989 | 0.179275579 | 0.347397896 | TP53BP2 | protein_coding | 39.79 | 0.154254406 | 0.154254406 | ENSG00000143514 | n.s. |
| ENSG00000091317 | 606.4991089 | 0.374548871 | 0.29876 | 1.253678 | 0.2099591 | 0.387597526 | CMTM6 | protein_coding | 38.68 | 0.154171433 | 0.154171433 | ENSG00000091317 | n.s. |
| ENSG00000273004 | 41.3135409 | 0.365510159 | 0.2868187 | 1.27436 | 0.202535989 | 0.378765423 | | lincRNA | 46.54 | 0.154109929 | 0.154109929 | ENSG00000273004 | n.s. |
| ENSG00000239942 | 10.07594609 | -0.457414288 | 0.4170777 | -1.096712 | 0.272767156 | 0.460666802 | RN7SL394P | misc_RNA | 55.67 | -0.153971634 | 0.153971634 | ENSG00000239942 | n.s. |

| | | | | | | | | | | | | | |
|---|---|---|---|---|---|---|---|---|---|---|---|---|---|
| ENSG00000215021 | 6075.384651 | 0.387351369 | 0.3165444 | 1.223687 | 0.221070175 | 0.400463186 | PHB2 | protein_coding | 53.17 | 0.153947922 | 0.153947922 | ENSG00000215021 | n.s. |
| ENSG00000164211 | 2896.803451 | 0.341244517 | 0.2568766 | 1.328437 | 0.184033625 | 0.354006653 | STARD4 | protein_coding | 36.15 | 0.153897379 | 0.153897379 | ENSG00000164211 | n.s. |
| ENSG00000166855 | 1204.34051 | -0.344349444 | 0.2606591 | -1.321072 | 0.18647739 | 0.357472767 | CLPX | protein_coding | 40 | -0.153840538 | 0.153840538 | ENSG00000166855 | n.s. |
| ENSG00000180747 | 370.2316028 | -0.324291559 | 0.2367387 | -1.369829 | 0.170740344 | 0.335765227 | SMG1P3 | transcribed_unprocessed_pseud | 43.71 | -0.153702616 | 0.153702616 | ENSG00000180747 | n.s. |
| ENSG00000258359 | 47.78460172 | -0.385491362 | 0.3142464 | -1.226717 | 0.219929 | 0.399303693 | PCNPP1 | processed_pseudogene | 41.29 | -0.153694124 | 0.153694124 | ENSG00000258359 | n.s. |
| ENSG00000138138 | 1652.241188 | -0.357786124 | 0.2775083 | -1.289281 | 0.197300529 | 0.372173861 | ATAD1 | protein_coding | 36.12 | -0.153581172 | 0.153581172 | ENSG00000138138 | n.s. |
| ENSG00000213326 | 120.2800064 | 0.500455248 | 0.486347 | 1.029009 | 0.303475608 | 0.493520981 | RPS7P11 | processed_pseudogene | 47.78 | 0.153486812 | 0.153486812 | ENSG00000213326 | n.s. |
| ENSG00000172757 | 4959.514453 | 0.349415058 | 0.2670994 | 1.308184 | 0.190811074 | 0.363727079 | CFL1 | protein_coding | 50.18 | 0.153471607 | 0.153471607 | ENSG00000172757 | n.s. |
| ENSG00000266531 | 2.087029204 | -0.600920677 | 0.6605832 | -0.909682 | 0.362990209 | 0.555497107 | MIR4706 | miRNA | 64.63 | -0.153425985 | 0.153425985 | ENSG00000266531 | n.s. |
| ENSG00000197121 | 3784.589334 | -0.338912275 | 0.254909 | -1.329542 | 0.183669094 | 0.353568693 | PGAP1 | protein_coding | 36 | -0.153027771 | 0.153027771 | ENSG00000197121 | n.s. |
| ENSG00000207633 | 10.0939005 | 0.450118854 | 0.4070798 | 1.105726 | 0.268845038 | 0.457128253 | MIR505 | miRNA | 52.38 | 0.153023277 | 0.153023277 | ENSG00000207633 | n.s. |
| ENSG00000102924 | 474.6160738 | 0.345769645 | 0.2633081 | 1.313175 | 0.189123864 | 0.361053087 | CBLN1 | protein_coding | 56.83 | 0.152978497 | 0.152978497 | ENSG00000102924 | n.s. |
| ENSG00000213189 | 30.99705886 | -0.397657325 | 0.3312952 | -1.200311 | 0.230018505 | 0.412593101 | BTF3L4P2 | processed_pseudogene | 39.31 | -0.152890508 | 0.152890508 | ENSG00000213189 | n.s. |
| ENSG00000232969 | 5.698626365 | 0.511215028 | 0.5061818 | 1.009944 | 0.312522341 | 0.502618881 | | antisense | 49.68 | 0.152731216 | 0.152731216 | ENSG00000232969 | n.s. |
| ENSG00000242173 | 110.4604729 | 0.361148161 | 0.2829651 | 1.276299 | 0.201849729 | 0.377684959 | ARHGDIG | protein_coding | 63.5 | 0.152718835 | 0.152718835 | ENSG00000242173 | n.s. |
| ENSG00000035499 | 749.0548934 | -0.357410096 | 0.2779317 | -1.285964 | 0.198455738 | 0.373897056 | DEPDC1B | protein_coding | 38.63 | -0.152702732 | 0.152702732 | ENSG00000035499 | n.s. |
| ENSG00000179409 | 721.3527042 | -0.328584659 | 0.2427038 | -1.353851 | 0.175783997 | 0.342988047 | GEMIN4 | protein_coding | 56.12 | -0.152700196 | 0.152700196 | ENSG00000179409 | n.s. |
| ENSG00000132677 | 33.04564417 | 0.388805888 | 0.3199796 | 1.215096 | 0.224329488 | 0.405049859 | RHBG | protein_coding | 51.82 | 0.152603012 | 0.152603012 | ENSG00000132677 | n.s. |
| ENSG00000145388 | 1328.931643 | -0.327486178 | 0.2415173 | -1.355954 | 0.175113958 | 0.342003722 | METTL14 | protein_coding | 37 | -0.152598462 | 0.152598462 | ENSG00000145388 | n.s. |
| ENSG00000104131 | 2102.521587 | -0.357384312 | 0.2780345 | -1.285395 | 0.198654124 | 0.374180764 | EIF3J | protein_coding | 42.09 | -0.15257399 | 0.15257399 | ENSG00000104131 | n.s. |
| ENSG00000172171 | 168.1836271 | -0.360095955 | 0.2817178 | -1.278215 | 0.201173548 | 0.377078528 | TEFM | protein_coding | 43.17 | -0.152525194 | 0.152525194 | ENSG00000172171 | n.s. |
| ENSG00000261485 | 77.82064795 | 0.331458385 | 0.2464691 | 1.344828 | 0.178680939 | 0.346728182 | PAN3-AS1 | antisense | 43.15 | 0.152474456 | 0.152474456 | ENSG00000261485 | n.s. |
| ENSG00000105953 | 509.4567173 | -0.357425505 | 0.2782596 | -1.284504 | 0.198965702 | 0.374464508 | OGDH | protein_coding | 46.23 | -0.15247391 | 0.15247391 | ENSG00000105953 | n.s. |
| ENSG00000105655 | 145.4484906 | 0.362935379 | 0.2854551 | 1.271427 | 0.20357672 | 0.38020103 | ISYNA1 | protein_coding | 66.58 | 0.152428036 | 0.152428036 | ENSG00000105655 | n.s. |
| ENSG00000226823 | 54.11707705 | 0.372036538 | 0.297771 | 1.249405 | 0.211516967 | 0.389337749 | SUGT1P1 | unprocessed_pseudogene | 38.61 | 0.152413505 | 0.152413505 | ENSG00000226823 | n.s. |
| ENSG00000070761 | 1004.564266 | 0.373163605 | 0.2992776 | 1.246881 | 0.212441102 | 0.390470953 | CFAP20 | protein_coding | 44.93 | 0.152404221 | 0.152404221 | ENSG00000070761 | n.s. |
| ENSG00000178035 | 2989.236663 | 0.348565549 | 0.2675132 | 1.302984 | 0.192580122 | 0.365622391 | IMPDH2 | protein_coding | 53.38 | 0.152311717 | 0.152311717 | ENSG00000178035 | n.s. |
| ENSG00000249138 | 26.12238806 | 0.372739203 | 0.2989217 | 1.246946 | 0.212417305 | 0.390470953 | | transcribed_processed_pseudog | 48.47 | 0.152230891 | 0.152230891 | ENSG00000249138 | n.s. |
| ENSG00000124783 | 1881.485915 | -0.348343978 | 0.2672957 | -1.303216 | 0.192501093 | 0.365622391 | SSR1 | protein_coding | 43.56 | -0.152214898 | 0.152214898 | ENSG00000124783 | n.s. |
| ENSG00000275126 | 1218.534489 | 0.299932288 | 0.209869 | 1.429141 | 0.152963783 | 0.310816514 | HIST1H4L | protein_coding | 58.97 | 0.152214411 | 0.152214411 | ENSG00000275126 | n.s. |
| ENSG00000183735 | 971.8927073 | -0.33141975 | 0.2467371 | -1.34321 | 0.179203965 | 0.347397896 | TBK1 | protein_coding | 37.87 | -0.15217894 | 0.15217894 | ENSG00000183735 | n.s. |
| ENSG00000132510 | 46.34937193 | -0.374375147 | 0.301145 | -1.243172 | 0.21380433 | 0.392251638 | KDM6B | protein_coding | 60.63 | -0.152159251 | 0.152159251 | ENSG00000132510 | n.s. |

| | | | | | | | | | | | | | |
|---|---|---|---|---|---|---|---|---|---|---|---|---|---|
| ENSG00000239280 | 140.9941568 | 0.329131922 | 0.2440022 | 1.348889 | 0.17737257 | 0.345151403 | | processed_pseudogene | 37.27 | 0.152055774 | 0.152055774 | ENSG00000239280 | n.s. |
| ENSG00000070814 | 2187.690284 | -0.320320344 | 0.2337719 | -1.370226 | 0.170616309 | 0.335616117 | TCOF1 | protein_coding | 52.45 | -0.151882195 | 0.151882195 | ENSG00000070814 | n.s. |
| ENSG00000013503 | 912.7762718 | -0.332029962 | 0.2477385 | -1.340243 | 0.180166202 | 0.348811396 | POLR3B | protein_coding | 39.75 | -0.151873604 | 0.151873604 | ENSG00000013503 | n.s. |
| ENSG00000067064 | 2456.611483 | -0.34672115 | 0.2656973 | -1.304948 | 0.191910536 | 0.365023278 | IDI1 | protein_coding | 43.32 | -0.151752718 | 0.151752718 | ENSG00000067064 | n.s. |
| ENSG00000158863 | 72.96930935 | -0.383734757 | 0.3146433 | -1.219587 | 0.222621558 | 0.402319608 | FAM160B2 | protein_coding | 57.17 | -0.151739775 | 0.151739775 | ENSG00000158863 | n.s. |
| ENSG00000105393 | 734.5570344 | 0.36546944 | 0.2898113 | 1.26106 | 0.207287287 | 0.384500519 | BABAM1 | protein_coding | 52.45 | 0.151707486 | 0.151707486 | ENSG00000105393 | n.s. |
| ENSG00000147684 | 2460.534442 | 0.383152503 | 0.3138558 | 1.220792 | 0.222164888 | 0.401923303 | NDUFB9 | protein_coding | 45.62 | 0.151673529 | 0.151673529 | ENSG00000147684 | n.s. |
| ENSG00000083312 | 17178.58516 | -0.355401408 | 0.2766348 | -1.284731 | 0.198886155 | 0.374454623 | TNPO1 | protein_coding | 37.25 | -0.151614526 | 0.151614526 | ENSG00000083312 | n.s. |
| ENSG00000141279 | 1658.519091 | -0.332572289 | 0.2487458 | -1.336997 | 0.181223736 | 0.350246818 | NPEPPS | protein_coding | 41.25 | -0.151528516 | 0.151528516 | ENSG00000141279 | n.s. |
| ENSG00000201164 | 5.102815316 | -0.690255736 | 0.8401537 | -0.821583 | 0.411314447 | 0.603385423 | RNU4-36P | snRNA | 38.19 | -0.151445688 | 0.151445688 | ENSG00000201164 | n.s. |
| ENSG00000110321 | 21500.99565 | 0.326948829 | 0.2420064 | 1.350993 | 0.176697799 | 0.344271404 | EIF4G2 | protein_coding | 41.78 | 0.151409692 | 0.151409692 | ENSG00000110321 | n.s. |
| ENSG00000199165 | 43.04246642 | 0.373007894 | 0.3003339 | 1.241978 | 0.214244858 | 0.392955742 | MIRLET7A1 | miRNA | 42.5 | 0.151313025 | 0.151313025 | ENSG00000199165 | n.s. |
| ENSG00000219545 | 177.4223985 | -0.374695983 | 0.3025915 | -1.23829 | 0.21560862 | 0.394625846 | UMAD1 | protein_coding | 37.6 | -0.151307661 | 0.151307661 | ENSG00000219545 | n.s. |
| ENSG00000254122 | 165.9667169 | -0.315817263 | 0.2291077 | -1.378466 | 0.168059364 | 0.331993547 | PCDHGB7 | protein_coding | 45.57 | -0.151235525 | 0.151235525 | ENSG00000254122 | n.s. |
| ENSG00000115204 | 485.803793 | -0.346036906 | 0.2656259 | -1.302723 | 0.192669455 | 0.365667505 | MPV17 | protein_coding | 50.33 | -0.151188241 | 0.151188241 | ENSG00000115204 | n.s. |
| ENSG00000165714 | 353.3778562 | -0.307851646 | 0.219731 | -1.401039 | 0.16120247 | 0.322772972 | BORCS5 | protein_coding | 42.13 | -0.151186818 | 0.151186818 | ENSG00000165714 | n.s. |
| ENSG00000110075 | 3376.784759 | -0.345031011 | 0.2641915 | -1.305989 | 0.191556462 | 0.364648705 | PPP6R3 | protein_coding | 41.89 | -0.151166824 | 0.151166824 | ENSG00000110075 | n.s. |
| ENSG00000130699 | 255.2374472 | -0.31734732 | 0.2311185 | -1.373094 | 0.169723209 | 0.334426301 | TAF4 | protein_coding | 52.35 | -0.150961985 | 0.150961985 | ENSG00000130699 | n.s. |
| ENSG00000088808 | 39.23061341 | 0.347867672 | 0.268329 | 1.296422 | 0.194830116 | 0.36866264 | PPP1R13B | protein_coding | 44.51 | 0.150755716 | 0.150755716 | ENSG00000088808 | n.s. |
| ENSG00000144354 | 3334.025268 | 0.327368594 | 0.2432006 | 1.346085 | 0.17827524 | 0.346375408 | CDCA7 | protein_coding | 43.24 | 0.150737835 | 0.150737835 | ENSG00000144354 | n.s. |
| ENSG00000069011 | 347.8392929 | 0.335671828 | 0.2533477 | 1.324945 | 0.185189396 | 0.355604674 | PITX1 | protein_coding | 59.4 | 0.150727573 | 0.150727573 | ENSG00000069011 | n.s. |
| ENSG00000068745 | 2576.315715 | 0.328235348 | 0.2444031 | 1.343008 | 0.179269308 | 0.347397896 | IP6K2 | protein_coding | 47.61 | 0.150716749 | 0.150716749 | ENSG00000068745 | n.s. |
| ENSG00000111206 | 1708.361754 | -0.323084925 | 0.2383038 | -1.355769 | 0.175172633 | 0.342065719 | FOXM1 | protein_coding | 48.07 | -0.15052218 | 0.15052218 | ENSG00000111206 | n.s. |
| ENSG00000048162 | 1167.995767 | 0.290075934 | 0.1999787 | 1.450534 | 0.146909606 | 0.302801251 | NOP16 | protein_coding | 49.33 | 0.150503675 | 0.150503675 | ENSG00000048162 | n.s. |
| ENSG00000149792 | 523.6487288 | -0.373123719 | 0.3016551 | -1.236921 | 0.216116248 | 0.395087516 | MRPL49 | protein_coding | 50.97 | -0.150483293 | 0.150483293 | ENSG00000149792 | n.s. |
| ENSG00000272677 | 73.84152513 | 0.33312942 | 0.2507157 | 1.328714 | 0.18394241 | 0.353880085 | | antisense | 45.19 | 0.150289298 | 0.150289298 | ENSG00000272677 | n.s. |
| ENSG00000104524 | 164.6861594 | -0.348504259 | 0.2696467 | -1.292448 | 0.196202121 | 0.370522683 | PYCR3 | protein_coding | 60.74 | -0.150269879 | 0.150269879 | ENSG00000104524 | n.s. |
| ENSG00000266228 | 5.797977045 | -0.505962818 | 0.5027499 | -1.006391 | 0.314227699 | 0.504670925 | MIR3611 | miRNA | 26.51 | -0.150266765 | 0.150266765 | ENSG00000266228 | n.s. |
| ENSG00000105875 | 406.0422337 | -0.341999338 | 0.2614785 | -1.307945 | 0.190892129 | 0.363831774 | WDR91 | protein_coding | 47.38 | -0.150171696 | 0.150171696 | ENSG00000105875 | n.s. |
| ENSG00000169976 | 1329.799712 | 0.383506853 | 0.3162642 | 1.212615 | 0.225276942 | 0.406128558 | SF3B5 | protein_coding | 56.57 | 0.150080218 | 0.150080218 | ENSG00000169976 | n.s. |
| ENSG00000124486 | 4060.794419 | -0.339038088 | 0.2581965 | -1.313101 | 0.189148978 | 0.361053087 | USP9X | protein_coding | 38.5 | -0.150000261 | 0.150000261 | ENSG00000124486 | n.s. |
| ENSG00000170035 | 2152.756856 | 0.341732277 | 0.2613657 | 1.307487 | 0.19104736 | 0.36397816 | UBE2E3 | protein_coding | 35.61 | 0.149994728 | 0.149994728 | ENSG00000170035 | n.s. |

| | | | | | | | | | | | | | |
|---|---|---|---|---|---|---|---|---|---|---|---|---|---|
| ENSG00000138867 | 543.5996689 | -0.342657496 | 0.262608 | -1.304825 | 0.191952354 | 0.365052948 | GUCD1 | protein_coding | 55.14 | -0.149962045 | 0.149962045 | ENSG00000138867 | n.s. |
| ENSG00000184368 | 515.0219623 | -0.351789892 | 0.2740792 | -1.283533 | 0.199305238 | 0.37481728 | MAP7D2 | protein_coding | 44.44 | -0.149925955 | 0.149925955 | ENSG00000184368 | n.s. |
| ENSG00000247077 | 823.3669365 | 0.359707095 | 0.2843055 | 1.265213 | 0.205794871 | 0.383010661 | PGAM5 | protein_coding | 55.09 | 0.14992201 | 0.14992201 | ENSG00000247077 | n.s. |
| ENSG00000167397 | 418.6390855 | 0.370219663 | 0.2984229 | 1.240588 | 0.214758166 | 0.393638487 | VKORC1 | protein_coding | 55.01 | 0.149902847 | 0.149902847 | ENSG00000167397 | n.s. |
| ENSG00000231795 | 27.02545978 | -0.375498195 | 0.3064938 | -1.225141 | 0.220522033 | 0.399782327 | ITCH-IT1 | sense_intronic | 44.02 | -0.149514523 | 0.149514523 | ENSG00000231795 | n.s. |
| ENSG00000157800 | 948.1777954 | -0.354393402 | 0.2781381 | -1.274163 | 0.202605529 | 0.378793712 | SLC37A3 | protein_coding | 45.22 | -0.14941128 | 0.14941128 | ENSG00000157800 | n.s. |
| ENSG00000226328 | 99.8014021 | -0.391477608 | 0.3276641 | -1.194753 | 0.232183574 | 0.415279727 | NUP50-AS1 | lincRNA | 41.42 | -0.149411058 | 0.149411058 | ENSG00000226328 | n.s. |
| ENSG00000162373 | 192.5274404 | -0.357655242 | 0.2823018 | -1.266925 | 0.205182012 | 0.382224933 | BEND5 | protein_coding | 44.07 | -0.149385794 | 0.149385794 | ENSG00000162373 | n.s. |
| ENSG00000198888 | 1653.335649 | -0.294754953 | 0.2064639 | -1.427634 | 0.153397105 | 0.311386031 | MT-ND1 | protein_coding | 47.7 | -0.149352593 | 0.149352593 | ENSG00000198888 | n.s. |
| ENSG00000274805 | 4.156653056 | 0.526295003 | 0.5386162 | 0.977124 | 0.328507634 | 0.520308138 | MIR6768 | miRNA | 62.5 | 0.149330618 | 0.149330618 | ENSG00000274805 | n.s. |
| ENSG00000110906 | 773.536976 | -0.35433165 | 0.2782271 | -1.273534 | 0.202828597 | 0.379007188 | KCTD10 | protein_coding | 47.69 | -0.149298545 | 0.149298545 | ENSG00000110906 | n.s. |
| ENSG00000093000 | 2379.556145 | -0.323000369 | 0.2394176 | -1.349109 | 0.177302067 | 0.345062438 | NUP50 | protein_coding | 43.62 | -0.149259218 | 0.149259218 | ENSG00000093000 | n.s. |
| ENSG00000104643 | 1431.287028 | -0.339486198 | 0.2593968 | -1.308752 | 0.19061823 | 0.363458999 | MTMR9 | protein_coding | 37.85 | -0.149219316 | 0.149219316 | ENSG00000104643 | n.s. |
| ENSG00000129596 | 151.6889292 | 0.329408759 | 0.2472262 | 1.332418 | 0.182722796 | 0.352459059 | CDO1 | protein_coding | 40.48 | 0.149186369 | 0.149186369 | ENSG00000129596 | n.s. |
| ENSG00000125844 | 421.6057061 | -0.322979958 | 0.2394758 | -1.348695 | 0.177434848 | 0.345224341 | RRBP1 | protein_coding | 52.51 | -0.149183988 | 0.149183988 | ENSG00000125844 | n.s. |
| ENSG00000140464 | 197.2226969 | 0.323876706 | 0.2408307 | 1.344831 | 0.178679749 | 0.346728182 | PML | protein_coding | 47.82 | 0.148986801 | 0.148986801 | ENSG00000140464 | n.s. |
| ENSG00000115325 | 299.4357455 | -0.306742169 | 0.2205914 | -1.390545 | 0.164363555 | 0.326976534 | DOK1 | protein_coding | 58.42 | -0.148918236 | 0.148918236 | ENSG00000115325 | n.s. |
| ENSG00000258984 | 280.5590089 | -0.386781921 | 0.3221081 | -1.200783 | 0.229835422 | 0.412370952 | UBE2F-SCLY | protein_coding | 45.35 | -0.148799621 | 0.148799621 | ENSG00000258984 | n.s. |
| ENSG00000104915 | 375.5911616 | 0.387260763 | 0.3228511 | 1.199503 | 0.230332481 | 0.41310307 | STX10 | protein_coding | 55.36 | 0.148685508 | 0.148685508 | ENSG00000104915 | n.s. |
| ENSG00000027644 | 520.7028135 | 0.328236521 | 0.2463807 | 1.332233 | 0.182783663 | 0.352527626 | INSRR | protein_coding | 55.51 | 0.148627743 | 0.148627743 | ENSG00000027644 | n.s. |
| ENSG00000138641 | 39.60591875 | 0.354011631 | 0.2785516 | 1.270901 | 0.20376372 | 0.380448205 | HERC3 | protein_coding | 39.32 | 0.148580261 | 0.148580261 | ENSG00000138641 | n.s. |
| ENSG00000164038 | 282.0128744 | -0.333213287 | 0.2527041 | -1.31859 | 0.187306052 | 0.358469025 | SLC9B2 | protein_coding | 37.26 | -0.148462635 | 0.148462635 | ENSG00000164038 | n.s. |
| ENSG00000078674 | 5537.075599 | -0.332922466 | 0.2524786 | -1.318617 | 0.187297351 | 0.358469025 | PCM1 | protein_coding | 36.54 | -0.148333061 | 0.148333061 | ENSG00000078674 | n.s. |
| ENSG00000182108 | 373.9524511 | -0.329400737 | 0.2482424 | -1.326932 | 0.184531249 | 0.354640458 | DEXI | protein_coding | 51.35 | -0.148300073 | 0.148300073 | ENSG00000182108 | n.s. |
| ENSG00000197381 | 17.07739116 | 0.407147093 | 0.3512449 | 1.159154 | 0.246393295 | 0.432395551 | ADARB1 | protein_coding | 45.43 | 0.148249904 | 0.148249904 | ENSG00000197381 | n.s. |
| ENSG00000143977 | 2259.990724 | 0.359432348 | 0.2864644 | 1.254719 | 0.209580828 | 0.387001839 | SNRPG | protein_coding | 41.09 | 0.148189274 | 0.148189274 | ENSG00000143977 | n.s. |
| ENSG00000142252 | 204.6259999 | -0.371684767 | 0.3033163 | -1.225403 | 0.220423334 | 0.399707486 | GEMIN7 | protein_coding | 50.45 | -0.148026327 | 0.148026327 | ENSG00000142252 | n.s. |
| ENSG00000141431 | 58.85003124 | -0.360217751 | 0.287749 | -1.251847 | 0.210625507 | 0.388210078 | ASXL3 | protein_coding | 36.09 | -0.148025431 | 0.148025431 | ENSG00000141431 | n.s. |
| ENSG00000128591 | 948.383752 | -0.306567688 | 0.2214082 | -1.384627 | 0.166166657 | 0.329376192 | FLNC | protein_coding | 60.86 | -0.147859986 | 0.147859986 | ENSG00000128591 | n.s. |
| ENSG00000090238 | 152.192184 | 0.365772957 | 0.2952066 | 1.23904 | 0.215330514 | 0.394324181 | YPEL3 | protein_coding | 58.41 | 0.147825893 | 0.147825893 | ENSG00000090238 | n.s. |
| ENSG00000178184 | 199.6924223 | -0.371007008 | 0.3026573 | -1.225832 | 0.220261864 | 0.399570805 | PARD6G | protein_coding | 47.5 | -0.147811511 | 0.147811511 | ENSG00000178184 | n.s. |
| ENSG00000240695 | 146.4978867 | 0.340063451 | 0.2618813 | 1.29854 | 0.194101722 | 0.367895694 | | processed_pseudogene | 43.42 | 0.147681156 | 0.147681156 | ENSG00000240695 | n.s. |

| | | | | | | | | | | | | | |
|---|---|---|---|---|---|---|---|---|---|---|---|---|---|
| ENSG00000170248 | 2090.93943 | -0.344652831 | 0.2677431 | -1.287252 | 0.198006406 | 0.373202002 | PDCD6IP | protein_coding | 36.93 | -0.147530724 | 0.147530724 | ENSG00000170248 | n.s. |
| ENSG00000213735 | 32.32166985 | -0.36703558 | 0.2976143 | -1.233259 | 0.217478992 | 0.396329843 | ANAPC10P1 | processed_pseudogene | 39.21 | -0.147527464 | 0.147527464 | ENSG00000213735 | n.s. |
| ENSG00000064932 | 255.4118796 | -0.322321187 | 0.2405895 | -1.339714 | 0.180338253 | 0.349019946 | SBNO2 | protein_coding | 63.18 | -0.147349051 | 0.147349051 | ENSG00000064932 | n.s. |
| ENSG00000170468 | 369.4618855 | -0.295190695 | 0.2088941 | -1.413112 | 0.157622851 | 0.317101338 | RIOX1 | protein_coding | 58.62 | -0.147241687 | 0.147241687 | ENSG00000170468 | n.s. |
| ENSG00000196782 | 1223.34644 | -0.305693547 | 0.2209921 | -1.383278 | 0.166579571 | 0.329912855 | MAML3 | protein_coding | 41.6 | -0.147222245 | 0.147222245 | ENSG00000196782 | n.s. |
| ENSG00000160051 | 79.32617384 | -0.345893033 | 0.2698497 | -1.281799 | 0.199913168 | 0.375626322 | IQCC | protein_coding | 55.4 | -0.147088931 | 0.147088931 | ENSG00000160051 | n.s. |
| ENSG00000165630 | 669.6713914 | 0.344896802 | 0.2686005 | 1.284051 | 0.199124022 | 0.374668254 | PRPF18 | protein_coding | 38.95 | 0.147047822 | 0.147047822 | ENSG00000165630 | n.s. |
| ENSG00000198825 | 682.3437425 | -0.332336699 | 0.2531542 | -1.312783 | 0.189255932 | 0.361158216 | INPP5F | protein_coding | 41.2 | -0.146993353 | 0.146993353 | ENSG00000198825 | n.s. |
| ENSG00000100201 | 9667.448269 | -0.323564796 | 0.2425424 | -1.334055 | 0.182185885 | 0.351569518 | DDX17 | protein_coding | 43.04 | -0.146894789 | 0.146894789 | ENSG00000100201 | n.s. |
| ENSG00000182400 | 677.7174894 | -0.353185332 | 0.2796136 | -1.263119 | 0.2065463 | 0.383841982 | TRAPPC6B | protein_coding | 40.78 | -0.146871247 | 0.146871247 | ENSG00000182400 | n.s. |
| ENSG00000010361 | 280.6587814 | 0.354233424 | 0.281414 | 1.258762 | 0.20811615 | 0.385165646 | FUZ | protein_coding | 52.29 | 0.146777489 | 0.146777489 | ENSG00000010361 | n.s. |
| ENSG00000148841 | 298.184381 | -0.322392736 | 0.2413997 | -1.335514 | 0.181708137 | 0.350793455 | ITPRIP | protein_coding | 52.49 | -0.146672098 | 0.146672098 | ENSG00000148841 | n.s. |
| ENSG00000159023 | 1460.890207 | -0.337414735 | 0.2596961 | -1.299267 | 0.193852154 | 0.367561424 | EPB41 | protein_coding | 41.87 | -0.146664088 | 0.146664088 | ENSG00000159023 | n.s. |
| ENSG00000015153 | 2185.567533 | 0.322146081 | 0.2411189 | 1.336047 | 0.181534074 | 0.350603262 | YAF2 | protein_coding | 39.02 | 0.146635757 | 0.146635757 | ENSG00000015153 | n.s. |
| ENSG00000116251 | 3888.29407 | 0.356295969 | 0.2843466 | 1.253034 | 0.210193318 | 0.387824221 | RPL22 | protein_coding | 50.61 | 0.146567716 | 0.146567716 | ENSG00000116251 | n.s. |
| ENSG00000137411 | 462.3888281 | 0.321851302 | 0.2410438 | 1.33524 | 0.181797876 | 0.35091804 | VARS2 | protein_coding | 53.81 | 0.146376139 | 0.146376139 | ENSG00000137411 | n.s. |
| ENSG00000130287 | 280.9501948 | -0.337690054 | 0.2605465 | -1.296084 | 0.194946681 | 0.368733506 | NCAN | protein_coding | 51.55 | -0.146316845 | 0.146316845 | ENSG00000130287 | n.s. |
| ENSG00000119203 | 2120.809079 | -0.358360566 | 0.2875934 | -1.246067 | 0.212739938 | 0.390865421 | CPSF3 | protein_coding | 42.1 | -0.146201346 | 0.146201346 | ENSG00000119203 | n.s. |
| ENSG00000158623 | 466.9055882 | -0.351306195 | 0.2781188 | -1.263151 | 0.206534771 | 0.383841982 | COPG2 | protein_coding | 40.91 | -0.146089812 | 0.146089812 | ENSG00000158623 | n.s. |
| ENSG00000157216 | 1291.204026 | -0.327378486 | 0.2481599 | -1.319224 | 0.187094336 | 0.358253346 | SSBP3 | protein_coding | 50.93 | -0.14594852 | 0.14594852 | ENSG00000157216 | n.s. |
| ENSG00000258289 | 1162.432328 | -0.36274189 | 0.293837 | -1.2345 | 0.217016575 | 0.395990382 | CHURC1 | protein_coding | 41.09 | -0.145936635 | 0.145936635 | ENSG00000258289 | n.s. |
| ENSG00000116016 | 202.1375423 | 0.34503367 | 0.2702766 | 1.276595 | 0.201745382 | 0.377649 | EPAS1 | protein_coding | 46.71 | 0.145918762 | 0.145918762 | ENSG00000116016 | n.s. |
| ENSG00000207744 | 15.56331094 | -0.425531575 | 0.3830225 | -1.110983 | 0.26657553 | 0.454383852 | MIR10B | miRNA | 40 | -0.145777377 | 0.145777377 | ENSG00000207744 | n.s. |
| ENSG00000155893 | 880.525788 | -0.35119124 | 0.2785621 | -1.260729 | 0.207406616 | 0.384523202 | PXYLP1 | protein_coding | 44.59 | -0.145771565 | 0.145771565 | ENSG00000155893 | n.s. |
| ENSG00000176396 | 293.641643 | -0.342765802 | 0.2674836 | -1.281446 | 0.200037002 | 0.375656215 | EID2 | protein_coding | 51.5 | -0.145747248 | 0.145747248 | ENSG00000176396 | n.s. |
| ENSG00000219665 | 246.2554768 | -0.323452018 | 0.2438938 | -1.3262 | 0.184773353 | 0.355037178 | | processed_transcript | 44.43 | -0.145464836 | 0.145464836 | ENSG00000219665 | n.s. |
| ENSG00000247137 | 494.9571817 | -0.280124638 | 0.1931493 | -1.450301 | 0.14697454 | 0.302845422 | | processed_transcript | 40.91 | -0.145322776 | 0.145322776 | ENSG00000247137 | n.s. |
| ENSG00000178917 | 268.295852 | -0.317918349 | 0.2374254 | -1.339024 | 0.180562828 | 0.349406012 | ZNF852 | protein_coding | 42.79 | -0.145183654 | 0.145183654 | ENSG00000178917 | n.s. |
| ENSG00000186111 | 187.1344449 | -0.320944063 | 0.2411453 | -1.330916 | 0.183216728 | 0.353069413 | PIP5K1C | protein_coding | 60.55 | -0.145111618 | 0.145111618 | ENSG00000186111 | n.s. |
| ENSG00000072778 | 775.2087661 | 0.3413705 | 0.266475 | 1.28106 | 0.20017246 | 0.375790487 | ACADVL | protein_coding | 58.5 | 0.145100971 | 0.145100971 | ENSG00000072778 | n.s. |
| ENSG00000136824 | 3457.218015 | -0.341199835 | 0.2662069 | -1.281709 | 0.199944723 | 0.375634918 | SMC2 | protein_coding | 35.98 | -0.145089785 | 0.145089785 | ENSG00000136824 | n.s. |
| ENSG00000175931 | 1490.030656 | -0.319284714 | 0.2392366 | -1.334598 | 0.18200794 | 0.35127482 | UBE2O | protein_coding | 49.87 | -0.145067962 | 0.145067962 | ENSG00000175931 | n.s. |

| | | | | | | | | | | | | | |
|---|---|---|---|---|---|---|---|---|---|---|---|---|---|
| ENSG00000111727 | 739.1857742 | -0.321499066 | 0.2419569 | -1.328745 | 0.18393201 | 0.353880085 | HCFC2 | protein_coding | 37.29 | -0.145042335 | 0.145042335 | ENSG00000111727 | n.s. |
| ENSG00000095059 | 845.5307612 | 0.356848745 | 0.2870311 | 1.243241 | 0.213779206 | 0.392251638 | DHPS | protein_coding | 54.74 | 0.145035903 | 0.145035903 | ENSG00000095059 | n.s. |
| ENSG00000183580 | 667.8449639 | -0.304411182 | 0.221695 | -1.373108 | 0.169718759 | 0.334426301 | FBXL7 | protein_coding | 39.82 | -0.14480827 | 0.14480827 | ENSG00000183580 | n.s. |
| ENSG00000168283 | 1944.202045 | 0.349559354 | 0.2777605 | 1.258492 | 0.208213862 | 0.385295268 | BMI1 | protein_coding | 41.6 | 0.144789695 | 0.144789695 | ENSG00000168283 | n.s. |
| ENSG00000025770 | 1022.690697 | 0.341422526 | 0.2669445 | 1.279002 | 0.200896435 | 0.376761996 | NCAPH2 | protein_coding | 55.08 | 0.144740245 | 0.144740245 | ENSG00000025770 | n.s. |
| ENSG00000165359 | 547.7966723 | -0.344200765 | 0.2706775 | -1.271627 | 0.20350574 | 0.380119456 | INTS6L | protein_coding | 38.59 | -0.144591823 | 0.144591823 | ENSG00000165359 | n.s. |
| ENSG00000128908 | 1154.814994 | -0.31707535 | 0.2370319 | -1.337691 | 0.18099727 | 0.349954866 | INO80 | protein_coding | 42.26 | -0.144582543 | 0.144582543 | ENSG00000128908 | n.s. |
| ENSG00000160326 | 183.3759566 | -0.330539945 | 0.2534591 | -1.304115 | 0.192194231 | 0.365313348 | SLC2A6 | protein_coding | 61.28 | -0.144556508 | 0.144556508 | ENSG00000160326 | n.s. |
| ENSG00000179604 | 1949.178007 | 0.319677057 | 0.2402105 | 1.33082 | 0.183248121 | 0.35308104 | CDC42EP4 | protein_coding | 52.99 | 0.144534182 | 0.144534182 | ENSG00000179604 | n.s. |
| ENSG00000260796 | 44.190828 | 0.345848976 | 0.2729262 | 1.267189 | 0.205087859 | 0.382112501 | | antisense | 47.47 | 0.14449873 | 0.14449873 | ENSG00000260796 | n.s. |
| ENSG00000238423 | 6.083931568 | -0.48261132 | 0.4774207 | -1.010872 | 0.312077633 | 0.502318715 | SNORD42B | snoRNA | 40.3 | -0.144310746 | 0.144310746 | ENSG00000238423 | n.s. |
| ENSG00000250467 | 229.2686232 | -0.326662542 | 0.2491799 | -1.310951 | 0.189874377 | 0.3622391 | | antisense | 41.2 | -0.144059708 | 0.144059708 | ENSG00000250467 | n.s. |
| ENSG00000149346 | 254.3787896 | -0.36366343 | 0.2977319 | -1.221446 | 0.221917158 | 0.401683904 | SLX4IP | protein_coding | 40.51 | -0.144052747 | 0.144052747 | ENSG00000149346 | n.s. |
| ENSG00000136026 | 2133.668927 | -0.347857126 | 0.2765725 | -1.257743 | 0.208484751 | 0.385489136 | CKAP4 | protein_coding | 44.3 | -0.144008626 | 0.144008626 | ENSG00000136026 | n.s. |
| ENSG00000055208 | 1459.201413 | -0.334466253 | 0.259043 | -1.291161 | 0.196647852 | 0.371093526 | TAB2 | protein_coding | 38.87 | -0.143993281 | 0.143993281 | ENSG00000055208 | n.s. |
| ENSG00000101350 | 657.1929237 | -0.339247384 | 0.2651445 | -1.279481 | 0.200727627 | 0.376496128 | KIF3B | protein_coding | 43.82 | -0.143922136 | 0.143922136 | ENSG00000101350 | n.s. |
| ENSG00000178585 | 134.4147444 | -0.346870217 | 0.2753978 | -1.259524 | 0.207840966 | 0.384912181 | CTNNBIP1 | protein_coding | 51.39 | -0.143825693 | 0.143825693 | ENSG00000178585 | n.s. |
| ENSG00000100300 | 346.9711067 | 0.395919028 | 0.3423534 | 1.156463 | 0.247491866 | 0.433383128 | TSPO | protein_coding | 57.81 | 0.143769285 | 0.143769285 | ENSG00000100300 | n.s. |
| ENSG00000254522 | 12.92560759 | -0.407425667 | 0.3598145 | -1.132321 | 0.257499344 | 0.443818194 | | processed_pseudogene | 51.34 | -0.143737697 | 0.143737697 | ENSG00000254522 | n.s. |
| ENSG00000112695 | 3390.409218 | 0.38002074 | 0.3199413 | 1.187783 | 0.234919045 | 0.418579199 | COX7A2 | protein_coding | 40.34 | 0.14373234 | 0.14373234 | ENSG00000112695 | n.s. |
| ENSG00000121892 | 4136.898353 | -0.344745652 | 0.2724861 | -1.265186 | 0.205804671 | 0.383010661 | PDS5A | protein_coding | 40.35 | -0.143686243 | 0.143686243 | ENSG00000121892 | n.s. |
| ENSG00000104529 | 3972.955982 | 0.343800222 | 0.2712306 | 1.267557 | 0.204956157 | 0.382008644 | EEF1D | protein_coding | 61.99 | 0.14368333 | 0.14368333 | ENSG00000104529 | n.s. |
| ENSG00000100697 | 4126.489129 | -0.322620177 | 0.2448518 | -1.317614 | 0.187632815 | 0.358897058 | DICER1 | protein_coding | 38 | -0.14357569 | 0.14357569 | ENSG00000100697 | n.s. |
| ENSG00000256591 | 295.8268135 | 0.29027878 | 0.2063602 | 1.406661 | 0.159528005 | 0.320264977 | | protein_coding | 48.75 | 0.143540114 | 0.143540114 | ENSG00000256591 | n.s. |
| ENSG00000266315 | 4.538636606 | -0.502358858 | 0.5117538 | -0.981642 | 0.326276449 | 0.517928058 | MIR4668 | miRNA | 30 | -0.143539279 | 0.143539279 | ENSG00000266315 | n.s. |
| ENSG00000068650 | 64.12405002 | -0.357614083 | 0.2902663 | -1.232021 | 0.217941308 | 0.396860689 | ATP11A | protein_coding | 50.33 | -0.143532674 | 0.143532674 | ENSG00000068650 | n.s. |
| ENSG00000122085 | 865.8712865 | -0.312423822 | 0.2326109 | -1.343118 | 0.17923384 | 0.347397896 | MTERF4 | protein_coding | 47.67 | -0.143456526 | 0.143456526 | ENSG00000122085 | n.s. |
| ENSG00000154079 | 302.6895771 | 0.336723796 | 0.262475 | 1.28288 | 0.199534256 | 0.374988734 | SDHAF4 | protein_coding | 40.87 | 0.143438202 | 0.143438202 | ENSG00000154079 | n.s. |
| ENSG00000267191 | 27.56041487 | -0.392272259 | 0.3374434 | -1.162483 | 0.245039299 | 0.430985618 | | antisense | 40.01 | -0.143390112 | 0.143390112 | ENSG00000267191 | n.s. |
| ENSG00000111581 | 3851.389601 | -0.33876639 | 0.2653 | -1.276918 | 0.201631088 | 0.377631105 | NUP107 | protein_coding | 39.32 | -0.143275229 | 0.143275229 | ENSG00000111581 | n.s. |
| ENSG00000223705 | 175.4150286 | 0.320258227 | 0.2422028 | 1.322273 | 0.186077277 | 0.35700068 | NSUN5P1 | transcribed_unprocessed_pseud | 53.54 | 0.143261419 | 0.143261419 | ENSG00000223705 | n.s. |
| ENSG00000131375 | 1066.970923 | -0.35155825 | 0.2824238 | -1.24479 | 0.213209073 | 0.391507069 | CAPN7 | protein_coding | 37.11 | -0.143175751 | 0.143175751 | ENSG00000131375 | n.s. |

| | | | | | | | | | | | | | |
|---|---|---|---|---|---|---|---|---|---|---|---|---|---|
| ENSG00000168785 | 648.3564085 | -0.337924237 | 0.2645991 | -1.277118 | 0.201560591 | 0.377549859 | TSPAN5 | protein_coding | 41.93 | -0.142950633 | 0.142950633 | ENSG00000168785 | n.s. |
| ENSG00000114125 | 1215.561267 | 0.365915597 | 0.3023107 | 1.210396 | 0.226127 | 0.407186527 | RNF7 | protein_coding | 39.34 | 0.142782681 | 0.142782681 | ENSG00000114125 | n.s. |
| ENSG00000166033 | 333.6293836 | -0.357296686 | 0.2908161 | -1.2286 | 0.219221755 | 0.398514754 | HTRA1 | protein_coding | 49.2 | -0.14275989 | 0.14275989 | ENSG00000166033 | n.s. |
| ENSG00000231607 | 2214.818945 | -0.321159982 | 0.243936 | -1.316575 | 0.187981151 | 0.359464574 | DLEU2 | antisense | 38.45 | -0.14270548 | 0.14270548 | ENSG00000231607 | n.s. |
| ENSG00000234616 | 1610.71133 | -0.308104686 | 0.2280511 | -1.351034 | 0.176684603 | 0.344271404 | JRK | protein_coding | 56.58 | -0.142682987 | 0.142682987 | ENSG00000234616 | n.s. |
| ENSG00000196205 | 1881.561028 | 0.380529732 | 0.3220753 | 1.181493 | 0.237406974 | 0.421930848 | EEF1A1P5 | processed_pseudogene | 48.31 | 0.142606836 | 0.142606836 | ENSG00000196205 | n.s. |
| ENSG00000131094 | 197.3854066 | -0.355608209 | 0.2888192 | -1.231248 | 0.218229942 | 0.397178495 | C1QL1 | protein_coding | 56.53 | -0.142603968 | 0.142603968 | ENSG00000131094 | n.s. |
| ENSG00000140694 | 845.0898441 | -0.320204828 | 0.242992 | -1.317759 | 0.187584416 | 0.358853783 | PARN | protein_coding | 41.91 | -0.142517556 | 0.142517556 | ENSG00000140694 | n.s. |
| ENSG00000164758 | 295.1518166 | 0.373435867 | 0.3131353 | 1.19257 | 0.233037716 | 0.416232469 | MED30 | protein_coding | 34.58 | 0.142153608 | 0.142153608 | ENSG00000164758 | n.s. |
| ENSG00000279069 | 308.7880149 | -0.296694837 | 0.2151646 | -1.37892 | 0.16791942 | 0.331905464 | | TEC | 57.27 | -0.142112554 | 0.142112554 | ENSG00000279069 | n.s. |
| ENSG00000273237 | 55.40532928 | -0.37302443 | 0.3126812 | -1.192986 | 0.2328747 | 0.416054722 | | antisense | 39.99 | -0.142066184 | 0.142066184 | ENSG00000273237 | n.s. |
| ENSG00000229809 | 115.9709056 | 0.326981378 | 0.2517508 | 1.298829 | 0.194002512 | 0.367796429 | ZNF688 | protein_coding | 56.77 | 0.142038255 | 0.142038255 | ENSG00000229809 | n.s. |
| ENSG00000134030 | 103.8456086 | -0.319961785 | 0.2433211 | -1.314978 | 0.18851745 | 0.360193283 | CTIF | protein_coding | 50.34 | -0.141891658 | 0.141891658 | ENSG00000134030 | n.s. |
| ENSG00000154640 | 1925.627745 | 0.329612067 | 0.2553237 | 1.290958 | 0.196718312 | 0.371125889 | BTG3 | protein_coding | 37.74 | 0.14189099 | 0.14189099 | ENSG00000154640 | n.s. |
| ENSG00000116001 | 4385.754927 | -0.361626529 | 0.2978349 | -1.214184 | 0.224677317 | 0.405309961 | TIA1 | protein_coding | 39.67 | -0.141834525 | 0.141834525 | ENSG00000116001 | n.s. |
| ENSG00000278135 | 4.968387076 | 0.547191355 | 0.5954278 | 0.918989 | 0.358101538 | 0.550983281 | | snRNA | 36.84 | 0.141646818 | 0.141646818 | ENSG00000278135 | n.s. |
| ENSG00000249087 | 89.31721947 | 0.317660651 | 0.2407281 | 1.319583 | 0.186974447 | 0.358220411 | ZNF436-AS1 | antisense | 51.57 | 0.141628897 | 0.141628897 | ENSG00000249087 | n.s. |
| ENSG00000219747 | 33.54213252 | -0.528778971 | 0.5621361 | -0.94066 | 0.34687909 | 0.539921472 | | processed_pseudogene | 51.11 | -0.141537951 | 0.141537951 | ENSG00000219747 | n.s. |
| ENSG00000165644 | 64.80503666 | 0.378692258 | 0.3214246 | 1.178168 | 0.238729583 | 0.423525992 | COMTD1 | protein_coding | 70.61 | 0.141297629 | 0.141297629 | ENSG00000165644 | n.s. |
| ENSG00000201700 | 18.37957835 | -0.418181928 | 0.3801085 | -1.100165 | 0.271260373 | 0.459330616 | SNORD113-3 | snoRNA | 38.89 | -0.141293054 | 0.141293054 | ENSG00000201700 | n.s. |
| ENSG00000156709 | 443.7757402 | -0.35332727 | 0.2874835 | -1.229035 | 0.219058629 | 0.39828691 | AIFM1 | protein_coding | 42.98 | -0.141261644 | 0.141261644 | ENSG00000156709 | n.s. |
| ENSG00000275379 | 1013.948294 | -0.305284815 | 0.2261928 | -1.349666 | 0.177122993 | 0.344858542 | HIST1H3I | protein_coding | 62.53 | -0.141151186 | 0.141151186 | ENSG00000275379 | n.s. |
| ENSG00000233276 | 1171.75554 | 0.374483196 | 0.3160438 | 1.184909 | 0.236053312 | 0.419955311 | GPX1 | protein_coding | 63.79 | 0.141104116 | 0.141104116 | ENSG00000233276 | n.s. |
| ENSG00000175215 | 1379.103531 | -0.322034289 | 0.2465817 | -1.305994 | 0.191554498 | 0.364648705 | CTDSP2 | protein_coding | 51.21 | -0.141091378 | 0.141091378 | ENSG00000175215 | n.s. |
| ENSG00000181523 | 25.6721045 | -0.376703197 | 0.3191221 | -1.180436 | 0.23782683 | 0.422325879 | SGSH | protein_coding | 58.83 | -0.14101971 | 0.14101971 | ENSG00000181523 | n.s. |
| ENSG00000058063 | 896.5854114 | -0.32226544 | 0.247336 | -1.302946 | 0.192593164 | 0.365622391 | ATP11B | protein_coding | 36.02 | -0.140819432 | 0.140819432 | ENSG00000058063 | n.s. |
| ENSG00000105323 | 2063.561693 | -0.321097238 | 0.2459037 | -1.305784 | 0.191625869 | 0.364681106 | HNRNPUL1 | protein_coding | 48.03 | -0.140668441 | 0.140668441 | ENSG00000105323 | n.s. |
| ENSG00000254333 | 14.25856984 | 0.399880676 | 0.3539478 | 1.129773 | 0.258571826 | 0.444950628 | NDST1-AS1 | lincRNA | 50.59 | 0.140633306 | 0.140633306 | ENSG00000254333 | n.s. |
| ENSG00000120697 | 960.8893723 | -0.339439457 | 0.2697988 | -1.258121 | 0.208348034 | 0.385411927 | ALG5 | protein_coding | 39.26 | -0.140553342 | 0.140553342 | ENSG00000120697 | n.s. |
| ENSG00000160767 | 216.7126576 | -0.305175083 | 0.2267704 | -1.345745 | 0.17838488 | 0.34653506 | FAM189B | protein_coding | 57.4 | -0.140457692 | 0.140457692 | ENSG00000160767 | n.s. |
| ENSG00000225648 | 418.7478292 | 0.328108777 | 0.254928 | 1.287064 | 0.198071859 | 0.373224319 | SBDSP1 | transcribed_unprocessed_pseud | 43.81 | 0.140440422 | 0.140440422 | ENSG00000225648 | n.s. |
| ENSG00000072133 | 428.6739787 | -0.340687213 | 0.2716015 | -1.254364 | 0.20970963 | 0.387188327 | RPS6KA6 | protein_coding | 35.17 | -0.140389618 | 0.140389618 | ENSG00000072133 | n.s. |

| | | | | | | | | | | | | | |
|---|---|---|---|---|---|---|---|---|---|---|---|---|---|
| ENSG00000215301 | 11496.86041 | 0.311788532 | 0.2349502 | 1.327041 | 0.184495146 | 0.354640458 | DDX3X | protein_coding | 42.68 | 0.140370852 | 0.140370852 | ENSG00000215301 | n.s. |
| ENSG00000117519 | 5256.236132 | 0.346691805 | 0.2796492 | 1.239738 | 0.215072249 | 0.3940067 | CNN3 | protein_coding | 43.03 | 0.140235585 | 0.140235585 | ENSG00000117519 | n.s. |
| ENSG00000163788 | 918.2895025 | -0.328991682 | 0.2563822 | -1.283208 | 0.199419241 | 0.374899025 | SNRK | protein_coding | 41.92 | -0.140178647 | 0.140178647 | ENSG00000163788 | n.s. |
| ENSG00000248275 | 162.4874253 | -0.341204656 | 0.2726267 | -1.251545 | 0.210735559 | 0.388323674 | TRIM52-AS1 | processed_transcript | 49.04 | -0.140168965 | 0.140168965 | ENSG00000248275 | n.s. |
| ENSG00000067836 | 82.69618205 | 0.35172653 | 0.2869595 | 1.225701 | 0.220311263 | 0.399608353 | ROGDI | protein_coding | 63.26 | 0.140115694 | 0.140115694 | ENSG00000067836 | n.s. |
| ENSG00000169122 | 43.18980063 | -0.337154215 | 0.2672416 | -1.261608 | 0.207089711 | 0.384338848 | FAM110B | protein_coding | 40.5 | -0.140015329 | 0.140015329 | ENSG00000169122 | n.s. |
| ENSG00000087095 | 680.4450112 | -0.309836418 | 0.2329902 | -1.329826 | 0.183575545 | 0.353467337 | NLK | protein_coding | 37.3 | -0.139937839 | 0.139937839 | ENSG00000087095 | n.s. |
| ENSG00000159086 | 4361.820223 | 0.322441716 | 0.2484803 | 1.297655 | 0.194405846 | 0.36821012 | PAXBP1 | protein_coding | 38.15 | 0.13990884 | 0.13990884 | ENSG00000159086 | n.s. |
| ENSG00000172765 | 261.9573147 | -0.324468204 | 0.2510441 | -1.292475 | 0.196192669 | 0.370522683 | TMCC1 | protein_coding | 39.37 | -0.139905888 | 0.139905888 | ENSG00000172765 | n.s. |
| ENSG00000111640 | 48156.04114 | 0.350944463 | 0.2861847 | 1.226286 | 0.220090914 | 0.399416813 | GAPDH | protein_coding | 60.61 | 0.139877218 | 0.139877218 | ENSG00000111640 | n.s. |
| ENSG00000154237 | 56.14532967 | -0.340862424 | 0.2725295 | -1.250736 | 0.21103093 | 0.38874967 | LRRK1 | protein_coding | 46.01 | -0.139866067 | 0.139866067 | ENSG00000154237 | n.s. |
| ENSG00000198919 | 1314.826902 | -0.356198712 | 0.2932108 | -1.214821 | 0.224434241 | 0.405127573 | DZIP3 | protein_coding | 35.7 | -0.139775295 | 0.139775295 | ENSG00000198919 | n.s. |
| ENSG00000103507 | 167.678454 | -0.319800677 | 0.2454385 | -1.302977 | 0.192582749 | 0.365622391 | BCKDK | protein_coding | 57.82 | -0.139742411 | 0.139742411 | ENSG00000103507 | n.s. |
| ENSG00000234327 | 83.44117766 | 0.358606159 | 0.2966249 | 1.208955 | 0.226680089 | 0.407918687 | | processed_transcript | 48.26 | 0.139650705 | 0.139650705 | ENSG00000234327 | n.s. |
| ENSG00000109184 | 1719.393372 | -0.320460133 | 0.2467882 | -1.298523 | 0.194107719 | 0.367895694 | DCUN1D4 | protein_coding | 40.07 | -0.139167919 | 0.139167919 | ENSG00000109184 | n.s. |
| ENSG00000179889 | 1168.825968 | -0.332790892 | 0.2625202 | -1.267677 | 0.204913304 | 0.381979908 | PDXDC1 | protein_coding | 46.49 | -0.139093108 | 0.139093108 | ENSG00000179889 | n.s. |
| ENSG00000175322 | 873.606505 | -0.327472632 | 0.2558759 | -1.27981 | 0.200611861 | 0.376329687 | ZNF519 | protein_coding | 39.05 | -0.138989707 | 0.138989707 | ENSG00000175322 | n.s. |
| ENSG00000117724 | 27259.65455 | -0.331399888 | 0.2608798 | -1.270316 | 0.203972033 | 0.380786083 | CENPF | protein_coding | 40.92 | -0.138962247 | 0.138962247 | ENSG00000117724 | n.s. |
| ENSG00000164620 | 278.1230033 | -0.339382524 | 0.2717545 | -1.248857 | 0.211717371 | 0.389552128 | RELL2 | protein_coding | 60 | -0.138954887 | 0.138954887 | ENSG00000164620 | n.s. |
| ENSG00000214578 | 55.54874657 | -0.376158037 | 0.3214191 | -1.170304 | 0.241878668 | 0.427207497 | HMGN2P15 | processed_pseudogene | 50 | -0.138938159 | 0.138938159 | ENSG00000214578 | n.s. |
| ENSG00000162992 | 3840.551967 | 0.282631134 | 0.2015696 | 1.402152 | 0.160869902 | 0.322457766 | NEUROD1 | protein_coding | 39.48 | 0.138920879 | 0.138920879 | ENSG00000162992 | n.s. |
| ENSG00000089009 | 30092.74046 | 0.358259587 | 0.2972307 | 1.205325 | 0.228077913 | 0.409745649 | RPL6 | protein_coding | 45.67 | 0.138820449 | 0.138820449 | ENSG00000089009 | n.s. |
| ENSG00000234690 | 81.33463921 | 0.326345675 | 0.2546637 | 1.281477 | 0.200026178 | 0.375656215 | | lincRNA | 46.05 | 0.138765256 | 0.138765256 | ENSG00000234690 | n.s. |
| ENSG00000143437 | 911.3285907 | -0.316900181 | 0.2429148 | -1.304573 | 0.192038163 | 0.365120229 | ARNT | protein_coding | 41.21 | -0.138664144 | 0.138664144 | ENSG00000143437 | n.s. |
| ENSG00000163605 | 2749.890108 | -0.343725096 | 0.2780005 | -1.236419 | 0.216302849 | 0.395324831 | PPP4R2 | protein_coding | 38.82 | -0.138536993 | 0.138536993 | ENSG00000163605 | n.s. |
| ENSG00000196498 | 874.8757733 | -0.306817799 | 0.2307868 | -1.329443 | 0.18370202 | 0.353568693 | NCOR2 | protein_coding | 57.37 | -0.138536274 | 0.138536274 | ENSG00000196498 | n.s. |
| ENSG00000102125 | 105.2580544 | 0.345288794 | 0.2805438 | 1.230784 | 0.218403688 | 0.39728698 | TAZ | protein_coding | 56.12 | 0.138424782 | 0.138424782 | ENSG00000102125 | n.s. |
| ENSG00000083444 | 214.682561 | -0.344667534 | 0.2796843 | -1.232345 | 0.217820219 | 0.396692074 | PLOD1 | protein_coding | 53.54 | -0.138400034 | 0.138400034 | ENSG00000083444 | n.s. |
| ENSG00000105248 | 315.2230973 | 0.332850575 | 0.2635942 | 1.262738 | 0.206683193 | 0.383891422 | CCDC94 | protein_coding | 52.8 | 0.13839647 | 0.13839647 | ENSG00000105248 | n.s. |
| ENSG00000123136 | 1918.95886 | 0.352636248 | 0.290485 | 1.213957 | 0.224764153 | 0.405414083 | DDX39A | protein_coding | 55.52 | 0.138269085 | 0.138269085 | ENSG00000123136 | n.s. |
| ENSG00000241553 | 430.9943762 | -0.332233235 | 0.2630834 | -1.262844 | 0.206645307 | 0.383872262 | ARPC4 | protein_coding | 48.44 | -0.138146987 | 0.138146987 | ENSG00000241553 | n.s. |
| ENSG00000115350 | 480.9269199 | 0.337508741 | 0.270373 | 1.248308 | 0.211918374 | 0.389668403 | POLE4 | protein_coding | 39.19 | 0.138143951 | 0.138143951 | ENSG00000115350 | n.s. |

| | | | | | | | | | | | | | |
|---|---|---|---|---|---|---|---|---|---|---|---|---|---|
| ENSG00000075131 | 608.9643488 | -0.336473342 | 0.2690362 | -1.250662 | 0.211057872 | 0.38874967 | TIPIN | protein_coding | 44.05 | -0.138065095 | 0.138065095 | ENSG00000075131 | n.s. |
| ENSG00000143315 | 390.360115 | -0.302901101 | 0.2266618 | -1.336357 | 0.181432583 | 0.350478572 | PIGM | protein_coding | 41.28 | -0.137922542 | 0.137922542 | ENSG00000143315 | n.s. |
| ENSG00000155438 | 2004.52181 | -0.348193104 | 0.2853724 | -1.220136 | 0.222413391 | 0.402059419 | NIFK | protein_coding | 43.01 | -0.137783409 | 0.137783409 | ENSG00000155438 | n.s. |
| ENSG00000278463 | 904.0797364 | 0.282866809 | 0.2029637 | 1.393682 | 0.163413721 | 0.325774519 | HIST1H2AB | protein_coding | 60.05 | 0.137779582 | 0.137779582 | ENSG00000278463 | n.s. |
| ENSG00000186141 | 995.7722061 | -0.325397147 | 0.2546748 | -1.277697 | 0.201356337 | 0.37727551 | POLR3C | protein_coding | 43.11 | -0.137754079 | 0.137754079 | ENSG00000186141 | n.s. |
| ENSG00000138768 | 1235.547988 | -0.345193095 | 0.2812293 | -1.227443 | 0.219655955 | 0.399095636 | USO1 | protein_coding | 37.52 | -0.137705472 | 0.137705472 | ENSG00000138768 | n.s. |
| ENSG00000263389 | 11.54333515 | -0.429181119 | 0.4047659 | -1.060319 | 0.2889993 | 0.477838278 | MIR3973 | miRNA | 46.73 | -0.137646567 | 0.137646567 | ENSG00000263389 | n.s. |
| ENSG00000181396 | 547.9094841 | -0.334362008 | 0.2667516 | -1.253458 | 0.210038899 | 0.387642045 | OGFOD3 | protein_coding | 54.32 | -0.137613079 | 0.137613079 | ENSG00000181396 | n.s. |
| ENSG00000102030 | 509.6868927 | 0.364291831 | 0.3069946 | 1.186639 | 0.235370006 | 0.41916815 | NAA10 | protein_coding | 58.17 | 0.137560866 | 0.137560866 | ENSG00000102030 | n.s. |
| ENSG00000116521 | 487.7060933 | 0.322508441 | 0.2512659 | 1.283535 | 0.199304819 | 0.37481728 | SCAMP3 | protein_coding | 51.95 | 0.137446775 | 0.137446775 | ENSG00000116521 | n.s. |
| ENSG00000197647 | 111.4687166 | -0.341335086 | 0.2766216 | -1.233942 | 0.217224452 | 0.396032547 | ZNF433 | protein_coding | 43.27 | -0.137308565 | 0.137308565 | ENSG00000197647 | n.s. |
| ENSG00000177542 | 236.9157856 | -0.329316798 | 0.2602388 | -1.265441 | 0.205713349 | 0.383009756 | SLC25A22 | protein_coding | 65.66 | -0.137256002 | 0.137256002 | ENSG00000177542 | n.s. |
| ENSG00000003249 | 132.6729038 | -0.331290976 | 0.2633938 | -1.257778 | 0.208472041 | 0.385489136 | DBNDD1 | protein_coding | 59.39 | -0.137150441 | 0.137150441 | ENSG00000003249 | n.s. |
| ENSG00000265808 | 1451.989294 | -0.343187196 | 0.2795091 | -1.227821 | 0.219514086 | 0.398889927 | SEC22B | protein_coding | 36.13 | -0.136982116 | 0.136982116 | ENSG00000265808 | n.s. |
| ENSG00000166822 | 522.6370324 | -0.339065532 | 0.2737719 | -1.238496 | 0.215532085 | 0.394537631 | TMEM170A | protein_coding | 44.64 | -0.136952491 | 0.136952491 | ENSG00000166822 | n.s. |
| ENSG00000167645 | 286.6747311 | 0.36553479 | 0.3095821 | 1.180736 | 0.237707508 | 0.422196918 | YIF1B | protein_coding | 55.37 | 0.136887275 | 0.136887275 | ENSG00000167645 | n.s. |
| ENSG00000264539 | 2.973899232 | 0.551424968 | 0.6184912 | 0.891565 | 0.372626265 | 0.564719221 | MIR548AR | miRNA | 33.33 | 0.136845717 | 0.136845717 | ENSG00000264539 | n.s. |
| ENSG00000150779 | 871.6893141 | 0.344728503 | 0.281971 | 1.222567 | 0.221493188 | 0.401020762 | TIMM8B | protein_coding | 46.22 | 0.136799695 | 0.136799695 | ENSG00000150779 | n.s. |
| ENSG00000213463 | 1077.936608 | -0.329313386 | 0.2609453 | -1.262001 | 0.206948289 | 0.384230035 | SYNJ2BP | protein_coding | 39.54 | -0.136799642 | 0.136799642 | ENSG00000213463 | n.s. |
| ENSG00000242853 | 36.76848856 | 0.362277807 | 0.3053835 | 1.186304 | 0.235502083 | 0.419324551 | RN7SL749P | misc_RNA | 55.86 | 0.136741653 | 0.136741653 | ENSG00000242853 | n.s. |
| ENSG00000198208 | 202.4142521 | -0.329114824 | 0.2609606 | -1.261167 | 0.207248742 | 0.384480245 | RPS6KL1 | protein_coding | 55.36 | -0.13662411 | 0.13662411 | ENSG00000198208 | n.s. |
| ENSG00000218426 | 280.4424616 | 0.379075487 | 0.3294757 | 1.150542 | 0.249920904 | 0.436210198 | | processed_pseudogene | 54.14 | 0.136582484 | 0.136582484 | ENSG00000218426 | n.s. |
| ENSG00000155393 | 665.3249417 | -0.326154726 | 0.2572045 | -1.268075 | 0.204770994 | 0.381816867 | HEATR3 | protein_coding | 42.6 | -0.136379932 | 0.136379932 | ENSG00000155393 | n.s. |
| ENSG00000164048 | 371.330163 | -0.315651598 | 0.2439599 | -1.293867 | 0.195711386 | 0.369827582 | ZNF589 | protein_coding | 45.4 | -0.136361712 | 0.136361712 | ENSG00000164048 | n.s. |
| ENSG00000123144 | 2028.232819 | 0.366576407 | 0.3120315 | 1.174806 | 0.240072412 | 0.424933663 | C19orf43 | protein_coding | 54.86 | 0.136248702 | 0.136248702 | ENSG00000123144 | n.s. |
| ENSG00000110011 | 78.4742097 | 0.337473729 | 0.272731 | 1.237387 | 0.215943513 | 0.39487543 | DNAJC4 | protein_coding | 58.22 | 0.136184111 | 0.136184111 | ENSG00000110011 | n.s. |
| ENSG00000103275 | 2507.990087 | 0.314610011 | 0.242985 | 1.294771 | 0.195399197 | 0.369388054 | UBE2I | protein_coding | 59.46 | 0.136074227 | 0.136074227 | ENSG00000103275 | n.s. |
| ENSG00000244038 | 944.6902171 | 0.358702115 | 0.3015518 | 1.189521 | 0.234234837 | 0.417894879 | DDOST | protein_coding | 50.61 | 0.135924051 | 0.135924051 | ENSG00000244038 | n.s. |
| ENSG00000169567 | 4172.242885 | 0.354174924 | 0.2954253 | 1.198865 | 0.230580572 | 0.413390021 | HINT1 | protein_coding | 43.5 | 0.135875672 | 0.135875672 | ENSG00000169567 | n.s. |
| ENSG00000250802 | 228.2793714 | -0.301660234 | 0.2272696 | -1.327324 | 0.184401672 | 0.354567666 | ZBED3-AS1 | antisense | 41.05 | -0.135837866 | 0.135837866 | ENSG00000250802 | n.s. |
| ENSG00000282381 | 192.343592 | 0.321694878 | 0.252348 | 1.274807 | 0.2023777 | 0.378577735 | | lincRNA | 39.38 | 0.135705354 | 0.135705354 | ENSG00000282381 | n.s. |
| ENSG00000099284 | 1690.978167 | -0.340880318 | 0.278295 | -1.224888 | 0.220617337 | 0.399903032 | H2AFY2 | protein_coding | 45.79 | -0.135685809 | 0.135685809 | ENSG00000099284 | n.s. |

| | | | | | | | | | | | | | |
|---|---|---|---|---|---|---|---|---|---|---|---|---|---|
| ENSG00000242818 | 8.442785332 | -0.428504832 | 0.4074524 | -1.051669 | 0.292951634 | 0.482351656 | RN7SL846P | misc_RNA | 54.11 | -0.135680153 | 0.135680153 | ENSG00000242818 | n.s. |
| ENSG00000230068 | 23.08309402 | 0.39954768 | 0.3618995 | 1.104029 | 0.269580381 | 0.457839429 | CDC42-IT1 | sense_intronic | 41.02 | 0.135561257 | 0.135561257 | ENSG00000230068 | n.s. |
| ENSG00000275038 | 106.8872093 | -0.33581126 | 0.2713564 | -1.237528 | 0.215891048 | 0.39487543 | | protein_coding | 53.55 | -0.135513238 | 0.135513238 | ENSG00000275038 | n.s. |
| ENSG00000137434 | 29.65332191 | 0.364509113 | 0.3103206 | 1.174621 | 0.240146345 | 0.42495734 | C6orf52 | protein_coding | 41.8 | 0.135471512 | 0.135471512 | ENSG00000137434 | n.s. |
| ENSG00000163444 | 1511.186556 | 0.339828709 | 0.2771836 | 1.226006 | 0.220196523 | 0.399556392 | TMEM183A | protein_coding | 41.5 | 0.135395206 | 0.135395206 | ENSG00000163444 | n.s. |
| ENSG00000279356 | 100.3638996 | -0.311659188 | 0.2401479 | -1.29778 | 0.194362877 | 0.368178822 | | TEC | 40.75 | -0.135241761 | 0.135241761 | ENSG00000279356 | n.s. |
| ENSG00000230126 | 22.05036845 | 0.397512989 | 0.3593038 | 1.106342 | 0.26857837 | 0.456862144 | FGF12-AS2 | antisense | 52.64 | 0.135239813 | 0.135239813 | ENSG00000230126 | n.s. |
| ENSG00000259398 | 34.5389137 | 0.37255852 | 0.3222473 | 1.156126 | 0.2476296 | 0.433569783 | | sense_intronic | 47.21 | 0.135216759 | 0.135216759 | ENSG00000259398 | n.s. |
| ENSG00000137770 | 2872.191178 | -0.329804228 | 0.2640136 | -1.249194 | 0.211594097 | 0.389428236 | CTDSPL2 | protein_coding | 40.06 | -0.135078763 | 0.135078763 | ENSG00000137770 | n.s. |
| ENSG00000183444 | 104.2697349 | -0.328452299 | 0.2623242 | -1.252085 | 0.210538725 | 0.388152896 | OR7E38P | unprocessed_pseudogene | 49.34 | -0.134992965 | 0.134992965 | ENSG00000183444 | n.s. |
| ENSG00000176623 | 619.7590217 | -0.335399815 | 0.2716044 | -1.234884 | 0.216873751 | 0.395892425 | RMDN1 | protein_coding | 38.38 | -0.134972535 | 0.134972535 | ENSG00000176623 | n.s. |
| ENSG00000144445 | 709.7196263 | -0.340018405 | 0.278109 | -1.222609 | 0.221477583 | 0.401020762 | KANSL1L | protein_coding | 36.36 | -0.134930572 | 0.134930572 | ENSG00000144445 | n.s. |
| ENSG00000055147 | 87.0651939 | -0.337486538 | 0.2746002 | -1.229011 | 0.219067813 | 0.39828691 | FAM114A2 | protein_coding | 37.35 | -0.134928457 | 0.134928457 | ENSG00000055147 | n.s. |
| ENSG00000104112 | 2092.946212 | -0.333022598 | 0.2683347 | -1.241072 | 0.214579213 | 0.393414062 | SCG3 | protein_coding | 40.12 | -0.134924146 | 0.134924146 | ENSG00000104112 | n.s. |
| ENSG00000164919 | 2388.829471 | 0.379047737 | 0.3326205 | 1.13958 | 0.254461291 | 0.440599995 | COX6C | protein_coding | 40.79 | 0.134924132 | 0.134924132 | ENSG00000164919 | n.s. |
| ENSG00000165995 | 44.66303423 | -0.376515325 | 0.3286224 | -1.145738 | 0.25190341 | 0.438191822 | CACNB2 | protein_coding | 40.47 | -0.134918894 | 0.134918894 | ENSG00000165995 | n.s. |
| ENSG00000125995 | 635.3223894 | 0.356288544 | 0.2997907 | 1.188458 | 0.234653166 | 0.418254926 | ROMO1 | protein_coding | 50.26 | 0.134876212 | 0.134876212 | ENSG00000125995 | n.s. |
| ENSG00000179965 | 253.6645066 | -0.335198161 | 0.2715406 | -1.234431 | 0.217042334 | 0.395990382 | ZNF771 | protein_coding | 51.85 | -0.13485537 | 0.13485537 | ENSG00000179965 | n.s. |
| ENSG00000158528 | 513.4306019 | -0.328026608 | 0.2620112 | -1.251956 | 0.210585717 | 0.388188128 | PPP1R9A | protein_coding | 36.45 | -0.134805077 | 0.134805077 | ENSG00000158528 | n.s. |
| ENSG00000105397 | 204.0080646 | 0.322125111 | 0.2539145 | 1.268636 | 0.204570913 | 0.381545992 | TYK2 | protein_coding | 54.34 | 0.134794253 | 0.134794253 | ENSG00000105397 | n.s. |
| ENSG00000166341 | 1549.317909 | -0.304570317 | 0.2319166 | -1.313275 | 0.189090265 | 0.361044702 | DCHS1 | protein_coding | 52.66 | -0.134753794 | 0.134753794 | ENSG00000166341 | n.s. |
| ENSG00000095906 | 632.4005213 | 0.340365019 | 0.2787636 | 1.220981 | 0.22209325 | 0.401898143 | NUBP2 | protein_coding | 62.18 | 0.134745065 | 0.134745065 | ENSG00000095906 | n.s. |
| ENSG00000169682 | 315.4336366 | 0.308107338 | 0.236278 | 1.304004 | 0.192232351 | 0.365335929 | SPNS1 | protein_coding | 55.79 | 0.134737685 | 0.134737685 | ENSG00000169682 | n.s. |
| ENSG00000008405 | 1495.560286 | 0.345296206 | 0.2853151 | 1.210227 | 0.226191631 | 0.407249961 | CRY1 | protein_coding | 37.62 | 0.134713498 | 0.134713498 | ENSG00000008405 | n.s. |
| ENSG00000111647 | 1463.329771 | -0.328016828 | 0.262302 | -1.250531 | 0.211105551 | 0.388786056 | UHRF1BP1L | protein_coding | 38.58 | -0.134581801 | 0.134581801 | ENSG00000111647 | n.s. |
| ENSG00000149541 | 315.618751 | 0.330241921 | 0.2653114 | 1.244733 | 0.213229771 | 0.391507069 | B3GAT3 | protein_coding | 53.95 | 0.134494454 | 0.134494454 | ENSG00000149541 | n.s. |
| ENSG00000109083 | 229.0281787 | 0.351398106 | 0.2935793 | 1.196944 | 0.231328295 | 0.414315534 | IFT20 | protein_coding | 47.79 | 0.134469086 | 0.134469086 | ENSG00000109083 | n.s. |
| ENSG00000103978 | 1442.640887 | 0.311828593 | 0.2412673 | 1.292461 | 0.196197463 | 0.370522683 | TMEM87A | protein_coding | 38.93 | 0.134455874 | 0.134455874 | ENSG00000103978 | n.s. |
| ENSG00000181315 | 342.052246 | -0.327538359 | 0.2619864 | -1.250211 | 0.211222423 | 0.388949847 | ZNF322 | protein_coding | 37.34 | -0.134325576 | 0.134325576 | ENSG00000181315 | n.s. |
| ENSG00000084731 | 838.8509599 | -0.306714436 | 0.2349734 | -1.305315 | 0.191785509 | 0.364881656 | KIF3C | protein_coding | 48.25 | -0.134294293 | 0.134294293 | ENSG00000084731 | n.s. |
| ENSG00000055163 | 48.43679638 | 0.334896042 | 0.2720436 | 1.231038 | 0.21830863 | 0.397217859 | CYFIP2 | protein_coding | 44.29 | 0.134283679 | 0.134283679 | ENSG00000055163 | n.s. |
| ENSG00000164332 | 724.0594691 | -0.345934713 | 0.2867715 | -1.206308 | 0.227698803 | 0.409170165 | UBLCP1 | protein_coding | 36.18 | -0.134255892 | 0.134255892 | ENSG00000164332 | n.s. |

| | | | | | | | | | | | | | |
|---|---|---|---|---|---|---|---|---|---|---|---|---|---|
| ENSG00000168256 | 566.1743518 | -0.30974968 | 0.2389254 | -1.296428 | 0.194827983 | 0.36866264 | NKIRAS2 | protein_coding | 49.4 | -0.134236489 | 0.134236489 | ENSG00000168256 | n.s. |
| ENSG00000167658 | 23069.12792 | 0.336079271 | 0.2736549 | 1.228113 | 0.219404384 | 0.398742623 | EEF2 | protein_coding | 59.53 | 0.134198919 | 0.134198919 | ENSG00000167658 | n.s. |
| ENSG00000198522 | 1242.635322 | 0.333505609 | 0.2705531 | 1.232681 | 0.217694885 | 0.396671363 | GPN1 | protein_coding | 41.23 | 0.133925564 | 0.133925564 | ENSG00000198522 | n.s. |
| ENSG00000075785 | 1070.030473 | -0.330987339 | 0.2670583 | -1.239382 | 0.215203935 | 0.394196078 | RAB7A | protein_coding | 44.06 | -0.13381411 | 0.13381411 | ENSG00000075785 | n.s. |
| ENSG00000145335 | 176.878845 | 0.348838047 | 0.291119 | 1.198266 | 0.230813415 | 0.413592688 | SNCA | protein_coding | 36.12 | 0.133753978 | 0.133753978 | ENSG00000145335 | n.s. |
| ENSG00000130726 | 4461.401805 | 0.316093482 | 0.247658 | 1.276331 | 0.201838619 | 0.377684959 | TRIM28 | protein_coding | 59.3 | 0.133666549 | 0.133666549 | ENSG00000130726 | n.s. |
| ENSG00000165806 | 168.3578055 | -0.343790069 | 0.2848109 | -1.207082 | 0.227400595 | 0.408792574 | CASP7 | protein_coding | 42.96 | -0.133561411 | 0.133561411 | ENSG00000165806 | n.s. |
| ENSG00000157483 | 68.06993194 | 0.329054145 | 0.2649965 | 1.24173 | 0.214336083 | 0.393020055 | MYO1E | protein_coding | 43.52 | 0.13345952 | 0.13345952 | ENSG00000157483 | n.s. |
| ENSG00000131116 | 1050.954662 | 0.343155387 | 0.2842451 | 1.207252 | 0.227335148 | 0.408792574 | ZNF428 | protein_coding | 52.67 | 0.133314839 | 0.133314839 | ENSG00000131116 | n.s. |
| ENSG00000120696 | 637.5584961 | -0.28103783 | 0.2050774 | -1.370399 | 0.170562464 | 0.335557607 | KBTBD7 | protein_coding | 42.21 | -0.133277388 | 0.133277388 | ENSG00000120696 | n.s. |
| ENSG00000166484 | 678.9460275 | -0.283592225 | 0.2081279 | -1.362586 | 0.173012985 | 0.339084994 | MAPK7 | protein_coding | 59.22 | -0.133200838 | 0.133200838 | ENSG00000166484 | n.s. |
| ENSG00000100413 | 734.7722709 | -0.330545087 | 0.2675752 | -1.235335 | 0.216705826 | 0.395791794 | POLR3H | protein_coding | 55.16 | -0.133055377 | 0.133055377 | ENSG00000100413 | n.s. |
| ENSG00000144043 | 550.398636 | -0.32703598 | 0.2629929 | -1.243516 | 0.213677605 | 0.392122486 | TEX261 | protein_coding | 52.63 | -0.132965717 | 0.132965717 | ENSG00000144043 | n.s. |
| ENSG00000253716 | 122.221478 | 0.303478085 | 0.2324049 | 1.305816 | 0.191615022 | 0.364681106 | MINCR | antisense | 49.93 | 0.132949724 | 0.132949724 | ENSG00000253716 | n.s. |
| ENSG00000145592 | 22321.16031 | 0.346773728 | 0.2894604 | 1.198001 | 0.230916747 | 0.4137246 | RPL37 | protein_coding | 42.83 | 0.132914436 | 0.132914436 | ENSG00000145592 | n.s. |
| ENSG00000237550 | 965.5054804 | 0.468681761 | 0.480066 | 0.976286 | 0.328922678 | 0.520526307 | RPL9P9 | transcribed_processed_pseudog | 41.84 | 0.132898143 | 0.132898143 | ENSG00000237550 | n.s. |
| ENSG00000240303 | 514.0582359 | -0.339857017 | 0.2804298 | -1.211915 | 0.225544979 | 0.406453888 | ACAD11 | protein_coding | 37.61 | -0.132880259 | 0.132880259 | ENSG00000240303 | n.s. |
| ENSG00000115318 | 459.4730726 | -0.338457211 | 0.2786256 | -1.214739 | 0.224465825 | 0.405127573 | LOXL3 | protein_coding | 50.8 | -0.132813384 | 0.132813384 | ENSG00000115318 | n.s. |
| ENSG00000080603 | 627.2270085 | -0.313593143 | 0.245453 | -1.27761 | 0.201387028 | 0.37727551 | SRCAP | protein_coding | 49.63 | -0.132756955 | 0.132756955 | ENSG00000080603 | n.s. |
| ENSG00000141994 | 764.3779787 | 0.335785535 | 0.2755006 | 1.21882 | 0.222912655 | 0.402752777 | DUS3L | protein_coding | 61.93 | 0.132622344 | 0.132622344 | ENSG00000141994 | n.s. |
| ENSG00000132964 | 639.2280192 | -0.329455663 | 0.2667361 | -1.235137 | 0.21677944 | 0.395791794 | CDK8 | protein_coding | 36.99 | -0.132616847 | 0.132616847 | ENSG00000132964 | n.s. |
| ENSG00000267871 | 29.48552833 | -0.371741167 | 0.3259232 | -1.140579 | 0.254045117 | 0.440427115 | | antisense | 45.15 | -0.132386677 | 0.132386677 | ENSG00000267871 | n.s. |
| ENSG00000074842 | 412.3628593 | 0.360491982 | 0.3090896 | 1.166302 | 0.243492217 | 0.42940369 | MYDGF | protein_coding | 54.74 | 0.132348945 | 0.132348945 | ENSG00000074842 | n.s. |
| ENSG00000172336 | 649.2651163 | 0.349610383 | 0.2941256 | 1.188643 | 0.234580093 | 0.418254926 | POP7 | protein_coding | 59.81 | 0.132348134 | 0.132348134 | ENSG00000172336 | n.s. |
| ENSG00000167972 | 386.2445249 | -0.318370859 | 0.2522439 | -1.262155 | 0.206893102 | 0.384178801 | ABCA3 | protein_coding | 53.03 | -0.132272459 | 0.132272459 | ENSG00000167972 | n.s. |
| ENSG00000164975 | 2451.857348 | -0.319060539 | 0.2534573 | -1.258834 | 0.208090433 | 0.385165646 | SNAPC3 | protein_coding | 39.78 | -0.132203518 | 0.132203518 | ENSG00000164975 | n.s. |
| ENSG00000146112 | 344.716374 | -0.304846905 | 0.2350237 | -1.29709 | 0.194600162 | 0.368477909 | PPP1R18 | protein_coding | 56.1 | -0.132178126 | 0.132178126 | ENSG00000146112 | n.s. |
| ENSG00000070961 | 2162.303574 | -0.313306877 | 0.2458495 | -1.274385 | 0.202527138 | 0.378765423 | ATP2B1 | protein_coding | 36.37 | -0.132099476 | 0.132099476 | ENSG00000070961 | n.s. |
| ENSG00000163597 | 11417.98982 | 0.284745578 | 0.2105539 | 1.352365 | 0.176258707 | 0.343656347 | SNHG16 | processed_transcript | 45.22 | 0.132086535 | 0.132086535 | ENSG00000163597 | n.s. |
| ENSG00000167552 | 13711.69115 | 0.368515623 | 0.3217419 | 1.145376 | 0.25205327 | 0.438338455 | TUBA1A | protein_coding | 48.62 | 0.131998768 | 0.131998768 | ENSG00000167552 | n.s. |
| ENSG00000241837 | 2907.239239 | 0.359392743 | 0.3081047 | 1.166463 | 0.243427299 | 0.429343588 | ATP5O | protein_coding | 43.51 | 0.131967225 | 0.131967225 | ENSG00000241837 | n.s. |
| ENSG00000096063 | 2468.062038 | -0.327163511 | 0.2644743 | -1.237033 | 0.216074693 | 0.395063421 | SRPK1 | protein_coding | 40.48 | -0.131955899 | 0.131955899 | ENSG00000096063 | n.s. |

| | | | | | | | | | | | | | |
|---|---|---|---|---|---|---|---|---|---|---|---|---|---|
| ENSG00000124562 | 1838.229702 | 0.360914963 | 0.3103999 | 1.162742 | 0.244934296 | 0.430909806 | SNRPC | protein_coding | 42.71 | 0.131955427 | 0.131955427 | ENSG00000124562 | n.s. |
| ENSG00000144619 | 53.99876861 | 0.327600617 | 0.265463 | 1.234073 | 0.217175865 | 0.396032547 | CNTN4 | protein_coding | 37.97 | 0.131783612 | 0.131783612 | ENSG00000144619 | n.s. |
| ENSG00000121940 | 1272.154992 | -0.328504798 | 0.2667608 | -1.231458 | 0.218151512 | 0.397087658 | CLCC1 | protein_coding | 41.67 | -0.131767746 | 0.131767746 | ENSG00000121940 | n.s. |
| ENSG00000267322 | 306.3468729 | -0.286213774 | 0.2127198 | -1.345497 | 0.178464819 | 0.346571869 | SNHG22 | antisense | 43.78 | -0.131717494 | 0.131717494 | ENSG00000267322 | n.s. |
| ENSG00000143158 | 527.9543479 | 0.348018945 | 0.2929589 | 1.187945 | 0.234855221 | 0.418519038 | MPC2 | protein_coding | 37.87 | 0.131650271 | 0.131650271 | ENSG00000143158 | n.s. |
| ENSG00000071082 | 22791.60031 | 0.358479252 | 0.3074648 | 1.16592 | 0.243646894 | 0.429471856 | RPL31 | protein_coding | 42.84 | 0.131585291 | 0.131585291 | ENSG00000071082 | n.s. |
| ENSG00000230898 | 96.27906082 | 0.31599228 | 0.2499784 | 1.264078 | 0.206201983 | 0.383611734 | | sense_intronic | 46.04 | 0.131486953 | 0.131486953 | ENSG00000230898 | n.s. |
| ENSG00000087995 | 959.5019666 | -0.320834158 | 0.2568857 | -1.248937 | 0.211687988 | 0.389549545 | METTL2A | protein_coding | 45.2 | -0.131361471 | 0.131361471 | ENSG00000087995 | n.s. |
| ENSG00000202048 | 23.08262634 | -0.598074 | 0.7278099 | -0.821745 | 0.411222172 | 0.603313625 | SNORD114-20 | snoRNA | 47.22 | -0.131251446 | 0.131251446 | ENSG00000202048 | n.s. |
| ENSG00000171960 | 437.1944759 | 0.349180153 | 0.2952328 | 1.182728 | 0.236916847 | 0.421222483 | PPIH | protein_coding | 43.8 | 0.131113116 | 0.131113116 | ENSG00000171960 | n.s. |
| ENSG00000165915 | 157.5884226 | 0.331253323 | 0.2714065 | 1.220506 | 0.222273067 | 0.401992549 | SLC39A13 | protein_coding | 60.6 | 0.131104101 | 0.131104101 | ENSG00000165915 | n.s. |
| ENSG00000205208 | 2014.972222 | 0.297671348 | 0.2272569 | 1.309845 | 0.190248272 | 0.362803272 | C4orf46 | protein_coding | 41.13 | 0.131073269 | 0.131073269 | ENSG00000205208 | n.s. |
| ENSG00000157110 | 5.569658343 | -0.479971318 | 0.5038617 | -0.952585 | 0.340800205 | 0.533305818 | RBPMS | protein_coding | 42.19 | -0.131043535 | 0.131043535 | ENSG00000157110 | n.s. |
| ENSG00000142156 | 376.6112433 | 0.318773975 | 0.2545452 | 1.252328 | 0.210450438 | 0.388085178 | COL6A1 | protein_coding | 64.67 | 0.131039356 | 0.131039356 | ENSG00000142156 | n.s. |
| ENSG00000256031 | 293.4772204 | -0.315858618 | 0.2506941 | -1.259936 | 0.207692281 | 0.384739173 | | sense_intronic | 44.24 | -0.131028765 | 0.131028765 | ENSG00000256031 | n.s. |
| ENSG00000056050 | 980.4271913 | -0.342390365 | 0.2860949 | -1.196772 | 0.231395359 | 0.414315534 | HPF1 | protein_coding | 38.6 | -0.131022105 | 0.131022105 | ENSG00000056050 | n.s. |
| ENSG00000105404 | 282.4802185 | 0.365894787 | 0.3196142 | 1.144801 | 0.252291443 | 0.438533553 | RABAC1 | protein_coding | 61.62 | 0.130989299 | 0.130989299 | ENSG00000105404 | n.s. |
| ENSG00000171533 | 693.3618622 | -0.326611514 | 0.2653686 | -1.230784 | 0.218403519 | 0.39728698 | MAP6 | protein_coding | 44.55 | -0.130937141 | 0.130937141 | ENSG00000171533 | n.s. |
| ENSG00000095066 | 7.957368518 | -0.451964777 | 0.4569483 | -0.989094 | 0.32261715 | 0.513289747 | HOOK2 | protein_coding | 53.64 | -0.130905908 | 0.130905908 | ENSG00000095066 | n.s. |
| ENSG00000127616 | 3266.192304 | -0.292599255 | 0.2213148 | -1.322095 | 0.186136513 | 0.357065124 | SMARCA4 | protein_coding | 52.73 | -0.130865768 | 0.130865768 | ENSG00000127616 | n.s. |
| ENSG00000120088 | 523.7276162 | -0.340854779 | 0.2841872 | -1.199402 | 0.230371556 | 0.413119935 | CRHR1 | protein_coding | 54.73 | -0.130862278 | 0.130862278 | ENSG00000120088 | n.s. |
| ENSG00000105245 | 339.5860582 | -0.336259158 | 0.2782671 | -1.208404 | 0.226891782 | 0.408194118 | NUMBL | protein_coding | 52.55 | -0.130849622 | 0.130849622 | ENSG00000105245 | n.s. |
| ENSG00000253250 | 480.0627846 | 0.351635241 | 0.2991476 | 1.175457 | 0.239811817 | 0.424580357 | C8orf88 | protein_coding | 32.98 | 0.130822411 | 0.130822411 | ENSG00000253250 | n.s. |
| ENSG00000148484 | 678.8321306 | -0.334839233 | 0.2766246 | -1.210446 | 0.226107618 | 0.407186527 | RSU1 | protein_coding | 41.15 | -0.130656479 | 0.130656479 | ENSG00000148484 | n.s. |
| ENSG00000244716 | 91.44816562 | 0.379732774 | 0.3411114 | 1.113222 | 0.265613055 | 0.453091223 | | processed_pseudogene | 46.13 | 0.130557577 | 0.130557577 | ENSG00000244716 | n.s. |
| ENSG00000257727 | 624.0464623 | -0.334172026 | 0.2758513 | -1.211421 | 0.225734185 | 0.40674221 | CNPY2 | protein_coding | 49.44 | -0.13055458 | 0.13055458 | ENSG00000257727 | n.s. |
| ENSG00000118096 | 223.1104265 | -0.35815721 | 0.3087261 | -1.160113 | 0.24600267 | 0.432025122 | IFT46 | protein_coding | 45 | -0.130545079 | 0.130545079 | ENSG00000118096 | n.s. |
| ENSG00000125676 | 2637.491411 | -0.333961238 | 0.2757465 | -1.211117 | 0.225850533 | 0.406846549 | THOC2 | protein_coding | 38.09 | -0.130435028 | 0.130435028 | ENSG00000125676 | n.s. |
| ENSG00000100836 | 1524.202688 | 0.313903366 | 0.2489803 | 1.260756 | 0.207396813 | 0.384523202 | PABPN1 | protein_coding | 50.48 | 0.130294209 | 0.130294209 | ENSG00000100836 | n.s. |
| ENSG00000165219 | 57.88156172 | 0.324439676 | 0.2632452 | 1.232462 | 0.217776635 | 0.396682343 | GAPVD1 | protein_coding | 42.11 | 0.130281065 | 0.130281065 | ENSG00000165219 | n.s. |
| ENSG00000182154 | 549.4481692 | 0.366340454 | 0.3217472 | 1.138597 | 0.254871224 | 0.440980744 | MRPL41 | protein_coding | 72.44 | 0.130263476 | 0.130263476 | ENSG00000182154 | n.s. |
| ENSG00000224080 | 38.48445498 | 0.371000421 | 0.3287817 | 1.12841 | 0.259146999 | 0.445664978 | UBE2FP1 | processed_pseudogene | 40.72 | 0.130217992 | 0.130217992 | ENSG00000224080 | n.s. |

| | | | | | | | | | | | | | |
|---|---|---|---|---|---|---|---|---|---|---|---|---|---|
| ENSG00000284329 | 27.28594271 | 0.3531792 | 0.3020717 | 1.16919 | 0.242327124 | 0.427857819 | MIR9-3 | miRNA | 44.44 | 0.130217357 | 0.130217357 | ENSG00000284329 | n.s. |
| ENSG00000257800 | 441.9802989 | 0.27929329 | 0.2058985 | 1.356461 | 0.174952474 | 0.34187555 | FNBP1P1 | processed_pseudogene | 41.55 | 0.130187528 | 0.130187528 | ENSG00000257800 | n.s. |
| ENSG00000169230 | 2007.692113 | 0.339360562 | 0.2832049 | 1.198286 | 0.230805545 | 0.413592688 | PRELID1 | protein_coding | 54.24 | 0.130120053 | 0.130120053 | ENSG00000169230 | n.s. |
| ENSG00000104635 | 967.4042417 | -0.347660745 | 0.2945954 | -1.180129 | 0.23794875 | 0.422409849 | SLC39A14 | protein_coding | 48.28 | -0.130117584 | 0.130117584 | ENSG00000104635 | n.s. |
| ENSG00000122490 | 296.2246413 | -0.340026279 | 0.284334 | -1.195869 | 0.23174758 | 0.414786162 | PQLC1 | protein_coding | 52.94 | -0.129949795 | 0.129949795 | ENSG00000122490 | n.s. |
| ENSG00000159131 | 1986.04775 | -0.325842729 | 0.2656661 | -1.226512 | 0.220006003 | 0.39934579 | GART | protein_coding | 41.63 | -0.129897494 | 0.129897494 | ENSG00000159131 | n.s. |
| ENSG00000180992 | 702.5686218 | 0.377184705 | 0.3383593 | 1.114746 | 0.264959204 | 0.452554959 | MRPL14 | protein_coding | 46.16 | 0.129875508 | 0.129875508 | ENSG00000180992 | n.s. |
| ENSG00000198816 | 297.2978951 | -0.314438775 | 0.250328 | -1.256107 | 0.209077068 | 0.386374341 | ZNF358 | protein_coding | 60.98 | -0.129860612 | 0.129860612 | ENSG00000198816 | n.s. |
| ENSG00000131584 | 568.0260916 | 0.321114107 | 0.2591435 | 1.239136 | 0.215295088 | 0.39431117 | ACAP3 | protein_coding | 67.3 | 0.129781773 | 0.129781773 | ENSG00000131584 | n.s. |
| ENSG00000103550 | 1723.86955 | -0.316994374 | 0.253885 | -1.248575 | 0.211820699 | 0.389668403 | KNOP1 | protein_coding | 51.83 | -0.129747322 | 0.129747322 | ENSG00000103550 | n.s. |
| ENSG00000178149 | 140.6643686 | 0.321435244 | 0.2597587 | 1.237438 | 0.215924633 | 0.39487543 | DALRD3 | protein_coding | 60.06 | 0.129711942 | 0.129711942 | ENSG00000178149 | n.s. |
| ENSG00000128185 | 296.051411 | 0.356567472 | 0.3079826 | 1.157752 | 0.246965276 | 0.432823448 | DGCR6L | protein_coding | 57.73 | 0.129679746 | 0.129679746 | ENSG00000128185 | n.s. |
| ENSG00000222489 | 175.5052856 | 0.345148403 | 0.2917692 | 1.18295 | 0.236828972 | 0.421120022 | SNORA79B | snoRNA | 45.95 | 0.129635707 | 0.129635707 | ENSG00000222489 | n.s. |
| ENSG00000110435 | 872.5172724 | -0.341392666 | 0.2866822 | -1.19084 | 0.233716284 | 0.417156799 | PDHX | protein_coding | 36.68 | -0.129627025 | 0.129627025 | ENSG00000110435 | n.s. |
| ENSG00000187837 | 3606.957519 | 0.281818002 | 0.2096934 | 1.343953 | 0.17896366 | 0.347045353 | HIST1H1C | protein_coding | 57.48 | 0.129527433 | 0.129527433 | ENSG00000187837 | n.s. |
| ENSG00000070610 | 735.5679335 | -0.325163432 | 0.2653413 | -1.225453 | 0.220404465 | 0.399707486 | GBA2 | protein_coding | 50.7 | -0.129498846 | 0.129498846 | ENSG00000070610 | n.s. |
| ENSG00000044115 | 2229.195597 | 0.333022122 | 0.2757598 | 1.207653 | 0.227180832 | 0.408661334 | CTNNA1 | protein_coding | 44.15 | 0.129424536 | 0.129424536 | ENSG00000044115 | n.s. |
| ENSG00000184863 | 1392.441435 | -0.304328742 | 0.2374869 | -1.281455 | 0.200033909 | 0.375656215 | RBM33 | protein_coding | 40.78 | -0.129403448 | 0.129403448 | ENSG00000184863 | n.s. |
| ENSG00000182117 | 985.0143098 | 0.36923147 | 0.3276871 | 1.126781 | 0.259835219 | 0.446462515 | NOP10 | protein_coding | 49.56 | 0.129310399 | 0.129310399 | ENSG00000182117 | n.s. |
| ENSG00000138061 | 410.8924687 | 0.314407665 | 0.2510716 | 1.252263 | 0.210474121 | 0.388085178 | CYP1B1 | protein_coding | 44.41 | 0.129244484 | 0.129244484 | ENSG00000138061 | n.s. |
| ENSG00000167641 | 365.6303637 | 0.358805687 | 0.3122339 | 1.149157 | 0.250491411 | 0.436511212 | PPP1R14A | protein_coding | 58.39 | 0.129171697 | 0.129171697 | ENSG00000167641 | n.s. |
| ENSG00000085840 | 1612.497086 | -0.323592228 | 0.2637563 | -1.226861 | 0.219874941 | 0.399303693 | ORC1 | protein_coding | 44.76 | -0.129015145 | 0.129015145 | ENSG00000085840 | n.s. |
| ENSG00000198720 | 733.7214516 | 0.294300463 | 0.2253102 | 1.306201 | 0.191484055 | 0.364610572 | ANKRD13B | protein_coding | 55.54 | 0.128953853 | 0.128953853 | ENSG00000198720 | n.s. |
| ENSG00000163013 | 375.7202134 | -0.31055382 | 0.246549 | -1.259603 | 0.207812733 | 0.384911093 | FBXO41 | protein_coding | 52.56 | -0.128767903 | 0.128767903 | ENSG00000163013 | n.s. |
| ENSG00000119414 | 964.4630151 | -0.334534516 | 0.278533 | -1.201059 | 0.22972846 | 0.412232162 | PPP6C | protein_coding | 42.35 | -0.128748327 | 0.128748327 | ENSG00000119414 | n.s. |
| ENSG00000149328 | 105.4978534 | -0.328184342 | 0.2702899 | -1.214194 | 0.224673763 | 0.405309961 | GLB1L2 | protein_coding | 51.13 | -0.128718074 | 0.128718074 | ENSG00000149328 | n.s. |
| ENSG00000207757 | 17.23103664 | 0.390264338 | 0.3614225 | 1.079801 | 0.280230851 | 0.467936452 | MIR93 | miRNA | 60 | 0.1287143 | 0.1287143 | ENSG00000207757 | n.s. |
| ENSG00000161647 | 444.2191015 | -0.322112902 | 0.2622645 | -1.228199 | 0.219372342 | 0.398736438 | MPP3 | protein_coding | 48.95 | -0.128624216 | 0.128624216 | ENSG00000161647 | n.s. |
| ENSG00000141337 | 94.88666701 | -0.337249743 | 0.2826082 | -1.193347 | 0.232733355 | 0.415855558 | ARSG | protein_coding | 45.96 | -0.128511541 | 0.128511541 | ENSG00000141337 | n.s. |
| ENSG00000052802 | 1476.746776 | -0.318085377 | 0.2570387 | -1.2375 | 0.215901548 | 0.39487543 | MSMO1 | protein_coding | 37.1 | -0.128360137 | 0.128360137 | ENSG00000052802 | n.s. |
| ENSG00000166575 | 95.37151608 | -0.322153612 | 0.2628005 | -1.225848 | 0.220255695 | 0.399570805 | TMEM135 | protein_coding | 36.32 | -0.128348013 | 0.128348013 | ENSG00000166575 | n.s. |
| ENSG00000185386 | 166.6843589 | 0.333715257 | 0.278004 | 1.200397 | 0.22998513 | 0.412586388 | MAPK11 | protein_coding | 65.39 | 0.128308545 | 0.128308545 | ENSG00000185386 | n.s. |

| | | | | | | | | | | | | | |
|---|---|---|---|---|---|---|---|---|---|---|---|---|---|
| ENSG00000107679 | 987.4926867 | -0.351042748 | 0.3020831 | -1.162073 | 0.24520568 | 0.431114874 | PLEKHA1 | protein_coding | 37.12 | -0.128273475 | 0.128273475 | ENSG00000107679 | n.s. |
| ENSG00000160299 | 1414.433893 | -0.303199084 | 0.2375166 | -1.276538 | 0.201765282 | 0.377649 | PCNT | protein_coding | 51.58 | -0.128226428 | 0.128226428 | ENSG00000160299 | n.s. |
| ENSG00000174938 | 282.5243788 | -0.311680675 | 0.2487934 | -1.252769 | 0.210289755 | 0.387899346 | SEZ6L2 | protein_coding | 52.94 | -0.128188325 | 0.128188325 | ENSG00000174938 | n.s. |
| ENSG00000235437 | 48.99959468 | -0.337801507 | 0.2838053 | -1.190258 | 0.233945109 | 0.417484973 | LINC01278 | processed_transcript | 38.18 | -0.128148093 | 0.128148093 | ENSG00000235437 | n.s. |
| ENSG00000148358 | 808.7903044 | -0.314926948 | 0.2531884 | -1.243844 | 0.213556936 | 0.391952705 | GPR107 | protein_coding | 46.73 | -0.12810168 | 0.12810168 | ENSG00000148358 | n.s. |
| ENSG00000175611 | 471.2463428 | 0.27998922 | 0.208916 | 1.3402 | 0.180180389 | 0.348811396 | LINC00476 | processed_transcript | 42.23 | 0.128069683 | 0.128069683 | ENSG00000175611 | n.s. |
| ENSG00000117054 | 2071.11525 | -0.343055835 | 0.291322 | -1.177583 | 0.238962888 | 0.423781259 | ACADM | protein_coding | 40.45 | -0.127911199 | 0.127911199 | ENSG00000117054 | n.s. |
| ENSG00000260442 | 63.81992049 | 0.320992062 | 0.2619982 | 1.225169 | 0.220511518 | 0.399782327 | ATP2A1-AS1 | antisense | 59.34 | 0.127811467 | 0.127811467 | ENSG00000260442 | n.s. |
| ENSG00000111554 | 1056.0851 | -0.321401257 | 0.2625327 | -1.224233 | 0.220864239 | 0.400295742 | MDM1 | protein_coding | 38.66 | -0.127795256 | 0.127795256 | ENSG00000111554 | n.s. |
| ENSG00000153989 | 1055.256949 | -0.332924428 | 0.2777824 | -1.198508 | 0.230719319 | 0.413569621 | NUS1 | protein_coding | 37.38 | -0.127660328 | 0.127660328 | ENSG00000153989 | n.s. |
| ENSG00000222588 | 25.71475349 | -0.378181895 | 0.3443197 | -1.098345 | 0.272053805 | 0.459908344 | | snoRNA | 36.43 | -0.127571611 | 0.127571611 | ENSG00000222588 | n.s. |
| ENSG00000227615 | 173.2392392 | 0.377020854 | 0.3422682 | 1.101536 | 0.270663281 | 0.458839018 | | processed_pseudogene | 46.87 | 0.127561107 | 0.127561107 | ENSG00000227615 | n.s. |
| ENSG00000196418 | 2179.409363 | 0.348121927 | 0.2990798 | 1.163977 | 0.244433445 | 0.430300525 | ZNF124 | protein_coding | 42.19 | 0.12749204 | 0.12749204 | ENSG00000196418 | n.s. |
| ENSG00000089220 | 7283.418701 | 0.349726635 | 0.301544 | 1.159786 | 0.24613579 | 0.432041872 | PEBP1 | protein_coding | 49.19 | 0.127466322 | 0.127466322 | ENSG00000089220 | n.s. |
| ENSG00000162819 | 953.7369544 | -0.326484626 | 0.2697882 | -1.210152 | 0.226220728 | 0.407249961 | BROX | protein_coding | 36.81 | -0.127374368 | 0.127374368 | ENSG00000162819 | n.s. |
| ENSG00000216775 | 1004.517775 | 0.293717361 | 0.2264379 | 1.297121 | 0.194589558 | 0.368477909 | | transcribed_unprocessed_pseud | 41.39 | 0.127352483 | 0.127352483 | ENSG00000216775 | n.s. |
| ENSG00000265396 | 27.17055585 | -0.357880116 | 0.3141664 | -1.139142 | 0.254643924 | 0.440806584 | MIR3128 | miRNA | 30.3 | -0.127316541 | 0.127316541 | ENSG00000265396 | n.s. |
| ENSG00000151576 | 1209.897098 | -0.315851475 | 0.2555654 | -1.235893 | 0.21649839 | 0.395526451 | QTRT2 | protein_coding | 38.31 | -0.127232702 | 0.127232702 | ENSG00000151576 | n.s. |
| ENSG00000116774 | 78.33909333 | 0.314461648 | 0.2536516 | 1.239738 | 0.215072161 | 0.3940067 | OLFML3 | protein_coding | 45.3 | 0.127198602 | 0.127198602 | ENSG00000116774 | n.s. |
| ENSG00000214293 | 147.8863675 | -0.351977647 | 0.3053906 | -1.152549 | 0.249095594 | 0.435151562 | APTR | lincRNA | 43.28 | -0.12719045 | 0.12719045 | ENSG00000214293 | n.s. |
| ENSG00000160208 | 1809.599936 | -0.316254967 | 0.2563572 | -1.233649 | 0.217333544 | 0.39611663 | RRP1B | protein_coding | 47.51 | -0.12719045 | 0.12719045 | ENSG00000160208 | n.s. |
| ENSG00000106628 | 1632.909855 | 0.315713362 | 0.2554722 | 1.235803 | 0.216531666 | 0.395535343 | POLD2 | protein_coding | 55.2 | 0.127173985 | 0.127173985 | ENSG00000106628 | n.s. |
| ENSG00000184678 | 1858.015898 | 0.282397979 | 0.2128252 | 1.326901 | 0.184541415 | 0.354640458 | HIST2H2BE | protein_coding | 47.99 | 0.12713888 | 0.12713888 | ENSG00000184678 | n.s. |
| ENSG00000239672 | 5616.513581 | 0.346346966 | 0.2970636 | 1.165902 | 0.243654255 | 0.429471856 | NME1 | protein_coding | 46.45 | 0.12713195 | 0.12713195 | ENSG00000239672 | n.s. |
| ENSG00000198931 | 497.5488013 | 0.354516236 | 0.3094233 | 1.145732 | 0.251906008 | 0.438191822 | APRT | protein_coding | 64.35 | 0.127035835 | 0.127035835 | ENSG00000198931 | n.s. |
| ENSG00000170638 | 348.2055843 | 0.350997204 | 0.3043192 | 1.153385 | 0.248752384 | 0.434770204 | TRABD | protein_coding | 62.8 | 0.126969808 | 0.126969808 | ENSG00000170638 | n.s. |
| ENSG00000213145 | 18.03146901 | 0.506430196 | 0.5642104 | 0.897591 | 0.369403586 | 0.561660132 | CRIP1 | protein_coding | 68.07 | 0.126874135 | 0.126874135 | ENSG00000213145 | n.s. |
| ENSG00000259377 | 145.5909516 | -0.285899958 | 0.2173392 | -1.315455 | 0.1883571 | 0.35998571 | | sense_intronic | 41.42 | -0.126858025 | 0.126858025 | ENSG00000259377 | n.s. |
| ENSG00000071243 | 673.2069018 | 0.323034431 | 0.2658354 | 1.215167 | 0.224302287 | 0.405049859 | ING3 | protein_coding | 35.55 | 0.126788273 | 0.126788273 | ENSG00000071243 | n.s. |
| ENSG00000161847 | 296.8334152 | -0.290377382 | 0.2229938 | -1.302177 | 0.192855916 | 0.365971503 | RAVER1 | protein_coding | 56.86 | -0.126765039 | 0.126765039 | ENSG00000161847 | n.s. |
| ENSG00000220867 | 28.94033393 | -0.349602491 | 0.3027512 | -1.154752 | 0.248192144 | 0.434172538 | HSPE1P26 | processed_pseudogene | 38.91 | -0.126674145 | 0.126674145 | ENSG00000220867 | n.s. |
| ENSG00000180573 | 900.3556032 | 0.28574931 | 0.2174905 | 1.313847 | 0.188897648 | 0.360771187 | HIST1H2AC | protein_coding | 41.82 | 0.126520695 | 0.126520695 | ENSG00000180573 | n.s. |

| | | | | | | | | | | | | | |
|---|---|---|---|---|---|---|---|---|---|---|---|---|---|
| ENSG00000168216 | 410.315767 | -0.340666216 | 0.2902173 | -1.173831 | 0.240462546 | 0.425222787 | LMBRD1 | protein_coding | 35 | -0.126517801 | 0.126517801 | ENSG00000168216 | n.s. |
| ENSG00000141458 | 64.61049177 | -0.331052687 | 0.2768603 | -1.195739 | 0.23179836 | 0.414823724 | NPC1 | protein_coding | 44.75 | -0.126507286 | 0.126507286 | ENSG00000141458 | n.s. |
| ENSG00000113296 | 33.2778017 | -0.350796638 | 0.304983 | -1.150217 | 0.250054573 | 0.436334104 | THBS4 | protein_coding | 42.04 | -0.126350228 | 0.126350228 | ENSG00000113296 | n.s. |
| ENSG00000100926 | 231.6732996 | -0.310262386 | 0.2493014 | -1.244527 | 0.213305623 | 0.39154307 | TM9SF1 | protein_coding | 51 | -0.126345191 | 0.126345191 | ENSG00000100926 | n.s. |
| ENSG00000110429 | 965.66455 | 0.330833396 | 0.2771046 | 1.193894 | 0.232519617 | 0.41552697 | FBXO3 | protein_coding | 37.03 | 0.126180116 | 0.126180116 | ENSG00000110429 | n.s. |
| ENSG00000244313 | 464.33196 | 0.375997712 | 0.343499 | 1.094611 | 0.273687249 | 0.461772609 | | processed_pseudogene | 54.22 | 0.126174241 | 0.126174241 | ENSG00000244313 | n.s. |
| ENSG00000125835 | 5279.67695 | 0.351030597 | 0.3058156 | 1.147851 | 0.251030255 | 0.437111142 | SNRPB | protein_coding | 46.19 | 0.126163248 | 0.126163248 | ENSG00000125835 | n.s. |
| ENSG00000185480 | 1116.570549 | -0.349030693 | 0.3030857 | -1.151591 | 0.249489204 | 0.43562054 | PARPBP | protein_coding | 36.45 | -0.125962263 | 0.125962263 | ENSG00000185480 | n.s. |
| ENSG00000006530 | 476.8796786 | -0.323355498 | 0.2675 | -1.208805 | 0.226737602 | 0.407969454 | AGK | protein_coding | 39.59 | -0.125905691 | 0.125905691 | ENSG00000006530 | n.s. |
| ENSG00000160124 | 634.8771492 | -0.344413369 | 0.2963096 | -1.162343 | 0.245096216 | 0.431031276 | CCDC58 | protein_coding | 39.16 | -0.125880061 | 0.125880061 | ENSG00000160124 | n.s. |
| ENSG00000112742 | 2443.316363 | -0.322087973 | 0.2659711 | -1.210989 | 0.225899724 | 0.406882518 | TTK | protein_coding | 34.98 | -0.125785329 | 0.125785329 | ENSG00000112742 | n.s. |
| ENSG00000094880 | 1237.000474 | -0.335848571 | 0.2845842 | -1.180138 | 0.237945495 | 0.422409849 | CDC23 | protein_coding | 41.21 | -0.125696689 | 0.125696689 | ENSG00000094880 | n.s. |
| ENSG00000165416 | 2466.485498 | -0.303320674 | 0.2409608 | -1.258797 | 0.208103747 | 0.385165646 | SUGT1 | protein_coding | 36.99 | -0.125681666 | 0.125681666 | ENSG00000165416 | n.s. |
| ENSG00000183020 | 894.3097419 | -0.30294786 | 0.2404456 | -1.259943 | 0.207689801 | 0.384739173 | AP2A2 | protein_coding | 53.1 | -0.125672949 | 0.125672949 | ENSG00000183020 | n.s. |
| ENSG00000151233 | 799.9237085 | -0.294872464 | 0.2297592 | -1.283398 | 0.199352821 | 0.374826377 | GXYLT1 | protein_coding | 37.61 | -0.125665755 | 0.125665755 | ENSG00000151233 | n.s. |
| ENSG00000109576 | 833.8369903 | -0.311006643 | 0.2511145 | -1.238505 | 0.215528834 | 0.394537631 | AADAT | protein_coding | 38.78 | -0.125619181 | 0.125619181 | ENSG00000109576 | n.s. |
| ENSG00000281842 | 3.277568064 | 0.545133648 | 0.643148 | 0.847602 | 0.396659555 | 0.588522058 | MIR1291 | miRNA | 51.72 | 0.125510075 | 0.125510075 | ENSG00000281842 | n.s. |
| ENSG00000126432 | 1201.276972 | 0.355969736 | 0.3144824 | 1.131923 | 0.257666922 | 0.444052056 | PRDX5 | protein_coding | 57.55 | 0.125502867 | 0.125502867 | ENSG00000126432 | n.s. |
| ENSG00000235954 | 150.1719841 | 0.304191194 | 0.242392 | 1.254955 | 0.209494976 | 0.386894622 | TTC28-AS1 | processed_transcript | 43.33 | 0.125450671 | 0.125450671 | ENSG00000235954 | n.s. |
| ENSG00000246089 | 153.8966789 | 0.301995897 | 0.2395632 | 1.260611 | 0.20744915 | 0.384544604 | | antisense | 49.69 | 0.125344387 | 0.125344387 | ENSG00000246089 | n.s. |
| ENSG00000197102 | 7663.814925 | -0.305620565 | 0.2444941 | -1.250012 | 0.211295217 | 0.38903244 | DYNC1H1 | protein_coding | 46.94 | -0.125308766 | 0.125308766 | ENSG00000197102 | n.s. |
| ENSG00000198874 | 623.4474662 | -0.303767134 | 0.2420533 | -1.25496 | 0.209493452 | 0.386894622 | TYW1 | protein_coding | 41.67 | -0.125275786 | 0.125275786 | ENSG00000198874 | n.s. |
| ENSG00000103043 | 269.0997662 | -0.300539528 | 0.237544 | -1.265195 | 0.205801301 | 0.383010661 | VAC14 | protein_coding | 54 | -0.125261611 | 0.125261611 | ENSG00000103043 | n.s. |
| ENSG00000108639 | 238.0344271 | 0.335914161 | 0.2852585 | 1.177578 | 0.238964782 | 0.423781259 | SYNGR2 | protein_coding | 62.13 | 0.125248367 | 0.125248367 | ENSG00000108639 | n.s. |
| ENSG00000122965 | 1271.641953 | -0.293841446 | 0.2289397 | -1.283488 | 0.199321062 | 0.37481728 | RBM19 | protein_coding | 50.43 | -0.125229464 | 0.125229464 | ENSG00000122965 | n.s. |
| ENSG00000152942 | 1075.80968 | -0.330085168 | 0.2773881 | -1.189976 | 0.234055784 | 0.417628949 | RAD17 | protein_coding | 39.7 | -0.1251714 | 0.1251714 | ENSG00000152942 | n.s. |
| ENSG00000279968 | 21.6848317 | 0.373625187 | 0.3419861 | 1.092516 | 0.274606455 | 0.462818747 | GVQW2 | protein_coding | 41.79 | 0.125010899 | 0.125010899 | ENSG00000279968 | n.s. |
| ENSG00000186020 | 923.9951633 | 0.316051377 | 0.2591589 | 1.219528 | 0.222644012 | 0.402319608 | ZNF529 | protein_coding | 40.96 | 0.124975817 | 0.124975817 | ENSG00000186020 | n.s. |
| ENSG00000162641 | 179.1995597 | -0.300963588 | 0.2387146 | -1.260767 | 0.207392719 | 0.384523202 | AKNAD1 | protein_coding | 43.48 | -0.124923199 | 0.124923199 | ENSG00000162641 | n.s. |
| ENSG00000137210 | 1227.526608 | 0.348675338 | 0.304656 | 1.144489 | 0.252421073 | 0.438656728 | TMEM14B | protein_coding | 42.53 | 0.124782258 | 0.124782258 | ENSG00000137210 | n.s. |
| ENSG00000183098 | 931.7911988 | -0.309374469 | 0.2505423 | -1.234819 | 0.216897753 | 0.395892425 | GPC6 | protein_coding | 37.15 | -0.124499343 | 0.124499343 | ENSG00000183098 | n.s. |
| ENSG00000100319 | 172.3454715 | 0.473757483 | 0.5102249 | 0.928527 | 0.353134417 | 0.546203422 | ZMAT5 | protein_coding | 52.77 | 0.12443031 | 0.12443031 | ENSG00000100319 | n.s. |

| | | | | | | | | | | | | | |
|---|---|---|---|---|---|---|---|---|---|---|---|---|---|
| ENSG00000270885 | 293.8423741 | -0.314191477 | 0.2572746 | -1.22123 | 0.221998926 | 0.401779676 | RASL10B | protein_coding | 54.98 | -0.124423606 | 0.124423606 | ENSG00000270885 | n.s. |
| ENSG00000259932 | 50.69174581 | -0.314219071 | 0.2574838 | -1.220345 | 0.222334166 | 0.402020597 | | sense_intronic | 56.18 | -0.124352731 | 0.124352731 | ENSG00000259932 | n.s. |
| ENSG00000160867 | 96.92068996 | 0.31050075 | 0.2524687 | 1.229858 | 0.218750201 | 0.397813355 | FGFR4 | protein_coding | 60.18 | 0.12429986 | 0.12429986 | ENSG00000160867 | n.s. |
| ENSG00000182957 | 507.015168 | -0.292285665 | 0.2281483 | -1.281121 | 0.200151132 | 0.375790487 | SPATA13 | protein_coding | 44.13 | -0.124237255 | 0.124237255 | ENSG00000182957 | n.s. |
| ENSG00000252620 | 21.4790299 | 0.373628998 | 0.3443371 | 1.085068 | 0.27789169 | 0.465952991 | RNU6ATAC24P | snRNA | 51.11 | 0.123917007 | 0.123917007 | ENSG00000252620 | n.s. |
| ENSG00000242251 | 27.01721105 | 0.350685805 | 0.3096864 | 1.13239 | 0.257470433 | 0.443818194 | RN7SL20P | misc_RNA | 52.19 | 0.123720163 | 0.123720163 | ENSG00000242251 | n.s. |
| ENSG00000260035 | 219.6616011 | -0.288531494 | 0.2240977 | -1.287526 | 0.197911181 | 0.373073026 | | sense_intronic | 50 | -0.12355096 | 0.12355096 | ENSG00000260035 | n.s. |
| ENSG00000113734 | 200.0499778 | 0.363384764 | 0.3287222 | 1.105446 | 0.268966275 | 0.457224948 | BNIP1 | protein_coding | 45.87 | 0.123503609 | 0.123503609 | ENSG00000113734 | n.s. |
| ENSG00000113384 | 948.9072367 | -0.341330634 | 0.2958505 | -1.153727 | 0.24861226 | 0.434688998 | GOLPH3 | protein_coding | 41.12 | -0.123500711 | 0.123500711 | ENSG00000113384 | n.s. |
| ENSG00000196890 | 474.4773216 | 0.29044454 | 0.2268041 | 1.280597 | 0.200335426 | 0.375912418 | HIST3H2BB | protein_coding | 52.96 | 0.123413756 | 0.123413756 | ENSG00000196890 | n.s. |
| ENSG00000129055 | 1238.766023 | -0.319259157 | 0.2652797 | -1.203481 | 0.228790048 | 0.410812979 | ANAPC13 | protein_coding | 45.33 | -0.123347641 | 0.123347641 | ENSG00000129055 | n.s. |
| ENSG00000273015 | 2721.125759 | 0.289474488 | 0.2256507 | 1.282844 | 0.199546903 | 0.374988734 | | lincRNA | 38.04 | 0.123310858 | 0.123310858 | ENSG00000273015 | n.s. |
| ENSG00000163602 | 1775.033478 | 0.323611742 | 0.2713717 | 1.192504 | 0.233063875 | 0.416232469 | RYBP | protein_coding | 38.33 | 0.123187355 | 0.123187355 | ENSG00000163602 | n.s. |
| ENSG00000282977 | 86.21563529 | -0.299785437 | 0.2395361 | -1.251525 | 0.21074292 | 0.388323674 | PCBP2-OT1 | non_coding | 50 | -0.123153696 | 0.123153696 | ENSG00000282977 | n.s. |
| ENSG00000133935 | 290.1959795 | -0.332673466 | 0.2840198 | -1.171304 | 0.241476649 | 0.426659821 | C14orf1 | protein_coding | 44.19 | -0.123061987 | 0.123061987 | ENSG00000133935 | n.s. |
| ENSG00000178718 | 360.2085344 | 0.293882514 | 0.2315446 | 1.269226 | 0.204360421 | 0.381408866 | RPP25 | protein_coding | 62.84 | 0.123021929 | 0.123021929 | ENSG00000178718 | n.s. |
| ENSG00000117525 | 580.7501457 | 0.292266708 | 0.2296453 | 1.272688 | 0.203128935 | 0.379466545 | F3 | protein_coding | 43.52 | 0.122993578 | 0.122993578 | ENSG00000117525 | n.s. |
| ENSG00000048740 | 229.4493171 | -0.328169424 | 0.2779023 | -1.180881 | 0.237650109 | 0.422148822 | CELF2 | protein_coding | 41.91 | -0.122910744 | 0.122910744 | ENSG00000048740 | n.s. |
| ENSG00000103056 | 323.1762211 | -0.300884641 | 0.241555 | -1.245616 | 0.212905556 | 0.391118096 | SMPD3 | protein_coding | 51.06 | -0.122668288 | 0.122668288 | ENSG00000103056 | n.s. |
| ENSG00000132017 | 405.0841659 | 0.303796981 | 0.2454211 | 1.23786 | 0.215767899 | 0.39481357 | DCAF15 | protein_coding | 59.89 | 0.12261487 | 0.12261487 | ENSG00000132017 | n.s. |
| ENSG00000165389 | 616.7301 | -0.354613252 | 0.317359 | -1.117388 | 0.263828278 | 0.451096785 | SPTSSA | protein_coding | 41.92 | -0.122600535 | 0.122600535 | ENSG00000165389 | n.s. |
| ENSG00000269425 | 12.41662104 | 0.395681397 | 0.3819395 | 1.035979 | 0.300211846 | 0.48996009 | | antisense | 51.49 | 0.122597645 | 0.122597645 | ENSG00000269425 | n.s. |
| ENSG00000136237 | 1184.97796 | -0.279624747 | 0.2140378 | -1.306427 | 0.191407483 | 0.364564485 | RAPGEF5 | protein_coding | 39.42 | -0.122538734 | 0.122538734 | ENSG00000136237 | n.s. |
| ENSG00000130723 | 4094.047853 | -0.295035571 | 0.2339162 | -1.261288 | 0.207205216 | 0.384476015 | PRRC2B | protein_coding | 49.02 | -0.122478338 | 0.122478338 | ENSG00000130723 | n.s. |
| ENSG00000131725 | 249.4277505 | -0.303701605 | 0.2457105 | -1.236014 | 0.216453396 | 0.395496147 | WDR44 | protein_coding | 39.07 | -0.122348543 | 0.122348543 | ENSG00000131725 | n.s. |
| ENSG00000146433 | 2843.661788 | -0.320496809 | 0.2681753 | -1.195102 | 0.232047144 | 0.415215575 | TMEM181 | protein_coding | 47.38 | -0.122342082 | 0.122342082 | ENSG00000146433 | n.s. |
| ENSG00000183856 | 668.0691981 | -0.32025237 | 0.2680836 | -1.194599 | 0.232243802 | 0.415300598 | IQGAP3 | protein_coding | 50.86 | -0.122220296 | 0.122220296 | ENSG00000183856 | n.s. |
| ENSG00000260293 | 477.9603895 | -0.292377159 | 0.2307097 | -1.267295 | 0.205049983 | 0.382112501 | | sense_intronic | 51.91 | -0.122157737 | 0.122157737 | ENSG00000260293 | n.s. |
| ENSG00000178935 | 175.3120062 | -0.316149624 | 0.2627662 | -1.203159 | 0.228914695 | 0.410930803 | ZNF552 | protein_coding | 47.58 | -0.122106881 | 0.122106881 | ENSG00000178935 | n.s. |
| ENSG00000143889 | 2047.077228 | -0.310626953 | 0.2557278 | -1.214678 | 0.22448892 | 0.405127573 | HNRNPLL | protein_coding | 37.07 | -0.121892563 | 0.121892563 | ENSG00000143889 | n.s. |
| ENSG00000101654 | 1700.061413 | -0.307670878 | 0.2520082 | -1.220876 | 0.222132853 | 0.401917571 | RNMT | protein_coding | 39.02 | -0.121795519 | 0.121795519 | ENSG00000101654 | n.s. |
| ENSG00000120915 | 35.45189909 | 0.342738971 | 0.301285 | 1.137591 | 0.255291421 | 0.441378674 | EPHX2 | protein_coding | 47.54 | 0.121736994 | 0.121736994 | ENSG00000120915 | n.s. |

| | | | | | | | | | | | | | |
|---|---|---|---|---|---|---|---|---|---|---|---|---|---|
| ENSG00000148229 | 2286.111964 | 0.332904775 | 0.2863337 | 1.162646 | 0.244973043 | 0.430923523 | POLE3 | protein_coding | 49.33 | 0.121709919 | 0.121709919 | ENSG00000148229 | n.s. |
| ENSG00000067992 | 113.8287039 | -0.321453761 | 0.2704465 | -1.188604 | 0.23459551 | 0.418254926 | PDK3 | protein_coding | 40.95 | -0.121689193 | 0.121689193 | ENSG00000067992 | n.s. |
| ENSG00000158470 | 894.2721524 | -0.303478177 | 0.2465159 | -1.23107 | 0.218296867 | 0.397217859 | B4GALT5 | protein_coding | 42.32 | -0.121686019 | 0.121686019 | ENSG00000158470 | n.s. |
| ENSG00000039123 | 3072.179776 | -0.324982497 | 0.2753153 | -1.180401 | 0.237840782 | 0.422325879 | SKIV2L2 | protein_coding | 36.85 | -0.121657947 | 0.121657947 | ENSG00000039123 | n.s. |
| ENSG00000127663 | 141.1400184 | -0.298175512 | 0.2395167 | -1.244905 | 0.213166641 | 0.391494413 | KDM4B | protein_coding | 57.26 | -0.121439262 | 0.121439262 | ENSG00000127663 | n.s. |
| ENSG00000073712 | 1501.382846 | -0.318334325 | 0.2665908 | -1.194093 | 0.232441482 | 0.41552697 | FERMT2 | protein_coding | 39.3 | -0.121412961 | 0.121412961 | ENSG00000073712 | n.s. |
| ENSG00000188917 | 174.2832412 | -0.337972147 | 0.2949577 | -1.145833 | 0.251864379 | 0.438191822 | TRMT2B | protein_coding | 44.9 | -0.121107497 | 0.121107497 | ENSG00000188917 | n.s. |
| ENSG00000261692 | 27.50063961 | 0.363007869 | 0.3329102 | 1.090408 | 0.275533544 | 0.463931978 | | sense_intronic | 52.6 | 0.121079711 | 0.121079711 | ENSG00000261692 | n.s. |
| ENSG00000157796 | 732.0957638 | -0.338239774 | 0.2955602 | -1.144402 | 0.252456824 | 0.438656728 | WDR19 | protein_coding | 39.3 | -0.121047629 | 0.121047629 | ENSG00000157796 | n.s. |
| ENSG00000174405 | 328.8019074 | -0.335076593 | 0.2907861 | -1.152313 | 0.249192488 | 0.435266217 | LIG4 | protein_coding | 40.58 | -0.121044757 | 0.121044757 | ENSG00000174405 | n.s. |
| ENSG00000092201 | 6973.033327 | -0.317233968 | 0.2657102 | -1.19391 | 0.232513273 | 0.41552697 | SUPT16H | protein_coding | 40.41 | -0.120993284 | 0.120993284 | ENSG00000092201 | n.s. |
| ENSG00000187186 | 44.1017795 | 0.367050661 | 0.3400447 | 1.079419 | 0.280401091 | 0.468164575 | | protein_coding | 61.23 | 0.120980431 | 0.120980431 | ENSG00000187186 | n.s. |
| ENSG00000166444 | 19.4983394 | -0.367787751 | 0.3412697 | -1.077704 | 0.281165825 | 0.468991467 | ST5 | protein_coding | 45.03 | -0.120941507 | 0.120941507 | ENSG00000166444 | n.s. |
| ENSG00000124172 | 4817.024034 | 0.339125192 | 0.2971181 | 1.141382 | 0.253711054 | 0.43995753 | ATP5E | protein_coding | 39.46 | 0.120928396 | 0.120928396 | ENSG00000124172 | n.s. |
| ENSG00000007545 | 416.6179385 | -0.299114222 | 0.2414378 | -1.238887 | 0.215387188 | 0.394376093 | CRAMP1 | protein_coding | 53.11 | -0.120868892 | 0.120868892 | ENSG00000007545 | n.s. |
| ENSG00000110931 | 863.4391882 | -0.326361928 | 0.2784811 | -1.171936 | 0.241222833 | 0.426319561 | CAMKK2 | protein_coding | 50.83 | -0.120840314 | 0.120840314 | ENSG00000110931 | n.s. |
| ENSG00000100139 | 372.6161719 | -0.30097889 | 0.2443937 | -1.231533 | 0.218123652 | 0.397087658 | MICALL1 | protein_coding | 55.92 | -0.120726729 | 0.120726729 | ENSG00000100139 | n.s. |
| ENSG00000086189 | 959.2000272 | -0.310670734 | 0.2574869 | -1.20655 | 0.227605654 | 0.409055573 | DIMT1 | protein_coding | 39.95 | -0.120607873 | 0.120607873 | ENSG00000086189 | n.s. |
| ENSG00000125484 | 2253.658581 | -0.300497668 | 0.2439306 | -1.231898 | 0.217987079 | 0.396892127 | GTF3C4 | protein_coding | 42.28 | -0.120597983 | 0.120597983 | ENSG00000125484 | n.s. |
| ENSG00000171552 | 672.7375991 | 0.281692217 | 0.2188581 | 1.2871 | 0.198059464 | 0.373224319 | BCL2L1 | protein_coding | 48.2 | 0.120572738 | 0.120572738 | ENSG00000171552 | n.s. |
| ENSG00000158201 | 249.034672 | 0.321972817 | 0.2728613 | 1.179987 | 0.238005256 | 0.422456294 | ABHD3 | protein_coding | 42.45 | 0.120488093 | 0.120488093 | ENSG00000158201 | n.s. |
| ENSG00000150768 | 885.8158093 | -0.332817234 | 0.2884269 | -1.153905 | 0.248539091 | 0.434615646 | DLAT | protein_coding | 41.27 | -0.120444774 | 0.120444774 | ENSG00000150768 | n.s. |
| ENSG00000105171 | 365.2499016 | -0.356435047 | 0.3239988 | -1.100112 | 0.271283176 | 0.459330616 | POP4 | protein_coding | 49.76 | -0.120430351 | 0.120430351 | ENSG00000105171 | n.s. |
| ENSG00000147854 | 2376.123598 | -0.297650183 | 0.2400494 | -1.239954 | 0.214992535 | 0.393964341 | UHRF2 | protein_coding | 38.99 | -0.120412324 | 0.120412324 | ENSG00000147854 | n.s. |
| ENSG00000199400 | 7.899871958 | 0.433911779 | 0.4506363 | 0.962887 | 0.335604334 | 0.527860273 | RNY4P19 | misc_RNA | 37.5 | 0.120402284 | 0.120402284 | ENSG00000199400 | n.s. |
| ENSG00000162873 | 141.1353371 | -0.329891414 | 0.2842845 | -1.160427 | 0.245875041 | 0.431855456 | KLHDC8A | protein_coding | 50.88 | -0.120298727 | 0.120298727 | ENSG00000162873 | n.s. |
| ENSG00000250742 | 154.2995205 | 0.339516003 | 0.2990302 | 1.13539 | 0.256211813 | 0.442310858 | LINC02381 | lincRNA | 51.73 | 0.120281149 | 0.120281149 | ENSG00000250742 | n.s. |
| ENSG00000230828 | 279.9693043 | -0.298907831 | 0.2421772 | -1.234253 | 0.217108727 | 0.39599795 | | processed_pseudogene | 42.49 | -0.120252732 | 0.120252732 | ENSG00000230828 | n.s. |
| ENSG00000230042 | 38.73530544 | 0.32994743 | 0.2847125 | 1.158879 | 0.246505374 | 0.432526065 | AK3P3 | processed_pseudogene | 46.54 | 0.120096811 | 0.120096811 | ENSG00000230042 | n.s. |
| ENSG00000185963 | 1234.667199 | -0.284737931 | 0.2235262 | -1.273846 | 0.202718037 | 0.378851441 | BICD2 | protein_coding | 51.35 | -0.120025881 | 0.120025881 | ENSG00000185963 | n.s. |
| ENSG00000134297 | 30.92069527 | -0.333373191 | 0.2900179 | -1.149492 | 0.250353189 | 0.436511212 | PLEKHA8P1 | transcribed_processed_pseudog | 41.34 | -0.120015881 | 0.120015881 | ENSG00000134297 | n.s. |
| ENSG00000005882 | 90.90712438 | 0.341053744 | 0.3018388 | 1.12992 | 0.258509893 | 0.44489904 | PDK2 | protein_coding | 56.67 | 0.119961743 | 0.119961743 | ENSG00000005882 | n.s. |

| | | | | | | | | | | | | | |
|---|---|---|---|---|---|---|---|---|---|---|---|---|---|
| ENSG00000164715 | 590.7066032 | -0.293065083 | 0.2347472 | -1.248429 | 0.211874168 | 0.389668403 | LMTK2 | protein_coding | 43.73 | -0.119952948 | 0.119952948 | ENSG00000164715 | n.s. |
| ENSG00000117620 | 1210.258727 | -0.310188615 | 0.2577092 | -1.203638 | 0.228729533 | 0.410757293 | SLC35A3 | protein_coding | 37.43 | -0.119861446 | 0.119861446 | ENSG00000117620 | n.s. |
| ENSG00000183087 | 200.7803435 | 0.332922956 | 0.2895859 | 1.149652 | 0.250287299 | 0.436511212 | GAS6 | protein_coding | 57.59 | 0.119853795 | 0.119853795 | ENSG00000183087 | n.s. |
| ENSG00000275651 | 2.388181517 | 0.553512422 | 0.6806107 | 0.813258 | 0.416069868 | 0.607417508 | MIR6851 | miRNA | 59.7 | 0.119842466 | 0.119842466 | ENSG00000275651 | n.s. |
| ENSG00000121988 | 286.8181216 | -0.336461599 | 0.2951089 | -1.140127 | 0.254233474 | 0.440444039 | ZRANB3 | protein_coding | 38.97 | -0.119817092 | 0.119817092 | ENSG00000121988 | n.s. |
| ENSG00000163798 | 1285.450896 | -0.317241375 | 0.267429 | -1.186264 | 0.235518063 | 0.419324551 | SLC4A1AP | protein_coding | 40.08 | -0.119742665 | 0.119742665 | ENSG00000163798 | n.s. |
| ENSG00000171109 | 1212.384502 | -0.326371241 | 0.2802914 | -1.1644 | 0.244262073 | 0.430053217 | MFN1 | protein_coding | 38.56 | -0.119607814 | 0.119607814 | ENSG00000171109 | n.s. |
| ENSG00000155974 | 477.8289668 | -0.332653996 | 0.2898649 | -1.147617 | 0.251126541 | 0.437218311 | GRIP1 | protein_coding | 38.92 | -0.119523132 | 0.119523132 | ENSG00000155974 | n.s. |
| ENSG00000151748 | 420.1720302 | 0.292814959 | 0.2349327 | 1.246378 | 0.212625571 | 0.390706853 | SAV1 | protein_coding | 37.89 | 0.119512124 | 0.119512124 | ENSG00000151748 | n.s. |
| ENSG00000226084 | 184.4816918 | 0.350869057 | 0.3168747 | 1.10728 | 0.26817286 | 0.45645119 | | processed_pseudogene | 45.77 | 0.119507985 | 0.119507985 | ENSG00000226084 | n.s. |
| ENSG00000163394 | 638.9235597 | -0.294440412 | 0.2370883 | -1.241902 | 0.214272786 | 0.392955742 | CCKAR | protein_coding | 42.88 | -0.119441626 | 0.119441626 | ENSG00000163394 | n.s. |
| ENSG00000103249 | 72.07874615 | 0.313755798 | 0.2630864 | 1.192596 | 0.233027546 | 0.416232469 | CLCN7 | protein_coding | 60.03 | 0.119435551 | 0.119435551 | ENSG00000103249 | n.s. |
| ENSG00000177788 | 17.37374085 | -0.387457694 | 0.3755296 | -1.031763 | 0.302182945 | 0.49190847 | | lincRNA | 44.3 | -0.119381798 | 0.119381798 | ENSG00000177788 | n.s. |
| ENSG00000173915 | 1761.278213 | 0.366618521 | 0.342158 | 1.071489 | 0.283949676 | 0.47250285 | USMG5 | protein_coding | 43.23 | 0.119369365 | 0.119369365 | ENSG00000173915 | n.s. |
| ENSG00000166441 | 14591.3561 | 0.337231894 | 0.2972343 | 1.134566 | 0.25655719 | 0.442742409 | RPL27A | protein_coding | 50.57 | 0.119329127 | 0.119329127 | ENSG00000166441 | n.s. |
| ENSG00000283514 | 20.0966925 | 0.370958527 | 0.3492225 | 1.062241 | 0.288126155 | 0.477341796 | MIR3975 | miRNA | 40 | 0.119140949 | 0.119140949 | ENSG00000283514 | n.s. |
| ENSG00000149557 | 152.6205774 | 0.342181432 | 0.304905 | 1.122256 | 0.261753665 | 0.448753527 | FEZ1 | protein_coding | 43.19 | 0.119076444 | 0.119076444 | ENSG00000149557 | n.s. |
| ENSG00000135521 | 1984.224595 | -0.317736583 | 0.2689623 | -1.181342 | 0.237466791 | 0.421930848 | LTV1 | protein_coding | 39.74 | -0.119074556 | 0.119074556 | ENSG00000135521 | n.s. |
| ENSG00000129197 | 2476.986985 | -0.312655717 | 0.2622273 | -1.192308 | 0.23314046 | 0.416315842 | RPAIN | protein_coding | 45.35 | -0.118989594 | 0.118989594 | ENSG00000129197 | n.s. |
| ENSG00000164111 | 1268.954504 | 0.335802298 | 0.2957348 | 1.135484 | 0.25617238 | 0.442297624 | ANXA5 | protein_coding | 39.13 | 0.118969849 | 0.118969849 | ENSG00000164111 | n.s. |
| ENSG00000184076 | 1143.398585 | 0.342410491 | 0.3053818 | 1.121254 | 0.262179903 | 0.449326784 | UQCR10 | protein_coding | 47.62 | 0.118966311 | 0.118966311 | ENSG00000184076 | n.s. |
| ENSG00000251136 | 85.87497465 | 0.308921382 | 0.2571508 | 1.201324 | 0.229625555 | 0.41210062 | | lincRNA | 38.48 | 0.118933722 | 0.118933722 | ENSG00000251136 | n.s. |
| ENSG00000172531 | 1100.525466 | 0.33553892 | 0.2953916 | 1.135912 | 0.255993372 | 0.442153048 | PPP1CA | protein_coding | 61.16 | 0.118924179 | 0.118924179 | ENSG00000172531 | n.s. |
| ENSG00000130560 | 590.4561352 | 0.325594995 | 0.2803752 | 1.161283 | 0.245526753 | 0.431355622 | UBAC1 | protein_coding | 51.82 | 0.118895746 | 0.118895746 | ENSG00000130560 | n.s. |
| ENSG00000201988 | 5.720835542 | -0.461864523 | 0.504552 | -0.915395 | 0.359984194 | 0.552936594 | | misc_RNA | 37.84 | -0.118849134 | 0.118849134 | ENSG00000201988 | n.s. |
| ENSG00000144597 | 875.4397507 | 0.307119045 | 0.2549981 | 1.204397 | 0.228436034 | 0.410336073 | EAF1 | protein_coding | 43.63 | 0.118812166 | 0.118812166 | ENSG00000144597 | n.s. |
| ENSG00000174276 | 94.51938686 | -0.303270898 | 0.24997 | -1.213229 | 0.225042264 | 0.405810574 | ZNHIT2 | protein_coding | 67.92 | -0.118784127 | 0.118784127 | ENSG00000174276 | n.s. |
| ENSG00000121281 | 295.2574707 | -0.329671107 | 0.2868199 | -1.149401 | 0.250390689 | 0.436511212 | ADCY7 | protein_coding | 54.43 | -0.118683114 | 0.118683114 | ENSG00000121281 | n.s. |
| ENSG00000251602 | 67.73043886 | 0.318580764 | 0.2707842 | 1.176512 | 0.239390408 | 0.424103913 | | antisense | 59.49 | 0.118680166 | 0.118680166 | ENSG00000251602 | n.s. |
| ENSG00000175221 | 244.3688119 | -0.298988605 | 0.2444922 | -1.222896 | 0.221368885 | 0.400952125 | MED16 | protein_coding | 62.67 | -0.118670814 | 0.118670814 | ENSG00000175221 | n.s. |
| ENSG00000184014 | 1298.847964 | -0.318292947 | 0.270662 | -1.175979 | 0.239603111 | 0.424318763 | DENND5A | protein_coding | 43.01 | -0.118502935 | 0.118502935 | ENSG00000184014 | n.s. |
| ENSG00000187522 | 789.1200472 | -0.325127797 | 0.2803277 | -1.159813 | 0.246124855 | 0.432041872 | HSPA14 | protein_coding | 39.55 | -0.118500681 | 0.118500681 | ENSG00000187522 | n.s. |

| | | | | | | | | | | | | | |
|---|---|---|---|---|---|---|---|---|---|---|---|---|---|
| ENSG00000147642 | 132.1555301 | -0.290295335 | 0.2328947 | -1.246466 | 0.212593404 | 0.390699309 | SYBU | protein_coding | 39.55 | -0.118486177 | 0.118486177 | ENSG00000147642 | n.s. |
| ENSG00000163596 | 32.9464441 | -0.333967787 | 0.2938483 | -1.136531 | 0.255734227 | 0.441803893 | ICA1L | protein_coding | 39.94 | -0.118481906 | 0.118481906 | ENSG00000163596 | n.s. |
| ENSG00000100353 | 4209.245898 | 0.317651351 | 0.2702761 | 1.175285 | 0.239880794 | 0.42464848 | EIF3D | protein_coding | 47.04 | 0.118156908 | 0.118156908 | ENSG00000100353 | n.s. |
| ENSG00000145191 | 1814.110531 | 0.299663532 | 0.2460859 | 1.217719 | 0.223330595 | 0.403405759 | EIF2B5 | protein_coding | 48.78 | 0.118144717 | 0.118144717 | ENSG00000145191 | n.s. |
| ENSG00000047230 | 773.9885886 | -0.31252041 | 0.2632565 | -1.187133 | 0.235175284 | 0.418874949 | CTPS2 | protein_coding | 43.51 | -0.118106342 | 0.118106342 | ENSG00000047230 | n.s. |
| ENSG00000170837 | 371.1644201 | 0.313440333 | 0.2645075 | 1.184996 | 0.236018977 | 0.419947887 | GPR27 | protein_coding | 54.84 | 0.118105759 | 0.118105759 | ENSG00000170837 | n.s. |
| ENSG00000119640 | 424.0847426 | -0.293531766 | 0.2377989 | -1.23437 | 0.217065187 | 0.395990382 | ACYP1 | protein_coding | 41.66 | -0.118092339 | 0.118092339 | ENSG00000119640 | n.s. |
| ENSG00000100075 | 409.9624271 | 0.330233442 | 0.2888804 | 1.143149 | 0.252976603 | 0.439230982 | SLC25A1 | protein_coding | 67.54 | 0.117994732 | 0.117994732 | ENSG00000100075 | n.s. |
| ENSG00000144746 | 791.3883949 | 0.356390549 | 0.3289451 | 1.083435 | 0.278615553 | 0.466582192 | ARL6IP5 | protein_coding | 42.5 | 0.117990875 | 0.117990875 | ENSG00000144746 | n.s. |
| ENSG00000118162 | 88.46769582 | 0.344076944 | 0.3095541 | 1.111525 | 0.266342628 | 0.454057599 | KPTN | protein_coding | 53.48 | 0.117980217 | 0.117980217 | ENSG00000118162 | n.s. |
| ENSG00000138326 | 27540.06382 | 0.343765597 | 0.3091272 | 1.112052 | 0.26611567 | 0.453768039 | RPS24 | protein_coding | 47.43 | 0.117968698 | 0.117968698 | ENSG00000138326 | n.s. |
| ENSG00000152784 | 971.2281058 | 0.283505279 | 0.2246177 | 1.262168 | 0.206888382 | 0.384178801 | PRDM8 | protein_coding | 47.9 | 0.117786975 | 0.117786975 | ENSG00000152784 | n.s. |
| ENSG00000170855 | 1009.738879 | 0.289212232 | 0.2323745 | 1.244595 | 0.213280575 | 0.39154307 | TRIAP1 | protein_coding | 44.54 | 0.117773138 | 0.117773138 | ENSG00000170855 | n.s. |
| ENSG00000134825 | 1225.847536 | 0.320758011 | 0.2751713 | 1.165667 | 0.243749171 | 0.429530402 | TMEM258 | protein_coding | 58.49 | 0.117720144 | 0.117720144 | ENSG00000134825 | n.s. |
| ENSG00000184281 | 596.7281854 | -0.310570143 | 0.261132 | -1.189322 | 0.234312826 | 0.417926913 | TSSC4 | protein_coding | 62.64 | -0.117674922 | 0.117674922 | ENSG00000184281 | n.s. |
| ENSG00000167671 | 371.2056021 | 0.338335058 | 0.3017998 | 1.121058 | 0.262263152 | 0.449383124 | UBXN6 | protein_coding | 59.86 | 0.117531929 | 0.117531929 | ENSG00000167671 | n.s. |
| ENSG00000133028 | 1286.247122 | -0.315410043 | 0.2680248 | -1.176794 | 0.239277601 | 0.424065941 | SCO1 | protein_coding | 41.24 | -0.11751125 | 0.11751125 | ENSG00000133028 | n.s. |
| ENSG00000109670 | 1708.092381 | -0.281087464 | 0.2217247 | -1.267732 | 0.204893822 | 0.381979908 | FBXW7 | protein_coding | 35.42 | -0.11748317 | 0.11748317 | ENSG00000109670 | n.s. |
| ENSG00000070756 | 6205.588501 | 0.3060683 | 0.2553864 | 1.198452 | 0.230741135 | 0.413569621 | PABPC1 | protein_coding | 40.75 | 0.117362309 | 0.117362309 | ENSG00000070756 | n.s. |
| ENSG00000088727 | 64.92745453 | 0.318997852 | 0.2731553 | 1.167826 | 0.242876943 | 0.4286444 | KIF9 | protein_coding | 48.45 | 0.117360217 | 0.117360217 | ENSG00000088727 | n.s. |
| ENSG00000166986 | 3633.941764 | -0.326617824 | 0.284682 | -1.147307 | 0.251254587 | 0.437386548 | MARS | protein_coding | 49.33 | -0.117299754 | 0.117299754 | ENSG00000166986 | n.s. |
| ENSG00000239305 | 1021.52966 | 0.304369362 | 0.2532581 | 1.201815 | 0.229435252 | 0.411812172 | RNF103 | protein_coding | 39.04 | 0.117273765 | 0.117273765 | ENSG00000239305 | n.s. |
| ENSG00000135862 | 1143.219694 | -0.311372466 | 0.2628514 | -1.184595 | 0.236177617 | 0.420122776 | LAMC1 | protein_coding | 40.08 | -0.117270273 | 0.117270273 | ENSG00000135862 | n.s. |
| ENSG00000278447 | 6.933407376 | 0.438088137 | 0.4662043 | 0.939691 | 0.347375878 | 0.540271933 | MIR6781 | miRNA | 62.5 | 0.117139333 | 0.117139333 | ENSG00000278447 | n.s. |
| ENSG00000277687 | 86.79092817 | -0.295535453 | 0.2419162 | -1.221644 | 0.221842275 | 0.401600572 | | sense_intronic | 34.21 | -0.117092824 | 0.117092824 | ENSG00000277687 | n.s. |
| ENSG00000138031 | 589.1374613 | -0.301167464 | 0.2494905 | -1.20713 | 0.227382029 | 0.408792574 | ADCY3 | protein_coding | 50.99 | -0.117002657 | 0.117002657 | ENSG00000138031 | n.s. |
| ENSG00000166266 | 1219.465484 | -0.321135587 | 0.2770469 | -1.159138 | 0.246399935 | 0.432395551 | CUL5 | protein_coding | 39.41 | -0.116931499 | 0.116931499 | ENSG00000166266 | n.s. |
| ENSG00000259630 | 37.73920387 | 0.329790733 | 0.290436 | 1.135502 | 0.25616492 | 0.442297624 | | transcribed_processed_pseudog | 43.6 | 0.116840039 | 0.116840039 | ENSG00000259630 | n.s. |
| ENSG00000090266 | 1709.749029 | 0.343938252 | 0.3113348 | 1.104721 | 0.269280353 | 0.457441479 | NDUFB2 | protein_coding | 43.57 | 0.1168236 | 0.1168236 | ENSG00000090266 | n.s. |
| ENSG00000244879 | 5500.798626 | -0.273445906 | 0.2127357 | -1.285378 | 0.198660074 | 0.374180764 | GABPB1-AS1 | antisense | 42.62 | -0.116739128 | 0.116739128 | ENSG00000244879 | n.s. |
| ENSG00000100147 | 77.94771651 | -0.313877352 | 0.267274 | -1.174365 | 0.240248691 | 0.425029546 | CCDC134 | protein_coding | 49.16 | -0.116630822 | 0.116630822 | ENSG00000100147 | n.s. |
| ENSG00000121152 | 1975.836895 | -0.324468864 | 0.2828597 | -1.147102 | 0.251339572 | 0.437425108 | NCAPH | protein_coding | 45.2 | -0.116515566 | 0.116515566 | ENSG00000121152 | n.s. |

| | | | | | | | | | | | | | |
|---|---|---|---|---|---|---|---|---|---|---|---|---|---|
| ENSG00000175756 | 420.7783255 | 0.324401736 | 0.2828964 | 1.146716 | 0.251499127 | 0.437648088 | AURKAIP1 | protein_coding | 66.87 | 0.116419661 | 0.116419661 | ENSG00000175756 | n.s. |
| ENSG00000101489 | 267.52226 | -0.312332674 | 0.2652867 | -1.17734 | 0.239059719 | 0.423894767 | CELF4 | protein_coding | 52.35 | -0.116419488 | 0.116419488 | ENSG00000101489 | n.s. |
| ENSG00000234771 | 255.0496261 | -0.297637795 | 0.2455212 | -1.212269 | 0.225409311 | 0.406314583 | SLC25A25-AS1 | antisense | 53.53 | -0.11641733 | 0.11641733 | ENSG00000234771 | n.s. |
| ENSG00000036448 | 56.25735529 | 0.326896562 | 0.2866913 | 1.140239 | 0.254186775 | 0.440444039 | MYOM2 | protein_coding | 46.64 | 0.116410894 | 0.116410894 | ENSG00000036448 | n.s. |
| ENSG00000212163 | 19.15138191 | 0.355437402 | 0.3305734 | 1.075215 | 0.282278521 | 0.470452938 | SNORD91A | snoRNA | 43.98 | 0.116399996 | 0.116399996 | ENSG00000212163 | n.s. |
| ENSG00000101132 | 886.5122378 | 0.355983179 | 0.3315679 | 1.073636 | 0.282985939 | 0.471180726 | PFDN4 | protein_coding | 41.84 | 0.116339747 | 0.116339747 | ENSG00000101132 | n.s. |
| ENSG00000182287 | 382.061216 | 0.324621613 | 0.2835413 | 1.144883 | 0.252257667 | 0.438529592 | AP1S2 | protein_coding | 36.77 | 0.116214893 | 0.116214893 | ENSG00000182287 | n.s. |
| ENSG00000133112 | 18099.62015 | 0.337739589 | 0.3032641 | 1.113681 | 0.265415944 | 0.45292146 | TPT1 | protein_coding | 44.18 | 0.116174685 | 0.116174685 | ENSG00000133112 | n.s. |
| ENSG00000006715 | 1223.173656 | -0.329268634 | 0.2907265 | -1.132572 | 0.257394179 | 0.443815802 | VPS41 | protein_coding | 38.35 | -0.116165065 | 0.116165065 | ENSG00000006715 | n.s. |
| ENSG00000130508 | 1345.964158 | -0.286814742 | 0.2312547 | -1.240255 | 0.214881174 | 0.393812109 | PXDN | protein_coding | 49.9 | -0.116077062 | 0.116077062 | ENSG00000130508 | n.s. |
| ENSG00000135404 | 1804.276279 | 0.34947517 | 0.3215913 | 1.086706 | 0.277166742 | 0.465523702 | CD63 | protein_coding | 59.18 | 0.116046097 | 0.116046097 | ENSG00000135404 | n.s. |
| ENSG00000184277 | 1290.866303 | 0.293128612 | 0.2402199 | 1.220251 | 0.222369585 | 0.402032428 | TM2D3 | protein_coding | 40.91 | 0.116002399 | 0.116002399 | ENSG00000184277 | n.s. |
| ENSG00000148843 | 2677.369928 | -0.290876519 | 0.2371107 | -1.226754 | 0.219914993 | 0.399303693 | PDCD11 | protein_coding | 46.81 | -0.115971501 | 0.115971501 | ENSG00000148843 | n.s. |
| ENSG00000082512 | 1071.26796 | -0.291624092 | 0.2382385 | -1.224085 | 0.220920238 | 0.400295742 | TRAF5 | protein_coding | 44.65 | -0.115955288 | 0.115955288 | ENSG00000082512 | n.s. |
| ENSG00000165443 | 1004.271006 | -0.300213849 | 0.2495057 | -1.203234 | 0.228885654 | 0.410930803 | PHYHIPL | protein_coding | 35.18 | -0.115951986 | 0.115951986 | ENSG00000165443 | n.s. |
| ENSG00000092470 | 1499.871254 | -0.304731986 | 0.2556937 | -1.191785 | 0.23334551 | 0.416628563 | WDR76 | protein_coding | 41.87 | -0.11587463 | 0.11587463 | ENSG00000092470 | n.s. |
| ENSG00000197879 | 277.4447312 | -0.304651997 | 0.255717 | -1.191364 | 0.233510716 | 0.416870074 | MYO1C | protein_coding | 60.07 | -0.11576754 | 0.11576754 | ENSG00000197879 | n.s. |
| ENSG00000170365 | 495.9601524 | -0.302470361 | 0.2527355 | -1.196786 | 0.231389901 | 0.414315534 | SMAD1 | protein_coding | 38.45 | -0.115745965 | 0.115745965 | ENSG00000170365 | n.s. |
| ENSG00000100505 | 359.5078087 | -0.312040324 | 0.2661026 | -1.172632 | 0.240943603 | 0.42588013 | TRIM9 | protein_coding | 40.47 | -0.115677287 | 0.115677287 | ENSG00000100505 | n.s. |
| ENSG00000235701 | 43.32528063 | -0.321796155 | 0.2805091 | -1.147186 | 0.251304658 | 0.437419021 | PCBP2P1 | processed_pseudogene | 49.54 | -0.115557751 | 0.115557751 | ENSG00000235701 | n.s. |
| ENSG00000100979 | 124.8871039 | 0.316412737 | 0.2724692 | 1.161279 | 0.2455285 | 0.431355622 | PLTP | protein_coding | 51.86 | 0.115542711 | 0.115542711 | ENSG00000100979 | n.s. |
| ENSG00000167549 | 398.2376607 | 0.287688928 | 0.2334312 | 1.232436 | 0.217786384 | 0.396682343 | CORO6 | protein_coding | 60.29 | 0.11552354 | 0.11552354 | ENSG00000167549 | n.s. |
| ENSG00000119523 | 661.102865 | 0.289733984 | 0.2361934 | 1.226681 | 0.219942543 | 0.399303693 | ALG2 | protein_coding | 42.69 | 0.115515976 | 0.115515976 | ENSG00000119523 | n.s. |
| ENSG00000168646 | 673.4698619 | 0.277686271 | 0.2197988 | 1.263365 | 0.206457874 | 0.383841982 | AXIN2 | protein_coding | 51.32 | 0.115475149 | 0.115475149 | ENSG00000168646 | n.s. |
| ENSG00000075391 | 469.7712399 | -0.309934475 | 0.2635994 | -1.175779 | 0.239683373 | 0.424406919 | RASAL2 | protein_coding | 36.99 | -0.115363049 | 0.115363049 | ENSG00000075391 | n.s. |
| ENSG00000134243 | 59.45728004 | 0.302446818 | 0.2533193 | 1.193935 | 0.232503361 | 0.41552697 | SORT1 | protein_coding | 42.27 | 0.115353453 | 0.115353453 | ENSG00000134243 | n.s. |
| ENSG00000275014 | 62.33949151 | -0.31713953 | 0.2740045 | -1.157424 | 0.247099041 | 0.432967524 | RN7SL166P | misc_RNA | 52.14 | -0.115294388 | 0.115294388 | ENSG00000275014 | n.s. |
| ENSG00000056487 | 1016.426201 | -0.309619868 | 0.263285 | -1.175987 | 0.239599932 | 0.424318763 | PHF21B | protein_coding | 52.73 | -0.115273881 | 0.115273881 | ENSG00000056487 | n.s. |
| ENSG00000104231 | 1275.701254 | -0.326140152 | 0.2877531 | -1.133403 | 0.257045143 | 0.443419592 | ZFAND1 | protein_coding | 37.45 | -0.115187849 | 0.115187849 | ENSG00000104231 | n.s. |
| ENSG00000119673 | 349.9705954 | 0.325089527 | 0.2862637 | 1.13563 | 0.256111558 | 0.442297624 | ACOT2 | protein_coding | 48.61 | 0.115174471 | 0.115174471 | ENSG00000119673 | n.s. |
| ENSG00000169180 | 2359.15876 | -0.288750114 | 0.2353059 | -1.227126 | 0.219775079 | 0.399259973 | XPO6 | protein_coding | 43.8 | -0.115137441 | 0.115137441 | ENSG00000169180 | n.s. |
| ENSG00000197857 | 262.9379398 | -0.303861461 | 0.2554809 | -1.189371 | 0.234293791 | 0.417926913 | ZNF44 | protein_coding | 43.35 | -0.115133005 | 0.115133005 | ENSG00000197857 | n.s. |

| | | | | | | | | | | | | | |
|---|---|---|---|---|---|---|---|---|---|---|---|---|---|
| ENSG00000226243 | 50.25407612 | 0.485604136 | 0.5613074 | 0.86513 | 0.386967211 | 0.579514303 | RPL37AP1 | processed_pseudogene | 53.05 | 0.115057024 | 0.115057024 | ENSG00000226243 | n.s. |
| ENSG00000109189 | 2340.11424 | -0.316467555 | 0.2735418 | -1.156926 | 0.247302674 | 0.433131075 | USP46 | protein_coding | 41.39 | -0.114998188 | 0.114998188 | ENSG00000109189 | n.s. |
| ENSG00000198258 | 1842.215436 | 0.340539087 | 0.3098403 | 1.099079 | 0.271733416 | 0.459615298 | UBL5 | protein_coding | 52.38 | 0.114967877 | 0.114967877 | ENSG00000198258 | n.s. |
| ENSG00000099203 | 198.8753033 | 0.324469007 | 0.2858145 | 1.135243 | 0.256273393 | 0.442362316 | TMED1 | protein_coding | 56.25 | 0.114934021 | 0.114934021 | ENSG00000099203 | n.s. |
| ENSG00000263790 | 7.186672577 | -0.458370239 | 0.5103618 | -0.898128 | 0.369117317 | 0.56139556 | MIR4473 | miRNA | 47.25 | -0.114927641 | 0.114927641 | ENSG00000263790 | n.s. |
| ENSG00000165238 | 185.4332814 | 0.284918476 | 0.2304141 | 1.23655 | 0.216254283 | 0.395287958 | WNK2 | protein_coding | 52.37 | 0.114846767 | 0.114846767 | ENSG00000165238 | n.s. |
| ENSG00000206768 | 10.51882571 | 0.405783474 | 0.4161646 | 0.975055 | 0.329532907 | 0.521314244 | | misc_RNA | 41.18 | 0.114796311 | 0.114796311 | ENSG00000206768 | n.s. |
| ENSG00000127511 | 267.1744126 | -0.304169663 | 0.2565303 | -1.185706 | 0.235738246 | 0.419555617 | SIN3B | protein_coding | 52.98 | -0.114735978 | 0.114735978 | ENSG00000127511 | n.s. |
| ENSG00000123684 | 3436.756081 | -0.307593581 | 0.2611696 | -1.177754 | 0.238894651 | 0.42376484 | LPGAT1 | protein_coding | 38.3 | -0.114693981 | 0.114693981 | ENSG00000123684 | n.s. |
| ENSG00000105612 | 111.8624563 | 0.315532804 | 0.2726802 | 1.157153 | 0.247209698 | 0.433106911 | DNASE2 | protein_coding | 52.62 | 0.114666163 | 0.114666163 | ENSG00000105612 | n.s. |
| ENSG00000248115 | 114.3111264 | 0.313632016 | 0.2699087 | 1.161993 | 0.245238399 | 0.431117958 | | antisense | 42.94 | 0.114602357 | 0.114602357 | ENSG00000248115 | n.s. |
| ENSG00000127989 | 727.1083003 | -0.311686433 | 0.2670891 | -1.166975 | 0.24322031 | 0.429137207 | MTERF1 | protein_coding | 38.41 | -0.114514789 | 0.114514789 | ENSG00000127989 | n.s. |
| ENSG00000240322 | 55.90541914 | -0.310570073 | 0.2656206 | -1.169224 | 0.242313291 | 0.427857819 | RN7SL481P | misc_RNA | 50.5 | -0.11450735 | 0.11450735 | ENSG00000240322 | n.s. |
| ENSG00000092820 | 5943.766803 | -0.299122899 | 0.2499636 | -1.196666 | 0.231436775 | 0.414336405 | EZR | protein_coding | 46.17 | -0.114458452 | 0.114458452 | ENSG00000092820 | n.s. |
| ENSG00000145782 | 2002.200468 | -0.310732722 | 0.2659162 | -1.168536 | 0.242590498 | 0.428247433 | ATG12 | protein_coding | 36.61 | -0.114444487 | 0.114444487 | ENSG00000145782 | n.s. |
| ENSG00000114200 | 525.1369433 | 0.329986076 | 0.2947292 | 1.119625 | 0.262873672 | 0.45001723 | BCHE | protein_coding | 32.3 | 0.114429555 | 0.114429555 | ENSG00000114200 | n.s. |
| ENSG00000000003 | 790.6962084 | 0.326833125 | 0.2902212 | 1.126152 | 0.260101263 | 0.446809362 | TSPAN6 | protein_coding | 40.4 | 0.114351633 | 0.114351633 | ENSG00000000003 | n.s. |
| ENSG00000198839 | 496.3150306 | -0.302426268 | 0.2546712 | -1.187517 | 0.235023931 | 0.418712501 | ZNF277 | protein_coding | 37.36 | -0.114342555 | 0.114342555 | ENSG00000198839 | n.s. |
| ENSG00000263597 | 6.518336625 | -0.440744501 | 0.4788914 | -0.920343 | 0.357393411 | 0.55031945 | MIR3936 | miRNA | 51.82 | -0.114322573 | 0.114322573 | ENSG00000263597 | n.s. |
| ENSG00000106591 | 2393.587487 | 0.311624713 | 0.2674998 | 1.164953 | 0.24403794 | 0.429767296 | MRPL32 | protein_coding | 40.85 | 0.114293548 | 0.114293548 | ENSG00000106591 | n.s. |
| ENSG00000221978 | 1770.015187 | 0.306570795 | 0.2605446 | 1.176654 | 0.239333767 | 0.424084559 | CCNL2 | protein_coding | 51.37 | 0.114212195 | 0.114212195 | ENSG00000221978 | n.s. |
| ENSG00000233382 | 85.94856458 | 0.340281795 | 0.3110831 | 1.093861 | 0.274015788 | 0.461991017 | NKAPP1 | transcribed_processed_pseudog | 42.11 | 0.114119104 | 0.114119104 | ENSG00000233382 | n.s. |
| ENSG00000224699 | 732.0303291 | -0.270225812 | 0.2120745 | -1.274202 | 0.202591773 | 0.378793712 | LAMTOR5-AS1 | processed_transcript | 43.14 | -0.113926456 | 0.113926456 | ENSG00000224699 | n.s. |
| ENSG00000004779 | 1400.00944 | 0.361077994 | 0.3443571 | 1.048557 | 0.294382187 | 0.483674844 | NDUFAB1 | protein_coding | 48.8 | 0.113900789 | 0.113900789 | ENSG00000004779 | n.s. |
| ENSG00000185825 | 637.8016591 | 0.342160345 | 0.3144217 | 1.088221 | 0.276497513 | 0.465129063 | BCAP31 | protein_coding | 54.34 | 0.113743174 | 0.113743174 | ENSG00000185825 | n.s. |
| ENSG00000248124 | 622.707041 | -0.288423973 | 0.2368566 | -1.217716 | 0.223332011 | 0.403405759 | RRN3P1 | transcribed_unprocessed_pseud | 41.35 | -0.113713432 | 0.113713432 | ENSG00000248124 | n.s. |
| ENSG00000005448 | 393.413725 | 0.345999881 | 0.3212444 | 1.077061 | 0.281452916 | 0.469357881 | WDR54 | protein_coding | 54.68 | 0.113659537 | 0.113659537 | ENSG00000005448 | n.s. |
| ENSG00000141447 | 1084.694776 | -0.319681226 | 0.2808459 | -1.13828 | 0.255003727 | 0.441134031 | OSBPL1A | protein_coding | 40.87 | -0.113624118 | 0.113624118 | ENSG00000141447 | n.s. |
| ENSG00000272779 | 348.337826 | 0.304453083 | 0.2583568 | 1.178421 | 0.23862876 | 0.423455038 | | unprocessed_pseudogene | 42.89 | 0.113619667 | 0.113619667 | ENSG00000272779 | n.s. |
| ENSG00000163673 | 219.0800789 | -0.297788757 | 0.2493374 | -1.194321 | 0.232352603 | 0.415441799 | DCLK3 | protein_coding | 43.39 | -0.113603376 | 0.113603376 | ENSG00000163673 | n.s. |
| ENSG00000254165 | 174.7481777 | 0.288816594 | 0.2374611 | 1.216269 | 0.223882462 | 0.404347584 | | antisense | 31.96 | 0.113575724 | 0.113575724 | ENSG00000254165 | n.s. |
| ENSG00000156253 | 316.4319884 | -0.344009078 | 0.3183194 | -1.080704 | 0.279828756 | 0.46760115 | RWDD2B | protein_coding | 45.42 | -0.113565801 | 0.113565801 | ENSG00000156253 | n.s. |

| | | | | | | | | | | | | | |
|---|---|---|---|---|---|---|---|---|---|---|---|---|---|
| ENSG00000131473 | 7741.457112 | -0.312035127 | 0.2694343 | -1.158112 | 0.246818252 | 0.432712562 | ACLY | protein_coding | 49.89 | -0.113518534 | 0.113518534 | ENSG00000131473 | n.s. |
| ENSG00000243199 | 268.0409757 | 0.353533364 | 0.3333319 | 1.060605 | 0.288869546 | 0.477838278 | | processed_pseudogene | 46.44 | 0.113384889 | 0.113384889 | ENSG00000243199 | n.s. |
| ENSG00000009830 | 70.44854041 | 0.313259527 | 0.2715763 | 1.153486 | 0.248710881 | 0.43475224 | POMT2 | protein_coding | 47.08 | 0.113324194 | 0.113324194 | ENSG00000009830 | n.s. |
| ENSG00000242299 | 286.4702058 | 0.351507578 | 0.3302497 | 1.064369 | 0.287161518 | 0.476310437 | | processed_pseudogene | 52.51 | 0.113224071 | 0.113224071 | ENSG00000242299 | n.s. |
| ENSG00000280389 | 34.09507637 | -0.329088281 | 0.2955624 | -1.113431 | 0.26552341 | 0.453022581 | | TEC | 43.68 | -0.113166927 | 0.113166927 | ENSG00000280389 | n.s. |
| ENSG00000189306 | 988.2762773 | -0.307342616 | 0.2630829 | -1.168235 | 0.242712079 | 0.428407744 | RRP7A | protein_coding | 57.96 | -0.113145937 | 0.113145937 | ENSG00000189306 | n.s. |
| ENSG00000100030 | 1783.618827 | -0.29982831 | 0.2528199 | -1.185936 | 0.235647496 | 0.419447724 | MAPK1 | protein_coding | 43.8 | -0.113131865 | 0.113131865 | ENSG00000100030 | n.s. |
| ENSG00000131470 | 1042.906372 | -0.274869262 | 0.219286 | -1.253474 | 0.210033362 | 0.387642045 | PSMC3IP | protein_coding | 48.67 | -0.113127701 | 0.113127701 | ENSG00000131470 | n.s. |
| ENSG00000112146 | 282.9681073 | -0.310716876 | 0.2682614 | -1.158262 | 0.246757287 | 0.432712562 | FBXO9 | protein_coding | 39.22 | -0.113038954 | 0.113038954 | ENSG00000112146 | n.s. |
| ENSG00000123124 | 76.25846565 | -0.310598854 | 0.2681937 | -1.158114 | 0.246817603 | 0.432712562 | WWP1 | protein_coding | 37.46 | -0.112996018 | 0.112996018 | ENSG00000123124 | n.s. |
| ENSG00000113312 | 630.5479818 | -0.333410597 | 0.3024034 | -1.102536 | 0.270228726 | 0.458549001 | TTC1 | protein_coding | 41.36 | -0.112897579 | 0.112897579 | ENSG00000113312 | n.s. |
| ENSG00000251474 | 364.6135847 | -0.283177917 | 0.2308892 | -1.226467 | 0.220023095 | 0.39934579 | RPL32P3 | transcribed_unprocessed_pseud | 50.11 | -0.112889129 | 0.112889129 | ENSG00000251474 | n.s. |
| ENSG00000138821 | 272.6258664 | 0.307507467 | 0.2637256 | 1.166013 | 0.243609169 | 0.429471856 | SLC39A8 | protein_coding | 38.15 | 0.112875318 | 0.112875318 | ENSG00000138821 | n.s. |
| ENSG00000103005 | 51.14147022 | 0.306971718 | 0.2630769 | 1.166852 | 0.243270326 | 0.429137207 | USB1 | protein_coding | 49.63 | 0.112782585 | 0.112782585 | ENSG00000103005 | n.s. |
| ENSG00000085999 | 291.8068232 | -0.271740881 | 0.2156309 | -1.260213 | 0.207592424 | 0.384662305 | RAD54L | protein_coding | 46.5 | -0.112750827 | 0.112750827 | ENSG00000085999 | n.s. |
| ENSG00000105669 | 1086.534607 | 0.339942706 | 0.3133653 | 1.084813 | 0.278004506 | 0.466007312 | COPE | protein_coding | 57.06 | 0.112727472 | 0.112727472 | ENSG00000105669 | n.s. |
| ENSG00000252481 | 419.0620916 | 0.287053549 | 0.2363512 | 1.214521 | 0.224548676 | 0.405182895 | SCARNA13 | scaRNA | 47.64 | 0.112625143 | 0.112625143 | ENSG00000252481 | n.s. |
| ENSG00000198924 | 592.6522892 | -0.27963624 | 0.2264033 | -1.235124 | 0.216784224 | 0.395791794 | DCLRE1A | protein_coding | 36.75 | -0.112562875 | 0.112562875 | ENSG00000198924 | n.s. |
| ENSG00000139505 | 1132.527505 | -0.323412691 | 0.2881418 | -1.122408 | 0.261689042 | 0.448753527 | MTMR6 | protein_coding | 37.84 | -0.11254507 | 0.11254507 | ENSG00000139505 | n.s. |
| ENSG00000108479 | 339.5054822 | 0.349224233 | 0.3281309 | 1.064283 | 0.287200421 | 0.47631822 | GALK1 | protein_coding | 60.24 | 0.112486105 | 0.112486105 | ENSG00000108479 | n.s. |
| ENSG00000207741 | 41.59795697 | 0.32763078 | 0.2946333 | 1.111995 | 0.266140186 | 0.453768039 | MIR590 | miRNA | 35.05 | 0.112431776 | 0.112431776 | ENSG00000207741 | n.s. |
| ENSG00000110344 | 670.2939007 | -0.306502502 | 0.2630747 | -1.165078 | 0.243987493 | 0.429747919 | UBE4A | protein_coding | 40.54 | -0.112420893 | 0.112420893 | ENSG00000110344 | n.s. |
| ENSG00000158270 | 405.269034 | -0.32487304 | 0.2906011 | -1.117934 | 0.263594978 | 0.450853003 | COLEC12 | protein_coding | 43.13 | -0.112394712 | 0.112394712 | ENSG00000158270 | n.s. |
| ENSG00000143149 | 1069.157722 | 0.318250954 | 0.2808695 | 1.133092 | 0.257175695 | 0.443589841 | ALDH9A1 | protein_coding | 40.34 | 0.112348445 | 0.112348445 | ENSG00000143149 | n.s. |
| ENSG00000167460 | 4065.55976 | -0.305822645 | 0.2622565 | -1.16612 | 0.243565832 | 0.429471856 | TPM4 | protein_coding | 50.92 | -0.112256878 | 0.112256878 | ENSG00000167460 | n.s. |
| ENSG00000243232 | 237.2320267 | -0.26980273 | 0.2135808 | -1.263235 | 0.206504669 | 0.383841982 | PCDHAC2 | protein_coding | 41.87 | -0.112196798 | 0.112196798 | ENSG00000243232 | n.s. |
| ENSG00000177042 | 128.6997008 | -0.314298886 | 0.2751154 | -1.142426 | 0.25327709 | 0.439599961 | TMEM80 | protein_coding | 55.94 | -0.112186585 | 0.112186585 | ENSG00000177042 | n.s. |
| ENSG00000153774 | 773.6127676 | -0.306861905 | 0.2640282 | -1.162232 | 0.245141436 | 0.431056353 | CFDP1 | protein_coding | 42.3 | -0.112147577 | 0.112147577 | ENSG00000153774 | n.s. |
| ENSG00000134594 | 112.7211499 | -0.323695337 | 0.2893874 | -1.118554 | 0.263330594 | 0.450633218 | RAB33A | protein_coding | 47.75 | -0.112055815 | 0.112055815 | ENSG00000134594 | n.s. |
| ENSG00000114446 | 957.9197936 | -0.313190809 | 0.2738635 | -1.143602 | 0.252788817 | 0.439014432 | IFT57 | protein_coding | 35.29 | -0.111972355 | 0.111972355 | ENSG00000114446 | n.s. |
| ENSG00000130734 | 329.2350702 | 0.300464095 | 0.2553125 | 1.176849 | 0.239255962 | 0.424065941 | ATG4D | protein_coding | 55.3 | 0.111942888 | 0.111942888 | ENSG00000130734 | n.s. |
| ENSG00000084623 | 1335.216591 | 0.336806814 | 0.3098606 | 1.086962 | 0.277053394 | 0.465523702 | EIF3I | protein_coding | 50.5 | 0.111839465 | 0.111839465 | ENSG00000084623 | n.s. |

| | | | | | | | | | | | | | |
|---|---|---|---|---|---|---|---|---|---|---|---|---|---|
| ENSG00000003147 | 1273.558615 | -0.309941742 | 0.2692643 | -1.151069 | 0.24970388 | 0.435886049 | ICA1 | protein_coding | 40.3 | -0.11177337 | 0.11177337 | ENSG00000003147 | n.s. |
| ENSG00000205664 | 267.8761026 | -0.300349391 | 0.2556992 | -1.17462 | 0.240146833 | 0.42495734 | | lincRNA | 49.39 | -0.111626252 | 0.111626252 | ENSG00000205664 | n.s. |
| ENSG00000082515 | 450.9973988 | 0.334463075 | 0.3067637 | 1.090296 | 0.275582955 | 0.463959066 | MRPL22 | protein_coding | 38.83 | 0.111550237 | 0.111550237 | ENSG00000082515 | n.s. |
| ENSG00000103852 | 271.234936 | -0.267782752 | 0.2118965 | -1.263743 | 0.206322315 | 0.383784314 | TTC23 | protein_coding | 41.8 | -0.111374269 | 0.111374269 | ENSG00000103852 | n.s. |
| ENSG00000092841 | 4441.354694 | 0.329658677 | 0.2997384 | 1.099821 | 0.271409992 | 0.459364831 | MYL6 | protein_coding | 51.56 | 0.111372631 | 0.111372631 | ENSG00000092841 | n.s. |
| ENSG00000233954 | 120.7246045 | 0.352438045 | 0.3356352 | 1.050063 | 0.293689289 | 0.483052093 | UQCRHL | protein_coding | 46.93 | 0.111372548 | 0.111372548 | ENSG00000233954 | n.s. |
| ENSG00000228222 | 23.12670274 | -0.342558325 | 0.320133 | -1.07005 | 0.284596832 | 0.473183866 | | lincRNA | 36.54 | -0.111321203 | 0.111321203 | ENSG00000228222 | n.s. |
| ENSG00000164535 | 280.4897743 | -0.295109732 | 0.2489116 | -1.185601 | 0.23578002 | 0.419576331 | DAGLB | protein_coding | 48.19 | -0.11131215 | 0.11131215 | ENSG00000164535 | n.s. |
| ENSG00000079785 | 174414.5998 | -0.306873497 | 0.2655971 | -1.15541 | 0.247922684 | 0.433864697 | DDX1 | protein_coding | 39.49 | -0.11128635 | 0.11128635 | ENSG00000079785 | n.s. |
| ENSG00000151092 | 355.2801087 | -0.319634729 | 0.2847242 | -1.122612 | 0.261602511 | 0.448723882 | NGLY1 | protein_coding | 37.55 | -0.111239539 | 0.111239539 | ENSG00000151092 | n.s. |
| ENSG00000278730 | 923.4271872 | -0.303219457 | 0.2602616 | -1.165057 | 0.243996071 | 0.429747919 | | lincRNA | 43.51 | -0.111216718 | 0.111216718 | ENSG00000278730 | n.s. |
| ENSG00000210156 | 7.548762114 | 0.419512753 | 0.4491662 | 0.933981 | 0.350313585 | 0.543323609 | MT-TK | Mt_tRNA | 32.86 | 0.111146306 | 0.111146306 | ENSG00000210156 | n.s. |
| ENSG00000142657 | 1031.220457 | 0.312157026 | 0.2740625 | 1.138999 | 0.254703468 | 0.440807521 | PGD | protein_coding | 47.69 | 0.111050176 | 0.111050176 | ENSG00000142657 | n.s. |
| ENSG00000204310 | 436.0001689 | -0.29075011 | 0.2433142 | -1.194957 | 0.232103747 | 0.415263489 | AGPAT1 | protein_coding | 50.57 | -0.110972412 | 0.110972412 | ENSG00000204310 | n.s. |
| ENSG00000196465 | 894.7445077 | 0.353180174 | 0.3379717 | 1.044999 | 0.296023311 | 0.485285831 | MYL6B | protein_coding | 53.91 | 0.110899418 | 0.110899418 | ENSG00000196465 | n.s. |
| ENSG00000048544 | 640.7610976 | -0.325288536 | 0.2941984 | -1.105677 | 0.26886614 | 0.457128253 | MRPS10 | protein_coding | 41.04 | -0.110585721 | 0.110585721 | ENSG00000048544 | n.s. |
| ENSG00000199104 | 3.735733935 | -0.471634303 | 0.5497368 | -0.857928 | 0.390932484 | 0.582819679 | MIR346 | miRNA | 67.37 | -0.110582111 | 0.110582111 | ENSG00000199104 | n.s. |
| ENSG00000131773 | 710.3383841 | 0.303317556 | 0.2615002 | 1.159913 | 0.246084027 | 0.432041872 | KHDRBS3 | protein_coding | 38.22 | 0.110551412 | 0.110551412 | ENSG00000131773 | n.s. |
| ENSG00000178449 | 316.7574217 | 0.361539861 | 0.3521158 | 1.026764 | 0.304531539 | 0.494649892 | COX14 | protein_coding | 49.49 | 0.110523488 | 0.110523488 | ENSG00000178449 | n.s. |
| ENSG00000124587 | 270.4337948 | -0.304499583 | 0.2634966 | -1.155611 | 0.247840321 | 0.433775083 | PEX6 | protein_coding | 52.03 | -0.110452777 | 0.110452777 | ENSG00000124587 | n.s. |
| ENSG00000179361 | 243.9490588 | -0.275816938 | 0.2242225 | -1.230104 | 0.21865824 | 0.397698063 | ARID3B | protein_coding | 46.67 | -0.110449934 | 0.110449934 | ENSG00000179361 | n.s. |
| ENSG00000107560 | 1173.549448 | -0.303924137 | 0.2626995 | -1.156927 | 0.247302235 | 0.433131075 | RAB11FIP2 | protein_coding | 35.43 | -0.110440152 | 0.110440152 | ENSG00000107560 | n.s. |
| ENSG00000240694 | 1326.470696 | -0.312890295 | 0.2761798 | -1.132922 | 0.257246826 | 0.44365757 | PNMA2 | protein_coding | 48.66 | -0.110435288 | 0.110435288 | ENSG00000240694 | n.s. |
| ENSG00000117143 | 1162.548728 | 0.304528878 | 0.263639 | 1.155098 | 0.248050345 | 0.43403355 | UAP1 | protein_coding | 39.68 | 0.110384623 | 0.110384623 | ENSG00000117143 | n.s. |
| ENSG00000047188 | 1876.108549 | -0.304991345 | 0.2643933 | -1.153552 | 0.248684057 | 0.43475224 | YTHDC2 | protein_coding | 34.67 | -0.110333112 | 0.110333112 | ENSG00000047188 | n.s. |
| ENSG00000263857 | 5.182115275 | -0.450391986 | 0.5095093 | -0.883972 | 0.376711304 | 0.569051913 | MIR4729 | miRNA | 43.06 | -0.110277628 | 0.110277628 | ENSG00000263857 | n.s. |
| ENSG00000162032 | 228.3492405 | 0.304821215 | 0.2643604 | 1.153052 | 0.248889123 | 0.434954596 | SPSB3 | protein_coding | 64.01 | 0.110209962 | 0.110209962 | ENSG00000162032 | n.s. |
| ENSG00000177853 | 195.5920514 | 0.27844727 | 0.2281491 | 1.220462 | 0.222289782 | 0.401992549 | ZNF518A | protein_coding | 38.02 | 0.110204416 | 0.110204416 | ENSG00000177853 | n.s. |
| ENSG00000019995 | 1120.442045 | -0.306420175 | 0.2669532 | -1.147842 | 0.251033591 | 0.437111142 | ZRANB1 | protein_coding | 39.68 | -0.1101299 | 0.1101299 | ENSG00000019995 | n.s. |
| ENSG00000051596 | 445.4937178 | 0.313744389 | 0.2782529 | 1.127551 | 0.259509477 | 0.446123034 | THOC3 | protein_coding | 43.01 | 0.109981638 | 0.109981638 | ENSG00000051596 | n.s. |
| ENSG00000122550 | 804.2281989 | -0.317329209 | 0.2836943 | -1.11856 | 0.263327685 | 0.450633218 | KLHL7 | protein_coding | 38.72 | -0.109852009 | 0.109852009 | ENSG00000122550 | n.s. |
| ENSG00000177932 | 1297.427109 | -0.306811929 | 0.2680915 | -1.14443 | 0.252445333 | 0.438656728 | ZNF354C | protein_coding | 42.43 | -0.109800382 | 0.109800382 | ENSG00000177932 | n.s. |

| | | | | | | | | | | | | | |
|---|---|---|---|---|---|---|---|---|---|---|---|---|---|
| ENSG00000104863 | 70.29388304 | 0.330458422 | 0.3038281 | 1.087649 | 0.276749991 | 0.465330464 | LIN7B | protein_coding | 58.86 | 0.109791015 | 0.109791015 | ENSG00000104863 | n.s. |
| ENSG00000100802 | 95.36221723 | 0.306916382 | 0.2683173 | 1.143856 | 0.252683392 | 0.438886086 | C14orf93 | protein_coding | 45.67 | 0.109768088 | 0.109768088 | ENSG00000100802 | n.s. |
| ENSG00000204334 | 24.69948986 | -0.357715975 | 0.3475555 | -1.029234 | 0.303369618 | 0.493494006 | ERICH2 | protein_coding | 37.22 | -0.109717971 | 0.109717971 | ENSG00000204334 | n.s. |
| ENSG00000071054 | 10711.84275 | -0.285873193 | 0.2384538 | -1.198862 | 0.230581549 | 0.413390021 | MAP4K4 | protein_coding | 41.04 | -0.109672395 | 0.109672395 | ENSG00000071054 | n.s. |
| ENSG00000278469 | 9.775076733 | -0.391114298 | 0.4034297 | -0.969473 | 0.332309172 | 0.52445484 | | misc_RNA | 52.26 | -0.109626171 | 0.109626171 | ENSG00000278469 | n.s. |
| ENSG00000075856 | 3435.233485 | -0.288488187 | 0.2422637 | -1.190802 | 0.23373125 | 0.417156799 | SART3 | protein_coding | 42.65 | -0.109539159 | 0.109539159 | ENSG00000075856 | n.s. |
| ENSG00000128463 | 1712.072412 | 0.309872688 | 0.2732932 | 1.133847 | 0.256858783 | 0.443206241 | EMC4 | protein_coding | 43.89 | 0.10950719 | 0.10950719 | ENSG00000128463 | n.s. |
| ENSG00000213625 | 1345.787914 | -0.303368447 | 0.263411 | -1.151692 | 0.249447564 | 0.435602463 | LEPROT | protein_coding | 40.78 | -0.109488607 | 0.109488607 | ENSG00000213625 | n.s. |
| ENSG00000158887 | 72.71996996 | 0.300482033 | 0.2593623 | 1.158542 | 0.246643034 | 0.432704491 | MPZ | protein_coding | 50.52 | 0.109317947 | 0.109317947 | ENSG00000158887 | n.s. |
| ENSG00000229833 | 744.5869803 | 0.326893118 | 0.2994371 | 1.091692 | 0.274968384 | 0.46328196 | PET100 | protein_coding | 55.95 | 0.109232843 | 0.109232843 | ENSG00000229833 | n.s. |
| ENSG00000267298 | 48.92387936 | -0.310857428 | 0.2753243 | -1.129059 | 0.258872827 | 0.445358503 | | sense_intronic | 31.13 | -0.109201184 | 0.109201184 | ENSG00000267298 | n.s. |
| ENSG00000140750 | 196.0575804 | -0.279542874 | 0.230801 | -1.211186 | 0.225824207 | 0.406846549 | ARHGAP17 | protein_coding | 44.36 | -0.109180883 | 0.109180883 | ENSG00000140750 | n.s. |
| ENSG00000244380 | 48.34525214 | 0.316900793 | 0.284286 | 1.114725 | 0.264968117 | 0.452554959 | | antisense | 48.45 | 0.109118029 | 0.109118029 | ENSG00000244380 | n.s. |
| ENSG00000154727 | 1120.407512 | -0.316623869 | 0.2841997 | -1.114089 | 0.265240921 | 0.452678273 | GABPA | protein_coding | 36.67 | -0.108985213 | 0.108985213 | ENSG00000154727 | n.s. |
| ENSG00000228502 | 78.52611985 | 0.351968205 | 0.3400255 | 1.035123 | 0.30061146 | 0.49038235 | EEF1A1P11 | processed_pseudogene | 46.72 | 0.1089219 | 0.1089219 | ENSG00000228502 | n.s. |
| ENSG00000114098 | 1808.763235 | -0.308706956 | 0.272613 | -1.1324 | 0.2574663 | 0.443818194 | ARMC8 | protein_coding | 39.28 | -0.108910239 | 0.108910239 | ENSG00000114098 | n.s. |
| ENSG00000177733 | 5879.336125 | 0.266925874 | 0.2143169 | 1.245473 | 0.212957927 | 0.391162694 | HNRNPA0 | protein_coding | 41.07 | 0.10881035 | 0.10881035 | ENSG00000177733 | n.s. |
| ENSG00000115944 | 1560.001011 | 0.344793318 | 0.3286227 | 1.049207 | 0.294082679 | 0.483527768 | COX7A2L | protein_coding | 44 | 0.108809386 | 0.108809386 | ENSG00000115944 | n.s. |
| ENSG00000063180 | 632.7342604 | 0.31997626 | 0.2895354 | 1.105137 | 0.26910022 | 0.457302863 | CA11 | protein_coding | 53.83 | 0.108726678 | 0.108726678 | ENSG00000063180 | n.s. |
| ENSG00000213780 | 211.0439497 | 0.29374193 | 0.2507746 | 1.171339 | 0.24146272 | 0.426659821 | GTF2H4 | protein_coding | 51.16 | 0.108660502 | 0.108660502 | ENSG00000213780 | n.s. |
| ENSG00000100911 | 474.7941795 | 0.321440319 | 0.2921125 | 1.100399 | 0.271158269 | 0.459230821 | PSME2 | protein_coding | 52.29 | 0.108636854 | 0.108636854 | ENSG00000100911 | n.s. |
| ENSG00000168066 | 2991.573372 | 0.265342903 | 0.2125631 | 1.248302 | 0.211920502 | 0.389668403 | SF1 | protein_coding | 49.33 | 0.108606126 | 0.108606126 | ENSG00000168066 | n.s. |
| ENSG00000270557 | 31.14223044 | 0.327391484 | 0.301733 | 1.085037 | 0.277905167 | 0.465952991 | | lincRNA | 40.43 | 0.108581971 | 0.108581971 | ENSG00000270557 | n.s. |
| ENSG00000186073 | 44.42252038 | 0.33175879 | 0.3088627 | 1.07413 | 0.282764267 | 0.470924269 | C15orf41 | protein_coding | 37.44 | 0.108501357 | 0.108501357 | ENSG00000186073 | n.s. |
| ENSG00000147099 | 305.375986 | -0.317992908 | 0.2869097 | -1.108338 | 0.267715892 | 0.455840573 | HDAC8 | protein_coding | 40.05 | -0.108495053 | 0.108495053 | ENSG00000147099 | n.s. |
| ENSG00000173457 | 1706.171141 | 0.313087085 | 0.28001 | 1.118128 | 0.263512263 | 0.450833281 | PPP1R14B | protein_coding | 65.56 | 0.108323132 | 0.108323132 | ENSG00000173457 | n.s. |
| ENSG00000185904 | 919.1317483 | -0.310459918 | 0.2763022 | -1.123625 | 0.261172308 | 0.448151596 | LINC00839 | lincRNA | 43.47 | -0.108218582 | 0.108218582 | ENSG00000185904 | n.s. |
| ENSG00000235162 | 659.485759 | 0.330572654 | 0.3075578 | 1.074831 | 0.282450328 | 0.470626606 | C12orf75 | protein_coding | 41.95 | 0.108204206 | 0.108204206 | ENSG00000235162 | n.s. |
| ENSG00000129824 | 4861.806808 | 0.320000536 | 0.2906245 | 1.101079 | 0.270862234 | 0.459062653 | RPS4Y1 | protein_coding | 40.56 | 0.108201153 | 0.108201153 | ENSG00000129824 | n.s. |
| ENSG00000130023 | 255.8592883 | 0.268018071 | 0.2165525 | 1.237659 | 0.215842695 | 0.39487543 | ERMARD | protein_coding | 43.29 | 0.108155982 | 0.108155982 | ENSG00000130023 | n.s. |
| ENSG00000266965 | 142.0369798 | 0.310633563 | 0.2767464 | 1.122448 | 0.261671855 | 0.448753527 | | sense_intronic | 40.97 | 0.108098034 | 0.108098034 | ENSG00000266965 | n.s. |
| ENSG00000100129 | 3391.411484 | 0.316782142 | 0.2859513 | 1.107819 | 0.267940168 | 0.456110889 | EIF3L | protein_coding | 46 | 0.108000396 | 0.108000396 | ENSG00000100129 | n.s. |

| | | | | | | | | | | | | | |
|---|---|---|---|---|---|---|---|---|---|---|---|---|---|
| ENSG00000011304 | 3832.440822 | 0.291807287 | 0.2492063 | 1.170947 | 0.241620126 | 0.426859156 | PTBP1 | protein_coding | 61.17 | 0.107885647 | 0.107885647 | ENSG00000011304 | n.s. |
| ENSG00000015676 | 1218.272778 | -0.287367118 | 0.2430187 | -1.18249 | 0.237011469 | 0.42128312 | NUDCD3 | protein_coding | 44.52 | -0.107885071 | 0.107885071 | ENSG00000015676 | n.s. |
| ENSG00000226396 | 90.54745434 | 0.321495246 | 0.2937812 | 1.094336 | 0.273807759 | 0.461888579 | | processed_pseudogene | 55.38 | 0.107849688 | 0.107849688 | ENSG00000226396 | n.s. |
| ENSG00000159593 | 2851.490109 | -0.307648432 | 0.272873 | -1.127442 | 0.259555654 | 0.446147329 | NAE1 | protein_coding | 46.31 | -0.107837452 | 0.107837452 | ENSG00000159593 | n.s. |
| ENSG00000080189 | 320.9022416 | -0.320074074 | 0.291801 | -1.096892 | 0.272688732 | 0.460590224 | SLC35C2 | protein_coding | 53.97 | -0.107764231 | 0.107764231 | ENSG00000080189 | n.s. |
| ENSG00000027001 | 189.8709972 | -0.319393905 | 0.2910904 | -1.097233 | 0.272539746 | 0.460394429 | MIPEP | protein_coding | 39.76 | -0.107594206 | 0.107594206 | ENSG00000027001 | n.s. |
| ENSG00000117592 | 2497.821483 | 0.321908385 | 0.2948788 | 1.091663 | 0.274981119 | 0.46328196 | PRDX6 | protein_coding | 40.72 | 0.107567171 | 0.107567171 | ENSG00000117592 | n.s. |
| ENSG00000061938 | 48.27976911 | -0.321800319 | 0.2948343 | -1.091462 | 0.275069754 | 0.463319162 | TNK2 | protein_coding | 58.27 | -0.107519838 | 0.107519838 | ENSG00000061938 | n.s. |
| ENSG00000135916 | 419.6587018 | -0.319076443 | 0.2906921 | -1.097644 | 0.272360039 | 0.460365842 | ITM2C | protein_coding | 58.43 | -0.107495867 | 0.107495867 | ENSG00000135916 | n.s. |
| ENSG00000204130 | 326.6858306 | 0.305399209 | 0.2702079 | 1.130238 | 0.258375989 | 0.444833547 | RUFY2 | protein_coding | 40.87 | 0.107440196 | 0.107440196 | ENSG00000204130 | n.s. |
| ENSG00000143294 | 693.4834266 | -0.291965008 | 0.250027 | -1.167734 | 0.24291411 | 0.428655659 | PRCC | protein_coding | 46.54 | -0.107411427 | 0.107411427 | ENSG00000143294 | n.s. |
| ENSG00000111669 | 10675.78389 | 0.317530205 | 0.2889146 | 1.099045 | 0.271748412 | 0.459615298 | TPI1 | protein_coding | 59.16 | 0.107199951 | 0.107199951 | ENSG00000111669 | n.s. |
| ENSG00000163319 | 1211.149238 | 0.286011723 | 0.242097 | 1.181393 | 0.237446573 | 0.421930848 | MRPS18C | protein_coding | 38.68 | 0.107185388 | 0.107185388 | ENSG00000163319 | n.s. |
| ENSG00000164347 | 1211.370123 | -0.324474592 | 0.3000764 | -1.081307 | 0.279560733 | 0.467427974 | GFM2 | protein_coding | 36.42 | -0.107169188 | 0.107169188 | ENSG00000164347 | n.s. |
| ENSG00000207370 | 77.36239 | 0.314822932 | 0.2844817 | 1.106654 | 0.268443415 | 0.456744186 | | misc_RNA | 38.94 | 0.107142735 | 0.107142735 | ENSG00000207370 | n.s. |
| ENSG00000163535 | 1208.446855 | -0.310271416 | 0.2778405 | -1.116725 | 0.264111867 | 0.451526219 | SGO2 | protein_coding | 37.75 | -0.107142003 | 0.107142003 | ENSG00000163535 | n.s. |
| ENSG00000107372 | 7894.033363 | 0.269953815 | 0.2207976 | 1.22263 | 0.221469314 | 0.401020762 | ZFAND5 | protein_coding | 39.89 | 0.10712662 | 0.10712662 | ENSG00000107372 | n.s. |
| ENSG00000146143 | 1090.253331 | -0.303009491 | 0.2672576 | -1.133773 | 0.256889633 | 0.443206241 | PRIM2 | protein_coding | 37.46 | -0.107081776 | 0.107081776 | ENSG00000146143 | n.s. |
| ENSG00000079462 | 665.0394481 | 0.316619475 | 0.2875927 | 1.10093 | 0.270927086 | 0.459062653 | PAFAH1B3 | protein_coding | 53.73 | 0.107057922 | 0.107057922 | ENSG00000079462 | n.s. |
| ENSG00000134744 | 2482.162867 | -0.303993921 | 0.2688465 | -1.130734 | 0.258167087 | 0.444638836 | ZCCHC11 | protein_coding | 38.59 | -0.107003613 | 0.107003613 | ENSG00000134744 | n.s. |
| ENSG00000165264 | 824.1094741 | 0.340179147 | 0.3251519 | 1.046216 | 0.295461309 | 0.484764059 | NDUFB6 | protein_coding | 41.42 | 0.106975996 | 0.106975996 | ENSG00000165264 | n.s. |
| ENSG00000200279 | 13.44253067 | -0.390513505 | 0.4092195 | -0.954289 | 0.33993759 | 0.532510245 | SNORD114-10 | snoRNA | 36.11 | -0.106872624 | 0.106872624 | ENSG00000200279 | n.s. |
| ENSG00000228716 | 2165.031713 | -0.307057123 | 0.2735994 | -1.122287 | 0.261740387 | 0.448753527 | DHFR | protein_coding | 41.24 | -0.106853461 | 0.106853461 | ENSG00000228716 | n.s. |
| ENSG00000170448 | 667.2643726 | -0.286659523 | 0.2434961 | -1.177265 | 0.239089679 | 0.423894767 | NFXL1 | protein_coding | 35.55 | -0.106850028 | 0.106850028 | ENSG00000170448 | n.s. |
| ENSG00000128563 | 386.9836963 | -0.294535795 | 0.2548553 | -1.155698 | 0.247804688 | 0.433767246 | PRKRIP1 | protein_coding | 46.23 | -0.10684087 | 0.10684087 | ENSG00000128563 | n.s. |
| ENSG00000070831 | 1013.630326 | 0.314208662 | 0.2841636 | 1.105732 | 0.268842644 | 0.457128253 | CDC42 | protein_coding | 41.82 | 0.106818985 | 0.106818985 | ENSG00000070831 | n.s. |
| ENSG00000106069 | 57.24403026 | 0.298732727 | 0.2612631 | 1.143417 | 0.252865468 | 0.439092781 | CHN2 | protein_coding | 42.02 | 0.106780132 | 0.106780132 | ENSG00000106069 | n.s. |
| ENSG00000103037 | 1111.80066 | 0.272029611 | 0.2239432 | 1.214726 | 0.22447056 | 0.405127573 | SETD6 | protein_coding | 48.83 | 0.106746649 | 0.106746649 | ENSG00000103037 | n.s. |
| ENSG00000104884 | 257.0863444 | -0.307523528 | 0.2745232 | -1.12021 | 0.262624362 | 0.449662977 | ERCC2 | protein_coding | 56.56 | -0.106745374 | 0.106745374 | ENSG00000104884 | n.s. |
| ENSG00000130731 | 356.5537618 | 0.324168512 | 0.3005096 | 1.078729 | 0.280708489 | 0.468509265 | METTL26 | protein_coding | 67.84 | 0.106742797 | 0.106742797 | ENSG00000130731 | n.s. |
| ENSG00000101363 | 275.2647009 | -0.31689177 | 0.2889755 | -1.096604 | 0.272814418 | 0.460670167 | MANBAL | protein_coding | 47.37 | -0.106668911 | 0.106668911 | ENSG00000101363 | n.s. |
| ENSG00000170906 | 410.121001 | 0.339098271 | 0.3240697 | 1.046375 | 0.295388118 | 0.484701093 | NDUFA3 | protein_coding | 54.99 | 0.106655223 | 0.106655223 | ENSG00000170906 | n.s. |

| | | | | | | | | | | | | | |
|---|---|---|---|---|---|---|---|---|---|---|---|---|---|
| ENSG00000185130 | 512.8387233 | 0.256896796 | 0.2038532 | 1.260205 | 0.20759553 | 0.384662305 | HIST1H2BL | protein_coding | 57.17 | 0.106591714 | 0.106591714 | ENSG00000185130 | n.s. |
| ENSG00000155755 | 3835.901744 | -0.304466158 | 0.2703225 | -1.126307 | 0.260035609 | 0.446751701 | TMEM237 | protein_coding | 38.83 | -0.106542995 | 0.106542995 | ENSG00000155755 | n.s. |
| ENSG00000130724 | 372.9188187 | 0.329193913 | 0.3084049 | 1.067408 | 0.28578751 | 0.474737728 | CHMP2A | protein_coding | 54.26 | 0.106509457 | 0.106509457 | ENSG00000130724 | n.s. |
| ENSG00000049759 | 139.2750189 | 0.288631655 | 0.2468709 | 1.16916 | 0.242339062 | 0.427857819 | NEDD4L | protein_coding | 43.36 | 0.106418643 | 0.106418643 | ENSG00000049759 | n.s. |
| ENSG00000158158 | 207.9756145 | 0.278285747 | 0.2326493 | 1.19616 | 0.231634147 | 0.41463644 | CNNM4 | protein_coding | 50.16 | 0.106397694 | 0.106397694 | ENSG00000158158 | n.s. |
| ENSG00000174243 | 2727.290914 | -0.314140002 | 0.2848936 | -1.102657 | 0.27017599 | 0.458515396 | DDX23 | protein_coding | 49.66 | -0.106382279 | 0.106382279 | ENSG00000174243 | n.s. |
| ENSG00000159063 | 1044.729907 | -0.282133463 | 0.2381912 | -1.184483 | 0.23622186 | 0.420147797 | ALG8 | protein_coding | 42.15 | -0.106250873 | 0.106250873 | ENSG00000159063 | n.s. |
| ENSG00000184260 | 1877.32997 | 0.255778966 | 0.2027054 | 1.261826 | 0.20701147 | 0.38424486 | HIST2H2AC | protein_coding | 62.82 | 0.106248519 | 0.106248519 | ENSG00000184260 | n.s. |
| ENSG00000064655 | 804.1316763 | -0.305763617 | 0.2727802 | -1.120916 | 0.262323722 | 0.449383124 | EYA2 | protein_coding | 45.44 | -0.106217156 | 0.106217156 | ENSG00000064655 | n.s. |
| ENSG00000178695 | 2656.210901 | 0.259335044 | 0.2077392 | 1.248368 | 0.21189624 | 0.389668403 | KCTD12 | protein_coding | 43.57 | 0.106147081 | 0.106147081 | ENSG00000178695 | n.s. |
| ENSG00000176597 | 500.2188184 | -0.319519887 | 0.2939823 | -1.086868 | 0.277095259 | 0.465523702 | B3GNT5 | protein_coding | 42.15 | -0.106099199 | 0.106099199 | ENSG00000176597 | n.s. |
| ENSG00000140319 | 2573.449998 | 0.312018697 | 0.2821843 | 1.105726 | 0.268844927 | 0.457128253 | SRP14 | protein_coding | 43.51 | 0.106074481 | 0.106074481 | ENSG00000140319 | n.s. |
| ENSG00000143393 | 692.9882868 | -0.300996563 | 0.2660379 | -1.131405 | 0.257884662 | 0.444372019 | PI4KB | protein_coding | 48.46 | -0.106027031 | 0.106027031 | ENSG00000143393 | n.s. |
| ENSG00000119487 | 1161.638907 | -0.318929286 | 0.2931761 | -1.087842 | 0.276664918 | 0.465330464 | MAPKAP1 | protein_coding | 41.82 | -0.105960592 | 0.105960592 | ENSG00000119487 | n.s. |
| ENSG00000119328 | 237.9366183 | -0.307520033 | 0.2758428 | -1.114838 | 0.264919839 | 0.452554959 | FAM206A | protein_coding | 39.01 | -0.105887964 | 0.105887964 | ENSG00000119328 | n.s. |
| ENSG00000158321 | 2196.127537 | -0.283788589 | 0.2411906 | -1.176615 | 0.239349024 | 0.424084559 | AUTS2 | protein_coding | 41.05 | -0.105724741 | 0.105724741 | ENSG00000158321 | n.s. |
| ENSG00000075240 | 355.7183791 | -0.297293266 | 0.2610996 | -1.13862 | 0.25486155 | 0.440980744 | GRAMD4 | protein_coding | 57.68 | -0.105711651 | 0.105711651 | ENSG00000075240 | n.s. |
| ENSG00000187535 | 274.9821122 | -0.312078263 | 0.2830932 | -1.102387 | 0.270293396 | 0.458550035 | IFT140 | protein_coding | 54.44 | -0.105673841 | 0.105673841 | ENSG00000187535 | n.s. |
| ENSG00000171735 | 208.8551726 | 0.341954172 | 0.3309126 | 1.033367 | 0.301432198 | 0.490915954 | CAMTA1 | protein_coding | 48 | 0.105661396 | 0.105661396 | ENSG00000171735 | n.s. |
| ENSG00000132326 | 435.4599911 | 0.27124942 | 0.22433 | 1.209154 | 0.226603843 | 0.407834192 | PER2 | protein_coding | 52.1 | 0.105656089 | 0.105656089 | ENSG00000132326 | n.s. |
| ENSG00000214026 | 655.0039988 | 0.332414806 | 0.3153579 | 1.054087 | 0.291842904 | 0.481408223 | MRPL23 | protein_coding | 62.94 | 0.105537212 | 0.105537212 | ENSG00000214026 | n.s. |
| ENSG00000156469 | 653.131106 | -0.28106085 | 0.2376607 | -1.182614 | 0.236962279 | 0.421249466 | MTERF3 | protein_coding | 38.81 | -0.105527286 | 0.105527286 | ENSG00000156469 | n.s. |
| ENSG00000196132 | 422.768313 | -0.31922177 | 0.2950597 | -1.081889 | 0.279301983 | 0.467282085 | MYT1 | protein_coding | 54.58 | -0.105477535 | 0.105477535 | ENSG00000196132 | n.s. |
| ENSG00000116584 | 1138.992055 | -0.29283271 | 0.2547079 | -1.149681 | 0.250275415 | 0.436511212 | ARHGEF2 | protein_coding | 50.95 | -0.105421122 | 0.105421122 | ENSG00000116584 | n.s. |
| ENSG00000085832 | 1070.911449 | -0.292746457 | 0.2550406 | -1.147843 | 0.251033556 | 0.437111142 | EPS15 | protein_coding | 38.05 | -0.105215455 | 0.105215455 | ENSG00000085832 | n.s. |
| ENSG00000113456 | 1254.302723 | -0.289249812 | 0.2500234 | -1.156891 | 0.247316817 | 0.433131075 | RAD1 | protein_coding | 40.32 | -0.105107787 | 0.105107787 | ENSG00000113456 | n.s. |
| ENSG00000245958 | 745.5115469 | -0.279789431 | 0.2365107 | -1.182989 | 0.236813648 | 0.421120022 | | transcribed_unprocessed_pseud | 36.55 | -0.105087262 | 0.105087262 | ENSG00000245958 | n.s. |
| ENSG00000119285 | 4807.573036 | -0.289969843 | 0.2511018 | -1.15479 | 0.24817645 | 0.434172538 | HEATR1 | protein_coding | 38.51 | -0.105066992 | 0.105066992 | ENSG00000119285 | n.s. |
| ENSG00000161996 | 39.68369245 | 0.311262352 | 0.2832522 | 1.098888 | 0.271817061 | 0.459675567 | WDR90 | protein_coding | 64.6 | 0.105066165 | 0.105066165 | ENSG00000161996 | n.s. |
| ENSG00000206560 | 1741.728774 | -0.291661865 | 0.2537523 | -1.149396 | 0.25039279 | 0.436511212 | ANKRD28 | protein_coding | 37.11 | -0.104999612 | 0.104999612 | ENSG00000206560 | n.s. |
| ENSG00000146872 | 2840.735803 | -0.274361177 | 0.2292274 | -1.196895 | 0.231347485 | 0.414315534 | TLK2 | protein_coding | 42.17 | -0.104989458 | 0.104989458 | ENSG00000146872 | n.s. |
| ENSG00000186166 | 284.1130666 | -0.293281113 | 0.2563324 | -1.144144 | 0.252563963 | 0.438788127 | CCDC84 | protein_coding | 45.88 | -0.10491989 | 0.10491989 | ENSG00000186166 | n.s. |

| | | | | | | | | | | | | | |
|---|---|---|---|---|---|---|---|---|---|---|---|---|---|
| ENSG00000101444 | 2847.256013 | 0.317583127 | 0.2939675 | 1.080334 | 0.279993347 | 0.467652035 | AHCY | protein_coding | 50.38 | 0.10482694 | 0.10482694 | ENSG00000101444 | n.s. |
| ENSG00000087206 | 952.9035368 | -0.281199643 | 0.2390689 | -1.176229 | 0.239503531 | 0.424250339 | UIMC1 | protein_coding | 42.38 | -0.104712505 | 0.104712505 | ENSG00000087206 | n.s. |
| ENSG00000189159 | 5966.744287 | 0.287834041 | 0.2485721 | 1.15795 | 0.246884487 | 0.432754968 | JPT1 | protein_coding | 46.27 | 0.104701917 | 0.104701917 | ENSG00000189159 | n.s. |
| ENSG00000170745 | 152.7542913 | 0.315555889 | 0.2907427 | 1.085344 | 0.277769201 | 0.465901588 | KCNS3 | protein_coding | 39.79 | 0.104671721 | 0.104671721 | ENSG00000170745 | n.s. |
| ENSG00000166925 | 130.6941135 | -0.315262718 | 0.2902489 | -1.086181 | 0.277399141 | 0.46572148 | TSC22D4 | protein_coding | 58.03 | -0.104627414 | 0.104627414 | ENSG00000166925 | n.s. |
| ENSG00000128581 | 171.0166918 | -0.326107868 | 0.3073204 | -1.061133 | 0.288629397 | 0.477725511 | IFT22 | protein_coding | 49.56 | -0.104622437 | 0.104622437 | ENSG00000128581 | n.s. |
| ENSG00000218227 | 135.1705694 | 0.42405067 | 0.4776507 | 0.887784 | 0.374656896 | 0.566686362 | | processed_pseudogene | 55.16 | 0.104595167 | 0.104595167 | ENSG00000218227 | n.s. |
| ENSG00000163430 | 1021.21245 | -0.307745595 | 0.2784225 | -1.105319 | 0.269021531 | 0.457224948 | FSTL1 | protein_coding | 43.92 | -0.10459352 | 0.10459352 | ENSG00000163430 | n.s. |
| ENSG00000100522 | 800.1323104 | -0.310310274 | 0.2828019 | -1.097271 | 0.272523021 | 0.460394429 | GNPNAT1 | protein_coding | 37.88 | -0.104534203 | 0.104534203 | ENSG00000100522 | n.s. |
| ENSG00000167186 | 609.2363674 | -0.298206672 | 0.2645304 | -1.127306 | 0.259613269 | 0.446168741 | COQ7 | protein_coding | 46.2 | -0.104521695 | 0.104521695 | ENSG00000167186 | n.s. |
| ENSG00000080200 | 4667.850701 | -0.300558357 | 0.2680247 | -1.121383 | 0.262124919 | 0.449287856 | CRYBG3 | protein_coding | 37.11 | -0.104436611 | 0.104436611 | ENSG00000080200 | n.s. |
| ENSG00000118894 | 103.7298226 | -0.287556775 | 0.2487795 | -1.15587 | 0.247734385 | 0.43369871 | EEF2KMT | protein_coding | 53.82 | -0.104329011 | 0.104329011 | ENSG00000118894 | n.s. |
| ENSG00000143320 | 450.8265595 | -0.321353903 | 0.3004807 | -1.069466 | 0.284859772 | 0.47356449 | CRABP2 | protein_coding | 55.42 | -0.104318184 | 0.104318184 | ENSG00000143320 | n.s. |
| ENSG00000178982 | 1653.707821 | 0.335214903 | 0.3226331 | 1.038997 | 0.298805957 | 0.488524582 | EIF3K | protein_coding | 50.37 | 0.104289908 | 0.104289908 | ENSG00000178982 | n.s. |
| ENSG00000257242 | 103.8131703 | -0.308467823 | 0.2802663 | -1.100624 | 0.271060298 | 0.459120746 | LINC01619 | processed_transcript | 38.41 | -0.104284668 | 0.104284668 | ENSG00000257242 | n.s. |
| ENSG00000159210 | 1818.755703 | -0.288374271 | 0.2501628 | -1.152746 | 0.249014529 | 0.435064536 | SNF8 | protein_coding | 47.13 | -0.104231818 | 0.104231818 | ENSG00000159210 | n.s. |
| ENSG00000275052 | 2960.633203 | -0.298879776 | 0.2660626 | -1.123344 | 0.261291494 | 0.448245622 | PPP4R3B | protein_coding | 38.14 | -0.104154803 | 0.104154803 | ENSG00000275052 | n.s. |
| ENSG00000140632 | 1458.659191 | -0.28744962 | 0.2490642 | -1.154119 | 0.248451555 | 0.434517146 | GLYR1 | protein_coding | 47.65 | -0.104054774 | 0.104054774 | ENSG00000140632 | n.s. |
| ENSG00000114554 | 2090.694193 | -0.267576038 | 0.2218346 | -1.206196 | 0.227741967 | 0.409194917 | PLXNA1 | protein_coding | 61.75 | -0.103838171 | 0.103838171 | ENSG00000114554 | n.s. |
| ENSG00000146425 | 4368.702408 | -0.312656191 | 0.2875841 | -1.087182 | 0.2769565 | 0.465483503 | DYNLT1 | protein_coding | 45.31 | -0.103831778 | 0.103831778 | ENSG00000146425 | n.s. |
| ENSG00000115307 | 2054.046857 | 0.29679321 | 0.2639477 | 1.124439 | 0.260826715 | 0.447834548 | AUP1 | protein_coding | 55.6 | 0.103545929 | 0.103545929 | ENSG00000115307 | n.s. |
| ENSG00000110711 | 446.4785042 | 0.339931511 | 0.332134 | 1.023477 | 0.306082297 | 0.495936671 | AIP | protein_coding | 56.24 | 0.103534224 | 0.103534224 | ENSG00000110711 | n.s. |
| ENSG00000068885 | 1212.970214 | -0.320374678 | 0.3004604 | -1.066279 | 0.286297481 | 0.475330291 | IFT80 | protein_coding | 36.52 | -0.103482465 | 0.103482465 | ENSG00000068885 | n.s. |
| ENSG00000140990 | 891.8150303 | 0.32936475 | 0.3150418 | 1.045464 | 0.295808671 | 0.485105315 | NDUFB10 | protein_coding | 58.95 | 0.103474536 | 0.103474536 | ENSG00000140990 | n.s. |
| ENSG00000272034 | 156.7273642 | 0.266375119 | 0.2207562 | 1.206648 | 0.227567678 | 0.409040121 | SNORD14A | snoRNA | 42.86 | 0.103415902 | 0.103415902 | ENSG00000272034 | n.s. |
| ENSG00000175166 | 3188.787913 | 0.289895513 | 0.2542676 | 1.14012 | 0.254236389 | 0.440444039 | PSMD2 | protein_coding | 49 | 0.103234477 | 0.103234477 | ENSG00000175166 | n.s. |
| ENSG00000173744 | 1747.673375 | -0.292589982 | 0.2583457 | -1.132552 | 0.257402327 | 0.443815802 | AGFG1 | protein_coding | 37.09 | -0.103224938 | 0.103224938 | ENSG00000173744 | n.s. |
| ENSG00000023516 | 2366.568932 | -0.297248259 | 0.2652914 | -1.120459 | 0.262518073 | 0.449629614 | AKAP11 | protein_coding | 36.45 | -0.103188275 | 0.103188275 | ENSG00000023516 | n.s. |
| ENSG00000196642 | 1411.195329 | -0.295851763 | 0.2632006 | -1.124054 | 0.260990007 | 0.447949194 | RABL6 | protein_coding | 57.06 | -0.103184586 | 0.103184586 | ENSG00000196642 | n.s. |
| ENSG00000116991 | 1328.123163 | -0.277730632 | 0.2365525 | -1.174076 | 0.240364623 | 0.425126606 | SIPA1L2 | protein_coding | 41.11 | -0.103171851 | 0.103171851 | ENSG00000116991 | n.s. |
| ENSG00000253966 | 84.49702694 | -0.308633016 | 0.2827659 | -1.091479 | 0.275062165 | 0.463319162 | | processed_pseudogene | 40.85 | -0.103120383 | 0.103120383 | ENSG00000253966 | n.s. |
| ENSG00000260563 | 45.25846157 | 0.31171528 | 0.2879516 | 1.082526 | 0.279018658 | 0.466991944 | | lincRNA | 47.36 | 0.103081317 | 0.103081317 | ENSG00000260563 | n.s. |

| | | | | | | | | | | | | | |
|---|---|---|---|---|---|---|---|---|---|---|---|---|---|
| ENSG00000099800 | 1175.209584 | 0.301465857 | 0.2717581 | 1.109317 | 0.267293403 | 0.455288238 | TIMM13 | protein_coding | 64.99 | 0.103014974 | 0.103014974 | ENSG00000099800 | n.s. |
| ENSG00000215908 | 278.0656921 | -0.292346852 | 0.2584273 | -1.131254 | 0.257948334 | 0.444372019 | CROCCP2 | transcribed_unprocessed_pseud | 52.04 | -0.102980142 | 0.102980142 | ENSG00000215908 | n.s. |
| ENSG00000196683 | 1318.8998 | 0.325064101 | 0.309019 | 1.051923 | 0.292835009 | 0.482216656 | TOMM7 | protein_coding | 41.61 | 0.102966587 | 0.102966587 | ENSG00000196683 | n.s. |
| ENSG00000196730 | 191.5321011 | 0.282914416 | 0.244295 | 1.158085 | 0.246829215 | 0.432712562 | DAPK1 | protein_coding | 45.13 | 0.102924405 | 0.102924405 | ENSG00000196730 | n.s. |
| ENSG00000173540 | 321.8691655 | -0.269612061 | 0.2256725 | -1.194705 | 0.232202303 | 0.415279727 | GMPPB | protein_coding | 61.13 | -0.102899942 | 0.102899942 | ENSG00000173540 | n.s. |
| ENSG00000198561 | 1468.103828 | -0.2889343 | 0.2534037 | -1.140214 | 0.254197348 | 0.440444039 | CTNND1 | protein_coding | 42.81 | -0.102892181 | 0.102892181 | ENSG00000198561 | n.s. |
| ENSG00000164134 | 3754.177014 | -0.285607062 | 0.2485412 | -1.149133 | 0.25050094 | 0.436511212 | NAA15 | protein_coding | 39.06 | -0.102819855 | 0.102819855 | ENSG00000164134 | n.s. |
| ENSG00000138617 | 429.3584166 | -0.263664517 | 0.217944 | -1.209781 | 0.22636302 | 0.407453437 | PARP16 | protein_coding | 46.92 | -0.1028086 | 0.1028086 | ENSG00000138617 | n.s. |
| ENSG00000172845 | 5027.826674 | -0.285435582 | 0.2484441 | -1.148892 | 0.250600311 | 0.436629724 | SP3 | protein_coding | 37.7 | -0.10272447 | 0.10272447 | ENSG00000172845 | n.s. |
| ENSG00000100564 | 298.8960627 | 0.323208525 | 0.3065017 | 1.054508 | 0.291650308 | 0.481176352 | PIGH | protein_coding | 43.92 | 0.102681967 | 0.102681967 | ENSG00000100564 | n.s. |
| ENSG00000122033 | 291.2081238 | -0.320593168 | 0.3027024 | -1.059104 | 0.289552646 | 0.478452586 | MTIF3 | protein_coding | 42.6 | -0.102641459 | 0.102641459 | ENSG00000122033 | n.s. |
| ENSG00000181544 | 436.152979 | -0.296799287 | 0.2655739 | -1.117577 | 0.263747617 | 0.451014255 | FANCB | protein_coding | 34.57 | -0.102636082 | 0.102636082 | ENSG00000181544 | n.s. |
| ENSG00000202347 | 22.4522267 | -0.351006855 | 0.3524765 | -0.99583 | 0.319332533 | 0.510045526 | RNU1-16P | snRNA | 48.78 | -0.102631265 | 0.102631265 | ENSG00000202347 | n.s. |
| ENSG00000213928 | 61.76522785 | -0.303096582 | 0.2750148 | -1.10211 | 0.270413739 | 0.458695236 | IRF9 | protein_coding | 53.95 | -0.102590849 | 0.102590849 | ENSG00000213928 | n.s. |
| ENSG00000172530 | 58.81258989 | 0.301116977 | 0.2720873 | 1.106692 | 0.268426932 | 0.456744186 | BANP | protein_coding | 51.17 | 0.102478229 | 0.102478229 | ENSG00000172530 | n.s. |
| ENSG00000127423 | 245.0063867 | 0.295808838 | 0.2646158 | 1.117881 | 0.263618013 | 0.450853003 | AUNIP | protein_coding | 42.54 | 0.102339515 | 0.102339515 | ENSG00000127423 | n.s. |
| ENSG00000214063 | 157.1204222 | 0.322300675 | 0.3058762 | 1.053696 | 0.292021947 | 0.481408223 | TSPAN4 | protein_coding | 65.01 | 0.102326112 | 0.102326112 | ENSG00000214063 | n.s. |
| ENSG00000111843 | 1068.605218 | 0.329017299 | 0.3168128 | 1.038523 | 0.299026726 | 0.488655998 | TMEM14C | protein_coding | 47.49 | 0.102323316 | 0.102323316 | ENSG00000111843 | n.s. |
| ENSG00000201570 | 18.20414847 | -0.363334999 | 0.37391 | -0.971718 | 0.331191008 | 0.523045871 | RNU4-56P | snRNA | 38.73 | -0.102264349 | 0.102264349 | ENSG00000201570 | n.s. |
| ENSG00000267321 | 233.1420284 | -0.307489208 | 0.2825697 | -1.088189 | 0.276511762 | 0.465129063 | LINC02001 | lincRNA | 44.42 | -0.102217568 | 0.102217568 | ENSG00000267321 | n.s. |
| ENSG00000144199 | 146.1249825 | -0.330587303 | 0.3197014 | -1.03405 | 0.301112715 | 0.49079751 | FAHD2B | protein_coding | 48.77 | -0.102183758 | 0.102183758 | ENSG00000144199 | n.s. |
| ENSG00000199024 | 15.21335372 | 0.358979004 | 0.3665774 | 0.979272 | 0.327445636 | 0.519279881 | MIR103A2 | miRNA | 44.87 | 0.102164887 | 0.102164887 | ENSG00000199024 | n.s. |
| ENSG00000277443 | 9436.812724 | 0.279365532 | 0.2402507 | 1.162808 | 0.244907304 | 0.430909806 | MARCKS | protein_coding | 48.49 | 0.102139844 | 0.102139844 | ENSG00000277443 | n.s. |
| ENSG00000188938 | 461.974137 | -0.293837301 | 0.2621251 | -1.120981 | 0.262295835 | 0.449383124 | FAM120AOS | protein_coding | 50.78 | -0.102074154 | 0.102074154 | ENSG00000188938 | n.s. |
| ENSG00000101557 | 1527.058518 | -0.291705535 | 0.2590527 | -1.126047 | 0.260145524 | 0.446830266 | USP14 | protein_coding | 38.05 | -0.102055345 | 0.102055345 | ENSG00000101557 | n.s. |
| ENSG00000144642 | 4100.975859 | -0.287185016 | 0.2523805 | -1.137905 | 0.255160272 | 0.441242533 | RBMS3 | protein_coding | 36.91 | -0.102043335 | 0.102043335 | ENSG00000144642 | n.s. |
| ENSG00000171150 | 877.7738061 | 0.284677877 | 0.2485367 | 1.145416 | 0.252037 | 0.438338455 | SOCS5 | protein_coding | 37.21 | 0.10196889 | 0.10196889 | ENSG00000171150 | n.s. |
| ENSG00000222000 | 114.3233487 | 0.274035074 | 0.233319 | 1.174508 | 0.240191527 | 0.424982416 | | antisense | 43.35 | 0.101839391 | 0.101839391 | ENSG00000222000 | n.s. |
| ENSG00000271380 | 93.70477885 | 0.271581654 | 0.229928 | 1.18116 | 0.23753931 | 0.422005845 | | antisense | 64.83 | 0.101756631 | 0.101756631 | ENSG00000271380 | n.s. |
| ENSG00000137941 | 475.4038032 | -0.285971846 | 0.2510752 | -1.138989 | 0.254707786 | 0.440807521 | TTLL7 | protein_coding | 35.53 | -0.101734772 | 0.101734772 | ENSG00000137941 | n.s. |
| ENSG00000157500 | 1520.683252 | -0.293568382 | 0.2624855 | -1.118417 | 0.263388789 | 0.450677414 | APPL1 | protein_coding | 37.38 | -0.101614059 | 0.101614059 | ENSG00000157500 | n.s. |
| ENSG00000147130 | 438.0793354 | -0.285656849 | 0.2509652 | -1.138233 | 0.255023186 | 0.441134031 | ZMYM3 | protein_coding | 53.76 | -0.101530853 | 0.101530853 | ENSG00000147130 | n.s. |

| | | | | | | | | | | | | | |
|---|---|---|---|---|---|---|---|---|---|---|---|---|---|
| ENSG00000166848 | 1791.715037 | 0.306152167 | 0.2824216 | 1.084025 | 0.278353535 | 0.466311668 | TERF2IP | protein_coding | 42.42 | 0.101435474 | 0.101435474 | ENSG00000166848 | n.s. |
| ENSG00000279117 | 113.5817499 | -0.289340823 | 0.256676 | -1.127261 | 0.259632201 | 0.446168741 | | TEC | 51.75 | -0.101414207 | 0.101414207 | ENSG00000279117 | n.s. |
| ENSG00000167992 | 114.2559872 | -0.275955614 | 0.2365005 | -1.166829 | 0.243279464 | 0.429137207 | VWCE | protein_coding | 51.58 | -0.101387149 | 0.101387149 | ENSG00000167992 | n.s. |
| ENSG00000237190 | 624.8681186 | 0.306855037 | 0.2837968 | 1.081249 | 0.279586295 | 0.467427974 | CDKN2AIPNL | protein_coding | 45.21 | 0.101349708 | 0.101349708 | ENSG00000237190 | n.s. |
| ENSG00000149499 | 215.8194519 | 0.284663746 | 0.2497661 | 1.139721 | 0.25440242 | 0.440552847 | EML3 | protein_coding | 57.83 | 0.101340859 | 0.101340859 | ENSG00000149499 | n.s. |
| ENSG00000105677 | 566.1849533 | 0.314360617 | 0.2953417 | 1.064396 | 0.287149205 | 0.476310437 | TMEM147 | protein_coding | 58.59 | 0.101258667 | 0.101258667 | ENSG00000105677 | n.s. |
| ENSG00000160803 | 646.9803486 | -0.295631731 | 0.266224 | -1.110462 | 0.266799832 | 0.454614376 | UBQLN4 | protein_coding | 50.71 | -0.101211544 | 0.101211544 | ENSG00000160803 | n.s. |
| ENSG00000072042 | 1705.83586 | -0.304571792 | 0.2800243 | -1.087662 | 0.276744332 | 0.465330464 | RDH11 | protein_coding | 42.66 | -0.10119048 | 0.10119048 | ENSG00000072042 | n.s. |
| ENSG00000200897 | 7.339127113 | -0.400543428 | 0.4438523 | -0.902425 | 0.366831079 | 0.558978779 | | snoRNA | 45.68 | -0.101179144 | 0.101179144 | ENSG00000200897 | n.s. |
| ENSG00000171729 | 100.540366 | 0.342585538 | 0.3418314 | 1.002206 | 0.316243983 | 0.506740128 | TMEM51 | protein_coding | 48.41 | 0.101136288 | 0.101136288 | ENSG00000171729 | n.s. |
| ENSG00000121413 | 463.1353623 | -0.314135976 | 0.2953842 | -1.063483 | 0.287563129 | 0.476749398 | ZSCAN18 | protein_coding | 50.18 | -0.101060637 | 0.101060637 | ENSG00000121413 | n.s. |
| ENSG00000178188 | 447.2652505 | 0.277769735 | 0.2397714 | 1.158478 | 0.246669219 | 0.432704491 | SH2B1 | protein_coding | 51.11 | 0.101055018 | 0.101055018 | ENSG00000178188 | n.s. |
| ENSG00000011105 | 523.8080846 | -0.293228906 | 0.2631639 | -1.114244 | 0.265174413 | 0.452675743 | TSPAN9 | protein_coding | 52.13 | -0.100933136 | 0.100933136 | ENSG00000011105 | n.s. |
| ENSG00000163875 | 422.3786168 | 0.314892658 | 0.2971877 | 1.059575 | 0.289338071 | 0.478211534 | MEAF6 | protein_coding | 42.74 | 0.100885295 | 0.100885295 | ENSG00000163875 | n.s. |
| ENSG00000158089 | 550.6547556 | -0.274735732 | 0.2357964 | -1.16514 | 0.243962417 | 0.429747919 | GALNT14 | protein_coding | 46.29 | -0.10076928 | 0.10076928 | ENSG00000158089 | n.s. |
| ENSG00000160710 | 3973.997826 | -0.275145518 | 0.2364494 | -1.163655 | 0.244563912 | 0.430421356 | ADAR | protein_coding | 45.35 | -0.100732477 | 0.100732477 | ENSG00000160710 | n.s. |
| ENSG00000197961 | 4084.146378 | -0.283604005 | 0.2494273 | -1.137021 | 0.255529681 | 0.441680912 | ZNF121 | protein_coding | 41.98 | -0.100648614 | 0.100648614 | ENSG00000197961 | n.s. |
| ENSG00000159199 | 1561.707921 | 0.303603584 | 0.2804833 | 1.08243 | 0.279061479 | 0.466991944 | ATP5G1 | protein_coding | 50.08 | 0.100398855 | 0.100398855 | ENSG00000159199 | n.s. |
| ENSG00000117000 | 664.0756354 | -0.27909061 | 0.2430227 | -1.148414 | 0.250797838 | 0.436919207 | RLF | protein_coding | 39.69 | -0.100360666 | 0.100360666 | ENSG00000117000 | n.s. |
| ENSG00000250329 | 76.99994042 | 0.327384916 | 0.3183013 | 1.028538 | 0.30369691 | 0.493737982 | KDELC1P1 | processed_pseudogene | 42.61 | 0.10034461 | 0.10034461 | ENSG00000250329 | n.s. |
| ENSG00000168005 | 914.2682134 | -0.301896479 | 0.2775877 | -1.087572 | 0.276784313 | 0.465330464 | C11orf84 | protein_coding | 52.9 | -0.100301638 | 0.100301638 | ENSG00000168005 | n.s. |
| ENSG00000158552 | 182.6192505 | 0.291535083 | 0.2619491 | 1.112946 | 0.265731734 | 0.45323814 | ZFAND2B | protein_coding | 50.41 | 0.100192897 | 0.100192897 | ENSG00000158552 | n.s. |
| ENSG00000100280 | 605.414932 | -0.279502016 | 0.2440429 | -1.145299 | 0.25208536 | 0.438339511 | AP1B1 | protein_coding | 49.33 | -0.100114654 | 0.100114654 | ENSG00000100280 | n.s. |
| ENSG00000245849 | 124.1556955 | -0.290342696 | 0.2602367 | -1.115687 | 0.264556129 | 0.452119182 | RAD51-AS1 | processed_transcript | 45.04 | -0.100094793 | 0.100094793 | ENSG00000245849 | n.s. |
| ENSG00000134109 | 404.2648903 | -0.280814178 | 0.2463131 | -1.14007 | 0.254257215 | 0.440444039 | EDEM1 | protein_coding | 43.38 | -0.10000053 | 0.10000053 | ENSG00000134109 | n.s. |
| ENSG00000111300 | 3073.677448 | -0.28000504 | 0.2450714 | -1.142545 | 0.253227594 | 0.439599961 | NAA25 | protein_coding | 41.42 | -0.099945658 | 0.099945658 | ENSG00000111300 | n.s. |
| ENSG00000092330 | 545.1405174 | -0.287294191 | 0.2563129 | -1.120873 | 0.262341883 | 0.449383124 | TINF2 | protein_coding | 53.3 | -0.099801187 | 0.099801187 | ENSG00000092330 | n.s. |
| ENSG00000139428 | 524.2809125 | -0.313451936 | 0.2968383 | -1.055969 | 0.290982526 | 0.480416187 | MMAB | protein_coding | 52.27 | -0.09979757 | 0.09979757 | ENSG00000139428 | n.s. |
| ENSG00000108582 | 1832.824048 | -0.287401967 | 0.2565514 | -1.120251 | 0.262606895 | 0.449662977 | CPD | protein_coding | 38.53 | -0.099760921 | 0.099760921 | ENSG00000108582 | n.s. |
| ENSG00000163904 | 993.1945915 | -0.295098159 | 0.2683511 | -1.099672 | 0.271475023 | 0.459364831 | SENP2 | protein_coding | 42.86 | -0.099696628 | 0.099696628 | ENSG00000163904 | n.s. |
| ENSG00000267519 | 66.30796803 | 0.319132051 | 0.3065704 | 1.040975 | 0.297887242 | 0.487308667 | | lincRNA | 55.64 | 0.099631707 | 0.099631707 | ENSG00000267519 | n.s. |
| ENSG00000224470 | 609.0656389 | -0.271452375 | 0.2328543 | -1.16576 | 0.243711299 | 0.429518027 | ATXN1L | protein_coding | 47.18 | -0.099628072 | 0.099628072 | ENSG00000224470 | n.s. |

| | | | | | | | | | | | | | |
|---|---|---|---|---|---|---|---|---|---|---|---|---|---|
| ENSG00000164117 | 1627.391396 | -0.254177612 | 0.2094519 | -1.213537 | 0.224924706 | 0.405651125 | FBXO8 | protein_coding | 33.92 | -0.099598815 | 0.099598815 | ENSG00000164117 | n.s. |
| ENSG00000121211 | 398.4811133 | -0.27577707 | 0.2393913 | -1.151993 | 0.249324063 | 0.435441412 | MND1 | protein_coding | 39.03 | -0.099574901 | 0.099574901 | ENSG00000121211 | n.s. |
| ENSG00000178409 | 499.7716574 | -0.2792117 | 0.2445678 | -1.141653 | 0.253598114 | 0.43995753 | BEND3 | protein_coding | 47.45 | -0.099563889 | 0.099563889 | ENSG00000178409 | n.s. |
| ENSG00000033011 | 452.7909373 | -0.280010243 | 0.246086 | -1.137855 | 0.255180987 | 0.441242533 | ALG1 | protein_coding | 49.28 | -0.099493976 | 0.099493976 | ENSG00000033011 | n.s. |
| ENSG00000047579 | 157.4571231 | -0.300337186 | 0.2773719 | -1.082796 | 0.278898976 | 0.466939084 | DTNBP1 | protein_coding | 39.93 | -0.099333453 | 0.099333453 | ENSG00000047579 | n.s. |
| ENSG00000204580 | 315.2935666 | -0.271767099 | 0.2339496 | -1.161648 | 0.245378461 | 0.431231836 | DDR1 | protein_coding | 55.25 | -0.099273582 | 0.099273582 | ENSG00000204580 | n.s. |
| ENSG00000238754 | 104.6502839 | 0.272116912 | 0.2344657 | 1.160583 | 0.245811513 | 0.431798347 | SCARNA18B | snoRNA | 40 | 0.099246215 | 0.099246215 | ENSG00000238754 | n.s. |
| ENSG00000196313 | 543.3552765 | -0.275514137 | 0.2395126 | -1.150312 | 0.250015538 | 0.436320674 | POM121 | protein_coding | 46.19 | -0.099238596 | 0.099238596 | ENSG00000196313 | n.s. |
| ENSG00000113161 | 3969.769263 | -0.288615366 | 0.2594291 | -1.112502 | 0.265922404 | 0.453507795 | HMGCR | protein_coding | 37.88 | -0.099114916 | 0.099114916 | ENSG00000113161 | n.s. |
| ENSG00000166736 | 116.1917125 | -0.320679467 | 0.3102074 | -1.033758 | 0.301249231 | 0.490839806 | HTR3A | protein_coding | 50.68 | -0.099109268 | 0.099109268 | ENSG00000166736 | n.s. |
| ENSG00000144320 | 835.6431285 | -0.308841694 | 0.2910391 | -1.061169 | 0.288613007 | 0.477725511 | LNPK | protein_coding | 35.26 | -0.099083076 | 0.099083076 | ENSG00000144320 | n.s. |
| ENSG00000127184 | 3079.968922 | 0.329303046 | 0.324711 | 1.014142 | 0.310515005 | 0.500497849 | COX7C | protein_coding | 42.27 | 0.098987766 | 0.098987766 | ENSG00000127184 | n.s. |
| ENSG00000111530 | 9888.151359 | -0.292268959 | 0.265042 | -1.102727 | 0.270145678 | 0.458515396 | CAND1 | protein_coding | 36.92 | -0.098975737 | 0.098975737 | ENSG00000111530 | n.s. |
| ENSG00000059377 | 49.66417857 | -0.30355238 | 0.2831224 | -1.072159 | 0.283648466 | 0.472058043 | TBXAS1 | protein_coding | 44.8 | -0.098959472 | 0.098959472 | ENSG00000059377 | n.s. |
| ENSG00000100056 | 288.632141 | -0.281298116 | 0.2489505 | -1.129936 | 0.258503263 | 0.44489904 | DGCR14 | protein_coding | 55.92 | -0.098943386 | 0.098943386 | ENSG00000100056 | n.s. |
| ENSG00000168282 | 678.7123536 | 0.258526816 | 0.2160089 | 1.196834 | 0.231371122 | 0.414315534 | MGAT2 | protein_coding | 47.6 | 0.098930142 | 0.098930142 | ENSG00000168282 | n.s. |
| ENSG00000047410 | 14805.56317 | -0.285935921 | 0.2557855 | -1.117874 | 0.263620936 | 0.450853003 | TPR | protein_coding | 34.16 | -0.09892383 | 0.09892383 | ENSG00000047410 | n.s. |
| ENSG00000144136 | 4003.059278 | -0.266390864 | 0.2269817 | -1.173623 | 0.240546204 | 0.425285689 | SLC20A1 | protein_coding | 42.45 | -0.098916051 | 0.098916051 | ENSG00000144136 | n.s. |
| ENSG00000134313 | 3513.563412 | -0.292526191 | 0.2660268 | -1.099612 | 0.271501343 | 0.459364831 | KIDINS220 | protein_coding | 38.39 | -0.098827709 | 0.098827709 | ENSG00000134313 | n.s. |
| ENSG00000250479 | 185.8061137 | 0.289435262 | 0.2611032 | 1.108509 | 0.267642083 | 0.455770636 | CHCHD10 | protein_coding | 64.1 | 0.098770842 | 0.098770842 | ENSG00000250479 | n.s. |
| ENSG00000129347 | 683.1185816 | -0.290205732 | 0.2622111 | -1.106764 | 0.268396102 | 0.456744186 | KRI1 | protein_coding | 55.09 | -0.098764838 | 0.098764838 | ENSG00000129347 | n.s. |
| ENSG00000135392 | 903.5201827 | -0.274295402 | 0.2386673 | -1.149279 | 0.250440789 | 0.436511212 | DNAJC14 | protein_coding | 49.07 | -0.098747605 | 0.098747605 | ENSG00000135392 | n.s. |
| ENSG00000225190 | 135.6517358 | -0.277365381 | 0.243437 | -1.139372 | 0.254547907 | 0.440695164 | PLEKHM1 | protein_coding | 50.52 | -0.09870372 | 0.09870372 | ENSG00000225190 | n.s. |
| ENSG00000177359 | 155.0118187 | 0.318607147 | 0.3080051 | 1.034422 | 0.30093906 | 0.490571892 | | protein_coding | 38.21 | 0.098544343 | 0.098544343 | ENSG00000177359 | n.s. |
| ENSG00000107404 | 278.5365209 | 0.302521056 | 0.2824046 | 1.071233 | 0.284064722 | 0.472637801 | DVL1 | protein_coding | 66.25 | 0.098461996 | 0.098461996 | ENSG00000107404 | n.s. |
| ENSG00000135052 | 1899.577493 | -0.29352039 | 0.2682743 | -1.094105 | 0.27390879 | 0.46192249 | GOLM1 | protein_coding | 45.43 | -0.098455806 | 0.098455806 | ENSG00000135052 | n.s. |
| ENSG00000155330 | 372.9719909 | 0.306768501 | 0.2892191 | 1.060678 | 0.288836038 | 0.477838278 | C16orf87 | protein_coding | 37.09 | 0.098386506 | 0.098386506 | ENSG00000155330 | n.s. |
| ENSG00000132128 | 428.9990881 | -0.272813186 | 0.2370041 | -1.151091 | 0.249694894 | 0.435886049 | LRRC41 | protein_coding | 47.41 | -0.098383809 | 0.098383809 | ENSG00000132128 | n.s. |
| ENSG00000088543 | 121.2717749 | -0.31358538 | 0.300424 | -1.043809 | 0.296573615 | 0.485844739 | C3orf18 | protein_coding | 55.98 | -0.098309799 | 0.098309799 | ENSG00000088543 | n.s. |
| ENSG00000168301 | 600.3747429 | 0.275637276 | 0.2414872 | 1.141415 | 0.253697076 | 0.43995753 | KCTD6 | protein_coding | 40.78 | 0.098289288 | 0.098289288 | ENSG00000168301 | n.s. |
| ENSG00000075624 | 14448.7142 | 0.264462717 | 0.2253071 | 1.173788 | 0.240480085 | 0.425222787 | ACTB | protein_coding | 56.7 | 0.098217081 | 0.098217081 | ENSG00000075624 | n.s. |
| ENSG00000100603 | 1636.934355 | -0.296083163 | 0.2727623 | -1.085499 | 0.277700727 | 0.465901588 | SNW1 | protein_coding | 41.12 | -0.098212505 | 0.098212505 | ENSG00000100603 | n.s. |

| | | | | | | | | | | | | | |
|---|---|---|---|---|---|---|---|---|---|---|---|---|---|
| ENSG00000175048 | 113.2340132 | -0.275760105 | 0.2418897 | -1.140024 | 0.254276319 | 0.440444039 | ZDHHC14 | protein_coding | 45.59 | -0.098200728 | 0.098200728 | ENSG00000175048 | n.s. |
| ENSG00000164597 | 738.5894711 | -0.300487249 | 0.2799103 | -1.073513 | 0.283041183 | 0.471216357 | COG5 | protein_coding | 37.1 | -0.098193116 | 0.098193116 | ENSG00000164597 | n.s. |
| ENSG00000120533 | 3013.853075 | 0.286559005 | 0.2579632 | 1.110852 | 0.266631909 | 0.454383852 | ENY2 | protein_coding | 37.11 | 0.098168556 | 0.098168556 | ENSG00000120533 | n.s. |
| ENSG00000223768 | 71.38365723 | -0.309406852 | 0.2939644 | -1.052532 | 0.292555731 | 0.481936263 | LINC00205 | lincRNA | 51.62 | -0.098085188 | 0.098085188 | ENSG00000223768 | n.s. |
| ENSG00000197223 | 1022.544965 | 0.283255449 | 0.2531985 | 1.118709 | 0.263264345 | 0.450630621 | C1D | protein_coding | 38.27 | 0.098057173 | 0.098057173 | ENSG00000197223 | n.s. |
| ENSG00000096654 | 541.7989416 | -0.284096903 | 0.2544982 | -1.116302 | 0.264292769 | 0.451724554 | ZNF184 | protein_coding | 38.79 | -0.098049314 | 0.098049314 | ENSG00000096654 | n.s. |
| ENSG00000161888 | 723.9125996 | -0.279344086 | 0.247616 | -1.128134 | 0.259263224 | 0.44580979 | SPC24 | protein_coding | 53.53 | -0.098007984 | 0.098007984 | ENSG00000161888 | n.s. |
| ENSG00000183807 | 756.6709985 | 0.310650802 | 0.2962515 | 1.048605 | 0.294359894 | 0.483674844 | FAM162B | protein_coding | 37.88 | 0.097993708 | 0.097993708 | ENSG00000183807 | n.s. |
| ENSG00000126247 | 663.0494158 | 0.315258836 | 0.3037335 | 1.037945 | 0.299295432 | 0.48892295 | CAPNS1 | protein_coding | 54.14 | 0.097969702 | 0.097969702 | ENSG00000126247 | n.s. |
| ENSG00000256725 | 52.60666254 | -0.304511065 | 0.2863961 | -1.063251 | 0.287667945 | 0.476809619 | | lincRNA | 50.12 | -0.097947504 | 0.097947504 | ENSG00000256725 | n.s. |
| ENSG00000142684 | 184.409318 | 0.319147693 | 0.3101587 | 1.028982 | 0.303488167 | 0.493520981 | ZNF593 | protein_coding | 62.91 | 0.097880804 | 0.097880804 | ENSG00000142684 | n.s. |
| ENSG00000158406 | 1338.087739 | 0.251916309 | 0.2086652 | 1.207275 | 0.227326248 | 0.408792574 | HIST1H4H | protein_coding | 41.76 | 0.097868731 | 0.097868731 | ENSG00000158406 | n.s. |
| ENSG00000051180 | 614.7182578 | -0.286083971 | 0.2577302 | -1.110013 | 0.266993226 | 0.454888239 | RAD51 | protein_coding | 43.47 | -0.097867979 | 0.097867979 | ENSG00000051180 | n.s. |
| ENSG00000252349 | 5.262498071 | -0.438464067 | 0.52798 | -0.830456 | 0.406281122 | 0.598129014 | | snoRNA | 41.76 | -0.097867429 | 0.097867429 | ENSG00000252349 | n.s. |
| ENSG00000110987 | 1026.955698 | -0.273812064 | 0.2396794 | -1.14241 | 0.253283839 | 0.439599961 | BCL7A | protein_coding | 52.98 | -0.097735123 | 0.097735123 | ENSG00000110987 | n.s. |
| ENSG00000201654 | 7.273816417 | 0.404284878 | 0.4611239 | 0.876738 | 0.380628927 | 0.573140901 | RNU6-7 | snRNA | 46.73 | 0.097731259 | 0.097731259 | ENSG00000201654 | n.s. |
| ENSG00000139220 | 134.9243237 | -0.278177769 | 0.2464158 | -1.128896 | 0.258941747 | 0.445422034 | PPFIA2 | protein_coding | 34.5 | -0.097703906 | 0.097703906 | ENSG00000139220 | n.s. |
| ENSG00000262728 | 42.91468694 | 0.308194517 | 0.2928157 | 1.05252 | 0.292560891 | 0.481936263 | | antisense | 45.14 | 0.097700865 | 0.097700865 | ENSG00000262728 | n.s. |
| ENSG00000077080 | 338.8294614 | 0.30347287 | 0.2853834 | 1.063387 | 0.287606649 | 0.476764778 | ACTL6B | protein_coding | 53.08 | 0.097625959 | 0.097625959 | ENSG00000077080 | n.s. |
| ENSG00000170419 | 619.012052 | -0.294389731 | 0.2714782 | -1.084396 | 0.278189457 | 0.466198989 | VSTM2A | protein_coding | 39.92 | -0.097569198 | 0.097569198 | ENSG00000170419 | n.s. |
| ENSG00000129657 | 159.79591 | 0.27250559 | 0.2382064 | 1.143989 | 0.252628019 | 0.438844655 | SEC14L1 | protein_coding | 46.75 | 0.097472303 | 0.097472303 | ENSG00000129657 | n.s. |
| ENSG00000175063 | 1648.0243 | 0.285298444 | 0.2572889 | 1.108864 | 0.26748872 | 0.455565192 | UBE2C | protein_coding | 51.32 | 0.097415001 | 0.097415001 | ENSG00000175063 | n.s. |
| ENSG00000133706 | 5953.55643 | -0.295814672 | 0.2742076 | -1.078798 | 0.280677761 | 0.468509265 | LARS | protein_coding | 38.69 | -0.097406393 | 0.097406393 | ENSG00000133706 | n.s. |
| ENSG00000201793 | 25.84817701 | -0.335394332 | 0.338426 | -0.991042 | 0.32166509 | 0.512419113 | RN7SKP9 | misc_RNA | 54.52 | -0.097390022 | 0.097390022 | ENSG00000201793 | n.s. |
| ENSG00000015479 | 2376.633409 | 0.266600825 | 0.229493 | 1.161695 | 0.245359372 | 0.431231836 | MATR3 | protein_coding | 38.8 | 0.097386398 | 0.097386398 | ENSG00000015479 | n.s. |
| ENSG00000089094 | 1036.822907 | -0.29347285 | 0.2703185 | -1.085656 | 0.277631295 | 0.465901588 | KDM2B | protein_coding | 50.79 | -0.097346649 | 0.097346649 | ENSG00000089094 | n.s. |
| ENSG00000126088 | 416.2199014 | 0.300679589 | 0.2817198 | 1.0673 | 0.285836324 | 0.474737728 | UROD | protein_coding | 52.29 | 0.097283754 | 0.097283754 | ENSG00000126088 | n.s. |
| ENSG00000272842 | 41.8965904 | -0.320663084 | 0.3143652 | -1.020034 | 0.307712491 | 0.49736255 | | antisense | 26.79 | -0.097265745 | 0.097265745 | ENSG00000272842 | n.s. |
| ENSG00000116062 | 6407.18953 | -0.288673425 | 0.2630863 | -1.097258 | 0.272528805 | 0.460394429 | MSH6 | protein_coding | 44.65 | -0.0972454 | 0.0972454 | ENSG00000116062 | n.s. |
| ENSG00000188243 | 859.6076789 | 0.409783474 | 0.4734388 | 0.865547 | 0.38673869 | 0.579415258 | COMMD6 | protein_coding | 40 | 0.097122811 | 0.097122811 | ENSG00000188243 | n.s. |
| ENSG00000263843 | 114.3720791 | -0.281685925 | 0.2525056 | -1.115563 | 0.264609173 | 0.452154335 | | antisense | 60.78 | -0.097100885 | 0.097100885 | ENSG00000263843 | n.s. |
| ENSG00000100330 | 114.5101534 | 0.28666201 | 0.2599341 | 1.102826 | 0.270102879 | 0.458503095 | MTMR3 | protein_coding | 40.65 | 0.097080305 | 0.097080305 | ENSG00000100330 | n.s. |

| | | | | | | | | | | | | | |
|---|---|---|---|---|---|---|---|---|---|---|---|---|---|
| ENSG00000223305 | 26.75991935 | -0.320643522 | 0.3147365 | -1.018768 | 0.30831304 | 0.498044142 | RN7SKP30 | misc_RNA | 50.6 | -0.097069107 | 0.097069107 | ENSG00000223305 | n.s. |
| ENSG00000164292 | 3349.114867 | -0.302606053 | 0.2853367 | -1.060523 | 0.288906846 | 0.477838278 | RHOBTB3 | protein_coding | 40.56 | -0.09705153 | 0.09705153 | ENSG00000164292 | n.s. |
| ENSG00000111707 | 970.3778467 | -0.277931997 | 0.2472017 | -1.124313 | 0.260880327 | 0.447871368 | SUDS3 | protein_coding | 44.49 | -0.09695566 | 0.09695566 | ENSG00000111707 | n.s. |
| ENSG00000086061 | 7833.861545 | 0.287104017 | 0.2613066 | 1.098725 | 0.271888092 | 0.459718939 | DNAJA1 | protein_coding | 41.22 | 0.096899789 | 0.096899789 | ENSG00000086061 | n.s. |
| ENSG00000163975 | 60.15987846 | -0.291741639 | 0.2684928 | -1.08659 | 0.277217906 | 0.465529479 | MELTF | protein_coding | 56.21 | -0.096873632 | 0.096873632 | ENSG00000163975 | n.s. |
| ENSG00000230043 | 6.856651755 | 0.426335468 | 0.5069459 | 0.840988 | 0.400354586 | 0.592678033 | TMSB4XP6 | processed_pseudogene | 42.96 | 0.09685539 | 0.09685539 | ENSG00000230043 | n.s. |
| ENSG00000272438 | 28.90447807 | -0.307014356 | 0.2927987 | -1.048551 | 0.294384792 | 0.483674844 | | lincRNA | 61.47 | -0.096846604 | 0.096846604 | ENSG00000272438 | n.s. |
| ENSG00000215256 | 172.1032962 | -0.272737327 | 0.2399227 | -1.136772 | 0.255633757 | 0.441696301 | DHRS4-AS1 | antisense | 43.99 | -0.096787997 | 0.096787997 | ENSG00000215256 | n.s. |
| ENSG00000185880 | 202.5756044 | -0.320090303 | 0.314434 | -1.017989 | 0.308683269 | 0.498526523 | TRIM69 | protein_coding | 39.92 | -0.096767054 | 0.096767054 | ENSG00000185880 | n.s. |
| ENSG00000213760 | 163.1924835 | 0.282896308 | 0.2548957 | 1.109851 | 0.267063069 | 0.454951561 | ATP6V1G2 | protein_coding | 49.67 | 0.096760394 | 0.096760394 | ENSG00000213760 | n.s. |
| ENSG00000258405 | 660.0264567 | 0.255563614 | 0.2150368 | 1.188465 | 0.234650422 | 0.418254926 | ZNF578 | protein_coding | 43.95 | 0.09674589 | 0.09674589 | ENSG00000258405 | n.s. |
| ENSG00000214182 | 165.6157836 | 0.409391061 | 0.4745971 | 0.862608 | 0.388353253 | 0.580590823 | PTMAP5 | transcribed_processed_pseudog | 44.58 | 0.096669443 | 0.096669443 | ENSG00000214182 | n.s. |
| ENSG00000139433 | 236.7557939 | -0.305884943 | 0.2912838 | -1.050127 | 0.293659809 | 0.483052093 | GLTP | protein_coding | 49.06 | -0.096661487 | 0.096661487 | ENSG00000139433 | n.s. |
| ENSG00000136811 | 588.7296014 | -0.291039936 | 0.2678185 | -1.086706 | 0.277166733 | 0.465523702 | ODF2 | protein_coding | 49.1 | -0.096642198 | 0.096642198 | ENSG00000136811 | n.s. |
| ENSG00000163378 | 129.6506662 | 0.263471201 | 0.2261 | 1.165286 | 0.243903207 | 0.42974745 | EOGT | protein_coding | 41.82 | 0.09663773 | 0.09663773 | ENSG00000163378 | n.s. |
| ENSG00000215447 | 76.58306747 | -0.29122359 | 0.2682441 | -1.085666 | 0.277626724 | 0.465901588 | | processed_transcript | 53.84 | -0.096600557 | 0.096600557 | ENSG00000215447 | n.s. |
| ENSG00000149582 | 77.03886032 | -0.295048888 | 0.2744997 | -1.07486 | 0.282437235 | 0.470626606 | TMEM25 | protein_coding | 49.48 | -0.096576442 | 0.096576442 | ENSG00000149582 | n.s. |
| ENSG00000196372 | 367.8807491 | -0.324741446 | 0.3224768 | -1.007023 | 0.313923818 | 0.504241039 | ASB13 | protein_coding | 50.97 | -0.096565705 | 0.096565705 | ENSG00000196372 | n.s. |
| ENSG00000236859 | 250.5844747 | 0.258679456 | 0.2198065 | 1.176851 | 0.239255152 | 0.424065941 | NIFK-AS1 | antisense | 44.09 | 0.096375327 | 0.096375327 | ENSG00000236859 | n.s. |
| ENSG00000113649 | 4898.56764 | -0.284820731 | 0.2585475 | -1.101619 | 0.270627434 | 0.458839018 | TCERG1 | protein_coding | 36.35 | -0.096366149 | 0.096366149 | ENSG00000113649 | n.s. |
| ENSG00000248527 | 180.3123279 | -0.270671404 | 0.2374857 | -1.139738 | 0.254395568 | 0.440552847 | MTATP6P1 | unprocessed_pseudogene | 43.91 | -0.096359558 | 0.096359558 | ENSG00000248527 | n.s. |
| ENSG00000230989 | 3206.40252 | 0.314619863 | 0.3061335 | 1.027721 | 0.304081031 | 0.49407401 | HSBP1 | protein_coding | 44.98 | 0.096339116 | 0.096339116 | ENSG00000230989 | n.s. |
| ENSG00000201229 | 6.06143797 | 0.413072795 | 0.4830561 | 0.855124 | 0.392482586 | 0.584504765 | SNORA63D | snoRNA | 43.18 | 0.096333509 | 0.096333509 | ENSG00000201229 | n.s. |
| ENSG00000138160 | 3196.059402 | -0.286903428 | 0.2620483 | -1.09485 | 0.27358257 | 0.461651936 | KIF11 | protein_coding | 39.18 | -0.096309275 | 0.096309275 | ENSG00000138160 | n.s. |
| ENSG00000141664 | 101.0573008 | -0.271335457 | 0.2386766 | -1.136833 | 0.255608144 | 0.441696301 | ZCCHC2 | protein_coding | 42.38 | -0.096290506 | 0.096290506 | ENSG00000141664 | n.s. |
| ENSG00000110756 | 153.2434286 | 0.270108287 | 0.2368944 | 1.140206 | 0.254200633 | 0.440444039 | HPS5 | protein_coding | 39.75 | 0.096188063 | 0.096188063 | ENSG00000110756 | n.s. |
| ENSG00000173120 | 707.861418 | 0.257745486 | 0.2187468 | 1.178282 | 0.238684033 | 0.423499144 | KDM2A | protein_coding | 42.99 | 0.096177074 | 0.096177074 | ENSG00000173120 | n.s. |
| ENSG00000184967 | 317.6664591 | 0.284488967 | 0.2586347 | 1.099964 | 0.271347622 | 0.459364831 | NOC4L | protein_coding | 63.79 | 0.096112395 | 0.096112395 | ENSG00000184967 | n.s. |
| ENSG00000110651 | 939.4422262 | 0.290637623 | 0.2684954 | 1.082468 | 0.279044777 | 0.466991944 | CD81 | protein_coding | 63.44 | 0.096111133 | 0.096111133 | ENSG00000110651 | n.s. |
| ENSG00000268603 | 23.60345852 | 0.325525543 | 0.3247374 | 1.002427 | 0.31613738 | 0.506702304 | | antisense | 66.04 | 0.096110479 | 0.096110479 | ENSG00000268603 | n.s. |
| ENSG00000153140 | 454.0409036 | -0.317649083 | 0.311912 | -1.018393 | 0.308491126 | 0.498274008 | CETN3 | protein_coding | 33.64 | -0.096098938 | 0.096098938 | ENSG00000153140 | n.s. |
| ENSG00000280063 | 90.00040821 | 0.273175947 | 0.2417313 | 1.130081 | 0.25844214 | 0.444892422 | | TEC | 53.91 | 0.096088271 | 0.096088271 | ENSG00000280063 | n.s. |

| | | | | | | | | | | | | | |
|---|---|---|---|---|---|---|---|---|---|---|---|---|---|
| ENSG00000118363 | 653.554281 | -0.283323226 | 0.2567156 | -1.103646 | 0.269746547 | 0.458009899 | SPCS2 | protein_coding | 39.25 | -0.096082028 | 0.096082028 | ENSG00000118363 | n.s. |
| ENSG00000227081 | 278.5398968 | 0.33906194 | 0.3474692 | 0.975804 | 0.329161485 | 0.520845022 | | processed_pseudogene | 49.02 | 0.096053359 | 0.096053359 | ENSG00000227081 | n.s. |
| ENSG00000035681 | 910.4177229 | -0.296539038 | 0.2776769 | -1.067929 | 0.285552755 | 0.474545142 | NSMAF | protein_coding | 38.41 | -0.095996349 | 0.095996349 | ENSG00000035681 | n.s. |
| ENSG00000158286 | 127.5082981 | 0.291705459 | 0.270599 | 1.077999 | 0.281034161 | 0.468828016 | RNF207 | protein_coding | 57.32 | 0.095967142 | 0.095967142 | ENSG00000158286 | n.s. |
| ENSG00000181163 | 42805.96381 | 0.305008726 | 0.2914463 | 1.046535 | 0.295314064 | 0.484636694 | NPM1 | protein_coding | 43.6 | 0.095950776 | 0.095950776 | ENSG00000181163 | n.s. |
| ENSG00000123989 | 171.7572352 | 0.268760341 | 0.2353302 | 1.142056 | 0.253430617 | 0.439745075 | CHPF | protein_coding | 61.17 | 0.095893425 | 0.095893425 | ENSG00000123989 | n.s. |
| ENSG00000170191 | 429.9733936 | -0.285831376 | 0.2611996 | -1.094302 | 0.273822332 | 0.461888579 | NANP | protein_coding | 43.71 | -0.095885787 | 0.095885787 | ENSG00000170191 | n.s. |
| ENSG00000106526 | 29.3823985 | 0.312752791 | 0.3040464 | 1.028635 | 0.303651192 | 0.493721307 | ACTR3C | protein_coding | 43.31 | 0.095864401 | 0.095864401 | ENSG00000106526 | n.s. |
| ENSG00000118680 | 1816.066556 | 0.314480833 | 0.3070585 | 1.024172 | 0.30575389 | 0.495808457 | MYL12B | protein_coding | 40.39 | 0.095817929 | 0.095817929 | ENSG00000118680 | n.s. |
| ENSG00000104221 | 358.1293022 | -0.257907705 | 0.2196522 | -1.174164 | 0.240329397 | 0.425118307 | BRF2 | protein_coding | 49.89 | -0.095810183 | 0.095810183 | ENSG00000104221 | n.s. |
| ENSG00000282826 | 188.8724846 | 0.292679806 | 0.27247 | 1.074173 | 0.282745254 | 0.470924269 | FRG1CP | unprocessed_pseudogene | 35.24 | 0.095720617 | 0.095720617 | ENSG00000282826 | n.s. |
| ENSG00000084636 | 64.34281743 | -0.301695693 | 0.2864477 | -1.053231 | 0.292235046 | 0.48168445 | COL16A1 | protein_coding | 56 | -0.095709149 | 0.095709149 | ENSG00000084636 | n.s. |
| ENSG00000210196 | 282.4087248 | -0.274489738 | 0.2442812 | -1.123663 | 0.261155937 | 0.448151596 | MT-TP | Mt_tRNA | 33.82 | -0.095680275 | 0.095680275 | ENSG00000210196 | n.s. |
| ENSG00000083544 | 462.1611158 | -0.289183573 | 0.267088 | -1.082728 | 0.27892928 | 0.466939084 | TDRD3 | protein_coding | 36.41 | -0.095644509 | 0.095644509 | ENSG00000083544 | n.s. |
| ENSG00000145494 | 775.4918547 | 0.307349091 | 0.2957827 | 1.039104 | 0.298756166 | 0.48850054 | NDUFS6 | protein_coding | 48.37 | 0.095627044 | 0.095627044 | ENSG00000145494 | n.s. |
| ENSG00000111832 | 1525.915625 | -0.289449927 | 0.2676346 | -1.081511 | 0.279469685 | 0.467427974 | RWDD1 | protein_coding | 36.42 | -0.095601056 | 0.095601056 | ENSG00000111832 | n.s. |
| ENSG00000148459 | 520.2365685 | -0.268954412 | 0.2363306 | -1.138043 | 0.255102561 | 0.441216517 | PDSS1 | protein_coding | 44.19 | -0.095572476 | 0.095572476 | ENSG00000148459 | n.s. |
| ENSG00000123179 | 785.5322892 | 0.32784197 | 0.3298933 | 0.993782 | 0.320329061 | 0.511117395 | EBPL | protein_coding | 45.56 | 0.095559161 | 0.095559161 | ENSG00000123179 | n.s. |
| ENSG00000244187 | 124.2353746 | 0.316034028 | 0.3104802 | 1.017888 | 0.308731178 | 0.498546068 | TMEM141 | protein_coding | 62.59 | 0.095535413 | 0.095535413 | ENSG00000244187 | n.s. |
| ENSG00000164978 | 126.2280229 | -0.297902146 | 0.2810003 | -1.060149 | 0.289076849 | 0.477893253 | NUDT2 | protein_coding | 46.38 | -0.095528013 | 0.095528013 | ENSG00000164978 | n.s. |
| ENSG00000120837 | 2417.736666 | -0.289790224 | 0.2686206 | -1.078809 | 0.280672965 | 0.468509265 | NFYB | protein_coding | 39.19 | -0.095422651 | 0.095422651 | ENSG00000120837 | n.s. |
| ENSG00000167792 | 1879.797497 | 0.287874437 | 0.2654864 | 1.084328 | 0.278219306 | 0.466198989 | NDUFV1 | protein_coding | 55.96 | 0.095409843 | 0.095409843 | ENSG00000167792 | n.s. |
| ENSG00000115207 | 1636.741944 | -0.281871249 | 0.2558767 | -1.10159 | 0.270639903 | 0.458839018 | GTF3C2 | protein_coding | 46.39 | -0.095368222 | 0.095368222 | ENSG00000115207 | n.s. |
| ENSG00000259713 | 44.24510776 | -0.291753199 | 0.271739 | -1.073652 | 0.282978626 | 0.471180726 | | sense_overlapping | 44.09 | -0.095348587 | 0.095348587 | ENSG00000259713 | n.s. |
| ENSG00000275056 | 99.04225672 | -0.280529035 | 0.2539068 | -1.10485 | 0.269224411 | 0.457439216 | | sense_intronic | 45.74 | -0.095286345 | 0.095286345 | ENSG00000275056 | n.s. |
| ENSG00000100321 | 317.1255679 | -0.301789734 | 0.2875629 | -1.049474 | 0.293960141 | 0.483383376 | SYNGR1 | protein_coding | 55.76 | -0.095277521 | 0.095277521 | ENSG00000100321 | n.s. |
| ENSG00000107223 | 2114.574302 | 0.309486475 | 0.3002945 | 1.03061 | 0.302723794 | 0.492616104 | EDF1 | protein_coding | 57.77 | 0.095164428 | 0.095164428 | ENSG00000107223 | n.s. |
| ENSG00000144401 | 440.1493842 | -0.312376886 | 0.3051994 | -1.023517 | 0.306063366 | 0.495936671 | METTL21A | protein_coding | 40.9 | -0.095141808 | 0.095141808 | ENSG00000144401 | n.s. |
| ENSG00000109133 | 3788.04867 | -0.292505394 | 0.2732203 | -1.070584 | 0.284356422 | 0.472897093 | TMEM33 | protein_coding | 35.29 | -0.095132511 | 0.095132511 | ENSG00000109133 | n.s. |
| ENSG00000177169 | 374.9899896 | -0.287554779 | 0.2661378 | -1.080473 | 0.279931516 | 0.467604859 | ULK1 | protein_coding | 62.39 | -0.094927867 | 0.094927867 | ENSG00000177169 | n.s. |
| ENSG00000172817 | 249.7656538 | -0.278909222 | 0.2520666 | -1.10649 | 0.26851445 | 0.456809224 | CYP7B1 | protein_coding | 37.55 | -0.094903084 | 0.094903084 | ENSG00000172817 | n.s. |
| ENSG00000175893 | 1002.415051 | -0.288068894 | 0.2670714 | -1.078621 | 0.280756645 | 0.468511216 | ZDHHC21 | protein_coding | 35.18 | -0.094855328 | 0.094855328 | ENSG00000175893 | n.s. |

| | | | | | | | | | | | | | |
|---|---|---|---|---|---|---|---|---|---|---|---|---|---|
| ENSG00000169679 | 2100.841775 | -0.301222212 | 0.2876196 | -1.047294 | 0.294964122 | 0.484290739 | BUB1 | protein_coding | 38.24 | -0.094853018 | 0.094853018 | ENSG00000169679 | n.s. |
| ENSG00000122696 | 318.6440678 | -0.271570367 | 0.2413975 | -1.124993 | 0.260592132 | 0.447486958 | SLC25A51 | protein_coding | 42.85 | -0.0948377 | 0.0948377 | ENSG00000122696 | n.s. |
| ENSG00000106263 | 4791.544601 | 0.28539658 | 0.2626241 | 1.086711 | 0.277164405 | 0.465523702 | EIF3B | protein_coding | 51.61 | 0.094768275 | 0.094768275 | ENSG00000106263 | n.s. |
| ENSG00000204536 | 189.7470514 | -0.284976613 | 0.2620645 | -1.087429 | 0.276847197 | 0.465355973 | CCHCR1 | protein_coding | 51.86 | -0.094673422 | 0.094673422 | ENSG00000204536 | n.s. |
| ENSG00000135617 | 165.6393383 | -0.32271619 | 0.3235087 | -0.99755 | 0.318497549 | 0.508906432 | PRADC1 | protein_coding | 52.11 | -0.094672686 | 0.094672686 | ENSG00000135617 | n.s. |
| ENSG00000163320 | 3449.917705 | -0.282493406 | 0.2584111 | -1.093194 | 0.274308795 | 0.462429029 | CGGBP1 | protein_coding | 35.99 | -0.094622554 | 0.094622554 | ENSG00000163320 | n.s. |
| ENSG00000178074 | 953.9947329 | -0.278425985 | 0.2522121 | -1.103936 | 0.269620993 | 0.457852553 | C2orf69 | protein_coding | 38 | -0.094462798 | 0.094462798 | ENSG00000178074 | n.s. |
| ENSG00000162545 | 90.35092718 | 0.281024254 | 0.2566133 | 1.095128 | 0.273460751 | 0.461502308 | CAMK2N1 | protein_coding | 57.6 | 0.094375286 | 0.094375286 | ENSG00000162545 | n.s. |
| ENSG00000163629 | 2525.045748 | -0.294543043 | 0.2780694 | -1.059243 | 0.289489248 | 0.478404605 | PTPN13 | protein_coding | 36.16 | -0.094314051 | 0.094314051 | ENSG00000163629 | n.s. |
| ENSG00000161204 | 488.3093273 | 0.27982397 | 0.2549939 | 1.097375 | 0.272477413 | 0.460394429 | ABCF3 | protein_coding | 51.95 | 0.094264284 | 0.094264284 | ENSG00000161204 | n.s. |
| ENSG00000130635 | 498.454763 | -0.265619396 | 0.233723 | -1.136471 | 0.25575949 | 0.441803893 | COL5A1 | protein_coding | 56 | -0.094233916 | 0.094233916 | ENSG00000130635 | n.s. |
| ENSG00000221445 | 2.812056486 | -0.467817347 | 0.6063681 | -0.771507 | 0.440406398 | 0.62896057 | MIR1301 | miRNA | 62.2 | -0.094207457 | 0.094207457 | ENSG00000221445 | n.s. |
| ENSG00000139133 | 690.7462457 | -0.258853749 | 0.2235928 | -1.157702 | 0.246985728 | 0.432823448 | ALG10 | protein_coding | 36.5 | -0.094142318 | 0.094142318 | ENSG00000139133 | n.s. |
| ENSG00000269032 | 13.60428787 | -0.37863864 | 0.424561 | -0.891836 | 0.372481067 | 0.564560625 | | transcribed_processed_pseudog | 58.79 | -0.094011966 | 0.094011966 | ENSG00000269032 | n.s. |
| ENSG00000183655 | 781.4489329 | -0.275750972 | 0.2489015 | -1.107872 | 0.267917197 | 0.456110889 | KLHL25 | protein_coding | 53.6 | -0.094011657 | 0.094011657 | ENSG00000183655 | n.s. |
| ENSG00000141101 | 576.9835601 | 0.279384511 | 0.2548688 | 1.09619 | 0.272995791 | 0.46082934 | NOB1 | protein_coding | 48.27 | 0.094001679 | 0.094001679 | ENSG00000141101 | n.s. |
| ENSG00000013563 | 136.7849726 | -0.278167387 | 0.2529661 | -1.099623 | 0.271496281 | 0.459364831 | DNASE1L1 | protein_coding | 54.47 | -0.093976698 | 0.093976698 | ENSG00000013563 | n.s. |
| ENSG00000172465 | 230.2152544 | 0.314382635 | 0.3111598 | 1.010358 | 0.312323981 | 0.502482874 | TCEAL1 | protein_coding | 50.67 | 0.093962284 | 0.093962284 | ENSG00000172465 | n.s. |
| ENSG00000143198 | 685.0591261 | 0.335998448 | 0.3471442 | 0.967893 | 0.333097813 | 0.525282687 | MGST3 | protein_coding | 43.51 | 0.093947489 | 0.093947489 | ENSG00000143198 | n.s. |
| ENSG00000135931 | 96.24148615 | -0.311510945 | 0.3065359 | -1.01623 | 0.309519901 | 0.499530031 | ARMC9 | protein_coding | 45.69 | -0.09390136 | 0.09390136 | ENSG00000135931 | n.s. |
| ENSG00000213033 | 47.54621356 | 0.307290755 | 0.2994224 | 1.026278 | 0.30476035 | 0.494800816 | AURKAPS1 | transcribed_processed_pseudog | 44.14 | 0.093898711 | 0.093898711 | ENSG00000213033 | n.s. |
| ENSG00000115561 | 1467.801456 | -0.29710836 | 0.2830019 | -1.049846 | 0.293789077 | 0.483159145 | CHMP3 | protein_coding | 40.96 | -0.093859441 | 0.093859441 | ENSG00000115561 | n.s. |
| ENSG00000173171 | 468.2275222 | 0.272354632 | 0.2443705 | 1.114515 | 0.265058166 | 0.452611464 | MTX1 | protein_coding | 55.87 | 0.093764741 | 0.093764741 | ENSG00000173171 | n.s. |
| ENSG00000277142 | 123.8038103 | 0.266223063 | 0.2354152 | 1.130866 | 0.258111439 | 0.444597992 | | lincRNA | 61.7 | 0.093719171 | 0.093719171 | ENSG00000277142 | n.s. |
| ENSG00000126012 | 371.1021048 | 0.264804787 | 0.2333926 | 1.134589 | 0.256547407 | 0.442742409 | KDM5C | protein_coding | 48.11 | 0.093700876 | 0.093700876 | ENSG00000126012 | n.s. |
| ENSG00000176125 | 50.30172007 | 0.282410649 | 0.2605871 | 1.083747 | 0.278476796 | 0.466462035 | UFSP1 | protein_coding | 66.77 | 0.093529803 | 0.093529803 | ENSG00000176125 | n.s. |
| ENSG00000105726 | 140.6158148 | 0.263417344 | 0.2316099 | 1.137332 | 0.255399538 | 0.441510773 | ATP13A1 | protein_coding | 57.98 | 0.093528618 | 0.093528618 | ENSG00000105726 | n.s. |
| ENSG00000197713 | 2073.362037 | -0.282857561 | 0.2614904 | -1.081713 | 0.279380046 | 0.467356528 | RPE | protein_coding | 38.52 | -0.093442472 | 0.093442472 | ENSG00000197713 | n.s. |
| ENSG00000214199 | 60.36139488 | 0.329304223 | 0.3372921 | 0.976318 | 0.328907111 | 0.520526307 | EEF1A1P12 | processed_pseudogene | 46.58 | 0.093376622 | 0.093376622 | ENSG00000214199 | n.s. |
| ENSG00000187678 | 136.3447687 | 0.282617835 | 0.2615533 | 1.080536 | 0.279903522 | 0.467604859 | SPRY4 | protein_coding | 51.21 | 0.093298078 | 0.093298078 | ENSG00000187678 | n.s. |
| ENSG00000275560 | 615.9304178 | 0.238212651 | 0.1965484 | 1.21198 | 0.225520151 | 0.406453888 | | sense_intronic | 43.17 | 0.093138459 | 0.093138459 | ENSG00000275560 | n.s. |
| ENSG00000186162 | 135.3651872 | -0.270456031 | 0.2426738 | -1.114484 | 0.265071743 | 0.452611464 | CIDECP | transcribed_unprocessed_pseud | 47.2 | -0.093111102 | 0.093111102 | ENSG00000186162 | n.s. |

| | | | | | | | | | | | | | |
|---|---|---|---|---|---|---|---|---|---|---|---|---|---|
| ENSG00000014123 | 1319.252817 | -0.270603701 | 0.2430445 | -1.113392 | 0.265540278 | 0.453022581 | UFL1 | protein_coding | 33.38 | -0.093055241 | 0.093055241 | ENSG00000014123 | n.s. |
| ENSG00000172270 | 1969.318988 | 0.29307093 | 0.278124 | 1.053742 | 0.292001144 | 0.481408223 | BSG | protein_coding | 62.58 | 0.093046063 | 0.093046063 | ENSG00000172270 | n.s. |
| ENSG00000241913 | 23.05383623 | -0.334337647 | 0.3466334 | -0.964528 | 0.334781265 | 0.527101612 | RPS29P21 | processed_pseudogene | 42.94 | -0.09298119 | 0.09298119 | ENSG00000241913 | n.s. |
| ENSG00000168395 | 1553.367258 | -0.260430991 | 0.2280253 | -1.142115 | 0.253406413 | 0.439745075 | ING5 | protein_coding | 51.69 | -0.092921521 | 0.092921521 | ENSG00000168395 | n.s. |
| ENSG00000156411 | 1787.274981 | 0.316659424 | 0.3174451 | 0.997525 | 0.318509707 | 0.508906432 | C14orf2 | protein_coding | 47.77 | 0.092895861 | 0.092895861 | ENSG00000156411 | n.s. |
| ENSG00000126787 | 1462.419091 | -0.29741721 | 0.2855861 | -1.041427 | 0.297677247 | 0.48717954 | DLGAP5 | protein_coding | 39.11 | -0.092886654 | 0.092886654 | ENSG00000126787 | n.s. |
| ENSG00000177707 | 552.6729098 | 0.269817833 | 0.2421383 | 1.114313 | 0.265144931 | 0.452675743 | NECTIN3 | protein_coding | 36.67 | 0.092874746 | 0.092874746 | ENSG00000177707 | n.s. |
| ENSG00000165943 | 956.6717247 | 0.283114535 | 0.2630938 | 1.076097 | 0.281883817 | 0.469907614 | MOAP1 | protein_coding | 51.56 | 0.092858034 | 0.092858034 | ENSG00000165943 | n.s. |
| ENSG00000054523 | 975.4843848 | -0.25774103 | 0.2242717 | -1.149236 | 0.250458831 | 0.436511212 | KIF1B | protein_coding | 41.99 | -0.092787955 | 0.092787955 | ENSG00000054523 | n.s. |
| ENSG00000166262 | 55.02782501 | 0.296840422 | 0.2851251 | 1.041088 | 0.297834487 | 0.487279619 | FAM227B | protein_coding | 36.14 | 0.092680037 | 0.092680037 | ENSG00000166262 | n.s. |
| ENSG00000164318 | 451.7519669 | -0.288239544 | 0.2712467 | -1.062647 | 0.287942061 | 0.477150359 | EGFLAM | protein_coding | 42.6 | -0.092624263 | 0.092624263 | ENSG00000164318 | n.s. |
| ENSG00000169189 | 401.0103017 | 0.310025212 | 0.3070584 | 1.009662 | 0.312657216 | 0.502728539 | NSMCE1 | protein_coding | 47.29 | 0.092594133 | 0.092594133 | ENSG00000169189 | n.s. |
| ENSG00000170275 | 1337.409287 | -0.290439537 | 0.2748532 | -1.056708 | 0.290645003 | 0.479972764 | CRTAP | protein_coding | 44.28 | -0.092587305 | 0.092587305 | ENSG00000170275 | n.s. |
| ENSG00000149231 | 672.1251658 | -0.288915857 | 0.2726338 | -1.059721 | 0.28927132 | 0.478157971 | CCDC82 | protein_coding | 34.71 | -0.092576904 | 0.092576904 | ENSG00000149231 | n.s. |
| ENSG00000109099 | 313.3366157 | -0.28041648 | 0.2594574 | -1.080781 | 0.279794709 | 0.46760115 | PMP22 | protein_coding | 45.52 | -0.092572331 | 0.092572331 | ENSG00000109099 | n.s. |
| ENSG00000148334 | 440.8889371 | 0.289880003 | 0.2740474 | 1.057773 | 0.290158757 | 0.479375189 | PTGES2 | protein_coding | 59.55 | 0.092565772 | 0.092565772 | ENSG00000148334 | n.s. |
| ENSG00000177548 | 27.5812691 | 0.328354529 | 0.3375399 | 0.972787 | 0.330659027 | 0.522561366 | RABEP2 | protein_coding | 52.63 | 0.092550895 | 0.092550895 | ENSG00000177548 | n.s. |
| ENSG00000177156 | 725.1289111 | 0.314735775 | 0.3150925 | 0.998868 | 0.317858668 | 0.508254468 | TALDO1 | protein_coding | 52.03 | 0.09250676 | 0.09250676 | ENSG00000177156 | n.s. |
| ENSG00000165997 | 4662.424425 | 0.288317377 | 0.2716838 | 1.061224 | 0.288588145 | 0.477725511 | ARL5B | protein_coding | 37.23 | 0.092498432 | 0.092498432 | ENSG00000165997 | n.s. |
| ENSG00000264834 | 6.219368679 | 0.401905646 | 0.4746176 | 0.846799 | 0.397107195 | 0.58899792 | MIR1273F | miRNA | 58.59 | 0.092392578 | 0.092392578 | ENSG00000264834 | n.s. |
| ENSG00000179598 | 632.2993203 | 0.264886188 | 0.2356475 | 1.124078 | 0.260979838 | 0.447949194 | PLD6 | protein_coding | 55.87 | 0.092384685 | 0.092384685 | ENSG00000179598 | n.s. |
| ENSG00000129084 | 2544.406542 | 0.303124384 | 0.296281 | 1.023098 | 0.306261625 | 0.495938661 | PSMA1 | protein_coding | 40.21 | 0.092323211 | 0.092323211 | ENSG00000129084 | n.s. |
| ENSG00000207008 | 35.47152194 | -0.3031183 | 0.2962581 | -1.023156 | 0.306234054 | 0.495938661 | SNORA54 | snoRNA | 49.59 | -0.092321358 | 0.092321358 | ENSG00000207008 | n.s. |
| ENSG00000167733 | 89.16533536 | 0.294430062 | 0.2820454 | 1.04391 | 0.296527034 | 0.485825593 | HSD11B1L | protein_coding | 58.24 | 0.092309599 | 0.092309599 | ENSG00000167733 | n.s. |
| ENSG00000197785 | 223.3799033 | -0.286890521 | 0.2697433 | -1.063569 | 0.287524108 | 0.476741473 | ATAD3A | protein_coding | 59.87 | -0.092297577 | 0.092297577 | ENSG00000197785 | n.s. |
| ENSG00000105364 | 346.8901539 | 0.304200738 | 0.2981652 | 1.020242 | 0.307613604 | 0.497322118 | MRPL4 | protein_coding | 56.13 | 0.092283013 | 0.092283013 | ENSG00000105364 | n.s. |
| ENSG00000255737 | 100.5706365 | -0.297678746 | 0.2874891 | -1.035444 | 0.300461834 | 0.490195701 | AGAP2-AS1 | antisense | 66.06 | -0.09217042 | 0.09217042 | ENSG00000255737 | n.s. |
| ENSG00000072274 | 6404.726228 | 0.29879575 | 0.2894542 | 1.032273 | 0.301944295 | 0.491577457 | TFRC | protein_coding | 42.47 | 0.092151012 | 0.092151012 | ENSG00000072274 | n.s. |
| ENSG00000111237 | 1225.125187 | -0.295629259 | 0.2843181 | -1.039783 | 0.298440593 | 0.488099187 | VPS29 | protein_coding | 40.72 | -0.092086124 | 0.092086124 | ENSG00000111237 | n.s. |
| ENSG00000159335 | 1809.907837 | 0.290754506 | 0.276578 | 1.051257 | 0.293140712 | 0.482434767 | PTMS | protein_coding | 61.84 | 0.092041654 | 0.092041654 | ENSG00000159335 | n.s. |
| ENSG00000164182 | 476.0758598 | 0.315211755 | 0.3168183 | 0.994929 | 0.319770743 | 0.510511187 | NDUFAF2 | protein_coding | 36.3 | 0.092040174 | 0.092040174 | ENSG00000164182 | n.s. |
| ENSG00000132570 | 139.2053453 | -0.301635095 | 0.2946508 | -1.023704 | 0.305975331 | 0.495878772 | PCBD2 | protein_coding | 42.21 | -0.091885435 | 0.091885435 | ENSG00000132570 | n.s. |

| | | | | | | | | | | | | | |
|---|---|---|---|---|---|---|---|---|---|---|---|---|---|
| ENSG00000235284 | 10.86087322 | -0.379818993 | 0.4330887 | -0.877 | 0.380486436 | 0.573013807 | SNORD62A | snoRNA | 46.51 | -0.091853493 | 0.091853493 | ENSG00000235284 | n.s. |
| ENSG00000249592 | 294.5542182 | -0.256318552 | 0.223858 | -1.145005 | 0.252207147 | 0.438496517 | | antisense | 50.65 | -0.091770718 | 0.091770718 | ENSG00000249592 | n.s. |
| ENSG00000064652 | 160.3941771 | -0.296565037 | 0.2865349 | -1.035005 | 0.300666534 | 0.490414733 | SNX24 | protein_coding | 40.32 | -0.091768044 | 0.091768044 | ENSG00000064652 | n.s. |
| ENSG00000204351 | 700.7309134 | 0.271849257 | 0.2474332 | 1.098678 | 0.271908745 | 0.459718939 | SKIV2L | protein_coding | 54.18 | 0.091751191 | 0.091751191 | ENSG00000204351 | n.s. |
| ENSG00000157927 | 283.8183054 | -0.2762775 | 0.2542414 | -1.086674 | 0.27718103 | 0.465523702 | RADIL | protein_coding | 53.13 | -0.09174021 | 0.09174021 | ENSG00000157927 | n.s. |
| ENSG00000130703 | 306.5667456 | -0.300000176 | 0.2923286 | -1.026243 | 0.304777117 | 0.494800816 | OSBPL2 | protein_coding | 52.75 | -0.091670932 | 0.091670932 | ENSG00000130703 | n.s. |
| ENSG00000163703 | 225.9074359 | 0.29218781 | 0.2797748 | 1.044368 | 0.296315187 | 0.48559279 | CRELD1 | protein_coding | 51.58 | 0.09166743 | 0.09166743 | ENSG00000163703 | n.s. |
| ENSG00000162585 | 105.199769 | 0.315405934 | 0.3180727 | 0.991616 | 0.321384912 | 0.512148578 | FAAP20 | protein_coding | 60.96 | 0.091658235 | 0.091658235 | ENSG00000162585 | n.s. |
| ENSG00000169018 | 3319.356769 | -0.285666615 | 0.2692106 | -1.061127 | 0.288632263 | 0.477725511 | FEM1B | protein_coding | 39.07 | -0.091648011 | 0.091648011 | ENSG00000169018 | n.s. |
| ENSG00000099617 | 32.01808852 | -0.300057507 | 0.2925946 | -1.025506 | 0.305124444 | 0.495133756 | EFNA2 | protein_coding | 64.19 | -0.091600795 | 0.091600795 | ENSG00000099617 | n.s. |
| ENSG00000132600 | 108.1497636 | -0.288881604 | 0.274591 | -1.052043 | 0.292779664 | 0.482182548 | PRMT7 | protein_coding | 48.59 | -0.091514374 | 0.091514374 | ENSG00000132600 | n.s. |
| ENSG00000128272 | 9918.05669 | 0.283349118 | 0.2658003 | 1.066022 | 0.286413522 | 0.475415093 | ATF4 | protein_coding | 58.94 | 0.091501091 | 0.091501091 | ENSG00000128272 | n.s. |
| ENSG00000258445 | 39.94128287 | -0.316496314 | 0.3204563 | -0.987643 | 0.323327685 | 0.513950377 | | processed_pseudogene | 41.59 | -0.091492378 | 0.091492378 | ENSG00000258445 | n.s. |
| ENSG00000115825 | 3002.207831 | -0.285093776 | 0.2686035 | -1.061393 | 0.288511514 | 0.477725511 | PRKD3 | protein_coding | 36.9 | -0.091464232 | 0.091464232 | ENSG00000115825 | n.s. |
| ENSG00000276410 | 556.5665295 | 0.274655464 | 0.2521985 | 1.089045 | 0.276134219 | 0.46471857 | HIST1H2BB | protein_coding | 54.07 | 0.091408077 | 0.091408077 | ENSG00000276410 | n.s. |
| ENSG00000254328 | 46.48952066 | 0.28497009 | 0.2686182 | 1.060874 | 0.288747192 | 0.477838278 | | processed_pseudogene | 50.1 | 0.09139534 | 0.09139534 | ENSG00000254328 | n.s. |
| ENSG00000196923 | 172.151775 | 0.275637297 | 0.2543749 | 1.083587 | 0.278547947 | 0.466525089 | PDLIM7 | protein_coding | 60.73 | 0.091270399 | 0.091270399 | ENSG00000196923 | n.s. |
| ENSG00000121350 | 570.6293203 | 0.261728034 | 0.2329733 | 1.123425 | 0.261257007 | 0.448241689 | PYROXD1 | protein_coding | 36.31 | 0.091209015 | 0.091209015 | ENSG00000121350 | n.s. |
| ENSG00000034533 | 442.1488693 | -0.246776839 | 0.2108615 | -1.170327 | 0.241869497 | 0.427207497 | ASTE1 | protein_coding | 39.7 | -0.091149773 | 0.091149773 | ENSG00000034533 | n.s. |
| ENSG00000183506 | 172.77616 | 0.268064956 | 0.2425162 | 1.105349 | 0.269008472 | 0.457224948 | PI4KAP2 | transcribed_unprocessed_pseud | 55.77 | 0.091107258 | 0.091107258 | ENSG00000183506 | n.s. |
| ENSG00000203930 | 153.4393513 | 0.265887599 | 0.2393384 | 1.110927 | 0.266599675 | 0.454383852 | LINC00632 | lincRNA | 40.81 | 0.091087005 | 0.091087005 | ENSG00000203930 | n.s. |
| ENSG00000196422 | 234.4290701 | -0.269582484 | 0.2451486 | -1.099669 | 0.271476159 | 0.459364831 | PPP1R26 | protein_coding | 59.96 | -0.091076355 | 0.091076355 | ENSG00000196422 | n.s. |
| ENSG00000112339 | 2995.373383 | -0.274507125 | 0.2529111 | -1.08539 | 0.277749125 | 0.465901588 | HBS1L | protein_coding | 36.66 | -0.091055608 | 0.091055608 | ENSG00000112339 | n.s. |
| ENSG00000120992 | 1583.279235 | 0.308833602 | 0.3086749 | 1.000514 | 0.317061826 | 0.507444026 | LYPLA1 | protein_coding | 41.54 | 0.09098604 | 0.09098604 | ENSG00000120992 | n.s. |
| ENSG00000067560 | 3063.549216 | 0.310562728 | 0.3115852 | 0.996718 | 0.318901191 | 0.509415016 | RHOA | protein_coding | 46.57 | 0.090972599 | 0.090972599 | ENSG00000067560 | n.s. |
| ENSG00000143924 | 3290.690463 | -0.276641927 | 0.2569462 | -1.076653 | 0.281635281 | 0.469549517 | EML4 | protein_coding | 38.02 | -0.090826691 | 0.090826691 | ENSG00000143924 | n.s. |
| ENSG00000205356 | 421.2616849 | -0.281564146 | 0.2643542 | -1.065102 | 0.286829891 | 0.475930471 | TECPR1 | protein_coding | 56.22 | -0.090792186 | 0.090792186 | ENSG00000205356 | n.s. |
| ENSG00000130164 | 1399.338588 | -0.25873276 | 0.2293663 | -1.128033 | 0.259305876 | 0.445828071 | LDLR | protein_coding | 51.08 | -0.090771885 | 0.090771885 | ENSG00000130164 | n.s. |
| ENSG00000130830 | 178.3183472 | 0.28555081 | 0.2709505 | 1.053885 | 0.291935384 | 0.481408223 | MPP1 | protein_coding | 41.68 | 0.090658526 | 0.090658526 | ENSG00000130830 | n.s. |
| ENSG00000134285 | 233.195229 | -0.270147754 | 0.2468869 | -1.094217 | 0.273859899 | 0.461895988 | FKBP11 | protein_coding | 51.54 | -0.090622634 | 0.090622634 | ENSG00000134285 | n.s. |
| ENSG00000126756 | 607.4076326 | 0.31951772 | 0.3272168 | 0.976471 | 0.328831107 | 0.520526307 | UXT | protein_coding | 44.96 | 0.090601587 | 0.090601587 | ENSG00000126756 | n.s. |
| ENSG00000177125 | 465.5333164 | 0.267746934 | 0.2430135 | 1.101778 | 0.270558193 | 0.458839018 | ZBTB34 | protein_coding | 43.23 | 0.090589406 | 0.090589406 | ENSG00000177125 | n.s. |

| | | | | | | | | | | | | | |
|---|---|---|---|---|---|---|---|---|---|---|---|---|---|
| ENSG00000244687 | 992.2854111 | -0.258943981 | 0.2300533 | -1.125583 | 0.260342264 | 0.447113031 | UBE2V1 | protein_coding | 44.49 | -0.090522329 | 0.090522329 | ENSG00000244687 | n.s. |
| ENSG00000100296 | 1289.288975 | -0.277534319 | 0.2587834 | -1.072458 | 0.283514441 | 0.4718914 | THOC5 | protein_coding | 47.17 | -0.090520021 | 0.090520021 | ENSG00000100296 | n.s. |
| ENSG00000242259 | 948.0686978 | 0.272867038 | 0.2516312 | 1.084393 | 0.278190702 | 0.466198989 | C22orf39 | protein_coding | 47.03 | 0.09043596 | 0.09043596 | ENSG00000242259 | n.s. |
| ENSG00000167107 | 254.7223465 | -0.290702312 | 0.2798984 | -1.038599 | 0.298991138 | 0.488655196 | ACSF2 | protein_coding | 49.96 | -0.090407686 | 0.090407686 | ENSG00000167107 | n.s. |
| ENSG00000077152 | 2433.779725 | -0.317961156 | 0.3249284 | -0.978558 | 0.327798577 | 0.519626772 | UBE2T | protein_coding | 40.71 | -0.090399053 | 0.090399053 | ENSG00000077152 | n.s. |
| ENSG00000117360 | 2525.122594 | -0.272562465 | 0.2512191 | -1.084959 | 0.277939823 | 0.465954986 | PRPF3 | protein_coding | 44.24 | -0.090396987 | 0.090396987 | ENSG00000117360 | n.s. |
| ENSG00000145439 | 901.7523283 | -0.277400154 | 0.2590396 | -1.070879 | 0.284223746 | 0.472732913 | CBR4 | protein_coding | 38.87 | -0.090261616 | 0.090261616 | ENSG00000145439 | n.s. |
| ENSG00000160446 | 88.79111862 | -0.29203499 | 0.2825852 | -1.033441 | 0.301397721 | 0.490915954 | ZDHHC12 | protein_coding | 61.4 | -0.090236725 | 0.090236725 | ENSG00000160446 | n.s. |
| ENSG00000103876 | 137.0782869 | -0.294816165 | 0.28714 | -1.026733 | 0.304546058 | 0.494649892 | FAH | protein_coding | 49.96 | -0.090125915 | 0.090125915 | ENSG00000103876 | n.s. |
| ENSG00000221227 | 13.05837098 | -0.356666166 | 0.3951849 | -0.90253 | 0.366775518 | 0.558955291 | MIR1305 | miRNA | 36.05 | -0.090102051 | 0.090102051 | ENSG00000221227 | n.s. |
| ENSG00000168884 | 214.0256454 | 0.283001627 | 0.2680999 | 1.055583 | 0.291158852 | 0.480593325 | TNIP2 | protein_coding | 54.04 | 0.090057419 | 0.090057419 | ENSG00000168884 | n.s. |
| ENSG00000104964 | 1574.405456 | 0.290453671 | 0.2804146 | 1.035801 | 0.300295135 | 0.489981151 | AES | protein_coding | 60.37 | 0.089988538 | 0.089988538 | ENSG00000104964 | n.s. |
| ENSG00000086589 | 1330.461449 | 0.259234053 | 0.2314205 | 1.120186 | 0.262634442 | 0.449662977 | RBM22 | protein_coding | 43.27 | 0.089983476 | 0.089983476 | ENSG00000086589 | n.s. |
| ENSG00000133226 | 1395.181231 | 0.252329754 | 0.2210366 | 1.141575 | 0.253630808 | 0.43995753 | SRRM1 | protein_coding | 41.76 | 0.089978076 | 0.089978076 | ENSG00000133226 | n.s. |
| ENSG00000090889 | 1365.082541 | -0.283362579 | 0.2688946 | -1.053806 | 0.291971951 | 0.481408223 | KIF4A | protein_coding | 40.48 | -0.089963792 | 0.089963792 | ENSG00000090889 | n.s. |
| ENSG00000176884 | 240.8625919 | 0.26874803 | 0.2459506 | 1.092691 | 0.27452942 | 0.462744929 | GRIN1 | protein_coding | 62.1 | 0.089938768 | 0.089938768 | ENSG00000176884 | n.s. |
| ENSG00000266019 | 144.7041694 | -0.261157838 | 0.2344046 | -1.114133 | 0.265222307 | 0.452678273 | MIR3609 | miRNA | 38.75 | -0.089893231 | 0.089893231 | ENSG00000266019 | n.s. |
| ENSG00000114770 | 1446.898218 | 0.278030342 | 0.2606152 | 1.066823 | 0.286051678 | 0.475035486 | ABCC5 | protein_coding | 44.7 | 0.089879966 | 0.089879966 | ENSG00000114770 | n.s. |
| ENSG00000198400 | 1898.131652 | 0.279217146 | 0.2625175 | 1.063614 | 0.287503771 | 0.476741473 | NTRK1 | protein_coding | 53.11 | 0.089828921 | 0.089828921 | ENSG00000198400 | n.s. |
| ENSG00000164885 | 233.0705958 | 0.303697399 | 0.3025434 | 1.003814 | 0.315468064 | 0.50609579 | CDK5 | protein_coding | 58.08 | 0.089823752 | 0.089823752 | ENSG00000164885 | n.s. |
| ENSG00000259956 | 2560.281759 | -0.238020956 | 0.2006872 | -1.18603 | 0.235610515 | 0.419435521 | RBM15B | protein_coding | 55.91 | -0.089813588 | 0.089813588 | ENSG00000259956 | n.s. |
| ENSG00000229638 | 547.0467982 | 0.322791628 | 0.3346373 | 0.964601 | 0.334744467 | 0.527101612 | RPL4P4 | processed_pseudogene | 48.29 | 0.089770177 | 0.089770177 | ENSG00000229638 | n.s. |
| ENSG00000151743 | 329.4685309 | -0.309947771 | 0.3137643 | -0.987836 | 0.323232758 | 0.513858152 | AMN1 | protein_coding | 40.55 | -0.08962349 | 0.08962349 | ENSG00000151743 | n.s. |
| ENSG00000090674 | 83.39034976 | 0.272298494 | 0.2525678 | 1.07812 | 0.280980072 | 0.468793953 | MCOLN1 | protein_coding | 57.01 | 0.089591105 | 0.089591105 | ENSG00000090674 | n.s. |
| ENSG00000122034 | 1680.976381 | 0.310202854 | 0.31434 | 0.986839 | 0.32372168 | 0.514400473 | GTF3A | protein_coding | 44.92 | 0.089555142 | 0.089555142 | ENSG00000122034 | n.s. |
| ENSG00000144579 | 387.2694364 | -0.26930453 | 0.2474499 | -1.088319 | 0.276454108 | 0.465129063 | CTDSP1 | protein_coding | 63.19 | -0.089523969 | 0.089523969 | ENSG00000144579 | n.s. |
| ENSG00000140263 | 1034.830691 | -0.293604241 | 0.286804 | -1.02371 | 0.305972047 | 0.495878772 | SORD | protein_coding | 45.28 | -0.08943904 | 0.08943904 | ENSG00000140263 | n.s. |
| ENSG00000100949 | 126.2334475 | 0.278765828 | 0.2628188 | 1.060677 | 0.288836686 | 0.477838278 | RABGGTA | protein_coding | 57.92 | 0.089405515 | 0.089405515 | ENSG00000100949 | n.s. |
| ENSG00000277075 | 2287.70117 | 0.244054928 | 0.2097006 | 1.163826 | 0.244494592 | 0.430353755 | HIST1H2AE | protein_coding | 56.19 | 0.089366668 | 0.089366668 | ENSG00000277075 | n.s. |
| ENSG00000100591 | 2250.591286 | 0.284019626 | 0.2713604 | 1.046651 | 0.295260657 | 0.484606168 | AHSA1 | protein_coding | 46.04 | 0.089355716 | 0.089355716 | ENSG00000100591 | n.s. |
| ENSG00000232112 | 2398.973637 | 0.301971842 | 0.3007052 | 1.004212 | 0.315276326 | 0.506004874 | TMA7 | protein_coding | 47.75 | 0.08933695 | 0.08933695 | ENSG00000232112 | n.s. |
| ENSG00000182674 | 1164.193001 | -0.265113366 | 0.2419097 | -1.095919 | 0.273114342 | 0.460973571 | KCNB2 | protein_coding | 38.88 | -0.089163981 | 0.089163981 | ENSG00000182674 | n.s. |

| | | | | | | | | | | | | | |
|---|---|---|---|---|---|---|---|---|---|---|---|---|---|
| ENSG00000113966 | 350.4646134 | -0.29160643 | 0.2840323 | -1.026666 | 0.304577571 | 0.494649892 | ARL6 | protein_coding | 34.36 | -0.089144693 | 0.089144693 | ENSG00000113966 | n.s. |
| ENSG00000169288 | 686.9939231 | -0.306111177 | 0.3082803 | -0.992964 | 0.320727585 | 0.511457021 | MRPL1 | protein_coding | 37.82 | -0.089136776 | 0.089136776 | ENSG00000169288 | n.s. |
| ENSG00000231940 | 41.32863551 | 0.301620892 | 0.3008759 | 1.002476 | 0.316113737 | 0.506702304 | RPS7P3 | processed_pseudogene | 46.03 | 0.0890527 | 0.0890527 | ENSG00000231940 | n.s. |
| ENSG00000234358 | 37.83505128 | -0.312371222 | 0.3186332 | -0.980348 | 0.326914612 | 0.518704518 | | processed_pseudogene | 48.76 | -0.089050778 | 0.089050778 | ENSG00000234358 | n.s. |
| ENSG00000096384 | 29179.81505 | 0.282988485 | 0.2705241 | 1.046075 | 0.295526499 | 0.484813886 | HSP90AB1 | protein_coding | 49.1 | 0.08897864 | 0.08897864 | ENSG00000096384 | n.s. |
| ENSG00000181588 | 513.9677156 | -0.269212851 | 0.2490505 | -1.080957 | 0.279716253 | 0.467581869 | MEX3D | protein_coding | 60.48 | -0.08887856 | 0.08887856 | ENSG00000181588 | n.s. |
| ENSG00000274605 | 99.85068553 | -0.286446672 | 0.276185 | -1.037155 | 0.299663662 | 0.489467054 | | lincRNA | 45.07 | -0.088877681 | 0.088877681 | ENSG00000274605 | n.s. |
| ENSG00000168795 | 800.873818 | -0.283105101 | 0.270964 | -1.044807 | 0.296112166 | 0.485374344 | ZBTB5 | protein_coding | 44.23 | -0.088873254 | 0.088873254 | ENSG00000168795 | n.s. |
| ENSG00000165338 | 928.336691 | -0.291971922 | 0.2855382 | -1.022532 | 0.306529168 | 0.496308404 | HECTD2 | protein_coding | 36.7 | -0.088831982 | 0.088831982 | ENSG00000165338 | n.s. |
| ENSG00000248174 | 1430.300498 | -0.244120486 | 0.2107945 | -1.158097 | 0.246824364 | 0.432712562 | LINC02268 | lincRNA | 36.76 | -0.088811154 | 0.088811154 | ENSG00000248174 | n.s. |
| ENSG00000170921 | 2505.625859 | -0.249282344 | 0.2185384 | -1.14068 | 0.254003151 | 0.440409199 | TANC2 | protein_coding | 37.68 | -0.088780316 | 0.088780316 | ENSG00000170921 | n.s. |
| ENSG00000160211 | 194.3363055 | -0.286250612 | 0.2761308 | -1.036649 | 0.299899615 | 0.48967457 | G6PD | protein_coding | 56.32 | -0.088764153 | 0.088764153 | ENSG00000160211 | n.s. |
| ENSG00000171456 | 1755.090077 | -0.267425693 | 0.2461755 | -1.086321 | 0.277336945 | 0.465673213 | ASXL1 | protein_coding | 42.17 | -0.0887636 | 0.0887636 | ENSG00000171456 | n.s. |
| ENSG00000059758 | 2060.305828 | 0.270222277 | 0.2509452 | 1.076818 | 0.281561687 | 0.469483038 | CDK17 | protein_coding | 37.82 | 0.088735617 | 0.088735617 | ENSG00000059758 | n.s. |
| ENSG00000144406 | 243.1465627 | -0.276262587 | 0.2601199 | -1.062059 | 0.288209015 | 0.477386636 | UNC80 | protein_coding | 37.57 | -0.088716133 | 0.088716133 | ENSG00000144406 | n.s. |
| ENSG00000232388 | 518.6254327 | 0.310144807 | 0.3158069 | 0.982071 | 0.326064817 | 0.517699599 | SMIM26 | protein_coding | 44.25 | 0.088677277 | 0.088677277 | ENSG00000232388 | n.s. |
| ENSG00000171314 | 1206.618534 | 0.269958622 | 0.2508993 | 1.075964 | 0.281943389 | 0.469950653 | PGAM1 | protein_coding | 47.47 | 0.088532321 | 0.088532321 | ENSG00000171314 | n.s. |
| ENSG00000082701 | 1808.539939 | -0.261335028 | 0.2369652 | -1.102842 | 0.270095968 | 0.458503095 | GSK3B | protein_coding | 37.15 | -0.088503127 | 0.088503127 | ENSG00000082701 | n.s. |
| ENSG00000100473 | 1100.440302 | -0.281214216 | 0.2689214 | -1.045712 | 0.295694247 | 0.484974792 | COCH | protein_coding | 39.92 | -0.088380239 | 0.088380239 | ENSG00000100473 | n.s. |
| ENSG00000165526 | 772.7433113 | 0.270769429 | 0.2524065 | 1.072752 | 0.283382593 | 0.471728341 | RPUSD4 | protein_coding | 43.35 | 0.088354239 | 0.088354239 | ENSG00000165526 | n.s. |
| ENSG00000102753 | 1330.3501 | -0.278733497 | 0.264855 | -1.0524 | 0.292615895 | 0.481969846 | KPNA3 | protein_coding | 38.71 | -0.088352981 | 0.088352981 | ENSG00000102753 | n.s. |
| ENSG00000149131 | 187.2722781 | -0.274731725 | 0.2584279 | -1.063088 | 0.287741845 | 0.476875337 | SERPING1 | protein_coding | 47.51 | -0.088352387 | 0.088352387 | ENSG00000149131 | n.s. |
| ENSG00000272888 | 10145.00153 | 0.250712332 | 0.2216209 | 1.131267 | 0.257942854 | 0.444372019 | LINC01578 | processed_transcript | 36.16 | 0.088314245 | 0.088314245 | ENSG00000272888 | n.s. |
| ENSG00000102362 | 2084.864463 | -0.275236228 | 0.2595784 | -1.06032 | 0.288998867 | 0.477838278 | SYTL4 | protein_coding | 41.39 | -0.088273505 | 0.088273505 | ENSG00000102362 | n.s. |
| ENSG00000144909 | 477.7253194 | -0.260666625 | 0.2366351 | -1.101555 | 0.27065507 | 0.458839018 | OSBPL11 | protein_coding | 39.18 | -0.088193857 | 0.088193857 | ENSG00000144909 | n.s. |
| ENSG00000101190 | 264.8003466 | -0.277295035 | 0.2628478 | -1.054964 | 0.291441622 | 0.480946035 | TCFL5 | protein_coding | 48.88 | -0.088153104 | 0.088153104 | ENSG00000101190 | n.s. |
| ENSG00000137413 | 209.8486442 | 0.250428848 | 0.2215404 | 1.130398 | 0.258308589 | 0.444772507 | TAF8 | protein_coding | 45.88 | 0.088116412 | 0.088116412 | ENSG00000137413 | n.s. |
| ENSG00000176788 | 2277.613458 | -0.290600107 | 0.2851142 | -1.019241 | 0.308088541 | 0.497739238 | BASP1 | protein_coding | 43.15 | -0.088051286 | 0.088051286 | ENSG00000176788 | n.s. |
| ENSG00000047849 | 1300.584066 | -0.275710661 | 0.2607488 | -1.05738 | 0.290338046 | 0.479522731 | MAP4 | protein_coding | 43.43 | -0.088004307 | 0.088004307 | ENSG00000047849 | n.s. |
| ENSG00000101911 | 531.6513386 | -0.298697681 | 0.2984397 | -1.000864 | 0.316892387 | 0.507444026 | PRPS2 | protein_coding | 42.76 | -0.087999878 | 0.087999878 | ENSG00000101911 | n.s. |
| ENSG00000176171 | 767.4270663 | 0.294393689 | 0.291316 | 1.010565 | 0.312224786 | 0.502482874 | BNIP3 | protein_coding | 49.42 | 0.087988013 | 0.087988013 | ENSG00000176171 | n.s. |
| ENSG00000260671 | 10.48163649 | 0.354587509 | 0.3977703 | 0.891438 | 0.372694309 | 0.564760869 | | sense_intronic | 50.67 | 0.087985714 | 0.087985714 | ENSG00000260671 | n.s. |

| | | | | | | | | | | | | | |
|---|---|---|---|---|---|---|---|---|---|---|---|---|---|
| ENSG00000117632 | 8917.98065 | 0.299072377 | 0.2992639 | 0.99936 | 0.317620288 | 0.507951673 | STMN1 | protein_coding | 48.24 | 0.087980396 | 0.087980396 | ENSG00000117632 | n.s. |
| ENSG00000101596 | 4704.430656 | -0.270291916 | 0.2523424 | -1.071131 | 0.284110307 | 0.472657164 | SMCHD1 | protein_coding | 36.8 | -0.087967519 | 0.087967519 | ENSG00000101596 | n.s. |
| ENSG00000108443 | 2552.033948 | -0.266225196 | 0.2461951 | -1.081359 | 0.279537523 | 0.467427974 | RPS6KB1 | protein_coding | 41.68 | -0.087930269 | 0.087930269 | ENSG00000108443 | n.s. |
| ENSG00000005075 | 567.4965993 | 0.303672615 | 0.3071429 | 0.988701 | 0.32280928 | 0.513536746 | POLR2J | protein_coding | 55.04 | 0.087891502 | 0.087891502 | ENSG00000005075 | n.s. |
| ENSG00000232442 | 42.46037718 | 0.294882634 | 0.2926021 | 1.007794 | 0.313553317 | 0.503762155 | MHENCR | antisense | 62.33 | 0.087808518 | 0.087808518 | ENSG00000232442 | n.s. |
| ENSG00000072571 | 1119.432903 | -0.302857342 | 0.305939 | -0.989927 | 0.322209767 | 0.512994356 | HMMR | protein_coding | 36.97 | -0.087794531 | 0.087794531 | ENSG00000072571 | n.s. |
| ENSG00000090924 | 136.3731694 | -0.274526269 | 0.2595436 | -1.057727 | 0.290179853 | 0.479375189 | PLEKHG2 | protein_coding | 56.77 | -0.087662949 | 0.087662949 | ENSG00000090924 | n.s. |
| ENSG00000126456 | 132.55989 | 0.29948294 | 0.3008039 | 0.995608 | 0.319440435 | 0.510159345 | IRF3 | protein_coding | 57.17 | 0.087537113 | 0.087537113 | ENSG00000126456 | n.s. |
| ENSG00000012983 | 1918.514012 | -0.270482735 | 0.2534281 | -1.067296 | 0.28583828 | 0.474737728 | MAP4K5 | protein_coding | 37.27 | -0.087513675 | 0.087513675 | ENSG00000012983 | n.s. |
| ENSG00000070047 | 903.824705 | -0.248770912 | 0.2202195 | -1.12965 | 0.258623743 | 0.444984969 | PHRF1 | protein_coding | 56.31 | -0.08748145 | 0.08748145 | ENSG00000070047 | n.s. |
| ENSG00000172409 | 371.2643685 | 0.284391055 | 0.2758866 | 1.030826 | 0.302622515 | 0.492508858 | CLP1 | protein_coding | 47.28 | 0.087474694 | 0.087474694 | ENSG00000172409 | n.s. |
| ENSG00000141627 | 832.0383904 | -0.281725152 | 0.2717246 | -1.036804 | 0.299827221 | 0.48967457 | DYM | protein_coding | 38.58 | -0.087360843 | 0.087360843 | ENSG00000141627 | n.s. |
| ENSG00000119471 | 400.7249473 | -0.296191215 | 0.2957818 | -1.001384 | 0.316641158 | 0.507236189 | HSDL2 | protein_coding | 40.81 | -0.08731414 | 0.08731414 | ENSG00000119471 | n.s. |
| ENSG00000136888 | 2051.794207 | 0.307413256 | 0.3146965 | 0.976856 | 0.328640361 | 0.520369344 | ATP6V1G1 | protein_coding | 44.34 | 0.087209543 | 0.087209543 | ENSG00000136888 | n.s. |
| ENSG00000235145 | 65.0968919 | -0.286090918 | 0.2793842 | -1.024005 | 0.305832783 | 0.495878635 | RPSAP16 | processed_pseudogene | 49.82 | -0.087150332 | 0.087150332 | ENSG00000235145 | n.s. |
| ENSG00000109046 | 1260.45525 | -0.281107468 | 0.2712412 | -1.036375 | 0.300027386 | 0.489773866 | WSB1 | protein_coding | 39.73 | -0.08714455 | 0.08714455 | ENSG00000109046 | n.s. |
| ENSG00000214046 | 1214.290885 | -0.290218324 | 0.2864512 | -1.013151 | 0.310987992 | 0.501023853 | SMIM7 | protein_coding | 47.85 | -0.087106591 | 0.087106591 | ENSG00000214046 | n.s. |
| ENSG00000267100 | 424.2786841 | 0.242339703 | 0.211113 | 1.147915 | 0.25100374 | 0.437111142 | ILF3-AS1 | lincRNA | 51.64 | 0.087098858 | 0.087098858 | ENSG00000267100 | n.s. |
| ENSG00000252933 | 5.154912758 | -0.40946558 | 0.5110682 | -0.801196 | 0.423018472 | 0.612998145 | RNU6-1223P | snRNA | 42.31 | -0.087028158 | 0.087028158 | ENSG00000252933 | n.s. |
| ENSG00000141556 | 2415.362663 | -0.267782536 | 0.2502266 | -1.07016 | 0.284547269 | 0.473157964 | TBCD | protein_coding | 52.84 | -0.087027676 | 0.087027676 | ENSG00000141556 | n.s. |
| ENSG00000013573 | 2244.70669 | -0.25272637 | 0.2267189 | -1.114713 | 0.264973643 | 0.452554959 | DDX11 | protein_coding | 52.28 | -0.087020935 | 0.087020935 | ENSG00000013573 | n.s. |
| ENSG00000248932 | 113.6830132 | -0.257172354 | 0.2336204 | -1.100813 | 0.270978109 | 0.459064834 | | antisense | 42.28 | -0.086956653 | 0.086956653 | ENSG00000248932 | n.s. |
| ENSG00000070087 | 5755.995814 | 0.269171365 | 0.2524294 | 1.066323 | 0.286277449 | 0.475330291 | PFN2 | protein_coding | 39.59 | 0.086943565 | 0.086943565 | ENSG00000070087 | n.s. |
| ENSG00000178952 | 1814.492649 | 0.291755037 | 0.2893919 | 1.008166 | 0.313374869 | 0.50353356 | TUFM | protein_coding | 55.48 | 0.086934709 | 0.086934709 | ENSG00000178952 | n.s. |
| ENSG00000235245 | 34.5356454 | 0.313547263 | 0.326074 | 0.961583 | 0.336259078 | 0.52837731 | | antisense | 58.53 | 0.086870101 | 0.086870101 | ENSG00000235245 | n.s. |
| ENSG00000169727 | 1864.055597 | 0.288133383 | 0.2835667 | 1.016104 | 0.309579684 | 0.499568606 | GPS1 | protein_coding | 66.61 | 0.086844802 | 0.086844802 | ENSG00000169727 | n.s. |
| ENSG00000243789 | 37.04413275 | 0.299545775 | 0.3027488 | 0.98942 | 0.322457592 | 0.51309452 | JMJD7 | protein_coding | 54.81 | 0.086809152 | 0.086809152 | ENSG00000243789 | n.s. |
| ENSG00000283172 | 34.09459521 | -0.305309026 | 0.3120137 | -0.978512 | 0.327821374 | 0.519626772 | | miRNA | 32.99 | -0.086801946 | 0.086801946 | ENSG00000283172 | n.s. |
| ENSG00000164609 | 1248.778652 | -0.277254058 | 0.2658798 | -1.04278 | 0.297050277 | 0.486413548 | SLU7 | protein_coding | 36.94 | -0.08677895 | 0.08677895 | ENSG00000164609 | n.s. |
| ENSG00000175548 | 1368.33498 | -0.260204981 | 0.2386856 | -1.090158 | 0.275643574 | 0.464005015 | ALG10B | protein_coding | 34.07 | -0.086772461 | 0.086772461 | ENSG00000175548 | n.s. |
| ENSG00000186106 | 660.8443612 | -0.304676499 | 0.3109623 | -0.979786 | 0.327191716 | 0.519085035 | ANKRD46 | protein_coding | 39.74 | -0.086760134 | 0.086760134 | ENSG00000186106 | n.s. |
| ENSG00000204442 | 475.6151603 | 0.261458658 | 0.2409017 | 1.085333 | 0.277774121 | 0.465901588 | FAM155A | protein_coding | 37.4 | 0.086727356 | 0.086727356 | ENSG00000204442 | n.s. |

| | | | | | | | | | | | | | |
|---|---|---|---|---|---|---|---|---|---|---|---|---|---|
| ENSG00000216895 | 112.7322364 | -0.2724857 | 0.2581368 | -1.055587 | 0.291157156 | 0.480593325 | | antisense | 42.24 | -0.086711017 | 0.086711017 | ENSG00000216895 | n.s. |
| ENSG00000149089 | 478.2664749 | -0.308326799 | 0.3174926 | -0.971131 | 0.331483276 | 0.523329363 | APIP | protein_coding | 40.49 | -0.086709172 | 0.086709172 | ENSG00000149089 | n.s. |
| ENSG00000105755 | 104.9415732 | 0.329719703 | 0.3548212 | 0.929256 | 0.352756445 | 0.545956869 | ETHE1 | protein_coding | 48.58 | 0.086664076 | 0.086664076 | ENSG00000105755 | n.s. |
| ENSG00000160679 | 3553.971331 | -0.257209959 | 0.2343441 | -1.097574 | 0.272390563 | 0.460365842 | CHTOP | protein_coding | 44.38 | -0.08665324 | 0.08665324 | ENSG00000160679 | n.s. |
| ENSG00000279570 | 41.11992636 | -0.28203449 | 0.2738975 | -1.029708 | 0.30314704 | 0.493189558 | | TEC | 41.6 | -0.086580676 | 0.086580676 | ENSG00000279570 | n.s. |
| ENSG00000214455 | 44.57329987 | -0.310178824 | 0.3208226 | -0.966823 | 0.333632315 | 0.526006423 | RCN1P2 | processed_pseudogene | 48.63 | -0.086542669 | 0.086542669 | ENSG00000214455 | n.s. |
| ENSG00000068137 | 302.1364066 | -0.279242424 | 0.2695814 | -1.035837 | 0.300278081 | 0.489981151 | PLEKHH3 | protein_coding | 62 | -0.086515063 | 0.086515063 | ENSG00000068137 | n.s. |
| ENSG00000137760 | 500.7899241 | -0.275246141 | 0.2633841 | -1.045037 | 0.296005856 | 0.485285831 | ALKBH8 | protein_coding | 36.18 | -0.086427946 | 0.086427946 | ENSG00000137760 | n.s. |
| ENSG00000253982 | 122.1518837 | 0.278809016 | 0.269057 | 1.036245 | 0.300087694 | 0.489814885 | | antisense | 64.54 | 0.086421879 | 0.086421879 | ENSG00000253982 | n.s. |
| ENSG00000109016 | 229.8558951 | -0.286673459 | 0.282286 | -1.015543 | 0.309847204 | 0.499826511 | DHRS7B | protein_coding | 44.22 | -0.086340516 | 0.086340516 | ENSG00000109016 | n.s. |
| ENSG00000179085 | 128.9378617 | 0.306638463 | 0.3154819 | 0.971968 | 0.331066226 | 0.52296345 | DPM3 | protein_coding | 60.71 | 0.086327516 | 0.086327516 | ENSG00000179085 | n.s. |
| ENSG00000171132 | 378.9934309 | -0.255212054 | 0.2318475 | -1.100776 | 0.270994317 | 0.459064834 | PRKCE | protein_coding | 44.43 | -0.086293825 | 0.086293825 | ENSG00000171132 | n.s. |
| ENSG00000117399 | 438.0703327 | 0.271326528 | 0.2572904 | 1.054554 | 0.291629438 | 0.481176352 | CDC20 | protein_coding | 52.2 | 0.086199278 | 0.086199278 | ENSG00000117399 | n.s. |
| ENSG00000119431 | 87.649292 | -0.287278184 | 0.283772 | -1.012356 | 0.31136805 | 0.501367422 | HDHD3 | protein_coding | 54.87 | -0.086138608 | 0.086138608 | ENSG00000119431 | n.s. |
| ENSG00000181722 | 28.89948254 | -0.305828602 | 0.3146715 | -0.971898 | 0.331101258 | 0.52296345 | ZBTB20 | protein_coding | 36.07 | -0.086099517 | 0.086099517 | ENSG00000181722 | n.s. |
| ENSG00000159259 | 813.3337231 | -0.272784387 | 0.2600845 | -1.04883 | 0.29425644 | 0.483674844 | CHAF1B | protein_coding | 46.73 | -0.08604888 | 0.08604888 | ENSG00000159259 | n.s. |
| ENSG00000112305 | 761.6361762 | -0.278084015 | 0.2687477 | -1.03474 | 0.300790283 | 0.490501652 | SMAP1 | protein_coding | 37.73 | -0.086027939 | 0.086027939 | ENSG00000112305 | n.s. |
| ENSG00000173757 | 835.5825089 | -0.285656017 | 0.2815853 | -1.014457 | 0.310364942 | 0.500371826 | STAT5B | protein_coding | 43.63 | -0.085898807 | 0.085898807 | ENSG00000173757 | n.s. |
| ENSG00000150459 | 2114.995596 | 0.281775696 | 0.2753866 | 1.0232 | 0.306213142 | 0.495938661 | SAP18 | protein_coding | 42.74 | 0.085820997 | 0.085820997 | ENSG00000150459 | n.s. |
| ENSG00000048392 | 458.7042173 | -0.294632177 | 0.2966947 | -0.993048 | 0.320686363 | 0.511457021 | RRM2B | protein_coding | 37.99 | -0.085794196 | 0.085794196 | ENSG00000048392 | n.s. |
| ENSG00000250565 | 150.9116311 | -0.271105811 | 0.2579946 | -1.05082 | 0.293341451 | 0.48265103 | ATP6V1E2 | protein_coding | 44.72 | -0.085768868 | 0.085768868 | ENSG00000250565 | n.s. |
| ENSG00000106333 | 1425.242274 | 0.270073544 | 0.2564876 | 1.052969 | 0.292355066 | 0.481793112 | PCOLCE | protein_coding | 60.91 | 0.085650965 | 0.085650965 | ENSG00000106333 | n.s. |
| ENSG00000143569 | 3082.184212 | -0.264413494 | 0.2476118 | -1.067855 | 0.285585991 | 0.474545142 | UBAP2L | protein_coding | 43.58 | -0.085596589 | 0.085596589 | ENSG00000143569 | n.s. |
| ENSG00000214176 | 463.0540774 | -0.271809157 | 0.2596052 | -1.04701 | 0.29509515 | 0.484391626 | PLEKHM1P1 | transcribed_unprocessed_pseud | 51.2 | -0.08556644 | 0.08556644 | ENSG00000214176 | n.s. |
| ENSG00000162521 | 2757.911121 | -0.273078083 | 0.261891 | -1.042717 | 0.297079514 | 0.486413548 | RBBP4 | protein_coding | 42.94 | -0.085471894 | 0.085471894 | ENSG00000162521 | n.s. |
| ENSG00000177225 | 615.7744877 | -0.273000531 | 0.2618253 | -1.042682 | 0.297095512 | 0.486413548 | GATD1 | protein_coding | 60.25 | -0.08544762 | 0.08544762 | ENSG00000177225 | n.s. |
| ENSG00000175928 | 993.5165952 | -0.258189452 | 0.2383623 | -1.083181 | 0.278728268 | 0.466714813 | LRRN1 | protein_coding | 40.88 | -0.08544739 | 0.08544739 | ENSG00000175928 | n.s. |
| ENSG00000202273 | 10.69732066 | 0.360927272 | 0.4178046 | 0.863866 | 0.387661436 | 0.580117173 | | misc_RNA | 40.78 | 0.085353625 | 0.085353625 | ENSG00000202273 | n.s. |
| ENSG00000065457 | 576.5648324 | -0.244910434 | 0.218243 | -1.122192 | 0.261780948 | 0.448753527 | ADAT1 | protein_coding | 44.65 | -0.085226902 | 0.085226902 | ENSG00000065457 | n.s. |
| ENSG00000168653 | 1568.308118 | 0.301070216 | 0.3088536 | 0.974799 | 0.329659908 | 0.521455908 | NDUFS5 | protein_coding | 46.54 | 0.085137361 | 0.085137361 | ENSG00000168653 | n.s. |
| ENSG00000115963 | 2229.438612 | -0.258506056 | 0.2396736 | -1.078576 | 0.280776958 | 0.468511216 | RND3 | protein_coding | 38.66 | -0.085120876 | 0.085120876 | ENSG00000115963 | n.s. |
| ENSG00000087191 | 2366.88217 | 0.259794731 | 0.2417749 | 1.074532 | 0.282584514 | 0.470737519 | PSMC5 | protein_coding | 50.95 | 0.08501034 | 0.08501034 | ENSG00000087191 | n.s. |

| | | | | | | | | | | | | | |
|---|---|---|---|---|---|---|---|---|---|---|---|---|---|
| ENSG00000142945 | 767.5870346 | -0.287296726 | 0.286141 | -1.004039 | 0.315359744 | 0.506022095 | KIF2C | protein_coding | 49.08 | -0.08499114 | 0.08499114 | ENSG00000142945 | n.s. |
| ENSG00000136816 | 351.0186465 | -0.279025892 | 0.272711 | -1.023156 | 0.306234098 | 0.495938661 | TOR1B | protein_coding | 50.46 | -0.084983484 | 0.084983484 | ENSG00000136816 | n.s. |
| ENSG00000264480 | 8.762448923 | 0.356694248 | 0.4104259 | 0.869083 | 0.384801581 | 0.57776258 | MIR4714 | miRNA | 46.75 | 0.084982615 | 0.084982615 | ENSG00000264480 | n.s. |
| ENSG00000129968 | 556.672782 | 0.274466292 | 0.2653015 | 1.034545 | 0.300881463 | 0.490571892 | ABHD17A | protein_coding | 65.16 | 0.084891694 | 0.084891694 | ENSG00000129968 | n.s. |
| ENSG00000176915 | 3435.120206 | -0.269539951 | 0.2574032 | -1.047151 | 0.295030004 | 0.484341793 | ANKLE2 | protein_coding | 51.03 | -0.084864129 | 0.084864129 | ENSG00000176915 | n.s. |
| ENSG00000198873 | 325.8163464 | -0.269024849 | 0.2567362 | -1.047865 | 0.294700861 | 0.483972646 | GRK5 | protein_coding | 50.47 | -0.084791032 | 0.084791032 | ENSG00000198873 | n.s. |
| ENSG00000182481 | 11945.37971 | 0.262557545 | 0.2463433 | 1.06582 | 0.286505144 | 0.475504962 | KPNA2 | protein_coding | 43.84 | 0.084765377 | 0.084765377 | ENSG00000182481 | n.s. |
| ENSG00000162616 | 605.7711069 | 0.260334003 | 0.243088 | 1.070945 | 0.284193939 | 0.472732913 | DNAJB4 | protein_coding | 37.07 | 0.084708561 | 0.084708561 | ENSG00000162616 | n.s. |
| ENSG00000172733 | 197.1224833 | -0.269734312 | 0.2581009 | -1.045073 | 0.295989114 | 0.485285831 | PURG | protein_coding | 36.48 | -0.084697218 | 0.084697218 | ENSG00000172733 | n.s. |
| ENSG00000221333 | 3.929073038 | 0.420650346 | 0.5455919 | 0.770998 | 0.440708046 | 0.629009177 | MIR548K | miRNA | 34.48 | 0.08469501 | 0.08469501 | ENSG00000221333 | n.s. |
| ENSG00000138660 | 781.6230549 | -0.29222854 | 0.2953343 | -0.989484 | 0.322426448 | 0.51309452 | AP1AR | protein_coding | 36.49 | -0.084688598 | 0.084688598 | ENSG00000138660 | n.s. |
| ENSG00000161179 | 307.1716629 | -0.271037038 | 0.2602435 | -1.041475 | 0.297655309 | 0.48717954 | YDJC | protein_coding | 67.56 | -0.084647837 | 0.084647837 | ENSG00000161179 | n.s. |
| ENSG00000113240 | 487.8169708 | -0.254677899 | 0.2341781 | -1.087539 | 0.2767986 | 0.465330464 | CLK4 | protein_coding | 39.49 | -0.084613807 | 0.084613807 | ENSG00000113240 | n.s. |
| ENSG00000266594 | 9.526758418 | -0.368933187 | 0.4364734 | -0.845259 | 0.397966128 | 0.589769281 | MIR4766 | miRNA | 36.84 | -0.084602966 | 0.084602966 | ENSG00000266594 | n.s. |
| ENSG00000105401 | 1708.968772 | 0.286313663 | 0.285518 | 1.002787 | 0.315963776 | 0.506582641 | CDC37 | protein_coding | 54.04 | 0.084562653 | 0.084562653 | ENSG00000105401 | n.s. |
| ENSG00000144224 | 3288.962217 | -0.281174657 | 0.2771653 | -1.014466 | 0.310360596 | 0.500371826 | UBXN4 | protein_coding | 39.83 | -0.08455123 | 0.08455123 | ENSG00000144224 | n.s. |
| ENSG00000252035 | 5.480372454 | -0.428712871 | 0.5650209 | -0.758756 | 0.447998703 | 0.635117192 | RNU6-397P | snRNA | 44.23 | -0.084519084 | 0.084519084 | ENSG00000252035 | n.s. |
| ENSG00000097007 | 1236.243126 | -0.254727188 | 0.2349716 | -1.084077 | 0.278330859 | 0.466311668 | ABL1 | protein_coding | 44.47 | -0.084397159 | 0.084397159 | ENSG00000097007 | n.s. |
| ENSG00000118197 | 544.5477953 | -0.275314773 | 0.2677575 | -1.028225 | 0.303844235 | 0.493828498 | DDX59 | protein_coding | 42 | -0.084363013 | 0.084363013 | ENSG00000118197 | n.s. |
| ENSG00000260167 | 44.84368419 | 0.290293591 | 0.292809 | 0.991409 | 0.321485774 | 0.512192063 | | antisense | 46.69 | 0.084349783 | 0.084349783 | ENSG00000260167 | n.s. |
| ENSG00000204519 | 1736.366915 | -0.252475617 | 0.2313781 | -1.091182 | 0.275192884 | 0.463470488 | ZNF551 | protein_coding | 45.44 | -0.08432128 | 0.08432128 | ENSG00000204519 | n.s. |
| ENSG00000244398 | 474.9556405 | 0.298390566 | 0.3063969 | 0.973869 | 0.330121429 | 0.52188948 | | processed_pseudogene | 46.73 | 0.084271899 | 0.084271899 | ENSG00000244398 | n.s. |
| ENSG00000275155 | 49.4092209 | 0.277760445 | 0.2722509 | 1.020237 | 0.307616037 | 0.497322118 | | sense_intronic | 49.11 | 0.084262027 | 0.084262027 | ENSG00000275155 | n.s. |
| ENSG00000188021 | 805.437165 | 0.250285065 | 0.2283153 | 1.096225 | 0.272980131 | 0.46082934 | UBQLN2 | protein_coding | 44.3 | 0.084210883 | 0.084210883 | ENSG00000188021 | n.s. |
| ENSG00000101452 | 668.7079363 | -0.281935889 | 0.279266 | -1.00956 | 0.312706024 | 0.502748957 | DHX35 | protein_coding | 41.33 | -0.084199821 | 0.084199821 | ENSG00000101452 | n.s. |
| ENSG00000179295 | 3470.307851 | -0.275546289 | 0.2685653 | -1.025994 | 0.304894512 | 0.494933694 | PTPN11 | protein_coding | 43.16 | -0.084166435 | 0.084166435 | ENSG00000179295 | n.s. |
| ENSG00000255468 | 49.7227076 | -0.276056834 | 0.2695087 | -1.024297 | 0.305695249 | 0.495771107 | | antisense | 48.63 | -0.084119704 | 0.084119704 | ENSG00000255468 | n.s. |
| ENSG00000252464 | 25.23569763 | -0.308073711 | 0.3233999 | -0.952609 | 0.340788131 | 0.533305818 | RN7SKP70 | misc_RNA | 51.29 | -0.084111418 | 0.084111418 | ENSG00000252464 | n.s. |
| ENSG00000244501 | 48.0384639 | -0.284306314 | 0.2831588 | -1.004053 | 0.315353259 | 0.506022095 | RN7SL623P | misc_RNA | 52.58 | -0.084106485 | 0.084106485 | ENSG00000244501 | n.s. |
| ENSG00000165688 | 899.400198 | 0.28122648 | 0.2784752 | 1.00988 | 0.312552918 | 0.502618881 | PMPCA | protein_coding | 53.93 | 0.084019561 | 0.084019561 | ENSG00000165688 | n.s. |
| ENSG00000177885 | 1956.069203 | 0.276899209 | 0.2714042 | 1.020247 | 0.307611569 | 0.497322118 | GRB2 | protein_coding | 44.22 | 0.08400076 | 0.08400076 | ENSG00000177885 | n.s. |
| ENSG00000166896 | 140.9868369 | -0.301616375 | 0.312647 | -0.964719 | 0.334685811 | 0.527101612 | ATP23 | protein_coding | 40.87 | -0.08388122 | 0.08388122 | ENSG00000166896 | n.s. |

| | | | | | | | | | | | | | |
|---|---|---|---|---|---|---|---|---|---|---|---|---|---|
| ENSG00000230844 | 192.199103 | 0.258639733 | 0.2419764 | 1.068863 | 0.28513124 | 0.473902622 | ZNF674-AS1 | lincRNA | 47.02 | 0.083879682 | 0.083879682 | ENSG00000230844 | n.s. |
| ENSG00000131051 | 13672.53362 | 0.247525716 | 0.224537 | 1.102383 | 0.270295205 | 0.458550035 | RBM39 | protein_coding | 40.28 | 0.083815492 | 0.083815492 | ENSG00000131051 | n.s. |
| ENSG00000113282 | 1967.28021 | -0.269408695 | 0.2593869 | -1.038636 | 0.298973928 | 0.488655196 | CLINT1 | protein_coding | 37.77 | -0.083785425 | 0.083785425 | ENSG00000113282 | n.s. |
| ENSG00000164548 | 1767.35289 | 0.251102249 | 0.2303549 | 1.090067 | 0.275683607 | 0.464016303 | TRA2A | protein_coding | 41.09 | 0.083734254 | 0.083734254 | ENSG00000164548 | n.s. |
| ENSG00000163050 | 1376.251336 | 0.275866767 | 0.2705196 | 1.019766 | 0.307839397 | 0.497484598 | COQ8A | protein_coding | 49.11 | 0.083648421 | 0.083648421 | ENSG00000163050 | n.s. |
| ENSG00000154277 | 7609.04747 | 0.28773843 | 0.2900945 | 0.991878 | 0.321256893 | 0.512120417 | UCHL1 | protein_coding | 44.57 | 0.083624818 | 0.083624818 | ENSG00000154277 | n.s. |
| ENSG00000062598 | 425.7548382 | -0.277572319 | 0.2734533 | -1.015063 | 0.310075787 | 0.500079368 | ELMO2 | protein_coding | 44.15 | -0.08353846 | 0.08353846 | ENSG00000062598 | n.s. |
| ENSG00000206601 | 21.39215 | 0.313395984 | 0.3342242 | 0.937682 | 0.348407977 | 0.541332916 | RNU6-431P | snRNA | 43.93 | 0.083531176 | 0.083531176 | ENSG00000206601 | n.s. |
| ENSG00000165632 | 401.7568831 | -0.26003664 | 0.2447996 | -1.062243 | 0.288125506 | 0.477341796 | TAF3 | protein_coding | 39.51 | -0.083516107 | 0.083516107 | ENSG00000165632 | n.s. |
| ENSG00000166887 | 830.9830247 | -0.265973132 | 0.2546065 | -1.044644 | 0.296187599 | 0.485440841 | VPS39 | protein_coding | 42.61 | -0.083479308 | 0.083479308 | ENSG00000166887 | n.s. |
| ENSG00000115226 | 105.5648312 | 0.296231883 | 0.3046999 | 0.972209 | 0.330946793 | 0.522838102 | FNDC4 | protein_coding | 57.84 | 0.083428606 | 0.083428606 | ENSG00000115226 | n.s. |
| ENSG00000087266 | 634.3307089 | -0.251201103 | 0.2314404 | -1.085381 | 0.277752745 | 0.465901588 | SH3BP2 | protein_coding | 59.23 | -0.083324865 | 0.083324865 | ENSG00000087266 | n.s. |
| ENSG00000208036 | 12.36349111 | 0.352046466 | 0.4077471 | 0.863394 | 0.387920755 | 0.580120522 | MIR106B | miRNA | 60.98 | 0.083252572 | 0.083252572 | ENSG00000208036 | n.s. |
| ENSG00000137414 | 750.0800094 | -0.285834294 | 0.2877819 | -0.993232 | 0.320596714 | 0.511419346 | FAM8A1 | protein_coding | 40.23 | -0.083241477 | 0.083241477 | ENSG00000137414 | n.s. |
| ENSG00000204438 | 338.3962746 | -0.257816571 | 0.2419171 | -1.065723 | 0.286548948 | 0.475520978 | GPANK1 | protein_coding | 53.99 | -0.083231006 | 0.083231006 | ENSG00000204438 | n.s. |
| ENSG00000167081 | 1179.021659 | -0.261543004 | 0.2478523 | -1.055238 | 0.291316697 | 0.480796867 | PBX3 | protein_coding | 37.29 | -0.083180711 | 0.083180711 | ENSG00000167081 | n.s. |
| ENSG00000100714 | 2403.28214 | -0.272911613 | 0.2666998 | -1.023292 | 0.306170021 | 0.495938661 | MTHFD1 | protein_coding | 43.75 | -0.083121245 | 0.083121245 | ENSG00000100714 | n.s. |
| ENSG00000109536 | 1146.668685 | 0.273928424 | 0.2684718 | 1.020325 | 0.307574479 | 0.497322118 | FRG1 | protein_coding | 35.99 | 0.083099536 | 0.083099536 | ENSG00000109536 | n.s. |
| ENSG00000224631 | 409.3882172 | 0.319178084 | 0.3458758 | 0.922811 | 0.356105548 | 0.549125877 | RPS27AP16 | transcribed_processed_pseudog | 42.05 | 0.083091021 | 0.083091021 | ENSG00000224631 | n.s. |
| ENSG00000196588 | 99.92362215 | 0.263303473 | 0.2511307 | 1.048472 | 0.294421255 | 0.483674844 | MKL1 | protein_coding | 43.58 | 0.083058159 | 0.083058159 | ENSG00000196588 | n.s. |
| ENSG00000101150 | 1081.7442 | 0.299393997 | 0.311039 | 0.962561 | 0.335767897 | 0.528057883 | TPD52L2 | protein_coding | 54.36 | 0.083027485 | 0.083027485 | ENSG00000101150 | n.s. |
| ENSG00000198171 | 536.7424082 | -0.270239389 | 0.2623827 | -1.029943 | 0.30303654 | 0.4930674 | DDRGK1 | protein_coding | 54.35 | -0.082988817 | 0.082988817 | ENSG00000198171 | n.s. |
| ENSG00000160226 | 197.9936483 | -0.27650599 | 0.2731372 | -1.012334 | 0.311378589 | 0.501367422 | C21orf2 | protein_coding | 60.43 | -0.082908631 | 0.082908631 | ENSG00000160226 | n.s. |
| ENSG00000143772 | 1091.113021 | -0.262549328 | 0.2506082 | -1.047648 | 0.29480061 | 0.484079359 | ITPKB | protein_coding | 49.82 | -0.082724944 | 0.082724944 | ENSG00000143772 | n.s. |
| ENSG00000135097 | 502.4004953 | -0.245756262 | 0.2241164 | -1.096556 | 0.272835323 | 0.460670167 | MSI1 | protein_coding | 54.01 | -0.082723994 | 0.082723994 | ENSG00000135097 | n.s. |
| ENSG00000204177 | 272.2280137 | -0.252282042 | 0.2347647 | -1.074616 | 0.282546508 | 0.470730529 | BMS1P1 | transcribed_unprocessed_pseud | 41.22 | -0.082553656 | 0.082553656 | ENSG00000204177 | n.s. |
| ENSG00000028203 | 3164.770572 | -0.275267676 | 0.271958 | -1.01217 | 0.311456828 | 0.501399307 | VEZT | protein_coding | 38.6 | -0.082529728 | 0.082529728 | ENSG00000028203 | n.s. |
| ENSG00000033050 | 1363.755487 | -0.278728567 | 0.2777877 | -1.003387 | 0.315674095 | 0.506260984 | ABCF2 | protein_coding | 49.19 | -0.082399284 | 0.082399284 | ENSG00000033050 | n.s. |
| ENSG00000278467 | 107.2408184 | 0.259958876 | 0.2472702 | 1.051315 | 0.293113953 | 0.482434767 | | lincRNA | 44.92 | 0.082292946 | 0.082292946 | ENSG00000278467 | n.s. |
| ENSG00000133134 | 729.0681094 | 0.300724185 | 0.3153236 | 0.9537 | 0.340235386 | 0.532916782 | BEX2 | protein_coding | 57.03 | 0.08220013 | 0.08220013 | ENSG00000133134 | n.s. |
| ENSG00000198056 | 1092.761795 | -0.278007638 | 0.2773559 | -1.00235 | 0.316174745 | 0.506702304 | PRIM1 | protein_coding | 42.82 | -0.082080955 | 0.082080955 | ENSG00000198056 | n.s. |
| ENSG00000166200 | 4424.903537 | -0.283788143 | 0.2872397 | -0.987984 | 0.323160552 | 0.513858152 | COPS2 | protein_coding | 35.44 | -0.082059257 | 0.082059257 | ENSG00000166200 | n.s. |

| | | | | | | | | | | | | | |
|---|---|---|---|---|---|---|---|---|---|---|---|---|---|
| ENSG00000177822 | 243.1003743 | -0.237586537 | 0.2128001 | -1.116478 | 0.264217778 | 0.451651826 | | antisense | 39.36 | -0.082013977 | 0.082013977 | ENSG00000177822 | n.s. |
| ENSG00000197111 | 5415.544732 | 0.24953695 | 0.2316244 | 1.077334 | 0.281330887 | 0.469210582 | PCBP2 | protein_coding | 43.79 | 0.082005878 | 0.082005878 | ENSG00000197111 | n.s. |
| ENSG00000069482 | 1037.859043 | 0.295297058 | 0.3064556 | 0.963588 | 0.335252376 | 0.527604706 | GAL | protein_coding | 57.14 | 0.082001436 | 0.082001436 | ENSG00000069482 | n.s. |
| ENSG00000142494 | 181.3430465 | -0.26573009 | 0.2573443 | -1.032586 | 0.301797674 | 0.491396212 | SLC47A1 | protein_coding | 47.07 | -0.081995855 | 0.081995855 | ENSG00000142494 | n.s. |
| ENSG00000263606 | 72.00377217 | -0.304287473 | 0.3225434 | -0.9434 | 0.345476268 | 0.538400054 | | transcribed_processed_pseudog | 40.49 | -0.081821351 | 0.081821351 | ENSG00000263606 | n.s. |
| ENSG00000119969 | 3639.256149 | -0.262558605 | 0.2524637 | -1.039985 | 0.298346676 | 0.48800291 | HELLS | protein_coding | 38.96 | -0.081807377 | 0.081807377 | ENSG00000119969 | n.s. |
| ENSG00000143797 | 1300.111041 | -0.272140538 | 0.2683281 | -1.014208 | 0.310483541 | 0.500497849 | MBOAT2 | protein_coding | 40.17 | -0.081804843 | 0.081804843 | ENSG00000143797 | n.s. |
| ENSG00000083168 | 2861.583081 | -0.241884996 | 0.2197053 | -1.100952 | 0.270917509 | 0.459062653 | KAT6A | protein_coding | 38.89 | -0.081788099 | 0.081788099 | ENSG00000083168 | n.s. |
| ENSG00000102007 | 262.5503836 | 0.269621456 | 0.2643242 | 1.020041 | 0.307709073 | 0.49736255 | PLP2 | protein_coding | 53.95 | 0.081783446 | 0.081783446 | ENSG00000102007 | n.s. |
| ENSG00000154845 | 1523.003723 | -0.268624841 | 0.2627904 | -1.022202 | 0.306685442 | 0.496336325 | PPP4R1 | protein_coding | 39.14 | -0.081722107 | 0.081722107 | ENSG00000154845 | n.s. |
| ENSG00000239559 | 47.92424004 | 0.39842303 | 0.5087635 | 0.78312 | 0.433556427 | 0.623767104 | | processed_pseudogene | 48.78 | 0.08166777 | 0.08166777 | ENSG00000239559 | n.s. |
| ENSG00000215769 | 179.8743795 | -0.28718432 | 0.2935533 | -0.978304 | 0.327924073 | 0.519654777 | ARHGAP27P1-BPTF | processed_transcript | 47.76 | -0.081642216 | 0.081642216 | ENSG00000215769 | n.s. |
| ENSG00000182952 | 933.213761 | 0.29881542 | 0.3135542 | 0.952994 | 0.340592903 | 0.533166099 | HMGN4 | protein_coding | 42.28 | 0.081617689 | 0.081617689 | ENSG00000182952 | n.s. |
| ENSG00000151892 | 1218.395224 | -0.256893456 | 0.2438077 | -1.053672 | 0.292032884 | 0.481408223 | GFRA1 | protein_coding | 44 | -0.081560203 | 0.081560203 | ENSG00000151892 | n.s. |
| ENSG00000054983 | 124.870758 | -0.297945338 | 0.3122102 | -0.95431 | 0.339926702 | 0.532510245 | GALC | protein_coding | 39.12 | -0.081539306 | 0.081539306 | ENSG00000054983 | n.s. |
| ENSG00000125482 | 953.7536357 | -0.265467407 | 0.2579927 | -1.028973 | 0.303492541 | 0.493520981 | TTF1 | protein_coding | 44.83 | -0.081417362 | 0.081417362 | ENSG00000125482 | n.s. |
| ENSG00000114439 | 1644.63189 | -0.278758879 | 0.2803207 | -0.994428 | 0.320014342 | 0.51078295 | BBX | protein_coding | 36.95 | -0.081331696 | 0.081331696 | ENSG00000114439 | n.s. |
| ENSG00000102054 | 2407.743011 | 0.275697905 | 0.2755406 | 1.000571 | 0.317034318 | 0.507444026 | RBBP7 | protein_coding | 42.25 | 0.081223871 | 0.081223871 | ENSG00000102054 | n.s. |
| ENSG00000110717 | 641.1990103 | 0.291813923 | 0.3025308 | 0.964576 | 0.334757311 | 0.527101612 | NDUFS8 | protein_coding | 57.63 | 0.081155102 | 0.081155102 | ENSG00000110717 | n.s. |
| ENSG00000164953 | 173.8697232 | 0.262532242 | 0.2539498 | 1.033796 | 0.301231658 | 0.490839806 | TMEM67 | protein_coding | 37.38 | 0.081138273 | 0.081138273 | ENSG00000164953 | n.s. |
| ENSG00000145912 | 1860.103565 | 0.296804699 | 0.3117538 | 0.952049 | 0.341072384 | 0.533349039 | NHP2 | protein_coding | 52.15 | 0.081024265 | 0.081024265 | ENSG00000145912 | n.s. |
| ENSG00000016082 | 2921.53959 | 0.244134796 | 0.2249305 | 1.085379 | 0.277753975 | 0.465901588 | ISL1 | protein_coding | 44.32 | 0.08098093 | 0.08098093 | ENSG00000016082 | n.s. |
| ENSG00000248487 | 154.9805587 | 0.28683477 | 0.2945989 | 0.973645 | 0.330232722 | 0.522006153 | ABHD14A | protein_coding | 54.78 | 0.080980448 | 0.080980448 | ENSG00000248487 | n.s. |
| ENSG00000135144 | 145.9668038 | -0.265743472 | 0.2595893 | -1.023708 | 0.305973438 | 0.495878772 | DTX1 | protein_coding | 53.96 | -0.080951968 | 0.080951968 | ENSG00000135144 | n.s. |
| ENSG00000117395 | 750.7837181 | 0.280078009 | 0.2836437 | 0.987429 | 0.323432393 | 0.514058128 | EBNA1BP2 | protein_coding | 44.38 | 0.080939119 | 0.080939119 | ENSG00000117395 | n.s. |
| ENSG00000174233 | 369.1780362 | -0.275334436 | 0.2756601 | -0.998818 | 0.317882634 | 0.508254468 | ADCY6 | protein_coding | 54.36 | -0.080925965 | 0.080925965 | ENSG00000174233 | n.s. |
| ENSG00000212232 | 482.5866368 | -0.245107882 | 0.2267968 | -1.080738 | 0.279813664 | 0.46760115 | SNORD17 | snoRNA | 49.79 | -0.0809161 | 0.0809161 | ENSG00000212232 | n.s. |
| ENSG00000073792 | 45.30874879 | 0.283504892 | 0.2891574 | 0.980452 | 0.326863164 | 0.518681996 | IGF2BP2 | protein_coding | 42.54 | 0.080826912 | 0.080826912 | ENSG00000073792 | n.s. |
| ENSG00000126524 | 1086.900463 | 0.278734699 | 0.2814979 | 0.990184 | 0.322084154 | 0.512910634 | SBDS | protein_coding | 42.81 | 0.080821439 | 0.080821439 | ENSG00000126524 | n.s. |
| ENSG00000205981 | 940.2994365 | 0.272919786 | 0.2716583 | 1.004644 | 0.315068507 | 0.505729632 | DNAJC19 | protein_coding | 39.17 | 0.080806527 | 0.080806527 | ENSG00000205981 | n.s. |
| ENSG00000259865 | 131.2923385 | -0.310407652 | 0.3364801 | -0.922514 | 0.356260388 | 0.549182172 | | lincRNA | 27.85 | -0.080794012 | 0.080794012 | ENSG00000259865 | n.s. |
| ENSG00000136273 | 540.3487334 | -0.24951477 | 0.2336783 | -1.06777 | 0.285624119 | 0.474551863 | HUS1 | protein_coding | 44.35 | -0.080772002 | 0.080772002 | ENSG00000136273 | n.s. |

| | | | | | | | | | | | | | |
|---|---|---|---|---|---|---|---|---|---|---|---|---|---|
| ENSG00000206885 | 103.1753943 | 0.254775761 | 0.2420463 | 1.052591 | 0.292528533 | 0.481936263 | SNORA75 | snoRNA | 38.69 | 0.080766564 | 0.080766564 | ENSG00000206885 | n.s. |
| ENSG00000265735 | 21.66491371 | -0.298624568 | 0.3154971 | -0.946521 | 0.343883065 | 0.53670032 | RN7SL5P | misc_RNA | 57.63 | -0.080708708 | 0.080708708 | ENSG00000265735 | n.s. |
| ENSG00000187653 | 61.9002322 | 0.310001997 | 0.3360332 | 0.922534 | 0.356250256 | 0.549182172 | TMSB4XP8 | processed_pseudogene | 44.44 | 0.080688426 | 0.080688426 | ENSG00000187653 | n.s. |
| ENSG00000119682 | 1069.177318 | -0.248502117 | 0.2324272 | -1.069161 | 0.284997071 | 0.473736177 | AREL1 | protein_coding | 42.31 | -0.080629854 | 0.080629854 | ENSG00000119682 | n.s. |
| ENSG00000198168 | 547.2574596 | -0.275948277 | 0.2774273 | -0.994669 | 0.319897381 | 0.510654808 | SVIP | protein_coding | 36.62 | -0.080541734 | 0.080541734 | ENSG00000198168 | n.s. |
| ENSG00000087086 | 4143.989841 | 0.272421811 | 0.2715114 | 1.003353 | 0.31569043 | 0.506260984 | FTL | protein_coding | 55.96 | 0.080534845 | 0.080534845 | ENSG00000087086 | n.s. |
| ENSG00000125458 | 935.05097 | -0.259080122 | 0.2495769 | -1.038078 | 0.299233964 | 0.488879897 | NT5C | protein_coding | 65.69 | -0.080521537 | 0.080521537 | ENSG00000125458 | n.s. |
| ENSG00000164532 | 210.5913251 | 0.252557748 | 0.2390415 | 1.056544 | 0.290719877 | 0.480039474 | TBX20 | protein_coding | 36.93 | 0.080495975 | 0.080495975 | ENSG00000164532 | n.s. |
| ENSG00000152253 | 536.0900625 | -0.255225649 | 0.2434803 | -1.04824 | 0.294528224 | 0.48374619 | SPC25 | protein_coding | 39.39 | -0.080493689 | 0.080493689 | ENSG00000152253 | n.s. |
| ENSG00000129484 | 902.9796703 | -0.263730117 | 0.2573701 | -1.024712 | 0.305499282 | 0.495568741 | PARP2 | protein_coding | 42.36 | -0.080410283 | 0.080410283 | ENSG00000129484 | n.s. |
| ENSG00000196850 | 617.0170193 | 0.261338096 | 0.2535177 | 1.030847 | 0.302612354 | 0.492508858 | PPTC7 | protein_coding | 42.74 | 0.080383928 | 0.080383928 | ENSG00000196850 | n.s. |
| ENSG00000252010 | 190.036639 | -0.263134783 | 0.2565041 | -1.02585 | 0.304962179 | 0.494985826 | SCARNA5 | scaRNA | 47.46 | -0.080363267 | 0.080363267 | ENSG00000252010 | n.s. |
| ENSG00000137656 | 499.9879061 | 0.266167246 | 0.2617234 | 1.016979 | 0.309163303 | 0.499070223 | BUD13 | protein_coding | 41.56 | 0.08033948 | 0.08033948 | ENSG00000137656 | n.s. |
| ENSG00000232593 | 365.4425627 | -0.284103795 | 0.2915729 | -0.974383 | 0.329866249 | 0.521663764 | KANTR | protein_coding | 42.45 | -0.080290383 | 0.080290383 | ENSG00000232593 | n.s. |
| ENSG00000131558 | 1795.858837 | -0.268947587 | 0.2664997 | -1.009185 | 0.312885764 | 0.502884552 | EXOC4 | protein_coding | 38.3 | -0.080289382 | 0.080289382 | ENSG00000131558 | n.s. |
| ENSG00000169857 | 360.6646442 | -0.268262802 | 0.2654756 | -1.010499 | 0.312256412 | 0.502482874 | AVEN | protein_coding | 39.85 | -0.080178047 | 0.080178047 | ENSG00000169857 | n.s. |
| ENSG00000143162 | 364.939947 | 0.292399372 | 0.3061825 | 0.954984 | 0.339585748 | 0.532225095 | CREG1 | protein_coding | 44.46 | 0.080089548 | 0.080089548 | ENSG00000143162 | n.s. |
| ENSG00000111358 | 872.062399 | -0.289640664 | 0.3017605 | -0.959836 | 0.33713756 | 0.529374789 | GTF2H3 | protein_coding | 42.84 | -0.080009394 | 0.080009394 | ENSG00000111358 | n.s. |
| ENSG00000162105 | 119.8126743 | 0.249257272 | 0.2348292 | 1.061441 | 0.288489649 | 0.477725511 | SHANK2 | protein_coding | 51.53 | 0.079967108 | 0.079967108 | ENSG00000162105 | n.s. |
| ENSG00000111885 | 615.4598432 | -0.260624049 | 0.2534948 | -1.028124 | 0.303891492 | 0.493828498 | MAN1A1 | protein_coding | 37.6 | -0.079861425 | 0.079861425 | ENSG00000111885 | n.s. |
| ENSG00000101608 | 1296.989194 | 0.274902932 | 0.2775255 | 0.99055 | 0.321905295 | 0.512684446 | MYL12A | protein_coding | 38.81 | 0.079763047 | 0.079763047 | ENSG00000101608 | n.s. |
| ENSG00000238297 | 78.28842423 | 0.260275186 | 0.253157 | 1.028118 | 0.303894461 | 0.493828498 | | snoRNA | 42.71 | 0.079754525 | 0.079754525 | ENSG00000238297 | n.s. |
| ENSG00000090372 | 1090.389348 | 0.253705151 | 0.2425906 | 1.045816 | 0.295646016 | 0.484952814 | STRN4 | protein_coding | 57.35 | 0.07973966 | 0.07973966 | ENSG00000090372 | n.s. |
| ENSG00000168894 | 528.2550606 | 0.290739445 | 0.30493 | 0.953463 | 0.340355654 | 0.533045193 | RNF181 | protein_coding | 54.37 | 0.079440474 | 0.079440474 | ENSG00000168894 | n.s. |
| ENSG00000166987 | 156.5942578 | 0.255320222 | 0.2458252 | 1.038625 | 0.298979054 | 0.488655196 | MBD6 | protein_coding | 57.85 | 0.079403946 | 0.079403946 | ENSG00000166987 | n.s. |
| ENSG00000241112 | 162.698242 | 0.255384622 | 0.2459873 | 1.038202 | 0.299175901 | 0.488842398 | RPL29P14 | processed_pseudogene | 55.14 | 0.079381492 | 0.079381492 | ENSG00000241112 | n.s. |
| ENSG00000103591 | 1386.014106 | -0.276215812 | 0.2806386 | -0.98424 | 0.324997415 | 0.516133114 | AAGAB | protein_coding | 39.49 | -0.079339754 | 0.079339754 | ENSG00000103591 | n.s. |
| ENSG00000123268 | 838.7626173 | -0.269398295 | 0.2694092 | -0.99996 | 0.317330099 | 0.507588226 | ATF1 | protein_coding | 41.93 | -0.079334689 | 0.079334689 | ENSG00000123268 | n.s. |
| ENSG00000132313 | 1392.55047 | -0.271804956 | 0.2735084 | -0.993772 | 0.320334008 | 0.511117395 | MRPL35 | protein_coding | 40.87 | -0.079225529 | 0.079225529 | ENSG00000132313 | n.s. |
| ENSG00000146918 | 1740.298193 | -0.260397485 | 0.2547232 | -1.022276 | 0.306650215 | 0.496336325 | NCAPG2 | protein_coding | 41.79 | -0.079219149 | 0.079219149 | ENSG00000146918 | n.s. |
| ENSG00000106012 | 39.63422842 | 0.291392548 | 0.3067522 | 0.949928 | 0.342148772 | 0.534770696 | IQCE | protein_coding | 52.81 | 0.079209935 | 0.079209935 | ENSG00000106012 | n.s. |
| ENSG00000206527 | 628.6567199 | -0.259084462 | 0.252318 | -1.026817 | 0.304506609 | 0.494649892 | HACD2 | protein_coding | 40.56 | -0.079202659 | 0.079202659 | ENSG00000206527 | n.s. |

| | | | | | | | | | | | | | |
|---|---|---|---|---|---|---|---|---|---|---|---|---|---|
| ENSG00000072195 | 600.7532174 | -0.271994394 | 0.2739234 | -0.992958 | 0.320730536 | 0.511457021 | SPEG | protein_coding | 57.68 | -0.07920228 | 0.07920228 | ENSG00000072195 | n.s. |
| ENSG00000136891 | 1461.726886 | 0.24995116 | 0.2377144 | 1.051477 | 0.293039748 | 0.482382645 | TEX10 | protein_coding | 37.95 | 0.079136618 | 0.079136618 | ENSG00000136891 | n.s. |
| ENSG00000138078 | 2172.736579 | -0.273387624 | 0.276706 | -0.988008 | 0.323148921 | 0.513858152 | PREPL | protein_coding | 37.28 | -0.079051877 | 0.079051877 | ENSG00000138078 | n.s. |
| ENSG00000263742 | 5.021130596 | 0.395727793 | 0.5165455 | 0.766104 | 0.443614212 | 0.631363278 | MIR3165 | miRNA | 53.33 | 0.079035025 | 0.079035025 | ENSG00000263742 | n.s. |
| ENSG00000143248 | 46.31624554 | -0.289237533 | 0.3034139 | -0.953277 | 0.340449643 | 0.533072468 | RGS5 | protein_coding | 37.67 | -0.07902367 | 0.07902367 | ENSG00000143248 | n.s. |
| ENSG00000179241 | 589.4190138 | -0.254767793 | 0.2457505 | -1.036693 | 0.299878946 | 0.48967457 | LDLRAD3 | protein_coding | 44.12 | -0.079001569 | 0.079001569 | ENSG00000179241 | n.s. |
| ENSG00000278588 | 639.195219 | 0.232715025 | 0.2107643 | 1.104148 | 0.26952876 | 0.457807604 | HIST1H2BI | protein_coding | 58.79 | 0.078964164 | 0.078964164 | ENSG00000278588 | n.s. |
| ENSG00000147679 | 955.3595285 | -0.24714726 | 0.2336842 | -1.057612 | 0.290232343 | 0.47940502 | UTP23 | protein_coding | 40.04 | -0.078913483 | 0.078913483 | ENSG00000147679 | n.s. |
| ENSG00000130396 | 4940.185757 | -0.259985901 | 0.2548215 | -1.020267 | 0.307602019 | 0.497322118 | AFDN | protein_coding | 38.79 | -0.078869902 | 0.078869902 | ENSG00000130396 | n.s. |
| ENSG00000174428 | 84.77213004 | -0.26979567 | 0.2711142 | -0.995137 | 0.319669773 | 0.510408515 | GTF2IRD2B | protein_coding | 46.64 | -0.078802483 | 0.078802483 | ENSG00000174428 | n.s. |
| ENSG00000072506 | 683.0759728 | 0.297835782 | 0.319392 | 0.932509 | 0.351073689 | 0.544082698 | HSD17B10 | protein_coding | 52.02 | 0.078728447 | 0.078728447 | ENSG00000072506 | n.s. |
| ENSG00000111361 | 1869.502805 | -0.26864091 | 0.2695184 | -0.996744 | 0.318888706 | 0.509415016 | EIF2B1 | protein_coding | 45.42 | -0.078692514 | 0.078692514 | ENSG00000111361 | n.s. |
| ENSG00000214765 | 814.9021101 | -0.262182889 | 0.2587803 | -1.013149 | 0.310989221 | 0.501023853 | SEPT7P2 | transcribed_unprocessed_pseud | 37.6 | -0.078691991 | 0.078691991 | ENSG00000214765 | n.s. |
| ENSG00000171246 | 900.9927723 | -0.254647145 | 0.246397 | -1.033483 | 0.301377782 | 0.490915954 | NPTX1 | protein_coding | 60.76 | -0.078684148 | 0.078684148 | ENSG00000171246 | n.s. |
| ENSG00000101898 | 191.2660465 | 0.235217421 | 0.2153973 | 1.092017 | 0.274825787 | 0.463132345 | MCTS2P | processed_pseudogene | 40.48 | 0.078631981 | 0.078631981 | ENSG00000101898 | n.s. |
| ENSG00000106028 | 3060.997187 | 0.258673133 | 0.2532898 | 1.021254 | 0.307134194 | 0.496889307 | SSBP1 | protein_coding | 40.78 | 0.078569468 | 0.078569468 | ENSG00000106028 | n.s. |
| ENSG00000225663 | 1182.062391 | 0.281551844 | 0.2913659 | 0.966317 | 0.333885596 | 0.526226982 | MCRIP1 | protein_coding | 59.8 | 0.078504224 | 0.078504224 | ENSG00000225663 | n.s. |
| ENSG00000142082 | 131.1207994 | -0.278380145 | 0.2862692 | -0.972442 | 0.330830844 | 0.522773561 | SIRT3 | protein_coding | 48.61 | -0.078415896 | 0.078415896 | ENSG00000142082 | n.s. |
| ENSG00000267606 | 40.70401831 | 0.275213138 | 0.2809568 | 0.979557 | 0.32730488 | 0.519151568 | | lincRNA | 40.14 | 0.078354785 | 0.078354785 | ENSG00000267606 | n.s. |
| ENSG00000185158 | 44.54108718 | 0.292754132 | 0.3113566 | 0.940254 | 0.347087518 | 0.539970485 | LRRC37B | protein_coding | 42.83 | 0.078349782 | 0.078349782 | ENSG00000185158 | n.s. |
| ENSG00000135801 | 1192.781282 | 0.261213926 | 0.2580804 | 1.012141 | 0.311470416 | 0.501399307 | TAF5L | protein_coding | 41.9 | 0.078316185 | 0.078316185 | ENSG00000135801 | n.s. |
| ENSG00000039560 | 5144.41551 | -0.269449617 | 0.2716264 | -0.991986 | 0.321204302 | 0.512095214 | RAI14 | protein_coding | 42.13 | -0.078315338 | 0.078315338 | ENSG00000039560 | n.s. |
| ENSG00000179632 | 549.5081575 | 0.281951851 | 0.2925531 | 0.963763 | 0.335164826 | 0.527526552 | MAF1 | protein_coding | 63.19 | 0.078313727 | 0.078313727 | ENSG00000179632 | n.s. |
| ENSG00000102974 | 1697.246615 | -0.262031029 | 0.2594084 | -1.01011 | 0.31244252 | 0.50255744 | CTCF | protein_coding | 45.32 | -0.078298617 | 0.078298617 | ENSG00000102974 | n.s. |
| ENSG00000234219 | 117.8113094 | 0.26574822 | 0.2657553 | 0.999973 | 0.317323389 | 0.507588226 | CDCA4P4 | unprocessed_pseudogene | 58.53 | 0.078259784 | 0.078259784 | ENSG00000234219 | n.s. |
| ENSG00000283761 | 213.9629988 | -0.261897683 | 0.2595933 | -1.008877 | 0.31303363 | 0.503043309 | | protein_coding | 38.29 | -0.07814886 | 0.07814886 | ENSG00000283761 | n.s. |
| ENSG00000106853 | 143.0450534 | -0.275321642 | 0.2817764 | -0.977093 | 0.328523224 | 0.520308138 | PTGR1 | protein_coding | 43.23 | -0.078119592 | 0.078119592 | ENSG00000106853 | n.s. |
| ENSG00000182325 | 213.7147404 | 0.264852672 | 0.264709 | 1.000543 | 0.317047926 | 0.507444026 | FBXL6 | protein_coding | 66.45 | 0.078028737 | 0.078028737 | ENSG00000182325 | n.s. |
| ENSG00000213942 | 15.72675074 | 0.363017957 | 0.4488313 | 0.808807 | 0.418626064 | 0.609615175 | | processed_pseudogene | 46.15 | 0.078028615 | 0.078028615 | ENSG00000213942 | n.s. |
| ENSG00000158850 | 626.8115799 | 0.27288118 | 0.2778805 | 0.982009 | 0.326095344 | 0.517699599 | B4GALT3 | protein_coding | 52.28 | 0.07802278 | 0.07802278 | ENSG00000158850 | n.s. |
| ENSG00000169223 | 820.095679 | 0.278998641 | 0.288231 | 0.967969 | 0.333059917 | 0.525282422 | LMAN2 | protein_coding | 50.49 | 0.07801001 | 0.07801001 | ENSG00000169223 | n.s. |
| ENSG00000144554 | 1616.05645 | -0.256708752 | 0.2515412 | -1.020544 | 0.307470763 | 0.497322118 | FANCD2 | protein_coding | 42.77 | -0.077875738 | 0.077875738 | ENSG00000144554 | n.s. |

| | | | | | | | | | | | | | |
|---|---|---|---|---|---|---|---|---|---|---|---|---|---|
| ENSG00000087053 | 765.8136543 | -0.263482714 | 0.2627054 | -1.002959 | 0.315880709 | 0.506507787 | MTMR2 | protein_coding | 36.56 | -0.077836449 | 0.077836449 | ENSG00000087053 | n.s. |
| ENSG00000178896 | 298.1591561 | -0.263842536 | 0.2635446 | -1.00113 | 0.31676377 | 0.507339464 | EXOSC4 | protein_coding | 58.65 | -0.077754753 | 0.077754753 | ENSG00000178896 | n.s. |
| ENSG00000199672 | 5.11901341 | -0.39289022 | 0.5163308 | -0.760927 | 0.446700491 | 0.634100082 | RNU4-21P | snRNA | 42.86 | -0.077730262 | 0.077730262 | ENSG00000199672 | n.s. |
| ENSG00000240849 | 276.5545514 | -0.248003738 | 0.2377838 | -1.04298 | 0.296957607 | 0.486359342 | TMEM189 | protein_coding | 52.37 | -0.077635769 | 0.077635769 | ENSG00000240849 | n.s. |
| ENSG00000265693 | 8.378229574 | -0.354879187 | 0.4329266 | -0.819721 | 0.412374947 | 0.604431665 | | transcribed_processed_pseudog | 46.92 | -0.077595325 | 0.077595325 | ENSG00000265693 | n.s. |
| ENSG00000103202 | 1479.970418 | 0.28737465 | 0.3038896 | 0.945655 | 0.344324656 | 0.537144194 | NME4 | protein_coding | 58.71 | 0.077565037 | 0.077565037 | ENSG00000103202 | n.s. |
| ENSG00000115129 | 726.9189359 | -0.250434538 | 0.2420969 | -1.034439 | 0.300930828 | 0.490571892 | TP53I3 | protein_coding | 49.47 | -0.077458736 | 0.077458736 | ENSG00000115129 | n.s. |
| ENSG00000013306 | 2244.962807 | 0.267188684 | 0.2699565 | 0.989747 | 0.322297643 | 0.513044479 | SLC25A39 | protein_coding | 61.11 | 0.077443299 | 0.077443299 | ENSG00000013306 | n.s. |
| ENSG00000083635 | 514.4909315 | -0.256943146 | 0.2529003 | -1.015986 | 0.309636181 | 0.499601871 | NUFIP1 | protein_coding | 38.71 | -0.077436483 | 0.077436483 | ENSG00000083635 | n.s. |
| ENSG00000134490 | 161.4707878 | -0.265283491 | 0.2669065 | -0.993919 | 0.320262171 | 0.511117395 | TMEM241 | protein_coding | 45.61 | -0.077324657 | 0.077324657 | ENSG00000134490 | n.s. |
| ENSG00000170836 | 1985.21985 | -0.239430916 | 0.2246037 | -1.066015 | 0.28641685 | 0.475415093 | PPM1D | protein_coding | 41 | -0.077318716 | 0.077318716 | ENSG00000170836 | n.s. |
| ENSG00000184465 | 690.9115461 | -0.254735252 | 0.2498196 | -1.019677 | 0.307881753 | 0.497484598 | WDR27 | protein_coding | 42.01 | -0.077240915 | 0.077240915 | ENSG00000184465 | n.s. |
| ENSG00000103335 | 448.1357847 | -0.249859155 | 0.241715 | -1.033693 | 0.30127968 | 0.490839806 | PIEZO1 | protein_coding | 61.99 | -0.077221526 | 0.077221526 | ENSG00000103335 | n.s. |
| ENSG00000178233 | 219.4343837 | -0.248128375 | 0.2387517 | -1.039274 | 0.298677406 | 0.488429119 | TMEM151B | protein_coding | 54.71 | -0.077217166 | 0.077217166 | ENSG00000178233 | n.s. |
| ENSG00000134644 | 2144.140345 | -0.251986048 | 0.2450676 | -1.028231 | 0.303841311 | 0.493828498 | PUM1 | protein_coding | 41.05 | -0.077214535 | 0.077214535 | ENSG00000134644 | n.s. |
| ENSG00000117122 | 126.7631373 | 0.301792755 | 0.3314246 | 0.910593 | 0.362510054 | 0.554884288 | MFAP2 | protein_coding | 62.61 | 0.077197854 | 0.077197854 | ENSG00000117122 | n.s. |
| ENSG00000123739 | 426.4774981 | -0.28568233 | 0.3018996 | -0.946283 | 0.344004516 | 0.536827491 | PLA2G12A | protein_coding | 39 | -0.077181439 | 0.077181439 | ENSG00000123739 | n.s. |
| ENSG00000155980 | 445.6122065 | -0.25602253 | 0.2523649 | -1.014493 | 0.310347438 | 0.500371826 | KIF5A | protein_coding | 47.7 | -0.076987806 | 0.076987806 | ENSG00000155980 | n.s. |
| ENSG00000250321 | 49.33016174 | 0.460291704 | 0.6879726 | 0.669055 | 0.503460228 | 0.680419022 | | processed_pseudogene | 47.18 | 0.076971615 | 0.076971615 | ENSG00000250321 | n.s. |
| ENSG00000143061 | 2445.476071 | -0.257817319 | 0.2554776 | -1.009158 | 0.31289872 | 0.502884552 | IGSF3 | protein_coding | 46.14 | -0.076966644 | 0.076966644 | ENSG00000143061 | n.s. |
| ENSG00000166333 | 381.5538811 | 0.275496734 | 0.2848718 | 0.96709 | 0.333498878 | 0.525855592 | ILK | protein_coding | 49.78 | 0.076900369 | 0.076900369 | ENSG00000166333 | n.s. |
| ENSG00000159228 | 509.5588045 | -0.282687266 | 0.2974719 | -0.950299 | 0.341960273 | 0.534596102 | CBR1 | protein_coding | 50.62 | -0.076883646 | 0.076883646 | ENSG00000159228 | n.s. |
| ENSG00000254402 | 63.40291444 | 0.283583936 | 0.2989825 | 0.948497 | 0.342876684 | 0.535667872 | LRRC24 | protein_coding | 63.08 | 0.076880853 | 0.076880853 | ENSG00000254402 | n.s. |
| ENSG00000138036 | 328.4086455 | -0.285534124 | 0.302495 | -0.94393 | 0.345205415 | 0.53809841 | DYNC2LI1 | protein_coding | 37.57 | -0.076848166 | 0.076848166 | ENSG00000138036 | n.s. |
| ENSG00000133056 | 272.2371822 | -0.238204663 | 0.2237423 | -1.064639 | 0.287039543 | 0.476221587 | PIK3C2B | protein_coding | 49.35 | -0.07674738 | 0.07674738 | ENSG00000133056 | n.s. |
| ENSG00000100852 | 2666.192176 | -0.271248652 | 0.2784853 | -0.974014 | 0.33004951 | 0.521835037 | ARHGAP5 | protein_coding | 36.95 | -0.076618729 | 0.076618729 | ENSG00000100852 | n.s. |
| ENSG00000252892 | 26.21161178 | -0.297465401 | 0.3251419 | -0.914879 | 0.360255354 | 0.552936594 | RNU6-548P | snRNA | 46.96 | -0.076545185 | 0.076545185 | ENSG00000252892 | n.s. |
| ENSG00000113643 | 2034.519339 | -0.264466789 | 0.2677213 | -0.987844 | 0.323229138 | 0.513858152 | RARS | protein_coding | 39.43 | -0.076472357 | 0.076472357 | ENSG00000113643 | n.s. |
| ENSG00000101400 | 150.253755 | -0.288796142 | 0.3094888 | -0.933139 | 0.350748051 | 0.543755163 | SNTA1 | protein_coding | 50.41 | -0.076414479 | 0.076414479 | ENSG00000101400 | n.s. |
| ENSG00000139146 | 6302.099087 | 0.275088517 | 0.2855244 | 0.96345 | 0.335321757 | 0.527612047 | FAM60A | protein_coding | 41.46 | 0.076388036 | 0.076388036 | ENSG00000139146 | n.s. |
| ENSG00000151929 | 103.7911325 | 0.262830374 | 0.2651041 | 0.991423 | 0.32147887 | 0.512192063 | BAG3 | protein_coding | 49.77 | 0.076369874 | 0.076369874 | ENSG00000151929 | n.s. |
| ENSG00000055950 | 409.0142751 | 0.288359927 | 0.3088813 | 0.933562 | 0.350529745 | 0.543598312 | MRPL43 | protein_coding | 53.18 | 0.076335188 | 0.076335188 | ENSG00000055950 | n.s. |

| | | | | | | | | | | | | | |
|---|---|---|---|---|---|---|---|---|---|---|---|---|---|
| ENSG00000272033 | 152.9023207 | -0.264807585 | 0.2685626 | -0.986018 | 0.324124319 | 0.51492274 | | sense_intronic | 36.08 | -0.076332886 | 0.076332886 | ENSG00000272033 | n.s. |
| ENSG00000186395 | 85.80359462 | 0.287261491 | 0.3071738 | 0.935176 | 0.349697717 | 0.542670676 | KRT10 | protein_coding | 42.64 | 0.076257483 | 0.076257483 | ENSG00000186395 | n.s. |
| ENSG00000136044 | 888.7897572 | -0.261649782 | 0.2633978 | -0.993363 | 0.320532874 | 0.511376098 | APPL2 | protein_coding | 43.65 | -0.076208004 | 0.076208004 | ENSG00000136044 | n.s. |
| ENSG00000113811 | 789.5349137 | 0.277943605 | 0.2909929 | 0.955156 | 0.339498803 | 0.532215916 | SELENOK | protein_coding | 40.23 | 0.076132128 | 0.076132128 | ENSG00000113811 | n.s. |
| ENSG00000075945 | 500.7374631 | -0.275082246 | 0.2862986 | -0.960823 | 0.336641148 | 0.528774222 | KIFAP3 | protein_coding | 34.27 | -0.076123434 | 0.076123434 | ENSG00000075945 | n.s. |
| ENSG00000263753 | 1567.017669 | -0.240604402 | 0.2290122 | -1.050618 | 0.293433968 | 0.482746205 | LINC00667 | lincRNA | 41.91 | -0.076098633 | 0.076098633 | ENSG00000263753 | n.s. |
| ENSG00000107105 | 3083.120859 | -0.258223903 | 0.2582161 | -1.00003 | 0.317295818 | 0.507588226 | ELAVL2 | protein_coding | 38.63 | -0.07604396 | 0.07604396 | ENSG00000107105 | n.s. |
| ENSG00000141098 | 458.0993608 | -0.253515285 | 0.2503551 | -1.012623 | 0.311240277 | 0.501260656 | GFOD2 | protein_coding | 46.49 | -0.076038457 | 0.076038457 | ENSG00000141098 | n.s. |
| ENSG00000183666 | 315.7613939 | -0.236748527 | 0.2229206 | -1.062031 | 0.288221796 | 0.477386636 | GUSBP1 | transcribed_unprocessed_pseud | 37.03 | -0.076026994 | 0.076026994 | ENSG00000183666 | n.s. |
| ENSG00000184056 | 365.267927 | -0.252870514 | 0.2495304 | -1.013386 | 0.310875922 | 0.501021584 | VPS33B | protein_coding | 46.83 | -0.075897457 | 0.075897457 | ENSG00000184056 | n.s. |
| ENSG00000267248 | 68.29942532 | -0.251223836 | 0.2468975 | -1.017523 | 0.308904834 | 0.498767888 | | processed_transcript | 47.27 | -0.075895102 | 0.075895102 | ENSG00000267248 | n.s. |
| ENSG00000166352 | 323.0755801 | -0.302525342 | 0.3369181 | -0.89792 | 0.369228436 | 0.561464342 | C11orf74 | protein_coding | 36.82 | -0.075836393 | 0.075836393 | ENSG00000166352 | n.s. |
| ENSG00000112759 | 556.1156213 | -0.274925005 | 0.2868034 | -0.958583 | 0.337768666 | 0.530007115 | SLC29A1 | protein_coding | 56.86 | -0.075801855 | 0.075801855 | ENSG00000112759 | n.s. |
| ENSG00000163382 | 439.8536302 | 0.283536272 | 0.301834 | 0.939378 | 0.347536544 | 0.540401083 | NAXE | protein_coding | 56.38 | 0.075784649 | 0.075784649 | ENSG00000163382 | n.s. |
| ENSG00000101084 | 331.241821 | 0.261063524 | 0.2634778 | 0.990837 | 0.321765313 | 0.51252013 | C20orf24 | protein_coding | 49.21 | 0.075783888 | 0.075783888 | ENSG00000101084 | n.s. |
| ENSG00000230629 | 193.820922 | 0.301616983 | 0.3356547 | 0.898593 | 0.368869418 | 0.561102251 | RPS23P8 | processed_pseudogene | 43.64 | 0.075693191 | 0.075693191 | ENSG00000230629 | n.s. |
| ENSG00000245910 | 2436.868765 | 0.26489488 | 0.2698994 | 0.981458 | 0.326367018 | 0.518012766 | SNHG6 | processed_transcript | 44.54 | 0.075669749 | 0.075669749 | ENSG00000245910 | n.s. |
| ENSG00000253230 | 1030.975825 | -0.239863343 | 0.2286714 | -1.048943 | 0.294204135 | 0.483670348 | LINC00599 | lincRNA | 53.37 | -0.07566502 | 0.07566502 | ENSG00000253230 | n.s. |
| ENSG00000010292 | 2980.775338 | -0.257384423 | 0.2578401 | -0.998233 | 0.318166441 | 0.50859143 | NCAPD2 | protein_coding | 46.34 | -0.075576036 | 0.075576036 | ENSG00000010292 | n.s. |
| ENSG00000101558 | 2902.421489 | -0.265792712 | 0.2716723 | -0.978358 | 0.327897423 | 0.519654777 | VAPA | protein_coding | 37.13 | -0.075560901 | 0.075560901 | ENSG00000101558 | n.s. |
| ENSG00000121064 | 612.7793822 | 0.27125988 | 0.2808975 | 0.96569 | 0.334199407 | 0.526602348 | SCPEP1 | protein_coding | 45.31 | 0.075550545 | 0.075550545 | ENSG00000121064 | n.s. |
| ENSG00000116586 | 391.1690252 | 0.290046324 | 0.3142698 | 0.922921 | 0.356048182 | 0.549125877 | LAMTOR2 | protein_coding | 50.94 | 0.075507205 | 0.075507205 | ENSG00000116586 | n.s. |
| ENSG00000117593 | 1318.14181 | -0.263301137 | 0.2678933 | -0.982858 | 0.325677161 | 0.51715364 | DARS2 | protein_coding | 40.56 | -0.075404289 | 0.075404289 | ENSG00000117593 | n.s. |
| ENSG00000124942 | 4958.068624 | -0.240326672 | 0.2302114 | -1.043939 | 0.296513494 | 0.485825593 | AHNAK | protein_coding | 50.05 | -0.075347125 | 0.075347125 | ENSG00000124942 | n.s. |
| ENSG00000198498 | 1371.077683 | -0.269371273 | 0.278256 | -0.96807 | 0.333009505 | 0.525282422 | TMA16 | protein_coding | 36.88 | -0.075318129 | 0.075318129 | ENSG00000198498 | n.s. |
| ENSG00000168214 | 2374.721108 | -0.255577822 | 0.2555706 | -1.000028 | 0.317296838 | 0.507588226 | RBPJ | protein_coding | 40.79 | -0.075264719 | 0.075264719 | ENSG00000168214 | n.s. |
| ENSG00000252496 | 6.52806591 | -0.461756504 | 0.7013619 | -0.658371 | 0.510299629 | 0.687123662 | RNA5SP33 | rRNA | 44.9 | -0.075250193 | 0.075250193 | ENSG00000252496 | n.s. |
| ENSG00000187838 | 87.52029304 | -0.261638599 | 0.2656928 | -0.984741 | 0.324751332 | 0.51580114 | PLSCR3 | protein_coding | 58.45 | -0.075225729 | 0.075225729 | ENSG00000187838 | n.s. |
| ENSG00000168056 | 110.7364973 | 0.27518467 | 0.2890468 | 0.952042 | 0.341075751 | 0.533349039 | LTBP3 | protein_coding | 60.22 | 0.075122246 | 0.075122246 | ENSG00000168056 | n.s. |
| ENSG00000141258 | 876.8481276 | 0.259758449 | 0.2628502 | 0.988237 | 0.32303636 | 0.513837368 | SGSM2 | protein_coding | 54.99 | 0.075115472 | 0.075115472 | ENSG00000141258 | n.s. |
| ENSG00000105926 | 400.7523564 | 0.284832075 | 0.306029 | 0.930736 | 0.351990321 | 0.544959861 | MPP6 | protein_coding | 35.88 | 0.075091842 | 0.075091842 | ENSG00000105926 | n.s. |
| ENSG00000187954 | 121.8101648 | -0.280385052 | 0.2981786 | -0.940326 | 0.347050478 | 0.539970485 | CYHR1 | protein_coding | 61.94 | -0.075039445 | 0.075039445 | ENSG00000187954 | n.s. |

| | | | | | | | | | | | | | |
|---|---|---|---|---|---|---|---|---|---|---|---|---|---|
| ENSG00000117411 | 466.434318 | -0.260234109 | 0.2639049 | -0.986091 | 0.324088696 | 0.51492274 | B4GALT2 | protein_coding | 56.94 | -0.075014545 | 0.075014545 | ENSG00000117411 | n.s. |
| ENSG00000108465 | 4443.920766 | 0.252646139 | 0.2511349 | 1.006018 | 0.314407099 | 0.504900813 | CDK5RAP3 | protein_coding | 51.51 | 0.074983841 | 0.074983841 | ENSG00000108465 | n.s. |
| ENSG00000136938 | 1216.379991 | -0.240067592 | 0.2305883 | -1.041109 | 0.297824976 | 0.487279619 | ANP32B | protein_coding | 43.52 | -0.074954324 | 0.074954324 | ENSG00000136938 | n.s. |
| ENSG00000080546 | 540.8967783 | -0.226977876 | 0.2100423 | -1.080629 | 0.279862126 | 0.46760115 | SESN1 | protein_coding | 37.66 | -0.074930942 | 0.074930942 | ENSG00000080546 | n.s. |
| ENSG00000260774 | 91.4564419 | 0.248881845 | 0.245165 | 1.015161 | 0.310029316 | 0.500062346 | | processed_transcript | 46.87 | 0.074907424 | 0.074907424 | ENSG00000260774 | n.s. |
| ENSG00000283045 | 156.2262921 | -0.258393762 | 0.2610801 | -0.989711 | 0.322315597 | 0.513044479 | | sense_intronic | 33.63 | -0.074894135 | 0.074894135 | ENSG00000283045 | n.s. |
| ENSG00000138081 | 2038.192443 | -0.254044089 | 0.2536706 | -1.001472 | 0.316598599 | 0.507236189 | FBXO11 | protein_coding | 39.46 | -0.074889598 | 0.074889598 | ENSG00000138081 | n.s. |
| ENSG00000175106 | 29.99444715 | 0.284706719 | 0.3067458 | 0.928152 | 0.353328676 | 0.546356637 | TVP23C | protein_coding | 42.63 | 0.074742283 | 0.074742283 | ENSG00000175106 | n.s. |
| ENSG00000146677 | 139.942018 | 0.298311387 | 0.3323704 | 0.897527 | 0.369437872 | 0.561660132 | | processed_pseudogene | 50.49 | 0.074734879 | 0.074734879 | ENSG00000146677 | n.s. |
| ENSG00000067177 | 292.6832639 | -0.25012953 | 0.2478283 | -1.009286 | 0.312837627 | 0.502884552 | PHKA1 | protein_coding | 37.87 | -0.074671595 | 0.074671595 | ENSG00000067177 | n.s. |
| ENSG00000132589 | 306.2630337 | -0.276056508 | 0.2914429 | -0.947206 | 0.343533688 | 0.536430627 | FLOT2 | protein_coding | 54.35 | -0.074669541 | 0.074669541 | ENSG00000132589 | n.s. |
| ENSG00000178921 | 1578.419339 | -0.244723319 | 0.2388866 | -1.024433 | 0.305630775 | 0.495724288 | PFAS | protein_coding | 51.84 | -0.074581831 | 0.074581831 | ENSG00000178921 | n.s. |
| ENSG00000117394 | 618.1161991 | 0.284533406 | 0.3069372 | 0.927008 | 0.353922129 | 0.546880191 | SLC2A1 | protein_coding | 52.89 | 0.074578427 | 0.074578427 | ENSG00000117394 | n.s. |
| ENSG00000167136 | 373.7857575 | 0.289492154 | 0.3160251 | 0.916042 | 0.359645035 | 0.552848354 | ENDOG | protein_coding | 61.54 | 0.074513537 | 0.074513537 | ENSG00000167136 | n.s. |
| ENSG00000105576 | 1737.600972 | -0.251625685 | 0.2506843 | -1.003755 | 0.315496646 | 0.50609579 | TNPO2 | protein_coding | 54.01 | -0.074422643 | 0.074422643 | ENSG00000105576 | n.s. |
| ENSG00000030110 | 176.4641154 | -0.252403972 | 0.2520664 | -1.001339 | 0.316662856 | 0.507236189 | BAK1 | protein_coding | 57.18 | -0.074406109 | 0.074406109 | ENSG00000030110 | n.s. |
| ENSG00000186908 | 2645.71035 | -0.267797767 | 0.2784724 | -0.961667 | 0.336216823 | 0.52837731 | ZDHHC17 | protein_coding | 36.11 | -0.074194935 | 0.074194935 | ENSG00000186908 | n.s. |
| ENSG00000160408 | 568.7553024 | -0.263226825 | 0.2705686 | -0.972865 | 0.330620286 | 0.522559455 | ST6GALNAC6 | protein_coding | 53.52 | -0.074194242 | 0.074194242 | ENSG00000160408 | n.s. |
| ENSG00000160796 | 342.3730768 | 0.244034843 | 0.2388672 | 1.021634 | 0.306954161 | 0.496713481 | NBEAL2 | protein_coding | 59.9 | 0.074160737 | 0.074160737 | ENSG00000160796 | n.s. |
| ENSG00000071462 | 799.4885606 | 0.274743384 | 0.2905271 | 0.945672 | 0.344315861 | 0.537144194 | BUD23 | protein_coding | 54.07 | 0.074155743 | 0.074155743 | ENSG00000071462 | n.s. |
| ENSG00000277534 | 429.3138308 | -0.237117366 | 0.2275421 | -1.042081 | 0.297374052 | 0.486812337 | | sense_intronic | 37.54 | -0.074131999 | 0.074131999 | ENSG00000277534 | n.s. |
| ENSG00000188610 | 120.3514805 | -0.271068313 | 0.2846167 | -0.952398 | 0.340895267 | 0.533305818 | FAM72B | protein_coding | 41.53 | -0.074008068 | 0.074008068 | ENSG00000188610 | n.s. |
| ENSG00000233476 | 221.8370343 | 0.288921696 | 0.3170859 | 0.911178 | 0.362201661 | 0.554599493 | EEF1A1P6 | processed_pseudogene | 48.27 | 0.073969886 | 0.073969886 | ENSG00000233476 | n.s. |
| ENSG00000199436 | 7.259912399 | -0.367263699 | 0.4762154 | -0.771213 | 0.440580491 | 0.62896057 | SNORD9 | snoRNA | 47.12 | -0.073958308 | 0.073958308 | ENSG00000199436 | n.s. |
| ENSG00000138413 | 2019.163536 | -0.269807618 | 0.2824566 | -0.955218 | 0.339467442 | 0.532215916 | IDH1 | protein_coding | 41.27 | -0.073903582 | 0.073903582 | ENSG00000138413 | n.s. |
| ENSG00000164054 | 209.1697029 | -0.268953182 | 0.2811386 | -0.956657 | 0.338740519 | 0.531292582 | SHISA5 | protein_coding | 50.15 | -0.073872361 | 0.073872361 | ENSG00000164054 | n.s. |
| ENSG00000143418 | 1486.27022 | 0.259804266 | 0.2655154 | 0.978491 | 0.327831758 | 0.519626772 | CERS2 | protein_coding | 52.38 | 0.073864556 | 0.073864556 | ENSG00000143418 | n.s. |
| ENSG00000166965 | 641.9411826 | -0.242787955 | 0.2376889 | -1.021453 | 0.307039993 | 0.496794633 | RCCD1 | protein_coding | 54.47 | -0.07376459 | 0.07376459 | ENSG00000166965 | n.s. |
| ENSG00000221923 | 593.1855945 | -0.237664666 | 0.2292776 | -1.036581 | 0.299931389 | 0.48967457 | ZNF880 | protein_coding | 41.86 | -0.073698019 | 0.073698019 | ENSG00000221923 | n.s. |
| ENSG00000064763 | 534.9891575 | -0.272621565 | 0.2879907 | -0.946633 | 0.343825697 | 0.53670032 | FAR2 | protein_coding | 38.9 | -0.073680925 | 0.073680925 | ENSG00000064763 | n.s. |
| ENSG00000184611 | 879.0089904 | -0.241194226 | 0.2352209 | -1.025395 | 0.305177067 | 0.495161438 | KCNH7 | protein_coding | 35.03 | -0.073625305 | 0.073625305 | ENSG00000184611 | n.s. |
| ENSG00000107140 | 136.9392406 | -0.248223535 | 0.2468704 | -1.005481 | 0.314665202 | 0.505198755 | TESK1 | protein_coding | 59.15 | -0.073607643 | 0.073607643 | ENSG00000107140 | n.s. |

| | | | | | | | | | | | | | | |
|---|---|---|---|---|---|---|---|---|---|---|---|---|---|---|
| ENSG00000104979 | 610.1996229 | 0.29275245 | 0.3254393 | 0.899561 | 0.368354024 | 0.560664776 | C19orf53 | | protein_coding | 54.88 | 0.073567733 | 0.073567733 | ENSG00000104979 | n.s. |
| ENSG00000182810 | 435.9964901 | 0.217641289 | 0.1979956 | 1.099223 | 0.271670955 | 0.459595955 | DDX28 | | protein_coding | 58.41 | 0.073480879 | 0.073480879 | ENSG00000182810 | n.s. |
| ENSG00000138758 | 3115.711893 | -0.249218137 | 0.2492419 | -0.999905 | 0.317356569 | 0.507588226 | | 11-Sep | protein_coding | 40.66 | -0.073391865 | 0.073391865 | ENSG00000138758 | n.s. |
| ENSG00000163349 | 1825.535775 | -0.239414621 | 0.2332147 | -1.026585 | 0.304616124 | 0.494654805 | HIPK1 | | protein_coding | 39.82 | -0.073188515 | 0.073188515 | ENSG00000163349 | n.s. |
| ENSG00000122729 | 386.6269731 | -0.243793629 | 0.2406214 | -1.013183 | 0.310972598 | 0.501023853 | ACO1 | | protein_coding | 41.71 | -0.073172609 | 0.073172609 | ENSG00000122729 | n.s. |
| ENSG00000180817 | 3685.329775 | 0.285478396 | 0.313062 | 0.911891 | 0.361826085 | 0.554446905 | PPA1 | | protein_coding | 39.81 | 0.073122447 | 0.073122447 | ENSG00000180817 | n.s. |
| ENSG00000134283 | 2311.757173 | -0.263795835 | 0.2743438 | -0.961552 | 0.336274606 | 0.52837731 | PPHLN1 | | protein_coding | 39.77 | -0.073086177 | 0.073086177 | ENSG00000134283 | n.s. |
| ENSG00000172315 | 1486.072518 | 0.233128375 | 0.2234012 | 1.043541 | 0.296697772 | 0.48599095 | TP53RK | | protein_coding | 45.1 | 0.073055863 | 0.073055863 | ENSG00000172315 | n.s. |
| ENSG00000047634 | 650.9912456 | 0.256737203 | 0.2623252 | 0.978698 | 0.327729171 | 0.519626772 | SCML1 | | protein_coding | 38.89 | 0.072992564 | 0.072992564 | ENSG00000047634 | n.s. |
| ENSG00000127838 | 408.8759336 | 0.285041532 | 0.3127737 | 0.911335 | 0.362119098 | 0.554599493 | PNKD | | protein_coding | 51.32 | 0.072976485 | 0.072976485 | ENSG00000127838 | n.s. |
| ENSG00000105185 | 1692.570832 | 0.264172249 | 0.2752203 | 0.959857 | 0.337126968 | 0.529374789 | PDCD5 | | protein_coding | 45.84 | 0.072974082 | 0.072974082 | ENSG00000105185 | n.s. |
| ENSG00000223508 | 105.63986 | 0.265080371 | 0.2769722 | 0.957065 | 0.338534494 | 0.531029266 | RPL23AP53 | | transcribed_processed_pseudog | 43.7 | 0.072865702 | 0.072865702 | ENSG00000223508 | n.s. |
| ENSG00000145741 | 8433.532315 | 0.272259357 | 0.2895434 | 0.940306 | 0.347060731 | 0.539970485 | BTF3 | | protein_coding | 38.63 | 0.072864766 | 0.072864766 | ENSG00000145741 | n.s. |
| ENSG00000116871 | 541.6592965 | -0.245608653 | 0.2442309 | -1.005641 | 0.31458816 | 0.505133312 | MAP7D1 | | protein_coding | 54.66 | -0.07284605 | 0.07284605 | ENSG00000116871 | n.s. |
| ENSG00000106554 | 1551.235356 | -0.270912299 | 0.2872422 | -0.943149 | 0.345604464 | 0.538479292 | CHCHD3 | | protein_coding | 39.02 | -0.072829622 | 0.072829622 | ENSG00000106554 | n.s. |
| ENSG00000100813 | 3259.843704 | -0.242237682 | 0.2391798 | -1.012785 | 0.311162986 | 0.501194152 | ACIN1 | | protein_coding | 45.36 | -0.072669852 | 0.072669852 | ENSG00000100813 | n.s. |
| ENSG00000143870 | 4842.023993 | 0.271132632 | 0.287993 | 0.941456 | 0.346471373 | 0.53952812 | PDIA6 | | protein_coding | 46.27 | 0.072659727 | 0.072659727 | ENSG00000143870 | n.s. |
| ENSG00000110911 | 1381.020091 | -0.25541235 | 0.2611732 | -0.977942 | 0.328102807 | 0.51976054 | SLC11A2 | | protein_coding | 43.54 | -0.072587346 | 0.072587346 | ENSG00000110911 | n.s. |
| ENSG00000203485 | 85.98805201 | -0.261092951 | 0.271122 | -0.963009 | 0.335542942 | 0.52782334 | INF2 | | protein_coding | 63.24 | -0.072456273 | 0.072456273 | ENSG00000203485 | n.s. |
| ENSG00000164163 | 5300.44683 | -0.272810035 | 0.2917696 | -0.935019 | 0.349778611 | 0.542735718 | ABCE1 | | protein_coding | 35.08 | -0.072406947 | 0.072406947 | ENSG00000164163 | n.s. |
| ENSG00000175130 | 2135.025797 | 0.237608804 | 0.2320676 | 1.023878 | 0.305893079 | 0.495878772 | MARCKSL1 | | protein_coding | 64.21 | 0.072381459 | 0.072381459 | ENSG00000175130 | n.s. |
| ENSG00000034713 | 1033.444977 | 0.282046347 | 0.3091429 | 0.912349 | 0.361584857 | 0.554199247 | GABARAPL2 | | protein_coding | 47.18 | 0.072298088 | 0.072298088 | ENSG00000034713 | n.s. |
| ENSG00000200525 | 16.63944381 | -0.303765169 | 0.3501206 | -0.867601 | 0.385612535 | 0.578408131 | RNU6-1209P | | snRNA | 38.32 | -0.072224911 | 0.072224911 | ENSG00000200525 | n.s. |
| ENSG00000170293 | 52.31569214 | 0.285283981 | 0.3156761 | 0.903724 | 0.366141935 | 0.558484609 | CMTM8 | | protein_coding | 43.06 | 0.072173649 | 0.072173649 | ENSG00000170293 | n.s. |
| ENSG00000258337 | 42.26700511 | 0.289673442 | 0.3240455 | 0.893928 | 0.371360184 | 0.563475136 | | | sense_intronic | 37.7 | 0.072164967 | 0.072164967 | ENSG00000258337 | n.s. |
| ENSG00000116809 | 250.9887907 | 0.248869524 | 0.2514022 | 0.989926 | 0.322210418 | 0.512994356 | ZBTB17 | | protein_coding | 51.22 | 0.072144143 | 0.072144143 | ENSG00000116809 | n.s. |
| ENSG00000165409 | 46.74811984 | 0.269544448 | 0.2868921 | 0.939533 | 0.347457402 | 0.540338369 | TSHR | | protein_coding | 38.85 | 0.072058447 | 0.072058447 | ENSG00000165409 | n.s. |
| ENSG00000149761 | 131.3730454 | -0.267030393 | 0.282452 | -0.945401 | 0.344454244 | 0.537228233 | NUDT22 | | protein_coding | 59.06 | -0.072055794 | 0.072055794 | ENSG00000149761 | n.s. |
| ENSG00000004455 | 1354.47154 | -0.272032429 | 0.2917962 | -0.932269 | 0.351197742 | 0.544149361 | AK2 | | protein_coding | 46.32 | -0.071893241 | 0.071893241 | ENSG00000004455 | n.s. |
| ENSG00000115866 | 2683.069211 | -0.275187854 | 0.2975996 | -0.924692 | 0.355126356 | 0.548162335 | DARS | | protein_coding | 37.04 | -0.071849019 | 0.071849019 | ENSG00000115866 | n.s. |
| ENSG00000084652 | 958.8985451 | -0.242544922 | 0.2416466 | -1.003718 | 0.315514721 | 0.50609579 | TXLNA | | protein_coding | 50.3 | -0.071736851 | 0.071736851 | ENSG00000084652 | n.s. |
| ENSG00000150456 | 96.59009328 | -0.2736732 | 0.2952425 | -0.926944 | 0.353955737 | 0.546880191 | EEF1AKMT1 | | protein_coding | 41.53 | -0.071731882 | 0.071731882 | ENSG00000150456 | n.s. |

| | | | | | | | | | | | | | |
|---|---|---|---|---|---|---|---|---|---|---|---|---|---|
| ENSG00000167325 | 3806.774475 | -0.264469656 | 0.2788707 | -0.94836 | 0.342946457 | 0.535716765 | RRM1 | protein_coding | 40.39 | -0.071688405 | 0.071688405 | ENSG00000167325 | n.s. |
| ENSG00000104960 | 1377.482252 | 0.249311509 | 0.25315 | 0.984837 | 0.324704155 | 0.515785049 | PTOV1 | protein_coding | 61.87 | 0.07168485 | 0.07168485 | ENSG00000104960 | n.s. |
| ENSG00000182923 | 549.083 | -0.23465737 | 0.2287792 | -1.025694 | 0.305035953 | 0.495047858 | CEP63 | protein_coding | 38.49 | -0.071653288 | 0.071653288 | ENSG00000182923 | n.s. |
| ENSG00000158859 | 393.5872526 | -0.227111919 | 0.2166252 | -1.048409 | 0.294450045 | 0.483674844 | ADAMTS4 | protein_coding | 51.48 | -0.07164166 | 0.07164166 | ENSG00000158859 | n.s. |
| ENSG00000203805 | 120.5873188 | -0.280707312 | 0.3086612 | -0.909435 | 0.363120508 | 0.555635436 | PLPP4 | protein_coding | 42.31 | -0.071639331 | 0.071639331 | ENSG00000203805 | n.s. |
| ENSG00000149182 | 1483.750258 | 0.245994295 | 0.2477676 | 0.992843 | 0.320786623 | 0.511487879 | ARFGAP2 | protein_coding | 53.19 | 0.071624844 | 0.071624844 | ENSG00000149182 | n.s. |
| ENSG00000108448 | 31.19221218 | 0.275965639 | 0.2997781 | 0.920566 | 0.35727682 | 0.550200771 | TRIM16L | protein_coding | 44.38 | 0.071607234 | 0.071607234 | ENSG00000108448 | n.s. |
| ENSG00000104442 | 873.1632553 | -0.275851629 | 0.2995692 | -0.920828 | 0.357140434 | 0.55011243 | ARMC1 | protein_coding | 42.6 | -0.071596888 | 0.071596888 | ENSG00000104442 | n.s. |
| ENSG00000139239 | 246.4074759 | 0.275974412 | 0.2999989 | 0.919918 | 0.357615528 | 0.550417975 | | processed_pseudogene | 49.46 | 0.071562205 | 0.071562205 | ENSG00000139239 | n.s. |
| ENSG00000162302 | 127.8579069 | -0.258289901 | 0.2685501 | -0.961794 | 0.336153085 | 0.52837731 | RPS6KA4 | protein_coding | 56.99 | -0.071560726 | 0.071560726 | ENSG00000162302 | n.s. |
| ENSG00000178922 | 75.89204565 | 0.261800396 | 0.2749941 | 0.952022 | 0.341085961 | 0.533349039 | HYI | protein_coding | 59.36 | 0.071468493 | 0.071468493 | ENSG00000178922 | n.s. |
| ENSG00000071894 | 714.3625981 | 0.244540472 | 0.2457132 | 0.995227 | 0.319625746 | 0.510396749 | CPSF1 | protein_coding | 59.98 | 0.07142834 | 0.07142834 | ENSG00000071894 | n.s. |
| ENSG00000085433 | 844.0753785 | 0.235538791 | 0.231 | 1.019649 | 0.307895193 | 0.497484598 | WDR47 | protein_coding | 41.17 | 0.071420157 | 0.071420157 | ENSG00000085433 | n.s. |
| ENSG00000081721 | 1578.819961 | 0.251652406 | 0.2576325 | 0.976788 | 0.328673994 | 0.520369344 | DUSP12 | protein_coding | 39.87 | 0.071390842 | 0.071390842 | ENSG00000081721 | n.s. |
| ENSG00000068697 | 1015.151894 | 0.272178818 | 0.2936461 | 0.926894 | 0.353981489 | 0.546880191 | LAPTM4A | protein_coding | 41.61 | 0.071340193 | 0.071340193 | ENSG00000068697 | n.s. |
| ENSG00000221464 | 4.783428542 | -0.383467323 | 0.5271889 | -0.727381 | 0.466992409 | 0.65175775 | MIR1271 | miRNA | 51.16 | -0.071291866 | 0.071291866 | ENSG00000221464 | n.s. |
| ENSG00000142949 | 693.504952 | -0.245186897 | 0.2472282 | -0.991743 | 0.32132286 | 0.512148578 | PTPRF | protein_coding | 55.69 | -0.0712523 | 0.0712523 | ENSG00000142949 | n.s. |
| ENSG00000136371 | 67.53411754 | -0.271179519 | 0.2920637 | -0.928494 | 0.353151132 | 0.546203422 | MTHFS | protein_coding | 40.35 | -0.071224103 | 0.071224103 | ENSG00000136371 | n.s. |
| ENSG00000126522 | 137.3231612 | 0.265284661 | 0.2815359 | 0.942276 | 0.346051138 | 0.539054597 | ASL | protein_coding | 54.6 | 0.071193714 | 0.071193714 | ENSG00000126522 | n.s. |
| ENSG00000281183 | 207.8372704 | 0.23063091 | 0.2233054 | 1.032805 | 0.30169532 | 0.49128701 | NPTN-IT1 | sense_intronic | 44.81 | 0.071187625 | 0.071187625 | ENSG00000281183 | n.s. |
| ENSG00000140474 | 876.9906956 | 0.249340586 | 0.2541859 | 0.980938 | 0.326623302 | 0.51836045 | ULK3 | protein_coding | 59.87 | 0.07115386 | 0.07115386 | ENSG00000140474 | n.s. |
| ENSG00000180176 | 535.2252494 | 0.258502116 | 0.2700049 | 0.957398 | 0.338366434 | 0.530825449 | TH | protein_coding | 65.25 | 0.071100558 | 0.071100558 | ENSG00000180176 | n.s. |
| ENSG00000184602 | 724.8733382 | -0.273754113 | 0.2973355 | -0.920691 | 0.357211711 | 0.550161355 | SNN | protein_coding | 59.43 | -0.071041908 | 0.071041908 | ENSG00000184602 | n.s. |
| ENSG00000206603 | 69.24713175 | 0.251176144 | 0.2577011 | 0.97468 | 0.329718891 | 0.521489961 | SNORA22B | snoRNA | 40.58 | 0.071021072 | 0.071021072 | ENSG00000206603 | n.s. |
| ENSG00000223482 | 279.0231248 | -0.266783556 | 0.2851487 | -0.935594 | 0.34948202 | 0.542517356 | NUTM2A-AS1 | antisense | 41.04 | -0.070854075 | 0.070854075 | ENSG00000223482 | n.s. |
| ENSG00000103479 | 1352.608649 | -0.26047893 | 0.274099 | -0.95031 | 0.34195491 | 0.534596102 | RBL2 | protein_coding | 38.71 | -0.070843552 | 0.070843552 | ENSG00000103479 | n.s. |
| ENSG00000004478 | 5720.256343 | 0.264528914 | 0.2811675 | 0.940823 | 0.346795497 | 0.539921472 | FKBP4 | protein_coding | 51.56 | 0.070806296 | 0.070806296 | ENSG00000004478 | n.s. |
| ENSG00000152932 | 573.0864049 | -0.269192988 | 0.2897642 | -0.929007 | 0.352885387 | 0.546035035 | RAB3C | protein_coding | 37.99 | -0.070738396 | 0.070738396 | ENSG00000152932 | n.s. |
| ENSG00000139233 | 1074.737946 | -0.256259291 | 0.2671823 | -0.959118 | 0.337499443 | 0.529763782 | LLPH | protein_coding | 39.72 | -0.070706483 | 0.070706483 | ENSG00000139233 | n.s. |
| ENSG00000196812 | 192.654457 | -0.234760907 | 0.231155 | -1.015599 | 0.309820207 | 0.499826511 | ZSCAN16 | protein_coding | 41.5 | -0.070705457 | 0.070705457 | ENSG00000196812 | n.s. |
| ENSG00000224597 | 310.3166754 | 0.248711048 | 0.2542568 | 0.978189 | 0.327981072 | 0.519683129 | SVIL-AS1 | antisense | 42.95 | 0.070698946 | 0.070698946 | ENSG00000224597 | n.s. |
| ENSG00000123505 | 6142.792029 | 0.244231643 | 0.2471569 | 0.988165 | 0.323072062 | 0.513837368 | AMD1 | protein_coding | 41.53 | 0.070625518 | 0.070625518 | ENSG00000123505 | n.s. |

| | | | | | | | | | | | | | |
|---|---|---|---|---|---|---|---|---|---|---|---|---|---|
| ENSG00000101972 | 4453.549545 | -0.250469464 | 0.2575933 | -0.972345 | 0.330879174 | 0.522790597 | STAG2 | protein_coding | 39.19 | -0.070550293 | 0.070550293 | ENSG00000101972 | n.s. |
| ENSG00000143409 | 130.6575025 | -0.252243471 | 0.2605799 | -0.968008 | 0.333040262 | 0.525282422 | MINDY1 | protein_coding | 49.22 | -0.070529074 | 0.070529074 | ENSG00000143409 | n.s. |
| ENSG00000174851 | 445.0771327 | 0.267625282 | 0.2876255 | 0.930465 | 0.352130628 | 0.545109488 | YIF1A | protein_coding | 58.9 | 0.070523613 | 0.070523613 | ENSG00000174851 | n.s. |
| ENSG00000224094 | 30.0852509 | 0.323981201 | 0.3968423 | 0.816398 | 0.414272583 | 0.605937302 | RPS24P8 | processed_pseudogene | 42.29 | 0.070489339 | 0.070489339 | ENSG00000224094 | n.s. |
| ENSG00000196967 | 120.057463 | 0.286487237 | 0.3232411 | 0.886296 | 0.375458142 | 0.56758998 | ZNF585A | protein_coding | 40.77 | 0.070465914 | 0.070465914 | ENSG00000196967 | n.s. |
| ENSG00000137497 | 4692.430039 | -0.236610451 | 0.2348533 | -1.007482 | 0.313703317 | 0.503944996 | NUMA1 | protein_coding | 46.62 | -0.07041926 | 0.07041926 | ENSG00000137497 | n.s. |
| ENSG00000117868 | 578.1838527 | -0.260798862 | 0.2757052 | -0.945934 | 0.344182286 | 0.537044712 | ESYT2 | protein_coding | 43.39 | -0.070412969 | 0.070412969 | ENSG00000117868 | n.s. |
| ENSG00000106524 | 201.9922666 | 0.266076985 | 0.2853594 | 0.932427 | 0.351115639 | 0.544082698 | ANKMY2 | protein_coding | 38.01 | 0.070333482 | 0.070333482 | ENSG00000106524 | n.s. |
| ENSG00000196655 | 366.7034808 | 0.26385734 | 0.281483 | 0.937383 | 0.348561569 | 0.541450711 | TRAPPC4 | protein_coding | 49.51 | 0.070302433 | 0.070302433 | ENSG00000196655 | n.s. |
| ENSG00000161981 | 516.9939437 | 0.279912432 | 0.3114057 | 0.898867 | 0.368723266 | 0.561005698 | SNRNP25 | protein_coding | 56.7 | 0.070267181 | 0.070267181 | ENSG00000161981 | n.s. |
| ENSG00000165280 | 5737.098689 | 0.252708883 | 0.2621283 | 0.964066 | 0.335012939 | 0.527347106 | VCP | protein_coding | 47.81 | 0.070228666 | 0.070228666 | ENSG00000165280 | n.s. |
| ENSG00000104765 | 1295.231285 | 0.264009917 | 0.2818996 | 0.936539 | 0.348995777 | 0.542004261 | BNIP3L | protein_coding | 43.64 | 0.070225925 | 0.070225925 | ENSG00000104765 | n.s. |
| ENSG00000116237 | 679.146633 | -0.253323817 | 0.2633609 | -0.961888 | 0.336105617 | 0.52837731 | ICMT | protein_coding | 49.59 | -0.070184843 | 0.070184843 | ENSG00000116237 | n.s. |
| ENSG00000154174 | 2366.902266 | -0.259151301 | 0.2742382 | -0.944986 | 0.344666 | 0.537438092 | TOMM70 | protein_coding | 38.61 | -0.069885734 | 0.069885734 | ENSG00000154174 | n.s. |
| ENSG00000207016 | 3.635028635 | -0.386616589 | 0.5430503 | -0.711935 | 0.47650495 | 0.659546517 | SNORA36C | snoRNA | 41.67 | -0.069882715 | 0.069882715 | ENSG00000207016 | n.s. |
| ENSG00000196284 | 137.2105663 | -0.264698751 | 0.2843876 | -0.930768 | 0.351973734 | 0.544959861 | SUPT3H | protein_coding | 36.3 | -0.069783984 | 0.069783984 | ENSG00000196284 | n.s. |
| ENSG00000169249 | 184.0608332 | -0.252346625 | 0.2629544 | -0.959659 | 0.337226722 | 0.529455081 | ZRSR2 | protein_coding | 41.5 | -0.069690789 | 0.069690789 | ENSG00000169249 | n.s. |
| ENSG00000112118 | 7438.996248 | -0.250342194 | 0.2596682 | -0.964085 | 0.335003277 | 0.527347106 | MCM3 | protein_coding | 43.17 | -0.069570955 | 0.069570955 | ENSG00000112118 | n.s. |
| ENSG00000131848 | 85.40493736 | -0.250862376 | 0.2609893 | -0.961198 | 0.336452596 | 0.528537685 | ZSCAN5A | protein_coding | 44.09 | -0.069469823 | 0.069469823 | ENSG00000131848 | n.s. |
| ENSG00000102316 | 1387.202387 | 0.27275558 | 0.3004393 | 0.907856 | 0.363954347 | 0.556307399 | MAGED2 | protein_coding | 52.37 | 0.0694668 | 0.0694668 | ENSG00000102316 | n.s. |
| ENSG00000216813 | 29.75374203 | 0.284228603 | 0.3219968 | 0.882706 | 0.377394974 | 0.569652019 | | processed_pseudogene | 44.58 | 0.06946273 | 0.06946273 | ENSG00000216813 | n.s. |
| ENSG00000247596 | 323.8302334 | 0.278271099 | 0.3109634 | 0.894868 | 0.370857862 | 0.563019737 | TWF2 | protein_coding | 59.9 | 0.069422067 | 0.069422067 | ENSG00000247596 | n.s. |
| ENSG00000116667 | 221.2544026 | -0.266958682 | 0.2896203 | -0.921754 | 0.356656879 | 0.549599094 | C1orf21 | protein_coding | 40.26 | -0.069396975 | 0.069396975 | ENSG00000116667 | n.s. |
| ENSG00000138035 | 2742.669069 | -0.257156639 | 0.2720819 | -0.945144 | 0.344585267 | 0.537372388 | PNPT1 | protein_coding | 39.78 | -0.069361485 | 0.069361485 | ENSG00000138035 | n.s. |
| ENSG00000146006 | 969.4236778 | 0.236226394 | 0.2367141 | 0.99794 | 0.31830856 | 0.508760198 | LRRTM2 | protein_coding | 38.27 | 0.069329346 | 0.069329346 | ENSG00000146006 | n.s. |
| ENSG00000100526 | 592.5290369 | -0.267779672 | 0.2915169 | -0.918573 | 0.358318805 | 0.551256655 | CDKN3 | protein_coding | 40.35 | -0.069260182 | 0.069260182 | ENSG00000100526 | n.s. |
| ENSG00000197965 | 1446.664183 | -0.247393491 | 0.2554993 | -0.968275 | 0.332907228 | 0.525220094 | MPZL1 | protein_coding | 41.24 | -0.069185734 | 0.069185734 | ENSG00000197965 | n.s. |
| ENSG00000198689 | 77.65928424 | 0.252490955 | 0.2644625 | 0.954733 | 0.339712945 | 0.532313459 | SLC9A6 | protein_coding | 40.97 | 0.069140243 | 0.069140243 | ENSG00000198689 | n.s. |
| ENSG00000112282 | 1903.724393 | -0.247830996 | 0.2564566 | -0.966366 | 0.33386096 | 0.526226982 | MED23 | protein_coding | 36.17 | -0.069101945 | 0.069101945 | ENSG00000112282 | n.s. |
| ENSG00000283928 | 4.646612327 | -0.378595208 | 0.5278153 | -0.717287 | 0.473196841 | 0.65702243 | MIR637 | miRNA | 61.62 | -0.069063264 | 0.069063264 | ENSG00000283928 | n.s. |
| ENSG00000023287 | 1918.247024 | -0.260190005 | 0.2783578 | -0.934732 | 0.34992625 | 0.542904299 | RB1CC1 | protein_coding | 37.73 | -0.069022351 | 0.069022351 | ENSG00000023287 | n.s. |
| ENSG00000131143 | 2162.940015 | 0.271068905 | 0.29872 | 0.907435 | 0.364176967 | 0.55645697 | COX4I1 | protein_coding | 50.84 | 0.069005581 | 0.069005581 | ENSG00000131143 | n.s. |

| | | | | | | | | | | | | | |
|---|---|---|---|---|---|---|---|---|---|---|---|---|---|
| ENSG00000099341 | 1546.276737 | 0.280689534 | 0.3168171 | 0.885967 | 0.375635312 | 0.567796163 | PSMD8 | protein_coding | 53.6 | 0.068995606 | 0.068995606 | ENSG00000099341 | n.s. |
| ENSG00000223745 | 378.0744692 | -0.248918503 | 0.2587147 | -0.962135 | 0.335981717 | 0.528274811 | CCDC18-AS1 | processed_transcript | 39.07 | -0.068985298 | 0.068985298 | ENSG00000223745 | n.s. |
| ENSG00000085978 | 402.9259225 | 0.242268222 | 0.2474055 | 0.979235 | 0.327463719 | 0.519279881 | ATG16L1 | protein_coding | 45.33 | 0.068949173 | 0.068949173 | ENSG00000085978 | n.s. |
| ENSG00000258748 | 25.29770281 | -0.285613367 | 0.3260452 | -0.875993 | 0.381033783 | 0.573651964 | | antisense | 65.82 | -0.068933219 | 0.068933219 | ENSG00000258748 | n.s. |
| ENSG00000099817 | 872.9417213 | 0.264683289 | 0.2868009 | 0.922882 | 0.356068802 | 0.549125877 | POLR2E | protein_coding | 61.47 | 0.068904495 | 0.068904495 | ENSG00000099817 | n.s. |
| ENSG00000160948 | 285.4256231 | 0.269092297 | 0.2953023 | 0.911243 | 0.362167159 | 0.554599493 | VPS28 | protein_coding | 63.23 | 0.068893153 | 0.068893153 | ENSG00000160948 | n.s. |
| ENSG00000136104 | 4587.946993 | -0.261135897 | 0.2804733 | -0.931054 | 0.351825437 | 0.544879372 | RNASEH2B | protein_coding | 38.33 | -0.06886144 | 0.06886144 | ENSG00000136104 | n.s. |
| ENSG00000143933 | 16341.25954 | 0.263215097 | 0.284355 | 0.925657 | 0.35462441 | 0.547508945 | CALM2 | protein_coding | 39.71 | 0.068859381 | 0.068859381 | ENSG00000143933 | n.s. |
| ENSG00000104886 | 413.6362778 | -0.267182642 | 0.2916866 | -0.915992 | 0.359671076 | 0.552848354 | PLEKHJ1 | protein_coding | 61.63 | -0.068771203 | 0.068771203 | ENSG00000104886 | n.s. |
| ENSG00000132507 | 3611.258447 | 0.259859676 | 0.2784395 | 0.933272 | 0.350679689 | 0.543709723 | EIF5A | protein_coding | 55.3 | 0.06876742 | 0.06876742 | ENSG00000132507 | n.s. |
| ENSG00000073803 | 469.4485011 | 0.27161866 | 0.3003626 | 0.904303 | 0.365835021 | 0.55825526 | MAP3K13 | protein_coding | 40.07 | 0.068764929 | 0.068764929 | ENSG00000073803 | n.s. |
| ENSG00000135924 | 183.884024 | -0.25607597 | 0.2719195 | -0.941734 | 0.346328682 | 0.539426585 | DNAJB2 | protein_coding | 56.92 | -0.068645684 | 0.068645684 | ENSG00000135924 | n.s. |
| ENSG00000197702 | 302.8890302 | -0.244314348 | 0.2517542 | -0.970448 | 0.331823172 | 0.523747197 | PARVA | protein_coding | 43.51 | -0.068622596 | 0.068622596 | ENSG00000197702 | n.s. |
| ENSG00000241120 | 49.95265635 | -0.29061468 | 0.337053 | -0.862223 | 0.388565022 | 0.580782695 | HMGN1P8 | processed_pseudogene | 43.06 | -0.068581093 | 0.068581093 | ENSG00000241120 | n.s. |
| ENSG00000177728 | 62.38428758 | 0.248402586 | 0.2590203 | 0.959008 | 0.33755468 | 0.529790757 | TMEM94 | protein_coding | 51.96 | 0.068533186 | 0.068533186 | ENSG00000177728 | n.s. |
| ENSG00000104756 | 441.8015069 | -0.254571106 | 0.2698053 | -0.943536 | 0.345406646 | 0.538351811 | KCTD9 | protein_coding | 37.3 | -0.068462781 | 0.068462781 | ENSG00000104756 | n.s. |
| ENSG00000169710 | 4859.121357 | 0.224814904 | 0.2198774 | 1.022456 | 0.306565251 | 0.496308404 | FASN | protein_coding | 66.65 | 0.068399568 | 0.068399568 | ENSG00000169710 | n.s. |
| ENSG00000070081 | 297.5590805 | -0.265864353 | 0.2908809 | -0.913997 | 0.360718278 | 0.55323646 | NUCB2 | protein_coding | 41.44 | -0.068350855 | 0.068350855 | ENSG00000070081 | n.s. |
| ENSG00000256683 | 355.1775007 | -0.241353672 | 0.2474445 | -0.975385 | 0.329369337 | 0.52111469 | ZNF350 | protein_coding | 42.15 | -0.068319184 | 0.068319184 | ENSG00000256683 | n.s. |
| ENSG00000011009 | 194.5650188 | 0.253661841 | 0.2686017 | 0.944379 | 0.34497599 | 0.537861222 | LYPLA2 | protein_coding | 62.39 | 0.068318684 | 0.068318684 | ENSG00000011009 | n.s. |
| ENSG00000205250 | 1063.77888 | 0.242568395 | 0.2497074 | 0.971411 | 0.331343804 | 0.523227831 | E2F4 | protein_coding | 56.55 | 0.068236712 | 0.068236712 | ENSG00000205250 | n.s. |
| ENSG00000243449 | 130.0251622 | 0.272127388 | 0.3033219 | 0.897157 | 0.369635116 | 0.561771595 | C4orf48 | protein_coding | 69.21 | 0.068151644 | 0.068151644 | ENSG00000243449 | n.s. |
| ENSG00000104805 | 337.664852 | 0.266869557 | 0.2935041 | 0.909253 | 0.363216478 | 0.555721211 | NUCB1 | protein_coding | 52.81 | 0.068089907 | 0.068089907 | ENSG00000104805 | n.s. |
| ENSG00000274578 | 122.6539996 | -0.253123157 | 0.2684762 | -0.942814 | 0.345776026 | 0.538686316 | | sense_intronic | 32.7 | -0.068005094 | 0.068005094 | ENSG00000274578 | n.s. |
| ENSG00000111728 | 166.4323986 | 0.250763101 | 0.2642984 | 0.948788 | 0.342728569 | 0.535572727 | ST8SIA1 | protein_coding | 38.38 | 0.068002326 | 0.068002326 | ENSG00000111728 | n.s. |
| ENSG00000100151 | 54.0795216 | -0.26120825 | 0.2835843 | -0.921096 | 0.357000467 | 0.549957678 | PICK1 | protein_coding | 54.9 | -0.067828137 | 0.067828137 | ENSG00000100151 | n.s. |
| ENSG00000104522 | 332.2997701 | 0.26458159 | 0.2900281 | 0.912262 | 0.3616309 | 0.554208809 | TSTA3 | protein_coding | 65.11 | 0.067819293 | 0.067819293 | ENSG00000104522 | n.s. |
| ENSG00000167654 | 737.5587308 | -0.252927646 | 0.2686072 | -0.941626 | 0.34638397 | 0.539452351 | ATCAY | protein_coding | 51.6 | -0.067796474 | 0.067796474 | ENSG00000167654 | n.s. |
| ENSG00000137274 | 287.0591074 | -0.256607957 | 0.2753541 | -0.93192 | 0.351377923 | 0.544367957 | BPHL | protein_coding | 47.11 | -0.067772072 | 0.067772072 | ENSG00000137274 | n.s. |
| ENSG00000263828 | 5.551453066 | -0.374643744 | 0.5260306 | -0.712209 | 0.476335322 | 0.659377281 | MIR4439 | miRNA | 42.5 | -0.067760323 | 0.067760323 | ENSG00000263828 | n.s. |
| ENSG00000161800 | 2032.433984 | -0.267920571 | 0.2967777 | -0.902765 | 0.366650619 | 0.558826116 | RACGAP1 | protein_coding | 43.65 | -0.067709772 | 0.067709772 | ENSG00000161800 | n.s. |
| ENSG00000182149 | 1379.476746 | 0.259654083 | 0.2815305 | 0.922295 | 0.356374826 | 0.549297763 | IST1 | protein_coding | 45.18 | 0.067559959 | 0.067559959 | ENSG00000182149 | n.s. |

| | | | | | | | | | | | | | |
|---|---|---|---|---|---|---|---|---|---|---|---|---|---|
| ENSG00000141480 | 460.4803362 | 0.24972257 | 0.2638329 | 0.946518 | 0.343884475 | 0.53670032 | ARRB2 | protein_coding | 56.76 | 0.067492056 | 0.067492056 | ENSG00000141480 | n.s. |
| ENSG00000171262 | 1074.1261 | -0.247157257 | 0.259578 | -0.95215 | 0.341020837 | 0.533349039 | FAM98B | protein_coding | 36.34 | -0.067471085 | 0.067471085 | ENSG00000171262 | n.s. |
| ENSG00000198162 | 2935.912841 | -0.253742142 | 0.2710994 | -0.935975 | 0.349286315 | 0.542394978 | MAN1A2 | protein_coding | 35.34 | -0.067415314 | 0.067415314 | ENSG00000198162 | n.s. |
| ENSG00000159461 | 462.1718782 | 0.261380426 | 0.285048 | 0.91697 | 0.359158493 | 0.552243368 | AMFR | protein_coding | 42.2 | 0.067402037 | 0.067402037 | ENSG00000159461 | n.s. |
| ENSG00000143190 | 323.9937621 | -0.224537344 | 0.221638 | -1.013081 | 0.311021271 | 0.501023853 | POU2F1 | protein_coding | 37.96 | -0.067392997 | 0.067392997 | ENSG00000143190 | n.s. |
| ENSG00000253729 | 10908.29906 | -0.239822297 | 0.247076 | -0.970642 | 0.331726622 | 0.523654168 | PRKDC | protein_coding | 41.93 | -0.067379378 | 0.067379378 | ENSG00000253729 | n.s. |
| ENSG00000101974 | 1807.711827 | -0.252776847 | 0.2695458 | -0.937788 | 0.348353335 | 0.541308424 | ATP11C | protein_coding | 39.51 | -0.067378987 | 0.067378987 | ENSG00000101974 | n.s. |
| ENSG00000279632 | 130.1816071 | -0.238329303 | 0.2446442 | -0.974188 | 0.32996342 | 0.521758172 | | TEC | 42.26 | -0.067335354 | 0.067335354 | ENSG00000279632 | n.s. |
| ENSG00000101210 | 793.2045981 | 0.253519762 | 0.2709658 | 0.935615 | 0.349471332 | 0.542517356 | EEF1A2 | protein_coding | 63.5 | 0.067331392 | 0.067331392 | ENSG00000101210 | n.s. |
| ENSG00000204946 | 717.2438605 | -0.240943933 | 0.2493286 | -0.966371 | 0.333858632 | 0.526226982 | ZNF783 | protein_coding | 51.5 | -0.067181647 | 0.067181647 | ENSG00000204946 | n.s. |
| ENSG00000157557 | 1443.120984 | -0.242425758 | 0.2520737 | -0.961726 | 0.336187446 | 0.52837731 | ETS2 | protein_coding | 46.04 | -0.067165472 | 0.067165472 | ENSG00000157557 | n.s. |
| ENSG00000101220 | 465.5955652 | 0.25237444 | 0.2694313 | 0.936693 | 0.348916436 | 0.541941492 | C20orf27 | protein_coding | 56.32 | 0.067143613 | 0.067143613 | ENSG00000101220 | n.s. |
| ENSG00000180834 | 87.94784691 | 0.263329198 | 0.2898232 | 0.908585 | 0.363568968 | 0.555955052 | MAP6D1 | protein_coding | 54.58 | 0.067138496 | 0.067138496 | ENSG00000180834 | n.s. |
| ENSG00000164674 | 148.7617704 | -0.255572092 | 0.2751649 | -0.928796 | 0.352994692 | 0.546143472 | SYTL3 | protein_coding | 46.02 | -0.067137064 | 0.067137064 | ENSG00000164674 | n.s. |
| ENSG00000196365 | 1430.218994 | -0.239589961 | 0.2472473 | -0.96903 | 0.332530403 | 0.524744508 | LONP1 | protein_coding | 57.82 | -0.067097671 | 0.067097671 | ENSG00000196365 | n.s. |
| ENSG00000164306 | 138.6480158 | -0.244292287 | 0.2554221 | -0.956426 | 0.33885714 | 0.531415629 | PRIMPOL | protein_coding | 41.31 | -0.067074277 | 0.067074277 | ENSG00000164306 | n.s. |
| ENSG00000188191 | 358.5869031 | -0.26046877 | 0.285015 | -0.913877 | 0.360781245 | 0.553254508 | PRKAR1B | protein_coding | 54.13 | -0.066960019 | 0.066960019 | ENSG00000188191 | n.s. |
| ENSG00000139668 | 2474.919848 | 0.221683295 | 0.2181407 | 1.01624 | 0.309515239 | 0.499530031 | WDFY2 | protein_coding | 39.68 | 0.066823857 | 0.066823857 | ENSG00000139668 | n.s. |
| ENSG00000107719 | 317.3677746 | -0.253367945 | 0.2721078 | -0.931131 | 0.351785901 | 0.544878751 | PALD1 | protein_coding | 52.2 | -0.06681316 | 0.06681316 | ENSG00000107719 | n.s. |
| ENSG00000238446 | 6.690066024 | 0.363823371 | 0.5049803 | 0.72047 | 0.471235428 | 0.655380168 | RNU7-38P | snRNA | 38.33 | 0.066764028 | 0.066764028 | ENSG00000238446 | n.s. |
| ENSG00000152240 | 759.8934356 | -0.2685323 | 0.3008448 | -0.892594 | 0.372074494 | 0.564169435 | HAUS1 | protein_coding | 41.32 | -0.066754564 | 0.066754564 | ENSG00000152240 | n.s. |
| ENSG00000182087 | 1239.492752 | 0.244311926 | 0.2562803 | 0.9533 | 0.340438225 | 0.533072468 | TMEM259 | protein_coding | 65.7 | 0.066749377 | 0.066749377 | ENSG00000182087 | n.s. |
| ENSG00000244112 | 58.47883056 | -0.257119721 | 0.2794749 | -0.92001 | 0.357567513 | 0.550417975 | RN7SL508P | misc_RNA | 53.47 | -0.066673044 | 0.066673044 | ENSG00000244112 | n.s. |
| ENSG00000170166 | 56.94528257 | -0.275335288 | 0.3145787 | -0.875251 | 0.381437409 | 0.574135464 | HOXD4 | protein_coding | 59.55 | -0.066351847 | 0.066351847 | ENSG00000170166 | n.s. |
| ENSG00000198356 | 573.8877029 | 0.278469229 | 0.3203958 | 0.869141 | 0.384769788 | 0.57776258 | ASNA1 | protein_coding | 53.33 | 0.066345458 | 0.066345458 | ENSG00000198356 | n.s. |
| ENSG00000260865 | 13.73756981 | -0.308970234 | 0.3828444 | -0.807039 | 0.419644222 | 0.610424303 | | transcribed_processed_pseudog | 52.1 | -0.066233388 | 0.066233388 | ENSG00000260865 | n.s. |
| ENSG00000130511 | 570.8299835 | 0.25474731 | 0.2765492 | 0.921164 | 0.356964561 | 0.549957678 | SSBP4 | protein_coding | 64.53 | 0.06615042 | 0.06615042 | ENSG00000130511 | n.s. |
| ENSG00000166557 | 939.8353218 | 0.262252129 | 0.2908877 | 0.901558 | 0.367291692 | 0.559496962 | TMED3 | protein_coding | 42.58 | 0.066140582 | 0.066140582 | ENSG00000166557 | n.s. |
| ENSG00000121766 | 292.5417406 | -0.272173069 | 0.3095341 | -0.879299 | 0.379239052 | 0.571691785 | ZCCHC17 | protein_coding | 41.86 | -0.066093977 | 0.066093977 | ENSG00000121766 | n.s. |
| ENSG00000010072 | 812.4651365 | -0.246593016 | 0.2623315 | -0.940005 | 0.347214752 | 0.540081666 | SPRTN | protein_coding | 41.57 | -0.065973635 | 0.065973635 | ENSG00000010072 | n.s. |
| ENSG00000075618 | 1760.350049 | 0.227021744 | 0.2289264 | 0.99168 | 0.321353526 | 0.512148578 | FSCN1 | protein_coding | 59.62 | 0.065973434 | 0.065973434 | ENSG00000075618 | n.s. |
| ENSG00000276291 | 445.0795689 | 0.236892543 | 0.2454879 | 0.964986 | 0.3345516 | 0.526978383 | FRG1HP | transcribed_unprocessed_pseud | 39.23 | 0.065905211 | 0.065905211 | ENSG00000276291 | n.s. |

| | | | | | | | | | | | | | |
|---|---|---|---|---|---|---|---|---|---|---|---|---|---|
| ENSG00000078967 | 36.76127978 | -0.262844756 | 0.2928967 | -0.897397 | 0.369506898 | 0.561703738 | UBE2D4 | protein_coding | 45.79 | -0.065840689 | 0.065840689 | ENSG00000078967 | n.s. |
| ENSG00000067248 | 1353.417862 | -0.257118332 | 0.28214 | -0.911315 | 0.362129536 | 0.554599493 | DHX29 | protein_coding | 37.08 | -0.065827572 | 0.065827572 | ENSG00000067248 | n.s. |
| ENSG00000105767 | 127.338225 | -0.25552734 | 0.2789035 | -0.916186 | 0.359569587 | 0.552814416 | CADM4 | protein_coding | 53.28 | -0.065778011 | 0.065778011 | ENSG00000105767 | n.s. |
| ENSG00000272145 | 348.6986377 | -0.212749408 | 0.2057603 | -1.033967 | 0.301151459 | 0.490803202 | NFYC-AS1 | antisense | 49.61 | -0.06575927 | 0.06575927 | ENSG00000272145 | n.s. |
| ENSG00000239953 | 13.00517859 | -0.308416465 | 0.3839934 | -0.803182 | 0.421869743 | 0.612098002 | RN7SL273P | misc_RNA | 52.84 | -0.065747925 | 0.065747925 | ENSG00000239953 | n.s. |
| ENSG00000114062 | 4258.516378 | -0.255440424 | 0.2793965 | -0.914258 | 0.360581385 | 0.553148371 | UBE3A | protein_coding | 36.21 | -0.065688641 | 0.065688641 | ENSG00000114062 | n.s. |
| ENSG00000064225 | 76.9655694 | -0.250117053 | 0.2696392 | -0.927599 | 0.35361564 | 0.546557566 | ST3GAL6 | protein_coding | 38.88 | -0.065621733 | 0.065621733 | ENSG00000064225 | n.s. |
| ENSG00000233327 | 334.5032754 | -0.265910986 | 0.2993633 | -0.888255 | 0.37440351 | 0.56654308 | USP32P2 | transcribed_unprocessed_pseud | 42.09 | -0.065618072 | 0.065618072 | ENSG00000233327 | n.s. |
| ENSG00000185896 | 898.9197694 | 0.244823813 | 0.2602424 | 0.940753 | 0.346831501 | 0.539921472 | LAMP1 | protein_coding | 50.64 | 0.065531843 | 0.065531843 | ENSG00000185896 | n.s. |
| ENSG00000213347 | 353.659753 | 0.244289624 | 0.2598216 | 0.94022 | 0.347104492 | 0.539970485 | MXD3 | protein_coding | 58.63 | 0.065379227 | 0.065379227 | ENSG00000213347 | n.s. |
| ENSG00000115392 | 1479.995462 | -0.253900183 | 0.2772497 | -0.915782 | 0.359781488 | 0.552936594 | FANCL | protein_coding | 34.76 | -0.06533478 | 0.06533478 | ENSG00000115392 | n.s. |
| ENSG00000104381 | 2132.058778 | -0.249519016 | 0.2693714 | -0.926301 | 0.354289493 | 0.547295309 | GDAP1 | protein_coding | 38 | -0.065318657 | 0.065318657 | ENSG00000104381 | n.s. |
| ENSG00000106460 | 1152.589698 | -0.253829337 | 0.2773099 | -0.915327 | 0.360019866 | 0.552936594 | TMEM106B | protein_coding | 34.61 | -0.06531655 | 0.06531655 | ENSG00000106460 | n.s. |
| ENSG00000185917 | 495.9645422 | -0.226000081 | 0.2288969 | -0.987344 | 0.323473851 | 0.514065338 | SETD4 | protein_coding | 44.84 | -0.065309883 | 0.065309883 | ENSG00000185917 | n.s. |
| ENSG00000157823 | 220.5150029 | 0.229025903 | 0.2338072 | 0.97955 | 0.327308228 | 0.519151568 | AP3S2 | protein_coding | 44.42 | 0.06520501 | 0.06520501 | ENSG00000157823 | n.s. |
| ENSG00000140153 | 91.74834339 | -0.264186561 | 0.2973976 | -0.888328 | 0.374364467 | 0.56654308 | WDR20 | protein_coding | 45.46 | -0.065192541 | 0.065192541 | ENSG00000140153 | n.s. |
| ENSG00000162738 | 420.0162268 | -0.219992888 | 0.218893 | -1.005025 | 0.314884904 | 0.5054932 | VANGL2 | protein_coding | 51.65 | -0.065180523 | 0.065180523 | ENSG00000162738 | n.s. |
| ENSG00000137409 | 1662.437703 | 0.257793268 | 0.2854807 | 0.903015 | 0.36651811 | 0.558746484 | MTCH1 | protein_coding | 52.18 | 0.065166322 | 0.065166322 | ENSG00000137409 | n.s. |
| ENSG00000170876 | 977.9996477 | -0.255706534 | 0.2815466 | -0.908221 | 0.363761448 | 0.556127226 | TMEM43 | protein_coding | 52.73 | -0.065160633 | 0.065160633 | ENSG00000170876 | n.s. |
| ENSG00000196268 | 1246.567195 | -0.228758029 | 0.233876 | -0.978117 | 0.328016616 | 0.519683129 | ZNF493 | protein_coding | 38.43 | -0.065027073 | 0.065027073 | ENSG00000196268 | n.s. |
| ENSG00000112078 | 2324.648524 | -0.256119067 | 0.2826687 | -0.906075 | 0.364896171 | 0.557372396 | KCTD20 | protein_coding | 42.39 | -0.065016989 | 0.065016989 | ENSG00000112078 | n.s. |
| ENSG00000136379 | 234.1700355 | -0.237057143 | 0.2481153 | -0.955431 | 0.339359552 | 0.532143603 | ABHD17C | protein_coding | 42.95 | -0.064946819 | 0.064946819 | ENSG00000136379 | n.s. |
| ENSG00000109775 | 459.6390232 | -0.275239766 | 0.319325 | -0.861942 | 0.388719206 | 0.580950784 | UFSP2 | protein_coding | 38.42 | -0.064918232 | 0.064918232 | ENSG00000109775 | n.s. |
| ENSG00000178338 | 503.1549453 | -0.25499757 | 0.2810046 | -0.90745 | 0.364168926 | 0.55645697 | ZNF354B | protein_coding | 44.33 | -0.064914327 | 0.064914327 | ENSG00000178338 | n.s. |
| ENSG00000108018 | 1153.418922 | -0.247008735 | 0.2662123 | -0.927864 | 0.353478179 | 0.546405761 | SORCS1 | protein_coding | 40.11 | -0.064836021 | 0.064836021 | ENSG00000108018 | n.s. |
| ENSG00000275901 | 10.83445744 | 0.31669625 | 0.405263 | 0.781459 | 0.43453283 | 0.624293171 | | misc_RNA | 60.63 | 0.064799668 | 0.064799668 | ENSG00000275901 | n.s. |
| ENSG00000207721 | 83.34343497 | -0.259275904 | 0.2895396 | -0.895476 | 0.370532505 | 0.562587139 | MIR186 | miRNA | 36.05 | -0.064769765 | 0.064769765 | ENSG00000207721 | n.s. |
| ENSG00000270006 | 51.01723268 | 0.255279321 | 0.2821739 | 0.904688 | 0.365630672 | 0.558126943 | | antisense | 59.44 | 0.064653831 | 0.064653831 | ENSG00000270006 | n.s. |
| ENSG00000207631 | 9.975402749 | 0.321106335 | 0.4167366 | 0.770526 | 0.440987992 | 0.629085636 | MIR641 | miRNA | 47.47 | 0.064635568 | 0.064635568 | ENSG00000207631 | n.s. |
| ENSG00000146842 | 597.8873101 | -0.220204245 | 0.2207434 | -0.997558 | 0.318493875 | 0.508906432 | TMEM209 | protein_coding | 37.56 | -0.064599571 | 0.064599571 | ENSG00000146842 | n.s. |
| ENSG00000207525 | 5.742565515 | -0.35276413 | 0.4905775 | -0.719079 | 0.472092017 | 0.656094752 | | misc_RNA | 40.59 | -0.064567631 | 0.064567631 | ENSG00000207525 | n.s. |
| ENSG00000159348 | 394.8278373 | -0.247833645 | 0.268452 | -0.923195 | 0.355905405 | 0.549076588 | CYB5R1 | protein_coding | 50.33 | -0.064527721 | 0.064527721 | ENSG00000159348 | n.s. |

| | | | | | | | | | | | | | |
|---|---|---|---|---|---|---|---|---|---|---|---|---|---|
| ENSG00000035928 | 3180.177127 | -0.239618966 | 0.2537685 | -0.944242 | 0.3450458 | 0.537909828 | RFC1 | protein_coding | 39.26 | -0.064527116 | 0.064527116 | ENSG00000035928 | n.s. |
| ENSG00000136908 | 280.695023 | -0.250756389 | 0.2742097 | -0.914469 | 0.360470257 | 0.553038831 | DPM2 | protein_coding | 56.79 | -0.06450567 | 0.06450567 | ENSG00000136908 | n.s. |
| ENSG00000205155 | 248.2279428 | 0.265248078 | 0.3013523 | 0.880193 | 0.378754983 | 0.57128353 | PSENEN | protein_coding | 52 | 0.064494618 | 0.064494618 | ENSG00000205155 | n.s. |
| ENSG00000143222 | 1057.347423 | 0.253429994 | 0.279362 | 0.907174 | 0.364314651 | 0.556606264 | UFC1 | protein_coding | 46.11 | 0.064485746 | 0.064485746 | ENSG00000143222 | n.s. |
| ENSG00000266538 | 19.29532094 | -0.293705515 | 0.3577683 | -0.820938 | 0.411681664 | 0.603733284 | | lincRNA | 50.16 | -0.064366997 | 0.064366997 | ENSG00000266538 | n.s. |
| ENSG00000198899 | 1859.593089 | -0.219101851 | 0.2194318 | -0.998496 | 0.318038728 | 0.508445654 | MT-ATP6 | protein_coding | 44.2 | -0.064362365 | 0.064362365 | ENSG00000198899 | n.s. |
| ENSG00000160932 | 987.5663957 | 0.251539954 | 0.2762978 | 0.910394 | 0.362614639 | 0.554983359 | LY6E | protein_coding | 63.58 | 0.064323807 | 0.064323807 | ENSG00000160932 | n.s. |
| ENSG00000273345 | 71.56955755 | 0.243201992 | 0.2607626 | 0.932657 | 0.350997153 | 0.544080757 | | processed_transcript | 41.3 | 0.064287196 | 0.064287196 | ENSG00000273345 | n.s. |
| ENSG00000122642 | 558.8939682 | -0.24201447 | 0.2586961 | -0.935516 | 0.349522244 | 0.542519309 | FKBP9 | protein_coding | 45.26 | -0.064275365 | 0.064275365 | ENSG00000122642 | n.s. |
| ENSG00000227379 | 25.99648672 | -0.269526402 | 0.3099801 | -0.869496 | 0.384576045 | 0.57760033 | PPIAP2 | processed_pseudogene | 47.97 | -0.064247701 | 0.064247701 | ENSG00000227379 | n.s. |
| ENSG00000205707 | 485.1149741 | 0.237218869 | 0.2505769 | 0.946691 | 0.343796302 | 0.53670032 | ETFRF1 | protein_coding | 38.05 | 0.064112704 | 0.064112704 | ENSG00000205707 | n.s. |
| ENSG00000179271 | 1493.692447 | 0.259257471 | 0.291484 | 0.88944 | 0.373766829 | 0.565953476 | GADD45GIP1 | protein_coding | 55.03 | 0.064093442 | 0.064093442 | ENSG00000179271 | n.s. |
| ENSG00000149212 | 1070.655128 | 0.263562952 | 0.2994143 | 0.880262 | 0.378717523 | 0.57128353 | SESN3 | protein_coding | 35.27 | 0.064084883 | 0.064084883 | ENSG00000149212 | n.s. |
| ENSG00000067365 | 429.483149 | -0.230737322 | 0.2394962 | -0.963428 | 0.335332831 | 0.527612047 | METTL22 | protein_coding | 49.56 | -0.064072362 | 0.064072362 | ENSG00000067365 | n.s. |
| ENSG00000100027 | 289.5968029 | 0.246093836 | 0.2667379 | 0.922606 | 0.356212779 | 0.549182172 | YPEL1 | protein_coding | 49.29 | 0.064054182 | 0.064054182 | ENSG00000100027 | n.s. |
| ENSG00000182670 | 7901.750377 | -0.241123954 | 0.2578333 | -0.935193 | 0.349688818 | 0.542670676 | TTC3 | protein_coding | 38.7 | -0.064009644 | 0.064009644 | ENSG00000182670 | n.s. |
| ENSG00000248643 | 163.0069539 | 0.22757822 | 0.2343187 | 0.971234 | 0.33143192 | 0.523307623 | RBM14-RBM4 | protein_coding | 47.43 | 0.064004767 | 0.064004767 | ENSG00000248643 | n.s. |
| ENSG00000120885 | 263.9774707 | 0.263279768 | 0.2992928 | 0.879673 | 0.379036485 | 0.571455166 | CLU | protein_coding | 49.9 | 0.06398168 | 0.06398168 | ENSG00000120885 | n.s. |
| ENSG00000125630 | 1802.049567 | -0.236101392 | 0.2490438 | -0.948032 | 0.343113343 | 0.535917329 | POLR1B | protein_coding | 42.17 | -0.063960387 | 0.063960387 | ENSG00000125630 | n.s. |
| ENSG00000261716 | 450.6375381 | 0.209639557 | 0.2045136 | 1.025064 | 0.305332963 | 0.495356659 | | sense_overlapping | 38.15 | 0.063957254 | 0.063957254 | ENSG00000261716 | n.s. |
| ENSG00000180228 | 1785.566457 | -0.24821644 | 0.2713464 | -0.914759 | 0.360318336 | 0.552936594 | PRKRA | protein_coding | 39.65 | -0.063872213 | 0.063872213 | ENSG00000180228 | n.s. |
| ENSG00000212413 | 13.52802225 | -0.298293553 | 0.3701671 | -0.805835 | 0.42033805 | 0.610830464 | RNU11-3P | snRNA | 45.86 | -0.063858479 | 0.063858479 | ENSG00000212413 | n.s. |
| ENSG00000137500 | 775.4293986 | -0.255055264 | 0.2844604 | -0.896628 | 0.369917192 | 0.561959308 | CCDC90B | protein_coding | 37.59 | -0.063839091 | 0.063839091 | ENSG00000137500 | n.s. |
| ENSG00000102898 | 1148.600696 | 0.252192369 | 0.2791199 | 0.903527 | 0.366246247 | 0.558484609 | NUTF2 | protein_coding | 51.22 | 0.063801842 | 0.063801842 | ENSG00000102898 | n.s. |
| ENSG00000188554 | 1918.698659 | -0.258536132 | 0.2910829 | -0.888187 | 0.374440094 | 0.56654308 | NBR1 | protein_coding | 44.16 | -0.063798201 | 0.063798201 | ENSG00000188554 | n.s. |
| ENSG00000263278 | 857.8798874 | 0.216456244 | 0.216355 | 1.000468 | 0.317084179 | 0.507444026 | | sense_intronic | 53.41 | 0.063770576 | 0.063770576 | ENSG00000263278 | n.s. |
| ENSG00000182247 | 321.6978008 | 0.247808349 | 0.270746 | 0.91528 | 0.360044816 | 0.552936594 | UBE2E2 | protein_coding | 37.68 | 0.063767201 | 0.063767201 | ENSG00000182247 | n.s. |
| ENSG00000135298 | 1244.721484 | -0.232750164 | 0.24373 | -0.954951 | 0.339602653 | 0.532225095 | ADGRB3 | protein_coding | 35.14 | -0.063751353 | 0.063751353 | ENSG00000135298 | n.s. |
| ENSG00000112893 | 933.0445613 | -0.228883907 | 0.237 | -0.965755 | 0.334166861 | 0.526602348 | MAN2A1 | protein_coding | 36.58 | -0.063748107 | 0.063748107 | ENSG00000112893 | n.s. |
| ENSG00000056998 | 202.2077767 | -0.251904398 | 0.2789581 | -0.903019 | 0.36651589 | 0.558746484 | GYG2 | protein_coding | 43.67 | -0.063677703 | 0.063677703 | ENSG00000056998 | n.s. |
| ENSG00000013583 | 87.27833586 | -0.270009197 | 0.3137132 | -0.860688 | 0.38940986 | 0.581366175 | HEBP1 | protein_coding | 43.52 | -0.063600731 | 0.063600731 | ENSG00000013583 | n.s. |
| ENSG00000238141 | 25.05531729 | -0.269937879 | 0.3135688 | -0.860857 | 0.389316898 | 0.581344827 | BRWD1-AS1 | antisense | 45.53 | -0.063588237 | 0.063588237 | ENSG00000238141 | n.s. |

| | | | | | | | | | | | | | |
|---|---|---|---|---|---|---|---|---|---|---|---|---|---|
| ENSG00000182534 | 1398.60233 | 0.260186449 | 0.2947101 | 0.882855 | 0.377314381 | 0.569652019 | MXRA7 | protein_coding | 52.39 | 0.06358706 | 0.06358706 | ENSG00000182534 | n.s. |
| ENSG00000214530 | 127.4451714 | 0.253407633 | 0.2824784 | 0.897087 | 0.369672581 | 0.561771595 | STARD10 | protein_coding | 54.42 | 0.063463464 | 0.063463464 | ENSG00000214530 | n.s. |
| ENSG00000235552 | 439.6078618 | 0.267697878 | 0.3092897 | 0.865525 | 0.386750925 | 0.579415258 | | transcribed_processed_pseudog | 46.79 | 0.063447093 | 0.063447093 | ENSG00000235552 | n.s. |
| ENSG00000089154 | 3304.723251 | -0.233266411 | 0.2454972 | -0.950179 | 0.342021067 | 0.534631111 | GCN1 | protein_coding | 49.67 | -0.063435814 | 0.063435814 | ENSG00000089154 | n.s. |
| ENSG00000164253 | 959.8358607 | -0.247729404 | 0.2718476 | -0.91128 | 0.362147691 | 0.554599493 | WDR41 | protein_coding | 37.31 | -0.063423814 | 0.063423814 | ENSG00000164253 | n.s. |
| ENSG00000130332 | 1175.414243 | 0.265595648 | 0.3051639 | 0.870338 | 0.384115822 | 0.577104221 | LSM7 | protein_coding | 57.71 | 0.063409832 | 0.063409832 | ENSG00000130332 | n.s. |
| ENSG00000152894 | 92.47284284 | 0.240709292 | 0.2588385 | 0.929959 | 0.35239213 | 0.545453655 | PTPRK | protein_coding | 35.41 | 0.06336483 | 0.06336483 | ENSG00000152894 | n.s. |
| ENSG00000258355 | 91.77016576 | 0.234092581 | 0.2471523 | 0.947159 | 0.343557559 | 0.536430627 | | sense_intronic | 52.08 | 0.063318867 | 0.063318867 | ENSG00000258355 | n.s. |
| ENSG00000273254 | 25.56987263 | 0.274404993 | 0.3232029 | 0.849018 | 0.39587146 | 0.587916623 | | antisense | 53.46 | 0.063300913 | 0.063300913 | ENSG00000273254 | n.s. |
| ENSG00000108960 | 3254.015333 | -0.249978604 | 0.2764029 | -0.904399 | 0.365783729 | 0.558238174 | MMD | protein_coding | 41.06 | -0.0632897 | 0.0632897 | ENSG00000108960 | n.s. |
| ENSG00000147601 | 1611.7012 | 0.236459228 | 0.2514901 | 0.940233 | 0.347098093 | 0.539970485 | TERF1 | protein_coding | 36.98 | 0.063283578 | 0.063283578 | ENSG00000147601 | n.s. |
| ENSG00000167771 | 1323.184882 | 0.229398707 | 0.2393077 | 0.958593 | 0.337763723 | 0.530007115 | RCOR2 | protein_coding | 60.76 | 0.063249422 | 0.063249422 | ENSG00000167771 | n.s. |
| ENSG00000144455 | 280.9038638 | -0.242325551 | 0.2621627 | -0.924333 | 0.355313192 | 0.548372243 | SUMF1 | protein_coding | 39.53 | -0.063228682 | 0.063228682 | ENSG00000144455 | n.s. |
| ENSG00000131148 | 390.5987502 | -0.245656517 | 0.2685734 | -0.914672 | 0.360363906 | 0.552936594 | EMC8 | protein_coding | 48.89 | -0.063213481 | 0.063213481 | ENSG00000131148 | n.s. |
| ENSG00000168036 | 10905.17791 | 0.245092274 | 0.2677399 | 0.915412 | 0.359975458 | 0.552936594 | CTNNB1 | protein_coding | 39.91 | 0.063068288 | 0.063068288 | ENSG00000168036 | n.s. |
| ENSG00000204178 | 493.9806844 | -0.252829439 | 0.2825816 | -0.894713 | 0.370940488 | 0.563083779 | TMEM57 | protein_coding | 40.69 | -0.063062484 | 0.063062484 | ENSG00000204178 | n.s. |
| ENSG00000239040 | 13.633771 | -0.30240353 | 0.3820525 | -0.791524 | 0.428638535 | 0.61875939 | | misc_RNA | 42.72 | -0.063044543 | 0.063044543 | ENSG00000239040 | n.s. |
| ENSG00000201581 | 12.2313508 | -0.29600006 | 0.3687917 | -0.802621 | 0.422193706 | 0.612376598 | RN7SKP78 | misc_RNA | 50.94 | -0.063042511 | 0.063042511 | ENSG00000201581 | n.s. |
| ENSG00000206605 | 6.513371788 | -0.34638678 | 0.4839426 | -0.71576 | 0.474139494 | 0.657665315 | RNU6-946P | snRNA | 42.99 | -0.063040683 | 0.063040683 | ENSG00000206605 | n.s. |
| ENSG00000160785 | 772.4116148 | 0.221964809 | 0.227329 | 0.976403 | 0.32886458 | 0.520526307 | SLC25A44 | protein_coding | 48.29 | 0.062939746 | 0.062939746 | ENSG00000160785 | n.s. |
| ENSG00000171700 | 103.2897548 | -0.265235666 | 0.3064969 | -0.865378 | 0.386831357 | 0.579415258 | RGS19 | protein_coding | 63.53 | -0.062863524 | 0.062863524 | ENSG00000171700 | n.s. |
| ENSG00000108953 | 5283.692942 | 0.249266686 | 0.2767191 | 0.900793 | 0.367698275 | 0.559993774 | YWHAE | protein_coding | 44.81 | 0.06276954 | 0.06276954 | ENSG00000108953 | n.s. |
| ENSG00000137185 | 904.0519662 | -0.224143068 | 0.2314435 | -0.968457 | 0.332816273 | 0.525136103 | ZSCAN9 | protein_coding | 43.42 | -0.06269912 | 0.06269912 | ENSG00000137185 | n.s. |
| ENSG00000105939 | 606.2702451 | -0.231079045 | 0.2435571 | -0.948768 | 0.342738849 | 0.535572727 | ZC3HAV1 | protein_coding | 43.4 | -0.062664373 | 0.062664373 | ENSG00000105939 | n.s. |
| ENSG00000087338 | 799.0529811 | -0.238195782 | 0.2563655 | -0.929126 | 0.352823887 | 0.546000554 | GMCL1 | protein_coding | 37.77 | -0.062599499 | 0.062599499 | ENSG00000087338 | n.s. |
| ENSG00000178999 | 979.6301682 | 0.227005293 | 0.2369102 | 0.958191 | 0.337966378 | 0.530257593 | AURKB | protein_coding | 52.69 | 0.062542933 | 0.062542933 | ENSG00000178999 | n.s. |
| ENSG00000255358 | 29.20332838 | -0.282259984 | 0.3416245 | -0.826229 | 0.408674353 | 0.600524856 | | processed_pseudogene | 44.64 | -0.06251184 | 0.06251184 | ENSG00000255358 | n.s. |
| ENSG00000224032 | 3198.189927 | 0.209152409 | 0.2069949 | 1.010423 | 0.312292725 | 0.502482874 | EPB41L4A-AS1 | lincRNA | 43.19 | 0.062511207 | 0.062511207 | ENSG00000224032 | n.s. |
| ENSG00000253218 | 51.80029321 | -0.257155452 | 0.2923335 | -0.879665 | 0.379041044 | 0.571455166 | KLF3P1 | processed_pseudogene | 48.92 | -0.062493362 | 0.062493362 | ENSG00000253218 | n.s. |
| ENSG00000276182 | 184.637823 | -0.228857433 | 0.2403009 | -0.952378 | 0.340905105 | 0.533305818 | | sense_intronic | 32.65 | -0.062483498 | 0.062483498 | ENSG00000276182 | n.s. |
| ENSG00000279527 | 72.82089904 | 0.250878252 | 0.2807203 | 0.893695 | 0.371485272 | 0.563603514 | | TEC | 37.48 | 0.062475285 | 0.062475285 | ENSG00000279527 | n.s. |
| ENSG00000175213 | 217.7901211 | 0.219911069 | 0.2250913 | 0.976986 | 0.328575989 | 0.520332517 | ZNF408 | protein_coding | 54.13 | 0.062392956 | 0.062392956 | ENSG00000175213 | n.s. |

| | | | | | | | | | | | | | |
|---|---|---|---|---|---|---|---|---|---|---|---|---|---|
| ENSG00000099901 | 3979.430879 | 0.254717013 | 0.2882059 | 0.883802 | 0.376802967 | 0.569128629 | RANBP1 | protein_coding | 57.69 | 0.062352067 | 0.062352067 | ENSG00000099901 | n.s. |
| ENSG00000221545 | 3.00059837 | -0.447774513 | 0.7669138 | -0.583866 | 0.559310778 | 0.725748738 | MIR1255B2 | miRNA | 43.28 | -0.062336351 | 0.062336351 | ENSG00000221545 | n.s. |
| ENSG00000131979 | 520.1847953 | 0.258679022 | 0.2958346 | 0.874404 | 0.381898234 | 0.574704833 | GCH1 | protein_coding | 42.22 | 0.062226571 | 0.062226571 | ENSG00000131979 | n.s. |
| ENSG00000167987 | 160.4865781 | -0.237585961 | 0.2565217 | -0.926183 | 0.354351087 | 0.54732973 | VPS37C | protein_coding | 49.88 | -0.062188353 | 0.062188353 | ENSG00000167987 | n.s. |
| ENSG00000172340 | 220.806314 | -0.261450351 | 0.3012407 | -0.867912 | 0.385442589 | 0.578350811 | SUCLG2 | protein_coding | 39.38 | -0.062175155 | 0.062175155 | ENSG00000172340 | n.s. |
| ENSG00000204685 | 117.6101252 | -0.232866723 | 0.2482114 | -0.938179 | 0.348152399 | 0.541056574 | STARD7-AS1 | processed_transcript | 46.6 | -0.062118903 | 0.062118903 | ENSG00000204685 | n.s. |
| ENSG00000100243 | 416.4275671 | -0.238585629 | 0.25849 | -0.922997 | 0.356008511 | 0.549125877 | CYB5R3 | protein_coding | 55.99 | -0.062110541 | 0.062110541 | ENSG00000100243 | n.s. |
| ENSG00000130764 | 360.2214779 | -0.241173111 | 0.2634954 | -0.915284 | 0.360042672 | 0.552936594 | LRRC47 | protein_coding | 54.71 | -0.06205979 | 0.06205979 | ENSG00000130764 | n.s. |
| ENSG00000085491 | 3057.975229 | -0.241172101 | 0.2634216 | -0.915536 | 0.359910092 | 0.552936594 | SLC25A24 | protein_coding | 39.44 | -0.062059531 | 0.062059531 | ENSG00000085491 | n.s. |
| ENSG00000140943 | 1042.932086 | -0.241081823 | 0.2635636 | -0.914701 | 0.360348727 | 0.552936594 | MBTPS1 | protein_coding | 45.14 | -0.0620363 | 0.0620363 | ENSG00000140943 | n.s. |
| ENSG00000198898 | 1744.76233 | 0.26220049 | 0.3037048 | 0.86334 | 0.38795056 | 0.580120522 | CAPZA2 | protein_coding | 38.99 | 0.062005636 | 0.062005636 | ENSG00000198898 | n.s. |
| ENSG00000196381 | 321.0551201 | -0.281471485 | 0.3420016 | -0.823012 | 0.410501111 | 0.602319208 | ZNF781 | protein_coding | 41.97 | -0.061972502 | 0.061972502 | ENSG00000196381 | n.s. |
| ENSG00000225206 | 1621.375822 | 0.226738959 | 0.2380722 | 0.952396 | 0.340896398 | 0.533305818 | MIR137HG | lincRNA | 34.52 | 0.061905105 | 0.061905105 | ENSG00000225206 | n.s. |
| ENSG00000183624 | 410.2717997 | -0.242694675 | 0.267091 | -0.908659 | 0.363530112 | 0.555955052 | HMCES | protein_coding | 44.53 | -0.061877512 | 0.061877512 | ENSG00000183624 | n.s. |
| ENSG00000150551 | 497.0633112 | -0.248505219 | 0.2781232 | -0.893508 | 0.371585407 | 0.563668197 | LYPD1 | protein_coding | 45.33 | -0.061871952 | 0.061871952 | ENSG00000150551 | n.s. |
| ENSG00000130600 | 441.9230936 | 0.234449326 | 0.2517891 | 0.931134 | 0.351784378 | 0.544878751 | H19 | processed_transcript | 62.11 | 0.061824317 | 0.061824317 | ENSG00000130600 | n.s. |
| ENSG00000182985 | 851.7022258 | -0.249418577 | 0.2800073 | -0.890758 | 0.373059269 | 0.565190882 | CADM1 | protein_coding | 38.54 | -0.061807134 | 0.061807134 | ENSG00000182985 | n.s. |
| ENSG00000070423 | 393.3151942 | 0.247796032 | 0.2771734 | 0.894011 | 0.371316135 | 0.563475136 | RNF126 | protein_coding | 63.55 | 0.061732247 | 0.061732247 | ENSG00000070423 | n.s. |
| ENSG00000100116 | 373.7192451 | 0.252300402 | 0.2856103 | 0.883373 | 0.377034897 | 0.569417168 | GCAT | protein_coding | 54.21 | 0.061704969 | 0.061704969 | ENSG00000100116 | n.s. |
| ENSG00000143761 | 3039.771555 | 0.250068023 | 0.2815972 | 0.888034 | 0.374522241 | 0.566605798 | ARF1 | protein_coding | 52.33 | 0.061696529 | 0.061696529 | ENSG00000143761 | n.s. |
| ENSG00000133104 | 1645.763249 | -0.252162817 | 0.2856651 | -0.882722 | 0.377386601 | 0.569652019 | SPG20 | protein_coding | 39.04 | -0.061626161 | 0.061626161 | ENSG00000133104 | n.s. |
| ENSG00000137076 | 1248.946807 | -0.232034851 | 0.2480069 | -0.935598 | 0.349480077 | 0.542517356 | TLN1 | protein_coding | 51.19 | -0.061625292 | 0.061625292 | ENSG00000137076 | n.s. |
| ENSG00000182944 | 12000.51645 | 0.222261763 | 0.2309848 | 0.962235 | 0.3359314 | 0.528255351 | EWSR1 | protein_coding | 44.02 | 0.061601202 | 0.061601202 | ENSG00000182944 | n.s. |
| ENSG00000283023 | 139.4229533 | 0.246073304 | 0.2744326 | 0.896662 | 0.369899282 | 0.561959308 | | unprocessed_pseudogene | 35.84 | 0.06159095 | 0.06159095 | ENSG00000283023 | n.s. |
| ENSG00000152465 | 464.9425822 | 0.241469209 | 0.2656946 | 0.908822 | 0.363443852 | 0.555955052 | NMT2 | protein_coding | 43.09 | 0.061565067 | 0.061565067 | ENSG00000152465 | n.s. |
| ENSG00000106462 | 2758.393272 | 0.230510618 | 0.2455991 | 0.938565 | 0.347954333 | 0.540837078 | EZH2 | protein_coding | 39.21 | 0.061531016 | 0.061531016 | ENSG00000106462 | n.s. |
| ENSG00000164808 | 338.4458336 | -0.228661609 | 0.2430719 | -0.940716 | 0.346850531 | 0.539921472 | SPIDR | protein_coding | 40.78 | -0.061205716 | 0.061205716 | ENSG00000164808 | n.s. |
| ENSG00000156587 | 214.7192102 | 0.258137401 | 0.2982148 | 0.865609 | 0.386704597 | 0.579415258 | UBE2L6 | protein_coding | 48.66 | 0.061181164 | 0.061181164 | ENSG00000156587 | n.s. |
| ENSG00000017260 | 2643.530239 | -0.235806274 | 0.2562453 | -0.920237 | 0.357449101 | 0.550344337 | ATP2C1 | protein_coding | 37.56 | -0.061160012 | 0.061160012 | ENSG00000017260 | n.s. |
| ENSG00000124702 | 1217.206541 | 0.266607139 | 0.3153308 | 0.845484 | 0.397840716 | 0.589682587 | KLHDC3 | protein_coding | 52.39 | 0.061154798 | 0.061154798 | ENSG00000124702 | n.s. |
| ENSG00000198763 | 1162.189879 | -0.207569177 | 0.2074397 | -1.000624 | 0.317008487 | 0.507444026 | MT-ND2 | protein_coding | 42.99 | -0.06115234 | 0.06115234 | ENSG00000198763 | n.s. |
| ENSG00000164151 | 2999.085255 | -0.231317451 | 0.2480626 | -0.932496 | 0.351080094 | 0.544082698 | ICE1 | protein_coding | 38.9 | -0.061145318 | 0.061145318 | ENSG00000164151 | n.s. |

| | | | | | | | | | | | | | |
|---|---|---|---|---|---|---|---|---|---|---|---|---|---|
| ENSG00000203879 | 1402.736624 | 0.24157925 | 0.2673317 | 0.903669 | 0.366171198 | 0.558484609 | GDI1 | protein_coding | 59.79 | 0.061116842 | 0.061116842 | ENSG00000203879 | n.s. |
| ENSG00000145817 | 901.7506165 | -0.254574582 | 0.2916595 | -0.872848 | 0.382745667 | 0.575622591 | YIPF5 | protein_coding | 35.8 | -0.061062812 | 0.061062812 | ENSG00000145817 | n.s. |
| ENSG00000177981 | 291.2551218 | -0.236775616 | 0.2590291 | -0.914089 | 0.360670114 | 0.553223531 | ASB8 | protein_coding | 42.11 | -0.060874858 | 0.060874858 | ENSG00000177981 | n.s. |
| ENSG00000122786 | 2167.112603 | -0.241487394 | 0.2679836 | -0.901128 | 0.367520496 | 0.559784253 | CALD1 | protein_coding | 40.05 | -0.06084983 | 0.06084983 | ENSG00000122786 | n.s. |
| ENSG00000250432 | 28.4425925 | 0.263475715 | 0.3101281 | 0.84957 | 0.395563921 | 0.587647937 | | lincRNA | 48.26 | 0.060832007 | 0.060832007 | ENSG00000250432 | n.s. |
| ENSG00000057935 | 292.1156426 | -0.227773024 | 0.2426914 | -0.93853 | 0.347972315 | 0.540837078 | MTA3 | protein_coding | 43.08 | -0.06080026 | 0.06080026 | ENSG00000057935 | n.s. |
| ENSG00000132153 | 2083.409552 | -0.234864513 | 0.2555594 | -0.919021 | 0.358084471 | 0.550983281 | DHX30 | protein_coding | 50.03 | -0.060797399 | 0.060797399 | ENSG00000132153 | n.s. |
| ENSG00000183060 | 506.7634986 | -0.240547741 | 0.2667755 | -0.901686 | 0.36722363 | 0.559496962 | LYSMD4 | protein_coding | 47.75 | -0.060666686 | 0.060666686 | ENSG00000183060 | n.s. |
| ENSG00000163590 | 557.3787021 | -0.236866841 | 0.260041 | -0.910883 | 0.362357176 | 0.554711266 | PPM1L | protein_coding | 38.44 | -0.060622044 | 0.060622044 | ENSG00000163590 | n.s. |
| ENSG00000274104 | 25.75388108 | 0.271485425 | 0.3266745 | 0.831058 | 0.405940955 | 0.598023057 | | lincRNA | 49.81 | 0.060617828 | 0.060617828 | ENSG00000274104 | n.s. |
| ENSG00000103160 | 423.6326018 | -0.253685317 | 0.2913447 | -0.870739 | 0.383896494 | 0.576901337 | HSDL1 | protein_coding | 45.07 | -0.06060503 | 0.06060503 | ENSG00000103160 | n.s. |
| ENSG00000170558 | 2548.706718 | -0.238874239 | 0.2639478 | -0.905006 | 0.365462448 | 0.557931323 | CDH2 | protein_coding | 37.22 | -0.060535332 | 0.060535332 | ENSG00000170558 | n.s. |
| ENSG00000165572 | 1400.630263 | 0.217410282 | 0.2251868 | 0.965466 | 0.334311313 | 0.526719069 | KBTBD6 | protein_coding | 42.89 | 0.060531579 | 0.060531579 | ENSG00000165572 | n.s. |
| ENSG00000120896 | 98.50779065 | 0.243757993 | 0.2734782 | 0.891325 | 0.37275475 | 0.564790988 | SORBS3 | protein_coding | 55.84 | 0.060479342 | 0.060479342 | ENSG00000120896 | n.s. |
| ENSG00000274943 | 106.2516023 | 0.256527445 | 0.2978646 | 0.861222 | 0.389115996 | 0.581169493 | | sense_intronic | 34.32 | 0.060462798 | 0.060462798 | ENSG00000274943 | n.s. |
| ENSG00000066468 | 323.434761 | -0.228081622 | 0.2441336 | -0.934249 | 0.350175448 | 0.543169871 | FGFR2 | protein_coding | 45.41 | -0.060456302 | 0.060456302 | ENSG00000066468 | n.s. |
| ENSG00000138032 | 1060.383568 | -0.240297859 | 0.2670467 | -0.899835 | 0.368208279 | 0.560525243 | PPM1B | protein_coding | 38.72 | -0.060412041 | 0.060412041 | ENSG00000138032 | n.s. |
| ENSG00000175390 | 2827.799552 | -0.233207917 | 0.2536761 | -0.919314 | 0.357931503 | 0.550843408 | EIF3F | protein_coding | 41.38 | -0.060394284 | 0.060394284 | ENSG00000175390 | n.s. |
| ENSG00000120053 | 694.0033242 | 0.254964441 | 0.2948024 | 0.864866 | 0.387112543 | 0.5796696 | GOT1 | protein_coding | 42.9 | 0.060380545 | 0.060380545 | ENSG00000120053 | n.s. |
| ENSG00000164221 | 1151.381726 | -0.238790475 | 0.2648471 | -0.901616 | 0.367260688 | 0.559496962 | CCDC112 | protein_coding | 36.27 | -0.060223499 | 0.060223499 | ENSG00000164221 | n.s. |
| ENSG00000116288 | 1201.166062 | 0.265714982 | 0.316519 | 0.839491 | 0.401193652 | 0.593502669 | PARK7 | protein_coding | 45.41 | 0.060204989 | 0.060204989 | ENSG00000116288 | n.s. |
| ENSG00000181610 | 2285.984194 | -0.252117813 | 0.2897025 | -0.870265 | 0.384155851 | 0.577104221 | MRPS23 | protein_coding | 44.44 | -0.060192056 | 0.060192056 | ENSG00000181610 | n.s. |
| ENSG00000214226 | 120.8490955 | -0.225748169 | 0.2407735 | -0.937596 | 0.348452208 | 0.541341228 | C17orf67 | protein_coding | 44.82 | -0.06016841 | 0.06016841 | ENSG00000214226 | n.s. |
| ENSG00000085377 | 1975.111419 | -0.229951081 | 0.248366 | -0.925856 | 0.354521016 | 0.547464454 | PREP | protein_coding | 40.95 | -0.060165337 | 0.060165337 | ENSG00000085377 | n.s. |
| ENSG00000132341 | 17364.34106 | 0.239434209 | 0.2664523 | 0.898601 | 0.368865345 | 0.561102251 | RAN | protein_coding | 45.79 | 0.060087928 | 0.060087928 | ENSG00000132341 | n.s. |
| ENSG00000112335 | 890.9113243 | 0.268254759 | 0.3221187 | 0.832782 | 0.404967447 | 0.59709433 | SNX3 | protein_coding | 41.16 | 0.060077545 | 0.060077545 | ENSG00000112335 | n.s. |
| ENSG00000122756 | 384.7064841 | -0.217051332 | 0.225967 | -0.960544 | 0.336781375 | 0.528934809 | CNTFR | protein_coding | 56.26 | -0.060035932 | 0.060035932 | ENSG00000122756 | n.s. |
| ENSG00000141433 | 257.9127296 | 0.229208868 | 0.2475234 | 0.926009 | 0.35444127 | 0.547408298 | ADCYAP1 | protein_coding | 52.82 | 0.059981353 | 0.059981353 | ENSG00000141433 | n.s. |
| ENSG00000249122 | 288.592678 | -0.217214668 | 0.2264024 | -0.959418 | 0.33734798 | 0.52958574 | | lincRNA | 45.08 | -0.059965089 | 0.059965089 | ENSG00000249122 | n.s. |
| ENSG00000239521 | 218.8800682 | -0.240605185 | 0.269151 | -0.893941 | 0.371353307 | 0.563475136 | GATS | protein_coding | 51.53 | -0.059940825 | 0.059940825 | ENSG00000239521 | n.s. |
| ENSG00000113712 | 3362.564211 | -0.228045512 | 0.2456746 | -0.928242 | 0.353282013 | 0.546345159 | CSNK1A1 | protein_coding | 39.26 | -0.05986945 | 0.05986945 | ENSG00000113712 | n.s. |
| ENSG00000105373 | 1955.007749 | 0.233054011 | 0.255146 | 0.913414 | 0.36102474 | 0.55352356 | NOP53 | protein_coding | 57.2 | 0.05986316 | 0.05986316 | ENSG00000105373 | n.s. |

| | | | | | | | | | | | | | |
|---|---|---|---|---|---|---|---|---|---|---|---|---|---|
| ENSG00000176244 | 157.7594059 | 0.265291956 | 0.3170816 | 0.836668 | 0.402779356 | 0.594813001 | ACBD7 | protein_coding | 46.04 | 0.059855051 | 0.059855051 | ENSG00000176244 | n.s. |
| ENSG00000121966 | 231.7490182 | 0.223927199 | 0.2384057 | 0.93927 | 0.347592369 | 0.540427532 | CXCR4 | protein_coding | 52.06 | 0.059847351 | 0.059847351 | ENSG00000121966 | n.s. |
| ENSG00000179915 | 664.5713217 | -0.242497047 | 0.2732643 | -0.887409 | 0.374858991 | 0.566745763 | NRXN1 | protein_coding | 35.81 | -0.059802613 | 0.059802613 | ENSG00000179915 | n.s. |
| ENSG00000138613 | 276.4278361 | -0.246017896 | 0.2796581 | -0.879709 | 0.379016761 | 0.571455166 | APH1B | protein_coding | 40.76 | -0.059786737 | 0.059786737 | ENSG00000138613 | n.s. |
| ENSG00000114120 | 3273.247978 | 0.240097837 | 0.2689386 | 0.892761 | 0.371985374 | 0.564116367 | SLC25A36 | protein_coding | 35.47 | 0.059695837 | 0.059695837 | ENSG00000114120 | n.s. |
| ENSG00000156273 | 2023.442062 | 0.231870852 | 0.2534936 | 0.914701 | 0.360348596 | 0.552936594 | BACH1 | protein_coding | 39.61 | 0.05966609 | 0.05966609 | ENSG00000156273 | n.s. |
| ENSG00000185883 | 250.5762278 | 0.247797741 | 0.2833025 | 0.874675 | 0.381750544 | 0.574544679 | ATP6V0C | protein_coding | 60.18 | 0.059639017 | 0.059639017 | ENSG00000185883 | n.s. |
| ENSG00000231566 | 32.19419084 | 0.255488338 | 0.2988302 | 0.854962 | 0.392572419 | 0.584513505 | | lincRNA | 48.32 | 0.059581273 | 0.059581273 | ENSG00000231566 | n.s. |
| ENSG00000004059 | 687.7238796 | 0.254265239 | 0.2966549 | 0.857108 | 0.391385304 | 0.583182533 | ARF5 | protein_coding | 56.5 | 0.059547773 | 0.059547773 | ENSG00000004059 | n.s. |
| ENSG00000172613 | 160.05227 | -0.234427427 | 0.2586366 | -0.906397 | 0.364725793 | 0.557173274 | RAD9A | protein_coding | 52.03 | -0.059546846 | 0.059546846 | ENSG00000172613 | n.s. |
| ENSG00000197774 | 251.3578334 | 0.23549955 | 0.2608411 | 0.902847 | 0.366607138 | 0.558821018 | EME2 | protein_coding | 64.06 | 0.059517158 | 0.059517158 | ENSG00000197774 | n.s. |
| ENSG00000167881 | 1207.628624 | -0.234698219 | 0.2592974 | -0.905132 | 0.365395715 | 0.557890618 | SRP68 | protein_coding | 46.03 | -0.059484485 | 0.059484485 | ENSG00000167881 | n.s. |
| ENSG00000152234 | 8300.915441 | 0.249977814 | 0.2884596 | 0.866596 | 0.386163565 | 0.57893349 | ATP5A1 | protein_coding | 44.44 | 0.059337565 | 0.059337565 | ENSG00000152234 | n.s. |
| ENSG00000108654 | 42540.12499 | 0.219311775 | 0.2315001 | 0.947351 | 0.343460106 | 0.536398772 | DDX5 | protein_coding | 45.11 | 0.059326516 | 0.059326516 | ENSG00000108654 | n.s. |
| ENSG00000263916 | 31.39650134 | 0.254746627 | 0.2982621 | 0.854103 | 0.393047813 | 0.585033644 | | sense_intronic | 46.34 | 0.059309895 | 0.059309895 | ENSG00000263916 | n.s. |
| ENSG00000102466 | 136.3825624 | 0.220457329 | 0.2337182 | 0.943261 | 0.345547194 | 0.538450317 | FGF14 | protein_coding | 37.27 | 0.059270915 | 0.059270915 | ENSG00000102466 | n.s. |
| ENSG00000160781 | 169.7745267 | 0.243315838 | 0.2762139 | 0.880896 | 0.378373872 | 0.57089904 | PAQR6 | protein_coding | 60.95 | 0.059232975 | 0.059232975 | ENSG00000160781 | n.s. |
| ENSG00000119403 | 1179.753155 | -0.223777451 | 0.239743 | -0.933406 | 0.350610574 | 0.543663105 | PHF19 | protein_coding | 51.61 | -0.059227209 | 0.059227209 | ENSG00000119403 | n.s. |
| ENSG00000162931 | 115.8084176 | 0.227108617 | 0.2457916 | 0.923989 | 0.35549228 | 0.548544721 | TRIM17 | protein_coding | 56.75 | 0.059227193 | 0.059227193 | ENSG00000162931 | n.s. |
| ENSG00000109519 | 1014.358711 | -0.236894429 | 0.2645141 | -0.895584 | 0.370475244 | 0.562561547 | GRPEL1 | protein_coding | 47.5 | -0.059183325 | 0.059183325 | ENSG00000109519 | n.s. |
| ENSG00000115652 | 698.0257643 | -0.23347252 | 0.25793 | -0.905178 | 0.365371111 | 0.557890618 | UXS1 | protein_coding | 45.03 | -0.05917383 | 0.05917383 | ENSG00000115652 | n.s. |
| ENSG00000135624 | 8665.711447 | 0.246649719 | 0.2825865 | 0.872829 | 0.382756155 | 0.575622591 | CCT7 | protein_coding | 46.07 | 0.059161937 | 0.059161937 | ENSG00000135624 | n.s. |
| ENSG00000200350 | 23.29118596 | 0.274996988 | 0.3409518 | 0.806557 | 0.419921844 | 0.610652258 | RNU6-1285P | snRNA | 42.86 | 0.058906014 | 0.058906014 | ENSG00000200350 | n.s. |
| ENSG00000167967 | 459.609646 | 0.237334903 | 0.2664939 | 0.890583 | 0.373152967 | 0.565271326 | E4F1 | protein_coding | 61.58 | 0.058798071 | 0.058798071 | ENSG00000167967 | n.s. |
| ENSG00000257954 | 48.7873708 | -0.252280937 | 0.2952791 | -0.854381 | 0.392893705 | 0.584866786 | | processed_pseudogene | 40.19 | -0.058767089 | 0.058767089 | ENSG00000257954 | n.s. |
| ENSG00000100814 | 3091.267922 | 0.236255639 | 0.2646991 | 0.892544 | 0.37210141 | 0.564169435 | CCNB1IP1 | protein_coding | 39.52 | 0.058730895 | 0.058730895 | ENSG00000100814 | n.s. |
| ENSG00000052723 | 1879.392563 | -0.238119232 | 0.2683286 | -0.887417 | 0.374854639 | 0.566745763 | SIKE1 | protein_coding | 36.9 | -0.058722993 | 0.058722993 | ENSG00000052723 | n.s. |
| ENSG00000207973 | 6.399459068 | 0.335143663 | 0.4818242 | 0.695572 | 0.486696609 | 0.668074228 | MIR589 | miRNA | 62.63 | 0.058708886 | 0.058708886 | ENSG00000207973 | n.s. |
| ENSG00000164751 | 586.4192502 | -0.249867167 | 0.2909486 | -0.858802 | 0.390449941 | 0.582287332 | PEX2 | protein_coding | 38.37 | -0.058684467 | 0.058684467 | ENSG00000164751 | n.s. |
| ENSG00000136802 | 617.412254 | -0.22630835 | 0.2459908 | -0.919987 | 0.357579528 | 0.550417975 | LRRC8A | protein_coding | 53.83 | -0.058683428 | 0.058683428 | ENSG00000136802 | n.s. |
| ENSG00000148985 | 134.6025174 | -0.254219623 | 0.2996063 | -0.848512 | 0.396152738 | 0.588134144 | PGAP2 | protein_coding | 50.46 | -0.058603624 | 0.058603624 | ENSG00000148985 | n.s. |
| ENSG00000163961 | 1364.395276 | -0.210846245 | 0.2189164 | -0.963136 | 0.335479317 | 0.527782885 | RNF168 | protein_coding | 42.36 | -0.058519257 | 0.058519257 | ENSG00000163961 | n.s. |

| | | | | | | | | | | | | | |
|---|---|---|---|---|---|---|---|---|---|---|---|---|---|
| ENSG00000174231 | 4014.10063 | -0.230652217 | 0.2547433 | -0.90543 | 0.365237717 | 0.557832901 | PRPF8 | protein_coding | 48.87 | -0.058469386 | 0.058469386 | ENSG00000174231 | n.s. |
| ENSG00000128039 | 422.7861451 | -0.247078639 | 0.2858823 | -0.864267 | 0.387441181 | 0.57997458 | SRD5A3 | protein_coding | 44.77 | -0.058456593 | 0.058456593 | ENSG00000128039 | n.s. |
| ENSG00000224415 | 39.41643445 | -0.270129853 | 0.3324107 | -0.812639 | 0.416425159 | 0.607617636 | | processed_pseudogene | 51.1 | -0.058447896 | 0.058447896 | ENSG00000224415 | n.s. |
| ENSG00000008018 | 4153.312376 | 0.253642828 | 0.2992117 | 0.847704 | 0.396603073 | 0.588503302 | PSMB1 | protein_coding | 39.81 | 0.058401539 | 0.058401539 | ENSG00000008018 | n.s. |
| ENSG00000207952 | 7.117193777 | 0.382668504 | 0.6095981 | 0.627739 | 0.530174946 | 0.703721171 | MIR624 | miRNA | 40.21 | 0.058394978 | 0.058394978 | ENSG00000207952 | n.s. |
| ENSG00000177963 | 753.5654471 | -0.239814748 | 0.2722459 | -0.880876 | 0.378385142 | 0.57089904 | RIC8A | protein_coding | 57.49 | -0.058380667 | 0.058380667 | ENSG00000177963 | n.s. |
| ENSG00000204370 | 601.7999414 | 0.246854981 | 0.2859291 | 0.863343 | 0.387948654 | 0.580120522 | SDHD | protein_coding | 41.79 | 0.058376703 | 0.058376703 | ENSG00000204370 | n.s. |
| ENSG00000123933 | 644.7462746 | 0.2358455 | 0.2649693 | 0.890086 | 0.373419712 | 0.565613865 | MXD4 | protein_coding | 62.91 | 0.058367033 | 0.058367033 | ENSG00000123933 | n.s. |
| ENSG00000054277 | 81.06303745 | 0.259220406 | 0.3103127 | 0.835352 | 0.403519413 | 0.595568498 | OPN3 | protein_coding | 39.62 | 0.058342291 | 0.058342291 | ENSG00000054277 | n.s. |
| ENSG00000171045 | 171.93104 | 0.230455144 | 0.2548864 | 0.904149 | 0.36591662 | 0.558318586 | TSNARE1 | protein_coding | 55.7 | 0.058332325 | 0.058332325 | ENSG00000171045 | n.s. |
| ENSG00000213047 | 227.4535911 | 0.220066103 | 0.2355506 | 0.934262 | 0.350168505 | 0.543169871 | DENND1B | protein_coding | 34.32 | 0.058331674 | 0.058331674 | ENSG00000213047 | n.s. |
| ENSG00000169684 | 854.0808397 | 0.233068868 | 0.2599753 | 0.896504 | 0.369983614 | 0.561998893 | CHRNA5 | protein_coding | 40.43 | 0.058328873 | 0.058328873 | ENSG00000169684 | n.s. |
| ENSG00000135269 | 463.1113366 | 0.243181806 | 0.2786677 | 0.872659 | 0.382849122 | 0.575638049 | TES | protein_coding | 37.73 | 0.05832728 | 0.05832728 | ENSG00000135269 | n.s. |
| ENSG00000131437 | 751.0436983 | -0.235869442 | 0.2652 | -0.889402 | 0.37378695 | 0.565953476 | KIF3A | protein_coding | 36.68 | -0.058311471 | 0.058311471 | ENSG00000131437 | n.s. |
| ENSG00000173011 | 260.7612094 | -0.231703139 | 0.2573378 | -0.900385 | 0.367915269 | 0.560201694 | TADA2B | protein_coding | 56.73 | -0.058309388 | 0.058309388 | ENSG00000173011 | n.s. |
| ENSG00000184270 | 411.3059748 | 0.215023025 | 0.2266839 | 0.948559 | 0.34284512 | 0.535667872 | HIST2H2AB | protein_coding | 62.85 | 0.058293688 | 0.058293688 | ENSG00000184270 | n.s. |
| ENSG00000168502 | 108.2434857 | -0.240226972 | 0.2735486 | -0.878188 | 0.379841893 | 0.572272939 | MTCL1 | protein_coding | 45.45 | -0.058230247 | 0.058230247 | ENSG00000168502 | n.s. |
| ENSG00000198155 | 135.294538 | -0.253758384 | 0.3001495 | -0.84544 | 0.397865275 | 0.589682587 | ZNF876P | transcribed_unprocessed_pseud | 38.73 | -0.058207529 | 0.058207529 | ENSG00000198155 | n.s. |
| ENSG00000116478 | 414.851609 | 0.24813152 | 0.2892546 | 0.857831 | 0.390985951 | 0.58283698 | HDAC1 | protein_coding | 46.79 | 0.058175155 | 0.058175155 | ENSG00000116478 | n.s. |
| ENSG00000231889 | 705.177292 | 0.212796398 | 0.2232741 | 0.953073 | 0.340553292 | 0.533166099 | TRAF3IP2-AS1 | antisense | 41.83 | 0.05812267 | 0.05812267 | ENSG00000231889 | n.s. |
| ENSG00000145996 | 1032.507878 | -0.221699389 | 0.2391562 | -0.927007 | 0.353923131 | 0.546880191 | CDKAL1 | protein_coding | 38.04 | -0.058109141 | 0.058109141 | ENSG00000145996 | n.s. |
| ENSG00000010256 | 1487.359578 | 0.244315063 | 0.2815906 | 0.867625 | 0.385599548 | 0.578408131 | UQCRC1 | protein_coding | 53.04 | 0.05808972 | 0.05808972 | ENSG00000010256 | n.s. |
| ENSG00000189050 | 257.4930384 | -0.239409845 | 0.2725295 | -0.878473 | 0.379687089 | 0.572181334 | RNFT1 | protein_coding | 38.09 | -0.058048823 | 0.058048823 | ENSG00000189050 | n.s. |
| ENSG00000252376 | 6.896172386 | -0.321878521 | 0.4530324 | -0.710498 | 0.477395466 | 0.660319598 | RNA5SP395 | rRNA | 45.76 | -0.058017256 | 0.058017256 | ENSG00000252376 | n.s. |
| ENSG00000105607 | 375.5629013 | 0.242047031 | 0.2776772 | 0.871685 | 0.383380265 | 0.576249965 | GCDH | protein_coding | 51.4 | 0.057943418 | 0.057943418 | ENSG00000105607 | n.s. |
| ENSG00000133256 | 42.44726588 | 0.244103305 | 0.2816479 | 0.866697 | 0.386108171 | 0.57893349 | PDE6B | protein_coding | 56.84 | 0.057943125 | 0.057943125 | ENSG00000133256 | n.s. |
| ENSG00000065183 | 3933.483506 | -0.22897817 | 0.2534291 | -0.90352 | 0.366250079 | 0.558484609 | WDR3 | protein_coding | 36.95 | -0.05792891 | 0.05792891 | ENSG00000065183 | n.s. |
| ENSG00000204922 | 302.5372961 | 0.238803057 | 0.2717953 | 0.878614 | 0.379610818 | 0.57212832 | UQCC3 | protein_coding | 52.86 | 0.057911306 | 0.057911306 | ENSG00000204922 | n.s. |
| ENSG00000100632 | 3055.264746 | 0.254821265 | 0.3029575 | 0.841112 | 0.400285004 | 0.592638038 | ERH | protein_coding | 40.33 | 0.057898061 | 0.057898061 | ENSG00000100632 | n.s. |
| ENSG00000175970 | 921.3176339 | -0.230390477 | 0.2563066 | -0.898886 | 0.36871327 | 0.561005698 | UNC119B | protein_coding | 48.18 | -0.05783555 | 0.05783555 | ENSG00000175970 | n.s. |
| ENSG00000152944 | 1205.214315 | 0.230184905 | 0.2560844 | 0.898864 | 0.368725357 | 0.561005698 | MED21 | protein_coding | 39.33 | 0.057783944 | 0.057783944 | ENSG00000152944 | n.s. |
| ENSG00000099624 | 267.9139979 | 0.243577877 | 0.2819018 | 0.864052 | 0.387559255 | 0.580026605 | ATP5D | protein_coding | 62.97 | 0.057618855 | 0.057618855 | ENSG00000099624 | n.s. |

| | | | | | | | | | | | | | |
|---|---|---|---|---|---|---|---|---|---|---|---|---|---|
| ENSG00000070476 | 704.4252822 | -0.224997566 | 0.24683 | -0.911549 | 0.362006402 | 0.554599493 | ZXDC | protein_coding | 48.68 | -0.057603996 | 0.057603996 | ENSG00000070476 | n.s. |
| ENSG00000113387 | 5518.697548 | 0.246515458 | 0.2880891 | 0.855692 | 0.392168293 | 0.58411326 | SUB1 | protein_coding | 42.87 | 0.057562083 | 0.057562083 | ENSG00000113387 | n.s. |
| ENSG00000116688 | 730.8385578 | -0.237828327 | 0.2712788 | -0.876693 | 0.380653158 | 0.573140901 | MFN2 | protein_coding | 51.12 | -0.057492286 | 0.057492286 | ENSG00000116688 | n.s. |
| ENSG00000115355 | 7057.932072 | -0.242478345 | 0.2802133 | -0.865335 | 0.386854945 | 0.579415258 | CCDC88A | protein_coding | 36.45 | -0.05746981 | 0.05746981 | ENSG00000115355 | n.s. |
| ENSG00000141858 | 574.8768052 | -0.22097351 | 0.2397335 | -0.921746 | 0.356660895 | 0.549599094 | SAMD1 | protein_coding | 64.37 | -0.057442946 | 0.057442946 | ENSG00000141858 | n.s. |
| ENSG00000135587 | 139.259603 | 0.224250692 | 0.246125 | 0.911125 | 0.362229488 | 0.554599493 | SMPD2 | protein_coding | 58.66 | 0.057412781 | 0.057412781 | ENSG00000135587 | n.s. |
| ENSG00000130313 | 303.9392874 | -0.250854372 | 0.297204 | -0.844048 | 0.398642638 | 0.590520423 | PGLS | protein_coding | 53.32 | -0.05738672 | 0.05738672 | ENSG00000130313 | n.s. |
| ENSG00000113441 | 1318.166565 | -0.222297526 | 0.2423233 | -0.917359 | 0.358954343 | 0.551990426 | LNPEP | protein_coding | 37.19 | -0.057367979 | 0.057367979 | ENSG00000113441 | n.s. |
| ENSG00000162604 | 989.5254316 | -0.240114335 | 0.2760686 | -0.869763 | 0.38442981 | 0.577453473 | TM2D1 | protein_coding | 39.63 | -0.057263188 | 0.057263188 | ENSG00000162604 | n.s. |
| ENSG00000112852 | 141.6612399 | 0.226051594 | 0.2501983 | 0.90349 | 0.366266104 | 0.558484609 | PCDHB2 | protein_coding | 48.38 | 0.057188519 | 0.057188519 | ENSG00000112852 | n.s. |
| ENSG00000138134 | 188.6804604 | 0.221271883 | 0.2417543 | 0.915276 | 0.360046818 | 0.552936594 | STAMBPL1 | protein_coding | 39.34 | 0.056938714 | 0.056938714 | ENSG00000138134 | n.s. |
| ENSG00000162972 | 589.8444848 | 0.232439476 | 0.2628622 | 0.884264 | 0.3765539 | 0.568937596 | MAIP1 | protein_coding | 39.93 | 0.056932648 | 0.056932648 | ENSG00000162972 | n.s. |
| ENSG00000129951 | 403.5186636 | -0.239969628 | 0.2770645 | -0.866115 | 0.386427214 | 0.579204016 | PLPPR3 | protein_coding | 61.48 | -0.056913221 | 0.056913221 | ENSG00000129951 | n.s. |
| ENSG00000122042 | 434.8110287 | 0.240443393 | 0.2781883 | 0.864319 | 0.387412751 | 0.57997458 | UBL3 | protein_coding | 36.43 | 0.056886753 | 0.056886753 | ENSG00000122042 | n.s. |
| ENSG00000103257 | 1141.9897 | -0.207712235 | 0.2175708 | -0.954688 | 0.339735502 | 0.532313459 | SLC7A5 | protein_coding | 58.41 | -0.056878372 | 0.056878372 | ENSG00000103257 | n.s. |
| ENSG00000179918 | 302.8256512 | 0.217767572 | 0.2358897 | 0.923175 | 0.355915837 | 0.549076588 | SEPHS2 | protein_coding | 51.31 | 0.056699505 | 0.056699505 | ENSG00000179918 | n.s. |
| ENSG00000261114 | 43.44783541 | -0.245671271 | 0.2896844 | -0.848065 | 0.396401592 | 0.588327402 | | sense_intronic | 38.63 | -0.056597974 | 0.056597974 | ENSG00000261114 | n.s. |
| ENSG00000090060 | 4356.263817 | -0.224798644 | 0.2497332 | -0.900155 | 0.36803773 | 0.56032688 | PAPOLA | protein_coding | 36.57 | -0.056550019 | 0.056550019 | ENSG00000090060 | n.s. |
| ENSG00000128536 | 161.8478407 | -0.236495242 | 0.2715206 | -0.871003 | 0.383752495 | 0.576747192 | CDHR3 | protein_coding | 43.76 | -0.056525795 | 0.056525795 | ENSG00000128536 | n.s. |
| ENSG00000168958 | 1548.908711 | -0.243651399 | 0.2859886 | -0.851962 | 0.394235284 | 0.586359754 | MFF | protein_coding | 36.79 | -0.056487128 | 0.056487128 | ENSG00000168958 | n.s. |
| ENSG00000273899 | 346.8319599 | 0.235279102 | 0.269357 | 0.873484 | 0.382399102 | 0.575334201 | NOL12 | protein_coding | 54.42 | 0.056485762 | 0.056485762 | ENSG00000273899 | n.s. |
| ENSG00000184640 | 4371.814329 | -0.225357209 | 0.2511584 | -0.897271 | 0.369574147 | 0.561744634 | 9-Sep | protein_coding | 54.75 | -0.056443207 | 0.056443207 | ENSG00000184640 | n.s. |
| ENSG00000116273 | 145.0190999 | 0.227941369 | 0.2562205 | 0.88963 | 0.373664767 | 0.56592348 | PHF13 | protein_coding | 54.99 | 0.056356745 | 0.056356745 | ENSG00000116273 | n.s. |
| ENSG00000113732 | 556.7950055 | -0.258581879 | 0.3167129 | -0.816455 | 0.414239905 | 0.605937302 | ATP6V0E1 | protein_coding | 44.09 | -0.056260257 | 0.056260257 | ENSG00000113732 | n.s. |
| ENSG00000236662 | 23.7272966 | -0.254566808 | 0.3089034 | -0.824098 | 0.409883664 | 0.601603443 | | antisense | 53.25 | -0.056180269 | 0.056180269 | ENSG00000236662 | n.s. |
| ENSG00000213465 | 283.5381274 | -0.265928444 | 0.3332723 | -0.797932 | 0.424910239 | 0.61509932 | ARL2 | protein_coding | 55.34 | -0.056125462 | 0.056125462 | ENSG00000213465 | n.s. |
| ENSG00000263409 | 1.993499029 | 0.431747655 | 0.777522 | 0.555287 | 0.578698543 | 0.741528472 | MIR4747 | miRNA | 62.96 | 0.056072005 | 0.056072005 | ENSG00000263409 | n.s. |
| ENSG00000108854 | 2137.626642 | -0.224128955 | 0.2501453 | -0.895995 | 0.370255474 | 0.562289155 | SMURF2 | protein_coding | 39.4 | -0.05604127 | 0.05604127 | ENSG00000108854 | n.s. |
| ENSG00000136444 | 955.4652093 | 0.21695959 | 0.2363904 | 0.917802 | 0.358722611 | 0.551755956 | RSAD1 | protein_coding | 50.31 | 0.056030459 | 0.056030459 | ENSG00000136444 | n.s. |
| ENSG00000237310 | 33.89523878 | 0.260515741 | 0.322366 | 0.808136 | 0.419012017 | 0.609927068 | | lincRNA | 54.44 | 0.055938485 | 0.055938485 | ENSG00000237310 | n.s. |
| ENSG00000163964 | 535.344224 | -0.23774102 | 0.276752 | -0.85904 | 0.390318459 | 0.582193879 | PIGX | protein_coding | 45.36 | -0.05585306 | 0.05585306 | ENSG00000163964 | n.s. |
| ENSG00000116957 | 867.9465514 | -0.239170822 | 0.2795114 | -0.855675 | 0.392177746 | 0.58411326 | TBCE | protein_coding | 44.4 | -0.055847089 | 0.055847089 | ENSG00000116957 | n.s. |

| | | | | | | | | | | | | | |
|---|---|---|---|---|---|---|---|---|---|---|---|---|---|
| ENSG00000206417 | 194.1388017 | 0.233130585 | 0.2673892 | 0.871877 | 0.38327526 | 0.576154335 | H1FX-AS1 | antisense | 57.61 | 0.055825722 | 0.055825722 | ENSG00000206417 | n.s. |
| ENSG00000167863 | 2687.737079 | 0.259013786 | 0.3196535 | 0.810296 | 0.41777031 | 0.608878471 | ATP5H | protein_coding | 47.61 | 0.05580954 | 0.05580954 | ENSG00000167863 | n.s. |
| ENSG00000226688 | 376.2475832 | -0.219123831 | 0.2413666 | -0.907847 | 0.363959211 | 0.556307399 | ENTPD1-AS1 | antisense | 39.46 | -0.055807589 | 0.055807589 | ENSG00000226688 | n.s. |
| ENSG00000278828 | 960.8096679 | 0.216768286 | 0.2369126 | 0.914972 | 0.360206555 | 0.552936594 | HIST1H3H | protein_coding | 49.07 | 0.055779827 | 0.055779827 | ENSG00000278828 | n.s. |
| ENSG00000103995 | 3030.041637 | -0.217474806 | 0.2383194 | -0.912535 | 0.361487068 | 0.554110365 | CEP152 | protein_coding | 36.6 | -0.055761351 | 0.055761351 | ENSG00000103995 | n.s. |
| ENSG00000139641 | 942.8953957 | -0.235478954 | 0.2727181 | -0.863452 | 0.387889086 | 0.580120522 | ESYT1 | protein_coding | 50.49 | -0.05568648 | 0.05568648 | ENSG00000139641 | n.s. |
| ENSG00000149782 | 482.1675077 | 0.217497297 | 0.2387203 | 0.911097 | 0.362244328 | 0.554599493 | PLCB3 | protein_coding | 60.32 | 0.055683774 | 0.055683774 | ENSG00000149782 | n.s. |
| ENSG00000106066 | 238.4833564 | 0.271700419 | 0.3471659 | 0.782624 | 0.433847851 | 0.623880263 | CPVL | protein_coding | 41 | 0.055671077 | 0.055671077 | ENSG00000106066 | n.s. |
| ENSG00000273035 | 91.38230201 | 0.23311046 | 0.2677989 | 0.870468 | 0.384044573 | 0.577061579 | | sense_intronic | 31.45 | 0.055661612 | 0.055661612 | ENSG00000273035 | n.s. |
| ENSG00000108344 | 1468.194677 | 0.25165633 | 0.304918 | 0.825325 | 0.40918726 | 0.601088228 | PSMD3 | protein_coding | 49.75 | 0.055631596 | 0.055631596 | ENSG00000108344 | n.s. |
| ENSG00000087269 | 1509.379804 | -0.223831636 | 0.2509131 | -0.892068 | 0.372356234 | 0.564432864 | NOP14 | protein_coding | 49.99 | -0.055597027 | 0.055597027 | ENSG00000087269 | n.s. |
| ENSG00000133983 | 753.8498153 | 0.241256492 | 0.284601 | 0.847701 | 0.396604646 | 0.588503302 | COX16 | protein_coding | 39.38 | 0.055549572 | 0.055549572 | ENSG00000133983 | n.s. |
| ENSG00000120725 | 57.19186578 | 0.22956652 | 0.2617182 | 0.877152 | 0.380404354 | 0.572952166 | SIL1 | protein_coding | 44.78 | 0.055527925 | 0.055527925 | ENSG00000120725 | n.s. |
| ENSG00000198752 | 1188.04717 | -0.215556759 | 0.2356382 | -0.914778 | 0.360308006 | 0.552936594 | CDC42BPB | protein_coding | 48.97 | -0.055468071 | 0.055468071 | ENSG00000198752 | n.s. |
| ENSG00000139269 | 90.12322163 | 0.226340831 | 0.2558981 | 0.884496 | 0.376428479 | 0.568809823 | INHBE | protein_coding | 53.48 | 0.055460953 | 0.055460953 | ENSG00000139269 | n.s. |
| ENSG00000138231 | 1005.393064 | 0.217452159 | 0.2393075 | 0.908672 | 0.36352309 | 0.555955052 | DBR1 | protein_coding | 39.44 | 0.055441672 | 0.055441672 | ENSG00000138231 | n.s. |
| ENSG00000185803 | 202.5304478 | 0.235354875 | 0.2735705 | 0.860308 | 0.389619356 | 0.58148465 | SLC52A2 | protein_coding | 65.97 | 0.05541707 | 0.05541707 | ENSG00000185803 | n.s. |
| ENSG00000235501 | 114.9421325 | 0.239538014 | 0.281619 | 0.850575 | 0.395005562 | 0.587228662 | | antisense | 41.99 | 0.055379453 | 0.055379453 | ENSG00000235501 | n.s. |
| ENSG00000038219 | 4099.543283 | -0.21740304 | 0.2395084 | -0.907705 | 0.364033926 | 0.556360522 | BOD1L1 | protein_coding | 38.88 | -0.055360313 | 0.055360313 | ENSG00000038219 | n.s. |
| ENSG00000167553 | 173.9422307 | 0.268577277 | 0.3417046 | 0.785993 | 0.431871777 | 0.622329823 | TUBA1C | protein_coding | 45.2 | 0.055321383 | 0.055321383 | ENSG00000167553 | n.s. |
| ENSG00000167088 | 4175.510132 | -0.239801908 | 0.2826219 | -0.84849 | 0.396165004 | 0.588134144 | SNRPD1 | protein_coding | 42.32 | -0.055280001 | 0.055280001 | ENSG00000167088 | n.s. |
| ENSG00000117650 | 1125.402518 | 0.225991898 | 0.2558624 | 0.883255 | 0.377098248 | 0.569451075 | NEK2 | protein_coding | 39.5 | 0.055264868 | 0.055264868 | ENSG00000117650 | n.s. |
| ENSG00000174028 | 26.60326351 | -0.262909601 | 0.3304477 | -0.795616 | 0.426255052 | 0.616533252 | FAM3C2 | processed_pseudogene | 46.24 | -0.055222451 | 0.055222451 | ENSG00000174028 | n.s. |
| ENSG00000037042 | 264.6593982 | -0.209242252 | 0.2248159 | -0.930727 | 0.351994831 | 0.544959861 | TUBG2 | protein_coding | 51.9 | -0.055163682 | 0.055163682 | ENSG00000037042 | n.s. |
| ENSG00000198380 | 1558.416766 | -0.214352839 | 0.2342751 | -0.914962 | 0.36021161 | 0.552936594 | GFPT1 | protein_coding | 39.5 | -0.055158273 | 0.055158273 | ENSG00000198380 | n.s. |
| ENSG00000088970 | 184.3400486 | -0.232269986 | 0.2680029 | -0.86667 | 0.386122943 | 0.57893349 | KIZ | protein_coding | 41.17 | -0.055134235 | 0.055134235 | ENSG00000088970 | n.s. |
| ENSG00000068383 | 276.9940407 | -0.240241951 | 0.2841095 | -0.845596 | 0.39777797 | 0.589678735 | INPP5A | protein_coding | 54.13 | -0.055107788 | 0.055107788 | ENSG00000068383 | n.s. |
| ENSG00000139719 | 861.2603336 | -0.237037002 | 0.2777522 | -0.853412 | 0.393430958 | 0.585541339 | VPS33A | protein_coding | 44.63 | -0.055097459 | 0.055097459 | ENSG00000139719 | n.s. |
| ENSG00000272028 | 7.227491744 | -0.327432091 | 0.4877744 | -0.671278 | 0.502043622 | 0.6790318 | RNU6-87P | snRNA | 41.67 | -0.055044572 | 0.055044572 | ENSG00000272028 | n.s. |
| ENSG00000255248 | 392.7814549 | 0.221217287 | 0.2477015 | 0.89308 | 0.37181432 | 0.563918386 | MIR100HG | processed_transcript | 39.21 | 0.055035265 | 0.055035265 | ENSG00000255248 | n.s. |
| ENSG00000128641 | 6801.469079 | -0.21980605 | 0.2451143 | -0.896749 | 0.369852754 | 0.561959308 | MYO1B | protein_coding | 39.67 | -0.055016384 | 0.055016384 | ENSG00000128641 | n.s. |
| ENSG00000174840 | 1462.641682 | 0.209567352 | 0.2258169 | 0.928041 | 0.353386325 | 0.546385097 | PDE12 | protein_coding | 41.24 | 0.055011672 | 0.055011672 | ENSG00000174840 | n.s. |

| | | | | | | | | | | | | | |
|---|---|---|---|---|---|---|---|---|---|---|---|---|---|
| ENSG00000270362 | 147.4674429 | 0.208981403 | 0.2250224 | 0.928714 | 0.353037494 | 0.546149004 | HMGN3-AS1 | antisense | 42.24 | 0.054897085 | 0.054897085 | ENSG00000270362 | n.s. |
| ENSG00000112367 | 355.8426481 | -0.246499057 | 0.2974488 | -0.828711 | 0.40726799 | 0.598900742 | FIG4 | protein_coding | 38.91 | -0.054881819 | 0.054881819 | ENSG00000112367 | n.s. |
| ENSG00000167657 | 283.0014545 | -0.228861302 | 0.2623379 | -0.872391 | 0.382994893 | 0.575795043 | DAPK3 | protein_coding | 58.44 | -0.054865396 | 0.054865396 | ENSG00000167657 | n.s. |
| ENSG00000125386 | 756.1096681 | -0.211838001 | 0.2306518 | -0.918432 | 0.35839282 | 0.551309605 | FAM193A | protein_coding | 45.62 | -0.054782248 | 0.054782248 | ENSG00000125386 | n.s. |
| ENSG00000156313 | 47.82396393 | -0.244907786 | 0.2943607 | -0.831999 | 0.40540961 | 0.597551897 | RPGR | protein_coding | 37.62 | -0.054767349 | 0.054767349 | ENSG00000156313 | n.s. |
| ENSG00000102543 | 157.6506465 | -0.226997476 | 0.2592428 | -0.875617 | 0.381238127 | 0.573897551 | CDADC1 | protein_coding | 38.43 | -0.054743989 | 0.054743989 | ENSG00000102543 | n.s. |
| ENSG00000107815 | 721.6748117 | -0.204837391 | 0.2182118 | -0.938709 | 0.347880192 | 0.54081464 | TWNK | protein_coding | 52.35 | -0.054681661 | 0.054681661 | ENSG00000107815 | n.s. |
| ENSG00000161835 | 86.94983299 | 0.236556481 | 0.2782018 | 0.850305 | 0.395155358 | 0.587291636 | GRASP | protein_coding | 58.66 | 0.054679128 | 0.054679128 | ENSG00000161835 | n.s. |
| ENSG00000166181 | 3373.880863 | -0.231063377 | 0.2673187 | -0.864374 | 0.387382304 | 0.57997458 | API5 | protein_coding | 37.75 | -0.054667525 | 0.054667525 | ENSG00000166181 | n.s. |
| ENSG00000125170 | 2262.301119 | -0.231977835 | 0.2695649 | -0.860564 | 0.389478205 | 0.581366175 | DOK4 | protein_coding | 58.39 | -0.054642435 | 0.054642435 | ENSG00000125170 | n.s. |
| ENSG00000110171 | 118.2692348 | -0.224672447 | 0.2553043 | -0.880018 | 0.378849507 | 0.571352003 | TRIM3 | protein_coding | 48.68 | -0.054617028 | 0.054617028 | ENSG00000110171 | n.s. |
| ENSG00000101367 | 2236.21073 | 0.235513975 | 0.276381 | 0.852135 | 0.394139128 | 0.586281953 | MAPRE1 | protein_coding | 44.44 | 0.054614154 | 0.054614154 | ENSG00000101367 | n.s. |
| ENSG00000157005 | 25.02151101 | 0.249641144 | 0.3048138 | 0.818996 | 0.41278889 | 0.604596579 | SST | protein_coding | 48.46 | 0.054555156 | 0.054555156 | ENSG00000157005 | n.s. |
| ENSG00000115947 | 1757.55019 | -0.232426829 | 0.2710001 | -0.857663 | 0.391078392 | 0.58291237 | ORC4 | protein_coding | 36.15 | -0.054480088 | 0.054480088 | ENSG00000115947 | n.s. |
| ENSG00000180879 | 1286.438062 | 0.25102124 | 0.3083646 | 0.81404 | 0.415621853 | 0.607118867 | SSR4 | protein_coding | 61.54 | 0.054402897 | 0.054402897 | ENSG00000180879 | n.s. |
| ENSG00000114302 | 979.2370647 | -0.228463756 | 0.263008 | -0.868657 | 0.385034691 | 0.577945263 | PRKAR2A | protein_coding | 43.41 | -0.054400257 | 0.054400257 | ENSG00000114302 | n.s. |
| ENSG00000284024 | 211.429899 | -0.222599554 | 0.2522119 | -0.882589 | 0.37745823 | 0.569685738 | | protein_coding | 46.24 | -0.054395463 | 0.054395463 | ENSG00000284024 | n.s. |
| ENSG00000178974 | 473.9227362 | -0.230804429 | 0.2682127 | -0.860528 | 0.389498217 | 0.581366175 | FBXO34 | protein_coding | 40.98 | -0.054366039 | 0.054366039 | ENSG00000178974 | n.s. |
| ENSG00000009844 | 1861.961754 | -0.248048703 | 0.3026642 | -0.819551 | 0.41247216 | 0.604446888 | VTA1 | protein_coding | 35.53 | -0.054233828 | 0.054233828 | ENSG00000009844 | n.s. |
| ENSG00000175040 | 987.1625502 | 0.214153132 | 0.2367717 | 0.904471 | 0.365745782 | 0.558238174 | CHST2 | protein_coding | 58.88 | 0.05421939 | 0.05421939 | ENSG00000175040 | n.s. |
| ENSG00000160284 | 40.18972874 | -0.253491365 | 0.3148774 | -0.805048 | 0.420792072 | 0.61115632 | SPATC1L | protein_coding | 50.77 | -0.054208544 | 0.054208544 | ENSG00000160284 | n.s. |
| ENSG00000196453 | 261.9147874 | -0.206219781 | 0.2222508 | -0.92787 | 0.353475243 | 0.546405761 | ZNF777 | protein_coding | 47.33 | -0.054129543 | 0.054129543 | ENSG00000196453 | n.s. |
| ENSG00000272330 | 6.427775835 | -0.31408203 | 0.4593971 | -0.683683 | 0.494175284 | 0.672841832 | | sense_intronic | 28.48 | -0.054049439 | 0.054049439 | ENSG00000272330 | n.s. |
| ENSG00000266751 | 7.852971552 | 0.310044429 | 0.4486993 | 0.690985 | 0.489575089 | 0.669780286 | MIR3661 | miRNA | 58.33 | 0.053968702 | 0.053968702 | ENSG00000266751 | n.s. |
| ENSG00000119559 | 68.86647893 | -0.228827631 | 0.2655854 | -0.861597 | 0.388909286 | 0.581051271 | C19orf25 | protein_coding | 62.65 | -0.053954248 | 0.053954248 | ENSG00000119559 | n.s. |
| ENSG00000198901 | 5948.082491 | 0.232717163 | 0.2734885 | 0.850921 | 0.394813176 | 0.587033729 | PRC1 | protein_coding | 45.15 | 0.053836078 | 0.053836078 | ENSG00000198901 | n.s. |
| ENSG00000158019 | 376.9086987 | -0.233305151 | 0.2747705 | -0.849091 | 0.395830682 | 0.587916623 | BABAM2 | protein_coding | 39.38 | -0.053819826 | 0.053819826 | ENSG00000158019 | n.s. |
| ENSG00000261544 | 27.27150223 | 0.257043936 | 0.3238342 | 0.793752 | 0.42734006 | 0.617589335 | | sense_intronic | 38.66 | 0.05379935 | 0.05379935 | ENSG00000261544 | n.s. |
| ENSG00000184178 | 692.1742488 | -0.217812122 | 0.2450727 | -0.888765 | 0.374129139 | 0.566318765 | SCFD2 | protein_coding | 39.78 | -0.053786318 | 0.053786318 | ENSG00000184178 | n.s. |
| ENSG00000068903 | 65.225482 | 0.247547265 | 0.3036466 | 0.815248 | 0.414930441 | 0.606517221 | SIRT2 | protein_coding | 53.55 | 0.053756588 | 0.053756588 | ENSG00000068903 | n.s. |
| ENSG00000197061 | 2125.547981 | 0.196748735 | 0.2064666 | 0.952932 | 0.340624323 | 0.533166099 | HIST1H4C | protein_coding | 49.66 | 0.053739452 | 0.053739452 | ENSG00000197061 | n.s. |
| ENSG00000152413 | 883.4475301 | -0.228434496 | 0.2658767 | -0.859175 | 0.390244219 | 0.582167606 | HOMER1 | protein_coding | 37.23 | -0.053671133 | 0.053671133 | ENSG00000152413 | n.s. |

| | | | | | | | | | | | | | |
|---|---|---|---|---|---|---|---|---|---|---|---|---|---|
| ENSG00000105254 | 982.6871129 | 0.246394894 | 0.3017895 | 0.816446 | 0.414245008 | 0.605937302 | TBCB | protein_coding | 55.29 | 0.053608707 | 0.053608707 | ENSG00000105254 | n.s. |
| ENSG00000266235 | 5.925343472 | -0.316012671 | 0.4674369 | -0.676054 | 0.499006213 | 0.6766985 | MIR3176 | miRNA | 74.44 | -0.053597261 | 0.053597261 | ENSG00000266235 | n.s. |
| ENSG00000206053 | 1902.068943 | -0.232895036 | 0.2750017 | -0.846886 | 0.397058818 | 0.588988911 | JPT2 | protein_coding | 47.78 | -0.053540911 | 0.053540911 | ENSG00000206053 | n.s. |
| ENSG00000169689 | 627.5693276 | 0.260648986 | 0.3323815 | 0.784186 | 0.43293089 | 0.623146572 | CENPX | protein_coding | 65.43 | 0.053539853 | 0.053539853 | ENSG00000169689 | n.s. |
| ENSG00000148450 | 172.9880841 | -0.240974805 | 0.291104 | -0.827796 | 0.407785853 | 0.599598947 | MSRB2 | protein_coding | 43.06 | -0.053529936 | 0.053529936 | ENSG00000148450 | n.s. |
| ENSG00000181524 | 148.8372306 | 0.27056617 | 0.3557201 | 0.760615 | 0.446886898 | 0.634100082 | RPL24P4 | processed_pseudogene | 48.69 | 0.053529404 | 0.053529404 | ENSG00000181524 | n.s. |
| ENSG00000126945 | 1826.974884 | -0.226115875 | 0.2618103 | -0.863663 | 0.387772998 | 0.580120522 | HNRNPH2 | protein_coding | 39.49 | -0.053472283 | 0.053472283 | ENSG00000126945 | n.s. |
| ENSG00000108651 | 1274.128212 | -0.228024943 | 0.2657493 | -0.858045 | 0.390867442 | 0.582785114 | UTP6 | protein_coding | 44.12 | -0.053469922 | 0.053469922 | ENSG00000108651 | n.s. |
| ENSG00000185414 | 1419.632163 | -0.249587259 | 0.3094969 | -0.806429 | 0.419995596 | 0.610652258 | MRPL30 | protein_coding | 40.26 | -0.053463097 | 0.053463097 | ENSG00000185414 | n.s. |
| ENSG00000140577 | 44.75899001 | 0.233664256 | 0.2768117 | 0.844127 | 0.39859839 | 0.590517704 | CRTC3 | protein_coding | 44.02 | 0.053454689 | 0.053454689 | ENSG00000140577 | n.s. |
| ENSG00000131899 | 495.7076663 | 0.225731509 | 0.2612156 | 0.864158 | 0.387501109 | 0.580001928 | LLGL1 | protein_coding | 59.89 | 0.05340143 | 0.05340143 | ENSG00000131899 | n.s. |
| ENSG00000182628 | 4133.454285 | -0.235955116 | 0.281445 | -0.83837 | 0.401822718 | 0.593995717 | SKA2 | protein_coding | 41.24 | -0.053376985 | 0.053376985 | ENSG00000182628 | n.s. |
| ENSG00000174721 | 161.2096113 | 0.207150805 | 0.2266858 | 0.913824 | 0.360809519 | 0.553254508 | FGFBP3 | protein_coding | 52.5 | 0.053253301 | 0.053253301 | ENSG00000174721 | n.s. |
| ENSG00000187953 | 385.6990333 | -0.213598941 | 0.2388154 | -0.89441 | 0.371102415 | 0.56326817 | PMS2CL | transcribed_unprocessed_pseud | 48.28 | -0.053246967 | 0.053246967 | ENSG00000187953 | n.s. |
| ENSG00000213626 | 150.124176 | 0.226727962 | 0.2640355 | 0.858702 | 0.390504705 | 0.582306631 | LBH | protein_coding | 46.97 | 0.053246668 | 0.053246668 | ENSG00000213626 | n.s. |
| ENSG00000126768 | 371.5458262 | 0.260079908 | 0.3329983 | 0.781025 | 0.434788003 | 0.624463933 | TIMM17B | protein_coding | 52.95 | 0.053184427 | 0.053184427 | ENSG00000126768 | n.s. |
| ENSG00000206612 | 30.52864591 | -0.244463915 | 0.299514 | -0.816202 | 0.414384694 | 0.606000609 | SNORA2A | snoRNA | 43.7 | -0.053177487 | 0.053177487 | ENSG00000206612 | n.s. |
| ENSG00000215184 | 38.36947111 | -0.243037271 | 0.2963992 | -0.819966 | 0.412235437 | 0.604290797 | RPS12P16 | processed_pseudogene | 48.16 | -0.05316538 | 0.05316538 | ENSG00000215184 | n.s. |
| ENSG00000165733 | 2736.12615 | -0.213290872 | 0.2387236 | -0.893464 | 0.371608876 | 0.563668197 | BMS1 | protein_coding | 40.62 | -0.053104408 | 0.053104408 | ENSG00000165733 | n.s. |
| ENSG00000111275 | 637.7533673 | -0.237465165 | 0.2854395 | -0.831928 | 0.405449456 | 0.597551897 | ALDH2 | protein_coding | 48.8 | -0.053102998 | 0.053102998 | ENSG00000111275 | n.s. |
| ENSG00000232677 | 580.7947916 | -0.222870476 | 0.2567125 | -0.868171 | 0.385300506 | 0.578262256 | LINC00665 | lincRNA | 45.72 | -0.053015349 | 0.053015349 | ENSG00000232677 | n.s. |
| ENSG00000086619 | 184.4897032 | 0.233684766 | 0.2782837 | 0.839736 | 0.401056548 | 0.593464801 | ERO1B | protein_coding | 37.47 | 0.052954144 | 0.052954144 | ENSG00000086619 | n.s. |
| ENSG00000074410 | 261.4543319 | 0.234634867 | 0.2804206 | 0.836725 | 0.402747359 | 0.594813001 | CA12 | protein_coding | 48.45 | 0.052938213 | 0.052938213 | ENSG00000074410 | n.s. |
| ENSG00000010438 | 49.7603387 | -0.321684843 | 0.4857932 | -0.662185 | 0.507852849 | 0.684892988 | PRSS3 | protein_coding | 42.84 | -0.052877679 | 0.052877679 | ENSG00000010438 | n.s. |
| ENSG00000179528 | 165.1883372 | 0.243180319 | 0.2985095 | 0.814649 | 0.415273522 | 0.606891282 | LBX2 | protein_coding | 60.07 | 0.052743161 | 0.052743161 | ENSG00000179528 | n.s. |
| ENSG00000065978 | 7345.724549 | 0.22363416 | 0.2595687 | 0.861561 | 0.388929408 | 0.581051271 | YBX1 | protein_coding | 40.41 | 0.052729702 | 0.052729702 | ENSG00000065978 | n.s. |
| ENSG00000137133 | 335.6982239 | 0.246146771 | 0.3055368 | 0.805621 | 0.420461641 | 0.610882545 | HINT2 | protein_coding | 54.66 | 0.052685817 | 0.052685817 | ENSG00000137133 | n.s. |
| ENSG00000137968 | 1007.679166 | -0.233298495 | 0.2786505 | -0.837244 | 0.402455284 | 0.59465671 | SLC44A5 | protein_coding | 36.99 | -0.052663327 | 0.052663327 | ENSG00000137968 | n.s. |
| ENSG00000105854 | 228.3628719 | -0.249802666 | 0.3134619 | -0.796916 | 0.425500066 | 0.615633111 | PON2 | protein_coding | 38.28 | -0.052627934 | 0.052627934 | ENSG00000105854 | n.s. |
| ENSG00000277283 | 164.9568083 | 0.209742397 | 0.2335453 | 0.89808 | 0.369142882 | 0.56139556 | | antisense | 52.48 | 0.052588927 | 0.052588927 | ENSG00000277283 | n.s. |
| ENSG00000137404 | 184.003807 | -0.20677033 | 0.2283972 | -0.90531 | 0.365301115 | 0.557868534 | NRM | protein_coding | 55.1 | -0.05240969 | 0.05240969 | ENSG00000137404 | n.s. |
| ENSG00000145354 | 973.1408314 | -0.243683887 | 0.3011832 | -0.809089 | 0.418464196 | 0.609506956 | CISD2 | protein_coding | 39.45 | -0.052397234 | 0.052397234 | ENSG00000145354 | n.s. |

| | | | | | | | | | | | | | |
|---|---|---|---|---|---|---|---|---|---|---|---|---|---|
| ENSG00000141298 | 55.59589073 | 0.2317985 | 0.2766744 | 0.837802 | 0.402141707 | 0.594312598 | SSH2 | protein_coding | 40.66 | 0.052382999 | 0.052382999 | ENSG00000141298 | n.s. |
| ENSG00000116704 | 925.7637365 | 0.2108463 | 0.236333 | 0.892158 | 0.372308357 | 0.56442174 | SLC35D1 | protein_coding | 37.83 | 0.052373434 | 0.052373434 | ENSG00000116704 | n.s. |
| ENSG00000114395 | 140.5643065 | 0.233436637 | 0.2801451 | 0.833271 | 0.40469215 | 0.59696375 | CYB561D2 | protein_coding | 55 | 0.052301955 | 0.052301955 | ENSG00000114395 | n.s. |
| ENSG00000198586 | 1704.221281 | -0.220387204 | 0.2546867 | -0.865327 | 0.386859458 | 0.579415258 | TLK1 | protein_coding | 38.08 | -0.052233987 | 0.052233987 | ENSG00000198586 | n.s. |
| ENSG00000185053 | 307.5216536 | 0.221980761 | 0.2582842 | 0.859444 | 0.390095755 | 0.582008486 | SGCZ | protein_coding | 35.48 | 0.052181169 | 0.052181169 | ENSG00000185053 | n.s. |
| ENSG00000075413 | 1499.611788 | -0.223464403 | 0.2611074 | -0.855833 | 0.392090005 | 0.584107547 | MARK3 | protein_coding | 42.15 | -0.052180543 | 0.052180543 | ENSG00000075413 | n.s. |
| ENSG00000179958 | 859.1023208 | 0.245773449 | 0.3070483 | 0.800439 | 0.423456472 | 0.613505147 | DCTPP1 | protein_coding | 53.23 | 0.05214865 | 0.05214865 | ENSG00000179958 | n.s. |
| ENSG00000134138 | 247.7230805 | 0.201792056 | 0.2199618 | 0.917396 | 0.358935161 | 0.551990426 | MEIS2 | protein_coding | 38.26 | 0.052076164 | 0.052076164 | ENSG00000134138 | n.s. |
| ENSG00000170873 | 1048.134445 | -0.214647334 | 0.2445059 | -0.877882 | 0.380007834 | 0.572416865 | MTSS1 | protein_coding | 45.65 | -0.052006383 | 0.052006383 | ENSG00000170873 | n.s. |
| ENSG00000198826 | 2568.047095 | -0.218107012 | 0.2508613 | -0.869433 | 0.384610548 | 0.57760033 | ARHGAP11A | protein_coding | 37.89 | -0.051990728 | 0.051990728 | ENSG00000198826 | n.s. |
| ENSG00000127580 | 142.4316207 | -0.210479886 | 0.2368861 | -0.888528 | 0.374256934 | 0.566450624 | WDR24 | protein_coding | 63.87 | -0.051954421 | 0.051954421 | ENSG00000127580 | n.s. |
| ENSG00000271793 | 149.0392322 | -0.238405648 | 0.2919395 | -0.816627 | 0.414141644 | 0.605937302 | | protein_coding | 38.3 | -0.051870468 | 0.051870468 | ENSG00000271793 | n.s. |
| ENSG00000142676 | 10053.26661 | 0.232093162 | 0.2793439 | 0.830851 | 0.406057767 | 0.598079974 | RPL11 | protein_coding | 46.18 | 0.051812649 | 0.051812649 | ENSG00000142676 | n.s. |
| ENSG00000111674 | 632.1706023 | 0.22954588 | 0.2744247 | 0.836462 | 0.402895018 | 0.594920706 | ENO2 | protein_coding | 55.33 | 0.051771988 | 0.051771988 | ENSG00000111674 | n.s. |
| ENSG00000179240 | 360.8950541 | -0.19816239 | 0.2142307 | -0.924996 | 0.354968245 | 0.547979031 | | protein_coding | 43.61 | -0.051767162 | 0.051767162 | ENSG00000179240 | n.s. |
| ENSG00000145216 | 2099.683498 | -0.213495941 | 0.2431247 | -0.878134 | 0.379871184 | 0.572272939 | FIP1L1 | protein_coding | 35.71 | -0.051750731 | 0.051750731 | ENSG00000145216 | n.s. |
| ENSG00000177370 | 392.5565382 | -0.224288047 | 0.2641685 | -0.849034 | 0.395862344 | 0.587916623 | TIMM22 | protein_coding | 48.68 | -0.051739722 | 0.051739722 | ENSG00000177370 | n.s. |
| ENSG00000177150 | 367.2025617 | -0.222964372 | 0.2615665 | -0.852419 | 0.393981321 | 0.586172466 | FAM210A | protein_coding | 40.65 | -0.051722068 | 0.051722068 | ENSG00000177150 | n.s. |
| ENSG00000153147 | 6401.128431 | -0.219342492 | 0.2546082 | -0.86149 | 0.388968089 | 0.581051271 | SMARCA5 | protein_coding | 35.51 | -0.051717789 | 0.051717789 | ENSG00000153147 | n.s. |
| ENSG00000186625 | 474.8536694 | -0.212676272 | 0.241909 | -0.879158 | 0.379315527 | 0.571745165 | KATNA1 | protein_coding | 42.14 | -0.051637267 | 0.051637267 | ENSG00000186625 | n.s. |
| ENSG00000188283 | 26.22775006 | -0.258389097 | 0.3371917 | -0.766297 | 0.443499466 | 0.631309995 | ZNF383 | protein_coding | 41.4 | -0.051615117 | 0.051615117 | ENSG00000188283 | n.s. |
| ENSG00000170502 | 354.9748997 | -0.222499185 | 0.261047 | -0.852334 | 0.394028899 | 0.586180613 | NUDT9 | protein_coding | 38.92 | -0.051612813 | 0.051612813 | ENSG00000170502 | n.s. |
| ENSG00000183723 | 504.6250902 | -0.227600237 | 0.2708799 | -0.840226 | 0.400781841 | 0.593247456 | CMTM4 | protein_coding | 45.26 | -0.051611566 | 0.051611566 | ENSG00000183723 | n.s. |
| ENSG00000124795 | 8619.25811 | -0.230857134 | 0.2777455 | -0.831182 | 0.405870713 | 0.59798285 | DEK | protein_coding | 39.43 | -0.051553 | 0.051553 | ENSG00000124795 | n.s. |
| ENSG00000272196 | 13.52151141 | 0.265656605 | 0.354616 | 0.749139 | 0.453773516 | 0.640240035 | HIST2H2AA4 | protein_coding | 61.7 | 0.051446307 | 0.051446307 | ENSG00000272196 | n.s. |
| ENSG00000256377 | 106.8612432 | 0.237655112 | 0.2923666 | 0.812867 | 0.416294434 | 0.607490558 | | antisense | 42.77 | 0.051442937 | 0.051442937 | ENSG00000256377 | n.s. |
| ENSG00000124380 | 789.2777992 | 0.227066551 | 0.2706308 | 0.839027 | 0.401454136 | 0.593674556 | SNRNP27 | protein_coding | 40.55 | 0.051419576 | 0.051419576 | ENSG00000124380 | n.s. |
| ENSG00000113648 | 6822.349838 | -0.22387775 | 0.2646291 | -0.846006 | 0.397549545 | 0.589402866 | H2AFY | protein_coding | 47.07 | -0.05139959 | 0.05139959 | ENSG00000113648 | n.s. |
| ENSG00000100445 | 470.7719833 | 0.203083691 | 0.2246562 | 0.903976 | 0.366008342 | 0.558397342 | SDR39U1 | protein_coding | 53.54 | 0.05139168 | 0.05139168 | ENSG00000100445 | n.s. |
| ENSG00000050438 | 95.54835779 | 0.220422889 | 0.2578776 | 0.854758 | 0.392685184 | 0.584618886 | SLC4A8 | protein_coding | 42.72 | 0.051386562 | 0.051386562 | ENSG00000050438 | n.s. |
| ENSG00000106080 | 404.6820887 | -0.22598922 | 0.2685513 | -0.841512 | 0.400060985 | 0.592432365 | FKBP14 | protein_coding | 38.7 | -0.051381186 | 0.051381186 | ENSG00000106080 | n.s. |
| ENSG00000079999 | 972.3174733 | -0.222256427 | 0.2613554 | -0.850399 | 0.395103257 | 0.587276892 | KEAP1 | protein_coding | 54.01 | -0.051376148 | 0.051376148 | ENSG00000079999 | n.s. |

| | | | | | | | | | | | | | |
|---|---|---|---|---|---|---|---|---|---|---|---|---|---|
| ENSG00000137547 | 1037.297033 | -0.239205212 | 0.2960261 | -0.808054 | 0.419059279 | 0.609927068 | MRPL15 | protein_coding | 43.26 | -0.051362644 | 0.051362644 | ENSG00000137547 | n.s. |
| ENSG00000151640 | 1100.229432 | -0.229142856 | 0.2750215 | -0.833182 | 0.404742328 | 0.59696375 | DPYSL4 | protein_coding | 61.67 | -0.051339924 | 0.051339924 | ENSG00000151640 | n.s. |
| ENSG00000241261 | 121.4355058 | 0.209158506 | 0.2368275 | 0.883168 | 0.377145417 | 0.56946054 | RPL17P19 | processed_pseudogene | 38.95 | 0.051146861 | 0.051146861 | ENSG00000241261 | n.s. |
| ENSG00000172428 | 846.6294511 | 0.25360272 | 0.3285707 | 0.771836 | 0.440211545 | 0.628944621 | COPS9 | protein_coding | 54.66 | 0.051072441 | 0.051072441 | ENSG00000172428 | n.s. |
| ENSG00000163902 | 2170.382652 | 0.229028951 | 0.2761071 | 0.829493 | 0.406825383 | 0.598565974 | RPN1 | protein_coding | 46.98 | 0.051047799 | 0.051047799 | ENSG00000163902 | n.s. |
| ENSG00000151665 | 411.579214 | -0.250508391 | 0.3218544 | -0.778328 | 0.436375438 | 0.625567375 | PIGF | protein_coding | 36.13 | -0.05103505 | 0.05103505 | ENSG00000151665 | n.s. |
| ENSG00000256537 | 2093.168793 | 0.209504782 | 0.2378341 | 0.880886 | 0.378379382 | 0.57089904 | SMIM10L1 | protein_coding | 38.73 | 0.051001988 | 0.051001988 | ENSG00000256537 | n.s. |
| ENSG00000102103 | 572.4643107 | 0.233137131 | 0.2845401 | 0.819347 | 0.412588327 | 0.604489874 | PQBP1 | protein_coding | 52.66 | 0.050966334 | 0.050966334 | ENSG00000102103 | n.s. |
| ENSG00000181817 | 56.41125117 | -0.238366031 | 0.2961906 | -0.804772 | 0.420951083 | 0.611226184 | LSM10 | protein_coding | 50.65 | -0.050962193 | 0.050962193 | ENSG00000181817 | n.s. |
| ENSG00000144228 | 1175.038697 | -0.220958577 | 0.2604492 | -0.848375 | 0.396229137 | 0.588134144 | SPOPL | protein_coding | 36.68 | -0.050936168 | 0.050936168 | ENSG00000144228 | n.s. |
| ENSG00000074356 | 1551.701139 | -0.21225001 | 0.2432033 | -0.872727 | 0.382812008 | 0.575638049 | NCBP3 | protein_coding | 44.57 | -0.050908273 | 0.050908273 | ENSG00000074356 | n.s. |
| ENSG00000141376 | 245.012219 | -0.228123051 | 0.2751511 | -0.829083 | 0.40705752 | 0.598723884 | BCAS3 | protein_coding | 39.85 | -0.050819751 | 0.050819751 | ENSG00000141376 | n.s. |
| ENSG00000067829 | 386.8845194 | 0.219706264 | 0.2584333 | 0.850147 | 0.395243505 | 0.587359944 | IDH3G | protein_coding | 58.77 | 0.050773167 | 0.050773167 | ENSG00000067829 | n.s. |
| ENSG00000235217 | 145.1574056 | -0.206901459 | 0.2337381 | -0.885185 | 0.376056887 | 0.568371693 | TSPY26P | transcribed_processed_pseudog | 55.99 | -0.050766906 | 0.050766906 | ENSG00000235217 | n.s. |
| ENSG00000110492 | 4615.567603 | 0.224802243 | 0.268677 | 0.836701 | 0.402760806 | 0.594813001 | MDK | protein_coding | 66.26 | 0.05071978 | 0.05071978 | ENSG00000110492 | n.s. |
| ENSG00000244657 | 4.371490149 | -0.326192987 | 0.5123336 | -0.636681 | 0.524332779 | 0.69912711 | | transcribed_unprocessed_pseud | 41.24 | -0.050704696 | 0.050704696 | ENSG00000244657 | n.s. |
| ENSG00000184428 | 282.1659396 | 0.228744624 | 0.2766763 | 0.826759 | 0.408373679 | 0.600273093 | TOP1MT | protein_coding | 54.3 | 0.050701503 | 0.050701503 | ENSG00000184428 | n.s. |
| ENSG00000169972 | 107.5790071 | 0.218467355 | 0.256226 | 0.852635 | 0.393861495 | 0.586056813 | PUSL1 | protein_coding | 66.76 | 0.050697596 | 0.050697596 | ENSG00000169972 | n.s. |
| ENSG00000162981 | 27257.75647 | 0.194172833 | 0.2100803 | 0.924279 | 0.355341118 | 0.548372243 | FAM84A | protein_coding | 41.56 | 0.050664456 | 0.050664456 | ENSG00000162981 | n.s. |
| ENSG00000113810 | 12544.88399 | -0.215052229 | 0.249878 | -0.860629 | 0.389442425 | 0.581366175 | SMC4 | protein_coding | 36.23 | -0.050655604 | 0.050655604 | ENSG00000113810 | n.s. |
| ENSG00000149289 | 1140.893131 | -0.206402375 | 0.2332392 | -0.884939 | 0.376189639 | 0.56851062 | ZC3H12C | protein_coding | 37.25 | -0.050622539 | 0.050622539 | ENSG00000149289 | n.s. |
| ENSG00000166974 | 654.5848168 | -0.202396679 | 0.2258832 | -0.896023 | 0.370240262 | 0.562289155 | MAPRE2 | protein_coding | 39.48 | -0.050607325 | 0.050607325 | ENSG00000166974 | n.s. |
| ENSG00000252712 | 21.15397096 | -0.266391411 | 0.360173 | -0.739621 | 0.459530159 | 0.645737461 | SCARNA14 | scaRNA | 42.65 | -0.050599455 | 0.050599455 | ENSG00000252712 | n.s. |
| ENSG00000198301 | 1726.377597 | -0.223646083 | 0.2667738 | -0.838336 | 0.401841963 | 0.593995717 | SDAD1 | protein_coding | 40.57 | -0.050592476 | 0.050592476 | ENSG00000198301 | n.s. |
| ENSG00000283041 | 599.9727466 | 0.246763045 | 0.3155185 | 0.782087 | 0.434163163 | 0.624056109 | | processed_pseudogene | 54.34 | 0.050531238 | 0.050531238 | ENSG00000283041 | n.s. |
| ENSG00000116133 | 3169.715819 | -0.216268247 | 0.2529555 | -0.854966 | 0.392570195 | 0.584513505 | DHCR24 | protein_coding | 47.68 | -0.050434934 | 0.050434934 | ENSG00000116133 | n.s. |
| ENSG00000138663 | 893.2742047 | 0.232326874 | 0.2853415 | 0.814206 | 0.41552669 | 0.607070513 | COPS4 | protein_coding | 40.06 | 0.050359373 | 0.050359373 | ENSG00000138663 | n.s. |
| ENSG00000271882 | 14.11913983 | 0.270842758 | 0.3726252 | 0.72685 | 0.467317663 | 0.651884964 | | lincRNA | 33.33 | 0.050330449 | 0.050330449 | ENSG00000271882 | n.s. |
| ENSG00000112514 | 1365.208047 | 0.230007926 | 0.2804981 | 0.819998 | 0.412217079 | 0.604290797 | CUTA | protein_coding | 54.9 | 0.050315158 | 0.050315158 | ENSG00000112514 | n.s. |
| ENSG00000115816 | 4487.090003 | -0.222803918 | 0.2661426 | -0.83716 | 0.402502681 | 0.59465671 | CEBPZ | protein_coding | 38.07 | -0.050294348 | 0.050294348 | ENSG00000115816 | n.s. |
| ENSG00000166451 | 732.1361864 | -0.239487669 | 0.3011612 | -0.795214 | 0.426488964 | 0.61674344 | CENPN | protein_coding | 43.26 | -0.050267375 | 0.050267375 | ENSG00000166451 | n.s. |
| ENSG00000171302 | 604.1000517 | -0.226761282 | 0.2742444 | -0.826858 | 0.408317411 | 0.600253755 | CANT1 | protein_coding | 57.12 | -0.050265066 | 0.050265066 | ENSG00000171302 | n.s. |

| | | | | | | | | | | | | | |
|---|---|---|---|---|---|---|---|---|---|---|---|---|---|
| ENSG00000063244 | 2679.987119 | 0.224626471 | 0.2699044 | 0.832245 | 0.405270831 | 0.597478375 | U2AF2 | protein_coding | 56.03 | 0.050243957 | 0.050243957 | ENSG00000063244 | n.s. |
| ENSG00000230470 | 39.44951525 | -0.238313816 | 0.298905 | -0.797289 | 0.425282923 | 0.615446849 | | sense_intronic | 39.79 | -0.050238804 | 0.050238804 | ENSG00000230470 | n.s. |
| ENSG00000075043 | 632.9982431 | 0.222262852 | 0.265267 | 0.837883 | 0.402096158 | 0.594308366 | KCNQ2 | protein_coding | 62.3 | 0.050228772 | 0.050228772 | ENSG00000075043 | n.s. |
| ENSG00000250251 | 132.7542073 | 0.219057667 | 0.259205 | 0.845114 | 0.398047406 | 0.589826951 | PKD1P6 | transcribed_unprocessed_pseud | 61.72 | 0.050224531 | 0.050224531 | ENSG00000250251 | n.s. |
| ENSG00000198677 | 4243.021264 | -0.223832106 | 0.2687187 | -0.832961 | 0.404866847 | 0.597009226 | TTC37 | protein_coding | 35.56 | -0.050142635 | 0.050142635 | ENSG00000198677 | n.s. |
| ENSG00000140688 | 449.8430369 | -0.234512987 | 0.2914415 | -0.804666 | 0.421012619 | 0.611236569 | C16orf58 | protein_coding | 51.18 | -0.050136689 | 0.050136689 | ENSG00000140688 | n.s. |
| ENSG00000087842 | 198.8400153 | 0.245971237 | 0.3160056 | 0.778376 | 0.436347254 | 0.625567375 | PIR | protein_coding | 38.99 | 0.050110714 | 0.050110714 | ENSG00000087842 | n.s. |
| ENSG00000186222 | 301.4427697 | 0.203171086 | 0.2289322 | 0.887473 | 0.37482447 | 0.566745763 | BLOC1S4 | protein_coding | 54.05 | 0.050104371 | 0.050104371 | ENSG00000186222 | n.s. |
| ENSG00000084463 | 3126.612871 | 0.210611031 | 0.2427153 | 0.867729 | 0.385542785 | 0.578408131 | WBP11 | protein_coding | 37.17 | 0.05007606 | 0.05007606 | ENSG00000084463 | n.s. |
| ENSG00000130725 | 474.6731299 | 0.236160895 | 0.2951656 | 0.800096 | 0.423655011 | 0.613728927 | UBE2M | protein_coding | 61.71 | 0.050071638 | 0.050071638 | ENSG00000130725 | n.s. |
| ENSG00000158435 | 646.8753294 | -0.221879209 | 0.265312 | -0.836295 | 0.402988703 | 0.594995939 | CNOT11 | protein_coding | 43.29 | -0.050030655 | 0.050030655 | ENSG00000158435 | n.s. |
| ENSG00000158805 | 268.9499578 | -0.223622328 | 0.2689732 | -0.831393 | 0.405751885 | 0.597871045 | ZNF276 | protein_coding | 56.58 | -0.049955546 | 0.049955546 | ENSG00000158805 | n.s. |
| ENSG00000167461 | 540.9222616 | -0.212578727 | 0.2474749 | -0.858991 | 0.390345461 | 0.582193879 | RAB8A | protein_coding | 53.79 | -0.049941623 | 0.049941623 | ENSG00000167461 | n.s. |
| ENSG00000184381 | 14.49886026 | -0.272824555 | 0.3792965 | -0.719291 | 0.471961665 | 0.656094752 | PLA2G6 | protein_coding | 51.54 | -0.049936016 | 0.049936016 | ENSG00000184381 | n.s. |
| ENSG00000213178 | 53.78537262 | 0.2682099 | 0.3684642 | 0.727913 | 0.46666687 | 0.651368704 | | processed_pseudogene | 41.46 | 0.049933472 | 0.049933472 | ENSG00000213178 | n.s. |
| ENSG00000276757 | 28.38763167 | -0.24821808 | 0.3224889 | -0.769695 | 0.441480807 | 0.62947723 | RN7SL192P | misc_RNA | 56.19 | -0.049896792 | 0.049896792 | ENSG00000276757 | n.s. |
| ENSG00000100865 | 339.0949089 | -0.238861181 | 0.3016089 | -0.791957 | 0.428385935 | 0.618523009 | CINP | protein_coding | 46.49 | -0.049836986 | 0.049836986 | ENSG00000100865 | n.s. |
| ENSG00000213533 | 378.4170705 | -0.211234401 | 0.2451545 | -0.861638 | 0.38888679 | 0.581051271 | TMEM110 | protein_coding | 48.39 | -0.049806018 | 0.049806018 | ENSG00000213533 | n.s. |
| ENSG00000123983 | 2895.625944 | 0.215509434 | 0.2536583 | 0.849605 | 0.395544606 | 0.587647937 | ACSL3 | protein_coding | 39.46 | 0.049757418 | 0.049757418 | ENSG00000123983 | n.s. |
| ENSG00000063660 | 810.9884204 | -0.220938575 | 0.2645014 | -0.835302 | 0.403547598 | 0.595568498 | GPC1 | protein_coding | 64.63 | -0.049726266 | 0.049726266 | ENSG00000063660 | n.s. |
| ENSG00000180198 | 676.5385476 | -0.226749061 | 0.2762021 | -0.820953 | 0.411672815 | 0.603733284 | RCC1 | protein_coding | 50 | -0.049693163 | 0.049693163 | ENSG00000180198 | n.s. |
| ENSG00000130669 | 190.2894802 | -0.228061655 | 0.27896 | -0.817543 | 0.41361846 | 0.605489414 | PAK4 | protein_coding | 56.17 | -0.04969314 | 0.04969314 | ENSG00000130669 | n.s. |
| ENSG00000178235 | 215.0053321 | 0.19740888 | 0.2192073 | 0.900558 | 0.367823459 | 0.560123157 | SLITRK1 | protein_coding | 47.31 | 0.049691067 | 0.049691067 | ENSG00000178235 | n.s. |
| ENSG00000148399 | 522.5081027 | 0.224760893 | 0.2723855 | 0.825157 | 0.409282259 | 0.601100938 | DPH7 | protein_coding | 52.97 | 0.049683978 | 0.049683978 | ENSG00000148399 | n.s. |
| ENSG00000167258 | 1789.917627 | -0.197305091 | 0.219347 | -0.899511 | 0.368380476 | 0.560664776 | CDK12 | protein_coding | 44.39 | -0.049582124 | 0.049582124 | ENSG00000167258 | n.s. |
| ENSG00000148660 | 640.0667232 | 0.215071776 | 0.2535085 | 0.848381 | 0.396225944 | 0.588134144 | CAMK2G | protein_coding | 51.65 | 0.049579122 | 0.049579122 | ENSG00000148660 | n.s. |
| ENSG00000255529 | 1691.826336 | 0.215525194 | 0.2543974 | 0.847199 | 0.396884306 | 0.588792775 | POLR2M | protein_coding | 42.66 | 0.049578883 | 0.049578883 | ENSG00000255529 | n.s. |
| ENSG00000171159 | 217.4434916 | 0.228860981 | 0.2812912 | 0.813609 | 0.415869121 | 0.607315478 | C9orf16 | protein_coding | 56.88 | 0.049568004 | 0.049568004 | ENSG00000171159 | n.s. |
| ENSG00000143321 | 6324.030932 | 0.231230138 | 0.2866865 | 0.806561 | 0.419919468 | 0.610652258 | HDGF | protein_coding | 51.35 | 0.04953089 | 0.04953089 | ENSG00000143321 | n.s. |
| ENSG00000184154 | 10.04494141 | 0.288372385 | 0.4226858 | 0.682238 | 0.49508832 | 0.673449188 | LRTOMT | protein_coding | 48.88 | 0.049512145 | 0.049512145 | ENSG00000184154 | n.s. |
| ENSG00000205808 | 263.8701848 | -0.202159816 | 0.2286647 | -0.884089 | 0.376648386 | 0.569018607 | PLPP6 | protein_coding | 47.55 | -0.049503588 | 0.049503588 | ENSG00000205808 | n.s. |
| ENSG00000111737 | 630.385337 | -0.227317977 | 0.2781989 | -0.817106 | 0.413867933 | 0.605720492 | RAB35 | protein_coding | 55.75 | -0.049493428 | 0.049493428 | ENSG00000111737 | n.s. |

| | | | | | | | | | | | | | |
|---|---|---|---|---|---|---|---|---|---|---|---|---|---|
| ENSG00000049860 | 342.1468517 | -0.232387508 | 0.2896395 | -0.802334 | 0.422360028 | 0.612554028 | HEXB | protein_coding | 42.11 | -0.049464982 | 0.049464982 | ENSG00000049860 | n.s. |
| ENSG00000114021 | 494.9826742 | -0.240717602 | 0.3068584 | -0.784458 | 0.432771291 | 0.623053394 | NIT2 | protein_coding | 43.01 | -0.049461385 | 0.049461385 | ENSG00000114021 | n.s. |
| ENSG00000134153 | 465.448517 | 0.230915833 | 0.2865408 | 0.805874 | 0.420315477 | 0.610830464 | EMC7 | protein_coding | 39.72 | 0.049434303 | 0.049434303 | ENSG00000134153 | n.s. |
| ENSG00000089199 | 1703.238434 | -0.208939367 | 0.2420368 | -0.863255 | 0.387997526 | 0.580121302 | CHGB | protein_coding | 42.87 | -0.04941023 | 0.04941023 | ENSG00000089199 | n.s. |
| ENSG00000096092 | 447.3247593 | 0.248812699 | 0.3260164 | 0.763191 | 0.445349623 | 0.633071582 | TMEM14A | protein_coding | 41.73 | 0.04940106 | 0.04940106 | ENSG00000096092 | n.s. |
| ENSG00000176619 | 2298.787256 | -0.207621803 | 0.2391935 | -0.868008 | 0.385390124 | 0.578334414 | LMNB2 | protein_coding | 59.78 | -0.049376817 | 0.049376817 | ENSG00000176619 | n.s. |
| ENSG00000147604 | 18745.18027 | 0.228072759 | 0.2805147 | 0.813051 | 0.416188771 | 0.607490298 | RPL7 | protein_coding | 50.72 | 0.049368778 | 0.049368778 | ENSG00000147604 | n.s. |
| ENSG00000172939 | 1256.522581 | -0.218663781 | 0.2611862 | -0.837195 | 0.402482914 | 0.59465671 | OXSR1 | protein_coding | 38.67 | -0.049359779 | 0.049359779 | ENSG00000172939 | n.s. |
| ENSG00000112130 | 615.8244883 | -0.223595612 | 0.2712849 | -0.82421 | 0.409820529 | 0.601574194 | RNF8 | protein_coding | 43.66 | -0.049349967 | 0.049349967 | ENSG00000112130 | n.s. |
| ENSG00000185359 | 1986.39495 | 0.214758736 | 0.2537094 | 0.846475 | 0.397287601 | 0.58913998 | HGS | protein_coding | 60.6 | 0.049347586 | 0.049347586 | ENSG00000185359 | n.s. |
| ENSG00000198727 | 2356.448201 | -0.20812634 | 0.2404437 | -0.865593 | 0.386713493 | 0.579415258 | MT-CYB | protein_coding | 46.28 | -0.049328039 | 0.049328039 | ENSG00000198727 | n.s. |
| ENSG00000158042 | 1158.695578 | 0.188443303 | 0.2035491 | 0.925788 | 0.354556272 | 0.547464454 | MRPL17 | protein_coding | 48.63 | 0.049305074 | 0.049305074 | ENSG00000158042 | n.s. |
| ENSG00000071564 | 1520.218745 | -0.208810822 | 0.2421089 | -0.862467 | 0.388430751 | 0.580644336 | TCF3 | protein_coding | 59.57 | -0.049298107 | 0.049298107 | ENSG00000071564 | n.s. |
| ENSG00000176108 | 223.4652634 | -0.22131674 | 0.2670572 | -0.828724 | 0.407260492 | 0.598900742 | CHMP6 | protein_coding | 58.37 | -0.049275098 | 0.049275098 | ENSG00000176108 | n.s. |
| ENSG00000179526 | 223.0379519 | -0.216992456 | 0.2584493 | -0.839594 | 0.401136119 | 0.593502669 | SHARPIN | protein_coding | 60.75 | -0.04916557 | 0.04916557 | ENSG00000179526 | n.s. |
| ENSG00000100403 | 1583.310142 | -0.215013158 | 0.2548156 | -0.843799 | 0.398781781 | 0.590663695 | ZC3H7B | protein_coding | 54.29 | -0.049164849 | 0.049164849 | ENSG00000100403 | n.s. |
| ENSG00000130305 | 376.1135945 | 0.218493611 | 0.2616311 | 0.835121 | 0.403649587 | 0.595655877 | NSUN5 | protein_coding | 53.93 | 0.049162061 | 0.049162061 | ENSG00000130305 | n.s. |
| ENSG00000148572 | 388.665724 | -0.243038046 | 0.3136372 | -0.774902 | 0.438397683 | 0.627683831 | NRBF2 | protein_coding | 42.23 | -0.049156646 | 0.049156646 | ENSG00000148572 | n.s. |
| ENSG00000144591 | 455.1285743 | 0.220123339 | 0.2651715 | 0.830117 | 0.406472744 | 0.598173555 | GMPPA | protein_coding | 54.38 | 0.04912554 | 0.04912554 | ENSG00000144591 | n.s. |
| ENSG00000186205 | 767.3814043 | -0.231400552 | 0.2890039 | -0.800683 | 0.423315076 | 0.613364118 | 1-Mar | protein_coding | 45.28 | -0.049122087 | 0.049122087 | ENSG00000186205 | n.s. |
| ENSG00000167113 | 594.8573719 | -0.217463057 | 0.2598997 | -0.836719 | 0.402750424 | 0.594813001 | COQ4 | protein_coding | 51.72 | -0.049063916 | 0.049063916 | ENSG00000167113 | n.s. |
| ENSG00000197818 | 54.35464254 | -0.22759083 | 0.2809768 | -0.809999 | 0.417940905 | 0.608943545 | SLC9A8 | protein_coding | 45.56 | -0.049028292 | 0.049028292 | ENSG00000197818 | n.s. |
| ENSG00000183527 | 1348.753139 | 0.216319105 | 0.2577456 | 0.839274 | 0.401315735 | 0.593532927 | PSMG1 | protein_coding | 41.56 | 0.049008215 | 0.049008215 | ENSG00000183527 | n.s. |
| ENSG00000107164 | 1459.136441 | -0.218574589 | 0.2623417 | -0.833168 | 0.404750255 | 0.59696375 | FUBP3 | protein_coding | 47.12 | -0.048972083 | 0.048972083 | ENSG00000107164 | n.s. |
| ENSG00000146066 | 487.4496937 | 0.225881629 | 0.277702 | 0.813396 | 0.415991272 | 0.607366452 | HIGD2A | protein_coding | 57.37 | 0.048914486 | 0.048914486 | ENSG00000146066 | n.s. |
| ENSG00000101182 | 3283.387438 | 0.230679101 | 0.2886415 | 0.799189 | 0.424180792 | 0.614234963 | PSMA7 | protein_coding | 52.28 | 0.0488268 | 0.0488268 | ENSG00000101182 | n.s. |
| ENSG00000176087 | 560.6870021 | -0.191463467 | 0.2107599 | -0.908443 | 0.363643977 | 0.556008686 | SLC35A4 | protein_coding | 54.5 | -0.048807565 | 0.048807565 | ENSG00000176087 | n.s. |
| ENSG00000105221 | 368.2381267 | 0.210723149 | 0.2475059 | 0.851386 | 0.394554807 | 0.586712227 | AKT2 | protein_coding | 53.61 | 0.048798184 | 0.048798184 | ENSG00000105221 | n.s. |
| ENSG00000227057 | 852.0657608 | 0.22554226 | 0.2782217 | 0.810657 | 0.417562842 | 0.60865543 | WDR46 | protein_coding | 51.88 | 0.048633339 | 0.048633339 | ENSG00000227057 | n.s. |
| ENSG00000139637 | 525.0408722 | 0.237172533 | 0.3028753 | 0.78307 | 0.433586011 | 0.623767104 | C12orf10 | protein_coding | 51.21 | 0.048615041 | 0.048615041 | ENSG00000139637 | n.s. |
| ENSG00000266589 | 13.02462675 | 0.261384983 | 0.3598826 | 0.726306 | 0.467650997 | 0.651988717 | MIR4512 | miRNA | 53.25 | 0.048554854 | 0.048554854 | ENSG00000266589 | n.s. |
| ENSG00000138111 | 118.8894529 | -0.217601769 | 0.2622234 | -0.829834 | 0.406632927 | 0.598346041 | MFSD13A | protein_coding | 53.46 | -0.048535548 | 0.048535548 | ENSG00000138111 | n.s. |

| | | | | | | | | | | | | | |
|---|---|---|---|---|---|---|---|---|---|---|---|---|---|
| ENSG00000138448 | 1553.700821 | -0.233894436 | 0.2964466 | -0.788993 | 0.430115818 | 0.620410218 | ITGAV | protein_coding | 35.78 | -0.048491242 | 0.048491242 | ENSG00000138448 | n.s. |
| ENSG00000171044 | 184.835816 | 0.204279208 | 0.2358473 | 0.86615 | 0.386407748 | 0.579204016 | XKR6 | protein_coding | 46.08 | 0.04844858 | 0.04844858 | ENSG00000171044 | n.s. |
| ENSG00000131652 | 502.6316977 | 0.230563981 | 0.2899979 | 0.795054 | 0.426582154 | 0.616763089 | THOC6 | protein_coding | 57.25 | 0.048391143 | 0.048391143 | ENSG00000131652 | n.s. |
| ENSG00000108773 | 588.992523 | 0.223423677 | 0.2746517 | 0.81348 | 0.415942869 | 0.607359472 | KAT2A | protein_coding | 58.67 | 0.048383334 | 0.048383334 | ENSG00000108773 | n.s. |
| ENSG00000186364 | 63.28922993 | -0.235564243 | 0.3004554 | -0.784024 | 0.433026082 | 0.623219159 | NUDT17 | protein_coding | 54.02 | -0.048375285 | 0.048375285 | ENSG00000186364 | n.s. |
| ENSG00000105467 | 61.62381975 | 0.234891091 | 0.2989744 | 0.785656 | 0.432068933 | 0.622380245 | SYNGR4 | protein_coding | 54.82 | 0.048374458 | 0.048374458 | ENSG00000105467 | n.s. |
| ENSG00000104626 | 853.8170299 | -0.212829455 | 0.2529834 | -0.841278 | 0.400192093 | 0.592563492 | ERI1 | protein_coding | 41.64 | -0.048368709 | 0.048368709 | ENSG00000104626 | n.s. |
| ENSG00000107938 | 1838.465984 | -0.205386692 | 0.2388406 | -0.859932 | 0.389826463 | 0.581669043 | EDRF1 | protein_coding | 39.09 | -0.048332428 | 0.048332428 | ENSG00000107938 | n.s. |
| ENSG00000078328 | 224.8246346 | -0.195657561 | 0.2200944 | -0.888971 | 0.374018695 | 0.566213144 | RBFOX1 | protein_coding | 41.37 | -0.048331341 | 0.048331341 | ENSG00000078328 | n.s. |
| ENSG00000149269 | 131.3222566 | -0.225866532 | 0.2804861 | -0.805268 | 0.420664886 | 0.611114066 | PAK1 | protein_coding | 39.88 | -0.048307819 | 0.048307819 | ENSG00000149269 | n.s. |
| ENSG00000053108 | 559.2335907 | -0.211953109 | 0.2516762 | -0.842166 | 0.399695032 | 0.591953401 | FSTL4 | protein_coding | 46.69 | -0.048264368 | 0.048264368 | ENSG00000053108 | n.s. |
| ENSG00000157540 | 2471.420616 | -0.205545247 | 0.2397348 | -0.857386 | 0.391231676 | 0.583078423 | DYRK1A | protein_coding | 38.86 | -0.048153708 | 0.048153708 | ENSG00000157540 | n.s. |
| ENSG00000198937 | 297.4964375 | 0.247062164 | 0.3283444 | 0.752448 | 0.45178159 | 0.638529598 | CCDC167 | protein_coding | 52.31 | 0.048132396 | 0.048132396 | ENSG00000198937 | n.s. |
| ENSG00000075790 | 258.0340526 | -0.225028427 | 0.2797175 | -0.804485 | 0.421117107 | 0.611260788 | BCAP29 | protein_coding | 37.42 | -0.048105106 | 0.048105106 | ENSG00000075790 | n.s. |
| ENSG00000140939 | 223.8114774 | -0.213055477 | 0.254993 | -0.835534 | 0.403416851 | 0.595564926 | NOL3 | protein_coding | 58.13 | -0.047952585 | 0.047952585 | ENSG00000140939 | n.s. |
| ENSG00000255121 | 96.49269897 | -0.211605066 | 0.252095 | -0.839386 | 0.401252648 | 0.593502669 | | lincRNA | 51.59 | -0.04794491 | 0.04794491 | ENSG00000255121 | n.s. |
| ENSG00000228107 | 33.34594092 | -0.237492389 | 0.3070143 | -0.773555 | 0.439194174 | 0.628393843 | | sense_overlapping | 47.84 | -0.047918383 | 0.047918383 | ENSG00000228107 | n.s. |
| ENSG00000233270 | 48.07771291 | 0.261486568 | 0.3632457 | 0.719861 | 0.471610284 | 0.655770497 | SNRPEP4 | processed_pseudogene | 43.01 | 0.047916924 | 0.047916924 | ENSG00000233270 | n.s. |
| ENSG00000120833 | 1755.547245 | 0.203090742 | 0.2357569 | 0.861441 | 0.388995108 | 0.581051271 | SOCS2 | protein_coding | 47.34 | 0.047885861 | 0.047885861 | ENSG00000120833 | n.s. |
| ENSG00000276259 | 49.74201554 | -0.247427863 | 0.3305952 | -0.748432 | 0.454199901 | 0.64054368 | | antisense | 42.84 | -0.047865229 | 0.047865229 | ENSG00000276259 | n.s. |
| ENSG00000132382 | 727.0526103 | -0.193998047 | 0.2185459 | -0.887676 | 0.374714873 | 0.566712489 | MYBBP1A | protein_coding | 59.89 | -0.047847138 | 0.047847138 | ENSG00000132382 | n.s. |
| ENSG00000254999 | 1182.171458 | 0.237691684 | 0.3084708 | 0.770548 | 0.440974637 | 0.629085636 | BRK1 | protein_coding | 45.07 | 0.047845014 | 0.047845014 | ENSG00000254999 | n.s. |
| ENSG00000177830 | 278.3137913 | 0.222643628 | 0.2757004 | 0.807557 | 0.419345913 | 0.610089148 | CHID1 | protein_coding | 59.37 | 0.047780814 | 0.047780814 | ENSG00000177830 | n.s. |
| ENSG00000213058 | 128.6816286 | 0.255426766 | 0.3500152 | 0.729759 | 0.465537447 | 0.650450548 | | processed_pseudogene | 53.78 | 0.047710071 | 0.047710071 | ENSG00000213058 | n.s. |
| ENSG00000108424 | 12762.98171 | -0.210460061 | 0.2506063 | -0.839803 | 0.401018634 | 0.593464801 | KPNB1 | protein_coding | 43.15 | -0.047691309 | 0.047691309 | ENSG00000108424 | n.s. |
| ENSG00000214548 | 1748.724566 | 0.206168748 | 0.2423994 | 0.850533 | 0.395028632 | 0.587228662 | MEG3 | lincRNA | 51.68 | 0.047664721 | 0.047664721 | ENSG00000214548 | n.s. |
| ENSG00000119953 | 1865.96162 | -0.20038402 | 0.2309849 | -0.86752 | 0.385657082 | 0.578408131 | SMNDC1 | protein_coding | 37.32 | -0.047644429 | 0.047644429 | ENSG00000119953 | n.s. |
| ENSG00000177602 | 368.5599186 | 0.192226924 | 0.2161393 | 0.889366 | 0.373806519 | 0.565953476 | GSG2 | protein_coding | 53.91 | 0.047522199 | 0.047522199 | ENSG00000177602 | n.s. |
| ENSG00000138757 | 3959.076797 | 0.221985064 | 0.2755155 | 0.805708 | 0.420411202 | 0.610873008 | G3BP2 | protein_coding | 41.77 | 0.047515695 | 0.047515695 | ENSG00000138757 | n.s. |
| ENSG00000179119 | 667.1440976 | 0.220127488 | 0.2715484 | 0.810638 | 0.41757356 | 0.60865543 | SPTY2D1 | protein_coding | 41.92 | 0.04746576 | 0.04746576 | ENSG00000179119 | n.s. |
| ENSG00000160014 | 1423.544539 | 0.219242026 | 0.2701641 | 0.811514 | 0.417070339 | 0.608193772 | CALM3 | protein_coding | 57.51 | 0.047347076 | 0.047347076 | ENSG00000160014 | n.s. |
| ENSG00000164616 | 217.1493206 | 0.231039104 | 0.2952021 | 0.782647 | 0.433834314 | 0.623880263 | FBXL21 | transcribed_unitary_pseudogene | 37.89 | 0.047339624 | 0.047339624 | ENSG00000164616 | n.s. |

| | | | | | | | | | | | | | |
|---|---|---|---|---|---|---|---|---|---|---|---|---|---|
| ENSG00000102471 | 629.3518348 | 0.222030616 | 0.276493 | 0.803024 | 0.42196066 | 0.612166122 | NDFIP2 | protein_coding | 35.47 | 0.047321543 | 0.047321543 | ENSG00000102471 | n.s. |
| ENSG00000239246 | 21.78168807 | 0.272186149 | 0.3944676 | 0.690009 | 0.490188625 | 0.670128457 | | processed_pseudogene | 52.35 | 0.047317368 | 0.047317368 | ENSG00000239246 | n.s. |
| ENSG00000115419 | 3357.652743 | -0.216690548 | 0.2647446 | -0.818489 | 0.413078079 | 0.604889156 | GLS | protein_coding | 35.11 | -0.047308791 | 0.047308791 | ENSG00000115419 | n.s. |
| ENSG00000124571 | 2879.258264 | -0.217757787 | 0.2671737 | -0.815042 | 0.415048329 | 0.606625854 | XPO5 | protein_coding | 43.8 | -0.047270661 | 0.047270661 | ENSG00000124571 | n.s. |
| ENSG00000266370 | 19.26730794 | 0.253091581 | 0.3469722 | 0.729429 | 0.465739341 | 0.650661133 | MIR3657 | miRNA | 29.91 | 0.047238312 | 0.047238312 | ENSG00000266370 | n.s. |
| ENSG00000221669 | 4.099827739 | -0.40351622 | 0.7830424 | -0.515319 | 0.606330466 | 0.763907255 | MIR548N | miRNA | 36 | -0.047195001 | 0.047195001 | ENSG00000221669 | n.s. |
| ENSG00000155313 | 1532.607578 | -0.211381336 | 0.2544529 | -0.830729 | 0.406126799 | 0.598079974 | USP25 | protein_coding | 35.41 | -0.047188926 | 0.047188926 | ENSG00000155313 | n.s. |
| ENSG00000221241 | 24.1534668 | 0.248283585 | 0.335552 | 0.739926 | 0.459344973 | 0.645671434 | SNORD88A | snoRNA | 70.33 | 0.047171008 | 0.047171008 | ENSG00000221241 | n.s. |
| ENSG00000264792 | 8.401279053 | -0.351517313 | 0.6173738 | -0.569375 | 0.569101567 | 0.73421063 | MIR4637 | miRNA | 30.95 | -0.047166358 | 0.047166358 | ENSG00000264792 | n.s. |
| ENSG00000261574 | 39.28976731 | 0.234437986 | 0.3045575 | 0.769766 | 0.441438735 | 0.62947723 | | sense_intronic | 52.59 | 0.047126718 | 0.047126718 | ENSG00000261574 | n.s. |
| ENSG00000115685 | 633.566481 | -0.226055497 | 0.2856791 | -0.791292 | 0.428773795 | 0.618826323 | PPP1R7 | protein_coding | 48.38 | -0.047117023 | 0.047117023 | ENSG00000115685 | n.s. |
| ENSG00000125447 | 140.7527574 | -0.215456669 | 0.262941 | -0.819411 | 0.412552132 | 0.604489874 | GGA3 | protein_coding | 51.82 | -0.047101191 | 0.047101191 | ENSG00000125447 | n.s. |
| ENSG00000100297 | 3250.480619 | -0.208089583 | 0.2481306 | -0.838629 | 0.40167733 | 0.593878461 | MCM5 | protein_coding | 52.51 | -0.047091177 | 0.047091177 | ENSG00000100297 | n.s. |
| ENSG00000140262 | 6211.225119 | -0.210036316 | 0.252127 | -0.833058 | 0.404812202 | 0.596991875 | TCF12 | protein_coding | 37.17 | -0.047054768 | 0.047054768 | ENSG00000140262 | n.s. |
| ENSG00000212978 | 109.0978803 | 0.209073232 | 0.2504303 | 0.834856 | 0.403798874 | 0.595813027 | | processed_transcript | 45.77 | 0.04701848 | 0.04701848 | ENSG00000212978 | n.s. |
| ENSG00000227248 | 98.56169734 | -0.235527819 | 0.3076698 | -0.765521 | 0.443961092 | 0.631579675 | FAM155A-IT1 | sense_intronic | 35.94 | -0.047004724 | 0.047004724 | ENSG00000227248 | n.s. |
| ENSG00000122705 | 2102.029645 | 0.238566866 | 0.3146345 | 0.758235 | 0.448310377 | 0.635300293 | CLTA | protein_coding | 44.48 | 0.047002669 | 0.047002669 | ENSG00000122705 | n.s. |
| ENSG00000139722 | 488.5179135 | 0.207625697 | 0.2475655 | 0.83867 | 0.401654578 | 0.593878461 | VPS37B | protein_coding | 47.3 | 0.046986199 | 0.046986199 | ENSG00000139722 | n.s. |
| ENSG00000135454 | 436.5553086 | -0.212103562 | 0.2566523 | -0.826424 | 0.408563653 | 0.600488942 | B4GALNT1 | protein_coding | 56.73 | -0.046979876 | 0.046979876 | ENSG00000135454 | n.s. |
| ENSG00000140463 | 420.9571965 | -0.230184807 | 0.2951649 | -0.779852 | 0.435478228 | 0.625068389 | BBS4 | protein_coding | 40.76 | -0.046974381 | 0.046974381 | ENSG00000140463 | n.s. |
| ENSG00000101343 | 1278.193866 | -0.215751429 | 0.2642381 | -0.816504 | 0.414212023 | 0.605937302 | CRNKL1 | protein_coding | 39.08 | -0.046941537 | 0.046941537 | ENSG00000101343 | n.s. |
| ENSG00000175348 | 329.2321907 | 0.21683351 | 0.2667388 | 0.812906 | 0.416272167 | 0.607490558 | TMEM9B | protein_coding | 42.7 | 0.046935883 | 0.046935883 | ENSG00000175348 | n.s. |
| ENSG00000155096 | 3595.195426 | 0.218518933 | 0.2705596 | 0.807655 | 0.419288996 | 0.610070915 | AZIN1 | protein_coding | 40.82 | 0.046898463 | 0.046898463 | ENSG00000155096 | n.s. |
| ENSG00000115514 | 1155.887034 | -0.238359082 | 0.3147981 | -0.757181 | 0.448941554 | 0.635741733 | TXNDC9 | protein_coding | 42.12 | -0.046889827 | 0.046889827 | ENSG00000115514 | n.s. |
| ENSG00000140600 | 282.6251388 | -0.228894826 | 0.2927249 | -0.781945 | 0.434246821 | 0.624056109 | SH3GL3 | protein_coding | 40.8 | -0.04687225 | 0.04687225 | ENSG00000140600 | n.s. |
| ENSG00000168495 | 1406.680532 | -0.197912737 | 0.2292024 | -0.863485 | 0.387871021 | 0.580120522 | POLR3D | protein_coding | 49.5 | -0.046802755 | 0.046802755 | ENSG00000168495 | n.s. |
| ENSG00000186648 | 206.3322382 | 0.21128209 | 0.2557876 | 0.826006 | 0.408800749 | 0.600647196 | CARMIL3 | protein_coding | 57.83 | 0.046773744 | 0.046773744 | ENSG00000186648 | n.s. |
| ENSG00000114054 | 922.3504949 | 0.223044121 | 0.2809056 | 0.794018 | 0.427184901 | 0.617429189 | PCCB | protein_coding | 41.76 | 0.046708304 | 0.046708304 | ENSG00000114054 | n.s. |
| ENSG00000090621 | 1370.13293 | 0.19839698 | 0.2307105 | 0.859939 | 0.389822696 | 0.581669043 | PABPC4 | protein_coding | 47.27 | 0.046687581 | 0.046687581 | ENSG00000090621 | n.s. |
| ENSG00000149925 | 3573.195965 | 0.217925884 | 0.2702979 | 0.806243 | 0.420102617 | 0.610743316 | ALDOA | protein_coding | 54.43 | 0.046666927 | 0.046666927 | ENSG00000149925 | n.s. |
| ENSG00000102401 | 727.7772405 | -0.221119283 | 0.2771765 | -0.797756 | 0.425011979 | 0.615178794 | ARMCX3 | protein_coding | 45.41 | -0.046655869 | 0.046655869 | ENSG00000102401 | n.s. |
| ENSG00000146858 | 332.454942 | -0.19174817 | 0.2178517 | -0.880178 | 0.378763073 | 0.57128353 | ZC3HAV1L | protein_coding | 46.48 | -0.046623241 | 0.046623241 | ENSG00000146858 | n.s. |

| | | | | | | | | | | | | | |
|---|---|---|---|---|---|---|---|---|---|---|---|---|---|
| ENSG00000141959 | 290.0255523 | 0.215757758 | 0.2659911 | 0.811147 | 0.417281542 | 0.608420873 | PFKL | protein_coding | 63.48 | 0.046559638 | 0.046559638 | ENSG00000141959 | n.s. |
| ENSG00000168734 | 156.4242127 | 0.245554399 | 0.3324983 | 0.738513 | 0.460202667 | 0.646363536 | PKIG | protein_coding | 44.32 | 0.046538244 | 0.046538244 | ENSG00000168734 | n.s. |
| ENSG00000134897 | 619.8158593 | -0.217542576 | 0.2702323 | -0.805021 | 0.420807759 | 0.61115632 | BIVM | protein_coding | 40.15 | -0.046520978 | 0.046520978 | ENSG00000134897 | n.s. |
| ENSG00000074319 | 746.0396823 | -0.23685407 | 0.3131987 | -0.756242 | 0.449504017 | 0.636214743 | TSG101 | protein_coding | 39.7 | -0.046517256 | 0.046517256 | ENSG00000074319 | n.s. |
| ENSG00000266237 | 40.06134454 | 0.231061573 | 0.2998145 | 0.770682 | 0.440895557 | 0.629082685 | | sense_intronic | 40.4 | 0.046510908 | 0.046510908 | ENSG00000266237 | n.s. |
| ENSG00000188985 | 47.70417054 | -0.255678493 | 0.3579511 | -0.714283 | 0.475052048 | 0.658233153 | DHFRP1 | processed_pseudogene | 42.04 | -0.046436391 | 0.046436391 | ENSG00000188985 | n.s. |
| ENSG00000160563 | 351.8999969 | -0.213198066 | 0.2608588 | -0.817293 | 0.413761094 | 0.605634536 | MED27 | protein_coding | 44 | -0.046432272 | 0.046432272 | ENSG00000160563 | n.s. |
| ENSG00000136628 | 5613.472655 | -0.218186092 | 0.2720436 | -0.802026 | 0.422537878 | 0.612620522 | EPRS | protein_coding | 38.15 | -0.046431846 | 0.046431846 | ENSG00000136628 | n.s. |
| ENSG00000131153 | 1222.39398 | 0.212535174 | 0.2598307 | 0.817976 | 0.413371167 | 0.605191037 | GINS2 | protein_coding | 49.77 | 0.046355518 | 0.046355518 | ENSG00000131153 | n.s. |
| ENSG00000254469 | 76.49336623 | 0.207811654 | 0.2506547 | 0.829075 | 0.407061717 | 0.598723884 | | protein_coding | 42.82 | 0.046294912 | 0.046294912 | ENSG00000254469 | n.s. |
| ENSG00000066629 | 132.1654713 | -0.231343947 | 0.3015638 | -0.767148 | 0.442993814 | 0.630987857 | EML1 | protein_coding | 45.19 | -0.046263931 | 0.046263931 | ENSG00000066629 | n.s. |
| ENSG00000145907 | 6993.119162 | -0.213820874 | 0.2633593 | -0.811898 | 0.416850128 | 0.607982855 | G3BP1 | protein_coding | 39.44 | -0.046208544 | 0.046208544 | ENSG00000145907 | n.s. |
| ENSG00000110697 | 11.9371492 | -0.266144285 | 0.3864103 | -0.688761 | 0.490973832 | 0.670468885 | PITPNM1 | protein_coding | 62.46 | -0.046208336 | 0.046208336 | ENSG00000110697 | n.s. |
| ENSG00000137944 | 928.7058031 | -0.235759231 | 0.3122541 | -0.755024 | 0.450234798 | 0.636980467 | KYAT3 | protein_coding | 38.59 | -0.046179077 | 0.046179077 | ENSG00000137944 | n.s. |
| ENSG00000099942 | 2379.824161 | 0.20333462 | 0.2422257 | 0.839443 | 0.401220823 | 0.593502669 | CRKL | protein_coding | 45.76 | 0.046071014 | 0.046071014 | ENSG00000099942 | n.s. |
| ENSG00000279074 | 17.18465348 | 0.251406911 | 0.3496805 | 0.718962 | 0.472164432 | 0.656094752 | | TEC | 44.73 | 0.046015871 | 0.046015871 | ENSG00000279074 | n.s. |
| ENSG00000132437 | 196.3187421 | -0.223343873 | 0.2842871 | -0.785628 | 0.432085574 | 0.622380245 | DDC | protein_coding | 44.42 | -0.045996376 | 0.045996376 | ENSG00000132437 | n.s. |
| ENSG00000178741 | 1407.862903 | 0.228267579 | 0.2956883 | 0.771987 | 0.440121945 | 0.628944621 | COX5A | protein_coding | 44.93 | 0.045970258 | 0.045970258 | ENSG00000178741 | n.s. |
| ENSG00000124784 | 1503.856347 | 0.208305101 | 0.2528255 | 0.823909 | 0.409991542 | 0.601698349 | RIOK1 | protein_coding | 40.11 | 0.045956517 | 0.045956517 | ENSG00000124784 | n.s. |
| ENSG00000226508 | 144.8671042 | 0.196615444 | 0.2296184 | 0.85627 | 0.391848318 | 0.583809967 | LINC01918 | lincRNA | 57.47 | 0.045954632 | 0.045954632 | ENSG00000226508 | n.s. |
| ENSG00000160695 | 461.6457093 | 0.205151732 | 0.2465743 | 0.832008 | 0.405404575 | 0.597551897 | VPS11 | protein_coding | 48.94 | 0.045876927 | 0.045876927 | ENSG00000160695 | n.s. |
| ENSG00000171425 | 195.58781 | -0.217889044 | 0.27355 | -0.796524 | 0.425727817 | 0.615898628 | ZNF581 | protein_coding | 58.42 | -0.045863631 | 0.045863631 | ENSG00000171425 | n.s. |
| ENSG00000279623 | 16.78981869 | 0.255799358 | 0.3619187 | 0.706787 | 0.479698998 | 0.661861971 | | sense_intronic | 32.71 | 0.045847577 | 0.045847577 | ENSG00000279623 | n.s. |
| ENSG00000160285 | 1062.783389 | 0.205244945 | 0.2471677 | 0.830387 | 0.406319737 | 0.598129014 | LSS | protein_coding | 53.33 | 0.045811725 | 0.045811725 | ENSG00000160285 | n.s. |
| ENSG00000121964 | 115.7450213 | -0.21700895 | 0.2720485 | -0.797685 | 0.425053508 | 0.615178794 | GTDC1 | protein_coding | 36.87 | -0.045788594 | 0.045788594 | ENSG00000121964 | n.s. |
| ENSG00000120008 | 2059.630866 | -0.211478434 | 0.2601149 | -0.813019 | 0.416206992 | 0.607490298 | WDR11 | protein_coding | 36.82 | -0.04577676 | 0.04577676 | ENSG00000120008 | n.s. |
| ENSG00000242338 | 81.50491505 | -0.229207042 | 0.2993068 | -0.765793 | 0.443799475 | 0.631543345 | BMS1P4 | transcribed_unprocessed_pseud | 39.67 | -0.045749001 | 0.045749001 | ENSG00000242338 | n.s. |
| ENSG00000099904 | 188.4188323 | 0.190678274 | 0.2183808 | 0.873146 | 0.38258361 | 0.575549606 | ZDHHC8 | protein_coding | 63.57 | 0.045747005 | 0.045747005 | ENSG00000099904 | n.s. |
| ENSG00000181915 | 1033.689581 | 0.188609508 | 0.2147362 | 0.878331 | 0.379763964 | 0.572235246 | ADO | protein_coding | 47.48 | 0.045723735 | 0.045723735 | ENSG00000181915 | n.s. |
| ENSG00000183722 | 82.67850076 | 0.21087315 | 0.2590592 | 0.813996 | 0.415647278 | 0.607118867 | LHFP | protein_coding | 39.43 | 0.045701751 | 0.045701751 | ENSG00000183722 | n.s. |
| ENSG00000111271 | 76.84709799 | -0.20473304 | 0.2465931 | -0.830246 | 0.406399517 | 0.598129014 | ACAD10 | protein_coding | 47.12 | -0.045697465 | 0.045697465 | ENSG00000111271 | n.s. |
| ENSG00000140400 | 1154.322934 | 0.193180367 | 0.2237616 | 0.863331 | 0.387955339 | 0.580120522 | MAN2C1 | protein_coding | 57.24 | 0.045683635 | 0.045683635 | ENSG00000140400 | n.s. |

| | | | | | | | | | | | | | |
|---|---|---|---|---|---|---|---|---|---|---|---|---|---|
| ENSG00000124217 | 404.6973793 | -0.185168686 | 0.2085394 | -0.887931 | 0.374577731 | 0.566628178 | MOCS3 | protein_coding | 45.61 | -0.045681454 | 0.045681454 | ENSG00000124217 | n.s. |
| ENSG00000146701 | 1972.161043 | 0.217873092 | 0.2747623 | 0.792951 | 0.42780627 | 0.61794239 | MDH2 | protein_coding | 50.93 | 0.045546808 | 0.045546808 | ENSG00000146701 | n.s. |
| ENSG00000134308 | 9242.309199 | 0.222169109 | 0.2841441 | 0.781889 | 0.434279943 | 0.624056109 | YWHAQ | protein_coding | 40.33 | 0.045494982 | 0.045494982 | ENSG00000134308 | n.s. |
| ENSG00000165699 | 75.10921716 | 0.212002832 | 0.2633609 | 0.80499 | 0.420825658 | 0.61115632 | TSC1 | protein_coding | 42.78 | 0.045336316 | 0.045336316 | ENSG00000165699 | n.s. |
| ENSG00000170231 | 151.6693946 | 0.23962703 | 0.3248831 | 0.737579 | 0.460770236 | 0.646899667 | FABP6 | protein_coding | 48.54 | 0.045328585 | 0.045328585 | ENSG00000170231 | n.s. |
| ENSG00000111328 | 2260.600543 | 0.224904323 | 0.2914265 | 0.771736 | 0.440270779 | 0.62896057 | CDK2AP1 | protein_coding | 58.95 | 0.045290463 | 0.045290463 | ENSG00000111328 | n.s. |
| ENSG00000274390 | 2.77108632 | -0.344082882 | 0.612919 | -0.561384 | 0.574535805 | 0.7386046 | MIR6885 | miRNA | 65.15 | -0.045277175 | 0.045277175 | ENSG00000274390 | n.s. |
| ENSG00000074855 | 143.210031 | 0.21039024 | 0.260316 | 0.808211 | 0.418969203 | 0.609927068 | ANO8 | protein_coding | 62.67 | 0.045175433 | 0.045175433 | ENSG00000074855 | n.s. |
| ENSG00000023697 | 1069.163009 | -0.220280597 | 0.281575 | -0.782316 | 0.43402895 | 0.623953027 | DERA | protein_coding | 39.16 | -0.045124063 | 0.045124063 | ENSG00000023697 | n.s. |
| ENSG00000117461 | 1245.307209 | -0.200969553 | 0.2409749 | -0.833986 | 0.404289108 | 0.596473165 | PIK3R3 | protein_coding | 40.27 | -0.045099395 | 0.045099395 | ENSG00000117461 | n.s. |
| ENSG00000013392 | 196.8391712 | -0.204124054 | 0.2476106 | -0.824375 | 0.409726405 | 0.601562874 | RWDD2A | protein_coding | 44.76 | -0.045054052 | 0.045054052 | ENSG00000013392 | n.s. |
| ENSG00000263740 | 128.2439273 | -0.20220599 | 0.2438453 | -0.829239 | 0.406969368 | 0.598714551 | RN7SL4P | misc_RNA | 58.64 | -0.045047488 | 0.045047488 | ENSG00000263740 | n.s. |
| ENSG00000105738 | 66.2020191 | 0.215263031 | 0.2712437 | 0.793615 | 0.427419657 | 0.617640258 | SIPA1L3 | protein_coding | 49.76 | 0.04504689 | 0.04504689 | ENSG00000105738 | n.s. |
| ENSG00000180233 | 422.1147601 | 0.219272317 | 0.2795807 | 0.78429 | 0.432870087 | 0.623123473 | ZNRF2 | protein_coding | 36.75 | 0.045044211 | 0.045044211 | ENSG00000180233 | n.s. |
| ENSG00000169635 | 616.1836857 | 0.193579128 | 0.2270284 | 0.852665 | 0.393845312 | 0.586056813 | HIC2 | protein_coding | 56.16 | 0.044922027 | 0.044922027 | ENSG00000169635 | n.s. |
| ENSG00000198618 | 445.6295729 | 0.237431582 | 0.3218574 | 0.737692 | 0.460701736 | 0.646868724 | PPIAP22 | processed_pseudogene | 49 | 0.04491822 | 0.04491822 | ENSG00000198618 | n.s. |
| ENSG00000148803 | 160.3273118 | -0.202961063 | 0.2460152 | -0.824994 | 0.409375005 | 0.601173736 | FUOM | protein_coding | 67.07 | -0.044854395 | 0.044854395 | ENSG00000148803 | n.s. |
| ENSG00000079134 | 2224.769554 | -0.208321389 | 0.257449 | -0.809175 | 0.418414294 | 0.609498035 | THOC1 | protein_coding | 37.31 | -0.044794867 | 0.044794867 | ENSG00000079134 | n.s. |
| ENSG00000171951 | 2267.564942 | 0.206588387 | 0.2536995 | 0.814303 | 0.415471123 | 0.607070513 | SCG2 | protein_coding | 36.25 | 0.044780277 | 0.044780277 | ENSG00000171951 | n.s. |
| ENSG00000274618 | 278.3223385 | 0.188304856 | 0.2170754 | 0.867463 | 0.385688507 | 0.578408131 | HIST1H4F | protein_coding | 57.37 | 0.044772419 | 0.044772419 | ENSG00000274618 | n.s. |
| ENSG00000100528 | 1959.056455 | 0.21399365 | 0.269847 | 0.793018 | 0.427767139 | 0.61794239 | CNIH1 | protein_coding | 38.26 | 0.044735803 | 0.044735803 | ENSG00000100528 | n.s. |
| ENSG00000183161 | 761.7095975 | -0.186435271 | 0.2135715 | -0.872941 | 0.382695309 | 0.575622591 | FANCF | protein_coding | 42.42 | -0.044718769 | 0.044718769 | ENSG00000183161 | n.s. |
| ENSG00000136854 | 52.30152145 | 0.225684131 | 0.2962931 | 0.761692 | 0.446243699 | 0.633811136 | STXBP1 | protein_coding | 47.08 | 0.044694516 | 0.044694516 | ENSG00000136854 | n.s. |
| ENSG00000139624 | 705.4991161 | -0.218209516 | 0.279262 | -0.781379 | 0.434579431 | 0.624293171 | CERS5 | protein_coding | 43.86 | -0.044648158 | 0.044648158 | ENSG00000139624 | n.s. |
| ENSG00000145365 | 635.1957004 | 0.206070885 | 0.2532611 | 0.81367 | 0.415834198 | 0.607315478 | TIFA | protein_coding | 40.42 | 0.044631997 | 0.044631997 | ENSG00000145365 | n.s. |
| ENSG00000174695 | 2276.812925 | -0.229203327 | 0.3050302 | -0.751412 | 0.452404791 | 0.638956372 | TMEM167A | protein_coding | 35.95 | -0.044586647 | 0.044586647 | ENSG00000174695 | n.s. |
| ENSG00000134253 | 271.9937901 | -0.187242736 | 0.2155234 | -0.868781 | 0.384966748 | 0.577945263 | TRIM45 | protein_coding | 46.89 | -0.044584984 | 0.044584984 | ENSG00000134253 | n.s. |
| ENSG00000166348 | 29.27455354 | -0.236247829 | 0.3208812 | -0.736247 | 0.461580357 | 0.647579938 | USP54 | protein_coding | 42.16 | -0.044581528 | 0.044581528 | ENSG00000166348 | n.s. |
| ENSG00000161682 | 615.6438359 | -0.208422588 | 0.2589891 | -0.804754 | 0.420961565 | 0.611226184 | FAM171A2 | protein_coding | 60.61 | -0.044560343 | 0.044560343 | ENSG00000161682 | n.s. |
| ENSG00000110330 | 630.5017344 | -0.209184903 | 0.2607788 | -0.802154 | 0.422463646 | 0.612576687 | BIRC2 | protein_coding | 38.86 | -0.044522818 | 0.044522818 | ENSG00000110330 | n.s. |
| ENSG00000091542 | 514.4068618 | 0.185117048 | 0.2117643 | 0.874166 | 0.382028122 | 0.574838166 | ALKBH5 | protein_coding | 50.24 | 0.04451221 | 0.04451221 | ENSG00000091542 | n.s. |
| ENSG00000270170 | 611.4607711 | 0.216309875 | 0.2754691 | 0.785242 | 0.432311743 | 0.622641606 | NCBP2-AS2 | protein_coding | 60.89 | 0.044508323 | 0.044508323 | ENSG00000270170 | n.s. |

| | | | | | | | | | | | | | |
|---|---|---|---|---|---|---|---|---|---|---|---|---|---|
| ENSG00000183828 | 81.27413786 | 0.237796681 | 0.3253509 | 0.730893 | 0.464844593 | 0.64992388 | NUDT14 | protein_coding | 64.58 | 0.044500678 | 0.044500678 | ENSG00000183828 | n.s. |
| ENSG00000210144 | 16.79065969 | -0.254136694 | 0.3658105 | -0.694722 | 0.487229334 | 0.668544293 | MT-TY | Mt_tRNA | 42.42 | -0.044440836 | 0.044440836 | ENSG00000210144 | n.s. |
| ENSG00000123358 | 231.3748044 | -0.208363503 | 0.2595974 | -0.802641 | 0.422182294 | 0.612376598 | NR4A1 | protein_coding | 56.01 | -0.044377554 | 0.044377554 | ENSG00000123358 | n.s. |
| ENSG00000147100 | 147.6965315 | -0.220206766 | 0.2852967 | -0.771852 | 0.440202249 | 0.628944621 | SLC16A2 | protein_coding | 43.6 | -0.044346911 | 0.044346911 | ENSG00000147100 | n.s. |
| ENSG00000283108 | 56.09454468 | 0.214625753 | 0.2725208 | 0.787557 | 0.43095569 | 0.621459977 | | processed_transcript | 42.44 | 0.044338855 | 0.044338855 | ENSG00000283108 | n.s. |
| ENSG00000228474 | 1203.578752 | 0.224037836 | 0.2945123 | 0.760708 | 0.446831615 | 0.634100082 | OST4 | protein_coding | 56.45 | 0.044324136 | 0.044324136 | ENSG00000228474 | n.s. |
| ENSG00000212296 | 118.4783105 | 0.200121976 | 0.2424131 | 0.825541 | 0.409064376 | 0.60097112 | SNORD72 | snoRNA | 31.25 | 0.044256254 | 0.044256254 | ENSG00000212296 | n.s. |
| ENSG00000110218 | 969.8579246 | 0.207975024 | 0.2593656 | 0.80186 | 0.422633671 | 0.612695606 | PANX1 | protein_coding | 43.1 | 0.044247775 | 0.044247775 | ENSG00000110218 | n.s. |
| ENSG00000274642 | 43.89845928 | -0.222774982 | 0.2922289 | -0.76233 | 0.445862825 | 0.633637655 | | unprocessed_pseudogene | 40.62 | -0.044144872 | 0.044144872 | ENSG00000274642 | n.s. |
| ENSG00000198954 | 186.759164 | -0.212913917 | 0.2698684 | -0.788954 | 0.430138607 | 0.620410218 | KIF1BP | protein_coding | 43.19 | -0.044141539 | 0.044141539 | ENSG00000198954 | n.s. |
| ENSG00000125970 | 1871.405547 | 0.210684642 | 0.2652766 | 0.794207 | 0.427074732 | 0.617334042 | RALY | protein_coding | 44.86 | 0.044134171 | 0.044134171 | ENSG00000125970 | n.s. |
| ENSG00000149257 | 714.7575694 | -0.200446559 | 0.2435519 | -0.823014 | 0.410500247 | 0.602319208 | SERPINH1 | protein_coding | 55.42 | -0.044132978 | 0.044132978 | ENSG00000149257 | n.s. |
| ENSG00000155254 | 759.1714181 | -0.194719393 | 0.231831 | -0.83992 | 0.400953389 | 0.593438295 | MARVELD1 | protein_coding | 57.63 | -0.044128172 | 0.044128172 | ENSG00000155254 | n.s. |
| ENSG00000264607 | 6.005123994 | -0.301400648 | 0.4970604 | -0.606366 | 0.54427155 | 0.714019861 | MIR3173 | miRNA | 48.53 | -0.044091813 | 0.044091813 | ENSG00000264607 | n.s. |
| ENSG00000170619 | 291.9861043 | 0.220097821 | 0.2866192 | 0.76791 | 0.442540573 | 0.630525215 | COMMD5 | protein_coding | 53.13 | 0.044085052 | 0.044085052 | ENSG00000170619 | n.s. |
| ENSG00000145248 | 819.4003745 | 0.199679553 | 0.2421969 | 0.824451 | 0.409683108 | 0.601562741 | SLC10A4 | protein_coding | 44.38 | 0.044073085 | 0.044073085 | ENSG00000145248 | n.s. |
| ENSG00000278637 | 414.5621022 | -0.197349379 | 0.2376923 | -0.830273 | 0.406384665 | 0.598129014 | HIST1H4A | protein_coding | 60.58 | -0.044049394 | 0.044049394 | ENSG00000278637 | n.s. |
| ENSG00000129636 | 555.5765531 | -0.219731644 | 0.2859829 | -0.768338 | 0.442286191 | 0.630356294 | ITFG1 | protein_coding | 37.51 | -0.044037277 | 0.044037277 | ENSG00000129636 | n.s. |
| ENSG00000266619 | 13.44463402 | 0.257010465 | 0.3776936 | 0.680473 | 0.496204786 | 0.674130097 | MIR4642 | miRNA | 58.54 | 0.044014657 | 0.044014657 | ENSG00000266619 | n.s. |
| ENSG00000166794 | 3775.450199 | 0.225954884 | 0.3004513 | 0.752052 | 0.452019962 | 0.638669167 | PPIB | protein_coding | 52.92 | 0.04399885 | 0.04399885 | ENSG00000166794 | n.s. |
| ENSG00000144895 | 3148.681433 | 0.218291099 | 0.2825034 | 0.772703 | 0.439698465 | 0.628727711 | EIF2A | protein_coding | 37.58 | 0.04399382 | 0.04399382 | ENSG00000144895 | n.s. |
| ENSG00000244005 | 58.21999084 | 0.225169827 | 0.2989453 | 0.753214 | 0.451321196 | 0.638008428 | NFS1 | protein_coding | 46.45 | 0.043947202 | 0.043947202 | ENSG00000244005 | n.s. |
| ENSG00000240616 | 62.71559313 | 0.229242245 | 0.3080882 | 0.74408 | 0.456828169 | 0.643320355 | RPS6P25 | processed_pseudogene | 45.86 | 0.043916557 | 0.043916557 | ENSG00000240616 | n.s. |
| ENSG00000184007 | 986.6916481 | 0.200166142 | 0.2436623 | 0.82149 | 0.41136726 | 0.603399322 | PTP4A2 | protein_coding | 41.84 | 0.043915487 | 0.043915487 | ENSG00000184007 | n.s. |
| ENSG00000064666 | 142.7594652 | -0.202503783 | 0.2487132 | -0.814206 | 0.415526969 | 0.607070513 | CNN2 | protein_coding | 59.73 | -0.043894895 | 0.043894895 | ENSG00000064666 | n.s. |
| ENSG00000117155 | 1416.626928 | -0.216989377 | 0.2801389 | -0.774578 | 0.438589231 | 0.627850901 | SSX2IP | protein_coding | 37.23 | -0.043862987 | 0.043862987 | ENSG00000117155 | n.s. |
| ENSG00000100422 | 3394.748313 | -0.214945612 | 0.2757607 | -0.779464 | 0.43570626 | 0.625202336 | CERK | protein_coding | 51.44 | -0.043844479 | 0.043844479 | ENSG00000100422 | n.s. |
| ENSG00000133731 | 892.3569925 | 0.218645874 | 0.2844843 | 0.768569 | 0.442149154 | 0.630225499 | IMPA1 | protein_coding | 37.59 | 0.043839379 | 0.043839379 | ENSG00000133731 | n.s. |
| ENSG00000129245 | 786.3015375 | -0.217457744 | 0.2816466 | -0.772094 | 0.440058562 | 0.628944621 | FXR2 | protein_coding | 48.07 | -0.043793291 | 0.043793291 | ENSG00000129245 | n.s. |
| ENSG00000226819 | 219.4844126 | -0.201129561 | 0.2461692 | -0.817038 | 0.413906827 | 0.605720492 | MEIS1-AS3 | antisense | 42.16 | -0.043791483 | 0.043791483 | ENSG00000226819 | n.s. |
| ENSG00000141753 | 415.7946672 | -0.214559589 | 0.2753132 | -0.779329 | 0.435785956 | 0.625252254 | IGFBP4 | protein_coding | 53 | -0.043758298 | 0.043758298 | ENSG00000141753 | n.s. |
| ENSG00000252614 | 42.26445579 | 0.21515312 | 0.2767206 | 0.77751 | 0.436857802 | 0.62614486 | RNU6-807P | snRNA | 35.51 | 0.043746047 | 0.043746047 | ENSG00000252614 | n.s. |

| | | | | | | | | | | | | | |
|---|---|---|---|---|---|---|---|---|---|---|---|---|---|
| ENSG00000207313 | 90.59602442 | 0.221704818 | 0.292311 | 0.758455 | 0.448178483 | 0.635212533 | SNORA2B | snoRNA | 40.15 | 0.043693794 | 0.043693794 | ENSG00000207313 | n.s. |
| ENSG00000103485 | 408.179229 | 0.224935644 | 0.2998745 | 0.750099 | 0.453194887 | 0.639747786 | QPRT | protein_coding | 51.92 | 0.043635537 | 0.043635537 | ENSG00000103485 | n.s. |
| ENSG00000143742 | 7935.656519 | 0.231582348 | 0.3149572 | 0.735282 | 0.462167816 | 0.648077601 | SRP9 | protein_coding | 40.6 | 0.043623859 | 0.043623859 | ENSG00000143742 | n.s. |
| ENSG00000204308 | 542.8810739 | 0.230669453 | 0.3131124 | 0.736698 | 0.461305747 | 0.647265842 | RNF5 | protein_coding | 51.43 | 0.043577453 | 0.043577453 | ENSG00000204308 | n.s. |
| ENSG00000125967 | 101.4960293 | -0.199230778 | 0.2431891 | -0.819242 | 0.412648149 | 0.604513908 | NECAB3 | protein_coding | 59.54 | -0.043550594 | 0.043550594 | ENSG00000125967 | n.s. |
| ENSG00000283050 | 56.44872353 | 0.215674703 | 0.2786924 | 0.773881 | 0.439001314 | 0.628247024 | GTF2IP12 | transcribed_unprocessed_pseud | 45.66 | 0.043538157 | 0.043538157 | ENSG00000283050 | n.s. |
| ENSG00000136874 | 227.2777902 | -0.202210863 | 0.2497548 | -0.809638 | 0.418148488 | 0.609174573 | STX17 | protein_coding | 36.32 | -0.043527553 | 0.043527553 | ENSG00000136874 | n.s. |
| ENSG00000059122 | 29.47060244 | -0.227451391 | 0.3058934 | -0.743564 | 0.457140193 | 0.643620479 | FLYWCH1 | protein_coding | 52.33 | -0.043527406 | 0.043527406 | ENSG00000059122 | n.s. |
| ENSG00000165283 | 1590.520268 | 0.223415347 | 0.2970064 | 0.752224 | 0.451916404 | 0.638655308 | STOML2 | protein_coding | 53.1 | 0.043506447 | 0.043506447 | ENSG00000165283 | n.s. |
| ENSG00000185347 | 177.8668715 | -0.211186276 | 0.2688031 | -0.785654 | 0.432070177 | 0.622380245 | C14orf80 | protein_coding | 66 | -0.043492589 | 0.043492589 | ENSG00000185347 | n.s. |
| ENSG00000204856 | 915.5143712 | -0.22402326 | 0.2987192 | -0.749946 | 0.453287311 | 0.639748528 | FAM216A | protein_coding | 43.77 | -0.04345843 | 0.04345843 | ENSG00000204856 | n.s. |
| ENSG00000138002 | 606.7159519 | -0.204006934 | 0.2543127 | -0.802189 | 0.422443452 | 0.612576687 | IFT172 | protein_coding | 45.51 | -0.043420741 | 0.043420741 | ENSG00000138002 | n.s. |
| ENSG00000077585 | 158.2033843 | -0.213040939 | 0.2736462 | -0.778527 | 0.436258378 | 0.625567375 | GPR137B | protein_coding | 43.63 | -0.043401959 | 0.043401959 | ENSG00000077585 | n.s. |
| ENSG00000171202 | 294.9322896 | 0.220392651 | 0.2909752 | 0.757428 | 0.448793701 | 0.635741733 | TMEM126A | protein_coding | 37.69 | 0.043355483 | 0.043355483 | ENSG00000171202 | n.s. |
| ENSG00000248774 | 239.5357888 | -0.183904296 | 0.2135847 | -0.861037 | 0.389217723 | 0.581259076 | | antisense | 35.99 | -0.043333416 | 0.043333416 | ENSG00000248774 | n.s. |
| ENSG00000226479 | 266.6354814 | 0.188493534 | 0.2226464 | 0.846605 | 0.397215429 | 0.589095698 | TMEM185B | protein_coding | 53.17 | 0.043318481 | 0.043318481 | ENSG00000226479 | n.s. |
| ENSG00000130175 | 811.2565083 | 0.200701297 | 0.2475839 | 0.81064 | 0.417572619 | 0.60865543 | PRKCSH | protein_coding | 56.85 | 0.043276919 | 0.043276919 | ENSG00000130175 | n.s. |
| ENSG00000149428 | 1020.669262 | 0.20227967 | 0.2514857 | 0.804339 | 0.421201458 | 0.611319493 | HYOU1 | protein_coding | 53.98 | 0.043233589 | 0.043233589 | ENSG00000149428 | n.s. |
| ENSG00000165948 | 173.0848187 | 0.219722596 | 0.2902437 | 0.757028 | 0.449033064 | 0.635742053 | IFI27L1 | protein_coding | 45.17 | 0.043223622 | 0.043223622 | ENSG00000165948 | n.s. |
| ENSG00000151532 | 622.4712959 | 0.195780444 | 0.2375289 | 0.824238 | 0.409804057 | 0.601574194 | VTI1A | protein_coding | 39.1 | 0.043210859 | 0.043210859 | ENSG00000151532 | n.s. |
| ENSG00000144233 | 998.7577038 | -0.19800177 | 0.2419793 | -0.818259 | 0.413209225 | 0.605017567 | AMMECR1L | protein_coding | 45.69 | -0.043210325 | 0.043210325 | ENSG00000144233 | n.s. |
| ENSG00000100568 | 936.8348824 | -0.210872402 | 0.270204 | -0.780419 | 0.435144211 | 0.624717761 | VTI1B | protein_coding | 42.97 | -0.043084637 | 0.043084637 | ENSG00000100568 | n.s. |
| ENSG00000259673 | 69.32964219 | 0.210245674 | 0.268692 | 0.782478 | 0.433933525 | 0.623880263 | IQCH-AS1 | lincRNA | 41.68 | 0.043079076 | 0.043079076 | ENSG00000259673 | n.s. |
| ENSG00000035141 | 1452.757744 | -0.215455744 | 0.280991 | -0.766771 | 0.44321764 | 0.631166943 | FAM136A | protein_coding | 48.46 | -0.043060071 | 0.043060071 | ENSG00000035141 | n.s. |
| ENSG00000171970 | 119.4709894 | -0.218881128 | 0.2891029 | -0.757105 | 0.448987177 | 0.635741733 | ZNF57 | protein_coding | 46.26 | -0.043058138 | 0.043058138 | ENSG00000171970 | n.s. |
| ENSG00000174442 | 1569.542622 | -0.219571913 | 0.2909817 | -0.75459 | 0.450494953 | 0.637184479 | ZWILCH | protein_coding | 40.27 | -0.042977867 | 0.042977867 | ENSG00000174442 | n.s. |
| ENSG00000138095 | 7265.379503 | -0.213284284 | 0.2760984 | -0.772494 | 0.439822131 | 0.628839959 | LRPPRC | protein_coding | 37.13 | -0.042968224 | 0.042968224 | ENSG00000138095 | n.s. |
| ENSG00000141985 | 442.7647802 | -0.209673962 | 0.2678623 | -0.782768 | 0.433763464 | 0.623880263 | SH3GL1 | protein_coding | 57.17 | -0.042961933 | 0.042961933 | ENSG00000141985 | n.s. |
| ENSG00000169895 | 940.5891476 | -0.211826421 | 0.2728581 | -0.776325 | 0.437557352 | 0.627018425 | SYAP1 | protein_coding | 43.74 | -0.042941388 | 0.042941388 | ENSG00000169895 | n.s. |
| ENSG00000168487 | 28.33000885 | -0.225690744 | 0.3050838 | -0.739766 | 0.459441739 | 0.645685609 | BMP1 | protein_coding | 54.06 | -0.042876478 | 0.042876478 | ENSG00000168487 | n.s. |
| ENSG00000111639 | 2628.215807 | 0.219996749 | 0.2923126 | 0.752608 | 0.451685664 | 0.638458832 | MRPL51 | protein_coding | 48.72 | 0.042870129 | 0.042870129 | ENSG00000111639 | n.s. |
| ENSG00000176101 | 645.6084845 | 0.229997176 | 0.3156465 | 0.728654 | 0.46621316 | 0.651061768 | SSNA1 | protein_coding | 65.89 | 0.042866369 | 0.042866369 | ENSG00000176101 | n.s. |

| | | | | | | | | | | | | | |
|---|---|---|---|---|---|---|---|---|---|---|---|---|---|
| ENSG00000128699 | 2531.598055 | -0.211618611 | 0.272819 | -0.775674 | 0.437941521 | 0.627258244 | ORMDL1 | protein_coding | 37.58 | -0.042864116 | 0.042864116 | ENSG00000128699 | n.s. |
| ENSG00000198353 | 163.9126118 | 0.209833565 | 0.2690259 | 0.779975 | 0.435405442 | 0.62502835 | HOXC4 | protein_coding | 51.68 | 0.04282709 | 0.04282709 | ENSG00000198353 | n.s. |
| ENSG00000272841 | 144.2174059 | 0.1994062 | 0.2466686 | 0.808397 | 0.418862016 | 0.609894985 | | antisense | 53.38 | 0.042821472 | 0.042821472 | ENSG00000272841 | n.s. |
| ENSG00000147121 | 89.09846118 | -0.201357803 | 0.2512399 | -0.801456 | 0.422867477 | 0.612906921 | KRBOX4 | protein_coding | 41.38 | -0.042809771 | 0.042809771 | ENSG00000147121 | n.s. |
| ENSG00000116815 | 138.3944163 | -0.226336044 | 0.3070538 | -0.737122 | 0.461048277 | 0.647029045 | CD58 | protein_coding | 41.62 | -0.042794764 | 0.042794764 | ENSG00000116815 | n.s. |
| ENSG00000196747 | 1090.668464 | 0.182718561 | 0.2131708 | 0.857146 | 0.391364063 | 0.583182533 | HIST1H2AI | protein_coding | 59.64 | 0.042791863 | 0.042791863 | ENSG00000196747 | n.s. |
| ENSG00000148341 | 382.6948248 | 0.212339236 | 0.2752588 | 0.771417 | 0.440459911 | 0.62896057 | SH3GLB2 | protein_coding | 55.75 | 0.042760149 | 0.042760149 | ENSG00000148341 | n.s. |
| ENSG00000124532 | 727.6107671 | -0.206332946 | 0.2616846 | -0.78848 | 0.430416201 | 0.620746299 | MRS2 | protein_coding | 39.3 | -0.042728636 | 0.042728636 | ENSG00000124532 | n.s. |
| ENSG00000133997 | 1209.455837 | 0.204246319 | 0.2575383 | 0.793072 | 0.427736084 | 0.61794239 | MED6 | protein_coding | 38.94 | 0.042698104 | 0.042698104 | ENSG00000133997 | n.s. |
| ENSG00000102580 | 663.3298024 | 0.195194753 | 0.2381457 | 0.819644 | 0.412418992 | 0.604432593 | DNAJC3 | protein_coding | 39.79 | 0.042679748 | 0.042679748 | ENSG00000102580 | n.s. |
| ENSG00000082212 | 1818.650573 | -0.217846251 | 0.2885539 | -0.754959 | 0.450273774 | 0.636980467 | ME2 | protein_coding | 38.73 | -0.042670392 | 0.042670392 | ENSG00000082212 | n.s. |
| ENSG00000174498 | 189.5186397 | -0.192037487 | 0.2320636 | -0.827521 | 0.407941812 | 0.599764926 | IGDCC3 | protein_coding | 51.86 | -0.042635959 | 0.042635959 | ENSG00000174498 | n.s. |
| ENSG00000169902 | 230.6095718 | 0.217105714 | 0.2871688 | 0.756021 | 0.449636513 | 0.636272911 | TPST1 | protein_coding | 42.23 | 0.042630133 | 0.042630133 | ENSG00000169902 | n.s. |
| ENSG00000212493 | 64.15110687 | 0.207562958 | 0.2648373 | 0.783738 | 0.433194068 | 0.623396487 | SNORD19 | snoRNA | 45.59 | 0.042599319 | 0.042599319 | ENSG00000212493 | n.s. |
| ENSG00000180185 | 605.1120542 | 0.198070085 | 0.2451545 | 0.80794 | 0.41912528 | 0.609959368 | FAHD1 | protein_coding | 45.47 | 0.042525468 | 0.042525468 | ENSG00000180185 | n.s. |
| ENSG00000172795 | 2238.422222 | -0.19777828 | 0.2447522 | -0.808075 | 0.419047141 | 0.609927068 | DCP2 | protein_coding | 37.79 | -0.042467366 | 0.042467366 | ENSG00000172795 | n.s. |
| ENSG00000166405 | 243.2068142 | 0.207197726 | 0.264779 | 0.782531 | 0.433902744 | 0.623880263 | RIC3 | protein_coding | 39.93 | 0.042454556 | 0.042454556 | ENSG00000166405 | n.s. |
| ENSG00000074800 | 10622.4655 | 0.225435985 | 0.3069518 | 0.734435 | 0.462683944 | 0.648279013 | ENO1 | protein_coding | 50.23 | 0.042435628 | 0.042435628 | ENSG00000074800 | n.s. |
| ENSG00000143147 | 59.08492204 | -0.214192219 | 0.2812991 | -0.76144 | 0.446394599 | 0.633811136 | GPR161 | protein_coding | 46.83 | -0.042418656 | 0.042418656 | ENSG00000143147 | n.s. |
| ENSG00000107581 | 9702.904355 | -0.194113715 | 0.2370678 | -0.818811 | 0.412894267 | 0.604683589 | EIF3A | protein_coding | 41.68 | -0.042408376 | 0.042408376 | ENSG00000107581 | n.s. |
| ENSG00000197608 | 667.2177491 | -0.211064474 | 0.2743942 | -0.769202 | 0.441773629 | 0.629754708 | ZNF841 | protein_coding | 41.11 | -0.042387778 | 0.042387778 | ENSG00000197608 | n.s. |
| ENSG00000258199 | 44.53826961 | -0.243825324 | 0.3534927 | -0.689761 | 0.490344801 | 0.670128457 | | sense_overlapping | 49.39 | -0.042387067 | 0.042387067 | ENSG00000258199 | n.s. |
| ENSG00000205485 | 236.5276836 | -0.201294823 | 0.252745 | -0.796434 | 0.42577959 | 0.615909531 | | transcribed_unprocessed_pseud | 41.83 | -0.042369153 | 0.042369153 | ENSG00000205485 | n.s. |
| ENSG00000163428 | 2668.319621 | 0.200663386 | 0.251443 | 0.798047 | 0.424843068 | 0.615066032 | LRRC58 | protein_coding | 38.94 | 0.042355678 | 0.042355678 | ENSG00000163428 | n.s. |
| ENSG00000233223 | 92.30181104 | -0.207576139 | 0.2663852 | -0.779233 | 0.43584246 | 0.625268889 | | antisense | 61.02 | -0.042331662 | 0.042331662 | ENSG00000233223 | n.s. |
| ENSG00000108819 | 1054.754877 | -0.19648675 | 0.2425794 | -0.80999 | 0.417946169 | 0.608943545 | PPP1R9B | protein_coding | 59.45 | -0.042327759 | 0.042327759 | ENSG00000108819 | n.s. |
| ENSG00000134480 | 1371.463762 | -0.205126897 | 0.2606796 | -0.786893 | 0.431344582 | 0.621827564 | CCNH | protein_coding | 36.69 | -0.042323839 | 0.042323839 | ENSG00000134480 | n.s. |
| ENSG00000235363 | 127.9404742 | 0.304343242 | 0.521746 | 0.583317 | 0.559680044 | 0.726024901 | SNRPGP10 | processed_pseudogene | 39.11 | 0.042318466 | 0.042318466 | ENSG00000235363 | n.s. |
| ENSG00000068438 | 556.7912253 | 0.226103341 | 0.309444 | 0.730676 | 0.464976982 | 0.64992388 | FTSJ1 | protein_coding | 50.6 | 0.042312415 | 0.042312415 | ENSG00000068438 | n.s. |
| ENSG00000202459 | 6.301459998 | -0.287507436 | 0.4721517 | -0.60893 | 0.542570722 | 0.712661525 | | misc_RNA | 46.3 | -0.042297141 | 0.042297141 | ENSG00000202459 | n.s. |
| ENSG00000242612 | 136.5164931 | 0.21643706 | 0.2870735 | 0.753943 | 0.450883512 | 0.637648667 | DECR2 | protein_coding | 56.95 | 0.042295815 | 0.042295815 | ENSG00000242612 | n.s. |
| ENSG00000185274 | 367.6073103 | -0.197056054 | 0.2439856 | -0.807654 | 0.419289563 | 0.610070915 | WBSCR17 | protein_coding | 44.5 | -0.042292107 | 0.042292107 | ENSG00000185274 | n.s. |

| | | | | | | | | | | | | | |
|---|---|---|---|---|---|---|---|---|---|---|---|---|---|
| ENSG00000181652 | 116.34704 | 0.199358426 | 0.2492489 | 0.799837 | 0.423805409 | 0.613882929 | ATG9B | protein_coding | 58.12 | 0.04224693 | 0.04224693 | ENSG00000181652 | n.s. |
| ENSG00000111412 | 968.6223402 | 0.209601295 | 0.2712365 | 0.772762 | 0.439663154 | 0.628727711 | C12orf49 | protein_coding | 44.28 | 0.0422425 | 0.0422425 | ENSG00000111412 | n.s. |
| ENSG00000228065 | 104.1229312 | -0.198718049 | 0.2479925 | -0.801307 | 0.422954198 | 0.612968804 | LINC01515 | lincRNA | 34.38 | -0.042239832 | 0.042239832 | ENSG00000228065 | n.s. |
| ENSG00000134910 | 966.2874759 | -0.205950089 | 0.2630777 | -0.782849 | 0.433715847 | 0.623880263 | STT3A | protein_coding | 41.03 | -0.042198917 | 0.042198917 | ENSG00000134910 | n.s. |
| ENSG00000112855 | 904.7657465 | -0.196932746 | 0.2443673 | -0.805888 | 0.420307222 | 0.610830464 | HARS2 | protein_coding | 45.91 | -0.042159227 | 0.042159227 | ENSG00000112855 | n.s. |
| ENSG00000276550 | 774.5862091 | -0.195527733 | 0.2413921 | -0.81 | 0.417939955 | 0.608943545 | HERC2P2 | transcribed_unprocessed_pseud | 42.65 | -0.042121165 | 0.042121165 | ENSG00000276550 | n.s. |
| ENSG00000136238 | 1444.810809 | 0.201485265 | 0.2540371 | 0.793133 | 0.427700279 | 0.61794239 | RAC1 | protein_coding | 45.86 | 0.0421209 | 0.0421209 | ENSG00000136238 | n.s. |
| ENSG00000088256 | 424.9170381 | -0.183685194 | 0.2173825 | -0.844986 | 0.398118511 | 0.589869536 | GNA11 | protein_coding | 61.66 | -0.042108735 | 0.042108735 | ENSG00000088256 | n.s. |
| ENSG00000160691 | 808.9691126 | 0.208229564 | 0.2690117 | 0.774054 | 0.438898864 | 0.628183283 | SHC1 | protein_coding | 53.79 | 0.042044386 | 0.042044386 | ENSG00000160691 | n.s. |
| ENSG00000142444 | 240.9548546 | 0.201935606 | 0.2554198 | 0.790603 | 0.429175856 | 0.619221478 | TIMM29 | protein_coding | 52.55 | 0.042033702 | 0.042033702 | ENSG00000142444 | n.s. |
| ENSG00000120860 | 247.3566412 | -0.222240248 | 0.3014672 | -0.737195 | 0.461003546 | 0.647029045 | WASHC3 | protein_coding | 36.36 | -0.042020347 | 0.042020347 | ENSG00000120860 | n.s. |
| ENSG00000163161 | 1628.93318 | -0.205565615 | 0.2633998 | -0.780432 | 0.435136603 | 0.624717761 | ERCC3 | protein_coding | 46.18 | -0.042000374 | 0.042000374 | ENSG00000163161 | n.s. |
| ENSG00000199121 | 7.031030619 | 0.28379144 | 0.4640129 | 0.611602 | 0.540800799 | 0.711273748 | MIR26B | miRNA | 55.84 | 0.041990695 | 0.041990695 | ENSG00000199121 | n.s. |
| ENSG00000148154 | 746.7575368 | 0.207270268 | 0.2671509 | 0.775855 | 0.437834744 | 0.627258244 | UGCG | protein_coding | 40.22 | 0.041983343 | 0.041983343 | ENSG00000148154 | n.s. |
| ENSG00000102144 | 4746.194289 | 0.219033861 | 0.294373 | 0.744069 | 0.456834615 | 0.643320355 | PGK1 | protein_coding | 41.35 | 0.04196091 | 0.04196091 | ENSG00000102144 | n.s. |
| ENSG00000185565 | 1335.549439 | -0.200859569 | 0.2536621 | -0.791839 | 0.428454545 | 0.618557925 | LSAMP | protein_coding | 37.2 | -0.041903248 | 0.041903248 | ENSG00000185565 | n.s. |
| ENSG00000092531 | 320.6136338 | -0.208211501 | 0.2704779 | -0.769791 | 0.441423752 | 0.62947723 | SNAP23 | protein_coding | 42.25 | -0.041854671 | 0.041854671 | ENSG00000092531 | n.s. |
| ENSG00000236535 | 120.7641642 | 0.209989649 | 0.2746026 | 0.764704 | 0.444447885 | 0.632018923 | RC3H1-IT1 | sense_intronic | 39.39 | 0.04184462 | 0.04184462 | ENSG00000236535 | n.s. |
| ENSG00000106723 | 2172.5478 | -0.211130601 | 0.2770328 | -0.762114 | 0.445991972 | 0.633688027 | SPIN1 | protein_coding | 41.25 | -0.041830144 | 0.041830144 | ENSG00000106723 | n.s. |
| ENSG00000168872 | 600.0556531 | -0.205190619 | 0.2634658 | -0.778813 | 0.436089863 | 0.625494918 | DDX19A | protein_coding | 47.09 | -0.041812968 | 0.041812968 | ENSG00000168872 | n.s. |
| ENSG00000077157 | 314.177839 | -0.191327166 | 0.2336132 | -0.818991 | 0.41279143 | 0.604596579 | PPP1R12B | protein_coding | 39.63 | -0.041811551 | 0.041811551 | ENSG00000077157 | n.s. |
| ENSG00000116898 | 580.2458825 | 0.227149793 | 0.3147659 | 0.721647 | 0.470511699 | 0.65465398 | MRPS15 | protein_coding | 49.28 | 0.041792879 | 0.041792879 | ENSG00000116898 | n.s. |
| ENSG00000104957 | 281.3941392 | -0.204408802 | 0.2618101 | -0.780752 | 0.434948239 | 0.624565216 | CCDC130 | protein_coding | 50.36 | -0.041785698 | 0.041785698 | ENSG00000104957 | n.s. |
| ENSG00000118600 | 828.2683325 | -0.190844371 | 0.2326166 | -0.820424 | 0.411974186 | 0.604098641 | TMEM5 | protein_coding | 38.5 | -0.041774333 | 0.041774333 | ENSG00000118600 | n.s. |
| ENSG00000163159 | 1030.958553 | -0.240225504 | 0.3483333 | -0.689643 | 0.490418978 | 0.670128457 | VPS72 | protein_coding | 47.36 | -0.041761268 | 0.041761268 | ENSG00000163159 | n.s. |
| ENSG00000072849 | 838.8568216 | -0.202452565 | 0.2573686 | -0.786625 | 0.431501443 | 0.621924906 | DERL2 | protein_coding | 41.38 | -0.041758281 | 0.041758281 | ENSG00000072849 | n.s. |
| ENSG00000163539 | 2642.617481 | -0.209547371 | 0.2740263 | -0.764698 | 0.444451431 | 0.632018923 | CLASP2 | protein_coding | 36.82 | -0.041756488 | 0.041756488 | ENSG00000163539 | n.s. |
| ENSG00000165606 | 430.8966166 | -0.207614458 | 0.2698371 | -0.769407 | 0.441651949 | 0.629645718 | DRGX | protein_coding | 48.29 | -0.041710522 | 0.041710522 | ENSG00000165606 | n.s. |
| ENSG00000111445 | 1853.556945 | -0.204575912 | 0.2626087 | -0.779014 | 0.435971334 | 0.625389335 | RFC5 | protein_coding | 45.74 | -0.041702703 | 0.041702703 | ENSG00000111445 | n.s. |
| ENSG00000103496 | 59.28884746 | -0.214067556 | 0.2847012 | -0.751902 | 0.452109697 | 0.638669167 | STX4 | protein_coding | 55.52 | -0.041684101 | 0.041684101 | ENSG00000103496 | n.s. |
| ENSG00000196961 | 264.2853756 | -0.203889486 | 0.2611203 | -0.780826 | 0.434904995 | 0.624565216 | AP2A1 | protein_coding | 54.57 | -0.041679539 | 0.041679539 | ENSG00000196961 | n.s. |
| ENSG00000050426 | 1251.858295 | -0.191777149 | 0.2351957 | -0.815394 | 0.414846878 | 0.606458745 | LETMD1 | protein_coding | 45.34 | -0.041653754 | 0.041653754 | ENSG00000050426 | n.s. |

| | | | | | | | | | | | | | |
|---|---|---|---|---|---|---|---|---|---|---|---|---|---|
| ENSG00000131669 | 78.64763006 | -0.213066674 | 0.2826515 | -0.753814 | 0.450960892 | 0.637693328 | NINJ1 | protein_coding | 59.68 | -0.041630698 | 0.041630698 | ENSG00000131669 | n.s. |
| ENSG00000202474 | 4.466185643 | -0.310553194 | 0.5459931 | -0.568786 | 0.569501458 | 0.734522353 | RNA5SP283 | rRNA | 58.82 | -0.04161257 | 0.04161257 | ENSG00000202474 | n.s. |
| ENSG00000136243 | 1139.505398 | -0.192497484 | 0.2370803 | -0.811951 | 0.416819971 | 0.607982855 | NUPL2 | protein_coding | 40.52 | -0.041600375 | 0.041600375 | ENSG00000136243 | n.s. |
| ENSG00000124214 | 1146.418111 | -0.199516077 | 0.2521249 | -0.791338 | 0.428746672 | 0.618826323 | STAU1 | protein_coding | 43.2 | -0.041585379 | 0.041585379 | ENSG00000124214 | n.s. |
| ENSG00000105993 | 1639.128276 | 0.206585535 | 0.268041 | 0.770724 | 0.440870687 | 0.629082685 | DNAJB6 | protein_coding | 48.15 | 0.041584071 | 0.041584071 | ENSG00000105993 | n.s. |
| ENSG00000261094 | 94.54033066 | -0.2237213 | 0.3080737 | -0.726194 | 0.467719838 | 0.651988717 | | sense_overlapping | 37.03 | -0.041558451 | 0.041558451 | ENSG00000261094 | n.s. |
| ENSG00000116685 | 321.2474189 | 0.206699784 | 0.2685412 | 0.769714 | 0.441469831 | 0.62947723 | KIAA2013 | protein_coding | 55.79 | 0.041550785 | 0.041550785 | ENSG00000116685 | n.s. |
| ENSG00000176410 | 207.6833476 | -0.180148338 | 0.21234 | -0.848396 | 0.396217593 | 0.588134144 | DNAJC30 | protein_coding | 54.45 | -0.041528445 | 0.041528445 | ENSG00000176410 | n.s. |
| ENSG00000262919 | 163.0217378 | 0.22294638 | 0.3061245 | 0.728287 | 0.466438207 | 0.651228628 | FAM58A | protein_coding | 56.23 | 0.041527445 | 0.041527445 | ENSG00000262919 | n.s. |
| ENSG00000076382 | 2602.269977 | -0.207774276 | 0.2712168 | -0.766082 | 0.443627591 | 0.631363278 | SPAG5 | protein_coding | 46.5 | -0.04149682 | 0.04149682 | ENSG00000076382 | n.s. |
| ENSG00000129625 | 737.0932403 | -0.205938597 | 0.2671332 | -0.770921 | 0.440753683 | 0.629009177 | REEP5 | protein_coding | 41.72 | -0.041464299 | 0.041464299 | ENSG00000129625 | n.s. |
| ENSG00000159917 | 382.9112972 | -0.193566494 | 0.2400295 | -0.806428 | 0.419996123 | 0.610652258 | ZNF235 | protein_coding | 39.47 | -0.041463111 | 0.041463111 | ENSG00000159917 | n.s. |
| ENSG00000118733 | 135.331813 | 0.221191752 | 0.3022434 | 0.731833 | 0.464270352 | 0.649521262 | OLFM3 | protein_coding | 35.12 | 0.0414528 | 0.0414528 | ENSG00000118733 | n.s. |
| ENSG00000006695 | 494.1687877 | -0.196257338 | 0.245844 | -0.7983 | 0.424696258 | 0.61491743 | COX10 | protein_coding | 38.47 | -0.041446252 | 0.041446252 | ENSG00000006695 | n.s. |
| ENSG00000075914 | 1005.429208 | -0.209248562 | 0.2747656 | -0.761553 | 0.446326787 | 0.633811136 | EXOSC7 | protein_coding | 46.87 | -0.041439613 | 0.041439613 | ENSG00000075914 | n.s. |
| ENSG00000104973 | 196.1729672 | -0.193809412 | 0.2410226 | -0.804113 | 0.421331646 | 0.611444706 | MED25 | protein_coding | 56.01 | -0.041405987 | 0.041405987 | ENSG00000104973 | n.s. |
| ENSG00000198231 | 6369.940682 | -0.20443377 | 0.2636399 | -0.775428 | 0.438086813 | 0.627389703 | DDX42 | protein_coding | 42 | -0.041390195 | 0.041390195 | ENSG00000198231 | n.s. |
| ENSG00000187634 | 1985.607795 | -0.203096832 | 0.2609594 | -0.77827 | 0.436409963 | 0.625567375 | SAMD11 | protein_coding | 66.58 | -0.041376087 | 0.041376087 | ENSG00000187634 | n.s. |
| ENSG00000166707 | 185.7170249 | 0.203960267 | 0.2626595 | 0.77652 | 0.437442157 | 0.626917875 | ZCCHC18 | protein_coding | 52.61 | 0.041360969 | 0.041360969 | ENSG00000166707 | n.s. |
| ENSG00000141933 | 173.7687803 | 0.216595455 | 0.2918953 | 0.742031 | 0.458068384 | 0.644340889 | TPGS1 | protein_coding | 61.72 | 0.041344674 | 0.041344674 | ENSG00000141933 | n.s. |
| ENSG00000198612 | 2246.438451 | 0.20215134 | 0.2587896 | 0.781142 | 0.434719165 | 0.624429478 | COPS8 | protein_coding | 39.13 | 0.041343305 | 0.041343305 | ENSG00000198612 | n.s. |
| ENSG00000125648 | 197.8162928 | -0.188885686 | 0.2303058 | -0.820152 | 0.412129602 | 0.604262894 | SLC25A23 | protein_coding | 52.6 | -0.041323291 | 0.041323291 | ENSG00000125648 | n.s. |
| ENSG00000135845 | 1309.777974 | 0.186581863 | 0.2258112 | 0.826274 | 0.408648748 | 0.600524856 | PIGC | protein_coding | 38.22 | 0.041322101 | 0.041322101 | ENSG00000135845 | n.s. |
| ENSG00000113763 | 192.434611 | -0.197806561 | 0.2497171 | -0.792123 | 0.428289222 | 0.618447505 | UNC5A | protein_coding | 58.17 | -0.041281667 | 0.041281667 | ENSG00000113763 | n.s. |
| ENSG00000186193 | 561.2523529 | -0.205829475 | 0.2680314 | -0.76793 | 0.442528566 | 0.630525215 | SAPCD2 | protein_coding | 63.61 | -0.041227138 | 0.041227138 | ENSG00000186193 | n.s. |
| ENSG00000073921 | 952.989759 | -0.207893028 | 0.2728104 | -0.762042 | 0.44603481 | 0.633688027 | PICALM | protein_coding | 37.33 | -0.041188702 | 0.041188702 | ENSG00000073921 | n.s. |
| ENSG00000204186 | 1645.483682 | 0.201139456 | 0.2572213 | 0.78197 | 0.434232018 | 0.624056109 | ZDBF2 | protein_coding | 37.53 | 0.041188606 | 0.041188606 | ENSG00000204186 | n.s. |
| ENSG00000185149 | 268.2047353 | -0.20604125 | 0.2686681 | -0.766899 | 0.443141595 | 0.631123203 | NPY2R | protein_coding | 39.63 | -0.041184731 | 0.041184731 | ENSG00000185149 | n.s. |
| ENSG00000168288 | 2369.880983 | 0.224305988 | 0.311296 | 0.720555 | 0.47118311 | 0.655372872 | MMADHC | protein_coding | 35.15 | 0.041162737 | 0.041162737 | ENSG00000168288 | n.s. |
| ENSG00000165119 | 17780.56554 | 0.195400858 | 0.2457754 | 0.795038 | 0.426591148 | 0.616763089 | HNRNPK | protein_coding | 39.12 | 0.04101105 | 0.04101105 | ENSG00000165119 | n.s. |
| ENSG00000100483 | 620.0011698 | -0.188449587 | 0.2309829 | -0.815859 | 0.41458055 | 0.606196702 | VCPKMT | protein_coding | 39.75 | -0.040966383 | 0.040966383 | ENSG00000100483 | n.s. |
| ENSG00000171094 | 1885.628916 | -0.183224702 | 0.2206222 | -0.830491 | 0.406261429 | 0.598129014 | ALK | protein_coding | 43.51 | -0.040896694 | 0.040896694 | ENSG00000171094 | n.s. |

| | | | | | | | | | | | | | |
|---|---|---|---|---|---|---|---|---|---|---|---|---|---|
| ENSG00000134769 | 20.51667489 | -0.236398218 | 0.3442172 | -0.686771 | 0.492227329 | 0.671561107 | DTNA | protein_coding | 37.99 | -0.040876671 | 0.040876671 | ENSG00000134769 | n.s. |
| ENSG00000115365 | 3363.014789 | -0.215656746 | 0.2919825 | -0.738595 | 0.460153133 | 0.646359168 | LANCL1 | protein_coding | 38.12 | -0.040872579 | 0.040872579 | ENSG00000115365 | n.s. |
| ENSG00000231213 | 239.4370472 | 0.213034139 | 0.2861426 | 0.744503 | 0.456572051 | 0.643140373 | PLSCR5 | protein_coding | 33.23 | 0.040837414 | 0.040837414 | ENSG00000231213 | n.s. |
| ENSG00000080608 | 2702.296466 | -0.196171 | 0.2481312 | -0.790594 | 0.429181015 | 0.619221478 | PUM3 | protein_coding | 39.61 | -0.040833776 | 0.040833776 | ENSG00000080608 | n.s. |
| ENSG00000085063 | 504.2295052 | 0.218158449 | 0.2985271 | 0.730783 | 0.464911895 | 0.64992388 | CD59 | protein_coding | 44.03 | 0.040825628 | 0.040825628 | ENSG00000085063 | n.s. |
| ENSG00000232303 | 81.91850222 | 0.206453866 | 0.2716796 | 0.759917 | 0.447304442 | 0.634336766 | DFFBP1 | processed_pseudogene | 56.69 | 0.040811824 | 0.040811824 | ENSG00000232303 | n.s. |
| ENSG00000214367 | 1051.248586 | 0.192563466 | 0.2408203 | 0.799615 | 0.42393397 | 0.614005271 | HAUS3 | protein_coding | 37.25 | 0.040790315 | 0.040790315 | ENSG00000214367 | n.s. |
| ENSG00000117758 | 251.9167987 | -0.219417813 | 0.3018073 | -0.727013 | 0.467218053 | 0.651845961 | STX12 | protein_coding | 41.41 | -0.040779902 | 0.040779902 | ENSG00000117758 | n.s. |
| ENSG00000151304 | 899.4548323 | -0.195464664 | 0.2470539 | -0.791182 | 0.428837701 | 0.618854406 | SRFBP1 | protein_coding | 36.16 | -0.040737086 | 0.040737086 | ENSG00000151304 | n.s. |
| ENSG00000074266 | 529.5374207 | 0.194944671 | 0.2459947 | 0.792475 | 0.428083662 | 0.618214793 | EED | protein_coding | 37.1 | 0.040716262 | 0.040716262 | ENSG00000074266 | n.s. |
| ENSG00000131115 | 870.2970255 | -0.197361426 | 0.2508624 | -0.786732 | 0.431438842 | 0.621899058 | ZNF227 | protein_coding | 41.09 | -0.040711735 | 0.040711735 | ENSG00000131115 | n.s. |
| ENSG00000186184 | 1249.801265 | 0.215868371 | 0.2934887 | 0.735525 | 0.46201957 | 0.648000249 | POLR1D | protein_coding | 40.03 | 0.040674961 | 0.040674961 | ENSG00000186184 | n.s. |
| ENSG00000171867 | 692.005007 | -0.216785558 | 0.2961015 | -0.732133 | 0.464087697 | 0.649396281 | PRNP | protein_coding | 41.96 | -0.040645168 | 0.040645168 | ENSG00000171867 | n.s. |
| ENSG00000100226 | 1006.924423 | 0.186447077 | 0.2284415 | 0.81617 | 0.414402916 | 0.606000609 | GTPBP1 | protein_coding | 51.1 | 0.040557262 | 0.040557262 | ENSG00000100226 | n.s. |
| ENSG00000143033 | 4218.830353 | -0.198632304 | 0.2551724 | -0.778424 | 0.436319166 | 0.625567375 | MTF2 | protein_coding | 37.37 | -0.040466547 | 0.040466547 | ENSG00000143033 | n.s. |
| ENSG00000157077 | 1631.616598 | -0.194971792 | 0.2470509 | -0.789197 | 0.429997041 | 0.620334557 | ZFYVE9 | protein_coding | 40.84 | -0.040432085 | 0.040432085 | ENSG00000157077 | n.s. |
| ENSG00000109586 | 739.7957088 | -0.204522849 | 0.2691199 | -0.759969 | 0.447272996 | 0.634336766 | GALNT7 | protein_coding | 38.98 | -0.040430101 | 0.040430101 | ENSG00000109586 | n.s. |
| ENSG00000132423 | 359.4094114 | -0.227017861 | 0.3224878 | -0.703958 | 0.481458932 | 0.663618499 | COQ3 | protein_coding | 39.36 | -0.040427685 | 0.040427685 | ENSG00000132423 | n.s. |
| ENSG00000029993 | 3186.267887 | -0.196191926 | 0.2498768 | -0.785155 | 0.43236298 | 0.62265099 | HMGB3 | protein_coding | 51.16 | -0.040367532 | 0.040367532 | ENSG00000029993 | n.s. |
| ENSG00000118705 | 3240.011509 | 0.205172787 | 0.2710936 | 0.756834 | 0.44914948 | 0.635792639 | RPN2 | protein_coding | 44.36 | 0.040354308 | 0.040354308 | ENSG00000118705 | n.s. |
| ENSG00000103266 | 492.7156471 | 0.210986803 | 0.283946 | 0.743053 | 0.457449847 | 0.643880016 | STUB1 | protein_coding | 64.94 | 0.040339633 | 0.040339633 | ENSG00000103266 | n.s. |
| ENSG00000199082 | 19.21657817 | -0.233341145 | 0.3398843 | -0.686531 | 0.492378316 | 0.671635475 | MIR342 | miRNA | 49.49 | -0.040336837 | 0.040336837 | ENSG00000199082 | n.s. |
| ENSG00000197045 | 3448.383755 | 0.203859204 | 0.2679717 | 0.760749 | 0.446807057 | 0.634100082 | GMFB | protein_coding | 36.79 | 0.040331952 | 0.040331952 | ENSG00000197045 | n.s. |
| ENSG00000168397 | 1013.101555 | 0.203061535 | 0.2660079 | 0.763367 | 0.445244736 | 0.633017712 | ATG4B | protein_coding | 52.33 | 0.0403248 | 0.0403248 | ENSG00000168397 | n.s. |
| ENSG00000229180 | 317.5185671 | 0.202348834 | 0.2645889 | 0.764767 | 0.444410404 | 0.632018923 | | transcribed_unprocessed_pseud | 47.61 | 0.040322036 | 0.040322036 | ENSG00000229180 | n.s. |
| ENSG00000126261 | 6006.940045 | 0.214149448 | 0.2915199 | 0.734596 | 0.462585353 | 0.648206106 | UBA2 | protein_coding | 42.17 | 0.040321532 | 0.040321532 | ENSG00000126261 | n.s. |
| ENSG00000157456 | 2217.634609 | 0.215383099 | 0.2947301 | 0.730781 | 0.464913008 | 0.64992388 | CCNB2 | protein_coding | 42.64 | 0.040306256 | 0.040306256 | ENSG00000157456 | n.s. |
| ENSG00000181029 | 306.6224719 | 0.223511103 | 0.3146221 | 0.710411 | 0.477449047 | 0.66032811 | TRAPPC5 | protein_coding | 58.05 | 0.040285689 | 0.040285689 | ENSG00000181029 | n.s. |
| ENSG00000217128 | 462.1075286 | -0.187454649 | 0.2322846 | -0.807004 | 0.419663968 | 0.610424303 | FNIP1 | protein_coding | 37.77 | -0.040184313 | 0.040184313 | ENSG00000217128 | n.s. |
| ENSG00000135249 | 629.1536987 | -0.198275573 | 0.2556219 | -0.77566 | 0.437949967 | 0.627258244 | RINT1 | protein_coding | 41.55 | -0.040161435 | 0.040161435 | ENSG00000135249 | n.s. |
| ENSG00000131018 | 115.2910566 | 0.185716957 | 0.2287067 | 0.812031 | 0.416773851 | 0.607982855 | SYNE1 | protein_coding | 38.41 | 0.040135044 | 0.040135044 | ENSG00000131018 | n.s. |
| ENSG00000089050 | 453.1088483 | -0.208558941 | 0.2795124 | -0.746153 | 0.455575133 | 0.64213126 | RBBP9 | protein_coding | 41.68 | -0.040121774 | 0.040121774 | ENSG00000089050 | n.s. |

| | | | | | | | | | | | | | |
|---|---|---|---|---|---|---|---|---|---|---|---|---|---|
| ENSG00000134684 | 1295.14512 | -0.20454628 | 0.2707625 | -0.755445 | 0.449981772 | 0.63669677 | YARS | protein_coding | 46.35 | -0.040104848 | 0.040104848 | ENSG00000134684 | n.s. |
| ENSG00000136710 | 221.0390916 | 0.220228145 | 0.3075809 | 0.716001 | 0.47399092 | 0.657665315 | CCDC115 | protein_coding | 51.35 | 0.040080435 | 0.040080435 | ENSG00000136710 | n.s. |
| ENSG00000267368 | 50.98008987 | -0.215404445 | 0.2966034 | -0.726237 | 0.467693396 | 0.651988717 | UPK3BL1 | protein_coding | 57.17 | -0.040013512 | 0.040013512 | ENSG00000267368 | n.s. |
| ENSG00000054967 | 287.5360123 | 0.190929966 | 0.240352 | 0.794376 | 0.426976317 | 0.617255868 | RELT | protein_coding | 59.84 | 0.040006467 | 0.040006467 | ENSG00000054967 | n.s. |
| ENSG00000120306 | 76.81926098 | -0.232888193 | 0.3411612 | -0.682634 | 0.49483828 | 0.673407621 | CYSTM1 | protein_coding | 44.09 | -0.03999202 | 0.03999202 | ENSG00000120306 | n.s. |
| ENSG00000145050 | 474.9580448 | 0.198580524 | 0.2574414 | 0.771362 | 0.44049236 | 0.62896057 | MANF | protein_coding | 52.06 | 0.039989467 | 0.039989467 | ENSG00000145050 | n.s. |
| ENSG00000112245 | 3437.878176 | 0.174084114 | 0.2057612 | 0.846049 | 0.397525162 | 0.589402866 | PTP4A1 | protein_coding | 39.41 | 0.039967581 | 0.039967581 | ENSG00000112245 | n.s. |
| ENSG00000114779 | 111.048089 | -0.195993362 | 0.2517683 | -0.778467 | 0.436293719 | 0.625567375 | ABHD14B | protein_coding | 55.05 | -0.039928926 | 0.039928926 | ENSG00000114779 | n.s. |
| ENSG00000138028 | 241.4265338 | -0.217425493 | 0.3017098 | -0.720645 | 0.471128244 | 0.655362028 | CGREF1 | protein_coding | 50.24 | -0.039901649 | 0.039901649 | ENSG00000138028 | n.s. |
| ENSG00000118690 | 63.65916125 | -0.219912366 | 0.3082127 | -0.713508 | 0.475531161 | 0.6585915 | ARMC2 | protein_coding | 41.62 | -0.039888558 | 0.039888558 | ENSG00000118690 | n.s. |
| ENSG00000114757 | 177.5710935 | 0.190771368 | 0.2406583 | 0.792706 | 0.427948873 | 0.618084249 | PEX5L | protein_coding | 38.14 | 0.039862121 | 0.039862121 | ENSG00000114757 | n.s. |
| ENSG00000267002 | 607.508017 | -0.197707601 | 0.2557824 | -0.772952 | 0.439550611 | 0.628645419 | | lincRNA | 44.38 | -0.039856715 | 0.039856715 | ENSG00000267002 | n.s. |
| ENSG00000087302 | 3737.587221 | -0.206930404 | 0.2771762 | -0.746566 | 0.45532549 | 0.641844366 | C14orf166 | protein_coding | 36.73 | -0.039848643 | 0.039848643 | ENSG00000087302 | n.s. |
| ENSG00000151413 | 162.2863117 | 0.193441592 | 0.2464059 | 0.785053 | 0.432422671 | 0.622672546 | NUBPL | protein_coding | 38.49 | 0.039798728 | 0.039798728 | ENSG00000151413 | n.s. |
| ENSG00000141965 | 1127.087394 | -0.171554989 | 0.2013818 | -0.851889 | 0.39427566 | 0.586359754 | FEM1A | protein_coding | 55.19 | -0.039772596 | 0.039772596 | ENSG00000141965 | n.s. |
| ENSG00000133657 | 3524.673023 | -0.196683827 | 0.2541028 | -0.774032 | 0.438911655 | 0.628183283 | ATP13A3 | protein_coding | 38.74 | -0.039713145 | 0.039713145 | ENSG00000133657 | n.s. |
| ENSG00000165617 | 304.5170304 | 0.192394747 | 0.2444135 | 0.787169 | 0.43118297 | 0.621723332 | DACT1 | protein_coding | 45.52 | 0.039710821 | 0.039710821 | ENSG00000165617 | n.s. |
| ENSG00000129472 | 401.514403 | 0.203893929 | 0.271162 | 0.751926 | 0.452095265 | 0.638669167 | RAB2B | protein_coding | 41.19 | 0.039703052 | 0.039703052 | ENSG00000129472 | n.s. |
| ENSG00000240682 | 646.3175572 | -0.205450921 | 0.2747511 | -0.747771 | 0.454598213 | 0.640884055 | ISY1 | protein_coding | 45.22 | -0.039697337 | 0.039697337 | ENSG00000240682 | n.s. |
| ENSG00000223756 | 470.0407598 | -0.18564335 | 0.2307532 | -0.80451 | 0.421102268 | 0.611260788 | TSSC2 | transcribed_unprocessed_pseud | 48.64 | -0.039685622 | 0.039685622 | ENSG00000223756 | n.s. |
| ENSG00000112667 | 370.7458171 | 0.218040454 | 0.3046468 | 0.715716 | 0.474166953 | 0.657665315 | DNPH1 | protein_coding | 55.32 | 0.039682286 | 0.039682286 | ENSG00000112667 | n.s. |
| ENSG00000267587 | 57.66199514 | -0.222828697 | 0.3167656 | -0.70345 | 0.48177555 | 0.663712596 | | processed_pseudogene | 48.68 | -0.039667951 | 0.039667951 | ENSG00000267587 | n.s. |
| ENSG00000174775 | 347.4405865 | 0.201591633 | 0.2666065 | 0.756139 | 0.449565778 | 0.63623748 | HRAS | protein_coding | 69.01 | 0.039588716 | 0.039588716 | ENSG00000174775 | n.s. |
| ENSG00000177303 | 343.1660558 | -0.1829071 | 0.2251097 | -0.812524 | 0.416490836 | 0.607649786 | CASKIN2 | protein_coding | 61.19 | -0.039571338 | 0.039571338 | ENSG00000177303 | n.s. |
| ENSG00000178913 | 4322.191687 | 0.198572058 | 0.2596471 | 0.764777 | 0.4444045 | 0.632018923 | TAF7 | protein_coding | 42.08 | 0.039569438 | 0.039569438 | ENSG00000178913 | n.s. |
| ENSG00000120438 | 7728.526638 | 0.196423418 | 0.254696 | 0.771207 | 0.440584101 | 0.62896057 | TCP1 | protein_coding | 40.93 | 0.039555076 | 0.039555076 | ENSG00000120438 | n.s. |
| ENSG00000104341 | 3219.064994 | 0.220001437 | 0.3103318 | 0.708923 | 0.478372092 | 0.66101376 | LAPTM4B | protein_coding | 44 | 0.039553948 | 0.039553948 | ENSG00000104341 | n.s. |
| ENSG00000253719 | 7400.631072 | 0.178885899 | 0.2167762 | 0.82521 | 0.409252265 | 0.601100938 | ATXN7L3B | protein_coding | 38.7 | 0.039543192 | 0.039543192 | ENSG00000253719 | n.s. |
| ENSG00000156931 | 309.507448 | -0.196033572 | 0.2535657 | -0.773108 | 0.43945868 | 0.62864309 | VPS8 | protein_coding | 38.45 | -0.039519555 | 0.039519555 | ENSG00000156931 | n.s. |
| ENSG00000114019 | 513.2988087 | -0.19928171 | 0.2617136 | -0.76145 | 0.446388554 | 0.633811136 | AMOTL2 | protein_coding | 55.04 | -0.039465777 | 0.039465777 | ENSG00000114019 | n.s. |
| ENSG00000108947 | 116.8807734 | -0.224449368 | 0.3220468 | -0.696947 | 0.485836258 | 0.667287976 | EFNB3 | protein_coding | 58.09 | -0.039432769 | 0.039432769 | ENSG00000108947 | n.s. |
| ENSG00000174579 | 1319.028568 | 0.184434213 | 0.2293904 | 0.804019 | 0.421386055 | 0.611459932 | MSL2 | protein_coding | 42.35 | 0.039401049 | 0.039401049 | ENSG00000174579 | n.s. |

| | | | | | | | | | | | | | |
|---|---|---|---|---|---|---|---|---|---|---|---|---|---|
| ENSG00000185900 | 541.3380206 | -0.182355316 | 0.2247156 | -0.811494 | 0.417082104 | 0.608193772 | POMK | protein_coding | 45.29 | -0.039381095 | 0.039381095 | ENSG00000185900 | n.s. |
| ENSG00000010322 | 1927.967608 | -0.198699318 | 0.2607263 | -0.762099 | 0.446000715 | 0.633688027 | NISCH | protein_coding | 53.28 | -0.039367203 | 0.039367203 | ENSG00000010322 | n.s. |
| ENSG00000113851 | 1077.936183 | -0.2113211 | 0.2902072 | -0.728173 | 0.466507592 | 0.651228628 | CRBN | protein_coding | 37.25 | -0.039362045 | 0.039362045 | ENSG00000113851 | n.s. |
| ENSG00000142867 | 473.8100556 | 0.196947155 | 0.2572224 | 0.765669 | 0.443873433 | 0.631569673 | BCL10 | protein_coding | 40.57 | 0.039306464 | 0.039306464 | ENSG00000142867 | n.s. |
| ENSG00000169032 | 657.099578 | -0.195011898 | 0.252778 | -0.771475 | 0.440425521 | 0.62896057 | MAP2K1 | protein_coding | 44.72 | -0.039270829 | 0.039270829 | ENSG00000169032 | n.s. |
| ENSG00000130520 | 2065.047739 | 0.225630466 | 0.3269319 | 0.690145 | 0.490102884 | 0.670128457 | LSM4 | protein_coding | 56.37 | 0.039224038 | 0.039224038 | ENSG00000130520 | n.s. |
| ENSG00000169221 | 416.6304509 | -0.184435851 | 0.230075 | -0.801634 | 0.422764973 | 0.612822147 | TBC1D10B | protein_coding | 54.1 | -0.039223151 | 0.039223151 | ENSG00000169221 | n.s. |
| ENSG00000233527 | 52.27487708 | -0.225898788 | 0.3280387 | -0.688635 | 0.491053202 | 0.670468885 | ZNF529-AS1 | antisense | 42.2 | -0.039220858 | 0.039220858 | ENSG00000233527 | n.s. |
| ENSG00000125753 | 136.7989383 | 0.204558162 | 0.2747838 | 0.744433 | 0.456614422 | 0.643140373 | VASP | protein_coding | 54.67 | 0.039212619 | 0.039212619 | ENSG00000125753 | n.s. |
| ENSG00000186522 | 821.5083655 | -0.20568525 | 0.2775 | -0.741208 | 0.458567298 | 0.644912372 | 10-Sep | protein_coding | 40.38 | -0.039182894 | 0.039182894 | ENSG00000186522 | n.s. |
| ENSG00000005469 | 286.8909664 | -0.209033655 | 0.28566 | -0.731757 | 0.464316966 | 0.649521262 | CROT | protein_coding | 37.43 | -0.039174292 | 0.039174292 | ENSG00000005469 | n.s. |
| ENSG00000142686 | 364.1098613 | 0.201893465 | 0.2691198 | 0.750199 | 0.453134703 | 0.639727688 | C1orf216 | protein_coding | 53.73 | 0.039168312 | 0.039168312 | ENSG00000142686 | n.s. |
| ENSG00000115641 | 507.036928 | -0.19800451 | 0.2605649 | -0.759905 | 0.447311525 | 0.634336766 | FHL2 | protein_coding | 44.54 | -0.039141555 | 0.039141555 | ENSG00000115641 | n.s. |
| ENSG00000277998 | 31.02134351 | -0.252862821 | 0.3984527 | -0.634612 | 0.525681605 | 0.700254974 | | unprocessed_pseudogene | 38.22 | -0.039128953 | 0.039128953 | ENSG00000277998 | n.s. |
| ENSG00000008300 | 968.0096445 | -0.184819696 | 0.2312219 | -0.799317 | 0.42410636 | 0.61419106 | CELSR3 | protein_coding | 59.89 | -0.039125684 | 0.039125684 | ENSG00000008300 | n.s. |
| ENSG00000107443 | 1289.136691 | 0.197918128 | 0.2605924 | 0.759493 | 0.447557604 | 0.634621094 | CCNJ | protein_coding | 39.76 | 0.03908596 | 0.03908596 | ENSG00000107443 | n.s. |
| ENSG00000161671 | 735.4438393 | 0.215087148 | 0.3011655 | 0.714182 | 0.475114363 | 0.658233153 | EMC10 | protein_coding | 57.11 | 0.039064181 | 0.039064181 | ENSG00000161671 | n.s. |
| ENSG00000176894 | 976.904021 | 0.221282984 | 0.3166285 | 0.698873 | 0.484631666 | 0.666224989 | PXMP2 | protein_coding | 52.39 | 0.03902969 | 0.03902969 | ENSG00000176894 | n.s. |
| ENSG00000163468 | 11580.37898 | 0.216674375 | 0.3052494 | 0.709827 | 0.477811249 | 0.660516096 | CCT3 | protein_coding | 45.93 | 0.03902665 | 0.03902665 | ENSG00000163468 | n.s. |
| ENSG00000116096 | 252.7437251 | 0.207812921 | 0.2834836 | 0.733068 | 0.463516679 | 0.648988608 | SPR | protein_coding | 54.78 | 0.039019562 | 0.039019562 | ENSG00000116096 | n.s. |
| ENSG00000160305 | 145.875988 | -0.221590878 | 0.3173819 | -0.698184 | 0.485062276 | 0.666685298 | DIP2A | protein_coding | 44.91 | -0.039017527 | 0.039017527 | ENSG00000160305 | n.s. |
| ENSG00000188599 | 80.81837246 | -0.205731482 | 0.2784408 | -0.73887 | 0.45998604 | 0.646203586 | NPIPP1 | transcribed_unprocessed_pseud | 45.29 | -0.039012992 | 0.039012992 | ENSG00000188599 | n.s. |
| ENSG00000182208 | 79.69693823 | -0.211379486 | 0.2923662 | -0.722996 | 0.469682512 | 0.653938962 | MOB2 | protein_coding | 54.17 | -0.038991649 | 0.038991649 | ENSG00000182208 | n.s. |
| ENSG00000111011 | 7780.405504 | 0.179388939 | 0.2199724 | 0.815507 | 0.414782364 | 0.606428106 | RSRC2 | protein_coding | 39.85 | 0.038966987 | 0.038966987 | ENSG00000111011 | n.s. |
| ENSG00000235545 | 54.83479691 | -0.207120987 | 0.2822468 | -0.733829 | 0.463052722 | 0.648665163 | | antisense | 42.23 | -0.038934484 | 0.038934484 | ENSG00000235545 | n.s. |
| ENSG00000079246 | 19637.02608 | -0.203583678 | 0.2740774 | -0.742796 | 0.457604942 | 0.643988195 | XRCC5 | protein_coding | 39.46 | -0.038909339 | 0.038909339 | ENSG00000079246 | n.s. |
| ENSG00000148090 | 321.1944557 | -0.184517448 | 0.2314657 | -0.79717 | 0.425352399 | 0.615483418 | AUH | protein_coding | 38.91 | -0.03889326 | 0.03889326 | ENSG00000148090 | n.s. |
| ENSG00000201573 | 27.79028936 | -0.230152358 | 0.3414538 | -0.674037 | 0.500288065 | 0.677749957 | | misc_RNA | 39.22 | -0.038879753 | 0.038879753 | ENSG00000201573 | n.s. |
| ENSG00000086065 | 1098.237832 | 0.22617757 | 0.3314751 | 0.682336 | 0.495026229 | 0.673449188 | CHMP5 | protein_coding | 41.65 | 0.038833596 | 0.038833596 | ENSG00000086065 | n.s. |
| ENSG00000133083 | 429.2362275 | -0.2074684 | 0.2839452 | -0.730664 | 0.464984684 | 0.64992388 | DCLK1 | protein_coding | 40.16 | -0.038825119 | 0.038825119 | ENSG00000133083 | n.s. |
| ENSG00000176386 | 125.6969238 | -0.213296327 | 0.2980413 | -0.71566 | 0.474201113 | 0.657665315 | CDC26 | protein_coding | 43.07 | -0.038818878 | 0.038818878 | ENSG00000176386 | n.s. |
| ENSG00000127419 | 121.9350578 | 0.204158249 | 0.2759126 | 0.739938 | 0.459337553 | 0.645671434 | TMEM175 | protein_coding | 51.83 | 0.038787705 | 0.038787705 | ENSG00000127419 | n.s. |

| | | | | | | | | | | | | | |
|---|---|---|---|---|---|---|---|---|---|---|---|---|---|
| ENSG00000022267 | 1315.204497 | 0.208219668 | 0.2859558 | 0.728153 | 0.466519739 | 0.651228628 | FHL1 | protein_coding | 42.07 | 0.038784352 | 0.038784352 | ENSG00000022267 | n.s. |
| ENSG00000105821 | 1935.848534 | -0.195312891 | 0.2559552 | -0.763074 | 0.445419035 | 0.633071582 | DNAJC2 | protein_coding | 40.56 | -0.038778824 | 0.038778824 | ENSG00000105821 | n.s. |
| ENSG00000197894 | 3070.487129 | -0.211866997 | 0.2949004 | -0.718436 | 0.472488565 | 0.656301346 | ADH5 | protein_coding | 40.37 | -0.038749776 | 0.038749776 | ENSG00000197894 | n.s. |
| ENSG00000229152 | 445.4989624 | 0.19389069 | 0.2529057 | 0.766652 | 0.443288412 | 0.631196898 | ANKRD10-IT1 | sense_intronic | 38.6 | 0.038746174 | 0.038746174 | ENSG00000229152 | n.s. |
| ENSG00000078804 | 163.8063371 | 0.180788572 | 0.2242676 | 0.806129 | 0.420168606 | 0.610775475 | TP53INP2 | protein_coding | 55.18 | 0.038710161 | 0.038710161 | ENSG00000078804 | n.s. |
| ENSG00000176641 | 307.3818975 | 0.192201421 | 0.2489837 | 0.771944 | 0.440147808 | 0.628944621 | RNF152 | protein_coding | 42.43 | 0.038706981 | 0.038706981 | ENSG00000176641 | n.s. |
| ENSG00000115761 | 1490.642143 | -0.197697196 | 0.2620012 | -0.754566 | 0.450509518 | 0.637184479 | NOL10 | protein_coding | 40.73 | -0.038696223 | 0.038696223 | ENSG00000115761 | n.s. |
| ENSG00000207821 | 13.29377933 | -0.246688015 | 0.3847681 | -0.641134 | 0.521435408 | 0.696865539 | MIR640 | miRNA | 52.08 | -0.038693265 | 0.038693265 | ENSG00000207821 | n.s. |
| ENSG00000173960 | 617.1909584 | -0.205738513 | 0.2804941 | -0.733486 | 0.463262017 | 0.648762529 | UBXN2A | protein_coding | 42.94 | -0.038661197 | 0.038661197 | ENSG00000173960 | n.s. |
| ENSG00000170089 | 295.4242949 | 0.216364204 | 0.3066601 | 0.705551 | 0.480467612 | 0.662725436 | | transcribed_unprocessed_pseud | 44.8 | 0.038657005 | 0.038657005 | ENSG00000170089 | n.s. |
| ENSG00000091947 | 476.5032637 | -0.188161428 | 0.2401156 | -0.783628 | 0.43325819 | 0.623424326 | TMEM101 | protein_coding | 49.94 | -0.038613784 | 0.038613784 | ENSG00000091947 | n.s. |
| ENSG00000170515 | 6061.372976 | 0.205549727 | 0.2803525 | 0.733183 | 0.463446701 | 0.64895589 | PA2G4 | protein_coding | 46.6 | 0.038599119 | 0.038599119 | ENSG00000170515 | n.s. |
| ENSG00000260230 | 378.2809236 | -0.212254483 | 0.2969536 | -0.714773 | 0.474749144 | 0.658032089 | FRRS1L | protein_coding | 42.27 | -0.038577874 | 0.038577874 | ENSG00000260230 | n.s. |
| ENSG00000060971 | 89.08874633 | 0.192658044 | 0.2511397 | 0.767135 | 0.443001243 | 0.630987857 | ACAA1 | protein_coding | 51.46 | 0.038527563 | 0.038527563 | ENSG00000060971 | n.s. |
| ENSG00000182224 | 228.9187701 | 0.186746819 | 0.2374477 | 0.786476 | 0.431588949 | 0.621986641 | CYB5D1 | protein_coding | 48.18 | 0.038510732 | 0.038510732 | ENSG00000182224 | n.s. |
| ENSG00000107175 | 217.0164373 | 0.204857778 | 0.2792354 | 0.733638 | 0.463169259 | 0.648762529 | CREB3 | protein_coding | 49.01 | 0.038495694 | 0.038495694 | ENSG00000107175 | n.s. |
| ENSG00000111696 | 1244.403708 | -0.187277238 | 0.2387373 | -0.784449 | 0.432776656 | 0.623053394 | NT5DC3 | protein_coding | 43.1 | -0.038480741 | 0.038480741 | ENSG00000111696 | n.s. |
| ENSG00000185619 | 1471.474693 | -0.20122559 | 0.2708174 | -0.74303 | 0.457463271 | 0.643880016 | PCGF3 | protein_coding | 56.09 | -0.038473338 | 0.038473338 | ENSG00000185619 | n.s. |
| ENSG00000249115 | 424.2688639 | -0.198278969 | 0.2643732 | -0.749996 | 0.453256828 | 0.639748528 | HAUS5 | protein_coding | 54.43 | -0.038464277 | 0.038464277 | ENSG00000249115 | n.s. |
| ENSG00000174672 | 359.0583897 | -0.189728706 | 0.2444604 | -0.776112 | 0.437682796 | 0.627133639 | BRSK2 | protein_coding | 63.85 | -0.038446607 | 0.038446607 | ENSG00000174672 | n.s. |
| ENSG00000213719 | 1040.90652 | 0.220950733 | 0.3198862 | 0.690717 | 0.489743656 | 0.66991916 | CLIC1 | protein_coding | 51 | 0.038440478 | 0.038440478 | ENSG00000213719 | n.s. |
| ENSG00000103549 | 1530.879529 | -0.192319094 | 0.2509032 | -0.766507 | 0.443374679 | 0.631196898 | RNF40 | protein_coding | 55.81 | -0.038432114 | 0.038432114 | ENSG00000103549 | n.s. |
| ENSG00000169258 | 1468.677251 | -0.194189709 | 0.2551811 | -0.760988 | 0.446664282 | 0.634100082 | GPRIN1 | protein_coding | 55.36 | -0.038418917 | 0.038418917 | ENSG00000169258 | n.s. |
| ENSG00000125124 | 329.4303764 | -0.211896943 | 0.2971136 | -0.713185 | 0.475731368 | 0.658803246 | BBS2 | protein_coding | 40.21 | -0.038405107 | 0.038405107 | ENSG00000125124 | n.s. |
| ENSG00000066654 | 2362.092566 | -0.189792195 | 0.2449313 | -0.774879 | 0.43841095 | 0.627683831 | THUMPD1 | protein_coding | 40.19 | -0.038387191 | 0.038387191 | ENSG00000066654 | n.s. |
| ENSG00000165169 | 423.8065711 | 0.21565469 | 0.3067998 | 0.702917 | 0.482107748 | 0.66388887 | DYNLT3 | protein_coding | 37.62 | 0.038365964 | 0.038365964 | ENSG00000165169 | n.s. |
| ENSG00000188612 | 7174.634599 | 0.218179937 | 0.3128413 | 0.697414 | 0.485543736 | 0.667083624 | SUMO2 | protein_coding | 46.32 | 0.038360336 | 0.038360336 | ENSG00000188612 | n.s. |
| ENSG00000198373 | 42.07970539 | -0.20954495 | 0.2916828 | -0.7184 | 0.472510572 | 0.656301346 | WWP2 | protein_coding | 45.6 | -0.038325081 | 0.038325081 | ENSG00000198373 | n.s. |
| ENSG00000163882 | 847.704476 | 0.210337699 | 0.2940201 | 0.715385 | 0.474370918 | 0.657704272 | POLR2H | protein_coding | 52.77 | 0.038275012 | 0.038275012 | ENSG00000163882 | n.s. |
| ENSG00000115946 | 1272.394046 | 0.203044014 | 0.2762949 | 0.734881 | 0.4624117 | 0.648137988 | PNO1 | protein_coding | 41.05 | 0.038239792 | 0.038239792 | ENSG00000115946 | n.s. |
| ENSG00000234268 | 23.33400598 | 0.221360882 | 0.3228843 | 0.685573 | 0.492982164 | 0.671937241 | | processed_pseudogene | 49.02 | 0.038222669 | 0.038222669 | ENSG00000234268 | n.s. |
| ENSG00000228201 | 21.97896895 | -0.220037375 | 0.3195279 | -0.688633 | 0.491054342 | 0.670468885 | | antisense | 66.35 | -0.038203191 | 0.038203191 | ENSG00000228201 | n.s. |

| | | | | | | | | | | | | | |
|---|---|---|---|---|---|---|---|---|---|---|---|---|---|
| ENSG00000185664 | 401.4626127 | -0.21170994 | 0.2977177 | -0.71111 | 0.477016239 | 0.660057354 | PMEL | protein_coding | 48.7 | -0.038196353 | 0.038196353 | ENSG00000185664 | n.s. |
| ENSG00000105894 | 105.6176491 | -0.205357229 | 0.282925 | -0.725836 | 0.467939144 | 0.652229122 | PTN | protein_coding | 38.25 | -0.038114263 | 0.038114263 | ENSG00000105894 | n.s. |
| ENSG00000176714 | 132.6262037 | 0.188356009 | 0.2430866 | 0.774851 | 0.438427441 | 0.627683831 | CCDC121 | protein_coding | 42.38 | 0.038096709 | 0.038096709 | ENSG00000176714 | n.s. |
| ENSG00000136986 | 977.8608529 | 0.204976519 | 0.2819091 | 0.727102 | 0.467163725 | 0.651845961 | DERL1 | protein_coding | 41.26 | 0.038095915 | 0.038095915 | ENSG00000136986 | n.s. |
| ENSG00000134352 | 1260.149242 | -0.188916548 | 0.2443959 | -0.772994 | 0.439525972 | 0.628645419 | IL6ST | protein_coding | 37.41 | -0.038084489 | 0.038084489 | ENSG00000134352 | n.s. |
| ENSG00000075142 | 1098.661199 | 0.201923251 | 0.2748484 | 0.734671 | 0.462539585 | 0.648206106 | SRI | protein_coding | 39.53 | 0.038019499 | 0.038019499 | ENSG00000075142 | n.s. |
| ENSG00000197114 | 98.39978104 | 0.195869185 | 0.2615794 | 0.748794 | 0.453981211 | 0.640401597 | ZGPAT | protein_coding | 49.61 | 0.037910009 | 0.037910009 | ENSG00000197114 | n.s. |
| ENSG00000160961 | 181.5639375 | 0.187307464 | 0.2421875 | 0.773398 | 0.439286579 | 0.628461471 | ZNF333 | protein_coding | 46.91 | 0.037783913 | 0.037783913 | ENSG00000160961 | n.s. |
| ENSG00000132694 | 828.2309661 | -0.179833441 | 0.2260612 | -0.795508 | 0.426318128 | 0.616560434 | ARHGEF11 | protein_coding | 44.51 | -0.037769401 | 0.037769401 | ENSG00000132694 | n.s. |
| ENSG00000103034 | 1410.416957 | 0.191724032 | 0.2530321 | 0.757706 | 0.44862683 | 0.63561935 | NDRG4 | protein_coding | 53.85 | 0.037731844 | 0.037731844 | ENSG00000103034 | n.s. |
| ENSG00000006118 | 2416.218337 | 0.186254325 | 0.2401117 | 0.775699 | 0.437926927 | 0.627258244 | TMEM132A | protein_coding | 58.71 | 0.037726488 | 0.037726488 | ENSG00000006118 | n.s. |
| ENSG00000134987 | 3246.531847 | -0.19170523 | 0.253152 | -0.757273 | 0.448886158 | 0.635741733 | WDR36 | protein_coding | 33.98 | -0.037712115 | 0.037712115 | ENSG00000134987 | n.s. |
| ENSG00000167978 | 3436.792862 | 0.190485147 | 0.2504712 | 0.760507 | 0.446951404 | 0.634100082 | SRRM2 | protein_coding | 52.86 | 0.037685999 | 0.037685999 | ENSG00000167978 | n.s. |
| ENSG00000138071 | 2451.706887 | -0.203125762 | 0.2800147 | -0.725411 | 0.468199817 | 0.652396522 | ACTR2 | protein_coding | 39.48 | -0.037677465 | 0.037677465 | ENSG00000138071 | n.s. |
| ENSG00000226174 | 41.39686796 | -0.20647133 | 0.2878999 | -0.717164 | 0.473273099 | 0.65702243 | TEX22 | protein_coding | 55.6 | -0.03766446 | 0.03766446 | ENSG00000226174 | n.s. |
| ENSG00000121864 | 2217.503556 | -0.190852374 | 0.2515829 | -0.758606 | 0.448088126 | 0.635179289 | ZNF639 | protein_coding | 41.32 | -0.037617704 | 0.037617704 | ENSG00000121864 | n.s. |
| ENSG00000050393 | 874.8960442 | -0.200343968 | 0.2734433 | -0.732671 | 0.463759202 | 0.649176514 | MCUR1 | protein_coding | 41.24 | -0.037591981 | 0.037591981 | ENSG00000050393 | n.s. |
| ENSG00000141380 | 2684.418179 | -0.19399175 | 0.258949 | -0.749151 | 0.453766475 | 0.640240035 | SS18 | protein_coding | 35.21 | -0.037567894 | 0.037567894 | ENSG00000141380 | n.s. |
| ENSG00000165629 | 2306.805405 | 0.223366229 | 0.3326252 | 0.671525 | 0.501885938 | 0.679019385 | ATP5C1 | protein_coding | 40.13 | 0.037551845 | 0.037551845 | ENSG00000165629 | n.s. |
| ENSG00000082146 | 549.8790408 | -0.203735785 | 0.2819665 | -0.722553 | 0.469954503 | 0.654234965 | STRADB | protein_coding | 38.23 | -0.037541629 | 0.037541629 | ENSG00000082146 | n.s. |
| ENSG00000279416 | 46.40021514 | -0.20504993 | 0.2851349 | -0.719133 | 0.472058896 | 0.656094752 | | TEC | 40.09 | -0.037530993 | 0.037530993 | ENSG00000279416 | n.s. |
| ENSG00000124787 | 201.8763641 | 0.198155714 | 0.268484 | 0.738054 | 0.460481665 | 0.646624934 | RPP40 | protein_coding | 43.41 | 0.037520299 | 0.037520299 | ENSG00000124787 | n.s. |
| ENSG00000236439 | 50.40616506 | 0.218897629 | 0.3213444 | 0.681193 | 0.495749237 | 0.673922732 | | processed_pseudogene | 49.61 | 0.037516841 | 0.037516841 | ENSG00000236439 | n.s. |
| ENSG00000274677 | 23.14585041 | -0.218436641 | 0.3201253 | -0.682347 | 0.495019301 | 0.673449188 | | sense_intronic | 52.12 | -0.037504515 | 0.037504515 | ENSG00000274677 | n.s. |
| ENSG00000181404 | 107.048747 | -0.208164658 | 0.2931373 | -0.710127 | 0.477625475 | 0.660440907 | WASHC1 | protein_coding | 54.03 | -0.037504201 | 0.037504201 | ENSG00000181404 | n.s. |
| ENSG00000161513 | 210.6637722 | 0.19697672 | 0.2658768 | 0.740857 | 0.458780041 | 0.645081248 | FDXR | protein_coding | 58.11 | 0.037501527 | 0.037501527 | ENSG00000161513 | n.s. |
| ENSG00000151287 | 496.1933111 | -0.206244427 | 0.2886895 | -0.714416 | 0.474969968 | 0.658207114 | TEX30 | protein_coding | 41.14 | -0.037461707 | 0.037461707 | ENSG00000151287 | n.s. |
| ENSG00000174891 | 1050.089699 | -0.209777305 | 0.2974511 | -0.70525 | 0.480654822 | 0.662917987 | RSRC1 | protein_coding | 35.48 | -0.037453681 | 0.037453681 | ENSG00000174891 | n.s. |
| ENSG00000129071 | 895.5850798 | 0.181411381 | 0.2305008 | 0.787031 | 0.431263439 | 0.621774968 | MBD4 | protein_coding | 39.9 | 0.03743728 | 0.03743728 | ENSG00000129071 | n.s. |
| ENSG00000132781 | 111.5832117 | 0.185640869 | 0.2403733 | 0.772302 | 0.439935287 | 0.628872587 | MUTYH | protein_coding | 51.03 | 0.037395002 | 0.037395002 | ENSG00000132781 | n.s. |
| ENSG00000083123 | 682.9838798 | -0.205890234 | 0.2882988 | -0.714156 | 0.475130964 | 0.658233153 | BCKDHB | protein_coding | 36.82 | -0.037393835 | 0.037393835 | ENSG00000083123 | n.s. |
| ENSG00000043355 | 1831.42301 | 0.161763927 | 0.1903909 | 0.849641 | 0.395524695 | 0.587647937 | ZIC2 | protein_coding | 56.6 | 0.037348506 | 0.037348506 | ENSG00000043355 | n.s. |

| | | | | | | | | | | | | | |
|---|---|---|---|---|---|---|---|---|---|---|---|---|---|
| ENSG00000116754 | 7614.298916 | -0.201221062 | 0.2773398 | -0.72554 | 0.468120894 | 0.65239371 | SRSF11 | protein_coding | 37.01 | -0.037324542 | 0.037324542 | ENSG00000116754 | n.s. |
| ENSG00000029364 | 1235.691866 | -0.191667787 | 0.255028 | -0.751556 | 0.452318278 | 0.638898994 | SLC39A9 | protein_coding | 40.2 | -0.037292379 | 0.037292379 | ENSG00000029364 | n.s. |
| ENSG00000233325 | 29.34804431 | 0.208046645 | 0.2942748 | 0.706981 | 0.479578361 | 0.661836342 | MIPEPP3 | transcribed_unprocessed_pseud | 39.33 | 0.037292234 | 0.037292234 | ENSG00000233325 | n.s. |
| ENSG00000155827 | 2286.76906 | -0.188788955 | 0.2486101 | -0.759378 | 0.447626729 | 0.634654475 | RNF20 | protein_coding | 38.17 | -0.037278768 | 0.037278768 | ENSG00000155827 | n.s. |
| ENSG00000148481 | 764.2781859 | -0.199618676 | 0.2736932 | -0.729352 | 0.465786353 | 0.650661522 | MINDY3 | protein_coding | 35.88 | -0.037257803 | 0.037257803 | ENSG00000148481 | n.s. |
| ENSG00000221944 | 1060.424962 | 0.166372287 | 0.2000098 | 0.831821 | 0.405510099 | 0.597578018 | TIGD1 | protein_coding | 43.77 | 0.037201739 | 0.037201739 | ENSG00000221944 | n.s. |
| ENSG00000168612 | 168.707002 | -0.187617278 | 0.2463894 | -0.761466 | 0.446378537 | 0.633811136 | ZSWIM1 | protein_coding | 50.32 | -0.037155751 | 0.037155751 | ENSG00000168612 | n.s. |
| ENSG00000132305 | 2974.803238 | -0.195435453 | 0.2642617 | -0.739553 | 0.459571393 | 0.645737461 | IMMT | protein_coding | 41.41 | -0.037121795 | 0.037121795 | ENSG00000132305 | n.s. |
| ENSG00000166170 | 1487.487599 | 0.184659103 | 0.2399161 | 0.769682 | 0.441488555 | 0.62947723 | BAG5 | protein_coding | 46.41 | 0.037120168 | 0.037120168 | ENSG00000166170 | n.s. |
| ENSG00000173599 | 355.4460072 | -0.201396101 | 0.2790077 | -0.72183 | 0.47039914 | 0.65465398 | PC | protein_coding | 51.92 | -0.037054504 | 0.037054504 | ENSG00000173599 | n.s. |
| ENSG00000233922 | 271.9038757 | -0.193283912 | 0.2596071 | -0.744525 | 0.45655902 | 0.643140373 | LINC01694 | lincRNA | 52.25 | -0.03705141 | 0.03705141 | ENSG00000233922 | n.s. |
| ENSG00000054356 | 594.787295 | -0.188151897 | 0.2486098 | -0.756816 | 0.449160124 | 0.635792639 | PTPRN | protein_coding | 53.59 | -0.037006563 | 0.037006563 | ENSG00000054356 | n.s. |
| ENSG00000083099 | 1938.801503 | -0.18372768 | 0.2382308 | -0.771217 | 0.440578317 | 0.62896057 | LYRM2 | protein_coding | 44.37 | -0.036998452 | 0.036998452 | ENSG00000083099 | n.s. |
| ENSG00000101057 | 1761.146081 | -0.184921401 | 0.2409576 | -0.767444 | 0.442817703 | 0.630855508 | MYBL2 | protein_coding | 49.88 | -0.036997243 | 0.036997243 | ENSG00000101057 | n.s. |
| ENSG00000077147 | 3598.292413 | -0.194677228 | 0.2631273 | -0.739859 | 0.459385278 | 0.645671434 | TM9SF3 | protein_coding | 37.03 | -0.03698642 | 0.03698642 | ENSG00000077147 | n.s. |
| ENSG00000155506 | 3866.069093 | -0.180721361 | 0.2312657 | -0.781445 | 0.434541017 | 0.624293171 | LARP1 | protein_coding | 46.52 | -0.036977654 | 0.036977654 | ENSG00000155506 | n.s. |
| ENSG00000143543 | 1518.202918 | 0.216376107 | 0.3185065 | 0.679346 | 0.496918635 | 0.67472188 | JTB | protein_coding | 50.96 | 0.036973312 | 0.036973312 | ENSG00000143543 | n.s. |
| ENSG00000196262 | 12356.80728 | 0.208536482 | 0.2974602 | 0.701057 | 0.483267564 | 0.664880582 | PPIA | protein_coding | 47.06 | 0.036964416 | 0.036964416 | ENSG00000196262 | n.s. |
| ENSG00000148337 | 644.0884787 | -0.193225359 | 0.2599407 | -0.743344 | 0.457273497 | 0.643743064 | CIZ1 | protein_coding | 52.19 | -0.036961584 | 0.036961584 | ENSG00000148337 | n.s. |
| ENSG00000108671 | 1488.149079 | 0.179367591 | 0.2282603 | 0.785803 | 0.431983034 | 0.622380245 | PSMD11 | protein_coding | 43.11 | 0.036939715 | 0.036939715 | ENSG00000108671 | n.s. |
| ENSG00000271427 | 25.82630187 | -0.214224093 | 0.3140144 | -0.682211 | 0.495105545 | 0.673449188 | | lincRNA | 29.39 | -0.036781241 | 0.036781241 | ENSG00000271427 | n.s. |
| ENSG00000101417 | 255.6510306 | -0.210285673 | 0.3028908 | -0.694262 | 0.487517682 | 0.668694533 | PXMP4 | protein_coding | 49.45 | -0.036752095 | 0.036752095 | ENSG00000101417 | n.s. |
| ENSG00000101856 | 2228.157777 | 0.206011184 | 0.292327 | 0.704729 | 0.480979195 | 0.663168284 | PGRMC1 | protein_coding | 42.58 | 0.036747503 | 0.036747503 | ENSG00000101856 | n.s. |
| ENSG00000149823 | 876.2617179 | 0.192864947 | 0.2602308 | 0.74113 | 0.458614377 | 0.644913441 | VPS51 | protein_coding | 50.69 | 0.036740497 | 0.036740497 | ENSG00000149823 | n.s. |
| ENSG00000213551 | 2817.91293 | 0.18399989 | 0.2403313 | 0.765609 | 0.443908699 | 0.631569673 | DNAJC9 | protein_coding | 40 | 0.036722465 | 0.036722465 | ENSG00000213551 | n.s. |
| ENSG00000106089 | 660.2669816 | 0.194984401 | 0.2652958 | 0.73497 | 0.462357846 | 0.648137988 | STX1A | protein_coding | 58.84 | 0.036721904 | 0.036721904 | ENSG00000106089 | n.s. |
| ENSG00000172500 | 581.8052967 | 0.218160758 | 0.3247703 | 0.671739 | 0.501750124 | 0.67896462 | FIBP | protein_coding | 55.12 | 0.036684355 | 0.036684355 | ENSG00000172500 | n.s. |
| ENSG00000123131 | 1551.192455 | 0.217327539 | 0.3229274 | 0.672992 | 0.500952422 | 0.678148816 | PRDX4 | protein_coding | 40.65 | 0.036657721 | 0.036657721 | ENSG00000123131 | n.s. |
| ENSG00000075426 | 603.2602774 | 0.170466641 | 0.2107404 | 0.808894 | 0.418576101 | 0.609606178 | FOSL2 | protein_coding | 51.8 | 0.036641914 | 0.036641914 | ENSG00000075426 | n.s. |
| ENSG00000233515 | 43.61495898 | 0.212077213 | 0.3094144 | 0.685415 | 0.493082151 | 0.671937241 | LINC01518 | lincRNA | 41.67 | 0.036619645 | 0.036619645 | ENSG00000233515 | n.s. |
| ENSG00000124570 | 125.0544246 | 0.21512241 | 0.317388 | 0.67779 | 0.497904921 | 0.675731573 | SERPINB6 | protein_coding | 47.23 | 0.036619382 | 0.036619382 | ENSG00000124570 | n.s. |
| ENSG00000168101 | 204.2701753 | 0.21073487 | 0.3060441 | 0.688577 | 0.491089685 | 0.670468885 | NUDT16L1 | protein_coding | 65.05 | 0.036588077 | 0.036588077 | ENSG00000168101 | n.s. |

| | | | | | | | | | | | | | |
|---|---|---|---|---|---|---|---|---|---|---|---|---|---|
| ENSG00000135469 | 322.9555201 | 0.194232985 | 0.2643236 | 0.73483 | 0.462442858 | 0.648137988 | COQ10A | protein_coding | 50.99 | 0.036580388 | 0.036580388 | ENSG00000135469 | n.s. |
| ENSG00000257390 | 64.54756829 | 0.195037446 | 0.2665771 | 0.731636 | 0.464390639 | 0.649559058 | | protein_coding | 42.96 | 0.036546381 | 0.036546381 | ENSG00000257390 | n.s. |
| ENSG00000106624 | 310.8100479 | 0.180940624 | 0.2338554 | 0.773729 | 0.439091295 | 0.628311213 | AEBP1 | protein_coding | 62.7 | 0.036518376 | 0.036518376 | ENSG00000106624 | n.s. |
| ENSG00000138600 | 853.667734 | -0.193005155 | 0.2619891 | -0.736691 | 0.461309987 | 0.647265842 | SPPL2A | protein_coding | 40.43 | -0.036462015 | 0.036462015 | ENSG00000138600 | n.s. |
| ENSG00000135736 | 137.6556835 | -0.198798104 | 0.2760393 | -0.72018 | 0.47141388 | 0.655562869 | CCDC102A | protein_coding | 56.06 | -0.03645672 | 0.03645672 | ENSG00000135736 | n.s. |
| ENSG00000049656 | 1035.734868 | 0.203157873 | 0.2871281 | 0.707551 | 0.479223893 | 0.661599828 | CLPTM1L | protein_coding | 56.17 | 0.03644746 | 0.03644746 | ENSG00000049656 | n.s. |
| ENSG00000135049 | 2058.595991 | -0.203333646 | 0.2876441 | -0.706893 | 0.479632887 | 0.661836342 | AGTPBP1 | protein_coding | 36.66 | -0.036447432 | 0.036447432 | ENSG00000135049 | n.s. |
| ENSG00000203668 | 5687.02887 | -0.197627532 | 0.2738647 | -0.721625 | 0.470525201 | 0.65465398 | CHML | protein_coding | 37.83 | -0.036361131 | 0.036361131 | ENSG00000203668 | n.s. |
| ENSG00000200142 | 20.27155127 | -0.216264092 | 0.3221375 | -0.671341 | 0.502003401 | 0.6790318 | | misc_RNA | 44.9 | -0.036356132 | 0.036356132 | ENSG00000200142 | n.s. |
| ENSG00000164687 | 3557.815378 | 0.178099041 | 0.2284238 | 0.779687 | 0.435575168 | 0.625110666 | FABP5 | protein_coding | 45.55 | 0.036339879 | 0.036339879 | ENSG00000164687 | n.s. |
| ENSG00000101266 | 2612.3993 | -0.191615033 | 0.259341 | -0.738854 | 0.45999596 | 0.646203586 | CSNK2A1 | protein_coding | 41.23 | -0.036336081 | 0.036336081 | ENSG00000101266 | n.s. |
| ENSG00000143549 | 3267.271138 | 0.198460663 | 0.2759007 | 0.719319 | 0.47194423 | 0.656094752 | TPM3 | protein_coding | 45.14 | 0.036324937 | 0.036324937 | ENSG00000143549 | n.s. |
| ENSG00000110237 | 433.4142954 | -0.183614927 | 0.2415485 | -0.760158 | 0.44716042 | 0.634316304 | ARHGEF17 | protein_coding | 55.27 | -0.036299592 | 0.036299592 | ENSG00000110237 | n.s. |
| ENSG00000138386 | 1406.827349 | -0.19884511 | 0.2773501 | -0.716946 | 0.473407241 | 0.65702243 | NAB1 | protein_coding | 37.7 | -0.036273286 | 0.036273286 | ENSG00000138386 | n.s. |
| ENSG00000071553 | 500.0601093 | 0.203044252 | 0.2879524 | 0.705131 | 0.48072848 | 0.662953906 | ATP6AP1 | protein_coding | 56.78 | 0.036246783 | 0.036246783 | ENSG00000071553 | n.s. |
| ENSG00000135452 | 152.0396223 | -0.193882399 | 0.2656826 | -0.729752 | 0.465541889 | 0.650450548 | TSPAN31 | protein_coding | 51.32 | -0.036214463 | 0.036214463 | ENSG00000135452 | n.s. |
| ENSG00000115540 | 1165.281378 | -0.207088833 | 0.2983952 | -0.694009 | 0.487676814 | 0.668694533 | MOB4 | protein_coding | 40.5 | -0.036193377 | 0.036193377 | ENSG00000115540 | n.s. |
| ENSG00000105483 | 86.17191812 | -0.221979805 | 0.3372211 | -0.658262 | 0.510369811 | 0.687123662 | CARD8 | protein_coding | 44.39 | -0.03617496 | 0.03617496 | ENSG00000105483 | n.s. |
| ENSG00000155368 | 2360.501422 | 0.208087194 | 0.3018242 | 0.689432 | 0.49055155 | 0.670128457 | DBI | protein_coding | 50.37 | 0.036174281 | 0.036174281 | ENSG00000155368 | n.s. |
| ENSG00000272993 | 61.21792888 | 0.190858225 | 0.2585846 | 0.738088 | 0.460460953 | 0.646624934 | | lincRNA | 33.98 | 0.036138538 | 0.036138538 | ENSG00000272993 | n.s. |
| ENSG00000005194 | 1059.700545 | -0.213985197 | 0.3177265 | -0.673489 | 0.500636421 | 0.677945841 | CIAPIN1 | protein_coding | 44.41 | -0.036121771 | 0.036121771 | ENSG00000005194 | n.s. |
| ENSG00000136738 | 1040.823094 | 0.188415297 | 0.253031 | 0.744633 | 0.456493388 | 0.643140373 | STAM | protein_coding | 37.21 | 0.036118125 | 0.036118125 | ENSG00000136738 | n.s. |
| ENSG00000146830 | 944.4528118 | 0.183006797 | 0.2413018 | 0.758415 | 0.448202824 | 0.635212533 | GIGYF1 | protein_coding | 62.01 | 0.036067152 | 0.036067152 | ENSG00000146830 | n.s. |
| ENSG00000182165 | 71.57770162 | 0.205653794 | 0.2955766 | 0.695772 | 0.486571903 | 0.667968902 | TP53TG1 | lincRNA | 38.94 | 0.036039543 | 0.036039543 | ENSG00000182165 | n.s. |
| ENSG00000168806 | 1437.271317 | -0.159971222 | 0.191495 | -0.835381 | 0.403503313 | 0.595568498 | LCMT2 | protein_coding | 46.08 | -0.036004448 | 0.036004448 | ENSG00000168806 | n.s. |
| ENSG00000134532 | 58.42903379 | -0.198672784 | 0.278791 | -0.712623 | 0.47607924 | 0.659219418 | SOX5 | protein_coding | 34.5 | -0.035953815 | 0.035953815 | ENSG00000134532 | n.s. |
| ENSG00000276404 | 13.81486928 | 0.242310621 | 0.3950612 | 0.61335 | 0.539645203 | 0.710628976 | MIR6835 | miRNA | 46.88 | 0.035948498 | 0.035948498 | ENSG00000276404 | n.s. |
| ENSG00000183207 | 1225.80528 | 0.2088105 | 0.3053584 | 0.683821 | 0.494088137 | 0.672841832 | RUVBL2 | protein_coding | 53.06 | 0.035933576 | 0.035933576 | ENSG00000183207 | n.s. |
| ENSG00000243279 | 57.04458729 | -0.197423931 | 0.275928 | -0.715491 | 0.474305935 | 0.657679667 | PRAF2 | protein_coding | 60.94 | -0.03592831 | 0.03592831 | ENSG00000243279 | n.s. |
| ENSG00000144021 | 1733.771322 | -0.198635721 | 0.278899 | -0.712214 | 0.47633236 | 0.659377281 | CIAO1 | protein_coding | 49.71 | -0.035926452 | 0.035926452 | ENSG00000144021 | n.s. |
| ENSG00000283103 | 194.9381928 | -0.206610492 | 0.2994846 | -0.689887 | 0.49026534 | 0.670128457 | | antisense | 48.25 | -0.035917569 | 0.035917569 | ENSG00000283103 | n.s. |
| ENSG00000182195 | 64.00323554 | 0.197373474 | 0.2760027 | 0.715114 | 0.474538499 | 0.657871106 | LDOC1 | protein_coding | 57.35 | 0.03589418 | 0.03589418 | ENSG00000182195 | n.s. |

| | | | | | | | | | | | | | |
|---|---|---|---|---|---|---|---|---|---|---|---|---|---|
| ENSG00000169193 | 147.2697416 | 0.199901079 | 0.2823552 | 0.707977 | 0.478959335 | 0.66139348 | CCDC126 | protein_coding | 41.98 | 0.035890258 | 0.035890258 | ENSG00000169193 | n.s. |
| ENSG00000141429 | 1730.32453 | -0.199859243 | 0.2822433 | -0.70811 | 0.478877066 | 0.66139348 | GALNT1 | protein_coding | 38.5 | -0.035882747 | 0.035882747 | ENSG00000141429 | n.s. |
| ENSG00000127445 | 1073.196786 | 0.196435998 | 0.2737381 | 0.717606 | 0.473000506 | 0.656850793 | PIN1 | protein_coding | 56.31 | 0.035856105 | 0.035856105 | ENSG00000127445 | n.s. |
| ENSG00000205423 | 336.8298127 | 0.180569681 | 0.236611 | 0.76315 | 0.445373961 | 0.633071582 | CNEP1R1 | protein_coding | 40.53 | 0.035851601 | 0.035851601 | ENSG00000205423 | n.s. |
| ENSG00000130640 | 229.0906151 | 0.195667514 | 0.2721583 | 0.718947 | 0.47217334 | 0.656094752 | TUBGCP2 | protein_coding | 55.72 | 0.035813698 | 0.035813698 | ENSG00000130640 | n.s. |
| ENSG00000140829 | 777.043355 | 0.17776117 | 0.2301596 | 0.772339 | 0.439913922 | 0.628872587 | DHX38 | protein_coding | 51.51 | 0.035807737 | 0.035807737 | ENSG00000140829 | n.s. |
| ENSG00000038002 | 135.346336 | -0.196521338 | 0.2746479 | -0.715539 | 0.474275901 | 0.657679667 | AGA | protein_coding | 39.89 | -0.035764051 | 0.035764051 | ENSG00000038002 | n.s. |
| ENSG00000232499 | 66.15524105 | -0.200823699 | 0.285328 | -0.703834 | 0.481535855 | 0.663618499 | | processed_pseudogene | 44.31 | -0.035762989 | 0.035762989 | ENSG00000232499 | n.s. |
| ENSG00000129562 | 829.5616508 | 0.220837887 | 0.3371437 | 0.655026 | 0.512450979 | 0.688892052 | DAD1 | protein_coding | 41.95 | 0.035742353 | 0.035742353 | ENSG00000129562 | n.s. |
| ENSG00000213366 | 168.4660642 | 0.19537589 | 0.2719149 | 0.718518 | 0.472437701 | 0.656301346 | GSTM2 | protein_coding | 46.7 | 0.035733607 | 0.035733607 | ENSG00000213366 | n.s. |
| ENSG00000184470 | 98.95612125 | -0.207059893 | 0.3024029 | -0.684715 | 0.493523591 | 0.672461266 | TXNRD2 | protein_coding | 54.14 | -0.035683196 | 0.035683196 | ENSG00000184470 | n.s. |
| ENSG00000185842 | 695.846753 | 0.198735859 | 0.2806977 | 0.708007 | 0.478941134 | 0.66139348 | DNAH14 | protein_coding | 37.4 | 0.035681054 | 0.035681054 | ENSG00000185842 | n.s. |
| ENSG00000167766 | 713.2714103 | -0.196032629 | 0.2739069 | -0.715691 | 0.474182324 | 0.657665315 | ZNF83 | protein_coding | 44.66 | -0.03567697 | 0.03567697 | ENSG00000167766 | n.s. |
| ENSG00000130340 | 1087.042799 | -0.18304607 | 0.2433793 | -0.752102 | 0.451989733 | 0.638669167 | SNX9 | protein_coding | 42.45 | -0.035643472 | 0.035643472 | ENSG00000130340 | n.s. |
| ENSG00000188811 | 643.4617236 | -0.179964478 | 0.2363427 | -0.761456 | 0.446384976 | 0.633811136 | NHLRC3 | protein_coding | 37.72 | -0.035640189 | 0.035640189 | ENSG00000188811 | n.s. |
| ENSG00000002549 | 995.4812082 | 0.192197638 | 0.2650584 | 0.725114 | 0.468381837 | 0.652519543 | LAP3 | protein_coding | 44.17 | 0.035634687 | 0.035634687 | ENSG00000002549 | n.s. |
| ENSG00000230185 | 68.36252954 | 0.184059091 | 0.2458317 | 0.74872 | 0.454026012 | 0.640401597 | C9orf147 | antisense | 40.99 | 0.035624194 | 0.035624194 | ENSG00000230185 | n.s. |
| ENSG00000146576 | 283.4788022 | -0.182282101 | 0.2418467 | -0.753709 | 0.451023757 | 0.637717453 | C7orf26 | protein_coding | 53.11 | -0.035612763 | 0.035612763 | ENSG00000146576 | n.s. |
| ENSG00000255076 | 13.05824502 | -0.242679005 | 0.3995045 | -0.60745 | 0.543552331 | 0.713277955 | | sense_intronic | 40.37 | -0.035611008 | 0.035611008 | ENSG00000255076 | n.s. |
| ENSG00000124257 | 55.5830296 | 0.192094085 | 0.2651345 | 0.724516 | 0.468749199 | 0.65290067 | NEURL2 | protein_coding | 56.82 | 0.035566774 | 0.035566774 | ENSG00000124257 | n.s. |
| ENSG00000050165 | 130.9959057 | 0.196374883 | 0.2756361 | 0.712443 | 0.476190774 | 0.659308288 | DKK3 | protein_coding | 49.18 | 0.035526468 | 0.035526468 | ENSG00000050165 | n.s. |
| ENSG00000265123 | 9.885165928 | 0.244583557 | 0.4057479 | 0.602797 | 0.546643813 | 0.71578311 | RN7SL200P | misc_RNA | 54.17 | 0.03551807 | 0.03551807 | ENSG00000265123 | n.s. |
| ENSG00000104435 | 1758.762651 | 0.205013626 | 0.2981446 | 0.687632 | 0.491684848 | 0.671084026 | STMN2 | protein_coding | 38.21 | 0.035513096 | 0.035513096 | ENSG00000104435 | n.s. |
| ENSG00000175826 | 263.6551186 | -0.185839577 | 0.2503741 | -0.742247 | 0.457937406 | 0.644286834 | CTDNEP1 | protein_coding | 52.53 | -0.03548063 | 0.03548063 | ENSG00000175826 | n.s. |
| ENSG00000272106 | 62.73998323 | 0.187646433 | 0.2545561 | 0.737152 | 0.461030185 | 0.647029045 | | antisense | 57.36 | 0.035479479 | 0.035479479 | ENSG00000272106 | n.s. |
| ENSG00000101146 | 1432.370683 | 0.190555973 | 0.2621227 | 0.726972 | 0.467242885 | 0.651845961 | RAE1 | protein_coding | 44.61 | 0.035415784 | 0.035415784 | ENSG00000101146 | n.s. |
| ENSG00000125812 | 979.9549091 | -0.188655976 | 0.2576618 | -0.732184 | 0.464056047 | 0.649396281 | GZF1 | protein_coding | 44.84 | -0.035371147 | 0.035371147 | ENSG00000125812 | n.s. |
| ENSG00000177595 | 37.17561142 | 0.197124831 | 0.2786463 | 0.707437 | 0.479294695 | 0.661631969 | PIDD1 | protein_coding | 62.59 | 0.035360946 | 0.035360946 | ENSG00000177595 | n.s. |
| ENSG00000106610 | 350.5965634 | -0.202142472 | 0.291166 | -0.694252 | 0.487524352 | 0.668694533 | STAG3L4 | transcribed_unprocessed_pseud | 44.75 | -0.03532889 | 0.03532889 | ENSG00000106610 | n.s. |
| ENSG00000134508 | 45.88289639 | 0.198010865 | 0.2813368 | 0.703821 | 0.481544042 | 0.663618499 | CABLES1 | protein_coding | 48.32 | 0.035262075 | 0.035262075 | ENSG00000134508 | n.s. |
| ENSG00000119812 | 800.3223416 | -0.188925753 | 0.2591001 | -0.729161 | 0.465903075 | 0.650694415 | FAM98A | protein_coding | 37.55 | -0.035257876 | 0.035257876 | ENSG00000119812 | n.s. |
| ENSG00000149179 | 46.36201906 | -0.205284697 | 0.3006158 | -0.682881 | 0.494682329 | 0.673390328 | C11orf49 | protein_coding | 41.37 | -0.035254183 | 0.035254183 | ENSG00000149179 | n.s. |

| | | | | | | | | | | | | | |
|---|---|---|---|---|---|---|---|---|---|---|---|---|---|
| ENSG00000148943 | 1921.691392 | 0.189593235 | 0.2607938 | 0.726985 | 0.467235065 | 0.651845961 | LIN7C | protein_coding | 37.11 | 0.035236855 | 0.035236855 | ENSG00000148943 | n.s. |
| ENSG00000252637 | 10.25425598 | 0.24892341 | 0.4216784 | 0.590316 | 0.554978991 | 0.722237406 | RNA5SP268 | rRNA | 41.13 | 0.035177862 | 0.035177862 | ENSG00000252637 | n.s. |
| ENSG00000271830 | 15.46813544 | 0.236249349 | 0.3839591 | 0.615298 | 0.538357751 | 0.709739131 | | lincRNA | 31.92 | 0.035177823 | 0.035177823 | ENSG00000271830 | n.s. |
| ENSG00000100068 | 337.4975139 | -0.174505216 | 0.2263391 | -0.77099 | 0.440712823 | 0.629009177 | LRP5L | protein_coding | 54.66 | -0.035135407 | 0.035135407 | ENSG00000100068 | n.s. |
| ENSG00000128739 | 1137.715881 | 0.201367469 | 0.2908273 | 0.692395 | 0.48868915 | 0.669199599 | SNRPN | protein_coding | 41.92 | 0.035127413 | 0.035127413 | ENSG00000128739 | n.s. |
| ENSG00000115520 | 262.6178365 | 0.203085795 | 0.2957998 | 0.686565 | 0.492356939 | 0.671635475 | COQ10B | protein_coding | 41.73 | 0.035106705 | 0.035106705 | ENSG00000115520 | n.s. |
| ENSG00000251247 | 30.67421966 | 0.206570911 | 0.3053249 | 0.676561 | 0.498684538 | 0.676559451 | ZNF345 | protein_coding | 39.29 | 0.035053851 | 0.035053851 | ENSG00000251247 | n.s. |
| ENSG00000143153 | 4523.213658 | 0.188597624 | 0.2595897 | 0.726522 | 0.467518841 | 0.651988717 | ATP1B1 | protein_coding | 43.48 | 0.035033879 | 0.035033879 | ENSG00000143153 | n.s. |
| ENSG00000258056 | 154.7042387 | 0.177768786 | 0.2347578 | 0.757243 | 0.448904165 | 0.635741733 | | antisense | 56.95 | 0.034970547 | 0.034970547 | ENSG00000258056 | n.s. |
| ENSG00000100941 | 8023.895882 | -0.177389252 | 0.2340114 | -0.758037 | 0.448428874 | 0.635403543 | PNN | protein_coding | 40.65 | -0.034936878 | 0.034936878 | ENSG00000100941 | n.s. |
| ENSG00000062194 | 2635.258775 | -0.185799284 | 0.2536361 | -0.732543 | 0.463837388 | 0.649176514 | GPBP1 | protein_coding | 40.02 | -0.034862858 | 0.034862858 | ENSG00000062194 | n.s. |
| ENSG00000105251 | 825.9080236 | -0.187734154 | 0.2588662 | -0.725217 | 0.468318903 | 0.652497156 | SHD | protein_coding | 54.76 | -0.034809925 | 0.034809925 | ENSG00000105251 | n.s. |
| ENSG00000124535 | 1040.948977 | 0.185904369 | 0.2544552 | 0.730598 | 0.465024901 | 0.64992388 | WRNIP1 | protein_coding | 44.69 | 0.03478968 | 0.03478968 | ENSG00000124535 | n.s. |
| ENSG00000233264 | 283.2767247 | -0.179715753 | 0.2401839 | -0.748242 | 0.454314002 | 0.640548238 | | processed_pseudogene | 38.76 | -0.034765681 | 0.034765681 | ENSG00000233264 | n.s. |
| ENSG00000275832 | 277.0977203 | 0.179344075 | 0.2393027 | 0.749445 | 0.45358931 | 0.640109869 | ARHGAP23 | protein_coding | 55.44 | 0.034747103 | 0.034747103 | ENSG00000275832 | n.s. |
| ENSG00000200840 | 9.485041905 | -0.375964406 | 0.883689 | -0.425449 | 0.670509553 | 0.808314048 | RNU6-82P | snRNA | 39.25 | -0.034746583 | 0.034746583 | ENSG00000200840 | n.s. |
| ENSG00000064115 | 829.1765903 | -0.191266369 | 0.2677074 | -0.714461 | 0.474942428 | 0.658207114 | TM7SF3 | protein_coding | 40.51 | -0.034741131 | 0.034741131 | ENSG00000064115 | n.s. |
| ENSG00000153914 | 7094.169626 | 0.185306727 | 0.2531815 | 0.731912 | 0.464222011 | 0.649518948 | SREK1 | protein_coding | 35.5 | 0.034727995 | 0.034727995 | ENSG00000153914 | n.s. |
| ENSG00000100216 | 1045.077669 | 0.195569523 | 0.2790781 | 0.70077 | 0.483446686 | 0.665055648 | TOMM22 | protein_coding | 48.7 | 0.034643579 | 0.034643579 | ENSG00000100216 | n.s. |
| ENSG00000105325 | 300.9243983 | -0.189153858 | 0.2630159 | -0.719173 | 0.472034464 | 0.656094752 | FZR1 | protein_coding | 60.71 | -0.034621481 | 0.034621481 | ENSG00000105325 | n.s. |
| ENSG00000005189 | 89.24577812 | 0.180798933 | 0.2431048 | 0.743708 | 0.457053346 | 0.643563283 | REXO5 | protein_coding | 39.89 | 0.034606496 | 0.034606496 | ENSG00000005189 | n.s. |
| ENSG00000232119 | 656.9680178 | 0.198944071 | 0.2884278 | 0.689753 | 0.49034932 | 0.670128457 | MCTS1 | protein_coding | 39.24 | 0.034584823 | 0.034584823 | ENSG00000232119 | n.s. |
| ENSG00000177697 | 495.6484334 | -0.195907038 | 0.2801888 | -0.699197 | 0.484429226 | 0.666012453 | CD151 | protein_coding | 63.43 | -0.03458105 | 0.03458105 | ENSG00000177697 | n.s. |
| ENSG00000112984 | 2494.512722 | 0.178646378 | 0.2385896 | 0.74876 | 0.454001839 | 0.640401597 | KIF20A | protein_coding | 45.27 | 0.034576577 | 0.034576577 | ENSG00000112984 | n.s. |
| ENSG00000104907 | 1203.69934 | 0.189892597 | 0.265278 | 0.715825 | 0.474099439 | 0.657665315 | TRMT1 | protein_coding | 53.66 | 0.034559515 | 0.034559515 | ENSG00000104907 | n.s. |
| ENSG00000183654 | 243.2445605 | -0.184419565 | 0.2522935 | -0.730972 | 0.464796066 | 0.64992388 | 11-Mar | protein_coding | 39.41 | -0.034511817 | 0.034511817 | ENSG00000183654 | n.s. |
| ENSG00000135047 | 586.5587045 | 0.186168653 | 0.2570883 | 0.724143 | 0.468978068 | 0.653154111 | CTSL | protein_coding | 46.28 | 0.034438285 | 0.034438285 | ENSG00000135047 | n.s. |
| ENSG00000122140 | 573.110042 | 0.193328777 | 0.2748185 | 0.703478 | 0.481757853 | 0.663712596 | MRPS2 | protein_coding | 63.07 | 0.034416377 | 0.034416377 | ENSG00000122140 | n.s. |
| ENSG00000130311 | 396.7129508 | 0.191655518 | 0.2707502 | 0.707868 | 0.479026923 | 0.66139348 | DDA1 | protein_coding | 56.15 | 0.034409849 | 0.034409849 | ENSG00000130311 | n.s. |
| ENSG00000140937 | 44.75492267 | 0.197034505 | 0.2841852 | 0.693331 | 0.488101715 | 0.669050361 | CDH11 | protein_coding | 40.38 | 0.034390637 | 0.034390637 | ENSG00000140937 | n.s. |
| ENSG00000147162 | 4509.990366 | 0.182578116 | 0.2484634 | 0.734829 | 0.462443638 | 0.648137988 | OGT | protein_coding | 40.6 | 0.034385397 | 0.034385397 | ENSG00000147162 | n.s. |
| ENSG00000256061 | 73.36745752 | -0.200193279 | 0.2931916 | -0.682807 | 0.494728842 | 0.673390328 | DNAAF4 | protein_coding | 42.67 | -0.034379818 | 0.034379818 | ENSG00000256061 | n.s. |

| | | | | | | | | | | | | | |
|---|---|---|---|---|---|---|---|---|---|---|---|---|---|
| ENSG00000131508 | 1278.183066 | -0.196950253 | 0.2843616 | -0.692605 | 0.488557565 | 0.669177272 | UBE2D2 | protein_coding | 44.35 | -0.034359708 | 0.034359708 | ENSG00000131508 | n.s. |
| ENSG00000197217 | 1343.375689 | -0.18774953 | 0.2613075 | -0.7185 | 0.472448846 | 0.656301346 | ENTPD4 | protein_coding | 44.43 | -0.03433877 | 0.03433877 | ENSG00000197217 | n.s. |
| ENSG00000170296 | 1742.748664 | 0.19243546 | 0.2735867 | 0.70338 | 0.481818841 | 0.663712596 | GABARAP | protein_coding | 50.38 | 0.034257349 | 0.034257349 | ENSG00000170296 | n.s. |
| ENSG00000183665 | 455.6632261 | 0.173140604 | 0.2275825 | 0.760782 | 0.44678748 | 0.634100082 | TRMT12 | protein_coding | 43.63 | 0.034254516 | 0.034254516 | ENSG00000183665 | n.s. |
| ENSG00000221838 | 44.64137077 | 0.199822975 | 0.2934029 | 0.681053 | 0.495837879 | 0.673977427 | AP4M1 | protein_coding | 56.13 | 0.034240596 | 0.034240596 | ENSG00000221838 | n.s. |
| ENSG00000172893 | 314.2125794 | -0.183416788 | 0.2515451 | -0.729161 | 0.46590337 | 0.650694415 | DHCR7 | protein_coding | 51.71 | -0.034229777 | 0.034229777 | ENSG00000172893 | n.s. |
| ENSG00000103326 | 651.4783608 | 0.17509419 | 0.2323583 | 0.753553 | 0.451117953 | 0.637785871 | CAPN15 | protein_coding | 65.01 | 0.034200291 | 0.034200291 | ENSG00000103326 | n.s. |
| ENSG00000166295 | 929.9347747 | -0.201878601 | 0.2988944 | -0.675418 | 0.499410313 | 0.677114596 | ANAPC16 | protein_coding | 43.46 | -0.034185683 | 0.034185683 | ENSG00000166295 | n.s. |
| ENSG00000131462 | 984.7017295 | -0.178988088 | 0.2411908 | -0.742102 | 0.458025799 | 0.644340889 | TUBG1 | protein_coding | 53.1 | -0.034166017 | 0.034166017 | ENSG00000131462 | n.s. |
| ENSG00000239553 | 19.96760194 | 0.219329002 | 0.3438282 | 0.637903 | 0.523536936 | 0.698601184 | RN7SL12P | misc_RNA | 54.14 | 0.034165028 | 0.034165028 | ENSG00000239553 | n.s. |
| ENSG00000154639 | 903.1272177 | -0.191737844 | 0.2729118 | -0.702563 | 0.482327889 | 0.664041452 | CXADR | protein_coding | 40.2 | -0.034091911 | 0.034091911 | ENSG00000154639 | n.s. |
| ENSG00000132963 | 1694.301782 | 0.20164751 | 0.2992051 | 0.673944 | 0.500346871 | 0.677749957 | POMP | protein_coding | 39.25 | 0.034064415 | 0.034064415 | ENSG00000132963 | n.s. |
| ENSG00000229589 | 89.21192866 | 0.178195612 | 0.2402747 | 0.741633 | 0.458309749 | 0.644615278 | ACVR2B-AS1 | antisense | 56.48 | 0.033981797 | 0.033981797 | ENSG00000229589 | n.s. |
| ENSG00000172380 | 992.4229343 | -0.194497741 | 0.2803631 | -0.693735 | 0.487848326 | 0.668863835 | GNG12 | protein_coding | 41.55 | -0.03397142 | 0.03397142 | ENSG00000172380 | n.s. |
| ENSG00000198176 | 1746.025745 | -0.190547211 | 0.2709218 | -0.703329 | 0.481850673 | 0.663712596 | TFDP1 | protein_coding | 53.33 | -0.033921203 | 0.033921203 | ENSG00000198176 | n.s. |
| ENSG00000145721 | 343.0742931 | 0.190925292 | 0.2720063 | 0.701915 | 0.482732198 | 0.664466672 | LIX1 | protein_coding | 39.28 | 0.033894356 | 0.033894356 | ENSG00000145721 | n.s. |
| ENSG00000180340 | 218.8001285 | 0.171185583 | 0.2250994 | 0.760489 | 0.446962451 | 0.634100082 | FZD2 | protein_coding | 64.91 | 0.033867731 | 0.033867731 | ENSG00000180340 | n.s. |
| ENSG00000132912 | 2703.631194 | 0.185631534 | 0.2588773 | 0.717064 | 0.47333475 | 0.65702243 | DCTN4 | protein_coding | 38.73 | 0.033862868 | 0.033862868 | ENSG00000132912 | n.s. |
| ENSG00000173726 | 11235.51927 | 0.19082978 | 0.272141 | 0.701216 | 0.483167943 | 0.66486927 | TOMM20 | protein_coding | 41.67 | 0.0338272 | 0.0338272 | ENSG00000173726 | n.s. |
| ENSG00000215039 | 620.784231 | 0.189182292 | 0.2680277 | 0.705831 | 0.480293142 | 0.662550423 | CD27-AS1 | processed_transcript | 49.5 | 0.033822211 | 0.033822211 | ENSG00000215039 | n.s. |
| ENSG00000162104 | 563.8785515 | -0.173836526 | 0.2313894 | -0.751272 | 0.452488683 | 0.639010034 | ADCY9 | protein_coding | 47.22 | -0.03380987 | 0.03380987 | ENSG00000162104 | n.s. |
| ENSG00000108179 | 1891.767287 | 0.173470106 | 0.2310947 | 0.750645 | 0.452866285 | 0.639413577 | PPIF | protein_coding | 55.59 | 0.033691042 | 0.033691042 | ENSG00000108179 | n.s. |
| ENSG00000130559 | 2646.827338 | -0.178554464 | 0.2428202 | -0.735336 | 0.462134725 | 0.648077601 | CAMSAP1 | protein_coding | 46.91 | -0.033634838 | 0.033634838 | ENSG00000130559 | n.s. |
| ENSG00000127955 | 807.5481052 | 0.18639737 | 0.2622386 | 0.710793 | 0.477212446 | 0.66013203 | GNAI1 | protein_coding | 36.74 | 0.033620344 | 0.033620344 | ENSG00000127955 | n.s. |
| ENSG00000228873 | 16.09412136 | 0.217702856 | 0.3435346 | 0.633715 | 0.526267133 | 0.700833792 | | lincRNA | 61.16 | 0.033610048 | 0.033610048 | ENSG00000228873 | n.s. |
| ENSG00000201228 | 11.98294877 | -0.28849829 | 0.5625594 | -0.512832 | 0.608069079 | 0.764782817 | | misc_RNA | 44.12 | -0.033599053 | 0.033599053 | ENSG00000201228 | n.s. |
| ENSG00000113971 | 122.3540475 | -0.186495863 | 0.2629129 | -0.709345 | 0.478110561 | 0.66071795 | NPHP3 | protein_coding | 36.77 | -0.033566252 | 0.033566252 | ENSG00000113971 | n.s. |
| ENSG00000249915 | 1153.661537 | 0.200073585 | 0.2983703 | 0.670555 | 0.502504248 | 0.679588795 | PDCD6 | protein_coding | 50.84 | 0.033563102 | 0.033563102 | ENSG00000249915 | n.s. |
| ENSG00000207344 | 6.558923092 | -0.268766483 | 0.4978417 | -0.539863 | 0.589291294 | 0.750109955 | SNORA22C | snoRNA | 42.54 | -0.033562234 | 0.033562234 | ENSG00000207344 | n.s. |
| ENSG00000178028 | 212.967342 | 0.174662028 | 0.234248 | 0.745629 | 0.455891799 | 0.642512554 | DMAP1 | protein_coding | 52.33 | 0.033555786 | 0.033555786 | ENSG00000178028 | n.s. |
| ENSG00000100883 | 867.9465837 | -0.192650669 | 0.278689 | -0.691275 | 0.489392946 | 0.669697666 | SRP54 | protein_coding | 39.06 | -0.033544569 | 0.033544569 | ENSG00000100883 | n.s. |
| ENSG00000167986 | 5334.718771 | 0.189702512 | 0.2710153 | 0.69997 | 0.48394615 | 0.665545461 | DDB1 | protein_coding | 47.67 | 0.03354363 | 0.03354363 | ENSG00000167986 | n.s. |

| | | | | | | | | | | | | | |
|---|---|---|---|---|---|---|---|---|---|---|---|---|---|
| ENSG00000108176 | 594.0448675 | 0.205544404 | 0.3118203 | 0.659176 | 0.509782821 | 0.686799793 | DNAJC12 | protein_coding | 39.63 | 0.033538648 | 0.033538648 | ENSG00000108176 | n.s. |
| ENSG00000100028 | 1927.175408 | 0.188919499 | 0.2692185 | 0.701733 | 0.48284576 | 0.664497525 | SNRPD3 | protein_coding | 50.71 | 0.033534464 | 0.033534464 | ENSG00000100028 | n.s. |
| ENSG00000152642 | 513.1672232 | 0.193363559 | 0.2810038 | 0.688117 | 0.49137898 | 0.670798073 | GPD1L | protein_coding | 44.49 | 0.033530826 | 0.033530826 | ENSG00000152642 | n.s. |
| ENSG00000122218 | 3704.998978 | -0.184546214 | 0.2584646 | -0.71401 | 0.475221214 | 0.658233153 | COPA | protein_coding | 40.67 | -0.033517329 | 0.033517329 | ENSG00000122218 | n.s. |
| ENSG00000212158 | 4.105864092 | 0.376093856 | 0.9132688 | 0.411811 | 0.680478197 | 0.814642987 | SNORD66 | snoRNA | 51.32 | 0.033484643 | 0.033484643 | ENSG00000212158 | n.s. |
| ENSG00000020426 | 686.4400248 | -0.193754477 | 0.2826663 | -0.685453 | 0.493058099 | 0.671937241 | MNAT1 | protein_coding | 36.62 | -0.033455835 | 0.033455835 | ENSG00000020426 | n.s. |
| ENSG00000104388 | 992.5475719 | -0.193567726 | 0.2823626 | -0.685529 | 0.493010212 | 0.671937241 | RAB2A | protein_coding | 39.18 | -0.033423589 | 0.033423589 | ENSG00000104388 | n.s. |
| ENSG00000270175 | 82.86962576 | -0.176765463 | 0.2397436 | -0.737311 | 0.460933545 | 0.647029045 | | antisense | 53.16 | -0.033422146 | 0.033422146 | ENSG00000270175 | n.s. |
| ENSG00000168556 | 140.1243592 | 0.194774903 | 0.2860826 | 0.680835 | 0.495976194 | 0.67399356 | ING2 | protein_coding | 46.91 | 0.03337356 | 0.03337356 | ENSG00000168556 | n.s. |
| ENSG00000103599 | 35.44561754 | -0.201934165 | 0.3041574 | -0.663913 | 0.506745691 | 0.683598513 | IQCH | protein_coding | 39.22 | -0.0333593 | 0.0333593 | ENSG00000103599 | n.s. |
| ENSG00000137486 | 201.9538814 | -0.187469781 | 0.266799 | -0.702663 | 0.482265762 | 0.664021578 | ARRB1 | protein_coding | 51.96 | -0.033335465 | 0.033335465 | ENSG00000137486 | n.s. |
| ENSG00000025800 | 661.2613621 | -0.175257622 | 0.2367505 | -0.740263 | 0.459140489 | 0.645522874 | KPNA6 | protein_coding | 44.21 | -0.033314435 | 0.033314435 | ENSG00000025800 | n.s. |
| ENSG00000187475 | 46.63926027 | 0.192914723 | 0.2814037 | 0.685544 | 0.493000465 | 0.671937241 | HIST1H1T | protein_coding | 48.33 | 0.033310834 | 0.033310834 | ENSG00000187475 | n.s. |
| ENSG00000111666 | 956.0766211 | -0.190848928 | 0.2753856 | -0.693024 | 0.488294206 | 0.669121135 | CHPT1 | protein_coding | 38.42 | -0.033302232 | 0.033302232 | ENSG00000111666 | n.s. |
| ENSG00000101745 | 331.69844 | -0.18385278 | 0.2582775 | -0.711842 | 0.476562618 | 0.659560768 | ANKRD12 | protein_coding | 37.33 | -0.033230505 | 0.033230505 | ENSG00000101745 | n.s. |
| ENSG00000081320 | 662.9686753 | 0.176119447 | 0.2392525 | 0.736124 | 0.46165536 | 0.647619907 | STK17B | protein_coding | 37.67 | 0.033230184 | 0.033230184 | ENSG00000081320 | n.s. |
| ENSG00000127948 | 283.1877776 | -0.195696466 | 0.2892378 | -0.676594 | 0.498663756 | 0.676559451 | POR | protein_coding | 50.98 | -0.033208523 | 0.033208523 | ENSG00000127948 | n.s. |
| ENSG00000167110 | 166.2414015 | -0.197507007 | 0.294152 | -0.671445 | 0.501936904 | 0.679019385 | GOLGA2 | protein_coding | 52.34 | -0.033204449 | 0.033204449 | ENSG00000167110 | n.s. |
| ENSG00000156639 | 130.8907377 | -0.182364149 | 0.2546095 | -0.71625 | 0.473836723 | 0.657552944 | ZFAND3 | protein_coding | 39.23 | -0.033202908 | 0.033202908 | ENSG00000156639 | n.s. |
| ENSG00000231312 | 59.76126749 | -0.206805764 | 0.317266 | -0.651837 | 0.514506206 | 0.690987741 | | antisense | 41.25 | -0.033198459 | 0.033198459 | ENSG00000231312 | n.s. |
| ENSG00000100083 | 139.0414304 | -0.181834792 | 0.2536099 | -0.716986 | 0.473382607 | 0.65702243 | GGA1 | protein_coding | 56.62 | -0.033170267 | 0.033170267 | ENSG00000100083 | n.s. |
| ENSG00000068001 | 829.9463514 | 0.183748955 | 0.2584656 | 0.710922 | 0.47713247 | 0.66013203 | HYAL2 | protein_coding | 61.05 | 0.033142651 | 0.033142651 | ENSG00000068001 | n.s. |
| ENSG00000145781 | 455.0636389 | 0.189362404 | 0.272413 | 0.69513 | 0.48697393 | 0.668389001 | COMMD10 | protein_coding | 37.46 | 0.033132874 | 0.033132874 | ENSG00000145781 | n.s. |
| ENSG00000132383 | 1041.044328 | -0.190459072 | 0.2758622 | -0.690414 | 0.489934031 | 0.670008007 | RPA1 | protein_coding | 44.49 | -0.033124643 | 0.033124643 | ENSG00000132383 | n.s. |
| ENSG00000249669 | 531.0197904 | 0.19125164 | 0.2782219 | 0.687407 | 0.49182637 | 0.6711456 | CARMN | lincRNA | 51.97 | 0.033121581 | 0.033121581 | ENSG00000249669 | n.s. |
| ENSG00000158373 | 1380.96566 | 0.165323884 | 0.2152572 | 0.76803 | 0.442469574 | 0.630525215 | HIST1H2BD | protein_coding | 42.24 | 0.033113967 | 0.033113967 | ENSG00000158373 | n.s. |
| ENSG00000142632 | 77.54898552 | 0.18205599 | 0.2546405 | 0.714953 | 0.474638129 | 0.657943715 | ARHGEF19 | protein_coding | 57.33 | 0.033099829 | 0.033099829 | ENSG00000142632 | n.s. |
| ENSG00000107937 | 3146.159914 | -0.187935964 | 0.269521 | -0.697296 | 0.485617516 | 0.667119159 | GTPBP4 | protein_coding | 47.14 | -0.033038501 | 0.033038501 | ENSG00000107937 | n.s. |
| ENSG00000210127 | 25.19876369 | 0.209698213 | 0.3264658 | 0.642328 | 0.520660054 | 0.696029758 | MT-TA | Mt_tRNA | 43.48 | 0.033000667 | 0.033000667 | ENSG00000210127 | n.s. |
| ENSG00000133872 | 2311.429393 | 0.193069523 | 0.284127 | 0.679518 | 0.49680942 | 0.67472188 | SARAF | protein_coding | 40.19 | 0.032990795 | 0.032990795 | ENSG00000133872 | n.s. |
| ENSG00000169762 | 237.0905564 | 0.198108862 | 0.2968557 | 0.667357 | 0.504543827 | 0.681552702 | TAPT1 | protein_coding | 38.85 | 0.032985236 | 0.032985236 | ENSG00000169762 | n.s. |
| ENSG00000150593 | 1570.036802 | -0.189072546 | 0.2731215 | -0.692265 | 0.488770771 | 0.669205963 | PDCD4 | protein_coding | 34.69 | -0.032981852 | 0.032981852 | ENSG00000150593 | n.s. |

| | | | | | | | | | | | | | |
|---|---|---|---|---|---|---|---|---|---|---|---|---|---|
| ENSG00000110328 | 151.2042022 | -0.184845999 | 0.2625359 | -0.704079 | 0.48138369 | 0.663594566 | GALNT18 | protein_coding | 46.35 | -0.032920551 | 0.032920551 | ENSG00000110328 | n.s. |
| ENSG00000107959 | 770.0456242 | -0.188323109 | 0.2713252 | -0.694086 | 0.487628102 | 0.668694533 | PITRM1 | protein_coding | 46.17 | -0.032913649 | 0.032913649 | ENSG00000107959 | n.s. |
| ENSG00000146409 | 179.5404547 | -0.194928984 | 0.2894781 | -0.673381 | 0.500705098 | 0.677945841 | SLC18B1 | protein_coding | 39.58 | -0.032904987 | 0.032904987 | ENSG00000146409 | n.s. |
| ENSG00000112658 | 743.3725643 | 0.186785173 | 0.2676489 | 0.697874 | 0.485256131 | 0.66675428 | SRF | protein_coding | 55.7 | 0.032880576 | 0.032880576 | ENSG00000112658 | n.s. |
| ENSG00000271819 | 25.63531494 | 0.213610408 | 0.3379918 | 0.631999 | 0.527387613 | 0.701856025 | RNU6-94P | snRNA | 37.38 | 0.032843021 | 0.032843021 | ENSG00000271819 | n.s. |
| ENSG00000213516 | 2012.800598 | -0.19324769 | 0.2858346 | -0.676082 | 0.498988406 | 0.6766985 | RBMXL1 | protein_coding | 41.18 | -0.032775733 | 0.032775733 | ENSG00000213516 | n.s. |
| ENSG00000140543 | 49.74027434 | -0.191706346 | 0.2821576 | -0.67943 | 0.496865388 | 0.67472188 | DET1 | protein_coding | 43.27 | -0.032757861 | 0.032757861 | ENSG00000140543 | n.s. |
| ENSG00000140044 | 191.3275985 | -0.190069578 | 0.2777078 | -0.684423 | 0.493708185 | 0.672461266 | JDP2 | protein_coding | 50 | -0.032755209 | 0.032755209 | ENSG00000140044 | n.s. |
| ENSG00000143258 | 429.8337394 | -0.168452102 | 0.2243105 | -0.750977 | 0.452666246 | 0.639195958 | USP21 | protein_coding | 53.43 | -0.032741358 | 0.032741358 | ENSG00000143258 | n.s. |
| ENSG00000166275 | 633.6052353 | 0.200964245 | 0.3055099 | 0.657799 | 0.510667058 | 0.687223801 | BORCS7 | protein_coding | 41.69 | 0.032737439 | 0.032737439 | ENSG00000166275 | n.s. |
| ENSG00000153015 | 843.0462899 | -0.1838518 | 0.2613628 | -0.703435 | 0.481784587 | 0.663712596 | CWC27 | protein_coding | 35.58 | -0.032729287 | 0.032729287 | ENSG00000153015 | n.s. |
| ENSG00000180385 | 578.4584395 | -0.173669485 | 0.2365172 | -0.734278 | 0.46277902 | 0.64834698 | EMC3-AS1 | transcribed_unprocessed_pseud | 43.98 | -0.032683296 | 0.032683296 | ENSG00000180385 | n.s. |
| ENSG00000100614 | 2201.613026 | -0.187935474 | 0.2724107 | -0.689897 | 0.490258666 | 0.670128457 | PPM1A | protein_coding | 36.08 | -0.032671067 | 0.032671067 | ENSG00000100614 | n.s. |
| ENSG00000152409 | 1151.788771 | -0.170730841 | 0.2298754 | -0.74271 | 0.45765711 | 0.643988195 | JMY | protein_coding | 40.2 | -0.032630436 | 0.032630436 | ENSG00000152409 | n.s. |
| ENSG00000157020 | 1267.926804 | 0.194645346 | 0.2905491 | 0.669922 | 0.502907319 | 0.679803747 | SEC13 | protein_coding | 46.85 | 0.032625761 | 0.032625761 | ENSG00000157020 | n.s. |
| ENSG00000059691 | 329.5674529 | -0.187517654 | 0.271501 | -0.69067 | 0.489772831 | 0.66991916 | GATB | protein_coding | 43.81 | -0.032623872 | 0.032623872 | ENSG00000059691 | n.s. |
| ENSG00000160193 | 315.8287753 | -0.168564901 | 0.2252336 | -0.7484 | 0.454218757 | 0.64054368 | WDR4 | protein_coding | 53.71 | -0.032609091 | 0.032609091 | ENSG00000160193 | n.s. |
| ENSG00000132676 | 1691.979998 | -0.192202188 | 0.2842369 | -0.676204 | 0.498910983 | 0.6766985 | DAP3 | protein_coding | 44.89 | -0.032598411 | 0.032598411 | ENSG00000132676 | n.s. |
| ENSG00000089248 | 1150.262218 | 0.195735647 | 0.293352 | 0.667238 | 0.504619986 | 0.681556024 | ERP29 | protein_coding | 47.22 | 0.03258968 | 0.03258968 | ENSG00000089248 | n.s. |
| ENSG00000091527 | 3402.132464 | -0.178646847 | 0.2491255 | -0.717096 | 0.473315068 | 0.65702243 | CDV3 | protein_coding | 42.4 | -0.032588723 | 0.032588723 | ENSG00000091527 | n.s. |
| ENSG00000172292 | 2417.13245 | -0.186297303 | 0.2683148 | -0.694324 | 0.487479269 | 0.668694533 | CERS6 | protein_coding | 39.32 | -0.032559595 | 0.032559595 | ENSG00000172292 | n.s. |
| ENSG00000197724 | 634.4079589 | -0.176337674 | 0.243832 | -0.723193 | 0.469561056 | 0.653835242 | PHF2 | protein_coding | 51.51 | -0.032539886 | 0.032539886 | ENSG00000197724 | n.s. |
| ENSG00000111602 | 2225.051492 | -0.186385896 | 0.2688729 | -0.693212 | 0.488176587 | 0.669050361 | TIMELESS | protein_coding | 46.62 | -0.032532016 | 0.032532016 | ENSG00000111602 | n.s. |
| ENSG00000157349 | 293.50247 | -0.184498699 | 0.2643136 | -0.698029 | 0.485158785 | 0.666686317 | DDX19B | protein_coding | 45.56 | -0.032486247 | 0.032486247 | ENSG00000157349 | n.s. |
| ENSG00000126214 | 622.08516 | -0.180421599 | 0.2548696 | -0.707898 | 0.479008769 | 0.66139348 | KLC1 | protein_coding | 49.58 | -0.03239291 | 0.03239291 | ENSG00000126214 | n.s. |
| ENSG00000109819 | 172.6581581 | 0.186327744 | 0.2703829 | 0.689125 | 0.490744309 | 0.670260252 | PPARGC1A | protein_coding | 38.32 | 0.032375663 | 0.032375663 | ENSG00000109819 | n.s. |
| ENSG00000189227 | 655.9798548 | 0.168646475 | 0.2263894 | 0.74494 | 0.456308152 | 0.642969173 | C15orf61 | protein_coding | 39.21 | 0.032348055 | 0.032348055 | ENSG00000189227 | n.s. |
| ENSG00000171097 | 92.99940414 | 0.213399784 | 0.3417009 | 0.624522 | 0.53228473 | 0.705406462 | KYAT1 | protein_coding | 49.05 | 0.032342992 | 0.032342992 | ENSG00000171097 | n.s. |
| ENSG00000087087 | 1826.469484 | 0.168445496 | 0.2260435 | 0.745191 | 0.456156461 | 0.642820486 | SRRT | protein_coding | 53.17 | 0.032326424 | 0.032326424 | ENSG00000087087 | n.s. |
| ENSG00000139438 | 259.2859348 | -0.162781393 | 0.2132384 | -0.763377 | 0.445238346 | 0.633017712 | FAM222A | protein_coding | 52.74 | -0.032325803 | 0.032325803 | ENSG00000139438 | n.s. |
| ENSG00000186019 | 314.3969483 | 0.158323052 | 0.2030696 | 0.779649 | 0.435597478 | 0.625110666 | | antisense | 39.85 | 0.032304725 | 0.032304725 | ENSG00000186019 | n.s. |
| ENSG00000188846 | 10844.13778 | 0.185136142 | 0.2672406 | 0.69277 | 0.488454199 | 0.669167354 | RPL14 | protein_coding | 44.36 | 0.032299824 | 0.032299824 | ENSG00000188846 | n.s. |

| | | | | | | | | | | | | | |
|---|---|---|---|---|---|---|---|---|---|---|---|---|---|
| ENSG00000069966 | 265.5444548 | -0.191096477 | 0.2833849 | -0.674336 | 0.500098031 | 0.677717041 | GNB5 | protein_coding | 45.39 | -0.032286054 | 0.032286054 | ENSG00000069966 | n.s. |
| ENSG00000141519 | 100.7592428 | 0.174046957 | 0.2399023 | 0.725491 | 0.468150941 | 0.65239371 | CCDC40 | protein_coding | 50.67 | 0.032284011 | 0.032284011 | ENSG00000141519 | n.s. |
| ENSG00000168924 | 658.3241728 | -0.171383443 | 0.2331258 | -0.735154 | 0.462245425 | 0.648121153 | LETM1 | protein_coding | 53.83 | -0.03227901 | 0.03227901 | ENSG00000168924 | n.s. |
| ENSG00000183765 | 110.9922523 | 0.202562028 | 0.3126892 | 0.647806 | 0.517110251 | 0.693081156 | CHEK2 | protein_coding | 43.89 | 0.032251097 | 0.032251097 | ENSG00000183765 | n.s. |
| ENSG00000187239 | 309.0285967 | -0.173051933 | 0.2376227 | -0.728263 | 0.466452397 | 0.651228628 | FNBP1 | protein_coding | 43.76 | -0.032233781 | 0.032233781 | ENSG00000187239 | n.s. |
| ENSG00000064545 | 211.4242415 | 0.197477588 | 0.2996013 | 0.659135 | 0.50980937 | 0.686799793 | TMEM161A | protein_coding | 51.6 | 0.032222387 | 0.032222387 | ENSG00000064545 | n.s. |
| ENSG00000074695 | 2965.459807 | -0.185948059 | 0.2704786 | -0.687478 | 0.491781657 | 0.6711456 | LMAN1 | protein_coding | 37.33 | -0.03220309 | 0.03220309 | ENSG00000074695 | n.s. |
| ENSG00000182004 | 4538.934041 | 0.200397206 | 0.3078156 | 0.65103 | 0.515027048 | 0.691301521 | SNRPE | protein_coding | 42.59 | 0.032130183 | 0.032130183 | ENSG00000182004 | n.s. |
| ENSG00000132405 | 1668.056362 | -0.187909904 | 0.2769231 | -0.678563 | 0.497414477 | 0.675282592 | TBC1D14 | protein_coding | 49.5 | -0.032041353 | 0.032041353 | ENSG00000132405 | n.s. |
| ENSG00000092931 | 108.6083968 | 0.17033176 | 0.2324066 | 0.732904 | 0.463616904 | 0.649063665 | MFSD11 | protein_coding | 44.16 | 0.031973435 | 0.031973435 | ENSG00000092931 | n.s. |
| ENSG00000269951 | 763.1721652 | 0.162516569 | 0.2146425 | 0.75715 | 0.448959931 | 0.635741733 | | sense_intronic | 37.65 | 0.031970142 | 0.031970142 | ENSG00000269951 | n.s. |
| ENSG00000064995 | 968.005438 | 0.188760916 | 0.2794506 | 0.675471 | 0.499376299 | 0.677114596 | TAF11 | protein_coding | 41.49 | 0.031964363 | 0.031964363 | ENSG00000064995 | n.s. |
| ENSG00000007202 | 2945.532094 | -0.180575897 | 0.2578232 | -0.700386 | 0.483686009 | 0.665319137 | KIAA0100 | protein_coding | 46.75 | -0.031956514 | 0.031956514 | ENSG00000007202 | n.s. |
| ENSG00000135677 | 1037.033671 | -0.186209901 | 0.2735144 | -0.680805 | 0.495994973 | 0.67399356 | GNS | protein_coding | 42.2 | -0.031905996 | 0.031905996 | ENSG00000135677 | n.s. |
| ENSG00000115762 | 1556.453858 | 0.181797159 | 0.2610142 | 0.696503 | 0.486113906 | 0.667603463 | PLEKHB2 | protein_coding | 42.42 | 0.031902024 | 0.031902024 | ENSG00000115762 | n.s. |
| ENSG00000132670 | 793.2566974 | -0.184077381 | 0.2676793 | -0.687679 | 0.491655082 | 0.671084026 | PTPRA | protein_coding | 43.04 | -0.031886454 | 0.031886454 | ENSG00000132670 | n.s. |
| ENSG00000137055 | 1236.349106 | -0.177801315 | 0.2514648 | -0.707063 | 0.479527631 | 0.661836342 | PLAA | protein_coding | 38.2 | -0.031870777 | 0.031870777 | ENSG00000137055 | n.s. |
| ENSG00000168090 | 658.4862098 | 0.191213687 | 0.2868008 | 0.666712 | 0.504955836 | 0.681778521 | COPS6 | protein_coding | 52.76 | 0.031809675 | 0.031809675 | ENSG00000168090 | n.s. |
| ENSG00000116560 | 8567.424003 | 0.170048667 | 0.2329698 | 0.729917 | 0.465440733 | 0.650439759 | SFPQ | protein_coding | 41.83 | 0.031763887 | 0.031763887 | ENSG00000116560 | n.s. |
| ENSG00000163463 | 516.4769323 | 0.194829707 | 0.2958605 | 0.658519 | 0.510204821 | 0.687123662 | KRTCAP2 | protein_coding | 50.64 | 0.031750442 | 0.031750442 | ENSG00000163463 | n.s. |
| ENSG00000138629 | 616.8563168 | 0.197274709 | 0.3022942 | 0.652592 | 0.514019453 | 0.690533852 | UBL7 | protein_coding | 47.95 | 0.031724737 | 0.031724737 | ENSG00000138629 | n.s. |
| ENSG00000058804 | 2651.32625 | -0.190657456 | 0.2862831 | -0.665975 | 0.505426983 | 0.682083928 | NDC1 | protein_coding | 40.26 | -0.031680059 | 0.031680059 | ENSG00000058804 | n.s. |
| ENSG00000182831 | 5387.179263 | -0.171363492 | 0.2367851 | -0.723709 | 0.46924451 | 0.653459823 | C16orf72 | protein_coding | 42.03 | -0.031664736 | 0.031664736 | ENSG00000182831 | n.s. |
| ENSG00000184117 | 2169.320625 | 0.196812947 | 0.3016312 | 0.652495 | 0.514081731 | 0.690550886 | NIPSNAP1 | protein_coding | 50.23 | 0.031648371 | 0.031648371 | ENSG00000184117 | n.s. |
| ENSG00000102181 | 294.8467694 | -0.191108956 | 0.2876687 | -0.664337 | 0.506474614 | 0.683299036 | CD99L2 | protein_coding | 43.02 | -0.031607356 | 0.031607356 | ENSG00000102181 | n.s. |
| ENSG00000154310 | 624.1822138 | -0.183949395 | 0.2694528 | -0.682678 | 0.494810501 | 0.673407621 | TNIK | protein_coding | 39.55 | -0.031588153 | 0.031588153 | ENSG00000154310 | n.s. |
| ENSG00000068323 | 140.5671548 | -0.175470005 | 0.2473359 | -0.70944 | 0.47805133 | 0.660701674 | TFE3 | protein_coding | 52.58 | -0.031583652 | 0.031583652 | ENSG00000068323 | n.s. |
| ENSG00000123815 | 196.6767032 | -0.178459535 | 0.2549832 | -0.699887 | 0.48399758 | 0.665550449 | COQ8B | protein_coding | 52.66 | -0.03155504 | 0.03155504 | ENSG00000123815 | n.s. |
| ENSG00000117036 | 870.7420387 | -0.16849776 | 0.2306101 | -0.730661 | 0.464986346 | 0.64992388 | ETV3 | protein_coding | 43.29 | -0.03153225 | 0.03153225 | ENSG00000117036 | n.s. |
| ENSG00000077522 | 39.69016539 | -0.211992238 | 0.345003 | -0.614465 | 0.538908203 | 0.709999258 | ACTN2 | protein_coding | 43.06 | -0.031532172 | 0.031532172 | ENSG00000077522 | n.s. |
| ENSG00000138594 | 871.1517889 | -0.177194884 | 0.2520933 | -0.702894 | 0.482121695 | 0.66388887 | TMOD3 | protein_coding | 41.02 | -0.031523787 | 0.031523787 | ENSG00000138594 | n.s. |
| ENSG00000185122 | 452.0911932 | 0.173556186 | 0.2430748 | 0.714003 | 0.475225142 | 0.658233153 | HSF1 | protein_coding | 56.87 | 0.031521318 | 0.031521318 | ENSG00000185122 | n.s. |

| | | | | | | | | | | | | | |
|---|---|---|---|---|---|---|---|---|---|---|---|---|---|
| ENSG00000210154 | 9.900470501 | -0.231746303 | 0.4036544 | -0.574121 | 0.565886182 | 0.731146373 | MT-TD | Mt_tRNA | 23.53 | -0.031516494 | 0.031516494 | ENSG00000210154 | n.s. |
| ENSG00000146938 | 836.4262269 | -0.180709091 | 0.261212 | -0.69181 | 0.489056676 | 0.669348494 | NLGN4X | protein_coding | 38.89 | -0.031506216 | 0.031506216 | ENSG00000146938 | n.s. |
| ENSG00000168993 | 183.0765549 | -0.175579859 | 0.2483627 | -0.706949 | 0.479597966 | 0.661836342 | CPLX1 | protein_coding | 55.28 | -0.031472583 | 0.031472583 | ENSG00000168993 | n.s. |
| ENSG00000275221 | 252.9858466 | -0.164575453 | 0.2215984 | -0.742674 | 0.45767889 | 0.643988195 | HIST1H2AK | protein_coding | 62.85 | -0.031454005 | 0.031454005 | ENSG00000275221 | n.s. |
| ENSG00000181555 | 2519.444801 | -0.171759616 | 0.2391619 | -0.718173 | 0.472650701 | 0.656430494 | SETD2 | protein_coding | 40.23 | -0.03139959 | 0.03139959 | ENSG00000181555 | n.s. |
| ENSG00000082068 | 834.8549358 | -0.188468002 | 0.2824759 | -0.6672 | 0.504644189 | 0.681556024 | WDR70 | protein_coding | 39 | -0.031379628 | 0.031379628 | ENSG00000082068 | n.s. |
| ENSG00000140830 | 169.2255738 | -0.178921229 | 0.2571235 | -0.695857 | 0.486518349 | 0.667961244 | TXNL4B | protein_coding | 44.45 | -0.031355719 | 0.031355719 | ENSG00000140830 | n.s. |
| ENSG00000126698 | 1125.324869 | 0.207061709 | 0.331868 | 0.623928 | 0.532674961 | 0.70575564 | DNAJC8 | protein_coding | 43.16 | 0.031337887 | 0.031337887 | ENSG00000126698 | n.s. |
| ENSG00000168827 | 2887.861195 | 0.170298258 | 0.2358929 | 0.72193 | 0.470337303 | 0.65465398 | GFM1 | protein_coding | 36.9 | 0.031332868 | 0.031332868 | ENSG00000168827 | n.s. |
| ENSG00000170270 | 357.0797865 | -0.196536198 | 0.3033874 | -0.647806 | 0.517110458 | 0.693081156 | GON7 | protein_coding | 43.35 | -0.031291689 | 0.031291689 | ENSG00000170270 | n.s. |
| ENSG00000154305 | 3235.570065 | -0.173437042 | 0.244343 | -0.70981 | 0.477822173 | 0.660516096 | MIA3 | protein_coding | 39.16 | -0.031238889 | 0.031238889 | ENSG00000154305 | n.s. |
| ENSG00000162236 | 184.3234583 | -0.178101088 | 0.2558222 | -0.696191 | 0.486309286 | 0.667740058 | STX5 | protein_coding | 47.05 | -0.031237607 | 0.031237607 | ENSG00000162236 | n.s. |
| ENSG00000102921 | 611.4353816 | 0.172876105 | 0.2439647 | 0.708611 | 0.478565752 | 0.66116372 | N4BP1 | protein_coding | 42.34 | 0.031064278 | 0.031064278 | ENSG00000102921 | n.s. |
| ENSG00000266338 | 334.2515776 | -0.189433508 | 0.2865557 | -0.66107 | 0.508567153 | 0.685590674 | NBPF15 | protein_coding | 42.75 | -0.031054801 | 0.031054801 | ENSG00000266338 | n.s. |
| ENSG00000279766 | 40.22857681 | 0.189912557 | 0.2878445 | 0.659775 | 0.509398345 | 0.686511776 | | TEC | 44.86 | 0.031022598 | 0.031022598 | ENSG00000279766 | n.s. |
| ENSG00000236698 | 26.22025713 | -0.20905183 | 0.3409954 | -0.613063 | 0.539834356 | 0.710687467 | EIF1AXP1 | processed_pseudogene | 38.62 | -0.031006848 | 0.031006848 | ENSG00000236698 | n.s. |
| ENSG00000115446 | 855.8870159 | -0.17150475 | 0.2410022 | -0.711632 | 0.476692963 | 0.659675591 | UNC50 | protein_coding | 42.81 | -0.030985692 | 0.030985692 | ENSG00000115446 | n.s. |
| ENSG00000152377 | 1027.859694 | -0.171770293 | 0.2416469 | -0.710832 | 0.477188469 | 0.66013203 | SPOCK1 | protein_coding | 42.8 | -0.030982069 | 0.030982069 | ENSG00000152377 | n.s. |
| ENSG00000122203 | 1437.042606 | 0.180529361 | 0.2648704 | 0.681576 | 0.495507046 | 0.673659272 | KIAA1191 | protein_coding | 47.1 | 0.030971565 | 0.030971565 | ENSG00000122203 | n.s. |
| ENSG00000072364 | 3077.913678 | -0.177381836 | 0.255971 | -0.692976 | 0.488324345 | 0.669121135 | AFF4 | protein_coding | 40.42 | -0.030952288 | 0.030952288 | ENSG00000072364 | n.s. |
| ENSG00000106588 | 653.9727878 | -0.180349366 | 0.2645036 | -0.681841 | 0.495339563 | 0.673628906 | PSMA2 | protein_coding | 38.3 | -0.030944215 | 0.030944215 | ENSG00000106588 | n.s. |
| ENSG00000114573 | 935.4015802 | -0.181684683 | 0.2679412 | -0.678077 | 0.497723079 | 0.675616499 | ATP6V1A | protein_coding | 38.22 | -0.030940857 | 0.030940857 | ENSG00000114573 | n.s. |
| ENSG00000221988 | 216.3972012 | 0.188981296 | 0.286075 | 0.660601 | 0.508868546 | 0.685930562 | PPT2 | protein_coding | 50.59 | 0.030939989 | 0.030939989 | ENSG00000221988 | n.s. |
| ENSG00000183520 | 798.2309349 | 0.181363509 | 0.2672844 | 0.678541 | 0.49742859 | 0.675282592 | UTP11 | protein_coding | 44.58 | 0.030925098 | 0.030925098 | ENSG00000183520 | n.s. |
| ENSG00000139180 | 1426.417473 | -0.179586788 | 0.2626789 | -0.683674 | 0.494180893 | 0.672841832 | NDUFA9 | protein_coding | 39.57 | -0.030904554 | 0.030904554 | ENSG00000139180 | n.s. |
| ENSG00000067601 | 56.74128947 | 0.204668462 | 0.3286449 | 0.622765 | 0.533439109 | 0.706331214 | PMS2P4 | transcribed_unprocessed_pseud | 43.32 | 0.030903219 | 0.030903219 | ENSG00000067601 | n.s. |
| ENSG00000255097 | 33.35466307 | -0.195990644 | 0.3047633 | -0.643091 | 0.520164792 | 0.695568037 | | sense_intronic | 45.92 | -0.03089996 | 0.03089996 | ENSG00000255097 | n.s. |
| ENSG00000157916 | 416.9475118 | -0.189532824 | 0.2879699 | -0.658169 | 0.510429552 | 0.687123662 | RER1 | protein_coding | 53.95 | -0.030887235 | 0.030887235 | ENSG00000157916 | n.s. |
| ENSG00000138385 | 5618.671703 | 0.181751455 | 0.2687777 | 0.676215 | 0.498904237 | 0.6766985 | SSB | protein_coding | 39.88 | 0.030825917 | 0.030825917 | ENSG00000138385 | n.s. |
| ENSG00000154719 | 729.4076003 | 0.183163879 | 0.272787 | 0.671454 | 0.501931416 | 0.679019385 | MRPL39 | protein_coding | 37.94 | 0.030793113 | 0.030793113 | ENSG00000154719 | n.s. |
| ENSG00000130119 | 413.1082497 | -0.170920055 | 0.2407759 | -0.709872 | 0.477783533 | 0.660516096 | GNL3L | protein_coding | 47.94 | -0.030785538 | 0.030785538 | ENSG00000130119 | n.s. |
| ENSG00000149554 | 1401.244633 | -0.176473452 | 0.2548739 | -0.692395 | 0.488689297 | 0.669199599 | CHEK1 | protein_coding | 37.72 | -0.030784793 | 0.030784793 | ENSG00000149554 | n.s. |

| | | | | | | | | | | | | | |
|---|---|---|---|---|---|---|---|---|---|---|---|---|---|
| ENSG00000106144 | 971.1855273 | 0.165700123 | 0.2280365 | 0.726638 | 0.46744745 | 0.651988717 | CASP2 | protein_coding | 43.43 | 0.030780442 | 0.030780442 | ENSG00000106144 | n.s. |
| ENSG00000110801 | 620.1479732 | 0.190058546 | 0.2902565 | 0.654795 | 0.512599596 | 0.689025311 | PSMD9 | protein_coding | 47.48 | 0.030744787 | 0.030744787 | ENSG00000110801 | n.s. |
| ENSG00000166716 | 859.0120522 | -0.16359997 | 0.2230358 | -0.733514 | 0.463244777 | 0.648762529 | ZNF592 | protein_coding | 45.9 | -0.030742764 | 0.030742764 | ENSG00000166716 | n.s. |
| ENSG00000150471 | 131.5579683 | -0.180643989 | 0.2666201 | -0.677533 | 0.498067687 | 0.675886589 | ADGRL3 | protein_coding | 35.25 | -0.03073227 | 0.03073227 | ENSG00000150471 | n.s. |
| ENSG00000171823 | 236.0083127 | 0.163056387 | 0.2216436 | 0.735669 | 0.461931954 | 0.647942635 | FBXL14 | protein_coding | 46.18 | 0.030730168 | 0.030730168 | ENSG00000171823 | n.s. |
| ENSG00000141698 | 1119.95421 | 0.191535101 | 0.2941882 | 0.651063 | 0.515005703 | 0.691301521 | NT5C3B | protein_coding | 48.83 | 0.030709299 | 0.030709299 | ENSG00000141698 | n.s. |
| ENSG00000151923 | 1828.933209 | 0.177785445 | 0.2592638 | 0.685732 | 0.492882143 | 0.671937241 | TIAL1 | protein_coding | 38.16 | 0.030698442 | 0.030698442 | ENSG00000151923 | n.s. |
| ENSG00000104660 | 1330.535245 | 0.183143592 | 0.2733753 | 0.669935 | 0.502899404 | 0.679803747 | LEPROTL1 | protein_coding | 42.61 | 0.030697878 | 0.030697878 | ENSG00000104660 | n.s. |
| ENSG00000264757 | 9.506886149 | 0.234928814 | 0.4213434 | 0.557571 | 0.577137477 | 0.740546777 | MIR3198-1 | miRNA | 56.25 | 0.030645877 | 0.030645877 | ENSG00000264757 | n.s. |
| ENSG00000146223 | 4533.881019 | -0.177746387 | 0.2596786 | -0.684486 | 0.493668259 | 0.672461266 | RPL7L1 | protein_coding | 46.45 | -0.030631519 | 0.030631519 | ENSG00000146223 | n.s. |
| ENSG00000128512 | 603.6556505 | -0.166405039 | 0.2306283 | -0.721529 | 0.47058405 | 0.654670436 | DOCK4 | protein_coding | 39.18 | -0.030614745 | 0.030614745 | ENSG00000128512 | n.s. |
| ENSG00000162613 | 13239.99972 | 0.176103851 | 0.2554324 | 0.689434 | 0.490549947 | 0.670128457 | FUBP1 | protein_coding | 36.32 | 0.030614235 | 0.030614235 | ENSG00000162613 | n.s. |
| ENSG00000139131 | 901.5174155 | 0.1754777 | 0.2533449 | 0.692643 | 0.488533321 | 0.669177272 | YARS2 | protein_coding | 39.51 | 0.030613632 | 0.030613632 | ENSG00000139131 | n.s. |
| ENSG00000074071 | 1064.412099 | 0.20161812 | 0.3225728 | 0.625031 | 0.531950427 | 0.705097657 | MRPS34 | protein_coding | 65.4 | 0.030595697 | 0.030595697 | ENSG00000074071 | n.s. |
| ENSG00000283419 | 3.490278879 | 0.279870948 | 0.5714861 | 0.489725 | 0.624328589 | 0.777831587 | MIR6874 | miRNA | 45.07 | 0.030537957 | 0.030537957 | ENSG00000283419 | n.s. |
| ENSG00000116906 | 1597.769957 | -0.194879885 | 0.3042919 | -0.640437 | 0.521888372 | 0.697203205 | GNPAT | protein_coding | 41.78 | -0.030526107 | 0.030526107 | ENSG00000116906 | n.s. |
| ENSG00000120158 | 882.5383963 | 0.180788638 | 0.2684529 | 0.673446 | 0.500663352 | 0.677945841 | RCL1 | protein_coding | 42.83 | 0.030518026 | 0.030518026 | ENSG00000120158 | n.s. |
| ENSG00000147044 | 221.3743817 | -0.169456707 | 0.2388319 | -0.709523 | 0.478000076 | 0.660696422 | CASK | protein_coding | 38.73 | -0.030501876 | 0.030501876 | ENSG00000147044 | n.s. |
| ENSG00000185909 | 106.6279563 | 0.176847787 | 0.2583858 | 0.684433 | 0.493701765 | 0.672461266 | KLHDC8B | protein_coding | 58.06 | 0.030476661 | 0.030476661 | ENSG00000185909 | n.s. |
| ENSG00000115310 | 4999.448678 | 0.174971652 | 0.2533678 | 0.690584 | 0.489827306 | 0.669927857 | RTN4 | protein_coding | 38.6 | 0.030440162 | 0.030440162 | ENSG00000115310 | n.s. |
| ENSG00000157014 | 747.7341343 | -0.173470308 | 0.2490919 | -0.696411 | 0.486171601 | 0.667616846 | TATDN2 | protein_coding | 46.98 | -0.030439306 | 0.030439306 | ENSG00000157014 | n.s. |
| ENSG00000151376 | 216.7843162 | -0.171708432 | 0.2449303 | -0.70105 | 0.483271672 | 0.664880582 | ME3 | protein_coding | 42.88 | -0.03043641 | 0.03043641 | ENSG00000151376 | n.s. |
| ENSG00000136451 | 3256.747062 | -0.171015687 | 0.2432134 | -0.703151 | 0.481961886 | 0.663800112 | VEZF1 | protein_coding | 40.94 | -0.03043441 | 0.03043441 | ENSG00000136451 | n.s. |
| ENSG00000101544 | 1315.003919 | 0.177902072 | 0.2616841 | 0.679835 | 0.496608846 | 0.674564386 | ADNP2 | protein_coding | 41.81 | 0.03041709 | 0.03041709 | ENSG00000101544 | n.s. |
| ENSG00000264346 | 40.66979799 | 0.205647068 | 0.3366116 | 0.610933 | 0.541244067 | 0.711590288 | SNORA77B | snoRNA | 55.2 | 0.030388465 | 0.030388465 | ENSG00000264346 | n.s. |
| ENSG00000134809 | 422.1082097 | -0.182654933 | 0.2740221 | -0.66657 | 0.505046837 | 0.681822569 | TIMM10 | protein_coding | 48.01 | -0.030380744 | 0.030380744 | ENSG00000134809 | n.s. |
| ENSG00000171298 | 452.2804014 | 0.169187453 | 0.238871 | 0.70828 | 0.478771704 | 0.661369028 | GAA | protein_coding | 58.33 | 0.030378647 | 0.030378647 | ENSG00000171298 | n.s. |
| ENSG00000283891 | 6.759405396 | 0.249420244 | 0.469896 | 0.530799 | 0.595558173 | 0.755522643 | MIR628 | miRNA | 41.05 | 0.030367542 | 0.030367542 | ENSG00000283891 | n.s. |
| ENSG00000187118 | 157.5017397 | 0.206716767 | 0.3402425 | 0.607557 | 0.543481244 | 0.713251896 | CMC1 | protein_coding | 35.93 | 0.030337146 | 0.030337146 | ENSG00000187118 | n.s. |
| ENSG00000231508 | 15.443973 | -0.212096487 | 0.3566373 | -0.594712 | 0.552035939 | 0.719662583 | RPL34P20 | processed_pseudogene | 44.73 | -0.030302453 | 0.030302453 | ENSG00000231508 | n.s. |
| ENSG00000090273 | 826.468031 | -0.173757095 | 0.2510695 | -0.692068 | 0.488894811 | 0.669309975 | NUDC | protein_coding | 48.43 | -0.030298494 | 0.030298494 | ENSG00000090273 | n.s. |
| ENSG00000105732 | 320.6704618 | -0.170597159 | 0.2429744 | -0.70212 | 0.482604317 | 0.664356328 | ZNF574 | protein_coding | 57 | -0.030297872 | 0.030297872 | ENSG00000105732 | n.s. |

| | | | | | | | | | | | | | |
|---|---|---|---|---|---|---|---|---|---|---|---|---|---|
| ENSG00000201592 | 103.9282277 | 0.174089603 | 0.252426 | 0.689666 | 0.490404352 | 0.670128457 | | snoRNA | 43.75 | 0.030264074 | 0.030264074 | ENSG00000201592 | n.s. |
| ENSG00000234545 | 1521.486906 | 0.169816413 | 0.2410915 | 0.704365 | 0.481205579 | 0.663414722 | FAM133B | protein_coding | 37.43 | 0.030263814 | 0.030263814 | ENSG00000234545 | n.s. |
| ENSG00000146729 | 986.9326067 | -0.192915682 | 0.3011109 | -0.64068 | 0.521730678 | 0.697059422 | NIPSNAP2 | protein_coding | 45.28 | -0.030235713 | 0.030235713 | ENSG00000146729 | n.s. |
| ENSG00000149658 | 795.72749 | -0.176523335 | 0.2593879 | -0.680538 | 0.496163798 | 0.674130097 | YTHDF1 | protein_coding | 49.38 | -0.03023073 | 0.03023073 | ENSG00000149658 | n.s. |
| ENSG00000077721 | 507.2272357 | -0.190119867 | 0.2939416 | -0.646795 | 0.517764785 | 0.693424307 | UBE2A | protein_coding | 40.14 | -0.030229237 | 0.030229237 | ENSG00000077721 | n.s. |
| ENSG00000103197 | 46.26603963 | 0.193139708 | 0.3019327 | 0.639678 | 0.522382002 | 0.697594917 | TSC2 | protein_coding | 60.06 | 0.030206411 | 0.030206411 | ENSG00000103197 | n.s. |
| ENSG00000169696 | 778.7478963 | 0.17829881 | 0.2645332 | 0.674013 | 0.50030312 | 0.677749957 | ASPSCR1 | protein_coding | 60.76 | 0.030120107 | 0.030120107 | ENSG00000169696 | n.s. |
| ENSG00000100938 | 137.4597036 | 0.186295117 | 0.2846353 | 0.654505 | 0.512786769 | 0.689210367 | GMPR2 | protein_coding | 46.62 | 0.03011427 | 0.03011427 | ENSG00000100938 | n.s. |
| ENSG00000004399 | 489.5864331 | -0.163568577 | 0.2266585 | -0.721652 | 0.470508441 | 0.65465398 | PLXND1 | protein_coding | 58.39 | -0.030094686 | 0.030094686 | ENSG00000004399 | n.s. |
| ENSG00000276855 | 24.75419262 | -0.214085607 | 0.3644313 | -0.587451 | 0.556900664 | 0.723497989 | | lincRNA | 33.62 | -0.030092445 | 0.030092445 | ENSG00000276855 | n.s. |
| ENSG00000173145 | 1768.868529 | 0.177728761 | 0.2633121 | 0.674974 | 0.49969247 | 0.677365228 | NOC3L | protein_coding | 37.01 | 0.030067637 | 0.030067637 | ENSG00000173145 | n.s. |
| ENSG00000168243 | 1576.997138 | -0.183275137 | 0.2771571 | -0.661268 | 0.508440387 | 0.685552538 | GNG4 | protein_coding | 48.07 | -0.030049655 | 0.030049655 | ENSG00000168243 | n.s. |
| ENSG00000224464 | 26.81965914 | -0.196602171 | 0.3129565 | -0.628209 | 0.529866883 | 0.703474455 | PGAM1P6 | processed_pseudogene | 52.21 | -0.03003131 | 0.03003131 | ENSG00000224464 | n.s. |
| ENSG00000187049 | 184.2967737 | -0.184228457 | 0.279989 | -0.657985 | 0.510548 | 0.687129995 | TMEM216 | protein_coding | 48.57 | -0.030022071 | 0.030022071 | ENSG00000187049 | n.s. |
| ENSG00000221829 | 692.0818094 | 0.171864916 | 0.2478973 | 0.693291 | 0.488127075 | 0.669050361 | FANCG | protein_coding | 53.25 | 0.029997507 | 0.029997507 | ENSG00000221829 | n.s. |
| ENSG00000158711 | 1571.32343 | -0.180578191 | 0.2712158 | -0.66581 | 0.505532546 | 0.682160268 | ELK4 | protein_coding | 41.85 | -0.02999649 | 0.02999649 | ENSG00000158711 | n.s. |
| ENSG00000144589 | 179.3904503 | 0.173552959 | 0.2529249 | 0.686184 | 0.492597256 | 0.671802488 | STK11IP | protein_coding | 53.34 | 0.02998273 | 0.02998273 | ENSG00000144589 | n.s. |
| ENSG00000182118 | 430.6120313 | -0.203517482 | 0.3336816 | -0.609915 | 0.541918038 | 0.712341847 | FAM89A | protein_coding | 47.02 | -0.029980474 | 0.029980474 | ENSG00000182118 | n.s. |
| ENSG00000171806 | 287.131003 | -0.162576259 | 0.22502 | -0.722497 | 0.469989091 | 0.654234965 | METTL18 | protein_coding | 40.77 | -0.029957318 | 0.029957318 | ENSG00000171806 | n.s. |
| ENSG00000038274 | 1749.671333 | -0.186738239 | 0.2867896 | -0.651133 | 0.514960453 | 0.691301521 | MAT2B | protein_coding | 38.25 | -0.029940206 | 0.029940206 | ENSG00000038274 | n.s. |
| ENSG00000110047 | 140.4659613 | -0.171717699 | 0.2482197 | -0.691797 | 0.489064659 | 0.669348494 | EHD1 | protein_coding | 54.84 | -0.029938588 | 0.029938588 | ENSG00000110047 | n.s. |
| ENSG00000105486 | 184.7169007 | 0.171449443 | 0.2473053 | 0.693271 | 0.488139807 | 0.669050361 | LIG1 | protein_coding | 51.48 | 0.02992499 | 0.02992499 | ENSG00000105486 | n.s. |
| ENSG00000109084 | 2220.349437 | -0.169669519 | 0.2428749 | -0.698588 | 0.484809472 | 0.666403621 | TMEM97 | protein_coding | 44.52 | -0.029906399 | 0.029906399 | ENSG00000109084 | n.s. |
| ENSG00000235244 | 109.7462948 | -0.174539605 | 0.2563432 | -0.680882 | 0.495945898 | 0.67399356 | DANT2 | lincRNA | 42.01 | -0.029906358 | 0.029906358 | ENSG00000235244 | n.s. |
| ENSG00000160633 | 1652.088715 | -0.159276415 | 0.2174195 | -0.732576 | 0.463816809 | 0.649176514 | SAFB | protein_coding | 47.35 | -0.029886181 | 0.029886181 | ENSG00000160633 | n.s. |
| ENSG00000105617 | 124.2317203 | -0.179349732 | 0.2686019 | -0.667716 | 0.504314978 | 0.681375783 | LENG1 | protein_coding | 56.01 | -0.029882052 | 0.029882052 | ENSG00000105617 | n.s. |
| ENSG00000198855 | 139.7553673 | 0.165558832 | 0.2331281 | 0.710163 | 0.477603327 | 0.660440907 | FICD | protein_coding | 50.41 | 0.029828078 | 0.029828078 | ENSG00000198855 | n.s. |
| ENSG00000155959 | 2054.682858 | 0.193897124 | 0.3079922 | 0.629552 | 0.528987711 | 0.702790513 | VBP1 | protein_coding | 40.41 | 0.029700019 | 0.029700019 | ENSG00000155959 | n.s. |
| ENSG00000167815 | 3175.013909 | 0.185446722 | 0.2851432 | 0.650364 | 0.515457448 | 0.691731475 | PRDX2 | protein_coding | 54.5 | 0.029683059 | 0.029683059 | ENSG00000167815 | n.s. |
| ENSG00000119777 | 299.9842094 | -0.169799066 | 0.2446434 | -0.694068 | 0.487639821 | 0.668694533 | TMEM214 | protein_coding | 55.01 | -0.029676161 | 0.029676161 | ENSG00000119777 | n.s. |
| ENSG00000100410 | 917.7389321 | -0.181677787 | 0.275402 | -0.659682 | 0.509457809 | 0.686525462 | PHF5A | protein_coding | 46.26 | -0.029675859 | 0.029675859 | ENSG00000100410 | n.s. |
| ENSG00000176422 | 990.6589024 | -0.148449717 | 0.1936679 | -0.766517 | 0.443368916 | 0.631196898 | SPRYD4 | protein_coding | 48.14 | -0.029665471 | 0.029665471 | ENSG00000176422 | n.s. |

| | | | | | | | | | | | | | |
|---|---|---|---|---|---|---|---|---|---|---|---|---|---|
| ENSG00000148290 | 307.5833086 | 0.189964071 | 0.2973067 | 0.63895 | 0.522855511 | 0.698026396 | SURF1 | protein_coding | 53.96 | 0.02965874 | 0.02965874 | ENSG00000148290 | n.s. |
| ENSG00000086504 | 483.3507057 | 0.196709362 | 0.3165539 | 0.621409 | 0.534330674 | 0.707041054 | MRPL28 | protein_coding | 64.85 | 0.02961565 | 0.02961565 | ENSG00000086504 | n.s. |
| ENSG00000108010 | 1685.515048 | 0.184899814 | 0.2844523 | 0.650021 | 0.515678941 | 0.691762009 | GLRX3 | protein_coding | 42.15 | 0.029591975 | 0.029591975 | ENSG00000108010 | n.s. |
| ENSG00000169964 | 189.504179 | -0.159225994 | 0.2192564 | -0.726209 | 0.467710673 | 0.651988717 | TMEM42 | protein_coding | 52.21 | -0.029577808 | 0.029577808 | ENSG00000169964 | n.s. |
| ENSG00000197226 | 1006.632142 | -0.177599209 | 0.266227 | -0.667097 | 0.504710186 | 0.681579041 | TBC1D9B | protein_coding | 54.22 | -0.029567386 | 0.029567386 | ENSG00000197226 | n.s. |
| ENSG00000186642 | 785.6475582 | 0.182544422 | 0.2786623 | 0.655074 | 0.512420102 | 0.688892052 | PDE2A | protein_coding | 55.35 | 0.0295446 | 0.0295446 | ENSG00000186642 | n.s. |
| ENSG00000092108 | 1318.275891 | -0.18119359 | 0.2753179 | -0.658125 | 0.510457838 | 0.687123662 | SCFD1 | protein_coding | 36.17 | -0.029528231 | 0.029528231 | ENSG00000092108 | n.s. |
| ENSG00000252914 | 11.47292 | 0.241990593 | 0.4548986 | 0.531966 | 0.594749543 | 0.755124739 | RNU6-789P | snRNA | 34.31 | 0.029518328 | 0.029518328 | ENSG00000252914 | n.s. |
| ENSG00000095752 | 110.2909568 | -0.192690738 | 0.3062339 | -0.629227 | 0.529200297 | 0.702790513 | IL11 | protein_coding | 60.87 | -0.029515232 | 0.029515232 | ENSG00000095752 | n.s. |
| ENSG00000125814 | 599.4232845 | -0.171249156 | 0.250241 | -0.684337 | 0.493762394 | 0.672469316 | NAPB | protein_coding | 40.72 | -0.029510944 | 0.029510944 | ENSG00000125814 | n.s. |
| ENSG00000196233 | 1882.794293 | -0.169118252 | 0.2442673 | -0.692349 | 0.488718058 | 0.669199599 | LCOR | protein_coding | 39.58 | -0.02950172 | 0.02950172 | ENSG00000196233 | n.s. |
| ENSG00000054965 | 1119.691845 | -0.170660478 | 0.2486359 | -0.686387 | 0.492469013 | 0.671693385 | FAM168A | protein_coding | 41.55 | -0.029495068 | 0.029495068 | ENSG00000054965 | n.s. |
| ENSG00000174238 | 508.7649663 | -0.18543746 | 0.2866687 | -0.64687 | 0.517715945 | 0.693424307 | PITPNA | protein_coding | 49.88 | -0.02948473 | 0.02948473 | ENSG00000174238 | n.s. |
| ENSG00000124275 | 1468.234962 | -0.164398171 | 0.2326775 | -0.70655 | 0.47984641 | 0.661999759 | MTRR | protein_coding | 39.8 | -0.029450645 | 0.029450645 | ENSG00000124275 | n.s. |
| ENSG00000137880 | 250.5268241 | 0.203821886 | 0.3397754 | 0.599872 | 0.548591265 | 0.717229859 | GCHFR | protein_coding | 57.52 | 0.029419985 | 0.029419985 | ENSG00000137880 | n.s. |
| ENSG00000115275 | 428.5311988 | 0.174098422 | 0.2581486 | 0.674412 | 0.500049619 | 0.677717041 | MOGS | protein_coding | 57.9 | 0.029414206 | 0.029414206 | ENSG00000115275 | n.s. |
| ENSG00000187790 | 360.7867366 | -0.165445475 | 0.2357693 | -0.701726 | 0.482850065 | 0.664497525 | FANCM | protein_coding | 37.51 | -0.029367669 | 0.029367669 | ENSG00000187790 | n.s. |
| ENSG00000156504 | 521.0845297 | -0.18754027 | 0.2931726 | -0.639692 | 0.522372688 | 0.697594917 | FAM122B | protein_coding | 40.17 | -0.029330678 | 0.029330678 | ENSG00000156504 | n.s. |
| ENSG00000133398 | 512.2473678 | 0.167998872 | 0.2424699 | 0.692865 | 0.488394379 | 0.669151243 | MED10 | protein_coding | 45.03 | 0.029311723 | 0.029311723 | ENSG00000133398 | n.s. |
| ENSG00000100109 | 471.2166497 | 0.166305741 | 0.2382279 | 0.698095 | 0.48511761 | 0.666686317 | TFIP11 | protein_coding | 44.89 | 0.029282858 | 0.029282858 | ENSG00000100109 | n.s. |
| ENSG00000145220 | 664.3216886 | -0.173946119 | 0.2588824 | -0.671912 | 0.501639927 | 0.678881484 | LYAR | protein_coding | 44.02 | -0.029258787 | 0.029258787 | ENSG00000145220 | n.s. |
| ENSG00000157881 | 191.1436407 | 0.183759122 | 0.2837317 | 0.647651 | 0.517210716 | 0.693081156 | PANK4 | protein_coding | 60.69 | 0.029257376 | 0.029257376 | ENSG00000157881 | n.s. |
| ENSG00000178053 | 271.971973 | 0.196410026 | 0.3189056 | 0.615888 | 0.537968663 | 0.70969473 | MLF1 | protein_coding | 37.51 | 0.029251035 | 0.029251035 | ENSG00000178053 | n.s. |
| ENSG00000240230 | 443.7706967 | -0.166121744 | 0.2382984 | -0.697117 | 0.485729836 | 0.667207627 | COX19 | protein_coding | 58.98 | -0.029194069 | 0.029194069 | ENSG00000240230 | n.s. |
| ENSG00000129925 | 166.1597409 | -0.175256218 | 0.2626032 | -0.66738 | 0.504529156 | 0.681552702 | TMEM8A | protein_coding | 61.61 | -0.029180258 | 0.029180258 | ENSG00000129925 | n.s. |
| ENSG00000146109 | 854.4960163 | -0.167590133 | 0.2431742 | -0.689177 | 0.490711731 | 0.670260252 | ABT1 | protein_coding | 47.69 | -0.029119881 | 0.029119881 | ENSG00000146109 | n.s. |
| ENSG00000107819 | 298.3758749 | -0.170923259 | 0.2521106 | -0.677969 | 0.497791128 | 0.675642999 | SFXN3 | protein_coding | 54.03 | -0.029105277 | 0.029105277 | ENSG00000107819 | n.s. |
| ENSG00000204568 | 600.1243678 | -0.189477739 | 0.3002489 | -0.631069 | 0.527995439 | 0.702329272 | MRPS18B | protein_coding | 46.36 | -0.029077108 | 0.029077108 | ENSG00000204568 | n.s. |
| ENSG00000100532 | 303.7786611 | 0.192078667 | 0.3080157 | 0.6236 | 0.532890165 | 0.705872873 | CGRRF1 | protein_coding | 37.44 | 0.029056414 | 0.029056414 | ENSG00000100532 | n.s. |
| ENSG00000003509 | 702.751132 | -0.18060265 | 0.2769426 | -0.65213 | 0.514317097 | 0.6908004 | NDUFAF7 | protein_coding | 38.97 | -0.02901335 | 0.02901335 | ENSG00000003509 | n.s. |
| ENSG00000223501 | 574.9566132 | -0.168889988 | 0.2477769 | -0.681621 | 0.495478502 | 0.673659272 | VPS52 | protein_coding | 45.57 | -0.028974717 | 0.028974717 | ENSG00000223501 | n.s. |
| ENSG00000119638 | 381.5668823 | -0.155964857 | 0.2147551 | -0.726245 | 0.467688548 | 0.651988717 | NEK9 | protein_coding | 42.1 | -0.028972019 | 0.028972019 | ENSG00000119638 | n.s. |

| | | | | | | | | | | | | | |
|---|---|---|---|---|---|---|---|---|---|---|---|---|---|
| ENSG00000164076 | 2318.5584 | 0.181853559 | 0.2808556 | 0.647498 | 0.517309443 | 0.693081156 | CAMKV | protein_coding | 56.12 | 0.02895398 | 0.02895398 | ENSG00000164076 | n.s. |
| ENSG00000276043 | 1625.333944 | -0.160880912 | 0.227042 | -0.708595 | 0.478575593 | 0.66116372 | UHRF1 | protein_coding | 52.99 | -0.02890885 | 0.02890885 | ENSG00000276043 | n.s. |
| ENSG00000105419 | 759.6513249 | 0.190015282 | 0.3035205 | 0.626038 | 0.531290177 | 0.704490774 | MEIS3 | protein_coding | 56.51 | 0.028906016 | 0.028906016 | ENSG00000105419 | n.s. |
| ENSG00000151150 | 1250.551805 | -0.165928656 | 0.2401106 | -0.691051 | 0.489533472 | 0.669780286 | ANK3 | protein_coding | 37.75 | -0.028882809 | 0.028882809 | ENSG00000151150 | n.s. |
| ENSG00000106245 | 812.3490911 | 0.181592136 | 0.2807525 | 0.646805 | 0.517758087 | 0.693424307 | BUD31 | protein_coding | 48.46 | 0.02887332 | 0.02887332 | ENSG00000106245 | n.s. |
| ENSG00000278600 | 471.8683043 | -0.161644375 | 0.2292938 | -0.704966 | 0.480831202 | 0.663029893 | | sense_intronic | 41.18 | -0.028848169 | 0.028848169 | ENSG00000278600 | n.s. |
| ENSG00000105618 | 1017.942717 | 0.18499684 | 0.2898459 | 0.638259 | 0.523304908 | 0.698425444 | PRPF31 | protein_coding | 57.08 | 0.028837298 | 0.028837298 | ENSG00000105618 | n.s. |
| ENSG00000167004 | 3682.408031 | 0.177315486 | 0.2700011 | 0.656721 | 0.511360034 | 0.687890411 | PDIA3 | protein_coding | 43.02 | 0.028810352 | 0.028810352 | ENSG00000167004 | n.s. |
| ENSG00000204070 | 301.9133859 | -0.16274989 | 0.2324908 | -0.700027 | 0.483910324 | 0.665545461 | SYS1 | protein_coding | 49.54 | -0.028777806 | 0.028777806 | ENSG00000204070 | n.s. |
| ENSG00000244462 | 3805.941805 | -0.184639971 | 0.290122 | -0.636422 | 0.524501522 | 0.699131025 | RBM12 | protein_coding | 39.8 | -0.0287007 | 0.0287007 | ENSG00000244462 | n.s. |
| ENSG00000115524 | 15870.93142 | 0.176078705 | 0.2675554 | 0.658102 | 0.51047273 | 0.687123662 | SF3B1 | protein_coding | 39.41 | 0.028694683 | 0.028694683 | ENSG00000115524 | n.s. |
| ENSG00000198815 | 383.4151473 | -0.166413917 | 0.2430721 | -0.684628 | 0.493578895 | 0.672461266 | FOXJ3 | protein_coding | 38.57 | -0.028678564 | 0.028678564 | ENSG00000198815 | n.s. |
| ENSG00000242114 | 238.8153406 | -0.184148504 | 0.2887983 | -0.637637 | 0.523709907 | 0.698698068 | MTFP1 | protein_coding | 56.98 | -0.028673848 | 0.028673848 | ENSG00000242114 | n.s. |
| ENSG00000154328 | 185.3260648 | -0.172962441 | 0.2601859 | -0.664765 | 0.506200969 | 0.682996035 | NEIL2 | protein_coding | 48.93 | -0.028639435 | 0.028639435 | ENSG00000154328 | n.s. |
| ENSG00000138495 | 1179.646987 | 0.180186225 | 0.2787336 | 0.646446 | 0.517990518 | 0.693593189 | COX17 | protein_coding | 41.28 | 0.028630723 | 0.028630723 | ENSG00000138495 | n.s. |
| ENSG00000264994 | 46.69217282 | -0.186847125 | 0.2968305 | -0.629474 | 0.529038751 | 0.702790513 | SNORD92 | snoRNA | 44.71 | -0.028620142 | 0.028620142 | ENSG00000264994 | n.s. |
| ENSG00000146963 | 1724.461126 | -0.17550528 | 0.2666019 | -0.658305 | 0.510342386 | 0.687123662 | LUC7L2 | protein_coding | 40.36 | -0.028601235 | 0.028601235 | ENSG00000146963 | n.s. |
| ENSG00000120899 | 24.60541045 | 0.189990914 | 0.306265 | 0.620348 | 0.535028615 | 0.707314789 | PTK2B | protein_coding | 46.91 | 0.028572213 | 0.028572213 | ENSG00000120899 | n.s. |
| ENSG00000142546 | 479.3844531 | 0.184439391 | 0.2904747 | 0.634959 | 0.525455442 | 0.700087667 | NOSIP | protein_coding | 50.75 | 0.028559992 | 0.028559992 | ENSG00000142546 | n.s. |
| ENSG00000147650 | 1424.315714 | 0.183034351 | 0.2871655 | 0.637383 | 0.523875535 | 0.698801298 | LRP12 | protein_coding | 35.69 | 0.028488619 | 0.028488619 | ENSG00000147650 | n.s. |
| ENSG00000133265 | 293.4473995 | 0.188339743 | 0.3022217 | 0.623184 | 0.533163636 | 0.706167932 | HSPBP1 | protein_coding | 51.91 | 0.02845663 | 0.02845663 | ENSG00000133265 | n.s. |
| ENSG00000088205 | 4838.20074 | -0.177664989 | 0.2732677 | -0.65015 | 0.515595406 | 0.691762009 | DDX18 | protein_coding | 39.26 | -0.02843409 | 0.02843409 | ENSG00000088205 | n.s. |
| ENSG00000235748 | 221.0678802 | -0.170467443 | 0.2552661 | -0.667803 | 0.504259467 | 0.681366903 | SEPT14P12 | processed_pseudogene | 40.36 | -0.028403109 | 0.028403109 | ENSG00000235748 | n.s. |
| ENSG00000127837 | 1245.732918 | 0.175198041 | 0.2673958 | 0.655201 | 0.512338267 | 0.688892052 | AAMP | protein_coding | 56.53 | 0.028355597 | 0.028355597 | ENSG00000127837 | n.s. |
| ENSG00000123353 | 248.5756208 | -0.184059115 | 0.2909197 | -0.63268 | 0.526942498 | 0.701397744 | ORMDL2 | protein_coding | 46.4 | -0.028351663 | 0.028351663 | ENSG00000123353 | n.s. |
| ENSG00000131844 | 762.194107 | -0.16914585 | 0.2524016 | -0.670146 | 0.502764917 | 0.679803747 | MCCC2 | protein_coding | 41.2 | -0.028351626 | 0.028351626 | ENSG00000131844 | n.s. |
| ENSG00000203808 | 40.01269045 | 0.184311547 | 0.2917349 | 0.631777 | 0.527532297 | 0.701981475 | BVES-AS1 | antisense | 39.74 | 0.028323957 | 0.028323957 | ENSG00000203808 | n.s. |
| ENSG00000127564 | 591.8661804 | -0.169861945 | 0.2542575 | -0.668071 | 0.504088508 | 0.681202008 | PKMYT1 | protein_coding | 62.22 | -0.028320077 | 0.028320077 | ENSG00000127564 | n.s. |
| ENSG00000055483 | 1605.050657 | 0.152859279 | 0.2109054 | 0.724777 | 0.468589046 | 0.6527429 | USP36 | protein_coding | 50.03 | 0.02831838 | 0.02831838 | ENSG00000055483 | n.s. |
| ENSG00000174996 | 222.7468976 | -0.181781207 | 0.2849448 | -0.637952 | 0.523504725 | 0.698601184 | KLC2 | protein_coding | 60.47 | -0.028316183 | 0.028316183 | ENSG00000174996 | n.s. |
| ENSG00000167112 | 741.1698218 | -0.188679577 | 0.3048365 | -0.618953 | 0.535947087 | 0.70817038 | TRUB2 | protein_coding | 51.14 | -0.028275944 | 0.028275944 | ENSG00000167112 | n.s. |
| ENSG00000117682 | 58.57043566 | 0.177441383 | 0.2739771 | 0.64765 | 0.51721112 | 0.693081156 | DHDDS | protein_coding | 45.47 | 0.028251491 | 0.028251491 | ENSG00000117682 | n.s. |

| | | | | | | | | | | | | | |
|---|---|---|---|---|---|---|---|---|---|---|---|---|---|
| ENSG00000186660 | 1383.886574 | -0.175140762 | 0.2681755 | -0.653083 | 0.513703033 | 0.690278796 | ZFP91 | protein_coding | 36.47 | -0.028193366 | 0.028193366 | ENSG00000186660 | n.s. |
| ENSG00000134602 | 527.0057577 | 0.189655637 | 0.3087946 | 0.614181 | 0.539095966 | 0.71017439 | STK26 | protein_coding | 36.62 | 0.028189464 | 0.028189464 | ENSG00000134602 | n.s. |
| ENSG00000227376 | 33.40309296 | 0.18933719 | 0.3078447 | 0.615041 | 0.538527444 | 0.709895627 | FTH1P16 | processed_pseudogene | 47.28 | 0.028174415 | 0.028174415 | ENSG00000227376 | n.s. |
| ENSG00000155876 | 721.4504421 | 0.162401973 | 0.2363206 | 0.68721 | 0.491950111 | 0.671248667 | RRAGA | protein_coding | 52.17 | 0.028114472 | 0.028114472 | ENSG00000155876 | n.s. |
| ENSG00000198833 | 1394.771108 | 0.178504323 | 0.2778325 | 0.642489 | 0.520555734 | 0.695957124 | UBE2J1 | protein_coding | 40.13 | 0.028099707 | 0.028099707 | ENSG00000198833 | n.s. |
| ENSG00000175175 | 823.412027 | -0.18469306 | 0.2951908 | -0.625673 | 0.531529148 | 0.704673427 | PPM1E | protein_coding | 39.61 | -0.028075581 | 0.028075581 | ENSG00000175175 | n.s. |
| ENSG00000105568 | 2480.364529 | -0.173106708 | 0.2638884 | -0.655985 | 0.511834028 | 0.688395012 | PPP2R1A | protein_coding | 48.38 | -0.028071379 | 0.028071379 | ENSG00000105568 | n.s. |
| ENSG00000001630 | 3677.893444 | 0.17096452 | 0.2586023 | 0.66111 | 0.50854186 | 0.685590674 | CYP51A1 | protein_coding | 39.91 | 0.028027085 | 0.028027085 | ENSG00000001630 | n.s. |
| ENSG00000163636 | 1642.007955 | -0.174767746 | 0.2687163 | -0.65038 | 0.515446667 | 0.691731475 | PSMD6 | protein_coding | 39.12 | -0.027973756 | 0.027973756 | ENSG00000163636 | n.s. |
| ENSG00000084234 | 2258.402976 | -0.177575525 | 0.2763378 | -0.642603 | 0.520481739 | 0.695925022 | APLP2 | protein_coding | 43.6 | -0.027957055 | 0.027957055 | ENSG00000084234 | n.s. |
| ENSG00000264538 | 1118.091921 | -0.16546795 | 0.2454744 | -0.674074 | 0.500264083 | 0.677749957 | SUZ12P1 | transcribed_unprocessed_pseud | 45.35 | -0.027952584 | 0.027952584 | ENSG00000264538 | n.s. |
| ENSG00000144827 | 935.6498568 | -0.175335502 | 0.2704902 | -0.648214 | 0.516846674 | 0.692861182 | ABHD10 | protein_coding | 35.84 | -0.027940373 | 0.027940373 | ENSG00000144827 | n.s. |
| ENSG00000171448 | 442.8711946 | -0.179739528 | 0.2823047 | -0.636686 | 0.52432915 | 0.69912711 | ZBTB26 | protein_coding | 39.97 | -0.027939405 | 0.027939405 | ENSG00000171448 | n.s. |
| ENSG00000205336 | 12.48939845 | -0.219347449 | 0.4010446 | -0.54694 | 0.584419794 | 0.745818221 | ADGRG1 | protein_coding | 55.51 | -0.027937629 | 0.027937629 | ENSG00000205336 | n.s. |
| ENSG00000180596 | 1151.140415 | 0.144234352 | 0.1927639 | 0.748244 | 0.45431319 | 0.640548238 | HIST1H2BC | protein_coding | 37.87 | 0.02790187 | 0.02790187 | ENSG00000180596 | n.s. |
| ENSG00000158604 | 1098.363871 | 0.17564984 | 0.2720163 | 0.645733 | 0.518452506 | 0.693944841 | TMED4 | protein_coding | 48.93 | 0.027871247 | 0.027871247 | ENSG00000158604 | n.s. |
| ENSG00000269556 | 70.59663446 | 0.181895067 | 0.289061 | 0.629262 | 0.529177668 | 0.702790513 | TMEM185A | protein_coding | 42.64 | 0.027861615 | 0.027861615 | ENSG00000269556 | n.s. |
| ENSG00000249264 | 67.05032525 | 0.186004591 | 0.3008488 | 0.618266 | 0.536399985 | 0.708435841 | EEF1A1P9 | processed_pseudogene | 47.65 | 0.02784479 | 0.02784479 | ENSG00000249264 | n.s. |
| ENSG00000174437 | 5073.591519 | -0.16240086 | 0.2386786 | -0.680417 | 0.496240707 | 0.674130097 | ATP2A2 | protein_coding | 43.42 | -0.027812168 | 0.027812168 | ENSG00000174437 | n.s. |
| ENSG00000165097 | 434.6968788 | -0.180555643 | 0.2853685 | -0.632711 | 0.526922694 | 0.701397744 | KDM1B | protein_coding | 42.05 | -0.027812003 | 0.027812003 | ENSG00000165097 | n.s. |
| ENSG00000183955 | 1002.477532 | -0.173435672 | 0.2668736 | -0.649879 | 0.515770105 | 0.691817647 | KMT5A | protein_coding | 51.73 | -0.027751158 | 0.027751158 | ENSG00000183955 | n.s. |
| ENSG00000175634 | 452.0734599 | 0.180764919 | 0.286744 | 0.630405 | 0.528429497 | 0.702412663 | RPS6KB2 | protein_coding | 58.79 | 0.027730725 | 0.027730725 | ENSG00000175634 | n.s. |
| ENSG00000155366 | 173.6144065 | 0.188242238 | 0.3092323 | 0.608741 | 0.542696427 | 0.712759396 | RHOC | protein_coding | 57.18 | 0.027682348 | 0.027682348 | ENSG00000155366 | n.s. |
| ENSG00000120727 | 2347.21068 | 0.17331685 | 0.2669984 | 0.649131 | 0.516253919 | 0.692333203 | PAIP2 | protein_coding | 43.58 | 0.027676073 | 0.027676073 | ENSG00000120727 | n.s. |
| ENSG00000128789 | 746.5674354 | -0.18581076 | 0.3019219 | -0.615426 | 0.538273099 | 0.70969473 | PSMG2 | protein_coding | 42.01 | -0.027672503 | 0.027672503 | ENSG00000128789 | n.s. |
| ENSG00000167747 | 2161.384615 | -0.158579706 | 0.2292299 | -0.691793 | 0.489067172 | 0.669348494 | C19orf48 | protein_coding | 54.79 | -0.027648008 | 0.027648008 | ENSG00000167747 | n.s. |
| ENSG00000008294 | 8708.030136 | -0.169960178 | 0.2586812 | -0.657026 | 0.511164365 | 0.687760094 | SPAG9 | protein_coding | 40 | -0.027629241 | 0.027629241 | ENSG00000008294 | n.s. |
| ENSG00000091164 | 1387.479638 | 0.174882518 | 0.2718664 | 0.643267 | 0.520051183 | 0.695508579 | TXNL1 | protein_coding | 38.58 | 0.027578538 | 0.027578538 | ENSG00000091164 | n.s. |
| ENSG00000137124 | 688.7150057 | -0.172522938 | 0.265968 | -0.648661 | 0.516557828 | 0.692645993 | ALDH1B1 | protein_coding | 51.48 | -0.027515455 | 0.027515455 | ENSG00000137124 | n.s. |
| ENSG00000116791 | 611.130018 | 0.18698155 | 0.3074455 | 0.608178 | 0.543069484 | 0.713114819 | CRYZ | protein_coding | 39.61 | 0.027456472 | 0.027456472 | ENSG00000116791 | n.s. |
| ENSG00000177432 | 420.6348853 | 0.173095221 | 0.2685103 | 0.64465 | 0.519153896 | 0.694616539 | NAP1L5 | protein_coding | 45.8 | 0.027393163 | 0.027393163 | ENSG00000177432 | n.s. |
| ENSG00000214753 | 434.046714 | -0.163420748 | 0.2438865 | -0.670069 | 0.502813919 | 0.679803747 | HNRNPUL2 | protein_coding | 46.07 | -0.027392005 | 0.027392005 | ENSG00000214753 | n.s. |

| | | | | | | | | | | | | | |
|---|---|---|---|---|---|---|---|---|---|---|---|---|---|
| ENSG00000128656 | 1601.931724 | -0.171928963 | 0.2655259 | -0.647504 | 0.517306022 | 0.693081156 | CHN1 | protein_coding | 37.22 | -0.027373826 | 0.027373826 | ENSG00000128656 | n.s. |
| ENSG00000101457 | 300.0040407 | -0.184000237 | 0.2992997 | -0.614769 | 0.538707148 | 0.709991464 | DNTTIP1 | protein_coding | 46 | -0.02736946 | 0.02736946 | ENSG00000101457 | n.s. |
| ENSG00000049449 | 2060.748945 | -0.179466779 | 0.2861728 | -0.627127 | 0.530575779 | 0.704079925 | RCN1 | protein_coding | 43.38 | -0.027346796 | 0.027346796 | ENSG00000049449 | n.s. |
| ENSG00000061794 | 1644.114809 | 0.179503644 | 0.2864193 | 0.626716 | 0.530845163 | 0.704207095 | MRPS35 | protein_coding | 38.64 | 0.027338334 | 0.027338334 | ENSG00000061794 | n.s. |
| ENSG00000228300 | 107.4685696 | 0.176903263 | 0.2789644 | 0.634143 | 0.525987583 | 0.700595535 | C19orf24 | protein_coding | 63.69 | 0.027337327 | 0.027337327 | ENSG00000228300 | n.s. |
| ENSG00000100387 | 1180.024033 | 0.191221459 | 0.3214824 | 0.594812 | 0.551969404 | 0.719643226 | RBX1 | protein_coding | 44.92 | 0.02732225 | 0.02732225 | ENSG00000100387 | n.s. |
| ENSG00000111252 | 184.5880185 | -0.161764474 | 0.2403288 | -0.673096 | 0.500885968 | 0.678124789 | SH2B3 | protein_coding | 54.02 | -0.02728811 | 0.02728811 | ENSG00000111252 | n.s. |
| ENSG00000137817 | 2428.363049 | -0.165992658 | 0.2507427 | -0.662004 | 0.507968616 | 0.684982764 | PARP6 | protein_coding | 43.94 | -0.027275973 | 0.027275973 | ENSG00000137817 | n.s. |
| ENSG00000117448 | 511.3846927 | 0.186585873 | 0.3081432 | 0.605517 | 0.544835653 | 0.714356041 | AKR1A1 | protein_coding | 49.1 | 0.027257449 | 0.027257449 | ENSG00000117448 | n.s. |
| ENSG00000163479 | 2719.107472 | 0.178916695 | 0.2855261 | 0.626621 | 0.530907631 | 0.704207095 | SSR2 | protein_coding | 46.99 | 0.027248942 | 0.027248942 | ENSG00000163479 | n.s. |
| ENSG00000128524 | 409.6327819 | 0.190625979 | 0.3203887 | 0.594983 | 0.551854539 | 0.719579658 | ATP6V1F | protein_coding | 52.57 | 0.027244479 | 0.027244479 | ENSG00000128524 | n.s. |
| ENSG00000228253 | 397.541678 | -0.161232943 | 0.2394034 | -0.673478 | 0.500643229 | 0.677945841 | MT-ATP8 | protein_coding | 39.61 | -0.027216927 | 0.027216927 | ENSG00000228253 | n.s. |
| ENSG00000122068 | 1673.450155 | 0.17353263 | 0.2707889 | 0.640841 | 0.521625944 | 0.696986374 | FYTTD1 | protein_coding | 42.18 | 0.027205702 | 0.027205702 | ENSG00000122068 | n.s. |
| ENSG00000167565 | 120.6856033 | 0.173997815 | 0.2722147 | 0.639193 | 0.52269712 | 0.697881857 | SERTAD3 | protein_coding | 55.9 | 0.027181607 | 0.027181607 | ENSG00000167565 | n.s. |
| ENSG00000198890 | 1473.73645 | -0.155275557 | 0.2235083 | -0.694719 | 0.487231124 | 0.668544293 | PRMT6 | protein_coding | 50.31 | -0.027153008 | 0.027153008 | ENSG00000198890 | n.s. |
| ENSG00000171444 | 4615.464277 | -0.161375215 | 0.240134 | -0.672021 | 0.501570074 | 0.678852935 | MCC | protein_coding | 41.55 | -0.027147233 | 0.027147233 | ENSG00000171444 | n.s. |
| ENSG00000051620 | 665.8481407 | 0.189883614 | 0.3190469 | 0.595159 | 0.551737229 | 0.719579658 | HEBP2 | protein_coding | 47.12 | 0.027138379 | 0.027138379 | ENSG00000051620 | n.s. |
| ENSG00000100266 | 795.5712703 | -0.166991449 | 0.2542533 | -0.656792 | 0.511314879 | 0.687890411 | PACSIN2 | protein_coding | 48.59 | -0.027132895 | 0.027132895 | ENSG00000100266 | n.s. |
| ENSG00000241878 | 303.710874 | -0.177473687 | 0.2827391 | -0.627694 | 0.530204344 | 0.703721171 | PISD | protein_coding | 51.17 | -0.027082375 | 0.027082375 | ENSG00000241878 | n.s. |
| ENSG00000207712 | 3.832860848 | -0.298492086 | 0.7136237 | -0.418277 | 0.675744908 | 0.811698417 | MIR627 | miRNA | 35.05 | -0.027044965 | 0.027044965 | ENSG00000207712 | n.s. |
| ENSG00000163900 | 819.6177831 | -0.152876493 | 0.2184761 | -0.69974 | 0.484089633 | 0.665611294 | TMEM41A | protein_coding | 44.99 | -0.027025402 | 0.027025402 | ENSG00000163900 | n.s. |
| ENSG00000168003 | 2529.166015 | 0.15443388 | 0.2224625 | 0.694202 | 0.487555695 | 0.668694533 | SLC3A2 | protein_coding | 48.27 | 0.026990753 | 0.026990753 | ENSG00000168003 | n.s. |
| ENSG00000116786 | 262.3520474 | -0.176168886 | 0.2798434 | -0.629527 | 0.529004322 | 0.702790513 | PLEKHM2 | protein_coding | 52 | -0.026984512 | 0.026984512 | ENSG00000116786 | n.s. |
| ENSG00000023734 | 3676.636813 | 0.175699039 | 0.278706 | 0.63041 | 0.528426361 | 0.702412663 | STRAP | protein_coding | 36.48 | 0.02695358 | 0.02695358 | ENSG00000023734 | n.s. |
| ENSG00000064300 | 915.2437554 | 0.171414273 | 0.2670482 | 0.641885 | 0.520947689 | 0.696347415 | NGFR | protein_coding | 56.74 | 0.026941873 | 0.026941873 | ENSG00000064300 | n.s. |
| ENSG00000146072 | 344.201687 | 0.160797008 | 0.2401014 | 0.669705 | 0.503046105 | 0.679925339 | TNFRSF21 | protein_coding | 41.38 | 0.026939734 | 0.026939734 | ENSG00000146072 | n.s. |
| ENSG00000182809 | 68.41505171 | 0.187220148 | 0.3131353 | 0.597889 | 0.549913992 | 0.717971916 | CRIP2 | protein_coding | 69.57 | 0.026939583 | 0.026939583 | ENSG00000182809 | n.s. |
| ENSG00000157350 | 425.7988966 | -0.155854888 | 0.2272743 | -0.685757 | 0.492866524 | 0.671937241 | ST3GAL2 | protein_coding | 51.87 | -0.026911664 | 0.026911664 | ENSG00000157350 | n.s. |
| ENSG00000089693 | 1409.413292 | 0.172949581 | 0.2715214 | 0.636965 | 0.524147735 | 0.699014264 | MLF2 | protein_coding | 52.27 | 0.026896075 | 0.026896075 | ENSG00000089693 | n.s. |
| ENSG00000231770 | 146.9461923 | 0.154169968 | 0.2230359 | 0.691234 | 0.489418498 | 0.669697666 | TMEM44-AS1 | antisense | 48.38 | 0.026844262 | 0.026844262 | ENSG00000231770 | n.s. |
| ENSG00000137269 | 214.4387355 | 0.165812939 | 0.2531372 | 0.655032 | 0.512447154 | 0.688892052 | LRRC1 | protein_coding | 39.72 | 0.026836629 | 0.026836629 | ENSG00000137269 | n.s. |
| ENSG00000282855 | 15.90099605 | 0.205235726 | 0.3685166 | 0.556924 | 0.577579411 | 0.740618727 | | processed_pseudogene | 44.48 | 0.026763829 | 0.026763829 | ENSG00000282855 | n.s. |

| | | | | | | | | | | | | | |
|---|---|---|---|---|---|---|---|---|---|---|---|---|---|
| ENSG00000206941 | 10.38895037 | 0.211500597 | 0.3887479 | 0.544056 | 0.586403015 | 0.747458499 | SNORD15A | snoRNA | 47.3 | 0.026736407 | 0.026736407 | ENSG00000206941 | n.s. |
| ENSG00000167842 | 1453.404837 | -0.15856347 | 0.2357552 | -0.672577 | 0.501216532 | 0.678440383 | MIS12 | protein_coding | 44.08 | -0.026716091 | 0.026716091 | ENSG00000167842 | n.s. |
| ENSG00000152242 | 1020.162746 | -0.166921336 | 0.2567816 | -0.650052 | 0.515658796 | 0.691762009 | C18orf25 | protein_coding | 40.58 | -0.02671464 | 0.02671464 | ENSG00000152242 | n.s. |
| ENSG00000160131 | 2027.586347 | 0.168203663 | 0.2603845 | 0.645982 | 0.518291098 | 0.693862207 | VMA21 | protein_coding | 41.5 | 0.026698423 | 0.026698423 | ENSG00000160131 | n.s. |
| ENSG00000179195 | 4090.398094 | -0.170471255 | 0.266261 | -0.640241 | 0.522015753 | 0.697306469 | ZNF664 | protein_coding | 44.11 | -0.026691759 | 0.026691759 | ENSG00000179195 | n.s. |
| ENSG00000127481 | 2746.225934 | -0.160474986 | 0.2408251 | -0.666355 | 0.505184344 | 0.681822569 | UBR4 | protein_coding | 43.74 | -0.026691584 | 0.026691584 | ENSG00000127481 | n.s. |
| ENSG00000166411 | 617.8796356 | 0.16897386 | 0.2624858 | 0.643745 | 0.519740907 | 0.695335125 | IDH3A | protein_coding | 44.46 | 0.026665061 | 0.026665061 | ENSG00000166411 | n.s. |
| ENSG00000249962 | 74.38768664 | 0.167159528 | 0.2578304 | 0.648331 | 0.516770602 | 0.692825911 | | processed_pseudogene | 75.2 | 0.026641197 | 0.026641197 | ENSG00000249962 | n.s. |
| ENSG00000134901 | 698.3074539 | -0.177994021 | 0.2881417 | -0.617731 | 0.536752754 | 0.708562491 | KDELC1 | protein_coding | 40.94 | -0.026631793 | 0.026631793 | ENSG00000134901 | n.s. |
| ENSG00000088247 | 1492.8521 | 0.152888109 | 0.2217272 | 0.689533 | 0.490488156 | 0.670128457 | KHSRP | protein_coding | 59.03 | 0.026578365 | 0.026578365 | ENSG00000088247 | n.s. |
| ENSG00000239268 | 33.97267299 | -0.176303934 | 0.2833572 | -0.622197 | 0.533812472 | 0.706758373 | | lincRNA | 36.82 | -0.026574122 | 0.026574122 | ENSG00000239268 | n.s. |
| ENSG00000182362 | 210.7060556 | 0.183164323 | 0.3042355 | 0.602048 | 0.547142328 | 0.716031829 | YBEY | protein_coding | 53.1 | 0.026571222 | 0.026571222 | ENSG00000182362 | n.s. |
| ENSG00000251022 | 3470.250975 | 0.154453796 | 0.2264045 | 0.682203 | 0.495110672 | 0.673449188 | THAP9-AS1 | antisense | 39.31 | 0.02651897 | 0.02651897 | ENSG00000251022 | n.s. |
| ENSG00000152484 | 1169.408489 | -0.164504445 | 0.2519056 | -0.65304 | 0.513730438 | 0.690278796 | USP12 | protein_coding | 38.86 | -0.02648118 | 0.02648118 | ENSG00000152484 | n.s. |
| ENSG00000207359 | 8.742638608 | 0.219076829 | 0.4150154 | 0.527876 | 0.597585098 | 0.757068 | RNU6-925P | snRNA | 36.54 | 0.026478745 | 0.026478745 | ENSG00000207359 | n.s. |
| ENSG00000184182 | 356.8306618 | -0.173811956 | 0.2773911 | -0.626595 | 0.530924504 | 0.704207095 | UBE2F | protein_coding | 44.85 | -0.026471492 | 0.026471492 | ENSG00000184182 | n.s. |
| ENSG00000204524 | 278.3185013 | -0.164985297 | 0.2532276 | -0.65153 | 0.514704556 | 0.691120796 | ZNF805 | protein_coding | 46.41 | -0.026471237 | 0.026471237 | ENSG00000204524 | n.s. |
| ENSG00000134057 | 2867.177418 | -0.179828934 | 0.2955396 | -0.608477 | 0.542871426 | 0.712921983 | CCNB1 | protein_coding | 43.63 | -0.026427299 | 0.026427299 | ENSG00000134057 | n.s. |
| ENSG00000008282 | 383.4929048 | 0.172211199 | 0.2730424 | 0.630712 | 0.528228584 | 0.702412663 | SYPL1 | protein_coding | 38.18 | 0.026418518 | 0.026418518 | ENSG00000008282 | n.s. |
| ENSG00000245870 | 7128.429978 | 0.166045894 | 0.2568908 | 0.646368 | 0.518041357 | 0.693594558 | LINC00682 | lincRNA | 51.34 | 0.02638375 | 0.02638375 | ENSG00000245870 | n.s. |
| ENSG00000133065 | 411.8972807 | -0.153715236 | 0.2255257 | -0.681586 | 0.495500534 | 0.673659272 | SLC41A1 | protein_coding | 50.84 | -0.026371341 | 0.026371341 | ENSG00000133065 | n.s. |
| ENSG00000161395 | 53.89036153 | -0.180300213 | 0.297465 | -0.606122 | 0.544433422 | 0.714097647 | PGAP3 | protein_coding | 51.61 | -0.026367536 | 0.026367536 | ENSG00000161395 | n.s. |
| ENSG00000042753 | 487.0165161 | 0.180693975 | 0.2988464 | 0.604638 | 0.545419304 | 0.714852017 | AP2S1 | protein_coding | 52.87 | 0.026342264 | 0.026342264 | ENSG00000042753 | n.s. |
| ENSG00000224568 | 55.89351312 | 0.166325388 | 0.2579003 | 0.644921 | 0.518978243 | 0.694515002 | LINC01886 | lincRNA | 38.69 | 0.026332364 | 0.026332364 | ENSG00000224568 | n.s. |
| ENSG00000152127 | 2097.156919 | -0.169357967 | 0.2661627 | -0.636295 | 0.524584225 | 0.699131025 | MGAT5 | protein_coding | 42.88 | -0.026325244 | 0.026325244 | ENSG00000152127 | n.s. |
| ENSG00000086232 | 1597.640655 | -0.173038077 | 0.2765185 | -0.625774 | 0.531463289 | 0.704653211 | EIF2AK1 | protein_coding | 44.7 | -0.026306038 | 0.026306038 | ENSG00000086232 | n.s. |
| ENSG00000169251 | 2341.004668 | 0.171357917 | 0.2718431 | 0.630356 | 0.528461728 | 0.702412663 | NMD3 | protein_coding | 37.99 | 0.026287618 | 0.026287618 | ENSG00000169251 | n.s. |
| ENSG00000214413 | 341.5685102 | -0.183918638 | 0.3090157 | -0.595176 | 0.551726054 | 0.719579658 | BBIP1 | protein_coding | 38.64 | -0.026285858 | 0.026285858 | ENSG00000214413 | n.s. |
| ENSG00000108852 | 368.3543248 | -0.167567939 | 0.2614573 | -0.6409 | 0.521587733 | 0.696986374 | MPP2 | protein_coding | 52.29 | -0.026270583 | 0.026270583 | ENSG00000108852 | n.s. |
| ENSG00000271551 | 82.74299556 | 0.16278235 | 0.2489397 | 0.653903 | 0.513174514 | 0.689664938 | | antisense | 61.08 | 0.026266862 | 0.026266862 | ENSG00000271551 | n.s. |
| ENSG00000134001 | 2998.942241 | -0.175998676 | 0.2854278 | -0.616614 | 0.537489562 | 0.709311771 | EIF2S1 | protein_coding | 37.7 | -0.026252461 | 0.026252461 | ENSG00000134001 | n.s. |
| ENSG00000130733 | 128.5746266 | 0.188829451 | 0.3240023 | 0.582803 | 0.560026033 | 0.726406042 | YIPF2 | protein_coding | 58.8 | 0.026213409 | 0.026213409 | ENSG00000130733 | n.s. |

| | | | | | | | | | | | | | |
|---|---|---|---|---|---|---|---|---|---|---|---|---|---|
| ENSG00000104835 | 346.2405015 | 0.165637633 | 0.2569116 | 0.644726 | 0.519104616 | 0.694616539 | SARS2 | protein_coding | 53.66 | 0.026212964 | 0.026212964 | ENSG00000104835 | n.s. |
| ENSG00000177879 | 992.6952356 | 0.178724199 | 0.2943681 | 0.607145 | 0.543754448 | 0.713475938 | AP3S1 | protein_coding | 36.62 | 0.026204662 | 0.026204662 | ENSG00000177879 | n.s. |
| ENSG00000140157 | 1182.957644 | -0.174128579 | 0.2804905 | -0.6208 | 0.534730973 | 0.707216152 | NIPA2 | protein_coding | 43.73 | -0.026197267 | 0.026197267 | ENSG00000140157 | n.s. |
| ENSG00000213995 | 848.5856944 | 0.167240942 | 0.2614336 | 0.639707 | 0.522363059 | 0.697594917 | NAXD | protein_coding | 50.13 | 0.02615593 | 0.02615593 | ENSG00000213995 | n.s. |
| ENSG00000125871 | 947.9295538 | -0.153965068 | 0.2275833 | -0.676522 | 0.498709304 | 0.676559451 | MGME1 | protein_coding | 42.82 | -0.026126954 | 0.026126954 | ENSG00000125871 | n.s. |
| ENSG00000111859 | 288.1614625 | -0.159899129 | 0.2427797 | -0.658618 | 0.510141038 | 0.687123662 | NEDD9 | protein_coding | 43.09 | -0.026057977 | 0.026057977 | ENSG00000111859 | n.s. |
| ENSG00000276180 | 924.7386192 | -0.154116697 | 0.2284518 | -0.674614 | 0.499921282 | 0.677609425 | HIST1H4I | protein_coding | 46.75 | -0.026048891 | 0.026048891 | ENSG00000276180 | n.s. |
| ENSG00000092096 | 275.0481074 | 0.170024807 | 0.2702015 | 0.629252 | 0.529184191 | 0.702790513 | SLC22A17 | protein_coding | 57.86 | 0.026043398 | 0.026043398 | ENSG00000092096 | n.s. |
| ENSG00000126746 | 373.3169231 | 0.15630114 | 0.2343629 | 0.666919 | 0.504823721 | 0.681666247 | ZNF384 | protein_coding | 46.35 | 0.026012919 | 0.026012919 | ENSG00000126746 | n.s. |
| ENSG00000091483 | 2676.947883 | 0.17636222 | 0.2893278 | 0.609558 | 0.54215435 | 0.712461375 | FH | protein_coding | 36.87 | 0.02596734 | 0.02596734 | ENSG00000091483 | n.s. |
| ENSG00000196368 | 816.405732 | 0.174793151 | 0.2847624 | 0.613821 | 0.539333561 | 0.710420168 | NUDT11 | protein_coding | 39.05 | 0.025954112 | 0.025954112 | ENSG00000196368 | n.s. |
| ENSG00000105063 | 593.6293367 | 0.154636514 | 0.230747 | 0.670156 | 0.502758281 | 0.679803747 | PPP6R1 | protein_coding | 56.47 | 0.025919623 | 0.025919623 | ENSG00000105063 | n.s. |
| ENSG00000114331 | 1667.744493 | -0.172894598 | 0.2797171 | -0.618105 | 0.536505919 | 0.708435841 | ACAP2 | protein_coding | 38.17 | -0.025882231 | 0.025882231 | ENSG00000114331 | n.s. |
| ENSG00000088682 | 532.1262286 | -0.18405354 | 0.3132221 | -0.587614 | 0.556791729 | 0.723491483 | COQ9 | protein_coding | 45.83 | -0.025871776 | 0.025871776 | ENSG00000088682 | n.s. |
| ENSG00000160957 | 780.5301218 | 0.159315406 | 0.2427108 | 0.6564 | 0.511566641 | 0.688101859 | RECQL4 | protein_coding | 65.51 | 0.025864421 | 0.025864421 | ENSG00000160957 | n.s. |
| ENSG00000248098 | 367.1030507 | 0.180093654 | 0.3017429 | 0.596845 | 0.550611098 | 0.718545161 | BCKDHA | protein_coding | 52.32 | 0.025851711 | 0.025851711 | ENSG00000248098 | n.s. |
| ENSG00000104695 | 782.0404535 | -0.166277428 | 0.2612463 | -0.636478 | 0.524465167 | 0.699131025 | PPP2CB | protein_coding | 41.95 | -0.0258464 | 0.0258464 | ENSG00000104695 | n.s. |
| ENSG00000081870 | 856.8979741 | -0.178690614 | 0.2978762 | -0.599882 | 0.548584815 | 0.717229859 | HSPB11 | protein_coding | 40.17 | -0.025792496 | 0.025792496 | ENSG00000081870 | n.s. |
| ENSG00000204852 | 251.5522483 | -0.175104934 | 0.287182 | -0.609735 | 0.542037266 | 0.712431309 | TCTN1 | protein_coding | 44.42 | -0.025785428 | 0.025785428 | ENSG00000204852 | n.s. |
| ENSG00000125962 | 639.994702 | 0.16142952 | 0.2486241 | 0.649292 | 0.516149928 | 0.692260423 | ARMCX5 | protein_coding | 45.27 | 0.025785217 | 0.025785217 | ENSG00000125962 | n.s. |
| ENSG00000134697 | 1541.985498 | -0.162413164 | 0.2514639 | -0.645871 | 0.518363026 | 0.693891781 | GNL2 | protein_coding | 42.77 | -0.02577631 | 0.02577631 | ENSG00000134697 | n.s. |
| ENSG00000132356 | 1321.621145 | -0.169228106 | 0.2699409 | -0.626908 | 0.530719604 | 0.704203664 | PRKAA1 | protein_coding | 37.74 | -0.025773731 | 0.025773731 | ENSG00000132356 | n.s. |
| ENSG00000143740 | 544.0662078 | -0.180298583 | 0.3029224 | -0.595197 | 0.551711566 | 0.719579658 | SNAP47 | protein_coding | 53.99 | -0.025768476 | 0.025768476 | ENSG00000143740 | n.s. |
| ENSG00000184517 | 684.826814 | -0.16675455 | 0.2634623 | -0.632935 | 0.526775995 | 0.701310211 | ZFP1 | protein_coding | 42.26 | -0.025695182 | 0.025695182 | ENSG00000184517 | n.s. |
| ENSG00000204065 | 205.899621 | -0.168706924 | 0.2692954 | -0.626475 | 0.531003214 | 0.704244403 | TCEAL5 | protein_coding | 50.63 | -0.025690116 | 0.025690116 | ENSG00000204065 | n.s. |
| ENSG00000114867 | 7555.634251 | 0.153956922 | 0.2310308 | 0.666391 | 0.505160967 | 0.681822569 | EIF4G1 | protein_coding | 52.77 | 0.025607444 | 0.025607444 | ENSG00000114867 | n.s. |
| ENSG00000134049 | 1204.966689 | -0.180612712 | 0.3063316 | -0.589599 | 0.555459674 | 0.722637362 | IER3IP1 | protein_coding | 38.68 | -0.025480767 | 0.025480767 | ENSG00000134049 | n.s. |
| ENSG00000099219 | 179.3557497 | 0.16029451 | 0.247896 | 0.64662 | 0.517877954 | 0.693509162 | ERMP1 | protein_coding | 39 | 0.025478459 | 0.025478459 | ENSG00000099219 | n.s. |
| ENSG00000175197 | 254.3961783 | 0.16674885 | 0.2653692 | 0.628366 | 0.529764489 | 0.703405586 | DDIT3 | protein_coding | 49.44 | 0.025478255 | 0.025478255 | ENSG00000175197 | n.s. |
| ENSG00000000419 | 1158.390196 | 0.176938939 | 0.2957705 | 0.59823 | 0.549686167 | 0.717943762 | DPM1 | protein_coding | 39.85 | 0.025463207 | 0.025463207 | ENSG00000000419 | n.s. |
| ENSG00000064601 | 86.04745651 | 0.173471514 | 0.2854156 | 0.607786 | 0.543329646 | 0.713251896 | CTSA | protein_coding | 56.04 | 0.02545817 | 0.02545817 | ENSG00000064601 | n.s. |
| ENSG00000134186 | 5142.148106 | 0.148009135 | 0.21664 | 0.683203 | 0.494478409 | 0.673181078 | PRPF38B | protein_coding | 37.93 | 0.025438049 | 0.025438049 | ENSG00000134186 | n.s. |

| | | | | | | | | | | | | | |
|---|---|---|---|---|---|---|---|---|---|---|---|---|---|
| ENSG00000159200 | 262.8301654 | -0.17654668 | 0.2950278 | -0.598407 | 0.54956847 | 0.717857379 | RCAN1 | protein_coding | 44.24 | -0.025415983 | 0.025415983 | ENSG00000159200 | n.s. |
| ENSG00000127990 | 699.7416985 | -0.171726591 | 0.2807263 | -0.611723 | 0.540721348 | 0.711238859 | SGCE | protein_coding | 35.25 | -0.025412878 | 0.025412878 | ENSG00000127990 | n.s. |
| ENSG00000108592 | 1870.088176 | -0.159327789 | 0.2456427 | -0.648616 | 0.516586654 | 0.692645993 | FTSJ3 | protein_coding | 50.46 | -0.025410978 | 0.025410978 | ENSG00000108592 | n.s. |
| ENSG00000119318 | 4439.681384 | 0.16879941 | 0.2717838 | 0.62108 | 0.534547201 | 0.707125955 | RAD23B | protein_coding | 38 | 0.025404855 | 0.025404855 | ENSG00000119318 | n.s. |
| ENSG00000271122 | 247.3113141 | 0.144999518 | 0.2086351 | 0.694991 | 0.487060955 | 0.668442551 | | antisense | 46 | 0.025365623 | 0.025365623 | ENSG00000271122 | n.s. |
| ENSG00000131370 | 189.3604339 | -0.170223331 | 0.2765547 | -0.615514 | 0.538215284 | 0.70969473 | SH3BP5 | protein_coding | 46.74 | -0.025351092 | 0.025351092 | ENSG00000131370 | n.s. |
| ENSG00000242588 | 76.79593066 | -0.159304518 | 0.2461542 | -0.647174 | 0.51751951 | 0.693295885 | | processed_transcript | 41.32 | -0.025342382 | 0.025342382 | ENSG00000242588 | n.s. |
| ENSG00000154473 | 7945.575535 | 0.162728999 | 0.255197 | 0.63766 | 0.523694905 | 0.698698068 | BUB3 | protein_coding | 39.33 | 0.025338607 | 0.025338607 | ENSG00000154473 | n.s. |
| ENSG00000185305 | 99.57921468 | 0.170302314 | 0.2770497 | 0.6147 | 0.538753129 | 0.709991464 | ARL15 | protein_coding | 38.5 | 0.025331937 | 0.025331937 | ENSG00000185305 | n.s. |
| ENSG00000155508 | 1055.020795 | 0.16823518 | 0.2708423 | 0.621155 | 0.534497371 | 0.707125955 | CNOT8 | protein_coding | 44.08 | 0.025319936 | 0.025319936 | ENSG00000155508 | n.s. |
| ENSG00000101391 | 523.3030273 | -0.161744327 | 0.252914 | -0.639523 | 0.522482675 | 0.697662442 | CDK5RAP1 | protein_coding | 44.4 | -0.025289479 | 0.025289479 | ENSG00000101391 | n.s. |
| ENSG00000134184 | 377.4123845 | 0.163360616 | 0.2573846 | 0.634695 | 0.525627662 | 0.700250114 | GSTM1 | protein_coding | 46.17 | 0.025279535 | 0.025279535 | ENSG00000134184 | n.s. |
| ENSG00000066926 | 578.7218786 | -0.174778884 | 0.2909954 | -0.600624 | 0.548090279 | 0.716953695 | FECH | protein_coding | 42.66 | -0.025257103 | 0.025257103 | ENSG00000066926 | n.s. |
| ENSG00000170430 | 283.8387051 | -0.170655319 | 0.2792048 | -0.611219 | 0.541054443 | 0.71147535 | MGMT | protein_coding | 45.49 | -0.025229707 | 0.025229707 | ENSG00000170430 | n.s. |
| ENSG00000091039 | 4320.528382 | -0.168923386 | 0.273907 | -0.616718 | 0.537420686 | 0.709311771 | OSBPL8 | protein_coding | 36.48 | -0.025197091 | 0.025197091 | ENSG00000091039 | n.s. |
| ENSG00000118900 | 1215.643925 | -0.154452024 | 0.234705 | -0.658069 | 0.510493942 | 0.687123662 | UBN1 | protein_coding | 47.93 | -0.025170289 | 0.025170289 | ENSG00000118900 | n.s. |
| ENSG00000163950 | 1692.435471 | 0.172086545 | 0.2839864 | 0.605968 | 0.544536316 | 0.714098062 | SLBP | protein_coding | 47.01 | 0.025166306 | 0.025166306 | ENSG00000163950 | n.s. |
| ENSG00000166377 | 187.1766883 | -0.174353905 | 0.2909741 | -0.599208 | 0.549034442 | 0.717429051 | ATP9B | protein_coding | 41.82 | -0.025145502 | 0.025145502 | ENSG00000166377 | n.s. |
| ENSG00000253251 | 307.4844658 | 0.161751941 | 0.2542414 | 0.636214 | 0.524637 | 0.699131025 | | protein_coding | 38.83 | 0.025142952 | 0.025142952 | ENSG00000253251 | n.s. |
| ENSG00000278053 | 1506.799214 | -0.166832659 | 0.2682057 | -0.622033 | 0.533920467 | 0.706834141 | DDX52 | protein_coding | 39.57 | -0.025138759 | 0.025138759 | ENSG00000278053 | n.s. |
| ENSG00000163565 | 51.61305816 | 0.165865624 | 0.2657878 | 0.624053 | 0.53259289 | 0.705747674 | IFI16 | protein_coding | 38.34 | 0.025103852 | 0.025103852 | ENSG00000163565 | n.s. |
| ENSG00000120690 | 117.7870412 | 0.170869461 | 0.28123 | 0.607579 | 0.54346673 | 0.713251896 | ELF1 | protein_coding | 38.8 | 0.025076301 | 0.025076301 | ENSG00000120690 | n.s. |
| ENSG00000241939 | 26.89382343 | -0.191692707 | 0.3431154 | -0.558683 | 0.576378152 | 0.739970725 | RN7SL517P | misc_RNA | 51.48 | -0.025070619 | 0.025070619 | ENSG00000241939 | n.s. |
| ENSG00000134905 | 744.4124624 | -0.163988576 | 0.2609244 | -0.628491 | 0.529682434 | 0.703363713 | CARS2 | protein_coding | 48.05 | -0.025060741 | 0.025060741 | ENSG00000134905 | n.s. |
| ENSG00000129351 | 7871.83073 | -0.163342681 | 0.2590256 | -0.630605 | 0.528299155 | 0.702412663 | ILF3 | protein_coding | 50.9 | -0.02505802 | 0.02505802 | ENSG00000129351 | n.s. |
| ENSG00000112208 | 911.2399549 | 0.159285649 | 0.2481895 | 0.64179 | 0.5210093 | 0.696362914 | BAG2 | protein_coding | 38.77 | 0.025034028 | 0.025034028 | ENSG00000112208 | n.s. |
| ENSG00000110172 | 2447.670611 | 0.153961153 | 0.2342911 | 0.657136 | 0.511093455 | 0.687731146 | CHORDC1 | protein_coding | 35.9 | 0.025031205 | 0.025031205 | ENSG00000110172 | n.s. |
| ENSG00000126001 | 82.90342865 | 0.15847082 | 0.2463252 | 0.64334 | 0.520003681 | 0.695508579 | CEP250 | protein_coding | 47.9 | 0.024990453 | 0.024990453 | ENSG00000126001 | n.s. |
| ENSG00000122692 | 1689.723733 | -0.162050347 | 0.2559084 | -0.633236 | 0.526579706 | 0.701182981 | SMU1 | protein_coding | 41.17 | -0.024983081 | 0.024983081 | ENSG00000122692 | n.s. |
| ENSG00000069248 | 3099.359085 | -0.167277125 | 0.2713066 | -0.616561 | 0.537524132 | 0.709311771 | NUP133 | protein_coding | 40.25 | -0.02495153 | 0.02495153 | ENSG00000069248 | n.s. |
| ENSG00000199248 | 11.50042408 | -0.207865456 | 0.396039 | -0.524861 | 0.599679746 | 0.758568275 | RNU6-28P | snRNA | 41.12 | -0.024944961 | 0.024944961 | ENSG00000199248 | n.s. |
| ENSG00000094631 | 683.487799 | 0.166458155 | 0.2690994 | 0.618575 | 0.536196423 | 0.708365385 | HDAC6 | protein_coding | 50.55 | 0.024925888 | 0.024925888 | ENSG00000094631 | n.s. |

| | | | | | | | | | | | | | |
|---|---|---|---|---|---|---|---|---|---|---|---|---|---|
| ENSG00000111144 | 2264.229775 | 0.171900201 | 0.2856312 | 0.601826 | 0.547290205 | 0.716158038 | LTA4H | protein_coding | 39.1 | 0.024924004 | 0.024924004 | ENSG00000111144 | n.s. |
| ENSG00000204237 | 470.5464973 | -0.154734668 | 0.2369868 | -0.652925 | 0.513804531 | 0.690311732 | OXLD1 | protein_coding | 62.12 | -0.024905279 | 0.024905279 | ENSG00000204237 | n.s. |
| ENSG00000196924 | 3744.262134 | -0.149466702 | 0.2242936 | -0.666389 | 0.505162726 | 0.681822569 | FLNA | protein_coding | 62.28 | -0.024860591 | 0.024860591 | ENSG00000196924 | n.s. |
| ENSG00000072803 | 1459.536075 | -0.167725574 | 0.2738188 | -0.612542 | 0.540179059 | 0.710861345 | FBXW11 | protein_coding | 39.72 | -0.024859463 | 0.024859463 | ENSG00000072803 | n.s. |
| ENSG00000159322 | 1423.548412 | -0.167487156 | 0.2729991 | -0.613508 | 0.539540433 | 0.710625433 | ADPGK | protein_coding | 43.53 | -0.024848269 | 0.024848269 | ENSG00000159322 | n.s. |
| ENSG00000160949 | 690.3851907 | 0.145309384 | 0.2138879 | 0.679372 | 0.496902427 | 0.67472188 | TONSL | protein_coding | 62.9 | 0.024829771 | 0.024829771 | ENSG00000160949 | n.s. |
| ENSG00000143256 | 910.2576108 | 0.165661893 | 0.2679566 | 0.618241 | 0.536416207 | 0.708435841 | PFDN2 | protein_coding | 44.14 | 0.024799498 | 0.024799498 | ENSG00000143256 | n.s. |
| ENSG00000100106 | 266.5180978 | 0.161800742 | 0.2568303 | 0.629991 | 0.528700533 | 0.702662993 | TRIOBP | protein_coding | 54.35 | 0.024796436 | 0.024796436 | ENSG00000100106 | n.s. |
| ENSG00000136261 | 2509.933118 | 0.168902727 | 0.2779475 | 0.607679 | 0.543400638 | 0.713251896 | BZW2 | protein_coding | 37.77 | 0.024787668 | 0.024787668 | ENSG00000136261 | n.s. |
| ENSG00000149809 | 107.2209815 | 0.178947682 | 0.3077217 | 0.581524 | 0.560887036 | 0.726981028 | TM7SF2 | protein_coding | 60.18 | 0.024780124 | 0.024780124 | ENSG00000149809 | n.s. |
| ENSG00000126804 | 1070.939797 | 0.168850882 | 0.2778957 | 0.607605 | 0.54344939 | 0.713251896 | ZBTB1 | protein_coding | 39.55 | 0.02478006 | 0.02478006 | ENSG00000126804 | n.s. |
| ENSG00000172301 | 772.3105263 | 0.168319778 | 0.2762937 | 0.609206 | 0.542387875 | 0.712570675 | COPRS | protein_coding | 49.04 | 0.024771968 | 0.024771968 | ENSG00000172301 | n.s. |
| ENSG00000134575 | 231.7011656 | -0.150365541 | 0.2268267 | -0.662909 | 0.507388538 | 0.684399401 | ACP2 | protein_coding | 52.75 | -0.024763758 | 0.024763758 | ENSG00000134575 | n.s. |
| ENSG00000177479 | 1691.645138 | -0.166344795 | 0.270682 | -0.61454 | 0.538858801 | 0.709999258 | ARIH2 | protein_coding | 44.51 | -0.024742475 | 0.024742475 | ENSG00000177479 | n.s. |
| ENSG00000160218 | 445.8126262 | -0.156878791 | 0.2438896 | -0.643237 | 0.520070374 | 0.695508579 | TRAPPC10 | protein_coding | 46.02 | -0.024739395 | 0.024739395 | ENSG00000160218 | n.s. |
| ENSG00000188295 | 1000.90382 | 0.144160101 | 0.2114012 | 0.681927 | 0.495285281 | 0.673620881 | ZNF669 | protein_coding | 43.07 | 0.024735633 | 0.024735633 | ENSG00000188295 | n.s. |
| ENSG00000103145 | 98.5025317 | 0.171292673 | 0.2854511 | 0.600077 | 0.548454782 | 0.717210099 | HCFC1R1 | protein_coding | 63.71 | 0.024726715 | 0.024726715 | ENSG00000103145 | n.s. |
| ENSG00000095319 | 1608.798466 | -0.153968056 | 0.2365052 | -0.651014 | 0.515037756 | 0.691301521 | NUP188 | protein_coding | 46.07 | -0.024686081 | 0.024686081 | ENSG00000095319 | n.s. |
| ENSG00000182768 | 6157.160013 | 0.17500672 | 0.2970438 | 0.589161 | 0.555753081 | 0.722859016 | NGRN | protein_coding | 45.25 | 0.024666567 | 0.024666567 | ENSG00000182768 | n.s. |
| ENSG00000204899 | 1178.20822 | -0.179680496 | 0.3109849 | -0.577779 | 0.563413407 | 0.72903391 | MZT1 | protein_coding | 38.9 | -0.024661556 | 0.024661556 | ENSG00000204899 | n.s. |
| ENSG00000202071 | 4.68262325 | -0.247701273 | 0.5442997 | -0.455082 | 0.649049896 | 0.795346039 | | misc_RNA | 39.22 | -0.024632375 | 0.024632375 | ENSG00000202071 | n.s. |
| ENSG00000184162 | 351.5341674 | -0.162711599 | 0.260802 | -0.623889 | 0.532700282 | 0.70575564 | NR2C2AP | protein_coding | 56.79 | -0.024625691 | 0.024625691 | ENSG00000184162 | n.s. |
| ENSG00000130159 | 504.2687496 | 0.174163957 | 0.2949354 | 0.590516 | 0.554844972 | 0.722175054 | ECSIT | protein_coding | 50.87 | 0.024619385 | 0.024619385 | ENSG00000130159 | n.s. |
| ENSG00000144659 | 673.0884609 | 0.167096379 | 0.2742651 | 0.609251 | 0.542357838 | 0.712570675 | SLC25A38 | protein_coding | 46.41 | 0.024591917 | 0.024591917 | ENSG00000144659 | n.s. |
| ENSG00000149100 | 4168.548721 | -0.166867413 | 0.2739173 | -0.609189 | 0.542399197 | 0.712570675 | EIF3M | protein_coding | 37.78 | -0.02455822 | 0.02455822 | ENSG00000149100 | n.s. |
| ENSG00000160799 | 561.1084054 | 0.155076507 | 0.2404044 | 0.645065 | 0.518884888 | 0.69445682 | CCDC12 | protein_coding | 51.44 | 0.024557102 | 0.024557102 | ENSG00000160799 | n.s. |
| ENSG00000084444 | 704.956808 | 0.151321567 | 0.2310014 | 0.655068 | 0.512424071 | 0.688892052 | FAM234B | protein_coding | 43.23 | 0.024491218 | 0.024491218 | ENSG00000084444 | n.s. |
| ENSG00000112406 | 484.4647052 | -0.15701216 | 0.2463461 | -0.637364 | 0.523887664 | 0.698801298 | HECA | protein_coding | 41.79 | -0.024438361 | 0.024438361 | ENSG00000112406 | n.s. |
| ENSG00000169021 | 2131.48349 | 0.162393719 | 0.2617892 | 0.620322 | 0.535045562 | 0.707314789 | UQCRFS1 | protein_coding | 42.39 | 0.024421947 | 0.024421947 | ENSG00000169021 | n.s. |
| ENSG00000105220 | 2049.891117 | 0.159392217 | 0.2531995 | 0.629512 | 0.529013666 | 0.702790513 | GPI | protein_coding | 52.45 | 0.024414761 | 0.024414761 | ENSG00000105220 | n.s. |
| ENSG00000172009 | 659.8214205 | -0.164566061 | 0.2684379 | -0.613051 | 0.539842753 | 0.710687467 | THOP1 | protein_coding | 59.11 | -0.024408659 | 0.024408659 | ENSG00000172009 | n.s. |
| ENSG00000115841 | 377.6753749 | -0.167893544 | 0.2782429 | -0.603406 | 0.546238432 | 0.715529532 | RMDN2 | protein_coding | 39.02 | -0.024407094 | 0.024407094 | ENSG00000115841 | n.s. |

| | | | | | | | | | | | | | |
|---|---|---|---|---|---|---|---|---|---|---|---|---|---|
| ENSG00000166971 | 100.1502226 | -0.161247289 | 0.2584984 | -0.623784 | 0.532769209 | 0.705779801 | AKTIP | protein_coding | 45.45 | -0.024401677 | 0.024401677 | ENSG00000166971 | n.s. |
| ENSG00000105135 | 435.2957389 | 0.174336039 | 0.2978465 | 0.585322 | 0.558331403 | 0.72474179 | ILVBL | protein_coding | 57.25 | 0.024375089 | 0.024375089 | ENSG00000105135 | n.s. |
| ENSG00000085719 | 1158.767017 | -0.177432703 | 0.3069308 | -0.578087 | 0.563205451 | 0.728832558 | CPNE3 | protein_coding | 37.62 | -0.024374327 | 0.024374327 | ENSG00000085719 | n.s. |
| ENSG00000200376 | 7.457585237 | -0.222945892 | 0.4546777 | -0.490338 | 0.62389452 | 0.777592284 | RNU5E-10P | snRNA | 36.44 | -0.024356406 | 0.024356406 | ENSG00000200376 | n.s. |
| ENSG00000124762 | 642.4523796 | 0.154236265 | 0.239668 | 0.643541 | 0.519872921 | 0.695444916 | CDKN1A | protein_coding | 54.4 | 0.024328806 | 0.024328806 | ENSG00000124762 | n.s. |
| ENSG00000106400 | 345.3887127 | 0.172021765 | 0.2914194 | 0.590289 | 0.554996625 | 0.722237406 | ZNHIT1 | protein_coding | 54.13 | 0.02431012 | 0.02431012 | ENSG00000106400 | n.s. |
| ENSG00000099204 | 1625.081966 | -0.156607619 | 0.246477 | -0.635384 | 0.525177842 | 0.699784774 | ABLIM1 | protein_coding | 42.59 | -0.024279742 | 0.024279742 | ENSG00000099204 | n.s. |
| ENSG00000169992 | 192.7392574 | 0.159263054 | 0.2538812 | 0.627313 | 0.530453905 | 0.703985294 | NLGN2 | protein_coding | 59.54 | 0.024277489 | 0.024277489 | ENSG00000169992 | n.s. |
| ENSG00000163714 | 9783.256989 | -0.163185786 | 0.2654598 | -0.614729 | 0.538733837 | 0.709991464 | U2SURP | protein_coding | 38.45 | -0.024273375 | 0.024273375 | ENSG00000163714 | n.s. |
| ENSG00000171824 | 997.8247615 | -0.158071956 | 0.2504743 | -0.63109 | 0.527981357 | 0.702329272 | EXOSC10 | protein_coding | 46.38 | -0.024257601 | 0.024257601 | ENSG00000171824 | n.s. |
| ENSG00000165802 | 843.8151786 | 0.163929742 | 0.2680658 | 0.611528 | 0.540850044 | 0.711273748 | NSMF | protein_coding | 65.22 | 0.024255573 | 0.024255573 | ENSG00000165802 | n.s. |
| ENSG00000120948 | 2133.963823 | -0.158765331 | 0.2528242 | -0.627967 | 0.53002538 | 0.703617789 | TARDBP | protein_coding | 43.58 | -0.024237622 | 0.024237622 | ENSG00000120948 | n.s. |
| ENSG00000164828 | 1863.613552 | 0.168363103 | 0.2815306 | 0.598028 | 0.549821402 | 0.717965759 | SUN1 | protein_coding | 52.69 | 0.024226822 | 0.024226822 | ENSG00000164828 | n.s. |
| ENSG00000116691 | 141.7517523 | 0.170401529 | 0.2874916 | 0.592718 | 0.553369721 | 0.720996316 | MIIP | protein_coding | 55.62 | 0.024208426 | 0.024208426 | ENSG00000116691 | n.s. |
| ENSG00000119231 | 1711.0963 | -0.159458254 | 0.2552471 | -0.624721 | 0.532154068 | 0.705300436 | SENP5 | protein_coding | 42.03 | -0.024177993 | 0.024177993 | ENSG00000119231 | n.s. |
| ENSG00000167900 | 1127.098518 | -0.169990863 | 0.2866703 | -0.592984 | 0.553191929 | 0.720899591 | TK1 | protein_coding | 53.2 | -0.024159989 | 0.024159989 | ENSG00000167900 | n.s. |
| ENSG00000158669 | 1861.014662 | -0.159850587 | 0.2566329 | -0.622876 | 0.53336574 | 0.706301237 | GPAT4 | protein_coding | 45.7 | -0.024139043 | 0.024139043 | ENSG00000158669 | n.s. |
| ENSG00000149657 | 841.2966814 | -0.162893153 | 0.266029 | -0.612313 | 0.540330412 | 0.710993321 | LSM14B | protein_coding | 54.14 | -0.024130092 | 0.024130092 | ENSG00000149657 | n.s. |
| ENSG00000175606 | 734.6785241 | 0.162577934 | 0.2652375 | 0.612952 | 0.539907909 | 0.710706043 | TMEM70 | protein_coding | 43.8 | 0.024111932 | 0.024111932 | ENSG00000175606 | n.s. |
| ENSG00000123080 | 2674.788712 | 0.157113944 | 0.2492055 | 0.630459 | 0.528394055 | 0.702412663 | CDKN2C | protein_coding | 40.97 | 0.024102484 | 0.024102484 | ENSG00000123080 | n.s. |
| ENSG00000167447 | 1726.262179 | 0.139520499 | 0.2034923 | 0.68563 | 0.492946313 | 0.671937241 | SMG8 | protein_coding | 44.25 | 0.024091184 | 0.024091184 | ENSG00000167447 | n.s. |
| ENSG00000204344 | 135.4663161 | -0.162251834 | 0.2647473 | -0.612856 | 0.53997187 | 0.710723042 | STK19 | protein_coding | 53.1 | -0.024061883 | 0.024061883 | ENSG00000204344 | n.s. |
| ENSG00000159164 | 657.2389507 | -0.154771502 | 0.243256 | -0.636249 | 0.524613842 | 0.699131025 | SV2A | protein_coding | 52.21 | -0.024057903 | 0.024057903 | ENSG00000159164 | n.s. |
| ENSG00000151502 | 412.3384151 | -0.159797399 | 0.2574852 | -0.620608 | 0.534857532 | 0.707216152 | VPS26B | protein_coding | 48.41 | -0.024041172 | 0.024041172 | ENSG00000151502 | n.s. |
| ENSG00000012660 | 3446.493696 | -0.165228653 | 0.2739263 | -0.603187 | 0.546384597 | 0.715567289 | ELOVL5 | protein_coding | 41.88 | -0.024015905 | 0.024015905 | ENSG00000012660 | n.s. |
| ENSG00000168002 | 850.7381443 | 0.182297101 | 0.3248516 | 0.56117 | 0.574681436 | 0.738671495 | POLR2G | protein_coding | 50.57 | 0.023980939 | 0.023980939 | ENSG00000168002 | n.s. |
| ENSG00000022277 | 1200.805327 | 0.166766936 | 0.2791879 | 0.597329 | 0.550287962 | 0.718190786 | RTFDC1 | protein_coding | 43.71 | 0.023974443 | 0.023974443 | ENSG00000022277 | n.s. |
| ENSG00000166228 | 321.1945109 | 0.167012839 | 0.2799255 | 0.596633 | 0.550752248 | 0.718594654 | PCBD1 | protein_coding | 51.19 | 0.023969018 | 0.023969018 | ENSG00000166228 | n.s. |
| ENSG00000169607 | 1265.563935 | -0.153768938 | 0.2413845 | -0.637029 | 0.524106005 | 0.699014264 | CKAP2L | protein_coding | 39.66 | -0.023913217 | 0.023913217 | ENSG00000169607 | n.s. |
| ENSG00000178127 | 2474.95 | 0.158895774 | 0.2560398 | 0.62059 | 0.534869358 | 0.707216152 | NDUFV2 | protein_coding | 37.35 | 0.023905524 | 0.023905524 | ENSG00000178127 | n.s. |
| ENSG00000059769 | 284.4785486 | -0.161032888 | 0.263136 | -0.611976 | 0.540553639 | 0.711085442 | DNAJC25 | protein_coding | 42.74 | -0.023845462 | 0.023845462 | ENSG00000059769 | n.s. |
| ENSG00000168291 | 1811.515009 | 0.15440577 | 0.243592 | 0.63387 | 0.526165387 | 0.700765323 | PDHB | protein_coding | 43.56 | 0.023844481 | 0.023844481 | ENSG00000168291 | n.s. |

| | | | | | | | | | | | | | |
|---|---|---|---|---|---|---|---|---|---|---|---|---|---|
| ENSG00000166171 | 138.2372618 | -0.169124994 | 0.2872474 | -0.588778 | 0.556010082 | 0.723015632 | DPCD | protein_coding | 50.19 | -0.023821647 | 0.023821647 | ENSG00000166171 | n.s. |
| ENSG00000225880 | 53.35085321 | 0.172149753 | 0.2963986 | 0.580805 | 0.561372024 | 0.727295193 | LINC00115 | lincRNA | 52.7 | 0.023806466 | 0.023806466 | ENSG00000225880 | n.s. |
| ENSG00000123473 | 104.6295484 | -0.157870065 | 0.2538621 | -0.621873 | 0.53402519 | 0.706905564 | STIL | protein_coding | 40.02 | -0.023781326 | 0.023781326 | ENSG00000123473 | n.s. |
| ENSG00000196981 | 398.2449644 | -0.140465174 | 0.208052 | -0.675145 | 0.499583847 | 0.677283925 | WDR5B | protein_coding | 40.54 | -0.023770814 | 0.023770814 | ENSG00000196981 | n.s. |
| ENSG00000237649 | 1253.252421 | -0.160359399 | 0.2624561 | -0.610995 | 0.541202784 | 0.711590288 | KIFC1 | protein_coding | 47.68 | -0.023696306 | 0.023696306 | ENSG00000237649 | n.s. |
| ENSG00000135390 | 3188.529712 | 0.165746064 | 0.2785105 | 0.595116 | 0.551765922 | 0.719579658 | ATP5G2 | protein_coding | 44.44 | 0.023688613 | 0.023688613 | ENSG00000135390 | n.s. |
| ENSG00000135241 | 515.4256087 | 0.162904176 | 0.2698472 | 0.60369 | 0.546049522 | 0.715475954 | PNPLA8 | protein_coding | 37.75 | 0.023687075 | 0.023687075 | ENSG00000135241 | n.s. |
| ENSG00000235262 | 23.59224945 | 0.179155638 | 0.3180084 | 0.563368 | 0.573184575 | 0.737632761 | KDM5C-IT1 | sense_intronic | 45.91 | 0.023677174 | 0.023677174 | ENSG00000235262 | n.s. |
| ENSG00000165501 | 642.2659686 | -0.151804108 | 0.2377633 | -0.638467 | 0.523169535 | 0.698378662 | LRR1 | protein_coding | 45.06 | -0.023667632 | 0.023667632 | ENSG00000165501 | n.s. |
| ENSG00000275700 | 728.0410637 | -0.157909978 | 0.2554907 | -0.618066 | 0.53653214 | 0.708435841 | AATF | protein_coding | 40.51 | -0.023639041 | 0.023639041 | ENSG00000275700 | n.s. |
| ENSG00000113391 | 538.8605523 | -0.158589141 | 0.2575745 | -0.615702 | 0.538091198 | 0.70969473 | FAM172A | protein_coding | 36.4 | -0.02361843 | 0.02361843 | ENSG00000113391 | n.s. |
| ENSG00000107798 | 506.0284325 | 0.177546781 | 0.3136901 | 0.565994 | 0.57139775 | 0.736286059 | LIPA | protein_coding | 41.43 | 0.023605452 | 0.023605452 | ENSG00000107798 | n.s. |
| ENSG00000142864 | 19828.12019 | -0.156793223 | 0.2526041 | -0.620707 | 0.534792192 | 0.707216152 | SERBP1 | protein_coding | 38.87 | -0.0235892 | 0.0235892 | ENSG00000142864 | n.s. |
| ENSG00000170899 | 1613.984681 | 0.173712229 | 0.3030236 | 0.573263 | 0.566466661 | 0.731692673 | GSTA4 | protein_coding | 41.01 | 0.023567763 | 0.023567763 | ENSG00000170899 | n.s. |
| ENSG00000213246 | 1691.645531 | 0.175308457 | 0.3076318 | 0.569865 | 0.568769543 | 0.733952029 | SUPT4H1 | protein_coding | 47.01 | 0.023549593 | 0.023549593 | ENSG00000213246 | n.s. |
| ENSG00000139354 | 767.7321276 | 0.163291772 | 0.27265 | 0.598906 | 0.549235477 | 0.717557056 | GAS2L3 | protein_coding | 36.61 | 0.023537457 | 0.023537457 | ENSG00000139354 | n.s. |
| ENSG00000114353 | 526.9667633 | 0.167301491 | 0.2844515 | 0.588155 | 0.556428396 | 0.723381378 | GNAI2 | protein_coding | 57.36 | 0.023528056 | 0.023528056 | ENSG00000114353 | n.s. |
| ENSG00000169740 | 593.0421638 | -0.178595945 | 0.3179696 | -0.561676 | 0.57433669 | 0.738432887 | ZNF32 | protein_coding | 46.34 | -0.023519116 | 0.023519116 | ENSG00000169740 | n.s. |
| ENSG00000149187 | 2547.49548 | -0.151293567 | 0.2377983 | -0.636226 | 0.524628817 | 0.699131025 | CELF1 | protein_coding | 43.94 | -0.023517288 | 0.023517288 | ENSG00000149187 | n.s. |
| ENSG00000276071 | 96.47562655 | -0.152390073 | 0.2406833 | -0.633156 | 0.526631833 | 0.701185331 | | lincRNA | 45.43 | -0.023493548 | 0.023493548 | ENSG00000276071 | n.s. |
| ENSG00000273703 | 715.0978002 | -0.138964737 | 0.2062053 | -0.673914 | 0.500365712 | 0.677749957 | HIST1H2BM | protein_coding | 55.38 | -0.023475383 | 0.023475383 | ENSG00000273703 | n.s. |
| ENSG00000225349 | 24.12408797 | -0.248258641 | 0.5720339 | -0.433993 | 0.664293601 | 0.804731603 | | processed_pseudogene | 47.09 | -0.02342294 | 0.02342294 | ENSG00000225349 | n.s. |
| ENSG00000077463 | 142.2115828 | 0.170459845 | 0.2947916 | 0.578238 | 0.563103126 | 0.728832558 | SIRT6 | protein_coding | 57.91 | 0.02341645 | 0.02341645 | ENSG00000077463 | n.s. |
| ENSG00000147140 | 9184.222112 | 0.157142312 | 0.2557409 | 0.614459 | 0.538912027 | 0.709999258 | NONO | protein_coding | 44.11 | 0.023373678 | 0.023373678 | ENSG00000147140 | n.s. |
| ENSG00000235301 | 62.41908961 | 0.156685369 | 0.2541192 | 0.616582 | 0.537510296 | 0.709311771 | HLA-Z | unprocessed_pseudogene | 53.33 | 0.023371634 | 0.023371634 | ENSG00000235301 | n.s. |
| ENSG00000176261 | 234.7083906 | -0.173697927 | 0.3044059 | -0.570613 | 0.56826214 | 0.733603502 | ZBTB8OS | protein_coding | 44.37 | -0.023369077 | 0.023369077 | ENSG00000176261 | n.s. |
| ENSG00000272627 | 21.1953809 | -0.195682793 | 0.3741824 | -0.522961 | 0.601001369 | 0.759805872 | | sense_intronic | 30.43 | -0.02334444 | 0.02334444 | ENSG00000272627 | n.s. |
| ENSG00000156873 | 221.3113859 | 0.171172981 | 0.2975928 | 0.575192 | 0.565161516 | 0.730639679 | PHKG2 | protein_coding | 52.63 | 0.02333032 | 0.02333032 | ENSG00000156873 | n.s. |
| ENSG00000170779 | 745.9214426 | -0.162194175 | 0.2713695 | -0.597688 | 0.550048424 | 0.718080095 | CDCA4 | protein_coding | 52.06 | -0.02332792 | 0.02332792 | ENSG00000170779 | n.s. |
| ENSG00000162664 | 4477.241009 | -0.157192145 | 0.2562798 | -0.613361 | 0.539637338 | 0.710628976 | ZNF326 | protein_coding | 35.67 | -0.023320569 | 0.023320569 | ENSG00000162664 | n.s. |
| ENSG00000110066 | 1191.561912 | 0.153404412 | 0.2453582 | 0.625226 | 0.531822589 | 0.704995326 | KMT5B | protein_coding | 42.48 | 0.023288901 | 0.023288901 | ENSG00000110066 | n.s. |
| ENSG00000228782 | 286.9743704 | -0.145387735 | 0.223616 | -0.650167 | 0.515584428 | 0.691762009 | MRPL45P2 | transcribed_unprocessed_pseud | 42.96 | -0.023268332 | 0.023268332 | ENSG00000228782 | n.s. |

| | | | | | | | | | | | | | |
|---|---|---|---|---|---|---|---|---|---|---|---|---|---|
| ENSG00000137947 | 1170.36859 | 0.164904421 | 0.2799201 | 0.589113 | 0.555785805 | 0.722859016 | GTF2B | protein_coding | 39.88 | 0.023242684 | 0.023242684 | ENSG00000137947 | n.s. |
| ENSG00000004534 | 2227.30272 | -0.152348158 | 0.2430853 | -0.626727 | 0.530838169 | 0.704207095 | RBM6 | protein_coding | 44.14 | -0.023202564 | 0.023202564 | ENSG00000004534 | n.s. |
| ENSG00000065613 | 2219.244274 | -0.155695326 | 0.2528335 | -0.615802 | 0.538025261 | 0.70969473 | SLK | protein_coding | 36.55 | -0.023187459 | 0.023187459 | ENSG00000065613 | n.s. |
| ENSG00000111667 | 2257.573643 | 0.161123522 | 0.2697121 | 0.597391 | 0.550246525 | 0.718190786 | USP5 | protein_coding | 52.77 | 0.023163145 | 0.023163145 | ENSG00000111667 | n.s. |
| ENSG00000272344 | 12.9181195 | -0.204295064 | 0.4062703 | -0.502855 | 0.615066184 | 0.770364212 | SNORD114-21 | snoRNA | 47.22 | -0.023147428 | 0.023147428 | ENSG00000272344 | n.s. |
| ENSG00000081181 | 129.7526185 | 0.172316581 | 0.3026935 | 0.569277 | 0.569167835 | 0.734228089 | ARG2 | protein_coding | 42.2 | 0.023119544 | 0.023119544 | ENSG00000081181 | n.s. |
| ENSG00000105127 | 601.4684874 | 0.150443776 | 0.2383836 | 0.631099 | 0.527975493 | 0.702329272 | AKAP8 | protein_coding | 53.24 | 0.023086986 | 0.023086986 | ENSG00000105127 | n.s. |
| ENSG00000165669 | 731.6116593 | 0.160139872 | 0.2673791 | 0.598924 | 0.5492233 | 0.717557056 | FAM204A | protein_coding | 38.05 | 0.023083131 | 0.023083131 | ENSG00000165669 | n.s. |
| ENSG00000136982 | 780.3501699 | -0.164164642 | 0.2793131 | -0.587744 | 0.556703996 | 0.723491483 | DSCC1 | protein_coding | 42.08 | -0.023076062 | 0.023076062 | ENSG00000136982 | n.s. |
| ENSG00000130803 | 766.5046196 | -0.154920962 | 0.2516684 | -0.615576 | 0.538174594 | 0.70969473 | ZNF317 | protein_coding | 44.48 | -0.023072134 | 0.023072134 | ENSG00000130803 | n.s. |
| ENSG00000237349 | 12.22273732 | 0.193077679 | 0.3689108 | 0.523372 | 0.60071525 | 0.759513115 | | processed_pseudogene | 40.28 | 0.023065971 | 0.023065971 | ENSG00000237349 | n.s. |
| ENSG00000201184 | 6.777469792 | -0.215566578 | 0.446425 | -0.482873 | 0.629185887 | 0.781705085 | RNU4-68P | snRNA | 40.43 | -0.023056368 | 0.023056368 | ENSG00000201184 | n.s. |
| ENSG00000156374 | 842.266714 | -0.155406229 | 0.2534076 | -0.613266 | 0.539700507 | 0.71063459 | PCGF6 | protein_coding | 43.89 | -0.023055082 | 0.023055082 | ENSG00000156374 | n.s. |
| ENSG00000136940 | 769.4880529 | 0.164128092 | 0.2796243 | 0.586959 | 0.557231081 | 0.723657153 | PDCL | protein_coding | 45.28 | 0.023054604 | 0.023054604 | ENSG00000136940 | n.s. |
| ENSG00000203814 | 1818.885881 | -0.140059898 | 0.2114117 | -0.662498 | 0.507651963 | 0.684688393 | HIST2H2BF | protein_coding | 41.91 | -0.023040839 | 0.023040839 | ENSG00000203814 | n.s. |
| ENSG00000207280 | 109.7975864 | 0.150051086 | 0.2376969 | 0.631271 | 0.527863581 | 0.702329272 | SNORD20 | snoRNA | 33.75 | 0.023026724 | 0.023026724 | ENSG00000207280 | n.s. |
| ENSG00000140406 | 728.2092442 | -0.140868277 | 0.2135839 | -0.659545 | 0.509545597 | 0.68657731 | MESDC1 | protein_coding | 55.46 | -0.023005276 | 0.023005276 | ENSG00000140406 | n.s. |
| ENSG00000109472 | 1551.846399 | 0.167912914 | 0.2909822 | 0.577056 | 0.56390189 | 0.729530386 | CPE | protein_coding | 38.23 | 0.022996784 | 0.022996784 | ENSG00000109472 | n.s. |
| ENSG00000237749 | 13.15507798 | 0.20572492 | 0.4128803 | 0.498268 | 0.61829537 | 0.773221727 | | processed_pseudogene | 49.8 | 0.022978641 | 0.022978641 | ENSG00000237749 | n.s. |
| ENSG00000135406 | 1287.554576 | 0.156967482 | 0.258927 | 0.606223 | 0.544366794 | 0.714077527 | PRPH | protein_coding | 62.39 | 0.022957222 | 0.022957222 | ENSG00000135406 | n.s. |
| ENSG00000158796 | 401.0906195 | -0.159080597 | 0.265373 | -0.59946 | 0.54886591 | 0.717343478 | DEDD | protein_coding | 46.36 | -0.02295101 | 0.02295101 | ENSG00000158796 | n.s. |
| ENSG00000143155 | 1550.437463 | -0.175848087 | 0.3154234 | -0.557499 | 0.577186853 | 0.740546777 | TIPRL | protein_coding | 38.59 | -0.022938944 | 0.022938944 | ENSG00000143155 | n.s. |
| ENSG00000172992 | 687.0701051 | -0.150418766 | 0.2401378 | -0.626385 | 0.53106228 | 0.704255655 | DCAKD | protein_coding | 51.25 | -0.022904214 | 0.022904214 | ENSG00000172992 | n.s. |
| ENSG00000182220 | 736.6938787 | 0.173111488 | 0.3074175 | 0.563115 | 0.573356413 | 0.737755431 | ATP6AP2 | protein_coding | 41.77 | 0.022865878 | 0.022865878 | ENSG00000182220 | n.s. |
| ENSG00000101189 | 343.5352898 | -0.165924394 | 0.2863726 | -0.5794 | 0.56231909 | 0.728294762 | MRGBP | protein_coding | 59.23 | -0.022846597 | 0.022846597 | ENSG00000101189 | n.s. |
| ENSG00000066855 | 484.1794585 | -0.169877458 | 0.2979338 | -0.570185 | 0.568552088 | 0.733773704 | MTFR1 | protein_coding | 40.22 | -0.022837961 | 0.022837961 | ENSG00000066855 | n.s. |
| ENSG00000080822 | 1401.324493 | 0.157176383 | 0.2609103 | 0.602415 | 0.546897719 | 0.715880764 | CLDND1 | protein_coding | 38.91 | 0.022815615 | 0.022815615 | ENSG00000080822 | n.s. |
| ENSG00000198858 | 201.7622281 | -0.1712921 | 0.302408 | -0.566427 | 0.571103436 | 0.73597493 | R3HDM4 | protein_coding | 58.93 | -0.022805313 | 0.022805313 | ENSG00000198858 | n.s. |
| ENSG00000003756 | 2684.875664 | -0.156105021 | 0.2579342 | -0.605213 | 0.545037716 | 0.714419142 | RBM5 | protein_coding | 44.45 | -0.022798657 | 0.022798657 | ENSG00000003756 | n.s. |
| ENSG00000160917 | 830.6085007 | -0.161357224 | 0.2734635 | -0.59005 | 0.555156866 | 0.72231091 | CPSF4 | protein_coding | 53.24 | -0.022795875 | 0.022795875 | ENSG00000160917 | n.s. |
| ENSG00000101474 | 953.1896633 | 0.165585853 | 0.2860659 | 0.578838 | 0.562698381 | 0.728514929 | APMAP | protein_coding | 46.54 | 0.022778246 | 0.022778246 | ENSG00000101474 | n.s. |
| ENSG00000136193 | 1860.887939 | -0.162254134 | 0.276592 | -0.586619 | 0.5574596 | 0.723818895 | SCRN1 | protein_coding | 43.91 | -0.022775626 | 0.022775626 | ENSG00000136193 | n.s. |

| | | | | | | | | | | | | | |
|---|---|---|---|---|---|---|---|---|---|---|---|---|---|
| ENSG00000132635 | 642.33281 | 0.15130326 | 0.2437439 | 0.620747 | 0.534766124 | 0.707216152 | PCED1A | protein_coding | 57.7 | 0.022763247 | 0.022763247 | ENSG00000132635 | n.s. |
| ENSG00000257315 | 4210.425872 | 0.139092348 | 0.2106828 | 0.660198 | 0.50912682 | 0.686212267 | ZBED6 | protein_coding | 40.71 | 0.022747374 | 0.022747374 | ENSG00000257315 | n.s. |
| ENSG00000136856 | 31.19024631 | -0.177034721 | 0.3216533 | -0.55039 | 0.582051948 | 0.743955685 | SLC2A8 | protein_coding | 57.79 | -0.022740629 | 0.022740629 | ENSG00000136856 | n.s. |
| ENSG00000068308 | 348.027913 | -0.151382512 | 0.2445252 | -0.619088 | 0.53585864 | 0.708120714 | OTUD5 | protein_coding | 49.93 | -0.022691134 | 0.022691134 | ENSG00000068308 | n.s. |
| ENSG00000238344 | 60.97355147 | -0.158730641 | 0.2670054 | -0.594485 | 0.552187974 | 0.719726003 | SNORD126 | snoRNA | 42.86 | -0.022671943 | 0.022671943 | ENSG00000238344 | n.s. |
| ENSG00000135597 | 1972.58586 | 0.156064353 | 0.2590131 | 0.602535 | 0.546818394 | 0.715880764 | REPS1 | protein_coding | 37.32 | 0.022654193 | 0.022654193 | ENSG00000135597 | n.s. |
| ENSG00000274303 | 15.12846231 | -0.20194311 | 0.4043866 | -0.499381 | 0.61751079 | 0.772401647 | | misc_RNA | 38.26 | -0.022649295 | 0.022649295 | ENSG00000274303 | n.s. |
| ENSG00000254325 | 62.36415239 | 0.161893803 | 0.2766226 | 0.585252 | 0.558378595 | 0.72474179 | | sense_intronic | 45.55 | 0.022635457 | 0.022635457 | ENSG00000254325 | n.s. |
| ENSG00000197694 | 3326.797526 | -0.154033307 | 0.2535071 | -0.60761 | 0.543446463 | 0.713251896 | SPTAN1 | protein_coding | 46.82 | -0.022605476 | 0.022605476 | ENSG00000197694 | n.s. |
| ENSG00000255256 | 29.25538685 | -0.173244909 | 0.3106066 | -0.557763 | 0.577006101 | 0.740546777 | | sense_intronic | 43.76 | -0.022599366 | 0.022599366 | ENSG00000255256 | n.s. |
| ENSG00000139974 | 41.50322188 | -0.162892523 | 0.2796599 | -0.582467 | 0.560252461 | 0.726564364 | SLC38A6 | protein_coding | 37.62 | -0.022597413 | 0.022597413 | ENSG00000139974 | n.s. |
| ENSG00000100219 | 1243.863402 | -0.154439259 | 0.2548316 | -0.606044 | 0.544485252 | 0.714098062 | XBP1 | protein_coding | 44.93 | -0.022585529 | 0.022585529 | ENSG00000100219 | n.s. |
| ENSG00000079332 | 1514.04699 | 0.159179832 | 0.2689111 | 0.591942 | 0.553889218 | 0.721470632 | SAR1A | protein_coding | 40.56 | 0.02256873 | 0.02256873 | ENSG00000079332 | n.s. |
| ENSG00000143862 | 462.5332204 | 0.160315407 | 0.2724312 | 0.588462 | 0.556222299 | 0.723224045 | ARL8A | protein_coding | 53.33 | 0.022560729 | 0.022560729 | ENSG00000143862 | n.s. |
| ENSG00000233061 | 231.3859958 | 0.149792846 | 0.2409394 | 0.621703 | 0.534136924 | 0.706986253 | TTLL7-IT1 | sense_intronic | 37.82 | 0.02255716 | 0.02255716 | ENSG00000233061 | n.s. |
| ENSG00000143553 | 365.7081184 | -0.173410149 | 0.3123467 | -0.555185 | 0.578768218 | 0.741528472 | SNAPIN | protein_coding | 51.31 | -0.022521152 | 0.022521152 | ENSG00000143553 | n.s. |
| ENSG00000223764 | 238.9658435 | -0.160041222 | 0.2721203 | -0.588127 | 0.556447214 | 0.723381378 | | lincRNA | 61.68 | -0.022507025 | 0.022507025 | ENSG00000223764 | n.s. |
| ENSG00000199872 | 34.27511021 | 0.169379319 | 0.2994816 | 0.565575 | 0.571682712 | 0.736448779 | RNU6-942P | snRNA | 38.32 | 0.022503305 | 0.022503305 | ENSG00000199872 | n.s. |
| ENSG00000107862 | 1052.055941 | -0.144176781 | 0.2258597 | -0.638347 | 0.523248012 | 0.698416458 | GBF1 | protein_coding | 43.61 | -0.022475075 | 0.022475075 | ENSG00000107862 | n.s. |
| ENSG00000116489 | 2587.109459 | -0.154505611 | 0.2560924 | -0.60332 | 0.54629598 | 0.715529532 | CAPZA1 | protein_coding | 40.34 | -0.022460857 | 0.022460857 | ENSG00000116489 | n.s. |
| ENSG00000227671 | 573.750554 | -0.160609265 | 0.2744939 | -0.585111 | 0.558473392 | 0.724797263 | | transcribed_unprocessed_pseud | 40.91 | -0.022450518 | 0.022450518 | ENSG00000227671 | n.s. |
| ENSG00000102978 | 1407.491629 | 0.158756512 | 0.2690124 | 0.590146 | 0.55509298 | 0.722295286 | POLR2C | protein_coding | 48.07 | 0.022429949 | 0.022429949 | ENSG00000102978 | n.s. |
| ENSG00000068120 | 728.1660056 | 0.155323493 | 0.258949 | 0.599823 | 0.548624434 | 0.717229859 | COASY | protein_coding | 57.56 | 0.022419647 | 0.022419647 | ENSG00000068120 | n.s. |
| ENSG00000186130 | 996.3702239 | -0.146068505 | 0.231529 | -0.630887 | 0.528114717 | 0.702353731 | ZBTB6 | protein_coding | 36.76 | -0.022413351 | 0.022413351 | ENSG00000186130 | n.s. |
| ENSG00000152102 | 3362.453924 | -0.158023816 | 0.2670606 | -0.591715 | 0.55404126 | 0.721601166 | FAM168B | protein_coding | 45.79 | -0.022392413 | 0.022392413 | ENSG00000152102 | n.s. |
| ENSG00000230590 | 436.6369622 | -0.149561426 | 0.2419967 | -0.618031 | 0.536555049 | 0.708435841 | FTX | lincRNA | 39.53 | -0.022389267 | 0.022389267 | ENSG00000230590 | n.s. |
| ENSG00000225265 | 61.10793817 | 0.159329942 | 0.2714096 | 0.587046 | 0.557172769 | 0.723648923 | TAF1A-AS1 | antisense | 41.4 | 0.022381408 | 0.022381408 | ENSG00000225265 | n.s. |
| ENSG00000199168 | 13.66160623 | 0.19253818 | 0.3760354 | 0.512021 | 0.608636018 | 0.765274285 | MIR374A | miRNA | 31.94 | 0.02236964 | 0.02236964 | ENSG00000199168 | n.s. |
| ENSG00000196990 | 73.76868596 | -0.172512847 | 0.3114858 | -0.553839 | 0.579689276 | 0.741907339 | FAM163B | protein_coding | 57.82 | -0.022366348 | 0.022366348 | ENSG00000196990 | n.s. |
| ENSG00000064703 | 2254.492298 | -0.153815975 | 0.2547611 | -0.603766 | 0.545999453 | 0.715475954 | DDX20 | protein_coding | 40.24 | -0.022365605 | 0.022365605 | ENSG00000064703 | n.s. |
| ENSG00000137812 | 2199.675592 | -0.157289357 | 0.2653352 | -0.592795 | 0.553318384 | 0.720996316 | KNL1 | protein_coding | 41.23 | -0.02234562 | 0.02234562 | ENSG00000137812 | n.s. |
| ENSG00000204392 | 513.0504492 | 0.172650578 | 0.3122934 | 0.552847 | 0.5803679 | 0.7424166 | LSM2 | protein_coding | 48.13 | 0.022332754 | 0.022332754 | ENSG00000204392 | n.s. |

| | | | | | | | | | | | | | |
|---|---|---|---|---|---|---|---|---|---|---|---|---|---|
| ENSG00000213420 | 477.1442006 | 0.158355053 | 0.2689376 | 0.588817 | 0.555983938 | 0.723015632 | GPC2 | protein_coding | 55.45 | 0.022304675 | 0.022304675 | ENSG00000213420 | n.s. |
| ENSG00000182685 | 109.629711 | 0.1476178 | 0.2369183 | 0.623075 | 0.533235357 | 0.706195745 | BRICD5 | protein_coding | 63.38 | 0.022301345 | 0.022301345 | ENSG00000182685 | n.s. |
| ENSG00000198406 | 73.48573882 | 0.185480779 | 0.3528385 | 0.525682 | 0.599109279 | 0.758240023 | BZW1P2 | processed_pseudogene | 40.19 | 0.022293546 | 0.022293546 | ENSG00000198406 | n.s. |
| ENSG00000113621 | 657.2518015 | 0.160936651 | 0.2769681 | 0.581066 | 0.561196222 | 0.72728024 | TXNDC15 | protein_coding | 46.17 | 0.022257251 | 0.022257251 | ENSG00000113621 | n.s. |
| ENSG00000237441 | 292.560985 | 0.162163743 | 0.2811983 | 0.576688 | 0.564150096 | 0.729783684 | RGL2 | protein_coding | 57.12 | 0.022184949 | 0.022184949 | ENSG00000237441 | n.s. |
| ENSG00000151276 | 170.5832374 | -0.1488727 | 0.2418987 | -0.615434 | 0.538268033 | 0.70969473 | MAGI1 | protein_coding | 40.3 | -0.022171376 | 0.022171376 | ENSG00000151276 | n.s. |
| ENSG00000144677 | 94.70069216 | -0.153361262 | 0.2553453 | -0.600603 | 0.548104225 | 0.716953695 | CTDSPL | protein_coding | 42.29 | -0.022162066 | 0.022162066 | ENSG00000144677 | n.s. |
| ENSG00000008513 | 74.11792927 | -0.156498845 | 0.2648171 | -0.590969 | 0.554540883 | 0.722075346 | ST3GAL1 | protein_coding | 49.4 | -0.022131673 | 0.022131673 | ENSG00000008513 | n.s. |
| ENSG00000143179 | 2993.90992 | -0.152799947 | 0.2540587 | -0.601436 | 0.547549865 | 0.716430483 | UCK2 | protein_coding | 44.22 | -0.022129397 | 0.022129397 | ENSG00000143179 | n.s. |
| ENSG00000173209 | 2746.3882 | -0.149296136 | 0.243704 | -0.612613 | 0.540132562 | 0.710861345 | AHSA2 | protein_coding | 44.61 | -0.022127942 | 0.022127942 | ENSG00000173209 | n.s. |
| ENSG00000198431 | 5802.492713 | 0.154744439 | 0.2602691 | 0.594556 | 0.552140617 | 0.719726003 | TXNRD1 | protein_coding | 42.37 | 0.022102583 | 0.022102583 | ENSG00000198431 | n.s. |
| ENSG00000262879 | 257.3974184 | -0.151746054 | 0.2515091 | -0.603342 | 0.546281025 | 0.715529532 | | processed_transcript | 43.56 | -0.022059694 | 0.022059694 | ENSG00000262879 | n.s. |
| ENSG00000163832 | 952.1662299 | 0.146469564 | 0.2356953 | 0.621436 | 0.534312713 | 0.707041054 | ELP6 | protein_coding | 48.73 | 0.022051779 | 0.022051779 | ENSG00000163832 | n.s. |
| ENSG00000087470 | 4320.72025 | -0.15412173 | 0.2590447 | -0.594962 | 0.551868964 | 0.719579658 | DNM1L | protein_coding | 38.21 | -0.022027251 | 0.022027251 | ENSG00000087470 | n.s. |
| ENSG00000170027 | 4742.101714 | -0.158757489 | 0.2726644 | -0.582245 | 0.560401587 | 0.726622397 | YWHAG | protein_coding | 44.56 | -0.02201827 | 0.02201827 | ENSG00000170027 | n.s. |
| ENSG00000156030 | 118.352583 | 0.1428696 | 0.2258627 | 0.632551 | 0.527027183 | 0.701443406 | ELMSAN1 | protein_coding | 51.27 | 0.02200297 | 0.02200297 | ENSG00000156030 | n.s. |
| ENSG00000260572 | 117.3165874 | 0.143645297 | 0.2282876 | 0.62923 | 0.529198772 | 0.702790513 | | antisense | 45.71 | 0.02200274 | 0.02200274 | ENSG00000260572 | n.s. |
| ENSG00000198692 | 707.0185723 | -0.164180996 | 0.2887151 | -0.568661 | 0.569586289 | 0.734563719 | EIF1AY | protein_coding | 35.64 | -0.021995414 | 0.021995414 | ENSG00000198692 | n.s. |
| ENSG00000166946 | 531.6034646 | 0.169547199 | 0.3060307 | 0.55402 | 0.579565008 | 0.741907339 | CCNDBP1 | protein_coding | 40.91 | 0.021981851 | 0.021981851 | ENSG00000166946 | n.s. |
| ENSG00000230081 | 11.15791321 | -0.20080323 | 0.4091245 | -0.490812 | 0.623559384 | 0.777244164 | HSPE1P28 | processed_pseudogene | 39.11 | -0.021976414 | 0.021976414 | ENSG00000230081 | n.s. |
| ENSG00000160752 | 3503.508448 | 0.163664258 | 0.2875373 | 0.569193 | 0.569225041 | 0.734233857 | FDPS | protein_coding | 50.48 | 0.021958112 | 0.021958112 | ENSG00000160752 | n.s. |
| ENSG00000205189 | 3823.472562 | 0.146690559 | 0.2374312 | 0.617824 | 0.53669168 | 0.708549048 | ZBTB10 | protein_coding | 38.89 | 0.02194932 | 0.02194932 | ENSG00000205189 | n.s. |
| ENSG00000204272 | 210.2230094 | 0.163825395 | 0.2883981 | 0.568053 | 0.569999068 | 0.734891844 | NBDY | protein_coding | 45.2 | 0.021916 | 0.021916 | ENSG00000204272 | n.s. |
| ENSG00000088833 | 417.2498595 | 0.149601549 | 0.2469196 | 0.605872 | 0.544600086 | 0.714114428 | NSFL1C | protein_coding | 45.86 | 0.021876562 | 0.021876562 | ENSG00000088833 | n.s. |
| ENSG00000165609 | 1666.019361 | -0.164796255 | 0.2915806 | -0.565183 | 0.571949635 | 0.736656315 | NUDT5 | protein_coding | 44.24 | -0.021874245 | 0.021874245 | ENSG00000165609 | n.s. |
| ENSG00000141385 | 2081.593813 | -0.153750035 | 0.2597062 | -0.592015 | 0.553840425 | 0.721470632 | AFG3L2 | protein_coding | 44.7 | -0.021798886 | 0.021798886 | ENSG00000141385 | n.s. |
| ENSG00000167085 | 5195.186484 | 0.16636817 | 0.2972152 | 0.559757 | 0.575645451 | 0.739569218 | PHB | protein_coding | 48.42 | 0.021797753 | 0.021797753 | ENSG00000167085 | n.s. |
| ENSG00000197969 | 127.8878136 | -0.165529147 | 0.2951136 | -0.5609 | 0.57486583 | 0.738840299 | VPS13A | protein_coding | 35.89 | -0.021758711 | 0.021758711 | ENSG00000197969 | n.s. |
| ENSG00000108389 | 3782.873119 | -0.149872063 | 0.2488016 | -0.602376 | 0.54692406 | 0.715880764 | MTMR4 | protein_coding | 47.85 | -0.021755325 | 0.021755325 | ENSG00000108389 | n.s. |
| ENSG00000182197 | 498.9766009 | -0.141747737 | 0.224665 | -0.630929 | 0.528086665 | 0.702353731 | EXT1 | protein_coding | 41.16 | -0.021750355 | 0.021750355 | ENSG00000182197 | n.s. |
| ENSG00000198546 | 433.2178857 | 0.16385002 | 0.2900619 | 0.56488 | 0.572155676 | 0.736853526 | ZNF511 | protein_coding | 57.36 | 0.021729599 | 0.021729599 | ENSG00000198546 | n.s. |
| ENSG00000116199 | 1843.808726 | -0.154208825 | 0.2624209 | -0.587639 | 0.556774388 | 0.723491483 | FAM20B | protein_coding | 43.17 | -0.021676607 | 0.021676607 | ENSG00000116199 | n.s. |

| | | | | | | | | | | | | | |
|---|---|---|---|---|---|---|---|---|---|---|---|---|---|
| ENSG00000170266 | 62.75199873 | 0.160405335 | 0.28055 | 0.571753 | 0.567489236 | 0.732741596 | GLB1 | protein_coding | 44.03 | 0.021662605 | 0.021662605 | ENSG00000170266 | n.s. |
| ENSG00000168936 | 366.6357478 | -0.158901532 | 0.2762956 | -0.575114 | 0.565214048 | 0.730639679 | TMEM129 | protein_coding | 62.46 | -0.021657762 | 0.021657762 | ENSG00000168936 | n.s. |
| ENSG00000228393 | 163.6981267 | 0.147865936 | 0.2437688 | 0.606583 | 0.5441279 | 0.713898676 | LINC01004 | antisense | 42 | 0.021642165 | 0.021642165 | ENSG00000228393 | n.s. |
| ENSG00000163344 | 184.8090138 | -0.163235946 | 0.289127 | -0.564582 | 0.572357978 | 0.737045887 | PMVK | protein_coding | 49.94 | -0.021629656 | 0.021629656 | ENSG00000163344 | n.s. |
| ENSG00000154144 | 571.2429638 | -0.153435353 | 0.2604152 | -0.589195 | 0.555730397 | 0.722859016 | TBRG1 | protein_coding | 40.05 | -0.02162616 | 0.02162616 | ENSG00000154144 | n.s. |
| ENSG00000179523 | 167.9514132 | 0.148934224 | 0.2472095 | 0.602462 | 0.546866891 | 0.715880764 | EIF3J-AS1 | lincRNA | 44.04 | 0.021619189 | 0.021619189 | ENSG00000179523 | n.s. |
| ENSG00000117069 | 1128.382466 | -0.158625367 | 0.2761744 | -0.574367 | 0.565719708 | 0.731134827 | ST6GALNAC5 | protein_coding | 39.93 | -0.021573451 | 0.021573451 | ENSG00000117069 | n.s. |
| ENSG00000076108 | 77.86525185 | -0.152001337 | 0.2566046 | -0.592356 | 0.553612007 | 0.7212445 | BAZ2A | protein_coding | 48.64 | -0.021571648 | 0.021571648 | ENSG00000076108 | n.s. |
| ENSG00000156858 | 137.5868986 | 0.145647224 | 0.2379584 | 0.61207 | 0.540491393 | 0.711070742 | PRR14 | protein_coding | 57.98 | 0.021568488 | 0.021568488 | ENSG00000156858 | n.s. |
| ENSG00000128513 | 253.8638026 | -0.143418579 | 0.2314655 | -0.619611 | 0.535513816 | 0.707799398 | POT1 | protein_coding | 34.37 | -0.021525667 | 0.021525667 | ENSG00000128513 | n.s. |
| ENSG00000145526 | 36.45361499 | -0.227560025 | 0.5232782 | -0.434874 | 0.663653959 | 0.804311909 | CDH18 | protein_coding | 34.89 | -0.021521603 | 0.021521603 | ENSG00000145526 | n.s. |
| ENSG00000157224 | 721.1181214 | -0.154907331 | 0.2658825 | -0.582616 | 0.560151948 | 0.726501684 | CLDN12 | protein_coding | 39.02 | -0.021495464 | 0.021495464 | ENSG00000157224 | n.s. |
| ENSG00000204316 | 136.0818985 | 0.147835366 | 0.2449522 | 0.603527 | 0.546157853 | 0.715529532 | MRPL38 | protein_coding | 57.66 | 0.021491187 | 0.021491187 | ENSG00000204316 | n.s. |
| ENSG00000241697 | 172.7520477 | 0.155385502 | 0.2674377 | 0.581016 | 0.561229903 | 0.72728024 | TMEFF1 | protein_coding | 39.38 | 0.021489537 | 0.021489537 | ENSG00000241697 | n.s. |
| ENSG00000138668 | 5647.178051 | 0.148440728 | 0.2468887 | 0.601246 | 0.547676388 | 0.716528692 | HNRNPD | protein_coding | 38.75 | 0.021489232 | 0.021489232 | ENSG00000138668 | n.s. |
| ENSG00000186376 | 350.8685207 | -0.154484167 | 0.2648105 | -0.583376 | 0.559640054 | 0.726024901 | ZNF75D | protein_coding | 41.67 | -0.021480789 | 0.021480789 | ENSG00000186376 | n.s. |
| ENSG00000137343 | 180.317222 | 0.150250309 | 0.2523105 | 0.595498 | 0.551510842 | 0.719517003 | ATAT1 | protein_coding | 48.87 | 0.021479625 | 0.021479625 | ENSG00000137343 | n.s. |
| ENSG00000242737 | 28.70204351 | -0.16465432 | 0.2954675 | -0.557267 | 0.577344937 | 0.740546777 | | processed_pseudogene | 42.62 | -0.021478745 | 0.021478745 | ENSG00000242737 | n.s. |
| ENSG00000099622 | 2588.368445 | 0.149197935 | 0.249506 | 0.597973 | 0.54985771 | 0.717965759 | CIRBP | protein_coding | 62.55 | 0.021469026 | 0.021469026 | ENSG00000099622 | n.s. |
| ENSG00000166912 | 161.6997632 | 0.149206174 | 0.249687 | 0.597573 | 0.550125015 | 0.71811275 | MTMR10 | protein_coding | 39.99 | 0.021456946 | 0.021456946 | ENSG00000166912 | n.s. |
| ENSG00000253771 | 33.78226488 | 0.167392498 | 0.30479 | 0.549206 | 0.582864049 | 0.74471172 | TPTE2P1 | transcribed_unprocessed_pseud | 41.61 | 0.021428217 | 0.021428217 | ENSG00000253771 | n.s. |
| ENSG00000178761 | 1367.684612 | -0.151331536 | 0.2560863 | -0.59094 | 0.554560919 | 0.722075346 | FAM219B | protein_coding | 51.13 | -0.021400925 | 0.021400925 | ENSG00000178761 | n.s. |
| ENSG00000265057 | 6.129845737 | 0.228757347 | 0.53254 | 0.429559 | 0.6675165 | 0.806336303 | MIR4765 | miRNA | 41.56 | 0.021385103 | 0.021385103 | ENSG00000265057 | n.s. |
| ENSG00000261864 | 101.8942084 | -0.154062066 | 0.2648609 | -0.581672 | 0.560787915 | 0.726957289 | | processed_transcript | 43.16 | -0.021336226 | 0.021336226 | ENSG00000261864 | n.s. |
| ENSG00000116698 | 1336.514629 | -0.143988209 | 0.2352013 | -0.612191 | 0.540411126 | 0.711032329 | SMG7 | protein_coding | 40.28 | -0.021326188 | 0.021326188 | ENSG00000116698 | n.s. |
| ENSG00000166508 | 7757.804578 | 0.153708344 | 0.2639404 | 0.58236 | 0.560324164 | 0.726589674 | MCM7 | protein_coding | 54.21 | 0.021321004 | 0.021321004 | ENSG00000166508 | n.s. |
| ENSG00000284324 | 42.10482492 | -0.164391892 | 0.296768 | -0.553941 | 0.579619368 | 0.741907339 | MIR3175 | miRNA | 71.43 | -0.021313463 | 0.021313463 | ENSG00000284324 | n.s. |
| ENSG00000111229 | 2137.898296 | 0.175047234 | 0.329739 | 0.530866 | 0.59551165 | 0.755522643 | ARPC3 | protein_coding | 45.85 | 0.021312441 | 0.021312441 | ENSG00000111229 | n.s. |
| ENSG00000145414 | 737.9361112 | 0.144526328 | 0.2371048 | 0.609546 | 0.542162484 | 0.712461375 | NAF1 | protein_coding | 36.36 | 0.021279865 | 0.021279865 | ENSG00000145414 | n.s. |
| ENSG00000179008 | 54.57757934 | -0.161376765 | 0.2871314 | -0.562031 | 0.574094827 | 0.73836766 | C14orf39 | protein_coding | 37.01 | -0.021257731 | 0.021257731 | ENSG00000179008 | n.s. |
| ENSG00000114902 | 671.2864259 | 0.164594278 | 0.2981014 | 0.552142 | 0.580851133 | 0.742761786 | SPCS1 | protein_coding | 47.37 | 0.021257425 | 0.021257425 | ENSG00000114902 | n.s. |
| ENSG00000078808 | 312.6975797 | 0.159618156 | 0.2820764 | 0.565869 | 0.571483164 | 0.736300218 | SDF4 | protein_coding | 61.35 | 0.021220447 | 0.021220447 | ENSG00000078808 | n.s. |

| | | | | | | | | | | | | | |
|---|---|---|---|---|---|---|---|---|---|---|---|---|---|
| ENSG00000157778 | 546.137243 | 0.16107575 | 0.2866198 | 0.561984 | 0.574126861 | 0.73836766 | PSMG3 | protein_coding | 57.7 | 0.021218079 | 0.021218079 | ENSG00000157778 | n.s. |
| ENSG00000127081 | 425.2634493 | -0.150963589 | 0.2570848 | -0.587213 | 0.557060484 | 0.723570587 | ZNF484 | protein_coding | 39.28 | -0.021213267 | 0.021213267 | ENSG00000127081 | n.s. |
| ENSG00000015475 | 1084.724529 | -0.163263576 | 0.2944701 | -0.554432 | 0.579283449 | 0.741830534 | BID | protein_coding | 52.51 | -0.021174518 | 0.021174518 | ENSG00000015475 | n.s. |
| ENSG00000125912 | 577.6239406 | -0.149421597 | 0.2529606 | -0.590691 | 0.554727343 | 0.722109313 | NCLN | protein_coding | 60.96 | -0.021127774 | 0.021127774 | ENSG00000125912 | n.s. |
| ENSG00000172922 | 812.1012699 | 0.150339964 | 0.2560201 | 0.587219 | 0.55705637 | 0.723570587 | RNASEH2C | protein_coding | 55.83 | 0.021125636 | 0.021125636 | ENSG00000172922 | n.s. |
| ENSG00000114450 | 1414.486722 | -0.156115818 | 0.2727593 | -0.572358 | 0.567079793 | 0.732348758 | GNB4 | protein_coding | 39.16 | -0.021119668 | 0.021119668 | ENSG00000114450 | n.s. |
| ENSG00000250132 | 193.5177035 | -0.140495284 | 0.2265634 | -0.620115 | 0.53518235 | 0.707428449 | | antisense | 45.76 | -0.021118897 | 0.021118897 | ENSG00000250132 | n.s. |
| ENSG00000138442 | 2409.108549 | -0.151701165 | 0.2600343 | -0.583389 | 0.559631435 | 0.726024901 | WDR12 | protein_coding | 39.21 | -0.021093817 | 0.021093817 | ENSG00000138442 | n.s. |
| ENSG00000213152 | 34.11438798 | 0.161204252 | 0.288495 | 0.558777 | 0.576314205 | 0.739970725 | | processed_pseudogene | 51.26 | 0.021083173 | 0.021083173 | ENSG00000213152 | n.s. |
| ENSG00000113580 | 512.3570095 | 0.146058694 | 0.2435925 | 0.599603 | 0.548771122 | 0.717313933 | NR3C1 | protein_coding | 37.82 | 0.021074916 | 0.021074916 | ENSG00000113580 | n.s. |
| ENSG00000141580 | 2817.144351 | 0.159164148 | 0.2822181 | 0.563976 | 0.572770686 | 0.737334773 | WDR45B | protein_coding | 49.34 | 0.021063033 | 0.021063033 | ENSG00000141580 | n.s. |
| ENSG00000134333 | 9052.848015 | 0.155097436 | 0.270622 | 0.573115 | 0.566567131 | 0.73175456 | LDHA | protein_coding | 44.3 | 0.021036573 | 0.021036573 | ENSG00000134333 | n.s. |
| ENSG00000089053 | 3568.285989 | 0.154324204 | 0.2682118 | 0.575382 | 0.565033141 | 0.730639679 | ANAPC5 | protein_coding | 44.55 | 0.021033887 | 0.021033887 | ENSG00000089053 | n.s. |
| ENSG00000041802 | 1966.442546 | 0.146140536 | 0.2443221 | 0.598147 | 0.549741857 | 0.717949149 | LSG1 | protein_coding | 41.69 | 0.021030546 | 0.021030546 | ENSG00000041802 | n.s. |
| ENSG00000135919 | 606.6699305 | -0.155527833 | 0.2719765 | -0.571843 | 0.567428398 | 0.732730995 | SERPINE2 | protein_coding | 43.21 | -0.021004879 | 0.021004879 | ENSG00000135919 | n.s. |
| ENSG00000112308 | 1833.482339 | -0.140045708 | 0.2261264 | -0.619325 | 0.535702422 | 0.707981473 | C6orf62 | protein_coding | 40.31 | -0.02100379 | 0.02100379 | ENSG00000112308 | n.s. |
| ENSG00000148824 | 492.6488488 | 0.157422059 | 0.2776423 | 0.566996 | 0.570716881 | 0.735544861 | MTG1 | protein_coding | 47.71 | 0.02099866 | 0.02099866 | ENSG00000148824 | n.s. |
| ENSG00000204209 | 1083.066093 | 0.149347797 | 0.2541027 | 0.587746 | 0.556702851 | 0.723491483 | DAXX | protein_coding | 51.32 | 0.020993309 | 0.020993309 | ENSG00000204209 | n.s. |
| ENSG00000163482 | 388.9638521 | -0.14609495 | 0.2448583 | -0.596651 | 0.550740358 | 0.718594654 | STK36 | protein_coding | 46.05 | -0.020966965 | 0.020966965 | ENSG00000163482 | n.s. |
| ENSG00000185238 | 1104.613663 | -0.159951708 | 0.2859656 | -0.559339 | 0.575930402 | 0.739682939 | PRMT3 | protein_coding | 36.02 | -0.02094638 | 0.02094638 | ENSG00000185238 | n.s. |
| ENSG00000037757 | 216.9107663 | -0.157364938 | 0.2781172 | -0.565822 | 0.571514505 | 0.736300218 | MRI1 | protein_coding | 52.16 | -0.020920893 | 0.020920893 | ENSG00000037757 | n.s. |
| ENSG00000146386 | 402.7555594 | 0.171243072 | 0.3215443 | 0.532564 | 0.594335154 | 0.754830464 | ABRACL | protein_coding | 44.2 | 0.020917441 | 0.020917441 | ENSG00000146386 | n.s. |
| ENSG00000165678 | 2421.784754 | 0.161006056 | 0.2898477 | 0.555485 | 0.578562993 | 0.741470228 | GHITM | protein_coding | 37.88 | 0.020915698 | 0.020915698 | ENSG00000165678 | n.s. |
| ENSG00000136521 | 1014.872985 | 0.211393295 | 0.4680341 | 0.451662 | 0.651512347 | 0.796400914 | NDUFB5 | protein_coding | 39.72 | 0.020900085 | 0.020900085 | ENSG00000136521 | n.s. |
| ENSG00000159479 | 214.6291026 | -0.156458049 | 0.2757798 | -0.56733 | 0.570490249 | 0.735388921 | MED8 | protein_coding | 46.78 | -0.020884477 | 0.020884477 | ENSG00000159479 | n.s. |
| ENSG00000125459 | 439.3484012 | -0.14450242 | 0.2406587 | -0.600445 | 0.548209406 | 0.717023913 | MSTO1 | protein_coding | 52.49 | -0.020875739 | 0.020875739 | ENSG00000125459 | n.s. |
| ENSG00000185829 | 210.2449775 | -0.151528291 | 0.2616727 | -0.579076 | 0.562538135 | 0.728375193 | ARL17A | protein_coding | 43.34 | -0.020857089 | 0.020857089 | ENSG00000185829 | n.s. |
| ENSG00000222858 | 9.00751344 | 0.201155673 | 0.4267379 | 0.47138 | 0.637369473 | 0.787618281 | RNU6-920P | snRNA | 41.3 | 0.020856667 | 0.020856667 | ENSG00000222858 | n.s. |
| ENSG00000075234 | 211.9106978 | -0.146406437 | 0.2466408 | -0.593602 | 0.552778395 | 0.720428119 | TTC38 | protein_coding | 54.16 | -0.020849645 | 0.020849645 | ENSG00000075234 | n.s. |
| ENSG00000125878 | 50.12985204 | 0.151344844 | 0.2611859 | 0.579453 | 0.562283767 | 0.728294762 | TCF15 | protein_coding | 56.76 | 0.020839097 | 0.020839097 | ENSG00000125878 | n.s. |
| ENSG00000146457 | 3358.719499 | 0.155123374 | 0.2722294 | 0.569826 | 0.56879569 | 0.733952029 | WTAP | protein_coding | 38.63 | 0.020838083 | 0.020838083 | ENSG00000146457 | n.s. |
| ENSG00000272636 | 662.4376453 | -0.148133068 | 0.2520045 | -0.587819 | 0.55665377 | 0.723491483 | DOC2B | protein_coding | 56.44 | -0.020822558 | 0.020822558 | ENSG00000272636 | n.s. |

| | | | | | | | | | | | | | |
|---|---|---|---|---|---|---|---|---|---|---|---|---|---|
| ENSG00000123064 | 1357.741966 | 0.139090972 | 0.2249983 | 0.618187 | 0.536452257 | 0.708435841 | DDX54 | protein_coding | 54.08 | 0.020821846 | 0.020821846 | ENSG00000123064 | n.s. |
| ENSG00000158985 | 534.0548282 | -0.149320605 | 0.2555506 | -0.584309 | 0.559012162 | 0.725428869 | CDC42SE2 | protein_coding | 39.71 | -0.020816064 | 0.020816064 | ENSG00000158985 | n.s. |
| ENSG00000138029 | 977.8878014 | 0.158830627 | 0.283996 | 0.559271 | 0.575977008 | 0.739682939 | HADHB | protein_coding | 41.22 | 0.020799569 | 0.020799569 | ENSG00000138029 | n.s. |
| ENSG00000102109 | 137.9190885 | 0.162249127 | 0.2960649 | 0.548019 | 0.583678974 | 0.745146008 | PCSK1N | protein_coding | 55.55 | 0.020728725 | 0.020728725 | ENSG00000102109 | n.s. |
| ENSG00000092094 | 639.0625618 | -0.146544167 | 0.2480829 | -0.590707 | 0.554717073 | 0.722109313 | OSGEP | protein_coding | 45.15 | -0.020720914 | 0.020720914 | ENSG00000092094 | n.s. |
| ENSG00000126947 | 276.0583223 | -0.159685422 | 0.2883355 | -0.553818 | 0.579703364 | 0.741907339 | ARMCX1 | protein_coding | 48.99 | -0.020703268 | 0.020703268 | ENSG00000126947 | n.s. |
| ENSG00000138085 | 1030.654066 | 0.15931692 | 0.2869237 | 0.555259 | 0.578717673 | 0.741528472 | ATRAID | protein_coding | 45.56 | 0.020690834 | 0.020690834 | ENSG00000138085 | n.s. |
| ENSG00000126249 | 153.1047691 | 0.167537685 | 0.3124121 | 0.536271 | 0.591770978 | 0.752509575 | PDCD2L | protein_coding | 45.07 | 0.020688889 | 0.020688889 | ENSG00000126249 | n.s. |
| ENSG00000011295 | 1877.247686 | 0.154457512 | 0.2717911 | 0.568295 | 0.569834637 | 0.734747884 | TTC19 | protein_coding | 41.07 | 0.020675938 | 0.020675938 | ENSG00000011295 | n.s. |
| ENSG00000121022 | 1200.603148 | -0.149954161 | 0.2588976 | -0.579203 | 0.562452415 | 0.728374265 | COPS5 | protein_coding | 39.82 | -0.020640501 | 0.020640501 | ENSG00000121022 | n.s. |
| ENSG00000092140 | 1927.935108 | -0.143309328 | 0.2394606 | -0.598467 | 0.549528265 | 0.717857379 | G2E3 | protein_coding | 35.48 | -0.020631073 | 0.020631073 | ENSG00000092140 | n.s. |
| ENSG00000174652 | 879.2103923 | -0.149053935 | 0.2566456 | -0.580777 | 0.561390608 | 0.727295193 | ZNF266 | protein_coding | 45 | -0.020612562 | 0.020612562 | ENSG00000174652 | n.s. |
| ENSG00000074219 | 460.5257559 | -0.158129363 | 0.2840551 | -0.556686 | 0.577742289 | 0.740759361 | TEAD2 | protein_coding | 53.57 | -0.02060787 | 0.02060787 | ENSG00000074219 | n.s. |
| ENSG00000147180 | 1903.4615 | -0.155871254 | 0.276923 | -0.562869 | 0.573524369 | 0.737797265 | ZNF711 | protein_coding | 34.34 | -0.020584819 | 0.020584819 | ENSG00000147180 | n.s. |
| ENSG00000250182 | 155.9130991 | 0.174979239 | 0.3384061 | 0.517069 | 0.605108145 | 0.762781412 | EEF1A1P13 | processed_pseudogene | 47.73 | 0.020577541 | 0.020577541 | ENSG00000250182 | n.s. |
| ENSG00000276966 | 3175.777228 | 0.128178434 | 0.1966771 | 0.65172 | 0.514581702 | 0.691022477 | HIST1H4E | protein_coding | 45.8 | 0.020573642 | 0.020573642 | ENSG00000276966 | n.s. |
| ENSG00000108469 | 406.6648981 | -0.141092475 | 0.233414 | -0.604473 | 0.545529224 | 0.714928782 | RECQL5 | protein_coding | 53.18 | -0.020562425 | 0.020562425 | ENSG00000108469 | n.s. |
| ENSG00000136156 | 1923.803302 | -0.167671142 | 0.3141003 | -0.533814 | 0.593470207 | 0.753992863 | ITM2B | protein_coding | 37.31 | -0.020561976 | 0.020561976 | ENSG00000136156 | n.s. |
| ENSG00000115457 | 923.0308925 | 0.158504277 | 0.2863314 | 0.553569 | 0.579873653 | 0.741940571 | IGFBP2 | protein_coding | 50.33 | 0.020547049 | 0.020547049 | ENSG00000115457 | n.s. |
| ENSG00000165410 | 1412.748567 | 0.145992396 | 0.2488387 | 0.586695 | 0.557408683 | 0.723818895 | CFL2 | protein_coding | 39.68 | 0.020492965 | 0.020492965 | ENSG00000165410 | n.s. |
| ENSG00000170310 | 214.8436012 | 0.174219469 | 0.3371689 | 0.516713 | 0.605356685 | 0.763025632 | STX8 | protein_coding | 43.43 | 0.020463971 | 0.020463971 | ENSG00000170310 | n.s. |
| ENSG00000163348 | 625.2165853 | -0.135738784 | 0.2183917 | -0.621538 | 0.53424549 | 0.707041054 | PYGO2 | protein_coding | 54.77 | -0.020436203 | 0.020436203 | ENSG00000163348 | n.s. |
| ENSG00000179820 | 121.217129 | -0.149697476 | 0.260311 | -0.575072 | 0.56524285 | 0.730639679 | MYADM | protein_coding | 58.37 | -0.020403279 | 0.020403279 | ENSG00000179820 | n.s. |
| ENSG00000206989 | 32.3100882 | -0.166950164 | 0.3135005 | -0.532536 | 0.594355123 | 0.754830464 | SNORD63 | snoRNA | 35.29 | -0.02039306 | 0.02039306 | ENSG00000206989 | n.s. |
| ENSG00000048140 | 528.9045577 | 0.160266838 | 0.2933676 | 0.5463 | 0.584859483 | 0.746310932 | TSPAN17 | protein_coding | 58.7 | 0.020366741 | 0.020366741 | ENSG00000048140 | n.s. |
| ENSG00000162702 | 3012.421804 | -0.136159967 | 0.2212357 | -0.615452 | 0.538256173 | 0.70969473 | ZNF281 | protein_coding | 43.03 | -0.020278089 | 0.020278089 | ENSG00000162702 | n.s. |
| ENSG00000224578 | 395.2139266 | -0.150181732 | 0.2633517 | -0.570271 | 0.568494244 | 0.733773704 | HNRNPA1P48 | transcribed_processed_pseudog | 45.03 | -0.02019011 | 0.02019011 | ENSG00000224578 | n.s. |
| ENSG00000115271 | 322.5412419 | 0.155245334 | 0.2793742 | 0.55569 | 0.57842307 | 0.741470228 | GCA | protein_coding | 36.25 | 0.020167344 | 0.020167344 | ENSG00000115271 | n.s. |
| ENSG00000208892 | 60.35433721 | -0.168740405 | 0.3223383 | -0.523489 | 0.600634287 | 0.759479718 | SNORA49 | snoRNA | 48.53 | -0.020161748 | 0.020161748 | ENSG00000208892 | n.s. |
| ENSG00000131504 | 873.3691063 | -0.137758061 | 0.2276048 | -0.605251 | 0.545012136 | 0.714419142 | DIAPH1 | protein_coding | 41.24 | -0.02011914 | 0.02011914 | ENSG00000131504 | n.s. |
| ENSG00000100902 | 844.8785774 | 0.145256434 | 0.2497406 | 0.581629 | 0.560816508 | 0.726957289 | PSMA6 | protein_coding | 42.27 | 0.020116724 | 0.020116724 | ENSG00000100902 | n.s. |
| ENSG00000160113 | 312.3768653 | 0.153190256 | 0.2733194 | 0.560481 | 0.575151522 | 0.73913925 | NR2F6 | protein_coding | 58.96 | 0.020109857 | 0.020109857 | ENSG00000160113 | n.s. |

| | | | | | | | | | | | | | |
|---|---|---|---|---|---|---|---|---|---|---|---|---|---|
| ENSG00000183023 | 465.7344059 | -0.139221423 | 0.231943 | -0.60024 | 0.548346502 | 0.717135858 | SLC8A1 | protein_coding | 37.43 | -0.020103373 | 0.020103373 | ENSG00000183023 | n.s. |
| ENSG00000119242 | 92.87307658 | -0.151856955 | 0.2692969 | -0.563902 | 0.57282111 | 0.737334773 | CCDC92 | protein_coding | 47.49 | -0.020096033 | 0.020096033 | ENSG00000119242 | n.s. |
| ENSG00000081026 | 2930.848718 | -0.150001425 | 0.2641506 | -0.567863 | 0.570127858 | 0.734989831 | MAGI3 | protein_coding | 37.11 | -0.020057991 | 0.020057991 | ENSG00000081026 | n.s. |
| ENSG00000183291 | 2388.574048 | 0.169599457 | 0.3266334 | 0.519235 | 0.603596952 | 0.761664412 | SELENOF | protein_coding | 37.87 | 0.020052818 | 0.020052818 | ENSG00000183291 | n.s. |
| ENSG00000273691 | 35.69450917 | -0.165357755 | 0.3125142 | -0.529121 | 0.596721717 | 0.756501939 | | sense_intronic | 39.08 | -0.020039699 | 0.020039699 | ENSG00000273691 | n.s. |
| ENSG00000162194 | 195.9674897 | -0.133772997 | 0.216231 | -0.618658 | 0.536141787 | 0.708360419 | LBHD1 | protein_coding | 52.45 | -0.020031933 | 0.020031933 | ENSG00000162194 | n.s. |
| ENSG00000122515 | 77.84854069 | 0.145554501 | 0.2515035 | 0.578737 | 0.56276637 | 0.728535209 | ZMIZ2 | protein_coding | 56.51 | 0.020020942 | 0.020020942 | ENSG00000122515 | n.s. |
| ENSG00000011478 | 85.99763792 | 0.146303036 | 0.2538468 | 0.576344 | 0.564382784 | 0.730016861 | QPCTL | protein_coding | 52.56 | 0.019994813 | 0.019994813 | ENSG00000011478 | n.s. |
| ENSG00000116670 | 274.3920672 | -0.151405315 | 0.2690596 | -0.56272 | 0.573625342 | 0.737858984 | MAD2L2 | protein_coding | 53.91 | -0.019989534 | 0.019989534 | ENSG00000116670 | n.s. |
| ENSG00000173486 | 231.2275693 | 0.157515773 | 0.2887192 | 0.545567 | 0.585363378 | 0.746712673 | FKBP2 | protein_coding | 63.61 | 0.019980321 | 0.019980321 | ENSG00000173486 | n.s. |
| ENSG00000277224 | 586.9564911 | 0.138465857 | 0.2309413 | 0.599572 | 0.548791784 | 0.717313933 | HIST1H2BF | protein_coding | 42.73 | 0.01997934 | 0.01997934 | ENSG00000277224 | n.s. |
| ENSG00000067113 | 219.9973168 | -0.155104117 | 0.2817378 | -0.550526 | 0.581958378 | 0.743904379 | PLPP1 | protein_coding | 39.62 | -0.019928224 | 0.019928224 | ENSG00000067113 | n.s. |
| ENSG00000114115 | 126.9310572 | 0.163978292 | 0.3093816 | 0.53002 | 0.59609838 | 0.756009997 | RBP1 | protein_coding | 48.01 | 0.019918847 | 0.019918847 | ENSG00000114115 | n.s. |
| ENSG00000163719 | 416.7576988 | -0.152610267 | 0.2739155 | -0.557144 | 0.577429294 | 0.740562644 | MTMR14 | protein_coding | 49.36 | -0.019906208 | 0.019906208 | ENSG00000163719 | n.s. |
| ENSG00000177030 | 403.3031195 | 0.158763424 | 0.2933362 | 0.541234 | 0.588346498 | 0.749249821 | DEAF1 | protein_coding | 53.21 | 0.019904703 | 0.019904703 | ENSG00000177030 | n.s. |
| ENSG00000204231 | 415.0831295 | -0.152138438 | 0.2723578 | -0.558598 | 0.576436291 | 0.739970725 | RXRB | protein_coding | 53.95 | -0.019897496 | 0.019897496 | ENSG00000204231 | n.s. |
| ENSG00000135720 | 3318.985508 | 0.142154958 | 0.242696 | 0.585733 | 0.558055239 | 0.724524719 | DYNC1LI2 | protein_coding | 45.09 | 0.01989413 | 0.01989413 | ENSG00000135720 | n.s. |
| ENSG00000186716 | 39.18006487 | 0.155355687 | 0.2829817 | 0.548996 | 0.583008471 | 0.74471172 | BCR | protein_coding | 53.07 | 0.019887363 | 0.019887363 | ENSG00000186716 | n.s. |
| ENSG00000205352 | 304.6868854 | -0.152031835 | 0.2722082 | -0.558513 | 0.57649414 | 0.739976759 | PRR13 | protein_coding | 48.6 | -0.019883015 | 0.019883015 | ENSG00000205352 | n.s. |
| ENSG00000071575 | 1021.602801 | -0.135992696 | 0.2246963 | -0.605229 | 0.545026927 | 0.714419142 | TRIB2 | protein_coding | 47.68 | -0.019861314 | 0.019861314 | ENSG00000071575 | n.s. |
| ENSG00000168092 | 1585.166829 | 0.150036462 | 0.2659923 | 0.564063 | 0.572711176 | 0.737334773 | PAFAH1B2 | protein_coding | 43.38 | 0.019855118 | 0.019855118 | ENSG00000168092 | n.s. |
| ENSG00000274001 | 49.26272046 | 0.152789914 | 0.2749994 | 0.555601 | 0.578483688 | 0.741470228 | | sense_intronic | 34.3 | 0.01984837 | 0.01984837 | ENSG00000274001 | n.s. |
| ENSG00000259494 | 1095.236212 | -0.133254436 | 0.2164194 | -0.615723 | 0.538077204 | 0.70969473 | MRPL46 | protein_coding | 41.02 | -0.019845373 | 0.019845373 | ENSG00000259494 | n.s. |
| ENSG00000171720 | 733.8902952 | 0.152128877 | 0.272936 | 0.557379 | 0.577268306 | 0.740546777 | HDAC3 | protein_coding | 45.27 | 0.019844832 | 0.019844832 | ENSG00000171720 | n.s. |
| ENSG00000279140 | 31.42771366 | 0.16747311 | 0.322376 | 0.519496 | 0.603414789 | 0.761663349 | | TEC | 65.57 | 0.019801508 | 0.019801508 | ENSG00000279140 | n.s. |
| ENSG00000278126 | 32.3411141 | 0.163967461 | 0.3109767 | 0.527266 | 0.598008873 | 0.757398121 | | sense_intronic | 43.43 | 0.019786901 | 0.019786901 | ENSG00000278126 | n.s. |
| ENSG00000143815 | 3691.060153 | 0.151690829 | 0.2723386 | 0.556993 | 0.577531886 | 0.740618727 | LBR | protein_coding | 41.69 | 0.01978129 | 0.01978129 | ENSG00000143815 | n.s. |
| ENSG00000096401 | 3468.066156 | 0.144901075 | 0.2519807 | 0.575048 | 0.565258644 | 0.730639679 | CDC5L | protein_coding | 38.5 | 0.019749545 | 0.019749545 | ENSG00000096401 | n.s. |
| ENSG00000144048 | 584.310678 | 0.156582674 | 0.2884934 | 0.54276 | 0.587295038 | 0.748321481 | DUSP11 | protein_coding | 40.29 | 0.019715604 | 0.019715604 | ENSG00000144048 | n.s. |
| ENSG00000196235 | 861.8108796 | 0.13663729 | 0.2280164 | 0.599243 | 0.549010768 | 0.717429051 | SUPT5H | protein_coding | 51.42 | 0.019705972 | 0.019705972 | ENSG00000196235 | n.s. |
| ENSG00000109180 | 3142.652719 | -0.15830246 | 0.2939602 | -0.538517 | 0.590220434 | 0.750812161 | OCIAD1 | protein_coding | 39.5 | -0.019703702 | 0.019703702 | ENSG00000109180 | n.s. |
| ENSG00000158457 | 101.1992682 | -0.152303304 | 0.2755844 | -0.552656 | 0.580499224 | 0.742509568 | TSPAN33 | protein_coding | 48.14 | -0.019692504 | 0.019692504 | ENSG00000158457 | n.s. |

| | | | | | | | | | | | | | |
|---|---|---|---|---|---|---|---|---|---|---|---|---|---|
| ENSG00000102218 | 208.4454486 | -0.155846554 | 0.2869637 | -0.543088 | 0.587069233 | 0.748102227 | RP2 | protein_coding | 39.34 | -0.019642752 | 0.019642752 | ENSG00000102218 | n.s. |
| ENSG00000197451 | 6182.080344 | 0.141542974 | 0.2431317 | 0.582166 | 0.560454896 | 0.726623849 | HNRNPAB | protein_coding | 53.64 | 0.019630645 | 0.019630645 | ENSG00000197451 | n.s. |
| ENSG00000121989 | 803.2917469 | 0.14434043 | 0.2517232 | 0.573409 | 0.566367551 | 0.73163253 | ACVR2A | protein_coding | 35.46 | 0.019588006 | 0.019588006 | ENSG00000121989 | n.s. |
| ENSG00000107518 | 323.5245463 | -0.153035603 | 0.2789124 | -0.548687 | 0.583220324 | 0.744838204 | ATRNL1 | protein_coding | 35.41 | -0.019579078 | 0.019579078 | ENSG00000107518 | n.s. |
| ENSG00000091140 | 1689.606108 | -0.150974896 | 0.2725331 | -0.553969 | 0.579600075 | 0.741907339 | DLD | protein_coding | 36.67 | -0.019573946 | 0.019573946 | ENSG00000091140 | n.s. |
| ENSG00000276931 | 26.34183644 | -0.165451342 | 0.3189164 | -0.518792 | 0.603905641 | 0.761955436 | | sense_intronic | 44.83 | -0.019534911 | 0.019534911 | ENSG00000276931 | n.s. |
| ENSG00000100220 | 2115.177187 | -0.153838337 | 0.2821283 | -0.545278 | 0.585562409 | 0.746728836 | RTCB | protein_coding | 41.29 | -0.019512405 | 0.019512405 | ENSG00000100220 | n.s. |
| ENSG00000134853 | 2474.660475 | -0.152239151 | 0.2770973 | -0.549407 | 0.582726273 | 0.744612509 | PDGFRA | protein_coding | 43.65 | -0.019497218 | 0.019497218 | ENSG00000134853 | n.s. |
| ENSG00000090054 | 854.1671896 | 0.152961096 | 0.2794903 | 0.547286 | 0.584182286 | 0.745651812 | SPTLC1 | protein_coding | 41.6 | 0.019497021 | 0.019497021 | ENSG00000090054 | n.s. |
| ENSG00000127527 | 60.50778951 | 0.162040877 | 0.3084539 | 0.525333 | 0.599352023 | 0.758409362 | EPS15L1 | protein_coding | 50.79 | 0.019460512 | 0.019460512 | ENSG00000127527 | n.s. |
| ENSG00000169564 | 1294.495565 | 0.133875638 | 0.2219711 | 0.603122 | 0.546427596 | 0.715567289 | PCBP1 | protein_coding | 55.2 | 0.01945876 | 0.01945876 | ENSG00000169564 | n.s. |
| ENSG00000151876 | 105.0331454 | -0.144668164 | 0.2539199 | -0.569739 | 0.568854477 | 0.733959863 | FBXO4 | protein_coding | 38.88 | -0.01943294 | 0.01943294 | ENSG00000151876 | n.s. |
| ENSG00000163608 | 1281.918642 | 0.142834759 | 0.2487195 | 0.57428 | 0.565778077 | 0.731142395 | NEPRO | protein_coding | 39.07 | 0.019425246 | 0.019425246 | ENSG00000163608 | n.s. |
| ENSG00000103260 | 28.97458836 | 0.174997539 | 0.3531447 | 0.495541 | 0.620218615 | 0.774536171 | METRN | protein_coding | 66.79 | 0.019417429 | 0.019417429 | ENSG00000103260 | n.s. |
| ENSG00000177556 | 230.8336197 | 0.159016166 | 0.2988111 | 0.532163 | 0.594613189 | 0.755038999 | ATOX1 | protein_coding | 46.3 | 0.019404841 | 0.019404841 | ENSG00000177556 | n.s. |
| ENSG00000201098 | 63.05455383 | -0.157451594 | 0.294038 | -0.53548 | 0.592317709 | 0.75299849 | RNY1 | misc_RNA | 44.25 | -0.019398964 | 0.019398964 | ENSG00000201098 | n.s. |
| ENSG00000102572 | 895.0037425 | 0.141762113 | 0.246531 | 0.575028 | 0.565272657 | 0.730639679 | STK24 | protein_coding | 47.42 | 0.019321715 | 0.019321715 | ENSG00000102572 | n.s. |
| ENSG00000173141 | 1023.361301 | 0.142015253 | 0.2473345 | 0.574183 | 0.565843944 | 0.731146373 | MRPL57 | protein_coding | 50 | 0.019313459 | 0.019313459 | ENSG00000173141 | n.s. |
| ENSG00000143436 | 1413.673086 | 0.154692885 | 0.2867209 | 0.539524 | 0.589525114 | 0.750270399 | MRPL9 | protein_coding | 48.42 | 0.019302917 | 0.019302917 | ENSG00000143436 | n.s. |
| ENSG00000164082 | 186.6499127 | 0.137819562 | 0.2354686 | 0.585299 | 0.558346681 | 0.72474179 | GRM2 | protein_coding | 55.81 | 0.019269476 | 0.019269476 | ENSG00000164082 | n.s. |
| ENSG00000139579 | 928.5210824 | 0.158216522 | 0.2978881 | 0.531127 | 0.595330573 | 0.755469937 | NABP2 | protein_coding | 51.22 | 0.019268053 | 0.019268053 | ENSG00000139579 | n.s. |
| ENSG00000171155 | 185.9785025 | 0.163248521 | 0.3148871 | 0.518435 | 0.604154722 | 0.762131576 | C1GALT1C1 | protein_coding | 40.73 | 0.019258435 | 0.019258435 | ENSG00000171155 | n.s. |
| ENSG00000201076 | 8.175219711 | -0.245679138 | 0.6679078 | -0.367834 | 0.712997085 | 0.835259438 | RNU4-51P | snRNA | 37.14 | -0.019206853 | 0.019206853 | ENSG00000201076 | n.s. |
| ENSG00000166407 | 72.27454674 | 0.155752977 | 0.2907201 | 0.535749 | 0.592132079 | 0.752831242 | LMO1 | protein_coding | 52.88 | 0.019204711 | 0.019204711 | ENSG00000166407 | n.s. |
| ENSG00000100994 | 1677.661243 | -0.14339377 | 0.2522684 | -0.568418 | 0.569751466 | 0.734708684 | PYGB | protein_coding | 52.17 | -0.01919825 | 0.01919825 | ENSG00000100994 | n.s. |
| ENSG00000236824 | 41.38296631 | 0.164446787 | 0.3199806 | 0.513927 | 0.607302755 | 0.764302292 | BCYRN1 | scRNA | 53 | 0.019196669 | 0.019196669 | ENSG00000236824 | n.s. |
| ENSG00000136933 | 1021.157363 | -0.163878195 | 0.3177324 | -0.515774 | 0.606012094 | 0.763644347 | RABEPK | protein_coding | 46.49 | -0.019191588 | 0.019191588 | ENSG00000136933 | n.s. |
| ENSG00000166164 | 2273.28494 | -0.13966111 | 0.2415878 | -0.578097 | 0.563198881 | 0.728832558 | BRD7 | protein_coding | 39.72 | -0.019185559 | 0.019185559 | ENSG00000166164 | n.s. |
| ENSG00000168917 | 1036.822921 | -0.140316993 | 0.2435736 | -0.576076 | 0.564563586 | 0.73018289 | SLC35G2 | protein_coding | 41.28 | -0.01916286 | 0.01916286 | ENSG00000168917 | n.s. |
| ENSG00000166265 | 435.1611535 | -0.145451332 | 0.2591217 | -0.561324 | 0.574576343 | 0.7386046 | CYYR1 | protein_coding | 37.29 | -0.019139649 | 0.019139649 | ENSG00000166265 | n.s. |
| ENSG00000123965 | 33.31244055 | -0.162329123 | 0.3137389 | -0.517402 | 0.604875558 | 0.762695381 | PMS2P5 | unprocessed_pseudogene | 38.92 | -0.01909784 | 0.01909784 | ENSG00000123965 | n.s. |
| ENSG00000147155 | 492.7834122 | 0.159989651 | 0.3060344 | 0.522783 | 0.601125098 | 0.759824307 | EBP | protein_coding | 49.69 | 0.019084656 | 0.019084656 | ENSG00000147155 | n.s. |

| | | | | | | | | | | | | | |
|---|---|---|---|---|---|---|---|---|---|---|---|---|---|
| ENSG00000114796 | 1380.662609 | -0.156623071 | 0.2947746 | -0.531332 | 0.595188966 | 0.755469937 | KLHL24 | protein_coding | 39.02 | -0.019073998 | 0.019073998 | ENSG00000114796 | n.s. |
| ENSG00000063241 | 223.3773437 | 0.169898544 | 0.3396517 | 0.500214 | 0.616924368 | 0.77222314 | ISOC2 | protein_coding | 56.69 | 0.019072333 | 0.019072333 | ENSG00000063241 | n.s. |
| ENSG00000256341 | 37.7804703 | 0.152785105 | 0.2832685 | 0.539365 | 0.589635104 | 0.750341793 | | antisense | 49.31 | 0.019058546 | 0.019058546 | ENSG00000256341 | n.s. |
| ENSG00000214174 | 405.461404 | -0.145316374 | 0.2597902 | -0.55936 | 0.575915787 | 0.739682939 | AMZ2P1 | transcribed_unprocessed_pseud | 41.33 | -0.019029818 | 0.019029818 | ENSG00000214174 | n.s. |
| ENSG00000241258 | 642.4814167 | -0.155719303 | 0.2929854 | -0.531492 | 0.595078079 | 0.755469937 | CRCP | protein_coding | 45 | -0.018963935 | 0.018963935 | ENSG00000241258 | n.s. |
| ENSG00000267649 | 46.20207403 | -0.150103501 | 0.2760802 | -0.543695 | 0.586651204 | 0.7477064 | | lincRNA | 48.38 | -0.018953404 | 0.018953404 | ENSG00000267649 | n.s. |
| ENSG00000178567 | 3133.627013 | 0.125861498 | 0.2026132 | 0.621191 | 0.534473888 | 0.707125955 | EPM2AIP1 | protein_coding | 38.52 | 0.018942561 | 0.018942561 | ENSG00000178567 | n.s. |
| ENSG00000050030 | 533.5133045 | -0.140605059 | 0.2465827 | -0.570215 | 0.568532123 | 0.733773704 | KIAA2022 | protein_coding | 39.49 | -0.018902643 | 0.018902643 | ENSG00000050030 | n.s. |
| ENSG00000128311 | 158.6424582 | 0.151417393 | 0.2808042 | 0.539228 | 0.589729831 | 0.750393752 | TST | protein_coding | 57.04 | 0.018883383 | 0.018883383 | ENSG00000128311 | n.s. |
| ENSG00000116213 | 127.1673033 | -0.153955178 | 0.2884349 | -0.533761 | 0.593507199 | 0.753992863 | WRAP73 | protein_coding | 55.25 | -0.018879949 | 0.018879949 | ENSG00000116213 | n.s. |
| ENSG00000137106 | 740.9263509 | 0.153781768 | 0.2879049 | 0.534141 | 0.593244086 | 0.75396964 | GRHPR | protein_coding | 53.09 | 0.01886074 | 0.01886074 | ENSG00000137106 | n.s. |
| ENSG00000115504 | 149.0316633 | -0.146027994 | 0.2644697 | -0.552154 | 0.580842863 | 0.742761786 | EHBP1 | protein_coding | 36.7 | -0.018859581 | 0.018859581 | ENSG00000115504 | n.s. |
| ENSG00000147274 | 11161.93182 | -0.148666522 | 0.2725258 | -0.545514 | 0.585400317 | 0.746712673 | RBMX | protein_coding | 43.01 | -0.018857825 | 0.018857825 | ENSG00000147274 | n.s. |
| ENSG00000141198 | 1435.975633 | -0.148478282 | 0.2722213 | -0.545432 | 0.585456223 | 0.746712673 | TOM1L1 | protein_coding | 40.86 | -0.018833947 | 0.018833947 | ENSG00000141198 | n.s. |
| ENSG00000103042 | 59.16103777 | -0.144492004 | 0.2596108 | -0.556572 | 0.577820091 | 0.740790896 | SLC38A7 | protein_coding | 52.78 | -0.018827939 | 0.018827939 | ENSG00000103042 | n.s. |
| ENSG00000057468 | 130.2111994 | -0.142096722 | 0.2519969 | -0.563883 | 0.572833875 | 0.737334773 | MSH4 | protein_coding | 37.01 | -0.01880441 | 0.01880441 | ENSG00000057468 | n.s. |
| ENSG00000145990 | 393.9852841 | -0.135949826 | 0.2340132 | -0.580949 | 0.561274594 | 0.72728024 | GFOD1 | protein_coding | 44.97 | -0.018801618 | 0.018801618 | ENSG00000145990 | n.s. |
| ENSG00000102309 | 608.8852699 | -0.155549516 | 0.294874 | -0.527512 | 0.597838267 | 0.757250927 | PIN4 | protein_coding | 44.57 | -0.018784189 | 0.018784189 | ENSG00000102309 | n.s. |
| ENSG00000184508 | 240.7059306 | 0.144921617 | 0.2618561 | 0.55344 | 0.579962165 | 0.741965747 | HDDC3 | protein_coding | 56.14 | 0.018784181 | 0.018784181 | ENSG00000184508 | n.s. |
| ENSG00000102119 | 286.7335421 | 0.145629904 | 0.2640065 | 0.551615 | 0.581212287 | 0.743147373 | EMD | protein_coding | 62.14 | 0.018775343 | 0.018775343 | ENSG00000102119 | n.s. |
| ENSG00000230082 | 56.33945755 | 0.1444115 | 0.260321 | 0.554744 | 0.579069751 | 0.741778282 | PRRT3-AS1 | antisense | 54.05 | 0.01873391 | 0.01873391 | ENSG00000230082 | n.s. |
| ENSG00000198393 | 1518.079379 | -0.142743752 | 0.254972 | -0.559841 | 0.575587994 | 0.739563644 | ZNF26 | protein_coding | 43.05 | -0.018702921 | 0.018702921 | ENSG00000198393 | n.s. |
| ENSG00000213585 | 3591.974812 | 0.157390615 | 0.301818 | 0.521475 | 0.602035703 | 0.760677787 | VDAC1 | protein_coding | 47.42 | 0.01869789 | 0.01869789 | ENSG00000213585 | n.s. |
| ENSG00000265056 | 8.453820315 | -0.189337042 | 0.4198785 | -0.450933 | 0.652037916 | 0.796633653 | MIR548S | miRNA | 26.83 | -0.018695395 | 0.018695395 | ENSG00000265056 | n.s. |
| ENSG00000167363 | 96.62939772 | 0.141392518 | 0.2509097 | 0.56352 | 0.573081128 | 0.737567808 | FN3K | protein_coding | 51.33 | 0.018691815 | 0.018691815 | ENSG00000167363 | n.s. |
| ENSG00000120756 | 39.74333208 | -0.161335673 | 0.3160531 | -0.51047 | 0.609722114 | 0.766031492 | PLS1 | protein_coding | 38.52 | -0.018675149 | 0.018675149 | ENSG00000120756 | n.s. |
| ENSG00000169683 | 487.2006551 | 0.143824383 | 0.2596349 | 0.553949 | 0.579614086 | 0.741907339 | LRRC45 | protein_coding | 64.94 | 0.018646879 | 0.018646879 | ENSG00000169683 | n.s. |
| ENSG00000112531 | 2593.495687 | -0.135459236 | 0.2338915 | -0.579154 | 0.562485104 | 0.728374265 | QKI | protein_coding | 36.39 | -0.018645342 | 0.018645342 | ENSG00000112531 | n.s. |
| ENSG00000166426 | 238.5539729 | 0.152035914 | 0.2848592 | 0.533723 | 0.593533131 | 0.753992863 | CRABP1 | protein_coding | 48.64 | 0.018644585 | 0.018644585 | ENSG00000166426 | n.s. |
| ENSG00000179222 | 817.228869 | 0.146931663 | 0.2694143 | 0.545374 | 0.585496104 | 0.746712673 | MAGED1 | protein_coding | 39 | 0.018637764 | 0.018637764 | ENSG00000179222 | n.s. |
| ENSG00000065268 | 439.8949881 | 0.152897734 | 0.2879063 | 0.531068 | 0.595371882 | 0.755469937 | WDR18 | protein_coding | 62.29 | 0.018620316 | 0.018620316 | ENSG00000065268 | n.s. |
| ENSG00000083457 | 274.1006941 | -0.159423925 | 0.3100944 | -0.514114 | 0.607172135 | 0.764302292 | ITGAE | protein_coding | 50.88 | -0.018610326 | 0.018610326 | ENSG00000083457 | n.s. |

| | | | | | | | | | | | | | |
|---|---|---|---|---|---|---|---|---|---|---|---|---|---|
| ENSG00000171723 | 361.8467585 | -0.147392794 | 0.2711864 | -0.543511 | 0.586778055 | 0.747799621 | GPHN | protein_coding | 36.43 | -0.018603146 | 0.018603146 | ENSG00000171723 | n.s. |
| ENSG00000197308 | 626.7433004 | 0.128060013 | 0.2126331 | 0.602258 | 0.547002243 | 0.715915795 | GATA3-AS1 | lincRNA | 60.72 | 0.018586381 | 0.018586381 | ENSG00000197308 | n.s. |
| ENSG00000141456 | 545.6838334 | -0.136291174 | 0.2370237 | -0.575011 | 0.565284107 | 0.730639679 | PELP1 | protein_coding | 48.03 | -0.018576044 | 0.018576044 | ENSG00000141456 | n.s. |
| ENSG00000165209 | 1030.926833 | -0.146830313 | 0.2697611 | -0.544298 | 0.58623668 | 0.747314896 | STRBP | protein_coding | 39.12 | -0.0185735 | 0.0185735 | ENSG00000165209 | n.s. |
| ENSG00000150527 | 42.43588107 | 0.153141178 | 0.2894544 | 0.529068 | 0.596757939 | 0.756501939 | CTAGE5 | protein_coding | 38.28 | 0.018559172 | 0.018559172 | ENSG00000150527 | n.s. |
| ENSG00000114742 | 1512.2669 | -0.148380012 | 0.2748468 | -0.539864 | 0.589290554 | 0.750109955 | WDR48 | protein_coding | 39.95 | -0.018528964 | 0.018528964 | ENSG00000114742 | n.s. |
| ENSG00000135829 | 16772.16425 | 0.143171379 | 0.2590936 | 0.552586 | 0.580547235 | 0.742509568 | DHX9 | protein_coding | 38.24 | 0.018511765 | 0.018511765 | ENSG00000135829 | n.s. |
| ENSG00000002330 | 220.7472976 | 0.158243955 | 0.3072867 | 0.514972 | 0.606572765 | 0.764004486 | BAD | protein_coding | 53.92 | 0.018499366 | 0.018499366 | ENSG00000002330 | n.s. |
| ENSG00000142192 | 4129.762938 | -0.14552704 | 0.2664261 | -0.546219 | 0.584915331 | 0.746313797 | APP | protein_coding | 39.73 | -0.018493362 | 0.018493362 | ENSG00000142192 | n.s. |
| ENSG00000099381 | 300.9233971 | -0.134520686 | 0.2325268 | -0.578517 | 0.562915194 | 0.728660119 | SETD1A | protein_coding | 55.71 | -0.018493231 | 0.018493231 | ENSG00000099381 | n.s. |
| ENSG00000115687 | 815.4914368 | -0.141192027 | 0.2527556 | -0.558611 | 0.576427261 | 0.739970725 | PASK | protein_coding | 50.98 | -0.018465864 | 0.018465864 | ENSG00000115687 | n.s. |
| ENSG00000197386 | 73.832658 | 0.16208566 | 0.3212813 | 0.504498 | 0.613911747 | 0.769489849 | HTT | protein_coding | 45.52 | 0.018444879 | 0.018444879 | ENSG00000197386 | n.s. |
| ENSG00000125744 | 353.1619736 | -0.146246215 | 0.2695129 | -0.542632 | 0.587383528 | 0.748349663 | RTN2 | protein_coding | 54.15 | -0.01841173 | 0.01841173 | ENSG00000125744 | n.s. |
| ENSG00000224167 | 32.2543352 | 0.157904245 | 0.3077284 | 0.513129 | 0.607861357 | 0.764659711 | | lincRNA | 45.99 | 0.018400865 | 0.018400865 | ENSG00000224167 | n.s. |
| ENSG00000111788 | 301.1280102 | -0.144275346 | 0.2636791 | -0.547163 | 0.584267025 | 0.745691611 | | unprocessed_pseudogene | 48.72 | -0.018386557 | 0.018386557 | ENSG00000111788 | n.s. |
| ENSG00000267197 | 32.40710256 | -0.162202474 | 0.3225065 | -0.502943 | 0.615004102 | 0.770364212 | | sense_intronic | 49.4 | -0.018378173 | 0.018378173 | ENSG00000267197 | n.s. |
| ENSG00000260751 | 49.14117376 | -0.15167622 | 0.2869646 | -0.528554 | 0.597115034 | 0.756747917 | | sense_intronic | 43.65 | -0.018360219 | 0.018360219 | ENSG00000260751 | n.s. |
| ENSG00000111877 | 1087.223119 | 0.141343255 | 0.255257 | 0.553729 | 0.579764172 | 0.741916964 | MCM9 | protein_coding | 39.07 | 0.018324404 | 0.018324404 | ENSG00000111877 | n.s. |
| ENSG00000230383 | 51.79199795 | 0.170102569 | 0.35085 | 0.48483 | 0.62779706 | 0.78043022 | | processed_pseudogene | 46.64 | 0.01831425 | 0.01831425 | ENSG00000230383 | n.s. |
| ENSG00000125870 | 3028.568855 | 0.155624471 | 0.3006382 | 0.517647 | 0.604704566 | 0.762683405 | SNRPB2 | protein_coding | 38.18 | 0.018310107 | 0.018310107 | ENSG00000125870 | n.s. |
| ENSG00000161203 | 2086.355719 | 0.148722723 | 0.2781645 | 0.534658 | 0.592886703 | 0.753653024 | AP2M1 | protein_coding | 54.13 | 0.018267397 | 0.018267397 | ENSG00000161203 | n.s. |
| ENSG00000170390 | 219.1867553 | -0.142720749 | 0.260216 | -0.54847 | 0.583369064 | 0.744838204 | DCLK2 | protein_coding | 40.72 | -0.018259415 | 0.018259415 | ENSG00000170390 | n.s. |
| ENSG00000159069 | 325.8424339 | 0.1488588 | 0.2790178 | 0.53351 | 0.593680604 | 0.754111399 | FBXW5 | protein_coding | 68.96 | 0.018244804 | 0.018244804 | ENSG00000159069 | n.s. |
| ENSG00000184209 | 628.7460348 | -0.128729328 | 0.2176598 | -0.591424 | 0.554236051 | 0.721787348 | SNRNP35 | protein_coding | 47.6 | -0.01822688 | 0.01822688 | ENSG00000184209 | n.s. |
| ENSG00000219200 | 222.5594725 | 0.147485812 | 0.2750718 | 0.536172 | 0.591839698 | 0.752528229 | RNASEK | protein_coding | 57.04 | 0.018211136 | 0.018211136 | ENSG00000219200 | n.s. |
| ENSG00000277157 | 1435.058808 | 0.123663947 | 0.2030784 | 0.608947 | 0.542559644 | 0.712661525 | HIST1H4D | protein_coding | 57.37 | 0.01819303 | 0.01819303 | ENSG00000277157 | n.s. |
| ENSG00000196357 | 91.3121724 | -0.139889637 | 0.2520485 | -0.555011 | 0.578887237 | 0.741612716 | ZNF565 | protein_coding | 45.12 | -0.018160869 | 0.018160869 | ENSG00000196357 | n.s. |
| ENSG00000166483 | 3203.529704 | 0.123059901 | 0.2015868 | 0.610456 | 0.54155967 | 0.711937993 | WEE1 | protein_coding | 40.54 | 0.018158452 | 0.018158452 | ENSG00000166483 | n.s. |
| ENSG00000145349 | 854.9264597 | -0.139997546 | 0.2525332 | -0.554373 | 0.579323665 | 0.741830534 | CAMK2D | protein_coding | 35.99 | -0.018157023 | 0.018157023 | ENSG00000145349 | n.s. |
| ENSG00000196968 | 228.0310751 | 0.136031866 | 0.2398781 | 0.567087 | 0.570654779 | 0.735532909 | FUT11 | protein_coding | 47.49 | 0.018146364 | 0.018146364 | ENSG00000196968 | n.s. |
| ENSG00000117862 | 1088.444662 | 0.152706426 | 0.2929669 | 0.521241 | 0.602198794 | 0.760767064 | TXNDC12 | protein_coding | 43.03 | 0.018133628 | 0.018133628 | ENSG00000117862 | n.s. |
| ENSG00000207484 | 9.237372835 | -0.180219815 | 0.3927338 | -0.458885 | 0.646316408 | 0.793440055 | | misc_RNA | 43.88 | -0.018109547 | 0.018109547 | ENSG00000207484 | n.s. |

| | | | | | | | | | | | | | |
|---|---|---|---|---|---|---|---|---|---|---|---|---|---|
| ENSG00000147586 | 508.4509897 | -0.150041045 | 0.2849317 | -0.526586 | 0.598481119 | 0.757720525 | MRPS28 | protein_coding | 37.19 | -0.018078588 | 0.018078588 | ENSG00000147586 | n.s. |
| ENSG00000216866 | 57.64735258 | 0.165909812 | 0.3395553 | 0.488609 | 0.625118499 | 0.77842097 | RPS2P55 | processed_pseudogene | 60.82 | 0.018048577 | 0.018048577 | ENSG00000216866 | n.s. |
| ENSG00000127054 | 351.8134171 | 0.148826507 | 0.2811419 | 0.529364 | 0.596552691 | 0.756448371 | INTS11 | protein_coding | 60.46 | 0.018040855 | 0.018040855 | ENSG00000127054 | n.s. |
| ENSG00000148331 | 612.4351187 | -0.137751914 | 0.2463209 | -0.559238 | 0.5759996 | 0.739682939 | ASB6 | protein_coding | 58.84 | -0.018039219 | 0.018039219 | ENSG00000148331 | n.s. |
| ENSG00000141540 | 318.902849 | -0.138270014 | 0.2481336 | -0.55724 | 0.5773634 | 0.740546777 | TTYH2 | protein_coding | 54.47 | -0.018036978 | 0.018036978 | ENSG00000141540 | n.s. |
| ENSG00000196155 | 499.2679193 | 0.144389984 | 0.2675067 | 0.539762 | 0.589361037 | 0.750130151 | PLEKHG4 | protein_coding | 59.77 | 0.018029021 | 0.018029021 | ENSG00000196155 | n.s. |
| ENSG00000124422 | 5135.883674 | -0.137374019 | 0.2452606 | -0.560114 | 0.575401372 | 0.739392091 | USP22 | protein_coding | 47.95 | -0.018013196 | 0.018013196 | ENSG00000124422 | n.s. |
| ENSG00000265345 | 4.736588736 | -0.206219186 | 0.5090231 | -0.405127 | 0.685383908 | 0.817983898 | MIR5188 | miRNA | 39.82 | -0.017993706 | 0.017993706 | ENSG00000265345 | n.s. |
| ENSG00000128228 | 105.2126695 | 0.155130846 | 0.3037124 | 0.510782 | 0.609503663 | 0.765964392 | SDF2L1 | protein_coding | 62.46 | 0.017962821 | 0.017962821 | ENSG00000128228 | n.s. |
| ENSG00000104147 | 288.2446136 | 0.15531713 | 0.30466 | 0.509805 | 0.610188208 | 0.766478747 | OIP5 | protein_coding | 45.46 | 0.01793911 | 0.01793911 | ENSG00000104147 | n.s. |
| ENSG00000237137 | 6.761093339 | 0.198279934 | 0.4746646 | 0.417726 | 0.676147192 | 0.811971481 | | processed_pseudogene | 44.88 | 0.017936249 | 0.017936249 | ENSG00000237137 | n.s. |
| ENSG00000269997 | 61.39592558 | 0.148039235 | 0.2803149 | 0.528118 | 0.597417751 | 0.756924866 | | sense_intronic | 41.59 | 0.017904935 | 0.017904935 | ENSG00000269997 | n.s. |
| ENSG00000106330 | 67.96024026 | 0.151013122 | 0.2900291 | 0.520683 | 0.602587863 | 0.7611097 | MOSPD3 | protein_coding | 57.23 | 0.01790302 | 0.01790302 | ENSG00000106330 | n.s. |
| ENSG00000149136 | 4664.475033 | 0.139832796 | 0.2549609 | 0.548448 | 0.583384372 | 0.744838204 | SSRP1 | protein_coding | 50.54 | 0.017889936 | 0.017889936 | ENSG00000149136 | n.s. |
| ENSG00000266507 | 7.163255492 | 0.18731114 | 0.4268656 | 0.438806 | 0.660802221 | 0.802752755 | MIR4479 | miRNA | 70.42 | 0.017872891 | 0.017872891 | ENSG00000266507 | n.s. |
| ENSG00000049883 | 400.951384 | -0.126399241 | 0.2139936 | -0.590668 | 0.5547426 | 0.722109313 | PTCD2 | protein_coding | 40.38 | -0.01787248 | 0.01787248 | ENSG00000049883 | n.s. |
| ENSG00000163364 | 180.2650889 | 0.148399596 | 0.2822505 | 0.525773 | 0.599046121 | 0.758240023 | LINC01116 | lincRNA | 38.98 | 0.017836636 | 0.017836636 | ENSG00000163364 | n.s. |
| ENSG00000048540 | 21.87365521 | -0.15952448 | 0.320237 | -0.498145 | 0.618381754 | 0.773221727 | LMO3 | protein_coding | 35.63 | -0.017818238 | 0.017818238 | ENSG00000048540 | n.s. |
| ENSG00000085449 | 966.0964722 | -0.143455201 | 0.2668808 | -0.537525 | 0.590904752 | 0.751476695 | WDFY1 | protein_coding | 42.72 | -0.017800564 | 0.017800564 | ENSG00000085449 | n.s. |
| ENSG00000076555 | 55.63548764 | 0.146821259 | 0.2774371 | 0.529206 | 0.596662848 | 0.756501939 | ACACB | protein_coding | 46.78 | 0.017793262 | 0.017793262 | ENSG00000076555 | n.s. |
| ENSG00000164022 | 1330.25044 | -0.150236105 | 0.2893526 | -0.519215 | 0.603611034 | 0.761664412 | AIMP1 | protein_coding | 36.71 | -0.017763367 | 0.017763367 | ENSG00000164022 | n.s. |
| ENSG00000023191 | 498.9984518 | 0.141543842 | 0.2613985 | 0.541487 | 0.588172029 | 0.749096155 | RNH1 | protein_coding | 60.36 | 0.017758435 | 0.017758435 | ENSG00000023191 | n.s. |
| ENSG00000037897 | 417.1413198 | -0.126270642 | 0.2149209 | -0.587521 | 0.556853626 | 0.723497989 | METTL1 | protein_coding | 54.22 | -0.017748939 | 0.017748939 | ENSG00000037897 | n.s. |
| ENSG00000148834 | 863.4739458 | 0.162476524 | 0.3318602 | 0.489593 | 0.624421723 | 0.777831587 | GSTO1 | protein_coding | 41.43 | 0.017728532 | 0.017728532 | ENSG00000148834 | n.s. |
| ENSG00000127415 | 13.70831478 | 0.174604476 | 0.3759334 | 0.464456 | 0.64232114 | 0.791643267 | IDUA | protein_coding | 65.45 | 0.0177172 | 0.0177172 | ENSG00000127415 | n.s. |
| ENSG00000182199 | 3104.11154 | 0.146115763 | 0.2762196 | 0.528984 | 0.596816488 | 0.756507279 | SHMT2 | protein_coding | 56.68 | 0.017707315 | 0.017707315 | ENSG00000182199 | n.s. |
| ENSG00000122417 | 1332.581771 | -0.142043149 | 0.2638072 | -0.538435 | 0.590276494 | 0.750814875 | ODF2L | protein_coding | 34.88 | -0.017679703 | 0.017679703 | ENSG00000122417 | n.s. |
| ENSG00000240211 | 10.44870794 | -0.178555603 | 0.3943195 | -0.45282 | 0.650678622 | 0.796220641 | | antisense | 53.54 | -0.017671035 | 0.017671035 | ENSG00000240211 | n.s. |
| ENSG00000273247 | 260.5057315 | -0.128742993 | 0.2230541 | -0.577183 | 0.563815977 | 0.729487023 | | antisense | 41.5 | -0.017635528 | 0.017635528 | ENSG00000273247 | n.s. |
| ENSG00000132463 | 2756.162553 | -0.142005548 | 0.2641077 | -0.537681 | 0.590797609 | 0.751409072 | GRSF1 | protein_coding | 43.53 | -0.017626235 | 0.017626235 | ENSG00000132463 | n.s. |
| ENSG00000066027 | 213.2533572 | 0.135710403 | 0.2447121 | 0.554572 | 0.579187627 | 0.741830534 | PPP2R5A | protein_coding | 39.28 | 0.017601001 | 0.017601001 | ENSG00000066027 | n.s. |
| ENSG00000224914 | 169.8929993 | 0.129380137 | 0.2254039 | 0.573993 | 0.565972859 | 0.731190509 | LINC00863 | lincRNA | 39.66 | 0.017591747 | 0.017591747 | ENSG00000224914 | n.s. |

| | | | | | | | | | | | | | |
|---|---|---|---|---|---|---|---|---|---|---|---|---|---|
| ENSG00000151468 | 568.6268661 | 0.138741638 | 0.2546722 | 0.544785 | 0.585901315 | 0.747024177 | CCDC3 | protein_coding | 43.73 | 0.017573757 | 0.017573757 | ENSG00000151468 | n.s. |
| ENSG00000178996 | 2086.509089 | 0.133022079 | 0.2362502 | 0.563056 | 0.573396834 | 0.737755431 | SNX18 | protein_coding | 42.68 | 0.017570565 | 0.017570565 | ENSG00000178996 | n.s. |
| ENSG00000251791 | 34.47432083 | -0.151190915 | 0.2958226 | -0.511086 | 0.609290508 | 0.76583477 | SCARNA6 | scaRNA | 53.21 | -0.017517723 | 0.017517723 | ENSG00000251791 | n.s. |
| ENSG00000169919 | 619.7680644 | -0.131952861 | 0.2334574 | -0.565212 | 0.571929789 | 0.736656315 | GUSB | protein_coding | 52.03 | -0.017514774 | 0.017514774 | ENSG00000169919 | n.s. |
| ENSG00000072110 | 510.7172442 | -0.135037646 | 0.2439499 | -0.553547 | 0.579889197 | 0.741940571 | ACTN1 | protein_coding | 50.81 | -0.017505049 | 0.017505049 | ENSG00000072110 | n.s. |
| ENSG00000189077 | 82.01316637 | 0.162157901 | 0.3339661 | 0.485552 | 0.62728488 | 0.779991215 | TMEM120A | protein_coding | 61.59 | 0.017498506 | 0.017498506 | ENSG00000189077 | n.s. |
| ENSG00000176022 | 238.5792108 | -0.126608557 | 0.2180859 | -0.580545 | 0.561547478 | 0.727430734 | B3GALT6 | protein_coding | 64.03 | -0.01749836 | 0.01749836 | ENSG00000176022 | n.s. |
| ENSG00000138303 | 267.475699 | -0.148687256 | 0.2874661 | -0.517234 | 0.604992841 | 0.762774186 | ASCC1 | protein_coding | 41.33 | -0.017486218 | 0.017486218 | ENSG00000138303 | n.s. |
| ENSG00000067840 | 903.1741157 | 0.134590609 | 0.2427171 | 0.554516 | 0.579225544 | 0.741830534 | PDZD4 | protein_coding | 56.35 | 0.017455769 | 0.017455769 | ENSG00000067840 | n.s. |
| ENSG00000173914 | 674.977883 | -0.135393626 | 0.2454802 | -0.551546 | 0.581259419 | 0.743147373 | RBM4B | protein_coding | 43.9 | -0.01745563 | 0.01745563 | ENSG00000173914 | n.s. |
| ENSG00000143442 | 2329.078748 | -0.133178294 | 0.2381005 | -0.559337 | 0.575932082 | 0.739682939 | POGZ | protein_coding | 41.79 | -0.017440283 | 0.017440283 | ENSG00000143442 | n.s. |
| ENSG00000014216 | 283.1251222 | 0.146339487 | 0.2801756 | 0.522313 | 0.601452109 | 0.760168638 | CAPN1 | protein_coding | 51.18 | 0.017427577 | 0.017427577 | ENSG00000014216 | n.s. |
| ENSG00000024862 | 251.297056 | -0.145144964 | 0.2767891 | -0.524388 | 0.600008442 | 0.758895125 | CCDC28A | protein_coding | 39.18 | -0.017391014 | 0.017391014 | ENSG00000024862 | n.s. |
| ENSG00000136240 | 1257.601211 | -0.147607744 | 0.2854643 | -0.51708 | 0.605100649 | 0.762781412 | KDELR2 | protein_coding | 46.8 | -0.017358656 | 0.017358656 | ENSG00000136240 | n.s. |
| ENSG00000162341 | 192.8136298 | 0.13304658 | 0.2386809 | 0.557424 | 0.577237439 | 0.740546777 | TPCN2 | protein_coding | 54.96 | 0.017355594 | 0.017355594 | ENSG00000162341 | n.s. |
| ENSG00000142396 | 862.8721935 | -0.135351698 | 0.2462474 | -0.549657 | 0.582554429 | 0.74446125 | | protein_coding | 45.5 | -0.01734639 | 0.01734639 | ENSG00000142396 | n.s. |
| ENSG00000102265 | 105.1524725 | 0.155290096 | 0.3117427 | 0.498135 | 0.618388561 | 0.773221727 | TIMP1 | protein_coding | 53.67 | 0.017345275 | 0.017345275 | ENSG00000102265 | n.s. |
| ENSG00000212452 | 37.81933156 | -0.151270584 | 0.2986056 | -0.50659 | 0.612442541 | 0.768409254 | SNORD69 | snoRNA | 41.56 | -0.017306476 | 0.017306476 | ENSG00000212452 | n.s. |
| ENSG00000167118 | 487.3856634 | 0.148748201 | 0.2905066 | 0.51203 | 0.608629733 | 0.765274285 | URM1 | protein_coding | 52.12 | 0.017281994 | 0.017281994 | ENSG00000167118 | n.s. |
| ENSG00000134371 | 1708.027151 | 0.130950847 | 0.233086 | 0.561814 | 0.574243057 | 0.738380687 | CDC73 | protein_coding | 35.71 | 0.017248803 | 0.017248803 | ENSG00000134371 | n.s. |
| ENSG00000143612 | 2749.298552 | -0.154305832 | 0.3099882 | -0.49778 | 0.618639383 | 0.773327 | C1orf43 | protein_coding | 45.88 | -0.017226213 | 0.017226213 | ENSG00000143612 | n.s. |
| ENSG00000071539 | 997.9164317 | -0.144800581 | 0.2778449 | -0.521156 | 0.602257969 | 0.76077279 | TRIP13 | protein_coding | 50.49 | -0.017194349 | 0.017194349 | ENSG00000071539 | n.s. |
| ENSG00000124575 | 3097.557444 | 0.127712763 | 0.2237576 | 0.570764 | 0.568159736 | 0.733539318 | HIST1H1D | protein_coding | 54.35 | 0.01718715 | 0.01718715 | ENSG00000124575 | n.s. |
| ENSG00000175305 | 1212.472216 | -0.134677213 | 0.2460089 | -0.547449 | 0.584070626 | 0.745577641 | CCNE2 | protein_coding | 41.61 | -0.017172304 | 0.017172304 | ENSG00000175305 | n.s. |
| ENSG00000157600 | 640.560232 | -0.136563216 | 0.2519361 | -0.542055 | 0.587780566 | 0.748666071 | TMEM164 | protein_coding | 42.06 | -0.017167614 | 0.017167614 | ENSG00000157600 | n.s. |
| ENSG00000080824 | 23790.69878 | 0.14484692 | 0.2784546 | 0.520182 | 0.602937057 | 0.761285267 | HSP90AA1 | protein_coding | 47.25 | 0.017157491 | 0.017157491 | ENSG00000080824 | n.s. |
| ENSG00000134709 | 253.6252184 | 0.141930267 | 0.2689317 | 0.527756 | 0.597668781 | 0.757105126 | HOOK1 | protein_coding | 34.37 | 0.017151395 | 0.017151395 | ENSG00000134709 | n.s. |
| ENSG00000147533 | 776.1426071 | -0.142921525 | 0.2722587 | -0.524948 | 0.599619661 | 0.758568275 | GOLGA7 | protein_coding | 39.89 | -0.017151344 | 0.017151344 | ENSG00000147533 | n.s. |
| ENSG00000169299 | 846.2354978 | -0.144621435 | 0.2783412 | -0.519583 | 0.603354078 | 0.761663349 | PGM2 | protein_coding | 39.92 | -0.017099596 | 0.017099596 | ENSG00000169299 | n.s. |
| ENSG00000167775 | 407.2298166 | 0.149125209 | 0.2940142 | 0.507204 | 0.612011568 | 0.768006945 | CD320 | protein_coding | 58.35 | 0.017094946 | 0.017094946 | ENSG00000167775 | n.s. |
| ENSG00000163171 | 2876.812718 | -0.134673331 | 0.2469261 | -0.545399 | 0.585478931 | 0.746712673 | CDC42EP3 | protein_coding | 41.38 | -0.017082838 | 0.017082838 | ENSG00000163171 | n.s. |
| ENSG00000136068 | 2280.552948 | -0.131463326 | 0.236659 | -0.555497 | 0.578554794 | 0.741470228 | FLNB | protein_coding | 46.61 | -0.017077912 | 0.017077912 | ENSG00000136068 | n.s. |

| | | | | | | | | | | | | | |
|---|---|---|---|---|---|---|---|---|---|---|---|---|---|
| ENSG00000159788 | 326.3052336 | -0.12915141 | 0.229401 | -0.562994 | 0.573438862 | 0.737755431 | RGS12 | protein_coding | 50.96 | -0.017059298 | 0.017059298 | ENSG00000159788 | n.s. |
| ENSG00000180900 | 759.6083854 | 0.136110545 | 0.2517889 | 0.540574 | 0.588801179 | 0.749760273 | SCRIB | protein_coding | 62.5 | 0.017024377 | 0.017024377 | ENSG00000180900 | n.s. |
| ENSG00000008311 | 328.4010042 | -0.135102013 | 0.249003 | -0.542572 | 0.587424655 | 0.748349663 | AASS | protein_coding | 36.48 | -0.017008726 | 0.017008726 | ENSG00000008311 | n.s. |
| ENSG00000105671 | 708.5415694 | 0.14384295 | 0.2768441 | 0.519581 | 0.60335562 | 0.761663349 | DDX49 | protein_coding | 57.12 | 0.017007551 | 0.017007551 | ENSG00000105671 | n.s. |
| ENSG00000110811 | 397.8181485 | 0.136005164 | 0.2516807 | 0.540388 | 0.588929689 | 0.749826453 | P3H3 | protein_coding | 56.26 | 0.017005983 | 0.017005983 | ENSG00000110811 | n.s. |
| ENSG00000112096 | 1813.337195 | 0.12883135 | 0.2293108 | 0.56182 | 0.574238861 | 0.738380687 | SOD2 | protein_coding | 42.3 | 0.016969623 | 0.016969623 | ENSG00000112096 | n.s. |
| ENSG00000153561 | 2734.095373 | -0.138000362 | 0.2583993 | -0.534059 | 0.593300948 | 0.753973083 | RMND5A | protein_coding | 40.38 | -0.016924938 | 0.016924938 | ENSG00000153561 | n.s. |
| ENSG00000173692 | 3290.987024 | -0.13869506 | 0.2610589 | -0.531279 | 0.595225593 | 0.755469937 | PSMD1 | protein_coding | 37.87 | -0.016890675 | 0.016890675 | ENSG00000173692 | n.s. |
| ENSG00000188342 | 1291.497538 | -0.146219169 | 0.2870031 | -0.509469 | 0.610423508 | 0.766705143 | GTF2F2 | protein_coding | 39.12 | -0.016869543 | 0.016869543 | ENSG00000188342 | n.s. |
| ENSG00000160294 | 1336.52754 | -0.134190196 | 0.2475531 | -0.542066 | 0.587772778 | 0.748666071 | MCM3AP | protein_coding | 47.4 | -0.016869298 | 0.016869298 | ENSG00000160294 | n.s. |
| ENSG00000140564 | 144.1875471 | 0.138069893 | 0.2599943 | 0.53105 | 0.595384384 | 0.755469937 | FURIN | protein_coding | 58.47 | 0.01681454 | 0.01681454 | ENSG00000140564 | n.s. |
| ENSG00000099331 | 431.0610089 | -0.132891811 | 0.2441 | -0.544415 | 0.586155637 | 0.747280005 | MYO9B | protein_coding | 52.36 | -0.016813025 | 0.016813025 | ENSG00000099331 | n.s. |
| ENSG00000126950 | 223.6242206 | 0.149759415 | 0.2995701 | 0.499914 | 0.617135323 | 0.772278906 | TMEM35A | protein_coding | 44.18 | 0.016806874 | 0.016806874 | ENSG00000126950 | n.s. |
| ENSG00000198856 | 1413.267416 | 0.152616278 | 0.3096106 | 0.49293 | 0.622062268 | 0.776142392 | OSTC | protein_coding | 39.57 | 0.016796733 | 0.016796733 | ENSG00000198856 | n.s. |
| ENSG00000125611 | 165.2934111 | 0.160800787 | 0.3393193 | 0.473892 | 0.635576738 | 0.786310195 | CHCHD5 | protein_coding | 53.69 | 0.016788582 | 0.016788582 | ENSG00000125611 | n.s. |
| ENSG00000244218 | 45.60830553 | -0.141670236 | 0.2723383 | -0.520199 | 0.602924605 | 0.761285267 | RN7SL81P | misc_RNA | 52.11 | -0.016781204 | 0.016781204 | ENSG00000244218 | n.s. |
| ENSG00000155660 | 775.6953702 | -0.13974518 | 0.2662266 | -0.524911 | 0.599645256 | 0.758568275 | PDIA4 | protein_coding | 50.87 | -0.016770166 | 0.016770166 | ENSG00000155660 | n.s. |
| ENSG00000070785 | 455.3314247 | -0.145587573 | 0.2861061 | -0.508859 | 0.610851285 | 0.767173231 | EIF2B3 | protein_coding | 42.55 | -0.016758085 | 0.016758085 | ENSG00000070785 | n.s. |
| ENSG00000135387 | 8793.636012 | -0.139441574 | 0.2654288 | -0.525345 | 0.599343649 | 0.758409362 | CAPRIN1 | protein_coding | 40.22 | -0.016746419 | 0.016746419 | ENSG00000135387 | n.s. |
| ENSG00000111711 | 1505.342984 | -0.146352085 | 0.2890882 | -0.506254 | 0.612678229 | 0.768427977 | GOLT1B | protein_coding | 35.99 | -0.016742215 | 0.016742215 | ENSG00000111711 | n.s. |
| ENSG00000164898 | 125.2352054 | -0.164799735 | 0.3549486 | -0.464292 | 0.64243864 | 0.791643267 | FMC1 | protein_coding | 46.51 | -0.016722308 | 0.016722308 | ENSG00000164898 | n.s. |
| ENSG00000256712 | 224.3627402 | -0.129548247 | 0.2351067 | -0.551019 | 0.581620741 | 0.74354105 | | sense_intronic | 41.28 | -0.016672218 | 0.016672218 | ENSG00000256712 | n.s. |
| ENSG00000265539 | 2.784473553 | -0.2217453 | 0.6235196 | -0.355635 | 0.72211406 | 0.84105689 | MIR3164 | miRNA | 50.6 | -0.01666962 | 0.01666962 | ENSG00000265539 | n.s. |
| ENSG00000164494 | 210.4916265 | -0.145206379 | 0.2861522 | -0.507444 | 0.611842976 | 0.76793381 | PDSS2 | protein_coding | 39.82 | -0.016651717 | 0.016651717 | ENSG00000164494 | n.s. |
| ENSG00000104408 | 8634.371768 | 0.155665182 | 0.3225362 | 0.482629 | 0.629359515 | 0.781758101 | EIF3E | protein_coding | 37.25 | 0.016644906 | 0.016644906 | ENSG00000104408 | n.s. |
| ENSG00000156795 | 397.0408493 | 0.148119468 | 0.2964817 | 0.499591 | 0.617363408 | 0.772376582 | WDYHV1 | protein_coding | 45.17 | 0.016614694 | 0.016614694 | ENSG00000156795 | n.s. |
| ENSG00000162851 | 767.9600712 | 0.141555404 | 0.2740934 | 0.516449 | 0.605540552 | 0.763188295 | TFB2M | protein_coding | 41.95 | 0.016614117 | 0.016614117 | ENSG00000162851 | n.s. |
| ENSG00000212418 | 8.375446382 | -0.178231043 | 0.4161642 | -0.428271 | 0.668453845 | 0.806885719 | RNY4P36 | misc_RNA | 38.54 | -0.016608989 | 0.016608989 | ENSG00000212418 | n.s. |
| ENSG00000213186 | 811.9191778 | 0.138616747 | 0.2645747 | 0.523923 | 0.600332148 | 0.75923558 | TRIM59 | protein_coding | 42.27 | 0.016581813 | 0.016581813 | ENSG00000213186 | n.s. |
| ENSG00000201672 | 17.49167947 | -0.170887999 | 0.3847257 | -0.444181 | 0.656911459 | 0.799840685 | SNORD113-4 | snoRNA | 40 | -0.016575539 | 0.016575539 | ENSG00000201672 | n.s. |
| ENSG00000218739 | 748.8069915 | -0.143114124 | 0.2801174 | -0.510908 | 0.609415771 | 0.765923071 | CEBPZOS | protein_coding | 38.06 | -0.016574741 | 0.016574741 | ENSG00000218739 | n.s. |
| ENSG00000200312 | 15.03501071 | 0.16490774 | 0.3592108 | 0.459083 | 0.646174225 | 0.793440055 | RN7SKP255 | misc_RNA | 56.23 | 0.016570899 | 0.016570899 | ENSG00000200312 | n.s. |

| | | | | | | | | | | | | | |
|---|---|---|---|---|---|---|---|---|---|---|---|---|---|
| ENSG00000202515 | 8.886920006 | 0.176743519 | 0.4110075 | 0.430025 | 0.667177425 | 0.806194128 | VTRNA1-3 | misc_RNA | 60.67 | 0.016536189 | 0.016536189 | ENSG00000202515 | n.s. |
| ENSG00000117500 | 1492.441713 | -0.142828081 | 0.2797325 | -0.510588 | 0.60963952 | 0.765996845 | TMED5 | protein_coding | 38.24 | -0.016535638 | 0.016535638 | ENSG00000117500 | n.s. |
| ENSG00000069345 | 1722.510051 | 0.142556609 | 0.2787092 | 0.511489 | 0.609008871 | 0.765619008 | DNAJA2 | protein_coding | 42.72 | 0.016534755 | 0.016534755 | ENSG00000069345 | n.s. |
| ENSG00000185129 | 560.4571992 | -0.129154819 | 0.2352424 | -0.549029 | 0.582985819 | 0.74471172 | PURA | protein_coding | 41.49 | -0.016533342 | 0.016533342 | ENSG00000185129 | n.s. |
| ENSG00000119335 | 14035.74887 | 0.135407128 | 0.2544568 | 0.532142 | 0.594627783 | 0.755038999 | SET | protein_coding | 49.47 | 0.016523815 | 0.016523815 | ENSG00000119335 | n.s. |
| ENSG00000170091 | 1011.554377 | 0.149238346 | 0.3016783 | 0.494694 | 0.62081638 | 0.77506923 | | protein_coding | 45.73 | 0.016514641 | 0.016514641 | ENSG00000170091 | n.s. |
| ENSG00000106344 | 1843.926085 | -0.128739968 | 0.2345068 | -0.548982 | 0.583017866 | 0.74471172 | RBM28 | protein_coding | 43.22 | -0.016480236 | 0.016480236 | ENSG00000106344 | n.s. |
| ENSG00000237819 | 353.2828289 | 0.155777045 | 0.3256314 | 0.478385 | 0.632376503 | 0.78410582 | | antisense | 37.79 | 0.016454001 | 0.016454001 | ENSG00000237819 | n.s. |
| ENSG00000100804 | 2255.297122 | 0.151613043 | 0.3109554 | 0.487572 | 0.625853223 | 0.778917888 | PSMB5 | protein_coding | 46.64 | 0.016451277 | 0.016451277 | ENSG00000100804 | n.s. |
| ENSG00000077097 | 8130.779359 | -0.135447849 | 0.2556586 | -0.5298 | 0.596250753 | 0.756134368 | TOP2B | protein_coding | 35.14 | -0.016443507 | 0.016443507 | ENSG00000077097 | n.s. |
| ENSG00000163558 | 1499.320401 | -0.140273695 | 0.2720547 | -0.515609 | 0.606127865 | 0.76372111 | PRKCI | protein_coding | 39.82 | -0.016421169 | 0.016421169 | ENSG00000163558 | n.s. |
| ENSG00000112787 | 132.5218458 | 0.127857712 | 0.2325716 | 0.549756 | 0.582486565 | 0.744442853 | FBRSL1 | protein_coding | 62.75 | 0.016387348 | 0.016387348 | ENSG00000112787 | n.s. |
| ENSG00000160447 | 254.6045276 | 0.142909021 | 0.2817547 | 0.507211 | 0.61200692 | 0.768006945 | PKN3 | protein_coding | 57.66 | 0.016382355 | 0.016382355 | ENSG00000160447 | n.s. |
| ENSG00000277462 | 557.2766326 | -0.145914759 | 0.2921284 | -0.499489 | 0.617435277 | 0.772376582 | ZNF670 | protein_coding | 41.24 | -0.01636739 | 0.01636739 | ENSG00000277462 | n.s. |
| ENSG00000157036 | 75.39722067 | 0.138112376 | 0.2654845 | 0.520228 | 0.602904898 | 0.761285267 | EXOG | protein_coding | 42.1 | 0.016359767 | 0.016359767 | ENSG00000157036 | n.s. |
| ENSG00000130826 | 2993.242713 | 0.133192246 | 0.2492297 | 0.534416 | 0.593054059 | 0.753796938 | DKC1 | protein_coding | 45.62 | 0.016348766 | 0.016348766 | ENSG00000130826 | n.s. |
| ENSG00000108559 | 1619.801878 | -0.13835509 | 0.2667294 | -0.51871 | 0.603963232 | 0.76195905 | NUP88 | protein_coding | 42.72 | -0.01633536 | 0.01633536 | ENSG00000108559 | n.s. |
| ENSG00000241685 | 261.8074955 | 0.147401919 | 0.2977929 | 0.494981 | 0.620613285 | 0.774903269 | ARPC1A | protein_coding | 45.22 | 0.016325132 | 0.016325132 | ENSG00000241685 | n.s. |
| ENSG00000182901 | 396.2621193 | -0.139823353 | 0.2719231 | -0.514202 | 0.607110951 | 0.764302292 | RGS7 | protein_coding | 39.15 | -0.016322256 | 0.016322256 | ENSG00000182901 | n.s. |
| ENSG00000183283 | 1686.198867 | 0.137883927 | 0.2654483 | 0.519438 | 0.603455318 | 0.761663349 | DAZAP2 | protein_coding | 46.84 | 0.016302974 | 0.016302974 | ENSG00000183283 | n.s. |
| ENSG00000145945 | 245.6460067 | -0.133805387 | 0.2519755 | -0.531025 | 0.595401251 | 0.755469937 | FAM50B | protein_coding | 58.75 | -0.016295196 | 0.016295196 | ENSG00000145945 | n.s. |
| ENSG00000102241 | 2969.898961 | -0.13515999 | 0.2570959 | -0.525718 | 0.599084056 | 0.758240023 | HTATSF1 | protein_coding | 37.88 | -0.016245325 | 0.016245325 | ENSG00000102241 | n.s. |
| ENSG00000119865 | 1507.68476 | 0.136150906 | 0.2603907 | 0.522872 | 0.601063612 | 0.759815569 | CNRIP1 | protein_coding | 40.04 | 0.016241688 | 0.016241688 | ENSG00000119865 | n.s. |
| ENSG00000233184 | 240.0625081 | -0.148047774 | 0.3013151 | -0.491339 | 0.623186895 | 0.777009606 | | antisense | 37.11 | -0.016222129 | 0.016222129 | ENSG00000233184 | n.s. |
| ENSG00000101161 | 1378.861571 | -0.132761495 | 0.2501855 | -0.530652 | 0.595659848 | 0.755522643 | PRPF6 | protein_coding | 50.53 | -0.016164046 | 0.016164046 | ENSG00000101161 | n.s. |
| ENSG00000089820 | 57.93237607 | 0.135171163 | 0.2580981 | 0.52372 | 0.60047323 | 0.759345031 | ARHGAP4 | protein_coding | 58.14 | 0.016161179 | 0.016161179 | ENSG00000089820 | n.s. |
| ENSG00000107771 | 538.9815885 | -0.13356478 | 0.2529067 | -0.528119 | 0.59741682 | 0.756924866 | CCSER2 | protein_coding | 37.56 | -0.01615429 | 0.01615429 | ENSG00000107771 | n.s. |
| ENSG00000207751 | 136.898315 | 0.123772171 | 0.2221166 | 0.55724 | 0.577363734 | 0.740546777 | MIR130B | processed_transcript | 53.58 | 0.016145771 | 0.016145771 | ENSG00000207751 | n.s. |
| ENSG00000143575 | 1603.945738 | 0.150891137 | 0.3123078 | 0.483149 | 0.628990136 | 0.781634208 | HAX1 | protein_coding | 49.78 | 0.016144815 | 0.016144815 | ENSG00000143575 | n.s. |
| ENSG00000206634 | 46.73845816 | -0.148673569 | 0.3054981 | -0.48666 | 0.62649957 | 0.779243046 | SNORA22 | snoRNA | 41.04 | -0.016105371 | 0.016105371 | ENSG00000206634 | n.s. |
| ENSG00000162368 | 537.6133451 | -0.150107167 | 0.3104144 | -0.48357 | 0.628690862 | 0.781332042 | CMPK1 | protein_coding | 42.2 | -0.016086139 | 0.016086139 | ENSG00000162368 | n.s. |
| ENSG00000130787 | 465.5384959 | 0.131088391 | 0.2454994 | 0.533966 | 0.59336492 | 0.75398556 | HIP1R | protein_coding | 59.16 | 0.016076283 | 0.016076283 | ENSG00000130787 | n.s. |

| | | | | | | | | | | | | | |
|---|---|---|---|---|---|---|---|---|---|---|---|---|---|
| ENSG00000109814 | 733.4713188 | -0.13191355 | 0.2485558 | -0.53072 | 0.595612843 | 0.755522643 | UGDH | protein_coding | 41.86 | -0.016060807 | 0.016060807 | ENSG00000109814 | n.s. |
| ENSG00000167508 | 253.1883618 | 0.135034074 | 0.2589732 | 0.521421 | 0.602073491 | 0.760677787 | MVD | protein_coding | 59.29 | 0.016041949 | 0.016041949 | ENSG00000167508 | n.s. |
| ENSG00000105520 | 105.5634071 | 0.127790204 | 0.2364985 | 0.540342 | 0.588960863 | 0.749826453 | PLPPR2 | protein_coding | 57.31 | 0.01597879 | 0.01597879 | ENSG00000105520 | n.s. |
| ENSG00000112651 | 293.8822112 | 0.136294624 | 0.2647043 | 0.514894 | 0.606627134 | 0.764004486 | MRPL2 | protein_coding | 49.64 | 0.015933399 | 0.015933399 | ENSG00000112651 | n.s. |
| ENSG00000142733 | 11.03475261 | -0.170840101 | 0.3989565 | -0.428217 | 0.668492909 | 0.806885719 | MAP3K6 | protein_coding | 60.79 | -0.015920242 | 0.015920242 | ENSG00000142733 | n.s. |
| ENSG00000156860 | 50.85936859 | 0.138253204 | 0.2719392 | 0.508397 | 0.611174612 | 0.767440849 | FBRS | protein_coding | 59.46 | 0.015892909 | 0.015892909 | ENSG00000156860 | n.s. |
| ENSG00000166337 | 361.6551751 | 0.131116176 | 0.248121 | 0.528436 | 0.597196504 | 0.756782282 | TAF10 | protein_coding | 52.56 | 0.015868866 | 0.015868866 | ENSG00000166337 | n.s. |
| ENSG00000158864 | 1811.556573 | -0.138315583 | 0.2724619 | -0.507651 | 0.611698011 | 0.767852603 | NDUFS2 | protein_coding | 49.73 | -0.01586786 | 0.01586786 | ENSG00000158864 | n.s. |
| ENSG00000072609 | 46.18502944 | 0.146063778 | 0.3001161 | 0.486691 | 0.626477363 | 0.779243046 | CHFR | protein_coding | 48.69 | 0.01582266 | 0.01582266 | ENSG00000072609 | n.s. |
| ENSG00000165672 | 3180.717707 | 0.150622432 | 0.3162435 | 0.476286 | 0.633870508 | 0.785176075 | PRDX3 | protein_coding | 46.03 | 0.015820317 | 0.015820317 | ENSG00000165672 | n.s. |
| ENSG00000265694 | 20.15852207 | -0.156725249 | 0.3397594 | -0.461283 | 0.644595573 | 0.792637386 | MIR4774 | miRNA | 34.21 | -0.015817566 | 0.015817566 | ENSG00000265694 | n.s. |
| ENSG00000068394 | 311.4287625 | -0.140959005 | 0.2821871 | -0.499523 | 0.617410807 | 0.772376582 | GPKOW | protein_coding | 50.87 | -0.015811499 | 0.015811499 | ENSG00000068394 | n.s. |
| ENSG00000135632 | 611.4747009 | 0.126574953 | 0.2349518 | 0.538727 | 0.590075032 | 0.750764386 | SMYD5 | protein_coding | 50.97 | 0.015758118 | 0.015758118 | ENSG00000135632 | n.s. |
| ENSG00000144834 | 1143.509665 | 0.150435904 | 0.3167371 | 0.474955 | 0.63481894 | 0.785892941 | TAGLN3 | protein_coding | 45.75 | 0.015741103 | 0.015741103 | ENSG00000144834 | n.s. |
| ENSG00000173465 | 256.6219903 | 0.139678065 | 0.2790129 | 0.500615 | 0.616642096 | 0.77193921 | SSSCA1 | protein_coding | 61.32 | 0.015702175 | 0.015702175 | ENSG00000173465 | n.s. |
| ENSG00000111786 | 3226.907938 | -0.134738749 | 0.2625666 | -0.51316 | 0.607839155 | 0.764659711 | SRSF9 | protein_coding | 46.53 | -0.015701348 | 0.015701348 | ENSG00000111786 | n.s. |
| ENSG00000141391 | 137.6545097 | -0.126094292 | 0.2341088 | -0.538614 | 0.590153228 | 0.750795266 | PRELID3A | protein_coding | 50.59 | -0.015696025 | 0.015696025 | ENSG00000141391 | n.s. |
| ENSG00000151503 | 1728.18217 | -0.132031517 | 0.2530317 | -0.521798 | 0.601810805 | 0.760552949 | NCAPD3 | protein_coding | 43.33 | -0.015694658 | 0.015694658 | ENSG00000151503 | n.s. |
| ENSG00000116729 | 1093.655213 | -0.137444145 | 0.2719341 | -0.505432 | 0.613255546 | 0.768947377 | WLS | protein_coding | 41.57 | -0.015682841 | 0.015682841 | ENSG00000116729 | n.s. |
| ENSG00000143845 | 108.5635822 | -0.13670249 | 0.2693033 | -0.507615 | 0.611723125 | 0.767852603 | ETNK2 | protein_coding | 52.5 | -0.015682802 | 0.015682802 | ENSG00000143845 | n.s. |
| ENSG00000101811 | 668.7545236 | -0.134510071 | 0.2621809 | -0.513043 | 0.607921304 | 0.764666033 | CSTF2 | protein_coding | 38.87 | -0.015674217 | 0.015674217 | ENSG00000101811 | n.s. |
| ENSG00000118939 | 107.4537337 | -0.147294823 | 0.3063871 | -0.480747 | 0.630696037 | 0.782845763 | UCHL3 | protein_coding | 35.78 | -0.015660945 | 0.015660945 | ENSG00000118939 | n.s. |
| ENSG00000119013 | 1191.809969 | 0.154360452 | 0.3330644 | 0.463455 | 0.643038081 | 0.791907764 | NDUFB3 | protein_coding | 41.49 | 0.015640634 | 0.015640634 | ENSG00000119013 | n.s. |
| ENSG00000149313 | 2113.865651 | -0.132706851 | 0.256414 | -0.517549 | 0.604772885 | 0.762683405 | AASDHPPT | protein_coding | 36.57 | -0.015613718 | 0.015613718 | ENSG00000149313 | n.s. |
| ENSG00000137821 | 405.3483822 | -0.131522146 | 0.2526277 | -0.520616 | 0.602634011 | 0.7611097 | LRRC49 | protein_coding | 38.17 | -0.015592311 | 0.015592311 | ENSG00000137821 | n.s. |
| ENSG00000137135 | 227.1040806 | -0.121845487 | 0.2221079 | -0.548587 | 0.583288999 | 0.744838204 | ARHGEF39 | protein_coding | 49.95 | -0.015588675 | 0.015588675 | ENSG00000137135 | n.s. |
| ENSG00000182919 | 161.2993942 | -0.149254089 | 0.3152241 | -0.473486 | 0.635866692 | 0.786319563 | C11orf54 | protein_coding | 41.12 | -0.015582264 | 0.015582264 | ENSG00000182919 | n.s. |
| ENSG00000209645 | 6.777917018 | 0.181304069 | 0.454049 | 0.399305 | 0.689668459 | 0.820601814 | SNORD105 | snoRNA | 47.06 | 0.015568132 | 0.015568132 | ENSG00000209645 | n.s. |
| ENSG00000096093 | 55.30142171 | -0.155149894 | 0.3385603 | -0.458264 | 0.646762976 | 0.793757825 | EFHC1 | protein_coding | 41.58 | -0.015563393 | 0.015563393 | ENSG00000096093 | n.s. |
| ENSG00000143420 | 3794.827619 | -0.138891364 | 0.2795254 | -0.496883 | 0.619271759 | 0.773909048 | ENSA | protein_coding | 45.32 | -0.015460008 | 0.015460008 | ENSG00000143420 | n.s. |
| ENSG00000137207 | 544.3558712 | 0.136556809 | 0.2718191 | 0.502381 | 0.615399397 | 0.770591402 | YIPF3 | protein_coding | 52.46 | 0.015454932 | 0.015454932 | ENSG00000137207 | n.s. |
| ENSG00000186501 | 220.6993981 | -0.141026269 | 0.286864 | -0.491614 | 0.62299246 | 0.777009606 | TMEM222 | protein_coding | 51.87 | -0.015452758 | 0.015452758 | ENSG00000186501 | n.s. |

| | | | | | | | | | | | | | |
|---|---|---|---|---|---|---|---|---|---|---|---|---|---|
| ENSG00000183255 | 1479.392344 | 0.138778206 | 0.279368 | 0.496758 | 0.619359889 | 0.773949716 | PTTG1IP | protein_coding | 51 | 0.015444246 | 0.015444246 | ENSG00000183255 | n.s. |
| ENSG00000123545 | 1139.386796 | -0.13761784 | 0.2753855 | -0.499728 | 0.617266575 | 0.772373733 | NDUFAF4 | protein_coding | 38.11 | -0.015436938 | 0.015436938 | ENSG00000123545 | n.s. |
| ENSG00000145284 | 431.0905543 | -0.140177425 | 0.2845206 | -0.492679 | 0.622239179 | 0.776293557 | SCD5 | protein_coding | 43.57 | -0.015415875 | 0.015415875 | ENSG00000145284 | n.s. |
| ENSG00000233369 | 252.0387492 | 0.146297974 | 0.3064102 | 0.477458 | 0.633036041 | 0.78463107 | GTF2IP4 | transcribed_processed_pseudog | 46.79 | 0.015410224 | 0.015410224 | ENSG00000233369 | n.s. |
| ENSG00000273014 | 22.28344907 | 0.157171202 | 0.3512284 | 0.44749 | 0.65452128 | 0.798047096 | | lincRNA | 28.6 | 0.015398295 | 0.015398295 | ENSG00000273014 | n.s. |
| ENSG00000244486 | 304.286383 | -0.131551379 | 0.2559385 | -0.513996 | 0.607254745 | 0.764302292 | SCARF2 | protein_coding | 63.11 | -0.015356629 | 0.015356629 | ENSG00000244486 | n.s. |
| ENSG00000061273 | 48.53005756 | 0.136807133 | 0.2735941 | 0.500037 | 0.617049143 | 0.772278906 | HDAC7 | protein_coding | 54.88 | 0.015353294 | 0.015353294 | ENSG00000061273 | n.s. |
| ENSG00000136159 | 1294.794278 | 0.140055266 | 0.285227 | 0.491031 | 0.623404571 | 0.777120767 | NUDT15 | protein_coding | 41.59 | 0.01533766 | 0.01533766 | ENSG00000136159 | n.s. |
| ENSG00000119280 | 948.7735035 | 0.134304417 | 0.2657895 | 0.505304 | 0.613345512 | 0.768987804 | C1orf198 | protein_coding | 47.8 | 0.015321522 | 0.015321522 | ENSG00000119280 | n.s. |
| ENSG00000130414 | 1770.254814 | 0.134745092 | 0.267281 | 0.504133 | 0.614168111 | 0.769672603 | NDUFA10 | protein_coding | 48.44 | 0.015319705 | 0.015319705 | ENSG00000130414 | n.s. |
| ENSG00000196591 | 9455.880269 | -0.134238866 | 0.2655943 | -0.505428 | 0.613258039 | 0.768947377 | HDAC2 | protein_coding | 38.55 | -0.015317108 | 0.015317108 | ENSG00000196591 | n.s. |
| ENSG00000198736 | 56.12556228 | -0.152425482 | 0.3320903 | -0.458988 | 0.646242761 | 0.793440055 | MSRB1 | protein_coding | 58.61 | -0.015316609 | 0.015316609 | ENSG00000198736 | n.s. |
| ENSG00000204574 | 2263.534444 | -0.130594445 | 0.2531076 | -0.515964 | 0.605879457 | 0.763546314 | ABCF1 | protein_coding | 47.97 | -0.015301048 | 0.015301048 | ENSG00000204574 | n.s. |
| ENSG00000100644 | 4194.943549 | 0.130564506 | 0.2540208 | 0.513991 | 0.607257979 | 0.764302292 | HIF1A | protein_coding | 37.25 | 0.015241426 | 0.015241426 | ENSG00000100644 | n.s. |
| ENSG00000270036 | 26.98576353 | -0.143097993 | 0.2971708 | -0.481535 | 0.630136631 | 0.782570004 | | lincRNA | 42.27 | -0.015236617 | 0.015236617 | ENSG00000270036 | n.s. |
| ENSG00000167700 | 144.1484403 | 0.136056276 | 0.2736836 | 0.49713 | 0.619097551 | 0.773830255 | MFSD3 | protein_coding | 66.96 | 0.015150451 | 0.015150451 | ENSG00000167700 | n.s. |
| ENSG00000108641 | 94.53176714 | -0.149389822 | 0.3221805 | -0.463684 | 0.642874464 | 0.791820697 | B9D1 | protein_coding | 51.57 | -0.015144117 | 0.015144117 | ENSG00000108641 | n.s. |
| ENSG00000128595 | 2423.894546 | 0.136207262 | 0.274498 | 0.496205 | 0.619749845 | 0.774159079 | CALU | protein_coding | 39.99 | 0.015142133 | 0.015142133 | ENSG00000128595 | n.s. |
| ENSG00000108797 | 435.9341736 | -0.133631427 | 0.2657763 | -0.502797 | 0.6151073 | 0.770364212 | CNTNAP1 | protein_coding | 54.63 | -0.015140962 | 0.015140962 | ENSG00000108797 | n.s. |
| ENSG00000234882 | 64.11476516 | -0.157285604 | 0.3562391 | -0.441517 | 0.658838761 | 0.801346385 | EIF3EP1 | processed_pseudogene | 39.97 | -0.015127685 | 0.015127685 | ENSG00000234882 | n.s. |
| ENSG00000109832 | 124.2096229 | 0.133188665 | 0.2644057 | 0.503728 | 0.614452185 | 0.769895805 | DDX25 | protein_coding | 44.92 | 0.015125976 | 0.015125976 | ENSG00000109832 | n.s. |
| ENSG00000136930 | 2655.822627 | 0.135987979 | 0.2742754 | 0.495808 | 0.620029814 | 0.774369851 | PSMB7 | protein_coding | 42.8 | 0.015101678 | 0.015101678 | ENSG00000136930 | n.s. |
| ENSG00000065911 | 7845.304561 | 0.139795871 | 0.287768 | 0.485794 | 0.627113453 | 0.779858933 | MTHFD2 | protein_coding | 43.3 | 0.01509571 | 0.01509571 | ENSG00000065911 | n.s. |
| ENSG00000087245 | 620.3614373 | -0.129487485 | 0.2523293 | -0.513169 | 0.607833362 | 0.764659711 | MMP2 | protein_coding | 45.67 | -0.015089409 | 0.015089409 | ENSG00000087245 | n.s. |
| ENSG00000201998 | 52.42635194 | 0.14657685 | 0.3125727 | 0.468937 | 0.639114809 | 0.789074708 | SNORA23 | snoRNA | 46.15 | 0.015080101 | 0.015080101 | ENSG00000201998 | n.s. |
| ENSG00000175600 | 110.780531 | -0.153256596 | 0.3417366 | -0.448464 | 0.653818304 | 0.797310413 | SUGCT | protein_coding | 38.71 | -0.015076244 | 0.015076244 | ENSG00000175600 | n.s. |
| ENSG00000115364 | 1745.825317 | -0.129129306 | 0.2511779 | -0.514095 | 0.607185633 | 0.764302292 | MRPL19 | protein_coding | 36.97 | -0.015073889 | 0.015073889 | ENSG00000115364 | n.s. |
| ENSG00000132002 | 931.1193492 | -0.128932172 | 0.2503505 | -0.515007 | 0.606548355 | 0.764004486 | DNAJB1 | protein_coding | 52.02 | -0.015072698 | 0.015072698 | ENSG00000132002 | n.s. |
| ENSG00000105849 | 1190.629965 | 0.131511504 | 0.259736 | 0.506328 | 0.612626684 | 0.768427977 | TWISTNB | protein_coding | 36.3 | 0.015044499 | 0.015044499 | ENSG00000105849 | n.s. |
| ENSG00000170144 | 9595.367728 | -0.124185931 | 0.2348688 | -0.528746 | 0.596981624 | 0.756647715 | HNRNPA3 | protein_coding | 40 | -0.015039696 | 0.015039696 | ENSG00000170144 | n.s. |
| ENSG00000161091 | 205.1894457 | 0.128761383 | 0.2505439 | 0.513927 | 0.607302735 | 0.764302292 | MFSD12 | protein_coding | 57.07 | 0.015030939 | 0.015030939 | ENSG00000161091 | n.s. |
| ENSG00000182872 | 1271.302718 | -0.131333797 | 0.2593343 | -0.506427 | 0.612557139 | 0.768427977 | RBM10 | protein_coding | 50.32 | -0.01502417 | 0.01502417 | ENSG00000182872 | n.s. |

| | | | | | | | | | | | | | |
|---|---|---|---|---|---|---|---|---|---|---|---|---|---|
| ENSG00000170525 | 642.9982467 | 0.124960127 | 0.2377048 | 0.525695 | 0.59910042 | 0.758240023 | PFKFB3 | protein_coding | 50.91 | 0.015019369 | 0.015019369 | ENSG00000170525 | n.s. |
| ENSG00000187260 | 92.1711303 | 0.13435871 | 0.2697864 | 0.498019 | 0.618470798 | 0.773255111 | WDR86 | protein_coding | 57.01 | 0.015004805 | 0.015004805 | ENSG00000187260 | n.s. |
| ENSG00000164118 | 6825.420498 | -0.132626415 | 0.2641291 | -0.502127 | 0.615577975 | 0.770699949 | CEP44 | protein_coding | 35.66 | -0.015001993 | 0.015001993 | ENSG00000164118 | n.s. |
| ENSG00000106268 | 631.6453517 | 0.136709179 | 0.277873 | 0.491984 | 0.622730363 | 0.776836739 | NUDT1 | protein_coding | 56.77 | 0.014992929 | 0.014992929 | ENSG00000106268 | n.s. |
| ENSG00000188242 | 553.0936612 | -0.124420019 | 0.2362674 | -0.526607 | 0.598466646 | 0.757720525 | | antisense | 58.18 | -0.014991486 | 0.014991486 | ENSG00000188242 | n.s. |
| ENSG00000280152 | 186.8057284 | -0.118269611 | 0.2169967 | -0.54503 | 0.585733153 | 0.746878166 | | TEC | 59.43 | -0.014990701 | 0.014990701 | ENSG00000280152 | n.s. |
| ENSG00000272398 | 12776.23099 | 0.131698231 | 0.2609963 | 0.504598 | 0.613841129 | 0.769470608 | CD24 | protein_coding | 50.84 | 0.014988308 | 0.014988308 | ENSG00000272398 | n.s. |
| ENSG00000161542 | 509.1942319 | 0.133763433 | 0.268044 | 0.499035 | 0.617754503 | 0.772637077 | PRPSAP1 | protein_coding | 48.2 | 0.014984776 | 0.014984776 | ENSG00000161542 | n.s. |
| ENSG00000086598 | 4258.914243 | 0.13585219 | 0.2763895 | 0.491524 | 0.623055632 | 0.777009606 | TMED2 | protein_coding | 42.27 | 0.014885815 | 0.014885815 | ENSG00000086598 | n.s. |
| ENSG00000219928 | 60.80709246 | -0.154911584 | 0.3513389 | -0.440918 | 0.659272482 | 0.801663823 | | processed_pseudogene | 52.3 | -0.014872707 | 0.014872707 | ENSG00000219928 | n.s. |
| ENSG00000140395 | 1399.367221 | -0.141245452 | 0.2960067 | -0.47717 | 0.633241208 | 0.784700146 | WDR61 | protein_coding | 43.39 | -0.014872619 | 0.014872619 | ENSG00000140395 | n.s. |
| ENSG00000129158 | 52.80251926 | 0.141074955 | 0.2956665 | 0.477142 | 0.633260852 | 0.784700146 | SERGEF | protein_coding | 42.67 | 0.014854666 | 0.014854666 | ENSG00000129158 | n.s. |
| ENSG00000185127 | 1154.216341 | 0.116039889 | 0.2115565 | 0.548505 | 0.583344866 | 0.744838204 | C6orf120 | protein_coding | 45.78 | 0.014845918 | 0.014845918 | ENSG00000185127 | n.s. |
| ENSG00000108669 | 344.4213636 | -0.132207273 | 0.2644588 | -0.499916 | 0.617134029 | 0.772278906 | CYTH1 | protein_coding | 45.26 | -0.014837071 | 0.014837071 | ENSG00000108669 | n.s. |
| ENSG00000166562 | 667.4921214 | 0.146652076 | 0.3175876 | 0.461769 | 0.644247089 | 0.792459777 | SEC11C | protein_coding | 43.41 | 0.014815199 | 0.014815199 | ENSG00000166562 | n.s. |
| ENSG00000137309 | 2647.248234 | 0.124475533 | 0.2386727 | 0.521532 | 0.601995936 | 0.760677787 | HMGA1 | protein_coding | 57.83 | 0.014787602 | 0.014787602 | ENSG00000137309 | n.s. |
| ENSG00000267751 | 23.31548703 | -0.145969078 | 0.3165019 | -0.461195 | 0.644658754 | 0.792637386 | | antisense | 53.95 | -0.014731994 | 0.014731994 | ENSG00000267751 | n.s. |
| ENSG00000176731 | 793.6149971 | 0.138015842 | 0.2866905 | 0.481411 | 0.63022467 | 0.782605516 | C8orf59 | protein_coding | 38.73 | 0.014692766 | 0.014692766 | ENSG00000176731 | n.s. |
| ENSG00000269967 | 34.06552298 | 0.144781409 | 0.3117483 | 0.464418 | 0.642348504 | 0.791643267 | | lincRNA | 43 | 0.014691039 | 0.014691039 | ENSG00000269967 | n.s. |
| ENSG00000137563 | 794.4805962 | 0.14267986 | 0.3041134 | 0.469167 | 0.63895058 | 0.789074708 | GGH | protein_coding | 38.86 | 0.014679172 | 0.014679172 | ENSG00000137563 | n.s. |
| ENSG00000139631 | 27.97877943 | 0.148372974 | 0.3284124 | 0.451789 | 0.651421252 | 0.796400914 | CSAD | protein_coding | 47.44 | 0.014669376 | 0.014669376 | ENSG00000139631 | n.s. |
| ENSG00000253833 | 16.70987021 | 0.147185329 | 0.3227818 | 0.45599 | 0.648397079 | 0.79496588 | | processed_pseudogene | 38.87 | 0.01466724 | 0.01466724 | ENSG00000253833 | n.s. |
| ENSG00000197808 | 110.594376 | -0.13489753 | 0.2763321 | -0.488172 | 0.625428184 | 0.778597694 | ZNF461 | protein_coding | 40.3 | -0.014661593 | 0.014661593 | ENSG00000197808 | n.s. |
| ENSG00000153113 | 959.3613233 | 0.128940241 | 0.2559751 | 0.503722 | 0.614456812 | 0.769895805 | CAST | protein_coding | 39.41 | 0.014643491 | 0.014643491 | ENSG00000153113 | n.s. |
| ENSG00000139644 | 4244.504891 | 0.132397082 | 0.2678972 | 0.494209 | 0.621158862 | 0.775293127 | TMBIM6 | protein_coding | 43.42 | 0.014634388 | 0.014634388 | ENSG00000139644 | n.s. |
| ENSG00000279369 | 653.505661 | 0.110458541 | 0.1958959 | 0.563863 | 0.572847104 | 0.737334773 | | TEC | 59.16 | 0.014617562 | 0.014617562 | ENSG00000279369 | n.s. |
| ENSG00000115234 | 1142.222597 | -0.13130325 | 0.2642501 | -0.49689 | 0.619266579 | 0.773909048 | SNX17 | protein_coding | 53.11 | -0.014615375 | 0.014615375 | ENSG00000115234 | n.s. |
| ENSG00000114423 | 216.6177528 | 0.132049303 | 0.2670968 | 0.494387 | 0.621032639 | 0.775205078 | CBLB | protein_coding | 35.4 | 0.01460246 | 0.01460246 | ENSG00000114423 | n.s. |
| ENSG00000130429 | 115.9965976 | -0.139941902 | 0.2957861 | -0.473119 | 0.636128545 | 0.786433825 | ARPC1B | protein_coding | 56.01 | -0.014601232 | 0.014601232 | ENSG00000130429 | n.s. |
| ENSG00000123562 | 4626.14468 | 0.144227354 | 0.3115733 | 0.4629 | 0.643435896 | 0.792091776 | MORF4L2 | protein_coding | 39.39 | 0.014599341 | 0.014599341 | ENSG00000123562 | n.s. |
| ENSG00000090932 | 322.3623691 | 0.133974025 | 0.2739133 | 0.489111 | 0.624763014 | 0.778115668 | DLL3 | protein_coding | 56.84 | 0.014597252 | 0.014597252 | ENSG00000090932 | n.s. |
| ENSG00000092199 | 15017.96445 | 0.142467205 | 0.3048341 | 0.46736 | 0.640242522 | 0.789890594 | HNRNPC | protein_coding | 39.78 | 0.014593351 | 0.014593351 | ENSG00000092199 | n.s. |

| | | | | | | | | | | | | | |
|---|---|---|---|---|---|---|---|---|---|---|---|---|---|
| ENSG00000183172 | 276.3537798 | 0.138060635 | 0.2885134 | 0.478524 | 0.632277183 | 0.78410582 | SMDT1 | protein_coding | 48.41 | 0.014582699 | 0.014582699 | ENSG00000183172 | n.s. |
| ENSG00000090565 | 453.383724 | -0.12868507 | 0.2558893 | -0.502893 | 0.615039174 | 0.770364212 | RAB11FIP3 | protein_coding | 52.85 | -0.01458052 | 0.01458052 | ENSG00000090565 | n.s. |
| ENSG00000252130 | 40.69974877 | 0.140250144 | 0.2971334 | 0.472011 | 0.636919157 | 0.787201547 | RNU6-1045P | snRNA | 31.78 | 0.014573962 | 0.014573962 | ENSG00000252130 | n.s. |
| ENSG00000102225 | 813.4158279 | -0.125845369 | 0.2464181 | -0.510699 | 0.609562102 | 0.765968695 | CDK16 | protein_coding | 54.42 | -0.014571507 | 0.014571507 | ENSG00000102225 | n.s. |
| ENSG00000129038 | 609.2724251 | 0.12443945 | 0.2417927 | 0.514653 | 0.606795231 | 0.76414707 | LOXL1 | protein_coding | 55.02 | 0.014537396 | 0.014537396 | ENSG00000129038 | n.s. |
| ENSG00000112972 | 3915.770334 | -0.128070168 | 0.2546659 | -0.502895 | 0.615038217 | 0.770364212 | HMGCS1 | protein_coding | 38.81 | -0.01451085 | 0.01451085 | ENSG00000112972 | n.s. |
| ENSG00000121905 | 323.2288489 | 0.145625908 | 0.3199119 | 0.455206 | 0.648960824 | 0.795346039 | HPCA | protein_coding | 57.12 | 0.014481605 | 0.014481605 | ENSG00000121905 | n.s. |
| ENSG00000115091 | 3039.249234 | -0.133518638 | 0.2740037 | -0.487288 | 0.626054498 | 0.779016478 | ACTR3 | protein_coding | 36.61 | -0.014480544 | 0.014480544 | ENSG00000115091 | n.s. |
| ENSG00000275479 | 9.304629747 | -0.175272139 | 0.4546374 | -0.385521 | 0.699851662 | 0.82681669 | | lincRNA | 34.3 | -0.01447586 | 0.01447586 | ENSG00000275479 | n.s. |
| ENSG00000119685 | 288.608707 | -0.120576562 | 0.2297404 | -0.524838 | 0.59969554 | 0.758568275 | TTLL5 | protein_coding | 40.03 | -0.014469829 | 0.014469829 | ENSG00000119685 | n.s. |
| ENSG00000185065 | 92.91518474 | 0.126487499 | 0.2497941 | 0.506367 | 0.612599006 | 0.768427977 | | antisense | 49.84 | 0.014469769 | 0.014469769 | ENSG00000185065 | n.s. |
| ENSG00000213064 | 1704.135847 | -0.127101114 | 0.2518183 | -0.504733 | 0.613746092 | 0.769420749 | SFT2D2 | protein_coding | 43.37 | -0.014468696 | 0.014468696 | ENSG00000213064 | n.s. |
| ENSG00000100605 | 353.5075072 | -0.131762046 | 0.2681825 | -0.491315 | 0.623203783 | 0.777009606 | ITPK1 | protein_coding | 53.85 | -0.014437643 | 0.014437643 | ENSG00000100605 | n.s. |
| ENSG00000137070 | 753.5704242 | 0.130530108 | 0.26426 | 0.493946 | 0.621344524 | 0.775455343 | IL11RA | protein_coding | 54.2 | 0.014416164 | 0.014416164 | ENSG00000137070 | n.s. |
| ENSG00000176834 | 32.18391129 | -0.150139316 | 0.3413188 | -0.43988 | 0.660023995 | 0.802157304 | VSIG10 | protein_coding | 47.55 | -0.014374407 | 0.014374407 | ENSG00000176834 | n.s. |
| ENSG00000173085 | 275.3967413 | -0.132541928 | 0.2723557 | -0.48665 | 0.62650626 | 0.779243046 | COQ2 | protein_coding | 39.35 | -0.014357878 | 0.014357878 | ENSG00000173085 | n.s. |
| ENSG00000177868 | 158.3617398 | 0.135728353 | 0.283731 | 0.47837 | 0.632387003 | 0.78410582 | SVBP | protein_coding | 41.39 | 0.014336351 | 0.014336351 | ENSG00000177868 | n.s. |
| ENSG00000181904 | 1900.118331 | -0.134431112 | 0.2790722 | -0.481707 | 0.630013939 | 0.782487429 | C5orf24 | protein_coding | 41.93 | -0.014319956 | 0.014319956 | ENSG00000181904 | n.s. |
| ENSG00000251867 | 34.97120082 | 0.141465405 | 0.306212 | 0.461985 | 0.644091925 | 0.792349946 | | antisense | 33.52 | 0.014299743 | 0.014299743 | ENSG00000251867 | n.s. |
| ENSG00000123728 | 347.9597253 | 0.134414399 | 0.2793969 | 0.481088 | 0.630454179 | 0.782754907 | RAP2C | protein_coding | 37.77 | 0.014298224 | 0.014298224 | ENSG00000123728 | n.s. |
| ENSG00000041880 | 123.6045775 | 0.122621566 | 0.2387702 | 0.513555 | 0.607563396 | 0.764561204 | PARP3 | protein_coding | 59.21 | 0.014296172 | 0.014296172 | ENSG00000041880 | n.s. |
| ENSG00000218596 | 105.1692384 | -0.133489724 | 0.2764729 | -0.482831 | 0.629215606 | 0.781705085 | | processed_pseudogene | 47.63 | -0.014277669 | 0.014277669 | ENSG00000218596 | n.s. |
| ENSG00000167100 | 36.56756717 | -0.142047714 | 0.3095128 | -0.45894 | 0.646277455 | 0.793440055 | SAMD14 | protein_coding | 55.35 | -0.014273789 | 0.014273789 | ENSG00000167100 | n.s. |
| ENSG00000116209 | 2053.209144 | 0.141749734 | 0.3084648 | 0.459533 | 0.645851515 | 0.793440055 | TMEM59 | protein_coding | 38.88 | 0.014243847 | 0.014243847 | ENSG00000116209 | n.s. |
| ENSG00000111341 | 2083.756145 | 0.130929074 | 0.2681514 | 0.488265 | 0.625361819 | 0.778584691 | MGP | protein_coding | 37.99 | 0.014231223 | 0.014231223 | ENSG00000111341 | n.s. |
| ENSG00000133874 | 225.1682761 | 0.135075199 | 0.2832867 | 0.476814 | 0.633494297 | 0.784919561 | RNF122 | protein_coding | 49.07 | 0.014206513 | 0.014206513 | ENSG00000133874 | n.s. |
| ENSG00000166734 | 1274.947863 | -0.134336307 | 0.2804757 | -0.478959 | 0.631967969 | 0.783935321 | CASC4 | protein_coding | 40.59 | -0.014202003 | 0.014202003 | ENSG00000166734 | n.s. |
| ENSG00000179152 | 437.9296909 | 0.122222543 | 0.238735 | 0.511959 | 0.608679698 | 0.765274285 | TCAIM | protein_coding | 39.47 | 0.014200167 | 0.014200167 | ENSG00000179152 | n.s. |
| ENSG00000164638 | 1001.951244 | -0.12533214 | 0.2493716 | -0.502592 | 0.615251176 | 0.770475096 | SLC29A4 | protein_coding | 57.52 | -0.014192786 | 0.014192786 | ENSG00000164638 | n.s. |
| ENSG00000140905 | 218.3835381 | -0.12481852 | 0.2475402 | -0.504235 | 0.614095971 | 0.769651472 | GCSH | protein_coding | 43.92 | -0.014192602 | 0.014192602 | ENSG00000140905 | n.s. |
| ENSG00000156500 | 22.35531587 | -0.142920204 | 0.3156196 | -0.452824 | 0.650675302 | 0.796220641 | FAM122C | protein_coding | 41.56 | -0.014144322 | 0.014144322 | ENSG00000156500 | n.s. |
| ENSG00000067057 | 1488.674542 | -0.130734438 | 0.2693297 | -0.485407 | 0.627387892 | 0.779991215 | PFKP | protein_coding | 51.45 | -0.014107591 | 0.014107591 | ENSG00000067057 | n.s. |

| | | | | | | | | | | | | | |
|---|---|---|---|---|---|---|---|---|---|---|---|---|---|
| ENSG00000164924 | 5500.02388 | -0.129974102 | 0.2669825 | -0.486826 | 0.626381476 | 0.779243046 | YWHAZ | protein_coding | 40.87 | -0.014079713 | 0.014079713 | ENSG00000164924 | n.s. |
| ENSG00000175216 | 6140.77173 | -0.128927137 | 0.2634288 | -0.489419 | 0.624544846 | 0.777915359 | CKAP5 | protein_coding | 40.42 | -0.01406178 | 0.01406178 | ENSG00000175216 | n.s. |
| ENSG00000136425 | 170.7694708 | 0.13606468 | 0.289463 | 0.470059 | 0.638312905 | 0.788434224 | CIB2 | protein_coding | 51.28 | 0.014046574 | 0.014046574 | ENSG00000136425 | n.s. |
| ENSG00000134759 | 2592.382975 | -0.128020301 | 0.2606932 | -0.491077 | 0.623372337 | 0.777120767 | ELP2 | protein_coding | 40.55 | -0.014019693 | 0.014019693 | ENSG00000134759 | n.s. |
| ENSG00000166471 | 967.2862421 | 0.126642134 | 0.2560391 | 0.49462 | 0.62086814 | 0.77506923 | TMEM41B | protein_coding | 42.5 | 0.014014156 | 0.014014156 | ENSG00000166471 | n.s. |
| ENSG00000176986 | 1083.175812 | -0.122614435 | 0.2423728 | -0.505892 | 0.612932513 | 0.76867766 | SEC24C | protein_coding | 47.03 | -0.014009403 | 0.014009403 | ENSG00000176986 | n.s. |
| ENSG00000100417 | 112.0154775 | 0.138238631 | 0.2985331 | 0.46306 | 0.643321583 | 0.792021081 | PMM1 | protein_coding | 54.84 | 0.013998494 | 0.013998494 | ENSG00000100417 | n.s. |
| ENSG00000169641 | 459.4374175 | -0.11893914 | 0.2297961 | -0.517586 | 0.604747444 | 0.762683405 | LUZP1 | protein_coding | 43.28 | -0.013993868 | 0.013993868 | ENSG00000169641 | n.s. |
| ENSG00000176401 | 187.9651105 | 0.116037498 | 0.2203159 | 0.526687 | 0.598410988 | 0.757720525 | EID2B | protein_coding | 48.95 | 0.013981468 | 0.013981468 | ENSG00000176401 | n.s. |
| ENSG00000107960 | 263.4898396 | -0.128855917 | 0.2644701 | -0.487223 | 0.626100293 | 0.779016478 | STN1 | protein_coding | 41.91 | -0.013974857 | 0.013974857 | ENSG00000107960 | n.s. |
| ENSG00000114544 | 322.431944 | -0.137329286 | 0.2948818 | -0.46571 | 0.641423355 | 0.791153205 | SLC41A3 | protein_coding | 45.99 | -0.0139718 | 0.0139718 | ENSG00000114544 | n.s. |
| ENSG00000133243 | 433.7480837 | -0.126875326 | 0.2573848 | -0.49294 | 0.622054851 | 0.776142392 | BTBD2 | protein_coding | 55.3 | -0.01396372 | 0.01396372 | ENSG00000133243 | n.s. |
| ENSG00000170860 | 1681.484646 | 0.140933755 | 0.3111708 | 0.452914 | 0.650610382 | 0.796220641 | LSM3 | protein_coding | 41.66 | 0.01394773 | 0.01394773 | ENSG00000170860 | n.s. |
| ENSG00000139970 | 1072.018633 | -0.138153674 | 0.2997663 | -0.460871 | 0.644890927 | 0.792685753 | RTN1 | protein_coding | 39.2 | -0.01393956 | 0.01393956 | ENSG00000139970 | n.s. |
| ENSG00000233906 | 25.60225901 | 0.146242306 | 0.3337841 | 0.438134 | 0.661288802 | 0.803063584 | | lincRNA | 40.52 | 0.01392959 | 0.01392959 | ENSG00000233906 | n.s. |
| ENSG00000183978 | 1108.909614 | 0.140857399 | 0.3119731 | 0.451505 | 0.651625702 | 0.796469551 | COA3 | protein_coding | 54.78 | 0.013921053 | 0.013921053 | ENSG00000183978 | n.s. |
| ENSG00000129083 | 2242.197415 | -0.137009555 | 0.294798 | -0.464757 | 0.64210517 | 0.791596249 | COPB1 | protein_coding | 38.27 | -0.013905959 | 0.013905959 | ENSG00000129083 | n.s. |
| ENSG00000263634 | 27.49166639 | 0.146974636 | 0.338133 | 0.434665 | 0.663805473 | 0.804365849 | MIR3919 | miRNA | 43.82 | 0.013895919 | 0.013895919 | ENSG00000263634 | n.s. |
| ENSG00000166477 | 2191.503036 | -0.120871412 | 0.2379853 | -0.507894 | 0.611527352 | 0.767772654 | LEO1 | protein_coding | 43.13 | -0.013872093 | 0.013872093 | ENSG00000166477 | n.s. |
| ENSG00000088298 | 98.80923912 | -0.137768772 | 0.3002597 | -0.458832 | 0.646354744 | 0.793440055 | EDEM2 | protein_coding | 45.82 | -0.013843816 | 0.013843816 | ENSG00000088298 | n.s. |
| ENSG00000259924 | 56.00677194 | 0.141643752 | 0.3173036 | 0.446398 | 0.655309687 | 0.798588776 | | processed_pseudogene | 40.46 | 0.013835308 | 0.013835308 | ENSG00000259924 | n.s. |
| ENSG00000130413 | 195.6088014 | -0.129899552 | 0.2700708 | -0.480983 | 0.630528383 | 0.782777233 | STK33 | protein_coding | 37.65 | -0.013816352 | 0.013816352 | ENSG00000130413 | n.s. |
| ENSG00000173511 | 135.3820804 | -0.130700745 | 0.2730066 | -0.478746 | 0.632119466 | 0.784053392 | VEGFB | protein_coding | 61.25 | -0.013809104 | 0.013809104 | ENSG00000173511 | n.s. |
| ENSG00000175224 | 224.9850415 | -0.125997036 | 0.2564112 | -0.491387 | 0.623153048 | 0.777009606 | ATG13 | protein_coding | 44.43 | -0.01380595 | 0.01380595 | ENSG00000175224 | n.s. |
| ENSG00000278986 | 123.596212 | 0.120170806 | 0.2366203 | 0.507863 | 0.611549144 | 0.767772654 | | TEC | 54.02 | 0.013791686 | 0.013791686 | ENSG00000278986 | n.s. |
| ENSG00000164933 | 992.0988677 | -0.136362385 | 0.2951634 | -0.461989 | 0.644088852 | 0.792349946 | SLC25A32 | protein_coding | 36.96 | -0.013783915 | 0.013783915 | ENSG00000164933 | n.s. |
| ENSG00000125827 | 1388.850789 | -0.134120147 | 0.2864603 | -0.468198 | 0.639642961 | 0.789446765 | TMX4 | protein_coding | 37.75 | -0.013771075 | 0.013771075 | ENSG00000125827 | n.s. |
| ENSG00000185420 | 479.4137775 | 0.135878646 | 0.2932904 | 0.46329 | 0.643156165 | 0.791957485 | SMYD3 | protein_coding | 40.6 | 0.013764253 | 0.013764253 | ENSG00000185420 | n.s. |
| ENSG00000263307 | 22.89410453 | 0.145767925 | 0.33571 | 0.434208 | 0.664137517 | 0.804698168 | | antisense | 41.96 | 0.01375568 | 0.01375568 | ENSG00000263307 | n.s. |
| ENSG00000129691 | 741.9651058 | -0.126713177 | 0.2597759 | -0.487779 | 0.62570652 | 0.778874554 | ASH2L | protein_coding | 42.99 | -0.013752496 | 0.013752496 | ENSG00000129691 | n.s. |
| ENSG00000130165 | 373.9093735 | 0.13492138 | 0.2894962 | 0.466056 | 0.641175518 | 0.79091758 | ELOF1 | protein_coding | 55.91 | 0.013744275 | 0.013744275 | ENSG00000130165 | n.s. |
| ENSG00000130204 | 1249.30999 | -0.131215677 | 0.2759618 | -0.475485 | 0.634441397 | 0.785728035 | TOMM40 | protein_coding | 56.25 | -0.013741923 | 0.013741923 | ENSG00000130204 | n.s. |

| | | | | | | | | | | | | | |
|---|---|---|---|---|---|---|---|---|---|---|---|---|---|
| ENSG00000168591 | 463.052766 | -0.123481014 | 0.2488034 | -0.496299 | 0.619683145 | 0.774145216 | TMUB2 | protein_coding | 52.96 | -0.013728319 | 0.013728319 | ENSG00000168591 | n.s. |
| ENSG00000147224 | 933.2471206 | 0.135833094 | 0.2945904 | 0.461091 | 0.64473306 | 0.792637386 | PRPS1 | protein_coding | 41.83 | 0.013709016 | 0.013709016 | ENSG00000147224 | n.s. |
| ENSG00000026025 | 2581.526412 | 0.118382036 | 0.23202 | 0.510223 | 0.609894902 | 0.766179439 | VIM | protein_coding | 46.25 | 0.013693192 | 0.013693192 | ENSG00000026025 | n.s. |
| ENSG00000241058 | 657.8966791 | -0.113527732 | 0.2154785 | -0.526863 | 0.598288472 | 0.757683318 | NSUN6 | protein_coding | 40.26 | -0.013681485 | 0.013681485 | ENSG00000241058 | n.s. |
| ENSG00000129103 | 767.6988372 | -0.127516176 | 0.26358 | -0.483785 | 0.62853812 | 0.781211949 | SUMF2 | protein_coding | 49.29 | -0.013673703 | 0.013673703 | ENSG00000129103 | n.s. |
| ENSG00000255496 | 10.16355883 | -0.159152398 | 0.398585 | -0.399294 | 0.689676951 | 0.820601814 | | sense_overlapping | 40.94 | -0.013666023 | 0.013666023 | ENSG00000255496 | n.s. |
| ENSG00000212529 | 13.23479118 | 0.214506014 | 0.6915621 | 0.310176 | 0.756427076 | 0.863752495 | | snoRNA | 53.1 | 0.013644874 | 0.013644874 | ENSG00000212529 | n.s. |
| ENSG00000004487 | 3614.858665 | 0.12853243 | 0.2677967 | 0.479963 | 0.631253848 | 0.78325883 | KDM1A | protein_coding | 39.16 | 0.01363661 | 0.01363661 | ENSG00000004487 | n.s. |
| ENSG00000115252 | 634.9155159 | 0.13511291 | 0.2929962 | 0.461142 | 0.644696542 | 0.792637386 | PDE1A | protein_coding | 36.57 | 0.013636331 | 0.013636331 | ENSG00000115252 | n.s. |
| ENSG00000135763 | 2056.182902 | 0.116521865 | 0.2262376 | 0.515042 | 0.606523714 | 0.764004486 | URB2 | protein_coding | 43.59 | 0.013621883 | 0.013621883 | ENSG00000135763 | n.s. |
| ENSG00000104613 | 1485.673042 | -0.131946095 | 0.2806054 | -0.470219 | 0.638198347 | 0.788432616 | INTS10 | protein_coding | 41.11 | -0.01362151 | 0.01362151 | ENSG00000104613 | n.s. |
| ENSG00000136144 | 998.5936316 | -0.137741121 | 0.3046851 | -0.452077 | 0.651213527 | 0.796385239 | RCBTB1 | protein_coding | 42.24 | -0.013619401 | 0.013619401 | ENSG00000136144 | n.s. |
| ENSG00000128951 | 3680.82522 | 0.132693869 | 0.2835627 | 0.467953 | 0.639818528 | 0.789593456 | DUT | protein_coding | 39.96 | 0.013613921 | 0.013613921 | ENSG00000128951 | n.s. |
| ENSG00000101255 | 202.4225528 | -0.137355212 | 0.302911 | -0.453451 | 0.650224246 | 0.796014436 | TRIB3 | protein_coding | 50.47 | -0.013609025 | 0.013609025 | ENSG00000101255 | n.s. |
| ENSG00000275342 | 243.6839786 | 0.115503937 | 0.2231986 | 0.517494 | 0.604811282 | 0.762683405 | PRAG1 | protein_coding | 43.93 | 0.013589697 | 0.013589697 | ENSG00000275342 | n.s. |
| ENSG00000162517 | 274.419621 | -0.127824533 | 0.2663033 | -0.479996 | 0.631230188 | 0.78325883 | PEF1 | protein_coding | 48.36 | -0.013561505 | 0.013561505 | ENSG00000162517 | n.s. |
| ENSG00000232437 | 16.46579412 | 0.150335356 | 0.3608164 | 0.416653 | 0.67693196 | 0.81263353 | RPS26P42 | processed_pseudogene | 47.21 | 0.013546007 | 0.013546007 | ENSG00000232437 | n.s. |
| ENSG00000206954 | 37.31729958 | -0.133919108 | 0.2896199 | -0.462396 | 0.64379732 | 0.792256504 | RNU6-1201P | snRNA | 40.19 | -0.0135438 | 0.0135438 | ENSG00000206954 | n.s. |
| ENSG00000174744 | 510.5792066 | 0.13723067 | 0.3045862 | 0.450548 | 0.652315386 | 0.796822874 | BRMS1 | protein_coding | 55.88 | 0.013536187 | 0.013536187 | ENSG00000174744 | n.s. |
| ENSG00000101966 | 44.58941345 | 0.134843661 | 0.2942684 | 0.458234 | 0.646784639 | 0.793757825 | XIAP | protein_coding | 41.98 | 0.013526435 | 0.013526435 | ENSG00000101966 | n.s. |
| ENSG00000109445 | 1372.805872 | 0.124621868 | 0.2555911 | 0.487583 | 0.625845242 | 0.778917888 | ZNF330 | protein_coding | 34.91 | 0.01352251 | 0.01352251 | ENSG00000109445 | n.s. |
| ENSG00000206120 | 37.54193127 | 0.133984744 | 0.2907665 | 0.460798 | 0.644943202 | 0.792685753 | EGFEM1P | transcribed_unitary_pseudogene | 36.09 | 0.01351892 | 0.01351892 | ENSG00000206120 | n.s. |
| ENSG00000164091 | 3268.508208 | -0.127096312 | 0.2643163 | -0.480849 | 0.630623596 | 0.782825636 | WDR82 | protein_coding | 44.88 | -0.013514781 | 0.013514781 | ENSG00000164091 | n.s. |
| ENSG00000264364 | 2183.923152 | 0.114046843 | 0.2196582 | 0.519201 | 0.603620277 | 0.761664412 | DYNLL2 | protein_coding | 49.02 | 0.013484481 | 0.013484481 | ENSG00000264364 | n.s. |
| ENSG00000137040 | 3511.962833 | 0.11712319 | 0.2302896 | 0.508591 | 0.611039063 | 0.767339846 | RANBP6 | protein_coding | 39.01 | 0.013470602 | 0.013470602 | ENSG00000137040 | n.s. |
| ENSG00000114978 | 2705.449919 | 0.128677572 | 0.2711284 | 0.4746 | 0.635072076 | 0.786015631 | MOB1A | protein_coding | 41.02 | 0.013455661 | 0.013455661 | ENSG00000114978 | n.s. |
| ENSG00000144635 | 1361.951629 | 0.132272583 | 0.284383 | 0.465121 | 0.641844611 | 0.79146245 | DYNC1LI1 | protein_coding | 39.16 | 0.013434885 | 0.013434885 | ENSG00000144635 | n.s. |
| ENSG00000106266 | 104.4507798 | 0.135799859 | 0.3006428 | 0.451698 | 0.651486303 | 0.796400914 | SNX8 | protein_coding | 51.46 | 0.013426294 | 0.013426294 | ENSG00000106266 | n.s. |
| ENSG00000132591 | 441.4836282 | 0.122926511 | 0.2508152 | 0.490108 | 0.624057548 | 0.777656247 | ERAL1 | protein_coding | 50.7 | 0.013425091 | 0.013425091 | ENSG00000132591 | n.s. |
| ENSG00000074590 | 873.2528092 | -0.115555403 | 0.2260702 | -0.511148 | 0.609247124 | 0.76583477 | NUAK1 | protein_coding | 45.97 | -0.013388818 | 0.013388818 | ENSG00000074590 | n.s. |
| ENSG00000143314 | 607.8730195 | -0.126852727 | 0.2653448 | -0.478067 | 0.632602218 | 0.784302822 | MRPL24 | protein_coding | 48.93 | -0.013385021 | 0.013385021 | ENSG00000143314 | n.s. |
| ENSG00000154813 | 653.0639536 | -0.135929345 | 0.3024874 | -0.449372 | 0.653163364 | 0.797221535 | DPH3 | protein_coding | 40.8 | -0.013378299 | 0.013378299 | ENSG00000154813 | n.s. |

| | | | | | | | | | | | | | | |
|---|---|---|---|---|---|---|---|---|---|---|---|---|---|---|
| ENSG00000104889 | 940.6215298 | 0.135202775 | 0.2989526 | 0.452255 | 0.651085405 | 0.796369304 | RNASEH2A | | protein_coding | 52.97 | 0.013369592 | 0.013369592 | ENSG00000104889 | n.s. |
| ENSG00000074582 | 577.0857963 | 0.129681459 | 0.2761092 | 0.469675 | 0.638587578 | 0.788703526 | BCS1L | | protein_coding | 54.42 | 0.013368371 | 0.013368371 | ENSG00000074582 | n.s. |
| ENSG00000243927 | 361.6060492 | -0.14066937 | 0.3222287 | -0.436551 | 0.662436787 | 0.803686597 | MRPS6 | | protein_coding | 40.3 | -0.01335139 | 0.01335139 | ENSG00000243927 | n.s. |
| ENSG00000168385 | 7386.476059 | -0.13157019 | 0.2835219 | -0.464056 | 0.642607284 | 0.791643267 | | 2-Sep | protein_coding | 41.89 | -0.01335049 | 0.01335049 | ENSG00000168385 | n.s. |
| ENSG00000181104 | 833.4423898 | -0.134302808 | 0.295768 | -0.454082 | 0.649770053 | 0.795668289 | F2R | | protein_coding | 42.79 | -0.013331964 | 0.013331964 | ENSG00000181104 | n.s. |
| ENSG00000267595 | 16.10376447 | -0.145541746 | 0.345557 | -0.42118 | 0.673623562 | 0.809935045 | | | unprocessed_pseudogene | 35 | -0.013324319 | 0.013324319 | ENSG00000267595 | n.s. |
| ENSG00000121089 | 45.88849573 | 0.137358736 | 0.3093886 | 0.443968 | 0.657065456 | 0.799914306 | NACA3P | | processed_pseudogene | 45.22 | 0.013317827 | 0.013317827 | ENSG00000121089 | n.s. |
| ENSG00000172775 | 1667.100304 | -0.128126763 | 0.2714267 | -0.472049 | 0.636891713 | 0.787201547 | FAM192A | | protein_coding | 41.93 | -0.013314172 | 0.013314172 | ENSG00000172775 | n.s. |
| ENSG00000198730 | 1198.118627 | 0.119496529 | 0.2407054 | 0.496443 | 0.619581829 | 0.774088102 | CTR9 | | protein_coding | 39.07 | 0.013289162 | 0.013289162 | ENSG00000198730 | n.s. |
| ENSG00000148019 | 2724.236175 | -0.124810282 | 0.2592996 | -0.481336 | 0.630277645 | 0.782605516 | CEP78 | | protein_coding | 37.2 | -0.01328694 | 0.01328694 | ENSG00000148019 | n.s. |
| ENSG00000152778 | 1370.440793 | -0.115852724 | 0.2284601 | -0.507103 | 0.612082861 | 0.768027188 | IFIT5 | | protein_coding | 41.44 | -0.013279434 | 0.013279434 | ENSG00000152778 | n.s. |
| ENSG00000120509 | 317.8730205 | 0.132392026 | 0.2894144 | 0.457448 | 0.647349122 | 0.794184333 | PDZD11 | | protein_coding | 49.29 | 0.01324962 | 0.01324962 | ENSG00000120509 | n.s. |
| ENSG00000163931 | 2784.241301 | 0.126901557 | 0.2681673 | 0.473218 | 0.636057735 | 0.786433825 | TKT | | protein_coding | 54.36 | 0.013240631 | 0.013240631 | ENSG00000163931 | n.s. |
| ENSG00000135185 | 601.2505774 | -0.118866785 | 0.239435 | -0.496447 | 0.619579078 | 0.774088102 | TMEM243 | | protein_coding | 43.9 | -0.013219129 | 0.013219129 | ENSG00000135185 | n.s. |
| ENSG00000283645 | 1.35703551 | -0.234481639 | 0.8445409 | -0.277644 | 0.781285731 | 0.878298727 | MIR5583-2 | | miRNA | 33.9 | -0.013214857 | 0.013214857 | ENSG00000283645 | n.s. |
| ENSG00000240964 | 31.12914525 | -0.143270247 | 0.3376446 | -0.424323 | 0.671330554 | 0.808693667 | RN7SL751P | | misc_RNA | 50.35 | -0.013211803 | 0.013211803 | ENSG00000240964 | n.s. |
| ENSG00000214113 | 560.4800627 | 0.133763548 | 0.2967318 | 0.450789 | 0.652141379 | 0.796680215 | LYRM4 | | protein_coding | 43.41 | 0.013204598 | 0.013204598 | ENSG00000214113 | n.s. |
| ENSG00000068912 | 739.7779775 | -0.133895192 | 0.2973778 | -0.450253 | 0.652528119 | 0.796942895 | ERLEC1 | | protein_coding | 38.1 | -0.013198424 | 0.013198424 | ENSG00000068912 | n.s. |
| ENSG00000282206 | 15.61838603 | -0.148983014 | 0.3635746 | -0.409773 | 0.681972562 | 0.815483027 | | | unprocessed_pseudogene | 65.35 | -0.013197671 | 0.013197671 | ENSG00000282206 | n.s. |
| ENSG00000166598 | 10802.80439 | 0.124590002 | 0.2598226 | 0.47952 | 0.631569069 | 0.783580124 | HSP90B1 | | protein_coding | 39.78 | 0.013196148 | 0.013196148 | ENSG00000166598 | n.s. |
| ENSG00000254207 | 19.40288147 | -0.146999882 | 0.3546346 | -0.414511 | 0.678500036 | 0.813323806 | | | sense_intronic | 61.74 | -0.013191257 | 0.013191257 | ENSG00000254207 | n.s. |
| ENSG00000037241 | 441.5751237 | 0.140966684 | 0.3278548 | 0.429967 | 0.667219807 | 0.806194128 | RPL26L1 | | protein_coding | 46.12 | 0.013188895 | 0.013188895 | ENSG00000037241 | n.s. |
| ENSG00000163812 | 503.7902123 | -0.1255442 | 0.26368 | -0.476123 | 0.633986483 | 0.785249872 | ZDHHC3 | | protein_coding | 47.26 | -0.013181152 | 0.013181152 | ENSG00000163812 | n.s. |
| ENSG00000136161 | 392.7354063 | -0.127841062 | 0.2729877 | -0.468303 | 0.639567645 | 0.789423788 | RCBTB2 | | protein_coding | 40.27 | -0.013127972 | 0.013127972 | ENSG00000136161 | n.s. |
| ENSG00000186635 | 318.1718287 | -0.123573069 | 0.2573535 | -0.480169 | 0.631107563 | 0.783216917 | ARAP1 | | protein_coding | 54.91 | -0.01311332 | 0.01311332 | ENSG00000186635 | n.s. |
| ENSG00000105402 | 642.6860191 | 0.125389767 | 0.2642222 | 0.474562 | 0.635099329 | 0.786015631 | NAPA | | protein_coding | 55.52 | 0.01311186 | 0.01311186 | ENSG00000105402 | n.s. |
| ENSG00000111676 | 277.8094433 | -0.122228194 | 0.252603 | -0.483875 | 0.628474794 | 0.781202978 | ATN1 | | protein_coding | 55.53 | -0.013107276 | 0.013107276 | ENSG00000111676 | n.s. |
| ENSG00000139971 | 142.5111714 | -0.132284887 | 0.2923595 | -0.452473 | 0.650927967 | 0.796331337 | C14orf37 | | protein_coding | 40.15 | -0.013083795 | 0.013083795 | ENSG00000139971 | n.s. |
| ENSG00000141622 | 661.3139511 | 0.111864062 | 0.2179322 | 0.513298 | 0.607743227 | 0.764659711 | RNF165 | | protein_coding | 49.03 | 0.01303572 | 0.01303572 | ENSG00000141622 | n.s. |
| ENSG00000103365 | 1631.919842 | 0.118208371 | 0.2395269 | 0.493508 | 0.621653882 | 0.775771893 | GGA2 | | protein_coding | 47.76 | 0.013034359 | 0.013034359 | ENSG00000103365 | n.s. |
| ENSG00000006740 | 53.81613748 | 0.124116292 | 0.2610985 | 0.475362 | 0.634529008 | 0.785728035 | ARHGAP44 | | protein_coding | 42.68 | 0.01299842 | 0.01299842 | ENSG00000006740 | n.s. |
| ENSG00000125868 | 1670.8252 | 0.136164293 | 0.3103753 | 0.438708 | 0.660872786 | 0.802768434 | DSTN | | protein_coding | 39.56 | 0.012991396 | 0.012991396 | ENSG00000125868 | n.s. |

| | | | | | | | | | | | | | |
|---|---|---|---|---|---|---|---|---|---|---|---|---|---|
| ENSG00000253586 | 19.56075723 | 0.14514434 | 0.3517447 | 0.412641 | 0.679869509 | 0.81418507 | | sense_intronic | 38.09 | 0.012958032 | 0.012958032 | ENSG00000253586 | n.s. |
| ENSG00000125656 | 317.6181036 | 0.131411241 | 0.2921366 | 0.449828 | 0.652834398 | 0.797170566 | CLPP | protein_coding | 54.14 | 0.012937272 | 0.012937272 | ENSG00000125656 | n.s. |
| ENSG00000132196 | 428.9090248 | 0.114322705 | 0.2276889 | 0.502101 | 0.615596792 | 0.770699949 | HSD17B7 | protein_coding | 39.34 | 0.012931575 | 0.012931575 | ENSG00000132196 | n.s. |
| ENSG00000150316 | 1105.949421 | 0.12774052 | 0.27608 | 0.462694 | 0.643583796 | 0.792112559 | CWC15 | protein_coding | 37.4 | 0.012929014 | 0.012929014 | ENSG00000150316 | n.s. |
| ENSG00000196943 | 970.2194097 | -0.118248301 | 0.2415204 | -0.4896 | 0.624417297 | 0.777831587 | NOP9 | protein_coding | 50.61 | -0.012902595 | 0.012902595 | ENSG00000196943 | n.s. |
| ENSG00000271601 | 567.3195505 | -0.122293006 | 0.255873 | -0.477944 | 0.632689934 | 0.784341728 | LIX1L | protein_coding | 43.6 | -0.012901261 | 0.012901261 | ENSG00000271601 | n.s. |
| ENSG00000186889 | 122.9642627 | -0.122365731 | 0.2561562 | -0.4777 | 0.632863969 | 0.784487628 | TMEM17 | protein_coding | 39.29 | -0.012899049 | 0.012899049 | ENSG00000186889 | n.s. |
| ENSG00000243989 | 263.498296 | 0.136099771 | 0.3120819 | 0.436103 | 0.662762212 | 0.803948385 | ACY1 | protein_coding | 57.5 | 0.012898425 | 0.012898425 | ENSG00000243989 | n.s. |
| ENSG00000279962 | 15.32047086 | -0.143930765 | 0.3480341 | -0.413554 | 0.679201051 | 0.813603737 | | TEC | 54.24 | -0.012894335 | 0.012894335 | ENSG00000279962 | n.s. |
| ENSG00000165502 | 1775.619551 | 0.127022802 | 0.2733067 | 0.464763 | 0.642101234 | 0.791596249 | RPL36AL | protein_coding | 45.59 | 0.012892341 | 0.012892341 | ENSG00000165502 | n.s. |
| ENSG00000111846 | 174.9607295 | 0.118855619 | 0.244313 | 0.486489 | 0.626620347 | 0.779315327 | GCNT2 | protein_coding | 43.51 | 0.012870492 | 0.012870492 | ENSG00000111846 | n.s. |
| ENSG00000271852 | 66.54279704 | 0.132206975 | 0.2969548 | 0.445209 | 0.65616867 | 0.799215851 | SNORD11B | snoRNA | 36.67 | 0.012868486 | 0.012868486 | ENSG00000271852 | n.s. |
| ENSG00000204427 | 355.6616775 | 0.12919042 | 0.2836134 | 0.455516 | 0.648738133 | 0.795243971 | ABHD16A | protein_coding | 50.35 | 0.012854397 | 0.012854397 | ENSG00000204427 | n.s. |
| ENSG00000143622 | 433.8698045 | -0.121389331 | 0.2533386 | -0.479158 | 0.631825962 | 0.783829 | RIT1 | protein_coding | 43.8 | -0.012840402 | 0.012840402 | ENSG00000143622 | n.s. |
| ENSG00000139370 | 643.5691888 | 0.110244336 | 0.2151266 | 0.512463 | 0.608327258 | 0.765038426 | SLC15A4 | protein_coding | 45.87 | 0.012823263 | 0.012823263 | ENSG00000139370 | n.s. |
| ENSG00000207468 | 71.80865836 | -0.124065301 | 0.2638323 | -0.470243 | 0.638181348 | 0.788432616 | SNORA19 | snoRNA | 32.81 | -0.012807933 | 0.012807933 | ENSG00000207468 | n.s. |
| ENSG00000162889 | 484.5810954 | 0.12261968 | 0.258746 | 0.4739 | 0.635571347 | 0.786310195 | MAPKAPK2 | protein_coding | 48.7 | 0.012802242 | 0.012802242 | ENSG00000162889 | n.s. |
| ENSG00000113141 | 2024.141133 | -0.122636215 | 0.2591811 | -0.473168 | 0.63609327 | 0.786433825 | IK | protein_coding | 45.21 | -0.012795595 | 0.012795595 | ENSG00000113141 | n.s. |
| ENSG00000136450 | 14939.95871 | -0.114491821 | 0.229938 | -0.497925 | 0.618537105 | 0.773268575 | SRSF1 | protein_coding | 43.01 | -0.01278526 | 0.01278526 | ENSG00000136450 | n.s. |
| ENSG00000255302 | 7891.409009 | 0.127098276 | 0.2768824 | 0.459033 | 0.646210134 | 0.793440055 | EID1 | protein_coding | 41.64 | 0.012771582 | 0.012771582 | ENSG00000255302 | n.s. |
| ENSG00000113048 | 1463.638871 | -0.122165483 | 0.2579912 | -0.473526 | 0.635838172 | 0.786319563 | MRPS27 | protein_coding | 38.95 | -0.012754189 | 0.012754189 | ENSG00000113048 | n.s. |
| ENSG00000135250 | 966.2506437 | -0.121283697 | 0.2544739 | -0.476606 | 0.633642962 | 0.784999875 | SRPK2 | protein_coding | 41.06 | -0.012750605 | 0.012750605 | ENSG00000135250 | n.s. |
| ENSG00000157637 | 415.9221495 | 0.129609671 | 0.289045 | 0.448407 | 0.653859822 | 0.797310413 | SLC38A10 | protein_coding | 56.73 | 0.012750036 | 0.012750036 | ENSG00000157637 | n.s. |
| ENSG00000101193 | 840.7658854 | -0.117487883 | 0.2410119 | -0.487478 | 0.625919941 | 0.778931296 | GID8 | protein_coding | 48.86 | -0.012747535 | 0.012747535 | ENSG00000101193 | n.s. |
| ENSG00000103264 | 82.02996008 | 0.123979886 | 0.2651323 | 0.467615 | 0.640059841 | 0.789751256 | FBXO31 | protein_coding | 51.99 | 0.012709138 | 0.012709138 | ENSG00000103264 | n.s. |
| ENSG00000206448 | 20.5482359 | 0.14365164 | 0.3511977 | 0.409034 | 0.682515057 | 0.81576698 | PPIAP30 | processed_pseudogene | 48.55 | 0.012703672 | 0.012703672 | ENSG00000206448 | n.s. |
| ENSG00000006744 | 1215.24841 | -0.126300166 | 0.2751895 | -0.458957 | 0.646264957 | 0.793440055 | ELAC2 | protein_coding | 48.08 | -0.012691383 | 0.012691383 | ENSG00000006744 | n.s. |
| ENSG00000143751 | 1212.876914 | 0.121082383 | 0.2547857 | 0.475232 | 0.634621427 | 0.785756703 | SDE2 | protein_coding | 40.61 | 0.012678767 | 0.012678767 | ENSG00000143751 | n.s. |
| ENSG00000134375 | 2456.73897 | 0.126685993 | 0.2770464 | 0.457274 | 0.647474421 | 0.794184333 | TIMM17A | protein_coding | 42.32 | 0.012678568 | 0.012678568 | ENSG00000134375 | n.s. |
| ENSG00000117697 | 1288.963871 | -0.132455074 | 0.3013423 | -0.43955 | 0.660262914 | 0.802287784 | NSL1 | protein_coding | 39.75 | -0.012671953 | 0.012671953 | ENSG00000117697 | n.s. |
| ENSG00000160714 | 1135.570563 | 0.125194286 | 0.2708343 | 0.462254 | 0.643899097 | 0.792311722 | UBE2Q1 | protein_coding | 50.99 | 0.012657633 | 0.012657633 | ENSG00000160714 | n.s. |
| ENSG00000162735 | 1101.111661 | -0.130851452 | 0.2954648 | -0.442866 | 0.657862314 | 0.800438433 | PEX19 | protein_coding | 46.35 | -0.01264968 | 0.01264968 | ENSG00000162735 | n.s. |

| | | | | | | | | | | | | | |
|---|---|---|---|---|---|---|---|---|---|---|---|---|---|
| ENSG00000143224 | 223.0433818 | 0.127237563 | 0.2797416 | 0.45484 | 0.649224628 | 0.795429212 | PPOX | protein_coding | 53.01 | 0.012647218 | 0.012647218 | ENSG00000143224 | n.s. |
| ENSG00000123143 | 740.3570008 | -0.127264919 | 0.2801127 | -0.454335 | 0.649587976 | 0.795585273 | PKN1 | protein_coding | 53.14 | -0.012639095 | 0.012639095 | ENSG00000123143 | n.s. |
| ENSG00000030419 | 27.1902038 | 0.13752767 | 0.3250433 | 0.423106 | 0.672218157 | 0.809347406 | IKZF2 | protein_coding | 34.84 | 0.012633982 | 0.012633982 | ENSG00000030419 | n.s. |
| ENSG00000170322 | 629.5300856 | -0.120507309 | 0.2540121 | -0.474416 | 0.635203532 | 0.786015631 | NFRKB | protein_coding | 45.4 | -0.012601307 | 0.012601307 | ENSG00000170322 | n.s. |
| ENSG00000139697 | 2717.977696 | -0.115852735 | 0.2372396 | -0.488336 | 0.625311597 | 0.778584691 | SBNO1 | protein_coding | 43.26 | -0.012592513 | 0.012592513 | ENSG00000139697 | n.s. |
| ENSG00000058600 | 484.5443422 | -0.127147444 | 0.2818036 | -0.451192 | 0.651851404 | 0.796633653 | POLR3E | protein_coding | 51.67 | -0.01255471 | 0.01255471 | ENSG00000058600 | n.s. |
| ENSG00000179262 | 862.4151366 | -0.120053743 | 0.2530643 | -0.4744 | 0.635214557 | 0.786015631 | RAD23A | protein_coding | 54.44 | -0.012553878 | 0.012553878 | ENSG00000179262 | n.s. |
| ENSG00000171603 | 387.0569707 | -0.121958891 | 0.2600145 | -0.469046 | 0.639036423 | 0.789074708 | CLSTN1 | protein_coding | 47.37 | -0.01254736 | 0.01254736 | ENSG00000171603 | n.s. |
| ENSG00000164880 | 767.1000384 | 0.112500672 | 0.2267823 | 0.496073 | 0.619842597 | 0.774205479 | INTS1 | protein_coding | 63.54 | 0.012503748 | 0.012503748 | ENSG00000164880 | n.s. |
| ENSG00000011260 | 2828.635899 | -0.123093296 | 0.2649537 | -0.464584 | 0.642229296 | 0.791643267 | UTP18 | protein_coding | 40.38 | -0.012490335 | 0.012490335 | ENSG00000011260 | n.s. |
| ENSG00000232732 | 49.50404125 | 0.126197703 | 0.2790654 | 0.452216 | 0.651113706 | 0.796369304 | | processed_transcript | 38.64 | 0.012479121 | 0.012479121 | ENSG00000232732 | n.s. |
| ENSG00000175283 | 240.9181886 | -0.116621782 | 0.2415129 | -0.48288 | 0.629180858 | 0.781705085 | DOLK | protein_coding | 56.89 | -0.012473523 | 0.012473523 | ENSG00000175283 | n.s. |
| ENSG00000148335 | 617.379573 | -0.122630316 | 0.2642648 | -0.464043 | 0.642616692 | 0.791643267 | NTMT1 | protein_coding | 55.42 | -0.012443357 | 0.012443357 | ENSG00000148335 | n.s. |
| ENSG00000131069 | 216.0637607 | -0.132801705 | 0.3083052 | -0.430748 | 0.666651944 | 0.806182257 | ACSS2 | protein_coding | 45.42 | -0.012425826 | 0.012425826 | ENSG00000131069 | n.s. |
| ENSG00000099256 | 1072.19467 | -0.131164416 | 0.3011483 | -0.435548 | 0.663165005 | 0.804079627 | PRTFDC1 | protein_coding | 41.87 | -0.012421394 | 0.012421394 | ENSG00000099256 | n.s. |
| ENSG00000184702 | 395.3447739 | 0.113455565 | 0.2314801 | 0.490131 | 0.624041174 | 0.777656247 | 5-Sep | protein_coding | 66.81 | 0.012390747 | 0.012390747 | ENSG00000184702 | n.s. |
| ENSG00000197619 | 340.2340024 | -0.123698236 | 0.2704359 | -0.457403 | 0.647381284 | 0.794184333 | ZNF615 | protein_coding | 39.06 | -0.012379557 | 0.012379557 | ENSG00000197619 | n.s. |
| ENSG00000105223 | 570.7453406 | 0.123865833 | 0.2714926 | 0.45624 | 0.648217273 | 0.79481542 | PLD3 | protein_coding | 52.33 | 0.0123536 | 0.0123536 | ENSG00000105223 | n.s. |
| ENSG00000183077 | 705.4948952 | -0.122882264 | 0.2676557 | -0.459106 | 0.646158296 | 0.793440055 | AFMID | protein_coding | 51.08 | -0.012347932 | 0.012347932 | ENSG00000183077 | n.s. |
| ENSG00000207524 | 2.932437063 | -0.284004441 | 1.2632294 | -0.224824 | 0.822116096 | 0.904752863 | RNU6-33P | snRNA | 44.86 | -0.012345683 | 0.012345683 | ENSG00000207524 | n.s. |
| ENSG00000164985 | 5511.069874 | -0.121864762 | 0.2631378 | -0.463122 | 0.643277256 | 0.792021081 | PSIP1 | protein_coding | 37.88 | -0.012340423 | 0.012340423 | ENSG00000164985 | n.s. |
| ENSG00000100023 | 28.51973773 | -0.134693258 | 0.3188191 | -0.422476 | 0.672677972 | 0.80983099 | PPIL2 | protein_coding | 54.06 | -0.012338658 | 0.012338658 | ENSG00000100023 | n.s. |
| ENSG00000183598 | 899.9099204 | 0.11139372 | 0.2250531 | 0.494966 | 0.620623885 | 0.774903269 | HIST2H3D | protein_coding | 67.4 | 0.012337134 | 0.012337134 | ENSG00000183598 | n.s. |
| ENSG00000138107 | 1371.776854 | -0.125902557 | 0.2815896 | -0.447114 | 0.654793022 | 0.798168731 | ACTR1A | protein_coding | 50.48 | -0.012326526 | 0.012326526 | ENSG00000138107 | n.s. |
| ENSG00000198728 | 2168.204132 | 0.120329294 | 0.2574998 | 0.467299 | 0.640286234 | 0.789890594 | LDB1 | protein_coding | 55.59 | 0.012325698 | 0.012325698 | ENSG00000198728 | n.s. |
| ENSG00000131408 | 199.6118245 | -0.124500065 | 0.2747267 | -0.453178 | 0.650420597 | 0.796184805 | NR1H2 | protein_coding | 51.27 | -0.012323778 | 0.012323778 | ENSG00000131408 | n.s. |
| ENSG00000103495 | 1923.253204 | 0.11793995 | 0.2490495 | 0.47356 | 0.635813577 | 0.786319563 | MAZ | protein_coding | 66.53 | 0.012313039 | 0.012313039 | ENSG00000103495 | n.s. |
| ENSG00000070748 | 1022.030041 | 0.113975532 | 0.2347912 | 0.485433 | 0.62736888 | 0.779991215 | CHAT | protein_coding | 51.04 | 0.012299133 | 0.012299133 | ENSG00000070748 | n.s. |
| ENSG00000106105 | 3968.097109 | 0.119524339 | 0.2548716 | 0.468959 | 0.639098882 | 0.789074708 | GARS | protein_coding | 38.71 | 0.012296888 | 0.012296888 | ENSG00000106105 | n.s. |
| ENSG00000272655 | 159.210223 | 0.112806656 | 0.2306723 | 0.489034 | 0.624817436 | 0.778115668 | POLR2J4 | transcribed_unprocessed_pseud | 54.06 | 0.012290944 | 0.012290944 | ENSG00000272655 | n.s. |
| ENSG00000213123 | 221.39369 | 0.136730508 | 0.3295167 | 0.414943 | 0.678183946 | 0.813086294 | TCTEX1D2 | protein_coding | 45.15 | 0.012287063 | 0.012287063 | ENSG00000213123 | n.s. |
| ENSG00000272574 | 31.72866612 | 0.130379541 | 0.3008936 | 0.433308 | 0.664791226 | 0.804958837 | | sense_intronic | 42.18 | 0.012285185 | 0.012285185 | ENSG00000272574 | n.s. |

| | | | | | | | | | | | | | |
|---|---|---|---|---|---|---|---|---|---|---|---|---|---|
| ENSG00000125952 | 472.8967602 | -0.118987167 | 0.2531211 | -0.47008 | 0.638297814 | 0.788434224 | MAX | protein_coding | 44.69 | -0.012283585 | 0.012283585 | ENSG00000125952 | n.s. |
| ENSG00000109689 | 287.5483972 | -0.116222223 | 0.2428167 | -0.478642 | 0.63219351 | 0.784075382 | STIM2 | protein_coding | 36.58 | -0.012277969 | 0.012277969 | ENSG00000109689 | n.s. |
| ENSG00000187713 | 427.4193538 | -0.124075711 | 0.2742273 | -0.452456 | 0.65094067 | 0.796331337 | TMEM203 | protein_coding | 56.71 | -0.012271856 | 0.012271856 | ENSG00000187713 | n.s. |
| ENSG00000114023 | 270.2959506 | 0.139031379 | 0.340406 | 0.408428 | 0.682959409 | 0.816134228 | FAM162A | protein_coding | 38.78 | 0.012267908 | 0.012267908 | ENSG00000114023 | n.s. |
| ENSG00000111790 | 1350.912824 | 0.122544798 | 0.2679523 | 0.457338 | 0.647428051 | 0.794184333 | FGFR1OP2 | protein_coding | 35.97 | 0.012264122 | 0.012264122 | ENSG00000111790 | n.s. |
| ENSG00000162300 | 277.4302605 | 0.117395523 | 0.24782 | 0.473713 | 0.635704614 | 0.786319563 | ZFPL1 | protein_coding | 56.15 | 0.012256201 | 0.012256201 | ENSG00000162300 | n.s. |
| ENSG00000072210 | 558.9353048 | 0.120518626 | 0.2589524 | 0.465408 | 0.641639044 | 0.791279045 | ALDH3A2 | protein_coding | 44.22 | 0.012253169 | 0.012253169 | ENSG00000072210 | n.s. |
| ENSG00000144468 | 68.37104795 | -0.129314012 | 0.2968161 | -0.43567 | 0.663075837 | 0.804041532 | RHBDD1 | protein_coding | 39.57 | -0.012248819 | 0.012248819 | ENSG00000144468 | n.s. |
| ENSG00000113658 | 2910.191018 | -0.119204075 | 0.2544439 | -0.468489 | 0.639435229 | 0.789330321 | SMAD5 | protein_coding | 37.17 | -0.012247171 | 0.012247171 | ENSG00000113658 | n.s. |
| ENSG00000172977 | 599.4539326 | 0.124384393 | 0.2765596 | 0.449756 | 0.6528863 | 0.797170566 | KAT5 | protein_coding | 55.16 | 0.012245487 | 0.012245487 | ENSG00000172977 | n.s. |
| ENSG00000137965 | 2631.059147 | -0.121333519 | 0.2632692 | -0.460872 | 0.644890182 | 0.792685753 | IFI44 | protein_coding | 35.31 | -0.012242424 | 0.012242424 | ENSG00000137965 | n.s. |
| ENSG00000182180 | 1425.154545 | -0.117384884 | 0.2482386 | -0.472871 | 0.636305014 | 0.786582146 | MRPS16 | protein_coding | 47.47 | -0.012238069 | 0.012238069 | ENSG00000182180 | n.s. |
| ENSG00000100811 | 2372.151446 | -0.121911063 | 0.2661589 | -0.458039 | 0.64692468 | 0.793859715 | YY1 | protein_coding | 45.55 | -0.012222344 | 0.012222344 | ENSG00000100811 | n.s. |
| ENSG00000280077 | 224.9410862 | -0.123625207 | 0.2741497 | -0.450941 | 0.652032378 | 0.796633653 | | TEC | 33.16 | -0.01220692 | 0.01220692 | ENSG00000280077 | n.s. |
| ENSG00000124207 | 5787.582664 | -0.123582428 | 0.2740298 | -0.450982 | 0.652002718 | 0.796633653 | CSE1L | protein_coding | 41.33 | -0.012202696 | 0.012202696 | ENSG00000124207 | n.s. |
| ENSG00000176912 | 58.92465305 | -0.122675693 | 0.2695471 | -0.455118 | 0.649024506 | 0.795346039 | TYMSOS | processed_transcript | 42.36 | -0.012199347 | 0.012199347 | ENSG00000176912 | n.s. |
| ENSG00000183048 | 234.4074535 | -0.121453571 | 0.2648488 | -0.458577 | 0.646537916 | 0.79359494 | SLC25A10 | protein_coding | 63.39 | -0.012194073 | 0.012194073 | ENSG00000183048 | n.s. |
| ENSG00000003056 | 1251.754374 | -0.123567704 | 0.2751468 | -0.449097 | 0.653361469 | 0.797221535 | M6PR | protein_coding | 42.71 | -0.012161654 | 0.012161654 | ENSG00000003056 | n.s. |
| ENSG00000188735 | 986.8368872 | 0.116420872 | 0.2457399 | 0.473757 | 0.635673526 | 0.786319563 | TMEM120B | protein_coding | 51.94 | 0.012154446 | 0.012154446 | ENSG00000188735 | n.s. |
| ENSG00000102805 | 163.7908687 | -0.119885 | 0.2586928 | -0.463426 | 0.643058908 | 0.791907764 | CLN5 | protein_coding | 38.81 | -0.012147396 | 0.012147396 | ENSG00000102805 | n.s. |
| ENSG00000175792 | 2123.108849 | -0.129633415 | 0.3009904 | -0.43069 | 0.666694139 | 0.806182257 | RUVBL1 | protein_coding | 45.86 | -0.012129379 | 0.012129379 | ENSG00000175792 | n.s. |
| ENSG00000183684 | 3689.921932 | 0.118294588 | 0.2529537 | 0.467653 | 0.640032611 | 0.789751256 | ALYREF | protein_coding | 57.57 | 0.01212634 | 0.01212634 | ENSG00000183684 | n.s. |
| ENSG00000278886 | 65.44645608 | -0.119506194 | 0.2583144 | -0.462639 | 0.643623456 | 0.792112559 | | TEC | 38.27 | -0.012095592 | 0.012095592 | ENSG00000278886 | n.s. |
| ENSG00000073111 | 4985.515957 | 0.116795552 | 0.2479874 | 0.470974 | 0.637659528 | 0.787906781 | MCM2 | protein_coding | 50.73 | 0.012091278 | 0.012091278 | ENSG00000073111 | n.s. |
| ENSG00000196951 | 43.86432803 | 0.129775969 | 0.3032709 | 0.427921 | 0.668708691 | 0.807076205 | SCOC-AS1 | antisense | 40.85 | 0.012080255 | 0.012080255 | ENSG00000196951 | n.s. |
| ENSG00000196700 | 631.834774 | -0.112930534 | 0.2339981 | -0.482613 | 0.629370578 | 0.781758101 | ZNF512B | protein_coding | 63.43 | -0.012075392 | 0.012075392 | ENSG00000196700 | n.s. |
| ENSG00000134986 | 6719.603312 | 0.121560017 | 0.2675061 | 0.45442 | 0.649526874 | 0.795585273 | NREP | protein_coding | 38.48 | 0.012072522 | 0.012072522 | ENSG00000134986 | n.s. |
| ENSG00000252361 | 6.131892634 | -0.172499917 | 0.5157997 | -0.334432 | 0.738053603 | 0.851299448 | RNU6-118P | snRNA | 36.27 | -0.012060788 | 0.012060788 | ENSG00000252361 | n.s. |
| ENSG00000075239 | 697.6066785 | 0.127105582 | 0.2922132 | 0.434976 | 0.663580199 | 0.804307547 | ACAT1 | protein_coding | 42.49 | 0.012021373 | 0.012021373 | ENSG00000075239 | n.s. |
| ENSG00000088832 | 839.5681588 | 0.122055812 | 0.271801 | 0.449063 | 0.653386119 | 0.797221535 | FKBP1A | protein_coding | 45.47 | 0.012012853 | 0.012012853 | ENSG00000088832 | n.s. |
| ENSG00000221296 | 4.907802477 | 0.170286595 | 0.5047479 | 0.33737 | 0.735838294 | 0.850072732 | MIR548T | miRNA | 36.49 | 0.012012683 | 0.012012683 | ENSG00000221296 | n.s. |
| ENSG00000079819 | 2452.061948 | -0.114678585 | 0.2414695 | -0.47492 | 0.634844352 | 0.785892941 | EPB41L2 | protein_coding | 38.6 | -0.011999579 | 0.011999579 | ENSG00000079819 | n.s. |

| | | | | | | | | | | | | | |
|---|---|---|---|---|---|---|---|---|---|---|---|---|---|
| ENSG00000267673 | 134.092846 | -0.130531616 | 0.3091207 | -0.422268 | 0.672829763 | 0.809846304 | FDX2 | protein_coding | 52.77 | -0.011956356 | 0.011956356 | ENSG00000267673 | n.s. |
| ENSG00000207967 | 6.733797466 | 0.162763136 | 0.4662881 | 0.349061 | 0.727043275 | 0.844466799 | MIR620 | miRNA | 21.05 | 0.01194965 | 0.01194965 | ENSG00000207967 | n.s. |
| ENSG00000188070 | 1321.132982 | -0.11393654 | 0.2396928 | -0.475344 | 0.634541839 | 0.785728035 | C11orf95 | protein_coding | 61.54 | -0.011932318 | 0.011932318 | ENSG00000188070 | n.s. |
| ENSG00000128059 | 2229.615859 | -0.119930513 | 0.2633515 | -0.455401 | 0.648820822 | 0.795275315 | PPAT | protein_coding | 37.44 | -0.011930987 | 0.011930987 | ENSG00000128059 | n.s. |
| ENSG00000100968 | 56.44701856 | 0.130404792 | 0.309843 | 0.420874 | 0.673847264 | 0.810117905 | NFATC4 | protein_coding | 56.84 | 0.011925749 | 0.011925749 | ENSG00000100968 | n.s. |
| ENSG00000253516 | 76.07150871 | -0.125326322 | 0.2868748 | -0.436868 | 0.66220732 | 0.803478215 | HMGB1P41 | processed_pseudogene | 42.27 | -0.011909246 | 0.011909246 | ENSG00000253516 | n.s. |
| ENSG00000222255 | 4.719862807 | -0.168135773 | 0.4965277 | -0.338623 | 0.734893638 | 0.849656869 | RNU6-101P | snRNA | 41.35 | -0.011896687 | 0.011896687 | ENSG00000222255 | n.s. |
| ENSG00000105197 | 1412.620673 | 0.119673128 | 0.2631166 | 0.454829 | 0.649232031 | 0.795429212 | TIMM50 | protein_coding | 53.07 | 0.011895325 | 0.011895325 | ENSG00000105197 | n.s. |
| ENSG00000229512 | 8.86907164 | -0.157471531 | 0.4407218 | -0.357304 | 0.720864445 | 0.840561437 | | sense_intronic | 67.61 | -0.011878162 | 0.011878162 | ENSG00000229512 | n.s. |
| ENSG00000110104 | 287.4562003 | -0.117658554 | 0.2556989 | -0.460145 | 0.645412186 | 0.793052146 | CCDC86 | protein_coding | 54.76 | -0.011848011 | 0.011848011 | ENSG00000110104 | n.s. |
| ENSG00000148356 | 142.3702193 | -0.12558573 | 0.2899057 | -0.433195 | 0.664872989 | 0.804958837 | LRSAM1 | protein_coding | 51.44 | -0.011833482 | 0.011833482 | ENSG00000148356 | n.s. |
| ENSG00000141425 | 3193.494538 | 0.125112271 | 0.2876341 | 0.43497 | 0.663584055 | 0.804307547 | RPRD1A | protein_coding | 38.63 | 0.01183285 | 0.01183285 | ENSG00000141425 | n.s. |
| ENSG00000068400 | 548.645448 | -0.112357407 | 0.2354311 | -0.477241 | 0.633190378 | 0.784700146 | GRIPAP1 | protein_coding | 50.22 | -0.011830815 | 0.011830815 | ENSG00000068400 | n.s. |
| ENSG00000163170 | 499.4335531 | 0.132679061 | 0.3218348 | 0.412258 | 0.680150122 | 0.814390278 | BOLA3 | protein_coding | 50.59 | 0.01183065 | 0.01183065 | ENSG00000163170 | n.s. |
| ENSG00000156136 | 817.3778822 | 0.126192241 | 0.2933404 | 0.430191 | 0.667057052 | 0.806194128 | DCK | protein_coding | 38.98 | 0.011806593 | 0.011806593 | ENSG00000156136 | n.s. |
| ENSG00000105355 | 109.565324 | 0.128678539 | 0.3061722 | 0.420282 | 0.674279758 | 0.810427924 | PLIN3 | protein_coding | 52.37 | 0.011746498 | 0.011746498 | ENSG00000105355 | n.s. |
| ENSG00000101844 | 115.2860309 | -0.123404227 | 0.2819204 | -0.437727 | 0.661584013 | 0.803281958 | ATG4A | protein_coding | 39.97 | -0.011739689 | 0.011739689 | ENSG00000101844 | n.s. |
| ENSG00000165895 | 88.17520915 | -0.122160742 | 0.2769286 | -0.441127 | 0.659120847 | 0.801553838 | ARHGAP42 | protein_coding | 38.03 | -0.011735653 | 0.011735653 | ENSG00000165895 | n.s. |
| ENSG00000135018 | 2674.235722 | -0.11741032 | 0.2569855 | -0.456875 | 0.647760672 | 0.794465454 | UBQLN1 | protein_coding | 38.75 | -0.011732224 | 0.011732224 | ENSG00000135018 | n.s. |
| ENSG00000114520 | 1362.181102 | -0.125276372 | 0.2914218 | -0.42988 | 0.667282995 | 0.806194128 | SNX4 | protein_coding | 40.09 | -0.011720904 | 0.011720904 | ENSG00000114520 | n.s. |
| ENSG00000196396 | 937.3939429 | -0.120892303 | 0.2724803 | -0.443674 | 0.657278601 | 0.799957861 | PTPN1 | protein_coding | 43.56 | -0.01171844 | 0.01171844 | ENSG00000196396 | n.s. |
| ENSG00000157326 | 85.17427461 | -0.122331207 | 0.2783905 | -0.439423 | 0.660355031 | 0.802287784 | DHRS4 | protein_coding | 45.27 | -0.011703405 | 0.011703405 | ENSG00000157326 | n.s. |
| ENSG00000130226 | 225.3648896 | -0.114900597 | 0.2467853 | -0.465589 | 0.641509502 | 0.79118937 | DPP6 | protein_coding | 43.87 | -0.011687638 | 0.011687638 | ENSG00000130226 | n.s. |
| ENSG00000132950 | 355.1989596 | 0.120445089 | 0.2714862 | 0.443651 | 0.657294974 | 0.799957861 | ZMYM5 | protein_coding | 42.61 | 0.011675091 | 0.011675091 | ENSG00000132950 | n.s. |
| ENSG00000166128 | 819.2284729 | 0.117521497 | 0.2587906 | 0.454118 | 0.649743835 | 0.795668289 | RAB8B | protein_coding | 39.83 | 0.011666118 | 0.011666118 | ENSG00000166128 | n.s. |
| ENSG00000189046 | 920.4938265 | -0.118178358 | 0.2623517 | -0.450458 | 0.652380346 | 0.796832315 | ALKBH2 | protein_coding | 51.05 | -0.011656293 | 0.011656293 | ENSG00000189046 | n.s. |
| ENSG00000280092 | 36.82196262 | 0.140178182 | 0.3603211 | 0.389037 | 0.697248861 | 0.825773724 | | TEC | 41.59 | 0.011654266 | 0.011654266 | ENSG00000280092 | n.s. |
| ENSG00000123737 | 2378.327311 | -0.115492649 | 0.2508297 | -0.460442 | 0.64519871 | 0.79285981 | EXOSC9 | protein_coding | 37.71 | -0.011642074 | 0.011642074 | ENSG00000123737 | n.s. |
| ENSG00000261236 | 1043.431191 | 0.122934849 | 0.2827349 | 0.434806 | 0.663703191 | 0.804311909 | BOP1 | protein_coding | 62.27 | 0.011626625 | 0.011626625 | ENSG00000261236 | n.s. |
| ENSG00000275227 | 50.48433851 | 0.121417104 | 0.2760215 | 0.439883 | 0.660022067 | 0.802157304 | | misc_RNA | 44.2 | 0.011624529 | 0.011624529 | ENSG00000275227 | n.s. |
| ENSG00000105619 | 158.4612286 | -0.118010688 | 0.262753 | -0.449132 | 0.653336672 | 0.797221535 | TFPT | protein_coding | 54.83 | -0.011614727 | 0.011614727 | ENSG00000105619 | n.s. |
| ENSG00000082996 | 1066.195559 | -0.118821073 | 0.266007 | -0.446684 | 0.655103199 | 0.798467659 | RNF13 | protein_coding | 36.39 | -0.011613889 | 0.011613889 | ENSG00000082996 | n.s. |

| | | | | | | | | | | | | | |
|---|---|---|---|---|---|---|---|---|---|---|---|---|---|
| ENSG00000094914 | 814.5634074 | -0.122638763 | 0.2816656 | -0.435406 | 0.663268093 | 0.804134593 | AAAS | protein_coding | 49.39 | -0.011610366 | 0.011610366 | ENSG00000094914 | n.s. |
| ENSG00000185627 | 1213.891322 | 0.130780943 | 0.3191837 | 0.409736 | 0.681999883 | 0.815483027 | PSMD13 | protein_coding | 44.35 | 0.011585239 | 0.011585239 | ENSG00000185627 | n.s. |
| ENSG00000010017 | 823.6438651 | 0.116997232 | 0.2589727 | 0.451774 | 0.651431501 | 0.796400914 | RANBP9 | protein_coding | 36.94 | 0.011567312 | 0.011567312 | ENSG00000010017 | n.s. |
| ENSG00000011332 | 77.5348396 | -0.124887526 | 0.2931967 | -0.425951 | 0.670143327 | 0.8080372 | DPF1 | protein_coding | 57.15 | -0.011560669 | 0.011560669 | ENSG00000011332 | n.s. |
| ENSG00000153130 | 1217.485077 | -0.128015434 | 0.3070192 | -0.416962 | 0.67670599 | 0.81243231 | SCOC | protein_coding | 40.94 | -0.011548632 | 0.011548632 | ENSG00000153130 | n.s. |
| ENSG00000102024 | 976.6646943 | 0.129736882 | 0.3153545 | 0.4114 | 0.680779193 | 0.814863197 | PLS3 | protein_coding | 38.9 | 0.011535593 | 0.011535593 | ENSG00000102024 | n.s. |
| ENSG00000134884 | 3154.858793 | 0.11701894 | 0.2605589 | 0.449107 | 0.653354169 | 0.797221535 | ARGLU1 | protein_coding | 37.66 | 0.011517119 | 0.011517119 | ENSG00000134884 | n.s. |
| ENSG00000162910 | 414.271455 | 0.122994357 | 0.2858603 | 0.43026 | 0.667006277 | 0.806194128 | MRPL55 | protein_coding | 56.68 | 0.011507397 | 0.011507397 | ENSG00000162910 | n.s. |
| ENSG00000188725 | 1424.461138 | 0.113815353 | 0.2463681 | 0.461973 | 0.64410089 | 0.792349946 | SMIM15 | protein_coding | 36.57 | 0.011504794 | 0.011504794 | ENSG00000188725 | n.s. |
| ENSG00000112640 | 817.2463103 | 0.122925462 | 0.2855197 | 0.430532 | 0.666808404 | 0.806194128 | PPP2R5D | protein_coding | 49.89 | 0.011500952 | 0.011500952 | ENSG00000112640 | n.s. |
| ENSG00000137700 | 307.4779479 | -0.123118712 | 0.2871258 | -0.428797 | 0.668070858 | 0.806516133 | SLC37A4 | protein_coding | 55.91 | -0.01149768 | 0.01149768 | ENSG00000137700 | n.s. |
| ENSG00000121691 | 999.7539961 | 0.122875178 | 0.2857839 | 0.429958 | 0.66722596 | 0.806194128 | CAT | protein_coding | 40.6 | 0.011496247 | 0.011496247 | ENSG00000121691 | n.s. |
| ENSG00000121753 | 325.1522576 | -0.116525901 | 0.2598398 | -0.448453 | 0.653826375 | 0.797310413 | ADGRB2 | protein_coding | 58.75 | -0.011462952 | 0.011462952 | ENSG00000121753 | n.s. |
| ENSG00000223959 | 123.5687947 | -0.112681667 | 0.2424661 | -0.464732 | 0.642123683 | 0.791596249 | AFG3L1P | transcribed_unitary_pseudogene | 50.98 | -0.011436769 | 0.011436769 | ENSG00000223959 | n.s. |
| ENSG00000100711 | 204.0363396 | -0.120574159 | 0.2766377 | -0.435856 | 0.662941277 | 0.803948385 | ZFYVE21 | protein_coding | 56.98 | -0.011427034 | 0.011427034 | ENSG00000100711 | n.s. |
| ENSG00000174939 | 274.6143352 | -0.124664562 | 0.2957069 | -0.421581 | 0.673330519 | 0.809846304 | ASPHD1 | protein_coding | 50.19 | -0.011418949 | 0.011418949 | ENSG00000174939 | n.s. |
| ENSG00000265241 | 4143.58699 | 0.112604795 | 0.2433122 | 0.4628 | 0.643508055 | 0.79211057 | RBM8A | protein_coding | 46.78 | 0.011397203 | 0.011397203 | ENSG00000265241 | n.s. |
| ENSG00000124006 | 703.6982579 | -0.114469238 | 0.2512929 | -0.455521 | 0.648734338 | 0.795243971 | OBSL1 | protein_coding | 58.29 | -0.011389645 | 0.011389645 | ENSG00000124006 | n.s. |
| ENSG00000274885 | 116.9889177 | -0.124339175 | 0.2946994 | -0.421919 | 0.673084346 | 0.809846304 | | sense_intronic | 33.33 | -0.011389145 | 0.011389145 | ENSG00000274885 | n.s. |
| ENSG00000090971 | 156.6708147 | 0.120803047 | 0.2787646 | 0.433351 | 0.664759461 | 0.804958837 | NAT14 | protein_coding | 66.08 | 0.011382827 | 0.011382827 | ENSG00000090971 | n.s. |
| ENSG00000113140 | 999.3946501 | 0.120081754 | 0.2760043 | 0.435072 | 0.663510144 | 0.804307547 | SPARC | protein_coding | 47.91 | 0.011357075 | 0.011357075 | ENSG00000113140 | n.s. |
| ENSG00000180304 | 650.366481 | 0.126086241 | 0.3031458 | 0.415926 | 0.677464144 | 0.812781842 | OAZ2 | protein_coding | 47.83 | 0.011351041 | 0.011351041 | ENSG00000180304 | n.s. |
| ENSG00000164284 | 1336.436754 | -0.11177958 | 0.2408252 | -0.464152 | 0.642538573 | 0.791643267 | GRPEL2 | protein_coding | 41.28 | -0.011342327 | 0.011342327 | ENSG00000164284 | n.s. |
| ENSG00000077942 | 864.5824119 | -0.115112961 | 0.2564686 | -0.448838 | 0.653548198 | 0.797299375 | FBLN1 | protein_coding | 53.13 | -0.011324649 | 0.011324649 | ENSG00000077942 | n.s. |
| ENSG00000243680 | 22.35474272 | 0.149306857 | 0.4163899 | 0.358575 | 0.719913317 | 0.83989887 | RPL37P23 | processed_pseudogene | 47.62 | 0.011313429 | 0.011313429 | ENSG00000243680 | n.s. |
| ENSG00000114650 | 738.380628 | 0.109861842 | 0.2343795 | 0.468735 | 0.63925919 | 0.789182984 | SCAP | protein_coding | 48.04 | 0.011296246 | 0.011296246 | ENSG00000114650 | n.s. |
| ENSG00000185324 | 563.1452785 | 0.112872415 | 0.2468136 | 0.457318 | 0.647442228 | 0.794184333 | CDK10 | protein_coding | 55.36 | 0.011296123 | 0.011296123 | ENSG00000185324 | n.s. |
| ENSG00000153574 | 394.7671941 | -0.114131288 | 0.2520275 | -0.452852 | 0.650654965 | 0.796220641 | RPIA | protein_coding | 41.25 | -0.011295182 | 0.011295182 | ENSG00000153574 | n.s. |
| ENSG00000163818 | 363.944368 | -0.122434328 | 0.288524 | -0.424347 | 0.671312708 | 0.808693667 | LZTFL1 | protein_coding | 42.03 | -0.011290399 | 0.011290399 | ENSG00000163818 | n.s. |
| ENSG00000184058 | 148.5155615 | -0.112321455 | 0.2443507 | -0.459673 | 0.645750847 | 0.793398257 | TBX1 | protein_coding | 57.77 | -0.01128929 | 0.01128929 | ENSG00000184058 | n.s. |
| ENSG00000134690 | 463.6736661 | -0.11329109 | 0.2493336 | -0.454376 | 0.649558505 | 0.795585273 | CDCA8 | protein_coding | 45.82 | -0.011251308 | 0.011251308 | ENSG00000134690 | n.s. |
| ENSG00000054392 | 186.4239197 | -0.117569772 | 0.2675606 | -0.439414 | 0.660361848 | 0.802287784 | HHAT | protein_coding | 42.26 | -0.011247879 | 0.011247879 | ENSG00000054392 | n.s. |

| | | | | | | | | | | | | | |
|---|---|---|---|---|---|---|---|---|---|---|---|---|---|
| ENSG00000104691 | 403.8152571 | -0.122738226 | 0.2907046 | -0.422209 | 0.672872181 | 0.809846304 | UBXN8 | protein_coding | 44.01 | -0.011242502 | 0.011242502 | ENSG00000104691 | n.s. |
| ENSG00000132406 | 519.8299294 | 0.11705787 | 0.2655916 | 0.440744 | 0.659398482 | 0.801747015 | TMEM128 | protein_coding | 41.69 | 0.011233183 | 0.011233183 | ENSG00000132406 | n.s. |
| ENSG00000160570 | 143.8249042 | 0.11959522 | 0.2767839 | 0.432089 | 0.66567692 | 0.805580891 | DEDD2 | protein_coding | 52.69 | 0.011228896 | 0.011228896 | ENSG00000160570 | n.s. |
| ENSG00000206659 | 10.49214069 | 0.141832782 | 0.3805977 | 0.372658 | 0.709402991 | 0.833411025 | | misc_RNA | 42.16 | 0.011224754 | 0.011224754 | ENSG00000206659 | n.s. |
| ENSG00000163346 | 185.728509 | -0.125246468 | 0.3025998 | -0.413901 | 0.678946356 | 0.813590918 | PBXIP1 | protein_coding | 54.24 | -0.011221321 | 0.011221321 | ENSG00000163346 | n.s. |
| ENSG00000174173 | 1339.067319 | -0.110989815 | 0.2406651 | -0.46118 | 0.644669747 | 0.792637386 | TRMT10C | protein_coding | 41.24 | -0.011201697 | 0.011201697 | ENSG00000174173 | n.s. |
| ENSG00000149929 | 452.7470823 | 0.105431329 | 0.2194249 | 0.480489 | 0.630879402 | 0.783003558 | HIRIP3 | protein_coding | 58.04 | 0.011200631 | 0.011200631 | ENSG00000149929 | n.s. |
| ENSG00000174109 | 247.5966125 | 0.113849763 | 0.2538 | 0.448581 | 0.653734179 | 0.797310413 | C16orf91 | protein_coding | 56.95 | 0.011199693 | 0.011199693 | ENSG00000174109 | n.s. |
| ENSG00000138796 | 566.4148762 | -0.122043596 | 0.2893986 | -0.421715 | 0.673233387 | 0.809846304 | HADH | protein_coding | 43.13 | -0.011178875 | 0.011178875 | ENSG00000138796 | n.s. |
| ENSG00000100604 | 3214.629135 | -0.116726755 | 0.2652696 | -0.440031 | 0.65991489 | 0.802157304 | CHGA | protein_coding | 56.35 | -0.011175473 | 0.011175473 | ENSG00000100604 | n.s. |
| ENSG00000106609 | 1225.20968 | -0.110170979 | 0.2374942 | -0.463889 | 0.642727203 | 0.791709355 | TMEM248 | protein_coding | 44.8 | -0.011175108 | 0.011175108 | ENSG00000106609 | n.s. |
| ENSG00000207067 | 161.732062 | -0.110637326 | 0.2397408 | -0.461487 | 0.644449077 | 0.792637386 | SNORA72 | snoRNA | 40.15 | -0.011166122 | 0.011166122 | ENSG00000207067 | n.s. |
| ENSG00000127540 | 651.3790613 | 0.128125089 | 0.3165685 | 0.404731 | 0.685675275 | 0.818191365 | UQCR11 | protein_coding | 56.38 | 0.011165475 | 0.011165475 | ENSG00000127540 | n.s. |
| ENSG00000152601 | 1858.712004 | -0.11490826 | 0.2583912 | -0.444707 | 0.656531809 | 0.799562477 | MBNL1 | protein_coding | 35.71 | -0.011163061 | 0.011163061 | ENSG00000152601 | n.s. |
| ENSG00000117877 | 288.6746402 | -0.112380601 | 0.2472289 | -0.454561 | 0.649425173 | 0.795585273 | CD3EAP | protein_coding | 53.31 | -0.011160884 | 0.011160884 | ENSG00000117877 | n.s. |
| ENSG00000078319 | 191.4774912 | 0.124108687 | 0.2986485 | 0.415568 | 0.677726292 | 0.812989277 | PMS2P1 | unprocessed_pseudogene | 44.3 | 0.011159255 | 0.011159255 | ENSG00000078319 | n.s. |
| ENSG00000141030 | 1847.251469 | 0.116897013 | 0.2672536 | 0.437401 | 0.66182054 | 0.803364895 | COPS3 | protein_coding | 44.79 | 0.011115404 | 0.011115404 | ENSG00000141030 | n.s. |
| ENSG00000146670 | 1348.479392 | 0.114622182 | 0.258277 | 0.443796 | 0.657190389 | 0.799957861 | CDCA5 | protein_coding | 52.67 | 0.011110659 | 0.011110659 | ENSG00000146670 | n.s. |
| ENSG00000113368 | 3136.480191 | -0.10949411 | 0.2358666 | -0.464221 | 0.642489739 | 0.791643267 | LMNB1 | protein_coding | 42.33 | -0.01111042 | 0.01111042 | ENSG00000113368 | n.s. |
| ENSG00000137492 | 2378.176388 | 0.11961853 | 0.2803382 | 0.426694 | 0.669602422 | 0.807759127 | THAP12 | protein_coding | 40.11 | 0.011090806 | 0.011090806 | ENSG00000137492 | n.s. |
| ENSG00000089597 | 1558.021881 | -0.117937208 | 0.2730466 | -0.431931 | 0.665791806 | 0.805580891 | GANAB | protein_coding | 50.02 | -0.011073224 | 0.011073224 | ENSG00000089597 | n.s. |
| ENSG00000160410 | 240.2263791 | 0.11842198 | 0.2758025 | 0.429372 | 0.667652251 | 0.80636342 | SHKBP1 | protein_coding | 53.53 | 0.011068806 | 0.011068806 | ENSG00000160410 | n.s. |
| ENSG00000196476 | 89.28348173 | -0.116266247 | 0.2655277 | -0.437869 | 0.661481578 | 0.803227631 | C20orf96 | protein_coding | 48.38 | -0.011064054 | 0.011064054 | ENSG00000196476 | n.s. |
| ENSG00000263013 | 231.9475958 | -0.100874331 | 0.2051931 | -0.491607 | 0.622997375 | 0.777009606 | | sense_intronic | 49.09 | -0.011053165 | 0.011053165 | ENSG00000263013 | n.s. |
| ENSG00000103035 | 1568.650242 | -0.11655889 | 0.2674205 | -0.435864 | 0.662935566 | 0.803948385 | PSMD7 | protein_coding | 43.55 | -0.0110465 | 0.0110465 | ENSG00000103035 | n.s. |
| ENSG00000185437 | 51.53957516 | 0.120484562 | 0.2856284 | 0.421823 | 0.673154366 | 0.809846304 | SH3BGR | protein_coding | 40.49 | 0.011036072 | 0.011036072 | ENSG00000185437 | n.s. |
| ENSG00000212123 | 81.68238008 | 0.122280078 | 0.2931658 | 0.417102 | 0.676603729 | 0.81243231 | PRR22 | protein_coding | 66.72 | 0.01103123 | 0.01103123 | ENSG00000212123 | n.s. |
| ENSG00000263786 | 551.7200375 | -0.104881759 | 0.2200786 | -0.476565 | 0.633671881 | 0.784999875 | | sense_intronic | 44.17 | -0.011026263 | 0.011026263 | ENSG00000263786 | n.s. |
| ENSG00000161011 | 850.2728069 | 0.113706453 | 0.2561309 | 0.443939 | 0.657086829 | 0.799914306 | SQSTM1 | protein_coding | 54.99 | 0.011024584 | 0.011024584 | ENSG00000161011 | n.s. |
| ENSG00000188002 | 153.3982599 | -0.110292037 | 0.2414923 | -0.45671 | 0.647879201 | 0.794540837 | | transcribed_unprocessed_pseud | 50.36 | -0.011016385 | 0.011016385 | ENSG00000188002 | n.s. |
| ENSG00000101577 | 778.1777044 | -0.111023277 | 0.2446489 | -0.453807 | 0.649968011 | 0.795840702 | LPIN2 | protein_coding | 41.63 | -0.011010606 | 0.011010606 | ENSG00000101577 | n.s. |
| ENSG00000158773 | 915.0095505 | 0.118779125 | 0.2785718 | 0.426386 | 0.669826532 | 0.807896365 | USF1 | protein_coding | 50.71 | 0.011004214 | 0.011004214 | ENSG00000158773 | n.s. |

| | | | | | | | | | | | | | |
|---|---|---|---|---|---|---|---|---|---|---|---|---|---|
| ENSG00000087301 | 616.1459863 | -0.117132868 | 0.2711636 | -0.431964 | 0.665767657 | 0.805580891 | TXNDC16 | protein_coding | 36.83 | -0.010997704 | 0.010997704 | ENSG00000087301 | n.s. |
| ENSG00000067704 | 2467.814318 | -0.119035325 | 0.2800364 | -0.425071 | 0.670784979 | 0.808462714 | IARS2 | protein_coding | 39.21 | -0.010991723 | 0.010991723 | ENSG00000067704 | n.s. |
| ENSG00000111231 | 649.3922739 | -0.119952957 | 0.2848145 | -0.421162 | 0.673636991 | 0.809935045 | GPN3 | protein_coding | 45.03 | -0.01098167 | 0.01098167 | ENSG00000111231 | n.s. |
| ENSG00000143545 | 277.5321976 | -0.117268812 | 0.2727094 | -0.430014 | 0.667185559 | 0.806194128 | RAB13 | protein_coding | 48.71 | -0.010971713 | 0.010971713 | ENSG00000143545 | n.s. |
| ENSG00000102893 | 72.84912265 | -0.114301939 | 0.2594268 | -0.440594 | 0.659506828 | 0.801808729 | PHKB | protein_coding | 38.19 | -0.010964896 | 0.010964896 | ENSG00000102893 | n.s. |
| ENSG00000145604 | 3369.861239 | -0.113421152 | 0.2564882 | -0.442208 | 0.658338627 | 0.800878 | SKP2 | protein_coding | 39.76 | -0.010937614 | 0.010937614 | ENSG00000145604 | n.s. |
| ENSG00000100823 | 2061.662307 | 0.116753911 | 0.2716966 | 0.429722 | 0.667398083 | 0.806263211 | APEX1 | protein_coding | 47.06 | 0.010919194 | 0.010919194 | ENSG00000100823 | n.s. |
| ENSG00000169446 | 584.893603 | 0.117850376 | 0.2765 | 0.426222 | 0.669946089 | 0.807901942 | MMGT1 | protein_coding | 39.98 | 0.010917818 | 0.010917818 | ENSG00000169446 | n.s. |
| ENSG00000197714 | 1712.167416 | 0.110361643 | 0.2440687 | 0.452174 | 0.6511433 | 0.796369304 | ZNF460 | protein_coding | 46.11 | 0.010913165 | 0.010913165 | ENSG00000197714 | n.s. |
| ENSG00000105701 | 1132.859793 | 0.116162263 | 0.2690229 | 0.431793 | 0.665891738 | 0.805631793 | FKBP8 | protein_coding | 56.64 | 0.010903385 | 0.010903385 | ENSG00000105701 | n.s. |
| ENSG00000257704 | 52.98122646 | -0.115521045 | 0.2662687 | -0.433851 | 0.664396307 | 0.804731603 | INAFM1 | protein_coding | 71.55 | -0.010899289 | 0.010899289 | ENSG00000257704 | n.s. |
| ENSG00000109794 | 50.90755786 | 0.122696557 | 0.2995902 | 0.409548 | 0.682137549 | 0.815577595 | FAM149A | protein_coding | 44.16 | 0.010862905 | 0.010862905 | ENSG00000109794 | n.s. |
| ENSG00000127774 | 312.2139319 | 0.121224122 | 0.2928916 | 0.413887 | 0.678956609 | 0.813590918 | EMC6 | protein_coding | 59.6 | 0.010860943 | 0.010860943 | ENSG00000127774 | n.s. |
| ENSG00000170004 | 6405.467669 | -0.107860889 | 0.2348285 | -0.459318 | 0.646006003 | 0.793440055 | CHD3 | protein_coding | 53.17 | -0.010838496 | 0.010838496 | ENSG00000170004 | n.s. |
| ENSG00000104731 | 77.55217878 | 0.114154116 | 0.2618875 | 0.43589 | 0.66291661 | 0.803948385 | KLHDC4 | protein_coding | 53.01 | 0.010818595 | 0.010818595 | ENSG00000104731 | n.s. |
| ENSG00000134453 | 1693.318617 | 0.111782155 | 0.2521951 | 0.443237 | 0.657594491 | 0.800252434 | RBM17 | protein_coding | 41.27 | 0.010817494 | 0.010817494 | ENSG00000134453 | n.s. |
| ENSG00000197170 | 3244.596671 | -0.11245491 | 0.2549291 | -0.441122 | 0.659124463 | 0.801553838 | PSMD12 | protein_coding | 40.05 | -0.01080324 | 0.01080324 | ENSG00000197170 | n.s. |
| ENSG00000204227 | 522.763655 | -0.119168737 | 0.2848663 | -0.418332 | 0.675704231 | 0.811698417 | RING1 | protein_coding | 53.03 | -0.010797319 | 0.010797319 | ENSG00000204227 | n.s. |
| ENSG00000165916 | 2030.16268 | 0.11787774 | 0.2795905 | 0.421609 | 0.673310799 | 0.809846304 | PSMC3 | protein_coding | 54.93 | 0.010797294 | 0.010797294 | ENSG00000165916 | n.s. |
| ENSG00000072864 | 275.2707243 | -0.111651578 | 0.2521041 | -0.442879 | 0.657853295 | 0.800438433 | NDE1 | protein_coding | 48.54 | -0.010793589 | 0.010793589 | ENSG00000072864 | n.s. |
| ENSG00000116731 | 507.3347116 | 0.108090575 | 0.2368893 | 0.456291 | 0.64818039 | 0.79481542 | PRDM2 | protein_coding | 43.44 | 0.010780275 | 0.010780275 | ENSG00000116731 | n.s. |
| ENSG00000182552 | 519.6929132 | 0.123974747 | 0.3069569 | 0.403883 | 0.686298583 | 0.818632266 | RWDD4 | protein_coding | 40.27 | 0.010774787 | 0.010774787 | ENSG00000182552 | n.s. |
| ENSG00000164258 | 982.125735 | 0.126320509 | 0.3187667 | 0.396279 | 0.691899265 | 0.821714898 | NDUFS4 | protein_coding | 36.57 | 0.010772466 | 0.010772466 | ENSG00000164258 | n.s. |
| ENSG00000199127 | 2.405744577 | -0.231170916 | 0.9691545 | -0.238528 | 0.811471256 | 0.898301652 | MIR383 | miRNA | 49.32 | -0.010767433 | 0.010767433 | ENSG00000199127 | n.s. |
| ENSG00000265354 | 1299.658097 | 0.118239365 | 0.2819141 | 0.419416 | 0.674911975 | 0.810977771 | TIMM23 | protein_coding | 40.16 | 0.010758724 | 0.010758724 | ENSG00000265354 | n.s. |
| ENSG00000170581 | 301.4812223 | -0.114823542 | 0.2676189 | -0.429056 | 0.667882378 | 0.806516133 | STAT2 | protein_coding | 47.77 | -0.010723019 | 0.010723019 | ENSG00000170581 | n.s. |
| ENSG00000108406 | 4187.24204 | -0.119373122 | 0.2877992 | -0.414779 | 0.678303497 | 0.813158222 | DHX40 | protein_coding | 39.64 | -0.010722684 | 0.010722684 | ENSG00000108406 | n.s. |
| ENSG00000164743 | 58.3729453 | 0.111946288 | 0.2545978 | 0.439699 | 0.660155394 | 0.802246971 | C8orf48 | protein_coding | 42.28 | 0.010712355 | 0.010712355 | ENSG00000164743 | n.s. |
| ENSG00000105556 | 67.96687666 | -0.113374259 | 0.2611812 | -0.434083 | 0.664228362 | 0.804731603 | MIER2 | protein_coding | 53.11 | -0.010696742 | 0.010696742 | ENSG00000105556 | n.s. |
| ENSG00000169718 | 1776.440446 | 0.113798641 | 0.2631491 | 0.432449 | 0.665414942 | 0.805474894 | DUS1L | protein_coding | 64.45 | 0.010691154 | 0.010691154 | ENSG00000169718 | n.s. |
| ENSG00000165973 | 421.2499477 | -0.118859712 | 0.2862879 | -0.415175 | 0.678013475 | 0.813086294 | NELL1 | protein_coding | 38.29 | -0.010681133 | 0.010681133 | ENSG00000165973 | n.s. |
| ENSG00000075213 | 68.15978282 | -0.115397624 | 0.2718903 | -0.424427 | 0.671254303 | 0.808693667 | SEMA3A | protein_coding | 34.35 | -0.010641502 | 0.010641502 | ENSG00000075213 | n.s. |

| | | | | | | | | | | | | | |
|---|---|---|---|---|---|---|---|---|---|---|---|---|---|
| ENSG00000151414 | 1649.066069 | -0.1160453 | 0.2750863 | -0.421851 | 0.673134139 | 0.809846304 | NEK7 | protein_coding | 35.84 | -0.010629447 | 0.010629447 | ENSG00000151414 | n.s. |
| ENSG00000168610 | 2015.442352 | -0.108662772 | 0.2433029 | -0.446615 | 0.655152951 | 0.798467659 | STAT3 | protein_coding | 45.02 | -0.010620989 | 0.010620989 | ENSG00000168610 | n.s. |
| ENSG00000109911 | 402.7604792 | 0.11348974 | 0.2643176 | 0.429369 | 0.667654781 | 0.80636342 | ELP4 | protein_coding | 35.79 | 0.010607793 | 0.010607793 | ENSG00000109911 | n.s. |
| ENSG00000151470 | 225.0890078 | 0.114426118 | 0.2684894 | 0.426185 | 0.669973125 | 0.807901942 | C4orf33 | protein_coding | 35.63 | 0.01060059 | 0.01060059 | ENSG00000151470 | n.s. |
| ENSG00000138382 | 780.310381 | -0.123435704 | 0.3092836 | -0.399102 | 0.689818021 | 0.820602898 | METTL5 | protein_coding | 39.58 | -0.010599048 | 0.010599048 | ENSG00000138382 | n.s. |
| ENSG00000238578 | 62.13305068 | 0.111812409 | 0.2564568 | 0.435989 | 0.662844503 | 0.803948385 | SNORD4A | snoRNA | 47.22 | 0.010596667 | 0.010596667 | ENSG00000238578 | n.s. |
| ENSG00000139263 | 1957.36059 | -0.101190997 | 0.2133101 | -0.474385 | 0.635225733 | 0.786015631 | LRIG3 | protein_coding | 39.12 | -0.010581423 | 0.010581423 | ENSG00000139263 | n.s. |
| ENSG00000238719 | 32.93610992 | 0.119376099 | 0.2913037 | 0.409799 | 0.681953057 | 0.815483027 | RNU7-96P | snRNA | 31.25 | 0.010574941 | 0.010574941 | ENSG00000238719 | n.s. |
| ENSG00000275023 | 1385.256286 | -0.107009931 | 0.2373138 | -0.450922 | 0.652046046 | 0.796633653 | MLLT6 | protein_coding | 56.24 | -0.010566305 | 0.010566305 | ENSG00000275023 | n.s. |
| ENSG00000137575 | 907.1776827 | -0.11725238 | 0.2816775 | -0.416265 | 0.67721641 | 0.812694714 | SDCBP | protein_coding | 35.86 | -0.010561223 | 0.010561223 | ENSG00000137575 | n.s. |
| ENSG00000116455 | 1182.615445 | 0.112629364 | 0.2613183 | 0.431004 | 0.666465107 | 0.806115341 | WDR77 | protein_coding | 44.57 | 0.010542425 | 0.010542425 | ENSG00000116455 | n.s. |
| ENSG00000226696 | 70.42514386 | -0.112270283 | 0.2599001 | -0.431975 | 0.665759812 | 0.805580891 | LENG8-AS1 | antisense | 48.1 | -0.010541152 | 0.010541152 | ENSG00000226696 | n.s. |
| ENSG00000263813 | 18.18277984 | 0.130702229 | 0.346313 | 0.377411 | 0.705868449 | 0.830814932 | MIR3679 | miRNA | 52.94 | 0.010520968 | 0.010520968 | ENSG00000263813 | n.s. |
| ENSG00000165752 | 97.48209414 | 0.116199298 | 0.2780179 | 0.417956 | 0.675979167 | 0.811839725 | STK32C | protein_coding | 55.1 | 0.010519488 | 0.010519488 | ENSG00000165752 | n.s. |
| ENSG00000160753 | 231.2434655 | 0.112023985 | 0.2597225 | 0.431322 | 0.666234355 | 0.805907732 | RUSC1 | protein_coding | 58.03 | 0.010498292 | 0.010498292 | ENSG00000160753 | n.s. |
| ENSG00000215190 | 430.7550552 | -0.115315021 | 0.2748323 | -0.419583 | 0.674789946 | 0.81097112 | LINC00680 | transcribed_unprocessed_pseud | 45.68 | -0.010493046 | 0.010493046 | ENSG00000215190 | n.s. |
| ENSG00000078668 | 2113.454534 | 0.118179926 | 0.2878054 | 0.410624 | 0.681348003 | 0.815125872 | VDAC3 | protein_coding | 40.42 | 0.010491461 | 0.010491461 | ENSG00000078668 | n.s. |
| ENSG00000164002 | 281.523242 | -0.112747569 | 0.2636264 | -0.427679 | 0.668884584 | 0.807148558 | EXO5 | protein_coding | 43.67 | -0.01049077 | 0.01049077 | ENSG00000164002 | n.s. |
| ENSG00000204348 | 208.8520712 | 0.107405386 | 0.2407803 | 0.446072 | 0.655545141 | 0.798735888 | DXO | protein_coding | 58.72 | 0.010482422 | 0.010482422 | ENSG00000204348 | n.s. |
| ENSG00000148935 | 66.52356488 | -0.121763048 | 0.3040489 | -0.400472 | 0.688808923 | 0.820294688 | GAS2 | protein_coding | 36.04 | -0.010475287 | 0.010475287 | ENSG00000148935 | n.s. |
| ENSG00000207523 | 123.0350741 | 0.106449972 | 0.2371796 | 0.448816 | 0.653564445 | 0.797299375 | SNORA66 | snoRNA | 47.37 | 0.010472397 | 0.010472397 | ENSG00000207523 | n.s. |
| ENSG00000259153 | 37.75616082 | 0.122234451 | 0.3070168 | 0.398136 | 0.690529955 | 0.821195719 | | lincRNA | 42.18 | 0.010457564 | 0.010457564 | ENSG00000259153 | n.s. |
| ENSG00000168575 | 400.143786 | 0.103623395 | 0.2249201 | 0.460712 | 0.645005227 | 0.792692009 | SLC20A2 | protein_coding | 44.17 | 0.010455136 | 0.010455136 | ENSG00000168575 | n.s. |
| ENSG00000254510 | 60.29194754 | -0.125641536 | 0.3231781 | -0.388769 | 0.697447211 | 0.825797901 | | processed_transcript | 53.5 | -0.010444107 | 0.010444107 | ENSG00000254510 | n.s. |
| ENSG00000139372 | 2753.288965 | -0.118076728 | 0.2886258 | -0.4091 | 0.682466532 | 0.81576698 | TDG | protein_coding | 40.65 | -0.010441983 | 0.010441983 | ENSG00000139372 | n.s. |
| ENSG00000162385 | 1317.729237 | -0.115886604 | 0.2783097 | -0.416394 | 0.677121432 | 0.812694714 | MAGOH | protein_coding | 41.63 | -0.010438204 | 0.010438204 | ENSG00000162385 | n.s. |
| ENSG00000138785 | 413.5685692 | -0.110621421 | 0.2549708 | -0.433859 | 0.66439072 | 0.804731603 | INTS12 | protein_coding | 36.56 | -0.010437014 | 0.010437014 | ENSG00000138785 | n.s. |
| ENSG00000065518 | 1367.378767 | 0.120078338 | 0.2975205 | 0.403597 | 0.686509219 | 0.818643628 | NDUFB4 | protein_coding | 39.39 | 0.010435422 | 0.010435422 | ENSG00000065518 | n.s. |
| ENSG00000135766 | 567.1426619 | 0.112291222 | 0.2628298 | 0.427239 | 0.669204997 | 0.80746522 | EGLN1 | protein_coding | 39.65 | 0.010429179 | 0.010429179 | ENSG00000135766 | n.s. |
| ENSG00000187135 | 111.6668309 | 0.11920283 | 0.2935545 | 0.406067 | 0.684693264 | 0.81758013 | VSTM2B | protein_coding | 51.2 | 0.010426632 | 0.010426632 | ENSG00000187135 | n.s. |
| ENSG00000122643 | 1106.901504 | 0.113391326 | 0.2679736 | 0.423144 | 0.672190498 | 0.809347406 | NT5C3A | protein_coding | 37.98 | 0.010416697 | 0.010416697 | ENSG00000122643 | n.s. |
| ENSG00000261324 | 249.8686473 | 0.100219772 | 0.212369 | 0.471913 | 0.636988672 | 0.787217582 | | sense_overlapping | 36.81 | 0.010413357 | 0.010413357 | ENSG00000261324 | n.s. |

| | | | | | | | | | | | | | |
|---|---|---|---|---|---|---|---|---|---|---|---|---|---|
| ENSG00000013374 | 864.3865583 | -0.116828767 | 0.2835777 | -0.411982 | 0.680352964 | 0.814563103 | NUB1 | protein_coding | 44.96 | -0.010406553 | 0.010406553 | ENSG00000013374 | n.s. |
| ENSG00000187735 | 1580.510311 | 0.116132697 | 0.280784 | 0.413601 | 0.679165979 | 0.813603737 | TCEA1 | protein_coding | 39.44 | 0.010403988 | 0.010403988 | ENSG00000187735 | n.s. |
| ENSG00000198843 | 1211.632019 | -0.1188637 | 0.2925748 | -0.406268 | 0.68454589 | 0.817474263 | SELENOT | protein_coding | 37.86 | -0.010403654 | 0.010403654 | ENSG00000198843 | n.s. |
| ENSG00000062716 | 1592.963401 | -0.112159415 | 0.262874 | -0.426666 | 0.669622628 | 0.807759127 | VMP1 | protein_coding | 39.75 | -0.010399211 | 0.010399211 | ENSG00000062716 | n.s. |
| ENSG00000171862 | 2968.978909 | -0.105623923 | 0.2352029 | -0.449076 | 0.653376934 | 0.797221535 | PTEN | protein_coding | 35.77 | -0.01039561 | 0.01039561 | ENSG00000171862 | n.s. |
| ENSG00000120742 | 2679.478377 | 0.108945321 | 0.2485983 | 0.438238 | 0.661213481 | 0.803042158 | SERP1 | protein_coding | 38.84 | 0.010378312 | 0.010378312 | ENSG00000120742 | n.s. |
| ENSG00000011275 | 1305.232154 | -0.110099517 | 0.2540888 | -0.433311 | 0.664788735 | 0.804958837 | RNF216 | protein_coding | 45.52 | -0.010374273 | 0.010374273 | ENSG00000011275 | n.s. |
| ENSG00000092621 | 4440.481671 | -0.114017794 | 0.2722452 | -0.418806 | 0.675358251 | 0.811443988 | PHGDH | protein_coding | 44.18 | -0.01034614 | 0.01034614 | ENSG00000092621 | n.s. |
| ENSG00000123130 | 411.3512486 | 0.121127259 | 0.3055224 | 0.396459 | 0.691766101 | 0.821656665 | ACOT9 | protein_coding | 44.61 | 0.01033332 | 0.01033332 | ENSG00000123130 | n.s. |
| ENSG00000201470 | 32.01293993 | -0.122148747 | 0.3102098 | -0.393762 | 0.693757004 | 0.823135405 | RNY4P7 | misc_RNA | 41.67 | -0.010325077 | 0.010325077 | ENSG00000201470 | n.s. |
| ENSG00000125901 | 720.5682154 | -0.112688306 | 0.2674901 | -0.42128 | 0.673550444 | 0.809935045 | MRPS26 | protein_coding | 61.08 | -0.010316593 | 0.010316593 | ENSG00000125901 | n.s. |
| ENSG00000114850 | 3498.299113 | 0.111259952 | 0.2609589 | 0.426351 | 0.669852448 | 0.807896365 | SSR3 | protein_coding | 38.72 | 0.010307606 | 0.010307606 | ENSG00000114850 | n.s. |
| ENSG00000213722 | 514.8133476 | 0.118551472 | 0.2938691 | 0.403416 | 0.686642267 | 0.818643628 | DDAH2 | protein_coding | 59.86 | 0.010302729 | 0.010302729 | ENSG00000213722 | n.s. |
| ENSG00000265706 | 37.19751016 | -0.122863788 | 0.3140105 | -0.391273 | 0.695595549 | 0.824446823 | SNORD53B | snoRNA | 46.15 | -0.010300575 | 0.010300575 | ENSG00000265706 | n.s. |
| ENSG00000162191 | 1178.688187 | 0.11601644 | 0.2825084 | 0.410665 | 0.681317884 | 0.815125872 | UBXN1 | protein_coding | 52.86 | 0.010299397 | 0.010299397 | ENSG00000162191 | n.s. |
| ENSG00000142186 | 512.9778113 | -0.11593572 | 0.2822501 | -0.410755 | 0.68125195 | 0.815125872 | SCYL1 | protein_coding | 57.59 | -0.010292231 | 0.010292231 | ENSG00000142186 | n.s. |
| ENSG00000065809 | 2399.993199 | 0.120457197 | 0.3032916 | 0.397166 | 0.691244867 | 0.82152815 | FAM107B | protein_coding | 43.19 | 0.01028434 | 0.01028434 | ENSG00000065809 | n.s. |
| ENSG00000263934 | 10909.30445 | -0.094159131 | 0.1921534 | -0.490021 | 0.624119296 | 0.77766359 | SNORD3A | snoRNA | 54.22 | -0.010282952 | 0.010282952 | ENSG00000263934 | n.s. |
| ENSG00000230002 | 306.9853569 | 0.103394191 | 0.2284148 | 0.45266 | 0.650793757 | 0.796291544 | ALMS1-IT1 | sense_intronic | 37.16 | 0.01022857 | 0.01022857 | ENSG00000230002 | n.s. |
| ENSG00000204371 | 1033.250477 | 0.104800098 | 0.2356878 | 0.444656 | 0.65656811 | 0.799562477 | EHMT2 | protein_coding | 53.82 | 0.010181078 | 0.010181078 | ENSG00000204371 | n.s. |
| ENSG00000106077 | 108.2992116 | -0.111061988 | 0.2633749 | -0.421688 | 0.673252908 | 0.809846304 | ABHD11 | protein_coding | 59.95 | -0.010172989 | 0.010172989 | ENSG00000106077 | n.s. |
| ENSG00000151746 | 3514.666277 | 0.105096049 | 0.2375157 | 0.44248 | 0.658141619 | 0.800708298 | BICD1 | protein_coding | 40.45 | 0.010144467 | 0.010144467 | ENSG00000151746 | n.s. |
| ENSG00000030582 | 323.912853 | 0.106671118 | 0.2439284 | 0.437305 | 0.661890152 | 0.803364895 | GRN | protein_coding | 57.64 | 0.010143053 | 0.010143053 | ENSG00000030582 | n.s. |
| ENSG00000230648 | 11.94327787 | -0.137595705 | 0.3928087 | -0.350287 | 0.726123476 | 0.84388858 | | antisense | 43.62 | -0.010142852 | 0.010142852 | ENSG00000230648 | n.s. |
| ENSG00000198712 | 10975.16467 | -0.102273048 | 0.2255342 | -0.45347 | 0.65021012 | 0.796014436 | MT-CO2 | protein_coding | 46.2 | -0.010133117 | 0.010133117 | ENSG00000198712 | n.s. |
| ENSG00000106305 | 865.6773763 | -0.116679085 | 0.2896442 | -0.402836 | 0.687068949 | 0.818801761 | AIMP2 | protein_coding | 46.93 | -0.010130222 | 0.010130222 | ENSG00000106305 | n.s. |
| ENSG00000146733 | 422.6243982 | 0.114060254 | 0.2777745 | 0.410622 | 0.681349917 | 0.815125872 | PSPH | protein_coding | 46.35 | 0.010125736 | 0.010125736 | ENSG00000146733 | n.s. |
| ENSG00000090097 | 1386.640274 | 0.111737147 | 0.2672791 | 0.418054 | 0.675907492 | 0.811823673 | PCBP4 | protein_coding | 58.75 | 0.01011649 | 0.01011649 | ENSG00000090097 | n.s. |
| ENSG00000170653 | 45.64220958 | -0.118129847 | 0.2970942 | -0.397618 | 0.690912141 | 0.821272922 | ATF7 | protein_coding | 42.93 | -0.010101578 | 0.010101578 | ENSG00000170653 | n.s. |
| ENSG00000143183 | 2383.3408 | 0.118070903 | 0.2967704 | 0.397853 | 0.690738779 | 0.821195719 | TMCO1 | protein_coding | 39.88 | 0.010101358 | 0.010101358 | ENSG00000143183 | n.s. |
| ENSG00000169714 | 8906.438899 | 0.118626514 | 0.2998709 | 0.395592 | 0.692406043 | 0.822066116 | CNBP | protein_coding | 42.41 | 0.010094316 | 0.010094316 | ENSG00000169714 | n.s. |
| ENSG00000164967 | 148.3660442 | -0.112239045 | 0.2701105 | -0.41553 | 0.677753827 | 0.812989277 | RPP25L | protein_coding | 58.74 | -0.010091994 | 0.010091994 | ENSG00000164967 | n.s. |

| | | | | | | | | | | | | | |
|---|---|---|---|---|---|---|---|---|---|---|---|---|---|
| ENSG00000135451 | 957.180456 | -0.10613404 | 0.242787 | -0.437149 | 0.662003431 | 0.803364895 | TROAP | protein_coding | 53.1 | -0.010091983 | 0.010091983 | ENSG00000135451 | n.s. |
| ENSG00000008256 | 966.5365463 | 0.114362425 | 0.2800567 | 0.408355 | 0.683013433 | 0.816134228 | CYTH3 | protein_coding | 44.78 | 0.010091158 | 0.010091158 | ENSG00000008256 | n.s. |
| ENSG00000168615 | 847.7140801 | 0.109195928 | 0.25689 | 0.425069 | 0.67078656 | 0.808462714 | ADAM9 | protein_coding | 37.5 | 0.010083153 | 0.010083153 | ENSG00000168615 | n.s. |
| ENSG00000112699 | 153.1244517 | -0.118201084 | 0.2983121 | -0.396233 | 0.691933166 | 0.821714898 | GMDS | protein_coding | 40.78 | -0.010080051 | 0.010080051 | ENSG00000112699 | n.s. |
| ENSG00000147316 | 901.6577235 | -0.10697029 | 0.2469209 | -0.433217 | 0.664857303 | 0.804958837 | MCPH1 | protein_coding | 42.19 | -0.010079418 | 0.010079418 | ENSG00000147316 | n.s. |
| ENSG00000265137 | 30.96416629 | 0.120279075 | 0.3075945 | 0.391031 | 0.695774075 | 0.82451804 | MIR3192 | miRNA | 48.05 | 0.010079367 | 0.010079367 | ENSG00000265137 | n.s. |
| ENSG00000124767 | 4512.862209 | 0.122011558 | 0.3162705 | 0.385782 | 0.699657926 | 0.82681669 | GLO1 | protein_coding | 39.8 | 0.010077028 | 0.010077028 | ENSG00000124767 | n.s. |
| ENSG00000108395 | 2263.656658 | -0.105934196 | 0.2422538 | -0.437286 | 0.661903988 | 0.803364895 | TRIM37 | protein_coding | 39.71 | -0.010072981 | 0.010072981 | ENSG00000108395 | n.s. |
| ENSG00000166136 | 1768.866108 | 0.118405402 | 0.2995799 | 0.395238 | 0.692667113 | 0.822235844 | NDUFB8 | protein_coding | 47.71 | 0.010064885 | 0.010064885 | ENSG00000166136 | n.s. |
| ENSG00000273238 | 397.4601061 | -0.107461579 | 0.2499168 | -0.429989 | 0.667203341 | 0.806194128 | | lincRNA | 33.02 | -0.010054145 | 0.010054145 | ENSG00000273238 | n.s. |
| ENSG00000137200 | 1184.895664 | 0.109182478 | 0.2577549 | 0.42359 | 0.671864666 | 0.809131714 | CMTR1 | protein_coding | 45.88 | 0.010042689 | 0.010042689 | ENSG00000137200 | n.s. |
| ENSG00000140350 | 6596.538261 | -0.109500333 | 0.2595525 | -0.421881 | 0.673111699 | 0.809846304 | ANP32A | protein_coding | 49.39 | -0.010029945 | 0.010029945 | ENSG00000140350 | n.s. |
| ENSG00000182916 | 617.4824992 | 0.121548008 | 0.3155643 | 0.385177 | 0.700106489 | 0.826977403 | TCEAL7 | protein_coding | 44.3 | 0.010028483 | 0.010028483 | ENSG00000182916 | n.s. |
| ENSG00000229588 | 18.72796663 | 0.166174973 | 0.5611842 | 0.296115 | 0.767142361 | 0.870349013 | | lincRNA | 44.6 | 0.010021441 | 0.010021441 | ENSG00000229588 | n.s. |
| ENSG00000076554 | 841.4173756 | 0.117175579 | 0.2958727 | 0.396034 | 0.692080128 | 0.821749243 | TPD52 | protein_coding | 41.15 | 0.00999047 | 0.00999047 | ENSG00000076554 | n.s. |
| ENSG00000201457 | 23.58131901 | 0.120450134 | 0.3103704 | 0.388085 | 0.697953065 | 0.826256316 | SNORA55 | snoRNA | 51.85 | 0.009983534 | 0.009983534 | ENSG00000201457 | n.s. |
| ENSG00000269925 | 22.29072281 | -0.128024884 | 0.3492206 | -0.366602 | 0.713916073 | 0.835843717 | | sense_intronic | 36.64 | -0.009969927 | 0.009969927 | ENSG00000269925 | n.s. |
| ENSG00000166889 | 440.7735084 | -0.101257396 | 0.2254382 | -0.449158 | 0.653317663 | 0.797221535 | PATL1 | protein_coding | 39.82 | -0.009965852 | 0.009965852 | ENSG00000166889 | n.s. |
| ENSG00000122359 | 279.2805509 | 0.105855359 | 0.2445046 | 0.432938 | 0.665059678 | 0.805114851 | ANXA11 | protein_coding | 50.39 | 0.009965452 | 0.009965452 | ENSG00000122359 | n.s. |
| ENSG00000133315 | 177.1977657 | 0.115990282 | 0.2901765 | 0.399723 | 0.689360444 | 0.820546125 | MACROD1 | protein_coding | 58.06 | 0.009963217 | 0.009963217 | ENSG00000133315 | n.s. |
| ENSG00000141002 | 742.0712561 | 0.11290536 | 0.2764707 | 0.408381 | 0.682994046 | 0.816134228 | TCF25 | protein_coding | 55.04 | 0.009962589 | 0.009962589 | ENSG00000141002 | n.s. |
| ENSG00000108468 | 9504.243973 | 0.112304404 | 0.2736646 | 0.410372 | 0.681532823 | 0.815274637 | CBX1 | protein_coding | 45.96 | 0.009960959 | 0.009960959 | ENSG00000108468 | n.s. |
| ENSG00000144395 | 136.7456028 | -0.106234977 | 0.2463019 | -0.43132 | 0.666235581 | 0.805907732 | CCDC150 | protein_coding | 38.16 | -0.009955777 | 0.009955777 | ENSG00000144395 | n.s. |
| ENSG00000179051 | 1550.060736 | -0.104450873 | 0.2382729 | -0.438367 | 0.6611206 | 0.8029994 | RCC2 | protein_coding | 51.68 | -0.009952578 | 0.009952578 | ENSG00000179051 | n.s. |
| ENSG00000143498 | 754.7560425 | 0.101185602 | 0.2261661 | 0.447395 | 0.654589757 | 0.798060699 | TAF1A | protein_coding | 36.34 | 0.009912554 | 0.009912554 | ENSG00000143498 | n.s. |
| ENSG00000129353 | 137.9018095 | -0.113353251 | 0.2797858 | -0.405143 | 0.685372413 | 0.817983898 | SLC44A2 | protein_coding | 51.73 | -0.009890666 | 0.009890666 | ENSG00000129353 | n.s. |
| ENSG00000175354 | 567.6422991 | -0.109727714 | 0.2637831 | -0.415977 | 0.677426761 | 0.812781842 | PTPN2 | protein_coding | 42.19 | -0.009878348 | 0.009878348 | ENSG00000175354 | n.s. |
| ENSG00000171786 | 266.0171151 | -0.101136419 | 0.2266589 | -0.446205 | 0.655448893 | 0.798688512 | NHLH1 | protein_coding | 50.12 | -0.009873196 | 0.009873196 | ENSG00000171786 | n.s. |
| ENSG00000002834 | 230.7320343 | 0.105020484 | 0.2429183 | 0.432328 | 0.665502794 | 0.805511206 | LASP1 | protein_coding | 53.69 | 0.009864407 | 0.009864407 | ENSG00000002834 | n.s. |
| ENSG00000125388 | 259.0790872 | -0.105255695 | 0.244308 | -0.430832 | 0.666590597 | 0.806182257 | GRK4 | protein_coding | 46.38 | -0.009848435 | 0.009848435 | ENSG00000125388 | n.s. |
| ENSG00000122547 | 921.2580057 | -0.100854672 | 0.2262551 | -0.445756 | 0.655773247 | 0.798804104 | EEPD1 | protein_coding | 45.46 | -0.009839353 | 0.009839353 | ENSG00000122547 | n.s. |
| ENSG00000164172 | 376.0629594 | -0.114223822 | 0.2856105 | -0.399929 | 0.689209119 | 0.820439344 | MOCS2 | protein_coding | 38.21 | -0.00981794 | 0.00981794 | ENSG00000164172 | n.s. |

| | | | | | | | | | | | | | |
|---|---|---|---|---|---|---|---|---|---|---|---|---|---|
| ENSG00000077458 | 1076.0909 | 0.108935921 | 0.2616178 | 0.416393 | 0.677122221 | 0.812694714 | FAM76B | protein_coding | 34.6 | 0.009812138 | 0.009812138 | ENSG00000077458 | n.s. |
| ENSG00000163938 | 5876.462635 | 0.106301488 | 0.2507022 | 0.424015 | 0.671554844 | 0.808828555 | GNL3 | protein_coding | 43.2 | 0.009794994 | 0.009794994 | ENSG00000163938 | n.s. |
| ENSG00000083896 | 2898.182765 | -0.107588147 | 0.256466 | -0.419502 | 0.674848931 | 0.810972006 | YTHDC1 | protein_coding | 35.4 | -0.009789891 | 0.009789891 | ENSG00000083896 | n.s. |
| ENSG00000132819 | 350.2183128 | 0.106156433 | 0.249988 | 0.424646 | 0.671094645 | 0.808693667 | RBM38 | protein_coding | 59.97 | 0.009789317 | 0.009789317 | ENSG00000132819 | n.s. |
| ENSG00000105647 | 426.664457 | -0.104811848 | 0.2443173 | -0.428999 | 0.667924025 | 0.806516133 | PIK3R2 | protein_coding | 57.77 | -0.009788058 | 0.009788058 | ENSG00000105647 | n.s. |
| ENSG00000197063 | 688.1815944 | 0.109249564 | 0.2641331 | 0.413616 | 0.679155685 | 0.813603737 | MAFG | protein_coding | 60.3 | 0.009787348 | 0.009787348 | ENSG00000197063 | n.s. |
| ENSG00000102931 | 651.4004711 | 0.118396899 | 0.306901 | 0.385782 | 0.699658074 | 0.82681669 | ARL2BP | protein_coding | 45.49 | 0.009778491 | 0.009778491 | ENSG00000102931 | n.s. |
| ENSG00000100376 | 98.01540625 | 0.110942084 | 0.2719959 | 0.407881 | 0.683360717 | 0.816339018 | FAM118A | protein_coding | 49.58 | 0.009777264 | 0.009777264 | ENSG00000100376 | n.s. |
| ENSG00000136003 | 1591.137058 | 0.109193396 | 0.2641578 | 0.413364 | 0.679339744 | 0.813699867 | ISCU | protein_coding | 45.76 | 0.009776713 | 0.009776713 | ENSG00000136003 | n.s. |
| ENSG00000100239 | 501.3303793 | 0.11215633 | 0.277029 | 0.404854 | 0.685584702 | 0.818153407 | PPP6R2 | protein_coding | 51.13 | 0.009776135 | 0.009776135 | ENSG00000100239 | n.s. |
| ENSG00000085733 | 1695.330584 | -0.106938949 | 0.2544178 | -0.420328 | 0.674245749 | 0.810427924 | CTTN | protein_coding | 52.81 | -0.009761987 | 0.009761987 | ENSG00000085733 | n.s. |
| ENSG00000172590 | 348.8556697 | 0.118949406 | 0.3100554 | 0.383639 | 0.701245853 | 0.82783161 | MRPL52 | protein_coding | 47.12 | 0.00976075 | 0.00976075 | ENSG00000172590 | n.s. |
| ENSG00000137842 | 161.3724558 | 0.105453084 | 0.2477019 | 0.425726 | 0.670307694 | 0.808165399 | TMEM62 | protein_coding | 40.55 | 0.009754384 | 0.009754384 | ENSG00000137842 | n.s. |
| ENSG00000183688 | 294.7794836 | -0.112550584 | 0.2797532 | -0.402321 | 0.687447828 | 0.819183166 | RFLNB | protein_coding | 48.81 | -0.009749017 | 0.009749017 | ENSG00000183688 | n.s. |
| ENSG00000110721 | 530.1900851 | -0.104386111 | 0.2434166 | -0.428837 | 0.668041712 | 0.806516133 | CHKA | protein_coding | 44.1 | -0.0097483 | 0.0097483 | ENSG00000110721 | n.s. |
| ENSG00000128191 | 1568.22947 | 0.099838219 | 0.2239191 | 0.445867 | 0.655693109 | 0.798776372 | DGCR8 | protein_coding | 52.21 | 0.009741693 | 0.009741693 | ENSG00000128191 | n.s. |
| ENSG00000168439 | 3083.815755 | 0.113996805 | 0.2874407 | 0.396592 | 0.691668087 | 0.821656665 | STIP1 | protein_coding | 50.06 | 0.009725024 | 0.009725024 | ENSG00000168439 | n.s. |
| ENSG00000124228 | 986.0865365 | -0.106355487 | 0.2528105 | -0.420692 | 0.673979645 | 0.810207098 | DDX27 | protein_coding | 49.05 | -0.009721312 | 0.009721312 | ENSG00000124228 | n.s. |
| ENSG00000172671 | 368.692983 | 0.108302437 | 0.261668 | 0.413892 | 0.678952846 | 0.813590918 | ZFAND4 | protein_coding | 37.95 | 0.009703239 | 0.009703239 | ENSG00000172671 | n.s. |
| ENSG00000160584 | 103.2321246 | -0.111117687 | 0.2740945 | -0.405399 | 0.685184077 | 0.817885622 | SIK3 | protein_coding | 40.48 | -0.009701399 | 0.009701399 | ENSG00000160584 | n.s. |
| ENSG00000166797 | 1150.020692 | 0.123662705 | 0.3351568 | 0.36897 | 0.712150341 | 0.83487542 | FAM96A | protein_coding | 41.73 | 0.009692476 | 0.009692476 | ENSG00000166797 | n.s. |
| ENSG00000131788 | 184.0280092 | 0.105057803 | 0.2474127 | 0.424626 | 0.671109579 | 0.808693667 | PIAS3 | protein_coding | 52.28 | 0.009688006 | 0.009688006 | ENSG00000131788 | n.s. |
| ENSG00000167969 | 409.2561038 | 0.119114276 | 0.3135353 | 0.379907 | 0.704014363 | 0.829429483 | ECI1 | protein_coding | 55.22 | 0.009674525 | 0.009674525 | ENSG00000167969 | n.s. |
| ENSG00000188706 | 330.5646578 | -0.104658096 | 0.2466907 | -0.424248 | 0.671384766 | 0.808693667 | ZDHHC9 | protein_coding | 43.4 | -0.009651147 | 0.009651147 | ENSG00000188706 | n.s. |
| ENSG00000223203 | 4.193047957 | -0.156690597 | 0.5207117 | -0.300916 | 0.763478367 | 0.868172926 | RNA5SP221 | rRNA | 52.43 | -0.009619825 | 0.009619825 | ENSG00000223203 | n.s. |
| ENSG00000102796 | 77.62827129 | -0.108747407 | 0.2657829 | -0.409159 | 0.682423102 | 0.81576698 | DHRS12 | protein_coding | 47.24 | -0.009616955 | 0.009616955 | ENSG00000102796 | n.s. |
| ENSG00000100285 | 1745.271838 | 0.098395817 | 0.2206588 | 0.445918 | 0.655656143 | 0.798776372 | NEFH | protein_coding | 52.82 | 0.009600951 | 0.009600951 | ENSG00000100285 | n.s. |
| ENSG00000224877 | 1149.881218 | 0.111492674 | 0.2781859 | 0.400785 | 0.688578633 | 0.820179682 | NDUFAF8 | protein_coding | 58.49 | 0.009598515 | 0.009598515 | ENSG00000224877 | n.s. |
| ENSG00000127952 | 184.4497646 | 0.119170022 | 0.3161349 | 0.376959 | 0.706203804 | 0.830936834 | STYXL1 | protein_coding | 48.42 | 0.009585082 | 0.009585082 | ENSG00000127952 | n.s. |
| ENSG00000102098 | 631.9805909 | -0.110283725 | 0.2730473 | -0.4039 | 0.686286445 | 0.818632266 | SCML2 | protein_coding | 39.57 | -0.009584884 | 0.009584884 | ENSG00000102098 | n.s. |
| ENSG00000257524 | 66.41518071 | 0.124562971 | 0.3432206 | 0.362924 | 0.716661731 | 0.837719862 | | processed_transcript | 56.77 | 0.00957904 | 0.00957904 | ENSG00000257524 | n.s. |
| ENSG00000132541 | 435.3720328 | 0.120866947 | 0.3244365 | 0.372544 | 0.709487718 | 0.833411025 | RIDA | protein_coding | 39.14 | 0.009565502 | 0.009565502 | ENSG00000132541 | n.s. |

| | | | | | | | | | | | | | |
|---|---|---|---|---|---|---|---|---|---|---|---|---|---|
| ENSG00000094841 | 175.2188999 | -0.115028778 | 0.2957488 | -0.388941 | 0.697319931 | 0.825787649 | UPRT | protein_coding | 37.52 | -0.009562528 | 0.009562528 | ENSG00000094841 | n.s. |
| ENSG00000196531 | 17813.3896 | 0.113092181 | 0.2872042 | 0.393769 | 0.693751438 | 0.823135405 | NACA | protein_coding | 47.57 | 0.009559537 | 0.009559537 | ENSG00000196531 | n.s. |
| ENSG00000161654 | 402.5908078 | 0.105151992 | 0.251195 | 0.418607 | 0.675503349 | 0.81154829 | LSM12 | protein_coding | 46.77 | 0.009535775 | 0.009535775 | ENSG00000161654 | n.s. |
| ENSG00000250957 | 460.1335939 | 0.10027246 | 0.2294165 | 0.437076 | 0.662056224 | 0.803364895 | | lincRNA | 34.31 | 0.009534622 | 0.009534622 | ENSG00000250957 | n.s. |
| ENSG00000199072 | 58.07612841 | 0.11081542 | 0.2768203 | 0.400315 | 0.688924286 | 0.820294688 | MIRLET7F1 | miRNA | 40.23 | 0.009533462 | 0.009533462 | ENSG00000199072 | n.s. |
| ENSG00000214736 | 1604.889243 | 0.115893091 | 0.3019416 | 0.383826 | 0.701107266 | 0.827738189 | TOMM6 | protein_coding | 50.87 | 0.009515635 | 0.009515635 | ENSG00000214736 | n.s. |
| ENSG00000173272 | 1133.16086 | 0.113299274 | 0.2890803 | 0.39193 | 0.695109837 | 0.82422196 | MZT2A | protein_coding | 51.73 | 0.009512133 | 0.009512133 | ENSG00000173272 | n.s. |
| ENSG00000169231 | 365.2479048 | 0.109386303 | 0.2712244 | 0.403305 | 0.686723505 | 0.818670379 | THBS3 | protein_coding | 55.3 | 0.009504677 | 0.009504677 | ENSG00000169231 | n.s. |
| ENSG00000196227 | 735.8285499 | -0.09642263 | 0.214286 | -0.449972 | 0.652730795 | 0.797120504 | FAM217B | protein_coding | 42.7 | -0.009495317 | 0.009495317 | ENSG00000196227 | n.s. |
| ENSG00000164438 | 113.5444793 | -0.111250721 | 0.2804826 | -0.39664 | 0.691632729 | 0.821656665 | TLX3 | protein_coding | 66.78 | -0.009490756 | 0.009490756 | ENSG00000164438 | n.s. |
| ENSG00000157870 | 99.33994875 | 0.114440286 | 0.2955277 | 0.38724 | 0.698578228 | 0.826664873 | FAM213B | protein_coding | 64.59 | 0.009460838 | 0.009460838 | ENSG00000157870 | n.s. |
| ENSG00000174013 | 1044.465431 | -0.11128869 | 0.2816906 | -0.395074 | 0.692788107 | 0.822309362 | FBXO45 | protein_coding | 41.25 | -0.009455617 | 0.009455617 | ENSG00000174013 | n.s. |
| ENSG00000244363 | 28.7179974 | -0.121848189 | 0.3334233 | -0.365446 | 0.714778534 | 0.836431461 | RPL7P23 | processed_pseudogene | 43.24 | -0.00945172 | 0.00945172 | ENSG00000244363 | n.s. |
| ENSG00000121957 | 1296.265214 | -0.116335333 | 0.3063955 | -0.37969 | 0.70417548 | 0.829429483 | GPSM2 | protein_coding | 38.78 | -0.009448818 | 0.009448818 | ENSG00000121957 | n.s. |
| ENSG00000136045 | 2249.46102 | -0.112561739 | 0.2874801 | -0.391546 | 0.695393601 | 0.824325748 | PWP1 | protein_coding | 40.59 | -0.009444058 | 0.009444058 | ENSG00000136045 | n.s. |
| ENSG00000214135 | 605.6381512 | -0.101048404 | 0.2356168 | -0.428868 | 0.668019537 | 0.806516133 | | transcribed_unprocessed_pseud | 45.89 | -0.009436601 | 0.009436601 | ENSG00000214135 | n.s. |
| ENSG00000199459 | 9.222994213 | 0.184899036 | 0.7210437 | 0.256433 | 0.797616887 | 0.889128896 | | misc_RNA | 43.64 | 0.009436373 | 0.009436373 | ENSG00000199459 | n.s. |
| ENSG00000252451 | 3.393953215 | -0.164105586 | 0.5806445 | -0.282627 | 0.777463097 | 0.876003824 | RNA5SP168 | rRNA | 44.64 | -0.009435086 | 0.009435086 | ENSG00000252451 | n.s. |
| ENSG00000108828 | 384.8539715 | 0.111670043 | 0.2838241 | 0.393448 | 0.693988582 | 0.823242994 | VAT1 | protein_coding | 54.7 | 0.009432987 | 0.009432987 | ENSG00000108828 | n.s. |
| ENSG00000108604 | 3035.288107 | -0.111529381 | 0.283296 | -0.393685 | 0.693813606 | 0.823135405 | SMARCD2 | protein_coding | 55.99 | -0.009427436 | 0.009427436 | ENSG00000108604 | n.s. |
| ENSG00000115756 | 221.4684432 | 0.105541921 | 0.2556987 | 0.412759 | 0.679783287 | 0.814161092 | HPCAL1 | protein_coding | 52.12 | 0.009423802 | 0.009423802 | ENSG00000115756 | n.s. |
| ENSG00000209082 | 3.267501149 | -0.16667018 | 0.5977099 | -0.278848 | 0.780361506 | 0.877937651 | MT-TL1 | Mt_tRNA | 38.67 | -0.00942292 | 0.00942292 | ENSG00000209082 | n.s. |
| ENSG00000140451 | 572.245042 | 0.104408526 | 0.2503584 | 0.417036 | 0.676651911 | 0.81243231 | PIF1 | protein_coding | 55.24 | 0.009418987 | 0.009418987 | ENSG00000140451 | n.s. |
| ENSG00000264419 | 4.054385052 | -0.159018541 | 0.5480536 | -0.290151 | 0.77170037 | 0.872517203 | MIR548AC | miRNA | 32.95 | -0.009418032 | 0.009418032 | ENSG00000264419 | n.s. |
| ENSG00000214022 | 1453.300564 | 0.105423246 | 0.2555269 | 0.412572 | 0.679920262 | 0.81418507 | REPIN1 | protein_coding | 62.28 | 0.009411857 | 0.009411857 | ENSG00000214022 | n.s. |
| ENSG00000276903 | 774.9108655 | -0.096075292 | 0.2148632 | -0.447146 | 0.65476943 | 0.798168731 | HIST1H2AL | protein_coding | 63.87 | -0.009406279 | 0.009406279 | ENSG00000276903 | n.s. |
| ENSG00000136436 | 104.7050474 | -0.113679687 | 0.2938473 | -0.386867 | 0.698854997 | 0.826664873 | CALCOCO2 | protein_coding | 43.98 | -0.009397958 | 0.009397958 | ENSG00000136436 | n.s. |
| ENSG00000188573 | 352.7002924 | 0.113756034 | 0.2950454 | 0.385554 | 0.699826755 | 0.82681669 | FBLL1 | protein_coding | 69.39 | 0.009395198 | 0.009395198 | ENSG00000188573 | n.s. |
| ENSG00000198026 | 420.7368339 | 0.094837618 | 0.2099604 | 0.451693 | 0.651490237 | 0.796400914 | ZNF335 | protein_coding | 53.47 | 0.009376429 | 0.009376429 | ENSG00000198026 | n.s. |
| ENSG00000259402 | 16.65827524 | 0.121507466 | 0.3336645 | 0.364161 | 0.715738117 | 0.837272879 | | sense_intronic | 43.06 | 0.009372233 | 0.009372233 | ENSG00000259402 | n.s. |
| ENSG00000101413 | 1002.361116 | -0.104037511 | 0.2506348 | -0.415096 | 0.678071581 | 0.813086294 | RPRD1B | protein_coding | 42.38 | -0.00934916 | 0.00934916 | ENSG00000101413 | n.s. |
| ENSG00000263666 | 5.740423919 | -0.149029487 | 0.4857276 | -0.306817 | 0.758982654 | 0.865676701 | SNORA70D | snoRNA | 53.33 | -0.009335844 | 0.009335844 | ENSG00000263666 | n.s. |

| | | | | | | | | | | | | | |
|---|---|---|---|---|---|---|---|---|---|---|---|---|---|
| ENSG00000180104 | 333.556069 | -0.105485078 | 0.2579017 | -0.409013 | 0.682530313 | 0.81576698 | EXOC3 | protein_coding | 51.24 | -0.009328455 | 0.009328455 | ENSG00000180104 | n.s. |
| ENSG00000112237 | 2130.305229 | -0.110440383 | 0.2810157 | -0.393004 | 0.694316248 | 0.823421219 | CCNC | protein_coding | 35.29 | -0.009318733 | 0.009318733 | ENSG00000112237 | n.s. |
| ENSG00000277258 | 212.9720213 | -0.101703384 | 0.2408393 | -0.422287 | 0.672815294 | 0.809846304 | PCGF2 | protein_coding | 57.06 | -0.009315765 | 0.009315765 | ENSG00000277258 | n.s. |
| ENSG00000107566 | 723.1741514 | -0.103661734 | 0.249805 | -0.414971 | 0.67816346 | 0.813086294 | ERLIN1 | protein_coding | 38.56 | -0.009315392 | 0.009315392 | ENSG00000107566 | n.s. |
| ENSG00000107874 | 847.1364807 | 0.115503288 | 0.306099 | 0.37734 | 0.705921184 | 0.830814932 | CUEDC2 | protein_coding | 52.94 | 0.009297519 | 0.009297519 | ENSG00000107874 | n.s. |
| ENSG00000148672 | 1917.787154 | 0.108663505 | 0.2728889 | 0.398197 | 0.690485029 | 0.821195719 | GLUD1 | protein_coding | 42.11 | 0.009296524 | 0.009296524 | ENSG00000148672 | n.s. |
| ENSG00000130741 | 2293.491525 | 0.104447235 | 0.2538219 | 0.411498 | 0.68070726 | 0.814847148 | EIF2S3 | protein_coding | 41.58 | 0.00928785 | 0.00928785 | ENSG00000130741 | n.s. |
| ENSG00000012061 | 557.5683831 | 0.112450795 | 0.2915746 | 0.385667 | 0.699743101 | 0.82681669 | ERCC1 | protein_coding | 52.1 | 0.009287397 | 0.009287397 | ENSG00000012061 | n.s. |
| ENSG00000176393 | 283.8766448 | 0.10812351 | 0.2709168 | 0.399102 | 0.689817865 | 0.820602898 | RNPEP | protein_coding | 48.75 | 0.009284236 | 0.009284236 | ENSG00000176393 | n.s. |
| ENSG00000221200 | 10.18511101 | 0.135670906 | 0.4120255 | 0.329278 | 0.741945593 | 0.854222153 | MIR1253 | miRNA | 45.71 | 0.009283847 | 0.009283847 | ENSG00000221200 | n.s. |
| ENSG00000148308 | 1058.547346 | -0.109664343 | 0.2784775 | -0.3938 | 0.69372897 | 0.823135405 | GTF3C5 | protein_coding | 51.81 | -0.009269786 | 0.009269786 | ENSG00000148308 | n.s. |
| ENSG00000130985 | 1808.608158 | 0.109660632 | 0.2785733 | 0.393651 | 0.693838764 | 0.823135405 | UBA1 | protein_coding | 51.31 | 0.009269473 | 0.009269473 | ENSG00000130985 | n.s. |
| ENSG00000138674 | 2162.701835 | -0.104598383 | 0.2551292 | -0.409982 | 0.681819114 | 0.81547698 | SEC31A | protein_coding | 38.78 | -0.009266192 | 0.009266192 | ENSG00000138674 | n.s. |
| ENSG00000064726 | 1872.767135 | -0.108297672 | 0.2721344 | -0.397957 | 0.690662248 | 0.821195719 | BTBD1 | protein_coding | 39.53 | -0.009265226 | 0.009265226 | ENSG00000064726 | n.s. |
| ENSG00000213676 | 351.1629043 | 0.106677792 | 0.2646988 | 0.403016 | 0.686936614 | 0.81878422 | ATF6B | protein_coding | 50.77 | 0.00926289 | 0.00926289 | ENSG00000213676 | n.s. |
| ENSG00000137574 | 1258.818555 | -0.104516785 | 0.2548855 | -0.410054 | 0.681766384 | 0.81547698 | TGS1 | protein_coding | 40.02 | -0.009258964 | 0.009258964 | ENSG00000137574 | n.s. |
| ENSG00000100258 | 294.2476454 | 0.105608067 | 0.2597949 | 0.406505 | 0.684371226 | 0.817335785 | LMF2 | protein_coding | 67.15 | 0.009251212 | 0.009251212 | ENSG00000100258 | n.s. |
| ENSG00000109919 | 1674.528856 | -0.11519978 | 0.3059998 | -0.37647 | 0.706567434 | 0.831224199 | MTCH2 | protein_coding | 43.47 | -0.009248448 | 0.009248448 | ENSG00000109919 | n.s. |
| ENSG00000185942 | 58.38549748 | 0.104088253 | 0.2532385 | 0.411028 | 0.681051658 | 0.815049186 | NKAIN3 | protein_coding | 37.47 | 0.009244721 | 0.009244721 | ENSG00000185942 | n.s. |
| ENSG00000221265 | 27.57600185 | -0.120019408 | 0.3301374 | -0.363544 | 0.716198719 | 0.83755976 | MIR1255A | miRNA | 33.63 | -0.009239598 | 0.009239598 | ENSG00000221265 | n.s. |
| ENSG00000162885 | 965.1200947 | -0.104872358 | 0.2573016 | -0.407585 | 0.683578157 | 0.816528713 | B3GALNT2 | protein_coding | 40.36 | -0.00923176 | 0.00923176 | ENSG00000162885 | n.s. |
| ENSG00000245937 | 95.98992015 | -0.102464297 | 0.2461131 | -0.41633 | 0.67716844 | 0.812694714 | LINC01184 | lincRNA | 37.74 | -0.009229222 | 0.009229222 | ENSG00000245937 | n.s. |
| ENSG00000169926 | 3075.397497 | 0.105514791 | 0.25996 | 0.405889 | 0.684824483 | 0.81766669 | KLF13 | protein_coding | 49.37 | 0.009224493 | 0.009224493 | ENSG00000169926 | n.s. |
| ENSG00000049541 | 1018.776576 | -0.108025159 | 0.2722973 | -0.396718 | 0.691575611 | 0.821656665 | RFC2 | protein_coding | 48.96 | -0.009215585 | 0.009215585 | ENSG00000049541 | n.s. |
| ENSG00000117640 | 252.2433532 | -0.113454257 | 0.2989385 | -0.379524 | 0.704299022 | 0.829429483 | MTFR1L | protein_coding | 49.43 | -0.009214815 | 0.009214815 | ENSG00000117640 | n.s. |
| ENSG00000174136 | 1643.908528 | 0.096079287 | 0.2182417 | 0.440243 | 0.659761472 | 0.802048282 | RGMB | protein_coding | 43.29 | 0.009204346 | 0.009204346 | ENSG00000174136 | n.s. |
| ENSG00000106617 | 106.125664 | -0.105910026 | 0.2628626 | -0.40291 | 0.68701426 | 0.818801761 | PRKAG2 | protein_coding | 48.91 | -0.009195239 | 0.009195239 | ENSG00000106617 | n.s. |
| ENSG00000242485 | 485.4000799 | -0.113142736 | 0.2987909 | -0.378669 | 0.70493397 | 0.829863504 | MRPL20 | protein_coding | 53.09 | -0.009163807 | 0.009163807 | ENSG00000242485 | n.s. |
| ENSG00000113448 | 67.00573527 | -0.111612243 | 0.2917031 | -0.382623 | 0.70199952 | 0.828019258 | PDE4D | protein_coding | 37.37 | -0.009147691 | 0.009147691 | ENSG00000113448 | n.s. |
| ENSG00000108528 | 543.9018387 | 0.105121653 | 0.2600146 | 0.404291 | 0.685998505 | 0.818436777 | SLC25A11 | protein_coding | 58.68 | 0.009147146 | 0.009147146 | ENSG00000108528 | n.s. |
| ENSG00000125454 | 321.3257851 | -0.095988754 | 0.2195727 | -0.437162 | 0.661994084 | 0.803364895 | SLC25A19 | protein_coding | 50.68 | -0.009127297 | 0.009127297 | ENSG00000125454 | n.s. |
| ENSG00000099821 | 408.6925602 | 0.110312376 | 0.2850896 | 0.386939 | 0.698801081 | 0.826664873 | POLRMT | protein_coding | 61.15 | 0.009119581 | 0.009119581 | ENSG00000099821 | n.s. |

| | | | | | | | | | | | | | |
|---|---|---|---|---|---|---|---|---|---|---|---|---|---|
| ENSG00000257365 | 379.7747479 | -0.105887637 | 0.2641332 | -0.400887 | 0.688503094 | 0.820159872 | FNTB | protein_coding | 43.84 | -0.009117083 | 0.009117083 | ENSG00000257365 | n.s. |
| ENSG00000272674 | 396.7050024 | -0.093713539 | 0.21086 | -0.444435 | 0.656728236 | 0.799687532 | PCDHB16 | protein_coding | 44.75 | -0.00909768 | 0.00909768 | ENSG00000272674 | n.s. |
| ENSG00000100519 | 3168.99593 | 0.102343863 | 0.2488633 | 0.411245 | 0.680892616 | 0.8149289 | PSMC6 | protein_coding | 37.9 | 0.009096351 | 0.009096351 | ENSG00000100519 | n.s. |
| ENSG00000261889 | 96.59394848 | -0.106617607 | 0.2688302 | -0.396598 | 0.691663666 | 0.821656665 | | lincRNA | 51.2 | -0.009095507 | 0.009095507 | ENSG00000261889 | n.s. |
| ENSG00000145777 | 82.69549709 | 0.101086218 | 0.2439302 | 0.414406 | 0.678576533 | 0.813345478 | TSLP | protein_coding | 43.11 | 0.009069955 | 0.009069955 | ENSG00000145777 | n.s. |
| ENSG00000154124 | 1192.581729 | -0.097966234 | 0.2302935 | -0.425397 | 0.670547099 | 0.808314048 | OTULIN | protein_coding | 41.68 | -0.009054027 | 0.009054027 | ENSG00000154124 | n.s. |
| ENSG00000176014 | 924.2623283 | -0.109769957 | 0.2852539 | -0.384815 | 0.700374515 | 0.827166021 | TUBB6 | protein_coding | 45.6 | -0.009045847 | 0.009045847 | ENSG00000176014 | n.s. |
| ENSG00000153560 | 1443.336355 | -0.103744739 | 0.2570974 | -0.403523 | 0.686563475 | 0.818643628 | UBP1 | protein_coding | 40.23 | -0.009015948 | 0.009015948 | ENSG00000153560 | n.s. |
| ENSG00000198933 | 1055.39393 | 0.102696381 | 0.2525097 | 0.406703 | 0.684226392 | 0.817232913 | TBKBP1 | protein_coding | 58.66 | 0.009001764 | 0.009001764 | ENSG00000198933 | n.s. |
| ENSG00000055070 | 575.6551537 | 0.109626201 | 0.2862132 | 0.383023 | 0.701702777 | 0.827946733 | SZRD1 | protein_coding | 48.1 | 0.008989086 | 0.008989086 | ENSG00000055070 | n.s. |
| ENSG00000249353 | 797.484305 | 0.112555238 | 0.2999274 | 0.375275 | 0.707456004 | 0.83205862 | NPM1P27 | processed_pseudogene | 41.69 | 0.008987094 | 0.008987094 | ENSG00000249353 | n.s. |
| ENSG00000244615 | 16.28052582 | -0.123487169 | 0.3563902 | -0.346494 | 0.728971249 | 0.845718415 | PSPC1P2 | transcribed_unprocessed_pseud | 33.35 | -0.008986681 | 0.008986681 | ENSG00000244615 | n.s. |
| ENSG00000031698 | 4932.16974 | -0.108697665 | 0.2811222 | -0.386656 | 0.69901067 | 0.826664873 | SARS | protein_coding | 43.12 | -0.008986092 | 0.008986092 | ENSG00000031698 | n.s. |
| ENSG00000129518 | 468.7320454 | -0.114563686 | 0.3104432 | -0.369033 | 0.712103322 | 0.83487542 | EAPP | protein_coding | 44.09 | -0.00897931 | 0.00897931 | ENSG00000129518 | n.s. |
| ENSG00000221676 | 9.594480901 | -0.137650198 | 0.433607 | -0.317454 | 0.750899292 | 0.860545793 | RNU6ATAC | snRNA | 50 | -0.008978374 | 0.008978374 | ENSG00000221676 | n.s. |
| ENSG00000137942 | 5459.924678 | -0.103958625 | 0.2588911 | -0.401553 | 0.688012672 | 0.819715937 | FNBP1L | protein_coding | 37.34 | -0.008975436 | 0.008975436 | ENSG00000137942 | n.s. |
| ENSG00000185262 | 413.0972525 | -0.103347722 | 0.2563801 | -0.403104 | 0.686872071 | 0.818777384 | UBALD2 | protein_coding | 58.45 | -0.008974113 | 0.008974113 | ENSG00000185262 | n.s. |
| ENSG00000250959 | 73.5168631 | -0.10844328 | 0.2803855 | -0.386765 | 0.698930187 | 0.826664873 | GLUD1P3 | transcribed_unprocessed_pseud | 52.12 | -0.008965062 | 0.008965062 | ENSG00000250959 | n.s. |
| ENSG00000234773 | 97.30622216 | 0.105789321 | 0.2690225 | 0.393236 | 0.694145237 | 0.823358676 | | transcribed_unprocessed_pseud | 42.34 | 0.008929775 | 0.008929775 | ENSG00000234773 | n.s. |
| ENSG00000171148 | 538.837806 | -0.101226176 | 0.2479705 | -0.408219 | 0.683113222 | 0.816183419 | TADA3 | protein_coding | 49.41 | -0.008929387 | 0.008929387 | ENSG00000171148 | n.s. |
| ENSG00000180329 | 1802.024606 | 0.103866793 | 0.2599485 | 0.399567 | 0.689475624 | 0.820546125 | CCDC43 | protein_coding | 39.85 | 0.008921846 | 0.008921846 | ENSG00000180329 | n.s. |
| ENSG00000198598 | 137.8675082 | -0.103522265 | 0.2586934 | -0.400174 | 0.689028685 | 0.820294688 | MMP17 | protein_coding | 61.93 | -0.008906031 | 0.008906031 | ENSG00000198598 | n.s. |
| ENSG00000278133 | 118.2014431 | -0.106134394 | 0.2709385 | -0.391729 | 0.695258644 | 0.824325748 | | sense_intronic | 46.39 | -0.008904796 | 0.008904796 | ENSG00000278133 | n.s. |
| ENSG00000140455 | 2678.056034 | -0.10756517 | 0.2781459 | -0.386722 | 0.69896193 | 0.826664873 | USP3 | protein_coding | 38.44 | -0.008892468 | 0.008892468 | ENSG00000140455 | n.s. |
| ENSG00000125037 | 557.9286555 | -0.109047005 | 0.2863625 | -0.380801 | 0.703351262 | 0.828935858 | EMC3 | protein_coding | 44.9 | -0.008885049 | 0.008885049 | ENSG00000125037 | n.s. |
| ENSG00000017797 | 1095.923556 | -0.10069101 | 0.2467091 | -0.408137 | 0.683173408 | 0.816185289 | RALBP1 | protein_coding | 41.38 | -0.008882079 | 0.008882079 | ENSG00000017797 | n.s. |
| ENSG00000241362 | 14.97085985 | 0.125796589 | 0.3725858 | 0.337631 | 0.735641115 | 0.850056539 | | processed_pseudogene | 44.63 | 0.008875225 | 0.008875225 | ENSG00000241362 | n.s. |
| ENSG00000139842 | 1353.327302 | -0.103276653 | 0.2595251 | -0.397945 | 0.690670861 | 0.821195719 | CUL4A | protein_coding | 45.97 | -0.008835661 | 0.008835661 | ENSG00000139842 | n.s. |
| ENSG00000203809 | 231.5068069 | 0.094348555 | 0.219253 | 0.430318 | 0.666964175 | 0.806194128 | LIN28B-AS1 | lincRNA | 40.64 | 0.008827286 | 0.008827286 | ENSG00000203809 | n.s. |
| ENSG00000112319 | 3236.915462 | -0.102497843 | 0.2561166 | -0.4002 | 0.689009254 | 0.820294688 | EYA4 | protein_coding | 36.95 | -0.0088179 | 0.0088179 | ENSG00000112319 | n.s. |
| ENSG00000152291 | 1089.284578 | -0.09560645 | 0.2252898 | -0.424371 | 0.671295228 | 0.808693667 | TGOLN2 | protein_coding | 47.66 | -0.00881644 | 0.00881644 | ENSG00000152291 | n.s. |
| ENSG00000213614 | 870.5877874 | 0.105178971 | 0.268909 | 0.391132 | 0.695699542 | 0.824499892 | HEXA | protein_coding | 45.73 | 0.008814986 | 0.008814986 | ENSG00000213614 | n.s. |

| | | | | | | | | | | | | | |
|---|---|---|---|---|---|---|---|---|---|---|---|---|---|
| ENSG00000166770 | 511.7944922 | 0.108528593 | 0.2859884 | 0.379486 | 0.704326997 | 0.829429483 | ZNF667-AS1 | lincRNA | 43.34 | 0.00881475 | 0.00881475 | ENSG00000166770 | n.s. |
| ENSG00000104679 | 452.3391768 | 0.1058869 | 0.2721034 | 0.389142 | 0.697170984 | 0.825751732 | R3HCC1 | protein_coding | 47.98 | 0.008804549 | 0.008804549 | ENSG00000104679 | n.s. |
| ENSG00000109270 | 1220.665307 | -0.104818593 | 0.2677419 | -0.391491 | 0.695434191 | 0.824325748 | LAMTOR3 | protein_coding | 34.97 | -0.008794399 | 0.008794399 | ENSG00000109270 | n.s. |
| ENSG00000134590 | 385.2290258 | 0.101959027 | 0.2541266 | 0.401214 | 0.688262903 | 0.819943903 | RTL8C | protein_coding | 60.29 | 0.008790485 | 0.008790485 | ENSG00000134590 | n.s. |
| ENSG00000174915 | 324.0403319 | 0.105971237 | 0.2732438 | 0.387827 | 0.698144246 | 0.826342119 | PTDSS2 | protein_coding | 56.05 | 0.008778669 | 0.008778669 | ENSG00000174915 | n.s. |
| ENSG00000167536 | 277.4201294 | -0.106486043 | 0.276961 | -0.38448 | 0.700622561 | 0.827236085 | DHRS13 | protein_coding | 54.96 | -0.008771312 | 0.008771312 | ENSG00000167536 | n.s. |
| ENSG00000163634 | 717.1296195 | -0.113690807 | 0.3127584 | -0.36351 | 0.716223982 | 0.83755976 | THOC7 | protein_coding | 39.89 | -0.008752395 | 0.008752395 | ENSG00000163634 | n.s. |
| ENSG00000259658 | 27.59506389 | -0.1120864 | 0.3050862 | -0.367393 | 0.713326182 | 0.8354878 | | transcribed_unprocessed_pseud | 44.33 | -0.008749452 | 0.008749452 | ENSG00000259658 | n.s. |
| ENSG00000204220 | 562.6710999 | 0.110373041 | 0.2959079 | 0.372998 | 0.709149955 | 0.833411025 | PFDN6 | protein_coding | 54.13 | 0.008735006 | 0.008735006 | ENSG00000204220 | n.s. |
| ENSG00000104774 | 283.6011056 | 0.100432921 | 0.248933 | 0.403454 | 0.686614557 | 0.818643628 | MAN2B1 | protein_coding | 53.98 | 0.008728134 | 0.008728134 | ENSG00000104774 | n.s. |
| ENSG00000160469 | 391.523803 | -0.106375966 | 0.2774008 | -0.383474 | 0.701368464 | 0.827906157 | BRSK1 | protein_coding | 54.48 | -0.008724838 | 0.008724838 | ENSG00000160469 | n.s. |
| ENSG00000255769 | 544.9362782 | 0.101407302 | 0.2537661 | 0.399609 | 0.689444296 | 0.820546125 | GOLGA2P10 | transcribed_unprocessed_pseud | 42.97 | 0.008710583 | 0.008710583 | ENSG00000255769 | n.s. |
| ENSG00000166704 | 594.4895798 | -0.090428294 | 0.2046363 | -0.441898 | 0.658563256 | 0.80108127 | ZNF606 | protein_coding | 45 | -0.008710363 | 0.008710363 | ENSG00000166704 | n.s. |
| ENSG00000146281 | 1160.035362 | -0.105456673 | 0.2734803 | -0.38561 | 0.699785676 | 0.82681669 | PM20D2 | protein_coding | 39.61 | -0.008709747 | 0.008709747 | ENSG00000146281 | n.s. |
| ENSG00000125877 | 432.2762008 | -0.110756699 | 0.2992687 | -0.370091 | 0.711314558 | 0.834542721 | ITPA | protein_coding | 50.89 | -0.008700097 | 0.008700097 | ENSG00000125877 | n.s. |
| ENSG00000151553 | 721.4349394 | -0.101014258 | 0.2523738 | -0.400256 | 0.688967641 | 0.820294688 | FAM160B1 | protein_coding | 39.9 | -0.008690267 | 0.008690267 | ENSG00000151553 | n.s. |
| ENSG00000165156 | 1910.364598 | -0.106005705 | 0.2769062 | -0.382822 | 0.701851929 | 0.827985461 | ZHX1 | protein_coding | 38.83 | -0.00869006 | 0.00869006 | ENSG00000165156 | n.s. |
| ENSG00000167182 | 132.674747 | -0.096493439 | 0.2325474 | -0.414941 | 0.678185099 | 0.813086294 | SP2 | protein_coding | 47.6 | -0.008671225 | 0.008671225 | ENSG00000167182 | n.s. |
| ENSG00000109320 | 77.62482657 | 0.101407391 | 0.2554029 | 0.397049 | 0.691331524 | 0.821561011 | NFKB1 | protein_coding | 37.86 | 0.008656153 | 0.008656153 | ENSG00000109320 | n.s. |
| ENSG00000180346 | 635.5394187 | 0.087882062 | 0.1954771 | 0.449577 | 0.653015281 | 0.797221535 | TIGD2 | protein_coding | 43.15 | 0.008649439 | 0.008649439 | ENSG00000180346 | n.s. |
| ENSG00000137720 | 222.6504655 | 0.112083523 | 0.3076843 | 0.364281 | 0.715648281 | 0.837238133 | C11orf1 | protein_coding | 42.92 | 0.008647356 | 0.008647356 | ENSG00000137720 | n.s. |
| ENSG00000205629 | 285.9901906 | -0.110678782 | 0.3013052 | -0.367331 | 0.713372044 | 0.8354878 | LCMT1 | protein_coding | 45.35 | -0.008639573 | 0.008639573 | ENSG00000205629 | n.s. |
| ENSG00000164649 | 951.2089167 | -0.102756948 | 0.2621344 | -0.392001 | 0.69505747 | 0.82422196 | CDCA7L | protein_coding | 41.49 | -0.008627044 | 0.008627044 | ENSG00000164649 | n.s. |
| ENSG00000099804 | 437.4177137 | 0.108085472 | 0.2885509 | 0.37458 | 0.707972693 | 0.832537372 | CDC34 | protein_coding | 64.75 | 0.0086032 | 0.0086032 | ENSG00000099804 | n.s. |
| ENSG00000135070 | 1296.218659 | 0.108794957 | 0.2927329 | 0.371653 | 0.710151486 | 0.833679523 | ISCA1 | protein_coding | 41.36 | 0.008594896 | 0.008594896 | ENSG00000135070 | n.s. |
| ENSG00000106477 | 49.33229945 | -0.10849271 | 0.2906859 | -0.37323 | 0.70897728 | 0.83328471 | CEP41 | protein_coding | 39.96 | -0.008593337 | 0.008593337 | ENSG00000106477 | n.s. |
| ENSG00000180537 | 327.8782196 | 0.108369592 | 0.2900968 | 0.373564 | 0.708729058 | 0.833203974 | RNF182 | protein_coding | 38.51 | 0.008588146 | 0.008588146 | ENSG00000180537 | n.s. |
| ENSG00000120805 | 1002.638208 | -0.110454995 | 0.3019008 | -0.365865 | 0.714465696 | 0.836135655 | ARL1 | protein_coding | 39.51 | -0.008584922 | 0.008584922 | ENSG00000120805 | n.s. |
| ENSG00000171497 | 1247.525114 | 0.108286088 | 0.2900428 | 0.373345 | 0.708891531 | 0.833254267 | PPID | protein_coding | 38.97 | 0.00857869 | 0.00857869 | ENSG00000171497 | n.s. |
| ENSG00000162636 | 697.8728527 | -0.100613447 | 0.2540107 | -0.396099 | 0.692031868 | 0.821749243 | FAM102B | protein_coding | 39.64 | -0.008578372 | 0.008578372 | ENSG00000162636 | n.s. |
| ENSG00000173456 | 205.9108513 | -0.098597729 | 0.2441574 | -0.403829 | 0.686338752 | 0.818632266 | RNF26 | protein_coding | 59.17 | -0.008569241 | 0.008569241 | ENSG00000173456 | n.s. |
| ENSG00000167693 | 453.0906299 | 0.102490393 | 0.2625369 | 0.390385 | 0.696252073 | 0.824944057 | NXN | protein_coding | 50.99 | 0.008565686 | 0.008565686 | ENSG00000167693 | n.s. |

| | | | | | | | | | | | | | |
|---|---|---|---|---|---|---|---|---|---|---|---|---|---|
| ENSG00000172732 | 362.2895038 | -0.103668739 | 0.2684788 | -0.386134 | 0.699397585 | 0.826771317 | MUS81 | protein_coding | 60.2 | -0.008564551 | 0.008564551 | ENSG00000172732 | n.s. |
| ENSG00000196419 | 8743.541339 | 0.110879205 | 0.3041653 | 0.364536 | 0.715457782 | 0.837085605 | XRCC6 | protein_coding | 46.22 | 0.008563215 | 0.008563215 | ENSG00000196419 | n.s. |
| ENSG00000133619 | 64.9315739 | -0.106465872 | 0.2826029 | -0.376733 | 0.70637199 | 0.831064493 | KRBA1 | protein_coding | 57.32 | -0.008556159 | 0.008556159 | ENSG00000133619 | n.s. |
| ENSG00000105887 | 2560.099483 | 0.105339769 | 0.2774182 | 0.379715 | 0.704157199 | 0.829429483 | MTPN | protein_coding | 36.4 | 0.008555752 | 0.008555752 | ENSG00000105887 | n.s. |
| ENSG00000130856 | 44.33501172 | -0.110976995 | 0.3052872 | -0.363517 | 0.716218927 | 0.83755976 | ZNF236 | protein_coding | 42.43 | -0.008543475 | 0.008543475 | ENSG00000130856 | n.s. |
| ENSG00000013275 | 1236.810574 | 0.104287988 | 0.2730545 | 0.381931 | 0.702512509 | 0.828553261 | PSMC4 | protein_coding | 50.25 | 0.008518198 | 0.008518198 | ENSG00000013275 | n.s. |
| ENSG00000153157 | 216.8989553 | 0.102948766 | 0.2664396 | 0.386387 | 0.699210203 | 0.826760396 | SYCP2L | protein_coding | 40.47 | 0.008505662 | 0.008505662 | ENSG00000153157 | n.s. |
| ENSG00000184613 | 4181.899913 | 0.103711464 | 0.2706388 | 0.38321 | 0.701564096 | 0.827940174 | NELL2 | protein_coding | 36.23 | 0.008504448 | 0.008504448 | ENSG00000184613 | n.s. |
| ENSG00000242028 | 328.7642889 | -0.08943632 | 0.2046222 | -0.43708 | 0.662053161 | 0.803364895 | HYPK | protein_coding | 49.58 | -0.008504245 | 0.008504245 | ENSG00000242028 | n.s. |
| ENSG00000132254 | 547.2594909 | -0.104806432 | 0.2766497 | -0.378842 | 0.704805422 | 0.829852506 | ARFIP2 | protein_coding | 52.8 | -0.008489226 | 0.008489226 | ENSG00000132254 | n.s. |
| ENSG00000214338 | 842.1676877 | 0.086270405 | 0.1923908 | 0.448412 | 0.653855606 | 0.797310413 | SOGA3 | protein_coding | 39.44 | 0.008486641 | 0.008486641 | ENSG00000214338 | n.s. |
| ENSG00000115694 | 1671.230275 | 0.104423429 | 0.2749727 | 0.379759 | 0.704124159 | 0.829429483 | STK25 | protein_coding | 59.31 | 0.008481326 | 0.008481326 | ENSG00000115694 | n.s. |
| ENSG00000154723 | 1294.56332 | 0.110382583 | 0.3043588 | 0.362673 | 0.716849524 | 0.837856033 | ATP5J | protein_coding | 41.96 | 0.00848076 | 0.00848076 | ENSG00000154723 | n.s. |
| ENSG00000198088 | 85.48020375 | 0.103798963 | 0.272137 | 0.381422 | 0.702890336 | 0.828718955 | NUP62CL | protein_coding | 37.06 | 0.00846924 | 0.00846924 | ENSG00000198088 | n.s. |
| ENSG00000063854 | 440.626102 | 0.112902692 | 0.318534 | 0.354445 | 0.723005658 | 0.841391489 | HAGH | protein_coding | 53.33 | 0.008467915 | 0.008467915 | ENSG00000063854 | n.s. |
| ENSG00000273742 | 4.858475746 | -0.14266366 | 0.4902605 | -0.290996 | 0.771054653 | 0.872370708 | MIR6075 | miRNA | 64.21 | -0.008459802 | 0.008459802 | ENSG00000273742 | n.s. |
| ENSG00000170802 | 1178.456769 | -0.09810521 | 0.2449512 | -0.400509 | 0.688781481 | 0.820294688 | FOXN2 | protein_coding | 36.31 | -0.008440001 | 0.008440001 | ENSG00000170802 | n.s. |
| ENSG00000126970 | 347.6152328 | -0.109789397 | 0.3029727 | -0.362374 | 0.717072631 | 0.837918778 | ZC4H2 | protein_coding | 39.24 | -0.008431615 | 0.008431615 | ENSG00000126970 | n.s. |
| ENSG00000100380 | 4603.335284 | 0.107832424 | 0.2934524 | 0.367461 | 0.71327492 | 0.8354878 | ST13 | protein_coding | 38.45 | 0.008417387 | 0.008417387 | ENSG00000100380 | n.s. |
| ENSG00000104823 | 268.5115426 | 0.111935044 | 0.3151165 | 0.355218 | 0.72242632 | 0.841187406 | ECH1 | protein_coding | 51.47 | 0.008407132 | 0.008407132 | ENSG00000104823 | n.s. |
| ENSG00000196363 | 1012.131189 | -0.103000596 | 0.2700384 | -0.381429 | 0.702884643 | 0.828718955 | WDR5 | protein_coding | 55.32 | -0.0084041 | 0.0084041 | ENSG00000196363 | n.s. |
| ENSG00000184220 | 1115.224379 | -0.106636328 | 0.2883325 | -0.369838 | 0.711503178 | 0.834542721 | CMSS1 | protein_coding | 38.77 | -0.008376436 | 0.008376436 | ENSG00000184220 | n.s. |
| ENSG00000084754 | 2585.216508 | 0.106245619 | 0.2864513 | 0.370903 | 0.71070983 | 0.834053688 | HADHA | protein_coding | 41.93 | 0.008372792 | 0.008372792 | ENSG00000084754 | n.s. |
| ENSG00000189221 | 208.8568763 | 0.100878117 | 0.260575 | 0.387137 | 0.698655077 | 0.826664873 | MAOA | protein_coding | 40.27 | 0.008339646 | 0.008339646 | ENSG00000189221 | n.s. |
| ENSG00000213160 | 2992.887319 | -0.10193724 | 0.2670111 | -0.381772 | 0.70263078 | 0.828553261 | KLHL23 | protein_coding | 42.29 | -0.00832619 | 0.00832619 | ENSG00000213160 | n.s. |
| ENSG00000177426 | 622.1289558 | -0.09090387 | 0.2157228 | -0.421392 | 0.673468824 | 0.809935045 | TGIF1 | protein_coding | 43.85 | -0.008322232 | 0.008322232 | ENSG00000177426 | n.s. |
| ENSG00000136819 | 808.5561196 | 0.096892488 | 0.2426782 | 0.399263 | 0.689699233 | 0.820601814 | C9orf78 | protein_coding | 48.01 | 0.008319918 | 0.008319918 | ENSG00000136819 | n.s. |
| ENSG00000163626 | 325.9854449 | -0.092535329 | 0.2229527 | -0.415045 | 0.678109136 | 0.813086294 | COX18 | protein_coding | 40.47 | -0.008315536 | 0.008315536 | ENSG00000163626 | n.s. |
| ENSG00000178988 | 2487.678542 | -0.097377568 | 0.2451726 | -0.39718 | 0.691234983 | 0.82152815 | MRFAP1L1 | protein_coding | 44.91 | -0.008313858 | 0.008313858 | ENSG00000178988 | n.s. |
| ENSG00000116752 | 1760.465292 | -0.104817887 | 0.2817301 | -0.372051 | 0.70985512 | 0.833559744 | BCAS2 | protein_coding | 40.33 | -0.008287245 | 0.008287245 | ENSG00000116752 | n.s. |
| ENSG00000143093 | 1543.539388 | 0.096825244 | 0.2433264 | 0.397923 | 0.690686722 | 0.821195719 | STRIP1 | protein_coding | 51.41 | 0.008283721 | 0.008283721 | ENSG00000143093 | n.s. |
| ENSG00000266967 | 1295.375046 | -0.10019778 | 0.2597706 | -0.385716 | 0.69970671 | 0.82681669 | AARSD1 | protein_coding | 48.21 | -0.008275411 | 0.008275411 | ENSG00000266967 | n.s. |

| | | | | | | | | | | | | | |
|---|---|---|---|---|---|---|---|---|---|---|---|---|---|
| ENSG00000168298 | 2845.832995 | 0.088929128 | 0.2079213 | 0.427706 | 0.668865344 | 0.807148558 | HIST1H1E | protein_coding | 60.08 | 0.008274546 | 0.008274546 | ENSG00000168298 | n.s. |
| ENSG00000171877 | 188.6525722 | -0.094834605 | 0.2343782 | -0.404622 | 0.685755346 | 0.818216786 | FRMD5 | protein_coding | 41.23 | -0.008263092 | 0.008263092 | ENSG00000171877 | n.s. |
| ENSG00000200120 | 74.84396933 | -0.11633844 | 0.3422303 | -0.339942 | 0.733900222 | 0.849243084 | | misc_RNA | 42.16 | -0.008256304 | 0.008256304 | ENSG00000200120 | n.s. |
| ENSG00000199150 | 12.63299993 | 0.122786536 | 0.377566 | 0.325205 | 0.745025563 | 0.856564072 | MIRLET7G | miRNA | 50 | 0.008256185 | 0.008256185 | ENSG00000199150 | n.s. |
| ENSG00000235238 | 53.12249183 | 0.112433679 | 0.3227494 | 0.348362 | 0.727568209 | 0.844758501 | SUMO2P1 | processed_pseudogene | 45.42 | 0.008237727 | 0.008237727 | ENSG00000235238 | n.s. |
| ENSG00000140931 | 188.0523187 | 0.099645811 | 0.2580238 | 0.386188 | 0.69935712 | 0.826771317 | CMTM3 | protein_coding | 58.84 | 0.008232199 | 0.008232199 | ENSG00000140931 | n.s. |
| ENSG00000196998 | 279.4075119 | 0.107886587 | 0.2994992 | 0.360223 | 0.718680163 | 0.838952285 | WDR45 | protein_coding | 50.92 | 0.008227727 | 0.008227727 | ENSG00000196998 | n.s. |
| ENSG00000169504 | 2152.760181 | 0.109174708 | 0.3066317 | 0.356045 | 0.721806771 | 0.840962945 | CLIC4 | protein_coding | 41.67 | 0.008212464 | 0.008212464 | ENSG00000169504 | n.s. |
| ENSG00000152952 | 874.3619438 | 0.1045365 | 0.2827604 | 0.3697 | 0.711606061 | 0.834542721 | PLOD2 | protein_coding | 36.22 | 0.008211491 | 0.008211491 | ENSG00000152952 | n.s. |
| ENSG00000164237 | 289.1348547 | 0.104552444 | 0.2834226 | 0.368892 | 0.712207934 | 0.83487542 | CMBL | protein_coding | 46.43 | 0.008194645 | 0.008194645 | ENSG00000164237 | n.s. |
| ENSG00000104980 | 674.0986434 | -0.099374156 | 0.2584169 | -0.38455 | 0.700570971 | 0.827236085 | TIMM44 | protein_coding | 56.41 | -0.008185502 | 0.008185502 | ENSG00000104980 | n.s. |
| ENSG00000168374 | 1276.118637 | 0.101699166 | 0.2695891 | 0.377238 | 0.705996954 | 0.830833876 | ARF4 | protein_coding | 41.71 | 0.008185339 | 0.008185339 | ENSG00000168374 | n.s. |
| ENSG00000181045 | 139.9948556 | -0.100972388 | 0.2665883 | -0.378758 | 0.70486771 | 0.829855667 | SLC26A11 | protein_coding | 53.49 | -0.008178504 | 0.008178504 | ENSG00000181045 | n.s. |
| ENSG00000041357 | 4396.902805 | 0.109820476 | 0.311687 | 0.352342 | 0.72458163 | 0.842760323 | PSMA4 | protein_coding | 42.32 | 0.008159213 | 0.008159213 | ENSG00000041357 | n.s. |
| ENSG00000169733 | 211.0556689 | 0.103064737 | 0.276667 | 0.372523 | 0.709503762 | 0.833411025 | RFNG | protein_coding | 67.12 | 0.008156621 | 0.008156621 | ENSG00000169733 | n.s. |
| ENSG00000161526 | 2589.091666 | -0.095929405 | 0.2426502 | -0.39534 | 0.692591779 | 0.822216519 | SAP30BP | protein_coding | 48.61 | -0.008155323 | 0.008155323 | ENSG00000161526 | n.s. |
| ENSG00000101439 | 242.5747813 | 0.102811958 | 0.2759293 | 0.372602 | 0.709444315 | 0.833411025 | CST3 | protein_coding | 51.65 | 0.008136616 | 0.008136616 | ENSG00000101439 | n.s. |
| ENSG00000196810 | 241.6356759 | -0.09072231 | 0.219252 | -0.413781 | 0.679034535 | 0.813603737 | CTBP1-AS2 | antisense | 46.78 | -0.008127546 | 0.008127546 | ENSG00000196810 | n.s. |
| ENSG00000244332 | 60.60137992 | -0.102953179 | 0.2773913 | -0.371148 | 0.710527403 | 0.833909891 | | sense_overlapping | 37.13 | -0.008121037 | 0.008121037 | ENSG00000244332 | n.s. |
| ENSG00000164162 | 537.7515847 | -0.097944921 | 0.2524569 | -0.387967 | 0.698040566 | 0.826289645 | ANAPC10 | protein_coding | 37.19 | -0.008116469 | 0.008116469 | ENSG00000164162 | n.s. |
| ENSG00000131748 | 226.77519 | 0.107874718 | 0.3029432 | 0.356089 | 0.721773933 | 0.840962945 | STARD3 | protein_coding | 54.9 | 0.008114675 | 0.008114675 | ENSG00000131748 | n.s. |
| ENSG00000116459 | 4131.470868 | 0.10806546 | 0.3046045 | 0.354773 | 0.722759641 | 0.841339606 | ATP5F1 | protein_coding | 39.38 | 0.008108007 | 0.008108007 | ENSG00000116459 | n.s. |
| ENSG00000214941 | 193.4272509 | -0.114772486 | 0.3397532 | -0.337811 | 0.735505322 | 0.849970169 | ZSWIM7 | protein_coding | 44.09 | -0.008102515 | 0.008102515 | ENSG00000214941 | n.s. |
| ENSG00000184216 | 1096.007823 | -0.098678245 | 0.2574264 | -0.383326 | 0.701478045 | 0.827940174 | IRAK1 | protein_coding | 61.78 | -0.008091719 | 0.008091719 | ENSG00000184216 | n.s. |
| ENSG00000108963 | 241.3565851 | 0.102311027 | 0.2750068 | 0.372031 | 0.709869847 | 0.833559744 | DPH1 | protein_coding | 56.54 | 0.008089044 | 0.008089044 | ENSG00000108963 | n.s. |
| ENSG00000065427 | 2885.675867 | 0.105903002 | 0.293783 | 0.36048 | 0.718487962 | 0.838798248 | KARS | protein_coding | 45.57 | 0.008084898 | 0.008084898 | ENSG00000065427 | n.s. |
| ENSG00000154096 | 575.892355 | -0.099183785 | 0.2604804 | -0.380773 | 0.703371991 | 0.828935858 | THY1 | protein_coding | 55.78 | -0.008081403 | 0.008081403 | ENSG00000154096 | n.s. |
| ENSG00000128245 | 1747.338207 | -0.10039349 | 0.2662013 | -0.377134 | 0.706074249 | 0.830854612 | YWHAH | protein_coding | 45.04 | -0.008079163 | 0.008079163 | ENSG00000128245 | n.s. |
| ENSG00000041353 | 377.3436694 | -0.109743028 | 0.3141712 | -0.34931 | 0.72685687 | 0.844391172 | RAB27B | protein_coding | 37.74 | -0.008061318 | 0.008061318 | ENSG00000041353 | n.s. |
| ENSG00000125826 | 171.8288033 | -0.103701771 | 0.2833012 | -0.366048 | 0.714329452 | 0.836116769 | RBCK1 | protein_coding | 51.12 | -0.008061057 | 0.008061057 | ENSG00000125826 | n.s. |
| ENSG00000137726 | 1973.750891 | -0.101362225 | 0.2708996 | -0.374169 | 0.708278583 | 0.832744698 | FXYD6 | protein_coding | 48.09 | -0.008057093 | 0.008057093 | ENSG00000137726 | n.s. |
| ENSG00000132436 | 1872.030523 | -0.094310455 | 0.2378651 | -0.396487 | 0.691745694 | 0.821656665 | FIGNL1 | protein_coding | 43.84 | -0.008045589 | 0.008045589 | ENSG00000132436 | n.s. |

| | | | | | | | | | | | | | |
|---|---|---|---|---|---|---|---|---|---|---|---|---|---|
| ENSG00000164086 | 263.8044859 | -0.08751899 | 0.2067218 | -0.423366 | 0.672028256 | 0.809258728 | DUSP7 | protein_coding | 59.41 | -0.008044099 | 0.008044099 | ENSG00000164086 | n.s. |
| ENSG00000079459 | 2477.985224 | 0.101668269 | 0.2732661 | 0.372049 | 0.709856623 | 0.833559744 | FDFT1 | protein_coding | 44.36 | 0.008038226 | 0.008038226 | ENSG00000079459 | n.s. |
| ENSG00000133119 | 3529.41791 | 0.09854216 | 0.2584489 | 0.381283 | 0.702993315 | 0.828770189 | RFC3 | protein_coding | 37.82 | 0.008037678 | 0.008037678 | ENSG00000133119 | n.s. |
| ENSG00000115661 | 406.4186043 | 0.092326133 | 0.2287872 | 0.403546 | 0.68654671 | 0.818643628 | STK16 | protein_coding | 53.12 | 0.008023613 | 0.008023613 | ENSG00000115661 | n.s. |
| ENSG00000099246 | 1295.584585 | -0.100061122 | 0.2664316 | -0.37556 | 0.707243852 | 0.831879374 | RAB18 | protein_coding | 36.66 | -0.00799885 | 0.00799885 | ENSG00000099246 | n.s. |
| ENSG00000069956 | 3159.180499 | 0.101276138 | 0.2726718 | 0.371421 | 0.710323707 | 0.833786049 | MAPK6 | protein_coding | 43.14 | 0.007995283 | 0.007995283 | ENSG00000069956 | n.s. |
| ENSG00000067596 | 3187.186592 | 0.095383115 | 0.2440641 | 0.390812 | 0.695936405 | 0.824640218 | DHX8 | protein_coding | 46.36 | 0.007986952 | 0.007986952 | ENSG00000067596 | n.s. |
| ENSG00000204843 | 3992.26676 | -0.095652539 | 0.2452554 | -0.390012 | 0.696527692 | 0.825130184 | DCTN1 | protein_coding | 49.96 | -0.007984837 | 0.007984837 | ENSG00000204843 | n.s. |
| ENSG00000200086 | 30.44944715 | 0.112982636 | 0.3358979 | 0.33636 | 0.736599341 | 0.850538274 | RNU6-433P | snRNA | 36.54 | 0.007943373 | 0.007943373 | ENSG00000200086 | n.s. |
| ENSG00000117479 | 650.9151904 | -0.099037948 | 0.2632658 | -0.37619 | 0.706775653 | 0.831398903 | SLC19A2 | protein_coding | 38.72 | -0.007941908 | 0.007941908 | ENSG00000117479 | n.s. |
| ENSG00000239857 | 375.4283178 | 0.105044179 | 0.2940346 | 0.357251 | 0.720903836 | 0.840561437 | GET4 | protein_coding | 59.75 | 0.007923539 | 0.007923539 | ENSG00000239857 | n.s. |
| ENSG00000182307 | 1523.336262 | -0.095805209 | 0.247558 | -0.387001 | 0.698755453 | 0.826664873 | C8orf33 | protein_coding | 51.6 | -0.007920266 | 0.007920266 | ENSG00000182307 | n.s. |
| ENSG00000165533 | 1993.068903 | 0.096300725 | 0.251363 | 0.383114 | 0.701635144 | 0.827940174 | TTC8 | protein_coding | 36.61 | 0.00789676 | 0.00789676 | ENSG00000165533 | n.s. |
| ENSG00000162630 | 350.2222259 | -0.090354489 | 0.222708 | -0.405708 | 0.684956977 | 0.817754758 | B3GALT2 | protein_coding | 32.61 | -0.007894897 | 0.007894897 | ENSG00000162630 | n.s. |
| ENSG00000164068 | 323.2823501 | 0.095494564 | 0.2473 | 0.386149 | 0.699386584 | 0.826771317 | RNF123 | protein_coding | 56.42 | 0.007889245 | 0.007889245 | ENSG00000164068 | n.s. |
| ENSG00000100348 | 307.6182457 | -0.100324788 | 0.2713245 | -0.369759 | 0.711561794 | 0.834542721 | TXN2 | protein_coding | 51.41 | -0.007880655 | 0.007880655 | ENSG00000100348 | n.s. |
| ENSG00000137171 | 203.5687722 | 0.101906056 | 0.2793192 | 0.364837 | 0.7152329 | 0.83689282 | KLC4 | protein_coding | 52 | 0.007880411 | 0.007880411 | ENSG00000137171 | n.s. |
| ENSG00000207559 | 19.27900771 | 0.114917791 | 0.3485984 | 0.329657 | 0.741659407 | 0.854163806 | MIR578 | miRNA | 34.38 | 0.007867138 | 0.007867138 | ENSG00000207559 | n.s. |
| ENSG00000115827 | 365.670378 | 0.090007436 | 0.2219808 | 0.405474 | 0.685129135 | 0.817885622 | DCAF17 | protein_coding | 37.36 | 0.007858318 | 0.007858318 | ENSG00000115827 | n.s. |
| ENSG00000137364 | 181.9220411 | -0.105646204 | 0.2997175 | -0.352486 | 0.72447384 | 0.842747872 | TPMT | protein_coding | 41.46 | -0.00784976 | 0.00784976 | ENSG00000137364 | n.s. |
| ENSG00000196507 | 267.9832332 | -0.100232206 | 0.2724731 | -0.367861 | 0.712976938 | 0.835259438 | TCEAL3 | protein_coding | 41.11 | -0.007836014 | 0.007836014 | ENSG00000196507 | n.s. |
| ENSG00000223803 | 26.27918433 | 0.168034618 | 0.7046617 | 0.238461 | 0.811523247 | 0.898301652 | RPS20P14 | processed_pseudogene | 46.11 | 0.007826683 | 0.007826683 | ENSG00000223803 | n.s. |
| ENSG00000244045 | 962.6797362 | -0.093700895 | 0.2402022 | -0.390092 | 0.696468652 | 0.825130184 | TMEM199 | protein_coding | 47.27 | -0.007821919 | 0.007821919 | ENSG00000244045 | n.s. |
| ENSG00000166908 | 191.7922376 | 0.094908905 | 0.2466956 | 0.384721 | 0.7004444 | 0.827166021 | PIP4K2C | protein_coding | 46.79 | 0.007821188 | 0.007821188 | ENSG00000166908 | n.s. |
| ENSG00000018610 | 459.4819756 | 0.105125213 | 0.298525 | 0.352149 | 0.724726664 | 0.842760323 | CXorf56 | protein_coding | 44.29 | 0.007810374 | 0.007810374 | ENSG00000018610 | n.s. |
| ENSG00000204673 | 309.3823693 | -0.091122687 | 0.2290749 | -0.397786 | 0.690788211 | 0.821195719 | AKT1S1 | protein_coding | 59.67 | -0.007795849 | 0.007795849 | ENSG00000204673 | n.s. |
| ENSG00000279665 | 38.35983011 | -0.108082296 | 0.3143649 | -0.343812 | 0.730987952 | 0.847006221 | | TEC | 46.43 | -0.007794182 | 0.007794182 | ENSG00000279665 | n.s. |
| ENSG00000129460 | 1459.81136 | -0.09560027 | 0.2510056 | -0.380869 | 0.703300397 | 0.828935858 | NGDN | protein_coding | 42.99 | -0.007789421 | 0.007789421 | ENSG00000129460 | n.s. |
| ENSG00000225630 | 42.91138247 | -0.103130191 | 0.2889496 | -0.356914 | 0.721156089 | 0.840644359 | MTND2P28 | unprocessed_pseudogene | 43.1 | -0.007774748 | 0.007774748 | ENSG00000225630 | n.s. |
| ENSG00000088451 | 338.7142209 | -0.10000596 | 0.2731593 | -0.366109 | 0.714284031 | 0.836116769 | TGDS | protein_coding | 36.98 | -0.00777377 | 0.00777377 | ENSG00000088451 | n.s. |
| ENSG00000207980 | 4.081512233 | -0.189508832 | 0.8839128 | -0.214398 | 0.830236978 | 0.910044832 | MIR23A | miRNA | 57.53 | -0.007757963 | 0.007757963 | ENSG00000207980 | n.s. |
| ENSG00000260032 | 5501.56765 | 0.091708682 | 0.2328716 | 0.393816 | 0.693716549 | 0.823135405 | NORAD | lincRNA | 41.56 | 0.007752017 | 0.007752017 | ENSG00000260032 | n.s. |

| | | | | | | | | | | | | | |
|---|---|---|---|---|---|---|---|---|---|---|---|---|---|
| ENSG00000189376 | 193.3193805 | -0.098797513 | 0.2678626 | -0.368837 | 0.712249532 | 0.83487542 | C8orf76 | protein_coding | 43.75 | -0.007743584 | 0.007743584 | ENSG00000189376 | n.s. |
| ENSG00000221740 | 32.18038369 | 0.103148233 | 0.2909494 | 0.354523 | 0.722947042 | 0.841391489 | SNORD93 | snoRNA | 41.89 | 0.007736312 | 0.007736312 | ENSG00000221740 | n.s. |
| ENSG00000182473 | 2758.557501 | -0.099408401 | 0.2716668 | -0.36592 | 0.714424584 | 0.836135655 | EXOC7 | protein_coding | 51.59 | -0.007726345 | 0.007726345 | ENSG00000182473 | n.s. |
| ENSG00000103811 | 67.48030841 | 0.11267793 | 0.3419955 | 0.329472 | 0.741798982 | 0.85418624 | CTSH | protein_coding | 52.42 | 0.007712515 | 0.007712515 | ENSG00000103811 | n.s. |
| ENSG00000225067 | 19.01914485 | 0.121474739 | 0.3921808 | 0.309742 | 0.756757432 | 0.864058859 | RPL23AP2 | processed_pseudogene | 48.93 | 0.007708383 | 0.007708383 | ENSG00000225067 | n.s. |
| ENSG00000256673 | 48.07382981 | 0.102508467 | 0.2884055 | 0.355432 | 0.72226617 | 0.841093332 | | unprocessed_pseudogene | 51.28 | 0.007704107 | 0.007704107 | ENSG00000256673 | n.s. |
| ENSG00000163093 | 288.9841274 | -0.092619438 | 0.2382239 | -0.388792 | 0.697430314 | 0.825797901 | BBS5 | protein_coding | 40.4 | -0.007699104 | 0.007699104 | ENSG00000163093 | n.s. |
| ENSG00000163257 | 2459.382157 | -0.092206781 | 0.2365199 | -0.389848 | 0.696649084 | 0.825203778 | DCAF16 | protein_coding | 39.37 | -0.007693622 | 0.007693622 | ENSG00000163257 | n.s. |
| ENSG00000163322 | 126.6067295 | 0.10345804 | 0.2937037 | 0.352253 | 0.724648428 | 0.842760323 | FAM175A | protein_coding | 42.23 | 0.00768651 | 0.00768651 | ENSG00000163322 | n.s. |
| ENSG00000161202 | 947.1450107 | -0.094496145 | 0.2484394 | -0.380359 | 0.703678951 | 0.829219094 | DVL3 | protein_coding | 52.91 | -0.007685438 | 0.007685438 | ENSG00000161202 | n.s. |
| ENSG00000117859 | 2425.838114 | -0.097877188 | 0.2649636 | -0.369399 | 0.711830582 | 0.83466542 | OSBPL9 | protein_coding | 40.08 | -0.007682144 | 0.007682144 | ENSG00000117859 | n.s. |
| ENSG00000234022 | 29.28359212 | -0.105282476 | 0.3034482 | -0.346954 | 0.728626076 | 0.84545886 | | antisense | 39.59 | -0.007675884 | 0.007675884 | ENSG00000234022 | n.s. |
| ENSG00000072832 | 4528.155202 | 0.099950017 | 0.2760737 | 0.362041 | 0.717321346 | 0.837976355 | CRMP1 | protein_coding | 48.98 | 0.007672986 | 0.007672986 | ENSG00000072832 | n.s. |
| ENSG00000100124 | 361.5051681 | 0.089642096 | 0.2251376 | 0.398166 | 0.69050791 | 0.821195719 | ANKRD54 | protein_coding | 50.74 | 0.007669179 | 0.007669179 | ENSG00000100124 | n.s. |
| ENSG00000131981 | 48.56034966 | 0.113311287 | 0.3466354 | 0.326889 | 0.743751879 | 0.855753868 | LGALS3 | protein_coding | 44.62 | 0.007665637 | 0.007665637 | ENSG00000131981 | n.s. |
| ENSG00000123154 | 225.5647486 | 0.096967357 | 0.2608258 | 0.371771 | 0.710063677 | 0.833679523 | WDR83 | protein_coding | 55.04 | 0.007660505 | 0.007660505 | ENSG00000123154 | n.s. |
| ENSG00000122008 | 251.828371 | -0.092704468 | 0.2399033 | -0.386424 | 0.699182444 | 0.826760396 | POLK | protein_coding | 36.45 | -0.007659274 | 0.007659274 | ENSG00000122008 | n.s. |
| ENSG00000162434 | 1438.943591 | 0.088499657 | 0.2201369 | 0.402021 | 0.687668608 | 0.819376126 | JAK1 | protein_coding | 42.69 | 0.007656698 | 0.007656698 | ENSG00000162434 | n.s. |
| ENSG00000092098 | 97.37579605 | -0.093398148 | 0.2440494 | -0.382702 | 0.701940902 | 0.828019258 | RNF31 | protein_coding | 52.92 | -0.007654871 | 0.007654871 | ENSG00000092098 | n.s. |
| ENSG00000154059 | 1102.42386 | 0.096703202 | 0.2593725 | 0.372835 | 0.709271101 | 0.833411025 | IMPACT | protein_coding | 38.35 | 0.007653165 | 0.007653165 | ENSG00000154059 | n.s. |
| ENSG00000177675 | 73.37811075 | -0.099884859 | 0.276716 | -0.360965 | 0.718125399 | 0.838532995 | CD163L1 | protein_coding | 36.25 | -0.007639178 | 0.007639178 | ENSG00000177675 | n.s. |
| ENSG00000099864 | 71.20119002 | -0.099863178 | 0.2766712 | -0.360945 | 0.718140302 | 0.838532995 | PALM | protein_coding | 59.66 | -0.00763752 | 0.00763752 | ENSG00000099864 | n.s. |
| ENSG00000254876 | 116.2924399 | 0.092166767 | 0.2378842 | 0.387444 | 0.698427694 | 0.826607343 | | processed_transcript | 40.69 | 0.00762226 | 0.00762226 | ENSG00000254876 | n.s. |
| ENSG00000149483 | 927.3166757 | -0.082057427 | 0.1923213 | -0.426668 | 0.669620886 | 0.807759127 | TMEM138 | protein_coding | 48.08 | -0.007608211 | 0.007608211 | ENSG00000149483 | n.s. |
| ENSG00000086475 | 1859.135507 | -0.098628673 | 0.2711268 | -0.363773 | 0.71602737 | 0.837540879 | SEPHS1 | protein_coding | 45.78 | -0.007593815 | 0.007593815 | ENSG00000086475 | n.s. |
| ENSG00000261013 | 28.76769635 | -0.107660851 | 0.3193829 | -0.33709 | 0.73604888 | 0.850174926 | | lincRNA | 36.34 | -0.007589197 | 0.007589197 | ENSG00000261013 | n.s. |
| ENSG00000241494 | 17.79626908 | 0.16494169 | 0.7022747 | 0.234868 | 0.814311358 | 0.899528486 | | processed_pseudogene | 43.84 | 0.007584857 | 0.007584857 | ENSG00000241494 | n.s. |
| ENSG00000143390 | 1436.274926 | -0.097391957 | 0.2656371 | -0.366635 | 0.713891022 | 0.835843717 | RFX5 | protein_coding | 49.78 | -0.007584391 | 0.007584391 | ENSG00000143390 | n.s. |
| ENSG00000053372 | 653.1839328 | 0.096902674 | 0.2631529 | 0.368237 | 0.71269637 | 0.835120738 | MRTO4 | protein_coding | 53.49 | 0.007582705 | 0.007582705 | ENSG00000053372 | n.s. |
| ENSG00000163466 | 2173.812094 | -0.099749197 | 0.2780119 | -0.358795 | 0.719748712 | 0.839889496 | ARPC2 | protein_coding | 44.32 | -0.00755878 | 0.00755878 | ENSG00000163466 | n.s. |
| ENSG00000126767 | 271.0568488 | -0.102047267 | 0.2911467 | -0.350501 | 0.725962597 | 0.843774709 | ELK1 | protein_coding | 49.22 | -0.007528384 | 0.007528384 | ENSG00000126767 | n.s. |
| ENSG00000102038 | 1698.440246 | -0.098751504 | 0.2743934 | -0.35989 | 0.718929197 | 0.839172637 | SMARCA1 | protein_coding | 36.87 | -0.007519797 | 0.007519797 | ENSG00000102038 | n.s. |

| | | | | | | | | | | | | | |
|---|---|---|---|---|---|---|---|---|---|---|---|---|---|
| ENSG00000279744 | 121.8039847 | 0.09108615 | 0.2364531 | 0.385219 | 0.700075384 | 0.826977403 | | TEC | 54.26 | 0.007515186 | 0.007515186 | ENSG00000279744 | n.s. |
| ENSG00000168118 | 1385.840655 | 0.103250776 | 0.2982221 | 0.346221 | 0.729176602 | 0.845815699 | RAB4A | protein_coding | 41.32 | 0.007508836 | 0.007508836 | ENSG00000168118 | n.s. |
| ENSG00000118242 | 403.1273629 | 0.098769533 | 0.2747255 | 0.359521 | 0.719205508 | 0.839424787 | MREG | protein_coding | 42.57 | 0.007508283 | 0.007508283 | ENSG00000118242 | n.s. |
| ENSG00000206687 | 15.49183477 | 0.115592109 | 0.3660248 | 0.315804 | 0.752151252 | 0.861129996 | RNU1-109P | snRNA | 48.98 | 0.007505544 | 0.007505544 | ENSG00000206687 | n.s. |
| ENSG00000165512 | 2384.690061 | 0.095941178 | 0.2607898 | 0.367887 | 0.712957497 | 0.835259438 | ZNF22 | protein_coding | 41.86 | 0.007500548 | 0.007500548 | ENSG00000165512 | n.s. |
| ENSG00000169760 | 29.87562544 | -0.10666861 | 0.3172087 | -0.336273 | 0.736665267 | 0.850538274 | NLGN1 | protein_coding | 35.49 | -0.007499458 | 0.007499458 | ENSG00000169760 | n.s. |
| ENSG00000196151 | 157.7902916 | 0.096162012 | 0.2619679 | 0.367076 | 0.713562682 | 0.835640779 | WDSUB1 | protein_coding | 39.11 | 0.00749875 | 0.00749875 | ENSG00000196151 | n.s. |
| ENSG00000187736 | 587.4084179 | -0.095647468 | 0.2593886 | -0.368742 | 0.712320035 | 0.834887773 | NHEJ1 | protein_coding | 43.12 | -0.007496074 | 0.007496074 | ENSG00000187736 | n.s. |
| ENSG00000064692 | 27.77995209 | -0.105864604 | 0.3126759 | -0.338576 | 0.734929087 | 0.849656869 | SNCAIP | protein_coding | 39.26 | -0.007490601 | 0.007490601 | ENSG00000064692 | n.s. |
| ENSG00000100426 | 1269.125673 | -0.094704037 | 0.2557038 | -0.370366 | 0.71110969 | 0.83445261 | ZBED4 | protein_coding | 47.49 | -0.007443578 | 0.007443578 | ENSG00000100426 | n.s. |
| ENSG00000041988 | 104.8206773 | 0.109890564 | 0.336254 | 0.326808 | 0.74381296 | 0.855753868 | THAP3 | protein_coding | 56.77 | 0.007434221 | 0.007434221 | ENSG00000041988 | n.s. |
| ENSG00000264314 | 12.80469847 | 0.11884309 | 0.3893301 | 0.30525 | 0.760175626 | 0.865917832 | MIR548AT | miRNA | 39.66 | 0.007430464 | 0.007430464 | ENSG00000264314 | n.s. |
| ENSG00000276368 | 1056.158708 | 0.08249516 | 0.198318 | 0.415974 | 0.677428902 | 0.812781842 | HIST1H2AJ | protein_coding | 62.53 | 0.00742671 | 0.00742671 | ENSG00000276368 | n.s. |
| ENSG00000175602 | 292.6786185 | -0.094802355 | 0.2574514 | -0.368234 | 0.712698743 | 0.835120738 | CCDC85B | protein_coding | 67.65 | -0.007418354 | 0.007418354 | ENSG00000175602 | n.s. |
| ENSG00000266412 | 2203.010472 | -0.099456329 | 0.2814565 | -0.353363 | 0.723816247 | 0.842194017 | NCOA4 | protein_coding | 41.24 | -0.007418234 | 0.007418234 | ENSG00000266412 | n.s. |
| ENSG00000186787 | 66.26174911 | -0.097641565 | 0.271977 | -0.359007 | 0.719590059 | 0.839803218 | SPIN2B | protein_coding | 51.44 | -0.007403424 | 0.007403424 | ENSG00000186787 | n.s. |
| ENSG00000185246 | 1668.725265 | -0.096151321 | 0.2647815 | -0.363135 | 0.716504303 | 0.837676498 | PRPF39 | protein_coding | 36.34 | -0.007396312 | 0.007396312 | ENSG00000185246 | n.s. |
| ENSG00000188529 | 6815.245327 | 0.09010429 | 0.2352923 | 0.382946 | 0.701759635 | 0.827946733 | SRSF10 | protein_coding | 39.02 | 0.007388336 | 0.007388336 | ENSG00000188529 | n.s. |
| ENSG00000010244 | 9037.056024 | 0.094012284 | 0.2543816 | 0.369572 | 0.711701539 | 0.834584395 | ZNF207 | protein_coding | 38.47 | 0.00738276 | 0.00738276 | ENSG00000010244 | n.s. |
| ENSG00000205571 | 65.83928242 | 0.104274557 | 0.307718 | 0.338864 | 0.734712154 | 0.849656869 | SMN2 | protein_coding | 42.22 | 0.007378095 | 0.007378095 | ENSG00000205571 | n.s. |
| ENSG00000187555 | 3875.849327 | 0.089963606 | 0.2347818 | 0.38318 | 0.701586532 | 0.827940174 | USP7 | protein_coding | 46.24 | 0.00737711 | 0.00737711 | ENSG00000187555 | n.s. |
| ENSG00000113716 | 1185.648789 | -0.089185324 | 0.2303496 | -0.387174 | 0.698627513 | 0.826664873 | HMGXB3 | protein_coding | 43.45 | -0.007372997 | 0.007372997 | ENSG00000113716 | n.s. |
| ENSG00000239779 | 517.7449367 | 0.102932358 | 0.3007976 | 0.342198 | 0.732201848 | 0.847982559 | WBP1 | protein_coding | 56.07 | 0.007371303 | 0.007371303 | ENSG00000239779 | n.s. |
| ENSG00000163001 | 605.8560437 | -0.100213355 | 0.2876247 | -0.348417 | 0.727527012 | 0.844758501 | CFAP36 | protein_coding | 40.44 | -0.007342375 | 0.007342375 | ENSG00000163001 | n.s. |
| ENSG00000153786 | 419.581386 | -0.095381698 | 0.2625509 | -0.363288 | 0.716389444 | 0.837676498 | ZDHHC7 | protein_coding | 48.57 | -0.00733711 | 0.00733711 | ENSG00000153786 | n.s. |
| ENSG00000116212 | 788.5130639 | -0.10224039 | 0.2994175 | -0.341464 | 0.732754068 | 0.848256103 | LRRC42 | protein_coding | 43.06 | -0.007307428 | 0.007307428 | ENSG00000116212 | n.s. |
| ENSG00000152556 | 2615.390642 | 0.098434841 | 0.2801084 | 0.351417 | 0.725275599 | 0.843052803 | PFKM | protein_coding | 44.84 | 0.007298473 | 0.007298473 | ENSG00000152556 | n.s. |
| ENSG00000229235 | 17.08413701 | 0.113156749 | 0.3601508 | 0.314193 | 0.753374663 | 0.86210532 | RPL37P18 | processed_pseudogene | 56.79 | 0.007291784 | 0.007291784 | ENSG00000229235 | n.s. |
| ENSG00000164164 | 1705.266575 | 0.089669374 | 0.2361989 | 0.379635 | 0.704216348 | 0.829429483 | OTUD4 | protein_coding | 37.41 | 0.007282994 | 0.007282994 | ENSG00000164164 | n.s. |
| ENSG00000135966 | 802.7392451 | 0.089104565 | 0.2333539 | 0.381843 | 0.702577776 | 0.828553261 | TGFBRAP1 | protein_coding | 44.1 | 0.007278022 | 0.007278022 | ENSG00000135966 | n.s. |
| ENSG00000233864 | 578.4095272 | -0.098098786 | 0.2790251 | -0.351577 | 0.725155546 | 0.843047817 | TTTY15 | lincRNA | 39.24 | -0.007273808 | 0.007273808 | ENSG00000233864 | n.s. |
| ENSG00000236618 | 134.0437727 | 0.103276858 | 0.306659 | 0.336781 | 0.736282146 | 0.850303278 | PITPNA-AS1 | antisense | 51.63 | 0.007273391 | 0.007273391 | ENSG00000236618 | n.s. |

| | | | | | | | | | | | | | |
|---|---|---|---|---|---|---|---|---|---|---|---|---|---|
| ENSG00000165868 | 648.6344126 | -0.100687748 | 0.2934568 | -0.343109 | 0.73151623 | 0.84732949 | HSPA12A | protein_coding | 47.75 | -0.00724425 | 0.00724425 | ENSG00000165868 | n.s. |
| ENSG00000164615 | 635.3757214 | 0.102956642 | 0.3063221 | 0.336106 | 0.736791074 | 0.850538274 | CAMLG | protein_coding | 43.3 | 0.007238484 | 0.007238484 | ENSG00000164615 | n.s. |
| ENSG00000166478 | 640.8670956 | 0.087147651 | 0.2245564 | 0.388088 | 0.697950938 | 0.826256316 | ZNF143 | protein_coding | 42.79 | 0.007223251 | 0.007223251 | ENSG00000166478 | n.s. |
| ENSG00000160271 | 2860.964097 | 0.091914552 | 0.2485786 | 0.369761 | 0.711560912 | 0.834542721 | RALGDS | protein_coding | 58.92 | 0.007220019 | 0.007220019 | ENSG00000160271 | n.s. |
| ENSG00000230699 | 166.5764319 | -0.091850931 | 0.2481484 | -0.370145 | 0.711274347 | 0.834542721 | | lincRNA | 58.54 | -0.007215022 | 0.007215022 | ENSG00000230699 | n.s. |
| ENSG00000166261 | 230.5110611 | 0.085497288 | 0.2174981 | 0.393094 | 0.694249696 | 0.82341243 | ZNF202 | protein_coding | 44.55 | 0.007214482 | 0.007214482 | ENSG00000166261 | n.s. |
| ENSG00000101158 | 1618.19446 | 0.095189251 | 0.2660219 | 0.357825 | 0.720474404 | 0.840201451 | NELFCD | protein_coding | 48.57 | 0.007197885 | 0.007197885 | ENSG00000101158 | n.s. |
| ENSG00000018236 | 2831.27564 | -0.101666221 | 0.2997905 | -0.339124 | 0.734516162 | 0.849602768 | CNTN1 | protein_coding | 35.21 | -0.00719635 | 0.00719635 | ENSG00000018236 | n.s. |
| ENSG00000115183 | 945.7776137 | -0.093494358 | 0.2574508 | -0.363154 | 0.716489627 | 0.837676498 | TANC1 | protein_coding | 43.42 | -0.007191929 | 0.007191929 | ENSG00000115183 | n.s. |
| ENSG00000166780 | 459.9526321 | -0.103035016 | 0.3089163 | -0.333537 | 0.73872894 | 0.851716051 | C16orf45 | protein_coding | 45.91 | -0.007182073 | 0.007182073 | ENSG00000166780 | n.s. |
| ENSG00000175416 | 362.4608365 | 0.099459269 | 0.2887421 | 0.344457 | 0.730502521 | 0.846859833 | CLTB | protein_coding | 53.09 | 0.007179812 | 0.007179812 | ENSG00000175416 | n.s. |
| ENSG00000101310 | 1117.713055 | -0.097602243 | 0.2791052 | -0.349697 | 0.726566153 | 0.844264776 | SEC23B | protein_coding | 42.98 | -0.007175847 | 0.007175847 | ENSG00000101310 | n.s. |
| ENSG00000100225 | 2201.465197 | 0.091770872 | 0.2506963 | 0.366064 | 0.714317365 | 0.836116769 | FBXO7 | protein_coding | 38.81 | 0.007133631 | 0.007133631 | ENSG00000100225 | n.s. |
| ENSG00000125821 | 870.8817096 | 0.100811639 | 0.297632 | 0.338712 | 0.734826413 | 0.849656869 | DTD1 | protein_coding | 43.21 | 0.007133071 | 0.007133071 | ENSG00000125821 | n.s. |
| ENSG00000176248 | 804.0327055 | -0.089809313 | 0.2405229 | -0.373392 | 0.7088567 | 0.833254267 | ANAPC2 | protein_coding | 61.65 | -0.007114914 | 0.007114914 | ENSG00000176248 | n.s. |
| ENSG00000271200 | 17.42968743 | 0.10730696 | 0.333369 | 0.321886 | 0.747538757 | 0.858461078 | | lincRNA | 61.12 | 0.00711224 | 0.00711224 | ENSG00000271200 | n.s. |
| ENSG00000128309 | 873.7560583 | -0.102201358 | 0.3070666 | -0.332831 | 0.73926169 | 0.852137523 | MPST | protein_coding | 55.28 | -0.007102004 | 0.007102004 | ENSG00000128309 | n.s. |
| ENSG00000157593 | 368.2999906 | -0.098711468 | 0.2878237 | -0.342958 | 0.731629945 | 0.84739071 | SLC35B2 | protein_coding | 58.08 | -0.007098964 | 0.007098964 | ENSG00000157593 | n.s. |
| ENSG00000057608 | 3918.48732 | 0.099673305 | 0.2927674 | 0.340452 | 0.73351605 | 0.848869085 | GDI2 | protein_coding | 43.74 | 0.007092681 | 0.007092681 | ENSG00000057608 | n.s. |
| ENSG00000212747 | 474.5096221 | -0.08601659 | 0.2235426 | -0.384788 | 0.700394248 | 0.827166021 | RTL8B | protein_coding | 51.18 | -0.007088396 | 0.007088396 | ENSG00000212747 | n.s. |
| ENSG00000099364 | 421.8016838 | -0.096198534 | 0.2756962 | -0.348929 | 0.727142263 | 0.844511321 | FBXL19 | protein_coding | 55.33 | -0.007060446 | 0.007060446 | ENSG00000099364 | n.s. |
| ENSG00000227896 | 118.865078 | 0.086760076 | 0.2281429 | 0.380288 | 0.703731438 | 0.829219094 | | antisense | 47.2 | 0.007056258 | 0.007056258 | ENSG00000227896 | n.s. |
| ENSG00000141503 | 243.3320174 | -0.093716549 | 0.2631516 | -0.356131 | 0.721742153 | 0.840962945 | MINK1 | protein_coding | 51.93 | -0.007049653 | 0.007049653 | ENSG00000141503 | n.s. |
| ENSG00000085741 | 119.492588 | -0.092057367 | 0.2548353 | -0.361243 | 0.717918096 | 0.838414163 | WNT11 | protein_coding | 57.84 | -0.007046199 | 0.007046199 | ENSG00000085741 | n.s. |
| ENSG00000165724 | 739.8861892 | 0.092946238 | 0.2591573 | 0.358648 | 0.719858482 | 0.83989887 | ZMYND19 | protein_coding | 57.96 | 0.007042816 | 0.007042816 | ENSG00000165724 | n.s. |
| ENSG00000259138 | 19.58930236 | -0.105122725 | 0.324218 | -0.324235 | 0.745760361 | 0.857055015 | | antisense | 40.33 | -0.007042308 | 0.007042308 | ENSG00000259138 | n.s. |
| ENSG00000130770 | 469.7847078 | 0.091668181 | 0.2531584 | 0.362098 | 0.717278656 | 0.837976355 | ATPIF1 | protein_coding | 44.37 | 0.007037205 | 0.007037205 | ENSG00000130770 | n.s. |
| ENSG00000133318 | 935.4587874 | 0.097249745 | 0.281861 | 0.345027 | 0.73007385 | 0.846574349 | RTN3 | protein_coding | 42.12 | 0.00703455 | 0.00703455 | ENSG00000133318 | n.s. |
| ENSG00000262766 | 38.01967975 | -0.101450947 | 0.3055044 | -0.332077 | 0.739831185 | 0.852492724 | | sense_intronic | 47.31 | -0.007031496 | 0.007031496 | ENSG00000262766 | n.s. |
| ENSG00000152492 | 2490.446924 | -0.094372406 | 0.2673468 | -0.352996 | 0.7240913 | 0.842443651 | CCDC50 | protein_coding | 36.59 | -0.007026888 | 0.007026888 | ENSG00000152492 | n.s. |
| ENSG00000131351 | 368.8015969 | 0.086048246 | 0.225876 | 0.380954 | 0.703237744 | 0.828935858 | HAUS8 | protein_coding | 49.64 | 0.007011131 | 0.007011131 | ENSG00000131351 | n.s. |
| ENSG00000130821 | 208.1466381 | 0.088527007 | 0.2375969 | 0.372593 | 0.709451147 | 0.833411025 | SLC6A8 | protein_coding | 64.7 | 0.007006094 | 0.007006094 | ENSG00000130821 | n.s. |

| | | | | | | | | | | | | | |
|---|---|---|---|---|---|---|---|---|---|---|---|---|---|
| ENSG00000224451 | 90.45182343 | -0.095657549 | 0.2750059 | -0.347838 | 0.727961758 | 0.84504015 | ATP5F1P1 | processed_pseudogene | 44.31 | -0.006994734 | 0.006994734 | ENSG00000224451 | n.s. |
| ENSG00000109332 | 7264.370977 | 0.093166978 | 0.2625233 | 0.35489 | 0.722671797 | 0.841339606 | UBE2D3 | protein_coding | 38.48 | 0.006990194 | 0.006990194 | ENSG00000109332 | n.s. |
| ENSG00000130309 | 605.926903 | -0.091454542 | 0.2535801 | -0.360653 | 0.718358562 | 0.838717505 | COLGALT1 | protein_coding | 52.86 | -0.00698569 | 0.00698569 | ENSG00000130309 | n.s. |
| ENSG00000117335 | 3183.401583 | 0.096051342 | 0.2774221 | 0.346228 | 0.729171224 | 0.845815699 | CD46 | protein_coding | 38.21 | 0.006985263 | 0.006985263 | ENSG00000117335 | n.s. |
| ENSG00000175581 | 738.0829314 | 0.100489987 | 0.3019445 | 0.332809 | 0.739278111 | 0.852137523 | MRPL48 | protein_coding | 43.35 | 0.00698308 | 0.00698308 | ENSG00000175581 | n.s. |
| ENSG00000206195 | 61.10524166 | -0.093972399 | 0.266805 | -0.352214 | 0.724677957 | 0.842760323 | | processed_transcript | 41.82 | -0.006981766 | 0.006981766 | ENSG00000206195 | n.s. |
| ENSG00000172936 | 278.1668818 | -0.085988185 | 0.2266966 | -0.379309 | 0.704458054 | 0.829513657 | MYD88 | protein_coding | 54.39 | -0.006980216 | 0.006980216 | ENSG00000172936 | n.s. |
| ENSG00000125149 | 369.8760156 | -0.088261369 | 0.2376643 | -0.37137 | 0.710361999 | 0.833786049 | C16orf70 | protein_coding | 47.85 | -0.006967827 | 0.006967827 | ENSG00000125149 | n.s. |
| ENSG00000108064 | 2727.456006 | 0.092590264 | 0.2598799 | 0.356281 | 0.721630175 | 0.840962945 | TFAM | protein_coding | 38.21 | 0.00696493 | 0.00696493 | ENSG00000108064 | n.s. |
| ENSG00000078061 | 327.6353738 | 0.092975416 | 0.2628474 | 0.353724 | 0.723545834 | 0.84194974 | ARAF | protein_coding | 51.23 | 0.00694655 | 0.00694655 | ENSG00000078061 | n.s. |
| ENSG00000131236 | 893.2626337 | 0.096250144 | 0.2799566 | 0.343804 | 0.730993805 | 0.847006221 | CAP1 | protein_coding | 43.27 | 0.006940925 | 0.006940925 | ENSG00000131236 | n.s. |
| ENSG00000101442 | 302.7489227 | -0.092303256 | 0.2594878 | -0.355713 | 0.722055252 | 0.84105689 | ACTR5 | protein_coding | 44.84 | -0.006938863 | 0.006938863 | ENSG00000101442 | n.s. |
| ENSG00000069998 | 790.4968142 | 0.09598126 | 0.2783915 | 0.344771 | 0.730266689 | 0.846674426 | HDHD5 | protein_coding | 50.12 | 0.006937867 | 0.006937867 | ENSG00000069998 | n.s. |
| ENSG00000203618 | 99.69303711 | 0.094034494 | 0.2685096 | 0.350209 | 0.726181791 | 0.84388858 | GP1BB | protein_coding | 70.83 | 0.006931742 | 0.006931742 | ENSG00000203618 | n.s. |
| ENSG00000162076 | 224.6137062 | -0.106018997 | 0.3333791 | -0.318013 | 0.750474801 | 0.860271748 | FLYWCH2 | protein_coding | 51.49 | -0.006929862 | 0.006929862 | ENSG00000162076 | n.s. |
| ENSG00000213977 | 78.3943172 | 0.102335207 | 0.3127282 | 0.327234 | 0.743491131 | 0.855718526 | TAX1BP3 | protein_coding | 57.28 | 0.006924928 | 0.006924928 | ENSG00000213977 | n.s. |
| ENSG00000166681 | 3547.161442 | 0.099157507 | 0.297138 | 0.333709 | 0.738599434 | 0.85163726 | BEX3 | protein_coding | 57.48 | 0.006915775 | 0.006915775 | ENSG00000166681 | n.s. |
| ENSG00000006634 | 2032.594393 | -0.091736713 | 0.2574008 | -0.356396 | 0.721543769 | 0.840955458 | DBF4 | protein_coding | 36.72 | -0.006901078 | 0.006901078 | ENSG00000006634 | n.s. |
| ENSG00000174903 | 454.0746305 | 0.095302481 | 0.2764388 | 0.344751 | 0.730281777 | 0.846674426 | RAB1B | protein_coding | 55.61 | 0.006888802 | 0.006888802 | ENSG00000174903 | n.s. |
| ENSG00000133142 | 806.1131555 | -0.094592947 | 0.2729384 | -0.346573 | 0.728912481 | 0.845718415 | TCEAL4 | protein_coding | 39.32 | -0.006883927 | 0.006883927 | ENSG00000133142 | n.s. |
| ENSG00000207416 | 10.92591244 | -0.120127901 | 0.4263283 | -0.281773 | 0.778117416 | 0.876515273 | | misc_RNA | 37.17 | -0.006876183 | 0.006876183 | ENSG00000207416 | n.s. |
| ENSG00000163728 | 2686.558975 | 0.086374691 | 0.2305989 | 0.374567 | 0.707982654 | 0.832537372 | TTC14 | protein_coding | 35.13 | 0.006875103 | 0.006875103 | ENSG00000163728 | n.s. |
| ENSG00000184672 | 101.7116066 | 0.089476994 | 0.2467603 | 0.362607 | 0.716898579 | 0.837856033 | RALYL | protein_coding | 35.32 | 0.006874571 | 0.006874571 | ENSG00000184672 | n.s. |
| ENSG00000240265 | 80.17796001 | 0.089429891 | 0.2470536 | 0.361986 | 0.71736265 | 0.837976355 | | processed_pseudogene | 60.68 | 0.006865375 | 0.006865375 | ENSG00000240265 | n.s. |
| ENSG00000129255 | 797.369386 | 0.090610021 | 0.2527677 | 0.358472 | 0.719990453 | 0.839918481 | MPDU1 | protein_coding | 56.21 | 0.006864875 | 0.006864875 | ENSG00000129255 | n.s. |
| ENSG00000187189 | 2447.429057 | -0.084885309 | 0.2244074 | -0.378264 | 0.705234263 | 0.830146825 | TSPYL4 | protein_coding | 47.75 | -0.00686256 | 0.00686256 | ENSG00000187189 | n.s. |
| ENSG00000173621 | 415.6668694 | -0.090544251 | 0.2523905 | -0.358747 | 0.719784634 | 0.839889496 | LRFN4 | protein_coding | 70.38 | -0.006861249 | 0.006861249 | ENSG00000173621 | n.s. |
| ENSG00000164241 | 108.4163092 | 0.086832013 | 0.233609 | 0.371698 | 0.71011766 | 0.833679523 | C5orf63 | protein_coding | 39.03 | 0.006859804 | 0.006859804 | ENSG00000164241 | n.s. |
| ENSG00000066379 | 245.0297026 | 0.097516268 | 0.290077 | 0.336174 | 0.736739844 | 0.850538274 | ZNRD1 | protein_coding | 41.39 | 0.006855992 | 0.006855992 | ENSG00000066379 | n.s. |
| ENSG00000158717 | 211.8016743 | 0.101580129 | 0.3118723 | 0.325711 | 0.744643288 | 0.856265982 | RNF166 | protein_coding | 62.53 | 0.006845618 | 0.006845618 | ENSG00000158717 | n.s. |
| ENSG00000129932 | 134.5374974 | -0.090711444 | 0.2539824 | -0.357156 | 0.720974732 | 0.840573706 | DOHH | protein_coding | 56.58 | -0.006841838 | 0.006841838 | ENSG00000129932 | n.s. |
| ENSG00000171055 | 609.1351402 | -0.097611814 | 0.2914943 | -0.334867 | 0.737725457 | 0.85105241 | FEZ2 | protein_coding | 40.54 | -0.006837092 | 0.006837092 | ENSG00000171055 | n.s. |

| | | | | | | | | | | | | | |
|---|---|---|---|---|---|---|---|---|---|---|---|---|---|
| ENSG00000183648 | 694.1999348 | 0.09825339 | 0.2955283 | 0.332467 | 0.739536677 | 0.852223899 | NDUFB1 | protein_coding | 44.2 | 0.006823333 | 0.006823333 | ENSG00000183648 | n.s. |
| ENSG00000128534 | 1734.205265 | -0.088649005 | 0.244537 | -0.362518 | 0.71696512 | 0.837863468 | LSM8 | protein_coding | 36.89 | -0.006810614 | 0.006810614 | ENSG00000128534 | n.s. |
| ENSG00000158290 | 711.8074348 | -0.095903753 | 0.2825453 | -0.339428 | 0.734287419 | 0.849408752 | CUL4B | protein_coding | 39.25 | -0.006797971 | 0.006797971 | ENSG00000158290 | n.s. |
| ENSG00000109065 | 655.7815681 | -0.092932056 | 0.2675119 | -0.347394 | 0.728295252 | 0.84523375 | NAT9 | protein_coding | 54.06 | -0.006786193 | 0.006786193 | ENSG00000109065 | n.s. |
| ENSG00000169855 | 1359.006817 | -0.088333174 | 0.2439084 | -0.362157 | 0.717234544 | 0.837976355 | ROBO1 | protein_coding | 35.1 | -0.006781182 | 0.006781182 | ENSG00000169855 | n.s. |
| ENSG00000180694 | 646.7570335 | -0.092166559 | 0.2637483 | -0.349449 | 0.726752288 | 0.844340129 | TMEM64 | protein_coding | 38.54 | -0.006772636 | 0.006772636 | ENSG00000180694 | n.s. |
| ENSG00000164081 | 183.575184 | 0.093285898 | 0.2697683 | 0.3458 | 0.729492952 | 0.84611216 | TEX264 | protein_coding | 54.18 | 0.00676995 | 0.00676995 | ENSG00000164081 | n.s. |
| ENSG00000273587 | 6.937935313 | 0.145075048 | 0.6080246 | 0.238601 | 0.811415289 | 0.898301652 | SNORA78 | snoRNA | 72.44 | 0.006757277 | 0.006757277 | ENSG00000273587 | n.s. |
| ENSG00000274460 | 66.29421383 | 0.093119051 | 0.2695281 | 0.345489 | 0.729726642 | 0.846242216 | | processed_transcript | 43.59 | 0.006751626 | 0.006751626 | ENSG00000274460 | n.s. |
| ENSG00000178467 | 186.4706564 | 0.093692817 | 0.272743 | 0.34352 | 0.731206949 | 0.847151728 | P4HTM | protein_coding | 54.55 | 0.006749518 | 0.006749518 | ENSG00000178467 | n.s. |
| ENSG00000121417 | 270.7131598 | 0.08987358 | 0.2533528 | 0.354737 | 0.722786694 | 0.841339606 | ZNF211 | protein_coding | 41.9 | 0.006743095 | 0.006743095 | ENSG00000121417 | n.s. |
| ENSG00000237973 | 141.8985317 | -0.094789388 | 0.2792316 | -0.339465 | 0.734259402 | 0.849408752 | MTCO1P12 | unprocessed_pseudogene | 45.76 | -0.006718982 | 0.006718982 | ENSG00000237973 | n.s. |
| ENSG00000113719 | 770.3807355 | -0.094907667 | 0.2801721 | -0.338748 | 0.734799732 | 0.849656869 | ERGIC1 | protein_coding | 50.91 | -0.006715327 | 0.006715327 | ENSG00000113719 | n.s. |
| ENSG00000119729 | 774.7118161 | -0.095171828 | 0.282091 | -0.33738 | 0.735830529 | 0.850072732 | RHOQ | protein_coding | 44.36 | -0.006713793 | 0.006713793 | ENSG00000119729 | n.s. |
| ENSG00000144724 | 816.2103345 | -0.084915375 | 0.2282308 | -0.372059 | 0.709848701 | 0.833559744 | PTPRG | protein_coding | 40.79 | -0.006713687 | 0.006713687 | ENSG00000144724 | n.s. |
| ENSG00000105447 | 746.0233831 | 0.093867882 | 0.2745994 | 0.341836 | 0.732474544 | 0.848218542 | GRWD1 | protein_coding | 55.81 | 0.006710825 | 0.006710825 | ENSG00000105447 | n.s. |
| ENSG00000145687 | 41.78125031 | 0.10875298 | 0.3603541 | 0.301795 | 0.762808503 | 0.867627587 | SSBP2 | protein_coding | 35.97 | 0.006706432 | 0.006706432 | ENSG00000145687 | n.s. |
| ENSG00000274523 | 657.663728 | -0.090007563 | 0.2559145 | -0.35171 | 0.725056129 | 0.843047817 | RCC1L | protein_coding | 51.13 | -0.006673862 | 0.006673862 | ENSG00000274523 | n.s. |
| ENSG00000141759 | 1576.334617 | 0.095842919 | 0.2876185 | 0.333229 | 0.738961193 | 0.851913281 | TXNL4A | protein_coding | 45.6 | 0.006671109 | 0.006671109 | ENSG00000141759 | n.s. |
| ENSG00000213281 | 2426.132005 | -0.094910214 | 0.2827489 | -0.33567 | 0.737120065 | 0.850706455 | NRAS | protein_coding | 39 | -0.006664621 | 0.006664621 | ENSG00000213281 | n.s. |
| ENSG00000270647 | 2285.668187 | -0.079319672 | 0.2025457 | -0.391614 | 0.695343633 | 0.824325748 | TAF15 | protein_coding | 46.76 | -0.00665501 | 0.00665501 | ENSG00000270647 | n.s. |
| ENSG00000142453 | 472.4956093 | -0.08802679 | 0.2472771 | -0.355984 | 0.721852198 | 0.840962945 | CARM1 | protein_coding | 54.32 | -0.006621652 | 0.006621652 | ENSG00000142453 | n.s. |
| ENSG00000114405 | 950.3536404 | -0.095820443 | 0.2893454 | -0.331163 | 0.740521515 | 0.853005796 | C3orf14 | protein_coding | 35.7 | -0.006616212 | 0.006616212 | ENSG00000114405 | n.s. |
| ENSG00000243156 | 92.27172631 | 0.086377208 | 0.2390558 | 0.361327 | 0.717855337 | 0.838414163 | MICAL3 | protein_coding | 49.78 | 0.006611432 | 0.006611432 | ENSG00000243156 | n.s. |
| ENSG00000108349 | 1465.068374 | -0.087819311 | 0.2463511 | -0.35648 | 0.72148088 | 0.840952557 | CASC3 | protein_coding | 45.51 | -0.006606515 | 0.006606515 | ENSG00000108349 | n.s. |
| ENSG00000103415 | 344.1690899 | -0.094504879 | 0.2827549 | -0.334229 | 0.738206747 | 0.851325481 | HMOX2 | protein_coding | 47.62 | -0.006606304 | 0.006606304 | ENSG00000103415 | n.s. |
| ENSG00000198055 | 465.8350975 | -0.09199499 | 0.2692995 | -0.341609 | 0.732645534 | 0.848256103 | GRK6 | protein_coding | 54.65 | -0.006575159 | 0.006575159 | ENSG00000198055 | n.s. |
| ENSG00000125772 | 1188.377839 | 0.091094627 | 0.2649055 | 0.343876 | 0.73093963 | 0.847006221 | GPCPD1 | protein_coding | 39.24 | 0.006569143 | 0.006569143 | ENSG00000125772 | n.s. |
| ENSG00000149930 | 586.1755745 | 0.083615723 | 0.2259864 | 0.370003 | 0.711379955 | 0.834542721 | TAOK2 | protein_coding | 55.16 | 0.006568134 | 0.006568134 | ENSG00000149930 | n.s. |
| ENSG00000258947 | 1440.876422 | 0.096477852 | 0.2940229 | 0.32813 | 0.742813038 | 0.855008758 | TUBB3 | protein_coding | 58.5 | 0.006563334 | 0.006563334 | ENSG00000258947 | n.s. |
| ENSG00000169241 | 168.422972 | 0.099850103 | 0.3124688 | 0.319552 | 0.749307768 | 0.859641732 | SLC50A1 | protein_coding | 55.69 | 0.006558405 | 0.006558405 | ENSG00000169241 | n.s. |
| ENSG00000005812 | 1019.974868 | 0.092084258 | 0.2703722 | 0.340583 | 0.733417315 | 0.848825377 | FBXL3 | protein_coding | 38.36 | 0.006554709 | 0.006554709 | ENSG00000005812 | n.s. |

| | | | | | | | | | | | | | |
|---|---|---|---|---|---|---|---|---|---|---|---|---|---|
| ENSG00000174780 | 2748.522572 | 0.092087249 | 0.2716274 | 0.33902 | 0.734594328 | 0.849622598 | SRP72 | protein_coding | 38.04 | 0.006517378 | 0.006517378 | ENSG00000174780 | n.s. |
| ENSG00000163249 | 199.5899122 | -0.088674842 | 0.2537542 | -0.349452 | 0.726750232 | 0.844340129 | CCNYL1 | protein_coding | 41.55 | -0.006516055 | 0.006516055 | ENSG00000163249 | n.s. |
| ENSG00000237289 | 347.8655332 | 0.087871052 | 0.2500529 | 0.35141 | 0.725280936 | 0.843052803 | CKMT1B | protein_coding | 49.85 | 0.006515219 | 0.006515219 | ENSG00000237289 | n.s. |
| ENSG00000076248 | 2663.662842 | -0.089079186 | 0.2564363 | -0.347374 | 0.728310659 | 0.84523375 | UNG | protein_coding | 47.32 | -0.006504845 | 0.006504845 | ENSG00000076248 | n.s. |
| ENSG00000131269 | 540.0254228 | -0.092424353 | 0.2748896 | -0.336224 | 0.736702333 | 0.850538274 | ABCB7 | protein_coding | 35.98 | -0.006497999 | 0.006497999 | ENSG00000131269 | n.s. |
| ENSG00000065665 | 220.2843552 | -0.09166335 | 0.2715935 | -0.337502 | 0.735738534 | 0.850072732 | SEC61A2 | protein_coding | 43.14 | -0.006466292 | 0.006466292 | ENSG00000065665 | n.s. |
| ENSG00000106052 | 1367.530811 | -0.093037224 | 0.2796979 | -0.332635 | 0.739410071 | 0.852174608 | TAX1BP1 | protein_coding | 36.81 | -0.006463427 | 0.006463427 | ENSG00000106052 | n.s. |
| ENSG00000159140 | 6366.070866 | -0.086053291 | 0.2422927 | -0.355163 | 0.722467787 | 0.841187406 | SON | protein_coding | 38.94 | -0.006463225 | 0.006463225 | ENSG00000159140 | n.s. |
| ENSG00000168874 | 3721.324444 | -0.081111188 | 0.216664 | -0.374364 | 0.708133631 | 0.832644586 | ATOH8 | protein_coding | 55.02 | -0.006451611 | 0.006451611 | ENSG00000168874 | n.s. |
| ENSG00000171224 | 135.3782289 | 0.08890363 | 0.2572065 | 0.345651 | 0.729605181 | 0.846171839 | C10orf35 | protein_coding | 56.17 | 0.006449197 | 0.006449197 | ENSG00000171224 | n.s. |
| ENSG00000120875 | 968.7233083 | -0.08232172 | 0.2235485 | -0.36825 | 0.712686853 | 0.835120738 | DUSP4 | protein_coding | 51.08 | -0.006441735 | 0.006441735 | ENSG00000120875 | n.s. |
| ENSG00000176209 | 504.0894001 | -0.098227847 | 0.3081084 | -0.318809 | 0.749871078 | 0.859931379 | SMIM19 | protein_coding | 44.11 | -0.00643748 | 0.00643748 | ENSG00000176209 | n.s. |
| ENSG00000166482 | 2230.056162 | -0.086720926 | 0.2466162 | -0.351643 | 0.725105823 | 0.843047817 | MFAP4 | protein_coding | 57.09 | -0.006430165 | 0.006430165 | ENSG00000166482 | n.s. |
| ENSG00000048405 | 923.7190662 | 0.092431113 | 0.2779008 | 0.332605 | 0.739432697 | 0.852174608 | ZNF800 | protein_coding | 37.8 | 0.00642132 | 0.00642132 | ENSG00000048405 | n.s. |
| ENSG00000206693 | 9.577473375 | 0.113929269 | 0.4106923 | 0.277408 | 0.781466971 | 0.878298727 | SNORA56 | snoRNA | 48.84 | 0.006420797 | 0.006420797 | ENSG00000206693 | n.s. |
| ENSG00000059145 | 37.7034353 | 0.097646305 | 0.3053946 | 0.319738 | 0.749166803 | 0.859550836 | UNKL | protein_coding | 57.82 | 0.006418139 | 0.006418139 | ENSG00000059145 | n.s. |
| ENSG00000279757 | 27.24606516 | -0.098374647 | 0.3106772 | -0.316646 | 0.751512363 | 0.860752438 | | TEC | 49.02 | -0.006406328 | 0.006406328 | ENSG00000279757 | n.s. |
| ENSG00000137713 | 632.1719263 | 0.088625643 | 0.2575566 | 0.344102 | 0.730769879 | 0.847006221 | PPP2R1B | protein_coding | 44.57 | 0.006391096 | 0.006391096 | ENSG00000137713 | n.s. |
| ENSG00000256060 | 125.2144031 | -0.092299395 | 0.2787908 | -0.33107 | 0.740591236 | 0.853015534 | | protein_coding | 50.95 | -0.006372633 | 0.006372633 | ENSG00000256060 | n.s. |
| ENSG00000159409 | 603.381632 | -0.084595708 | 0.2379268 | -0.355554 | 0.722174922 | 0.841057418 | CELF3 | protein_coding | 57.78 | -0.006359428 | 0.006359428 | ENSG00000159409 | n.s. |
| ENSG00000205339 | 6887.466146 | 0.090936374 | 0.2719877 | 0.33434 | 0.738123029 | 0.851299448 | IPO7 | protein_coding | 40.4 | 0.006358057 | 0.006358057 | ENSG00000205339 | n.s. |
| ENSG00000166851 | 632.7136352 | -0.083894175 | 0.2342668 | -0.358114 | 0.720258098 | 0.84015346 | PLK1 | protein_coding | 50.69 | -0.006345871 | 0.006345871 | ENSG00000166851 | n.s. |
| ENSG00000179041 | 513.1357841 | -0.084164551 | 0.2366112 | -0.355708 | 0.722059094 | 0.84105689 | RRS1 | protein_coding | 56.33 | -0.006327039 | 0.006327039 | ENSG00000179041 | n.s. |
| ENSG00000150753 | 9434.08387 | 0.093838899 | 0.2879066 | 0.325935 | 0.7444734 | 0.856233313 | CCT5 | protein_coding | 43.56 | 0.006325481 | 0.006325481 | ENSG00000150753 | n.s. |
| ENSG00000161653 | 92.63926233 | -0.086613969 | 0.2492501 | -0.347498 | 0.728217069 | 0.84523375 | NAGS | protein_coding | 63.61 | -0.006324827 | 0.006324827 | ENSG00000161653 | n.s. |
| ENSG00000182796 | 23.45469915 | 0.103792897 | 0.3475186 | 0.298669 | 0.765192903 | 0.869199706 | TMEM198B | transcribed_unitary_pseudogene | 54.51 | 0.006318956 | 0.006318956 | ENSG00000182796 | n.s. |
| ENSG00000130299 | 208.9976831 | 0.089421879 | 0.2645467 | 0.338019 | 0.735348703 | 0.849859716 | GTPBP3 | protein_coding | 56.97 | 0.006317902 | 0.006317902 | ENSG00000130299 | n.s. |
| ENSG00000214013 | 56.24985259 | -0.089392679 | 0.26435 | -0.33816 | 0.735242416 | 0.849859716 | GANC | protein_coding | 40.5 | -0.006315839 | 0.006315839 | ENSG00000214013 | n.s. |
| ENSG00000199366 | 8.982888792 | -0.1147247 | 0.4210275 | -0.272487 | 0.785247246 | 0.881124871 | | misc_RNA | 43.4 | -0.006305561 | 0.006305561 | ENSG00000199366 | n.s. |
| ENSG00000134086 | 1334.132264 | -0.089350486 | 0.2650573 | -0.337099 | 0.736042458 | 0.850174926 | VHL | protein_coding | 47.55 | -0.006298468 | 0.006298468 | ENSG00000134086 | n.s. |
| ENSG00000272872 | 24.17519232 | 0.100500003 | 0.3274202 | 0.306945 | 0.758885241 | 0.865676701 | | sense_intronic | 48.56 | 0.006295749 | 0.006295749 | ENSG00000272872 | n.s. |
| ENSG00000092051 | 267.8782044 | -0.089594967 | 0.2667822 | -0.335836 | 0.736994851 | 0.850683802 | JPH4 | protein_coding | 56.69 | -0.006292419 | 0.006292419 | ENSG00000092051 | n.s. |

| | | | | | | | | | | | | | |
|---|---|---|---|---|---|---|---|---|---|---|---|---|---|
| ENSG00000214194 | 571.3965239 | 0.100199244 | 0.3260119 | 0.307348 | 0.758578161 | 0.865569885 | LINC00998 | protein_coding | 37.39 | 0.006282278 | 0.006282278 | ENSG00000214194 | n.s. |
| ENSG00000151575 | 50.91371952 | 0.092091041 | 0.2803241 | 0.328516 | 0.742521336 | 0.854743659 | TEX9 | protein_coding | 36.87 | 0.006277304 | 0.006277304 | ENSG00000151575 | n.s. |
| ENSG00000263511 | 8.800897032 | -0.114867279 | 0.4233678 | -0.271318 | 0.78614649 | 0.881778585 | MIR5699 | miRNA | 57.78 | -0.0062764 | 0.0062764 | ENSG00000263511 | n.s. |
| ENSG00000247516 | 802.1470629 | -0.082972925 | 0.2317414 | -0.358041 | 0.720312566 | 0.84015346 | MIR4458HG | lincRNA | 40.34 | -0.006276187 | 0.006276187 | ENSG00000247516 | n.s. |
| ENSG00000022840 | 1043.287633 | 0.081987605 | 0.2269113 | 0.36132 | 0.717860172 | 0.838414163 | RNF10 | protein_coding | 45.25 | 0.006275446 | 0.006275446 | ENSG00000022840 | n.s. |
| ENSG00000223692 | 25.3978363 | 0.100016177 | 0.325978 | 0.306819 | 0.758981326 | 0.865676701 | DIP2A-IT1 | sense_intronic | 40.16 | 0.00626544 | 0.00626544 | ENSG00000223692 | n.s. |
| ENSG00000089063 | 1618.102956 | 0.094978921 | 0.2965565 | 0.320273 | 0.74876171 | 0.859298491 | TMEM230 | protein_coding | 44.98 | 0.006254927 | 0.006254927 | ENSG00000089063 | n.s. |
| ENSG00000134748 | 2263.419237 | -0.084951179 | 0.2438864 | -0.348323 | 0.727597784 | 0.844758501 | PRPF38A | protein_coding | 41.39 | -0.006224154 | 0.006224154 | ENSG00000134748 | n.s. |
| ENSG00000114784 | 465.2603469 | 0.086129594 | 0.2504735 | 0.343867 | 0.730946266 | 0.847006221 | EIF1B | protein_coding | 41.77 | 0.006211098 | 0.006211098 | ENSG00000114784 | n.s. |
| ENSG00000208772 | 89.54417752 | -0.087386412 | 0.2574484 | -0.339433 | 0.734283811 | 0.849408752 | SNORD94 | snoRNA | 54.01 | -0.006194235 | 0.006194235 | ENSG00000208772 | n.s. |
| ENSG00000161980 | 360.6342317 | -0.092435323 | 0.2852652 | -0.324033 | 0.745913111 | 0.857159809 | POLR3K | protein_coding | 48.85 | -0.006187454 | 0.006187454 | ENSG00000161980 | n.s. |
| ENSG00000252396 | 21.91927841 | 0.102164354 | 0.3437885 | 0.297172 | 0.766335128 | 0.870007109 | RN7SKP195 | misc_RNA | 53.47 | 0.006178614 | 0.006178614 | ENSG00000252396 | n.s. |
| ENSG00000128602 | 351.1767404 | -0.085731411 | 0.2496001 | -0.343475 | 0.731241074 | 0.847151728 | SMO | protein_coding | 49.17 | -0.006175988 | 0.006175988 | ENSG00000128602 | n.s. |
| ENSG00000137312 | 923.7233908 | -0.095104696 | 0.3016113 | -0.315322 | 0.752517163 | 0.861412189 | FLOT1 | protein_coding | 49.72 | -0.006161737 | 0.006161737 | ENSG00000137312 | n.s. |
| ENSG00000147955 | 1163.787747 | -0.083810085 | 0.240003 | -0.349204 | 0.726935907 | 0.844412534 | SIGMAR1 | protein_coding | 57.25 | -0.006155459 | 0.006155459 | ENSG00000147955 | n.s. |
| ENSG00000104738 | 5136.470715 | -0.085197596 | 0.2475334 | -0.344186 | 0.730706215 | 0.847006221 | MCM4 | protein_coding | 45.8 | -0.006143888 | 0.006143888 | ENSG00000104738 | n.s. |
| ENSG00000103254 | 205.091126 | 0.093309259 | 0.2916423 | 0.319944 | 0.74901067 | 0.859513357 | FAM173A | protein_coding | 71.94 | 0.006134838 | 0.006134838 | ENSG00000103254 | n.s. |
| ENSG00000148248 | 1342.216621 | 0.090625249 | 0.2773223 | 0.326787 | 0.743829154 | 0.855753868 | SURF4 | protein_coding | 52.38 | 0.0061309 | 0.0061309 | ENSG00000148248 | n.s. |
| ENSG00000085872 | 456.9059242 | 0.081184847 | 0.2274486 | 0.356937 | 0.721138924 | 0.840644359 | CHERP | protein_coding | 57.3 | 0.006120339 | 0.006120339 | ENSG00000085872 | n.s. |
| ENSG00000268266 | 56.10773036 | -0.094410707 | 0.2994158 | -0.315316 | 0.752521472 | 0.861412189 | | lincRNA | 42.65 | -0.006116774 | 0.006116774 | ENSG00000268266 | n.s. |
| ENSG00000130816 | 3045.586978 | -0.083464754 | 0.2397245 | -0.348169 | 0.727712961 | 0.844821776 | DNMT1 | protein_coding | 50.43 | -0.006112533 | 0.006112533 | ENSG00000130816 | n.s. |
| ENSG00000202150 | 6.307965703 | 0.151719363 | 0.7187927 | 0.211075 | 0.832828535 | 0.911467305 | RNU6-407P | snRNA | 40.38 | 0.006108055 | 0.006108055 | ENSG00000202150 | n.s. |
| ENSG00000175279 | 64.62038427 | 0.093138716 | 0.2923028 | 0.318638 | 0.750001235 | 0.859941299 | CENPS | protein_coding | 46.89 | 0.006103491 | 0.006103491 | ENSG00000175279 | n.s. |
| ENSG00000100271 | 115.5159895 | -0.095588666 | 0.3073319 | -0.311028 | 0.755779695 | 0.863354436 | TTLL1 | protein_coding | 51.32 | -0.006099596 | 0.006099596 | ENSG00000100271 | n.s. |
| ENSG00000129480 | 428.5952549 | -0.095266687 | 0.3055942 | -0.311742 | 0.75523628 | 0.862962972 | DTD2 | protein_coding | 38.52 | -0.006097815 | 0.006097815 | ENSG00000129480 | n.s. |
| ENSG00000014641 | 3568.99664 | 0.093426091 | 0.2941058 | 0.317661 | 0.750741724 | 0.860506876 | MDH1 | protein_coding | 38.05 | 0.006095647 | 0.006095647 | ENSG00000014641 | n.s. |
| ENSG00000158402 | 353.1792146 | -0.086947902 | 0.2600209 | -0.334388 | 0.73808665 | 0.851299448 | CDC25C | protein_coding | 44.09 | -0.006079193 | 0.006079193 | ENSG00000158402 | n.s. |
| ENSG00000141499 | 325.849037 | 0.090482284 | 0.2784722 | 0.324924 | 0.745238603 | 0.85673461 | WRAP53 | protein_coding | 50.13 | 0.006076219 | 0.006076219 | ENSG00000141499 | n.s. |
| ENSG00000198746 | 142.5008266 | 0.086475333 | 0.2575383 | 0.335777 | 0.737039338 | 0.850683802 | GPATCH3 | protein_coding | 50.12 | 0.006073321 | 0.006073321 | ENSG00000198746 | n.s. |
| ENSG00000124226 | 1183.364171 | 0.093387133 | 0.2953527 | 0.316189 | 0.751859404 | 0.861079089 | RNF114 | protein_coding | 46.63 | 0.006066144 | 0.006066144 | ENSG00000124226 | n.s. |
| ENSG00000129128 | 3216.117906 | 0.085697128 | 0.2534864 | 0.338074 | 0.735307549 | 0.849859716 | SPCS3 | protein_coding | 36.77 | 0.006054738 | 0.006054738 | ENSG00000129128 | n.s. |
| ENSG00000114388 | 167.2173561 | 0.097010646 | 0.3183705 | 0.30471 | 0.760587118 | 0.866206187 | NPRL2 | protein_coding | 56.73 | 0.0060514 | 0.0060514 | ENSG00000114388 | n.s. |

| | | | | | | | | | | | | | |
|---|---|---|---|---|---|---|---|---|---|---|---|---|---|
| ENSG00000235859 | 122.9263288 | -0.094375518 | 0.3026961 | -0.311783 | 0.755205386 | 0.862962972 | | processed_pseudogene | 47.57 | -0.006040773 | 0.006040773 | ENSG00000235859 | n.s. |
| ENSG00000204435 | 836.0969989 | 0.090135539 | 0.277806 | 0.324455 | 0.745593573 | 0.857004814 | CSNK2B | protein_coding | 52.7 | 0.00604059 | 0.00604059 | ENSG00000204435 | n.s. |
| ENSG00000183751 | 337.5250824 | 0.090774121 | 0.2820057 | 0.321888 | 0.747537869 | 0.858461078 | TBL3 | protein_coding | 64.29 | 0.006016453 | 0.006016453 | ENSG00000183751 | n.s. |
| ENSG00000143641 | 1887.202293 | -0.085405842 | 0.2535592 | -0.336828 | 0.736246591 | 0.850303278 | GALNT2 | protein_coding | 45.94 | -0.006014804 | 0.006014804 | ENSG00000143641 | n.s. |
| ENSG00000207109 | 19.64509472 | -0.096823329 | 0.3195333 | -0.303015 | 0.761878583 | 0.866849351 | SNORD38C | snoRNA | 34.78 | -0.006008504 | 0.006008504 | ENSG00000207109 | n.s. |
| ENSG00000165912 | 137.8757019 | 0.089062251 | 0.273323 | 0.32585 | 0.744537983 | 0.856233313 | PACSIN3 | protein_coding | 59.66 | 0.006003497 | 0.006003497 | ENSG00000165912 | n.s. |
| ENSG00000130772 | 102.9669972 | -0.084292019 | 0.2474523 | -0.340639 | 0.733375041 | 0.848825377 | MED18 | protein_coding | 46.05 | -0.006000045 | 0.006000045 | ENSG00000130772 | n.s. |
| ENSG00000176273 | 604.0416625 | -0.085553134 | 0.2552866 | -0.335126 | 0.737530151 | 0.850897605 | SLC35G1 | protein_coding | 40.3 | -0.005999217 | 0.005999217 | ENSG00000176273 | n.s. |
| ENSG00000213753 | 337.3818301 | -0.091940641 | 0.289908 | -0.317137 | 0.751139462 | 0.860548536 | CENPBD1P1 | transcribed_processed_pseudog | 45.66 | -0.005996794 | 0.005996794 | ENSG00000213753 | n.s. |
| ENSG00000105518 | 233.4676309 | 0.093820626 | 0.301331 | 0.311354 | 0.755531455 | 0.863225377 | TMEM205 | protein_coding | 54.21 | 0.005992868 | 0.005992868 | ENSG00000105518 | n.s. |
| ENSG00000134597 | 545.4843288 | -0.095849409 | 0.3139392 | -0.305312 | 0.760128563 | 0.865917832 | RBMX2 | protein_coding | 43.77 | -0.005992823 | 0.005992823 | ENSG00000134597 | n.s. |
| ENSG00000100442 | 1963.21032 | -0.092118808 | 0.2916485 | -0.315856 | 0.752112162 | 0.861129996 | FKBP3 | protein_coding | 40.63 | -0.005981392 | 0.005981392 | ENSG00000100442 | n.s. |
| ENSG00000120253 | 1638.824472 | -0.086468239 | 0.2608974 | -0.331426 | 0.740322566 | 0.852847186 | NUP43 | protein_coding | 43.95 | -0.005977444 | 0.005977444 | ENSG00000120253 | n.s. |
| ENSG00000154839 | 400.9266381 | -0.083471811 | 0.2442372 | -0.341765 | 0.732527455 | 0.848218542 | SKA1 | protein_coding | 39.41 | -0.005967587 | 0.005967587 | ENSG00000154839 | n.s. |
| ENSG00000109738 | 130.4977763 | -0.095194576 | 0.3110601 | -0.306033 | 0.759579742 | 0.865692775 | GLRB | protein_coding | 35.21 | -0.005962627 | 0.005962627 | ENSG00000109738 | n.s. |
| ENSG00000185624 | 4051.294228 | 0.089884586 | 0.2790722 | 0.322084 | 0.747389337 | 0.858461078 | P4HB | protein_coding | 55.79 | 0.005957496 | 0.005957496 | ENSG00000185624 | n.s. |
| ENSG00000170632 | 343.9480098 | -0.092997923 | 0.2980318 | -0.31204 | 0.755009894 | 0.862962972 | ARMC10 | protein_coding | 40.09 | -0.005952596 | 0.005952596 | ENSG00000170632 | n.s. |
| ENSG00000188603 | 47.00492517 | -0.09483332 | 0.310014 | -0.3059 | 0.759680696 | 0.865692775 | CLN3 | protein_coding | 50.61 | -0.005939999 | 0.005939999 | ENSG00000188603 | n.s. |
| ENSG00000184831 | 259.309844 | 0.094817189 | 0.3096978 | 0.30616 | 0.759482554 | 0.865692775 | APOO | protein_coding | 43.27 | 0.005938989 | 0.005938989 | ENSG00000184831 | n.s. |
| ENSG00000131871 | 888.9014291 | -0.086257076 | 0.2618609 | -0.3294 | 0.74185305 | 0.85418624 | SELENOS | protein_coding | 45.06 | -0.005904075 | 0.005904075 | ENSG00000131871 | n.s. |
| ENSG00000143368 | 652.1259046 | 0.08091631 | 0.2331531 | 0.347052 | 0.728552071 | 0.845443448 | SF3B4 | protein_coding | 48.41 | 0.005900048 | 0.005900048 | ENSG00000143368 | n.s. |
| ENSG00000200874 | 6.090399003 | 0.113971381 | 0.441053 | 0.258407 | 0.796092469 | 0.887998353 | | misc_RNA | 42.16 | 0.005879537 | 0.005879537 | ENSG00000200874 | n.s. |
| ENSG00000079739 | 659.3389671 | -0.091623672 | 0.2925087 | -0.313234 | 0.75410287 | 0.862640004 | PGM1 | protein_coding | 42.42 | -0.005879528 | 0.005879528 | ENSG00000079739 | n.s. |
| ENSG00000111652 | 491.1103032 | -0.087213636 | 0.2681132 | -0.325287 | 0.744964129 | 0.856564072 | COPS7A | protein_coding | 50.55 | -0.005864258 | 0.005864258 | ENSG00000111652 | n.s. |
| ENSG00000276704 | 78.67385648 | 0.088470479 | 0.2746391 | 0.322134 | 0.747351472 | 0.858461078 | | sense_intronic | 33.88 | 0.005863769 | 0.005863769 | ENSG00000276704 | n.s. |
| ENSG00000138750 | 1700.938522 | -0.089176234 | 0.2788043 | -0.319852 | 0.749080216 | 0.859522321 | NUP54 | protein_coding | 36.8 | -0.005862699 | 0.005862699 | ENSG00000138750 | n.s. |
| ENSG00000182934 | 529.9850128 | 0.078728445 | 0.2231344 | 0.35283 | 0.724216105 | 0.842518452 | SRPRA | protein_coding | 49.55 | 0.005859016 | 0.005859016 | ENSG00000182934 | n.s. |
| ENSG00000113269 | 924.2523092 | -0.091417366 | 0.2932508 | -0.311738 | 0.755239805 | 0.862962972 | RNF130 | protein_coding | 42.83 | -0.005851428 | 0.005851428 | ENSG00000113269 | n.s. |
| ENSG00000197557 | 336.2867583 | 0.075009631 | 0.2044932 | 0.366807 | 0.713762661 | 0.83580467 | TTC30A | protein_coding | 39.41 | 0.005842891 | 0.005842891 | ENSG00000197557 | n.s. |
| ENSG00000164588 | 11.44902224 | -0.107581637 | 0.3990047 | -0.269625 | 0.787448769 | 0.882599357 | HCN1 | protein_coding | 34.76 | -0.00583484 | 0.00583484 | ENSG00000164588 | n.s. |
| ENSG00000141452 | 328.2599224 | 0.084339679 | 0.2543828 | 0.331546 | 0.740231823 | 0.852813213 | C18orf8 | protein_coding | 45.14 | 0.005831758 | 0.005831758 | ENSG00000141452 | n.s. |
| ENSG00000274620 | 15.25246228 | -0.103121911 | 0.3701084 | -0.278626 | 0.7805317 | 0.877937651 | MIR378J | miRNA | 42.2 | -0.005830134 | 0.005830134 | ENSG00000274620 | n.s. |

| | | | | | | | | | | | | | |
|---|---|---|---|---|---|---|---|---|---|---|---|---|---|
| ENSG00000214485 | 197.5820543 | 0.09514925 | 0.3167477 | 0.300394 | 0.763876287 | 0.86848355 | RPL7P1 | processed_pseudogene | 43.51 | 0.005826788 | 0.005826788 | ENSG00000214485 | n.s. |
| ENSG00000111364 | 2194.235955 | -0.082341029 | 0.2435941 | -0.338025 | 0.735343979 | 0.849859716 | DDX55 | protein_coding | 45.83 | -0.005817621 | 0.005817621 | ENSG00000111364 | n.s. |
| ENSG00000204962 | 90.63490607 | -0.084513834 | 0.2568645 | -0.329021 | 0.742139781 | 0.854375076 | PCDHA8 | protein_coding | 40.02 | -0.005776641 | 0.005776641 | ENSG00000204962 | n.s. |
| ENSG00000136936 | 555.37367 | 0.088103541 | 0.2764201 | 0.318731 | 0.74993082 | 0.859931379 | XPA | protein_coding | 38.79 | 0.005773972 | 0.005773972 | ENSG00000136936 | n.s. |
| ENSG00000256338 | 42.1731149 | 0.094723574 | 0.3169754 | 0.298836 | 0.765065378 | 0.869148157 | RPL41P2 | processed_pseudogene | 50.61 | 0.005769252 | 0.005769252 | ENSG00000256338 | n.s. |
| ENSG00000260267 | 337.168387 | 0.074993086 | 0.2066038 | 0.36298 | 0.716619677 | 0.837719862 | | antisense | 59.88 | 0.005767057 | 0.005767057 | ENSG00000260267 | n.s. |
| ENSG00000171130 | 315.8849579 | -0.088399939 | 0.2786409 | -0.317254 | 0.751050847 | 0.860548536 | ATP6V0E2 | protein_coding | 54.93 | -0.005765853 | 0.005765853 | ENSG00000171130 | n.s. |
| ENSG00000077312 | 1291.450723 | -0.086691283 | 0.2685708 | -0.322787 | 0.746856197 | 0.858101901 | SNRPA | protein_coding | 50.77 | -0.005761601 | 0.005761601 | ENSG00000077312 | n.s. |
| ENSG00000078304 | 1310.228095 | -0.088586207 | 0.2803452 | -0.31599 | 0.752010256 | 0.861129996 | PPP2R5C | protein_coding | 44.63 | -0.005752016 | 0.005752016 | ENSG00000078304 | n.s. |
| ENSG00000067167 | 970.6022526 | 0.087044209 | 0.2711637 | 0.321002 | 0.748208586 | 0.858947882 | TRAM1 | protein_coding | 38.27 | 0.005747806 | 0.005747806 | ENSG00000067167 | n.s. |
| ENSG00000166173 | 196.3873345 | 0.081957564 | 0.2445422 | 0.335147 | 0.737514279 | 0.850897605 | LARP6 | protein_coding | 44.62 | 0.005747086 | 0.005747086 | ENSG00000166173 | n.s. |
| ENSG00000095574 | 468.7492023 | 0.081896611 | 0.2443028 | 0.335226 | 0.737454722 | 0.850897605 | IKZF5 | protein_coding | 38.33 | 0.005742811 | 0.005742811 | ENSG00000095574 | n.s. |
| ENSG00000107672 | 1369.298929 | -0.089398041 | 0.2860936 | -0.312478 | 0.754677017 | 0.862804362 | NSMCE4A | protein_coding | 43.84 | -0.005729312 | 0.005729312 | ENSG00000107672 | n.s. |
| ENSG00000148700 | 824.8691626 | -0.088465662 | 0.2808499 | -0.314993 | 0.752767233 | 0.861551845 | ADD3 | protein_coding | 38.65 | -0.005725372 | 0.005725372 | ENSG00000148700 | n.s. |
| ENSG00000122545 | 5485.865188 | -0.087712048 | 0.2763851 | -0.317354 | 0.750974673 | 0.860548536 | 7-Sep | protein_coding | 36.13 | -0.005720986 | 0.005720986 | ENSG00000122545 | n.s. |
| ENSG00000231181 | 26.75882862 | -0.091938655 | 0.3027448 | -0.303684 | 0.761368929 | 0.866533092 | | processed_pseudogene | 44.54 | -0.005719949 | 0.005719949 | ENSG00000231181 | n.s. |
| ENSG00000105176 | 2415.764842 | -0.089583965 | 0.2881633 | -0.310879 | 0.755892449 | 0.863354436 | URI1 | protein_coding | 37.88 | -0.005716431 | 0.005716431 | ENSG00000105176 | n.s. |
| ENSG00000140307 | 1613.557379 | 0.093842102 | 0.3133748 | 0.299456 | 0.764591818 | 0.869148157 | GTF2A2 | protein_coding | 39.73 | 0.005715565 | 0.005715565 | ENSG00000140307 | n.s. |
| ENSG00000205903 | 471.080346 | -0.081936802 | 0.2455153 | -0.333734 | 0.738580341 | 0.85163726 | ZNF316 | protein_coding | 61.59 | -0.005714711 | 0.005714711 | ENSG00000205903 | n.s. |
| ENSG00000079616 | 2317.589704 | 0.086132598 | 0.267531 | 0.321954 | 0.747487771 | 0.858461078 | KIF22 | protein_coding | 50.19 | 0.005708816 | 0.005708816 | ENSG00000079616 | n.s. |
| ENSG00000129559 | 2076.332117 | 0.090625813 | 0.2940285 | 0.308221 | 0.757914004 | 0.865194459 | NEDD8 | protein_coding | 43.63 | 0.005699119 | 0.005699119 | ENSG00000129559 | n.s. |
| ENSG00000185721 | 1135.690371 | 0.088581781 | 0.2831515 | 0.312842 | 0.754400438 | 0.862782758 | DRG1 | protein_coding | 44.41 | 0.005677963 | 0.005677963 | ENSG00000185721 | n.s. |
| ENSG00000038210 | 673.3179277 | 0.08388979 | 0.2564627 | 0.327103 | 0.743589753 | 0.855753868 | PI4K2B | protein_coding | 41.82 | 0.005675239 | 0.005675239 | ENSG00000038210 | n.s. |
| ENSG00000110958 | 4817.041572 | 0.088406305 | 0.2822952 | 0.31317 | 0.7541517 | 0.862640004 | PTGES3 | protein_coding | 44.13 | 0.005673068 | 0.005673068 | ENSG00000110958 | n.s. |
| ENSG00000254377 | 771.7137979 | -0.079919072 | 0.2353842 | -0.339526 | 0.734213408 | 0.849408752 | MIR124-2HG | lincRNA | 44.12 | -0.005664925 | 0.005664925 | ENSG00000254377 | n.s. |
| ENSG00000160404 | 166.3176063 | -0.079225635 | 0.232062 | -0.341399 | 0.732803592 | 0.848256103 | TOR2A | protein_coding | 59.86 | -0.005662494 | 0.005662494 | ENSG00000160404 | n.s. |
| ENSG00000162227 | 321.0552467 | -0.088163833 | 0.283533 | -0.310947 | 0.755840649 | 0.863354436 | TAF6L | protein_coding | 51.76 | -0.005625811 | 0.005625811 | ENSG00000162227 | n.s. |
| ENSG00000116157 | 758.3889686 | -0.089542782 | 0.2925576 | -0.306069 | 0.759552218 | 0.865692775 | GPX7 | protein_coding | 49.67 | -0.00560862 | 0.00560862 | ENSG00000116157 | n.s. |
| ENSG00000121900 | 44.46756831 | 0.095275727 | 0.3323144 | 0.286704 | 0.774339294 | 0.873885547 | TMEM54 | protein_coding | 57.33 | 0.00557796 | 0.00557796 | ENSG00000121900 | n.s. |
| ENSG00000155115 | 1368.390213 | 0.08588028 | 0.2718649 | 0.315893 | 0.75208358 | 0.861129996 | GTF3C6 | protein_coding | 45.43 | 0.005576317 | 0.005576317 | ENSG00000155115 | n.s. |
| ENSG00000212607 | 84.96062046 | -0.079384244 | 0.2366442 | -0.335458 | 0.737279462 | 0.850819889 | SNORA3B | snoRNA | 52.67 | -0.005569786 | 0.005569786 | ENSG00000212607 | n.s. |
| ENSG00000135549 | 815.8773586 | 0.094386892 | 0.3270683 | 0.288585 | 0.772899215 | 0.873217822 | PKIB | protein_coding | 37.28 | 0.005557256 | 0.005557256 | ENSG00000135549 | n.s. |

| | | | | | | | | | | | | | |
|---|---|---|---|---|---|---|---|---|---|---|---|---|---|
| ENSG00000263712 | 40.98832396 | -0.093859406 | 0.3245227 | -0.289223 | 0.772410752 | 0.872841077 | MIR4639 | miRNA | 46.38 | -0.005543789 | 0.005543789 | ENSG00000263712 | n.s. |
| ENSG00000004142 | 892.7066979 | -0.086340935 | 0.2759658 | -0.312868 | 0.754380758 | 0.862782758 | POLDIP2 | protein_coding | 50.16 | -0.005534328 | 0.005534328 | ENSG00000004142 | n.s. |
| ENSG00000165985 | 352.2535131 | 0.084201873 | 0.2636447 | 0.319376 | 0.749441202 | 0.859723973 | C1QL3 | protein_coding | 44.9 | 0.005527092 | 0.005527092 | ENSG00000165985 | n.s. |
| ENSG00000277149 | 303.0603726 | -0.081588756 | 0.249549 | -0.326945 | 0.743709606 | 0.855753868 | TYW1B | protein_coding | 41.66 | -0.005519571 | 0.005519571 | ENSG00000277149 | n.s. |
| ENSG00000221926 | 266.3751818 | 0.076524982 | 0.2228791 | 0.343348 | 0.731337045 | 0.847192418 | TRIM16 | protein_coding | 45.66 | 0.005511172 | 0.005511172 | ENSG00000221926 | n.s. |
| ENSG00000117419 | 311.2509563 | 0.091166276 | 0.3071561 | 0.296808 | 0.766613349 | 0.870103338 | ERI3 | protein_coding | 47.73 | 0.005509102 | 0.005509102 | ENSG00000117419 | n.s. |
| ENSG00000132467 | 886.2790247 | -0.079555596 | 0.2397857 | -0.331778 | 0.740056966 | 0.852682318 | UTP3 | protein_coding | 44.81 | -0.005506261 | 0.005506261 | ENSG00000132467 | n.s. |
| ENSG00000156298 | 473.2436709 | 0.090213894 | 0.3019107 | 0.29881 | 0.765085097 | 0.869148157 | TSPAN7 | protein_coding | 39.6 | 0.005494584 | 0.005494584 | ENSG00000156298 | n.s. |
| ENSG00000107438 | 456.198028 | 0.084997452 | 0.2700727 | 0.314721 | 0.752973785 | 0.86171741 | PDLIM1 | protein_coding | 43.74 | 0.005493822 | 0.005493822 | ENSG00000107438 | n.s. |
| ENSG00000162607 | 6739.693421 | 0.083771198 | 0.2625264 | 0.319096 | 0.749653547 | 0.859896715 | USP1 | protein_coding | 39.48 | 0.005491513 | 0.005491513 | ENSG00000162607 | n.s. |
| ENSG00000108039 | 975.585769 | 0.085451795 | 0.2734771 | 0.312464 | 0.754687773 | 0.862804362 | XPNPEP1 | protein_coding | 44.62 | 0.005476406 | 0.005476406 | ENSG00000108039 | n.s. |
| ENSG00000129534 | 2296.406722 | -0.081484327 | 0.2508392 | -0.324847 | 0.745296962 | 0.85673461 | MIS18BP1 | protein_coding | 37.65 | -0.005471973 | 0.005471973 | ENSG00000129534 | n.s. |
| ENSG00000174307 | 387.4783639 | 0.072138499 | 0.2015385 | 0.357939 | 0.72038887 | 0.840172075 | PHLDA3 | protein_coding | 57.71 | 0.005455961 | 0.005455961 | ENSG00000174307 | n.s. |
| ENSG00000103502 | 325.2608295 | 0.086915551 | 0.2831067 | 0.307006 | 0.758838565 | 0.865676701 | CDIPT | protein_coding | 56.47 | 0.005444761 | 0.005444761 | ENSG00000103502 | n.s. |
| ENSG00000171490 | 5798.638258 | 0.083083075 | 0.2619886 | 0.317125 | 0.751148916 | 0.860548536 | RSL1D1 | protein_coding | 45.24 | 0.005419063 | 0.005419063 | ENSG00000171490 | n.s. |
| ENSG00000120251 | 1699.113381 | -0.086276375 | 0.2813439 | -0.306658 | 0.759103678 | 0.865686399 | GRIA2 | protein_coding | 34.59 | -0.005404301 | 0.005404301 | ENSG00000120251 | n.s. |
| ENSG00000213190 | 4154.388431 | 0.081686293 | 0.2546029 | 0.320838 | 0.748333121 | 0.859003814 | MLLT11 | protein_coding | 46.4 | 0.005391696 | 0.005391696 | ENSG00000213190 | n.s. |
| ENSG00000176973 | 484.3352555 | 0.086446941 | 0.2844968 | 0.303859 | 0.761235244 | 0.866471411 | FAM89B | protein_coding | 64.22 | 0.005380956 | 0.005380956 | ENSG00000176973 | n.s. |
| ENSG00000173848 | 2391.498089 | 0.081310093 | 0.25288 | 0.321536 | 0.747804042 | 0.858694902 | NET1 | protein_coding | 37.02 | 0.005379566 | 0.005379566 | ENSG00000173848 | n.s. |
| ENSG00000108774 | 595.7008179 | 0.085789685 | 0.2799797 | 0.306414 | 0.759289481 | 0.865692775 | RAB5C | protein_coding | 49.38 | 0.00537354 | 0.00537354 | ENSG00000108774 | n.s. |
| ENSG00000249572 | 39.11882615 | -0.085781689 | 0.2801759 | -0.306171 | 0.759474558 | 0.865692775 | | antisense | 40.72 | -0.00537304 | 0.00537304 | ENSG00000249572 | n.s. |
| ENSG00000246695 | 131.9467428 | -0.082198349 | 0.2583702 | -0.318142 | 0.750377413 | 0.860271748 | RASSF8-AS1 | antisense | 43.94 | -0.005372841 | 0.005372841 | ENSG00000246695 | n.s. |
| ENSG00000136271 | 1372.178092 | -0.082079661 | 0.2580792 | -0.318041 | 0.750454172 | 0.860271748 | DDX56 | protein_coding | 53.74 | -0.005365083 | 0.005365083 | ENSG00000136271 | n.s. |
| ENSG00000135446 | 5140.169876 | 0.084071185 | 0.2705421 | 0.310751 | 0.755990002 | 0.863395029 | CDK4 | protein_coding | 53.24 | 0.005362939 | 0.005362939 | ENSG00000135446 | n.s. |
| ENSG00000152404 | 640.9672983 | -0.081168513 | 0.2531605 | -0.320621 | 0.748497795 | 0.859066425 | CWF19L2 | protein_coding | 36.3 | -0.005354951 | 0.005354951 | ENSG00000152404 | n.s. |
| ENSG00000132323 | 1101.470199 | -0.081652228 | 0.2560785 | -0.318856 | 0.749835516 | 0.859931379 | ILKAP | protein_coding | 44.85 | -0.005351177 | 0.005351177 | ENSG00000132323 | n.s. |
| ENSG00000162627 | 579.6865253 | 0.085035083 | 0.2759185 | 0.308189 | 0.757938434 | 0.865194459 | SNX7 | protein_coding | 36.05 | 0.005347539 | 0.005347539 | ENSG00000162627 | n.s. |
| ENSG00000229539 | 22.03505991 | 0.092669616 | 0.3272606 | 0.283168 | 0.777048335 | 0.87583137 | | antisense | 53.17 | 0.00533587 | 0.00533587 | ENSG00000229539 | n.s. |
| ENSG00000133895 | 238.425797 | -0.085483645 | 0.2806229 | -0.304621 | 0.760654793 | 0.866206187 | MEN1 | protein_coding | 57.78 | -0.00533236 | 0.00533236 | ENSG00000133895 | n.s. |
| ENSG00000043514 | 402.0084004 | -0.0849999 | 0.2764373 | -0.307483 | 0.758475432 | 0.865534077 | TRIT1 | protein_coding | 42.46 | -0.005330839 | 0.005330839 | ENSG00000043514 | n.s. |
| ENSG00000106009 | 535.0389721 | 0.07952106 | 0.2452313 | 0.32427 | 0.745733918 | 0.857055015 | BRAT1 | protein_coding | 55.18 | 0.005327219 | 0.005327219 | ENSG00000106009 | n.s. |
| ENSG00000164062 | 859.1786481 | 0.087460155 | 0.2926442 | 0.298862 | 0.765045515 | 0.869148157 | APEH | protein_coding | 58.08 | 0.005326865 | 0.005326865 | ENSG00000164062 | n.s. |

| | | | | | | | | | | | | | |
|---|---|---|---|---|---|---|---|---|---|---|---|---|---|
| ENSG00000269439 | 51.69157867 | -0.089916212 | 0.3095771 | -0.290449 | 0.771473117 | 0.872517203 | | lincRNA | 52.93 | -0.005325378 | 0.005325378 | ENSG00000269439 | n.s. |
| ENSG00000227535 | 17.8493192 | 0.092484736 | 0.3264577 | 0.283298 | 0.77694857 | 0.87583137 | | sense_overlapping | 39.23 | 0.005325225 | 0.005325225 | ENSG00000227535 | n.s. |
| ENSG00000184164 | 157.3539494 | 0.083605284 | 0.2694401 | 0.310293 | 0.756338444 | 0.863722125 | CRELD2 | protein_coding | 59.36 | 0.005319466 | 0.005319466 | ENSG00000184164 | n.s. |
| ENSG00000268471 | 570.6514743 | -0.072510202 | 0.2079168 | -0.348746 | 0.727279855 | 0.844600669 | | lincRNA | 43.57 | -0.00531852 | 0.00531852 | ENSG00000268471 | n.s. |
| ENSG00000274641 | 702.0101353 | 0.070886406 | 0.1998684 | 0.354665 | 0.722840219 | 0.841339606 | HIST1H2BO | protein_coding | 59.32 | 0.005318513 | 0.005318513 | ENSG00000274641 | n.s. |
| ENSG00000117751 | 757.7343276 | 0.079817936 | 0.2469286 | 0.323243 | 0.746511267 | 0.857776378 | PPP1R8 | protein_coding | 44.44 | 0.005317943 | 0.005317943 | ENSG00000117751 | n.s. |
| ENSG00000145241 | 1652.010462 | -0.081040145 | 0.254232 | -0.318765 | 0.749905093 | 0.859931379 | CENPC | protein_coding | 36.85 | -0.005311063 | 0.005311063 | ENSG00000145241 | n.s. |
| ENSG00000272277 | 62.08252941 | 0.084902249 | 0.279629 | 0.303625 | 0.761413908 | 0.866533092 | | antisense | 65.06 | 0.00528218 | 0.00528218 | ENSG00000272277 | n.s. |
| ENSG00000105341 | 346.1458627 | -0.080939165 | 0.2553944 | -0.316918 | 0.75130558 | 0.860586391 | ATP5SL | protein_coding | 51.01 | -0.005277681 | 0.005277681 | ENSG00000105341 | n.s. |
| ENSG00000004975 | 595.6828551 | -0.08413858 | 0.2736469 | -0.307471 | 0.758484613 | 0.865534077 | DVL2 | protein_coding | 55.14 | -0.005276821 | 0.005276821 | ENSG00000004975 | n.s. |
| ENSG00000248152 | 45.25656949 | 0.087688561 | 0.2974225 | 0.294828 | 0.768125074 | 0.870825292 | | antisense | 37.21 | 0.005267361 | 0.005267361 | ENSG00000248152 | n.s. |
| ENSG00000006007 | 665.4488228 | 0.090796606 | 0.3188271 | 0.284783 | 0.775810246 | 0.875049098 | GDE1 | protein_coding | 40.17 | 0.005263259 | 0.005263259 | ENSG00000006007 | n.s. |
| ENSG00000243554 | 109.4128204 | -0.082177994 | 0.2635858 | -0.311769 | 0.755215761 | 0.862962972 | | unprocessed_pseudogene | 46.12 | -0.005260036 | 0.005260036 | ENSG00000243554 | n.s. |
| ENSG00000168438 | 1612.710228 | -0.079339171 | 0.2463945 | -0.322001 | 0.747452296 | 0.858461078 | CDC40 | protein_coding | 37.34 | -0.005258552 | 0.005258552 | ENSG00000168438 | n.s. |
| ENSG00000122952 | 2383.771553 | 0.077462522 | 0.2372509 | 0.3265 | 0.744045711 | 0.855932284 | ZWINT | protein_coding | 48.07 | 0.005233414 | 0.005233414 | ENSG00000122952 | n.s. |
| ENSG00000254682 | 65.06487749 | 0.083665904 | 0.2740436 | 0.305301 | 0.760136589 | 0.865917832 | | antisense | 45.01 | 0.00523107 | 0.00523107 | ENSG00000254682 | n.s. |
| ENSG00000066117 | 2195.89716 | -0.077507106 | 0.2378462 | -0.325871 | 0.744522186 | 0.856233313 | SMARCD1 | protein_coding | 48.68 | -0.00522459 | 0.00522459 | ENSG00000066117 | n.s. |
| ENSG00000064547 | 151.2396163 | -0.079948406 | 0.2517823 | -0.31753 | 0.750841585 | 0.860545793 | LPAR2 | protein_coding | 58.96 | -0.005214716 | 0.005214716 | ENSG00000064547 | n.s. |
| ENSG00000050344 | 176.7466019 | 0.081265027 | 0.260043 | 0.312506 | 0.754655958 | 0.862804362 | NFE2L3 | protein_coding | 44.64 | 0.005208086 | 0.005208086 | ENSG00000050344 | n.s. |
| ENSG00000156973 | 297.4518238 | 0.092169955 | 0.3313702 | 0.278148 | 0.780898757 | 0.87815632 | PDE6D | protein_coding | 42.51 | 0.005200982 | 0.005200982 | ENSG00000156973 | n.s. |
| ENSG00000165732 | 7726.150837 | 0.082932949 | 0.2701736 | 0.306962 | 0.758872557 | 0.865676701 | DDX21 | protein_coding | 41.17 | 0.005195274 | 0.005195274 | ENSG00000165732 | n.s. |
| ENSG00000089006 | 4467.922502 | 0.075849793 | 0.2301348 | 0.329589 | 0.741710868 | 0.854163806 | SNX5 | protein_coding | 42.13 | 0.005192588 | 0.005192588 | ENSG00000089006 | n.s. |
| ENSG00000168259 | 2375.660455 | 0.072594825 | 0.2126022 | 0.341458 | 0.732758451 | 0.848256103 | DNAJC7 | protein_coding | 45.79 | 0.00518857 | 0.00518857 | ENSG00000168259 | n.s. |
| ENSG00000164713 | 218.3760826 | 0.087152316 | 0.3001571 | 0.290356 | 0.77154418 | 0.872517203 | BRI3 | protein_coding | 51.97 | 0.005161683 | 0.005161683 | ENSG00000164713 | n.s. |
| ENSG00000154978 | 365.274327 | -0.085866581 | 0.2917121 | -0.294354 | 0.768487503 | 0.870869029 | VOPP1 | protein_coding | 46.42 | -0.005156044 | 0.005156044 | ENSG00000154978 | n.s. |
| ENSG00000206651 | 8.349454039 | -0.108591726 | 0.4479443 | -0.242422 | 0.808452884 | 0.896439772 | | misc_RNA | 41.59 | -0.005155814 | 0.005155814 | ENSG00000206651 | n.s. |
| ENSG00000132388 | 1133.671602 | -0.082162312 | 0.2679417 | -0.306642 | 0.75911551 | 0.865686399 | UBE2G1 | protein_coding | 41.77 | -0.005146598 | 0.005146598 | ENSG00000132388 | n.s. |
| ENSG00000112739 | 3358.300366 | -0.080326793 | 0.2572299 | -0.312276 | 0.754830614 | 0.862849478 | PRPF4B | protein_coding | 38.36 | -0.005146133 | 0.005146133 | ENSG00000112739 | n.s. |
| ENSG00000069943 | 120.5533273 | 0.077829303 | 0.2424579 | 0.321001 | 0.748209432 | 0.858947882 | PIGB | protein_coding | 39.99 | 0.005139317 | 0.005139317 | ENSG00000069943 | n.s. |
| ENSG00000180902 | 226.8524348 | -0.077817426 | 0.2425926 | -0.320774 | 0.748381546 | 0.859003814 | D2HGDH | protein_coding | 56.88 | -0.005136332 | 0.005136332 | ENSG00000180902 | n.s. |
| ENSG00000100314 | 396.3635709 | 0.085341903 | 0.2898527 | 0.294432 | 0.768427839 | 0.870869029 | CABP7 | protein_coding | 62.85 | 0.005124538 | 0.005124538 | ENSG00000100314 | n.s. |
| ENSG00000211460 | 3494.279616 | 0.08215443 | 0.2698145 | 0.304485 | 0.760758562 | 0.866212091 | TSN | protein_coding | 44 | 0.005124445 | 0.005124445 | ENSG00000211460 | n.s. |

| | | | | | | | | | | | | | |
|---|---|---|---|---|---|---|---|---|---|---|---|---|---|
| ENSG00000113758 | 3503.495701 | -0.08201455 | 0.2690427 | -0.304838 | 0.760489193 | 0.86618873 | DBN1 | protein_coding | 59.54 | -0.00511668 | 0.00511668 | ENSG00000113758 | n.s. |
| ENSG00000110074 | 349.6873749 | -0.086029957 | 0.2974017 | -0.289272 | 0.772373328 | 0.872841077 | FOXRED1 | protein_coding | 51.05 | -0.005081344 | 0.005081344 | ENSG00000110074 | n.s. |
| ENSG00000161057 | 1547.924334 | -0.084436588 | 0.2866288 | -0.294585 | 0.768310841 | 0.870869029 | PSMC2 | protein_coding | 38.99 | -0.005070177 | 0.005070177 | ENSG00000161057 | n.s. |
| ENSG00000060656 | 225.159201 | -0.078512091 | 0.2499591 | -0.3141 | 0.753445263 | 0.862115253 | PTPRU | protein_coding | 51.58 | -0.005058901 | 0.005058901 | ENSG00000060656 | n.s. |
| ENSG00000184983 | 1740.736882 | 0.085430893 | 0.2952884 | 0.289313 | 0.772341545 | 0.872841077 | NDUFA6 | protein_coding | 48.79 | 0.005045961 | 0.005045961 | ENSG00000184983 | n.s. |
| ENSG00000164631 | 1312.898667 | -0.080666554 | 0.2642791 | -0.305232 | 0.760189159 | 0.865917832 | ZNF12 | protein_coding | 43.43 | -0.005043541 | 0.005043541 | ENSG00000164631 | n.s. |
| ENSG00000132780 | 4327.58091 | -0.077171704 | 0.2434278 | -0.317021 | 0.751227778 | 0.860568073 | NASP | protein_coding | 42.9 | -0.005032735 | 0.005032735 | ENSG00000132780 | n.s. |
| ENSG00000112110 | 1153.889531 | -0.086069908 | 0.3010013 | -0.285945 | 0.774920011 | 0.87425746 | MRPL18 | protein_coding | 43.85 | -0.005023096 | 0.005023096 | ENSG00000112110 | n.s. |
| ENSG00000241506 | 78.74810236 | -0.083205756 | 0.2818999 | -0.295161 | 0.767871174 | 0.870709774 | PSMC1P1 | processed_pseudogene | 45.76 | -0.005002878 | 0.005002878 | ENSG00000241506 | n.s. |
| ENSG00000197586 | 484.4073732 | -0.081506791 | 0.2711257 | -0.300624 | 0.763701491 | 0.868355725 | ENTPD6 | protein_coding | 50.79 | -0.004996556 | 0.004996556 | ENSG00000197586 | n.s. |
| ENSG00000143653 | 1334.418726 | -0.08498875 | 0.2954388 | -0.28767 | 0.773599674 | 0.8735015 | SCCPDH | protein_coding | 41.94 | -0.004991929 | 0.004991929 | ENSG00000143653 | n.s. |
| ENSG00000117906 | 4979.433951 | -0.086549357 | 0.3057661 | -0.283057 | 0.777132856 | 0.87583137 | RCN2 | protein_coding | 38.97 | -0.004983469 | 0.004983469 | ENSG00000117906 | n.s. |
| ENSG00000175193 | 681.6021564 | 0.084345489 | 0.291823 | 0.28903 | 0.772558701 | 0.872937412 | PARL | protein_coding | 41.83 | 0.004977809 | 0.004977809 | ENSG00000175193 | n.s. |
| ENSG00000146416 | 289.7894582 | 0.082430974 | 0.2783433 | 0.296149 | 0.767116624 | 0.870349013 | AIG1 | protein_coding | 39.67 | 0.004971128 | 0.004971128 | ENSG00000146416 | n.s. |
| ENSG00000107537 | 226.6576963 | 0.087538569 | 0.3127372 | 0.279911 | 0.779545809 | 0.877487475 | PHYH | protein_coding | 46.07 | 0.004968608 | 0.004968608 | ENSG00000107537 | n.s. |
| ENSG00000213588 | 237.6643945 | 0.073700552 | 0.2261936 | 0.325829 | 0.74455338 | 0.856233313 | ZBTB9 | protein_coding | 53.65 | 0.004967998 | 0.004967998 | ENSG00000213588 | n.s. |
| ENSG00000113845 | 1109.860537 | -0.084602881 | 0.2944378 | -0.287337 | 0.773854248 | 0.873621397 | TIMMDC1 | protein_coding | 38.48 | -0.004964221 | 0.004964221 | ENSG00000113845 | n.s. |
| ENSG00000273353 | 31.03746199 | -0.089363125 | 0.3258778 | -0.274223 | 0.783913414 | 0.880101421 | | sense_intronic | 33.02 | -0.004956729 | 0.004956729 | ENSG00000273353 | n.s. |
| ENSG00000213523 | 506.8415765 | 0.081623123 | 0.2740905 | 0.297796 | 0.765858655 | 0.869743072 | SRA1 | protein_coding | 45.85 | 0.004947098 | 0.004947098 | ENSG00000213523 | n.s. |
| ENSG00000132718 | 1572.084188 | 0.083656124 | 0.2888355 | 0.289632 | 0.772097486 | 0.87269957 | SYT11 | protein_coding | 46.5 | 0.004947025 | 0.004947025 | ENSG00000132718 | n.s. |
| ENSG00000085998 | 334.7216553 | 0.082446926 | 0.2804774 | 0.293952 | 0.768794486 | 0.871087698 | POMGNT1 | protein_coding | 49.39 | 0.004941714 | 0.004941714 | ENSG00000085998 | n.s. |
| ENSG00000117528 | 2033.528007 | -0.079968953 | 0.264859 | -0.30193 | 0.762705167 | 0.867627587 | ABCD3 | protein_coding | 37.04 | -0.004931417 | 0.004931417 | ENSG00000117528 | n.s. |
| ENSG00000165813 | 457.4841274 | -0.079314194 | 0.2615297 | -0.30327 | 0.761683828 | 0.866769426 | CCDC186 | protein_coding | 37.11 | -0.004925127 | 0.004925127 | ENSG00000165813 | n.s. |
| ENSG00000164104 | 12201.26925 | 0.081626612 | 0.2753402 | 0.296457 | 0.766880897 | 0.870292962 | HMGB2 | protein_coding | 50.22 | 0.004924903 | 0.004924903 | ENSG00000164104 | n.s. |
| ENSG00000140521 | 1544.018578 | 0.073840652 | 0.2299919 | 0.321058 | 0.748166675 | 0.858947882 | POLG | protein_coding | 49.14 | 0.004875934 | 0.004875934 | ENSG00000140521 | n.s. |
| ENSG00000117543 | 1156.146697 | -0.075326343 | 0.2390897 | -0.315055 | 0.752720142 | 0.861551845 | DPH5 | protein_coding | 37.71 | -0.004875014 | 0.004875014 | ENSG00000117543 | n.s. |
| ENSG00000132872 | 1904.142984 | 0.075790179 | 0.2416345 | 0.313656 | 0.753782143 | 0.862429844 | SYT4 | protein_coding | 34.83 | 0.004871506 | 0.004871506 | ENSG00000132872 | n.s. |
| ENSG00000128626 | 339.1661131 | 0.081798675 | 0.2819778 | 0.290089 | 0.77174814 | 0.872517203 | MRPS12 | protein_coding | 55.6 | 0.004844608 | 0.004844608 | ENSG00000128626 | n.s. |
| ENSG00000134265 | 717.3434051 | 0.076969708 | 0.2498453 | 0.308069 | 0.758029467 | 0.865227437 | NAPG | protein_coding | 39.34 | 0.004839064 | 0.004839064 | ENSG00000134265 | n.s. |
| ENSG00000125445 | 2320.477168 | 0.080504058 | 0.2740194 | 0.29379 | 0.768918699 | 0.871157544 | MRPS7 | protein_coding | 50.49 | 0.004822459 | 0.004822459 | ENSG00000125445 | n.s. |
| ENSG00000100918 | 951.2021728 | 0.077236078 | 0.2535878 | 0.304573 | 0.760691162 | 0.866206187 | REC8 | protein_coding | 55.21 | 0.004817888 | 0.004817888 | ENSG00000100918 | n.s. |
| ENSG00000140848 | 233.9937575 | -0.079069769 | 0.2648194 | -0.29858 | 0.765260536 | 0.869205616 | CPNE2 | protein_coding | 54.06 | -0.004813568 | 0.004813568 | ENSG00000140848 | n.s. |

| | | | | | | | | | | | | | |
|---|---|---|---|---|---|---|---|---|---|---|---|---|---|
| ENSG00000178057 | 336.5386471 | 0.075076529 | 0.2400682 | 0.31273 | 0.754485761 | 0.862804362 | NDUFAF3 | protein_coding | 64.14 | 0.00481148 | 0.00481148 | ENSG00000178057 | n.s. |
| ENSG00000205476 | 589.8610109 | -0.081292605 | 0.2804815 | -0.289832 | 0.771944515 | 0.872668356 | CCDC85C | protein_coding | 56.04 | -0.00480852 | 0.00480852 | ENSG00000205476 | n.s. |
| ENSG00000102172 | 1778.987802 | 0.079940279 | 0.2708689 | 0.295125 | 0.767898105 | 0.870709774 | SMS | protein_coding | 43.48 | 0.004806536 | 0.004806536 | ENSG00000102172 | n.s. |
| ENSG00000111641 | 1259.306639 | 0.077109603 | 0.2536279 | 0.304027 | 0.761107692 | 0.86644493 | NOP2 | protein_coding | 49.45 | 0.00480077 | 0.00480077 | ENSG00000111641 | n.s. |
| ENSG00000281091 | 21.25712291 | -0.087336818 | 0.3206781 | -0.27235 | 0.785352601 | 0.881172074 | | lincRNA | 41.04 | -0.004798221 | 0.004798221 | ENSG00000281091 | n.s. |
| ENSG00000163002 | 1923.428758 | 0.077418835 | 0.2557462 | 0.302717 | 0.762105198 | 0.867036335 | NUP35 | protein_coding | 38.01 | 0.00479708 | 0.00479708 | ENSG00000163002 | n.s. |
| ENSG00000164610 | 365.0625136 | -0.080870128 | 0.2785802 | -0.290294 | 0.771591423 | 0.872517203 | RP9 | protein_coding | 42.71 | -0.004789614 | 0.004789614 | ENSG00000164610 | n.s. |
| ENSG00000011677 | 289.5917718 | 0.081909054 | 0.2861141 | 0.286281 | 0.77466283 | 0.874038135 | GABRA3 | protein_coding | 37.36 | 0.004789191 | 0.004789191 | ENSG00000011677 | n.s. |
| ENSG00000120265 | 1654.015418 | -0.085649305 | 0.3109989 | -0.275401 | 0.783008406 | 0.879463217 | PCMT1 | protein_coding | 41.51 | -0.004777717 | 0.004777717 | ENSG00000120265 | n.s. |
| ENSG00000167699 | 1237.158632 | -0.085209342 | 0.3082847 | -0.276398 | 0.782242206 | 0.878886236 | GLOD4 | protein_coding | 45.54 | -0.004777461 | 0.004777461 | ENSG00000167699 | n.s. |
| ENSG00000136813 | 3037.319974 | 0.076043314 | 0.2482178 | 0.306357 | 0.75933272 | 0.865692775 | KIAA0368 | protein_coding | 39.33 | 0.004763065 | 0.004763065 | ENSG00000136813 | n.s. |
| ENSG00000144118 | 252.9204851 | 0.081174503 | 0.2827743 | 0.287065 | 0.774062824 | 0.873778366 | RALB | protein_coding | 46.37 | 0.004756721 | 0.004756721 | ENSG00000144118 | n.s. |
| ENSG00000115317 | 485.7574089 | -0.080748375 | 0.2799297 | -0.28846 | 0.772995019 | 0.873217822 | HTRA2 | protein_coding | 54.73 | -0.004754255 | 0.004754255 | ENSG00000115317 | n.s. |
| ENSG00000082898 | 9472.186249 | 0.079132416 | 0.2688618 | 0.294324 | 0.768510525 | 0.870869029 | XPO1 | protein_coding | 40.9 | 0.004751676 | 0.004751676 | ENSG00000082898 | n.s. |
| ENSG00000212664 | 42.9920745 | 0.114892883 | 0.5319586 | 0.215981 | 0.829002663 | 0.909192917 | | processed_pseudogene | 46.13 | 0.004750126 | 0.004750126 | ENSG00000212664 | n.s. |
| ENSG00000120705 | 3072.910949 | -0.080772035 | 0.2803733 | -0.288087 | 0.773279767 | 0.87336617 | ETF1 | protein_coding | 43.04 | -0.004749689 | 0.004749689 | ENSG00000120705 | n.s. |
| ENSG00000188010 | 212.0386405 | 0.089681355 | 0.3391741 | 0.264411 | 0.791463232 | 0.885317152 | MORN2 | protein_coding | 39.59 | 0.004744244 | 0.004744244 | ENSG00000188010 | n.s. |
| ENSG00000143727 | 4876.402005 | 0.08221775 | 0.2899 | 0.283607 | 0.776711386 | 0.875710797 | ACP1 | protein_coding | 42.67 | 0.004738973 | 0.004738973 | ENSG00000143727 | n.s. |
| ENSG00000151726 | 968.5963129 | -0.079152593 | 0.2704379 | -0.292683 | 0.769764447 | 0.871573602 | ACSL1 | protein_coding | 44.71 | -0.004725088 | 0.004725088 | ENSG00000151726 | n.s. |
| ENSG00000092758 | 157.2643404 | -0.077488359 | 0.25892 | -0.299275 | 0.764730014 | 0.869148157 | COL9A3 | protein_coding | 61.84 | -0.00471952 | 0.00471952 | ENSG00000092758 | n.s. |
| ENSG00000145029 | 90.87264186 | 0.082933915 | 0.2957392 | 0.280429 | 0.779148252 | 0.877321562 | NICN1 | protein_coding | 52.25 | 0.004714063 | 0.004714063 | ENSG00000145029 | n.s. |
| ENSG00000166086 | 1043.015179 | 0.080117561 | 0.2781207 | 0.288068 | 0.773294996 | 0.87336617 | JAM3 | protein_coding | 41.98 | 0.004711204 | 0.004711204 | ENSG00000166086 | n.s. |
| ENSG00000168496 | 3292.130454 | 0.075207027 | 0.2454587 | 0.306394 | 0.759304844 | 0.865692775 | FEN1 | protein_coding | 49.78 | 0.004710683 | 0.004710683 | ENSG00000168496 | n.s. |
| ENSG00000189403 | 18007.43037 | 0.077316126 | 0.2586521 | 0.298919 | 0.765001518 | 0.869148157 | HMGB1 | protein_coding | 41.99 | 0.00470903 | 0.00470903 | ENSG00000189403 | n.s. |
| ENSG00000127022 | 9675.760897 | -0.078797998 | 0.2689719 | -0.29296 | 0.769552736 | 0.87155587 | CANX | protein_coding | 45.8 | -0.004704616 | 0.004704616 | ENSG00000127022 | n.s. |
| ENSG00000196776 | 2289.845818 | -0.081276223 | 0.2863095 | -0.283875 | 0.776505906 | 0.875550027 | CD47 | protein_coding | 36.88 | -0.004691185 | 0.004691185 | ENSG00000196776 | n.s. |
| ENSG00000110367 | 3212.907316 | -0.073189286 | 0.2342734 | -0.31241 | 0.754729177 | 0.862804362 | DDX6 | protein_coding | 41.56 | -0.004690531 | 0.004690531 | ENSG00000110367 | n.s. |
| ENSG00000065548 | 5145.25937 | -0.077723061 | 0.2624574 | -0.296136 | 0.76712626 | 0.870349013 | ZC3H15 | protein_coding | 37.22 | -0.00468721 | 0.00468721 | ENSG00000065548 | n.s. |
| ENSG00000262814 | 862.8079402 | -0.078902965 | 0.2710051 | -0.291149 | 0.770937117 | 0.872370708 | MRPL12 | protein_coding | 57.67 | -0.004678861 | 0.004678861 | ENSG00000262814 | n.s. |
| ENSG00000197930 | 958.0949358 | -0.075228078 | 0.2482569 | -0.303025 | 0.761870675 | 0.866849351 | ERO1A | protein_coding | 37.59 | -0.004668382 | 0.004668382 | ENSG00000197930 | n.s. |
| ENSG00000126821 | 286.0866082 | -0.082544668 | 0.2958724 | -0.278987 | 0.780254477 | 0.877928163 | SGPP1 | protein_coding | 39.66 | -0.00466716 | 0.00466716 | ENSG00000126821 | n.s. |
| ENSG00000052749 | 711.0345163 | -0.07658632 | 0.2562609 | -0.298861 | 0.765046352 | 0.869148157 | RRP12 | protein_coding | 50.49 | -0.004664581 | 0.004664581 | ENSG00000052749 | n.s. |

| | | | | | | | | | | | | | |
|---|---|---|---|---|---|---|---|---|---|---|---|---|---|
| ENSG00000177917 | 1174.760162 | 0.080261271 | 0.2815175 | 0.285102 | 0.775565835 | 0.874915177 | ARL6IP6 | protein_coding | 36.91 | 0.004657887 | 0.004657887 | ENSG00000177917 | n.s. |
| ENSG00000258920 | 47.97752256 | 0.080165701 | 0.2812832 | 0.285 | 0.77564418 | 0.874932668 | FOXN3-AS1 | antisense | 54.67 | 0.004651644 | 0.004651644 | ENSG00000258920 | n.s. |
| ENSG00000117477 | 206.8347268 | 0.077438574 | 0.262956 | 0.294493 | 0.76838157 | 0.870869029 | CCDC181 | protein_coding | 36.8 | 0.004649966 | 0.004649966 | ENSG00000117477 | n.s. |
| ENSG00000185049 | 401.3485393 | 0.076655483 | 0.2572838 | 0.297941 | 0.765747949 | 0.869688286 | NELFA | protein_coding | 48.3 | 0.004648111 | 0.004648111 | ENSG00000185049 | n.s. |
| ENSG00000196312 | 387.7620068 | 0.074483636 | 0.2448558 | 0.304194 | 0.760980171 | 0.866393566 | MFSD14C | protein_coding | 44.07 | 0.004639197 | 0.004639197 | ENSG00000196312 | n.s. |
| ENSG00000100764 | 1635.723381 | -0.075164755 | 0.2497737 | -0.300931 | 0.763466803 | 0.868172926 | PSMC1 | protein_coding | 44.79 | -0.004614647 | 0.004614647 | ENSG00000100764 | n.s. |
| ENSG00000151176 | 173.5325229 | 0.077185864 | 0.2636009 | 0.292813 | 0.769664808 | 0.871573602 | PLBD2 | protein_coding | 51.13 | 0.004607682 | 0.004607682 | ENSG00000151176 | n.s. |
| ENSG00000198804 | 38049.13877 | -0.073978945 | 0.2433748 | -0.303971 | 0.761149748 | 0.86644493 | MT-CO1 | protein_coding | 46.24 | -0.004605858 | 0.004605858 | ENSG00000198804 | n.s. |
| ENSG00000196504 | 8182.122886 | -0.076896097 | 0.2620686 | -0.29342 | 0.769201355 | 0.871406873 | PRPF40A | protein_coding | 36.31 | -0.004596773 | 0.004596773 | ENSG00000196504 | n.s. |
| ENSG00000152382 | 1143.919718 | -0.081276892 | 0.2917165 | -0.278616 | 0.780539501 | 0.877937651 | TADA1 | protein_coding | 40.95 | -0.004595097 | 0.004595097 | ENSG00000152382 | n.s. |
| ENSG00000076356 | 60.64848898 | 0.07517334 | 0.2515218 | 0.298874 | 0.765036188 | 0.869148157 | PLXNA2 | protein_coding | 47.98 | 0.004578521 | 0.004578521 | ENSG00000076356 | n.s. |
| ENSG00000236194 | 276.4404641 | 0.072418332 | 0.234399 | 0.308953 | 0.757357076 | 0.864672618 | | sense_intronic | 36.73 | 0.004573094 | 0.004573094 | ENSG00000236194 | n.s. |
| ENSG00000129515 | 1107.628406 | -0.085613043 | 0.3215001 | -0.266292 | 0.790014019 | 0.884368353 | SNX6 | protein_coding | 44.34 | -0.004568895 | 0.004568895 | ENSG00000129515 | n.s. |
| ENSG00000198910 | 659.8143817 | -0.074592356 | 0.2483984 | -0.300293 | 0.763953541 | 0.868500461 | L1CAM | protein_coding | 56.65 | -0.004567286 | 0.004567286 | ENSG00000198910 | n.s. |
| ENSG00000136146 | 1319.048703 | 0.077954041 | 0.2718825 | 0.28672 | 0.774327039 | 0.873885547 | MED4 | protein_coding | 40.77 | 0.004563854 | 0.004563854 | ENSG00000136146 | n.s. |
| ENSG00000162704 | 1417.078464 | 0.084900443 | 0.3179973 | 0.266985 | 0.789480882 | 0.883880855 | ARPC5 | protein_coding | 38.38 | 0.004551196 | 0.004551196 | ENSG00000162704 | n.s. |
| ENSG00000144034 | 946.4027333 | -0.084651156 | 0.3164459 | -0.267506 | 0.78907964 | 0.883573736 | TPRKB | protein_coding | 40.52 | -0.004550609 | 0.004550609 | ENSG00000144034 | n.s. |
| ENSG00000173786 | 434.9852041 | 0.075378261 | 0.2542911 | 0.296425 | 0.766905434 | 0.870292962 | CNP | protein_coding | 54.63 | 0.004547912 | 0.004547912 | ENSG00000173786 | n.s. |
| ENSG00000102882 | 192.8681543 | -0.080186028 | 0.2872133 | -0.279186 | 0.780101861 | 0.877827383 | MAPK3 | protein_coding | 56.36 | -0.004537798 | 0.004537798 | ENSG00000102882 | n.s. |
| ENSG00000164902 | 1443.074687 | -0.074485065 | 0.249257 | -0.298828 | 0.765071004 | 0.869148157 | PHAX | protein_coding | 43.51 | -0.004536601 | 0.004536601 | ENSG00000164902 | n.s. |
| ENSG00000187778 | 1509.619789 | 0.077021221 | 0.2673957 | 0.288042 | 0.773314526 | 0.87336617 | MCRS1 | protein_coding | 55.12 | 0.004529128 | 0.004529128 | ENSG00000187778 | n.s. |
| ENSG00000278040 | 7.917116504 | -0.10603496 | 0.4787939 | -0.221463 | 0.824732238 | 0.906365691 | | misc_RNA | 50.6 | -0.004527326 | 0.004527326 | ENSG00000278040 | n.s. |
| ENSG00000159111 | 628.8631563 | -0.078722524 | 0.2786347 | -0.282529 | 0.777537546 | 0.876003824 | MRPL10 | protein_coding | 49.17 | -0.004526073 | 0.004526073 | ENSG00000159111 | n.s. |
| ENSG00000064607 | 2528.192388 | -0.075783198 | 0.259107 | -0.292478 | 0.769920867 | 0.871583725 | SUGP2 | protein_coding | 50 | -0.004523566 | 0.004523566 | ENSG00000064607 | n.s. |
| ENSG00000104331 | 1559.136398 | 0.076267013 | 0.2619209 | 0.291183 | 0.770911035 | 0.872370708 | IMPAD1 | protein_coding | 38.21 | 0.004522552 | 0.004522552 | ENSG00000104331 | n.s. |
| ENSG00000181666 | 382.6085667 | -0.072063987 | 0.2342289 | -0.307665 | 0.75833741 | 0.865507973 | HKR1 | protein_coding | 43.83 | -0.004520497 | 0.004520497 | ENSG00000181666 | n.s. |
| ENSG00000119396 | 1031.743457 | -0.076257902 | 0.2624403 | -0.290572 | 0.771378427 | 0.872517203 | RAB14 | protein_coding | 39.61 | -0.004516451 | 0.004516451 | ENSG00000119396 | n.s. |
| ENSG00000185513 | 34.37633052 | -0.082935129 | 0.3066155 | -0.270486 | 0.78678659 | 0.882212265 | L3MBTL1 | protein_coding | 49.19 | -0.004513903 | 0.004513903 | ENSG00000185513 | n.s. |
| ENSG00000172020 | 328.695839 | -0.083065617 | 0.3075846 | -0.270058 | 0.787115808 | 0.882439278 | GAP43 | protein_coding | 38.61 | -0.004511723 | 0.004511723 | ENSG00000172020 | n.s. |
| ENSG00000132300 | 5458.868062 | -0.074680486 | 0.2525811 | -0.295669 | 0.767482608 | 0.870522228 | PTCD3 | protein_coding | 41.22 | -0.004497269 | 0.004497269 | ENSG00000132300 | n.s. |
| ENSG00000137513 | 873.7318021 | -0.074781053 | 0.2532336 | -0.295305 | 0.767761175 | 0.87069633 | NARS2 | protein_coding | 38.77 | -0.00449683 | 0.00449683 | ENSG00000137513 | n.s. |
| ENSG00000154957 | 262.3877267 | -0.075296839 | 0.2570218 | -0.292959 | 0.769553516 | 0.87155587 | ZNF18 | protein_coding | 41.67 | -0.00449558 | 0.00449558 | ENSG00000154957 | n.s. |

| | | | | | | | | | | | | | |
|---|---|---|---|---|---|---|---|---|---|---|---|---|---|
| ENSG00000141562 | 1622.012992 | 0.075303426 | 0.2573843 | 0.292572 | 0.769849325 | 0.871573602 | NARF | protein_coding | 51.72 | 0.004495308 | 0.004495308 | ENSG00000141562 | n.s. |
| ENSG00000108523 | 421.84547 | 0.078158082 | 0.2763472 | 0.282826 | 0.777310497 | 0.875960668 | RNF167 | protein_coding | 52.54 | 0.004495293 | 0.004495293 | ENSG00000108523 | n.s. |
| ENSG00000164024 | 1428.884958 | 0.077540548 | 0.2727119 | 0.284331 | 0.776156447 | 0.875297766 | METAP1 | protein_coding | 39.1 | 0.004485269 | 0.004485269 | ENSG00000164024 | n.s. |
| ENSG00000117984 | 314.450224 | 0.07450653 | 0.2526951 | 0.294848 | 0.768110374 | 0.870825292 | CTSD | protein_coding | 64.73 | 0.00447553 | 0.00447553 | ENSG00000117984 | n.s. |
| ENSG00000169062 | 1346.102522 | 0.071421826 | 0.2334638 | 0.305922 | 0.759663697 | 0.865692775 | UPF3A | protein_coding | 47.58 | 0.004473592 | 0.004473592 | ENSG00000169062 | n.s. |
| ENSG00000139946 | 2875.2027 | -0.074497786 | 0.2531471 | -0.294287 | 0.768538947 | 0.870869029 | PELI2 | protein_coding | 39.91 | -0.00447338 | 0.00447338 | ENSG00000139946 | n.s. |
| ENSG00000120314 | 362.847519 | -0.0743345 | 0.251642 | -0.295398 | 0.767689985 | 0.870686515 | WDR55 | protein_coding | 53.17 | -0.004470342 | 0.004470342 | ENSG00000120314 | n.s. |
| ENSG00000217527 | 88.11017676 | -0.077127354 | 0.2710377 | -0.284563 | 0.775978786 | 0.875168298 | RPS16P5 | transcribed_processed_pseudog | 50 | -0.004466323 | 0.004466323 | ENSG00000217527 | n.s. |
| ENSG00000230922 | 193.5107961 | -0.073586097 | 0.2477642 | -0.297001 | 0.766466077 | 0.870007109 | | sense_intronic | 51.69 | -0.004450281 | 0.004450281 | ENSG00000230922 | n.s. |
| ENSG00000111906 | 2067.699587 | -0.080108169 | 0.2927693 | -0.273622 | 0.784375014 | 0.880372696 | HDDC2 | protein_coding | 40.11 | -0.00443266 | 0.00443266 | ENSG00000111906 | n.s. |
| ENSG00000085365 | 1669.923462 | -0.080570492 | 0.2959666 | -0.272228 | 0.785446482 | 0.881206395 | SCAMP1 | protein_coding | 37.59 | -0.004425122 | 0.004425122 | ENSG00000085365 | n.s. |
| ENSG00000059573 | 1726.071493 | -0.077962752 | 0.2789422 | -0.279494 | 0.77986555 | 0.877684408 | ALDH18A1 | protein_coding | 43.16 | -0.004417496 | 0.004417496 | ENSG00000059573 | n.s. |
| ENSG00000160058 | 387.9715249 | -0.079165235 | 0.2876898 | -0.275176 | 0.783181274 | 0.879586422 | BSDC1 | protein_coding | 47.99 | -0.004411204 | 0.004411204 | ENSG00000160058 | n.s. |
| ENSG00000159399 | 3173.957141 | 0.071576559 | 0.2376108 | 0.301234 | 0.763235756 | 0.868038836 | HK2 | protein_coding | 47.62 | 0.004399156 | 0.004399156 | ENSG00000159399 | n.s. |
| ENSG00000161013 | 339.582304 | 0.074234413 | 0.2557903 | 0.290216 | 0.771651043 | 0.872517203 | MGAT4B | protein_coding | 64.04 | 0.004396607 | 0.004396607 | ENSG00000161013 | n.s. |
| ENSG00000114346 | 3805.114773 | -0.076381872 | 0.2702853 | -0.282597 | 0.777485651 | 0.876003824 | ECT2 | protein_coding | 34.96 | -0.004391499 | 0.004391499 | ENSG00000114346 | n.s. |
| ENSG00000200105 | 4.44939933 | 0.112585657 | 0.5521407 | 0.203908 | 0.838425757 | 0.914127002 | RNU6-251P | snRNA | 39.42 | 0.004390105 | 0.004390105 | ENSG00000200105 | n.s. |
| ENSG00000134899 | 86.01972407 | -0.07352842 | 0.2510187 | -0.29292 | 0.76958327 | 0.87155587 | ERCC5 | protein_coding | 39.83 | -0.004389997 | 0.004389997 | ENSG00000134899 | n.s. |
| ENSG00000077549 | 317.5424526 | -0.077927604 | 0.2810545 | -0.277269 | 0.781573846 | 0.878347942 | CAPZB | protein_coding | 49.21 | -0.004389928 | 0.004389928 | ENSG00000077549 | n.s. |
| ENSG00000151292 | 2194.16433 | -0.077308841 | 0.2762743 | -0.279826 | 0.77961072 | 0.877487475 | CSNK1G3 | protein_coding | 34.48 | -0.004387978 | 0.004387978 | ENSG00000151292 | n.s. |
| ENSG00000203880 | 1425.293787 | 0.07816758 | 0.2827385 | 0.276466 | 0.782190213 | 0.878886236 | PCMTD2 | protein_coding | 45.7 | 0.004382648 | 0.004382648 | ENSG00000203880 | n.s. |
| ENSG00000065485 | 66.90689235 | 0.072294941 | 0.243064 | 0.297432 | 0.766136954 | 0.869932466 | PDIA5 | protein_coding | 45.21 | 0.004374889 | 0.004374889 | ENSG00000065485 | n.s. |
| ENSG00000253092 | 8.510161937 | -0.096276124 | 0.4126748 | -0.233298 | 0.815530176 | 0.900660478 | | snoRNA | 37.88 | -0.004374681 | 0.004374681 | ENSG00000253092 | n.s. |
| ENSG00000239437 | 13.61744685 | -0.093347331 | 0.3891487 | -0.239876 | 0.810426582 | 0.897730074 | RN7SL752P | misc_RNA | 55.4 | -0.004373717 | 0.004373717 | ENSG00000239437 | n.s. |
| ENSG00000242110 | 235.2465446 | -0.07316237 | 0.2500026 | -0.292646 | 0.76979242 | 0.871573602 | AMACR | protein_coding | 41.36 | -0.004367496 | 0.004367496 | ENSG00000242110 | n.s. |
| ENSG00000139173 | 131.0029901 | -0.08069929 | 0.2998147 | -0.269164 | 0.787803552 | 0.882854866 | TMEM117 | protein_coding | 37.35 | -0.004366694 | 0.004366694 | ENSG00000139173 | n.s. |
| ENSG00000103353 | 1631.834143 | 0.073661961 | 0.2542594 | 0.289712 | 0.772036651 | 0.87269957 | UBFD1 | protein_coding | 47.48 | 0.004356018 | 0.004356018 | ENSG00000103353 | n.s. |
| ENSG00000222561 | 8.395749073 | -0.10018352 | 0.4457202 | -0.224768 | 0.822159967 | 0.904752863 | RNU6-1025P | snRNA | 36.54 | -0.004354981 | 0.004354981 | ENSG00000222561 | n.s. |
| ENSG00000238713 | 38.091455 | -0.083127787 | 0.3181687 | -0.26127 | 0.793884682 | 0.886695569 | | misc_RNA | 43.36 | -0.004341387 | 0.004341387 | ENSG00000238713 | n.s. |
| ENSG00000146776 | 66.78949273 | 0.072920653 | 0.2497839 | 0.291935 | 0.770336344 | 0.871953515 | ATXN7L1 | protein_coding | 45.57 | 0.004339265 | 0.004339265 | ENSG00000146776 | n.s. |
| ENSG00000128606 | 129.5989674 | -0.079805019 | 0.2957713 | -0.26982 | 0.787298698 | 0.882508391 | LRRC17 | protein_coding | 39.14 | -0.004331909 | 0.004331909 | ENSG00000128606 | n.s. |
| ENSG00000167550 | 52.36712943 | 0.080553606 | 0.3008807 | 0.267726 | 0.788910196 | 0.883573736 | RHEBL1 | protein_coding | 49.73 | 0.004330336 | 0.004330336 | ENSG00000167550 | n.s. |

| | | | | | | | | | | | | | |
|---|---|---|---|---|---|---|---|---|---|---|---|---|---|
| ENSG00000228106 | 159.4013401 | 0.081248834 | 0.3056411 | 0.265831 | 0.790369476 | 0.884555268 | | lincRNA | 37.92 | 0.004328534 | 0.004328534 | ENSG00000228106 | n.s. |
| ENSG00000247708 | 63.36036651 | -0.075059295 | 0.2651268 | -0.283107 | 0.777094679 | 0.87583137 | STX18-AS1 | antisense | 45.62 | -0.004321877 | 0.004321877 | ENSG00000247708 | n.s. |
| ENSG00000165996 | 167.4383573 | -0.081433191 | 0.3077577 | -0.264602 | 0.791316368 | 0.885223991 | HACD1 | protein_coding | 41.97 | -0.004311629 | 0.004311629 | ENSG00000165996 | n.s. |
| ENSG00000106976 | 30.81777785 | -0.083313511 | 0.3217709 | -0.258922 | 0.795695544 | 0.887767529 | DNM1 | protein_coding | 54.26 | -0.004307371 | 0.004307371 | ENSG00000106976 | n.s. |
| ENSG00000071626 | 1697.376083 | 0.078925845 | 0.2912266 | 0.271012 | 0.786381952 | 0.881971638 | DAZAP1 | protein_coding | 58.75 | 0.00430504 | 0.00430504 | ENSG00000071626 | n.s. |
| ENSG00000198042 | 1380.527192 | -0.07099066 | 0.2390157 | -0.297013 | 0.766456953 | 0.870007109 | MAK16 | protein_coding | 40.72 | -0.004293316 | 0.004293316 | ENSG00000198042 | n.s. |
| ENSG00000228223 | 635.5603303 | 0.068510746 | 0.2241256 | 0.30568 | 0.759848149 | 0.865812732 | HCG11 | lincRNA | 39.3 | 0.004287131 | 0.004287131 | ENSG00000228223 | n.s. |
| ENSG00000173852 | 590.9237176 | -0.075384231 | 0.2692199 | -0.28001 | 0.779469893 | 0.877487475 | DPY19L1 | protein_coding | 37.49 | -0.004278739 | 0.004278739 | ENSG00000173852 | n.s. |
| ENSG00000142330 | 94.60450027 | 0.076910149 | 0.2799979 | 0.274681 | 0.783561202 | 0.87980021 | CAPN10 | protein_coding | 54.59 | 0.00427743 | 0.00427743 | ENSG00000142330 | n.s. |
| ENSG00000181929 | 346.17223 | -0.078261752 | 0.2889575 | -0.270842 | 0.786512767 | 0.881976261 | PRKAG1 | protein_coding | 45.41 | -0.004268639 | 0.004268639 | ENSG00000181929 | n.s. |
| ENSG00000233593 | 31.16911352 | -0.087133608 | 0.3508262 | -0.248367 | 0.803850555 | 0.893359776 | | processed_transcript | 41.8 | -0.004267247 | 0.004267247 | ENSG00000233593 | n.s. |
| ENSG00000066697 | 601.4867016 | -0.079362137 | 0.2966388 | -0.267538 | 0.789054998 | 0.883573736 | MSANTD3 | protein_coding | 43.76 | -0.004266286 | 0.004266286 | ENSG00000066697 | n.s. |
| ENSG00000178927 | 833.885415 | -0.071438144 | 0.2447456 | -0.291887 | 0.770372778 | 0.871953515 | C17orf62 | protein_coding | 60.43 | -0.004251046 | 0.004251046 | ENSG00000178927 | n.s. |
| ENSG00000171004 | 189.2731926 | -0.081421241 | 0.3119915 | -0.260973 | 0.794113615 | 0.88678568 | HS6ST2 | protein_coding | 41.21 | -0.004248668 | 0.004248668 | ENSG00000171004 | n.s. |
| ENSG00000201863 | 13.12207254 | 0.092066864 | 0.3911234 | 0.235391 | 0.813905383 | 0.899282662 | | snoRNA | 53.23 | 0.00424463 | 0.00424463 | ENSG00000201863 | n.s. |
| ENSG00000141542 | 311.0900082 | -0.081110375 | 0.3106088 | -0.261134 | 0.793989527 | 0.886718231 | RAB40B | protein_coding | 53.03 | -0.004235126 | 0.004235126 | ENSG00000141542 | n.s. |
| ENSG00000131943 | 287.4841474 | -0.0820185 | 0.3177576 | -0.258117 | 0.796316969 | 0.888106469 | C19orf12 | protein_coding | 50.38 | -0.004226821 | 0.004226821 | ENSG00000131943 | n.s. |
| ENSG00000105058 | 1550.33025 | 0.078614339 | 0.2940607 | 0.267341 | 0.789207 | 0.88364528 | FAM32A | protein_coding | 52.93 | 0.004223322 | 0.004223322 | ENSG00000105058 | n.s. |
| ENSG00000100664 | 7816.294214 | -0.072104845 | 0.2517235 | -0.286445 | 0.774537569 | 0.874016492 | EIF5 | protein_coding | 44.94 | -0.004216718 | 0.004216718 | ENSG00000100664 | n.s. |
| ENSG00000028528 | 1704.535829 | -0.073518701 | 0.2606859 | -0.28202 | 0.777927995 | 0.876372803 | SNX1 | protein_coding | 43.68 | -0.004213438 | 0.004213438 | ENSG00000028528 | n.s. |
| ENSG00000093009 | 1196.36303 | -0.075494852 | 0.2744449 | -0.275082 | 0.783253234 | 0.879596288 | CDC45 | protein_coding | 48.03 | -0.004206317 | 0.004206317 | ENSG00000093009 | n.s. |
| ENSG00000127483 | 2103.900432 | -0.074489633 | 0.2676894 | -0.278269 | 0.780805934 | 0.878122861 | HP1BP3 | protein_coding | 41.17 | -0.004204546 | 0.004204546 | ENSG00000127483 | n.s. |
| ENSG00000168883 | 2195.135411 | 0.073942536 | 0.2636122 | 0.280497 | 0.779095893 | 0.877321562 | USP39 | protein_coding | 46.55 | 0.004202982 | 0.004202982 | ENSG00000168883 | n.s. |
| ENSG00000171843 | 335.3722124 | 0.075532604 | 0.274891 | 0.274773 | 0.783490687 | 0.879791986 | MLLT3 | protein_coding | 37.12 | 0.004201123 | 0.004201123 | ENSG00000171843 | n.s. |
| ENSG00000248333 | 218.3255534 | -0.071692959 | 0.2498086 | -0.286992 | 0.774118807 | 0.873778366 | CDK11B | protein_coding | 51.91 | -0.004201115 | 0.004201115 | ENSG00000248333 | n.s. |
| ENSG00000101940 | 328.0357315 | 0.074487481 | 0.2682794 | 0.277649 | 0.781281892 | 0.878298727 | WDR13 | protein_coding | 51.72 | 0.004197947 | 0.004197947 | ENSG00000101940 | n.s. |
| ENSG00000235316 | 62.20621872 | -0.07392961 | 0.2638862 | -0.280157 | 0.779356892 | 0.877485525 | DUSP8P5 | processed_pseudogene | 67.6 | -0.004196248 | 0.004196248 | ENSG00000235316 | n.s. |
| ENSG00000134108 | 1047.529859 | -0.079294011 | 0.3008944 | -0.263528 | 0.792143864 | 0.885691491 | ARL8B | protein_coding | 40.82 | -0.004180184 | 0.004180184 | ENSG00000134108 | n.s. |
| ENSG00000185495 | 181.012018 | -0.075470043 | 0.2758345 | -0.273606 | 0.784387222 | 0.880372696 | | transcribed_unprocessed_pseud | 36.59 | -0.004176017 | 0.004176017 | ENSG00000185495 | n.s. |
| ENSG00000262652 | 53.28177137 | -0.079008158 | 0.2991279 | -0.264128 | 0.79168099 | 0.885424735 | | sense_intronic | 45.29 | -0.004175451 | 0.004175451 | ENSG00000262652 | n.s. |
| ENSG00000173436 | 310.4228356 | 0.074341558 | 0.269387 | 0.275966 | 0.782574411 | 0.879089755 | MINOS1 | protein_coding | 42.96 | 0.004160658 | 0.004160658 | ENSG00000173436 | n.s. |
| ENSG00000160075 | 621.2425507 | -0.073787116 | 0.2658448 | -0.277557 | 0.781352364 | 0.878298727 | SSU72 | protein_coding | 53.87 | -0.004158476 | 0.004158476 | ENSG00000160075 | n.s. |

| | | | | | | | | | | | | | |
|---|---|---|---|---|---|---|---|---|---|---|---|---|---|
| ENSG00000153037 | 1091.434252 | -0.07423133 | 0.2689585 | -0.275995 | 0.78255154 | 0.879089755 | SRP19 | protein_coding | 41.11 | -0.004154489 | 0.004154489 | ENSG00000153037 | n.s. |
| ENSG00000180964 | 915.7534801 | -0.081889157 | 0.3212601 | -0.2549 | 0.798800431 | 0.889878253 | TCEAL8 | protein_coding | 46.4 | -0.004149275 | 0.004149275 | ENSG00000180964 | n.s. |
| ENSG00000166402 | 2251.848836 | 0.068440387 | 0.2303891 | 0.297064 | 0.766417429 | 0.870007109 | TUB | protein_coding | 49.43 | 0.004139083 | 0.004139083 | ENSG00000166402 | n.s. |
| ENSG00000206728 | 11.88998794 | 0.092492242 | 0.4021879 | 0.229973 | 0.818112995 | 0.90236768 | | misc_RNA | 44.23 | 0.004126677 | 0.004126677 | ENSG00000206728 | n.s. |
| ENSG00000277775 | 929.0567629 | 0.0642652 | 0.2050683 | 0.313384 | 0.753988659 | 0.862595242 | HIST1H3F | protein_coding | 60.83 | 0.004125372 | 0.004125372 | ENSG00000277775 | n.s. |
| ENSG00000115806 | 1194.243492 | 0.072355024 | 0.2575109 | 0.280978 | 0.778726924 | 0.877059939 | GORASP2 | protein_coding | 41.04 | 0.004122118 | 0.004122118 | ENSG00000115806 | n.s. |
| ENSG00000010270 | 806.5723111 | -0.079835962 | 0.3094416 | -0.258 | 0.796406839 | 0.888135556 | STARD3NL | protein_coding | 38.4 | -0.004113208 | 0.004113208 | ENSG00000010270 | n.s. |
| ENSG00000259488 | 711.8108893 | -0.064415276 | 0.2069772 | -0.311219 | 0.755633965 | 0.863271658 | | antisense | 43.49 | -0.004113077 | 0.004113077 | ENSG00000259488 | n.s. |
| ENSG00000146410 | 95.53793713 | -0.069437001 | 0.239171 | -0.290324 | 0.771568643 | 0.872517203 | MTFR2 | protein_coding | 39.81 | -0.004112476 | 0.004112476 | ENSG00000146410 | n.s. |
| ENSG00000140022 | 141.7501966 | 0.071914263 | 0.2555111 | 0.281453 | 0.778363247 | 0.87672126 | STON2 | protein_coding | 41.03 | 0.00410907 | 0.00410907 | ENSG00000140022 | n.s. |
| ENSG00000101246 | 388.2729243 | -0.067901586 | 0.2283066 | -0.297414 | 0.766150391 | 0.869932466 | ARFRP1 | protein_coding | 62.43 | -0.004109028 | 0.004109028 | ENSG00000101246 | n.s. |
| ENSG00000089737 | 5094.143403 | 0.070231941 | 0.2452334 | 0.286388 | 0.774580873 | 0.874016492 | DDX24 | protein_coding | 43.37 | 0.00410719 | 0.00410719 | ENSG00000089737 | n.s. |
| ENSG00000126216 | 1152.973279 | -0.068060537 | 0.2299292 | -0.296007 | 0.767225105 | 0.870371966 | TUBGCP3 | protein_coding | 42.87 | -0.004103717 | 0.004103717 | ENSG00000126216 | n.s. |
| ENSG00000051009 | 255.1394899 | 0.07229352 | 0.2583385 | 0.27984 | 0.779600006 | 0.877487475 | FAM160A2 | protein_coding | 44.55 | 0.004103313 | 0.004103313 | ENSG00000051009 | n.s. |
| ENSG00000171204 | 508.4191257 | -0.078927741 | 0.3039939 | -0.259636 | 0.795144637 | 0.88743878 | TMEM126B | protein_coding | 37.32 | -0.004093319 | 0.004093319 | ENSG00000171204 | n.s. |
| ENSG00000196535 | 273.4010321 | -0.072180923 | 0.2583111 | -0.279434 | 0.779911764 | 0.877684408 | MYO18A | protein_coding | 53.74 | -0.004089888 | 0.004089888 | ENSG00000196535 | n.s. |
| ENSG00000277544 | 60.28953966 | -0.081518186 | 0.3228295 | -0.252512 | 0.800645672 | 0.891007089 | | unprocessed_pseudogene | 40.59 | -0.004085597 | 0.004085597 | ENSG00000277544 | n.s. |
| ENSG00000108588 | 3131.233908 | -0.072226136 | 0.2591247 | -0.278731 | 0.780451141 | 0.877937651 | CCDC47 | protein_coding | 42.02 | -0.0040834 | 0.0040834 | ENSG00000108588 | n.s. |
| ENSG00000152223 | 619.6737229 | -0.068259914 | 0.2328161 | -0.293192 | 0.769375067 | 0.871532753 | EPG5 | protein_coding | 40.87 | -0.004076229 | 0.004076229 | ENSG00000152223 | n.s. |
| ENSG00000118507 | 91.85549766 | -0.074795851 | 0.2767291 | -0.270285 | 0.786940687 | 0.882313994 | AKAP7 | protein_coding | 38.22 | -0.004067161 | 0.004067161 | ENSG00000118507 | n.s. |
| ENSG00000105472 | 52.29322544 | 0.077093507 | 0.2926873 | 0.263399 | 0.792243183 | 0.885691491 | CLEC11A | protein_coding | 66.4 | 0.004064179 | 0.004064179 | ENSG00000105472 | n.s. |
| ENSG00000011114 | 420.8150495 | 0.070207955 | 0.2471287 | 0.284095 | 0.776337815 | 0.875431392 | BTBD7 | protein_coding | 40.08 | 0.004056467 | 0.004056467 | ENSG00000011114 | n.s. |
| ENSG00000084093 | 725.1254568 | -0.068405277 | 0.23505 | -0.291024 | 0.771032677 | 0.872370708 | REST | protein_coding | 43.58 | -0.004056359 | 0.004056359 | ENSG00000084093 | n.s. |
| ENSG00000260614 | 52.97872665 | -0.079568939 | 0.3113565 | -0.255556 | 0.79829389 | 0.889456292 | | antisense | 37.67 | -0.004048101 | 0.004048101 | ENSG00000260614 | n.s. |
| ENSG00000153044 | 1201.201774 | -0.078610337 | 0.3060976 | -0.256815 | 0.7973219 | 0.88894241 | CENPH | protein_coding | 42.65 | -0.004019061 | 0.004019061 | ENSG00000153044 | n.s. |
| ENSG00000145293 | 1450.022624 | 0.077732902 | 0.3001331 | 0.258995 | 0.795639263 | 0.887767529 | ENOPH1 | protein_coding | 43.17 | 0.004018849 | 0.004018849 | ENSG00000145293 | n.s. |
| ENSG00000168765 | 94.54287411 | 0.076440852 | 0.2910965 | 0.262596 | 0.792861796 | 0.886089894 | GSTM4 | protein_coding | 49.98 | 0.004014843 | 0.004014843 | ENSG00000168765 | n.s. |
| ENSG00000047249 | 701.2710434 | -0.073472041 | 0.2712289 | -0.270886 | 0.786478952 | 0.881976261 | ATP6V1H | protein_coding | 39.2 | -0.004007393 | 0.004007393 | ENSG00000047249 | n.s. |
| ENSG00000139197 | 1220.841814 | -0.071019884 | 0.2557212 | -0.277724 | 0.781224333 | 0.878298727 | PEX5 | protein_coding | 42.58 | -0.004002521 | 0.004002521 | ENSG00000139197 | n.s. |
| ENSG00000083937 | 457.4578981 | -0.075844742 | 0.2879232 | -0.26342 | 0.79222685 | 0.885691491 | CHMP2B | protein_coding | 34.78 | -0.003998347 | 0.003998347 | ENSG00000083937 | n.s. |
| ENSG00000093010 | 595.589137 | 0.081760764 | 0.3297932 | 0.247915 | 0.804200007 | 0.89357122 | COMT | protein_coding | 53.41 | 0.003995716 | 0.003995716 | ENSG00000093010 | n.s. |
| ENSG00000107745 | 736.7486017 | -0.079053057 | 0.3112463 | -0.253989 | 0.79950428 | 0.890168997 | MICU1 | protein_coding | 40.17 | -0.003994356 | 0.003994356 | ENSG00000107745 | n.s. |

| | | | | | | | | | | | | | |
|---|---|---|---|---|---|---|---|---|---|---|---|---|---|
| ENSG00000132294 | 913.389367 | -0.072856164 | 0.2679588 | -0.271893 | 0.785704147 | 0.881424449 | EFR3A | protein_coding | 35.32 | -0.003993604 | 0.003993604 | ENSG00000132294 | n.s. |
| ENSG00000111897 | 4857.618127 | -0.074772885 | 0.2808326 | -0.266254 | 0.790043351 | 0.884368353 | SERINC1 | protein_coding | 35.93 | -0.00399039 | 0.00399039 | ENSG00000111897 | n.s. |
| ENSG00000198087 | 988.1910749 | 0.06745617 | 0.2338215 | 0.288494 | 0.772968409 | 0.873217822 | CD2AP | protein_coding | 36.38 | 0.003971644 | 0.003971644 | ENSG00000198087 | n.s. |
| ENSG00000145194 | 138.8692627 | -0.077592629 | 0.3030164 | -0.256067 | 0.797898781 | 0.88919258 | ECE2 | protein_coding | 51.09 | -0.003957548 | 0.003957548 | ENSG00000145194 | n.s. |
| ENSG00000225439 | 229.2959475 | -0.06710543 | 0.2332968 | -0.28764 | 0.773622566 | 0.8735015 | BOLA3-AS1 | antisense | 49.2 | -0.003941528 | 0.003941528 | ENSG00000225439 | n.s. |
| ENSG00000122565 | 10861.77994 | -0.073994397 | 0.2787335 | -0.265466 | 0.790650116 | 0.88469192 | CBX3 | protein_coding | 41.57 | -0.003937089 | 0.003937089 | ENSG00000122565 | n.s. |
| ENSG00000249509 | 122.2972943 | 0.0682863 | 0.2410506 | 0.283286 | 0.776957472 | 0.87583137 | | antisense | 40.94 | 0.003931891 | 0.003931891 | ENSG00000249509 | n.s. |
| ENSG00000138780 | 347.2633068 | 0.069428824 | 0.2502601 | 0.277427 | 0.781452503 | 0.878298727 | GSTCD | protein_coding | 35.68 | 0.003912852 | 0.003912852 | ENSG00000138780 | n.s. |
| ENSG00000274828 | 178.9555071 | 0.064590388 | 0.2182839 | 0.295901 | 0.76730586 | 0.870392658 | | lincRNA | 49.81 | 0.003893817 | 0.003893817 | ENSG00000274828 | n.s. |
| ENSG00000179010 | 3285.680147 | 0.074906679 | 0.288038 | 0.260058 | 0.794818796 | 0.887222432 | MRFAP1 | protein_coding | 53.11 | 0.003892712 | 0.003892712 | ENSG00000179010 | n.s. |
| ENSG00000242707 | 22.80525568 | 0.083037323 | 0.3460551 | 0.239954 | 0.810365854 | 0.897730074 | RN7SL362P | misc_RNA | 51.69 | 0.003890649 | 0.003890649 | ENSG00000242707 | n.s. |
| ENSG00000108262 | 404.4973767 | 0.065670326 | 0.2263205 | 0.290165 | 0.771689895 | 0.872517203 | GIT1 | protein_coding | 60.72 | 0.003889391 | 0.003889391 | ENSG00000108262 | n.s. |
| ENSG00000264477 | 2.73764771 | -0.115277724 | 0.6407812 | -0.179902 | 0.857229632 | 0.925281663 | MIR5682 | miRNA | 59.21 | -0.003887862 | 0.003887862 | ENSG00000264477 | n.s. |
| ENSG00000081154 | 2166.244739 | -0.076852577 | 0.3026885 | -0.2539 | 0.7995729 | 0.890168997 | PCNP | protein_coding | 39.82 | -0.003883171 | 0.003883171 | ENSG00000081154 | n.s. |
| ENSG00000159685 | 268.7098303 | 0.073323988 | 0.2777492 | 0.263994 | 0.791784882 | 0.885453552 | CHCHD6 | protein_coding | 46.24 | 0.003874015 | 0.003874015 | ENSG00000159685 | n.s. |
| ENSG00000068878 | 3934.783277 | -0.071886408 | 0.2683614 | -0.267872 | 0.788798099 | 0.883573736 | PSME4 | protein_coding | 37.98 | -0.003864412 | 0.003864412 | ENSG00000068878 | n.s. |
| ENSG00000230199 | 2.672952596 | 0.120918909 | 0.7101284 | 0.170278 | 0.864791885 | 0.92917891 | | processed_pseudogene | 53.75 | 0.003857393 | 0.003857393 | ENSG00000230199 | n.s. |
| ENSG00000248866 | 232.2444638 | 0.062530734 | 0.2071992 | 0.30179 | 0.762811843 | 0.867627587 | USP46-AS1 | lincRNA | 49.93 | 0.003856061 | 0.003856061 | ENSG00000248866 | n.s. |
| ENSG00000234031 | 57.65205287 | 0.073607444 | 0.2813167 | 0.261653 | 0.793588777 | 0.886483997 | RPS3AP44 | processed_pseudogene | 41.01 | 0.003851811 | 0.003851811 | ENSG00000234031 | n.s. |
| ENSG00000168014 | 122.5665043 | 0.06784767 | 0.2435608 | 0.278566 | 0.780578193 | 0.877937651 | C2CD3 | protein_coding | 40.35 | 0.003835858 | 0.003835858 | ENSG00000168014 | n.s. |
| ENSG00000174807 | 159.7610025 | 0.070626049 | 0.2617594 | 0.269813 | 0.787304225 | 0.882508391 | CD248 | protein_coding | 65.4 | 0.003833664 | 0.003833664 | ENSG00000174807 | n.s. |
| ENSG00000184357 | 4252.451663 | 0.065244929 | 0.2269643 | 0.287468 | 0.773754128 | 0.873579202 | HIST1H1B | protein_coding | 59.32 | 0.003829728 | 0.003829728 | ENSG00000184357 | n.s. |
| ENSG00000007080 | 452.250935 | 0.072404638 | 0.2741342 | 0.264121 | 0.791686599 | 0.885424735 | CCDC124 | protein_coding | 55.84 | 0.003826466 | 0.003826466 | ENSG00000007080 | n.s. |
| ENSG00000112659 | 33.47758517 | -0.082369272 | 0.3467861 | -0.237522 | 0.812251956 | 0.898562507 | CUL9 | protein_coding | 49.09 | -0.003826193 | 0.003826193 | ENSG00000112659 | n.s. |
| ENSG00000130024 | 2681.166606 | -0.072916283 | 0.2783519 | -0.261957 | 0.79335448 | 0.886364502 | PHF10 | protein_coding | 39.5 | -0.003819912 | 0.003819912 | ENSG00000130024 | n.s. |
| ENSG00000133027 | 117.3100415 | -0.075542776 | 0.2974734 | -0.253948 | 0.799535745 | 0.890168997 | PEMT | protein_coding | 55.22 | -0.00381699 | 0.00381699 | ENSG00000133027 | n.s. |
| ENSG00000184207 | 426.8679192 | -0.068636966 | 0.2504591 | -0.274045 | 0.784050374 | 0.880136528 | PGP | protein_coding | 61.69 | -0.003805917 | 0.003805917 | ENSG00000184207 | n.s. |
| ENSG00000137504 | 4351.921611 | 0.064047608 | 0.2200577 | 0.291049 | 0.771013753 | 0.872370708 | CREBZF | protein_coding | 40.72 | 0.003797954 | 0.003797954 | ENSG00000137504 | n.s. |
| ENSG00000165689 | 782.5981332 | -0.075569413 | 0.2989689 | -0.252767 | 0.80044844 | 0.890858803 | SDCCAG3 | protein_coding | 56.51 | -0.003792914 | 0.003792914 | ENSG00000165689 | n.s. |
| ENSG00000164402 | 815.8678224 | -0.070514854 | 0.2633267 | -0.267785 | 0.788865097 | 0.883573736 | 8-Sep | protein_coding | 48.12 | -0.003790681 | 0.003790681 | ENSG00000164402 | n.s. |
| ENSG00000254087 | 958.6299555 | 0.072800085 | 0.2800328 | 0.25997 | 0.794887066 | 0.887222432 | LYN | protein_coding | 43.2 | 0.003783237 | 0.003783237 | ENSG00000254087 | n.s. |
| ENSG00000107949 | 2511.749988 | -0.071009913 | 0.2671655 | -0.26579 | 0.790400925 | 0.884555268 | BCCIP | protein_coding | 42.86 | -0.003783055 | 0.003783055 | ENSG00000107949 | n.s. |

| | | | | | | | | | | | | | |
|---|---|---|---|---|---|---|---|---|---|---|---|---|---|
| ENSG00000101052 | 627.0869587 | 0.075297017 | 0.2984145 | 0.252324 | 0.800790981 | 0.891097573 | IFT52 | protein_coding | 44.51 | 0.003770479 | 0.003770479 | ENSG00000101052 | n.s. |
| ENSG00000099282 | 72.63772156 | -0.077034168 | 0.3115515 | -0.24726 | 0.80470715 | 0.893812338 | TSPAN15 | protein_coding | 51.44 | -0.003755697 | 0.003755697 | ENSG00000099282 | n.s. |
| ENSG00000239082 | 3.848215454 | -0.106907 | 0.5747336 | -0.186011 | 0.852435822 | 0.922333766 | RNU7-170P | snRNA | 42.62 | -0.003753707 | 0.003753707 | ENSG00000239082 | n.s. |
| ENSG00000100567 | 1824.65604 | 0.072318475 | 0.2792489 | 0.258975 | 0.795654536 | 0.887767529 | PSMA3 | protein_coding | 40.29 | 0.003738919 | 0.003738919 | ENSG00000100567 | n.s. |
| ENSG00000128609 | 2287.756353 | 0.074686407 | 0.2966202 | 0.251791 | 0.801202351 | 0.891270401 | NDUFA5 | protein_coding | 35.76 | 0.003733612 | 0.003733612 | ENSG00000128609 | n.s. |
| ENSG00000196704 | 1289.823297 | -0.071068557 | 0.2707981 | -0.262441 | 0.792981403 | 0.886089894 | AMZ2 | protein_coding | 43.73 | -0.003732678 | 0.003732678 | ENSG00000196704 | n.s. |
| ENSG00000175567 | 113.6761465 | 0.067275082 | 0.2461968 | 0.273257 | 0.784655407 | 0.880602711 | UCP2 | protein_coding | 53.18 | 0.003714929 | 0.003714929 | ENSG00000175567 | n.s. |
| ENSG00000271002 | 10.25697502 | -0.087461349 | 0.3965376 | -0.220563 | 0.825433054 | 0.906845899 | | processed_pseudogene | 42.54 | -0.003714178 | 0.003714178 | ENSG00000271002 | n.s. |
| ENSG00000128891 | 203.0537551 | -0.073551196 | 0.289976 | -0.253646 | 0.799769174 | 0.89024514 | C15orf57 | protein_coding | 45.99 | -0.003713628 | 0.003713628 | ENSG00000128891 | n.s. |
| ENSG00000248266 | 60.48531065 | -0.074364649 | 0.2956146 | -0.251559 | 0.801381625 | 0.891398607 | | sense_intronic | 36.47 | -0.003712882 | 0.003712882 | ENSG00000248266 | n.s. |
| ENSG00000225032 | 108.64688 | -0.070529158 | 0.2682335 | -0.262939 | 0.792597294 | 0.885873976 | | antisense | 55.15 | -0.003711812 | 0.003711812 | ENSG00000225032 | n.s. |
| ENSG00000153179 | 489.8854709 | -0.069039032 | 0.2579702 | -0.267624 | 0.788988711 | 0.883573736 | RASSF3 | protein_coding | 43.87 | -0.003711345 | 0.003711345 | ENSG00000153179 | n.s. |
| ENSG00000123374 | 1485.009499 | -0.068995979 | 0.2579201 | -0.267509 | 0.789077158 | 0.883573736 | CDK2 | protein_coding | 47.74 | -0.003709031 | 0.003709031 | ENSG00000123374 | n.s. |
| ENSG00000137962 | 1518.774007 | -0.069778455 | 0.2634661 | -0.264848 | 0.791126573 | 0.885082787 | ARHGAP29 | protein_coding | 38.07 | -0.003699382 | 0.003699382 | ENSG00000137962 | n.s. |
| ENSG00000100897 | 827.0143192 | 0.07009 | 0.2662373 | 0.263261 | 0.79234911 | 0.885738793 | DCAF11 | protein_coding | 50.99 | 0.003693346 | 0.003693346 | ENSG00000100897 | n.s. |
| ENSG00000055609 | 553.6353532 | -0.069625228 | 0.2627356 | -0.265001 | 0.791008593 | 0.885021909 | KMT2C | protein_coding | 38.65 | -0.003693338 | 0.003693338 | ENSG00000055609 | n.s. |
| ENSG00000140538 | 107.3059226 | -0.068489719 | 0.2550726 | -0.268511 | 0.788306234 | 0.883347098 | NTRK3 | protein_coding | 45.3 | -0.003689446 | 0.003689446 | ENSG00000140538 | n.s. |
| ENSG00000168237 | 171.7632123 | -0.066974551 | 0.2457495 | -0.272532 | 0.785213168 | 0.881124871 | GLYCTK | protein_coding | 56.59 | -0.003681091 | 0.003681091 | ENSG00000168237 | n.s. |
| ENSG00000101773 | 1541.047776 | -0.067928756 | 0.2523263 | -0.26921 | 0.787768113 | 0.882854866 | RBBP8 | protein_coding | 37.83 | -0.003675672 | 0.003675672 | ENSG00000101773 | n.s. |
| ENSG00000145824 | 351.7385993 | -0.075936428 | 0.3089861 | -0.24576 | 0.805867999 | 0.894724579 | CXCL14 | protein_coding | 52.05 | -0.003668537 | 0.003668537 | ENSG00000145824 | n.s. |
| ENSG00000090863 | 807.205774 | -0.068380689 | 0.2570764 | -0.265994 | 0.790244064 | 0.884521915 | GLG1 | protein_coding | 42.38 | -0.003644103 | 0.003644103 | ENSG00000090863 | n.s. |
| ENSG00000121289 | 272.4538156 | 0.064642235 | 0.2329372 | 0.277509 | 0.781389097 | 0.878298727 | CEP89 | protein_coding | 45.48 | 0.003643091 | 0.003643091 | ENSG00000121289 | n.s. |
| ENSG00000187714 | 212.9366811 | 0.064773298 | 0.2340756 | 0.27672 | 0.781995465 | 0.878750837 | SLC18A3 | protein_coding | 67.19 | 0.003636 | 0.003636 | ENSG00000187714 | n.s. |
| ENSG00000174738 | 1419.977531 | -0.070931339 | 0.2761619 | -0.256847 | 0.797296935 | 0.88894241 | NR1D2 | protein_coding | 40.69 | -0.003626462 | 0.003626462 | ENSG00000174738 | n.s. |
| ENSG00000125166 | 1469.886249 | -0.071548734 | 0.2808245 | -0.254781 | 0.798892285 | 0.889891999 | GOT2 | protein_coding | 44.18 | -0.003624852 | 0.003624852 | ENSG00000125166 | n.s. |
| ENSG00000106682 | 3070.749262 | 0.069899257 | 0.2699223 | 0.258961 | 0.795665613 | 0.887767529 | EIF4H | protein_coding | 45.4 | 0.003613844 | 0.003613844 | ENSG00000106682 | n.s. |
| ENSG00000254858 | 129.7386915 | -0.071356665 | 0.280648 | -0.254257 | 0.799297152 | 0.890075521 | MPV17L2 | protein_coding | 59.62 | -0.003608731 | 0.003608731 | ENSG00000254858 | n.s. |
| ENSG00000175376 | 375.7346983 | 0.061235933 | 0.2128563 | 0.287687 | 0.773586587 | 0.8735015 | EIF1AD | protein_coding | 48.54 | 0.003596775 | 0.003596775 | ENSG00000175376 | n.s. |
| ENSG00000069329 | 2149.74046 | -0.069499018 | 0.2685817 | -0.258763 | 0.795818069 | 0.887767529 | VPS35 | protein_coding | 38.86 | -0.003593151 | 0.003593151 | ENSG00000069329 | n.s. |
| ENSG00000050130 | 1293.737872 | -0.068110198 | 0.2596207 | -0.262345 | 0.793055424 | 0.88610149 | JKAMP | protein_coding | 36.78 | -0.003576911 | 0.003576911 | ENSG00000050130 | n.s. |
| ENSG00000090316 | 929.8734031 | -0.076839239 | 0.3260212 | -0.235688 | 0.81367487 | 0.89925347 | MAEA | protein_coding | 55.03 | -0.003543663 | 0.003543663 | ENSG00000090316 | n.s. |
| ENSG00000116863 | 125.3400778 | -0.069454645 | 0.2719536 | -0.255391 | 0.798420749 | 0.889526453 | ADPRHL2 | protein_coding | 55.16 | -0.003531153 | 0.003531153 | ENSG00000116863 | n.s. |

| | | | | | | | | | | | | | |
|---|---|---|---|---|---|---|---|---|---|---|---|---|---|
| ENSG00000102897 | 398.108762 | 0.069755882 | 0.2742169 | 0.254382 | 0.799200314 | 0.890038872 | LYRM1 | protein_coding | 44.36 | 0.003529022 | 0.003529022 | ENSG00000102897 | n.s. |
| ENSG00000173418 | 712.6079689 | -0.07339198 | 0.3003461 | -0.244358 | 0.806953568 | 0.895274636 | NAA20 | protein_coding | 41.08 | -0.003526024 | 0.003526024 | ENSG00000173418 | n.s. |
| ENSG00000123472 | 457.6067066 | -0.072956191 | 0.2966885 | -0.245902 | 0.805758347 | 0.894706475 | ATPAF1 | protein_coding | 41.86 | -0.003525201 | 0.003525201 | ENSG00000123472 | n.s. |
| ENSG00000110442 | 559.3898824 | 0.069110593 | 0.2699341 | 0.256028 | 0.797929476 | 0.88919258 | COMMD9 | protein_coding | 42.17 | 0.003524928 | 0.003524928 | ENSG00000110442 | n.s. |
| ENSG00000130881 | 828.654126 | 0.065385034 | 0.2442214 | 0.267729 | 0.788908298 | 0.883573736 | LRP3 | protein_coding | 54.69 | 0.003514916 | 0.003514916 | ENSG00000130881 | n.s. |
| ENSG00000178537 | 188.469078 | -0.072182981 | 0.292199 | -0.247034 | 0.804882212 | 0.893935469 | SLC25A20 | protein_coding | 47.4 | -0.003514866 | 0.003514866 | ENSG00000178537 | n.s. |
| ENSG00000084112 | 233.4322237 | 0.063099829 | 0.2301506 | 0.274168 | 0.783955888 | 0.880101421 | SSH1 | protein_coding | 48.43 | 0.003499976 | 0.003499976 | ENSG00000084112 | n.s. |
| ENSG00000105643 | 95.21122844 | 0.066823085 | 0.2557895 | 0.261242 | 0.793905549 | 0.886695569 | ARRDC2 | protein_coding | 58.01 | 0.003489866 | 0.003489866 | ENSG00000105643 | n.s. |
| ENSG00000171793 | 2236.380421 | 0.064692017 | 0.2411895 | 0.268221 | 0.788529469 | 0.883526139 | CTPS1 | protein_coding | 45.27 | 0.003479175 | 0.003479175 | ENSG00000171793 | n.s. |
| ENSG00000136937 | 2453.390317 | -0.06676854 | 0.2563137 | -0.260495 | 0.794481672 | 0.887125537 | NCBP1 | protein_coding | 37.78 | -0.00347296 | 0.00347296 | ENSG00000136937 | n.s. |
| ENSG00000167394 | 288.1224398 | 0.066246993 | 0.2531603 | 0.26168 | 0.79356817 | 0.886483997 | ZNF668 | protein_coding | 54.5 | 0.003466646 | 0.003466646 | ENSG00000167394 | n.s. |
| ENSG00000113552 | 1019.409667 | 0.074581739 | 0.3141491 | 0.237409 | 0.812339745 | 0.898562507 | GNPDA1 | protein_coding | 44.88 | 0.003464449 | 0.003464449 | ENSG00000113552 | n.s. |
| ENSG00000223131 | 26.06642999 | -0.074537892 | 0.3141356 | -0.237279 | 0.812440063 | 0.898602081 | RNA5SP304 | rRNA | 57.69 | -0.003460987 | 0.003460987 | ENSG00000223131 | n.s. |
| ENSG00000108219 | 376.8713377 | -0.066598108 | 0.2561378 | -0.260009 | 0.7948569 | 0.887222432 | TSPAN14 | protein_coding | 49.31 | -0.003460936 | 0.003460936 | ENSG00000108219 | n.s. |
| ENSG00000261766 | 21.95111671 | 0.077937468 | 0.3421177 | 0.227809 | 0.81979479 | 0.903363899 | | antisense | 53.53 | 0.003439947 | 0.003439947 | ENSG00000261766 | n.s. |
| ENSG00000115461 | 6923.469932 | 0.065478159 | 0.2494342 | 0.262507 | 0.792930802 | 0.886089894 | IGFBP5 | protein_coding | 50.77 | 0.003439058 | 0.003439058 | ENSG00000115461 | n.s. |
| ENSG00000124802 | 1101.477452 | -0.071414468 | 0.2923069 | -0.244313 | 0.806988198 | 0.895274636 | EEF1E1 | protein_coding | 39.63 | -0.003431017 | 0.003431017 | ENSG00000124802 | n.s. |
| ENSG00000132429 | 1089.064733 | -0.072194631 | 0.2979648 | -0.242293 | 0.80855352 | 0.896439772 | POPDC3 | protein_coding | 39.7 | -0.00342772 | 0.00342772 | ENSG00000132429 | n.s. |
| ENSG00000265996 | 54.77015786 | 0.07648035 | 0.332537 | 0.22999 | 0.818099173 | 0.90236768 | MIR3671 | miRNA | 27.27 | 0.003412283 | 0.003412283 | ENSG00000265996 | n.s. |
| ENSG00000121680 | 177.9253867 | 0.069688076 | 0.2812394 | 0.247789 | 0.804297531 | 0.89357122 | PEX16 | protein_coding | 58.04 | 0.003405714 | 0.003405714 | ENSG00000121680 | n.s. |
| ENSG00000282870 | 107.0687099 | 0.068058729 | 0.2700261 | 0.252045 | 0.801006247 | 0.89119466 | FRG1DP | unprocessed_pseudogene | 35.11 | 0.003404803 | 0.003404803 | ENSG00000282870 | n.s. |
| ENSG00000066032 | 593.1199917 | -0.068702514 | 0.2748817 | -0.249935 | 0.802637735 | 0.892368057 | CTNNA2 | protein_coding | 38.65 | -0.00339775 | 0.00339775 | ENSG00000066032 | n.s. |
| ENSG00000134058 | 480.5197238 | 0.068417249 | 0.2729829 | 0.250628 | 0.802101482 | 0.892056788 | CDK7 | protein_coding | 40.47 | 0.003394008 | 0.003394008 | ENSG00000134058 | n.s. |
| ENSG00000187109 | 17505.22807 | -0.066933959 | 0.2627997 | -0.254696 | 0.798958107 | 0.889891999 | NAP1L1 | protein_coding | 37.2 | -0.003391055 | 0.003391055 | ENSG00000187109 | n.s. |
| ENSG00000175701 | 300.5665564 | 0.06414916 | 0.2430605 | 0.263923 | 0.791839558 | 0.885453552 | LINC00116 | protein_coding | 40.57 | 0.003389271 | 0.003389271 | ENSG00000175701 | n.s. |
| ENSG00000196204 | 222.7569104 | 0.067103855 | 0.2644416 | 0.253757 | 0.79968346 | 0.890220901 | RNF216P1 | transcribed_unprocessed_pseud | 45.46 | 0.003388893 | 0.003388893 | ENSG00000196204 | n.s. |
| ENSG00000162174 | 580.9910465 | -0.069305033 | 0.2795116 | -0.24795 | 0.804172707 | 0.89357122 | ASRGL1 | protein_coding | 44.11 | -0.003386994 | 0.003386994 | ENSG00000162174 | n.s. |
| ENSG00000226465 | 15.8046904 | 0.081500966 | 0.3764493 | 0.216499 | 0.828598642 | 0.908821403 | | antisense | 44.49 | 0.003384039 | 0.003384039 | ENSG00000226465 | n.s. |
| ENSG00000196371 | 330.4361966 | 0.061655574 | 0.2269252 | 0.2717 | 0.785852675 | 0.881520045 | FUT4 | protein_coding | 49.18 | 0.003376741 | 0.003376741 | ENSG00000196371 | n.s. |
| ENSG00000105708 | 359.5355765 | -0.0687299 | 0.2776712 | -0.247523 | 0.804503793 | 0.893657756 | ZNF14 | protein_coding | 42.02 | -0.003355997 | 0.003355997 | ENSG00000105708 | n.s. |
| ENSG00000143570 | 1271.10514 | 0.067707781 | 0.2706286 | 0.250187 | 0.802442685 | 0.892222449 | SLC39A1 | protein_coding | 52.3 | 0.003353353 | 0.003353353 | ENSG00000143570 | n.s. |
| ENSG00000114956 | 1263.193893 | -0.072477329 | 0.3077343 | -0.235519 | 0.813805814 | 0.899282662 | DGUOK | protein_coding | 42.81 | -0.003341479 | 0.003341479 | ENSG00000114956 | n.s. |

| | | | | | | | | | | | | | |
|---|---|---|---|---|---|---|---|---|---|---|---|---|---|
| ENSG00000163866 | 317.683401 | 0.065500865 | 0.2556666 | 0.256196 | 0.797799211 | 0.88919258 | SMIM12 | protein_coding | 47.95 | 0.003340817 | 0.003340817 | ENSG00000163866 | n.s. |
| ENSG00000185475 | 149.9076914 | -0.071867433 | 0.3030923 | -0.237114 | 0.812568358 | 0.898672591 | TMEM179B | protein_coding | 52.76 | -0.003334541 | 0.003334541 | ENSG00000185475 | n.s. |
| ENSG00000199636 | 9.797505496 | -0.100457194 | 0.5715892 | -0.175751 | 0.860489819 | 0.926587593 | | misc_RNA | 36.19 | -0.003326491 | 0.003326491 | ENSG00000199636 | n.s. |
| ENSG00000102081 | 1484.774024 | 0.064694083 | 0.2510698 | 0.257674 | 0.796658737 | 0.888345315 | FMR1 | protein_coding | 36.35 | 0.003326452 | 0.003326452 | ENSG00000102081 | n.s. |
| ENSG00000113196 | 1583.786043 | 0.05941094 | 0.2153227 | 0.275916 | 0.782612764 | 0.879089755 | HAND1 | protein_coding | 56.03 | 0.003325039 | 0.003325039 | ENSG00000113196 | n.s. |
| ENSG00000087365 | 2991.568906 | 0.065091764 | 0.2540765 | 0.25619 | 0.797804456 | 0.88919258 | SF3B2 | protein_coding | 48.41 | 0.003319951 | 0.003319951 | ENSG00000087365 | n.s. |
| ENSG00000101198 | 81.25171757 | 0.074114828 | 0.3239427 | 0.22879 | 0.819032217 | 0.902850135 | NKAIN4 | protein_coding | 59.79 | 0.003289537 | 0.003289537 | ENSG00000101198 | n.s. |
| ENSG00000130558 | 2750.269091 | -0.069237081 | 0.2859635 | -0.242119 | 0.808688283 | 0.896446414 | OLFM1 | protein_coding | 55.24 | -0.003287077 | 0.003287077 | ENSG00000130558 | n.s. |
| ENSG00000148384 | 273.7385664 | -0.063644894 | 0.2464348 | -0.258263 | 0.796204279 | 0.888051925 | INPP5E | protein_coding | 63.05 | -0.003281635 | 0.003281635 | ENSG00000148384 | n.s. |
| ENSG00000132773 | 518.2859541 | -0.057557982 | 0.2049886 | -0.280786 | 0.778874354 | 0.877155029 | TOE1 | protein_coding | 53.53 | -0.00327641 | 0.00327641 | ENSG00000132773 | n.s. |
| ENSG00000197771 | 3775.47579 | 0.065792019 | 0.2633568 | 0.249821 | 0.802725922 | 0.892391999 | MCMBP | protein_coding | 41.74 | 0.003253042 | 0.003253042 | ENSG00000197771 | n.s. |
| ENSG00000069849 | 1365.615125 | 0.069896189 | 0.2940671 | 0.237688 | 0.812123165 | 0.898537096 | ATP1B3 | protein_coding | 40.61 | 0.003247656 | 0.003247656 | ENSG00000069849 | n.s. |
| ENSG00000263272 | 88.76769903 | 0.063829842 | 0.2506713 | 0.254636 | 0.799004516 | 0.889891999 | | antisense | 45.52 | 0.003233792 | 0.003233792 | ENSG00000263272 | n.s. |
| ENSG00000171606 | 132.4999 | 0.060434847 | 0.2276021 | 0.265529 | 0.79060232 | 0.88469192 | ZNF274 | protein_coding | 47.52 | 0.003215614 | 0.003215614 | ENSG00000171606 | n.s. |
| ENSG00000197780 | 535.7326781 | 0.064420469 | 0.2564285 | 0.251222 | 0.801642539 | 0.891617596 | TAF13 | protein_coding | 43.54 | 0.003209516 | 0.003209516 | ENSG00000197780 | n.s. |
| ENSG00000106636 | 942.8094913 | 0.0665103 | 0.2712478 | 0.245201 | 0.80630062 | 0.894939695 | YKT6 | protein_coding | 48.11 | 0.003206211 | 0.003206211 | ENSG00000106636 | n.s. |
| ENSG00000063978 | 1427.696057 | -0.061559415 | 0.2366065 | -0.260176 | 0.794727769 | 0.887222432 | RNF4 | protein_coding | 47.38 | -0.003199088 | 0.003199088 | ENSG00000063978 | n.s. |
| ENSG00000152332 | 2733.612278 | 0.063796462 | 0.2530165 | 0.252143 | 0.800930188 | 0.891181252 | UHMK1 | protein_coding | 38.32 | 0.00319199 | 0.00319199 | ENSG00000152332 | n.s. |
| ENSG00000166347 | 467.4959459 | 0.071153062 | 0.3078612 | 0.231121 | 0.817221096 | 0.901884061 | CYB5A | protein_coding | 44.31 | 0.003191164 | 0.003191164 | ENSG00000166347 | n.s. |
| ENSG00000077254 | 3405.524586 | 0.066357602 | 0.2714781 | 0.244431 | 0.80689722 | 0.895274636 | USP33 | protein_coding | 38.35 | 0.003188066 | 0.003188066 | ENSG00000077254 | n.s. |
| ENSG00000090487 | 910.3987755 | -0.066668749 | 0.2737005 | -0.243583 | 0.807553891 | 0.895830836 | SPG21 | protein_coding | 44.04 | -0.003185032 | 0.003185032 | ENSG00000090487 | n.s. |
| ENSG00000147872 | 1132.361737 | 0.065983069 | 0.2689665 | 0.245321 | 0.8062081 | 0.894908346 | PLIN2 | protein_coding | 44.99 | 0.003181799 | 0.003181799 | ENSG00000147872 | n.s. |
| ENSG00000006042 | 1000.579359 | 0.070524361 | 0.3039943 | 0.231992 | 0.816543926 | 0.901334392 | TMEM98 | protein_coding | 48.47 | 0.00318164 | 0.00318164 | ENSG00000006042 | n.s. |
| ENSG00000087274 | 1453.888116 | -0.066789273 | 0.2748891 | -0.242968 | 0.808030161 | 0.896216357 | ADD1 | protein_coding | 44.84 | -0.00317831 | 0.00317831 | ENSG00000087274 | n.s. |
| ENSG00000233237 | 112.7529443 | 0.060307373 | 0.229206 | 0.263114 | 0.792462451 | 0.885794373 | LINC00472 | lincRNA | 37.07 | 0.003176213 | 0.003176213 | ENSG00000233237 | n.s. |
| ENSG00000170037 | 667.1419833 | 0.061376287 | 0.2371955 | 0.258758 | 0.795821772 | 0.887767529 | CNTROB | protein_coding | 48.77 | 0.0031732 | 0.0031732 | ENSG00000170037 | n.s. |
| ENSG00000131435 | 723.780307 | 0.068682768 | 0.2909849 | 0.236035 | 0.813405138 | 0.899098106 | PDLIM4 | protein_coding | 56.99 | 0.003172657 | 0.003172657 | ENSG00000131435 | n.s. |
| ENSG00000100612 | 559.4610029 | 0.068466793 | 0.2899962 | 0.236095 | 0.813358602 | 0.899098106 | DHRS7 | protein_coding | 39.65 | 0.003162681 | 0.003162681 | ENSG00000100612 | n.s. |
| ENSG00000183726 | 353.262656 | 0.067739637 | 0.2843429 | 0.238232 | 0.811700966 | 0.898426946 | TMEM50A | protein_coding | 42.04 | 0.00315106 | 0.00315106 | ENSG00000183726 | n.s. |
| ENSG00000134970 | 1153.510515 | 0.067788715 | 0.2848117 | 0.238012 | 0.811871494 | 0.898498787 | TMED7 | protein_coding | 37.91 | 0.003150989 | 0.003150989 | ENSG00000134970 | n.s. |
| ENSG00000084676 | 238.4827989 | 0.064991378 | 0.264282 | 0.245917 | 0.805746673 | 0.894706475 | NCOA1 | protein_coding | 37.1 | 0.003140346 | 0.003140346 | ENSG00000084676 | n.s. |
| ENSG00000144029 | 1420.465507 | -0.065120801 | 0.2654403 | -0.245331 | 0.80619996 | 0.894908346 | MRPS5 | protein_coding | 42.25 | -0.003140219 | 0.003140219 | ENSG00000144029 | n.s. |

| | | | | | | | | | | | | | |
|---|---|---|---|---|---|---|---|---|---|---|---|---|---|
| ENSG00000121644 | 1829.59067 | -0.063377351 | 0.2533024 | -0.250204 | 0.802429332 | 0.892222449 | DESI2 | protein_coding | 40.41 | -0.003138881 | 0.003138881 | ENSG00000121644 | n.s. |
| ENSG00000186952 | 35.39067193 | 0.069148672 | 0.2969574 | 0.232857 | 0.815872281 | 0.900823931 | TMEM232 | protein_coding | 36.01 | 0.00313659 | 0.00313659 | ENSG00000186952 | n.s. |
| ENSG00000263628 | 5.50366544 | -0.094283192 | 0.5355052 | -0.176064 | 0.860243653 | 0.926523739 | MIR3155A | miRNA | 65.85 | -0.00312487 | 0.00312487 | ENSG00000263628 | n.s. |
| ENSG00000259781 | 389.5997378 | 0.068926412 | 0.2963361 | 0.232595 | 0.816075579 | 0.900976947 | HMGB1P6 | processed_pseudogene | 43.65 | 0.003121424 | 0.003121424 | ENSG00000259781 | n.s. |
| ENSG00000164105 | 738.2379251 | -0.066370462 | 0.2761758 | -0.24032 | 0.810082466 | 0.897498545 | SAP30 | protein_coding | 42.24 | -0.003117172 | 0.003117172 | ENSG00000164105 | n.s. |
| ENSG00000137509 | 414.1493294 | 0.070765566 | 0.31167 | 0.227053 | 0.820382614 | 0.90381826 | PRCP | protein_coding | 39.37 | 0.003107945 | 0.003107945 | ENSG00000137509 | n.s. |
| ENSG00000120327 | 322.7649715 | 0.060541435 | 0.2360242 | 0.256505 | 0.797560759 | 0.889128896 | PCDHB14 | protein_coding | 44.93 | 0.003089749 | 0.003089749 | ENSG00000120327 | n.s. |
| ENSG00000149600 | 202.9970298 | 0.069687707 | 0.3069563 | 0.227028 | 0.820401866 | 0.90381826 | COMMD7 | protein_coding | 48.22 | 0.003060607 | 0.003060607 | ENSG00000149600 | n.s. |
| ENSG00000005893 | 666.4884721 | 0.067894058 | 0.2942485 | 0.230737 | 0.817518987 | 0.902082335 | LAMP2 | protein_coding | 38.79 | 0.003038519 | 0.003038519 | ENSG00000005893 | n.s. |
| ENSG00000264853 | 258.8679911 | -0.064591033 | 0.2683118 | -0.240731 | 0.809763456 | 0.897352483 | | sense_intronic | 41.47 | -0.003038164 | 0.003038164 | ENSG00000264853 | n.s. |
| ENSG00000165716 | 422.2838792 | 0.065163957 | 0.2729942 | 0.238701 | 0.811337544 | 0.898301652 | FAM69B | protein_coding | 63.98 | 0.003035194 | 0.003035194 | ENSG00000165716 | n.s. |
| ENSG00000111186 | 183.0087852 | -0.060583107 | 0.2404691 | -0.251937 | 0.801089589 | 0.891216168 | WNT5B | protein_coding | 47.53 | -0.003030182 | 0.003030182 | ENSG00000111186 | n.s. |
| ENSG00000137216 | 137.4236726 | 0.065264128 | 0.2756974 | 0.236724 | 0.812871083 | 0.898793209 | TMEM63B | protein_coding | 54.28 | 0.003024354 | 0.003024354 | ENSG00000137216 | n.s. |
| ENSG00000269974 | 109.9503986 | -0.065411359 | 0.2767945 | -0.236317 | 0.813186359 | 0.899032954 | | lincRNA | 29.47 | -0.0030236 | 0.0030236 | ENSG00000269974 | n.s. |
| ENSG00000009307 | 20393.80362 | 0.064193388 | 0.2663986 | 0.240967 | 0.809580361 | 0.897220996 | CSDE1 | protein_coding | 38.28 | 0.003023545 | 0.003023545 | ENSG00000009307 | n.s. |
| ENSG00000165271 | 1161.116033 | -0.058081068 | 0.2233903 | -0.259998 | 0.794865148 | 0.887222432 | NOL6 | protein_coding | 54.48 | -0.003018327 | 0.003018327 | ENSG00000165271 | n.s. |
| ENSG00000213689 | 86.68668466 | -0.061042115 | 0.2444214 | -0.249741 | 0.802787459 | 0.892391999 | TREX1 | protein_coding | 62.72 | -0.003018187 | 0.003018187 | ENSG00000213689 | n.s. |
| ENSG00000121440 | 767.189643 | -0.062620316 | 0.2557069 | -0.244891 | 0.80654081 | 0.894992245 | PDZRN3 | protein_coding | 41.51 | -0.003017092 | 0.003017092 | ENSG00000121440 | n.s. |
| ENSG00000136718 | 1431.315942 | -0.063224571 | 0.2608611 | -0.242369 | 0.808494456 | 0.896439772 | IMP4 | protein_coding | 53.58 | -0.003001832 | 0.003001832 | ENSG00000136718 | n.s. |
| ENSG00000140575 | 2080.838959 | -0.058451643 | 0.2284982 | -0.255808 | 0.798099163 | 0.889310496 | IQGAP1 | protein_coding | 40.77 | -0.002977911 | 0.002977911 | ENSG00000140575 | n.s. |
| ENSG00000278845 | 557.1666098 | -0.069176461 | 0.3106891 | -0.222655 | 0.823804055 | 0.905989246 | MRPL45 | protein_coding | 45.16 | -0.002966076 | 0.002966076 | ENSG00000278845 | n.s. |
| ENSG00000132970 | 913.6072187 | -0.063621458 | 0.2667783 | -0.238481 | 0.811508319 | 0.898301652 | WASF3 | protein_coding | 39.78 | -0.002963348 | 0.002963348 | ENSG00000132970 | n.s. |
| ENSG00000140391 | 7964.736692 | -0.066934975 | 0.2937689 | -0.227849 | 0.819763568 | 0.903363899 | TSPAN3 | protein_coding | 41.92 | -0.002954327 | 0.002954327 | ENSG00000140391 | n.s. |
| ENSG00000166803 | 1719.476353 | -0.06897 | 0.3114293 | -0.221463 | 0.824732127 | 0.906365691 | PCLAF | protein_coding | 43.34 | -0.002944781 | 0.002944781 | ENSG00000166803 | n.s. |
| ENSG00000270195 | 69.93536804 | -0.060997364 | 0.2488942 | -0.245073 | 0.806399531 | 0.894978132 | | lincRNA | 60.47 | -0.002939315 | 0.002939315 | ENSG00000270195 | n.s. |
| ENSG00000139684 | 2621.299424 | -0.067299911 | 0.2981123 | -0.225754 | 0.821393104 | 0.904409347 | ESD | protein_coding | 34.63 | -0.002936629 | 0.002936629 | ENSG00000139684 | n.s. |
| ENSG00000089289 | 486.9610261 | 0.065255309 | 0.282024 | 0.231382 | 0.817017976 | 0.901731374 | IGBP1 | protein_coding | 42.89 | 0.002931452 | 0.002931452 | ENSG00000089289 | n.s. |
| ENSG00000173163 | 301.2304969 | -0.07015894 | 0.3223167 | -0.217671 | 0.827685616 | 0.908320866 | COMMD1 | protein_coding | 41.05 | -0.002929887 | 0.002929887 | ENSG00000173163 | n.s. |
| ENSG00000124098 | 279.372129 | -0.063167537 | 0.2666336 | -0.236908 | 0.812728418 | 0.898734076 | FAM210B | protein_coding | 49.44 | -0.002929002 | 0.002929002 | ENSG00000124098 | n.s. |
| ENSG00000201182 | 9.773178271 | 0.079756682 | 0.4105564 | 0.194265 | 0.845968512 | 0.918923716 | RNU6-670P | snRNA | 41.35 | 0.002928708 | 0.002928708 | ENSG00000201182 | n.s. |
| ENSG00000181019 | 140.5379912 | 0.068144894 | 0.305315 | 0.223195 | 0.823383452 | 0.905812879 | NQO1 | protein_coding | 47.09 | 0.002927607 | 0.002927607 | ENSG00000181019 | n.s. |
| ENSG00000151694 | 906.8161835 | -0.062985921 | 0.2647998 | -0.237862 | 0.811987837 | 0.898530175 | ADAM17 | protein_coding | 41 | -0.002926788 | 0.002926788 | ENSG00000151694 | n.s. |

| | | | | | | | | | | | | | |
|---|---|---|---|---|---|---|---|---|---|---|---|---|---|
| ENSG00000069667 | 48.77042255 | -0.072499491 | 0.3431135 | -0.211299 | 0.832654082 | 0.911467305 | RORA | protein_coding | 41.64 | -0.00291875 | 0.00291875 | ENSG00000069667 | n.s. |
| ENSG00000180190 | 189.3440552 | -0.066828944 | 0.2959223 | -0.225833 | 0.821331533 | 0.904409347 | TDRP | protein_coding | 42.86 | -0.002916078 | 0.002916078 | ENSG00000180190 | n.s. |
| ENSG00000175582 | 845.235352 | -0.061850888 | 0.2564934 | -0.24114 | 0.809446379 | 0.89714392 | RAB6A | protein_coding | 41.4 | -0.00291552 | 0.00291552 | ENSG00000175582 | n.s. |
| ENSG00000167280 | 558.2138459 | -0.062674726 | 0.2633589 | -0.237982 | 0.811894939 | 0.898498787 | ENGASE | protein_coding | 58.4 | -0.002913278 | 0.002913278 | ENSG00000167280 | n.s. |
| ENSG00000135930 | 1128.553812 | 0.063669686 | 0.2722699 | 0.233848 | 0.815103207 | 0.900260348 | EIF4E2 | protein_coding | 44.37 | 0.002905367 | 0.002905367 | ENSG00000135930 | n.s. |
| ENSG00000207189 | 9.784040551 | 0.07935826 | 0.4127622 | 0.192261 | 0.847537416 | 0.919453284 | | misc_RNA | 45.1 | 0.002894222 | 0.002894222 | ENSG00000207189 | n.s. |
| ENSG00000138439 | 1304.782679 | 0.062199076 | 0.2625808 | 0.236876 | 0.812753052 | 0.898734076 | FAM117B | protein_coding | 39.53 | 0.002884096 | 0.002884096 | ENSG00000138439 | n.s. |
| ENSG00000145386 | 3548.796193 | 0.058807409 | 0.2372659 | 0.247854 | 0.80424706 | 0.89357122 | CCNA2 | protein_coding | 39.44 | 0.002873967 | 0.002873967 | ENSG00000145386 | n.s. |
| ENSG00000197006 | 1762.025495 | 0.067695602 | 0.3073368 | 0.220265 | 0.825664604 | 0.906960811 | METTL9 | protein_coding | 42.83 | 0.002871071 | 0.002871071 | ENSG00000197006 | n.s. |
| ENSG00000127554 | 351.0284119 | -0.059544247 | 0.2430943 | -0.244943 | 0.806500537 | 0.894992245 | GFER | protein_coding | 60.99 | -0.002868885 | 0.002868885 | ENSG00000127554 | n.s. |
| ENSG00000204634 | 458.9964735 | 0.058707479 | 0.2371723 | 0.247531 | 0.804497319 | 0.893657756 | TBC1D8 | protein_coding | 44.32 | 0.002866614 | 0.002866614 | ENSG00000204634 | n.s. |
| ENSG00000144567 | 668.7349558 | -0.064208517 | 0.2793669 | -0.229836 | 0.818219356 | 0.902413505 | RETREG2 | protein_coding | 53.85 | -0.002863341 | 0.002863341 | ENSG00000144567 | n.s. |
| ENSG00000115241 | 2311.641447 | 0.063914807 | 0.278368 | 0.229605 | 0.818398356 | 0.902467948 | PPM1G | protein_coding | 45.48 | 0.002848569 | 0.002848569 | ENSG00000115241 | n.s. |
| ENSG00000104320 | 3901.994558 | -0.061101267 | 0.2570644 | -0.237689 | 0.812122644 | 0.898537096 | NBN | protein_coding | 37.15 | -0.002839009 | 0.002839009 | ENSG00000104320 | n.s. |
| ENSG00000110660 | 402.7808081 | 0.061525308 | 0.2618567 | 0.234958 | 0.814241363 | 0.899522534 | SLC35F2 | protein_coding | 43.51 | 0.002829423 | 0.002829423 | ENSG00000110660 | n.s. |
| ENSG00000124688 | 665.384178 | 0.058534451 | 0.2382349 | 0.245701 | 0.805914023 | 0.894724579 | MAD2L1BP | protein_coding | 49.25 | 0.002827836 | 0.002827836 | ENSG00000124688 | n.s. |
| ENSG00000128708 | 1770.46689 | -0.062263813 | 0.2670995 | -0.233111 | 0.815675261 | 0.900749259 | HAT1 | protein_coding | 39.93 | -0.002826533 | 0.002826533 | ENSG00000128708 | n.s. |
| ENSG00000276728 | 139.7002514 | 0.05700218 | 0.2277473 | 0.250287 | 0.80236545 | 0.892222449 | | lincRNA | 54.45 | 0.002823139 | 0.002823139 | ENSG00000276728 | n.s. |
| ENSG00000261377 | 181.0228849 | -0.063350514 | 0.276264 | -0.229312 | 0.818626796 | 0.902648363 | PDCD6IPP2 | transcribed_unprocessed_pseud | 41.42 | -0.00281792 | 0.00281792 | ENSG00000261377 | n.s. |
| ENSG00000101079 | 459.562275 | 0.065804879 | 0.2962734 | 0.222109 | 0.824229295 | 0.906170599 | NDRG3 | protein_coding | 44.49 | 0.002815793 | 0.002815793 | ENSG00000101079 | n.s. |
| ENSG00000225159 | 58.46886004 | 0.068390108 | 0.3182398 | 0.214901 | 0.82984437 | 0.9096861 | NPM1P39 | processed_pseudogene | 41.41 | 0.002811411 | 0.002811411 | ENSG00000225159 | n.s. |
| ENSG00000201643 | 4.559752414 | -0.085367102 | 0.4896435 | -0.174345 | 0.861594006 | 0.927045854 | SNORA14A | snoRNA | 45.19 | -0.002808474 | 0.002808474 | ENSG00000201643 | n.s. |
| ENSG00000204304 | 346.9029037 | -0.061840529 | 0.2654229 | -0.232989 | 0.815770197 | 0.900782652 | PBX2 | protein_coding | 54.92 | -0.002806322 | 0.002806322 | ENSG00000204304 | n.s. |
| ENSG00000082258 | 4150.337462 | -0.059776698 | 0.2493061 | -0.239772 | 0.810506812 | 0.897747521 | CCNT2 | protein_coding | 37.48 | -0.002800286 | 0.002800286 | ENSG00000082258 | n.s. |
| ENSG00000131747 | 12862.28419 | -0.063881898 | 0.2824634 | -0.22616 | 0.821077033 | 0.904204344 | TOP2A | protein_coding | 39.83 | -0.002793773 | 0.002793773 | ENSG00000131747 | n.s. |
| ENSG00000183431 | 1978.674849 | -0.062358105 | 0.2702731 | -0.230723 | 0.817530338 | 0.902082335 | SF3A3 | protein_coding | 44.59 | -0.002790764 | 0.002790764 | ENSG00000183431 | n.s. |
| ENSG00000164929 | 44.68897404 | 0.070499544 | 0.342363 | 0.20592 | 0.836853032 | 0.913127332 | BAALC | protein_coding | 42.75 | 0.002782522 | 0.002782522 | ENSG00000164929 | n.s. |
| ENSG00000162073 | 739.8643785 | -0.060224242 | 0.2554832 | -0.235727 | 0.813644634 | 0.89925347 | PAQR4 | protein_coding | 64.48 | -0.002777414 | 0.002777414 | ENSG00000162073 | n.s. |
| ENSG00000139636 | 53.60915686 | -0.064878841 | 0.2931831 | -0.221291 | 0.824865675 | 0.906369252 | LMBR1L | protein_coding | 51.62 | -0.002769991 | 0.002769991 | ENSG00000139636 | n.s. |
| ENSG00000136783 | 222.1467328 | -0.068612391 | 0.3274408 | -0.209541 | 0.834025657 | 0.911685579 | NIPSNAP3A | protein_coding | 36.49 | -0.002755125 | 0.002755125 | ENSG00000136783 | n.s. |
| ENSG00000250151 | 29.12938425 | 0.068577743 | 0.3269272 | 0.209765 | 0.833851453 | 0.911652929 | ARPC4-TTLL3 | protein_coding | 49.96 | 0.0027548 | 0.0027548 | ENSG00000250151 | n.s. |
| ENSG00000042429 | 2769.0866 | -0.057730218 | 0.2377146 | -0.242855 | 0.808117567 | 0.896241906 | MED17 | protein_coding | 39.79 | -0.002746501 | 0.002746501 | ENSG00000042429 | n.s. |

| | | | | | | | | | | | | | |
|---|---|---|---|---|---|---|---|---|---|---|---|---|---|
| ENSG00000169813 | 3007.226553 | -0.063135433 | 0.2814212 | -0.224345 | 0.822488873 | 0.904971754 | HNRNPF | protein_coding | 47.66 | -0.002737866 | 0.002737866 | ENSG00000169813 | n.s. |
| ENSG00000158941 | 1795.011413 | 0.058721919 | 0.2461808 | 0.238532 | 0.811468733 | 0.898301652 | CCAR2 | protein_coding | 50.83 | 0.002735138 | 0.002735138 | ENSG00000158941 | n.s. |
| ENSG00000171566 | 1625.887864 | 0.06281728 | 0.2793125 | 0.2249 | 0.822057375 | 0.904752863 | PLRG1 | protein_coding | 34.71 | 0.002730669 | 0.002730669 | ENSG00000171566 | n.s. |
| ENSG00000155099 | 240.9516374 | -0.064976577 | 0.2971219 | -0.218687 | 0.826894181 | 0.907881688 | TMEM55A | protein_coding | 35.6 | -0.002727115 | 0.002727115 | ENSG00000155099 | n.s. |
| ENSG00000187866 | 384.7564257 | -0.054272122 | 0.2146337 | -0.252859 | 0.800376921 | 0.890850418 | FAM122A | protein_coding | 40.85 | -0.002724201 | 0.002724201 | ENSG00000187866 | n.s. |
| ENSG00000072401 | 811.6380595 | -0.060681055 | 0.2634304 | -0.230349 | 0.817820216 | 0.902187693 | UBE2D1 | protein_coding | 36.05 | -0.002712631 | 0.002712631 | ENSG00000072401 | n.s. |
| ENSG00000111911 | 583.0148581 | -0.065985255 | 0.3070235 | -0.214919 | 0.829830279 | 0.9096861 | HINT3 | protein_coding | 38.08 | -0.002712551 | 0.002712551 | ENSG00000111911 | n.s. |
| ENSG00000116221 | 1409.378797 | -0.06414755 | 0.2918191 | -0.21982 | 0.826011656 | 0.907270455 | MRPL37 | protein_coding | 48.9 | -0.002711083 | 0.002711083 | ENSG00000116221 | n.s. |
| ENSG00000270808 | 15.99091339 | 0.07057976 | 0.3497707 | 0.201789 | 0.840081948 | 0.915359286 | | processed_pseudogene | 58.04 | 0.002710856 | 0.002710856 | ENSG00000270808 | n.s. |
| ENSG00000175467 | 773.9203229 | -0.058147467 | 0.2449193 | -0.237415 | 0.812335006 | 0.898562507 | SART1 | protein_coding | 55.3 | -0.002701049 | 0.002701049 | ENSG00000175467 | n.s. |
| ENSG00000201882 | 61.3387354 | 0.059669177 | 0.2570107 | 0.232166 | 0.816409 | 0.90127359 | | snoRNA | 40 | 0.002693667 | 0.002693667 | ENSG00000201882 | n.s. |
| ENSG00000213462 | 652.6539773 | -0.054648639 | 0.2194378 | -0.249039 | 0.803330347 | 0.892852912 | ERV3-1 | protein_coding | 45.75 | -0.00268981 | 0.00268981 | ENSG00000213462 | n.s. |
| ENSG00000182173 | 610.6162533 | -0.061681649 | 0.2734564 | -0.225563 | 0.821541333 | 0.904488092 | TSEN54 | protein_coding | 54.69 | -0.002689144 | 0.002689144 | ENSG00000182173 | n.s. |
| ENSG00000077616 | 564.4224992 | -0.0602661 | 0.2623488 | -0.229717 | 0.818311346 | 0.902443474 | NAALAD2 | protein_coding | 34.98 | -0.002686662 | 0.002686662 | ENSG00000077616 | n.s. |
| ENSG00000253738 | 1177.088137 | 0.055550281 | 0.2259173 | 0.245888 | 0.805769194 | 0.894706475 | OTUD6B-AS1 | antisense | 35.41 | 0.002684157 | 0.002684157 | ENSG00000253738 | n.s. |
| ENSG00000198300 | 969.6172411 | 0.056324029 | 0.2320201 | 0.242755 | 0.80819526 | 0.896256679 | PEG3 | protein_coding | 45.75 | 0.002679199 | 0.002679199 | ENSG00000198300 | n.s. |
| ENSG00000156110 | 735.3232596 | -0.063668884 | 0.2907374 | -0.218991 | 0.826657037 | 0.907728254 | ADK | protein_coding | 37.42 | -0.002676904 | 0.002676904 | ENSG00000156110 | n.s. |
| ENSG00000101347 | 446.8885856 | 0.06506092 | 0.3024871 | 0.215087 | 0.829699781 | 0.909675692 | SAMHD1 | protein_coding | 43.82 | 0.002674876 | 0.002674876 | ENSG00000101347 | n.s. |
| ENSG00000106605 | 495.3147083 | -0.061243623 | 0.2715969 | -0.225495 | 0.821594549 | 0.904488092 | BLVRA | protein_coding | 44.31 | -0.002670047 | 0.002670047 | ENSG00000106605 | n.s. |
| ENSG00000252274 | 12.77571858 | 0.075566964 | 0.4064256 | 0.185931 | 0.852499162 | 0.922333766 | | scaRNA | 31.39 | 0.002653299 | 0.002653299 | ENSG00000252274 | n.s. |
| ENSG00000116649 | 266.4640693 | 0.056921897 | 0.2384357 | 0.238731 | 0.811314525 | 0.898301652 | SRM | protein_coding | 59.88 | 0.002651297 | 0.002651297 | ENSG00000116649 | n.s. |
| ENSG00000259330 | 776.2275897 | -0.057475182 | 0.2442417 | -0.235321 | 0.813959643 | 0.899282662 | INAFM2 | protein_coding | 56.16 | -0.002649823 | 0.002649823 | ENSG00000259330 | n.s. |
| ENSG00000140326 | 520.9054893 | -0.057737239 | 0.2462332 | -0.234482 | 0.814610861 | 0.899787943 | CDAN1 | protein_coding | 54.46 | -0.00264782 | 0.00264782 | ENSG00000140326 | n.s. |
| ENSG00000144744 | 1679.473133 | -0.061904592 | 0.2791419 | -0.221768 | 0.82449488 | 0.906287556 | UBA3 | protein_coding | 37.06 | -0.00264543 | 0.00264543 | ENSG00000144744 | n.s. |
| ENSG00000102879 | 94.25803668 | -0.066560589 | 0.3206229 | -0.207598 | 0.83554309 | 0.912553886 | CORO1A | protein_coding | 64.42 | -0.002645216 | 0.002645216 | ENSG00000102879 | n.s. |
| ENSG00000166582 | 3567.02291 | 0.063161222 | 0.2894708 | 0.218195 | 0.827276803 | 0.908086955 | CENPV | protein_coding | 48.01 | 0.002644722 | 0.002644722 | ENSG00000166582 | n.s. |
| ENSG00000106244 | 1377.299842 | -0.060171702 | 0.2652358 | -0.226861 | 0.820531709 | 0.903889784 | PDAP1 | protein_coding | 54.05 | -0.002640606 | 0.002640606 | ENSG00000106244 | n.s. |
| ENSG00000119878 | 393.3307022 | 0.06551023 | 0.3101722 | 0.211206 | 0.832726509 | 0.911467305 | CRIPT | protein_coding | 39.14 | 0.00263737 | 0.00263737 | ENSG00000119878 | n.s. |
| ENSG00000132680 | 3168.556037 | -0.055486466 | 0.2291054 | -0.242188 | 0.80863488 | 0.896446414 | KIAA0907 | protein_coding | 42.57 | -0.002634257 | 0.002634257 | ENSG00000132680 | n.s. |
| ENSG00000176058 | 262.1814489 | 0.058825241 | 0.2552149 | 0.230493 | 0.817708728 | 0.902136183 | TPRN | protein_coding | 62.56 | 0.002631129 | 0.002631129 | ENSG00000176058 | n.s. |
| ENSG00000142173 | 531.3195946 | 0.059165411 | 0.2585307 | 0.228853 | 0.818983522 | 0.902850135 | COL6A2 | protein_coding | 55.88 | 0.002626017 | 0.002626017 | ENSG00000142173 | n.s. |
| ENSG00000005206 | 49.00908931 | 0.060833138 | 0.2735753 | 0.222363 | 0.824031001 | 0.906024135 | SPPL2B | protein_coding | 61.06 | 0.002607322 | 0.002607322 | ENSG00000005206 | n.s. |

| | | | | | | | | | | | | | |
|---|---|---|---|---|---|---|---|---|---|---|---|---|---|
| ENSG00000094975 | 2387.368461 | 0.059804858 | 0.2656393 | 0.225136 | 0.82187381 | 0.904723993 | SUCO | protein_coding | 36.42 | 0.002600548 | 0.002600548 | ENSG00000094975 | n.s. |
| ENSG00000121653 | 487.0324444 | -0.059201166 | 0.2605933 | -0.227178 | 0.820285032 | 0.90381826 | MAPK8IP1 | protein_coding | 58.04 | -0.002600049 | 0.002600049 | ENSG00000121653 | n.s. |
| ENSG00000163399 | 8857.825502 | 0.058488087 | 0.2556942 | 0.228742 | 0.81906917 | 0.902850135 | ATP1A1 | protein_coding | 44.46 | 0.002595955 | 0.002595955 | ENSG00000163399 | n.s. |
| ENSG00000164542 | 129.9703293 | 0.063478608 | 0.2967963 | 0.213879 | 0.830641135 | 0.910253357 | KIAA0895 | protein_coding | 39.56 | 0.002592321 | 0.002592321 | ENSG00000164542 | n.s. |
| ENSG00000183873 | 171.0830318 | 0.055654787 | 0.2330306 | 0.23883 | 0.811237079 | 0.898301652 | SCN5A | protein_coding | 50.93 | 0.002592278 | 0.002592278 | ENSG00000183873 | n.s. |
| ENSG00000196636 | 258.4816192 | 0.065396808 | 0.3165904 | 0.206566 | 0.836348845 | 0.912791798 | SDHAF3 | protein_coding | 38.69 | 0.002591562 | 0.002591562 | ENSG00000196636 | n.s. |
| ENSG00000012817 | 368.6120137 | 0.054972591 | 0.2279462 | 0.241165 | 0.809427415 | 0.89714392 | KDM5D | protein_coding | 38.66 | 0.002591291 | 0.002591291 | ENSG00000012817 | n.s. |
| ENSG00000160310 | 340.7924253 | 0.064678736 | 0.310407 | 0.208367 | 0.834942043 | 0.912208166 | PRMT2 | protein_coding | 45.52 | 0.002581073 | 0.002581073 | ENSG00000160310 | n.s. |
| ENSG00000202354 | 3.000720771 | 0.095149608 | 0.6469514 | 0.147074 | 0.883073764 | 0.939484681 | RNY3 | misc_RNA | 45.1 | 0.002579534 | 0.002579534 | ENSG00000202354 | n.s. |
| ENSG00000108946 | 4869.882947 | -0.058938251 | 0.2603634 | -0.226369 | 0.820914275 | 0.90416814 | PRKAR1A | protein_coding | 41.97 | -0.002578596 | 0.002578596 | ENSG00000108946 | n.s. |
| ENSG00000142208 | 1691.721727 | 0.059827595 | 0.2678247 | 0.223383 | 0.823237086 | 0.905723426 | AKT1 | protein_coding | 64.08 | 0.00257285 | 0.00257285 | ENSG00000142208 | n.s. |
| ENSG00000166902 | 391.8321546 | 0.063807306 | 0.3037926 | 0.210036 | 0.833639767 | 0.911652929 | MRPL16 | protein_coding | 44.6 | 0.002563169 | 0.002563169 | ENSG00000166902 | n.s. |
| ENSG00000131165 | 400.8924278 | -0.060777712 | 0.277511 | -0.21901 | 0.826642189 | 0.907728254 | CHMP1A | protein_coding | 58.66 | -0.002555347 | 0.002555347 | ENSG00000131165 | n.s. |
| ENSG00000182718 | 7595.171678 | 0.060965019 | 0.279125 | 0.218415 | 0.827105943 | 0.908042584 | ANXA2 | protein_coding | 44.95 | 0.002554055 | 0.002554055 | ENSG00000182718 | n.s. |
| ENSG00000130479 | 243.160287 | -0.056572913 | 0.2439239 | -0.231928 | 0.816593552 | 0.901334392 | MAP1S | protein_coding | 57.53 | -0.002552234 | 0.002552234 | ENSG00000130479 | n.s. |
| ENSG00000132329 | 206.7711858 | 0.063247066 | 0.3004366 | 0.210517 | 0.833264016 | 0.911604157 | RAMP1 | protein_coding | 52.41 | 0.002542134 | 0.002542134 | ENSG00000132329 | n.s. |
| ENSG00000273270 | 279.2746912 | -0.051521885 | 0.2065783 | -0.249406 | 0.803046687 | 0.892608895 | | lincRNA | 51.96 | -0.002542027 | 0.002542027 | ENSG00000273270 | n.s. |
| ENSG00000196576 | 796.093833 | -0.057199352 | 0.2499034 | -0.228886 | 0.818957612 | 0.902850135 | PLXNB2 | protein_coding | 64.97 | -0.002538755 | 0.002538755 | ENSG00000196576 | n.s. |
| ENSG00000185432 | 334.2629707 | -0.057328439 | 0.2511301 | -0.228282 | 0.819427178 | 0.90310349 | METTL7A | protein_coding | 42.85 | -0.002537499 | 0.002537499 | ENSG00000185432 | n.s. |
| ENSG00000282995 | 102.9723699 | 0.060478162 | 0.2770872 | 0.218264 | 0.827223399 | 0.908086955 | FRG1EP | unprocessed_pseudogene | 34.33 | 0.002532376 | 0.002532376 | ENSG00000282995 | n.s. |
| ENSG00000138777 | 1244.406581 | -0.065818278 | 0.3260614 | -0.201859 | 0.840027294 | 0.915359286 | PPA2 | protein_coding | 36.65 | -0.002527975 | 0.002527975 | ENSG00000138777 | n.s. |
| ENSG00000162384 | 1384.391084 | 0.052940849 | 0.2175755 | 0.243322 | 0.807756144 | 0.895983811 | C1orf123 | protein_coding | 46.92 | 0.00252527 | 0.00252527 | ENSG00000162384 | n.s. |
| ENSG00000164039 | 371.8371027 | -0.064535568 | 0.3149172 | -0.204929 | 0.837627853 | 0.913901152 | BDH2 | protein_coding | 38.56 | -0.002523391 | 0.002523391 | ENSG00000164039 | n.s. |
| ENSG00000186297 | 310.5580482 | 0.058844104 | 0.2658409 | 0.221351 | 0.824819266 | 0.906369252 | GABRA5 | protein_coding | 43.55 | 0.002512339 | 0.002512339 | ENSG00000186297 | n.s. |
| ENSG00000138430 | 2897.719337 | -0.060053964 | 0.2757881 | -0.217754 | 0.827620779 | 0.908320866 | OLA1 | protein_coding | 36.5 | -0.002507896 | 0.002507896 | ENSG00000138430 | n.s. |
| ENSG00000074696 | 7635.453953 | 0.058299483 | 0.2614812 | 0.222959 | 0.82356774 | 0.905872464 | HACD3 | protein_coding | 42.55 | 0.002502968 | 0.002502968 | ENSG00000074696 | n.s. |
| ENSG00000134287 | 932.6964852 | -0.057710126 | 0.2572134 | -0.224367 | 0.822471938 | 0.904971754 | ARF3 | protein_coding | 47.76 | -0.002502598 | 0.002502598 | ENSG00000134287 | n.s. |
| ENSG00000163138 | 624.7915972 | -0.052936054 | 0.2200719 | -0.24054 | 0.809911769 | 0.897445409 | PACRGL | protein_coding | 38.27 | -0.002487569 | 0.002487569 | ENSG00000163138 | n.s. |
| ENSG00000172725 | 244.3505989 | 0.058374481 | 0.2647571 | 0.220483 | 0.825494859 | 0.906845899 | CORO1B | protein_coding | 64.53 | 0.00247896 | 0.00247896 | ENSG00000172725 | n.s. |
| ENSG00000104067 | 1039.406798 | -0.052616512 | 0.2189511 | -0.240312 | 0.810088646 | 0.897498545 | TJP1 | protein_coding | 40.21 | -0.0024712 | 0.0024712 | ENSG00000104067 | n.s. |
| ENSG00000200520 | 21.80767097 | 0.065885641 | 0.3334253 | 0.197602 | 0.843356133 | 0.917409811 | RNU6-1214P | snRNA | 34.86 | 0.002466536 | 0.002466536 | ENSG00000200520 | n.s. |
| ENSG00000230679 | 18.6855536 | 0.067098406 | 0.3465244 | 0.193633 | 0.846463585 | 0.918932748 | ENO1-AS1 | antisense | 59.53 | 0.002463603 | 0.002463603 | ENSG00000230679 | n.s. |

| | | | | | | | | | | | | | |
|---|---|---|---|---|---|---|---|---|---|---|---|---|---|
| ENSG00000088930 | 4854.806668 | 0.060513713 | 0.2839669 | 0.213101 | 0.831247953 | 0.910651134 | XRN2 | protein_coding | 37.22 | 0.002459759 | 0.002459759 | ENSG00000088930 | n.s. |
| ENSG00000169105 | 265.5351285 | 0.054841313 | 0.2378754 | 0.230546 | 0.817667225 | 0.902136183 | CHST14 | protein_coding | 57.11 | 0.002452937 | 0.002452937 | ENSG00000169105 | n.s. |
| ENSG00000136807 | 499.5758199 | 0.05315703 | 0.2258173 | 0.235398 | 0.813899546 | 0.899282662 | CDK9 | protein_coding | 57.43 | 0.00245074 | 0.00245074 | ENSG00000136807 | n.s. |
| ENSG00000177034 | 3342.242332 | -0.055821356 | 0.2467894 | -0.22619 | 0.821053417 | 0.904204344 | MTX3 | protein_coding | 37.49 | -0.002441258 | 0.002441258 | ENSG00000177034 | n.s. |
| ENSG00000138279 | 530.7931127 | 0.060396649 | 0.2848992 | 0.211993 | 0.832112473 | 0.911239732 | ANXA7 | protein_coding | 40.79 | 0.002438053 | 0.002438053 | ENSG00000138279 | n.s. |
| ENSG00000253457 | 136.3948234 | 0.066357557 | 0.3441331 | 0.192825 | 0.847095826 | 0.919189088 | SMIM18 | protein_coding | 36.33 | 0.002428364 | 0.002428364 | ENSG00000253457 | n.s. |
| ENSG00000133169 | 4262.653615 | 0.059195858 | 0.2763798 | 0.214183 | 0.830404348 | 0.910156635 | BEX1 | protein_coding | 56.79 | 0.002420155 | 0.002420155 | ENSG00000133169 | n.s. |
| ENSG00000169599 | 421.5423382 | -0.064563427 | 0.326674 | -0.197639 | 0.843327749 | 0.917409811 | NFU1 | protein_coding | 41.47 | -0.002417036 | 0.002417036 | ENSG00000169599 | n.s. |
| ENSG00000117614 | 548.7485086 | -0.057448068 | 0.2644204 | -0.21726 | 0.828005463 | 0.908481446 | SYF2 | protein_coding | 43.61 | -0.002394662 | 0.002394662 | ENSG00000117614 | n.s. |
| ENSG00000099251 | 104.7454756 | 0.058201377 | 0.2704855 | 0.215174 | 0.829631872 | 0.909675692 | HSD17B7P2 | transcribed_unprocessed_pseud | 39.3 | 0.002392857 | 0.002392857 | ENSG00000099251 | n.s. |
| ENSG00000259349 | 31.94240642 | 0.059938082 | 0.2874964 | 0.208483 | 0.834851956 | 0.912208166 | | antisense | 43.04 | 0.002391892 | 0.002391892 | ENSG00000259349 | n.s. |
| ENSG00000122958 | 843.0669441 | -0.059494582 | 0.283765 | -0.209661 | 0.833931942 | 0.911654713 | VPS26A | protein_coding | 39.68 | -0.002389875 | 0.002389875 | ENSG00000122958 | n.s. |
| ENSG00000153904 | 582.7287256 | -0.055308703 | 0.2479726 | -0.223044 | 0.823501581 | 0.905871258 | DDAH1 | protein_coding | 39.19 | -0.002374597 | 0.002374597 | ENSG00000153904 | n.s. |
| ENSG00000101104 | 230.8584624 | -0.054530753 | 0.2424841 | -0.224884 | 0.822069681 | 0.904752863 | PABPC1L | protein_coding | 47.22 | -0.002370454 | 0.002370454 | ENSG00000101104 | n.s. |
| ENSG00000274963 | 35.81327849 | -0.06212831 | 0.3107707 | -0.199917 | 0.841545562 | 0.91635782 | RN7SL600P | misc_RNA | 50.84 | -0.002356832 | 0.002356832 | ENSG00000274963 | n.s. |
| ENSG00000135506 | 1921.127062 | -0.058767162 | 0.2807496 | -0.209322 | 0.834196593 | 0.911800845 | OS9 | protein_coding | 43.99 | -0.002356564 | 0.002356564 | ENSG00000135506 | n.s. |
| ENSG00000232818 | 32.3701813 | 0.062377926 | 0.3136681 | 0.198866 | 0.842367576 | 0.916797802 | RPS2P32 | processed_pseudogene | 61.32 | 0.002353297 | 0.002353297 | ENSG00000232818 | n.s. |
| ENSG00000153485 | 214.399305 | 0.057126687 | 0.265212 | 0.2154 | 0.829455424 | 0.909617822 | TMEM251 | protein_coding | 48.11 | 0.002350251 | 0.002350251 | ENSG00000153485 | n.s. |
| ENSG00000134717 | 2483.641659 | -0.058483974 | 0.2785329 | -0.209972 | 0.833689909 | 0.911652929 | BTF3L4 | protein_coding | 41.19 | -0.002349329 | 0.002349329 | ENSG00000134717 | n.s. |
| ENSG00000206888 | 8.429002841 | 0.07299744 | 0.4258409 | 0.17142 | 0.863893913 | 0.928730982 | RNU6-48P | snRNA | 45.79 | 0.002343953 | 0.002343953 | ENSG00000206888 | n.s. |
| ENSG00000207088 | 20.10802016 | 0.064313418 | 0.3361023 | 0.191351 | 0.848250823 | 0.919868853 | SNORA7B | snoRNA | 56.83 | 0.00233291 | 0.00233291 | ENSG00000207088 | n.s. |
| ENSG00000126602 | 2418.28964 | 0.059927702 | 0.2947266 | 0.203333 | 0.838874627 | 0.914473179 | TRAP1 | protein_coding | 50.88 | 0.002326934 | 0.002326934 | ENSG00000126602 | n.s. |
| ENSG00000170522 | 71.15345043 | -0.055745952 | 0.2566196 | -0.217232 | 0.828027692 | 0.908481446 | ELOVL6 | protein_coding | 39.87 | -0.002323711 | 0.002323711 | ENSG00000170522 | n.s. |
| ENSG00000187630 | 82.70991267 | -0.060526263 | 0.3004258 | -0.201468 | 0.84033243 | 0.915408802 | DHRS4L2 | protein_coding | 43.65 | -0.002323296 | 0.002323296 | ENSG00000187630 | n.s. |
| ENSG00000178307 | 465.3470491 | 0.052488108 | 0.2299289 | 0.22828 | 0.819428743 | 0.90310349 | TMEM11 | protein_coding | 51.96 | 0.002323254 | 0.002323254 | ENSG00000178307 | n.s. |
| ENSG00000113013 | 9218.984446 | 0.057159988 | 0.2684777 | 0.212904 | 0.831401814 | 0.91067638 | HSPA9 | protein_coding | 42.74 | 0.002322749 | 0.002322749 | ENSG00000113013 | n.s. |
| ENSG00000251456 | 77.83214796 | 0.058143947 | 0.27908 | 0.208341 | 0.834962355 | 0.912208166 | | sense_intronic | 32.97 | 0.002320295 | 0.002320295 | ENSG00000251456 | n.s. |
| ENSG00000199005 | 2.686673717 | 0.116080203 | 1.038084 | 0.111822 | 0.910964869 | 0.955092628 | MIR370 | miRNA | 57.33 | 0.002316323 | 0.002316323 | ENSG00000199005 | n.s. |
| ENSG00000136378 | 373.9086641 | -0.054418099 | 0.2464248 | -0.22083 | 0.825224441 | 0.906691911 | ADAMTS7 | protein_coding | 56.03 | -0.00231496 | 0.00231496 | ENSG00000136378 | n.s. |
| ENSG00000210140 | 38.84123345 | -0.061353965 | 0.30852 | -0.198865 | 0.842368009 | 0.916797802 | MT-TC | Mt_tRNA | 45.45 | -0.002314666 | 0.002314666 | ENSG00000210140 | n.s. |
| ENSG00000091640 | 662.3751957 | 0.058178699 | 0.2800787 | 0.207723 | 0.835445499 | 0.912553886 | SPAG7 | protein_coding | 51.65 | 0.002312108 | 0.002312108 | ENSG00000091640 | n.s. |
| ENSG00000076513 | 762.0646965 | -0.057492081 | 0.273371 | -0.210308 | 0.833427373 | 0.911604157 | ANKRD13A | protein_coding | 44 | -0.00231082 | 0.00231082 | ENSG00000076513 | n.s. |

| | | | | | | | | | | | | | |
|---|---|---|---|---|---|---|---|---|---|---|---|---|---|
| ENSG00000118508 | 88.44881262 | -0.058121134 | 0.2801937 | -0.207432 | 0.835672504 | 0.912553886 | RAB32 | protein_coding | 39.47 | -0.00230982 | 0.00230982 | ENSG00000118508 | n.s. |
| ENSG00000100982 | 483.9269952 | 0.054026214 | 0.2436058 | 0.221777 | 0.824487332 | 0.906287556 | PCIF1 | protein_coding | 53.89 | 0.002308755 | 0.002308755 | ENSG00000100982 | n.s. |
| ENSG00000113407 | 7065.919143 | 0.057008762 | 0.2689063 | 0.212002 | 0.832105189 | 0.911239732 | TARS | protein_coding | 38.63 | 0.002301293 | 0.002301293 | ENSG00000113407 | n.s. |
| ENSG00000111247 | 2031.974838 | -0.057011222 | 0.2697377 | -0.211358 | 0.832607925 | 0.911467305 | RAD51AP1 | protein_coding | 36.4 | -0.002295209 | 0.002295209 | ENSG00000111247 | n.s. |
| ENSG00000260942 | 383.8816271 | -0.049599085 | 0.2099184 | -0.236278 | 0.813217052 | 0.899032954 | CAPN10-AS1 | antisense | 54.57 | -0.002292687 | 0.002292687 | ENSG00000260942 | n.s. |
| ENSG00000184840 | 338.9580882 | -0.056442757 | 0.2682431 | -0.210416 | 0.833342695 | 0.911604157 | TMED9 | protein_coding | 53.16 | -0.002268643 | 0.002268643 | ENSG00000184840 | n.s. |
| ENSG00000147649 | 2481.479882 | -0.056744039 | 0.2721626 | -0.208493 | 0.834843907 | 0.912208166 | MTDH | protein_coding | 40.6 | -0.00226443 | 0.00226443 | ENSG00000147649 | n.s. |
| ENSG00000179967 | 67.30815122 | 0.058106439 | 0.2851718 | 0.203759 | 0.838541525 | 0.914181634 | PPP1R14BP3 | processed_pseudogene | 65.17 | 0.002264263 | 0.002264263 | ENSG00000179967 | n.s. |
| ENSG00000275084 | 25.58802047 | -0.061760265 | 0.3209967 | -0.192402 | 0.847427664 | 0.919405857 | SNORD91B | snoRNA | 40.99 | -0.002253801 | 0.002253801 | ENSG00000275084 | n.s. |
| ENSG00000133597 | 164.110314 | -0.056072536 | 0.2666025 | -0.210323 | 0.833415859 | 0.911604157 | ADCK2 | protein_coding | 50.51 | -0.002253763 | 0.002253763 | ENSG00000133597 | n.s. |
| ENSG00000207820 | 21.7840473 | -0.064169532 | 0.3451291 | -0.185929 | 0.852500376 | 0.922333766 | MIR545 | miRNA | 41.51 | -0.002253114 | 0.002253114 | ENSG00000207820 | n.s. |
| ENSG00000111850 | 716.1653847 | -0.056485052 | 0.2723301 | -0.207414 | 0.835686645 | 0.912553886 | SMIM8 | protein_coding | 40.32 | -0.0022448 | 0.0022448 | ENSG00000111850 | n.s. |
| ENSG00000162068 | 95.47110337 | -0.053551159 | 0.2462849 | -0.217436 | 0.827868718 | 0.908450202 | NTN3 | protein_coding | 70.83 | -0.002233023 | 0.002233023 | ENSG00000162068 | n.s. |
| ENSG00000214078 | 578.7384389 | 0.054764029 | 0.2567747 | 0.213277 | 0.831111253 | 0.910644682 | CPNE1 | protein_coding | 43.12 | 0.002226215 | 0.002226215 | ENSG00000214078 | n.s. |
| ENSG00000108826 | 2362.994853 | 0.055274346 | 0.2612631 | 0.211566 | 0.832445778 | 0.911461353 | MRPL27 | protein_coding | 47.14 | 0.002225441 | 0.002225441 | ENSG00000108826 | n.s. |
| ENSG00000107951 | 1437.333578 | -0.055278628 | 0.2634493 | -0.209826 | 0.833803146 | 0.911652929 | MTPAP | protein_coding | 43.36 | -0.002220568 | 0.002220568 | ENSG00000107951 | n.s. |
| ENSG00000145088 | 72.14402296 | -0.05934408 | 0.3008445 | -0.197258 | 0.843625383 | 0.917568644 | EAF2 | protein_coding | 36.14 | -0.00221718 | 0.00221718 | ENSG00000145088 | n.s. |
| ENSG00000143793 | 381.8797032 | 0.056592885 | 0.2762858 | 0.204835 | 0.837701365 | 0.91390974 | C1orf35 | protein_coding | 60.86 | 0.002212595 | 0.002212595 | ENSG00000143793 | n.s. |
| ENSG00000100227 | 1199.094175 | 0.055815766 | 0.2710382 | 0.205933 | 0.836843096 | 0.913127332 | POLDIP3 | protein_coding | 49.1 | 0.002202973 | 0.002202973 | ENSG00000100227 | n.s. |
| ENSG00000160214 | 742.7677818 | -0.052108907 | 0.2375039 | -0.219402 | 0.826336682 | 0.907555859 | RRP1 | protein_coding | 57.27 | -0.002195173 | 0.002195173 | ENSG00000160214 | n.s. |
| ENSG00000255152 | 191.7792126 | -0.056232436 | 0.2753565 | -0.204217 | 0.838184035 | 0.914034657 | MSH5-SAPCD1 | protein_coding | 48.78 | -0.002195165 | 0.002195165 | ENSG00000255152 | n.s. |
| ENSG00000105723 | 518.3137145 | 0.053345611 | 0.2480257 | 0.215081 | 0.829704203 | 0.909675692 | GSK3A | protein_coding | 53.29 | 0.00219322 | 0.00219322 | ENSG00000105723 | n.s. |
| ENSG00000173020 | 701.9702942 | -0.054434855 | 0.2585652 | -0.210527 | 0.83325667 | 0.911604157 | GRK2 | protein_coding | 61.02 | -0.002187938 | 0.002187938 | ENSG00000173020 | n.s. |
| ENSG00000111665 | 224.8625413 | 0.054945239 | 0.2655036 | 0.206947 | 0.83605105 | 0.912681415 | CDCA3 | protein_coding | 52.08 | 0.002180271 | 0.002180271 | ENSG00000111665 | n.s. |
| ENSG00000213639 | 3130.082069 | -0.056783829 | 0.2818542 | -0.201465 | 0.84033478 | 0.915408802 | PPP1CB | protein_coding | 38.98 | -0.002179642 | 0.002179642 | ENSG00000213639 | n.s. |
| ENSG00000272030 | 38.67354419 | -0.057483507 | 0.2879102 | -0.199658 | 0.841748262 | 0.916457699 | | antisense | 53.92 | -0.002177911 | 0.002177911 | ENSG00000272030 | n.s. |
| ENSG00000142188 | 208.2626598 | 0.056706164 | 0.2816256 | 0.201353 | 0.840422556 | 0.915408802 | TMEM50B | protein_coding | 43.36 | 0.002176661 | 0.002176661 | ENSG00000142188 | n.s. |
| ENSG00000257698 | 210.0927585 | 0.06161546 | 0.3286029 | 0.187507 | 0.851262847 | 0.921914485 | | lincRNA | 40.3 | 0.002175603 | 0.002175603 | ENSG00000257698 | n.s. |
| ENSG00000144381 | 23304.92888 | 0.054555939 | 0.2621236 | 0.208131 | 0.835127011 | 0.912316468 | HSPD1 | protein_coding | 42.78 | 0.002174299 | 0.002174299 | ENSG00000144381 | n.s. |
| ENSG00000156603 | 345.4717323 | -0.055658452 | 0.2725119 | -0.204242 | 0.838164151 | 0.914034657 | MED19 | protein_coding | 45.7 | -0.002172758 | 0.002172758 | ENSG00000156603 | n.s. |
| ENSG00000128567 | 263.3100679 | 0.053338955 | 0.2502396 | 0.213152 | 0.831208746 | 0.910651134 | PODXL | protein_coding | 50.47 | 0.00216812 | 0.00216812 | ENSG00000128567 | n.s. |
| ENSG00000134758 | 1639.14068 | -0.057845494 | 0.2936077 | -0.197016 | 0.843814825 | 0.917631321 | RNF138 | protein_coding | 40.11 | -0.002159475 | 0.002159475 | ENSG00000134758 | n.s. |

| | | | | | | | | | | | | | |
|---|---|---|---|---|---|---|---|---|---|---|---|---|---|
| ENSG00000132155 | 3021.34643 | 0.051891328 | 0.2393852 | 0.216769 | 0.828388265 | 0.90866224 | RAF1 | protein_coding | 42.69 | 0.002158551 | 0.002158551 | ENSG00000132155 | n.s. |
| ENSG00000168275 | 388.9187501 | 0.057944591 | 0.2947242 | 0.196606 | 0.84413576 | 0.917908637 | COA6 | protein_coding | 44.99 | 0.00215557 | 0.00215557 | ENSG00000168275 | n.s. |
| ENSG00000273749 | 1476.152092 | 0.055053703 | 0.2693533 | 0.204392 | 0.838047051 | 0.914034657 | CYFIP1 | protein_coding | 48.83 | 0.00214915 | 0.00214915 | ENSG00000273749 | n.s. |
| ENSG00000169217 | 1361.722546 | -0.053497968 | 0.255059 | -0.209747 | 0.833864831 | 0.911652929 | CD2BP2 | protein_coding | 57.88 | -0.002149038 | 0.002149038 | ENSG00000169217 | n.s. |
| ENSG00000197860 | 665.2589291 | -0.054128664 | 0.2617591 | -0.206788 | 0.836175347 | 0.912745539 | SGTB | protein_coding | 39.79 | -0.002146217 | 0.002146217 | ENSG00000197860 | n.s. |
| ENSG00000105298 | 551.8866632 | -0.051288885 | 0.2352402 | -0.218028 | 0.82740749 | 0.908158808 | CACTIN | protein_coding | 61.58 | -0.002145835 | 0.002145835 | ENSG00000105298 | n.s. |
| ENSG00000160606 | 42.70197012 | -0.055766613 | 0.2767085 | -0.201536 | 0.84027981 | 0.915408802 | TLCD1 | protein_coding | 60.34 | -0.002140597 | 0.002140597 | ENSG00000160606 | n.s. |
| ENSG00000168566 | 712.0534093 | -0.05361257 | 0.2569766 | -0.208628 | 0.834738498 | 0.912208166 | SNRNP48 | protein_coding | 37.33 | -0.002139466 | 0.002139466 | ENSG00000168566 | n.s. |
| ENSG00000120685 | 935.044652 | 0.052086751 | 0.2434231 | 0.213976 | 0.830565629 | 0.910253357 | PROSER1 | protein_coding | 37.33 | 0.002127104 | 0.002127104 | ENSG00000120685 | n.s. |
| ENSG00000269929 | 923.7826689 | 0.049554165 | 0.2228265 | 0.222389 | 0.824011067 | 0.906024135 | | lincRNA | 36.45 | 0.002123903 | 0.002123903 | ENSG00000269929 | n.s. |
| ENSG00000198189 | 461.356345 | 0.060484185 | 0.3248257 | 0.186205 | 0.852283983 | 0.922333766 | HSD17B11 | protein_coding | 39.92 | 0.002123714 | 0.002123714 | ENSG00000198189 | n.s. |
| ENSG00000234964 | 64.07956892 | -0.075292929 | 0.492485 | -0.152884 | 0.878489991 | 0.937154913 | FABP5P7 | processed_pseudogene | 44.44 | -0.002122403 | 0.002122403 | ENSG00000234964 | n.s. |
| ENSG00000269688 | 149.8710624 | 0.050364917 | 0.2300296 | 0.21895 | 0.826689238 | 0.907728254 | | sense_intronic | 48.12 | 0.00211755 | 0.00211755 | ENSG00000269688 | n.s. |
| ENSG00000233757 | 467.1939486 | -0.049315024 | 0.2213855 | -0.222756 | 0.823725167 | 0.90597405 | | protein_coding | 41.52 | -0.002114837 | 0.002114837 | ENSG00000233757 | n.s. |
| ENSG00000265982 | 14.30637266 | 0.06383093 | 0.3632238 | 0.175734 | 0.860502591 | 0.926587593 | | unprocessed_pseudogene | 72.92 | 0.002113667 | 0.002113667 | ENSG00000265982 | n.s. |
| ENSG00000200075 | 17.91303038 | 0.062424453 | 0.3456101 | 0.180621 | 0.856665067 | 0.925266075 | | snoRNA | 41.73 | 0.002105787 | 0.002105787 | ENSG00000200075 | n.s. |
| ENSG00000173208 | 466.4949077 | -0.0479727 | 0.2117128 | -0.226593 | 0.820740012 | 0.90404772 | ABCD2 | protein_coding | 34.95 | -0.002101619 | 0.002101619 | ENSG00000173208 | n.s. |
| ENSG00000260682 | 9.928181166 | 0.069678675 | 0.4310243 | 0.161658 | 0.871574912 | 0.933026104 | RN7SKP176 | misc_RNA | 55.66 | 0.002097761 | 0.002097761 | ENSG00000260682 | n.s. |
| ENSG00000075089 | 979.7660487 | -0.05957774 | 0.3188786 | -0.186835 | 0.85178985 | 0.922317592 | ACTR6 | protein_coding | 41 | -0.002092341 | 0.002092341 | ENSG00000075089 | n.s. |
| ENSG00000106608 | 86.03649476 | -0.054808806 | 0.2730833 | -0.200704 | 0.840930332 | 0.915853646 | URGCP | protein_coding | 45.69 | -0.002092267 | 0.002092267 | ENSG00000106608 | n.s. |
| ENSG00000018699 | 474.0953293 | -0.05278377 | 0.2557664 | -0.206375 | 0.836498054 | 0.912883086 | TTC27 | protein_coding | 39.46 | -0.002089437 | 0.002089437 | ENSG00000018699 | n.s. |
| ENSG00000120334 | 919.8315495 | -0.050002925 | 0.2305951 | -0.216843 | 0.828330715 | 0.90866224 | CENPL | protein_coding | 40.71 | -0.002079998 | 0.002079998 | ENSG00000120334 | n.s. |
| ENSG00000103051 | 747.0666324 | -0.055691568 | 0.2825867 | -0.197078 | 0.843766669 | 0.917631321 | COG4 | protein_coding | 47.24 | -0.002079065 | 0.002079065 | ENSG00000103051 | n.s. |
| ENSG00000117118 | 442.4607775 | 0.05702784 | 0.2968643 | 0.192101 | 0.847663308 | 0.919518211 | SDHB | protein_coding | 43.51 | 0.002078075 | 0.002078075 | ENSG00000117118 | n.s. |
| ENSG00000103245 | 189.5381364 | -0.057148813 | 0.2981609 | -0.191671 | 0.847999887 | 0.919685642 | NARFL | protein_coding | 60.82 | -0.002077965 | 0.002077965 | ENSG00000103245 | n.s. |
| ENSG00000163956 | 536.4050784 | -0.051693351 | 0.2457494 | -0.21035 | 0.833394638 | 0.911604157 | LRPAP1 | protein_coding | 57.65 | -0.002077747 | 0.002077747 | ENSG00000163956 | n.s. |
| ENSG00000151465 | 1548.068356 | -0.054602057 | 0.27277 | -0.200176 | 0.841342768 | 0.916231158 | CDC123 | protein_coding | 42.91 | -0.002074602 | 0.002074602 | ENSG00000151465 | n.s. |
| ENSG00000224531 | 1093.977169 | 0.056247079 | 0.2882411 | 0.195139 | 0.845284126 | 0.918762803 | SMIM13 | protein_coding | 42.19 | 0.002069701 | 0.002069701 | ENSG00000224531 | n.s. |
| ENSG00000091651 | 2142.674408 | -0.051907665 | 0.2502853 | -0.207394 | 0.835702203 | 0.912553886 | ORC6 | protein_coding | 44.16 | -0.002062888 | 0.002062888 | ENSG00000091651 | n.s. |
| ENSG00000109534 | 372.2426778 | 0.0511329 | 0.2413878 | 0.211829 | 0.832240575 | 0.911308339 | GAR1 | protein_coding | 39.37 | 0.002062428 | 0.002062428 | ENSG00000109534 | n.s. |
| ENSG00000111325 | 86.17331122 | -0.053560721 | 0.2648938 | -0.202197 | 0.839762732 | 0.915226342 | OGFOD2 | protein_coding | 59.54 | -0.002060561 | 0.002060561 | ENSG00000111325 | n.s. |
| ENSG00000227698 | 44.36308762 | 0.054605246 | 0.275819 | 0.197975 | 0.843064708 | 0.917317125 | | sense_intronic | 51.92 | 0.002046632 | 0.002046632 | ENSG00000227698 | n.s. |

| | | | | | | | | | | | | | |
|---|---|---|---|---|---|---|---|---|---|---|---|---|---|
| ENSG00000163376 | 398.7942051 | 0.054173592 | 0.2724099 | 0.198868 | 0.842366085 | 0.916797802 | KBTBD8 | protein_coding | 36.69 | 0.002043777 | 0.002043777 | ENSG00000163376 | n.s. |
| ENSG00000198642 | 1697.034882 | -0.050639459 | 0.239965 | -0.211028 | 0.832865038 | 0.911467305 | KLHL9 | protein_coding | 39.32 | -0.002038689 | 0.002038689 | ENSG00000198642 | n.s. |
| ENSG00000100577 | 227.454914 | -0.058910704 | 0.3202483 | -0.183953 | 0.854050147 | 0.923453667 | GSTZ1 | protein_coding | 54.52 | -0.00203742 | 0.00203742 | ENSG00000100577 | n.s. |
| ENSG00000012822 | 349.7080807 | 0.055336507 | 0.2861178 | 0.193405 | 0.846642077 | 0.91905486 | CALCOCO1 | protein_coding | 49.2 | 0.002028557 | 0.002028557 | ENSG00000012822 | n.s. |
| ENSG00000162062 | 535.9664015 | -0.05103973 | 0.2463536 | -0.207181 | 0.835868685 | 0.912553886 | C16orf59 | protein_coding | 62.04 | -0.002028394 | 0.002028394 | ENSG00000162062 | n.s. |
| ENSG00000254577 | 34.81041022 | -0.060120508 | 0.3339936 | -0.180005 | 0.85714867 | 0.925266075 | | sense_overlapping | 39.9 | -0.002028067 | 0.002028067 | ENSG00000254577 | n.s. |
| ENSG00000162998 | 787.0710497 | -0.054643297 | 0.2787955 | -0.195998 | 0.844611933 | 0.918139596 | FRZB | protein_coding | 36.85 | -0.00202679 | 0.00202679 | ENSG00000162998 | n.s. |
| ENSG00000170892 | 427.5531203 | 0.057619904 | 0.3094345 | 0.18621 | 0.852279802 | 0.922333766 | TSEN34 | protein_coding | 55.95 | 0.002023144 | 0.002023144 | ENSG00000170892 | n.s. |
| ENSG00000099991 | 238.1418927 | 0.050873724 | 0.2455523 | 0.207181 | 0.835868624 | 0.912553886 | CABIN1 | protein_coding | 50.24 | 0.002021797 | 0.002021797 | ENSG00000099991 | n.s. |
| ENSG00000172932 | 69.59432368 | -0.054397794 | 0.2769681 | -0.196405 | 0.844293553 | 0.918008523 | ANKRD13D | protein_coding | 53.51 | -0.002021057 | 0.002021057 | ENSG00000172932 | n.s. |
| ENSG00000262803 | 43.46966338 | 0.056002351 | 0.2945854 | 0.190106 | 0.849226358 | 0.920424925 | | processed_pseudogene | 38.8 | 0.002016736 | 0.002016736 | ENSG00000262803 | n.s. |
| ENSG00000131089 | 128.1245327 | 0.050784509 | 0.2457532 | 0.206648 | 0.836284472 | 0.912791798 | ARHGEF9 | protein_coding | 40.8 | 0.002012502 | 0.002012502 | ENSG00000131089 | n.s. |
| ENSG00000154582 | 1031.912289 | 0.054789028 | 0.2819734 | 0.194306 | 0.845936562 | 0.918923716 | ELOC | protein_coding | 41.19 | 0.002011883 | 0.002011883 | ENSG00000154582 | n.s. |
| ENSG00000138074 | 1078.502274 | -0.051472104 | 0.2521907 | -0.2041 | 0.83827541 | 0.914034657 | SLC5A6 | protein_coding | 53.48 | -0.002009334 | 0.002009334 | ENSG00000138074 | n.s. |
| ENSG00000170606 | 4483.915992 | -0.054800721 | 0.2837901 | -0.193103 | 0.846878287 | 0.919178013 | HSPA4 | protein_coding | 41.31 | -0.002005727 | 0.002005727 | ENSG00000170606 | n.s. |
| ENSG00000257621 | 2195.373632 | 0.049916796 | 0.239499 | 0.208422 | 0.834899703 | 0.912208166 | PSMA3-AS1 | antisense | 38.57 | 0.001991982 | 0.001991982 | ENSG00000257621 | n.s. |
| ENSG00000079950 | 207.9935169 | -0.054057137 | 0.2787713 | -0.193912 | 0.846244667 | 0.918923716 | STX7 | protein_coding | 37.41 | -0.001985007 | 0.001985007 | ENSG00000079950 | n.s. |
| ENSG00000079150 | 900.3009102 | -0.050638033 | 0.2474134 | -0.20467 | 0.837830165 | 0.913950189 | FKBP7 | protein_coding | 37.58 | -0.001978806 | 0.001978806 | ENSG00000079150 | n.s. |
| ENSG00000186480 | 1354.336764 | -0.050676378 | 0.2482704 | -0.204118 | 0.83826156 | 0.914034657 | INSIG1 | protein_coding | 44.41 | -0.001978271 | 0.001978271 | ENSG00000186480 | n.s. |
| ENSG00000198938 | 15083.61606 | 0.049044845 | 0.2336821 | 0.209878 | 0.833762532 | 0.911652929 | MT-CO3 | protein_coding | 46.56 | 0.001970154 | 0.001970154 | ENSG00000198938 | n.s. |
| ENSG00000118855 | 673.4939929 | -0.054618466 | 0.286715 | -0.190497 | 0.848919395 | 0.92030715 | MFSD1 | protein_coding | 40.83 | -0.001969935 | 0.001969935 | ENSG00000118855 | n.s. |
| ENSG00000196123 | 663.1509621 | 0.051164049 | 0.2532245 | 0.20205 | 0.839877535 | 0.915279815 | KIAA0895L | protein_coding | 56.58 | 0.001967059 | 0.001967059 | ENSG00000196123 | n.s. |
| ENSG00000182704 | 741.3467782 | 0.048703141 | 0.2296553 | 0.212071 | 0.832051931 | 0.911239732 | TSKU | protein_coding | 55.35 | 0.001966017 | 0.001966017 | ENSG00000182704 | n.s. |
| ENSG00000280069 | 360.7276797 | -0.050304988 | 0.245847 | -0.204619 | 0.837869728 | 0.913950189 | | TEC | 37.97 | -0.001965792 | 0.001965792 | ENSG00000280069 | n.s. |
| ENSG00000143401 | 5383.718676 | -0.051177905 | 0.2542243 | -0.20131 | 0.840456131 | 0.915408802 | ANP32E | protein_coding | 42.62 | -0.00196446 | 0.00196446 | ENSG00000143401 | n.s. |
| ENSG00000005486 | 50.35243922 | 0.05312597 | 0.2728668 | 0.194696 | 0.845631267 | 0.918762803 | RHBDD2 | protein_coding | 47.79 | 0.001954855 | 0.001954855 | ENSG00000005486 | n.s. |
| ENSG00000171766 | 219.1832111 | 0.05259993 | 0.2683123 | 0.19604 | 0.844578895 | 0.918139596 | GATM | protein_coding | 40.25 | 0.001950999 | 0.001950999 | ENSG00000171766 | n.s. |
| ENSG00000167315 | 1607.365456 | -0.052958888 | 0.2717372 | -0.19489 | 0.845478991 | 0.918762803 | ACAA2 | protein_coding | 39.54 | -0.001948707 | 0.001948707 | ENSG00000167315 | n.s. |
| ENSG00000114744 | 1231.952365 | -0.05414099 | 0.2850242 | -0.189952 | 0.849346559 | 0.920483548 | COMMD2 | protein_coding | 37.91 | -0.001948208 | 0.001948208 | ENSG00000114744 | n.s. |
| ENSG00000197157 | 2748.612189 | -0.053030442 | 0.2734504 | -0.193931 | 0.846230126 | 0.918923716 | SND1 | protein_coding | 42.44 | -0.001947306 | 0.001947306 | ENSG00000197157 | n.s. |
| ENSG00000120694 | 5461.966318 | 0.050247991 | 0.2476285 | 0.202917 | 0.839200041 | 0.914756296 | HSPH1 | protein_coding | 36.82 | 0.001944326 | 0.001944326 | ENSG00000120694 | n.s. |
| ENSG00000165516 | 1153.614323 | 0.053971279 | 0.2836311 | 0.190287 | 0.849084346 | 0.920414307 | KLHDC2 | protein_coding | 41.17 | 0.001943864 | 0.001943864 | ENSG00000165516 | n.s. |

| | | | | | | | | | | | | | |
|---|---|---|---|---|---|---|---|---|---|---|---|---|---|
| ENSG00000269386 | 69.71969181 | 0.05189736 | 0.2627326 | 0.197529 | 0.843413458 | 0.917409811 | RAB11B-AS1 | antisense | 48.55 | 0.001942862 | 0.001942862 | ENSG00000269386 | n.s. |
| ENSG00000108100 | 1059.576826 | -0.051363859 | 0.2583208 | -0.198837 | 0.842389879 | 0.916797802 | CCNY | protein_coding | 43.24 | -0.001937775 | 0.001937775 | ENSG00000108100 | n.s. |
| ENSG00000112304 | 166.7587863 | -0.055073198 | 0.2951594 | -0.186588 | 0.851983702 | 0.922333766 | ACOT13 | protein_coding | 41.43 | -0.001933724 | 0.001933724 | ENSG00000112304 | n.s. |
| ENSG00000175879 | 411.1654616 | 0.048004509 | 0.2275402 | 0.210972 | 0.832909468 | 0.911467305 | HOXD8 | protein_coding | 52.17 | 0.001932609 | 0.001932609 | ENSG00000175879 | n.s. |
| ENSG00000160072 | 300.9875504 | 0.049738314 | 0.2453291 | 0.202741 | 0.839337358 | 0.914834353 | ATAD3B | protein_coding | 58.83 | 0.001922761 | 0.001922761 | ENSG00000160072 | n.s. |
| ENSG00000115211 | 600.3791286 | -0.047661628 | 0.2259001 | -0.210985 | 0.832898638 | 0.911467305 | EIF2B4 | protein_coding | 50.64 | -0.001918805 | 0.001918805 | ENSG00000115211 | n.s. |
| ENSG00000100462 | 1616.079521 | 0.052719523 | 0.2750813 | 0.191651 | 0.848015821 | 0.919685642 | PRMT5 | protein_coding | 47.75 | 0.001916913 | 0.001916913 | ENSG00000100462 | n.s. |
| ENSG00000136699 | 615.3778502 | 0.045971027 | 0.2120119 | 0.216832 | 0.828339014 | 0.90866224 | SMPD4 | protein_coding | 53.96 | 0.001912281 | 0.001912281 | ENSG00000136699 | n.s. |
| ENSG00000185222 | 992.0850627 | -0.051855881 | 0.2660009 | -0.194946 | 0.84543504 | 0.918762803 | TCEAL9 | protein_coding | 44.15 | -0.00190812 | 0.00190812 | ENSG00000185222 | n.s. |
| ENSG00000026297 | 109.9213535 | -0.071710041 | 0.4999497 | -0.143435 | 0.885947027 | 0.94067265 | RNASET2 | protein_coding | 48.7 | -0.001904725 | 0.001904725 | ENSG00000026297 | n.s. |
| ENSG00000104671 | 782.2519064 | -0.054075768 | 0.2904095 | -0.186205 | 0.852283842 | 0.922333766 | DCTN6 | protein_coding | 41.56 | -0.001898703 | 0.001898703 | ENSG00000104671 | n.s. |
| ENSG00000156261 | 6017.96906 | 0.054198606 | 0.2927255 | 0.185152 | 0.853110076 | 0.922694911 | CCT8 | protein_coding | 38.32 | 0.001893801 | 0.001893801 | ENSG00000156261 | n.s. |
| ENSG00000196305 | 7762.746903 | -0.051724975 | 0.2681126 | -0.192923 | 0.847019601 | 0.919178013 | IARS | protein_coding | 41.23 | -0.001893153 | 0.001893153 | ENSG00000196305 | n.s. |
| ENSG00000034053 | 1712.480672 | -0.051497335 | 0.2673208 | -0.192642 | 0.847239 | 0.919272803 | APBA2 | protein_coding | 48.63 | -0.001882515 | 0.001882515 | ENSG00000034053 | n.s. |
| ENSG00000215252 | 950.6303879 | 0.051067004 | 0.2623074 | 0.194684 | 0.845640485 | 0.918762803 | GOLGA8B | protein_coding | 47.09 | 0.001879092 | 0.001879092 | ENSG00000215252 | n.s. |
| ENSG00000197903 | 1472.965764 | -0.0436284 | 0.196183 | -0.222386 | 0.824013255 | 0.906024135 | HIST1H2BK | protein_coding | 61.15 | -0.001869923 | 0.001869923 | ENSG00000197903 | n.s. |
| ENSG00000251580 | 20.99112914 | -0.05999665 | 0.3599894 | -0.166662 | 0.867635813 | 0.93127488 | | lincRNA | 53.54 | -0.001855223 | 0.001855223 | ENSG00000251580 | n.s. |
| ENSG00000251562 | 44817.14079 | -0.045379064 | 0.2122319 | -0.213818 | 0.830688727 | 0.910253357 | MALAT1 | lincRNA | 40.27 | -0.001853177 | 0.001853177 | ENSG00000251562 | n.s. |
| ENSG00000240429 | 222.6268049 | -0.045504359 | 0.2137125 | -0.212923 | 0.831386817 | 0.91067638 | LRRFIP1P1 | processed_pseudogene | 47.15 | -0.001849112 | 0.001849112 | ENSG00000240429 | n.s. |
| ENSG00000263675 | 8.733350572 | 0.063222444 | 0.4002676 | 0.15795 | 0.874495836 | 0.935290023 | MIR5581 | miRNA | 48.33 | 0.001836846 | 0.001836846 | ENSG00000263675 | n.s. |
| ENSG00000115484 | 9346.273729 | 0.052231797 | 0.2799692 | 0.186563 | 0.852003568 | 0.922333766 | CCT4 | protein_coding | 41.18 | 0.001833957 | 0.001833957 | ENSG00000115484 | n.s. |
| ENSG00000159840 | 263.0726457 | -0.046140979 | 0.2227012 | -0.207188 | 0.835863148 | 0.912553886 | ZYX | protein_coding | 58.32 | -0.001833711 | 0.001833711 | ENSG00000159840 | n.s. |
| ENSG00000204590 | 343.8618633 | -0.05222053 | 0.2808834 | -0.185915 | 0.852511186 | 0.922333766 | GNL1 | protein_coding | 49.56 | -0.001833562 | 0.001833562 | ENSG00000204590 | n.s. |
| ENSG00000111752 | 487.2885923 | 0.04887748 | 0.2467991 | 0.198046 | 0.843009343 | 0.917317125 | PHC1 | protein_coding | 44.13 | 0.001831952 | 0.001831952 | ENSG00000111752 | n.s. |
| ENSG00000186416 | 503.444344 | -0.050653647 | 0.2654442 | -0.190826 | 0.848661966 | 0.920171369 | NKRF | protein_coding | 42.08 | -0.001830181 | 0.001830181 | ENSG00000186416 | n.s. |
| ENSG00000067082 | 1244.426891 | 0.042777767 | 0.192935 | 0.221721 | 0.824530954 | 0.906287556 | KLF6 | protein_coding | 48.08 | 0.001828064 | 0.001828064 | ENSG00000067082 | n.s. |
| ENSG00000172062 | 170.7470707 | 0.055842106 | 0.3215775 | 0.173651 | 0.862140112 | 0.927418434 | SMN1 | protein_coding | 42.22 | 0.001827393 | 0.001827393 | ENSG00000172062 | n.s. |
| ENSG00000101997 | 125.7384421 | -0.054309335 | 0.3050994 | -0.178005 | 0.858718748 | 0.925994426 | CCDC22 | protein_coding | 53.55 | -0.001813477 | 0.001813477 | ENSG00000101997 | n.s. |
| ENSG00000175324 | 623.8434258 | -0.053826167 | 0.3000232 | -0.179407 | 0.85761837 | 0.925557753 | LSM1 | protein_coding | 42.96 | -0.00180837 | 0.00180837 | ENSG00000175324 | n.s. |
| ENSG00000070501 | 557.7823431 | 0.054130633 | 0.3039045 | 0.178117 | 0.858630909 | 0.925994426 | POLB | protein_coding | 40.44 | 0.00180751 | 0.00180751 | ENSG00000070501 | n.s. |
| ENSG00000166788 | 1085.102506 | 0.050086341 | 0.2627396 | 0.190631 | 0.848814626 | 0.920265226 | SAAL1 | protein_coding | 39.77 | 0.001807465 | 0.001807465 | ENSG00000166788 | n.s. |
| ENSG00000110536 | 673.70585 | 0.053522939 | 0.2973416 | 0.180005 | 0.857148734 | 0.925266075 | PTPMT1 | protein_coding | 47.83 | 0.001805509 | 0.001805509 | ENSG00000110536 | n.s. |

| | | | | | | | | | | | | | |
|---|---|---|---|---|---|---|---|---|---|---|---|---|---|
| ENSG00000204463 | 1124.667539 | -0.049011592 | 0.2514608 | -0.194907 | 0.845465363 | 0.918762803 | BAG6 | protein_coding | 52.36 | -0.00180346 | 0.00180346 | ENSG00000204463 | n.s. |
| ENSG00000189067 | 337.1997909 | -0.051013339 | 0.2718252 | -0.18767 | 0.851135644 | 0.92184843 | LITAF | protein_coding | 49.14 | -0.001802836 | 0.001802836 | ENSG00000189067 | n.s. |
| ENSG00000143228 | 3025.550272 | -0.053070189 | 0.2940634 | -0.180472 | 0.856782084 | 0.925266075 | NUF2 | protein_coding | 36.44 | -0.001790236 | 0.001790236 | ENSG00000143228 | n.s. |
| ENSG00000235529 | 56.74894173 | 0.05219064 | 0.2860238 | 0.18247 | 0.855214229 | 0.92422673 | AGAP1-IT1 | sense_intronic | 50.66 | 0.001786041 | 0.001786041 | ENSG00000235529 | n.s. |
| ENSG00000151353 | 1258.155402 | 0.052196896 | 0.2865631 | 0.182148 | 0.855466598 | 0.924310216 | TMEM18 | protein_coding | 53.39 | 0.001784207 | 0.001784207 | ENSG00000151353 | n.s. |
| ENSG00000140548 | 229.0220177 | -0.048869836 | 0.2547327 | -0.191848 | 0.847861658 | 0.919661722 | ZNF710 | protein_coding | 50.7 | -0.001777488 | 0.001777488 | ENSG00000140548 | n.s. |
| ENSG00000008735 | 543.1205624 | -0.048413372 | 0.2508481 | -0.192999 | 0.846959968 | 0.919178013 | MAPK8IP2 | protein_coding | 63.07 | -0.001771947 | 0.001771947 | ENSG00000008735 | n.s. |
| ENSG00000142669 | 146.0643281 | 0.051555033 | 0.2816822 | 0.183026 | 0.854777995 | 0.923924697 | SH3BGRL3 | protein_coding | 61.74 | 0.001771607 | 0.001771607 | ENSG00000142669 | n.s. |
| ENSG00000068724 | 127.2357065 | -0.048023917 | 0.2478794 | -0.193739 | 0.846380232 | 0.918923716 | TTC7A | protein_coding | 50.88 | -0.001763464 | 0.001763464 | ENSG00000068724 | n.s. |
| ENSG00000062485 | 2717.328143 | 0.049198304 | 0.2598837 | 0.189309 | 0.849850716 | 0.920958244 | CS | protein_coding | 46.16 | 0.001759334 | 0.001759334 | ENSG00000062485 | n.s. |
| ENSG00000156304 | 1308.463279 | -0.046357831 | 0.2328223 | -0.199113 | 0.84217474 | 0.916797802 | SCAF4 | protein_coding | 38.12 | -0.001748916 | 0.001748916 | ENSG00000156304 | n.s. |
| ENSG00000137449 | 106.2324536 | -0.048522097 | 0.2551951 | -0.190137 | 0.849201562 | 0.920424925 | CPEB2 | protein_coding | 34.97 | -0.00174736 | 0.00174736 | ENSG00000137449 | n.s. |
| ENSG00000060762 | 882.0539604 | 0.05493335 | 0.3243043 | 0.169388 | 0.865491253 | 0.929515174 | MPC1 | protein_coding | 38.93 | 0.001743778 | 0.001743778 | ENSG00000060762 | n.s. |
| ENSG00000253475 | 23.76969513 | 0.053871862 | 0.3122053 | 0.172553 | 0.863003055 | 0.928274995 | | sense_intronic | 47.61 | 0.001741319 | 0.001741319 | ENSG00000253475 | n.s. |
| ENSG00000131759 | 115.2142506 | 0.046933627 | 0.2392876 | 0.196139 | 0.844501366 | 0.918139596 | RARA | protein_coding | 56.99 | 0.001740828 | 0.001740828 | ENSG00000131759 | n.s. |
| ENSG00000177565 | 5917.4129 | -0.04918102 | 0.2618516 | -0.18782 | 0.851017596 | 0.921792282 | TBL1XR1 | protein_coding | 39.31 | -0.001739382 | 0.001739382 | ENSG00000177565 | n.s. |
| ENSG00000186132 | 144.5714741 | -0.052255569 | 0.2943948 | -0.177502 | 0.859114349 | 0.926238894 | C2orf76 | protein_coding | 39.21 | -0.001738908 | 0.001738908 | ENSG00000186132 | n.s. |
| ENSG00000124444 | 254.9220995 | -0.046063383 | 0.2319493 | -0.198592 | 0.842581562 | 0.916934742 | ZNF576 | protein_coding | 50.29 | -0.001734819 | 0.001734819 | ENSG00000124444 | n.s. |
| ENSG00000163374 | 1066.567406 | 0.047172054 | 0.2429621 | 0.194154 | 0.846055331 | 0.918923716 | YY1AP1 | protein_coding | 44.63 | 0.001732183 | 0.001732183 | ENSG00000163374 | n.s. |
| ENSG00000240344 | 558.36649 | 0.049349561 | 0.2656379 | 0.185778 | 0.852619223 | 0.922378977 | PPIL3 | protein_coding | 41.39 | 0.001731706 | 0.001731706 | ENSG00000240344 | n.s. |
| ENSG00000008324 | 380.5288933 | 0.051690667 | 0.2903468 | 0.178031 | 0.858698786 | 0.925994426 | SS18L2 | protein_coding | 47.77 | 0.001726036 | 0.001726036 | ENSG00000008324 | n.s. |
| ENSG00000134440 | 4916.568683 | -0.050689213 | 0.2797163 | -0.181217 | 0.856197642 | 0.924956531 | NARS | protein_coding | 42.81 | -0.001717283 | 0.001717283 | ENSG00000134440 | n.s. |
| ENSG00000095380 | 418.2080816 | -0.046753169 | 0.2413347 | -0.193727 | 0.846389265 | 0.918923716 | NANS | protein_coding | 46.23 | -0.001716802 | 0.001716802 | ENSG00000095380 | n.s. |
| ENSG00000143374 | 828.8253179 | -0.048462924 | 0.2578563 | -0.187945 | 0.850919423 | 0.921757654 | TARS2 | protein_coding | 48.52 | -0.001714776 | 0.001714776 | ENSG00000143374 | n.s. |
| ENSG00000151500 | 338.9284807 | -0.049515919 | 0.2689981 | -0.184075 | 0.853954345 | 0.923453667 | THYN1 | protein_coding | 43.95 | -0.001712503 | 0.001712503 | ENSG00000151500 | n.s. |
| ENSG00000051128 | 198.5124857 | -0.051557312 | 0.2919429 | -0.176601 | 0.859822056 | 0.926439893 | HOMER3 | protein_coding | 56.77 | -0.001710814 | 0.001710814 | ENSG00000051128 | n.s. |
| ENSG00000211574 | 10.44670843 | -0.061616925 | 0.4084484 | -0.150856 | 0.880089246 | 0.938101559 | MIR770 | miRNA | 65.31 | -0.001709878 | 0.001709878 | ENSG00000211574 | n.s. |
| ENSG00000144867 | 1251.062263 | 0.049936571 | 0.2740178 | 0.182238 | 0.855395631 | 0.924305267 | SRPRB | protein_coding | 41.15 | 0.00170706 | 0.00170706 | ENSG00000144867 | n.s. |
| ENSG00000011426 | 2708.072984 | 0.045307369 | 0.2291375 | 0.19773 | 0.843256288 | 0.917409811 | ANLN | protein_coding | 37.88 | 0.001696155 | 0.001696155 | ENSG00000011426 | n.s. |
| ENSG00000057252 | 819.5594245 | 0.046235396 | 0.2396258 | 0.192948 | 0.846999444 | 0.919178013 | SOAT1 | protein_coding | 39.63 | 0.001692233 | 0.001692233 | ENSG00000057252 | n.s. |
| ENSG00000155229 | 1140.793966 | -0.047379378 | 0.25159 | -0.18832 | 0.850625956 | 0.92158316 | MMS19 | protein_coding | 44.91 | -0.001680332 | 0.001680332 | ENSG00000155229 | n.s. |
| ENSG00000232024 | 29.386954 | -0.057497411 | 0.3627029 | -0.158525 | 0.874043251 | 0.935021451 | LSM12P1 | processed_pseudogene | 48.98 | -0.001677684 | 0.001677684 | ENSG00000232024 | n.s. |

| | | | | | | | | | | | | | | |
|---|---|---|---|---|---|---|---|---|---|---|---|---|---|---|
| ENSG00000100554 | 1151.094306 | -0.050056735 | 0.279621 | -0.179016 | 0.857924851 | 0.925804766 | ATP6V1D | protein_coding | | 38.54 | -0.001675929 | 0.001675929 | ENSG00000100554 | n.s. |
| ENSG00000147677 | 4064.703085 | 0.050006999 | 0.279863 | 0.178684 | 0.858185962 | 0.925831073 | EIF3H | protein_coding | | 37.79 | 0.001673647 | 0.001673647 | ENSG00000147677 | n.s. |
| ENSG00000163930 | 1197.839701 | -0.047570337 | 0.255776 | -0.185984 | 0.852457035 | 0.922333766 | BAP1 | protein_coding | | 56.55 | -0.001670285 | 0.001670285 | ENSG00000163930 | n.s. |
| ENSG00000197291 | 395.2888429 | -0.045267928 | 0.2325304 | -0.194675 | 0.845647153 | 0.918762803 | RAMP2-AS1 | lincRNA | | 49.8 | -0.001665706 | 0.001665706 | ENSG00000197291 | n.s. |
| ENSG00000085415 | 1069.9812 | 0.047395591 | 0.253752 | 0.186779 | 0.851833794 | 0.922317592 | SEH1L | protein_coding | | 43.55 | 0.00166451 | 0.00166451 | ENSG00000085415 | n.s. |
| ENSG00000132386 | 1756.209841 | -0.050093009 | 0.2835457 | -0.176666 | 0.859770418 | 0.926439893 | SERPINF1 | protein_coding | | 53.48 | -0.001662224 | 0.001662224 | ENSG00000132386 | n.s. |
| ENSG00000188312 | 233.2980515 | -0.049026086 | 0.2720042 | -0.18024 | 0.856964019 | 0.925266075 | CENPP | protein_coding | | 40.03 | -0.001653815 | 0.001653815 | ENSG00000188312 | n.s. |
| ENSG00000086848 | 256.3088262 | 0.04901303 | 0.2716966 | 0.180396 | 0.856841539 | 0.925266075 | ALG9 | protein_coding | | 40.69 | 0.001653374 | 0.001653374 | ENSG00000086848 | n.s. |
| ENSG00000070061 | 1964.465974 | 0.047694761 | 0.2593247 | 0.183919 | 0.854076914 | 0.923453667 | IKBKAP | protein_coding | | 39.91 | 0.001649518 | 0.001649518 | ENSG00000070061 | n.s. |
| ENSG00000213904 | 20.86877699 | -0.055900761 | 0.3504401 | -0.159516 | 0.873262437 | 0.934525601 | LIPE-AS1 | antisense | | 46.03 | -0.001643974 | 0.001643974 | ENSG00000213904 | n.s. |
| ENSG00000125534 | 164.3973751 | 0.04464722 | 0.2303441 | 0.193828 | 0.846310251 | 0.918923716 | PPDPF | protein_coding | | 71.07 | 0.00163947 | 0.00163947 | ENSG00000125534 | n.s. |
| ENSG00000104969 | 787.4718214 | 0.049433133 | 0.2823496 | 0.175078 | 0.861018529 | 0.926784921 | SGTA | protein_coding | | 57.8 | 0.001632334 | 0.001632334 | ENSG00000104969 | n.s. |
| ENSG00000122691 | 3420.511928 | 0.042994916 | 0.215126 | 0.199859 | 0.841590709 | 0.91635782 | TWIST1 | protein_coding | | 37.36 | 0.001631008 | 0.001631008 | ENSG00000122691 | n.s. |
| ENSG00000196839 | 114.5134843 | -0.05286379 | 0.3189425 | -0.165747 | 0.868355988 | 0.931657581 | ADA | protein_coding | | 53.46 | -0.001625227 | 0.001625227 | ENSG00000196839 | n.s. |
| ENSG00000143486 | 697.5674243 | 0.048069243 | 0.2662107 | 0.180568 | 0.856706348 | 0.925266075 | EIF2D | protein_coding | | 46.17 | 0.001621537 | 0.001621537 | ENSG00000143486 | n.s. |
| ENSG00000215717 | 758.5573116 | 0.048833056 | 0.2769695 | 0.176312 | 0.860048819 | 0.926439893 | TMEM167B | protein_coding | | 43.73 | 0.001620415 | 0.001620415 | ENSG00000215717 | n.s. |
| ENSG00000115415 | 1224.282526 | -0.04627642 | 0.2497315 | -0.185305 | 0.852990031 | 0.922636747 | STAT1 | protein_coding | | 41.75 | -0.001618252 | 0.001618252 | ENSG00000115415 | n.s. |
| ENSG00000134291 | 876.2659975 | -0.05055282 | 0.2968596 | -0.170292 | 0.864780499 | 0.92917891 | TMEM106C | protein_coding | | 46.67 | -0.001612668 | 0.001612668 | ENSG00000134291 | n.s. |
| ENSG00000170759 | 6124.859864 | 0.045447909 | 0.2412277 | 0.188403 | 0.85056113 | 0.92158316 | KIF5B | protein_coding | | 37.58 | 0.001611831 | 0.001611831 | ENSG00000170759 | n.s. |
| ENSG00000165060 | 314.9141267 | 0.048459218 | 0.2754869 | 0.175904 | 0.860369428 | 0.926587527 | FXN | protein_coding | | 45.22 | 0.001604657 | 0.001604657 | ENSG00000165060 | n.s. |
| ENSG00000124785 | 296.3359335 | -0.048333776 | 0.2740816 | -0.176348 | 0.860020441 | 0.926439893 | NRN1 | protein_coding | | 52.74 | -0.001603848 | 0.001603848 | ENSG00000124785 | n.s. |
| ENSG00000277972 | 142.9538683 | 0.048481314 | 0.2785815 | 0.174029 | 0.861842533 | 0.927169957 | CISD3 | protein_coding | | 56.94 | 0.001592158 | 0.001592158 | ENSG00000277972 | n.s. |
| ENSG00000158526 | 369.287953 | 0.047421502 | 0.2654277 | 0.178661 | 0.858204111 | 0.925831073 | TSR2 | protein_coding | | 47.57 | 0.001587115 | 0.001587115 | ENSG00000158526 | n.s. |
| ENSG00000108883 | 4687.647222 | 0.046768682 | 0.2597864 | 0.180027 | 0.857130996 | 0.925266075 | EFTUD2 | protein_coding | | 46.25 | 0.001577665 | 0.001577665 | ENSG00000108883 | n.s. |
| ENSG00000235408 | 107.6333965 | -0.045223651 | 0.2447364 | -0.184785 | 0.85339757 | 0.922934159 | SNORA71B | snoRNA | | 51.84 | -0.001575107 | 0.001575107 | ENSG00000235408 | n.s. |
| ENSG00000127870 | 1243.492413 | -0.044896867 | 0.2421489 | -0.18541 | 0.852907319 | 0.922618957 | RNF6 | protein_coding | | 40.18 | -0.001570386 | 0.001570386 | ENSG00000127870 | n.s. |
| ENSG00000103512 | 575.9223482 | -0.046982797 | 0.264008 | -0.17796 | 0.858754581 | 0.925994426 | NOMO1 | protein_coding | | 45.56 | -0.001568832 | 0.001568832 | ENSG00000103512 | n.s. |
| ENSG00000173992 | 165.4273734 | -0.047103229 | 0.2684136 | -0.175488 | 0.860696574 | 0.926724801 | CCS | protein_coding | | 51.72 | -0.001556725 | 0.001556725 | ENSG00000173992 | n.s. |
| ENSG00000054116 | 218.167359 | -0.063691705 | 0.4780462 | -0.133233 | 0.894008819 | 0.945356024 | TRAPPC3 | protein_coding | | 49.8 | -0.001554371 | 0.001554371 | ENSG00000054116 | n.s. |
| ENSG00000001084 | 1371.658895 | 0.046747401 | 0.2636844 | 0.177285 | 0.859284225 | 0.926350311 | GCLC | protein_coding | | 40.16 | 0.001553171 | 0.001553171 | ENSG00000001084 | n.s. |
| ENSG00000115274 | 160.1073986 | -0.046670706 | 0.2644755 | -0.176465 | 0.859928518 | 0.926439893 | INO80B | protein_coding | | 53.1 | -0.001548662 | 0.001548662 | ENSG00000115274 | n.s. |
| ENSG00000264616 | 25.66987453 | 0.053678718 | 0.3439094 | 0.156084 | 0.875966911 | 0.935856914 | MIR4755 | miRNA | | 47.22 | 0.00154544 | 0.00154544 | ENSG00000264616 | n.s. |

| | | | | | | | | | | | | | |
|---|---|---|---|---|---|---|---|---|---|---|---|---|---|
| ENSG00000246350 | 222.7180143 | -0.04363601 | 0.2321403 | -0.187973 | 0.850898131 | 0.921757654 | | antisense | 43.62 | -0.001543984 | 0.001543984 | ENSG00000246350 | n.s. |
| ENSG00000118418 | 2595.866034 | 0.050015925 | 0.3008997 | 0.166221 | 0.867982851 | 0.931519532 | HMGN3 | protein_coding | 38.62 | 0.001540892 | 0.001540892 | ENSG00000118418 | n.s. |
| ENSG00000172766 | 2198.131233 | 0.045207896 | 0.2487098 | 0.18177 | 0.855763476 | 0.924559236 | NAA16 | protein_coding | 37.34 | 0.001540019 | 0.001540019 | ENSG00000172766 | n.s. |
| ENSG00000126953 | 338.5703739 | 0.041884589 | 0.2161845 | 0.193745 | 0.846375823 | 0.918923716 | TIMM8A | protein_coding | 45.87 | 0.001538025 | 0.001538025 | ENSG00000126953 | n.s. |
| ENSG00000266931 | 10.77509614 | -0.059038933 | 0.4208568 | -0.140283 | 0.888436631 | 0.942149933 | | unprocessed_pseudogene | 41.96 | -0.001527926 | 0.001527926 | ENSG00000266931 | n.s. |
| ENSG00000158301 | 709.6162618 | -0.047552729 | 0.2772688 | -0.171504 | 0.863827428 | 0.928730982 | GPRASP2 | protein_coding | 51.49 | -0.001526921 | 0.001526921 | ENSG00000158301 | n.s. |
| ENSG00000157827 | 2652.564225 | -0.045804122 | 0.2595406 | -0.176481 | 0.859915682 | 0.926439893 | FMNL2 | protein_coding | 40.57 | -0.001519907 | 0.001519907 | ENSG00000157827 | n.s. |
| ENSG00000164983 | 499.2980829 | -0.045799874 | 0.2595812 | -0.176438 | 0.859950209 | 0.926439893 | TMEM65 | protein_coding | 35.86 | -0.001519766 | 0.001519766 | ENSG00000164983 | n.s. |
| ENSG00000067334 | 5254.591488 | -0.044185302 | 0.2422232 | -0.182416 | 0.855256568 | 0.92422673 | DNTTIP2 | protein_coding | 39.28 | -0.001512086 | 0.001512086 | ENSG00000067334 | n.s. |
| ENSG00000165804 | 236.4094742 | -0.046845375 | 0.2718563 | -0.172317 | 0.863188587 | 0.928402834 | ZNF219 | protein_coding | 55.48 | -0.001511398 | 0.001511398 | ENSG00000165804 | n.s. |
| ENSG00000143499 | 1205.216853 | -0.045759572 | 0.262251 | -0.174488 | 0.861482217 | 0.926997211 | SMYD2 | protein_coding | 43.58 | -0.001506477 | 0.001506477 | ENSG00000143499 | n.s. |
| ENSG00000154358 | 213.0773367 | -0.04478308 | 0.2502605 | -0.178946 | 0.857980241 | 0.925804766 | OBSCN | protein_coding | 53.87 | -0.001499364 | 0.001499364 | ENSG00000154358 | n.s. |
| ENSG00000178719 | 215.4494158 | -0.045604765 | 0.2618352 | -0.174174 | 0.861729068 | 0.927119528 | GRINA | protein_coding | 66.68 | -0.001498767 | 0.001498767 | ENSG00000178719 | n.s. |
| ENSG00000131828 | 1566.849683 | 0.046988911 | 0.2766766 | 0.169833 | 0.865141202 | 0.929382012 | PDHA1 | protein_coding | 44.45 | 0.001494517 | 0.001494517 | ENSG00000131828 | n.s. |
| ENSG00000119139 | 377.4626348 | -0.044829795 | 0.2532395 | -0.177025 | 0.85948851 | 0.926427075 | TJP2 | protein_coding | 43.45 | -0.001487845 | 0.001487845 | ENSG00000119139 | n.s. |
| ENSG00000213888 | 166.1093867 | 0.04112948 | 0.2153733 | 0.190968 | 0.848550401 | 0.920122059 | LINC01521 | lincRNA | 59.44 | 0.001487018 | 0.001487018 | ENSG00000213888 | n.s. |
| ENSG00000180667 | 1672.543811 | -0.043271821 | 0.2362364 | -0.183172 | 0.854663311 | 0.923924697 | YOD1 | protein_coding | 39.45 | -0.001486968 | 0.001486968 | ENSG00000180667 | n.s. |
| ENSG00000170852 | 1391.806491 | -0.044764495 | 0.2539379 | -0.176281 | 0.860072959 | 0.926439893 | KBTBD2 | protein_coding | 41.11 | -0.001485409 | 0.001485409 | ENSG00000170852 | n.s. |
| ENSG00000079805 | 691.6813162 | -0.044983682 | 0.257137 | -0.174941 | 0.861126361 | 0.926790009 | DNM2 | protein_coding | 53.07 | -0.001485301 | 0.001485301 | ENSG00000079805 | n.s. |
| ENSG00000085721 | 2382.201572 | -0.047006656 | 0.2786702 | -0.168682 | 0.866046739 | 0.929968283 | RRN3 | protein_coding | 42.09 | -0.001482207 | 0.001482207 | ENSG00000085721 | n.s. |
| ENSG00000277161 | 655.3768848 | -0.041950325 | 0.2238669 | -0.18739 | 0.851355164 | 0.921942751 | PIGW | protein_coding | 44.58 | -0.001480681 | 0.001480681 | ENSG00000277161 | n.s. |
| ENSG00000239467 | 66.58545465 | 0.049129881 | 0.3030439 | 0.162121 | 0.871210302 | 0.933016882 | | lincRNA | 40.11 | 0.001479325 | 0.001479325 | ENSG00000239467 | n.s. |
| ENSG00000112697 | 2486.241778 | 0.045927467 | 0.2672081 | 0.171879 | 0.863532663 | 0.928629433 | TMEM30A | protein_coding | 37.37 | 0.001476915 | 0.001476915 | ENSG00000112697 | n.s. |
| ENSG00000066651 | 1803.828255 | -0.04470887 | 0.2557597 | -0.174808 | 0.861230433 | 0.926797906 | TRMT11 | protein_coding | 36.02 | -0.001476061 | 0.001476061 | ENSG00000066651 | n.s. |
| ENSG00000163754 | 243.6144006 | -0.047023368 | 0.279961 | -0.167964 | 0.866611604 | 0.930431321 | GYG1 | protein_coding | 42.09 | -0.001472569 | 0.001472569 | ENSG00000163754 | n.s. |
| ENSG00000137168 | 1084.705899 | -0.05080894 | 0.3230103 | -0.157298 | 0.875009824 | 0.935480441 | PPIL1 | protein_coding | 44.26 | -0.001471696 | 0.001471696 | ENSG00000137168 | n.s. |
| ENSG00000168393 | 1293.000383 | 0.048140018 | 0.2918225 | 0.164963 | 0.868972808 | 0.932070341 | DTYMK | protein_coding | 51.17 | 0.001470741 | 0.001470741 | ENSG00000168393 | n.s. |
| ENSG00000166197 | 5477.114523 | 0.044300896 | 0.2513557 | 0.176248 | 0.860099264 | 0.926439893 | NOLC1 | protein_coding | 47.22 | 0.001470026 | 0.001470026 | ENSG00000166197 | n.s. |
| ENSG00000163156 | 484.3239838 | -0.04565932 | 0.2665723 | -0.171283 | 0.864001213 | 0.928774621 | SCNM1 | protein_coding | 48.32 | -0.001465192 | 0.001465192 | ENSG00000163156 | n.s. |
| ENSG00000061936 | 2688.131583 | 0.040849434 | 0.2159817 | 0.189134 | 0.849987967 | 0.921035292 | SFSWAP | protein_coding | 46.45 | 0.001459294 | 0.001459294 | ENSG00000061936 | n.s. |
| ENSG00000178252 | 2387.289497 | -0.0457276 | 0.2682448 | -0.17047 | 0.864640793 | 0.92917891 | WDR6 | protein_coding | 56.28 | -0.00145874 | 0.00145874 | ENSG00000178252 | n.s. |
| ENSG00000255409 | 45.58217937 | -0.047567778 | 0.2879828 | -0.165176 | 0.868805661 | 0.931996396 | RSF1-IT1 | sense_intronic | 43.19 | -0.001454897 | 0.001454897 | ENSG00000255409 | n.s. |

| | | | | | | | | | | | | | |
|---|---|---|---|---|---|---|---|---|---|---|---|---|---|
| ENSG00000256248 | 14.13849777 | -0.055304658 | 0.3878332 | -0.142599 | 0.886606816 | 0.941229638 | | lincRNA | 38.25 | -0.001454756 | 0.001454756 | ENSG00000256248 | n.s. |
| ENSG00000279591 | 41.67075467 | -0.050617299 | 0.3250023 | -0.155744 | 0.876234481 | 0.935999131 | | TEC | 51.96 | -0.001453959 | 0.001453959 | ENSG00000279591 | n.s. |
| ENSG00000187446 | 371.5063717 | -0.043681776 | 0.2460694 | -0.177518 | 0.859101465 | 0.926238894 | CHP1 | protein_coding | 44.02 | -0.001453598 | 0.001453598 | ENSG00000187446 | n.s. |
| ENSG00000176105 | 1596.099025 | -0.045491101 | 0.2669779 | -0.170393 | 0.864701251 | 0.92917891 | YES1 | protein_coding | 40.01 | -0.001451196 | 0.001451196 | ENSG00000176105 | n.s. |
| ENSG00000253910 | 88.47907092 | -0.043488093 | 0.2483933 | -0.175078 | 0.861018671 | 0.926784921 | PCDHGB2 | protein_coding | 45.22 | -0.001436022 | 0.001436022 | ENSG00000253910 | n.s. |
| ENSG00000183775 | 1134.22857 | -0.043423802 | 0.2477426 | -0.175278 | 0.860861296 | 0.926758821 | KCTD16 | protein_coding | 37.85 | -0.00143443 | 0.00143443 | ENSG00000183775 | n.s. |
| ENSG00000106459 | 249.4350779 | -0.042849111 | 0.2402312 | -0.178366 | 0.858435447 | 0.925944716 | NRF1 | protein_coding | 41.15 | -0.001431801 | 0.001431801 | ENSG00000106459 | n.s. |
| ENSG00000104825 | 197.7133568 | -0.048629376 | 0.3045097 | -0.159697 | 0.873119548 | 0.934525601 | NFKBIB | protein_coding | 50.76 | -0.001430131 | 0.001430131 | ENSG00000104825 | n.s. |
| ENSG00000204822 | 433.359294 | 0.049227015 | 0.3119356 | 0.157811 | 0.874605371 | 0.935322942 | MRPL53 | protein_coding | 53.92 | 0.001429474 | 0.001429474 | ENSG00000204822 | n.s. |
| ENSG00000132485 | 5433.574953 | -0.04571236 | 0.2726632 | -0.167651 | 0.866857573 | 0.93062364 | ZRANB2 | protein_coding | 34.21 | -0.00142741 | 0.00142741 | ENSG00000132485 | n.s. |
| ENSG00000101166 | 1812.68563 | 0.04710645 | 0.2892842 | 0.162838 | 0.870646036 | 0.932820682 | PRELID3B | protein_coding | 40.89 | 0.001422701 | 0.001422701 | ENSG00000101166 | n.s. |
| ENSG00000141522 | 1621.14503 | -0.042412186 | 0.2371472 | -0.178843 | 0.858060766 | 0.925819904 | ARHGDIA | protein_coding | 68.12 | -0.001419684 | 0.001419684 | ENSG00000141522 | n.s. |
| ENSG00000197037 | 259.0717626 | -0.041275722 | 0.2254519 | -0.18308 | 0.854735306 | 0.923924697 | ZSCAN25 | protein_coding | 48.45 | -0.001418375 | 0.001418375 | ENSG00000197037 | n.s. |
| ENSG00000238723 | 36.9409389 | 0.049170744 | 0.3145948 | 0.156299 | 0.875797597 | 0.935799494 | | misc_RNA | 40.2 | 0.001416963 | 0.001416963 | ENSG00000238723 | n.s. |
| ENSG00000147400 | 317.9245562 | 0.050865671 | 0.3363334 | 0.151236 | 0.879789647 | 0.937925824 | CETN2 | protein_coding | 43.47 | 0.001415668 | 0.001415668 | ENSG00000147400 | n.s. |
| ENSG00000101843 | 408.3648929 | 0.049241073 | 0.3177972 | 0.154945 | 0.876864715 | 0.936169575 | PSMD10 | protein_coding | 37.94 | 0.001410534 | 0.001410534 | ENSG00000101843 | n.s. |
| ENSG00000088882 | 1276.843783 | 0.043744196 | 0.2541171 | 0.172142 | 0.863325992 | 0.928478896 | CPXM1 | protein_coding | 59.72 | 0.001409786 | 0.001409786 | ENSG00000088882 | n.s. |
| ENSG00000225973 | 141.2490821 | -0.046383888 | 0.2840008 | -0.163323 | 0.870264036 | 0.932698497 | PIGBOS1 | protein_coding | 47.12 | -0.001403517 | 0.001403517 | ENSG00000225973 | n.s. |
| ENSG00000097021 | 262.911797 | -0.047811319 | 0.300519 | -0.159096 | 0.873593391 | 0.934683838 | ACOT7 | protein_coding | 54 | -0.001402558 | 0.001402558 | ENSG00000097021 | n.s. |
| ENSG00000099326 | 300.34565 | 0.043509379 | 0.2554958 | 0.170294 | 0.864779001 | 0.92917891 | MZF1 | protein_coding | 55.33 | 0.001387978 | 0.001387978 | ENSG00000099326 | n.s. |
| ENSG00000269834 | 510.1669511 | 0.040021007 | 0.2178266 | 0.183729 | 0.854226251 | 0.923543414 | ZNF528-AS1 | processed_transcript | 46.27 | 0.001382433 | 0.001382433 | ENSG00000269834 | n.s. |
| ENSG00000197858 | 333.162575 | -0.043526256 | 0.2568195 | -0.169482 | 0.865417606 | 0.929507778 | GPAA1 | protein_coding | 64.3 | -0.001381827 | 0.001381827 | ENSG00000197858 | n.s. |
| ENSG00000143106 | 2292.872236 | 0.047546329 | 0.3007124 | 0.158112 | 0.874368292 | 0.935290023 | PSMA5 | protein_coding | 40.74 | 0.001381397 | 0.001381397 | ENSG00000143106 | n.s. |
| ENSG00000265641 | 4.345080367 | 0.064406374 | 0.5423088 | 0.118763 | 0.905462905 | 0.9519635 | MIR3929 | miRNA | 52.73 | 0.001376989 | 0.001376989 | ENSG00000265641 | n.s. |
| ENSG00000137824 | 598.1050051 | -0.042647278 | 0.2486881 | -0.171489 | 0.863839274 | 0.928730982 | RMDN3 | protein_coding | 49.18 | -0.001369407 | 0.001369407 | ENSG00000137824 | n.s. |
| ENSG00000178234 | 794.5196062 | -0.044529443 | 0.2684724 | -0.165862 | 0.868265323 | 0.931657581 | GALNT11 | protein_coding | 41.23 | -0.001368999 | 0.001368999 | ENSG00000178234 | n.s. |
| ENSG00000186591 | 506.8187006 | -0.042875856 | 0.2518988 | -0.170211 | 0.864844465 | 0.92917891 | UBE2H | protein_coding | 41.85 | -0.001367768 | 0.001367768 | ENSG00000186591 | n.s. |
| ENSG00000166454 | 1399.641237 | -0.045114785 | 0.2766469 | -0.163077 | 0.870457712 | 0.93277033 | ATMIN | protein_coding | 44.63 | -0.001363607 | 0.001363607 | ENSG00000166454 | n.s. |
| ENSG00000095209 | 126.9796434 | 0.045244554 | 0.2792408 | 0.162027 | 0.871284568 | 0.933016882 | TMEM38B | protein_coding | 37.52 | 0.001362336 | 0.001362336 | ENSG00000095209 | n.s. |
| ENSG00000159055 | 527.4838063 | 0.044826395 | 0.2731917 | 0.164084 | 0.869664995 | 0.93241535 | MIS18A | protein_coding | 40.72 | 0.0013623 | 0.0013623 | ENSG00000159055 | n.s. |
| ENSG00000155256 | 421.6652148 | -0.044793434 | 0.2728451 | -0.164172 | 0.869596013 | 0.932413191 | ZFYVE27 | protein_coding | 48.53 | -0.001361344 | 0.001361344 | ENSG00000155256 | n.s. |
| ENSG00000135974 | 926.7158907 | -0.042214602 | 0.2458204 | -0.171729 | 0.863650207 | 0.928684108 | C2orf49 | protein_coding | 38.53 | -0.001356439 | 0.001356439 | ENSG00000135974 | n.s. |

| | | | | | | | | | | | | | | |
|---|---|---|---|---|---|---|---|---|---|---|---|---|---|---|
| ENSG00000114383 | 234.0251396 | -0.046603342 | 0.294955 | -0.158002 | 0.874455581 | 0.935290023 | TUSC2 | | protein_coding | 59.39 | -0.001353999 | 0.001353999 | ENSG00000114383 | n.s. |
| ENSG00000090263 | 591.7910536 | 0.047260077 | 0.3045202 | 0.155195 | 0.876667418 | 0.936169575 | MRPS33 | | protein_coding | 41.83 | 0.001353787 | 0.001353787 | ENSG00000090263 | n.s. |
| ENSG00000239002 | 303.2900671 | -0.042427215 | 0.2488712 | -0.170479 | 0.864633757 | 0.92917891 | SCARNA10 | | snoRNA | 45.45 | -0.001353456 | 0.001353456 | ENSG00000239002 | n.s. |
| ENSG00000138069 | 1674.251296 | 0.043764827 | 0.2633807 | 0.166166 | 0.868026604 | 0.931519532 | RAB1A | | protein_coding | 42.71 | 0.001348308 | 0.001348308 | ENSG00000138069 | n.s. |
| ENSG00000025156 | 2289.213141 | -0.045798201 | 0.2874539 | -0.159324 | 0.873413892 | 0.934563604 | HSF2 | | protein_coding | 35.55 | -0.001346061 | 0.001346061 | ENSG00000025156 | n.s. |
| ENSG00000165487 | 606.4305328 | 0.045687496 | 0.2865247 | 0.159454 | 0.873311254 | 0.934525601 | MICU2 | | protein_coding | 38.3 | 0.001343614 | 0.001343614 | ENSG00000165487 | n.s. |
| ENSG00000174227 | 337.966039 | 0.040145615 | 0.2250855 | 0.178357 | 0.858442462 | 0.925944716 | PIGG | | protein_coding | 49.98 | 0.001341463 | 0.001341463 | ENSG00000174227 | n.s. |
| ENSG00000109606 | 9621.992108 | -0.044344478 | 0.2720357 | -0.16301 | 0.870510719 | 0.93277033 | DHX15 | | protein_coding | 37.26 | -0.001340324 | 0.001340324 | ENSG00000109606 | n.s. |
| ENSG00000239523 | 52.13770643 | 0.043269124 | 0.2592689 | 0.166889 | 0.867457413 | 0.931195802 | MYLK-AS1 | | antisense | 44.17 | 0.001339568 | 0.001339568 | ENSG00000239523 | n.s. |
| ENSG00000273989 | 29.68844849 | -0.048599627 | 0.3259323 | -0.14911 | 0.881467164 | 0.938612008 | | | sense_intronic | 25.75 | -0.001337165 | 0.001337165 | ENSG00000273989 | n.s. |
| ENSG00000163870 | 353.0636077 | -0.043557091 | 0.2641135 | -0.164918 | 0.869008482 | 0.932070341 | TPRA1 | | protein_coding | 55.15 | -0.001330726 | 0.001330726 | ENSG00000163870 | n.s. |
| ENSG00000032389 | 605.4322522 | -0.0463336 | 0.2978057 | -0.155583 | 0.87636149 | 0.936028319 | TSSC1 | | protein_coding | 48.27 | -0.001330285 | 0.001330285 | ENSG00000032389 | n.s. |
| ENSG00000125450 | 2361.049976 | -0.043741234 | 0.2663069 | -0.164251 | 0.869533378 | 0.932413191 | NUP85 | | protein_coding | 48.14 | -0.001329366 | 0.001329366 | ENSG00000125450 | n.s. |
| ENSG00000173559 | 278.9047066 | 0.039208968 | 0.2181437 | 0.179739 | 0.857357347 | 0.925347779 | NABP1 | | protein_coding | 38.76 | 0.001321147 | 0.001321147 | ENSG00000173559 | n.s. |
| ENSG00000106355 | 1710.452303 | -0.043531976 | 0.2661303 | -0.163574 | 0.870066562 | 0.932630437 | LSM5 | | protein_coding | 39.73 | -0.001318602 | 0.001318602 | ENSG00000106355 | n.s. |
| ENSG00000114126 | 1528.981888 | -0.043734065 | 0.2701803 | -0.16187 | 0.871408267 | 0.933016882 | TFDP2 | | protein_coding | 39.89 | -0.001316854 | 0.001316854 | ENSG00000114126 | n.s. |
| ENSG00000117505 | 3110.123106 | 0.042574972 | 0.2555132 | 0.166625 | 0.867664853 | 0.93127488 | DR1 | | protein_coding | 36.47 | 0.001316508 | 0.001316508 | ENSG00000117505 | n.s. |
| ENSG00000270116 | 105.4103449 | 0.042489381 | 0.2563309 | 0.16576 | 0.868345927 | 0.931657581 | | | sense_intronic | 36.04 | 0.00130628 | 0.00130628 | ENSG00000270116 | n.s. |
| ENSG00000189266 | 1397.136217 | 0.039354093 | 0.2222031 | 0.177109 | 0.859423071 | 0.926427075 | PNRC2 | | protein_coding | 43.71 | 0.001306114 | 0.001306114 | ENSG00000189266 | n.s. |
| ENSG00000108578 | 1878.260445 | -0.042573748 | 0.2575042 | -0.165332 | 0.868682499 | 0.931936079 | BLMH | | protein_coding | 40.18 | -0.001303347 | 0.001303347 | ENSG00000108578 | n.s. |
| ENSG00000198783 | 870.8832612 | -0.038618671 | 0.2142444 | -0.180255 | 0.856952198 | 0.925266075 | ZNF830 | | protein_coding | 42.79 | -0.001302738 | 0.001302738 | ENSG00000198783 | n.s. |
| ENSG00000240024 | 197.0378825 | 0.044602725 | 0.2827594 | 0.157741 | 0.874660971 | 0.935322942 | LINC00888 | | transcribed_unprocessed_pseud | 39.85 | 0.001295192 | 0.001295192 | ENSG00000240024 | n.s. |
| ENSG00000274290 | 807.2543032 | -0.037685093 | 0.206116 | -0.182834 | 0.85492795 | 0.924015048 | HIST1H2BE | | protein_coding | 41.16 | -0.001293388 | 0.001293388 | ENSG00000274290 | n.s. |
| ENSG00000184489 | 91.68590072 | 0.044808416 | 0.2874507 | 0.155882 | 0.87612596 | 0.935955017 | PTP4A3 | | protein_coding | 62.4 | 0.001288019 | 0.001288019 | ENSG00000184489 | n.s. |
| ENSG00000087111 | 576.4545815 | 0.044607481 | 0.2846386 | 0.156716 | 0.875468525 | 0.93575528 | PIGS | | protein_coding | 47.27 | 0.001286378 | 0.001286378 | ENSG00000087111 | n.s. |
| ENSG00000145495 | 4239.907439 | -0.042591096 | 0.2623637 | -0.162336 | 0.871041196 | 0.933016882 | | 6-Mar | protein_coding | 39.32 | -0.001282439 | 0.001282439 | ENSG00000145495 | n.s. |
| ENSG00000136518 | 2166.934496 | 0.045440772 | 0.2969456 | 0.153027 | 0.878376771 | 0.937154913 | ACTL6A | | protein_coding | 39.25 | 0.001280912 | 0.001280912 | ENSG00000136518 | n.s. |
| ENSG00000177380 | 337.4496898 | 0.0424985 | 0.2624145 | 0.161952 | 0.871343794 | 0.933016882 | PPFIA3 | | protein_coding | 55.2 | 0.001279651 | 0.001279651 | ENSG00000177380 | n.s. |
| ENSG00000147853 | 1093.623888 | 0.045304626 | 0.2963327 | 0.152884 | 0.878489494 | 0.937154913 | AK3 | | protein_coding | 41.47 | 0.001277075 | 0.001277075 | ENSG00000147853 | n.s. |
| ENSG00000121390 | 2092.736791 | -0.041848655 | 0.2540703 | -0.164713 | 0.869169949 | 0.932171715 | PSPC1 | | protein_coding | 40.29 | -0.001276555 | 0.001276555 | ENSG00000121390 | n.s. |
| ENSG00000144848 | 603.4290421 | -0.042344595 | 0.260613 | -0.162481 | 0.870927293 | 0.933001127 | ATG3 | | protein_coding | 36.32 | -0.001275327 | 0.001275327 | ENSG00000144848 | n.s. |
| ENSG00000121060 | 655.4065902 | -0.042173375 | 0.2587598 | -0.162983 | 0.870532046 | 0.93277033 | TRIM25 | | protein_coding | 49.18 | -0.001274702 | 0.001274702 | ENSG00000121060 | n.s. |

| | | | | | | | | | | | | | |
|---|---|---|---|---|---|---|---|---|---|---|---|---|---|
| ENSG00000198742 | 556.9025324 | 0.041998275 | 0.2570894 | 0.163361 | 0.870234552 | 0.932698497 | SMURF1 | protein_coding | 42.72 | 0.001270814 | 0.001270814 | ENSG00000198742 | n.s. |
| ENSG00000048707 | 67.19012482 | -0.044097009 | 0.2821341 | -0.156298 | 0.875798087 | 0.935799494 | VPS13D | protein_coding | 42.34 | -0.001270752 | 0.001270752 | ENSG00000048707 | n.s. |
| ENSG00000171848 | 3847.124279 | -0.037579774 | 0.2085507 | -0.180195 | 0.856999564 | 0.925266075 | RRM2 | protein_coding | 45.64 | -0.001267692 | 0.001267692 | ENSG00000171848 | n.s. |
| ENSG00000144369 | 1327.511875 | 0.041926201 | 0.2591967 | 0.161754 | 0.871499286 | 0.933016882 | FAM171B | protein_coding | 37.17 | 0.001262419 | 0.001262419 | ENSG00000144369 | n.s. |
| ENSG00000120656 | 145.5679967 | -0.044230428 | 0.2866493 | -0.154302 | 0.877371973 | 0.936423916 | TAF12 | protein_coding | 47.17 | -0.001261784 | 0.001261784 | ENSG00000120656 | n.s. |
| ENSG00000181938 | 1646.304573 | 0.043928971 | 0.2834485 | 0.15498 | 0.876836748 | 0.936169575 | GINS3 | protein_coding | 41.93 | 0.001258366 | 0.001258366 | ENSG00000181938 | n.s. |
| ENSG00000198478 | 531.3225053 | 0.043319889 | 0.2755746 | 0.157198 | 0.875088479 | 0.935492698 | SH3BGRL2 | protein_coding | 38.84 | 0.001254527 | 0.001254527 | ENSG00000198478 | n.s. |
| ENSG00000143319 | 1044.820135 | 0.03783149 | 0.2163009 | 0.174902 | 0.861156529 | 0.926790009 | ISG20L2 | protein_coding | 50.3 | 0.001249145 | 0.001249145 | ENSG00000143319 | n.s. |
| ENSG00000137807 | 2687.527886 | 0.03945381 | 0.2334499 | 0.169003 | 0.865794025 | 0.929768624 | KIF23 | protein_coding | 39.36 | 0.001247731 | 0.001247731 | ENSG00000137807 | n.s. |
| ENSG00000179562 | 646.9036747 | 0.037641561 | 0.2146183 | 0.175388 | 0.860774467 | 0.926737002 | GCC1 | protein_coding | 52.07 | 0.001243809 | 0.001243809 | ENSG00000179562 | n.s. |
| ENSG00000144736 | 687.1917413 | -0.042856261 | 0.2721371 | -0.15748 | 0.874866254 | 0.935398775 | SHQ1 | protein_coding | 39.56 | -0.001242969 | 0.001242969 | ENSG00000144736 | n.s. |
| ENSG00000164754 | 6868.938928 | -0.04208603 | 0.2637384 | -0.159575 | 0.873215945 | 0.934525601 | RAD21 | protein_coding | 36.02 | -0.0012377 | 0.0012377 | ENSG00000164754 | n.s. |
| ENSG00000119421 | 399.8633733 | -0.045976813 | 0.3158079 | -0.145585 | 0.884249214 | 0.939936396 | NDUFA8 | protein_coding | 41.5 | -0.001236847 | 0.001236847 | ENSG00000119421 | n.s. |
| ENSG00000125520 | 99.90752263 | -0.042666557 | 0.2717138 | -0.157028 | 0.875223118 | 0.935564796 | SLC2A4RG | protein_coding | 69.47 | -0.001234178 | 0.001234178 | ENSG00000125520 | n.s. |
| ENSG00000083857 | 1411.771273 | 0.038852888 | 0.2291057 | 0.169585 | 0.865336536 | 0.929492408 | FAT1 | protein_coding | 41.53 | 0.001233741 | 0.001233741 | ENSG00000083857 | n.s. |
| ENSG00000186918 | 577.831906 | -0.04087667 | 0.2516202 | -0.162454 | 0.870948477 | 0.933001127 | ZNF395 | protein_coding | 48.07 | -0.001231117 | 0.001231117 | ENSG00000186918 | n.s. |
| ENSG00000027697 | 607.8978477 | -0.044997869 | 0.3033859 | -0.148319 | 0.882091087 | 0.939013243 | IFNGR1 | protein_coding | 38.81 | -0.001229715 | 0.001229715 | ENSG00000027697 | n.s. |
| ENSG00000118402 | 326.980324 | 0.042853579 | 0.2774853 | 0.154435 | 0.877266377 | 0.936382994 | ELOVL4 | protein_coding | 36.66 | 0.001223319 | 0.001223319 | ENSG00000118402 | n.s. |
| ENSG00000234604 | 34.52353234 | -0.044222914 | 0.2946064 | -0.150108 | 0.880679057 | 0.938390482 | | processed_pseudogene | 35 | -0.001221278 | 0.001221278 | ENSG00000234604 | n.s. |
| ENSG00000105676 | 678.4021761 | 0.045326544 | 0.3113526 | 0.145579 | 0.884253395 | 0.939936396 | ARMC6 | protein_coding | 51.45 | 0.001219354 | 0.001219354 | ENSG00000105676 | n.s. |
| ENSG00000126067 | 763.837453 | -0.042239797 | 0.2698851 | -0.15651 | 0.87563082 | 0.935785071 | PSMB2 | protein_coding | 41.1 | -0.001217515 | 0.001217515 | ENSG00000126067 | n.s. |
| ENSG00000248773 | 114.3449439 | -0.043160632 | 0.283624 | -0.152176 | 0.879048516 | 0.937494637 | | sense_intronic | 33.4 | -0.001209845 | 0.001209845 | ENSG00000248773 | n.s. |
| ENSG00000106772 | 70.81704723 | 0.041634932 | 0.2642397 | 0.157565 | 0.874799583 | 0.935398775 | PRUNE2 | protein_coding | 41.04 | 0.001207546 | 0.001207546 | ENSG00000106772 | n.s. |
| ENSG00000231259 | 87.41887589 | -0.04185077 | 0.2678666 | -0.156237 | 0.875845953 | 0.935799494 | | unprocessed_pseudogene | 38.72 | -0.001206022 | 0.001206022 | ENSG00000231259 | n.s. |
| ENSG00000141570 | 657.0368812 | 0.039814254 | 0.2432938 | 0.163647 | 0.870009171 | 0.932630437 | CBX8 | protein_coding | 60.55 | 0.00120599 | 0.00120599 | ENSG00000141570 | n.s. |
| ENSG00000198722 | 980.6869219 | 0.041624668 | 0.2676153 | 0.155539 | 0.876396263 | 0.936028319 | UNC13B | protein_coding | 41.63 | 0.001195086 | 0.001195086 | ENSG00000198722 | n.s. |
| ENSG00000163864 | 38.72315524 | 0.046061546 | 0.329307 | 0.139874 | 0.888759407 | 0.942149933 | NMNAT3 | protein_coding | 41.66 | 0.001192072 | 0.001192072 | ENSG00000163864 | n.s. |
| ENSG00000102317 | 4211.583007 | 0.040081444 | 0.2496445 | 0.160554 | 0.87244463 | 0.933885338 | RBM3 | protein_coding | 48.05 | 0.001190677 | 0.001190677 | ENSG00000102317 | n.s. |
| ENSG00000111276 | 3768.630159 | 0.039225735 | 0.2393743 | 0.163868 | 0.869835221 | 0.932526047 | CDKN1B | protein_coding | 45.43 | 0.001190071 | 0.001190071 | ENSG00000111276 | n.s. |
| ENSG00000078369 | 1984.02542 | -0.041217127 | 0.2631541 | -0.156627 | 0.87553856 | 0.9357583 | GNB1 | protein_coding | 46.7 | -0.00118855 | 0.00118855 | ENSG00000078369 | n.s. |
| ENSG00000138801 | 1436.729164 | 0.042929939 | 0.2860369 | 0.150085 | 0.880697295 | 0.938390482 | PAPSS1 | protein_coding | 38.61 | 0.001185571 | 0.001185571 | ENSG00000138801 | n.s. |
| ENSG00000107362 | 777.227068 | 0.042542572 | 0.2814986 | 0.151129 | 0.879874036 | 0.937943971 | ABHD17B | protein_coding | 37.98 | 0.001183666 | 0.001183666 | ENSG00000107362 | n.s. |

| | | | | | | | | | | | | | |
|---|---|---|---|---|---|---|---|---|---|---|---|---|---|
| ENSG00000266472 | 2189.140244 | -0.043132968 | 0.2923927 | -0.147517 | 0.882723746 | 0.939212574 | MRPS21 | protein_coding | 46.58 | -0.001174774 | 0.001174774 | ENSG00000266472 | n.s. |
| ENSG00000277450 | 42.88512508 | 0.042495709 | 0.2834498 | 0.149923 | 0.880825201 | 0.938435851 | | sense_intronic | 45.6 | 0.001172686 | 0.001172686 | ENSG00000277450 | n.s. |
| ENSG00000058799 | 392.4043452 | -0.042872447 | 0.2894393 | -0.148122 | 0.882246162 | 0.939016105 | YIPF1 | protein_coding | 43.46 | -0.001171574 | 0.001171574 | ENSG00000058799 | n.s. |
| ENSG00000197444 | 754.5495317 | -0.041851959 | 0.2757879 | -0.151754 | 0.879380841 | 0.937633592 | OGDHL | protein_coding | 55.46 | -0.001170467 | 0.001170467 | ENSG00000197444 | n.s. |
| ENSG00000105865 | 110.036597 | 0.040844887 | 0.2633869 | 0.155076 | 0.876761681 | 0.936169575 | DUS4L | protein_coding | 37.93 | 0.001170021 | 0.001170021 | ENSG00000105865 | n.s. |
| ENSG00000214114 | 205.5246127 | -0.045427136 | 0.3265668 | -0.139105 | 0.889367024 | 0.942578753 | MYCBP | protein_coding | 44.31 | -0.001166676 | 0.001166676 | ENSG00000214114 | n.s. |
| ENSG00000200792 | 22.05108416 | -0.045187685 | 0.3236516 | -0.139618 | 0.8889616 | 0.942292528 | SNORA80A | snoRNA | 55.88 | -0.001166486 | 0.001166486 | ENSG00000200792 | n.s. |
| ENSG00000140740 | 2483.024166 | 0.042752623 | 0.2895367 | 0.147659 | 0.882612081 | 0.939208805 | UQCRC2 | protein_coding | 40.33 | 0.001164489 | 0.001164489 | ENSG00000140740 | n.s. |
| ENSG00000213672 | 262.6061353 | -0.040503604 | 0.2613856 | -0.154957 | 0.87685497 | 0.936169575 | NCKIPSD | protein_coding | 56.62 | -0.001160245 | 0.001160245 | ENSG00000213672 | n.s. |
| ENSG00000210100 | 53.03901604 | 0.042959642 | 0.2938402 | 0.146201 | 0.883762963 | 0.939790948 | MT-TI | Mt_tRNA | 28.99 | 0.001158568 | 0.001158568 | ENSG00000210100 | n.s. |
| ENSG00000134330 | 740.6598285 | -0.043113477 | 0.2973449 | -0.144995 | 0.884714939 | 0.940009623 | IAH1 | protein_coding | 45.86 | -0.00115836 | 0.00115836 | ENSG00000134330 | n.s. |
| ENSG00000185630 | 150.7089524 | -0.038038537 | 0.2313614 | -0.164412 | 0.869407 | 0.932354129 | PBX1 | protein_coding | 41.67 | -0.001157098 | 0.001157098 | ENSG00000185630 | n.s. |
| ENSG00000204954 | 289.6940586 | 0.036776702 | 0.2184383 | 0.168362 | 0.866298495 | 0.930166881 | C12orf73 | protein_coding | 41.85 | 0.001156227 | 0.001156227 | ENSG00000204954 | n.s. |
| ENSG00000179348 | 2091.297525 | -0.036326001 | 0.2139334 | -0.169801 | 0.865167008 | 0.929382012 | GATA2 | protein_coding | 60.45 | -0.001155375 | 0.001155375 | ENSG00000179348 | n.s. |
| ENSG00000163689 | 38.5380601 | 0.042562865 | 0.2898695 | 0.146835 | 0.883262583 | 0.939613764 | C3orf67 | protein_coding | 38.65 | 0.001151352 | 0.001151352 | ENSG00000163689 | n.s. |
| ENSG00000105968 | 2702.069449 | -0.040765784 | 0.2667299 | -0.152835 | 0.878528041 | 0.937154913 | H2AFV | protein_coding | 41.41 | -0.001149131 | 0.001149131 | ENSG00000105968 | n.s. |
| ENSG00000172057 | 311.2135779 | 0.041882969 | 0.2826491 | 0.14818 | 0.882200654 | 0.939016105 | ORMDL3 | protein_coding | 57.09 | 0.001144534 | 0.001144534 | ENSG00000172057 | n.s. |
| ENSG00000032219 | 133.3725143 | 0.037922448 | 0.2344177 | 0.161773 | 0.871484628 | 0.933016882 | ARID4A | protein_coding | 37.48 | 0.001141864 | 0.001141864 | ENSG00000032219 | n.s. |
| ENSG00000126254 | 437.906419 | -0.04174288 | 0.2820967 | -0.147974 | 0.88236355 | 0.939016105 | RBM42 | protein_coding | 53.79 | -0.001140706 | 0.001140706 | ENSG00000126254 | n.s. |
| ENSG00000169957 | 703.2652862 | -0.040074373 | 0.26036 | -0.153919 | 0.877673528 | 0.936673964 | ZNF768 | protein_coding | 64.52 | -0.001138575 | 0.001138575 | ENSG00000169957 | n.s. |
| ENSG00000266111 | 38.83094965 | 0.042097963 | 0.2894839 | 0.145424 | 0.884375982 | 0.939936396 | | lincRNA | 44.26 | 0.0011325 | 0.0011325 | ENSG00000266111 | n.s. |
| ENSG00000213341 | 1372.703442 | -0.040180857 | 0.2632179 | -0.152652 | 0.878672364 | 0.937237058 | CHUK | protein_coding | 38.39 | -0.001131113 | 0.001131113 | ENSG00000213341 | n.s. |
| ENSG00000155100 | 943.6380595 | -0.041858029 | 0.286482 | -0.146111 | 0.883834136 | 0.939790948 | OTUD6B | protein_coding | 34.94 | -0.001128858 | 0.001128858 | ENSG00000155100 | n.s. |
| ENSG00000100360 | 208.4661887 | 0.043381377 | 0.3100369 | 0.139923 | 0.888720604 | 0.942149933 | IFT27 | protein_coding | 51.18 | 0.001122709 | 0.001122709 | ENSG00000100360 | n.s. |
| ENSG00000259429 | 277.6223079 | 0.040618306 | 0.2713251 | 0.149703 | 0.880998598 | 0.938440302 | UBE2Q2P2 | transcribed_unprocessed_pseud | 39.44 | 0.001120795 | 0.001120795 | ENSG00000259429 | n.s. |
| ENSG00000214706 | 617.2690079 | 0.039954831 | 0.2635879 | 0.151581 | 0.87951766 | 0.937707664 | IFRD2 | protein_coding | 59.19 | 0.00111604 | 0.00111604 | ENSG00000214706 | n.s. |
| ENSG00000136522 | 1157.373408 | 0.041657469 | 0.2890279 | 0.14413 | 0.885398143 | 0.940304984 | MRPL47 | protein_coding | 38.56 | 0.001113557 | 0.001113557 | ENSG00000136522 | n.s. |
| ENSG00000139168 | 1199.951738 | -0.043375991 | 0.3140927 | -0.138099 | 0.890161913 | 0.942914513 | ZCRB1 | protein_coding | 37.28 | -0.001107288 | 0.001107288 | ENSG00000139168 | n.s. |
| ENSG00000162909 | 1402.20113 | 0.04085824 | 0.2817479 | 0.145017 | 0.884697454 | 0.940009623 | CAPN2 | protein_coding | 48.73 | 0.001097767 | 0.001097767 | ENSG00000162909 | n.s. |
| ENSG00000131653 | 771.80565 | 0.038156366 | 0.2466853 | 0.154676 | 0.877076519 | 0.936323906 | TRAF7 | protein_coding | 63.76 | 0.001090276 | 0.001090276 | ENSG00000131653 | n.s. |
| ENSG00000104549 | 3782.848016 | 0.039481135 | 0.2637626 | 0.149684 | 0.88101366 | 0.938440302 | SQLE | protein_coding | 37.59 | 0.001089417 | 0.001089417 | ENSG00000104549 | n.s. |
| ENSG00000124074 | 158.2698767 | 0.042569585 | 0.3087428 | 0.13788 | 0.890334933 | 0.942954254 | ENKD1 | protein_coding | 60.38 | 0.001085924 | 0.001085924 | ENSG00000124074 | n.s. |

| | | | | | | | | | | | | | |
|---|---|---|---|---|---|---|---|---|---|---|---|---|---|
| ENSG00000070010 | 849.3113283 | -0.040244104 | 0.2749476 | -0.14637 | 0.883629229 | 0.939788386 | UFD1 | protein_coding | 46.94 | -0.001085381 | 0.001085381 | ENSG00000070010 | n.s. |
| ENSG00000160813 | 77.44221143 | 0.041348312 | 0.290511 | 0.14233 | 0.886819677 | 0.941383834 | PPP1R35 | protein_coding | 67.13 | 0.001084701 | 0.001084701 | ENSG00000160813 | n.s. |
| ENSG00000124194 | 766.5633625 | -0.041531263 | 0.293208 | -0.141644 | 0.887360922 | 0.941660457 | GDAP1L1 | protein_coding | 50.31 | -0.001084201 | 0.001084201 | ENSG00000124194 | n.s. |
| ENSG00000162066 | 148.9406586 | -0.039273345 | 0.262066 | -0.14986 | 0.880874678 | 0.938435851 | AMDHD2 | protein_coding | 59.17 | -0.001083764 | 0.001083764 | ENSG00000162066 | n.s. |
| ENSG00000001497 | 1087.978278 | 0.039304831 | 0.2642442 | 0.148744 | 0.881755378 | 0.938799428 | LAS1L | protein_coding | 50.45 | 0.001078021 | 0.001078021 | ENSG00000001497 | n.s. |
| ENSG00000166226 | 11211.82135 | -0.039148075 | 0.2623818 | -0.149203 | 0.881393662 | 0.938612008 | CCT2 | protein_coding | 39.53 | -0.001077116 | 0.001077116 | ENSG00000166226 | n.s. |
| ENSG00000182054 | 2688.723864 | -0.041343082 | 0.2927753 | -0.141211 | 0.887703276 | 0.941890924 | IDH2 | protein_coding | 52.54 | -0.001074895 | 0.001074895 | ENSG00000182054 | n.s. |
| ENSG00000181061 | 1050.559749 | 0.043928293 | 0.328346 | 0.133787 | 0.893571319 | 0.9452272 | HIGD1A | protein_coding | 42.66 | 0.001074653 | 0.001074653 | ENSG00000181061 | n.s. |
| ENSG00000270133 | 62.47302753 | 0.038815048 | 0.2580647 | 0.150408 | 0.880442565 | 0.938334494 | | sense_intronic | 36.21 | 0.001072938 | 0.001072938 | ENSG00000270133 | n.s. |
| ENSG00000092036 | 292.1574207 | 0.039908969 | 0.275177 | 0.14503 | 0.884687024 | 0.940009623 | HAUS4 | protein_coding | 45.59 | 0.001072262 | 0.001072262 | ENSG00000092036 | n.s. |
| ENSG00000252391 | 21.27950184 | -0.043984815 | 0.3318846 | -0.13253 | 0.894564754 | 0.945711851 | RNU6-638P | snRNA | 36.54 | -0.001066243 | 0.001066243 | ENSG00000252391 | n.s. |
| ENSG00000011485 | 1169.907809 | 0.039696718 | 0.2744776 | 0.144626 | 0.885005818 | 0.940114196 | PPP5C | protein_coding | 50.75 | 0.001064642 | 0.001064642 | ENSG00000011485 | n.s. |
| ENSG00000240137 | 35.05720715 | -0.055118425 | 0.5121932 | -0.107613 | 0.914303024 | 0.956627415 | ERICH6-AS1 | antisense | 42.97 | -0.001061425 | 0.001061425 | ENSG00000240137 | n.s. |
| ENSG00000165462 | 2438.95386 | -0.037122973 | 0.240223 | -0.154535 | 0.87718753 | 0.936370619 | PHOX2A | protein_coding | 61.28 | -0.001059943 | 0.001059943 | ENSG00000165462 | n.s. |
| ENSG00000239335 | 32.08946867 | -0.041478806 | 0.2996373 | -0.13843 | 0.889900564 | 0.942914513 | LLPH-AS1 | antisense | 44.42 | -0.001058858 | 0.001058858 | ENSG00000239335 | n.s. |
| ENSG00000112081 | 14183.72065 | 0.036142892 | 0.227466 | 0.158894 | 0.873752696 | 0.934782449 | SRSF3 | protein_coding | 40.22 | 0.001058605 | 0.001058605 | ENSG00000112081 | n.s. |
| ENSG00000187325 | 650.4532232 | -0.039460491 | 0.2734298 | -0.144317 | 0.885250393 | 0.94021979 | TAF9B | protein_coding | 38.54 | -0.001056381 | 0.001056381 | ENSG00000187325 | n.s. |
| ENSG00000104064 | 2332.336233 | -0.034975406 | 0.2157621 | -0.162102 | 0.871225727 | 0.933016882 | GABPB1 | protein_coding | 39.38 | -0.001053127 | 0.001053127 | ENSG00000104064 | n.s. |
| ENSG00000168701 | 477.4682325 | -0.040169223 | 0.2827199 | -0.142081 | 0.887015747 | 0.941485657 | TMEM208 | protein_coding | 55.38 | -0.001051883 | 0.001051883 | ENSG00000168701 | n.s. |
| ENSG00000073578 | 1211.440536 | 0.037230206 | 0.2435251 | 0.15288 | 0.878492644 | 0.937154913 | SDHA | protein_coding | 46.07 | 0.001049468 | 0.001049468 | ENSG00000073578 | n.s. |
| ENSG00000146731 | 8410.065279 | -0.038885363 | 0.2664623 | -0.145932 | 0.88397512 | 0.93986908 | CCT6A | protein_coding | 41.33 | -0.001047285 | 0.001047285 | ENSG00000146731 | n.s. |
| ENSG00000132522 | 392.8500592 | 0.037215506 | 0.2441406 | 0.152435 | 0.878844086 | 0.937348415 | GPS2 | protein_coding | 56.28 | 0.001045717 | 0.001045717 | ENSG00000132522 | n.s. |
| ENSG00000105698 | 385.4365335 | 0.038727822 | 0.2649575 | 0.146166 | 0.883790215 | 0.939790948 | USF2 | protein_coding | 57.36 | 0.001044441 | 0.001044441 | ENSG00000105698 | n.s. |
| ENSG00000136770 | 257.7922152 | 0.038240774 | 0.2592888 | 0.147483 | 0.882750538 | 0.939212574 | DNAJC1 | protein_coding | 37.73 | 0.00104153 | 0.00104153 | ENSG00000136770 | n.s. |
| ENSG00000158163 | 125.3200595 | 0.037759424 | 0.2529928 | 0.149251 | 0.881355573 | 0.938612008 | DZIP1L | protein_coding | 45.71 | 0.001038909 | 0.001038909 | ENSG00000158163 | n.s. |
| ENSG00000141127 | 1132.38249 | -0.040569428 | 0.2935783 | -0.138189 | 0.890090669 | 0.942914513 | PRPSAP2 | protein_coding | 44.41 | -0.001035643 | 0.001035643 | ENSG00000141127 | n.s. |
| ENSG00000241785 | 30.63574743 | -0.040224622 | 0.2919061 | -0.1378 | 0.89039859 | 0.942954254 | RN7SL390P | misc_RNA | 50.7 | -0.001026105 | 0.001026105 | ENSG00000241785 | n.s. |
| ENSG00000187147 | 563.19094 | -0.038021061 | 0.2629148 | -0.144614 | 0.885015928 | 0.940114196 | RNF220 | protein_coding | 47.21 | -0.001019702 | 0.001019702 | ENSG00000187147 | n.s. |
| ENSG00000053371 | 209.4635251 | 0.04177105 | 0.3130318 | 0.13344 | 0.89384518 | 0.945344709 | AKR7A2 | protein_coding | 52.87 | 0.001019623 | 0.001019623 | ENSG00000053371 | n.s. |
| ENSG00000107262 | 1882.241284 | 0.036399863 | 0.2397937 | 0.151797 | 0.879347369 | 0.937633592 | BAG1 | protein_coding | 46.54 | 0.001017989 | 0.001017989 | ENSG00000107262 | n.s. |
| ENSG00000037474 | 2256.421798 | 0.036824905 | 0.2453076 | 0.150117 | 0.880672118 | 0.938390482 | NSUN2 | protein_coding | 43.29 | 0.001016972 | 0.001016972 | ENSG00000037474 | n.s. |
| ENSG00000057757 | 407.098385 | -0.037143476 | 0.2509626 | -0.148004 | 0.882339571 | 0.939016105 | PITHD1 | protein_coding | 44.59 | -0.001015018 | 0.001015018 | ENSG00000057757 | n.s. |

| | | | | | | | | | | | | | |
|---|---|---|---|---|---|---|---|---|---|---|---|---|---|
| ENSG00000278535 | 179.9851568 | -0.037100092 | 0.2500418 | -0.148376 | 0.882046402 | 0.939013243 | DHRS11 | protein_coding | 54.49 | -0.001013882 | 0.001013882 | ENSG00000278535 | n.s. |
| ENSG00000250317 | 218.0263332 | -0.043727311 | 0.3424942 | -0.127673 | 0.898407636 | 0.948117437 | SMIM20 | protein_coding | 45.19 | -0.001011757 | 0.001011757 | ENSG00000250317 | n.s. |
| ENSG00000223572 | 259.6040145 | -0.03641416 | 0.2419762 | -0.150487 | 0.880380777 | 0.938334494 | CKMT1A | protein_coding | 50.24 | -0.001006572 | 0.001006572 | ENSG00000223572 | n.s. |
| ENSG00000171311 | 585.5020274 | -0.037190418 | 0.2540556 | -0.146387 | 0.883615967 | 0.939788386 | EXOSC1 | protein_coding | 46.3 | -0.001003023 | 0.001003023 | ENSG00000171311 | n.s. |
| ENSG00000113575 | 4360.106906 | -0.038280137 | 0.2695579 | -0.142011 | 0.887071445 | 0.941485657 | PPP2CA | protein_coding | 40.38 | -0.001002415 | 0.001002415 | ENSG00000113575 | n.s. |
| ENSG00000100029 | 1823.449057 | 0.038224235 | 0.2699994 | 0.141572 | 0.887418434 | 0.941660457 | PES1 | protein_coding | 50.48 | 0.000997869 | 0.000997869 | ENSG00000100029 | n.s. |
| ENSG00000210112 | 10.26410737 | 0.046562233 | 0.3910799 | 0.119061 | 0.905227296 | 0.951852552 | MT-TM | Mt_tRNA | 45.59 | 0.000997844 | 0.000997844 | ENSG00000210112 | n.s. |
| ENSG00000140320 | 572.3607415 | 0.03537869 | 0.2308035 | 0.153285 | 0.878173608 | 0.937135828 | BAHD1 | protein_coding | 53.77 | 0.000997589 | 0.000997589 | ENSG00000140320 | n.s. |
| ENSG00000251703 | 44.10961869 | 0.042792703 | 0.335849 | 0.127416 | 0.898610766 | 0.948117437 | RNU6-998P | snRNA | 38.61 | 0.000990132 | 0.000990132 | ENSG00000251703 | n.s. |
| ENSG00000101224 | 573.8524418 | -0.037134107 | 0.2585127 | -0.143645 | 0.885780661 | 0.940567735 | CDC25B | protein_coding | 53.03 | -0.000988136 | 0.000988136 | ENSG00000101224 | n.s. |
| ENSG00000159692 | 1659.300771 | -0.036649073 | 0.2522108 | -0.145311 | 0.884465108 | 0.93995937 | CTBP1 | protein_coding | 60.99 | -0.000985527 | 0.000985527 | ENSG00000159692 | n.s. |
| ENSG00000143799 | 8160.763876 | -0.037498721 | 0.2641661 | -0.141951 | 0.887118461 | 0.941485657 | PARP1 | protein_coding | 46.8 | -0.000981952 | 0.000981952 | ENSG00000143799 | n.s. |
| ENSG00000141295 | 378.2584485 | -0.038344449 | 0.2764079 | -0.138724 | 0.889668152 | 0.942805167 | SCRN2 | protein_coding | 59.28 | -0.000980776 | 0.000980776 | ENSG00000141295 | n.s. |
| ENSG00000198840 | 1751.665848 | 0.035057066 | 0.2308289 | 0.151875 | 0.879285762 | 0.937633592 | MT-ND3 | protein_coding | 40.17 | 0.000980435 | 0.000980435 | ENSG00000198840 | n.s. |
| ENSG00000119326 | 1221.189237 | 0.039223857 | 0.2885583 | 0.13593 | 0.89187627 | 0.944160075 | CTNNAL1 | protein_coding | 39.11 | 0.000978807 | 0.000978807 | ENSG00000119326 | n.s. |
| ENSG00000055044 | 7272.386632 | -0.03627613 | 0.2493633 | -0.145475 | 0.884335838 | 0.939936396 | NOP58 | protein_coding | 41.15 | -0.000975884 | 0.000975884 | ENSG00000055044 | n.s. |
| ENSG00000100908 | 161.5737555 | 0.037642873 | 0.2677045 | 0.140614 | 0.888175279 | 0.942104556 | EMC9 | protein_coding | 53.54 | 0.000974984 | 0.000974984 | ENSG00000100908 | n.s. |
| ENSG00000255624 | 75.58638416 | 0.037567489 | 0.268506 | 0.139913 | 0.888728713 | 0.942149933 | | transcribed_processed_pseudog | 37.42 | 0.000972246 | 0.000972246 | ENSG00000255624 | n.s. |
| ENSG00000118816 | 2033.688016 | -0.03603481 | 0.2496349 | -0.14435 | 0.885224065 | 0.94021979 | CCNI | protein_coding | 39.09 | -0.000964674 | 0.000964674 | ENSG00000118816 | n.s. |
| ENSG00000160325 | 187.2020906 | 0.036119777 | 0.2509314 | 0.143943 | 0.885545622 | 0.940389879 | CACFD1 | protein_coding | 58.04 | 0.000964111 | 0.000964111 | ENSG00000160325 | n.s. |
| ENSG00000139613 | 1329.030895 | 0.036179515 | 0.2534873 | 0.142727 | 0.886505669 | 0.941194024 | SMARCC2 | protein_coding | 48.99 | 0.000952275 | 0.000952275 | ENSG00000139613 | n.s. |
| ENSG00000140553 | 840.4468943 | -0.037419066 | 0.2721008 | -0.137519 | 0.890620454 | 0.943117477 | UNC45A | protein_coding | 53.82 | -0.000951724 | 0.000951724 | ENSG00000140553 | n.s. |
| ENSG00000108829 | 4241.550418 | 0.037190077 | 0.2691965 | 0.138152 | 0.89012018 | 0.942914513 | LRRC59 | protein_coding | 47.65 | 0.000949376 | 0.000949376 | ENSG00000108829 | n.s. |
| ENSG00000131067 | 370.078539 | -0.035123659 | 0.24148 | -0.145452 | 0.884354308 | 0.939936396 | GGT7 | protein_coding | 49.05 | -0.00094488 | 0.00094488 | ENSG00000131067 | n.s. |
| ENSG00000180979 | 451.5013405 | 0.036338408 | 0.2588972 | 0.140358 | 0.888376773 | 0.942149933 | LRRC57 | protein_coding | 43.17 | 0.000940437 | 0.000940437 | ENSG00000180979 | n.s. |
| ENSG00000167962 | 543.5690532 | -0.035023463 | 0.241947 | -0.144757 | 0.88490291 | 0.940114196 | ZNF598 | protein_coding | 64.25 | -0.000939308 | 0.000939308 | ENSG00000167962 | n.s. |
| ENSG00000272812 | 107.5703599 | -0.03469607 | 0.2367462 | -0.146554 | 0.883484177 | 0.939777691 | | sense_intronic | 36.24 | -0.000935922 | 0.000935922 | ENSG00000272812 | n.s. |
| ENSG00000127399 | 177.2494828 | -0.035714402 | 0.2531858 | -0.14106 | 0.887822496 | 0.941945638 | LRRC61 | protein_coding | 54.13 | -0.000927652 | 0.000927652 | ENSG00000127399 | n.s. |
| ENSG00000146007 | 1440.597179 | -0.036932763 | 0.2721084 | -0.135728 | 0.892036202 | 0.944218831 | ZMAT2 | protein_coding | 43.7 | -0.000920636 | 0.000920636 | ENSG00000146007 | n.s. |
| ENSG00000163512 | 1221.08918 | -0.037076476 | 0.2760017 | -0.134334 | 0.893138281 | 0.944921298 | AZI2 | protein_coding | 34.24 | -0.000912243 | 0.000912243 | ENSG00000163512 | n.s. |
| ENSG00000187840 | 1763.860551 | -0.038141471 | 0.2919384 | -0.130649 | 0.896052942 | 0.946781901 | EIF4EBP1 | protein_coding | 49.34 | -0.000905862 | 0.000905862 | ENSG00000187840 | n.s. |
| ENSG00000166997 | 142.5847044 | -0.039137968 | 0.3059905 | -0.127906 | 0.898223494 | 0.948117437 | CNPY4 | protein_coding | 50.62 | -0.000905569 | 0.000905569 | ENSG00000166997 | n.s. |

| | | | | | | | | | | | | | |
|---|---|---|---|---|---|---|---|---|---|---|---|---|---|
| ENSG00000130758 | 134.113825 | -0.03873363 | 0.303062 | -0.127808 | 0.898301229 | 0.948117437 | MAP3K10 | protein_coding | 52.96 | -0.000896214 | 0.000896214 | ENSG00000130758 | n.s. |
| ENSG00000065600 | 722.7492199 | 0.039201 | 0.3100512 | 0.126434 | 0.899388421 | 0.948769819 | TMEM206 | protein_coding | 44.57 | 0.000895317 | 0.000895317 | ENSG00000065600 | n.s. |
| ENSG00000132768 | 351.8425651 | -0.035887406 | 0.2644834 | -0.135689 | 0.892067406 | 0.944218831 | DPH2 | protein_coding | 56.05 | -0.000894578 | 0.000894578 | ENSG00000132768 | n.s. |
| ENSG00000104472 | 249.4842202 | 0.034360061 | 0.2455913 | 0.139907 | 0.888733107 | 0.942149933 | CHRAC1 | protein_coding | 47.65 | 0.000889238 | 0.000889238 | ENSG00000104472 | n.s. |
| ENSG00000167005 | 4928.826901 | 0.036807978 | 0.2783708 | 0.132226 | 0.894805176 | 0.945894196 | NUDT21 | protein_coding | 39.54 | 0.000889186 | 0.000889186 | ENSG00000167005 | n.s. |
| ENSG00000122390 | 80.84567522 | -0.034906851 | 0.2541917 | -0.137325 | 0.890774 | 0.943208336 | NAA60 | protein_coding | 50.69 | -0.000886368 | 0.000886368 | ENSG00000122390 | n.s. |
| ENSG00000259943 | 226.3584261 | 0.032123459 | 0.2155172 | 0.149053 | 0.881511931 | 0.938612008 | | antisense | 40.43 | 0.000883841 | 0.000883841 | ENSG00000259943 | n.s. |
| ENSG00000104824 | 4118.873267 | 0.03398059 | 0.2411797 | 0.140893 | 0.887954292 | 0.942013685 | HNRNPL | protein_coding | 53.73 | 0.000881551 | 0.000881551 | ENSG00000104824 | n.s. |
| ENSG00000072135 | 288.0571647 | -0.033851062 | 0.2406557 | -0.140662 | 0.888137118 | 0.942104556 | PTPN18 | protein_coding | 47.65 | -0.000876773 | 0.000876773 | ENSG00000072135 | n.s. |
| ENSG00000020129 | 205.4679482 | -0.034133508 | 0.2453367 | -0.139129 | 0.88934801 | 0.942578753 | NCDN | protein_coding | 57.71 | -0.000876629 | 0.000876629 | ENSG00000020129 | n.s. |
| ENSG00000174799 | 447.5308664 | 0.033870818 | 0.2414353 | 0.140289 | 0.888431308 | 0.942149933 | CEP135 | protein_coding | 37.81 | 0.000876576 | 0.000876576 | ENSG00000174799 | n.s. |
| ENSG00000279432 | 303.1113545 | -0.034098136 | 0.246564 | -0.138293 | 0.890008646 | 0.942914513 | | TEC | 39.69 | -0.000870446 | 0.000870446 | ENSG00000279432 | n.s. |
| ENSG00000248489 | 21.17177116 | -0.040426051 | 0.3371839 | -0.119893 | 0.904567759 | 0.951658271 | LINC02062 | lincRNA | 39.81 | -0.000869927 | 0.000869927 | ENSG00000248489 | n.s. |
| ENSG00000136754 | 245.854954 | 0.034884001 | 0.2580571 | 0.135179 | 0.892470058 | 0.944501415 | ABI1 | protein_coding | 38.44 | 0.000865032 | 0.000865032 | ENSG00000136754 | n.s. |
| ENSG00000224745 | 153.6412839 | 0.035249642 | 0.2630525 | 0.134002 | 0.893400746 | 0.945127163 | | sense_intronic | 34.91 | 0.00086396 | 0.00086396 | ENSG00000224745 | n.s. |
| ENSG00000115568 | 414.8667107 | -0.031349378 | 0.2097132 | -0.149487 | 0.881169451 | 0.938534445 | ZNF142 | protein_coding | 49.52 | -0.000863669 | 0.000863669 | ENSG00000115568 | n.s. |
| ENSG00000150093 | 4163.213804 | 0.035776339 | 0.2720308 | 0.131516 | 0.895367329 | 0.94627291 | ITGB1 | protein_coding | 39.15 | 0.000858045 | 0.000858045 | ENSG00000150093 | n.s. |
| ENSG00000159256 | 930.4084183 | -0.034617972 | 0.2567063 | -0.134854 | 0.892726998 | 0.944629731 | MORC3 | protein_coding | 42.3 | -0.000856393 | 0.000856393 | ENSG00000159256 | n.s. |
| ENSG00000130844 | 455.6479359 | -0.034522615 | 0.2552279 | -0.135262 | 0.892404793 | 0.944501415 | ZNF331 | protein_coding | 44.28 | -0.000856071 | 0.000856071 | ENSG00000130844 | n.s. |
| ENSG00000142687 | 36.42514024 | 0.036959474 | 0.288173 | 0.128254 | 0.897947611 | 0.948117437 | KIAA0319L | protein_coding | 41.89 | 0.000855163 | 0.000855163 | ENSG00000142687 | n.s. |
| ENSG00000140374 | 2386.507789 | 0.036645024 | 0.2837634 | 0.129139 | 0.897247385 | 0.947745415 | ETFA | protein_coding | 40.14 | 0.000854134 | 0.000854134 | ENSG00000140374 | n.s. |
| ENSG00000165704 | 316.293359 | 0.039340934 | 0.325053 | 0.121029 | 0.903667831 | 0.951290429 | HPRT1 | protein_coding | 40.13 | 0.000853182 | 0.000853182 | ENSG00000165704 | n.s. |
| ENSG00000253559 | 56.88598989 | -0.035497926 | 0.2697756 | -0.131583 | 0.895314021 | 0.94627291 | OSGEPL1-AS1 | antisense | 40.66 | -0.000851368 | 0.000851368 | ENSG00000253559 | n.s. |
| ENSG00000156471 | 1030.574625 | 0.034586239 | 0.2592695 | 0.133399 | 0.893877978 | 0.945344709 | PTDSS1 | protein_coding | 43.71 | 0.000844243 | 0.000844243 | ENSG00000156471 | n.s. |
| ENSG00000267719 | 290.6367944 | 0.032526708 | 0.232539 | 0.139876 | 0.888757696 | 0.942149933 | LINC02078 | lincRNA | 56.79 | 0.00084179 | 0.00084179 | ENSG00000267719 | n.s. |
| ENSG00000111716 | 28524.4253 | 0.036091669 | 0.2832422 | 0.127423 | 0.898605332 | 0.948117437 | LDHB | protein_coding | 38.01 | 0.000835084 | 0.000835084 | ENSG00000111716 | n.s. |
| ENSG00000127884 | 620.7047759 | 0.03880586 | 0.3255098 | 0.119216 | 0.905104521 | 0.951852552 | ECHS1 | protein_coding | 56.3 | 0.000831622 | 0.000831622 | ENSG00000127884 | n.s. |
| ENSG00000104853 | 362.4879468 | 0.034628667 | 0.2640305 | 0.131154 | 0.895653455 | 0.946431622 | CLPTM1 | protein_coding | 52.83 | 0.000827998 | 0.000827998 | ENSG00000104853 | n.s. |
| ENSG00000128989 | 6595.244879 | 0.034302886 | 0.2598272 | 0.132022 | 0.894966961 | 0.945993395 | ARPP19 | protein_coding | 38.46 | 0.000827108 | 0.000827108 | ENSG00000128989 | n.s. |
| ENSG00000177889 | 3279.313805 | -0.035653539 | 0.2795803 | -0.127525 | 0.898524714 | 0.948117437 | UBE2N | protein_coding | 40.11 | -0.000824947 | 0.000824947 | ENSG00000177889 | n.s. |
| ENSG00000151348 | 634.1020678 | -0.035165253 | 0.2711241 | -0.129702 | 0.896802434 | 0.947430024 | EXT2 | protein_coding | 40.36 | -0.000824726 | 0.000824726 | ENSG00000151348 | n.s. |
| ENSG00000213015 | 114.4598035 | 0.033191267 | 0.2460008 | 0.134923 | 0.892672435 | 0.944629731 | ZNF580 | protein_coding | 57.45 | 0.000821098 | 0.000821098 | ENSG00000213015 | n.s. |

| | | | | | | | | | | | | | |
|---|---|---|---|---|---|---|---|---|---|---|---|---|---|
| ENSG00000185651 | 1176.404107 | 0.035464375 | 0.276594 | 0.128218 | 0.897976327 | 0.948117437 | UBE2L3 | protein_coding | 49.73 | 0.00082057 | 0.00082057 | ENSG00000185651 | n.s. |
| ENSG00000096746 | 3674.464993 | 0.032063639 | 0.2312335 | 0.138663 | 0.889716088 | 0.942805167 | HNRNPH3 | protein_coding | 39.82 | 0.000820125 | 0.000820125 | ENSG00000096746 | n.s. |
| ENSG00000169862 | 136.170398 | 0.03382688 | 0.2550612 | 0.132623 | 0.894491867 | 0.945706604 | CTNND2 | protein_coding | 39.43 | 0.000820085 | 0.000820085 | ENSG00000169862 | n.s. |
| ENSG00000143753 | 1985.365872 | 0.035422962 | 0.2769497 | 0.127904 | 0.898224999 | 0.948117437 | DEGS1 | protein_coding | 46.06 | 0.000819612 | 0.000819612 | ENSG00000143753 | n.s. |
| ENSG00000055211 | 439.6313819 | 0.035395878 | 0.2775546 | 0.127528 | 0.898522817 | 0.948117437 | GINM1 | protein_coding | 42.05 | 0.000818985 | 0.000818985 | ENSG00000055211 | n.s. |
| ENSG00000201499 | 15.08454073 | 0.040217799 | 0.3561385 | 0.112927 | 0.910088101 | 0.954462039 | RNU6-312P | snRNA | 38.32 | 0.000814062 | 0.000814062 | ENSG00000201499 | n.s. |
| ENSG00000160256 | 279.9201518 | 0.034822845 | 0.2698059 | 0.129066 | 0.897305183 | 0.947745415 | FAM207A | protein_coding | 56.7 | 0.000811662 | 0.000811662 | ENSG00000160256 | n.s. |
| ENSG00000168904 | 36.22726327 | -0.03736533 | 0.3150056 | -0.118618 | 0.905578024 | 0.9519635 | LRRC28 | protein_coding | 38.12 | -0.00079886 | 0.00079886 | ENSG00000168904 | n.s. |
| ENSG00000144028 | 5892.856929 | -0.032293065 | 0.2397582 | -0.13469 | 0.892856881 | 0.944695368 | SNRNP200 | protein_coding | 47.58 | -0.000797904 | 0.000797904 | ENSG00000144028 | n.s. |
| ENSG00000181789 | 1418.688576 | -0.034261202 | 0.2675542 | -0.128053 | 0.898106776 | 0.948117437 | COPG1 | protein_coding | 48.18 | -0.000792731 | 0.000792731 | ENSG00000181789 | n.s. |
| ENSG00000087884 | 114.9309915 | -0.035301596 | 0.2829515 | -0.124762 | 0.900711983 | 0.949614792 | AAMDC | protein_coding | 41.82 | -0.00079261 | 0.00079261 | ENSG00000087884 | n.s. |
| ENSG00000108840 | 1279.833231 | -0.032336666 | 0.2418399 | -0.133711 | 0.893631088 | 0.9452272 | HDAC5 | protein_coding | 55.66 | -0.000791077 | 0.000791077 | ENSG00000108840 | n.s. |
| ENSG00000165914 | 41.81886167 | -0.03516189 | 0.2815227 | -0.124899 | 0.900603537 | 0.949614792 | TTC7B | protein_coding | 46.46 | -0.000789473 | 0.000789473 | ENSG00000165914 | n.s. |
| ENSG00000136950 | 611.5288123 | 0.033590529 | 0.2585547 | 0.129917 | 0.896632485 | 0.947322362 | ARPC5L | protein_coding | 52.55 | 0.000789452 | 0.000789452 | ENSG00000136950 | n.s. |
| ENSG00000139726 | 1902.596299 | 0.034840516 | 0.2771111 | 0.125728 | 0.899947555 | 0.949240137 | DENR | protein_coding | 42.12 | 0.000788229 | 0.000788229 | ENSG00000139726 | n.s. |
| ENSG00000222346 | 4.436350807 | -0.049790145 | 0.5558894 | -0.089568 | 0.928630169 | 0.964226851 | RNA5SP237 | rRNA | 45.13 | -0.000787719 | 0.000787719 | ENSG00000222346 | n.s. |
| ENSG00000131467 | 2480.613155 | -0.032138295 | 0.2415546 | -0.133048 | 0.894155603 | 0.945422878 | PSME3 | protein_coding | 46.04 | -0.000783335 | 0.000783335 | ENSG00000131467 | n.s. |
| ENSG00000063245 | 325.3888437 | -0.032642049 | 0.248579 | -0.131315 | 0.895526467 | 0.94636926 | EPN1 | protein_coding | 54.85 | -0.00078143 | 0.00078143 | ENSG00000063245 | n.s. |
| ENSG00000136490 | 2258.658953 | 0.031238069 | 0.2294623 | 0.136136 | 0.891713805 | 0.944113697 | LIMD2 | protein_coding | 66.27 | 0.000780193 | 0.000780193 | ENSG00000136490 | n.s. |
| ENSG00000135486 | 33241.04955 | 0.031201526 | 0.2293022 | 0.136072 | 0.89176465 | 0.944113697 | HNRNPA1 | protein_coding | 42.13 | 0.00077928 | 0.00077928 | ENSG00000135486 | n.s. |
| ENSG00000272931 | 40.86290572 | 0.035767739 | 0.2951839 | 0.121171 | 0.903555557 | 0.95124416 | | lincRNA | 49.93 | 0.000776446 | 0.000776446 | ENSG00000272931 | n.s. |
| ENSG00000167965 | 380.1211032 | 0.034760356 | 0.2805296 | 0.12391 | 0.901386712 | 0.950110318 | MLST8 | protein_coding | 61.98 | 0.000772582 | 0.000772582 | ENSG00000167965 | n.s. |
| ENSG00000170469 | 33.44402557 | -0.038941281 | 0.35055 | -0.111086 | 0.91154797 | 0.955396145 | SPATA24 | protein_coding | 51.17 | -0.000771681 | 0.000771681 | ENSG00000170469 | n.s. |
| ENSG00000203644 | 154.2704259 | 0.036872402 | 0.3178041 | 0.116022 | 0.907634776 | 0.953172575 | | sense_intronic | 33.96 | 0.000767995 | 0.000767995 | ENSG00000203644 | n.s. |
| ENSG00000140612 | 1505.686974 | 0.035661695 | 0.2986032 | 0.119428 | 0.904935986 | 0.951761878 | SEC11A | protein_coding | 42.41 | 0.000765717 | 0.000765717 | ENSG00000140612 | n.s. |
| ENSG00000112576 | 150.0783596 | -0.033915005 | 0.2718288 | -0.124766 | 0.900708755 | 0.949614792 | CCND3 | protein_coding | 47.06 | -0.000761478 | 0.000761478 | ENSG00000112576 | n.s. |
| ENSG00000220868 | 22.89479153 | -0.037319007 | 0.3290198 | -0.113425 | 0.909693783 | 0.95424294 | MRPL35P1 | processed_pseudogene | 41.96 | -0.000759108 | 0.000759108 | ENSG00000220868 | n.s. |
| ENSG00000184005 | 697.1242018 | 0.035215286 | 0.2933638 | 0.12004 | 0.904451752 | 0.951658271 | ST6GALNAC3 | protein_coding | 38.78 | 0.000757797 | 0.000757797 | ENSG00000184005 | n.s. |
| ENSG00000122507 | 39.32332132 | -0.038959524 | 0.3594423 | -0.108389 | 0.913687278 | 0.956270613 | BBS9 | protein_coding | 37.38 | -0.000756562 | 0.000756562 | ENSG00000122507 | n.s. |
| ENSG00000177732 | 831.2962108 | -0.030987691 | 0.2326166 | -0.133214 | 0.894024475 | 0.945356024 | SOX12 | protein_coding | 61.99 | -0.000756242 | 0.000756242 | ENSG00000177732 | n.s. |
| ENSG00000131446 | 398.5692452 | 0.032635755 | 0.2556992 | 0.127633 | 0.898439116 | 0.948117437 | MGAT1 | protein_coding | 50.9 | 0.000755122 | 0.000755122 | ENSG00000131446 | n.s. |
| ENSG00000141696 | 393.3498691 | 0.033909467 | 0.2735001 | 0.123983 | 0.901328425 | 0.950110318 | P3H4 | protein_coding | 56.65 | 0.000753671 | 0.000753671 | ENSG00000141696 | n.s. |

| | | | | | | | | | | | | | |
|---|---|---|---|---|---|---|---|---|---|---|---|---|---|
| ENSG00000168273 | 298.704546 | 0.032495378 | 0.2549682 | 0.127449 | 0.898585255 | 0.948117437 | SMIM4 | protein_coding | 43.07 | 0.000751874 | 0.000751874 | ENSG00000168273 | n.s. |
| ENSG00000166313 | 692.4154955 | 0.034989216 | 0.2926995 | 0.11954 | 0.90484776 | 0.951740981 | APBB1 | protein_coding | 51.87 | 0.000751611 | 0.000751611 | ENSG00000166313 | n.s. |
| ENSG00000141560 | 548.4736429 | -0.034057651 | 0.2765962 | -0.123131 | 0.902003119 | 0.950541495 | FN3KRP | protein_coding | 53.24 | -0.000750253 | 0.000750253 | ENSG00000141560 | n.s. |
| ENSG00000166839 | 190.5854221 | -0.034446937 | 0.2840249 | -0.121281 | 0.903468151 | 0.951224067 | ANKDD1A | protein_coding | 46.75 | -0.00074809 | 0.00074809 | ENSG00000166839 | n.s. |
| ENSG00000196072 | 766.9695479 | -0.034439349 | 0.2837199 | -0.121385 | 0.903386083 | 0.951221687 | BLOC1S2 | protein_coding | 47.38 | -0.000747963 | 0.000747963 | ENSG00000196072 | n.s. |
| ENSG00000127463 | 481.4726047 | -0.032732467 | 0.2590065 | -0.126377 | 0.899433516 | 0.948769819 | EMC1 | protein_coding | 46.63 | -0.000747581 | 0.000747581 | ENSG00000127463 | n.s. |
| ENSG00000125304 | 1733.208602 | 0.03461545 | 0.2890377 | 0.119761 | 0.904672472 | 0.951658271 | TM9SF2 | protein_coding | 38.93 | 0.000744889 | 0.000744889 | ENSG00000125304 | n.s. |
| ENSG00000184867 | 381.8281527 | -0.035577993 | 0.3041911 | -0.116959 | 0.906892274 | 0.952955557 | ARMCX2 | protein_coding | 51.79 | -0.000744553 | 0.000744553 | ENSG00000184867 | n.s. |
| ENSG00000129315 | 3021.809387 | 0.031824665 | 0.2458544 | 0.129445 | 0.897005391 | 0.947572538 | CCNT1 | protein_coding | 40.95 | 0.0007443 | 0.0007443 | ENSG00000129315 | n.s. |
| ENSG00000243770 | 75.62155457 | 0.032829186 | 0.261491 | 0.125546 | 0.900091214 | 0.949247839 | RN7SL65P | misc_RNA | 49.84 | 0.000742609 | 0.000742609 | ENSG00000243770 | n.s. |
| ENSG00000106603 | 1876.984854 | -0.033560838 | 0.2734226 | -0.122743 | 0.902310256 | 0.950580031 | COA1 | protein_coding | 40.39 | -0.000738718 | 0.000738718 | ENSG00000106603 | n.s. |
| ENSG00000114686 | 3553.011924 | 0.033885148 | 0.2779806 | 0.121898 | 0.902980168 | 0.95106989 | MRPL3 | protein_coding | 35.94 | 0.000738275 | 0.000738275 | ENSG00000114686 | n.s. |
| ENSG00000125352 | 193.4996368 | -0.033493228 | 0.2725357 | -0.122895 | 0.90219039 | 0.950541495 | RNF113A | protein_coding | 54.55 | -0.00073782 | 0.00073782 | ENSG00000125352 | n.s. |
| ENSG00000129219 | 34.24701095 | 0.035370089 | 0.3040332 | 0.116336 | 0.907386049 | 0.953172575 | PLD2 | protein_coding | 54.27 | 0.000736705 | 0.000736705 | ENSG00000129219 | n.s. |
| ENSG00000176946 | 598.9106191 | -0.033052995 | 0.2677853 | -0.123431 | 0.901765846 | 0.950419544 | THAP4 | protein_coding | 49.81 | -0.000729964 | 0.000729964 | ENSG00000176946 | n.s. |
| ENSG00000114209 | 1191.495915 | 0.03387119 | 0.2816012 | 0.120281 | 0.904260789 | 0.951658271 | PDCD10 | protein_coding | 35.73 | 0.000728873 | 0.000728873 | ENSG00000114209 | n.s. |
| ENSG00000184436 | 489.2340661 | -0.028527123 | 0.2066902 | -0.138019 | 0.890225618 | 0.942914513 | THAP7 | protein_coding | 62.2 | -0.000728231 | 0.000728231 | ENSG00000184436 | n.s. |
| ENSG00000166579 | 162.849777 | 0.032020621 | 0.256487 | 0.124843 | 0.900647782 | 0.949614792 | NDEL1 | protein_coding | 44.62 | 0.000718944 | 0.000718944 | ENSG00000166579 | n.s. |
| ENSG00000225377 | 53.79765083 | 0.033314449 | 0.2779374 | 0.119863 | 0.904591568 | 0.951658271 | NRSN2-AS1 | antisense | 52 | 0.000716893 | 0.000716893 | ENSG00000225377 | n.s. |
| ENSG00000103769 | 2547.888662 | 0.034297511 | 0.2956049 | 0.116025 | 0.907632859 | 0.953172575 | RAB11A | protein_coding | 40.1 | 0.000714364 | 0.000714364 | ENSG00000103769 | n.s. |
| ENSG00000130402 | 195.3038101 | 0.031147841 | 0.2461833 | 0.126523 | 0.899317959 | 0.948769819 | ACTN4 | protein_coding | 53.01 | 0.00071139 | 0.00071139 | ENSG00000130402 | n.s. |
| ENSG00000159128 | 551.4089515 | 0.033782975 | 0.288825 | 0.116967 | 0.90688624 | 0.952955557 | IFNGR2 | protein_coding | 44.83 | 0.000706988 | 0.000706988 | ENSG00000159128 | n.s. |
| ENSG00000120802 | 15018.38052 | -0.033584464 | 0.2863641 | -0.117279 | 0.906639059 | 0.952833304 | TMPO | protein_coding | 39.75 | -0.000704705 | 0.000704705 | ENSG00000120802 | n.s. |
| ENSG00000028310 | 1040.687718 | 0.030409604 | 0.2364904 | 0.128587 | 0.897684424 | 0.948074053 | BRD9 | protein_coding | 55.14 | 0.000704218 | 0.000704218 | ENSG00000028310 | n.s. |
| ENSG00000206732 | 17.26181546 | -0.050248193 | 0.6260785 | -0.080259 | 0.936031578 | 0.968301335 | RNU6-936P | snRNA | 33.64 | -0.000702946 | 0.000702946 | ENSG00000206732 | n.s. |
| ENSG00000083520 | 3423.192272 | 0.032304212 | 0.2657099 | 0.121577 | 0.903234043 | 0.951193373 | DIS3 | protein_coding | 36.46 | 0.000702009 | 0.000702009 | ENSG00000083520 | n.s. |
| ENSG00000170412 | 119.194614 | 0.033653908 | 0.2886972 | 0.116572 | 0.907199497 | 0.953136309 | GPRC5C | protein_coding | 54.5 | 0.000701515 | 0.000701515 | ENSG00000170412 | n.s. |
| ENSG00000157191 | 211.6391192 | -0.031724918 | 0.2581868 | -0.122876 | 0.902205405 | 0.950541495 | NECAP2 | protein_coding | 49.4 | -0.000698866 | 0.000698866 | ENSG00000157191 | n.s. |
| ENSG00000207696 | 9.84696544 | 0.039226632 | 0.3984045 | 0.098459 | 0.921567585 | 0.960403105 | MIR659 | miRNA | 53.61 | 0.000688288 | 0.000688288 | ENSG00000207696 | n.s. |
| ENSG00000156170 | 393.7833958 | -0.03192576 | 0.2655091 | -0.120244 | 0.904290224 | 0.951658271 | NDUFAF6 | protein_coding | 42.88 | -0.00068701 | 0.00068701 | ENSG00000156170 | n.s. |
| ENSG00000119900 | 665.2390193 | -0.031060305 | 0.2526605 | -0.122933 | 0.902160186 | 0.950541495 | OGFRL1 | protein_coding | 36.22 | -0.000684225 | 0.000684225 | ENSG00000119900 | n.s. |
| ENSG00000132563 | 30.2594385 | -0.033411125 | 0.2929865 | -0.114036 | 0.909208947 | 0.954020102 | REEP2 | protein_coding | 58.31 | -0.000683006 | 0.000683006 | ENSG00000132563 | n.s. |

| | | | | | | | | | | | | | |
|---|---|---|---|---|---|---|---|---|---|---|---|---|---|
| ENSG00000197928 | 565.1923687 | -0.031301488 | 0.2579003 | -0.121371 | 0.90339757 | 0.951221687 | ZNF677 | protein_coding | 38.84 | -0.000679814 | 0.000679814 | ENSG00000197928 | n.s. |
| ENSG00000125503 | 402.3351605 | 0.032687406 | 0.2830545 | 0.115481 | 0.908063924 | 0.95339522 | PPP1R12C | protein_coding | 57.6 | 0.000677513 | 0.000677513 | ENSG00000125503 | n.s. |
| ENSG00000125651 | 1030.261743 | -0.031260682 | 0.2624869 | -0.119094 | 0.905200695 | 0.951852552 | GTF2F1 | protein_coding | 54.75 | -0.000669926 | 0.000669926 | ENSG00000125651 | n.s. |
| ENSG00000230180 | 4.145055651 | -0.060113972 | 0.9510336 | -0.063209 | 0.949600002 | 0.974765398 | RPL12P49 | processed_pseudogene | 46.28 | -0.000667259 | 0.000667259 | ENSG00000230180 | n.s. |
| ENSG00000204394 | 1346.758072 | -0.030171432 | 0.2445668 | -0.123367 | 0.901816606 | 0.950419544 | VARS | protein_coding | 53.81 | -0.000666325 | 0.000666325 | ENSG00000204394 | n.s. |
| ENSG00000158169 | 206.6350539 | -0.030902293 | 0.2580492 | -0.119753 | 0.904678436 | 0.951658271 | FANCC | protein_coding | 41.78 | -0.000664986 | 0.000664986 | ENSG00000158169 | n.s. |
| ENSG00000145247 | 709.1496431 | 0.033497168 | 0.3011665 | 0.111225 | 0.911438136 | 0.955352945 | OCIAD2 | protein_coding | 41.87 | 0.000664455 | 0.000664455 | ENSG00000145247 | n.s. |
| ENSG00000071859 | 599.3400678 | 0.030926525 | 0.2604788 | 0.11873 | 0.905489629 | 0.9519635 | FAM50A | protein_coding | 60.48 | 0.0006612 | 0.0006612 | ENSG00000071859 | n.s. |
| ENSG00000100997 | 397.1555486 | -0.031628527 | 0.2733749 | -0.115697 | 0.907893046 | 0.953338791 | ABHD12 | protein_coding | 47.42 | -0.000656378 | 0.000656378 | ENSG00000100997 | n.s. |
| ENSG00000131188 | 67.64466525 | 0.032817934 | 0.2926332 | 0.112147 | 0.910706847 | 0.955017808 | PRR7 | protein_coding | 62.31 | 0.000655982 | 0.000655982 | ENSG00000131188 | n.s. |
| ENSG00000135631 | 240.7464185 | 0.028991138 | 0.2307987 | 0.125612 | 0.900038904 | 0.949247839 | RAB11FIP5 | protein_coding | 48.58 | 0.000655791 | 0.000655791 | ENSG00000135631 | n.s. |
| ENSG00000143434 | 213.2943351 | 0.029920181 | 0.2445792 | 0.122333 | 0.902635053 | 0.950850256 | SEMA6C | protein_coding | 57.48 | 0.000654889 | 0.000654889 | ENSG00000143434 | n.s. |
| ENSG00000151725 | 3729.087184 | -0.031796999 | 0.2791839 | -0.113893 | 0.909322882 | 0.954020102 | CENPU | protein_coding | 40.02 | -0.000650009 | 0.000650009 | ENSG00000151725 | n.s. |
| ENSG00000201435 | 39.82966555 | 0.033303362 | 0.3032336 | 0.109827 | 0.912546256 | 0.956082589 | RNU4-24P | snRNA | 38.85 | 0.000649568 | 0.000649568 | ENSG00000201435 | n.s. |
| ENSG00000205302 | 1153.945589 | 0.030145699 | 0.2514638 | 0.119881 | 0.904577529 | 0.951658271 | SNX2 | protein_coding | 36.08 | 0.000648705 | 0.000648705 | ENSG00000205302 | n.s. |
| ENSG00000102977 | 220.7946655 | 0.031930844 | 0.2828193 | 0.112902 | 0.910108312 | 0.954462039 | ACD | protein_coding | 62.02 | 0.000646323 | 0.000646323 | ENSG00000102977 | n.s. |
| ENSG00000129946 | 256.0548599 | 0.032352115 | 0.2896056 | 0.111711 | 0.911052589 | 0.955092628 | SHC2 | protein_coding | 58.69 | 0.000645571 | 0.000645571 | ENSG00000129946 | n.s. |
| ENSG00000205531 | 3063.587135 | -0.029608524 | 0.2427424 | -0.121975 | 0.90291877 | 0.95106989 | NAP1L4 | protein_coding | 44.27 | -0.000645098 | 0.000645098 | ENSG00000205531 | n.s. |
| ENSG00000164045 | 1906.3917 | -0.029643374 | 0.2438234 | -0.121577 | 0.903233866 | 0.951193373 | CDC25A | protein_coding | 45.04 | -0.000644186 | 0.000644186 | ENSG00000164045 | n.s. |
| ENSG00000115539 | 555.3728253 | -0.030111319 | 0.2539276 | -0.118582 | 0.905606303 | 0.9519635 | PDCL3 | protein_coding | 46.99 | -0.000643771 | 0.000643771 | ENSG00000115539 | n.s. |
| ENSG00000068305 | 286.9124093 | -0.030292169 | 0.2565564 | -0.118072 | 0.906010462 | 0.952244559 | MEF2A | protein_coding | 36.86 | -0.000643754 | 0.000643754 | ENSG00000068305 | n.s. |
| ENSG00000132967 | 583.5178122 | 0.033465517 | 0.3140185 | 0.106572 | 0.915128698 | 0.95670047 | HMGB1P5 | transcribed_processed_pseudog | 39.42 | 0.000643342 | 0.000643342 | ENSG00000132967 | n.s. |
| ENSG00000183837 | 140.4122497 | 0.03086718 | 0.2676873 | 0.115311 | 0.908198942 | 0.953454521 | PNMA3 | protein_coding | 60.7 | 0.000638951 | 0.000638951 | ENSG00000183837 | n.s. |
| ENSG00000106799 | 2937.959135 | -0.031010505 | 0.273887 | -0.113224 | 0.909853189 | 0.954338252 | TGFBR1 | protein_coding | 39.03 | -0.000629441 | 0.000629441 | ENSG00000106799 | n.s. |
| ENSG00000116005 | 451.7015762 | -0.030730556 | 0.2696588 | -0.113961 | 0.909268812 | 0.954020102 | PCYOX1 | protein_coding | 45.23 | -0.000628209 | 0.000628209 | ENSG00000116005 | n.s. |
| ENSG00000228624 | 40.97089026 | 0.04464378 | 0.5535759 | 0.080646 | 0.935723338 | 0.968126331 | HDAC2-AS2 | antisense | 37.94 | 0.000628047 | 0.000628047 | ENSG00000228624 | n.s. |
| ENSG00000181191 | 555.5409505 | -0.03199556 | 0.2951249 | -0.108414 | 0.913667607 | 0.956270613 | PJA1 | protein_coding | 51.45 | -0.000621328 | 0.000621328 | ENSG00000181191 | n.s. |
| ENSG00000198130 | 590.5631428 | 0.030954616 | 0.2770058 | 0.111747 | 0.911023863 | 0.955092628 | HIBCH | protein_coding | 38.94 | 0.000617684 | 0.000617684 | ENSG00000198130 | n.s. |
| ENSG00000069509 | 139.9118792 | -0.031608123 | 0.2886808 | -0.109492 | 0.912812586 | 0.956091452 | FUNDC1 | protein_coding | 41.18 | -0.000616376 | 0.000616376 | ENSG00000069509 | n.s. |
| ENSG00000184887 | 345.278001 | -0.030123772 | 0.2645026 | -0.113888 | 0.909326294 | 0.954020102 | BTBD6 | protein_coding | 64.36 | -0.000615804 | 0.000615804 | ENSG00000184887 | n.s. |
| ENSG00000008838 | 828.9733942 | 0.028760571 | 0.2431189 | 0.118298 | 0.905831245 | 0.95212807 | MED24 | protein_coding | 50.63 | 0.000612733 | 0.000612733 | ENSG00000008838 | n.s. |
| ENSG00000204469 | 2621.730277 | -0.027524359 | 0.2216654 | -0.124171 | 0.901180068 | 0.950036351 | PRRC2A | protein_coding | 53.44 | -0.000612686 | 0.000612686 | ENSG00000204469 | n.s. |

| | | | | | | | | | | | | | |
|---|---|---|---|---|---|---|---|---|---|---|---|---|---|
| ENSG00000105290 | 83.56273698 | 0.031456024 | 0.2898533 | 0.108524 | 0.913580072 | 0.956270613 | APLP1 | protein_coding | 57.02 | 0.000610851 | 0.000610851 | ENSG00000105290 | n.s. |
| ENSG00000148362 | 537.1696971 | 0.031305188 | 0.2858474 | 0.109517 | 0.91279234 | 0.956091452 | C9orf142 | protein_coding | 64.9 | 0.000610469 | 0.000610469 | ENSG00000148362 | n.s. |
| ENSG00000196459 | 332.0277761 | -0.02969116 | 0.2588092 | -0.114722 | 0.908665318 | 0.953810858 | TRAPPC2 | protein_coding | 42.18 | -0.000609789 | 0.000609789 | ENSG00000196459 | n.s. |
| ENSG00000131966 | 1046.342678 | -0.032851373 | 0.3167824 | -0.103703 | 0.917404824 | 0.958216607 | ACTR10 | protein_coding | 38.67 | -0.000608943 | 0.000608943 | ENSG00000131966 | n.s. |
| ENSG00000142230 | 2551.805559 | -0.03112919 | 0.2877829 | -0.108169 | 0.913861623 | 0.956381192 | SAE1 | protein_coding | 47.18 | -0.00060294 | 0.00060294 | ENSG00000142230 | n.s. |
| ENSG00000172331 | 380.1446546 | 0.031745551 | 0.3003516 | 0.105695 | 0.915824649 | 0.957284279 | BPGM | protein_coding | 39.66 | 0.000601866 | 0.000601866 | ENSG00000172331 | n.s. |
| ENSG00000105819 | 2149.4805 | 0.029393945 | 0.2576219 | 0.114097 | 0.909160705 | 0.954020102 | PMPCB | protein_coding | 38.52 | 0.000600885 | 0.000600885 | ENSG00000105819 | n.s. |
| ENSG00000108239 | 423.6682257 | 0.029345835 | 0.2577234 | 0.113866 | 0.909344306 | 0.954020102 | TBC1D12 | protein_coding | 38.55 | 0.000599901 | 0.000599901 | ENSG00000108239 | n.s. |
| ENSG00000213453 | 42.78983409 | 0.030650254 | 0.284003 | 0.107922 | 0.91405735 | 0.956442243 | FTH1P3 | processed_pseudogene | 47.95 | 0.000592814 | 0.000592814 | ENSG00000213453 | n.s. |
| ENSG00000263764 | 2.694357593 | -0.056409598 | 0.938497 | -0.060106 | 0.952070959 | 0.976118113 | SNORD43 | snoRNA | 45.31 | -0.000592167 | 0.000592167 | ENSG00000263764 | n.s. |
| ENSG00000132603 | 989.1623309 | 0.028380842 | 0.2446463 | 0.116008 | 0.907646484 | 0.953172575 | NIP7 | protein_coding | 45.71 | 0.000591129 | 0.000591129 | ENSG00000132603 | n.s. |
| ENSG00000141424 | 3527.728647 | -0.030297839 | 0.2757458 | -0.109876 | 0.912507726 | 0.956082589 | SLC39A6 | protein_coding | 39.45 | -0.000590947 | 0.000590947 | ENSG00000141424 | n.s. |
| ENSG00000121774 | 2238.635298 | 0.030143161 | 0.2751704 | 0.109544 | 0.912771315 | 0.956091452 | KHDRBS1 | protein_coding | 43.63 | 0.000587809 | 0.000587809 | ENSG00000121774 | n.s. |
| ENSG00000108984 | 396.5263884 | -0.030448651 | 0.2847443 | -0.106933 | 0.914841885 | 0.95670047 | MAP2K6 | protein_coding | 41 | -0.000585345 | 0.000585345 | ENSG00000108984 | n.s. |
| ENSG00000104687 | 982.924547 | 0.029446839 | 0.2643527 | 0.111392 | 0.911305306 | 0.955285633 | GSR | protein_coding | 44.72 | 0.000585013 | 0.000585013 | ENSG00000104687 | n.s. |
| ENSG00000055118 | 682.7714071 | -0.028692552 | 0.2525017 | -0.113633 | 0.909528628 | 0.954141583 | KCNH2 | protein_coding | 59.57 | -0.00058496 | 0.00058496 | ENSG00000055118 | n.s. |
| ENSG00000263603 | 56.7203609 | 0.030401567 | 0.2836614 | 0.107176 | 0.914649696 | 0.95670047 | | lincRNA | 54.04 | 0.00058444 | 0.00058444 | ENSG00000263603 | n.s. |
| ENSG00000102383 | 47.34983605 | -0.030185526 | 0.2814751 | -0.10724 | 0.914598203 | 0.95670047 | ZDHHC15 | protein_coding | 38.32 | -0.000580287 | 0.000580287 | ENSG00000102383 | n.s. |
| ENSG00000263697 | 3.377201409 | 0.044603828 | 0.6059241 | 0.073613 | 0.941318406 | 0.970635872 | MIR3621 | miRNA | 72.94 | 0.000577337 | 0.000577337 | ENSG00000263697 | n.s. |
| ENSG00000167862 | 866.4923917 | 0.032594818 | 0.3248436 | 0.10034 | 0.920074377 | 0.960095128 | MRPL58 | protein_coding | 48.66 | 0.000576463 | 0.000576463 | ENSG00000167862 | n.s. |
| ENSG00000153310 | 1581.91751 | 0.031008834 | 0.2986143 | 0.103842 | 0.91729441 | 0.958173161 | FAM49B | protein_coding | 42.04 | 0.0005754 | 0.0005754 | ENSG00000153310 | n.s. |
| ENSG00000033030 | 1130.023424 | 0.027700601 | 0.2403797 | 0.115237 | 0.908257367 | 0.953454521 | ZCCHC8 | protein_coding | 42.68 | 0.000573403 | 0.000573403 | ENSG00000033030 | n.s. |
| ENSG00000175344 | 852.6839534 | -0.027517875 | 0.2367571 | -0.116228 | 0.907471593 | 0.953172575 | CHRNA7 | protein_coding | 41.21 | -0.000573155 | 0.000573155 | ENSG00000175344 | n.s. |
| ENSG00000213563 | 269.2895456 | 0.029573353 | 0.2737074 | 0.108047 | 0.91395815 | 0.956410321 | C8orf82 | protein_coding | 67.88 | 0.000572414 | 0.000572414 | ENSG00000213563 | n.s. |
| ENSG00000138101 | 250.6529992 | 0.027442219 | 0.2364139 | 0.116077 | 0.90759149 | 0.953172575 | DTNB | protein_coding | 41.46 | 0.000571579 | 0.000571579 | ENSG00000138101 | n.s. |
| ENSG00000196208 | 300.2734722 | -0.02786449 | 0.244609 | -0.113914 | 0.909305633 | 0.954020102 | GREB1 | protein_coding | 47.32 | -0.000569619 | 0.000569619 | ENSG00000196208 | n.s. |
| ENSG00000239789 | 491.1576427 | 0.031767677 | 0.3136938 | 0.10127 | 0.919336359 | 0.959611634 | MRPS17 | protein_coding | 49.9 | 0.000568784 | 0.000568784 | ENSG00000239789 | n.s. |
| ENSG00000124635 | 889.05331 | 0.026236376 | 0.2191382 | 0.119725 | 0.904700774 | 0.951658271 | HIST1H2BJ | protein_coding | 41.7 | 0.00056458 | 0.00056458 | ENSG00000124635 | n.s. |
| ENSG00000185559 | 71.93833544 | 0.031776972 | 0.3211641 | 0.098943 | 0.921183435 | 0.960134566 | DLK1 | protein_coding | 55.89 | 0.000561432 | 0.000561432 | ENSG00000185559 | n.s. |
| ENSG00000187051 | 614.6034895 | 0.03133141 | 0.3100114 | 0.101065 | 0.919498586 | 0.959611634 | RPS19BP1 | protein_coding | 54.82 | 0.000560973 | 0.000560973 | ENSG00000187051 | n.s. |
| ENSG00000084090 | 4403.755747 | -0.02877348 | 0.2638898 | -0.109036 | 0.913173967 | 0.956270613 | STARD7 | protein_coding | 42.97 | -0.000558758 | 0.000558758 | ENSG00000084090 | n.s. |
| ENSG00000088325 | 2630.231716 | 0.028524527 | 0.2589029 | 0.110175 | 0.912270883 | 0.955937947 | TPX2 | protein_coding | 41.52 | 0.000558234 | 0.000558234 | ENSG00000088325 | n.s. |

| | | | | | | | | | | | | | |
|---|---|---|---|---|---|---|---|---|---|---|---|---|---|
| ENSG00000117362 | 1453.597021 | -0.028291034 | 0.2589892 | -0.109236 | 0.913015049 | 0.956213971 | APH1A | protein_coding | 53.83 | -0.000550117 | 0.000550117 | ENSG00000117362 | n.s. |
| ENSG00000099899 | 440.1730902 | -0.028245515 | 0.259518 | -0.108838 | 0.913330699 | 0.956270613 | TRMT2A | protein_coding | 63.27 | -0.000548505 | 0.000548505 | ENSG00000099899 | n.s. |
| ENSG00000166900 | 515.6567783 | -0.028512843 | 0.2670241 | -0.10678 | 0.914963472 | 0.95670047 | STX3 | protein_coding | 44.87 | -0.000548131 | 0.000548131 | ENSG00000166900 | n.s. |
| ENSG00000170348 | 1576.322489 | -0.029358841 | 0.2817043 | -0.104219 | 0.916995828 | 0.958005021 | TMED10 | protein_coding | 42.21 | -0.00054702 | 0.00054702 | ENSG00000170348 | n.s. |
| ENSG00000164818 | 704.6966884 | 0.025420309 | 0.2119742 | 0.119922 | 0.904545155 | 0.951658271 | DNAAF5 | protein_coding | 53.29 | 0.000547019 | 0.000547019 | ENSG00000164818 | n.s. |
| ENSG00000114857 | 5406.584274 | 0.027889117 | 0.254764 | 0.10947 | 0.912829395 | 0.956091452 | NKTR | protein_coding | 37.54 | 0.000543854 | 0.000543854 | ENSG00000114857 | n.s. |
| ENSG00000273559 | 401.6708448 | -0.027603305 | 0.2498287 | -0.110489 | 0.912021639 | 0.955748703 | CWC25 | protein_coding | 47.75 | -0.000542578 | 0.000542578 | ENSG00000273559 | n.s. |
| ENSG00000198721 | 553.6154769 | -0.030584518 | 0.3070317 | -0.099614 | 0.920651146 | 0.960095128 | ECI2 | protein_coding | 41.7 | -0.00054091 | 0.00054091 | ENSG00000198721 | n.s. |
| ENSG00000146083 | 598.1196292 | -0.02577895 | 0.2211466 | -0.11657 | 0.907201204 | 0.953136309 | RNF44 | protein_coding | 60.11 | -0.000537362 | 0.000537362 | ENSG00000146083 | n.s. |
| ENSG00000001167 | 1634.899806 | 0.025812777 | 0.2232261 | 0.115635 | 0.907941706 | 0.953338791 | NFYA | protein_coding | 40 | 0.000535686 | 0.000535686 | ENSG00000001167 | n.s. |
| ENSG00000114354 | 1695.406523 | 0.027846339 | 0.2611738 | 0.10662 | 0.915090503 | 0.95670047 | TFG | protein_coding | 36.38 | 0.000535318 | 0.000535318 | ENSG00000114354 | n.s. |
| ENSG00000130224 | 543.8784426 | -0.028797152 | 0.2779363 | -0.103611 | 0.917478361 | 0.95822153 | LRCH2 | protein_coding | 37.13 | -0.000533729 | 0.000533729 | ENSG00000130224 | n.s. |
| ENSG00000198952 | 2373.594963 | -0.027455839 | 0.2529447 | -0.108545 | 0.91356352 | 0.956270613 | SMG5 | protein_coding | 49.82 | -0.00053317 | 0.00053317 | ENSG00000198952 | n.s. |
| ENSG00000205323 | 1705.72478 | 0.028208849 | 0.2677232 | 0.105366 | 0.916085603 | 0.95741329 | SARNP | protein_coding | 42.48 | 0.000533163 | 0.000533163 | ENSG00000205323 | n.s. |
| ENSG00000119333 | 601.5554528 | -0.02988616 | 0.2980531 | -0.100271 | 0.920128976 | 0.960095128 | WDR34 | protein_coding | 53.82 | -0.000528559 | 0.000528559 | ENSG00000119333 | n.s. |
| ENSG00000063046 | 10534.88526 | 0.028520775 | 0.2761075 | 0.103296 | 0.917728109 | 0.958272671 | EIF4B | protein_coding | 42.8 | 0.000527945 | 0.000527945 | ENSG00000063046 | n.s. |
| ENSG00000104164 | 2466.813744 | -0.028455537 | 0.275495 | -0.103289 | 0.917733807 | 0.958272671 | BLOC1S6 | protein_coding | 39.68 | -0.000526738 | 0.000526738 | ENSG00000104164 | n.s. |
| ENSG00000124299 | 420.9538167 | 0.029127531 | 0.2877418 | 0.101228 | 0.919369465 | 0.959611634 | PEPD | protein_coding | 53.89 | 0.000521514 | 0.000521514 | ENSG00000124299 | n.s. |
| ENSG00000119314 | 897.3001634 | 0.026987548 | 0.2515135 | 0.107301 | 0.914550529 | 0.95670047 | PTBP3 | protein_coding | 37.7 | 0.000518809 | 0.000518809 | ENSG00000119314 | n.s. |
| ENSG00000173276 | 188.1501397 | -0.025933362 | 0.2319874 | -0.111788 | 0.910991664 | 0.955092628 | ZBTB21 | protein_coding | 42.8 | -0.000517487 | 0.000517487 | ENSG00000173276 | n.s. |
| ENSG00000144741 | 309.4769463 | -0.028766865 | 0.2827626 | -0.101735 | 0.918966998 | 0.959461067 | SLC25A26 | protein_coding | 40.15 | -0.000517017 | 0.000517017 | ENSG00000144741 | n.s. |
| ENSG00000107020 | 322.0601069 | 0.029904102 | 0.3084036 | 0.096964 | 0.922754843 | 0.961071477 | PLGRKT | protein_coding | 39.88 | 0.000515676 | 0.000515676 | ENSG00000107020 | n.s. |
| ENSG00000137804 | 3657.566657 | 0.029042506 | 0.2931022 | 0.099087 | 0.921069498 | 0.960134566 | NUSAP1 | protein_coding | 44.52 | 0.00051312 | 0.00051312 | ENSG00000137804 | n.s. |
| ENSG00000100284 | 207.9655638 | 0.027538653 | 0.2660247 | 0.103519 | 0.917550948 | 0.958225462 | TOM1 | protein_coding | 53.51 | 0.000510354 | 0.000510354 | ENSG00000100284 | n.s. |
| ENSG00000112031 | 1088.914126 | 0.026211126 | 0.2415142 | 0.108528 | 0.913576623 | 0.956270613 | MTRF1L | protein_coding | 37.68 | 0.000508999 | 0.000508999 | ENSG00000112031 | n.s. |
| ENSG00000228423 | 15.93012855 | 0.031635536 | 0.3496654 | 0.090474 | 0.927910752 | 0.963695464 | | lincRNA | 44.71 | 0.000508073 | 0.000508073 | ENSG00000228423 | n.s. |
| ENSG00000265683 | 110.714378 | 0.026406065 | 0.2457809 | 0.107437 | 0.914441975 | 0.95670047 | | processed_pseudogene | 41.68 | 0.000507631 | 0.000507631 | ENSG00000265683 | n.s. |
| ENSG00000229492 | 76.17242932 | -0.028571325 | 0.2863428 | -0.09978 | 0.920518866 | 0.960095128 | | processed_pseudogene | 56.54 | -0.000505305 | 0.000505305 | ENSG00000229492 | n.s. |
| ENSG00000131100 | 972.7551742 | -0.029907026 | 0.3128499 | -0.095595 | 0.923841882 | 0.961916375 | ATP6V1E1 | protein_coding | 45.05 | -0.000504313 | 0.000504313 | ENSG00000131100 | n.s. |
| ENSG00000178531 | 148.8857502 | 0.025876807 | 0.2381507 | 0.108657 | 0.913474324 | 0.956270613 | CTXN1 | protein_coding | 74.79 | 0.000502507 | 0.000502507 | ENSG00000178531 | n.s. |
| ENSG00000207546 | 9.590270463 | -0.035153223 | 0.4331864 | -0.08115 | 0.935322391 | 0.967927282 | MIR548C | miRNA | 34.02 | -0.000497674 | 0.000497674 | ENSG00000207546 | n.s. |
| ENSG00000222937 | 45.75680524 | -0.02880408 | 0.298076 | -0.096633 | 0.923017588 | 0.961194755 | SNORD63B | snoRNA | 42.86 | -0.000495102 | 0.000495102 | ENSG00000222937 | n.s. |

| | | | | | | | | | | | | | |
|---|---|---|---|---|---|---|---|---|---|---|---|---|---|
| ENSG00000110880 | 2445.492093 | 0.025896145 | 0.2440404 | 0.106114 | 0.915491777 | 0.957008184 | CORO1C | protein_coding | 41.35 | 0.000494211 | 0.000494211 | ENSG00000110880 | n.s. |
| ENSG00000234841 | 181.1129524 | -0.026016885 | 0.2468288 | -0.105405 | 0.9160548 | 0.95741329 | | processed_pseudogene | 41.2 | -0.000491733 | 0.000491733 | ENSG00000234841 | n.s. |
| ENSG00000196405 | 2160.861427 | -0.025198806 | 0.231815 | -0.108702 | 0.913438665 | 0.956270613 | EVL | protein_coding | 45.48 | -0.00048934 | 0.00048934 | ENSG00000196405 | n.s. |
| ENSG00000264354 | 6.649447744 | -0.036191403 | 0.4701951 | -0.076971 | 0.938646585 | 0.969369489 | MIR3134 | miRNA | 33.78 | -0.00048897 | 0.00048897 | ENSG00000264354 | n.s. |
| ENSG00000252835 | 497.7312485 | -0.024680012 | 0.2230625 | -0.110642 | 0.911900495 | 0.955693684 | SCARNA21 | scaRNA | 64.03 | -0.000485734 | 0.000485734 | ENSG00000252835 | n.s. |
| ENSG00000089902 | 801.3365889 | 0.025223005 | 0.2360685 | 0.106846 | 0.914911063 | 0.95670047 | RCOR1 | protein_coding | 43.48 | 0.000484887 | 0.000484887 | ENSG00000089902 | n.s. |
| ENSG00000101782 | 943.674195 | -0.026888558 | 0.2646659 | -0.101594 | 0.919078665 | 0.959461067 | RIOK3 | protein_coding | 41.88 | -0.000483258 | 0.000483258 | ENSG00000101782 | n.s. |
| ENSG00000109805 | 4943.458093 | 0.025031234 | 0.2345865 | 0.106704 | 0.915024093 | 0.95670047 | NCAPG | protein_coding | 35.41 | 0.000481201 | 0.000481201 | ENSG00000109805 | n.s. |
| ENSG00000266916 | 112.643247 | -0.025739926 | 0.2466718 | -0.104349 | 0.916892484 | 0.958005021 | ZNF793-AS1 | antisense | 44.46 | -0.000479592 | 0.000479592 | ENSG00000266916 | n.s. |
| ENSG00000136143 | 1050.982462 | -0.027140237 | 0.2764018 | -0.098191 | 0.921780441 | 0.960486468 | SUCLA2 | protein_coding | 38.08 | -0.000475192 | 0.000475192 | ENSG00000136143 | n.s. |
| ENSG00000156256 | 1918.532459 | -0.026857089 | 0.2696242 | -0.099609 | 0.920654465 | 0.960095128 | USP16 | protein_coding | 38.76 | -0.000474987 | 0.000474987 | ENSG00000156256 | n.s. |
| ENSG00000186298 | 5053.759151 | 0.026766202 | 0.2689359 | 0.099526 | 0.920720401 | 0.960095128 | PPP1CC | protein_coding | 41.52 | 0.00047338 | 0.00047338 | ENSG00000186298 | n.s. |
| ENSG00000172354 | 689.2440739 | 0.026770764 | 0.2707397 | 0.09888 | 0.921233472 | 0.960134566 | GNB2 | protein_coding | 65.91 | 0.000472983 | 0.000472983 | ENSG00000172354 | n.s. |
| ENSG00000131507 | 1144.183504 | 0.02824915 | 0.2978338 | 0.094849 | 0.924434995 | 0.962188236 | NDFIP1 | protein_coding | 39.29 | 0.00047289 | 0.00047289 | ENSG00000131507 | n.s. |
| ENSG00000176142 | 501.1394055 | -0.02490456 | 0.2365677 | -0.105275 | 0.916157962 | 0.957417046 | TMEM39A | protein_coding | 39.09 | -0.000470667 | 0.000470667 | ENSG00000176142 | n.s. |
| ENSG00000127922 | 1676.131119 | -0.026498424 | 0.2680109 | -0.098871 | 0.921240944 | 0.960134566 | SEM1 | protein_coding | 37.66 | -0.000468171 | 0.000468171 | ENSG00000127922 | n.s. |
| ENSG00000179115 | 834.2657524 | -0.029532444 | 0.3303343 | -0.089402 | 0.928762672 | 0.964292519 | FARSA | protein_coding | 53.91 | -0.000466353 | 0.000466353 | ENSG00000179115 | n.s. |
| ENSG00000246174 | 80.44542206 | 0.024872682 | 0.2385763 | 0.104255 | 0.916967274 | 0.958005021 | KCTD21-AS1 | antisense | 47.55 | 0.000463433 | 0.000463433 | ENSG00000246174 | n.s. |
| ENSG00000137161 | 538.3287152 | 0.026546359 | 0.2722583 | 0.097504 | 0.922325908 | 0.960787344 | CNPY3 | protein_coding | 50.57 | 0.000461183 | 0.000461183 | ENSG00000137161 | n.s. |
| ENSG00000105928 | 168.7266169 | -0.027270268 | 0.2881548 | -0.094638 | 0.924602724 | 0.962197753 | DFNA5 | protein_coding | 45.73 | -0.000456386 | 0.000456386 | ENSG00000105928 | n.s. |
| ENSG00000144560 | 763.1804267 | -0.026062548 | 0.2670586 | -0.097591 | 0.92225699 | 0.960787344 | VGLL4 | protein_coding | 44.93 | -0.000452778 | 0.000452778 | ENSG00000144560 | n.s. |
| ENSG00000138175 | 447.5470129 | -0.027963547 | 0.3061923 | -0.091327 | 0.927232972 | 0.963527976 | ARL3 | protein_coding | 44.57 | -0.000451211 | 0.000451211 | ENSG00000138175 | n.s. |
| ENSG00000105379 | 295.9144914 | -0.027247209 | 0.2906249 | -0.093754 | 0.925304679 | 0.962721954 | ETFB | protein_coding | 51.5 | -0.000449555 | 0.000449555 | ENSG00000105379 | n.s. |
| ENSG00000267374 | 144.0030149 | -0.025986926 | 0.2682679 | -0.096869 | 0.922830179 | 0.961071477 | | lincRNA | 37.1 | -0.000448127 | 0.000448127 | ENSG00000267374 | n.s. |
| ENSG00000118513 | 183.1759204 | 0.026641475 | 0.2808369 | 0.094865 | 0.924422389 | 0.962188236 | MYB | protein_coding | 39.74 | 0.000445977 | 0.000445977 | ENSG00000118513 | n.s. |
| ENSG00000102393 | 256.369771 | -0.026597861 | 0.2804999 | -0.094823 | 0.924455364 | 0.962188236 | GLA | protein_coding | 43.7 | -0.000445247 | 0.000445247 | ENSG00000102393 | n.s. |
| ENSG00000163541 | 1361.002208 | 0.027533252 | 0.3011522 | 0.091426 | 0.927153816 | 0.963527976 | SUCLG1 | protein_coding | 39.25 | 0.000444268 | 0.000444268 | ENSG00000163541 | n.s. |
| ENSG00000196678 | 182.4752655 | -0.023035563 | 0.21567 | -0.106809 | 0.914940246 | 0.95670047 | ERI2 | protein_coding | 40.84 | -0.000442836 | 0.000442836 | ENSG00000196678 | n.s. |
| ENSG00000182979 | 1117.748547 | -0.02494229 | 0.2505204 | -0.099562 | 0.920692142 | 0.960095128 | MTA1 | protein_coding | 59.42 | -0.000441123 | 0.000441123 | ENSG00000182979 | n.s. |
| ENSG00000096717 | 1171.556848 | 0.024528943 | 0.2412933 | 0.101656 | 0.919029621 | 0.959461067 | SIRT1 | protein_coding | 40.87 | 0.00044085 | 0.00044085 | ENSG00000096717 | n.s. |
| ENSG00000006712 | 534.0013709 | -0.025254765 | 0.2586616 | -0.097636 | 0.922221092 | 0.960787344 | PAF1 | protein_coding | 52.73 | -0.000438744 | 0.000438744 | ENSG00000006712 | n.s. |
| ENSG00000188343 | 995.2558334 | 0.024449821 | 0.2467493 | 0.099088 | 0.921068629 | 0.960134566 | FAM92A | protein_coding | 38.54 | 0.000431977 | 0.000431977 | ENSG00000188343 | n.s. |

| | | | | | | | | | | | | | |
|---|---|---|---|---|---|---|---|---|---|---|---|---|---|
| ENSG00000156959 | 455.3270166 | -0.024579456 | 0.2503387 | -0.098185 | 0.921785548 | 0.960486468 | LHFPL4 | protein_coding | 46.67 | -0.000430356 | 0.000430356 | ENSG00000156959 | n.s. |
| ENSG00000132792 | 906.4700152 | -0.024356407 | 0.245378 | -0.099261 | 0.920931232 | 0.960134566 | CTNNBL1 | protein_coding | 42.04 | -0.000430326 | 0.000430326 | ENSG00000132792 | n.s. |
| ENSG00000132361 | 820.6122679 | -0.024124234 | 0.2421112 | -0.099641 | 0.920629252 | 0.960095128 | CLUH | protein_coding | 62.54 | -0.000426655 | 0.000426655 | ENSG00000132361 | n.s. |
| ENSG00000117308 | 76.46785894 | -0.028088483 | 0.3227661 | -0.087024 | 0.930652232 | 0.965634737 | GALE | protein_coding | 60.1 | -0.000426583 | 0.000426583 | ENSG00000117308 | n.s. |
| ENSG00000213593 | 1371.857262 | -0.027645804 | 0.3137205 | -0.088122 | 0.929779395 | 0.965100848 | TMX2 | protein_coding | 45.19 | -0.0004265 | 0.0004265 | ENSG00000213593 | n.s. |
| ENSG00000138381 | 2300.192512 | -0.025188333 | 0.2653327 | -0.094931 | 0.924369524 | 0.962188236 | ASNSD1 | protein_coding | 37.75 | -0.000421652 | 0.000421652 | ENSG00000138381 | n.s. |
| ENSG00000172046 | 939.5166798 | 0.02499882 | 0.26153 | 0.095587 | 0.923848723 | 0.961916375 | USP19 | protein_coding | 56.09 | 0.000421547 | 0.000421547 | ENSG00000172046 | n.s. |
| ENSG00000108384 | 2116.831119 | -0.022554784 | 0.2157599 | -0.104536 | 0.916743616 | 0.957957173 | RAD51C | protein_coding | 41.71 | -0.000420735 | 0.000420735 | ENSG00000108384 | n.s. |
| ENSG00000169228 | 289.4476124 | -0.024799813 | 0.2602235 | -0.095302 | 0.924074975 | 0.962080003 | RAB24 | protein_coding | 57.83 | -0.000416359 | 0.000416359 | ENSG00000169228 | n.s. |
| ENSG00000117139 | 1995.431289 | -0.023486064 | 0.237325 | -0.098962 | 0.921168769 | 0.960134566 | KDM5B | protein_coding | 41.09 | -0.000414949 | 0.000414949 | ENSG00000117139 | n.s. |
| ENSG00000083750 | 334.0228975 | 0.0255364 | 0.2770082 | 0.092186 | 0.92654992 | 0.963298327 | RRAGB | protein_coding | 40.87 | 0.000414691 | 0.000414691 | ENSG00000083750 | n.s. |
| ENSG00000171206 | 276.6451325 | -0.024769299 | 0.2616309 | -0.094673 | 0.924574817 | 0.962197753 | TRIM8 | protein_coding | 57.48 | -0.000414531 | 0.000414531 | ENSG00000171206 | n.s. |
| ENSG00000170579 | 63.07382908 | -0.027294277 | 0.3156744 | -0.086463 | 0.931098073 | 0.965709064 | DLGAP1 | protein_coding | 39.67 | -0.000413609 | 0.000413609 | ENSG00000170579 | n.s. |
| ENSG00000177084 | 3731.715334 | 0.023101507 | 0.2305439 | 0.100204 | 0.92018208 | 0.960095128 | POLE | protein_coding | 50.84 | 0.000408567 | 0.000408567 | ENSG00000177084 | n.s. |
| ENSG00000198089 | 22.07699316 | 0.029071498 | 0.3662907 | 0.079367 | 0.936740497 | 0.968465636 | SFI1 | protein_coding | 46.58 | 0.000404553 | 0.000404553 | ENSG00000198089 | n.s. |
| ENSG00000168389 | 146.0084457 | 0.024400308 | 0.2623252 | 0.093016 | 0.925891243 | 0.96289093 | MFSD2A | protein_coding | 53.7 | 0.000400724 | 0.000400724 | ENSG00000168389 | n.s. |
| ENSG00000261886 | 11.99808259 | -0.028759597 | 0.3623975 | -0.079359 | 0.936746873 | 0.968465636 | | lincRNA | 42.81 | -0.000400212 | 0.000400212 | ENSG00000261886 | n.s. |
| ENSG00000262049 | 403.6864279 | -0.021334216 | 0.2054311 | -0.103851 | 0.917287626 | 0.958173161 | | antisense | 61.96 | -0.000395878 | 0.000395878 | ENSG00000262049 | n.s. |
| ENSG00000114503 | 3072.304043 | 0.024416616 | 0.2691945 | 0.090703 | 0.927728982 | 0.963578561 | NCBP2 | protein_coding | 44 | 0.000393422 | 0.000393422 | ENSG00000114503 | n.s. |
| ENSG00000131778 | 1974.551731 | -0.023708511 | 0.2529352 | -0.093734 | 0.925320818 | 0.962721954 | CHD1L | protein_coding | 40.27 | -0.00039117 | 0.00039117 | ENSG00000131778 | n.s. |
| ENSG00000156925 | 886.8767021 | 0.022113657 | 0.2210225 | 0.100052 | 0.92030334 | 0.960095128 | ZIC3 | protein_coding | 48.02 | 0.000391096 | 0.000391096 | ENSG00000156925 | n.s. |
| ENSG00000250197 | 50.95131846 | -0.026751598 | 0.3209112 | -0.083361 | 0.933564209 | 0.966898347 | HMGN1P15 | processed_pseudogene | 43.05 | -0.000391086 | 0.000391086 | ENSG00000250197 | n.s. |
| ENSG00000167272 | 514.0398907 | 0.024248722 | 0.2667182 | 0.090915 | 0.92756003 | 0.96354684 | POP5 | protein_coding | 48.65 | 0.000391063 | 0.000391063 | ENSG00000167272 | n.s. |
| ENSG00000201223 | 98.37175891 | 0.022454894 | 0.2303691 | 0.097474 | 0.92235033 | 0.960787344 | RNU6-1305P | snRNA | 46.15 | 0.000390103 | 0.000390103 | ENSG00000201223 | n.s. |
| ENSG00000205309 | 209.6178257 | 0.023634984 | 0.2537719 | 0.093135 | 0.925796517 | 0.962864339 | NT5M | protein_coding | 49.17 | 0.000388439 | 0.000388439 | ENSG00000205309 | n.s. |
| ENSG00000204682 | 244.1244456 | -0.021583373 | 0.2134287 | -0.101127 | 0.919449754 | 0.959611634 | CASC10 | protein_coding | 57.39 | -0.000386439 | 0.000386439 | ENSG00000204682 | n.s. |
| ENSG00000203950 | 216.5350636 | -0.023327507 | 0.2480749 | -0.094034 | 0.925082043 | 0.96262461 | RTL8A | protein_coding | 60.16 | -0.000385908 | 0.000385908 | ENSG00000203950 | n.s. |
| ENSG00000161999 | 359.2008625 | -0.023883751 | 0.2609197 | -0.091537 | 0.927066063 | 0.963527976 | JMJD8 | protein_coding | 65.22 | -0.00038538 | 0.00038538 | ENSG00000161999 | n.s. |
| ENSG00000242779 | 74.01171629 | -0.024051928 | 0.2664097 | -0.090282 | 0.92806333 | 0.963782035 | ZNF702P | transcribed_processed_pseudog | 45.63 | -0.00038534 | 0.00038534 | ENSG00000242779 | n.s. |
| ENSG00000127337 | 1537.749749 | 0.027078225 | 0.3312773 | 0.081739 | 0.934854381 | 0.967802628 | YEATS4 | protein_coding | 39.36 | 0.000384868 | 0.000384868 | ENSG00000127337 | n.s. |
| ENSG00000133275 | 1008.170242 | 0.025102892 | 0.2870372 | 0.087455 | 0.930309693 | 0.965466746 | CSNK1G2 | protein_coding | 61.03 | 0.000383137 | 0.000383137 | ENSG00000133275 | n.s. |
| ENSG00000258732 | 488.0522795 | -0.023660445 | 0.2582196 | -0.091629 | 0.926992671 | 0.963527976 | | unprocessed_pseudogene | 40.36 | -0.000381777 | 0.000381777 | ENSG00000258732 | n.s. |

| | | | | | | | | | | | | | |
|---|---|---|---|---|---|---|---|---|---|---|---|---|---|
| ENSG00000179933 | 566.8504728 | -0.025946238 | 0.3092101 | -0.083911 | 0.933126912 | 0.96673309 | C14orf119 | protein_coding | 47.09 | -0.000381239 | 0.000381239 | ENSG00000179933 | n.s. |
| ENSG00000122694 | 283.5802577 | -0.025764053 | 0.30316 | -0.084985 | 0.93227332 | 0.966567982 | GLIPR2 | protein_coding | 51.06 | -0.000380473 | 0.000380473 | ENSG00000122694 | n.s. |
| ENSG00000116874 | 435.1428507 | -0.023526919 | 0.2587009 | -0.090943 | 0.927538234 | 0.96354684 | WARS2 | protein_coding | 38.1 | -0.000379423 | 0.000379423 | ENSG00000116874 | n.s. |
| ENSG00000102003 | 933.2263614 | 0.024979464 | 0.2867635 | 0.087108 | 0.93058547 | 0.965634737 | SYP | protein_coding | 52.69 | 0.000379366 | 0.000379366 | ENSG00000102003 | n.s. |
| ENSG00000272994 | 665.0299948 | -0.021374666 | 0.2146772 | -0.099567 | 0.920688446 | 0.960095128 | | lincRNA | 41.14 | -0.000378027 | 0.000378027 | ENSG00000272994 | n.s. |
| ENSG00000167522 | 108.4669777 | -0.023306683 | 0.256187 | -0.090975 | 0.927512245 | 0.96354684 | ANKRD11 | protein_coding | 51.19 | -0.000375871 | 0.000375871 | ENSG00000167522 | n.s. |
| ENSG00000276102 | 11.86176776 | -0.029728148 | 0.4176819 | -0.071174 | 0.943259163 | 0.971734912 | MIR6747 | miRNA | 52.46 | -0.000370181 | 0.000370181 | ENSG00000276102 | n.s. |
| ENSG00000122176 | 253.9203384 | -0.023858837 | 0.2693931 | -0.088565 | 0.929427515 | 0.964910841 | FMOD | protein_coding | 50.17 | -0.000370118 | 0.000370118 | ENSG00000122176 | n.s. |
| ENSG00000198492 | 1786.399399 | -0.022407001 | 0.2399839 | -0.093369 | 0.925610595 | 0.962814817 | YTHDF2 | protein_coding | 43.57 | -0.000368757 | 0.000368757 | ENSG00000198492 | n.s. |
| ENSG00000132286 | 678.6838972 | -0.022171066 | 0.2367307 | -0.093655 | 0.925383029 | 0.962721954 | TIMM10B | protein_coding | 47.6 | -0.000365803 | 0.000365803 | ENSG00000132286 | n.s. |
| ENSG00000182093 | 391.1945002 | 0.026249841 | 0.3302628 | 0.079482 | 0.936649508 | 0.968465636 | WRB | protein_coding | 44.24 | 0.000365287 | 0.000365287 | ENSG00000182093 | n.s. |
| ENSG00000269176 | 38.24925468 | 0.024758227 | 0.2943942 | 0.084099 | 0.9329778 | 0.966650537 | | antisense | 52.69 | 0.000364701 | 0.000364701 | ENSG00000269176 | n.s. |
| ENSG00000099783 | 4909.97838 | -0.022456987 | 0.2440979 | -0.092 | 0.92669811 | 0.963298327 | HNRNPM | protein_coding | 46.68 | -0.000364683 | 0.000364683 | ENSG00000099783 | n.s. |
| ENSG00000217555 | 129.6314934 | -0.026317875 | 0.334267 | -0.078733 | 0.937244916 | 0.968620915 | CKLF | protein_coding | 41.69 | -0.000364401 | 0.000364401 | ENSG00000217555 | n.s. |
| ENSG00000114107 | 746.1910862 | -0.024580948 | 0.2900849 | -0.084737 | 0.932470423 | 0.966650537 | CEP70 | protein_coding | 39.55 | -0.00036209 | 0.00036209 | ENSG00000114107 | n.s. |
| ENSG00000141873 | 283.2182998 | -0.024532215 | 0.2914305 | -0.084179 | 0.932914415 | 0.966650537 | SLC39A3 | protein_coding | 57.96 | -0.000361372 | 0.000361372 | ENSG00000141873 | n.s. |
| ENSG00000008083 | 1009.918555 | -0.021965051 | 0.2356078 | -0.093227 | 0.925723088 | 0.962859895 | JARID2 | protein_coding | 43.92 | -0.000361037 | 0.000361037 | ENSG00000008083 | n.s. |
| ENSG00000111670 | 465.8755607 | -0.021750427 | 0.2328375 | -0.093415 | 0.925574169 | 0.962814817 | GNPTAB | protein_coding | 40.94 | -0.000357952 | 0.000357952 | ENSG00000111670 | n.s. |
| ENSG00000159079 | 165.563774 | 0.025139505 | 0.3086052 | 0.081462 | 0.93507478 | 0.967896477 | C21orf59 | protein_coding | 47.5 | 0.000356254 | 0.000356254 | ENSG00000159079 | n.s. |
| ENSG00000196743 | 205.0477555 | -0.024473954 | 0.2948592 | -0.083002 | 0.933849808 | 0.96705027 | GM2A | protein_coding | 47.41 | -0.000356119 | 0.000356119 | ENSG00000196743 | n.s. |
| ENSG00000002822 | 533.1199355 | -0.021826247 | 0.2358263 | -0.092552 | 0.926259315 | 0.963129831 | MAD1L1 | protein_coding | 55.65 | -0.000356099 | 0.000356099 | ENSG00000002822 | n.s. |
| ENSG00000239119 | 29.62832999 | -0.025119631 | 0.3105223 | -0.080895 | 0.935525647 | 0.967993727 | RNU7-119P | snRNA | 36.51 | -0.000354877 | 0.000354877 | ENSG00000239119 | n.s. |
| ENSG00000240053 | 240.6098537 | -0.021966211 | 0.2406297 | -0.091286 | 0.927265054 | 0.963527976 | LY6G5B | protein_coding | 52.19 | -0.00035444 | 0.00035444 | ENSG00000240053 | n.s. |
| ENSG00000124383 | 2573.048591 | -0.021964166 | 0.2412765 | -0.091033 | 0.927466224 | 0.96354684 | MPHOSPH10 | protein_coding | 40.5 | -0.00035422 | 0.00035422 | ENSG00000124383 | n.s. |
| ENSG00000121068 | 3923.456624 | -0.021945051 | 0.2419253 | -0.09071 | 0.927723014 | 0.963578561 | TBX2 | protein_coding | 61.32 | -0.000353598 | 0.000353598 | ENSG00000121068 | n.s. |
| ENSG00000166450 | 228.0100314 | -0.021587443 | 0.2329664 | -0.092663 | 0.926171031 | 0.963109961 | PRTG | protein_coding | 38.58 | -0.000352396 | 0.000352396 | ENSG00000166450 | n.s. |
| ENSG00000101365 | 1148.99873 | 0.022814362 | 0.2594255 | 0.087942 | 0.929922874 | 0.965137247 | IDH3B | protein_coding | 51.38 | 0.00035159 | 0.00035159 | ENSG00000101365 | n.s. |
| ENSG00000177239 | 282.551167 | 0.023201222 | 0.2682352 | 0.086496 | 0.93107229 | 0.965709064 | MAN1B1 | protein_coding | 54.87 | 0.000351584 | 0.000351584 | ENSG00000177239 | n.s. |
| ENSG00000166073 | 264.2967288 | -0.02297747 | 0.2646708 | -0.086815 | 0.930818344 | 0.965634737 | GPR176 | protein_coding | 40.8 | -0.000348962 | 0.000348962 | ENSG00000166073 | n.s. |
| ENSG00000006194 | 1535.228569 | -0.021245893 | 0.230723 | -0.092084 | 0.92663131 | 0.963298327 | ZNF263 | protein_coding | 47.09 | -0.000345016 | 0.000345016 | ENSG00000006194 | n.s. |
| ENSG00000167674 | 927.9724315 | -0.023192991 | 0.2749077 | -0.084366 | 0.932765077 | 0.966650537 | | protein_coding | 57.07 | -0.000341644 | 0.000341644 | ENSG00000167674 | n.s. |
| ENSG00000221823 | 1297.487357 | -0.023081763 | 0.2739411 | -0.084258 | 0.932851204 | 0.966650537 | PPP3R1 | protein_coding | 37.41 | -0.000340006 | 0.000340006 | ENSG00000221823 | n.s. |

| | | | | | | | | | | | | | |
|---|---|---|---|---|---|---|---|---|---|---|---|---|---|
| ENSG00000114737 | 187.0134647 | -0.020869096 | 0.2268169 | -0.092009 | 0.926691251 | 0.963298327 | CISH | protein_coding | 57 | -0.000338897 | 0.000338897 | ENSG00000114737 | n.s. |
| ENSG00000163527 | 2795.417666 | 0.022846061 | 0.2702511 | 0.084536 | 0.932629949 | 0.966650537 | STT3B | protein_coding | 35.96 | 0.000336534 | 0.000336534 | ENSG00000163527 | n.s. |
| ENSG00000243317 | 1239.52994 | -0.024258207 | 0.3109947 | -0.078002 | 0.937826464 | 0.968790463 | C7orf73 | protein_coding | 43.1 | -0.000334039 | 0.000334039 | ENSG00000243317 | n.s. |
| ENSG00000280238 | 14.6591945 | -0.027255335 | 0.390385 | -0.069817 | 0.944339675 | 0.972200628 | | TEC | 59.34 | -0.000333717 | 0.000333717 | ENSG00000280238 | n.s. |
| ENSG00000139826 | 953.8329024 | -0.023144283 | 0.2809187 | -0.082388 | 0.934338317 | 0.96741224 | ABHD13 | protein_coding | 35.43 | -0.00033301 | 0.00033301 | ENSG00000139826 | n.s. |
| ENSG00000197457 | 1460.684736 | 0.022508182 | 0.2674502 | 0.084158 | 0.932930491 | 0.966650537 | STMN3 | protein_coding | 59.38 | 0.000331557 | 0.000331557 | ENSG00000197457 | n.s. |
| ENSG00000172687 | 1128.462918 | 0.02141361 | 0.2426119 | 0.088263 | 0.929667797 | 0.965088334 | ZNF738 | protein_coding | 40.08 | 0.000330475 | 0.000330475 | ENSG00000172687 | n.s. |
| ENSG00000221514 | 24.04870694 | 0.023985116 | 0.3120633 | 0.07686 | 0.938735101 | 0.969369489 | SNORD111B | snoRNA | 37.5 | 0.000324055 | 0.000324055 | ENSG00000221514 | n.s. |
| ENSG00000234171 | 355.3846225 | -0.021358637 | 0.246859 | -0.086522 | 0.931051787 | 0.965709064 | RNASEH1-AS1 | antisense | 47.42 | -0.000323662 | 0.000323662 | ENSG00000234171 | n.s. |
| ENSG00000100241 | 1341.795001 | -0.021179359 | 0.2438333 | -0.08686 | 0.930782806 | 0.965634737 | SBF1 | protein_coding | 60.67 | -0.000321653 | 0.000321653 | ENSG00000100241 | n.s. |
| ENSG00000159720 | 439.5767553 | 0.023018698 | 0.288994 | 0.079651 | 0.936514721 | 0.968465636 | ATP6V0D1 | protein_coding | 53.78 | 0.000320323 | 0.000320323 | ENSG00000159720 | n.s. |
| ENSG00000199933 | 31.08856714 | 0.025512818 | 0.3549696 | 0.071873 | 0.942702778 | 0.97158315 | RNY1P16 | misc_RNA | 45.05 | 0.000319421 | 0.000319421 | ENSG00000199933 | n.s. |
| ENSG00000103855 | 521.9577458 | -0.021099921 | 0.2446756 | -0.086236 | 0.931278574 | 0.965824327 | CD276 | protein_coding | 52.51 | -0.000318648 | 0.000318648 | ENSG00000103855 | n.s. |
| ENSG00000204569 | 2370.204419 | -0.018440439 | 0.1903602 | -0.096871 | 0.922828591 | 0.961071477 | PPP1R10 | protein_coding | 46.34 | -0.000317993 | 0.000317993 | ENSG00000204569 | n.s. |
| ENSG00000162129 | 1025.959969 | -0.021560761 | 0.2545297 | -0.084708 | 0.932493354 | 0.966650537 | CLPB | protein_coding | 45.92 | -0.000317601 | 0.000317601 | ENSG00000162129 | n.s. |
| ENSG00000134294 | 10359.43669 | 0.020854496 | 0.240082 | 0.086864 | 0.930779569 | 0.965634737 | SLC38A2 | protein_coding | 38.54 | 0.00031672 | 0.00031672 | ENSG00000134294 | n.s. |
| ENSG00000119048 | 943.8241756 | -0.022550339 | 0.2841188 | -0.079369 | 0.936738805 | 0.968465636 | UBE2B | protein_coding | 41.55 | -0.000313806 | 0.000313806 | ENSG00000119048 | n.s. |
| ENSG00000131043 | 759.4171806 | 0.021282869 | 0.2542797 | 0.083699 | 0.933296034 | 0.966764429 | AAR2 | protein_coding | 47.41 | 0.000312418 | 0.000312418 | ENSG00000131043 | n.s. |
| ENSG00000140854 | 664.7290055 | -0.021926411 | 0.2692008 | -0.08145 | 0.935084071 | 0.967896477 | KATNB1 | protein_coding | 56.59 | -0.000310721 | 0.000310721 | ENSG00000140854 | n.s. |
| ENSG00000101421 | 286.1904975 | -0.022562803 | 0.2883275 | -0.078254 | 0.937625935 | 0.968790463 | CHMP4B | protein_coding | 46.43 | -0.000310693 | 0.000310693 | ENSG00000101421 | n.s. |
| ENSG00000133193 | 1008.061027 | -0.021775625 | 0.2686394 | -0.081059 | 0.935395076 | 0.967930557 | FAM104A | protein_coding | 43.89 | -0.000308251 | 0.000308251 | ENSG00000133193 | n.s. |
| ENSG00000047315 | 4940.805738 | -0.022141729 | 0.2797019 | -0.079162 | 0.93690386 | 0.968484107 | POLR2B | protein_coding | 40.19 | -0.000307936 | 0.000307936 | ENSG00000047315 | n.s. |
| ENSG00000177383 | 1819.919421 | -0.022137893 | 0.2793284 | -0.079254 | 0.936830602 | 0.968480285 | MAGEF1 | protein_coding | 52.87 | -0.000307921 | 0.000307921 | ENSG00000177383 | n.s. |
| ENSG00000265566 | 13.48790515 | -0.025291063 | 0.3656573 | -0.069166 | 0.94485747 | 0.9723742 | RN7SL605P | misc_RNA | 55.16 | -0.000307706 | 0.000307706 | ENSG00000265566 | n.s. |
| ENSG00000129993 | 21.76741903 | 0.023514811 | 0.3173205 | 0.074104 | 0.940927396 | 0.970409788 | CBFA2T3 | protein_coding | 60.67 | 0.000306747 | 0.000306747 | ENSG00000129993 | n.s. |
| ENSG00000258904 | 60.76739858 | -0.021033786 | 0.2529393 | -0.083157 | 0.933726353 | 0.966994347 | | sense_intronic | 50.52 | -0.00030659 | 0.00030659 | ENSG00000258904 | n.s. |
| ENSG00000188785 | 470.3029161 | -0.021221423 | 0.2570566 | -0.082555 | 0.934205035 | 0.967346177 | ZNF548 | protein_coding | 45.35 | -0.000305972 | 0.000305972 | ENSG00000188785 | n.s. |
| ENSG00000203778 | 316.2423146 | -0.023584944 | 0.3203593 | -0.07362 | 0.94131253 | 0.970635872 | FAM229B | protein_coding | 36.12 | -0.000305276 | 0.000305276 | ENSG00000203778 | n.s. |
| ENSG00000185504 | 601.8834722 | 0.020761336 | 0.2477084 | 0.083814 | 0.933204633 | 0.966741674 | FAAP100 | protein_coding | 61.6 | 0.000304975 | 0.000304975 | ENSG00000185504 | n.s. |
| ENSG00000197153 | 1072.863325 | -0.019693928 | 0.2236086 | -0.088073 | 0.929818486 | 0.965100848 | HIST1H3J | protein_coding | 48.84 | -0.000303824 | 0.000303824 | ENSG00000197153 | n.s. |
| ENSG00000143801 | 401.093105 | -0.022982643 | 0.3065506 | -0.074972 | 0.940237174 | 0.970129107 | PSEN2 | protein_coding | 52.42 | -0.000302692 | 0.000302692 | ENSG00000143801 | n.s. |
| ENSG00000159346 | 1216.244541 | -0.022180446 | 0.2885713 | -0.076863 | 0.938732558 | 0.969369489 | ADIPOR1 | protein_coding | 44.28 | -0.000299672 | 0.000299672 | ENSG00000159346 | n.s. |

| | | | | | | | | | | | | | |
|---|---|---|---|---|---|---|---|---|---|---|---|---|---|
| ENSG00000202125 | 11.79061486 | -0.025470747 | 0.3850578 | -0.066148 | 0.947260115 | 0.973391392 | RNU1-100P | snRNA | 48.48 | -0.000298326 | 0.000298326 | ENSG00000202125 | n.s. |
| ENSG00000146540 | 363.5560675 | 0.022779805 | 0.306997 | 0.074202 | 0.94084961 | 0.970409788 | C7orf50 | protein_coding | 57.69 | 0.000297159 | 0.000297159 | ENSG00000146540 | n.s. |
| ENSG00000111481 | 1376.208496 | 0.022474653 | 0.2985501 | 0.075279 | 0.939992464 | 0.970020388 | COPZ1 | protein_coding | 46.59 | 0.000297096 | 0.000297096 | ENSG00000111481 | n.s. |
| ENSG00000176593 | 257.8435054 | -0.018671367 | 0.2080279 | -0.089754 | 0.928482586 | 0.964145514 | | processed_transcript | 50.45 | -0.00029608 | 0.00029608 | ENSG00000176593 | n.s. |
| ENSG00000113083 | 315.5631647 | -0.02147972 | 0.2752579 | -0.078035 | 0.937800292 | 0.968790463 | LOX | protein_coding | 37.93 | -0.000295779 | 0.000295779 | ENSG00000113083 | n.s. |
| ENSG00000143669 | 33.07267246 | 0.024237214 | 0.3513108 | 0.068991 | 0.944996936 | 0.97244582 | LYST | protein_coding | 37.77 | 0.000294109 | 0.000294109 | ENSG00000143669 | n.s. |
| ENSG00000102409 | 1342.143425 | -0.02115675 | 0.2706885 | -0.078159 | 0.937701558 | 0.968790463 | BEX4 | protein_coding | 52.02 | -0.000291332 | 0.000291332 | ENSG00000102409 | n.s. |
| ENSG00000175768 | 1412.93152 | 0.019774632 | 0.2349003 | 0.084183 | 0.932910856 | 0.966650537 | TOMM5 | protein_coding | 40.98 | 0.00029129 | 0.00029129 | ENSG00000175768 | n.s. |
| ENSG00000242247 | 442.8158285 | -0.021825316 | 0.2879488 | -0.075796 | 0.939581544 | 0.969811983 | ARFGAP3 | protein_coding | 45 | -0.000290549 | 0.000290549 | ENSG00000242247 | n.s. |
| ENSG00000185485 | 418.4823022 | -0.019857244 | 0.2379975 | -0.083435 | 0.933505919 | 0.966898347 | SDHAP1 | transcribed_unprocessed_pseud | 48.24 | -0.000290297 | 0.000290297 | ENSG00000185485 | n.s. |
| ENSG00000154447 | 494.2302907 | 0.019560817 | 0.2296132 | 0.08519 | 0.932110111 | 0.966484672 | SH3RF1 | protein_coding | 39.27 | 0.000289598 | 0.000289598 | ENSG00000154447 | n.s. |
| ENSG00000119185 | 659.1662567 | 0.022636693 | 0.3101255 | 0.072992 | 0.941812453 | 0.971034862 | ITGB1BP1 | protein_coding | 45.09 | 0.000288961 | 0.000288961 | ENSG00000119185 | n.s. |
| ENSG00000131373 | 136.1628448 | 0.020600202 | 0.2562985 | 0.080376 | 0.935938349 | 0.968276834 | HACL1 | protein_coding | 39.06 | 0.000288412 | 0.000288412 | ENSG00000131373 | n.s. |
| ENSG00000274627 | 358.1161284 | -0.019214808 | 0.2252133 | -0.085318 | 0.932008357 | 0.966484672 | | sense_intronic | 50.38 | -0.000284476 | 0.000284476 | ENSG00000274627 | n.s. |
| ENSG00000134056 | 494.4597641 | -0.023228664 | 0.3307494 | -0.07023 | 0.944010271 | 0.972200628 | MRPS36 | protein_coding | 42.53 | -0.000284414 | 0.000284414 | ENSG00000134056 | n.s. |
| ENSG00000082153 | 5692.892174 | -0.020444322 | 0.2614635 | -0.078192 | 0.937675425 | 0.968790463 | BZW1 | protein_coding | 41.12 | -0.000281521 | 0.000281521 | ENSG00000082153 | n.s. |
| ENSG00000132424 | 9207.209041 | 0.020144604 | 0.2520693 | 0.079917 | 0.936303333 | 0.968391356 | PNISR | protein_coding | 36 | 0.000280999 | 0.000280999 | ENSG00000132424 | n.s. |
| ENSG00000114631 | 2074.261462 | 0.020680733 | 0.2698966 | 0.076625 | 0.938922145 | 0.969418837 | PODXL2 | protein_coding | 48.64 | 0.000278953 | 0.000278953 | ENSG00000114631 | n.s. |
| ENSG00000122674 | 473.1037151 | -0.020676416 | 0.2697232 | -0.076658 | 0.938895695 | 0.969418837 | CCZ1 | protein_coding | 46.24 | -0.000278895 | 0.000278895 | ENSG00000122674 | n.s. |
| ENSG00000149926 | 146.765816 | -0.020113384 | 0.2551714 | -0.078823 | 0.93717339 | 0.968618893 | FAM57B | protein_coding | 52 | -0.000278511 | 0.000278511 | ENSG00000149926 | n.s. |
| ENSG00000064490 | 117.6623747 | -0.023005207 | 0.335113 | -0.068649 | 0.945268933 | 0.972581981 | RFXANK | protein_coding | 54.66 | -0.00027776 | 0.00027776 | ENSG00000064490 | n.s. |
| ENSG00000126775 | 517.1358762 | -0.020804403 | 0.274036 | -0.075919 | 0.939483925 | 0.969783118 | ATG14 | protein_coding | 41.79 | -0.000277227 | 0.000277227 | ENSG00000126775 | n.s. |
| ENSG00000272333 | 361.8609034 | 0.019348186 | 0.235909 | 0.082015 | 0.934634418 | 0.967646862 | KMT2B | protein_coding | 59.77 | 0.000276352 | 0.000276352 | ENSG00000272333 | n.s. |
| ENSG00000126934 | 607.6353375 | 0.021156977 | 0.2846604 | 0.074324 | 0.940752934 | 0.970409788 | MAP2K2 | protein_coding | 57.42 | 0.000275989 | 0.000275989 | ENSG00000126934 | n.s. |
| ENSG00000176853 | 1202.944041 | -0.02095404 | 0.279942 | -0.074851 | 0.940332985 | 0.970156069 | FAM91A1 | protein_coding | 36.91 | -0.000275722 | 0.000275722 | ENSG00000176853 | n.s. |
| ENSG00000025772 | 424.0571894 | -0.020801183 | 0.2766521 | -0.075189 | 0.940064378 | 0.970022703 | TOMM34 | protein_coding | 45.11 | -0.000274952 | 0.000274952 | ENSG00000025772 | n.s. |
| ENSG00000153066 | 206.3400902 | -0.019420316 | 0.2390922 | -0.081225 | 0.935262849 | 0.967927282 | TXNDC11 | protein_coding | 43.55 | -0.000274939 | 0.000274939 | ENSG00000153066 | n.s. |
| ENSG00000123159 | 322.0359052 | 0.022397673 | 0.3223953 | 0.069473 | 0.944613363 | 0.972338596 | GIPC1 | protein_coding | 53.7 | 0.000272859 | 0.000272859 | ENSG00000123159 | n.s. |
| ENSG00000115657 | 981.6699712 | -0.019496354 | 0.2440496 | -0.079887 | 0.936327259 | 0.968391356 | ABCB6 | protein_coding | 54.46 | -0.000271957 | 0.000271957 | ENSG00000115657 | n.s. |
| ENSG00000151014 | 289.7544571 | -0.019192787 | 0.2362091 | -0.081253 | 0.935240451 | 0.967927282 | NOCT | protein_coding | 44.81 | -0.000271717 | 0.000271717 | ENSG00000151014 | n.s. |
| ENSG00000141577 | 577.7794269 | 0.019251398 | 0.2401485 | 0.080165 | 0.936106376 | 0.968306766 | CEP131 | protein_coding | 59.72 | 0.00026927 | 0.00026927 | ENSG00000141577 | n.s. |
| ENSG00000005249 | 561.5449192 | 0.0197768 | 0.2568232 | 0.077005 | 0.93861917 | 0.969369489 | PRKAR2B | protein_coding | 38.63 | 0.000267198 | 0.000267198 | ENSG00000005249 | n.s. |

| | | | | | | | | | | | | | |
|---|---|---|---|---|---|---|---|---|---|---|---|---|---|
| ENSG00000113583 | 909.8287381 | 0.022657934 | 0.3386114 | 0.066914 | 0.946649957 | 0.973355539 | C5orf15 | protein_coding | 39.91 | 0.000265743 | 0.000265743 | ENSG00000113583 | n.s. |
| ENSG00000225091 | 106.7869304 | 0.019368549 | 0.2494245 | 0.077653 | 0.938104121 | 0.969005392 | SNORA71A | snoRNA | 53.73 | 0.000264842 | 0.000264842 | ENSG00000225091 | n.s. |
| ENSG00000265855 | 2.448313936 | -0.035040028 | 0.821084 | -0.042675 | 0.965960347 | 0.982831475 | MIR4514 | miRNA | 45.61 | -0.000263534 | 0.000263534 | ENSG00000265855 | n.s. |
| ENSG00000186469 | 383.8053884 | 0.020881887 | 0.2891954 | 0.072207 | 0.942437298 | 0.971391361 | GNG2 | protein_coding | 40.23 | 0.000263232 | 0.000263232 | ENSG00000186469 | n.s. |
| ENSG00000200488 | 29.58849352 | -0.021613971 | 0.3121003 | -0.069253 | 0.944788014 | 0.9723742 | RN7SKP203 | misc_RNA | 58.43 | -0.000262968 | 0.000262968 | ENSG00000200488 | n.s. |
| ENSG00000062822 | 908.8692519 | 0.020049652 | 0.2704611 | 0.074131 | 0.940905862 | 0.970409788 | POLD1 | protein_coding | 55.06 | 0.000261544 | 0.000261544 | ENSG00000062822 | n.s. |
| ENSG00000169756 | 1625.458547 | 0.020201286 | 0.274601 | 0.073566 | 0.941355756 | 0.970635872 | LIMS1 | protein_coding | 41.38 | 0.000261479 | 0.000261479 | ENSG00000169756 | n.s. |
| ENSG00000179603 | 389.2621959 | -0.019494342 | 0.2577007 | -0.075647 | 0.939699759 | 0.969862102 | GRM8 | protein_coding | 37.51 | -0.00025908 | 0.00025908 | ENSG00000179603 | n.s. |
| ENSG00000101746 | 336.545203 | -0.018528834 | 0.2346088 | -0.078978 | 0.937050454 | 0.968563732 | NOL4 | protein_coding | 34.88 | -0.000257028 | 0.000257028 | ENSG00000101746 | n.s. |
| ENSG00000196756 | 1492.809453 | 0.01729043 | 0.2030027 | 0.085173 | 0.932123549 | 0.966484672 | SNHG17 | processed_transcript | 51.31 | 0.000255985 | 0.000255985 | ENSG00000196756 | n.s. |
| ENSG00000222467 | 15.0787318 | -0.022302727 | 0.3448361 | -0.064676 | 0.94843173 | 0.973964816 | | misc_RNA | 36.96 | -0.000255516 | 0.000255516 | ENSG00000222467 | n.s. |
| ENSG00000204564 | 91.25494217 | -0.019044702 | 0.2497651 | -0.07625 | 0.939219855 | 0.969616742 | C6orf136 | protein_coding | 52.14 | -0.000255197 | 0.000255197 | ENSG00000204564 | n.s. |
| ENSG00000224358 | 120.2972183 | -0.018449615 | 0.2348249 | -0.078568 | 0.937376605 | 0.968685109 | | antisense | 42.24 | -0.000254925 | 0.000254925 | ENSG00000224358 | n.s. |
| ENSG00000099797 | 287.1039888 | 0.020286415 | 0.287379 | 0.070591 | 0.943723145 | 0.97206905 | TECR | protein_coding | 53.54 | 0.000249581 | 0.000249581 | ENSG00000099797 | n.s. |
| ENSG00000188986 | 556.7530603 | -0.019668033 | 0.270701 | -0.072656 | 0.942079914 | 0.9712387 | NELFB | protein_coding | 56.65 | -0.000249273 | 0.000249273 | ENSG00000188986 | n.s. |
| ENSG00000105552 | 200.924123 | -0.020786623 | 0.3047769 | -0.068203 | 0.945624232 | 0.972875651 | BCAT2 | protein_coding | 54.54 | -0.000248248 | 0.000248248 | ENSG00000105552 | n.s. |
| ENSG00000067225 | 18592.45678 | -0.020172205 | 0.2886966 | -0.069873 | 0.944294441 | 0.972200628 | PKM | protein_coding | 48.38 | -0.000246991 | 0.000246991 | ENSG00000067225 | n.s. |
| ENSG00000258429 | 75.87622243 | -0.019219052 | 0.2657306 | -0.072325 | 0.94234301 | 0.971381701 | PDF | protein_coding | 61.27 | -0.000242354 | 0.000242354 | ENSG00000258429 | n.s. |
| ENSG00000081307 | 1698.521245 | 0.018462034 | 0.249106 | 0.074113 | 0.940920344 | 0.970409788 | UBA5 | protein_coding | 36.76 | 0.000240834 | 0.000240834 | ENSG00000081307 | n.s. |
| ENSG00000131475 | 1421.342082 | -0.020071408 | 0.296396 | -0.067718 | 0.946009949 | 0.973045663 | VPS25 | protein_coding | 49.25 | -0.000238183 | 0.000238183 | ENSG00000131475 | n.s. |
| ENSG00000100412 | 815.0140731 | -0.019389926 | 0.2766408 | -0.070091 | 0.944121523 | 0.972200628 | ACO2 | protein_coding | 50.27 | -0.000237412 | 0.000237412 | ENSG00000100412 | n.s. |
| ENSG00000168286 | 563.0332855 | -0.019110328 | 0.2761534 | -0.069202 | 0.944828942 | 0.9723742 | THAP11 | protein_coding | 60.31 | -0.000232507 | 0.000232507 | ENSG00000168286 | n.s. |
| ENSG00000210176 | 4.128175419 | -0.027215846 | 0.5687832 | -0.047849 | 0.961836393 | 0.98064207 | MT-TH | Mt_tRNA | 36.23 | -0.000231048 | 0.000231048 | ENSG00000210176 | n.s. |
| ENSG00000172534 | 1240.468506 | 0.017205167 | 0.2282659 | 0.075373 | 0.939917652 | 0.970015082 | HCFC1 | protein_coding | 59.51 | 0.000227478 | 0.000227478 | ENSG00000172534 | n.s. |
| ENSG00000088992 | 1215.94014 | 0.018707903 | 0.2715103 | 0.068903 | 0.945066753 | 0.97244582 | TESC | protein_coding | 51.4 | 0.000227013 | 0.000227013 | ENSG00000088992 | n.s. |
| ENSG00000135111 | 3022.26737 | 0.01693367 | 0.2222014 | 0.076209 | 0.939253106 | 0.969616742 | TBX3 | protein_coding | 47.27 | 0.000226909 | 0.000226909 | ENSG00000135111 | n.s. |
| ENSG00000071994 | 2936.810416 | -0.01798727 | 0.2487649 | -0.072306 | 0.942358158 | 0.971381701 | PDCD2 | protein_coding | 41.64 | -0.000226821 | 0.000226821 | ENSG00000071994 | n.s. |
| ENSG00000177576 | 70.89832056 | 0.019611709 | 0.3026118 | 0.064808 | 0.948326765 | 0.973928859 | C18orf32 | protein_coding | 43.19 | 0.000225001 | 0.000225001 | ENSG00000177576 | n.s. |
| ENSG00000033327 | 414.7435957 | 0.017853077 | 0.2491678 | 0.071651 | 0.942879796 | 0.97158315 | GAB2 | protein_coding | 41.71 | 0.000223521 | 0.000223521 | ENSG00000033327 | n.s. |
| ENSG00000183605 | 463.0035688 | 0.019693839 | 0.3077217 | 0.063999 | 0.948971128 | 0.97423131 | SFXN4 | protein_coding | 47.78 | 0.000223287 | 0.000223287 | ENSG00000183605 | n.s. |
| ENSG00000102226 | 932.8801143 | 0.018894644 | 0.281119 | 0.067212 | 0.946412731 | 0.97318349 | USP11 | protein_coding | 48.33 | 0.000223056 | 0.000223056 | ENSG00000102226 | n.s. |
| ENSG00000119707 | 5678.01371 | 0.017773885 | 0.2484667 | 0.071534 | 0.942972552 | 0.97158315 | RBM25 | protein_coding | 40.7 | 0.000222529 | 0.000222529 | ENSG00000119707 | n.s. |

| | | | | | | | | | | | | | |
|---|---|---|---|---|---|---|---|---|---|---|---|---|---|
| ENSG00000142409 | 399.7544485 | -0.018771565 | 0.2781172 | -0.067495 | 0.946187509 | 0.973095634 | ZNF787 | protein_coding | 58.61 | -0.000222339 | 0.000222339 | ENSG00000142409 | n.s. |
| ENSG00000173039 | 260.3291582 | -0.017708433 | 0.2478544 | -0.071447 | 0.943042066 | 0.97158315 | RELA | protein_coding | 55.62 | -0.00022171 | 0.00022171 | ENSG00000173039 | n.s. |
| ENSG00000090520 | 1183.722606 | -0.018424105 | 0.270878 | -0.068016 | 0.945772701 | 0.9729094 | DNAJB11 | protein_coding | 40.07 | -0.000219755 | 0.000219755 | ENSG00000090520 | n.s. |
| ENSG00000124406 | 2975.956861 | -0.018415201 | 0.272237 | -0.067644 | 0.946069032 | 0.973045663 | ATP8A1 | protein_coding | 38.15 | -0.000218529 | 0.000218529 | ENSG00000124406 | n.s. |
| ENSG00000119650 | 200.7820928 | 0.020560563 | 0.3408165 | 0.060327 | 0.951894898 | 0.976081357 | IFT43 | protein_coding | 43.54 | 0.000216173 | 0.000216173 | ENSG00000119650 | n.s. |
| ENSG00000105700 | 481.7715565 | 0.018408917 | 0.2785848 | 0.06608 | 0.947314038 | 0.973391392 | KXD1 | protein_coding | 53.42 | 0.000215614 | 0.000215614 | ENSG00000105700 | n.s. |
| ENSG00000110497 | 176.9215507 | 0.017100959 | 0.2391048 | 0.071521 | 0.942983293 | 0.97158315 | AMBRA1 | protein_coding | 43.63 | 0.000214104 | 0.000214104 | ENSG00000110497 | n.s. |
| ENSG00000184634 | 536.5962185 | -0.017502107 | 0.2514497 | -0.069605 | 0.944508218 | 0.972302249 | MED12 | protein_coding | 51.08 | -0.000213503 | 0.000213503 | ENSG00000184634 | n.s. |
| ENSG00000151552 | 290.5771291 | -0.017927425 | 0.2659572 | -0.067407 | 0.946257554 | 0.973095791 | QDPR | protein_coding | 45.09 | -0.00021234 | 0.00021234 | ENSG00000151552 | n.s. |
| ENSG00000272821 | 109.5489498 | -0.017985366 | 0.2709242 | -0.066385 | 0.947071124 | 0.973378571 | | antisense | 61.4 | -0.000210756 | 0.000210756 | ENSG00000272821 | n.s. |
| ENSG00000168061 | 288.444971 | 0.016974593 | 0.2403924 | 0.070612 | 0.943706544 | 0.97206905 | SAC3D1 | protein_coding | 58.58 | 0.000208836 | 0.000208836 | ENSG00000168061 | n.s. |
| ENSG00000060491 | 357.9946405 | -0.016749931 | 0.239811 | -0.069846 | 0.944315935 | 0.972200628 | OGFR | protein_coding | 63.45 | -0.000205088 | 0.000205088 | ENSG00000060491 | n.s. |
| ENSG00000145569 | 336.5485666 | 0.018452111 | 0.2934822 | 0.062873 | 0.949867628 | 0.974792289 | FAM105A | protein_coding | 42.96 | 0.000204595 | 0.000204595 | ENSG00000145569 | n.s. |
| ENSG00000278272 | 1733.580894 | 0.016166173 | 0.2255601 | 0.071671 | 0.942863532 | 0.97158315 | HIST1H3C | protein_coding | 59.85 | 0.000202401 | 0.000202401 | ENSG00000278272 | n.s. |
| ENSG00000230483 | 16.62142326 | 0.019923745 | 0.351071 | 0.056751 | 0.954743289 | 0.977059229 | | antisense | 64.79 | 0.000200814 | 0.000200814 | ENSG00000230483 | n.s. |
| ENSG00000166913 | 2918.401948 | 0.017067057 | 0.2591353 | 0.065862 | 0.947488038 | 0.973426502 | YWHAB | protein_coding | 40.42 | 0.000199631 | 0.000199631 | ENSG00000166913 | n.s. |
| ENSG00000170677 | 1235.247083 | -0.017519848 | 0.2732399 | -0.064119 | 0.948875537 | 0.974205007 | SOCS6 | protein_coding | 39.2 | -0.000198844 | 0.000198844 | ENSG00000170677 | n.s. |
| ENSG00000263584 | 3.535088503 | -0.030324596 | 0.8196242 | -0.036998 | 0.970486465 | 0.985132915 | MIR4480 | miRNA | 53.52 | -0.000197267 | 0.000197267 | ENSG00000263584 | n.s. |
| ENSG00000224593 | 45.0830391 | -0.019046783 | 0.3254713 | -0.058521 | 0.953333941 | 0.976581921 | | transcribed_processed_pseudog | 41.23 | -0.000196017 | 0.000196017 | ENSG00000224593 | n.s. |
| ENSG00000117152 | 8771.022817 | -0.016695862 | 0.2518823 | -0.066284 | 0.947151417 | 0.973378571 | RGS4 | protein_coding | 39.47 | -0.000195646 | 0.000195646 | ENSG00000117152 | n.s. |
| ENSG00000100425 | 1701.353073 | -0.015515339 | 0.2200906 | -0.070495 | 0.943799479 | 0.972075761 | BRD1 | protein_coding | 54 | -0.000190837 | 0.000190837 | ENSG00000100425 | n.s. |
| ENSG00000197021 | 150.684377 | -0.017113847 | 0.271675 | -0.062994 | 0.949771414 | 0.974765398 | CXorf40B | protein_coding | 52.2 | -0.000189962 | 0.000189962 | ENSG00000197021 | n.s. |
| ENSG00000169744 | 61.04878857 | 0.016696619 | 0.259448 | 0.064354 | 0.948688049 | 0.974156185 | LDB2 | protein_coding | 40.21 | 0.000189864 | 0.000189864 | ENSG00000169744 | n.s. |
| ENSG00000188707 | 626.5208378 | 0.016466705 | 0.253131 | 0.065052 | 0.948132516 | 0.973801196 | ZBED6CL | protein_coding | 55.68 | 0.000189856 | 0.000189856 | ENSG00000188707 | n.s. |
| ENSG00000110917 | 2284.333429 | 0.015909303 | 0.2340086 | 0.067986 | 0.945796791 | 0.9729094 | MLEC | protein_coding | 47.71 | 0.00018976 | 0.00018976 | ENSG00000110917 | n.s. |
| ENSG00000168758 | 897.0510595 | 0.016111227 | 0.2421633 | 0.06653 | 0.946955534 | 0.973378571 | SEMA4C | protein_coding | 62.17 | 0.000188795 | 0.000188795 | ENSG00000168758 | n.s. |
| ENSG00000110063 | 117.098875 | -0.016567948 | 0.2623462 | -0.063153 | 0.949644667 | 0.974765398 | DCPS | protein_coding | 49.42 | -0.000183902 | 0.000183902 | ENSG00000110063 | n.s. |
| ENSG00000226471 | 102.2814712 | 0.015891584 | 0.2435003 | 0.065263 | 0.947964533 | 0.973772332 | | antisense | 45.98 | 0.00018343 | 0.00018343 | ENSG00000226471 | n.s. |
| ENSG00000185163 | 677.2299878 | 0.015630136 | 0.2355519 | 0.066355 | 0.947094899 | 0.973378571 | DDX51 | protein_coding | 60.51 | 0.000183157 | 0.000183157 | ENSG00000185163 | n.s. |
| ENSG00000198961 | 2420.025756 | 0.016613998 | 0.2672116 | 0.062175 | 0.950423129 | 0.975110944 | PJA2 | protein_coding | 37.66 | 0.000181856 | 0.000181856 | ENSG00000198961 | n.s. |
| ENSG00000099949 | 573.9915707 | -0.016586546 | 0.2659295 | -0.062372 | 0.950266614 | 0.975110944 | LZTR1 | protein_coding | 58.44 | -0.000181556 | 0.000181556 | ENSG00000099949 | n.s. |
| ENSG00000011007 | 874.0407347 | -0.015990878 | 0.2493111 | -0.06414 | 0.948858542 | 0.974205007 | ELOA | protein_coding | 45.7 | -0.000181491 | 0.000181491 | ENSG00000011007 | n.s. |

| | | | | | | | | | | | | | |
|---|---|---|---|---|---|---|---|---|---|---|---|---|---|
| ENSG00000087088 | 395.1823473 | 0.016317627 | 0.2587983 | 0.063052 | 0.949725483 | 0.974765398 | BAX | protein_coding | 53.57 | 0.000181124 | 0.000181124 | ENSG00000087088 | n.s. |
| ENSG00000155275 | 137.5325268 | 0.015627454 | 0.2399285 | 0.065134 | 0.948067466 | 0.973801196 | TRMT44 | protein_coding | 49.1 | 0.00018018 | 0.00018018 | ENSG00000155275 | n.s. |
| ENSG00000181472 | 728.7212775 | 0.016330172 | 0.2635795 | 0.061955 | 0.950598351 | 0.975110944 | ZBTB2 | protein_coding | 44.33 | 0.00017875 | 0.00017875 | ENSG00000181472 | n.s. |
| ENSG00000248383 | 178.1182322 | -0.015144984 | 0.2274418 | -0.066588 | 0.946909385 | 0.973378571 | PCDHAC1 | protein_coding | 39.4 | -0.000177472 | 0.000177472 | ENSG00000248383 | n.s. |
| ENSG00000147164 | 353.3878677 | 0.017101448 | 0.2904501 | 0.058879 | 0.953048387 | 0.976544943 | SNX12 | protein_coding | 42.55 | 0.000176278 | 0.000176278 | ENSG00000147164 | n.s. |
| ENSG00000183160 | 129.1925445 | 0.016612938 | 0.2741875 | 0.06059 | 0.951685975 | 0.976010889 | TMEM119 | protein_coding | 56.18 | 0.000175189 | 0.000175189 | ENSG00000183160 | n.s. |
| ENSG00000075702 | 295.1191032 | 0.014303626 | 0.2041568 | 0.070062 | 0.94414434 | 0.972200628 | WDR62 | protein_coding | 51.45 | 0.000175135 | 0.000175135 | ENSG00000075702 | n.s. |
| ENSG00000200118 | 13.99583235 | -0.018936851 | 0.3638403 | -0.052047 | 0.958491127 | 0.979083804 | | misc_RNA | 47 | -0.000173843 | 0.000173843 | ENSG00000200118 | n.s. |
| ENSG00000162961 | 835.0159696 | 0.017202922 | 0.3025641 | 0.056857 | 0.954659017 | 0.977044802 | DPY30 | protein_coding | 39 | 0.0001735 | 0.0001735 | ENSG00000162961 | n.s. |
| ENSG00000172594 | 260.0074457 | -0.016309202 | 0.267766 | -0.060908 | 0.951432155 | 0.97582246 | SMPDL3A | protein_coding | 40.47 | -0.000173354 | 0.000173354 | ENSG00000172594 | n.s. |
| ENSG00000122566 | 20932.41935 | 0.014732786 | 0.2211535 | 0.066618 | 0.946885875 | 0.973378571 | HNRNPA2B1 | protein_coding | 40.07 | 0.000172642 | 0.000172642 | ENSG00000122566 | n.s. |
| ENSG00000116030 | 3594.432115 | 0.016866458 | 0.2937391 | 0.05742 | 0.954210741 | 0.976801422 | SUMO1 | protein_coding | 41.66 | 0.000171932 | 0.000171932 | ENSG00000116030 | n.s. |
| ENSG00000008382 | 117.5672399 | 0.016353637 | 0.2714929 | 0.060236 | 0.951967697 | 0.976084118 | MPND | protein_coding | 55.95 | 0.000171922 | 0.000171922 | ENSG00000008382 | n.s. |
| ENSG00000137714 | 258.1315352 | 0.016776928 | 0.290773 | 0.057698 | 0.953989438 | 0.9767579 | FDX1 | protein_coding | 39.82 | 0.000171344 | 0.000171344 | ENSG00000137714 | n.s. |
| ENSG00000136504 | 4074.853775 | -0.01474454 | 0.2251878 | -0.065477 | 0.947794508 | 0.973669515 | KAT7 | protein_coding | 42.42 | -0.000170866 | 0.000170866 | ENSG00000136504 | n.s. |
| ENSG00000175265 | 940.2378983 | 0.016552154 | 0.2792955 | 0.059264 | 0.952741875 | 0.976528772 | GOLGA8A | protein_coding | 47.08 | 0.000170735 | 0.000170735 | ENSG00000175265 | n.s. |
| ENSG00000176887 | 8902.069349 | -0.014590664 | 0.2214735 | -0.06588 | 0.947473398 | 0.973426502 | SOX11 | protein_coding | 47.75 | -0.000170665 | 0.000170665 | ENSG00000176887 | n.s. |
| ENSG00000149196 | 433.5204765 | -0.016993514 | 0.3024253 | -0.056191 | 0.955189821 | 0.977300697 | HIKESHI | protein_coding | 37.85 | -0.000169456 | 0.000169456 | ENSG00000149196 | n.s. |
| ENSG00000108309 | 878.0920171 | 0.016801592 | 0.2951384 | 0.056928 | 0.95460267 | 0.977044802 | RUNDC3A | protein_coding | 59.03 | 0.000169453 | 0.000169453 | ENSG00000108309 | n.s. |
| ENSG00000171530 | 2018.629197 | -0.01771035 | 0.3282547 | -0.053953 | 0.956972554 | 0.978262032 | TBCA | protein_coding | 39.32 | -0.000169042 | 0.000169042 | ENSG00000171530 | n.s. |
| ENSG00000106236 | 371.190306 | -0.015412347 | 0.2485154 | -0.062018 | 0.950548763 | 0.975110944 | NPTX2 | protein_coding | 54.62 | -0.000168703 | 0.000168703 | ENSG00000106236 | n.s. |
| ENSG00000119383 | 544.7986872 | -0.016218459 | 0.2742889 | -0.059129 | 0.952849265 | 0.976528772 | PTPA | protein_coding | 52.59 | -0.000167293 | 0.000167293 | ENSG00000119383 | n.s. |
| ENSG00000121073 | 1667.223199 | -0.016108563 | 0.2730867 | -0.058987 | 0.952962467 | 0.976528772 | SLC35B1 | protein_coding | 47.79 | -0.000166159 | 0.000166159 | ENSG00000121073 | n.s. |
| ENSG00000076928 | 51.81829093 | 0.016022736 | 0.2735429 | 0.058575 | 0.953290735 | 0.976581921 | ARHGEF1 | protein_coding | 58.25 | 0.000164895 | 0.000164895 | ENSG00000076928 | n.s. |
| ENSG00000128050 | 7855.832816 | -0.016082289 | 0.2771731 | -0.058023 | 0.953730674 | 0.976716643 | PAICS | protein_coding | 39.99 | -0.000164545 | 0.000164545 | ENSG00000128050 | n.s. |
| ENSG00000281398 | 512.2818135 | -0.013975957 | 0.2108897 | -0.066271 | 0.947161738 | 0.973378571 | SNHG4 | processed_transcript | 43.8 | -0.000163773 | 0.000163773 | ENSG00000281398 | n.s. |
| ENSG00000113594 | 1288.647665 | 0.015567264 | 0.2577342 | 0.0604 | 0.951836697 | 0.976081357 | LIFR | protein_coding | 39.45 | 0.000163674 | 0.000163674 | ENSG00000113594 | n.s. |
| ENSG00000007520 | 501.715994 | -0.01577483 | 0.2697402 | -0.058482 | 0.953365042 | 0.976581921 | TSR3 | protein_coding | 62.57 | -0.000162344 | 0.000162344 | ENSG00000007520 | n.s. |
| ENSG00000087460 | 13232.98143 | -0.015572028 | 0.2633131 | -0.059139 | 0.952841529 | 0.976528772 | GNAS | protein_coding | 47.09 | -0.000160625 | 0.000160625 | ENSG00000087460 | n.s. |
| ENSG00000170043 | 189.8697999 | 0.016265953 | 0.2935521 | 0.055411 | 0.955811204 | 0.977649088 | TRAPPC1 | protein_coding | 57.62 | 0.000159683 | 0.000159683 | ENSG00000170043 | n.s. |
| ENSG00000123297 | 836.391952 | 0.016381965 | 0.3000608 | 0.054595 | 0.956460732 | 0.978026055 | TSFM | protein_coding | 44.22 | 0.000158079 | 0.000158079 | ENSG00000123297 | n.s. |
| ENSG00000164338 | 1381.087222 | -0.014430863 | 0.2327932 | -0.06199 | 0.950570743 | 0.975110944 | UTP15 | protein_coding | 37.9 | -0.00015796 | 0.00015796 | ENSG00000164338 | n.s. |

| | | | | | | | | | | | | | |
|---|---|---|---|---|---|---|---|---|---|---|---|---|---|
| ENSG00000180773 | 238.3434571 | -0.015635843 | 0.2741561 | -0.057033 | 0.954519195 | 0.977044802 | SLC36A4 | protein_coding | 35.08 | -0.000157696 | 0.000157696 | ENSG00000180773 | n.s. |
| ENSG00000152795 | 19932.19553 | -0.014348503 | 0.2308979 | -0.062142 | 0.950449595 | 0.975110944 | HNRNPDL | protein_coding | 42.24 | -0.000157058 | 0.000157058 | ENSG00000152795 | n.s. |
| ENSG00000198960 | 151.0901959 | 0.015321132 | 0.2665261 | 0.057485 | 0.954159213 | 0.976801422 | ARMCX6 | protein_coding | 51.56 | 0.000156179 | 0.000156179 | ENSG00000198960 | n.s. |
| ENSG00000002919 | 401.8431121 | -0.015284405 | 0.265127 | -0.057649 | 0.954027916 | 0.9767579 | SNX11 | protein_coding | 48 | -0.000156101 | 0.000156101 | ENSG00000002919 | n.s. |
| ENSG00000237380 | 175.1906238 | 0.016316548 | 0.3041392 | 0.053648 | 0.957215387 | 0.978438428 | HOXD-AS2 | antisense | 51.9 | 0.000154461 | 0.000154461 | ENSG00000237380 | n.s. |
| ENSG00000005100 | 689.5065109 | -0.014741198 | 0.2498772 | -0.058994 | 0.952957075 | 0.976528772 | DHX33 | protein_coding | 45.81 | -0.000152055 | 0.000152055 | ENSG00000005100 | n.s. |
| ENSG00000132275 | 887.5479961 | -0.013763072 | 0.2229722 | -0.061725 | 0.950781437 | 0.9752269 | RRP8 | protein_coding | 47.4 | -0.000149939 | 0.000149939 | ENSG00000132275 | n.s. |
| ENSG00000112473 | 775.1494487 | 0.014496968 | 0.2473104 | 0.058619 | 0.953255955 | 0.976581921 | SLC39A7 | protein_coding | 52.92 | 0.000149193 | 0.000149193 | ENSG00000112473 | n.s. |
| ENSG00000169490 | 460.8873972 | 0.01494585 | 0.2657785 | 0.056234 | 0.95515521 | 0.977300697 | TM2D2 | protein_coding | 42.34 | 0.000149037 | 0.000149037 | ENSG00000169490 | n.s. |
| ENSG00000168078 | 1308.887155 | 0.016225921 | 0.3121973 | 0.051973 | 0.958549977 | 0.979083804 | PBK | protein_coding | 37.25 | 0.000148956 | 0.000148956 | ENSG00000168078 | n.s. |
| ENSG00000180776 | 1039.882303 | -0.015233858 | 0.276513 | -0.055093 | 0.956064584 | 0.977836421 | ZDHHC20 | protein_coding | 39.65 | -0.000148283 | 0.000148283 | ENSG00000180776 | n.s. |
| ENSG00000044090 | 391.208976 | 0.014697065 | 0.2596688 | 0.056599 | 0.954864424 | 0.977111375 | CUL7 | protein_coding | 54.7 | 0.000147793 | 0.000147793 | ENSG00000044090 | n.s. |
| ENSG00000149547 | 872.6844679 | 0.015759689 | 0.2991572 | 0.05268 | 0.957986649 | 0.978689024 | EI24 | protein_coding | 43.49 | 0.000147436 | 0.000147436 | ENSG00000149547 | n.s. |
| ENSG00000180543 | 185.2260559 | -0.014390737 | 0.2486473 | -0.057876 | 0.953847317 | 0.976716643 | TSPYL5 | protein_coding | 49.7 | -0.000147238 | 0.000147238 | ENSG00000180543 | n.s. |
| ENSG00000104866 | 373.1011525 | 0.014831845 | 0.2658512 | 0.05579 | 0.955509086 | 0.977555516 | PPP1R37 | protein_coding | 55.56 | 0.000146221 | 0.000146221 | ENSG00000104866 | n.s. |
| ENSG00000125779 | 1762.223657 | 0.01400748 | 0.2357269 | 0.059422 | 0.952615604 | 0.976528772 | PANK2 | protein_coding | 45.01 | 0.000144487 | 0.000144487 | ENSG00000125779 | n.s. |
| ENSG00000149743 | 110.091259 | -0.015382522 | 0.2899322 | -0.053056 | 0.957687616 | 0.978620328 | TRPT1 | protein_coding | 60.14 | -0.000144377 | 0.000144377 | ENSG00000149743 | n.s. |
| ENSG00000050405 | 2323.198514 | -0.014652683 | 0.2695145 | -0.054367 | 0.956642808 | 0.978068572 | LIMA1 | protein_coding | 42.32 | -0.000141116 | 0.000141116 | ENSG00000050405 | n.s. |
| ENSG00000267254 | 249.9260204 | 0.013715357 | 0.2354551 | 0.05825 | 0.953549168 | 0.976698674 | | antisense | 40.52 | 0.000140437 | 0.000140437 | ENSG00000267254 | n.s. |
| ENSG00000107562 | 190.8896569 | -0.014739454 | 0.2765367 | -0.0533 | 0.957492745 | 0.978506422 | CXCL12 | protein_coding | 52.37 | -0.000139086 | 0.000139086 | ENSG00000107562 | n.s. |
| ENSG00000213397 | 286.4546382 | -0.014107993 | 0.2539578 | -0.055553 | 0.955698295 | 0.977649088 | HAUS7 | protein_coding | 58.62 | -0.000138498 | 0.000138498 | ENSG00000213397 | n.s. |
| ENSG00000112320 | 240.1328348 | -0.014418078 | 0.2662703 | -0.054148 | 0.956817037 | 0.978174874 | SOBP | protein_coding | 41.63 | -0.000138176 | 0.000138176 | ENSG00000112320 | n.s. |
| ENSG00000153551 | 212.6988095 | -0.014606347 | 0.2728797 | -0.053527 | 0.957312263 | 0.978465618 | CMTM7 | protein_coding | 45.15 | -0.000138095 | 0.000138095 | ENSG00000153551 | n.s. |
| ENSG00000171865 | 1167.247849 | 0.013078915 | 0.2183686 | 0.059894 | 0.952240255 | 0.976219797 | RNASEH1 | protein_coding | 45.37 | 0.000136706 | 0.000136706 | ENSG00000171865 | n.s. |
| ENSG00000163923 | 427.6997538 | 0.015185288 | 0.2970288 | 0.051124 | 0.959226745 | 0.97948764 | RPL39L | protein_coding | 41.84 | 0.000136683 | 0.000136683 | ENSG00000163923 | n.s. |
| ENSG00000278705 | 1389.391244 | 0.013217064 | 0.228283 | 0.057898 | 0.95383011 | 0.976716643 | HIST1H4B | protein_coding | 53.42 | 0.000135229 | 0.000135229 | ENSG00000278705 | n.s. |
| ENSG00000163785 | 1399.641899 | 0.014232197 | 0.2703952 | 0.052635 | 0.958022891 | 0.978689024 | RYK | protein_coding | 40.6 | 0.000133146 | 0.000133146 | ENSG00000163785 | n.s. |
| ENSG00000181924 | 948.4529223 | 0.01407585 | 0.2656631 | 0.052984 | 0.957744781 | 0.978620328 | COA4 | protein_coding | 46.92 | 0.000132113 | 0.000132113 | ENSG00000181924 | n.s. |
| ENSG00000160789 | 53.66401072 | 0.014770052 | 0.2923292 | 0.050525 | 0.959703702 | 0.979759103 | LMNA | protein_coding | 53.04 | 0.000131168 | 0.000131168 | ENSG00000160789 | n.s. |
| ENSG00000167721 | 2627.434758 | -0.013340781 | 0.2406354 | -0.05544 | 0.955788078 | 0.977649088 | TSR1 | protein_coding | 45.82 | -0.000130966 | 0.000130966 | ENSG00000167721 | n.s. |
| ENSG00000170584 | 965.3792938 | -0.013359574 | 0.2444124 | -0.05466 | 0.956409364 | 0.978026055 | NUDCD2 | protein_coding | 37.65 | -0.000128914 | 0.000128914 | ENSG00000170584 | n.s. |
| ENSG00000283341 | 173.209461 | 0.012576822 | 0.216245 | 0.05816 | 0.953621128 | 0.976700527 | | processed_transcript | 38.5 | 0.000128769 | 0.000128769 | ENSG00000283341 | n.s. |

| | | | | | | | | | | | | | |
|---|---|---|---|---|---|---|---|---|---|---|---|---|---|
| ENSG00000155463 | 808.0061853 | -0.013339175 | 0.2437597 | -0.054723 | 0.956359434 | 0.978026055 | OXA1L | protein_coding | 46.66 | -0.000128717 | 0.000128717 | ENSG00000155463 | n.s. |
| ENSG00000006625 | 1705.850801 | 0.014579267 | 0.292272 | 0.049883 | 0.960215995 | 0.97992284 | GGCT | protein_coding | 40.55 | 0.000128416 | 0.000128416 | ENSG00000006625 | n.s. |
| ENSG00000172458 | 60.57560756 | 0.015037703 | 0.3136585 | 0.047943 | 0.961761733 | 0.980637759 | IL17D | protein_coding | 52.71 | 0.000127691 | 0.000127691 | ENSG00000172458 | n.s. |
| ENSG00000206663 | 3.207671423 | -0.024019702 | 0.7891964 | -0.030436 | 0.975719618 | 0.987852257 | | misc_RNA | 44.12 | -0.000127497 | 0.000127497 | ENSG00000206663 | n.s. |
| ENSG00000254986 | 365.0312472 | -0.013774731 | 0.2661765 | -0.05175 | 0.958727603 | 0.9791475 | DPP3 | protein_coding | 52.26 | -0.000126065 | 0.000126065 | ENSG00000254986 | n.s. |
| ENSG00000164949 | 237.9509515 | 0.012992541 | 0.238912 | 0.054382 | 0.956630719 | 0.978068572 | GEM | protein_coding | 44.2 | 0.000125127 | 0.000125127 | ENSG00000164949 | n.s. |
| ENSG00000215154 | 115.8157455 | -0.013006577 | 0.243651 | -0.053382 | 0.957427544 | 0.978506422 | | transcribed_unprocessed_pseud | 54.23 | -0.000122734 | 0.000122734 | ENSG00000215154 | n.s. |
| ENSG00000149357 | 446.6381445 | 0.014043597 | 0.2873142 | 0.048879 | 0.961015819 | 0.980307952 | LAMTOR1 | protein_coding | 48.06 | 0.000121301 | 0.000121301 | ENSG00000149357 | n.s. |
| ENSG00000136319 | 490.6237465 | 0.013530082 | 0.2660818 | 0.050849 | 0.959445575 | 0.979576016 | TTC5 | protein_coding | 39.84 | 0.000121255 | 0.000121255 | ENSG00000136319 | n.s. |
| ENSG00000263001 | 2565.324309 | 0.013408379 | 0.2637432 | 0.050839 | 0.959454006 | 0.979576016 | GTF2I | protein_coding | 42.79 | 0.000120164 | 0.000120164 | ENSG00000263001 | n.s. |
| ENSG00000115649 | 235.9630439 | 0.014252481 | 0.3009941 | 0.047351 | 0.962233186 | 0.980833299 | CNPPD1 | protein_coding | 54.28 | 0.000119789 | 0.000119789 | ENSG00000115649 | n.s. |
| ENSG00000111845 | 982.5050298 | 0.013464699 | 0.2676221 | 0.050312 | 0.959873472 | 0.979772018 | PAK1IP1 | protein_coding | 40.83 | 0.000119499 | 0.000119499 | ENSG00000111845 | n.s. |
| ENSG00000104325 | 814.1775911 | -0.014244562 | 0.3086518 | -0.046151 | 0.963189964 | 0.981088146 | DECR1 | protein_coding | 38.15 | -0.000118116 | 0.000118116 | ENSG00000104325 | n.s. |
| ENSG00000159352 | 2824.100077 | 0.014144969 | 0.3049349 | 0.046387 | 0.963001921 | 0.981040151 | PSMD4 | protein_coding | 47.67 | 0.00011759 | 0.00011759 | ENSG00000159352 | n.s. |
| ENSG00000145919 | 2079.221217 | 0.013332193 | 0.2683273 | 0.049686 | 0.960372373 | 0.979938772 | BOD1 | protein_coding | 44.51 | 0.000117337 | 0.000117337 | ENSG00000145919 | n.s. |
| ENSG00000171612 | 340.6064703 | -0.013985767 | 0.2963923 | -0.047187 | 0.962364449 | 0.980893135 | SLC25A33 | protein_coding | 45.5 | -0.000117177 | 0.000117177 | ENSG00000171612 | n.s. |
| ENSG00000148110 | 796.4803827 | -0.012996278 | 0.2541606 | -0.051134 | 0.959218659 | 0.97948764 | MFSD14B | protein_coding | 40.66 | -0.00011698 | 0.00011698 | ENSG00000148110 | n.s. |
| ENSG00000185236 | 173.7395535 | 0.013385949 | 0.2724382 | 0.049134 | 0.96081259 | 0.980172456 | RAB11B | protein_coding | 59.05 | 0.000116424 | 0.000116424 | ENSG00000185236 | n.s. |
| ENSG00000215105 | 278.4940286 | 0.013200677 | 0.2653477 | 0.049749 | 0.960322734 | 0.979938772 | TTC3P1 | processed_pseudogene | 40.31 | 0.00011618 | 0.00011618 | ENSG00000215105 | n.s. |
| ENSG00000100983 | 507.4992885 | -0.013809729 | 0.29166 | -0.047349 | 0.962235296 | 0.980833299 | GSS | protein_coding | 46.29 | -0.000116068 | 0.000116068 | ENSG00000100983 | n.s. |
| ENSG00000221630 | 3.564729323 | 0.025248388 | 0.9235699 | 0.027338 | 0.978190291 | 0.989490222 | MIR1179 | miRNA | 41.76 | 0.000115852 | 0.000115852 | ENSG00000221630 | n.s. |
| ENSG00000136758 | 4039.001158 | -0.01297511 | 0.2585972 | -0.050175 | 0.959982946 | 0.979772018 | YME1L1 | protein_coding | 38.36 | -0.000115154 | 0.000115154 | ENSG00000136758 | n.s. |
| ENSG00000169826 | 529.9195869 | 0.012817986 | 0.2555608 | 0.050156 | 0.959997836 | 0.979772018 | CSGALNACT2 | protein_coding | 38.88 | 0.000113759 | 0.000113759 | ENSG00000169826 | n.s. |
| ENSG00000197535 | 125.9076467 | -0.01261799 | 0.2506017 | -0.050351 | 0.95984287 | 0.979772018 | MYO5A | protein_coding | 39.92 | -0.000111984 | 0.000111984 | ENSG00000197535 | n.s. |
| ENSG00000111142 | 4813.677609 | -0.013037457 | 0.2692145 | -0.048428 | 0.961375331 | 0.980531004 | METAP2 | protein_coding | 39.23 | -0.000111323 | 0.000111323 | ENSG00000111142 | n.s. |
| ENSG00000121481 | 998.7768031 | -0.013078459 | 0.2727326 | -0.047953 | 0.961753373 | 0.980637759 | RNF2 | protein_coding | 39.39 | -0.000111054 | 0.000111054 | ENSG00000121481 | n.s. |
| ENSG00000263399 | 3.057874839 | -0.019950489 | 0.6285274 | -0.031742 | 0.974678091 | 0.987300332 | MIR3170 | miRNA | 49.35 | -0.00011074 | 0.00011074 | ENSG00000263399 | n.s. |
| ENSG00000139343 | 2316.092888 | -0.012668611 | 0.2572672 | -0.049243 | 0.960725645 | 0.980155576 | SNRPF | protein_coding | 39.09 | -0.00011028 | 0.00011028 | ENSG00000139343 | n.s. |
| ENSG00000166342 | 80.77941222 | -0.012791524 | 0.2625465 | -0.048721 | 0.961141655 | 0.980364489 | NETO1 | protein_coding | 36.85 | -0.000110166 | 0.000110166 | ENSG00000166342 | n.s. |
| ENSG00000114738 | 247.1468034 | 0.013183758 | 0.2824324 | 0.046679 | 0.962768797 | 0.981030744 | MAPKAPK3 | protein_coding | 53.69 | 0.000109654 | 0.000109654 | ENSG00000114738 | n.s. |
| ENSG00000146574 | 367.293039 | 0.012790312 | 0.2649435 | 0.048276 | 0.961496578 | 0.980571751 | CCZ1B | protein_coding | 46.69 | 0.000108981 | 0.000108981 | ENSG00000146574 | n.s. |
| ENSG00000167513 | 478.1179609 | -0.01155788 | 0.2187086 | -0.052846 | 0.9578546 | 0.978660717 | CDT1 | protein_coding | 65.55 | -0.000108272 | 0.000108272 | ENSG00000167513 | n.s. |

| | | | | | | | | | | | | | |
|---|---|---|---|---|---|---|---|---|---|---|---|---|---|
| ENSG00000256185 | 8.081095482 | -0.022777967 | 0.810453 | -0.028105 | 0.977578224 | 0.989158547 | | 3prime_overlapping_ncRNA | 36.43 | -0.000107833 | 0.000107833 | ENSG00000256185 | n.s. |
| ENSG00000151498 | 164.0141478 | 0.012997715 | 0.2813345 | 0.0462 | 0.963150665 | 0.981088146 | ACAD8 | protein_coding | 47.88 | 0.000107777 | 0.000107777 | ENSG00000151498 | n.s. |
| ENSG00000148291 | 192.3083367 | 0.013977091 | 0.3215799 | 0.043464 | 0.965331806 | 0.982417293 | SURF2 | protein_coding | 59.53 | 0.00010768 | 0.00010768 | ENSG00000148291 | n.s. |
| ENSG00000214826 | 337.4830976 | 0.011721175 | 0.226634 | 0.051719 | 0.958752989 | 0.9791475 | DDX12P | unprocessed_pseudogene | 52.17 | 0.000107271 | 0.000107271 | ENSG00000214826 | n.s. |
| ENSG00000148688 | 1297.661711 | -0.012715011 | 0.2712018 | -0.046884 | 0.962605722 | 0.981030744 | RPP30 | protein_coding | 37.93 | -0.000105756 | 0.000105756 | ENSG00000148688 | n.s. |
| ENSG00000236519 | 42.75711641 | -0.012673802 | 0.2726321 | -0.046487 | 0.962922226 | 0.981030744 | LINC01424 | antisense | 45.59 | -0.000105413 | 0.000105413 | ENSG00000236519 | n.s. |
| ENSG00000023572 | 265.8111812 | -0.013355294 | 0.3026495 | -0.044128 | 0.96480243 | 0.982083948 | GLRX2 | protein_coding | 41.92 | -0.000104858 | 0.000104858 | ENSG00000023572 | n.s. |
| ENSG00000103942 | 356.0381767 | -0.011923752 | 0.2417598 | -0.049321 | 0.960663753 | 0.980155576 | HOMER2 | protein_coding | 45.38 | -0.000103796 | 0.000103796 | ENSG00000103942 | n.s. |
| ENSG00000125971 | 422.3195292 | 0.013234821 | 0.3045881 | 0.043452 | 0.965341592 | 0.982417293 | DYNLRB1 | protein_coding | 48.33 | 0.000101961 | 0.000101961 | ENSG00000125971 | n.s. |
| ENSG00000102078 | 212.6221543 | -0.012871646 | 0.2893737 | -0.044481 | 0.964520958 | 0.982012819 | SLC25A14 | protein_coding | 37.86 | -0.000101465 | 0.000101465 | ENSG00000102078 | n.s. |
| ENSG00000173905 | 3385.795805 | 0.011664651 | 0.2420007 | 0.048201 | 0.961556139 | 0.980571751 | GOLIM4 | protein_coding | 38.33 | 9.94E-05 | 9.94E-05 | ENSG00000173905 | n.s. |
| ENSG00000138760 | 2679.532633 | 0.012242385 | 0.2764828 | 0.044279 | 0.964682005 | 0.982083948 | SCARB2 | protein_coding | 41.79 | 9.61E-05 | 9.61E-05 | ENSG00000138760 | n.s. |
| ENSG00000088766 | 824.5030152 | -0.012227209 | 0.2766933 | -0.04419 | 0.964752574 | 0.982083948 | CRLS1 | protein_coding | 39.36 | -9.60E-05 | 9.60E-05 | ENSG00000088766 | n.s. |
| ENSG00000119801 | 476.1292109 | 0.012800985 | 0.3026211 | 0.0423 | 0.966259246 | 0.982920105 | YPEL5 | protein_coding | 41.9 | 9.58E-05 | 9.58E-05 | ENSG00000119801 | n.s. |
| ENSG00000100347 | 652.812548 | 0.012715041 | 0.3005862 | 0.042301 | 0.966258894 | 0.982920105 | SAMM50 | protein_coding | 47.18 | 9.51E-05 | 9.51E-05 | ENSG00000100347 | n.s. |
| ENSG00000177519 | 239.3062383 | 0.011935332 | 0.2661299 | 0.044848 | 0.964228657 | 0.981858816 | RPRM | protein_coding | 59.62 | 9.49E-05 | 9.49E-05 | ENSG00000177519 | n.s. |
| ENSG00000111605 | 8412.359836 | -0.011806952 | 0.2629927 | -0.044895 | 0.964191326 | 0.981858816 | CPSF6 | protein_coding | 35.78 | -9.39E-05 | 9.39E-05 | ENSG00000111605 | n.s. |
| ENSG00000207003 | 11.39715395 | -0.014125429 | 0.3716323 | -0.038009 | 0.969680384 | 0.984817273 | RNU6-611P | snRNA | 42.31 | -9.39E-05 | 9.39E-05 | ENSG00000207003 | n.s. |
| ENSG00000126822 | 73.27704169 | -0.011762643 | 0.2620411 | -0.044889 | 0.96419615 | 0.981858816 | PLEKHG3 | protein_coding | 52.94 | -9.35E-05 | 9.35E-05 | ENSG00000126822 | n.s. |
| ENSG00000196696 | 187.8708302 | -0.011241519 | 0.2412613 | -0.046595 | 0.962836187 | 0.981030744 | | processed_transcript | 44.73 | -9.35E-05 | 9.35E-05 | ENSG00000196696 | n.s. |
| ENSG00000204356 | 1020.828584 | 0.012420408 | 0.2901731 | 0.042803 | 0.96585822 | 0.982818755 | NELFE | protein_coding | 49.59 | 9.35E-05 | 9.35E-05 | ENSG00000204356 | n.s. |
| ENSG00000087152 | 1330.390739 | -0.011452587 | 0.255955 | -0.044745 | 0.964310938 | 0.981870791 | ATXN7L3 | protein_coding | 59.15 | -9.10E-05 | 9.10E-05 | ENSG00000087152 | n.s. |
| ENSG00000123600 | 113.8680567 | 0.010849824 | 0.2331844 | 0.046529 | 0.962888663 | 0.981030744 | METTL8 | protein_coding | 39.39 | 9.02E-05 | 9.02E-05 | ENSG00000123600 | n.s. |
| ENSG00000149503 | 1127.588349 | 0.010541995 | 0.2220925 | 0.047467 | 0.962141279 | 0.980833299 | INCENP | protein_coding | 53.88 | 8.86E-05 | 8.86E-05 | ENSG00000149503 | n.s. |
| ENSG00000070366 | 583.9439007 | -0.010532728 | 0.2262046 | -0.046563 | 0.962861657 | 0.981030744 | SMG6 | protein_coding | 45.75 | -8.76E-05 | 8.76E-05 | ENSG00000070366 | n.s. |
| ENSG00000273611 | 725.6444875 | 0.012159165 | 0.3072814 | 0.03957 | 0.96843584 | 0.984036005 | ZNHIT3 | protein_coding | 45.77 | 8.50E-05 | 8.50E-05 | ENSG00000273611 | n.s. |
| ENSG00000149531 | 134.977344 | -0.011770148 | 0.2928886 | -0.040186 | 0.967944489 | 0.983700082 | FRG1BP | unprocessed_pseudogene | 35.24 | -8.40E-05 | 8.40E-05 | ENSG00000149531 | n.s. |
| ENSG00000132581 | 444.5277372 | 0.01113748 | 0.2603458 | 0.04278 | 0.965877255 | 0.982818755 | SDF2 | protein_coding | 47.11 | 8.38E-05 | 8.38E-05 | ENSG00000132581 | n.s. |
| ENSG00000168268 | 2061.843072 | 0.011112453 | 0.2625688 | 0.042322 | 0.966241953 | 0.982920105 | NT5DC2 | protein_coding | 61.51 | 8.31E-05 | 8.31E-05 | ENSG00000168268 | n.s. |
| ENSG00000155252 | 661.9455034 | 0.011090766 | 0.2663487 | 0.04164 | 0.966785666 | 0.983069002 | PI4K2A | protein_coding | 44.74 | 8.22E-05 | 8.22E-05 | ENSG00000155252 | n.s. |
| ENSG00000197978 | 381.2111603 | -0.010504392 | 0.2402175 | -0.043729 | 0.965120691 | 0.98233609 | GOLGA6L9 | protein_coding | 47.43 | -8.13E-05 | 8.13E-05 | ENSG00000197978 | n.s. |
| ENSG00000186314 | 91.5035418 | 0.010707112 | 0.2563667 | 0.041765 | 0.966686174 | 0.983069002 | PRELID2 | protein_coding | 38.53 | 7.94E-05 | 7.94E-05 | ENSG00000186314 | n.s. |

| | | | | | | | | | | | | | |
|---|---|---|---|---|---|---|---|---|---|---|---|---|---|
| ENSG00000156735 | 696.3124251 | 0.010624099 | 0.2555927 | 0.041567 | 0.966844265 | 0.983069002 | BAG4 | protein_coding | 41.83 | 7.88E-05 | 7.88E-05 | ENSG00000156735 | n.s. |
| ENSG00000104885 | 633.1721457 | -0.009863282 | 0.2193542 | -0.044965 | 0.964135136 | 0.981858816 | DOT1L | protein_coding | 59.45 | -7.84E-05 | 7.84E-05 | ENSG00000104885 | n.s. |
| ENSG00000225733 | 2870.771701 | 0.010573007 | 0.2530605 | 0.041781 | 0.966673646 | 0.983069002 | FGD5-AS1 | antisense | 45.41 | 7.84E-05 | 7.84E-05 | ENSG00000225733 | n.s. |
| ENSG00000223189 | 8.173591908 | -0.014553885 | 0.4707005 | -0.03092 | 0.975333633 | 0.987693882 | RNU6-177P | snRNA | 40.38 | -7.83E-05 | 7.83E-05 | ENSG00000223189 | n.s. |
| ENSG00000064102 | 1080.15338 | -0.010515136 | 0.2511833 | -0.041862 | 0.966608386 | 0.983069002 | INTS13 | protein_coding | 36.38 | -7.80E-05 | 7.80E-05 | ENSG00000064102 | n.s. |
| ENSG00000164346 | 2044.778201 | -0.010997551 | 0.2783776 | -0.039506 | 0.968487063 | 0.984036005 | NSA2 | protein_coding | 37.93 | -7.69E-05 | 7.69E-05 | ENSG00000164346 | n.s. |
| ENSG00000084092 | 479.1336785 | -0.011080232 | 0.2826272 | -0.039204 | 0.968727415 | 0.984208391 | NOA1 | protein_coding | 45.66 | -7.66E-05 | 7.66E-05 | ENSG00000084092 | n.s. |
| ENSG00000265470 | 2.557142789 | 0.020626354 | 0.9008494 | 0.022897 | 0.981732782 | 0.991488868 | MIR548AQ | miRNA | 34.48 | 7.66E-05 | 7.66E-05 | ENSG00000265470 | n.s. |
| ENSG00000105835 | 988.2474102 | -0.010617937 | 0.2617964 | -0.040558 | 0.967648273 | 0.983614435 | NAMPT | protein_coding | 37.12 | -7.62E-05 | 7.62E-05 | ENSG00000105835 | n.s. |
| ENSG00000160360 | 786.224528 | -0.010989482 | 0.2815749 | -0.039029 | 0.968867566 | 0.984278958 | GPSM1 | protein_coding | 66.05 | -7.56E-05 | 7.56E-05 | ENSG00000160360 | n.s. |
| ENSG00000132613 | 287.0445923 | -0.010193151 | 0.2456373 | -0.041497 | 0.966899872 | 0.983069002 | MTSS1L | protein_coding | 56.24 | -7.56E-05 | 7.56E-05 | ENSG00000132613 | n.s. |
| ENSG00000111642 | 5643.756431 | 0.009865728 | 0.2369446 | 0.041637 | 0.966787861 | 0.983069002 | CHD4 | protein_coding | 48.35 | 7.32E-05 | 7.32E-05 | ENSG00000111642 | n.s. |
| ENSG00000263793 | 4.354091389 | -0.015675656 | 0.5634993 | -0.027818 | 0.977806983 | 0.989318118 | MIR3115 | miRNA | 41.18 | -7.31E-05 | 7.31E-05 | ENSG00000263793 | n.s. |
| ENSG00000141384 | 800.907406 | 0.009855031 | 0.2399651 | 0.041069 | 0.967241201 | 0.983344231 | TAF4B | protein_coding | 38.65 | 7.19E-05 | 7.19E-05 | ENSG00000141384 | n.s. |
| ENSG00000135596 | 1580.245212 | -0.010016441 | 0.2468483 | -0.040577 | 0.967632871 | 0.983614435 | MICAL1 | protein_coding | 50.36 | -7.19E-05 | 7.19E-05 | ENSG00000135596 | n.s. |
| ENSG00000162694 | 1099.453499 | -0.010842181 | 0.2905668 | -0.037314 | 0.970234718 | 0.984949178 | EXTL2 | protein_coding | 38.7 | -7.14E-05 | 7.14E-05 | ENSG00000162694 | n.s. |
| ENSG00000175573 | 155.192988 | -0.009920021 | 0.2455827 | -0.040394 | 0.967779166 | 0.98361631 | C11orf68 | protein_coding | 61.82 | -7.12E-05 | 7.12E-05 | ENSG00000175573 | n.s. |
| ENSG00000176890 | 1908.449741 | -0.010100127 | 0.2540879 | -0.039751 | 0.968292017 | 0.983981443 | TYMS | protein_coding | 45.77 | -7.08E-05 | 7.08E-05 | ENSG00000176890 | n.s. |
| ENSG00000046653 | 514.9448487 | -0.01062398 | 0.2813797 | -0.037757 | 0.969881636 | 0.984877983 | GPM6B | protein_coding | 41.5 | -7.03E-05 | 7.03E-05 | ENSG00000046653 | n.s. |
| ENSG00000161914 | 45.04618542 | -0.010507422 | 0.2790856 | -0.037649 | 0.969967174 | 0.984893011 | ZNF653 | protein_coding | 55.67 | -6.95E-05 | 6.95E-05 | ENSG00000161914 | n.s. |
| ENSG00000163811 | 5259.184908 | 0.00954986 | 0.2365089 | 0.040378 | 0.967791412 | 0.98361631 | WDR43 | protein_coding | 39.29 | 6.85E-05 | 6.85E-05 | ENSG00000163811 | n.s. |
| ENSG00000105443 | 477.9390351 | 0.010161124 | 0.2644054 | 0.03843 | 0.969344769 | 0.984620065 | CYTH2 | protein_coding | 58.29 | 6.84E-05 | 6.84E-05 | ENSG00000105443 | n.s. |
| ENSG00000148798 | 2126.590048 | -0.009828118 | 0.2564742 | -0.03832 | 0.969432461 | 0.984637303 | INA | protein_coding | 45.88 | -6.61E-05 | 6.61E-05 | ENSG00000148798 | n.s. |
| ENSG00000010318 | 41.10966827 | -0.010243417 | 0.280606 | -0.036505 | 0.970879985 | 0.985291345 | PHF7 | protein_coding | 45.18 | -6.59E-05 | 6.59E-05 | ENSG00000010318 | n.s. |
| ENSG00000104723 | 2143.105212 | -0.010165891 | 0.2796003 | -0.036359 | 0.970996377 | 0.985291345 | TUSC3 | protein_coding | 36.98 | -6.54E-05 | 6.54E-05 | ENSG00000104723 | n.s. |
| ENSG00000136247 | 170.7060465 | 0.010742835 | 0.3091887 | 0.034745 | 0.972282889 | 0.986165562 | ZDHHC4 | protein_coding | 47.01 | 6.50E-05 | 6.50E-05 | ENSG00000136247 | n.s. |
| ENSG00000176390 | 357.760495 | 0.010001608 | 0.2746601 | 0.036414 | 0.970951857 | 0.985291345 | CRLF3 | protein_coding | 42.46 | 6.44E-05 | 6.44E-05 | ENSG00000176390 | n.s. |
| ENSG00000281394 | 73.19930766 | -0.009890303 | 0.2715668 | -0.036419 | 0.970947939 | 0.985291345 | SCARNA4 | scaRNA | 46.51 | -6.36E-05 | 6.36E-05 | ENSG00000281394 | n.s. |
| ENSG00000076924 | 485.7773356 | 0.009616753 | 0.2544156 | 0.037799 | 0.969847634 | 0.984877983 | XAB2 | protein_coding | 60.82 | 6.36E-05 | 6.36E-05 | ENSG00000076924 | n.s. |
| ENSG00000137970 | 275.0320723 | 0.01096102 | 0.3316988 | 0.033045 | 0.973638621 | 0.986837714 | RPL7P9 | processed_pseudogene | 42.97 | 6.31E-05 | 6.31E-05 | ENSG00000137970 | n.s. |
| ENSG00000223509 | 90.61419569 | 0.010582466 | 0.3129442 | 0.033816 | 0.97302402 | 0.986414259 | | transcribed_processed_pseudog | 39.96 | 6.29E-05 | 6.29E-05 | ENSG00000223509 | n.s. |
| ENSG00000197265 | 625.007926 | 0.010302743 | 0.2981762 | 0.034553 | 0.97243655 | 0.986215306 | GTF2E2 | protein_coding | 39.44 | 6.21E-05 | 6.21E-05 | ENSG00000197265 | n.s. |

| | | | | | | | | | | | | | |
|---|---|---|---|---|---|---|---|---|---|---|---|---|---|
| ENSG00000172239 | 2047.750983 | 0.00990577 | 0.2784257 | 0.035578 | 0.971619022 | 0.985707639 | PAIP1 | protein_coding | 38.36 | 6.19E-05 | 6.19E-05 | ENSG00000172239 | n.s. |
| ENSG00000154217 | 249.5707169 | 0.008913817 | 0.229515 | 0.038838 | 0.96901985 | 0.984361839 | PITPNC1 | protein_coding | 44.62 | 6.10E-05 | 6.10E-05 | ENSG00000154217 | n.s. |
| ENSG00000129195 | 1270.543792 | -0.009922312 | 0.2900771 | -0.034206 | 0.972713057 | 0.986314463 | PIMREG | protein_coding | 53.2 | -5.94E-05 | 5.94E-05 | ENSG00000129195 | n.s. |
| ENSG00000256546 | 107.1519959 | 0.008952134 | 0.2398004 | 0.037332 | 0.970220602 | 0.984949178 | | processed_transcript | 50.95 | 5.90E-05 | 5.90E-05 | ENSG00000256546 | n.s. |
| ENSG00000088356 | 493.8959115 | -0.009449541 | 0.2667176 | -0.035429 | 0.971737656 | 0.985756167 | PDRG1 | protein_coding | 49.41 | -5.89E-05 | 5.89E-05 | ENSG00000088356 | n.s. |
| ENSG00000124733 | 759.1425719 | 0.009843984 | 0.2899893 | 0.033946 | 0.972920191 | 0.986414259 | MEA1 | protein_coding | 56.59 | 5.85E-05 | 5.85E-05 | ENSG00000124733 | n.s. |
| ENSG00000158186 | 156.9193115 | -0.008824844 | 0.2349886 | -0.037554 | 0.97004301 | 0.984898187 | MRAS | protein_coding | 48.52 | -5.83E-05 | 5.83E-05 | ENSG00000158186 | n.s. |
| ENSG00000107643 | 2490.685235 | 0.009108213 | 0.2551316 | 0.0357 | 0.971521524 | 0.985680549 | MAPK8 | protein_coding | 37.34 | 5.71E-05 | 5.71E-05 | ENSG00000107643 | n.s. |
| ENSG00000241973 | 1598.479352 | 0.008508383 | 0.2333581 | 0.036461 | 0.970915076 | 0.985291345 | PI4KA | protein_coding | 47.81 | 5.48E-05 | 5.48E-05 | ENSG00000241973 | n.s. |
| ENSG00000115073 | 896.068225 | -0.009375884 | 0.2838995 | -0.033025 | 0.973654362 | 0.986837714 | ACTR1B | protein_coding | 58.67 | -5.40E-05 | 5.40E-05 | ENSG00000115073 | n.s. |
| ENSG00000115677 | 6233.150164 | -0.008893241 | 0.2555008 | -0.034807 | 0.972233566 | 0.986165562 | HDLBP | protein_coding | 46.99 | -5.38E-05 | 5.38E-05 | ENSG00000115677 | n.s. |
| ENSG00000162413 | 311.956848 | -0.008348388 | 0.231539 | -0.036056 | 0.971237644 | 0.985464343 | KLHL21 | protein_coding | 54.46 | -5.31E-05 | 5.31E-05 | ENSG00000162413 | n.s. |
| ENSG00000088305 | 235.4319568 | -0.008723337 | 0.2528058 | -0.034506 | 0.9724736 | 0.986215306 | DNMT3B | protein_coding | 49.4 | -5.26E-05 | 5.26E-05 | ENSG00000088305 | n.s. |
| ENSG00000131791 | 937.6370679 | -0.008712516 | 0.2571882 | -0.033876 | 0.972976003 | 0.986414259 | PRKAB2 | protein_coding | 39.83 | -5.18E-05 | 5.18E-05 | ENSG00000131791 | n.s. |
| ENSG00000225761 | 33.43521147 | 0.010327823 | 0.3566401 | 0.028959 | 0.976897554 | 0.988613508 | | sense_intronic | 40.99 | 5.14E-05 | 5.14E-05 | ENSG00000225761 | n.s. |
| ENSG00000260608 | 36.72836724 | 0.0091105 | 0.2862734 | 0.031824 | 0.974612034 | 0.987300332 | | antisense | 39.41 | 5.06E-05 | 5.06E-05 | ENSG00000260608 | n.s. |
| ENSG00000173674 | 2588.276752 | -0.008743462 | 0.2749622 | -0.031799 | 0.974632524 | 0.987300332 | EIF1AX | protein_coding | 38.58 | -4.85E-05 | 4.85E-05 | ENSG00000173674 | n.s. |
| ENSG00000125457 | 86.74199587 | -0.009194151 | 0.3075209 | -0.029898 | 0.976148684 | 0.98807111 | MIF4GD | protein_coding | 55.74 | -4.79E-05 | 4.79E-05 | ENSG00000125457 | n.s. |
| ENSG00000163661 | 327.8432156 | -0.008485092 | 0.2616883 | -0.032424 | 0.974133581 | 0.987201989 | PTX3 | protein_coding | 42.75 | -4.75E-05 | 4.75E-05 | ENSG00000163661 | n.s. |
| ENSG00000261430 | 13.31524555 | -0.010469453 | 0.3912845 | -0.026757 | 0.978653847 | 0.98968448 | | sense_intronic | 53.35 | -4.71E-05 | 4.71E-05 | ENSG00000261430 | n.s. |
| ENSG00000101180 | 104.07236 | 0.00847163 | 0.2659892 | 0.03185 | 0.974592049 | 0.987300332 | HRH3 | protein_coding | 66.12 | 4.70E-05 | 4.70E-05 | ENSG00000101180 | n.s. |
| ENSG00000160766 | 184.2116932 | -0.007820085 | 0.2275545 | -0.034366 | 0.972585485 | 0.986256935 | GBAP1 | transcribed_unprocessed_pseud | 49.29 | -4.70E-05 | 4.70E-05 | ENSG00000160766 | n.s. |
| ENSG00000223837 | 55.6138168 | -0.008359643 | 0.2580388 | -0.032397 | 0.974155579 | 0.987201989 | | sense_intronic | 58.47 | -4.68E-05 | 4.68E-05 | ENSG00000223837 | n.s. |
| ENSG00000065150 | 5995.231755 | -0.008585763 | 0.2796797 | -0.030699 | 0.975509943 | 0.987711798 | IPO5 | protein_coding | 40.27 | -4.61E-05 | 4.61E-05 | ENSG00000065150 | n.s. |
| ENSG00000235875 | 92.03229127 | 0.007932862 | 0.2377964 | 0.03336 | 0.973387595 | 0.986710993 | ARHGEF7-AS2 | antisense | 60.85 | 4.61E-05 | 4.61E-05 | ENSG00000235875 | n.s. |
| ENSG00000129187 | 925.3143376 | -0.008217837 | 0.2598143 | -0.03163 | 0.974767394 | 0.98731896 | DCTD | protein_coding | 46.1 | -4.55E-05 | 4.55E-05 | ENSG00000129187 | n.s. |
| ENSG00000161956 | 373.6364906 | 0.008889951 | 0.302029 | 0.029434 | 0.976518377 | 0.988301619 | SENP3 | protein_coding | 51.55 | 4.54E-05 | 4.54E-05 | ENSG00000161956 | n.s. |
| ENSG00000174151 | 196.0760568 | 0.008146677 | 0.255195 | 0.031923 | 0.974533187 | 0.987300332 | CYB561D1 | protein_coding | 52.44 | 4.52E-05 | 4.52E-05 | ENSG00000174151 | n.s. |
| ENSG00000066557 | 2099.279389 | 0.008388679 | 0.2729527 | 0.030733 | 0.975482401 | 0.987711798 | LRRC40 | protein_coding | 35.14 | 4.50E-05 | 4.50E-05 | ENSG00000066557 | n.s. |
| ENSG00000154429 | 1361.849294 | 0.00832144 | 0.2762659 | 0.030121 | 0.975970455 | 0.988034367 | CCSAP | protein_coding | 43.58 | 4.35E-05 | 4.35E-05 | ENSG00000154429 | n.s. |
| ENSG00000131477 | 320.968716 | 0.007852874 | 0.2541495 | 0.030899 | 0.975350369 | 0.987693882 | RAMP2 | protein_coding | 55.13 | 4.22E-05 | 4.22E-05 | ENSG00000131477 | n.s. |
| ENSG00000100991 | 1090.946612 | -0.008052332 | 0.2688263 | -0.029954 | 0.976104005 | 0.98807111 | TRPC4AP | protein_coding | 42.95 | -4.20E-05 | 4.20E-05 | ENSG00000100991 | n.s. |

| | | | | | | | | | | | | | |
|---|---|---|---|---|---|---|---|---|---|---|---|---|---|
| ENSG00000151778 | 86.87426727 | 0.009365532 | 0.3577253 | 0.026181 | 0.979113136 | 0.989920282 | SERP2 | protein_coding | 44.49 | 4.12E-05 | 4.12E-05 | ENSG00000151778 | n.s. |
| ENSG00000163291 | 1632.646395 | 0.008365257 | 0.2943713 | 0.028417 | 0.977329274 | 0.988978521 | PAQR3 | protein_coding | 35.92 | 4.03E-05 | 4.03E-05 | ENSG00000163291 | n.s. |
| ENSG00000112996 | 2664.601427 | -0.007440156 | 0.2387313 | -0.031165 | 0.975137636 | 0.987622122 | MRPS30 | protein_coding | 36.6 | -4.02E-05 | 4.02E-05 | ENSG00000112996 | n.s. |
| ENSG00000182518 | 186.4591648 | -0.008456275 | 0.3052487 | -0.027703 | 0.977899109 | 0.989339435 | FAM104B | protein_coding | 36.9 | -3.94E-05 | 3.94E-05 | ENSG00000182518 | n.s. |
| ENSG00000100242 | 497.6430747 | 0.007649135 | 0.259241 | 0.029506 | 0.976461129 | 0.988301619 | SUN2 | protein_coding | 54.74 | 3.91E-05 | 3.91E-05 | ENSG00000100242 | n.s. |
| ENSG00000274267 | 2867.945714 | 0.006827516 | 0.2116663 | 0.032256 | 0.974267873 | 0.98724393 | HIST1H3B | protein_coding | 59.12 | 3.81E-05 | 3.81E-05 | ENSG00000274267 | n.s. |
| ENSG00000252283 | 8.939735706 | 0.009237428 | 0.4092547 | 0.022571 | 0.981992205 | 0.991491591 | | misc_RNA | 49.02 | 3.43E-05 | 3.43E-05 | ENSG00000252283 | n.s. |
| ENSG00000168159 | 1049.738048 | 0.007557444 | 0.2905283 | 0.026013 | 0.97924716 | 0.989983895 | RNF187 | protein_coding | 56.59 | 3.30E-05 | 3.30E-05 | ENSG00000168159 | n.s. |
| ENSG00000170312 | 4740.015868 | 0.007170389 | 0.2690432 | 0.026651 | 0.978737743 | 0.98968448 | CDK1 | protein_coding | 35.07 | 3.23E-05 | 3.23E-05 | ENSG00000170312 | n.s. |
| ENSG00000133606 | 1122.495233 | -0.006991741 | 0.2554186 | -0.027374 | 0.978161708 | 0.989490222 | MKRN1 | protein_coding | 45.78 | -3.21E-05 | 3.21E-05 | ENSG00000133606 | n.s. |
| ENSG00000185728 | 2187.783997 | -0.007113232 | 0.2684729 | -0.026495 | 0.978862392 | 0.989738641 | YTHDF3 | protein_coding | 37.56 | -3.19E-05 | 3.19E-05 | ENSG00000185728 | n.s. |
| ENSG00000119720 | 350.0451711 | -0.007044946 | 0.2730805 | -0.025798 | 0.979418415 | 0.990085137 | NRDE2 | protein_coding | 44.18 | -3.05E-05 | 3.05E-05 | ENSG00000119720 | n.s. |
| ENSG00000196230 | 19398.5813 | -0.006733458 | 0.2522437 | -0.026694 | 0.978703592 | 0.98968448 | TUBB | protein_coding | 53.06 | -3.03E-05 | 3.03E-05 | ENSG00000196230 | n.s. |
| ENSG00000279529 | 114.2774835 | 0.006367827 | 0.2344909 | 0.027156 | 0.978335333 | 0.989565044 | | antisense | 50.53 | 2.90E-05 | 2.90E-05 | ENSG00000279529 | n.s. |
| ENSG00000106348 | 617.2683477 | -0.006579215 | 0.2608125 | -0.025226 | 0.979874826 | 0.9904027 | IMPDH1 | protein_coding | 57.06 | -2.76E-05 | 2.76E-05 | ENSG00000106348 | n.s. |
| ENSG00000074201 | 2744.363077 | 0.00710164 | 0.2977989 | 0.023847 | 0.980974572 | 0.991360305 | CLNS1A | protein_coding | 41.76 | 2.68E-05 | 2.68E-05 | ENSG00000074201 | n.s. |
| ENSG00000140025 | 80.55380961 | -0.007077938 | 0.3073554 | -0.023029 | 0.981627527 | 0.991488868 | EFCAB11 | protein_coding | 40 | -2.63E-05 | 2.63E-05 | ENSG00000140025 | n.s. |
| ENSG00000243406 | 744.3514876 | -0.005800697 | 0.2172436 | -0.026701 | 0.978697941 | 0.98968448 | MRPS31P5 | transcribed_unprocessed_pseud | 39.42 | -2.61E-05 | 2.61E-05 | ENSG00000243406 | n.s. |
| ENSG00000252242 | 1.799781568 | -0.010784119 | 0.7206718 | -0.014964 | 0.988060916 | 0.994489419 | RNU7-115P | snRNA | 36.36 | -2.59E-05 | 2.59E-05 | ENSG00000252242 | n.s. |
| ENSG00000102312 | 76.20505759 | -0.006829373 | 0.297671 | -0.022943 | 0.981695991 | 0.991488868 | PORCN | protein_coding | 51.68 | -2.54E-05 | 2.54E-05 | ENSG00000102312 | n.s. |
| ENSG00000100354 | 274.8758835 | 0.005892522 | 0.2314116 | 0.025463 | 0.97968536 | 0.990283089 | TNRC6B | protein_coding | 42.15 | 2.50E-05 | 2.50E-05 | ENSG00000100354 | n.s. |
| ENSG00000164109 | 4912.871104 | -0.006680757 | 0.2869318 | -0.023283 | 0.981424187 | 0.991484724 | MAD2L1 | protein_coding | 38.16 | -2.48E-05 | 2.48E-05 | ENSG00000164109 | n.s. |
| ENSG00000145730 | 282.4071164 | 0.006456343 | 0.2864892 | 0.022536 | 0.982020329 | 0.991491591 | PAM | protein_coding | 35.92 | 2.40E-05 | 2.40E-05 | ENSG00000145730 | n.s. |
| ENSG00000089685 | 3859.675824 | 0.006347572 | 0.2670916 | 0.023766 | 0.981039637 | 0.991360305 | BIRC5 | protein_coding | 49.27 | 2.39E-05 | 2.39E-05 | ENSG00000089685 | n.s. |
| ENSG00000108679 | 652.0146738 | -0.006337024 | 0.278852 | -0.022725 | 0.981869315 | 0.991491591 | LGALS3BP | protein_coding | 58.85 | -2.35E-05 | 2.35E-05 | ENSG00000108679 | n.s. |
| ENSG00000167740 | 138.8318014 | 0.006193734 | 0.2655728 | 0.023322 | 0.981393287 | 0.991484724 | CYB5D2 | protein_coding | 49.97 | 2.30E-05 | 2.30E-05 | ENSG00000167740 | n.s. |
| ENSG00000138709 | 402.4230597 | -0.006191716 | 0.2664001 | -0.023242 | 0.981457098 | 0.991484724 | LARP1B | protein_coding | 39.11 | -2.30E-05 | 2.30E-05 | ENSG00000138709 | n.s. |
| ENSG00000152117 | 817.5821734 | -0.005684313 | 0.2298524 | -0.02473 | 0.980270109 | 0.990730309 | | transcribed_processed_pseudog | 53.93 | -2.30E-05 | 2.30E-05 | ENSG00000152117 | n.s. |
| ENSG00000134152 | 915.945158 | -0.006192228 | 0.2728242 | -0.022697 | 0.981892155 | 0.991491591 | KATNBL1 | protein_coding | 40.54 | -2.30E-05 | 2.30E-05 | ENSG00000134152 | n.s. |
| ENSG00000086666 | 1385.114695 | -0.006312679 | 0.2966232 | -0.021282 | 0.983020852 | 0.992142031 | ZFAND6 | protein_coding | 38.17 | -2.16E-05 | 2.16E-05 | ENSG00000086666 | n.s. |
| ENSG00000105438 | 707.0427341 | 0.00586801 | 0.2652877 | 0.022119 | 0.982352695 | 0.991683337 | KDELR1 | protein_coding | 53.35 | 2.13E-05 | 2.13E-05 | ENSG00000105438 | n.s. |
| ENSG00000119446 | 931.0352839 | -0.006007058 | 0.2762588 | -0.021744 | 0.982651916 | 0.99191348 | RBM18 | protein_coding | 40.59 | -2.12E-05 | 2.12E-05 | ENSG00000119446 | n.s. |

| | | | | | | | | | | | | | |
|---|---|---|---|---|---|---|---|---|---|---|---|---|---|
| ENSG00000207789 | 3.841350595 | 0.008651955 | 0.5864141 | 0.014754 | 0.988228437 | 0.994537862 | MIR26A2 | miRNA | 47.62 | 2.06E-05 | 2.06E-05 | ENSG00000207789 | n.s. |
| ENSG00000116171 | 1142.086185 | 0.0060249 | 0.2881462 | 0.020909 | 0.983318107 | 0.992271006 | SCP2 | protein_coding | 40.01 | 2.03E-05 | 2.03E-05 | ENSG00000116171 | n.s. |
| ENSG00000095015 | 722.0945879 | 0.00537807 | 0.2271049 | 0.023681 | 0.981107063 | 0.991360305 | MAP3K1 | protein_coding | 38.07 | 2.03E-05 | 2.03E-05 | ENSG00000095015 | n.s. |
| ENSG00000183840 | 125.7162574 | 0.005434678 | 0.2345614 | 0.02317 | 0.981515043 | 0.991484724 | GPR39 | protein_coding | 42.93 | 2.02E-05 | 2.02E-05 | ENSG00000183840 | n.s. |
| ENSG00000174456 | 151.7003517 | -0.005928782 | 0.284305 | -0.020854 | 0.983362446 | 0.992271006 | C12orf76 | protein_coding | 45.73 | -2.00E-05 | 2.00E-05 | ENSG00000174456 | n.s. |
| ENSG00000113328 | 2132.625346 | 0.005694398 | 0.2661404 | 0.021396 | 0.98292959 | 0.992121841 | CCNG1 | protein_coding | 37.5 | 1.96E-05 | 1.96E-05 | ENSG00000113328 | n.s. |
| ENSG00000122378 | 2606.265074 | -0.005698522 | 0.2851207 | -0.019986 | 0.984054264 | 0.992614422 | FAM213A | protein_coding | 47.5 | -1.83E-05 | 1.83E-05 | ENSG00000122378 | n.s. |
| ENSG00000161618 | 304.1707417 | 0.005488341 | 0.2705539 | 0.020286 | 0.983815566 | 0.992584357 | ALDH16A1 | protein_coding | 56.83 | 1.77E-05 | 1.77E-05 | ENSG00000161618 | n.s. |
| ENSG00000049245 | 353.7913188 | 0.005233305 | 0.2502331 | 0.020914 | 0.983314479 | 0.992271006 | VAMP3 | protein_coding | 42.77 | 1.76E-05 | 1.76E-05 | ENSG00000049245 | n.s. |
| ENSG00000077380 | 2870.286113 | 0.005476263 | 0.2765599 | 0.019801 | 0.984201831 | 0.992614422 | DYNC1I2 | protein_coding | 36.49 | 1.76E-05 | 1.76E-05 | ENSG00000077380 | n.s. |
| ENSG00000145743 | 220.4055493 | -0.005336937 | 0.2618052 | -0.020385 | 0.983736135 | 0.992576144 | FBXL17 | protein_coding | 35.81 | -1.73E-05 | 1.73E-05 | ENSG00000145743 | n.s. |
| ENSG00000084693 | 444.775148 | 0.005360264 | 0.2698663 | 0.019863 | 0.984152927 | 0.992614422 | AGBL5 | protein_coding | 48.47 | 1.73E-05 | 1.73E-05 | ENSG00000084693 | n.s. |
| ENSG00000148840 | 2129.725177 | 0.004690005 | 0.2109625 | 0.022231 | 0.98226332 | 0.991665014 | PPRC1 | protein_coding | 51.99 | 1.70E-05 | 1.70E-05 | ENSG00000148840 | n.s. |
| ENSG00000215957 | 4.083542452 | -0.008632522 | 0.7170592 | -0.012039 | 0.990394671 | 0.995471051 | MIR300 | miRNA | 43.37 | -1.70E-05 | 1.70E-05 | ENSG00000215957 | n.s. |
| ENSG00000099956 | 1209.798443 | -0.005493258 | 0.2858075 | -0.01922 | 0.984665498 | 0.9929382 | SMARCB1 | protein_coding | 50.77 | -1.69E-05 | 1.69E-05 | ENSG00000099956 | n.s. |
| ENSG00000198513 | 615.899928 | -0.005052745 | 0.2539296 | -0.019898 | 0.984124571 | 0.992614422 | ATL1 | protein_coding | 37.9 | -1.63E-05 | 1.63E-05 | ENSG00000198513 | n.s. |
| ENSG00000164096 | 1079.194146 | -0.00541189 | 0.2929599 | -0.018473 | 0.985261402 | 0.99332328 | C4orf3 | protein_coding | 42.89 | -1.57E-05 | 1.57E-05 | ENSG00000164096 | n.s. |
| ENSG00000174574 | 952.8137744 | 0.005561343 | 0.3089139 | 0.018003 | 0.98563655 | 0.993512695 | AKIRIN1 | protein_coding | 43.9 | 1.57E-05 | 1.57E-05 | ENSG00000174574 | n.s. |
| ENSG00000029363 | 15172.72971 | -0.004879804 | 0.244816 | -0.019933 | 0.984097188 | 0.992614422 | BCLAF1 | protein_coding | 35.59 | -1.57E-05 | 1.57E-05 | ENSG00000029363 | n.s. |
| ENSG00000091157 | 373.3287967 | 0.005078146 | 0.2635519 | 0.019268 | 0.984627225 | 0.9929382 | WDR7 | protein_coding | 36.37 | 1.56E-05 | 1.56E-05 | ENSG00000091157 | n.s. |
| ENSG00000023909 | 737.8110108 | -0.004826506 | 0.2551148 | -0.018919 | 0.984905758 | 0.993108551 | GCLM | protein_coding | 36.37 | -1.45E-05 | 1.45E-05 | ENSG00000023909 | n.s. |
| ENSG00000180581 | 51.57746032 | -0.005373612 | 0.3433592 | -0.01565 | 0.987513521 | 0.994082189 | SRP9P1 | processed_pseudogene | 45.21 | -1.39E-05 | 1.39E-05 | ENSG00000180581 | n.s. |
| ENSG00000116857 | 812.43268 | 0.004842851 | 0.2708553 | 0.01788 | 0.98573471 | 0.993512695 | TMEM9 | protein_coding | 48.53 | 1.37E-05 | 1.37E-05 | ENSG00000116857 | n.s. |
| ENSG00000113068 | 892.962081 | 0.004879713 | 0.2788327 | 0.017501 | 0.986037329 | 0.993601921 | PFDN1 | protein_coding | 40.78 | 1.36E-05 | 1.36E-05 | ENSG00000113068 | n.s. |
| ENSG00000197043 | 2515.595001 | 0.0047881 | 0.2652415 | 0.018052 | 0.98559749 | 0.993512695 | ANXA6 | protein_coding | 49.02 | 1.35E-05 | 1.35E-05 | ENSG00000197043 | n.s. |
| ENSG00000137710 | 1187.823127 | -0.004716089 | 0.2694484 | -0.017503 | 0.986035534 | 0.993601921 | RDX | protein_coding | 39.96 | -1.31E-05 | 1.31E-05 | ENSG00000137710 | n.s. |
| ENSG00000164934 | 1997.883872 | 0.004747472 | 0.2772439 | 0.017124 | 0.986337842 | 0.993688985 | DCAF13 | protein_coding | 37.09 | 1.31E-05 | 1.31E-05 | ENSG00000164934 | n.s. |
| ENSG00000280206 | 75.89499181 | 0.004937802 | 0.3045534 | 0.016213 | 0.987064258 | 0.993989271 | | lincRNA | 40.82 | 1.29E-05 | 1.29E-05 | ENSG00000280206 | n.s. |
| ENSG00000184160 | 177.3368522 | 0.004563072 | 0.2629899 | 0.017351 | 0.986156799 | 0.993650392 | ADRA2C | protein_coding | 69.62 | 1.26E-05 | 1.26E-05 | ENSG00000184160 | n.s. |
| ENSG00000198408 | 4293.081391 | -0.004467803 | 0.2530878 | -0.017653 | 0.985915536 | 0.993601921 | MGEA5 | protein_coding | 41.92 | -1.25E-05 | 1.25E-05 | ENSG00000198408 | n.s. |
| ENSG00000106290 | 288.4235744 | -0.004372259 | 0.244164 | -0.017907 | 0.985712995 | 0.993512695 | TAF6 | protein_coding | 53.95 | -1.24E-05 | 1.24E-05 | ENSG00000106290 | n.s. |
| ENSG00000199133 | 71.04617134 | 0.00471539 | 0.2945326 | 0.01601 | 0.987226625 | 0.994004133 | MIRLET7D | miRNA | 48.28 | 1.23E-05 | 1.23E-05 | ENSG00000199133 | n.s. |

| | | | | | | | | | | | | | |
|---|---|---|---|---|---|---|---|---|---|---|---|---|---|
| ENSG00000133835 | 1594.692134 | 0.004660447 | 0.2794655 | 0.016676 | 0.986694867 | 0.993933861 | HSD17B4 | protein_coding | 37.34 | 1.23E-05 | 1.23E-05 | ENSG00000133835 | n.s. |
| ENSG00000162757 | 236.3819327 | -0.004135598 | 0.2212945 | -0.018688 | 0.985089831 | 0.993222225 | C1orf74 | protein_coding | 44.56 | -1.22E-05 | 1.22E-05 | ENSG00000162757 | n.s. |
| ENSG00000135999 | 2361.665579 | 0.004348688 | 0.2535493 | 0.017151 | 0.986315952 | 0.993688985 | EPC2 | protein_coding | 37.09 | 1.20E-05 | 1.20E-05 | ENSG00000135999 | n.s. |
| ENSG00000222071 | 6.754872803 | -0.006965195 | 0.6453092 | -0.010794 | 0.991388138 | 0.996140876 | MIR1915 | miRNA | 78.75 | -1.17E-05 | 1.17E-05 | ENSG00000222071 | n.s. |
| ENSG00000234788 | 80.01207281 | 0.004453936 | 0.2796593 | 0.015926 | 0.987293195 | 0.994004133 | HSPA8P3 | processed_pseudogene | 45.25 | 1.16E-05 | 1.16E-05 | ENSG00000234788 | n.s. |
| ENSG00000178802 | 83.22639202 | 0.004235272 | 0.2569881 | 0.01648 | 0.986851122 | 0.993933861 | MPI | protein_coding | 51.86 | 1.12E-05 | 1.12E-05 | ENSG00000178802 | n.s. |
| ENSG00000261476 | 48.18617543 | 0.004223102 | 0.2651125 | 0.015929 | 0.987290657 | 0.994004133 | | sense_intronic | 46.6 | 1.10E-05 | 1.10E-05 | ENSG00000261476 | n.s. |
| ENSG00000250299 | 96.52115352 | 0.004023612 | 0.2546437 | 0.015801 | 0.987393193 | 0.99403293 | MRPS31P4 | transcribed_unprocessed_pseud | 41.83 | 1.05E-05 | 1.05E-05 | ENSG00000250299 | n.s. |
| ENSG00000100934 | 1204.288501 | 0.004384917 | 0.3010284 | 0.014566 | 0.988378062 | 0.994537862 | SEC23A | protein_coding | 38.8 | 1.04E-05 | 1.04E-05 | ENSG00000100934 | n.s. |
| ENSG00000112655 | 1893.507338 | -0.003910835 | 0.2364506 | -0.01654 | 0.986803785 | 0.993933861 | PTK7 | protein_coding | 49.42 | -1.03E-05 | 1.03E-05 | ENSG00000112655 | n.s. |
| ENSG00000115053 | 22239.15257 | 0.003775582 | 0.2293625 | 0.016461 | 0.986866458 | 0.993933861 | NCL | protein_coding | 47.89 | 9.98E-06 | 9.98E-06 | ENSG00000115053 | n.s. |
| ENSG00000254505 | 191.4130202 | -0.003758368 | 0.2312717 | -0.016251 | 0.987034246 | 0.993989271 | CHMP4A | protein_coding | 50.64 | -9.84E-06 | 9.84E-06 | ENSG00000254505 | n.s. |
| ENSG00000260924 | 133.9422913 | 0.003792916 | 0.2460897 | 0.015413 | 0.9877029 | 0.994200945 | LINC01311 | antisense | 63.11 | 9.58E-06 | 9.58E-06 | ENSG00000260924 | n.s. |
| ENSG00000111961 | 66.09303238 | 0.003989194 | 0.2715227 | 0.014692 | 0.988277951 | 0.994537862 | SASH1 | protein_coding | 43.29 | 9.49E-06 | 9.49E-06 | ENSG00000111961 | n.s. |
| ENSG00000198860 | 1354.335938 | 0.004067083 | 0.2896065 | 0.014043 | 0.988795292 | 0.9948079 | TSEN15 | protein_coding | 35.04 | 9.19E-06 | 9.19E-06 | ENSG00000198860 | n.s. |
| ENSG00000143771 | 1437.640481 | -0.004034769 | 0.2920572 | -0.013815 | 0.98897758 | 0.9948079 | CNIH4 | protein_coding | 43.22 | -9.12E-06 | 9.12E-06 | ENSG00000143771 | n.s. |
| ENSG00000120686 | 1626.419741 | -0.003999705 | 0.2875902 | -0.013908 | 0.988903654 | 0.9948079 | UFM1 | protein_coding | 33.9 | -9.04E-06 | 9.04E-06 | ENSG00000120686 | n.s. |
| ENSG00000198556 | 492.085827 | 0.003649875 | 0.2509279 | 0.014546 | 0.988394771 | 0.994537862 | ZNF789 | protein_coding | 47.78 | 8.68E-06 | 8.68E-06 | ENSG00000198556 | n.s. |
| ENSG00000225921 | 1150.149774 | 0.003725851 | 0.2772721 | 0.013438 | 0.98927873 | 0.994923993 | NOL7 | protein_coding | 38.69 | 8.23E-06 | 8.23E-06 | ENSG00000225921 | n.s. |
| ENSG00000102900 | 1493.553818 | 0.003641985 | 0.2747262 | 0.013257 | 0.989422927 | 0.994925279 | NUP93 | protein_coding | 42.43 | 8.05E-06 | 8.05E-06 | ENSG00000102900 | n.s. |
| ENSG00000281896 | 59.24688761 | -0.003676078 | 0.2836229 | -0.012961 | 0.989658791 | 0.995018728 | | transcribed_unprocessed_pseud | 48.09 | -7.97E-06 | 7.97E-06 | ENSG00000281896 | n.s. |
| ENSG00000239719 | 9.198933782 | -0.004737059 | 0.4409869 | -0.010742 | 0.991429328 | 0.996140876 | | processed_pseudogene | 50.83 | -7.95E-06 | 7.95E-06 | ENSG00000239719 | n.s. |
| ENSG00000185164 | 158.1014138 | 0.003572703 | 0.2693366 | 0.013265 | 0.989416509 | 0.994925279 | NOMO2 | protein_coding | 46.04 | 7.89E-06 | 7.89E-06 | ENSG00000185164 | n.s. |
| ENSG00000100726 | 350.1866817 | -0.00344567 | 0.2503879 | -0.013761 | 0.989020395 | 0.9948079 | TELO2 | protein_coding | 63.09 | -7.79E-06 | 7.79E-06 | ENSG00000100726 | n.s. |
| ENSG00000244389 | 62.00125965 | 0.00345359 | 0.2540382 | 0.013595 | 0.989153282 | 0.994869692 | RN7SL242P | misc_RNA | 53.54 | 7.71E-06 | 7.71E-06 | ENSG00000244389 | n.s. |
| ENSG00000215305 | 546.888015 | 0.003508943 | 0.284979 | 0.012313 | 0.990175904 | 0.995323017 | VPS16 | protein_coding | 48.8 | 7.14E-06 | 7.14E-06 | ENSG00000215305 | n.s. |
| ENSG00000239264 | 135.2238289 | 0.003150522 | 0.2262074 | 0.013928 | 0.988887755 | 0.9948079 | TXNDC5 | protein_coding | 47.02 | 7.12E-06 | 7.12E-06 | ENSG00000239264 | n.s. |
| ENSG00000100058 | 485.1900823 | -0.003167313 | 0.2405828 | -0.013165 | 0.989496019 | 0.994926921 | CRYBB2P1 | transcribed_unprocessed_pseud | 45.32 | -7.00E-06 | 7.00E-06 | ENSG00000100058 | n.s. |
| ENSG00000242802 | 240.2777916 | -0.003282962 | 0.2590546 | -0.012673 | 0.989888794 | 0.995178112 | AP5Z1 | protein_coding | 58.29 | -6.89E-06 | 6.89E-06 | ENSG00000242802 | n.s. |
| ENSG00000073050 | 526.619769 | 0.003214938 | 0.2593355 | 0.012397 | 0.990109016 | 0.995323017 | XRCC1 | protein_coding | 50.94 | 6.55E-06 | 6.55E-06 | ENSG00000073050 | n.s. |
| ENSG00000242798 | 18.26101761 | 0.003676434 | 0.3311781 | 0.011101 | 0.991142804 | 0.996007309 | | antisense | 51.6 | 6.39E-06 | 6.39E-06 | ENSG00000242798 | n.s. |
| ENSG00000186998 | 170.2912833 | -0.003245903 | 0.2799278 | -0.011596 | 0.990748335 | 0.995754643 | EMID1 | protein_coding | 52.65 | -6.00E-06 | 6.00E-06 | ENSG00000186998 | n.s. |

| | | | | | | | | | | | | | |
|---|---|---|---|---|---|---|---|---|---|---|---|---|---|
| ENSG00000171421 | 300.5281504 | 0.003541999 | 0.3321 | 0.010665 | 0.991490358 | 0.996140876 | MRPL36 | protein_coding | 55.79 | 5.95E-06 | 5.95E-06 | ENSG00000171421 | n.s. |
| ENSG00000201367 | 12.30934331 | -0.003802698 | 0.40167 | -0.009467 | 0.992446364 | 0.996549647 | RNU6-522P | snRNA | 35.51 | -5.71E-06 | 5.71E-06 | ENSG00000201367 | n.s. |
| ENSG00000149091 | 384.8660638 | -0.002911681 | 0.2610858 | -0.011152 | 0.991102019 | 0.996007309 | DGKZ | protein_coding | 56.98 | -5.06E-06 | 5.06E-06 | ENSG00000149091 | n.s. |
| ENSG00000160783 | 345.3274492 | 0.003140185 | 0.3122426 | 0.010057 | 0.991975908 | 0.996412998 | PMF1 | protein_coding | 48.91 | 4.90E-06 | 4.90E-06 | ENSG00000160783 | n.s. |
| ENSG00000182013 | 504.9856576 | 0.002841742 | 0.279086 | 0.010182 | 0.991875825 | 0.996412998 | PNMA8A | protein_coding | 52.55 | 4.43E-06 | 4.43E-06 | ENSG00000182013 | n.s. |
| ENSG00000186577 | 158.2212613 | -0.002740689 | 0.2710226 | -0.010112 | 0.991931607 | 0.996412998 | SMIM29 | protein_coding | 61.63 | -4.28E-06 | 4.28E-06 | ENSG00000186577 | n.s. |
| ENSG00000103510 | 579.8766388 | 0.002770949 | 0.2882491 | 0.009613 | 0.992330026 | 0.996549647 | KAT8 | protein_coding | 52.56 | 4.16E-06 | 4.16E-06 | ENSG00000103510 | n.s. |
| ENSG00000067445 | 857.8688461 | 0.002746333 | 0.2826428 | 0.009717 | 0.992247379 | 0.996549647 | TRO | protein_coding | 51.41 | 4.12E-06 | 4.12E-06 | ENSG00000067445 | n.s. |
| ENSG00000139921 | 1831.326356 | -0.002608753 | 0.2688705 | -0.009703 | 0.992258536 | 0.996549647 | TMX1 | protein_coding | 36.96 | -3.92E-06 | 3.92E-06 | ENSG00000139921 | n.s. |
| ENSG00000101019 | 631.1010725 | 0.002553338 | 0.2731397 | 0.009348 | 0.992541403 | 0.996549647 | UQCC1 | protein_coding | 43.02 | 3.83E-06 | 3.83E-06 | ENSG00000101019 | n.s. |
| ENSG00000156642 | 978.5384076 | 0.002537249 | 0.2707587 | 0.009371 | 0.992523223 | 0.996549647 | NPTN | protein_coding | 41.2 | 3.81E-06 | 3.81E-06 | ENSG00000156642 | n.s. |
| ENSG00000132004 | 76.71211992 | 0.002523051 | 0.2859875 | 0.008822 | 0.992960962 | 0.996755261 | FBXW9 | protein_coding | 56.55 | 3.56E-06 | 3.56E-06 | ENSG00000132004 | n.s. |
| ENSG00000130702 | 39.75099029 | -0.002813222 | 0.3687982 | -0.007628 | 0.993913733 | 0.997280261 | LAMA5 | protein_coding | 61.89 | -3.33E-06 | 3.33E-06 | ENSG00000130702 | n.s. |
| ENSG00000196557 | 540.3409983 | -0.002311132 | 0.2571126 | -0.008989 | 0.992828079 | 0.99669373 | CACNA1H | protein_coding | 65.12 | -3.32E-06 | 3.32E-06 | ENSG00000196557 | n.s. |
| ENSG00000008130 | 298.0123924 | 0.002209332 | 0.2413802 | 0.009153 | 0.992697137 | 0.996634139 | NADK | protein_coding | 53.7 | 3.23E-06 | 3.23E-06 | ENSG00000008130 | n.s. |
| ENSG00000188690 | 615.7422133 | 0.002344996 | 0.2908869 | 0.008062 | 0.993567892 | 0.997146071 | UROS | protein_coding | 46.1 | 2.91E-06 | 2.91E-06 | ENSG00000188690 | n.s. |
| ENSG00000214331 | 262.3746102 | 0.002046619 | 0.2370887 | 0.008632 | 0.993112515 | 0.996835523 | | transcribed_unprocessed_pseud | 46.88 | 2.82E-06 | 2.82E-06 | ENSG00000214331 | n.s. |
| ENSG00000140365 | 1328.738684 | -0.00213396 | 0.2675737 | -0.007975 | 0.993636759 | 0.997146071 | COMMD4 | protein_coding | 53.48 | -2.65E-06 | 2.65E-06 | ENSG00000140365 | n.s. |
| ENSG00000198945 | 192.1708277 | -0.001940865 | 0.2333106 | -0.008319 | 0.993362632 | 0.9970147 | L3MBTL3 | protein_coding | 38.41 | -2.52E-06 | 2.52E-06 | ENSG00000198945 | n.s. |
| ENSG00000120063 | 2970.281143 | 0.002059699 | 0.2653256 | 0.007763 | 0.993806155 | 0.997244187 | GNA13 | protein_coding | 40.35 | 2.47E-06 | 2.47E-06 | ENSG00000120063 | n.s. |
| ENSG00000175203 | 1662.047579 | -0.002164714 | 0.2994708 | -0.007228 | 0.99423257 | 0.997528289 | DCTN2 | protein_coding | 48.08 | -2.33E-06 | 2.33E-06 | ENSG00000175203 | n.s. |
| ENSG00000204619 | 230.1133133 | 0.001743857 | 0.2478913 | 0.007035 | 0.994387116 | 0.997611459 | PPP1R11 | protein_coding | 49.16 | 1.81E-06 | 1.81E-06 | ENSG00000204619 | n.s. |
| ENSG00000188933 | 600.1424052 | -0.001811003 | 0.28307 | -0.006398 | 0.994895392 | 0.997905665 | USP32P1 | transcribed_unprocessed_pseud | 42.86 | -1.65E-06 | 1.65E-06 | ENSG00000188933 | n.s. |
| ENSG00000058262 | 2638.440118 | 0.001693074 | 0.2513061 | 0.006737 | 0.994624614 | 0.997777829 | SEC61A1 | protein_coding | 47.3 | 1.64E-06 | 1.64E-06 | ENSG00000058262 | n.s. |
| ENSG00000207392 | 28.26365011 | 0.001980486 | 0.3497235 | 0.005663 | 0.9954816 | 0.998134115 | SNORA20 | snoRNA | 40.15 | 1.61E-06 | 1.61E-06 | ENSG00000207392 | n.s. |
| ENSG00000186866 | 27.7922198 | 0.001855262 | 0.3200689 | 0.005796 | 0.995375133 | 0.998099242 | POFUT2 | protein_coding | 52.95 | 1.53E-06 | 1.53E-06 | ENSG00000186866 | n.s. |
| ENSG00000273619 | 54.77481478 | 0.001640402 | 0.2540858 | 0.006456 | 0.994848819 | 0.997905665 | | antisense | 65.72 | 1.49E-06 | 1.49E-06 | ENSG00000273619 | n.s. |
| ENSG00000206635 | 6.610094114 | -0.002190162 | 0.4613885 | -0.004747 | 0.996212541 | 0.99843559 | RNU6-1062P | snRNA | 40.19 | -1.49E-06 | 1.49E-06 | ENSG00000206635 | n.s. |
| ENSG00000240853 | 17.97433162 | -0.001835955 | 0.3586301 | -0.005119 | 0.995915363 | 0.998300097 | RN7SL328P | misc_RNA | 54 | -1.36E-06 | 1.36E-06 | ENSG00000240853 | n.s. |
| ENSG00000033627 | 877.2426578 | -0.00144809 | 0.2407324 | -0.006015 | 0.995200473 | 0.998067866 | ATP6V0A1 | protein_coding | 45.47 | -1.22E-06 | 1.22E-06 | ENSG00000033627 | n.s. |
| ENSG00000184787 | 1853.527699 | 0.001418402 | 0.2439042 | 0.005815 | 0.995360004 | 0.998099242 | UBE2G2 | protein_coding | 45 | 1.17E-06 | 1.17E-06 | ENSG00000184787 | n.s. |
| ENSG00000169592 | 498.5730307 | -0.001480134 | 0.2738698 | -0.005405 | 0.995687841 | 0.998197135 | INO80E | protein_coding | 57.4 | -1.16E-06 | 1.16E-06 | ENSG00000169592 | n.s. |

| | | | | | | | | | | | | | |
|---|---|---|---|---|---|---|---|---|---|---|---|---|---|
| ENSG00000145675 | 1658.700667 | 0.001377188 | 0.2286074 | 0.006024 | 0.995193374 | 0.998067866 | PIK3R1 | protein_coding | 39.22 | 1.16E-06 | 1.16E-06 | ENSG00000145675 | n.s. |
| ENSG00000160908 | 383.4625887 | -0.001405783 | 0.2534651 | -0.005546 | 0.995574749 | 0.99815563 | ZNF394 | protein_coding | 47.6 | -1.13E-06 | 1.13E-06 | ENSG00000160908 | n.s. |
| ENSG00000124181 | 2887.970078 | -0.001249672 | 0.245223 | -0.005096 | 0.995933947 | 0.998300097 | PLCG1 | protein_coding | 49.84 | -9.23E-07 | 9.23E-07 | ENSG00000124181 | n.s. |
| ENSG00000244165 | 163.1434775 | 0.001197568 | 0.2608358 | 0.004591 | 0.996336709 | 0.998488159 | P2RY11 | protein_coding | 62.43 | 7.87E-07 | 7.87E-07 | ENSG00000244165 | n.s. |
| ENSG00000101447 | 1210.940865 | -0.001087288 | 0.2288681 | -0.004751 | 0.996209488 | 0.99843559 | FAM83D | protein_coding | 46.5 | -7.39E-07 | 7.39E-07 | ENSG00000101447 | n.s. |
| ENSG00000108848 | 18186.4933 | -0.001057302 | 0.2522249 | -0.004192 | 0.996655355 | 0.998735606 | LUC7L3 | protein_coding | 39.68 | -5.81E-07 | 5.81E-07 | ENSG00000108848 | n.s. |
| ENSG00000241420 | 29.6761043 | -0.001134489 | 0.3248581 | -0.003492 | 0.997213584 | 0.998866441 | RN7SL505P | misc_RNA | 54.61 | -5.59E-07 | 5.59E-07 | ENSG00000241420 | n.s. |
| ENSG00000214253 | 527.8328663 | 0.001035454 | 0.3101014 | 0.003339 | 0.997335803 | 0.998866441 | FIS1 | protein_coding | 51.92 | 5.10E-07 | 5.10E-07 | ENSG00000214253 | n.s. |
| ENSG00000175550 | 460.2763787 | 0.001030654 | 0.2937099 | 0.003509 | 0.997200159 | 0.998866441 | DRAP1 | protein_coding | 65.51 | 5.08E-07 | 5.08E-07 | ENSG00000175550 | n.s. |
| ENSG00000137142 | 2372.127404 | -0.001008212 | 0.2650199 | -0.003804 | 0.996964624 | 0.998866441 | IGFBPL1 | protein_coding | 49.74 | -4.97E-07 | 4.97E-07 | ENSG00000137142 | n.s. |
| ENSG00000119689 | 1464.528179 | -0.00094977 | 0.2466984 | -0.00385 | 0.996928211 | 0.998866441 | DLST | protein_coding | 44.92 | -4.68E-07 | 4.68E-07 | ENSG00000119689 | n.s. |
| ENSG00000210195 | 116.3651325 | 0.000884765 | 0.2812612 | 0.003146 | 0.997490094 | 0.998866441 | MT-TT | Mt_tRNA | 37.88 | 4.36E-07 | 4.36E-07 | ENSG00000210195 | n.s. |
| ENSG00000137996 | 1030.469463 | -0.000883909 | 0.2441149 | -0.003621 | 0.997110968 | 0.998866441 | RTCA | protein_coding | 39.07 | -4.35E-07 | 4.35E-07 | ENSG00000137996 | n.s. |
| ENSG00000102178 | 513.449296 | -0.000883213 | 0.2822574 | -0.003129 | 0.99750334 | 0.998866441 | UBL4A | protein_coding | 61.17 | -4.35E-07 | 4.35E-07 | ENSG00000102178 | n.s. |
| ENSG00000164896 | 547.9041572 | 0.000851281 | 0.2441951 | 0.003486 | 0.997218526 | 0.998866441 | FASTK | protein_coding | 61.75 | 4.19E-07 | 4.19E-07 | ENSG00000164896 | n.s. |
| ENSG00000183010 | 2041.042593 | -0.000858337 | 0.2868395 | -0.002992 | 0.997612419 | 0.998903827 | PYCR1 | protein_coding | 62.3 | -4.09E-07 | 4.09E-07 | ENSG00000183010 | n.s. |
| ENSG00000240457 | 8.831078295 | 0.001034367 | 0.4045781 | 0.002557 | 0.997960085 | 0.999093601 | RN7SL472P | misc_RNA | 53.24 | 4.07E-07 | 4.07E-07 | ENSG00000240457 | n.s. |
| ENSG00000105281 | 978.7273531 | -0.000756634 | 0.2386746 | -0.00317 | 0.997470591 | 0.998866441 | SLC1A5 | protein_coding | 54.4 | -3.73E-07 | 3.73E-07 | ENSG00000105281 | n.s. |
| ENSG00000212482 | 3.461166586 | 0.001203204 | 0.6339749 | 0.001898 | 0.998485717 | 0.999347038 | RNU6-530P | snRNA | 34.58 | 3.41E-07 | 3.41E-07 | ENSG00000212482 | n.s. |
| ENSG00000156486 | 204.4355833 | 0.000642595 | 0.2274728 | 0.002825 | 0.997746033 | 0.998965771 | KCNS2 | protein_coding | 51.66 | 2.89E-07 | 2.89E-07 | ENSG00000156486 | n.s. |
| ENSG00000184047 | 1411.781875 | 0.000645262 | 0.2596581 | 0.002485 | 0.998017223 | 0.999093601 | DIABLO | protein_coding | 47.05 | 2.54E-07 | 2.54E-07 | ENSG00000184047 | n.s. |
| ENSG00000163040 | 125.0747946 | -0.000630362 | 0.2882358 | -0.002187 | 0.998255055 | 0.999188003 | CCDC74A | protein_coding | 61.5 | -2.22E-07 | 2.22E-07 | ENSG00000163040 | n.s. |
| ENSG00000184194 | 278.4798911 | 0.000576769 | 0.2601856 | 0.002217 | 0.998231283 | 0.999188003 | GPR173 | protein_coding | 45.9 | 2.03E-07 | 2.03E-07 | ENSG00000184194 | n.s. |
| ENSG00000197448 | 275.0819123 | 0.000408788 | 0.2608887 | 0.001567 | 0.99874979 | 0.999539486 | GSTK1 | protein_coding | 41.49 | 8.18E-08 | 8.18E-08 | ENSG00000197448 | n.s. |
| ENSG00000131779 | 308.4209464 | 0.000313858 | 0.2800148 | 0.001121 | 0.999105681 | 0.999626144 | PEX11B | protein_coding | 45.5 | 5.10E-08 | 5.10E-08 | ENSG00000131779 | n.s. |
| ENSG00000226632 | 16.58118468 | 0.000279981 | 0.3379393 | 0.000828 | 0.999338957 | 0.999626144 | UBE2V1P1 | processed_pseudogene | 40.9 | 4.55E-08 | 4.55E-08 | ENSG00000226632 | n.s. |
| ENSG00000123595 | 241.6147844 | 0.000268789 | 0.2677941 | 0.001004 | 0.99919915 | 0.999626144 | RAB9A | protein_coding | 41.6 | 4.36E-08 | 4.36E-08 | ENSG00000123595 | n.s. |
| ENSG00000119862 | 500.0468287 | -0.000249108 | 0.2541645 | -0.00098 | 0.999217988 | 0.999626144 | LGALSL | protein_coding | 44.8 | -4.05E-08 | 4.05E-08 | ENSG00000119862 | n.s. |
| ENSG00000243943 | 2170.773805 | 0.000235555 | 0.2681725 | 0.000878 | 0.999299161 | 0.999626144 | ZNF512 | protein_coding | 41.21 | 3.83E-08 | 3.83E-08 | ENSG00000243943 | n.s. |
| ENSG00000136717 | 604.978465 | 0.000230048 | 0.2709271 | 0.000849 | 0.999322506 | 0.999626144 | BIN1 | protein_coding | 59.66 | 3.74E-08 | 3.74E-08 | ENSG00000136717 | n.s. |
| ENSG00000117222 | 926.4672921 | -0.000228837 | 0.2373503 | -0.000964 | 0.999230735 | 0.999626144 | RBBP5 | protein_coding | 40.77 | -3.72E-08 | 3.72E-08 | ENSG00000117222 | n.s. |
| ENSG00000138622 | 451.8883324 | -0.000153621 | 0.2272851 | -0.000676 | 0.999460715 | 0.999676116 | HCN4 | protein_coding | 55.08 | -2.16E-08 | 2.16E-08 | ENSG00000138622 | n.s. |

| | | | | | | | | | | | | | |
|---|---|---|---|---|---|---|---|---|---|---|---|---|---|
| ENSG00000102158 | 640.0325865 | -0.000111212 | 0.2864628 | -0.000388 | 0.99969024 | 0.999833864 | MAGT1 | protein_coding | 40.93 | -8.02E-09 | 8.02E-09 | ENSG00000102158 | n.s. |
| ENSG00000239382 | 169.1935968 | -7.96E-05 | 0.2827015 | -0.000282 | 0.999775206 | 0.999847018 | ALKBH6 | protein_coding | 56.33 | -5.29E-09 | 5.29E-09 | ENSG00000239382 | n.s. |
| ENSG00000278311 | 1613.281286 | -1.78E-05 | 0.2282511 | -7.78E-05 | 0.999937941 | 0.999937941 | GGNBP2 | protein_coding | 42.1 | -4.78E-10 | 4.78E-10 | ENSG00000278311 | n.s. |
