## Supplementary material for "CDK12 loss in cancer cells affects DNA damage response genes through premature cleavage and polyadenylation"

**Supplementary Table 2: TT-seq expression changes with 30min THZ531 treatment**

| gene_id | baseMean | log2FoldChange | lfcSE | stat | pvalue | padj | hgnc_symbol | gene_biotype | gene_gc(%) | PI | absPI | ensembl_gene_id | significant |
|---|---|---|---|---|---|---|---|---|---|---|---|---|---|
| ENSG00000170345 | 387.0462 | 3.937465127 | 0.260997 | 15.086257 | 1.99E-51 | 2.78E-47 | FOS | protein_coding | 53.92 | 183.3144128 | 183.3144128 | ENSG00000170345 | up |
| ENSG00000120738 | 379.4916 | 3.567818445 | 0.271229 | 13.154253 | 1.61E-39 | 1.12E-35 | EGR1 | protein_coding | 56.56 | 124.6982343 | 124.6982343 | ENSG00000120738 | up |
| ENSG00000142871 | 216.8154 | 2.446829491 | 0.247273 | 9.8952364 | 4.37E-23 | 2.03E-19 | CYR61 | protein_coding | 50.25 | 45.73938337 | 45.73938337 | ENSG00000142871 | up |
| ENSG00000244405 | 458.9768 | -2.322103353 | 0.280048 | -8.291813 | 1.12E-16 | 3.88E-13 | ETV5 | protein_coding | 42.69 | -28.81942954 | 28.81942954 | ENSG00000244405 | down |
| ENSG00000248540 | 200.7061 | -2.235505486 | 0.277464 | -8.056933 | 7.82E-16 | 2.18E-12 | | lincRNA | 56.18 | -26.07013306 | 26.07013306 | ENSG00000248540 | down |
| ENSG00000236824 | 41.38297 | -3.0476498 | 0.507991 | -5.999412 | 1.98E-09 | 1.84E-06 | BCYRN1 | scRNA | 53 | -17.48016145 | 17.48016145 | ENSG00000236824 | down |
| ENSG00000187678 | 136.3448 | -2.140641491 | 0.308924 | -6.929339 | 4.23E-12 | 8.41E-09 | SPRY4 | protein_coding | 51.21 | -17.286145 | 17.286145 | ENSG00000187678 | down |
| ENSG00000119630 | 2207.669 | -1.628045406 | 0.229587 | -7.091199 | 1.33E-12 | 3.09E-09 | PGF | protein_coding | 56.15 | -13.85582959 | 13.85582959 | ENSG00000119630 | down |
| ENSG00000198576 | 404.9207 | 1.917399358 | 0.296453 | 6.4678127 | 9.94E-11 | 1.54E-07 | ARC | protein_coding | 66.75 | 13.06321318 | 13.06321318 | ENSG00000198576 | up |
| ENSG00000261326 | 163.7239 | -1.713296944 | 0.253784 | -6.750999 | 1.47E-11 | 2.56E-08 | LINC01355 | lincRNA | 36 | -13.00827661 | 13.00827661 | ENSG00000261326 | down |
| ENSG00000137868 | 67.54105 | -1.985129595 | 0.314068 | -6.320699 | 2.60E-10 | 3.30E-07 | STRA6 | protein_coding | 55.21 | -12.86770292 | 12.86770292 | ENSG00000137868 | down |
| ENSG00000114315 | 53.41176 | -3.00560512 | 0.573205 | -5.243509 | 1.58E-07 | 7.08E-05 | HES1 | protein_coding | 53.32 | -12.47388168 | 12.47388168 | ENSG00000114315 | down |
| ENSG00000257531 | 120.8225 | 1.595886577 | 0.252435 | 6.321968 | 2.58E-10 | 3.30E-07 | | transcribed_processed_pseudog | 51.61 | 10.34461147 | 10.34461147 | ENSG00000257531 | up |
| ENSG00000271815 | 23.95374 | -2.221877396 | 0.40636 | -5.467751 | 4.56E-08 | 2.64E-05 | | lincRNA | 48.24 | -10.17113637 | 10.17113637 | ENSG00000271815 | down |
| ENSG00000274995 | 73.23911 | -1.709621698 | 0.287955 | -5.937121 | 2.90E-09 | 2.38E-06 | | sense_intronic | 47.1 | -9.615276872 | 9.615276872 | ENSG00000274995 | down |
| ENSG00000172183 | 20.58612 | -2.242967536 | 0.421568 | -5.320534 | 1.03E-07 | 5.34E-05 | ISG20 | protein_coding | 52.19 | -9.583852649 | 9.583852649 | ENSG00000172183 | down |
| ENSG00000279598 | 265.7687 | -1.524146141 | 0.244601 | -6.231153 | 4.63E-10 | 5.37E-07 | | TEC | 39.95 | -9.556172787 | 9.556172787 | ENSG00000279598 | down |
| ENSG00000151012 | 402.6646 | -1.668115552 | 0.284726 | -5.85867 | 4.67E-09 | 3.61E-06 | SLC7A11 | protein_coding | 35.52 | -9.078894295 | 9.078894295 | ENSG00000151012 | down |
| ENSG00000277039 | 16.32999 | -2.364722378 | 0.468147 | -5.051238 | 4.39E-07 | 0.000165973 | | misc_RNA | 55.7 | -8.938564417 | 8.938564417 | ENSG00000277039 | down |
| ENSG00000107338 | 274.07 | -1.487975668 | 0.244575 | -6.083923 | 1.17E-09 | 1.25E-06 | SHB | protein_coding | 50.89 | -8.782195396 | 8.782195396 | ENSG00000107338 | down |
| ENSG00000230734 | 29.42651 | -2.10577584 | 0.406795 | -5.176504 | 2.26E-07 | 8.99E-05 | RPL10P3 | processed_pseudogene | 50.85 | -8.52011392 | 8.52011392 | ENSG00000230734 | down |
| ENSG00000251745 | 17.93117 | -2.351312642 | 0.485909 | -4.838998 | 1.30E-06 | 0.000394976 | RNU7-124P | snRNA | 37.1 | -8.002526881 | 8.002526881 | ENSG00000251745 | down |
| ENSG00000228950 | 51.69443 | 1.743665271 | 0.32278 | 5.4020306 | 6.59E-08 | 3.53E-05 | | lincRNA | 46.07 | 7.763523625 | 7.763523625 | ENSG00000228950 | up |
| ENSG00000147459 | 988.4931 | -1.357740835 | 0.228252 | -5.94844 | 2.71E-09 | 2.36E-06 | DOCK5 | protein_coding | 42.37 | -7.641207902 | 7.641207902 | ENSG00000147459 | down |
| ENSG00000214535 | 92.15451 | 1.875990649 | 0.362332 | 5.1775432 | 2.25E-07 | 8.99E-05 | RPS15AP1 | processed_pseudogene | 46.29 | 7.590387229 | 7.590387229 | ENSG00000214535 | up |
| ENSG00000162772 | 495.5344 | 1.548904969 | 0.274739 | 5.6377356 | 1.72E-08 | 1.26E-05 | ATF3 | protein_coding | 43.16 | 7.587672527 | 7.587672527 | ENSG00000162772 | up |
| ENSG00000225761 | 33.43521 | 1.815589566 | 0.34544 | 5.2558734 | 1.47E-07 | 6.84E-05 | | sense_intronic | 40.99 | 7.562141098 | 7.562141098 | ENSG00000225761 | up |
| ENSG00000235655 | 286.3312 | -1.766210752 | 0.337345 | -5.235626 | 1.64E-07 | 7.15E-05 | H3F3AP4 | processed_pseudogene | 51.58 | -7.321728533 | 7.321728533 | ENSG00000235655 | down |
| ENSG00000171604 | 375.8457 | -1.53949529 | 0.281268 | -5.473416 | 4.41E-08 | 2.64E-05 | CXXC5 | protein_coding | 58.2 | -7.047381 | 7.047381 | ENSG00000171604 | down |
| ENSG00000177606 | 2056.877 | 1.169789761 | 0.192639 | 6.0724474 | 1.26E-09 | 1.25E-06 | JUN | protein_coding | 56.67 | 6.904227319 | 6.904227319 | ENSG00000177606 | up |

| | | | | | | | | | | | | | |
|---|---|---|---|---|---|---|---|---|---|---|---|---|---|
| ENSG00000260198 | 65.40523 | -1.621665239 | 0.30854 | -5.255939 | 1.47E-07 | 6.84E-05 | | sense_intronic | 49.45 | -6.754423787 | 6.754423787 | ENSG00000260198 | down |
| ENSG00000213942 | 15.72675 | 2.062226834 | 0.434269 | 4.7487338 | 2.05E-06 | 0.000558815 | | processed_pseudogene | 46.15 | 6.707871163 | 6.707871163 | ENSG00000213942 | up |
| ENSG00000165507 | 161.607 | 1.422290822 | 0.2576 | 5.5213228 | 3.36E-08 | 2.13E-05 | C10orf10 | protein_coding | 57.22 | 6.644601621 | 6.644601621 | ENSG00000165507 | up |
| ENSG00000225401 | 16.1875 | -2.106508226 | 0.452114 | -4.659238 | 3.17E-06 | 0.000803439 | TGIF2P1 | processed_pseudogene | 59.42 | -6.519742495 | 6.519742495 | ENSG00000225401 | down |
| ENSG00000259707 | 17.81024 | -2.110018845 | 0.456475 | -4.622417 | 3.79E-06 | 0.000895072 | | processed_pseudogene | 52.2 | -6.431637502 | 6.431637502 | ENSG00000259707 | down |
| ENSG00000178404 | 274.9042 | -1.311021814 | 0.233891 | -5.605269 | 2.08E-08 | 1.45E-05 | CEP295NL | protein_coding | 53.83 | -6.344528528 | 6.344528528 | ENSG00000178404 | down |
| ENSG00000242999 | 11.04143 | -2.48334638 | 0.576416 | -4.308255 | 1.65E-05 | 0.002899999 | RN7SL239P | misc_RNA | 53.67 | -6.301745019 | 6.301745019 | ENSG00000242999 | down |
| ENSG00000242539 | 54.7383 | -1.50706957 | 0.289986 | -5.197035 | 2.02E-07 | 8.54E-05 | | antisense | 38.14 | -6.131319404 | 6.131319404 | ENSG00000242539 | down |
| ENSG00000227008 | 131.6455 | 1.581757001 | 0.313199 | 5.0503212 | 4.41E-07 | 0.000165973 | | processed_pseudogene | 42.31 | 5.978983824 | 5.978983824 | ENSG00000227008 | up |
| ENSG00000100234 | 1749.022 | -1.29699477 | 0.244643 | -5.30158 | 1.15E-07 | 5.71E-05 | TIMP3 | protein_coding | 47.18 | -5.503751524 | 5.503751524 | ENSG00000100234 | down |
| ENSG00000272345 | 26.82769 | -1.773444007 | 0.382272 | -4.63922 | 3.50E-06 | 0.000855042 | | antisense | 35.29 | -5.440948094 | 5.440948094 | ENSG00000272345 | down |
| ENSG00000207134 | 4.850406 | -3.674883572 | 1.075677 | -3.416346 | 0.000635 | 0.034134195 | RNU6-106P | snRNA | 37.38 | -5.390357194 | 5.390357194 | ENSG00000207134 | down |
| ENSG00000159388 | 493.4199 | 1.177801183 | 0.215781 | 5.4583076 | 4.81E-08 | 2.68E-05 | BTG2 | protein_coding | 53.33 | 5.385305282 | 5.385305282 | ENSG00000159388 | up |
| ENSG00000279616 | 46.10231 | -1.5349239 | 0.312544 | -4.911065 | 9.06E-07 | 0.000323382 | | TEC | 42.03 | -5.357320513 | 5.357320513 | ENSG00000279616 | down |
| ENSG00000187536 | 21.32062 | -1.791730934 | 0.393246 | -4.556256 | 5.21E-06 | 0.001169384 | TPM3P7 | processed_pseudogene | 48.5 | -5.25343163 | 5.25343163 | ENSG00000187536 | down |
| ENSG00000254180 | 14.32834 | -2.0136305 | 0.466663 | -4.314952 | 1.60E-05 | 0.002865686 | | lincRNA | 38.39 | -5.120202157 | 5.120202157 | ENSG00000254180 | down |
| ENSG00000254783 | 21.45059 | -1.75707977 | 0.387116 | -4.5389 | 5.65E-06 | 0.001230192 | | processed_pseudogene | 48.11 | -5.113149816 | 5.113149816 | ENSG00000254783 | down |
| ENSG00000230778 | 58.21863 | -1.460144801 | 0.298523 | -4.891224 | 1.00E-06 | 0.000348809 | ANKRD63 | protein_coding | 73.84 | -5.048323367 | 5.048323367 | ENSG00000230778 | down |
| ENSG00000113369 | 4404.683 | 1.060633462 | 0.190606 | 5.5645467 | 2.63E-08 | 1.74E-05 | ARRDC3 | protein_coding | 35.81 | 5.047346333 | 5.047346333 | ENSG00000113369 | up |
| ENSG00000279962 | 15.32047 | -2.375624274 | 0.598789 | -3.967384 | 7.27E-05 | 0.00815907 | | TEC | 54.24 | -4.961157126 | 4.961157126 | ENSG00000279962 | down |
| ENSG00000269939 | 63.74911 | -1.409806014 | 0.290501 | -4.853014 | 1.22E-06 | 0.000384779 | PCF11-AS1 | lincRNA | 35.62 | -4.814190251 | 4.814190251 | ENSG00000269939 | down |
| ENSG00000225342 | 72.7787 | 1.460431494 | 0.309182 | 4.7235329 | 2.32E-06 | 0.000597612 | | antisense | 36.31 | 4.707818319 | 4.707818319 | ENSG00000225342 | up |
| ENSG00000127364 | 66.05128 | -1.372241053 | 0.283113 | -4.846981 | 1.25E-06 | 0.000387846 | TAS2R4 | protein_coding | 41.46 | -4.681181384 | 4.681181384 | ENSG00000127364 | down |
| ENSG00000224324 | 13.67144 | -1.960597393 | 0.472179 | -4.152234 | 3.29E-05 | 0.004284193 | THAP5P1 | processed_pseudogene | 40.08 | -4.64295147 | 4.64295147 | ENSG00000224324 | down |
| ENSG00000252636 | 25.48403 | -1.771152119 | 0.404818 | -4.375186 | 1.21E-05 | 0.002448209 | RNU6-826P | snRNA | 47.12 | -4.624746548 | 4.624746548 | ENSG00000252636 | down |
| ENSG00000255330 | 317.1729 | -1.417865633 | 0.297931 | -4.759044 | 1.95E-06 | 0.000552693 | SOGA3 | protein_coding | 38.15 | -4.618720251 | 4.618720251 | ENSG00000255330 | down |
| ENSG00000224967 | 9.542305 | -2.503095929 | 0.666635 | -3.754821 | 0.000173 | 0.014290854 | | processed_pseudogene | 46.41 | -4.618066374 | 4.618066374 | ENSG00000224967 | down |
| ENSG00000222714 | 9.993773 | -2.273425702 | 0.581935 | -3.906663 | 9.36E-05 | 0.009580195 | RN7SKP38 | misc_RNA | 49.51 | -4.589195484 | 4.589195484 | ENSG00000222714 | down |
| ENSG00000272159 | 22.82392 | -1.763924428 | 0.406116 | -4.343399 | 1.40E-05 | 0.002665434 | | antisense | 42.59 | -4.54075085 | 4.54075085 | ENSG00000272159 | down |
| ENSG00000273786 | 16.8608 | -1.822978315 | 0.427145 | -4.267823 | 1.97E-05 | 0.003271743 | | sense_intronic | 33.48 | -4.530503582 | 4.530503582 | ENSG00000273786 | down |
| ENSG00000143333 | 75.60933 | 1.472406762 | 0.317395 | 4.6390291 | 3.50E-06 | 0.000855042 | RGS16 | protein_coding | 50.88 | 4.517362112 | 4.517362112 | ENSG00000143333 | up |
| ENSG00000207739 | 60.0268 | -1.383759677 | 0.292076 | -4.737675 | 2.16E-06 | 0.000578833 | MIR218-2 | miRNA | 55.45 | -4.479848123 | 4.479848123 | ENSG00000207739 | down |

| | | | | | | | | | | | | | |
|---|---|---|---|---|---|---|---|---|---|---|---|---|---|
| ENSG00000189367 | 377.1393 | -1.374458392 | 0.289083 | -4.754543 | 1.99E-06 | 0.000553845 | KIAA0408 | protein_coding | 35.14 | -4.476077085 | 4.476077085 | ENSG00000189367 | down |
| ENSG00000278576 | 118.9103 | -1.314849119 | 0.273556 | -4.806516 | 1.54E-06 | 0.000454965 | | sense_intronic | 43.2 | -4.394254179 | 4.394254179 | ENSG00000278576 | down |
| ENSG00000253833 | 16.70987 | -2.180699938 | 0.562004 | -3.880222 | 0.000104 | 0.010152591 | | processed_pseudogene | 38.87 | -4.34705763 | 4.34705763 | ENSG00000253833 | down |
| ENSG00000277681 | 13.13238 | -2.201510226 | 0.570756 | -3.857186 | 0.000115 | 0.010646463 | MIR6739 | miRNA | 38.67 | -4.343127593 | 4.343127593 | ENSG00000277681 | down |
| ENSG00000143507 | 436.0326 | 1.221356301 | 0.24816 | 4.9216524 | 8.58E-07 | 0.000314427 | DUSP10 | protein_coding | 41.35 | 4.277775659 | 4.277775659 | ENSG00000143507 | up |
| ENSG00000280347 | 127.4656 | -1.220650143 | 0.25049 | -4.873043 | 1.10E-06 | 0.00035582 | | TEC | 37.71 | -4.209740746 | 4.209740746 | ENSG00000280347 | down |
| ENSG00000259630 | 37.7392 | -1.686962906 | 0.395635 | -4.263932 | 2.01E-05 | 0.003290095 | | transcribed_processed_pseudog | 43.6 | -4.188377312 | 4.188377312 | ENSG00000259630 | down |
| ENSG00000188483 | 561.8934 | -1.207934847 | 0.247265 | -4.885188 | 1.03E-06 | 0.000349363 | IER5L | protein_coding | 64.99 | -4.175496502 | 4.175496502 | ENSG00000188483 | down |
| ENSG00000258904 | 60.7674 | -1.409749288 | 0.309835 | -4.549998 | 5.36E-06 | 0.001185587 | | sense_intronic | 50.52 | -4.125020436 | 4.125020436 | ENSG00000258904 | down |
| ENSG00000272196 | 13.52151 | -2.42748893 | 0.671796 | -3.613432 | 0.000302 | 0.020522559 | HIST2H2AA4 | protein_coding | 61.7 | -4.097039306 | 4.097039306 | ENSG00000272196 | down |
| ENSG00000279764 | 57.66792 | -1.337990855 | 0.289227 | -4.626088 | 3.73E-06 | 0.000894523 | | TEC | 55.67 | -4.078742984 | 4.078742984 | ENSG00000279764 | down |
| ENSG00000159307 | 72.47561 | -1.440147795 | 0.321005 | -4.48637 | 7.24E-06 | 0.001551814 | SCUBE1 | protein_coding | 55.48 | -4.045606056 | 4.045606056 | ENSG00000159307 | down |
| ENSG00000154319 | 348.8334 | -1.328036577 | 0.287667 | -4.616573 | 3.90E-06 | 0.000905294 | FAM167A | protein_coding | 52.59 | -4.041494851 | 4.041494851 | ENSG00000154319 | down |
| ENSG00000272746 | 22.83034 | -1.817402185 | 0.45042 | -4.034901 | 5.46E-05 | 0.006285512 | | sense_intronic | 38.95 | -4.001300434 | 4.001300434 | ENSG00000272746 | down |
| ENSG00000279638 | 8.241201 | -2.699941248 | 0.795068 | -3.39586 | 0.000684 | 0.035944079 | | TEC | 32.61 | -3.899721275 | 3.899721275 | ENSG00000279638 | down |
| ENSG00000263675 | 8.733351 | -2.944963677 | 0.90706 | -3.246715 | 0.001167 | 0.048550416 | MIR5581 | miRNA | 48.33 | -3.869114025 | 3.869114025 | ENSG00000263675 | down |
| ENSG00000151136 | 875.6979 | -1.095580589 | 0.224444 | -4.881301 | 1.05E-06 | 0.000349363 | BTBD11 | protein_coding | 46.12 | -3.787118922 | 3.787118922 | ENSG00000151136 | down |
| ENSG00000229330 | 31.65608 | -1.460932163 | 0.334913 | -4.36212 | 1.29E-05 | 0.002561997 | | sense_intronic | 51.89 | -3.785890775 | 3.785890775 | ENSG00000229330 | down |
| ENSG00000233133 | 4.804095 | -2.891603599 | 0.903491 | -3.200479 | 0.001372 | 0.053209578 | | unprocessed_pseudogene | 68.18 | -3.683932433 | 3.683932433 | ENSG00000233133 | down |
| ENSG00000267543 | 19.43561 | -1.659380279 | 0.410983 | -4.03759 | 5.40E-05 | 0.006265703 | | sense_intronic | 46.92 | -3.65566482 | 3.65566482 | ENSG00000267543 | down |
| ENSG00000248559 | 14.4921 | -1.775973515 | 0.451772 | -3.93113 | 8.45E-05 | 0.008985934 | | antisense | 44.13 | -3.634417502 | 3.634417502 | ENSG00000248559 | down |
| ENSG00000270137 | 38.04456 | -1.394519431 | 0.320577 | -4.350032 | 1.36E-05 | 0.002665434 | | sense_intronic | 36.55 | -3.589816655 | 3.589816655 | ENSG00000270137 | down |
| ENSG00000226221 | 169.6477 | 1.429297883 | 0.333518 | 4.2855156 | 1.82E-05 | 0.00309559 | | processed_pseudogene | 43.84 | 3.586475177 | 3.586475177 | ENSG00000226221 | up |
| ENSG00000185641 | 137.9017 | 1.42328424 | 0.331997 | 4.2870359 | 1.81E-05 | 0.00309559 | | processed_pseudogene | 47.72 | 3.571385404 | 3.571385404 | ENSG00000185641 | up |
| ENSG00000272788 | 18.3547 | -1.690453829 | 0.425143 | -3.976204 | 7.00E-05 | 0.007926385 | | sense_intronic | 45.54 | -3.551516424 | 3.551516424 | ENSG00000272788 | down |
| ENSG00000187653 | 61.90023 | 1.429127006 | 0.337682 | 4.2321646 | 2.31E-05 | 0.003582447 | TMSB4XP8 | processed_pseudogene | 44.44 | 3.495387795 | 3.495387795 | ENSG00000187653 | up |
| ENSG00000157240 | 291.5722 | -1.084185629 | 0.229467 | -4.724795 | 2.30E-06 | 0.000597612 | FZD1 | protein_coding | 47.74 | -3.494959526 | 3.494959526 | ENSG00000157240 | down |
| ENSG00000271789 | 58.08649 | -1.346297642 | 0.310327 | -4.338313 | 1.44E-05 | 0.002665434 | | lincRNA | 39.49 | -3.465682579 | 3.465682579 | ENSG00000271789 | down |
| ENSG00000224631 | 409.3882 | 1.440830126 | 0.346125 | 4.1627438 | 3.14E-05 | 0.004284193 | RPS27AP16 | nscribed_processed_pseudog | 42.05 | 3.41207449 | 3.41207449 | ENSG00000224631 | up |
| ENSG00000231767 | 90.93392 | 1.43043174 | 0.343942 | 4.158928 | 3.20E-05 | 0.004284193 | | processed_pseudogene | 42.92 | 3.387449751 | 3.387449751 | ENSG00000231767 | up |
| ENSG00000201643 | 4.559752 | -3.539367277 | 1.246527 | -2.839382 | 0.00452 | 0.110509521 | SNORA14A | snoRNA | 45.19 | -3.385759812 | 3.385759812 | ENSG00000201643 | n.s. |
| ENSG00000272440 | 13.14367 | -1.82658124 | 0.487499 | -3.746841 | 0.000179 | 0.014495721 | | lincRNA | 26.98 | -3.358644865 | 3.358644865 | ENSG00000272440 | down |

| | | | | | | | | | | | | | |
|---|---|---|---|---|---|---|---|---|---|---|---|---|---|
| ENSG00000262380 | 16.49198 | -1.808331779 | 0.480466 | -3.763704 | 0.000167 | 0.013957589 | | sense_intronic | 48.01 | -3.354798323 | 3.354798323 | ENSG00000262380 | down |
| ENSG00000225159 | 58.46886 | 1.352461633 | 0.318064 | 4.2521737 | 2.12E-05 | 0.003427414 | NPM1P39 | processed_pseudogene | 41.41 | 3.333863181 | 3.333863181 | ENSG00000225159 | up |
| ENSG00000266533 | 31.14586 | -1.378187854 | 0.331818 | -4.15344 | 3.28E-05 | 0.004284193 | MIR3619 | miRNA | 65.06 | -3.263729384 | 3.263729384 | ENSG00000266533 | down |
| ENSG00000264470 | 5.669703 | -2.799691212 | 0.908173 | -3.082773 | 0.002051 | 0.06830421 | MIR4794 | miRNA | 38.96 | -3.263187176 | 3.263187176 | ENSG00000264470 | down |
| ENSG00000240589 | 27.44189 | -1.45843941 | 0.358908 | -4.063547 | 4.83E-05 | 0.005801164 | RN7SL258P | misc_RNA | 53.33 | -3.261777633 | 3.261777633 | ENSG00000240589 | down |
| ENSG00000229891 | 102.0747 | -1.347326172 | 0.321708 | -4.18804 | 2.81E-05 | 0.004088489 | LINC01315 | lincRNA | 55.65 | -3.218003954 | 3.218003954 | ENSG00000229891 | down |
| ENSG00000064195 | 81.91217 | -1.303310167 | 0.308322 | -4.227102 | 2.37E-05 | 0.003582447 | DLX3 | protein_coding | 57.41 | -3.187662419 | 3.187662419 | ENSG00000064195 | down |
| ENSG00000237260 | 35.2575 | -1.337192128 | 0.321755 | -4.155934 | 3.24E-05 | 0.004284193 | | processed_pseudogene | 45.35 | -3.166646133 | 3.166646133 | ENSG00000237260 | down |
| ENSG00000272248 | 9.973832 | -3.127062629 | 1.080769 | -2.893368 | 0.003811 | 0.101076701 | | lincRNA | 47.76 | -3.112518475 | 3.112518475 | ENSG00000272248 | n.s. |
| ENSG00000137834 | 4497.287 | -0.922402864 | 0.193201 | -4.774329 | 1.80E-06 | 0.000523003 | SMAD6 | protein_coding | 50.83 | -3.026860923 | 3.026860923 | ENSG00000137834 | n.s. |
| ENSG00000206846 | 16.84665 | -1.626313578 | 0.434008 | -3.747193 | 0.000179 | 0.014495721 | | misc_RNA | 43.62 | -2.990400661 | 2.990400661 | ENSG00000206846 | down |
| ENSG00000234233 | 423.173 | 0.980779239 | 0.212987 | 4.6048771 | 4.13E-06 | 0.000941992 | KCNH1-IT1 | sense_intronic | 47.01 | 2.967791481 | 2.967791481 | ENSG00000234233 | n.s. |
| ENSG00000279233 | 92.09456 | -1.150476976 | 0.265023 | -4.341054 | 1.42E-05 | 0.002665434 | | TEC | 53.04 | -2.961594739 | 2.961594739 | ENSG00000279233 | down |
| ENSG00000266680 | 39.05017 | 1.45365878 | 0.376065 | 3.8654445 | 0.000111 | 0.010574525 | | antisense | 45.23 | 2.872050551 | 2.872050551 | ENSG00000266680 | up |
| ENSG00000107731 | 2201.251 | -1.044979376 | 0.235324 | -4.440605 | 8.97E-06 | 0.001892392 | UNC5B | protein_coding | 54.37 | -2.845467181 | 2.845467181 | ENSG00000107731 | down |
| ENSG00000140557 | 403.8329 | -1.15224809 | 0.272019 | -4.235903 | 2.28E-05 | 0.003582447 | ST8SIA2 | protein_coding | 48 | -2.818191729 | 2.818191729 | ENSG00000140557 | down |
| ENSG00000207417 | 7.308151 | -2.013478811 | 0.605506 | -3.325284 | 0.000883 | 0.040189476 | | misc_RNA | 41.12 | -2.810590209 | 2.810590209 | ENSG00000207417 | down |
| ENSG00000279587 | 32.3253 | -1.333474013 | 0.337519 | -3.95081 | 7.79E-05 | 0.008538755 | | TEC | 30.42 | -2.758431632 | 2.758431632 | ENSG00000279587 | down |
| ENSG00000242086 | 742.7367 | -1.049663737 | 0.239291 | -4.386559 | 1.15E-05 | 0.002357861 | LINC00969 | lincRNA | 48.33 | -2.757972386 | 2.757972386 | ENSG00000242086 | down |
| ENSG00000095015 | 722.0946 | -1.024342897 | 0.231705 | -4.420885 | 9.83E-06 | 0.002042682 | MAP3K1 | protein_coding | 38.07 | -2.755276695 | 2.755276695 | ENSG00000095015 | down |
| ENSG00000166886 | 553.7551 | -1.155083941 | 0.276419 | -4.178749 | 2.93E-05 | 0.004122295 | NAB2 | protein_coding | 60.22 | -2.754714606 | 2.754714606 | ENSG00000166886 | down |
| ENSG00000214389 | 248.8257 | 1.385592254 | 0.357786 | 3.8726877 | 0.000108 | 0.010335831 | RPS3AP26 | processed_pseudogene | 43.11 | 2.751307619 | 2.751307619 | ENSG00000214389 | up |
| ENSG00000213935 | 26.63029 | -1.486030547 | 0.395631 | -3.756105 | 0.000173 | 0.014290854 | | processed_pseudogene | 43.16 | -2.741639911 | 2.741639911 | ENSG00000213935 | down |
| ENSG00000146592 | 133.8404 | 1.175826867 | 0.284968 | 4.1261764 | 3.69E-05 | 0.004668564 | CREB5 | protein_coding | 40.7 | 2.740636915 | 2.740636915 | ENSG00000146592 | up |
| ENSG00000253168 | 21.87786 | -2.250362802 | 0.718914 | -3.130225 | 0.001747 | 0.062358007 | | antisense | 38.68 | -2.711929708 | 2.711929708 | ENSG00000253168 | down |
| ENSG00000232623 | 17.41952 | -1.471190333 | 0.393134 | -3.742206 | 0.000182 | 0.014596081 | | antisense | 48.46 | -2.700757844 | 2.700757844 | ENSG00000232623 | down |
| ENSG00000230325 | 15.76393 | -1.553728078 | 0.425257 | -3.653624 | 0.000259 | 0.018556694 | | antisense | 43.55 | -2.690279217 | 2.690279217 | ENSG00000230325 | down |
| ENSG00000280033 | 41.80334 | -1.369139152 | 0.356386 | -3.841729 | 0.000122 | 0.010974075 | | TEC | 61.64 | -2.683009004 | 2.683009004 | ENSG00000280033 | down |
| ENSG00000135116 | 467.646 | -1.10121958 | 0.261165 | -4.216569 | 2.48E-05 | 0.003713512 | HRK | protein_coding | 49 | -2.676200481 | 2.676200481 | ENSG00000135116 | down |
| ENSG00000259183 | 10.09101 | -1.822877657 | 0.534142 | -3.41272 | 0.000643 | 0.034310298 | MED28P6 | processed_pseudogene | 40.69 | -2.669741976 | 2.669741976 | ENSG00000259183 | down |
| ENSG00000168675 | 69.03122 | -1.193392969 | 0.293908 | -4.060432 | 4.90E-05 | 0.005828861 | LDLRAD4 | protein_coding | 46.44 | -2.666536737 | 2.666536737 | ENSG00000168675 | down |
| ENSG00000271011 | 41.65509 | -1.314975511 | 0.336361 | -3.909422 | 9.25E-05 | 0.009541591 | | antisense | 39.81 | -2.656749147 | 2.656749147 | ENSG00000271011 | down |

| | | | | | | | | | | | | | |
|---|---|---|---|---|---|---|---|---|---|---|---|---|---|
| ENSG00000206734 | 8.007412 | -2.13818819 | 0.676075 | -3.162651 | 0.001563 | 0.057891374 | | misc_RNA | 43.14 | -2.645764435 | 2.645764435 | ENSG00000206734 | down |
| ENSG00000257180 | 44.55671 | -1.30253158 | 0.333793 | -3.902216 | 9.53E-05 | 0.009663101 | | sense_intronic | 41.44 | -2.624449383 | 2.624449383 | ENSG00000257180 | down |
| ENSG00000280176 | 13.65182 | -1.619455683 | 0.455117 | -3.558329 | 0.000373 | 0.024057275 | | TEC | 47.22 | -2.62149967 | 2.62149967 | ENSG00000280176 | down |
| ENSG00000008853 | 202.4666 | -1.033551901 | 0.240371 | -4.299819 | 1.71E-05 | 0.002974955 | RHOBTB2 | protein_coding | 53.51 | -2.611289147 | 2.611289147 | ENSG00000008853 | down |
| ENSG00000198618 | 445.6296 | 1.268980447 | 0.32225 | 3.9378785 | 8.22E-05 | 0.008804176 | PPIAP22 | processed_pseudogene | 49 | 2.608149802 | 2.608149802 | ENSG00000198618 | up |
| ENSG00000270956 | 52.02539 | -1.290935823 | 0.331069 | -3.8993 | 9.65E-05 | 0.009663101 | | antisense | 36.15 | -2.601085284 | 2.601085284 | ENSG00000270956 | down |
| ENSG00000165606 | 430.8966 | -1.131564641 | 0.276206 | -4.096815 | 4.19E-05 | 0.005115776 | DRGX | protein_coding | 48.29 | -2.592514735 | 2.592514735 | ENSG00000165606 | down |
| ENSG00000250170 | 31.76624 | -1.489350693 | 0.407003 | -3.659309 | 0.000253 | 0.018415263 | RASA2-IT1 | sense_intronic | 40.44 | -2.583758457 | 2.583758457 | ENSG00000250170 | down |
| ENSG00000141905 | 16.44307 | -1.60249355 | 0.451643 | -3.548141 | 0.000388 | 0.024664733 | NFIC | protein_coding | 55.9 | -2.576687155 | 2.576687155 | ENSG00000141905 | down |
| ENSG00000237481 | 23.93056 | -1.362888108 | 0.359892 | -3.786938 | 0.000153 | 0.012869504 | | antisense | 38.05 | -2.576455719 | 2.576455719 | ENSG00000237481 | down |
| ENSG00000127362 | 36.96033 | -1.298733662 | 0.336535 | -3.859132 | 0.000114 | 0.010646463 | TAS2R3 | protein_coding | 45.69 | -2.562134819 | 2.562134819 | ENSG00000127362 | down |
| ENSG00000281881 | 16.96573 | -2.046393265 | 0.644162 | -3.17683 | 0.001489 | 0.056028529 | | lincRNA | 47.65 | -2.561246937 | 2.561246937 | ENSG00000281881 | down |
| ENSG00000141639 | 250.2079 | -1.068038622 | 0.255519 | -4.179877 | 2.92E-05 | 0.004122295 | MAPK4 | protein_coding | 45.75 | -2.547123622 | 2.547123622 | ENSG00000141639 | down |
| ENSG00000259548 | 15.90035 | -1.519646019 | 0.424275 | -3.581747 | 0.000341 | 0.022482711 | | lincRNA | 46.83 | -2.504606596 | 2.504606596 | ENSG00000259548 | down |
| ENSG00000229087 | 236.8498 | -1.018664206 | 0.240847 | -4.2295 | 2.34E-05 | 0.003582447 | | processed_pseudogene | 48.85 | -2.491469559 | 2.491469559 | ENSG00000229087 | down |
| ENSG00000115884 | 373.0587 | -1.071151365 | 0.261026 | -4.103615 | 4.07E-05 | 0.005011598 | SDC1 | protein_coding | 57.56 | -2.463673621 | 2.463673621 | ENSG00000115884 | down |
| ENSG00000259581 | 126.5533 | -1.030624975 | 0.247965 | -4.156334 | 3.23E-05 | 0.004284193 | TYRO3P | processed_pseudogene | 57.26 | -2.440654954 | 2.440654954 | ENSG00000259581 | down |
| ENSG00000275632 | 70.67399 | -1.221287995 | 0.314434 | -3.884081 | 0.000103 | 0.010071383 | | lincRNA | 40.65 | -2.438803303 | 2.438803303 | ENSG00000275632 | down |
| ENSG00000207042 | 7.154694 | -2.079692704 | 0.67337 | -3.088483 | 0.002012 | 0.067495056 | RNU6-196P | snRNA | 39.25 | -2.434754554 | 2.434754554 | ENSG00000207042 | down |
| ENSG00000108312 | 5261.092 | -1.0124345 | 0.241803 | -4.187025 | 2.83E-05 | 0.004088489 | UBTF | protein_coding | 59.52 | -2.418136224 | 2.418136224 | ENSG00000108312 | down |
| ENSG00000213090 | 75.8163 | -1.188686066 | 0.303319 | -3.918925 | 8.89E-05 | 0.009241637 | | processed_pseudogene | 38.19 | -2.418085924 | 2.418085924 | ENSG00000213090 | down |
| ENSG00000095951 | 955.7391 | -1.005108429 | 0.23997 | -4.188475 | 2.81E-05 | 0.004088489 | HIVEP1 | protein_coding | 39.37 | -2.400638365 | 2.400638365 | ENSG00000095951 | down |
| ENSG00000198369 | 542.8321 | -1.039331586 | 0.252861 | -4.110286 | 3.95E-05 | 0.004956703 | SPRED2 | protein_coding | 43.76 | -2.39545886 | 2.39545886 | ENSG00000198369 | down |
| ENSG00000249353 | 797.4843 | 1.176342881 | 0.300136 | 3.9193624 | 8.88E-05 | 0.009241637 | NPM1P27 | processed_pseudogene | 41.69 | 2.392976786 | 2.392976786 | ENSG00000249353 | up |
| ENSG00000166833 | 1727.013 | -0.939879458 | 0.217596 | -4.319384 | 1.56E-05 | 0.002865686 | NAV2 | protein_coding | 44.51 | -2.389898657 | 2.389898657 | ENSG00000166833 | n.s. |
| ENSG00000207305 | 3.665837 | -2.564602745 | 0.91642 | -2.798503 | 0.005134 | 0.117181746 | | misc_RNA | 38.89 | -2.388004292 | 2.388004292 | ENSG00000207305 | n.s. |
| ENSG00000155760 | 299.017 | -0.96783869 | 0.227886 | -4.247033 | 2.17E-05 | 0.003466675 | FZD7 | protein_coding | 52.76 | -2.380967283 | 2.380967283 | ENSG00000155760 | n.s. |
| ENSG00000267397 | 41.51963 | -1.217278288 | 0.31805 | -3.827319 | 0.00013 | 0.011415674 | | lincRNA | 37.35 | -2.364561161 | 2.364561161 | ENSG00000267397 | down |
| ENSG00000207484 | 9.237373 | -2.076143843 | 0.681687 | -3.045597 | 0.002322 | 0.073314736 | | misc_RNA | 43.88 | -2.356026144 | 2.356026144 | ENSG00000207484 | down |
| ENSG00000197744 | 189.9928 | 1.304950106 | 0.351039 | 3.7173902 | 0.000201 | 0.015833788 | PTMAP2 | processed_pseudogene | 51.99 | 2.349451965 | 2.349451965 | ENSG00000197744 | up |
| ENSG00000087258 | 377.7388 | -0.942866663 | 0.220665 | -4.27284 | 1.93E-05 | 0.0032375 | GNAO1 | protein_coding | 49.48 | -2.347540252 | 2.347540252 | ENSG00000087258 | n.s. |
| ENSG00000212333 | 2.452781 | -3.171215749 | 1.25789 | -2.521059 | 0.0117 | 0.182089199 | RNA5SP213 | rRNA | 46.23 | -2.345798439 | 2.345798439 | ENSG00000212333 | n.s. |

| | | | | | | | | | | | | | |
|---|---|---|---|---|---|---|---|---|---|---|---|---|---|
| ENSG00000196498 | 874.8758 | -0.975711121 | 0.233914 | -4.171237 | 3.03E-05 | 0.004217973 | NCOR2 | protein_coding | 57.37 | -2.317212663 | 2.317212663 | ENSG00000196498 | n.s. |
| ENSG00000264966 | 3.538604 | -2.868988758 | 1.092057 | -2.627142 | 0.008611 | 0.157676123 | MIR5094 | miRNA | 38.82 | -2.301600524 | 2.301600524 | ENSG00000264966 | n.s. |
| ENSG00000223013 | 3.603497 | -2.866681923 | 1.092018 | -2.625123 | 0.008662 | 0.157676123 | RNA5SP344 | rRNA | 47.12 | -2.299749902 | 2.299749902 | ENSG00000223013 | n.s. |
| ENSG00000243680 | 22.35474 | 1.460229027 | 0.415259 | 3.5164284 | 0.000437 | 0.026946218 | RPL37P23 | processed_pseudogene | 47.62 | 2.291832652 | 2.291832652 | ENSG00000243680 | up |
| ENSG00000242318 | 22.64577 | -1.340707138 | 0.369095 | -3.632415 | 0.000281 | 0.019744027 | | processed_pseudogene | 40.72 | -2.285321467 | 2.285321467 | ENSG00000242318 | down |
| ENSG00000207248 | 12.55145 | -1.628268571 | 0.488285 | -3.334665 | 0.000854 | 0.039650653 | RNU6-1005P | snRNA | 43.93 | -2.282424913 | 2.282424913 | ENSG00000207248 | down |
| ENSG00000280165 | 173.0616 | 0.974767798 | 0.236217 | 4.1265699 | 3.68E-05 | 0.004668564 | PCDH20 | protein_coding | 44.3 | 2.272005076 | 2.272005076 | ENSG00000280165 | n.s. |
| ENSG00000276137 | 3.735289 | -2.514920792 | 0.911379 | -2.759467 | 0.00579 | 0.126147369 | MIR6133 | miRNA | 50.93 | -2.261220117 | 2.261220117 | ENSG00000276137 | n.s. |
| ENSG00000121089 | 45.8885 | 1.187162474 | 0.312912 | 3.793917 | 0.000148 | 0.012666449 | NACA3P | processed_pseudogene | 45.22 | 2.252456941 | 2.252456941 | ENSG00000121089 | up |
| ENSG00000207737 | 6.554938 | -1.893333906 | 0.609341 | -3.107185 | 0.001889 | 0.064772181 | MIR181B2 | miRNA | 46.07 | -2.25043842 | 2.25043842 | ENSG00000207737 | down |
| ENSG00000135747 | 117.6833 | -1.125450059 | 0.289677 | -3.885185 | 0.000102 | 0.010071383 | ZNF670-ZNF695 | protein_coding | 42.2 | -2.247423485 | 2.247423485 | ENSG00000135747 | down |
| ENSG00000227615 | 173.2392 | 1.265850958 | 0.34373 | 3.6826939 | 0.000231 | 0.017284945 | | processed_pseudogene | 46.87 | 2.230849657 | 2.230849657 | ENSG00000227615 | up |
| ENSG00000218823 | 183.9297 | -1.078816713 | 0.273273 | -3.94776 | 7.89E-05 | 0.00857042 | PAPOLB | protein_coding | 40.47 | -2.229911871 | 2.229911871 | ENSG00000218823 | down |
| ENSG00000277118 | 18.24661 | -1.556997458 | 0.46211 | -3.369319 | 0.000754 | 0.037280859 | | misc_RNA | 54.55 | -2.224192805 | 2.224192805 | ENSG00000277118 | down |
| ENSG00000266262 | 4.767531 | -2.505246252 | 0.912686 | -2.744915 | 0.006053 | 0.130381308 | MIR4428 | miRNA | 58.9 | -2.216603469 | 2.216603469 | ENSG00000266262 | n.s. |
| ENSG00000143878 | 4358.802 | 0.870277798 | 0.201747 | 4.313703 | 1.61E-05 | 0.002865686 | RHOB | protein_coding | 57.34 | 2.212917543 | 2.212917543 | ENSG00000143878 | n.s. |
| ENSG00000121068 | 3923.457 | -0.987229913 | 0.242699 | -4.067716 | 4.75E-05 | 0.005747924 | TBX2 | protein_coding | 61.32 | -2.211877759 | 2.211877759 | ENSG00000121068 | n.s. |
| ENSG00000222724 | 321.8994 | 0.931614385 | 0.224307 | 4.1533037 | 3.28E-05 | 0.004284193 | RNU2-63P | snRNA | 38.66 | 2.206184906 | 2.206184906 | ENSG00000222724 | n.s. |
| ENSG00000272367 | 53.49799 | -1.230147207 | 0.331848 | -3.706964 | 0.00021 | 0.016224866 | | antisense | 32.76 | -2.201740684 | 2.201740684 | ENSG00000272367 | down |
| ENSG00000280152 | 186.8057 | -0.95695008 | 0.233093 | -4.105451 | 4.04E-05 | 0.005011598 | | TEC | 59.43 | -2.20100795 | 2.20100795 | ENSG00000280152 | n.s. |
| ENSG00000165801 | 228.8646 | -1.077884995 | 0.274528 | -3.926321 | 8.63E-05 | 0.009097956 | ARHGEF40 | protein_coding | 53.84 | -2.200023835 | 2.200023835 | ENSG00000165801 | down |
| ENSG00000166068 | 783.2143 | -1.016762744 | 0.253451 | -4.011674 | 6.03E-05 | 0.006880443 | SPRED1 | protein_coding | 35.39 | -2.198631094 | 2.198631094 | ENSG00000166068 | down |
| ENSG00000212126 | 43.39919 | -1.155744282 | 0.304618 | -3.794079 | 0.000148 | 0.012666449 | TAS2R50 | protein_coding | 38.5 | -2.192845788 | 2.192845788 | ENSG00000212126 | down |
| ENSG00000274093 | 32.46154 | -1.300870105 | 0.360271 | -3.610809 | 0.000305 | 0.02063064 | | sense_intronic | 31.79 | -2.192600027 | 2.192600027 | ENSG00000274093 | down |
| ENSG00000202408 | 3.882003 | 3.252292567 | 1.341459 | 2.4244442 | 0.015332 | 0.213678848 | RNU1-122P | snRNA | 48.72 | 2.179811582 | 2.179811582 | ENSG00000202408 | n.s. |
| ENSG00000229638 | 547.0468 | 1.233984702 | 0.33509 | 3.6825497 | 0.000231 | 0.017284945 | RPL4P4 | processed_pseudogene | 48.29 | 2.174690734 | 2.174690734 | ENSG00000229638 | up |
| ENSG00000136099 | 1511.683 | 0.84454372 | 0.194398 | 4.3444065 | 1.40E-05 | 0.002665434 | PCDH8 | protein_coding | 56.58 | 2.174051538 | 2.174051538 | ENSG00000136099 | n.s. |
| ENSG00000272386 | 13.65124 | -1.501116169 | 0.443415 | -3.385356 | 0.000711 | 0.03643289 | | sense_intronic | 38.35 | -2.159365183 | 2.159365183 | ENSG00000272386 | down |
| ENSG00000213139 | 54.93819 | -1.089877017 | 0.282449 | -3.858671 | 0.000114 | 0.010646463 | CRYGS | protein_coding | 37.09 | -2.150103547 | 2.150103547 | ENSG00000213139 | down |
| ENSG00000103257 | 1141.99 | -0.911804911 | 0.220112 | -4.142465 | 3.44E-05 | 0.004429482 | SLC7A5 | protein_coding | 58.41 | -2.146066927 | 2.146066927 | ENSG00000103257 | n.s. |
| ENSG00000100345 | 1422.964 | -0.873406641 | 0.206578 | -4.227977 | 2.36E-05 | 0.003582447 | MYH9 | protein_coding | 52.37 | -2.136195663 | 2.136195663 | ENSG00000100345 | n.s. |
| ENSG00000139239 | 246.4075 | 1.138111115 | 0.301272 | 3.7776878 | 0.000158 | 0.013276429 | | processed_pseudogene | 49.46 | 2.136142047 | 2.136142047 | ENSG00000139239 | up |

| | | | | | | | | | | | | | |
|---|---|---|---|---|---|---|---|---|---|---|---|---|---|
| ENSG00000267280 | 346.684 | -0.969298458 | 0.239986 | -4.038978 | 5.37E-05 | 0.006265703 | TBX2-AS1 | antisense | 62.75 | -2.135393749 | 2.135393749 | ENSG00000267280 | n.s. |
| ENSG00000101463 | 239.9842 | -1.05760714 | 0.271213 | -3.89955 | 9.64E-05 | 0.009663101 | SYNDIG1 | protein_coding | 46.16 | -2.130955171 | 2.130955171 | ENSG00000101463 | down |
| ENSG00000266124 | 9.985842 | -2.258416245 | 0.804763 | -2.806311 | 0.005011 | 0.115515592 | MIR5587 | miRNA | 64.15 | -2.116947674 | 2.116947674 | ENSG00000266124 | n.s. |
| ENSG00000064042 | 4094.003 | -1.01300637 | 0.255538 | -3.964204 | 7.36E-05 | 0.00820246 | LIMCH1 | protein_coding | 40.5 | -2.113187874 | 2.113187874 | ENSG00000064042 | down |
| ENSG00000258739 | 91.7235 | -1.101517045 | 0.289159 | -3.809381 | 0.000139 | 0.012199277 | ENO1P2 | ıscribed_processed_pseudog | 43.23 | -2.107935609 | 2.107935609 | ENSG00000258739 | down |
| ENSG00000283615 | 4.231863 | -2.261942757 | 0.807958 | -2.799579 | 0.005117 | 0.116983481 | MIR924 | miRNA | 43.4 | -2.107848947 | 2.107848947 | ENSG00000283615 | n.s. |
| ENSG00000181524 | 148.8372 | 1.277949385 | 0.356873 | 3.580959 | 0.000342 | 0.022482711 | RPL24P4 | processed_pseudogene | 48.69 | 2.106253968 | 2.106253968 | ENSG00000181524 | up |
| ENSG00000279951 | 51.74315 | -1.074428617 | 0.279624 | -3.842399 | 0.000122 | 0.010974075 | | TEC | 40.65 | -2.105484785 | 2.105484785 | ENSG00000279951 | down |
| ENSG00000214944 | 81.15456 | -1.047633588 | 0.269506 | -3.887231 | 0.000101 | 0.010071383 | ARHGEF28 | protein_coding | 39.36 | -2.092030925 | 2.092030925 | ENSG00000214944 | down |
| ENSG00000123870 | 52.17075 | -1.173027041 | 0.317068 | -3.699609 | 0.000216 | 0.016518788 | ZNF137P | cribed_unprocessed_pseudc | 46.4 | -2.090359769 | 2.090359769 | ENSG00000123870 | down |
| ENSG00000184160 | 177.3369 | -1.069951165 | 0.279239 | -3.831669 | 0.000127 | 0.011287091 | ADRA2C | protein_coding | 69.62 | -2.08364214 | 2.08364214 | ENSG00000184160 | down |
| ENSG00000239494 | 10.12802 | -1.623199499 | 0.505405 | -3.211681 | 0.00132 | 0.052047939 | RN7SL333P | misc_RNA | 54.88 | -2.083533129 | 2.083533129 | ENSG00000239494 | down |
| ENSG00000218682 | 14.10467 | -1.465072234 | 0.436125 | -3.359292 | 0.000781 | 0.038444384 | | processed_pseudogene | 53.56 | -2.073322017 | 2.073322017 | ENSG00000218682 | down |
| ENSG00000268598 | 10.1236 | -1.609620858 | 0.500434 | -3.21645 | 0.001298 | 0.051629384 | VN1R80P | unprocessed_pseudogene | 40.83 | -2.071747923 | 2.071747923 | ENSG00000268598 | down |
| ENSG00000198719 | 133.3135 | -1.050304138 | 0.272949 | -3.84799 | 0.000119 | 0.010857493 | DLL1 | protein_coding | 53.53 | -2.063081393 | 2.063081393 | ENSG00000198719 | down |
| ENSG00000222160 | 4.29007 | -2.244921241 | 0.806918 | -2.782093 | 0.005401 | 0.120702132 | | misc_RNA | 45.19 | -2.061477634 | 2.061477634 | ENSG00000222160 | n.s. |
| ENSG00000272910 | 14.93177 | -1.454196036 | 0.435155 | -3.341786 | 0.000832 | 0.039650653 | | antisense | 33.73 | -2.038418796 | 2.038418796 | ENSG00000272910 | down |
| ENSG00000239218 | 66.72055 | -1.200169982 | 0.331878 | -3.616298 | 0.000299 | 0.020468836 | RPS20P22 | ıscribed_processed_pseudog | 47.12 | -2.026975302 | 2.026975302 | ENSG00000239218 | down |
| ENSG00000275908 | 2.76264 | -2.758397557 | 1.099394 | -2.509017 | 0.012107 | 0.185127237 | | snRNA | 38.95 | -2.020608084 | 2.020608084 | ENSG00000275908 | n.s. |
| ENSG00000270135 | 11.86729 | -1.524914262 | 0.468619 | -3.254061 | 0.001138 | 0.047710603 | | antisense | 29.45 | -2.01499898 | 2.01499898 | ENSG00000270135 | down |
| ENSG00000264814 | 7.975241 | -1.711534634 | 0.555527 | -3.080919 | 0.002064 | 0.06830421 | MIR1273C | miRNA | 49.35 | -1.994883523 | 1.994883523 | ENSG00000264814 | down |
| ENSG00000270277 | 163.6683 | -1.006077942 | 0.260686 | -3.859353 | 0.000114 | 0.010646463 | | antisense | 39.77 | -1.984785181 | 1.984785181 | ENSG00000270277 | down |
| ENSG00000235518 | 47.00196 | -1.218832599 | 0.342968 | -3.553776 | 0.00038 | 0.024364755 | | lincRNA | 41.75 | -1.96626701 | 1.96626701 | ENSG00000235518 | down |
| ENSG00000200291 | 6.031477 | -1.983144041 | 0.687217 | -2.885761 | 0.003905 | 0.102382029 | | misc_RNA | 45.1 | -1.96286891 | 1.96286891 | ENSG00000200291 | n.s. |
| ENSG00000222627 | 8.288038 | -1.63645679 | 0.52576 | -3.112556 | 0.001855 | 0.063919891 | RNU2-37P | snRNA | 45.2 | -1.954525036 | 1.954525036 | ENSG00000222627 | down |
| ENSG00000214842 | 78.84556 | 1.12599526 | 0.308048 | 3.6552629 | 0.000257 | 0.018534076 | RAD51AP2 | protein_coding | 34.45 | 1.95025652 | 1.95025652 | ENSG00000214842 | up |
| ENSG00000275291 | 132.2251 | -1.027834359 | 0.271167 | -3.790416 | 0.00015 | 0.012768007 | | snRNA | 46.91 | -1.94659173 | 1.94659173 | ENSG00000275291 | down |
| ENSG00000240399 | 51.54914 | -1.124541008 | 0.308178 | -3.649001 | 0.000263 | 0.018700992 | | transcribed_processed_pseudog | 43.65 | -1.943359063 | 1.943359063 | ENSG00000240399 | down |
| ENSG00000276702 | 16.75788 | -1.386084649 | 0.41429 | -3.345688 | 0.000821 | 0.039650653 | | antisense | 27.46 | -1.942943682 | 1.942943682 | ENSG00000276702 | down |
| ENSG00000252112 | 9.683195 | -1.616559423 | 0.518241 | -3.119321 | 0.001813 | 0.062968329 | | snoRNA | 43.75 | -1.941290369 | 1.941290369 | ENSG00000252112 | down |
| ENSG00000244398 | 474.9556 | 1.121043403 | 0.307106 | 3.6503459 | 0.000262 | 0.01869876 | | processed_pseudogene | 46.73 | 1.937372846 | 1.937372846 | ENSG00000244398 | up |
| ENSG00000251805 | 4.151919 | -2.41209131 | 0.918691 | -2.625573 | 0.00865 | 0.157676123 | | scaRNA | 47.66 | -1.935061825 | 1.935061825 | ENSG00000251805 | n.s. |

| | | | | | | | | | | | | | |
|---|---|---|---|---|---|---|---|---|---|---|---|---|---|
| ENSG00000067082 | 1244.427 | 0.809595177 | 0.19344 | 4.1852595 | 2.85E-05 | 0.004088489 | KLF6 | protein_coding | 48.08 | 1.93366724 | 1.93366724 | ENSG00000067082 | n.s. |
| ENSG00000277998 | 31.02134 | 1.331434309 | 0.391938 | 3.3970516 | 0.000681 | 0.035923511 | | unprocessed_pseudogene | 38.22 | 1.923418262 | 1.923418262 | ENSG00000277998 | up |
| ENSG00000263755 | 11.51244 | -1.497151451 | 0.466356 | -3.210321 | 0.001326 | 0.052147157 | RN7SL498P | misc_RNA | 54.36 | -1.920500009 | 1.920500009 | ENSG00000263755 | down |
| ENSG00000202187 | 3.450333 | -2.402935723 | 0.917452 | -2.619141 | 0.008815 | 0.159007629 | RNA5SP355 | rRNA | 47.41 | -1.918941309 | 1.918941309 | ENSG00000202187 | n.s. |
| ENSG00000223476 | 23.98237 | -1.274003219 | 0.368666 | -3.455715 | 0.000549 | 0.031189581 | VN1R42P | processed_pseudogene | 38.37 | -1.918636691 | 1.918636691 | ENSG00000223476 | down |
| ENSG00000181619 | 187.0888 | -0.958899411 | 0.24722 | -3.878728 | 0.000105 | 0.010152591 | GPR135 | protein_coding | 40.34 | -1.911492237 | 1.911492237 | ENSG00000181619 | n.s. |
| ENSG00000239627 | 90.17264 | -1.232889418 | 0.352228 | -3.500257 | 0.000465 | 0.02826005 | RPL12P20 | processed_pseudogene | 46.35 | -1.909532509 | 1.909532509 | ENSG00000239627 | down |
| ENSG00000231066 | 42.23991 | -1.123225576 | 0.310322 | -3.619548 | 0.000295 | 0.020341233 | NPM1P9 | processed_pseudogene | 41.35 | -1.900073908 | 1.900073908 | ENSG00000231066 | down |
| ENSG00000278281 | 2.186165 | -2.995026593 | 1.262272 | -2.372727 | 0.017657 | 0.232108003 | MIR7856 | miRNA | 42.86 | -1.899774972 | 1.899774972 | ENSG00000278281 | n.s. |
| ENSG00000252685 | 8.234135 | -2.135700971 | 0.777284 | -2.747646 | 0.006002 | 0.129972657 | RNU6-928P | snRNA | 40.38 | -1.892547147 | 1.892547147 | ENSG00000252685 | n.s. |
| ENSG00000163565 | 51.61306 | -1.154125587 | 0.323281 | -3.570035 | 0.000357 | 0.023331418 | IFI16 | protein_coding | 38.34 | -1.883600893 | 1.883600893 | ENSG00000163565 | down |
| ENSG00000268375 | 21.23805 | -1.315623349 | 0.39006 | -3.372878 | 0.000744 | 0.037254979 | | antisense | 49.56 | -1.879783268 | 1.879783268 | ENSG00000268375 | down |
| ENSG00000251213 | 108.9384 | -1.065119273 | 0.289538 | -3.678686 | 0.000234 | 0.017362167 | | lincRNA | 35.61 | -1.875031796 | 1.875031796 | ENSG00000251213 | down |
| ENSG00000252034 | 7.120954 | -1.738453416 | 0.583307 | -2.980342 | 0.002879 | 0.083706059 | RNY4P37 | misc_RNA | 40.62 | -1.872736955 | 1.872736955 | ENSG00000252034 | down |
| ENSG00000174307 | 387.4784 | -0.849266829 | 0.210156 | -4.041132 | 5.32E-05 | 0.006265703 | PHLDA3 | protein_coding | 57.71 | -1.870960449 | 1.870960449 | ENSG00000174307 | n.s. |
| ENSG00000276136 | 33.48764 | -1.164336816 | 0.328767 | -3.541523 | 0.000398 | 0.025062965 | | lincRNA | 42.69 | -1.864065465 | 1.864065465 | ENSG00000276136 | down |
| ENSG00000279900 | 35.89458 | -1.142772277 | 0.320537 | -3.565186 | 0.000364 | 0.023545975 | | TEC | 40.27 | -1.860528492 | 1.860528492 | ENSG00000279900 | down |
| ENSG00000264931 | 6.445777 | -1.944440247 | 0.686471 | -2.832515 | 0.004618 | 0.111247803 | MIR3138 | miRNA | 67.07 | -1.854429304 | 1.854429304 | ENSG00000264931 | n.s. |
| ENSG00000225213 | 27.84137 | -1.254531282 | 0.36582 | -3.429369 | 0.000605 | 0.033277115 | | sense_intronic | 39.83 | -1.854014489 | 1.854014489 | ENSG00000225213 | down |
| ENSG00000251666 | 31.78643 | -1.351139437 | 0.409247 | -3.301523 | 0.000962 | 0.042690184 | ZNF346-IT1 | sense_intronic | 44.4 | -1.850617824 | 1.850617824 | ENSG00000251666 | down |
| ENSG00000199630 | 17.19111 | -1.3740312 | 0.419319 | -3.276817 | 0.00105 | 0.045009285 | | misc_RNA | 40.62 | -1.850404911 | 1.850404911 | ENSG00000199630 | down |
| ENSG00000122863 | 264.9944 | -0.938218259 | 0.243403 | -3.85458 | 0.000116 | 0.010689213 | CHST3 | protein_coding | 51.39 | -1.849279114 | 1.849279114 | ENSG00000122863 | n.s. |
| ENSG00000207268 | 11.78797 | -1.605866877 | 0.524749 | -3.060259 | 0.002211 | 0.071230366 | SNORA70C | snoRNA | 50.37 | -1.84246699 | 1.84246699 | ENSG00000207268 | down |
| ENSG00000064666 | 142.7595 | -0.999304803 | 0.26696 | -3.743277 | 0.000182 | 0.014596081 | CNN2 | protein_coding | 59.73 | -1.834487507 | 1.834487507 | ENSG00000064666 | n.s. |
| ENSG00000200522 | 6.215984 | -1.863700054 | 0.648255 | -2.874947 | 0.004041 | 0.103804772 | RNU6-957P | snRNA | 37.37 | -1.833475835 | 1.833475835 | ENSG00000200522 | n.s. |
| ENSG00000202347 | 22.45223 | -1.432720684 | 0.448241 | -3.196315 | 0.001392 | 0.053388725 | RNU1-16P | snRNA | 48.78 | -1.823209356 | 1.823209356 | ENSG00000202347 | down |
| ENSG00000229447 | 17.9972 | -1.369242166 | 0.420048 | -3.259724 | 0.001115 | 0.046909423 | | processed_pseudogene | 39.13 | -1.819366705 | 1.819366705 | ENSG00000229447 | down |
| ENSG00000279499 | 16.34761 | -1.309292207 | 0.397192 | -3.296375 | 0.000979 | 0.043100984 | | TEC | 29.16 | -1.787855292 | 1.787855292 | ENSG00000279499 | down |
| ENSG00000259407 | 32.55434 | -1.225835088 | 0.359864 | -3.406387 | 0.000658 | 0.034849263 | | antisense | 48.91 | -1.787030234 | 1.787030234 | ENSG00000259407 | down |
| ENSG00000200257 | 5.188204 | -2.022902931 | 0.739896 | -2.734037 | 0.006256 | 0.132179666 | RNU6-97P | snRNA | 41.35 | -1.777798607 | 1.777798607 | ENSG00000200257 | n.s. |
| ENSG00000253180 | 15.58801 | -1.37245432 | 0.425195 | -3.227824 | 0.001247 | 0.050780579 | | processed_pseudogene | 41.97 | -1.776370852 | 1.776370852 | ENSG00000253180 | down |
| ENSG00000278492 | 15.53029 | -1.358342489 | 0.419461 | -3.238305 | 0.001202 | 0.049530614 | | sense_intronic | 32.64 | -1.772808491 | 1.772808491 | ENSG00000278492 | down |

| | | | | | | | | | | | | | |
|---|---|---|---|---|---|---|---|---|---|---|---|---|---|
| ENSG00000200874 | 6.090399 | -2.340593921 | 0.913526 | -2.562155 | 0.010402 | 0.174827325 | | misc_RNA | 42.16 | -1.77274404 | 1.77274404 | ENSG00000200874 | n.s. |
| ENSG00000242931 | 86.36592 | -0.992912622 | 0.268308 | -3.700645 | 0.000215 | 0.016518788 | RPL7P49 | processed_pseudogene | 43.64 | -1.769391946 | 1.769391946 | ENSG00000242931 | n.s. |
| ENSG00000199899 | 3.942939 | -2.342967449 | 0.916513 | -2.556394 | 0.010576 | 0.175750036 | | misc_RNA | 37.37 | -1.769185444 | 1.769185444 | ENSG00000199899 | n.s. |
| ENSG00000225900 | 30.28744 | -1.22977902 | 0.362836 | -3.389356 | 0.000701 | 0.03643289 | HSPE1P13 | processed_pseudogene | 38.38 | -1.769044964 | 1.769044964 | ENSG00000225900 | down |
| ENSG00000279838 | 913.8272 | -0.855082967 | 0.216374 | -3.951867 | 7.75E-05 | 0.008538755 | | TEC | 41.03 | -1.768829299 | 1.768829299 | ENSG00000279838 | n.s. |
| ENSG00000228412 | 63.49496 | 1.099126644 | 0.309698 | 3.5490256 | 0.000387 | 0.024664733 | | antisense | 38.79 | 1.767311642 | 1.767311642 | ENSG00000228412 | up |
| ENSG00000260583 | 30.4413 | -1.17002325 | 0.338193 | -3.459634 | 0.000541 | 0.03099004 | | antisense | 44.17 | -1.765305183 | 1.765305183 | ENSG00000260583 | down |
| ENSG00000146677 | 139.942 | 1.159120979 | 0.334384 | 3.4664349 | 0.000527 | 0.030432353 | | processed_pseudogene | 50.49 | 1.757997596 | 1.757997596 | ENSG00000146677 | up |
| ENSG00000260452 | 39.10098 | -1.133166761 | 0.323949 | -3.497979 | 0.000469 | 0.028378547 | TPRKBP2 | processed_pseudogene | 33.68 | -1.75302013 | 1.75302013 | ENSG00000260452 | down |
| ENSG00000234709 | 28.42508 | -1.295419711 | 0.394491 | -3.283778 | 0.001024 | 0.044425871 | UPF3AP3 | ıscribed_processed_pseudog | 52.28 | -1.751879043 | 1.751879043 | ENSG00000234709 | down |
| ENSG00000277840 | 38.85082 | -1.122338711 | 0.319881 | -3.508609 | 0.00045 | 0.027507512 | | sense_intronic | 38.84 | -1.751464209 | 1.751464209 | ENSG00000277840 | down |
| ENSG00000261474 | 24.83301 | -1.249459503 | 0.37454 | -3.335982 | 0.00085 | 0.039650653 | | lincRNA | 48.37 | -1.751429431 | 1.751429431 | ENSG00000261474 | down |
| ENSG00000217767 | 20.39148 | -1.381875242 | 0.433581 | -3.18712 | 0.001437 | 0.054514828 | NDUFAB1P1 | processed_pseudogene | 47.39 | -1.745979132 | 1.745979132 | ENSG00000217767 | down |
| ENSG00000168811 | 64.49115 | 1.040323275 | 0.288852 | 3.60158 | 0.000316 | 0.02127388 | IL12A | protein_coding | 43.49 | 1.739579989 | 1.739579989 | ENSG00000168811 | up |
| ENSG00000280037 | 26.38781 | -1.239706573 | 0.371484 | -3.337171 | 0.000846 | 0.039650653 | | TEC | 45.03 | -1.737758265 | 1.737758265 | ENSG00000280037 | down |
| ENSG00000270574 | 158.1868 | -1.021384317 | 0.282005 | -3.621872 | 0.000292 | 0.020259632 | | antisense | 40.39 | -1.729579968 | 1.729579968 | ENSG00000270574 | down |
| ENSG00000235369 | 8.743061 | -1.562937038 | 0.519481 | -3.00865 | 0.002624 | 0.078827723 | RPL36AP15 | processed_pseudogene | 44.74 | -1.724421284 | 1.724421284 | ENSG00000235369 | down |
| ENSG00000265010 | 10.23897 | -1.475134511 | 0.479198 | -3.078343 | 0.002082 | 0.068607507 | | sense_intronic | 44.73 | -1.716508354 | 1.716508354 | ENSG00000265010 | down |
| ENSG00000226084 | 184.4817 | 1.113467521 | 0.318849 | 3.4921455 | 0.000479 | 0.028755623 | | processed_pseudogene | 45.77 | 1.716162127 | 1.716162127 | ENSG00000226084 | up |
| ENSG00000236670 | 38.74586 | -1.13440852 | 0.328793 | -3.450221 | 0.00056 | 0.031516313 | KRT18P5 | processed_pseudogene | 54.69 | -1.703274228 | 1.703274228 | ENSG00000236670 | down |
| ENSG00000232346 | 98.69908 | 1.214822931 | 0.363694 | 3.3402301 | 0.000837 | 0.039650653 | | processed_pseudogene | 50.61 | 1.702877628 | 1.702877628 | ENSG00000232346 | up |
| ENSG00000239272 | 20.55712 | -1.312730526 | 0.407613 | -3.220528 | 0.00128 | 0.051192934 | RPL21P10 | processed_pseudogene | 47.08 | -1.694459413 | 1.694459413 | ENSG00000239272 | down |
| ENSG00000081377 | 351.5189 | -0.958974624 | 0.259984 | -3.688596 | 0.000225 | 0.017156128 | CDC14B | protein_coding | 42.84 | -1.693147109 | 1.693147109 | ENSG00000081377 | n.s. |
| ENSG00000261229 | 26.86731 | -1.183888874 | 0.351204 | -3.370943 | 0.000749 | 0.037274749 | | lincRNA | 40.63 | -1.691286205 | 1.691286205 | ENSG00000261229 | down |
| ENSG00000258913 | 44.69886 | -1.103395596 | 0.317315 | -3.477286 | 0.000507 | 0.029882384 | | lincRNA | 53.02 | -1.682220108 | 1.682220108 | ENSG00000258913 | down |
| ENSG00000244313 | 464.332 | 1.165577468 | 0.344186 | 3.3864729 | 0.000708 | 0.03643289 | | processed_pseudogene | 54.22 | 1.676690621 | 1.676690621 | ENSG00000244313 | up |
| ENSG00000270878 | 40.37156 | -1.10224263 | 0.318048 | -3.465649 | 0.000529 | 0.030432353 | | processed_pseudogene | 60.03 | -1.671732225 | 1.671732225 | ENSG00000270878 | down |
| ENSG00000140464 | 197.2227 | -0.948106974 | 0.257623 | -3.680212 | 0.000233 | 0.017350904 | PML | protein_coding | 47.82 | -1.669310993 | 1.669310993 | ENSG00000140464 | n.s. |
| ENSG00000124635 | 889.0533 | -0.853433166 | 0.222593 | -3.834049 | 0.000126 | 0.011250075 | HIST1H2BJ | protein_coding | 41.7 | -1.663208591 | 1.663208591 | ENSG00000124635 | n.s. |
| ENSG00000259713 | 44.24511 | -1.114261557 | 0.324173 | -3.437246 | 0.000588 | 0.032748711 | | sense_overlapping | 44.09 | -1.654462011 | 1.654462011 | ENSG00000259713 | down |
| ENSG00000204172 | 108.6033 | -0.949040354 | 0.259425 | -3.658244 | 0.000254 | 0.018415263 | AGAP9 | protein_coding | 40.48 | -1.646416155 | 1.646416155 | ENSG00000204172 | n.s. |
| ENSG00000260060 | 94.04481 | -0.973379317 | 0.269233 | -3.615378 | 0.0003 | 0.020468836 | | antisense | 50.06 | -1.643946995 | 1.643946995 | ENSG00000260060 | n.s. |

| | | | | | | | | | | | | | |
|---|---|---|---|---|---|---|---|---|---|---|---|---|---|
| ENSG00000271907 | 3.58109 | -2.291143331 | 0.920584 | -2.488792 | 0.012818 | 0.191814442 | SNORA35B | snoRNA | 43.41 | -1.643021723 | 1.643021723 | ENSG00000271907 | n.s. |
| ENSG00000237887 | 43.62714 | -1.080489161 | 0.311555 | -3.468049 | 0.000524 | 0.030413132 | RPL23AP32 | processed_pseudogene | 48.8 | -1.639035988 | 1.639035988 | ENSG00000237887 | down |
| ENSG00000224635 | 132.6636 | -0.896542113 | 0.240558 | -3.726921 | 0.000194 | 0.015421387 | | lincRNA | 48.32 | -1.624423649 | 1.624423649 | ENSG00000224635 | n.s. |
| ENSG00000227535 | 17.84932 | -1.428514885 | 0.468631 | -3.048272 | 0.002302 | 0.072996398 | | sense_overlapping | 39.23 | -1.623790823 | 1.623790823 | ENSG00000227535 | down |
| ENSG00000139438 | 259.2859 | -0.851076036 | 0.22406 | -3.798423 | 0.000146 | 0.012592912 | FAM222A | protein_coding | 52.74 | -1.616937084 | 1.616937084 | ENSG00000139438 | n.s. |
| ENSG00000254665 | 37.87312 | -1.053974465 | 0.302404 | -3.485323 | 0.000492 | 0.029238493 | | antisense | 47.55 | -1.61684428 | 1.61684428 | ENSG00000254665 | down |
| ENSG00000166123 | 829.9534 | -0.919381557 | 0.250163 | -3.675127 | 0.000238 | 0.017512841 | GPT2 | protein_coding | 51.96 | -1.615025546 | 1.615025546 | ENSG00000166123 | n.s. |
| ENSG00000154734 | 930.1911 | 0.780852617 | 0.197875 | 3.9461827 | 7.94E-05 | 0.00857042 | ADAMTS1 | protein_coding | 45.12 | 1.614020712 | 1.614020712 | ENSG00000154734 | n.s. |
| ENSG00000200755 | 4.466846 | -2.270761704 | 0.915982 | -2.479047 | 0.013173 | 0.194706159 | RNA5SP68 | rRNA | 50.42 | -1.613649387 | 1.613649387 | ENSG00000200755 | n.s. |
| ENSG00000272192 | 41.06789 | 1.153959615 | 0.346985 | 3.3256727 | 0.000882 | 0.040189476 | | lincRNA | 39.66 | 1.610797977 | 1.610797977 | ENSG00000272192 | up |
| ENSG00000264395 | 159.2402 | 0.934023948 | 0.257428 | 3.6282895 | 0.000285 | 0.01996133 | MIR3193 | miRNA | 56.36 | 1.587663733 | 1.587663733 | ENSG00000264395 | n.s. |
| ENSG00000066735 | 2757.427 | -0.879203209 | 0.236256 | -3.721395 | 0.000198 | 0.015673305 | KIF26A | protein_coding | 63.77 | -1.586820613 | 1.586820613 | ENSG00000066735 | n.s. |
| ENSG00000222898 | 13.19561 | -1.351464821 | 0.438074 | -3.085014 | 0.002035 | 0.067959896 | RN7SKP97 | misc_RNA | 49.85 | -1.578169386 | 1.578169386 | ENSG00000222898 | down |
| ENSG00000263361 | 3.752405 | -2.093325141 | 0.820118 | -2.552467 | 0.010696 | 0.176242214 | MIR378H | miRNA | 51.81 | -1.578137019 | 1.578137019 | ENSG00000263361 | n.s. |
| ENSG00000243911 | 9.665854 | -1.654599504 | 0.585374 | -2.82657 | 0.004705 | 0.111596326 | RN7SL430P | misc_RNA | 52.89 | -1.575758006 | 1.575758006 | ENSG00000243911 | n.s. |
| ENSG00000165197 | 176.3806 | 1.072431763 | 0.314138 | 3.4138836 | 0.00064 | 0.034295524 | VEGFD | protein_coding | 39.3 | 1.570857899 | 1.570857899 | ENSG00000165197 | up |
| ENSG00000265713 | 52.17684 | -1.023801129 | 0.293836 | -3.484258 | 0.000494 | 0.029238493 | | transcribed_processed_pseudog | 42.83 | -1.570557024 | 1.570557024 | ENSG00000265713 | down |
| ENSG00000183691 | 61.23458 | -1.12150982 | 0.336626 | -3.331622 | 0.000863 | 0.039805646 | NOG | protein_coding | 59.46 | -1.570175789 | 1.570175789 | ENSG00000183691 | down |
| ENSG00000078725 | 138.5118 | -0.926099924 | 0.255672 | -3.622213 | 0.000292 | 0.020259632 | BRINP1 | protein_coding | 40.76 | -1.568228384 | 1.568228384 | ENSG00000078725 | n.s. |
| ENSG00000166963 | 1657.109 | -0.820824115 | 0.215699 | -3.805414 | 0.000142 | 0.012319021 | MAP1A | protein_coding | 52.57 | -1.567301103 | 1.567301103 | ENSG00000166963 | n.s. |
| ENSG00000149212 | 1070.655 | 1.03678416 | 0.299775 | 3.4585462 | 0.000543 | 0.03099004 | SESN3 | protein_coding | 35.27 | 1.564276993 | 1.564276993 | ENSG00000149212 | up |
| ENSG00000278463 | 904.0797 | -0.796152855 | 0.206926 | -3.847533 | 0.000119 | 0.010857493 | HIST1H2AB | protein_coding | 60.05 | -1.563859535 | 1.563859535 | ENSG00000278463 | n.s. |
| ENSG00000136848 | 625.5376 | -0.849458056 | 0.226485 | -3.750612 | 0.000176 | 0.01444747 | DAB2IP | protein_coding | 53.14 | -1.563179677 | 1.563179677 | ENSG00000136848 | n.s. |
| ENSG00000256973 | 54.40198 | -1.086095596 | 0.32095 | -3.384004 | 0.000714 | 0.03643289 | | lincRNA | 42.08 | -1.562355442 | 1.562355442 | ENSG00000256973 | down |
| ENSG00000221164 | 37.6031 | -1.11305824 | 0.333093 | -3.341584 | 0.000833 | 0.039650653 | SNORA11F | snoRNA | 50.78 | -1.560229008 | 1.560229008 | ENSG00000221164 | down |
| ENSG00000213553 | 843.8021 | 1.088933796 | 0.323246 | 3.368751 | 0.000755 | 0.037280859 | RPLP0P6 | processed_pseudogene | 53.56 | 1.555557269 | 1.555557269 | ENSG00000213553 | up |
| ENSG00000235552 | 439.6079 | 1.059693699 | 0.310104 | 3.4172157 | 0.000633 | 0.034134195 | | transcribed_processed_pseudog | 46.79 | 1.554369667 | 1.554369667 | ENSG00000235552 | up |
| ENSG00000107130 | 2006.873 | -0.878989741 | 0.238613 | -3.683748 | 0.00023 | 0.017284945 | NCS1 | protein_coding | 54.7 | -1.549071752 | 1.549071752 | ENSG00000107130 | n.s. |
| ENSG00000226243 | 50.25408 | 1.602953003 | 0.562548 | 2.8494513 | 0.004379 | 0.108407336 | RPL37AP1 | processed_pseudogene | 53.05 | 1.546755601 | 1.546755601 | ENSG00000226243 | n.s. |
| ENSG00000239899 | 32.0358 | -1.131834692 | 0.343767 | -3.292452 | 0.000993 | 0.043348054 | RN7SL674P | misc_RNA | 54.18 | -1.542725091 | 1.542725091 | ENSG00000239899 | down |
| ENSG00000199530 | 7.956461 | -2.036622716 | 0.79496 | -2.56192 | 0.01041 | 0.174827325 | | misc_RNA | 38.89 | -1.54251908 | 1.54251908 | ENSG00000199530 | n.s. |
| ENSG00000226686 | 131.0089 | -0.902376997 | 0.248142 | -3.636537 | 0.000276 | 0.019529526 | LINC01535 | lincRNA | 44.46 | -1.542440479 | 1.542440479 | ENSG00000226686 | n.s. |

| | | | | | | | | | | | | | |
|---|---|---|---|---|---|---|---|---|---|---|---|---|---|
| ENSG00000130956 | 248.0711 | -0.925472147 | 0.257369 | -3.595897 | 0.000323 | 0.02163929 | HABP4 | protein_coding | 43.78 | -1.540686224 | 1.540686224 | ENSG00000130956 | n.s. |
| ENSG00000167528 | 54.07167 | -1.075154351 | 0.318502 | -3.375665 | 0.000736 | 0.037146987 | ZNF641 | protein_coding | 46.15 | -1.537552875 | 1.537552875 | ENSG00000167528 | down |
| ENSG00000279573 | 57.77104 | -1.035746661 | 0.302207 | -3.427271 | 0.00061 | 0.033288525 | | TEC | 50.46 | -1.530528488 | 1.530528488 | ENSG00000279573 | down |
| ENSG00000235843 | 41.10706 | -1.112326309 | 0.336913 | -3.30152 | 0.000962 | 0.042690184 | | antisense | 42.48 | -1.523522174 | 1.523522174 | ENSG00000235843 | down |
| ENSG00000213197 | 46.66883 | -1.090575545 | 0.327927 | -3.325671 | 0.000882 | 0.040189476 | | processed_pseudogene | 52.88 | -1.522320937 | 1.522320937 | ENSG00000213197 | down |
| ENSG00000278770 | 9.802025 | -1.511292307 | 0.520633 | -2.9028 | 0.003698 | 0.098747222 | | misc_RNA | 53.02 | -1.519566801 | 1.519566801 | ENSG00000278770 | down |
| ENSG00000201581 | 12.23135 | -1.550485275 | 0.540452 | -2.86887 | 0.004119 | 0.104852945 | RN7SKP78 | misc_RNA | 50.94 | -1.518575309 | 1.518575309 | ENSG00000201581 | n.s. |
| ENSG00000252546 | 3.752787 | 2.048293709 | 0.811638 | 2.5236554 | 0.011614 | 0.181889957 | RNA5SP466 | rRNA | 43.52 | 1.516129143 | 1.516129143 | ENSG00000252546 | n.s. |
| ENSG00000213178 | 53.78537 | 1.19092549 | 0.372441 | 3.1976232 | 0.001386 | 0.053388725 | | processed_pseudogene | 41.46 | 1.515512773 | 1.515512773 | ENSG00000213178 | up |
| ENSG00000250569 | 34.26071 | -1.171885652 | 0.363481 | -3.224059 | 0.001264 | 0.050948982 | NTAN1P2 | processed_pseudogene | 48.2 | -1.515089349 | 1.515089349 | ENSG00000250569 | down |
| ENSG00000206633 | 205.6271 | 0.844370109 | 0.22751 | 3.711361 | 0.000206 | 0.016124703 | SNORA80B | snoRNA | 58.82 | 1.513540413 | 1.513540413 | ENSG00000206633 | n.s. |
| ENSG00000249140 | 27.41193 | -1.133961139 | 0.347233 | -3.265705 | 0.001092 | 0.046349923 | PRDX2P3 | processed_pseudogene | 54.61 | -1.512648576 | 1.512648576 | ENSG00000249140 | down |
| ENSG00000206638 | 2.617607 | -2.544274159 | 1.100647 | -2.311617 | 0.020799 | 0.254913212 | RNU6-672P | snRNA | 39.25 | -1.510300616 | 1.510300616 | ENSG00000206638 | n.s. |
| ENSG00000185338 | 36.46492 | 1.254818878 | 0.401626 | 3.1243463 | 0.001782 | 0.062653718 | SOCS1 | protein_coding | 68.56 | 1.509613805 | 1.509613805 | ENSG00000185338 | up |
| ENSG00000272807 | 119.1181 | -1.015532925 | 0.295464 | -3.437075 | 0.000588 | 0.032748711 | | antisense | 31.62 | -1.507869166 | 1.507869166 | ENSG00000272807 | down |
| ENSG00000269867 | 38.82369 | -1.046721267 | 0.30886 | -3.388984 | 0.000702 | 0.03643289 | | sense_intronic | 46.52 | -1.505715218 | 1.505715218 | ENSG00000269867 | down |
| ENSG00000184347 | 4846.85 | -0.83889803 | 0.226293 | -3.707136 | 0.00021 | 0.016224866 | SLIT3 | protein_coding | 45.01 | -1.501475524 | 1.501475524 | ENSG00000184347 | n.s. |
| ENSG00000270062 | 26.50707 | -1.146291951 | 0.353509 | -3.242614 | 0.001184 | 0.049077962 | | sense_intronic | 32.06 | -1.500626249 | 1.500626249 | ENSG00000270062 | down |
| ENSG00000236552 | 659.6026 | 1.112730008 | 0.339599 | 3.2766038 | 0.001051 | 0.045009285 | RPL13AP5 | processed_pseudogene | 57.35 | 1.498511149 | 1.498511149 | ENSG00000236552 | up |
| ENSG00000235672 | 17.28103 | -1.274057527 | 0.412138 | -3.09134 | 0.001993 | 0.067010372 | | transcribed_processed_pseudog | 47.87 | -1.495562587 | 1.495562587 | ENSG00000235672 | down |
| ENSG00000161149 | 102.7753 | -0.955679982 | 0.272128 | -3.511882 | 0.000445 | 0.027290678 | TUBA3FP | processed_transcript | 54.66 | -1.494669805 | 1.494669805 | ENSG00000161149 | n.s. |
| ENSG00000236439 | 50.40617 | 1.082295806 | 0.327017 | 3.309598 | 0.000934 | 0.041827218 | | processed_pseudogene | 49.61 | 1.491989167 | 1.491989167 | ENSG00000236439 | up |
| ENSG00000279957 | 69.73197 | 1.035485294 | 0.305916 | 3.3848642 | 0.000712 | 0.03643289 | | TEC | 35.72 | 1.489552199 | 1.489552199 | ENSG00000279957 | up |
| ENSG00000207943 | 3.693346 | -2.047728899 | 0.818265 | -2.502526 | 0.012331 | 0.187429218 | MIR634 | miRNA | 52.58 | -1.489032091 | 1.489032091 | ENSG00000207943 | n.s. |
| ENSG00000234268 | 23.33401 | -1.322307137 | 0.436273 | -3.030914 | 0.002438 | 0.074968112 | | processed_pseudogene | 49.02 | -1.487758739 | 1.487758739 | ENSG00000234268 | down |
| ENSG00000035403 | 859.5011 | -0.970518875 | 0.279335 | -3.474394 | 0.000512 | 0.030078912 | VCL | protein_coding | 41.45 | -1.476875334 | 1.476875334 | ENSG00000035403 | n.s. |
| ENSG00000274825 | 25.32644 | -1.124725602 | 0.347575 | -3.235919 | 0.001213 | 0.049750266 | | lincRNA | 46.08 | -1.465747573 | 1.465747573 | ENSG00000274825 | down |
| ENSG00000267519 | 66.30797 | -1.128748502 | 0.3493 | -3.231455 | 0.001232 | 0.050286744 | | lincRNA | 55.64 | -1.465732403 | 1.465732403 | ENSG00000267519 | down |
| ENSG00000207766 | 4.286617 | -2.213661991 | 0.917406 | -2.412959 | 0.015824 | 0.217740811 | MIR626 | miRNA | 31.91 | -1.465577421 | 1.465577421 | ENSG00000207766 | n.s. |
| ENSG00000275673 | 38.81046 | -1.069516726 | 0.324219 | -3.298746 | 0.000971 | 0.04292603 | | sense_intronic | 52.38 | -1.462328057 | 1.462328057 | ENSG00000275673 | down |
| ENSG00000200651 | 12.87729 | -1.338799426 | 0.447142 | -2.994129 | 0.002752 | 0.0814896 | | misc_RNA | 42.86 | -1.457815765 | 1.457815765 | ENSG00000200651 | down |
| ENSG00000271002 | 10.25698 | -1.671382982 | 0.61557 | -2.715179 | 0.006624 | 0.135426965 | | processed_pseudogene | 42.54 | -1.451253243 | 1.451253243 | ENSG00000271002 | n.s. |

| | | | | | | | | | | | | | |
|---|---|---|---|---|---|---|---|---|---|---|---|---|---|
| ENSG00000224094 | 30.08525 | 1.24324423 | 0.40338 | 3.0820674 | 0.002056 | 0.06830421 | RPS24P8 | processed_pseudogene | 42.29 | 1.449066458 | 1.449066458 | ENSG00000224094 | up |
| ENSG00000252062 | 8.328129 | -1.728304684 | 0.644407 | -2.682008 | 0.007318 | 0.145143654 | RNU6-469P | snRNA | 42.57 | -1.448668353 | 1.448668353 | ENSG00000252062 | n.s. |
| ENSG00000213131 | 55.21168 | -1.045639158 | 0.31518 | -3.317596 | 0.000908 | 0.04117752 | YWHAZP4 | processed_pseudogene | 46.71 | -1.448565551 | 1.448565551 | ENSG00000213131 | down |
| ENSG00000197275 | 235.0341 | -0.936238007 | 0.268218 | -3.49059 | 0.000482 | 0.028799362 | RAD54B | protein_coding | 36.72 | -1.442384314 | 1.442384314 | ENSG00000197275 | n.s. |
| ENSG00000171488 | 308.7818 | -0.900465091 | 0.254143 | -3.543148 | 0.000395 | 0.025022235 | LRRC8C | protein_coding | 39.69 | -1.442251432 | 1.442251432 | ENSG00000171488 | n.s. |
| ENSG00000279348 | 750.5157 | -0.901652628 | 0.254825 | -3.538321 | 0.000403 | 0.025254562 | | TEC | 40.08 | -1.44053448 | 1.44053448 | ENSG00000279348 | n.s. |
| ENSG00000218426 | 280.4425 | 1.080009252 | 0.330876 | 3.2640924 | 0.001098 | 0.046472913 | | processed_pseudogene | 54.14 | 1.439436447 | 1.439436447 | ENSG00000218426 | up |
| ENSG00000258441 | 1144.384 | -0.882555196 | 0.247498 | -3.565914 | 0.000363 | 0.023545975 | LINC00641 | processed_transcript | 41.26 | -1.436873401 | 1.436873401 | ENSG00000258441 | n.s. |
| ENSG00000129993 | 21.76742 | -1.291617275 | 0.428259 | -3.015971 | 0.002562 | 0.077701305 | CBFA2T3 | protein_coding | 60.67 | -1.433141961 | 1.433141961 | ENSG00000129993 | down |
| ENSG00000259469 | 26.01171 | -1.122395752 | 0.350489 | -3.20237 | 0.001363 | 0.053157763 | | lincRNA | 35.16 | -1.43041853 | 1.43041853 | ENSG00000259469 | down |
| ENSG00000264572 | 11.39483 | -1.381693429 | 0.471099 | -2.932918 | 0.003358 | 0.092447097 | MIR4296 | miRNA | 56.82 | -1.428818462 | 1.428818462 | ENSG00000264572 | down |
| ENSG00000272707 | 7.406907 | -1.659345355 | 0.612983 | -2.706999 | 0.006789 | 0.138200935 | | lincRNA | 41.25 | -1.426189111 | 1.426189111 | ENSG00000272707 | n.s. |
| ENSG00000137936 | 47.91365 | -1.051033969 | 0.320616 | -3.278172 | 0.001045 | 0.045009285 | BCAR3 | protein_coding | 45.08 | -1.415425224 | 1.415425224 | ENSG00000137936 | down |
| ENSG00000242699 | 50.2921 | -1.092329429 | 0.338762 | -3.224471 | 0.001262 | 0.050948982 | RN7SL516P | misc_RNA | 49.16 | -1.412233932 | 1.412233932 | ENSG00000242699 | down |
| ENSG00000267632 | 117.9785 | -0.882961846 | 0.249759 | -3.535251 | 0.000407 | 0.025435227 | | sense_intronic | 50.58 | -1.407939505 | 1.407939505 | ENSG00000267632 | n.s. |
| ENSG00000163352 | 20.0417 | -1.172239774 | 0.376023 | -3.11747 | 0.001824 | 0.063176664 | LENEP | protein_coding | 54.66 | -1.406035154 | 1.406035154 | ENSG00000163352 | down |
| ENSG00000166444 | 19.49834 | -1.318160062 | 0.444018 | -2.968705 | 0.002991 | 0.086028362 | ST5 | protein_coding | 45.03 | -1.404312822 | 1.404312822 | ENSG00000166444 | down |
| ENSG00000237406 | 68.14397 | -0.989783107 | 0.294899 | -3.356346 | 0.00079 | 0.038623611 | NDUFA9P1 | processed_pseudogene | 47.05 | -1.39870915 | 1.39870915 | ENSG00000237406 | n.s. |
| ENSG00000200170 | 5.210873 | -1.803129246 | 0.697124 | -2.586526 | 0.009695 | 0.167712137 | | misc_RNA | 47.52 | -1.398210441 | 1.398210441 | ENSG00000200170 | n.s. |
| ENSG00000214485 | 197.5821 | 1.040104831 | 0.317983 | 3.2709413 | 0.001072 | 0.045639412 | RPL7P1 | processed_pseudogene | 43.51 | 1.394426896 | 1.394426896 | ENSG00000214485 | up |
| ENSG00000197182 | 1238.254 | -0.792265924 | 0.216231 | -3.663978 | 0.000248 | 0.018197252 | MIRLET7BHG | lincRNA | 56.82 | -1.378538113 | 1.378538113 | ENSG00000197182 | n.s. |
| ENSG00000200869 | 3.930956 | 1.895633348 | 0.75924 | 2.4967524 | 0.012534 | 0.189886728 | RNU6-457P | snRNA | 40.19 | 1.367709677 | 1.367709677 | ENSG00000200869 | n.s. |
| ENSG00000238658 | 13.86915 | -1.297021214 | 0.438778 | -2.955982 | 0.003117 | 0.088405426 | RNU6-625P | snRNA | 38.32 | -1.366439191 | 1.366439191 | ENSG00000238658 | down |
| ENSG00000074590 | 873.2528 | -0.823097384 | 0.229189 | -3.591345 | 0.000329 | 0.021915438 | NUAK1 | protein_coding | 45.97 | -1.365724202 | 1.365724202 | ENSG00000074590 | n.s. |
| ENSG00000207563 | 25.98121 | -1.105023294 | 0.349769 | -3.159298 | 0.001581 | 0.058251827 | MIR23B | miRNA | 50.52 | -1.364361704 | 1.364361704 | ENSG00000207563 | down |
| ENSG00000178882 | 164.9261 | -0.92312006 | 0.269254 | -3.428431 | 0.000607 | 0.033277115 | RFLNA | protein_coding | 47.18 | -1.36423698 | 1.36423698 | ENSG00000178882 | n.s. |
| ENSG00000229816 | 91.91605 | -0.95130194 | 0.282022 | -3.373152 | 0.000743 | 0.037254979 | DDX50P1 | processed_pseudogene | 42.67 | -1.359235127 | 1.359235127 | ENSG00000229816 | n.s. |
| ENSG00000230832 | 29.25846 | 1.537184045 | 0.562186 | 2.7342978 | 0.006251 | 0.132179666 | | processed_pseudogene | 35.29 | 1.350931679 | 1.350931679 | ENSG00000230832 | n.s. |
| ENSG00000252021 | 3.027931 | -2.458229571 | 1.100933 | -2.232861 | 0.025558 | 0.282417274 | | misc_RNA | 49.53 | -1.349835352 | 1.349835352 | ENSG00000252021 | n.s. |
| ENSG00000103546 | 882.1609 | -0.909466005 | 0.26489 | -3.433372 | 0.000596 | 0.033066993 | SLC6A2 | protein_coding | 48.47 | -1.346560184 | 1.346560184 | ENSG00000103546 | n.s. |
| ENSG00000100003 | 8.460541 | -1.860805774 | 0.744914 | -2.498015 | 0.012489 | 0.189624386 | SEC14L2 | protein_coding | 50.43 | -1.343698665 | 1.343698665 | ENSG00000100003 | n.s. |
| ENSG00000234797 | 472.7711 | 1.107529234 | 0.353001 | 3.1374677 | 0.001704 | 0.061309434 | RPS3AP6 | processed_pseudogene | 42.26 | 1.342848949 | 1.342848949 | ENSG00000234797 | up |

| | | | | | | | | | | | | | |
|---|---|---|---|---|---|---|---|---|---|---|---|---|---|
| ENSG00000158089 | 550.6548 | -0.846827682 | 0.240241 | -3.524909 | 0.000424 | 0.02621414 | GALNT14 | protein_coding | 46.29 | -1.339227821 | 1.339227821 | ENSG00000158089 | n.s. |
| ENSG00000259932 | 50.69175 | -0.989157695 | 0.301369 | -3.282215 | 0.00103 | 0.04453414 | | sense_intronic | 56.18 | -1.336655646 | 1.336655646 | ENSG00000259932 | n.s. |
| ENSG00000260955 | 31.91739 | -1.099837647 | 0.351328 | -3.130516 | 0.001745 | 0.062358007 | | lincRNA | 43.05 | -1.325422899 | 1.325422899 | ENSG00000260955 | down |
| ENSG00000202382 | 4.797181 | -1.752432557 | 0.685155 | -2.557716 | 0.010536 | 0.17570541 | | misc_RNA | 44.12 | -1.323463121 | 1.323463121 | ENSG00000202382 | n.s. |
| ENSG00000249274 | 38.60441 | -1.031772813 | 0.321698 | -3.20727 | 0.00134 | 0.052407271 | PDLIM1P4 | processed_pseudogene | 45.03 | -1.321296988 | 1.321296988 | ENSG00000249274 | down |
| ENSG00000264853 | 258.868 | -0.933123523 | 0.278041 | -3.35606 | 0.000791 | 0.038623611 | | sense_intronic | 41.47 | -1.318640822 | 1.318640822 | ENSG00000264853 | n.s. |
| ENSG00000072071 | 778.407 | -0.798873658 | 0.223065 | -3.581346 | 0.000342 | 0.022482711 | ADGRL1 | protein_coding | 56.42 | -1.316664675 | 1.316664675 | ENSG00000072071 | n.s. |
| ENSG00000213860 | 32.89212 | -1.359692324 | 0.476719 | -2.852186 | 0.004342 | 0.107997903 | RPL21P75 | processed_pseudogene | 43.06 | -1.314257766 | 1.314257766 | ENSG00000213860 | n.s. |
| ENSG00000234624 | 20.20075 | -1.217293639 | 0.40811 | -2.982757 | 0.002857 | 0.083706059 | | processed_pseudogene | 54.98 | -1.311321179 | 1.311321179 | ENSG00000234624 | down |
| ENSG00000147437 | 47.01563 | -0.971168627 | 0.296533 | -3.275081 | 0.001056 | 0.045113847 | GNRH1 | protein_coding | 39.78 | -1.306892041 | 1.306892041 | ENSG00000147437 | n.s. |
| ENSG00000278765 | 72.19942 | -0.932116117 | 0.279472 | -3.335272 | 0.000852 | 0.039650653 | | lincRNA | 38.47 | -1.306593448 | 1.306593448 | ENSG00000278765 | n.s. |
| ENSG00000235238 | 53.12249 | 1.040835831 | 0.327322 | 3.1798527 | 0.001473 | 0.055597647 | SUMO2P1 | processed_pseudogene | 45.42 | 1.306190253 | 1.306190253 | ENSG00000235238 | up |
| ENSG00000272777 | 74.59139 | -0.950502202 | 0.287928 | -3.301185 | 0.000963 | 0.042690184 | | lincRNA | 35.4 | -1.30187623 | 1.30187623 | ENSG00000272777 | n.s. |
| ENSG00000252751 | 4.395732 | -1.96982698 | 0.817671 | -2.409069 | 0.015993 | 0.219167644 | RNU6-143P | snRNA | 41.94 | -1.298556356 | 1.298556356 | ENSG00000252751 | n.s. |
| ENSG00000197859 | 138.0209 | -0.922863193 | 0.276764 | -3.334482 | 0.000855 | 0.039650653 | ADAMTSL2 | protein_coding | 59.43 | -1.293623166 | 1.293623166 | ENSG00000197859 | n.s. |
| ENSG00000249493 | 21.4142 | -1.15830053 | 0.383481 | -3.020488 | 0.002524 | 0.076718623 | ANKRD20A18P | unprocessed_pseudogene | 35.73 | -1.291619998 | 1.291619998 | ENSG00000249493 | down |
| ENSG00000237172 | 116.117 | -0.875556181 | 0.255918 | -3.421237 | 0.000623 | 0.033902989 | B3GNT9 | protein_coding | 62.77 | -1.286859214 | 1.286859214 | ENSG00000237172 | n.s. |
| ENSG00000128564 | 165.7651 | -0.966222509 | 0.296235 | -3.261676 | 0.001108 | 0.046728867 | VGF | protein_coding | 68.27 | -1.285476675 | 1.285476675 | ENSG00000128564 | n.s. |
| ENSG00000221783 | 4.456583 | -1.824093233 | 0.738635 | -2.469546 | 0.013528 | 0.197974563 | MIR1183 | miRNA | 48.31 | -1.283050045 | 1.283050045 | ENSG00000221783 | n.s. |
| ENSG00000266258 | 36.70402 | -0.999128649 | 0.311041 | -3.212208 | 0.001317 | 0.052047939 | LINC01909 | lincRNA | 39.71 | -1.282477997 | 1.282477997 | ENSG00000266258 | n.s. |
| ENSG00000239559 | 47.92424 | 1.414753621 | 0.511122 | 2.7679382 | 0.005641 | 0.124079983 | | processed_pseudogene | 48.78 | 1.282188764 | 1.282188764 | ENSG00000239559 | n.s. |
| ENSG00000273338 | 182.3303 | 0.914487584 | 0.273818 | 3.3397663 | 0.000838 | 0.039650653 | | antisense | 28.71 | 1.281882658 | 1.281882658 | ENSG00000273338 | n.s. |
| ENSG00000244018 | 20.94376 | -1.187663308 | 0.398506 | -2.980287 | 0.00288 | 0.083706059 | RPL35P6 | processed_pseudogene | 48.39 | -1.27940211 | 1.27940211 | ENSG00000244018 | down |
| ENSG00000103528 | 528.0853 | -0.863146286 | 0.251483 | -3.432222 | 0.000599 | 0.03307576 | SYT17 | protein_coding | 45.9 | -1.277879587 | 1.277879587 | ENSG00000103528 | n.s. |
| ENSG00000071127 | 1059.6 | -0.885597061 | 0.260965 | -3.393542 | 0.00069 | 0.036113461 | WDR1 | protein_coding | 54.98 | -1.277323998 | 1.277323998 | ENSG00000071127 | n.s. |
| ENSG00000175105 | 743.5033 | -0.901374947 | 0.268676 | -3.354873 | 0.000794 | 0.038653993 | ZNF654 | protein_coding | 35.51 | -1.273467605 | 1.273467605 | ENSG00000175105 | n.s. |
| ENSG00000276653 | 8.389379 | -1.548420656 | 0.584159 | -2.650682 | 0.008033 | 0.152397234 | RN7SL856P | misc_RNA | 52.68 | -1.265095159 | 1.265095159 | ENSG00000276653 | n.s. |
| ENSG00000259408 | 65.6212 | -0.985106033 | 0.307066 | -3.20812 | 0.001336 | 0.052399788 | | antisense | 42.23 | -1.261596207 | 1.261596207 | ENSG00000259408 | n.s. |
| ENSG00000125945 | 426.8667 | -0.827526358 | 0.238328 | -3.47222 | 0.000516 | 0.030196123 | ZNF436 | protein_coding | 46 | -1.257880461 | 1.257880461 | ENSG00000125945 | n.s. |
| ENSG00000101445 | 304.979 | -0.893644915 | 0.26724 | -3.343973 | 0.000826 | 0.039650653 | PPP1R16B | protein_coding | 47.52 | -1.252666454 | 1.252666454 | ENSG00000101445 | n.s. |
| ENSG00000226360 | 84.52727 | 1.062698171 | 0.343592 | 3.0929037 | 0.001982 | 0.066819761 | | processed_pseudogene | 51.69 | 1.248771393 | 1.248771393 | ENSG00000226360 | up |
| ENSG00000071242 | 950.2332 | -0.829990052 | 0.240441 | -3.451955 | 0.000557 | 0.031498886 | RPS6KA2 | protein_coding | 46.48 | -1.246400045 | 1.246400045 | ENSG00000071242 | n.s. |

| | | | | | | | | | | | | | |
|---|---|---|---|---|---|---|---|---|---|---|---|---|---|
| ENSG00000146373 | 120.5179 | -0.900065325 | 0.27141 | -3.316255 | 0.000912 | 0.04124125 | RNF217 | protein_coding | 36.66 | -1.246291814 | 1.246291814 | ENSG00000146373 | n.s. |
| ENSG00000249264 | 67.05033 | 0.978971952 | 0.306255 | 3.1965921 | 0.001391 | 0.053388725 | EEF1A1P9 | processed_pseudogene | 47.65 | 1.245791201 | 1.245791201 | ENSG00000249264 | n.s. |
| ENSG00000260743 | 53.70397 | -0.9741462 | 0.304365 | -3.200584 | 0.001371 | 0.053209578 | | lincRNA | 34.74 | -1.24107218 | 1.24107218 | ENSG00000260743 | n.s. |
| ENSG00000164683 | 110.0871 | -0.958116527 | 0.297234 | -3.223441 | 0.001267 | 0.050948982 | HEY1 | protein_coding | 46.45 | -1.238714837 | 1.238714837 | ENSG00000164683 | n.s. |
| ENSG00000229065 | 19.1709 | -1.134922715 | 0.378623 | -2.9975 | 0.002722 | 0.08115396 | | antisense | 47.48 | -1.237849179 | 1.237849179 | ENSG00000229065 | down |
| ENSG00000233895 | 44.39235 | -0.990265727 | 0.312325 | -3.170624 | 0.001521 | 0.056779015 | | lincRNA | 44.35 | -1.233685067 | 1.233685067 | ENSG00000233895 | n.s. |
| ENSG00000266531 | 2.087029 | -2.664981763 | 1.277492 | -2.086105 | 0.036969 | 0.344525344 | MIR4706 | miRNA | 64.63 | -1.233297127 | 1.233297127 | ENSG00000266531 | n.s. |
| ENSG00000280123 | 339.3956 | -0.873707079 | 0.262378 | -3.329951 | 0.000869 | 0.039913228 | | TEC | 30.55 | -1.222214107 | 1.222214107 | ENSG00000280123 | n.s. |
| ENSG00000207450 | 4.616658 | -1.80539164 | 0.743214 | -2.429167 | 0.015134 | 0.211662698 | | misc_RNA | 35.64 | -1.217476092 | 1.217476092 | ENSG00000207450 | n.s. |
| ENSG00000254905 | 24.86789 | -1.07637813 | 0.354114 | -3.039637 | 0.002369 | 0.07427583 | | antisense | 50.41 | -1.215395044 | 1.215395044 | ENSG00000254905 | down |
| ENSG00000124762 | 642.4524 | -0.83210041 | 0.244233 | -3.406988 | 0.000657 | 0.034849263 | CDKN1A | protein_coding | 54.4 | -1.213041302 | 1.213041302 | ENSG00000124762 | n.s. |
| ENSG00000101665 | 877.6073 | -0.797728218 | 0.229889 | -3.470059 | 0.00052 | 0.030312718 | SMAD7 | protein_coding | 52.06 | -1.211250679 | 1.211250679 | ENSG00000101665 | n.s. |
| ENSG00000279161 | 99.15677 | -1.07479095 | 0.354553 | -3.031397 | 0.002434 | 0.074968112 | | TEC | 56.04 | -1.209272478 | 1.209272478 | ENSG00000279161 | down |
| ENSG00000138696 | 304.5048 | -0.845233335 | 0.250755 | -3.370758 | 0.00075 | 0.037274749 | BMPR1B | protein_coding | 36.7 | -1.207487891 | 1.207487891 | ENSG00000138696 | n.s. |
| ENSG00000239183 | 33.92261 | 1.053364163 | 0.344561 | 3.0571236 | 0.002235 | 0.071526477 | SNORA84 | snoRNA | 55.64 | 1.206663587 | 1.206663587 | ENSG00000239183 | up |
| ENSG00000226761 | 98.39988 | -0.951055348 | 0.298044 | -3.190992 | 0.001418 | 0.054084184 | TAS2R46 | protein_coding | 38.71 | -1.204920283 | 1.204920283 | ENSG00000226761 | n.s. |
| ENSG00000207047 | 13.17666 | 1.260438614 | 0.445628 | 2.8284553 | 0.004677 | 0.111596326 | SNORD51 | snoRNA | 38.75 | 1.200378843 | 1.200378843 | ENSG00000207047 | n.s. |
| ENSG00000203814 | 1818.886 | -0.752435511 | 0.212916 | -3.533962 | 0.000409 | 0.02544553 | HIST2H2BF | protein_coding | 41.91 | -1.199674522 | 1.199674522 | ENSG00000203814 | n.s. |
| ENSG00000271924 | 26.01934 | -1.06306931 | 0.350446 | -3.033471 | 0.002418 | 0.074799793 | RNA5SP108 | rRNA | 47.86 | -1.197121928 | 1.197121928 | ENSG00000271924 | down |
| ENSG00000221176 | 8.13428 | -1.490398964 | 0.567759 | -2.625054 | 0.008664 | 0.157676123 | MIR1207 | miRNA | 62.07 | -1.195648825 | 1.195648825 | ENSG00000221176 | n.s. |
| ENSG00000206987 | 5.608192 | 1.925787554 | 0.818425 | 2.3530417 | 0.018621 | 0.239827812 | | snoRNA | 44.36 | 1.194181739 | 1.194181739 | ENSG00000206987 | n.s. |
| ENSG00000186815 | 50.07171 | -1.00677113 | 0.324372 | -3.103754 | 0.001911 | 0.065206718 | TPCN1 | protein_coding | 50.56 | -1.193736239 | 1.193736239 | ENSG00000186815 | down |
| ENSG00000223111 | 20.96147 | 1.166971006 | 0.39997 | 2.9176445 | 0.003527 | 0.095163761 | | snoRNA | 52.24 | 1.192094027 | 1.192094027 | ENSG00000223111 | up |
| ENSG00000278716 | 172.5057 | -0.924907426 | 0.28743 | -3.21785 | 0.001292 | 0.051525136 | | lincRNA | 57.9 | -1.191263054 | 1.191263054 | ENSG00000278716 | n.s. |
| ENSG00000164532 | 210.5913 | -0.846989349 | 0.253407 | -3.342413 | 0.000831 | 0.039650653 | TBX20 | protein_coding | 36.93 | -1.187267031 | 1.187267031 | ENSG00000164532 | n.s. |
| ENSG00000213976 | 88.24768 | -0.860339171 | 0.259851 | -3.31089 | 0.00093 | 0.041768834 | | unprocessed_pseudogene | 37.56 | -1.186534744 | 1.186534744 | ENSG00000213976 | n.s. |
| ENSG00000231333 | 29.26773 | -1.019873497 | 0.331596 | -3.07565 | 0.0021 | 0.068810475 | RPL34P6 | processed_pseudogene | 42.61 | -1.18544531 | 1.18544531 | ENSG00000231333 | down |
| ENSG00000200683 | 2.828865 | -2.097693745 | 0.928159 | -2.260058 | 0.023818 | 0.272248483 | RNU6-379P | snRNA | 38.46 | -1.185269403 | 1.185269403 | ENSG00000200683 | n.s. |
| ENSG00000283041 | 599.9727 | 0.98630099 | 0.316171 | 3.1195166 | 0.001811 | 0.062968329 | | processed_pseudogene | 54.34 | 1.184426991 | 1.184426991 | ENSG00000283041 | n.s. |
| ENSG00000231307 | 34.90794 | -1.010854098 | 0.328128 | -3.080668 | 0.002065 | 0.06830421 | RPS3P2 | processed_pseudogene | 46.93 | -1.178203551 | 1.178203551 | ENSG00000231307 | down |
| ENSG00000164086 | 263.8045 | -0.759781079 | 0.217321 | -3.496119 | 0.000472 | 0.028453472 | DUSP7 | protein_coding | 59.41 | -1.174518772 | 1.174518772 | ENSG00000164086 | n.s. |
| ENSG00000264676 | 8.61265 | -1.418502716 | 0.532247 | -2.665123 | 0.007696 | 0.148615313 | RN7SL204P | misc_RNA | 58.4 | -1.174430087 | 1.174430087 | ENSG00000264676 | n.s. |

| | | | | | | | | | | | | | |
|---|---|---|---|---|---|---|---|---|---|---|---|---|---|
| ENSG00000252305 | 10.05049 | -1.343675227 | 0.494373 | -2.717938 | 0.006569 | 0.134873528 | | snoRNA | 53.89 | -1.169095917 | 1.169095917 | ENSG00000252305 | n.s. |
| ENSG00000281026 | 341.0772 | -0.832928771 | 0.249856 | -3.333631 | 0.000857 | 0.039650653 | N4BP2L2-IT2 | sense_intronic | 38.1 | -1.16755762 | 1.16755762 | ENSG00000281026 | n.s. |
| ENSG00000238311 | 6.472953 | -1.562438652 | 0.615658 | -2.537835 | 0.011154 | 0.179742753 | SNORD13E | snoRNA | 42.31 | -1.164561479 | 1.164561479 | ENSG00000238311 | n.s. |
| ENSG00000255115 | 11.57289 | -1.250842399 | 0.44785 | -2.792996 | 0.005222 | 0.118034481 | | processed_pseudogene | 41.26 | -1.16077062 | 1.16077062 | ENSG00000255115 | n.s. |
| ENSG00000128591 | 948.3838 | -0.7726024 | 0.223967 | -3.44962 | 0.000561 | 0.031516313 | FLNC | protein_coding | 60.86 | -1.160035149 | 1.160035149 | ENSG00000128591 | n.s. |
| ENSG00000223225 | 6.387061 | -1.552098837 | 0.612392 | -2.534486 | 0.011261 | 0.180009224 | RNU6-1054P | snRNA | 40.38 | -1.155856136 | 1.155856136 | ENSG00000223225 | n.s. |
| ENSG00000266649 | 393.7753 | -0.823735233 | 0.24649 | -3.341865 | 0.000832 | 0.039650653 | MIR3126 | miRNA | 40.54 | -1.154670582 | 1.154670582 | ENSG00000266649 | n.s. |
| ENSG00000266288 | 21.20042 | -1.093896984 | 0.369994 | -2.956527 | 0.003111 | 0.088405426 | | lincRNA | 37.89 | -1.152443533 | 1.152443533 | ENSG00000266288 | down |
| ENSG00000143320 | 450.8266 | -0.954208175 | 0.305026 | -3.12828 | 0.001758 | 0.062452009 | CRABP2 | protein_coding | 55.42 | -1.149299456 | 1.149299456 | ENSG00000143320 | n.s. |
| ENSG00000278700 | 13.25986 | -1.274191059 | 0.461568 | -2.760571 | 0.00577 | 0.126116552 | | misc_RNA | 54.82 | -1.145788164 | 1.145788164 | ENSG00000278700 | n.s. |
| ENSG00000264370 | 9.132719 | -1.339306275 | 0.496181 | -2.699232 | 0.00695 | 0.140238403 | MIR3125 | miRNA | 57.69 | -1.142606436 | 1.142606436 | ENSG00000264370 | n.s. |
| ENSG00000280378 | 53.84317 | -0.95132468 | 0.305292 | -3.116114 | 0.001833 | 0.063310472 | | TEC | 47.81 | -1.140185884 | 1.140185884 | ENSG00000280378 | n.s. |
| ENSG00000242299 | 286.4702 | 1.007517479 | 0.331733 | 3.0371292 | 0.002388 | 0.074429557 | | processed_pseudogene | 52.51 | 1.136736198 | 1.136736198 | ENSG00000242299 | up |
| ENSG00000261758 | 86.5988 | 0.897911782 | 0.281568 | 3.1889693 | 0.001428 | 0.054315315 | | antisense | 43.73 | 1.135928173 | 1.135928173 | ENSG00000261758 | n.s. |
| ENSG00000243951 | 18.96431 | -1.160972742 | 0.405686 | -2.861754 | 0.004213 | 0.106457698 | RN7SL308P | misc_RNA | 52.53 | -1.129420904 | 1.129420904 | ENSG00000243951 | n.s. |
| ENSG00000212370 | 4.770392 | -1.771229436 | 0.745571 | -2.375668 | 0.017517 | 0.230963869 | RNU6-482P | snRNA | 42.99 | -1.127309521 | 1.127309521 | ENSG00000212370 | n.s. |
| ENSG00000226525 | 80.64006 | 1.002342014 | 0.330995 | 3.0282715 | 0.00246 | 0.075262844 | RPS7P10 | processed_pseudogene | 45.77 | 1.126050437 | 1.126050437 | ENSG00000226525 | up |
| ENSG00000260951 | 146.3368 | -0.943378381 | 0.303709 | -3.106193 | 0.001895 | 0.064830103 | | lincRNA | 40.38 | -1.120944161 | 1.120944161 | ENSG00000260951 | n.s. |
| ENSG00000279948 | 119.3438 | -0.857858633 | 0.265151 | -3.235358 | 0.001215 | 0.049750266 | | TEC | 46.34 | -1.117965312 | 1.117965312 | ENSG00000279948 | n.s. |
| ENSG00000215012 | 454.4097 | -0.776403145 | 0.229421 | -3.384192 | 0.000714 | 0.03643289 | C22orf29 | protein_coding | 55.11 | -1.116860876 | 1.116860876 | ENSG00000215012 | n.s. |
| ENSG00000263883 | 77.44665 | -1.011369856 | 0.335987 | -3.010149 | 0.002611 | 0.078692029 | EEF1DP7 | ıscribed_processed_pseudog | 55.75 | -1.116622366 | 1.116622366 | ENSG00000263883 | down |
| ENSG00000252326 | 8.338623 | -1.382088221 | 0.524431 | -2.635404 | 0.008404 | 0.156104918 | SNORD116-25 | snoRNA | 43.48 | -1.114769438 | 1.114769438 | ENSG00000252326 | n.s. |
| ENSG00000273284 | 40.44267 | -1.055034502 | 0.35641 | -2.960172 | 0.003075 | 0.087902938 | | sense_intronic | 32.4 | -1.114112859 | 1.114112859 | ENSG00000273284 | down |
| ENSG00000270072 | 28.65134 | -1.098977657 | 0.377849 | -2.908512 | 0.003632 | 0.097421639 | | sense_intronic | 43.27 | -1.111445083 | 1.111445083 | ENSG00000270072 | down |
| ENSG00000200563 | 4.349129 | -1.757765945 | 0.743575 | -2.36394 | 0.018082 | 0.235607368 | RNU6-640P | snRNA | 36.19 | -1.103545027 | 1.103545027 | ENSG00000200563 | n.s. |
| ENSG00000173065 | 42.77232 | -0.972619832 | 0.319826 | -3.041093 | 0.002357 | 0.074252089 | FAM222B | protein_coding | 45.72 | -1.098371139 | 1.098371139 | ENSG00000173065 | n.s. |
| ENSG00000201003 | 9.614583 | -1.353060199 | 0.513722 | -2.633835 | 0.008443 | 0.156104918 | | snoRNA | 47.79 | -1.091355917 | 1.091355917 | ENSG00000201003 | n.s. |
| ENSG00000238015 | 13.64114 | -1.20258221 | 0.434269 | -2.769213 | 0.005619 | 0.123987433 | | processed_pseudogene | 42.92 | -1.090287888 | 1.090287888 | ENSG00000238015 | n.s. |
| ENSG00000265093 | 20.4054 | -1.122240312 | 0.394178 | -2.847039 | 0.004413 | 0.108742185 | RN7SL246P | misc_RNA | 50 | -1.081392946 | 1.081392946 | ENSG00000265093 | n.s. |
| ENSG00000136826 | 78.81473 | 0.955724487 | 0.314345 | 3.0403637 | 0.002363 | 0.074264146 | KLF4 | protein_coding | 56.93 | 1.079223979 | 1.079223979 | ENSG00000136826 | n.s. |
| ENSG00000278886 | 65.44646 | -0.910809027 | 0.29408 | -3.097149 | 0.001954 | 0.066190691 | | TEC | 38.27 | -1.074028817 | 1.074028817 | ENSG00000278886 | n.s. |
| ENSG00000257818 | 65.32771 | -0.878955651 | 0.279489 | -3.144864 | 0.001662 | 0.060247415 | C1GALT1P1 | processed_pseudogene | 37.83 | -1.07238002 | 1.07238002 | ENSG00000257818 | n.s. |

| | | | | | | | | | | | | | |
|---|---|---|---|---|---|---|---|---|---|---|---|---|---|
| ENSG00000225750 | 16.31977 | -1.119413807 | 0.394363 | -2.838533 | 0.004532 | 0.110509521 | | lincRNA | 47.46 | -1.070831587 | 1.070831587 | ENSG00000225750 | n.s. |
| ENSG00000236723 | 75.00366 | -0.864036822 | 0.273068 | -3.164182 | 0.001555 | 0.057741321 | | antisense | 47.26 | -1.070121084 | 1.070121084 | ENSG00000236723 | n.s. |
| ENSG00000163263 | 10.24205 | -1.284694084 | 0.480243 | -2.675094 | 0.007471 | 0.147123666 | C1orf189 | protein_coding | 46.57 | -1.06927332 | 1.06927332 | ENSG00000163263 | n.s. |
| ENSG00000122644 | 660.4771 | 0.728443836 | 0.213231 | 3.4162175 | 0.000635 | 0.034134195 | ARL4A | protein_coding | 42.68 | 1.068488946 | 1.068488946 | ENSG00000122644 | n.s. |
| ENSG00000163738 | 548.7699 | 0.74256462 | 0.219563 | 3.3820151 | 0.00072 | 0.036563728 | MTHFD2L | protein_coding | 36.15 | 1.067027884 | 1.067027884 | ENSG00000163738 | n.s. |
| ENSG00000200997 | 91.77368 | 0.88624232 | 0.284127 | 3.1191774 | 0.001814 | 0.062968329 | RNU1-85P | snRNA | 41.46 | 1.064268753 | 1.064268753 | ENSG00000200997 | n.s. |
| ENSG00000211491 | 5.892627 | -1.522622767 | 0.618687 | -2.461054 | 0.013853 | 0.200076043 | MIR320D1 | miRNA | 47.92 | -1.064016267 | 1.064016267 | ENSG00000211491 | n.s. |
| ENSG00000183570 | 77.86466 | -0.856973323 | 0.271106 | -3.16103 | 0.001572 | 0.058060074 | PCBP3 | protein_coding | 46.92 | -1.059323932 | 1.059323932 | ENSG00000183570 | n.s. |
| ENSG00000278743 | 87.9798 | 0.892739861 | 0.287906 | 3.1008039 | 0.00193 | 0.065538642 | | lincRNA | 60.14 | 1.056559911 | 1.056559911 | ENSG00000278743 | n.s. |
| ENSG00000254678 | 10.23413 | 1.414669815 | 0.556602 | 2.5416204 | 0.011034 | 0.179260564 | | processed_pseudogene | 58.8 | 1.056072579 | 1.056072579 | ENSG00000254678 | n.s. |
| ENSG00000246863 | 135.475 | 0.845811505 | 0.266493 | 3.1738658 | 0.001504 | 0.056451276 | | lincRNA | 41.23 | 1.055848692 | 1.055848692 | ENSG00000246863 | n.s. |
| ENSG00000197362 | 132.1357 | -0.857259392 | 0.271991 | -3.151794 | 0.001623 | 0.059180368 | ZNF786 | protein_coding | 46.13 | -1.052562233 | 1.052562233 | ENSG00000197362 | n.s. |
| ENSG00000166435 | 30.97216 | -1.044734521 | 0.359946 | -2.902479 | 0.003702 | 0.098747222 | XRRA1 | protein_coding | 41.82 | -1.050454559 | 1.050454559 | ENSG00000166435 | down |
| ENSG00000266852 | 39.26893 | 0.974362131 | 0.32711 | 2.9787 | 0.002895 | 0.083791014 | MIR4482 | miRNA | 48.57 | 1.049195632 | 1.049195632 | ENSG00000266852 | n.s. |
| ENSG00000138606 | 1810.031 | -0.754306624 | 0.227592 | -3.314299 | 0.000919 | 0.041396465 | SHF | protein_coding | 50.88 | -1.04323378 | 1.04323378 | ENSG00000138606 | n.s. |
| ENSG00000242182 | 15.40929 | -1.198559236 | 0.44081 | -2.718994 | 0.006548 | 0.134873528 | RN7SL745P | misc_RNA | 50.84 | -1.04283437 | 1.04283437 | ENSG00000242182 | n.s. |
| ENSG00000258650 | 7.801902 | -1.396371916 | 0.550359 | -2.537202 | 0.011174 | 0.179859901 | HSBP1P1 | processed_pseudogene | 46.39 | -1.040388748 | 1.040388748 | ENSG00000258650 | n.s. |
| ENSG00000230629 | 193.8209 | 0.994639821 | 0.337692 | 2.9454037 | 0.003225 | 0.089992658 | RPS23P8 | processed_pseudogene | 43.64 | 1.040187283 | 1.040187283 | ENSG00000230629 | n.s. |
| ENSG00000237082 | 26.68818 | -1.047646889 | 0.362723 | -2.888286 | 0.003873 | 0.101755569 | COX5BP6 | processed_pseudogene | 52.25 | -1.039728579 | 1.039728579 | ENSG00000237082 | n.s. |
| ENSG00000229127 | 188.9909 | -0.849580125 | 0.270047 | -3.146042 | 0.001655 | 0.060161893 | | antisense | 37.9 | -1.0370642 | 1.0370642 | ENSG00000229127 | n.s. |
| ENSG00000213442 | 322.006 | 1.004591784 | 0.343362 | 2.9257499 | 0.003436 | 0.093626613 | RPL18AP3 | processed_pseudogene | 60.23 | 1.033323799 | 1.033323799 | ENSG00000213442 | up |
| ENSG00000234851 | 270.6344 | 0.94680485 | 0.316063 | 2.9956195 | 0.002739 | 0.081307947 | RPL23AP42 | processed_pseudogene | 49.26 | 1.031891368 | 1.031891368 | ENSG00000234851 | n.s. |
| ENSG00000199460 | 6.062684 | -1.511346523 | 0.618969 | -2.441717 | 0.014618 | 0.207886636 | RNU6-1216P | snRNA | 40.19 | -1.03100044 | 1.03100044 | ENSG00000199460 | n.s. |
| ENSG00000245059 | 30.30232 | 1.172608043 | 0.429462 | 2.7304132 | 0.006325 | 0.132435959 | | lincRNA | 48.28 | 1.029542918 | 1.029542918 | ENSG00000245059 | n.s. |
| ENSG00000227034 | 20.7054 | 1.046283456 | 0.363884 | 2.8753251 | 0.004036 | 0.103804772 | | processed_pseudogene | 43.97 | 1.029315543 | 1.029315543 | ENSG00000227034 | n.s. |
| ENSG00000223825 | 14.20265 | -1.13654169 | 0.41081 | -2.766589 | 0.005665 | 0.124398086 | DAZAP2P1 | processed_pseudogene | 54.87 | -1.028781968 | 1.028781968 | ENSG00000223825 | n.s. |
| ENSG00000111252 | 184.588 | -0.809337115 | 0.253373 | -3.194253 | 0.001402 | 0.053623963 | SH2B3 | protein_coding | 54.02 | -1.028376996 | 1.028376996 | ENSG00000111252 | n.s. |
| ENSG00000140534 | 2096.263 | -0.733551899 | 0.219969 | -3.334794 | 0.000854 | 0.039650653 | TICRR | protein_coding | 42.67 | -1.028256124 | 1.028256124 | ENSG00000140534 | n.s. |
| ENSG00000163110 | 148.733 | 0.828018952 | 0.261591 | 3.1653167 | 0.001549 | 0.05767034 | PDLIM5 | protein_coding | 36.89 | 1.025954804 | 1.025954804 | ENSG00000163110 | n.s. |
| ENSG00000201136 | 5.842873 | -1.558126219 | 0.647536 | -2.40624 | 0.016118 | 0.219790693 | RNU6-353P | snRNA | 38.89 | -1.025232563 | 1.025232563 | ENSG00000201136 | n.s. |
| ENSG00000141858 | 574.8768 | -0.785819763 | 0.243837 | -3.222721 | 0.00127 | 0.050948982 | SAMD1 | protein_coding | 64.37 | -1.015958468 | 1.015958468 | ENSG00000141858 | n.s. |
| ENSG00000254409 | 17.20247 | -1.093067428 | 0.390957 | -2.795879 | 0.005176 | 0.117751086 | | sense_overlapping | 48.33 | -1.015497986 | 1.015497986 | ENSG00000254409 | n.s. |

| | | | | | | | | | | | | | |
|---|---|---|---|---|---|---|---|---|---|---|---|---|---|
| ENSG00000238731 | 3.880168 | -1.729303615 | 0.752625 | -2.297697 | 0.021579 | 0.258781015 | RNU7-90P | snRNA | 43.55 | -1.015218103 | 1.015218103 | ENSG00000238731 | n.s. |
| ENSG00000260576 | 17.41506 | -1.151498864 | 0.421078 | -2.734644 | 0.006245 | 0.132179666 | EIF5A2P1 | processed_pseudogene | 40.87 | -1.011977908 | 1.011977908 | ENSG00000260576 | n.s. |
| ENSG00000229623 | 34.83063 | -0.958438368 | 0.324247 | -2.955893 | 0.003118 | 0.088405426 | METTL21AP1 | processed_pseudogene | 46.93 | -1.009735025 | 1.009735025 | ENSG00000229623 | n.s. |
| ENSG00000212371 | 28.55538 | -1.009429388 | 0.34919 | -2.890776 | 0.003843 | 0.101335127 | | snoRNA | 41.83 | -1.003615058 | 1.003615058 | ENSG00000212371 | n.s. |
| ENSG00000266289 | 20.47166 | -1.031654448 | 0.361118 | -2.856837 | 0.004279 | 0.10714823 | | lincRNA | 31.92 | -1.000720289 | 1.000720289 | ENSG00000266289 | n.s. |
| ENSG00000204196 | 279.232 | 0.998964799 | 0.344628 | 2.8986728 | 0.003747 | 0.099762615 | RPL12P16 | processed_pseudogene | 51.81 | 0.999995906 | 0.999995906 | ENSG00000204196 | n.s. |
| ENSG00000167992 | 114.256 | -0.809068705 | 0.256278 | -3.156996 | 0.001594 | 0.058558673 | VWCE | protein_coding | 51.58 | -0.997103374 | 0.997103374 | ENSG00000167992 | n.s. |
| ENSG00000261291 | 39.13316 | -0.97507297 | 0.334006 | -2.919327 | 0.003508 | 0.095020015 | | sense_intronic | 44.24 | -0.996704871 | 0.996704871 | ENSG00000261291 | n.s. |
| ENSG00000275745 | 7.732027 | 1.612678023 | 0.686318 | 2.3497541 | 0.018786 | 0.241064423 | | misc_RNA | 41.44 | 0.996420339 | 0.996420339 | ENSG00000275745 | n.s. |
| ENSG00000272148 | 25.51996 | -1.030192035 | 0.361352 | -2.850935 | 0.004359 | 0.108184704 | | antisense | 42.5 | -0.994994637 | 0.994994637 | ENSG00000272148 | n.s. |
| ENSG00000230732 | 137.9459 | -0.809705704 | 0.256918 | -3.151614 | 0.001624 | 0.059180368 | | sense_intronic | 39.19 | -0.994174752 | 0.994174752 | ENSG00000230732 | n.s. |
| ENSG00000102786 | 4400.68 | -0.690657252 | 0.204364 | -3.379543 | 0.000726 | 0.036759926 | INTS6 | protein_coding | 36.13 | -0.99083442 | 0.99083442 | ENSG00000102786 | n.s. |
| ENSG00000107929 | 869.5081 | -0.788821706 | 0.248016 | -3.18053 | 0.00147 | 0.055597647 | LARP4B | protein_coding | 42.7 | -0.989926742 | 0.989926742 | ENSG00000107929 | n.s. |
| ENSG00000236184 | 30.62085 | -1.024955271 | 0.359741 | -2.849149 | 0.004384 | 0.108407336 | TCEA1P4 | processed_pseudogene | 41.59 | -0.989021702 | 0.989021702 | ENSG00000236184 | n.s. |
| ENSG00000279792 | 214.7222 | -0.722229176 | 0.219197 | -3.294886 | 0.000985 | 0.043109503 | | TEC | 56.49 | -0.98615121 | 0.98615121 | ENSG00000279792 | n.s. |
| ENSG00000248476 | 35.6394 | -0.936695788 | 0.317335 | -2.951756 | 0.00316 | 0.089147501 | BACH1-IT1 | sense_intronic | 43.74 | -0.983428315 | 0.983428315 | ENSG00000248476 | n.s. |
| ENSG00000207109 | 19.64509 | -1.150312543 | 0.425749 | -2.701856 | 0.006895 | 0.139674846 | SNORD38C | snoRNA | 34.78 | -0.983381256 | 0.983381256 | ENSG00000207109 | n.s. |
| ENSG00000276043 | 1625.334 | -0.743403658 | 0.228583 | -3.252232 | 0.001145 | 0.047874483 | UHRF1 | protein_coding | 52.99 | -0.981215442 | 0.981215442 | ENSG00000276043 | n.s. |
| ENSG00000233072 | 14.81811 | -1.126034397 | 0.414344 | -2.717631 | 0.006575 | 0.134873528 | RPS15AP6 | processed_pseudogene | 41.3 | -0.979732445 | 0.979732445 | ENSG00000233072 | n.s. |
| ENSG00000230002 | 306.9854 | -0.763219164 | 0.23758 | -3.212475 | 0.001316 | 0.052047939 | ALMS1-IT1 | sense_intronic | 37.16 | -0.979665417 | 0.979665417 | ENSG00000230002 | n.s. |
| ENSG00000180611 | 229.5247 | -0.841361912 | 0.273371 | -3.07773 | 0.002086 | 0.068607507 | MB21D2 | protein_coding | 42.73 | -0.979032583 | 0.979032583 | ENSG00000180611 | n.s. |
| ENSG00000240233 | 10.98899 | -1.280386249 | 0.497459 | -2.573855 | 0.010057 | 0.171947811 | RN7SL587P | misc_RNA | 55.87 | -0.978987615 | 0.978987615 | ENSG00000240233 | n.s. |
| ENSG00000239291 | 113.0272 | -0.840222568 | 0.273045 | -3.077233 | 0.002089 | 0.068607507 | | processed_pseudogene | 58.28 | -0.97770681 | 0.97770681 | ENSG00000239291 | n.s. |
| ENSG00000204842 | 570.5495 | -0.74882053 | 0.231177 | -3.239162 | 0.001199 | 0.049528509 | ATXN2 | protein_coding | 41.99 | -0.977319182 | 0.977319182 | ENSG00000204842 | n.s. |
| ENSG00000277382 | 10.89332 | 1.280450642 | 0.498255 | 2.5698681 | 0.010174 | 0.172742391 | | antisense | 58.74 | 0.976473034 | 0.976473034 | ENSG00000277382 | n.s. |
| ENSG00000199331 | 16.64885 | -1.104690593 | 0.402997 | -2.741188 | 0.006122 | 0.130768951 | | misc_RNA | 44.12 | -0.975989014 | 0.975989014 | ENSG00000199331 | n.s. |
| ENSG00000252985 | 11.98094 | -1.21790811 | 0.464243 | -2.623427 | 0.008705 | 0.158017967 | | snoRNA | 43.59 | -0.975901887 | 0.975901887 | ENSG00000252985 | n.s. |
| ENSG00000281910 | 25.28392 | -1.026235408 | 0.364066 | -2.818819 | 0.00482 | 0.11285058 | SNORA50A | snoRNA | 43.38 | -0.972354153 | 0.972354153 | ENSG00000281910 | n.s. |
| ENSG00000078549 | 1486.378 | -0.718647736 | 0.218776 | -3.284861 | 0.00102 | 0.044393851 | ADCYAP1R1 | protein_coding | 51.89 | -0.972098391 | 0.972098391 | ENSG00000078549 | n.s. |
| ENSG00000219433 | 43.59676 | -0.929566735 | 0.315819 | -2.943353 | 0.003247 | 0.09040971 | BTBD10P2 | processed_pseudogene | 44.41 | -0.970267745 | 0.970267745 | ENSG00000219433 | n.s. |
| ENSG00000137496 | 145.4473 | -0.777721631 | 0.24523 | -3.171393 | 0.001517 | 0.056779015 | IL18BP | protein_coding | 57.03 | -0.968895052 | 0.968895052 | ENSG00000137496 | n.s. |
| ENSG00000232082 | 44.61335 | -1.01719921 | 0.359801 | -2.827119 | 0.004697 | 0.111596326 | RPS6KA2-IT1 | sense_intronic | 53.76 | -0.968729772 | 0.968729772 | ENSG00000232082 | n.s. |

| | | | | | | | | | | | | | |
|---|---|---|---|---|---|---|---|---|---|---|---|---|---|
| ENSG00000103248 | 67.7074 | -0.845447116 | 0.276676 | -3.055735 | 0.002245 | 0.071693796 | MTHFSD | protein_coding | 49.14 | -0.967629799 | 0.967629799 | ENSG00000103248 | n.s. |
| ENSG00000252947 | 45.0547 | 0.920082681 | 0.311969 | 2.9492789 | 0.003185 | 0.089409345 | SCARNA1 | scaRNA | 40.36 | 0.964814419 | 0.964814419 | ENSG00000252947 | n.s. |
| ENSG00000207290 | 4.195205 | -1.692316313 | 0.745743 | -2.269302 | 0.02325 | 0.269182866 | | misc_RNA | 40.35 | -0.964540061 | 0.964540061 | ENSG00000207290 | n.s. |
| ENSG00000274124 | 10.66551 | -1.272925136 | 0.496809 | -2.562202 | 0.010401 | 0.174827325 | | antisense | 43.27 | -0.964101645 | 0.964101645 | ENSG00000274124 | n.s. |
| ENSG00000207864 | 51.11139 | -0.932051696 | 0.317635 | -2.934348 | 0.003342 | 0.092447097 | MIR27B | miRNA | 47.42 | -0.963840923 | 0.963840923 | ENSG00000207864 | n.s. |
| ENSG00000199787 | 6.163091 | -1.473174018 | 0.614485 | -2.397412 | 0.016511 | 0.223191469 | SNORA80C | snoRNA | 54.81 | -0.959511252 | 0.959511252 | ENSG00000199787 | n.s. |
| ENSG00000264292 | 5.855943 | -1.484639965 | 0.621705 | -2.388012 | 0.01694 | 0.226129282 | MIR2467 | miRNA | 60.49 | -0.958547691 | 0.958547691 | ENSG00000264292 | n.s. |
| ENSG00000116652 | 105.5091 | -0.830405983 | 0.270699 | -3.067634 | 0.002158 | 0.070118383 | DLEU2L | sense_intronic | 35.86 | -0.958428103 | 0.958428103 | ENSG00000116652 | n.s. |
| ENSG00000219470 | 61.74371 | -0.867112033 | 0.288255 | -3.008141 | 0.002629 | 0.078827723 | | processed_pseudogene | 60.77 | -0.956702931 | 0.956702931 | ENSG00000219470 | n.s. |
| ENSG00000280010 | 61.16568 | -0.857358272 | 0.283703 | -3.02203 | 0.002511 | 0.076663636 | | TEC | 38.67 | -0.956306496 | 0.956306496 | ENSG00000280010 | n.s. |
| ENSG00000260035 | 219.6616 | -0.750690004 | 0.23462 | -3.199605 | 0.001376 | 0.053223087 | | sense_intronic | 50 | -0.956303952 | 0.956303952 | ENSG00000260035 | n.s. |
| ENSG00000267248 | 68.29943 | -0.847025321 | 0.278977 | -3.036181 | 0.002396 | 0.074461754 | | processed_transcript | 47.27 | -0.955501092 | 0.955501092 | ENSG00000267248 | n.s. |
| ENSG00000127366 | 32.21228 | -0.9743662 | 0.339494 | -2.870058 | 0.004104 | 0.104803968 | TAS2R5 | protein_coding | 48.39 | -0.954510831 | 0.954510831 | ENSG00000127366 | n.s. |
| ENSG00000148331 | 612.4351 | -0.785603848 | 0.250235 | -3.139468 | 0.001693 | 0.061050122 | ASB6 | protein_coding | 58.84 | -0.95396933 | 0.95396933 | ENSG00000148331 | n.s. |
| ENSG00000242876 | 9.110902 | -1.308622812 | 0.522436 | -2.504846 | 0.01225 | 0.186772923 | RN7SL812P | misc_RNA | 52.69 | -0.953575232 | 0.953575232 | ENSG00000242876 | n.s. |
| ENSG00000264448 | 25.70776 | 1.140583011 | 0.425934 | 2.6778398 | 0.00741 | 0.146129673 | | lincRNA | 52.48 | 0.952685175 | 0.952685175 | ENSG00000264448 | n.s. |
| ENSG00000279265 | 13.5882 | -1.141851064 | 0.426935 | -2.67453 | 0.007483 | 0.147163073 | | TEC | 39.48 | -0.950249772 | 0.950249772 | ENSG00000279265 | n.s. |
| ENSG00000153714 | 71.34725 | 0.828137166 | 0.270547 | 3.0609724 | 0.002206 | 0.071230366 | LURAP1L | protein_coding | 36.85 | 0.950150609 | 0.950150609 | ENSG00000153714 | n.s. |
| ENSG00000259891 | 77.37864 | -0.806202064 | 0.26043 | -3.09566 | 0.001964 | 0.06636238 | | antisense | 48.41 | -0.949768954 | 0.949768954 | ENSG00000259891 | n.s. |
| ENSG00000265727 | 36.58038 | -0.952699632 | 0.330402 | -2.883452 | 0.003933 | 0.102598935 | RN7SL648P | misc_RNA | 53.16 | -0.942083842 | 0.942083842 | ENSG00000265727 | n.s. |
| ENSG00000238123 | 20.92633 | -1.074299794 | 0.393895 | -2.727375 | 0.006384 | 0.132862373 | MID1IP1-AS1 | antisense | 46.07 | -0.941729047 | 0.941729047 | ENSG00000238123 | n.s. |
| ENSG00000215957 | 4.083542 | 1.587367433 | 0.688202 | 2.3065424 | 0.02108 | 0.255337128 | MIR300 | miRNA | 43.37 | 0.941127976 | 0.941127976 | ENSG00000215957 | n.s. |
| ENSG00000129173 | 368.3021 | -0.714059733 | 0.219945 | -3.246541 | 0.001168 | 0.048550416 | E2F8 | protein_coding | 41.92 | -0.938136708 | 0.938136708 | ENSG00000129173 | n.s. |
| ENSG00000168646 | 673.4699 | -0.724278666 | 0.224449 | -3.226918 | 0.001251 | 0.050793086 | AXIN2 | protein_coding | 51.32 | -0.937358119 | 0.937358119 | ENSG00000168646 | n.s. |
| ENSG00000240445 | 161.1899 | -0.811560264 | 0.264507 | -3.068198 | 0.002154 | 0.070118383 | FOXO3B | processed_pseudogene | 60.32 | -0.936676976 | 0.936676976 | ENSG00000240445 | n.s. |
| ENSG00000203804 | 404.0739 | 0.685401646 | 0.20796 | 3.2958263 | 0.000981 | 0.043100984 | ADAMTSL4-AS1 | processed_transcript | 48.94 | 0.935924733 | 0.935924733 | ENSG00000203804 | n.s. |
| ENSG00000185630 | 150.709 | -0.776244046 | 0.248283 | -3.126453 | 0.001769 | 0.062653718 | PBX1 | protein_coding | 41.67 | -0.933862846 | 0.933862846 | ENSG00000185630 | n.s. |
| ENSG00000280184 | 139.3177 | -0.84824265 | 0.282351 | -3.004209 | 0.002663 | 0.079555916 | | TEC | 45.01 | -0.93249649 | 0.93249649 | ENSG00000280184 | n.s. |
| ENSG00000207978 | 2.634238 | -1.96404383 | 0.932264 | -2.106747 | 0.03514 | 0.335560808 | MIR154 | miRNA | 40.48 | -0.931406086 | 0.931406086 | ENSG00000207978 | n.s. |
| ENSG00000267698 | 38.5902 | -0.957050196 | 0.334601 | -2.860273 | 0.004233 | 0.106466113 | | antisense | 51.74 | -0.93100752 | 0.93100752 | ENSG00000267698 | n.s. |
| ENSG00000259330 | 776.2276 | -0.773305319 | 0.24746 | -3.124971 | 0.001778 | 0.062653718 | INAFM2 | protein_coding | 56.16 | -0.930327402 | 0.930327402 | ENSG00000259330 | n.s. |
| ENSG00000178951 | 252.4711 | -0.77311532 | 0.247333 | -3.125804 | 0.001773 | 0.062653718 | ZBTB7A | protein_coding | 62.12 | -0.930098823 | 0.930098823 | ENSG00000178951 | n.s. |

| | | | | | | | | | | | | | |
|---|---|---|---|---|---|---|---|---|---|---|---|---|---|
| ENSG00000078687 | 334.908 | -0.768956943 | 0.246179 | -3.123564 | 0.001787 | 0.06266224 | TNRC6C | protein_coding | 42.22 | -0.925050655 | 0.925050655 | ENSG00000078687 | n.s. |
| ENSG00000168826 | 106.0674 | -0.857492603 | 0.287856 | -2.978899 | 0.002893 | 0.083791014 | ZBTB49 | protein_coding | 43.97 | -0.923350226 | 0.923350226 | ENSG00000168826 | n.s. |
| ENSG00000076650 | 281.0924 | -0.833616298 | 0.277165 | -3.007659 | 0.002633 | 0.078827723 | GPATCH1 | protein_coding | 45.71 | -0.919746382 | 0.919746382 | ENSG00000076650 | n.s. |
| ENSG00000242071 | 266.7724 | 0.962876578 | 0.339766 | 2.8339411 | 0.004598 | 0.111136994 | RPL7AP6 | processed_pseudogene | 53.06 | 0.918720362 | 0.918720362 | ENSG00000242071 | n.s. |
| ENSG00000224877 | 1149.881 | -0.840203137 | 0.28037 | -2.996761 | 0.002729 | 0.081177199 | NDUFAF8 | protein_coding | 58.49 | -0.916296921 | 0.916296921 | ENSG00000224877 | n.s. |
| ENSG00000272077 | 160.4845 | -0.801459726 | 0.262556 | -3.052526 | 0.002269 | 0.072133889 | | lincRNA | 41.53 | -0.915155328 | 0.915155328 | ENSG00000272077 | n.s. |
| ENSG00000267649 | 46.20207 | -0.928005263 | 0.322649 | -2.876203 | 0.004025 | 0.103804772 | | lincRNA | 48.38 | -0.912955505 | 0.912955505 | ENSG00000267649 | n.s. |
| ENSG00000155363 | 63.07681 | -0.85080064 | 0.286254 | -2.972187 | 0.002957 | 0.085411807 | MOV10 | protein_coding | 48.61 | -0.909065246 | 0.909065246 | ENSG00000155363 | n.s. |
| ENSG00000074047 | 59.27588 | -0.896347259 | 0.307925 | -2.910924 | 0.003604 | 0.09685932 | GLI2 | protein_coding | 52.42 | -0.908769363 | 0.908769363 | ENSG00000074047 | n.s. |
| ENSG00000070159 | 86.14746 | -0.922116599 | 0.320299 | -2.878925 | 0.00399 | 0.103651976 | PTPN3 | protein_coding | 45.34 | -0.90775225 | 0.90775225 | ENSG00000070159 | n.s. |
| ENSG00000267222 | 19.45961 | -1.155462655 | 0.444422 | -2.59992 | 0.009325 | 0.163921434 | | sense_intronic | 38.89 | -0.907459069 | 0.907459069 | ENSG00000267222 | n.s. |
| ENSG00000266146 | 9.673693 | -1.262155296 | 0.506824 | -2.490321 | 0.012763 | 0.191689721 | MIR5190 | miRNA | 61.25 | -0.905471693 | 0.905471693 | ENSG00000266146 | n.s. |
| ENSG00000240457 | 8.831078 | -1.462006127 | 0.621787 | -2.351297 | 0.018708 | 0.240510243 | RN7SL472P | misc_RNA | 53.24 | -0.904786501 | 0.904786501 | ENSG00000240457 | n.s. |
| ENSG00000215030 | 240.7614 | 0.945684484 | 0.33336 | 2.8368291 | 0.004556 | 0.110713338 | | processed_pseudogene | 61.32 | 0.903885281 | 0.903885281 | ENSG00000215030 | n.s. |
| ENSG00000255142 | 13.30303 | -1.183069472 | 0.459945 | -2.572197 | 0.010106 | 0.17218396 | | lincRNA | 59.44 | -0.903873724 | 0.903873724 | ENSG00000255142 | n.s. |
| ENSG00000073712 | 1501.383 | -0.809530514 | 0.267973 | -3.020939 | 0.00252 | 0.076718623 | FERMT2 | protein_coding | 39.3 | -0.902706831 | 0.902706831 | ENSG00000073712 | n.s. |
| ENSG00000200086 | 30.44945 | -1.089491568 | 0.408593 | -2.66645 | 0.007666 | 0.148580262 | RNU6-433P | snRNA | 36.54 | -0.902141376 | 0.902141376 | ENSG00000200086 | n.s. |
| ENSG00000259326 | 76.20241 | -0.862619664 | 0.292838 | -2.945719 | 0.003222 | 0.089992658 | | antisense | 36.31 | -0.902121538 | 0.902121538 | ENSG00000259326 | n.s. |
| ENSG00000111249 | 470.3195 | -0.747155079 | 0.238749 | -3.129456 | 0.001751 | 0.062361556 | CUX2 | protein_coding | 49.28 | -0.900383927 | 0.900383927 | ENSG00000111249 | n.s. |
| ENSG00000222267 | 6.150232 | -1.407467908 | 0.592698 | -2.374679 | 0.017564 | 0.231140499 | RNU6-892P | snRNA | 34.34 | -0.89532405 | 0.89532405 | ENSG00000222267 | n.s. |
| ENSG00000219150 | 8.951505 | -1.89519571 | 0.900892 | -2.103688 | 0.035406 | 0.337177445 | | processed_pseudogene | 47.6 | -0.894800517 | 0.894800517 | ENSG00000219150 | n.s. |
| ENSG00000231697 | 28.50422 | -0.963943671 | 0.345053 | -2.793608 | 0.005212 | 0.118034481 | NANOGP5 | ıscribed_processed_pseudog | 46.86 | -0.894531152 | 0.894531152 | ENSG00000231697 | n.s. |
| ENSG00000251456 | 77.83215 | 0.841240524 | 0.284308 | 2.9589015 | 0.003087 | 0.088085155 | | sense_intronic | 32.97 | 0.887590587 | 0.887590587 | ENSG00000251456 | n.s. |
| ENSG00000159082 | 217.3889 | -0.777115306 | 0.254527 | -3.05317 | 0.002264 | 0.072133889 | SYNJ1 | protein_coding | 38.08 | -0.887357393 | 0.887357393 | ENSG00000159082 | n.s. |
| ENSG00000207034 | 16.83958 | -1.104207002 | 0.420473 | -2.626106 | 0.008637 | 0.157676123 | | misc_RNA | 41.03 | -0.885832475 | 0.885832475 | ENSG00000207034 | n.s. |
| ENSG00000263266 | 477.6557 | 0.966435113 | 0.348102 | 2.7762951 | 0.005498 | 0.121897799 | RPS7P1 | processed_pseudogene | 47.52 | 0.88332569 | 0.88332569 | ENSG00000263266 | n.s. |
| ENSG00000143344 | 199.7719 | -0.732917429 | 0.234046 | -3.131505 | 0.001739 | 0.062358007 | RGL1 | protein_coding | 39.98 | -0.883244491 | 0.883244491 | ENSG00000143344 | n.s. |
| ENSG00000200090 | 22.15802 | -1.030409498 | 0.381416 | -2.701537 | 0.006902 | 0.139674846 | | misc_RNA | 44.44 | -0.880878325 | 0.880878325 | ENSG00000200090 | n.s. |
| ENSG00000140470 | 215.1435 | -0.743950974 | 0.239903 | -3.10105 | 0.001928 | 0.065538642 | ADAMTS17 | protein_coding | 46.62 | -0.880467883 | 0.880467883 | ENSG00000140470 | n.s. |
| ENSG00000109118 | 402.7417 | -0.722161929 | 0.229796 | -3.142627 | 0.001674 | 0.060551982 | PHF12 | protein_coding | 46.18 | -0.879500529 | 0.879500529 | ENSG00000109118 | n.s. |
| ENSG00000127418 | 118.3804 | -0.8157782 | 0.273675 | -2.980828 | 0.002875 | 0.083706059 | FGFRL1 | protein_coding | 66.74 | -0.878791441 | 0.878791441 | ENSG00000127418 | n.s. |
| ENSG00000272533 | 100.6271 | 0.779657643 | 0.257246 | 3.0307874 | 0.002439 | 0.074968112 | SNORA28 | snoRNA | 41.27 | 0.877211081 | 0.877211081 | ENSG00000272533 | n.s. |

| | | | | | | | | | | | | | |
|---|---|---|---|---|---|---|---|---|---|---|---|---|---|
| ENSG00000207493 | 15.29446 | -1.120259767 | 0.431968 | -2.593386 | 0.009504 | 0.165191859 | SNORA46 | snoRNA | 43.7 | -0.876055865 | 0.876055865 | ENSG00000207493 | n.s. |
| ENSG00000142733 | 11.03475 | -1.345099285 | 0.561237 | -2.396667 | 0.016545 | 0.223428884 | MAP3K6 | protein_coding | 60.79 | -0.875472242 | 0.875472242 | ENSG00000142733 | n.s. |
| ENSG00000223086 | 7.904165 | 1.318726827 | 0.545661 | 2.4167501 | 0.01566 | 0.216946191 | RNA5SP155 | rRNA | 51.16 | 0.875170383 | 0.875170383 | ENSG00000223086 | n.s. |
| ENSG00000228415 | 14.77277 | -1.186011481 | 0.471213 | -2.516935 | 0.011838 | 0.183252281 | PTMAP1 | processed_pseudogene | 50.46 | -0.874031884 | 0.874031884 | ENSG00000228415 | n.s. |
| ENSG00000236153 | 37.34534 | 1.054524817 | 0.395213 | 2.668243 | 0.007625 | 0.148477748 | | antisense | 46.54 | 0.87350364 | 0.87350364 | ENSG00000236153 | n.s. |
| ENSG00000144802 | 130.1778 | 0.776258004 | 0.25632 | 3.0284756 | 0.002458 | 0.075262844 | NFKBIZ | protein_coding | 40.23 | 0.87206328 | 0.87206328 | ENSG00000144802 | n.s. |
| ENSG00000170385 | 903.8353 | 0.726067261 | 0.232731 | 3.1197745 | 0.00181 | 0.062968329 | SLC30A1 | protein_coding | 38.51 | 0.871918075 | 0.871918075 | ENSG00000170385 | n.s. |
| ENSG00000279555 | 513.6193 | -0.757632548 | 0.247614 | -3.059737 | 0.002215 | 0.071230366 | | TEC | 49.86 | -0.869258206 | 0.869258206 | ENSG00000279555 | n.s. |
| ENSG00000236255 | 12.24604 | -1.164265503 | 0.457893 | -2.542657 | 0.011001 | 0.179260564 | | lincRNA | 48.08 | -0.869141944 | 0.869141944 | ENSG00000236255 | n.s. |
| ENSG00000265800 | 117.8723 | -0.784173392 | 0.260187 | -3.013879 | 0.002579 | 0.078004891 | | sense_intronic | 43.68 | -0.868768581 | 0.868768581 | ENSG00000265800 | n.s. |
| ENSG00000199855 | 5.987223 | -1.396937502 | 0.59295 | -2.355912 | 0.018477 | 0.238865111 | RNU6-490P | snRNA | 39.25 | -0.868681785 | 0.868681785 | ENSG00000199855 | n.s. |
| ENSG00000234202 | 7.782649 | -1.648652734 | 0.751514 | -2.193776 | 0.028252 | 0.297290527 | | processed_pseudogene | 50.64 | -0.868541468 | 0.868541468 | ENSG00000234202 | n.s. |
| ENSG00000229692 | 60.73241 | -0.946524956 | 0.340425 | -2.780418 | 0.005429 | 0.120938432 | SOS1-IT1 | sense_intronic | 38.13 | -0.868375753 | 0.868375753 | ENSG00000229692 | n.s. |
| ENSG00000253509 | 8.532193 | -1.260250042 | 0.51515 | -2.446373 | 0.01443 | 0.206062749 | | sense_intronic | 48.86 | -0.864532172 | 0.864532172 | ENSG00000253509 | n.s. |
| ENSG00000235313 | 20.2624 | -1.012721292 | 0.375089 | -2.69995 | 0.006935 | 0.140139316 | HM13-IT1 | sense_intronic | 52.69 | -0.864296867 | 0.864296867 | ENSG00000235313 | n.s. |
| ENSG00000228079 | 14.86303 | -1.425651726 | 0.612371 | -2.328087 | 0.019908 | 0.248850741 | | lincRNA | 39.55 | -0.861180696 | 0.861180696 | ENSG00000228079 | n.s. |
| ENSG00000149115 | 844.9648 | -0.723215077 | 0.232738 | -3.107419 | 0.001887 | 0.064772181 | TNKS1BP1 | protein_coding | 55.55 | -0.859621744 | 0.859621744 | ENSG00000149115 | n.s. |
| ENSG00000223764 | 238.9658 | -0.825583533 | 0.281272 | -2.935182 | 0.003334 | 0.092447097 | | lincRNA | 61.68 | -0.853741481 | 0.853741481 | ENSG00000223764 | n.s. |
| ENSG00000162396 | 93.74959 | -0.881947591 | 0.309077 | -2.853493 | 0.004324 | 0.107997903 | PARS2 | protein_coding | 49.77 | -0.852477027 | 0.852477027 | ENSG00000162396 | n.s. |
| ENSG00000169955 | 251.4317 | -0.692520109 | 0.219606 | -3.153467 | 0.001613 | 0.059115429 | ZNF747 | protein_coding | 57.48 | -0.850621864 | 0.850621864 | ENSG00000169955 | n.s. |
| ENSG00000229054 | 22.28322 | -1.049320997 | 0.397352 | -2.640786 | 0.008271 | 0.154788381 | RPS29P14 | processed_pseudogene | 52.63 | -0.850224553 | 0.850224553 | ENSG00000229054 | n.s. |
| ENSG00000127152 | 708.1999 | -0.740172267 | 0.241827 | -3.06075 | 0.002208 | 0.071230366 | BCL11B | protein_coding | 50.77 | -0.849225417 | 0.849225417 | ENSG00000127152 | n.s. |
| ENSG00000183864 | 317.2561 | -0.739010333 | 0.241581 | -3.059056 | 0.00222 | 0.071230366 | TOB2 | protein_coding | 51.46 | -0.84789229 | 0.84789229 | ENSG00000183864 | n.s. |
| ENSG00000279801 | 74.15234 | -0.777225117 | 0.259863 | -2.990907 | 0.002782 | 0.082048502 | | TEC | 53.69 | -0.844011563 | 0.844011563 | ENSG00000279801 | n.s. |
| ENSG00000267144 | 19.50567 | 1.117087572 | 0.436854 | 2.5571172 | 0.010554 | 0.175750036 | | lincRNA | 39.12 | 0.843517938 | 0.843517938 | ENSG00000267144 | n.s. |
| ENSG00000234383 | 35.75791 | -0.869834438 | 0.304986 | -2.852051 | 0.004344 | 0.107997903 | CTBP2P8 | processed_pseudogene | 55.86 | -0.84076864 | 0.84076864 | ENSG00000234383 | n.s. |
| ENSG00000249471 | 113.154 | -0.812641538 | 0.277342 | -2.93011 | 0.003388 | 0.092685471 | ZNF324B | protein_coding | 58.99 | -0.839449233 | 0.839449233 | ENSG00000249471 | n.s. |
| ENSG00000254522 | 12.92561 | -1.195362154 | 0.485278 | -2.463254 | 0.013768 | 0.199953179 | | processed_pseudogene | 51.34 | -0.835643838 | 0.835643838 | ENSG00000254522 | n.s. |
| ENSG00000252452 | 12.75138 | -1.105769585 | 0.432334 | -2.557672 | 0.010538 | 0.17570541 | RNU6-107P | snRNA | 44.12 | -0.83509363 | 0.83509363 | ENSG00000252452 | n.s. |
| ENSG00000265213 | 32.49314 | 0.942808359 | 0.344594 | 2.7359939 | 0.006219 | 0.131997212 | MIR3684 | miRNA | 36.49 | 0.829138897 | 0.829138897 | ENSG00000265213 | n.s. |
| ENSG00000264160 | 11.14939 | -1.151654101 | 0.462792 | -2.488492 | 0.012829 | 0.191814442 | MIR4762 | miRNA | 44 | -0.825872689 | 0.825872689 | ENSG00000264160 | n.s. |
| ENSG00000199886 | 9.672514 | -1.264378847 | 0.527185 | -2.398358 | 0.016469 | 0.223049174 | RNU6-249P | snRNA | 43.14 | -0.823868474 | 0.823868474 | ENSG00000199886 | n.s. |

| | | | | | | | | | | | | | |
|---|---|---|---|---|---|---|---|---|---|---|---|---|---|
| ENSG00000170604 | 260.6528 | -0.742970871 | 0.24655 | -3.013469 | 0.002583 | 0.078004891 | IRF2BP1 | protein_coding | 68.49 | -0.823121207 | 0.823121207 | ENSG00000170604 | n.s. |
| ENSG00000264916 | 19.71435 | -1.06117584 | 0.410281 | -2.586459 | 0.009697 | 0.167712137 | RN7SL230P | misc_RNA | 55.12 | -0.822873426 | 0.822873426 | ENSG00000264916 | n.s. |
| ENSG00000255837 | 75.40858 | -0.874936177 | 0.31145 | -2.809238 | 0.004966 | 0.114850754 | TAS2R20 | protein_coding | 34.76 | -0.822322898 | 0.822322898 | ENSG00000255837 | n.s. |
| ENSG00000221287 | 3.503791 | -1.729800711 | 0.820693 | -2.107732 | 0.035054 | 0.335533508 | MIR1289-2 | miRNA | 53.15 | -0.820382381 | 0.820382381 | ENSG00000221287 | n.s. |
| ENSG00000225224 | 76.86827 | -0.783143741 | 0.265833 | -2.945999 | 0.003219 | 0.089992658 | RPS27AP12 | processed_pseudogene | 44.09 | -0.819006179 | 0.819006179 | ENSG00000225224 | n.s. |
| ENSG00000207003 | 11.39715 | -1.286614959 | 0.542397 | -2.372091 | 0.017688 | 0.232108003 | RNU6-611P | snRNA | 42.31 | -0.816112586 | 0.816112586 | ENSG00000207003 | n.s. |
| ENSG00000145685 | 259.5192 | -0.77708296 | 0.26329 | -2.951433 | 0.003163 | 0.089147501 | LHFPL2 | protein_coding | 43.31 | -0.815852272 | 0.815852272 | ENSG00000145685 | n.s. |
| ENSG00000206965 | 4.24005 | -1.531495355 | 0.696148 | -2.199956 | 0.02781 | 0.293497624 | RNU6-5P | snRNA | 45.79 | -0.815361097 | 0.815361097 | ENSG00000206965 | n.s. |
| ENSG00000275004 | 183.2271 | -0.756882614 | 0.253802 | -2.982179 | 0.002862 | 0.083706059 | ZNF280B | protein_coding | 41.55 | -0.815346577 | 0.815346577 | ENSG00000275004 | n.s. |
| ENSG00000173705 | 698.1254 | -0.788257013 | 0.2688 | -2.932507 | 0.003362 | 0.092447097 | SUSD5 | protein_coding | 41.61 | -0.815141875 | 0.815141875 | ENSG00000173705 | n.s. |
| ENSG00000258748 | 25.2977 | -1.035177134 | 0.397763 | -2.602495 | 0.009255 | 0.16331424 | | antisense | 65.82 | -0.814659505 | 0.814659505 | ENSG00000258748 | n.s. |
| ENSG00000257023 | 36.08891 | 0.981916298 | 0.368576 | 2.6640809 | 0.00772 | 0.148870041 | | antisense | 38.09 | 0.812233984 | 0.812233984 | ENSG00000257023 | n.s. |
| ENSG00000116273 | 145.0191 | -0.799872087 | 0.274658 | -2.912252 | 0.003588 | 0.096635009 | PHF13 | protein_coding | 54.99 | -0.811762592 | 0.811762592 | ENSG00000116273 | n.s. |
| ENSG00000267672 | 55.7886 | -0.903008417 | 0.327438 | -2.757797 | 0.005819 | 0.126595758 | | sense_intronic | 41.34 | -0.81052306 | 0.81052306 | ENSG00000267672 | n.s. |
| ENSG00000108557 | 948.6977 | -0.697472877 | 0.226977 | -3.072885 | 0.00212 | 0.069288322 | RAI1 | protein_coding | 56.45 | -0.808608176 | 0.808608176 | ENSG00000108557 | n.s. |
| ENSG00000092820 | 5943.767 | -0.747509306 | 0.250322 | -2.986189 | 0.002825 | 0.083148981 | EZR | protein_coding | 46.17 | -0.807416996 | 0.807416996 | ENSG00000092820 | n.s. |
| ENSG00000273014 | 22.28345 | -1.118338418 | 0.447794 | -2.49744 | 0.012509 | 0.189725185 | | lincRNA | 28.6 | -0.807300563 | 0.807300563 | ENSG00000273014 | n.s. |
| ENSG00000278928 | 390.4078 | -0.848602126 | 0.300642 | -2.822632 | 0.004763 | 0.112211468 | | TEC | 41.49 | -0.806140415 | 0.806140415 | ENSG00000278928 | n.s. |
| ENSG00000198406 | 73.48574 | 0.959346972 | 0.357262 | 2.685274 | 0.007247 | 0.144556561 | BZW1P2 | processed_pseudogene | 40.19 | 0.805815187 | 0.805815187 | ENSG00000198406 | n.s. |
| ENSG00000131149 | 3205.992 | -0.713440053 | 0.23511 | -3.034493 | 0.002409 | 0.074712853 | GSE1 | protein_coding | 56.58 | -0.8037649 | 0.8037649 | ENSG00000131149 | n.s. |
| ENSG00000251550 | 122.93 | -0.773228546 | 0.263646 | -2.932828 | 0.003359 | 0.092447097 | | processed_pseudogene | 38.31 | -0.799600837 | 0.799600837 | ENSG00000251550 | n.s. |
| ENSG00000265195 | 24.24199 | -0.972945614 | 0.366104 | -2.657565 | 0.007871 | 0.150725964 | MIR4312 | miRNA | 59.21 | -0.799578312 | 0.799578312 | ENSG00000265195 | n.s. |
| ENSG00000222276 | 7.466166 | -1.314889594 | 0.563447 | -2.333652 | 0.019614 | 0.247135688 | RNU2-33P | snRNA | 38.02 | -0.79822284 | 0.79822284 | ENSG00000222276 | n.s. |
| ENSG00000266276 | 2.732574 | -1.876177683 | 0.932475 | -2.012042 | 0.044216 | 0.375547181 | MIR4743 | miRNA | 53.62 | -0.798004962 | 0.798004962 | ENSG00000266276 | n.s. |
| ENSG00000130707 | 4611.76 | -0.841420097 | 0.298496 | -2.818862 | 0.004819 | 0.11285058 | ASS1 | protein_coding | 55.65 | -0.797242348 | 0.797242348 | ENSG00000130707 | n.s. |
| ENSG00000279583 | 34.54418 | -0.980751503 | 0.371024 | -2.643367 | 0.008209 | 0.154088881 | | TEC | 63.49 | -0.796594516 | 0.796594516 | ENSG00000279583 | n.s. |
| ENSG00000138756 | 328.1603 | -0.775420661 | 0.265525 | -2.920335 | 0.003497 | 0.09489754 | BMP2K | protein_coding | 35.68 | -0.793057639 | 0.793057639 | ENSG00000138756 | n.s. |
| ENSG00000275038 | 106.8872 | -0.823783037 | 0.289607 | -2.844488 | 0.004448 | 0.109423042 | | protein_coding | 53.55 | -0.791565885 | 0.791565885 | ENSG00000275038 | n.s. |
| ENSG00000260273 | 39.00475 | 0.943647786 | 0.351707 | 2.6830507 | 0.007295 | 0.145105366 | | antisense | 47.71 | 0.791075534 | 0.791075534 | ENSG00000260273 | n.s. |
| ENSG00000102935 | 355.1604 | -0.700554144 | 0.230674 | -3.036985 | 0.00239 | 0.074429557 | ZNF423 | protein_coding | 50.74 | -0.790403413 | 0.790403413 | ENSG00000102935 | n.s. |
| ENSG00000159753 | 333.4214 | -0.740501252 | 0.249266 | -2.970722 | 0.002971 | 0.0856425 | CARMIL2 | protein_coding | 62.9 | -0.790344882 | 0.790344882 | ENSG00000159753 | n.s. |
| ENSG00000218175 | 110.766 | 1.186131854 | 0.490288 | 2.4192568 | 0.015552 | 0.215671391 | | processed_pseudogene | 53.91 | 0.79020989 | 0.79020989 | ENSG00000218175 | n.s. |

| | | | | | | | | | | | | | |
|---|---|---|---|---|---|---|---|---|---|---|---|---|---|
| ENSG00000215952 | 13.75371 | -1.072265941 | 0.426283 | -2.515387 | 0.01189 | 0.18333021 | MIR921 | miRNA | 55.36 | -0.790009048 | 0.790009048 | ENSG00000215952 | n.s. |
| ENSG00000179134 | 132.0336 | -0.828505525 | 0.292869 | -2.828933 | 0.00467 | 0.111596326 | SAMD4B | protein_coding | 49.82 | -0.789027322 | 0.789027322 | ENSG00000179134 | n.s. |
| ENSG00000238832 | 7.44103 | -1.788539681 | 0.878005 | -2.037051 | 0.041645 | 0.36238932 | | snoRNA | 32.41 | -0.788432308 | 0.788432308 | ENSG00000238832 | n.s. |
| ENSG00000218227 | 135.1706 | 1.16469842 | 0.479502 | 2.4289752 | 0.015142 | 0.211662698 | | processed_pseudogene | 55.16 | 0.785420985 | 0.785420985 | ENSG00000218227 | n.s. |
| ENSG00000273320 | 51.81978 | 0.897128889 | 0.329198 | 2.7251928 | 0.006426 | 0.133345298 | | antisense | 47.11 | 0.785007779 | 0.785007779 | ENSG00000273320 | n.s. |
| ENSG00000151490 | 549.7632 | -0.746201685 | 0.252994 | -2.949487 | 0.003183 | 0.089409345 | PTPRO | protein_coding | 37.5 | -0.782479835 | 0.782479835 | ENSG00000151490 | n.s. |
| ENSG00000212418 | 8.375446 | -1.397004623 | 0.621853 | -2.246518 | 0.024671 | 0.276340982 | RNY4P36 | misc_RNA | 38.54 | -0.780303503 | 0.780303503 | ENSG00000212418 | n.s. |
| ENSG00000103199 | 249.4561 | -0.755905569 | 0.258214 | -2.927438 | 0.003418 | 0.09330237 | ZNF500 | protein_coding | 54.32 | -0.778663864 | 0.778663864 | ENSG00000103199 | n.s. |
| ENSG00000140332 | 2417.415 | -0.672453018 | 0.219238 | -3.067232 | 0.002161 | 0.070118383 | TLE3 | protein_coding | 52.15 | -0.776123829 | 0.776123829 | ENSG00000140332 | n.s. |
| ENSG00000261886 | 11.99808 | -1.233693355 | 0.52207 | -2.363082 | 0.018124 | 0.235607368 | | lincRNA | 42.81 | -0.774526421 | 0.774526421 | ENSG00000261886 | n.s. |
| ENSG00000131378 | 207.0154 | -0.748754831 | 0.255433 | -2.931321 | 0.003375 | 0.092506687 | RFTN1 | protein_coding | 43.5 | -0.774082866 | 0.774082866 | ENSG00000131378 | n.s. |
| ENSG00000241506 | 78.7481 | 0.810689217 | 0.285953 | 2.8350399 | 0.004582 | 0.111136994 | PSMC1P1 | processed_pseudogene | 45.76 | 0.773512107 | 0.773512107 | ENSG00000241506 | n.s. |
| ENSG00000272936 | 59.19138 | -0.809268596 | 0.285245 | -2.837098 | 0.004553 | 0.110713338 | | antisense | 26.74 | -0.773498968 | 0.773498968 | ENSG00000272936 | n.s. |
| ENSG00000263553 | 23.63959 | 0.99928315 | 0.386646 | 2.5844903 | 0.009752 | 0.168254454 | | lincRNA | 42.41 | 0.773478564 | 0.773478564 | ENSG00000263553 | n.s. |
| ENSG00000202400 | 40.3654 | 0.874394051 | 0.319056 | 2.7405678 | 0.006133 | 0.130768951 | SNORD82 | snoRNA | 37.14 | 0.772523087 | 0.772523087 | ENSG00000202400 | n.s. |
| ENSG00000072609 | 46.18503 | -0.9328283 | 0.349827 | -2.666545 | 0.007664 | 0.148580262 | CHFR | protein_coding | 48.69 | -0.7724181 | 0.7724181 | ENSG00000072609 | n.s. |
| ENSG00000157450 | 687.4105 | -0.775362655 | 0.26808 | -2.892285 | 0.003825 | 0.101168416 | RNF111 | protein_coding | 39.52 | -0.771450989 | 0.771450989 | ENSG00000157450 | n.s. |
| ENSG00000269973 | 110.1799 | -0.808975468 | 0.285724 | -2.831314 | 0.004636 | 0.111473467 | | lincRNA | 42.62 | -0.770814873 | 0.770814873 | ENSG00000269973 | n.s. |
| ENSG00000197256 | 220.3838 | -0.785414945 | 0.273433 | -2.872423 | 0.004073 | 0.104433016 | KANK2 | protein_coding | 52.98 | -0.77061944 | 0.77061944 | ENSG00000197256 | n.s. |
| ENSG00000242818 | 8.442785 | -1.339485662 | 0.588537 | -2.275958 | 0.022849 | 0.265986481 | RN7SL846P | misc_RNA | 54.11 | -0.770392367 | 0.770392367 | ENSG00000242818 | n.s. |
| ENSG00000281469 | 24.89694 | -0.968604843 | 0.371321 | -2.608536 | 0.009093 | 0.161482502 | | TEC | 29.71 | -0.767013505 | 0.767013505 | ENSG00000281469 | n.s. |
| ENSG00000259402 | 16.65828 | -1.123859367 | 0.460352 | -2.441304 | 0.014634 | 0.207911968 | | sense_intronic | 43.06 | -0.766607526 | 0.766607526 | ENSG00000259402 | n.s. |
| ENSG00000279146 | 92.30481 | -0.802370422 | 0.283678 | -2.828458 | 0.004677 | 0.111596326 | | TEC | 55.12 | -0.764137553 | 0.764137553 | ENSG00000279146 | n.s. |
| ENSG00000188394 | 34.71239 | -0.899660635 | 0.333793 | -2.695269 | 0.007033 | 0.141610603 | GPR21 | protein_coding | 45.81 | -0.763725717 | 0.763725717 | ENSG00000188394 | n.s. |
| ENSG00000180573 | 900.3556 | -0.672886159 | 0.2209 | -3.046118 | 0.002318 | 0.073314736 | HIST1H2AC | protein_coding | 41.82 | -0.763597083 | 0.763597083 | ENSG00000180573 | n.s. |
| ENSG00000279306 | 28.75907 | -1.008071095 | 0.393306 | -2.563073 | 0.010375 | 0.174827325 | | TEC | 58.41 | -0.76350366 | 0.76350366 | ENSG00000279306 | n.s. |
| ENSG00000233875 | 15.89623 | -0.995064145 | 0.386283 | -2.576001 | 0.009995 | 0.171380592 | | antisense | 53.59 | -0.762257306 | 0.762257306 | ENSG00000233875 | n.s. |
| ENSG00000267376 | 9.620441 | -1.187696519 | 0.498609 | -2.382022 | 0.017218 | 0.228309094 | | processed_pseudogene | 47.06 | -0.76187975 | 0.76187975 | ENSG00000267376 | n.s. |
| ENSG00000218730 | 18.77108 | -1.028437555 | 0.409003 | -2.514498 | 0.01192 | 0.183386434 | | processed_pseudogene | 44.33 | -0.75758083 | 0.75758083 | ENSG00000218730 | n.s. |
| ENSG00000207194 | 2.565147 | -2.459308672 | 1.435876 | -1.712759 | 0.086757 | 0.492652156 | RNU6-1026P | snRNA | 37.96 | -0.756138091 | 0.756138091 | ENSG00000207194 | n.s. |
| ENSG00000184489 | 91.6859 | -0.854793429 | 0.312024 | -2.739512 | 0.006153 | 0.130792064 | PTP4A3 | protein_coding | 62.4 | -0.75514042 | 0.75514042 | ENSG00000184489 | n.s. |
| ENSG00000199788 | 7.845722 | -1.235396577 | 0.528084 | -2.339395 | 0.019315 | 0.245525197 | RNY3P2 | misc_RNA | 44.12 | -0.753473231 | 0.753473231 | ENSG00000199788 | n.s. |

| | | | | | | | | | | | | | |
|---|---|---|---|---|---|---|---|---|---|---|---|---|---|
| ENSG00000272689 | 12.61895 | -1.155059623 | 0.482204 | -2.395373 | 0.016603 | 0.22376458 | | sense_intronic | 35.95 | -0.751029767 | 0.751029767 | ENSG00000272689 | n.s. |
| ENSG00000212526 | 4.617297 | -1.583043537 | 0.751206 | -2.107336 | 0.035088 | 0.335533508 | RNU6-466P | snRNA | 39.81 | -0.750780722 | 0.750780722 | ENSG00000212526 | n.s. |
| ENSG00000156097 | 22.56285 | 1.018063847 | 0.404648 | 2.515927 | 0.011872 | 0.183252281 | GPR61 | protein_coding | 49.33 | 0.750262773 | 0.750262773 | ENSG00000156097 | n.s. |
| ENSG00000212499 | 3.40563 | -1.687056852 | 0.825867 | -2.042772 | 0.041075 | 0.359904219 | RNA5SP300 | rRNA | 54.21 | -0.748737869 | 0.748737869 | ENSG00000212499 | n.s. |
| ENSG00000184068 | 86.33993 | 0.817651407 | 0.294398 | 2.77737 | 0.00548 | 0.121689136 | | antisense | 56.07 | 0.747945146 | 0.747945146 | ENSG00000184068 | n.s. |
| ENSG00000265201 | 7.795742 | -1.232840129 | 0.528638 | -2.332108 | 0.019695 | 0.247485201 | MIR4677 | miRNA | 46.25 | -0.74765684 | 0.74765684 | ENSG00000265201 | n.s. |
| ENSG00000187634 | 1985.608 | -0.755388974 | 0.262046 | -2.882662 | 0.003943 | 0.10262154 | SAMD11 | protein_coding | 66.58 | -0.746899514 | 0.746899514 | ENSG00000187634 | n.s. |
| ENSG00000207234 | 17.81746 | -1.005618003 | 0.397989 | -2.526749 | 0.011512 | 0.181823456 | RNU6-125P | snRNA | 41.12 | -0.744509382 | 0.744509382 | ENSG00000207234 | n.s. |
| ENSG00000185485 | 418.4823 | -0.716774887 | 0.243925 | -2.93851 | 0.003298 | 0.091651249 | SDHAP1 | cribed_unprocessed_pseudc | 48.24 | -0.743913139 | 0.743913139 | ENSG00000185485 | n.s. |
| ENSG00000145198 | 236.0454 | -0.751054859 | 0.260483 | -2.883321 | 0.003935 | 0.102598935 | VWA5B2 | protein_coding | 58.09 | -0.742685967 | 0.742685967 | ENSG00000145198 | n.s. |
| ENSG00000135953 | 374.9439 | -0.754230581 | 0.262245 | -2.876052 | 0.004027 | 0.103804772 | MFSD9 | protein_coding | 41.84 | -0.741998983 | 0.741998983 | ENSG00000135953 | n.s. |
| ENSG00000227081 | 278.5399 | 0.91989071 | 0.349105 | 2.6349955 | 0.008414 | 0.156104918 | | processed_pseudogene | 49.02 | 0.741968591 | 0.741968591 | ENSG00000227081 | n.s. |
| ENSG00000226216 | 72.03732 | -0.848809624 | 0.312348 | -2.717509 | 0.006578 | 0.134873528 | RPS12P5 | processed_pseudogene | 44.74 | -0.738526577 | 0.738526577 | ENSG00000226216 | n.s. |
| ENSG00000272102 | 25.01696 | -0.939191409 | 0.361344 | -2.59916 | 0.009345 | 0.163964393 | | lincRNA | 39.93 | -0.737500482 | 0.737500482 | ENSG00000272102 | n.s. |
| ENSG00000232693 | 88.91236 | -0.87769288 | 0.327192 | -2.682501 | 0.007307 | 0.145136682 | | sense_intronic | 43.37 | -0.735702192 | 0.735702192 | ENSG00000232693 | n.s. |
| ENSG00000179029 | 876.9073 | 0.629385951 | 0.204005 | 3.0851502 | 0.002034 | 0.067959896 | TMEM107 | protein_coding | 49.51 | 0.73496374 | 0.73496374 | ENSG00000179029 | n.s. |
| ENSG00000167157 | 270.1954 | -0.89664344 | 0.337918 | -2.653433 | 0.007968 | 0.151550608 | PRRX2 | protein_coding | 54.69 | -0.734747577 | 0.734747577 | ENSG00000167157 | n.s. |
| ENSG00000064393 | 897.2576 | -0.739072464 | 0.255768 | -2.889615 | 0.003857 | 0.101517698 | HIPK2 | protein_coding | 45.62 | -0.734237627 | 0.734237627 | ENSG00000064393 | n.s. |
| ENSG00000151883 | 193.5926 | 0.709461094 | 0.241961 | 2.9321334 | 0.003366 | 0.092447097 | PARP8 | protein_coding | 35.7 | 0.733658486 | 0.733658486 | ENSG00000151883 | n.s. |
| ENSG00000237162 | 14.49375 | -1.0038589 | 0.400412 | -2.507063 | 0.012174 | 0.185852348 | | processed_pseudogene | 42.78 | -0.733652155 | 0.733652155 | ENSG00000237162 | n.s. |
| ENSG00000169379 | 285.1126 | -0.77442152 | 0.274824 | -2.817883 | 0.004834 | 0.112918712 | ARL13B | protein_coding | 36.2 | -0.73355846 | 0.73355846 | ENSG00000169379 | n.s. |
| ENSG00000169136 | 440.4963 | -0.744206738 | 0.258695 | -2.876772 | 0.004018 | 0.103804772 | ATF5 | protein_coding | 55.56 | -0.7321377 | 0.7321377 | ENSG00000169136 | n.s. |
| ENSG00000138744 | 67.50406 | -0.786870213 | 0.281301 | -2.797251 | 0.005154 | 0.117444319 | NAAA | protein_coding | 44.41 | -0.731921482 | 0.731921482 | ENSG00000138744 | n.s. |
| ENSG00000199781 | 11.45967 | -1.077941276 | 0.442747 | -2.434664 | 0.014906 | 0.209839048 | | misc_RNA | 36.36 | -0.730966738 | 0.730966738 | ENSG00000199781 | n.s. |
| ENSG00000249655 | 36.94557 | -0.825349812 | 0.301215 | -2.740073 | 0.006143 | 0.130768951 | | antisense | 43.27 | -0.72919273 | 0.72919273 | ENSG00000249655 | n.s. |
| ENSG00000270012 | 87.67422 | -0.742027412 | 0.258377 | -2.871878 | 0.00408 | 0.104433016 | | lincRNA | 49.5 | -0.728049233 | 0.728049233 | ENSG00000270012 | n.s. |
| ENSG00000137764 | 190.2217 | -0.728031393 | 0.251435 | -2.895506 | 0.003785 | 0.100582346 | MAP2K5 | protein_coding | 39.55 | -0.726195471 | 0.726195471 | ENSG00000137764 | n.s. |
| ENSG00000269038 | 21.36956 | -1.006992164 | 0.403882 | -2.493285 | 0.012657 | 0.190920497 | | lincRNA | 48.67 | -0.72417584 | 0.72417584 | ENSG00000269038 | n.s. |
| ENSG00000273449 | 164.1207 | -0.740739642 | 0.258426 | -2.866356 | 0.004152 | 0.105304418 | | lincRNA | 33.87 | -0.724112561 | 0.724112561 | ENSG00000273449 | n.s. |
| ENSG00000107984 | 1498.683 | -0.640960652 | 0.211041 | -3.037144 | 0.002388 | 0.074429557 | DKK1 | protein_coding | 43.26 | -0.723166784 | 0.723166784 | ENSG00000107984 | n.s. |
| ENSG00000180828 | 257.9177 | -0.681482333 | 0.229888 | -2.964411 | 0.003033 | 0.08687934 | BHLHE22 | protein_coding | 53.59 | -0.723109651 | 0.723109651 | ENSG00000180828 | n.s. |
| ENSG00000276103 | 5.502146 | -1.331836927 | 0.600712 | -2.217098 | 0.026616 | 0.287074936 | | snRNA | 44.44 | -0.721861906 | 0.721861906 | ENSG00000276103 | n.s. |

| | | | | | | | | | | | | | |
|---|---|---|---|---|---|---|---|---|---|---|---|---|---|
| ENSG00000273742 | 4.858476 | -1.676192643 | 0.829274 | -2.021278 | 0.043251 | 0.371106396 | MIR6075 | miRNA | 64.21 | -0.72160355 | 0.72160355 | ENSG00000273742 | n.s. |
| ENSG00000177614 | 358.9189 | -0.783613684 | 0.281407 | -2.784629 | 0.005359 | 0.120342054 | PGBD5 | protein_coding | 48.34 | -0.720597495 | 0.720597495 | ENSG00000177614 | n.s. |
| ENSG00000241549 | 91.59085 | -0.819185684 | 0.299845 | -2.732027 | 0.006295 | 0.132435959 | GUSBP2 | cribed_unprocessed_pseudc | 39.56 | -0.71924018 | 0.71924018 | ENSG00000241549 | n.s. |
| ENSG00000220744 | 39.82749 | -0.895517496 | 0.340694 | -2.628507 | 0.008576 | 0.157512796 | RPL5P18 | processed_pseudogene | 40.52 | -0.718817707 | 0.718817707 | ENSG00000220744 | n.s. |
| ENSG00000276564 | 137.7297 | -0.771127516 | 0.275393 | -2.800098 | 0.005109 | 0.116983481 | | lincRNA | 32.83 | -0.718594809 | 0.718594809 | ENSG00000276564 | n.s. |
| ENSG00000274934 | 54.37316 | 0.791659609 | 0.285962 | 2.7684044 | 0.005633 | 0.124079983 | | misc_RNA | 42.51 | 0.717479736 | 0.717479736 | ENSG00000274934 | n.s. |
| ENSG00000276116 | 54.05623 | 0.887811133 | 0.336791 | 2.636093 | 0.008387 | 0.156104918 | FUT8-AS1 | antisense | 56.67 | 0.716093736 | 0.716093736 | ENSG00000276116 | n.s. |
| ENSG00000223375 | 19.52546 | -0.930338608 | 0.36065 | -2.579619 | 0.009891 | 0.170014443 | | processed_pseudogene | 44.91 | -0.715908754 | 0.715908754 | ENSG00000223375 | n.s. |
| ENSG00000262265 | 86.89279 | -0.765282315 | 0.272972 | -2.803522 | 0.005055 | 0.11613472 | | sense_intronic | 43.33 | -0.715567985 | 0.715567985 | ENSG00000262265 | n.s. |
| ENSG00000088808 | 39.23061 | -0.881552593 | 0.33411 | -2.638509 | 0.008327 | 0.155351425 | PPP1R13B | protein_coding | 44.51 | -0.712898145 | 0.712898145 | ENSG00000088808 | n.s. |
| ENSG00000207016 | 3.635029 | -1.82404165 | 0.942334 | -1.935664 | 0.052909 | 0.406621815 | SNORA36C | snoRNA | 41.67 | -0.712852486 | 0.712852486 | ENSG00000207016 | n.s. |
| ENSG00000272335 | 165.9493 | -0.811457406 | 0.297067 | -2.731566 | 0.006303 | 0.132435959 | | lincRNA | 35.2 | -0.712454797 | 0.712454797 | ENSG00000272335 | n.s. |
| ENSG00000060709 | 700.5522 | -0.731866745 | 0.255897 | -2.860007 | 0.004236 | 0.106466113 | RIMBP2 | protein_coding | 50.38 | -0.711951626 | 0.711951626 | ENSG00000060709 | n.s. |
| ENSG00000238365 | 8.243648 | -1.560829779 | 0.753108 | -2.072518 | 0.038217 | 0.350082264 | RNU7-57P | snRNA | 40 | -0.711472869 | 0.711472869 | ENSG00000238365 | n.s. |
| ENSG00000266315 | 4.538637 | -1.666854816 | 0.826789 | -2.016059 | 0.043794 | 0.374378906 | MIR4668 | miRNA | 30 | -0.711227997 | 0.711227997 | ENSG00000266315 | n.s. |
| ENSG00000259659 | 69.54784 | -0.874105096 | 0.330678 | -2.643373 | 0.008208 | 0.154088881 | | sense_intronic | 46.69 | -0.709973243 | 0.709973243 | ENSG00000259659 | n.s. |
| ENSG00000171246 | 900.9928 | -0.717548967 | 0.248779 | -2.884277 | 0.003923 | 0.102598935 | NPTX1 | protein_coding | 60.76 | -0.709553425 | 0.709553425 | ENSG00000171246 | n.s. |
| ENSG00000261061 | 76.47043 | -0.827575837 | 0.306124 | -2.703397 | 0.006863 | 0.139300314 | | sense_intronic | 43.44 | -0.70844456 | 0.70844456 | ENSG00000261061 | n.s. |
| ENSG00000268713 | 52.84346 | -0.856239742 | 0.321728 | -2.661378 | 0.007782 | 0.149575275 | | lincRNA | 40.14 | -0.706517823 | 0.706517823 | ENSG00000268713 | n.s. |
| ENSG00000284031 | 1.703143 | -2.040584618 | 1.125435 | -1.813151 | 0.069809 | 0.451018079 | MIR4761 | miRNA | 62.2 | -0.705646505 | 0.705646505 | ENSG00000284031 | n.s. |
| ENSG00000276696 | 7.239095 | -1.221207142 | 0.535571 | -2.280196 | 0.022596 | 0.264374115 | | misc_RNA | 35.38 | -0.705590367 | 0.705590367 | ENSG00000276696 | n.s. |
| ENSG00000252317 | 5.231885 | -1.356391644 | 0.622094 | -2.180365 | 0.02923 | 0.302134081 | | misc_RNA | 37.63 | -0.705052761 | 0.705052761 | ENSG00000252317 | n.s. |
| ENSG00000227773 | 26.63112 | -0.871799006 | 0.330956 | -2.634186 | 0.008434 | 0.156104918 | ASH1L-IT1 | sense_intronic | 43.24 | -0.703178619 | 0.703178619 | ENSG00000227773 | n.s. |
| ENSG00000233476 | 221.837 | 0.849849584 | 0.319417 | 2.6606308 | 0.007799 | 0.149575275 | EEF1A1P6 | processed_pseudogene | 48.27 | 0.701245047 | 0.701245047 | ENSG00000233476 | n.s. |
| ENSG00000252485 | 2.528898 | -2.033002352 | 1.12254 | -1.811073 | 0.07013 | 0.452404539 | | misc_RNA | 48.42 | -0.700314514 | 0.700314514 | ENSG00000252485 | n.s. |
| ENSG00000202361 | 10.53538 | -1.115314243 | 0.471877 | -2.363572 | 0.0181 | 0.235607368 | | misc_RNA | 38.38 | -0.700206697 | 0.700206697 | ENSG00000202361 | n.s. |
| ENSG00000222932 | 3.304636 | -2.245697084 | 1.303538 | -1.722771 | 0.08493 | 0.487906764 | RNU6-172P | snRNA | 39.78 | -0.699901053 | 0.699901053 | ENSG00000222932 | n.s. |
| ENSG00000230409 | 51.10064 | 0.846497533 | 0.318142 | 2.6607523 | 0.007797 | 0.149575275 | TCEA1P2 | processed_pseudogene | 41.72 | 0.698479135 | 0.698479135 | ENSG00000230409 | n.s. |
| ENSG00000280551 | 212.5012 | -0.722534315 | 0.253252 | -2.853028 | 0.00433 | 0.107997903 | | TEC | 38.4 | -0.698390598 | 0.698390598 | ENSG00000280551 | n.s. |
| ENSG00000138166 | 153.4619 | 0.763229517 | 0.274753 | 2.777878 | 0.005472 | 0.121689136 | DUSP5 | protein_coding | 49.09 | 0.69816282 | 0.69816282 | ENSG00000138166 | n.s. |
| ENSG00000163673 | 219.0801 | -0.731517236 | 0.258717 | -2.827479 | 0.004692 | 0.111596326 | DCLK3 | protein_coding | 43.39 | -0.696660515 | 0.696660515 | ENSG00000163673 | n.s. |
| ENSG00000198496 | 190.4607 | -0.71565687 | 0.250162 | -2.860779 | 0.004226 | 0.106466113 | NBR2 | cribed_unprocessed_pseudc | 47.53 | -0.696182844 | 0.696182844 | ENSG00000198496 | n.s. |

| | | | | | | | | | | | | | |
|---|---|---|---|---|---|---|---|---|---|---|---|---|---|
| ENSG00000276410 | 556.5665 | -0.728411625 | 0.257108 | -2.833093 | 0.00461 | 0.111239262 | HIST1H2BB | protein_coding | 54.07 | -0.694716689 | 0.694716689 | ENSG00000276410 | n.s. |
| ENSG00000070748 | 1022.03 | -0.689294623 | 0.237391 | -2.90362 | 0.003689 | 0.098747222 | CHAT | protein_coding | 51.04 | -0.693068588 | 0.693068588 | ENSG00000070748 | n.s. |
| ENSG00000199104 | 3.735734 | -1.7975382 | 0.935784 | -1.920891 | 0.054745 | 0.412398929 | MIR346 | miRNA | 67.37 | -0.691481438 | 0.691481438 | ENSG00000199104 | n.s. |
| ENSG00000265688 | 74.54119 | -0.816761733 | 0.303473 | -2.691382 | 0.007116 | 0.142548954 | MAFG-AS1 | antisense | 65.54 | -0.691009802 | 0.691009802 | ENSG00000265688 | n.s. |
| ENSG00000268362 | 281.8118 | -0.705469038 | 0.24602 | -2.867526 | 0.004137 | 0.105107004 | | lincRNA | 45.45 | -0.690208574 | 0.690208574 | ENSG00000268362 | n.s. |
| ENSG00000257246 | 39.78429 | -0.847297569 | 0.32014 | -2.646644 | 0.008129 | 0.15357772 | PHBP19 | processed_pseudogene | 51.59 | -0.689422126 | 0.689422126 | ENSG00000257246 | n.s. |
| ENSG00000254676 | 59.76503 | -0.777518607 | 0.283169 | -2.745772 | 0.006037 | 0.130381308 | | sense_overlapping | 35.34 | -0.687936541 | 0.687936541 | ENSG00000254676 | n.s. |
| ENSG00000256006 | 54.50804 | -0.776361696 | 0.282945 | -2.743861 | 0.006072 | 0.130381308 | | sense_intronic | 43.46 | -0.686912924 | 0.686912924 | ENSG00000256006 | n.s. |
| ENSG00000201012 | 4.364326 | -1.40400867 | 0.658551 | -2.131965 | 0.03301 | 0.324413031 | | misc_RNA | 36.79 | -0.686422238 | 0.686422238 | ENSG00000201012 | n.s. |
| ENSG00000139514 | 1631.932 | -0.697534787 | 0.242439 | -2.877152 | 0.004013 | 0.103804772 | SLC7A1 | protein_coding | 47.44 | -0.686222643 | 0.686222643 | ENSG00000139514 | n.s. |
| ENSG00000106070 | 156.8724 | -0.792539099 | 0.29216 | -2.712685 | 0.006674 | 0.136250333 | GRB10 | protein_coding | 45.52 | -0.68607132 | 0.68607132 | ENSG00000106070 | n.s. |
| ENSG00000221445 | 2.812056 | -2.016243064 | 1.119645 | -1.800788 | 0.071736 | 0.457319201 | MIR1301 | miRNA | 62.2 | -0.685080205 | 0.685080205 | ENSG00000221445 | n.s. |
| ENSG00000104756 | 441.8015 | -0.758050605 | 0.274369 | -2.76289 | 0.005729 | 0.125420964 | KCTD9 | protein_coding | 37.3 | -0.683481064 | 0.683481064 | ENSG00000104756 | n.s. |
| ENSG00000238812 | 3.931102 | -1.53489397 | 0.75238 | -2.040052 | 0.041345 | 0.360907877 | RNU7-127P | snRNA | 46.77 | -0.679349657 | 0.679349657 | ENSG00000238812 | n.s. |
| ENSG00000110925 | 655.189 | -0.692777894 | 0.24142 | -2.869596 | 0.00411 | 0.104803968 | CSRNP2 | protein_coding | 46.64 | -0.678660655 | 0.678660655 | ENSG00000110925 | n.s. |
| ENSG00000115170 | 243.213 | -0.709310487 | 0.249842 | -2.839031 | 0.004525 | 0.110509521 | ACVR1 | protein_coding | 40.31 | -0.678526627 | 0.678526627 | ENSG00000115170 | n.s. |
| ENSG00000238250 | 20.75893 | -0.98234211 | 0.400989 | -2.449799 | 0.014294 | 0.20432188 | ST6GAL2-IT1 | sense_intronic | 39.64 | -0.67750674 | 0.67750674 | ENSG00000238250 | n.s. |
| ENSG00000040199 | 388.8424 | -0.712025524 | 0.252126 | -2.824085 | 0.004742 | 0.111902762 | PHLPP2 | protein_coding | 41.82 | -0.677249623 | 0.677249623 | ENSG00000040199 | n.s. |
| ENSG00000276365 | 3.711824 | -1.540342172 | 0.756818 | -2.035288 | 0.041822 | 0.363474721 | MIR145 | miRNA | 47.73 | -0.677020108 | 0.677020108 | ENSG00000276365 | n.s. |
| ENSG00000264885 | 200.6267 | -0.695773287 | 0.243372 | -2.858885 | 0.004251 | 0.106650879 | | sense_intronic | 46.1 | -0.676316378 | 0.676316378 | ENSG00000264885 | n.s. |
| ENSG00000261056 | 167.2258 | -0.728499158 | 0.260918 | -2.792067 | 0.005237 | 0.11804641 | | transcribed_processed_pseudog | 45.32 | -0.676008767 | 0.676008767 | ENSG00000261056 | n.s. |
| ENSG00000232906 | 82.79604 | -0.846559481 | 0.323492 | -2.616939 | 0.008872 | 0.159803295 | | processed_pseudogene | 33.95 | -0.67421205 | 0.67421205 | ENSG00000232906 | n.s. |
| ENSG00000229235 | 17.08414 | -1.110180508 | 0.475715 | -2.333711 | 0.019611 | 0.247135688 | RPL37P18 | processed_pseudogene | 56.79 | -0.673951214 | 0.673951214 | ENSG00000229235 | n.s. |
| ENSG00000255571 | 758.1176 | -0.718489105 | 0.256124 | -2.805236 | 0.005028 | 0.115709901 | MIR9-3HG | lincRNA | 53.82 | -0.672958077 | 0.672958077 | ENSG00000255571 | n.s. |
| ENSG00000225030 | 26.31439 | -0.888215732 | 0.347649 | -2.554924 | 0.010621 | 0.175835251 | | sense_intronic | 52.3 | -0.670508776 | 0.670508776 | ENSG00000225030 | n.s. |
| ENSG00000119737 | 114.4329 | -0.853781101 | 0.328508 | -2.598964 | 0.009351 | 0.163964393 | GPR75 | protein_coding | 44.92 | -0.670431998 | 0.670431998 | ENSG00000119737 | n.s. |
| ENSG00000184260 | 1877.33 | -0.61180586 | 0.204369 | -2.993641 | 0.002757 | 0.0814896 | HIST2H2AC | protein_coding | 62.82 | -0.666194062 | 0.666194062 | ENSG00000184260 | n.s. |
| ENSG00000208308 | 19.26275 | -0.980078263 | 0.402299 | -2.436196 | 0.014843 | 0.209375753 | SNORA40B | snoRNA | 32.03 | -0.665545287 | 0.665545287 | ENSG00000208308 | n.s. |
| ENSG00000266297 | 10.72983 | -1.176807066 | 0.520276 | -2.261892 | 0.023704 | 0.27210013 | MIR744 | miRNA | 54.08 | -0.665215202 | 0.665215202 | ENSG00000266297 | n.s. |
| ENSG00000198121 | 634.0187 | -0.700429014 | 0.248443 | -2.819273 | 0.004813 | 0.11285058 | LPAR1 | protein_coding | 38.26 | -0.663653832 | 0.663653832 | ENSG00000198121 | n.s. |
| ENSG00000222522 | 8.307713 | -1.141099365 | 0.499685 | -2.283637 | 0.022393 | 0.262880506 | RNU6-519P | snRNA | 39.81 | -0.662113343 | 0.662113343 | ENSG00000222522 | n.s. |
| ENSG00000222266 | 4.174381 | -1.452388264 | 0.701538 | -2.070291 | 0.038425 | 0.350082264 | RNU6-757P | snRNA | 38 | -0.662041985 | 0.662041985 | ENSG00000222266 | n.s. |

| | | | | | | | | | | | | | |
|---|---|---|---|---|---|---|---|---|---|---|---|---|---|
| ENSG00000131873 | 1299.412 | -0.716714951 | 0.257082 | -2.787882 | 0.005305 | 0.11933258 | CHSY1 | protein_coding | 43.36 | -0.661700607 | 0.661700607 | ENSG00000131873 | n.s. |
| ENSG00000234788 | 80.01207 | -0.811735238 | 0.3071 | -2.643232 | 0.008212 | 0.154088881 | HSPA8P3 | processed_pseudogene | 45.25 | -0.659314656 | 0.659314656 | ENSG00000234788 | n.s. |
| ENSG00000167632 | 246.9614 | -0.696094581 | 0.247146 | -2.816536 | 0.004854 | 0.113024587 | TRAPPC9 | protein_coding | 47.51 | -0.659081193 | 0.659081193 | ENSG00000167632 | n.s. |
| ENSG00000265981 | 62.81255 | 0.793630268 | 0.29712 | 2.6710727 | 0.007561 | 0.147918794 | MIR544B | miRNA | 39.74 | 0.658694586 | 0.658694586 | ENSG00000265981 | n.s. |
| ENSG00000188315 | 186.9402 | -0.709560265 | 0.254044 | -2.793062 | 0.005221 | 0.118034481 | C3orf62 | protein_coding | 48.07 | -0.658465614 | 0.658465614 | ENSG00000188315 | n.s. |
| ENSG00000272438 | 28.90448 | -0.891973022 | 0.354361 | -2.517128 | 0.011832 | 0.183252281 | | lincRNA | 61.47 | -0.657340062 | 0.657340062 | ENSG00000272438 | n.s. |
| ENSG00000182841 | 539.4686 | -0.642154352 | 0.220076 | -2.917875 | 0.003524 | 0.095163761 | RRP7BP | cribed_unprocessed_pseudc | 54.88 | -0.655978909 | 0.655978909 | ENSG00000182841 | n.s. |
| ENSG00000279672 | 65.04436 | -0.772777265 | 0.286838 | -2.694128 | 0.007057 | 0.141787631 | | TEC | 62.07 | -0.655594598 | 0.655594598 | ENSG00000279672 | n.s. |
| ENSG00000132688 | 3572.082 | -0.65887085 | 0.227834 | -2.891889 | 0.003829 | 0.101168416 | NES | protein_coding | 59 | -0.655546879 | 0.655546879 | ENSG00000132688 | n.s. |
| ENSG00000260350 | 35.83307 | -0.865643144 | 0.338013 | -2.560976 | 0.010438 | 0.174908566 | | antisense | 57.21 | -0.655455397 | 0.655455397 | ENSG00000260350 | n.s. |
| ENSG00000183779 | 181.9959 | -0.686733821 | 0.242323 | -2.833964 | 0.004597 | 0.111136994 | ZNF703 | protein_coding | 65.89 | -0.655241138 | 0.655241138 | ENSG00000183779 | n.s. |
| ENSG00000075461 | 251.8407 | -0.718356694 | 0.25898 | -2.77379 | 0.005541 | 0.122645009 | CACNG4 | protein_coding | 52.37 | -0.654674459 | 0.654674459 | ENSG00000075461 | n.s. |
| ENSG00000253816 | 47.28265 | -0.823903944 | 0.315504 | -2.61139 | 0.009018 | 0.160756371 | | unprocessed_pseudogene | 40.49 | -0.654041157 | 0.654041157 | ENSG00000253816 | n.s. |
| ENSG00000131797 | 308.9444 | -0.703592359 | 0.252011 | -2.791914 | 0.00524 | 0.11804641 | CLUHP3 | cribed_unprocessed_pseudc | 55.17 | -0.652896572 | 0.652896572 | ENSG00000131797 | n.s. |
| ENSG00000201247 | 13.05811 | 1.023862601 | 0.43093 | 2.3759364 | 0.017504 | 0.230963869 | SNORD114-13 | snoRNA | 45.95 | 0.651643448 | 0.651643448 | ENSG00000201247 | n.s. |
| ENSG00000196787 | 1879.403 | -0.620624477 | 0.210197 | -2.95258 | 0.003151 | 0.089147501 | HIST1H2AG | protein_coding | 46.8 | -0.651587945 | 0.651587945 | ENSG00000196787 | n.s. |
| ENSG00000253476 | 17.02039 | -0.958075414 | 0.393208 | -2.436559 | 0.014828 | 0.209375753 | | lincRNA | 40.57 | -0.650603733 | 0.650603733 | ENSG00000253476 | n.s. |
| ENSG00000271265 | 26.97047 | -0.878246025 | 0.347202 | -2.529496 | 0.011423 | 0.181687298 | | lincRNA | 36.1 | -0.650495256 | 0.650495256 | ENSG00000271265 | n.s. |
| ENSG00000232450 | 25.71539 | -0.881368959 | 0.350178 | -2.516919 | 0.011839 | 0.183252281 | | transcribed_processed_pseudog | 54.14 | -0.649525392 | 0.649525392 | ENSG00000232450 | n.s. |
| ENSG00000160293 | 490.1683 | -0.69013464 | 0.245766 | -2.808099 | 0.004983 | 0.115066504 | VAV2 | protein_coding | 56.31 | -0.648071681 | 0.648071681 | ENSG00000160293 | n.s. |
| ENSG00000241395 | 21.3064 | -0.930432253 | 0.378838 | -2.456019 | 0.014049 | 0.201855602 | RN7SL344P | misc_RNA | 55.07 | -0.646612448 | 0.646612448 | ENSG00000241395 | n.s. |
| ENSG00000272330 | 6.427776 | -1.436161339 | 0.698697 | -2.055484 | 0.039832 | 0.355503043 | | sense_intronic | 28.48 | -0.645061455 | 0.645061455 | ENSG00000272330 | n.s. |
| ENSG00000243173 | 28.92566 | -0.843341714 | 0.327692 | -2.573582 | 0.010065 | 0.171947811 | RN7SL861P | misc_RNA | 52.28 | -0.644821899 | 0.644821899 | ENSG00000243173 | n.s. |
| ENSG00000222255 | 4.719863 | -1.61739483 | 0.828881 | -1.951299 | 0.051022 | 0.399465302 | RNU6-101P | snRNA | 41.35 | -0.644565706 | 0.644565706 | ENSG00000222255 | n.s. |
| ENSG00000231729 | 12.63167 | -1.058446095 | 0.452961 | -2.336729 | 0.019453 | 0.246449406 | ARHGEF9-IT1 | sense_intronic | 43.53 | -0.64382336 | 0.64382336 | ENSG00000231729 | n.s. |
| ENSG00000274641 | 702.0101 | -0.604556161 | 0.203918 | -2.964701 | 0.00303 | 0.08687934 | HIST1H2BO | protein_coding | 59.32 | -0.64148456 | 0.64148456 | ENSG00000274641 | n.s. |
| ENSG00000151687 | 79.26155 | -0.774505257 | 0.290845 | -2.662945 | 0.007746 | 0.14916674 | ANKAR | protein_coding | 37.84 | -0.63999538 | 0.63999538 | ENSG00000151687 | n.s. |
| ENSG00000243609 | 55.78519 | -0.795644904 | 0.302462 | -2.630563 | 0.008524 | 0.156990291 | | processed_pseudogene | 55.79 | -0.639799713 | 0.639799713 | ENSG00000243609 | n.s. |
| ENSG00000115902 | 619.249 | -0.702524988 | 0.253385 | -2.772559 | 0.005562 | 0.122914497 | SLC1A4 | protein_coding | 45.52 | -0.639576565 | 0.639576565 | ENSG00000115902 | n.s. |
| ENSG00000252782 | 50.01767 | 0.809241041 | 0.310534 | 2.6059686 | 0.009161 | 0.162077988 | RNU6-341P | snRNA | 43.75 | 0.639523743 | 0.639523743 | ENSG00000252782 | n.s. |
| ENSG00000267766 | 11.29945 | -1.04737899 | 0.447514 | -2.340439 | 0.019261 | 0.245220122 | | sense_intronic | 45.41 | -0.639366111 | 0.639366111 | ENSG00000267766 | n.s. |
| ENSG00000201610 | 4.270781 | -1.608715028 | 0.825553 | -1.948652 | 0.051337 | 0.400783303 | RNA5SP84 | rRNA | 57.94 | -0.638805261 | 0.638805261 | ENSG00000201610 | n.s. |

| | | | | | | | | | | | | | |
|---|---|---|---|---|---|---|---|---|---|---|---|---|---|
| ENSG00000131849 | 292.8114 | -0.676501804 | 0.240801 | -2.809375 | 0.004964 | 0.114850754 | ZNF132 | protein_coding | 48.36 | -0.635821147 | 0.635821147 | ENSG00000131849 | n.s. |
| ENSG00000267419 | 46.57069 | -0.802214115 | 0.307708 | -2.60706 | 0.009132 | 0.161767835 | | transcribed_unprocessed_pseudc | 41.25 | -0.634637866 | 0.634637866 | ENSG00000267419 | n.s. |
| ENSG00000144191 | 1671.918 | -0.723786649 | 0.26534 | -2.727774 | 0.006376 | 0.132862373 | CNGA3 | protein_coding | 47.34 | -0.634469927 | 0.634469927 | ENSG00000144191 | n.s. |
| ENSG00000256019 | 68.1885 | -0.79684466 | 0.304987 | -2.612715 | 0.008983 | 0.160685001 | | processed_pseudogene | 39.68 | -0.632714311 | 0.632714311 | ENSG00000256019 | n.s. |
| ENSG00000253106 | 205.0493 | 0.713744985 | 0.260152 | 2.7435688 | 0.006078 | 0.130381308 | | antisense | 40.87 | 0.631510619 | 0.631510619 | ENSG00000253106 | n.s. |
| ENSG00000274012 | 22225.71 | -0.58412203 | 0.195674 | -2.985187 | 0.002834 | 0.083245819 | RN7SL2 | misc_RNA | 62.67 | -0.630640089 | 0.630640089 | ENSG00000274012 | n.s. |
| ENSG00000162490 | 260.5677 | -0.722080859 | 0.265373 | -2.720999 | 0.006508 | 0.134505157 | DRAXIN | protein_coding | 53.39 | -0.629120938 | 0.629120938 | ENSG00000162490 | n.s. |
| ENSG00000198799 | 1049.526 | -0.715966107 | 0.262096 | -2.731696 | 0.006301 | 0.132435959 | LRIG2 | protein_coding | 41.19 | -0.628614003 | 0.628614003 | ENSG00000198799 | n.s. |
| ENSG00000231241 | 11.78932 | -1.048504901 | 0.452285 | -2.318241 | 0.020436 | 0.252377705 | RPS3AP3 | processed_pseudogene | 35.95 | -0.626952472 | 0.626952472 | ENSG00000231241 | n.s. |
| ENSG00000173621 | 415.6669 | -0.707437921 | 0.257792 | -2.74422 | 0.006065 | 0.130381308 | LRFN4 | protein_coding | 70.38 | -0.625930225 | 0.625930225 | ENSG00000173621 | n.s. |
| ENSG00000235529 | 56.74894 | -0.828747744 | 0.324194 | -2.556337 | 0.010578 | 0.175750036 | AGAP1-IT1 | sense_intronic | 50.66 | -0.625791214 | 0.625791214 | ENSG00000235529 | n.s. |
| ENSG00000269189 | 49.2324 | -0.789979942 | 0.302962 | -2.607519 | 0.00912 | 0.161757115 | | sense_intronic | 40.82 | -0.624982051 | 0.624982051 | ENSG00000269189 | n.s. |
| ENSG00000271200 | 17.42969 | -1.040713788 | 0.448031 | -2.322861 | 0.020187 | 0.251394113 | | lincRNA | 61.12 | -0.624058712 | 0.624058712 | ENSG00000271200 | n.s. |
| ENSG00000272469 | 16.66702 | -1.019442535 | 0.435603 | -2.340302 | 0.019268 | 0.245220122 | RN7SL760P | misc_RNA | 54.44 | -0.622312472 | 0.622312472 | ENSG00000272469 | n.s. |
| ENSG00000123094 | 562.013 | -0.709618249 | 0.260239 | -2.726791 | 0.006395 | 0.132899384 | RASSF8 | protein_coding | 38.09 | -0.621964096 | 0.621964096 | ENSG00000123094 | n.s. |
| ENSG00000146006 | 969.4237 | 0.665778009 | 0.23758 | 2.8023343 | 0.005073 | 0.116371214 | LRRTM2 | protein_coding | 38.27 | 0.621939481 | 0.621939481 | ENSG00000146006 | n.s. |
| ENSG00000272677 | 73.84153 | -0.76032917 | 0.286905 | -2.650107 | 0.008047 | 0.152426182 | | antisense | 45.19 | -0.621143638 | 0.621143638 | ENSG00000272677 | n.s. |
| ENSG00000274997 | 806.3903 | -0.635913726 | 0.222012 | -2.864319 | 0.004179 | 0.105791239 | HIST1H2AH | protein_coding | 61.76 | -0.620365823 | 0.620365823 | ENSG00000274997 | n.s. |
| ENSG00000166716 | 859.0121 | -0.642979212 | 0.225759 | -2.848083 | 0.004398 | 0.108578166 | ZNF592 | protein_coding | 45.9 | -0.619997527 | 0.619997527 | ENSG00000166716 | n.s. |
| ENSG00000123908 | 1578.321 | -0.652119759 | 0.231464 | -2.817374 | 0.004842 | 0.112918712 | AGO2 | protein_coding | 51.18 | -0.617710063 | 0.617710063 | ENSG00000123908 | n.s. |
| ENSG00000175274 | 1361.893 | -0.649315126 | 0.229923 | -2.824058 | 0.004742 | 0.111902762 | TP53I11 | protein_coding | 54.85 | -0.617602052 | 0.617602052 | ENSG00000175274 | n.s. |
| ENSG00000101680 | 395.8464 | -0.649204997 | 0.229808 | -2.824983 | 0.004728 | 0.111902762 | LAMA1 | protein_coding | 43.14 | -0.617497302 | 0.617497302 | ENSG00000101680 | n.s. |
| ENSG00000222000 | 114.3233 | -0.702799779 | 0.25732 | -2.731225 | 0.00631 | 0.132435959 | | antisense | 43.35 | -0.617054045 | 0.617054045 | ENSG00000222000 | n.s. |
| ENSG00000152292 | 27.43826 | -0.955152751 | 0.39963 | -2.390092 | 0.016844 | 0.225935655 | SH2D6 | protein_coding | 54.62 | -0.617043222 | 0.617043222 | ENSG00000152292 | n.s. |
| ENSG00000204580 | 315.2936 | -0.669749473 | 0.240678 | -2.782759 | 0.00539 | 0.120648285 | DDR1 | protein_coding | 55.25 | -0.615150725 | 0.615150725 | ENSG00000204580 | n.s. |
| ENSG00000264049 | 8.239763 | -1.202913925 | 0.555185 | -2.16669 | 0.030258 | 0.308894909 | MIR4737 | miRNA | 62.96 | -0.613713752 | 0.613713752 | ENSG00000264049 | n.s. |
| ENSG00000243373 | 19.71684 | -0.929392882 | 0.385564 | -2.410478 | 0.015932 | 0.218753659 | RN7SL173P | misc_RNA | 51.71 | -0.613440822 | 0.613440822 | ENSG00000243373 | n.s. |
| ENSG00000226819 | 219.4844 | -0.69867687 | 0.255854 | -2.73076 | 0.006319 | 0.132435959 | MEIS1-AS3 | antisense | 42.16 | -0.613434155 | 0.613434155 | ENSG00000226819 | n.s. |
| ENSG00000253284 | 377.1865 | -0.732779778 | 0.273482 | -2.679446 | 0.007374 | 0.145843776 | | sense_intronic | 38.08 | -0.612686039 | 0.612686039 | ENSG00000253284 | n.s. |
| ENSG00000221650 | 11.17149 | -1.100870881 | 0.490791 | -2.243055 | 0.024893 | 0.27771574 | MIR1267 | miRNA | 44.87 | -0.612524013 | 0.612524013 | ENSG00000221650 | n.s. |
| ENSG00000224003 | 218.4177 | -0.743363252 | 0.279763 | -2.657122 | 0.007881 | 0.150725964 | YES1P1 | processed_pseudogene | 42.44 | -0.610904788 | 0.610904788 | ENSG00000224003 | n.s. |
| ENSG00000222958 | 3.368032 | -1.496409659 | 0.758747 | -1.972211 | 0.048585 | 0.391129217 | MIR1913 | miRNA | 63.75 | -0.610055902 | 0.610055902 | ENSG00000222958 | n.s. |

| | | | | | | | | | | | | | |
|---|---|---|---|---|---|---|---|---|---|---|---|---|---|
| ENSG00000266017 | 15.46812 | -1.014410353 | 0.436371 | -2.324653 | 0.020091 | 0.25045014 | MIR4477B | miRNA | 41.98 | -0.609943361 | 0.609943361 | ENSG00000266017 | n.s. |
| ENSG00000124302 | 523.226 | -0.734371882 | 0.274856 | -2.671844 | 0.007544 | 0.147918794 | CHST8 | protein_coding | 50.64 | -0.609511509 | 0.609511509 | ENSG00000124302 | n.s. |
| ENSG00000130749 | 547.3335 | -0.636693766 | 0.224265 | -2.839021 | 0.004525 | 0.110509521 | ZC3H4 | protein_coding | 51.95 | -0.60906145 | 0.60906145 | ENSG00000130749 | n.s. |
| ENSG00000206622 | 16.61318 | 1.026122915 | 0.444038 | 2.3108908 | 0.020839 | 0.255099509 | SNORA69 | snoRNA | 36.36 | 0.608788851 | 0.608788851 | ENSG00000206622 | n.s. |
| ENSG00000214140 | 103.6433 | -0.848151974 | 0.340843 | -2.488395 | 0.012832 | 0.191814442 | PRCD | protein_coding | 55 | -0.608225639 | 0.608225639 | ENSG00000214140 | n.s. |
| ENSG00000165449 | 393.1317 | -0.688309342 | 0.25107 | -2.741503 | 0.006116 | 0.130768951 | SLC16A9 | protein_coding | 40.06 | -0.60811811 | 0.60811811 | ENSG00000165449 | n.s. |
| ENSG00000167625 | 216.1325 | -0.661614066 | 0.237736 | -2.78298 | 0.005386 | 0.120648285 | ZNF526 | protein_coding | 54.5 | -0.607678526 | 0.607678526 | ENSG00000167625 | n.s. |
| ENSG00000167548 | 235.0711 | -0.671901715 | 0.243097 | -2.763921 | 0.005711 | 0.125222476 | KMT2D | protein_coding | 54.13 | -0.606268818 | 0.606268818 | ENSG00000167548 | n.s. |
| ENSG00000196924 | 3744.262 | -0.636480352 | 0.224921 | -2.829798 | 0.004658 | 0.111596326 | FLNA | protein_coding | 62.28 | -0.606152128 | 0.606152128 | ENSG00000196924 | n.s. |
| ENSG00000178386 | 151.9602 | -0.755051002 | 0.287547 | -2.625832 | 0.008644 | 0.157676123 | ZNF223 | protein_coding | 43.75 | -0.605727637 | 0.605727637 | ENSG00000178386 | n.s. |
| ENSG00000185950 | 241.8011 | -0.694400813 | 0.255214 | -2.720858 | 0.006511 | 0.134505157 | IRS2 | protein_coding | 46.75 | -0.605004391 | 0.605004391 | ENSG00000185950 | n.s. |
| ENSG00000164438 | 113.5445 | -0.773665167 | 0.298505 | -2.5918 | 0.009548 | 0.165748533 | TLX3 | protein_coding | 66.78 | -0.603884584 | 0.603884584 | ENSG00000164438 | n.s. |
| ENSG00000183598 | 899.9099 | -0.641050222 | 0.228014 | -2.811457 | 0.004932 | 0.114441759 | HIST2H3D | protein_coding | 67.4 | -0.603494599 | 0.603494599 | ENSG00000183598 | n.s. |
| ENSG00000140479 | 43.53725 | -0.850525207 | 0.343408 | -2.47672 | 0.01326 | 0.1953577 | PCSK6 | protein_coding | 48.11 | -0.603166507 | 0.603166507 | ENSG00000140479 | n.s. |
| ENSG00000259767 | 53.43751 | -0.838764418 | 0.336536 | -2.492347 | 0.01269 | 0.191218231 | | sense_intronic | 42.08 | -0.602627661 | 0.602627661 | ENSG00000259767 | n.s. |
| ENSG00000201690 | 3.702575 | -1.485263094 | 0.755681 | -1.965463 | 0.049361 | 0.393439254 | RNY1P2 | misc_RNA | 42.74 | -0.601713219 | 0.601713219 | ENSG00000201690 | n.s. |
| ENSG00000277728 | 30.1637 | -0.808035544 | 0.319953 | -2.525479 | 0.011554 | 0.181859152 | | antisense | 44.61 | -0.598160301 | 0.598160301 | ENSG00000277728 | n.s. |
| ENSG00000263968 | 53.51775 | -0.761322922 | 0.292806 | -2.600092 | 0.00932 | 0.163921434 | RN7SL381P | misc_RNA | 56.71 | -0.597915811 | 0.597915811 | ENSG00000263968 | n.s. |
| ENSG00000279656 | 36.9589 | -0.801649346 | 0.315704 | -2.539242 | 0.011109 | 0.179645377 | | TEC | 46.87 | -0.597696889 | 0.597696889 | ENSG00000279656 | n.s. |
| ENSG00000149256 | 435.0787 | -0.623335549 | 0.219364 | -2.841557 | 0.004489 | 0.110239383 | TENM4 | protein_coding | 44.45 | -0.596945532 | 0.596945532 | ENSG00000149256 | n.s. |
| ENSG00000207759 | 8.916595 | -1.098012975 | 0.494476 | -2.22056 | 0.026381 | 0.286058985 | MIR181A1 | miRNA | 41.82 | -0.59681881 | 0.59681881 | ENSG00000207759 | n.s. |
| ENSG00000252427 | 8.35953 | -1.149861069 | 0.527652 | -2.179203 | 0.029317 | 0.302800461 | | snoRNA | 45.37 | -0.596597904 | 0.596597904 | ENSG00000252427 | n.s. |
| ENSG00000272574 | 31.72867 | -0.891938342 | 0.36857 | -2.419996 | 0.015521 | 0.21544805 | | sense_intronic | 42.18 | -0.594617325 | 0.594617325 | ENSG00000272574 | n.s. |
| ENSG00000267681 | 13.17859 | -1.054149089 | 0.466859 | -2.257962 | 0.023948 | 0.272897252 | | transcribed_processed_pseudog | 47 | -0.594540969 | 0.594540969 | ENSG00000267681 | n.s. |
| ENSG00000238259 | 24.34387 | -0.891624837 | 0.36839 | -2.420329 | 0.015506 | 0.21544805 | | processed_pseudogene | 46.67 | -0.594408324 | 0.594408324 | ENSG00000238259 | n.s. |
| ENSG00000241785 | 30.63575 | -0.870830147 | 0.356901 | -2.439975 | 0.014688 | 0.208099767 | RN7SL390P | misc_RNA | 50.7 | -0.593669649 | 0.593669649 | ENSG00000241785 | n.s. |
| ENSG00000280239 | 123.1104 | -0.706217019 | 0.262984 | -2.685399 | 0.007244 | 0.144556561 | | TEC | 54.27 | -0.593195596 | 0.593195596 | ENSG00000280239 | n.s. |
| ENSG00000197903 | 1472.966 | -0.578494526 | 0.198014 | -2.921481 | 0.003484 | 0.094734017 | HIST1H2BK | protein_coding | 61.15 | -0.592085703 | 0.592085703 | ENSG00000197903 | n.s. |
| ENSG00000265820 | 2.382168 | -1.924305521 | 1.124916 | -1.710622 | 0.087151 | 0.492652156 | MIR3177 | miRNA | 63.41 | -0.591646229 | 0.591646229 | ENSG00000265820 | n.s. |
| ENSG00000239197 | 19.62546 | -0.943547147 | 0.399605 | -2.361197 | 0.018216 | 0.236587468 | | snoRNA | 41.96 | -0.590668311 | 0.590668311 | ENSG00000239197 | n.s. |
| ENSG00000230735 | 16.52453 | -1.005255632 | 0.438052 | -2.29483 | 0.021743 | 0.259183371 | | sense_intronic | 38.58 | -0.589474731 | 0.589474731 | ENSG00000230735 | n.s. |
| ENSG00000234167 | 14.03456 | -1.032230726 | 0.454511 | -2.271077 | 0.023142 | 0.26850849 | ELOBP4 | processed_pseudogene | 58.45 | -0.589447075 | 0.589447075 | ENSG00000234167 | n.s. |

| | | | | | | | | | | | | | |
|---|---|---|---|---|---|---|---|---|---|---|---|---|---|
| ENSG00000186260 | 267.243 | -0.710644551 | 0.266591 | -2.665669 | 0.007684 | 0.148580262 | MKL2 | protein_coding | 38.53 | -0.588441318 | 0.588441318 | ENSG00000186260 | n.s. |
| ENSG00000227986 | 58.82462 | -0.800107894 | 0.318548 | -2.511736 | 0.012014 | 0.184217437 | TRIM60P18 | unprocessed_pseudogene | 40.73 | -0.587814679 | 0.587814679 | ENSG00000227986 | n.s. |
| ENSG00000146426 | 289.4563 | -0.67451756 | 0.247902 | -2.7209 | 0.00651 | 0.134505157 | TIAM2 | protein_coding | 43.49 | -0.587680888 | 0.587680888 | ENSG00000146426 | n.s. |
| ENSG00000252438 | 6.832492 | -1.288558845 | 0.621792 | -2.072331 | 0.038235 | 0.350082264 | | snoRNA | 39.73 | -0.587363639 | 0.587363639 | ENSG00000252438 | n.s. |
| ENSG00000271141 | 95.22506 | -0.788099925 | 0.310888 | -2.534993 | 0.011245 | 0.180009224 | | lincRNA | 38.31 | -0.586902143 | 0.586902143 | ENSG00000271141 | n.s. |
| ENSG00000212447 | 3.996866 | -1.401690706 | 0.701402 | -1.998414 | 0.045672 | 0.381456593 | SNORD90 | snoRNA | 33.33 | -0.586684478 | 0.586684478 | ENSG00000212447 | n.s. |
| ENSG00000109787 | 629.8989 | -0.669171507 | 0.245295 | -2.728024 | 0.006371 | 0.132862373 | KLF3 | protein_coding | 41.91 | -0.586594403 | 0.586594403 | ENSG00000109787 | n.s. |
| ENSG00000139174 | 225.2281 | 0.618018537 | 0.219623 | 2.8139969 | 0.004893 | 0.113730911 | PRICKLE1 | protein_coding | 41.36 | 0.583484578 | 0.583484578 | ENSG00000139174 | n.s. |
| ENSG00000254500 | 12.08379 | -0.991501597 | 0.431115 | -2.299856 | 0.021456 | 0.258017317 | RANP3 | processed_pseudogene | 41.67 | -0.583351099 | 0.583351099 | ENSG00000254500 | n.s. |
| ENSG00000253475 | 23.7697 | -0.933367058 | 0.395736 | -2.358561 | 0.018346 | 0.237609746 | | sense_intronic | 47.61 | -0.582547748 | 0.582547748 | ENSG00000253475 | n.s. |
| ENSG00000104728 | 120.2716 | -0.776653405 | 0.305064 | -2.54587 | 0.010901 | 0.177923733 | ARHGEF10 | protein_coding | 50 | -0.582308409 | 0.582308409 | ENSG00000104728 | n.s. |
| ENSG00000282100 | 131.1766 | -0.768778644 | 0.300196 | -2.560922 | 0.010439 | 0.174908566 | | processed_pseudogene | 45.94 | -0.582110671 | 0.582110671 | ENSG00000282100 | n.s. |
| ENSG00000049540 | 121.5072 | -0.731274536 | 0.279679 | -2.614694 | 0.008931 | 0.160029118 | ELN | protein_coding | 54.95 | -0.581948999 | 0.581948999 | ENSG00000049540 | n.s. |
| ENSG00000228886 | 13.39989 | -0.990741146 | 0.431275 | -2.297236 | 0.021605 | 0.258843947 | | lincRNA | 44.26 | -0.581527391 | 0.581527391 | ENSG00000228886 | n.s. |
| ENSG00000240616 | 62.71559 | 0.793809233 | 0.316406 | 2.5088318 | 0.012113 | 0.185127237 | RPS6P25 | processed_pseudogene | 45.86 | 0.581488825 | 0.581488825 | ENSG00000240616 | n.s. |
| ENSG00000200897 | 7.339127 | -1.297981497 | 0.631983 | -2.053824 | 0.039993 | 0.356478089 | | snoRNA | 45.68 | -0.581453084 | 0.581453084 | ENSG00000200897 | n.s. |
| ENSG00000199072 | 58.07613 | 0.741873176 | 0.285912 | 2.5947575 | 0.009466 | 0.164910296 | MIRLET7F1 | miRNA | 40.23 | 0.580702883 | 0.580702883 | ENSG00000199072 | n.s. |
| ENSG00000120149 | 216.8336 | -0.762209263 | 0.296903 | -2.567203 | 0.010252 | 0.173441096 | MSX2 | protein_coding | 47.07 | -0.579925386 | 0.579925386 | ENSG00000120149 | n.s. |
| ENSG00000183798 | 152.8614 | -0.746367403 | 0.288543 | -2.586676 | 0.009691 | 0.167712137 | EMILIN3 | protein_coding | 62.66 | -0.578759786 | 0.578759786 | ENSG00000183798 | n.s. |
| ENSG00000279281 | 302.4588 | -0.719245846 | 0.273371 | -2.63102 | 0.008513 | 0.156990291 | | TEC | 45.42 | -0.578365152 | 0.578365152 | ENSG00000279281 | n.s. |
| ENSG00000144791 | 502.682 | -0.610394928 | 0.216556 | -2.818646 | 0.004823 | 0.11285058 | LIMD1 | protein_coding | 46.55 | -0.578346877 | 0.578346877 | ENSG00000144791 | n.s. |
| ENSG00000135111 | 3022.267 | -0.62309697 | 0.223048 | -2.793562 | 0.005213 | 0.118034481 | TBX3 | protein_coding | 47.27 | -0.578228446 | 0.578228446 | ENSG00000135111 | n.s. |
| ENSG00000170017 | 347.5441 | -0.780744603 | 0.309696 | -2.521006 | 0.011702 | 0.182089199 | ALCAM | protein_coding | 35.29 | -0.577529129 | 0.577529129 | ENSG00000170017 | n.s. |
| ENSG00000177519 | 239.3062 | -0.721586385 | 0.275511 | -2.619084 | 0.008817 | 0.159007629 | RPRM | protein_coding | 59.62 | -0.576245927 | 0.576245927 | ENSG00000177519 | n.s. |
| ENSG00000259926 | 23.91056 | -0.903648746 | 0.380378 | -2.375659 | 0.017518 | 0.230963869 | | sense_intronic | 40.21 | -0.575132625 | 0.575132625 | ENSG00000259926 | n.s. |
| ENSG00000179988 | 338.0648 | 0.712469755 | 0.270492 | 2.6339791 | 0.008439 | 0.156104918 | PSTK | protein_coding | 40.38 | 0.574666288 | 0.574666288 | ENSG00000179988 | n.s. |
| ENSG00000273723 | 333.0081 | 0.602669839 | 0.21318 | 2.8270525 | 0.004698 | 0.111596326 | | lincRNA | 38.77 | 0.573952683 | 0.573952683 | ENSG00000273723 | n.s. |
| ENSG00000152495 | 2226.791 | -0.686093346 | 0.256011 | -2.679941 | 0.007364 | 0.145835185 | CAMK4 | protein_coding | 37.31 | -0.5736685 | 0.5736685 | ENSG00000152495 | n.s. |
| ENSG00000146094 | 26.13925 | -0.898494266 | 0.377718 | -2.378741 | 0.017372 | 0.229913168 | DOK3 | protein_coding | 58.13 | -0.573631223 | 0.573631223 | ENSG00000146094 | n.s. |
| ENSG00000196205 | 1881.561 | 0.801299539 | 0.322399 | 2.4854248 | 0.01294 | 0.192890069 | EEF1A1P5 | processed_pseudogene | 48.31 | 0.572680872 | 0.572680872 | ENSG00000196205 | n.s. |
| ENSG00000244265 | 20.6596 | -1.01190032 | 0.447375 | -2.261862 | 0.023706 | 0.27210013 | SIAH2-AS1 | antisense | 56.64 | -0.57199816 | 0.57199816 | ENSG00000244265 | n.s. |
| ENSG00000254876 | 116.2924 | -0.690481458 | 0.259013 | -2.665819 | 0.00768 | 0.148580262 | | processed_transcript | 40.69 | -0.571745492 | 0.571745492 | ENSG00000254876 | n.s. |

| | | | | | | | | | | | | | |
|---|---|---|---|---|---|---|---|---|---|---|---|---|---|
| ENSG00000239151 | 1.984665 | -1.90200489 | 1.123582 | -1.692804 | 0.090493 | 0.501165391 | RNU7-195P | snRNA | 38.71 | -0.570637467 | 0.570637467 | ENSG00000239151 | n.s. |
| ENSG00000265558 | 28.62198 | -0.871509969 | 0.362821 | -2.402039 | 0.016304 | 0.221694304 | MIR3918 | miRNA | 62.37 | -0.570181445 | 0.570181445 | ENSG00000265558 | n.s. |
| ENSG00000143479 | 202.82 | -0.673328943 | 0.250304 | -2.690048 | 0.007144 | 0.142914438 | DYRK3 | protein_coding | 44.46 | -0.568911713 | 0.568911713 | ENSG00000143479 | n.s. |
| ENSG00000185379 | 439.8785 | -0.685001778 | 0.256466 | -2.670922 | 0.007564 | 0.147918794 | RAD51D | protein_coding | 47.17 | -0.568535476 | 0.568535476 | ENSG00000185379 | n.s. |
| ENSG00000173275 | 173.6005 | -0.717183139 | 0.274886 | -2.60902 | 0.00908 | 0.161460238 | ZNF449 | protein_coding | 37.9 | -0.567962008 | 0.567962008 | ENSG00000173275 | n.s. |
| ENSG00000244657 | 4.37149 | -1.549484595 | 0.830741 | -1.865184 | 0.062156 | 0.430475522 | | transcribed_unprocessed_pseudc | 41.24 | -0.56719122 | 0.56719122 | ENSG00000244657 | n.s. |
| ENSG00000113657 | 820.5003 | -0.6825044 | 0.255755 | -2.668586 | 0.007617 | 0.148477748 | DPYSL3 | protein_coding | 41.43 | -0.565344758 | 0.565344758 | ENSG00000113657 | n.s. |
| ENSG00000198252 | 268.4776 | -0.74180209 | 0.288757 | -2.568953 | 0.010201 | 0.172988151 | STYX | protein_coding | 38.27 | -0.565241058 | 0.565241058 | ENSG00000198252 | n.s. |
| ENSG00000279623 | 16.78982 | -1.060649496 | 0.482171 | -2.199735 | 0.027826 | 0.293497624 | | sense_intronic | 32.71 | -0.564684923 | 0.564684923 | ENSG00000279623 | n.s. |
| ENSG00000265421 | 3.453025 | -1.544950647 | 0.832539 | -1.855709 | 0.063495 | 0.432716997 | MIR4459 | miRNA | 65.15 | -0.562046936 | 0.562046936 | ENSG00000265421 | n.s. |
| ENSG00000253784 | 79.34962 | 0.757616186 | 0.300804 | 2.5186371 | 0.011781 | 0.183065705 | | lincRNA | 42.76 | 0.558660879 | 0.558660879 | ENSG00000253784 | n.s. |
| ENSG00000197933 | 213.7703 | -0.743499843 | 0.291695 | -2.548894 | 0.010807 | 0.177428307 | ZNF823 | protein_coding | 44.81 | -0.558351349 | 0.558351349 | ENSG00000197933 | n.s. |
| ENSG00000226609 | 38.15428 | 0.817577812 | 0.334771 | 2.442197 | 0.014598 | 0.207822555 | | antisense | 43.09 | 0.557839326 | 0.557839326 | ENSG00000226609 | n.s. |
| ENSG00000136010 | 1217.673 | -0.664215664 | 0.247546 | -2.683206 | 0.007292 | 0.145105366 | ALDH1L2 | protein_coding | 39.68 | -0.556822968 | 0.556822968 | ENSG00000136010 | n.s. |
| ENSG00000276903 | 774.9109 | -0.606324109 | 0.218009 | -2.781193 | 0.005416 | 0.120843585 | HIST1H2AL | protein_coding | 63.87 | -0.556469957 | 0.556469957 | ENSG00000276903 | n.s. |
| ENSG00000256436 | 189.0177 | -0.75201747 | 0.298218 | -2.521706 | 0.011679 | 0.182089199 | TAS2R31 | protein_coding | 40.45 | -0.556279215 | 0.556279215 | ENSG00000256436 | n.s. |
| ENSG00000207637 | 5.24112 | -1.324239362 | 0.662357 | -1.999284 | 0.045578 | 0.381356862 | MIR595 | miRNA | 53.12 | -0.554417221 | 0.554417221 | ENSG00000207637 | n.s. |
| ENSG00000124126 | 264.6526 | -0.661584884 | 0.247029 | -2.678166 | 0.007403 | 0.146129673 | PREX1 | protein_coding | 50.46 | -0.55259644 | 0.55259644 | ENSG00000124126 | n.s. |
| ENSG00000196267 | 148.4559 | -0.739945077 | 0.291113 | -2.541783 | 0.011029 | 0.179260564 | ZNF836 | protein_coding | 43.18 | -0.552380278 | 0.552380278 | ENSG00000196267 | n.s. |
| ENSG00000284324 | 42.10482 | -0.826527646 | 0.341302 | -2.421693 | 0.015448 | 0.21487333 | MIR3175 | miRNA | 71.43 | -0.551969615 | 0.551969615 | ENSG00000284324 | n.s. |
| ENSG00000011114 | 420.815 | -0.671613707 | 0.252872 | -2.655939 | 0.007909 | 0.150841114 | BTBD7 | protein_coding | 40.08 | -0.551717409 | 0.551717409 | ENSG00000011114 | n.s. |
| ENSG00000152782 | 440.0717 | -0.721576041 | 0.280569 | -2.571835 | 0.010116 | 0.17218396 | PANK1 | protein_coding | 39.91 | -0.551289369 | 0.551289369 | ENSG00000152782 | n.s. |
| ENSG00000112941 | 1038.593 | -0.616714417 | 0.223909 | -2.754313 | 0.005882 | 0.127751781 | PAPD7 | protein_coding | 47.97 | -0.551116378 | 0.551116378 | ENSG00000112941 | n.s. |
| ENSG00000239763 | 23.70523 | -0.887415272 | 0.37691 | -2.354447 | 0.01855 | 0.239365741 | | processed_pseudogene | 50.13 | -0.551029871 | 0.551029871 | ENSG00000239763 | n.s. |
| ENSG00000183873 | 171.083 | -0.661253841 | 0.247379 | -2.673039 | 0.007517 | 0.147610278 | SCN5A | protein_coding | 50.93 | -0.549424841 | 0.549424841 | ENSG00000183873 | n.s. |
| ENSG00000188215 | 269.8357 | -0.647149803 | 0.240128 | -2.695026 | 0.007038 | 0.141610603 | DCUN1D3 | protein_coding | 43.13 | -0.549368204 | 0.549368204 | ENSG00000188215 | n.s. |
| ENSG00000275106 | 34.87125 | -0.765205787 | 0.307529 | -2.48824 | 0.012838 | 0.191814442 | | TEC | 40.38 | -0.548743377 | 0.548743377 | ENSG00000275106 | n.s. |
| ENSG00000272081 | 15.07104 | 1.010873198 | 0.45596 | 2.2170216 | 0.026622 | 0.287074936 | | lincRNA | 54.66 | 0.547898049 | 0.547898049 | ENSG00000272081 | n.s. |
| ENSG00000251348 | 145.2564 | -0.782639557 | 0.317539 | -2.464701 | 0.013713 | 0.199710058 | HSPD1P11 | processed_pseudogene | 40.36 | -0.547534683 | 0.547534683 | ENSG00000251348 | n.s. |
| ENSG00000265210 | 3.526453 | -1.538787481 | 0.837865 | -1.836557 | 0.066275 | 0.440875092 | MIR4486 | miRNA | 57.14 | -0.547322758 | 0.547322758 | ENSG00000265210 | n.s. |
| ENSG00000105327 | 77.10632 | -0.759379312 | 0.304417 | -2.494533 | 0.012612 | 0.190456473 | BBC3 | protein_coding | 59.72 | -0.546908218 | 0.546908218 | ENSG00000105327 | n.s. |
| ENSG00000278535 | 179.9852 | -0.686426747 | 0.262512 | -2.614836 | 0.008927 | 0.160029118 | DHRS11 | protein_coding | 54.49 | -0.546259085 | 0.546259085 | ENSG00000278535 | n.s. |

| | | | | | | | | | | | | | |
|---|---|---|---|---|---|---|---|---|---|---|---|---|---|
| ENSG00000264201 | 1.259779 | -2.090019049 | 1.310561 | -1.594752 | 0.110768 | 0.548299349 | MIR4701 | miRNA | 58.73 | -0.545457916 | 0.545457916 | ENSG00000264201 | n.s. |
| ENSG00000186684 | 207.7068 | -0.695999417 | 0.268172 | -2.595352 | 0.009449 | 0.164867498 | CYP27C1 | protein_coding | 45.93 | -0.544873551 | 0.544873551 | ENSG00000186684 | n.s. |
| ENSG00000150995 | 1536.897 | -0.634755816 | 0.234731 | -2.704188 | 0.006847 | 0.139171968 | ITPR1 | protein_coding | 42.9 | -0.543635495 | 0.543635495 | ENSG00000150995 | n.s. |
| ENSG00000010539 | 359.7874 | -0.613248166 | 0.223509 | -2.743731 | 0.006075 | 0.130381308 | ZNF200 | protein_coding | 43.67 | -0.542592574 | 0.542592574 | ENSG00000010539 | n.s. |
| ENSG00000257999 | 36.01902 | 0.890560071 | 0.381399 | 2.3349809 | 0.019544 | 0.247090218 | | antisense | 39.27 | 0.540698602 | 0.540698602 | ENSG00000257999 | n.s. |
| ENSG00000252549 | 11.76512 | -0.976319459 | 0.436075 | -2.238879 | 0.025164 | 0.279613138 | RNU6-759P | snRNA | 39.81 | -0.54033661 | 0.54033661 | ENSG00000252549 | n.s. |
| ENSG00000259732 | 52.42167 | -0.725706577 | 0.286441 | -2.53353 | 0.011292 | 0.180090288 | | sense_overlapping | 42.36 | -0.540295592 | 0.540295592 | ENSG00000259732 | n.s. |
| ENSG00000252242 | 1.799782 | -2.081636213 | 1.309924 | -1.589128 | 0.112032 | 0.550170175 | RNU7-115P | snRNA | 36.36 | -0.540190752 | 0.540190752 | ENSG00000252242 | n.s. |
| ENSG00000225183 | 6.559135 | -1.151947399 | 0.549468 | -2.096478 | 0.03604 | 0.340720178 | | processed_pseudogene | 44.54 | -0.538653076 | 0.538653076 | ENSG00000225183 | n.s. |
| ENSG00000266126 | 21.41007 | -0.915335437 | 0.398125 | -2.299118 | 0.021498 | 0.258034534 | | sense_intronic | 39.89 | -0.538512127 | 0.538512127 | ENSG00000266126 | n.s. |
| ENSG00000254936 | 61.55102 | -0.747219357 | 0.299526 | -2.494672 | 0.012607 | 0.190456473 | | sense_intronic | 43.59 | -0.538150566 | 0.538150566 | ENSG00000254936 | n.s. |
| ENSG00000222881 | 19.38442 | -0.886865371 | 0.380241 | -2.332378 | 0.019681 | 0.247485201 | | misc_RNA | 38.32 | -0.537840183 | 0.537840183 | ENSG00000222881 | n.s. |
| ENSG00000202169 | 7.013172 | 1.41596392 | 0.74576 | 1.8986867 | 0.057606 | 0.417869145 | | misc_RNA | 36.46 | 0.536593256 | 0.536593256 | ENSG00000202169 | n.s. |
| ENSG00000264229 | 321.6101 | 0.617367935 | 0.227355 | 2.7154375 | 0.006619 | 0.135426965 | RNU4ATAC | snRNA | 45.67 | 0.536057401 | 0.536057401 | ENSG00000264229 | n.s. |
| ENSG00000283196 | 46.92665 | -0.81115414 | 0.337215 | -2.40545 | 0.016153 | 0.220051277 | | lincRNA | 46.72 | -0.533314466 | 0.533314466 | ENSG00000283196 | n.s. |
| ENSG00000149289 | 1140.893 | 0.629173931 | 0.233652 | 2.6927827 | 0.007086 | 0.142155897 | ZC3H12C | protein_coding | 37.25 | 0.533058251 | 0.533058251 | ENSG00000149289 | n.s. |
| ENSG00000284059 | 5.25121 | -1.253169309 | 0.623072 | -2.011274 | 0.044297 | 0.375547181 | MIR5572 | miRNA | 59.12 | -0.533017387 | 0.533017387 | ENSG00000284059 | n.s. |
| ENSG00000275191 | 17.22621 | -1.185496179 | 0.57652 | -2.056296 | 0.039754 | 0.355381165 | | sense_intronic | 35.29 | -0.532650064 | 0.532650064 | ENSG00000275191 | n.s. |
| ENSG00000109819 | 172.6582 | 0.702666743 | 0.274164 | 2.5629476 | 0.010379 | 0.174827325 | PPARGC1A | protein_coding | 38.32 | 0.532193248 | 0.532193248 | ENSG00000109819 | n.s. |
| ENSG00000159917 | 382.9113 | -0.651005148 | 0.245604 | -2.650631 | 0.008034 | 0.152397234 | ZNF235 | protein_coding | 39.47 | -0.531886124 | 0.531886124 | ENSG00000159917 | n.s. |
| ENSG00000251595 | 179.5944 | -0.767183783 | 0.312612 | -2.454105 | 0.014124 | 0.202724384 | ABCA11P | ıscribed_processed_pseudog | 39.69 | -0.531730485 | 0.531730485 | ENSG00000251595 | n.s. |
| ENSG00000252768 | 10.6815 | -0.981906712 | 0.443186 | -2.215564 | 0.026721 | 0.287735335 | RNU6-856P | snRNA | 40.82 | -0.531218209 | 0.531218209 | ENSG00000252768 | n.s. |
| ENSG00000226419 | 815.4853 | -0.590557609 | 0.214008 | -2.759518 | 0.005789 | 0.126147369 | SLC16A1-AS1 | antisense | 43.76 | -0.530983222 | 0.530983222 | ENSG00000226419 | n.s. |
| ENSG00000156515 | 1375.944 | -0.655045549 | 0.248293 | -2.638191 | 0.008335 | 0.155351425 | HK1 | protein_coding | 47.64 | -0.529725351 | 0.529725351 | ENSG00000156515 | n.s. |
| ENSG00000262248 | 21.25814 | -0.912592552 | 0.399727 | -2.283041 | 0.022428 | 0.262893343 | | processed_pseudogene | 49.54 | -0.529504826 | 0.529504826 | ENSG00000262248 | n.s. |
| ENSG00000166603 | 37.38601 | 0.884359829 | 0.381018 | 2.3210449 | 0.020284 | 0.251935742 | MC4R | protein_coding | 43.7 | 0.529475263 | 0.529475263 | ENSG00000166603 | n.s. |
| ENSG00000168280 | 6674.787 | -0.658592232 | 0.250598 | -2.628086 | 0.008587 | 0.157512796 | KIF5C | protein_coding | 43.05 | -0.528641551 | 0.528641551 | ENSG00000168280 | n.s. |
| ENSG00000261770 | 232.6029 | -0.712933264 | 0.282174 | -2.52657 | 0.011518 | 0.181823456 | | lincRNA | 46.86 | -0.527820208 | 0.527820208 | ENSG00000261770 | n.s. |
| ENSG00000122678 | 57.59219 | -0.732392238 | 0.29355 | -2.49495 | 0.012597 | 0.190456473 | POLM | protein_coding | 56.76 | -0.527472012 | 0.527472012 | ENSG00000122678 | n.s. |
| ENSG00000181222 | 969.3574 | -0.652057149 | 0.247074 | -2.639118 | 0.008312 | 0.155343538 | POLR2A | protein_coding | 49.69 | -0.527323055 | 0.527323055 | ENSG00000181222 | n.s. |
| ENSG00000237298 | 1421.921 | -0.655388934 | 0.249127 | -2.630742 | 0.00852 | 0.156990291 | TTN-AS1 | antisense | 37.97 | -0.527016072 | 0.527016072 | ENSG00000237298 | n.s. |
| ENSG00000281468 | 35.19888 | -0.852850286 | 0.36323 | -2.347963 | 0.018876 | 0.241336844 | | sense_intronic | 30.71 | -0.526529615 | 0.526529615 | ENSG00000281468 | n.s. |

| | | | | | | | | | | | | | |
|---|---|---|---|---|---|---|---|---|---|---|---|---|---|
| ENSG00000109066 | 299.0206 | -0.658524583 | 0.25132 | -2.620259 | 0.008786 | 0.15887218 | TMEM104 | protein_coding | 54.36 | -0.526129628 | 0.526129628 | ENSG00000109066 | n.s. |
| ENSG00000054598 | 225.2434 | -0.662059094 | 0.25347 | -2.61198 | 0.009002 | 0.160685001 | FOXC1 | protein_coding | 56.32 | -0.525691248 | 0.525691248 | ENSG00000054598 | n.s. |
| ENSG00000220804 | 61.57148 | -0.770564234 | 0.315821 | -2.439874 | 0.014692 | 0.208099767 | LINC01881 | cribed_unprocessed_pseudc | 39.11 | -0.525315528 | 0.525315528 | ENSG00000220804 | n.s. |
| ENSG00000263278 | 857.8799 | -0.59914532 | 0.219652 | -2.727705 | 0.006378 | 0.132862373 | | sense_intronic | 53.41 | -0.525209587 | 0.525209587 | ENSG00000263278 | n.s. |
| ENSG00000273437 | 25.22193 | -0.84202277 | 0.357162 | -2.35754 | 0.018396 | 0.238042868 | | lincRNA | 40 | -0.524870536 | 0.524870536 | ENSG00000273437 | n.s. |
| ENSG00000261167 | 365.5399 | -0.709444539 | 0.281424 | -2.520913 | 0.011705 | 0.182089199 | | sense_overlapping | 39.27 | -0.524787345 | 0.524787345 | ENSG00000261167 | n.s. |
| ENSG00000100207 | 1223.293 | -0.610010205 | 0.225261 | -2.708009 | 0.006769 | 0.137982682 | TCF20 | protein_coding | 49.75 | -0.524715783 | 0.524715783 | ENSG00000100207 | n.s. |
| ENSG00000238386 | 7.377223 | -1.330355677 | 0.68419 | -1.944424 | 0.051844 | 0.403482109 | RNU7-48P | snRNA | 33.33 | -0.524393904 | 0.524393904 | ENSG00000238386 | n.s. |
| ENSG00000259946 | 88.09591 | -1.147427661 | 0.554297 | -2.070059 | 0.038447 | 0.350082264 | | lincRNA | 32.1 | -0.523031826 | 0.523031826 | ENSG00000259946 | n.s. |
| ENSG00000166343 | 85.68544 | -0.717714714 | 0.286565 | -2.504541 | 0.012261 | 0.186772923 | MSS51 | protein_coding | 42.14 | -0.522988724 | 0.522988724 | ENSG00000166343 | n.s. |
| ENSG00000215218 | 152.7403 | -0.711731581 | 0.283569 | -2.509909 | 0.012076 | 0.184969559 | UBE2QL1 | protein_coding | 47.32 | -0.521627889 | 0.521627889 | ENSG00000215218 | n.s. |
| ENSG00000241400 | 17.76736 | -0.879711602 | 0.381014 | -2.308869 | 0.020951 | 0.255337128 | | processed_pseudogene | 45.3 | -0.521568719 | 0.521568719 | ENSG00000241400 | n.s. |
| ENSG00000252965 | 3.903038 | -1.414651268 | 0.754836 | -1.874119 | 0.060914 | 0.42849416 | | misc_RNA | 49.06 | -0.520669603 | 0.520669603 | ENSG00000252965 | n.s. |
| ENSG00000253437 | 4.878856 | -1.413838073 | 0.754549 | -1.873752 | 0.060965 | 0.42849416 | RNU6-988P | snRNA | 35.51 | -0.520370303 | 0.520370303 | ENSG00000253437 | n.s. |
| ENSG00000157985 | 615.0521 | -0.628024367 | 0.235462 | -2.667198 | 0.007649 | 0.148580262 | AGAP1 | protein_coding | 47.33 | -0.520028593 | 0.520028593 | ENSG00000157985 | n.s. |
| ENSG00000175414 | 597.6163 | -0.701730038 | 0.278054 | -2.52372 | 0.011612 | 0.181889957 | ARL10 | protein_coding | 50.75 | -0.519414455 | 0.519414455 | ENSG00000175414 | n.s. |
| ENSG00000163684 | 532.0848 | -0.687123412 | 0.268639 | -2.55779 | 0.010534 | 0.17570541 | RPP14 | protein_coding | 44.14 | -0.518925816 | 0.518925816 | ENSG00000163684 | n.s. |
| ENSG00000279865 | 96.65491 | 0.700497392 | 0.277583 | 2.5235588 | 0.011617 | 0.181889957 | | TEC | 38.59 | 0.518502062 | 0.518502062 | ENSG00000279865 | n.s. |
| ENSG00000254463 | 40.62757 | -0.802993066 | 0.336241 | -2.388144 | 0.016934 | 0.226129282 | | processed_pseudogene | 45.21 | -0.518447009 | 0.518447009 | ENSG00000254463 | n.s. |
| ENSG00000142207 | 1410.087 | -0.595420467 | 0.219075 | -2.717879 | 0.00657 | 0.134873528 | URB1 | protein_coding | 46.41 | -0.518059441 | 0.518059441 | ENSG00000142207 | n.s. |
| ENSG00000008283 | 76.53464 | -0.73642119 | 0.298227 | -2.469328 | 0.013537 | 0.197974563 | CYB561 | protein_coding | 57.41 | -0.517991747 | 0.517991747 | ENSG00000008283 | n.s. |
| ENSG00000177051 | 144.6449 | -0.686707742 | 0.268919 | -2.553583 | 0.010662 | 0.176185284 | FBXO46 | protein_coding | 51.45 | -0.517798493 | 0.517798493 | ENSG00000177051 | n.s. |
| ENSG00000119685 | 288.6087 | -0.630745752 | 0.237578 | -2.654899 | 0.007933 | 0.151099897 | TTLL5 | protein_coding | 40.03 | -0.517675639 | 0.517675639 | ENSG00000119685 | n.s. |
| ENSG00000135643 | 161.5015 | -0.699436469 | 0.277447 | -2.520977 | 0.011703 | 0.182089199 | KCNMB4 | protein_coding | 39.69 | -0.517384217 | 0.517384217 | ENSG00000135643 | n.s. |
| ENSG00000270140 | 39.69049 | -0.81960512 | 0.346062 | -2.368377 | 0.017866 | 0.233789887 | | sense_intronic | 40.27 | -0.517313671 | 0.517313671 | ENSG00000270140 | n.s. |
| ENSG00000284138 | 23.99744 | 0.864756019 | 0.372807 | 2.3195789 | 0.020364 | 0.252244945 | | processed_pseudogene | 58.18 | 0.517277619 | 0.517277619 | ENSG00000284138 | n.s. |
| ENSG00000136205 | 535.6009 | -0.64989689 | 0.248492 | -2.615363 | 0.008913 | 0.160029118 | TNS3 | protein_coding | 48.46 | -0.517188589 | 0.517188589 | ENSG00000136205 | n.s. |
| ENSG00000218996 | 37.68814 | -0.78080057 | 0.323484 | -2.413721 | 0.015791 | 0.217674794 | ARL4AP5 | processed_pseudogene | 39.73 | -0.517039779 | 0.517039779 | ENSG00000218996 | n.s. |
| ENSG00000212248 | 27.92986 | 0.872068699 | 0.37795 | 2.3073674 | 0.021034 | 0.255337128 | RNU6-750P | snRNA | 43.93 | 0.517037349 | 0.517037349 | ENSG00000212248 | n.s. |
| ENSG00000251754 | 3.28931 | -1.411987479 | 0.75654 | -1.866376 | 0.061989 | 0.430475522 | RNU6-999P | snRNA | 44.76 | -0.51686019 | 0.51686019 | ENSG00000251754 | n.s. |
| ENSG00000198987 | 31.3742 | 0.827561655 | 0.350724 | 2.3595784 | 0.018296 | 0.237400972 | MIR16-2 | miRNA | 37.04 | 0.516826743 | 0.516826743 | ENSG00000198987 | n.s. |
| ENSG00000154229 | 1497.39 | -0.623221919 | 0.233482 | -2.669248 | 0.007602 | 0.148449585 | PRKCA | protein_coding | 43.87 | -0.516290133 | 0.516290133 | ENSG00000154229 | n.s. |

| | | | | | | | | | | | | | |
|---|---|---|---|---|---|---|---|---|---|---|---|---|---|
| ENSG00000261560 | 95.7044 | -0.730771923 | 0.295412 | -2.473735 | 0.013371 | 0.196598957 | | sense_intronic | 43.21 | -0.516231018 | 0.516231018 | ENSG00000261560 | n.s. |
| ENSG00000105281 | 978.7274 | -0.63670452 | 0.241099 | -2.640846 | 0.00827 | 0.154788381 | SLC1A5 | protein_coding | 54.4 | -0.515897249 | 0.515897249 | ENSG00000105281 | n.s. |
| ENSG00000167889 | 271.4707 | -0.655410578 | 0.251886 | -2.602008 | 0.009268 | 0.163338944 | MGAT5B | protein_coding | 54.13 | -0.515749306 | 0.515749306 | ENSG00000167889 | n.s. |
| ENSG00000196361 | 927.414 | -0.642051384 | 0.244297 | -2.62816 | 0.008585 | 0.157512796 | ELAVL3 | protein_coding | 53.1 | -0.515364474 | 0.515364474 | ENSG00000196361 | n.s. |
| ENSG00000079335 | 188.538 | -0.665214536 | 0.257343 | -2.584929 | 0.00974 | 0.168249203 | CDC14A | protein_coding | 38.39 | -0.514907306 | 0.514907306 | ENSG00000079335 | n.s. |
| ENSG00000179943 | 143.9022 | -0.672449506 | 0.26135 | -2.572985 | 0.010083 | 0.172033469 | FIZ1 | protein_coding | 57.1 | -0.514011697 | 0.514011697 | ENSG00000179943 | n.s. |
| ENSG00000171786 | 266.0171 | -0.625066646 | 0.235309 | -2.656367 | 0.007899 | 0.150841114 | NHLH1 | protein_coding | 50.12 | -0.513479916 | 0.513479916 | ENSG00000171786 | n.s. |
| ENSG00000203865 | 75.8333 | -0.696430983 | 0.276742 | -2.516538 | 0.011851 | 0.183252281 | ATP1A1-AS1 | processed_transcript | 45.35 | -0.513235237 | 0.513235237 | ENSG00000203865 | n.s. |
| ENSG00000196182 | 162.3544 | -0.679562621 | 0.265636 | -2.558251 | 0.01052 | 0.17570541 | STK40 | protein_coding | 52.28 | -0.513215794 | 0.513215794 | ENSG00000196182 | n.s. |
| ENSG00000170881 | 910.2221 | 0.692388747 | 0.273755 | 2.5292301 | 0.011431 | 0.181687298 | RNF139 | protein_coding | 39.17 | 0.512835335 | 0.512835335 | ENSG00000170881 | n.s. |
| ENSG00000134297 | 30.9207 | -0.813701021 | 0.344275 | -2.363523 | 0.018102 | 0.235607368 | PLEKHA8P1 | ıscribed_processed_pseudog | 41.34 | -0.510850559 | 0.510850559 | ENSG00000134297 | n.s. |
| ENSG00000233956 | 17.68298 | -0.869811144 | 0.378841 | -2.295976 | 0.021677 | 0.258843947 | BTF3P6 | processed_pseudogene | 40.28 | -0.510546076 | 0.510546076 | ENSG00000233956 | n.s. |
| ENSG00000256037 | 23.98719 | 0.872630641 | 0.380857 | 2.2912313 | 0.02195 | 0.260036752 | MRPL40P1 | processed_pseudogene | 50.99 | 0.510458617 | 0.510458617 | ENSG00000256037 | n.s. |
| ENSG00000207582 | 4.550919 | -1.344781267 | 0.705781 | -1.90538 | 0.056731 | 0.417346136 | MIR30B | miRNA | 43.18 | -0.510349336 | 0.510349336 | ENSG00000207582 | n.s. |
| ENSG00000267127 | 36.84574 | -0.784526284 | 0.327534 | -2.395252 | 0.016609 | 0.22376458 | | protein_coding | 46.26 | -0.510105782 | 0.510105782 | ENSG00000267127 | n.s. |
| ENSG00000277423 | 11.88626 | -0.947656308 | 0.428865 | -2.209685 | 0.027127 | 0.29009854 | | lincRNA | 37.98 | -0.509322106 | 0.509322106 | ENSG00000277423 | n.s. |
| ENSG00000253948 | 20.74829 | -0.911488599 | 0.405388 | -2.248437 | 0.024548 | 0.276340982 | | lincRNA | 38.08 | -0.509116245 | 0.509116245 | ENSG00000253948 | n.s. |
| ENSG00000268117 | 9.936984 | -1.009934454 | 0.46833 | -2.156458 | 0.031048 | 0.314156871 | VN1R84P | unprocessed_pseudogene | 41.28 | -0.507849012 | 0.507849012 | ENSG00000268117 | n.s. |
| ENSG00000206778 | 5.81117 | -1.193482354 | 0.593478 | -2.010996 | 0.044326 | 0.375547181 | RNU6-213P | snRNA | 41.35 | -0.507630406 | 0.507630406 | ENSG00000206778 | n.s. |
| ENSG00000232142 | 14.95911 | -0.924212551 | 0.413819 | -2.233371 | 0.025524 | 0.282417274 | RPS25P9 | processed_pseudogene | 45.63 | -0.507493193 | 0.507493193 | ENSG00000232142 | n.s. |
| ENSG00000261578 | 175.7835 | -0.647026145 | 0.249393 | -2.594402 | 0.009476 | 0.164910296 | | sense_overlapping | 47.28 | -0.506461157 | 0.506461157 | ENSG00000261578 | n.s. |
| ENSG00000199730 | 23.69924 | -0.867824115 | 0.379352 | -2.28765 | 0.022158 | 0.261111919 | RN7SKP95 | misc_RNA | 50.5 | -0.506091855 | 0.506091855 | ENSG00000199730 | n.s. |
| ENSG00000084710 | 310.4418 | -0.673591813 | 0.264347 | -2.548137 | 0.01083 | 0.177604223 | EFR3B | protein_coding | 48.38 | -0.505562122 | 0.505562122 | ENSG00000084710 | n.s. |
| ENSG00000207326 | 8.402547 | -1.105370732 | 0.532483 | -2.075879 | 0.037905 | 0.349043056 | | misc_RNA | 39.47 | -0.505288172 | 0.505288172 | ENSG00000207326 | n.s. |
| ENSG00000212124 | 183.2008 | -0.711414718 | 0.287124 | -2.477724 | 0.013222 | 0.195015287 | TAS2R19 | protein_coding | 40.52 | -0.505055607 | 0.505055607 | ENSG00000212124 | n.s. |
| ENSG00000261542 | 105.0129 | -0.745607291 | 0.306596 | -2.431891 | 0.01502 | 0.210813098 | | lincRNA | 32.38 | -0.504106886 | 0.504106886 | ENSG00000261542 | n.s. |
| ENSG00000199075 | 2.772314 | -1.626663857 | 0.946576 | -1.718471 | 0.085711 | 0.490059299 | MIR26A1 | miRNA | 55.84 | -0.503861351 | 0.503861351 | ENSG00000199075 | n.s. |
| ENSG00000272650 | 15.21147 | -0.894881741 | 0.397047 | -2.253841 | 0.024206 | 0.274225075 | | lincRNA | 28.52 | -0.50282764 | 0.50282764 | ENSG00000272650 | n.s. |
| ENSG00000262251 | 196.1508 | 0.618462302 | 0.233829 | 2.6449287 | 0.008171 | 0.153940789 | | sense_intronic | 49.82 | 0.502591096 | 0.502591096 | ENSG00000262251 | n.s. |
| ENSG00000265435 | 6.574597 | -1.154797221 | 0.569515 | -2.027684 | 0.042592 | 0.36833247 | MIR3121 | miRNA | 35.06 | -0.500904837 | 0.500904837 | ENSG00000265435 | n.s. |
| ENSG00000223023 | 6.794646 | -1.155951192 | 0.570569 | -2.025964 | 0.042769 | 0.368937997 | | misc_RNA | 37.96 | -0.50058075 | 0.50058075 | ENSG00000223023 | n.s. |
| ENSG00000206784 | 4.326781 | -1.281085679 | 0.662317 | -1.934248 | 0.053083 | 0.407272581 | | misc_RNA | 40.71 | -0.499770518 | 0.499770518 | ENSG00000206784 | n.s. |

| | | | | | | | | | | | | | |
|---|---|---|---|---|---|---|---|---|---|---|---|---|---|
| ENSG00000272721 | 23.15009 | -0.91095521 | 0.408819 | -2.228258 | 0.025863 | 0.283560235 | | antisense | 53.28 | -0.498615591 | 0.498615591 | ENSG00000272721 | n.s. |
| ENSG00000231888 | 26.11946 | -0.783345725 | 0.329665 | -2.376189 | 0.017493 | 0.230963869 | MTND5P15 | processed_pseudogene | 44.99 | -0.498565051 | 0.498565051 | ENSG00000231888 | n.s. |
| ENSG00000176058 | 262.1814 | -0.668990414 | 0.263996 | -2.534089 | 0.011274 | 0.180009224 | TPRN | protein_coding | 62.56 | -0.498200666 | 0.498200666 | ENSG00000176058 | n.s. |
| ENSG00000238829 | 2.754505 | -1.617076407 | 0.943965 | -1.713069 | 0.0867 | 0.492652156 | RNU7-45P | snRNA | 39.68 | -0.497185685 | 0.497185685 | ENSG00000238829 | n.s. |
| ENSG00000229419 | 325.7098 | -0.671538855 | 0.266138 | -2.523269 | 0.011627 | 0.181889957 | RALGAPA1P1 | processed_pseudogene | 40.13 | -0.497067205 | 0.497067205 | ENSG00000229419 | n.s. |
| ENSG00000122778 | 846.039 | -0.609670529 | 0.230496 | -2.645037 | 0.008168 | 0.153940789 | KIAA1549 | protein_coding | 44.25 | -0.495446495 | 0.495446495 | ENSG00000122778 | n.s. |
| ENSG00000141664 | 101.0573 | -0.657440695 | 0.258222 | -2.546031 | 0.010896 | 0.177923733 | ZCCHC2 | protein_coding | 42.38 | -0.492926759 | 0.492926759 | ENSG00000141664 | n.s. |
| ENSG00000222432 | 11.33611 | -1.080418733 | 0.521623 | -2.071263 | 0.038334 | 0.350082264 | | misc_RNA | 40.71 | -0.492487154 | 0.492487154 | ENSG00000222432 | n.s. |
| ENSG00000159110 | 167.9414 | -0.699642518 | 0.283287 | -2.469734 | 0.013521 | 0.197974563 | IFNAR2 | protein_coding | 44.16 | -0.492121975 | 0.492121975 | ENSG00000159110 | n.s. |
| ENSG00000283160 | 17.39207 | -0.918578784 | 0.416655 | -2.20465 | 0.027479 | 0.29160484 | MIR4521 | miRNA | 41.67 | -0.491628208 | 0.491628208 | ENSG00000283160 | n.s. |
| ENSG00000087116 | 431.6091 | -0.638293126 | 0.247563 | -2.578307 | 0.009929 | 0.17045049 | ADAMTS2 | protein_coding | 53.87 | -0.490465553 | 0.490465553 | ENSG00000087116 | n.s. |
| ENSG00000144857 | 62.97426 | -0.738507586 | 0.305639 | -2.416273 | 0.01568 | 0.217014788 | BOC | protein_coding | 47.04 | -0.490007664 | 0.490007664 | ENSG00000144857 | n.s. |
| ENSG00000181201 | 125.4412 | -0.692814653 | 0.280072 | -2.473704 | 0.013372 | 0.196598957 | HIST3H2BA | unitary_pseudogene | 61.07 | -0.48941729 | 0.48941729 | ENSG00000181201 | n.s. |
| ENSG00000201516 | 5.514059 | -1.319435191 | 0.702011 | -1.879508 | 0.060175 | 0.425937056 | | snoRNA | 46.34 | -0.48905469 | 0.48905469 | ENSG00000201516 | n.s. |
| ENSG00000272215 | 8.452778 | -1.135451737 | 0.561773 | -2.021193 | 0.04326 | 0.371106396 | | snRNA | 46.03 | -0.488813745 | 0.488813745 | ENSG00000272215 | n.s. |
| ENSG00000176595 | 560.0979 | -0.655334785 | 0.258187 | -2.538218 | 0.011142 | 0.179742753 | KBTBD11 | protein_coding | 50.18 | -0.488452872 | 0.488452872 | ENSG00000176595 | n.s. |
| ENSG00000146267 | 1024.406 | -0.638174409 | 0.247965 | -2.573646 | 0.010063 | 0.171947811 | FAXC | protein_coding | 41.71 | -0.487950291 | 0.487950291 | ENSG00000146267 | n.s. |
| ENSG00000035862 | 2305.102 | -0.614495327 | 0.235243 | -2.61217 | 0.008997 | 0.160685001 | TIMP2 | protein_coding | 52.6 | -0.487924443 | 0.487924443 | ENSG00000035862 | n.s. |
| ENSG00000252503 | 4.719327 | -1.267484721 | 0.659817 | -1.920965 | 0.054736 | 0.412398929 | RNU6-531P | snRNA | 46.15 | -0.487579155 | 0.487579155 | ENSG00000252503 | n.s. |
| ENSG00000271576 | 696.8178 | 0.551767129 | 0.20136 | 2.7401967 | 0.00614 | 0.130768951 | | lincRNA | 43.79 | 0.487483699 | 0.487483699 | ENSG00000271576 | n.s. |
| ENSG00000277142 | 123.8038 | -0.651864052 | 0.256603 | -2.540362 | 0.011074 | 0.179362555 | | lincRNA | 61.7 | -0.486465425 | 0.486465425 | ENSG00000277142 | n.s. |
| ENSG00000067955 | 901.6484 | -0.655855442 | 0.259365 | -2.528695 | 0.011449 | 0.181717699 | CBFB | protein_coding | 40.45 | -0.485728357 | 0.485728357 | ENSG00000067955 | n.s. |
| ENSG00000251703 | 44.10962 | -0.85786103 | 0.378983 | -2.263585 | 0.0236 | 0.271595762 | RNU6-998P | snRNA | 38.61 | -0.485615408 | 0.485615408 | ENSG00000251703 | n.s. |
| ENSG00000265986 | 9.66373 | -1.039425107 | 0.496268 | -2.094482 | 0.036217 | 0.341167655 | MIR4735 | miRNA | 37.68 | -0.485444943 | 0.485444943 | ENSG00000265986 | n.s. |
| ENSG00000168993 | 183.0766 | -0.655739057 | 0.259594 | -2.526016 | 0.011536 | 0.181859152 | CPLX1 | protein_coding | 55.28 | -0.485420567 | 0.485420567 | ENSG00000168993 | n.s. |
| ENSG00000124225 | 324.0416 | -0.640041628 | 0.249548 | -2.564801 | 0.010323 | 0.174434351 | PMEPA1 | protein_coding | 54.59 | -0.48538708 | 0.48538708 | ENSG00000124225 | n.s. |
| ENSG00000260903 | 1065.678 | -0.641975763 | 0.25146 | -2.552992 | 0.01068 | 0.176185284 | XKR7 | protein_coding | 51.87 | -0.484069222 | 0.484069222 | ENSG00000260903 | n.s. |
| ENSG00000259820 | 386.7622 | -0.616281965 | 0.237232 | -2.597806 | 0.009382 | 0.164161265 | | lincRNA | 36.21 | -0.483614521 | 0.483614521 | ENSG00000259820 | n.s. |
| ENSG00000267270 | 72.71776 | -0.655813591 | 0.26106 | -2.512114 | 0.012001 | 0.184217437 | PARD6G-AS1 | antisense | 48.33 | -0.481806089 | 0.481806089 | ENSG00000267270 | n.s. |
| ENSG00000110921 | 183.6285 | -0.731117721 | 0.303842 | -2.406242 | 0.016118 | 0.219790693 | MVK | protein_coding | 52.45 | -0.481068662 | 0.481068662 | ENSG00000110921 | n.s. |
| ENSG00000072840 | 318.9702 | -0.630475681 | 0.245278 | -2.570452 | 0.010157 | 0.172661866 | EVC | protein_coding | 47.4 | -0.4809291 | 0.4809291 | ENSG00000072840 | n.s. |
| ENSG00000263720 | 27.70127 | -0.804092739 | 0.346716 | -2.319166 | 0.020386 | 0.252297924 | | sense_intronic | 40.05 | -0.480916869 | 0.480916869 | ENSG00000263720 | n.s. |

| | | | | | | | | | | | | | |
|---|---|---|---|---|---|---|---|---|---|---|---|---|---|
| ENSG00000257870 | 135.5669 | 0.624363726 | 0.241962 | 2.5804164 | 0.009868 | 0.169831763 | | lincRNA | 42.38 | 0.480748257 | 0.480748257 | ENSG00000257870 | n.s. |
| ENSG00000256294 | 223.2486 | -0.642381144 | 0.252886 | -2.5402 | 0.011079 | 0.179362555 | ZNF225 | protein_coding | 41.4 | -0.479388632 | 0.479388632 | ENSG00000256294 | n.s. |
| ENSG00000103647 | 254.1325 | -0.636805287 | 0.249703 | -2.550253 | 0.010764 | 0.176946537 | CORO2B | protein_coding | 49.34 | -0.478978149 | 0.478978149 | ENSG00000103647 | n.s. |
| ENSG00000059122 | 29.4706 | -0.829379339 | 0.363954 | -2.278805 | 0.022679 | 0.264673416 | FLYWCH1 | protein_coding | 52.33 | -0.478792132 | 0.478792132 | ENSG00000059122 | n.s. |
| ENSG00000199135 | 17.92254 | 0.855869474 | 0.380889 | 2.2470303 | 0.024638 | 0.276340982 | MIR101-1 | miRNA | 45.33 | 0.478049921 | 0.478049921 | ENSG00000199135 | n.s. |
| ENSG00000236536 | 26.26852 | -0.842230147 | 0.371566 | -2.266702 | 0.023408 | 0.270693867 | | antisense | 37.7 | -0.477983787 | 0.477983787 | ENSG00000236536 | n.s. |
| ENSG00000279880 | 124.1806 | -0.66897626 | 0.269226 | -2.484817 | 0.012962 | 0.193012971 | | TEC | 40.21 | -0.477925675 | 0.477925675 | ENSG00000279880 | n.s. |
| ENSG00000151151 | 433.7273 | -0.639245385 | 0.251629 | -2.540429 | 0.011072 | 0.179362555 | IPMK | protein_coding | 38.62 | -0.477048514 | 0.477048514 | ENSG00000151151 | n.s. |
| ENSG00000202358 | 10.15406 | -1.03037219 | 0.493905 | -2.086175 | 0.036963 | 0.344525344 | RNU6-652P | snRNA | 36.45 | -0.476834431 | 0.476834431 | ENSG00000202358 | n.s. |
| ENSG00000199545 | 29.41616 | 0.851024808 | 0.378191 | 2.2502508 | 0.024433 | 0.275450235 | RNA5SP195 | rRNA | 50.43 | 0.476537173 | 0.476537173 | ENSG00000199545 | n.s. |
| ENSG00000115221 | 16.10963 | -0.882350978 | 0.398709 | -2.213022 | 0.026896 | 0.288501124 | ITGB6 | protein_coding | 40.66 | -0.476339373 | 0.476339373 | ENSG00000115221 | n.s. |
| ENSG00000202150 | 6.307966 | -1.594975241 | 0.943792 | -1.689964 | 0.091035 | 0.502967617 | RNU6-407P | snRNA | 40.38 | -0.476036271 | 0.476036271 | ENSG00000202150 | n.s. |
| ENSG00000270993 | 3.83856 | 1.586158494 | 0.937942 | 1.6911056 | 0.090817 | 0.501961137 | | processed_pseudogene | 60 | 0.474784672 | 0.474784672 | ENSG00000270993 | n.s. |
| ENSG00000276182 | 184.6378 | -0.637369477 | 0.251308 | -2.536209 | 0.011206 | 0.180009224 | | sense_intronic | 32.65 | -0.474652389 | 0.474652389 | ENSG00000276182 | n.s. |
| ENSG00000272669 | 35.66758 | -0.813864631 | 0.355718 | -2.287947 | 0.022141 | 0.261111919 | | antisense | 56.82 | -0.474624125 | 0.474624125 | ENSG00000272669 | n.s. |
| ENSG00000080854 | 68.81996 | -0.749020615 | 0.315929 | -2.370847 | 0.017747 | 0.23267083 | IGSF9B | protein_coding | 56.66 | -0.474323343 | 0.474323343 | ENSG00000080854 | n.s. |
| ENSG00000249459 | 468.8199 | -0.596011788 | 0.22791 | -2.61512 | 0.00892 | 0.160029118 | ZNF286B | protein_coding | 42.4 | -0.474306771 | 0.474306771 | ENSG00000249459 | n.s. |
| ENSG00000116741 | 1574.903 | 0.535008382 | 0.194686 | 2.7480598 | 0.005995 | 0.129972657 | RGS2 | protein_coding | 40.83 | 0.474096608 | 0.474096608 | ENSG00000116741 | n.s. |
| ENSG00000116514 | 123.3173 | -0.672518565 | 0.272085 | -2.471718 | 0.013447 | 0.197277396 | RNF19B | protein_coding | 44.34 | -0.474073584 | 0.474073584 | ENSG00000116514 | n.s. |
| ENSG00000177542 | 236.9158 | -0.664847364 | 0.26799 | -2.480867 | 0.013106 | 0.194127146 | SLC25A22 | protein_coding | 65.66 | -0.473313967 | 0.473313967 | ENSG00000177542 | n.s. |
| ENSG00000196152 | 172.2019 | -0.603356611 | 0.232363 | -2.596608 | 0.009415 | 0.164471885 | ZNF79 | protein_coding | 47.15 | -0.472976274 | 0.472976274 | ENSG00000196152 | n.s. |
| ENSG00000265817 | 328.5439 | -0.641033373 | 0.25478 | -2.516026 | 0.011869 | 0.183252281 | FSBP | protein_coding | 35.6 | -0.472409935 | 0.472409935 | ENSG00000265817 | n.s. |
| ENSG00000159882 | 211.3671 | -0.639340767 | 0.254245 | -2.514665 | 0.011915 | 0.183386434 | ZNF230 | protein_coding | 40.86 | -0.470959376 | 0.470959376 | ENSG00000159882 | n.s. |
| ENSG00000256588 | 31.70906 | -0.770350258 | 0.329389 | -2.338722 | 0.01935 | 0.245585944 | | sense_intronic | 41.14 | -0.469756886 | 0.469756886 | ENSG00000256588 | n.s. |
| ENSG00000182253 | 669.4622 | -0.598343788 | 0.230337 | -2.597688 | 0.009385 | 0.164161265 | SYNM | protein_coding | 49.91 | -0.469537908 | 0.469537908 | ENSG00000182253 | n.s. |
| ENSG00000265281 | 20.9707 | -0.829267896 | 0.366256 | -2.264177 | 0.023563 | 0.271581624 | MIR3935 | miRNA | 46.15 | -0.46944824 | 0.46944824 | ENSG00000265281 | n.s. |
| ENSG00000233012 | 114.4382 | -0.654346241 | 0.262982 | -2.488177 | 0.01284 | 0.191814442 | HDAC1P2 | ıscribed_processed_pseudog | 46.87 | -0.469243924 | 0.469243924 | ENSG00000233012 | n.s. |
| ENSG00000070047 | 903.8247 | -0.58421734 | 0.222603 | -2.624481 | 0.008678 | 0.157735679 | PHRF1 | protein_coding | 56.31 | -0.468583237 | 0.468583237 | ENSG00000070047 | n.s. |
| ENSG00000257954 | 48.78737 | -0.771990183 | 0.331005 | -2.332263 | 0.019687 | 0.247485201 | | processed_pseudogene | 40.19 | -0.468174037 | 0.468174037 | ENSG00000257954 | n.s. |
| ENSG00000161021 | 933.8078 | -0.626526913 | 0.246475 | -2.541953 | 0.011024 | 0.179260564 | MAML1 | protein_coding | 49.38 | -0.46771189 | 0.46771189 | ENSG00000161021 | n.s. |
| ENSG00000259682 | 26.90814 | -0.847681438 | 0.378985 | -2.236714 | 0.025305 | 0.280734627 | | sense_intronic | 45.14 | -0.467669254 | 0.467669254 | ENSG00000259682 | n.s. |
| ENSG00000204084 | 255.728 | -0.627630686 | 0.247487 | -2.536013 | 0.011212 | 0.180009224 | INPP5B | protein_coding | 46.64 | -0.46739986 | 0.46739986 | ENSG00000204084 | n.s. |

| | | | | | | | | | | | | | |
|---|---|---|---|---|---|---|---|---|---|---|---|---|---|
| ENSG00000267405 | 54.71966 | -0.728396137 | 0.305782 | -2.382075 | 0.017215 | 0.228309094 | | lincRNA | 47.65 | -0.467249216 | 0.467249216 | ENSG00000267405 | n.s. |
| ENSG00000223203 | 4.193048 | -1.468556011 | 0.8415 | -1.745164 | 0.080956 | 0.480663403 | RNA5SP221 | rRNA | 52.43 | -0.467234229 | 0.467234229 | ENSG00000223203 | n.s. |
| ENSG00000162407 | 603.4973 | 0.672074497 | 0.273626 | 2.4561806 | 0.014042 | 0.201855602 | PLPP3 | protein_coding | 41.42 | 0.46706435 | 0.46706435 | ENSG00000162407 | n.s. |
| ENSG00000158859 | 393.5873 | -0.582665068 | 0.222188 | -2.622396 | 0.008731 | 0.15829041 | ADAMTS4 | protein_coding | 51.48 | -0.466449838 | 0.466449838 | ENSG00000158859 | n.s. |
| ENSG00000257298 | 213.2951 | 0.617140874 | 0.241466 | 2.555806 | 0.010594 | 0.175765447 | | sense_intronic | 43.93 | 0.465982398 | 0.465982398 | ENSG00000257298 | n.s. |
| ENSG00000219438 | 1146.579 | -0.715116835 | 0.298581 | -2.395052 | 0.016618 | 0.22376458 | FAM19A5 | protein_coding | 54.37 | -0.464975158 | 0.464975158 | ENSG00000219438 | n.s. |
| ENSG00000153363 | 272.0443 | -0.672632664 | 0.274335 | -2.451869 | 0.014212 | 0.203778276 | LINC00467 | lincRNA | 39.05 | -0.464682972 | 0.464682972 | ENSG00000153363 | n.s. |
| ENSG00000133858 | 395.7816 | -0.701213427 | 0.290486 | -2.413933 | 0.015781 | 0.217674794 | ZFC3H1 | protein_coding | 36.14 | -0.464337822 | 0.464337822 | ENSG00000133858 | n.s. |
| ENSG00000236636 | 32.9203 | -0.850995454 | 0.382472 | -2.224987 | 0.026082 | 0.285167272 | | processed_pseudogene | 40.74 | -0.463707694 | 0.463707694 | ENSG00000236636 | n.s. |
| ENSG00000160145 | 147.2539 | -0.625108256 | 0.247537 | -2.525309 | 0.01156 | 0.181859152 | KALRN | protein_coding | 42.68 | -0.462745662 | 0.462745662 | ENSG00000160145 | n.s. |
| ENSG00000202125 | 11.79061 | -1.063739867 | 0.524805 | -2.026924 | 0.04267 | 0.368546519 | RNU1-100P | snRNA | 48.48 | -0.461139408 | 0.461139408 | ENSG00000202125 | n.s. |
| ENSG00000167470 | 407.4416 | -0.581520455 | 0.22286 | -2.609352 | 0.009071 | 0.161460238 | MIDN | protein_coding | 65.97 | -0.46052606 | 0.46052606 | ENSG00000167470 | n.s. |
| ENSG00000199798 | 5.641886 | 1.090546338 | 0.544199 | 2.0039488 | 0.045076 | 0.378748866 | SNORD114-5 | snoRNA | 44.29 | 0.459827401 | 0.459827401 | ENSG00000199798 | n.s. |
| ENSG00000176248 | 804.0327 | -0.616839466 | 0.243272 | -2.535596 | 0.011226 | 0.180009224 | ANAPC2 | protein_coding | 61.65 | -0.459363582 | 0.459363582 | ENSG00000176248 | n.s. |
| ENSG00000105072 | 91.81539 | -0.733653412 | 0.310989 | -2.359095 | 0.01832 | 0.23748915 | C19orf44 | protein_coding | 51.63 | -0.458061076 | 0.458061076 | ENSG00000105072 | n.s. |
| ENSG00000213693 | 109.0437 | -0.702477176 | 0.292868 | -2.39861 | 0.016457 | 0.223049174 | SEC14L1P1 | processed_pseudogene | 50.02 | -0.457733693 | 0.457733693 | ENSG00000213693 | n.s. |
| ENSG00000112182 | 157.3895 | -0.605613907 | 0.236987 | -2.555477 | 0.010604 | 0.175765447 | BACH2 | protein_coding | 40.5 | -0.457278771 | 0.457278771 | ENSG00000112182 | n.s. |
| ENSG00000207139 | 4.2059 | -1.363608653 | 0.763033 | -1.78709 | 0.073923 | 0.462574432 | | misc_RNA | 38.61 | -0.456561234 | 0.456561234 | ENSG00000207139 | n.s. |
| ENSG00000252941 | 3.745908 | -1.364058384 | 0.763534 | -1.786507 | 0.074017 | 0.462895871 | RNA5SP340 | rRNA | 41.12 | -0.4563003 | 0.4563003 | ENSG00000252941 | n.s. |
| ENSG00000257027 | 129.5985 | -0.705762579 | 0.295425 | -2.388975 | 0.016895 | 0.226129282 | | sense_intronic | 41.43 | -0.455670806 | 0.455670806 | ENSG00000257027 | n.s. |
| ENSG00000260804 | 861.1427 | -0.618244048 | 0.245935 | -2.513853 | 0.011942 | 0.18351933 | LINC01963 | lincRNA | 44.66 | -0.455224335 | 0.455224335 | ENSG00000260804 | n.s. |
| ENSG00000273373 | 742.6362 | -0.703139295 | 0.294384 | -2.388509 | 0.016917 | 0.226129282 | | antisense | 34.98 | -0.453977101 | 0.453977101 | ENSG00000273373 | n.s. |
| ENSG00000275902 | 36.59816 | -0.785496107 | 0.344651 | -2.279106 | 0.022661 | 0.264673416 | | antisense | 54.49 | -0.453458795 | 0.453458795 | ENSG00000275902 | n.s. |
| ENSG00000213699 | 478.1746 | -0.600268721 | 0.235088 | -2.553376 | 0.010668 | 0.176185284 | SLC35F6 | protein_coding | 52.66 | -0.452620846 | 0.452620846 | ENSG00000213699 | n.s. |
| ENSG00000239653 | 126.0462 | -0.606876265 | 0.239129 | -2.537866 | 0.011153 | 0.179742753 | PSMD6-AS2 | antisense | 38.82 | -0.452334381 | 0.452334381 | ENSG00000239653 | n.s. |
| ENSG00000207815 | 13.27646 | -1.169726873 | 0.607604 | -1.925148 | 0.054211 | 0.41109933 | MIR563 | miRNA | 44.3 | -0.451576839 | 0.451576839 | ENSG00000207815 | n.s. |
| ENSG00000228956 | 505.7685 | -0.646203693 | 0.262552 | -2.461238 | 0.013846 | 0.200076043 | SATB1-AS1 | processed_transcript | 37.1 | -0.451570314 | 0.451570314 | ENSG00000228956 | n.s. |
| ENSG00000185219 | 1356.042 | -0.605244425 | 0.238807 | -2.534452 | 0.011262 | 0.180009224 | ZNF445 | protein_coding | 44.45 | -0.450728695 | 0.450728695 | ENSG00000185219 | n.s. |
| ENSG00000107140 | 136.9392 | -0.643049675 | 0.261109 | -2.462762 | 0.013787 | 0.199953179 | TESK1 | protein_coding | 59.15 | -0.449537822 | 0.449537822 | ENSG00000107140 | n.s. |
| ENSG00000079156 | 73.60841 | -0.662785573 | 0.272403 | -2.43311 | 0.01497 | 0.210316903 | OSBPL6 | protein_coding | 39.34 | -0.448789226 | 0.448789226 | ENSG00000079156 | n.s. |
| ENSG00000108175 | 1705.787 | -0.605835757 | 0.239662 | -2.527878 | 0.011475 | 0.181717699 | ZMIZ1 | protein_coding | 55.32 | -0.44868364 | 0.44868364 | ENSG00000108175 | n.s. |
| ENSG00000235173 | 154.3836 | -0.642619669 | 0.261378 | -2.458586 | 0.013949 | 0.200832917 | HGH1 | protein_coding | 64.59 | -0.448012009 | 0.448012009 | ENSG00000235173 | n.s. |

| | | | | | | | | | | | | | |
|---|---|---|---|---|---|---|---|---|---|---|---|---|---|
| ENSG00000225793 | 83.00784 | -0.62864867 | 0.253324 | -2.481602 | 0.013079 | 0.193987598 | | lincRNA | 37.06 | -0.447739949 | 0.447739949 | ENSG00000225793 | n.s. |
| ENSG00000266507 | 7.163255 | -1.224806136 | 0.659043 | -1.85846 | 0.063104 | 0.431139465 | MIR4479 | miRNA | 70.42 | -0.447522387 | 0.447522387 | ENSG00000266507 | n.s. |
| ENSG00000215930 | 10.00649 | -0.998094682 | 0.485903 | -2.054103 | 0.039966 | 0.356465505 | MIR942 | miRNA | 44.19 | -0.447128941 | 0.447128941 | ENSG00000215930 | n.s. |
| ENSG00000163635 | 238.1973 | -0.596042706 | 0.234043 | -2.546725 | 0.010874 | 0.177905487 | ATXN7 | protein_coding | 40.77 | -0.446919172 | 0.446919172 | ENSG00000163635 | n.s. |
| ENSG00000200953 | 15.64821 | -0.925918807 | 0.436652 | -2.120496 | 0.033964 | 0.329535967 | | misc_RNA | 38.18 | -0.446382844 | 0.446382844 | ENSG00000200953 | n.s. |
| ENSG00000270361 | 20.49627 | 0.978618201 | 0.47202 | 2.0732577 | 0.038148 | 0.350082264 | | lincRNA | 45.8 | 0.446083428 | 0.446083428 | ENSG00000270361 | n.s. |
| ENSG00000173706 | 1161.646 | -0.625966616 | 0.252253 | -2.481502 | 0.013083 | 0.193987598 | HEG1 | protein_coding | 44.98 | -0.44582972 | 0.44582972 | ENSG00000173706 | n.s. |
| ENSG00000249055 | 30.12683 | -0.801092783 | 0.357041 | -2.243698 | 0.024852 | 0.27771574 | TBCAP3 | ıscribed_processed_pseudog | 40.71 | -0.445727628 | 0.445727628 | ENSG00000249055 | n.s. |
| ENSG00000204315 | 64.08202 | -0.685753631 | 0.286524 | -2.393357 | 0.016695 | 0.224150681 | FKBPL | protein_coding | 54.57 | -0.44536951 | 0.44536951 | ENSG00000204315 | n.s. |
| ENSG00000175309 | 165.4882 | -0.634221799 | 0.257031 | -2.467495 | 0.013606 | 0.198781797 | PHYKPL | protein_coding | 51.43 | -0.444984847 | 0.444984847 | ENSG00000175309 | n.s. |
| ENSG00000106683 | 205.4242 | -0.635977365 | 0.258023 | -2.464811 | 0.013709 | 0.199710058 | LIMK1 | protein_coding | 54.59 | -0.444929805 | 0.444929805 | ENSG00000106683 | n.s. |
| ENSG00000118420 | 361.8759 | -0.636152539 | 0.258352 | -2.462345 | 0.013803 | 0.199953179 | UBE3D | protein_coding | 38.49 | -0.444716229 | 0.444716229 | ENSG00000118420 | n.s. |
| ENSG00000200169 | 182.0134 | 0.618705818 | 0.248299 | 2.4917807 | 0.01271 | 0.191316306 | RNU5D-1 | snRNA | 37.93 | 0.444384224 | 0.444384224 | ENSG00000200169 | n.s. |
| ENSG00000198948 | 71.12989 | -0.695326102 | 0.29229 | -2.378894 | 0.017365 | 0.229913168 | MFAP3L | protein_coding | 42.91 | -0.443921322 | 0.443921322 | ENSG00000198948 | n.s. |
| ENSG00000159167 | 1343.519 | -0.554730679 | 0.211651 | -2.620971 | 0.008768 | 0.158746962 | STC1 | protein_coding | 43.81 | -0.443393224 | 0.443393224 | ENSG00000159167 | n.s. |
| ENSG00000264802 | 7.109952 | -1.092086078 | 0.555356 | -1.966461 | 0.049245 | 0.393340111 | MIR5191 | miRNA | 56.67 | -0.442547968 | 0.442547968 | ENSG00000264802 | n.s. |
| ENSG00000136770 | 257.7922 | -0.652216371 | 0.267882 | -2.434715 | 0.014904 | 0.209839048 | DNAJC1 | protein_coding | 37.73 | -0.442276852 | 0.442276852 | ENSG00000136770 | n.s. |
| ENSG00000181026 | 2967.791 | -0.539543719 | 0.203699 | -2.64873 | 0.008079 | 0.152840499 | AEN | protein_coding | 51.46 | -0.440139022 | 0.440139022 | ENSG00000181026 | n.s. |
| ENSG00000199165 | 43.04247 | 0.733451055 | 0.315961 | 2.3213339 | 0.020269 | 0.251935742 | MIRLET7A1 | miRNA | 42.5 | 0.439124638 | 0.439124638 | ENSG00000199165 | n.s. |
| ENSG00000252756 | 7.259923 | -1.140466488 | 0.593702 | -1.92094 | 0.054739 | 0.412398929 | RNU6-577P | snRNA | 40.82 | -0.438717468 | 0.438717468 | ENSG00000252756 | n.s. |
| ENSG00000275769 | 22.26598 | -0.808734734 | 0.364662 | -2.217767 | 0.026571 | 0.287074936 | | sense_intronic | 34.05 | -0.438338046 | 0.438338046 | ENSG00000275769 | n.s. |
| ENSG00000274799 | 10.63028 | -0.963231462 | 0.465949 | -2.067246 | 0.038711 | 0.350893753 | | misc_RNA | 55.32 | -0.438101136 | 0.438101136 | ENSG00000274799 | n.s. |
| ENSG00000238886 | 11.34433 | -0.970815864 | 0.471025 | -2.061072 | 0.039296 | 0.354033183 | SNORD121A | snoRNA | 39.02 | -0.437795268 | 0.437795268 | ENSG00000238886 | n.s. |
| ENSG00000200176 | 10.17012 | -0.98420677 | 0.4824 | -2.04023 | 0.041327 | 0.360907877 | RNU1-19P | snRNA | 52.73 | -0.435613498 | 0.435613498 | ENSG00000200176 | n.s. |
| ENSG00000283736 | 26.46515 | 0.854758583 | 0.394768 | 2.16522 | 0.030371 | 0.30955579 | MIR484 | miRNA | 59.49 | 0.435295269 | 0.435295269 | ENSG00000283736 | n.s. |
| ENSG00000272407 | 15.95736 | -0.920558206 | 0.437548 | -2.1039 | 0.035387 | 0.337177445 | | transcribed_processed_pseudog | 44.18 | -0.434633718 | 0.434633718 | ENSG00000272407 | n.s. |
| ENSG00000088992 | 1215.94 | -0.658500696 | 0.273277 | -2.409643 | 0.015968 | 0.21903854 | TESC | protein_coding | 51.4 | -0.434267686 | 0.434267686 | ENSG00000088992 | n.s. |
| ENSG00000141298 | 55.59589 | -0.731987196 | 0.317283 | -2.307047 | 0.021052 | 0.255337128 | SSH2 | protein_coding | 40.66 | -0.433984983 | 0.433984983 | ENSG00000141298 | n.s. |
| ENSG00000282527 | 11.70532 | 1.043526556 | 0.524369 | 1.9900613 | 0.046584 | 0.384553485 | | antisense | 36.18 | 0.433108652 | 0.433108652 | ENSG00000282527 | n.s. |
| ENSG00000157741 | 987.9845 | -0.609814637 | 0.246113 | -2.477784 | 0.01322 | 0.195015287 | UBN2 | protein_coding | 39.02 | -0.432926525 | 0.432926525 | ENSG00000157741 | n.s. |
| ENSG00000067798 | 46.12474 | 0.787578181 | 0.352451 | 2.2345752 | 0.025445 | 0.282065493 | NAV3 | protein_coding | 35.89 | 0.432892379 | 0.432892379 | ENSG00000067798 | n.s. |
| ENSG00000138080 | 1086.172 | -0.607491493 | 0.244681 | -2.482785 | 0.013036 | 0.193910185 | EMILIN1 | protein_coding | 61.63 | -0.432776562 | 0.432776562 | ENSG00000138080 | n.s. |

| | | | | | | | | | | | | | |
|---|---|---|---|---|---|---|---|---|---|---|---|---|---|
| ENSG00000278351 | 23.1858 | -0.830016871 | 0.380229 | -2.182942 | 0.02904 | 0.301509943 | | sense_intronic | 29.92 | -0.432188425 | 0.432188425 | ENSG00000278351 | n.s. |
| ENSG00000010318 | 41.10967 | -0.748811696 | 0.328673 | -2.278291 | 0.022709 | 0.264808346 | PHF7 | protein_coding | 45.18 | -0.432115516 | 0.432115516 | ENSG00000010318 | n.s. |
| ENSG00000220867 | 28.94033 | -0.79352112 | 0.357315 | -2.220787 | 0.026365 | 0.286058985 | HSPE1P26 | processed_pseudogene | 38.91 | -0.431313966 | 0.431313966 | ENSG00000220867 | n.s. |
| ENSG00000273703 | 715.0978 | -0.54582614 | 0.209504 | -2.605331 | 0.009179 | 0.162173762 | HIST1H2BM | protein_coding | 55.38 | -0.431213244 | 0.431213244 | ENSG00000273703 | n.s. |
| ENSG00000168807 | 442.4183 | -0.63690962 | 0.26198 | -2.431138 | 0.015051 | 0.210899376 | SNTB2 | protein_coding | 41.92 | -0.430502947 | 0.430502947 | ENSG00000168807 | n.s. |
| ENSG00000163660 | 10578.8 | 0.558738651 | 0.216514 | 2.5806093 | 0.009863 | 0.169831763 | CCNL1 | protein_coding | 37.3 | 0.430218191 | 0.430218191 | ENSG00000163660 | n.s. |
| ENSG00000228446 | 102.901 | -0.718500286 | 0.309474 | -2.321685 | 0.02025 | 0.251935742 | | processed_pseudogene | 41.88 | -0.43017346 | 0.43017346 | ENSG00000228446 | n.s. |
| ENSG00000275202 | 52.24875 | 0.725324543 | 0.314244 | 2.3081571 | 0.02099 | 0.255337128 | | antisense | 61.76 | 0.430034789 | 0.430034789 | ENSG00000275202 | n.s. |
| ENSG00000199591 | 4.751307 | -1.212786972 | 0.662892 | -1.82954 | 0.067319 | 0.442113119 | | misc_RNA | 42.99 | -0.429892472 | 0.429892472 | ENSG00000199591 | n.s. |
| ENSG00000206715 | 6.061492 | -1.173077069 | 0.628489 | -1.866503 | 0.061971 | 0.430475522 | RNU6-444P | snRNA | 41.67 | -0.429406666 | 0.429406666 | ENSG00000206715 | n.s. |
| ENSG00000145526 | 36.45361 | 1.038712496 | 0.523371 | 1.9846579 | 0.047183 | 0.387108156 | CDH18 | protein_coding | 34.89 | 0.428123718 | 0.428123718 | ENSG00000145526 | n.s. |
| ENSG00000263080 | 20.62274 | -0.915862483 | 0.437141 | -2.095119 | 0.03616 | 0.340867363 | | antisense | 46.49 | -0.428087477 | 0.428087477 | ENSG00000263080 | n.s. |
| ENSG00000275832 | 277.0977 | -0.611738282 | 0.248297 | -2.463741 | 0.01375 | 0.199953179 | ARHGAP23 | protein_coding | 55.44 | -0.427648913 | 0.427648913 | ENSG00000275832 | n.s. |
| ENSG00000239884 | 15.91245 | -0.884774466 | 0.417122 | -2.121142 | 0.03391 | 0.329262928 | RN7SL608P | misc_RNA | 55.2 | -0.426865779 | 0.426865779 | ENSG00000239884 | n.s. |
| ENSG00000155816 | 420.7399 | -0.586039396 | 0.234076 | -2.503629 | 0.012293 | 0.187050139 | FMN2 | protein_coding | 39.28 | -0.426661273 | 0.426661273 | ENSG00000155816 | n.s. |
| ENSG00000256274 | 170.447 | -0.718377881 | 0.311315 | -2.307557 | 0.021024 | 0.255337128 | | transcribed_processed_pseudog | 33.67 | -0.42591621 | 0.42591621 | ENSG00000256274 | n.s. |
| ENSG00000278223 | 4.633996 | -1.263379343 | 0.705478 | -1.790814 | 0.073323 | 0.460477385 | MIR6783 | miRNA | 59.38 | -0.425495669 | 0.425495669 | ENSG00000278223 | n.s. |
| ENSG00000237738 | 9.793656 | -0.96695341 | 0.475073 | -2.035378 | 0.041813 | 0.363474721 | RNF216-IT1 | sense_intronic | 48.47 | -0.42500096 | 0.42500096 | ENSG00000237738 | n.s. |
| ENSG00000267340 | 17.67745 | -0.882047877 | 0.416551 | -2.117503 | 0.034217 | 0.329831947 | | processed_pseudogene | 50.51 | -0.424888885 | 0.424888885 | ENSG00000267340 | n.s. |
| ENSG00000272644 | 29.29711 | -0.750555987 | 0.33182 | -2.261935 | 0.023701 | 0.27210013 | | lincRNA | 44.96 | -0.424267722 | 0.424267722 | ENSG00000272644 | n.s. |
| ENSG00000283575 | 4.37169 | -1.541520046 | 0.947206 | -1.627439 | 0.103644 | 0.531112578 | | snRNA | 42.71 | -0.423630386 | 0.423630386 | ENSG00000283575 | n.s. |
| ENSG00000116698 | 1336.515 | -0.589932845 | 0.236814 | -2.491118 | 0.012734 | 0.191466556 | SMG7 | protein_coding | 40.28 | -0.423516963 | 0.423516963 | ENSG00000116698 | n.s. |
| ENSG00000133624 | 383.6507 | -0.629649745 | 0.259457 | -2.426799 | 0.015233 | 0.212530539 | ZNF767P | cribed_unprocessed_pseudc | 46.36 | -0.423488979 | 0.423488979 | ENSG00000133624 | n.s. |
| ENSG00000212306 | 5.336784 | -1.114443063 | 0.586329 | -1.900712 | 0.05734 | 0.417869145 | | misc_RNA | 48.42 | -0.422329004 | 0.422329004 | ENSG00000212306 | n.s. |
| ENSG00000184058 | 148.5156 | -0.627227068 | 0.258462 | -2.426764 | 0.015234 | 0.212530539 | TBX1 | protein_coding | 57.77 | -0.421859539 | 0.421859539 | ENSG00000184058 | n.s. |
| ENSG00000173451 | 194.5158 | -0.566520324 | 0.223645 | -2.53312 | 0.011305 | 0.180094534 | THAP2 | protein_coding | 37.49 | -0.421774079 | 0.421774079 | ENSG00000173451 | n.s. |
| ENSG00000256667 | 210.552 | -0.690746316 | 0.295151 | -2.340315 | 0.019267 | 0.245220122 | KLRA1P | cribed_unprocessed_pseudc | 33.5 | -0.421661872 | 0.421661872 | ENSG00000256667 | n.s. |
| ENSG00000273156 | 64.63272 | -0.720964173 | 0.314787 | -2.290326 | 0.022002 | 0.260271766 | | lincRNA | 30.83 | -0.421456147 | 0.421456147 | ENSG00000273156 | n.s. |
| ENSG00000243403 | 14.7289 | -1.14353676 | 0.610235 | -1.873928 | 0.06094 | 0.42849416 | | processed_pseudogene | 52.59 | -0.420884528 | 0.420884528 | ENSG00000243403 | n.s. |
| ENSG00000227987 | 186.7802 | -0.612953136 | 0.250605 | -2.445894 | 0.014449 | 0.206125079 | | sense_intronic | 51.01 | -0.420405655 | 0.420405655 | ENSG00000227987 | n.s. |
| ENSG00000176542 | 633.4491 | -0.652088693 | 0.273228 | -2.386612 | 0.017004 | 0.226774472 | USF3 | protein_coding | 40.61 | -0.420209759 | 0.420209759 | ENSG00000176542 | n.s. |
| ENSG00000269929 | 923.7827 | 0.565461961 | 0.223717 | 2.5275721 | 0.011485 | 0.181717699 | | lincRNA | 36.45 | 0.418782695 | 0.418782695 | ENSG00000269929 | n.s. |

| | | | | | | | | | | | | | |
|---|---|---|---|---|---|---|---|---|---|---|---|---|---|
| ENSG00000142751 | 206.3968 | -0.598133295 | 0.242896 | -2.462507 | 0.013797 | 0.199953179 | GPN2 | protein_coding | 48.74 | -0.418138052 | 0.418138052 | ENSG00000142751 | n.s. |
| ENSG00000230648 | 11.94328 | -1.025033936 | 0.519846 | -1.971805 | 0.048632 | 0.39116228 | | antisense | 43.62 | -0.41784794 | 0.41784794 | ENSG00000230648 | n.s. |
| ENSG00000063587 | 325.3816 | -0.611916026 | 0.250998 | -2.437927 | 0.014772 | 0.209011298 | ZNF275 | protein_coding | 49.08 | -0.415999018 | 0.415999018 | ENSG00000063587 | n.s. |
| ENSG00000103264 | 82.02996 | -0.682882566 | 0.292182 | -2.337184 | 0.01943 | 0.246373825 | FBXO31 | protein_coding | 51.99 | -0.415469464 | 0.415469464 | ENSG00000103264 | n.s. |
| ENSG00000100058 | 485.1901 | -0.601184109 | 0.245262 | -2.451195 | 0.014238 | 0.203950388 | CRYBB2P1 | cribed_unprocessed_pseudc | 45.32 | -0.415102876 | 0.415102876 | ENSG00000100058 | n.s. |
| ENSG00000121101 | 103.5113 | 0.768602604 | 0.347278 | 2.213217 | 0.026883 | 0.288501124 | TEX14 | protein_coding | 45.25 | 0.41493203 | 0.41493203 | ENSG00000121101 | n.s. |
| ENSG00000116128 | 635.2698 | -0.578502244 | 0.232591 | -2.487212 | 0.012875 | 0.192129254 | BCL9 | protein_coding | 42.09 | -0.41444277 | 0.41444277 | ENSG00000116128 | n.s. |
| ENSG00000157693 | 260.4337 | -0.592035518 | 0.240186 | -2.464909 | 0.013705 | 0.199710058 | TMEM268 | protein_coding | 47.67 | -0.414188085 | 0.414188085 | ENSG00000157693 | n.s. |
| ENSG00000118491 | 36.75339 | -0.759507547 | 0.341427 | -2.224511 | 0.026114 | 0.285167272 | ZC2HC1B | protein_coding | 39.82 | -0.413855905 | 0.413855905 | ENSG00000118491 | n.s. |
| ENSG00000274667 | 174.7558 | 0.580892779 | 0.234051 | 2.4819048 | 0.013068 | 0.193987598 | | sense_intronic | 28.97 | 0.41372696 | 0.41372696 | ENSG00000274667 | n.s. |
| ENSG00000225447 | 15.72103 | -0.898159964 | 0.431378 | -2.082071 | 0.037336 | 0.346783837 | RPS15AP10 | processed_pseudogene | 43.19 | -0.41310073 | 0.41310073 | ENSG00000225447 | n.s. |
| ENSG00000200769 | 14.65197 | -0.872144793 | 0.415247 | -2.100305 | 0.035702 | 0.338379377 | | misc_RNA | 42.39 | -0.41042795 | 0.41042795 | ENSG00000200769 | n.s. |
| ENSG00000228532 | 31.59862 | 1.151685188 | 0.626314 | 1.8388316 | 0.06594 | 0.440875092 | | processed_pseudogene | 45.49 | 0.409636497 | 0.409636497 | ENSG00000228532 | n.s. |
| ENSG00000166845 | 475.2857 | -0.603725327 | 0.248087 | -2.433525 | 0.014953 | 0.210287972 | C18orf54 | protein_coding | 34.77 | -0.408834078 | 0.408834078 | ENSG00000166845 | n.s. |
| ENSG00000202515 | 8.88692 | -1.116856289 | 0.600151 | -1.860958 | 0.06275 | 0.430564731 | VTRNA1-3 | misc_RNA | 60.67 | -0.408726456 | 0.408726456 | ENSG00000202515 | n.s. |
| ENSG00000255496 | 10.16356 | -1.047249483 | 0.541856 | -1.932707 | 0.053272 | 0.407947312 | | sense_overlapping | 40.94 | -0.40779468 | 0.40779468 | ENSG00000255496 | n.s. |
| ENSG00000263603 | 56.72036 | -0.721332384 | 0.319063 | -2.260787 | 0.023772 | 0.272193206 | | lincRNA | 54.04 | -0.407641319 | 0.407641319 | ENSG00000263603 | n.s. |
| ENSG00000085741 | 119.4926 | 0.622702643 | 0.259242 | 2.4020152 | 0.016305 | 0.221694304 | WNT11 | protein_coding | 57.84 | 0.40740038 | 0.40740038 | ENSG00000085741 | n.s. |
| ENSG00000213842 | 37.93586 | -0.732771934 | 0.326997 | -2.240915 | 0.025032 | 0.278366304 | SUGT1P2 | processed_pseudogene | 38.41 | -0.406969321 | 0.406969321 | ENSG00000213842 | n.s. |
| ENSG00000251726 | 6.55886 | -1.06138594 | 0.554789 | -1.913136 | 0.055731 | 0.413612824 | RNU7-41P | snRNA | 44.26 | -0.406941742 | 0.406941742 | ENSG00000251726 | n.s. |
| ENSG00000264421 | 12.67819 | 0.975392051 | 0.489014 | 1.9946107 | 0.046085 | 0.382988464 | | antisense | 38.18 | 0.406557362 | 0.406557362 | ENSG00000264421 | n.s. |
| ENSG00000221886 | 294.6309 | -0.541943167 | 0.212759 | -2.547216 | 0.010859 | 0.177864178 | ZBED8 | protein_coding | 40 | -0.406409418 | 0.406409418 | ENSG00000221886 | n.s. |
| ENSG00000279059 | 203.1274 | -0.593242498 | 0.242661 | -2.444742 | 0.014496 | 0.206573024 | | TEC | 39.83 | -0.406327444 | 0.406327444 | ENSG00000279059 | n.s. |
| ENSG00000265458 | 89.41533 | 0.612242304 | 0.253738 | 2.41289 | 0.015827 | 0.217740811 | | antisense | 48.51 | 0.40534124 | 0.40534124 | ENSG00000265458 | n.s. |
| ENSG00000249002 | 18.72325 | -0.823513422 | 0.385335 | -2.137138 | 0.032587 | 0.322462993 | | processed_pseudogene | 41.62 | -0.404773415 | 0.404773415 | ENSG00000249002 | n.s. |
| ENSG00000230042 | 38.73531 | -0.756588732 | 0.343292 | -2.203919 | 0.02753 | 0.291927257 | AK3P3 | processed_pseudogene | 46.54 | -0.404567177 | 0.404567177 | ENSG00000230042 | n.s. |
| ENSG00000206878 | 8.830956 | -1.038389675 | 0.537526 | -1.931793 | 0.053385 | 0.407947312 | | snoRNA | 47.2 | -0.404344707 | 0.404344707 | ENSG00000206878 | n.s. |
| ENSG00000268912 | 125.2607 | -0.654533086 | 0.278718 | -2.34837 | 0.018856 | 0.241336844 | | lincRNA | 45.69 | -0.404093261 | 0.404093261 | ENSG00000268912 | n.s. |
| ENSG00000214067 | 45.26203 | -0.712905314 | 0.314954 | -2.263523 | 0.023603 | 0.271595762 | | processed_pseudogene | 43.08 | -0.403559309 | 0.403559309 | ENSG00000214067 | n.s. |
| ENSG00000088367 | 184.3581 | -0.66665492 | 0.286033 | -2.330696 | 0.019769 | 0.24819613 | EPB41L1 | protein_coding | 49.98 | -0.403462886 | 0.403462886 | ENSG00000088367 | n.s. |
| ENSG00000111215 | 140.1193 | -0.743135696 | 0.335235 | -2.216761 | 0.026639 | 0.287074936 | PRR4 | protein_coding | 36.64 | -0.402783058 | 0.402783058 | ENSG00000111215 | n.s. |
| ENSG00000260000 | 199.8334 | 0.567973211 | 0.229721 | 2.4724466 | 0.013419 | 0.197083545 | | antisense | 42.32 | 0.400619696 | 0.400619696 | ENSG00000260000 | n.s. |

| | | | | | | | | | | | | | |
|---|---|---|---|---|---|---|---|---|---|---|---|---|---|
| ENSG00000104332 | 890.719 | -0.614706922 | 0.256404 | -2.397415 | 0.016511 | 0.223191469 | SFRP1 | protein_coding | 48.62 | -0.400372394 | 0.400372394 | ENSG00000104332 | n.s. |
| ENSG00000268201 | 27.54486 | -0.760153153 | 0.347205 | -2.189352 | 0.028571 | 0.297752712 | | sense_intronic | 48.14 | -0.399950229 | 0.399950229 | ENSG00000268201 | n.s. |
| ENSG00000239280 | 140.9942 | 0.604596717 | 0.250661 | 2.4120124 | 0.015865 | 0.218050121 | | processed_pseudogene | 37.27 | 0.39990667 | 0.39990667 | ENSG00000239280 | n.s. |
| ENSG00000279070 | 81.76294 | 0.743239861 | 0.336469 | 2.2089407 | 0.027179 | 0.290191679 | | lincRNA | 44.39 | 0.399353959 | 0.399353959 | ENSG00000279070 | n.s. |
| ENSG00000225568 | 42.02158 | -0.734595912 | 0.33067 | -2.221537 | 0.026315 | 0.286058985 | | processed_pseudogene | 42.25 | -0.399285499 | 0.399285499 | ENSG00000225568 | n.s. |
| ENSG00000252658 | 13.05465 | -0.949949754 | 0.47453 | -2.001875 | 0.045298 | 0.380160717 | RNU6-786P | snRNA | 36.67 | -0.399010018 | 0.399010018 | ENSG00000252658 | n.s. |
| ENSG00000139517 | 200.1822 | -0.658240205 | 0.282477 | -2.330243 | 0.019793 | 0.24827217 | LNX2 | protein_coding | 37.54 | -0.398282691 | 0.398282691 | ENSG00000139517 | n.s. |
| ENSG00000225595 | 16.33209 | 1.353950458 | 0.806992 | 1.6777751 | 0.093391 | 0.508121556 | LINC01623 | lincRNA | 44.93 | 0.398105276 | 0.398105276 | ENSG00000225595 | n.s. |
| ENSG00000222078 | 18.52516 | -0.853644092 | 0.408129 | -2.091604 | 0.036474 | 0.341750791 | RN7SKP110 | misc_RNA | 50.88 | -0.398046107 | 0.398046107 | ENSG00000222078 | n.s. |
| ENSG00000270039 | 23.57664 | -0.807950283 | 0.377795 | -2.138593 | 0.032469 | 0.321971749 | | lincRNA | 37.89 | -0.397658775 | 0.397658775 | ENSG00000270039 | n.s. |
| ENSG00000265745 | 19.92521 | -0.797777956 | 0.371555 | -2.147134 | 0.031783 | 0.318122825 | RN7SL375P | misc_RNA | 52.68 | -0.396818879 | 0.396818879 | ENSG00000265745 | n.s. |
| ENSG00000240996 | 26.5351 | -0.772326159 | 0.35555 | -2.172199 | 0.029841 | 0.306529566 | | lincRNA | 36.1 | -0.396610821 | 0.396610821 | ENSG00000240996 | n.s. |
| ENSG00000225693 | 30.69493 | -0.703808287 | 0.312233 | -2.254109 | 0.024189 | 0.274225075 | LAGE3P1 | processed_pseudogene | 62.19 | -0.395464835 | 0.395464835 | ENSG00000225693 | n.s. |
| ENSG00000262766 | 38.01968 | 0.707197742 | 0.314779 | 2.2466456 | 0.024663 | 0.276340982 | | sense_intronic | 47.31 | 0.395008625 | 0.395008625 | ENSG00000262766 | n.s. |
| ENSG00000256064 | 25.38883 | -0.758753354 | 0.347928 | -2.180779 | 0.0292 | 0.302134081 | | lincRNA | 37.64 | -0.394400208 | 0.394400208 | ENSG00000256064 | n.s. |
| ENSG00000278619 | 92.08641 | -0.613051121 | 0.257074 | -2.384723 | 0.017092 | 0.227506482 | MRM1 | protein_coding | 52.69 | -0.394195687 | 0.394195687 | ENSG00000278619 | n.s. |
| ENSG00000260855 | 153.5032 | -0.668088134 | 0.290351 | -2.300966 | 0.021394 | 0.258017317 | | antisense | 38.91 | -0.393070418 | 0.393070418 | ENSG00000260855 | n.s. |
| ENSG00000239419 | 112.0479 | 0.604504952 | 0.252446 | 2.3945878 | 0.016639 | 0.223831511 | RN7SL535P | misc_RNA | 53.72 | 0.392975837 | 0.392975837 | ENSG00000239419 | n.s. |
| ENSG00000232536 | 21.85057 | 0.87123206 | 0.423069 | 2.0593141 | 0.039464 | 0.354033183 | | sense_intronic | 45.61 | 0.392887351 | 0.392887351 | ENSG00000232536 | n.s. |
| ENSG00000215184 | 38.36947 | -0.745244431 | 0.339922 | -2.1924 | 0.028351 | 0.297290527 | RPS12P16 | processed_pseudogene | 48.16 | -0.392608873 | 0.392608873 | ENSG00000215184 | n.s. |
| ENSG00000230359 | 72.7968 | 0.66881833 | 0.291252 | 2.2963533 | 0.021656 | 0.258843947 | TPI1P2 | ıscribed_processed_pseudog | 50.02 | 0.392570935 | 0.392570935 | ENSG00000230359 | n.s. |
| ENSG00000259959 | 242.4497 | -0.607155121 | 0.254074 | -2.38968 | 0.016863 | 0.225971734 | | lincRNA | 37.48 | -0.39218935 | 0.39218935 | ENSG00000259959 | n.s. |
| ENSG00000149328 | 105.4979 | -0.660882702 | 0.286649 | -2.305548 | 0.021136 | 0.255669083 | GLB1L2 | protein_coding | 51.13 | -0.391455222 | 0.391455222 | ENSG00000149328 | n.s. |
| ENSG00000100314 | 396.3636 | -0.673746083 | 0.295117 | -2.282977 | 0.022432 | 0.262893343 | CABP7 | protein_coding | 62.85 | -0.390921229 | 0.390921229 | ENSG00000100314 | n.s. |
| ENSG00000129932 | 134.5375 | -0.632405757 | 0.269193 | -2.349266 | 0.01881 | 0.241158369 | DOHH | protein_coding | 56.58 | -0.390635557 | 0.390635557 | ENSG00000129932 | n.s. |
| ENSG00000152932 | 573.0864 | 0.665765247 | 0.290281 | 2.2935222 | 0.021818 | 0.259855787 | RAB3C | protein_coding | 37.99 | 0.389650833 | 0.389650833 | ENSG00000152932 | n.s. |
| ENSG00000207023 | 48.31816 | 0.683264458 | 0.3011 | 2.2692309 | 0.023254 | 0.269182866 | RNU6-975P | snRNA | 38.1 | 0.389428345 | 0.389428345 | ENSG00000207023 | n.s. |
| ENSG00000213888 | 166.1094 | -0.563663637 | 0.230008 | -2.450623 | 0.014261 | 0.204064516 | LINC01521 | lincRNA | 59.44 | -0.389058964 | 0.389058964 | ENSG00000213888 | n.s. |
| ENSG00000173638 | 264.9776 | -0.605650352 | 0.254091 | -2.383597 | 0.017144 | 0.227985755 | SLC19A1 | protein_coding | 59.34 | -0.38888342 | 0.38888342 | ENSG00000173638 | n.s. |
| ENSG00000276326 | 3.630489 | -1.237409763 | 0.71467 | -1.731442 | 0.083373 | 0.485191567 | MIR2909 | miRNA | 52.17 | -0.388654018 | 0.388654018 | ENSG00000276326 | n.s. |
| ENSG00000223044 | 12.46616 | -1.108075068 | 0.608851 | -1.819944 | 0.068768 | 0.446156243 | RNU6-130P | snRNA | 39.81 | -0.388394745 | 0.388394745 | ENSG00000223044 | n.s. |
| ENSG00000103494 | 382.8559 | -0.654896009 | 0.283539 | -2.309724 | 0.020903 | 0.25529669 | RPGRIP1L | protein_coding | 37.27 | -0.388323742 | 0.388323742 | ENSG00000103494 | n.s. |

| | | | | | | | | | | | | | |
|---|---|---|---|---|---|---|---|---|---|---|---|---|---|
| ENSG00000197061 | 2125.548 | -0.524454796 | 0.20778 | -2.524084 | 0.0116 | 0.181889957 | HIST1H4C | protein_coding | 49.66 | -0.388196867 | 0.388196867 | ENSG00000197061 | n.s. |
| ENSG00000177125 | 465.5333 | -0.594965316 | 0.248517 | -2.394068 | 0.016663 | 0.223932851 | ZBTB34 | protein_coding | 43.23 | -0.386657363 | 0.386657363 | ENSG00000177125 | n.s. |
| ENSG00000260920 | 78.95614 | -0.65192618 | 0.282568 | -2.307146 | 0.021047 | 0.255337128 | | sense_overlapping | 37.89 | -0.386517925 | 0.386517925 | ENSG00000260920 | n.s. |
| ENSG00000124766 | 4640.244 | 0.507999112 | 0.197855 | 2.5675286 | 0.010243 | 0.173441096 | SOX4 | protein_coding | 50.73 | 0.386510103 | 0.386510103 | ENSG00000124766 | n.s. |
| ENSG00000272183 | 63.38944 | -0.655244328 | 0.284515 | -2.303023 | 0.021278 | 0.257159356 | | antisense | 57.94 | -0.386461578 | 0.386461578 | ENSG00000272183 | n.s. |
| ENSG00000276368 | 1056.159 | -0.512609984 | 0.200875 | -2.55188 | 0.010714 | 0.176330602 | HIST1H2AJ | protein_coding | 62.53 | -0.38633995 | 0.38633995 | ENSG00000276368 | n.s. |
| ENSG00000186063 | 414.2206 | -0.638830644 | 0.274281 | -2.329108 | 0.019853 | 0.248801 | AIDA | protein_coding | 38.46 | -0.385948178 | 0.385948178 | ENSG00000186063 | n.s. |
| ENSG00000100068 | 337.4975 | -0.567075866 | 0.232642 | -2.437551 | 0.014787 | 0.209016529 | LRP5L | protein_coding | 54.66 | -0.385509157 | 0.385509157 | ENSG00000100068 | n.s. |
| ENSG00000167196 | 2024.988 | -0.634845805 | 0.271847 | -2.335303 | 0.019528 | 0.247090218 | FBXO22 | protein_coding | 39.46 | -0.385443105 | 0.385443105 | ENSG00000167196 | n.s. |
| ENSG00000232454 | 21.44285 | -0.774532465 | 0.360937 | -2.145894 | 0.031881 | 0.31813913 | | processed_pseudogene | 37.61 | -0.385239212 | 0.385239212 | ENSG00000232454 | n.s. |
| ENSG00000106701 | 1510.377 | -0.62103346 | 0.263889 | -2.35339 | 0.018603 | 0.239825334 | FSD1L | protein_coding | 37.37 | -0.385105918 | 0.385105918 | ENSG00000106701 | n.s. |
| ENSG00000264857 | 6.557379 | 1.289627957 | 0.763928 | 1.6881542 | 0.091382 | 0.503484921 | MIR5696 | miRNA | 41.18 | 0.384326582 | 0.384326582 | ENSG00000264857 | n.s. |
| ENSG00000173894 | 3381.225 | -0.584028285 | 0.242646 | -2.406915 | 0.016088 | 0.219790693 | CBX2 | protein_coding | 58.13 | -0.384285181 | 0.384285181 | ENSG00000173894 | n.s. |
| ENSG00000138623 | 106.2915 | -0.646593466 | 0.279644 | -2.312206 | 0.020766 | 0.254739827 | SEMA7A | protein_coding | 57.93 | -0.384013897 | 0.384013897 | ENSG00000138623 | n.s. |
| ENSG00000150510 | 299.0932 | -0.563252811 | 0.230835 | -2.440065 | 0.014685 | 0.208099767 | FAM124A | protein_coding | 44.27 | -0.383985441 | 0.383985441 | ENSG00000150510 | n.s. |
| ENSG00000251839 | 4.80472 | -1.434680722 | 0.890589 | -1.610934 | 0.107194 | 0.540549786 | RNU7-12P | snRNA | 41.94 | -0.383295471 | 0.383295471 | ENSG00000251839 | n.s. |
| ENSG00000172209 | 481.9753 | 0.546880687 | 0.222127 | 2.4620218 | 0.013816 | 0.199953179 | GPR22 | protein_coding | 30.79 | 0.382308805 | 0.382308805 | ENSG00000172209 | n.s. |
| ENSG00000210191 | 20.0702 | 0.857550421 | 0.41939 | 2.0447581 | 0.040879 | 0.35907784 | MT-TL2 | Mt_tRNA | 35.21 | 0.381448199 | 0.381448199 | ENSG00000210191 | n.s. |
| ENSG00000279347 | 80.75665 | -0.723727498 | 0.330506 | -2.189754 | 0.028542 | 0.297671493 | | TEC | 44.12 | -0.380870826 | 0.380870826 | ENSG00000279347 | n.s. |
| ENSG00000117298 | 245.7465 | -0.577735045 | 0.239875 | -2.408489 | 0.016019 | 0.219300552 | ECE1 | protein_coding | 52.19 | -0.380704445 | 0.380704445 | ENSG00000117298 | n.s. |
| ENSG00000233250 | 10.66869 | -1.08895264 | 0.598863 | -1.818367 | 0.069008 | 0.447091133 | | antisense | 45.34 | -0.380702137 | 0.380702137 | ENSG00000233250 | n.s. |
| ENSG00000180221 | 29.36581 | -0.783188343 | 0.368243 | -2.126827 | 0.033434 | 0.326854919 | TPT1P10 | processed_pseudogene | 42.14 | -0.380351483 | 0.380351483 | ENSG00000180221 | n.s. |
| ENSG00000222164 | 52.88413 | 0.645287002 | 0.28098 | 2.2965585 | 0.021644 | 0.258843947 | RN7SKP266 | misc_RNA | 43.55 | 0.378758941 | 0.378758941 | ENSG00000222164 | n.s. |
| ENSG00000251374 | 9.770409 | -0.919696825 | 0.464226 | -1.98114 | 0.047576 | 0.387927349 | | processed_pseudogene | 39.81 | -0.378224953 | 0.378224953 | ENSG00000251374 | n.s. |
| ENSG00000207993 | 3.413267 | -1.284226865 | 0.765037 | -1.678647 | 0.093221 | 0.508003334 | MIR134 | miRNA | 60.27 | -0.377734062 | 0.377734062 | ENSG00000207993 | n.s. |
| ENSG00000151176 | 173.5325 | -0.637038864 | 0.276159 | -2.306785 | 0.021067 | 0.255337128 | PLBD2 | protein_coding | 51.13 | -0.377691443 | 0.377691443 | ENSG00000151176 | n.s. |
| ENSG00000280000 | 68.75941 | -0.693981908 | 0.312579 | -2.220179 | 0.026407 | 0.286116459 | | TEC | 38.91 | -0.377149436 | 0.377149436 | ENSG00000280000 | n.s. |
| ENSG00000132677 | 33.04564 | -0.807321507 | 0.385704 | -2.093111 | 0.036339 | 0.341167655 | RHBG | protein_coding | 51.82 | -0.377045099 | 0.377045099 | ENSG00000132677 | n.s. |
| ENSG00000244716 | 91.44817 | 0.749664474 | 0.347695 | 2.156099 | 0.031076 | 0.314212213 | | processed_pseudogene | 46.13 | 0.376914009 | 0.376914009 | ENSG00000244716 | n.s. |
| ENSG00000167491 | 1108.07 | -0.539260533 | 0.219151 | -2.460684 | 0.013867 | 0.200076043 | GATAD2A | protein_coding | 50.87 | -0.376837909 | 0.376837909 | ENSG00000167491 | n.s. |
| ENSG00000107821 | 67.14768 | -0.64026344 | 0.278383 | -2.299941 | 0.021452 | 0.258017317 | KAZALD1 | protein_coding | 58.78 | -0.376699728 | 0.376699728 | ENSG00000107821 | n.s. |
| ENSG00000207698 | 5.059014 | -1.122399844 | 0.627432 | -1.78888 | 0.073634 | 0.462013656 | MIR32 | miRNA | 38.57 | -0.376391386 | 0.376391386 | ENSG00000207698 | n.s. |

| | | | | | | | | | | | | | |
|---|---|---|---|---|---|---|---|---|---|---|---|---|---|
| ENSG00000206660 | 3.946916 | 1.279653118 | 0.762488 | 1.6782602 | 0.093296 | 0.508003334 | | misc_RNA | 50 | 0.37638877 | 0.37638877 | ENSG00000206660 | n.s. |
| ENSG00000264901 | 4.430056 | -1.214427495 | 0.706931 | -1.717886 | 0.085817 | 0.490059299 | MIR3616 | miRNA | 54.35 | -0.376170575 | 0.376170575 | ENSG00000264901 | n.s. |
| ENSG00000276030 | 12.83015 | -1.11223514 | 0.619595 | -1.795099 | 0.072638 | 0.459013748 | | lincRNA | 32.28 | -0.376129347 | 0.376129347 | ENSG00000276030 | n.s. |
| ENSG00000199122 | 5.361896 | -1.059247216 | 0.579048 | -1.829291 | 0.067356 | 0.442148543 | MIR148B | miRNA | 43.43 | -0.375430899 | 0.375430899 | ENSG00000199122 | n.s. |
| ENSG00000255284 | 37.22224 | -0.719083632 | 0.329647 | -2.181376 | 0.029156 | 0.302134081 | | lincRNA | 59.11 | -0.373779876 | 0.373779876 | ENSG00000255284 | n.s. |
| ENSG00000266245 | 12.88013 | -0.905824733 | 0.456182 | -1.985664 | 0.047071 | 0.387052655 | MIR4644 | miRNA | 59.52 | -0.373408083 | 0.373408083 | ENSG00000266245 | n.s. |
| ENSG00000262172 | 25.69953 | -0.769591749 | 0.362044 | -2.125685 | 0.03353 | 0.327371305 | | sense_intronic | 62.58 | -0.373220745 | 0.373220745 | ENSG00000262172 | n.s. |
| ENSG00000222511 | 6.1392 | -1.133940881 | 0.638545 | -1.775821 | 0.075762 | 0.468789069 | | misc_RNA | 33.98 | -0.373092091 | 0.373092091 | ENSG00000222511 | n.s. |
| ENSG00000120129 | 608.7003 | 0.502788497 | 0.198886 | 2.5280194 | 0.011471 | 0.181717699 | DUSP1 | protein_coding | 52.99 | 0.372366554 | 0.372366554 | ENSG00000120129 | n.s. |
| ENSG00000270558 | 32.53812 | -0.726928678 | 0.335054 | -2.169586 | 0.030038 | 0.307717153 | | processed_pseudogene | 36.11 | -0.372077204 | 0.372077204 | ENSG00000270558 | n.s. |
| ENSG00000208032 | 9.11939 | -0.929907558 | 0.475785 | -1.95447 | 0.050646 | 0.398383517 | MIR548A3 | miRNA | 36.08 | -0.371682785 | 0.371682785 | ENSG00000208032 | n.s. |
| ENSG00000280968 | 4.623634 | -1.21184373 | 0.709319 | -1.708462 | 0.087551 | 0.493646579 | MIR3653 | miRNA | 56.36 | -0.371531739 | 0.371531739 | ENSG00000280968 | n.s. |
| ENSG00000165891 | 1102.004 | -0.560971967 | 0.232383 | -2.413994 | 0.015779 | 0.217674794 | E2F7 | protein_coding | 40.18 | -0.371471068 | 0.371471068 | ENSG00000165891 | n.s. |
| ENSG00000207712 | 3.832861 | 1.191046191 | 0.691123 | 1.7233479 | 0.084826 | 0.48779901 | MIR627 | miRNA | 35.05 | 0.371319471 | 0.371319471 | ENSG00000207712 | n.s. |
| ENSG00000204388 | 269.6415 | 0.554793473 | 0.228972 | 2.4229782 | 0.015394 | 0.214328592 | HSPA1B | protein_coding | 58.75 | 0.371112389 | 0.371112389 | ENSG00000204388 | n.s. |
| ENSG00000260966 | 289.1682 | -0.6913713 | 0.31325 | -2.207094 | 0.027307 | 0.290674245 | | sense_overlapping | 34.44 | -0.370985309 | 0.370985309 | ENSG00000260966 | n.s. |
| ENSG00000277476 | 107.1927 | -0.610906731 | 0.261698 | -2.334399 | 0.019575 | 0.247090218 | | lincRNA | 59.74 | -0.370908629 | 0.370908629 | ENSG00000277476 | n.s. |
| ENSG00000123689 | 27.95577 | 0.768006218 | 0.362489 | 2.1187033 | 0.034116 | 0.329631296 | G0S2 | protein_coding | 57.59 | 0.370157156 | 0.370157156 | ENSG00000123689 | n.s. |
| ENSG00000112210 | 650.3252 | -0.605886172 | 0.258828 | -2.340887 | 0.019238 | 0.245220122 | RAB23 | protein_coding | 37.66 | -0.369859515 | 0.369859515 | ENSG00000112210 | n.s. |
| ENSG00000213799 | 432.3341 | -0.628112775 | 0.273165 | -2.299393 | 0.021483 | 0.258034534 | ZNF845 | protein_coding | 44.17 | -0.369532669 | 0.369532669 | ENSG00000213799 | n.s. |
| ENSG00000263235 | 41.98022 | -0.67422299 | 0.302736 | -2.227096 | 0.025941 | 0.283942182 | | sense_overlapping | 50.84 | -0.368644961 | 0.368644961 | ENSG00000263235 | n.s. |
| ENSG00000143147 | 59.08492 | -0.68597259 | 0.311043 | -2.205395 | 0.027426 | 0.291271934 | GPR161 | protein_coding | 46.83 | -0.36747645 | 0.36747645 | ENSG00000143147 | n.s. |
| ENSG00000244389 | 62.00126 | -0.649263676 | 0.287592 | -2.257588 | 0.023971 | 0.272897252 | RN7SL242P | misc_RNA | 53.54 | -0.366185257 | 0.366185257 | ENSG00000244389 | n.s. |
| ENSG00000280376 | 39.72556 | -0.751572412 | 0.352896 | -2.12973 | 0.033194 | 0.325902089 | | TEC | 43.92 | -0.36595027 | 0.36595027 | ENSG00000280376 | n.s. |
| ENSG00000283236 | 157.0694 | -0.645019113 | 0.284626 | -2.266201 | 0.023439 | 0.270823271 | | transcribed_unprocessed_pseudc | 38.76 | -0.365928387 | 0.365928387 | ENSG00000283236 | n.s. |
| ENSG00000130164 | 1399.339 | -0.555613211 | 0.230806 | -2.407277 | 0.016072 | 0.219790693 | LDLR | protein_coding | 51.08 | -0.365588326 | 0.365588326 | ENSG00000130164 | n.s. |
| ENSG00000242971 | 16.93061 | -0.858502501 | 0.426306 | -2.013819 | 0.044029 | 0.375283011 | RN7SL233P | misc_RNA | 52.71 | -0.365413945 | 0.365413945 | ENSG00000242971 | n.s. |
| ENSG00000200394 | 46.30863 | -0.651920874 | 0.289636 | -2.250826 | 0.024397 | 0.275291793 | SNORA38B | snoRNA | 48.09 | -0.365210464 | 0.365210464 | ENSG00000200394 | n.s. |
| ENSG00000249138 | 26.12239 | -0.795170443 | 0.382374 | -2.079564 | 0.037566 | 0.347525043 | | transcribed_processed_pseudog | 48.47 | -0.364994282 | 0.364994282 | ENSG00000249138 | n.s. |
| ENSG00000167395 | 640.7172 | -0.513477136 | 0.207108 | -2.479266 | 0.013165 | 0.194706159 | ZNF646 | protein_coding | 58.57 | -0.364887282 | 0.364887282 | ENSG00000167395 | n.s. |
| ENSG00000183688 | 294.7795 | -0.646535908 | 0.286223 | -2.258858 | 0.023892 | 0.272665375 | RFLNB | protein_coding | 48.81 | -0.364885475 | 0.364885475 | ENSG00000183688 | n.s. |
| ENSG00000178429 | 68.95758 | 0.977871355 | 0.518706 | 1.8852146 | 0.059401 | 0.423990171 | RPS3AP5 | processed_pseudogene | 42.39 | 0.364398099 | 0.364398099 | ENSG00000178429 | n.s. |

| | | | | | | | | | | | | | |
|---|---|---|---|---|---|---|---|---|---|---|---|---|---|
| ENSG00000213015 | 114.4598 | -0.612594989 | 0.265124 | -2.310594 | 0.020855 | 0.255099509 | ZNF580 | protein_coding | 57.45 | -0.363446712 | 0.363446712 | ENSG00000213015 | n.s. |
| ENSG00000279197 | 90.06404 | 0.71852132 | 0.33281 | 2.1589567 | 0.030854 | 0.31264457 | | TEC | 36.38 | 0.3628167 | 0.3628167 | ENSG00000279197 | n.s. |
| ENSG00000233237 | 112.7529 | -0.586967628 | 0.24977 | -2.350037 | 0.018772 | 0.241064423 | LINC00472 | lincRNA | 37.07 | -0.362667857 | 0.362667857 | ENSG00000233237 | n.s. |
| ENSG00000266501 | 17.40116 | -0.788817893 | 0.378996 | -2.081336 | 0.037403 | 0.346944726 | | processed_pseudogene | 47.49 | -0.362650909 | 0.362650909 | ENSG00000266501 | n.s. |
| ENSG00000143970 | 1083.881 | -0.640487753 | 0.283303 | -2.260783 | 0.023773 | 0.272193206 | ASXL2 | protein_coding | 39.53 | -0.361954181 | 0.361954181 | ENSG00000143970 | n.s. |
| ENSG00000243199 | 268.041 | 0.722346703 | 0.335672 | 2.1519437 | 0.031402 | 0.315901054 | | processed_pseudogene | 46.44 | 0.361497631 | 0.361497631 | ENSG00000243199 | n.s. |
| ENSG00000176155 | 102.3779 | -0.590971081 | 0.252268 | -2.34263 | 0.019148 | 0.244589652 | CCDC57 | protein_coding | 51.82 | -0.361415409 | 0.361415409 | ENSG00000176155 | n.s. |
| ENSG00000272263 | 93.4571 | -0.609413708 | 0.264231 | -2.306367 | 0.02109 | 0.255337128 | | antisense | 43.14 | -0.361312874 | 0.361312874 | ENSG00000272263 | n.s. |
| ENSG00000125629 | 182.986 | -0.632272279 | 0.27829 | -2.271991 | 0.023087 | 0.268314471 | INSIG2 | protein_coding | 37.96 | -0.361252499 | 0.361252499 | ENSG00000125629 | n.s. |
| ENSG00000225573 | 46.37262 | 0.986522114 | 0.529967 | 1.8614781 | 0.062677 | 0.430511912 | RPL35P5 | processed_pseudogene | 56.49 | 0.361081721 | 0.361081721 | ENSG00000225573 | n.s. |
| ENSG00000237731 | 48.24383 | 0.642959046 | 0.285391 | 2.252909 | 0.024265 | 0.2744889 | RNGTTP1 | processed_pseudogene | 41.35 | 0.361005567 | 0.361005567 | ENSG00000237731 | n.s. |
| ENSG00000243446 | 8.780257 | -1.002830988 | 0.542205 | -1.84954 | 0.06438 | 0.436711663 | RN7SL284P | misc_RNA | 53.82 | -0.360823814 | 0.360823814 | ENSG00000243446 | n.s. |
| ENSG00000235651 | 93.07037 | -0.671227768 | 0.304 | -2.207984 | 0.027245 | 0.29045754 | | processed_pseudogene | 47.42 | -0.360393831 | 0.360393831 | ENSG00000235651 | n.s. |
| ENSG00000207368 | 9.122327 | -0.968956536 | 0.514688 | -1.88261 | 0.059753 | 0.424678304 | | misc_RNA | 43.24 | -0.360393623 | 0.360393623 | ENSG00000207368 | n.s. |
| ENSG00000141040 | 306.1635 | -0.543233263 | 0.224864 | -2.415829 | 0.015699 | 0.217063736 | ZNF287 | protein_coding | 40.28 | -0.36038786 | 0.36038786 | ENSG00000141040 | n.s. |
| ENSG00000260949 | 132.5443 | -0.642069874 | 0.285108 | -2.252022 | 0.024321 | 0.274853743 | | sense_overlapping | 46.98 | -0.360135929 | 0.360135929 | ENSG00000260949 | n.s. |
| ENSG00000244044 | 39.61914 | -0.73759077 | 0.346438 | -2.129067 | 0.033249 | 0.325902089 | RN7SL735P | misc_RNA | 55.82 | -0.359142429 | 0.359142429 | ENSG00000244044 | n.s. |
| ENSG00000251369 | 351.5849 | -0.615742037 | 0.26913 | -2.287899 | 0.022143 | 0.261111919 | ZNF550 | protein_coding | 44.48 | -0.359084317 | 0.359084317 | ENSG00000251369 | n.s. |
| ENSG00000118762 | 977.6014 | -0.622158472 | 0.273129 | -2.277895 | 0.022733 | 0.264861281 | PKD2 | protein_coding | 40.51 | -0.358973944 | 0.358973944 | ENSG00000118762 | n.s. |
| ENSG00000140320 | 572.3607 | -0.559600266 | 0.234894 | -2.382353 | 0.017202 | 0.228309094 | BAHD1 | protein_coding | 53.77 | -0.358970582 | 0.358970582 | ENSG00000140320 | n.s. |
| ENSG00000182809 | 68.41505 | -0.73156714 | 0.342608 | -2.135286 | 0.032738 | 0.323186501 | CRIP2 | protein_coding | 69.57 | -0.35886791 | 0.35886791 | ENSG00000182809 | n.s. |
| ENSG00000184619 | 166.1973 | -0.549344543 | 0.228905 | -2.399877 | 0.016401 | 0.222558474 | KRBA2 | protein_coding | 48.93 | -0.358478003 | 0.358478003 | ENSG00000184619 | n.s. |
| ENSG00000227331 | 68.0205 | -0.654146205 | 0.293516 | -2.228659 | 0.025837 | 0.283560235 | | processed_pseudogene | 50.75 | -0.358049982 | 0.358049982 | ENSG00000227331 | n.s. |
| ENSG00000145014 | 98.50127 | -0.657260584 | 0.295662 | -2.22301 | 0.026215 | 0.285820844 | TMEM44 | protein_coding | 50.87 | -0.357488043 | 0.357488043 | ENSG00000145014 | n.s. |
| ENSG00000212385 | 9.050304 | -1.334759108 | 0.828024 | -1.611981 | 0.106966 | 0.539862594 | RNU6-817P | snRNA | 44.23 | -0.357337388 | 0.357337388 | ENSG00000212385 | n.s. |
| ENSG00000245857 | 49.72752 | -0.677567041 | 0.309224 | -2.191185 | 0.028438 | 0.297290527 | | lincRNA | 46.12 | -0.356955143 | 0.356955143 | ENSG00000245857 | n.s. |
| ENSG00000109674 | 496.3885 | -0.614289361 | 0.268959 | -2.283948 | 0.022375 | 0.262880506 | NEIL3 | protein_coding | 37.34 | -0.356436253 | 0.356436253 | ENSG00000109674 | n.s. |
| ENSG00000159399 | 3173.957 | -0.56433962 | 0.238352 | -2.367676 | 0.0179 | 0.234013871 | HK2 | protein_coding | 47.62 | -0.355961955 | 0.355961955 | ENSG00000159399 | n.s. |
| ENSG00000277806 | 35.93172 | -0.689163172 | 0.316701 | -2.176068 | 0.02955 | 0.304535394 | | sense_intronic | 29.65 | -0.355857829 | 0.355857829 | ENSG00000277806 | n.s. |
| ENSG00000270100 | 29.76893 | -0.737495363 | 0.347605 | -2.121645 | 0.033868 | 0.329262928 | | lincRNA | 40.06 | -0.355809921 | 0.355809921 | ENSG00000270100 | n.s. |
| ENSG00000199289 | 2.690311 | 1.773632543 | 1.278067 | 1.3877463 | 0.165214 | 0.630558849 | RNU6-502P | snRNA | 44.44 | 0.355213149 | 0.355213149 | ENSG00000199289 | n.s. |
| ENSG00000207263 | 5.767921 | -1.041421795 | 0.577701 | -1.8027 | 0.071435 | 0.456235439 | SNORD116-16 | snoRNA | 43.48 | -0.354927986 | 0.354927986 | ENSG00000207263 | n.s. |

| | | | | | | | | | | | | | |
|---|---|---|---|---|---|---|---|---|---|---|---|---|---|
| ENSG00000229591 | 24.77397 | -0.725383601 | 0.340141 | -2.132598 | 0.032958 | 0.324308154 | | antisense | 43.97 | -0.354743143 | 0.354743143 | ENSG00000229591 | n.s. |
| ENSG00000120370 | 323.1977 | -0.620898448 | 0.273338 | -2.271541 | 0.023114 | 0.268406769 | GORAB | protein_coding | 36.49 | -0.354661251 | 0.354661251 | ENSG00000120370 | n.s. |
| ENSG00000235613 | 53.1283 | -0.642602306 | 0.287237 | -2.237182 | 0.025274 | 0.280618873 | NSRP1P1 | processed_pseudogene | 41.35 | -0.354641368 | 0.354641368 | ENSG00000235613 | n.s. |
| ENSG00000187013 | 191.4083 | -0.556803229 | 0.234431 | -2.375121 | 0.017543 | 0.231082019 | C17orf82 | lincRNA | 70.85 | -0.35425706 | 0.35425706 | ENSG00000187013 | n.s. |
| ENSG00000125841 | 279.8166 | -0.590634903 | 0.254799 | -2.318045 | 0.020447 | 0.252377705 | NRSN2 | protein_coding | 51.94 | -0.353169558 | 0.353169558 | ENSG00000125841 | n.s. |
| ENSG00000233644 | 27.22624 | -0.755330394 | 0.360496 | -2.095254 | 0.036148 | 0.340867363 | ARHGEF7-IT1 | sense_intronic | 43.09 | -0.353052438 | 0.353052438 | ENSG00000233644 | n.s. |
| ENSG00000225026 | 10.70457 | 0.871530127 | 0.443691 | 1.9642719 | 0.049499 | 0.393585711 | | processed_pseudogene | 40.81 | 0.35293543 | 0.35293543 | ENSG00000225026 | n.s. |
| ENSG00000196505 | 884.7874 | -0.591506017 | 0.25549 | -2.31518 | 0.020603 | 0.253631062 | GDAP2 | protein_coding | 36.55 | -0.352417841 | 0.352417841 | ENSG00000196505 | n.s. |
| ENSG00000276216 | 751.9747 | 0.504353481 | 0.205084 | 2.4592562 | 0.013923 | 0.200665921 | | lincRNA | 49.57 | 0.351799857 | 0.351799857 | ENSG00000276216 | n.s. |
| ENSG00000181027 | 261.1413 | -0.621691828 | 0.275032 | -2.260436 | 0.023794 | 0.272216048 | FKRP | protein_coding | 51.13 | -0.351309519 | 0.351309519 | ENSG00000181027 | n.s. |
| ENSG00000082397 | 2098.765 | -0.575821288 | 0.246166 | -2.339155 | 0.019327 | 0.245525197 | EPB41L3 | protein_coding | 39.83 | -0.351195668 | 0.351195668 | ENSG00000082397 | n.s. |
| ENSG00000205485 | 236.5277 | -0.598706289 | 0.260867 | -2.295067 | 0.021729 | 0.259183371 | | transcribed_unprocessed_pseudc | 41.83 | -0.351077097 | 0.351077097 | ENSG00000205485 | n.s. |
| ENSG00000117877 | 288.6746 | -0.588944467 | 0.254332 | -2.315654 | 0.020577 | 0.253535977 | CD3EAP | protein_coding | 53.31 | -0.350987582 | 0.350987582 | ENSG00000117877 | n.s. |
| ENSG00000168763 | 218.4389 | -0.620729113 | 0.274932 | -2.257758 | 0.023961 | 0.272897252 | CNNM3 | protein_coding | 53.5 | -0.35009174 | 0.35009174 | ENSG00000168763 | n.s. |
| ENSG00000270231 | 193.3702 | -0.645855185 | 0.291682 | -2.214243 | 0.026812 | 0.288266097 | NBPF8 | unprocessed_pseudogene | 44.63 | -0.348895125 | 0.348895125 | ENSG00000270231 | n.s. |
| ENSG00000189343 | 285.9149 | 0.702166232 | 0.327469 | 2.1442208 | 0.032015 | 0.318846689 | | processed_pseudogene | 60.82 | 0.348568019 | 0.348568019 | ENSG00000189343 | n.s. |
| ENSG00000221333 | 3.929073 | -1.45901703 | 0.955681 | -1.526679 | 0.126841 | 0.576937759 | MIR548K | miRNA | 34.48 | -0.348516911 | 0.348516911 | ENSG00000221333 | n.s. |
| ENSG00000212664 | 42.99207 | 0.981488064 | 0.536115 | 1.8307429 | 0.067139 | 0.441892852 | | processed_pseudogene | 46.13 | 0.348117154 | 0.348117154 | ENSG00000212664 | n.s. |
| ENSG00000187987 | 352.9476 | -0.591632887 | 0.257254 | -2.299797 | 0.02146 | 0.258017317 | ZSCAN23 | protein_coding | 39.86 | -0.348087886 | 0.348087886 | ENSG00000187987 | n.s. |
| ENSG00000203506 | 193.7705 | 0.559620018 | 0.237573 | 2.3555696 | 0.018494 | 0.238865111 | RBMS3-AS2 | antisense | 36.04 | 0.347998186 | 0.347998186 | ENSG00000203506 | n.s. |
| ENSG00000202470 | 20.325 | -0.846141051 | 0.427171 | -1.980802 | 0.047613 | 0.387927349 | | misc_RNA | 40.78 | -0.34797517 | 0.34797517 | ENSG00000202470 | n.s. |
| ENSG00000181826 | 304.0954 | -0.578127409 | 0.248699 | -2.324609 | 0.020093 | 0.25045014 | RELL1 | protein_coding | 42.83 | -0.34761571 | 0.34761571 | ENSG00000181826 | n.s. |
| ENSG00000137970 | 275.0321 | 0.7107625 | 0.333213 | 2.1330606 | 0.03292 | 0.324308154 | RPL7P9 | processed_pseudogene | 42.97 | 0.347592808 | 0.347592808 | ENSG00000137970 | n.s. |
| ENSG00000158195 | 309.6492 | -0.539837646 | 0.226286 | -2.385639 | 0.017049 | 0.2271575 | WASF2 | protein_coding | 44 | -0.347478873 | 0.347478873 | ENSG00000158195 | n.s. |
| ENSG00000202222 | 27.51846 | -0.813979415 | 0.403827 | -2.015662 | 0.043835 | 0.374378906 | | misc_RNA | 41.59 | -0.347315761 | 0.347315761 | ENSG00000202222 | n.s. |
| ENSG00000065060 | 989.3244 | -0.594675712 | 0.259874 | -2.288325 | 0.022119 | 0.261111919 | UHRF1BP1 | protein_coding | 42.65 | -0.346798999 | 0.346798999 | ENSG00000065060 | n.s. |
| ENSG00000266111 | 38.83095 | -0.717816305 | 0.338846 | -2.118417 | 0.03414 | 0.329631296 | | lincRNA | 44.26 | -0.345967046 | 0.345967046 | ENSG00000266111 | n.s. |
| ENSG00000273243 | 27.89137 | -0.741525025 | 0.354461 | -2.091979 | 0.03644 | 0.341750791 | | lincRNA | 43.97 | -0.345766054 | 0.345766054 | ENSG00000273243 | n.s. |
| ENSG00000213185 | 51.15728 | -0.758080076 | 0.365929 | -2.071657 | 0.038297 | 0.350082264 | FAM24B | protein_coding | 41.03 | -0.345555559 | 0.345555559 | ENSG00000213185 | n.s. |
| ENSG00000278177 | 45.6207 | 0.664577591 | 0.304764 | 2.1806317 | 0.029211 | 0.302134081 | | antisense | 52.69 | 0.34544762 | 0.34544762 | ENSG00000278177 | n.s. |
| ENSG00000234287 | 125.0746 | 0.739522819 | 0.353299 | 2.0931943 | 0.036332 | 0.341167655 | | processed_pseudogene | 49.41 | 0.345380932 | 0.345380932 | ENSG00000234287 | n.s. |
| ENSG00000135637 | 276.2149 | -0.548806427 | 0.23195 | -2.36605 | 0.017979 | 0.234823323 | CCDC142 | protein_coding | 52.12 | -0.345341258 | 0.345341258 | ENSG00000135637 | n.s. |

| | | | | | | | | | | | | | |
|---|---|---|---|---|---|---|---|---|---|---|---|---|---|
| ENSG00000139800 | 1001.483 | -0.510856423 | 0.210141 | -2.431013 | 0.015057 | 0.210899376 | ZIC5 | protein_coding | 51.78 | -0.345300477 | 0.345300477 | ENSG00000139800 | n.s. |
| ENSG00000226276 | 16.07133 | -0.811566335 | 0.40311 | -2.013261 | 0.044087 | 0.375429044 | | lincRNA | 37.65 | -0.345298859 | 0.345298859 | ENSG00000226276 | n.s. |
| ENSG00000226396 | 90.54745 | 0.659899981 | 0.301821 | 2.1863935 | 0.028787 | 0.299774863 | | processed_pseudogene | 55.38 | 0.345262829 | 0.345262829 | ENSG00000226396 | n.s. |
| ENSG00000235990 | 19.83929 | -0.770236583 | 0.375438 | -2.051569 | 0.040212 | 0.35721239 | RPL23AP20 | processed_pseudogene | 49.64 | -0.344352354 | 0.344352354 | ENSG00000235990 | n.s. |
| ENSG00000007047 | 272.136 | -0.590290231 | 0.258129 | -2.286804 | 0.022207 | 0.261141374 | MARK4 | protein_coding | 53.57 | -0.344212587 | 0.344212587 | ENSG00000007047 | n.s. |
| ENSG00000183020 | 894.3097 | -0.566225074 | 0.242555 | -2.334422 | 0.019574 | 0.247090218 | AP2A2 | protein_coding | 53.1 | -0.34378041 | 0.34378041 | ENSG00000183020 | n.s. |
| ENSG00000212354 | 3.800172 | -1.196869967 | 0.720835 | -1.660394 | 0.096835 | 0.516736278 | RNU6-1242P | snRNA | 33.01 | -0.343179779 | 0.343179779 | ENSG00000212354 | n.s. |
| ENSG00000203327 | 29.50765 | -0.742147442 | 0.355919 | -2.085161 | 0.037055 | 0.34486157 | | antisense | 37.18 | -0.343135728 | 0.343135728 | ENSG00000203327 | n.s. |
| ENSG00000238778 | 10.55989 | -0.910495244 | 0.480677 | -1.894192 | 0.0582 | 0.420504599 | RNU7-80P | snRNA | 40.98 | -0.342554943 | 0.342554943 | ENSG00000238778 | n.s. |
| ENSG00000276449 | 123.5 | -0.628448562 | 0.282511 | -2.224509 | 0.026114 | 0.285167272 | | antisense | 41.89 | -0.342441822 | 0.342441822 | ENSG00000276449 | n.s. |
| ENSG00000139645 | 451.8951 | -0.540119563 | 0.227867 | -2.370325 | 0.017772 | 0.232780921 | ANKRD52 | protein_coding | 53.52 | -0.341924103 | 0.341924103 | ENSG00000139645 | n.s. |
| ENSG00000158636 | 263.1893 | -0.604862085 | 0.26748 | -2.261333 | 0.023739 | 0.272193206 | EMSY | protein_coding | 37.68 | -0.341821306 | 0.341821306 | ENSG00000158636 | n.s. |
| ENSG00000228873 | 16.09412 | -0.883406361 | 0.458422 | -1.92706 | 0.053972 | 0.410295828 | | lincRNA | 61.16 | -0.341792482 | 0.341792482 | ENSG00000228873 | n.s. |
| ENSG00000266589 | 13.02463 | -0.928096849 | 0.495673 | -1.872396 | 0.061152 | 0.42849416 | MIR4512 | miRNA | 53.25 | -0.341590772 | 0.341590772 | ENSG00000266589 | n.s. |
| ENSG00000229119 | 61.6117 | 0.723622465 | 0.344192 | 2.1023822 | 0.03552 | 0.337341243 | | processed_pseudogene | 55.25 | 0.341499554 | 0.341499554 | ENSG00000229119 | n.s. |
| ENSG00000227598 | 55.04894 | -0.634362013 | 0.287175 | -2.208977 | 0.027176 | 0.290191679 | | antisense | 43.79 | -0.340852253 | 0.340852253 | ENSG00000227598 | n.s. |
| ENSG00000068024 | 56.59484 | -0.674134695 | 0.312049 | -2.160346 | 0.030746 | 0.312237522 | HDAC4 | protein_coding | 51.33 | -0.340785139 | 0.340785139 | ENSG00000068024 | n.s. |
| ENSG00000148803 | 160.3273 | -0.588682093 | 0.258057 | -2.281209 | 0.022536 | 0.26389408 | FUOM | protein_coding | 67.07 | -0.340594006 | 0.340594006 | ENSG00000148803 | n.s. |
| ENSG00000151131 | 2292.365 | -0.578818742 | 0.251612 | -2.300446 | 0.021423 | 0.258017317 | C12orf45 | protein_coding | 39.89 | -0.340548669 | 0.340548669 | ENSG00000151131 | n.s. |
| ENSG00000207622 | 4.239024 | -1.57094924 | 1.07997 | -1.454623 | 0.145774 | 0.607109641 | MIR619 | miRNA | 56.57 | -0.340476338 | 0.340476338 | ENSG00000207622 | n.s. |
| ENSG00000138411 | 598.919 | -0.563499574 | 0.242014 | -2.328375 | 0.019892 | 0.248850741 | HECW2 | protein_coding | 39.16 | -0.340388151 | 0.340388151 | ENSG00000138411 | n.s. |
| ENSG00000200241 | 19.88688 | -0.780182064 | 0.384219 | -2.030568 | 0.042299 | 0.36624784 | | misc_RNA | 43.24 | -0.340334864 | 0.340334864 | ENSG00000200241 | n.s. |
| ENSG00000280040 | 23.04828 | 0.82958508 | 0.419208 | 1.9789351 | 0.047823 | 0.3891173 | | TEC | 34.38 | 0.340063069 | 0.340063069 | ENSG00000280040 | n.s. |
| ENSG00000156500 | 22.35532 | -0.787778997 | 0.389298 | -2.023588 | 0.043013 | 0.370584328 | FAM122C | protein_coding | 41.56 | -0.339621728 | 0.339621728 | ENSG00000156500 | n.s. |
| ENSG00000085978 | 402.9259 | -0.579849219 | 0.253495 | -2.287419 | 0.022171 | 0.261111919 | ATG16L1 | protein_coding | 45.33 | -0.338152584 | 0.338152584 | ENSG00000085978 | n.s. |
| ENSG00000228423 | 15.93013 | -0.874756902 | 0.454803 | -1.923377 | 0.054433 | 0.411605417 | | lincRNA | 44.71 | -0.337235335 | 0.337235335 | ENSG00000228423 | n.s. |
| ENSG00000183655 | 781.4489 | -0.575990583 | 0.251289 | -2.292143 | 0.021897 | 0.260036752 | KLHL25 | protein_coding | 53.6 | -0.336934486 | 0.336934486 | ENSG00000183655 | n.s. |
| ENSG00000278588 | 639.1952 | -0.51618966 | 0.215049 | -2.400339 | 0.01638 | 0.222495094 | HIST1H2BI | protein_coding | 58.79 | -0.336906441 | 0.336906441 | ENSG00000278588 | n.s. |
| ENSG00000266405 | 78.71391 | 0.627200131 | 0.284273 | 2.2063324 | 0.027361 | 0.29088361 | | transcribed_processed_pseudog | 40.82 | 0.336355356 | 0.336355356 | ENSG00000266405 | n.s. |
| ENSG00000206612 | 30.52865 | -0.732465067 | 0.352268 | -2.079285 | 0.037591 | 0.347531413 | SNORA2A | snoRNA | 43.7 | -0.336205812 | 0.336205812 | ENSG00000206612 | n.s. |
| ENSG00000251748 | 15.80065 | -0.82540623 | 0.418914 | -1.970346 | 0.048799 | 0.391599066 | RNU4ATAC11P | snRNA | 49.21 | -0.336071041 | 0.336071041 | ENSG00000251748 | n.s. |
| ENSG00000234160 | 99.28932 | -0.637726355 | 0.290742 | -2.193446 | 0.028275 | 0.297290527 | | antisense | 48.87 | -0.335966315 | 0.335966315 | ENSG00000234160 | n.s. |

| | | | | | | | | | | | | | |
|---|---|---|---|---|---|---|---|---|---|---|---|---|---|
| ENSG00000238523 | 1.876995 | -1.43914977 | 0.956728 | -1.504241 | 0.132519 | 0.584436416 | RNU7-107P | snRNA | 32.79 | -0.335700006 | 0.335700006 | ENSG00000238523 | n.s. |
| ENSG00000143013 | 1851.071 | -0.602051481 | 0.2682 | -2.244783 | 0.024782 | 0.277363624 | LMO4 | protein_coding | 42.55 | -0.335312871 | 0.335312871 | ENSG00000143013 | n.s. |
| ENSG00000235092 | 48.58451 | 0.673006318 | 0.313681 | 2.1455115 | 0.031912 | 0.31813913 | ID2-AS1 | antisense | 48.44 | 0.334741841 | 0.334741841 | ENSG00000235092 | n.s. |
| ENSG00000255270 | 104.5843 | -0.64386787 | 0.295257 | -2.1807 | 0.029206 | 0.302134081 | NAV2-IT1 | sense_intronic | 44.27 | -0.334682701 | 0.334682701 | ENSG00000255270 | n.s. |
| ENSG00000214425 | 571.3439 | -0.593437773 | 0.262904 | -2.257238 | 0.023993 | 0.272922535 | LRRC37A4P | cribed_unprocessed_pseudc | 42.83 | -0.334675524 | 0.334675524 | ENSG00000214425 | n.s. |
| ENSG00000244501 | 48.03846 | 0.634210816 | 0.289443 | 2.1911423 | 0.028441 | 0.297290527 | RN7SL623P | misc_RNA | 52.58 | 0.334114263 | 0.334114263 | ENSG00000244501 | n.s. |
| ENSG00000100485 | 978.4978 | -0.567617015 | 0.247215 | -2.29605 | 0.021673 | 0.258843947 | SOS2 | protein_coding | 38.27 | -0.333169611 | 0.333169611 | ENSG00000100485 | n.s. |
| ENSG00000197857 | 262.9379 | -0.591643065 | 0.262287 | -2.255712 | 0.024089 | 0.273561526 | ZNF44 | protein_coding | 43.35 | -0.333062494 | 0.333062494 | ENSG00000197857 | n.s. |
| ENSG00000277568 | 9.9432 | -0.869050274 | 0.454365 | -1.912669 | 0.05579 | 0.413827257 | | misc_RNA | 57.05 | -0.333003473 | 0.333003473 | ENSG00000277568 | n.s. |
| ENSG00000065802 | 829.1665 | -0.598675336 | 0.266981 | -2.24239 | 0.024936 | 0.277971334 | ASB1 | protein_coding | 47.31 | -0.33286348 | 0.33286348 | ENSG00000065802 | n.s. |
| ENSG00000165914 | 41.81886 | -0.690490509 | 0.326257 | -2.1164 | 0.034311 | 0.329909835 | TTC7B | protein_coding | 46.46 | -0.3325435 | 0.3325435 | ENSG00000165914 | n.s. |
| ENSG00000261799 | 214.9498 | -0.593173196 | 0.263541 | -2.250784 | 0.024399 | 0.275291793 | | sense_overlapping | 39.78 | -0.332299619 | 0.332299619 | ENSG00000261799 | n.s. |
| ENSG00000167562 | 326.5934 | -0.558963213 | 0.241939 | -2.310347 | 0.020869 | 0.255099509 | ZNF701 | protein_coding | 45.33 | -0.331627495 | 0.331627495 | ENSG00000167562 | n.s. |
| ENSG00000252914 | 11.47292 | 0.900274141 | 0.480638 | 1.8730807 | 0.061057 | 0.42849416 | RNU6-789P | snRNA | 34.31 | 0.331350482 | 0.331350482 | ENSG00000252914 | n.s. |
| ENSG00000145545 | 601.2293 | -0.609238793 | 0.274559 | -2.218972 | 0.026489 | 0.28655881 | SRD5A1 | protein_coding | 44.23 | -0.330686431 | 0.330686431 | ENSG00000145545 | n.s. |
| ENSG00000249685 | 12.83343 | -0.820921106 | 0.41876 | -1.960363 | 0.049953 | 0.395620546 | | lincRNA | 41.75 | -0.330602302 | 0.330602302 | ENSG00000249685 | n.s. |
| ENSG00000196150 | 624.8661 | -0.612283509 | 0.276609 | -2.213534 | 0.026861 | 0.288501124 | ZNF250 | protein_coding | 47.13 | -0.330542777 | 0.330542777 | ENSG00000196150 | n.s. |
| ENSG00000203872 | 204.4538 | -0.619901344 | 0.28162 | -2.2012 | 0.027722 | 0.293049415 | C6orf163 | protein_coding | 40.23 | -0.330444078 | 0.330444078 | ENSG00000203872 | n.s. |
| ENSG00000236698 | 26.22026 | 0.724148326 | 0.349817 | 2.0700795 | 0.038445 | 0.350082264 | EIF1AXP1 | processed_pseudogene | 38.62 | 0.330088453 | 0.330088453 | ENSG00000236698 | n.s. |
| ENSG00000270117 | 26.67302 | 0.805753776 | 0.408196 | 1.9739367 | 0.048389 | 0.390788403 | | lincRNA | 52.78 | 0.328794543 | 0.328794543 | ENSG00000270117 | n.s. |
| ENSG00000279250 | 26.49499 | -0.721501192 | 0.349071 | -2.066919 | 0.038742 | 0.350944672 | | TEC | 48.4 | -0.328110853 | 0.328110853 | ENSG00000279250 | n.s. |
| ENSG00000242798 | 18.26102 | -0.824952671 | 0.423376 | -1.948512 | 0.051354 | 0.400783303 | | antisense | 51.6 | -0.327580769 | 0.327580769 | ENSG00000242798 | n.s. |
| ENSG00000179862 | 250.2027 | -0.586538309 | 0.261462 | -2.243303 | 0.024877 | 0.27771574 | CITED4 | protein_coding | 73.48 | -0.326349625 | 0.326349625 | ENSG00000179862 | n.s. |
| ENSG00000232931 | 184.5923 | -0.619026767 | 0.282361 | -2.192324 | 0.028356 | 0.297290527 | LINC00342 | lincRNA | 39.66 | -0.326115019 | 0.326115019 | ENSG00000232931 | n.s. |
| ENSG00000162738 | 420.0162 | -0.526463713 | 0.223841 | -2.351958 | 0.018675 | 0.240305064 | VANGL2 | protein_coding | 51.65 | -0.326005851 | 0.326005851 | ENSG00000162738 | n.s. |
| ENSG00000200871 | 3.785251 | -1.164947798 | 0.709371 | -1.642227 | 0.100543 | 0.525174439 | RNU6-810P | snRNA | 38.32 | -0.325831729 | 0.325831729 | ENSG00000200871 | n.s. |
| ENSG00000224728 | 79.35774 | -0.639348908 | 0.295254 | -2.165422 | 0.030355 | 0.30955579 | IMPDH1P8 | processed_pseudogene | 57.11 | -0.325595508 | 0.325595508 | ENSG00000224728 | n.s. |
| ENSG00000183853 | 287.4707 | -0.553209842 | 0.240523 | -2.300027 | 0.021447 | 0.258017317 | KIRREL | protein_coding | 49.16 | -0.325481644 | 0.325481644 | ENSG00000183853 | n.s. |
| ENSG00000211520 | 3.496073 | -1.165520503 | 0.710174 | -1.641177 | 0.100761 | 0.525821245 | MIR216B | miRNA | 45.12 | -0.325368884 | 0.325368884 | ENSG00000211520 | n.s. |
| ENSG00000265056 | 8.45382 | -1.014715504 | 0.578454 | -1.754186 | 0.079399 | 0.478143199 | MIR548S | miRNA | 26.83 | -0.325157483 | 0.325157483 | ENSG00000265056 | n.s. |
| ENSG00000141582 | 631.581 | -0.581702124 | 0.258867 | -2.247109 | 0.024633 | 0.276340982 | CBX4 | protein_coding | 62.53 | -0.324912458 | 0.324912458 | ENSG00000141582 | n.s. |
| ENSG00000224660 | 224.6666 | -0.576086955 | 0.25526 | -2.256861 | 0.024017 | 0.272967872 | SH3BP5-AS1 | antisense | 45.6 | -0.324848789 | 0.324848789 | ENSG00000224660 | n.s. |

| | | | | | | | | | | | | | |
|---|---|---|---|---|---|---|---|---|---|---|---|---|---|
| ENSG00000109689 | 287.5484 | -0.567129424 | 0.249929 | -2.269163 | 0.023258 | 0.269182866 | STIM2 | protein_coding | 36.58 | -0.323236882 | 0.323236882 | ENSG00000109689 | n.s. |
| ENSG00000119640 | 424.0847 | -0.554039943 | 0.242238 | -2.287169 | 0.022186 | 0.261111919 | ACYP1 | protein_coding | 41.66 | -0.323101304 | 0.323101304 | ENSG00000119640 | n.s. |
| ENSG00000200550 | 9.668706 | -0.871341084 | 0.463867 | -1.87843 | 0.060322 | 0.426183381 | RNU6-137P | snRNA | 37.96 | -0.322747778 | 0.322747778 | ENSG00000200550 | n.s. |
| ENSG00000263648 | 27.7657 | -0.711410802 | 0.344501 | -2.06505 | 0.038918 | 0.35209789 | | processed_pseudogene | 43.95 | -0.322508538 | 0.322508538 | ENSG00000263648 | n.s. |
| ENSG00000130309 | 605.9269 | -0.577257529 | 0.256925 | -2.24679 | 0.024653 | 0.276340982 | COLGALT1 | protein_coding | 52.86 | -0.322429908 | 0.322429908 | ENSG00000130309 | n.s. |
| ENSG00000153885 | 1142.446 | -0.57719057 | 0.256714 | -2.24838 | 0.024552 | 0.276340982 | KCTD15 | protein_coding | 56 | -0.322392508 | 0.322392508 | ENSG00000153885 | n.s. |
| ENSG00000206659 | 10.49214 | -0.956748604 | 0.534009 | -1.791635 | 0.073191 | 0.460406637 | | misc_RNA | 42.16 | -0.322288827 | 0.322288827 | ENSG00000206659 | n.s. |
| ENSG00000267719 | 290.6368 | -0.550917583 | 0.240369 | -2.291965 | 0.021908 | 0.260036752 | LINC02078 | lincRNA | 56.79 | -0.322267651 | 0.322267651 | ENSG00000267719 | n.s. |
| ENSG00000252960 | 20.35287 | -0.730822678 | 0.358727 | -2.037269 | 0.041623 | 0.36238932 | RN7SKP258 | misc_RNA | 44.08 | -0.322164622 | 0.322164622 | ENSG00000252960 | n.s. |
| ENSG00000180891 | 631.7641 | -0.550680029 | 0.240358 | -2.291087 | 0.021958 | 0.260036752 | CUEDC1 | protein_coding | 51.38 | -0.32212869 | 0.32212869 | ENSG00000180891 | n.s. |
| ENSG00000186638 | 334.8159 | -0.591931083 | 0.266408 | -2.2219 | 0.02629 | 0.286058985 | KIF24 | protein_coding | 42.81 | -0.321740829 | 0.321740829 | ENSG00000186638 | n.s. |
| ENSG00000131323 | 273.7892 | -0.549252244 | 0.239744 | -2.290992 | 0.021964 | 0.260036752 | TRAF3 | protein_coding | 45.83 | -0.321293486 | 0.321293486 | ENSG00000131323 | n.s. |
| ENSG00000180423 | 74.94113 | -0.591671089 | 0.266583 | -2.219467 | 0.026455 | 0.28641715 | HARBI1 | protein_coding | 46.15 | -0.321277982 | 0.321277982 | ENSG00000180423 | n.s. |
| ENSG00000269974 | 109.9504 | -0.635204656 | 0.294102 | -2.159808 | 0.030787 | 0.312250224 | | lincRNA | 29.47 | -0.321094202 | 0.321094202 | ENSG00000269974 | n.s. |
| ENSG00000241868 | 16.49497 | -0.817317751 | 0.420943 | -1.941637 | 0.052181 | 0.404745128 | RN7SL434P | misc_RNA | 54.61 | -0.321057426 | 0.321057426 | ENSG00000241868 | n.s. |
| ENSG00000087448 | 2118.328 | -0.567198598 | 0.251 | -2.25976 | 0.023836 | 0.272248483 | KLHL42 | protein_coding | 43.03 | -0.320486794 | 0.320486794 | ENSG00000087448 | n.s. |
| ENSG00000197162 | 432.063 | -0.538993177 | 0.232963 | -2.313647 | 0.020687 | 0.25444002 | ZNF785 | protein_coding | 49.75 | -0.320385402 | 0.320385402 | ENSG00000197162 | n.s. |
| ENSG00000238141 | 25.05532 | -0.753221501 | 0.374496 | -2.011293 | 0.044294 | 0.375547181 | BRWD1-AS1 | antisense | 45.53 | -0.320371839 | 0.320371839 | ENSG00000238141 | n.s. |
| ENSG00000258986 | 355.8379 | -0.563542844 | 0.248799 | -2.265053 | 0.023509 | 0.271387839 | TMEM179 | protein_coding | 53.82 | -0.319196087 | 0.319196087 | ENSG00000258986 | n.s. |
| ENSG00000243508 | 52.76249 | -0.630532113 | 0.291766 | -2.161087 | 0.030689 | 0.312109223 | | processed_pseudogene | 41.37 | -0.318855927 | 0.318855927 | ENSG00000243508 | n.s. |
| ENSG00000184678 | 1858.016 | -0.506531995 | 0.214327 | -2.363365 | 0.01811 | 0.235607368 | HIST2H2BE | protein_coding | 47.99 | -0.318006425 | 0.318006425 | ENSG00000184678 | n.s. |
| ENSG00000214182 | 165.6158 | 0.882476768 | 0.476913 | 1.8503926 | 0.064257 | 0.436414668 | PTMAP5 | ıscribed_processed_pseudog | 44.58 | 0.317780467 | 0.317780467 | ENSG00000214182 | n.s. |
| ENSG00000279696 | 327.7591 | -0.532809467 | 0.230079 | -2.315771 | 0.020571 | 0.253535977 | | TEC | 37.34 | -0.317533345 | 0.317533345 | ENSG00000279696 | n.s. |
| ENSG00000155970 | 83.66288 | -0.657363425 | 0.309689 | -2.122659 | 0.033782 | 0.329031605 | MICU3 | protein_coding | 35.97 | -0.317350334 | 0.317350334 | ENSG00000155970 | n.s. |
| ENSG00000259377 | 145.591 | -0.533730469 | 0.230828 | -2.312247 | 0.020764 | 0.254739827 | | sense_intronic | 41.42 | -0.316984207 | 0.316984207 | ENSG00000259377 | n.s. |
| ENSG00000131080 | 118.1415 | -0.601291643 | 0.274346 | -2.191731 | 0.028399 | 0.297290527 | EDA2R | protein_coding | 39.73 | -0.31677182 | 0.31677182 | ENSG00000131080 | n.s. |
| ENSG00000256525 | 714.1144 | -0.582380498 | 0.262251 | -2.220699 | 0.026371 | 0.286058985 | POLG2 | protein_coding | 41.34 | -0.316549662 | 0.316549662 | ENSG00000256525 | n.s. |
| ENSG00000266235 | 5.925343 | -1.104114697 | 0.665739 | -1.65848 | 0.097221 | 0.516972948 | MIR3176 | miRNA | 74.44 | -0.316364393 | 0.316364393 | ENSG00000266235 | n.s. |
| ENSG00000279088 | 373.7903 | 0.512175714 | 0.218115 | 2.3481955 | 0.018865 | 0.241336844 | | TEC | 42.48 | 0.316205184 | 0.316205184 | ENSG00000279088 | n.s. |
| ENSG00000211514 | 43.22248 | -0.653854611 | 0.308519 | -2.119336 | 0.034062 | 0.329615114 | MIR454 | miRNA | 39.13 | -0.315153269 | 0.315153269 | ENSG00000211514 | n.s. |
| ENSG00000185130 | 512.8387 | -0.496571781 | 0.209333 | -2.372157 | 0.017685 | 0.232108003 | HIST1H2BL | protein_coding | 57.17 | -0.314980389 | 0.314980389 | ENSG00000185130 | n.s. |
| ENSG00000200879 | 116.0934 | -0.585943413 | 0.265091 | -2.210348 | 0.027081 | 0.29009854 | SNORD14E | snoRNA | 44.71 | -0.314917899 | 0.314917899 | ENSG00000200879 | n.s. |

| | | | | | | | | | | | | | |
|---|---|---|---|---|---|---|---|---|---|---|---|---|---|
| ENSG00000048707 | 67.19012 | -0.655311823 | 0.310053 | -2.113551 | 0.034554 | 0.33155762 | VPS13D | protein_coding | 42.34 | -0.314183347 | 0.314183347 | ENSG00000048707 | n.s. |
| ENSG00000197016 | 462.0326 | -0.557913165 | 0.247648 | -2.252845 | 0.024269 | 0.2744889 | ZNF470 | protein_coding | 40.11 | -0.313254413 | 0.313254413 | ENSG00000197016 | n.s. |
| ENSG00000103653 | 279.9154 | -0.571176599 | 0.256053 | -2.230701 | 0.025701 | 0.283320929 | CSK | protein_coding | 56.64 | -0.312845614 | 0.312845614 | ENSG00000103653 | n.s. |
| ENSG00000068305 | 286.9124 | -0.582196674 | 0.263717 | -2.207655 | 0.027268 | 0.290479939 | MEF2A | protein_coding | 36.86 | -0.312571995 | 0.312571995 | ENSG00000068305 | n.s. |
| ENSG00000271870 | 101.5837 | -0.648642277 | 0.306431 | -2.116764 | 0.03428 | 0.329909835 | | antisense | 33.81 | -0.312389193 | 0.312389193 | ENSG00000271870 | n.s. |
| ENSG00000077514 | 962.3944 | -0.561131193 | 0.2501 | -2.243629 | 0.024856 | 0.27771574 | POLD3 | protein_coding | 41.72 | -0.312213118 | 0.312213118 | ENSG00000077514 | n.s. |
| ENSG00000215158 | 778.6243 | -0.583511362 | 0.265008 | -2.201867 | 0.027675 | 0.293015122 | | transcribed_unprocessed_pseudc | 38.71 | -0.311075722 | 0.311075722 | ENSG00000215158 | n.s. |
| ENSG00000260565 | 620.0398 | -0.523112831 | 0.226673 | -2.307783 | 0.021011 | 0.255337128 | ERVK13-1 | processed_transcript | 47.26 | -0.31014629 | 0.31014629 | ENSG00000260565 | n.s. |
| ENSG00000248794 | 20.83291 | -0.774458256 | 0.395985 | -1.955779 | 0.050491 | 0.398128225 | | processed_pseudogene | 44.44 | -0.309765515 | 0.309765515 | ENSG00000248794 | n.s. |
| ENSG00000163328 | 62.09549 | 0.629641856 | 0.294877 | 2.135271 | 0.032739 | 0.323186501 | GPR155 | protein_coding | 39.18 | 0.30886879 | 0.30886879 | ENSG00000163328 | n.s. |
| ENSG00000140471 | 869.255 | -0.600661092 | 0.27687 | -2.169469 | 0.030047 | 0.307717153 | LINS1 | protein_coding | 40.18 | -0.307447355 | 0.307447355 | ENSG00000140471 | n.s. |
| ENSG00000105409 | 916.8302 | -0.560182716 | 0.251403 | -2.228229 | 0.025865 | 0.283560235 | ATP1A3 | protein_coding | 56.13 | -0.306618627 | 0.306618627 | ENSG00000105409 | n.s. |
| ENSG00000224687 | 72.96724 | 0.628767439 | 0.295426 | 2.128341 | 0.033309 | 0.325902089 | RASAL2-AS1 | lincRNA | 45.58 | 0.30615495 | 0.30615495 | ENSG00000224687 | n.s. |
| ENSG00000134109 | 404.2649 | -0.55909401 | 0.250866 | -2.228654 | 0.025837 | 0.283560235 | EDEM1 | protein_coding | 43.38 | -0.306022719 | 0.306022719 | ENSG00000134109 | n.s. |
| ENSG00000279140 | 31.42771 | -0.751776176 | 0.381674 | -1.96968 | 0.048875 | 0.391760224 | | TEC | 65.57 | -0.305957613 | 0.305957613 | ENSG00000279140 | n.s. |
| ENSG00000207136 | 6.837913 | -0.957795168 | 0.546916 | -1.751267 | 0.0799 | 0.479305998 | RNU6-769P | snRNA | 36.54 | -0.305907456 | 0.305907456 | ENSG00000207136 | n.s. |
| ENSG00000166135 | 826.535 | -0.54421727 | 0.241399 | -2.254435 | 0.024169 | 0.274225075 | HIF1AN | protein_coding | 45.24 | -0.305791785 | 0.305791785 | ENSG00000166135 | n.s. |
| ENSG00000235381 | 221.8503 | -0.585326309 | 0.26788 | -2.185031 | 0.028887 | 0.300364159 | | antisense | 41.83 | -0.305746304 | 0.305746304 | ENSG00000235381 | n.s. |
| ENSG00000207063 | 8.53904 | -0.893781712 | 0.494922 | -1.805904 | 0.070933 | 0.455117569 | SNORD116-1 | snoRNA | 42.11 | -0.305562873 | 0.305562873 | ENSG00000207063 | n.s. |
| ENSG00000135503 | 972.809 | -0.560853783 | 0.252269 | -2.223239 | 0.0262 | 0.285820844 | ACVR1B | protein_coding | 44.84 | -0.305051795 | 0.305051795 | ENSG00000135503 | n.s. |
| ENSG00000170011 | 767.0565 | -0.556115523 | 0.249219 | -2.231436 | 0.025652 | 0.283008461 | MYRIP | protein_coding | 40.16 | -0.304862852 | 0.304862852 | ENSG00000170011 | n.s. |
| ENSG00000230149 | 39.3289 | -0.664129057 | 0.319292 | -2.080006 | 0.037525 | 0.347525043 | | antisense | 57.97 | -0.304844465 | 0.304844465 | ENSG00000230149 | n.s. |
| ENSG00000282870 | 107.0687 | -0.618493849 | 0.289208 | -2.13858 | 0.03247 | 0.321971749 | FRG1DP | unprocessed_pseudogene | 35.11 | -0.304411684 | 0.304411684 | ENSG00000282870 | n.s. |
| ENSG00000260310 | 21.9112 | 0.809932091 | 0.428027 | 1.8922432 | 0.058459 | 0.42095504 | | sense_intronic | 56.74 | 0.304343555 | 0.304343555 | ENSG00000260310 | n.s. |
| ENSG00000265444 | 2.335306 | 1.51088636 | 1.084392 | 1.3933027 | 0.163528 | 0.629299052 | MIR4733 | miRNA | 52.63 | 0.303904098 | 0.303904098 | ENSG00000265444 | n.s. |
| ENSG00000125319 | 316.3874 | -0.58853023 | 0.270504 | -2.175679 | 0.029579 | 0.304609998 | C17orf53 | protein_coding | 50.23 | -0.303832172 | 0.303832172 | ENSG00000125319 | n.s. |
| ENSG00000120254 | 1362.001 | -0.568771053 | 0.258381 | -2.201287 | 0.027716 | 0.293049415 | MTHFD1L | protein_coding | 43.51 | -0.303188609 | 0.303188609 | ENSG00000120254 | n.s. |
| ENSG00000136014 | 39.7309 | 0.6638168 | 0.320757 | 2.0695336 | 0.038496 | 0.350082264 | USP44 | protein_coding | 41.64 | 0.30258754 | 0.30258754 | ENSG00000136014 | n.s. |
| ENSG00000254512 | 40.17365 | -0.917139497 | 0.516073 | -1.777151 | 0.075543 | 0.46814761 | PHBP2 | processed_pseudogene | 53.1 | -0.302304943 | 0.302304943 | ENSG00000254512 | n.s. |
| ENSG00000201221 | 8.423535 | -0.903120787 | 0.506087 | -1.784518 | 0.07434 | 0.463348535 | RNU4-40P | snRNA | 37.06 | -0.301725616 | 0.301725616 | ENSG00000201221 | n.s. |
| ENSG00000186185 | 1316.813 | -0.532683258 | 0.235204 | -2.264768 | 0.023527 | 0.271387839 | KIF18B | protein_coding | 49.25 | -0.30171692 | 0.30171692 | ENSG00000186185 | n.s. |
| ENSG00000237349 | 12.22274 | -0.901859313 | 0.505866 | -1.782803 | 0.074618 | 0.463799422 | | processed_pseudogene | 40.28 | -0.300923214 | 0.300923214 | ENSG00000237349 | n.s. |

| | | | | | | | | | | | | | |
|---|---|---|---|---|---|---|---|---|---|---|---|---|---|
| ENSG00000267088 | 37.57222 | -0.624021576 | 0.294561 | -2.118477 | 0.034135 | 0.329631296 | | processed_pseudogene | 32.88 | -0.300760654 | 0.300760654 | ENSG00000267088 | n.s. |
| ENSG00000227379 | 25.99649 | -0.731013444 | 0.368803 | -1.982127 | 0.047465 | 0.387927349 | PPIAP2 | processed_pseudogene | 47.97 | -0.300628988 | 0.300628988 | ENSG00000227379 | n.s. |
| ENSG00000273783 | 172.1532 | 0.55297072 | 0.249367 | 2.2174932 | 0.026589 | 0.287074936 | | antisense | 46.82 | 0.299712742 | 0.299712742 | ENSG00000273783 | n.s. |
| ENSG00000196912 | 586.9211 | -0.605819358 | 0.282801 | -2.14221 | 0.032177 | 0.320224581 | ANKRD36B | protein_coding | 35.8 | -0.299605137 | 0.299605137 | ENSG00000196912 | n.s. |
| ENSG00000182541 | 405.4371 | -0.572786158 | 0.26204 | -2.185873 | 0.028825 | 0.299946903 | LIMK2 | protein_coding | 47.35 | -0.299541739 | 0.299541739 | ENSG00000182541 | n.s. |
| ENSG00000264812 | 19.50526 | -0.851702484 | 0.467374 | -1.822314 | 0.068407 | 0.445487047 | | sense_intronic | 50.59 | -0.299088034 | 0.299088034 | ENSG00000264812 | n.s. |
| ENSG00000266579 | 42.82215 | -0.682122195 | 0.335598 | -2.032556 | 0.042097 | 0.36495768 | | lincRNA | 34.79 | -0.298604102 | 0.298604102 | ENSG00000266579 | n.s. |
| ENSG00000275092 | 42.56008 | 0.649471005 | 0.312276 | 2.0797968 | 0.037544 | 0.347525043 | | sense_intronic | 46.68 | 0.298116216 | 0.298116216 | ENSG00000275092 | n.s. |
| ENSG00000213152 | 34.11439 | -0.697617926 | 0.346177 | -2.015204 | 0.043883 | 0.374378906 | | processed_pseudogene | 51.26 | -0.297665637 | 0.297665637 | ENSG00000213152 | n.s. |
| ENSG00000275713 | 1844.203 | -0.496961854 | 0.214366 | -2.318289 | 0.020434 | 0.252377705 | HIST1H2BH | protein_coding | 41.55 | -0.297157851 | 0.297157851 | ENSG00000275713 | n.s. |
| ENSG00000207336 | 7.383754 | -0.924765306 | 0.525829 | -1.758682 | 0.078632 | 0.477239282 | RNU6-658P | snRNA | 38.68 | -0.297093631 | 0.297093631 | ENSG00000207336 | n.s. |
| ENSG00000105722 | 251.2084 | -0.573656389 | 0.263507 | -2.177003 | 0.02948 | 0.304103432 | ERF | protein_coding | 64.09 | -0.296568122 | 0.296568122 | ENSG00000105722 | n.s. |
| ENSG00000232640 | 16.72496 | 0.802092781 | 0.427388 | 1.87673 | 0.060555 | 0.426932326 | | antisense | 47.25 | 0.296486346 | 0.296486346 | ENSG00000232640 | n.s. |
| ENSG00000265520 | 11.24789 | -0.833840893 | 0.455522 | -1.830517 | 0.067173 | 0.441892852 | MIR548V | miRNA | 37.5 | -0.295749209 | 0.295749209 | ENSG00000265520 | n.s. |
| ENSG00000253092 | 8.510162 | -0.976202107 | 0.574729 | -1.698543 | 0.089405 | 0.498053895 | | snoRNA | 37.88 | -0.295519474 | 0.295519474 | ENSG00000253092 | n.s. |
| ENSG00000102804 | 2940.749 | 0.489626623 | 0.210422 | 2.3268802 | 0.019972 | 0.249385825 | TSC22D1 | protein_coding | 38.08 | 0.295307642 | 0.295307642 | ENSG00000102804 | n.s. |
| ENSG00000162929 | 366.5184 | -0.580612626 | 0.268388 | -2.163333 | 0.030516 | 0.31080316 | KIAA1841 | protein_coding | 39.67 | -0.29466937 | 0.29466937 | ENSG00000162929 | n.s. |
| ENSG00000206687 | 15.49183 | -0.853961027 | 0.471262 | -1.812074 | 0.069975 | 0.451882747 | RNU1-109P | snRNA | 48.98 | -0.294594556 | 0.294594556 | ENSG00000206687 | n.s. |
| ENSG00000273017 | 18.52064 | -0.767916874 | 0.400354 | -1.918095 | 0.055099 | 0.413454097 | | sense_intronic | 27.88 | -0.294551949 | 0.294551949 | ENSG00000273017 | n.s. |
| ENSG00000261072 | 126.9439 | -0.635531992 | 0.304525 | -2.086962 | 0.036892 | 0.344263918 | | processed_pseudogene | 44.95 | -0.294320263 | 0.294320263 | ENSG00000261072 | n.s. |
| ENSG00000241494 | 17.79627 | 1.127100016 | 0.70799 | 1.5919728 | 0.111391 | 0.548299349 | | processed_pseudogene | 43.84 | 0.294153121 | 0.294153121 | ENSG00000241494 | n.s. |
| ENSG00000229276 | 37.16479 | -0.657831929 | 0.320699 | -2.051243 | 0.040243 | 0.35721239 | REV3L-IT1 | sense_intronic | 37.95 | -0.294099214 | 0.294099214 | ENSG00000229276 | n.s. |
| ENSG00000226824 | 38.18228 | -0.661171955 | 0.323276 | -2.045222 | 0.040833 | 0.35907784 | | sense_intronic | 50.03 | -0.29409682 | 0.29409682 | ENSG00000226824 | n.s. |
| ENSG00000238723 | 36.94094 | -0.717221477 | 0.362771 | -1.977061 | 0.048035 | 0.389875966 | | misc_RNA | 40.2 | -0.293396325 | 0.293396325 | ENSG00000238723 | n.s. |
| ENSG00000136295 | 572.3679 | -0.53903016 | 0.242512 | -2.222693 | 0.026237 | 0.285830294 | TTYH3 | protein_coding | 62.29 | -0.293174054 | 0.293174054 | ENSG00000136295 | n.s. |
| ENSG00000104427 | 492.1828 | -0.607461987 | 0.286367 | -2.121269 | 0.033899 | 0.329262928 | ZC2HC1A | protein_coding | 34.79 | -0.293074387 | 0.293074387 | ENSG00000104427 | n.s. |
| ENSG00000130294 | 4625.845 | -0.535341799 | 0.240096 | -2.229699 | 0.025767 | 0.283560235 | KIF1A | protein_coding | 58.61 | -0.293021835 | 0.293021835 | ENSG00000130294 | n.s. |
| ENSG00000142166 | 460.5663 | -0.571079495 | 0.263043 | -2.171053 | 0.029927 | 0.307056519 | IFNAR1 | protein_coding | 41.3 | -0.292839102 | 0.292839102 | ENSG00000142166 | n.s. |
| ENSG00000212429 | 10.36406 | -0.832690431 | 0.456742 | -1.82311 | 0.068287 | 0.445110495 | RNU11-6P | snRNA | 43.61 | -0.29271747 | 0.29271747 | ENSG00000212429 | n.s. |
| ENSG00000213337 | 166.448 | -0.625951129 | 0.298577 | -2.096445 | 0.036043 | 0.340720178 | ANKRD39 | protein_coding | 51.61 | -0.292696091 | 0.292696091 | ENSG00000213337 | n.s. |
| ENSG00000255893 | 25.78609 | -0.731249661 | 0.373711 | -1.956724 | 0.05038 | 0.397866833 | | antisense | 39.22 | -0.292691644 | 0.292691644 | ENSG00000255893 | n.s. |
| ENSG00000221203 | 7.230674 | -0.918464698 | 0.526126 | -1.745714 | 0.080861 | 0.480505215 | MIR1262 | miRNA | 35.48 | -0.292349053 | 0.292349053 | ENSG00000221203 | n.s. |

| | | | | | | | | | | | | | |
|---|---|---|---|---|---|---|---|---|---|---|---|---|---|
| ENSG00000272418 | 13.48694 | -0.798190274 | 0.428583 | -1.862393 | 0.062548 | 0.430475522 | | sense_intronic | 43.51 | -0.292178778 | 0.292178778 | ENSG00000272418 | n.s. |
| ENSG00000147324 | 650.9938 | -0.490489274 | 0.212123 | -2.312287 | 0.020762 | 0.254739827 | MFHAS1 | protein_coding | 44.16 | -0.291303125 | 0.291303125 | ENSG00000147324 | n.s. |
| ENSG00000157152 | 2238.716 | -0.543156347 | 0.246194 | -2.206215 | 0.027369 | 0.29088361 | SYN2 | protein_coding | 40.96 | -0.291284292 | 0.291284292 | ENSG00000157152 | n.s. |
| ENSG00000163795 | 183.3892 | -0.583846312 | 0.271684 | -2.148989 | 0.031635 | 0.317332638 | ZNF513 | protein_coding | 57.87 | -0.291038779 | 0.291038779 | ENSG00000163795 | n.s. |
| ENSG00000259349 | 31.94241 | -0.6914259 | 0.345531 | -2.001053 | 0.045387 | 0.38020912 | | antisense | 43.04 | -0.2903833 | 0.2903833 | ENSG00000259349 | n.s. |
| ENSG00000224255 | 25.49061 | -0.714645057 | 0.36331 | -1.96704 | 0.049179 | 0.393287465 | PDCL3P6 | processed_pseudogene | 44.76 | -0.28963842 | 0.28963842 | ENSG00000224255 | n.s. |
| ENSG00000265137 | 30.96417 | 0.660531578 | 0.324797 | 2.0336781 | 0.041984 | 0.364429018 | MIR3192 | miRNA | 48.05 | 0.289568489 | 0.289568489 | ENSG00000265137 | n.s. |
| ENSG00000284231 | 3.175152 | -1.537740632 | 1.144793 | -1.343248 | 0.179192 | 0.64843101 | MIR424 | miRNA | 51.02 | -0.289304716 | 0.289304716 | ENSG00000284231 | n.s. |
| ENSG00000173404 | 1847.086 | 0.483496851 | 0.208424 | 2.3197756 | 0.020353 | 0.252244945 | INSM1 | protein_coding | 63.13 | 0.289216951 | 0.289216951 | ENSG00000173404 | n.s. |
| ENSG00000196747 | 1090.668 | -0.494062874 | 0.215567 | -2.291924 | 0.02191 | 0.260036752 | HIST1H2AI | protein_coding | 59.64 | -0.289009621 | 0.289009621 | ENSG00000196747 | n.s. |
| ENSG00000164080 | 1080.955 | -0.504773019 | 0.222085 | -2.272885 | 0.023033 | 0.267911188 | RAD54L2 | protein_coding | 44.38 | -0.288734785 | 0.288734785 | ENSG00000164080 | n.s. |
| ENSG00000160392 | 153.0546 | -0.579522433 | 0.270096 | -2.145618 | 0.031903 | 0.31813913 | C19orf47 | protein_coding | 50.5 | -0.288244555 | 0.288244555 | ENSG00000160392 | n.s. |
| ENSG00000196440 | 43.10553 | -0.632722898 | 0.306477 | -2.064501 | 0.03897 | 0.35209789 | ARMCX4 | protein_coding | 41.06 | -0.286836433 | 0.286836433 | ENSG00000196440 | n.s. |
| ENSG00000157168 | 70.11374 | -0.573673443 | 0.266773 | -2.150419 | 0.031522 | 0.316882126 | NRG1 | protein_coding | 37.9 | -0.28632171 | 0.28632171 | ENSG00000157168 | n.s. |
| ENSG00000206635 | 6.610094 | -1.075182427 | 0.668977 | -1.607205 | 0.108009 | 0.542338372 | RNU6-1062P | snRNA | 40.19 | -0.285707869 | 0.285707869 | ENSG00000206635 | n.s. |
| ENSG00000006062 | 369.3666 | -0.513269498 | 0.22898 | -2.24155 | 0.02499 | 0.278131598 | MAP3K14 | protein_coding | 49.97 | -0.285249353 | 0.285249353 | ENSG00000006062 | n.s. |
| ENSG00000249962 | 74.38769 | -0.604010129 | 0.287059 | -2.104134 | 0.035367 | 0.337177445 | | processed_pseudogene | 75.2 | -0.285178239 | 0.285178239 | ENSG00000249962 | n.s. |
| ENSG00000185829 | 210.245 | -0.578757542 | 0.270584 | -2.138917 | 0.032442 | 0.321971749 | ARL17A | protein_coding | 43.34 | -0.28485418 | 0.28485418 | ENSG00000185829 | n.s. |
| ENSG00000199568 | 740.6958 | 0.471003141 | 0.202319 | 2.3280223 | 0.019911 | 0.248850741 | RNU5A-1 | snRNA | 41.38 | 0.284514658 | 0.284514658 | ENSG00000199568 | n.s. |
| ENSG00000253257 | 12.54024 | -0.787966628 | 0.425399 | -1.8523 | 0.063983 | 0.435510653 | MTND4P7 | processed_pseudogene | 35.38 | -0.284456912 | 0.284456912 | ENSG00000253257 | n.s. |
| ENSG00000151240 | 592.9321 | -0.544409033 | 0.249274 | -2.183977 | 0.028964 | 0.30094404 | DIP2C | protein_coding | 48.43 | -0.28391707 | 0.28391707 | ENSG00000151240 | n.s. |
| ENSG00000225423 | 69.19149 | -0.607749915 | 0.290314 | -2.093423 | 0.036311 | 0.341167655 | TNPO1P1 | processed_pseudogene | 41.12 | -0.283838749 | 0.283838749 | ENSG00000225423 | n.s. |
| ENSG00000172379 | 1216.294 | -0.548968067 | 0.252176 | -2.176921 | 0.029486 | 0.304103432 | ARNT2 | protein_coding | 45.48 | -0.283804786 | 0.283804786 | ENSG00000172379 | n.s. |
| ENSG00000201457 | 23.58132 | -0.739978217 | 0.386423 | -1.914941 | 0.0555 | 0.413574865 | SNORA55 | snoRNA | 51.85 | -0.283741586 | 0.283741586 | ENSG00000201457 | n.s. |
| ENSG00000112242 | 1226.227 | -0.510551029 | 0.227749 | -2.241725 | 0.024979 | 0.278131598 | E2F3 | protein_coding | 42.72 | -0.283738565 | 0.283738565 | ENSG00000112242 | n.s. |
| ENSG00000251807 | 33.77093 | 0.681277506 | 0.341955 | 1.9923003 | 0.046338 | 0.383342621 | RNU6-202P | snRNA | 44.79 | 0.283692736 | 0.283692736 | ENSG00000251807 | n.s. |
| ENSG00000023902 | 1952.646 | -0.527579642 | 0.23876 | -2.209666 | 0.027128 | 0.29009854 | PLEKHO1 | protein_coding | 51.63 | -0.28355003 | 0.28355003 | ENSG00000023902 | n.s. |
| ENSG00000249859 | 449.99 | -0.507625391 | 0.225904 | -2.247083 | 0.024635 | 0.276340982 | PVT1 | lincRNA | 46.6 | -0.28353655 | 0.28353655 | ENSG00000249859 | n.s. |
| ENSG00000265355 | 12.09691 | 0.889089992 | 0.508912 | 1.7470424 | 0.08063 | 0.479877462 | MIR3136 | miRNA | 38.46 | 0.283503812 | 0.283503812 | ENSG00000265355 | n.s. |
| ENSG00000117262 | 212.1817 | -0.577826681 | 0.270628 | -2.13513 | 0.03275 | 0.323186501 | GPR89A | protein_coding | 38.91 | -0.283451022 | 0.283451022 | ENSG00000117262 | n.s. |
| ENSG00000277440 | 33.89799 | -0.636914024 | 0.311245 | -2.046342 | 0.040723 | 0.35907784 | | sense_intronic | 43.81 | -0.283306616 | 0.283306616 | ENSG00000277440 | n.s. |
| ENSG00000278899 | 47.63301 | 0.708873485 | 0.36277 | 1.9540586 | 0.050694 | 0.398541231 | | antisense | 59.54 | 0.283213914 | 0.283213914 | ENSG00000278899 | n.s. |

| | | | | | | | | | | | | | |
|---|---|---|---|---|---|---|---|---|---|---|---|---|---|
| ENSG00000239246 | 21.78169 | 0.773188158 | 0.414496 | 1.8653704 | 0.06213 | 0.430475522 | | processed_pseudogene | 52.35 | 0.283026715 | 0.283026715 | ENSG00000239246 | n.s. |
| ENSG00000280028 | 116.2159 | -0.584446472 | 0.275116 | -2.124365 | 0.03364 | 0.327986808 | | TEC | 58.77 | -0.282956033 | 0.282956033 | ENSG00000280028 | n.s. |
| ENSG00000171843 | 335.3722 | 0.586775297 | 0.276875 | 2.1192766 | 0.034067 | 0.329615114 | MLLT3 | protein_coding | 37.12 | 0.282821517 | 0.282821517 | ENSG00000171843 | n.s. |
| ENSG00000186566 | 2192.665 | -0.566879379 | 0.263718 | -2.14957 | 0.031589 | 0.317099477 | GPATCH8 | protein_coding | 40.95 | -0.28276197 | 0.28276197 | ENSG00000186566 | n.s. |
| ENSG00000225484 | 238.2907 | -0.573987042 | 0.26822 | -2.139984 | 0.032356 | 0.321780677 | NUTM2B-AS1 | antisense | 39.68 | -0.282654201 | 0.282654201 | ENSG00000225484 | n.s. |
| ENSG00000268543 | 21.20292 | -0.735474216 | 0.38328 | -1.918894 | 0.054998 | 0.413017377 | | TEC | 47.14 | -0.282445395 | 0.282445395 | ENSG00000268543 | n.s. |
| ENSG00000277610 | 23.90309 | 0.769013057 | 0.411923 | 1.8668861 | 0.061917 | 0.430475522 | RNVU1-4 | snRNA | 43.9 | 0.281498413 | 0.281498413 | ENSG00000277610 | n.s. |
| ENSG00000135956 | 305.578 | -0.517795548 | 0.23311 | -2.221251 | 0.026334 | 0.286058985 | TMEM127 | protein_coding | 51.14 | -0.281444874 | 0.281444874 | ENSG00000135956 | n.s. |
| ENSG00000247595 | 43.32306 | -0.650111404 | 0.32098 | -2.025393 | 0.042827 | 0.369213631 | SPTY2D1-AS1 | protein_coding | 43.98 | -0.281317684 | 0.281317684 | ENSG00000247595 | n.s. |
| ENSG00000223692 | 25.39784 | -0.748689771 | 0.3959 | -1.891108 | 0.05861 | 0.421486462 | DIP2A-IT1 | sense_intronic | 40.16 | -0.280920658 | 0.280920658 | ENSG00000223692 | n.s. |
| ENSG00000119596 | 1800.44 | -0.5226127 | 0.236677 | -2.208129 | 0.027235 | 0.29045754 | YLPM1 | protein_coding | 39.67 | -0.280599823 | 0.280599823 | ENSG00000119596 | n.s. |
| ENSG00000107485 | 1476.987 | -0.468417239 | 0.20192 | -2.319813 | 0.020351 | 0.252244945 | GATA3 | protein_coding | 48.73 | -0.280196667 | 0.280196667 | ENSG00000107485 | n.s. |
| ENSG00000172201 | 431.3673 | 0.554229197 | 0.256547 | 2.1603436 | 0.030746 | 0.312237522 | ID4 | protein_coding | 53.08 | 0.280171122 | 0.280171122 | ENSG00000172201 | n.s. |
| ENSG00000224977 | 22.13751 | 0.788846274 | 0.430061 | 1.8342654 | 0.066615 | 0.441444624 | | lincRNA | 54.51 | 0.280138061 | 0.280138061 | ENSG00000224977 | n.s. |
| ENSG00000172244 | 330.5349 | -0.558241452 | 0.2591 | -2.154541 | 0.031198 | 0.315037554 | C5orf34 | protein_coding | 37.92 | -0.280034943 | 0.280034943 | ENSG00000172244 | n.s. |
| ENSG00000246985 | 114.65 | -0.614252247 | 0.296691 | -2.070345 | 0.03842 | 0.350082264 | SOCS2-AS1 | processed_transcript | 42.92 | -0.279994535 | 0.279994535 | ENSG00000246985 | n.s. |
| ENSG00000197386 | 73.83266 | -0.685324078 | 0.347013 | -1.974925 | 0.048277 | 0.390382659 | HTT | protein_coding | 45.52 | -0.279961384 | 0.279961384 | ENSG00000197386 | n.s. |
| ENSG00000167550 | 52.36713 | -0.671552326 | 0.336582 | -1.995214 | 0.04602 | 0.382988464 | RHEBL1 | protein_coding | 49.73 | -0.279912617 | 0.279912617 | ENSG00000167550 | n.s. |
| ENSG00000099622 | 2588.368 | -0.544029137 | 0.250392 | -2.172707 | 0.029802 | 0.306453766 | CIRBP | protein_coding | 62.55 | -0.279432426 | 0.279432426 | ENSG00000099622 | n.s. |
| ENSG00000233060 | 34.62778 | 0.693404945 | 0.353674 | 1.9605747 | 0.049929 | 0.395620546 | | lincRNA | 46.88 | 0.279248846 | 0.279248846 | ENSG00000233060 | n.s. |
| ENSG00000237892 | 610.6932 | 0.546283472 | 0.252055 | 2.167321 | 0.03021 | 0.308824672 | KLF7-IT1 | sense_intronic | 42.55 | 0.278761907 | 0.278761907 | ENSG00000237892 | n.s. |
| ENSG00000177225 | 615.7745 | -0.56626707 | 0.264683 | -2.139413 | 0.032402 | 0.321971749 | GATD1 | protein_coding | 60.25 | -0.278706591 | 0.278706591 | ENSG00000177225 | n.s. |
| ENSG00000103404 | 888.5894 | -0.508808757 | 0.228259 | -2.229087 | 0.025808 | 0.283560235 | USP31 | protein_coding | 41.37 | -0.278498851 | 0.278498851 | ENSG00000103404 | n.s. |
| ENSG00000207004 | 24.95802 | -0.700992384 | 0.359703 | -1.948811 | 0.051318 | 0.400783303 | RNU6-301P | snRNA | 44.86 | -0.278357332 | 0.278357332 | ENSG00000207004 | n.s. |
| ENSG00000197563 | 366.8904 | -0.604478591 | 0.290247 | -2.082636 | 0.037284 | 0.346536178 | PIGN | protein_coding | 38.03 | -0.27821213 | 0.27821213 | ENSG00000197563 | n.s. |
| ENSG00000273175 | 166.5811 | 0.559116517 | 0.260612 | 2.1453946 | 0.031921 | 0.31813913 | | lincRNA | 45.05 | 0.278095001 | 0.278095001 | ENSG00000273175 | n.s. |
| ENSG00000185920 | 3381.779 | -0.507085741 | 0.227232 | -2.231575 | 0.025643 | 0.283008461 | PTCH1 | protein_coding | 47.49 | -0.277984697 | 0.277984697 | ENSG00000185920 | n.s. |
| ENSG00000093217 | 406.999 | -0.570513373 | 0.268036 | -2.128499 | 0.033296 | 0.325902089 | XYLB | protein_coding | 44.83 | -0.277790296 | 0.277790296 | ENSG00000093217 | n.s. |
| ENSG00000207975 | 15.61865 | -0.747836776 | 0.397879 | -1.879557 | 0.060169 | 0.425937056 | MIR181B1 | miRNA | 44.55 | -0.277189122 | 0.277189122 | ENSG00000207975 | n.s. |
| ENSG00000225215 | 42.70785 | -0.622879815 | 0.304687 | -2.044328 | 0.040921 | 0.35907784 | SMARCE1P1 | processed_pseudogene | 47.22 | -0.27706404 | 0.27706404 | ENSG00000225215 | n.s. |
| ENSG00000250510 | 123.2463 | 0.662556514 | 0.331962 | 1.9958826 | 0.045947 | 0.382837889 | GPR162 | protein_coding | 61.64 | 0.276276186 | 0.276276186 | ENSG00000250510 | n.s. |
| ENSG00000054967 | 287.536 | -0.539270524 | 0.248589 | -2.169327 | 0.030058 | 0.307717153 | RELT | protein_coding | 59.84 | -0.276024698 | 0.276024698 | ENSG00000054967 | n.s. |

| | | | | | | | | | | | | | |
|---|---|---|---|---|---|---|---|---|---|---|---|---|---|
| ENSG00000187801 | 167.1932 | -0.613332984 | 0.299061 | -2.050862 | 0.04028 | 0.35721239 | ZFP69B | protein_coding | 41.8 | -0.274204915 | 0.274204915 | ENSG00000187801 | n.s. |
| ENSG00000275377 | 6.031804 | -0.991183393 | 0.607478 | -1.631636 | 0.102756 | 0.528900013 | MIR1299 | miRNA | 50.6 | -0.274187516 | 0.274187516 | ENSG00000275377 | n.s. |
| ENSG00000181523 | 25.6721 | -0.71495842 | 0.373042 | -1.916563 | 0.055293 | 0.413559567 | SGSH | protein_coding | 58.83 | -0.274159334 | 0.274159334 | ENSG00000181523 | n.s. |
| ENSG00000237021 | 1189.086 | 0.46787338 | 0.20416 | 2.2916996 | 0.021923 | 0.260036752 | | lincRNA | 34.41 | 0.273689677 | 0.273689677 | ENSG00000237021 | n.s. |
| ENSG00000272145 | 348.6986 | 0.473553468 | 0.207804 | 2.2788414 | 0.022676 | 0.264673416 | NFYC-AS1 | antisense | 49.61 | 0.273377529 | 0.273377529 | ENSG00000272145 | n.s. |
| ENSG00000283204 | 11.55916 | 0.872350982 | 0.504493 | 1.7291647 | 0.08378 | 0.486229087 | MIR4434 | miRNA | 33.96 | 0.273184618 | 0.273184618 | ENSG00000283204 | n.s. |
| ENSG00000182095 | 784.7466 | -0.497190377 | 0.222644 | -2.233121 | 0.025541 | 0.282417274 | TNRC18 | protein_coding | 54.68 | -0.273011583 | 0.273011583 | ENSG00000182095 | n.s. |
| ENSG00000280088 | 127.6948 | -0.603857412 | 0.293128 | -2.060044 | 0.039394 | 0.354033183 | | TEC | 39.54 | -0.272313141 | 0.272313141 | ENSG00000280088 | n.s. |
| ENSG00000261351 | 79.66267 | -0.578365184 | 0.275317 | -2.100726 | 0.035665 | 0.338259327 | | antisense | 45.05 | -0.272265537 | 0.272265537 | ENSG00000261351 | n.s. |
| ENSG00000239043 | 10.38473 | -0.872448666 | 0.506446 | -1.722689 | 0.084945 | 0.487906764 | SNORD127 | snoRNA | 36.05 | -0.271910109 | 0.271910109 | ENSG00000239043 | n.s. |
| ENSG00000080802 | 841.7081 | -0.532527445 | 0.245662 | -2.167723 | 0.03018 | 0.308738089 | CNOT4 | protein_coding | 37.19 | -0.271807221 | 0.271807221 | ENSG00000080802 | n.s. |
| ENSG00000169914 | 630.1867 | -0.515797498 | 0.235498 | -2.190244 | 0.028507 | 0.297523446 | OTUD3 | protein_coding | 41.28 | -0.27155645 | 0.27155645 | ENSG00000169914 | n.s. |
| ENSG00000226174 | 41.39687 | -0.652071517 | 0.327258 | -1.992531 | 0.046313 | 0.383342621 | TEX22 | protein_coding | 55.6 | -0.271530986 | 0.271530986 | ENSG00000226174 | n.s. |
| ENSG00000274390 | 2.771086 | -1.34834648 | 0.964427 | -1.398081 | 0.162089 | 0.628957308 | MIR6885 | miRNA | 65.15 | -0.27152844 | 0.27152844 | ENSG00000274390 | n.s. |
| ENSG00000267274 | 25.46712 | -0.701602815 | 0.364102 | -1.92694 | 0.053987 | 0.410295828 | | sense_overlapping | 45.7 | -0.271452163 | 0.271452163 | ENSG00000267274 | n.s. |
| ENSG00000079102 | 2943.288 | 0.531629974 | 0.24537 | 2.166649 | 0.030262 | 0.308894909 | RUNX1T1 | protein_coding | 38.11 | 0.271231897 | 0.271231897 | ENSG00000079102 | n.s. |
| ENSG00000135976 | 1766.861 | -0.571473485 | 0.271257 | -2.106759 | 0.035138 | 0.335560808 | ANKRD36 | protein_coding | 35.92 | -0.271009167 | 0.271009167 | ENSG00000135976 | n.s. |
| ENSG00000259416 | 30.69895 | -0.66390855 | 0.337061 | -1.969698 | 0.048873 | 0.391760224 | | antisense | 55.74 | -0.270197277 | 0.270197277 | ENSG00000259416 | n.s. |
| ENSG00000254333 | 14.25857 | -0.845741656 | 0.483374 | -1.749662 | 0.080177 | 0.479305998 | NDST1-AS1 | lincRNA | 50.59 | -0.270119005 | 0.270119005 | ENSG00000254333 | n.s. |
| ENSG00000245904 | 28.90221 | -0.724768137 | 0.384513 | -1.8849 | 0.059443 | 0.423990171 | | antisense | 39.81 | -0.270080651 | 0.270080651 | ENSG00000245904 | n.s. |
| ENSG00000278771 | 266.1484 | -0.491769155 | 0.220148 | -2.23381 | 0.025496 | 0.28239823 | RN7SL3 | misc_RNA | 62.21 | -0.270049145 | 0.270049145 | ENSG00000278771 | n.s. |
| ENSG00000026025 | 2581.526 | -0.511330093 | 0.232937 | -2.195144 | 0.028153 | 0.296503687 | VIM | protein_coding | 46.25 | -0.2699669 | 0.2699669 | ENSG00000026025 | n.s. |
| ENSG00000157216 | 1291.204 | -0.537593482 | 0.249524 | -2.154477 | 0.031203 | 0.315037554 | SSBP3 | protein_coding | 50.93 | -0.269677143 | 0.269677143 | ENSG00000157216 | n.s. |
| ENSG00000214826 | 337.4831 | -0.511208268 | 0.233288 | -2.191318 | 0.028429 | 0.297290527 | DDX12P | unprocessed_pseudogene | 52.17 | -0.269314192 | 0.269314192 | ENSG00000214826 | n.s. |
| ENSG00000274051 | 19.65009 | -0.747101138 | 0.404197 | -1.84836 | 0.06455 | 0.436711663 | | misc_RNA | 60.22 | -0.268810882 | 0.268810882 | ENSG00000274051 | n.s. |
| ENSG00000117791 | 11.27725 | -0.848959357 | 0.48763 | -1.740989 | 0.081685 | 0.482429187 | 2-Mar | protein_coding | 44.02 | -0.268752029 | 0.268752029 | ENSG00000117791 | n.s. |
| ENSG00000250853 | 70.71751 | 0.597791394 | 0.29077 | 2.0558903 | 0.039793 | 0.355381165 | RNF138P1 | processed_pseudogene | 37.73 | 0.268591017 | 0.268591017 | ENSG00000250853 | n.s. |
| ENSG00000273173 | 69.38388 | -0.639385185 | 0.319472 | -2.001382 | 0.045351 | 0.38020912 | SNURF | protein_coding | 40.75 | -0.268527372 | 0.268527372 | ENSG00000273173 | n.s. |
| ENSG00000100147 | 77.94772 | -0.593577113 | 0.288225 | -2.059423 | 0.039454 | 0.354033183 | CCDC134 | protein_coding | 49.16 | -0.267677178 | 0.267677178 | ENSG00000100147 | n.s. |
| ENSG00000217331 | 62.427 | -0.620865451 | 0.306878 | -2.023165 | 0.043056 | 0.370730246 | | processed_pseudogene | 30.92 | -0.267556987 | 0.267556987 | ENSG00000217331 | n.s. |
| ENSG00000244486 | 304.2864 | -0.555278469 | 0.262228 | -2.117543 | 0.034214 | 0.329831947 | SCARF2 | protein_coding | 63.11 | -0.267481682 | 0.267481682 | ENSG00000244486 | n.s. |
| ENSG00000207181 | 234.007 | 0.501404255 | 0.227815 | 2.2009285 | 0.027741 | 0.293049415 | SNORA14B | snoRNA | 46.67 | 0.267278122 | 0.267278122 | ENSG00000207181 | n.s. |

| | | | | | | | | | | | | | |
|---|---|---|---|---|---|---|---|---|---|---|---|---|---|
| ENSG00000202078 | 17.67475 | -0.750674813 | 0.408257 | -1.83873 | 0.065955 | 0.440875092 | | misc_RNA | 40.71 | -0.267003348 | 0.267003348 | ENSG00000202078 | n.s. |
| ENSG00000259378 | 75.27112 | -0.627696068 | 0.312003 | -2.01183 | 0.044238 | 0.375547181 | DCAF13P3 | processed_pseudogene | 39.43 | -0.266981417 | 0.266981417 | ENSG00000259378 | n.s. |
| ENSG00000283645 | 1.357036 | -1.666102592 | 1.342963 | -1.240617 | 0.214747 | 0.691472186 | MIR5583-2 | miRNA | 33.9 | -0.266951761 | 0.266951761 | ENSG00000283645 | n.s. |
| ENSG00000252118 | 5.931497 | 1.005062016 | 0.626397 | 1.6045131 | 0.108601 | 0.543082569 | RNU6ATAC39P | snRNA | 37.58 | 0.26647625 | 0.26647625 | ENSG00000252118 | n.s. |
| ENSG00000160888 | 986.5843 | 0.503784253 | 0.229428 | 2.1958298 | 0.028104 | 0.296210251 | IER2 | protein_coding | 62.33 | 0.266199557 | 0.266199557 | ENSG00000160888 | n.s. |
| ENSG00000196313 | 543.3553 | -0.525269749 | 0.242916 | -2.162348 | 0.030591 | 0.311347454 | POM121 | protein_coding | 46.19 | -0.266182904 | 0.266182904 | ENSG00000196313 | n.s. |
| ENSG00000090924 | 136.3732 | -0.569935345 | 0.272223 | -2.093633 | 0.036293 | 0.341167655 | PLEKHG2 | protein_coding | 56.77 | -0.266178129 | 0.266178129 | ENSG00000090924 | n.s. |
| ENSG00000268439 | 73.19051 | -0.590159467 | 0.286624 | -2.059001 | 0.039494 | 0.354073606 | EMG1 | protein_coding | 45.52 | -0.266106708 | 0.266106708 | ENSG00000268439 | n.s. |
| ENSG00000207507 | 4.034995 | -1.329607756 | 0.959844 | -1.385233 | 0.165981 | 0.63130003 | RNU6-9 | snRNA | 45.79 | -0.265608017 | 0.265608017 | ENSG00000207507 | n.s. |
| ENSG00000004139 | 958.6822 | -0.539882156 | 0.252577 | -2.137495 | 0.032558 | 0.322405076 | SARM1 | protein_coding | 51.55 | -0.265405058 | 0.265405058 | ENSG00000004139 | n.s. |
| ENSG00000263511 | 8.800897 | -0.94784315 | 0.578168 | -1.63939 | 0.101132 | 0.526181024 | MIR5699 | miRNA | 57.78 | -0.264320108 | 0.264320108 | ENSG00000263511 | n.s. |
| ENSG00000245534 | 56.27095 | -0.591067631 | 0.288196 | -2.050923 | 0.040274 | 0.35721239 | RORA-AS1 | antisense | 40.18 | -0.264250666 | 0.264250666 | ENSG00000245534 | n.s. |
| ENSG00000130544 | 534.6384 | -0.549904838 | 0.260142 | -2.113863 | 0.034527 | 0.331530132 | ZNF557 | protein_coding | 45.81 | -0.263666718 | 0.263666718 | ENSG00000130544 | n.s. |
| ENSG00000201317 | 9.297995 | -0.85120791 | 0.495025 | -1.719526 | 0.085519 | 0.490059299 | RNU4-59P | snRNA | 37.07 | -0.263662812 | 0.263662812 | ENSG00000201317 | n.s. |
| ENSG00000279659 | 80.33979 | -0.592682382 | 0.289648 | -2.046215 | 0.040735 | 0.35907784 | | TEC | 36.03 | -0.263631877 | 0.263631877 | ENSG00000279659 | n.s. |
| ENSG00000108599 | 628.7021 | -0.558419591 | 0.265606 | -2.102434 | 0.035515 | 0.337341243 | AKAP10 | protein_coding | 41.26 | -0.26353527 | 0.26353527 | ENSG00000108599 | n.s. |
| ENSG00000265110 | 2.737684 | -1.32479045 | 0.958272 | -1.382478 | 0.166825 | 0.632544316 | MIR4731 | miRNA | 64.29 | -0.2635128 | 0.2635128 | ENSG00000265110 | n.s. |
| ENSG00000259291 | 122.0835 | -0.555614042 | 0.263599 | -2.107801 | 0.035048 | 0.335533508 | ZNF710-AS1 | antisense | 53.18 | -0.263507795 | 0.263507795 | ENSG00000259291 | n.s. |
| ENSG00000260686 | 90.52238 | -0.555977375 | 0.264131 | -2.104927 | 0.035298 | 0.336840039 | | sense_overlapping | 31.22 | -0.262741726 | 0.262741726 | ENSG00000260686 | n.s. |
| ENSG00000156486 | 204.4356 | 0.504927008 | 0.231452 | 2.1815589 | 0.029142 | 0.302134081 | KCNS2 | protein_coding | 51.66 | 0.262461202 | 0.262461202 | ENSG00000156486 | n.s. |
| ENSG00000278526 | 12.41213 | -0.809421333 | 0.458939 | -1.76368 | 0.077786 | 0.475213657 | | misc_RNA | 59.12 | -0.261533007 | 0.261533007 | ENSG00000278526 | n.s. |
| ENSG00000125834 | 449.9708 | -0.496548025 | 0.226673 | -2.190591 | 0.028481 | 0.297484595 | STK35 | protein_coding | 48.13 | -0.261450172 | 0.261450172 | ENSG00000125834 | n.s. |
| ENSG00000168546 | 2768.338 | -0.560517591 | 0.268021 | -2.091318 | 0.0365 | 0.341750791 | GFRA2 | protein_coding | 52.15 | -0.261364012 | 0.261364012 | ENSG00000168546 | n.s. |
| ENSG00000137275 | 218.2997 | -0.594346658 | 0.292084 | -2.03485 | 0.041866 | 0.36363053 | RIPK1 | protein_coding | 45.29 | -0.261120059 | 0.261120059 | ENSG00000137275 | n.s. |
| ENSG00000196937 | 587.8153 | -0.560152303 | 0.267978 | -2.090293 | 0.036591 | 0.342151034 | FAM3C | protein_coding | 35.92 | -0.26090894 | 0.26090894 | ENSG00000196937 | n.s. |
| ENSG00000237179 | 116.425 | 0.587372999 | 0.287529 | 2.0428312 | 0.041069 | 0.359904219 | LINC01797 | lincRNA | 36.28 | 0.26068381 | 0.26068381 | ENSG00000237179 | n.s. |
| ENSG00000278135 | 4.968387 | 1.022858805 | 0.64916 | 1.5756647 | 0.115103 | 0.556094484 | | snRNA | 36.84 | 0.260677011 | 0.260677011 | ENSG00000278135 | n.s. |
| ENSG00000198909 | 823.9553 | -0.519876087 | 0.24143 | -2.153321 | 0.031293 | 0.31549522 | MAP3K3 | protein_coding | 45.21 | -0.260461671 | 0.260461671 | ENSG00000198909 | n.s. |
| ENSG00000146555 | 301.4633 | -0.520421788 | 0.241837 | -2.151954 | 0.031401 | 0.315901054 | SDK1 | protein_coding | 43.12 | -0.260444525 | 0.260444525 | ENSG00000146555 | n.s. |
| ENSG00000196646 | 476.0878 | -0.522683455 | 0.243329 | -2.14805 | 0.03171 | 0.317622748 | ZNF136 | protein_coding | 44.4 | -0.260342567 | 0.260342567 | ENSG00000196646 | n.s. |
| ENSG00000264536 | 3.489372 | -1.087608463 | 0.711506 | -1.528601 | 0.126363 | 0.576460831 | MIR5706 | miRNA | 45 | -0.260188787 | 0.260188787 | ENSG00000264536 | n.s. |
| ENSG00000167281 | 403.5642 | -0.539392586 | 0.25485 | -2.116512 | 0.034301 | 0.329909835 | RBFOX3 | protein_coding | 55.28 | -0.259774024 | 0.259774024 | ENSG00000167281 | n.s. |

| | | | | | | | | | | | | | |
|---|---|---|---|---|---|---|---|---|---|---|---|---|---|
| ENSG00000233937 | 399.177 | -0.492571914 | 0.224758 | -2.191564 | 0.028411 | 0.297290527 | | processed_transcript | 48.01 | -0.259496208 | 0.259496208 | ENSG00000233937 | n.s. |
| ENSG00000283419 | 3.490279 | -1.319950108 | 0.958313 | -1.377368 | 0.168399 | 0.636128582 | MIR6874 | miRNA | 45.07 | -0.259310918 | 0.259310918 | ENSG00000283419 | n.s. |
| ENSG00000270036 | 26.98576 | -0.681582423 | 0.357218 | -1.908028 | 0.056388 | 0.416711332 | | lincRNA | 42.27 | -0.259113571 | 0.259113571 | ENSG00000270036 | n.s. |
| ENSG00000283785 | 7.702943 | -0.902666632 | 0.542458 | -1.66403 | 0.096107 | 0.516637382 | MIR15A | miRNA | 42.17 | -0.258897584 | 0.258897584 | ENSG00000283785 | n.s. |
| ENSG00000153404 | 485.3468 | -0.542145241 | 0.256781 | -2.111317 | 0.034745 | 0.333165317 | PLEKHG4B | protein_coding | 49.1 | -0.258787727 | 0.258787727 | ENSG00000153404 | n.s. |
| ENSG00000081760 | 709.3421 | -0.52842546 | 0.247688 | -2.13343 | 0.03289 | 0.324306352 | AACS | protein_coding | 51.22 | -0.258423589 | 0.258423589 | ENSG00000081760 | n.s. |
| ENSG00000177084 | 3731.715 | -0.50242435 | 0.231147 | -2.173611 | 0.029734 | 0.305980493 | POLE | protein_coding | 50.84 | -0.258399989 | 0.258399989 | ENSG00000177084 | n.s. |
| ENSG00000272854 | 13.92328 | 0.847253177 | 0.497263 | 1.7038329 | 0.088412 | 0.495479338 | | antisense | 49.1 | 0.258390573 | 0.258390573 | ENSG00000272854 | n.s. |
| ENSG00000276672 | 581.5485 | -0.519193718 | 0.241914 | -2.146194 | 0.031858 | 0.31813913 | | sense_intronic | 39.6 | -0.258238083 | 0.258238083 | ENSG00000276672 | n.s. |
| ENSG00000261115 | 495.43 | -0.517884802 | 0.241057 | -2.148395 | 0.031682 | 0.317576343 | TMEM178B | protein_coding | 43.27 | -0.257985276 | 0.257985276 | ENSG00000261115 | n.s. |
| ENSG00000242791 | 27.68916 | -0.658474059 | 0.340237 | -1.935339 | 0.052949 | 0.406621815 | | lincRNA | 37.87 | -0.257337802 | 0.257337802 | ENSG00000242791 | n.s. |
| ENSG00000163956 | 536.4051 | -0.530318298 | 0.249579 | -2.124853 | 0.033599 | 0.327818303 | LRPAP1 | protein_coding | 57.65 | -0.256868578 | 0.256868578 | ENSG00000163956 | n.s. |
| ENSG00000180581 | 51.57746 | 0.671221871 | 0.35125 | 1.9109506 | 0.056011 | 0.414367789 | SRP9P1 | processed_pseudogene | 45.21 | 0.256818893 | 0.256818893 | ENSG00000180581 | n.s. |
| ENSG00000222365 | 419.3768 | 0.474662029 | 0.214275 | 2.2152035 | 0.026746 | 0.287778928 | SNORD12B | snoRNA | 41.76 | 0.256764157 | 0.256764157 | ENSG00000222365 | n.s. |
| ENSG00000110092 | 10369.09 | -0.523308026 | 0.245097 | -2.135101 | 0.032753 | 0.323186501 | CCND1 | protein_coding | 58.58 | -0.256707071 | 0.256707071 | ENSG00000110092 | n.s. |
| ENSG00000163249 | 199.5899 | -0.550922965 | 0.263526 | -2.090581 | 0.036566 | 0.34213948 | CCNYL1 | protein_coding | 41.55 | -0.256618159 | 0.256618159 | ENSG00000163249 | n.s. |
| ENSG00000165417 | 1368.444 | -0.559360219 | 0.269244 | -2.077524 | 0.037753 | 0.348104442 | GTF2A1 | protein_coding | 37.75 | -0.256349438 | 0.256349438 | ENSG00000165417 | n.s. |
| ENSG00000029153 | 602.7281 | -0.561614623 | 0.271217 | -2.070723 | 0.038385 | 0.350082264 | ARNTL2 | protein_coding | 39.26 | -0.256000732 | 0.256000732 | ENSG00000029153 | n.s. |
| ENSG00000090857 | 874.901 | -0.523583702 | 0.245597 | -2.131879 | 0.033017 | 0.324413031 | PDPR | protein_coding | 47.32 | -0.255980967 | 0.255980967 | ENSG00000090857 | n.s. |
| ENSG00000230119 | 11.51864 | -0.814140379 | 0.469151 | -1.73535 | 0.082679 | 0.484982269 | | processed_pseudogene | 39.58 | -0.255863267 | 0.255863267 | ENSG00000230119 | n.s. |
| ENSG00000050438 | 95.54836 | -0.574746299 | 0.281153 | -2.044247 | 0.040929 | 0.35907784 | SLC4A8 | protein_coding | 42.72 | -0.255653703 | 0.255653703 | ENSG00000050438 | n.s. |
| ENSG00000272686 | 532.1744 | 0.484809226 | 0.221252 | 2.1912091 | 0.028437 | 0.297290527 | | antisense | 41.64 | 0.255406677 | 0.255406677 | ENSG00000272686 | n.s. |
| ENSG00000174871 | 116.0856 | -0.566660229 | 0.275359 | -2.057893 | 0.0396 | 0.354422376 | CNIH2 | protein_coding | 61.39 | -0.255268458 | 0.255268458 | ENSG00000174871 | n.s. |
| ENSG00000263731 | 639.6698 | -0.524196313 | 0.246239 | -2.128808 | 0.03327 | 0.325902089 | | lincRNA | 43.7 | -0.255237924 | 0.255237924 | ENSG00000263731 | n.s. |
| ENSG00000265778 | 8.329712 | -1.075643193 | 0.710393 | -1.514151 | 0.129987 | 0.5791453 | | antisense | 55.01 | -0.255155972 | 0.255155972 | ENSG00000265778 | n.s. |
| ENSG00000129292 | 3709.88 | -0.564912083 | 0.274275 | -2.059654 | 0.039432 | 0.354033183 | PHF20L1 | protein_coding | 34.77 | -0.25475051 | 0.25475051 | ENSG00000129292 | n.s. |
| ENSG00000106012 | 39.63423 | -0.675201105 | 0.356847 | -1.892132 | 0.058473 | 0.42095504 | IQCE | protein_coding | 52.81 | -0.253716462 | 0.253716462 | ENSG00000106012 | n.s. |
| ENSG00000277775 | 929.0568 | -0.462708808 | 0.207744 | -2.227301 | 0.025927 | 0.283942182 | HIST1H3F | protein_coding | 60.83 | -0.252995334 | 0.252995334 | ENSG00000277775 | n.s. |
| ENSG00000160298 | 405.3243 | -0.469825665 | 0.212584 | -2.210072 | 0.0271 | 0.29009854 | C21orf58 | protein_coding | 52.19 | -0.252509897 | 0.252509897 | ENSG00000160298 | n.s. |
| ENSG00000235912 | 11.37125 | 1.064321035 | 0.70252 | 1.5150049 | 0.129771 | 0.5791453 | | processed_pseudogene | 53.65 | 0.252470215 | 0.252470215 | ENSG00000235912 | n.s. |
| ENSG00000148773 | 8696.547 | -0.498844659 | 0.230974 | -2.159748 | 0.030792 | 0.312250224 | MKI67 | protein_coding | 43.51 | -0.252164599 | 0.252164599 | ENSG00000148773 | n.s. |
| ENSG00000244627 | 89.08334 | -0.584992217 | 0.289284 | -2.022205 | 0.043155 | 0.370895024 | | transcribed_unprocessed_pseudc | 43.54 | -0.251984809 | 0.251984809 | ENSG00000244627 | n.s. |

| | | | | | | | | | | | | | |
|---|---|---|---|---|---|---|---|---|---|---|---|---|---|
| ENSG00000131759 | 115.2143 | -0.539325426 | 0.25787 | -2.091464 | 0.036486 | 0.341750791 | RARA | protein_coding | 56.99 | -0.251482308 | 0.251482308 | ENSG00000131759 | n.s. |
| ENSG00000034152 | 41.98346 | -0.643265473 | 0.332363 | -1.935431 | 0.052937 | 0.406621815 | MAP2K3 | protein_coding | 56.97 | -0.251394145 | 0.251394145 | ENSG00000034152 | n.s. |
| ENSG00000207779 | 39.96102 | 0.625374505 | 0.319199 | 1.9592006 | 0.050089 | 0.396455218 | MIR15B | miRNA | 36.73 | 0.251279142 | 0.251279142 | ENSG00000207779 | n.s. |
| ENSG00000231416 | 58.97858 | -0.595590834 | 0.297021 | -2.005217 | 0.04494 | 0.378563498 | | processed_pseudogene | 44.38 | -0.251256702 | 0.251256702 | ENSG00000231416 | n.s. |
| ENSG00000271938 | 6.040458 | 1.057803584 | 0.697799 | 1.515915 | 0.129541 | 0.579048024 | | lincRNA | 39.73 | 0.251001363 | 0.251001363 | ENSG00000271938 | n.s. |
| ENSG00000151208 | 879.3019 | -0.488221353 | 0.224777 | -2.172025 | 0.029854 | 0.306529566 | DLG5 | protein_coding | 50 | -0.250715154 | 0.250715154 | ENSG00000151208 | n.s. |
| ENSG00000068137 | 302.1364 | -0.56349702 | 0.275163 | -2.047863 | 0.040573 | 0.35907784 | PLEKHH3 | protein_coding | 62 | -0.250649896 | 0.250649896 | ENSG00000068137 | n.s. |
| ENSG00000168661 | 109.0787 | -0.605448122 | 0.304583 | -1.987791 | 0.046835 | 0.385618457 | ZNF30 | protein_coding | 42.7 | -0.250559975 | 0.250559975 | ENSG00000168661 | n.s. |
| ENSG00000236577 | 6.195203 | -0.894741007 | 0.544704 | -1.642618 | 0.100462 | 0.525174439 | SNRPGP14 | processed_pseudogene | 41.27 | -0.250255857 | 0.250255857 | ENSG00000236577 | n.s. |
| ENSG00000160229 | 356.7281 | -0.566625372 | 0.277957 | -2.038533 | 0.041497 | 0.361758374 | ZNF66 | protein_coding | 38.46 | -0.250211229 | 0.250211229 | ENSG00000160229 | n.s. |
| ENSG00000130766 | 205.7674 | -0.586134239 | 0.29083 | -2.015384 | 0.043864 | 0.374378906 | SESN2 | protein_coding | 50.06 | -0.250096815 | 0.250096815 | ENSG00000130766 | n.s. |
| ENSG00000274349 | 141.1081 | -0.542025602 | 0.260118 | -2.083765 | 0.037182 | 0.345810463 | ZNF658 | protein_coding | 38.33 | -0.249961541 | 0.249961541 | ENSG00000274349 | n.s. |
| ENSG00000185298 | 882.4903 | -0.536336881 | 0.256641 | -2.089834 | 0.036633 | 0.342307043 | CCDC137 | protein_coding | 57.88 | -0.249709959 | 0.249709959 | ENSG00000185298 | n.s. |
| ENSG00000154359 | 592.758 | -0.555629278 | 0.270244 | -2.056031 | 0.03978 | 0.355381165 | LONRF1 | protein_coding | 38.66 | -0.249647342 | 0.249647342 | ENSG00000154359 | n.s. |
| ENSG00000197150 | 28.06669 | -0.669596891 | 0.354919 | -1.886618 | 0.059212 | 0.42398874 | ABCB8 | protein_coding | 57.04 | -0.249522387 | 0.249522387 | ENSG00000197150 | n.s. |
| ENSG00000264522 | 688.0559 | -0.529103067 | 0.251783 | -2.101428 | 0.035603 | 0.337904892 | OTUD7B | protein_coding | 42.46 | -0.249316286 | 0.249316286 | ENSG00000264522 | n.s. |
| ENSG00000173166 | 537.4411 | -0.559457357 | 0.273333 | -2.046798 | 0.040678 | 0.35907784 | RAPH1 | protein_coding | 39.16 | -0.248853008 | 0.248853008 | ENSG00000173166 | n.s. |
| ENSG00000166025 | 1698.233 | -0.496015447 | 0.23026 | -2.154153 | 0.031228 | 0.315065145 | AMOTL1 | protein_coding | 41.79 | -0.248801169 | 0.248801169 | ENSG00000166025 | n.s. |
| ENSG00000230470 | 39.44952 | -0.648786982 | 0.338481 | -1.916761 | 0.055268 | 0.413559567 | | sense_intronic | 39.79 | -0.248785107 | 0.248785107 | ENSG00000230470 | n.s. |
| ENSG00000187514 | 25294.64 | -0.609664719 | 0.309097 | -1.972408 | 0.048563 | 0.391129217 | PTMA | protein_coding | 61.71 | -0.248547955 | 0.248547955 | ENSG00000187514 | n.s. |
| ENSG00000225218 | 12.4431 | -0.768287144 | 0.43539 | -1.764596 | 0.077632 | 0.474896813 | | lincRNA | 51.1 | -0.248464634 | 0.248464634 | ENSG00000225218 | n.s. |
| ENSG00000200312 | 15.03501 | -0.802014407 | 0.466464 | -1.71935 | 0.085551 | 0.490059299 | RN7SKP255 | misc_RNA | 56.23 | -0.248425057 | 0.248425057 | ENSG00000200312 | n.s. |
| ENSG00000196968 | 228.0311 | -0.525304911 | 0.249842 | -2.102547 | 0.035505 | 0.337341243 | FUT11 | protein_coding | 47.49 | -0.24790744 | 0.24790744 | ENSG00000196968 | n.s. |
| ENSG00000225830 | 338.1015 | -0.536091623 | 0.257068 | -2.085411 | 0.037032 | 0.34486157 | ERCC6 | protein_coding | 40.54 | -0.247864749 | 0.247864749 | ENSG00000225830 | n.s. |
| ENSG00000101624 | 530.7861 | 0.49479237 | 0.229902 | 2.1521926 | 0.031382 | 0.315901054 | CEP76 | protein_coding | 40.84 | 0.24761831 | 0.24761831 | ENSG00000101624 | n.s. |
| ENSG00000234816 | 167.2876 | -0.556481481 | 0.271774 | -2.047591 | 0.0406 | 0.35907784 | HIST1H2APS5 | unprocessed_pseudogene | 54.47 | -0.247529304 | 0.247529304 | ENSG00000234816 | n.s. |
| ENSG00000188807 | 215.3965 | -0.542026616 | 0.261518 | -2.07262 | 0.038208 | 0.350082264 | TMEM201 | protein_coding | 59.34 | -0.247071934 | 0.247071934 | ENSG00000188807 | n.s. |
| ENSG00000279425 | 100.5735 | -0.54699594 | 0.265292 | -2.061866 | 0.039221 | 0.35389993 | | TEC | 43.76 | -0.246760548 | 0.246760548 | ENSG00000279425 | n.s. |
| ENSG00000125967 | 101.496 | -0.541204227 | 0.261386 | -2.070518 | 0.038404 | 0.350082264 | NECAB3 | protein_coding | 59.54 | -0.246697064 | 0.246697064 | ENSG00000125967 | n.s. |
| ENSG00000278472 | 19.04427 | -0.752383672 | 0.42485 | -1.770939 | 0.076571 | 0.471098423 | | sense_intronic | 35.67 | -0.245945456 | 0.245945456 | ENSG00000278472 | n.s. |
| ENSG00000267796 | 67.65546 | -0.641102233 | 0.334731 | -1.915273 | 0.055458 | 0.413574865 | LIN37 | protein_coding | 54.65 | -0.245827999 | 0.245827999 | ENSG00000267796 | n.s. |
| ENSG00000267080 | 308.5713 | -0.487349454 | 0.225915 | -2.157223 | 0.030988 | 0.313782097 | ASB16-AS1 | antisense | 56.18 | -0.245317991 | 0.245317991 | ENSG00000267080 | n.s. |

| | | | | | | | | | | | | | |
|---|---|---|---|---|---|---|---|---|---|---|---|---|---|
| ENSG00000148339 | 444.4895 | -0.547725082 | 0.267133 | -2.05038 | 0.040327 | 0.357401664 | SLC25A25 | protein_coding | 49.9 | -0.244747354 | 0.244747354 | ENSG00000148339 | n.s. |
| ENSG00000135916 | 419.6587 | -0.587085937 | 0.29443 | -1.993973 | 0.046155 | 0.383193955 | ITM2C | protein_coding | 58.43 | -0.244569053 | 0.244569053 | ENSG00000135916 | n.s. |
| ENSG00000253726 | 264.2581 | -0.533505251 | 0.256834 | -2.077237 | 0.03778 | 0.348118472 | | sense_intronic | 38 | -0.244491015 | 0.244491015 | ENSG00000253726 | n.s. |
| ENSG00000272953 | 57.43866 | -0.602668951 | 0.306694 | -1.96505 | 0.049408 | 0.393543473 | | lincRNA | 54.91 | -0.244085316 | 0.244085316 | ENSG00000272953 | n.s. |
| ENSG00000135406 | 1287.555 | -0.538179627 | 0.260639 | -2.064845 | 0.038938 | 0.35209789 | PRPH | protein_coding | 62.39 | -0.24397651 | 0.24397651 | ENSG00000135406 | n.s. |
| ENSG00000151065 | 444.3183 | -0.576905462 | 0.287531 | -2.006413 | 0.044812 | 0.377904228 | DCP1B | protein_coding | 38.72 | -0.243810776 | 0.243810776 | ENSG00000151065 | n.s. |
| ENSG00000275770 | 4.016826 | -1.066081234 | 0.716226 | -1.488471 | 0.136627 | 0.590925983 | MIR6505 | miRNA | 50.7 | -0.24356429 | 0.24356429 | ENSG00000275770 | n.s. |
| ENSG00000272812 | 107.5704 | 0.513236449 | 0.243526 | 2.1075255 | 0.035072 | 0.335533508 | | sense_intronic | 36.24 | 0.243409624 | 0.243409624 | ENSG00000272812 | n.s. |
| ENSG00000091844 | 239.2241 | -0.587446955 | 0.29558 | -1.987436 | 0.046874 | 0.38571389 | RGS17 | protein_coding | 37.43 | -0.243047202 | 0.243047202 | ENSG00000091844 | n.s. |
| ENSG00000187122 | 318.4477 | -0.54841618 | 0.268678 | -2.041169 | 0.041234 | 0.360843563 | SLIT1 | protein_coding | 50.45 | -0.242773443 | 0.242773443 | ENSG00000187122 | n.s. |
| ENSG00000119321 | 590.8495 | -0.502874395 | 0.236923 | -2.12252 | 0.033794 | 0.329031605 | FKBP15 | protein_coding | 41.19 | -0.242768842 | 0.242768842 | ENSG00000119321 | n.s. |
| ENSG00000222043 | 39.29219 | 0.663702777 | 0.356861 | 1.8598368 | 0.062909 | 0.430754942 | | antisense | 68.45 | 0.242762386 | 0.242762386 | ENSG00000222043 | n.s. |
| ENSG00000161551 | 332.3515 | -0.594121425 | 0.300896 | -1.974505 | 0.048324 | 0.390493645 | ZNF577 | protein_coding | 41 | -0.24263089 | 0.24263089 | ENSG00000161551 | n.s. |
| ENSG00000000460 | 861.6233 | -0.485717618 | 0.225912 | -2.150034 | 0.031553 | 0.316959535 | C1orf112 | protein_coding | 39.22 | -0.242371235 | 0.242371235 | ENSG00000000460 | n.s. |
| ENSG00000171574 | 160.5818 | -0.547451269 | 0.268288 | -2.040533 | 0.041297 | 0.360907877 | ZNF584 | protein_coding | 51.81 | -0.242303924 | 0.242303924 | ENSG00000171574 | n.s. |
| ENSG00000108219 | 376.8713 | -0.537843138 | 0.261339 | -2.058026 | 0.039588 | 0.354422376 | TSPAN14 | protein_coding | 49.31 | -0.242286967 | 0.242286967 | ENSG00000108219 | n.s. |
| ENSG00000267002 | 607.508 | -0.534337699 | 0.258794 | -2.064719 | 0.03895 | 0.35209789 | | lincRNA | 44.38 | -0.242234823 | 0.242234823 | ENSG00000267002 | n.s. |
| ENSG00000070269 | 57.08577 | -0.639053589 | 0.336334 | -1.900055 | 0.057426 | 0.417869145 | TMEM260 | protein_coding | 39.13 | -0.242175553 | 0.242175553 | ENSG00000070269 | n.s. |
| ENSG00000210112 | 10.26411 | -0.867099516 | 0.529326 | -1.63812 | 0.101397 | 0.526378134 | MT-TM | Mt_tRNA | 45.59 | -0.241662508 | 0.241662508 | ENSG00000210112 | n.s. |
| ENSG00000130699 | 255.2374 | -0.50359605 | 0.238208 | -2.114106 | 0.034506 | 0.331530132 | TAF4 | protein_coding | 52.35 | -0.24146272 | 0.24146272 | ENSG00000130699 | n.s. |
| ENSG00000249784 | 225.2636 | 0.840592998 | 0.506017 | 1.6611964 | 0.096674 | 0.516736278 | SCARNA22 | scaRNA | 56 | 0.241024111 | 0.241024111 | ENSG00000249784 | n.s. |
| ENSG00000180346 | 635.5394 | 0.439947663 | 0.197275 | 2.2301224 | 0.025739 | 0.283519382 | TIGD2 | protein_coding | 43.15 | 0.240834938 | 0.240834938 | ENSG00000180346 | n.s. |
| ENSG00000135913 | 1868.226 | -0.499209377 | 0.235773 | -2.117332 | 0.034232 | 0.329831947 | USP37 | protein_coding | 39.62 | -0.240472792 | 0.240472792 | ENSG00000135913 | n.s. |
| ENSG00000175662 | 28.73456 | -0.648455712 | 0.345139 | -1.878823 | 0.060269 | 0.426183381 | TOM1L2 | protein_coding | 48.26 | -0.240190259 | 0.240190259 | ENSG00000175662 | n.s. |
| ENSG00000222445 | 11.79319 | -0.752762613 | 0.43066 | -1.747929 | 0.080476 | 0.479859335 | RN7SKP56 | misc_RNA | 45.62 | -0.240045498 | 0.240045498 | ENSG00000222445 | n.s. |
| ENSG00000186174 | 28.55924 | -0.677633787 | 0.370752 | -1.827728 | 0.06759 | 0.44261708 | BCL9L | protein_coding | 59.25 | -0.239863273 | 0.239863273 | ENSG00000186174 | n.s. |
| ENSG00000244357 | 23.03395 | -0.656170237 | 0.353055 | -1.858552 | 0.063091 | 0.431139465 | RN7SL145P | misc_RNA | 50.68 | -0.239752939 | 0.239752939 | ENSG00000244357 | n.s. |
| ENSG00000252157 | 14.55142 | 0.757318727 | 0.434886 | 1.7414181 | 0.08161 | 0.482429187 | RNU6-479P | snRNA | 36.19 | 0.239741682 | 0.239741682 | ENSG00000252157 | n.s. |
| ENSG00000274364 | 14.13142 | -0.762446889 | 0.43966 | -1.734175 | 0.082887 | 0.485101957 | | sense_intronic | 41.76 | -0.239535632 | 0.239535632 | ENSG00000274364 | n.s. |
| ENSG00000167106 | 77.52983 | -0.59132684 | 0.301083 | -1.963997 | 0.04953 | 0.393614248 | FAM102A | protein_coding | 55.6 | -0.239445498 | 0.239445498 | ENSG00000167106 | n.s. |
| ENSG00000276292 | 14.06749 | -0.744227515 | 0.422584 | -1.761133 | 0.078216 | 0.477043998 | | antisense | 35.68 | -0.239225657 | 0.239225657 | ENSG00000276292 | n.s. |
| ENSG00000259924 | 56.00677 | -0.653211456 | 0.350042 | -1.866093 | 0.062028 | 0.430475522 | | processed_pseudogene | 40.46 | -0.239109059 | 0.239109059 | ENSG00000259924 | n.s. |

| | | | | | | | | | | | | | |
|---|---|---|---|---|---|---|---|---|---|---|---|---|---|
| ENSG00000227376 | 33.40309 | -0.668855852 | 0.363131 | -1.841912 | 0.065488 | 0.439628633 | FTH1P16 | processed_pseudogene | 47.28 | -0.238724037 | 0.238724037 | ENSG00000227376 | n.s. |
| ENSG00000278784 | 35.93924 | -0.606940433 | 0.312415 | -1.942739 | 0.052048 | 0.404612394 | | sense_intronic | 44.67 | -0.238503809 | 0.238503809 | ENSG00000278784 | n.s. |
| ENSG00000272333 | 361.8609 | -0.507772429 | 0.2419 | -2.099104 | 0.035808 | 0.33915039 | KMT2B | protein_coding | 59.77 | -0.238453832 | 0.238453832 | ENSG00000272333 | n.s. |
| ENSG00000175387 | 3669.024 | -0.522686486 | 0.252291 | -2.071763 | 0.038288 | 0.350082264 | SMAD2 | protein_coding | 37.52 | -0.238256124 | 0.238256124 | ENSG00000175387 | n.s. |
| ENSG00000181588 | 513.9677 | -0.522263125 | 0.252516 | -2.068236 | 0.038618 | 0.350505588 | MEX3D | protein_coding | 60.48 | -0.23778904 | 0.23778904 | ENSG00000181588 | n.s. |
| ENSG00000199477 | 31.18764 | 0.625681394 | 0.328108 | 1.906938 | 0.056529 | 0.416937262 | SNORA31 | snoRNA | 43.08 | 0.237714688 | 0.237714688 | ENSG00000199477 | n.s. |
| ENSG00000204104 | 330.8438 | -0.521276217 | 0.251913 | -2.069269 | 0.038521 | 0.350082264 | TRAF3IP1 | protein_coding | 44.49 | -0.237613281 | 0.237613281 | ENSG00000204104 | n.s. |
| ENSG00000134255 | 1215.127 | -0.545925772 | 0.269024 | -2.029285 | 0.042429 | 0.36714891 | CEPT1 | protein_coding | 36.54 | -0.237563834 | 0.237563834 | ENSG00000134255 | n.s. |
| ENSG00000079387 | 979.7556 | -0.492484831 | 0.23231 | -2.119949 | 0.03401 | 0.329588879 | SENP1 | protein_coding | 38.62 | -0.237391207 | 0.237391207 | ENSG00000079387 | n.s. |
| ENSG00000104327 | 149.1491 | -0.565869172 | 0.28311 | -1.998762 | 0.045634 | 0.381370674 | CALB1 | protein_coding | 36.25 | -0.23690266 | 0.23690266 | ENSG00000104327 | n.s. |
| ENSG00000171016 | 1435.297 | -0.556658583 | 0.276575 | -2.012684 | 0.044148 | 0.375486418 | PYGO1 | protein_coding | 39.31 | -0.236805772 | 0.236805772 | ENSG00000171016 | n.s. |
| ENSG00000140044 | 191.3276 | -0.570521682 | 0.2867 | -1.989963 | 0.046595 | 0.384553485 | JDP2 | protein_coding | 50 | -0.236791172 | 0.236791172 | ENSG00000140044 | n.s. |
| ENSG00000278272 | 1733.581 | -0.483794788 | 0.226859 | -2.132575 | 0.03296 | 0.324308154 | HIST1H3C | protein_coding | 59.85 | -0.236596035 | 0.236596035 | ENSG00000278272 | n.s. |
| ENSG00000203485 | 85.98805 | -0.574968332 | 0.290259 | -1.980878 | 0.047605 | 0.387927349 | INF2 | protein_coding | 63.24 | -0.236455498 | 0.236455498 | ENSG00000203485 | n.s. |
| ENSG00000135919 | 606.6699 | -0.554104422 | 0.274933 | -2.015415 | 0.043861 | 0.374378906 | SERPINE2 | protein_coding | 43.21 | -0.236430056 | 0.236430056 | ENSG00000135919 | n.s. |
| ENSG00000258808 | 15.72676 | 1.11724742 | 0.780795 | 1.4309101 | 0.152456 | 0.614654298 | LINC02310 | lincRNA | 44.19 | 0.236151556 | 0.236151556 | ENSG00000258808 | n.s. |
| ENSG00000279698 | 21.15367 | -0.665373101 | 0.363179 | -1.832081 | 0.066939 | 0.441705004 | | TEC | 41.33 | -0.236119409 | 0.236119409 | ENSG00000279698 | n.s. |
| ENSG00000206699 | 4.752244 | -1.240034621 | 0.919452 | -1.348666 | 0.177444 | 0.645396531 | | misc_RNA | 37.25 | -0.235821567 | 0.235821567 | ENSG00000206699 | n.s. |
| ENSG00000207385 | 7.863009 | -0.853092378 | 0.523176 | -1.630602 | 0.102974 | 0.529239659 | RNU6-310P | snRNA | 38.32 | -0.235750047 | 0.235750047 | ENSG00000207385 | n.s. |
| ENSG00000236337 | 25.8145 | -0.6545547 | 0.353719 | -1.850495 | 0.064242 | 0.436414668 | FMR1-IT1 | sense_intronic | 45.88 | -0.23570558 | 0.23570558 | ENSG00000236337 | n.s. |
| ENSG00000131981 | 48.56035 | -0.688536142 | 0.381678 | -1.80397 | 0.071236 | 0.455380982 | LGALS3 | protein_coding | 44.62 | -0.235221236 | 0.235221236 | ENSG00000131981 | n.s. |
| ENSG00000168140 | 74.11133 | -0.613862864 | 0.321079 | -1.911875 | 0.055892 | 0.414070243 | VASN | protein_coding | 63.46 | -0.235064038 | 0.235064038 | ENSG00000168140 | n.s. |
| ENSG00000102870 | 953.6613 | -0.515249753 | 0.249243 | -2.067261 | 0.03871 | 0.350893753 | ZNF629 | protein_coding | 57.75 | -0.234348141 | 0.234348141 | ENSG00000102870 | n.s. |
| ENSG00000166153 | 53.92521 | -0.574226224 | 0.291034 | -1.973055 | 0.048489 | 0.391129217 | DEPDC4 | protein_coding | 39.88 | -0.234100398 | 0.234100398 | ENSG00000166153 | n.s. |
| ENSG00000212615 | 41.37913 | 0.639096706 | 0.342473 | 1.8661233 | 0.062024 | 0.430475522 | | snoRNA | 33.33 | 0.233942333 | 0.233942333 | ENSG00000212615 | n.s. |
| ENSG00000236860 | 13.80314 | -0.749756396 | 0.435026 | -1.723474 | 0.084803 | 0.48779901 | RPL39P29 | processed_pseudogene | 40.13 | -0.233743369 | 0.233743369 | ENSG00000236860 | n.s. |
| ENSG00000060656 | 225.1592 | -0.527484953 | 0.258637 | -2.03948 | 0.041402 | 0.361178828 | PTPRU | protein_coding | 51.58 | -0.23329484 | 0.23329484 | ENSG00000060656 | n.s. |
| ENSG00000264638 | 7.36761 | 0.793411794 | 0.47289 | 1.677794 | 0.093387 | 0.508121556 | MIR3152 | miRNA | 45.95 | 0.233288759 | 0.233288759 | ENSG00000264638 | n.s. |
| ENSG00000177706 | 377.9621 | -0.517199443 | 0.251131 | -2.059482 | 0.039448 | 0.354033183 | FAM20C | protein_coding | 59.85 | -0.233234208 | 0.233234208 | ENSG00000177706 | n.s. |
| ENSG00000101004 | 1035.74 | -0.504181452 | 0.242633 | -2.077955 | 0.037713 | 0.347968518 | NINL | protein_coding | 49.07 | -0.231147052 | 0.231147052 | ENSG00000101004 | n.s. |
| ENSG00000204381 | 1114.433 | -0.538980598 | 0.26694 | -2.019109 | 0.043476 | 0.37264772 | LAYN | protein_coding | 44.3 | -0.231061808 | 0.231061808 | ENSG00000204381 | n.s. |
| ENSG00000230071 | 78.3885 | -0.566547148 | 0.28751 | -1.970531 | 0.048778 | 0.391599066 | RPL4P6 | processed_pseudogene | 46.63 | -0.230674403 | 0.230674403 | ENSG00000230071 | n.s. |

| | | | | | | | | | | | | | |
|---|---|---|---|---|---|---|---|---|---|---|---|---|---|
| ENSG00000197937 | 477.3217 | -0.577868599 | 0.296171 | -1.951132 | 0.051041 | 0.399465302 | ZNF347 | protein_coding | 44.35 | -0.230292737 | 0.230292737 | ENSG00000197937 | n.s. |
| ENSG00000165238 | 185.4333 | -0.504594476 | 0.243816 | -2.069572 | 0.038492 | 0.350082264 | WNK2 | protein_coding | 52.37 | -0.230009245 | 0.230009245 | ENSG00000165238 | n.s. |
| ENSG00000185963 | 1234.667 | -0.477107825 | 0.225065 | -2.11987 | 0.034017 | 0.329588879 | BICD2 | protein_coding | 51.35 | -0.229979068 | 0.229979068 | ENSG00000185963 | n.s. |
| ENSG00000115297 | 5176.768 | -0.509370262 | 0.24701 | -2.062147 | 0.039194 | 0.35388756 | TLX2 | protein_coding | 64.99 | -0.229794603 | 0.229794603 | ENSG00000115297 | n.s. |
| ENSG00000280077 | 224.9411 | -0.558760378 | 0.282127 | -1.980525 | 0.047645 | 0.387927349 | | TEC | 33.16 | -0.229789983 | 0.229789983 | ENSG00000280077 | n.s. |
| ENSG00000005206 | 49.00909 | -0.598856345 | 0.312249 | -1.917883 | 0.055126 | 0.413454097 | SPPL2B | protein_coding | 61.06 | -0.229704945 | 0.229704945 | ENSG00000005206 | n.s. |
| ENSG00000092140 | 1927.935 | -0.500258499 | 0.240483 | -2.080222 | 0.037505 | 0.347525043 | G2E3 | protein_coding | 35.48 | -0.229625602 | 0.229625602 | ENSG00000092140 | n.s. |
| ENSG00000091136 | 5684.583 | -0.515856754 | 0.252279 | -2.044787 | 0.040876 | 0.35907784 | LAMB1 | protein_coding | 42.42 | -0.229458963 | 0.229458963 | ENSG00000091136 | n.s. |
| ENSG00000139044 | 242.4056 | -0.508653628 | 0.246805 | -2.060957 | 0.039307 | 0.354033183 | B4GALNT3 | protein_coding | 49.06 | -0.22938042 | 0.22938042 | ENSG00000139044 | n.s. |
| ENSG00000264364 | 2183.923 | -0.470067633 | 0.220764 | -2.12928 | 0.033231 | 0.325902089 | DYNLL2 | protein_coding | 49.02 | -0.228881974 | 0.228881974 | ENSG00000264364 | n.s. |
| ENSG00000171533 | 693.3619 | -0.538094318 | 0.26774 | -2.009767 | 0.044456 | 0.375858958 | MAP6 | protein_coding | 44.55 | -0.228676683 | 0.228676683 | ENSG00000171533 | n.s. |
| ENSG00000251756 | 11.89445 | 0.733370504 | 0.425308 | 1.7243295 | 0.084648 | 0.48779901 | RNA5SP385 | rRNA | 48.18 | 0.228634918 | 0.228634918 | ENSG00000251756 | n.s. |
| ENSG00000066322 | 98.16207 | -0.551536083 | 0.277355 | -1.988559 | 0.04675 | 0.385375413 | ELOVL1 | protein_coding | 58.74 | -0.228399915 | 0.228399915 | ENSG00000066322 | n.s. |
| ENSG00000265491 | 1362.3 | -0.513409131 | 0.250874 | -2.046483 | 0.040709 | 0.35907784 | RNF115 | protein_coding | 41.91 | -0.228370232 | 0.228370232 | ENSG00000265491 | n.s. |
| ENSG00000197063 | 688.1816 | -0.537128044 | 0.267183 | -2.010338 | 0.044395 | 0.375755532 | MAFG | protein_coding | 60.3 | -0.22833024 | 0.22833024 | ENSG00000197063 | n.s. |
| ENSG00000265452 | 16.86615 | -0.683044631 | 0.382802 | -1.784327 | 0.074371 | 0.463348535 | MIR3682 | miRNA | 29.76 | -0.228199888 | 0.228199888 | ENSG00000265452 | n.s. |
| ENSG00000163125 | 1204.316 | -0.468937017 | 0.22051 | -2.126602 | 0.033453 | 0.326854919 | RPRD2 | protein_coding | 41.76 | -0.227736906 | 0.227736906 | ENSG00000163125 | n.s. |
| ENSG00000197535 | 125.9076 | -0.53551007 | 0.266461 | -2.009712 | 0.044462 | 0.375858958 | MYO5A | protein_coding | 39.92 | -0.227578442 | 0.227578442 | ENSG00000197535 | n.s. |
| ENSG00000154124 | 1192.582 | -0.486432442 | 0.232032 | -2.096398 | 0.036047 | 0.340720178 | OTULIN | protein_coding | 41.68 | -0.227456854 | 0.227456854 | ENSG00000154124 | n.s. |
| ENSG00000250959 | 73.51686 | -0.586347922 | 0.304042 | -1.928512 | 0.053792 | 0.409408975 | GLUD1P3 | cribed_unprocessed_pseudc | 52.12 | -0.227410727 | 0.227410727 | ENSG00000250959 | n.s. |
| ENSG00000164442 | 1338.993 | 0.425431994 | 0.19311 | 2.2030558 | 0.027591 | 0.292349299 | CITED2 | protein_coding | 50.32 | 0.227222353 | 0.227222353 | ENSG00000164442 | n.s. |
| ENSG00000240265 | 80.17796 | -0.544668981 | 0.273089 | -1.994473 | 0.0461 | 0.382988464 | | processed_pseudogene | 60.68 | -0.227025824 | 0.227025824 | ENSG00000240265 | n.s. |
| ENSG00000206621 | 10.15851 | -0.769244277 | 0.457777 | -1.680389 | 0.092882 | 0.507041328 | SNORD116-14 | snoRNA | 41.3 | -0.226893707 | 0.226893707 | ENSG00000206621 | n.s. |
| ENSG00000160570 | 143.8249 | 0.555911347 | 0.281762 | 1.9729811 | 0.048498 | 0.391129217 | DEDD2 | protein_coding | 52.69 | 0.226633794 | 0.226633794 | ENSG00000160570 | n.s. |
| ENSG00000176018 | 196.559 | -0.568328436 | 0.29109 | -1.952412 | 0.050889 | 0.399312737 | LYSMD3 | protein_coding | 35.92 | -0.226585066 | 0.226585066 | ENSG00000176018 | n.s. |
| ENSG00000184530 | 191.1304 | 0.496910087 | 0.239952 | 2.0708735 | 0.038371 | 0.350082264 | C6orf58 | protein_coding | 35.52 | 0.226506471 | 0.226506471 | ENSG00000184530 | n.s. |
| ENSG00000054793 | 1073.619 | -0.496131838 | 0.239814 | -2.068818 | 0.038563 | 0.350237668 | ATP9A | protein_coding | 46.69 | -0.226056096 | 0.226056096 | ENSG00000054793 | n.s. |
| ENSG00000182223 | 208.1276 | -0.495386787 | 0.23879 | -2.074568 | 0.038027 | 0.349698832 | ZAR1 | protein_coding | 53.24 | -0.226047873 | 0.226047873 | ENSG00000182223 | n.s. |
| ENSG00000179774 | 48.03791 | 0.617354616 | 0.331184 | 1.8640813 | 0.06231 | 0.430475522 | ATOH7 | protein_coding | 54.26 | 0.225983607 | 0.225983607 | ENSG00000179774 | n.s. |
| ENSG00000006025 | 86.48849 | -0.58177967 | 0.30164 | -1.928721 | 0.053766 | 0.409408975 | OSBPL7 | protein_coding | 57.86 | -0.225638964 | 0.225638964 | ENSG00000006025 | n.s. |
| ENSG00000047056 | 328.3336 | -0.506984448 | 0.247617 | -2.047455 | 0.040613 | 0.35907784 | WDR37 | protein_coding | 45.5 | -0.22551246 | 0.22551246 | ENSG00000047056 | n.s. |
| ENSG00000128482 | 339.9292 | -0.548173258 | 0.276494 | -1.982586 | 0.047414 | 0.387927349 | RNF112 | protein_coding | 60.03 | -0.225436034 | 0.225436034 | ENSG00000128482 | n.s. |

| | | | | | | | | | | | | | |
|---|---|---|---|---|---|---|---|---|---|---|---|---|---|
| ENSG00000167987 | 160.4866 | -0.535510676 | 0.267386 | -2.002766 | 0.045202 | 0.379585924 | VPS37C | protein_coding | 49.88 | -0.225283933 | 0.225283933 | ENSG00000167987 | n.s. |
| ENSG00000278434 | 103.2934 | -0.533904623 | 0.266264 | -2.005171 | 0.044945 | 0.378563498 | | sense_intronic | 52.78 | -0.225233679 | 0.225233679 | ENSG00000278434 | n.s. |
| ENSG00000261609 | 750.2298 | -0.531659352 | 0.264786 | -2.007884 | 0.044656 | 0.377040512 | GAN | protein_coding | 41.53 | -0.225217273 | 0.225217273 | ENSG00000261609 | n.s. |
| ENSG00000206732 | 17.26182 | -1.015334228 | 0.690605 | -1.470209 | 0.141505 | 0.600114276 | RNU6-936P | snRNA | 33.64 | -0.225166653 | 0.225166653 | ENSG00000206732 | n.s. |
| ENSG00000150712 | 530.4899 | -0.498978049 | 0.242273 | -2.059568 | 0.03944 | 0.354033183 | MTMR12 | protein_coding | 44.09 | -0.22501716 | 0.22501716 | ENSG00000150712 | n.s. |
| ENSG00000088756 | 254.2503 | -0.545676803 | 0.275041 | -1.983981 | 0.047258 | 0.387498315 | ARHGAP28 | protein_coding | 39.1 | -0.224671609 | 0.224671609 | ENSG00000088756 | n.s. |
| ENSG00000172493 | 1282.803 | -0.489456778 | 0.235467 | -2.078668 | 0.037648 | 0.347594073 | AFF1 | protein_coding | 41.19 | -0.224625243 | 0.224625243 | ENSG00000172493 | n.s. |
| ENSG00000090905 | 1980.243 | -0.498016586 | 0.241658 | -2.060828 | 0.039319 | 0.354033183 | TNRC6A | protein_coding | 42.59 | -0.224583582 | 0.224583582 | ENSG00000090905 | n.s. |
| ENSG00000279600 | 48.50048 | -0.624422188 | 0.337942 | -1.847721 | 0.064643 | 0.436903102 | | TEC | 41.15 | -0.224551505 | 0.224551505 | ENSG00000279600 | n.s. |
| ENSG00000096654 | 541.7989 | -0.521205516 | 0.257687 | -2.022631 | 0.043111 | 0.370895024 | ZNF184 | protein_coding | 38.79 | -0.224508751 | 0.224508751 | ENSG00000096654 | n.s. |
| ENSG00000275221 | 252.9858 | -0.480614015 | 0.229563 | -2.093602 | 0.036295 | 0.341167655 | HIST1H2AK | protein_coding | 62.85 | -0.224462197 | 0.224462197 | ENSG00000275221 | n.s. |
| ENSG00000242737 | 28.70204 | -0.638208399 | 0.350374 | -1.821508 | 0.06853 | 0.445699067 | | processed_pseudogene | 42.62 | -0.223984519 | 0.223984519 | ENSG00000242737 | n.s. |
| ENSG00000268357 | 65.90085 | -0.611692632 | 0.328568 | -1.861692 | 0.062647 | 0.430511912 | VN1R81P | unprocessed_pseudogene | 39.25 | -0.223888573 | 0.223888573 | ENSG00000268357 | n.s. |
| ENSG00000245975 | 41.21998 | 0.611398855 | 0.327995 | 1.8640495 | 0.062315 | 0.430475522 | | lincRNA | 45.68 | 0.223803492 | 0.223803492 | ENSG00000245975 | n.s. |
| ENSG00000106077 | 108.2992 | -0.551380065 | 0.280218 | -1.967683 | 0.049104 | 0.39292039 | ABHD11 | protein_coding | 59.95 | -0.223692375 | 0.223692375 | ENSG00000106077 | n.s. |
| ENSG00000090776 | 169.2298 | -0.52401238 | 0.259961 | -2.015734 | 0.043828 | 0.374378906 | EFNB1 | protein_coding | 56.43 | -0.223590124 | 0.223590124 | ENSG00000090776 | n.s. |
| ENSG00000084636 | 64.34282 | -0.589808921 | 0.310195 | -1.901413 | 0.057248 | 0.417869145 | COL16A1 | protein_coding | 56 | -0.223513809 | 0.223513809 | ENSG00000084636 | n.s. |
| ENSG00000127080 | 260.5878 | -0.529714915 | 0.264238 | -2.00469 | 0.044996 | 0.378748866 | IPPK | protein_coding | 47.93 | -0.223353584 | 0.223353584 | ENSG00000127080 | n.s. |
| ENSG00000256594 | 170.2365 | -0.48998527 | 0.236323 | -2.073373 | 0.038138 | 0.350082264 | | transcribed_unprocessed_pseudc | 39.19 | -0.223349933 | 0.223349933 | ENSG00000256594 | n.s. |
| ENSG00000140564 | 144.1875 | -0.543402738 | 0.274622 | -1.978728 | 0.047847 | 0.3891173 | FURIN | protein_coding | 58.47 | -0.222751357 | 0.222751357 | ENSG00000140564 | n.s. |
| ENSG00000061938 | 48.27977 | -0.608164708 | 0.325283 | -1.869647 | 0.061533 | 0.430280775 | TNK2 | protein_coding | 58.27 | -0.222739143 | 0.222739143 | ENSG00000061938 | n.s. |
| ENSG00000140538 | 107.3059 | -0.540626597 | 0.272806 | -1.981722 | 0.04751 | 0.387927349 | NTRK3 | protein_coding | 45.3 | -0.222332473 | 0.222332473 | ENSG00000140538 | n.s. |
| ENSG00000146826 | 130.4901 | -0.522556992 | 0.259613 | -2.012827 | 0.044133 | 0.375486418 | C7orf43 | protein_coding | 60.47 | -0.222298758 | 0.222298758 | ENSG00000146826 | n.s. |
| ENSG00000000457 | 418.1321 | -0.558100334 | 0.28617 | -1.95024 | 0.051148 | 0.400071404 | SCYL3 | protein_coding | 40.14 | -0.222047189 | 0.222047189 | ENSG00000000457 | n.s. |
| ENSG00000206609 | 3.842543 | -0.991683862 | 0.671724 | -1.476326 | 0.139856 | 0.597163054 | SNORD116-11 | snoRNA | 40.22 | -0.222045028 | 0.222045028 | ENSG00000206609 | n.s. |
| ENSG00000266072 | 5.929869 | -0.869123511 | 0.550935 | -1.577542 | 0.114671 | 0.55529213 | MIR5693 | miRNA | 46.58 | -0.222042355 | 0.222042355 | ENSG00000266072 | n.s. |
| ENSG00000180855 | 530.1419 | -0.542163329 | 0.274351 | -1.976166 | 0.048136 | 0.389875966 | ZNF443 | protein_coding | 41.62 | -0.22178467 | 0.22178467 | ENSG00000180855 | n.s. |
| ENSG00000105643 | 95.21123 | -0.545663783 | 0.277157 | -1.968787 | 0.048978 | 0.39235624 | ARRDC2 | protein_coding | 58.01 | -0.221713801 | 0.221713801 | ENSG00000105643 | n.s. |
| ENSG00000187187 | 32.41505 | -0.64663413 | 0.357568 | -1.808421 | 0.070541 | 0.45427456 | ZNF546 | protein_coding | 39.84 | -0.221589608 | 0.221589608 | ENSG00000187187 | n.s. |
| ENSG00000100523 | 1459.231 | -0.52031413 | 0.25839 | -2.01368 | 0.044043 | 0.375283011 | DDHD1 | protein_coding | 38.26 | -0.221467076 | 0.221467076 | ENSG00000100523 | n.s. |
| ENSG00000279529 | 114.2775 | 0.4953347 | 0.241443 | 2.0515588 | 0.040213 | 0.35721239 | | antisense | 50.53 | 0.221451011 | 0.221451011 | ENSG00000279529 | n.s. |
| ENSG00000118596 | 41.11445 | -0.590317252 | 0.31228 | -1.890343 | 0.058712 | 0.421582342 | SLC16A7 | protein_coding | 35.78 | -0.221438384 | 0.221438384 | ENSG00000118596 | n.s. |

| | | | | | | | | | | | | | |
|---|---|---|---|---|---|---|---|---|---|---|---|---|---|
| ENSG00000168040 | 190.293 | -0.497491334 | 0.2433 | -2.044762 | 0.040878 | 0.35907784 | FADD | protein_coding | 55.01 | -0.221289814 | 0.221289814 | ENSG00000168040 | n.s. |
| ENSG00000175727 | 565.0892 | -0.494388482 | 0.24082 | -2.052934 | 0.040079 | 0.356790345 | MLXIP | protein_coding | 50.38 | -0.221281811 | 0.221281811 | ENSG00000175727 | n.s. |
| ENSG00000197153 | 1072.863 | -0.473011528 | 0.225662 | -2.096111 | 0.036072 | 0.340729828 | HIST1H3J | protein_coding | 48.84 | -0.221175387 | 0.221175387 | ENSG00000197153 | n.s. |
| ENSG00000148110 | 796.4804 | -0.517564983 | 0.256692 | -2.016289 | 0.04377 | 0.374378906 | MFSD14B | protein_coding | 40.66 | -0.220839093 | 0.220839093 | ENSG00000148110 | n.s. |
| ENSG00000231074 | 1667.605 | -0.534993094 | 0.269481 | -1.98527 | 0.047114 | 0.387052655 | HCG18 | antisense | 42.72 | -0.220540176 | 0.220540176 | ENSG00000231074 | n.s. |
| ENSG00000267758 | 59.62473 | -0.593603657 | 0.315507 | -1.881429 | 0.059914 | 0.425223094 | | antisense | 38.82 | -0.2204544 | 0.2204544 | ENSG00000267758 | n.s. |
| ENSG00000159905 | 247.3797 | -0.538531433 | 0.272437 | -1.976719 | 0.048073 | 0.389875966 | ZNF221 | protein_coding | 41.33 | -0.220298958 | 0.220298958 | ENSG00000159905 | n.s. |
| ENSG00000242889 | 56.87654 | -0.581228716 | 0.305368 | -1.903374 | 0.056992 | 0.417869145 | RN7SL449P | misc_RNA | 55.94 | -0.220262257 | 0.220262257 | ENSG00000242889 | n.s. |
| ENSG00000238964 | 6.709943 | -0.857427399 | 0.542468 | -1.580604 | 0.113969 | 0.553658141 | RNU7-125P | snRNA | 35.48 | -0.22015161 | 0.22015161 | ENSG00000238964 | n.s. |
| ENSG00000206948 | 17.10609 | 0.705816657 | 0.409177 | 1.7249662 | 0.084534 | 0.48779901 | SNORA36A | snoRNA | 40.15 | 0.220044756 | 0.220044756 | ENSG00000206948 | n.s. |
| ENSG00000101019 | 631.1011 | -0.542835749 | 0.276195 | -1.965407 | 0.049367 | 0.393439254 | UQCC1 | protein_coding | 43.02 | -0.219914874 | 0.219914874 | ENSG00000101019 | n.s. |
| ENSG00000277453 | 17.2911 | -0.849414985 | 0.535809 | -1.585295 | 0.112899 | 0.551349488 | | lincRNA | 49.15 | -0.219635801 | 0.219635801 | ENSG00000277453 | n.s. |
| ENSG00000239263 | 55.84455 | -0.572401728 | 0.298719 | -1.91619 | 0.055341 | 0.413559567 | RBM43P1 | processed_pseudogene | 37.28 | -0.21949427 | 0.21949427 | ENSG00000239263 | n.s. |
| ENSG00000272137 | 30.16223 | -0.660647061 | 0.370995 | -1.780745 | 0.074954 | 0.465679039 | | lincRNA | 29.94 | -0.219277553 | 0.219277553 | ENSG00000272137 | n.s. |
| ENSG00000181638 | 174.2402 | -0.477486134 | 0.229704 | -2.078706 | 0.037644 | 0.347594073 | ZFP41 | protein_coding | 57.2 | -0.219131584 | 0.219131584 | ENSG00000181638 | n.s. |
| ENSG00000213058 | 128.6816 | 0.640692097 | 0.354612 | 1.8067395 | 0.070803 | 0.455117569 | | processed_pseudogene | 53.78 | 0.219037507 | 0.219037507 | ENSG00000213058 | n.s. |
| ENSG00000173681 | 284.8163 | -0.525815999 | 0.263884 | -1.992606 | 0.046305 | 0.383342621 | CXorf23 | protein_coding | 39.56 | -0.218956561 | 0.218956561 | ENSG00000173681 | n.s. |
| ENSG00000125449 | 125.3808 | -0.520390563 | 0.26007 | -2.000962 | 0.045397 | 0.38020912 | ARMC7 | protein_coding | 51.99 | -0.218552312 | 0.218552312 | ENSG00000125449 | n.s. |
| ENSG00000131711 | 12065.35 | -0.493389783 | 0.241667 | -2.04161 | 0.04119 | 0.360686468 | MAP1B | protein_coding | 42.29 | -0.218507608 | 0.218507608 | ENSG00000131711 | n.s. |
| ENSG00000237280 | 56.30261 | -0.559690214 | 0.289369 | -1.934172 | 0.053092 | 0.407272581 | | sense_intronic | 42.61 | -0.218343451 | 0.218343451 | ENSG00000237280 | n.s. |
| ENSG00000259562 | 67.55977 | -0.540511061 | 0.275457 | -1.962231 | 0.049736 | 0.394568837 | | transcribed_processed_pseudog | 41.58 | -0.218300104 | 0.218300104 | ENSG00000259562 | n.s. |
| ENSG00000278571 | 59.84347 | -0.57575531 | 0.302449 | -1.903642 | 0.056957 | 0.417869145 | MIR7161 | miRNA | 52.38 | -0.218188057 | 0.218188057 | ENSG00000278571 | n.s. |
| ENSG00000247363 | 31.82545 | 0.652731326 | 0.365583 | 1.7854544 | 0.074188 | 0.463232209 | | lincRNA | 42.44 | 0.218143626 | 0.218143626 | ENSG00000247363 | n.s. |
| ENSG00000169410 | 622.4953 | -0.53029222 | 0.267433 | -1.9829 | 0.047379 | 0.387927349 | PTPN9 | protein_coding | 45.12 | -0.218082464 | 0.218082464 | ENSG00000169410 | n.s. |
| ENSG00000088256 | 424.917 | -0.465513031 | 0.222159 | -2.095408 | 0.036135 | 0.340867363 | GNA11 | protein_coding | 61.66 | -0.217587578 | 0.217587578 | ENSG00000088256 | n.s. |
| ENSG00000010803 | 42.90679 | -0.611009702 | 0.332413 | -1.838105 | 0.066047 | 0.440875092 | SCMH1 | protein_coding | 39.5 | -0.217326641 | 0.217326641 | ENSG00000010803 | n.s. |
| ENSG00000237491 | 64.68419 | 0.541517692 | 0.276647 | 1.9574309 | 0.050297 | 0.397436283 | | lincRNA | 40.42 | 0.217003732 | 0.217003732 | ENSG00000237491 | n.s. |
| ENSG00000162695 | 1265.406 | -0.518313779 | 0.259298 | -1.998914 | 0.045618 | 0.381370674 | SLC30A7 | protein_coding | 35.85 | -0.216993466 | 0.216993466 | ENSG00000162695 | n.s. |
| ENSG00000279589 | 104.0631 | -0.585469634 | 0.311663 | -1.878537 | 0.060308 | 0.426183381 | | TEC | 36.46 | -0.216859995 | 0.216859995 | ENSG00000279589 | n.s. |
| ENSG00000241772 | 51.95689 | -0.61266271 | 0.335406 | -1.826628 | 0.067756 | 0.442685196 | | 3prime_overlapping_ncRNA | 32.74 | -0.216824397 | 0.216824397 | ENSG00000241772 | n.s. |
| ENSG00000264210 | 1.586539 | 1.767066502 | 1.686034 | 1.0480608 | 0.294611 | 0.754039228 | MIR4716 | miRNA | 45.24 | 0.216653062 | 0.216653062 | ENSG00000264210 | n.s. |
| ENSG00000257752 | 24.04999 | -0.61664215 | 0.33865 | -1.820885 | 0.068624 | 0.445830359 | | processed_pseudogene | 43.24 | -0.216336788 | 0.216336788 | ENSG00000257752 | n.s. |

| | | | | | | | | | | | | | |
|---|---|---|---|---|---|---|---|---|---|---|---|---|---|
| ENSG00000135801 | 1192.781 | -0.518569993 | 0.260006 | -1.994451 | 0.046103 | 0.382988464 | TAF5L | protein_coding | 41.9 | -0.216147392 | 0.216147392 | ENSG00000135801 | n.s. |
| ENSG00000241839 | 143.5896 | -0.548096384 | 0.281828 | -1.94479 | 0.0518 | 0.403363759 | PLEKHO2 | protein_coding | 53.96 | -0.216116116 | 0.216116116 | ENSG00000241839 | n.s. |
| ENSG00000235183 | 177.0304 | 0.483230145 | 0.235459 | 2.052289 | 0.040142 | 0.357119014 | | processed_pseudogene | 46.79 | 0.216094251 | 0.216094251 | ENSG00000235183 | n.s. |
| ENSG00000196417 | 406.1974 | -0.525444089 | 0.265131 | -1.981828 | 0.047499 | 0.387927349 | ZNF765 | protein_coding | 44.73 | -0.216088672 | 0.216088672 | ENSG00000196417 | n.s. |
| ENSG00000118733 | 135.3318 | 0.579847465 | 0.307491 | 1.8857362 | 0.059331 | 0.423990171 | OLFM3 | protein_coding | 35.12 | 0.216076801 | 0.216076801 | ENSG00000118733 | n.s. |
| ENSG00000221883 | 55.38766 | -0.579776165 | 0.307702 | -1.884215 | 0.059536 | 0.423999253 | ARIH2OS | protein_coding | 61.45 | -0.216044838 | 0.216044838 | ENSG00000221883 | n.s. |
| ENSG00000100077 | 3125.284 | -0.503799066 | 0.249536 | -2.018945 | 0.043493 | 0.37264772 | GRK3 | protein_coding | 40.55 | -0.215979431 | 0.215979431 | ENSG00000100077 | n.s. |
| ENSG00000159217 | 4913.173 | -0.487521202 | 0.238915 | -2.04056 | 0.041295 | 0.360907877 | IGF2BP1 | protein_coding | 47.35 | -0.215778658 | 0.215778658 | ENSG00000159217 | n.s. |
| ENSG00000261574 | 39.28977 | -0.63556605 | 0.35333 | -1.798789 | 0.072052 | 0.45772936 | | sense_intronic | 52.59 | -0.215705543 | 0.215705543 | ENSG00000261574 | n.s. |
| ENSG00000168824 | 452.459 | -0.552693885 | 0.28592 | -1.933036 | 0.053232 | 0.407895615 | | protein_coding | 50.8 | -0.215247166 | 0.215247166 | ENSG00000168824 | n.s. |
| ENSG00000242970 | 16.11401 | -0.705623736 | 0.414176 | -1.703683 | 0.08844 | 0.495479338 | | processed_pseudogene | 51.88 | -0.215197212 | 0.215197212 | ENSG00000242970 | n.s. |
| ENSG00000279133 | 514.9976 | -0.499386066 | 0.24694 | -2.022298 | 0.043146 | 0.370895024 | | TEC | 40.51 | -0.215110046 | 0.215110046 | ENSG00000279133 | n.s. |
| ENSG00000278642 | 7.04116 | -1.072996277 | 0.772905 | -1.388264 | 0.165057 | 0.63030432 | | lincRNA | 50.84 | -0.215081799 | 0.215081799 | ENSG00000278642 | n.s. |
| ENSG00000271869 | 17.325 | -0.777806412 | 0.476851 | -1.631132 | 0.102862 | 0.529239659 | | lincRNA | 54.09 | -0.21494495 | 0.21494495 | ENSG00000271869 | n.s. |
| ENSG00000267216 | 511.269 | 0.43199807 | 0.201298 | 2.1460662 | 0.031868 | 0.31813913 | | protein_coding | 47.15 | 0.214868458 | 0.214868458 | ENSG00000267216 | n.s. |
| ENSG00000206603 | 69.24713 | 0.530824405 | 0.270339 | 1.9635542 | 0.049582 | 0.393797785 | SNORA22B | snoRNA | 40.58 | 0.214838826 | 0.214838826 | ENSG00000206603 | n.s. |
| ENSG00000101413 | 1002.361 | -0.50703413 | 0.252562 | -2.007565 | 0.044689 | 0.37709809 | RPRD1B | protein_coding | 42.38 | -0.214752109 | 0.214752109 | ENSG00000101413 | n.s. |
| ENSG00000113273 | 307.8001 | -0.497465473 | 0.24606 | -2.021724 | 0.043205 | 0.371092907 | ARSB | protein_coding | 41.01 | -0.214167516 | 0.214167516 | ENSG00000113273 | n.s. |
| ENSG00000255468 | 49.72271 | -0.569631532 | 0.301313 | -1.8905 | 0.058691 | 0.421582342 | | antisense | 48.63 | -0.213678806 | 0.213678806 | ENSG00000255468 | n.s. |
| ENSG00000215154 | 115.8157 | -0.518185726 | 0.261022 | -1.985218 | 0.04712 | 0.387052655 | | transcribed_unprocessed_pseudc | 54.23 | -0.213611675 | 0.213611675 | ENSG00000215154 | n.s. |
| ENSG00000172534 | 1240.469 | -0.473385217 | 0.230044 | -2.0578 | 0.039609 | 0.354422376 | HCFC1 | protein_coding | 59.51 | -0.213250036 | 0.213250036 | ENSG00000172534 | n.s. |
| ENSG00000160062 | 281.4579 | -0.552195115 | 0.286882 | -1.924816 | 0.054252 | 0.41118975 | ZBTB8A | protein_coding | 43.45 | -0.213123968 | 0.213123968 | ENSG00000160062 | n.s. |
| ENSG00000256031 | 293.4772 | -0.511182738 | 0.256455 | -1.993262 | 0.046233 | 0.383206589 | | sense_intronic | 44.24 | -0.212941876 | 0.212941876 | ENSG00000256031 | n.s. |
| ENSG00000027847 | 132.6314 | -0.566414984 | 0.299373 | -1.892003 | 0.058491 | 0.42095504 | B4GALT7 | protein_coding | 57.92 | -0.212838522 | 0.212838522 | ENSG00000027847 | n.s. |
| ENSG00000253463 | 15.99814 | -0.661288796 | 0.375499 | -1.761094 | 0.078222 | 0.477043998 | HMGB1P19 | processed_pseudogene | 43.15 | -0.212565705 | 0.212565705 | ENSG00000253463 | n.s. |
| ENSG00000261131 | 17.22649 | -0.671846854 | 0.386387 | -1.738792 | 0.082071 | 0.482958503 | | sense_intronic | 43.42 | -0.212364195 | 0.212364195 | ENSG00000261131 | n.s. |
| ENSG00000080546 | 540.8968 | -0.449673226 | 0.213786 | -2.103378 | 0.035433 | 0.337204269 | SESN1 | protein_coding | 37.66 | -0.212293849 | 0.212293849 | ENSG00000080546 | n.s. |
| ENSG00000243353 | 9.362583 | -0.802681959 | 0.500757 | -1.602936 | 0.108949 | 0.544064048 | RPS29P19 | processed_pseudogene | 47.95 | -0.212188953 | 0.212188953 | ENSG00000243353 | n.s. |
| ENSG00000221821 | 36.0983 | -0.653161862 | 0.369676 | -1.766847 | 0.077254 | 0.473625887 | C6orf226 | protein_coding | 59.6 | -0.211993198 | 0.211993198 | ENSG00000221821 | n.s. |
| ENSG00000166736 | 116.1917 | -0.594045037 | 0.32268 | -1.840975 | 0.065625 | 0.44012549 | HTR3A | protein_coding | 50.68 | -0.211731599 | 0.211731599 | ENSG00000166736 | n.s. |
| ENSG00000272447 | 141.4446 | -0.522696386 | 0.266068 | -1.964518 | 0.04947 | 0.393585711 | | lincRNA | 63.73 | -0.211671482 | 0.211671482 | ENSG00000272447 | n.s. |
| ENSG00000207963 | 5.849511 | 0.858282619 | 0.551593 | 1.5560084 | 0.119706 | 0.566955067 | MIR569 | miRNA | 32.29 | 0.211524918 | 0.211524918 | ENSG00000207963 | n.s. |

| | | | | | | | | | | | | | |
|---|---|---|---|---|---|---|---|---|---|---|---|---|---|
| ENSG00000149639 | 786.3132 | -0.497542923 | 0.247618 | -2.009316 | 0.044504 | 0.375985676 | SOGA1 | protein_coding | 51.57 | -0.211370514 | 0.211370514 | ENSG00000149639 | n.s. |
| ENSG00000127580 | 142.4316 | -0.500712283 | 0.249802 | -2.004437 | 0.045023 | 0.378748866 | WDR24 | protein_coding | 63.87 | -0.211124663 | 0.211124663 | ENSG00000127580 | n.s. |
| ENSG00000132205 | 260.4774 | -0.515398144 | 0.260551 | -1.978109 | 0.047916 | 0.389457194 | EMILIN2 | protein_coding | 47.46 | -0.211076295 | 0.211076295 | ENSG00000132205 | n.s. |
| ENSG00000128578 | 220.4557 | -0.55682816 | 0.29331 | -1.898428 | 0.05764 | 0.417869145 | STRIP2 | protein_coding | 43.81 | -0.21101543 | 0.21101543 | ENSG00000128578 | n.s. |
| ENSG00000201558 | 254.3454 | 0.462250206 | 0.222806 | 2.0746765 | 0.038017 | 0.349698832 | RNVU1-6 | snRNA | 50.61 | 0.210927459 | 0.210927459 | ENSG00000201558 | n.s. |
| ENSG00000267481 | 97.49376 | -0.567884928 | 0.301792 | -1.881709 | 0.059875 | 0.425223094 | | sense_intronic | 40.74 | -0.210902898 | 0.210902898 | ENSG00000267481 | n.s. |
| ENSG00000187260 | 92.17113 | -0.555644251 | 0.29159 | -1.905564 | 0.056707 | 0.417346136 | WDR86 | protein_coding | 57.01 | -0.210868995 | 0.210868995 | ENSG00000187260 | n.s. |
| ENSG00000114648 | 411.7968 | -0.526864213 | 0.269498 | -1.954987 | 0.050585 | 0.398128225 | KLHL18 | protein_coding | 44.87 | -0.210733584 | 0.210733584 | ENSG00000114648 | n.s. |
| ENSG00000124831 | 1400.384 | -0.508766611 | 0.25594 | -1.987837 | 0.04683 | 0.385618457 | LRRFIP1 | protein_coding | 45.52 | -0.210549087 | 0.210549087 | ENSG00000124831 | n.s. |
| ENSG00000275155 | 49.40922 | -0.582545345 | 0.314517 | -1.852188 | 0.063999 | 0.435510653 | | sense_intronic | 49.11 | -0.210299579 | 0.210299579 | ENSG00000275155 | n.s. |
| ENSG00000239942 | 10.07595 | -0.825647262 | 0.524193 | -1.575083 | 0.115237 | 0.556520073 | RN7SL394P | misc_RNA | 55.67 | -0.210143053 | 0.210143053 | ENSG00000239942 | n.s. |
| ENSG00000261428 | 278.1891 | -0.573563196 | 0.307695 | -1.864064 | 0.062313 | 0.430475522 | | antisense | 40.23 | -0.209953691 | 0.209953691 | ENSG00000261428 | n.s. |
| ENSG00000259153 | 37.75616 | 0.591493111 | 0.322738 | 1.8327358 | 0.066842 | 0.441705004 | | lincRNA | 42.18 | 0.209901789 | 0.209901789 | ENSG00000259153 | n.s. |
| ENSG00000260257 | 143.7125 | -0.513035856 | 0.259469 | -1.977252 | 0.048013 | 0.389875966 | | lincRNA | 40.4 | -0.209869391 | 0.209869391 | ENSG00000260257 | n.s. |
| ENSG00000239908 | 19.08822 | -0.682490916 | 0.39884 | -1.71119 | 0.087046 | 0.492652156 | RN7SL75P | misc_RNA | 50.89 | -0.209838392 | 0.209838392 | ENSG00000239908 | n.s. |
| ENSG00000238363 | 142.4502 | 0.484033157 | 0.238798 | 2.026957 | 0.042667 | 0.368546519 | SNORA13 | snoRNA | 40.6 | 0.209832094 | 0.209832094 | ENSG00000238363 | n.s. |
| ENSG00000214188 | 103.5333 | 0.573049158 | 0.30758 | 1.8630919 | 0.062449 | 0.430475522 | ST7-OT4 | sense_intronic | 38.94 | 0.209765526 | 0.209765526 | ENSG00000214188 | n.s. |
| ENSG00000104936 | 19.89032 | -0.711119423 | 0.423131 | -1.680614 | 0.092838 | 0.507041328 | DMPK | protein_coding | 59.04 | -0.209749396 | 0.209749396 | ENSG00000104936 | n.s. |
| ENSG00000198380 | 1558.417 | -0.478763114 | 0.235492 | -2.033031 | 0.042049 | 0.364768459 | GFPT1 | protein_coding | 39.5 | -0.209689972 | 0.209689972 | ENSG00000198380 | n.s. |
| ENSG00000201649 | 4.063046 | -1.013216532 | 0.714337 | -1.418401 | 0.156074 | 0.621038953 | RNY4P34 | misc_RNA | 40 | -0.20961541 | 0.20961541 | ENSG00000201649 | n.s. |
| ENSG00000226632 | 16.58118 | -0.716752177 | 0.428221 | -1.673789 | 0.094172 | 0.509979371 | UBE2V1P1 | processed_pseudogene | 40.9 | -0.209612304 | 0.209612304 | ENSG00000226632 | n.s. |
| ENSG00000266643 | 6.387919 | -1.049823264 | 0.758278 | -1.384484 | 0.16621 | 0.631590186 | MIR3677 | miRNA | 71.67 | -0.209507588 | 0.209507588 | ENSG00000266643 | n.s. |
| ENSG00000178343 | 116.6805 | -0.542926948 | 0.282255 | -1.923532 | 0.054413 | 0.411605417 | SHISA3 | protein_coding | 50.29 | -0.20930861 | 0.20930861 | ENSG00000178343 | n.s. |
| ENSG00000067141 | 163.3835 | -0.524965798 | 0.268998 | -1.951557 | 0.050991 | 0.399465302 | NEO1 | protein_coding | 38.73 | -0.209209863 | 0.209209863 | ENSG00000067141 | n.s. |
| ENSG00000228218 | 52.74242 | -0.58770559 | 0.319437 | -1.839817 | 0.065795 | 0.440839489 | ATF4P3 | processed_pseudogene | 52.08 | -0.209058344 | 0.209058344 | ENSG00000228218 | n.s. |
| ENSG00000138767 | 680.2516 | -0.46924924 | 0.229199 | -2.047346 | 0.040624 | 0.35907784 | CNOT6L | protein_coding | 36.25 | -0.20872741 | 0.20872741 | ENSG00000138767 | n.s. |
| ENSG00000283414 | 4.250168 | 0.91286374 | 0.613349 | 1.4883272 | 0.136665 | 0.590925983 | | snRNA | 35.56 | 0.208559162 | 0.208559162 | ENSG00000283414 | n.s. |
| ENSG00000177688 | 159.0061 | -0.506799843 | 0.25583 | -1.981001 | 0.047591 | 0.387927349 | SUMO4 | protein_coding | 39.87 | -0.208421234 | 0.208421234 | ENSG00000177688 | n.s. |
| ENSG00000266922 | 19.30209 | -0.666875385 | 0.386132 | -1.727065 | 0.084156 | 0.48719542 | | sense_intronic | 41.07 | -0.208263051 | 0.208263051 | ENSG00000266922 | n.s. |
| ENSG00000232015 | 17.47508 | -0.660580821 | 0.38022 | -1.737365 | 0.082323 | 0.484029016 | HSPE1P25 | processed_pseudogene | 44.71 | -0.208167911 | 0.208167911 | ENSG00000232015 | n.s. |
| ENSG00000259797 | 29.32613 | -0.651131288 | 0.372146 | -1.749667 | 0.080176 | 0.479305998 | | sense_intronic | 48.39 | -0.207962957 | 0.207962957 | ENSG00000259797 | n.s. |
| ENSG00000221083 | 3.320784 | -1.065940342 | 0.776697 | -1.372401 | 0.169939 | 0.638266663 | | snoRNA | 55.2 | -0.207856063 | 0.207856063 | ENSG00000221083 | n.s. |

| | | | | | | | | | | | | | |
|---|---|---|---|---|---|---|---|---|---|---|---|---|---|
| ENSG00000283676 | 2.73129 | 1.669374164 | 1.58 | 1.0565657 | 0.29071 | 0.750798148 | MIR5087 | miRNA | 42.11 | 0.207798366 | 0.207798366 | ENSG00000283676 | n.s. |
| ENSG00000146278 | 1221.925 | 0.451705543 | 0.217011 | 2.0814882 | 0.037389 | 0.346944726 | PNRC1 | protein_coding | 48.53 | 0.207666975 | 0.207666975 | ENSG00000146278 | n.s. |
| ENSG00000281156 | 5.126393 | -1.130505153 | 0.851131 | -1.328238 | 0.184099 | 0.655193675 | MIR3651 | miRNA | 53.33 | -0.207595004 | 0.207595004 | ENSG00000281156 | n.s. |
| ENSG00000279089 | 376.274 | -0.539791855 | 0.281222 | -1.919448 | 0.054928 | 0.413017377 | | TEC | 37.78 | -0.207297171 | 0.207297171 | ENSG00000279089 | n.s. |
| ENSG00000215630 | 25.88938 | -0.620058491 | 0.347695 | -1.783342 | 0.074531 | 0.463461407 | GUSBP9 | unprocessed_pseudogene | 50.96 | -0.207091118 | 0.207091118 | ENSG00000215630 | n.s. |
| ENSG00000272337 | 9.772336 | -0.80421632 | 0.508273 | -1.582254 | 0.113592 | 0.552791606 | RNU6-90P | snRNA | 40.57 | -0.207036292 | 0.207036292 | ENSG00000272337 | n.s. |
| ENSG00000225190 | 135.6517 | -0.506162579 | 0.256234 | -1.97539 | 0.048224 | 0.390361031 | PLEKHM1 | protein_coding | 50.52 | -0.206784392 | 0.206784392 | ENSG00000225190 | n.s. |
| ENSG00000278963 | 1508.09 | 0.420052377 | 0.19644 | 2.138319 | 0.032491 | 0.321971749 | | TEC | 40.35 | 0.206742317 | 0.206742317 | ENSG00000278963 | n.s. |
| ENSG00000264678 | 14.96898 | -0.69464561 | 0.411941 | -1.686273 | 0.091743 | 0.504081279 | MIR3140 | miRNA | 36.67 | -0.206656674 | 0.206656674 | ENSG00000264678 | n.s. |
| ENSG00000111676 | 277.8094 | -0.509947869 | 0.259351 | -1.966248 | 0.04927 | 0.393340111 | ATN1 | protein_coding | 55.53 | -0.206647075 | 0.206647075 | ENSG00000111676 | n.s. |
| ENSG00000162992 | 3840.552 | 0.428295939 | 0.201921 | 2.1211111 | 0.033912 | 0.329262928 | NEUROD1 | protein_coding | 39.48 | 0.206634444 | 0.206634444 | ENSG00000162992 | n.s. |
| ENSG00000114019 | 513.2988 | -0.517865675 | 0.265143 | -1.953156 | 0.050801 | 0.399155977 | AMOTL2 | protein_coding | 55.04 | -0.206554537 | 0.206554537 | ENSG00000114019 | n.s. |
| ENSG00000251722 | 11.885 | 0.72296361 | 0.43683 | 1.6550243 | 0.09792 | 0.518435958 | RNU5E-3P | snRNA | 32.35 | 0.206265049 | 0.206265049 | ENSG00000251722 | n.s. |
| ENSG00000076356 | 60.64849 | -0.542753275 | 0.284512 | -1.907665 | 0.056435 | 0.416836968 | PLXNA2 | protein_coding | 47.98 | -0.206264574 | 0.206264574 | ENSG00000076356 | n.s. |
| ENSG00000207416 | 10.92591 | 0.734218906 | 0.446315 | 1.6450693 | 0.099956 | 0.524173499 | | misc_RNA | 37.17 | 0.205966715 | 0.205966715 | ENSG00000207416 | n.s. |
| ENSG00000265462 | 31.04785 | -0.65938791 | 0.382222 | -1.725145 | 0.084501 | 0.48779901 | MIR3680-1 | miRNA | 43.68 | -0.205570172 | 0.205570172 | ENSG00000265462 | n.s. |
| ENSG00000221296 | 4.907802 | -1.055632155 | 0.769819 | -1.371274 | 0.17029 | 0.638865917 | MIR548T | miRNA | 36.49 | -0.205415757 | 0.205415757 | ENSG00000221296 | n.s. |
| ENSG00000221230 | 3.561225 | -1.056150812 | 0.770812 | -1.370179 | 0.170631 | 0.639142968 | MIR548L | miRNA | 39.53 | -0.205317814 | 0.205317814 | ENSG00000221230 | n.s. |
| ENSG00000135069 | 1842.163 | -0.552841116 | 0.29385 | -1.88137 | 0.059922 | 0.425223094 | PSAT1 | protein_coding | 45.59 | -0.205315879 | 0.205315879 | ENSG00000135069 | n.s. |
| ENSG00000133027 | 117.31 | -0.574395843 | 0.31193 | -1.841423 | 0.06556 | 0.439897079 | PEMT | protein_coding | 55.22 | -0.204857658 | 0.204857658 | ENSG00000133027 | n.s. |
| ENSG00000177679 | 367.2321 | -0.521271553 | 0.268456 | -1.941743 | 0.052168 | 0.404745128 | SRRM3 | protein_coding | 52.93 | -0.204765042 | 0.204765042 | ENSG00000177679 | n.s. |
| ENSG00000101384 | 153.0444 | -0.507372945 | 0.258661 | -1.961538 | 0.049816 | 0.394984536 | JAG1 | protein_coding | 47.52 | -0.204684346 | 0.204684346 | ENSG00000101384 | n.s. |
| ENSG00000154027 | 105.08 | -0.559163759 | 0.300365 | -1.861616 | 0.062657 | 0.430511912 | AK5 | protein_coding | 39.61 | -0.204662226 | 0.204662226 | ENSG00000154027 | n.s. |
| ENSG00000137449 | 106.2325 | -0.526601794 | 0.273084 | -1.92835 | 0.053812 | 0.409408975 | CPEB2 | protein_coding | 34.97 | -0.204238631 | 0.204238631 | ENSG00000137449 | n.s. |
| ENSG00000224020 | 165.8832 | -0.479667628 | 0.238455 | -2.011563 | 0.044266 | 0.375547181 | MIR181A2HG | antisense | 38.12 | -0.204019668 | 0.204019668 | ENSG00000224020 | n.s. |
| ENSG00000106346 | 453.4506 | -0.439678547 | 0.210603 | -2.087714 | 0.036824 | 0.343859992 | USP42 | protein_coding | 45.58 | -0.203843043 | 0.203843043 | ENSG00000106346 | n.s. |
| ENSG00000168389 | 146.0084 | -0.529363139 | 0.275687 | -1.92016 | 0.054838 | 0.412705132 | MFSD2A | protein_coding | 53.7 | -0.203466085 | 0.203466085 | ENSG00000168389 | n.s. |
| ENSG00000259439 | 36.73174 | -0.601279527 | 0.334891 | -1.795448 | 0.072582 | 0.459013748 | LINC01833 | lincRNA | 52.5 | -0.203337287 | 0.203337287 | ENSG00000259439 | n.s. |
| ENSG00000189180 | 1724.835 | -0.522467254 | 0.270526 | -1.931304 | 0.053445 | 0.408185051 | ZNF33A | protein_coding | 39.26 | -0.203314425 | 0.203314425 | ENSG00000189180 | n.s. |
| ENSG00000101191 | 2338.241 | -0.497556279 | 0.252398 | -1.971314 | 0.048688 | 0.391387419 | DIDO1 | protein_coding | 49.19 | -0.202701014 | 0.202701014 | ENSG00000101191 | n.s. |
| ENSG00000271833 | 98.2526 | -0.528387988 | 0.276081 | -1.913885 | 0.055635 | 0.413574865 | | sense_intronic | 45.7 | -0.202608188 | 0.202608188 | ENSG00000271833 | n.s. |
| ENSG00000265539 | 2.784474 | -1.22119427 | 0.96871 | -1.26064 | 0.207438 | 0.682785213 | MIR3164 | miRNA | 50.6 | -0.202371299 | 0.202371299 | ENSG00000265539 | n.s. |

| | | | | | | | | | | | | | |
|---|---|---|---|---|---|---|---|---|---|---|---|---|---|
| ENSG00000007545 | 416.6179 | -0.488487857 | 0.245629 | -1.988722 | 0.046732 | 0.385375413 | CRAMP1 | protein_coding | 53.11 | -0.202290636 | 0.202290636 | ENSG00000007545 | n.s. |
| ENSG00000107829 | 202.4368 | -0.520896891 | 0.269857 | -1.930273 | 0.053573 | 0.40893514 | FBXW4 | protein_coding | 45.8 | -0.202287999 | 0.202287999 | ENSG00000107829 | n.s. |
| ENSG00000248383 | 178.1182 | -0.479626396 | 0.239313 | -2.004179 | 0.045051 | 0.378748866 | PCDHAC1 | protein_coding | 39.4 | -0.202233827 | 0.202233827 | ENSG00000248383 | n.s. |
| ENSG00000271862 | 80.45479 | -0.53813805 | 0.284443 | -1.891904 | 0.058504 | 0.42095504 | | lincRNA | 40.34 | -0.20221306 | 0.20221306 | ENSG00000271862 | n.s. |
| ENSG00000274159 | 26.34759 | -0.599666002 | 0.334728 | -1.791503 | 0.073213 | 0.460406637 | | sense_intronic | 38.52 | -0.202002545 | 0.202002545 | ENSG00000274159 | n.s. |
| ENSG00000167981 | 193.804 | -0.490870948 | 0.247772 | -1.981137 | 0.047576 | 0.387927349 | ZNF597 | protein_coding | 42.53 | -0.201870482 | 0.201870482 | ENSG00000167981 | n.s. |
| ENSG00000116016 | 202.1375 | 0.526375382 | 0.274948 | 1.9144512 | 0.055563 | 0.413574865 | EPAS1 | protein_coding | 46.71 | 0.201836463 | 0.201836463 | ENSG00000116016 | n.s. |
| ENSG00000239407 | 72.36306 | -0.59567448 | 0.331318 | -1.797896 | 0.072193 | 0.458344637 | | lincRNA | 38.6 | -0.201819187 | 0.201819187 | ENSG00000239407 | n.s. |
| ENSG00000113838 | 274.366 | -0.532213072 | 0.279568 | -1.903701 | 0.056949 | 0.417869145 | TBCCD1 | protein_coding | 40.74 | -0.201687304 | 0.201687304 | ENSG00000113838 | n.s. |
| ENSG00000140750 | 196.0576 | -0.480195913 | 0.240001 | -2.000807 | 0.045413 | 0.38020912 | ARHGAP17 | protein_coding | 44.36 | -0.201671465 | 0.201671465 | ENSG00000140750 | n.s. |
| ENSG00000272567 | 15.12664 | -0.727207679 | 0.444802 | -1.634902 | 0.10207 | 0.528171612 | | lincRNA | 29.8 | -0.201600108 | 0.201600108 | ENSG00000272567 | n.s. |
| ENSG00000121274 | 703.0714 | -0.503736168 | 0.257522 | -1.956092 | 0.050454 | 0.398128225 | PAPD5 | protein_coding | 41.51 | -0.201482898 | 0.201482898 | ENSG00000121274 | n.s. |
| ENSG00000275054 | 7.626967 | -1.201856076 | 0.947001 | -1.269119 | 0.204399 | 0.679838368 | | misc_RNA | 32.43 | -0.201424262 | 0.201424262 | ENSG00000275054 | n.s. |
| ENSG00000256044 | 6.27683 | -1.121978974 | 0.851606 | -1.317486 | 0.187676 | 0.661470368 | | processed_pseudogene | 50 | -0.201383565 | 0.201383565 | ENSG00000256044 | n.s. |
| ENSG00000251934 | 15.21244 | -0.680273803 | 0.404154 | -1.683206 | 0.092335 | 0.505936376 | RNU6-1143P | snRNA | 34.62 | -0.201295804 | 0.201295804 | ENSG00000251934 | n.s. |
| ENSG00000102189 | 1440.629 | -0.523257481 | 0.272357 | -1.921221 | 0.054704 | 0.412398929 | EEA1 | protein_coding | 36.99 | -0.201287981 | 0.201287981 | ENSG00000102189 | n.s. |
| ENSG00000170734 | 487.718 | -0.482931538 | 0.242289 | -1.993204 | 0.046239 | 0.383206589 | POLH | protein_coding | 43.39 | -0.201173358 | 0.201173358 | ENSG00000170734 | n.s. |
| ENSG00000250764 | 49.37963 | -0.548821371 | 0.29361 | -1.869218 | 0.061592 | 0.430281981 | | antisense | 41.29 | -0.201004093 | 0.201004093 | ENSG00000250764 | n.s. |
| ENSG00000199914 | 6.387237 | 0.785213796 | 0.49739 | 1.5786697 | 0.114412 | 0.554650439 | SNORD114-16 | snoRNA | 47.14 | 0.200999529 | 0.200999529 | ENSG00000199914 | n.s. |
| ENSG00000101098 | 265.9676 | -0.491100459 | 0.248504 | -1.976226 | 0.048129 | 0.389875966 | RIMS4 | protein_coding | 49.54 | -0.200896201 | 0.200896201 | ENSG00000101098 | n.s. |
| ENSG00000252414 | 5.068561 | -0.926869893 | 0.640572 | -1.446942 | 0.147913 | 0.607109641 | RNU6-100P | snRNA | 40.57 | -0.200883173 | 0.200883173 | ENSG00000252414 | n.s. |
| ENSG00000080371 | 1522.713 | -0.511638356 | 0.263707 | -1.940179 | 0.052358 | 0.405289476 | RAB21 | protein_coding | 37.14 | -0.200682304 | 0.200682304 | ENSG00000080371 | n.s. |
| ENSG00000272948 | 64.30182 | 0.548360165 | 0.29479 | 1.8601748 | 0.062861 | 0.430713982 | | antisense | 57.28 | 0.200596197 | 0.200596197 | ENSG00000272948 | n.s. |
| ENSG00000110619 | 1257.042 | -0.532419837 | 0.281151 | -1.893713 | 0.058263 | 0.420691467 | CARS | protein_coding | 52.02 | -0.200209184 | 0.200209184 | ENSG00000110619 | n.s. |
| ENSG00000168438 | 1612.71 | -0.489409515 | 0.247615 | -1.976497 | 0.048098 | 0.389875966 | CDC40 | protein_coding | 37.34 | -0.200204481 | 0.200204481 | ENSG00000168438 | n.s. |
| ENSG00000196456 | 115.7495 | -0.555367606 | 0.300434 | -1.848554 | 0.064522 | 0.436711663 | ZNF775 | protein_coding | 52.5 | -0.199824158 | 0.199824158 | ENSG00000196456 | n.s. |
| ENSG00000261889 | 96.59395 | -0.539064105 | 0.287021 | -1.878133 | 0.060363 | 0.426183381 | | lincRNA | 51.2 | -0.199671225 | 0.199671225 | ENSG00000261889 | n.s. |
| ENSG00000146587 | 1096.445 | -0.517157912 | 0.268882 | -1.923367 | 0.054434 | 0.411605417 | RBAK | protein_coding | 40.26 | -0.199374159 | 0.199374159 | ENSG00000146587 | n.s. |
| ENSG00000204611 | 667.1189 | -0.493037827 | 0.251197 | -1.962753 | 0.049675 | 0.394311879 | ZNF616 | protein_coding | 42.23 | -0.199266237 | 0.199266237 | ENSG00000204611 | n.s. |
| ENSG00000155858 | 389.1508 | -0.519582337 | 0.270998 | -1.917293 | 0.055201 | 0.413559567 | LSM11 | protein_coding | 43.31 | -0.199240044 | 0.199240044 | ENSG00000155858 | n.s. |
| ENSG00000267692 | 7.924907 | -0.791827008 | 0.504881 | -1.568343 | 0.116801 | 0.560379317 | | processed_pseudogene | 43.31 | -0.199158668 | 0.199158668 | ENSG00000267692 | n.s. |
| ENSG00000231622 | 14.936 | 0.705374687 | 0.427822 | 1.6487566 | 0.099197 | 0.521983162 | RPS29P7 | processed_pseudogene | 49.71 | 0.199157962 | 0.199157962 | ENSG00000231622 | n.s. |

| | | | | | | | | | | | | | |
|---|---|---|---|---|---|---|---|---|---|---|---|---|---|
| ENSG00000232442 | 42.46038 | -0.603753403 | 0.339643 | -1.777611 | 0.075468 | 0.468034287 | MHENCR | antisense | 62.33 | -0.199070979 | 0.199070979 | ENSG00000232442 | n.s. |
| ENSG00000227124 | 516.1928 | -0.47696813 | 0.239267 | -1.993459 | 0.046211 | 0.383206589 | ZNF717 | protein_coding | 41.48 | -0.198689198 | 0.198689198 | ENSG00000227124 | n.s. |
| ENSG00000275318 | 43.55079 | -0.558182772 | 0.30376 | -1.837581 | 0.066124 | 0.440875092 | | unprocessed_pseudogene | 35.56 | -0.198536924 | 0.198536924 | ENSG00000275318 | n.s. |
| ENSG00000176125 | 50.30172 | -0.557691523 | 0.303161 | -1.839589 | 0.065829 | 0.440852534 | UFSP1 | protein_coding | 66.77 | -0.198374589 | 0.198374589 | ENSG00000176125 | n.s. |
| ENSG00000269609 | 208.0781 | -0.522944223 | 0.274708 | -1.903636 | 0.056958 | 0.417869145 | RPARP-AS1 | processed_transcript | 51.6 | -0.198174783 | 0.198174783 | ENSG00000269609 | n.s. |
| ENSG00000205138 | 252.1606 | -0.505798363 | 0.261037 | -1.937648 | 0.052666 | 0.406112801 | SDHAF1 | protein_coding | 63.38 | -0.197945869 | 0.197945869 | ENSG00000205138 | n.s. |
| ENSG00000270130 | 27.89279 | -0.621450444 | 0.356195 | -1.744689 | 0.081039 | 0.480948486 | | sense_intronic | 42.05 | -0.19755999 | 0.19755999 | ENSG00000270130 | n.s. |
| ENSG00000064692 | 27.77995 | -0.633581919 | 0.367646 | -1.723348 | 0.084826 | 0.48779901 | SNCAIP | protein_coding | 39.26 | -0.19752492 | 0.19752492 | ENSG00000064692 | n.s. |
| ENSG00000135457 | 709.5828 | -0.503923519 | 0.25996 | -1.938463 | 0.052567 | 0.405702385 | TFCP2 | protein_coding | 41.94 | -0.197433425 | 0.197433425 | ENSG00000135457 | n.s. |
| ENSG00000177303 | 343.1661 | -0.464096162 | 0.230799 | -2.010826 | 0.044344 | 0.375547181 | CASKIN2 | protein_coding | 61.19 | -0.19739657 | 0.19739657 | ENSG00000177303 | n.s. |
| ENSG00000206828 | 529.8121 | 0.443329886 | 0.216729 | 2.0455449 | 0.040801 | 0.35907784 | | snRNA | 45.12 | 0.197198185 | 0.197198185 | ENSG00000206828 | n.s. |
| ENSG00000243449 | 130.0252 | -0.575416924 | 0.318617 | -1.805983 | 0.070921 | 0.455117569 | C4orf48 | protein_coding | 69.21 | -0.196721466 | 0.196721466 | ENSG00000243449 | n.s. |
| ENSG00000175348 | 329.2322 | -0.518902393 | 0.27326 | -1.898932 | 0.057573 | 0.417869145 | TMEM9B | protein_coding | 42.7 | -0.196643093 | 0.196643093 | ENSG00000175348 | n.s. |
| ENSG00000070778 | 462.2419 | -0.491382816 | 0.251329 | -1.955141 | 0.050566 | 0.398128225 | PTPN21 | protein_coding | 41.28 | -0.196541841 | 0.196541841 | ENSG00000070778 | n.s. |
| ENSG00000265333 | 7.054509 | -0.816135949 | 0.531474 | -1.535608 | 0.124635 | 0.574507637 | MIR3137 | miRNA | 54.67 | -0.196447345 | 0.196447345 | ENSG00000265333 | n.s. |
| ENSG00000264480 | 8.762449 | -0.86775181 | 0.584848 | -1.483722 | 0.137883 | 0.593795878 | MIR4714 | miRNA | 46.75 | -0.196426748 | 0.196426748 | ENSG00000264480 | n.s. |
| ENSG00000249212 | 69.72479 | -0.554777841 | 0.3036 | -1.827332 | 0.06765 | 0.44261708 | ATP1B1P1 | unprocessed_pseudogene | 43.17 | -0.196375729 | 0.196375729 | ENSG00000249212 | n.s. |
| ENSG00000160999 | 87.00954 | -0.526660287 | 0.279384 | -1.885077 | 0.059419 | 0.423990171 | SH2B2 | protein_coding | 56.99 | -0.196256907 | 0.196256907 | ENSG00000160999 | n.s. |
| ENSG00000115977 | 393.7825 | -0.500873562 | 0.258355 | -1.938705 | 0.052537 | 0.405702385 | AAK1 | protein_coding | 41.95 | -0.196238475 | 0.196238475 | ENSG00000115977 | n.s. |
| ENSG00000146834 | 572.2137 | -0.453123133 | 0.223642 | -2.026113 | 0.042753 | 0.368937997 | MEPCE | protein_coding | 58.8 | -0.196223439 | 0.196223439 | ENSG00000146834 | n.s. |
| ENSG00000169122 | 43.1898 | -0.552871731 | 0.30205 | -1.830401 | 0.06719 | 0.441892852 | FAM110B | protein_coding | 40.5 | -0.196094217 | 0.196094217 | ENSG00000169122 | n.s. |
| ENSG00000269837 | 313.5014 | -0.525836383 | 0.27892 | -1.885262 | 0.059395 | 0.423990171 | IPO5P1 | ıscribed_processed_pseudog | 46.62 | -0.195949884 | 0.195949884 | ENSG00000269837 | n.s. |
| ENSG00000207389 | 40.52027 | 0.564793166 | 0.311407 | 1.813683 | 0.069727 | 0.450697584 | RNU1-4 | snRNA | 54.27 | 0.195483257 | 0.195483257 | ENSG00000207389 | n.s. |
| ENSG00000170260 | 241.1088 | -0.530184631 | 0.283071 | -1.872973 | 0.061072 | 0.42849416 | ZNF212 | protein_coding | 45.75 | -0.195137153 | 0.195137153 | ENSG00000170260 | n.s. |
| ENSG00000180787 | 181.897 | -0.508102249 | 0.264766 | -1.919062 | 0.054976 | 0.413017377 | ZFP3 | protein_coding | 45.78 | -0.195127358 | 0.195127358 | ENSG00000180787 | n.s. |
| ENSG00000163393 | 327.3272 | -0.524133958 | 0.278323 | -1.883184 | 0.059675 | 0.424480483 | SLC22A15 | protein_coding | 39.5 | -0.195052402 | 0.195052402 | ENSG00000163393 | n.s. |
| ENSG00000243845 | 11.95627 | -0.871095581 | 0.589816 | -1.476894 | 0.139704 | 0.597163054 | RN7SL30P | misc_RNA | 55.56 | -0.195044459 | 0.195044459 | ENSG00000243845 | n.s. |
| ENSG00000272672 | 18.30846 | -0.628361696 | 0.365236 | -1.720424 | 0.085355 | 0.489659038 | | lincRNA | 44.06 | -0.194858873 | 0.194858873 | ENSG00000272672 | n.s. |
| ENSG00000180822 | 778.83 | -0.492309256 | 0.252893 | -1.946708 | 0.05157 | 0.402182089 | PSMG4 | protein_coding | 49.83 | -0.194746354 | 0.194746354 | ENSG00000180822 | n.s. |
| ENSG00000157657 | 667.8937 | -0.47348335 | 0.239063 | -1.98058 | 0.047638 | 0.387927349 | ZNF618 | protein_coding | 45.61 | -0.194719839 | 0.194719839 | ENSG00000157657 | n.s. |
| ENSG00000182771 | 544.5009 | -0.477273642 | 0.241977 | -1.972394 | 0.048565 | 0.391129217 | GRID1 | protein_coding | 45.04 | -0.194574795 | 0.194574795 | ENSG00000182771 | n.s. |
| ENSG00000185477 | 2474.557 | -0.513289141 | 0.270244 | -1.899351 | 0.057518 | 0.417869145 | GPRIN3 | protein_coding | 39.05 | -0.194515897 | 0.194515897 | ENSG00000185477 | n.s. |

| | | | | | | | | | | | | | |
|---|---|---|---|---|---|---|---|---|---|---|---|---|---|
| ENSG00000274591 | 67.78608 | -0.55624975 | 0.30588 | -1.818522 | 0.068984 | 0.447091133 | | sense_intronic | 35.05 | -0.194467106 | 0.194467106 | ENSG00000274591 | n.s. |
| ENSG00000230483 | 16.62142 | -0.713075948 | 0.43941 | -1.622805 | 0.104631 | 0.53381423 | | antisense | 64.79 | -0.19439153 | 0.19439153 | ENSG00000230483 | n.s. |
| ENSG00000255020 | 32.39787 | -0.620036812 | 0.358486 | -1.729599 | 0.083702 | 0.485981036 | | antisense | 53.9 | -0.194307556 | 0.194307556 | ENSG00000255020 | n.s. |
| ENSG00000157326 | 85.17427 | -0.547143582 | 0.298325 | -1.834052 | 0.066646 | 0.441444767 | DHRS4 | protein_coding | 45.27 | -0.194303613 | 0.194303613 | ENSG00000157326 | n.s. |
| ENSG00000152193 | 1021.765 | -0.506234971 | 0.264481 | -1.914071 | 0.055611 | 0.413574865 | RNF219 | protein_coding | 38.52 | -0.194113705 | 0.194113705 | ENSG00000152193 | n.s. |
| ENSG00000214875 | 27.75422 | -0.652399615 | 0.38676 | -1.686833 | 0.091635 | 0.504081279 | MED28P1 | processed_pseudogene | 47.99 | -0.194088515 | 0.194088515 | ENSG00000214875 | n.s. |
| ENSG00000274115 | 73.88855 | -0.529309424 | 0.284187 | -1.862539 | 0.062527 | 0.430475522 | MIR6081 | miRNA | 64.95 | -0.193754529 | 0.193754529 | ENSG00000274115 | n.s. |
| ENSG00000249087 | 89.31722 | -0.509568042 | 0.267028 | -1.908297 | 0.056353 | 0.416675859 | ZNF436-AS1 | antisense | 51.57 | -0.193738616 | 0.193738616 | ENSG00000249087 | n.s. |
| ENSG00000099889 | 99.23508 | -0.546898466 | 0.299266 | -1.827468 | 0.067629 | 0.44261708 | ARVCF | protein_coding | 58.03 | -0.193586652 | 0.193586652 | ENSG00000099889 | n.s. |
| ENSG00000164244 | 1324.912 | -0.527594122 | 0.282063 | -1.870486 | 0.061416 | 0.429914933 | PRRC1 | protein_coding | 36.29 | -0.193425222 | 0.193425222 | ENSG00000164244 | n.s. |
| ENSG00000106236 | 371.1903 | -0.492810042 | 0.253905 | -1.94092 | 0.052268 | 0.405192803 | NPTX2 | protein_coding | 54.62 | -0.193348243 | 0.193348243 | ENSG00000106236 | n.s. |
| ENSG00000172869 | 556.9112 | -0.493990612 | 0.254964 | -1.937492 | 0.052685 | 0.406112801 | DMXL1 | protein_coding | 37.58 | -0.193324867 | 0.193324867 | ENSG00000172869 | n.s. |
| ENSG00000118454 | 1199.915 | -0.498243885 | 0.258368 | -1.928428 | 0.053802 | 0.409408975 | ANKRD13C | protein_coding | 37.55 | -0.193240224 | 0.193240224 | ENSG00000118454 | n.s. |
| ENSG00000233264 | 283.2767 | 0.476575633 | 0.242443 | 1.9657222 | 0.049331 | 0.393439254 | | processed_pseudogene | 38.76 | 0.193071423 | 0.193071423 | ENSG00000233264 | n.s. |
| ENSG00000260007 | 146.236 | -0.497783651 | 0.258076 | -1.928823 | 0.053753 | 0.409408975 | | protein_coding | 50.05 | -0.193061726 | 0.193061726 | ENSG00000260007 | n.s. |
| ENSG00000225193 | 30.97618 | 0.615005607 | 0.355298 | 1.7309577 | 0.083459 | 0.485381733 | | transcribed_processed_pseudog | 47.47 | 0.193060452 | 0.193060452 | ENSG00000225193 | n.s. |
| ENSG00000256299 | 40.9402 | -0.569665483 | 0.317319 | -1.795243 | 0.072615 | 0.459013748 | | lincRNA | 43.4 | -0.19264623 | 0.19264623 | ENSG00000256299 | n.s. |
| ENSG00000240770 | 22.2669 | -0.642062223 | 0.379434 | -1.692157 | 0.090616 | 0.501449528 | C21orf91-OT1 | lincRNA | 39.23 | -0.192472773 | 0.192472773 | ENSG00000240770 | n.s. |
| ENSG00000005189 | 89.24578 | -0.507708424 | 0.267384 | -1.898795 | 0.057591 | 0.417869145 | REXO5 | protein_coding | 39.89 | -0.19240103 | 0.19240103 | ENSG00000005189 | n.s. |
| ENSG00000221439 | 43.29636 | -0.603223764 | 0.345036 | -1.748292 | 0.080413 | 0.479859335 | RNU4ATAC16P | snRNA | 43.65 | -0.192359645 | 0.192359645 | ENSG00000221439 | n.s. |
| ENSG00000224553 | 15.6773 | -0.699387462 | 0.429704 | -1.627603 | 0.103609 | 0.531112578 | | processed_pseudogene | 50.86 | -0.192201056 | 0.192201056 | ENSG00000224553 | n.s. |
| ENSG00000135297 | 988.5013 | -0.458877701 | 0.229719 | -1.997558 | 0.045765 | 0.381774093 | MTO1 | protein_coding | 42.61 | -0.191899693 | 0.191899693 | ENSG00000135297 | n.s. |
| ENSG00000276180 | 924.7386 | -0.459955205 | 0.230588 | -1.994708 | 0.046075 | 0.382988464 | HIST1H4I | protein_coding | 46.75 | -0.19171591 | 0.19171591 | ENSG00000276180 | n.s. |
| ENSG00000069011 | 347.8393 | -0.499194268 | 0.260139 | -1.918951 | 0.054991 | 0.413017377 | PITX1 | protein_coding | 59.4 | -0.191706411 | 0.191706411 | ENSG00000069011 | n.s. |
| ENSG00000183475 | 392.6668 | -0.504289951 | 0.26446 | -1.906866 | 0.056538 | 0.416937262 | ASB7 | protein_coding | 40.07 | -0.191594523 | 0.191594523 | ENSG00000183475 | n.s. |
| ENSG00000279665 | 38.35983 | -0.612968394 | 0.354899 | -1.727164 | 0.084138 | 0.48719542 | | TEC | 46.43 | -0.19142807 | 0.19142807 | ENSG00000279665 | n.s. |
| ENSG00000162105 | 119.8127 | -0.491347468 | 0.254269 | -1.932392 | 0.053311 | 0.407947312 | SHANK2 | protein_coding | 51.53 | -0.191328701 | 0.191328701 | ENSG00000162105 | n.s. |
| ENSG00000186868 | 65.10641 | -0.565354164 | 0.314691 | -1.796535 | 0.07241 | 0.458879248 | MAPT | protein_coding | 49.7 | -0.191260208 | 0.191260208 | ENSG00000186868 | n.s. |
| ENSG00000183495 | 1299.594 | -0.472263365 | 0.240408 | -1.964422 | 0.049481 | 0.393585711 | EP400 | protein_coding | 47.47 | -0.191248092 | 0.191248092 | ENSG00000183495 | n.s. |
| ENSG00000224167 | 32.25434 | -0.62200802 | 0.363057 | -1.713253 | 0.086666 | 0.492652156 | | lincRNA | 45.99 | -0.191242345 | 0.191242345 | ENSG00000224167 | n.s. |
| ENSG00000279207 | 213.0389 | -0.503645626 | 0.264337 | -1.905319 | 0.056739 | 0.417346136 | | TEC | 36.67 | -0.19113533 | 0.19113533 | ENSG00000279207 | n.s. |
| ENSG00000279528 | 25.05825 | -0.666454308 | 0.401424 | -1.660225 | 0.096869 | 0.516736278 | | TEC | 42.12 | -0.191093142 | 0.191093142 | ENSG00000279528 | n.s. |

| | | | | | | | | | | | | | |
|---|---|---|---|---|---|---|---|---|---|---|---|---|---|
| ENSG00000004399 | 489.5864 | -0.459333403 | 0.230677 | -1.991241 | 0.046454 | 0.383848684 | PLXND1 | protein_coding | 58.39 | -0.191009176 | 0.191009176 | ENSG00000004399 | n.s. |
| ENSG00000176641 | 307.3819 | -0.493578481 | 0.256209 | -1.926469 | 0.054046 | 0.410330693 | RNF152 | protein_coding | 42.43 | -0.19094873 | 0.19094873 | ENSG00000176641 | n.s. |
| ENSG00000270607 | 97.52601 | 0.536729344 | 0.292106 | 1.8374466 | 0.066144 | 0.440875092 | | lincRNA | 47.57 | 0.190906274 | 0.190906274 | ENSG00000270607 | n.s. |
| ENSG00000273444 | 5.537585 | -0.83703229 | 0.562737 | -1.48743 | 0.136901 | 0.591765387 | | unprocessed_pseudogene | 40.12 | -0.190718176 | 0.190718176 | ENSG00000273444 | n.s. |
| ENSG00000271335 | 120.2904 | -0.495646463 | 0.257917 | -1.921728 | 0.05464 | 0.412398929 | | antisense | 42.38 | -0.190666507 | 0.190666507 | ENSG00000271335 | n.s. |
| ENSG00000206712 | 16.47648 | -0.649295147 | 0.38737 | -1.676161 | 0.093707 | 0.508646797 | RNU6-26P | snRNA | 44.86 | -0.190622463 | 0.190622463 | ENSG00000206712 | n.s. |
| ENSG00000272688 | 35.58033 | 0.592417154 | 0.336633 | 1.7598303 | 0.078437 | 0.477221369 | | lincRNA | 46.94 | 0.190331853 | 0.190331853 | ENSG00000272688 | n.s. |
| ENSG00000080845 | 204.1282 | -0.531646689 | 0.288511 | -1.842723 | 0.065369 | 0.439256007 | DLGAP4 | protein_coding | 51.09 | -0.189947947 | 0.189947947 | ENSG00000080845 | n.s. |
| ENSG00000198863 | 490.8419 | -0.466043523 | 0.236447 | -1.971031 | 0.04872 | 0.391421328 | RUNDC1 | protein_coding | 47.47 | -0.189845399 | 0.189845399 | ENSG00000198863 | n.s. |
| ENSG00000150401 | 720.0526 | -0.490577758 | 0.254656 | -1.926431 | 0.054051 | 0.410330693 | DCUN1D2 | protein_coding | 44.89 | -0.189787853 | 0.189787853 | ENSG00000150401 | n.s. |
| ENSG00000186716 | 39.18006 | -0.584581644 | 0.330921 | -1.76653 | 0.077307 | 0.473643369 | BCR | protein_coding | 53.07 | -0.189725118 | 0.189725118 | ENSG00000186716 | n.s. |
| ENSG00000146833 | 372.6352 | -0.479496721 | 0.246334 | -1.946533 | 0.051591 | 0.402182089 | TRIM4 | protein_coding | 43.25 | -0.189678006 | 0.189678006 | ENSG00000146833 | n.s. |
| ENSG00000131094 | 197.3854 | -0.539805637 | 0.296255 | -1.8221 | 0.06844 | 0.445487047 | C1QL1 | protein_coding | 56.53 | -0.189560803 | 0.189560803 | ENSG00000131094 | n.s. |
| ENSG00000186283 | 321.5788 | -0.512789753 | 0.273242 | -1.876687 | 0.060561 | 0.426932326 | TOR3A | protein_coding | 52.61 | -0.189548097 | 0.189548097 | ENSG00000186283 | n.s. |
| ENSG00000065833 | 217.3464 | -0.512746222 | 0.273795 | -1.872735 | 0.061105 | 0.42849416 | ME1 | protein_coding | 36.91 | -0.188718858 | 0.188718858 | ENSG00000065833 | n.s. |
| ENSG00000265967 | 17.71905 | -0.87067877 | 0.600713 | -1.44941 | 0.147223 | 0.607109641 | | antisense | 42.96 | -0.188704709 | 0.188704709 | ENSG00000265967 | n.s. |
| ENSG00000215883 | 293.006 | -0.509384463 | 0.271206 | -1.878219 | 0.060351 | 0.426183381 | CYB5RL | protein_coding | 48.99 | -0.188677783 | 0.188677783 | ENSG00000215883 | n.s. |
| ENSG00000102760 | 62.74133 | 0.574666927 | 0.324045 | 1.7734163 | 0.07616 | 0.469606803 | RGCC | protein_coding | 45.66 | 0.188643395 | 0.188643395 | ENSG00000102760 | n.s. |
| ENSG00000064651 | 955.5603 | -0.457187335 | 0.230326 | -1.984956 | 0.047149 | 0.387063744 | SLC12A2 | protein_coding | 35.24 | -0.188460623 | 0.188460623 | ENSG00000064651 | n.s. |
| ENSG00000165355 | 215.1257 | -0.51441738 | 0.276832 | -1.858231 | 0.063136 | 0.431139465 | FBXO33 | protein_coding | 38.98 | -0.187958965 | 0.187958965 | ENSG00000165355 | n.s. |
| ENSG00000148296 | 801.338 | -0.47832994 | 0.2463 | -1.942059 | 0.05213 | 0.404745128 | SURF6 | protein_coding | 53.39 | -0.187896787 | 0.187896787 | ENSG00000148296 | n.s. |
| ENSG00000169116 | 752.7341 | -0.499835252 | 0.264105 | -1.892565 | 0.058416 | 0.42095504 | PARM1 | protein_coding | 40.47 | -0.187820237 | 0.187820237 | ENSG00000169116 | n.s. |
| ENSG00000112787 | 132.5218 | -0.481558375 | 0.249238 | -1.932124 | 0.053344 | 0.407947312 | FBRSL1 | protein_coding | 62.75 | -0.187516869 | 0.187516869 | ENSG00000112787 | n.s. |
| ENSG00000271380 | 93.70478 | -0.48871065 | 0.255089 | -1.915846 | 0.055385 | 0.413559567 | | antisense | 64.83 | -0.187401928 | 0.187401928 | ENSG00000271380 | n.s. |
| ENSG00000197121 | 3784.589 | -0.488689957 | 0.255341 | -1.913873 | 0.055636 | 0.413574865 | PGAP1 | protein_coding | 36 | -0.187386143 | 0.187386143 | ENSG00000197121 | n.s. |
| ENSG00000187951 | 966.3989 | -0.468382672 | 0.239509 | -1.955592 | 0.050513 | 0.398128225 | ARHGAP11B | protein_coding | 40.92 | -0.187342311 | 0.187342311 | ENSG00000187951 | n.s. |
| ENSG00000144677 | 94.70069 | -0.511715561 | 0.273988 | -1.86766 | 0.06181 | 0.430475522 | CTDSPL | protein_coding | 42.29 | -0.187314269 | 0.187314269 | ENSG00000144677 | n.s. |
| ENSG00000063015 | 320.1341 | -0.477559504 | 0.246288 | -1.93903 | 0.052498 | 0.405702385 | SEZ6 | protein_coding | 54.21 | -0.187104203 | 0.187104203 | ENSG00000063015 | n.s. |
| ENSG00000115107 | 195.8313 | -0.546913652 | 0.302763 | -1.806408 | 0.070855 | 0.455117569 | STEAP3 | protein_coding | 51.67 | -0.18697687 | 0.18697687 | ENSG00000115107 | n.s. |
| ENSG00000049130 | 462.5943 | -0.484927365 | 0.252016 | -1.92419 | 0.054331 | 0.411560102 | KITLG | protein_coding | 36.84 | -0.18697186 | 0.18697186 | ENSG00000049130 | n.s. |
| ENSG00000139220 | 134.9243 | -0.493138626 | 0.258997 | -1.904034 | 0.056906 | 0.417869145 | PPFIA2 | protein_coding | 34.5 | -0.186879663 | 0.186879663 | ENSG00000139220 | n.s. |
| ENSG00000271936 | 60.54148 | -0.546688238 | 0.302833 | -1.805249 | 0.071036 | 0.455354508 | | antisense | 53.25 | -0.186776233 | 0.186776233 | ENSG00000271936 | n.s. |

| | | | | | | | | | | | | | |
|---|---|---|---|---|---|---|---|---|---|---|---|---|---|
| ENSG00000201376 | 35.42814 | 0.568961341 | 0.320662 | 1.7743327 | 0.076008 | 0.469606803 | | snoRNA | 45.93 | 0.186770447 | 0.186770447 | ENSG00000201376 | n.s. |
| ENSG00000151612 | 247.2525 | -0.483048507 | 0.250875 | -1.925458 | 0.054172 | 0.411028382 | ZNF827 | protein_coding | 41.53 | -0.186518645 | 0.186518645 | ENSG00000151612 | n.s. |
| ENSG00000197312 | 471.1742 | -0.46913533 | 0.240619 | -1.949699 | 0.051212 | 0.400350899 | DDI2 | protein_coding | 42.4 | -0.186509063 | 0.186509063 | ENSG00000197312 | n.s. |
| ENSG00000174516 | 120.1634 | -0.523093767 | 0.284978 | -1.835557 | 0.066423 | 0.441084074 | PELI3 | protein_coding | 56.57 | -0.185948652 | 0.185948652 | ENSG00000174516 | n.s. |
| ENSG00000252338 | 10.93023 | 0.717926284 | 0.452199 | 1.5876319 | 0.11237 | 0.55089864 | RNU6-503P | snRNA | 37.14 | 0.185891432 | 0.185891432 | ENSG00000252338 | n.s. |
| ENSG00000231587 | 10.21537 | -0.765888919 | 0.496753 | -1.541789 | 0.123125 | 0.572027647 | SNORD62B | snoRNA | 46.51 | -0.185791617 | 0.185791617 | ENSG00000231587 | n.s. |
| ENSG00000236780 | 6973.332 | -0.523394017 | 0.286158 | -1.829038 | 0.067394 | 0.442188857 | LINC01829 | lincRNA | 36.25 | -0.18548676 | 0.18548676 | ENSG00000236780 | n.s. |
| ENSG00000256304 | 171.6967 | -0.522506553 | 0.285141 | -1.83245 | 0.066884 | 0.441705004 | CCDC150P1 | processed_pseudogene | 44.83 | -0.185420689 | 0.185420689 | ENSG00000256304 | n.s. |
| ENSG00000229659 | 18.94073 | -0.855351289 | 0.589795 | -1.450251 | 0.146989 | 0.607109641 | RPL26P6 | processed_pseudogene | 42.01 | -0.18538274 | 0.18538274 | ENSG00000229659 | n.s. |
| ENSG00000075336 | 730.6413 | -0.47240419 | 0.243435 | -1.940573 | 0.05231 | 0.405289476 | TIMM21 | protein_coding | 42.31 | -0.185293303 | 0.185293303 | ENSG00000075336 | n.s. |
| ENSG00000208036 | 12.36349 | 0.709949826 | 0.445968 | 1.5919295 | 0.111401 | 0.548299349 | MIR106B | miRNA | 60.98 | 0.185284317 | 0.185284317 | ENSG00000208036 | n.s. |
| ENSG00000138622 | 451.8883 | -0.456627466 | 0.232028 | -1.967987 | 0.04907 | 0.392866719 | HCN4 | protein_coding | 55.08 | -0.185278768 | 0.185278768 | ENSG00000138622 | n.s. |
| ENSG00000237729 | 11.75957 | 0.761738686 | 0.49373 | 1.5428258 | 0.122873 | 0.571396639 | | processed_pseudogene | 47.62 | 0.185149971 | 0.185149971 | ENSG00000237729 | n.s. |
| ENSG00000264070 | 32.51063 | -0.620814095 | 0.367634 | -1.688677 | 0.091281 | 0.503331271 | | processed_pseudogene | 65.34 | -0.185093293 | 0.185093293 | ENSG00000264070 | n.s. |
| ENSG00000131504 | 873.3691 | -0.453352257 | 0.229885 | -1.972087 | 0.0486 | 0.391129217 | DIAPH1 | protein_coding | 41.24 | -0.184822531 | 0.184822531 | ENSG00000131504 | n.s. |
| ENSG00000096060 | 1879.245 | -0.459827112 | 0.234728 | -1.958975 | 0.050116 | 0.396455218 | FKBP5 | protein_coding | 41.97 | -0.184761229 | 0.184761229 | ENSG00000096060 | n.s. |
| ENSG00000272168 | 1003.341 | 0.422089433 | 0.207724 | 2.0319717 | 0.042157 | 0.365242806 | CASC15 | lincRNA | 40.31 | 0.184629655 | 0.184629655 | ENSG00000272168 | n.s. |
| ENSG00000069482 | 1037.859 | -0.551285148 | 0.3084 | -1.787565 | 0.073846 | 0.462574432 | GAL | protein_coding | 57.14 | -0.184580398 | 0.184580398 | ENSG00000069482 | n.s. |
| ENSG00000221214 | 12.95027 | -0.690385907 | 0.42872 | -1.610341 | 0.107323 | 0.540616659 | MIR548E | miRNA | 35.23 | -0.184409377 | 0.184409377 | ENSG00000221214 | n.s. |
| ENSG00000116095 | 1003.46 | -0.500907002 | 0.267477 | -1.872712 | 0.061108 | 0.42849416 | PLEKHA3 | protein_coding | 37.26 | -0.184361373 | 0.184361373 | ENSG00000116095 | n.s. |
| ENSG00000157927 | 283.8183 | -0.491640145 | 0.260158 | -1.889772 | 0.058788 | 0.421913319 | RADIL | protein_coding | 53.13 | -0.184255303 | 0.184255303 | ENSG00000157927 | n.s. |
| ENSG00000279069 | 308.788 | -0.441469006 | 0.22119 | -1.995879 | 0.045947 | 0.382837889 | | TEC | 57.27 | -0.184085992 | 0.184085992 | ENSG00000279069 | n.s. |
| ENSG00000250220 | 61.48281 | -0.550737933 | 0.308429 | -1.785624 | 0.07416 | 0.463232209 | | antisense | 39.01 | -0.184057307 | 0.184057307 | ENSG00000250220 | n.s. |
| ENSG00000272662 | 44.36147 | 0.572805348 | 0.325586 | 1.7593048 | 0.078526 | 0.477221369 | | lincRNA | 42.86 | 0.18403097 | 0.18403097 | ENSG00000272662 | n.s. |
| ENSG00000109171 | 289.9933 | -0.479874249 | 0.25043 | -1.916198 | 0.05534 | 0.413559567 | SLAIN2 | protein_coding | 37.94 | -0.184013505 | 0.184013505 | ENSG00000109171 | n.s. |
| ENSG00000111647 | 1463.33 | -0.495454675 | 0.263399 | -1.881002 | 0.059972 | 0.425223094 | UHRF1BP1L | protein_coding | 38.58 | -0.184003521 | 0.184003521 | ENSG00000111647 | n.s. |
| ENSG00000071655 | 800.5978 | -0.468883647 | 0.24168 | -1.940099 | 0.052368 | 0.405289476 | MBD3 | protein_coding | 58.04 | -0.183912424 | 0.183912424 | ENSG00000071655 | n.s. |
| ENSG00000200913 | 47.77338 | -0.558883637 | 0.314724 | -1.775789 | 0.075768 | 0.468789069 | SNORD46 | snoRNA | 48.08 | -0.183885305 | 0.183885305 | ENSG00000200913 | n.s. |
| ENSG00000196597 | 422.011 | -0.478865466 | 0.249955 | -1.915809 | 0.055389 | 0.413559567 | ZNF782 | protein_coding | 38.77 | -0.183626675 | 0.183626675 | ENSG00000196597 | n.s. |
| ENSG00000269343 | 820.3314 | -0.498434147 | 0.266187 | -1.872496 | 0.061138 | 0.42849416 | ZNF587B | protein_coding | 47.01 | -0.183451226 | 0.183451226 | ENSG00000269343 | n.s. |
| ENSG00000182118 | 430.612 | -0.583804309 | 0.337072 | -1.731986 | 0.083276 | 0.485101957 | FAM89A | protein_coding | 47.02 | -0.183412033 | 0.183412033 | ENSG00000182118 | n.s. |
| ENSG00000164663 | 519.3595 | -0.459275317 | 0.235285 | -1.951995 | 0.050939 | 0.399336224 | USP49 | protein_coding | 41.51 | -0.183095292 | 0.183095292 | ENSG00000164663 | n.s. |

| | | | | | | | | | | | | | |
|---|---|---|---|---|---|---|---|---|---|---|---|---|---|
| ENSG00000274421 | 88.17296 | -0.499778142 | 0.268106 | -1.864109 | 0.062306 | 0.430475522 | | sense_intronic | 39.25 | -0.182944558 | 0.182944558 | ENSG00000274421 | n.s. |
| ENSG00000172817 | 249.7657 | -0.488724724 | 0.258798 | -1.888442 | 0.058967 | 0.422755861 | CYP7B1 | protein_coding | 37.55 | -0.182739239 | 0.182739239 | ENSG00000172817 | n.s. |
| ENSG00000165118 | 374.5229 | -0.508439846 | 0.275427 | -1.846009 | 0.064891 | 0.437274745 | C9orf64 | protein_coding | 44.72 | -0.18265478 | 0.18265478 | ENSG00000165118 | n.s. |
| ENSG00000255409 | 45.58218 | -0.569865853 | 0.324697 | -1.755071 | 0.079247 | 0.478143199 | RSF1-IT1 | sense_intronic | 43.19 | -0.182608964 | 0.182608964 | ENSG00000255409 | n.s. |
| ENSG00000282386 | 30.80579 | -0.641749044 | 0.388193 | -1.65317 | 0.098296 | 0.519362534 | | antisense | 45.63 | -0.18259646 | 0.18259646 | ENSG00000282386 | n.s. |
| ENSG00000199005 | 2.686674 | 1.233926097 | 1.049479 | 1.1757511 | 0.239694 | 0.711266989 | MIR370 | miRNA | 57.33 | 0.182580771 | 0.182580771 | ENSG00000199005 | n.s. |
| ENSG00000226555 | 60.62235 | -0.514712656 | 0.281043 | -1.83144 | 0.067035 | 0.441892852 | AGKP1 | processed_pseudogene | 44.81 | -0.182559841 | 0.182559841 | ENSG00000226555 | n.s. |
| ENSG00000180957 | 1095.799 | -0.47319411 | 0.246363 | -1.920717 | 0.054767 | 0.412398929 | PITPNB | protein_coding | 39.71 | -0.18202948 | 0.18202948 | ENSG00000180957 | n.s. |
| ENSG00000259287 | 35.55545 | -0.567535774 | 0.32316 | -1.756209 | 0.079053 | 0.477920232 | | antisense | 43.53 | -0.181977272 | 0.181977272 | ENSG00000259287 | n.s. |
| ENSG00000135778 | 1366.971 | -0.511274284 | 0.2782 | -1.837797 | 0.066092 | 0.440875092 | NTPCR | protein_coding | 42.72 | -0.181852306 | 0.181852306 | ENSG00000135778 | n.s. |
| ENSG00000189227 | 655.9799 | 0.448390916 | 0.228004 | 1.9665964 | 0.04923 | 0.393340111 | C15orf61 | protein_coding | 39.21 | 0.181702242 | 0.181702242 | ENSG00000189227 | n.s. |
| ENSG00000267793 | 38.83027 | -0.579398505 | 0.334955 | -1.729778 | 0.08367 | 0.485981036 | | transcribed_processed_pseudog | 35.26 | -0.181572296 | 0.181572296 | ENSG00000267793 | n.s. |
| ENSG00000147536 | 632.7045 | -0.466264477 | 0.241335 | -1.93202 | 0.053357 | 0.407947312 | GINS4 | protein_coding | 46.76 | -0.181561487 | 0.181561487 | ENSG00000147536 | n.s. |
| ENSG00000219928 | 60.80709 | 0.604647517 | 0.357187 | 1.6928059 | 0.090492 | 0.501165391 | | processed_pseudogene | 52.3 | 0.1814057 | 0.1814057 | ENSG00000219928 | n.s. |
| ENSG00000151116 | 180.6522 | -0.542605289 | 0.304048 | -1.784605 | 0.074325 | 0.463348535 | UEVLD | protein_coding | 40.83 | -0.181280198 | 0.181280198 | ENSG00000151116 | n.s. |
| ENSG00000084112 | 233.4322 | -0.463582484 | 0.239456 | -1.935978 | 0.05287 | 0.406621815 | SSH1 | protein_coding | 48.43 | -0.181172358 | 0.181172358 | ENSG00000084112 | n.s. |
| ENSG00000263790 | 7.186673 | 0.771538056 | 0.511789 | 1.5075327 | 0.131674 | 0.582369535 | MIR4473 | miRNA | 47.25 | 0.181158179 | 0.181158179 | ENSG00000263790 | n.s. |
| ENSG00000175073 | 831.142 | -0.470736518 | 0.244916 | -1.92203 | 0.054602 | 0.412398929 | VCPIP1 | protein_coding | 39.75 | -0.181084087 | 0.181084087 | ENSG00000175073 | n.s. |
| ENSG00000248115 | 114.3111 | 0.509291522 | 0.27798 | 1.8321163 | 0.066934 | 0.441705004 | | antisense | 42.94 | 0.180731101 | 0.180731101 | ENSG00000248115 | n.s. |
| ENSG00000178665 | 271.1651 | -0.514550009 | 0.282395 | -1.822096 | 0.06844 | 0.445487047 | ZNF713 | protein_coding | 43 | -0.180691912 | 0.180691912 | ENSG00000178665 | n.s. |
| ENSG00000263857 | 5.182115 | -0.934388493 | 0.684573 | -1.364921 | 0.172278 | 0.640837445 | MIR4729 | miRNA | 43.06 | -0.180572557 | 0.180572557 | ENSG00000263857 | n.s. |
| ENSG00000279766 | 40.22858 | -0.577553559 | 0.334315 | -1.727573 | 0.084065 | 0.48719542 | | TEC | 44.86 | -0.18036813 | 0.18036813 | ENSG00000279766 | n.s. |
| ENSG00000161888 | 723.9126 | -0.475431701 | 0.249984 | -1.90185 | 0.057191 | 0.417869145 | SPC24 | protein_coding | 53.53 | -0.180169452 | 0.180169452 | ENSG00000161888 | n.s. |
| ENSG00000119938 | 202.4875 | 0.469683786 | 0.244916 | 1.9177318 | 0.055145 | 0.413454097 | PPP1R3C | protein_coding | 40.41 | 0.180157878 | 0.180157878 | ENSG00000119938 | n.s. |
| ENSG00000102317 | 4211.583 | -0.475284803 | 0.250132 | -1.900132 | 0.057416 | 0.417869145 | RBM3 | protein_coding | 48.05 | -0.180113784 | 0.180113784 | ENSG00000102317 | n.s. |
| ENSG00000196312 | 387.762 | -0.475081339 | 0.250277 | -1.89822 | 0.057667 | 0.417869145 | MFSD14C | protein_coding | 44.07 | -0.18003668 | 0.18003668 | ENSG00000196312 | n.s. |
| ENSG00000130479 | 243.1603 | -0.477541823 | 0.251943 | -1.895435 | 0.058035 | 0.419811599 | MAP1S | protein_coding | 57.53 | -0.180007273 | 0.180007273 | ENSG00000130479 | n.s. |
| ENSG00000281195 | 389.8002 | -0.474654775 | 0.249949 | -1.899007 | 0.057563 | 0.417869145 | | processed_transcript | 36.8 | -0.179875029 | 0.179875029 | ENSG00000281195 | n.s. |
| ENSG00000180596 | 1151.14 | -0.400482629 | 0.195051 | -2.053222 | 0.040051 | 0.356769879 | HIST1H2BC | protein_coding | 37.87 | -0.179260758 | 0.179260758 | ENSG00000180596 | n.s. |
| ENSG00000270022 | 465.1173 | 0.427714794 | 0.214117 | 1.997571 | 0.045763 | 0.381774093 | | lincRNA | 49.25 | 0.178867566 | 0.178867566 | ENSG00000270022 | n.s. |
| ENSG00000163873 | 488.1455 | -0.476624217 | 0.252063 | -1.890896 | 0.058638 | 0.421486462 | GRIK3 | protein_coding | 49.57 | -0.178837209 | 0.178837209 | ENSG00000163873 | n.s. |
| ENSG00000092445 | 449.925 | -0.488472601 | 0.262274 | -1.862452 | 0.062539 | 0.430475522 | TYRO3 | protein_coding | 53.19 | -0.178806147 | 0.178806147 | ENSG00000092445 | n.s. |

| | | | | | | | | | | | | | |
|---|---|---|---|---|---|---|---|---|---|---|---|---|---|
| ENSG00000181396 | 547.9095 | -0.497615161 | 0.269613 | -1.845666 | 0.064941 | 0.437274745 | OGFOD3 | protein_coding | 54.32 | -0.17876606 | 0.17876606 | ENSG00000181396 | n.s. |
| ENSG00000262663 | 29.38054 | -0.582791448 | 0.340951 | -1.70931 | 0.087394 | 0.493622914 | | lincRNA | 52.98 | -0.17868659 | 0.17868659 | ENSG00000262663 | n.s. |
| ENSG00000149658 | 795.7275 | -0.48762107 | 0.2616 | -1.863991 | 0.062323 | 0.430475522 | YTHDF1 | protein_coding | 49.38 | -0.178494443 | 0.178494443 | ENSG00000149658 | n.s. |
| ENSG00000226085 | 43.93924 | 0.558791627 | 0.319128 | 1.7509936 | 0.079947 | 0.479305998 | UQCRFS1P1 | processed_pseudogene | 53.9 | 0.178470858 | 0.178470858 | ENSG00000226085 | n.s. |
| ENSG00000213904 | 20.86878 | -0.672418505 | 0.419027 | -1.604715 | 0.108557 | 0.543082569 | LIPE-AS1 | antisense | 46.03 | -0.1782811 | 0.1782811 | ENSG00000213904 | n.s. |
| ENSG00000073417 | 92.96603 | -0.502942898 | 0.274842 | -1.829936 | 0.06726 | 0.442113119 | PDE8A | protein_coding | 41.8 | -0.178276458 | 0.178276458 | ENSG00000073417 | n.s. |
| ENSG00000138829 | 743.024 | -0.477228505 | 0.253111 | -1.885451 | 0.059369 | 0.423990171 | FBN2 | protein_coding | 37.12 | -0.177836439 | 0.177836439 | ENSG00000138829 | n.s. |
| ENSG00000273270 | 279.2747 | -0.427212797 | 0.214479 | -1.991864 | 0.046386 | 0.38351048 | | lincRNA | 51.96 | -0.177815691 | 0.177815691 | ENSG00000273270 | n.s. |
| ENSG00000201164 | 5.102815 | -1.16620864 | 0.971724 | -1.200144 | 0.230084 | 0.703952151 | RNU4-36P | snRNA | 38.19 | -0.177796507 | 0.177796507 | ENSG00000201164 | n.s. |
| ENSG00000156475 | 231.1732 | -0.499870422 | 0.272041 | -1.837485 | 0.066138 | 0.440875092 | PPP2R2B | protein_coding | 39.61 | -0.177796129 | 0.177796129 | ENSG00000156475 | n.s. |
| ENSG00000130635 | 498.4548 | -0.457983511 | 0.237322 | -1.929797 | 0.053632 | 0.409160547 | COL5A1 | protein_coding | 56 | -0.177746263 | 0.177746263 | ENSG00000130635 | n.s. |
| ENSG00000272829 | 43.72846 | 0.58212659 | 0.341411 | 1.7050587 | 0.088183 | 0.495072019 | | lincRNA | 52.91 | 0.177741654 | 0.177741654 | ENSG00000272829 | n.s. |
| ENSG00000051341 | 1739.757 | -0.467088333 | 0.245099 | -1.905714 | 0.056687 | 0.417346136 | POLQ | protein_coding | 39.06 | -0.177261705 | 0.177261705 | ENSG00000051341 | n.s. |
| ENSG00000177595 | 37.17561 | -0.56828107 | 0.329675 | -1.723759 | 0.084751 | 0.48779901 | PIDD1 | protein_coding | 62.59 | -0.177166787 | 0.177166787 | ENSG00000177595 | n.s. |
| ENSG00000200089 | 4.521759 | 1.079810998 | 0.861197 | 1.2538485 | 0.209897 | 0.685525959 | SNORD114-31 | snoRNA | 50.67 | 0.17706319 | 0.17706319 | ENSG00000200089 | n.s. |
| ENSG00000006831 | 1851.246 | -0.474507164 | 0.251821 | -1.884302 | 0.059524 | 0.423999253 | ADIPOR2 | protein_coding | 39.4 | -0.176817934 | 0.176817934 | ENSG00000006831 | n.s. |
| ENSG00000107816 | 899.5196 | -0.470543605 | 0.248712 | -1.891924 | 0.058501 | 0.42095504 | LZTS2 | protein_coding | 60.49 | -0.176813482 | 0.176813482 | ENSG00000107816 | n.s. |
| ENSG00000157570 | 669.9952 | -0.466203969 | 0.245103 | -1.902073 | 0.057162 | 0.417869145 | TSPAN18 | protein_coding | 50.12 | -0.176672514 | 0.176672514 | ENSG00000157570 | n.s. |
| ENSG00000236778 | 199.6441 | 0.449701695 | 0.231887 | 1.9393148 | 0.052463 | 0.405702385 | INTS6-AS1 | antisense | 40.22 | 0.176189724 | 0.176189724 | ENSG00000236778 | n.s. |
| ENSG00000255121 | 96.4927 | -0.495351113 | 0.269716 | -1.836564 | 0.066274 | 0.440875092 | | lincRNA | 51.59 | -0.176188681 | 0.176188681 | ENSG00000255121 | n.s. |
| ENSG00000197122 | 298.9307 | -0.459135469 | 0.239881 | -1.914013 | 0.055619 | 0.413574865 | SRC | protein_coding | 54.1 | -0.176053596 | 0.176053596 | ENSG00000197122 | n.s. |
| ENSG00000160716 | 876.13 | -0.445836147 | 0.229164 | -1.945487 | 0.051716 | 0.402936503 | CHRNB2 | protein_coding | 54.69 | -0.175999788 | 0.175999788 | ENSG00000160716 | n.s. |
| ENSG00000265972 | 3713.492 | 0.464249167 | 0.24434 | 1.9000163 | 0.057431 | 0.417869145 | TXNIP | protein_coding | 42.49 | 0.175931723 | 0.175931723 | ENSG00000265972 | n.s. |
| ENSG00000243702 | 21.19259 | 0.645855123 | 0.398405 | 1.6211029 | 0.104996 | 0.534498535 | RN7SL638P | misc_RNA | 51.72 | 0.175707135 | 0.175707135 | ENSG00000243702 | n.s. |
| ENSG00000117971 | 598.7489 | -0.47529413 | 0.25333 | -1.876188 | 0.060629 | 0.42719796 | CHRNB4 | protein_coding | 46.53 | -0.175559787 | 0.175559787 | ENSG00000117971 | n.s. |
| ENSG00000119866 | 1102.246 | -0.438806799 | 0.224425 | -1.95525 | 0.050554 | 0.398128225 | BCL11A | protein_coding | 44.56 | -0.175512641 | 0.175512641 | ENSG00000119866 | n.s. |
| ENSG00000006607 | 659.8754 | -0.465517178 | 0.245637 | -1.895143 | 0.058073 | 0.419811599 | FARP2 | protein_coding | 45.6 | -0.175474636 | 0.175474636 | ENSG00000006607 | n.s. |
| ENSG00000196367 | 1774.814 | -0.439876562 | 0.225247 | -1.952859 | 0.050836 | 0.399206745 | TRRAP | protein_coding | 46.21 | -0.17542371 | 0.17542371 | ENSG00000196367 | n.s. |
| ENSG00000206149 | 514.3512 | -0.457632994 | 0.239292 | -1.912446 | 0.055819 | 0.413827257 | HERC2P9 | cribed_unprocessed_pseudc | 42.75 | -0.175356226 | 0.175356226 | ENSG00000206149 | n.s. |
| ENSG00000275227 | 50.48434 | -0.548987185 | 0.313305 | -1.752244 | 0.079732 | 0.479305998 | | misc_RNA | 44.2 | -0.175339445 | 0.175339445 | ENSG00000275227 | n.s. |
| ENSG00000052802 | 1476.747 | -0.479503368 | 0.258139 | -1.857539 | 0.063235 | 0.431364185 | MSMO1 | protein_coding | 37.1 | -0.175093491 | 0.175093491 | ENSG00000052802 | n.s. |
| ENSG00000180626 | 956.7204 | -0.488676288 | 0.264994 | -1.844103 | 0.065168 | 0.438326836 | ZNF594 | protein_coding | 43.95 | -0.175044794 | 0.175044794 | ENSG00000180626 | n.s. |

| | | | | | | | | | | | | | |
|---|---|---|---|---|---|---|---|---|---|---|---|---|---|
| ENSG00000172667 | 617.5334 | -0.465447727 | 0.245808 | -1.893542 | 0.058286 | 0.420691467 | ZMAT3 | protein_coding | 40.73 | -0.17502524 | 0.17502524 | ENSG00000172667 | n.s. |
| ENSG00000261242 | 46.99437 | 0.556784779 | 0.321504 | 1.7318141 | 0.083307 | 0.485101957 | | antisense | 47.72 | 0.174923389 | 0.174923389 | ENSG00000261242 | n.s. |
| ENSG00000161847 | 296.8334 | -0.444985722 | 0.229098 | -1.942336 | 0.052096 | 0.404745128 | RAVER1 | protein_coding | 56.86 | -0.174798566 | 0.174798566 | ENSG00000161847 | n.s. |
| ENSG00000231010 | 71.70142 | 0.551220646 | 0.3163 | 1.7427169 | 0.081383 | 0.48187821 | | lincRNA | 46.96 | 0.174771513 | 0.174771513 | ENSG00000231010 | n.s. |
| ENSG00000224738 | 67.81315 | 0.538497899 | 0.304852 | 1.7664244 | 0.077325 | 0.473643369 | | antisense | 42.19 | 0.174768706 | 0.174768706 | ENSG00000224738 | n.s. |
| ENSG00000273165 | 27.85164 | -0.586002832 | 0.34699 | -1.688816 | 0.091255 | 0.503331271 | | lincRNA | 34.15 | -0.174714451 | 0.174714451 | ENSG00000273165 | n.s. |
| ENSG00000201184 | 6.77747 | -0.856213926 | 0.60762 | -1.409128 | 0.158797 | 0.625264564 | RNU4-68P | snRNA | 40.43 | -0.174613001 | 0.174613001 | ENSG00000201184 | n.s. |
| ENSG00000235445 | 16.10616 | -0.719733515 | 0.466826 | -1.54176 | 0.123132 | 0.572027647 | | sense_intronic | 53.98 | -0.174595101 | 0.174595101 | ENSG00000235445 | n.s. |
| ENSG00000204335 | 69.78438 | 0.555333676 | 0.320327 | 1.7336463 | 0.082981 | 0.485101957 | SP5 | protein_coding | 66.42 | 0.1744675 | 0.1744675 | ENSG00000204335 | n.s. |
| ENSG00000111788 | 301.128 | -0.492667073 | 0.2695 | -1.828081 | 0.067537 | 0.44250464 | | unprocessed_pseudogene | 48.72 | -0.174444627 | 0.174444627 | ENSG00000111788 | n.s. |
| ENSG00000104228 | 370.2892 | -0.466508773 | 0.247075 | -1.888123 | 0.059009 | 0.422845397 | TRIM35 | protein_coding | 45.69 | -0.17438956 | 0.17438956 | ENSG00000104228 | n.s. |
| ENSG00000269951 | 763.1722 | 0.426834865 | 0.216133 | 1.9748725 | 0.048283 | 0.390382659 | | sense_intronic | 37.65 | 0.17436609 | 0.17436609 | ENSG00000269951 | n.s. |
| ENSG00000090470 | 680.3877 | -0.476052316 | 0.255144 | -1.865819 | 0.062067 | 0.430475522 | PDCD7 | protein_coding | 45.72 | -0.174259683 | 0.174259683 | ENSG00000090470 | n.s. |
| ENSG00000258365 | 52.63595 | -0.543386513 | 0.309734 | -1.754363 | 0.079368 | 0.478143199 | | antisense | 45.53 | -0.17412387 | 0.17412387 | ENSG00000258365 | n.s. |
| ENSG00000199962 | 4.187937 | -1.134175977 | 0.942913 | -1.202843 | 0.229037 | 0.702737638 | | misc_RNA | 39.45 | -0.173763455 | 0.173763455 | ENSG00000199962 | n.s. |
| ENSG00000197020 | 976.559 | -0.460977495 | 0.243236 | -1.895185 | 0.058068 | 0.419811599 | ZNF100 | protein_coding | 38.56 | -0.173763423 | 0.173763423 | ENSG00000197020 | n.s. |
| ENSG00000273238 | 397.4601 | -0.474688461 | 0.254613 | -1.864351 | 0.062272 | 0.430475522 | | lincRNA | 33.02 | -0.173760442 | 0.173760442 | ENSG00000273238 | n.s. |
| ENSG00000164506 | 514.4498 | -0.48355366 | 0.261878 | -1.846484 | 0.064822 | 0.437274745 | STXBP5 | protein_coding | 34.37 | -0.173714527 | 0.173714527 | ENSG00000164506 | n.s. |
| ENSG00000196453 | 261.9148 | -0.44428181 | 0.229537 | -1.93556 | 0.052922 | 0.406621815 | ZNF777 | protein_coding | 47.33 | -0.173629474 | 0.173629474 | ENSG00000196453 | n.s. |
| ENSG00000176953 | 618.7497 | -0.48839267 | 0.266099 | -1.835377 | 0.06645 | 0.441084074 | NFATC2IP | protein_coding | 49.47 | -0.173613154 | 0.173613154 | ENSG00000176953 | n.s. |
| ENSG00000052795 | 1001.254 | -0.443576769 | 0.228965 | -1.93731 | 0.052707 | 0.406112801 | FNIP2 | protein_coding | 40.56 | -0.173595242 | 0.173595242 | ENSG00000052795 | n.s. |
| ENSG00000253513 | 14.75131 | -0.855653566 | 0.608385 | -1.406434 | 0.159595 | 0.626988507 | | antisense | 49.42 | -0.173475563 | 0.173475563 | ENSG00000253513 | n.s. |
| ENSG00000141098 | 458.0994 | -0.473877217 | 0.254094 | -1.864968 | 0.062186 | 0.430475522 | GFOD2 | protein_coding | 46.49 | -0.173463484 | 0.173463484 | ENSG00000141098 | n.s. |
| ENSG00000104369 | 254.4214 | -0.475538535 | 0.256112 | -1.856761 | 0.063345 | 0.431907163 | JPH1 | protein_coding | 39.71 | -0.173385912 | 0.173385912 | ENSG00000104369 | n.s. |
| ENSG00000135185 | 601.2506 | -0.45985191 | 0.242626 | -1.895309 | 0.058051 | 0.419811599 | TMEM243 | protein_coding | 43.9 | -0.173339139 | 0.173339139 | ENSG00000135185 | n.s. |
| ENSG00000264566 | 4.096127 | -0.956506086 | 0.723528 | -1.322004 | 0.186167 | 0.658994794 | MIR23C | miRNA | 45 | -0.173240485 | 0.173240485 | ENSG00000264566 | n.s. |
| ENSG00000202189 | 6.115068 | -0.812947205 | 0.564895 | -1.439111 | 0.150119 | 0.612216404 | SNORA30B | snoRNA | 53.49 | -0.173235015 | 0.173235015 | ENSG00000202189 | n.s. |
| ENSG00000120693 | 3816.437 | -0.487185336 | 0.26538 | -1.835803 | 0.066387 | 0.441084074 | SMAD9 | protein_coding | 41.07 | -0.173183972 | 0.173183972 | ENSG00000120693 | n.s. |
| ENSG00000093000 | 2379.556 | -0.456740038 | 0.240134 | -1.902018 | 0.057169 | 0.417869145 | NUP50 | protein_coding | 43.62 | -0.173086066 | 0.173086066 | ENSG00000093000 | n.s. |
| ENSG00000256341 | 37.78047 | -0.566152506 | 0.331935 | -1.705612 | 0.08808 | 0.494891637 | | antisense | 49.31 | -0.172953864 | 0.172953864 | ENSG00000256341 | n.s. |
| ENSG00000233718 | 1544.368 | -0.388806129 | 0.190034 | -2.045983 | 0.040758 | 0.35907784 | MYCNOS | protein_coding | 59.15 | -0.172945397 | 0.172945397 | ENSG00000233718 | n.s. |
| ENSG00000103111 | 345.6301 | -0.509448982 | 0.283111 | -1.799464 | 0.071945 | 0.45772936 | MON1B | protein_coding | 49.31 | -0.172902516 | 0.172902516 | ENSG00000103111 | n.s. |

| | | | | | | | | | | | | | |
|---|---|---|---|---|---|---|---|---|---|---|---|---|---|
| ENSG00000197136 | 358.8154 | -0.433645288 | 0.222125 | -1.952262 | 0.050907 | 0.399312737 | PCNX3 | protein_coding | 60.5 | -0.172888668 | 0.172888668 | ENSG00000197136 | n.s. |
| ENSG00000230551 | 1143.937 | -0.456163678 | 0.239901 | -1.901464 | 0.057241 | 0.417869145 | | processed_transcript | 37.36 | -0.172867648 | 0.172867648 | ENSG00000230551 | n.s. |
| ENSG00000254884 | 71.03639 | 0.531714729 | 0.300875 | 1.7672271 | 0.07719 | 0.473444607 | PRR13P2 | processed_pseudogene | 56.65 | 0.172664163 | 0.172664163 | ENSG00000254884 | n.s. |
| ENSG00000252255 | 12.44898 | -0.701444852 | 0.451561 | -1.553378 | 0.120333 | 0.567746164 | RNU2-35P | snRNA | 35.21 | -0.172447265 | 0.172447265 | ENSG00000252255 | n.s. |
| ENSG00000185716 | 272.8967 | -0.539776211 | 0.308443 | -1.750005 | 0.080117 | 0.479305998 | C16orf52 | protein_coding | 39.3 | -0.172397578 | 0.172397578 | ENSG00000185716 | n.s. |
| ENSG00000183814 | 990.5543 | -0.504263618 | 0.279497 | -1.804185 | 0.071202 | 0.455380982 | LIN9 | protein_coding | 39.7 | -0.172269114 | 0.172269114 | ENSG00000183814 | n.s. |
| ENSG00000204569 | 2370.204 | -0.390097667 | 0.191384 | -2.038294 | 0.04152 | 0.361758374 | PPP1R10 | protein_coding | 46.34 | -0.172259877 | 0.172259877 | ENSG00000204569 | n.s. |
| ENSG00000130764 | 360.2215 | -0.48902878 | 0.2681 | -1.824052 | 0.068144 | 0.444389214 | LRRC47 | protein_coding | 54.71 | -0.172253781 | 0.172253781 | ENSG00000130764 | n.s. |
| ENSG00000278828 | 960.8097 | -0.45426943 | 0.23933 | -1.898086 | 0.057685 | 0.417869145 | HIST1H3H | protein_coding | 49.07 | -0.172149805 | 0.172149805 | ENSG00000278828 | n.s. |
| ENSG00000224914 | 169.893 | -0.454192174 | 0.238651 | -1.903164 | 0.057019 | 0.417869145 | LINC00863 | lincRNA | 39.66 | -0.172120528 | 0.172120528 | ENSG00000224914 | n.s. |
| ENSG00000201209 | 9.21365 | -0.739477988 | 0.49212 | -1.502636 | 0.132933 | 0.5851482 | | snoRNA | 30.88 | -0.172101764 | 0.172101764 | ENSG00000201209 | n.s. |
| ENSG00000236711 | 115.8942 | -0.501786056 | 0.277415 | -1.808795 | 0.070483 | 0.454110874 | SMAD9-IT1 | sense_intronic | 42.29 | -0.172031377 | 0.172031377 | ENSG00000236711 | n.s. |
| ENSG00000188322 | 280.4788 | -0.439844343 | 0.227116 | -1.93665 | 0.052788 | 0.406509778 | SBK1 | protein_coding | 55.29 | -0.171947911 | 0.171947911 | ENSG00000188322 | n.s. |
| ENSG00000261335 | 185.7248 | -0.469506264 | 0.251674 | -1.865531 | 0.062107 | 0.430475522 | | antisense | 46.77 | -0.17186349 | 0.17186349 | ENSG00000261335 | n.s. |
| ENSG00000201036 | 10.19016 | 0.708600178 | 0.459825 | 1.5410203 | 0.123312 | 0.572276225 | | snoRNA | 38.2 | 0.171760641 | 0.171760641 | ENSG00000201036 | n.s. |
| ENSG00000212123 | 81.68238 | -0.546654348 | 0.315651 | -1.731831 | 0.083304 | 0.485101957 | PRR22 | protein_coding | 66.72 | -0.171740742 | 0.171740742 | ENSG00000212123 | n.s. |
| ENSG00000207573 | 10.12587 | 0.791804605 | 0.544021 | 1.4554679 | 0.14554 | 0.607098916 | MIR550A2 | miRNA | 53.61 | 0.171616159 | 0.171616159 | ENSG00000207573 | n.s. |
| ENSG00000184208 | 363.7606 | -0.460509449 | 0.24411 | -1.886483 | 0.05923 | 0.42398874 | C22orf46 | protein_coding | 51.21 | -0.171606855 | 0.171606855 | ENSG00000184208 | n.s. |
| ENSG00000144218 | 437.3634 | -0.468589776 | 0.251421 | -1.863766 | 0.062355 | 0.430475522 | AFF3 | protein_coding | 41.63 | -0.171528009 | 0.171528009 | ENSG00000144218 | n.s. |
| ENSG00000174989 | 796.7715 | -0.457035665 | 0.241883 | -1.889489 | 0.058826 | 0.421967198 | FBXW8 | protein_coding | 43.26 | -0.171261002 | 0.171261002 | ENSG00000174989 | n.s. |
| ENSG00000213540 | 40.49926 | -0.539461173 | 0.309406 | -1.743536 | 0.08124 | 0.481731421 | | processed_pseudogene | 53.75 | -0.171114401 | 0.171114401 | ENSG00000213540 | n.s. |
| ENSG00000163935 | 208.1367 | -0.482068197 | 0.263164 | -1.831817 | 0.066979 | 0.441754845 | SFMBT1 | protein_coding | 40.71 | -0.171046801 | 0.171046801 | ENSG00000163935 | n.s. |
| ENSG00000115129 | 726.9189 | -0.45987443 | 0.244518 | -1.880738 | 0.060008 | 0.425223094 | TP53I3 | protein_coding | 49.47 | -0.170789618 | 0.170789618 | ENSG00000115129 | n.s. |
| ENSG00000105426 | 1302.488 | -0.45060817 | 0.237221 | -1.899526 | 0.057495 | 0.417869145 | PTPRS | protein_coding | 52.47 | -0.170762335 | 0.170762335 | ENSG00000105426 | n.s. |
| ENSG00000075618 | 1760.35 | -0.442899218 | 0.230293 | -1.923199 | 0.054455 | 0.411605417 | FSCN1 | protein_coding | 59.62 | -0.170746028 | 0.170746028 | ENSG00000075618 | n.s. |
| ENSG00000102109 | 137.9191 | -0.539167845 | 0.309718 | -1.740835 | 0.081712 | 0.482429187 | PCSK1N | protein_coding | 55.55 | -0.170682438 | 0.170682438 | ENSG00000102109 | n.s. |
| ENSG00000252376 | 6.896172 | -0.847375113 | 0.60613 | -1.398008 | 0.162111 | 0.628957308 | RNA5SP395 | rRNA | 45.76 | -0.170643411 | 0.170643411 | ENSG00000252376 | n.s. |
| ENSG00000201616 | 14.42745 | -0.69351398 | 0.446586 | -1.552924 | 0.120441 | 0.567746164 | RNU1-91P | snRNA | 46.01 | -0.170497493 | 0.170497493 | ENSG00000201616 | n.s. |
| ENSG00000196890 | 474.4773 | -0.444504737 | 0.232026 | -1.915752 | 0.055397 | 0.413559567 | HIST3H2BB | protein_coding | 52.96 | -0.170450644 | 0.170450644 | ENSG00000196890 | n.s. |
| ENSG00000265345 | 4.736589 | -0.949493733 | 0.720415 | -1.317981 | 0.18751 | 0.661470368 | MIR5188 | miRNA | 39.82 | -0.170424256 | 0.170424256 | ENSG00000265345 | n.s. |
| ENSG00000186001 | 918.1696 | -0.478643363 | 0.260507 | -1.837353 | 0.066158 | 0.440875092 | LRCH3 | protein_coding | 40.91 | -0.170245995 | 0.170245995 | ENSG00000186001 | n.s. |
| ENSG00000235423 | 134.0881 | -0.467155949 | 0.251783 | -1.855393 | 0.06354 | 0.432812928 | | antisense | 46.01 | -0.169904514 | 0.169904514 | ENSG00000235423 | n.s. |

| | | | | | | | | | | | | | |
|---|---|---|---|---|---|---|---|---|---|---|---|---|---|
| ENSG00000166987 | 156.5943 | -0.477759629 | 0.26018 | -1.836268 | 0.066318 | 0.440948881 | MBD6 | protein_coding | 57.85 | -0.16989694 | 0.16989694 | ENSG00000166987 | n.s. |
| ENSG00000197019 | 197.5825 | -0.52870714 | 0.300376 | -1.760153 | 0.078382 | 0.477221369 | SERTAD1 | protein_coding | 52.57 | -0.169863092 | 0.169863092 | ENSG00000197019 | n.s. |
| ENSG00000198886 | 3640.122 | -0.431042301 | 0.221732 | -1.943975 | 0.051898 | 0.403677294 | MT-ND4 | protein_coding | 44.27 | -0.169815872 | 0.169815872 | ENSG00000198886 | n.s. |
| ENSG00000063245 | 325.3888 | -0.471191668 | 0.254552 | -1.851064 | 0.06416 | 0.436183432 | EPN1 | protein_coding | 54.85 | -0.169784887 | 0.169784887 | ENSG00000063245 | n.s. |
| ENSG00000279819 | 38.40611 | -0.551951612 | 0.322617 | -1.710855 | 0.087108 | 0.492652156 | | TEC | 42.5 | -0.169702829 | 0.169702829 | ENSG00000279819 | n.s. |
| ENSG00000105732 | 320.6705 | -0.46324491 | 0.248666 | -1.862918 | 0.062474 | 0.430475522 | ZNF574 | protein_coding | 57 | -0.169571512 | 0.169571512 | ENSG00000105732 | n.s. |
| ENSG00000264402 | 10.83388 | 0.775356989 | 0.530733 | 1.4609178 | 0.144038 | 0.6044126 | MIR548AL | miRNA | 32.99 | 0.169544592 | 0.169544592 | ENSG00000264402 | n.s. |
| ENSG00000200731 | 12.83909 | -0.694856348 | 0.450159 | -1.543581 | 0.12269 | 0.570735495 | RNU1-124P | snRNA | 48.28 | -0.169242768 | 0.169242768 | ENSG00000200731 | n.s. |
| ENSG00000102100 | 105.7937 | -0.53773768 | 0.309712 | -1.736253 | 0.082519 | 0.484641218 | SLC35A2 | protein_coding | 52.04 | -0.169161332 | 0.169161332 | ENSG00000102100 | n.s. |
| ENSG00000110888 | 221.4596 | -0.506265157 | 0.283871 | -1.783432 | 0.074516 | 0.463461407 | CAPRIN2 | protein_coding | 37.58 | -0.16908569 | 0.16908569 | ENSG00000110888 | n.s. |
| ENSG00000049618 | 4906.878 | -0.460976918 | 0.246446 | -1.8705 | 0.061414 | 0.429914933 | ARID1B | protein_coding | 40.87 | -0.169002191 | 0.169002191 | ENSG00000049618 | n.s. |
| ENSG00000181827 | 2186.749 | -0.461411493 | 0.247006 | -1.868021 | 0.061759 | 0.430475522 | RFX7 | protein_coding | 37.13 | -0.168900387 | 0.168900387 | ENSG00000181827 | n.s. |
| ENSG00000136895 | 1154.828 | -0.440440537 | 0.230163 | -1.913604 | 0.055671 | 0.413584774 | GARNL3 | protein_coding | 43.3 | -0.168880519 | 0.168880519 | ENSG00000136895 | n.s. |
| ENSG00000262633 | 67.96406 | -0.533395317 | 0.306592 | -1.739757 | 0.081902 | 0.482429187 | | protein_coding | 45.82 | -0.168855049 | 0.168855049 | ENSG00000262633 | n.s. |
| ENSG00000105793 | 660.6668 | -0.519074233 | 0.293612 | -1.767893 | 0.077079 | 0.472970027 | GTPBP10 | protein_coding | 38.02 | -0.16878549 | 0.16878549 | ENSG00000105793 | n.s. |
| ENSG00000122912 | 520.4569 | -0.493514155 | 0.273188 | -1.806503 | 0.07084 | 0.455117569 | SLC25A16 | protein_coding | 43.34 | -0.168720842 | 0.168720842 | ENSG00000122912 | n.s. |
| ENSG00000241634 | 145.6323 | -0.534198404 | 0.307351 | -1.73807 | 0.082198 | 0.483501903 | | processed_pseudogene | 50.49 | -0.168593984 | 0.168593984 | ENSG00000241634 | n.s. |
| ENSG00000178233 | 219.4344 | -0.460112448 | 0.246743 | -1.864742 | 0.062218 | 0.430475522 | TMEM151B | protein_coding | 54.71 | -0.16842487 | 0.16842487 | ENSG00000178233 | n.s. |
| ENSG00000260278 | 94.59995 | 0.525378377 | 0.299315 | 1.7552719 | 0.079213 | 0.478143199 | | antisense | 45.09 | 0.168353307 | 0.168353307 | ENSG00000260278 | n.s. |
| ENSG00000276641 | 3.473183 | -0.994181327 | 0.779109 | -1.276049 | 0.201938 | 0.677624898 | MIR6769A | miRNA | 64.38 | -0.168027227 | 0.168027227 | ENSG00000276641 | n.s. |
| ENSG00000282977 | 86.21564 | -0.473069784 | 0.258711 | -1.828565 | 0.067465 | 0.442237706 | PCBP2-OT1 | non_coding | 50 | -0.167629549 | 0.167629549 | ENSG00000282977 | n.s. |
| ENSG00000214773 | 102.1455 | -0.46927433 | 0.254746 | -1.84213 | 0.065456 | 0.439626611 | | sense_intronic | 45.6 | -0.16749153 | 0.16749153 | ENSG00000214773 | n.s. |
| ENSG00000106993 | 534.4617 | -0.470714824 | 0.256512 | -1.83506 | 0.066497 | 0.441084074 | CDC37L1 | protein_coding | 36.8 | -0.167329058 | 0.167329058 | ENSG00000106993 | n.s. |
| ENSG00000283183 | 229.3683 | -0.469119228 | 0.254897 | -1.840423 | 0.065706 | 0.440455774 | | lincRNA | 45.49 | -0.167052275 | 0.167052275 | ENSG00000283183 | n.s. |
| ENSG00000167378 | 810.4923 | -0.456297691 | 0.245188 | -1.861011 | 0.062743 | 0.430564731 | IRGQ | protein_coding | 52.71 | -0.166987409 | 0.166987409 | ENSG00000167378 | n.s. |
| ENSG00000165650 | 1475.485 | -0.456061828 | 0.24494 | -1.861934 | 0.062612 | 0.430511912 | PDZD8 | protein_coding | 37.51 | -0.166925391 | 0.166925391 | ENSG00000165650 | n.s. |
| ENSG00000006704 | 553.222 | -0.463281058 | 0.250567 | -1.848933 | 0.064467 | 0.436711663 | GTF2IRD1 | protein_coding | 52.72 | -0.166690938 | 0.166690938 | ENSG00000006704 | n.s. |
| ENSG00000005889 | 216.7714 | 0.425374499 | 0.219427 | 1.9385662 | 0.052554 | 0.405702385 | ZFX | protein_coding | 40.77 | 0.166658513 | 0.166658513 | ENSG00000005889 | n.s. |
| ENSG00000214331 | 262.3746 | -0.454909353 | 0.244848 | -1.857927 | 0.063179 | 0.431199305 | | transcribed_unprocessed_pseudc | 46.88 | -0.166188371 | 0.166188371 | ENSG00000214331 | n.s. |
| ENSG00000168610 | 2015.442 | -0.454165271 | 0.244248 | -1.859442 | 0.062964 | 0.430788188 | STAT3 | protein_coding | 45.02 | -0.166104687 | 0.166104687 | ENSG00000168610 | n.s. |
| ENSG00000237749 | 13.15508 | 0.685184481 | 0.444694 | 1.5408006 | 0.123365 | 0.572276225 | | processed_pseudogene | 49.8 | 0.16608481 | 0.16608481 | ENSG00000237749 | n.s. |
| ENSG00000103260 | 28.97459 | -0.646518278 | 0.408965 | -1.580865 | 0.113909 | 0.553589858 | METRN | protein_coding | 66.79 | -0.166033571 | 0.166033571 | ENSG00000103260 | n.s. |

| | | | | | | | | | | | | | |
|---|---|---|---|---|---|---|---|---|---|---|---|---|---|
| ENSG00000185670 | 111.0888 | -0.473244425 | 0.259897 | -1.820894 | 0.068623 | 0.445830359 | ZBTB3 | protein_coding | 51.23 | -0.166028513 | 0.166028513 | ENSG00000185670 | n.s. |
| ENSG00000108091 | 837.1407 | -0.438085847 | 0.230615 | -1.89964 | 0.05748 | 0.417869145 | CCDC6 | protein_coding | 40.92 | -0.166016879 | 0.166016879 | ENSG00000108091 | n.s. |
| ENSG00000084070 | 161.0842 | -0.489124933 | 0.271821 | -1.799441 | 0.071949 | 0.45772936 | SMAP2 | protein_coding | 41.95 | -0.166004712 | 0.166004712 | ENSG00000084070 | n.s. |
| ENSG00000060237 | 4328.807 | -0.432727267 | 0.226156 | -1.913398 | 0.055697 | 0.413584774 | WNK1 | protein_coding | 38.72 | -0.165922978 | 0.165922978 | ENSG00000060237 | n.s. |
| ENSG00000166426 | 238.554 | -0.516023109 | 0.292966 | -1.761378 | 0.078174 | 0.477043998 | CRABP1 | protein_coding | 48.64 | -0.165871275 | 0.165871275 | ENSG00000166426 | n.s. |
| ENSG00000173898 | 2805.917 | -0.43247062 | 0.225858 | -1.914789 | 0.055519 | 0.413574865 | SPTBN2 | protein_coding | 55.04 | -0.16582907 | 0.16582907 | ENSG00000173898 | n.s. |
| ENSG00000277075 | 2287.701 | -0.412810256 | 0.210833 | -1.957997 | 0.05023 | 0.397136868 | HIST1H2AE | protein_coding | 56.19 | -0.165561596 | 0.165561596 | ENSG00000277075 | n.s. |
| ENSG00000277977 | 12.77363 | -0.695572937 | 0.456563 | -1.523499 | 0.127634 | 0.57808993 | | unprocessed_pseudogene | 45.36 | -0.165549556 | 0.165549556 | ENSG00000277977 | n.s. |
| ENSG00000197124 | 282.9167 | -0.484463463 | 0.268546 | -1.804025 | 0.071227 | 0.455380982 | ZNF682 | protein_coding | 41.4 | -0.165504884 | 0.165504884 | ENSG00000197124 | n.s. |
| ENSG00000144395 | 136.7456 | -0.472364205 | 0.259769 | -1.818401 | 0.069003 | 0.447091133 | CCDC150 | protein_coding | 38.16 | -0.165140389 | 0.165140389 | ENSG00000144395 | n.s. |
| ENSG00000049883 | 400.9514 | -0.423660124 | 0.21912 | -1.933461 | 0.053179 | 0.407718571 | PTCD2 | protein_coding | 40.38 | -0.165074722 | 0.165074722 | ENSG00000049883 | n.s. |
| ENSG00000080603 | 627.227 | -0.458678765 | 0.248109 | -1.848699 | 0.064501 | 0.436711663 | SRCAP | protein_coding | 49.63 | -0.165035009 | 0.165035009 | ENSG00000080603 | n.s. |
| ENSG00000199266 | 13.62962 | 0.687970377 | 0.449624 | 1.5301012 | 0.125992 | 0.575897021 | SNORA60 | snoRNA | 50.75 | 0.164875656 | 0.164875656 | ENSG00000199266 | n.s. |
| ENSG00000153814 | 132.9411 | -0.520668395 | 0.299289 | -1.739684 | 0.081915 | 0.482429187 | JAZF1 | protein_coding | 41.44 | -0.164826133 | 0.164826133 | ENSG00000153814 | n.s. |
| ENSG00000130338 | 2476.111 | -0.449323109 | 0.240418 | -1.868921 | 0.061634 | 0.430354829 | TULP4 | protein_coding | 43.15 | -0.164530134 | 0.164530134 | ENSG00000130338 | n.s. |
| ENSG00000164082 | 186.6499 | -0.457064243 | 0.247175 | -1.849153 | 0.064436 | 0.436711663 | GRM2 | protein_coding | 55.81 | -0.164454096 | 0.164454096 | ENSG00000164082 | n.s. |
| ENSG00000068654 | 2544.21 | -0.44914151 | 0.240463 | -1.867819 | 0.061787 | 0.430475522 | POLR1A | protein_coding | 46.69 | -0.164408941 | 0.164408941 | ENSG00000068654 | n.s. |
| ENSG00000173327 | 391.7176 | -0.433202229 | 0.227369 | -1.905283 | 0.056743 | 0.417346136 | MAP3K11 | protein_coding | 57.13 | -0.164401806 | 0.164401806 | ENSG00000173327 | n.s. |
| ENSG00000169951 | 185.6025 | -0.432290455 | 0.226636 | -1.907423 | 0.056466 | 0.416847613 | ZNF764 | protein_coding | 58.06 | -0.164280178 | 0.164280178 | ENSG00000169951 | n.s. |
| ENSG00000102595 | 1276.922 | -0.491645368 | 0.27567 | -1.783455 | 0.074512 | 0.463461407 | UGGT2 | protein_coding | 37.01 | -0.164202878 | 0.164202878 | ENSG00000102595 | n.s. |
| ENSG00000145555 | 527.1649 | -0.428607223 | 0.224199 | -1.911727 | 0.055911 | 0.414070243 | MYO10 | protein_coding | 44.1 | -0.164124841 | 0.164124841 | ENSG00000145555 | n.s. |
| ENSG00000111186 | 183.0088 | -0.461226279 | 0.251024 | -1.83738 | 0.066154 | 0.440875092 | WNT5B | protein_coding | 47.53 | -0.164051009 | 0.164051009 | ENSG00000111186 | n.s. |
| ENSG00000106868 | 501.6895 | -0.462519379 | 0.252582 | -1.831167 | 0.067076 | 0.441892852 | SUSD1 | protein_coding | 42.78 | -0.164047772 | 0.164047772 | ENSG00000106868 | n.s. |
| ENSG00000162073 | 739.8644 | -0.469078103 | 0.258023 | -1.817973 | 0.069068 | 0.447272801 | PAQR4 | protein_coding | 64.48 | -0.163908794 | 0.163908794 | ENSG00000162073 | n.s. |
| ENSG00000253276 | 375.917 | 0.43213745 | 0.227404 | 1.9003087 | 0.057393 | 0.417869145 | CCDC71L | protein_coding | 43.76 | 0.163762676 | 0.163762676 | ENSG00000253276 | n.s. |
| ENSG00000147813 | 69.37526 | -0.520576103 | 0.299976 | -1.73539 | 0.082672 | 0.484982269 | NAPRT | protein_coding | 67.89 | -0.163603607 | 0.163603607 | ENSG00000147813 | n.s. |
| ENSG00000110080 | 67.01157 | -0.50560587 | 0.286507 | -1.764723 | 0.07761 | 0.474896813 | ST3GAL4 | protein_coding | 52.01 | -0.163513315 | 0.163513315 | ENSG00000110080 | n.s. |
| ENSG00000167695 | 176.9833 | -0.481741041 | 0.268104 | -1.796847 | 0.07236 | 0.458773825 | FAM57A | protein_coding | 49.91 | -0.163021827 | 0.163021827 | ENSG00000167695 | n.s. |
| ENSG00000189308 | 1001.598 | -0.452130327 | 0.244248 | -1.851115 | 0.064153 | 0.436183432 | LIN54 | protein_coding | 39.03 | -0.162916498 | 0.162916498 | ENSG00000189308 | n.s. |
| ENSG00000067842 | 322.6125 | -0.461465263 | 0.252899 | -1.824698 | 0.068047 | 0.443960682 | ATP2B3 | protein_coding | 58.21 | -0.162738259 | 0.162738259 | ENSG00000067842 | n.s. |
| ENSG00000138101 | 250.653 | -0.452151077 | 0.24465 | -1.848156 | 0.06458 | 0.436711663 | DTNB | protein_coding | 41.46 | -0.162686313 | 0.162686313 | ENSG00000138101 | n.s. |
| ENSG00000278818 | 22.16953 | -0.650635008 | 0.416062 | -1.563792 | 0.117866 | 0.562372792 | | misc_RNA | 51.51 | -0.162642941 | 0.162642941 | ENSG00000278818 | n.s. |

| | | | | | | | | | | | | | |
|---|---|---|---|---|---|---|---|---|---|---|---|---|---|
| ENSG00000141522 | 1621.145 | -0.444224424 | 0.238387 | -1.863455 | 0.062398 | 0.430475522 | ARHGDIA | protein_coding | 68.12 | -0.162609034 | 0.162609034 | ENSG00000141522 | n.s. |
| ENSG00000101333 | 73.69367 | -0.506394215 | 0.288091 | -1.757758 | 0.078789 | 0.477489649 | PLCB4 | protein_coding | 39.14 | -0.162570793 | 0.162570793 | ENSG00000101333 | n.s. |
| ENSG00000141753 | 415.7947 | -0.495192517 | 0.279223 | -1.773465 | 0.076152 | 0.469606803 | IGFBP4 | protein_coding | 53 | -0.162554679 | 0.162554679 | ENSG00000141753 | n.s. |
| ENSG00000157014 | 747.7341 | -0.459304602 | 0.251487 | -1.826352 | 0.067797 | 0.442749255 | TATDN2 | protein_coding | 46.98 | -0.162521333 | 0.162521333 | ENSG00000157014 | n.s. |
| ENSG00000143416 | 174.5242 | 0.526196022 | 0.306641 | 1.7160001 | 0.086162 | 0.491092308 | SELENBP1 | protein_coding | 54.96 | 0.162508732 | 0.162508732 | ENSG00000143416 | n.s. |
| ENSG00000240729 | 33.15502 | -0.544727948 | 0.322736 | -1.687844 | 0.091441 | 0.503597002 | | processed_pseudogene | 42.23 | -0.162283642 | 0.162283642 | ENSG00000240729 | n.s. |
| ENSG00000127616 | 3266.192 | -0.424057904 | 0.221874 | -1.911254 | 0.055972 | 0.414299599 | SMARCA4 | protein_coding | 52.73 | -0.162280805 | 0.162280805 | ENSG00000127616 | n.s. |
| ENSG00000252585 | 20.59565 | -0.626905946 | 0.395081 | -1.586778 | 0.112563 | 0.551060121 | | misc_RNA | 43.36 | -0.162243896 | 0.162243896 | ENSG00000252585 | n.s. |
| ENSG00000152133 | 899.2456 | -0.508756761 | 0.291222 | -1.746972 | 0.080642 | 0.479877462 | GPATCH11 | protein_coding | 38.7 | -0.162227089 | 0.162227089 | ENSG00000152133 | n.s. |
| ENSG00000130449 | 1657.142 | -0.474863183 | 0.263133 | -1.804651 | 0.071129 | 0.455380982 | ZSWIM6 | protein_coding | 38.17 | -0.162225188 | 0.162225188 | ENSG00000130449 | n.s. |
| ENSG00000221545 | 3.000598 | 0.958605786 | 0.749078 | 1.279714 | 0.200646 | 0.677491543 | MIR1255B2 | miRNA | 43.28 | 0.162096521 | 0.162096521 | ENSG00000221545 | n.s. |
| ENSG00000200215 | 5.686913 | 0.769881617 | 0.539352 | 1.4274206 | 0.153459 | 0.615840022 | SNORD113-6 | snoRNA | 40 | 0.162084786 | 0.162084786 | ENSG00000200215 | n.s. |
| ENSG00000176124 | 150.6319 | 0.479159378 | 0.266926 | 1.7950996 | 0.072638 | 0.459013748 | DLEU1 | protein_coding | 39.72 | 0.16203939 | 0.16203939 | ENSG00000176124 | n.s. |
| ENSG00000160584 | 103.2321 | -0.507936437 | 0.290561 | -1.748126 | 0.080442 | 0.479859335 | SIK3 | protein_coding | 40.48 | -0.161973845 | 0.161973845 | ENSG00000160584 | n.s. |
| ENSG00000213533 | 378.4171 | -0.456839504 | 0.249809 | -1.828755 | 0.067436 | 0.442237706 | TMEM110 | protein_coding | 48.39 | -0.161878443 | 0.161878443 | ENSG00000213533 | n.s. |
| ENSG00000155506 | 3866.069 | -0.435862118 | 0.231756 | -1.880697 | 0.060013 | 0.425223094 | LARP1 | protein_coding | 46.52 | -0.161871849 | 0.161871849 | ENSG00000155506 | n.s. |
| ENSG00000174028 | 26.60326 | -0.610299452 | 0.38023 | -1.605081 | 0.108476 | 0.543082569 | FAM3C2 | processed_pseudogene | 46.24 | -0.161811218 | 0.161811218 | ENSG00000174028 | n.s. |
| ENSG00000269825 | 301.9558 | -0.476674992 | 0.264965 | -1.799012 | 0.072017 | 0.45772936 | | sense_intronic | 41.71 | -0.161779312 | 0.161779312 | ENSG00000269825 | n.s. |
| ENSG00000123124 | 76.25847 | -0.505146625 | 0.288088 | -1.753448 | 0.079525 | 0.478697507 | WWP1 | protein_coding | 37.46 | -0.161616023 | 0.161616023 | ENSG00000123124 | n.s. |
| ENSG00000087266 | 634.3307 | -0.438783369 | 0.234282 | -1.872887 | 0.061084 | 0.42849416 | SH3BP2 | protein_coding | 59.23 | -0.161496454 | 0.161496454 | ENSG00000087266 | n.s. |
| ENSG00000130382 | 530.6624 | -0.441496708 | 0.237403 | -1.859695 | 0.062929 | 0.430754942 | MLLT1 | protein_coding | 55.57 | -0.161486132 | 0.161486132 | ENSG00000130382 | n.s. |
| ENSG00000092098 | 97.3758 | -0.473890938 | 0.262959 | -1.802149 | 0.071522 | 0.456370327 | RNF31 | protein_coding | 52.92 | -0.161446397 | 0.161446397 | ENSG00000092098 | n.s. |
| ENSG00000119772 | 2331.441 | -0.449268938 | 0.243344 | -1.846231 | 0.064859 | 0.437274745 | DNMT3A | protein_coding | 53.43 | -0.161397891 | 0.161397891 | ENSG00000119772 | n.s. |
| ENSG00000198743 | 2080.335 | -0.491567656 | 0.277285 | -1.772788 | 0.076264 | 0.469832545 | SLC5A3 | protein_coding | 39.03 | -0.161262163 | 0.161262163 | ENSG00000198743 | n.s. |
| ENSG00000208012 | 2.763822 | -0.977318472 | 0.777048 | -1.257732 | 0.208489 | 0.683940032 | MIRLET7F2 | miRNA | 42.17 | -0.161239932 | 0.161239932 | ENSG00000208012 | n.s. |
| ENSG00000110697 | 11.93715 | -0.71729091 | 0.484488 | -1.480514 | 0.138736 | 0.595996411 | PITPNM1 | protein_coding | 62.46 | -0.161215691 | 0.161215691 | ENSG00000110697 | n.s. |
| ENSG00000116786 | 262.352 | -0.501865064 | 0.286075 | -1.754314 | 0.079377 | 0.478143199 | PLEKHM2 | protein_coding | 52 | -0.160818653 | 0.160818653 | ENSG00000116786 | n.s. |
| ENSG00000135315 | 721.8788 | -0.492847235 | 0.278482 | -1.769766 | 0.076766 | 0.471809602 | CEP162 | protein_coding | 37 | -0.160783143 | 0.160783143 | ENSG00000135315 | n.s. |
| ENSG00000166033 | 333.6294 | -0.511457744 | 0.295105 | -1.733137 | 0.083071 | 0.485101957 | HTRA1 | protein_coding | 49.2 | -0.160683132 | 0.160683132 | ENSG00000166033 | n.s. |
| ENSG00000134532 | 58.42903 | 0.502905698 | 0.287268 | 1.75065 | 0.080006 | 0.479305998 | SOX5 | protein_coding | 34.5 | 0.160621611 | 0.160621611 | ENSG00000134532 | n.s. |
| ENSG00000200556 | 20.50354 | 0.674249993 | 0.442257 | 1.5245652 | 0.127368 | 0.578027275 | RNU6-103P | snRNA | 45.79 | 0.160506336 | 0.160506336 | ENSG00000200556 | n.s. |
| ENSG00000244642 | 15.77966 | -0.641365737 | 0.409968 | -1.564427 | 0.117717 | 0.562372792 | RN7SL396P | misc_RNA | 56.27 | -0.160325849 | 0.160325849 | ENSG00000244642 | n.s. |

| | | | | | | | | | | | | | |
|---|---|---|---|---|---|---|---|---|---|---|---|---|---|
| ENSG00000178409 | 499.7717 | -0.452989077 | 0.247961 | -1.826857 | 0.067721 | 0.442685196 | BEND3 | protein_coding | 47.45 | -0.1603151 | 0.1603151 | ENSG00000178409 | n.s. |
| ENSG00000279355 | 92.77424 | -0.497408276 | 0.282797 | -1.758891 | 0.078596 | 0.477239282 | AGPAT4-IT1 | TEC | 51.33 | -0.159799281 | 0.159799281 | ENSG00000279355 | n.s. |
| ENSG00000179886 | 268.037 | -0.421560037 | 0.221739 | -1.901151 | 0.057282 | 0.417869145 | TIGD5 | protein_coding | 64.53 | -0.159754263 | 0.159754263 | ENSG00000179886 | n.s. |
| ENSG00000274954 | 19.31715 | 0.675188853 | 0.446922 | 1.5107532 | 0.130851 | 0.580204884 | | sense_intronic | 26.62 | 0.159627216 | 0.159627216 | ENSG00000274954 | n.s. |
| ENSG00000200350 | 23.29119 | 0.59382798 | 0.367541 | 1.6156798 | 0.106164 | 0.538671714 | RNU6-1285P | snRNA | 42.86 | 0.159547225 | 0.159547225 | ENSG00000200350 | n.s. |
| ENSG00000169155 | 430.6756 | -0.497591837 | 0.283579 | -1.754684 | 0.079313 | 0.478143199 | ZBTB43 | protein_coding | 41.82 | -0.159449332 | 0.159449332 | ENSG00000169155 | n.s. |
| ENSG00000212135 | 18.74305 | -0.610528735 | 0.383422 | -1.592315 | 0.111314 | 0.548299349 | SNORD67 | snoRNA | 51.35 | -0.159337175 | 0.159337175 | ENSG00000212135 | n.s. |
| ENSG00000256442 | 27.68759 | -0.671403344 | 0.442683 | -1.51667 | 0.12935 | 0.579048024 | | antisense | 50.9 | -0.159314222 | 0.159314222 | ENSG00000256442 | n.s. |
| ENSG00000124164 | 1343.575 | -0.483066488 | 0.271709 | -1.77788 | 0.075424 | 0.467968938 | VAPB | protein_coding | 42.86 | -0.159307103 | 0.159307103 | ENSG00000124164 | n.s. |
| ENSG00000243319 | 153.6068 | -0.483161063 | 0.271883 | -1.777091 | 0.075553 | 0.46814761 | FGF14-IT1 | sense_intronic | 37.7 | -0.159258192 | 0.159258192 | ENSG00000243319 | n.s. |
| ENSG00000155744 | 975.8445 | -0.475945266 | 0.266745 | -1.784273 | 0.074379 | 0.463348535 | FAM126B | protein_coding | 39.16 | -0.159009604 | 0.159009604 | ENSG00000155744 | n.s. |
| ENSG00000169710 | 4859.121 | -0.419084992 | 0.220384 | -1.901609 | 0.057222 | 0.417869145 | FASN | protein_coding | 66.65 | -0.158816321 | 0.158816321 | ENSG00000169710 | n.s. |
| ENSG00000269688 | 149.8711 | -0.447379037 | 0.244054 | -1.833116 | 0.066785 | 0.441705004 | | sense_intronic | 48.12 | -0.158760361 | 0.158760361 | ENSG00000269688 | n.s. |
| ENSG00000189144 | 98.84875 | -0.49705614 | 0.284087 | -1.74966 | 0.080177 | 0.479305998 | ZNF573 | protein_coding | 41.38 | -0.158753337 | 0.158753337 | ENSG00000189144 | n.s. |
| ENSG00000168077 | 278.6249 | -0.471339228 | 0.263141 | -1.791202 | 0.073261 | 0.460477385 | SCARA3 | protein_coding | 50.25 | -0.158743137 | 0.158743137 | ENSG00000168077 | n.s. |
| ENSG00000270883 | 21.38912 | -0.567265223 | 0.345104 | -1.64375 | 0.100228 | 0.525008902 | | lincRNA | 41.47 | -0.158739718 | 0.158739718 | ENSG00000270883 | n.s. |
| ENSG00000204604 | 335.2623 | -0.470642054 | 0.262597 | -1.792263 | 0.073091 | 0.460211177 | ZNF468 | protein_coding | 43.76 | -0.158626534 | 0.158626534 | ENSG00000204604 | n.s. |
| ENSG00000173218 | 392.094 | -0.425389594 | 0.225714 | -1.884643 | 0.059478 | 0.423999253 | VANGL1 | protein_coding | 45.32 | -0.158515012 | 0.158515012 | ENSG00000173218 | n.s. |
| ENSG00000243960 | 43.85723 | -0.535696519 | 0.318253 | -1.683243 | 0.092328 | 0.505936376 | | sense_overlapping | 41.95 | -0.158514794 | 0.158514794 | ENSG00000243960 | n.s. |
| ENSG00000274805 | 4.156653 | -1.034386681 | 0.858793 | -1.204465 | 0.22841 | 0.702737638 | MIR6768 | miRNA | 62.5 | -0.158475058 | 0.158475058 | ENSG00000274805 | n.s. |
| ENSG00000255374 | 64.71261 | -0.528360958 | 0.312193 | -1.692417 | 0.090566 | 0.501374421 | TAS2R43 | protein_coding | 38.95 | -0.158422601 | 0.158422601 | ENSG00000255374 | n.s. |
| ENSG00000165271 | 1161.116 | -0.424529058 | 0.225179 | -1.885298 | 0.05939 | 0.423990171 | NOL6 | protein_coding | 54.48 | -0.158198296 | 0.158198296 | ENSG00000165271 | n.s. |
| ENSG00000151233 | 799.9237 | -0.432156846 | 0.231959 | -1.863074 | 0.062452 | 0.430475522 | GXYLT1 | protein_coding | 37.61 | -0.158191679 | 0.158191679 | ENSG00000151233 | n.s. |
| ENSG00000260404 | 187.7693 | -0.452893433 | 0.249198 | -1.817401 | 0.069156 | 0.447423558 | | transcribed_unprocessed_pseudc | 46.61 | -0.15818714 | 0.15818714 | ENSG00000260404 | n.s. |
| ENSG00000169635 | 616.1837 | -0.431527013 | 0.230832 | -1.869442 | 0.061561 | 0.430280775 | HIC2 | protein_coding | 56.16 | -0.15804593 | 0.15804593 | ENSG00000169635 | n.s. |
| ENSG00000101181 | 226.7618 | -0.499197377 | 0.286807 | -1.740535 | 0.081765 | 0.482429187 | MTG2 | protein_coding | 49.97 | -0.158029129 | 0.158029129 | ENSG00000101181 | n.s. |
| ENSG00000251804 | 3.954144 | -1.2445022 | 1.167683 | -1.065788 | 0.28652 | 0.746611882 | RNU6-1294P | snRNA | 37.04 | -0.15793368 | 0.15793368 | ENSG00000251804 | n.s. |
| ENSG00000208037 | 11.06466 | 0.924256314 | 0.719118 | 1.2852632 | 0.1987 | 0.674743473 | MIR320A | miRNA | 59.76 | 0.157919642 | 0.157919642 | ENSG00000208037 | n.s. |
| ENSG00000182903 | 1076.114 | -0.465146485 | 0.258455 | -1.79972 | 0.071905 | 0.45772936 | ZNF721 | protein_coding | 41.11 | -0.157866638 | 0.157866638 | ENSG00000182903 | n.s. |
| ENSG00000148841 | 298.1844 | -0.449555381 | 0.246939 | -1.820512 | 0.068681 | 0.445830359 | ITPRIP | protein_coding | 52.49 | -0.157717677 | 0.157717677 | ENSG00000148841 | n.s. |
| ENSG00000221771 | 8.527647 | -0.993559309 | 0.806228 | -1.232355 | 0.217816 | 0.693853089 | MIR1205 | miRNA | 52.38 | -0.157710127 | 0.157710127 | ENSG00000221771 | n.s. |
| ENSG00000152128 | 90.58928 | -0.520648946 | 0.306476 | -1.698825 | 0.089352 | 0.498053895 | TMEM163 | protein_coding | 43.72 | -0.157612754 | 0.157612754 | ENSG00000152128 | n.s. |

| | | | | | | | | | | | | | |
|---|---|---|---|---|---|---|---|---|---|---|---|---|---|
| ENSG00000075568 | 2990.608 | -0.438382423 | 0.237514 | -1.845713 | 0.064934 | 0.437274745 | TMEM131 | protein_coding | 39.29 | -0.157486959 | 0.157486959 | ENSG00000075568 | n.s. |
| ENSG00000132394 | 156.2929 | -0.46625853 | 0.260006 | -1.79326 | 0.072931 | 0.459467368 | EEFSEC | protein_coding | 47.87 | -0.15747664 | 0.15747664 | ENSG00000132394 | n.s. |
| ENSG00000159069 | 325.8424 | -0.496661892 | 0.285012 | -1.742601 | 0.081403 | 0.48187821 | FBXW5 | protein_coding | 68.96 | -0.157472966 | 0.157472966 | ENSG00000159069 | n.s. |
| ENSG00000100889 | 205.4114 | -0.504683891 | 0.292356 | -1.726262 | 0.0843 | 0.487561582 | PCK2 | protein_coding | 49.2 | -0.157446497 | 0.157446497 | ENSG00000100889 | n.s. |
| ENSG00000167522 | 108.467 | -0.483273491 | 0.273268 | -1.768499 | 0.076978 | 0.472556476 | ANKRD11 | protein_coding | 51.19 | -0.157327888 | 0.157327888 | ENSG00000167522 | n.s. |
| ENSG00000168487 | 28.33001 | -0.575731797 | 0.3546 | -1.623609 | 0.104459 | 0.533328875 | BMP1 | protein_coding | 54.06 | -0.157177603 | 0.157177603 | ENSG00000168487 | n.s. |
| ENSG00000205181 | 329.3015 | -0.489197869 | 0.277978 | -1.759841 | 0.078435 | 0.477221369 | LINC00654 | lincRNA | 50.26 | -0.157169549 | 0.157169549 | ENSG00000205181 | n.s. |
| ENSG00000272599 | 139.6789 | -0.494846734 | 0.283761 | -1.743887 | 0.081179 | 0.481573667 | | antisense | 40.62 | -0.157033311 | 0.157033311 | ENSG00000272599 | n.s. |
| ENSG00000048991 | 804.6654 | -0.464660624 | 0.258997 | -1.794077 | 0.072801 | 0.459269401 | R3HDM1 | protein_coding | 37.98 | -0.157023921 | 0.157023921 | ENSG00000048991 | n.s. |
| ENSG00000184357 | 4252.452 | -0.426360718 | 0.227482 | -1.874258 | 0.060895 | 0.42849416 | HIST1H1B | protein_coding | 59.32 | -0.156924234 | 0.156924234 | ENSG00000184357 | n.s. |
| ENSG00000274315 | 68.27442 | -0.468847234 | 0.262599 | -1.785413 | 0.074194 | 0.463232209 | | antisense | 33.4 | -0.156689332 | 0.156689332 | ENSG00000274315 | n.s. |
| ENSG00000146263 | 4131.121 | -0.457794827 | 0.253454 | -1.806224 | 0.070883 | 0.455117569 | MMS22L | protein_coding | 35.58 | -0.156509247 | 0.156509247 | ENSG00000146263 | n.s. |
| ENSG00000252451 | 3.393953 | -1.026894574 | 0.859091 | -1.195326 | 0.23196 | 0.704043192 | RNA5SP168 | rRNA | 44.64 | -0.156499448 | 0.156499448 | ENSG00000252451 | n.s. |
| ENSG00000231113 | 70.9492 | -0.532056669 | 0.317009 | -1.678367 | 0.093276 | 0.508003334 | | antisense | 48.07 | -0.156495657 | 0.156495657 | ENSG00000231113 | n.s. |
| ENSG00000269961 | 30.19704 | 0.604221652 | 0.380533 | 1.5878309 | 0.112325 | 0.55089864 | | lincRNA | 48.32 | 0.156450085 | 0.156450085 | ENSG00000269961 | n.s. |
| ENSG00000100393 | 1664.346 | -0.441991456 | 0.241967 | -1.826664 | 0.06775 | 0.442685196 | EP300 | protein_coding | 42.33 | -0.156422987 | 0.156422987 | ENSG00000100393 | n.s. |
| ENSG00000151967 | 1064.906 | 0.445590609 | 0.244634 | 1.8214593 | 0.068537 | 0.445699067 | SCHIP1 | protein_coding | 39.91 | 0.156383712 | 0.156383712 | ENSG00000151967 | n.s. |
| ENSG00000196208 | 300.2735 | -0.454434044 | 0.251103 | -1.809751 | 0.070334 | 0.453364303 | GREB1 | protein_coding | 47.32 | -0.156122033 | 0.156122033 | ENSG00000196208 | n.s. |
| ENSG00000116127 | 4445.293 | -0.43573283 | 0.236259 | -1.844304 | 0.065139 | 0.438326836 | ALMS1 | protein_coding | 38.28 | -0.156080345 | 0.156080345 | ENSG00000116127 | n.s. |
| ENSG00000205763 | 154.2706 | -0.48931516 | 0.279915 | -1.748087 | 0.080449 | 0.479859335 | RP9P | cribed_unprocessed_pseudc | 43.81 | -0.15603578 | 0.15603578 | ENSG00000205763 | n.s. |
| ENSG00000256325 | 82.78935 | -0.487797911 | 0.278734 | -1.750049 | 0.08011 | 0.479305998 | | antisense | 37.61 | -0.155796378 | 0.155796378 | ENSG00000256325 | n.s. |
| ENSG00000148926 | 208.6266 | 0.432869519 | 0.233995 | 1.8499077 | 0.064327 | 0.436676128 | ADM | protein_coding | 60.01 | 0.155764005 | 0.155764005 | ENSG00000148926 | n.s. |
| ENSG00000251445 | 4.384105 | 1.06505363 | 0.910776 | 1.1693908 | 0.242246 | 0.714515282 | | antisense | 53.73 | 0.155485558 | 0.155485558 | ENSG00000251445 | n.s. |
| ENSG00000110063 | 117.0989 | -0.486785412 | 0.278017 | -1.750921 | 0.079959 | 0.479305998 | DCPS | protein_coding | 49.42 | -0.155472999 | 0.155472999 | ENSG00000110063 | n.s. |
| ENSG00000197013 | 283.5782 | -0.446659237 | 0.245998 | -1.815704 | 0.069416 | 0.448897065 | ZNF429 | protein_coding | 41.06 | -0.15537186 | 0.15537186 | ENSG00000197013 | n.s. |
| ENSG00000213928 | 61.76523 | -0.509182408 | 0.29895 | -1.703238 | 0.088523 | 0.495550072 | IRF9 | protein_coding | 53.95 | -0.155256059 | 0.155256059 | ENSG00000213928 | n.s. |
| ENSG00000148444 | 735.3238 | 0.431172356 | 0.233621 | 1.8456094 | 0.064949 | 0.437274745 | COMMD3 | protein_coding | 41.08 | 0.154896774 | 0.154896774 | ENSG00000148444 | n.s. |
| ENSG00000228801 | 87.2561 | -0.514556804 | 0.303616 | -1.69476 | 0.090121 | 0.500068613 | | antisense | 40.39 | -0.154866369 | 0.154866369 | ENSG00000228801 | n.s. |
| ENSG00000197147 | 1570.239 | -0.484831054 | 0.276696 | -1.752213 | 0.079737 | 0.479305998 | LRRC8B | protein_coding | 38.63 | -0.154848802 | 0.154848802 | ENSG00000197147 | n.s. |
| ENSG00000160087 | 71.27824 | -0.492478063 | 0.284264 | -1.732466 | 0.083191 | 0.485101957 | UBE2J2 | protein_coding | 56.33 | -0.154720342 | 0.154720342 | ENSG00000160087 | n.s. |
| ENSG00000212597 | 5.023141 | 0.800240677 | 0.58647 | 1.3645039 | 0.172409 | 0.640837445 | RNU6-876P | snRNA | 36.54 | 0.154648207 | 0.154648207 | ENSG00000212597 | n.s. |
| ENSG00000250329 | 76.99994 | -0.559436784 | 0.343014 | -1.630947 | 0.102902 | 0.529239659 | KDELC1P1 | processed_pseudogene | 42.61 | -0.154599023 | 0.154599023 | ENSG00000250329 | n.s. |

| | | | | | | | | | | | | | |
|---|---|---|---|---|---|---|---|---|---|---|---|---|---|
| ENSG00000114554 | 2090.694 | -0.417989157 | 0.222714 | -1.876794 | 0.060546 | 0.426932326 | PLXNA1 | protein_coding | 61.75 | -0.154505914 | 0.154505914 | ENSG00000114554 | n.s. |
| ENSG00000225178 | 817.7083 | 0.554777311 | 0.338924 | 1.6368798 | 0.101656 | 0.526936211 | RPSAP58 | processed_pseudogene | 52.36 | 0.154362324 | 0.154362324 | ENSG00000225178 | n.s. |
| ENSG00000188803 | 66.56579 | -0.485785918 | 0.278662 | -1.743281 | 0.081285 | 0.481790215 | SHISA6 | protein_coding | 43.68 | -0.15406313 | 0.15406313 | ENSG00000188803 | n.s. |
| ENSG00000263766 | 33.7769 | -0.521483972 | 0.310181 | -1.681222 | 0.09272 | 0.506647149 | | antisense | 43.3 | -0.153991295 | 0.153991295 | ENSG00000263766 | n.s. |
| ENSG00000208018 | 8.935318 | -0.709907135 | 0.490602 | -1.447013 | 0.147893 | 0.607109641 | MIR645 | miRNA | 56.38 | -0.153860211 | 0.153860211 | ENSG00000208018 | n.s. |
| ENSG00000127125 | 136.8526 | -0.479087648 | 0.272607 | -1.757432 | 0.078844 | 0.477489649 | PPCS | protein_coding | 42.3 | -0.1538044 | 0.1538044 | ENSG00000127125 | n.s. |
| ENSG00000143811 | 1051.674 | -0.433146574 | 0.236392 | -1.832325 | 0.066903 | 0.441705004 | PYCR2 | protein_coding | 54.97 | -0.15370972 | 0.15370972 | ENSG00000143811 | n.s. |
| ENSG00000182957 | 507.0152 | -0.427573487 | 0.231619 | -1.846025 | 0.064889 | 0.437274745 | SPATA13 | protein_coding | 44.13 | -0.153603896 | 0.153603896 | ENSG00000182957 | n.s. |
| ENSG00000231856 | 190.9373 | 0.431554288 | 0.235169 | 1.8350791 | 0.066494 | 0.441084074 | | antisense | 47.93 | 0.153408324 | 0.153408324 | ENSG00000231856 | n.s. |
| ENSG00000214176 | 463.0541 | -0.466875045 | 0.263114 | -1.774424 | 0.075993 | 0.469606803 | PLEKHM1P1 | cribed_unprocessed_pseudo | 51.2 | -0.153259026 | 0.153259026 | ENSG00000214176 | n.s. |
| ENSG00000234946 | 39.74675 | 0.547878965 | 0.333443 | 1.6430984 | 0.100363 | 0.525174439 | SDHCP3 | processed_pseudogene | 49.01 | 0.153239785 | 0.153239785 | ENSG00000234946 | n.s. |
| ENSG00000001167 | 1634.9 | -0.418553798 | 0.224555 | -1.863928 | 0.062332 | 0.430475522 | NFYA | protein_coding | 40 | -0.153212262 | 0.153212262 | ENSG00000001167 | n.s. |
| ENSG00000089057 | 554.8838 | -0.457265188 | 0.255724 | -1.788117 | 0.073757 | 0.462368337 | SLC23A2 | protein_coding | 44.93 | -0.153189286 | 0.153189286 | ENSG00000089057 | n.s. |
| ENSG00000170836 | 1985.22 | -0.419092636 | 0.225536 | -1.858207 | 0.06314 | 0.431139465 | PPM1D | protein_coding | 41 | -0.153128998 | 0.153128998 | ENSG00000170836 | n.s. |
| ENSG00000224593 | 45.08304 | -0.577495183 | 0.359741 | -1.605306 | 0.108426 | 0.543082569 | | transcribed_processed_pseudog | 41.23 | -0.153113687 | 0.153113687 | ENSG00000224593 | n.s. |
| ENSG00000197024 | 642.2392 | -0.45264767 | 0.252178 | -1.794951 | 0.072662 | 0.459013748 | ZNF398 | protein_coding | 45.92 | -0.153073812 | 0.153073812 | ENSG00000197024 | n.s. |
| ENSG00000197238 | 106.3893 | -0.473740082 | 0.268581 | -1.763863 | 0.077755 | 0.475213657 | HIST1H4J | protein_coding | 62.2 | -0.153070673 | 0.153070673 | ENSG00000197238 | n.s. |
| ENSG00000162337 | 406.1018 | -0.430189405 | 0.234461 | -1.834805 | 0.066535 | 0.441124412 | LRP5 | protein_coding | 54.62 | -0.152906052 | 0.152906052 | ENSG00000162337 | n.s. |
| ENSG00000151718 | 481.398 | -0.456284296 | 0.255167 | -1.788182 | 0.073747 | 0.462368337 | WWC2 | protein_coding | 39.31 | -0.152860675 | 0.152860675 | ENSG00000151718 | n.s. |
| ENSG00000279075 | 183.8112 | 0.42946182 | 0.233769 | 1.8371241 | 0.066192 | 0.440875092 | | TEC | 38.6 | 0.152752885 | 0.152752885 | ENSG00000279075 | n.s. |
| ENSG00000243302 | 71.44435 | -0.49672605 | 0.290183 | -1.71177 | 0.086939 | 0.492652156 | | processed_pseudogene | 58.96 | -0.152723199 | 0.152723199 | ENSG00000243302 | n.s. |
| ENSG00000075213 | 68.15978 | 0.485958096 | 0.280592 | 1.7319055 | 0.08329 | 0.485101957 | SEMA3A | protein_coding | 34.35 | 0.152671984 | 0.152671984 | ENSG00000075213 | n.s. |
| ENSG00000135749 | 751.5847 | -0.477577925 | 0.272746 | -1.751 | 0.079946 | 0.479305998 | PCNX2 | protein_coding | 40.87 | -0.152532246 | 0.152532246 | ENSG00000135749 | n.s. |
| ENSG00000065526 | 1070.185 | -0.446378739 | 0.247362 | -1.804554 | 0.071145 | 0.455380982 | SPEN | protein_coding | 42.99 | -0.152494186 | 0.152494186 | ENSG00000065526 | n.s. |
| ENSG00000129810 | 524.9466 | -0.450654028 | 0.251067 | -1.794953 | 0.072661 | 0.459013748 | SGO1 | protein_coding | 37.06 | -0.152399613 | 0.152399613 | ENSG00000129810 | n.s. |
| ENSG00000099954 | 1427.344 | -0.42833821 | 0.233061 | -1.837881 | 0.06608 | 0.440875092 | CECR2 | protein_coding | 44.03 | -0.152353235 | 0.152353235 | ENSG00000099954 | n.s. |
| ENSG00000077044 | 1130.451 | -0.445927606 | 0.247257 | -1.8035 | 0.07131 | 0.455642836 | DGKD | protein_coding | 47.76 | -0.152228739 | 0.152228739 | ENSG00000077044 | n.s. |
| ENSG00000170871 | 1265.915 | -0.441906572 | 0.243988 | -1.811182 | 0.070113 | 0.452404539 | KIAA0232 | protein_coding | 42.27 | -0.152224903 | 0.152224903 | ENSG00000170871 | n.s. |
| ENSG00000137404 | 184.0038 | -0.433803944 | 0.23829 | -1.820484 | 0.068685 | 0.445830359 | NRM | protein_coding | 55.1 | -0.152191594 | 0.152191594 | ENSG00000137404 | n.s. |
| ENSG00000204682 | 244.1244 | -0.415481777 | 0.22222 | -1.869687 | 0.061527 | 0.430280775 | CASC10 | protein_coding | 57.39 | -0.152169394 | 0.152169394 | ENSG00000204682 | n.s. |
| ENSG00000255067 | 30.77279 | -0.532782781 | 0.321929 | -1.65497 | 0.097931 | 0.518435958 | | sense_intronic | 47.1 | -0.15200553 | 0.15200553 | ENSG00000255067 | n.s. |
| ENSG00000078328 | 224.8246 | -0.42230887 | 0.228506 | -1.848134 | 0.064583 | 0.436711663 | RBFOX1 | protein_coding | 41.37 | -0.151948932 | 0.151948932 | ENSG00000078328 | n.s. |

| | | | | | | | | | | | | | |
|---|---|---|---|---|---|---|---|---|---|---|---|---|---|
| ENSG00000231795 | 27.02546 | -0.56937459 | 0.353775 | -1.609425 | 0.107523 | 0.541309679 | ITCH-IT1 | sense_intronic | 44.02 | -0.151769193 | 0.151769193 | ENSG00000231795 | n.s. |
| ENSG00000137992 | 768.9775 | -0.472265293 | 0.268394 | -1.759595 | 0.078477 | 0.477221369 | DBT | protein_coding | 39.15 | -0.151729449 | 0.151729449 | ENSG00000137992 | n.s. |
| ENSG00000265855 | 2.448314 | 1.005169127 | 0.84486 | 1.1897465 | 0.234146 | 0.706508138 | MIR4514 | miRNA | 45.61 | 0.151662764 | 0.151662764 | ENSG00000265855 | n.s. |
| ENSG00000186594 | 53.82664 | 0.551501009 | 0.338937 | 1.6271476 | 0.103706 | 0.531234393 | MIR22HG | lincRNA | 56.95 | 0.151504946 | 0.151504946 | ENSG00000186594 | n.s. |
| ENSG00000149548 | 194.284 | -0.474543664 | 0.27162 | -1.747086 | 0.080622 | 0.479877462 | CCDC15 | protein_coding | 37.57 | -0.151317571 | 0.151317571 | ENSG00000149548 | n.s. |
| ENSG00000184270 | 411.306 | -0.42611046 | 0.232391 | -1.833594 | 0.066714 | 0.441474604 | HIST2H2AB | protein_coding | 62.85 | -0.151309385 | 0.151309385 | ENSG00000184270 | n.s. |
| ENSG00000157483 | 68.06993 | -0.50201811 | 0.296105 | -1.695404 | 0.089999 | 0.499822278 | MYO1E | protein_coding | 43.52 | -0.151200018 | 0.151200018 | ENSG00000157483 | n.s. |
| ENSG00000148154 | 746.7575 | -0.472980902 | 0.269938 | -1.752184 | 0.079742 | 0.479305998 | UGCG | protein_coding | 40.22 | -0.151064016 | 0.151064016 | ENSG00000148154 | n.s. |
| ENSG00000180176 | 535.2252 | -0.477142989 | 0.273955 | -1.741685 | 0.081564 | 0.482429187 | TH | protein_coding | 65.25 | -0.15104745 | 0.15104745 | ENSG00000180176 | n.s. |
| ENSG00000259781 | 389.5997 | -0.507521115 | 0.300919 | -1.686572 | 0.091686 | 0.504081279 | HMGB1P6 | processed_pseudogene | 43.65 | -0.150987243 | 0.150987243 | ENSG00000259781 | n.s. |
| ENSG00000166619 | 721.5601 | -0.444794971 | 0.247283 | -1.798729 | 0.072062 | 0.45772936 | BLCAP | protein_coding | 47.98 | -0.150959512 | 0.150959512 | ENSG00000166619 | n.s. |
| ENSG00000074410 | 261.4543 | 0.487316412 | 0.283733 | 1.7175161 | 0.085885 | 0.490071988 | CA12 | protein_coding | 48.45 | 0.150941444 | 0.150941444 | ENSG00000074410 | n.s. |
| ENSG00000181896 | 283.2225 | -0.480261142 | 0.27723 | -1.732359 | 0.08321 | 0.485101957 | ZNF101 | protein_coding | 46.28 | -0.15088219 | 0.15088219 | ENSG00000181896 | n.s. |
| ENSG00000227698 | 44.36309 | 0.495659589 | 0.291379 | 1.7010808 | 0.088928 | 0.496448249 | | sense_intronic | 51.92 | 0.150742976 | 0.150742976 | ENSG00000227698 | n.s. |
| ENSG00000178585 | 134.4147 | -0.490033483 | 0.286274 | -1.711764 | 0.08694 | 0.492652156 | CTNNBIP1 | protein_coding | 51.39 | -0.150665505 | 0.150665505 | ENSG00000178585 | n.s. |
| ENSG00000180901 | 781.0757 | -0.442377557 | 0.245569 | -1.801439 | 0.071634 | 0.456874505 | KCTD2 | protein_coding | 49.14 | -0.150498206 | 0.150498206 | ENSG00000180901 | n.s. |
| ENSG00000186017 | 596.8881 | -0.468578511 | 0.266601 | -1.757604 | 0.078815 | 0.477489649 | ZNF566 | protein_coding | 40.96 | -0.150430589 | 0.150430589 | ENSG00000186017 | n.s. |
| ENSG00000207762 | 2.533197 | 0.933881808 | 0.749983 | 1.2452045 | 0.213057 | 0.69012437 | MIR329-2 | miRNA | 41.67 | 0.150422805 | 0.150422805 | ENSG00000207762 | n.s. |
| ENSG00000182372 | 333.2263 | -0.475088284 | 0.273099 | -1.739619 | 0.081926 | 0.482429187 | CLN8 | protein_coding | 48.06 | -0.150397 | 0.150397 | ENSG00000182372 | n.s. |
| ENSG00000272843 | 41.09284 | 0.544922879 | 0.334488 | 1.6291263 | 0.103286 | 0.530060844 | | antisense | 51.27 | 0.15022122 | 0.15022122 | ENSG00000272843 | n.s. |
| ENSG00000163961 | 1364.395 | -0.409909547 | 0.220319 | -1.860529 | 0.062811 | 0.430582973 | RNF168 | protein_coding | 42.36 | -0.15000359 | 0.15000359 | ENSG00000163961 | n.s. |
| ENSG00000278861 | 21.68597 | -0.563141526 | 0.350193 | -1.608091 | 0.107815 | 0.541917403 | | TEC | 37.44 | -0.149833321 | 0.149833321 | ENSG00000278861 | n.s. |
| ENSG00000160993 | 163.7218 | -0.5250385 | 0.31723 | -1.65507 | 0.09791 | 0.518435958 | ALKBH4 | protein_coding | 54.42 | -0.149796048 | 0.149796048 | ENSG00000160993 | n.s. |
| ENSG00000132326 | 435.46 | -0.421603233 | 0.229928 | -1.833633 | 0.066709 | 0.441474604 | PER2 | protein_coding | 52.1 | -0.149708894 | 0.149708894 | ENSG00000132326 | n.s. |
| ENSG00000180357 | 1562.372 | -0.448009468 | 0.251133 | -1.783952 | 0.074431 | 0.463461407 | ZNF609 | protein_coding | 42.67 | -0.14962908 | 0.14962908 | ENSG00000180357 | n.s. |
| ENSG00000174740 | 113.4529 | 0.441690462 | 0.245747 | 1.7973414 | 0.072281 | 0.458693923 | PABPC5 | protein_coding | 41.28 | 0.149502069 | 0.149502069 | ENSG00000174740 | n.s. |
| ENSG00000135870 | 1668.737 | -0.441350742 | 0.24594 | -1.794544 | 0.072726 | 0.459050639 | RC3H1 | protein_coding | 38.58 | -0.149238077 | 0.149238077 | ENSG00000135870 | n.s. |
| ENSG00000172020 | 328.6958 | -0.519440778 | 0.312449 | -1.662483 | 0.096416 | 0.516706636 | GAP43 | protein_coding | 38.61 | -0.148952739 | 0.148952739 | ENSG00000172020 | n.s. |
| ENSG00000010404 | 357.3692 | -0.473200159 | 0.272989 | -1.733402 | 0.083024 | 0.485101957 | IDS | protein_coding | 45.34 | -0.148663862 | 0.148663862 | ENSG00000010404 | n.s. |
| ENSG00000258284 | 7.590023 | -0.726658036 | 0.514865 | -1.411358 | 0.158139 | 0.6244391 | POLR2KP1 | processed_pseudogene | 41.07 | -0.14860877 | 0.14860877 | ENSG00000258284 | n.s. |
| ENSG00000070601 | 304.4219 | -0.434352176 | 0.240307 | -1.807485 | 0.070687 | 0.455002684 | FRMPD1 | protein_coding | 44.15 | -0.148542381 | 0.148542381 | ENSG00000070601 | n.s. |
| ENSG00000168564 | 1695.406 | 0.391671212 | 0.206237 | 1.8991297 | 0.057547 | 0.417869145 | CDKN2AIP | protein_coding | 43.85 | 0.148427603 | 0.148427603 | ENSG00000168564 | n.s. |

| | | | | | | | | | | | | | |
|---|---|---|---|---|---|---|---|---|---|---|---|---|---|
| ENSG00000134909 | 201.1384 | -0.438722688 | 0.244482 | -1.794501 | 0.072733 | 0.459050639 | ARHGAP32 | protein_coding | 36.86 | -0.148349428 | 0.148349428 | ENSG00000134909 | n.s. |
| ENSG00000202566 | 11.99783 | -0.655049563 | 0.441365 | -1.484144 | 0.137771 | 0.593774169 | MIR421 | miRNA | 38.82 | -0.148289268 | 0.148289268 | ENSG00000202566 | n.s. |
| ENSG00000270673 | 38.33686 | 0.531337199 | 0.323857 | 1.6406545 | 0.100869 | 0.525948113 | YTHDF3-AS1 | lincRNA | 55.29 | 0.148273417 | 0.148273417 | ENSG00000270673 | n.s. |
| ENSG00000138363 | 1004.859 | -0.433625791 | 0.240115 | -1.805909 | 0.070933 | 0.455117569 | ATIC | protein_coding | 41.16 | -0.148246424 | 0.148246424 | ENSG00000138363 | n.s. |
| ENSG00000251718 | 9.439856 | -0.709796897 | 0.498692 | -1.423317 | 0.154644 | 0.618354676 | RNU2-13P | snRNA | 38.05 | -0.148178869 | 0.148178869 | ENSG00000251718 | n.s. |
| ENSG00000114480 | 205.1782 | -0.481589993 | 0.281309 | -1.711958 | 0.086904 | 0.492652156 | GBE1 | protein_coding | 35.29 | -0.148069473 | 0.148069473 | ENSG00000114480 | n.s. |
| ENSG00000266144 | 3.401678 | 1.047032897 | 0.912886 | 1.1469485 | 0.251403 | 0.722279621 | MIR4654 | miRNA | 56.58 | 0.147940135 | 0.147940135 | ENSG00000266144 | n.s. |
| ENSG00000068650 | 64.12405 | -0.516727943 | 0.31175 | -1.657506 | 0.097417 | 0.517489317 | ATP11A | protein_coding | 50.33 | -0.147835147 | 0.147835147 | ENSG00000068650 | n.s. |
| ENSG00000267422 | 12.4866 | -0.723031482 | 0.512544 | -1.410672 | 0.158341 | 0.624627103 | | transcribed_processed_pseudog | 43.57 | -0.147772578 | 0.147772578 | ENSG00000267422 | n.s. |
| ENSG00000118473 | 162.7366 | -0.480507243 | 0.280664 | -1.712037 | 0.08689 | 0.492652156 | SGIP1 | protein_coding | 37.83 | -0.147736571 | 0.147736571 | ENSG00000118473 | n.s. |
| ENSG00000101577 | 778.1777 | -0.4408639 | 0.247042 | -1.784571 | 0.074331 | 0.463348535 | LPIN2 | protein_coding | 41.63 | -0.147289193 | 0.147289193 | ENSG00000101577 | n.s. |
| ENSG00000177888 | 1008.47 | -0.459515948 | 0.261917 | -1.754435 | 0.079356 | 0.478143199 | ZBTB41 | protein_coding | 33.4 | -0.147248217 | 0.147248217 | ENSG00000177888 | n.s. |
| ENSG00000048471 | 148.0752 | -0.460511259 | 0.262869 | -1.751864 | 0.079797 | 0.479305998 | SNX29 | protein_coding | 47.13 | -0.147081372 | 0.147081372 | ENSG00000048471 | n.s. |
| ENSG00000278040 | 7.917117 | 0.714627727 | 0.504983 | 1.4151521 | 0.157024 | 0.622581912 | | misc_RNA | 50.6 | 0.147072888 | 0.147072888 | ENSG00000278040 | n.s. |
| ENSG00000257135 | 60.43327 | 0.482803281 | 0.283763 | 1.7014311 | 0.088862 | 0.496280219 | | lincRNA | 54.18 | 0.14690402 | 0.14690402 | ENSG00000257135 | n.s. |
| ENSG00000167977 | 490.3779 | -0.479999751 | 0.281216 | -1.706871 | 0.087846 | 0.494318319 | KCTD5 | protein_coding | 59.23 | -0.146876705 | 0.146876705 | ENSG00000167977 | n.s. |
| ENSG00000101849 | 85.74761 | -0.478736989 | 0.280224 | -1.708409 | 0.08756 | 0.493646579 | TBL1X | protein_coding | 45.23 | -0.146773038 | 0.146773038 | ENSG00000101849 | n.s. |
| ENSG00000207650 | 10.92384 | -0.667374559 | 0.455312 | -1.465752 | 0.142716 | 0.602678881 | MIR570 | miRNA | 37.11 | -0.146765027 | 0.146765027 | ENSG00000207650 | n.s. |
| ENSG00000115970 | 1830.148 | -0.448007046 | 0.252882 | -1.771603 | 0.076461 | 0.470627671 | THADA | protein_coding | 40.47 | -0.146642805 | 0.146642805 | ENSG00000115970 | n.s. |
| ENSG00000274290 | 807.2543 | -0.393277616 | 0.208853 | -1.88304 | 0.059695 | 0.424480483 | HIST1H2BE | protein_coding | 41.16 | -0.146355226 | 0.146355226 | ENSG00000274290 | n.s. |
| ENSG00000160789 | 53.66401 | -0.528445839 | 0.323747 | -1.632283 | 0.10262 | 0.528723147 | LMNA | protein_coding | 53.04 | -0.146258841 | 0.146258841 | ENSG00000160789 | n.s. |
| ENSG00000201470 | 32.01294 | -0.565385145 | 0.356557 | -1.585678 | 0.112812 | 0.551349488 | RNY4P7 | misc_RNA | 41.67 | -0.146193347 | 0.146193347 | ENSG00000201470 | n.s. |
| ENSG00000122359 | 279.2806 | -0.446352094 | 0.251944 | -1.771629 | 0.076456 | 0.470627671 | ANXA11 | protein_coding | 50.39 | -0.146101102 | 0.146101102 | ENSG00000122359 | n.s. |
| ENSG00000109762 | 436.9294 | -0.457092548 | 0.261233 | -1.749747 | 0.080162 | 0.479305998 | SNX25 | protein_coding | 41.36 | -0.14598948 | 0.14598948 | ENSG00000109762 | n.s. |
| ENSG00000213066 | 419.3565 | -0.478605647 | 0.28095 | -1.703527 | 0.08847 | 0.495479338 | FGFR1OP | protein_coding | 41.03 | -0.145962495 | 0.145962495 | ENSG00000213066 | n.s. |
| ENSG00000119004 | 521.7407 | -0.460032623 | 0.263946 | -1.742902 | 0.081351 | 0.48187821 | CYP20A1 | protein_coding | 42.38 | -0.145859191 | 0.145859191 | ENSG00000119004 | n.s. |
| ENSG00000103723 | 1045.889 | -0.463920488 | 0.267349 | -1.735265 | 0.082694 | 0.484982269 | AP3B2 | protein_coding | 48.94 | -0.145798212 | 0.145798212 | ENSG00000103723 | n.s. |
| ENSG00000261455 | 237.3571 | 0.442200837 | 0.248864 | 1.776874 | 0.075589 | 0.468160757 | LINC01003 | lincRNA | 60.83 | 0.145751604 | 0.145751604 | ENSG00000261455 | n.s. |
| ENSG00000152767 | 1574.83 | -0.41695639 | 0.229388 | -1.81769 | 0.069112 | 0.447345712 | FARP1 | protein_coding | 44.52 | -0.145666517 | 0.145666517 | ENSG00000152767 | n.s. |
| ENSG00000222675 | 3.429057 | 1.102932452 | 1.003821 | 1.0987343 | 0.271884 | 0.737902688 | RNA5SP146 | rRNA | 55.28 | 0.145588085 | 0.145588085 | ENSG00000222675 | n.s. |
| ENSG00000231466 | 72.6334 | -0.497227188 | 0.29695 | -1.674446 | 0.094043 | 0.509648797 | | processed_pseudogene | 41.79 | -0.145552816 | 0.145552816 | ENSG00000231466 | n.s. |
| ENSG00000206859 | 6.982533 | -0.732595 | 0.530194 | -1.38175 | 0.167049 | 0.633047814 | RNU6-767P | snRNA | 33.64 | -0.145466617 | 0.145466617 | ENSG00000206859 | n.s. |

| | | | | | | | | | | | | | |
|---|---|---|---|---|---|---|---|---|---|---|---|---|---|
| ENSG00000244112 | 58.47883 | -0.507265341 | 0.305012 | -1.663097 | 0.096293 | 0.516706636 | RN7SL508P | misc_RNA | 53.47 | -0.14546136 | 0.14546136 | ENSG00000244112 | n.s. |
| ENSG00000253540 | 89.36168 | -0.445716458 | 0.251794 | -1.770161 | 0.0767 | 0.471686734 | FAM86HP | cribed_unprocessed_pseudc | 53.04 | -0.145457934 | 0.145457934 | ENSG00000253540 | n.s. |
| ENSG00000273271 | 31.44045 | -0.819932549 | 0.626128 | -1.309529 | 0.190355 | 0.665072813 | | antisense | 35.77 | -0.145235313 | 0.145235313 | ENSG00000273271 | n.s. |
| ENSG00000238735 | 17.80344 | -0.754708549 | 0.555522 | -1.358558 | 0.174287 | 0.642125589 | RNU7-113P | snRNA | 37.1 | -0.145190848 | 0.145190848 | ENSG00000238735 | n.s. |
| ENSG00000277258 | 212.972 | -0.442502067 | 0.249599 | -1.772849 | 0.076254 | 0.469832545 | PCGF2 | protein_coding | 57.06 | -0.145165858 | 0.145165858 | ENSG00000277258 | n.s. |
| ENSG00000144619 | 53.99877 | -0.505853646 | 0.30429 | -1.662404 | 0.096432 | 0.516706636 | CNTN4 | protein_coding | 37.97 | -0.145056548 | 0.145056548 | ENSG00000144619 | n.s. |
| ENSG00000183654 | 243.2446 | -0.454059422 | 0.259383 | -1.750539 | 0.080025 | 0.479305998 | 11-Mar | protein_coding | 39.41 | -0.145020739 | 0.145020739 | ENSG00000183654 | n.s. |
| ENSG00000085999 | 291.8068 | -0.407624519 | 0.221899 | -1.836984 | 0.066212 | 0.440875092 | RAD54L | protein_coding | 46.5 | -0.144985697 | 0.144985697 | ENSG00000085999 | n.s. |
| ENSG00000261801 | 126.8806 | -0.470323914 | 0.274252 | -1.714936 | 0.086357 | 0.491757974 | LOXL1-AS1 | antisense | 50.5 | -0.144976683 | 0.144976683 | ENSG00000261801 | n.s. |
| ENSG00000162545 | 90.35093 | -0.476544605 | 0.280239 | -1.700491 | 0.089039 | 0.496668574 | CAMK2N1 | protein_coding | 57.6 | -0.144837783 | 0.144837783 | ENSG00000162545 | n.s. |
| ENSG00000258640 | 22.08898 | -0.574336269 | 0.36577 | -1.57021 | 0.116366 | 0.559643895 | RPL21P5 | processed_pseudogene | 45.75 | -0.144783413 | 0.144783413 | ENSG00000258640 | n.s. |
| ENSG00000111412 | 968.6223 | -0.468919396 | 0.273353 | -1.715437 | 0.086265 | 0.491436255 | C12orf49 | protein_coding | 44.28 | -0.144677018 | 0.144677018 | ENSG00000111412 | n.s. |
| ENSG00000237631 | 52.78592 | -0.506331212 | 0.305864 | -1.655411 | 0.097841 | 0.518435958 | | processed_pseudogene | 44.82 | -0.144458768 | 0.144458768 | ENSG00000237631 | n.s. |
| ENSG00000180953 | 119.2775 | -0.544603397 | 0.339137 | -1.605849 | 0.108307 | 0.543082569 | ST20 | protein_coding | 41.13 | -0.144392951 | 0.144392951 | ENSG00000180953 | n.s. |
| ENSG00000258978 | 14.29229 | 0.607887566 | 0.39973 | 1.5207451 | 0.128324 | 0.579048024 | HIF1AP1 | processed_pseudogene | 36 | 0.144242854 | 0.144242854 | ENSG00000258978 | n.s. |
| ENSG00000139977 | 533.3837 | -0.471142371 | 0.276049 | -1.706733 | 0.087872 | 0.494318319 | NAA30 | protein_coding | 37.28 | -0.144166406 | 0.144166406 | ENSG00000139977 | n.s. |
| ENSG00000165195 | 248.1442 | 0.472297477 | 0.277057 | 1.7046926 | 0.088252 | 0.495195739 | PIGA | protein_coding | 39.56 | 0.1441561 | 0.1441561 | ENSG00000165195 | n.s. |
| ENSG00000100211 | 154.4622 | -0.484396975 | 0.287251 | -1.686321 | 0.091734 | 0.504081279 | CBY1 | protein_coding | 47.17 | -0.144107825 | 0.144107825 | ENSG00000100211 | n.s. |
| ENSG00000265735 | 21.66491 | -0.583738701 | 0.374788 | -1.557516 | 0.119348 | 0.56654653 | RN7SL5P | misc_RNA | 57.63 | -0.14404594 | 0.14404594 | ENSG00000265735 | n.s. |
| ENSG00000162994 | 75.82173 | -0.502236023 | 0.302399 | -1.660841 | 0.096745 | 0.516736278 | CLHC1 | protein_coding | 39.9 | -0.144006661 | 0.144006661 | ENSG00000162994 | n.s. |
| ENSG00000101353 | 86.88191 | -0.472781512 | 0.277729 | -1.702315 | 0.088696 | 0.495951531 | MROH8 | protein_coding | 45.33 | -0.143990699 | 0.143990699 | ENSG00000101353 | n.s. |
| ENSG00000265291 | 3.393087 | 1.037082109 | 0.91385 | 1.1348489 | 0.256439 | 0.726402587 | MIR4710 | miRNA | 64.29 | 0.143970453 | 0.143970453 | ENSG00000265291 | n.s. |
| ENSG00000252779 | 10.17901 | -0.689626389 | 0.484747 | -1.422652 | 0.154837 | 0.618415021 | RNU6-182P | snRNA | 33.33 | -0.143938799 | 0.143938799 | ENSG00000252779 | n.s. |
| ENSG00000204070 | 301.9134 | -0.427360836 | 0.238632 | -1.790882 | 0.073312 | 0.460477385 | SYS1 | protein_coding | 49.54 | -0.14393158 | 0.14393158 | ENSG00000204070 | n.s. |
| ENSG00000087299 | 368.822 | -0.447665606 | 0.25427 | -1.760595 | 0.078307 | 0.477221369 | L2HGDH | protein_coding | 40.78 | -0.143826059 | 0.143826059 | ENSG00000087299 | n.s. |
| ENSG00000227036 | 53.80252 | -0.584651868 | 0.37635 | -1.553481 | 0.120308 | 0.567746164 | LINC00511 | lincRNA | 47.33 | -0.143734201 | 0.143734201 | ENSG00000227036 | n.s. |
| ENSG00000274677 | 23.14585 | -0.586088256 | 0.378332 | -1.549136 | 0.121349 | 0.568737594 | | sense_intronic | 52.12 | -0.143643236 | 0.143643236 | ENSG00000274677 | n.s. |
| ENSG00000274270 | 89.69732 | 0.47088059 | 0.276368 | 1.7038167 | 0.088415 | 0.495479338 | | sense_intronic | 58.61 | 0.14360655 | 0.14360655 | ENSG00000274270 | n.s. |
| ENSG00000247572 | 82.82118 | -0.470087647 | 0.275674 | -1.705233 | 0.088151 | 0.495072019 | CKMT2-AS1 | antisense | 41.97 | -0.143532622 | 0.143532622 | ENSG00000247572 | n.s. |
| ENSG00000144579 | 387.2694 | -0.444059937 | 0.251758 | -1.763835 | 0.07776 | 0.475213657 | CTDSP1 | protein_coding | 63.19 | -0.143480689 | 0.143480689 | ENSG00000144579 | n.s. |
| ENSG00000244968 | 113.3441 | 0.446554653 | 0.253662 | 1.7604302 | 0.078335 | 0.477221369 | LIFR-AS1 | antisense | 41.55 | 0.143469132 | 0.143469132 | ENSG00000244968 | n.s. |
| ENSG00000223658 | 90.08568 | 0.499597644 | 0.300146 | 1.6645173 | 0.096009 | 0.516512586 | C1GALT1C1L | protein_coding | 38.99 | 0.143344102 | 0.143344102 | ENSG00000223658 | n.s. |

| | | | | | | | | | | | | | |
|---|---|---|---|---|---|---|---|---|---|---|---|---|---|
| ENSG00000279031 | 24.30495 | -0.55342291 | 0.348701 | -1.587098 | 0.11249 | 0.55089864 | | TEC | 47.62 | -0.143296853 | 0.143296853 | ENSG00000279031 | n.s. |
| ENSG00000275651 | 2.388182 | -1.203810744 | 1.170098 | -1.028812 | 0.303568 | 0.760781413 | MIR6851 | miRNA | 59.7 | -0.142940616 | 0.142940616 | ENSG00000275651 | n.s. |
| ENSG00000236144 | 292.5426 | -0.405077534 | 0.221984 | -1.824807 | 0.06803 | 0.443960682 | TMEM147-AS1 | antisense | 56.8 | -0.142852816 | 0.142852816 | ENSG00000236144 | n.s. |
| ENSG00000256628 | 107.5596 | -0.434997569 | 0.24527 | -1.773544 | 0.076139 | 0.469606803 | ZBTB11-AS1 | antisense | 52.17 | -0.142794747 | 0.142794747 | ENSG00000256628 | n.s. |
| ENSG00000095739 | 1233.457 | -0.414345802 | 0.228804 | -1.810921 | 0.070153 | 0.452404539 | BAMBI | protein_coding | 47.11 | -0.142730961 | 0.142730961 | ENSG00000095739 | n.s. |
| ENSG00000130222 | 805.6213 | 0.430710618 | 0.24201 | 1.7797251 | 0.075121 | 0.46629918 | GADD45G | protein_coding | 66.23 | 0.142709653 | 0.142709653 | ENSG00000130222 | n.s. |
| ENSG00000225930 | 270.8121 | -0.455330451 | 0.263227 | -1.729798 | 0.083666 | 0.485981036 | LINC02249 | lincRNA | 40.51 | -0.142691765 | 0.142691765 | ENSG00000225930 | n.s. |
| ENSG00000204713 | 2017.161 | -0.447174333 | 0.255924 | -1.747291 | 0.080587 | 0.479877462 | TRIM27 | protein_coding | 44.66 | -0.142590322 | 0.142590322 | ENSG00000204713 | n.s. |
| ENSG00000243854 | 16.19856 | -0.594905901 | 0.388703 | -1.53049 | 0.125896 | 0.575897021 | RN7SL67P | misc_RNA | 53.38 | -0.142572274 | 0.142572274 | ENSG00000243854 | n.s. |
| ENSG00000185088 | 2372.55 | -0.383042842 | 0.203395 | -1.883246 | 0.059667 | 0.424480483 | RPS27L | protein_coding | 39.85 | -0.142546434 | 0.142546434 | ENSG00000185088 | n.s. |
| ENSG00000166024 | 605.3448 | -0.450241477 | 0.258791 | -1.73979 | 0.081896 | 0.482429187 | R3HCC1L | protein_coding | 37.83 | -0.142531335 | 0.142531335 | ENSG00000166024 | n.s. |
| ENSG00000235301 | 62.41909 | -0.481414439 | 0.28589 | -1.683913 | 0.092199 | 0.505784153 | HLA-Z | unprocessed_pseudogene | 53.33 | -0.142515418 | 0.142515418 | ENSG00000235301 | n.s. |
| ENSG00000240695 | 146.4979 | 0.461429544 | 0.268905 | 1.715959 | 0.08617 | 0.491092308 | | processed_pseudogene | 43.42 | 0.142506455 | 0.142506455 | ENSG00000240695 | n.s. |
| ENSG00000171848 | 3847.124 | -0.389125852 | 0.209121 | -1.860766 | 0.062777 | 0.430564731 | RRM2 | protein_coding | 45.64 | -0.142405099 | 0.142405099 | ENSG00000171848 | n.s. |
| ENSG00000238959 | 11.95918 | -0.657006705 | 0.452458 | -1.452084 | 0.146478 | 0.607109641 | RNU7-19P | snRNA | 42.42 | -0.142394949 | 0.142394949 | ENSG00000238959 | n.s. |
| ENSG00000196757 | 582.7366 | -0.48027557 | 0.285084 | -1.684679 | 0.09205 | 0.505369926 | ZNF700 | protein_coding | 44.6 | -0.142349167 | 0.142349167 | ENSG00000196757 | n.s. |
| ENSG00000143376 | 726.0237 | -0.425066432 | 0.237843 | -1.78717 | 0.07391 | 0.462574432 | SNX27 | protein_coding | 40.41 | -0.142320052 | 0.142320052 | ENSG00000143376 | n.s. |
| ENSG00000132182 | 758.5574 | -0.417720245 | 0.231766 | -1.802334 | 0.071493 | 0.456370327 | NUP210 | protein_coding | 52.6 | -0.142310019 | 0.142310019 | ENSG00000132182 | n.s. |
| ENSG00000122965 | 1271.642 | -0.415856068 | 0.230313 | -1.805613 | 0.070979 | 0.455199058 | RBM19 | protein_coding | 50.43 | -0.14213904 | 0.14213904 | ENSG00000122965 | n.s. |
| ENSG00000076053 | 96.38651 | 0.471396537 | 0.278002 | 1.6956612 | 0.08995 | 0.499750469 | RBM7 | protein_coding | 38.27 | 0.142006694 | 0.142006694 | ENSG00000076053 | n.s. |
| ENSG00000223960 | 425.5806 | -0.427272405 | 0.239968 | -1.780539 | 0.074988 | 0.465679963 | | antisense | 39.56 | -0.14181703 | 0.14181703 | ENSG00000223960 | n.s. |
| ENSG00000185475 | 149.9077 | -0.512383494 | 0.313762 | -1.633033 | 0.102462 | 0.528723147 | TMEM179B | protein_coding | 52.76 | -0.141813239 | 0.141813239 | ENSG00000185475 | n.s. |
| ENSG00000197782 | 735.1619 | -0.441666625 | 0.251378 | -1.756983 | 0.078921 | 0.477537429 | ZNF780A | protein_coding | 38.52 | -0.141771711 | 0.141771711 | ENSG00000197782 | n.s. |
| ENSG00000038532 | 428.9989 | -0.447313958 | 0.256925 | -1.741031 | 0.081678 | 0.482429187 | CLEC16A | protein_coding | 48.06 | -0.141604581 | 0.141604581 | ENSG00000038532 | n.s. |
| ENSG00000174501 | 1646.966 | -0.463329404 | 0.271586 | -1.706012 | 0.088006 | 0.494760783 | ANKRD36C | protein_coding | 35.73 | -0.141595658 | 0.141595658 | ENSG00000174501 | n.s. |
| ENSG00000207632 | 4.387543 | -0.854409737 | 0.677825 | -1.260517 | 0.207483 | 0.682785213 | MIR588 | miRNA | 44.58 | -0.141589272 | 0.141589272 | ENSG00000207632 | n.s. |
| ENSG00000260727 | 174.7152 | -0.444020491 | 0.254188 | -1.74682 | 0.080668 | 0.479877462 | SLC7A5P1 | unprocessed_pseudogene | 69.27 | -0.141584657 | 0.141584657 | ENSG00000260727 | n.s. |
| ENSG00000013561 | 683.4934 | -0.476663285 | 0.28284 | -1.685277 | 0.091935 | 0.504935967 | RNF14 | protein_coding | 42.04 | -0.141456357 | 0.141456357 | ENSG00000013561 | n.s. |
| ENSG00000168569 | 246.1656 | -0.482390205 | 0.288276 | -1.673365 | 0.094256 | 0.510034879 | TMEM223 | protein_coding | 52.1 | -0.141050955 | 0.141050955 | ENSG00000168569 | n.s. |
| ENSG00000185753 | 167.8321 | -0.470786103 | 0.278359 | -1.691293 | 0.090781 | 0.501961137 | CXorf38 | protein_coding | 45.19 | -0.14092036 | 0.14092036 | ENSG00000185753 | n.s. |
| ENSG00000275401 | 109.9728 | -0.431206243 | 0.24367 | -1.769634 | 0.076788 | 0.471809602 | | lincRNA | 50.69 | -0.140673803 | 0.140673803 | ENSG00000275401 | n.s. |
| ENSG00000152439 | 219.1255 | -0.4511193 | 0.261741 | -1.723536 | 0.084792 | 0.48779901 | ZNF773 | protein_coding | 43.76 | -0.140640541 | 0.140640541 | ENSG00000152439 | n.s. |

| | | | | | | | | | | | | | |
|---|---|---|---|---|---|---|---|---|---|---|---|---|---|
| ENSG00000273802 | 2208.785 | -0.391252559 | 0.211806 | -1.847223 | 0.064715 | 0.437178558 | HIST1H2BG | protein_coding | 48.76 | -0.140593143 | 0.140593143 | ENSG00000273802 | n.s. |
| ENSG00000136856 | 31.19025 | -0.570468832 | 0.366506 | -1.556508 | 0.119587 | 0.566955067 | SLC2A8 | protein_coding | 57.79 | -0.140592819 | 0.140592819 | ENSG00000136856 | n.s. |
| ENSG00000263712 | 40.98832 | 0.536960992 | 0.336016 | 1.5980232 | 0.110038 | 0.54732402 | MIR4639 | miRNA | 46.38 | 0.140552489 | 0.140552489 | ENSG00000263712 | n.s. |
| ENSG00000197081 | 1221.331 | -0.411178863 | 0.227831 | -1.804755 | 0.071113 | 0.455380982 | IGF2R | protein_coding | 45.55 | -0.140469025 | 0.140469025 | ENSG00000197081 | n.s. |
| ENSG00000238713 | 38.09146 | -0.559963946 | 0.357359 | -1.566952 | 0.117126 | 0.561405206 | | misc_RNA | 43.36 | -0.140396156 | 0.140396156 | ENSG00000238713 | n.s. |
| ENSG00000081019 | 1543.292 | -0.437830431 | 0.249396 | -1.75556 | 0.079164 | 0.478143199 | RSBN1 | protein_coding | 37.22 | -0.140299266 | 0.140299266 | ENSG00000081019 | n.s. |
| ENSG00000207761 | 3.847069 | 0.808798896 | 0.624574 | 1.2949606 | 0.195334 | 0.670750289 | MIR329-1 | miRNA | 42.5 | 0.140277378 | 0.140277378 | ENSG00000207761 | n.s. |
| ENSG00000166855 | 1204.341 | -0.450915451 | 0.261933 | -1.721489 | 0.085162 | 0.488752167 | CLPX | protein_coding | 40 | -0.140194711 | 0.140194711 | ENSG00000166855 | n.s. |
| ENSG00000104957 | 281.3941 | -0.457142306 | 0.267683 | -1.707772 | 0.087679 | 0.493830606 | CCDC130 | protein_coding | 50.36 | -0.140078458 | 0.140078458 | ENSG00000104957 | n.s. |
| ENSG00000168710 | 2152.347 | -0.455491484 | 0.265901 | -1.713014 | 0.08671 | 0.492652156 | AHCYL1 | protein_coding | 40.97 | -0.140045235 | 0.140045235 | ENSG00000168710 | n.s. |
| ENSG00000277383 | 26.45763 | -0.595775372 | 0.395096 | -1.507927 | 0.131573 | 0.582107445 | | antisense | 65.46 | -0.140005333 | 0.140005333 | ENSG00000277383 | n.s. |
| ENSG00000181220 | 413.6609 | -0.429466075 | 0.242762 | -1.769082 | 0.07688 | 0.472167535 | ZNF746 | protein_coding | 46.99 | -0.139964659 | 0.139964659 | ENSG00000181220 | n.s. |
| ENSG00000242802 | 240.2778 | -0.45611492 | 0.266804 | -1.70955 | 0.087349 | 0.49357262 | AP5Z1 | protein_coding | 58.29 | -0.139867157 | 0.139867157 | ENSG00000242802 | n.s. |
| ENSG00000006756 | 219.9228 | -0.49736084 | 0.302329 | -1.645099 | 0.099949 | 0.524173499 | ARSD | protein_coding | 48.88 | -0.13952212 | 0.13952212 | ENSG00000006756 | n.s. |
| ENSG00000135119 | 242.1246 | -0.411007607 | 0.228501 | -1.798716 | 0.072064 | 0.45772936 | RNFT2 | protein_coding | 47.15 | -0.139492377 | 0.139492377 | ENSG00000135119 | n.s. |
| ENSG00000126216 | 1152.973 | -0.414699168 | 0.231656 | -1.790152 | 0.073429 | 0.460936835 | TUBGCP3 | protein_coding | 42.87 | -0.139487625 | 0.139487625 | ENSG00000126216 | n.s. |
| ENSG00000225032 | 108.6469 | -0.475952314 | 0.284132 | -1.675112 | 0.093912 | 0.509365634 | | antisense | 55.15 | -0.139439921 | 0.139439921 | ENSG00000225032 | n.s. |
| ENSG00000254087 | 958.63 | 0.472187813 | 0.280827 | 1.6814189 | 0.092682 | 0.506647149 | LYN | protein_coding | 43.2 | 0.139434416 | 0.139434416 | ENSG00000254087 | n.s. |
| ENSG00000262803 | 43.46966 | -0.532632645 | 0.333485 | -1.597172 | 0.110227 | 0.54732402 | | processed_pseudogene | 38.8 | -0.13941952 | 0.13941952 | ENSG00000262803 | n.s. |
| ENSG00000179348 | 2091.298 | -0.393311468 | 0.214959 | -1.829703 | 0.067294 | 0.442113119 | GATA2 | protein_coding | 60.45 | -0.139415778 | 0.139415778 | ENSG00000179348 | n.s. |
| ENSG00000124942 | 4958.069 | -0.413214182 | 0.230574 | -1.792109 | 0.073116 | 0.460211177 | AHNAK | protein_coding | 50.05 | -0.139270881 | 0.139270881 | ENSG00000124942 | n.s. |
| ENSG00000175322 | 873.6065 | -0.445044328 | 0.257672 | -1.727172 | 0.084137 | 0.48719542 | ZNF519 | protein_coding | 39.05 | -0.138985921 | 0.138985921 | ENSG00000175322 | n.s. |
| ENSG00000121753 | 325.1523 | -0.453085345 | 0.265221 | -1.708331 | 0.087575 | 0.493646579 | ADGRB2 | protein_coding | 58.75 | -0.138908658 | 0.138908658 | ENSG00000121753 | n.s. |
| ENSG00000141568 | 2375.443 | -0.431238494 | 0.244695 | -1.762354 | 0.07801 | 0.4763713 | FOXK2 | protein_coding | 49.72 | -0.13888226 | 0.13888226 | ENSG00000141568 | n.s. |
| ENSG00000222679 | 33.32015 | 0.544866777 | 0.34576 | 1.5758504 | 0.11506 | 0.556094484 | RNU6-1267P | snRNA | 41.12 | 0.138860068 | 0.138860068 | ENSG00000222679 | n.s. |
| ENSG00000005469 | 286.891 | -0.484133621 | 0.291068 | -1.663304 | 0.096252 | 0.516706636 | CROT | protein_coding | 37.43 | -0.138828202 | 0.138828202 | ENSG00000005469 | n.s. |
| ENSG00000228701 | 63.08969 | 0.487244734 | 0.29462 | 1.6538074 | 0.098167 | 0.519093939 | TNKS2-AS1 | antisense | 38.44 | 0.138744907 | 0.138744907 | ENSG00000228701 | n.s. |
| ENSG00000261596 | 94.14914 | -0.497253317 | 0.303278 | -1.639598 | 0.101089 | 0.526152672 | | sense_intronic | 39.37 | -0.138678092 | 0.138678092 | ENSG00000261596 | n.s. |
| ENSG00000171467 | 606.0734 | -0.431954535 | 0.245895 | -1.756666 | 0.078975 | 0.477656661 | ZNF318 | protein_coding | 42.92 | -0.138607369 | 0.138607369 | ENSG00000171467 | n.s. |
| ENSG00000169967 | 2581.441 | -0.440429077 | 0.253627 | -1.736522 | 0.082472 | 0.484641218 | MAP3K2 | protein_coding | 38.69 | -0.138550026 | 0.138550026 | ENSG00000169967 | n.s. |
| ENSG00000177576 | 70.89832 | -0.522188647 | 0.325827 | -1.602657 | 0.10901 | 0.544064048 | C18orf32 | protein_coding | 43.19 | -0.138040554 | 0.138040554 | ENSG00000177576 | n.s. |
| ENSG00000178691 | 1429.447 | -0.442691391 | 0.256782 | -1.723996 | 0.084709 | 0.48779901 | SUZ12 | protein_coding | 41 | -0.138013063 | 0.138013063 | ENSG00000178691 | n.s. |

| | | | | | | | | | | | | | |
|---|---|---|---|---|---|---|---|---|---|---|---|---|---|
| ENSG00000252212 | 14.5366 | -0.60112932 | 0.402928 | -1.491904 | 0.135724 | 0.589422885 | RNU2-58P | snRNA | 43.72 | -0.138003065 | 0.138003065 | ENSG00000252212 | n.s. |
| ENSG00000275441 | 28.7058 | -0.652294444 | 0.455608 | -1.4317 | 0.15223 | 0.614628915 | | antisense | 53.24 | -0.137886574 | 0.137886574 | ENSG00000275441 | n.s. |
| ENSG00000228170 | 27.21567 | -0.580717758 | 0.38246 | -1.518375 | 0.12892 | 0.579048024 | | antisense | 48.47 | -0.137795855 | 0.137795855 | ENSG00000228170 | n.s. |
| ENSG00000182199 | 3104.112 | -0.465893874 | 0.276848 | -1.682851 | 0.092404 | 0.506113336 | SHMT2 | protein_coding | 56.68 | -0.137789147 | 0.137789147 | ENSG00000182199 | n.s. |
| ENSG00000202142 | 4.062152 | 0.785349314 | 0.604086 | 1.3000623 | 0.19358 | 0.668123216 | SNORD114-18 | snoRNA | 37.5 | 0.137548778 | 0.137548778 | ENSG00000202142 | n.s. |
| ENSG00000206941 | 10.38895 | -0.714106517 | 0.524854 | -1.36058 | 0.173646 | 0.642002364 | SNORD15A | snoRNA | 47.3 | -0.137439349 | 0.137439349 | ENSG00000206941 | n.s. |
| ENSG00000189067 | 337.1998 | -0.465502054 | 0.277037 | -1.680289 | 0.092901 | 0.507041328 | LITAF | protein_coding | 49.14 | -0.137302922 | 0.137302922 | ENSG00000189067 | n.s. |
| ENSG00000226642 | 185.1782 | -0.48349374 | 0.292702 | -1.651832 | 0.098569 | 0.520035603 | ACTG1P12 | processed_pseudogene | 56.43 | -0.137296229 | 0.137296229 | ENSG00000226642 | n.s. |
| ENSG00000205683 | 81.52747 | -0.481490674 | 0.291078 | -1.654164 | 0.098094 | 0.518907679 | DPF3 | protein_coding | 45.75 | -0.137181461 | 0.137181461 | ENSG00000205683 | n.s. |
| ENSG00000187186 | 44.10178 | -0.577813099 | 0.380686 | -1.517822 | 0.129059 | 0.579048024 | | protein_coding | 61.23 | -0.137106622 | 0.137106622 | ENSG00000187186 | n.s. |
| ENSG00000118412 | 2693.776 | -0.442611163 | 0.257601 | -1.718202 | 0.08576 | 0.490059299 | CASP8AP2 | protein_coding | 42.57 | -0.137099412 | 0.137099412 | ENSG00000118412 | n.s. |
| ENSG00000265828 | 40.59648 | -0.519603646 | 0.324425 | -1.601615 | 0.109241 | 0.544920997 | MIR3939 | miRNA | 54.72 | -0.137002052 | 0.137002052 | ENSG00000265828 | n.s. |
| ENSG00000125534 | 164.3974 | -0.426793334 | 0.242888 | -1.757158 | 0.078891 | 0.477537429 | PPDPF | protein_coding | 71.07 | -0.136997495 | 0.136997495 | ENSG00000125534 | n.s. |
| ENSG00000159658 | 331.2835 | -0.430394187 | 0.246575 | -1.74549 | 0.0809 | 0.480530903 | EFCAB14 | protein_coding | 40.17 | -0.136985293 | 0.136985293 | ENSG00000159658 | n.s. |
| ENSG00000276730 | 4.282702 | -0.819408783 | 0.646304 | -1.267837 | 0.204856 | 0.680575501 | | misc_RNA | 46.75 | -0.136942618 | 0.136942618 | ENSG00000276730 | n.s. |
| ENSG00000061273 | 48.53006 | -0.504462697 | 0.311358 | -1.620201 | 0.105189 | 0.535288187 | HDAC7 | protein_coding | 54.88 | -0.136917401 | 0.136917401 | ENSG00000061273 | n.s. |
| ENSG00000239268 | 33.97267 | -0.522926827 | 0.327301 | -1.597695 | 0.110111 | 0.54732402 | | lincRNA | 36.82 | -0.136878969 | 0.136878969 | ENSG00000239268 | n.s. |
| ENSG00000065328 | 2118.994 | -0.446345806 | 0.261168 | -1.709038 | 0.087444 | 0.493646579 | MCM10 | protein_coding | 43.59 | -0.136842424 | 0.136842424 | ENSG00000065328 | n.s. |
| ENSG00000099617 | 32.01809 | -0.530777439 | 0.335352 | -1.582747 | 0.113479 | 0.552437472 | EFNA2 | protein_coding | 64.19 | -0.1367903 | 0.1367903 | ENSG00000099617 | n.s. |
| ENSG00000171161 | 329.6759 | -0.465564965 | 0.277711 | -1.67644 | 0.093652 | 0.508556034 | ZNF672 | protein_coding | 52.37 | -0.136718361 | 0.136718361 | ENSG00000171161 | n.s. |
| ENSG00000178171 | 2618.344 | -0.426352716 | 0.242834 | -1.755734 | 0.079134 | 0.478143199 | AMER3 | protein_coding | 55.57 | -0.136621324 | 0.136621324 | ENSG00000178171 | n.s. |
| ENSG00000161692 | 405.8437 | -0.444288509 | 0.259635 | -1.711206 | 0.087043 | 0.492652156 | DBF4B | protein_coding | 47.89 | -0.136600773 | 0.136600773 | ENSG00000161692 | n.s. |
| ENSG00000255256 | 29.25539 | -0.555992044 | 0.359177 | -1.547963 | 0.121631 | 0.568759707 | | sense_intronic | 43.76 | -0.136257625 | 0.136257625 | ENSG00000255256 | n.s. |
| ENSG00000149294 | 2454.231 | -0.439839565 | 0.255931 | -1.718583 | 0.08569 | 0.490059299 | NCAM1 | protein_coding | 41.71 | -0.136240906 | 0.136240906 | ENSG00000149294 | n.s. |
| ENSG00000202119 | 8.541143 | -0.676455385 | 0.482962 | -1.400639 | 0.161322 | 0.628931954 | RNU6-302P | snRNA | 45.79 | -0.136235639 | 0.136235639 | ENSG00000202119 | n.s. |
| ENSG00000206848 | 3.977707 | -0.878695226 | 0.722249 | -1.21661 | 0.223753 | 0.69988307 | RNU6-890P | snRNA | 36.45 | -0.136175364 | 0.136175364 | ENSG00000206848 | n.s. |
| ENSG00000266308 | 11.59311 | -0.676622654 | 0.485339 | -1.394123 | 0.163281 | 0.629215564 | RN7SL510P | misc_RNA | 57.41 | -0.136136845 | 0.136136845 | ENSG00000266308 | n.s. |
| ENSG00000170681 | 322.4471 | -0.43001165 | 0.247204 | -1.7395 | 0.081947 | 0.482429187 | CAVIN4 | protein_coding | 40.73 | -0.136127251 | 0.136127251 | ENSG00000170681 | n.s. |
| ENSG00000276722 | 9.750063 | 0.700367222 | 0.511719 | 1.3686552 | 0.171107 | 0.639550064 | | misc_RNA | 56.14 | 0.135959104 | 0.135959104 | ENSG00000276722 | n.s. |
| ENSG00000252283 | 8.939736 | -0.730418897 | 0.546315 | -1.336992 | 0.181225 | 0.651484256 | | misc_RNA | 49.02 | -0.13592809 | 0.13592809 | ENSG00000252283 | n.s. |
| ENSG00000283871 | 9.333523 | -0.764203354 | 0.582219 | -1.31257 | 0.189328 | 0.663959974 | MIR130B | miRNA | 57.32 | -0.135919757 | 0.135919757 | ENSG00000283871 | n.s. |
| ENSG00000185262 | 413.0973 | -0.444719078 | 0.26069 | -1.705928 | 0.088021 | 0.494760783 | UBALD2 | protein_coding | 58.45 | -0.135908254 | 0.135908254 | ENSG00000185262 | n.s. |

| | | | | | | | | | | | | | |
|---|---|---|---|---|---|---|---|---|---|---|---|---|---|
| ENSG00000199347 | 388.7089 | 0.383083252 | 0.209237 | 1.830862 | 0.067121 | 0.441892852 | RNU5E-1 | snRNA | 40 | 0.135873126 | 0.135873126 | ENSG00000199347 | n.s. |
| ENSG00000121653 | 487.0324 | -0.448817729 | 0.264303 | -1.69812 | 0.089485 | 0.498053895 | MAPK8IP1 | protein_coding | 58.04 | -0.135867745 | 0.135867745 | ENSG00000121653 | n.s. |
| ENSG00000225591 | 18.63241 | -0.58664616 | 0.391518 | -1.498389 | 0.134032 | 0.586819166 | | processed_pseudogene | 47.45 | -0.135806069 | 0.135806069 | ENSG00000225591 | n.s. |
| ENSG00000073711 | 551.6927 | -0.432116791 | 0.249274 | -1.7335 | 0.083007 | 0.485101957 | PPP2R3A | protein_coding | 39.07 | -0.135756824 | 0.135756824 | ENSG00000073711 | n.s. |
| ENSG00000271646 | 49.60297 | 0.52016175 | 0.326022 | 1.5954794 | 0.110605 | 0.548299349 | | lincRNA | 50.34 | 0.135752994 | 0.135752994 | ENSG00000271646 | n.s. |
| ENSG00000134852 | 822.3996 | -0.432003878 | 0.249385 | -1.732279 | 0.083224 | 0.485101957 | CLOCK | protein_coding | 37.44 | -0.135721351 | 0.135721351 | ENSG00000134852 | n.s. |
| ENSG00000278730 | 923.4272 | -0.445707986 | 0.26196 | -1.701434 | 0.088862 | 0.496280219 | | lincRNA | 43.51 | -0.135616921 | 0.135616921 | ENSG00000278730 | n.s. |
| ENSG00000221634 | 11.33211 | -0.67064591 | 0.477329 | -1.404998 | 0.160022 | 0.627825878 | MIR1276 | miRNA | 56.63 | -0.135578305 | 0.135578305 | ENSG00000221634 | n.s. |
| ENSG00000266402 | 244.3067 | 0.401090485 | 0.223633 | 1.7935199 | 0.07289 | 0.459467368 | SNHG25 | lincRNA | 63.71 | 0.135466437 | 0.135466437 | ENSG00000266402 | n.s. |
| ENSG00000155903 | 624.1657 | -0.440349181 | 0.257276 | -1.711581 | 0.086974 | 0.492652156 | RASA2 | protein_coding | 36.35 | -0.135389588 | 0.135389588 | ENSG00000155903 | n.s. |
| ENSG00000253251 | 307.4845 | 0.440164424 | 0.257302 | 1.7106899 | 0.087138 | 0.492652156 | | protein_coding | 38.83 | 0.135332783 | 0.135332783 | ENSG00000253251 | n.s. |
| ENSG00000102743 | 349.2724 | -0.467212932 | 0.279987 | -1.668698 | 0.095177 | 0.513341297 | SLC25A15 | protein_coding | 46.36 | -0.135301967 | 0.135301967 | ENSG00000102743 | n.s. |
| ENSG00000196850 | 617.017 | -0.440022694 | 0.257074 | -1.711657 | 0.08696 | 0.492652156 | PPTC7 | protein_coding | 42.74 | -0.135289207 | 0.135289207 | ENSG00000196850 | n.s. |
| ENSG00000200397 | 41.79838 | -0.521865274 | 0.329162 | -1.585437 | 0.112867 | 0.551349488 | | misc_RNA | 48.04 | -0.134940282 | 0.134940282 | ENSG00000200397 | n.s. |
| ENSG00000173153 | 245.9436 | -0.477229138 | 0.289274 | -1.649748 | 0.098995 | 0.521709512 | ESRRA | protein_coding | 57.05 | -0.134851232 | 0.134851232 | ENSG00000173153 | n.s. |
| ENSG00000151474 | 515.5196 | -0.432141808 | 0.250382 | -1.725933 | 0.084359 | 0.487561582 | FRMD4A | protein_coding | 43.78 | -0.134815506 | 0.134815506 | ENSG00000151474 | n.s. |
| ENSG00000269834 | 510.167 | -0.398306701 | 0.222107 | -1.793311 | 0.072923 | 0.459467368 | ZNF528-AS1 | processed_transcript | 46.27 | -0.134526227 | 0.134526227 | ENSG00000269834 | n.s. |
| ENSG00000221063 | 23.05117 | -0.58073976 | 0.387481 | -1.498756 | 0.133937 | 0.586819166 | MIR1296 | miRNA | 57.61 | -0.134438763 | 0.134438763 | ENSG00000221063 | n.s. |
| ENSG00000166398 | 410.9842 | -0.433970418 | 0.252639 | -1.717751 | 0.085842 | 0.490059299 | KIAA0355 | protein_coding | 44.95 | -0.13442293 | 0.13442293 | ENSG00000166398 | n.s. |
| ENSG00000230699 | 166.5764 | -0.440987962 | 0.25901 | -1.702591 | 0.088645 | 0.495861421 | | lincRNA | 58.54 | -0.134342431 | 0.134342431 | ENSG00000230699 | n.s. |
| ENSG00000183955 | 1002.478 | -0.452314266 | 0.268536 | -1.684371 | 0.09211 | 0.505497667 | KMT5A | protein_coding | 51.73 | -0.134012053 | 0.134012053 | ENSG00000183955 | n.s. |
| ENSG00000219085 | 44.27442 | -0.499830002 | 0.309857 | -1.613101 | 0.106722 | 0.539737388 | NPM1P37 | processed_pseudogene | 39.24 | -0.13386322 | 0.13386322 | ENSG00000219085 | n.s. |
| ENSG00000126062 | 368.5361 | -0.407641727 | 0.229768 | -1.774143 | 0.076039 | 0.469606803 | TMEM115 | protein_coding | 57.53 | -0.133814764 | 0.133814764 | ENSG00000126062 | n.s. |
| ENSG00000199390 | 10.69684 | 0.791795383 | 0.62157 | 1.2738637 | 0.202712 | 0.677959669 | SNORD114-7 | snoRNA | 40.26 | 0.133652005 | 0.133652005 | ENSG00000199390 | n.s. |
| ENSG00000203684 | 22.69265 | -0.552845038 | 0.359326 | -1.538563 | 0.123911 | 0.57343851 | IBA57-AS1 | lincRNA | 52.98 | -0.133519343 | 0.133519343 | ENSG00000203684 | n.s. |
| ENSG00000276083 | 6.04995 | 0.780390248 | 0.606266 | 1.2872085 | 0.198022 | 0.674426423 | MIR7976 | miRNA | 33.33 | 0.133497788 | 0.133497788 | ENSG00000276083 | n.s. |
| ENSG00000168758 | 897.0511 | -0.4243027 | 0.244388 | -1.736184 | 0.082531 | 0.484641218 | SEMA4C | protein_coding | 62.17 | -0.133476996 | 0.133476996 | ENSG00000168758 | n.s. |
| ENSG00000162302 | 127.8579 | -0.465592018 | 0.280603 | -1.659254 | 0.097065 | 0.516830893 | RPS6KA4 | protein_coding | 56.99 | -0.133462667 | 0.133462667 | ENSG00000162302 | n.s. |
| ENSG00000088247 | 1492.852 | -0.39884485 | 0.223272 | -1.786362 | 0.074041 | 0.462895871 | KHSRP | protein_coding | 59.03 | -0.13342026 | 0.13342026 | ENSG00000088247 | n.s. |
| ENSG00000132694 | 828.231 | -0.405425537 | 0.228326 | -1.775645 | 0.075791 | 0.468789069 | ARHGEF11 | protein_coding | 44.51 | -0.133394134 | 0.133394134 | ENSG00000132694 | n.s. |
| ENSG00000115145 | 497.3434 | -0.445628073 | 0.2635 | -1.691191 | 0.0908 | 0.501961137 | STAM2 | protein_coding | 39.09 | -0.133389809 | 0.133389809 | ENSG00000115145 | n.s. |
| ENSG00000144485 | 2316.264 | -0.438169023 | 0.257454 | -1.701929 | 0.088769 | 0.496157209 | HES6 | protein_coding | 70.08 | -0.133370191 | 0.133370191 | ENSG00000144485 | n.s. |

| | | | | | | | | | | | | | |
|---|---|---|---|---|---|---|---|---|---|---|---|---|---|
| ENSG00000232995 | 324.1361 | -0.406040133 | 0.228894 | -1.773921 | 0.076076 | 0.469606803 | | antisense | 37.85 | -0.133289016 | 0.133289016 | ENSG00000232995 | n.s. |
| ENSG00000227470 | 47.89386 | -0.533104735 | 0.340778 | -1.564377 | 0.117729 | 0.562372792 | | processed_pseudogene | 43.79 | -0.133263229 | 0.133263229 | ENSG00000227470 | n.s. |
| ENSG00000273363 | 25.98465 | 0.553697243 | 0.361129 | 1.5332395 | 0.125217 | 0.574749865 | | sense_intronic | 53.32 | 0.133175882 | 0.133175882 | ENSG00000273363 | n.s. |
| ENSG00000261544 | 27.2715 | -0.577028311 | 0.386102 | -1.494499 | 0.135045 | 0.587941172 | | sense_intronic | 38.66 | -0.133100885 | 0.133100885 | ENSG00000261544 | n.s. |
| ENSG00000185085 | 202.1293 | -0.426911923 | 0.24754 | -1.724619 | 0.084596 | 0.48779901 | INTS5 | protein_coding | 54.61 | -0.133093671 | 0.133093671 | ENSG00000185085 | n.s. |
| ENSG00000165868 | 648.6344 | 0.480750843 | 0.294361 | 1.6332037 | 0.102426 | 0.528723147 | HSPA12A | protein_coding | 47.75 | 0.133058217 | 0.133058217 | ENSG00000165868 | n.s. |
| ENSG00000142528 | 421.2985 | -0.450254647 | 0.267721 | -1.681806 | 0.092607 | 0.50662512 | ZNF473 | protein_coding | 48.48 | -0.132966177 | 0.132966177 | ENSG00000142528 | n.s. |
| ENSG00000141002 | 742.0713 | -0.463693108 | 0.279033 | -1.661784 | 0.096556 | 0.516736278 | TCF25 | protein_coding | 55.04 | -0.13295521 | 0.13295521 | ENSG00000141002 | n.s. |
| ENSG00000198742 | 556.9025 | -0.441528889 | 0.260552 | -1.694593 | 0.090153 | 0.500068613 | SMURF1 | protein_coding | 42.72 | -0.132887128 | 0.132887128 | ENSG00000198742 | n.s. |
| ENSG00000147439 | 115.1728 | -0.483409299 | 0.296963 | -1.627843 | 0.103558 | 0.531064946 | BIN3 | protein_coding | 49.55 | -0.132866189 | 0.132866189 | ENSG00000147439 | n.s. |
| ENSG00000230184 | 18.70004 | -0.605065826 | 0.412992 | -1.46508 | 0.142899 | 0.603270163 | SMYD3-IT1 | sense_intronic | 46.52 | -0.132804781 | 0.132804781 | ENSG00000230184 | n.s. |
| ENSG00000212195 | 1065.698 | 0.371457863 | 0.201574 | 1.8427873 | 0.06536 | 0.439256007 | | snoRNA | 43.37 | 0.132715316 | 0.132715316 | ENSG00000212195 | n.s. |
| ENSG00000201121 | 8.272153 | -0.70562562 | 0.525707 | -1.34224 | 0.179518 | 0.648529087 | RNY1P12 | misc_RNA | 43.81 | -0.132707392 | 0.132707392 | ENSG00000201121 | n.s. |
| ENSG00000159433 | 1446.897 | -0.422378595 | 0.243961 | -1.731337 | 0.083392 | 0.485191567 | STARD9 | protein_coding | 43.49 | -0.132663523 | 0.132663523 | ENSG00000159433 | n.s. |
| ENSG00000167962 | 543.5691 | -0.42400648 | 0.245501 | -1.727103 | 0.084149 | 0.48719542 | ZNF598 | protein_coding | 64.25 | -0.132415868 | 0.132415868 | ENSG00000167962 | n.s. |
| ENSG00000243232 | 237.232 | -0.395471994 | 0.221247 | -1.787466 | 0.073862 | 0.462574432 | PCDHAC2 | protein_coding | 41.87 | -0.132411291 | 0.132411291 | ENSG00000243232 | n.s. |
| ENSG00000255081 | 19.77174 | -0.571868952 | 0.381859 | -1.49759 | 0.13424 | 0.586819166 | | sense_intronic | 45.8 | -0.132385209 | 0.132385209 | ENSG00000255081 | n.s. |
| ENSG00000012174 | 319.8195 | -0.456797213 | 0.273736 | -1.668751 | 0.095167 | 0.513341297 | MBTPS2 | protein_coding | 38.49 | -0.132285639 | 0.132285639 | ENSG00000012174 | n.s. |
| ENSG00000109111 | 1521.521 | -0.427004392 | 0.248392 | -1.719071 | 0.085601 | 0.490059299 | SUPT6H | protein_coding | 48.02 | -0.132265193 | 0.132265193 | ENSG00000109111 | n.s. |
| ENSG00000146112 | 344.7164 | -0.417735139 | 0.239871 | -1.741496 | 0.081597 | 0.482429187 | PPP1R18 | protein_coding | 56.1 | -0.13224092 | 0.13224092 | ENSG00000146112 | n.s. |
| ENSG00000011105 | 523.8081 | -0.447626537 | 0.266137 | -1.681939 | 0.092581 | 0.50662512 | TSPAN9 | protein_coding | 52.13 | -0.132190061 | 0.132190061 | ENSG00000011105 | n.s. |
| ENSG00000205268 | 1716.36 | -0.430782754 | 0.25223 | -1.707893 | 0.087656 | 0.493830606 | PDE7A | protein_coding | 37.7 | -0.132001312 | 0.132001312 | ENSG00000205268 | n.s. |
| ENSG00000088876 | 349.2963 | -0.442608909 | 0.262079 | -1.688836 | 0.091251 | 0.503331271 | ZNF343 | protein_coding | 43.04 | -0.131962114 | 0.131962114 | ENSG00000088876 | n.s. |
| ENSG00000111145 | 379.2619 | -0.433692047 | 0.254994 | -1.700792 | 0.088982 | 0.496552408 | ELK3 | protein_coding | 45.11 | -0.131857521 | 0.131857521 | ENSG00000111145 | n.s. |
| ENSG00000141179 | 328.5862 | -0.460821537 | 0.278125 | -1.656887 | 0.097542 | 0.51775891 | PCTP | protein_coding | 40.32 | -0.131736168 | 0.131736168 | ENSG00000141179 | n.s. |
| ENSG00000130881 | 828.6541 | -0.424158122 | 0.246674 | -1.71951 | 0.085522 | 0.490059299 | LRP3 | protein_coding | 54.69 | -0.131383557 | 0.131383557 | ENSG00000130881 | n.s. |
| ENSG00000158158 | 207.9756 | -0.420787847 | 0.243801 | -1.725949 | 0.084357 | 0.487561582 | CNNM4 | protein_coding | 50.16 | -0.131273404 | 0.131273404 | ENSG00000158158 | n.s. |
| ENSG00000126461 | 294.9995 | -0.418282223 | 0.241701 | -1.730579 | 0.083527 | 0.485572115 | SCAF1 | protein_coding | 58.2 | -0.131234484 | 0.131234484 | ENSG00000126461 | n.s. |
| ENSG00000271474 | 366.5004 | 0.387447201 | 0.215596 | 1.7970995 | 0.07232 | 0.458728412 | | antisense | 44.58 | 0.13112932 | 0.13112932 | ENSG00000271474 | n.s. |
| ENSG00000042445 | 217.6281 | -0.419591984 | 0.242799 | -1.728145 | 0.083962 | 0.487085887 | RETSAT | protein_coding | 51.7 | -0.131078209 | 0.131078209 | ENSG00000042445 | n.s. |
| ENSG00000224418 | 67.04866 | 0.501957222 | 0.315185 | 1.5925787 | 0.111255 | 0.548299349 | STK24-AS1 | antisense | 55.01 | 0.131001935 | 0.131001935 | ENSG00000224418 | n.s. |
| ENSG00000185453 | 15.29429 | -0.597786337 | 0.408407 | -1.463701 | 0.143276 | 0.603872801 | C19orf68 | protein_coding | 50.04 | -0.130947806 | 0.130947806 | ENSG00000185453 | n.s. |

| | | | | | | | | | | | | | |
|---|---|---|---|---|---|---|---|---|---|---|---|---|---|
| ENSG00000238058 | 31.75964 | -0.546741823 | 0.357591 | -1.528959 | 0.126275 | 0.5764328 | | antisense | 51.51 | -0.1308087 | 0.1308087 | ENSG00000238058 | n.s. |
| ENSG00000008311 | 328.401 | -0.431989096 | 0.254406 | -1.698033 | 0.089502 | 0.498053895 | AASS | protein_coding | 36.48 | -0.13077332 | 0.13077332 | ENSG00000008311 | n.s. |
| ENSG00000263013 | 231.9476 | -0.385258472 | 0.214128 | -1.799194 | 0.071988 | 0.45772936 | | sense_intronic | 49.09 | -0.130753347 | 0.130753347 | ENSG00000263013 | n.s. |
| ENSG00000082269 | 391.856 | -0.426094634 | 0.249376 | -1.708641 | 0.087518 | 0.493646579 | FAM135A | protein_coding | 36.09 | -0.130633741 | 0.130633741 | ENSG00000082269 | n.s. |
| ENSG00000266490 | 72.454 | -0.517366915 | 0.32934 | -1.570919 | 0.116202 | 0.559237383 | | lincRNA | 37.01 | -0.130585378 | 0.130585378 | ENSG00000266490 | n.s. |
| ENSG00000115421 | 1131.416 | -0.423604841 | 0.247104 | -1.714278 | 0.086478 | 0.49224449 | PAPOLG | protein_coding | 39.6 | -0.130393676 | 0.130393676 | ENSG00000115421 | n.s. |
| ENSG00000137040 | 3511.963 | 0.405561919 | 0.230588 | 1.7588171 | 0.078609 | 0.477239282 | RANBP6 | protein_coding | 39.01 | 0.13029237 | 0.13029237 | ENSG00000137040 | n.s. |
| ENSG00000199030 | 65.24885 | 0.471808252 | 0.289506 | 1.6296988 | 0.103165 | 0.529634455 | MIRLET7C | miRNA | 47.62 | 0.130230292 | 0.130230292 | ENSG00000199030 | n.s. |
| ENSG00000125354 | 805.6566 | -0.437032381 | 0.258956 | -1.687671 | 0.091474 | 0.503597002 | 6-Sep | protein_coding | 47.21 | -0.130199316 | 0.130199316 | ENSG00000125354 | n.s. |
| ENSG00000211459 | 632.1567 | -0.442989386 | 0.264162 | -1.676963 | 0.09355 | 0.508462135 | MT-RNR1 | Mt_rRNA | 45.49 | -0.130124315 | 0.130124315 | ENSG00000211459 | n.s. |
| ENSG00000273989 | 29.68845 | -0.562017199 | 0.375228 | -1.4978 | 0.134185 | 0.586819166 | | sense_intronic | 25.75 | -0.130104571 | 0.130104571 | ENSG00000273989 | n.s. |
| ENSG00000005339 | 403.5719 | -0.41396097 | 0.239013 | -1.731962 | 0.08328 | 0.485101957 | CREBBP | protein_coding | 45.01 | -0.130052865 | 0.130052865 | ENSG00000005339 | n.s. |
| ENSG00000178338 | 503.1549 | -0.465871649 | 0.284023 | -1.640261 | 0.100951 | 0.526024946 | ZNF354B | protein_coding | 44.33 | -0.129975236 | 0.129975236 | ENSG00000178338 | n.s. |
| ENSG00000124120 | 375.5742 | -0.453245783 | 0.273265 | -1.658631 | 0.09719 | 0.516972948 | TTPAL | protein_coding | 46.55 | -0.129869503 | 0.129869503 | ENSG00000124120 | n.s. |
| ENSG00000156675 | 436.2912 | -0.416349527 | 0.241532 | -1.723789 | 0.084746 | 0.48779901 | RAB11FIP1 | protein_coding | 45.87 | -0.129800748 | 0.129800748 | ENSG00000156675 | n.s. |
| ENSG00000265123 | 9.885166 | -0.722968908 | 0.54819 | -1.318829 | 0.187226 | 0.661405956 | RN7SL200P | misc_RNA | 54.17 | -0.129795981 | 0.129795981 | ENSG00000265123 | n.s. |
| ENSG00000143494 | 609.1753 | -0.436262804 | 0.258707 | -1.686323 | 0.091734 | 0.504081279 | VASH2 | protein_coding | 48.23 | -0.129787936 | 0.129787936 | ENSG00000143494 | n.s. |
| ENSG00000160226 | 197.9936 | -0.462119527 | 0.280792 | -1.645771 | 0.099811 | 0.524007414 | C21orf2 | protein_coding | 60.43 | -0.129699653 | 0.129699653 | ENSG00000160226 | n.s. |
| ENSG00000266962 | 256.9967 | -0.406699329 | 0.23286 | -1.746544 | 0.080717 | 0.479877462 | | antisense | 63.34 | -0.129684072 | 0.129684072 | ENSG00000266962 | n.s. |
| ENSG00000112877 | 222.5862 | -0.452269817 | 0.272273 | -1.661089 | 0.096696 | 0.516736278 | CEP72 | protein_coding | 53.81 | -0.129679798 | 0.129679798 | ENSG00000112877 | n.s. |
| ENSG00000270522 | 73.91678 | 0.451262928 | 0.270912 | 1.6657164 | 0.09577 | 0.516023395 | | lincRNA | 47.71 | 0.129661652 | 0.129661652 | ENSG00000270522 | n.s. |
| ENSG00000139998 | 174.9146 | -0.451269409 | 0.271077 | -1.66473 | 0.095967 | 0.516512586 | RAB15 | protein_coding | 52.23 | -0.129477809 | 0.129477809 | ENSG00000139998 | n.s. |
| ENSG00000163611 | 468.9628 | -0.451460476 | 0.271616 | -1.66213 | 0.096487 | 0.516706636 | SPICE1 | protein_coding | 38.02 | -0.129458983 | 0.129458983 | ENSG00000163611 | n.s. |
| ENSG00000213326 | 120.28 | 0.669449387 | 0.49068 | 1.3643291 | 0.172464 | 0.640837445 | RPS7P11 | processed_pseudogene | 47.78 | 0.129372513 | 0.129372513 | ENSG00000213326 | n.s. |
| ENSG00000265641 | 4.34508 | -0.907181682 | 0.787841 | -1.151478 | 0.249536 | 0.720401565 | MIR3929 | miRNA | 52.73 | -0.129205671 | 0.129205671 | ENSG00000265641 | n.s. |
| ENSG00000236605 | 807.1931 | 0.399307469 | 0.226227 | 1.7650748 | 0.077551 | 0.474822032 | | lincRNA | 46.76 | 0.129163643 | 0.129163643 | ENSG00000236605 | n.s. |
| ENSG00000140526 | 773.0153 | -0.414901125 | 0.240942 | -1.721995 | 0.08507 | 0.488426713 | ABHD2 | protein_coding | 44.01 | -0.129117475 | 0.129117475 | ENSG00000140526 | n.s. |
| ENSG00000010278 | 334.0952 | -0.540442602 | 0.353967 | -1.526817 | 0.126807 | 0.576937759 | CD9 | protein_coding | 51.04 | -0.129096085 | 0.129096085 | ENSG00000010278 | n.s. |
| ENSG00000272428 | 11.99597 | -0.612954452 | 0.429421 | -1.427397 | 0.153466 | 0.615840022 | | lincRNA | 32.23 | -0.129046582 | 0.129046582 | ENSG00000272428 | n.s. |
| ENSG00000210184 | 3.227036 | -0.903432978 | 0.783656 | -1.152844 | 0.248975 | 0.720036883 | MT-TS2 | Mt_tRNA | 45.76 | -0.128870429 | 0.128870429 | ENSG00000210184 | n.s. |
| ENSG00000258900 | 44.37735 | -0.469371095 | 0.288626 | -1.626228 | 0.103901 | 0.53160968 | HNRNPCP1 | processed_pseudogene | 45.95 | -0.12879877 | 0.12879877 | ENSG00000258900 | n.s. |
| ENSG00000100095 | 1169.456 | -0.41884131 | 0.24445 | -1.713405 | 0.086638 | 0.492652156 | SEZ6L | protein_coding | 44.56 | -0.128776787 | 0.128776787 | ENSG00000100095 | n.s. |

| | | | | | | | | | | | | | |
|---|---|---|---|---|---|---|---|---|---|---|---|---|---|
| ENSG00000180488 | 700.4646 | -0.469203583 | 0.288548 | -1.626084 | 0.103932 | 0.53160968 | MIGA1 | protein_coding | 39.53 | -0.128752804 | 0.128752804 | ENSG00000180488 | n.s. |
| ENSG00000201882 | 61.33874 | 0.448061226 | 0.270043 | 1.6592247 | 0.097071 | 0.516830893 | | snoRNA | 40 | 0.128437438 | 0.128437438 | ENSG00000201882 | n.s. |
| ENSG00000228825 | 12.40363 | 0.776894568 | 0.617026 | 1.2590952 | 0.207996 | 0.683512059 | LAMTOR3P1 | processed_pseudogene | 39.46 | 0.128384794 | 0.128384794 | ENSG00000228825 | n.s. |
| ENSG00000120458 | 311.3008 | -0.454694956 | 0.275823 | -1.648505 | 0.099249 | 0.522042316 | MSANTD2 | protein_coding | 40.09 | -0.128357791 | 0.128357791 | ENSG00000120458 | n.s. |
| ENSG00000197457 | 1460.685 | -0.447353504 | 0.268719 | -1.664761 | 0.095961 | 0.516512586 | STMN3 | protein_coding | 59.38 | -0.128354261 | 0.128354261 | ENSG00000197457 | n.s. |
| ENSG00000253633 | 59.38751 | 0.494347766 | 0.311002 | 1.5895331 | 0.11194 | 0.550138341 | | lincRNA | 43.71 | 0.12829713 | 0.12829713 | ENSG00000253633 | n.s. |
| ENSG00000208009 | 10.34453 | -0.659182953 | 0.480858 | -1.370846 | 0.170423 | 0.638865917 | MIR130A | miRNA | 50.56 | -0.128270596 | 0.128270596 | ENSG00000208009 | n.s. |
| ENSG00000187605 | 1669.144 | -0.399026199 | 0.226785 | -1.759492 | 0.078494 | 0.477221369 | TET3 | protein_coding | 46.75 | -0.128199184 | 0.128199184 | ENSG00000187605 | n.s. |
| ENSG00000112167 | 262.4174 | -0.39900405 | 0.226922 | -1.758334 | 0.078691 | 0.477390225 | SAYSD1 | protein_coding | 41.97 | -0.128130765 | 0.128130765 | ENSG00000112167 | n.s. |
| ENSG00000186350 | 381.0325 | -0.401651043 | 0.229639 | -1.749054 | 0.080282 | 0.479725712 | RXRA | protein_coding | 62.16 | -0.128129496 | 0.128129496 | ENSG00000186350 | n.s. |
| ENSG00000225527 | 32.29399 | -0.502464501 | 0.318765 | -1.576284 | 0.11496 | 0.555954895 | | antisense | 37.37 | -0.128108571 | 0.128108571 | ENSG00000225527 | n.s. |
| ENSG00000159556 | 154.6685 | -0.453419755 | 0.275009 | -1.648742 | 0.0992 | 0.521983162 | ISL2 | protein_coding | 59.5 | -0.128020123 | 0.128020123 | ENSG00000159556 | n.s. |
| ENSG00000135108 | 1919.952 | -0.422699691 | 0.248907 | -1.698226 | 0.089465 | 0.498053895 | FBXO21 | protein_coding | 44.38 | -0.127961197 | 0.127961197 | ENSG00000135108 | n.s. |
| ENSG00000258525 | 39.61957 | -0.489417668 | 0.307389 | -1.592176 | 0.111345 | 0.548299349 | | antisense | 39.27 | -0.127729334 | 0.127729334 | ENSG00000258525 | n.s. |
| ENSG00000161202 | 947.145 | -0.424315766 | 0.25037 | -1.694753 | 0.090122 | 0.500068613 | DVL3 | protein_coding | 52.91 | -0.127706487 | 0.127706487 | ENSG00000161202 | n.s. |
| ENSG00000251730 | 3.48543 | -0.904158592 | 0.789057 | -1.145873 | 0.251848 | 0.722544863 | | snoRNA | 37.78 | -0.127608588 | 0.127608588 | ENSG00000251730 | n.s. |
| ENSG00000162981 | 27257.76 | 0.377235455 | 0.210127 | 1.7952746 | 0.07261 | 0.459013748 | FAM84A | protein_coding | 41.56 | 0.127571338 | 0.127571338 | ENSG00000162981 | n.s. |
| ENSG00000212420 | 12.86527 | -0.623514399 | 0.441873 | -1.411072 | 0.158223 | 0.624594454 | RNU6-1111P | snRNA | 39.25 | -0.127447513 | 0.127447513 | ENSG00000212420 | n.s. |
| ENSG00000207012 | 4.922801 | -0.923084742 | 0.816698 | -1.130264 | 0.258365 | 0.728120901 | RNU6-72P | snRNA | 39.24 | -0.127197849 | 0.127197849 | ENSG00000207012 | n.s. |
| ENSG00000137478 | 224.5563 | -0.430577598 | 0.256108 | -1.681232 | 0.092718 | 0.506647149 | FCHSD2 | protein_coding | 38.87 | -0.127147152 | 0.127147152 | ENSG00000137478 | n.s. |
| ENSG00000204149 | 259.7934 | -0.465624063 | 0.286714 | -1.623999 | 0.104376 | 0.533294163 | AGAP6 | protein_coding | 41.1 | -0.127130814 | 0.127130814 | ENSG00000204149 | n.s. |
| ENSG00000101294 | 1087.173 | -0.407510292 | 0.236487 | -1.72318 | 0.084856 | 0.48779901 | HM13 | protein_coding | 49.84 | -0.127045036 | 0.127045036 | ENSG00000101294 | n.s. |
| ENSG00000147852 | 2095.311 | 0.413207696 | 0.24144 | 1.7114289 | 0.087002 | 0.492652156 | VLDLR | protein_coding | 41.97 | 0.127044678 | 0.127044678 | ENSG00000147852 | n.s. |
| ENSG00000158545 | 572.3023 | -0.419526534 | 0.247132 | -1.69758 | 0.089587 | 0.498130861 | ZC3H18 | protein_coding | 51.75 | -0.126972454 | 0.126972454 | ENSG00000158545 | n.s. |
| ENSG00000183048 | 234.4075 | -0.448764174 | 0.272096 | -1.649288 | 0.099089 | 0.521983162 | SLC25A10 | protein_coding | 63.39 | -0.12670565 | 0.12670565 | ENSG00000183048 | n.s. |
| ENSG00000088854 | 437.8788 | -0.437007186 | 0.261902 | -1.668591 | 0.095198 | 0.513341297 | C20orf194 | protein_coding | 43.79 | -0.12655457 | 0.12655457 | ENSG00000088854 | n.s. |
| ENSG00000217512 | 124.1463 | -0.453377971 | 0.276496 | -1.639727 | 0.101062 | 0.526152672 | | processed_pseudogene | 36.91 | -0.126441775 | 0.126441775 | ENSG00000217512 | n.s. |
| ENSG00000136854 | 52.30152 | -0.513983033 | 0.331127 | -1.552222 | 0.120609 | 0.567746164 | STXBP1 | protein_coding | 47.08 | -0.126360566 | 0.126360566 | ENSG00000136854 | n.s. |
| ENSG00000270313 | 7.220586 | -0.901598075 | 0.791043 | -1.139758 | 0.254387 | 0.72418384 | COX6CP16 | processed_pseudogene | 49.07 | -0.126360025 | 0.126360025 | ENSG00000270313 | n.s. |
| ENSG00000058056 | 1268.862 | -0.433183372 | 0.259062 | -1.67212 | 0.094501 | 0.510964378 | USP13 | protein_coding | 42.73 | -0.126320333 | 0.126320333 | ENSG00000058056 | n.s. |
| ENSG00000115756 | 221.4684 | -0.440082732 | 0.2646 | -1.663197 | 0.096273 | 0.516706636 | HPCAL1 | protein_coding | 52.12 | -0.126196347 | 0.126196347 | ENSG00000115756 | n.s. |
| ENSG00000074755 | 1078.269 | -0.425733013 | 0.252929 | -1.683209 | 0.092335 | 0.505936376 | ZZEF1 | protein_coding | 44.49 | -0.125976142 | 0.125976142 | ENSG00000074755 | n.s. |

| | | | | | | | | | | | | | |
|---|---|---|---|---|---|---|---|---|---|---|---|---|---|
| ENSG00000198231 | 6369.941 | -0.439161512 | 0.263898 | -1.664133 | 0.096086 | 0.516637382 | DDX42 | protein_coding | 42 | -0.125957746 | 0.125957746 | ENSG00000198231 | n.s. |
| ENSG00000167513 | 478.118 | -0.392198496 | 0.223104 | -1.757917 | 0.078762 | 0.477489649 | CDT1 | protein_coding | 65.55 | -0.125909851 | 0.125909851 | ENSG00000167513 | n.s. |
| ENSG00000119283 | 442.3126 | -0.406431479 | 0.236611 | -1.717721 | 0.085847 | 0.490059299 | TRIM67 | protein_coding | 48.93 | -0.125892705 | 0.125892705 | ENSG00000119283 | n.s. |
| ENSG00000243279 | 57.04459 | -0.481476494 | 0.302596 | -1.591152 | 0.111575 | 0.548733438 | PRAF2 | protein_coding | 60.94 | -0.125491347 | 0.125491347 | ENSG00000243279 | n.s. |
| ENSG00000181450 | 1802.019 | -0.458329292 | 0.282067 | -1.624893 | 0.104185 | 0.532515969 | ZNF678 | protein_coding | 40.32 | -0.125429769 | 0.125429769 | ENSG00000181450 | n.s. |
| ENSG00000207971 | 34.19433 | 0.511383722 | 0.330141 | 1.5489835 | 0.121386 | 0.568737594 | MIR125B1 | miRNA | 53.41 | 0.125334046 | 0.125334046 | ENSG00000207971 | n.s. |
| ENSG00000266173 | 625.0196 | -0.420056428 | 0.248803 | -1.688312 | 0.091351 | 0.503484921 | STRADA | protein_coding | 48.88 | -0.1251825 | 0.1251825 | ENSG00000266173 | n.s. |
| ENSG00000234444 | 922.5887 | -0.42659106 | 0.254607 | -1.675491 | 0.093838 | 0.509160888 | ZNF736 | protein_coding | 38.55 | -0.125053021 | 0.125053021 | ENSG00000234444 | n.s. |
| ENSG00000257680 | 52.26548 | -0.536091 | 0.356519 | -1.503682 | 0.132663 | 0.584700682 | | processed_pseudogene | 41.86 | -0.124944799 | 0.124944799 | ENSG00000257680 | n.s. |
| ENSG00000276058 | 19.41404 | -0.581826772 | 0.403303 | -1.442656 | 0.149118 | 0.609918763 | STMN1P1 | processed_pseudogene | 43.86 | -0.124934503 | 0.124934503 | ENSG00000276058 | n.s. |
| ENSG00000198538 | 434.348 | -0.487680171 | 0.308913 | -1.578697 | 0.114406 | 0.554650439 | ZNF28 | protein_coding | 45.1 | -0.124836682 | 0.124836682 | ENSG00000198538 | n.s. |
| ENSG00000171295 | 396.4226 | -0.439142536 | 0.265776 | -1.652304 | 0.098473 | 0.519724994 | ZNF440 | protein_coding | 44.1 | -0.124815901 | 0.124815901 | ENSG00000171295 | n.s. |
| ENSG00000083097 | 1165.166 | -0.47537927 | 0.297199 | -1.59953 | 0.109703 | 0.546473088 | DOPEY1 | protein_coding | 35.96 | -0.124754362 | 0.124754362 | ENSG00000083097 | n.s. |
| ENSG00000224531 | 1093.977 | -0.466901668 | 0.289867 | -1.610745 | 0.107235 | 0.540549786 | SMIM13 | protein_coding | 42.19 | -0.124739457 | 0.124739457 | ENSG00000224531 | n.s. |
| ENSG00000196204 | 222.7569 | -0.447211358 | 0.272894 | -1.638771 | 0.101261 | 0.526261702 | RNF216P1 | cribed_unprocessed_pseudc | 45.46 | -0.124681737 | 0.124681737 | ENSG00000196204 | n.s. |
| ENSG00000140548 | 229.022 | -0.435243516 | 0.262663 | -1.657041 | 0.097511 | 0.51775891 | ZNF710 | protein_coding | 50.7 | -0.124424117 | 0.124424117 | ENSG00000140548 | n.s. |
| ENSG00000154447 | 494.2303 | -0.401445575 | 0.233774 | -1.717238 | 0.085936 | 0.490161172 | SH3RF1 | protein_coding | 39.27 | -0.124312076 | 0.124312076 | ENSG00000154447 | n.s. |
| ENSG00000247828 | 267.2749 | -0.453936767 | 0.279302 | -1.625254 | 0.104108 | 0.532317753 | TMEM161B-AS1 | antisense | 37.29 | -0.124301073 | 0.124301073 | ENSG00000247828 | n.s. |
| ENSG00000132872 | 1904.143 | 0.410917446 | 0.242135 | 1.6970577 | 0.089686 | 0.498480974 | SYT4 | protein_coding | 34.83 | 0.12424147 | 0.12424147 | ENSG00000132872 | n.s. |
| ENSG00000135837 | 3652.059 | -0.433330665 | 0.261146 | -1.659345 | 0.097046 | 0.516830893 | CEP350 | protein_coding | 36.47 | -0.1242149 | 0.1242149 | ENSG00000135837 | n.s. |
| ENSG00000166341 | 1549.318 | -0.400435487 | 0.233007 | -1.718554 | 0.085696 | 0.490059299 | DCHS1 | protein_coding | 52.66 | -0.124035439 | 0.124035439 | ENSG00000166341 | n.s. |
| ENSG00000164815 | 609.2011 | -0.455122809 | 0.280531 | -1.622364 | 0.104726 | 0.533987611 | ORC5 | protein_coding | 36.03 | -0.124006774 | 0.124006774 | ENSG00000164815 | n.s. |
| ENSG00000109458 | 54.47258 | -0.497138475 | 0.31819 | -1.562396 | 0.118195 | 0.563117176 | GAB1 | protein_coding | 38.61 | -0.123986945 | 0.123986945 | ENSG00000109458 | n.s. |
| ENSG00000037757 | 216.9108 | -0.461417559 | 0.285497 | -1.616192 | 0.106053 | 0.538671714 | MRI1 | protein_coding | 52.16 | -0.123971745 | 0.123971745 | ENSG00000037757 | n.s. |
| ENSG00000201942 | 6.398427 | 0.697221704 | 0.531721 | 1.3112539 | 0.189772 | 0.664231317 | RNA5SP363 | rRNA | 41.74 | 0.123882807 | 0.123882807 | ENSG00000201942 | n.s. |
| ENSG00000258469 | 44.20725 | -0.514984967 | 0.33584 | -1.533425 | 0.125171 | 0.574749865 | CHMP4BP1 | processed_pseudogene | 53.57 | -0.123864762 | 0.123864762 | ENSG00000258469 | n.s. |
| ENSG00000206652 | 157.7348 | 0.442700687 | 0.269533 | 1.6424734 | 0.100492 | 0.525174439 | RNU1-1 | snRNA | 54.27 | 0.123821797 | 0.123821797 | ENSG00000206652 | n.s. |
| ENSG00000197302 | 467.3882 | -0.442593671 | 0.269416 | -1.64279 | 0.100426 | 0.525174439 | ZNF720 | protein_coding | 38.78 | -0.123791865 | 0.123791865 | ENSG00000197302 | n.s. |
| ENSG00000283149 | 26.86859 | 0.56897281 | 0.390194 | 1.4581798 | 0.144791 | 0.606306192 | | protein_coding | 41.04 | 0.123642341 | 0.123642341 | ENSG00000283149 | n.s. |
| ENSG00000151327 | 423.6421 | -0.473599435 | 0.297182 | -1.593636 | 0.111018 | 0.548299349 | FAM177A1 | protein_coding | 41.39 | -0.123601055 | 0.123601055 | ENSG00000151327 | n.s. |
| ENSG00000271659 | 46.71162 | -0.475815285 | 0.299447 | -1.588979 | 0.112065 | 0.550170175 | | antisense | 41.18 | -0.123475473 | 0.123475473 | ENSG00000271659 | n.s. |
| ENSG00000268043 | 403.3967 | -0.432293297 | 0.261095 | -1.655693 | 0.097784 | 0.518435958 | NBPF12 | protein_coding | 43.89 | -0.123335389 | 0.123335389 | ENSG00000268043 | n.s. |

| | | | | | | | | | | | | | |
|---|---|---|---|---|---|---|---|---|---|---|---|---|---|
| ENSG00000130856 | 44.33501 | -0.516963554 | 0.338819 | -1.525783 | 0.127064 | 0.577520151 | ZNF236 | protein_coding | 42.43 | -0.123261097 | 0.123261097 | ENSG00000130856 | n.s. |
| ENSG00000160953 | 629.8501 | -0.446356283 | 0.273859 | -1.629877 | 0.103127 | 0.529634455 | MUM1 | protein_coding | 52.99 | -0.123204944 | 0.123204944 | ENSG00000160953 | n.s. |
| ENSG00000168228 | 334.5421 | -0.443864435 | 0.271448 | -1.635172 | 0.102013 | 0.528171612 | ZCCHC4 | protein_coding | 38.93 | -0.123050293 | 0.123050293 | ENSG00000168228 | n.s. |
| ENSG00000252122 | 5.923904 | -0.740534704 | 0.586531 | -1.262566 | 0.206745 | 0.682214558 | | snoRNA | 56.3 | -0.122987276 | 0.122987276 | ENSG00000252122 | n.s. |
| ENSG00000127526 | 651.1278 | -0.424642999 | 0.254399 | -1.669201 | 0.095078 | 0.513341297 | SLC35E1 | protein_coding | 50.12 | -0.122973978 | 0.122973978 | ENSG00000127526 | n.s. |
| ENSG00000111727 | 739.1858 | -0.412419072 | 0.24414 | -1.689275 | 0.091167 | 0.503296715 | HCFC2 | protein_coding | 37.29 | -0.12297343 | 0.12297343 | ENSG00000111727 | n.s. |
| ENSG00000078043 | 1760.417 | -0.416179932 | 0.247401 | -1.682207 | 0.092529 | 0.506597509 | PIAS2 | protein_coding | 38.59 | -0.122913312 | 0.122913312 | ENSG00000078043 | n.s. |
| ENSG00000248909 | 37.98754 | -0.521794076 | 0.345781 | -1.50903 | 0.131291 | 0.581414031 | HMGB1P21 | processed_pseudogene | 42.69 | -0.122890067 | 0.122890067 | ENSG00000248909 | n.s. |
| ENSG00000279421 | 16.24118 | -0.566667706 | 0.390226 | -1.452154 | 0.146459 | 0.607109641 | | TEC | 51.64 | -0.122815519 | 0.122815519 | ENSG00000279421 | n.s. |
| ENSG00000143322 | 1617.253 | -0.418027919 | 0.249356 | -1.676433 | 0.093653 | 0.508556034 | ABL2 | protein_coding | 41.31 | -0.122758576 | 0.122758576 | ENSG00000143322 | n.s. |
| ENSG00000144893 | 378.4461 | -0.393750253 | 0.228406 | -1.723905 | 0.084725 | 0.48779901 | MED12L | protein_coding | 38.81 | -0.122755219 | 0.122755219 | ENSG00000144893 | n.s. |
| ENSG00000104863 | 70.29388 | -0.508368336 | 0.330758 | -1.53698 | 0.124298 | 0.573617848 | LIN7B | protein_coding | 58.86 | -0.122708599 | 0.122708599 | ENSG00000104863 | n.s. |
| ENSG00000072422 | 634.9114 | -0.45282571 | 0.279604 | -1.619524 | 0.105335 | 0.535832676 | RHOBTB1 | protein_coding | 38.94 | -0.122702548 | 0.122702548 | ENSG00000072422 | n.s. |
| ENSG00000197417 | 306.9351 | -0.44619141 | 0.274047 | -1.628158 | 0.103491 | 0.530917502 | SHPK | protein_coding | 50.5 | -0.122690572 | 0.122690572 | ENSG00000197417 | n.s. |
| ENSG00000254324 | 6.82117 | -0.694780369 | 0.531787 | -1.306502 | 0.191382 | 0.666152146 | MIR151A | miRNA | 52.22 | -0.122577716 | 0.122577716 | ENSG00000254324 | n.s. |
| ENSG00000100139 | 372.6162 | -0.417086299 | 0.248725 | -1.676894 | 0.093563 | 0.508462135 | MICALL1 | protein_coding | 55.92 | -0.122515506 | 0.122515506 | ENSG00000100139 | n.s. |
| ENSG00000080839 | 944.1927 | -0.446359497 | 0.27448 | -1.626199 | 0.103907 | 0.53160968 | RBL1 | protein_coding | 43.07 | -0.122484224 | 0.122484224 | ENSG00000080839 | n.s. |
| ENSG00000114933 | 1203.094 | -0.417710629 | 0.249591 | -1.673579 | 0.094213 | 0.51000526 | INO80D | protein_coding | 40.81 | -0.122149175 | 0.122149175 | ENSG00000114933 | n.s. |
| ENSG00000197497 | 101.8094 | -0.441269128 | 0.270352 | -1.632205 | 0.102636 | 0.528723147 | ZNF665 | protein_coding | 45.26 | -0.122130796 | 0.122130796 | ENSG00000197497 | n.s. |
| ENSG00000162959 | 486.2399 | -0.456640365 | 0.283525 | -1.61058 | 0.107271 | 0.540549786 | MEMO1 | protein_coding | 38.06 | -0.121998003 | 0.121998003 | ENSG00000162959 | n.s. |
| ENSG00000178718 | 360.2085 | 0.400032128 | 0.234888 | 1.7030785 | 0.088553 | 0.495550072 | RPP25 | protein_coding | 62.84 | 0.121974779 | 0.121974779 | ENSG00000178718 | n.s. |
| ENSG00000167524 | 142.1519 | -0.447650301 | 0.275938 | -1.622287 | 0.104742 | 0.533987611 | | protein_coding | 45.98 | -0.121970749 | 0.121970749 | ENSG00000167524 | n.s. |
| ENSG00000082512 | 1071.268 | -0.406034562 | 0.239792 | -1.693277 | 0.090403 | 0.501066377 | TRAF5 | protein_coding | 44.65 | -0.121852896 | 0.121852896 | ENSG00000082512 | n.s. |
| ENSG00000268603 | 23.60346 | -0.571173819 | 0.396869 | -1.439199 | 0.150094 | 0.612216404 | | antisense | 66.04 | -0.121714307 | 0.121714307 | ENSG00000268603 | n.s. |
| ENSG00000168273 | 298.7045 | -0.430928445 | 0.261332 | -1.648969 | 0.099154 | 0.521983162 | SMIM4 | protein_coding | 43.07 | -0.121669848 | 0.121669848 | ENSG00000168273 | n.s. |
| ENSG00000150551 | 497.0633 | -0.45397909 | 0.28119 | -1.614491 | 0.106421 | 0.539538058 | LYPD1 | protein_coding | 45.33 | -0.12165637 | 0.12165637 | ENSG00000150551 | n.s. |
| ENSG00000273680 | 101.0827 | -0.419252809 | 0.251034 | -1.670103 | 0.094899 | 0.512720075 | | antisense | 36.71 | -0.12163349 | 0.12163349 | ENSG00000273680 | n.s. |
| ENSG00000198353 | 163.9126 | -0.453784583 | 0.281157 | -1.61399 | 0.10653 | 0.539562772 | HOXC4 | protein_coding | 51.68 | -0.12159522 | 0.12159522 | ENSG00000198353 | n.s. |
| ENSG00000207370 | 77.36239 | 0.469862967 | 0.296221 | 1.586192 | 0.112696 | 0.55132222 | | misc_RNA | 38.94 | 0.121503981 | 0.121503981 | ENSG00000207370 | n.s. |
| ENSG00000263627 | 20.36921 | 0.589802197 | 0.417415 | 1.4129881 | 0.157659 | 0.622955867 | PPP4R1-AS1 | lincRNA | 42.71 | 0.121229548 | 0.121229548 | ENSG00000263627 | n.s. |
| ENSG00000157778 | 546.1372 | -0.463138365 | 0.290057 | -1.596715 | 0.110329 | 0.547347748 | PSMG3 | protein_coding | 57.7 | -0.121220291 | 0.121220291 | ENSG00000157778 | n.s. |
| ENSG00000162065 | 390.3564 | -0.43772282 | 0.268107 | -1.632643 | 0.102544 | 0.528723147 | TBC1D24 | protein_coding | 55.83 | -0.121149279 | 0.121149279 | ENSG00000162065 | n.s. |

| | | | | | | | | | | | | | |
|---|---|---|---|---|---|---|---|---|---|---|---|---|---|
| ENSG00000144711 | 459.6279 | -0.435381344 | 0.265958 | -1.637031 | 0.101624 | 0.526936211 | IQSEC1 | protein_coding | 54.36 | -0.121141357 | 0.121141357 | ENSG00000144711 | n.s. |
| ENSG00000077238 | 43.22491 | -0.483144504 | 0.308342 | -1.56691 | 0.117136 | 0.561405206 | IL4R | protein_coding | 49.92 | -0.121135712 | 0.121135712 | ENSG00000077238 | n.s. |
| ENSG00000241956 | 47.84377 | 0.4925853 | 0.317149 | 1.5531655 | 0.120384 | 0.567746164 | | antisense | 35.56 | 0.121100023 | 0.121100023 | ENSG00000241956 | n.s. |
| ENSG00000168874 | 3721.324 | -0.379689032 | 0.217213 | -1.748001 | 0.080464 | 0.479859335 | ATOH8 | protein_coding | 55.02 | -0.121077537 | 0.121077537 | ENSG00000168874 | n.s. |
| ENSG00000270006 | 51.01723 | -0.493971445 | 0.318867 | -1.549145 | 0.121347 | 0.568737594 | | antisense | 59.44 | -0.121066505 | 0.121066505 | ENSG00000270006 | n.s. |
| ENSG00000160352 | 1735.285 | -0.43437927 | 0.26521 | -1.63787 | 0.101449 | 0.526451999 | ZNF714 | protein_coding | 38.78 | -0.12103597 | 0.12103597 | ENSG00000160352 | n.s. |
| ENSG00000174231 | 4014.101 | -0.422927864 | 0.255154 | -1.657537 | 0.097411 | 0.517489317 | PRPF8 | protein_coding | 48.87 | -0.120999075 | 0.120999075 | ENSG00000174231 | n.s. |
| ENSG00000259663 | 640.1431 | -0.394929329 | 0.231303 | -1.707408 | 0.087746 | 0.494011481 | | antisense | 41.79 | -0.120952224 | 0.120952224 | ENSG00000259663 | n.s. |
| ENSG00000226977 | 9.542898 | -0.614661056 | 0.446024 | -1.37809 | 0.168175 | 0.63576037 | HMGN1P24 | processed_pseudogene | 47.65 | -0.120907854 | 0.120907854 | ENSG00000226977 | n.s. |
| ENSG00000075340 | 1782.14 | -0.398898665 | 0.234964 | -1.697701 | 0.089564 | 0.498130861 | ADD2 | protein_coding | 44.81 | -0.120729294 | 0.120729294 | ENSG00000075340 | n.s. |
| ENSG00000213793 | 178.1436 | -0.450171437 | 0.278952 | -1.613796 | 0.106572 | 0.539562772 | ZNF888 | protein_coding | 45.68 | -0.120627048 | 0.120627048 | ENSG00000213793 | n.s. |
| ENSG00000110880 | 2445.492 | -0.41062722 | 0.244849 | -1.67706 | 0.093531 | 0.508462135 | CORO1C | protein_coding | 41.35 | -0.120618208 | 0.120618208 | ENSG00000110880 | n.s. |
| ENSG00000065135 | 4733.577 | -0.423217754 | 0.256029 | -1.653007 | 0.098329 | 0.519362534 | GNAI3 | protein_coding | 39.95 | -0.120417886 | 0.120417886 | ENSG00000065135 | n.s. |
| ENSG00000167460 | 4065.56 | -0.43087924 | 0.262635 | -1.640599 | 0.100881 | 0.525948113 | TPM4 | protein_coding | 50.92 | -0.120239911 | 0.120239911 | ENSG00000167460 | n.s. |
| ENSG00000175221 | 244.3688 | -0.418241985 | 0.251062 | -1.665894 | 0.095735 | 0.516023395 | MED16 | protein_coding | 62.67 | -0.120173724 | 0.120173724 | ENSG00000175221 | n.s. |
| ENSG00000274627 | 358.1161 | -0.393533961 | 0.230871 | -1.704559 | 0.088277 | 0.495195739 | | sense_intronic | 50.38 | -0.120115656 | 0.120115656 | ENSG00000274627 | n.s. |
| ENSG00000234689 | 11.49053 | 0.595974249 | 0.426459 | 1.3974945 | 0.162265 | 0.628957308 | LINC00444 | lincRNA | 41.89 | 0.120016598 | 0.120016598 | ENSG00000234689 | n.s. |
| ENSG00000274925 | 222.9676 | -0.376229652 | 0.215417 | -1.74652 | 0.080721 | 0.479877462 | | lincRNA | 48.73 | -0.119968216 | 0.119968216 | ENSG00000274925 | n.s. |
| ENSG00000101400 | 150.2538 | -0.491791383 | 0.318558 | -1.543805 | 0.122635 | 0.570672973 | SNTA1 | protein_coding | 50.41 | -0.119806624 | 0.119806624 | ENSG00000101400 | n.s. |
| ENSG00000273691 | 35.69451 | -0.523482621 | 0.351502 | -1.489273 | 0.136416 | 0.590399179 | | sense_intronic | 39.08 | -0.119801225 | 0.119801225 | ENSG00000273691 | n.s. |
| ENSG00000163877 | 221.5838 | -0.446916797 | 0.276996 | -1.613444 | 0.106648 | 0.539737388 | SNIP1 | protein_coding | 43.23 | -0.119692138 | 0.119692138 | ENSG00000163877 | n.s. |
| ENSG00000152969 | 208.2018 | -0.455886719 | 0.284961 | -1.599821 | 0.109638 | 0.546361779 | JAKMIP1 | protein_coding | 49.33 | -0.11967924 | 0.11967924 | ENSG00000152969 | n.s. |
| ENSG00000265693 | 8.37823 | -0.70557817 | 0.550697 | -1.281246 | 0.200107 | 0.67702417 | | transcribed_processed_pseudog | 46.92 | -0.119521997 | 0.119521997 | ENSG00000265693 | n.s. |
| ENSG00000110987 | 1026.956 | -0.405318897 | 0.241308 | -1.679674 | 0.093021 | 0.507296335 | BCL7A | protein_coding | 52.98 | -0.119462992 | 0.119462992 | ENSG00000110987 | n.s. |
| ENSG00000187764 | 47.00759 | -0.471123482 | 0.299531 | -1.572873 | 0.115748 | 0.558020731 | SEMA4D | protein_coding | 51.34 | -0.119358977 | 0.119358977 | ENSG00000187764 | n.s. |
| ENSG00000213033 | 47.54621 | 0.489155368 | 0.316845 | 1.5438324 | 0.122629 | 0.570672973 | AURKAPS1 | nscribed_processed_pseudog | 44.14 | 0.119164458 | 0.119164458 | ENSG00000213033 | n.s. |
| ENSG00000275591 | 231.0897 | -0.468201862 | 0.297373 | -1.574461 | 0.115381 | 0.556634437 | XKR5 | protein_coding | 49.65 | -0.119124566 | 0.119124566 | ENSG00000275591 | n.s. |
| ENSG00000182446 | 2442.06 | -0.410426236 | 0.245708 | -1.670385 | 0.094843 | 0.51261685 | NPLOC4 | protein_coding | 50.06 | -0.119108617 | 0.119108617 | ENSG00000182446 | n.s. |
| ENSG00000219608 | 13.48679 | -0.59160456 | 0.424663 | -1.393116 | 0.163585 | 0.629342315 | HIGD1AP16 | processed_pseudogene | 39.92 | -0.118979407 | 0.118979407 | ENSG00000219608 | n.s. |
| ENSG00000146858 | 332.4549 | -0.384078613 | 0.223482 | -1.718612 | 0.085685 | 0.490059299 | ZC3HAV1L | protein_coding | 46.48 | -0.118968875 | 0.118968875 | ENSG00000146858 | n.s. |
| ENSG00000168672 | 363.6463 | 0.378680054 | 0.218355 | 1.7342415 | 0.082875 | 0.485101957 | FAM84B | protein_coding | 50.4 | 0.118968766 | 0.118968766 | ENSG00000168672 | n.s. |
| ENSG00000188234 | 88.31168 | -0.505398951 | 0.335098 | -1.508211 | 0.1315 | 0.581970996 | AGAP4 | protein_coding | 41.11 | -0.118818616 | 0.118818616 | ENSG00000188234 | n.s. |

| | | | | | | | | | | | | | |
|---|---|---|---|---|---|---|---|---|---|---|---|---|---|
| ENSG00000198920 | 391.4371 | -0.44137909 | 0.272916 | -1.617269 | 0.10582 | 0.538106349 | KIAA0753 | protein_coding | 41.58 | -0.118789186 | 0.118789186 | ENSG00000198920 | n.s. |
| ENSG00000031691 | 598.0823 | -0.446837455 | 0.278081 | -1.606858 | 0.108086 | 0.542492677 | CENPQ | protein_coding | 37.99 | -0.118682763 | 0.118682763 | ENSG00000031691 | n.s. |
| ENSG00000265144 | 2.198486 | -0.928938328 | 0.866755 | -1.071742 | 0.283836 | 0.745300357 | MIR548AM | miRNA | 37.84 | -0.118596321 | 0.118596321 | ENSG00000265144 | n.s. |
| ENSG00000122966 | 1627.053 | -0.402255995 | 0.239473 | -1.679753 | 0.093005 | 0.507296335 | CIT | protein_coding | 43.74 | -0.118560238 | 0.118560238 | ENSG00000122966 | n.s. |
| ENSG00000275494 | 22.47791 | 0.558836415 | 0.389609 | 1.4343525 | 0.151472 | 0.613599141 | | sense_intronic | 64.04 | 0.11853773 | 0.11853773 | ENSG00000275494 | n.s. |
| ENSG00000114999 | 2643.365 | -0.417453771 | 0.252754 | -1.651619 | 0.098612 | 0.520067531 | TTL | protein_coding | 42.38 | -0.118531932 | 0.118531932 | ENSG00000114999 | n.s. |
| ENSG00000270638 | 500.8852 | 0.374264663 | 0.215141 | 1.7396269 | 0.081925 | 0.482429187 | | lincRNA | 41.96 | 0.118479627 | 0.118479627 | ENSG00000270638 | n.s. |
| ENSG00000183783 | 110.7027 | 0.441756918 | 0.273727 | 1.6138592 | 0.106558 | 0.539562772 | KCTD8 | protein_coding | 35.64 | 0.11837231 | 0.11837231 | ENSG00000183783 | n.s. |
| ENSG00000048028 | 101.5296 | -0.462914981 | 0.293383 | -1.577852 | 0.1146 | 0.555290912 | USP28 | protein_coding | 41.45 | -0.118265257 | 0.118265257 | ENSG00000048028 | n.s. |
| ENSG00000138092 | 1435.626 | -0.404723074 | 0.241922 | -1.672951 | 0.094337 | 0.510277244 | CENPO | protein_coding | 43.93 | -0.118257573 | 0.118257573 | ENSG00000138092 | n.s. |
| ENSG00000200597 | 13.99165 | -0.753685926 | 0.615916 | -1.223683 | 0.221072 | 0.696894187 | RNU1-87P | snRNA | 45.29 | -0.118202944 | 0.118202944 | ENSG00000200597 | n.s. |
| ENSG00000240211 | 10.44871 | -0.661181878 | 0.50287 | -1.314818 | 0.188571 | 0.662619911 | | antisense | 53.54 | -0.118176685 | 0.118176685 | ENSG00000240211 | n.s. |
| ENSG00000129317 | 1705.457 | -0.457328049 | 0.288689 | -1.584155 | 0.113158 | 0.551745164 | PUS7L | protein_coding | 35.9 | -0.118110212 | 0.118110212 | ENSG00000129317 | n.s. |
| ENSG00000239264 | 135.2238 | -0.403460481 | 0.240975 | -1.674286 | 0.094074 | 0.509648797 | TXNDC5 | protein_coding | 47.02 | -0.118104582 | 0.118104582 | ENSG00000239264 | n.s. |
| ENSG00000166881 | 1622.432 | -0.456237148 | 0.287776 | -1.585392 | 0.112877 | 0.551349488 | NEMP1 | protein_coding | 44.16 | -0.117970619 | 0.117970619 | ENSG00000166881 | n.s. |
| ENSG00000228265 | 43.99837 | 0.48265646 | 0.312194 | 1.5460123 | 0.122102 | 0.569616291 | RALY-AS1 | lincRNA | 49.5 | 0.117969733 | 0.117969733 | ENSG00000228265 | n.s. |
| ENSG00000100294 | 273.8484 | -0.451753162 | 0.283417 | -1.593954 | 0.110946 | 0.548299349 | MCAT | protein_coding | 52.52 | -0.117899566 | 0.117899566 | ENSG00000100294 | n.s. |
| ENSG00000125741 | 276.2776 | -0.431628298 | 0.265883 | -1.623375 | 0.104509 | 0.533387737 | OPA3 | protein_coding | 49.78 | -0.117815954 | 0.117815954 | ENSG00000125741 | n.s. |
| ENSG00000064932 | 255.4119 | -0.410393058 | 0.246835 | -1.662622 | 0.096388 | 0.516706636 | SBNO2 | protein_coding | 63.18 | -0.117682656 | 0.117682656 | ENSG00000064932 | n.s. |
| ENSG00000216099 | 2.33199 | -0.924754335 | 0.865707 | -1.068207 | 0.285427 | 0.746113205 | MIR889 | miRNA | 35.44 | -0.11762438 | 0.11762438 | ENSG00000216099 | n.s. |
| ENSG00000123485 | 1538.345 | -0.409883044 | 0.246377 | -1.663642 | 0.096184 | 0.516706636 | HJURP | protein_coding | 48.25 | -0.117536406 | 0.117536406 | ENSG00000123485 | n.s. |
| ENSG00000131389 | 459.8065 | -0.409691977 | 0.246657 | -1.660976 | 0.096718 | 0.516736278 | SLC6A6 | protein_coding | 52.21 | -0.117471409 | 0.117471409 | ENSG00000131389 | n.s. |
| ENSG00000254726 | 2367.487 | -0.40965443 | 0.246346 | -1.662922 | 0.096328 | 0.516706636 | MEX3A | protein_coding | 53.44 | -0.11747085 | 0.11747085 | ENSG00000254726 | n.s. |
| ENSG00000171970 | 119.471 | -0.470906607 | 0.301496 | -1.561899 | 0.118312 | 0.563356985 | ZNF57 | protein_coding | 46.26 | -0.11735761 | 0.11735761 | ENSG00000171970 | n.s. |
| ENSG00000176148 | 471.0855 | -0.393457759 | 0.2329 | -1.689387 | 0.091145 | 0.503296715 | TCP11L1 | protein_coding | 42.1 | -0.117319623 | 0.117319623 | ENSG00000176148 | n.s. |
| ENSG00000279283 | 74.74812 | -0.476504775 | 0.306886 | -1.55271 | 0.120492 | 0.567746164 | | TEC | 57.82 | -0.117146694 | 0.117146694 | ENSG00000279283 | n.s. |
| ENSG00000100151 | 54.07952 | -0.479136417 | 0.309981 | -1.545697 | 0.122178 | 0.569616291 | PICK1 | protein_coding | 54.9 | -0.117109372 | 0.117109372 | ENSG00000100151 | n.s. |
| ENSG00000143493 | 2158.005 | -0.441331592 | 0.274977 | -1.604976 | 0.108499 | 0.543082569 | INTS7 | protein_coding | 38.12 | -0.117012071 | 0.117012071 | ENSG00000143493 | n.s. |
| ENSG00000101331 | 127.485 | -0.462347744 | 0.294102 | -1.572065 | 0.115935 | 0.558535693 | CCM2L | protein_coding | 50.16 | -0.116950431 | 0.116950431 | ENSG00000101331 | n.s. |
| ENSG00000089902 | 801.3366 | -0.399485156 | 0.238582 | -1.674411 | 0.09405 | 0.509648797 | RCOR1 | protein_coding | 43.48 | -0.116940889 | 0.116940889 | ENSG00000089902 | n.s. |
| ENSG00000100842 | 354.4226 | -0.388417583 | 0.229234 | -1.694413 | 0.090187 | 0.500068613 | EFS | protein_coding | 54.28 | -0.116902197 | 0.116902197 | ENSG00000100842 | n.s. |
| ENSG00000130119 | 413.1082 | -0.407291347 | 0.245045 | -1.662111 | 0.096491 | 0.516706636 | GNL3L | protein_coding | 47.94 | -0.116793222 | 0.116793222 | ENSG00000130119 | n.s. |

| | | | | | | | | | | | | | |
|---|---|---|---|---|---|---|---|---|---|---|---|---|---|
| ENSG00000250366 | 119.7641 | -0.470961823 | 0.301881 | -1.560089 | 0.118739 | 0.565003283 | TUNAR | protein_coding | 47.34 | -0.116774527 | 0.116774527 | ENSG00000250366 | n.s. |
| ENSG00000225742 | 31.56493 | 0.554906957 | 0.389123 | 1.4260463 | 0.153855 | 0.61644154 | LINC02036 | lincRNA | 44.61 | 0.116590448 | 0.116590448 | ENSG00000225742 | n.s. |
| ENSG00000142632 | 77.54899 | -0.446673118 | 0.280187 | -1.594199 | 0.110892 | 0.548299349 | ARHGEF19 | protein_coding | 57.33 | -0.116573764 | 0.116573764 | ENSG00000142632 | n.s. |
| ENSG00000243107 | 107.2567 | -0.502795393 | 0.335323 | -1.499434 | 0.133761 | 0.586803051 | | sense_overlapping | 38.79 | -0.116400973 | 0.116400973 | ENSG00000243107 | n.s. |
| ENSG00000213977 | 78.39432 | -0.501137981 | 0.333871 | -1.500995 | 0.133357 | 0.585903557 | TAX1BP3 | protein_coding | 57.28 | -0.116351142 | 0.116351142 | ENSG00000213977 | n.s. |
| ENSG00000203497 | 44.32838 | 0.481253022 | 0.31305 | 1.5373028 | 0.124219 | 0.573519528 | PDCD4-AS1 | antisense | 55.62 | 0.116199404 | 0.116199404 | ENSG00000203497 | n.s. |
| ENSG00000198440 | 340.7708 | -0.43427568 | 0.269555 | -1.611083 | 0.107162 | 0.540549786 | ZNF583 | protein_coding | 38.95 | -0.116022958 | 0.116022958 | ENSG00000198440 | n.s. |
| ENSG00000205356 | 421.2617 | -0.432729672 | 0.268006 | -1.614626 | 0.106392 | 0.539538058 | TECPR1 | protein_coding | 56.22 | -0.115961995 | 0.115961995 | ENSG00000205356 | n.s. |
| ENSG00000197021 | 150.6844 | -0.448539499 | 0.283211 | -1.583765 | 0.113247 | 0.551745164 | CXorf40B | protein_coding | 52.2 | -0.115840468 | 0.115840468 | ENSG00000197021 | n.s. |
| ENSG00000239774 | 24.5938 | 0.541058963 | 0.37544 | 1.4411348 | 0.149547 | 0.610955836 | | lincRNA | 38.78 | 0.115781306 | 0.115781306 | ENSG00000239774 | n.s. |
| ENSG00000172986 | 159.2578 | -0.430612927 | 0.266344 | -1.616757 | 0.105931 | 0.538471109 | GXYLT2 | protein_coding | 43.26 | -0.115764943 | 0.115764943 | ENSG00000172986 | n.s. |
| ENSG00000213865 | 121.8632 | -0.459334491 | 0.292613 | -1.569768 | 0.116469 | 0.55975287 | C8orf44 | protein_coding | 40.76 | -0.115753978 | 0.115753978 | ENSG00000213865 | n.s. |
| ENSG00000132017 | 405.0842 | -0.41277295 | 0.250989 | -1.644583 | 0.100056 | 0.524304665 | DCAF15 | protein_coding | 59.89 | -0.115748254 | 0.115748254 | ENSG00000132017 | n.s. |
| ENSG00000274267 | 2867.946 | -0.368399614 | 0.212426 | -1.734248 | 0.082874 | 0.485101957 | HIST1H3B | protein_coding | 59.12 | -0.115738992 | 0.115738992 | ENSG00000274267 | n.s. |
| ENSG00000250420 | 562.5147 | -0.420106322 | 0.258249 | -1.62675 | 0.10379 | 0.531471845 | AACSP1 | cribed_unprocessed_pseudc | 48.81 | -0.115327477 | 0.115327477 | ENSG00000250420 | n.s. |
| ENSG00000207022 | 6.031849 | -0.715563921 | 0.575011 | -1.244434 | 0.21334 | 0.690152016 | | snoRNA | 54.84 | -0.115245318 | 0.115245318 | ENSG00000207022 | n.s. |
| ENSG00000235127 | 429.7877 | -0.413335607 | 0.252446 | -1.637323 | 0.101563 | 0.526848168 | | lincRNA | 41.51 | -0.115037303 | 0.115037303 | ENSG00000235127 | n.s. |
| ENSG00000166166 | 304.0636 | -0.414821142 | 0.253722 | -1.634941 | 0.102061 | 0.528171612 | TRMT61A | protein_coding | 64.22 | -0.114998768 | 0.114998768 | ENSG00000166166 | n.s. |
| ENSG00000118046 | 366.6827 | -0.429549449 | 0.266484 | -1.611918 | 0.10698 | 0.539862594 | STK11 | protein_coding | 57.49 | -0.114997588 | 0.114997588 | ENSG00000118046 | n.s. |
| ENSG00000186501 | 220.6994 | -0.459996997 | 0.293966 | -1.564799 | 0.11763 | 0.562372792 | TMEM222 | protein_coding | 51.87 | -0.114988071 | 0.114988071 | ENSG00000186501 | n.s. |
| ENSG00000257702 | 119.2228 | -0.43234758 | 0.268951 | -1.607533 | 0.107938 | 0.542336313 | LBX2-AS1 | antisense | 60.7 | -0.114888291 | 0.114888291 | ENSG00000257702 | n.s. |
| ENSG00000175931 | 1490.031 | -0.400065873 | 0.240325 | -1.664687 | 0.095975 | 0.516512586 | UBE2O | protein_coding | 49.87 | -0.114786537 | 0.114786537 | ENSG00000175931 | n.s. |
| ENSG00000211643 | 55.32629 | -0.467982311 | 0.301843 | -1.550414 | 0.121042 | 0.568618694 | IGLV5-52 | IG_V_gene | 53 | -0.114739372 | 0.114739372 | ENSG00000211643 | n.s. |
| ENSG00000232024 | 29.38695 | 0.541477117 | 0.377668 | 1.4337378 | 0.151647 | 0.614039101 | LSM12P1 | processed_pseudogene | 48.98 | 0.114687005 | 0.114687005 | ENSG00000232024 | n.s. |
| ENSG00000261824 | 674.4222 | -0.414075654 | 0.25345 | -1.633757 | 0.10231 | 0.528723147 | LINC00662 | lincRNA | 42.21 | -0.114604413 | 0.114604413 | ENSG00000261824 | n.s. |
| ENSG00000273063 | 20.56297 | 0.598110149 | 0.442177 | 1.3526492 | 0.176168 | 0.643404323 | | antisense | 55.92 | 0.114547678 | 0.114547678 | ENSG00000273063 | n.s. |
| ENSG00000132437 | 196.3187 | -0.456973522 | 0.291928 | -1.565362 | 0.117498 | 0.561980157 | DDC | protein_coding | 44.42 | -0.114370885 | 0.114370885 | ENSG00000132437 | n.s. |
| ENSG00000128973 | 1323.14 | -0.400431264 | 0.24176 | -1.656317 | 0.097658 | 0.518072194 | CLN6 | protein_coding | 47.26 | -0.11436706 | 0.11436706 | ENSG00000128973 | n.s. |
| ENSG00000202459 | 6.30146 | -0.746045694 | 0.619867 | -1.203558 | 0.228761 | 0.702737638 | | misc_RNA | 46.3 | -0.114299262 | 0.114299262 | ENSG00000202459 | n.s. |
| ENSG00000241462 | 21.70775 | -0.567382531 | 0.40485 | -1.401465 | 0.161075 | 0.628931954 | | processed_pseudogene | 43.35 | -0.114268765 | 0.114268765 | ENSG00000241462 | n.s. |
| ENSG00000103319 | 527.4893 | -0.407251998 | 0.247622 | -1.644654 | 0.100041 | 0.524304665 | EEF2K | protein_coding | 47.84 | -0.114200089 | 0.114200089 | ENSG00000103319 | n.s. |
| ENSG00000258215 | 36.48673 | -0.481154661 | 0.317506 | -1.515421 | 0.129666 | 0.5791453 | | processed_pseudogene | 51.99 | -0.114135883 | 0.114135883 | ENSG00000258215 | n.s. |

| | | | | | | | | | | | | | |
|---|---|---|---|---|---|---|---|---|---|---|---|---|---|
| ENSG00000109099 | 313.3366 | -0.426246187 | 0.26441 | -1.612064 | 0.106948 | 0.539862594 | PMP22 | protein_coding | 45.52 | -0.11411325 | 0.11411325 | ENSG00000109099 | n.s. |
| ENSG00000228106 | 159.4013 | -0.480848801 | 0.31621 | -1.520662 | 0.128345 | 0.579048024 | | lincRNA | 37.92 | -0.114098408 | 0.114098408 | ENSG00000228106 | n.s. |
| ENSG00000176624 | 4551.953 | 0.363024737 | 0.209618 | 1.7318426 | 0.083302 | 0.485101957 | MEX3C | protein_coding | 41.02 | 0.114050383 | 0.114050383 | ENSG00000176624 | n.s. |
| ENSG00000108187 | 230.6272 | -0.466545754 | 0.301854 | -1.5456 | 0.122201 | 0.569616291 | PBLD | protein_coding | 43.3 | -0.114031993 | 0.114031993 | ENSG00000108187 | n.s. |
| ENSG00000178385 | 730.4363 | -0.405798078 | 0.246532 | -1.646024 | 0.099759 | 0.523930874 | PLEKHM3 | protein_coding | 41.34 | -0.113918075 | 0.113918075 | ENSG00000178385 | n.s. |
| ENSG00000154814 | 559.0249 | -0.402891961 | 0.244217 | -1.649728 | 0.098999 | 0.521709512 | OXNAD1 | protein_coding | 42.08 | -0.113845683 | 0.113845683 | ENSG00000154814 | n.s. |
| ENSG00000148459 | 520.2366 | -0.397272946 | 0.239558 | -1.658356 | 0.097246 | 0.516972948 | PDSS1 | protein_coding | 44.19 | -0.113831484 | 0.113831484 | ENSG00000148459 | n.s. |
| ENSG00000268895 | 133.2292 | -0.436108184 | 0.273767 | -1.592988 | 0.111163 | 0.548299349 | A1BG-AS1 | antisense | 55.91 | -0.113816504 | 0.113816504 | ENSG00000268895 | n.s. |
| ENSG00000244232 | 13.59818 | -0.575201855 | 0.416961 | -1.379509 | 0.167738 | 0.634623865 | RN7SL698P | misc_RNA | 52.98 | -0.113592933 | 0.113592933 | ENSG00000244232 | n.s. |
| ENSG00000175093 | 519.6858 | -0.422688084 | 0.261612 | -1.615703 | 0.106158 | 0.538671714 | SPSB4 | protein_coding | 47.04 | -0.113566071 | 0.113566071 | ENSG00000175093 | n.s. |
| ENSG00000278236 | 43.58513 | -0.478515737 | 0.315318 | -1.517564 | 0.129124 | 0.579048024 | | misc_RNA | 52.33 | -0.113544806 | 0.113544806 | ENSG00000278236 | n.s. |
| ENSG00000256725 | 52.60666 | -0.476422777 | 0.312406 | -1.525011 | 0.127256 | 0.577882813 | | lincRNA | 50.12 | -0.113464954 | 0.113464954 | ENSG00000256725 | n.s. |
| ENSG00000222467 | 15.07873 | -0.588869862 | 0.432935 | -1.360181 | 0.173773 | 0.642002364 | | misc_RNA | 36.96 | -0.11333588 | 0.11333588 | ENSG00000222467 | n.s. |
| ENSG00000164715 | 590.7066 | -0.39461065 | 0.237562 | -1.661087 | 0.096696 | 0.516736278 | LMTK2 | protein_coding | 43.73 | -0.113147125 | 0.113147125 | ENSG00000164715 | n.s. |
| ENSG00000122435 | 1148.373 | -0.437856302 | 0.276474 | -1.583719 | 0.113258 | 0.551745164 | TRMT13 | protein_coding | 35.8 | -0.11308141 | 0.11308141 | ENSG00000122435 | n.s. |
| ENSG00000070614 | 1038.616 | -0.390388702 | 0.233891 | -1.669108 | 0.095096 | 0.513341297 | NDST1 | protein_coding | 52.04 | -0.113054146 | 0.113054146 | ENSG00000070614 | n.s. |
| ENSG00000204406 | 420.6614 | -0.414685552 | 0.255715 | -1.621671 | 0.104874 | 0.534173967 | MBD5 | protein_coding | 36.28 | -0.112926042 | 0.112926042 | ENSG00000204406 | n.s. |
| ENSG00000259744 | 31.1596 | 0.520735652 | 0.35961 | 1.448058 | 0.147601 | 0.607109641 | | sense_intronic | 37.84 | 0.112860533 | 0.112860533 | ENSG00000259744 | n.s. |
| ENSG00000143614 | 752.983 | -0.40280094 | 0.245332 | -1.641862 | 0.100619 | 0.525276988 | GATAD2B | protein_coding | 45.03 | -0.112627825 | 0.112627825 | ENSG00000143614 | n.s. |
| ENSG00000113522 | 215.1821 | -0.40008837 | 0.242969 | -1.646667 | 0.099627 | 0.523631644 | RAD50 | protein_coding | 38.81 | -0.112414476 | 0.112414476 | ENSG00000113522 | n.s. |
| ENSG00000207185 | 5.154095 | -0.736233758 | 0.613178 | -1.200686 | 0.229873 | 0.703699853 | RNU6-1157P | snRNA | 39.62 | -0.112358504 | 0.112358504 | ENSG00000207185 | n.s. |
| ENSG00000145569 | 336.5486 | -0.460837405 | 0.298502 | -1.543835 | 0.122628 | 0.570672973 | FAM105A | protein_coding | 42.96 | -0.112265842 | 0.112265842 | ENSG00000145569 | n.s. |
| ENSG00000204439 | 290.063 | -0.39175483 | 0.236181 | -1.658705 | 0.097175 | 0.516972948 | C6orf47 | protein_coding | 57.01 | -0.112250366 | 0.112250366 | ENSG00000204439 | n.s. |
| ENSG00000240668 | 57.69406 | -0.480852798 | 0.319585 | -1.504616 | 0.132423 | 0.584381334 | KRT8P36 | processed_pseudogene | 56.53 | -0.11218472 | 0.11218472 | ENSG00000240668 | n.s. |
| ENSG00000178295 | 2889.189 | -0.418631324 | 0.259318 | -1.614355 | 0.106451 | 0.539538058 | GEN1 | protein_coding | 35.87 | -0.11218395 | 0.11218395 | ENSG00000178295 | n.s. |
| ENSG00000271551 | 82.743 | -0.433161735 | 0.272905 | -1.587228 | 0.112461 | 0.55089864 | | antisense | 61.08 | -0.112157831 | 0.112157831 | ENSG00000271551 | n.s. |
| ENSG00000073578 | 1211.441 | -0.402016954 | 0.245156 | -1.639839 | 0.101039 | 0.526152672 | SDHA | protein_coding | 46.07 | -0.112117793 | 0.112117793 | ENSG00000073578 | n.s. |
| ENSG00000175467 | 773.9203 | -0.404287374 | 0.247323 | -1.634654 | 0.102122 | 0.528171612 | SART1 | protein_coding | 55.3 | -0.112078545 | 0.112078545 | ENSG00000175467 | n.s. |
| ENSG00000267108 | 7.480421 | -0.686929467 | 0.549014 | -1.251205 | 0.21086 | 0.687090943 | | lincRNA | 52.44 | -0.111959732 | 0.111959732 | ENSG00000267108 | n.s. |
| ENSG00000231728 | 57.87148 | -0.439747967 | 0.27923 | -1.57486 | 0.115289 | 0.556576156 | | antisense | 52.99 | -0.111905041 | 0.111905041 | ENSG00000231728 | n.s. |
| ENSG00000143067 | 393.2101 | -0.404246377 | 0.247488 | -1.633395 | 0.102386 | 0.528723147 | ZNF697 | protein_coding | 44.33 | -0.111883948 | 0.111883948 | ENSG00000143067 | n.s. |
| ENSG00000008196 | 288.789 | -0.40420939 | 0.247539 | -1.632912 | 0.102488 | 0.528723147 | TFAP2B | protein_coding | 44.14 | -0.111873711 | 0.111873711 | ENSG00000008196 | n.s. |

| | | | | | | | | | | | | | |
|---|---|---|---|---|---|---|---|---|---|---|---|---|---|
| ENSG00000163026 | 730.6143 | -0.447745682 | 0.28644 | -1.563141 | 0.11802 | 0.562542305 | WDCP | protein_coding | 43.29 | -0.111866936 | 0.111866936 | ENSG00000163026 | n.s. |
| ENSG00000238227 | 294.9751 | -0.379976104 | 0.226382 | -1.678472 | 0.093255 | 0.508003334 | TMEM250 | protein_coding | 65.35 | -0.111763678 | 0.111763678 | ENSG00000238227 | n.s. |
| ENSG00000083812 | 275.0462 | -0.399407505 | 0.243216 | -1.642195 | 0.10055 | 0.525174439 | ZNF324 | protein_coding | 57.75 | -0.111712849 | 0.111712849 | ENSG00000083812 | n.s. |
| ENSG00000236662 | 23.7273 | -0.523582059 | 0.363671 | -1.439715 | 0.149948 | 0.612057538 | | antisense | 53.25 | -0.111631752 | 0.111631752 | ENSG00000236662 | n.s. |
| ENSG00000183741 | 1183.991 | -0.35773295 | 0.207253 | -1.72607 | 0.084335 | 0.487561582 | CBX6 | protein_coding | 58.47 | -0.111602135 | 0.111602135 | ENSG00000183741 | n.s. |
| ENSG00000188070 | 1321.133 | -0.397050009 | 0.241081 | -1.646956 | 0.099567 | 0.523516907 | C11orf95 | protein_coding | 61.54 | -0.111598564 | 0.111598564 | ENSG00000188070 | n.s. |
| ENSG00000130717 | 404.539 | -0.42011093 | 0.261838 | -1.604468 | 0.108611 | 0.543082569 | UCK1 | protein_coding | 57.27 | -0.111385749 | 0.111385749 | ENSG00000130717 | n.s. |
| ENSG00000136492 | 3828.512 | -0.402414292 | 0.246535 | -1.632282 | 0.10262 | 0.528723147 | BRIP1 | protein_coding | 35.92 | -0.111376878 | 0.111376878 | ENSG00000136492 | n.s. |
| ENSG00000116731 | 507.3347 | -0.396723525 | 0.241002 | -1.64614 | 0.099735 | 0.523930874 | PRDM2 | protein_coding | 43.44 | -0.111370612 | 0.111370612 | ENSG00000116731 | n.s. |
| ENSG00000176473 | 108.8926 | -0.440693691 | 0.280422 | -1.57154 | 0.116057 | 0.5589301 | WDR25 | protein_coding | 46.07 | -0.111337957 | 0.111337957 | ENSG00000176473 | n.s. |
| ENSG00000246695 | 131.9467 | -0.430447865 | 0.271464 | -1.585655 | 0.112818 | 0.551349488 | RASSF8-AS1 | antisense | 43.94 | -0.111302206 | 0.111302206 | ENSG00000246695 | n.s. |
| ENSG00000127720 | 171.8158 | -0.485893477 | 0.325913 | -1.490871 | 0.135995 | 0.590213403 | METTL25 | protein_coding | 35.3 | -0.1112652 | 0.1112652 | ENSG00000127720 | n.s. |
| ENSG00000103335 | 448.1358 | -0.401225545 | 0.245431 | -1.634782 | 0.102095 | 0.528171612 | PIEZO1 | protein_coding | 61.99 | -0.111229729 | 0.111229729 | ENSG00000103335 | n.s. |
| ENSG00000116670 | 274.3921 | -0.434241652 | 0.274996 | -1.579083 | 0.114317 | 0.554576902 | MAD2L2 | protein_coding | 53.91 | -0.111182461 | 0.111182461 | ENSG00000116670 | n.s. |
| ENSG00000075420 | 86.80706 | -0.466651976 | 0.306091 | -1.524551 | 0.127371 | 0.578027275 | FNDC3B | protein_coding | 39.76 | -0.111087282 | 0.111087282 | ENSG00000075420 | n.s. |
| ENSG00000138083 | 312.8699 | -0.387416934 | 0.23337 | -1.660098 | 0.096895 | 0.516736278 | SIX3 | protein_coding | 57.2 | -0.111084463 | 0.111084463 | ENSG00000138083 | n.s. |
| ENSG00000197951 | 581.5101 | -0.432457935 | 0.273562 | -1.580839 | 0.113915 | 0.553589858 | ZNF71 | protein_coding | 46.22 | -0.111060333 | 0.111060333 | ENSG00000197951 | n.s. |
| ENSG00000108788 | 1109 | -0.424238646 | 0.265444 | -1.598222 | 0.109993 | 0.54732402 | MLX | protein_coding | 52.95 | -0.111046796 | 0.111046796 | ENSG00000108788 | n.s. |
| ENSG00000125846 | 323.3663 | -0.451081618 | 0.29023 | -1.554222 | 0.120132 | 0.567746164 | ZNF133 | protein_coding | 40.41 | -0.110896517 | 0.110896517 | ENSG00000125846 | n.s. |
| ENSG00000088305 | 235.432 | -0.41817922 | 0.26065 | -1.604372 | 0.108632 | 0.543082569 | DNMT3B | protein_coding | 49.4 | -0.110873586 | 0.110873586 | ENSG00000088305 | n.s. |
| ENSG00000163349 | 1825.536 | -0.387489288 | 0.234164 | -1.654777 | 0.09797 | 0.518446014 | HIPK1 | protein_coding | 39.82 | -0.110549322 | 0.110549322 | ENSG00000163349 | n.s. |
| ENSG00000109046 | 1260.455 | -0.430529305 | 0.272438 | -1.580285 | 0.114042 | 0.553662697 | WSB1 | protein_coding | 39.73 | -0.110540438 | 0.110540438 | ENSG00000109046 | n.s. |
| ENSG00000114200 | 525.1369 | 0.456391235 | 0.296497 | 1.5392795 | 0.123736 | 0.572922348 | BCHE | protein_coding | 32.3 | 0.110402973 | 0.110402973 | ENSG00000114200 | n.s. |
| ENSG00000172403 | 2526.613 | -0.422611505 | 0.264947 | -1.595081 | 0.110694 | 0.548299349 | SYNPO2 | protein_coding | 38.36 | -0.11029411 | 0.11029411 | ENSG00000172403 | n.s. |
| ENSG00000225067 | 19.01914 | 0.569940305 | 0.417256 | 1.3659256 | 0.171962 | 0.640837445 | RPL23AP2 | processed_pseudogene | 48.93 | 0.110142172 | 0.110142172 | ENSG00000225067 | n.s. |
| ENSG00000144233 | 998.7577 | -0.397724711 | 0.243707 | -1.631979 | 0.102684 | 0.528723147 | AMMECR1L | protein_coding | 45.69 | -0.110078935 | 0.110078935 | ENSG00000144233 | n.s. |
| ENSG00000213137 | 23.81987 | -0.549812786 | 0.396286 | -1.387415 | 0.165315 | 0.630679057 | ARF1P2 | processed_pseudogene | 55.2 | -0.110067896 | 0.110067896 | ENSG00000213137 | n.s. |
| ENSG00000175066 | 761.8386 | -0.424865075 | 0.267673 | -1.587255 | 0.112455 | 0.55089864 | GK5 | protein_coding | 38.79 | -0.110009591 | 0.110009591 | ENSG00000175066 | n.s. |
| ENSG00000275185 | 58.87908 | -0.464297306 | 0.307159 | -1.511588 | 0.130639 | 0.579675642 | | antisense | 37.27 | -0.109952542 | 0.109952542 | ENSG00000275185 | n.s. |
| ENSG00000068615 | 382.345 | -0.437132871 | 0.278722 | -1.568346 | 0.1168 | 0.560379317 | REEP1 | protein_coding | 46.05 | -0.109946743 | 0.109946743 | ENSG00000068615 | n.s. |
| ENSG00000261476 | 48.18618 | -0.459523152 | 0.300538 | -1.529001 | 0.126264 | 0.5764328 | | sense_intronic | 46.6 | -0.109941518 | 0.109941518 | ENSG00000261476 | n.s. |
| ENSG00000101350 | 657.1929 | -0.424442532 | 0.267386 | -1.58738 | 0.112427 | 0.55089864 | KIF3B | protein_coding | 43.82 | -0.109900183 | 0.109900183 | ENSG00000101350 | n.s. |

| | | | | | | | | | | | | | |
|---|---|---|---|---|---|---|---|---|---|---|---|---|---|
| ENSG00000159079 | 165.5638 | -0.476066331 | 0.318355 | -1.495393 | 0.134812 | 0.587845023 | C21orf59 | protein_coding | 47.5 | -0.10984619 | 0.10984619 | ENSG00000159079 | n.s. |
| ENSG00000106086 | 1193.163 | -0.392598453 | 0.239078 | -1.642138 | 0.100561 | 0.525174439 | PLEKHA8 | protein_coding | 39.62 | -0.109808382 | 0.109808382 | ENSG00000106086 | n.s. |
| ENSG00000257923 | 282.6386 | -0.393358109 | 0.239777 | -1.640514 | 0.100898 | 0.525948113 | CUX1 | protein_coding | 48.61 | -0.109769373 | 0.109769373 | ENSG00000257923 | n.s. |
| ENSG00000145868 | 1148.166 | -0.447517092 | 0.288845 | -1.549331 | 0.121302 | 0.568737594 | FBXO38 | protein_coding | 37.05 | -0.109681098 | 0.109681098 | ENSG00000145868 | n.s. |
| ENSG00000144043 | 550.3986 | -0.422386346 | 0.265703 | -1.589695 | 0.111904 | 0.550138341 | TEX261 | protein_coding | 52.63 | -0.10962112 | 0.10962112 | ENSG00000144043 | n.s. |
| ENSG00000259415 | 38.11007 | 0.505632308 | 0.347789 | 1.4538466 | 0.145989 | 0.607109641 | | sense_intronic | 45.47 | 0.109587141 | 0.109587141 | ENSG00000259415 | n.s. |
| ENSG00000254172 | 4.044169 | -0.816066076 | 0.731336 | -1.115857 | 0.264483 | 0.734131278 | RNU5A-3P | snRNA | 36.84 | -0.109537507 | 0.109537507 | ENSG00000254172 | n.s. |
| ENSG00000196378 | 87.98568 | -0.457777471 | 0.29931 | -1.529441 | 0.126155 | 0.5764328 | ZNF34 | protein_coding | 49.73 | -0.109523862 | 0.109523862 | ENSG00000196378 | n.s. |
| ENSG00000200525 | 16.63944 | -0.568643422 | 0.418504 | -1.358752 | 0.174225 | 0.64206882 | RNU6-1209P | snRNA | 38.32 | -0.109417469 | 0.109417469 | ENSG00000200525 | n.s. |
| ENSG00000252198 | 6.29225 | -0.710322483 | 0.587597 | -1.208859 | 0.226717 | 0.701406486 | | misc_RNA | 53.02 | -0.10941113 | 0.10941113 | ENSG00000252198 | n.s. |
| ENSG00000124459 | 571.7935 | -0.38147691 | 0.229798 | -1.660051 | 0.096904 | 0.516736278 | ZNF45 | protein_coding | 41.36 | -0.109381274 | 0.109381274 | ENSG00000124459 | n.s. |
| ENSG00000186272 | 309.9148 | -0.410578101 | 0.255228 | -1.608674 | 0.107688 | 0.541549688 | ZNF17 | protein_coding | 47.68 | -0.109362277 | 0.109362277 | ENSG00000186272 | n.s. |
| ENSG00000276740 | 35.36896 | -0.504250838 | 0.346224 | -1.45643 | 0.145274 | 0.607098916 | | sense_intronic | 44.14 | -0.1092916 | 0.1092916 | ENSG00000276740 | n.s. |
| ENSG00000170471 | 1658.995 | -0.409043782 | 0.254184 | -1.60924 | 0.107564 | 0.541309679 | RALGAPB | protein_coding | 39.37 | -0.109032341 | 0.109032341 | ENSG00000170471 | n.s. |
| ENSG00000280383 | 1356.657 | 0.346972858 | 0.200214 | 1.7330138 | 0.083093 | 0.485101957 | | sense_overlapping | 41.87 | 0.109007413 | 0.109007413 | ENSG00000280383 | n.s. |
| ENSG00000135144 | 145.9668 | -0.426196348 | 0.27013 | -1.577747 | 0.114624 | 0.555290912 | DTX1 | protein_coding | 53.96 | -0.108884402 | 0.108884402 | ENSG00000135144 | n.s. |
| ENSG00000241975 | 43.38371 | -0.511001164 | 0.355197 | -1.438643 | 0.150252 | 0.612366927 | ELOCP19 | processed_pseudogene | 42.14 | -0.108837255 | 0.108837255 | ENSG00000241975 | n.s. |
| ENSG00000135912 | 577.3603 | -0.392694482 | 0.240601 | -1.63214 | 0.10265 | 0.528723147 | TTLL4 | protein_coding | 44.96 | -0.10868671 | 0.10868671 | ENSG00000135912 | n.s. |
| ENSG00000106105 | 3968.097 | -0.409837676 | 0.255374 | -1.60485 | 0.108527 | 0.543082569 | GARS | protein_coding | 38.71 | -0.108661958 | 0.108661958 | ENSG00000106105 | n.s. |
| ENSG00000101421 | 286.1905 | -0.451264593 | 0.294139 | -1.534186 | 0.124984 | 0.574749865 | CHMP4B | protein_coding | 46.43 | -0.108538666 | 0.108538666 | ENSG00000101421 | n.s. |
| ENSG00000271851 | 60.34832 | -0.473317541 | 0.317638 | -1.490116 | 0.136194 | 0.590213403 | | sense_intronic | 40.23 | -0.108385425 | 0.108385425 | ENSG00000271851 | n.s. |
| ENSG00000243156 | 92.27173 | -0.414999111 | 0.260387 | -1.59378 | 0.110985 | 0.548299349 | MICAL3 | protein_coding | 49.78 | -0.10830741 | 0.10830741 | ENSG00000243156 | n.s. |
| ENSG00000258682 | 52.15593 | -0.472849008 | 0.317267 | -1.490384 | 0.136123 | 0.590213403 | | sense_overlapping | 37.85 | -0.108278135 | 0.108278135 | ENSG00000258682 | n.s. |
| ENSG00000076770 | 1082.368 | -0.421379583 | 0.266469 | -1.581347 | 0.113799 | 0.553411977 | MBNL3 | protein_coding | 36.96 | -0.108274093 | 0.108274093 | ENSG00000076770 | n.s. |
| ENSG00000163626 | 325.9854 | -0.379088006 | 0.228886 | -1.656228 | 0.097676 | 0.518072194 | COX18 | protein_coding | 40.47 | -0.108271218 | 0.108271218 | ENSG00000163626 | n.s. |
| ENSG00000131931 | 363.1401 | 0.38574077 | 0.234416 | 1.6455428 | 0.099858 | 0.524056489 | THAP1 | protein_coding | 41.03 | 0.108247307 | 0.108247307 | ENSG00000131931 | n.s. |
| ENSG00000176871 | 849.2861 | -0.432883695 | 0.276875 | -1.563464 | 0.117944 | 0.562372792 | WSB2 | protein_coding | 46.42 | -0.108210404 | 0.108210404 | ENSG00000176871 | n.s. |
| ENSG00000119888 | 55.45308 | 0.455670192 | 0.299764 | 1.5200954 | 0.128487 | 0.579048024 | EPCAM | protein_coding | 44.35 | 0.108123891 | 0.108123891 | ENSG00000119888 | n.s. |
| ENSG00000011451 | 470.0854 | -0.387292138 | 0.236414 | -1.638194 | 0.101381 | 0.526378134 | WIZ | protein_coding | 58.16 | -0.107939155 | 0.107939155 | ENSG00000011451 | n.s. |
| ENSG00000089820 | 57.93238 | -0.446505421 | 0.29046 | -1.537235 | 0.124236 | 0.573519528 | ARHGAP4 | protein_coding | 58.14 | -0.107809533 | 0.107809533 | ENSG00000089820 | n.s. |
| ENSG00000199691 | 21.94148 | -0.535179112 | 0.382711 | -1.39839 | 0.161996 | 0.628957308 | RN7SKP173 | misc_RNA | 48.25 | -0.107773745 | 0.107773745 | ENSG00000199691 | n.s. |
| ENSG00000145012 | 17.67407 | -0.530704148 | 0.377153 | -1.407132 | 0.159388 | 0.626527722 | LPP | protein_coding | 39.17 | -0.107764629 | 0.107764629 | ENSG00000145012 | n.s. |

| | | | | | | | | | | | | | |
|---|---|---|---|---|---|---|---|---|---|---|---|---|---|
| ENSG00000097021 | 262.9118 | -0.461363517 | 0.306553 | -1.505002 | 0.132324 | 0.584331244 | ACOT7 | protein_coding | 54 | -0.107654974 | 0.107654974 | ENSG00000097021 | n.s. |
| ENSG00000266993 | 15.18445 | -0.561292608 | 0.414861 | -1.352966 | 0.176067 | 0.643404323 | | antisense | 42.62 | -0.107496529 | 0.107496529 | ENSG00000266993 | n.s. |
| ENSG00000198824 | 625.4276 | -0.409323061 | 0.255858 | -1.599808 | 0.109641 | 0.546361779 | CHAMP1 | protein_coding | 42.93 | -0.107455363 | 0.107455363 | ENSG00000198824 | n.s. |
| ENSG00000274943 | 106.2516 | -0.469502664 | 0.315247 | -1.489315 | 0.136404 | 0.590399179 | | sense_intronic | 34.32 | -0.107447683 | 0.107447683 | ENSG00000274943 | n.s. |
| ENSG00000198853 | 329.0719 | -0.385222591 | 0.235129 | -1.638346 | 0.101349 | 0.526378134 | RUSC2 | protein_coding | 44.22 | -0.107362368 | 0.107362368 | ENSG00000198853 | n.s. |
| ENSG00000159692 | 1659.301 | -0.406119238 | 0.253319 | -1.60319 | 0.108893 | 0.544064048 | CTBP1 | protein_coding | 60.99 | -0.107357609 | 0.107357609 | ENSG00000159692 | n.s. |
| ENSG00000101104 | 230.8585 | -0.402937317 | 0.250489 | -1.608603 | 0.107703 | 0.541549688 | PABPC1L | protein_coding | 47.22 | -0.107327065 | 0.107327065 | ENSG00000101104 | n.s. |
| ENSG00000109089 | 252.3407 | -0.406878269 | 0.254066 | -1.601466 | 0.109274 | 0.544920997 | CDR2L | protein_coding | 53.8 | -0.107280152 | 0.107280152 | ENSG00000109089 | n.s. |
| ENSG00000180537 | 327.8782 | -0.450475072 | 0.295519 | -1.524354 | 0.12742 | 0.57806181 | RNF182 | protein_coding | 38.51 | -0.107224654 | 0.107224654 | ENSG00000180537 | n.s. |
| ENSG00000273306 | 76.36795 | -0.459248528 | 0.305184 | -1.504823 | 0.13237 | 0.584331244 | | antisense | 44.41 | -0.107161461 | 0.107161461 | ENSG00000273306 | n.s. |
| ENSG00000118515 | 273.9507 | 0.434555867 | 0.279347 | 1.5556117 | 0.1198 | 0.566955067 | SGK1 | protein_coding | 42.27 | 0.107096884 | 0.107096884 | ENSG00000118515 | n.s. |
| ENSG00000005100 | 689.5065 | -0.404873306 | 0.252588 | -1.6029 | 0.108957 | 0.544064048 | DHX33 | protein_coding | 45.81 | -0.107028247 | 0.107028247 | ENSG00000005100 | n.s. |
| ENSG00000227372 | 300.2269 | -0.412375217 | 0.259482 | -1.589226 | 0.112009 | 0.550170175 | TP73-AS1 | anscribed_unitary_pseudoge | 54.46 | -0.107012588 | 0.107012588 | ENSG00000227372 | n.s. |
| ENSG00000134343 | 675.1586 | 0.450838055 | 0.297395 | 1.5159576 | 0.12953 | 0.579048024 | ANO3 | protein_coding | 36.36 | 0.106977295 | 0.106977295 | ENSG00000134343 | n.s. |
| ENSG00000207359 | 8.742639 | -0.687606033 | 0.564078 | -1.218991 | 0.222847 | 0.698928583 | RNU6-925P | snRNA | 36.54 | -0.106968944 | 0.106968944 | ENSG00000207359 | n.s. |
| ENSG00000208024 | 79.89764 | 0.424565532 | 0.270635 | 1.5687751 | 0.1167 | 0.560282428 | MIR199A2 | miRNA | 53.64 | 0.106817715 | 0.106817715 | ENSG00000208024 | n.s. |
| ENSG00000002016 | 254.7032 | -0.42330723 | 0.269692 | -1.569594 | 0.11651 | 0.55975287 | RAD52 | protein_coding | 45.17 | -0.106674976 | 0.106674976 | ENSG00000002016 | n.s. |
| ENSG00000207525 | 5.742566 | -0.745538224 | 0.645486 | -1.155004 | 0.248089 | 0.719455242 | | misc_RNA | 40.59 | -0.106609147 | 0.106609147 | ENSG00000207525 | n.s. |
| ENSG00000136938 | 1216.38 | -0.38100879 | 0.232018 | -1.642154 | 0.100558 | 0.525174439 | ANP32B | protein_coding | 43.52 | -0.106566794 | 0.106566794 | ENSG00000136938 | n.s. |
| ENSG00000199082 | 19.21658 | -0.551039794 | 0.403712 | -1.364934 | 0.172274 | 0.640837445 | MIR342 | miRNA | 49.49 | -0.106489608 | 0.106489608 | ENSG00000199082 | n.s. |
| ENSG00000066739 | 1194.606 | -0.408032503 | 0.256036 | -1.59365 | 0.111014 | 0.548299349 | ATG2B | protein_coding | 37.31 | -0.106489249 | 0.106489249 | ENSG00000066739 | n.s. |
| ENSG00000138621 | 99.66918 | -0.485591994 | 0.331844 | -1.463312 | 0.143382 | 0.603872801 | PPCDC | protein_coding | 47.94 | -0.106371127 | 0.106371127 | ENSG00000138621 | n.s. |
| ENSG00000237953 | 25.54917 | 0.516759625 | 0.365278 | 1.414702 | 0.157156 | 0.622581912 | | processed_pseudogene | 40.45 | 0.10635094 | 0.10635094 | ENSG00000237953 | n.s. |
| ENSG00000179528 | 165.1883 | -0.462311294 | 0.309691 | -1.492817 | 0.135485 | 0.589119333 | LBX2 | protein_coding | 60.07 | -0.106237621 | 0.106237621 | ENSG00000179528 | n.s. |
| ENSG00000199015 | 3.036551 | -0.801586705 | 0.727966 | -1.101132 | 0.270839 | 0.737020734 | MIR377 | miRNA | 39.13 | -0.106226506 | 0.106226506 | ENSG00000199015 | n.s. |
| ENSG00000139971 | 142.5112 | -0.455758289 | 0.303078 | -1.503768 | 0.132641 | 0.584700682 | C14orf37 | protein_coding | 40.15 | -0.106221943 | 0.106221943 | ENSG00000139971 | n.s. |
| ENSG00000167130 | 373.8374 | -0.434382813 | 0.281064 | -1.545494 | 0.122227 | 0.569616291 | DOLPP1 | protein_coding | 57.76 | -0.106170804 | 0.106170804 | ENSG00000167130 | n.s. |
| ENSG00000135482 | 517.7139 | -0.350645644 | 0.206429 | -1.698627 | 0.089389 | 0.498053895 | ZC3H10 | protein_coding | 48.69 | -0.106148732 | 0.106148732 | ENSG00000135482 | n.s. |
| ENSG00000106771 | 1472.173 | -0.395879873 | 0.245515 | -1.612448 | 0.106864 | 0.539862594 | TMEM245 | protein_coding | 39.52 | -0.105983678 | 0.105983678 | ENSG00000106771 | n.s. |
| ENSG00000237765 | 1127.02 | -0.415733837 | 0.263851 | -1.575639 | 0.115109 | 0.556094484 | FAM200B | protein_coding | 40.86 | -0.105950356 | 0.105950356 | ENSG00000237765 | n.s. |
| ENSG00000227500 | 51.6376 | -0.457650709 | 0.30538 | -1.498625 | 0.133971 | 0.586819166 | SCAMP4 | protein_coding | 60.16 | -0.105944176 | 0.105944176 | ENSG00000227500 | n.s. |
| ENSG00000278876 | 46.02169 | -0.488540887 | 0.337107 | -1.449215 | 0.147278 | 0.607109641 | | TEC | 45.85 | -0.105882869 | 0.105882869 | ENSG00000278876 | n.s. |

| | | | | | | | | | | | | | |
|---|---|---|---|---|---|---|---|---|---|---|---|---|---|
| ENSG00000206656 | 5.914185 | -0.863509744 | 0.823879 | -1.048102 | 0.294591 | 0.754039228 | SNORD116-17 | snoRNA | 42.39 | -0.105871528 | 0.105871528 | ENSG00000206656 | n.s. |
| ENSG00000101040 | 68.52714 | -0.430461752 | 0.277073 | -1.553604 | 0.120279 | 0.567746164 | ZMYND8 | protein_coding | 46.13 | -0.10582721 | 0.10582721 | ENSG00000101040 | n.s. |
| ENSG00000213689 | 86.68668 | -0.416874442 | 0.26493 | -1.573526 | 0.115597 | 0.557484721 | TREX1 | protein_coding | 62.72 | -0.105788989 | 0.105788989 | ENSG00000213689 | n.s. |
| ENSG00000100056 | 288.6321 | -0.405025523 | 0.25443 | -1.591895 | 0.111408 | 0.548299349 | DGCR14 | protein_coding | 55.92 | -0.10570448 | 0.10570448 | ENSG00000100056 | n.s. |
| ENSG00000158234 | 596.3288 | -0.44527453 | 0.293432 | -1.51747 | 0.129148 | 0.579048024 | FAIM | protein_coding | 40.46 | -0.105657152 | 0.105657152 | ENSG00000158234 | n.s. |
| ENSG00000200488 | 29.58849 | -0.512199624 | 0.361808 | -1.415667 | 0.156873 | 0.622581912 | RN7SKP203 | misc_RNA | 58.43 | -0.105412476 | 0.105412476 | ENSG00000200488 | n.s. |
| ENSG00000100297 | 3250.481 | -0.398481593 | 0.248648 | -1.602596 | 0.109024 | 0.544064048 | MCM5 | protein_coding | 52.51 | -0.105338598 | 0.105338598 | ENSG00000100297 | n.s. |
| ENSG00000160072 | 300.9876 | -0.40205852 | 0.251792 | -1.596789 | 0.110313 | 0.547347748 | ATAD3B | protein_coding | 58.83 | -0.105233456 | 0.105233456 | ENSG00000160072 | n.s. |
| ENSG00000207231 | 5.713614 | -0.704587678 | 0.595739 | -1.182711 | 0.236924 | 0.709700242 | | misc_RNA | 43.86 | -0.104930753 | 0.104930753 | ENSG00000207231 | n.s. |
| ENSG00000144026 | 897.2898 | -0.391753539 | 0.242982 | -1.612273 | 0.106902 | 0.539862594 | ZNF514 | protein_coding | 46.72 | -0.104878989 | 0.104878989 | ENSG00000144026 | n.s. |
| ENSG00000168517 | 130.9608 | -0.394363851 | 0.245377 | -1.607176 | 0.108016 | 0.542338372 | HEXIM2 | protein_coding | 53.43 | -0.104794175 | 0.104794175 | ENSG00000168517 | n.s. |
| ENSG00000117569 | 4037.083 | -0.427566813 | 0.276102 | -1.54858 | 0.121483 | 0.568737594 | PTBP2 | protein_coding | 35.67 | -0.104791522 | 0.104791522 | ENSG00000117569 | n.s. |
| ENSG00000180998 | 247.8714 | -0.442004163 | 0.292451 | -1.511378 | 0.130692 | 0.579683575 | GPR137C | protein_coding | 37.31 | -0.104670566 | 0.104670566 | ENSG00000180998 | n.s. |
| ENSG00000164180 | 1048.84 | -0.403991445 | 0.254424 | -1.587865 | 0.112317 | 0.55089864 | TMEM161B | protein_coding | 35.47 | -0.104604818 | 0.104604818 | ENSG00000164180 | n.s. |
| ENSG00000047621 | 838.1216 | -0.436086719 | 0.284974 | -1.530271 | 0.12595 | 0.575897021 | C12orf4 | protein_coding | 36.55 | -0.104510436 | 0.104510436 | ENSG00000047621 | n.s. |
| ENSG00000250321 | 49.33016 | 0.775559837 | 0.694401 | 1.116876 | 0.264047 | 0.733524057 | | processed_pseudogene | 47.18 | 0.104379216 | 0.104379216 | ENSG00000250321 | n.s. |
| ENSG00000175550 | 460.2764 | -0.447284729 | 0.297314 | -1.504421 | 0.132473 | 0.58441765 | DRAP1 | protein_coding | 65.51 | -0.104341095 | 0.104341095 | ENSG00000175550 | n.s. |
| ENSG00000175567 | 113.6761 | -0.413102595 | 0.263013 | -1.570657 | 0.116262 | 0.559337018 | UCP2 | protein_coding | 53.18 | -0.104236706 | 0.104236706 | ENSG00000175567 | n.s. |
| ENSG00000113083 | 315.5632 | -0.431621304 | 0.280696 | -1.537684 | 0.124126 | 0.573519528 | LOX | protein_coding | 37.93 | -0.104215736 | 0.104215736 | ENSG00000113083 | n.s. |
| ENSG00000164088 | 129.573 | 0.436760264 | 0.28619 | 1.526121 | 0.12698 | 0.577380418 | PPM1M | protein_coding | 58.15 | 0.104183898 | 0.104183898 | ENSG00000164088 | n.s. |
| ENSG00000167565 | 120.6856 | 0.430432998 | 0.279561 | 1.539673 | 0.12364 | 0.572922348 | SERTAD3 | protein_coding | 55.9 | 0.104123566 | 0.104123566 | ENSG00000167565 | n.s. |
| ENSG00000092964 | 3107.27 | -0.365846181 | 0.221317 | -1.653043 | 0.098322 | 0.519362534 | DPYSL2 | protein_coding | 44.97 | -0.104093988 | 0.104093988 | ENSG00000092964 | n.s. |
| ENSG00000225526 | 11.69168 | 0.685170848 | 0.574367 | 1.1929153 | 0.232903 | 0.704958888 | MKRN2OS | protein_coding | 43.71 | 0.104033744 | 0.104033744 | ENSG00000225526 | n.s. |
| ENSG00000264539 | 2.973899 | -0.964766601 | 0.989157 | -0.975343 | 0.32939 | 0.780141891 | MIR548AR | miRNA | 33.33 | -0.104027311 | 0.104027311 | ENSG00000264539 | n.s. |
| ENSG00000184144 | 1149.695 | -0.398920553 | 0.250698 | -1.591241 | 0.111555 | 0.548733438 | CNTN2 | protein_coding | 54.18 | -0.103974084 | 0.103974084 | ENSG00000184144 | n.s. |
| ENSG00000118402 | 326.9803 | -0.433311313 | 0.28289 | -1.53173 | 0.125589 | 0.575724414 | ELOVL4 | protein_coding | 36.66 | -0.103901706 | 0.103901706 | ENSG00000118402 | n.s. |
| ENSG00000177192 | 575.7463 | -0.362154899 | 0.218095 | -1.660541 | 0.096806 | 0.516736278 | PUS1 | protein_coding | 51.99 | -0.103841053 | 0.103841053 | ENSG00000177192 | n.s. |
| ENSG00000096968 | 230.356 | -0.423688279 | 0.273609 | -1.548515 | 0.121498 | 0.568737594 | JAK2 | protein_coding | 37.67 | -0.10384094 | 0.10384094 | ENSG00000096968 | n.s. |
| ENSG00000200612 | 7.384098 | 0.61436649 | 0.481587 | 1.2757117 | 0.202057 | 0.677624898 | SNORD114-25 | snoRNA | 47.22 | 0.103834477 | 0.103834477 | ENSG00000200612 | n.s. |
| ENSG00000279786 | 58.52924 | -0.431120001 | 0.280877 | -1.534906 | 0.124807 | 0.574721329 | | TEC | 43.34 | -0.103702763 | 0.103702763 | ENSG00000279786 | n.s. |
| ENSG00000151689 | 89.29405 | -0.43685008 | 0.287422 | -1.519891 | 0.128538 | 0.579048024 | INPP1 | protein_coding | 40.14 | -0.103658153 | 0.103658153 | ENSG00000151689 | n.s. |
| ENSG00000121966 | 231.749 | -0.395603359 | 0.247775 | -1.596621 | 0.11035 | 0.547347748 | CXCR4 | protein_coding | 52.06 | -0.103543903 | 0.103543903 | ENSG00000121966 | n.s. |

| | | | | | | | | | | | | | |
|---|---|---|---|---|---|---|---|---|---|---|---|---|---|
| ENSG00000120071 | 7771.912 | -0.373914964 | 0.229038 | -1.632545 | 0.102565 | 0.528723147 | KANSL1 | protein_coding | 40.92 | -0.103489072 | 0.103489072 | ENSG00000120071 | n.s. |
| ENSG00000132356 | 1321.621 | -0.42092947 | 0.271159 | -1.552333 | 0.120582 | 0.567746164 | PRKAA1 | protein_coding | 37.74 | -0.103483739 | 0.103483739 | ENSG00000132356 | n.s. |
| ENSG00000251609 | 44.51253 | -0.494471284 | 0.34717 | -1.424294 | 0.154362 | 0.61775666 | SETP12 | processed_pseudogene | 40.75 | -0.103434771 | 0.103434771 | ENSG00000251609 | n.s. |
| ENSG00000119682 | 1069.177 | -0.379496015 | 0.234029 | -1.621579 | 0.104893 | 0.534173967 | AREL1 | protein_coding | 42.31 | -0.103343323 | 0.103343323 | ENSG00000119682 | n.s. |
| ENSG00000228629 | 16.07163 | -0.536552666 | 0.394237 | -1.360991 | 0.173516 | 0.642002364 | | antisense | 43.73 | -0.103266736 | 0.103266736 | ENSG00000228629 | n.s. |
| ENSG00000151640 | 1100.229 | -0.425865912 | 0.276412 | -1.540695 | 0.123391 | 0.572276225 | DPYSL4 | protein_coding | 61.67 | -0.103227468 | 0.103227468 | ENSG00000151640 | n.s. |
| ENSG00000196814 | 692.4453 | -0.39400191 | 0.246679 | -1.597223 | 0.110216 | 0.54732402 | MVB12B | protein_coding | 47.23 | -0.103132164 | 0.103132164 | ENSG00000196814 | n.s. |
| ENSG00000151748 | 420.172 | -0.386844088 | 0.240399 | -1.609177 | 0.107578 | 0.541309679 | SAV1 | protein_coding | 37.89 | -0.103114919 | 0.103114919 | ENSG00000151748 | n.s. |
| ENSG00000260318 | 47.25194 | -0.462519755 | 0.314017 | -1.472913 | 0.140775 | 0.598646262 | COX6CP1 | processed_pseudogene | 45.58 | -0.10306315 | 0.10306315 | ENSG00000260318 | n.s. |
| ENSG00000184635 | 983.0639 | -0.403495854 | 0.255907 | -1.576729 | 0.114858 | 0.555652001 | ZNF93 | protein_coding | 40 | -0.102970979 | 0.102970979 | ENSG00000184635 | n.s. |
| ENSG00000225746 | 292.8317 | 0.459793811 | 0.311115 | 1.4778925 | 0.139437 | 0.597163054 | MEG8 | lincRNA | 38.95 | 0.102951085 | 0.102951085 | ENSG00000225746 | n.s. |
| ENSG00000140406 | 728.2092 | -0.358957312 | 0.21626 | -1.659844 | 0.096946 | 0.516760143 | MESDC1 | protein_coding | 55.46 | -0.102917006 | 0.102917006 | ENSG00000140406 | n.s. |
| ENSG00000114279 | 232.2413 | -0.433667026 | 0.285843 | -1.517149 | 0.129229 | 0.579048024 | FGF12 | protein_coding | 37.5 | -0.102902861 | 0.102902861 | ENSG00000114279 | n.s. |
| ENSG00000272426 | 33.06696 | 0.515345349 | 0.372311 | 1.3841793 | 0.166304 | 0.631771973 | | lincRNA | 40.87 | 0.102780293 | 0.102780293 | ENSG00000272426 | n.s. |
| ENSG00000283162 | 81.95281 | -0.410507781 | 0.262145 | -1.565959 | 0.117358 | 0.561890629 | | transcribed_unprocessed_pseudc | 41.54 | -0.102769888 | 0.102769888 | ENSG00000283162 | n.s. |
| ENSG00000267751 | 23.31549 | -0.516260676 | 0.373346 | -1.382796 | 0.166727 | 0.632544316 | | antisense | 53.95 | -0.102688917 | 0.102688917 | ENSG00000267751 | n.s. |
| ENSG00000167978 | 3436.793 | -0.39753196 | 0.251079 | -1.583295 | 0.113354 | 0.552022176 | SRRM2 | protein_coding | 52.86 | -0.102580528 | 0.102580528 | ENSG00000167978 | n.s. |
| ENSG00000224358 | 120.2972 | -0.396723116 | 0.250483 | -1.583835 | 0.113231 | 0.551745164 | | antisense | 42.24 | -0.102458293 | 0.102458293 | ENSG00000224358 | n.s. |
| ENSG00000007944 | 2143.25 | 0.396514965 | 0.250332 | 1.5839549 | 0.113204 | 0.551745164 | MYLIP | protein_coding | 40.19 | 0.102404536 | 0.102404536 | ENSG00000007944 | n.s. |
| ENSG00000163975 | 60.15988 | -0.439293087 | 0.292212 | -1.503335 | 0.132753 | 0.584762676 | MELTF | protein_coding | 56.21 | -0.102364231 | 0.102364231 | ENSG00000163975 | n.s. |
| ENSG00000175470 | 462.0557 | -0.431506768 | 0.28496 | -1.514272 | 0.129957 | 0.5791453 | PPP2R2D | protein_coding | 48.11 | -0.102358783 | 0.102358783 | ENSG00000175470 | n.s. |
| ENSG00000183150 | 618.8086 | -0.368864689 | 0.225643 | -1.634728 | 0.102106 | 0.528171612 | GPR19 | protein_coding | 41.27 | -0.102258493 | 0.102258493 | ENSG00000183150 | n.s. |
| ENSG00000276853 | 180.7864 | 0.423325278 | 0.27546 | 1.5367966 | 0.124343 | 0.57363467 | | sense_intronic | 32.32 | 0.10217574 | 0.10217574 | ENSG00000276853 | n.s. |
| ENSG00000183309 | 522.3797 | -0.416647343 | 0.269215 | -1.547639 | 0.121709 | 0.568759707 | ZNF623 | protein_coding | 47.94 | -0.102108255 | 0.102108255 | ENSG00000183309 | n.s. |
| ENSG00000078246 | 528.2761 | -0.424432105 | 0.276611 | -1.534402 | 0.124931 | 0.574749865 | TULP3 | protein_coding | 45 | -0.102084886 | 0.102084886 | ENSG00000078246 | n.s. |
| ENSG00000039319 | 1186.273 | -0.409743662 | 0.262402 | -1.561513 | 0.118403 | 0.563596794 | ZFYVE16 | protein_coding | 37.65 | -0.102039073 | 0.102039073 | ENSG00000039319 | n.s. |
| ENSG00000164087 | 257.9825 | -0.424149477 | 0.276607 | -1.533402 | 0.125177 | 0.574749865 | POC1A | protein_coding | 51.67 | -0.102016908 | 0.102016908 | ENSG00000164087 | n.s. |
| ENSG00000279570 | 41.11993 | -0.454954787 | 0.30774 | -1.478374 | 0.139308 | 0.596977843 | | TEC | 41.6 | -0.101928884 | 0.101928884 | ENSG00000279570 | n.s. |
| ENSG00000112541 | 642.6138 | -0.389268405 | 0.243672 | -1.597512 | 0.110152 | 0.54732402 | PDE10A | protein_coding | 40.93 | -0.101893143 | 0.101893143 | ENSG00000112541 | n.s. |
| ENSG00000133773 | 3289.115 | 0.36507046 | 0.222739 | 1.6390076 | 0.101212 | 0.526202467 | CCDC59 | protein_coding | 35.35 | 0.101798846 | 0.101798846 | ENSG00000133773 | n.s. |
| ENSG00000281183 | 207.8373 | -0.378704685 | 0.234276 | -1.616487 | 0.105989 | 0.538571713 | NPTN-IT1 | sense_intronic | 44.81 | -0.101779331 | 0.101779331 | ENSG00000281183 | n.s. |
| ENSG00000254491 | 61.26771 | -0.428900937 | 0.282347 | -1.519058 | 0.128748 | 0.579048024 | | processed_pseudogene | 48.99 | -0.101771937 | 0.101771937 | ENSG00000254491 | n.s. |

| | | | | | | | | | | | | | |
|---|---|---|---|---|---|---|---|---|---|---|---|---|---|
| ENSG00000168803 | 516.0396 | -0.428761909 | 0.281874 | -1.521109 | 0.128232 | 0.579048024 | ADAL | protein_coding | 39.85 | -0.101738948 | 0.101738948 | ENSG00000168803 | n.s. |
| ENSG00000240808 | 25.73394 | -0.50000991 | 0.355257 | -1.40746 | 0.159291 | 0.626322307 | | processed_pseudogene | 43.1 | -0.101603074 | 0.101603074 | ENSG00000240808 | n.s. |
| ENSG00000089123 | 257.3659 | -0.403962991 | 0.257608 | -1.568128 | 0.116851 | 0.560427522 | TASP1 | protein_coding | 37.32 | -0.101588833 | 0.101588833 | ENSG00000089123 | n.s. |
| ENSG00000075303 | 550.058 | -0.43609121 | 0.290289 | -1.502264 | 0.133029 | 0.58538567 | SLC25A40 | protein_coding | 35.47 | -0.101416462 | 0.101416462 | ENSG00000075303 | n.s. |
| ENSG00000235688 | 14.47778 | 0.642936204 | 0.523316 | 1.228581 | 0.219229 | 0.695448706 | | antisense | 57.25 | 0.101413475 | 0.101413475 | ENSG00000235688 | n.s. |
| ENSG00000176273 | 604.0417 | -0.40450038 | 0.258191 | -1.566669 | 0.117192 | 0.561481501 | SLC35G1 | protein_coding | 40.3 | -0.101393905 | 0.101393905 | ENSG00000176273 | n.s. |
| ENSG00000204681 | 21.99739 | -0.546088141 | 0.408931 | -1.335404 | 0.181744 | 0.652131419 | GABBR1 | protein_coding | 45.01 | -0.101389387 | 0.101389387 | ENSG00000204681 | n.s. |
| ENSG00000073169 | 216.7374 | -0.413134859 | 0.266473 | -1.550379 | 0.12105 | 0.568618694 | SELENOO | protein_coding | 55.78 | -0.101291936 | 0.101291936 | ENSG00000073169 | n.s. |
| ENSG00000146648 | 455.9786 | -0.388097874 | 0.243607 | -1.593133 | 0.11113 | 0.548299349 | EGFR | protein_coding | 44.57 | -0.101286664 | 0.101286664 | ENSG00000146648 | n.s. |
| ENSG00000109572 | 1016.283 | -0.412466653 | 0.265862 | -1.551433 | 0.120798 | 0.568199666 | CLCN3 | protein_coding | 37.1 | -0.101260162 | 0.101260162 | ENSG00000109572 | n.s. |
| ENSG00000156735 | 696.3124 | -0.404294428 | 0.258254 | -1.565491 | 0.117468 | 0.561980157 | BAG4 | protein_coding | 41.83 | -0.101186413 | 0.101186413 | ENSG00000156735 | n.s. |
| ENSG00000112062 | 1376.068 | -0.393195393 | 0.24856 | -1.581895 | 0.113673 | 0.552996448 | MAPK14 | protein_coding | 39.88 | -0.10116039 | 0.10116039 | ENSG00000112062 | n.s. |
| ENSG00000154822 | 884.4752 | -0.38626773 | 0.241736 | -1.597889 | 0.110068 | 0.54732402 | PLCL2 | protein_coding | 39.07 | -0.1011077 | 0.1011077 | ENSG00000154822 | n.s. |
| ENSG00000134461 | 162.3746 | -0.387221255 | 0.242784 | -1.594921 | 0.11073 | 0.548299349 | ANKRD16 | protein_coding | 46.86 | -0.101057882 | 0.101057882 | ENSG00000134461 | n.s. |
| ENSG00000224415 | 39.41643 | 0.485818435 | 0.342064 | 1.4202556 | 0.155533 | 0.619951348 | | processed_pseudogene | 51.1 | 0.100876502 | 0.100876502 | ENSG00000224415 | n.s. |
| ENSG00000278987 | 122.8894 | -0.403493956 | 0.257961 | -1.564166 | 0.117779 | 0.562372792 | | TEC | 45 | -0.100863684 | 0.100863684 | ENSG00000278987 | n.s. |
| ENSG00000116260 | 447.4056 | -0.416913838 | 0.270843 | -1.539319 | 0.123726 | 0.572922348 | QSOX1 | protein_coding | 51.87 | -0.100853224 | 0.100853224 | ENSG00000116260 | n.s. |
| ENSG00000140153 | 91.74834 | -0.458077221 | 0.3124 | -1.466318 | 0.142562 | 0.602521359 | WDR20 | protein_coding | 45.46 | -0.10078961 | 0.10078961 | ENSG00000140153 | n.s. |
| ENSG00000253054 | 5.741793 | -0.785492044 | 0.728619 | -1.078055 | 0.281009 | 0.744242186 | RNU7-77P | snRNA | 46.67 | -0.10076741 | 0.10076741 | ENSG00000253054 | n.s. |
| ENSG00000070882 | 283.985 | -0.392613158 | 0.248503 | -1.579915 | 0.114126 | 0.553845288 | OSBPL3 | protein_coding | 40.72 | -0.100749065 | 0.100749065 | ENSG00000070882 | n.s. |
| ENSG00000085872 | 456.9059 | -0.374961401 | 0.232043 | -1.615914 | 0.106113 | 0.538671714 | CHERP | protein_coding | 57.3 | -0.100743065 | 0.100743065 | ENSG00000085872 | n.s. |
| ENSG00000213613 | 58.44041 | -0.436983011 | 0.292502 | -1.493947 | 0.135189 | 0.588384714 | RPL11P3 | processed_pseudogene | 46.27 | -0.100654063 | 0.100654063 | ENSG00000213613 | n.s. |
| ENSG00000162374 | 4826.2 | -0.408077764 | 0.262113 | -1.556876 | 0.1195 | 0.566955067 | ELAVL4 | protein_coding | 40.1 | -0.100571319 | 0.100571319 | ENSG00000162374 | n.s. |
| ENSG00000221680 | 20.02539 | 0.534363114 | 0.398013 | 1.3425759 | 0.179409 | 0.648472544 | MIR1278 | miRNA | 37.04 | 0.100518194 | 0.100518194 | ENSG00000221680 | n.s. |
| ENSG00000207980 | 4.081512 | -0.982980739 | 1.051178 | -0.935123 | 0.349725 | 0.7903292 | MIR23A | miRNA | 57.53 | -0.10045274 | 0.10045274 | ENSG00000207980 | n.s. |
| ENSG00000198429 | 284.2081 | -0.423889649 | 0.280251 | -1.512536 | 0.130397 | 0.579675642 | ZNF69 | protein_coding | 43.23 | -0.100383405 | 0.100383405 | ENSG00000198429 | n.s. |
| ENSG00000283982 | 296.1042 | -0.363016665 | 0.222624 | -1.630629 | 0.102969 | 0.529239659 | | lincRNA | 39.73 | -0.100318791 | 0.100318791 | ENSG00000283982 | n.s. |
| ENSG00000196693 | 1302.616 | -0.40530988 | 0.259944 | -1.559219 | 0.118945 | 0.565789092 | ZNF33B | protein_coding | 40.2 | -0.100251546 | 0.100251546 | ENSG00000196693 | n.s. |
| ENSG00000134871 | 1479.181 | -0.374239659 | 0.231979 | -1.613248 | 0.106691 | 0.539737388 | COL4A2 | protein_coding | 47.83 | -0.100227929 | 0.100227929 | ENSG00000134871 | n.s. |
| ENSG00000113360 | 1268.482 | -0.390226896 | 0.24694 | -1.580253 | 0.114049 | 0.553662697 | DROSHA | protein_coding | 39.75 | -0.100192604 | 0.100192604 | ENSG00000113360 | n.s. |
| ENSG00000154548 | 459.582 | -0.435992554 | 0.292197 | -1.492117 | 0.135668 | 0.589422885 | SRSF12 | protein_coding | 40.77 | -0.100092121 | 0.100092121 | ENSG00000154548 | n.s. |
| ENSG00000114030 | 1685.065 | -0.421347152 | 0.277643 | -1.517589 | 0.129118 | 0.579048024 | KPNA1 | protein_coding | 38.27 | -0.099979534 | 0.099979534 | ENSG00000114030 | n.s. |

| | | | | | | | | | | | | | |
|---|---|---|---|---|---|---|---|---|---|---|---|---|---|
| ENSG00000012232 | 582.2645 | -0.406628098 | 0.262002 | -1.552006 | 0.120661 | 0.567746164 | EXTL3 | protein_coding | 44.36 | -0.099967807 | 0.099967807 | ENSG00000012232 | n.s. |
| ENSG00000105835 | 988.2474 | -0.407772418 | 0.263615 | -1.546846 | 0.1219 | 0.569091093 | NAMPT | protein_coding | 37.12 | -0.099830114 | 0.099830114 | ENSG00000105835 | n.s. |
| ENSG00000130023 | 255.8593 | 0.36122545 | 0.221634 | 1.6298276 | 0.103138 | 0.529634455 | ERMARD | protein_coding | 43.29 | 0.09970681 | 0.09970681 | ENSG00000130023 | n.s. |
| ENSG00000207750 | 36.04718 | -0.49829841 | 0.35926 | -1.387014 | 0.165437 | 0.630850594 | MIR553 | miRNA | 32.35 | -0.099696299 | 0.099696299 | ENSG00000207750 | n.s. |
| ENSG00000163618 | 209.5711 | -0.420059915 | 0.276588 | -1.518723 | 0.128832 | 0.579048024 | CADPS | protein_coding | 40 | -0.099674092 | 0.099674092 | ENSG00000163618 | n.s. |
| ENSG00000140650 | 900.1497 | -0.380613297 | 0.238199 | -1.59788 | 0.11007 | 0.54732402 | PMM2 | protein_coding | 46.16 | -0.099627621 | 0.099627621 | ENSG00000140650 | n.s. |
| ENSG00000232795 | 33.16678 | -0.448744181 | 0.305177 | -1.470439 | 0.141443 | 0.600033044 | SCAND3P1 | processed_pseudogene | 36.22 | -0.099542603 | 0.099542603 | ENSG00000232795 | n.s. |
| ENSG00000160326 | 183.376 | -0.405695024 | 0.261585 | -1.550912 | 0.120923 | 0.568402184 | SLC2A6 | protein_coding | 61.28 | -0.099534946 | 0.099534946 | ENSG00000160326 | n.s. |
| ENSG00000177302 | 1513.374 | -0.389540079 | 0.246951 | -1.577399 | 0.114704 | 0.55529213 | TOP3A | protein_coding | 48.23 | -0.099519108 | 0.099519108 | ENSG00000177302 | n.s. |
| ENSG00000147454 | 897.3058 | -0.397410705 | 0.253842 | -1.565581 | 0.117447 | 0.561980157 | SLC25A37 | protein_coding | 48.45 | -0.099463561 | 0.099463561 | ENSG00000147454 | n.s. |
| ENSG00000082805 | 859.1675 | -0.381102188 | 0.239003 | -1.59455 | 0.110813 | 0.548299349 | ERC1 | protein_coding | 39.35 | -0.099460914 | 0.099460914 | ENSG00000082805 | n.s. |
| ENSG00000161682 | 615.6438 | -0.405328226 | 0.261602 | -1.549408 | 0.121284 | 0.568737594 | FAM171A2 | protein_coding | 60.61 | -0.09934111 | 0.09934111 | ENSG00000161682 | n.s. |
| ENSG00000233527 | 52.27488 | -0.493253505 | 0.352854 | -1.397896 | 0.162144 | 0.628957308 | ZNF529-AS1 | antisense | 42.2 | -0.099330815 | 0.099330815 | ENSG00000233527 | n.s. |
| ENSG00000100994 | 1677.661 | -0.396670006 | 0.25329 | -1.56607 | 0.117332 | 0.561890629 | PYGB | protein_coding | 52.17 | -0.099305626 | 0.099305626 | ENSG00000100994 | n.s. |
| ENSG00000273619 | 54.77481 | -0.428233294 | 0.285909 | -1.497794 | 0.134187 | 0.586819166 | | antisense | 65.72 | -0.09913417 | 0.09913417 | ENSG00000273619 | n.s. |
| ENSG00000133639 | 3984.676 | 0.347951097 | 0.210525 | 1.6527758 | 0.098377 | 0.519414517 | BTG1 | protein_coding | 42.46 | 0.098987187 | 0.098987187 | ENSG00000133639 | n.s. |
| ENSG00000168297 | 369.2663 | -0.400232451 | 0.256884 | -1.558025 | 0.119227 | 0.566347442 | PXK | protein_coding | 43.61 | -0.098824219 | 0.098824219 | ENSG00000168297 | n.s. |
| ENSG00000199172 | 11.221 | -0.564398161 | 0.434412 | -1.299224 | 0.193867 | 0.668296258 | MIR331 | miRNA | 51.06 | -0.098787158 | 0.098787158 | ENSG00000199172 | n.s. |
| ENSG00000254331 | 11.40736 | -0.569320789 | 0.439717 | -1.294745 | 0.195408 | 0.670750289 | CKS1BP7 | processed_pseudogene | 45.96 | -0.098742503 | 0.098742503 | ENSG00000254331 | n.s. |
| ENSG00000173295 | 162.809 | -0.42240564 | 0.280546 | -1.505656 | 0.132155 | 0.58394178 | FAM86B3P | cribed_unprocessed_pseudc | 52.58 | -0.09868682 | 0.09868682 | ENSG00000173295 | n.s. |
| ENSG00000166974 | 654.5848 | -0.368561782 | 0.228615 | -1.612151 | 0.106929 | 0.539862594 | MAPRE2 | protein_coding | 39.48 | -0.098670167 | 0.098670167 | ENSG00000166974 | n.s. |
| ENSG00000131473 | 7741.457 | -0.412263519 | 0.269624 | -1.529032 | 0.126257 | 0.5764328 | ACLY | protein_coding | 49.89 | -0.098634589 | 0.098634589 | ENSG00000131473 | n.s. |
| ENSG00000127564 | 591.8662 | -0.399954824 | 0.257077 | -1.555777 | 0.119761 | 0.566955067 | PKMYT1 | protein_coding | 62.22 | -0.09856941 | 0.09856941 | ENSG00000127564 | n.s. |
| ENSG00000255737 | 100.5706 | -0.443229637 | 0.301201 | -1.471542 | 0.141145 | 0.599316087 | AGAP2-AS1 | antisense | 66.06 | -0.098549479 | 0.098549479 | ENSG00000255737 | n.s. |
| ENSG00000168038 | 170.1144 | -0.41192909 | 0.269472 | -1.528653 | 0.126351 | 0.576460831 | ULK4 | protein_coding | 39.39 | -0.098545877 | 0.098545877 | ENSG00000168038 | n.s. |
| ENSG00000237036 | 203.6769 | -0.394185666 | 0.252042 | -1.563969 | 0.117825 | 0.562372792 | ZEB1-AS1 | antisense | 38.57 | -0.098536837 | 0.098536837 | ENSG00000237036 | n.s. |
| ENSG00000187790 | 360.7867 | -0.381269756 | 0.240649 | -1.584341 | 0.113116 | 0.551745164 | FANCM | protein_coding | 37.51 | -0.098467286 | 0.098467286 | ENSG00000187790 | n.s. |
| ENSG00000108176 | 594.0449 | 0.45417447 | 0.313154 | 1.4503237 | 0.146968 | 0.607109641 | DNAJC12 | protein_coding | 39.63 | 0.098434537 | 0.098434537 | ENSG00000108176 | n.s. |
| ENSG00000261071 | 94.51264 | 0.409184325 | 0.2666 | 1.5348269 | 0.124826 | 0.574721329 | | antisense | 54.16 | 0.098426296 | 0.098426296 | ENSG00000261071 | n.s. |
| ENSG00000131148 | 390.5988 | -0.414430472 | 0.272471 | -1.521008 | 0.128258 | 0.579048024 | EMC8 | protein_coding | 48.89 | -0.098338307 | 0.098338307 | ENSG00000131148 | n.s. |
| ENSG00000160877 | 421.2863 | -0.376775723 | 0.23631 | -1.594414 | 0.110843 | 0.548299349 | NACC1 | protein_coding | 55.78 | -0.098331783 | 0.098331783 | ENSG00000160877 | n.s. |
| ENSG00000092470 | 1499.871 | -0.399216581 | 0.256715 | -1.555097 | 0.119923 | 0.567149766 | WDR76 | protein_coding | 41.87 | -0.098327939 | 0.098327939 | ENSG00000092470 | n.s. |

| | | | | | | | | | | | | | |
|---|---|---|---|---|---|---|---|---|---|---|---|---|---|
| ENSG00000136715 | 518.5546 | -0.389378225 | 0.247839 | -1.571095 | 0.11616 | 0.559233228 | SAP130 | protein_coding | 43.39 | -0.098281802 | 0.098281802 | ENSG00000136715 | n.s. |
| ENSG00000168813 | 1232.387 | -0.389885684 | 0.248395 | -1.569618 | 0.116504 | 0.55975287 | ZNF507 | protein_coding | 39.82 | -0.098252623 | 0.098252623 | ENSG00000168813 | n.s. |
| ENSG00000165102 | 118.2485 | -0.401961702 | 0.260099 | -1.54542 | 0.122245 | 0.569616291 | HGSNAT | protein_coding | 44.1 | -0.098246514 | 0.098246514 | ENSG00000165102 | n.s. |
| ENSG00000168101 | 204.2702 | -0.454720567 | 0.31472 | -1.444841 | 0.148503 | 0.60842133 | NUDT16L1 | protein_coding | 65.05 | -0.098126683 | 0.098126683 | ENSG00000168101 | n.s. |
| ENSG00000198646 | 713.3679 | -0.379726269 | 0.23962 | -1.584701 | 0.113034 | 0.551745164 | NCOA6 | protein_coding | 43.48 | -0.098068662 | 0.098068662 | ENSG00000198646 | n.s. |
| ENSG00000160075 | 621.2426 | -0.410341593 | 0.268591 | -1.527756 | 0.126573 | 0.576883472 | SSU72 | protein_coding | 53.87 | -0.098035491 | 0.098035491 | ENSG00000160075 | n.s. |
| ENSG00000229809 | 115.9709 | -0.411803636 | 0.270225 | -1.523928 | 0.127527 | 0.57808993 | ZNF688 | protein_coding | 56.77 | -0.098011158 | 0.098011158 | ENSG00000229809 | n.s. |
| ENSG00000174276 | 94.51939 | -0.407544047 | 0.265864 | -1.532901 | 0.1253 | 0.574811915 | ZNHIT2 | protein_coding | 67.92 | -0.098003844 | 0.098003844 | ENSG00000174276 | n.s. |
| ENSG00000271843 | 34.16641 | -0.452009747 | 0.311957 | -1.448948 | 0.147352 | 0.607109641 | | lincRNA | 36.82 | -0.09796537 | 0.09796537 | ENSG00000271843 | n.s. |
| ENSG00000062370 | 615.3924 | -0.412175995 | 0.27137 | -1.518872 | 0.128795 | 0.579048024 | ZNF112 | protein_coding | 38.31 | -0.097803352 | 0.097803352 | ENSG00000062370 | n.s. |
| ENSG00000084676 | 238.4828 | -0.41217201 | 0.271973 | -1.515491 | 0.129648 | 0.5791453 | NCOA1 | protein_coding | 37.1 | -0.097772338 | 0.097772338 | ENSG00000084676 | n.s. |
| ENSG00000153558 | 170.3536 | -0.410461902 | 0.2694 | -1.523615 | 0.127605 | 0.57808993 | FBXL2 | protein_coding | 38.73 | -0.097691819 | 0.097691819 | ENSG00000153558 | n.s. |
| ENSG00000103148 | 103.8874 | -0.411651962 | 0.270489 | -1.521881 | 0.128039 | 0.579048024 | NPRL3 | protein_coding | 51.3 | -0.097679007 | 0.097679007 | ENSG00000103148 | n.s. |
| ENSG00000227097 | 299.7583 | 0.448903663 | 0.307682 | 1.4589839 | 0.14457 | 0.605912571 | RPS28P7 | processed_pseudogene | 61.9 | 0.097676965 | 0.097676965 | ENSG00000227097 | n.s. |
| ENSG00000256603 | 25.18015 | 0.504490624 | 0.368948 | 1.3673742 | 0.171508 | 0.640361041 | | antisense | 44.96 | 0.097656822 | 0.097656822 | ENSG00000256603 | n.s. |
| ENSG00000276603 | 116.6359 | -0.396051371 | 0.254646 | -1.555302 | 0.119874 | 0.567111082 | | sense_intronic | 43.68 | -0.097560073 | 0.097560073 | ENSG00000276603 | n.s. |
| ENSG00000256188 | 290.0232 | -0.435430815 | 0.294797 | -1.477053 | 0.139661 | 0.597163054 | TAS2R30 | protein_coding | 35.98 | -0.097496038 | 0.097496038 | ENSG00000256188 | n.s. |
| ENSG00000188985 | 47.70417 | 0.500994218 | 0.365411 | 1.3710423 | 0.170362 | 0.638865917 | DHFRP1 | processed_pseudogene | 42.04 | 0.097488605 | 0.097488605 | ENSG00000188985 | n.s. |
| ENSG00000173214 | 674.0041 | -0.38951243 | 0.24912 | -1.563554 | 0.117922 | 0.562372792 | MFSD4B | protein_coding | 43.27 | -0.097368642 | 0.097368642 | ENSG00000173214 | n.s. |
| ENSG00000206717 | 5.091238 | -0.6828559 | 0.593244 | -1.151054 | 0.24971 | 0.720412302 | | misc_RNA | 44.9 | -0.097251572 | 0.097251572 | ENSG00000206717 | n.s. |
| ENSG00000149262 | 563.5844 | -0.40873164 | 0.268412 | -1.522776 | 0.127815 | 0.57834404 | INTS4 | protein_coding | 40.73 | -0.097201998 | 0.097201998 | ENSG00000149262 | n.s. |
| ENSG00000225513 | 56.94318 | 0.433750115 | 0.293358 | 1.4785697 | 0.139255 | 0.596937344 | | processed_pseudogene | 54.22 | 0.097190931 | 0.097190931 | ENSG00000225513 | n.s. |
| ENSG00000168434 | 187.8087 | -0.41204098 | 0.272964 | -1.509508 | 0.131169 | 0.58124285 | COG7 | protein_coding | 45.97 | -0.097094316 | 0.097094316 | ENSG00000168434 | n.s. |
| ENSG00000164074 | 279.6089 | -0.418832612 | 0.279262 | -1.499785 | 0.13367 | 0.586724086 | ABHD18 | protein_coding | 39.82 | -0.096987427 | 0.096987427 | ENSG00000164074 | n.s. |
| ENSG00000248161 | 21.74244 | 0.530779599 | 0.400479 | 1.3253606 | 0.185052 | 0.656928407 | | lincRNA | 38.25 | 0.096857701 | 0.096857701 | ENSG00000248161 | n.s. |
| ENSG00000129951 | 403.5187 | -0.419705863 | 0.280756 | -1.494916 | 0.134936 | 0.587941172 | PLPPR3 | protein_coding | 61.48 | -0.096811926 | 0.096811926 | ENSG00000129951 | n.s. |
| ENSG00000207696 | 9.846965 | -0.628214054 | 0.519302 | -1.209728 | 0.226383 | 0.701289341 | MIR659 | miRNA | 53.61 | -0.096809521 | 0.096809521 | ENSG00000207696 | n.s. |
| ENSG00000152454 | 674.8549 | 0.364257809 | 0.226751 | 1.606425 | 0.108181 | 0.542774082 | ZNF256 | protein_coding | 48.53 | 0.096667065 | 0.096667065 | ENSG00000152454 | n.s. |
| ENSG00000083857 | 1411.771 | -0.368363501 | 0.230555 | -1.597723 | 0.110105 | 0.54732402 | FAT1 | protein_coding | 41.53 | -0.096421169 | 0.096421169 | ENSG00000083857 | n.s. |
| ENSG00000151657 | 613.3653 | -0.404940236 | 0.265858 | -1.523148 | 0.127722 | 0.578111285 | KIN | protein_coding | 39.94 | -0.096371141 | 0.096371141 | ENSG00000151657 | n.s. |
| ENSG00000113296 | 33.2778 | -0.47823997 | 0.341425 | -1.400716 | 0.161299 | 0.628931954 | THBS4 | protein_coding | 42.04 | -0.09631578 | 0.09631578 | ENSG00000113296 | n.s. |
| ENSG00000175906 | 784.9441 | 0.378519662 | 0.240394 | 1.5745776 | 0.115354 | 0.556634437 | ARL4D | protein_coding | 55.99 | 0.096306731 | 0.096306731 | ENSG00000175906 | n.s. |

| | | | | | | | | | | | | | |
|---|---|---|---|---|---|---|---|---|---|---|---|---|---|
| ENSG00000134318 | 2473.187 | -0.389782082 | 0.25007 | -1.55869 | 0.11907 | 0.566191443 | ROCK2 | protein_coding | 36.75 | -0.096290479 | 0.096290479 | ENSG00000134318 | n.s. |
| ENSG00000108443 | 2552.034 | -0.38561399 | 0.246827 | -1.562284 | 0.118221 | 0.563117176 | RPS6KB1 | protein_coding | 41.68 | -0.096172602 | 0.096172602 | ENSG00000108443 | n.s. |
| ENSG00000146733 | 422.6244 | -0.420116002 | 0.282084 | -1.489329 | 0.136401 | 0.590399179 | PSPH | protein_coding | 46.35 | -0.096145335 | 0.096145335 | ENSG00000146733 | n.s. |
| ENSG00000240531 | 17.91889 | -0.514267367 | 0.383888 | -1.33963 | 0.180366 | 0.650241161 | | transcribed_processed_pseudog | 41.28 | -0.096129707 | 0.096129707 | ENSG00000240531 | n.s. |
| ENSG00000236358 | 15.1349 | -0.54080965 | 0.412402 | -1.311367 | 0.189734 | 0.664231317 | | antisense | 40.86 | -0.096091411 | 0.096091411 | ENSG00000236358 | n.s. |
| ENSG00000165494 | 3336.204 | -0.344517706 | 0.210192 | -1.639063 | 0.1012 | 0.526202467 | PCF11 | protein_coding | 38.25 | -0.09606777 | 0.09606777 | ENSG00000165494 | n.s. |
| ENSG00000264881 | 2.690766 | -1.051456519 | 1.208527 | -0.870031 | 0.384283 | 0.810719585 | MIR1273D | miRNA | 54.65 | -0.095818533 | 0.095818533 | ENSG00000264881 | n.s. |
| ENSG00000048828 | 1260.048 | -0.37528936 | 0.237976 | -1.577005 | 0.114794 | 0.555537493 | FAM120A | protein_coding | 42.4 | -0.095806354 | 0.095806354 | ENSG00000048828 | n.s. |
| ENSG00000199377 | 29.50172 | 0.475713974 | 0.340979 | 1.3951405 | 0.162973 | 0.629036117 | RNU5F-1 | snRNA | 35.04 | 0.095772839 | 0.095772839 | ENSG00000199377 | n.s. |
| ENSG00000203993 | 226.3042 | -0.389538297 | 0.250909 | -1.552509 | 0.12054 | 0.567746164 | ARRDC1-AS1 | antisense | 56.78 | -0.095766351 | 0.095766351 | ENSG00000203993 | n.s. |
| ENSG00000264358 | 49.85999 | -0.477012714 | 0.343076 | -1.390401 | 0.164407 | 0.63030432 | MIR3122 | miRNA | 50.68 | -0.095617063 | 0.095617063 | ENSG00000264358 | n.s. |
| ENSG00000138463 | 376.5224 | -0.403309739 | 0.266572 | -1.512948 | 0.130293 | 0.579675642 | DIRC2 | protein_coding | 38.14 | -0.095509775 | 0.095509775 | ENSG00000138463 | n.s. |
| ENSG00000100722 | 1753.49 | -0.394012613 | 0.255653 | -1.541202 | 0.123268 | 0.572276225 | ZC3H14 | protein_coding | 40.2 | -0.09550641 | 0.09550641 | ENSG00000100722 | n.s. |
| ENSG00000231485 | 24.66181 | 0.486555474 | 0.353406 | 1.3767603 | 0.168586 | 0.636399681 | | lincRNA | 73.9 | 0.095496266 | 0.095496266 | ENSG00000231485 | n.s. |
| ENSG00000276573 | 71.32622 | 0.413781521 | 0.276841 | 1.4946525 | 0.135005 | 0.587941172 | | sense_intronic | 39.01 | 0.095445381 | 0.095445381 | ENSG00000276573 | n.s. |
| ENSG00000140992 | 472.8394 | -0.408128364 | 0.271034 | -1.50582 | 0.132113 | 0.583941213 | PDPK1 | protein_coding | 52.51 | -0.095351386 | 0.095351386 | ENSG00000140992 | n.s. |
| ENSG00000139636 | 53.60916 | -0.457259174 | 0.321576 | -1.421931 | 0.155046 | 0.618896168 | LMBR1L | protein_coding | 51.62 | -0.095284676 | 0.095284676 | ENSG00000139636 | n.s. |
| ENSG00000250539 | 117.1355 | -0.401445609 | 0.264608 | -1.517135 | 0.129233 | 0.579048024 | KRT8P33 | processed_pseudogene | 58.61 | -0.095257188 | 0.095257188 | ENSG00000250539 | n.s. |
| ENSG00000100836 | 1524.203 | -0.388133377 | 0.250437 | -1.549822 | 0.121184 | 0.568737594 | PABPN1 | protein_coding | 50.48 | -0.095126858 | 0.095126858 | ENSG00000100836 | n.s. |
| ENSG00000274642 | 43.89846 | -0.459170736 | 0.323661 | -1.418679 | 0.155993 | 0.620893263 | | unprocessed_pseudogene | 40.62 | -0.09504056 | 0.09504056 | ENSG00000274642 | n.s. |
| ENSG00000096070 | 167.3471 | -0.38478059 | 0.246992 | -1.557865 | 0.119265 | 0.566347442 | BRPF3 | protein_coding | 47.04 | -0.095008891 | 0.095008891 | ENSG00000096070 | n.s. |
| ENSG00000164818 | 704.6967 | -0.348751123 | 0.215019 | -1.621952 | 0.104814 | 0.534157909 | DNAAF5 | protein_coding | 53.29 | -0.094975511 | 0.094975511 | ENSG00000164818 | n.s. |
| ENSG00000175866 | 415.5844 | -0.363837405 | 0.228332 | -1.59346 | 0.111057 | 0.548299349 | BAIAP2 | protein_coding | 60.69 | -0.094955112 | 0.094955112 | ENSG00000175866 | n.s. |
| ENSG00000275029 | 35.2279 | -0.470024677 | 0.334954 | -1.40325 | 0.160542 | 0.62809251 | HMGB1P24 | ıscribed_processed_pseudog | 44.34 | -0.094933885 | 0.094933885 | ENSG00000275029 | n.s. |
| ENSG00000244218 | 45.60831 | -0.441112123 | 0.305512 | -1.443848 | 0.148782 | 0.609261602 | RN7SL81P | misc_RNA | 52.11 | -0.094925649 | 0.094925649 | ENSG00000244218 | n.s. |
| ENSG00000128731 | 3080.021 | -0.379515496 | 0.242685 | -1.563821 | 0.11786 | 0.562372792 | HERC2 | protein_coding | 43.74 | -0.094869651 | 0.094869651 | ENSG00000128731 | n.s. |
| ENSG00000188399 | 56.4326 | -0.460535306 | 0.325331 | -1.415591 | 0.156895 | 0.622581912 | ANKRD36P1 | unprocessed_pseudogene | 34.53 | -0.094779778 | 0.094779778 | ENSG00000188399 | n.s. |
| ENSG00000207105 | 6.856408 | -0.640195018 | 0.543891 | -1.177064 | 0.23917 | 0.711226106 | | misc_RNA | 39.64 | -0.09474394 | 0.09474394 | ENSG00000207105 | n.s. |
| ENSG00000145920 | 3031.288 | -0.39507199 | 0.258187 | -1.530177 | 0.125973 | 0.575897021 | CPLX2 | protein_coding | 50.92 | -0.094681044 | 0.094681044 | ENSG00000145920 | n.s. |
| ENSG00000124222 | 513.9251 | -0.390385571 | 0.253412 | -1.540518 | 0.123434 | 0.572285304 | STX16 | protein_coding | 41.01 | -0.094624546 | 0.094624546 | ENSG00000124222 | n.s. |
| ENSG00000265057 | 6.129846 | 0.671290458 | 0.586236 | 1.1450858 | 0.252174 | 0.722876965 | MIR4765 | miRNA | 41.56 | 0.094608733 | 0.094608733 | ENSG00000265057 | n.s. |
| ENSG00000265135 | 6.720378 | 0.629741823 | 0.5308 | 1.186402 | 0.235464 | 0.70779073 | MIR5687 | miRNA | 36.36 | 0.094521181 | 0.094521181 | ENSG00000265135 | n.s. |

| | | | | | | | | | | | | | |
|---|---|---|---|---|---|---|---|---|---|---|---|---|---|
| ENSG00000132613 | 287.0446 | -0.388776013 | 0.252113 | -1.542069 | 0.123057 | 0.572027647 | MTSS1L | protein_coding | 56.24 | -0.094310444 | 0.094310444 | ENSG00000132613 | n.s. |
| ENSG00000105486 | 184.7169 | -0.394568834 | 0.258272 | -1.527727 | 0.12658 | 0.576883472 | LIG1 | protein_coding | 51.48 | -0.094267191 | 0.094267191 | ENSG00000105486 | n.s. |
| ENSG00000159259 | 813.3337 | -0.397206101 | 0.261966 | -1.516249 | 0.129456 | 0.579048024 | CHAF1B | protein_coding | 46.73 | -0.094251215 | 0.094251215 | ENSG00000159259 | n.s. |
| ENSG00000181744 | 1672.36 | 0.381704874 | 0.244976 | 1.5581347 | 0.119201 | 0.566347442 | C3orf58 | protein_coding | 37.13 | 0.094249444 | 0.094249444 | ENSG00000181744 | n.s. |
| ENSG00000166889 | 440.7735 | -0.364408612 | 0.229727 | -1.586271 | 0.112678 | 0.55132222 | PATL1 | protein_coding | 39.82 | -0.094234064 | 0.094234064 | ENSG00000166889 | n.s. |
| ENSG00000260160 | 40.01809 | -0.449581779 | 0.31554 | -1.4248 | 0.154215 | 0.617347914 | | sense_overlapping | 42.96 | -0.094173902 | 0.094173902 | ENSG00000260160 | n.s. |
| ENSG00000082641 | 2436.631 | -0.39409229 | 0.257947 | -1.527804 | 0.126561 | 0.576883472 | NFE2L1 | protein_coding | 54.35 | -0.094153339 | 0.094153339 | ENSG00000082641 | n.s. |
| ENSG00000124370 | 222.6213 | -0.394402177 | 0.258512 | -1.525665 | 0.127093 | 0.577520151 | MCEE | protein_coding | 41.39 | -0.094038438 | 0.094038438 | ENSG00000124370 | n.s. |
| ENSG00000089022 | 4181.552 | -0.396317948 | 0.261613 | -1.514902 | 0.129797 | 0.5791453 | MAPKAPK5 | protein_coding | 43.78 | -0.094011557 | 0.094011557 | ENSG00000089022 | n.s. |
| ENSG00000111203 | 596.4412 | -0.396654251 | 0.262098 | -1.51338 | 0.130183 | 0.579456054 | ITFG2 | protein_coding | 50.2 | -0.093998924 | 0.093998924 | ENSG00000111203 | n.s. |
| ENSG00000172062 | 170.7471 | -0.464707574 | 0.33074 | -1.405055 | 0.160005 | 0.627825878 | SMN1 | protein_coding | 42.22 | -0.093945649 | 0.093945649 | ENSG00000172062 | n.s. |
| ENSG00000065413 | 9.278559 | -0.728733026 | 0.672307 | -1.083929 | 0.278396 | 0.743262409 | ANKRD44 | protein_coding | 40.24 | -0.093902957 | 0.093902957 | ENSG00000065413 | n.s. |
| ENSG00000080644 | 4898.926 | -0.383112595 | 0.247536 | -1.547704 | 0.121694 | 0.568759707 | CHRNA3 | protein_coding | 47.02 | -0.093889855 | 0.093889855 | ENSG00000080644 | n.s. |
| ENSG00000130669 | 190.2895 | -0.421923375 | 0.28672 | -1.471551 | 0.141142 | 0.599316087 | PAK4 | protein_coding | 56.17 | -0.093812158 | 0.093812158 | ENSG00000130669 | n.s. |
| ENSG00000276926 | 15.63808 | -0.529754472 | 0.404736 | -1.308887 | 0.190572 | 0.665181845 | MIR6797 | miRNA | 65.28 | -0.093798121 | 0.093798121 | ENSG00000276926 | n.s. |
| ENSG00000264384 | 32.49352 | -0.485041426 | 0.3555 | -1.364394 | 0.172444 | 0.640837445 | RN7SL431P | misc_RNA | 51.8 | -0.093735284 | 0.093735284 | ENSG00000264384 | n.s. |
| ENSG00000222297 | 6.111091 | -0.760691189 | 0.722697 | -1.052573 | 0.292537 | 0.752972779 | RNU6-1156P | snRNA | 37 | -0.093732919 | 0.093732919 | ENSG00000222297 | n.s. |
| ENSG00000130147 | 992.1299 | -0.381958042 | 0.246823 | -1.547496 | 0.121744 | 0.568759707 | SH3BP4 | protein_coding | 50.77 | -0.093606907 | 0.093606907 | ENSG00000130147 | n.s. |
| ENSG00000175595 | 196.6483 | -0.405316954 | 0.271044 | -1.495392 | 0.134812 | 0.587845023 | ERCC4 | protein_coding | 39.57 | -0.093521681 | 0.093521681 | ENSG00000175595 | n.s. |
| ENSG00000256229 | 639.0795 | -0.413265348 | 0.278586 | -1.483439 | 0.137958 | 0.593935213 | ZNF486 | protein_coding | 40.9 | -0.0935058 | 0.0935058 | ENSG00000256229 | n.s. |
| ENSG00000276524 | 63.26198 | 0.432703972 | 0.2994 | 1.4452381 | 0.148391 | 0.608376989 | | lincRNA | 58.04 | 0.093389296 | 0.093389296 | ENSG00000276524 | n.s. |
| ENSG00000138134 | 188.6805 | 0.38240896 | 0.247564 | 1.5446867 | 0.122422 | 0.570251713 | STAMBPL1 | protein_coding | 39.34 | 0.093282318 | 0.093282318 | ENSG00000138134 | n.s. |
| ENSG00000037637 | 397.3277 | -0.378254492 | 0.243016 | -1.556503 | 0.119589 | 0.566955067 | FBXO42 | protein_coding | 43.85 | -0.093221333 | 0.093221333 | ENSG00000037637 | n.s. |
| ENSG00000148985 | 134.6025 | -0.442525958 | 0.309807 | -1.42839 | 0.15318 | 0.615680787 | PGAP2 | protein_coding | 50.46 | -0.093215614 | 0.093215614 | ENSG00000148985 | n.s. |
| ENSG00000102466 | 136.3826 | -0.38441397 | 0.249433 | -1.541153 | 0.12328 | 0.572276225 | FGF14 | protein_coding | 37.27 | -0.093179754 | 0.093179754 | ENSG00000102466 | n.s. |
| ENSG00000242615 | 72.7635 | -0.429390613 | 0.295557 | -1.452816 | 0.146275 | 0.607109641 | | transcribed_processed_pseudog | 38.76 | -0.09306306 | 0.09306306 | ENSG00000242615 | n.s. |
| ENSG00000121671 | 148.288 | -0.387907115 | 0.253191 | -1.532072 | 0.125505 | 0.575560299 | CRY2 | protein_coding | 49.13 | -0.093062474 | 0.093062474 | ENSG00000121671 | n.s. |
| ENSG00000278469 | 9.775077 | -0.603008964 | 0.497786 | -1.211383 | 0.225749 | 0.701204614 | | misc_RNA | 52.26 | -0.092956989 | 0.092956989 | ENSG00000278469 | n.s. |
| ENSG00000130706 | 961.4722 | -0.390041981 | 0.255728 | -1.525224 | 0.127203 | 0.577830913 | ADRM1 | protein_coding | 64.09 | -0.092907699 | 0.092907699 | ENSG00000130706 | n.s. |
| ENSG00000181007 | 655.191 | -0.404207215 | 0.270768 | -1.492819 | 0.135485 | 0.589119333 | ZFP82 | protein_coding | 40.74 | -0.092885494 | 0.092885494 | ENSG00000181007 | n.s. |
| ENSG00000099725 | 528.5817 | -0.376218639 | 0.241763 | -1.556146 | 0.119673 | 0.566955067 | PRKY | cribed_unprocessed_pseudo | 44.3 | -0.092719595 | 0.092719595 | ENSG00000099725 | n.s. |
| ENSG00000074657 | 2679.184 | -0.383938576 | 0.249702 | -1.537588 | 0.124149 | 0.573519528 | ZNF532 | protein_coding | 43.34 | -0.092702657 | 0.092702657 | ENSG00000074657 | n.s. |

| | | | | | | | | | | | | | |
|---|---|---|---|---|---|---|---|---|---|---|---|---|---|
| ENSG00000162769 | 1326.495 | -0.391332216 | 0.258868 | -1.511708 | 0.130608 | 0.579675642 | FLVCR1 | protein_coding | 40.27 | -0.092673318 | 0.092673318 | ENSG00000162769 | n.s. |
| ENSG00000188878 | 62.47214 | -0.450227101 | 0.317915 | -1.416189 | 0.15672 | 0.622543961 | FBF1 | protein_coding | 53.45 | -0.092670233 | 0.092670233 | ENSG00000188878 | n.s. |
| ENSG00000122557 | 281.1831 | -0.390339665 | 0.257696 | -1.514726 | 0.129842 | 0.5791453 | HERPUD2 | protein_coding | 38.32 | -0.092593434 | 0.092593434 | ENSG00000122557 | n.s. |
| ENSG00000125875 | 506.5933 | -0.389693237 | 0.256996 | -1.516342 | 0.129433 | 0.579048024 | TBC1D20 | protein_coding | 46.39 | -0.092468522 | 0.092468522 | ENSG00000125875 | n.s. |
| ENSG00000176476 | 196.8177 | -0.426025796 | 0.294415 | -1.447023 | 0.147891 | 0.607109641 | SGF29 | protein_coding | 48.31 | -0.092333794 | 0.092333794 | ENSG00000176476 | n.s. |
| ENSG00000213703 | 56.31246 | -0.425748666 | 0.294091 | -1.447677 | 0.147707 | 0.607109641 | | processed_pseudogene | 50.73 | -0.09227373 | 0.09227373 | ENSG00000213703 | n.s. |
| ENSG00000261008 | 38.63563 | -0.457975652 | 0.32683 | -1.401266 | 0.161134 | 0.628931954 | LINC01572 | lincRNA | 37.41 | -0.09223462 | 0.09223462 | ENSG00000261008 | n.s. |
| ENSG00000198393 | 1518.079 | -0.388705924 | 0.256082 | -1.517898 | 0.12904 | 0.579048024 | ZNF26 | protein_coding | 43.05 | -0.092234247 | 0.092234247 | ENSG00000198393 | n.s. |
| ENSG00000216977 | 40.48634 | -0.457994882 | 0.328292 | -1.395083 | 0.162991 | 0.629036117 | RPL21P65 | processed_pseudogene | 45.49 | -0.092205553 | 0.092205553 | ENSG00000216977 | n.s. |
| ENSG00000215193 | 1227.02 | -0.398272672 | 0.265883 | -1.497927 | 0.134152 | 0.586819166 | PEX26 | protein_coding | 48.63 | -0.092198415 | 0.092198415 | ENSG00000215193 | n.s. |
| ENSG00000095209 | 126.9796 | -0.425215568 | 0.292723 | -1.452622 | 0.146329 | 0.607109641 | TMEM38B | protein_coding | 37.52 | -0.092158191 | 0.092158191 | ENSG00000095209 | n.s. |
| ENSG00000147475 | 624.7442 | -0.407080862 | 0.27431 | -1.484015 | 0.137805 | 0.593774169 | ERLIN2 | protein_coding | 44.02 | -0.092154436 | 0.092154436 | ENSG00000147475 | n.s. |
| ENSG00000198018 | 290.7497 | -0.388225174 | 0.255234 | -1.521058 | 0.128245 | 0.579048024 | ENTPD7 | protein_coding | 40.08 | -0.092120172 | 0.092120172 | ENSG00000198018 | n.s. |
| ENSG00000172725 | 244.3506 | -0.404646307 | 0.272177 | -1.486701 | 0.137094 | 0.592046125 | CORO1B | protein_coding | 64.53 | -0.092115489 | 0.092115489 | ENSG00000172725 | n.s. |
| ENSG00000033627 | 877.2427 | -0.37587158 | 0.242911 | -1.547366 | 0.121775 | 0.568759707 | ATP6V0A1 | protein_coding | 45.47 | -0.092115291 | 0.092115291 | ENSG00000033627 | n.s. |
| ENSG00000118058 | 2114.263 | -0.375822245 | 0.242673 | -1.54868 | 0.121459 | 0.568737594 | KMT2A | protein_coding | 41.25 | -0.092109546 | 0.092109546 | ENSG00000118058 | n.s. |
| ENSG00000007376 | 202.2166 | -0.425134989 | 0.293908 | -1.44649 | 0.14804 | 0.607294556 | RPUSD1 | protein_coding | 67.17 | -0.092084499 | 0.092084499 | ENSG00000007376 | n.s. |
| ENSG00000178607 | 175.5213 | -0.418323703 | 0.285222 | -1.46666 | 0.142468 | 0.60247798 | ERN1 | protein_coding | 45.1 | -0.092055821 | 0.092055821 | ENSG00000178607 | n.s. |
| ENSG00000166813 | 553.0734 | -0.39810702 | 0.266024 | -1.496506 | 0.134522 | 0.587308708 | KIF7 | protein_coding | 49.35 | -0.092015893 | 0.092015893 | ENSG00000166813 | n.s. |
| ENSG00000102221 | 357.3956 | -0.410847087 | 0.278105 | -1.47731 | 0.139593 | 0.597163054 | JADE3 | protein_coding | 40.39 | -0.091991567 | 0.091991567 | ENSG00000102221 | n.s. |
| ENSG00000071462 | 799.4886 | -0.424325367 | 0.29292 | -1.448603 | 0.147448 | 0.607109641 | BUD23 | protein_coding | 54.07 | -0.091965255 | 0.091965255 | ENSG00000071462 | n.s. |
| ENSG00000078902 | 26.94003 | -0.501116404 | 0.377611 | -1.32707 | 0.184485 | 0.65558715 | TOLLIP | protein_coding | 57.49 | -0.091889498 | 0.091889498 | ENSG00000078902 | n.s. |
| ENSG00000271991 | 187.1112 | 0.400708359 | 0.268946 | 1.4899192 | 0.136245 | 0.590213403 | | lincRNA | 46.06 | 0.091758581 | 0.091758581 | ENSG00000271991 | n.s. |
| ENSG00000135631 | 240.7464 | -0.371994496 | 0.239033 | -1.55625 | 0.119649 | 0.566955067 | RAB11FIP5 | protein_coding | 48.58 | -0.091678549 | 0.091678549 | ENSG00000135631 | n.s. |
| ENSG00000107771 | 538.9816 | -0.387079013 | 0.25605 | -1.511731 | 0.130602 | 0.579675642 | CCSER2 | protein_coding | 37.56 | -0.091666096 | 0.091666096 | ENSG00000107771 | n.s. |
| ENSG00000126749 | 1176.193 | -0.396926653 | 0.265334 | -1.495953 | 0.134666 | 0.587575478 | EMG1 | protein_coding | 48.3 | -0.091664787 | 0.091664787 | ENSG00000126749 | n.s. |
| ENSG00000012048 | 3778.135 | -0.397287644 | 0.265826 | -1.494539 | 0.135035 | 0.587941172 | BRCA1 | protein_coding | 44.09 | -0.091640802 | 0.091640802 | ENSG00000012048 | n.s. |
| ENSG00000234473 | 39.26156 | -0.422491569 | 0.291587 | -1.448937 | 0.147355 | 0.607109641 | ZNF101P2 | processed_pseudogene | 44.02 | -0.09156781 | 0.09156781 | ENSG00000234473 | n.s. |
| ENSG00000265185 | 645.7146 | -0.335224412 | 0.206446 | -1.623788 | 0.104421 | 0.533328575 | SNORD3B-1 | snoRNA | 55.28 | -0.09151799 | 0.09151799 | ENSG00000265185 | n.s. |
| ENSG00000279145 | 421.435 | -0.34442333 | 0.214658 | -1.604523 | 0.108599 | 0.543082569 | | TEC | 38.91 | -0.091318382 | 0.091318382 | ENSG00000279145 | n.s. |
| ENSG00000239553 | 19.9676 | -0.537441627 | 0.419254 | -1.2819 | 0.199878 | 0.676613539 | RN7SL12P | misc_RNA | 54.14 | -0.091181979 | 0.091181979 | ENSG00000239553 | n.s. |
| ENSG00000162241 | 16.88584 | -0.578621833 | 0.471181 | -1.228025 | 0.219438 | 0.695793572 | SLC25A45 | protein_coding | 59.38 | -0.091144273 | 0.091144273 | ENSG00000162241 | n.s. |

| | | | | | | | | | | | | | |
|---|---|---|---|---|---|---|---|---|---|---|---|---|---|
| ENSG00000236859 | 250.5845 | -0.360194498 | 0.229091 | -1.572275 | 0.115887 | 0.558494404 | NIFK-AS1 | antisense | 44.09 | -0.091122427 | 0.091122427 | ENSG00000236859 | n.s. |
| ENSG00000100744 | 420.4435 | -0.406907504 | 0.275671 | -1.476062 | 0.139927 | 0.597163054 | GSKIP | protein_coding | 38.94 | -0.091109467 | 0.091109467 | ENSG00000100744 | n.s. |
| ENSG00000178921 | 1578.419 | -0.371394008 | 0.239939 | -1.547869 | 0.121654 | 0.568759707 | PFAS | protein_coding | 51.84 | -0.091017967 | 0.091017967 | ENSG00000178921 | n.s. |
| ENSG00000254577 | 34.81041 | -0.495592361 | 0.373047 | -1.328499 | 0.184013 | 0.655193675 | | sense_overlapping | 39.9 | -0.091005776 | 0.091005776 | ENSG00000254577 | n.s. |
| ENSG00000140350 | 6596.538 | -0.390529521 | 0.25981 | -1.503138 | 0.132803 | 0.584763747 | ANP32A | protein_coding | 49.39 | -0.091001017 | 0.091001017 | ENSG00000140350 | n.s. |
| ENSG00000228175 | 60.88871 | 0.419833302 | 0.2896 | 1.4496982 | 0.147143 | 0.607109641 | GEMIN8P4 | processed_pseudogene | 53.77 | 0.090991677 | 0.090991677 | ENSG00000228175 | n.s. |
| ENSG00000147118 | 204.5313 | -0.402813218 | 0.271696 | -1.482587 | 0.138184 | 0.594542171 | ZNF182 | protein_coding | 40.39 | -0.090962208 | 0.090962208 | ENSG00000147118 | n.s. |
| ENSG00000152253 | 536.0901 | -0.378082675 | 0.246497 | -1.53382 | 0.125074 | 0.574749865 | SPC25 | protein_coding | 39.39 | -0.090936869 | 0.090936869 | ENSG00000152253 | n.s. |
| ENSG00000252153 | 7.998113 | -0.594113303 | 0.494194 | -1.202187 | 0.229291 | 0.702989524 | MIR2278 | miRNA | 58.33 | -0.090929722 | 0.090929722 | ENSG00000252153 | n.s. |
| ENSG00000145022 | 161.22 | -0.405007053 | 0.273805 | -1.479181 | 0.139092 | 0.596580707 | TCTA | protein_coding | 53.23 | -0.090855554 | 0.090855554 | ENSG00000145022 | n.s. |
| ENSG00000107290 | 2825.757 | -0.419139096 | 0.288683 | -1.4519 | 0.146529 | 0.607109641 | SETX | protein_coding | 41 | -0.090841219 | 0.090841219 | ENSG00000107290 | n.s. |
| ENSG00000163093 | 288.9841 | -0.376120696 | 0.244475 | -1.538486 | 0.12393 | 0.57343851 | BBS5 | protein_coding | 40.4 | -0.090838092 | 0.090838092 | ENSG00000163093 | n.s. |
| ENSG00000107566 | 723.1742 | -0.382532313 | 0.252216 | -1.516687 | 0.129346 | 0.579048024 | ERLIN1 | protein_coding | 38.56 | -0.090769339 | 0.090769339 | ENSG00000107566 | n.s. |
| ENSG00000260614 | 52.97873 | -0.464836351 | 0.338608 | -1.372788 | 0.169818 | 0.638037836 | | antisense | 37.67 | -0.090714472 | 0.090714472 | ENSG00000260614 | n.s. |
| ENSG00000040608 | 129.3257 | -0.38202658 | 0.251153 | -1.521092 | 0.128237 | 0.579048024 | RTN4R | protein_coding | 60.45 | -0.090649336 | 0.090649336 | ENSG00000040608 | n.s. |
| ENSG00000116649 | 266.4641 | -0.376865459 | 0.245801 | -1.533212 | 0.125224 | 0.574749865 | SRM | protein_coding | 59.88 | -0.090644103 | 0.090644103 | ENSG00000116649 | n.s. |
| ENSG00000077157 | 314.1778 | -0.369831079 | 0.239105 | -1.546733 | 0.121928 | 0.569091093 | PPP1R12B | protein_coding | 39.63 | -0.090541383 | 0.090541383 | ENSG00000077157 | n.s. |
| ENSG00000088826 | 117.3225 | -0.416601712 | 0.285828 | -1.457527 | 0.144971 | 0.606317657 | SMOX | protein_coding | 49.87 | -0.090527462 | 0.090527462 | ENSG00000088826 | n.s. |
| ENSG00000157106 | 3415.712 | -0.374342001 | 0.243351 | -1.538282 | 0.12398 | 0.573477749 | SMG1 | protein_coding | 39.95 | -0.09039739 | 0.09039739 | ENSG00000157106 | n.s. |
| ENSG00000196696 | 187.8708 | -0.380717163 | 0.251153 | -1.515876 | 0.129551 | 0.579048024 | | processed_transcript | 44.73 | -0.09033863 | 0.09033863 | ENSG00000196696 | n.s. |
| ENSG00000083814 | 142.1244 | -0.394242648 | 0.264478 | -1.490647 | 0.136054 | 0.590213403 | ZNF671 | protein_coding | 49.44 | -0.090277992 | 0.090277992 | ENSG00000083814 | n.s. |
| ENSG00000184007 | 986.6916 | -0.37647175 | 0.245804 | -1.531592 | 0.125623 | 0.575724414 | PTP4A2 | protein_coding | 41.84 | -0.090272412 | 0.090272412 | ENSG00000184007 | n.s. |
| ENSG00000146966 | 158.6065 | -0.426931659 | 0.298259 | -1.431413 | 0.152312 | 0.614654298 | DENND2A | protein_coding | 47.56 | -0.090240151 | 0.090240151 | ENSG00000146966 | n.s. |
| ENSG00000232648 | 22.91253 | -0.482360737 | 0.360113 | -1.339472 | 0.180417 | 0.650258454 | | antisense | 40.1 | -0.090159971 | 0.090159971 | ENSG00000232648 | n.s. |
| ENSG00000240207 | 158.1724 | -0.415867985 | 0.286758 | -1.450239 | 0.146992 | 0.607109641 | | antisense | 43.38 | -0.090132262 | 0.090132262 | ENSG00000240207 | n.s. |
| ENSG00000171853 | 725.2682 | -0.398021799 | 0.268216 | -1.483962 | 0.137819 | 0.593774169 | TRAPPC12 | protein_coding | 47.39 | -0.090103657 | 0.090103657 | ENSG00000171853 | n.s. |
| ENSG00000137337 | 620.3207 | -0.374434232 | 0.244166 | -1.53352 | 0.125148 | 0.574749865 | MDC1 | protein_coding | 48.84 | -0.090059342 | 0.090059342 | ENSG00000137337 | n.s. |
| ENSG00000137310 | 383.0133 | -0.38113943 | 0.25235 | -1.510359 | 0.130952 | 0.580465481 | TCF19 | protein_coding | 51.1 | -0.090034129 | 0.090034129 | ENSG00000137310 | n.s. |
| ENSG00000204764 | 889.9685 | -0.401132397 | 0.271211 | -1.479041 | 0.139129 | 0.596580707 | RANBP17 | protein_coding | 37.4 | -0.08998635 | 0.08998635 | ENSG00000204764 | n.s. |
| ENSG00000230749 | 48.12789 | -0.445036869 | 0.316762 | -1.404955 | 0.160035 | 0.627825878 | MEIS1-AS2 | antisense | 50.78 | -0.089969004 | 0.089969004 | ENSG00000230749 | n.s. |
| ENSG00000277872 | 2.238437 | -1.016937197 | 1.184338 | -0.858655 | 0.390531 | 0.815710819 | MIR6817 | miRNA | 54.55 | -0.089962106 | 0.089962106 | ENSG00000277872 | n.s. |
| ENSG00000125877 | 432.2762 | -0.430761276 | 0.302757 | -1.422796 | 0.154795 | 0.618415021 | ITPA | protein_coding | 50.89 | -0.089908481 | 0.089908481 | ENSG00000125877 | n.s. |

| | | | | | | | | | | | | | |
|---|---|---|---|---|---|---|---|---|---|---|---|---|---|
| ENSG00000214753 | 434.0467 | -0.377660969 | 0.247817 | -1.52395 | 0.127521 | 0.57808993 | HNRNPUL2 | protein_coding | 46.07 | -0.089885046 | 0.089885046 | ENSG00000214753 | n.s. |
| ENSG00000175826 | 263.6551 | -0.385731106 | 0.256589 | -1.503302 | 0.132761 | 0.584762676 | CTDNEP1 | protein_coding | 52.53 | -0.089883199 | 0.089883199 | ENSG00000175826 | n.s. |
| ENSG00000179967 | 67.30815 | 0.424604523 | 0.296358 | 1.4327435 | 0.151931 | 0.614510829 | PPP1R14BP3 | processed_pseudogene | 65.17 | 0.089791314 | 0.089791314 | ENSG00000179967 | n.s. |
| ENSG00000184661 | 1017.678 | -0.364030553 | 0.234003 | -1.555664 | 0.119788 | 0.566955067 | CDCA2 | protein_coding | 39.74 | -0.089715824 | 0.089715824 | ENSG00000184661 | n.s. |
| ENSG00000280269 | 66.55156 | -0.43331172 | 0.305397 | -1.418846 | 0.155944 | 0.620877176 | | TEC | 38.04 | -0.089693058 | 0.089693058 | ENSG00000280269 | n.s. |
| ENSG00000200063 | 9.089553 | -0.588474318 | 0.491483 | -1.197345 | 0.231172 | 0.704043192 | | snoRNA | 59.69 | -0.089683896 | 0.089683896 | ENSG00000200063 | n.s. |
| ENSG00000168884 | 214.0256 | -0.406983426 | 0.277508 | -1.466562 | 0.142495 | 0.60247798 | TNIP2 | protein_coding | 54.04 | -0.089560293 | 0.089560293 | ENSG00000168884 | n.s. |
| ENSG00000145362 | 5668.015 | -0.371980719 | 0.242251 | -1.53552 | 0.124656 | 0.574507637 | ANK2 | protein_coding | 37.08 | -0.089537319 | 0.089537319 | ENSG00000145362 | n.s. |
| ENSG00000239396 | 20.45608 | -0.48575629 | 0.364921 | -1.331126 | 0.183148 | 0.654498372 | RN7SL414P | misc_RNA | 53.06 | -0.08942357 | 0.08942357 | ENSG00000239396 | n.s. |
| ENSG00000223131 | 26.06643 | -0.486936924 | 0.366727 | -1.327791 | 0.184247 | 0.655193675 | RNA5SP304 | rRNA | 57.69 | -0.089416375 | 0.089416375 | ENSG00000223131 | n.s. |
| ENSG00000115993 | 975.0594 | -0.39913186 | 0.270147 | -1.477463 | 0.139552 | 0.597163054 | TRAK2 | protein_coding | 37.73 | -0.089368445 | 0.089368445 | ENSG00000115993 | n.s. |
| ENSG00000184640 | 4371.814 | -0.379484273 | 0.251528 | -1.508717 | 0.131371 | 0.581584033 | 9-Sep | protein_coding | 54.75 | -0.089325864 | 0.089325864 | ENSG00000184640 | n.s. |
| ENSG00000112531 | 2593.496 | -0.363891251 | 0.234588 | -1.551191 | 0.120856 | 0.568279648 | QKI | protein_coding | 36.39 | -0.089312704 | 0.089312704 | ENSG00000112531 | n.s. |
| ENSG00000152749 | 710.606 | -0.420908808 | 0.293438 | -1.434407 | 0.151456 | 0.613599141 | GPR180 | protein_coding | 36.49 | -0.08928118 | 0.08928118 | ENSG00000152749 | n.s. |
| ENSG00000177990 | 175.9216 | -0.37681021 | 0.249099 | -1.51269 | 0.130358 | 0.579675642 | DPY19L2 | protein_coding | 36.31 | -0.08923429 | 0.08923429 | ENSG00000177990 | n.s. |
| ENSG00000068724 | 127.2357 | -0.389761628 | 0.261773 | -1.488932 | 0.136505 | 0.590603821 | TTC7A | protein_coding | 50.88 | -0.089139945 | 0.089139945 | ENSG00000068724 | n.s. |
| ENSG00000090615 | 1453.637 | -0.373024334 | 0.244228 | -1.52736 | 0.126672 | 0.576921481 | GOLGA3 | protein_coding | 53.56 | -0.08910928 | 0.08910928 | ENSG00000090615 | n.s. |
| ENSG00000201644 | 8.985775 | 0.605543305 | 0.515802 | 1.1739845 | 0.240401 | 0.712634415 | | misc_RNA | 36.17 | 0.089095529 | 0.089095529 | ENSG00000201644 | n.s. |
| ENSG00000121075 | 99.33883 | 0.400920982 | 0.272516 | 1.4711848 | 0.141241 | 0.599542886 | TBX4 | protein_coding | 52.81 | 0.089076522 | 0.089076522 | ENSG00000121075 | n.s. |
| ENSG00000125952 | 472.8968 | -0.384723267 | 0.25672 | -1.49861 | 0.133975 | 0.586819166 | MAX | protein_coding | 44.69 | -0.089061786 | 0.089061786 | ENSG00000125952 | n.s. |
| ENSG00000260196 | 152.5494 | -0.419866203 | 0.292637 | -1.434767 | 0.151353 | 0.613599141 | | antisense | 48.42 | -0.089060027 | 0.089060027 | ENSG00000260196 | n.s. |
| ENSG00000133302 | 931.5459 | -0.398358955 | 0.270063 | -1.47506 | 0.140196 | 0.597636216 | SLF1 | protein_coding | 36.06 | -0.08905836 | 0.08905836 | ENSG00000133302 | n.s. |
| ENSG00000101003 | 779.5521 | -0.397958199 | 0.269719 | -1.475457 | 0.14009 | 0.597387275 | GINS1 | protein_coding | 41.38 | -0.089040772 | 0.089040772 | ENSG00000101003 | n.s. |
| ENSG00000197566 | 64.70253 | -0.398548219 | 0.270347 | -1.474212 | 0.140424 | 0.597898852 | ZNF624 | protein_coding | 37.68 | -0.089024625 | 0.089024625 | ENSG00000197566 | n.s. |
| ENSG00000183647 | 435.438 | -0.367818901 | 0.23893 | -1.539441 | 0.123697 | 0.572922348 | ZNF530 | protein_coding | 46.12 | -0.08897695 | 0.08897695 | ENSG00000183647 | n.s. |
| ENSG00000270395 | 4.089938 | -0.698827615 | 0.65432 | -1.06802 | 0.285511 | 0.746113205 | MREGP1 | processed_pseudogene | 51.76 | -0.08888757 | 0.08888757 | ENSG00000270395 | n.s. |
| ENSG00000113580 | 512.357 | -0.374735155 | 0.247571 | -1.513647 | 0.130115 | 0.579456054 | NR3C1 | protein_coding | 37.82 | -0.088804548 | 0.088804548 | ENSG00000113580 | n.s. |
| ENSG00000103091 | 22.77017 | -0.48049986 | 0.360579 | -1.332577 | 0.182671 | 0.65347449 | WDR59 | protein_coding | 44.73 | -0.088782614 | 0.088782614 | ENSG00000103091 | n.s. |
| ENSG00000227051 | 5108.552 | -0.387530847 | 0.260036 | -1.490299 | 0.136146 | 0.590213403 | C14orf132 | protein_coding | 47.76 | -0.08874105 | 0.08874105 | ENSG00000227051 | n.s. |
| ENSG00000078967 | 36.76128 | -0.452443371 | 0.32874 | -1.376297 | 0.16873 | 0.636645906 | UBE2D4 | protein_coding | 45.79 | -0.088725072 | 0.088725072 | ENSG00000078967 | n.s. |
| ENSG00000134802 | 373.5739 | -0.373882677 | 0.246143 | -1.518965 | 0.128771 | 0.579048024 | SLC43A3 | protein_coding | 48.17 | -0.088716906 | 0.088716906 | ENSG00000134802 | n.s. |
| ENSG00000198435 | 127.9195 | 0.395879238 | 0.26812 | 1.476501 | 0.139809 | 0.597163054 | NRARP | protein_coding | 67.29 | 0.08864016 | 0.08864016 | ENSG00000198435 | n.s. |

| | | | | | | | | | | | | | |
|---|---|---|---|---|---|---|---|---|---|---|---|---|---|
| ENSG00000205571 | 65.83928 | -0.455399931 | 0.332134 | -1.371132 | 0.170334 | 0.638865917 | SMN2 | protein_coding | 42.22 | -0.0886164 | 0.0886164 | ENSG00000205571 | n.s. |
| ENSG00000251768 | 8.077195 | -0.598709518 | 0.508994 | -1.17626 | 0.239491 | 0.71125159 | RNA5SP217 | rRNA | 49.45 | -0.088595088 | 0.088595088 | ENSG00000251768 | n.s. |
| ENSG00000214401 | 45.73692 | 0.435921754 | 0.30966 | 1.4077442 | 0.159207 | 0.626322307 | KANSL1-AS1 | antisense | 45.58 | 0.088580225 | 0.088580225 | ENSG00000214401 | n.s. |
| ENSG00000173320 | 857.9552 | -0.375267119 | 0.248649 | -1.509223 | 0.131242 | 0.581381012 | STOX2 | protein_coding | 42.71 | -0.088390101 | 0.088390101 | ENSG00000173320 | n.s. |
| ENSG00000162063 | 477.3487 | -0.372888654 | 0.246661 | -1.511744 | 0.130599 | 0.579675642 | CCNF | protein_coding | 52.76 | -0.088305607 | 0.088305607 | ENSG00000162063 | n.s. |
| ENSG00000269921 | 75.2681 | 0.400903373 | 0.273192 | 1.4674795 | 0.142246 | 0.602215775 | | lincRNA | 42.72 | 0.088298115 | 0.088298115 | ENSG00000269921 | n.s. |
| ENSG00000197724 | 634.408 | -0.372832009 | 0.246499 | -1.512506 | 0.130405 | 0.579675642 | PHF2 | protein_coding | 51.51 | -0.088292193 | 0.088292193 | ENSG00000197724 | n.s. |
| ENSG00000109501 | 130.9126 | -0.400838842 | 0.273158 | -1.467426 | 0.14226 | 0.602215775 | WFS1 | protein_coding | 57.04 | -0.088283902 | 0.088283902 | ENSG00000109501 | n.s. |
| ENSG00000091127 | 1464.872 | -0.403205034 | 0.275847 | -1.4617 | 0.143823 | 0.604031311 | PUS7 | protein_coding | 43 | -0.088277931 | 0.088277931 | ENSG00000091127 | n.s. |
| ENSG00000119547 | 1176.154 | -0.394187361 | 0.267099 | -1.475812 | 0.139994 | 0.597163054 | ONECUT2 | protein_coding | 44.08 | -0.088261337 | 0.088261337 | ENSG00000119547 | n.s. |
| ENSG00000160917 | 830.6085 | -0.402602168 | 0.275352 | -1.462135 | 0.143704 | 0.604031311 | CPSF4 | protein_coding | 53.24 | -0.088145939 | 0.088145939 | ENSG00000160917 | n.s. |
| ENSG00000164045 | 1906.392 | -0.37139011 | 0.244797 | -1.517132 | 0.129233 | 0.579048024 | CDC25A | protein_coding | 45.04 | -0.088125457 | 0.088125457 | ENSG00000164045 | n.s. |
| ENSG00000248174 | 1430.3 | -0.33760349 | 0.212069 | -1.591952 | 0.111396 | 0.548299349 | LINC02268 | lincRNA | 36.76 | -0.088108525 | 0.088108525 | ENSG00000248174 | n.s. |
| ENSG00000099810 | 1449.789 | -0.391973193 | 0.264852 | -1.479968 | 0.138882 | 0.596095432 | MTAP | protein_coding | 40.09 | -0.088070186 | 0.088070186 | ENSG00000099810 | n.s. |
| ENSG00000207997 | 15.92832 | -0.522306139 | 0.410354 | -1.272819 | 0.203082 | 0.678385894 | MIR644A | miRNA | 32.98 | -0.088020697 | 0.088020697 | ENSG00000207997 | n.s. |
| ENSG00000187626 | 182.7437 | -0.371443246 | 0.245737 | -1.511549 | 0.130649 | 0.579675642 | ZKSCAN4 | protein_coding | 44.69 | -0.087963313 | 0.087963313 | ENSG00000187626 | n.s. |
| ENSG00000212283 | 17.70822 | -0.498516379 | 0.381468 | -1.306837 | 0.191268 | 0.666152146 | SNORD89 | snoRNA | 47.37 | -0.087951534 | 0.087951534 | ENSG00000212283 | n.s. |
| ENSG00000116685 | 321.2474 | -0.401369165 | 0.274584 | -1.461737 | 0.143813 | 0.604031311 | KIAA2013 | protein_coding | 55.79 | -0.087875985 | 0.087875985 | ENSG00000116685 | n.s. |
| ENSG00000148516 | 4819.633 | -0.385882787 | 0.259594 | -1.486486 | 0.137151 | 0.592108152 | ZEB1 | protein_coding | 36.04 | -0.087826521 | 0.087826521 | ENSG00000148516 | n.s. |
| ENSG00000232545 | 38.97369 | -0.448018571 | 0.325777 | -1.375229 | 0.16906 | 0.636857974 | | sense_intronic | 55.52 | -0.087792558 | 0.087792558 | ENSG00000232545 | n.s. |
| ENSG00000273768 | 775.7121 | 0.336233967 | 0.21115 | 1.5923934 | 0.111296 | 0.548299349 | | snRNA | 54.27 | 0.087751104 | 0.087751104 | ENSG00000273768 | n.s. |
| ENSG00000160961 | 181.5639 | -0.379363002 | 0.253526 | -1.49635 | 0.134563 | 0.587308708 | ZNF333 | protein_coding | 46.91 | -0.087683521 | 0.087683521 | ENSG00000160961 | n.s. |
| ENSG00000278705 | 1389.391 | -0.356548073 | 0.229725 | -1.552066 | 0.120646 | 0.567746164 | HIST1H4B | protein_coding | 53.42 | -0.087655843 | 0.087655843 | ENSG00000278705 | n.s. |
| ENSG00000099251 | 104.7455 | -0.412506323 | 0.287124 | -1.436681 | 0.150809 | 0.613237549 | HSD17B7P2 | cribed_unprocessed_pseudc | 39.3 | -0.087604488 | 0.087604488 | ENSG00000099251 | n.s. |
| ENSG00000090447 | 456.4902 | -0.368840586 | 0.242475 | -1.52115 | 0.128222 | 0.579048024 | TFAP4 | protein_coding | 56.39 | -0.087520492 | 0.087520492 | ENSG00000090447 | n.s. |
| ENSG00000116580 | 1596.741 | -0.369283463 | 0.243993 | -1.513499 | 0.130153 | 0.579456054 | GON4L | protein_coding | 44.03 | -0.087512609 | 0.087512609 | ENSG00000116580 | n.s. |
| ENSG00000265625 | 35.592 | -0.439249623 | 0.317584 | -1.383096 | 0.166636 | 0.632544316 | | sense_intronic | 39.73 | -0.087370722 | 0.087370722 | ENSG00000265625 | n.s. |
| ENSG00000132199 | 76.84665 | -0.424795768 | 0.300665 | -1.412853 | 0.157699 | 0.622955867 | ENOSF1 | protein_coding | 46.1 | -0.087313677 | 0.087313677 | ENSG00000132199 | n.s. |
| ENSG00000201182 | 9.773178 | -0.61457728 | 0.534423 | -1.149983 | 0.250151 | 0.721235893 | RNU6-670P | snRNA | 41.35 | -0.087222447 | 0.087222447 | ENSG00000201182 | n.s. |
| ENSG00000065491 | 257.8198 | -0.373395805 | 0.248129 | -1.504843 | 0.132365 | 0.584331244 | TBC1D22B | protein_coding | 43.34 | -0.08712851 | 0.08712851 | ENSG00000065491 | n.s. |
| ENSG00000065559 | 893.8837 | -0.40058313 | 0.274767 | -1.457901 | 0.144868 | 0.606306192 | MAP2K4 | protein_coding | 37.09 | -0.087049917 | 0.087049917 | ENSG00000065559 | n.s. |
| ENSG00000196642 | 1411.195 | -0.389609508 | 0.264252 | -1.474388 | 0.140377 | 0.597880976 | RABL6 | protein_coding | 57.06 | -0.087033023 | 0.087033023 | ENSG00000196642 | n.s. |

| | | | | | | | | | | | | | |
|---|---|---|---|---|---|---|---|---|---|---|---|---|---|
| ENSG00000280389 | 34.09508 | -0.452200117 | 0.332429 | -1.360291 | 0.173738 | 0.642002364 | | TEC | 43.68 | -0.08703196 | 0.08703196 | ENSG00000280389 | n.s. |
| ENSG00000234936 | 29.5282 | -0.473902837 | 0.35696 | -1.327607 | 0.184308 | 0.655193675 | | antisense | 49.76 | -0.087022922 | 0.087022922 | ENSG00000234936 | n.s. |
| ENSG00000104447 | 83.07315 | 0.401503451 | 0.275968 | 1.4548935 | 0.145699 | 0.607109641 | TRPS1 | protein_coding | 35.81 | 0.087018995 | 0.087018995 | ENSG00000104447 | n.s. |
| ENSG00000250132 | 193.5177 | -0.361618664 | 0.23587 | -1.533124 | 0.125245 | 0.574749865 | | antisense | 45.76 | -0.086976927 | 0.086976927 | ENSG00000250132 | n.s. |
| ENSG00000168395 | 1553.367 | -0.354765619 | 0.229117 | -1.548404 | 0.121525 | 0.568737594 | ING5 | protein_coding | 51.69 | -0.086948819 | 0.086948819 | ENSG00000168395 | n.s. |
| ENSG00000213735 | 32.32167 | -0.453609148 | 0.335148 | -1.353458 | 0.175909 | 0.643168111 | ANAPC10P1 | processed_pseudogene | 39.21 | -0.086945759 | 0.086945759 | ENSG00000213735 | n.s. |
| ENSG00000033100 | 622.7799 | -0.359781123 | 0.234015 | -1.537424 | 0.124189 | 0.573519528 | CHPF2 | protein_coding | 56.82 | -0.086869796 | 0.086869796 | ENSG00000033100 | n.s. |
| ENSG00000231259 | 87.41888 | -0.410760818 | 0.286747 | -1.432483 | 0.152006 | 0.614510829 | | unprocessed_pseudogene | 38.72 | -0.086863779 | 0.086863779 | ENSG00000231259 | n.s. |
| ENSG00000201662 | 27.9413 | -0.493965366 | 0.378981 | -1.303404 | 0.192437 | 0.667270349 | RNU6-60P | snRNA | 41.75 | -0.086788812 | 0.086788812 | ENSG00000201662 | n.s. |
| ENSG00000201950 | 8.31242 | 0.571227687 | 0.478725 | 1.1932264 | 0.232781 | 0.704872993 | SNORD113-9 | snoRNA | 41.67 | 0.086763276 | 0.086763276 | ENSG00000201950 | n.s. |
| ENSG00000038382 | 2964.404 | -0.359551021 | 0.234015 | -1.536447 | 0.124429 | 0.573839353 | TRIO | protein_coding | 43.35 | -0.086727184 | 0.086727184 | ENSG00000038382 | n.s. |
| ENSG00000271840 | 25.03017 | -0.448651049 | 0.32851 | -1.365715 | 0.172028 | 0.640837445 | | lincRNA | 43.32 | -0.086702766 | 0.086702766 | ENSG00000271840 | n.s. |
| ENSG00000196511 | 43.19102 | 0.448594626 | 0.328899 | 1.3639267 | 0.172591 | 0.640837445 | TPK1 | protein_coding | 37.29 | 0.086691862 | 0.086691862 | ENSG00000196511 | n.s. |
| ENSG00000128191 | 1568.229 | -0.350997797 | 0.225282 | -1.558038 | 0.119224 | 0.566347442 | DGCR8 | protein_coding | 52.21 | -0.086667343 | 0.086667343 | ENSG00000128191 | n.s. |
| ENSG00000225783 | 1618.223 | -0.365015577 | 0.240443 | -1.518098 | 0.12899 | 0.579048024 | MIAT | lincRNA | 49.99 | -0.086612873 | 0.086612873 | ENSG00000225783 | n.s. |
| ENSG00000154217 | 249.5707 | -0.362559542 | 0.237355 | -1.527497 | 0.126637 | 0.576921481 | PITPNC1 | protein_coding | 44.62 | -0.086609416 | 0.086609416 | ENSG00000154217 | n.s. |
| ENSG00000213468 | 240.5441 | -0.365109959 | 0.24097 | -1.515169 | 0.12973 | 0.5791453 | FIRRE | lincRNA | 40.91 | -0.086608633 | 0.086608633 | ENSG00000213468 | n.s. |
| ENSG00000246350 | 222.718 | -0.364898103 | 0.240535 | -1.517028 | 0.12926 | 0.579048024 | | antisense | 43.62 | -0.086584998 | 0.086584998 | ENSG00000246350 | n.s. |
| ENSG00000213853 | 548.9622 | -0.36107884 | 0.235828 | -1.531108 | 0.125743 | 0.575893201 | EMP2 | protein_coding | 48.31 | -0.08653545 | 0.08653545 | ENSG00000213853 | n.s. |
| ENSG00000253925 | 78.60252 | 0.420997324 | 0.297714 | 1.4141005 | 0.157332 | 0.622955867 | | antisense | 44.09 | 0.086532935 | 0.086532935 | ENSG00000253925 | n.s. |
| ENSG00000176974 | 410.8938 | -0.392005819 | 0.266954 | -1.468439 | 0.141985 | 0.601734228 | SHMT1 | protein_coding | 49.9 | -0.086474635 | 0.086474635 | ENSG00000176974 | n.s. |
| ENSG00000272945 | 24.08698 | -0.469884487 | 0.353221 | -1.330286 | 0.183424 | 0.654656406 | | sense_intronic | 49.8 | -0.086452439 | 0.086452439 | ENSG00000272945 | n.s. |
| ENSG00000139624 | 705.4991 | -0.405175917 | 0.281368 | -1.440021 | 0.149861 | 0.61188309 | CERS5 | protein_coding | 43.86 | -0.086436806 | 0.086436806 | ENSG00000139624 | n.s. |
| ENSG00000204685 | 117.6101 | -0.385990649 | 0.261543 | -1.475822 | 0.139992 | 0.597163054 | STARD7-AS1 | processed_transcript | 46.6 | -0.086426035 | 0.086426035 | ENSG00000204685 | n.s. |
| ENSG00000136161 | 392.7354 | -0.400268493 | 0.277038 | -1.444815 | 0.14851 | 0.60842133 | RCBTB2 | protein_coding | 40.27 | -0.086376167 | 0.086376167 | ENSG00000136161 | n.s. |
| ENSG00000176912 | 58.92465 | -0.418978728 | 0.295702 | -1.416896 | 0.156513 | 0.622084796 | TYMSOS | processed_transcript | 42.36 | -0.086372638 | 0.086372638 | ENSG00000176912 | n.s. |
| ENSG00000169375 | 1636.001 | -0.371541957 | 0.247408 | -1.50174 | 0.133164 | 0.585639918 | SIN3A | protein_coding | 44.13 | -0.086334955 | 0.086334955 | ENSG00000169375 | n.s. |
| ENSG00000117713 | 962.8436 | -0.372345518 | 0.248422 | -1.498844 | 0.133914 | 0.586819166 | ARID1A | protein_coding | 46.19 | -0.08619639 | 0.08619639 | ENSG00000117713 | n.s. |
| ENSG00000140022 | 141.7502 | -0.39316619 | 0.268516 | -1.464218 | 0.143134 | 0.603629167 | STON2 | protein_coding | 41.03 | -0.086193739 | 0.086193739 | ENSG00000140022 | n.s. |
| ENSG00000136169 | 335.4551 | -0.397613641 | 0.274376 | -1.449157 | 0.147294 | 0.607109641 | SETDB2 | protein_coding | 37.41 | -0.086175946 | 0.086175946 | ENSG00000136169 | n.s. |
| ENSG00000207205 | 332.4423 | 0.342502993 | 0.218306 | 1.5689106 | 0.116669 | 0.560282428 | RNVU1-15 | snRNA | 52.73 | 0.086171355 | 0.086171355 | ENSG00000207205 | n.s. |
| ENSG00000202124 | 5.011647 | -0.668932379 | 0.618729 | -1.081139 | 0.279635 | 0.743390786 | RNY4P3 | misc_RNA | 43.16 | -0.086147002 | 0.086147002 | ENSG00000202124 | n.s. |

| | | | | | | | | | | | | | |
|---|---|---|---|---|---|---|---|---|---|---|---|---|---|
| ENSG00000126790 | 250.1037 | -0.350291683 | 0.225391 | -1.554152 | 0.120148 | 0.567746164 | L3HYPDH | protein_coding | 42.53 | -0.086117736 | 0.086117736 | ENSG00000126790 | n.s. |
| ENSG00000265401 | 28.01342 | 0.508772451 | 0.397422 | 1.2801832 | 0.200481 | 0.677328602 | | antisense | 54.44 | 0.086084596 | 0.086084596 | ENSG00000265401 | n.s. |
| ENSG00000228716 | 2165.032 | -0.396895076 | 0.274259 | -1.447154 | 0.147854 | 0.607109641 | DHFR | protein_coding | 41.24 | -0.086020209 | 0.086020209 | ENSG00000228716 | n.s. |
| ENSG00000132434 | 765.1785 | -0.399908587 | 0.277065 | -1.443374 | 0.148915 | 0.609468503 | LANCL2 | protein_coding | 42.91 | -0.085999835 | 0.085999835 | ENSG00000132434 | n.s. |
| ENSG00000137331 | 82.37704 | 0.432538404 | 0.312984 | 1.3819829 | 0.166977 | 0.632948522 | IER3 | protein_coding | 58.55 | 0.0859158 | 0.0859158 | ENSG00000137331 | n.s. |
| ENSG00000189007 | 390.8436 | -0.361627838 | 0.238373 | -1.517065 | 0.12925 | 0.579048024 | ADAT2 | protein_coding | 38.36 | -0.085809012 | 0.085809012 | ENSG00000189007 | n.s. |
| ENSG00000117318 | 1055.045 | -0.362297284 | 0.239652 | -1.511765 | 0.130594 | 0.579675642 | ID3 | protein_coding | 57.27 | -0.085797412 | 0.085797412 | ENSG00000117318 | n.s. |
| ENSG00000267264 | 17.62883 | -0.487478721 | 0.373609 | -1.304784 | 0.191966 | 0.66685299 | | processed_pseudogene | 43.98 | -0.08578158 | 0.08578158 | ENSG00000267264 | n.s. |
| ENSG00000264559 | 4.815665 | -0.794878583 | 0.816577 | -0.973427 | 0.330341 | 0.780141891 | MIR3162 | miRNA | 48.78 | -0.085708897 | 0.085708897 | ENSG00000264559 | n.s. |
| ENSG00000161647 | 444.2191 | -0.38943446 | 0.265534 | -1.466611 | 0.142482 | 0.60247798 | MPP3 | protein_coding | 48.95 | -0.085698488 | 0.085698488 | ENSG00000161647 | n.s. |
| ENSG00000064652 | 160.3942 | -0.416917104 | 0.295062 | -1.412982 | 0.157661 | 0.622955867 | SNX24 | protein_coding | 40.32 | -0.085694275 | 0.085694275 | ENSG00000064652 | n.s. |
| ENSG00000145990 | 393.9853 | -0.36116188 | 0.238525 | -1.514147 | 0.129988 | 0.5791453 | GFOD1 | protein_coding | 44.97 | -0.085672099 | 0.085672099 | ENSG00000145990 | n.s. |
| ENSG00000239344 | 38.62714 | -0.460211363 | 0.344092 | -1.337467 | 0.18107 | 0.65143233 | | processed_pseudogene | 43.26 | -0.085659459 | 0.085659459 | ENSG00000239344 | n.s. |
| ENSG00000126775 | 517.1359 | -0.399743189 | 0.277304 | -1.441534 | 0.149434 | 0.610673934 | ATG14 | protein_coding | 41.79 | -0.08562124 | 0.08562124 | ENSG00000126775 | n.s. |
| ENSG00000124406 | 2975.957 | 0.394843537 | 0.272501 | 1.4489615 | 0.147348 | 0.607109641 | ATP8A1 | protein_coding | 38.15 | 0.085575573 | 0.085575573 | ENSG00000124406 | n.s. |
| ENSG00000234665 | 84.74572 | -0.415715195 | 0.293786 | -1.415028 | 0.15706 | 0.622581912 | | lincRNA | 38.27 | -0.085555643 | 0.085555643 | ENSG00000234665 | n.s. |
| ENSG00000164970 | 322.6039 | -0.390400059 | 0.266797 | -1.463287 | 0.143389 | 0.603872801 | FAM219A | protein_coding | 47.16 | -0.085518902 | 0.085518902 | ENSG00000164970 | n.s. |
| ENSG00000100731 | 1482.246 | -0.369170274 | 0.246408 | -1.49821 | 0.134079 | 0.586819166 | PCNX1 | protein_coding | 36.87 | -0.085461335 | 0.085461335 | ENSG00000100731 | n.s. |
| ENSG00000282206 | 15.61839 | -0.53897118 | 0.437845 | -1.230964 | 0.218336 | 0.694198502 | | unprocessed_pseudogene | 65.35 | -0.085435732 | 0.085435732 | ENSG00000282206 | n.s. |
| ENSG00000084731 | 838.851 | -0.360052025 | 0.23688 | -1.519978 | 0.128517 | 0.579048024 | KIF3C | protein_coding | 48.25 | -0.085435095 | 0.085435095 | ENSG00000084731 | n.s. |
| ENSG00000138061 | 410.8925 | 0.377019952 | 0.25389 | 1.484972 | 0.137551 | 0.593469445 | CYP1B1 | protein_coding | 44.41 | 0.085433338 | 0.085433338 | ENSG00000138061 | n.s. |
| ENSG00000241261 | 121.4355 | 0.36891396 | 0.245954 | 1.499928 | 0.133633 | 0.586724086 | RPL17P19 | processed_pseudogene | 38.95 | 0.085427961 | 0.085427961 | ENSG00000241261 | n.s. |
| ENSG00000144182 | 83.40791 | 0.389633367 | 0.266352 | 1.4628538 | 0.143507 | 0.604031311 | LIPT1 | protein_coding | 41.3 | 0.085306543 | 0.085306543 | ENSG00000144182 | n.s. |
| ENSG00000156239 | 446.0419 | -0.410393008 | 0.289001 | -1.420039 | 0.155596 | 0.620025262 | N6AMT1 | protein_coding | 38.62 | -0.085193737 | 0.085193737 | ENSG00000156239 | n.s. |
| ENSG00000107651 | 1096.759 | -0.389102833 | 0.266011 | -1.462733 | 0.143541 | 0.604031311 | SEC23IP | protein_coding | 40.11 | -0.085190388 | 0.085190388 | ENSG00000107651 | n.s. |
| ENSG00000173542 | 960.9746 | -0.392511937 | 0.269939 | -1.454074 | 0.145926 | 0.607109641 | MOB1B | protein_coding | 40.16 | -0.085070239 | 0.085070239 | ENSG00000173542 | n.s. |
| ENSG00000258813 | 82.88749 | 0.392387927 | 0.271007 | 1.4478885 | 0.147648 | 0.607109641 | | antisense | 53.45 | 0.085043362 | 0.085043362 | ENSG00000258813 | n.s. |
| ENSG00000141994 | 764.378 | -0.399823907 | 0.278173 | -1.437321 | 0.150627 | 0.613029999 | DUS3L | protein_coding | 61.93 | -0.084969886 | 0.084969886 | ENSG00000141994 | n.s. |
| ENSG00000187231 | 1570.429 | -0.375270835 | 0.253011 | -1.483219 | 0.138016 | 0.594002688 | SESTD1 | protein_coding | 36.9 | -0.084890612 | 0.084890612 | ENSG00000187231 | n.s. |
| ENSG00000166402 | 2251.849 | -0.354093737 | 0.231299 | -1.530891 | 0.125796 | 0.575897021 | TUB | protein_coding | 49.43 | -0.084860394 | 0.084860394 | ENSG00000166402 | n.s. |
| ENSG00000136868 | 628.7563 | -0.401426391 | 0.280402 | -1.431609 | 0.152256 | 0.614628915 | SLC31A1 | protein_coding | 41.28 | -0.084856326 | 0.084856326 | ENSG00000136868 | n.s. |
| ENSG00000257727 | 624.0465 | -0.399541069 | 0.278076 | -1.436804 | 0.150774 | 0.613237549 | CNPY2 | protein_coding | 49.44 | -0.08485104 | 0.08485104 | ENSG00000257727 | n.s. |

| | | | | | | | | | | | | | |
|---|---|---|---|---|---|---|---|---|---|---|---|---|---|
| ENSG00000274248 | 20.12534 | 0.531612002 | 0.430154 | 1.2358651 | 0.216509 | 0.692499549 | | lincRNA | 63.23 | 0.084834912 | 0.084834912 | ENSG00000274248 | n.s. |
| ENSG00000241697 | 172.752 | -0.39938523 | 0.278257 | -1.435309 | 0.151199 | 0.613566131 | TMEFF1 | protein_coding | 39.38 | -0.084725032 | 0.084725032 | ENSG00000241697 | n.s. |
| ENSG00000259138 | 19.5893 | -0.500390792 | 0.390712 | -1.280714 | 0.200294 | 0.677196354 | | antisense | 40.33 | -0.08470885 | 0.08470885 | ENSG00000259138 | n.s. |
| ENSG00000106003 | 343.2641 | -0.374164425 | 0.252149 | -1.483901 | 0.137835 | 0.593774169 | LFNG | protein_coding | 62.83 | -0.084702856 | 0.084702856 | ENSG00000106003 | n.s. |
| ENSG00000128536 | 161.8478 | -0.401073482 | 0.28062 | -1.429238 | 0.152936 | 0.61505617 | CDHR3 | protein_coding | 43.76 | -0.084660684 | 0.084660684 | ENSG00000128536 | n.s. |
| ENSG00000135333 | 814.4331 | 0.365656911 | 0.244203 | 1.4973465 | 0.134303 | 0.586912197 | EPHA7 | protein_coding | 33.25 | 0.084622833 | 0.084622833 | ENSG00000135333 | n.s. |
| ENSG00000233184 | 240.0625 | 0.421892547 | 0.303745 | 1.3889687 | 0.164842 | 0.63030432 | | antisense | 37.11 | 0.084568241 | 0.084568241 | ENSG00000233184 | n.s. |
| ENSG00000235217 | 145.1574 | -0.366542755 | 0.245242 | -1.494619 | 0.135014 | 0.587941172 | TSPY26P | nscribed_processed_pseudog | 55.99 | -0.084548998 | 0.084548998 | ENSG00000235217 | n.s. |
| ENSG00000103423 | 876.3915 | -0.4183831 | 0.298161 | -1.403214 | 0.160553 | 0.62809251 | DNAJA3 | protein_coding | 49.02 | -0.084503506 | 0.084503506 | ENSG00000103423 | n.s. |
| ENSG00000237984 | 202.9897 | 0.355744549 | 0.234152 | 1.5192915 | 0.128689 | 0.579048024 | PTENP1 | nscribed_processed_pseudog | 41.99 | 0.084412993 | 0.084412993 | ENSG00000237984 | n.s. |
| ENSG00000228253 | 397.5417 | -0.364709447 | 0.243719 | -1.496432 | 0.134541 | 0.587308708 | MT-ATP8 | protein_coding | 39.61 | -0.084296593 | 0.084296593 | ENSG00000228253 | n.s. |
| ENSG00000173992 | 165.4274 | -0.39871771 | 0.278481 | -1.431759 | 0.152213 | 0.614628915 | CCS | protein_coding | 51.72 | -0.084283745 | 0.084283745 | ENSG00000173992 | n.s. |
| ENSG00000134698 | 286.276 | -0.364013596 | 0.243054 | -1.497666 | 0.13422 | 0.586819166 | AGO4 | protein_coding | 41.53 | -0.084267586 | 0.084267586 | ENSG00000134698 | n.s. |
| ENSG00000251791 | 34.47432 | -0.449989911 | 0.335724 | -1.340355 | 0.18013 | 0.649773904 | SCARNA6 | scaRNA | 53.21 | -0.084255092 | 0.084255092 | ENSG00000251791 | n.s. |
| ENSG00000186918 | 577.8319 | -0.376279571 | 0.254706 | -1.47731 | 0.139593 | 0.597163054 | ZNF395 | protein_coding | 48.07 | -0.084251656 | 0.084251656 | ENSG00000186918 | n.s. |
| ENSG00000173889 | 2056.388 | -0.366876089 | 0.246039 | -1.491131 | 0.135927 | 0.590119307 | PHC3 | protein_coding | 37.75 | -0.084036702 | 0.084036702 | ENSG00000173889 | n.s. |
| ENSG00000267368 | 50.98009 | -0.437813793 | 0.323257 | -1.354382 | 0.175614 | 0.642859735 | UPK3BL1 | protein_coding | 57.17 | -0.084009363 | 0.084009363 | ENSG00000267368 | n.s. |
| ENSG00000103550 | 1723.87 | -0.375741531 | 0.254754 | -1.47492 | 0.140234 | 0.597636216 | KNOP1 | protein_coding | 51.83 | -0.084001939 | 0.084001939 | ENSG00000103550 | n.s. |
| ENSG00000142599 | 173.7158 | -0.361425097 | 0.240744 | -1.501283 | 0.133282 | 0.5857901 | RERE | protein_coding | 41.9 | -0.08394386 | 0.08394386 | ENSG00000142599 | n.s. |
| ENSG00000263002 | 490.4866 | -0.366455965 | 0.245931 | -1.490076 | 0.136204 | 0.590213403 | ZNF234 | protein_coding | 41.72 | -0.083915093 | 0.083915093 | ENSG00000263002 | n.s. |
| ENSG00000251868 | 10.05624 | 0.544095916 | 0.448902 | 1.2120601 | 0.225489 | 0.701091738 | RNU7-71P | snRNA | 43.33 | 0.083913275 | 0.083913275 | ENSG00000251868 | n.s. |
| ENSG00000117523 | 7166.762 | -0.375256315 | 0.254464 | -1.474692 | 0.140295 | 0.597714988 | PRRC2C | protein_coding | 38.3 | -0.083871983 | 0.083871983 | ENSG00000117523 | n.s. |
| ENSG00000167615 | 1087.61 | -0.34101877 | 0.219724 | -1.552032 | 0.120655 | 0.567746164 | LENG8 | protein_coding | 58.96 | -0.083838029 | 0.083838029 | ENSG00000167615 | n.s. |
| ENSG00000136026 | 2133.669 | -0.396627115 | 0.277217 | -1.430747 | 0.152503 | 0.614654298 | CKAP4 | protein_coding | 44.3 | -0.083834707 | 0.083834707 | ENSG00000136026 | n.s. |
| ENSG00000181472 | 728.7213 | -0.386731215 | 0.266021 | -1.453763 | 0.146012 | 0.607109641 | ZBTB2 | protein_coding | 44.33 | -0.083817366 | 0.083817366 | ENSG00000181472 | n.s. |
| ENSG00000175764 | 109.2472 | -0.392097886 | 0.272157 | -1.440703 | 0.149669 | 0.611274515 | TTLL11 | protein_coding | 44.23 | -0.083816299 | 0.083816299 | ENSG00000175764 | n.s. |
| ENSG00000059377 | 49.66418 | -0.426529157 | 0.309757 | -1.376981 | 0.168518 | 0.636365056 | TBXAS1 | protein_coding | 44.8 | -0.083724976 | 0.083724976 | ENSG00000059377 | n.s. |
| ENSG00000151948 | 192.9851 | -0.407285266 | 0.288153 | -1.413435 | 0.157528 | 0.622955867 | GLT1D1 | protein_coding | 45.85 | -0.083714521 | 0.083714521 | ENSG00000151948 | n.s. |
| ENSG00000230043 | 6.856652 | 0.630954443 | 0.572476 | 1.1021506 | 0.270396 | 0.737020734 | TMSB4XP6 | processed_pseudogene | 42.96 | 0.083614268 | 0.083614268 | ENSG00000230043 | n.s. |
| ENSG00000278012 | 254.8973 | -0.371570917 | 0.25088 | -1.481071 | 0.138588 | 0.595725392 | | sense_intronic | 39.78 | -0.083586322 | 0.083586322 | ENSG00000278012 | n.s. |
| ENSG00000159055 | 527.4838 | -0.395497186 | 0.276514 | -1.430298 | 0.152632 | 0.614774803 | MIS18A | protein_coding | 40.72 | -0.083562204 | 0.083562204 | ENSG00000159055 | n.s. |
| ENSG00000154237 | 56.14533 | -0.414318089 | 0.296312 | -1.398248 | 0.162039 | 0.628957308 | LRRK1 | protein_coding | 46.01 | -0.083434893 | 0.083434893 | ENSG00000154237 | n.s. |

| | | | | | | | | | | | | | |
|---|---|---|---|---|---|---|---|---|---|---|---|---|---|
| ENSG00000172932 | 69.59432 | -0.415985444 | 0.299645 | -1.388263 | 0.165057 | 0.63030432 | ANKRD13D | protein_coding | 53.51 | -0.083384164 | 0.083384164 | ENSG00000172932 | n.s. |
| ENSG00000274904 | 19.22879 | -0.562660876 | 0.477499 | -1.17835 | 0.238657 | 0.710909549 | | antisense | 61.54 | -0.083378269 | 0.083378269 | ENSG00000274904 | n.s. |
| ENSG00000283327 | 4.136473 | 0.785232985 | 0.814481 | 0.9640903 | 0.335001 | 0.783111834 | MIR6841 | miRNA | 45.83 | 0.083373065 | 0.083373065 | ENSG00000283327 | n.s. |
| ENSG00000273893 | 55.83311 | -0.412781271 | 0.294121 | -1.403438 | 0.160486 | 0.62809251 | | sense_intronic | 46 | -0.083372069 | 0.083372069 | ENSG00000273893 | n.s. |
| ENSG00000143624 | 838.7731 | -0.372286467 | 0.252243 | -1.475904 | 0.13997 | 0.597163054 | INTS3 | protein_coding | 48.32 | -0.083357572 | 0.083357572 | ENSG00000143624 | n.s. |
| ENSG00000129007 | 52.05926 | -0.426532965 | 0.310679 | -1.372905 | 0.169782 | 0.638037836 | CALML4 | protein_coding | 50.69 | -0.083239429 | 0.083239429 | ENSG00000129007 | n.s. |
| ENSG00000130779 | 1402.885 | -0.384021666 | 0.265172 | -1.448201 | 0.147561 | 0.607109641 | CLIP1 | protein_coding | 43.88 | -0.083230118 | 0.083230118 | ENSG00000130779 | n.s. |
| ENSG00000140105 | 617.6984 | -0.393721146 | 0.275162 | -1.430872 | 0.152467 | 0.614654298 | WARS | protein_coding | 46.89 | -0.083220475 | 0.083220475 | ENSG00000140105 | n.s. |
| ENSG00000105619 | 158.4612 | -0.392045286 | 0.272957 | -1.436291 | 0.150919 | 0.613504028 | TFPT | protein_coding | 54.83 | -0.083185181 | 0.083185181 | ENSG00000105619 | n.s. |
| ENSG00000172465 | 230.2153 | 0.429698564 | 0.315013 | 1.3640664 | 0.172547 | 0.640837445 | TCEAL1 | protein_coding | 50.67 | 0.083040158 | 0.083040158 | ENSG00000172465 | n.s. |
| ENSG00000198876 | 616.5907 | -0.382378933 | 0.263268 | -1.452433 | 0.146381 | 0.607109641 | DCAF12 | protein_coding | 45.99 | -0.082874084 | 0.082874084 | ENSG00000198876 | n.s. |
| ENSG00000206954 | 37.3173 | -0.440030783 | 0.327625 | -1.343095 | 0.179241 | 0.64843101 | RNU6-1201P | snRNA | 40.19 | -0.08278573 | 0.08278573 | ENSG00000206954 | n.s. |
| ENSG00000198455 | 296.2873 | 0.336487166 | 0.216522 | 1.5540533 | 0.120172 | 0.567746164 | ZXDB | protein_coding | 46.1 | 0.082723954 | 0.082723954 | ENSG00000198455 | n.s. |
| ENSG00000118200 | 3465.604 | -0.38311613 | 0.265065 | -1.445369 | 0.148354 | 0.608376989 | CAMSAP2 | protein_coding | 37.69 | -0.08268689 | 0.08268689 | ENSG00000118200 | n.s. |
| ENSG00000198786 | 1757.292 | -0.347352841 | 0.228035 | -1.523245 | 0.127697 | 0.578111285 | MT-ND5 | protein_coding | 44.92 | -0.082666 | 0.082666 | ENSG00000198786 | n.s. |
| ENSG00000105894 | 105.6176 | -0.412018013 | 0.296586 | -1.389204 | 0.164771 | 0.63030432 | PTN | protein_coding | 38.25 | -0.082588894 | 0.082588894 | ENSG00000105894 | n.s. |
| ENSG00000144283 | 1060.297 | -0.387261819 | 0.269209 | -1.438515 | 0.150288 | 0.612366927 | PKP4 | protein_coding | 38.59 | -0.082482226 | 0.082482226 | ENSG00000144283 | n.s. |
| ENSG00000108819 | 1054.755 | -0.362790773 | 0.244163 | -1.485853 | 0.137318 | 0.592647019 | PPP1R9B | protein_coding | 59.45 | -0.082427477 | 0.082427477 | ENSG00000108819 | n.s. |
| ENSG00000166912 | 161.6998 | -0.380189199 | 0.261774 | -1.452356 | 0.146403 | 0.607109641 | MTMR10 | protein_coding | 39.99 | -0.082399497 | 0.082399497 | ENSG00000166912 | n.s. |
| ENSG00000196072 | 766.9695 | -0.403077747 | 0.285844 | -1.410131 | 0.158501 | 0.624627103 | BLOC1S2 | protein_coding | 47.38 | -0.082380698 | 0.082380698 | ENSG00000196072 | n.s. |
| ENSG00000198382 | 146.7564 | -0.390123258 | 0.272939 | -1.429343 | 0.152906 | 0.61505617 | UVRAG | protein_coding | 38.73 | -0.082349254 | 0.082349254 | ENSG00000198382 | n.s. |
| ENSG00000115252 | 634.9155 | 0.4097672 | 0.294304 | 1.3923243 | 0.163824 | 0.629718088 | PDE1A | protein_coding | 36.57 | 0.082303312 | 0.082303312 | ENSG00000115252 | n.s. |
| ENSG00000272578 | 275.3154 | -0.379713847 | 0.261841 | -1.450172 | 0.147011 | 0.607109641 | | transcribed_unprocessed_pseudc | 49.78 | -0.082296472 | 0.082296472 | ENSG00000272578 | n.s. |
| ENSG00000264701 | 15.68764 | -0.540046142 | 0.452207 | -1.194245 | 0.232382 | 0.704163047 | | antisense | 51.52 | -0.082263484 | 0.082263484 | ENSG00000264701 | n.s. |
| ENSG00000139433 | 236.7558 | -0.411333155 | 0.296953 | -1.385178 | 0.165998 | 0.63130003 | GLTP | protein_coding | 49.06 | -0.082169635 | 0.082169635 | ENSG00000139433 | n.s. |
| ENSG00000124160 | 777.5628 | -0.385567498 | 0.268013 | -1.438613 | 0.15026 | 0.612366927 | NCOA5 | protein_coding | 41.78 | -0.082121356 | 0.082121356 | ENSG00000124160 | n.s. |
| ENSG00000141576 | 1247.411 | -0.357422098 | 0.239572 | -1.49192 | 0.13572 | 0.589422885 | RNF157 | protein_coding | 44.99 | -0.082054465 | 0.082054465 | ENSG00000141576 | n.s. |
| ENSG00000178950 | 600.9922 | -0.367934675 | 0.249826 | -1.472763 | 0.140815 | 0.598646262 | GAK | protein_coding | 56.5 | -0.081986783 | 0.081986783 | ENSG00000178950 | n.s. |
| ENSG00000134954 | 248.4486 | -0.374095884 | 0.25561 | -1.463544 | 0.143318 | 0.603872801 | ETS1 | protein_coding | 42.5 | -0.081947398 | 0.081947398 | ENSG00000134954 | n.s. |
| ENSG00000204514 | 1295.641 | -0.356881062 | 0.23915 | -1.492293 | 0.135622 | 0.589422885 | ZNF814 | protein_coding | 46.12 | -0.081930258 | 0.081930258 | ENSG00000204514 | n.s. |
| ENSG00000234690 | 81.33464 | 0.385356867 | 0.268228 | 1.4366768 | 0.15081 | 0.613237549 | | lincRNA | 46.05 | 0.081838724 | 0.081838724 | ENSG00000234690 | n.s. |
| ENSG00000265694 | 20.15852 | -0.502096858 | 0.401299 | -1.251177 | 0.21087 | 0.687090943 | MIR4774 | miRNA | 34.21 | -0.081834646 | 0.081834646 | ENSG00000265694 | n.s. |

| | | | | | | | | | | | | | |
|---|---|---|---|---|---|---|---|---|---|---|---|---|---|
| ENSG00000143756 | 1030.314 | -0.406035953 | 0.290542 | -1.39751 | 0.16226 | 0.628957308 | FBXO28 | protein_coding | 40.97 | -0.081767046 | 0.081767046 | ENSG00000143756 | n.s. |
| ENSG00000105520 | 105.5634 | -0.373300702 | 0.255302 | -1.462193 | 0.143688 | 0.604031311 | PLPPR2 | protein_coding | 57.31 | -0.08173066 | 0.08173066 | ENSG00000105520 | n.s. |
| ENSG00000029639 | 388.1248 | -0.363351428 | 0.245384 | -1.480745 | 0.138674 | 0.595914867 | TFB1M | protein_coding | 38.35 | -0.081687135 | 0.081687135 | ENSG00000029639 | n.s. |
| ENSG00000196247 | 1482.964 | -0.384316873 | 0.267359 | -1.437457 | 0.150588 | 0.613029999 | ZNF107 | protein_coding | 39.91 | -0.081674358 | 0.081674358 | ENSG00000196247 | n.s. |
| ENSG00000206914 | 13.92823 | -0.52623003 | 0.432576 | -1.216502 | 0.223794 | 0.69988307 | | misc_RNA | 41.18 | -0.081552242 | 0.081552242 | ENSG00000206914 | n.s. |
| ENSG00000156860 | 50.85937 | -0.419495415 | 0.306334 | -1.369405 | 0.170873 | 0.639237118 | FBRS | protein_coding | 59.46 | -0.081523906 | 0.081523906 | ENSG00000156860 | n.s. |
| ENSG00000136527 | 25464.6 | 0.332400741 | 0.214539 | 1.5493718 | 0.121292 | 0.568737594 | TRA2B | protein_coding | 38.67 | 0.081467454 | 0.081467454 | ENSG00000136527 | n.s. |
| ENSG00000260442 | 63.81992 | -0.406705013 | 0.293153 | -1.387346 | 0.165336 | 0.630679057 | ATP2A1-AS1 | antisense | 59.34 | -0.081418923 | 0.081418923 | ENSG00000260442 | n.s. |
| ENSG00000100350 | 476.073 | -0.338239067 | 0.220305 | -1.535322 | 0.124705 | 0.574541601 | FOXRED2 | protein_coding | 52.7 | -0.081406878 | 0.081406878 | ENSG00000100350 | n.s. |
| ENSG00000187239 | 309.0286 | -0.360366228 | 0.243137 | -1.482155 | 0.138299 | 0.594852203 | FNBP1 | protein_coding | 43.76 | -0.081295351 | 0.081295351 | ENSG00000187239 | n.s. |
| ENSG00000084444 | 704.9568 | -0.350871387 | 0.234009 | -1.49939 | 0.133772 | 0.586803051 | FAM234B | protein_coding | 43.23 | -0.081229406 | 0.081229406 | ENSG00000084444 | n.s. |
| ENSG00000141622 | 661.314 | -0.338733541 | 0.221219 | -1.531217 | 0.125716 | 0.575893201 | RNF165 | protein_coding | 49.03 | -0.081180219 | 0.081180219 | ENSG00000141622 | n.s. |
| ENSG00000207973 | 6.399459 | -0.683676244 | 0.664089 | -1.029494 | 0.303247 | 0.760781413 | MIR589 | miRNA | 62.63 | -0.08117979 | 0.08117979 | ENSG00000207973 | n.s. |
| ENSG00000140948 | 481.8832 | -0.370289213 | 0.252862 | -1.46439 | 0.143087 | 0.603629167 | ZCCHC14 | protein_coding | 48.02 | -0.081178424 | 0.081178424 | ENSG00000140948 | n.s. |
| ENSG00000085449 | 966.0965 | -0.384074953 | 0.268537 | -1.430251 | 0.152645 | 0.614774803 | WDFY1 | protein_coding | 42.72 | -0.081148869 | 0.081148869 | ENSG00000085449 | n.s. |
| ENSG00000119778 | 1433.888 | -0.36808815 | 0.250779 | -1.467778 | 0.142165 | 0.602177752 | ATAD2B | protein_coding | 37.98 | -0.081080726 | 0.081080726 | ENSG00000119778 | n.s. |
| ENSG00000268392 | 24.77313 | -0.438289378 | 0.328851 | -1.33279 | 0.182601 | 0.653392268 | | lincRNA | 41.22 | -0.081007276 | 0.081007276 | ENSG00000268392 | n.s. |
| ENSG00000166908 | 191.7922 | -0.373709872 | 0.256699 | -1.455832 | 0.145439 | 0.607098916 | PIP4K2C | protein_coding | 46.79 | -0.08099808 | 0.08099808 | ENSG00000166908 | n.s. |
| ENSG00000281091 | 21.25712 | -0.484643592 | 0.38243 | -1.267274 | 0.205057 | 0.680575501 | | lincRNA | 41.04 | -0.080995425 | 0.080995425 | ENSG00000281091 | n.s. |
| ENSG00000253878 | 265.2774 | -0.348353369 | 0.23197 | -1.501715 | 0.133171 | 0.585639918 | | sense_intronic | 46.4 | -0.080946638 | 0.080946638 | ENSG00000253878 | n.s. |
| ENSG00000187257 | 1083.254 | -0.372249746 | 0.255289 | -1.458153 | 0.144798 | 0.606306192 | RSBN1L | protein_coding | 39.26 | -0.080892846 | 0.080892846 | ENSG00000187257 | n.s. |
| ENSG00000132563 | 30.25944 | -0.450189203 | 0.341567 | -1.318013 | 0.187499 | 0.661470368 | REEP2 | protein_coding | 58.31 | -0.080804283 | 0.080804283 | ENSG00000132563 | n.s. |
| ENSG00000222872 | 4.433131 | -0.672553953 | 0.64817 | -1.037619 | 0.299447 | 0.758335061 | RNU4-78P | snRNA | 42.06 | -0.080799868 | 0.080799868 | ENSG00000222872 | n.s. |
| ENSG00000100815 | 1367.316 | -0.38725653 | 0.272276 | -1.422296 | 0.15494 | 0.618650071 | TRIP11 | protein_coding | 37.97 | -0.080764259 | 0.080764259 | ENSG00000100815 | n.s. |
| ENSG00000170088 | 534.7844 | -0.383525987 | 0.268727 | -1.427195 | 0.153524 | 0.615840022 | TMEM192 | protein_coding | 41.86 | -0.080744528 | 0.080744528 | ENSG00000170088 | n.s. |
| ENSG00000166479 | 1889.134 | -0.381312547 | 0.266183 | -1.43252 | 0.151995 | 0.614510829 | TMX3 | protein_coding | 36.18 | -0.08063634 | 0.08063634 | ENSG00000166479 | n.s. |
| ENSG00000204634 | 458.9965 | -0.357760388 | 0.241452 | -1.481704 | 0.138419 | 0.595184611 | TBC1D8 | protein_coding | 44.32 | -0.080620698 | 0.080620698 | ENSG00000204634 | n.s. |
| ENSG00000221044 | 17.31638 | -0.477001798 | 0.373895 | -1.275765 | 0.202039 | 0.677624898 | | snoRNA | 50.49 | -0.080618382 | 0.080618382 | ENSG00000221044 | n.s. |
| ENSG00000157193 | 2421.56 | -0.399947327 | 0.285644 | -1.400161 | 0.161465 | 0.628931954 | LRP8 | protein_coding | 51.78 | -0.080547928 | 0.080547928 | ENSG00000157193 | n.s. |
| ENSG00000161533 | 1058.614 | -0.381511956 | 0.266904 | -1.429397 | 0.15289 | 0.61505617 | ACOX1 | protein_coding | 44.33 | -0.080531535 | 0.080531535 | ENSG00000161533 | n.s. |
| ENSG00000120334 | 919.8315 | -0.348145565 | 0.232655 | -1.496402 | 0.134549 | 0.587308708 | CENPL | protein_coding | 40.71 | -0.080468124 | 0.080468124 | ENSG00000120334 | n.s. |
| ENSG00000087245 | 620.3614 | -0.371199261 | 0.255039 | -1.455459 | 0.145542 | 0.607098916 | MMP2 | protein_coding | 45.67 | -0.080453929 | 0.080453929 | ENSG00000087245 | n.s. |

| | | | | | | | | | | | | | |
|---|---|---|---|---|---|---|---|---|---|---|---|---|---|
| ENSG00000204131 | 602.1967 | -0.358019081 | 0.241914 | -1.479946 | 0.138888 | 0.596095432 | NHSL2 | protein_coding | 45.33 | -0.080441233 | 0.080441233 | ENSG00000204131 | n.s. |
| ENSG00000167461 | 540.9223 | -0.366719458 | 0.250469 | -1.464133 | 0.143158 | 0.603629167 | RAB8A | protein_coding | 53.79 | -0.080395827 | 0.080395827 | ENSG00000167461 | n.s. |
| ENSG00000163625 | 2255.488 | -0.370878083 | 0.255747 | -1.450177 | 0.147009 | 0.607109641 | WDFY3 | protein_coding | 36.88 | -0.080381471 | 0.080381471 | ENSG00000163625 | n.s. |
| ENSG00000167549 | 398.2377 | -0.354770699 | 0.23907 | -1.483959 | 0.13782 | 0.593774169 | CORO6 | protein_coding | 60.29 | -0.080312529 | 0.080312529 | ENSG00000167549 | n.s. |
| ENSG00000112977 | 314.3214 | -0.4005375 | 0.288376 | -1.388941 | 0.164851 | 0.63030432 | DAP | protein_coding | 46.31 | -0.080287628 | 0.080287628 | ENSG00000112977 | n.s. |
| ENSG00000143384 | 9260.439 | 0.337128838 | 0.221243 | 1.5237973 | 0.127559 | 0.57808993 | MCL1 | protein_coding | 44.74 | 0.080238213 | 0.080238213 | ENSG00000143384 | n.s. |
| ENSG00000130921 | 399.694 | -0.377339304 | 0.262486 | -1.437559 | 0.150559 | 0.613029999 | C12orf65 | protein_coding | 47.41 | -0.080191497 | 0.080191497 | ENSG00000130921 | n.s. |
| ENSG00000274344 | 10.45695 | -0.525778434 | 0.440009 | -1.194927 | 0.232116 | 0.704043192 | | misc_RNA | 37.5 | -0.080129 | 0.080129 | ENSG00000274344 | n.s. |
| ENSG00000109381 | 367.9588 | -0.369709887 | 0.254509 | -1.452637 | 0.146324 | 0.607109641 | ELF2 | protein_coding | 39.88 | -0.080128285 | 0.080128285 | ENSG00000109381 | n.s. |
| ENSG00000133275 | 1008.17 | -0.400318013 | 0.288688 | -1.38668 | 0.165539 | 0.630850594 | CSNK1G2 | protein_coding | 61.03 | -0.08009302 | 0.08009302 | ENSG00000133275 | n.s. |
| ENSG00000111725 | 431.7376 | -0.359173522 | 0.243774 | -1.47339 | 0.140646 | 0.598476209 | PRKAB1 | protein_coding | 47.82 | -0.080078853 | 0.080078853 | ENSG00000111725 | n.s. |
| ENSG00000165495 | 29.29919 | -0.450583992 | 0.343556 | -1.311531 | 0.189678 | 0.664231317 | PKNOX2 | protein_coding | 49.03 | -0.080060057 | 0.080060057 | ENSG00000165495 | n.s. |
| ENSG00000163703 | 225.9074 | 0.397276668 | 0.28419 | 1.3979263 | 0.162135 | 0.628957308 | CRELD1 | protein_coding | 51.58 | 0.080003111 | 0.080003111 | ENSG00000163703 | n.s. |
| ENSG00000129968 | 556.6728 | -0.382831257 | 0.268934 | -1.423516 | 0.154587 | 0.61830251 | ABHD17A | protein_coding | 65.16 | -0.07993478 | 0.07993478 | ENSG00000129968 | n.s. |
| ENSG00000206644 | 5.213785 | -0.673019312 | 0.652188 | -1.031941 | 0.3021 | 0.760781413 | RNU6-1279P | snRNA | 49.51 | -0.079914385 | 0.079914385 | ENSG00000206644 | n.s. |
| ENSG00000232587 | 68.85632 | -0.400032928 | 0.28878 | -1.385249 | 0.165976 | 0.63130003 | EEF1A1P3 | processed_pseudogene | 45.49 | -0.079912254 | 0.079912254 | ENSG00000232587 | n.s. |
| ENSG00000077279 | 935.1812 | -0.368700589 | 0.254723 | -1.447459 | 0.147768 | 0.607109641 | DCX | protein_coding | 39.48 | -0.079909537 | 0.079909537 | ENSG00000077279 | n.s. |
| ENSG00000184083 | 117.8308 | -0.376595484 | 0.262234 | -1.436104 | 0.150973 | 0.613541057 | FAM120C | protein_coding | 42.69 | -0.079897131 | 0.079897131 | ENSG00000184083 | n.s. |
| ENSG00000262333 | 64.04649 | -0.411121155 | 0.30025 | -1.369262 | 0.170917 | 0.639237118 | HNRNPA1P16 | processed_pseudogene | 46.28 | -0.079896469 | 0.079896469 | ENSG00000262333 | n.s. |
| ENSG00000172071 | 727.4025 | -0.33668125 | 0.22189 | -1.517333 | 0.129183 | 0.579048024 | EIF2AK3 | protein_coding | 38.57 | -0.07988955 | 0.07988955 | ENSG00000172071 | n.s. |
| ENSG00000226766 | 59.87821 | -0.415066088 | 0.305432 | -1.358948 | 0.174163 | 0.642010653 | FABP7P1 | processed_pseudogene | 40.41 | -0.079882692 | 0.079882692 | ENSG00000226766 | n.s. |
| ENSG00000252236 | 16.8887 | 0.596352298 | 0.534962 | 1.1147558 | 0.264955 | 0.734601921 | | snoRNA | 47.52 | 0.079880162 | 0.079880162 | ENSG00000252236 | n.s. |
| ENSG00000168256 | 566.1744 | -0.356697789 | 0.24168 | -1.475911 | 0.139968 | 0.597163054 | NKIRAS2 | protein_coding | 49.4 | -0.079867157 | 0.079867157 | ENSG00000168256 | n.s. |
| ENSG00000173875 | 759.8908 | -0.367190275 | 0.251896 | -1.457708 | 0.144921 | 0.606306192 | ZNF791 | protein_coding | 45.22 | -0.079793383 | 0.079793383 | ENSG00000173875 | n.s. |
| ENSG00000139767 | 683.1049 | -0.336213834 | 0.222032 | -1.514256 | 0.129961 | 0.5791453 | SRRM4 | protein_coding | 44.14 | -0.079754112 | 0.079754112 | ENSG00000139767 | n.s. |
| ENSG00000263666 | 5.740424 | -0.669509263 | 0.647416 | -1.034126 | 0.301077 | 0.760228309 | SNORA70D | snoRNA | 53.33 | -0.07970907 | 0.07970907 | ENSG00000263666 | n.s. |
| ENSG00000119771 | 446.5778 | -0.367599441 | 0.254076 | -1.446807 | 0.147951 | 0.607109641 | KLHL29 | protein_coding | 50.5 | -0.079670882 | 0.079670882 | ENSG00000119771 | n.s. |
| ENSG00000124224 | 71.04004 | 0.395553748 | 0.282599 | 1.3996985 | 0.161604 | 0.628931954 | PPP4R1L | cribed_unprocessed_pseudc | 41.96 | 0.079663077 | 0.079663077 | ENSG00000124224 | n.s. |
| ENSG00000241112 | 162.6982 | 0.367373767 | 0.25288 | 1.4527606 | 0.14629 | 0.607109641 | RPL29P14 | processed_pseudogene | 55.14 | 0.079621971 | 0.079621971 | ENSG00000241112 | n.s. |
| ENSG00000198400 | 1898.132 | -0.376948012 | 0.2636 | -1.430003 | 0.152716 | 0.614883793 | NTRK1 | protein_coding | 53.11 | -0.079614041 | 0.079614041 | ENSG00000198400 | n.s. |
| ENSG00000271983 | 21.43394 | 0.49888434 | 0.403515 | 1.2363464 | 0.21633 | 0.692499549 | | lincRNA | 43.09 | 0.079612215 | 0.079612215 | ENSG00000271983 | n.s. |
| ENSG00000283023 | 139.423 | -0.39882301 | 0.288066 | -1.384485 | 0.16621 | 0.631590186 | | unprocessed_pseudogene | 35.84 | -0.079590965 | 0.079590965 | ENSG00000283023 | n.s. |

| | | | | | | | | | | | | | |
|---|---|---|---|---|---|---|---|---|---|---|---|---|---|
| ENSG00000254325 | 62.36415 | -0.413967546 | 0.304862 | -1.357884 | 0.174501 | 0.642377343 | | sense_intronic | 45.55 | -0.079568613 | 0.079568613 | ENSG00000254325 | n.s. |
| ENSG00000174607 | 365.9944 | -0.394930465 | 0.282539 | -1.397789 | 0.162176 | 0.628957308 | UGT8 | protein_coding | 36.21 | -0.079530636 | 0.079530636 | ENSG00000174607 | n.s. |
| ENSG00000188419 | 389.0645 | -0.386324266 | 0.273003 | -1.415092 | 0.157042 | 0.622581912 | CHM | protein_coding | 36.96 | -0.079506886 | 0.079506886 | ENSG00000188419 | n.s. |
| ENSG00000034693 | 529.814 | -0.394765638 | 0.283092 | -1.39448 | 0.163173 | 0.629147846 | PEX3 | protein_coding | 35.11 | -0.079445513 | 0.079445513 | ENSG00000034693 | n.s. |
| ENSG00000275014 | 62.33949 | -0.406146078 | 0.295587 | -1.374034 | 0.169431 | 0.637567981 | RN7SL166P | misc_RNA | 52.14 | -0.079390796 | 0.079390796 | ENSG00000275014 | n.s. |
| ENSG00000153250 | 2288.665 | -0.355564449 | 0.241277 | -1.473677 | 0.140569 | 0.598330163 | RBMS1 | protein_coding | 39.2 | -0.079311887 | 0.079311887 | ENSG00000153250 | n.s. |
| ENSG00000134262 | 781.3073 | -0.362006358 | 0.247663 | -1.461689 | 0.143826 | 0.604031311 | AP4B1 | protein_coding | 43.25 | -0.07925787 | 0.07925787 | ENSG00000134262 | n.s. |
| ENSG00000165312 | 102.7655 | 0.389086203 | 0.276218 | 1.4086185 | 0.158948 | 0.625601663 | OTUD1 | protein_coding | 49.91 | 0.079257679 | 0.079257679 | ENSG00000165312 | n.s. |
| ENSG00000270015 | 161.025 | -0.393613636 | 0.282334 | -1.39414 | 0.163275 | 0.629215564 | | sense_intronic | 51.16 | -0.079195277 | 0.079195277 | ENSG00000270015 | n.s. |
| ENSG00000254791 | 108.6385 | 0.387988752 | 0.275238 | 1.4096479 | 0.158644 | 0.625012983 | FAR1-IT1 | sense_intronic | 44.96 | 0.079192757 | 0.079192757 | ENSG00000254791 | n.s. |
| ENSG00000196741 | 120.254 | -0.365349654 | 0.252375 | -1.447645 | 0.147716 | 0.607109641 | LINC01560 | lincRNA | 39.98 | -0.079183279 | 0.079183279 | ENSG00000196741 | n.s. |
| ENSG00000177822 | 243.1004 | -0.333582225 | 0.220044 | -1.515977 | 0.129525 | 0.579048024 | | antisense | 39.36 | -0.079154197 | 0.079154197 | ENSG00000177822 | n.s. |
| ENSG00000238297 | 78.28842 | -0.391455001 | 0.278654 | -1.404805 | 0.160079 | 0.627825878 | | snoRNA | 42.71 | -0.079136851 | 0.079136851 | ENSG00000238297 | n.s. |
| ENSG00000204138 | 297.3741 | -0.365086826 | 0.25213 | -1.448009 | 0.147615 | 0.607109641 | PHACTR4 | protein_coding | 42.87 | -0.079126316 | 0.079126316 | ENSG00000204138 | n.s. |
| ENSG00000175602 | 292.6786 | -0.375639375 | 0.263168 | -1.427377 | 0.153471 | 0.615840022 | CCDC85B | protein_coding | 67.65 | -0.079084143 | 0.079084143 | ENSG00000175602 | n.s. |
| ENSG00000176095 | 674.2971 | -0.383403825 | 0.270609 | -1.416818 | 0.156536 | 0.622084796 | IP6K1 | protein_coding | 44.81 | -0.079038857 | 0.079038857 | ENSG00000176095 | n.s. |
| ENSG00000135900 | 809.8005 | 0.362056693 | 0.247929 | 1.4603214 | 0.144202 | 0.604917447 | MRPL44 | protein_coding | 37.45 | 0.079038384 | 0.079038384 | ENSG00000135900 | n.s. |
| ENSG00000186532 | 364.0577 | -0.364495872 | 0.25114 | -1.451364 | 0.146679 | 0.607109641 | SMYD4 | protein_coding | 46.16 | -0.078998237 | 0.078998237 | ENSG00000186532 | n.s. |
| ENSG00000111846 | 174.9607 | 0.364208839 | 0.250107 | 1.4562101 | 0.145335 | 0.607098916 | GCNT2 | protein_coding | 43.51 | 0.078938821 | 0.078938821 | ENSG00000111846 | n.s. |
| ENSG00000185504 | 601.8835 | -0.364131846 | 0.250795 | -1.451912 | 0.146526 | 0.607109641 | FAAP100 | protein_coding | 61.6 | -0.07891934 | 0.07891934 | ENSG00000185504 | n.s. |
| ENSG00000163138 | 624.7916 | -0.336371412 | 0.223199 | -1.507046 | 0.131799 | 0.582735568 | PACRGL | protein_coding | 38.27 | -0.078888674 | 0.078888674 | ENSG00000163138 | n.s. |
| ENSG00000270141 | 41.93188 | -0.513305561 | 0.425409 | -1.206617 | 0.22758 | 0.70225867 | TERC | lincRNA | 66.73 | -0.078793887 | 0.078793887 | ENSG00000270141 | n.s. |
| ENSG00000105229 | 235.6081 | -0.392097308 | 0.281528 | -1.392747 | 0.163696 | 0.62959761 | PIAS4 | protein_coding | 59.38 | -0.078786829 | 0.078786829 | ENSG00000105229 | n.s. |
| ENSG00000182054 | 2688.724 | -0.402711766 | 0.293363 | -1.372744 | 0.169832 | 0.638037836 | IDH2 | protein_coding | 52.54 | -0.078590638 | 0.078590638 | ENSG00000182054 | n.s. |
| ENSG00000205643 | 83.64814 | -0.409855342 | 0.303165 | -1.351921 | 0.1764 | 0.643565193 | CDPF1 | protein_coding | 57.88 | -0.078449367 | 0.078449367 | ENSG00000205643 | n.s. |
| ENSG00000254635 | 519.9529 | -0.369783562 | 0.257648 | -1.435226 | 0.151223 | 0.613566131 | WAC-AS1 | antisense | 43.31 | -0.078445375 | 0.078445375 | ENSG00000254635 | n.s. |
| ENSG00000105323 | 2063.562 | -0.359904989 | 0.24664 | -1.459231 | 0.144501 | 0.605862372 | HNRNPUL1 | protein_coding | 48.03 | -0.07832469 | 0.07832469 | ENSG00000105323 | n.s. |
| ENSG00000165060 | 314.9141 | -0.390695439 | 0.280966 | -1.390545 | 0.164363 | 0.63030432 | FXN | protein_coding | 45.22 | -0.07831479 | 0.07831479 | ENSG00000165060 | n.s. |
| ENSG00000200367 | 10.25138 | 0.515532808 | 0.432325 | 1.1924671 | 0.233078 | 0.705095593 | SNORD113-8 | snoRNA | 39.19 | 0.078233134 | 0.078233134 | ENSG00000200367 | n.s. |
| ENSG00000085511 | 1291.545 | -0.356986797 | 0.24425 | -1.461565 | 0.14386 | 0.604031311 | MAP3K4 | protein_coding | 38.79 | -0.078158885 | 0.078158885 | ENSG00000085511 | n.s. |
| ENSG00000125848 | 573.1138 | 0.32705548 | 0.214217 | 1.5267471 | 0.126824 | 0.576937759 | FLRT3 | protein_coding | 35.11 | 0.078124082 | 0.078124082 | ENSG00000125848 | n.s. |
| ENSG00000157212 | 935.6585 | -0.339022412 | 0.226957 | -1.493772 | 0.135235 | 0.588399638 | PAXIP1 | protein_coding | 40.81 | -0.078086219 | 0.078086219 | ENSG00000157212 | n.s. |

| | | | | | | | | | | | | | |
|---|---|---|---|---|---|---|---|---|---|---|---|---|---|
| ENSG00000275714 | 1484.034 | -0.348050679 | 0.235308 | -1.479127 | 0.139106 | 0.596580707 | HIST1H3A | protein_coding | 60.58 | -0.078078486 | 0.078078486 | ENSG00000275714 | n.s. |
| ENSG00000169180 | 2359.159 | -0.348678529 | 0.235987 | -1.477534 | 0.139532 | 0.597163054 | XPO6 | protein_coding | 43.8 | -0.078071588 | 0.078071588 | ENSG00000169180 | n.s. |
| ENSG00000157833 | 386.0586 | -0.356653646 | 0.244077 | -1.461233 | 0.143952 | 0.604232315 | GAREM2 | protein_coding | 58.42 | -0.07803441 | 0.07803441 | ENSG00000157833 | n.s. |
| ENSG00000261777 | 55.42097 | 0.405782884 | 0.299163 | 1.3563958 | 0.174973 | 0.642395686 | | processed_transcript | 46.95 | 0.077990409 | 0.077990409 | ENSG00000261777 | n.s. |
| ENSG00000268521 | 47.64211 | -0.441381797 | 0.337801 | -1.306633 | 0.191338 | 0.666152146 | VN1R83P | unprocessed_pseudogene | 35.96 | -0.077871476 | 0.077871476 | ENSG00000268521 | n.s. |
| ENSG00000169604 | 24650.15 | -0.359024968 | 0.247944 | -1.448006 | 0.147615 | 0.607109641 | ANTXR1 | protein_coding | 41.88 | -0.077812512 | 0.077812512 | ENSG00000169604 | n.s. |
| ENSG00000076321 | 810.6981 | -0.373928353 | 0.263125 | -1.421108 | 0.155285 | 0.619317868 | KLHL20 | protein_coding | 38.33 | -0.077809401 | 0.077809401 | ENSG00000076321 | n.s. |
| ENSG00000197785 | 223.3799 | -0.386374677 | 0.276134 | -1.39923 | 0.161744 | 0.628957308 | ATAD3A | protein_coding | 59.87 | -0.077807681 | 0.077807681 | ENSG00000197785 | n.s. |
| ENSG00000121289 | 272.4538 | -0.352689729 | 0.240199 | -1.468322 | 0.142017 | 0.601734228 | CEP89 | protein_coding | 45.48 | -0.077801691 | 0.077801691 | ENSG00000121289 | n.s. |
| ENSG00000205060 | 1298.22 | -0.377980053 | 0.267109 | -1.41508 | 0.157045 | 0.622581912 | SLC35B4 | protein_coding | 40.49 | -0.077789618 | 0.077789618 | ENSG00000205060 | n.s. |
| ENSG00000128590 | 600.5067 | 0.334634281 | 0.222903 | 1.5012573 | 0.133289 | 0.5857901 | DNAJB9 | protein_coding | 38.78 | 0.07772148 | 0.07772148 | ENSG00000128590 | n.s. |
| ENSG00000236194 | 276.4405 | 0.350205551 | 0.238145 | 1.470554 | 0.141412 | 0.600033044 | | sense_intronic | 36.73 | 0.077684288 | 0.077684288 | ENSG00000236194 | n.s. |
| ENSG00000252641 | 8.286399 | 0.554178478 | 0.486306 | 1.1395681 | 0.254466 | 0.72418384 | RNU6-678P | snRNA | 37.61 | 0.077668762 | 0.077668762 | ENSG00000252641 | n.s. |
| ENSG00000172936 | 278.1669 | -0.345376761 | 0.233345 | -1.48011 | 0.138844 | 0.596095432 | MYD88 | protein_coding | 54.39 | -0.077600703 | 0.077600703 | ENSG00000172936 | n.s. |
| ENSG00000116141 | 1265.227 | -0.36840434 | 0.258097 | -1.427389 | 0.153468 | 0.615840022 | MARK1 | protein_coding | 37.1 | -0.077560936 | 0.077560936 | ENSG00000116141 | n.s. |
| ENSG00000170448 | 667.2644 | -0.357649947 | 0.245832 | -1.454855 | 0.145709 | 0.607109641 | NFXL1 | protein_coding | 35.55 | -0.0775145 | 0.0775145 | ENSG00000170448 | n.s. |
| ENSG00000196876 | 344.8979 | -0.345922178 | 0.234191 | -1.477095 | 0.13965 | 0.597163054 | SCN8A | protein_coding | 41.03 | -0.077454421 | 0.077454421 | ENSG00000196876 | n.s. |
| ENSG00000119782 | 81.24635 | 0.437114264 | 0.333962 | 1.3088727 | 0.190577 | 0.665181845 | FKBP1B | protein_coding | 48.23 | 0.077395282 | 0.077395282 | ENSG00000119782 | n.s. |
| ENSG00000186487 | 1157.901 | -0.339856743 | 0.228544 | -1.487053 | 0.137001 | 0.592011337 | MYT1L | protein_coding | 44.5 | -0.077375177 | 0.077375177 | ENSG00000186487 | n.s. |
| ENSG00000198900 | 2033.223 | -0.33984661 | 0.228571 | -1.486831 | 0.137059 | 0.592046125 | TOP1 | protein_coding | 39.86 | -0.077364197 | 0.077364197 | ENSG00000198900 | n.s. |
| ENSG00000107282 | 644.3403 | -0.356871418 | 0.246446 | -1.448074 | 0.147596 | 0.607109641 | APBA1 | protein_coding | 41.44 | -0.077345767 | 0.077345767 | ENSG00000107282 | n.s. |
| ENSG00000148429 | 860.7813 | -0.359822778 | 0.249357 | -1.443 | 0.14902 | 0.609700915 | USP6NL | protein_coding | 38.22 | -0.077319853 | 0.077319853 | ENSG00000148429 | n.s. |
| ENSG00000277138 | 2.278812 | -0.793277464 | 0.871323 | -0.910428 | 0.362597 | 0.799058295 | MIR6509 | miRNA | 37.65 | -0.077282309 | 0.077282309 | ENSG00000277138 | n.s. |
| ENSG00000118655 | 509.5366 | -0.356513161 | 0.24492 | -1.455628 | 0.145495 | 0.607098916 | DCLRE1B | protein_coding | 45.65 | -0.077270856 | 0.077270856 | ENSG00000118655 | n.s. |
| ENSG00000270048 | 20.09089 | -0.506760788 | 0.422438 | -1.19961 | 0.230291 | 0.704043192 | | sense_intronic | 44.87 | -0.077230697 | 0.077230697 | ENSG00000270048 | n.s. |
| ENSG00000274340 | 90.51309 | 0.363966868 | 0.253678 | 1.4347618 | 0.151355 | 0.613599141 | | lincRNA | 37.51 | 0.077202926 | 0.077202926 | ENSG00000274340 | n.s. |
| ENSG00000101187 | 112.7156 | -0.402202369 | 0.296885 | -1.35474 | 0.1755 | 0.642859735 | SLCO4A1 | protein_coding | 58.22 | -0.077176109 | 0.077176109 | ENSG00000101187 | n.s. |
| ENSG00000183337 | 1919.969 | -0.350314217 | 0.238948 | -1.466069 | 0.142629 | 0.602521359 | BCOR | protein_coding | 53.23 | -0.07707878 | 0.07707878 | ENSG00000183337 | n.s. |
| ENSG00000051128 | 198.5125 | -0.404546976 | 0.299743 | -1.349644 | 0.17713 | 0.644922042 | HOMER3 | protein_coding | 56.77 | -0.077063278 | 0.077063278 | ENSG00000051128 | n.s. |
| ENSG00000038427 | 2136.372 | -0.366750795 | 0.257181 | -1.426043 | 0.153856 | 0.61644154 | VCAN | protein_coding | 37.22 | -0.077057314 | 0.077057314 | ENSG00000038427 | n.s. |
| ENSG00000109911 | 402.7605 | 0.376056498 | 0.266635 | 1.4103775 | 0.158428 | 0.624627103 | ELP4 | protein_coding | 35.79 | 0.076858117 | 0.076858117 | ENSG00000109911 | n.s. |
| ENSG00000110108 | 233.5844 | -0.387124359 | 0.280267 | -1.381271 | 0.167196 | 0.633309461 | TMEM109 | protein_coding | 51.1 | -0.076799288 | 0.076799288 | ENSG00000110108 | n.s. |

| | | | | | | | | | | | | | |
|---|---|---|---|---|---|---|---|---|---|---|---|---|---|
| ENSG00000178878 | 350.8039 | 0.323066683 | 0.21241 | 1.520961 | 0.12827 | 0.579048024 | APOLD1 | protein_coding | 42.82 | 0.076659012 | 0.076659012 | ENSG00000178878 | n.s. |
| ENSG00000124532 | 727.6108 | -0.372879013 | 0.263816 | -1.413403 | 0.157537 | 0.622955867 | MRS2 | protein_coding | 39.3 | -0.076642566 | 0.076642566 | ENSG00000124532 | n.s. |
| ENSG00000136425 | 170.7695 | -0.40361181 | 0.299546 | -1.34741 | 0.177848 | 0.646016851 | CIB2 | protein_coding | 51.28 | -0.076587825 | 0.076587825 | ENSG00000136425 | n.s. |
| ENSG00000198888 | 1653.336 | -0.31841272 | 0.207533 | -1.534275 | 0.124962 | 0.574749865 | MT-ND1 | protein_coding | 47.7 | -0.076584985 | 0.076584985 | ENSG00000198888 | n.s. |
| ENSG00000101882 | 317.048 | -0.397888991 | 0.292519 | -1.360214 | 0.173762 | 0.642002364 | NKAP | protein_coding | 41.82 | -0.076579057 | 0.076579057 | ENSG00000101882 | n.s. |
| ENSG00000078061 | 327.6354 | -0.377561604 | 0.268438 | -1.406512 | 0.159572 | 0.626988507 | ARAF | protein_coding | 51.23 | -0.076546998 | 0.076546998 | ENSG00000078061 | n.s. |
| ENSG00000070770 | 990.2442 | -0.349066369 | 0.238341 | -1.464565 | 0.14304 | 0.603629167 | CSNK2A2 | protein_coding | 41.31 | -0.076525745 | 0.076525745 | ENSG00000070770 | n.s. |
| ENSG00000198911 | 1454.354 | -0.36073194 | 0.251354 | -1.435154 | 0.151243 | 0.613566131 | SREBF2 | protein_coding | 48.21 | -0.076525177 | 0.076525177 | ENSG00000198911 | n.s. |
| ENSG00000141433 | 257.9127 | 0.361026853 | 0.251824 | 1.4336458 | 0.151673 | 0.614039101 | ADCYAP1 | protein_coding | 52.82 | 0.076466922 | 0.076466922 | ENSG00000141433 | n.s. |
| ENSG00000196295 | 162.8655 | -0.395552499 | 0.289608 | -1.36582 | 0.171996 | 0.640837445 | | processed_transcript | 42.17 | -0.076441359 | 0.076441359 | ENSG00000196295 | n.s. |
| ENSG00000206418 | 491.3821 | -0.385175662 | 0.278864 | -1.381229 | 0.167209 | 0.633309461 | RAB12 | protein_coding | 42.23 | -0.076412698 | 0.076412698 | ENSG00000206418 | n.s. |
| ENSG00000137073 | 1044.518 | -0.352167018 | 0.241968 | -1.45543 | 0.14555 | 0.607098916 | UBAP2 | protein_coding | 43.26 | -0.07632887 | 0.07632887 | ENSG00000137073 | n.s. |
| ENSG00000137824 | 598.105 | -0.360597255 | 0.251662 | -1.432865 | 0.151896 | 0.614510829 | RMDN3 | protein_coding | 49.18 | -0.076255667 | 0.076255667 | ENSG00000137824 | n.s. |
| ENSG00000057252 | 819.5594 | -0.351453903 | 0.241984 | -1.452386 | 0.146394 | 0.607109641 | SOAT1 | protein_coding | 39.63 | -0.076171613 | 0.076171613 | ENSG00000057252 | n.s. |
| ENSG00000074356 | 1551.701 | -0.352889305 | 0.244259 | -1.444733 | 0.148533 | 0.60842133 | NCBP3 | protein_coding | 44.57 | -0.076151948 | 0.076151948 | ENSG00000074356 | n.s. |
| ENSG00000153048 | 2612.693 | -0.372586892 | 0.264218 | -1.410152 | 0.158495 | 0.624627103 | CARHSP1 | protein_coding | 55.97 | -0.076149002 | 0.076149002 | ENSG00000153048 | n.s. |
| ENSG00000101346 | 634.5514 | -0.369340222 | 0.260698 | -1.416736 | 0.15656 | 0.622084796 | POFUT1 | protein_coding | 47.65 | -0.076139639 | 0.076139639 | ENSG00000101346 | n.s. |
| ENSG00000081320 | 662.9687 | -0.351289856 | 0.242383 | -1.449317 | 0.147249 | 0.607109641 | STK17B | protein_coding | 37.67 | -0.076136059 | 0.076136059 | ENSG00000081320 | n.s. |
| ENSG00000139618 | 2045.635 | -0.379781161 | 0.27313 | -1.390478 | 0.164384 | 0.63030432 | BRCA2 | protein_coding | 38.11 | -0.076127026 | 0.076127026 | ENSG00000139618 | n.s. |
| ENSG00000143079 | 727.8382 | -0.351242327 | 0.242494 | -1.448456 | 0.14749 | 0.607109641 | CTTNBP2NL | protein_coding | 39.63 | -0.076125758 | 0.076125758 | ENSG00000143079 | n.s. |
| ENSG00000104626 | 853.817 | -0.363211291 | 0.25484 | -1.425251 | 0.154085 | 0.617181083 | ERI1 | protein_coding | 41.64 | -0.076124508 | 0.076124508 | ENSG00000104626 | n.s. |
| ENSG00000141026 | 436.4171 | -0.318634889 | 0.208685 | -1.526867 | 0.126794 | 0.576937759 | MED9 | protein_coding | 47.9 | -0.076112646 | 0.076112646 | ENSG00000141026 | n.s. |
| ENSG00000155850 | 600.0089 | -0.383678497 | 0.277897 | -1.380652 | 0.167386 | 0.633636749 | SLC26A2 | protein_coding | 41.19 | -0.076029594 | 0.076029594 | ENSG00000155850 | n.s. |
| ENSG00000229676 | 510.6049 | -0.374068396 | 0.265766 | -1.407511 | 0.159276 | 0.626322307 | ZNF492 | protein_coding | 39.03 | -0.076011491 | 0.076011491 | ENSG00000229676 | n.s. |
| ENSG00000241157 | 28.21521 | -0.44916302 | 0.350809 | -1.280362 | 0.200418 | 0.677286179 | | processed_pseudogene | 46.39 | -0.076010864 | 0.076010864 | ENSG00000241157 | n.s. |
| ENSG00000108001 | 4194.072 | -0.354868797 | 0.246163 | -1.441603 | 0.149414 | 0.610673934 | EBF3 | protein_coding | 48.8 | -0.076009567 | 0.076009567 | ENSG00000108001 | n.s. |
| ENSG00000136478 | 799.3352 | -0.376211613 | 0.267998 | -1.403783 | 0.160383 | 0.62809251 | TEX2 | protein_coding | 43.58 | -0.075985862 | 0.075985862 | ENSG00000136478 | n.s. |
| ENSG00000125818 | 437.3961 | -0.377222592 | 0.270408 | -1.395012 | 0.163012 | 0.629036117 | PSMF1 | protein_coding | 45.44 | -0.075944119 | 0.075944119 | ENSG00000125818 | n.s. |
| ENSG00000189306 | 988.2763 | -0.372624909 | 0.264548 | -1.408535 | 0.158973 | 0.625601663 | RRP7A | protein_coding | 57.96 | -0.075904479 | 0.075904479 | ENSG00000189306 | n.s. |
| ENSG00000169087 | 257.45 | -0.394307428 | 0.290054 | -1.359428 | 0.174011 | 0.642002364 | HSPBAP1 | protein_coding | 39.97 | -0.075889737 | 0.075889737 | ENSG00000169087 | n.s. |
| ENSG00000203930 | 153.4394 | -0.361095469 | 0.253191 | -1.42618 | 0.153816 | 0.61644154 | LINC00632 | lincRNA | 40.81 | -0.075869084 | 0.075869084 | ENSG00000203930 | n.s. |
| ENSG00000104142 | 364.2276 | -0.350042395 | 0.240539 | -1.455242 | 0.145602 | 0.607109641 | VPS18 | protein_coding | 53.64 | -0.075865693 | 0.075865693 | ENSG00000104142 | n.s. |

| | | | | | | | | | | | | | |
|---|---|---|---|---|---|---|---|---|---|---|---|---|---|
| ENSG00000258429 | 75.87622 | -0.392186042 | 0.28739 | -1.364649 | 0.172363 | 0.640837445 | PDF | protein_coding | 61.27 | -0.075790784 | 0.075790784 | ENSG00000258429 | n.s. |
| ENSG00000207763 | 12.50455 | -0.491636677 | 0.405862 | -1.211339 | 0.225765 | 0.701204614 | MIR617 | miRNA | 45.36 | -0.075788367 | 0.075788367 | ENSG00000207763 | n.s. |
| ENSG00000100226 | 1006.924 | -0.339693163 | 0.230599 | -1.473091 | 0.140726 | 0.598635657 | GTPBP1 | protein_coding | 51.1 | -0.075696347 | 0.075696347 | ENSG00000100226 | n.s. |
| ENSG00000232228 | 9.901075 | -0.565973731 | 0.509156 | -1.111591 | 0.266314 | 0.734987182 | | processed_pseudogene | 46.86 | -0.07568214 | 0.07568214 | ENSG00000232228 | n.s. |
| ENSG00000249637 | 30.01932 | 0.464316079 | 0.371292 | 1.2505422 | 0.211102 | 0.687111024 | | antisense | 44.25 | 0.075671023 | 0.075671023 | ENSG00000249637 | n.s. |
| ENSG00000054611 | 193.6785 | -0.357870391 | 0.249944 | -1.431802 | 0.152201 | 0.614628915 | TBC1D22A | protein_coding | 49.42 | -0.075649153 | 0.075649153 | ENSG00000054611 | n.s. |
| ENSG00000223551 | 18.05029 | -0.508651439 | 0.430501 | -1.181534 | 0.237391 | 0.710029832 | TMSB4XP4 | processed_pseudogene | 43.04 | -0.075648373 | 0.075648373 | ENSG00000223551 | n.s. |
| ENSG00000224016 | 14.16286 | -0.701015766 | 0.718262 | -0.975989 | 0.32907 | 0.780141891 | | unprocessed_pseudogene | 39.96 | -0.075588007 | 0.075588007 | ENSG00000224016 | n.s. |
| ENSG00000241030 | 14.92493 | -0.489888533 | 0.404808 | -1.210175 | 0.226212 | 0.701289341 | | processed_pseudogene | 54.23 | -0.075493176 | 0.075493176 | ENSG00000241030 | n.s. |
| ENSG00000111275 | 637.7534 | -0.391429584 | 0.287663 | -1.360721 | 0.173602 | 0.642002364 | ALDH2 | protein_coding | 48.8 | -0.075335858 | 0.075335858 | ENSG00000111275 | n.s. |
| ENSG00000141349 | 686.5495 | -0.314100439 | 0.205262 | -1.530242 | 0.125957 | 0.575897021 | G6PC3 | protein_coding | 53.16 | -0.075275794 | 0.075275794 | ENSG00000141349 | n.s. |
| ENSG00000200142 | 20.27155 | -0.47170753 | 0.381397 | -1.23679 | 0.216165 | 0.692499549 | | misc_RNA | 44.9 | -0.075275326 | 0.075275326 | ENSG00000200142 | n.s. |
| ENSG00000136485 | 4325.358 | -0.373829244 | 0.267967 | -1.395056 | 0.162999 | 0.629036117 | DCAF7 | protein_coding | 43.96 | -0.075260955 | 0.075260955 | ENSG00000136485 | n.s. |
| ENSG00000076984 | 312.219 | -0.366156634 | 0.259055 | -1.41343 | 0.157529 | 0.622955867 | MAP2K7 | protein_coding | 63.14 | -0.07526083 | 0.07526083 | ENSG00000076984 | n.s. |
| ENSG00000275175 | 11.11897 | 0.611865944 | 0.582454 | 1.050497 | 0.29349 | 0.753629384 | | lincRNA | 36.71 | 0.075162944 | 0.075162944 | ENSG00000275175 | n.s. |
| ENSG00000174903 | 454.0746 | -0.384961606 | 0.280324 | -1.373275 | 0.169667 | 0.638037836 | RAB1B | protein_coding | 55.61 | -0.075126631 | 0.075126631 | ENSG00000174903 | n.s. |
| ENSG00000148730 | 1582.068 | -0.353758464 | 0.246368 | -1.435894 | 0.151032 | 0.613566131 | EIF4EBP2 | protein_coding | 43.66 | -0.075045833 | 0.075045833 | ENSG00000148730 | n.s. |
| ENSG00000170430 | 283.8387 | -0.388191564 | 0.284469 | -1.364618 | 0.172373 | 0.640837445 | MGMT | protein_coding | 45.49 | -0.075018843 | 0.075018843 | ENSG00000170430 | n.s. |
| ENSG00000247416 | 50.34718 | 0.426926103 | 0.327456 | 1.3037649 | 0.192314 | 0.667270349 | | antisense | 53.01 | 0.075010137 | 0.075010137 | ENSG00000247416 | n.s. |
| ENSG00000254122 | 165.9667 | -0.346074401 | 0.238516 | -1.450947 | 0.146795 | 0.607109641 | PCDHGB7 | protein_coding | 45.57 | -0.075005698 | 0.075005698 | ENSG00000254122 | n.s. |
| ENSG00000032219 | 133.3725 | -0.355306366 | 0.248649 | -1.428947 | 0.153019 | 0.615215178 | ARID4A | protein_coding | 37.48 | -0.074960035 | 0.074960035 | ENSG00000032219 | n.s. |
| ENSG00000143340 | 585.0951 | -0.375168518 | 0.270694 | -1.385952 | 0.165762 | 0.631263714 | FAM163A | protein_coding | 47.32 | -0.074954608 | 0.074954608 | ENSG00000143340 | n.s. |
| ENSG00000232546 | 13.02168 | -0.504787287 | 0.428077 | -1.179198 | 0.238319 | 0.71051867 | | unprocessed_pseudogene | 41.56 | -0.074922804 | 0.074922804 | ENSG00000232546 | n.s. |
| ENSG00000183751 | 337.5251 | -0.389475994 | 0.287121 | -1.356488 | 0.174944 | 0.642395686 | TBL3 | protein_coding | 64.29 | -0.074856267 | 0.074856267 | ENSG00000183751 | n.s. |
| ENSG00000274963 | 35.81328 | 0.426208533 | 0.327144 | 1.3028156 | 0.192638 | 0.667375338 | RN7SL600P | misc_RNA | 50.84 | 0.074854939 | 0.074854939 | ENSG00000274963 | n.s. |
| ENSG00000226989 | 23.94861 | 0.46456566 | 0.373233 | 1.2447075 | 0.213239 | 0.690152016 | | processed_pseudogene | 50.63 | 0.074820733 | 0.074820733 | ENSG00000226989 | n.s. |
| ENSG00000170776 | 107.441 | -0.371282299 | 0.266194 | -1.394779 | 0.163082 | 0.629036117 | AKAP13 | protein_coding | 39.92 | -0.074748193 | 0.074748193 | ENSG00000170776 | n.s. |
| ENSG00000201570 | 18.20415 | -0.505132919 | 0.429297 | -1.176652 | 0.239334 | 0.71125159 | RNU4-56P | snRNA | 38.73 | -0.074747927 | 0.074747927 | ENSG00000201570 | n.s. |
| ENSG00000166503 | 4647.608 | -0.379793779 | 0.275586 | -1.378131 | 0.168163 | 0.63576037 | | protein_coding | 39.61 | -0.074707923 | 0.074707923 | ENSG00000166503 | n.s. |
| ENSG00000121989 | 803.2917 | -0.360671852 | 0.254161 | -1.419067 | 0.155879 | 0.62079796 | ACVR2A | protein_coding | 35.46 | -0.074677006 | 0.074677006 | ENSG00000121989 | n.s. |
| ENSG00000272602 | 613.1663 | -0.387881763 | 0.285365 | -1.359246 | 0.174069 | 0.642002364 | ZNF595 | protein_coding | 38.31 | -0.074653032 | 0.074653032 | ENSG00000272602 | n.s. |
| ENSG00000248240 | 27.63556 | -0.474360977 | 0.386937 | -1.225938 | 0.220222 | 0.696377966 | | antisense | 45.83 | -0.074548189 | 0.074548189 | ENSG00000248240 | n.s. |

| | | | | | | | | | | | | | |
|---|---|---|---|---|---|---|---|---|---|---|---|---|---|
| ENSG00000244710 | 31.09259 | -0.44076445 | 0.344564 | -1.279195 | 0.200828 | 0.677491543 | RN7SL47P | misc_RNA | 54.85 | -0.074531559 | 0.074531559 | ENSG00000244710 | n.s. |
| ENSG00000113163 | 563.4962 | -0.357550788 | 0.251484 | -1.421766 | 0.155094 | 0.618910335 | COL4A3BP | protein_coding | 37.13 | -0.074503667 | 0.074503667 | ENSG00000113163 | n.s. |
| ENSG00000207460 | 44.29735 | -0.431643592 | 0.334374 | -1.290899 | 0.196739 | 0.672118672 | SNORD116-19 | snoRNA | 42.39 | -0.074481845 | 0.074481845 | ENSG00000207460 | n.s. |
| ENSG00000163162 | 928.6317 | -0.376101768 | 0.272487 | -1.380256 | 0.167508 | 0.633925508 | RNF149 | protein_coding | 41.19 | -0.074453773 | 0.074453773 | ENSG00000163162 | n.s. |
| ENSG00000154721 | 210.0607 | -0.368595048 | 0.262566 | -1.403819 | 0.160373 | 0.62809251 | JAM2 | protein_coding | 39.65 | -0.074447496 | 0.074447496 | ENSG00000154721 | n.s. |
| ENSG00000170085 | 111.1786 | -0.382701416 | 0.279487 | -1.369302 | 0.170905 | 0.639237118 | SIMC1 | protein_coding | 42.19 | -0.074373434 | 0.074373434 | ENSG00000170085 | n.s. |
| ENSG00000243701 | 165.3319 | -0.386947613 | 0.28531 | -1.356238 | 0.175024 | 0.642395686 | DUBR | lincRNA | 38.56 | -0.074370319 | 0.074370319 | ENSG00000243701 | n.s. |
| ENSG00000159267 | 385.6938 | -0.370937803 | 0.266777 | -1.390443 | 0.164394 | 0.63030432 | HLCS | protein_coding | 44.35 | -0.074354377 | 0.074354377 | ENSG00000159267 | n.s. |
| ENSG00000113615 | 681.3096 | -0.350486365 | 0.244325 | -1.434512 | 0.151426 | 0.613599141 | SEC24A | protein_coding | 42.26 | -0.074343505 | 0.074343505 | ENSG00000113615 | n.s. |
| ENSG00000259658 | 27.59506 | -0.44731073 | 0.35433 | -1.262413 | 0.2068 | 0.682214558 | | transcribed_unprocessed_pseudc | 44.33 | -0.074288926 | 0.074288926 | ENSG00000259658 | n.s. |
| ENSG00000166037 | 1172.559 | -0.386445412 | 0.285042 | -1.355751 | 0.175178 | 0.642395686 | CEP57 | protein_coding | 37.69 | -0.074273797 | 0.074273797 | ENSG00000166037 | n.s. |
| ENSG00000136935 | 197.4976 | -0.367863536 | 0.262241 | -1.402771 | 0.160685 | 0.628432732 | GOLGA1 | protein_coding | 44.12 | -0.074213232 | 0.074213232 | ENSG00000136935 | n.s. |
| ENSG00000240849 | 276.5546 | -0.349728499 | 0.243588 | -1.435741 | 0.151076 | 0.613566131 | TMEM189 | protein_coding | 52.37 | -0.074190922 | 0.074190922 | ENSG00000240849 | n.s. |
| ENSG00000097007 | 1236.243 | -0.34207641 | 0.236291 | -1.447689 | 0.147704 | 0.607109641 | ABL1 | protein_coding | 44.47 | -0.074139202 | 0.074139202 | ENSG00000097007 | n.s. |
| ENSG00000272906 | 34.46799 | 0.454252819 | 0.362884 | 1.2517848 | 0.210648 | 0.686934651 | | lincRNA | 54.1 | 0.074081629 | 0.074081629 | ENSG00000272906 | n.s. |
| ENSG00000100604 | 3214.629 | -0.369421692 | 0.265774 | -1.389985 | 0.164533 | 0.63030432 | CHGA | protein_coding | 56.35 | -0.074050473 | 0.074050473 | ENSG00000100604 | n.s. |
| ENSG00000212567 | 19.19454 | 0.482766163 | 0.401046 | 1.203767 | 0.22868 | 0.702737638 | | snoRNA | 54.69 | 0.073963052 | 0.073963052 | ENSG00000212567 | n.s. |
| ENSG00000267152 | 61.31358 | -0.421392647 | 0.323812 | -1.301348 | 0.193139 | 0.667684687 | | lincRNA | 35.52 | -0.073924316 | 0.073924316 | ENSG00000267152 | n.s. |
| ENSG00000198818 | 389.5123 | -0.376729754 | 0.273905 | -1.3754 | 0.169007 | 0.636857974 | SFT2D1 | protein_coding | 43.24 | -0.073822987 | 0.073822987 | ENSG00000198818 | n.s. |
| ENSG00000006652 | 3180.323 | 0.340580921 | 0.235329 | 1.4472563 | 0.147825 | 0.607109641 | IFRD1 | protein_coding | 39.45 | 0.073815081 | 0.073815081 | ENSG00000006652 | n.s. |
| ENSG00000103496 | 59.28885 | -0.40737038 | 0.308027 | -1.322514 | 0.185997 | 0.658994794 | STX4 | protein_coding | 55.52 | -0.073782115 | 0.073782115 | ENSG00000103496 | n.s. |
| ENSG00000209645 | 6.777917 | -0.628714002 | 0.618327 | -1.016799 | 0.309249 | 0.763298957 | SNORD105 | snoRNA | 47.06 | -0.073751504 | 0.073751504 | ENSG00000209645 | n.s. |
| ENSG00000207009 | 14.33119 | 0.562413411 | 0.514747 | 1.0926019 | 0.274569 | 0.739386355 | | misc_RNA | 46.36 | 0.073748465 | 0.073748465 | ENSG00000207009 | n.s. |
| ENSG00000149308 | 887.6463 | -0.370750091 | 0.26808 | -1.382981 | 0.166671 | 0.632544316 | NPAT | protein_coding | 40.27 | -0.073745546 | 0.073745546 | ENSG00000149308 | n.s. |
| ENSG00000199065 | 4.706632 | -0.775131654 | 0.860564 | -0.900725 | 0.367735 | 0.803384223 | MIR101-2 | miRNA | 40.51 | -0.073696961 | 0.073696961 | ENSG00000199065 | n.s. |
| ENSG00000149557 | 152.6206 | 0.409728562 | 0.311049 | 1.3172457 | 0.187756 | 0.661470368 | FEZ1 | protein_coding | 43.19 | 0.073542018 | 0.073542018 | ENSG00000149557 | n.s. |
| ENSG00000157895 | 421.4881 | -0.391664914 | 0.292011 | -1.341268 | 0.179833 | 0.649162475 | C12orf43 | protein_coding | 49.65 | -0.073494587 | 0.073494587 | ENSG00000157895 | n.s. |
| ENSG00000040275 | 1460.822 | -0.364822829 | 0.261006 | -1.397757 | 0.162186 | 0.628957308 | SPDL1 | protein_coding | 37.28 | -0.073467595 | 0.073467595 | ENSG00000040275 | n.s. |
| ENSG00000139734 | 1085.769 | -0.364838346 | 0.261377 | -1.395834 | 0.162764 | 0.629036117 | DIAPH3 | protein_coding | 35.91 | -0.073450868 | 0.073450868 | ENSG00000139734 | n.s. |
| ENSG00000088038 | 209.4694 | -0.36608267 | 0.263501 | -1.389303 | 0.164741 | 0.63030432 | CNOT3 | protein_coding | 57.91 | -0.073381167 | 0.073381167 | ENSG00000088038 | n.s. |
| ENSG00000094916 | 8814.108 | -0.345808522 | 0.240923 | -1.435348 | 0.151188 | 0.613566131 | CBX5 | protein_coding | 42.96 | -0.073359343 | 0.073359343 | ENSG00000094916 | n.s. |
| ENSG00000188566 | 248.7947 | -0.363206629 | 0.258674 | -1.404109 | 0.160286 | 0.62809251 | NDOR1 | protein_coding | 59.32 | -0.073359162 | 0.073359162 | ENSG00000188566 | n.s. |

| | | | | | | | | | | | | | |
|---|---|---|---|---|---|---|---|---|---|---|---|---|---|
| ENSG00000276966 | 3175.777 | -0.303635749 | 0.197428 | -1.537959 | 0.124059 | 0.573519528 | HIST1H4E | protein_coding | 45.8 | -0.073313395 | 0.073313395 | ENSG00000276966 | n.s. |
| ENSG00000101180 | 104.0724 | -0.382002311 | 0.282119 | -1.354046 | 0.175722 | 0.642859735 | HRH3 | protein_coding | 66.12 | -0.073300045 | 0.073300045 | ENSG00000101180 | n.s. |
| ENSG00000238423 | 6.083932 | -0.617227901 | 0.598385 | -1.03149 | 0.302311 | 0.760781413 | SNORD42B | snoRNA | 40.3 | -0.073289707 | 0.073289707 | ENSG00000238423 | n.s. |
| ENSG00000244026 | 153.393 | -0.365345036 | 0.262947 | -1.389425 | 0.164703 | 0.63030432 | FAM86DP | cribed_unprocessed_pseudc | 52.79 | -0.073233309 | 0.073233309 | ENSG00000244026 | n.s. |
| ENSG00000080345 | 8855.735 | -0.350856052 | 0.246619 | -1.422662 | 0.154834 | 0.618415021 | RIF1 | protein_coding | 38.04 | -0.073230665 | 0.073230665 | ENSG00000080345 | n.s. |
| ENSG00000188428 | 271.1241 | -0.411950385 | 0.314028 | -1.311826 | 0.189579 | 0.664193421 | BLOC1S5 | protein_coding | 40 | -0.07320582 | 0.07320582 | ENSG00000188428 | n.s. |
| ENSG00000101447 | 1210.941 | -0.336309035 | 0.230477 | -1.459186 | 0.144514 | 0.605862372 | FAM83D | protein_coding | 46.5 | -0.073189596 | 0.073189596 | ENSG00000101447 | n.s. |
| ENSG00000086102 | 1610.547 | -0.363387103 | 0.259746 | -1.399012 | 0.161809 | 0.628957308 | NFX1 | protein_coding | 42.57 | -0.07317847 | 0.07317847 | ENSG00000086102 | n.s. |
| ENSG00000166452 | 193.8953 | -0.398344644 | 0.299699 | -1.329151 | 0.183798 | 0.655151359 | AKIP1 | protein_coding | 46.42 | -0.073159321 | 0.073159321 | ENSG00000166452 | n.s. |
| ENSG00000177200 | 6241.501 | -0.363065809 | 0.259154 | -1.400965 | 0.161225 | 0.628931954 | CHD9 | protein_coding | 38.92 | -0.073120125 | 0.073120125 | ENSG00000177200 | n.s. |
| ENSG00000200816 | 31.4661 | 0.451171168 | 0.361497 | 1.2480639 | 0.212008 | 0.688575085 | SNORA38 | snoRNA | 44.7 | 0.073111699 | 0.073111699 | ENSG00000200816 | n.s. |
| ENSG00000028528 | 1704.536 | -0.364183087 | 0.261673 | -1.391749 | 0.163998 | 0.629889786 | SNX1 | protein_coding | 43.68 | -0.07310445 | 0.07310445 | ENSG00000028528 | n.s. |
| ENSG00000205559 | 41.25465 | -0.43185328 | 0.337664 | -1.278944 | 0.200917 | 0.677491543 | CHKB-AS1 | antisense | 63.15 | -0.073024715 | 0.073024715 | ENSG00000205559 | n.s. |
| ENSG00000221264 | 3.418453 | 0.780858873 | 0.872295 | 0.8951776 | 0.370692 | 0.806303209 | MIR1284 | miRNA | 30 | 0.073011563 | 0.073011563 | ENSG00000221264 | n.s. |
| ENSG00000146414 | 4471.13 | -0.379629362 | 0.279507 | -1.35821 | 0.174397 | 0.64236221 | SHPRH | protein_coding | 35.43 | -0.072972362 | 0.072972362 | ENSG00000146414 | n.s. |
| ENSG00000206755 | 9.651684 | -0.547800914 | 0.495892 | -1.104677 | 0.269299 | 0.735983077 | SNORA30 | snoRNA | 51.94 | -0.072929925 | 0.072929925 | ENSG00000206755 | n.s. |
| ENSG00000274220 | 56.85954 | 0.431481237 | 0.338429 | 1.2749524 | 0.202326 | 0.677624898 | | lincRNA | 46.42 | 0.072924922 | 0.072924922 | ENSG00000274220 | n.s. |
| ENSG00000270175 | 82.86963 | -0.362106712 | 0.259203 | -1.397002 | 0.162413 | 0.629027704 | | antisense | 53.16 | -0.072903026 | 0.072903026 | ENSG00000270175 | n.s. |
| ENSG00000166783 | 800.8585 | -0.36169312 | 0.258422 | -1.399624 | 0.161626 | 0.628931954 | MARF1 | protein_coding | 44.18 | -0.07284367 | 0.07284367 | ENSG00000166783 | n.s. |
| ENSG00000196584 | 1659.897 | -0.381649866 | 0.282677 | -1.350128 | 0.176975 | 0.64464925 | XRCC2 | protein_coding | 42.57 | -0.072771667 | 0.072771667 | ENSG00000196584 | n.s. |
| ENSG00000240869 | 15.86466 | 0.472740112 | 0.391477 | 1.207581 | 0.227208 | 0.70173545 | RN7SL128P | misc_RNA | 51.8 | 0.072719996 | 0.072719996 | ENSG00000240869 | n.s. |
| ENSG00000260274 | 60.57278 | 0.386505064 | 0.287814 | 1.3428989 | 0.179305 | 0.64843101 | | lincRNA | 54.55 | 0.072715603 | 0.072715603 | ENSG00000260274 | n.s. |
| ENSG00000139793 | 1995.955 | 0.370961853 | 0.269714 | 1.3753878 | 0.169011 | 0.636857974 | MBNL2 | protein_coding | 40.24 | 0.072692724 | 0.072692724 | ENSG00000139793 | n.s. |
| ENSG00000261366 | 284.8641 | 0.328753965 | 0.223764 | 1.469199 | 0.141779 | 0.601092151 | MANEA-AS1 | antisense | 32.72 | 0.072674004 | 0.072674004 | ENSG00000261366 | n.s. |
| ENSG00000117620 | 1210.259 | -0.360842772 | 0.258916 | -1.393669 | 0.163418 | 0.629252671 | SLC35A3 | protein_coding | 37.43 | -0.07259252 | 0.07259252 | ENSG00000117620 | n.s. |
| ENSG00000100263 | 483.2564 | -0.355741093 | 0.252454 | -1.409131 | 0.158796 | 0.625264564 | RHBDD3 | protein_coding | 56.01 | -0.072548481 | 0.072548481 | ENSG00000100263 | n.s. |
| ENSG00000199664 | 3.134663 | -0.671989112 | 0.689773 | -0.974217 | 0.329949 | 0.780141891 | RNU6-1266P | snRNA | 39.05 | -0.072458168 | 0.072458168 | ENSG00000199664 | n.s. |
| ENSG00000083799 | 688.3691 | -0.365408283 | 0.264622 | -1.380869 | 0.167319 | 0.633555789 | CYLD | protein_coding | 36.42 | -0.072429453 | 0.072429453 | ENSG00000083799 | n.s. |
| ENSG00000176407 | 1195.805 | -0.368930337 | 0.26799 | -1.376656 | 0.168619 | 0.636399681 | KCMF1 | protein_coding | 41.72 | -0.072409975 | 0.072409975 | ENSG00000176407 | n.s. |
| ENSG00000169756 | 1625.459 | -0.375418192 | 0.275648 | -1.361946 | 0.173215 | 0.641657113 | LIMS1 | protein_coding | 41.38 | -0.072341955 | 0.072341955 | ENSG00000169756 | n.s. |
| ENSG00000112218 | 410.6984 | -0.360674706 | 0.259513 | -1.389812 | 0.164586 | 0.63030432 | GPR63 | protein_coding | 36.26 | -0.072297142 | 0.072297142 | ENSG00000112218 | n.s. |
| ENSG00000104067 | 1039.407 | -0.325021829 | 0.220831 | -1.471811 | 0.141072 | 0.599316087 | TJP1 | protein_coding | 40.21 | -0.072266675 | 0.072266675 | ENSG00000104067 | n.s. |

| | | | | | | | | | | | | | |
|---|---|---|---|---|---|---|---|---|---|---|---|---|---|
| ENSG00000197763 | 236.3525 | -0.371244768 | 0.270853 | -1.370649 | 0.170484 | 0.638865917 | TXNRD3 | protein_coding | 39.52 | -0.072240624 | 0.072240624 | ENSG00000197763 | n.s. |
| ENSG00000139437 | 725.5753 | -0.340731737 | 0.237691 | -1.433508 | 0.151713 | 0.614039101 | TCHP | protein_coding | 46.02 | -0.072168336 | 0.072168336 | ENSG00000139437 | n.s. |
| ENSG00000185163 | 677.23 | -0.341323156 | 0.238375 | -1.431872 | 0.15218 | 0.614628915 | DDX51 | protein_coding | 60.51 | -0.072151282 | 0.072151282 | ENSG00000185163 | n.s. |
| ENSG00000186472 | 175.371 | -0.368915219 | 0.268393 | -1.374534 | 0.169276 | 0.637498207 | PCLO | protein_coding | 33.81 | -0.072130684 | 0.072130684 | ENSG00000186472 | n.s. |
| ENSG00000136816 | 351.0186 | -0.375286255 | 0.276744 | -1.356076 | 0.175075 | 0.642395686 | TOR1B | protein_coding | 50.46 | -0.072129037 | 0.072129037 | ENSG00000136816 | n.s. |
| ENSG00000078699 | 640.5405 | -0.359745711 | 0.258831 | -1.389888 | 0.164563 | 0.63030432 | CBFA2T2 | protein_coding | 42.36 | -0.072110926 | 0.072110926 | ENSG00000078699 | n.s. |
| ENSG00000142544 | 87.85771 | -0.387244071 | 0.289452 | -1.337855 | 0.180944 | 0.65131356 | CTU1 | protein_coding | 54.31 | -0.072108672 | 0.072108672 | ENSG00000142544 | n.s. |
| ENSG00000127554 | 351.0284 | -0.350695249 | 0.24815 | -1.413237 | 0.157586 | 0.622955867 | GFER | protein_coding | 60.99 | -0.072082855 | 0.072082855 | ENSG00000127554 | n.s. |
| ENSG00000264349 | 41.44224 | -0.426451509 | 0.334535 | -1.274761 | 0.202394 | 0.677624898 | MIR4258 | miRNA | 68.13 | -0.072074845 | 0.072074845 | ENSG00000264349 | n.s. |
| ENSG00000081014 | 949.6763 | -0.366381494 | 0.265909 | -1.377846 | 0.168251 | 0.635872961 | AP4E1 | protein_coding | 37.92 | -0.072041462 | 0.072041462 | ENSG00000081014 | n.s. |
| ENSG00000176024 | 186.416 | -0.382103601 | 0.28432 | -1.343921 | 0.178974 | 0.648076085 | ZNF613 | protein_coding | 41.55 | -0.071978385 | 0.071978385 | ENSG00000176024 | n.s. |
| ENSG00000134046 | 630.8449 | -0.348938663 | 0.246664 | -1.41463 | 0.157177 | 0.622581912 | MBD2 | protein_coding | 39.24 | -0.071812799 | 0.071812799 | ENSG00000134046 | n.s. |
| ENSG00000022840 | 1043.288 | -0.331321171 | 0.228877 | -1.447595 | 0.14773 | 0.607109641 | RNF10 | protein_coding | 45.25 | -0.071808188 | 0.071808188 | ENSG00000022840 | n.s. |
| ENSG00000233220 | 52.48999 | 0.416008074 | 0.322246 | 1.290965 | 0.196716 | 0.672118672 | LINC00167 | lincRNA | 54.19 | 0.071783874 | 0.071783874 | ENSG00000233220 | n.s. |
| ENSG00000175048 | 113.234 | -0.356424194 | 0.255343 | -1.395863 | 0.162756 | 0.629036117 | ZDHHC14 | protein_coding | 45.59 | -0.071756893 | 0.071756893 | ENSG00000175048 | n.s. |
| ENSG00000249492 | 72.49271 | 0.373312906 | 0.275241 | 1.3563119 | 0.175 | 0.642395686 | | processed_transcript | 36.71 | 0.071749764 | 0.071749764 | ENSG00000249492 | n.s. |
| ENSG00000123066 | 2639.573 | -0.33731832 | 0.23458 | -1.437965 | 0.150444 | 0.612823957 | MED13L | protein_coding | 37.29 | -0.071735553 | 0.071735553 | ENSG00000123066 | n.s. |
| ENSG00000152578 | 144.4711 | 0.348954488 | 0.246998 | 1.4127854 | 0.157719 | 0.622955867 | GRIA4 | protein_coding | 35.32 | 0.071725054 | 0.071725054 | ENSG00000152578 | n.s. |
| ENSG00000106772 | 70.81705 | -0.385409832 | 0.288211 | -1.337249 | 0.181141 | 0.651484256 | PRUNE2 | protein_coding | 41.04 | -0.071723257 | 0.071723257 | ENSG00000106772 | n.s. |
| ENSG00000278829 | 181.4669 | 0.347631911 | 0.245321 | 1.4170506 | 0.156468 | 0.622084796 | | sense_intronic | 36.65 | 0.071664462 | 0.071664462 | ENSG00000278829 | n.s. |
| ENSG00000046651 | 171.4437 | 0.330576984 | 0.227082 | 1.4557631 | 0.145458 | 0.607098916 | OFD1 | protein_coding | 39.8 | 0.071649435 | 0.071649435 | ENSG00000046651 | n.s. |
| ENSG00000198792 | 189.8836 | -0.37370583 | 0.276235 | -1.352856 | 0.176102 | 0.643404323 | TMEM184B | protein_coding | 53.51 | -0.071570655 | 0.071570655 | ENSG00000198792 | n.s. |
| ENSG00000167508 | 253.1884 | -0.366068408 | 0.266373 | -1.37427 | 0.169358 | 0.637567981 | MVD | protein_coding | 59.29 | -0.071556673 | 0.071556673 | ENSG00000167508 | n.s. |
| ENSG00000135679 | 4536.524 | -0.355568327 | 0.255077 | -1.393967 | 0.163328 | 0.629222471 | MDM2 | protein_coding | 39.81 | -0.071538845 | 0.071538845 | ENSG00000135679 | n.s. |
| ENSG00000241913 | 23.05384 | -0.469635406 | 0.393302 | -1.194083 | 0.232446 | 0.70416476 | RPS29P21 | processed_pseudogene | 42.94 | -0.071537548 | 0.071537548 | ENSG00000241913 | n.s. |
| ENSG00000119723 | 48.09713 | -0.406389682 | 0.311495 | -1.304641 | 0.192015 | 0.66685649 | COQ6 | protein_coding | 45.54 | -0.071511424 | 0.071511424 | ENSG00000119723 | n.s. |
| ENSG00000197603 | 1195.091 | -0.356533994 | 0.256769 | -1.388539 | 0.164973 | 0.63030432 | C5orf42 | protein_coding | 38.94 | -0.071467138 | 0.071467138 | ENSG00000197603 | n.s. |
| ENSG00000133114 | 834.9577 | -0.38403579 | 0.28728 | -1.3368 | 0.181288 | 0.651541877 | GPALPP1 | protein_coding | 38.86 | -0.071452802 | 0.071452802 | ENSG00000133114 | n.s. |
| ENSG00000151332 | 512.3756 | 0.354883329 | 0.254045 | 1.3969337 | 0.162434 | 0.629027704 | MBIP | protein_coding | 35.27 | 0.071448741 | 0.071448741 | ENSG00000151332 | n.s. |
| ENSG00000196466 | 282.9519 | -0.354599479 | 0.254484 | -1.393408 | 0.163497 | 0.629299052 | ZNF799 | protein_coding | 41.86 | -0.071325175 | 0.071325175 | ENSG00000196466 | n.s. |
| ENSG00000271270 | 212.857 | -0.388393181 | 0.292492 | -1.327877 | 0.184219 | 0.655193675 | TMCC1-AS1 | antisense | 44.04 | -0.071320758 | 0.071320758 | ENSG00000271270 | n.s. |
| ENSG00000261783 | 23.53153 | -0.449530042 | 0.365295 | -1.230594 | 0.218475 | 0.694480193 | | sense_intronic | 40.26 | -0.071178648 | 0.071178648 | ENSG00000261783 | n.s. |

| | | | | | | | | | | | | | |
|---|---|---|---|---|---|---|---|---|---|---|---|---|---|
| ENSG00000267100 | 424.2787 | -0.319036449 | 0.216659 | -1.472529 | 0.140878 | 0.598731586 | ILF3-AS1 | lincRNA | 51.64 | -0.071071057 | 0.071071057 | ENSG00000267100 | n.s. |
| ENSG00000077232 | 3518.147 | -0.367356683 | 0.269473 | -1.363244 | 0.172806 | 0.640837445 | DNAJC10 | protein_coding | 36.39 | -0.070992458 | 0.070992458 | ENSG00000077232 | n.s. |
| ENSG00000162733 | 4501.201 | -0.365287426 | 0.266647 | -1.36993 | 0.170709 | 0.639237118 | DDR2 | protein_coding | 40.47 | -0.070989233 | 0.070989233 | ENSG00000162733 | n.s. |
| ENSG00000115998 | 236.4955 | -0.327522453 | 0.225291 | -1.453773 | 0.146009 | 0.607109641 | C2orf42 | protein_coding | 43.21 | -0.070984881 | 0.070984881 | ENSG00000115998 | n.s. |
| ENSG00000165511 | 88.21265 | 0.372125707 | 0.275644 | 1.3500223 | 0.177009 | 0.64464925 | C10orf25 | antisense | 44.41 | 0.070955634 | 0.070955634 | ENSG00000165511 | n.s. |
| ENSG00000233270 | 48.07771 | 0.458570224 | 0.377574 | 1.2145176 | 0.22455 | 0.700517875 | SNRPEP4 | processed_pseudogene | 43.01 | 0.070886142 | 0.070886142 | ENSG00000233270 | n.s. |
| ENSG00000101161 | 1378.862 | -0.351883408 | 0.251398 | -1.399706 | 0.161602 | 0.628931954 | PRPF6 | protein_coding | 50.53 | -0.07086803 | 0.07086803 | ENSG00000101161 | n.s. |
| ENSG00000198142 | 612.9004 | 0.326955998 | 0.224555 | 1.4560202 | 0.145387 | 0.607098916 | SOWAHC | protein_coding | 47.58 | 0.07086462 | 0.07086462 | ENSG00000198142 | n.s. |
| ENSG00000107854 | 1374.363 | -0.350496332 | 0.249493 | -1.404836 | 0.16007 | 0.627825878 | TNKS2 | protein_coding | 37.63 | -0.070856614 | 0.070856614 | ENSG00000107854 | n.s. |
| ENSG00000249572 | 39.11883 | -0.410513763 | 0.317728 | -1.292029 | 0.196347 | 0.672118672 | | antisense | 40.72 | -0.070835808 | 0.070835808 | ENSG00000249572 | n.s. |
| ENSG00000198816 | 297.2979 | -0.353867567 | 0.255267 | -1.386263 | 0.165667 | 0.631074156 | ZNF358 | protein_coding | 60.98 | -0.070745064 | 0.070745064 | ENSG00000198816 | n.s. |
| ENSG00000225265 | 61.10794 | -0.394809068 | 0.300047 | -1.315824 | 0.188233 | 0.662144925 | TAF1A-AS1 | antisense | 41.4 | -0.070689358 | 0.070689358 | ENSG00000225265 | n.s. |
| ENSG00000111752 | 487.2886 | -0.350950187 | 0.250636 | -1.400236 | 0.161443 | 0.628931954 | PHC1 | protein_coding | 44.13 | -0.070680083 | 0.070680083 | ENSG00000111752 | n.s. |
| ENSG00000119965 | 351.1456 | -0.377983633 | 0.282143 | -1.33969 | 0.180346 | 0.650241161 | C10orf88 | protein_coding | 36.24 | -0.070654796 | 0.070654796 | ENSG00000119965 | n.s. |
| ENSG00000135341 | 2109.613 | -0.365584344 | 0.268135 | -1.363433 | 0.172746 | 0.640837445 | MAP3K7 | protein_coding | 35.81 | -0.070649949 | 0.070649949 | ENSG00000135341 | n.s. |
| ENSG00000129925 | 166.1597 | -0.368082789 | 0.271853 | -1.353975 | 0.175744 | 0.642859735 | TMEM8A | protein_coding | 61.61 | -0.070629115 | 0.070629115 | ENSG00000129925 | n.s. |
| ENSG00000104221 | 358.1293 | -0.325705149 | 0.224375 | -1.451611 | 0.14661 | 0.607109641 | BRF2 | protein_coding | 49.89 | -0.070591012 | 0.070591012 | ENSG00000104221 | n.s. |
| ENSG00000278520 | 8.536431 | -0.537975876 | 0.492293 | -1.092795 | 0.274484 | 0.739386355 | MIR7851 | miRNA | 59.38 | -0.070544006 | 0.070544006 | ENSG00000278520 | n.s. |
| ENSG00000112406 | 484.4647 | -0.349692007 | 0.249767 | -1.400074 | 0.161491 | 0.628931954 | HECA | protein_coding | 41.79 | -0.07042669 | 0.07042669 | ENSG00000112406 | n.s. |
| ENSG00000126005 | 90.74001 | -0.392921167 | 0.298562 | -1.316046 | 0.188159 | 0.662050319 | MMP24-AS1 | protein_coding | 47.16 | -0.070375718 | 0.070375718 | ENSG00000126005 | n.s. |
| ENSG00000223787 | 15.09699 | -0.469396143 | 0.395909 | -1.185617 | 0.235774 | 0.708083978 | | processed_pseudogene | 40.75 | -0.070369632 | 0.070369632 | ENSG00000223787 | n.s. |
| ENSG00000198951 | 202.7957 | -0.3659502 | 0.269703 | -1.356866 | 0.174824 | 0.642395686 | NAGA | protein_coding | 53.29 | -0.070334671 | 0.070334671 | ENSG00000198951 | n.s. |
| ENSG00000270829 | 14.24278 | 0.536150979 | 0.491276 | 1.0913444 | 0.275121 | 0.739442303 | | lincRNA | 47.19 | 0.070287092 | 0.070287092 | ENSG00000270829 | n.s. |
| ENSG00000106415 | 1144.286 | 0.364993125 | 0.268085 | 1.3614844 | 0.173361 | 0.641941562 | GLCCI1 | protein_coding | 36.7 | 0.070262821 | 0.070262821 | ENSG00000106415 | n.s. |
| ENSG00000137364 | 181.922 | -0.400422246 | 0.307716 | -1.301274 | 0.193165 | 0.667684687 | TPMT | protein_coding | 41.46 | -0.070245508 | 0.070245508 | ENSG00000137364 | n.s. |
| ENSG00000207744 | 15.56331 | -0.503348068 | 0.44252 | -1.137459 | 0.255346 | 0.725252428 | MIR10B | miRNA | 40 | -0.070222496 | 0.070222496 | ENSG00000207744 | n.s. |
| ENSG00000161267 | 570.2633 | -0.348504288 | 0.248682 | -1.401406 | 0.161093 | 0.628931954 | BDH1 | protein_coding | 48.52 | -0.070187488 | 0.070187488 | ENSG00000161267 | n.s. |
| ENSG00000185347 | 177.8669 | -0.372491922 | 0.277109 | -1.344205 | 0.178882 | 0.648076085 | C14orf80 | protein_coding | 66 | -0.070167795 | 0.070167795 | ENSG00000185347 | n.s. |
| ENSG00000169857 | 360.6646 | -0.365281543 | 0.2695 | -1.355407 | 0.175288 | 0.642584104 | AVEN | protein_coding | 39.85 | -0.070159634 | 0.070159634 | ENSG00000169857 | n.s. |
| ENSG00000012963 | 984.9564 | -0.379267204 | 0.284508 | -1.333061 | 0.182512 | 0.653245015 | UBR7 | protein_coding | 43.29 | -0.070135569 | 0.070135569 | ENSG00000012963 | n.s. |
| ENSG00000112139 | 7.358695 | -0.565247105 | 0.535505 | -1.055539 | 0.291179 | 0.751590786 | MDGA1 | protein_coding | 52.18 | -0.070101128 | 0.070101128 | ENSG00000112139 | n.s. |
| ENSG00000116688 | 730.8386 | -0.368605379 | 0.273279 | -1.348823 | 0.177394 | 0.645396531 | MFN2 | protein_coding | 51.12 | -0.070098928 | 0.070098928 | ENSG00000116688 | n.s. |

| | | | | | | | | | | | | | |
|---|---|---|---|---|---|---|---|---|---|---|---|---|---|
| ENSG00000158417 | 3020.051 | -0.334368376 | 0.234673 | -1.424826 | 0.154207 | 0.617347914 | EIF5B | protein_coding | 38.48 | -0.070040149 | 0.070040149 | ENSG00000158417 | n.s. |
| ENSG00000141873 | 283.2183 | -0.390877699 | 0.296956 | -1.316281 | 0.18808 | 0.661975752 | SLC39A3 | protein_coding | 57.96 | -0.070028835 | 0.070028835 | ENSG00000141873 | n.s. |
| ENSG00000212246 | 14.60762 | 0.520119267 | 0.465311 | 1.1177881 | 0.263658 | 0.733524057 | RNU6-360P | snRNA | 42.31 | 0.070000584 | 0.070000584 | ENSG00000212246 | n.s. |
| ENSG00000207092 | 5.147825 | -0.829636434 | 0.989496 | -0.838444 | 0.401782 | 0.823540971 | | misc_RNA | 40.2 | -0.069950622 | 0.069950622 | ENSG00000207092 | n.s. |
| ENSG00000084774 | 1624.445 | -0.33074299 | 0.231182 | -1.430663 | 0.152527 | 0.614654298 | CAD | protein_coding | 51.17 | -0.069908841 | 0.069908841 | ENSG00000084774 | n.s. |
| ENSG00000141570 | 657.0369 | -0.345462521 | 0.246161 | -1.403402 | 0.160497 | 0.62809251 | CBX8 | protein_coding | 60.55 | -0.069775271 | 0.069775271 | ENSG00000141570 | n.s. |
| ENSG00000164366 | 546.8643 | -0.359954462 | 0.263129 | -1.367977 | 0.171319 | 0.640170938 | CCDC127 | protein_coding | 51.76 | -0.069724636 | 0.069724636 | ENSG00000164366 | n.s. |
| ENSG00000110721 | 530.1901 | -0.345923694 | 0.246675 | -1.402345 | 0.160812 | 0.628753199 | CHKA | protein_coding | 44.1 | -0.069710471 | 0.069710471 | ENSG00000110721 | n.s. |
| ENSG00000158966 | 1079.667 | -0.338942014 | 0.239792 | -1.413483 | 0.157514 | 0.622955867 | CACHD1 | protein_coding | 39.92 | -0.069667063 | 0.069667063 | ENSG00000158966 | n.s. |
| ENSG00000119927 | 1082.03 | -0.361769262 | 0.266121 | -1.359416 | 0.174015 | 0.642002364 | GPAM | protein_coding | 39.93 | -0.069627332 | 0.069627332 | ENSG00000119927 | n.s. |
| ENSG00000089775 | 542.9851 | -0.345795386 | 0.247631 | -1.396413 | 0.16259 | 0.629027704 | ZBTB25 | protein_coding | 42.25 | -0.069619063 | 0.069619063 | ENSG00000089775 | n.s. |
| ENSG00000178966 | 447.8489 | -0.359670912 | 0.263082 | -1.367145 | 0.17158 | 0.640457684 | RMI1 | protein_coding | 38.66 | -0.06959976 | 0.06959976 | ENSG00000178966 | n.s. |
| ENSG00000196151 | 157.7903 | 0.362699419 | 0.267851 | 1.3541072 | 0.175702 | 0.642859735 | WDSUB1 | protein_coding | 39.11 | 0.069596134 | 0.069596134 | ENSG00000196151 | n.s. |
| ENSG00000171940 | 1097.781 | -0.317750907 | 0.217272 | -1.462455 | 0.143617 | 0.604031311 | ZNF217 | protein_coding | 44.04 | -0.069568558 | 0.069568558 | ENSG00000171940 | n.s. |
| ENSG00000030582 | 323.9129 | -0.34706217 | 0.249901 | -1.388797 | 0.164894 | 0.63030432 | GRN | protein_coding | 57.64 | -0.069568514 | 0.069568514 | ENSG00000030582 | n.s. |
| ENSG00000278099 | 202.4581 | 0.344090056 | 0.244885 | 1.4051111 | 0.159988 | 0.627825878 | | snRNA | 51.52 | 0.069561517 | 0.069561517 | ENSG00000278099 | n.s. |
| ENSG00000103495 | 1923.253 | -0.347080311 | 0.250057 | -1.388003 | 0.165136 | 0.630433369 | MAZ | protein_coding | 66.53 | -0.069541291 | 0.069541291 | ENSG00000103495 | n.s. |
| ENSG00000138600 | 853.6677 | -0.359718835 | 0.2638 | -1.363604 | 0.172692 | 0.640837445 | SPPL2A | protein_coding | 40.43 | -0.069516427 | 0.069516427 | ENSG00000138600 | n.s. |
| ENSG00000149761 | 131.373 | -0.386431425 | 0.292865 | -1.319484 | 0.187007 | 0.661005765 | NUDT22 | protein_coding | 59.06 | -0.069478343 | 0.069478343 | ENSG00000149761 | n.s. |
| ENSG00000115808 | 801.2342 | -0.344928272 | 0.24721 | -1.395286 | 0.162929 | 0.629036117 | STRN | protein_coding | 37.76 | -0.069442484 | 0.069442484 | ENSG00000115808 | n.s. |
| ENSG00000177675 | 73.37811 | -0.390124789 | 0.297311 | -1.312178 | 0.18946 | 0.663959974 | CD163L1 | protein_coding | 36.25 | -0.069386854 | 0.069386854 | ENSG00000177675 | n.s. |
| ENSG00000215417 | 8461.243 | 0.29884408 | 0.199159 | 1.5005312 | 0.133477 | 0.586245566 | MIR17HG | processed_transcript | 44.48 | 0.069308047 | 0.069308047 | ENSG00000215417 | n.s. |
| ENSG00000185664 | 401.4626 | -0.393289826 | 0.301156 | -1.305933 | 0.191575 | 0.666492649 | PMEL | protein_coding | 48.7 | -0.069299491 | 0.069299491 | ENSG00000185664 | n.s. |
| ENSG00000199872 | 34.27511 | -0.43045644 | 0.346278 | -1.243095 | 0.213833 | 0.690444009 | RNU6-942P | snRNA | 38.32 | -0.069248188 | 0.069248188 | ENSG00000199872 | n.s. |
| ENSG00000227726 | 20.11487 | -0.454335571 | 0.380144 | -1.195167 | 0.232022 | 0.704043192 | | sense_intronic | 48.1 | -0.069241058 | 0.069241058 | ENSG00000227726 | n.s. |
| ENSG00000125878 | 50.12985 | -0.389293083 | 0.296593 | -1.312548 | 0.189335 | 0.663959974 | TCF15 | protein_coding | 56.76 | -0.069238929 | 0.069238929 | ENSG00000125878 | n.s. |
| ENSG00000258001 | 19.27754 | 0.517290641 | 0.46517 | 1.1120462 | 0.266118 | 0.7348601 | | lincRNA | 43.48 | 0.069211073 | 0.069211073 | ENSG00000258001 | n.s. |
| ENSG00000274750 | 763.1374 | -0.321813098 | 0.222962 | -1.443357 | 0.14892 | 0.609468503 | HIST1H3E | protein_coding | 42.63 | -0.069205499 | 0.069205499 | ENSG00000274750 | n.s. |
| ENSG00000162601 | 1967.893 | -0.371879527 | 0.278137 | -1.337038 | 0.18121 | 0.651484256 | MYSM1 | protein_coding | 36.33 | -0.06920532 | 0.06920532 | ENSG00000162601 | n.s. |
| ENSG00000145675 | 1658.701 | 0.327708126 | 0.22923 | 1.4296015 | 0.152831 | 0.61505617 | PIK3R1 | protein_coding | 39.22 | 0.069174342 | 0.069174342 | ENSG00000145675 | n.s. |
| ENSG00000202082 | 7.488066 | -0.751458152 | 0.852104 | -0.881885 | 0.377839 | 0.809223062 | RNU6-290P | snRNA | 33.66 | -0.069082862 | 0.069082862 | ENSG00000202082 | n.s. |
| ENSG00000277151 | 246.9212 | -0.343020937 | 0.24565 | -1.396382 | 0.162599 | 0.629027704 | | lincRNA | 34.33 | -0.069060483 | 0.069060483 | ENSG00000277151 | n.s. |

| | | | | | | | | | | | | | |
|---|---|---|---|---|---|---|---|---|---|---|---|---|---|
| ENSG00000258732 | 488.0523 | -0.357067652 | 0.261754 | -1.364132 | 0.172526 | 0.640837445 | | unprocessed_pseudogene | 40.36 | -0.069004081 | 0.069004081 | ENSG00000258732 | n.s. |
| ENSG00000151692 | 1190.954 | -0.363317848 | 0.269543 | -1.347901 | 0.17769 | 0.645779714 | RNF144A | protein_coding | 44.89 | -0.068999728 | 0.068999728 | ENSG00000151692 | n.s. |
| ENSG00000186166 | 284.1131 | -0.357031127 | 0.261476 | -1.365447 | 0.172113 | 0.640837445 | CCDC84 | protein_coding | 45.88 | -0.068997022 | 0.068997022 | ENSG00000186166 | n.s. |
| ENSG00000127511 | 267.1744 | -0.35702138 | 0.261948 | -1.362949 | 0.172899 | 0.640912816 | SIN3B | protein_coding | 52.98 | -0.068976903 | 0.068976903 | ENSG00000127511 | n.s. |
| ENSG00000084234 | 2258.403 | -0.370523589 | 0.277002 | -1.337621 | 0.18102 | 0.651419475 | APLP2 | protein_coding | 43.6 | -0.068968987 | 0.068968987 | ENSG00000084234 | n.s. |
| ENSG00000168061 | 288.445 | -0.343535365 | 0.24683 | -1.39179 | 0.163986 | 0.629889786 | SAC3D1 | protein_coding | 58.58 | -0.068959721 | 0.068959721 | ENSG00000168061 | n.s. |
| ENSG00000144227 | 54.48544 | -0.388481913 | 0.29653 | -1.310095 | 0.190164 | 0.664737354 | NXPH2 | protein_coding | 38.25 | -0.068897235 | 0.068897235 | ENSG00000144227 | n.s. |
| ENSG00000186666 | 144.4485 | -0.334731453 | 0.236584 | -1.414851 | 0.157112 | 0.622581912 | BCDIN3D | protein_coding | 45.14 | -0.068888905 | 0.068888905 | ENSG00000186666 | n.s. |
| ENSG00000196081 | 1068.421 | -0.37743334 | 0.284852 | -1.325016 | 0.185166 | 0.656998505 | ZNF724 | protein_coding | 40.39 | -0.068857285 | 0.068857285 | ENSG00000196081 | n.s. |
| ENSG00000057935 | 292.1156 | -0.344053267 | 0.248127 | -1.386604 | 0.165563 | 0.630850594 | MTA3 | protein_coding | 43.08 | -0.068835936 | 0.068835936 | ENSG00000057935 | n.s. |
| ENSG00000251787 | 2.377733 | -0.7199636 | 0.797783 | -0.902455 | 0.366815 | 0.802563919 | RNU7-47P | snRNA | 45.16 | -0.068771189 | 0.068771189 | ENSG00000251787 | n.s. |
| ENSG00000112137 | 88.09597 | -0.369973958 | 0.276974 | -1.33577 | 0.181624 | 0.651909066 | PHACTR1 | protein_coding | 41.2 | -0.068745963 | 0.068745963 | ENSG00000112137 | n.s. |
| ENSG00000264911 | 28.4202 | -0.462873863 | 0.391987 | -1.180839 | 0.237667 | 0.710398214 | | processed_pseudogene | 39.06 | -0.068735907 | 0.068735907 | ENSG00000264911 | n.s. |
| ENSG00000240775 | 32.52566 | -0.421752095 | 0.337205 | -1.25073 | 0.211033 | 0.687111024 | | processed_pseudogene | 47.13 | -0.068734239 | 0.068734239 | ENSG00000240775 | n.s. |
| ENSG00000116984 | 3123.175 | -0.341377807 | 0.244954 | -1.393642 | 0.163426 | 0.629252671 | MTR | protein_coding | 40.73 | -0.068676657 | 0.068676657 | ENSG00000116984 | n.s. |
| ENSG00000105698 | 385.4365 | -0.362755931 | 0.269489 | -1.346089 | 0.178274 | 0.646887381 | USF2 | protein_coding | 57.36 | -0.068623019 | 0.068623019 | ENSG00000105698 | n.s. |
| ENSG00000155542 | 314.1162 | 0.330872127 | 0.233066 | 1.4196517 | 0.155709 | 0.620297005 | SETD9 | protein_coding | 38.6 | 0.068622984 | 0.068622984 | ENSG00000155542 | n.s. |
| ENSG00000087088 | 395.1823 | -0.356882187 | 0.263243 | -1.355713 | 0.17519 | 0.642395686 | BAX | protein_coding | 53.57 | -0.068591823 | 0.068591823 | ENSG00000087088 | n.s. |
| ENSG00000120437 | 2506.572 | -0.373223387 | 0.280774 | -1.329267 | 0.18376 | 0.655151359 | ACAT2 | protein_coding | 44.57 | -0.068545593 | 0.068545593 | ENSG00000120437 | n.s. |
| ENSG00000153767 | 251.0905 | -0.379743558 | 0.287584 | -1.320459 | 0.186682 | 0.660022735 | GTF2E1 | protein_coding | 38.04 | -0.068521347 | 0.068521347 | ENSG00000153767 | n.s. |
| ENSG00000095319 | 1608.798 | -0.335066077 | 0.237574 | -1.410362 | 0.158433 | 0.624627103 | NUP188 | protein_coding | 46.07 | -0.068480529 | 0.068480529 | ENSG00000095319 | n.s. |
| ENSG00000100601 | 317.3255 | -0.354289721 | 0.259792 | -1.363744 | 0.172648 | 0.640837445 | ALKBH1 | protein_coding | 42.35 | -0.06846724 | 0.06846724 | ENSG00000100601 | n.s. |
| ENSG00000125107 | 3723.63 | -0.339763722 | 0.242446 | -1.401399 | 0.161095 | 0.628931954 | CNOT1 | protein_coding | 41.44 | -0.06842717 | 0.06842717 | ENSG00000125107 | n.s. |
| ENSG00000276656 | 3.917634 | -0.656624657 | 0.691739 | -0.949238 | 0.3425 | 0.786682723 | MIR6083 | miRNA | 50 | -0.068420544 | 0.068420544 | ENSG00000276656 | n.s. |
| ENSG00000258405 | 660.0265 | 0.315484115 | 0.217117 | 1.4530608 | 0.146207 | 0.607109641 | ZNF578 | protein_coding | 43.95 | 0.068375778 | 0.068375778 | ENSG00000258405 | n.s. |
| ENSG00000183807 | 756.671 | 0.388344769 | 0.297538 | 1.305192 | 0.191827 | 0.666819478 | FAM162B | protein_coding | 37.88 | 0.068345465 | 0.068345465 | ENSG00000183807 | n.s. |
| ENSG00000035687 | 1412.072 | -0.360157682 | 0.267273 | -1.347529 | 0.17781 | 0.646016851 | ADSS | protein_coding | 38.3 | -0.068342137 | 0.068342137 | ENSG00000035687 | n.s. |
| ENSG00000128965 | 239.9765 | 0.406296328 | 0.319534 | 1.2715287 | 0.203541 | 0.679010816 | CHAC1 | protein_coding | 57.9 | 0.068307883 | 0.068307883 | ENSG00000128965 | n.s. |
| ENSG00000235748 | 221.0679 | -0.355580346 | 0.262396 | -1.355128 | 0.175377 | 0.64274023 | SEPT14P12 | processed_pseudogene | 40.36 | -0.068258809 | 0.068258809 | ENSG00000235748 | n.s. |
| ENSG00000072182 | 554.706 | -0.331623515 | 0.234397 | -1.414792 | 0.15713 | 0.622581912 | ASIC4 | protein_coding | 56.96 | -0.06824928 | 0.06824928 | ENSG00000072182 | n.s. |
| ENSG00000243679 | 64.74554 | -0.393439939 | 0.303951 | -1.294418 | 0.195521 | 0.670778798 | | processed_pseudogene | 67.24 | -0.068230619 | 0.068230619 | ENSG00000243679 | n.s. |
| ENSG00000165115 | 174.3995 | -0.355515424 | 0.262593 | -1.353867 | 0.175779 | 0.642859735 | KIF27 | protein_coding | 38.37 | -0.068217642 | 0.068217642 | ENSG00000165115 | n.s. |

| | | | | | | | | | | | | | |
|---|---|---|---|---|---|---|---|---|---|---|---|---|---|
| ENSG00000065615 | 1218.959 | -0.359583028 | 0.266926 | -1.347126 | 0.17794 | 0.646179893 | CYB5R4 | protein_coding | 37.46 | -0.068193684 | 0.068193684 | ENSG00000065615 | n.s. |
| ENSG00000164687 | 3557.815 | -0.326142359 | 0.229028 | -1.424026 | 0.154439 | 0.617889294 | FABP5 | protein_coding | 45.55 | -0.068192887 | 0.068192887 | ENSG00000164687 | n.s. |
| ENSG00000174599 | 293.1782 | 0.323838333 | 0.226933 | 1.427024 | 0.153573 | 0.615840022 | TRAM1L1 | protein_coding | 40.63 | 0.068178361 | 0.068178361 | ENSG00000174599 | n.s. |
| ENSG00000279453 | 133.4004 | -0.403775372 | 0.316994 | -1.273763 | 0.202747 | 0.677959669 | | TEC | 38.44 | -0.068155725 | 0.068155725 | ENSG00000279453 | n.s. |
| ENSG00000159885 | 203.4445 | -0.381983818 | 0.290759 | -1.313748 | 0.188931 | 0.663594149 | ZNF222 | protein_coding | 42.04 | -0.068030345 | 0.068030345 | ENSG00000159885 | n.s. |
| ENSG00000161217 | 659.1502 | -0.355209491 | 0.262793 | -1.351673 | 0.17648 | 0.643565193 | PCYT1A | protein_coding | 46.69 | -0.067989744 | 0.067989744 | ENSG00000161217 | n.s. |
| ENSG00000165272 | 64.09179 | -0.425924216 | 0.344565 | -1.236122 | 0.216413 | 0.692499549 | AQP3 | protein_coding | 55.65 | -0.067969202 | 0.067969202 | ENSG00000165272 | n.s. |
| ENSG00000163348 | 625.2166 | -0.318556352 | 0.221339 | -1.439224 | 0.150087 | 0.612216404 | PYGO2 | protein_coding | 54.77 | -0.067882778 | 0.067882778 | ENSG00000163348 | n.s. |
| ENSG00000132604 | 271.2505 | -0.350587626 | 0.256356 | -1.367581 | 0.171443 | 0.640290395 | TERF2 | protein_coding | 44.29 | -0.067881833 | 0.067881833 | ENSG00000132604 | n.s. |
| ENSG00000152457 | 330.4632 | -0.350355136 | 0.256153 | -1.367756 | 0.171388 | 0.6402576 | DCLRE1C | protein_coding | 41.86 | -0.067844611 | 0.067844611 | ENSG00000152457 | n.s. |
| ENSG00000060749 | 2690.66 | -0.350872436 | 0.256865 | -1.365977 | 0.171946 | 0.640837445 | QSER1 | protein_coding | 38.54 | -0.067806842 | 0.067806842 | ENSG00000060749 | n.s. |
| ENSG00000133678 | 142.8343 | -0.401188809 | 0.314325 | -1.27635 | 0.201832 | 0.677624898 | TMEM254 | protein_coding | 42.93 | -0.067805179 | 0.067805179 | ENSG00000133678 | n.s. |
| ENSG00000278991 | 706.4078 | 0.324860525 | 0.228328 | 1.4227794 | 0.1548 | 0.618415021 | | TEC | 37.19 | 0.067804879 | 0.067804879 | ENSG00000278991 | n.s. |
| ENSG00000138641 | 39.60592 | -0.409676374 | 0.325375 | -1.25909 | 0.207998 | 0.683512059 | HERC3 | protein_coding | 39.32 | -0.067700585 | 0.067700585 | ENSG00000138641 | n.s. |
| ENSG00000226465 | 15.80469 | -0.506234769 | 0.456423 | -1.109135 | 0.267372 | 0.734987182 | | antisense | 44.49 | -0.067693832 | 0.067693832 | ENSG00000226465 | n.s. |
| ENSG00000275576 | 34.10286 | 0.441645174 | 0.367181 | 1.2027998 | 0.229054 | 0.702737638 | | sense_intronic | 44.66 | 0.067663037 | 0.067663037 | ENSG00000275576 | n.s. |
| ENSG00000187824 | 29.46956 | -0.443957939 | 0.371537 | -1.194922 | 0.232118 | 0.704043192 | TMEM220 | protein_coding | 40.31 | -0.067659499 | 0.067659499 | ENSG00000187824 | n.s. |
| ENSG00000241764 | 26.80138 | 0.445280153 | 0.373351 | 1.1926588 | 0.233003 | 0.705086103 | | antisense | 56.77 | 0.067574757 | 0.067574757 | ENSG00000241764 | n.s. |
| ENSG00000107819 | 298.3759 | -0.350637434 | 0.257464 | -1.36189 | 0.173233 | 0.641657113 | SFXN3 | protein_coding | 54.03 | -0.067566777 | 0.067566777 | ENSG00000107819 | n.s. |
| ENSG00000088356 | 493.8959 | -0.36169108 | 0.270144 | -1.338885 | 0.180608 | 0.650636782 | PDRG1 | protein_coding | 49.41 | -0.067513759 | 0.067513759 | ENSG00000088356 | n.s. |
| ENSG00000276007 | 49.32111 | -0.397095306 | 0.309457 | -1.283199 | 0.199422 | 0.676054647 | | sense_intronic | 37.34 | -0.067513428 | 0.067513428 | ENSG00000276007 | n.s. |
| ENSG00000125149 | 369.876 | -0.336654862 | 0.242432 | -1.388658 | 0.164937 | 0.63030432 | C16orf70 | protein_coding | 47.85 | -0.067482371 | 0.067482371 | ENSG00000125149 | n.s. |
| ENSG00000133026 | 3578.107 | -0.334608691 | 0.239826 | -1.395214 | 0.162951 | 0.629036117 | MYH10 | protein_coding | 42 | -0.0673649 | 0.0673649 | ENSG00000133026 | n.s. |
| ENSG00000128928 | 778.4733 | -0.329484608 | 0.233564 | -1.410684 | 0.158338 | 0.624627103 | IVD | protein_coding | 50.9 | -0.067339792 | 0.067339792 | ENSG00000128928 | n.s. |
| ENSG00000269293 | 150.3168 | -0.363935395 | 0.272893 | -1.33362 | 0.182328 | 0.653089584 | ZSCAN16-AS1 | antisense | 40.71 | -0.067337963 | 0.067337963 | ENSG00000269293 | n.s. |
| ENSG00000095752 | 110.291 | -0.402833664 | 0.318385 | -1.265242 | 0.205785 | 0.680704186 | IL11 | protein_coding | 60.87 | -0.067289971 | 0.067289971 | ENSG00000095752 | n.s. |
| ENSG00000141642 | 332.1136 | 0.309625798 | 0.212389 | 1.4578275 | 0.144888 | 0.606306192 | ELAC1 | protein_coding | 41.78 | 0.067284162 | 0.067284162 | ENSG00000141642 | n.s. |
| ENSG00000221909 | 276.2065 | -0.319385778 | 0.223747 | -1.427443 | 0.153452 | 0.615840022 | FAM200A | protein_coding | 44.02 | -0.067240956 | 0.067240956 | ENSG00000221909 | n.s. |
| ENSG00000174371 | 3310.195 | -0.326144878 | 0.23018 | -1.416914 | 0.156508 | 0.622084796 | EXO1 | protein_coding | 39.58 | -0.067234901 | 0.067234901 | ENSG00000174371 | n.s. |
| ENSG00000266676 | 12.48651 | -0.471280183 | 0.409073 | -1.152068 | 0.249293 | 0.720036883 | MIR4297 | miRNA | 68.42 | -0.067225883 | 0.067225883 | ENSG00000266676 | n.s. |
| ENSG00000246174 | 80.44542 | -0.352461955 | 0.261003 | -1.350416 | 0.176883 | 0.64464925 | KCTD21-AS1 | antisense | 47.55 | -0.067206218 | 0.067206218 | ENSG00000246174 | n.s. |
| ENSG00000199575 | 14.64553 | 0.63674095 | 0.661982 | 0.961871 | 0.336114 | 0.784266959 | SNORD114-1 | snoRNA | 48.61 | 0.067199145 | 0.067199145 | ENSG00000199575 | n.s. |

| | | | | | | | | | | | | | |
|---|---|---|---|---|---|---|---|---|---|---|---|---|---|
| ENSG00000224114 | 222.512 | 0.428979371 | 0.350783 | 1.2229188 | 0.22136 | 0.697285172 | | processed_pseudogene | 54.27 | 0.067173695 | 0.067173695 | ENSG00000224114 | n.s. |
| ENSG00000144868 | 688.3029 | -0.33362372 | 0.238867 | -1.39669 | 0.162507 | 0.629027704 | TMEM108 | protein_coding | 40.21 | -0.067168539 | 0.067168539 | ENSG00000144868 | n.s. |
| ENSG00000132466 | 3022.775 | -0.350353156 | 0.25898 | -1.35282 | 0.176113 | 0.643404323 | ANKRD17 | protein_coding | 36.17 | -0.067098244 | 0.067098244 | ENSG00000132466 | n.s. |
| ENSG00000009335 | 1615.464 | -0.356024957 | 0.264893 | -1.344035 | 0.178937 | 0.648076085 | UBE3C | protein_coding | 42.12 | -0.067065847 | 0.067065847 | ENSG00000009335 | n.s. |
| ENSG00000200075 | 17.91303 | -0.478310047 | 0.419335 | -1.14064 | 0.25402 | 0.72418384 | | snoRNA | 41.73 | -0.067035713 | 0.067035713 | ENSG00000200075 | n.s. |
| ENSG00000248489 | 21.17177 | 0.437036316 | 0.362497 | 1.2056289 | 0.227961 | 0.702691634 | LINC02062 | lincRNA | 39.81 | 0.066969355 | 0.066969355 | ENSG00000248489 | n.s. |
| ENSG00000188707 | 626.5208 | -0.347905357 | 0.255989 | -1.359063 | 0.174127 | 0.642010653 | ZBED6CL | protein_coding | 55.68 | -0.066957088 | 0.066957088 | ENSG00000188707 | n.s. |
| ENSG00000264354 | 6.649448 | -0.609832655 | 0.619898 | -0.983763 | 0.325232 | 0.776688313 | MIR3134 | miRNA | 33.78 | -0.066931103 | 0.066931103 | ENSG00000264354 | n.s. |
| ENSG00000224032 | 3198.19 | 0.304106653 | 0.207433 | 1.4660479 | 0.142635 | 0.602521359 | EPB41L4A-AS1 | lincRNA | 43.19 | 0.066911843 | 0.066911843 | ENSG00000224032 | n.s. |
| ENSG00000123200 | 2513.885 | -0.348046206 | 0.256634 | -1.356199 | 0.175036 | 0.642395686 | ZC3H13 | protein_coding | 34.9 | -0.06689357 | 0.06689357 | ENSG00000123200 | n.s. |
| ENSG00000066933 | 110.0615 | -0.389602146 | 0.302416 | -1.288298 | 0.197642 | 0.673794115 | MYO9A | protein_coding | 37.93 | -0.066806166 | 0.066806166 | ENSG00000066933 | n.s. |
| ENSG00000138182 | 2810.932 | -0.343283377 | 0.250469 | -1.370563 | 0.170511 | 0.638865917 | KIF20B | protein_coding | 35.15 | -0.066799609 | 0.066799609 | ENSG00000138182 | n.s. |
| ENSG00000153944 | 8456.895 | -0.333786773 | 0.240694 | -1.38677 | 0.165512 | 0.630850594 | MSI2 | protein_coding | 47.5 | -0.066781883 | 0.066781883 | ENSG00000153944 | n.s. |
| ENSG00000095564 | 3365.914 | -0.354452738 | 0.263927 | -1.342997 | 0.179273 | 0.64843101 | BTAF1 | protein_coding | 37.37 | -0.0666854 | 0.0666854 | ENSG00000095564 | n.s. |
| ENSG00000244300 | 153.7278 | -0.371861626 | 0.282516 | -1.316251 | 0.18809 | 0.661975752 | GATA2-AS1 | antisense | 55.65 | -0.066621955 | 0.066621955 | ENSG00000244300 | n.s. |
| ENSG00000245556 | 142.6155 | -0.332271798 | 0.238963 | -1.390477 | 0.164384 | 0.63030432 | SCAMP1-AS1 | lincRNA | 42.36 | -0.066603788 | 0.066603788 | ENSG00000245556 | n.s. |
| ENSG00000271855 | 91.75151 | 0.351216517 | 0.260806 | 1.3466593 | 0.17809 | 0.646388678 | | lincRNA | 60.09 | 0.066557728 | 0.066557728 | ENSG00000271855 | n.s. |
| ENSG00000198814 | 211.9065 | -0.393648974 | 0.308623 | -1.275502 | 0.202132 | 0.677624898 | GK | protein_coding | 38.63 | -0.066530867 | 0.066530867 | ENSG00000198814 | n.s. |
| ENSG00000212163 | 19.15138 | -0.472063056 | 0.41266 | -1.143952 | 0.252644 | 0.723181779 | SNORD91A | snoRNA | 43.98 | -0.066444067 | 0.066444067 | ENSG00000212163 | n.s. |
| ENSG00000274211 | 902.0054 | -0.328835617 | 0.234246 | -1.403802 | 0.160378 | 0.62809251 | SOCS7 | protein_coding | 44.99 | -0.066417029 | 0.066417029 | ENSG00000274211 | n.s. |
| ENSG00000185986 | 234.5051 | -0.366589745 | 0.277325 | -1.32188 | 0.186208 | 0.658994794 | SDHAP3 | cribed_unprocessed_pseudc | 47.9 | -0.066396007 | 0.066396007 | ENSG00000185986 | n.s. |
| ENSG00000261173 | 37.07925 | 0.411303978 | 0.330178 | 1.2457048 | 0.212873 | 0.689743466 | | antisense | 43.26 | 0.066348434 | 0.066348434 | ENSG00000261173 | n.s. |
| ENSG00000061987 | 724.341 | -0.366216776 | 0.276932 | -1.322406 | 0.186033 | 0.658994794 | MON2 | protein_coding | 36.21 | -0.066328456 | 0.066328456 | ENSG00000061987 | n.s. |
| ENSG00000128563 | 386.9837 | -0.348722624 | 0.258635 | -1.348321 | 0.177555 | 0.645457024 | PRKRIP1 | protein_coding | 46.23 | -0.066303563 | 0.066303563 | ENSG00000128563 | n.s. |
| ENSG00000151881 | 174.3149 | -0.391989488 | 0.307783 | -1.273591 | 0.202808 | 0.677959669 | TMEM267 | protein_coding | 38.29 | -0.066166313 | 0.066166313 | ENSG00000151881 | n.s. |
| ENSG00000212413 | 13.52802 | -0.494178818 | 0.444604 | -1.111504 | 0.266351 | 0.734987182 | RNU11-3P | snRNA | 45.86 | -0.066081708 | 0.066081708 | ENSG00000212413 | n.s. |
| ENSG00000125686 | 1359.242 | -0.343785954 | 0.253452 | -1.356413 | 0.174968 | 0.642395686 | MED1 | protein_coding | 45.01 | -0.066074761 | 0.066074761 | ENSG00000125686 | n.s. |
| ENSG00000169221 | 416.6305 | -0.328065657 | 0.234167 | -1.400992 | 0.161217 | 0.628931954 | TBC1D10B | protein_coding | 54.1 | -0.066071223 | 0.066071223 | ENSG00000169221 | n.s. |
| ENSG00000162616 | 605.7711 | 0.337161904 | 0.245062 | 1.3758245 | 0.168876 | 0.636852728 | DNAJB4 | protein_coding | 37.07 | 0.066070579 | 0.066070579 | ENSG00000162616 | n.s. |
| ENSG00000184863 | 1392.441 | -0.330619388 | 0.238597 | -1.385682 | 0.165844 | 0.63130003 | RBM33 | protein_coding | 40.78 | -0.066045914 | 0.066045914 | ENSG00000184863 | n.s. |
| ENSG00000167967 | 459.6096 | -0.359596239 | 0.270704 | -1.328372 | 0.184055 | 0.655193675 | E4F1 | protein_coding | 61.58 | -0.066032766 | 0.066032766 | ENSG00000167967 | n.s. |
| ENSG00000252233 | 25.63269 | -0.422456359 | 0.345786 | -1.221727 | 0.221811 | 0.697757205 | RN7SKP253 | misc_RNA | 42.07 | -0.0660281 | 0.0660281 | ENSG00000252233 | n.s. |

| | | | | | | | | | | | | | | |
|---|---|---|---|---|---|---|---|---|---|---|---|---|---|---|
| ENSG00000152117 | 817.5822 | -0.326143568 | 0.232193 | -1.404621 | 0.160134 | 0.627863998 | | transcribed_processed_pseudog | 53.93 | -0.065924841 | 0.065924841 | ENSG00000152117 | | n.s. |
| ENSG00000166261 | 230.5111 | 0.316370348 | 0.22255 | 1.4215673 | 0.155152 | 0.618962761 | ZNF202 | protein_coding | 44.55 | 0.065911168 | 0.065911168 | ENSG00000166261 | | n.s. |
| ENSG00000278948 | 81.06409 | -0.363764121 | 0.275212 | -1.321757 | 0.186249 | 0.658994794 | | TEC | 52.24 | -0.065884236 | 0.065884236 | ENSG00000278948 | | n.s. |
| ENSG00000175305 | 1212.472 | -0.338812323 | 0.247393 | -1.369531 | 0.170833 | 0.639237118 | CCNE2 | protein_coding | 41.61 | -0.065844114 | 0.065844114 | ENSG00000175305 | | n.s. |
| ENSG00000177628 | 177.1404 | -0.326914955 | 0.233351 | -1.400955 | 0.161227 | 0.628931954 | GBA | protein_coding | 51.52 | -0.065839475 | 0.065839475 | ENSG00000177628 | | n.s. |
| ENSG00000231940 | 41.32864 | -0.419279282 | 0.342189 | -1.225286 | 0.220467 | 0.696655251 | RPS7P3 | processed_pseudogene | 46.03 | -0.065819336 | 0.065819336 | ENSG00000231940 | | n.s. |
| ENSG00000110768 | 814.6018 | -0.373583455 | 0.286398 | -1.304419 | 0.192091 | 0.666952386 | GTF2H1 | protein_coding | 40.71 | -0.065715261 | 0.065715261 | ENSG00000110768 | | n.s. |
| ENSG00000228065 | 104.1229 | -0.351931864 | 0.262859 | -1.338862 | 0.180616 | 0.650636782 | LINC01515 | lincRNA | 34.38 | -0.06569209 | 0.06569209 | ENSG00000228065 | | n.s. |
| ENSG00000162139 | 553.5394 | -0.34991981 | 0.260878 | -1.341317 | 0.179818 | 0.649162475 | NEU3 | protein_coding | 43.57 | -0.06566126 | 0.06566126 | ENSG00000162139 | | n.s. |
| ENSG00000266573 | 7.929917 | 0.593194529 | 0.601343 | 0.9864503 | 0.323912 | 0.775183815 | | lincRNA | 41.83 | 0.065604529 | 0.065604529 | ENSG00000266573 | | n.s. |
| ENSG00000207518 | 4.414636 | -0.644499638 | 0.690802 | -0.932973 | 0.350834 | 0.791294165 | RNU6-59P | snRNA | 39.25 | -0.065521146 | 0.065521146 | ENSG00000207518 | | n.s. |
| ENSG00000239039 | 207.5229 | 0.326870297 | 0.235172 | 1.3899228 | 0.164552 | 0.63030432 | SNORD13 | snoRNA | 49.04 | 0.065521058 | 0.065521058 | ENSG00000239039 | | n.s. |
| ENSG00000167113 | 594.8574 | -0.351252317 | 0.262453 | -1.338343 | 0.180785 | 0.650908532 | COQ4 | protein_coding | 51.72 | -0.065501544 | 0.065501544 | ENSG00000167113 | | n.s. |
| ENSG00000140743 | 298.3047 | -0.364743364 | 0.27676 | -1.317905 | 0.187535 | 0.661470368 | CDR2 | protein_coding | 43.33 | -0.065467643 | 0.065467643 | ENSG00000140743 | | n.s. |
| ENSG00000136960 | 507.3449 | 0.38305656 | 0.29793 | 1.285726 | 0.198539 | 0.674743473 | ENPP2 | protein_coding | 39.28 | 0.065449545 | 0.065449545 | ENSG00000136960 | | n.s. |
| ENSG00000277948 | 28.58634 | -0.436285832 | 0.367902 | -1.185876 | 0.235671 | 0.707931694 | | misc_RNA | 40.64 | -0.065446646 | 0.065446646 | ENSG00000277948 | | n.s. |
| ENSG00000135823 | 1575.611 | -0.340439281 | 0.251101 | -1.355785 | 0.175167 | 0.642395686 | STX6 | protein_coding | 40.71 | -0.065431539 | 0.065431539 | ENSG00000135823 | | n.s. |
| ENSG00000173334 | 831.1106 | 0.297104697 | 0.202492 | 1.4672448 | 0.142309 | 0.602241628 | TRIB1 | protein_coding | 51.37 | 0.065431138 | 0.065431138 | ENSG00000173334 | | n.s. |
| ENSG00000250565 | 150.9116 | -0.355893839 | 0.267609 | -1.329904 | 0.18355 | 0.654903931 | ATP6V1E2 | protein_coding | 44.72 | -0.065421261 | 0.065421261 | ENSG00000250565 | | n.s. |
| ENSG00000275479 | 9.30463 | 0.516452684 | 0.48651 | 1.0615465 | 0.288442 | 0.747018696 | | lincRNA | 34.3 | 0.065418302 | 0.065418302 | ENSG00000275479 | | n.s. |
| ENSG00000233010 | 36.11342 | -0.41231882 | 0.334889 | -1.231209 | 0.218245 | 0.694065444 | RPEP4 | processed_pseudogene | 47.34 | -0.065393591 | 0.065393591 | ENSG00000233010 | | n.s. |
| ENSG00000252933 | 5.154913 | -0.624155986 | 0.653922 | -0.95448 | 0.339841 | 0.785979841 | RNU6-1223P | snRNA | 42.31 | -0.065279596 | 0.065279596 | ENSG00000252933 | | n.s. |
| ENSG00000197291 | 395.2888 | -0.327916823 | 0.237177 | -1.382585 | 0.166792 | 0.632544316 | RAMP2-AS1 | lincRNA | 49.8 | -0.065225621 | 0.065225621 | ENSG00000197291 | | n.s. |
| ENSG00000117597 | 1845.692 | -0.333586687 | 0.24278 | -1.374031 | 0.169432 | 0.637567981 | DIEXF | protein_coding | 41.44 | -0.065207358 | 0.065207358 | ENSG00000117597 | | n.s. |
| ENSG00000199237 | 6.32791 | -0.552247424 | 0.541787 | -1.019306 | 0.308058 | 0.761997697 | RNU6-834P | snRNA | 40 | -0.065190788 | 0.065190788 | ENSG00000199237 | | n.s. |
| ENSG00000153815 | 175.6064 | -0.354834719 | 0.267095 | -1.328497 | 0.184014 | 0.655193675 | CMIP | protein_coding | 51.33 | -0.065158407 | 0.065158407 | ENSG00000153815 | | n.s. |
| ENSG00000173540 | 321.8692 | -0.323446403 | 0.230732 | -1.401826 | 0.160967 | 0.628931954 | GMPPB | protein_coding | 61.13 | -0.065140922 | 0.065140922 | ENSG00000173540 | | n.s. |
| ENSG00000264314 | 12.8047 | -0.515241507 | 0.485749 | -1.060715 | 0.28882 | 0.747460922 | MIR548AT | miRNA | 39.66 | -0.065132456 | 0.065132456 | ENSG00000264314 | | n.s. |
| ENSG00000189362 | 417.1244 | -0.340106826 | 0.251547 | -1.352063 | 0.176355 | 0.643565193 | NEMP2 | protein_coding | 40.26 | -0.065098981 | 0.065098981 | ENSG00000189362 | | n.s. |
| ENSG00000266709 | 122.8628 | 0.350557393 | 0.262532 | 1.3352946 | 0.18178 | 0.652131419 | | lincRNA | 52.3 | 0.065086195 | 0.065086195 | ENSG00000266709 | | n.s. |
| ENSG00000204977 | 1456.337 | -0.331253389 | 0.240504 | -1.377331 | 0.16841 | 0.636128582 | TRIM13 | protein_coding | 41.26 | -0.065076415 | 0.065076415 | ENSG00000204977 | | n.s. |
| ENSG00000207652 | 45.8893 | -0.419894876 | 0.345413 | -1.21563 | 0.224126 | 0.700099826 | MIR621 | miRNA | 55.21 | -0.065016535 | 0.065016535 | ENSG00000207652 | | n.s. |

| | | | | | | | | | | | | | |
|---|---|---|---|---|---|---|---|---|---|---|---|---|---|
| ENSG00000167554 | 218.7349 | -0.325775074 | 0.235256 | -1.384771 | 0.166123 | 0.631590186 | ZNF610 | protein_coding | 44.88 | -0.065013181 | 0.065013181 | ENSG00000167554 | n.s. |
| ENSG00000280798 | 210.5262 | -0.36694483 | 0.280323 | -1.309007 | 0.190532 | 0.665181845 | LINC00294 | lincRNA | 45.1 | -0.064971109 | 0.064971109 | ENSG00000280798 | n.s. |
| ENSG00000271798 | 9.637858 | 0.549895936 | 0.535987 | 1.0259509 | 0.304915 | 0.761997697 | SNORA51 | snoRNA | 50.76 | 0.064913203 | 0.064913203 | ENSG00000271798 | n.s. |
| ENSG00000265420 | 17.46912 | 0.464510233 | 0.40812 | 1.1381709 | 0.255049 | 0.72487977 | MIR4779 | miRNA | 36.14 | 0.064907882 | 0.064907882 | ENSG00000265420 | n.s. |
| ENSG00000265390 | 4.699456 | -0.705963012 | 0.800302 | -0.88212 | 0.377712 | 0.809223062 | MIR4999 | miRNA | 28.57 | -0.064900414 | 0.064900414 | ENSG00000265390 | n.s. |
| ENSG00000110497 | 176.9216 | -0.337970277 | 0.249478 | -1.354711 | 0.17551 | 0.642859735 | AMBRA1 | protein_coding | 43.63 | -0.064851012 | 0.064851012 | ENSG00000110497 | n.s. |
| ENSG00000229521 | 54.99002 | -0.384185957 | 0.301568 | -1.273961 | 0.202677 | 0.677959669 | MYCBP2-AS2 | sense_intronic | 42.35 | -0.064849107 | 0.064849107 | ENSG00000229521 | n.s. |
| ENSG00000252916 | 12.32097 | -0.494416823 | 0.45231 | -1.093093 | 0.274353 | 0.739386355 | RNU6-762P | snRNA | 40.2 | -0.06483217 | 0.06483217 | ENSG00000252916 | n.s. |
| ENSG00000100592 | 314.2354 | -0.332391063 | 0.242225 | -1.372242 | 0.169988 | 0.638281144 | DAAM1 | protein_coding | 39.75 | -0.064812264 | 0.064812264 | ENSG00000100592 | n.s. |
| ENSG00000275662 | 4.400465 | 0.593664441 | 0.604948 | 0.9813473 | 0.326422 | 0.778045194 | SNORD112 | snoRNA | 37.66 | 0.06470656 | 0.06470656 | ENSG00000275662 | n.s. |
| ENSG00000260475 | 38.70198 | 0.424562007 | 0.354397 | 1.1979844 | 0.230923 | 0.704043192 | | lincRNA | 59.61 | 0.064703546 | 0.064703546 | ENSG00000260475 | n.s. |
| ENSG00000197774 | 251.3578 | -0.355667014 | 0.268593 | -1.324183 | 0.185442 | 0.657811821 | EME2 | protein_coding | 64.06 | -0.064695234 | 0.064695234 | ENSG00000197774 | n.s. |
| ENSG00000207956 | 9.70667 | -0.504650478 | 0.469333 | -1.075251 | 0.282262 | 0.744567264 | MIR579 | miRNA | 32.65 | -0.064643739 | 0.064643739 | ENSG00000207956 | n.s. |
| ENSG00000273797 | 22.64197 | 0.451557576 | 0.390839 | 1.1553551 | 0.247945 | 0.719195543 | | sense_intronic | 36.05 | 0.064641828 | 0.064641828 | ENSG00000273797 | n.s. |
| ENSG00000198763 | 1162.19 | -0.301426989 | 0.209016 | -1.442121 | 0.149268 | 0.610355798 | MT-ND2 | protein_coding | 42.99 | -0.064631048 | 0.064631048 | ENSG00000198763 | n.s. |
| ENSG00000152443 | 954.9757 | -0.324877312 | 0.234867 | -1.38324 | 0.166591 | 0.632544316 | ZNF776 | protein_coding | 43.81 | -0.064621035 | 0.064621035 | ENSG00000152443 | n.s. |
| ENSG00000105875 | 406.0422 | -0.351656532 | 0.264892 | -1.327548 | 0.184328 | 0.655193675 | WDR91 | protein_coding | 47.38 | -0.064574796 | 0.064574796 | ENSG00000105875 | n.s. |
| ENSG00000100600 | 306.6647 | -0.363352485 | 0.277157 | -1.310998 | 0.189858 | 0.664260131 | LGMN | protein_coding | 47.28 | -0.064553861 | 0.064553861 | ENSG00000100600 | n.s. |
| ENSG00000145725 | 2572.605 | -0.367536055 | 0.282329 | -1.301803 | 0.192984 | 0.667577363 | PPIP5K2 | protein_coding | 34.29 | -0.064501989 | 0.064501989 | ENSG00000145725 | n.s. |
| ENSG00000105656 | 143.5555 | -0.336711872 | 0.248949 | -1.352533 | 0.176205 | 0.643404323 | ELL | protein_coding | 55.52 | -0.064485719 | 0.064485719 | ENSG00000105656 | n.s. |
| ENSG00000122729 | 386.627 | -0.333635277 | 0.2447 | -1.363446 | 0.172742 | 0.640837445 | ACO1 | protein_coding | 41.71 | -0.064475725 | 0.064475725 | ENSG00000122729 | n.s. |
| ENSG00000104312 | 255.1737 | 0.341564814 | 0.254091 | 1.3442637 | 0.178863 | 0.648076085 | RIPK2 | protein_coding | 36.01 | 0.064341932 | 0.064341932 | ENSG00000104312 | n.s. |
| ENSG00000130254 | 1023.597 | -0.318806453 | 0.228262 | -1.396666 | 0.162514 | 0.629027704 | SAFB2 | protein_coding | 49.34 | -0.064185375 | 0.064185375 | ENSG00000130254 | n.s. |
| ENSG00000139350 | 1443.875 | -0.33729293 | 0.250117 | -1.348539 | 0.177485 | 0.645396531 | NEDD1 | protein_coding | 35.02 | -0.064144134 | 0.064144134 | ENSG00000139350 | n.s. |
| ENSG00000163576 | 23.73677 | 0.44091699 | 0.378652 | 1.1644375 | 0.244247 | 0.715606581 | EFHB | protein_coding | 40.39 | 0.064076559 | 0.064076559 | ENSG00000163576 | n.s. |
| ENSG00000188895 | 1198.643 | -0.345623807 | 0.259092 | -1.333982 | 0.18221 | 0.652831893 | MSL1 | protein_coding | 45.42 | -0.064009058 | 0.064009058 | ENSG00000188895 | n.s. |
| ENSG00000141560 | 548.4736 | -0.364183553 | 0.27954 | -1.302797 | 0.192644 | 0.667375338 | FN3KRP | protein_coding | 53.24 | -0.063961501 | 0.063961501 | ENSG00000141560 | n.s. |
| ENSG00000237200 | 9.216408 | -0.500895115 | 0.46787 | -1.070586 | 0.284356 | 0.745309826 | ZBTB40-IT1 | sense_intronic | 44.07 | -0.06394585 | 0.06394585 | ENSG00000237200 | n.s. |
| ENSG00000186642 | 785.6476 | 0.36434274 | 0.279879 | 1.3017849 | 0.19299 | 0.667577363 | PDE2A | protein_coding | 55.35 | 0.063941567 | 0.063941567 | ENSG00000186642 | n.s. |
| ENSG00000163346 | 185.7285 | 0.385797201 | 0.305979 | 1.2608605 | 0.207359 | 0.682785213 | PBXIP1 | protein_coding | 54.24 | 0.063932728 | 0.063932728 | ENSG00000163346 | n.s. |
| ENSG00000135097 | 502.4005 | -0.317210435 | 0.227436 | -1.394726 | 0.163098 | 0.629036117 | MSI1 | protein_coding | 54.01 | -0.063862206 | 0.063862206 | ENSG00000135097 | n.s. |
| ENSG00000111670 | 465.8756 | -0.325722526 | 0.23683 | -1.375343 | 0.169025 | 0.636857974 | GNPTAB | protein_coding | 40.94 | -0.063827742 | 0.063827742 | ENSG00000111670 | n.s. |

| | | | | | | | | | | | | | |
|---|---|---|---|---|---|---|---|---|---|---|---|---|---|
| ENSG00000257315 | 4210.426 | 0.301955652 | 0.210993 | 1.4311191 | 0.152396 | 0.614654298 | ZBED6 | protein_coding | 40.71 | 0.063824088 | 0.063824088 | ENSG00000257315 | n.s. |
| ENSG00000279369 | 653.5057 | -0.29151028 | 0.199445 | -1.461607 | 0.143849 | 0.604031311 | | TEC | 59.16 | -0.06382342 | 0.06382342 | ENSG00000279369 | n.s. |
| ENSG00000130559 | 2646.827 | -0.331225093 | 0.243442 | -1.360589 | 0.173643 | 0.642002364 | CAMSAP1 | protein_coding | 46.91 | -0.063748699 | 0.063748699 | ENSG00000130559 | n.s. |
| ENSG00000184992 | 949.0986 | -0.363256464 | 0.279315 | -1.300528 | 0.19342 | 0.667903971 | BRI3BP | protein_coding | 48.32 | -0.063673763 | 0.063673763 | ENSG00000184992 | n.s. |
| ENSG00000227630 | 22.67648 | 0.501230119 | 0.470196 | 1.0660026 | 0.286422 | 0.746611882 | LINC01132 | lincRNA | 50.62 | 0.06360866 | 0.06360866 | ENSG00000227630 | n.s. |
| ENSG00000104524 | 164.6862 | -0.361926195 | 0.277728 | -1.303167 | 0.192518 | 0.667270349 | PYCR3 | protein_coding | 60.74 | -0.063589771 | 0.063589771 | ENSG00000104524 | n.s. |
| ENSG00000155016 | 311.7335 | -0.343244479 | 0.257272 | -1.334168 | 0.182149 | 0.652831893 | CYP2U1 | protein_coding | 40.45 | -0.06356841 | 0.06356841 | ENSG00000155016 | n.s. |
| ENSG00000266038 | 10.66339 | -0.497791964 | 0.464966 | -1.070598 | 0.28435 | 0.745309826 | MIR4659A | miRNA | 38.27 | -0.063549692 | 0.063549692 | ENSG00000266038 | n.s. |
| ENSG00000231125 | 137.4215 | -0.375617472 | 0.293642 | -1.279169 | 0.200838 | 0.677491543 | | sense_intronic | 40.43 | -0.063515458 | 0.063515458 | ENSG00000231125 | n.s. |
| ENSG00000164916 | 581.9869 | -0.317877319 | 0.229416 | -1.385593 | 0.165871 | 0.63130003 | FOXK1 | protein_coding | 52.09 | -0.063500505 | 0.063500505 | ENSG00000164916 | n.s. |
| ENSG00000177082 | 1133.917 | -0.336607065 | 0.25022 | -1.345244 | 0.178546 | 0.647707196 | WDR73 | protein_coding | 48.66 | -0.063491255 | 0.063491255 | ENSG00000177082 | n.s. |
| ENSG00000100926 | 231.6733 | -0.341490204 | 0.255583 | -1.33612 | 0.18151 | 0.651834927 | TM9SF1 | protein_coding | 51 | -0.063470179 | 0.063470179 | ENSG00000100926 | n.s. |
| ENSG00000198740 | 1232.796 | -0.32972533 | 0.242267 | -1.361001 | 0.173513 | 0.642002364 | ZNF652 | protein_coding | 41.52 | -0.063460049 | 0.063460049 | ENSG00000198740 | n.s. |
| ENSG00000239912 | 11.7357 | -0.49263087 | 0.455702 | -1.081037 | 0.279681 | 0.743390786 | RPL39P36 | processed_pseudogene | 37.82 | -0.063442396 | 0.063442396 | ENSG00000239912 | n.s. |
| ENSG00000177283 | 738.799 | -0.301274063 | 0.211112 | -1.427081 | 0.153557 | 0.615840022 | FZD8 | protein_coding | 64.57 | -0.063427858 | 0.063427858 | ENSG00000177283 | n.s. |
| ENSG00000180592 | 695.7631 | -0.291824404 | 0.200196 | -1.457695 | 0.144925 | 0.606306192 | SKIDA1 | protein_coding | 45.4 | -0.063415776 | 0.063415776 | ENSG00000180592 | n.s. |
| ENSG00000274460 | 66.29421 | 0.36581241 | 0.2827 | 1.2939958 | 0.195667 | 0.671002415 | | processed_transcript | 43.59 | 0.063386481 | 0.063386481 | ENSG00000274460 | n.s. |
| ENSG00000178235 | 215.0053 | 0.314734805 | 0.225103 | 1.398181 | 0.162059 | 0.628957308 | SLITRK1 | protein_coding | 47.31 | 0.063380927 | 0.063380927 | ENSG00000178235 | n.s. |
| ENSG00000123064 | 1357.742 | -0.315482087 | 0.226554 | -1.392527 | 0.163763 | 0.629679644 | DDX54 | protein_coding | 54.08 | -0.063374151 | 0.063374151 | ENSG00000123064 | n.s. |
| ENSG00000070214 | 1023.797 | -0.352989155 | 0.268047 | -1.316892 | 0.187875 | 0.661552858 | SLC44A1 | protein_coding | 38.86 | -0.063338768 | 0.063338768 | ENSG00000070214 | n.s. |
| ENSG00000168137 | 2657.241 | -0.33213964 | 0.245986 | -1.35024 | 0.176939 | 0.64464925 | SETD5 | protein_coding | 39.13 | -0.06333123 | 0.06333123 | ENSG00000168137 | n.s. |
| ENSG00000170234 | 944.9101 | -0.33664734 | 0.250731 | -1.342662 | 0.179382 | 0.648472544 | PWWP2A | protein_coding | 42.31 | -0.063326195 | 0.063326195 | ENSG00000170234 | n.s. |
| ENSG00000136720 | 414.5192 | -0.344797281 | 0.25971 | -1.327624 | 0.184302 | 0.655193675 | HS6ST1 | protein_coding | 53.08 | -0.063315229 | 0.063315229 | ENSG00000136720 | n.s. |
| ENSG00000282995 | 102.9724 | -0.374340321 | 0.293177 | -1.276842 | 0.201658 | 0.677624898 | FRG1EP | unprocessed_pseudogene | 34.33 | -0.063267499 | 0.063267499 | ENSG00000282995 | n.s. |
| ENSG00000185730 | 155.6791 | -0.326809267 | 0.239376 | -1.365256 | 0.172173 | 0.640837445 | ZNF696 | protein_coding | 58.48 | -0.063156584 | 0.063156584 | ENSG00000185730 | n.s. |
| ENSG00000156304 | 1308.463 | -0.322277313 | 0.234232 | -1.37589 | 0.168856 | 0.636852728 | SCAF4 | protein_coding | 38.12 | -0.06315378 | 0.06315378 | ENSG00000156304 | n.s. |
| ENSG00000244356 | 15.35257 | -0.448560753 | 0.392059 | -1.144115 | 0.252576 | 0.723136889 | RN7SL398P | misc_RNA | 53.36 | -0.063148151 | 0.063148151 | ENSG00000244356 | n.s. |
| ENSG00000165409 | 46.74812 | 0.38466304 | 0.306601 | 1.2546065 | 0.209622 | 0.685286106 | TSHR | protein_coding | 38.85 | 0.063134004 | 0.063134004 | ENSG00000165409 | n.s. |
| ENSG00000205730 | 812.1826 | -0.317098188 | 0.229328 | -1.382729 | 0.166748 | 0.632544316 | ITPRIPL2 | protein_coding | 49.31 | -0.063073697 | 0.063073697 | ENSG00000205730 | n.s. |
| ENSG00000160007 | 1500.021 | -0.329473402 | 0.243747 | -1.351701 | 0.176471 | 0.643565193 | ARHGAP35 | protein_coding | 47.96 | -0.063063665 | 0.063063665 | ENSG00000160007 | n.s. |
| ENSG00000092847 | 690.5579 | -0.313968834 | 0.225521 | -1.392195 | 0.163863 | 0.629718088 | AGO1 | protein_coding | 44.84 | -0.063061843 | 0.063061843 | ENSG00000092847 | n.s. |
| ENSG00000065485 | 66.90689 | -0.354213002 | 0.269915 | -1.312315 | 0.189414 | 0.663959974 | PDIA5 | protein_coding | 45.21 | -0.062999652 | 0.062999652 | ENSG00000065485 | n.s. |

| | | | | | | | | | | | | | |
|---|---|---|---|---|---|---|---|---|---|---|---|---|---|
| ENSG00000260796 | 44.19083 | -0.391802196 | 0.315279 | -1.242714 | 0.213973 | 0.690737299 | | antisense | 47.47 | -0.062957555 | 0.062957555 | ENSG00000260796 | n.s. |
| ENSG00000118260 | 1854.78 | -0.362982653 | 0.280346 | -1.294766 | 0.195401 | 0.670750289 | CREB1 | protein_coding | 37.47 | -0.062955396 | 0.062955396 | ENSG00000118260 | n.s. |
| ENSG00000101198 | 81.25172 | -0.411296896 | 0.342076 | -1.202354 | 0.229226 | 0.702977342 | NKAIN4 | protein_coding | 59.79 | -0.062952557 | 0.062952557 | ENSG00000101198 | n.s. |
| ENSG00000110060 | 299.0386 | -0.35911402 | 0.276126 | -1.300542 | 0.193415 | 0.667903971 | PUS3 | protein_coding | 40.24 | -0.062947651 | 0.062947651 | ENSG00000110060 | n.s. |
| ENSG00000207597 | 42.54256 | -0.396510417 | 0.321584 | -1.232991 | 0.217579 | 0.693853089 | MIR490 | miRNA | 51.56 | -0.062939079 | 0.062939079 | ENSG00000207597 | n.s. |
| ENSG00000284428 | 580.2908 | -0.303016776 | 0.213332 | -1.420398 | 0.155492 | 0.619951348 | | processed_transcript | 45.13 | -0.062919128 | 0.062919128 | ENSG00000284428 | n.s. |
| ENSG00000207771 | 14.11051 | 0.48477088 | 0.445681 | 1.0877089 | 0.276724 | 0.741896779 | MIR550A1 | miRNA | 50.52 | 0.062853703 | 0.062853703 | ENSG00000207771 | n.s. |
| ENSG00000164466 | 1476.807 | -0.359058409 | 0.276365 | -1.299216 | 0.19387 | 0.668296258 | SFXN1 | protein_coding | 42.43 | -0.062846342 | 0.062846342 | ENSG00000164466 | n.s. |
| ENSG00000168348 | 123.2669 | 0.323542432 | 0.236354 | 1.36889 | 0.171034 | 0.639447334 | INSM2 | protein_coding | 58.21 | 0.062830393 | 0.062830393 | ENSG00000168348 | n.s. |
| ENSG00000111696 | 1244.404 | -0.326312912 | 0.240067 | -1.359259 | 0.174064 | 0.642002364 | NT5DC3 | protein_coding | 43.1 | -0.062803284 | 0.062803284 | ENSG00000111696 | n.s. |
| ENSG00000121210 | 1240.399 | -0.322925979 | 0.235847 | -1.369217 | 0.170932 | 0.639237118 | KIAA0922 | protein_coding | 41.78 | -0.062756794 | 0.062756794 | ENSG00000121210 | n.s. |
| ENSG00000148143 | 2841.309 | -0.311713792 | 0.22349 | -1.394758 | 0.163089 | 0.629036117 | ZNF462 | protein_coding | 40.08 | -0.062755598 | 0.062755598 | ENSG00000148143 | n.s. |
| ENSG00000125485 | 662.1512 | -0.327485334 | 0.242219 | -1.352022 | 0.176368 | 0.643565193 | DDX31 | protein_coding | 44.73 | -0.062683134 | 0.062683134 | ENSG00000125485 | n.s. |
| ENSG00000109670 | 1708.092 | -0.311247054 | 0.222692 | -1.397659 | 0.162216 | 0.628957308 | FBXW7 | protein_coding | 35.42 | -0.062678568 | 0.062678568 | ENSG00000109670 | n.s. |
| ENSG00000157540 | 2471.421 | -0.326019022 | 0.240395 | -1.356181 | 0.175042 | 0.642395686 | DYRK1A | protein_coding | 38.86 | -0.062660003 | 0.062660003 | ENSG00000157540 | n.s. |
| ENSG00000152217 | 1477.942 | -0.329469422 | 0.24432 | -1.348517 | 0.177492 | 0.645396531 | SETBP1 | protein_coding | 42.22 | -0.062656311 | 0.062656311 | ENSG00000152217 | n.s. |
| ENSG00000140443 | 142.0168 | -0.325808517 | 0.23996 | -1.357761 | 0.17454 | 0.642377343 | IGF1R | protein_coding | 44.52 | -0.062623585 | 0.062623585 | ENSG00000140443 | n.s. |
| ENSG00000170852 | 1391.806 | -0.339796555 | 0.255173 | -1.33163 | 0.182982 | 0.65425144 | KBTBD2 | protein_coding | 41.11 | -0.062609321 | 0.062609321 | ENSG00000170852 | n.s. |
| ENSG00000164172 | 376.063 | -0.369966426 | 0.289584 | -1.277578 | 0.201398 | 0.677503939 | MOCS2 | protein_coding | 38.21 | -0.062556948 | 0.062556948 | ENSG00000164172 | n.s. |
| ENSG00000143434 | 213.2943 | -0.337705107 | 0.253105 | -1.334249 | 0.182122 | 0.652831893 | SEMA6C | protein_coding | 57.48 | -0.062542526 | 0.062542526 | ENSG00000143434 | n.s. |
| ENSG00000169258 | 1468.677 | -0.34050149 | 0.256245 | -1.328813 | 0.18391 | 0.655193675 | GPRIN1 | protein_coding | 55.36 | -0.062526392 | 0.062526392 | ENSG00000169258 | n.s. |
| ENSG00000185722 | 946.9706 | -0.336456362 | 0.251858 | -1.335899 | 0.181582 | 0.651909066 | ANKFY1 | protein_coding | 44.91 | -0.062517959 | 0.062517959 | ENSG00000185722 | n.s. |
| ENSG00000100106 | 266.5181 | -0.34807111 | 0.263931 | -1.318796 | 0.187237 | 0.661405956 | TRIOBP | protein_coding | 54.35 | -0.062489867 | 0.062489867 | ENSG00000100106 | n.s. |
| ENSG00000131018 | 115.2911 | -0.331393679 | 0.24646 | -1.344615 | 0.17875 | 0.647938638 | SYNE1 | protein_coding | 38.41 | -0.062456481 | 0.062456481 | ENSG00000131018 | n.s. |
| ENSG00000057019 | 1520.046 | -0.33254411 | 0.247852 | -1.341703 | 0.179692 | 0.648989471 | DCBLD2 | protein_coding | 38.58 | -0.062439263 | 0.062439263 | ENSG00000057019 | n.s. |
| ENSG00000198346 | 290.986 | -0.373736463 | 0.29536 | -1.265361 | 0.205742 | 0.680704186 | ZNF813 | protein_coding | 43.99 | -0.062429529 | 0.062429529 | ENSG00000198346 | n.s. |
| ENSG00000138246 | 1890.595 | -0.354125077 | 0.27136 | -1.305 | 0.191893 | 0.666819478 | DNAJC13 | protein_coding | 36.8 | -0.062323082 | 0.062323082 | ENSG00000138246 | n.s. |
| ENSG00000128039 | 422.7861 | -0.368616018 | 0.289134 | -1.274896 | 0.202346 | 0.677624898 | SRD5A3 | protein_coding | 44.77 | -0.062300031 | 0.062300031 | ENSG00000128039 | n.s. |
| ENSG00000259429 | 277.6223 | -0.359089398 | 0.277434 | -1.294324 | 0.195553 | 0.670778798 | UBE2Q2P2 | cribed_unprocessed_pseudc | 39.44 | -0.062273525 | 0.062273525 | ENSG00000259429 | n.s. |
| ENSG00000174606 | 2023.622 | -0.337251383 | 0.253147 | -1.332237 | 0.182782 | 0.653705624 | ANGEL2 | protein_coding | 38.68 | -0.062262601 | 0.062262601 | ENSG00000174606 | n.s. |
| ENSG00000161091 | 205.1894 | -0.343409306 | 0.259738 | -1.32214 | 0.186122 | 0.658994794 | MFSD12 | protein_coding | 57.07 | -0.062197612 | 0.062197612 | ENSG00000161091 | n.s. |
| ENSG00000134049 | 1204.967 | -0.383721588 | 0.307464 | -1.24802 | 0.212024 | 0.688575085 | IER3IP1 | protein_coding | 38.68 | -0.062181583 | 0.062181583 | ENSG00000134049 | n.s. |

| | | | | | | | | | | | | | |
|---|---|---|---|---|---|---|---|---|---|---|---|---|---|
| ENSG00000011198 | 385.2579 | -0.338378523 | 0.254529 | -1.329429 | 0.183706 | 0.655151359 | ABHD5 | protein_coding | 40.26 | -0.062146042 | 0.062146042 | ENSG00000011198 | n.s. |
| ENSG00000013392 | 196.8392 | 0.335382926 | 0.251392 | 1.334102 | 0.18217 | 0.652831893 | RWDD2A | protein_coding | 44.76 | 0.062112461 | 0.062112461 | ENSG00000013392 | n.s. |
| ENSG00000156374 | 842.2667 | -0.338696901 | 0.255314 | -1.32659 | 0.184644 | 0.655769671 | PCGF6 | protein_coding | 43.89 | -0.062065758 | 0.062065758 | ENSG00000156374 | n.s. |
| ENSG00000187840 | 1763.861 | -0.371226627 | 0.292814 | -1.267791 | 0.204873 | 0.680575501 | EIF4EBP1 | protein_coding | 49.34 | -0.062040763 | 0.062040763 | ENSG00000187840 | n.s. |
| ENSG00000266097 | 14.36413 | -0.538577012 | 0.535451 | -1.005838 | 0.314493 | 0.767300529 | MIR5192 | miRNA | 55.43 | -0.061954939 | 0.061954939 | ENSG00000266097 | n.s. |
| ENSG00000277224 | 586.9565 | -0.32032049 | 0.234432 | -1.36637 | 0.171823 | 0.640837445 | HIST1H2BF | protein_coding | 42.73 | -0.061902614 | 0.061902614 | ENSG00000277224 | n.s. |
| ENSG00000199357 | 11.09582 | 0.524321726 | 0.51412 | 1.0198436 | 0.307803 | 0.761997697 | | misc_RNA | 39.18 | 0.061894261 | 0.061894261 | ENSG00000199357 | n.s. |
| ENSG00000275764 | 194.7281 | -0.360427614 | 0.27967 | -1.288763 | 0.197481 | 0.673407427 | | lincRNA | 32.72 | -0.061893386 | 0.061893386 | ENSG00000275764 | n.s. |
| ENSG00000106125 | 222.7167 | -0.34473169 | 0.261414 | -1.318719 | 0.187263 | 0.661405956 | MINDY4 | protein_coding | 48.22 | -0.061890335 | 0.061890335 | ENSG00000106125 | n.s. |
| ENSG00000266228 | 5.797977 | -0.593403245 | 0.624937 | -0.949541 | 0.342346 | 0.786682723 | MIR3611 | miRNA | 26.51 | -0.061832848 | 0.061832848 | ENSG00000266228 | n.s. |
| ENSG00000124733 | 759.1426 | -0.36982191 | 0.292085 | -1.266144 | 0.205462 | 0.680704186 | MEA1 | protein_coding | 56.59 | -0.061775635 | 0.061775635 | ENSG00000124733 | n.s. |
| ENSG00000221406 | 13.88221 | -0.586400526 | 0.611208 | -0.959412 | 0.337351 | 0.78475552 | MIR320B2 | miRNA | 36.96 | -0.061727816 | 0.061727816 | ENSG00000221406 | n.s. |
| ENSG00000141084 | 233.8258 | -0.315555793 | 0.229683 | -1.373875 | 0.169481 | 0.637578225 | RANBP10 | protein_coding | 49.73 | -0.061680594 | 0.061680594 | ENSG00000141084 | n.s. |
| ENSG00000253716 | 122.2215 | -0.333426863 | 0.250056 | -1.333408 | 0.182398 | 0.653170746 | MINCR | antisense | 49.93 | -0.061675059 | 0.061675059 | ENSG00000253716 | n.s. |
| ENSG00000160445 | 200.347 | -0.360378056 | 0.280229 | -1.286011 | 0.198439 | 0.674743473 | ZER1 | protein_coding | 51.42 | -0.061574666 | 0.061574666 | ENSG00000160445 | n.s. |
| ENSG00000100280 | 605.4149 | -0.330027677 | 0.246556 | -1.338552 | 0.180717 | 0.650832322 | AP1B1 | protein_coding | 49.33 | -0.061560355 | 0.061560355 | ENSG00000100280 | n.s. |
| ENSG00000073536 | 732.1554 | -0.305642769 | 0.218747 | -1.397243 | 0.16234 | 0.629027704 | NLE1 | protein_coding | 52.13 | -0.061535128 | 0.061535128 | ENSG00000073536 | n.s. |
| ENSG00000201700 | 18.37958 | 0.446312051 | 0.394487 | 1.1313721 | 0.257899 | 0.728045648 | SNORD113-3 | snoRNA | 38.89 | 0.061520273 | 0.061520273 | ENSG00000201700 | n.s. |
| ENSG00000197451 | 6182.08 | -0.326962895 | 0.243453 | -1.343025 | 0.179264 | 0.64843101 | HNRNPAB | protein_coding | 53.64 | -0.061513565 | 0.061513565 | ENSG00000197451 | n.s. |
| ENSG00000100296 | 1289.289 | -0.342687263 | 0.259917 | -1.318449 | 0.187354 | 0.661470368 | THOC5 | protein_coding | 47.17 | -0.061508802 | 0.061508802 | ENSG00000100296 | n.s. |
| ENSG00000240053 | 240.6099 | -0.331126029 | 0.248068 | -1.334818 | 0.181936 | 0.652523039 | LY6G5B | protein_coding | 52.19 | -0.06139214 | 0.06139214 | ENSG00000240053 | n.s. |
| ENSG00000199248 | 11.50042 | -0.50445273 | 0.48292 | -1.044589 | 0.296213 | 0.755758758 | RNU6-28P | snRNA | 41.12 | -0.061349932 | 0.061349932 | ENSG00000199248 | n.s. |
| ENSG00000202523 | 11.9416 | -0.481301155 | 0.449941 | -1.069698 | 0.284755 | 0.745682152 | | misc_RNA | 34.31 | -0.061340028 | 0.061340028 | ENSG00000202523 | n.s. |
| ENSG00000200502 | 19.81113 | -0.433892036 | 0.378057 | -1.147689 | 0.251097 | 0.722169533 | | misc_RNA | 46.53 | -0.061335342 | 0.061335342 | ENSG00000200502 | n.s. |
| ENSG00000281128 | 121.3017 | 0.329845932 | 0.246833 | 1.3363132 | 0.181447 | 0.651776425 | | antisense | 36.98 | 0.061318805 | 0.061318805 | ENSG00000281128 | n.s. |
| ENSG00000164077 | 170.0511 | -0.362320891 | 0.284241 | -1.274698 | 0.202416 | 0.677624898 | MON1A | protein_coding | 51.16 | -0.061236088 | 0.061236088 | ENSG00000164077 | n.s. |
| ENSG00000113595 | 619.5035 | -0.344609815 | 0.262814 | -1.311229 | 0.18978 | 0.664231317 | TRIM23 | protein_coding | 37.19 | -0.061230497 | 0.061230497 | ENSG00000113595 | n.s. |
| ENSG00000074855 | 143.21 | -0.353757252 | 0.273532 | -1.293295 | 0.195909 | 0.671501886 | ANO8 | protein_coding | 62.67 | -0.061183294 | 0.061183294 | ENSG00000074855 | n.s. |
| ENSG00000141551 | 902.495 | -0.348314968 | 0.267508 | -1.302075 | 0.192891 | 0.667565484 | CSNK1D | protein_coding | 55.31 | -0.061131411 | 0.061131411 | ENSG00000141551 | n.s. |
| ENSG00000272627 | 21.19538 | -0.466130117 | 0.425625 | -1.095166 | 0.273444 | 0.739386355 | | sense_intronic | 30.43 | -0.061122975 | 0.061122975 | ENSG00000272627 | n.s. |
| ENSG00000151247 | 2129.908 | -0.347299177 | 0.266125 | -1.305025 | 0.191884 | 0.666819478 | EIF4E | protein_coding | 39.19 | -0.061121781 | 0.061121781 | ENSG00000151247 | n.s. |
| ENSG00000270116 | 105.4103 | -0.352937368 | 0.272813 | -1.293699 | 0.19577 | 0.671189316 | | sense_intronic | 36.04 | -0.061112858 | 0.061112858 | ENSG00000270116 | n.s. |

| | | | | | | | | | | | | | |
|---|---|---|---|---|---|---|---|---|---|---|---|---|---|
| ENSG00000250802 | 228.2794 | -0.317883704 | 0.234118 | -1.35779 | 0.17453 | 0.642377343 | ZBED3-AS1 | antisense | 41.05 | -0.061100358 | 0.061100358 | ENSG00000250802 | n.s. |
| ENSG00000111341 | 2083.756 | -0.349415667 | 0.269024 | -1.298828 | 0.194003 | 0.66858954 | MGP | protein_coding | 37.99 | -0.061091983 | 0.061091983 | ENSG00000111341 | n.s. |
| ENSG00000153317 | 953.166 | -0.320317751 | 0.237178 | -1.350538 | 0.176844 | 0.64464925 | ASAP1 | protein_coding | 42.15 | -0.061077073 | 0.061077073 | ENSG00000153317 | n.s. |
| ENSG00000162687 | 1331.951 | -0.365841386 | 0.28934 | -1.264399 | 0.206087 | 0.680908009 | KCNT2 | protein_coding | 33.86 | -0.061063156 | 0.061063156 | ENSG00000162687 | n.s. |
| ENSG00000169372 | 187.3212 | 0.350585972 | 0.270297 | 1.2970407 | 0.194617 | 0.669854827 | CRADD | protein_coding | 40.55 | 0.061008729 | 0.061008729 | ENSG00000169372 | n.s. |
| ENSG00000196922 | 723.8176 | -0.361616697 | 0.284089 | -1.272899 | 0.203054 | 0.678385894 | ZNF252P | anscribed_unitary_pseudoge | 41.65 | -0.0609408 | 0.0609408 | ENSG00000196922 | n.s. |
| ENSG00000169926 | 3075.397 | -0.342199082 | 0.260558 | -1.313331 | 0.189071 | 0.663694184 | KLF13 | protein_coding | 49.37 | -0.060922383 | 0.060922383 | ENSG00000169926 | n.s. |
| ENSG00000162889 | 484.5811 | -0.343844783 | 0.262565 | -1.30956 | 0.190345 | 0.665072813 | MAPKAPK2 | protein_coding | 48.7 | -0.060905503 | 0.060905503 | ENSG00000162889 | n.s. |
| ENSG00000201557 | 9.231047 | 0.500798025 | 0.479488 | 1.0444434 | 0.29628 | 0.755765348 | SNORD114-15 | snoRNA | 45.83 | 0.060903563 | 0.060903563 | ENSG00000201557 | n.s. |
| ENSG00000108423 | 313.5328 | -0.360792082 | 0.283257 | -1.273728 | 0.20276 | 0.677959669 | TUBD1 | protein_coding | 45.29 | -0.060900311 | 0.060900311 | ENSG00000108423 | n.s. |
| ENSG00000257704 | 52.98123 | -0.369789156 | 0.294548 | -1.255446 | 0.209317 | 0.68461467 | INAFM1 | protein_coding | 71.55 | -0.06085021 | 0.06085021 | ENSG00000257704 | n.s. |
| ENSG00000267334 | 106.5195 | 0.359770502 | 0.282139 | 1.2751531 | 0.202255 | 0.677624898 | | lincRNA | 49.61 | 0.060805045 | 0.060805045 | ENSG00000267334 | n.s. |
| ENSG00000267637 | 13.45184 | -0.478998701 | 0.449937 | -1.06459 | 0.287062 | 0.746611882 | | sense_intronic | 21.08 | -0.060787379 | 0.060787379 | ENSG00000267637 | n.s. |
| ENSG00000080200 | 4667.851 | 0.348051523 | 0.268154 | 1.297953 | 0.194303 | 0.669140798 | CRYBG3 | protein_coding | 37.11 | 0.060728896 | 0.060728896 | ENSG00000080200 | n.s. |
| ENSG00000280128 | 441.7884 | -0.365715849 | 0.289768 | -1.262099 | 0.206913 | 0.68234303 | | TEC | 41.69 | -0.060707822 | 0.060707822 | ENSG00000280128 | n.s. |
| ENSG00000198910 | 659.8144 | -0.332596794 | 0.250988 | -1.325151 | 0.185121 | 0.656998505 | L1CAM | protein_coding | 56.65 | -0.060677502 | 0.060677502 | ENSG00000198910 | n.s. |
| ENSG00000112851 | 1553.531 | -0.349832955 | 0.270219 | -1.294627 | 0.195449 | 0.670750289 | ERBIN | protein_coding | 37.27 | -0.060674724 | 0.060674724 | ENSG00000112851 | n.s. |
| ENSG00000132612 | 441.7586 | -0.35597401 | 0.277503 | -1.282778 | 0.19957 | 0.676374453 | VPS4A | protein_coding | 54.49 | -0.060448945 | 0.060448945 | ENSG00000132612 | n.s. |
| ENSG00000201801 | 91.90373 | 0.352399709 | 0.273593 | 1.2880442 | 0.197731 | 0.673929742 | RNU5E-4P | snRNA | 33.33 | 0.060396157 | 0.060396157 | ENSG00000201801 | n.s. |
| ENSG00000224905 | 80.08207 | -0.366839821 | 0.292133 | -1.255727 | 0.209215 | 0.68461467 | | antisense | 42.94 | -0.060364886 | 0.060364886 | ENSG00000224905 | n.s. |
| ENSG00000003987 | 193.7947 | -0.349938833 | 0.271215 | -1.290265 | 0.196959 | 0.672285905 | MTMR7 | protein_coding | 40.36 | -0.06034555 | 0.06034555 | ENSG00000003987 | n.s. |
| ENSG00000099250 | 505.7369 | -0.327846505 | 0.246406 | -1.330515 | 0.183349 | 0.654554818 | NRP1 | protein_coding | 41.34 | -0.060341453 | 0.060341453 | ENSG00000099250 | n.s. |
| ENSG00000229770 | 20.10675 | -0.451234338 | 0.40686 | -1.109065 | 0.267402 | 0.734987182 | | antisense | 44.21 | -0.060339162 | 0.060339162 | ENSG00000229770 | n.s. |
| ENSG00000103599 | 35.44562 | 0.388829917 | 0.319316 | 1.2176958 | 0.22334 | 0.699562905 | IQCH | protein_coding | 39.22 | 0.060335993 | 0.060335993 | ENSG00000103599 | n.s. |
| ENSG00000182010 | 727.7593 | -0.367576249 | 0.292985 | -1.254592 | 0.209627 | 0.685286106 | RTKN2 | protein_coding | 35.12 | -0.060329582 | 0.060329582 | ENSG00000182010 | n.s. |
| ENSG00000121390 | 2092.737 | -0.33576189 | 0.254891 | -1.317275 | 0.187746 | 0.661470368 | PSPC1 | protein_coding | 40.29 | -0.06026577 | 0.06026577 | ENSG00000121390 | n.s. |
| ENSG00000167107 | 254.7223 | -0.360753827 | 0.285186 | -1.264979 | 0.205879 | 0.680704186 | ACSF2 | protein_coding | 49.96 | -0.060260889 | 0.060260889 | ENSG00000167107 | n.s. |
| ENSG00000117625 | 1324.018 | -0.349075821 | 0.270336 | -1.291264 | 0.196612 | 0.672118672 | RCOR3 | protein_coding | 37.38 | -0.060234443 | 0.060234443 | ENSG00000117625 | n.s. |
| ENSG00000198793 | 1182.632 | -0.327183376 | 0.245846 | -1.330849 | 0.183239 | 0.654498372 | MTOR | protein_coding | 44.54 | -0.060231655 | 0.060231655 | ENSG00000198793 | n.s. |
| ENSG00000133193 | 1008.061 | -0.349008093 | 0.270285 | -1.291259 | 0.196614 | 0.672118672 | FAM104A | protein_coding | 43.89 | -0.060222756 | 0.060222756 | ENSG00000133193 | n.s. |
| ENSG00000261765 | 34.30084 | 0.392588077 | 0.3263 | 1.2031525 | 0.228917 | 0.702737638 | | sense_intronic | 49.38 | 0.060147157 | 0.060147157 | ENSG00000261765 | n.s. |
| ENSG00000117533 | 486.5144 | -0.3643456 | 0.289585 | -1.258165 | 0.208332 | 0.6837835 | VAMP4 | protein_coding | 36.68 | -0.060146676 | 0.060146676 | ENSG00000117533 | n.s. |

| | | | | | | | | | | | | | |
|---|---|---|---|---|---|---|---|---|---|---|---|---|---|
| ENSG00000171823 | 236.0083 | -0.312757571 | 0.230601 | -1.356271 | 0.175013 | 0.642395686 | FBXL14 | protein_coding | 46.18 | -0.060111187 | 0.060111187 | ENSG00000171823 | n.s. |
| ENSG00000131351 | 368.8016 | -0.313251131 | 0.231322 | -1.354177 | 0.17568 | 0.642859735 | HAUS8 | protein_coding | 49.64 | -0.060107809 | 0.060107809 | ENSG00000131351 | n.s. |
| ENSG00000204954 | 289.6941 | -0.308880412 | 0.225343 | -1.370714 | 0.170464 | 0.638865917 | C12orf73 | protein_coding | 41.85 | -0.060105126 | 0.060105126 | ENSG00000204954 | n.s. |
| ENSG00000131368 | 1990.358 | -0.342607903 | 0.263326 | -1.301078 | 0.193232 | 0.667750696 | MRPS25 | protein_coding | 47.97 | -0.06008851 | 0.06008851 | ENSG00000131368 | n.s. |
| ENSG00000203666 | 204.9015 | -0.37072123 | 0.296964 | -1.248372 | 0.211895 | 0.688575085 | EFCAB2 | protein_coding | 42.37 | -0.060074892 | 0.060074892 | ENSG00000203666 | n.s. |
| ENSG00000252637 | 10.25426 | -0.534997483 | 0.539183 | -0.992238 | 0.321082 | 0.772226289 | RNA5SP268 | rRNA | 41.13 | -0.060056366 | 0.060056366 | ENSG00000252637 | n.s. |
| ENSG00000269858 | 668.4146 | -0.332084055 | 0.251382 | -1.321036 | 0.186489 | 0.659510569 | EGLN2 | protein_coding | 59.14 | -0.060033571 | 0.060033571 | ENSG00000269858 | n.s. |
| ENSG00000262202 | 315.7603 | -0.31060268 | 0.227318 | -1.366382 | 0.171819 | 0.640837445 | | lincRNA | 56.2 | -0.060024626 | 0.060024626 | ENSG00000262202 | n.s. |
| ENSG00000197299 | 2049.703 | -0.337821312 | 0.257697 | -1.310922 | 0.189884 | 0.664260131 | BLM | protein_coding | 40.73 | -0.060017947 | 0.060017947 | ENSG00000197299 | n.s. |
| ENSG00000227896 | 118.8651 | -0.325486269 | 0.244465 | -1.331421 | 0.18305 | 0.654329035 | | antisense | 47.2 | -0.05995581 | 0.05995581 | ENSG00000227896 | n.s. |
| ENSG00000177335 | 81.99467 | -0.360898337 | 0.285785 | -1.262834 | 0.206649 | 0.682201186 | C8orf31 | processed_transcript | 51.12 | -0.059940713 | 0.059940713 | ENSG00000177335 | n.s. |
| ENSG00000048740 | 229.4493 | -0.358778778 | 0.283646 | -1.264881 | 0.205914 | 0.680704186 | CELF2 | protein_coding | 41.91 | -0.059930973 | 0.059930973 | ENSG00000048740 | n.s. |
| ENSG00000100023 | 28.51974 | -0.419179723 | 0.363126 | -1.154363 | 0.248351 | 0.719540199 | PPIL2 | protein_coding | 54.06 | -0.059919619 | 0.059919619 | ENSG00000100023 | n.s. |
| ENSG00000134285 | 233.1952 | -0.333777722 | 0.253313 | -1.317649 | 0.187621 | 0.661470368 | FKBP11 | protein_coding | 51.54 | -0.059909632 | 0.059909632 | ENSG00000134285 | n.s. |
| ENSG00000181192 | 1158.589 | -0.356185518 | 0.280318 | -1.270647 | 0.203854 | 0.679010816 | DHTKD1 | protein_coding | 46.21 | -0.059883088 | 0.059883088 | ENSG00000181192 | n.s. |
| ENSG00000127423 | 245.0064 | -0.350471181 | 0.272683 | -1.285268 | 0.198699 | 0.674743473 | AUNIP | protein_coding | 42.54 | -0.059881964 | 0.059881964 | ENSG00000127423 | n.s. |
| ENSG00000135040 | 801.4583 | -0.339238057 | 0.259689 | -1.306323 | 0.191443 | 0.666197179 | NAA35 | protein_coding | 37.88 | -0.059840647 | 0.059840647 | ENSG00000135040 | n.s. |
| ENSG00000276404 | 13.81487 | 0.468549284 | 0.436927 | 1.0723746 | 0.283552 | 0.745300357 | MIR6835 | miRNA | 46.88 | 0.059819064 | 0.059819064 | ENSG00000276404 | n.s. |
| ENSG00000197619 | 340.234 | -0.352576905 | 0.274969 | -1.282242 | 0.199758 | 0.676613539 | ZNF615 | protein_coding | 39.06 | -0.059817957 | 0.059817957 | ENSG00000197619 | n.s. |
| ENSG00000130723 | 4094.048 | -0.315538139 | 0.234297 | -1.346742 | 0.178063 | 0.646388678 | PRRC2B | protein_coding | 49.02 | -0.059796453 | 0.059796453 | ENSG00000130723 | n.s. |
| ENSG00000236519 | 42.75712 | -0.379093576 | 0.308532 | -1.228702 | 0.219183 | 0.695448706 | LINC01424 | antisense | 45.59 | -0.059796286 | 0.059796286 | ENSG00000236519 | n.s. |
| ENSG00000108433 | 1523.351 | -0.339694061 | 0.260228 | -1.305371 | 0.191767 | 0.666819478 | GOSR2 | protein_coding | 47.31 | -0.059783343 | 0.059783343 | ENSG00000108433 | n.s. |
| ENSG00000204623 | 210.4555 | -0.307074557 | 0.224001 | -1.370863 | 0.170418 | 0.638865917 | ZNRD1ASP | anscribed_unitary_pseudoge | 41.59 | -0.059753724 | 0.059753724 | ENSG00000204623 | n.s. |
| ENSG00000136371 | 67.53412 | -0.380642668 | 0.311075 | -1.223635 | 0.22109 | 0.696894187 | MTHFS | protein_coding | 40.35 | -0.059697392 | 0.059697392 | ENSG00000136371 | n.s. |
| ENSG00000274213 | 21.33479 | 0.496792537 | 0.47836 | 1.0385329 | 0.299022 | 0.758335061 | | lincRNA | 63.51 | 0.059684091 | 0.059684091 | ENSG00000274213 | n.s. |
| ENSG00000196437 | 560.6268 | -0.335114374 | 0.255182 | -1.313239 | 0.189103 | 0.663694184 | ZNF569 | protein_coding | 39.27 | -0.059661078 | 0.059661078 | ENSG00000196437 | n.s. |
| ENSG00000141699 | 826.2522 | -0.324083644 | 0.243505 | -1.330913 | 0.183218 | 0.654498372 | RETREG3 | protein_coding | 46.4 | -0.059661021 | 0.059661021 | ENSG00000141699 | n.s. |
| ENSG00000138592 | 1964.321 | -0.329339319 | 0.249063 | -1.322312 | 0.186064 | 0.658994794 | USP8 | protein_coding | 39.77 | -0.059649284 | 0.059649284 | ENSG00000138592 | n.s. |
| ENSG00000201998 | 52.42635 | 0.391173247 | 0.32738 | 1.1948612 | 0.232141 | 0.704043192 | SNORA23 | snoRNA | 46.15 | 0.059615075 | 0.059615075 | ENSG00000201998 | n.s. |
| ENSG00000252906 | 25.87341 | -0.408346028 | 0.349418 | -1.168646 | 0.242546 | 0.714515282 | SCARNA3 | scaRNA | 46.85 | -0.059613815 | 0.059613815 | ENSG00000252906 | n.s. |
| ENSG00000148925 | 379.7023 | -0.363314277 | 0.289635 | -1.254389 | 0.209701 | 0.685367285 | BTBD10 | protein_coding | 36.22 | -0.059611382 | 0.059611382 | ENSG00000148925 | n.s. |
| ENSG00000140937 | 44.75492 | -0.388269539 | 0.321759 | -1.206708 | 0.227545 | 0.70225867 | CDH11 | protein_coding | 40.38 | -0.059600496 | 0.059600496 | ENSG00000140937 | n.s. |

| | | | | | | | | | | | | | |
|---|---|---|---|---|---|---|---|---|---|---|---|---|---|
| ENSG00000204469 | 2621.73 | -0.305345158 | 0.222404 | -1.372933 | 0.169773 | 0.638037836 | PRRC2A | protein_coding | 53.44 | -0.059589197 | 0.059589197 | ENSG00000204469 | n.s. |
| ENSG00000093009 | 1196.363 | -0.352280346 | 0.275765 | -1.277468 | 0.201437 | 0.677503939 | CDC45 | protein_coding | 48.03 | -0.059566441 | 0.059566441 | ENSG00000093009 | n.s. |
| ENSG00000108306 | 456.308 | -0.328872623 | 0.24885 | -1.321568 | 0.186312 | 0.659050036 | FBXL20 | protein_coding | 43.28 | -0.059552785 | 0.059552785 | ENSG00000108306 | n.s. |
| ENSG00000265342 | 16.53985 | 0.462138788 | 0.427401 | 1.0812763 | 0.279574 | 0.743390786 | | antisense | 48.16 | 0.059515539 | 0.059515539 | ENSG00000265342 | n.s. |
| ENSG00000105576 | 1737.601 | -0.331469591 | 0.251559 | -1.31766 | 0.187617 | 0.661470368 | TNPO2 | protein_coding | 54.01 | -0.059495346 | 0.059495346 | ENSG00000105576 | n.s. |
| ENSG00000150961 | 50.54231 | -0.383245263 | 0.314891 | -1.217075 | 0.223576 | 0.699734492 | SEC24D | protein_coding | 38.92 | -0.059428585 | 0.059428585 | ENSG00000150961 | n.s. |
| ENSG00000143294 | 693.4834 | -0.331732798 | 0.252157 | -1.315578 | 0.188316 | 0.662268051 | PRCC | protein_coding | 46.54 | -0.059368957 | 0.059368957 | ENSG00000143294 | n.s. |
| ENSG00000145901 | 58.61619 | 0.378735215 | 0.309612 | 1.2232583 | 0.221232 | 0.697038953 | TNIP1 | protein_coding | 50.01 | 0.059364075 | 0.059364075 | ENSG00000145901 | n.s. |
| ENSG00000262652 | 53.28177 | -0.389510993 | 0.325525 | -1.196564 | 0.231477 | 0.704043192 | | sense_intronic | 45.29 | -0.059361747 | 0.059361747 | ENSG00000262652 | n.s. |
| ENSG00000224165 | 67.83987 | 0.371722056 | 0.30037 | 1.2375488 | 0.215883 | 0.692499549 | DNAJC27-AS1 | antisense | 45.18 | 0.059319593 | 0.059319593 | ENSG00000224165 | n.s. |
| ENSG00000112039 | 280.512 | -0.359504296 | 0.285945 | -1.257249 | 0.208664 | 0.683940032 | FANCE | protein_coding | 50.42 | -0.059311729 | 0.059311729 | ENSG00000112039 | n.s. |
| ENSG00000201500 | 8.414398 | 0.488138495 | 0.467962 | 1.0431164 | 0.296894 | 0.756014066 | | snoRNA | 39.74 | 0.059294244 | 0.059294244 | ENSG00000201500 | n.s. |
| ENSG00000116205 | 538.8103 | -0.314207118 | 0.233626 | -1.344914 | 0.178653 | 0.64775724 | TCEANC2 | protein_coding | 42.31 | -0.059255605 | 0.059255605 | ENSG00000116205 | n.s. |
| ENSG00000279203 | 37.92953 | -0.388808498 | 0.325 | -1.196335 | 0.231566 | 0.704043192 | | TEC | 39.46 | -0.059254686 | 0.059254686 | ENSG00000279203 | n.s. |
| ENSG00000213221 | 142.5678 | -0.359112162 | 0.285632 | -1.257255 | 0.208661 | 0.683940032 | DNLZ | protein_coding | 64.59 | -0.059247034 | 0.059247034 | ENSG00000213221 | n.s. |
| ENSG00000261423 | 99.89168 | 0.350104017 | 0.274142 | 1.2770908 | 0.20157 | 0.677624898 | TMEM202-AS1 | lincRNA | 40.16 | 0.059171306 | 0.059171306 | ENSG00000261423 | n.s. |
| ENSG00000268471 | 570.6515 | 0.293779575 | 0.209831 | 1.4000738 | 0.161491 | 0.628931954 | | lincRNA | 43.57 | 0.059166131 | 0.059166131 | ENSG00000268471 | n.s. |
| ENSG00000174206 | 224.66 | -0.306045274 | 0.224404 | -1.363814 | 0.172626 | 0.640837445 | C12orf66 | protein_coding | 39.91 | -0.059143898 | 0.059143898 | ENSG00000174206 | n.s. |
| ENSG00000115325 | 299.4357 | -0.307151062 | 0.225924 | -1.359531 | 0.173978 | 0.642002364 | DOK1 | protein_coding | 58.42 | -0.059115329 | 0.059115329 | ENSG00000115325 | n.s. |
| ENSG00000176108 | 223.4653 | -0.349537839 | 0.273569 | -1.277696 | 0.201356 | 0.677503939 | CHMP6 | protein_coding | 58.37 | -0.059102715 | 0.059102715 | ENSG00000176108 | n.s. |
| ENSG00000274191 | 28.50138 | -0.414840872 | 0.360287 | -1.151418 | 0.24956 | 0.720401565 | | lincRNA | 43.93 | -0.059083857 | 0.059083857 | ENSG00000274191 | n.s. |
| ENSG00000272478 | 19.57105 | -0.43134124 | 0.382803 | -1.126797 | 0.259828 | 0.729499422 | | antisense | 45.5 | -0.059082987 | 0.059082987 | ENSG00000272478 | n.s. |
| ENSG00000184162 | 351.5342 | -0.342251845 | 0.265199 | -1.290546 | 0.196861 | 0.672118672 | NR2C2AP | protein_coding | 56.79 | -0.059056938 | 0.059056938 | ENSG00000184162 | n.s. |
| ENSG00000253596 | 25.45352 | 0.425396983 | 0.374847 | 1.1348542 | 0.256436 | 0.726402587 | | antisense | 36.36 | 0.059054723 | 0.059054723 | ENSG00000253596 | n.s. |
| ENSG00000229536 | 14.78754 | -0.48534125 | 0.46495 | -1.043857 | 0.296552 | 0.755765348 | | lincRNA | 43.98 | -0.059023817 | 0.059023817 | ENSG00000229536 | n.s. |
| ENSG00000222011 | 133.7794 | -0.375884863 | 0.30707 | -1.2241 | 0.220914 | 0.696826406 | FAM185A | protein_coding | 37.7 | -0.058967088 | 0.058967088 | ENSG00000222011 | n.s. |
| ENSG00000143207 | 1519.051 | -0.334147685 | 0.255596 | -1.307327 | 0.191102 | 0.666152146 | RFWD2 | protein_coding | 36.32 | -0.058952529 | 0.058952529 | ENSG00000143207 | n.s. |
| ENSG00000267587 | 57.662 | -0.398284033 | 0.338295 | -1.177328 | 0.239064 | 0.711216825 | | processed_pseudogene | 48.68 | -0.058945232 | 0.058945232 | ENSG00000267587 | n.s. |
| ENSG00000264419 | 4.054385 | -0.645045723 | 0.737746 | -0.874347 | 0.38193 | 0.81032471 | MIR548AC | miRNA | 32.95 | -0.058919068 | 0.058919068 | ENSG00000264419 | n.s. |
| ENSG00000123374 | 1485.009 | -0.336606975 | 0.259041 | -1.299435 | 0.193795 | 0.668296258 | CDK2 | protein_coding | 47.74 | -0.058916645 | 0.058916645 | ENSG00000123374 | n.s. |
| ENSG00000283929 | 24.77013 | -0.416926372 | 0.363354 | -1.14744 | 0.2512 | 0.722279621 | MIR5010 | miRNA | 54.17 | -0.058909461 | 0.058909461 | ENSG00000283929 | n.s. |
| ENSG00000163755 | 659.1752 | -0.347635554 | 0.271283 | -1.281449 | 0.200036 | 0.676981321 | HPS3 | protein_coding | 38.69 | -0.058897567 | 0.058897567 | ENSG00000163755 | n.s. |

| | | | | | | | | | | | | | |
|---|---|---|---|---|---|---|---|---|---|---|---|---|---|
| ENSG00000168724 | 1372.284 | -0.3362543 | 0.258626 | -1.300156 | 0.193547 | 0.668123216 | DNAJC21 | protein_coding | 39.24 | -0.058892734 | 0.058892734 | ENSG00000168724 | n.s. |
| ENSG00000074621 | 36.97451 | -0.397871646 | 0.337987 | -1.177182 | 0.239123 | 0.711226106 | SLC24A1 | protein_coding | 44.73 | -0.058881944 | 0.058881944 | ENSG00000074621 | n.s. |
| ENSG00000257390 | 64.54757 | 0.355175912 | 0.281567 | 1.261427 | 0.207155 | 0.682685326 | | protein_coding | 42.96 | 0.058880861 | 0.058880861 | ENSG00000257390 | n.s. |
| ENSG00000263439 | 12.45127 | -0.458851367 | 0.426186 | -1.076646 | 0.281638 | 0.744567264 | MIR4753 | miRNA | 38.55 | -0.058777053 | 0.058777053 | ENSG00000263439 | n.s. |
| ENSG00000221036 | 2.988054 | -0.644974947 | 0.740612 | -0.870868 | 0.383826 | 0.810719585 | MIR1193 | miRNA | 55.13 | -0.058776138 | 0.058776138 | ENSG00000221036 | n.s. |
| ENSG00000272735 | 37.04163 | 0.410544031 | 0.355327 | 1.1553992 | 0.247927 | 0.719195543 | | lincRNA | 53.38 | 0.058770615 | 0.058770615 | ENSG00000272735 | n.s. |
| ENSG00000100167 | 647.2072 | -0.333102757 | 0.254821 | -1.3072 | 0.191145 | 0.666152146 | 3-Sep | protein_coding | 51.53 | -0.058768176 | 0.058768176 | ENSG00000100167 | n.s. |
| ENSG00000243260 | 34.64868 | -0.396536628 | 0.336497 | -1.178426 | 0.238627 | 0.710909549 | RN7SL558P | misc_RNA | 49.5 | -0.058761039 | 0.058761039 | ENSG00000243260 | n.s. |
| ENSG00000116199 | 1843.809 | -0.340247192 | 0.263267 | -1.292406 | 0.196217 | 0.672118672 | FAM20B | protein_coding | 43.17 | -0.058711027 | 0.058711027 | ENSG00000116199 | n.s. |
| ENSG00000229955 | 33.63795 | -0.403909923 | 0.346686 | -1.165061 | 0.243994 | 0.715606021 | | sense_intronic | 36.7 | -0.058698619 | 0.058698619 | ENSG00000229955 | n.s. |
| ENSG00000223482 | 279.0231 | -0.361982125 | 0.289976 | -1.248319 | 0.211914 | 0.688575085 | NUTM2A-AS1 | antisense | 41.04 | -0.058658731 | 0.058658731 | ENSG00000223482 | n.s. |
| ENSG00000010438 | 49.76034 | -0.507816361 | 0.50307 | -1.009435 | 0.312766 | 0.766641623 | PRSS3 | protein_coding | 42.84 | -0.05860587 | 0.05860587 | ENSG00000010438 | n.s. |
| ENSG00000160199 | 392.7281 | -0.34237743 | 0.265905 | -1.287592 | 0.197888 | 0.674264807 | PKNOX1 | protein_coding | 46.99 | -0.058604576 | 0.058604576 | ENSG00000160199 | n.s. |
| ENSG00000118894 | 103.7298 | -0.339497377 | 0.262701 | -1.292334 | 0.196241 | 0.672118672 | EEF2KMT | protein_coding | 53.82 | -0.058581644 | 0.058581644 | ENSG00000118894 | n.s. |
| ENSG00000253218 | 51.80029 | -0.382139312 | 0.317138 | -1.204964 | 0.228217 | 0.702737638 | KLF3P1 | processed_pseudogene | 48.92 | -0.058546336 | 0.058546336 | ENSG00000253218 | n.s. |
| ENSG00000223039 | 27.29349 | -0.41772345 | 0.366587 | -1.139493 | 0.254498 | 0.72418384 | RN7SKP268 | misc_RNA | 52.85 | -0.05854443 | 0.05854443 | ENSG00000223039 | n.s. |
| ENSG00000148297 | 518.5074 | -0.354425798 | 0.281913 | -1.257218 | 0.208675 | 0.683940032 | MED22 | protein_coding | 56.94 | -0.058473868 | 0.058473868 | ENSG00000148297 | n.s. |
| ENSG00000106133 | 180.8645 | -0.352773385 | 0.279898 | -1.260365 | 0.207538 | 0.682785213 | NSUN5P2 | cribed_unprocessed_pseudc | 52.77 | -0.058460157 | 0.058460157 | ENSG00000106133 | n.s. |
| ENSG00000100503 | 2545.585 | -0.31829992 | 0.239661 | -1.328126 | 0.184136 | 0.655193675 | NIN | protein_coding | 41.15 | -0.058449511 | 0.058449511 | ENSG00000100503 | n.s. |
| ENSG00000283200 | 8.60478 | -0.491316306 | 0.475283 | -1.033734 | 0.301261 | 0.760388234 | MIR1206 | miRNA | 42.37 | -0.058449254 | 0.058449254 | ENSG00000283200 | n.s. |
| ENSG00000101152 | 717.1837 | -0.336768715 | 0.259966 | -1.295433 | 0.195171 | 0.670750289 | DNAJC5 | protein_coding | 51.37 | -0.058408873 | 0.058408873 | ENSG00000101152 | n.s. |
| ENSG00000197119 | 902.2724 | -0.347336853 | 0.273213 | -1.271303 | 0.203621 | 0.679010816 | SLC25A29 | protein_coding | 57.04 | -0.058395421 | 0.058395421 | ENSG00000197119 | n.s. |
| ENSG00000105771 | 178.3128 | -0.349295201 | 0.27555 | -1.267627 | 0.204931 | 0.680575501 | SMG9 | protein_coding | 50.25 | -0.058375502 | 0.058375502 | ENSG00000105771 | n.s. |
| ENSG00000233924 | 50.23675 | -0.37898514 | 0.31375 | -1.207922 | 0.227077 | 0.701641249 | RPSAP13 | processed_pseudogene | 49.6 | -0.058320085 | 0.058320085 | ENSG00000233924 | n.s. |
| ENSG00000106608 | 86.03649 | -0.362051096 | 0.291064 | -1.243886 | 0.213541 | 0.690180176 | URGCP | protein_coding | 45.69 | -0.05830381 | 0.05830381 | ENSG00000106608 | n.s. |
| ENSG00000129680 | 530.4002 | -0.362007165 | 0.290936 | -1.244283 | 0.213395 | 0.690152016 | MAP7D3 | protein_coding | 40.66 | -0.05830315 | 0.05830315 | ENSG00000129680 | n.s. |
| ENSG00000148356 | 142.3702 | 0.364564815 | 0.294615 | 1.2374278 | 0.215928 | 0.692499549 | LRSAM1 | protein_coding | 51.44 | 0.058177437 | 0.058177437 | ENSG00000148356 | n.s. |
| ENSG00000105011 | 331.3869 | -0.370360695 | 0.302386 | -1.224795 | 0.220653 | 0.696655251 | ASF1B | protein_coding | 51.11 | -0.058139994 | 0.058139994 | ENSG00000105011 | n.s. |
| ENSG00000104853 | 362.4879 | -0.343820187 | 0.268779 | -1.279194 | 0.200829 | 0.677491543 | CLPTM1 | protein_coding | 52.83 | -0.05813866 | 0.05813866 | ENSG00000104853 | n.s. |
| ENSG00000188573 | 352.7003 | -0.368409208 | 0.29971 | -1.229217 | 0.21899 | 0.695448706 | FBLL1 | protein_coding | 69.39 | -0.058110988 | 0.058110988 | ENSG00000188573 | n.s. |
| ENSG00000092978 | 1301.2 | -0.328121221 | 0.250601 | -1.309335 | 0.190421 | 0.665135717 | GPATCH2 | protein_coding | 36.36 | -0.058106899 | 0.058106899 | ENSG00000092978 | n.s. |
| ENSG00000272030 | 38.67354 | -0.386185807 | 0.325093 | -1.187923 | 0.234864 | 0.707216633 | | antisense | 53.92 | -0.058100702 | 0.058100702 | ENSG00000272030 | n.s. |

| | | | | | | | | | | | | | |
|---|---|---|---|---|---|---|---|---|---|---|---|---|---|
| ENSG00000273559 | 401.6708 | -0.330985762 | 0.254149 | -1.302331 | 0.192803 | 0.667523235 | CWC25 | protein_coding | 47.75 | -0.058099127 | 0.058099127 | ENSG00000273559 | n.s. |
| ENSG00000198974 | 6.030316 | -0.555639465 | 0.583728 | -0.951881 | 0.341157 | 0.786054116 | MIR30E | miRNA | 51.09 | -0.058090747 | 0.058090747 | ENSG00000198974 | n.s. |
| ENSG00000130429 | 115.9966 | -0.374586824 | 0.307823 | -1.216891 | 0.223646 | 0.699734492 | ARPC1B | protein_coding | 56.01 | -0.058085949 | 0.058085949 | ENSG00000130429 | n.s. |
| ENSG00000250151 | 29.12938 | -0.422451611 | 0.374507 | -1.128021 | 0.259311 | 0.72892993 | ARPC4-TTLL3 | protein_coding | 49.96 | -0.058008612 | 0.058008612 | ENSG00000250151 | n.s. |
| ENSG00000258920 | 47.97752 | -0.378445206 | 0.314438 | -1.20356 | 0.22876 | 0.702737638 | FOXN3-AS1 | antisense | 54.67 | -0.057980373 | 0.057980373 | ENSG00000258920 | n.s. |
| ENSG00000013503 | 912.7763 | -0.326708914 | 0.249323 | -1.310386 | 0.190065 | 0.664559877 | POLR3B | protein_coding | 39.75 | -0.057979686 | 0.057979686 | ENSG00000013503 | n.s. |
| ENSG00000076043 | 423.0306 | 0.350465415 | 0.278232 | 1.2596161 | 0.207808 | 0.683351224 | REXO2 | protein_coding | 39.61 | 0.057951567 | 0.057951567 | ENSG00000076043 | n.s. |
| ENSG00000229492 | 76.17243 | -0.373018814 | 0.30624 | -1.21806 | 0.223201 | 0.699286493 | | processed_pseudogene | 56.54 | -0.057946556 | 0.057946556 | ENSG00000229492 | n.s. |
| ENSG00000231213 | 239.437 | 0.360450035 | 0.290113 | 1.2424478 | 0.214071 | 0.690827094 | PLSCR5 | protein_coding | 33.23 | 0.057899319 | 0.057899319 | ENSG00000231213 | n.s. |
| ENSG00000279602 | 2286.881 | 0.288295677 | 0.207331 | 1.3905107 | 0.164374 | 0.63030432 | | TEC | 46.4 | 0.057788787 | 0.057788787 | ENSG00000279602 | n.s. |
| ENSG00000204394 | 1346.758 | -0.323268375 | 0.245885 | -1.314714 | 0.188606 | 0.662619911 | VARS | protein_coding | 53.81 | -0.05777954 | 0.05777954 | ENSG00000204394 | n.s. |
| ENSG00000141985 | 442.7648 | -0.344371421 | 0.271179 | -1.269903 | 0.204119 | 0.679567798 | SH3GL1 | protein_coding | 57.17 | -0.057774232 | 0.057774232 | ENSG00000141985 | n.s. |
| ENSG00000137571 | 543.1902 | -0.329881504 | 0.253785 | -1.299846 | 0.193654 | 0.668213857 | SLCO5A1 | protein_coding | 40.04 | -0.057757146 | 0.057757146 | ENSG00000137571 | n.s. |
| ENSG00000134030 | 103.8456 | -0.332851316 | 0.2571 | -1.294638 | 0.195445 | 0.670750289 | CTIF | protein_coding | 50.34 | -0.057729443 | 0.057729443 | ENSG00000134030 | n.s. |
| ENSG00000257004 | 16.35537 | 0.478174157 | 0.459482 | 1.040682 | 0.298023 | 0.757323737 | | lincRNA | 41.37 | 0.057724434 | 0.057724434 | ENSG00000257004 | n.s. |
| ENSG00000197321 | 38.81381 | -0.39511847 | 0.337645 | -1.170217 | 0.241914 | 0.714350709 | SVIL | protein_coding | 43.61 | -0.057722272 | 0.057722272 | ENSG00000197321 | n.s. |
| ENSG00000115935 | 151.6133 | -0.341457712 | 0.267447 | -1.27673 | 0.201698 | 0.677624898 | WIPF1 | protein_coding | 41.82 | -0.057709988 | 0.057709988 | ENSG00000115935 | n.s. |
| ENSG00000103018 | 1497.987 | -0.367519793 | 0.300137 | -1.224506 | 0.220762 | 0.696659758 | CYB5B | protein_coding | 39.78 | -0.057692991 | 0.057692991 | ENSG00000103018 | n.s. |
| ENSG00000198300 | 969.6172 | -0.311971287 | 0.234027 | -1.333058 | 0.182513 | 0.653245015 | PEG3 | protein_coding | 45.75 | -0.057690946 | 0.057690946 | ENSG00000198300 | n.s. |
| ENSG00000264994 | 46.69217 | 0.374210578 | 0.309287 | 1.2099131 | 0.226312 | 0.701289341 | SNORD92 | snoRNA | 44.71 | 0.057666884 | 0.057666884 | ENSG00000264994 | n.s. |
| ENSG00000145945 | 245.646 | -0.334023386 | 0.258517 | -1.292075 | 0.196331 | 0.672118672 | FAM50B | protein_coding | 58.75 | -0.057637085 | 0.057637085 | ENSG00000145945 | n.s. |
| ENSG00000177463 | 1444.239 | -0.324253546 | 0.247405 | -1.310617 | 0.189987 | 0.66445359 | NR2C2 | protein_coding | 42.09 | -0.057566466 | 0.057566466 | ENSG00000177463 | n.s. |
| ENSG00000227248 | 98.5617 | -0.381519403 | 0.320589 | -1.190056 | 0.234024 | 0.706508138 | FAM155A-IT1 | sense_intronic | 35.94 | -0.057564728 | 0.057564728 | ENSG00000227248 | n.s. |
| ENSG00000235545 | 54.8348 | -0.372526254 | 0.306938 | -1.213685 | 0.224868 | 0.700612508 | | antisense | 42.23 | -0.057563544 | 0.057563544 | ENSG00000235545 | n.s. |
| ENSG00000251562 | 44817.14 | -0.292587786 | 0.212276 | -1.378336 | 0.1681 | 0.63576037 | MALAT1 | lincRNA | 40.27 | -0.057553933 | 0.057553933 | ENSG00000251562 | n.s. |
| ENSG00000125970 | 1871.406 | -0.340271166 | 0.266295 | -1.277799 | 0.20132 | 0.677503939 | RALY | protein_coding | 44.86 | -0.057535831 | 0.057535831 | ENSG00000125970 | n.s. |
| ENSG00000245888 | 96.77449 | 0.350510274 | 0.279475 | 1.2541725 | 0.209779 | 0.685462895 | | antisense | 44.09 | 0.05748931 | 0.05748931 | ENSG00000245888 | n.s. |
| ENSG00000132819 | 350.2183 | -0.330775373 | 0.255324 | -1.295514 | 0.195143 | 0.670750289 | RBM38 | protein_coding | 59.97 | -0.057369394 | 0.057369394 | ENSG00000132819 | n.s. |
| ENSG00000129946 | 256.0549 | -0.363580082 | 0.29581 | -1.229101 | 0.219034 | 0.695448706 | SHC2 | protein_coding | 58.69 | -0.057349267 | 0.057349267 | ENSG00000129946 | n.s. |
| ENSG00000239306 | 2039.615 | -0.285950266 | 0.205958 | -1.388391 | 0.165018 | 0.63030432 | RBM14 | protein_coding | 50.54 | -0.05731865 | 0.05731865 | ENSG00000239306 | n.s. |
| ENSG00000152223 | 619.6737 | -0.312671236 | 0.235714 | -1.326488 | 0.184678 | 0.655769671 | EPG5 | protein_coding | 40.87 | -0.057296589 | 0.057296589 | ENSG00000152223 | n.s. |
| ENSG00000167930 | 281.4757 | -0.342672789 | 0.270401 | -1.267275 | 0.205057 | 0.680575501 | FAM234A | protein_coding | 52.35 | -0.05726874 | 0.05726874 | ENSG00000167930 | n.s. |

| | | | | | | | | | | | | | |
|---|---|---|---|---|---|---|---|---|---|---|---|---|---|
| ENSG00000119401 | 362.5091 | -0.33509091 | 0.260573 | -1.285975 | 0.198452 | 0.674743473 | TRIM32 | protein_coding | 43.3 | -0.057254071 | 0.057254071 | ENSG00000119401 | n.s. |
| ENSG00000185803 | 202.5304 | -0.353050318 | 0.282707 | -1.248822 | 0.21173 | 0.688445029 | SLC52A2 | protein_coding | 65.97 | -0.057240306 | 0.057240306 | ENSG00000185803 | n.s. |
| ENSG00000210082 | 5352.061 | -0.342174649 | 0.269798 | -1.268263 | 0.204704 | 0.680538672 | MT-RNR2 | Mt_rRNA | 42.81 | -0.057193531 | 0.057193531 | ENSG00000210082 | n.s. |
| ENSG00000226964 | 14.80939 | 0.489124548 | 0.481591 | 1.0156438 | 0.309799 | 0.763980297 | RHEBP2 | processed_pseudogene | 40.67 | 0.057187387 | 0.057187387 | ENSG00000226964 | n.s. |
| ENSG00000143748 | 1613.163 | -0.33456872 | 0.260371 | -1.28497 | 0.198803 | 0.674775366 | NVL | protein_coding | 41.59 | -0.057157981 | 0.057157981 | ENSG00000143748 | n.s. |
| ENSG00000223837 | 55.61382 | -0.356596175 | 0.287244 | -1.241439 | 0.214444 | 0.691453912 | | sense_intronic | 58.47 | -0.057139816 | 0.057139816 | ENSG00000223837 | n.s. |
| ENSG00000132680 | 3168.556 | -0.307010952 | 0.229688 | -1.336645 | 0.181338 | 0.651554717 | KIAA0907 | protein_coding | 42.57 | -0.057119113 | 0.057119113 | ENSG00000132680 | n.s. |
| ENSG00000251774 | 11.14482 | -0.463095845 | 0.440127 | -1.052186 | 0.292714 | 0.752972779 | RNU6-377P | snRNA | 40.2 | -0.057063005 | 0.057063005 | ENSG00000251774 | n.s. |
| ENSG00000206698 | 18.46619 | -0.445071822 | 0.41449 | -1.073783 | 0.28292 | 0.744776963 | RNU1-73P | snRNA | 45.12 | -0.056957517 | 0.056957517 | ENSG00000206698 | n.s. |
| ENSG00000241015 | 302.1537 | -0.341316622 | 0.270009 | -1.264095 | 0.206196 | 0.681106972 | TPM3P9 | ıscribed_processed_pseudog | 45.51 | -0.056926382 | 0.056926382 | ENSG00000241015 | n.s. |
| ENSG00000109062 | 184.1829 | -0.336545443 | 0.263172 | -1.278805 | 0.200966 | 0.677493231 | SLC9A3R1 | protein_coding | 56.2 | -0.056908165 | 0.056908165 | ENSG00000109062 | n.s. |
| ENSG00000205664 | 267.8761 | -0.334802512 | 0.261024 | -1.282653 | 0.199614 | 0.676374453 | | lincRNA | 49.39 | -0.056853753 | 0.056853753 | ENSG00000205664 | n.s. |
| ENSG00000167900 | 1127.099 | -0.356118571 | 0.287941 | -1.236777 | 0.21617 | 0.692499549 | TK1 | protein_coding | 53.2 | -0.056829581 | 0.056829581 | ENSG00000167900 | n.s. |
| ENSG00000075785 | 1070.03 | -0.339885664 | 0.26833 | -1.266668 | 0.205274 | 0.680704186 | RAB7A | protein_coding | 44.06 | -0.056775038 | 0.056775038 | ENSG00000075785 | n.s. |
| ENSG00000105325 | 300.9244 | -0.33967668 | 0.268004 | -1.267431 | 0.205001 | 0.680575501 | FZR1 | protein_coding | 60.71 | -0.05676802 | 0.05676802 | ENSG00000105325 | n.s. |
| ENSG00000187676 | 193.9483 | -0.350490721 | 0.280982 | -1.24738 | 0.212258 | 0.688714317 | B3GLCT | protein_coding | 41.08 | -0.056765789 | 0.056765789 | ENSG00000187676 | n.s. |
| ENSG00000062725 | 1897.352 | -0.357865773 | 0.290658 | -1.231228 | 0.218238 | 0.694065444 | APPBP2 | protein_coding | 39.69 | -0.056757361 | 0.056757361 | ENSG00000062725 | n.s. |
| ENSG00000162631 | 165.1836 | 0.321394709 | 0.245695 | 1.3081035 | 0.190838 | 0.665924932 | NTNG1 | protein_coding | 37.35 | 0.056750182 | 0.056750182 | ENSG00000162631 | n.s. |
| ENSG00000153487 | 449.98 | -0.313286323 | 0.237018 | -1.321784 | 0.18624 | 0.658994794 | ING1 | protein_coding | 49.86 | -0.056741797 | 0.056741797 | ENSG00000153487 | n.s. |
| ENSG00000249690 | 152.5579 | -0.322317514 | 0.246995 | -1.304953 | 0.191909 | 0.666819478 | | antisense | 45.58 | -0.056725215 | 0.056725215 | ENSG00000249690 | n.s. |
| ENSG00000104866 | 373.1012 | -0.341482401 | 0.270374 | -1.263002 | 0.206588 | 0.682201186 | PPP1R37 | protein_coding | 55.56 | -0.056715968 | 0.056715968 | ENSG00000104866 | n.s. |
| ENSG00000279356 | 100.3639 | -0.328639526 | 0.254584 | -1.290887 | 0.196743 | 0.672118672 | | TEC | 40.75 | -0.056708078 | 0.056708078 | ENSG00000279356 | n.s. |
| ENSG00000100796 | 2359.04 | -0.31844602 | 0.242477 | -1.313303 | 0.189081 | 0.663694184 | PPP4R3A | protein_coding | 40.03 | -0.056693578 | 0.056693578 | ENSG00000100796 | n.s. |
| ENSG00000106089 | 660.267 | -0.339397815 | 0.268139 | -1.265754 | 0.205601 | 0.680704186 | STX1A | protein_coding | 58.84 | -0.056693547 | 0.056693547 | ENSG00000106089 | n.s. |
| ENSG00000162433 | 497.4348 | -0.339167353 | 0.268012 | -1.265494 | 0.205694 | 0.680704186 | AK4 | protein_coding | 42.62 | -0.05665505 | 0.05665505 | ENSG00000162433 | n.s. |
| ENSG00000196449 | 591.5784 | 0.334921302 | 0.261935 | 1.2786451 | 0.201022 | 0.677503939 | YRDC | protein_coding | 49.48 | 0.056631232 | 0.056631232 | ENSG00000196449 | n.s. |
| ENSG00000258982 | 22.94109 | -0.439556871 | 0.406531 | -1.081239 | 0.279591 | 0.743390786 | | antisense | 48.69 | -0.056607376 | 0.056607376 | ENSG00000258982 | n.s. |
| ENSG00000158161 | 279.911 | -0.32936432 | 0.255562 | -1.288784 | 0.197473 | 0.673407427 | EYA3 | protein_coding | 39.52 | -0.056559132 | 0.056559132 | ENSG00000158161 | n.s. |
| ENSG00000197308 | 626.7433 | -0.29385726 | 0.216088 | -1.359894 | 0.173864 | 0.642002364 | GATA3-AS1 | lincRNA | 60.72 | -0.056556759 | 0.056556759 | ENSG00000197308 | n.s. |
| ENSG00000196705 | 2082.385 | -0.33838601 | 0.267224 | -1.266303 | 0.205405 | 0.680704186 | ZNF431 | protein_coding | 39.12 | -0.056524533 | 0.056524533 | ENSG00000196705 | n.s. |
| ENSG00000242073 | 29.74829 | -0.413507869 | 0.367413 | -1.125459 | 0.260395 | 0.730043745 | | transcribed_unprocessed_pseudo | 41.4 | -0.056506312 | 0.056506312 | ENSG00000242073 | n.s. |
| ENSG00000272667 | 137.4672 | -0.315922257 | 0.240191 | -1.315297 | 0.18841 | 0.662432976 | | lincRNA | 51.9 | -0.056505241 | 0.056505241 | ENSG00000272667 | n.s. |

| | | | | | | | | | | | | | |
|---|---|---|---|---|---|---|---|---|---|---|---|---|---|
| ENSG00000226801 | 38.15905 | -0.379413954 | 0.320886 | -1.182395 | 0.237049 | 0.709700242 | OSTCP8 | processed_pseudogene | 44.2 | -0.056504241 | 0.056504241 | ENSG00000226801 | n.s. |
| ENSG00000235088 | 37.93138 | -0.381670659 | 0.324409 | -1.176512 | 0.23939 | 0.71125159 | | processed_pseudogene | 47.74 | -0.056478384 | 0.056478384 | ENSG00000235088 | n.s. |
| ENSG00000238300 | 10.41952 | -0.460377882 | 0.438779 | -1.049224 | 0.294075 | 0.753915388 | SNORD121B | snoRNA | 40.74 | -0.056477958 | 0.056477958 | ENSG00000238300 | n.s. |
| ENSG00000252192 | 26.2347 | 0.405781171 | 0.357429 | 1.1352775 | 0.256259 | 0.726402587 | | snoRNA | 46.51 | 0.056331604 | 0.056331604 | ENSG00000252192 | n.s. |
| ENSG00000108094 | 785.8006 | -0.337190146 | 0.266128 | -1.267021 | 0.205148 | 0.680704186 | CUL2 | protein_coding | 39.72 | -0.056324774 | 0.056324774 | ENSG00000108094 | n.s. |
| ENSG00000271966 | 52.51285 | 0.361843328 | 0.296882 | 1.2188105 | 0.222916 | 0.698928583 | | lincRNA | 42.32 | 0.056290953 | 0.056290953 | ENSG00000271966 | n.s. |
| ENSG00000242707 | 22.80526 | 0.417996799 | 0.373163 | 1.1201454 | 0.262652 | 0.733524057 | RN7SL362P | misc_RNA | 51.69 | 0.056256366 | 0.056256366 | ENSG00000242707 | n.s. |
| ENSG00000163364 | 180.2651 | 0.354176578 | 0.287326 | 1.2326667 | 0.2177 | 0.693853089 | LINC01116 | lincRNA | 38.98 | 0.056219324 | 0.056219324 | ENSG00000163364 | n.s. |
| ENSG00000207654 | 8.924134 | -0.472930463 | 0.45787 | -1.032891 | 0.301655 | 0.760565055 | MIR128-1 | miRNA | 50 | -0.056214233 | 0.056214233 | ENSG00000207654 | n.s. |
| ENSG00000143995 | 21763.09 | -0.290865549 | 0.213376 | -1.363159 | 0.172832 | 0.640837445 | MEIS1 | protein_coding | 39.2 | -0.056210384 | 0.056210384 | ENSG00000143995 | n.s. |
| ENSG00000259467 | 45.59871 | 0.364363333 | 0.300432 | 1.2127961 | 0.225208 | 0.701091738 | NDUFAF4P1 | processed_pseudogene | 42 | 0.05619399 | 0.05619399 | ENSG00000259467 | n.s. |
| ENSG00000172748 | 239.6962 | -0.348801441 | 0.280309 | -1.244347 | 0.213372 | 0.690152016 | ZNF596 | protein_coding | 40.95 | -0.056176299 | 0.056176299 | ENSG00000172748 | n.s. |
| ENSG00000164011 | 107.0841 | -0.338807197 | 0.268596 | -1.261402 | 0.207164 | 0.682685326 | ZNF691 | protein_coding | 47.25 | -0.056167265 | 0.056167265 | ENSG00000164011 | n.s. |
| ENSG00000170340 | 640.0353 | -0.348084286 | 0.279408 | -1.245794 | 0.21284 | 0.689743466 | B3GNT2 | protein_coding | 45.4 | -0.056150313 | 0.056150313 | ENSG00000170340 | n.s. |
| ENSG00000180938 | 213.8208 | -0.299863484 | 0.223696 | -1.340496 | 0.180084 | 0.649773904 | ZNF572 | protein_coding | 39.84 | -0.056145759 | 0.056145759 | ENSG00000180938 | n.s. |
| ENSG00000100908 | 161.5738 | -0.346811522 | 0.278092 | -1.247112 | 0.212357 | 0.688872544 | EMC9 | protein_coding | 53.54 | -0.056135303 | 0.056135303 | ENSG00000100908 | n.s. |
| ENSG00000230525 | 117.6642 | 0.339957768 | 0.270263 | 1.2578765 | 0.208436 | 0.683940032 | LINC01799 | lincRNA | 38.12 | 0.056086904 | 0.056086904 | ENSG00000230525 | n.s. |
| ENSG00000167394 | 288.1224 | -0.331557441 | 0.259409 | -1.278124 | 0.201206 | 0.677503939 | ZNF668 | protein_coding | 54.5 | -0.056062443 | 0.056062443 | ENSG00000167394 | n.s. |
| ENSG00000200455 | 12.24196 | 0.472063 | 0.458202 | 1.0302519 | 0.302892 | 0.760781413 | RNU6-1112P | snRNA | 41.41 | 0.056052811 | 0.056052811 | ENSG00000200455 | n.s. |
| ENSG00000015532 | 283.7771 | -0.33340239 | 0.262323 | -1.270961 | 0.203743 | 0.679010816 | XYLT2 | protein_coding | 56.54 | -0.056052713 | 0.056052713 | ENSG00000015532 | n.s. |
| ENSG00000124275 | 1468.235 | -0.309276345 | 0.233833 | -1.322637 | 0.185956 | 0.658994794 | MTRR | protein_coding | 39.8 | -0.056015518 | 0.056015518 | ENSG00000124275 | n.s. |
| ENSG00000278396 | 28.37594 | -0.438448741 | 0.408941 | -1.072155 | 0.28365 | 0.745300357 | | sense_intronic | 37.36 | -0.055976168 | 0.055976168 | ENSG00000278396 | n.s. |
| ENSG00000100241 | 1341.795 | -0.319032978 | 0.245163 | -1.301312 | 0.193152 | 0.667684687 | SBF1 | protein_coding | 60.67 | -0.055967504 | 0.055967504 | ENSG00000100241 | n.s. |
| ENSG00000115750 | 1013.871 | -0.334985456 | 0.264514 | -1.266419 | 0.205363 | 0.680704186 | TAF1B | protein_coding | 39.53 | -0.055956499 | 0.055956499 | ENSG00000115750 | n.s. |
| ENSG00000267680 | 306.8681 | -0.330533518 | 0.258129 | -1.280499 | 0.20037 | 0.677286179 | ZNF224 | protein_coding | 39.97 | -0.055935456 | 0.055935456 | ENSG00000267680 | n.s. |
| ENSG00000199762 | 7.929275 | 0.559837708 | 0.605219 | 0.9250162 | 0.354957 | 0.794524781 | | misc_RNA | 38.46 | 0.055923618 | 0.055923618 | ENSG00000199762 | n.s. |
| ENSG00000260924 | 133.9423 | -0.330651225 | 0.259128 | -1.276014 | 0.201951 | 0.677624898 | LINC01311 | antisense | 63.11 | -0.055883577 | 0.055883577 | ENSG00000260924 | n.s. |
| ENSG00000023909 | 737.811 | -0.329808969 | 0.257463 | -1.280998 | 0.200195 | 0.67702417 | GCLM | protein_coding | 36.37 | -0.055868263 | 0.055868263 | ENSG00000023909 | n.s. |
| ENSG00000159208 | 183.6306 | 0.34818564 | 0.280391 | 1.241786 | 0.214316 | 0.691201113 | CIART | protein_coding | 47.53 | 0.055847435 | 0.055847435 | ENSG00000159208 | n.s. |
| ENSG00000270750 | 48.22341 | -0.355836359 | 0.290679 | -1.224154 | 0.220894 | 0.696826406 | | lincRNA | 34.73 | -0.055821971 | 0.055821971 | ENSG00000270750 | n.s. |
| ENSG00000130165 | 373.9094 | -0.358538587 | 0.293999 | -1.219524 | 0.222645 | 0.698928583 | ELOF1 | protein_coding | 55.91 | -0.055776844 | 0.055776844 | ENSG00000130165 | n.s. |
| ENSG00000174483 | 76.35521 | -0.360226213 | 0.296492 | -1.21496 | 0.224381 | 0.700148043 | BBS1 | protein_coding | 49.91 | -0.055766664 | 0.055766664 | ENSG00000174483 | n.s. |

| | | | | | | | | | | | | | |
|---|---|---|---|---|---|---|---|---|---|---|---|---|---|
| ENSG00000114757 | 177.5711 | 0.32046188 | 0.247095 | 1.2969201 | 0.194659 | 0.669854827 | PEX5L | protein_coding | 38.14 | 0.055766555 | 0.055766555 | ENSG00000114757 | n.s. |
| ENSG00000283146 | 3.623768 | -0.630287011 | 0.73705 | -0.855148 | 0.392469 | 0.815710819 | MIR548AU | miRNA | 35.19 | -0.05575757 | 0.05575757 | ENSG00000283146 | n.s. |
| ENSG00000139974 | 41.50322 | -0.371231475 | 0.312934 | -1.186292 | 0.235507 | 0.70779073 | SLC38A6 | protein_coding | 37.62 | -0.055720036 | 0.055720036 | ENSG00000139974 | n.s. |
| ENSG00000169105 | 265.5351 | -0.31828465 | 0.244934 | -1.299469 | 0.193783 | 0.668296258 | CHST14 | protein_coding | 57.11 | -0.055709671 | 0.055709671 | ENSG00000169105 | n.s. |
| ENSG00000106785 | 189.0036 | -0.349013591 | 0.281842 | -1.23833 | 0.215594 | 0.692470486 | TRIM14 | protein_coding | 46.67 | -0.055702127 | 0.055702127 | ENSG00000106785 | n.s. |
| ENSG00000241362 | 14.97086 | 0.438087793 | 0.41055 | 1.0670742 | 0.285938 | 0.746239075 | | processed_pseudogene | 44.63 | 0.055690602 | 0.055690602 | ENSG00000241362 | n.s. |
| ENSG00000138758 | 3115.712 | -0.322626785 | 0.24973 | -1.291902 | 0.196391 | 0.672118672 | 11-Sep | protein_coding | 40.66 | -0.055670555 | 0.055670555 | ENSG00000138758 | n.s. |
| ENSG00000158691 | 1108.903 | -0.289648038 | 0.213621 | -1.355896 | 0.175132 | 0.642395686 | ZSCAN12 | protein_coding | 41.13 | -0.055669595 | 0.055669595 | ENSG00000158691 | n.s. |
| ENSG00000160781 | 169.7745 | -0.352832797 | 0.287026 | -1.229272 | 0.21897 | 0.695448706 | PAQR6 | protein_coding | 60.95 | -0.055654045 | 0.055654045 | ENSG00000160781 | n.s. |
| ENSG00000196482 | 1933.263 | -0.325453577 | 0.252898 | -1.286898 | 0.19813 | 0.674626594 | ESRRG | protein_coding | 38.06 | -0.055631909 | 0.055631909 | ENSG00000196482 | n.s. |
| ENSG00000196867 | 429.7812 | -0.337983679 | 0.269215 | -1.255442 | 0.209318 | 0.68461467 | ZFP28 | protein_coding | 40.78 | -0.055616498 | 0.055616498 | ENSG00000196867 | n.s. |
| ENSG00000275854 | 67.22735 | -0.377592493 | 0.321969 | -1.17276 | 0.240892 | 0.712813552 | | sense_intronic | 29.27 | -0.055515178 | 0.055515178 | ENSG00000275854 | n.s. |
| ENSG00000146433 | 2843.662 | -0.337285147 | 0.268656 | -1.255456 | 0.209313 | 0.68461467 | TMEM181 | protein_coding | 47.38 | -0.055501552 | 0.055501552 | ENSG00000146433 | n.s. |
| ENSG00000163888 | 109.1595 | -0.345937117 | 0.278777 | -1.240911 | 0.214639 | 0.691472186 | CAMK2N2 | protein_coding | 68.28 | -0.055427873 | 0.055427873 | ENSG00000163888 | n.s. |
| ENSG00000229587 | 45.31013 | -0.414504909 | 0.373126 | -1.110897 | 0.266613 | 0.734987182 | | sense_intronic | 38.86 | -0.055427694 | 0.055427694 | ENSG00000229587 | n.s. |
| ENSG00000165985 | 352.2535 | 0.335350024 | 0.266347 | 1.2590727 | 0.208004 | 0.683512059 | C1QL3 | protein_coding | 44.9 | 0.055417872 | 0.055417872 | ENSG00000165985 | n.s. |
| ENSG00000105063 | 593.6293 | -0.308701485 | 0.234205 | -1.318082 | 0.187476 | 0.661470368 | PPP6R1 | protein_coding | 56.47 | -0.055408708 | 0.055408708 | ENSG00000105063 | n.s. |
| ENSG00000261094 | 94.54033 | -0.37702643 | 0.321578 | -1.172425 | 0.241026 | 0.712940305 | | sense_overlapping | 37.03 | -0.05540284 | 0.05540284 | ENSG00000261094 | n.s. |
| ENSG00000065183 | 3933.484 | -0.32573283 | 0.253816 | -1.283344 | 0.199372 | 0.676054647 | WDR3 | protein_coding | 36.95 | -0.055380509 | 0.055380509 | ENSG00000065183 | n.s. |
| ENSG00000130311 | 396.713 | -0.343184964 | 0.275371 | -1.246266 | 0.212667 | 0.689718524 | DDA1 | protein_coding | 56.15 | -0.055365382 | 0.055365382 | ENSG00000130311 | n.s. |
| ENSG00000005175 | 1463.207 | -0.335189551 | 0.266327 | -1.258562 | 0.208189 | 0.683663723 | RPAP3 | protein_coding | 36.57 | -0.055359056 | 0.055359056 | ENSG00000005175 | n.s. |
| ENSG00000185164 | 158.1014 | -0.346485282 | 0.279667 | -1.238922 | 0.215374 | 0.692210945 | NOMO2 | protein_coding | 46.04 | -0.055355021 | 0.055355021 | ENSG00000185164 | n.s. |
| ENSG00000147874 | 2577.097 | -0.348521146 | 0.282488 | -1.233757 | 0.217294 | 0.693734586 | HAUS6 | protein_coding | 39.7 | -0.055347477 | 0.055347477 | ENSG00000147874 | n.s. |
| ENSG00000261529 | 126.1184 | -0.328266664 | 0.257937 | -1.272662 | 0.203138 | 0.678409233 | | sense_intronic | 37.99 | -0.055315641 | 0.055315641 | ENSG00000261529 | n.s. |
| ENSG00000144909 | 477.7253 | -0.313518174 | 0.239901 | -1.306864 | 0.191259 | 0.666152146 | OSBPL11 | protein_coding | 39.18 | -0.055312936 | 0.055312936 | ENSG00000144909 | n.s. |
| ENSG00000053372 | 653.1839 | -0.334706024 | 0.265887 | -1.258826 | 0.208093 | 0.683643799 | MRTO4 | protein_coding | 53.49 | -0.055283434 | 0.055283434 | ENSG00000053372 | n.s. |
| ENSG00000251474 | 364.6136 | -0.309254256 | 0.235171 | -1.31502 | 0.188503 | 0.662593286 | RPL32P3 | cribed_unprocessed_pseudc | 50.11 | -0.055280116 | 0.055280116 | ENSG00000251474 | n.s. |
| ENSG00000214013 | 56.24985 | -0.355127209 | 0.291337 | -1.218957 | 0.222861 | 0.698928583 | GANC | protein_coding | 40.5 | -0.055246145 | 0.055246145 | ENSG00000214013 | n.s. |
| ENSG00000186522 | 821.5084 | -0.345855251 | 0.279241 | -1.238553 | 0.215511 | 0.692470486 | 10-Sep | protein_coding | 40.38 | -0.055198059 | 0.055198059 | ENSG00000186522 | n.s. |
| ENSG00000136819 | 808.5561 | -0.317659192 | 0.245042 | -1.296347 | 0.194856 | 0.670367996 | C9orf78 | protein_coding | 48.01 | -0.055173185 | 0.055173185 | ENSG00000136819 | n.s. |
| ENSG00000255076 | 13.05825 | -0.478374479 | 0.474953 | -1.007203 | 0.313837 | 0.766827179 | | sense_intronic | 40.37 | -0.055157774 | 0.055157774 | ENSG00000255076 | n.s. |
| ENSG00000188612 | 7174.635 | -0.370332082 | 0.313073 | -1.182893 | 0.236852 | 0.709700242 | SUMO2 | protein_coding | 46.32 | -0.055151723 | 0.055151723 | ENSG00000188612 | n.s. |

| | | | | | | | | | | | | | |
|---|---|---|---|---|---|---|---|---|---|---|---|---|---|
| ENSG00000041353 | 377.3437 | 0.372170905 | 0.315844 | 1.178336 | 0.238663 | 0.710909549 | RAB27B | protein_coding | 37.74 | 0.055150388 | 0.055150388 | ENSG00000041353 | n.s. |
| ENSG00000117174 | 1235.925 | -0.322630979 | 0.251032 | -1.285221 | 0.198715 | 0.674743473 | ZNHIT6 | protein_coding | 36.46 | -0.055125151 | 0.055125151 | ENSG00000117174 | n.s. |
| ENSG00000235408 | 107.6334 | -0.329640033 | 0.260408 | -1.265858 | 0.205564 | 0.680704186 | SNORA71B | snoRNA | 51.84 | -0.055063591 | 0.055063591 | ENSG00000235408 | n.s. |
| ENSG00000139083 | 303.4576 | -0.313196282 | 0.24027 | -1.303516 | 0.192399 | 0.667270349 | ETV6 | protein_coding | 43.19 | -0.055028015 | 0.055028015 | ENSG00000139083 | n.s. |
| ENSG00000115687 | 815.4914 | -0.325387498 | 0.254723 | -1.277415 | 0.201456 | 0.677503939 | PASK | protein_coding | 50.98 | -0.055019178 | 0.055019178 | ENSG00000115687 | n.s. |
| ENSG00000074964 | 189.1521 | -0.335358266 | 0.267429 | -1.254008 | 0.209839 | 0.68549731 | ARHGEF10L | protein_coding | 54.36 | -0.054996826 | 0.054996826 | ENSG00000074964 | n.s. |
| ENSG00000264247 | 719.686 | -0.28531084 | 0.209499 | -1.361874 | 0.173238 | 0.641657113 | LINC00909 | lincRNA | 43.19 | -0.054978539 | 0.054978539 | ENSG00000264247 | n.s. |
| ENSG00000125755 | 147.3215 | -0.344420672 | 0.27814 | -1.238301 | 0.215604 | 0.692470486 | SYMPK | protein_coding | 51.56 | -0.054969103 | 0.054969103 | ENSG00000125755 | n.s. |
| ENSG00000213923 | 1567.553 | -0.306200742 | 0.232484 | -1.317082 | 0.187811 | 0.661496606 | CSNK1E | protein_coding | 55.79 | -0.054954576 | 0.054954576 | ENSG00000213923 | n.s. |
| ENSG00000260288 | 79.89204 | 0.349757901 | 0.285759 | 1.2239613 | 0.220967 | 0.696833726 | | processed_transcript | 49.1 | 0.054866815 | 0.054866815 | ENSG00000260288 | n.s. |
| ENSG00000151148 | 644.0973 | -0.317884876 | 0.246264 | -1.290829 | 0.196763 | 0.672118672 | UBE3B | protein_coding | 46.32 | -0.054852319 | 0.054852319 | ENSG00000151148 | n.s. |
| ENSG00000056487 | 1016.426 | -0.332636337 | 0.264654 | -1.256873 | 0.208799 | 0.684188053 | PHF21B | protein_coding | 52.73 | -0.054826622 | 0.054826622 | ENSG00000056487 | n.s. |
| ENSG00000238754 | 104.6503 | 0.317735843 | 0.24618 | 1.2906635 | 0.19682 | 0.672118672 | SCARNA18B | snoRNA | 40 | 0.054826603 | 0.054826603 | ENSG00000238754 | n.s. |
| ENSG00000075624 | 14448.71 | -0.299090457 | 0.225462 | -1.326565 | 0.184653 | 0.655769671 | ACTB | protein_coding | 56.7 | -0.054807929 | 0.054807929 | ENSG00000075624 | n.s. |
| ENSG00000140025 | 80.55381 | -0.378172022 | 0.325269 | -1.162644 | 0.244974 | 0.716280109 | EFCAB11 | protein_coding | 40 | -0.054803596 | 0.054803596 | ENSG00000140025 | n.s. |
| ENSG00000206885 | 103.1754 | 0.324064518 | 0.253381 | 1.2789612 | 0.200911 | 0.677491543 | SNORA75 | snoRNA | 38.69 | 0.054798053 | 0.054798053 | ENSG00000206885 | n.s. |
| ENSG00000112290 | 2792.688 | -0.325656416 | 0.256134 | -1.271431 | 0.203575 | 0.679010816 | WASF1 | protein_coding | 36.65 | -0.054750434 | 0.054750434 | ENSG00000112290 | n.s. |
| ENSG00000186787 | 66.26175 | -0.355677763 | 0.294445 | -1.20796 | 0.227063 | 0.701641249 | SPIN2B | protein_coding | 51.44 | -0.054733432 | 0.054733432 | ENSG00000186787 | n.s. |
| ENSG00000115232 | 1791.367 | 0.321438835 | 0.250364 | 1.2838871 | 0.199181 | 0.675731841 | ITGA4 | protein_coding | 36.12 | 0.054717124 | 0.054717124 | ENSG00000115232 | n.s. |
| ENSG00000119541 | 965.4511 | -0.354826724 | 0.292791 | -1.211877 | 0.225559 | 0.701152989 | VPS4B | protein_coding | 39.19 | -0.054709743 | 0.054709743 | ENSG00000119541 | n.s. |
| ENSG00000236095 | 25.81002 | -0.396734595 | 0.351224 | -1.129577 | 0.258654 | 0.728120901 | | sense_intronic | 42.14 | -0.054668639 | 0.054668639 | ENSG00000236095 | n.s. |
| ENSG00000140987 | 417.6916 | -0.344079235 | 0.278882 | -1.233782 | 0.217284 | 0.693734586 | ZSCAN32 | protein_coding | 45.48 | -0.054642072 | 0.054642072 | ENSG00000140987 | n.s. |
| ENSG00000178904 | 1965.699 | -0.342262573 | 0.276617 | -1.237315 | 0.21597 | 0.692499549 | DPY19L3 | protein_coding | 40.45 | -0.054618434 | 0.054618434 | ENSG00000178904 | n.s. |
| ENSG00000167202 | 583.7199 | -0.326728513 | 0.258344 | -1.264702 | 0.205978 | 0.680710383 | TBC1D2B | protein_coding | 47.99 | -0.054575955 | 0.054575955 | ENSG00000167202 | n.s. |
| ENSG00000028310 | 1040.688 | -0.31054455 | 0.238296 | -1.303188 | 0.192511 | 0.667270349 | BRD9 | protein_coding | 55.14 | -0.05456211 | 0.05456211 | ENSG00000028310 | n.s. |
| ENSG00000231992 | 13.14754 | 0.462153328 | 0.45237 | 1.0216263 | 0.306958 | 0.761997697 | | sense_intronic | 42.34 | 0.05455551 | 0.05455551 | ENSG00000231992 | n.s. |
| ENSG00000129521 | 493.5157 | -0.310395924 | 0.238124 | -1.303505 | 0.192402 | 0.667270349 | EGLN3 | protein_coding | 41.68 | -0.054535997 | 0.054535997 | ENSG00000129521 | n.s. |
| ENSG00000145777 | 82.6955 | -0.333170551 | 0.265962 | -1.252698 | 0.210315 | 0.686087755 | TSLP | protein_coding | 43.11 | -0.054513476 | 0.054513476 | ENSG00000145777 | n.s. |
| ENSG00000185112 | 1181.358 | -0.286493451 | 0.212415 | -1.348742 | 0.17742 | 0.645396531 | FAM43A | protein_coding | 60.99 | -0.054483426 | 0.054483426 | ENSG00000185112 | n.s. |
| ENSG00000100154 | 1551.468 | -0.288878305 | 0.214768 | -1.345074 | 0.178601 | 0.647737535 | TTC28 | protein_coding | 39.91 | -0.054482719 | 0.054482719 | ENSG00000100154 | n.s. |
| ENSG00000146576 | 283.4788 | -0.318473664 | 0.2475 | -1.286762 | 0.198177 | 0.674626594 | C7orf26 | protein_coding | 53.11 | -0.054438787 | 0.054438787 | ENSG00000146576 | n.s. |
| ENSG00000224046 | 147.8363 | 0.324686908 | 0.25586 | 1.2690015 | 0.204441 | 0.679838368 | | antisense | 47.9 | 0.054415684 | 0.054415684 | ENSG00000224046 | n.s. |

| | | | | | | | | | | | | | |
|---|---|---|---|---|---|---|---|---|---|---|---|---|---|
| ENSG00000109790 | 615.5415 | -0.347988801 | 0.284794 | -1.221898 | 0.221746 | 0.697710933 | KLHL5 | protein_coding | 36.93 | -0.054399166 | 0.054399166 | ENSG00000109790 | n.s. |
| ENSG00000101247 | 1079.267 | -0.328123679 | 0.260074 | -1.261657 | 0.207072 | 0.682685326 | NDUFAF5 | protein_coding | 39.88 | -0.054396157 | 0.054396157 | ENSG00000101247 | n.s. |
| ENSG00000149582 | 77.03886 | -0.352960638 | 0.291525 | -1.210738 | 0.225996 | 0.701289341 | TMEM25 | protein_coding | 49.48 | -0.054392209 | 0.054392209 | ENSG00000149582 | n.s. |
| ENSG00000131127 | 1442.044 | -0.334367573 | 0.26756 | -1.24969 | 0.211413 | 0.687733955 | ZNF141 | protein_coding | 40.31 | -0.054361324 | 0.054361324 | ENSG00000131127 | n.s. |
| ENSG00000210140 | 38.84123 | -0.389907385 | 0.342932 | -1.136983 | 0.255545 | 0.725521442 | MT-TC | Mt_tRNA | 45.45 | -0.054333496 | 0.054333496 | ENSG00000210140 | n.s. |
| ENSG00000124508 | 238.5536 | 0.336794019 | 0.270317 | 1.2459209 | 0.212793 | 0.689743466 | BTN2A2 | protein_coding | 45.78 | 0.054329053 | 0.054329053 | ENSG00000124508 | n.s. |
| ENSG00000261762 | 206.3247 | -0.344123554 | 0.280062 | -1.228739 | 0.21917 | 0.695448706 | | antisense | 39.46 | -0.054280293 | 0.054280293 | ENSG00000261762 | n.s. |
| ENSG00000116473 | 840.2461 | -0.336843489 | 0.270547 | -1.245046 | 0.213115 | 0.69012437 | RAP1A | protein_coding | 39.31 | -0.054256269 | 0.054256269 | ENSG00000116473 | n.s. |
| ENSG00000215612 | 705.8395 | -0.329711817 | 0.262629 | -1.255429 | 0.209323 | 0.68461467 | HMX1 | protein_coding | 59.36 | -0.054255332 | 0.054255332 | ENSG00000215612 | n.s. |
| ENSG00000088387 | 174.9697 | -0.35184663 | 0.290809 | -1.20989 | 0.226321 | 0.701289341 | DOCK9 | protein_coding | 41.78 | -0.054220537 | 0.054220537 | ENSG00000088387 | n.s. |
| ENSG00000264538 | 1118.092 | -0.317327805 | 0.246922 | -1.285136 | 0.198745 | 0.674743473 | SUZ12P1 | cribed_unprocessed_pseudc | 45.35 | -0.054219044 | 0.054219044 | ENSG00000264538 | n.s. |
| ENSG00000127586 | 597.7117 | -0.32336476 | 0.254732 | -1.26943 | 0.204288 | 0.679803349 | CHTF18 | protein_coding | 66.52 | -0.054201334 | 0.054201334 | ENSG00000127586 | n.s. |
| ENSG00000187475 | 46.63926 | 0.358225133 | 0.300732 | 1.1911773 | 0.233584 | 0.706076839 | HIST1H1T | protein_coding | 48.33 | 0.054145025 | 0.054145025 | ENSG00000187475 | n.s. |
| ENSG00000136279 | 723.1388 | -0.342828917 | 0.2788 | -1.229661 | 0.218824 | 0.695273403 | DBNL | protein_coding | 53.2 | -0.054113619 | 0.054113619 | ENSG00000136279 | n.s. |
| ENSG00000180747 | 370.2316 | -0.311950745 | 0.240753 | -1.295727 | 0.195069 | 0.670750289 | SMG1P3 | cribed_unprocessed_pseudc | 43.71 | -0.054104466 | 0.054104466 | ENSG00000180747 | n.s. |
| ENSG00000250182 | 155.9131 | 0.389730732 | 0.343247 | 1.1354237 | 0.256198 | 0.726402587 | EEF1A1P13 | processed_pseudogene | 47.73 | 0.05410344 | 0.05410344 | ENSG00000250182 | n.s. |
| ENSG00000100284 | 207.9656 | -0.338904438 | 0.274096 | -1.236445 | 0.216293 | 0.692499549 | TOM1 | protein_coding | 53.51 | -0.054082541 | 0.054082541 | ENSG00000100284 | n.s. |
| ENSG00000130787 | 465.5385 | -0.319525543 | 0.24962 | -1.280049 | 0.200528 | 0.677328602 | HIP1R | protein_coding | 59.16 | -0.054063909 | 0.054063909 | ENSG00000130787 | n.s. |
| ENSG00000131845 | 439.4487 | -0.319557204 | 0.249845 | -1.279024 | 0.200889 | 0.677491543 | ZNF304 | protein_coding | 47.4 | -0.054035884 | 0.054035884 | ENSG00000131845 | n.s. |
| ENSG00000198182 | 277.7976 | -0.3674811 | 0.313346 | -1.172763 | 0.240891 | 0.712813552 | ZNF607 | protein_coding | 41.55 | -0.05402856 | 0.05402856 | ENSG00000198182 | n.s. |
| ENSG00000136870 | 738.7199 | -0.338461682 | 0.273376 | -1.238082 | 0.215686 | 0.692499549 | ZNF189 | protein_coding | 39.9 | -0.054011886 | 0.054011886 | ENSG00000136870 | n.s. |
| ENSG00000132541 | 435.372 | 0.377086329 | 0.326199 | 1.1560013 | 0.247681 | 0.719177715 | RIDA | protein_coding | 39.14 | 0.053985104 | 0.053985104 | ENSG00000132541 | n.s. |
| ENSG00000275457 | 110.074 | 0.342685073 | 0.279185 | 1.2274477 | 0.219654 | 0.695847029 | | antisense | 63.04 | 0.053968178 | 0.053968178 | ENSG00000275457 | n.s. |
| ENSG00000178531 | 148.8858 | -0.319279388 | 0.25031 | -1.275537 | 0.202119 | 0.677624898 | CTXN1 | protein_coding | 74.79 | -0.053961615 | 0.053961615 | ENSG00000178531 | n.s. |
| ENSG00000140983 | 866.6796 | -0.325621498 | 0.258277 | -1.260745 | 0.207401 | 0.682785213 | RHOT2 | protein_coding | 66.37 | -0.053960657 | 0.053960657 | ENSG00000140983 | n.s. |
| ENSG00000161996 | 39.68369 | -0.377619216 | 0.327388 | -1.153429 | 0.248734 | 0.719685597 | WDR90 | protein_coding | 64.6 | -0.053945619 | 0.053945619 | ENSG00000161996 | n.s. |
| ENSG00000201785 | 35.17526 | 0.387152262 | 0.340551 | 1.1368407 | 0.255605 | 0.7255427 | SNORD117 | snoRNA | 44.74 | 0.053944644 | 0.053944644 | ENSG00000201785 | n.s. |
| ENSG00000108510 | 5489.947 | -0.330961983 | 0.264667 | -1.250484 | 0.211123 | 0.687111024 | MED13 | protein_coding | 38.73 | -0.053937895 | 0.053937895 | ENSG00000108510 | n.s. |
| ENSG00000198039 | 634.4332 | -0.353899421 | 0.296054 | -1.195389 | 0.231935 | 0.704043192 | ZNF273 | protein_coding | 44.96 | -0.053934518 | 0.053934518 | ENSG00000198039 | n.s. |
| ENSG00000274561 | 83.64639 | -0.353718067 | 0.295608 | -1.196578 | 0.231471 | 0.704043192 | | lincRNA | 33.41 | -0.05390688 | 0.05390688 | ENSG00000274561 | n.s. |
| ENSG00000259972 | 119.0606 | -0.34338472 | 0.280416 | -1.224556 | 0.220743 | 0.696659758 | | processed_transcript | 47.73 | -0.053904285 | 0.053904285 | ENSG00000259972 | n.s. |
| ENSG00000054118 | 1442.235 | -0.305489225 | 0.233797 | -1.306644 | 0.191334 | 0.666152146 | THRAP3 | protein_coding | 43.72 | -0.053896416 | 0.053896416 | ENSG00000054118 | n.s. |

| | | | | | | | | | | | | | |
|---|---|---|---|---|---|---|---|---|---|---|---|---|---|
| ENSG00000072864 | 275.2707 | -0.325158468 | 0.257986 | -1.260373 | 0.207535 | 0.682785213 | NDE1 | protein_coding | 48.54 | -0.053883926 | 0.053883926 | ENSG00000072864 | n.s. |
| ENSG00000160336 | 391.0824 | -0.339051915 | 0.275106 | -1.232439 | 0.217785 | 0.693853089 | ZNF761 | protein_coding | 44.23 | -0.053818549 | 0.053818549 | ENSG00000160336 | n.s. |
| ENSG00000168769 | 885.9728 | -0.30228192 | 0.230222 | -1.312999 | 0.189183 | 0.663809724 | TET2 | protein_coding | 36.29 | -0.053792999 | 0.053792999 | ENSG00000168769 | n.s. |
| ENSG00000134287 | 932.6965 | -0.325631598 | 0.258995 | -1.25729 | 0.208649 | 0.683940032 | ARF3 | protein_coding | 47.76 | -0.053723344 | 0.053723344 | ENSG00000134287 | n.s. |
| ENSG00000090861 | 1025.389 | -0.336638735 | 0.272059 | -1.237373 | 0.215949 | 0.692499549 | AARS | protein_coding | 49.47 | -0.053720979 | 0.053720979 | ENSG00000090861 | n.s. |
| ENSG00000264405 | 2.607396 | 0.786186908 | 1.052192 | 0.7471899 | 0.454949 | 0.854433778 | MIR3913-1 | miRNA | 33.33 | 0.053713541 | 0.053713541 | ENSG00000264405 | n.s. |
| ENSG00000207588 | 6.915881 | -0.512633191 | 0.536684 | -0.955187 | 0.339483 | 0.785717908 | MIR593 | miRNA | 61 | -0.053689791 | 0.053689791 | ENSG00000207588 | n.s. |
| ENSG00000229539 | 22.03506 | -0.418893975 | 0.389139 | -1.076464 | 0.28172 | 0.744567264 | | antisense | 53.17 | -0.053658669 | 0.053658669 | ENSG00000229539 | n.s. |
| ENSG00000060491 | 357.9946 | -0.31446388 | 0.244809 | -1.284527 | 0.198958 | 0.67513656 | OGFR | protein_coding | 63.45 | -0.053650171 | 0.053650171 | ENSG00000060491 | n.s. |
| ENSG00000141564 | 1464.135 | -0.2910767 | 0.218739 | -1.330705 | 0.183286 | 0.654498957 | RPTOR | protein_coding | 49.12 | -0.053584613 | 0.053584613 | ENSG00000141564 | n.s. |
| ENSG00000240877 | 13.79825 | -0.471954559 | 0.474083 | -0.995511 | 0.319488 | 0.770123065 | RN7SL521P | misc_RNA | 67.34 | -0.053538464 | 0.053538464 | ENSG00000240877 | n.s. |
| ENSG00000256087 | 446.6311 | -0.339822433 | 0.276816 | -1.227612 | 0.219593 | 0.695810058 | ZNF432 | protein_coding | 41.15 | -0.053525193 | 0.053525193 | ENSG00000256087 | n.s. |
| ENSG00000166265 | 435.1612 | 0.326959404 | 0.260914 | 1.2531321 | 0.210158 | 0.686023341 | CYYR1 | protein_coding | 37.29 | 0.053510538 | 0.053510538 | ENSG00000166265 | n.s. |
| ENSG00000143630 | 187.2369 | -0.334376611 | 0.269657 | -1.240007 | 0.214973 | 0.691877944 | HCN3 | protein_coding | 53.79 | -0.053490398 | 0.053490398 | ENSG00000143630 | n.s. |
| ENSG00000122783 | 731.7139 | -0.303627672 | 0.232795 | -1.304268 | 0.192142 | 0.666964456 | C7orf49 | protein_coding | 40.03 | -0.053407291 | 0.053407291 | ENSG00000122783 | n.s. |
| ENSG00000185127 | 1154.216 | 0.285053439 | 0.212676 | 1.3403157 | 0.180143 | 0.649773904 | C6orf120 | protein_coding | 45.78 | 0.05337276 | 0.05337276 | ENSG00000185127 | n.s. |
| ENSG00000152683 | 893.5063 | -0.349197538 | 0.290638 | -1.201487 | 0.229562 | 0.703335843 | SLC30A6 | protein_coding | 39.74 | -0.053370391 | 0.053370391 | ENSG00000152683 | n.s. |
| ENSG00000206675 | 11.939 | -0.456589035 | 0.44995 | -1.014755 | 0.310223 | 0.764194956 | RNU6-32P | snRNA | 44.86 | -0.053327697 | 0.053327697 | ENSG00000206675 | n.s. |
| ENSG00000155252 | 661.9455 | -0.333307256 | 0.268886 | -1.239586 | 0.215128 | 0.691899391 | PI4K2A | protein_coding | 44.74 | -0.053314846 | 0.053314846 | ENSG00000155252 | n.s. |
| ENSG00000139990 | 624.6756 | -0.316862914 | 0.249097 | -1.272048 | 0.203356 | 0.678974657 | DCAF5 | protein_coding | 40.67 | -0.05327937 | 0.05327937 | ENSG00000139990 | n.s. |
| ENSG00000185559 | 71.93834 | -0.385833116 | 0.340907 | -1.131784 | 0.257725 | 0.727758051 | DLK1 | protein_coding | 55.89 | -0.053249979 | 0.053249979 | ENSG00000185559 | n.s. |
| ENSG00000013563 | 136.785 | -0.328616757 | 0.263424 | -1.247484 | 0.21222 | 0.688714317 | DNASE1L1 | protein_coding | 54.47 | -0.053223062 | 0.053223062 | ENSG00000013563 | n.s. |
| ENSG00000117481 | 201.0115 | -0.318232318 | 0.251533 | -1.265172 | 0.20581 | 0.680704186 | NSUN4 | protein_coding | 47.35 | -0.053158029 | 0.053158029 | ENSG00000117481 | n.s. |
| ENSG00000159788 | 326.3052 | -0.304625822 | 0.234694 | -1.297969 | 0.194298 | 0.669140798 | RGS12 | protein_coding | 50.96 | -0.053151872 | 0.053151872 | ENSG00000159788 | n.s. |
| ENSG00000105298 | 551.8867 | -0.307942891 | 0.238477 | -1.291291 | 0.196603 | 0.672118672 | CACTIN | protein_coding | 61.58 | -0.05313679 | 0.05313679 | ENSG00000105298 | n.s. |
| ENSG00000162341 | 192.8136 | -0.316052664 | 0.248703 | -1.270803 | 0.203799 | 0.679010816 | TPCN2 | protein_coding | 54.96 | -0.053135819 | 0.053135819 | ENSG00000162341 | n.s. |
| ENSG00000101057 | 1761.146 | -0.310879334 | 0.241878 | -1.285273 | 0.198697 | 0.674743473 | MYBL2 | protein_coding | 49.88 | -0.053117249 | 0.053117249 | ENSG00000101057 | n.s. |
| ENSG00000175040 | 987.1626 | -0.308122469 | 0.238876 | -1.289883 | 0.197091 | 0.672408622 | CHST2 | protein_coding | 58.88 | -0.053110061 | 0.053110061 | ENSG00000175040 | n.s. |
| ENSG00000137094 | 230.7256 | -0.378672843 | 0.331911 | -1.140888 | 0.253917 | 0.72418384 | DNAJB5 | protein_coding | 55.15 | -0.053071442 | 0.053071442 | ENSG00000137094 | n.s. |
| ENSG00000168917 | 1036.823 | -0.313373803 | 0.245165 | -1.278214 | 0.201174 | 0.677503939 | SLC35G2 | protein_coding | 41.28 | -0.052987805 | 0.052987805 | ENSG00000168917 | n.s. |
| ENSG00000141577 | 577.7794 | -0.311911166 | 0.24332 | -1.281897 | 0.199879 | 0.676613539 | CEP131 | protein_coding | 59.72 | -0.052918635 | 0.052918635 | ENSG00000141577 | n.s. |
| ENSG00000067057 | 1488.675 | -0.333322067 | 0.270358 | -1.232889 | 0.217617 | 0.693853089 | PFKP | protein_coding | 51.45 | -0.052909036 | 0.052909036 | ENSG00000067057 | n.s. |

| | | | | | | | | | | | | | |
|---|---|---|---|---|---|---|---|---|---|---|---|---|---|
| ENSG00000259196 | 23.98115 | -0.39646351 | 0.357937 | -1.107634 | 0.26802 | 0.735486422 | HMBOX1-IT1 | sense_intronic | 38.15 | -0.052898279 | 0.052898279 | ENSG00000259196 | n.s. |
| ENSG00000179909 | 399.0971 | -0.326532642 | 0.261754 | -1.247479 | 0.212222 | 0.688714317 | ZNF154 | protein_coding | 46.08 | -0.052885517 | 0.052885517 | ENSG00000179909 | n.s. |
| ENSG00000252161 | 3.445859 | -0.623823042 | 0.742373 | -0.84031 | 0.400735 | 0.822682214 | RNA5SP318 | rRNA | 50 | -0.052880164 | 0.052880164 | ENSG00000252161 | n.s. |
| ENSG00000103160 | 423.6326 | -0.350389935 | 0.294464 | -1.189925 | 0.234076 | 0.706508138 | HSDL1 | protein_coding | 45.07 | -0.052867826 | 0.052867826 | ENSG00000103160 | n.s. |
| ENSG00000263593 | 54.12558 | -0.394545062 | 0.35462 | -1.112586 | 0.265886 | 0.734612586 | MIR4804 | miRNA | 45.21 | -0.052846011 | 0.052846011 | ENSG00000263593 | n.s. |
| ENSG00000210196 | 282.4087 | -0.316113338 | 0.249549 | -1.266738 | 0.205249 | 0.680704186 | MT-TP | Mt_tRNA | 33.82 | -0.052804071 | 0.052804071 | ENSG00000210196 | n.s. |
| ENSG00000167912 | 66.78405 | 0.340393685 | 0.279722 | 1.2168999 | 0.223642 | 0.699734492 | | antisense | 54.4 | 0.052783731 | 0.052783731 | ENSG00000167912 | n.s. |
| ENSG00000131725 | 249.4278 | -0.317626485 | 0.251533 | -1.262764 | 0.206674 | 0.682201186 | WDR44 | protein_coding | 39.07 | -0.052753798 | 0.052753798 | ENSG00000131725 | n.s. |
| ENSG00000164114 | 615.2195 | -0.341977082 | 0.282908 | -1.208795 | 0.226742 | 0.701406486 | MAP9 | protein_coding | 33.75 | -0.052674806 | 0.052674806 | ENSG00000164114 | n.s. |
| ENSG00000280186 | 110.7991 | 0.329406073 | 0.265804 | 1.2392804 | 0.215242 | 0.692103837 | | TEC | 42.04 | 0.052648559 | 0.052648559 | ENSG00000280186 | n.s. |
| ENSG00000120875 | 968.7233 | -0.295815543 | 0.225441 | -1.312165 | 0.189464 | 0.663959974 | DUSP4 | protein_coding | 51.08 | -0.052613191 | 0.052613191 | ENSG00000120875 | n.s. |
| ENSG00000072110 | 510.7172 | -0.313655584 | 0.247169 | -1.26899 | 0.204445 | 0.679838368 | ACTN1 | protein_coding | 50.81 | -0.052566897 | 0.052566897 | ENSG00000072110 | n.s. |
| ENSG00000230630 | 1655.845 | -0.273036517 | 0.200816 | -1.359634 | 0.173946 | 0.642002364 | DNM3OS | antisense | 33.12 | -0.052549529 | 0.052549529 | ENSG00000230630 | n.s. |
| ENSG00000125827 | 1388.851 | -0.344674829 | 0.287505 | -1.198846 | 0.230588 | 0.704043192 | TMX4 | protein_coding | 37.75 | -0.052528684 | 0.052528684 | ENSG00000125827 | n.s. |
| ENSG00000271819 | 25.63531 | 0.400361736 | 0.366344 | 1.0928585 | 0.274456 | 0.739386355 | RNU6-94P | snRNA | 37.38 | 0.052498861 | 0.052498861 | ENSG00000271819 | n.s. |
| ENSG00000143772 | 1091.113 | -0.317422391 | 0.251972 | -1.259755 | 0.207758 | 0.683347672 | ITPKB | protein_coding | 49.82 | -0.052488421 | 0.052488421 | ENSG00000143772 | n.s. |
| ENSG00000252498 | 94.17565 | -0.371186248 | 0.323403 | -1.147751 | 0.251071 | 0.722169533 | RNU6-1016P | snRNA | 36.17 | -0.052471199 | 0.052471199 | ENSG00000252498 | n.s. |
| ENSG00000198482 | 358.742 | -0.333744731 | 0.272157 | -1.226296 | 0.220087 | 0.696377966 | ZNF808 | protein_coding | 44.95 | -0.052449646 | 0.052449646 | ENSG00000198482 | n.s. |
| ENSG00000198055 | 465.8351 | -0.3339709 | 0.272645 | -1.224928 | 0.220602 | 0.696655251 | GRK6 | protein_coding | 54.65 | -0.052427448 | 0.052427448 | ENSG00000198055 | n.s. |
| ENSG00000141232 | 1879.745 | 0.285180765 | 0.214456 | 1.3297896 | 0.183588 | 0.654903931 | TOB1 | protein_coding | 50.17 | 0.052422613 | 0.052422613 | ENSG00000141232 | n.s. |
| ENSG00000160211 | 194.3363 | -0.341436274 | 0.283008 | -1.206453 | 0.227643 | 0.702297783 | G6PD | protein_coding | 56.32 | -0.052403196 | 0.052403196 | ENSG00000160211 | n.s. |
| ENSG00000196141 | 79.90904 | -0.365804392 | 0.315956 | -1.157771 | 0.246958 | 0.719028029 | SPATS2L | protein_coding | 40.19 | -0.052403008 | 0.052403008 | ENSG00000196141 | n.s. |
| ENSG00000127481 | 2746.226 | -0.309277109 | 0.241425 | -1.281049 | 0.200177 | 0.67702417 | UBR4 | protein_coding | 43.74 | -0.052390252 | 0.052390252 | ENSG00000127481 | n.s. |
| ENSG00000185158 | 44.54109 | -0.389074565 | 0.348261 | -1.117194 | 0.263911 | 0.733524057 | LRRC37B | protein_coding | 42.83 | -0.052363849 | 0.052363849 | ENSG00000185158 | n.s. |
| ENSG00000070413 | 528.8909 | -0.338127179 | 0.278231 | -1.215275 | 0.224261 | 0.700099826 | DGCR2 | protein_coding | 48.62 | -0.052355623 | 0.052355623 | ENSG00000070413 | n.s. |
| ENSG00000200087 | 398.4772 | 0.288014956 | 0.217561 | 1.3238365 | 0.185557 | 0.658052755 | SNORA73B | snoRNA | 54.9 | 0.052343633 | 0.052343633 | ENSG00000200087 | n.s. |
| ENSG00000014138 | 1116.558 | -0.317663987 | 0.252793 | -1.256617 | 0.208892 | 0.684330895 | POLA2 | protein_coding | 47.9 | -0.052330013 | 0.052330013 | ENSG00000014138 | n.s. |
| ENSG00000110906 | 773.537 | -0.339259482 | 0.279886 | -1.212134 | 0.225461 | 0.701091738 | KCTD10 | protein_coding | 47.69 | -0.052322345 | 0.052322345 | ENSG00000110906 | n.s. |
| ENSG00000161298 | 474.9475 | -0.32962327 | 0.267721 | -1.231218 | 0.218241 | 0.694065444 | ZNF382 | protein_coding | 41.92 | -0.052278112 | 0.052278112 | ENSG00000161298 | n.s. |
| ENSG00000122126 | 646.3081 | -0.317450811 | 0.252761 | -1.255932 | 0.209141 | 0.68461467 | OCRL | protein_coding | 39.83 | -0.052237737 | 0.052237737 | ENSG00000122126 | n.s. |
| ENSG00000147862 | 9121.784 | -0.308839266 | 0.241728 | -1.277633 | 0.201379 | 0.677503939 | NFIB | protein_coding | 38.29 | -0.052221068 | 0.052221068 | ENSG00000147862 | n.s. |
| ENSG00000143845 | 108.5636 | -0.34083327 | 0.282946 | -1.204586 | 0.228363 | 0.702737638 | ETNK2 | protein_coding | 52.5 | -0.05221797 | 0.05221797 | ENSG00000143845 | n.s. |

| | | | | | | | | | | | | | |
|---|---|---|---|---|---|---|---|---|---|---|---|---|---|
| ENSG00000127663 | 141.14 | -0.315102601 | 0.249919 | -1.260818 | 0.207374 | 0.682785213 | KDM4B | protein_coding | 57.26 | -0.052217509 | 0.052217509 | ENSG00000127663 | n.s. |
| ENSG00000135835 | 791.408 | -0.310517503 | 0.24425 | -1.271313 | 0.203617 | 0.679010816 | KIAA1614 | protein_coding | 54.17 | -0.05220523 | 0.05220523 | ENSG00000135835 | n.s. |
| ENSG00000138735 | 1124.091 | -0.332758988 | 0.27197 | -1.223513 | 0.221136 | 0.696894187 | PDE5A | protein_coding | 36.45 | -0.052187643 | 0.052187643 | ENSG00000138735 | n.s. |
| ENSG00000131779 | 308.4209 | -0.342103245 | 0.285162 | -1.199682 | 0.230263 | 0.704043192 | PEX11B | protein_coding | 45.5 | -0.052136773 | 0.052136773 | ENSG00000131779 | n.s. |
| ENSG00000250461 | 106.7288 | -0.363812108 | 0.314054 | -1.158437 | 0.246686 | 0.719028029 | | processed_pseudogene | 37.77 | -0.052117605 | 0.052117605 | ENSG00000250461 | n.s. |
| ENSG00000105085 | 66.2388 | 0.35087764 | 0.297508 | 1.1793894 | 0.238243 | 0.71051867 | MED26 | protein_coding | 52.52 | 0.05207884 | 0.05207884 | ENSG00000105085 | n.s. |
| ENSG00000132510 | 46.34937 | -0.371488009 | 0.325669 | -1.140693 | 0.253998 | 0.72418384 | KDM6B | protein_coding | 60.63 | -0.052064479 | 0.052064479 | ENSG00000132510 | n.s. |
| ENSG00000275379 | 1013.948 | -0.296594639 | 0.227736 | -1.302359 | 0.192794 | 0.667523235 | HIST1H3I | protein_coding | 62.53 | -0.052062329 | 0.052062329 | ENSG00000275379 | n.s. |
| ENSG00000228939 | 73.7878 | -0.386534725 | 0.345409 | -1.119065 | 0.263112 | 0.733524057 | AKT3-IT1 | sense_intronic | 43.31 | -0.052022023 | 0.052022023 | ENSG00000228939 | n.s. |
| ENSG00000115652 | 698.0258 | -0.323282526 | 0.260048 | -1.243166 | 0.213807 | 0.690444009 | UXS1 | protein_coding | 45.03 | -0.052006956 | 0.052006956 | ENSG00000115652 | n.s. |
| ENSG00000177697 | 495.6484 | -0.340137694 | 0.283041 | -1.201725 | 0.22947 | 0.703260264 | CD151 | protein_coding | 63.43 | -0.052001584 | 0.052001584 | ENSG00000177697 | n.s. |
| ENSG00000087008 | 148.5094 | -0.330853892 | 0.26977 | -1.226432 | 0.220036 | 0.696377966 | ACOX3 | protein_coding | 51.93 | -0.051995336 | 0.051995336 | ENSG00000087008 | n.s. |
| ENSG00000235194 | 142.9967 | -0.303682954 | 0.235873 | -1.287484 | 0.197926 | 0.674264807 | PPP1R3E | protein_coding | 51.79 | -0.051981262 | 0.051981262 | ENSG00000235194 | n.s. |
| ENSG00000138867 | 543.5997 | -0.32721548 | 0.265093 | -1.234344 | 0.217075 | 0.693680706 | GUCD1 | protein_coding | 55.14 | -0.051975033 | 0.051975033 | ENSG00000138867 | n.s. |
| ENSG00000079805 | 691.6813 | -0.32274437 | 0.259548 | -1.243488 | 0.213688 | 0.690296427 | DNM2 | protein_coding | 53.07 | -0.051950346 | 0.051950346 | ENSG00000079805 | n.s. |
| ENSG00000187714 | 212.9367 | -0.308803506 | 0.242953 | -1.271044 | 0.203713 | 0.679010816 | SLC18A3 | protein_coding | 67.19 | -0.051917067 | 0.051917067 | ENSG00000187714 | n.s. |
| ENSG00000113648 | 6822.35 | -0.326712317 | 0.264844 | -1.233604 | 0.21735 | 0.693734586 | H2AFY | protein_coding | 47.07 | -0.05188409 | 0.05188409 | ENSG00000113648 | n.s. |
| ENSG00000154380 | 7160.896 | -0.326554307 | 0.264617 | -1.234062 | 0.21718 | 0.693734586 | ENAH | protein_coding | 38.55 | -0.051858997 | 0.051858997 | ENSG00000154380 | n.s. |
| ENSG00000206824 | 22.54945 | -0.387120595 | 0.347607 | -1.113675 | 0.265419 | 0.734612586 | | misc_RNA | 44.12 | -0.051851566 | 0.051851566 | ENSG00000206824 | n.s. |
| ENSG00000269925 | 22.29072 | 0.402520961 | 0.371858 | 1.0824583 | 0.279049 | 0.743390786 | | sense_intronic | 36.64 | 0.051837787 | 0.051837787 | ENSG00000269925 | n.s. |
| ENSG00000264477 | 2.737648 | -0.695866571 | 0.888017 | -0.783619 | 0.433264 | 0.842381919 | MIR5682 | miRNA | 59.21 | -0.051835771 | 0.051835771 | ENSG00000264477 | n.s. |
| ENSG00000173486 | 231.2276 | -0.349204943 | 0.296008 | -1.179716 | 0.238113 | 0.71051867 | FKBP2 | protein_coding | 63.61 | -0.051830571 | 0.051830571 | ENSG00000173486 | n.s. |
| ENSG00000203791 | 629.5364 | -0.34445437 | 0.289882 | -1.188255 | 0.234733 | 0.707216633 | EEF1AKMT2 | protein_coding | 40.87 | -0.05182231 | 0.05182231 | ENSG00000203791 | n.s. |
| ENSG00000139324 | 1119.396 | -0.339972384 | 0.284433 | -1.195264 | 0.231984 | 0.704043192 | TMTC3 | protein_coding | 34.6 | -0.051812028 | 0.051812028 | ENSG00000139324 | n.s. |
| ENSG00000115963 | 2229.439 | -0.306550722 | 0.240367 | -1.275342 | 0.202188 | 0.677624898 | RND3 | protein_coding | 38.66 | -0.051810335 | 0.051810335 | ENSG00000115963 | n.s. |
| ENSG00000105447 | 746.0234 | -0.335775725 | 0.276895 | -1.212644 | 0.225266 | 0.701091738 | GRWD1 | protein_coding | 55.81 | -0.051785062 | 0.051785062 | ENSG00000105447 | n.s. |
| ENSG00000274494 | 3.655744 | -0.618402805 | 0.742017 | -0.833407 | 0.404615 | 0.824799944 | MIR6832 | miRNA | 52.78 | -0.051730246 | 0.051730246 | ENSG00000274494 | n.s. |
| ENSG00000103043 | 269.0998 | -0.308608172 | 0.243093 | -1.269508 | 0.20426 | 0.679803349 | VAC14 | protein_coding | 54 | -0.051727883 | 0.051727883 | ENSG00000103043 | n.s. |
| ENSG00000231249 | 24.03262 | 0.403772287 | 0.375425 | 1.0755063 | 0.282148 | 0.744567264 | ITPR1-AS1 | antisense | 47.6 | 0.05172164 | 0.05172164 | ENSG00000231249 | n.s. |
| ENSG00000139323 | 390.0482 | -0.36155062 | 0.313409 | -1.153607 | 0.248661 | 0.719646518 | POC1B | protein_coding | 39.79 | -0.051658631 | 0.051658631 | ENSG00000139323 | n.s. |
| ENSG00000068028 | 317.862 | -0.297663742 | 0.229691 | -1.295932 | 0.194999 | 0.67069365 | RASSF1 | protein_coding | 54.57 | -0.051637457 | 0.051637457 | ENSG00000068028 | n.s. |
| ENSG00000153827 | 3580.509 | -0.322266755 | 0.259708 | -1.240879 | 0.21465 | 0.691472186 | TRIP12 | protein_coding | 38.19 | -0.051635282 | 0.051635282 | ENSG00000153827 | n.s. |

| | | | | | | | | | | | | | |
|---|---|---|---|---|---|---|---|---|---|---|---|---|---|
| ENSG00000073008 | 201.2139 | -0.328468676 | 0.267904 | -1.226069 | 0.220173 | 0.696377966 | PVR | protein_coding | 50.36 | -0.051620488 | 0.051620488 | ENSG00000073008 | n.s. |
| ENSG00000143442 | 2329.079 | -0.305269274 | 0.238825 | -1.278211 | 0.201175 | 0.677503939 | POGZ | protein_coding | 41.79 | -0.051617425 | 0.051617425 | ENSG00000143442 | n.s. |
| ENSG00000070010 | 849.3113 | -0.334914343 | 0.276812 | -1.209898 | 0.226318 | 0.701289341 | UFD1 | protein_coding | 46.94 | -0.051611225 | 0.051611225 | ENSG00000070010 | n.s. |
| ENSG00000164219 | 1592.558 | -0.315348167 | 0.251629 | -1.253225 | 0.210124 | 0.686023341 | PGGT1B | protein_coding | 35.39 | -0.05161023 | 0.05161023 | ENSG00000164219 | n.s. |
| ENSG00000068097 | 1372.31 | -0.318157744 | 0.254954 | -1.247902 | 0.212067 | 0.688575085 | HEATR6 | protein_coding | 40.4 | -0.051557047 | 0.051557047 | ENSG00000068097 | n.s. |
| ENSG00000179604 | 1949.178 | -0.306659904 | 0.241325 | -1.270732 | 0.203824 | 0.679010816 | CDC42EP4 | protein_coding | 52.99 | -0.051556677 | 0.051556677 | ENSG00000179604 | n.s. |
| ENSG00000013573 | 2244.707 | -0.295207143 | 0.227442 | -1.297943 | 0.194307 | 0.669140798 | DDX11 | protein_coding | 52.28 | -0.051508477 | 0.051508477 | ENSG00000013573 | n.s. |
| ENSG00000122299 | 1308.263 | -0.317725283 | 0.25454 | -1.248234 | 0.211945 | 0.688575085 | ZC3H7A | protein_coding | 42.77 | -0.051486968 | 0.051486968 | ENSG00000122299 | n.s. |
| ENSG00000174579 | 1319.029 | -0.29809431 | 0.230979 | -1.290568 | 0.196853 | 0.672118672 | MSL2 | protein_coding | 42.35 | -0.051437377 | 0.051437377 | ENSG00000174579 | n.s. |
| ENSG00000175782 | 639.4653 | -0.332753581 | 0.274179 | -1.213635 | 0.224887 | 0.700612508 | SLC35E3 | protein_coding | 43.41 | -0.051417787 | 0.051417787 | ENSG00000175782 | n.s. |
| ENSG00000063322 | 430.7806 | -0.332596138 | 0.27404 | -1.213679 | 0.22487 | 0.700612508 | MED29 | protein_coding | 52.36 | -0.051393458 | 0.051393458 | ENSG00000063322 | n.s. |
| ENSG00000240291 | 102.2804 | -0.358822596 | 0.310324 | -1.156283 | 0.247565 | 0.7191431 | | antisense | 39.11 | -0.051377899 | 0.051377899 | ENSG00000240291 | n.s. |
| ENSG00000071564 | 1520.219 | -0.30754684 | 0.24315 | -1.264844 | 0.205927 | 0.680704186 | TCF3 | protein_coding | 59.57 | -0.05137311 | 0.05137311 | ENSG00000071564 | n.s. |
| ENSG00000126215 | 318.2281 | -0.314769027 | 0.251494 | -1.251597 | 0.210717 | 0.686934651 | XRCC3 | protein_coding | 59.79 | -0.051333973 | 0.051333973 | ENSG00000126215 | n.s. |
| ENSG00000104886 | 413.6363 | -0.346795178 | 0.294838 | -1.176221 | 0.239506 | 0.71125159 | PLEKHJ1 | protein_coding | 61.63 | -0.051317623 | 0.051317623 | ENSG00000104886 | n.s. |
| ENSG00000207082 | 13.03598 | -0.418271104 | 0.398768 | -1.048907 | 0.294221 | 0.753915388 | RNU6-171P | snRNA | 43.93 | -0.051312408 | 0.051312408 | ENSG00000207082 | n.s. |
| ENSG00000138604 | 1771.126 | -0.326508321 | 0.266299 | -1.226095 | 0.220163 | 0.696377966 | GLCE | protein_coding | 36.64 | -0.051312408 | 0.051312408 | ENSG00000138604 | n.s. |
| ENSG00000100726 | 350.1867 | -0.318091003 | 0.255341 | -1.245749 | 0.212857 | 0.689743466 | TELO2 | protein_coding | 63.09 | -0.051312025 | 0.051312025 | ENSG00000100726 | n.s. |
| ENSG00000113161 | 3969.769 | -0.321495283 | 0.259787 | -1.237533 | 0.215889 | 0.692499549 | HMGCR | protein_coding | 37.88 | -0.05130438 | 0.05130438 | ENSG00000113161 | n.s. |
| ENSG00000224464 | 26.81966 | -0.390917307 | 0.357584 | -1.093217 | 0.274299 | 0.739386355 | PGAM1P6 | processed_pseudogene | 52.21 | -0.051260427 | 0.051260427 | ENSG00000224464 | n.s. |
| ENSG00000068784 | 1082.916 | -0.318276958 | 0.25588 | -1.243853 | 0.213554 | 0.690180176 | SRBD1 | protein_coding | 36.55 | -0.051254532 | 0.051254532 | ENSG00000068784 | n.s. |
| ENSG00000086475 | 1859.136 | -0.330784692 | 0.27196 | -1.216298 | 0.223871 | 0.69996872 | SEPHS1 | protein_coding | 45.78 | -0.051245617 | 0.051245617 | ENSG00000086475 | n.s. |
| ENSG00000143476 | 3997.853 | -0.322788987 | 0.261936 | -1.232322 | 0.217829 | 0.693853089 | DTL | protein_coding | 38.44 | -0.051237094 | 0.051237094 | ENSG00000143476 | n.s. |
| ENSG00000211997 | 22.37419 | -0.402767684 | 0.377262 | -1.067609 | 0.285697 | 0.74621359 | MIR708 | miRNA | 48.86 | -0.051206613 | 0.051206613 | ENSG00000211997 | n.s. |
| ENSG00000269053 | 12.09425 | 0.486985095 | 0.508609 | 0.9574849 | 0.338323 | 0.784993146 | | antisense | 47.45 | 0.051198757 | 0.051198757 | ENSG00000269053 | n.s. |
| ENSG00000241990 | 357.8669 | -0.289896593 | 0.221821 | -1.306891 | 0.19125 | 0.666152146 | PRR34-AS1 | antisense | 55.48 | -0.051145461 | 0.051145461 | ENSG00000241990 | n.s. |
| ENSG00000197728 | 4511.2 | -0.338857143 | 0.284759 | -1.18998 | 0.234054 | 0.706508138 | RPS26 | protein_coding | 46.62 | -0.051127725 | 0.051127725 | ENSG00000197728 | n.s. |
| ENSG00000155093 | 163.6613 | -0.335420472 | 0.27997 | -1.198059 | 0.230894 | 0.704043192 | PTPRN2 | protein_coding | 51.39 | -0.051118314 | 0.051118314 | ENSG00000155093 | n.s. |
| ENSG00000103222 | 1006.089 | -0.302676791 | 0.23762 | -1.273784 | 0.20274 | 0.677959669 | ABCC1 | protein_coding | 48.48 | -0.051090674 | 0.051090674 | ENSG00000103222 | n.s. |
| ENSG00000182400 | 677.7175 | -0.335967009 | 0.281492 | -1.193522 | 0.232665 | 0.704676403 | TRAPPC6B | protein_coding | 40.78 | -0.051070436 | 0.051070436 | ENSG00000182400 | n.s. |
| ENSG00000277581 | 109.0188 | 0.307464351 | 0.243566 | 1.2623435 | 0.206825 | 0.682214558 | | sense_intronic | 57.47 | 0.051063377 | 0.051063377 | ENSG00000277581 | n.s. |
| ENSG00000162734 | 380.5999 | -0.335020905 | 0.279878 | -1.197026 | 0.231296 | 0.704043192 | PEA15 | protein_coding | 51.85 | -0.051057419 | 0.051057419 | ENSG00000162734 | n.s. |

| | | | | | | | | | | | | | |
|---|---|---|---|---|---|---|---|---|---|---|---|---|---|
| ENSG00000178184 | 199.6924 | -0.356518043 | 0.308532 | -1.155531 | 0.247873 | 0.719195543 | PARD6G | protein_coding | 47.5 | -0.051036632 | 0.051036632 | ENSG00000178184 | n.s. |
| ENSG00000203326 | 112.1602 | -0.344296399 | 0.292112 | -1.178645 | 0.23854 | 0.710909549 | ZNF525 | protein_coding | 43.64 | -0.051019786 | 0.051019786 | ENSG00000203326 | n.s. |
| ENSG00000278864 | 185.6107 | -0.313949297 | 0.251286 | -1.249371 | 0.211529 | 0.687952251 | | TEC | 53.36 | -0.050998459 | 0.050998459 | ENSG00000278864 | n.s. |
| ENSG00000143387 | 75.63171 | -0.349192217 | 0.298709 | -1.169007 | 0.242401 | 0.714515282 | CTSK | protein_coding | 43 | -0.05097804 | 0.05097804 | ENSG00000143387 | n.s. |
| ENSG00000258399 | 229.6885 | 0.338115099 | 0.28437 | 1.1889959 | 0.234441 | 0.706826753 | | lincRNA | 46.71 | 0.050949557 | 0.050949557 | ENSG00000258399 | n.s. |
| ENSG00000120784 | 999.2313 | -0.324327999 | 0.264806 | -1.224775 | 0.22066 | 0.696655251 | ZFP30 | protein_coding | 41.09 | -0.050913685 | 0.050913685 | ENSG00000120784 | n.s. |
| ENSG00000164327 | 1179.948 | -0.333970196 | 0.279272 | -1.195861 | 0.231751 | 0.704043192 | RICTOR | protein_coding | 35.53 | -0.05089729 | 0.05089729 | ENSG00000164327 | n.s. |
| ENSG00000104880 | 324.7553 | -0.333680472 | 0.279251 | -1.194914 | 0.232121 | 0.704043192 | ARHGEF18 | protein_coding | 51.44 | -0.050853136 | 0.050853136 | ENSG00000104880 | n.s. |
| ENSG00000184557 | 109.2447 | 0.341460638 | 0.288921 | 1.1818486 | 0.237266 | 0.709809046 | SOCS3 | protein_coding | 62.24 | 0.050829308 | 0.050829308 | ENSG00000184557 | n.s. |
| ENSG00000251821 | 28.03023 | -0.38804439 | 0.355764 | -1.090734 | 0.27539 | 0.739632414 | RNU6-583P | snRNA | 40.22 | -0.050827631 | 0.050827631 | ENSG00000251821 | n.s. |
| ENSG00000252627 | 7.296492 | -0.509352332 | 0.551791 | -0.923088 | 0.355961 | 0.794802462 | RNU6-122P | snRNA | 34.58 | -0.050803207 | 0.050803207 | ENSG00000252627 | n.s. |
| ENSG00000108262 | 404.4974 | -0.297309251 | 0.231168 | -1.286117 | 0.198402 | 0.674743473 | GIT1 | protein_coding | 60.72 | -0.050798647 | 0.050798647 | ENSG00000108262 | n.s. |
| ENSG00000124214 | 1146.418 | -0.315390879 | 0.253467 | -1.244306 | 0.213387 | 0.690152016 | STAU1 | protein_coding | 43.2 | -0.050795353 | 0.050795353 | ENSG00000124214 | n.s. |
| ENSG00000251862 | 8.703528 | 0.460231353 | 0.467058 | 0.9853832 | 0.324436 | 0.775604549 | MIR1343 | miRNA | 79.76 | 0.050790972 | 0.050790972 | ENSG00000251862 | n.s. |
| ENSG00000198464 | 1008.886 | -0.331407607 | 0.274987 | -1.205177 | 0.228135 | 0.702737638 | ZNF480 | protein_coding | 42.41 | -0.050773894 | 0.050773894 | ENSG00000198464 | n.s. |
| ENSG00000269887 | 37.23154 | -0.375853675 | 0.335139 | -1.121485 | 0.262082 | 0.732723331 | | sense_intronic | 33.15 | -0.05076279 | 0.05076279 | ENSG00000269887 | n.s. |
| ENSG00000166002 | 45.16062 | -0.394019587 | 0.363615 | -1.083618 | 0.278534 | 0.743390786 | SMCO4 | protein_coding | 45.2 | -0.050742956 | 0.050742956 | ENSG00000166002 | n.s. |
| ENSG00000174672 | 359.0584 | -0.311236315 | 0.248845 | -1.250723 | 0.211035 | 0.687111024 | BRSK2 | protein_coding | 63.85 | -0.050723142 | 0.050723142 | ENSG00000174672 | n.s. |
| ENSG00000272716 | 81.11627 | 0.354361468 | 0.306976 | 1.1543631 | 0.248351 | 0.719540199 | | lincRNA | 47.89 | 0.050654178 | 0.050654178 | ENSG00000272716 | n.s. |
| ENSG00000186153 | 123.1269 | -0.349193476 | 0.30028 | -1.162893 | 0.244873 | 0.716253458 | WWOX | protein_coding | 42.64 | -0.050609752 | 0.050609752 | ENSG00000186153 | n.s. |
| ENSG00000151445 | 347.5548 | -0.329571436 | 0.27309 | -1.206825 | 0.2275 | 0.70225867 | VIPAS39 | protein_coding | 41.62 | -0.050590168 | 0.050590168 | ENSG00000151445 | n.s. |
| ENSG00000122756 | 384.7065 | -0.296047382 | 0.230249 | -1.285769 | 0.198524 | 0.674743473 | CNTFR | protein_coding | 56.26 | -0.050583043 | 0.050583043 | ENSG00000122756 | n.s. |
| ENSG00000169217 | 1361.723 | -0.316855476 | 0.256285 | -1.236338 | 0.216333 | 0.692499549 | CD2BP2 | protein_coding | 57.88 | -0.050563957 | 0.050563957 | ENSG00000169217 | n.s. |
| ENSG00000153187 | 38646.08 | -0.295647254 | 0.229902 | -1.285971 | 0.198453 | 0.674743473 | HNRNPU | protein_coding | 38.67 | -0.050514676 | 0.050514676 | ENSG00000153187 | n.s. |
| ENSG00000146701 | 1972.161 | -0.330988699 | 0.275692 | -1.200573 | 0.229917 | 0.703699853 | MDH2 | protein_coding | 50.93 | -0.050513026 | 0.050513026 | ENSG00000146701 | n.s. |
| ENSG00000089050 | 453.1088 | -0.335699973 | 0.282601 | -1.187894 | 0.234875 | 0.707216633 | RBBP9 | protein_coding | 41.68 | -0.050505233 | 0.050505233 | ENSG00000089050 | n.s. |
| ENSG00000187838 | 87.52029 | -0.33460093 | 0.28126 | -1.189652 | 0.234183 | 0.706508138 | PLSCR3 | protein_coding | 58.45 | -0.050485536 | 0.050485536 | ENSG00000187838 | n.s. |
| ENSG00000271452 | 32.15388 | -0.380955457 | 0.345667 | -1.102087 | 0.270424 | 0.737020734 | | sense_intronic | 34.76 | -0.050484329 | 0.050484329 | ENSG00000271452 | n.s. |
| ENSG00000130024 | 2681.167 | -0.332588098 | 0.278925 | -1.192392 | 0.233107 | 0.705095593 | PHF10 | protein_coding | 39.5 | -0.050470909 | 0.050470909 | ENSG00000130024 | n.s. |
| ENSG00000142208 | 1691.722 | -0.326460041 | 0.268839 | -1.214334 | 0.22462 | 0.700529123 | AKT1 | protein_coding | 64.08 | -0.050462171 | 0.050462171 | ENSG00000142208 | n.s. |
| ENSG00000160818 | 1644.399 | -0.285691748 | 0.218657 | -1.306577 | 0.191356 | 0.666152146 | GPATCH4 | protein_coding | 48.64 | -0.050403615 | 0.050403615 | ENSG00000160818 | n.s. |
| ENSG00000105612 | 111.8625 | -0.340575521 | 0.289473 | -1.176536 | 0.239381 | 0.71125159 | DNASE2 | protein_coding | 52.62 | -0.050397259 | 0.050397259 | ENSG00000105612 | n.s. |

| | | | | | | | | | | | | | |
|---|---|---|---|---|---|---|---|---|---|---|---|---|---|
| ENSG00000164631 | 1312.899 | -0.323862472 | 0.265494 | -1.219847 | 0.222523 | 0.698891176 | ZNF12 | protein_coding | 43.43 | -0.050389905 | 0.050389905 | ENSG00000164631 | n.s. |
| ENSG00000214435 | 427.7788 | -0.338661467 | 0.286696 | -1.181255 | 0.237501 | 0.710056308 | AS3MT | protein_coding | 43.69 | -0.050361402 | 0.050361402 | ENSG00000214435 | n.s. |
| ENSG00000111300 | 3073.677 | -0.307689095 | 0.245557 | -1.253023 | 0.210197 | 0.686023341 | NAA25 | protein_coding | 41.42 | -0.050356738 | 0.050356738 | ENSG00000111300 | n.s. |
| ENSG00000130402 | 195.3038 | -0.31546711 | 0.255257 | -1.235881 | 0.216503 | 0.692499549 | ACTN4 | protein_coding | 53.01 | -0.050342401 | 0.050342401 | ENSG00000130402 | n.s. |
| ENSG00000104643 | 1431.287 | -0.31961176 | 0.260352 | -1.227613 | 0.219592 | 0.695810058 | MTMR9 | protein_coding | 37.85 | -0.050341824 | 0.050341824 | ENSG00000104643 | n.s. |
| ENSG00000101945 | 232.8527 | -0.326667611 | 0.270022 | -1.209781 | 0.226363 | 0.701289341 | SUV39H1 | protein_coding | 54.35 | -0.050340381 | 0.050340381 | ENSG00000101945 | n.s. |
| ENSG00000153976 | 155.1051 | 0.317022441 | 0.257174 | 1.2327159 | 0.217682 | 0.693853089 | HS3ST3A1 | protein_coding | 40.94 | 0.050321756 | 0.050321756 | ENSG00000153976 | n.s. |
| ENSG00000198917 | 576.3601 | -0.31235141 | 0.251143 | -1.243721 | 0.213602 | 0.690180176 | SPOUT1 | protein_coding | 57.61 | -0.050300296 | 0.050300296 | ENSG00000198917 | n.s. |
| ENSG00000249709 | 353.966 | -0.326206999 | 0.269682 | -1.2096 | 0.226432 | 0.701289341 | ZNF564 | protein_coding | 44.24 | -0.0502694 | 0.0502694 | ENSG00000249709 | n.s. |
| ENSG00000204147 | 294.7008 | -0.326314232 | 0.269867 | -1.209168 | 0.226598 | 0.701406486 | ASAH2B | protein_coding | 38.24 | -0.050262254 | 0.050262254 | ENSG00000204147 | n.s. |
| ENSG00000054523 | 975.4844 | -0.291479878 | 0.225937 | -1.290096 | 0.197017 | 0.67232198 | KIF1B | protein_coding | 41.99 | -0.050257745 | 0.050257745 | ENSG00000054523 | n.s. |
| ENSG00000228222 | 23.1267 | -0.393195662 | 0.366421 | -1.07307 | 0.28324 | 0.745049657 | | lincRNA | 36.54 | -0.050256217 | 0.050256217 | ENSG00000228222 | n.s. |
| ENSG00000125170 | 2262.301 | -0.326315421 | 0.270196 | -1.207701 | 0.227162 | 0.70173545 | DOK4 | protein_coding | 58.39 | -0.050195986 | 0.050195986 | ENSG00000125170 | n.s. |
| ENSG00000134253 | 271.9938 | -0.287909823 | 0.221897 | -1.297492 | 0.194462 | 0.669509439 | TRIM45 | protein_coding | 46.89 | -0.050166356 | 0.050166356 | ENSG00000134253 | n.s. |
| ENSG00000169021 | 2131.483 | 0.320605283 | 0.262282 | 1.2223701 | 0.221568 | 0.697595355 | UQCRFS1 | protein_coding | 42.39 | 0.050141518 | 0.050141518 | ENSG00000169021 | n.s. |
| ENSG00000111737 | 630.3853 | -0.333190354 | 0.280403 | -1.188256 | 0.234732 | 0.707216633 | RAB35 | protein_coding | 55.75 | -0.050127667 | 0.050127667 | ENSG00000111737 | n.s. |
| ENSG00000186998 | 170.2913 | -0.339654937 | 0.289112 | -1.174823 | 0.240066 | 0.712065189 | EMID1 | protein_coding | 52.65 | -0.050092393 | 0.050092393 | ENSG00000186998 | n.s. |
| ENSG00000173818 | 68.70473 | -0.3573738 | 0.313172 | -1.141142 | 0.253811 | 0.72418384 | ENDOV | protein_coding | 56.47 | -0.050086356 | 0.050086356 | ENSG00000173818 | n.s. |
| ENSG00000144815 | 905.1222 | -0.315203457 | 0.255363 | -1.234337 | 0.217077 | 0.693680706 | NXPE3 | protein_coding | 39.47 | -0.050067038 | 0.050067038 | ENSG00000144815 | n.s. |
| ENSG00000130477 | 993.311 | -0.311500579 | 0.250731 | -1.242369 | 0.2141 | 0.690827094 | UNC13A | protein_coding | 52.17 | -0.050036537 | 0.050036537 | ENSG00000130477 | n.s. |
| ENSG00000171132 | 378.9934 | -0.299242777 | 0.235994 | -1.268012 | 0.204793 | 0.680575501 | PRKCE | protein_coding | 44.43 | -0.050010557 | 0.050010557 | ENSG00000171132 | n.s. |
| ENSG00000277782 | 57.93485 | 0.349499781 | 0.302837 | 1.1540849 | 0.248465 | 0.719540199 | | lincRNA | 47.56 | 0.049959225 | 0.049959225 | ENSG00000277782 | n.s. |
| ENSG00000215966 | 4.508426 | -0.745305801 | 1.005761 | -0.741037 | 0.458671 | 0.856995387 | MIR876 | miRNA | 37.04 | -0.049951525 | 0.049951525 | ENSG00000215966 | n.s. |
| ENSG00000164741 | 3709.793 | -0.295422404 | 0.231349 | -1.276953 | 0.201619 | 0.677624898 | DLC1 | protein_coding | 38.54 | -0.049929531 | 0.049929531 | ENSG00000164741 | n.s. |
| ENSG00000104129 | 186.6419 | -0.335281759 | 0.283816 | -1.181334 | 0.23747 | 0.710056308 | DNAJC17 | protein_coding | 50.19 | -0.049858815 | 0.049858815 | ENSG00000104129 | n.s. |
| ENSG00000099194 | 6690.525 | -0.313841284 | 0.254333 | -1.233976 | 0.217212 | 0.693734586 | SCD | protein_coding | 47.04 | -0.049840084 | 0.049840084 | ENSG00000099194 | n.s. |
| ENSG00000243094 | 23.38298 | -0.396462395 | 0.374453 | -1.058778 | 0.289701 | 0.748759774 | | processed_pseudogene | 46.04 | -0.049818472 | 0.049818472 | ENSG00000243094 | n.s. |
| ENSG00000156017 | 407.3084 | -0.311761651 | 0.251632 | -1.238961 | 0.21536 | 0.692210945 | CARNMT1 | protein_coding | 37.59 | -0.049807521 | 0.049807521 | ENSG00000156017 | n.s. |
| ENSG00000272576 | 62.56177 | -0.325043852 | 0.269909 | -1.204274 | 0.228484 | 0.702737638 | | lincRNA | 35.08 | -0.049798924 | 0.049798924 | ENSG00000272576 | n.s. |
| ENSG00000283101 | 48.69666 | 0.355171074 | 0.311629 | 1.1397248 | 0.254401 | 0.72418384 | BMS1P21 | cribed_unprocessed_pseudc | 36.5 | 0.049777642 | 0.049777642 | ENSG00000283101 | n.s. |
| ENSG00000198046 | 976.5739 | -0.297897313 | 0.235423 | -1.265369 | 0.205739 | 0.680704186 | ZNF667 | protein_coding | 43.97 | -0.049761237 | 0.049761237 | ENSG00000198046 | n.s. |
| ENSG00000200840 | 9.485042 | -0.710916678 | 0.940665 | -0.75576 | 0.449793 | 0.85118703 | RNU6-82P | snRNA | 39.25 | -0.049746396 | 0.049746396 | ENSG00000200840 | n.s. |

| | | | | | | | | | | | | | |
|---|---|---|---|---|---|---|---|---|---|---|---|---|---|
| ENSG00000121900 | 44.46757 | -0.38958201 | 0.363651 | -1.071308 | 0.284031 | 0.745300357 | TMEM54 | protein_coding | 57.33 | -0.049737417 | 0.049737417 | ENSG00000121900 | n.s. |
| ENSG00000117152 | 8771.023 | 0.311511978 | 0.25199 | 1.2362094 | 0.216381 | 0.692499549 | RGS4 | protein_coding | 39.47 | 0.049711239 | 0.049711239 | ENSG00000117152 | n.s. |
| ENSG00000156853 | 411.2352 | -0.288074661 | 0.222908 | -1.292347 | 0.196237 | 0.672118672 | ZNF689 | protein_coding | 48.18 | -0.049708446 | 0.049708446 | ENSG00000156853 | n.s. |
| ENSG00000126254 | 437.9064 | -0.335881865 | 0.285606 | -1.176033 | 0.239582 | 0.71125159 | RBM42 | protein_coding | 53.79 | -0.049702707 | 0.049702707 | ENSG00000126254 | n.s. |
| ENSG00000171603 | 387.057 | -0.320859777 | 0.264053 | -1.215133 | 0.224316 | 0.700099826 | CLSTN1 | protein_coding | 47.37 | -0.049681938 | 0.049681938 | ENSG00000171603 | n.s. |
| ENSG00000164031 | 570.2021 | -0.346651274 | 0.299373 | -1.157923 | 0.246896 | 0.719028029 | DNAJB14 | protein_coding | 34.8 | -0.049659244 | 0.049659244 | ENSG00000164031 | n.s. |
| ENSG00000186889 | 122.9643 | -0.324013978 | 0.268764 | -1.20557 | 0.227983 | 0.702691634 | TMEM17 | protein_coding | 39.29 | -0.049650352 | 0.049650352 | ENSG00000186889 | n.s. |
| ENSG00000104331 | 1559.136 | -0.320155473 | 0.26305 | -1.217088 | 0.223571 | 0.699734492 | IMPAD1 | protein_coding | 38.21 | -0.049645458 | 0.049645458 | ENSG00000104331 | n.s. |
| ENSG00000093183 | 1376.783 | -0.323656831 | 0.268339 | -1.206149 | 0.22776 | 0.702503504 | SEC22C | protein_coding | 43.29 | -0.049633262 | 0.049633262 | ENSG00000093183 | n.s. |
| ENSG00000127957 | 227.3967 | -0.328394887 | 0.275846 | -1.190499 | 0.23385 | 0.706422166 | PMS2P3 | cribed_unprocessed_pseudc | 42.92 | -0.049566506 | 0.049566506 | ENSG00000127957 | n.s. |
| ENSG00000187391 | 9.421689 | -0.47142679 | 0.492647 | -0.956925 | 0.338605 | 0.784993146 | MAGI2 | protein_coding | 36.83 | -0.049563048 | 0.049563048 | ENSG00000187391 | n.s. |
| ENSG00000265442 | 4.013811 | -0.562090253 | 0.658926 | -0.85304 | 0.393637 | 0.816295599 | MIR3941 | miRNA | 40.78 | -0.049549686 | 0.049549686 | ENSG00000265442 | n.s. |
| ENSG00000086619 | 184.4897 | 0.334293206 | 0.283789 | 1.1779644 | 0.238811 | 0.710909549 | ERO1B | protein_coding | 37.47 | 0.049537457 | 0.049537457 | ENSG00000086619 | n.s. |
| ENSG00000237370 | 12.73484 | 0.607681549 | 0.742676 | 0.8182324 | 0.413224 | 0.829007487 | | antisense | 58.51 | 0.049490526 | 0.049490526 | ENSG00000237370 | n.s. |
| ENSG00000167291 | 2258.996 | -0.304253854 | 0.243423 | -1.249897 | 0.211337 | 0.687648811 | TBC1D16 | protein_coding | 52.48 | -0.049481809 | 0.049481809 | ENSG00000167291 | n.s. |
| ENSG00000185883 | 250.5762 | -0.33928148 | 0.290452 | -1.168116 | 0.24276 | 0.714766539 | ATP6V0C | protein_coding | 60.18 | -0.049479381 | 0.049479381 | ENSG00000185883 | n.s. |
| ENSG00000164983 | 499.2981 | -0.319443111 | 0.26287 | -1.215215 | 0.224284 | 0.700099826 | TMEM65 | protein_coding | 35.86 | -0.049462581 | 0.049462581 | ENSG00000164983 | n.s. |
| ENSG00000160207 | 31.85919 | 0.386853886 | 0.360568 | 1.0729022 | 0.283315 | 0.745106685 | HSF2BP | protein_coding | 43.01 | 0.049432785 | 0.049432785 | ENSG00000160207 | n.s. |
| ENSG00000138443 | 2065.139 | -0.315551655 | 0.258074 | -1.22272 | 0.221435 | 0.697363879 | ABI2 | protein_coding | 39.51 | -0.04939663 | 0.04939663 | ENSG00000138443 | n.s. |
| ENSG00000206195 | 61.10524 | -0.339498663 | 0.291175 | -1.16596 | 0.243631 | 0.715324966 | | processed_transcript | 41.82 | -0.049395907 | 0.049395907 | ENSG00000206195 | n.s. |
| ENSG00000165684 | 345.7078 | -0.290476692 | 0.226364 | -1.283226 | 0.199413 | 0.676054647 | SNAPC4 | protein_coding | 58.99 | -0.049386324 | 0.049386324 | ENSG00000165684 | n.s. |
| ENSG00000154781 | 644.3452 | -0.323951844 | 0.271156 | -1.194705 | 0.232202 | 0.704043192 | CCDC174 | protein_coding | 42.28 | -0.049370487 | 0.049370487 | ENSG00000154781 | n.s. |
| ENSG00000143952 | 1047.002 | -0.329409803 | 0.277889 | -1.185402 | 0.235858 | 0.7081858 | VPS54 | protein_coding | 36.99 | -0.049362977 | 0.049362977 | ENSG00000143952 | n.s. |
| ENSG00000101337 | 433.1061 | -0.309264918 | 0.250128 | -1.236426 | 0.2163 | 0.692499549 | TM9SF4 | protein_coding | 47.45 | -0.049352652 | 0.049352652 | ENSG00000101337 | n.s. |
| ENSG00000116285 | 247.8748 | -0.335497525 | 0.286042 | -1.172897 | 0.240837 | 0.712813552 | ERRFI1 | protein_coding | 42.8 | -0.049326205 | 0.049326205 | ENSG00000116285 | n.s. |
| ENSG00000104549 | 3782.848 | -0.319924611 | 0.264216 | -1.210845 | 0.225955 | 0.701289341 | SQLE | protein_coding | 37.59 | -0.049301266 | 0.049301266 | ENSG00000104549 | n.s. |
| ENSG00000242612 | 136.5165 | -0.345384613 | 0.299683 | -1.152501 | 0.249115 | 0.720036883 | DECR2 | protein_coding | 56.95 | -0.049267477 | 0.049267477 | ENSG00000242612 | n.s. |
| ENSG00000207189 | 9.784041 | -0.488186785 | 0.525314 | -0.929324 | 0.352721 | 0.79272638 | | misc_RNA | 45.1 | -0.049246667 | 0.049246667 | ENSG00000207189 | n.s. |
| ENSG00000106299 | 947.5056 | -0.311086539 | 0.252853 | -1.230306 | 0.218582 | 0.694664257 | WASL | protein_coding | 35.43 | -0.049221682 | 0.049221682 | ENSG00000106299 | n.s. |
| ENSG00000166004 | 1665.635 | -0.313334183 | 0.255788 | -1.224975 | 0.220584 | 0.696655251 | CEP295 | protein_coding | 38.58 | -0.049187853 | 0.049187853 | ENSG00000166004 | n.s. |
| ENSG00000183960 | 491.6447 | -0.319458118 | 0.264402 | -1.20823 | 0.226959 | 0.701641249 | KCNH8 | protein_coding | 36.03 | -0.049159776 | 0.049159776 | ENSG00000183960 | n.s. |
| ENSG00000101126 | 2848.115 | -0.309365646 | 0.250805 | -1.23349 | 0.217393 | 0.693734586 | ADNP | protein_coding | 41.52 | -0.049129323 | 0.049129323 | ENSG00000101126 | n.s. |

| | | | | | | | | | | | | | |
|---|---|---|---|---|---|---|---|---|---|---|---|---|---|
| ENSG00000204946 | 717.2439 | -0.309740569 | 0.251423 | -1.231948 | 0.217968 | 0.694065444 | ZNF783 | protein_coding | 51.5 | -0.049124724 | 0.049124724 | ENSG00000204946 | n.s. |
| ENSG00000207313 | 90.59602 | 0.347419537 | 0.302622 | 1.1480308 | 0.250956 | 0.722169533 | SNORA2B | snoRNA | 40.15 | 0.049111517 | 0.049111517 | ENSG00000207313 | n.s. |
| ENSG00000067992 | 113.8287 | -0.331683556 | 0.281884 | -1.176665 | 0.239329 | 0.71125159 | PDK3 | protein_coding | 40.95 | -0.049081454 | 0.049081454 | ENSG00000067992 | n.s. |
| ENSG00000255310 | 187.9891 | -0.33572292 | 0.287476 | -1.167829 | 0.242876 | 0.714766539 | | sense_intronic | 42.08 | -0.048960416 | 0.048960416 | ENSG00000255310 | n.s. |
| ENSG00000109016 | 229.8559 | -0.33598864 | 0.287968 | -1.166756 | 0.243309 | 0.715115708 | DHRS7B | protein_coding | 44.22 | -0.048927903 | 0.048927903 | ENSG00000109016 | n.s. |
| ENSG00000186575 | 879.2381 | -0.277289945 | 0.212097 | -1.307373 | 0.191086 | 0.666152146 | NF2 | protein_coding | 45.79 | -0.048921313 | 0.048921313 | ENSG00000186575 | n.s. |
| ENSG00000054654 | 5880.404 | -0.319271062 | 0.265 | -1.204798 | 0.228281 | 0.702737638 | SYNE2 | protein_coding | 40.76 | -0.048914493 | 0.048914493 | ENSG00000054654 | n.s. |
| ENSG00000131375 | 1066.971 | -0.332670685 | 0.283605 | -1.173005 | 0.240794 | 0.712813552 | CAPN7 | protein_coding | 37.11 | -0.048910592 | 0.048910592 | ENSG00000131375 | n.s. |
| ENSG00000176087 | 560.687 | -0.278582067 | 0.213923 | -1.302251 | 0.19283 | 0.667523235 | SLC35A4 | protein_coding | 54.5 | -0.048900517 | 0.048900517 | ENSG00000176087 | n.s. |
| ENSG00000134265 | 717.3434 | -0.310032962 | 0.252387 | -1.228401 | 0.219296 | 0.695503935 | NAPG | protein_coding | 39.34 | -0.048892325 | 0.048892325 | ENSG00000134265 | n.s. |
| ENSG00000168264 | 4556.08 | 0.259857755 | 0.193595 | 1.342276 | 0.179507 | 0.648529087 | IRF2BP2 | protein_coding | 48.01 | 0.048871589 | 0.048871589 | ENSG00000168264 | n.s. |
| ENSG00000178028 | 212.9673 | -0.303310956 | 0.243627 | -1.24498 | 0.213139 | 0.69012437 | DMAP1 | protein_coding | 52.33 | -0.048855095 | 0.048855095 | ENSG00000178028 | n.s. |
| ENSG00000170265 | 434.5219 | -0.310686007 | 0.253331 | -1.226402 | 0.220047 | 0.696377966 | ZNF282 | protein_coding | 51.85 | -0.048825853 | 0.048825853 | ENSG00000170265 | n.s. |
| ENSG00000100413 | 734.7723 | -0.321353055 | 0.269389 | -1.192895 | 0.23291 | 0.704958888 | POLR3H | protein_coding | 55.16 | -0.04879303 | 0.04879303 | ENSG00000100413 | n.s. |
| ENSG00000168701 | 477.4682 | -0.33415128 | 0.285932 | -1.168638 | 0.24255 | 0.714515282 | TMEM208 | protein_coding | 55.38 | -0.048782237 | 0.048782237 | ENSG00000168701 | n.s. |
| ENSG00000248527 | 180.3123 | -0.304867142 | 0.245822 | -1.240194 | 0.214904 | 0.691815763 | MTATP6P1 | unprocessed_pseudogene | 43.91 | -0.048781653 | 0.048781653 | ENSG00000248527 | n.s. |
| ENSG00000166507 | 217.6183 | -0.30433046 | 0.245296 | -1.240667 | 0.214729 | 0.691472186 | NDST2 | protein_coding | 53.04 | -0.048761435 | 0.048761435 | ENSG00000166507 | n.s. |
| ENSG00000173588 | 1224.279 | -0.316271967 | 0.261434 | -1.209758 | 0.226372 | 0.701289341 | CEP83 | protein_coding | 37.69 | -0.048738384 | 0.048738384 | ENSG00000173588 | n.s. |
| ENSG00000148950 | 314.9472 | -0.35101844 | 0.309337 | -1.134746 | 0.256482 | 0.726402587 | IMMP1L | protein_coding | 34.91 | -0.048729299 | 0.048729299 | ENSG00000148950 | n.s. |
| ENSG00000150403 | 567.4288 | -0.32115155 | 0.269379 | -1.192192 | 0.233186 | 0.705179682 | TMCO3 | protein_coding | 48.38 | -0.048718757 | 0.048718757 | ENSG00000150403 | n.s. |
| ENSG00000198551 | 537.9379 | -0.306479683 | 0.248242 | -1.2346 | 0.216979 | 0.693680706 | ZNF627 | protein_coding | 48.23 | -0.048681351 | 0.048681351 | ENSG00000198551 | n.s. |
| ENSG00000066044 | 865.0584 | -0.325757188 | 0.275199 | -1.183715 | 0.236526 | 0.709118937 | ELAVL1 | protein_coding | 52.21 | -0.048629332 | 0.048629332 | ENSG00000066044 | n.s. |
| ENSG00000102763 | 885.4804 | -0.317910695 | 0.26468 | -1.201112 | 0.229708 | 0.703369603 | VWA8 | protein_coding | 37.16 | -0.048581969 | 0.048581969 | ENSG00000102763 | n.s. |
| ENSG00000166199 | 493.69 | -0.302746104 | 0.243779 | -1.241885 | 0.214279 | 0.691201113 | ALKBH3 | protein_coding | 40.79 | -0.048559134 | 0.048559134 | ENSG00000166199 | n.s. |
| ENSG00000264720 | 6.756495 | -0.494362321 | 0.540942 | -0.913891 | 0.360774 | 0.797857548 | MIR3117 | miRNA | 46.15 | -0.048484408 | 0.048484408 | ENSG00000264720 | n.s. |
| ENSG00000108854 | 2137.627 | -0.307849957 | 0.250856 | -1.227198 | 0.219748 | 0.695985401 | SMURF2 | protein_coding | 39.4 | -0.048455543 | 0.048455543 | ENSG00000108854 | n.s. |
| ENSG00000207605 | 4.62063 | -0.823961868 | 1.207824 | -0.682187 | 0.495121 | 0.873485531 | MIR191 | miRNA | 59.78 | -0.04840305 | 0.04840305 | ENSG00000207605 | n.s. |
| ENSG00000257242 | 103.8132 | 0.332905443 | 0.285763 | 1.1649696 | 0.244031 | 0.715606021 | LINC01619 | processed_transcript | 38.41 | 0.048379821 | 0.048379821 | ENSG00000257242 | n.s. |
| ENSG00000204310 | 436.0002 | -0.304550576 | 0.246692 | -1.234536 | 0.217003 | 0.693680706 | AGPAT1 | protein_coding | 50.57 | -0.048374931 | 0.048374931 | ENSG00000204310 | n.s. |
| ENSG00000255647 | 132.7199 | 0.313404474 | 0.259068 | 1.2097397 | 0.226379 | 0.701289341 | | processed_transcript | 46.57 | 0.048296495 | 0.048296495 | ENSG00000255647 | n.s. |
| ENSG00000275126 | 1218.534 | -0.27552316 | 0.211832 | -1.300665 | 0.193373 | 0.667903971 | HIST1H4L | protein_coding | 58.97 | -0.048295346 | 0.048295346 | ENSG00000275126 | n.s. |
| ENSG00000135476 | 1137.094 | -0.286421542 | 0.224965 | -1.273181 | 0.202954 | 0.678282423 | ESPL1 | protein_coding | 51.69 | -0.048287647 | 0.048287647 | ENSG00000135476 | n.s. |

| | | | | | | | | | | | | | |
|---|---|---|---|---|---|---|---|---|---|---|---|---|---|
| ENSG00000178188 | 447.2653 | -0.302413479 | 0.244458 | -1.237078 | 0.216058 | 0.692499549 | SH2B1 | protein_coding | 51.11 | -0.048259296 | 0.048259296 | ENSG00000178188 | n.s. |
| ENSG00000165055 | 605.2922 | -0.313583 | 0.259558 | -1.208141 | 0.226993 | 0.701641249 | METTL2B | protein_coding | 45.29 | -0.048255684 | 0.048255684 | ENSG00000165055 | n.s. |
| ENSG00000138617 | 429.3584 | -0.284312863 | 0.221759 | -1.282081 | 0.199814 | 0.676613539 | PARP16 | protein_coding | 46.92 | -0.048236326 | 0.048236326 | ENSG00000138617 | n.s. |
| ENSG00000198466 | 576.0502 | -0.310064187 | 0.254438 | -1.218625 | 0.222986 | 0.698928583 | ZNF587 | protein_coding | 46.53 | -0.048235817 | 0.048235817 | ENSG00000198466 | n.s. |
| ENSG00000103707 | 375.532 | -0.327528388 | 0.278995 | -1.17396 | 0.240411 | 0.712634415 | MTFMT | protein_coding | 40 | -0.048190302 | 0.048190302 | ENSG00000103707 | n.s. |
| ENSG00000263465 | 725.0645 | 0.301267653 | 0.243025 | 1.2396567 | 0.215102 | 0.691899391 | SRSF8 | protein_coding | 43.87 | 0.048189885 | 0.048189885 | ENSG00000263465 | n.s. |
| ENSG00000134901 | 698.3075 | -0.335986181 | 0.290131 | -1.15805 | 0.246844 | 0.719028029 | KDELC1 | protein_coding | 40.94 | -0.048131425 | 0.048131425 | ENSG00000134901 | n.s. |
| ENSG00000254144 | 27.85741 | 0.404614129 | 0.393187 | 1.0290624 | 0.30345 | 0.760781413 | | antisense | 54.3 | 0.048043925 | 0.048043925 | ENSG00000254144 | n.s. |
| ENSG00000234964 | 64.07957 | 0.468328153 | 0.498173 | 0.9400911 | 0.347171 | 0.789748319 | FABP5P7 | processed_pseudogene | 44.44 | 0.048008923 | 0.048008923 | ENSG00000234964 | n.s. |
| ENSG00000132740 | 102.9653 | -0.326682829 | 0.27872 | -1.172081 | 0.241165 | 0.712994199 | IGHMBP2 | protein_coding | 50.65 | -0.047994282 | 0.047994282 | ENSG00000132740 | n.s. |
| ENSG00000015171 | 1549.795 | -0.331112879 | 0.284724 | -1.162925 | 0.24486 | 0.716253458 | ZMYND11 | protein_coding | 37.8 | -0.047989272 | 0.047989272 | ENSG00000015171 | n.s. |
| ENSG00000234219 | 117.8113 | -0.328891247 | 0.281596 | -1.167955 | 0.242825 | 0.714766539 | CDCA4P4 | unprocessed_pseudogene | 58.53 | -0.047964114 | 0.047964114 | ENSG00000234219 | n.s. |
| ENSG00000119969 | 3639.256 | -0.308296545 | 0.252867 | -1.219203 | 0.222767 | 0.698928583 | HELLS | protein_coding | 38.96 | -0.04796083 | 0.04796083 | ENSG00000119969 | n.s. |
| ENSG00000100105 | 1004.603 | -0.29037291 | 0.230724 | -1.258531 | 0.2082 | 0.683663723 | PATZ1 | protein_coding | 52.8 | -0.047957254 | 0.047957254 | ENSG00000100105 | n.s. |
| ENSG00000033011 | 452.7909 | -0.305469051 | 0.249333 | -1.225146 | 0.22052 | 0.696655251 | ALG1 | protein_coding | 49.28 | -0.047953169 | 0.047953169 | ENSG00000033011 | n.s. |
| ENSG00000199366 | 8.982889 | -0.483736512 | 0.526973 | -0.917952 | 0.358644 | 0.796009826 | | misc_RNA | 43.4 | -0.047929374 | 0.047929374 | ENSG00000199366 | n.s. |
| ENSG00000213189 | 30.99706 | 0.37166675 | 0.343459 | 1.0821283 | 0.279196 | 0.743390786 | BTF3L4P2 | processed_pseudogene | 39.31 | 0.047864294 | 0.047864294 | ENSG00000213189 | n.s. |
| ENSG00000251493 | 622.8315 | -0.307153888 | 0.252047 | -1.218636 | 0.222982 | 0.698928583 | FOXD1 | protein_coding | 53.9 | -0.04778307 | 0.04778307 | ENSG00000251493 | n.s. |
| ENSG00000232044 | 131.259 | 0.301192451 | 0.244566 | 1.2315366 | 0.218122 | 0.694065444 | LINC01105 | processed_transcript | 43.52 | 0.047768996 | 0.047768996 | ENSG00000232044 | n.s. |
| ENSG00000162885 | 965.1201 | -0.31167355 | 0.258954 | -1.203585 | 0.22875 | 0.702737638 | B3GALNT2 | protein_coding | 40.36 | -0.047750503 | 0.047750503 | ENSG00000162885 | n.s. |
| ENSG00000171208 | 2301.945 | -0.316598221 | 0.26621 | -1.189279 | 0.23433 | 0.706643631 | NETO2 | protein_coding | 38.7 | -0.04774287 | 0.04774287 | ENSG00000171208 | n.s. |
| ENSG00000113812 | 841.886 | -0.31303228 | 0.261959 | -1.194968 | 0.2321 | 0.704043192 | ACTR8 | protein_coding | 42.11 | -0.047706337 | 0.047706337 | ENSG00000113812 | n.s. |
| ENSG00000105127 | 601.4685 | -0.298698319 | 0.241657 | -1.236044 | 0.216442 | 0.692499549 | AKAP8 | protein_coding | 53.24 | -0.047666429 | 0.047666429 | ENSG00000105127 | n.s. |
| ENSG00000060069 | 136.7663 | 0.29797901 | 0.240365 | 1.2396923 | 0.215089 | 0.691899391 | CTDP1 | protein_coding | 53.52 | 0.047663844 | 0.047663844 | ENSG00000060069 | n.s. |
| ENSG00000161013 | 339.5823 | -0.312573563 | 0.260996 | -1.19762 | 0.231065 | 0.704043192 | MGAT4B | protein_coding | 64.04 | -0.047636429 | 0.047636429 | ENSG00000161013 | n.s. |
| ENSG00000167674 | 927.9724 | -0.324209531 | 0.276617 | -1.172052 | 0.241176 | 0.712994199 | | protein_coding | 57.07 | -0.04763092 | 0.04763092 | ENSG00000167674 | n.s. |
| ENSG00000132382 | 727.0526 | -0.281667655 | 0.22091 | -1.275033 | 0.202297 | 0.677624898 | MYBBP1A | protein_coding | 59.89 | -0.047604832 | 0.047604832 | ENSG00000132382 | n.s. |
| ENSG00000253696 | 63.59256 | -0.339439645 | 0.297485 | -1.141032 | 0.253856 | 0.72418384 | KBTBD11-OT1 | lincRNA | 54.8 | -0.047572864 | 0.047572864 | ENSG00000253696 | n.s. |
| ENSG00000206992 | 28.39801 | -0.412517155 | 0.409336 | -1.007772 | 0.313564 | 0.766827179 | RNU6-574P | snRNA | 42.99 | -0.04756426 | 0.04756426 | ENSG00000206992 | n.s. |
| ENSG00000124006 | 703.6983 | -0.307800311 | 0.253587 | -1.213786 | 0.22483 | 0.700612508 | OBSL1 | protein_coding | 58.29 | -0.047561955 | 0.047561955 | ENSG00000124006 | n.s. |
| ENSG00000170522 | 71.15345 | -0.325692833 | 0.278647 | -1.168838 | 0.242469 | 0.714515282 | ELOVL6 | protein_coding | 39.87 | -0.047547401 | 0.047547401 | ENSG00000170522 | n.s. |
| ENSG00000259865 | 131.2923 | -0.371000803 | 0.34494 | -1.075552 | 0.282128 | 0.744567264 | | lincRNA | 27.85 | -0.047523742 | 0.047523742 | ENSG00000259865 | n.s. |

| | | | | | | | | | | | | | |
|---|---|---|---|---|---|---|---|---|---|---|---|---|---|
| ENSG00000143324 | 1519.365 | -0.30881809 | 0.255603 | -1.208194 | 0.226973 | 0.701641249 | XPR1 | protein_coding | 36.95 | -0.047522437 | 0.047522437 | ENSG00000143324 | n.s. |
| ENSG00000066135 | 569.3453 | -0.301541106 | 0.245598 | -1.227785 | 0.219528 | 0.695810058 | KDM4A | protein_coding | 46.75 | -0.047495528 | 0.047495528 | ENSG00000066135 | n.s. |
| ENSG00000156639 | 130.8907 | -0.315901755 | 0.266038 | -1.187433 | 0.235057 | 0.707610349 | ZFAND3 | protein_coding | 39.23 | -0.047450283 | 0.047450283 | ENSG00000156639 | n.s. |
| ENSG00000143390 | 1436.275 | 0.315787575 | 0.266207 | 1.1862501 | 0.235524 | 0.70779073 | RFX5 | protein_coding | 49.78 | 0.047398177 | 0.047398177 | ENSG00000143390 | n.s. |
| ENSG00000255883 | 22.74221 | -0.401359062 | 0.393488 | -1.020004 | 0.307727 | 0.761997697 | FUNDC2P1 | processed_pseudogene | 51.05 | -0.047378969 | 0.047378969 | ENSG00000255883 | n.s. |
| ENSG00000257093 | 912.9604 | -0.318133744 | 0.26904 | -1.182477 | 0.237016 | 0.709700242 | KIAA1147 | protein_coding | 43.51 | -0.047378083 | 0.047378083 | ENSG00000257093 | n.s. |
| ENSG00000228107 | 33.34594 | -0.369260613 | 0.342753 | -1.077338 | 0.281329 | 0.744242186 | | sense_overlapping | 47.84 | -0.047370862 | 0.047370862 | ENSG00000228107 | n.s. |
| ENSG00000110047 | 140.466 | -0.31073932 | 0.259121 | -1.199206 | 0.230448 | 0.704043192 | EHD1 | protein_coding | 54.84 | -0.047356889 | 0.047356889 | ENSG00000110047 | n.s. |
| ENSG00000090621 | 1370.133 | -0.290466133 | 0.232234 | -1.250746 | 0.211027 | 0.687111024 | PABPC4 | protein_coding | 47.27 | -0.047338161 | 0.047338161 | ENSG00000090621 | n.s. |
| ENSG00000162852 | 907.3855 | -0.322313063 | 0.275057 | -1.171805 | 0.241275 | 0.713070766 | CNST | protein_coding | 41.32 | -0.047337271 | 0.047337271 | ENSG00000162852 | n.s. |
| ENSG00000197498 | 1343.45 | -0.318847774 | 0.270253 | -1.179812 | 0.238075 | 0.71051867 | RPF2 | protein_coding | 42.58 | -0.047324823 | 0.047324823 | ENSG00000197498 | n.s. |
| ENSG00000131242 | 52.87361 | -0.343336114 | 0.303868 | -1.129886 | 0.258524 | 0.728120901 | RAB11FIP4 | protein_coding | 51.38 | -0.047310516 | 0.047310516 | ENSG00000131242 | n.s. |
| ENSG00000160218 | 445.8126 | -0.302489467 | 0.247488 | -1.22224 | 0.221617 | 0.697595355 | TRAPPC10 | protein_coding | 46.02 | -0.04730827 | 0.04730827 | ENSG00000160218 | n.s. |
| ENSG00000184381 | 14.49886 | -0.437079629 | 0.446955 | -0.977906 | 0.328121 | 0.779459027 | PLA2G6 | protein_coding | 51.54 | -0.047294948 | 0.047294948 | ENSG00000184381 | n.s. |
| ENSG00000103194 | 1244.845 | -0.310314044 | 0.259668 | -1.195042 | 0.23207 | 0.704043192 | USP10 | protein_coding | 43.62 | -0.047292076 | 0.047292076 | ENSG00000103194 | n.s. |
| ENSG00000004700 | 1888.928 | -0.313965995 | 0.264113 | -1.188756 | 0.234536 | 0.706958548 | RECQL | protein_coding | 37.57 | -0.047285178 | 0.047285178 | ENSG00000004700 | n.s. |
| ENSG00000274523 | 657.6637 | -0.309647157 | 0.258364 | -1.198491 | 0.230726 | 0.704043192 | RCC1L | protein_coding | 51.13 | -0.047190442 | 0.047190442 | ENSG00000274523 | n.s. |
| ENSG00000142186 | 512.9778 | -0.329485805 | 0.285101 | -1.155681 | 0.247812 | 0.719195543 | SCYL1 | protein_coding | 57.59 | -0.047166886 | 0.047166886 | ENSG00000142186 | n.s. |
| ENSG00000152763 | 32.71347 | -0.35938159 | 0.329348 | -1.091192 | 0.275189 | 0.739442303 | WDR78 | protein_coding | 36.44 | -0.047113384 | 0.047113384 | ENSG00000152763 | n.s. |
| ENSG00000221869 | 145.0123 | 0.313645758 | 0.264369 | 1.1863945 | 0.235467 | 0.70779073 | CEBPD | protein_coding | 70.02 | 0.047076701 | 0.047076701 | ENSG00000221869 | n.s. |
| ENSG00000263399 | 3.057875 | -0.667492915 | 0.878398 | -0.759898 | 0.447315 | 0.850164607 | MIR3170 | miRNA | 49.35 | -0.047056234 | 0.047056234 | ENSG00000263399 | n.s. |
| ENSG00000126214 | 622.0852 | -0.308654453 | 0.257323 | -1.199481 | 0.230341 | 0.704043192 | KLC1 | protein_coding | 49.58 | -0.047039154 | 0.047039154 | ENSG00000126214 | n.s. |
| ENSG00000166579 | 162.8498 | -0.315539099 | 0.266909 | -1.182197 | 0.237127 | 0.709700242 | NDEL1 | protein_coding | 44.62 | -0.046991675 | 0.046991675 | ENSG00000166579 | n.s. |
| ENSG00000125447 | 140.7528 | -0.319948489 | 0.273071 | -1.171669 | 0.24133 | 0.71308035 | GGA3 | protein_coding | 51.82 | -0.046988125 | 0.046988125 | ENSG00000125447 | n.s. |
| ENSG00000138814 | 1474.288 | -0.306648346 | 0.254588 | -1.204488 | 0.228401 | 0.702737638 | PPP3CA | protein_coding | 35.79 | -0.046980607 | 0.046980607 | ENSG00000138814 | n.s. |
| ENSG00000206775 | 10.99318 | 0.473801074 | 0.515926 | 0.9183508 | 0.358435 | 0.796009826 | SNORD37 | snoRNA | 43.94 | 0.046944955 | 0.046944955 | ENSG00000206775 | n.s. |
| ENSG00000137726 | 1973.751 | -0.318879097 | 0.271676 | -1.173748 | 0.240496 | 0.712660043 | FXYD6 | protein_coding | 48.09 | -0.046912724 | 0.046912724 | ENSG00000137726 | n.s. |
| ENSG00000130226 | 225.3649 | -0.306130831 | 0.253952 | -1.205468 | 0.228023 | 0.702691634 | DPP6 | protein_coding | 43.87 | -0.046910024 | 0.046910024 | ENSG00000130226 | n.s. |
| ENSG00000116871 | 541.6593 | -0.301359942 | 0.247025 | -1.21996 | 0.22248 | 0.698891176 | MAP7D1 | protein_coding | 54.66 | -0.046888727 | 0.046888727 | ENSG00000116871 | n.s. |
| ENSG00000241685 | 261.8075 | -0.342347276 | 0.303983 | -1.126205 | 0.260079 | 0.729661551 | ARPC1A | protein_coding | 45.22 | -0.046859995 | 0.046859995 | ENSG00000241685 | n.s. |
| ENSG00000201208 | 31.34004 | -0.395656885 | 0.385153 | -1.027273 | 0.304292 | 0.76144055 | | misc_RNA | 42.27 | -0.046831531 | 0.046831531 | ENSG00000201208 | n.s. |
| ENSG00000070366 | 583.9439 | -0.287121136 | 0.229407 | -1.25158 | 0.210723 | 0.686934651 | SMG6 | protein_coding | 45.75 | -0.046825029 | 0.046825029 | ENSG00000070366 | n.s. |

| | | | | | | | | | | | | | |
|---|---|---|---|---|---|---|---|---|---|---|---|---|---|
| ENSG00000278133 | 118.2014 | -0.327426107 | 0.283514 | -1.154884 | 0.248138 | 0.719455242 | | sense_intronic | 46.39 | -0.046820695 | 0.046820695 | ENSG00000278133 | n.s. |
| ENSG00000133059 | 452.2418 | -0.294805677 | 0.239182 | -1.232556 | 0.217742 | 0.693853089 | DSTYK | protein_coding | 43.55 | -0.046795234 | 0.046795234 | ENSG00000133059 | n.s. |
| ENSG00000241472 | 152.6645 | -0.303391933 | 0.250272 | -1.212247 | 0.225418 | 0.701091738 | PTPRG-AS1 | processed_transcript | 37.73 | -0.046790667 | 0.046790667 | ENSG00000241472 | n.s. |
| ENSG00000160959 | 466.8205 | -0.279891865 | 0.220988 | -1.266546 | 0.205318 | 0.680704186 | LRRC14 | protein_coding | 63.14 | -0.046753579 | 0.046753579 | ENSG00000160959 | n.s. |
| ENSG00000122970 | 1159.472 | -0.315932666 | 0.268662 | -1.175949 | 0.239615 | 0.71125159 | IFT81 | protein_coding | 39.9 | -0.046750689 | 0.046750689 | ENSG00000122970 | n.s. |
| ENSG00000230922 | 193.5108 | -0.306692142 | 0.256276 | -1.196725 | 0.231414 | 0.704043192 | | sense_intronic | 51.69 | -0.046740096 | 0.046740096 | ENSG00000230922 | n.s. |
| ENSG00000141452 | 328.2599 | -0.30940609 | 0.25981 | -1.190893 | 0.233696 | 0.706261259 | C18orf8 | protein_coding | 45.14 | -0.04673103 | 0.04673103 | ENSG00000141452 | n.s. |
| ENSG00000212445 | 9.786545 | -0.448390347 | 0.47233 | -0.949316 | 0.34246 | 0.786682723 | | snoRNA | 46.67 | -0.046722448 | 0.046722448 | ENSG00000212445 | n.s. |
| ENSG00000162669 | 427.9975 | -0.33900242 | 0.299861 | -1.130532 | 0.258252 | 0.728120901 | HFM1 | protein_coding | 37.16 | -0.046713348 | 0.046713348 | ENSG00000162669 | n.s. |
| ENSG00000117707 | 866.8398 | 0.30219281 | 0.249023 | 1.2135158 | 0.224933 | 0.700612508 | PROX1 | protein_coding | 41.17 | 0.046695472 | 0.046695472 | ENSG00000117707 | n.s. |
| ENSG00000199127 | 2.405745 | -0.803829407 | 1.188555 | -0.676308 | 0.498845 | 0.87484798 | MIR383 | miRNA | 49.32 | -0.046676289 | 0.046676289 | ENSG00000199127 | n.s. |
| ENSG00000185053 | 307.5217 | 0.310776358 | 0.261908 | 1.1865855 | 0.235391 | 0.70779073 | SGCZ | protein_coding | 35.48 | 0.046646018 | 0.046646018 | ENSG00000185053 | n.s. |
| ENSG00000139318 | 294.0251 | 0.282432758 | 0.224438 | 1.2584014 | 0.208247 | 0.683663723 | DUSP6 | protein_coding | 46.57 | 0.046645878 | 0.046645878 | ENSG00000139318 | n.s. |
| ENSG00000143303 | 92.45495 | -0.330427678 | 0.28826 | -1.146282 | 0.251678 | 0.722498818 | RRNAD1 | protein_coding | 51 | -0.046644116 | 0.046644116 | ENSG00000143303 | n.s. |
| ENSG00000172456 | 52.76126 | -0.339638791 | 0.301159 | -1.127772 | 0.259416 | 0.728949992 | FGGY | protein_coding | 40.3 | -0.046633175 | 0.046633175 | ENSG00000172456 | n.s. |
| ENSG00000275523 | 3.028753 | -0.80897141 | 1.207434 | -0.669992 | 0.502863 | 0.875883784 | MIR6508 | miRNA | 48.33 | -0.046559147 | 0.046559147 | ENSG00000275523 | n.s. |
| ENSG00000212534 | 102.163 | 0.305193858 | 0.254424 | 1.1995461 | 0.230316 | 0.704043192 | SNORD70 | snoRNA | 37.5 | 0.046511756 | 0.046511756 | ENSG00000212534 | n.s. |
| ENSG00000011566 | 1715.021 | -0.301720083 | 0.249452 | -1.209529 | 0.22646 | 0.701289341 | MAP4K3 | protein_coding | 36.9 | -0.046495898 | 0.046495898 | ENSG00000011566 | n.s. |
| ENSG00000166225 | 1013.382 | -0.301808548 | 0.249681 | -1.208775 | 0.226749 | 0.701406486 | FRS2 | protein_coding | 37.91 | -0.046487638 | 0.046487638 | ENSG00000166225 | n.s. |
| ENSG00000088888 | 775.0346 | -0.301716445 | 0.2495 | -1.209286 | 0.226553 | 0.701406486 | MAVS | protein_coding | 52.52 | -0.046473451 | 0.046473451 | ENSG00000088888 | n.s. |
| ENSG00000111707 | 970.3778 | -0.30136 | 0.248715 | -1.211667 | 0.22564 | 0.701204614 | SUDS3 | protein_coding | 44.49 | -0.046456222 | 0.046456222 | ENSG00000111707 | n.s. |
| ENSG00000224078 | 2199.352 | -0.310496135 | 0.26213 | -1.184513 | 0.23621 | 0.708727536 | SNHG14 | processed_transcript | 42.05 | -0.046425597 | 0.046425597 | ENSG00000224078 | n.s. |
| ENSG00000103512 | 575.9223 | -0.313728269 | 0.2668 | -1.175892 | 0.239638 | 0.71125159 | NOMO1 | protein_coding | 45.56 | -0.046424489 | 0.046424489 | ENSG00000103512 | n.s. |
| ENSG00000152270 | 804.9352 | -0.309078579 | 0.260527 | -1.186359 | 0.23548 | 0.70779073 | PDE3B | protein_coding | 36.96 | -0.046391189 | 0.046391189 | ENSG00000152270 | n.s. |
| ENSG00000230376 | 8.927321 | 0.543007496 | 0.644367 | 0.8426986 | 0.399397 | 0.82151052 | MEMO1P4 | processed_pseudogene | 44.04 | 0.046365711 | 0.046365711 | ENSG00000230376 | n.s. |
| ENSG00000075234 | 211.9107 | -0.30410536 | 0.254083 | -1.196872 | 0.231356 | 0.704043192 | TTC38 | protein_coding | 54.16 | -0.046345869 | 0.046345869 | ENSG00000075234 | n.s. |
| ENSG00000020129 | 205.4679 | -0.304064744 | 0.253592 | -1.199029 | 0.230517 | 0.704043192 | NCDN | protein_coding | 57.71 | -0.046339679 | 0.046339679 | ENSG00000020129 | n.s. |
| ENSG00000196557 | 540.341 | -0.308713455 | 0.260234 | -1.18629 | 0.235508 | 0.70779073 | CACNA1H | protein_coding | 65.12 | -0.046336386 | 0.046336386 | ENSG00000196557 | n.s. |
| ENSG00000181090 | 239.6793 | -0.315205954 | 0.268948 | -1.171997 | 0.241198 | 0.712994199 | EHMT1 | protein_coding | 49.25 | -0.046308169 | 0.046308169 | ENSG00000181090 | n.s. |
| ENSG00000115415 | 1224.283 | -0.302097066 | 0.251112 | -1.203035 | 0.228963 | 0.702737638 | STAT1 | protein_coding | 41.75 | -0.04628332 | 0.04628332 | ENSG00000115415 | n.s. |
| ENSG00000201044 | 3.642522 | -0.702823843 | 0.958866 | -0.732974 | 0.463574 | 0.859318855 | RNU6-268P | snRNA | 42.99 | -0.046277899 | 0.046277899 | ENSG00000201044 | n.s. |
| ENSG00000164620 | 278.123 | -0.32085834 | 0.276397 | -1.160861 | 0.245698 | 0.717427972 | RELL2 | protein_coding | 60 | -0.046274733 | 0.046274733 | ENSG00000164620 | n.s. |

| | | | | | | | | | | | | | |
|---|---|---|---|---|---|---|---|---|---|---|---|---|---|
| ENSG00000138594 | 871.1518 | -0.303624984 | 0.253864 | -1.196012 | 0.231692 | 0.704043192 | TMOD3 | protein_coding | 41.02 | -0.046272659 | 0.046272659 | ENSG00000138594 | n.s. |
| ENSG00000166508 | 7757.805 | -0.311749451 | 0.264175 | -1.180088 | 0.237965 | 0.71051867 | MCM7 | protein_coding | 54.21 | -0.046271258 | 0.046271258 | ENSG00000166508 | n.s. |
| ENSG00000183763 | 384.3577 | -0.306422168 | 0.257556 | -1.189728 | 0.234153 | 0.706508138 | TRAIP | protein_coding | 49.84 | -0.046233844 | 0.046233844 | ENSG00000183763 | n.s. |
| ENSG00000184967 | 317.6665 | -0.311972672 | 0.264805 | -1.178124 | 0.238747 | 0.710909549 | NOC4L | protein_coding | 63.79 | -0.046229874 | 0.046229874 | ENSG00000184967 | n.s. |
| ENSG00000278840 | 44.56527 | -0.354337256 | 0.325159 | -1.089734 | 0.27583 | 0.740529536 | | sense_intronic | 29.41 | -0.046225993 | 0.046225993 | ENSG00000278840 | n.s. |
| ENSG00000268798 | 164.6682 | -0.303060399 | 0.253285 | -1.196521 | 0.231493 | 0.704043192 | | lincRNA | 57 | -0.046186616 | 0.046186616 | ENSG00000268798 | n.s. |
| ENSG00000144228 | 1175.039 | -0.30946868 | 0.261693 | -1.182563 | 0.236982 | 0.709700242 | SPOPL | protein_coding | 36.68 | -0.046087638 | 0.046087638 | ENSG00000144228 | n.s. |
| ENSG00000175691 | 168.0425 | -0.336900102 | 0.299315 | -1.125572 | 0.260347 | 0.730043745 | ZNF77 | protein_coding | 49.74 | -0.046037775 | 0.046037775 | ENSG00000175691 | n.s. |
| ENSG00000251865 | 8.646519 | -0.461383382 | 0.499622 | -0.923465 | 0.355765 | 0.794802373 | RNU6-518P | snRNA | 36.46 | -0.046018768 | 0.046018768 | ENSG00000251865 | n.s. |
| ENSG00000163029 | 2324.788 | -0.32218054 | 0.279671 | -1.152 | 0.249321 | 0.720036883 | SMC6 | protein_coding | 36.86 | -0.045957526 | 0.045957526 | ENSG00000163029 | n.s. |
| ENSG00000136754 | 245.855 | -0.311368942 | 0.264974 | -1.175094 | 0.239957 | 0.711894641 | ABI1 | protein_coding | 38.44 | -0.04595316 | 0.04595316 | ENSG00000136754 | n.s. |
| ENSG00000139154 | 1455.985 | -0.305396787 | 0.257079 | -1.187949 | 0.234853 | 0.707216633 | AEBP2 | protein_coding | 40.33 | -0.045946193 | 0.045946193 | ENSG00000139154 | n.s. |
| ENSG00000165895 | 88.17521 | 0.326354673 | 0.285212 | 1.1442532 | 0.252519 | 0.723128487 | ARHGAP42 | protein_coding | 38.03 | 0.045945689 | 0.045945689 | ENSG00000165895 | n.s. |
| ENSG00000252035 | 5.480372 | 0.511243032 | 0.589977 | 0.8665479 | 0.38619 | 0.813115862 | RNU6-397P | snRNA | 44.23 | 0.045933942 | 0.045933942 | ENSG00000252035 | n.s. |
| ENSG00000187531 | 328.3233 | -0.291078989 | 0.236895 | -1.228723 | 0.219176 | 0.695448706 | SIRT7 | protein_coding | 57.63 | -0.045913314 | 0.045913314 | ENSG00000187531 | n.s. |
| ENSG00000168234 | 96.16949 | -0.333116459 | 0.294615 | -1.130683 | 0.258189 | 0.728120901 | TTC39C | protein_coding | 41.64 | -0.045902283 | 0.045902283 | ENSG00000168234 | n.s. |
| ENSG00000162408 | 552.1234 | -0.297296481 | 0.245212 | -1.212404 | 0.225358 | 0.701091738 | NOL9 | protein_coding | 47.78 | -0.045850595 | 0.045850595 | ENSG00000162408 | n.s. |
| ENSG00000213047 | 227.4536 | 0.294081717 | 0.240959 | 1.2204645 | 0.222289 | 0.698471535 | DENND1B | protein_coding | 34.32 | 0.045833014 | 0.045833014 | ENSG00000213047 | n.s. |
| ENSG00000197969 | 127.8878 | -0.341074553 | 0.30561 | -1.116045 | 0.264403 | 0.734053608 | VPS13A | protein_coding | 35.89 | -0.045796838 | 0.045796838 | ENSG00000197969 | n.s. |
| ENSG00000080819 | 324.7135 | -0.325206122 | 0.284143 | -1.144516 | 0.25241 | 0.72310706 | CPOX | protein_coding | 38.07 | -0.045788176 | 0.045788176 | ENSG00000080819 | n.s. |
| ENSG00000179151 | 676.3464 | -0.290997326 | 0.237192 | -1.226844 | 0.219881 | 0.69624936 | EDC3 | protein_coding | 43.7 | -0.04575502 | 0.04575502 | ENSG00000179151 | n.s. |
| ENSG00000125633 | 621.3629 | -0.294054311 | 0.241248 | -1.218888 | 0.222887 | 0.698928583 | CCDC93 | protein_coding | 42.78 | -0.045745205 | 0.045745205 | ENSG00000125633 | n.s. |
| ENSG00000145248 | 819.4004 | -0.296574702 | 0.244631 | -1.212333 | 0.225385 | 0.701091738 | SLC10A4 | protein_coding | 44.38 | -0.045739278 | 0.045739278 | ENSG00000145248 | n.s. |
| ENSG00000174106 | 1066.22 | -0.311076055 | 0.265177 | -1.17309 | 0.24076 | 0.712813552 | LEMD3 | protein_coding | 35.54 | -0.045735662 | 0.045735662 | ENSG00000174106 | n.s. |
| ENSG00000153066 | 206.3401 | -0.298433109 | 0.247556 | -1.20552 | 0.228003 | 0.702691634 | TXNDC11 | protein_coding | 43.55 | -0.045730462 | 0.045730462 | ENSG00000153066 | n.s. |
| ENSG00000125844 | 421.6057 | -0.295207255 | 0.242914 | -1.215273 | 0.224262 | 0.700099826 | RRBP1 | protein_coding | 52.51 | -0.0457099 | 0.0457099 | ENSG00000125844 | n.s. |
| ENSG00000126391 | 175.1334 | -0.298262057 | 0.247925 | -1.203032 | 0.228964 | 0.702737638 | FRMD8 | protein_coding | 56.27 | -0.045695771 | 0.045695771 | ENSG00000126391 | n.s. |
| ENSG00000168209 | 1287.65 | 0.328877215 | 0.289724 | 1.1351402 | 0.256317 | 0.726402587 | DDIT4 | protein_coding | 58.72 | 0.045655596 | 0.045655596 | ENSG00000168209 | n.s. |
| ENSG00000131470 | 1042.906 | -0.277362503 | 0.220839 | -1.255947 | 0.209135 | 0.68461467 | PSMC3IP | protein_coding | 48.67 | -0.045641053 | 0.045641053 | ENSG00000131470 | n.s. |
| ENSG00000132600 | 108.1498 | -0.326332756 | 0.286635 | -1.138498 | 0.254913 | 0.724760001 | PRMT7 | protein_coding | 48.59 | -0.045623206 | 0.045623206 | ENSG00000132600 | n.s. |
| ENSG00000052126 | 975.3424 | -0.29910436 | 0.250302 | -1.194973 | 0.232098 | 0.704043192 | PLEKHA5 | protein_coding | 38.09 | -0.045583713 | 0.045583713 | ENSG00000052126 | n.s. |
| ENSG00000162222 | 197.6128 | -0.296926354 | 0.246886 | -1.202687 | 0.229098 | 0.702737638 | TTC9C | protein_coding | 47.12 | -0.045491132 | 0.045491132 | ENSG00000162222 | n.s. |

| | | | | | | | | | | | | | |
|---|---|---|---|---|---|---|---|---|---|---|---|---|---|
| ENSG00000166233 | 3130.345 | -0.305431044 | 0.258398 | -1.182016 | 0.237199 | 0.709762958 | ARIH1 | protein_coding | 38.73 | -0.045474611 | 0.045474611 | ENSG00000166233 | n.s. |
| ENSG00000090989 | 949.7714 | -0.322074665 | 0.281167 | -1.145491 | 0.252006 | 0.722544863 | EXOC1 | protein_coding | 34.99 | -0.045456067 | 0.045456067 | ENSG00000090989 | n.s. |
| ENSG00000188486 | 1003.378 | -0.277553202 | 0.221542 | -1.252822 | 0.21027 | 0.686087755 | H2AFX | protein_coding | 66.42 | -0.045413347 | 0.045413347 | ENSG00000188486 | n.s. |
| ENSG00000207425 | 4.877358 | -0.635593128 | 0.831858 | -0.764065 | 0.444829 | 0.848321918 | | misc_RNA | 46 | -0.045406336 | 0.045406336 | ENSG00000207425 | n.s. |
| ENSG00000262180 | 60.08482 | -0.354211328 | 0.329523 | -1.074921 | 0.28241 | 0.744597466 | OCLM | protein_coding | 35.52 | -0.045366836 | 0.045366836 | ENSG00000262180 | n.s. |
| ENSG00000184708 | 504.427 | -0.289990556 | 0.237283 | -1.222128 | 0.221659 | 0.697595355 | EIF4ENIF1 | protein_coding | 44.22 | -0.045353485 | 0.045353485 | ENSG00000184708 | n.s. |
| ENSG00000222629 | 8.6273 | -0.468302463 | 0.516512 | -0.906663 | 0.364585 | 0.800144301 | RNU2-42P | snRNA | 32.74 | -0.045346516 | 0.045346516 | ENSG00000222629 | n.s. |
| ENSG00000124789 | 3435.594 | -0.289954173 | 0.237391 | -1.221419 | 0.221928 | 0.697887666 | NUP153 | protein_coding | 40.18 | -0.04529504 | 0.04529504 | ENSG00000124789 | n.s. |
| ENSG00000123989 | 171.7572 | -0.296879424 | 0.247379 | -1.200101 | 0.2301 | 0.703952151 | CHPF | protein_coding | 61.17 | -0.045261305 | 0.045261305 | ENSG00000123989 | n.s. |
| ENSG00000172831 | 378.8207 | -0.301620446 | 0.254344 | -1.185874 | 0.235672 | 0.707931694 | CES2 | protein_coding | 55.49 | -0.045245674 | 0.045245674 | ENSG00000172831 | n.s. |
| ENSG00000226479 | 266.6355 | 0.282220145 | 0.22747 | 1.2406941 | 0.214719 | 0.691472186 | TMEM185B | protein_coding | 53.17 | 0.045218803 | 0.045218803 | ENSG00000226479 | n.s. |
| ENSG00000162402 | 4365.274 | -0.28977003 | 0.237333 | -1.220943 | 0.222107 | 0.698197164 | USP24 | protein_coding | 37.78 | -0.045210476 | 0.045210476 | ENSG00000162402 | n.s. |
| ENSG00000189369 | 132.3397 | 0.322569472 | 0.282491 | 1.1418743 | 0.253506 | 0.72418384 | GSPT2 | protein_coding | 44.83 | 0.045208489 | 0.045208489 | ENSG00000189369 | n.s. |
| ENSG00000112419 | 1038.015 | -0.322514027 | 0.282823 | -1.14034 | 0.254145 | 0.72418384 | PHACTR2 | protein_coding | 40.02 | -0.045200718 | 0.045200718 | ENSG00000112419 | n.s. |
| ENSG00000102879 | 94.25804 | -0.356145572 | 0.334649 | -1.064235 | 0.287222 | 0.746611882 | CORO1A | protein_coding | 64.42 | -0.04519669 | 0.04519669 | ENSG00000102879 | n.s. |
| ENSG00000200156 | 569.4784 | 0.260390697 | 0.201084 | 1.294934 | 0.195343 | 0.670750289 | RNU5B-1 | snRNA | 39.66 | 0.045161936 | 0.045161936 | ENSG00000200156 | n.s. |
| ENSG00000131023 | 1606.433 | -0.296436588 | 0.248228 | -1.194213 | 0.232395 | 0.704163047 | LATS1 | protein_coding | 41.22 | -0.045155228 | 0.045155228 | ENSG00000131023 | n.s. |
| ENSG00000169439 | 894.3769 | 0.335401854 | 0.300308 | 1.1168607 | 0.264054 | 0.733524057 | SDC2 | protein_coding | 41.4 | 0.045140273 | 0.045140273 | ENSG00000169439 | n.s. |
| ENSG00000160551 | 3276.971 | -0.292735737 | 0.241877 | -1.210268 | 0.226176 | 0.701289341 | TAOK1 | protein_coding | 41.03 | -0.045111386 | 0.045111386 | ENSG00000160551 | n.s. |
| ENSG00000143473 | 210.9584 | -0.319112475 | 0.278242 | -1.146887 | 0.251428 | 0.722279621 | KCNH1 | protein_coding | 41.06 | -0.045088882 | 0.045088882 | ENSG00000143473 | n.s. |
| ENSG00000252494 | 24.83355 | -0.368528924 | 0.352045 | -1.046823 | 0.295181 | 0.754508255 | RNU6-126P | snRNA | 36.89 | -0.045084356 | 0.045084356 | ENSG00000252494 | n.s. |
| ENSG00000069966 | 265.5445 | -0.32663499 | 0.288568 | -1.131918 | 0.257669 | 0.727758051 | GNB5 | protein_coding | 45.39 | -0.045079869 | 0.045079869 | ENSG00000069966 | n.s. |
| ENSG00000138709 | 402.4231 | -0.31366223 | 0.270445 | -1.1598 | 0.24613 | 0.718271454 | LARP1B | protein_coding | 39.11 | -0.045076836 | 0.045076836 | ENSG00000138709 | n.s. |
| ENSG00000184194 | 278.4799 | -0.310127878 | 0.26614 | -1.165279 | 0.243906 | 0.715606021 | GPR173 | protein_coding | 45.9 | -0.045069648 | 0.045069648 | ENSG00000184194 | n.s. |
| ENSG00000275667 | 3.893765 | -0.562084363 | 0.692867 | -0.811244 | 0.417225 | 0.831423508 | MIR6839 | miRNA | 49.56 | -0.045066632 | 0.045066632 | ENSG00000275667 | n.s. |
| ENSG00000137710 | 1187.823 | -0.313899804 | 0.270815 | -1.159092 | 0.246419 | 0.718522635 | RDX | protein_coding | 39.96 | -0.045063313 | 0.045063313 | ENSG00000137710 | n.s. |
| ENSG00000197912 | 680.0538 | -0.295641281 | 0.247168 | -1.196115 | 0.231652 | 0.704043192 | SPG7 | protein_coding | 52.29 | -0.045055937 | 0.045055937 | ENSG00000197912 | n.s. |
| ENSG00000179051 | 1550.061 | -0.290872305 | 0.239366 | -1.215179 | 0.224298 | 0.700099826 | RCC2 | protein_coding | 51.68 | -0.045038676 | 0.045038676 | ENSG00000179051 | n.s. |
| ENSG00000185278 | 1224.484 | -0.3117113 | 0.268349 | -1.161591 | 0.245402 | 0.717133775 | ZBTB37 | protein_coding | 40.41 | -0.045011057 | 0.045011057 | ENSG00000185278 | n.s. |
| ENSG00000162923 | 2287.193 | -0.294334683 | 0.244993 | -1.201401 | 0.229596 | 0.703335843 | WDR26 | protein_coding | 38.39 | -0.044985303 | 0.044985303 | ENSG00000162923 | n.s. |
| ENSG00000087053 | 765.8137 | -0.308754134 | 0.26454 | -1.167137 | 0.243155 | 0.715021889 | MTMR2 | protein_coding | 36.56 | -0.044979507 | 0.044979507 | ENSG00000087053 | n.s. |
| ENSG00000243770 | 75.62155 | 0.315244558 | 0.27348 | 1.1527167 | 0.249027 | 0.720036883 | RN7SL65P | misc_RNA | 49.84 | 0.044968141 | 0.044968141 | ENSG00000243770 | n.s. |

| | | | | | | | | | | | | | |
|---|---|---|---|---|---|---|---|---|---|---|---|---|---|
| ENSG00000179820 | 121.2171 | -0.314567032 | 0.272548 | -1.15417 | 0.24843 | 0.719540199 | MYADM | protein_coding | 58.37 | -0.044965765 | 0.044965765 | ENSG00000179820 | n.s. |
| ENSG00000184281 | 596.7282 | -0.307864119 | 0.263419 | -1.168724 | 0.242515 | 0.714515282 | TSSC4 | protein_coding | 62.64 | -0.044944614 | 0.044944614 | ENSG00000184281 | n.s. |
| ENSG00000157500 | 1520.683 | -0.307787754 | 0.263393 | -1.168549 | 0.242585 | 0.714515282 | APPL1 | protein_coding | 37.38 | -0.044933466 | 0.044933466 | ENSG00000157500 | n.s. |
| ENSG00000197149 | 345.4157 | -0.351836134 | 0.328205 | -1.072001 | 0.28372 | 0.745300357 | | processed_pseudogene | 60.02 | -0.044918452 | 0.044918452 | ENSG00000197149 | n.s. |
| ENSG00000120690 | 117.787 | 0.326508505 | 0.289339 | 1.1284646 | 0.259124 | 0.728550156 | ELF1 | protein_coding | 38.8 | 0.044908158 | 0.044908158 | ENSG00000120690 | n.s. |
| ENSG00000206786 | 4.618405 | -0.499885477 | 0.577481 | -0.865631 | 0.386692 | 0.813136725 | RNU6-701P | snRNA | 40.19 | -0.044907924 | 0.044907924 | ENSG00000206786 | n.s. |
| ENSG00000196865 | 1529.554 | -0.308889334 | 0.265155 | -1.164938 | 0.244044 | 0.715606021 | NHLRC2 | protein_coding | 35.52 | -0.044889656 | 0.044889656 | ENSG00000196865 | n.s. |
| ENSG00000164144 | 589.4737 | -0.335602789 | 0.302519 | -1.109362 | 0.267274 | 0.734987182 | ARFIP1 | protein_coding | 37.33 | -0.044876884 | 0.044876884 | ENSG00000164144 | n.s. |
| ENSG00000099956 | 1209.798 | -0.325088274 | 0.287085 | -1.132378 | 0.257476 | 0.727737514 | SMARCB1 | protein_coding | 50.77 | -0.044870386 | 0.044870386 | ENSG00000099956 | n.s. |
| ENSG00000160325 | 187.2021 | -0.305093795 | 0.260172 | -1.17266 | 0.240932 | 0.712813552 | CACFD1 | protein_coding | 58.04 | -0.044856126 | 0.044856126 | ENSG00000160325 | n.s. |
| ENSG00000272055 | 5.597383 | -0.490605155 | 0.559708 | -0.876537 | 0.380738 | 0.810319533 | RNU6-6P | snRNA | 45.79 | -0.044813686 | 0.044813686 | ENSG00000272055 | n.s. |
| ENSG00000085840 | 1612.497 | -0.308275707 | 0.264591 | -1.165103 | 0.243977 | 0.715606021 | ORC1 | protein_coding | 44.76 | -0.04480048 | 0.04480048 | ENSG00000085840 | n.s. |
| ENSG00000188321 | 578.3858 | 0.334016645 | 0.30024 | 1.1125003 | 0.265923 | 0.734612586 | ZNF559 | protein_coding | 40.31 | 0.044738736 | 0.044738736 | ENSG00000188321 | n.s. |
| ENSG00000154370 | 823.0271 | -0.279090965 | 0.224861 | -1.24117 | 0.214543 | 0.691472186 | TRIM11 | protein_coding | 56.97 | -0.044717429 | 0.044717429 | ENSG00000154370 | n.s. |
| ENSG00000134690 | 463.6737 | -0.299289165 | 0.252804 | -1.183877 | 0.236462 | 0.709079996 | CDCA8 | protein_coding | 45.82 | -0.044685299 | 0.044685299 | ENSG00000134690 | n.s. |
| ENSG00000071205 | 525.13 | -0.305877362 | 0.261667 | -1.168957 | 0.242421 | 0.714515282 | ARHGAP10 | protein_coding | 40.94 | -0.04465457 | 0.04465457 | ENSG00000071205 | n.s. |
| ENSG00000071537 | 969.3569 | -0.305663952 | 0.261391 | -1.169373 | 0.242253 | 0.714515282 | SEL1L | protein_coding | 38.48 | -0.044623415 | 0.044623415 | ENSG00000071537 | n.s. |
| ENSG00000200183 | 8.482831 | -0.44949656 | 0.489129 | -0.918973 | 0.35811 | 0.795806072 | RNU6-238P | snRNA | 41.51 | -0.0445868 | 0.0445868 | ENSG00000200183 | n.s. |
| ENSG00000118620 | 888.1897 | -0.303065466 | 0.258218 | -1.173683 | 0.240522 | 0.712660043 | ZNF430 | protein_coding | 39.05 | -0.044586261 | 0.044586261 | ENSG00000118620 | n.s. |
| ENSG00000243398 | 17.32961 | -0.389841694 | 0.389845 | -0.999991 | 0.317315 | 0.768744443 | RN7SL141P | misc_RNA | 54.51 | -0.044526943 | 0.044526943 | ENSG00000243398 | n.s. |
| ENSG00000204568 | 600.1244 | -0.334123572 | 0.30245 | -1.104722 | 0.26928 | 0.735983077 | MRPS18B | protein_coding | 46.36 | -0.044482597 | 0.044482597 | ENSG00000204568 | n.s. |
| ENSG00000177169 | 374.99 | -0.311194364 | 0.269762 | -1.153588 | 0.248669 | 0.719646518 | ULK1 | protein_coding | 62.39 | -0.044463691 | 0.044463691 | ENSG00000177169 | n.s. |
| ENSG00000125386 | 756.1097 | -0.284317106 | 0.232789 | -1.221352 | 0.221953 | 0.697887666 | FAM193A | protein_coding | 45.62 | -0.044414448 | 0.044414448 | ENSG00000125386 | n.s. |
| ENSG00000206634 | 46.73846 | 0.34482816 | 0.318731 | 1.0818797 | 0.279306 | 0.743390786 | SNORA22 | snoRNA | 41.04 | 0.044407945 | 0.044407945 | ENSG00000206634 | n.s. |
| ENSG00000070423 | 393.3152 | -0.319794565 | 0.281785 | -1.134888 | 0.256422 | 0.726402587 | RNF126 | protein_coding | 63.55 | -0.044394719 | 0.044394719 | ENSG00000070423 | n.s. |
| ENSG00000162813 | 504.3278 | -0.348963438 | 0.326589 | -1.068509 | 0.285291 | 0.746113205 | BPNT1 | protein_coding | 40.42 | -0.0443865 | 0.0443865 | ENSG00000162813 | n.s. |
| ENSG00000103942 | 356.0382 | -0.293719833 | 0.246672 | -1.190728 | 0.23376 | 0.706302782 | HOMER2 | protein_coding | 45.38 | -0.044354363 | 0.044354363 | ENSG00000103942 | n.s. |
| ENSG00000244332 | 60.60138 | -0.332932399 | 0.300909 | -1.106424 | 0.268543 | 0.735983077 | | sense_overlapping | 37.13 | -0.044324013 | 0.044324013 | ENSG00000244332 | n.s. |
| ENSG00000160767 | 216.7127 | -0.28476262 | 0.23374 | -1.218289 | 0.223114 | 0.699171362 | FAM189B | protein_coding | 57.4 | -0.044256772 | 0.044256772 | ENSG00000160767 | n.s. |
| ENSG00000198752 | 1188.047 | -0.287048259 | 0.236974 | -1.211308 | 0.225778 | 0.701204614 | CDC42BPB | protein_coding | 48.97 | -0.044249992 | 0.044249992 | ENSG00000198752 | n.s. |
| ENSG00000269968 | 77.06277 | 0.319813416 | 0.28224 | 1.1331276 | 0.257161 | 0.727284297 | | antisense | 56.5 | 0.044228849 | 0.044228849 | ENSG00000269968 | n.s. |
| ENSG00000114982 | 1682.413 | -0.309978401 | 0.268947 | -1.152565 | 0.249089 | 0.720036883 | KANSL3 | protein_coding | 44.51 | -0.044216949 | 0.044216949 | ENSG00000114982 | n.s. |

| | | | | | | | | | | | | | |
|---|---|---|---|---|---|---|---|---|---|---|---|---|---|
| ENSG00000127947 | 910.0926 | -0.302751221 | 0.259045 | -1.168722 | 0.242516 | 0.714515282 | PTPN12 | protein_coding | 38.16 | -0.04419819 | 0.04419819 | ENSG00000127947 | n.s. |
| ENSG00000271533 | 258.9678 | -0.34820183 | 0.327229 | -1.064093 | 0.287287 | 0.746611882 | | sense_intronic | 32.02 | -0.044188589 | 0.044188589 | ENSG00000271533 | n.s. |
| ENSG00000087502 | 2542.11 | -0.3085615 | 0.266802 | -1.156517 | 0.24747 | 0.7191431 | ERGIC2 | protein_coding | 34.04 | -0.04418128 | 0.04418128 | ENSG00000087502 | n.s. |
| ENSG00000214063 | 157.1204 | -0.342908381 | 0.31682 | -1.082345 | 0.279099 | 0.743390786 | TSPAN4 | protein_coding | 65.01 | -0.04416071 | 0.04416071 | ENSG00000214063 | n.s. |
| ENSG00000083168 | 2861.583 | -0.273994682 | 0.220284 | -1.243826 | 0.213564 | 0.690180176 | KAT6A | protein_coding | 38.89 | -0.044123424 | 0.044123424 | ENSG00000083168 | n.s. |
| ENSG00000272474 | 4.551947 | 0.498582472 | 0.581962 | 0.8567274 | 0.391596 | 0.815710819 | SNORD113-5 | snoRNA | 41.03 | 0.044106489 | 0.044106489 | ENSG00000272474 | n.s. |
| ENSG00000254911 | 449.313 | -0.285206636 | 0.234884 | -1.214245 | 0.224654 | 0.700529123 | SCARNA9 | antisense | 40.23 | -0.044085475 | 0.044085475 | ENSG00000254911 | n.s. |
| ENSG00000257151 | 379.4715 | -0.3422439 | 0.316322 | -1.081949 | 0.279275 | 0.743390786 | PWAR6 | lincRNA | 34.45 | -0.044075137 | 0.044075137 | ENSG00000257151 | n.s. |
| ENSG00000171492 | 688.1673 | -0.307558388 | 0.266046 | -1.156036 | 0.247667 | 0.719177715 | LRRC8D | protein_coding | 40.31 | -0.044031221 | 0.044031221 | ENSG00000171492 | n.s. |
| ENSG00000155849 | 39.98879 | -0.33070206 | 0.299448 | -1.104371 | 0.269432 | 0.735983077 | ELMO1 | protein_coding | 41.67 | -0.044027083 | 0.044027083 | ENSG00000155849 | n.s. |
| ENSG00000081692 | 652.9177 | -0.283583273 | 0.232974 | -1.217233 | 0.223516 | 0.699734492 | JMJD4 | protein_coding | 61.26 | -0.043974327 | 0.043974327 | ENSG00000081692 | n.s. |
| ENSG00000119523 | 661.1029 | 0.286834628 | 0.238189 | 1.2042316 | 0.2285 | 0.702737638 | ALG2 | protein_coding | 42.69 | 0.043945011 | 0.043945011 | ENSG00000119523 | n.s. |
| ENSG00000206630 | 45.75281 | -0.347510134 | 0.327601 | -1.060773 | 0.288793 | 0.747460922 | SNORD60 | snoRNA | 42.17 | -0.04392928 | 0.04392928 | ENSG00000206630 | n.s. |
| ENSG00000237190 | 624.8681 | -0.322384886 | 0.286743 | -1.124299 | 0.260886 | 0.730701535 | CDKN2AIPNL | protein_coding | 45.21 | -0.043928158 | 0.043928158 | ENSG00000237190 | n.s. |
| ENSG00000155966 | 74.7755 | -0.328140213 | 0.295492 | -1.110486 | 0.26679 | 0.734987182 | AFF2 | protein_coding | 38.84 | -0.043878986 | 0.043878986 | ENSG00000155966 | n.s. |
| ENSG00000252908 | 40.13201 | -0.327438215 | 0.293823 | -1.114405 | 0.265105 | 0.734612586 | RNU6-1003P | snRNA | 37.14 | -0.04385761 | 0.04385761 | ENSG00000252908 | n.s. |
| ENSG00000063978 | 1427.696 | -0.286161911 | 0.237828 | -1.203229 | 0.228888 | 0.702737638 | RNF4 | protein_coding | 47.38 | -0.043841946 | 0.043841946 | ENSG00000063978 | n.s. |
| ENSG00000204177 | 272.228 | -0.287612753 | 0.240389 | -1.196447 | 0.231522 | 0.704043192 | BMS1P1 | cribed_unprocessed_pseudc | 41.22 | -0.043832384 | 0.043832384 | ENSG00000204177 | n.s. |
| ENSG00000273882 | 14.3845 | -0.415569738 | 0.432292 | -0.961317 | 0.336393 | 0.78441387 | MIR8066 | miRNA | 39.74 | -0.043823797 | 0.043823797 | ENSG00000273882 | n.s. |
| ENSG00000250571 | 77.21149 | -0.327047171 | 0.293947 | -1.112606 | 0.265878 | 0.734612586 | GLI4 | protein_coding | 64.03 | -0.043805233 | 0.043805233 | ENSG00000250571 | n.s. |
| ENSG00000244270 | 61.26771 | -0.333840312 | 0.304958 | -1.094708 | 0.273645 | 0.739386355 | | processed_pseudogene | 46.9 | -0.043776002 | 0.043776002 | ENSG00000244270 | n.s. |
| ENSG00000170242 | 3086.114 | -0.30567925 | 0.264254 | -1.156765 | 0.247368 | 0.7191431 | USP47 | protein_coding | 35.32 | -0.043768586 | 0.043768586 | ENSG00000170242 | n.s. |
| ENSG00000275127 | 6.335523 | -0.46942953 | 0.52547 | -0.893352 | 0.371669 | 0.806899249 | SNORD116-22 | snoRNA | 43.48 | -0.043741767 | 0.043741767 | ENSG00000275127 | n.s. |
| ENSG00000100997 | 397.1555 | -0.315004791 | 0.277315 | -1.135911 | 0.255994 | 0.726402587 | ABHD12 | protein_coding | 47.42 | -0.04372979 | 0.04372979 | ENSG00000100997 | n.s. |
| ENSG00000130305 | 376.1136 | -0.306916378 | 0.266598 | -1.151233 | 0.249636 | 0.720401565 | NSUN5 | protein_coding | 53.93 | -0.043712673 | 0.043712673 | ENSG00000130305 | n.s. |
| ENSG00000167182 | 132.6747 | 0.286798913 | 0.239643 | 1.1967761 | 0.231394 | 0.704043192 | SP2 | protein_coding | 47.6 | 0.043708354 | 0.043708354 | ENSG00000167182 | n.s. |
| ENSG00000110048 | 637.5631 | -0.311818101 | 0.273349 | -1.140734 | 0.253981 | 0.72418384 | OSBP | protein_coding | 42.8 | -0.043701672 | 0.043701672 | ENSG00000110048 | n.s. |
| ENSG00000120800 | 3560.573 | -0.293428142 | 0.248187 | -1.182288 | 0.237091 | 0.709700242 | UTP20 | protein_coding | 39.31 | -0.0436988 | 0.0436988 | ENSG00000120800 | n.s. |
| ENSG00000164880 | 767.1 | -0.280042707 | 0.229388 | -1.220827 | 0.222151 | 0.698197164 | INTS1 | protein_coding | 63.54 | -0.043692801 | 0.043692801 | ENSG00000164880 | n.s. |
| ENSG00000118518 | 727.2403 | -0.32813025 | 0.297091 | -1.104479 | 0.269386 | 0.735983077 | RNF146 | protein_coding | 36.49 | -0.043684693 | 0.043684693 | ENSG00000118518 | n.s. |
| ENSG00000106723 | 2172.548 | -0.314952013 | 0.277674 | -1.134251 | 0.256689 | 0.726695143 | SPIN1 | protein_coding | 41.25 | -0.043667386 | 0.043667386 | ENSG00000106723 | n.s. |
| ENSG00000260339 | 116.6067 | 0.299245179 | 0.256191 | 1.1680557 | 0.242784 | 0.714766539 | HEXA-AS1 | antisense | 41.18 | 0.043640656 | 0.043640656 | ENSG00000260339 | n.s. |

| | | | | | | | | | | | | | |
|---|---|---|---|---|---|---|---|---|---|---|---|---|---|
| ENSG00000111879 | 733.8636 | -0.309941728 | 0.27078 | -1.144626 | 0.252364 | 0.72310706 | FAM184A | protein_coding | 39.55 | -0.043638989 | 0.043638989 | ENSG00000111879 | n.s. |
| ENSG00000244314 | 22.89057 | -0.3563591 | 0.34059 | -1.046299 | 0.295423 | 0.75457253 | RN7SL36P | misc_RNA | 54.71 | -0.043582365 | 0.043582365 | ENSG00000244314 | n.s. |
| ENSG00000220842 | 31.49426 | 0.478189914 | 0.549437 | 0.8703275 | 0.384121 | 0.810719585 | RPL21P16 | processed_pseudogene | 43.27 | 0.043577129 | 0.043577129 | ENSG00000220842 | n.s. |
| ENSG00000253966 | 84.49703 | -0.328281924 | 0.297475 | -1.10356 | 0.269784 | 0.736654834 | | processed_pseudogene | 40.85 | -0.043574815 | 0.043574815 | ENSG00000253966 | n.s. |
| ENSG00000123415 | 402.4407 | -0.304290256 | 0.263104 | -1.156538 | 0.247461 | 0.7191431 | SMUG1 | protein_coding | 50.76 | -0.043569704 | 0.043569704 | ENSG00000123415 | n.s. |
| ENSG00000267500 | 118.0281 | -0.315590726 | 0.278855 | -1.131739 | 0.257744 | 0.727758051 | ZNF887P | cribed_unprocessed_pseudc | 43.51 | -0.043555617 | 0.043555617 | ENSG00000267500 | n.s. |
| ENSG00000134504 | 166.5201 | -0.288718037 | 0.24275 | -1.189362 | 0.234297 | 0.706643631 | KCTD1 | protein_coding | 42.61 | -0.04353855 | 0.04353855 | ENSG00000134504 | n.s. |
| ENSG00000218336 | 206.0071 | -0.310545913 | 0.272059 | -1.141465 | 0.253676 | 0.72418384 | TENM3 | protein_coding | 38.92 | -0.043523373 | 0.043523373 | ENSG00000218336 | n.s. |
| ENSG00000179331 | 361.5054 | -0.310537979 | 0.272559 | -1.139341 | 0.254561 | 0.72418384 | RAB39A | protein_coding | 42.88 | -0.043522261 | 0.043522261 | ENSG00000179331 | n.s. |
| ENSG00000186448 | 961.2785 | -0.29945348 | 0.257218 | -1.164199 | 0.244343 | 0.715606581 | ZNF197 | protein_coding | 40.2 | -0.043518279 | 0.043518279 | ENSG00000186448 | n.s. |
| ENSG00000077549 | 317.5425 | -0.32087877 | 0.28573 | -1.123015 | 0.261431 | 0.731785945 | CAPZB | protein_coding | 49.21 | -0.043516275 | 0.043516275 | ENSG00000077549 | n.s. |
| ENSG00000135966 | 802.7392 | -0.283956934 | 0.235751 | -1.20448 | 0.228404 | 0.702737638 | TGFBRAP1 | protein_coding | 44.1 | -0.043504129 | 0.043504129 | ENSG00000135966 | n.s. |
| ENSG00000256944 | 10.00627 | 0.489067915 | 0.567203 | 0.8622449 | 0.388553 | 0.814823372 | | antisense | 48.7 | 0.043495999 | 0.043495999 | ENSG00000256944 | n.s. |
| ENSG00000149972 | 22.38354 | -0.415814413 | 0.435904 | -0.953913 | 0.340128 | 0.785979841 | CNTN5 | protein_coding | 35.04 | -0.043489444 | 0.043489444 | ENSG00000149972 | n.s. |
| ENSG00000168476 | 140.3066 | -0.299774785 | 0.257711 | -1.16322 | 0.24474 | 0.716165582 | REEP4 | protein_coding | 62.11 | -0.043463313 | 0.043463313 | ENSG00000168476 | n.s. |
| ENSG00000197779 | 336.8413 | -0.324371612 | 0.291421 | -1.113068 | 0.265679 | 0.734612586 | ZNF81 | protein_coding | 39.93 | -0.043446864 | 0.043446864 | ENSG00000197779 | n.s. |
| ENSG00000160299 | 1414.434 | -0.285012176 | 0.238563 | -1.194703 | 0.232203 | 0.704043192 | PCNT | protein_coding | 51.58 | -0.043436054 | 0.043436054 | ENSG00000160299 | n.s. |
| ENSG00000180263 | 576.9939 | 0.276306473 | 0.225416 | 1.2257635 | 0.220288 | 0.696427255 | FGD6 | protein_coding | 41.01 | 0.043414445 | 0.043414445 | ENSG00000180263 | n.s. |
| ENSG00000111877 | 1087.223 | 0.298565411 | 0.256251 | 1.1651299 | 0.243966 | 0.715606021 | MCM9 | protein_coding | 39.07 | 0.043389321 | 0.043389321 | ENSG00000111877 | n.s. |
| ENSG00000102924 | 474.6161 | -0.306882504 | 0.267508 | -1.14719 | 0.251303 | 0.722279621 | CBLN1 | protein_coding | 56.83 | -0.043360852 | 0.043360852 | ENSG00000102924 | n.s. |
| ENSG00000196743 | 205.0478 | -0.329964948 | 0.302007 | -1.092573 | 0.274581 | 0.739386355 | GM2A | protein_coding | 47.41 | -0.043267831 | 0.043267831 | ENSG00000196743 | n.s. |
| ENSG00000127419 | 121.9351 | -0.322956248 | 0.290152 | -1.113059 | 0.265683 | 0.734612586 | TMEM175 | protein_coding | 51.83 | -0.043257288 | 0.043257288 | ENSG00000127419 | n.s. |
| ENSG00000183506 | 172.7762 | -0.29658842 | 0.254151 | -1.166978 | 0.243219 | 0.715021889 | PI4KAP2 | cribed_unprocessed_pseudc | 55.77 | -0.043207198 | 0.043207198 | ENSG00000183506 | n.s. |
| ENSG00000253854 | 35.67915 | -0.336745433 | 0.312384 | -1.077984 | 0.281041 | 0.744242186 | | antisense | 40.97 | -0.043199629 | 0.043199629 | ENSG00000253854 | n.s. |
| ENSG00000108474 | 202.1405 | 0.281397674 | 0.233184 | 1.2067606 | 0.227524 | 0.70225867 | PIGL | protein_coding | 44.78 | 0.043195356 | 0.043195356 | ENSG00000108474 | n.s. |
| ENSG00000139926 | 2641.774 | 0.333899507 | 0.307368 | 1.0863175 | 0.277339 | 0.742417658 | FRMD6 | protein_coding | 39.42 | 0.043190471 | 0.043190471 | ENSG00000139926 | n.s. |
| ENSG00000232882 | 25.40162 | -0.359381203 | 0.346454 | -1.037313 | 0.29959 | 0.758335061 | PHKA1P1 | processed_pseudogene | 49.11 | -0.04317565 | 0.04317565 | ENSG00000232882 | n.s. |
| ENSG00000100490 | 30.46844 | -0.363538397 | 0.352588 | -1.031058 | 0.302514 | 0.760781413 | CDKL1 | protein_coding | 43.04 | -0.043166588 | 0.043166588 | ENSG00000100490 | n.s. |
| ENSG00000128602 | 351.1767 | -0.296515908 | 0.254231 | -1.166326 | 0.243483 | 0.715191409 | SMO | protein_coding | 49.17 | -0.043166107 | 0.043166107 | ENSG00000128602 | n.s. |
| ENSG00000187135 | 111.6668 | 0.329101096 | 0.301387 | 1.0919551 | 0.274853 | 0.739386355 | VSTM2B | protein_coding | 51.2 | 0.043154556 | 0.043154556 | ENSG00000187135 | n.s. |
| ENSG00000100811 | 2372.151 | -0.305671157 | 0.266801 | -1.145691 | 0.251923 | 0.722544863 | YY1 | protein_coding | 45.55 | -0.043140954 | 0.043140954 | ENSG00000100811 | n.s. |
| ENSG00000170185 | 1253.726 | -0.299896344 | 0.258536 | -1.159978 | 0.246058 | 0.718210323 | USP38 | protein_coding | 35.49 | -0.043109607 | 0.043109607 | ENSG00000170185 | n.s. |

| | | | | | | | | | | | | | |
|---|---|---|---|---|---|---|---|---|---|---|---|---|---|
| ENSG00000180667 | 1672.544 | 0.282840864 | 0.23684 | 1.1942256 | 0.23239 | 0.704163047 | YOD1 | protein_coding | 39.45 | 0.043084235 | 0.043084235 | ENSG00000180667 | n.s. |
| ENSG00000118965 | 1594.517 | -0.320117674 | 0.286398 | -1.117738 | 0.263679 | 0.733524057 | WDR35 | protein_coding | 36.9 | -0.043083241 | 0.043083241 | ENSG00000118965 | n.s. |
| ENSG00000202444 | 12.21963 | 0.453390306 | 0.503742 | 0.900045 | 0.368096 | 0.803511983 | RNU5E-6P | snRNA | 37.39 | 0.043075544 | 0.043075544 | ENSG00000202444 | n.s. |
| ENSG00000221944 | 1060.425 | 0.256133839 | 0.20139 | 1.2718305 | 0.203433 | 0.679010816 | TIGD1 | protein_coding | 43.77 | 0.043062068 | 0.043062068 | ENSG00000221944 | n.s. |
| ENSG00000257509 | 45.11478 | -0.339236674 | 0.318975 | -1.063522 | 0.287545 | 0.746611882 | | sense_intronic | 46.25 | -0.043050865 | 0.043050865 | ENSG00000257509 | n.s. |
| ENSG00000240370 | 65.21121 | -0.319749659 | 0.286065 | -1.117753 | 0.263672 | 0.733524057 | RPL13P5 | ıscribed_processed_pseudog | 48.71 | -0.043033712 | 0.043033712 | ENSG00000240370 | n.s. |
| ENSG00000171223 | 1388.7 | 0.286683487 | 0.241494 | 1.1871241 | 0.235179 | 0.70779073 | JUNB | protein_coding | 64.89 | 0.043029795 | 0.043029795 | ENSG00000171223 | n.s. |
| ENSG00000204599 | 235.433 | -0.299718918 | 0.258549 | -1.159234 | 0.246361 | 0.718522635 | TRIM39 | protein_coding | 44.36 | -0.043027512 | 0.043027512 | ENSG00000204599 | n.s. |
| ENSG00000206645 | 4.879367 | -0.505020557 | 0.599962 | -0.841754 | 0.399926 | 0.821869704 | | misc_RNA | 39.22 | -0.043026251 | 0.043026251 | ENSG00000206645 | n.s. |
| ENSG00000079432 | 421.5749 | -0.282297832 | 0.235587 | -1.198273 | 0.230811 | 0.704043192 | CIC | protein_coding | 61.52 | -0.043022386 | 0.043022386 | ENSG00000079432 | n.s. |
| ENSG00000157349 | 293.5025 | -0.306956359 | 0.269271 | -1.139952 | 0.254306 | 0.72418384 | DDX19B | protein_coding | 45.56 | -0.043020293 | 0.043020293 | ENSG00000157349 | n.s. |
| ENSG00000278318 | 414.7179 | -0.319600733 | 0.28574 | -1.1185 | 0.263353 | 0.733524057 | ZNF229 | protein_coding | 41.83 | -0.043013669 | 0.043013669 | ENSG00000278318 | n.s. |
| ENSG00000214530 | 127.4452 | 0.32199512 | 0.290499 | 1.1084207 | 0.26768 | 0.735235906 | STARD10 | protein_coding | 54.42 | 0.043009948 | 0.043009948 | ENSG00000214530 | n.s. |
| ENSG00000178913 | 4322.192 | 0.300339204 | 0.259905 | 1.1555748 | 0.247855 | 0.719195543 | TAF7 | protein_coding | 42.08 | 0.042994462 | 0.042994462 | ENSG00000178913 | n.s. |
| ENSG00000172785 | 323.3376 | -0.322679623 | 0.292174 | -1.10441 | 0.269415 | 0.735983077 | CBWD1 | protein_coding | 36.26 | -0.042959039 | 0.042959039 | ENSG00000172785 | n.s. |
| ENSG00000279235 | 100.134 | -0.318513676 | 0.284177 | -1.12083 | 0.26236 | 0.73306092 | | TEC | 44.07 | -0.042954733 | 0.042954733 | ENSG00000279235 | n.s. |
| ENSG00000128045 | 4563.076 | 0.254094406 | 0.19919 | 1.2756402 | 0.202083 | 0.677624898 | RASL11B | protein_coding | 47.73 | 0.04294466 | 0.04294466 | ENSG00000128045 | n.s. |
| ENSG00000101997 | 125.7384 | -0.337627204 | 0.31613 | -1.068002 | 0.28552 | 0.746113205 | CCDC22 | protein_coding | 53.55 | -0.042944585 | 0.042944585 | ENSG00000101997 | n.s. |
| ENSG00000272800 | 50.5045 | -0.363772877 | 0.354879 | -1.025061 | 0.305334 | 0.761997697 | | lincRNA | 30.68 | -0.042942057 | 0.042942057 | ENSG00000272800 | n.s. |
| ENSG00000163319 | 1211.149 | 0.287766173 | 0.24316 | 1.1834449 | 0.236633 | 0.709287251 | MRPS18C | protein_coding | 38.68 | 0.042928338 | 0.042928338 | ENSG00000163319 | n.s. |
| ENSG00000067900 | 1959.733 | -0.281660488 | 0.235319 | -1.19693 | 0.231334 | 0.704043192 | ROCK1 | protein_coding | 36.92 | -0.042925255 | 0.042925255 | ENSG00000067900 | n.s. |
| ENSG00000107758 | 625.0604 | -0.331828606 | 0.305611 | -1.085788 | 0.277573 | 0.742417658 | PPP3CB | protein_coding | 38.91 | -0.042922597 | 0.042922597 | ENSG00000107758 | n.s. |
| ENSG00000100605 | 353.5075 | -0.309151174 | 0.272397 | -1.134929 | 0.256405 | 0.726402587 | ITPK1 | protein_coding | 53.85 | -0.042917175 | 0.042917175 | ENSG00000100605 | n.s. |
| ENSG00000148153 | 564.939 | -0.318779306 | 0.284777 | -1.119399 | 0.26297 | 0.733524057 | INIP | protein_coding | 40.14 | -0.042903116 | 0.042903116 | ENSG00000148153 | n.s. |
| ENSG00000174151 | 196.0761 | -0.302941103 | 0.263735 | -1.148656 | 0.250698 | 0.721766875 | CYB561D1 | protein_coding | 52.44 | -0.042897385 | 0.042897385 | ENSG00000174151 | n.s. |
| ENSG00000198932 | 402.9134 | -0.289672161 | 0.246001 | -1.177525 | 0.238986 | 0.71113531 | GPRASP1 | protein_coding | 48.65 | -0.042885314 | 0.042885314 | ENSG00000198932 | n.s. |
| ENSG00000104218 | 736.9191 | -0.299439623 | 0.258949 | -1.156365 | 0.247532 | 0.7191431 | CSPP1 | protein_coding | 38.01 | -0.042875167 | 0.042875167 | ENSG00000104218 | n.s. |
| ENSG00000105865 | 110.0366 | 0.308496649 | 0.2718 | 1.1350151 | 0.256369 | 0.726402587 | DUS4L | protein_coding | 37.93 | 0.042826312 | 0.042826312 | ENSG00000105865 | n.s. |
| ENSG00000280238 | 14.65919 | 0.406515374 | 0.423298 | 0.9603521 | 0.336878 | 0.784604238 | | TEC | 59.34 | 0.042826131 | 0.042826131 | ENSG00000280238 | n.s. |
| ENSG00000196405 | 2160.861 | -0.279669258 | 0.232654 | -1.202083 | 0.229331 | 0.702989524 | EVL | protein_coding | 45.48 | -0.042803701 | 0.042803701 | ENSG00000196405 | n.s. |
| ENSG00000073614 | 4008.111 | -0.286252034 | 0.241684 | -1.184406 | 0.236252 | 0.708727536 | KDM5A | protein_coding | 39.46 | -0.042800603 | 0.042800603 | ENSG00000073614 | n.s. |
| ENSG00000137055 | 1236.349 | -0.294321247 | 0.252707 | -1.164674 | 0.244151 | 0.715606021 | PLAA | protein_coding | 38.2 | -0.042772534 | 0.042772534 | ENSG00000137055 | n.s. |

| | | | | | | | | | | | | | |
|---|---|---|---|---|---|---|---|---|---|---|---|---|---|
| ENSG00000100304 | 730.2784 | -0.305177629 | 0.267763 | -1.139732 | 0.254398 | 0.72418384 | TTLL12 | protein_coding | 58.31 | -0.042771002 | 0.042771002 | ENSG00000100304 | n.s. |
| ENSG00000176014 | 924.2623 | -0.319269503 | 0.286817 | -1.113145 | 0.265646 | 0.734612586 | TUBB6 | protein_coding | 45.6 | -0.042763479 | 0.042763479 | ENSG00000176014 | n.s. |
| ENSG00000149716 | 415.2108 | -0.280597542 | 0.234289 | -1.197654 | 0.231052 | 0.704043192 | ORAOV1 | protein_coding | 53.52 | -0.042763261 | 0.042763261 | ENSG00000149716 | n.s. |
| ENSG00000272842 | 41.89659 | -0.354632254 | 0.340948 | -1.040136 | 0.298277 | 0.757691034 | | antisense | 26.79 | -0.042735971 | 0.042735971 | ENSG00000272842 | n.s. |
| ENSG00000271830 | 15.46814 | 0.406371276 | 0.424742 | 0.9567494 | 0.338694 | 0.784993146 | | lincRNA | 31.92 | 0.042723493 | 0.042723493 | ENSG00000271830 | n.s. |
| ENSG00000215190 | 430.7551 | -0.313034168 | 0.278265 | -1.124952 | 0.26061 | 0.730367609 | LINC00680 | cribed_unprocessed_pseudc | 45.68 | -0.042716172 | 0.042716172 | ENSG00000215190 | n.s. |
| ENSG00000177511 | 1315.624 | -0.301465509 | 0.262403 | -1.148864 | 0.250612 | 0.721669788 | ST8SIA3 | protein_coding | 41.88 | -0.042706049 | 0.042706049 | ENSG00000177511 | n.s. |
| ENSG00000207162 | 7.603322 | -0.457855536 | 0.512884 | -0.892708 | 0.372013 | 0.806907815 | RNU6-549P | snRNA | 39.25 | -0.042661183 | 0.042661183 | ENSG00000207162 | n.s. |
| ENSG00000138459 | 638.7635 | -0.332374773 | 0.308227 | -1.078343 | 0.280881 | 0.744242186 | SLC35A5 | protein_coding | 38.43 | -0.042638936 | 0.042638936 | ENSG00000138459 | n.s. |
| ENSG00000176225 | 1053.051 | -0.284696905 | 0.240227 | -1.185118 | 0.235971 | 0.708370066 | RTTN | protein_coding | 36.79 | -0.042630457 | 0.042630457 | ENSG00000176225 | n.s. |
| ENSG00000206903 | 41.05689 | 0.342474235 | 0.324079 | 1.05676 | 0.290621 | 0.750798148 | SNORA24B | snoRNA | 43.51 | 0.042630099 | 0.042630099 | ENSG00000206903 | n.s. |
| ENSG00000145723 | 112.1275 | -0.297520606 | 0.257142 | -1.157029 | 0.24726 | 0.7191431 | GIN1 | protein_coding | 36.01 | -0.042600393 | 0.042600393 | ENSG00000145723 | n.s. |
| ENSG00000182022 | 260.6092 | -0.296638867 | 0.255903 | -1.159184 | 0.246381 | 0.718522635 | CHST15 | protein_coding | 50.24 | -0.042585341 | 0.042585341 | ENSG00000182022 | n.s. |
| ENSG00000186812 | 1878.72 | -0.253348399 | 0.199431 | -1.270358 | 0.203957 | 0.679190265 | ZNF397 | protein_coding | 41.49 | -0.042564696 | 0.042564696 | ENSG00000186812 | n.s. |
| ENSG00000197472 | 724.5656 | -0.308361286 | 0.272873 | -1.130053 | 0.258454 | 0.728120901 | ZNF695 | protein_coding | 42.51 | -0.042491107 | 0.042491107 | ENSG00000197472 | n.s. |
| ENSG00000136731 | 2893.071 | -0.292292348 | 0.25104 | -1.164324 | 0.244293 | 0.715606581 | UGGT1 | protein_coding | 40.84 | -0.042477583 | 0.042477583 | ENSG00000136731 | n.s. |
| ENSG00000133958 | 1082.046 | -0.289566174 | 0.247246 | -1.171166 | 0.241532 | 0.713526369 | UNC79 | protein_coding | 39.32 | -0.042447498 | 0.042447498 | ENSG00000133958 | n.s. |
| ENSG00000199646 | 6.694124 | -0.538063295 | 0.66966 | -0.803487 | 0.421693 | 0.834219429 | RNU6-1272P | snRNA | 43.93 | -0.042356181 | 0.042356181 | ENSG00000199646 | n.s. |
| ENSG00000255100 | 13.07046 | 0.46452748 | 0.5327 | 0.8720251 | 0.383195 | 0.810719585 | | antisense | 50.41 | 0.04233208 | 0.04233208 | ENSG00000255100 | n.s. |
| ENSG00000018189 | 1653.536 | -0.295619018 | 0.255473 | -1.157143 | 0.247214 | 0.7191431 | RUFY3 | protein_coding | 39.91 | -0.042328115 | 0.042328115 | ENSG00000018189 | n.s. |
| ENSG00000107957 | 1105.522 | -0.29191867 | 0.250945 | -1.16328 | 0.244716 | 0.716165582 | SH3PXD2A | protein_coding | 51.58 | -0.042324282 | 0.042324282 | ENSG00000107957 | n.s. |
| ENSG00000152422 | 182.1134 | -0.320587799 | 0.291904 | -1.098265 | 0.272089 | 0.73797827 | XRCC4 | protein_coding | 35.2 | -0.04230362 | 0.04230362 | ENSG00000152422 | n.s. |
| ENSG00000141431 | 58.85003 | -0.330453019 | 0.307517 | -1.074584 | 0.282561 | 0.744776963 | ASXL3 | protein_coding | 36.09 | -0.042289317 | 0.042289317 | ENSG00000141431 | n.s. |
| ENSG00000134247 | 1260.983 | -0.290159001 | 0.248616 | -1.167095 | 0.243172 | 0.715021889 | PTGFRN | protein_coding | 44.19 | -0.042270556 | 0.042270556 | ENSG00000134247 | n.s. |
| ENSG00000234912 | 513.7343 | -0.284571784 | 0.2413 | -1.17933 | 0.238267 | 0.71051867 | SNHG20 | processed_transcript | 49.6 | -0.042237426 | 0.042237426 | ENSG00000234912 | n.s. |
| ENSG00000122484 | 1491.762 | -0.307373742 | 0.272584 | -1.127631 | 0.259476 | 0.728949992 | RPAP2 | protein_coding | 38.47 | -0.042203111 | 0.042203111 | ENSG00000122484 | n.s. |
| ENSG00000139722 | 488.5179 | -0.292212866 | 0.251545 | -1.161672 | 0.245369 | 0.717133775 | VPS37B | protein_coding | 47.3 | -0.042195486 | 0.042195486 | ENSG00000139722 | n.s. |
| ENSG00000182963 | 3083.435 | -0.308083391 | 0.27351 | -1.126408 | 0.259993 | 0.729661551 | GJC1 | protein_coding | 44.78 | -0.042170004 | 0.042170004 | ENSG00000182963 | n.s. |
| ENSG00000163848 | 1602.25 | -0.301614976 | 0.264918 | -1.138523 | 0.254902 | 0.724760001 | ZNF148 | protein_coding | 36.86 | -0.042167518 | 0.042167518 | ENSG00000163848 | n.s. |
| ENSG00000254985 | 52.16525 | -0.328913067 | 0.305682 | -1.075997 | 0.281929 | 0.744567264 | RSF1-IT2 | sense_intronic | 39 | -0.042132469 | 0.042132469 | ENSG00000254985 | n.s. |
| ENSG00000102780 | 1266.655 | -0.314582407 | 0.282827 | -1.112278 | 0.266019 | 0.73473105 | DGKH | protein_coding | 38.24 | -0.042113652 | 0.042113652 | ENSG00000102780 | n.s. |
| ENSG00000104884 | 257.0863 | -0.312888243 | 0.279564 | -1.119199 | 0.263055 | 0.733524057 | ERCC2 | protein_coding | 56.56 | -0.042110264 | 0.042110264 | ENSG00000104884 | n.s. |

| | | | | | | | | | | | | | |
|---|---|---|---|---|---|---|---|---|---|---|---|---|---|
| ENSG00000147100 | 147.6965 | -0.321137093 | 0.294242 | -1.091405 | 0.275095 | 0.739442303 | SLC16A2 | protein_coding | 43.6 | -0.042099694 | 0.042099694 | ENSG00000147100 | n.s. |
| ENSG00000279354 | 55.83532 | 0.328544692 | 0.305517 | 1.0753719 | 0.282208 | 0.744567264 | | TEC | 41.63 | 0.042085281 | 0.042085281 | ENSG00000279354 | n.s. |
| ENSG00000274565 | 33.71762 | -0.376957502 | 0.380625 | -0.990364 | 0.321996 | 0.773444264 | | antisense | 59.01 | -0.042057517 | 0.042057517 | ENSG00000274565 | n.s. |
| ENSG00000154096 | 575.8924 | -0.299867501 | 0.263189 | -1.139361 | 0.254552 | 0.72418384 | THY1 | protein_coding | 55.78 | -0.042026781 | 0.042026781 | ENSG00000154096 | n.s. |
| ENSG00000272583 | 30.52857 | 0.355956528 | 0.348047 | 1.0227258 | 0.306438 | 0.761997697 | | lincRNA | 38.79 | 0.042019366 | 0.042019366 | ENSG00000272583 | n.s. |
| ENSG00000175416 | 362.4608 | -0.320442159 | 0.293173 | -1.093013 | 0.274388 | 0.739386355 | CLTB | protein_coding | 53.09 | -0.042019121 | 0.042019121 | ENSG00000175416 | n.s. |
| ENSG00000143149 | 1069.158 | -0.314031975 | 0.282614 | -1.111168 | 0.266496 | 0.734987182 | ALDH9A1 | protein_coding | 40.34 | -0.041992429 | 0.041992429 | ENSG00000143149 | n.s. |
| ENSG00000207821 | 13.29378 | -0.42098682 | 0.455862 | -0.923496 | 0.355749 | 0.794802373 | MIR640 | miRNA | 52.08 | -0.041989581 | 0.041989581 | ENSG00000207821 | n.s. |
| ENSG00000196670 | 1073.981 | -0.311490488 | 0.277979 | -1.120554 | 0.262478 | 0.733242258 | ZFP62 | protein_coding | 42.93 | -0.041974126 | 0.041974126 | ENSG00000196670 | n.s. |
| ENSG00000078747 | 1398.117 | -0.297329483 | 0.259374 | -1.146334 | 0.251657 | 0.722498818 | ITCH | protein_coding | 40.53 | -0.04197188 | 0.04197188 | ENSG00000078747 | n.s. |
| ENSG00000171634 | 6232.458 | -0.297317275 | 0.25943 | -1.146041 | 0.251778 | 0.722544863 | BPTF | protein_coding | 40.86 | -0.041961928 | 0.041961928 | ENSG00000171634 | n.s. |
| ENSG00000199047 | 2.459154 | 0.781560758 | 1.20702 | 0.6475126 | 0.5173 | 0.88372671 | MIR378A | miRNA | 53.03 | 0.041955759 | 0.041955759 | ENSG00000199047 | n.s. |
| ENSG00000283514 | 20.09669 | -0.401159236 | 0.421545 | -0.951641 | 0.341279 | 0.786054116 | MIR3975 | miRNA | 40 | -0.041940217 | 0.041940217 | ENSG00000283514 | n.s. |
| ENSG00000051180 | 614.7183 | -0.297644752 | 0.259996 | -1.144807 | 0.252289 | 0.723058869 | RAD51 | protein_coding | 43.47 | -0.041916222 | 0.041916222 | ENSG00000051180 | n.s. |
| ENSG00000143569 | 3082.184 | -0.288733336 | 0.248088 | -1.163834 | 0.244491 | 0.715888782 | UBAP2L | protein_coding | 43.58 | -0.041910927 | 0.041910927 | ENSG00000143569 | n.s. |
| ENSG00000228084 | 38.06585 | 0.348756696 | 0.336126 | 1.0375788 | 0.299466 | 0.758335061 | | antisense | 43.18 | 0.041899233 | 0.041899233 | ENSG00000228084 | n.s. |
| ENSG00000113716 | 1185.649 | -0.276312371 | 0.231821 | -1.191923 | 0.233292 | 0.705346165 | HMGXB3 | protein_coding | 43.45 | -0.041888317 | 0.041888317 | ENSG00000113716 | n.s. |
| ENSG00000119943 | 58.78502 | -0.324884141 | 0.300586 | -1.080837 | 0.27977 | 0.743390786 | PYROXD2 | protein_coding | 50.39 | -0.041839498 | 0.041839498 | ENSG00000119943 | n.s. |
| ENSG00000105939 | 606.2702 | -0.287056446 | 0.246053 | -1.166644 | 0.243354 | 0.715115708 | ZC3HAV1 | protein_coding | 43.4 | -0.041802217 | 0.041802217 | ENSG00000105939 | n.s. |
| ENSG00000159461 | 462.1719 | -0.316892511 | 0.288894 | -1.096915 | 0.272679 | 0.738276701 | AMFR | protein_coding | 42.2 | -0.04176036 | 0.04176036 | ENSG00000159461 | n.s. |
| ENSG00000153914 | 7094.17 | 0.292266856 | 0.253342 | 1.153645 | 0.248646 | 0.719646518 | SREK1 | protein_coding | 35.5 | 0.041759314 | 0.041759314 | ENSG00000153914 | n.s. |
| ENSG00000164823 | 659.9713 | -0.310766322 | 0.278357 | -1.11643 | 0.264238 | 0.733889699 | OSGIN2 | protein_coding | 38.2 | -0.041757425 | 0.041757425 | ENSG00000164823 | n.s. |
| ENSG00000151458 | 1534.208 | -0.283154611 | 0.24107 | -1.174575 | 0.240165 | 0.712207974 | ANKRD50 | protein_coding | 35.15 | -0.041735055 | 0.041735055 | ENSG00000151458 | n.s. |
| ENSG00000138593 | 1061.718 | -0.281039743 | 0.238578 | -1.177979 | 0.238805 | 0.710909549 | SECISBP2L | protein_coding | 36.79 | -0.041646058 | 0.041646058 | ENSG00000138593 | n.s. |
| ENSG00000250486 | 280.0769 | 0.262557141 | 0.213205 | 1.2314785 | 0.218144 | 0.694065444 | FAM218A | protein_coding | 39.77 | 0.041641452 | 0.041641452 | ENSG00000250486 | n.s. |
| ENSG00000096080 | 573.7555 | -0.312729666 | 0.282911 | -1.1054 | 0.268986 | 0.735983077 | MRPS18A | protein_coding | 49.03 | -0.04163438 | 0.04163438 | ENSG00000096080 | n.s. |
| ENSG00000154655 | 155.6311 | -0.321089859 | 0.295257 | -1.087493 | 0.276819 | 0.741896779 | L3MBTL4 | protein_coding | 39.96 | -0.041631392 | 0.041631392 | ENSG00000154655 | n.s. |
| ENSG00000169760 | 29.87563 | 0.349175236 | 0.33735 | 1.0350522 | 0.300645 | 0.759962588 | NLGN1 | protein_coding | 35.49 | 0.041624407 | 0.041624407 | ENSG00000169760 | n.s. |
| ENSG00000154114 | 367.2134 | -0.32428671 | 0.300719 | -1.078373 | 0.280867 | 0.744242186 | TBCEL | protein_coding | 36.59 | -0.041601353 | 0.041601353 | ENSG00000154114 | n.s. |
| ENSG00000264834 | 6.219369 | -0.525932117 | 0.65325 | -0.805101 | 0.420761 | 0.833560144 | MIR1273F | miRNA | 58.59 | -0.041581802 | 0.041581802 | ENSG00000264834 | n.s. |
| ENSG00000174446 | 342.2588 | -0.278006312 | 0.234737 | -1.184329 | 0.236283 | 0.708727536 | SNAPC5 | protein_coding | 44.54 | -0.041567697 | 0.041567697 | ENSG00000174446 | n.s. |
| ENSG00000134324 | 1759.243 | -0.290549887 | 0.251729 | -1.154215 | 0.248412 | 0.719540199 | LPIN1 | protein_coding | 46.76 | -0.041532636 | 0.041532636 | ENSG00000134324 | n.s. |

| | | | | | | | | | | | | | |
|---|---|---|---|---|---|---|---|---|---|---|---|---|---|
| ENSG00000161999 | 359.2009 | -0.300647766 | 0.265439 | -1.132645 | 0.257363 | 0.727567253 | JMJD8 | protein_coding | 65.22 | -0.041527531 | 0.041527531 | ENSG00000161999 | n.s. |
| ENSG00000255062 | 55.29209 | -0.31334101 | 0.284366 | -1.101893 | 0.270508 | 0.737020734 | | antisense | 45.33 | -0.041524043 | 0.041524043 | ENSG00000255062 | n.s. |
| ENSG00000136643 | 702.4987 | -0.296162106 | 0.259183 | -1.142675 | 0.253173 | 0.72418384 | RPS6KC1 | protein_coding | 37.51 | -0.041507466 | 0.041507466 | ENSG00000136643 | n.s. |
| ENSG00000173163 | 301.2305 | -0.341042259 | 0.326722 | -1.04383 | 0.296564 | 0.755765348 | COMMD1 | protein_coding | 41.05 | -0.041475181 | 0.041475181 | ENSG00000173163 | n.s. |
| ENSG00000133657 | 3524.673 | -0.292507013 | 0.254529 | -1.149208 | 0.25047 | 0.721473621 | ATP13A3 | protein_coding | 38.74 | -0.041471511 | 0.041471511 | ENSG00000133657 | n.s. |
| ENSG00000232499 | 66.15524 | -0.325963984 | 0.30502 | -1.068665 | 0.285221 | 0.746113205 | | processed_pseudogene | 44.31 | -0.041461078 | 0.041461078 | ENSG00000232499 | n.s. |
| ENSG00000168916 | 1362.033 | 0.277110666 | 0.233986 | 1.1843023 | 0.236293 | 0.708727536 | ZNF608 | protein_coding | 40.42 | 0.041433779 | 0.041433779 | ENSG00000168916 | n.s. |
| ENSG00000104497 | 167.8625 | -0.30930309 | 0.277987 | -1.112654 | 0.265857 | 0.734612586 | SNX16 | protein_coding | 35.17 | -0.041428562 | 0.041428562 | ENSG00000104497 | n.s. |
| ENSG00000153786 | 419.5814 | -0.300678217 | 0.266241 | -1.129346 | 0.258752 | 0.728144462 | ZDHHC7 | protein_coding | 48.57 | -0.041428181 | 0.041428181 | ENSG00000153786 | n.s. |
| ENSG00000274712 | 147.6389 | 0.295502741 | 0.259268 | 1.1397576 | 0.254387 | 0.72418384 | | lincRNA | 51.97 | 0.041415055 | 0.041415055 | ENSG00000274712 | n.s. |
| ENSG00000163466 | 2173.812 | -0.309565199 | 0.278694 | -1.11077 | 0.266667 | 0.734987182 | ARPC2 | protein_coding | 44.32 | -0.041395131 | 0.041395131 | ENSG00000163466 | n.s. |
| ENSG00000233954 | 120.7246 | 0.350622542 | 0.343141 | 1.0218021 | 0.306875 | 0.761997697 | UQCRHL | protein_coding | 46.93 | 0.041389708 | 0.041389708 | ENSG00000233954 | n.s. |
| ENSG00000077942 | 864.5824 | -0.295035919 | 0.25828 | -1.142312 | 0.253325 | 0.72418384 | FBLN1 | protein_coding | 53.13 | -0.041349629 | 0.041349629 | ENSG00000077942 | n.s. |
| ENSG00000130313 | 303.9393 | -0.32345838 | 0.301358 | -1.073336 | 0.283121 | 0.745017596 | PGLS | protein_coding | 53.32 | -0.041348807 | 0.041348807 | ENSG00000130313 | n.s. |
| ENSG00000155287 | 225.5921 | -0.310487579 | 0.281004 | -1.104924 | 0.269193 | 0.735983077 | SLC25A28 | protein_coding | 45.02 | -0.041335886 | 0.041335886 | ENSG00000155287 | n.s. |
| ENSG00000273951 | 52.58063 | -0.321888002 | 0.298303 | -1.079063 | 0.28056 | 0.744242186 | | sense_intronic | 33.74 | -0.041293633 | 0.041293633 | ENSG00000273951 | n.s. |
| ENSG00000238719 | 32.93611 | -0.347634672 | 0.33689 | -1.031893 | 0.302122 | 0.760781413 | RNU7-96P | snRNA | 31.25 | -0.041278178 | 0.041278178 | ENSG00000238719 | n.s. |
| ENSG00000166130 | 718.4178 | -0.321720572 | 0.298169 | -1.078987 | 0.280594 | 0.744242186 | IKBIP | protein_coding | 40.95 | -0.041272154 | 0.041272154 | ENSG00000166130 | n.s. |
| ENSG00000152242 | 1020.163 | -0.294470112 | 0.258266 | -1.140183 | 0.25421 | 0.72418384 | C18orf25 | protein_coding | 40.58 | -0.041270331 | 0.041270331 | ENSG00000152242 | n.s. |
| ENSG00000167637 | 169.6356 | -0.312737803 | 0.284949 | -1.097523 | 0.272413 | 0.738113065 | ZNF283 | protein_coding | 38.88 | -0.041242957 | 0.041242957 | ENSG00000167637 | n.s. |
| ENSG00000019549 | 808.727 | 0.283446759 | 0.243085 | 1.166038 | 0.243599 | 0.715324966 | SNAI2 | protein_coding | 41.13 | 0.041240545 | 0.041240545 | ENSG00000019549 | n.s. |
| ENSG00000164187 | 485.5511 | -0.335623918 | 0.31945 | -1.050632 | 0.293428 | 0.753623766 | LMBRD2 | protein_coding | 35.54 | -0.041229859 | 0.041229859 | ENSG00000164187 | n.s. |
| ENSG00000106009 | 535.039 | -0.28766609 | 0.248635 | -1.156984 | 0.247279 | 0.7191431 | BRAT1 | protein_coding | 55.18 | -0.041189378 | 0.041189378 | ENSG00000106009 | n.s. |
| ENSG00000261839 | 78.48817 | 0.306041993 | 0.273676 | 1.1182658 | 0.263454 | 0.733524057 | | lincRNA | 43.07 | 0.041188857 | 0.041188857 | ENSG00000261839 | n.s. |
| ENSG00000164961 | 924.946 | -0.305242455 | 0.272255 | -1.121163 | 0.262218 | 0.732958382 | WASHC5 | protein_coding | 40.15 | -0.041183521 | 0.041183521 | ENSG00000164961 | n.s. |
| ENSG00000047578 | 46.58539 | -0.342500009 | 0.329932 | -1.038093 | 0.299227 | 0.758335061 | KIAA0556 | protein_coding | 47.84 | -0.041147562 | 0.041147562 | ENSG00000047578 | n.s. |
| ENSG00000167264 | 137.5627 | -0.277228689 | 0.234972 | -1.179835 | 0.238066 | 0.71051867 | DUS2 | protein_coding | 48.49 | -0.041147531 | 0.041147531 | ENSG00000167264 | n.s. |
| ENSG00000253516 | 76.07151 | -0.324213908 | 0.304916 | -1.06329 | 0.287651 | 0.746611882 | HMGB1P41 | processed_pseudogene | 42.27 | -0.041144399 | 0.041144399 | ENSG00000253516 | n.s. |
| ENSG00000169071 | 380.672 | -0.301914012 | 0.268524 | -1.124348 | 0.260865 | 0.730701535 | ROR2 | protein_coding | 44.96 | -0.041138797 | 0.041138797 | ENSG00000169071 | n.s. |
| ENSG00000061337 | 629.6424 | -0.298526451 | 0.264044 | -1.130592 | 0.258227 | 0.728120901 | LZTS1 | protein_coding | 50.43 | -0.041135901 | 0.041135901 | ENSG00000061337 | n.s. |
| ENSG00000104812 | 200.5575 | -0.310243716 | 0.281683 | -1.101394 | 0.270725 | 0.737020734 | GYS1 | protein_coding | 52.54 | -0.041113589 | 0.041113589 | ENSG00000104812 | n.s. |
| ENSG00000158092 | 445.056 | -0.307288831 | 0.276807 | -1.110118 | 0.266948 | 0.734987182 | NCK1 | protein_coding | 39.43 | -0.041090735 | 0.041090735 | ENSG00000158092 | n.s. |

| | | | | | | | | | | | | | |
|---|---|---|---|---|---|---|---|---|---|---|---|---|---|
| ENSG00000249855 | 90.71446 | -0.320807554 | 0.298458 | -1.074885 | 0.282426 | 0.744597466 | EEF1A1P19 | processed_pseudogene | 47.68 | -0.041088533 | 0.041088533 | ENSG00000249855 | n.s. |
| ENSG00000137054 | 1569.82 | -0.281828068 | 0.24142 | -1.167378 | 0.243058 | 0.715021889 | POLR1E | protein_coding | 46.29 | -0.0410569 | 0.0410569 | ENSG00000137054 | n.s. |
| ENSG00000063601 | 474.1044 | -0.310222594 | 0.28202 | -1.1 | 0.271332 | 0.737359702 | MTMR1 | protein_coding | 42.48 | -0.04104884 | 0.04104884 | ENSG00000063601 | n.s. |
| ENSG00000252367 | 68.60533 | 0.308285703 | 0.279092 | 1.1046007 | 0.269333 | 0.735983077 | | misc_RNA | 37.36 | 0.041042745 | 0.041042745 | ENSG00000252367 | n.s. |
| ENSG00000175395 | 305.3799 | -0.309707502 | 0.28127 | -1.101105 | 0.270851 | 0.737020734 | ZNF25 | protein_coding | 40.15 | -0.041042529 | 0.041042529 | ENSG00000175395 | n.s. |
| ENSG00000209082 | 3.267501 | -0.599989936 | 0.803066 | -0.747124 | 0.454989 | 0.854433778 | MT-TL1 | Mt_tRNA | 38.67 | -0.040992267 | 0.040992267 | ENSG00000209082 | n.s. |
| ENSG00000253008 | 98.00531 | 0.306546278 | 0.276404 | 1.1090523 | 0.267408 | 0.734987182 | MIR2355 | miRNA | 41.38 | 0.04099144 | 0.04099144 | ENSG00000253008 | n.s. |
| ENSG00000042088 | 667.7927 | -0.306007783 | 0.274871 | -1.113279 | 0.265589 | 0.734612586 | TDP1 | protein_coding | 42.14 | -0.040987183 | 0.040987183 | ENSG00000042088 | n.s. |
| ENSG00000254028 | 22.13262 | 0.406303606 | 0.437182 | 0.929369 | 0.352698 | 0.79272638 | | lincRNA | 43.09 | 0.040986563 | 0.040986563 | ENSG00000254028 | n.s. |
| ENSG00000256223 | 717.9668 | -0.304494021 | 0.272617 | -1.11693 | 0.264024 | 0.733524057 | ZNF10 | protein_coding | 40.03 | -0.040980522 | 0.040980522 | ENSG00000256223 | n.s. |
| ENSG00000197702 | 302.889 | -0.292367712 | 0.256534 | -1.139686 | 0.254417 | 0.72418384 | PARVA | protein_coding | 43.51 | -0.040975677 | 0.040975677 | ENSG00000197702 | n.s. |
| ENSG00000092439 | 3435.524 | -0.299109468 | 0.265598 | -1.126173 | 0.260092 | 0.729661551 | TRPM7 | protein_coding | 38.89 | -0.040941667 | 0.040941667 | ENSG00000092439 | n.s. |
| ENSG00000137955 | 11571.31 | 0.258099953 | 0.209567 | 1.2315854 | 0.218104 | 0.694065444 | RABGGTB | protein_coding | 37.86 | 0.040934544 | 0.040934544 | ENSG00000137955 | n.s. |
| ENSG00000158321 | 2196.128 | -0.281668164 | 0.241862 | -1.16458 | 0.244189 | 0.715606021 | AUTS2 | protein_coding | 41.05 | -0.040933711 | 0.040933711 | ENSG00000158321 | n.s. |
| ENSG00000147642 | 132.1555 | -0.283397055 | 0.243989 | -1.161518 | 0.245431 | 0.717133775 | SYBU | protein_coding | 39.55 | -0.040922485 | 0.040922485 | ENSG00000147642 | n.s. |
| ENSG00000200999 | 6.543109 | -0.456231435 | 0.527432 | -0.865006 | 0.387036 | 0.813399117 | | snoRNA | 42.5 | -0.040922273 | 0.040922273 | ENSG00000200999 | n.s. |
| ENSG00000171055 | 609.1351 | -0.317705641 | 0.293826 | -1.081272 | 0.279576 | 0.743390786 | FEZ2 | protein_coding | 40.54 | -0.04091503 | 0.04091503 | ENSG00000171055 | n.s. |
| ENSG00000073060 | 1517.081 | -0.291879036 | 0.256178 | -1.139359 | 0.254553 | 0.72418384 | SCARB1 | protein_coding | 51.45 | -0.040907188 | 0.040907188 | ENSG00000073060 | n.s. |
| ENSG00000196372 | 367.8807 | -0.338800758 | 0.32554 | -1.040735 | 0.297998 | 0.757323737 | ASB13 | protein_coding | 50.97 | -0.040899496 | 0.040899496 | ENSG00000196372 | n.s. |
| ENSG00000119906 | 548.7382 | -0.267480338 | 0.222667 | -1.201259 | 0.229651 | 0.703349338 | SLF2 | protein_coding | 39.04 | -0.04087873 | 0.04087873 | ENSG00000119906 | n.s. |
| ENSG00000206922 | 16.33843 | -0.374821301 | 0.382133 | -0.980865 | 0.326659 | 0.778045194 | RNU6-80P | snRNA | 41.12 | -0.040853714 | 0.040853714 | ENSG00000206922 | n.s. |
| ENSG00000116497 | 55.18826 | 0.321158828 | 0.300642 | 1.068245 | 0.28541 | 0.746113205 | S100PBP | protein_coding | 41.44 | 0.040849885 | 0.040849885 | ENSG00000116497 | n.s. |
| ENSG00000148468 | 1302.951 | -0.305374525 | 0.275306 | -1.109218 | 0.267336 | 0.734987182 | FAM171A1 | protein_coding | 44.98 | -0.040834753 | 0.040834753 | ENSG00000148468 | n.s. |
| ENSG00000165392 | 1380.728 | -0.302334283 | 0.269558 | -1.121591 | 0.262036 | 0.732723331 | WRN | protein_coding | 35.97 | -0.040833262 | 0.040833262 | ENSG00000165392 | n.s. |
| ENSG00000116133 | 3169.716 | -0.289980226 | 0.253424 | -1.144247 | 0.252521 | 0.723128487 | DHCR24 | protein_coding | 47.68 | -0.04082473 | 0.04082473 | ENSG00000116133 | n.s. |
| ENSG00000065268 | 439.895 | -0.316360171 | 0.291673 | -1.08464 | 0.278081 | 0.742965563 | WDR18 | protein_coding | 62.29 | -0.040820369 | 0.040820369 | ENSG00000065268 | n.s. |
| ENSG00000261377 | 181.0229 | -0.311152176 | 0.284523 | -1.093593 | 0.274133 | 0.739386355 | PDCD6IPP2 | cribed_unprocessed_pseudc | 41.42 | -0.040800939 | 0.040800939 | ENSG00000261377 | n.s. |
| ENSG00000101246 | 388.2729 | -0.274766823 | 0.232839 | -1.180071 | 0.237972 | 0.71051867 | ARFRP1 | protein_coding | 62.43 | -0.04078213 | 0.04078213 | ENSG00000101246 | n.s. |
| ENSG00000100266 | 795.5713 | -0.291556046 | 0.256167 | -1.138147 | 0.255059 | 0.72487977 | PACSIN2 | protein_coding | 48.59 | -0.040740298 | 0.040740298 | ENSG00000100266 | n.s. |
| ENSG00000149257 | 714.7576 | -0.284268839 | 0.245709 | -1.156931 | 0.247301 | 0.7191431 | SERPINH1 | protein_coding | 55.42 | -0.040702943 | 0.040702943 | ENSG00000149257 | n.s. |
| ENSG00000100330 | 114.5102 | -0.30549355 | 0.276452 | -1.105049 | 0.269138 | 0.735983077 | MTMR3 | protein_coding | 40.65 | -0.04067102 | 0.04067102 | ENSG00000100330 | n.s. |
| ENSG00000159899 | 274.7749 | -0.288019861 | 0.251222 | -1.146476 | 0.251598 | 0.722498818 | NPR2 | protein_coding | 48.3 | -0.040657707 | 0.040657707 | ENSG00000159899 | n.s. |

| | | | | | | | | | | | | | |
|---|---|---|---|---|---|---|---|---|---|---|---|---|---|
| ENSG00000091542 | 514.4069 | -0.262497809 | 0.216009 | -1.215217 | 0.224283 | 0.700099826 | ALKBH5 | protein_coding | 50.24 | -0.040645169 | 0.040645169 | ENSG00000091542 | n.s. |
| ENSG00000132361 | 820.6123 | -0.283076651 | 0.244225 | -1.159082 | 0.246423 | 0.718522635 | CLUH | protein_coding | 62.54 | -0.040638356 | 0.040638356 | ENSG00000132361 | n.s. |
| ENSG00000011422 | 55.62185 | -0.313409217 | 0.288233 | -1.087348 | 0.276883 | 0.741925232 | PLAUR | protein_coding | 49.94 | -0.040630327 | 0.040630327 | ENSG00000011422 | n.s. |
| ENSG00000239900 | 507.8311 | -0.287492245 | 0.250668 | -1.146903 | 0.251422 | 0.722279621 | ADSL | protein_coding | 44.97 | -0.040621113 | 0.040621113 | ENSG00000239900 | n.s. |
| ENSG00000168243 | 1576.997 | -0.306222373 | 0.278048 | -1.101327 | 0.270754 | 0.737020734 | GNG4 | protein_coding | 48.07 | -0.040580679 | 0.040580679 | ENSG00000168243 | n.s. |
| ENSG00000186104 | 213.7605 | -0.30342449 | 0.273371 | -1.109935 | 0.267027 | 0.734987182 | CYP2R1 | protein_coding | 39.86 | -0.040573994 | 0.040573994 | ENSG00000186104 | n.s. |
| ENSG00000172086 | 265.5147 | -0.31625491 | 0.293179 | -1.07871 | 0.280717 | 0.744242186 | KRCC1 | protein_coding | 38.55 | -0.040570988 | 0.040570988 | ENSG00000172086 | n.s. |
| ENSG00000159842 | 457.7219 | -0.303369853 | 0.273497 | -1.109226 | 0.267333 | 0.734987182 | ABR | protein_coding | 55.2 | -0.040566688 | 0.040566688 | ENSG00000159842 | n.s. |
| ENSG00000115109 | 620.9683 | -0.289316506 | 0.253119 | -1.143005 | 0.253037 | 0.724157976 | EPB41L5 | protein_coding | 38.5 | -0.040552535 | 0.040552535 | ENSG00000115109 | n.s. |
| ENSG00000165568 | 43.92715 | -0.364609216 | 0.368586 | -0.98921 | 0.322561 | 0.774129273 | AKR1E2 | protein_coding | 42.59 | -0.040539626 | 0.040539626 | ENSG00000165568 | n.s. |
| ENSG00000184588 | 1557.962 | -0.301183163 | 0.269322 | -1.118301 | 0.263438 | 0.733524057 | PDE4B | protein_coding | 37.45 | -0.040534928 | 0.040534928 | ENSG00000184588 | n.s. |
| ENSG00000165752 | 97.48209 | -0.31650462 | 0.29471 | -1.073951 | 0.282845 | 0.744776963 | STK32C | protein_coding | 55.1 | -0.040504288 | 0.040504288 | ENSG00000165752 | n.s. |
| ENSG00000089639 | 161.5772 | -0.302349092 | 0.271316 | -1.11438 | 0.265116 | 0.734612586 | GMIP | protein_coding | 56.14 | -0.040497132 | 0.040497132 | ENSG00000089639 | n.s. |
| ENSG00000256338 | 42.17311 | 0.342949135 | 0.334932 | 1.0239353 | 0.305866 | 0.761997697 | RPL41P2 | processed_pseudogene | 50.61 | 0.040483891 | 0.040483891 | ENSG00000256338 | n.s. |
| ENSG00000135838 | 108.8495 | -0.305444919 | 0.276906 | -1.103064 | 0.269999 | 0.737020734 | NPL | protein_coding | 41.95 | -0.040477651 | 0.040477651 | ENSG00000135838 | n.s. |
| ENSG00000196715 | 808.9505 | -0.316287397 | 0.294512 | -1.073938 | 0.28285 | 0.744776963 | VKORC1L1 | protein_coding | 41.16 | -0.040476489 | 0.040476489 | ENSG00000196715 | n.s. |
| ENSG00000137274 | 287.0591 | -0.30711698 | 0.280002 | -1.09684 | 0.272711 | 0.738276701 | BPHL | protein_coding | 47.11 | -0.040472133 | 0.040472133 | ENSG00000137274 | n.s. |
| ENSG00000100867 | 173.5222 | -0.27253487 | 0.230853 | -1.180557 | 0.237779 | 0.71051867 | DHRS2 | protein_coding | 49.5 | -0.040450853 | 0.040450853 | ENSG00000100867 | n.s. |
| ENSG00000101109 | 927.1073 | -0.288381921 | 0.253029 | -1.139719 | 0.254403 | 0.72418384 | STK4 | protein_coding | 38.46 | -0.040417064 | 0.040417064 | ENSG00000101109 | n.s. |
| ENSG00000186687 | 630.3363 | -0.312632689 | 0.288119 | -1.085081 | 0.277886 | 0.74275391 | LYRM7 | protein_coding | 38.11 | -0.040378092 | 0.040378092 | ENSG00000186687 | n.s. |
| ENSG00000175215 | 1379.104 | -0.284638139 | 0.247602 | -1.149581 | 0.250317 | 0.721375365 | CTDSP2 | protein_coding | 51.21 | -0.040372702 | 0.040372702 | ENSG00000175215 | n.s. |
| ENSG00000118900 | 1215.644 | -0.276149383 | 0.236056 | -1.169848 | 0.242062 | 0.714515282 | UBN1 | protein_coding | 47.93 | -0.040314628 | 0.040314628 | ENSG00000118900 | n.s. |
| ENSG00000164066 | 751.7434 | -0.306866714 | 0.279965 | -1.096089 | 0.27304 | 0.739022549 | INTU | protein_coding | 37.11 | -0.040304583 | 0.040304583 | ENSG00000164066 | n.s. |
| ENSG00000119280 | 948.7735 | -0.299469371 | 0.267645 | -1.118906 | 0.26318 | 0.733524057 | C1orf198 | protein_coding | 47.8 | -0.040304276 | 0.040304276 | ENSG00000119280 | n.s. |
| ENSG00000076003 | 3556.998 | -0.301369723 | 0.271452 | -1.110212 | 0.266908 | 0.734987182 | MCM6 | protein_coding | 40.28 | -0.04029923 | 0.04029923 | ENSG00000076003 | n.s. |
| ENSG00000242689 | 114.9822 | -0.304055079 | 0.275808 | -1.102415 | 0.270281 | 0.737020734 | CNTF | protein_coding | 41.59 | -0.040293469 | 0.040293469 | ENSG00000242689 | n.s. |
| ENSG00000119392 | 488.0842 | -0.301926327 | 0.272596 | -1.107598 | 0.268035 | 0.735486422 | GLE1 | protein_coding | 45.27 | -0.040284623 | 0.040284623 | ENSG00000119392 | n.s. |
| ENSG00000158526 | 369.288 | -0.300721055 | 0.269937 | -1.114041 | 0.265262 | 0.734612586 | TSR2 | protein_coding | 47.57 | -0.04027907 | 0.04027907 | ENSG00000158526 | n.s. |
| ENSG00000196712 | 2041.088 | -0.295194602 | 0.262441 | -1.124804 | 0.260672 | 0.730396063 | NF1 | protein_coding | 37.62 | -0.040276817 | 0.040276817 | ENSG00000196712 | n.s. |
| ENSG00000156973 | 297.4518 | 0.340961524 | 0.333927 | 1.0210673 | 0.307223 | 0.761997697 | PDE6D | protein_coding | 42.51 | 0.040249261 | 0.040249261 | ENSG00000156973 | n.s. |
| ENSG00000069275 | 11688.09 | -0.301230906 | 0.271895 | -1.107893 | 0.267908 | 0.735446299 | NUCKS1 | protein_coding | 39.57 | -0.040198973 | 0.040198973 | ENSG00000069275 | n.s. |
| ENSG00000164684 | 2415.981 | -0.293582491 | 0.260692 | -1.126165 | 0.260096 | 0.729661551 | ZNF704 | protein_coding | 40.26 | -0.040185142 | 0.040185142 | ENSG00000164684 | n.s. |

| | | | | | | | | | | | | | |
|---|---|---|---|---|---|---|---|---|---|---|---|---|---|
| ENSG00000152520 | 674.1213 | -0.298461458 | 0.266885 | -1.118314 | 0.263433 | 0.733524057 | PAN3 | protein_coding | 37.49 | -0.040168626 | 0.040168626 | ENSG00000152520 | n.s. |
| ENSG00000074219 | 460.5258 | -0.311301602 | 0.287077 | -1.084384 | 0.278195 | 0.742965563 | TEAD2 | protein_coding | 53.57 | -0.040167655 | 0.040167655 | ENSG00000074219 | n.s. |
| ENSG00000119285 | 4807.573 | -0.286399807 | 0.251397 | -1.139233 | 0.254606 | 0.72418384 | HEATR1 | protein_coding | 38.51 | -0.040139268 | 0.040139268 | ENSG00000119285 | n.s. |
| ENSG00000164252 | 1001.59 | -0.299668438 | 0.269077 | -1.113689 | 0.265413 | 0.734612586 | AGGF1 | protein_coding | 39.02 | -0.040138081 | 0.040138081 | ENSG00000164252 | n.s. |
| ENSG00000220515 | 59.39515 | -0.337724522 | 0.32712 | -1.032418 | 0.301876 | 0.760781413 | PGAM1P10 | processed_pseudogene | 53.09 | -0.040101446 | 0.040101446 | ENSG00000220515 | n.s. |
| ENSG00000109971 | 27688.61 | 0.299842895 | 0.269736 | 1.1116154 | 0.266304 | 0.734987182 | HSPA8 | protein_coding | 49.25 | 0.040095062 | 0.040095062 | ENSG00000109971 | n.s. |
| ENSG00000206870 | 7.392674 | -0.437210251 | 0.497865 | -0.87817 | 0.379851 | 0.809760709 | RNU6-398P | snRNA | 42.2 | -0.04006739 | 0.04006739 | ENSG00000206870 | n.s. |
| ENSG00000147955 | 1163.788 | -0.279605076 | 0.241451 | -1.158021 | 0.246856 | 0.719028029 | SIGMAR1 | protein_coding | 57.25 | -0.040054596 | 0.040054596 | ENSG00000147955 | n.s. |
| ENSG00000138398 | 3197.567 | -0.295857581 | 0.263587 | -1.122429 | 0.26168 | 0.73233574 | PPIG | protein_coding | 37.96 | -0.040026504 | 0.040026504 | ENSG00000138398 | n.s. |
| ENSG00000239521 | 218.8801 | -0.302715683 | 0.275391 | -1.09922 | 0.271672 | 0.737615059 | GATS | protein_coding | 51.53 | -0.04001 | 0.04001 | ENSG00000239521 | n.s. |
| ENSG00000171827 | 427.9104 | -0.30182425 | 0.273886 | -1.102006 | 0.270459 | 0.737020734 | ZNF570 | protein_coding | 41.65 | -0.039997838 | 0.039997838 | ENSG00000171827 | n.s. |
| ENSG00000130540 | 375.825 | -0.281898147 | 0.245309 | -1.149155 | 0.250492 | 0.721473621 | SULT4A1 | protein_coding | 56.02 | -0.03996739 | 0.03996739 | ENSG00000130540 | n.s. |
| ENSG00000163584 | 5324.442 | 0.323517182 | 0.307192 | 1.0531422 | 0.292276 | 0.752608927 | RPL22L1 | protein_coding | 40.36 | 0.039931931 | 0.039931931 | ENSG00000163584 | n.s. |
| ENSG00000263584 | 3.535089 | -0.678946246 | 0.99144 | -0.684808 | 0.493465 | 0.873348456 | MIR4480 | miRNA | 53.52 | -0.039930488 | 0.039930488 | ENSG00000263584 | n.s. |
| ENSG00000208028 | 27.86257 | 0.391307588 | 0.419019 | 0.9338654 | 0.350373 | 0.790639816 | MIR616 | miRNA | 40.21 | 0.039921716 | 0.039921716 | ENSG00000208028 | n.s. |
| ENSG00000241258 | 642.4814 | -0.314383477 | 0.295101 | -1.065342 | 0.286721 | 0.746611882 | CRCP | protein_coding | 45 | -0.039896867 | 0.039896867 | ENSG00000241258 | n.s. |
| ENSG00000272371 | 73.32908 | 0.296324043 | 0.264576 | 1.1199958 | 0.262716 | 0.733524057 | | lincRNA | 43.42 | 0.03988096 | 0.03988096 | ENSG00000272371 | n.s. |
| ENSG00000137574 | 1258.819 | -0.289378278 | 0.256141 | -1.12976 | 0.258578 | 0.728120901 | TGS1 | protein_coding | 40.02 | -0.039875315 | 0.039875315 | ENSG00000137574 | n.s. |
| ENSG00000255857 | 133.9826 | -0.325921904 | 0.311429 | -1.046536 | 0.295314 | 0.75457253 | PXN-AS1 | antisense | 49.38 | -0.039859926 | 0.039859926 | ENSG00000255857 | n.s. |
| ENSG00000264575 | 167.8333 | 0.278088843 | 0.240121 | 1.1581205 | 0.246815 | 0.719028029 | LINC00526 | lincRNA | 54.56 | 0.039837389 | 0.039837389 | ENSG00000264575 | n.s. |
| ENSG00000254694 | 50.17326 | -0.337368835 | 0.330676 | -1.020241 | 0.307614 | 0.761997697 | | antisense | 35.49 | -0.039825157 | 0.039825157 | ENSG00000254694 | n.s. |
| ENSG00000173011 | 260.7612 | -0.294762244 | 0.262817 | -1.121547 | 0.262055 | 0.732723331 | TADA2B | protein_coding | 56.73 | -0.039810583 | 0.039810583 | ENSG00000173011 | n.s. |
| ENSG00000283734 | 7.789649 | 0.478738605 | 0.577269 | 0.8293157 | 0.406926 | 0.825802394 | MIR4785 | miRNA | 80.82 | 0.039794602 | 0.039794602 | ENSG00000283734 | n.s. |
| ENSG00000131626 | 1081.333 | -0.290254777 | 0.257531 | -1.127067 | 0.259714 | 0.729326218 | PPFIA1 | protein_coding | 45 | -0.039787595 | 0.039787595 | ENSG00000131626 | n.s. |
| ENSG00000101220 | 465.5956 | -0.30067982 | 0.273395 | -1.0998 | 0.271419 | 0.737359702 | C20orf27 | protein_coding | 56.32 | -0.039786134 | 0.039786134 | ENSG00000101220 | n.s. |
| ENSG00000234006 | 38.14424 | -0.360165636 | 0.365607 | -0.985117 | 0.324566 | 0.775648559 | DDX39B-AS1 | antisense | 55.21 | -0.039738879 | 0.039738879 | ENSG00000234006 | n.s. |
| ENSG00000140694 | 845.0898 | -0.280862395 | 0.244673 | -1.147909 | 0.251006 | 0.722169533 | PARN | protein_coding | 41.91 | -0.039702943 | 0.039702943 | ENSG00000140694 | n.s. |
| ENSG00000264616 | 25.66987 | -0.377096448 | 0.393219 | -0.959 | 0.337559 | 0.78475552 | MIR4755 | miRNA | 47.22 | -0.039695292 | 0.039695292 | ENSG00000264616 | n.s. |
| ENSG00000116539 | 3641.324 | -0.275170695 | 0.237047 | -1.160827 | 0.245712 | 0.717427972 | ASH1L | protein_coding | 41.38 | -0.039685584 | 0.039685584 | ENSG00000116539 | n.s. |
| ENSG00000135127 | 127.8628 | -0.294780766 | 0.26333 | -1.119433 | 0.262955 | 0.733524057 | BICDL1 | protein_coding | 43.81 | -0.039673258 | 0.039673258 | ENSG00000135127 | n.s. |
| ENSG00000257261 | 16.30347 | -0.37726398 | 0.394152 | -0.957154 | 0.33849 | 0.784993146 | | lincRNA | 38.48 | -0.039663322 | 0.039663322 | ENSG00000257261 | n.s. |
| ENSG00000283341 | 173.2095 | -0.267598541 | 0.227194 | -1.177843 | 0.238859 | 0.710909549 | | processed_transcript | 38.5 | -0.039654264 | 0.039654264 | ENSG00000283341 | n.s. |

| | | | | | | | | | | | | | |
|---|---|---|---|---|---|---|---|---|---|---|---|---|---|
| ENSG00000139190 | 200.0248 | -0.276876643 | 0.239435 | -1.156377 | 0.247527 | 0.7191431 | VAMP1 | protein_coding | 48.87 | -0.039644494 | 0.039644494 | ENSG00000139190 | n.s. |
| ENSG00000185090 | 490.0718 | -0.297687985 | 0.269286 | -1.10547 | 0.268956 | 0.735983077 | MANEAL | protein_coding | 54.73 | -0.039631848 | 0.039631848 | ENSG00000185090 | n.s. |
| ENSG00000235501 | 114.9421 | 0.310779809 | 0.290511 | 1.0697712 | 0.284722 | 0.745682152 | | antisense | 41.99 | 0.039607722 | 0.039607722 | ENSG00000235501 | n.s. |
| ENSG00000106246 | 350.201 | -0.307275058 | 0.284247 | -1.081013 | 0.279691 | 0.743390786 | PTCD1 | protein_coding | 53.4 | -0.03957175 | 0.03957175 | ENSG00000106246 | n.s. |
| ENSG00000107890 | 794.0273 | -0.307262095 | 0.283902 | -1.082283 | 0.279127 | 0.743390786 | ANKRD26 | protein_coding | 38.15 | -0.039570081 | 0.039570081 | ENSG00000107890 | n.s. |
| ENSG00000130810 | 642.7557 | -0.272104007 | 0.23361 | -1.164779 | 0.244108 | 0.715606021 | PPAN | protein_coding | 58.7 | -0.039543791 | 0.039543791 | ENSG00000130810 | n.s. |
| ENSG00000112425 | 82.72901 | -0.312776538 | 0.294786 | -1.061029 | 0.288677 | 0.747460922 | EPM2A | protein_coding | 37.55 | -0.039538554 | 0.039538554 | ENSG00000112425 | n.s. |
| ENSG00000164904 | 1988.785 | -0.270821595 | 0.231661 | -1.169043 | 0.242386 | 0.714515282 | ALDH7A1 | protein_coding | 43.05 | -0.039536832 | 0.039536832 | ENSG00000164904 | n.s. |
| ENSG00000100417 | 112.0155 | -0.325649684 | 0.31244 | -1.042278 | 0.297283 | 0.756138711 | PMM1 | protein_coding | 54.84 | -0.039533392 | 0.039533392 | ENSG00000100417 | n.s. |
| ENSG00000161618 | 304.1707 | -0.301275826 | 0.275787 | -1.092423 | 0.274647 | 0.739386355 | ALDH16A1 | protein_coding | 56.83 | -0.039505868 | 0.039505868 | ENSG00000161618 | n.s. |
| ENSG00000273204 | 21.75505 | -0.39193463 | 0.422184 | -0.928349 | 0.353226 | 0.792950339 | | antisense | 54.08 | -0.039488986 | 0.039488986 | ENSG00000273204 | n.s. |
| ENSG00000065000 | 1081.09 | -0.2865823 | 0.253769 | -1.129302 | 0.25877 | 0.728144462 | AP3D1 | protein_coding | 56.41 | -0.039486011 | 0.039486011 | ENSG00000065000 | n.s. |
| ENSG00000066777 | 1388.33 | -0.294990651 | 0.266017 | -1.108916 | 0.267466 | 0.734987182 | ARFGEF1 | protein_coding | 38.36 | -0.039446219 | 0.039446219 | ENSG00000066777 | n.s. |
| ENSG00000182141 | 882.9777 | -0.297628012 | 0.270333 | -1.100968 | 0.270911 | 0.737020734 | ZNF708 | protein_coding | 39.17 | -0.039441752 | 0.039441752 | ENSG00000182141 | n.s. |
| ENSG00000185515 | 513.2479 | -0.310064783 | 0.290718 | -1.06655 | 0.286175 | 0.746285571 | BRCC3 | protein_coding | 37.76 | -0.03940767 | 0.03940767 | ENSG00000185515 | n.s. |
| ENSG00000205808 | 263.8702 | 0.270629468 | 0.232024 | 1.1663873 | 0.243458 | 0.715191409 | PLPP6 | protein_coding | 47.55 | 0.039397618 | 0.039397618 | ENSG00000205808 | n.s. |
| ENSG00000166986 | 3633.942 | -0.307089467 | 0.285025 | -1.077414 | 0.281295 | 0.744242186 | MARS | protein_coding | 49.33 | -0.039395192 | 0.039395192 | ENSG00000166986 | n.s. |
| ENSG00000135537 | 56.83054 | -0.309510523 | 0.289606 | -1.068731 | 0.285191 | 0.746113205 | AFG1L | protein_coding | 39.15 | -0.039368276 | 0.039368276 | ENSG00000135537 | n.s. |
| ENSG00000132952 | 896.1439 | -0.29250332 | 0.261736 | -1.117549 | 0.26376 | 0.733524057 | USPL1 | protein_coding | 38.94 | -0.039366746 | 0.039366746 | ENSG00000132952 | n.s. |
| ENSG00000121005 | 383.8197 | -0.304263549 | 0.279991 | -1.08669 | 0.277174 | 0.742417658 | CRISPLD1 | protein_coding | 35.22 | -0.03935701 | 0.03935701 | ENSG00000121005 | n.s. |
| ENSG00000100084 | 866.8536 | -0.278507791 | 0.242847 | -1.146843 | 0.251446 | 0.722279621 | HIRA | protein_coding | 46.11 | -0.039351658 | 0.039351658 | ENSG00000100084 | n.s. |
| ENSG00000154001 | 1171.118 | -0.292242758 | 0.261305 | -1.118396 | 0.263398 | 0.733524057 | PPP2R5E | protein_coding | 40.18 | -0.039331678 | 0.039331678 | ENSG00000154001 | n.s. |
| ENSG00000147548 | 2849.502 | -0.285379809 | 0.252509 | -1.130179 | 0.258401 | 0.728120901 | NSD3 | protein_coding | 38.79 | -0.039324339 | 0.039324339 | ENSG00000147548 | n.s. |
| ENSG00000102265 | 105.1525 | -0.333066018 | 0.326165 | -1.021159 | 0.307179 | 0.761997697 | TIMP1 | protein_coding | 53.67 | -0.039317225 | 0.039317225 | ENSG00000102265 | n.s. |
| ENSG00000145740 | 1003.259 | -0.31812439 | 0.301815 | -1.054039 | 0.291865 | 0.75237733 | SLC30A5 | protein_coding | 39.35 | -0.039308817 | 0.039308817 | ENSG00000145740 | n.s. |
| ENSG00000010322 | 1927.968 | -0.292064985 | 0.261484 | -1.116953 | 0.264015 | 0.733524057 | NISCH | protein_coding | 53.28 | -0.039307752 | 0.039307752 | ENSG00000010322 | n.s. |
| ENSG00000252874 | 21.70943 | -0.360570487 | 0.367333 | -0.981589 | 0.326302 | 0.778045194 | | misc_RNA | 38.89 | -0.039300444 | 0.039300444 | ENSG00000252874 | n.s. |
| ENSG00000270722 | 281.5241 | 0.257844059 | 0.21557 | 1.1961061 | 0.231655 | 0.704043192 | | snRNA | 53.66 | 0.039295614 | 0.039295614 | ENSG00000270722 | n.s. |
| ENSG00000272990 | 29.24322 | -0.469637842 | 0.564083 | -0.832568 | 0.405088 | 0.824799944 | | antisense | 38.19 | -0.039285852 | 0.039285852 | ENSG00000272990 | n.s. |
| ENSG00000040933 | 1228.696 | -0.297451173 | 0.270764 | -1.098564 | 0.271958 | 0.737938074 | INPP4A | protein_coding | 47.33 | -0.03925763 | 0.03925763 | ENSG00000040933 | n.s. |
| ENSG00000265871 | 14.42696 | 0.405334265 | 0.446712 | 0.9073731 | 0.36421 | 0.800117344 | MIR3174 | miRNA | 45.98 | 0.03925513 | 0.03925513 | ENSG00000265871 | n.s. |
| ENSG00000051108 | 1624.744 | 0.254495989 | 0.209904 | 1.2124404 | 0.225344 | 0.701091738 | HERPUD1 | protein_coding | 43.16 | 0.039249682 | 0.039249682 | ENSG00000051108 | n.s. |

| | | | | | | | | | | | | | |
|---|---|---|---|---|---|---|---|---|---|---|---|---|---|
| ENSG00000196700 | 631.8348 | -0.27395596 | 0.236658 | -1.157603 | 0.247026 | 0.719076249 | ZNF512B | protein_coding | 63.43 | -0.039237358 | 0.039237358 | ENSG00000196700 | n.s. |
| ENSG00000269032 | 13.60429 | 0.403916646 | 0.444761 | 0.9081647 | 0.363791 | 0.799601391 | | transcribed_processed_pseudog | 58.79 | 0.039230994 | 0.039230994 | ENSG00000269032 | n.s. |
| ENSG00000165714 | 353.3779 | -0.264305286 | 0.224092 | -1.17945 | 0.238219 | 0.71051867 | BORCS5 | protein_coding | 42.13 | -0.039229381 | 0.039229381 | ENSG00000165714 | n.s. |
| ENSG00000079313 | 325.8378 | -0.271835136 | 0.234184 | -1.160777 | 0.245732 | 0.717427972 | REXO1 | protein_coding | 60.94 | -0.039204524 | 0.039204524 | ENSG00000079313 | n.s. |
| ENSG00000281394 | 73.19931 | 0.305510255 | 0.283198 | 1.0787867 | 0.280683 | 0.744242186 | SCARNA4 | scaRNA | 46.51 | 0.039192602 | 0.039192602 | ENSG00000281394 | n.s. |
| ENSG00000259642 | 138.6015 | 0.276850213 | 0.241079 | 1.148379 | 0.250812 | 0.72194701 | ST20-AS1 | antisense | 56.17 | 0.039172832 | 0.039172832 | ENSG00000259642 | n.s. |
| ENSG00000130349 | 318.9236 | -0.298653294 | 0.273609 | -1.091534 | 0.275038 | 0.739442303 | C6orf203 | protein_coding | 47.04 | -0.039152165 | 0.039152165 | ENSG00000130349 | n.s. |
| ENSG00000230177 | 75.25752 | -0.308406695 | 0.289646 | -1.064769 | 0.28698 | 0.746611882 | | antisense | 41.18 | -0.039138383 | 0.039138383 | ENSG00000230177 | n.s. |
| ENSG00000174996 | 222.7469 | -0.308952568 | 0.291014 | -1.06164 | 0.288399 | 0.747018696 | KLC2 | protein_coding | 60.47 | -0.039134567 | 0.039134567 | ENSG00000174996 | n.s. |
| ENSG00000086758 | 3850.476 | -0.274199953 | 0.23793 | -1.152438 | 0.249141 | 0.720036883 | HUWE1 | protein_coding | 41.2 | -0.039113323 | 0.039113323 | ENSG00000086758 | n.s. |
| ENSG00000200636 | 4.481362 | 0.665221442 | 0.974041 | 0.6829502 | 0.494638 | 0.873443478 | SNORD114-27 | snoRNA | 45.71 | 0.039091867 | 0.039091867 | ENSG00000200636 | n.s. |
| ENSG00000062822 | 908.8693 | -0.297909755 | 0.272241 | -1.094288 | 0.273829 | 0.739386355 | POLD1 | protein_coding | 55.06 | -0.03906448 | 0.03906448 | ENSG00000062822 | n.s. |
| ENSG00000027001 | 189.871 | -0.31416644 | 0.297471 | -1.056124 | 0.290912 | 0.751040718 | MIPEP | protein_coding | 39.76 | -0.039062361 | 0.039062361 | ENSG00000027001 | n.s. |
| ENSG00000177150 | 367.2026 | -0.293195531 | 0.265433 | -1.104594 | 0.269335 | 0.735983077 | FAM210A | protein_coding | 40.65 | -0.039033758 | 0.039033758 | ENSG00000177150 | n.s. |
| ENSG00000065609 | 32.19086 | -0.34860789 | 0.351625 | -0.991421 | 0.32148 | 0.772918048 | SNAP91 | protein_coding | 36.62 | -0.038997562 | 0.038997562 | ENSG00000065609 | n.s. |
| ENSG00000165617 | 304.517 | 0.28152369 | 0.248296 | 1.1338232 | 0.256869 | 0.726907198 | DACT1 | protein_coding | 45.52 | 0.038996952 | 0.038996952 | ENSG00000165617 | n.s. |
| ENSG00000265140 | 3.862594 | 0.584274433 | 0.791895 | 0.7378178 | 0.460625 | 0.857620257 | MIR4301 | miRNA | 56.06 | 0.038974008 | 0.038974008 | ENSG00000265140 | n.s. |
| ENSG00000235319 | 27.66853 | 0.37457209 | 0.395114 | 0.9480111 | 0.343124 | 0.787036737 | | lincRNA | 48.96 | 0.038957369 | 0.038957369 | ENSG00000235319 | n.s. |
| ENSG00000198131 | 998.4061 | -0.277919731 | 0.243924 | -1.13937 | 0.254549 | 0.72418384 | ZNF544 | protein_coding | 47.34 | -0.038950776 | 0.038950776 | ENSG00000198131 | n.s. |
| ENSG00000137814 | 824.176 | -0.306892566 | 0.288683 | -1.063078 | 0.287747 | 0.746611882 | HAUS2 | protein_coding | 42.11 | -0.038946233 | 0.038946233 | ENSG00000137814 | n.s. |
| ENSG00000141644 | 798.8983 | -0.302186655 | 0.278913 | -1.083445 | 0.278611 | 0.743390786 | MBD1 | protein_coding | 49.32 | -0.038916451 | 0.038916451 | ENSG00000141644 | n.s. |
| ENSG00000165948 | 173.0848 | -0.315675792 | 0.299944 | -1.052451 | 0.292593 | 0.752972779 | IFI27L1 | protein_coding | 45.17 | -0.0388978 | 0.0388978 | ENSG00000165948 | n.s. |
| ENSG00000089094 | 1036.823 | -0.296602842 | 0.271596 | -1.092074 | 0.2748 | 0.739386355 | KDM2B | protein_coding | 50.79 | -0.038893106 | 0.038893106 | ENSG00000089094 | n.s. |
| ENSG00000154642 | 1275.223 | -0.278631322 | 0.244941 | -1.137546 | 0.25531 | 0.725252428 | C21orf91 | protein_coding | 37.16 | -0.038872081 | 0.038872081 | ENSG00000154642 | n.s. |
| ENSG00000200571 | 8.823911 | -0.424159664 | 0.483083 | -0.878026 | 0.37993 | 0.809760709 | RNU6-1284P | snRNA | 38 | -0.038871391 | 0.038871391 | ENSG00000200571 | n.s. |
| ENSG00000117411 | 466.4343 | -0.293685126 | 0.266856 | -1.100537 | 0.271098 | 0.737350747 | B4GALT2 | protein_coding | 56.94 | -0.038862141 | 0.038862141 | ENSG00000117411 | n.s. |
| ENSG00000134444 | 1493.555 | -0.280352441 | 0.247258 | -1.133848 | 0.256858 | 0.726907198 | KIAA1468 | protein_coding | 34.63 | -0.038834709 | 0.038834709 | ENSG00000134444 | n.s. |
| ENSG00000133703 | 1290.689 | -0.299252874 | 0.275021 | -1.088108 | 0.276548 | 0.741780068 | KRAS | protein_coding | 36.37 | -0.038820531 | 0.038820531 | ENSG00000133703 | n.s. |
| ENSG00000173559 | 278.9047 | -0.263978663 | 0.225066 | -1.172897 | 0.240837 | 0.712813552 | NABP1 | protein_coding | 38.76 | -0.038811213 | 0.038811213 | ENSG00000173559 | n.s. |
| ENSG00000171241 | 695.382 | -0.264836326 | 0.226232 | -1.170639 | 0.241744 | 0.714000369 | SHCBP1 | protein_coding | 39.54 | -0.038745969 | 0.038745969 | ENSG00000171241 | n.s. |
| ENSG00000154874 | 991.9964 | -0.2953448 | 0.270446 | -1.092065 | 0.274805 | 0.739386355 | CCDC144B | cribed_unprocessed_pseudc | 38.34 | -0.038728141 | 0.038728141 | ENSG00000154874 | n.s. |
| ENSG00000087157 | 1008.912 | -0.280012678 | 0.247141 | -1.133008 | 0.257211 | 0.727284297 | PGS1 | protein_coding | 52.04 | -0.038724574 | 0.038724574 | ENSG00000087157 | n.s. |

| | | | | | | | | | | | | | |
|---|---|---|---|---|---|---|---|---|---|---|---|---|---|
| ENSG00000212907 | 788.4732 | -0.266689676 | 0.229173 | -1.163705 | 0.244544 | 0.71589159 | MT-ND4L | protein_coding | 43.1 | -0.038710737 | 0.038710737 | ENSG00000212907 | n.s. |
| ENSG00000163806 | 134.6766 | -0.290389366 | 0.262259 | -1.107264 | 0.26818 | 0.735737755 | SPDYA | protein_coding | 39.58 | -0.038702211 | 0.038702211 | ENSG00000163806 | n.s. |
| ENSG00000108840 | 1279.833 | -0.276748423 | 0.243186 | -1.138011 | 0.255116 | 0.724893845 | HDAC5 | protein_coding | 55.66 | -0.038668836 | 0.038668836 | ENSG00000108840 | n.s. |
| ENSG00000198171 | 536.7424 | -0.291720582 | 0.264943 | -1.101071 | 0.270866 | 0.737020734 | DDRGK1 | protein_coding | 54.35 | -0.038658897 | 0.038658897 | ENSG00000198171 | n.s. |
| ENSG00000248774 | 239.5358 | -0.260382975 | 0.220693 | -1.179843 | 0.238063 | 0.71051867 | | antisense | 35.99 | -0.038647214 | 0.038647214 | ENSG00000248774 | n.s. |
| ENSG00000200855 | 9.930363 | -0.40636667 | 0.451034 | -0.900966 | 0.367606 | 0.803356361 | | misc_RNA | 44.12 | -0.038642123 | 0.038642123 | ENSG00000200855 | n.s. |
| ENSG00000161204 | 488.3093 | -0.288433825 | 0.259004 | -1.113625 | 0.26544 | 0.734612586 | ABCF3 | protein_coding | 51.95 | -0.038633298 | 0.038633298 | ENSG00000161204 | n.s. |
| ENSG00000030110 | 176.4641 | -0.288889442 | 0.260081 | -1.110766 | 0.266669 | 0.734987182 | BAK1 | protein_coding | 57.18 | -0.038630364 | 0.038630364 | ENSG00000030110 | n.s. |
| ENSG00000105856 | 481.443 | 0.303794987 | 0.284641 | 1.0672919 | 0.28584 | 0.746239075 | HBP1 | protein_coding | 37.95 | 0.038619031 | 0.038619031 | ENSG00000105856 | n.s. |
| ENSG00000125931 | 147.8035 | -0.294124592 | 0.268461 | -1.095594 | 0.273256 | 0.739321311 | CITED1 | protein_coding | 56.77 | -0.038579374 | 0.038579374 | ENSG00000125931 | n.s. |
| ENSG00000157851 | 818.9527 | -0.28558583 | 0.254575 | -1.121814 | 0.261941 | 0.732723331 | DPYSL5 | protein_coding | 46.42 | -0.038571216 | 0.038571216 | ENSG00000157851 | n.s. |
| ENSG00000163510 | 1189.882 | -0.289017943 | 0.260841 | -1.108024 | 0.267852 | 0.735446299 | CWC22 | protein_coding | 35.03 | -0.038569165 | 0.038569165 | ENSG00000163510 | n.s. |
| ENSG00000165934 | 1832.367 | -0.298016029 | 0.274244 | -1.08668 | 0.277178 | 0.742417658 | CPSF2 | protein_coding | 39.6 | -0.038548882 | 0.038548882 | ENSG00000165934 | n.s. |
| ENSG00000158258 | 1683.346 | -0.279685098 | 0.247603 | -1.12957 | 0.258657 | 0.728120901 | CLSTN2 | protein_coding | 42.71 | -0.038539628 | 0.038539628 | ENSG00000158258 | n.s. |
| ENSG00000131979 | 520.1848 | -0.313993214 | 0.299119 | -1.049726 | 0.293844 | 0.753915388 | GCH1 | protein_coding | 42.22 | -0.038519869 | 0.038519869 | ENSG00000131979 | n.s. |
| ENSG00000127314 | 2241.149 | -0.303713333 | 0.285928 | -1.062201 | 0.288144 | 0.746804204 | RAP1B | protein_coding | 38.96 | -0.038508799 | 0.038508799 | ENSG00000127314 | n.s. |
| ENSG00000140939 | 223.8115 | -0.289188881 | 0.261463 | -1.106043 | 0.268708 | 0.735983077 | NOL3 | protein_coding | 58.13 | -0.038500344 | 0.038500344 | ENSG00000140939 | n.s. |
| ENSG00000277157 | 1435.059 | -0.248645771 | 0.204608 | -1.215231 | 0.224278 | 0.700099826 | HIST1H4D | protein_coding | 57.37 | -0.038500319 | 0.038500319 | ENSG00000277157 | n.s. |
| ENSG00000077235 | 1878.124 | -0.271180214 | 0.235779 | -1.150148 | 0.250083 | 0.721189828 | GTF3C1 | protein_coding | 49.42 | -0.038494142 | 0.038494142 | ENSG00000077235 | n.s. |
| ENSG00000181315 | 342.0522 | -0.291693829 | 0.265835 | -1.097276 | 0.272521 | 0.738113065 | ZNF322 | protein_coding | 37.34 | -0.038467738 | 0.038467738 | ENSG00000181315 | n.s. |
| ENSG00000174842 | 726.2889 | -0.278128735 | 0.245465 | -1.133068 | 0.257186 | 0.727284297 | GLMN | protein_coding | 38 | -0.038464033 | 0.038464033 | ENSG00000174842 | n.s. |
| ENSG00000273247 | 260.5057 | -0.266521783 | 0.229609 | -1.160763 | 0.245738 | 0.717427972 | | antisense | 41.5 | -0.038438223 | 0.038438223 | ENSG00000273247 | n.s. |
| ENSG00000182568 | 7736.578 | -0.263746668 | 0.226003 | -1.167003 | 0.243209 | 0.715021889 | SATB1 | protein_coding | 36.43 | -0.038422789 | 0.038422789 | ENSG00000182568 | n.s. |
| ENSG00000172888 | 901.0941 | -0.292783364 | 0.267405 | -1.094906 | 0.273558 | 0.739386355 | ZNF621 | protein_coding | 41.97 | -0.038392263 | 0.038392263 | ENSG00000172888 | n.s. |
| ENSG00000140497 | 273.6093 | -0.295875505 | 0.271926 | -1.088074 | 0.276563 | 0.741780068 | SCAMP2 | protein_coding | 51.05 | -0.038382403 | 0.038382403 | ENSG00000140497 | n.s. |
| ENSG00000008130 | 298.0124 | -0.279006978 | 0.247217 | -1.128591 | 0.259071 | 0.728547758 | NADK | protein_coding | 53.7 | -0.03837517 | 0.03837517 | ENSG00000008130 | n.s. |
| ENSG00000212309 | 11.40137 | 0.515373375 | 0.659006 | 0.7820461 | 0.434187 | 0.84244096 | | snoRNA | 36.47 | 0.038374972 | 0.038374972 | ENSG00000212309 | n.s. |
| ENSG00000251867 | 34.9712 | 0.33248798 | 0.329618 | 1.0087072 | 0.313115 | 0.766650888 | | antisense | 33.52 | 0.038369897 | 0.038369897 | ENSG00000251867 | n.s. |
| ENSG00000265584 | 3.115054 | -0.585637265 | 0.799994 | -0.732052 | 0.464137 | 0.860018452 | MIR3978 | miRNA | 48.51 | -0.038354691 | 0.038354691 | ENSG00000265584 | n.s. |
| ENSG00000172270 | 1969.319 | -0.299660872 | 0.279059 | -1.073825 | 0.282901 | 0.744776963 | BSG | protein_coding | 62.58 | -0.03834873 | 0.03834873 | ENSG00000172270 | n.s. |
| ENSG00000149970 | 404.2038 | -0.299269696 | 0.278099 | -1.076126 | 0.281871 | 0.744567264 | CNKSR2 | protein_coding | 34.96 | -0.03833527 | 0.03833527 | ENSG00000149970 | n.s. |
| ENSG00000119314 | 897.3002 | -0.284107891 | 0.253436 | -1.121024 | 0.262278 | 0.732977057 | PTBP3 | protein_coding | 37.7 | -0.038328887 | 0.038328887 | ENSG00000119314 | n.s. |

| | | | | | | | | | | | | | |
|---|---|---|---|---|---|---|---|---|---|---|---|---|---|
| ENSG00000127191 | 217.4607 | -0.292197192 | 0.266889 | -1.094826 | 0.273593 | 0.739386355 | TRAF2 | protein_coding | 55.12 | -0.038315399 | 0.038315399 | ENSG00000127191 | n.s. |
| ENSG00000167740 | 138.8318 | -0.298578922 | 0.277017 | -1.077836 | 0.281107 | 0.744242186 | CYB5D2 | protein_coding | 49.97 | -0.038303411 | 0.038303411 | ENSG00000167740 | n.s. |
| ENSG00000251580 | 20.99113 | 0.367172062 | 0.386316 | 0.9504456 | 0.341886 | 0.786560769 | | lincRNA | 53.54 | 0.038284193 | 0.038284193 | ENSG00000251580 | n.s. |
| ENSG00000110911 | 1381.02 | -0.288883491 | 0.262186 | -1.101828 | 0.270536 | 0.737020734 | SLC11A2 | protein_coding | 43.54 | -0.038282925 | 0.038282925 | ENSG00000110911 | n.s. |
| ENSG00000148660 | 640.0667 | -0.28575778 | 0.256482 | -1.114143 | 0.265218 | 0.734612586 | CAMK2G | protein_coding | 51.65 | -0.038274864 | 0.038274864 | ENSG00000148660 | n.s. |
| ENSG00000131446 | 398.5692 | -0.287485506 | 0.259952 | -1.10592 | 0.268761 | 0.735983077 | MGAT1 | protein_coding | 50.9 | -0.03827357 | 0.03827357 | ENSG00000131446 | n.s. |
| ENSG00000256053 | 135.2988 | -0.332597428 | 0.331143 | -1.004393 | 0.315189 | 0.767425784 | APOPT1 | protein_coding | 48.97 | -0.038236602 | 0.038236602 | ENSG00000256053 | n.s. |
| ENSG00000070814 | 2187.69 | -0.269487984 | 0.234442 | -1.149489 | 0.250355 | 0.721375365 | TCOF1 | protein_coding | 52.45 | -0.038223823 | 0.038223823 | ENSG00000070814 | n.s. |
| ENSG00000135776 | 577.6973 | -0.301021828 | 0.28319 | -1.062968 | 0.287797 | 0.746611882 | ABCB10 | protein_coding | 43.62 | -0.038201206 | 0.038201206 | ENSG00000135776 | n.s. |
| ENSG00000133315 | 177.1978 | -0.312762582 | 0.299114 | -1.045629 | 0.295732 | 0.754947373 | MACROD1 | protein_coding | 58.06 | -0.038183095 | 0.038183095 | ENSG00000133315 | n.s. |
| ENSG00000185220 | 196.6086 | -0.323310629 | 0.317134 | -1.019477 | 0.307977 | 0.761997697 | PGBD2 | protein_coding | 46.25 | -0.038165637 | 0.038165637 | ENSG00000185220 | n.s. |
| ENSG00000144589 | 179.3905 | -0.288955635 | 0.263121 | -1.098185 | 0.272124 | 0.73797827 | STK11IP | protein_coding | 53.34 | -0.038129553 | 0.038129553 | ENSG00000144589 | n.s. |
| ENSG00000146856 | 395.2764 | 0.25685399 | 0.217816 | 1.1792273 | 0.238308 | 0.71051867 | AGBL3 | protein_coding | 36.82 | 0.038123426 | 0.038123426 | ENSG00000146856 | n.s. |
| ENSG00000101558 | 2902.421 | -0.295108063 | 0.272136 | -1.084415 | 0.278181 | 0.742965563 | VAPA | protein_coding | 37.13 | -0.038078182 | 0.038078182 | ENSG00000101558 | n.s. |
| ENSG00000176444 | 1509.714 | -0.281654331 | 0.251 | -1.122128 | 0.261808 | 0.732546475 | CLK2 | protein_coding | 50.41 | -0.038069756 | 0.038069756 | ENSG00000176444 | n.s. |
| ENSG00000106344 | 1843.926 | -0.269659266 | 0.235409 | -1.145491 | 0.252006 | 0.722544863 | RBM28 | protein_coding | 43.22 | -0.03805841 | 0.03805841 | ENSG00000106344 | n.s. |
| ENSG00000177105 | 67.10044 | -0.307932665 | 0.292251 | -1.053657 | 0.29204 | 0.75237733 | RHOG | protein_coding | 51.22 | -0.038049484 | 0.038049484 | ENSG00000177105 | n.s. |
| ENSG00000085832 | 1070.911 | -0.284487177 | 0.256336 | -1.10982 | 0.267076 | 0.734987182 | EPS15 | protein_coding | 38.05 | -0.038041692 | 0.038041692 | ENSG00000085832 | n.s. |
| ENSG00000181991 | 1041.691 | -0.284011155 | 0.254974 | -1.113883 | 0.265329 | 0.734612586 | MRPS11 | protein_coding | 45.97 | -0.038040919 | 0.038040919 | ENSG00000181991 | n.s. |
| ENSG00000236296 | 142.7168 | 0.276969238 | 0.245635 | 1.1275664 | 0.259503 | 0.728949992 | GUSBP5 | ıscribed_processed_pseudog | 39.63 | 0.038028504 | 0.038028504 | ENSG00000236296 | n.s. |
| ENSG00000160193 | 315.8288 | -0.265980866 | 0.230475 | -1.154056 | 0.248477 | 0.719540199 | WDR4 | protein_coding | 53.71 | -0.038020618 | 0.038020618 | ENSG00000160193 | n.s. |
| ENSG00000121542 | 317.3922 | -0.321762566 | 0.314498 | -1.0231 | 0.30626 | 0.761997697 | SEC22A | protein_coding | 37.72 | -0.037982894 | 0.037982894 | ENSG00000121542 | n.s. |
| ENSG00000270704 | 50.46612 | -0.335121854 | 0.33676 | -0.995137 | 0.31967 | 0.770355344 | | sense_intronic | 37.74 | -0.037972289 | 0.037972289 | ENSG00000270704 | n.s. |
| ENSG00000234284 | 202.6182 | -0.275036514 | 0.242943 | -1.132102 | 0.257592 | 0.727758051 | ZNF879 | protein_coding | 43.57 | -0.037958609 | 0.037958609 | ENSG00000234284 | n.s. |
| ENSG00000121406 | 347.5926 | -0.293441306 | 0.270225 | -1.085916 | 0.277516 | 0.742417658 | ZNF549 | protein_coding | 43.95 | -0.037957134 | 0.037957134 | ENSG00000121406 | n.s. |
| ENSG00000100335 | 1034.223 | -0.276283499 | 0.245039 | -1.127508 | 0.259528 | 0.728949992 | MIEF1 | protein_coding | 49.85 | -0.03793435 | 0.03793435 | ENSG00000100335 | n.s. |
| ENSG00000221514 | 24.04871 | -0.354470046 | 0.366763 | -0.966484 | 0.333802 | 0.781622542 | SNORD111B | snoRNA | 37.5 | -0.037929331 | 0.037929331 | ENSG00000221514 | n.s. |
| ENSG00000148606 | 948.3886 | -0.283100828 | 0.254367 | -1.112963 | 0.265724 | 0.734612586 | POLR3A | protein_coding | 44.7 | -0.037918988 | 0.037918988 | ENSG00000148606 | n.s. |
| ENSG00000084093 | 725.1255 | -0.270490611 | 0.237419 | -1.139296 | 0.25458 | 0.72418384 | REST | protein_coding | 43.58 | -0.037909576 | 0.037909576 | ENSG00000084093 | n.s. |
| ENSG00000208022 | 43.73645 | -0.330355491 | 0.329422 | -1.002835 | 0.315941 | 0.767820213 | MIR618 | miRNA | 45.92 | -0.037905141 | 0.037905141 | ENSG00000208022 | n.s. |
| ENSG00000056586 | 1601.145 | -0.289067721 | 0.264953 | -1.091015 | 0.275266 | 0.739442303 | RC3H2 | protein_coding | 40.5 | -0.037895537 | 0.037895537 | ENSG00000056586 | n.s. |
| ENSG00000143924 | 3290.69 | -0.284633406 | 0.257371 | -1.105927 | 0.268758 | 0.735983077 | EML4 | protein_coding | 38.02 | -0.037893863 | 0.037893863 | ENSG00000143924 | n.s. |

| | | | | | | | | | | | | | |
|---|---|---|---|---|---|---|---|---|---|---|---|---|---|
| ENSG00000162585 | 105.1998 | -0.332722298 | 0.333474 | -0.997745 | 0.318403 | 0.769456789 | FAAP20 | protein_coding | 60.96 | -0.037869043 | 0.037869043 | ENSG00000162585 | n.s. |
| ENSG00000166851 | 632.7136 | -0.270184749 | 0.23696 | -1.140212 | 0.254198 | 0.72418384 | PLK1 | protein_coding | 50.69 | -0.037866709 | 0.037866709 | ENSG00000166851 | n.s. |
| ENSG00000133313 | 1008.28 | -0.292492561 | 0.269352 | -1.085914 | 0.277517 | 0.742417658 | CNDP2 | protein_coding | 51.18 | -0.037834412 | 0.037834412 | ENSG00000133313 | n.s. |
| ENSG00000234171 | 355.3846 | -0.28110248 | 0.251605 | -1.117238 | 0.263892 | 0.733524057 | RNASEH1-AS1 | antisense | 47.42 | -0.037832357 | 0.037832357 | ENSG00000234171 | n.s. |
| ENSG00000130204 | 1249.31 | -0.296560128 | 0.277119 | -1.070153 | 0.28455 | 0.745679202 | TOMM40 | protein_coding | 56.25 | -0.037795986 | 0.037795986 | ENSG00000130204 | n.s. |
| ENSG00000179833 | 648.9265 | -0.274256179 | 0.242776 | -1.129667 | 0.258617 | 0.728120901 | SERTAD2 | protein_coding | 40.14 | -0.037791542 | 0.037791542 | ENSG00000179833 | n.s. |
| ENSG00000109920 | 2381.686 | -0.285789539 | 0.259948 | -1.099409 | 0.27159 | 0.73753588 | FNBP4 | protein_coding | 43.42 | -0.037786192 | 0.037786192 | ENSG00000109920 | n.s. |
| ENSG00000164949 | 237.951 | 0.274641104 | 0.243299 | 1.1288219 | 0.258973 | 0.728529796 | GEM | protein_coding | 44.2 | 0.03777762 | 0.03777762 | ENSG00000164949 | n.s. |
| ENSG00000166822 | 522.637 | -0.295750566 | 0.276184 | -1.070848 | 0.284238 | 0.745309826 | TMEM170A | protein_coding | 44.64 | -0.03775645 | 0.03775645 | ENSG00000166822 | n.s. |
| ENSG00000131381 | 1019.303 | -0.287909141 | 0.263349 | -1.093259 | 0.27428 | 0.739386355 | RBSN | protein_coding | 44.4 | -0.037753113 | 0.037753113 | ENSG00000131381 | n.s. |
| ENSG00000253057 | 16.60001 | -0.362194372 | 0.381726 | -0.948834 | 0.342705 | 0.786723798 | RN7SKP57 | misc_RNA | 44.71 | -0.037732581 | 0.037732581 | ENSG00000253057 | n.s. |
| ENSG00000220323 | 25.8984 | -0.368376355 | 0.393278 | -0.936681 | 0.348923 | 0.790028419 | | processed_pseudogene | 63.29 | -0.037706004 | 0.037706004 | ENSG00000220323 | n.s. |
| ENSG00000140632 | 1458.659 | -0.280007453 | 0.250039 | -1.119857 | 0.262775 | 0.733524057 | GLYR1 | protein_coding | 47.65 | -0.037684982 | 0.037684982 | ENSG00000140632 | n.s. |
| ENSG00000235437 | 48.99959 | -0.317119776 | 0.307728 | -1.030521 | 0.302766 | 0.760781413 | LINC01278 | processed_transcript | 38.18 | -0.037654836 | 0.037654836 | ENSG00000235437 | n.s. |
| ENSG00000283050 | 56.44872 | -0.316639679 | 0.3079 | -1.028383 | 0.303769 | 0.760781413 | GTF2IP12 | cribed_unprocessed_pseudc | 45.66 | -0.037597829 | 0.037597829 | ENSG00000283050 | n.s. |
| ENSG00000154710 | 359.9176 | -0.286688303 | 0.261857 | -1.094828 | 0.273592 | 0.739386355 | RABGEF1 | protein_coding | 44.49 | -0.037593027 | 0.037593027 | ENSG00000154710 | n.s. |
| ENSG00000218510 | 169.6528 | -0.27227772 | 0.240498 | -1.132141 | 0.257575 | 0.727758051 | LINC00339 | lincRNA | 47.63 | -0.03757786 | 0.03757786 | ENSG00000218510 | n.s. |
| ENSG00000254996 | 720.6003 | -0.28483708 | 0.259573 | -1.097328 | 0.272498 | 0.738113065 | NKHD1-EIF4EBP | protein_coding | 38.97 | -0.03756349 | 0.03756349 | ENSG00000254996 | n.s. |
| ENSG00000120963 | 1198.536 | 0.272410722 | 0.241063 | 1.1300397 | 0.25846 | 0.728120901 | ZNF706 | protein_coding | 40.04 | 0.037537245 | 0.037537245 | ENSG00000120963 | n.s. |
| ENSG00000207741 | 41.59796 | 0.322501022 | 0.318361 | 1.0130046 | 0.311058 | 0.76510176 | MIR590 | miRNA | 35.05 | 0.037500676 | 0.037500676 | ENSG00000207741 | n.s. |
| ENSG00000089234 | 1153.108 | -0.28587056 | 0.26135 | -1.093824 | 0.274032 | 0.739386355 | BRAP | protein_coding | 44.41 | -0.037485797 | 0.037485797 | ENSG00000089234 | n.s. |
| ENSG00000105402 | 642.686 | -0.289754494 | 0.266935 | -1.085485 | 0.277707 | 0.742417658 | NAPA | protein_coding | 55.52 | -0.037480239 | 0.037480239 | ENSG00000105402 | n.s. |
| ENSG00000215769 | 179.8744 | -0.311749377 | 0.300359 | -1.037921 | 0.299307 | 0.758335061 | P27P1-BPTFP1-K | processed_transcript | 47.76 | -0.037453216 | 0.037453216 | ENSG00000215769 | n.s. |
| ENSG00000166224 | 851.837 | -0.280039886 | 0.252039 | -1.1111 | 0.266525 | 0.734987182 | SGPL1 | protein_coding | 41.07 | -0.037446999 | 0.037446999 | ENSG00000166224 | n.s. |
| ENSG00000144029 | 1420.466 | -0.289337955 | 0.266536 | -1.085547 | 0.277679 | 0.742417658 | MRPS5 | protein_coding | 42.25 | -0.037426358 | 0.037426358 | ENSG00000144029 | n.s. |
| ENSG00000184154 | 10.04494 | -0.447320758 | 0.538303 | -0.830984 | 0.405983 | 0.824799944 | LRTOMT | protein_coding | 48.88 | -0.037418998 | 0.037418998 | ENSG00000184154 | n.s. |
| ENSG00000108963 | 241.3566 | 0.295896791 | 0.279 | 1.0605617 | 0.288889 | 0.747482487 | DPH1 | protein_coding | 56.54 | 0.037401052 | 0.037401052 | ENSG00000108963 | n.s. |
| ENSG00000174938 | 282.5244 | -0.280816676 | 0.253677 | -1.106986 | 0.2683 | 0.735922141 | SEZ6L2 | protein_coding | 52.94 | -0.037395832 | 0.037395832 | ENSG00000174938 | n.s. |
| ENSG00000207757 | 17.23104 | 0.374893883 | 0.406802 | 0.9215625 | 0.356757 | 0.794806874 | MIR93 | miRNA | 60 | 0.037391311 | 0.037391311 | ENSG00000207757 | n.s. |
| ENSG00000161048 | 163.5475 | -0.264747533 | 0.231049 | -1.145851 | 0.251857 | 0.722544863 | NAPEPLD | protein_coding | 40.35 | -0.037365191 | 0.037365191 | ENSG00000161048 | n.s. |
| ENSG00000179041 | 513.1358 | -0.271009748 | 0.239895 | -1.129702 | 0.258602 | 0.728120901 | RRS1 | protein_coding | 56.33 | -0.037344195 | 0.037344195 | ENSG00000179041 | n.s. |
| ENSG00000272325 | 1832.638 | -0.281489449 | 0.255667 | -1.101 | 0.270897 | 0.737020734 | NUDT3 | protein_coding | 42.12 | -0.037303065 | 0.037303065 | ENSG00000272325 | n.s. |

| | | | | | | | | | | | | | |
|---|---|---|---|---|---|---|---|---|---|---|---|---|---|
| ENSG00000234127 | 375.4286 | -0.293846302 | 0.276513 | -1.062685 | 0.287925 | 0.746611882 | TRIM26 | protein_coding | 46.9 | -0.037290595 | 0.037290595 | ENSG00000234127 | n.s. |
| ENSG00000152684 | 1361.68 | 0.261813317 | 0.227429 | 1.1511893 | 0.249654 | 0.720401565 | PELO | protein_coding | 38.85 | 0.037288854 | 0.037288854 | ENSG00000152684 | n.s. |
| ENSG00000212127 | 110.0722 | -0.345791324 | 0.356017 | -0.971278 | 0.33141 | 0.780141891 | TAS2R14 | protein_coding | 36.14 | -0.037285434 | 0.037285434 | ENSG00000212127 | n.s. |
| ENSG00000250467 | 229.2686 | -0.281109612 | 0.255104 | -1.101942 | 0.270487 | 0.737020734 | | antisense | 41.2 | -0.037252728 | 0.037252728 | ENSG00000250467 | n.s. |
| ENSG00000222222 | 49.31725 | 0.345470418 | 0.356339 | 0.9694985 | 0.332297 | 0.780141891 | RNU2-17P | snRNA | 42.63 | 0.037250832 | 0.037250832 | ENSG00000222222 | n.s. |
| ENSG00000121774 | 2238.635 | -0.293499347 | 0.275884 | -1.063849 | 0.287397 | 0.746611882 | KHDRBS1 | protein_coding | 43.63 | -0.037246565 | 0.037246565 | ENSG00000121774 | n.s. |
| ENSG00000099624 | 267.914 | -0.303030988 | 0.288423 | -1.050649 | 0.29342 | 0.753623766 | ATP5D | protein_coding | 62.97 | -0.037225967 | 0.037225967 | ENSG00000099624 | n.s. |
| ENSG00000026297 | 109.9214 | -0.430535317 | 0.508325 | -0.846969 | 0.397012 | 0.819511304 | RNASET2 | protein_coding | 48.7 | -0.037217648 | 0.037217648 | ENSG00000026297 | n.s. |
| ENSG00000260865 | 13.73757 | -0.400384071 | 0.448961 | -0.891802 | 0.372499 | 0.807333164 | | transcribed_processed_pseudog | 52.1 | -0.037214581 | 0.037214581 | ENSG00000260865 | n.s. |
| ENSG00000137338 | 427.6678 | -0.290084763 | 0.26871 | -1.079545 | 0.280345 | 0.744242186 | PGBD1 | protein_coding | 39.87 | -0.037213732 | 0.037213732 | ENSG00000137338 | n.s. |
| ENSG00000122779 | 2965.637 | -0.277707815 | 0.24911 | -1.114802 | 0.264935 | 0.734601921 | TRIM24 | protein_coding | 39.19 | -0.03719839 | 0.03719839 | ENSG00000122779 | n.s. |
| ENSG00000083817 | 279.6409 | -0.279309183 | 0.252552 | -1.105948 | 0.268749 | 0.735983077 | ZNF416 | protein_coding | 48.61 | -0.037185038 | 0.037185038 | ENSG00000083817 | n.s. |
| ENSG00000163812 | 503.7902 | -0.288461774 | 0.266656 | -1.081777 | 0.279352 | 0.743390786 | ZDHHC3 | protein_coding | 47.26 | -0.037148922 | 0.037148922 | ENSG00000163812 | n.s. |
| ENSG00000143183 | 2383.341 | -0.308586335 | 0.297433 | -1.037498 | 0.299504 | 0.758335061 | TMCO1 | protein_coding | 39.88 | -0.037073212 | 0.037073212 | ENSG00000143183 | n.s. |
| ENSG00000146842 | 597.8873 | -0.258736194 | 0.223477 | -1.157775 | 0.246956 | 0.719028029 | TMEM209 | protein_coding | 37.56 | -0.037065041 | 0.037065041 | ENSG00000146842 | n.s. |
| ENSG00000121481 | 998.7768 | -0.291549876 | 0.2743 | -1.062887 | 0.287833 | 0.746611882 | RNF2 | protein_coding | 39.39 | -0.036999167 | 0.036999167 | ENSG00000121481 | n.s. |
| ENSG00000196586 | 479.6745 | -0.291456394 | 0.273836 | -1.064347 | 0.287172 | 0.746611882 | MYO6 | protein_coding | 37.66 | -0.036987304 | 0.036987304 | ENSG00000196586 | n.s. |
| ENSG00000258634 | 261.2695 | -0.288704469 | 0.268362 | -1.075802 | 0.282016 | 0.744567264 | | antisense | 53.13 | -0.036981906 | 0.036981906 | ENSG00000258634 | n.s. |
| ENSG00000114491 | 1594.642 | -0.289625865 | 0.270208 | -1.071864 | 0.283781 | 0.745300357 | UMPS | protein_coding | 42.81 | -0.036976149 | 0.036976149 | ENSG00000114491 | n.s. |
| ENSG00000267041 | 1628.871 | -0.287939908 | 0.266633 | -1.079913 | 0.280181 | 0.744032323 | ZNF850 | protein_coding | 41.15 | -0.036973844 | 0.036973844 | ENSG00000267041 | n.s. |
| ENSG00000178177 | 772.2798 | -0.288192106 | 0.26743 | -1.077636 | 0.281196 | 0.744242186 | LCORL | protein_coding | 36.02 | -0.036970931 | 0.036970931 | ENSG00000178177 | n.s. |
| ENSG00000275413 | 23.81627 | 0.342752135 | 0.352746 | 0.9716692 | 0.331215 | 0.780141891 | | antisense | 32.33 | 0.036957729 | 0.036957729 | ENSG00000275413 | n.s. |
| ENSG00000229388 | 49.28697 | 0.324686321 | 0.325499 | 0.9975044 | 0.31852 | 0.769456789 | LINC01715 | lincRNA | 47.96 | 0.036954422 | 0.036954422 | ENSG00000229388 | n.s. |
| ENSG00000214087 | 1296.75 | -0.278780501 | 0.252812 | -1.10272 | 0.270149 | 0.737020734 | ARL16 | protein_coding | 56.96 | -0.036944074 | 0.036944074 | ENSG00000214087 | n.s. |
| ENSG00000047346 | 289.7503 | -0.288643761 | 0.268739 | -1.074066 | 0.282793 | 0.744776963 | FAM214A | protein_coding | 37.21 | -0.036938829 | 0.036938829 | ENSG00000047346 | n.s. |
| ENSG00000180902 | 226.8524 | -0.276765546 | 0.249816 | -1.107876 | 0.267915 | 0.735446299 | D2HGDH | protein_coding | 56.88 | -0.036934095 | 0.036934095 | ENSG00000180902 | n.s. |
| ENSG00000230590 | 436.637 | -0.274422705 | 0.245623 | -1.117251 | 0.263887 | 0.733524057 | FTX | lincRNA | 39.53 | -0.036933355 | 0.036933355 | ENSG00000230590 | n.s. |
| ENSG00000227836 | 19.75982 | -0.354396515 | 0.373488 | -0.948883 | 0.34268 | 0.786723798 | | processed_pseudogene | 45.37 | -0.036920218 | 0.036920218 | ENSG00000227836 | n.s. |
| ENSG00000188859 | 398.2362 | -0.2815128 | 0.257658 | -1.092582 | 0.274577 | 0.739386355 | FAM78B | protein_coding | 45.08 | -0.03691437 | 0.03691437 | ENSG00000188859 | n.s. |
| ENSG00000101745 | 331.6984 | -0.285301067 | 0.262686 | -1.086092 | 0.277438 | 0.742417658 | ANKRD12 | protein_coding | 37.33 | -0.03690418 | 0.03690418 | ENSG00000101745 | n.s. |
| ENSG00000252311 | 7.205673 | -0.554301181 | 0.752237 | -0.73687 | 0.461201 | 0.857958626 | RNU1-103P | snRNA | 48.8 | -0.036879684 | 0.036879684 | ENSG00000252311 | n.s. |
| ENSG00000155097 | 785.6182 | -0.305333385 | 0.293311 | -1.040989 | 0.297881 | 0.757237831 | ATP6V1C1 | protein_coding | 38.39 | -0.03687441 | 0.03687441 | ENSG00000155097 | n.s. |

| | | | | | | | | | | | | | |
|---|---|---|---|---|---|---|---|---|---|---|---|---|---|
| ENSG00000113758 | 3503.496 | -0.288706879 | 0.269477 | -1.071359 | 0.284008 | 0.745300357 | DBN1 | protein_coding | 59.54 | -0.036858823 | 0.036858823 | ENSG00000113758 | n.s. |
| ENSG00000123975 | 1791.101 | 0.289887803 | 0.271759 | 1.0667098 | 0.286103 | 0.746239075 | CKS2 | protein_coding | 38.43 | 0.036851121 | 0.036851121 | ENSG00000123975 | n.s. |
| ENSG00000111490 | 867.1361 | -0.271609801 | 0.241823 | -1.123175 | 0.261363 | 0.731743443 | TBC1D30 | protein_coding | 41.2 | -0.036841469 | 0.036841469 | ENSG00000111490 | n.s. |
| ENSG00000160633 | 1652.089 | -0.25399515 | 0.218468 | -1.162617 | 0.244985 | 0.716280109 | SAFB | protein_coding | 47.35 | -0.036808243 | 0.036808243 | ENSG00000160633 | n.s. |
| ENSG00000256546 | 107.152 | -0.280163125 | 0.255686 | -1.095732 | 0.273196 | 0.739301676 | | processed_transcript | 50.95 | -0.036751325 | 0.036751325 | ENSG00000256546 | n.s. |
| ENSG00000170949 | 667.3923 | -0.302175547 | 0.289402 | -1.044137 | 0.296422 | 0.755765348 | ZNF160 | protein_coding | 45.26 | -0.036748482 | 0.036748482 | ENSG00000170949 | n.s. |
| ENSG00000101557 | 1527.059 | -0.282986527 | 0.259948 | -1.088628 | 0.276318 | 0.741552755 | USP14 | protein_coding | 38.05 | -0.03674805 | 0.03674805 | ENSG00000101557 | n.s. |
| ENSG00000272579 | 54.70078 | -0.35129563 | 0.367969 | -0.954689 | 0.339735 | 0.785979841 | | antisense | 29.38 | -0.036741516 | 0.036741516 | ENSG00000272579 | n.s. |
| ENSG00000117016 | 1799.845 | -0.280116912 | 0.256539 | -1.091908 | 0.274874 | 0.739386355 | RIMS3 | protein_coding | 51.39 | -0.03673133 | 0.03673133 | ENSG00000117016 | n.s. |
| ENSG00000106477 | 49.3323 | 0.31096955 | 0.305045 | 1.0194225 | 0.308002 | 0.761997697 | CEP41 | protein_coding | 39.96 | 0.036708818 | 0.036708818 | ENSG00000106477 | n.s. |
| ENSG00000115020 | 1952.512 | -0.276862432 | 0.251042 | -1.102853 | 0.270091 | 0.737020734 | PIKFYVE | protein_coding | 37.08 | -0.036689891 | 0.036689891 | ENSG00000115020 | n.s. |
| ENSG00000144357 | 2267.47 | -0.289069265 | 0.271358 | -1.06527 | 0.286754 | 0.746611882 | UBR3 | protein_coding | 36.88 | -0.036684365 | 0.036684365 | ENSG00000144357 | n.s. |
| ENSG00000120742 | 2679.478 | 0.27544905 | 0.249012 | 1.1061679 | 0.268654 | 0.735983077 | SERP1 | protein_coding | 38.84 | 0.03667113 | 0.03667113 | ENSG00000120742 | n.s. |
| ENSG00000163040 | 125.0748 | -0.308745437 | 0.300032 | -1.02904 | 0.303461 | 0.760781413 | CCDC74A | protein_coding | 61.5 | -0.036660466 | 0.036660466 | ENSG00000163040 | n.s. |
| ENSG00000283761 | 213.963 | -0.286157563 | 0.266014 | -1.075726 | 0.28205 | 0.744567264 | | protein_coding | 38.29 | -0.036655657 | 0.036655657 | ENSG00000283761 | n.s. |
| ENSG00000153574 | 394.7672 | -0.279534975 | 0.25598 | -1.092018 | 0.274825 | 0.739386355 | RPIA | protein_coding | 41.25 | -0.036655021 | 0.036655021 | ENSG00000153574 | n.s. |
| ENSG00000121067 | 1773.504 | -0.288815276 | 0.271767 | -1.06273 | 0.287904 | 0.746611882 | SPOP | protein_coding | 40.44 | -0.036652132 | 0.036652132 | ENSG00000121067 | n.s. |
| ENSG00000228505 | 1171.497 | -0.298836879 | 0.285102 | -1.048177 | 0.294557 | 0.754039228 | | lincRNA | 40.25 | -0.036639212 | 0.036639212 | ENSG00000228505 | n.s. |
| ENSG00000181722 | 28.89948 | 0.328393275 | 0.331689 | 0.9900627 | 0.322143 | 0.773444264 | ZBTB20 | protein_coding | 36.07 | 0.036639159 | 0.036639159 | ENSG00000181722 | n.s. |
| ENSG00000206562 | 425.0369 | -0.273468218 | 0.245324 | -1.114722 | 0.26497 | 0.734601921 | METTL6 | protein_coding | 42.52 | -0.036630505 | 0.036630505 | ENSG00000206562 | n.s. |
| ENSG00000177370 | 392.5565 | -0.286650397 | 0.267728 | -1.070676 | 0.284315 | 0.745309826 | TIMM22 | protein_coding | 48.68 | -0.036594694 | 0.036594694 | ENSG00000177370 | n.s. |
| ENSG00000103342 | 2874.889 | -0.282857289 | 0.260354 | -1.086435 | 0.277287 | 0.742417658 | GSPT1 | protein_coding | 42.29 | -0.036588073 | 0.036588073 | ENSG00000103342 | n.s. |
| ENSG00000226314 | 94.92709 | 0.27883592 | 0.255046 | 1.0932762 | 0.274273 | 0.739386355 | ZNF192P1 | cribed_unprocessed_pseudc | 42.26 | 0.036563355 | 0.036563355 | ENSG00000226314 | n.s. |
| ENSG00000207606 | 5.901692 | -0.547333834 | 0.740637 | -0.739004 | 0.459905 | 0.857465447 | MIR554 | miRNA | 48.96 | -0.036552799 | 0.036552799 | ENSG00000207606 | n.s. |
| ENSG00000104885 | 633.1721 | -0.256173866 | 0.222323 | -1.152259 | 0.249215 | 0.720036883 | DOT1L | protein_coding | 59.45 | -0.036541987 | 0.036541987 | ENSG00000104885 | n.s. |
| ENSG00000137962 | 1518.774 | 0.284751874 | 0.264046 | 1.0784166 | 0.280848 | 0.744242186 | ARHGAP29 | protein_coding | 38.07 | 0.036529598 | 0.036529598 | ENSG00000137962 | n.s. |
| ENSG00000100325 | 183.3364 | -0.287183734 | 0.26886 | -1.068155 | 0.285451 | 0.746113205 | ASCC2 | protein_coding | 48.94 | -0.036528414 | 0.036528414 | ENSG00000100325 | n.s. |
| ENSG00000121879 | 1113.633 | -0.2713878 | 0.242422 | -1.119486 | 0.262933 | 0.733524057 | PIK3CA | protein_coding | 35.78 | -0.0365249 | 0.0365249 | ENSG00000121879 | n.s. |
| ENSG00000143367 | 226.0565 | -0.278505116 | 0.254486 | -1.094385 | 0.273786 | 0.739386355 | TUFT1 | protein_coding | 45.93 | -0.036519977 | 0.036519977 | ENSG00000143367 | n.s. |
| ENSG00000102710 | 1529.135 | -0.284600639 | 0.264064 | -1.07777 | 0.281136 | 0.744242186 | SUPT20H | protein_coding | 35.69 | -0.036510197 | 0.036510197 | ENSG00000102710 | n.s. |
| ENSG00000006530 | 476.8797 | -0.287402038 | 0.27021 | -1.063626 | 0.287498 | 0.746611882 | AGK | protein_coding | 39.59 | -0.036472785 | 0.036472785 | ENSG00000006530 | n.s. |
| ENSG00000167114 | 205.749 | -0.281581222 | 0.259397 | -1.085522 | 0.277691 | 0.742417658 | SLC27A4 | protein_coding | 54.67 | -0.036423012 | 0.036423012 | ENSG00000167114 | n.s. |

| | | | | | | | | | | | | | |
|---|---|---|---|---|---|---|---|---|---|---|---|---|---|
| ENSG00000254858 | 129.7387 | -0.303150924 | 0.29177 | -1.039005 | 0.298802 | 0.758335061 | MPV17L2 | protein_coding | 59.62 | -0.036420208 | 0.036420208 | ENSG00000254858 | n.s. |
| ENSG00000103061 | 632.0726 | -0.282113767 | 0.260181 | -1.0843 | 0.278232 | 0.742965563 | SLC7A6OS | protein_coding | 47.19 | -0.03640151 | 0.03640151 | ENSG00000103061 | n.s. |
| ENSG00000161960 | 1312.508 | 0.233895474 | 0.191811 | 1.2194051 | 0.22269 | 0.698928583 | EIF4A1 | protein_coding | 52.92 | 0.036386464 | 0.036386464 | ENSG00000161960 | n.s. |
| ENSG00000186615 | 116.8816 | -0.293249356 | 0.277856 | -1.055399 | 0.291243 | 0.751617438 | KTN1-AS1 | antisense | 40.78 | -0.036363845 | 0.036363845 | ENSG00000186615 | n.s. |
| ENSG00000256464 | 31.11102 | -0.337138563 | 0.347514 | -0.970145 | 0.331974 | 0.780141891 | YWHABP2 | processed_pseudogene | 43.2 | -0.036352438 | 0.036352438 | ENSG00000256464 | n.s. |
| ENSG00000155636 | 386.3078 | -0.285836376 | 0.267893 | -1.066981 | 0.28598 | 0.746239075 | RBM45 | protein_coding | 36.95 | -0.036336096 | 0.036336096 | ENSG00000155636 | n.s. |
| ENSG00000065883 | 1296.728 | -0.271264823 | 0.243243 | -1.115199 | 0.264765 | 0.734601921 | CDK13 | protein_coding | 38.87 | -0.036335364 | 0.036335364 | ENSG00000065883 | n.s. |
| ENSG00000144580 | 2314.338 | -0.276883479 | 0.253474 | -1.092355 | 0.274677 | 0.739386355 | CNOT9 | protein_coding | 41.17 | -0.036307334 | 0.036307334 | ENSG00000144580 | n.s. |
| ENSG00000251988 | 20.53476 | -0.33654672 | 0.345467 | -0.974179 | 0.329968 | 0.780141891 | RNU4ATAC18P | snRNA | 47.62 | -0.036288622 | 0.036288622 | ENSG00000251988 | n.s. |
| ENSG00000136490 | 2258.659 | -0.261348983 | 0.230288 | -1.134877 | 0.256427 | 0.726402587 | LIMD2 | protein_coding | 66.27 | -0.03628115 | 0.03628115 | ENSG00000136490 | n.s. |
| ENSG00000076685 | 767.7763 | -0.307110445 | 0.299857 | -1.02419 | 0.305745 | 0.761997697 | NT5C2 | protein_coding | 40.09 | -0.036253264 | 0.036253264 | ENSG00000076685 | n.s. |
| ENSG00000263540 | 3.37408 | -0.528107428 | 0.70559 | -0.748463 | 0.454181 | 0.853840253 | MIR5582 | miRNA | 32.35 | -0.036240514 | 0.036240514 | ENSG00000263540 | n.s. |
| ENSG00000122687 | 668.0776 | -0.283687773 | 0.264785 | -1.071388 | 0.283995 | 0.745300357 | MRM2 | protein_coding | 49.95 | -0.036218041 | 0.036218041 | ENSG00000122687 | n.s. |
| ENSG00000147316 | 901.6577 | -0.273557672 | 0.2487 | -1.099952 | 0.271353 | 0.737359702 | MCPH1 | protein_coding | 42.19 | -0.036197315 | 0.036197315 | ENSG00000147316 | n.s. |
| ENSG00000135596 | 1580.245 | -0.2730484 | 0.247929 | -1.101318 | 0.270758 | 0.737020734 | MICAL1 | protein_coding | 50.36 | -0.036184454 | 0.036184454 | ENSG00000135596 | n.s. |
| ENSG00000162373 | 192.5274 | -0.300068882 | 0.288523 | -1.040018 | 0.298332 | 0.757692692 | BEND5 | protein_coding | 44.07 | -0.036160371 | 0.036160371 | ENSG00000162373 | n.s. |
| ENSG00000129351 | 7871.831 | -0.280769955 | 0.259215 | -1.083154 | 0.27874 | 0.743390786 | ILF3 | protein_coding | 50.9 | -0.036158348 | 0.036158348 | ENSG00000129351 | n.s. |
| ENSG00000119711 | 227.96 | -0.280765801 | 0.259194 | -1.083225 | 0.278709 | 0.743390786 | ALDH6A1 | protein_coding | 43.41 | -0.036157813 | 0.036157813 | ENSG00000119711 | n.s. |
| ENSG00000099256 | 1072.195 | -0.307654585 | 0.302397 | -1.017388 | 0.308969 | 0.762996547 | PRTFDC1 | protein_coding | 41.87 | -0.036142469 | 0.036142469 | ENSG00000099256 | n.s. |
| ENSG00000215105 | 278.494 | 0.285276946 | 0.268709 | 1.0616561 | 0.288392 | 0.747018696 | TTC3P1 | processed_pseudogene | 40.31 | 0.036135611 | 0.036135611 | ENSG00000215105 | n.s. |
| ENSG00000163312 | 300.6263 | -0.271363954 | 0.245525 | -1.105241 | 0.269055 | 0.735983077 | HELQ | protein_coding | 38.46 | -0.036127273 | 0.036127273 | ENSG00000163312 | n.s. |
| ENSG00000277688 | 19.30888 | 0.401907121 | 0.464267 | 0.8656812 | 0.386665 | 0.813136725 | | antisense | 39.22 | 0.036105898 | 0.036105898 | ENSG00000277688 | n.s. |
| ENSG00000130770 | 469.7847 | -0.279069297 | 0.25691 | -1.086251 | 0.277368 | 0.742417658 | ATPIF1 | protein_coding | 44.37 | -0.03609809 | 0.03609809 | ENSG00000130770 | n.s. |
| ENSG00000226944 | 19.69323 | -0.349581469 | 0.369853 | -0.945189 | 0.344562 | 0.788387641 | | antisense | 58.13 | -0.03609785 | 0.03609785 | ENSG00000226944 | n.s. |
| ENSG00000066697 | 601.4867 | -0.305746783 | 0.298962 | -1.022696 | 0.306452 | 0.761997697 | MSANTD3 | protein_coding | 43.76 | -0.036092289 | 0.036092289 | ENSG00000066697 | n.s. |
| ENSG00000139496 | 3726.204 | -0.278191024 | 0.255784 | -1.0876 | 0.276772 | 0.741896779 | NUP58 | protein_coding | 36.36 | -0.036069279 | 0.036069279 | ENSG00000139496 | n.s. |
| ENSG00000222020 | 65.82223 | 0.305544846 | 0.299341 | 1.0207248 | 0.307385 | 0.761997697 | | antisense | 64.05 | 0.036068451 | 0.036068451 | ENSG00000222020 | n.s. |
| ENSG00000234608 | 773.243 | 0.270003195 | 0.243582 | 1.1084697 | 0.267659 | 0.735235906 | MAPKAPK5-AS1 | lincRNA | 53.55 | 0.036065216 | 0.036065216 | ENSG00000234608 | n.s. |
| ENSG00000279765 | 593.8244 | -0.252678391 | 0.219303 | -1.15219 | 0.249243 | 0.720036883 | | protein_coding | 38.36 | -0.036043374 | 0.036043374 | ENSG00000279765 | n.s. |
| ENSG00000265942 | 54.21578 | 0.311333366 | 0.308182 | 1.0102251 | 0.312387 | 0.766004018 | RN7SL577P | misc_RNA | 52.15 | 0.036042738 | 0.036042738 | ENSG00000265942 | n.s. |
| ENSG00000106692 | 113.6427 | -0.303482308 | 0.294341 | -1.031058 | 0.302514 | 0.760781413 | FKTN | protein_coding | 37.44 | -0.036035522 | 0.036035522 | ENSG00000106692 | n.s. |
| ENSG00000055609 | 553.6354 | -0.283346263 | 0.26554 | -1.067058 | 0.285946 | 0.746239075 | KMT2C | protein_coding | 38.65 | -0.036019548 | 0.036019548 | ENSG00000055609 | n.s. |

| | | | | | | | | | | | | | |
|---|---|---|---|---|---|---|---|---|---|---|---|---|---|
| ENSG00000126773 | 1424.145 | -0.296620592 | 0.284535 | -1.042474 | 0.297192 | 0.756138711 | PCNX4 | protein_coding | 37.99 | -0.036009303 | 0.036009303 | ENSG00000126773 | n.s. |
| ENSG00000081665 | 1047.743 | -0.272926213 | 0.24872 | -1.097323 | 0.2725 | 0.738113065 | ZNF506 | protein_coding | 40.49 | -0.03599272 | 0.03599272 | ENSG00000081665 | n.s. |
| ENSG00000134014 | 1075.5 | -0.280528693 | 0.260255 | -1.077899 | 0.281079 | 0.744242186 | ELP3 | protein_coding | 40.16 | -0.035987824 | 0.035987824 | ENSG00000134014 | n.s. |
| ENSG00000156858 | 137.5869 | -0.274433884 | 0.251524 | -1.091085 | 0.275235 | 0.739442303 | PRR14 | protein_coding | 57.98 | -0.035977104 | 0.035977104 | ENSG00000156858 | n.s. |
| ENSG00000176410 | 207.6833 | -0.253540353 | 0.220521 | -1.149733 | 0.250254 | 0.721375365 | DNAJC30 | protein_coding | 54.45 | -0.035961832 | 0.035961832 | ENSG00000176410 | n.s. |
| ENSG00000251739 | 6.848215 | -0.44469179 | 0.546562 | -0.813616 | 0.415865 | 0.830119829 | RNU6-1053P | snRNA | 46.53 | -0.035957428 | 0.035957428 | ENSG00000251739 | n.s. |
| ENSG00000145721 | 343.0743 | 0.290170113 | 0.275087 | 1.0548292 | 0.291503 | 0.751776696 | LIX1 | protein_coding | 39.28 | 0.03595531 | 0.03595531 | ENSG00000145721 | n.s. |
| ENSG00000146909 | 1148.296 | -0.253250688 | 0.220161 | -1.150297 | 0.250022 | 0.721162708 | NOM1 | protein_coding | 47.02 | -0.035953174 | 0.035953174 | ENSG00000146909 | n.s. |
| ENSG00000269190 | 188.2982 | 0.33224947 | 0.339801 | 0.9777772 | 0.328185 | 0.779459027 | FBXO17 | protein_coding | 49.36 | 0.035951622 | 0.035951622 | ENSG00000269190 | n.s. |
| ENSG00000122512 | 434.7599 | -0.296132496 | 0.284132 | -1.042235 | 0.297303 | 0.756138711 | PMS2 | protein_coding | 45.05 | -0.035950049 | 0.035950049 | ENSG00000122512 | n.s. |
| ENSG00000104435 | 1758.763 | 0.30445989 | 0.298696 | 1.0192975 | 0.308062 | 0.761997697 | STMN2 | protein_coding | 38.21 | 0.035940376 | 0.035940376 | ENSG00000104435 | n.s. |
| ENSG00000102312 | 76.20506 | -0.315708104 | 0.31618 | -0.998508 | 0.318033 | 0.76941363 | PORCN | protein_coding | 51.68 | -0.03594025 | 0.03594025 | ENSG00000102312 | n.s. |
| ENSG00000164674 | 148.7618 | -0.295906044 | 0.283937 | -1.042155 | 0.29734 | 0.756138711 | SYTL3 | protein_coding | 46.02 | -0.035922558 | 0.035922558 | ENSG00000164674 | n.s. |
| ENSG00000142687 | 36.42514 | 0.312105349 | 0.310147 | 1.0063157 | 0.314264 | 0.767300529 | KIAA0319L | protein_coding | 41.89 | 0.035902884 | 0.035902884 | ENSG00000142687 | n.s. |
| ENSG00000252481 | 419.0621 | 0.267334345 | 0.239535 | 1.1160569 | 0.264398 | 0.734053608 | SCARNA13 | scaRNA | 47.64 | 0.035895577 | 0.035895577 | ENSG00000252481 | n.s. |
| ENSG00000136280 | 172.2237 | -0.303633148 | 0.296024 | -1.025704 | 0.305031 | 0.761997697 | CCM2 | protein_coding | 46.33 | -0.035842782 | 0.035842782 | ENSG00000136280 | n.s. |
| ENSG00000011007 | 874.0407 | -0.273932815 | 0.251234 | -1.090348 | 0.27556 | 0.739945777 | ELOA | protein_coding | 45.7 | -0.035830441 | 0.035830441 | ENSG00000011007 | n.s. |
| ENSG00000084652 | 958.8985 | -0.269084491 | 0.243203 | -1.106418 | 0.268546 | 0.735983077 | TXLNA | protein_coding | 50.3 | -0.035823803 | 0.035823803 | ENSG00000084652 | n.s. |
| ENSG00000154803 | 213.925 | -0.285022327 | 0.269168 | -1.058903 | 0.289644 | 0.748759774 | FLCN | protein_coding | 53.29 | -0.035815192 | 0.035815192 | ENSG00000154803 | n.s. |
| ENSG00000183666 | 315.7614 | -0.258409764 | 0.227941 | -1.133672 | 0.256932 | 0.726939222 | GUSBP1 | cribed_unprocessed_pseudc | 37.03 | -0.035790243 | 0.035790243 | ENSG00000183666 | n.s. |
| ENSG00000135622 | 340.6954 | -0.292449729 | 0.279352 | -1.046885 | 0.295152 | 0.754508255 | SEMA4F | protein_coding | 44.97 | -0.035777131 | 0.035777131 | ENSG00000135622 | n.s. |
| ENSG00000277072 | 173.5646 | -0.271191559 | 0.247148 | -1.097286 | 0.272517 | 0.738113065 | STAG3L2 | cribed_unprocessed_pseudc | 47.41 | -0.035763958 | 0.035763958 | ENSG00000277072 | n.s. |
| ENSG00000122696 | 318.6441 | -0.270318423 | 0.245949 | -1.099082 | 0.271732 | 0.737635112 | SLC25A51 | protein_coding | 42.85 | -0.035724855 | 0.035724855 | ENSG00000122696 | n.s. |
| ENSG00000196199 | 939.2849 | -0.281420561 | 0.2647 | -1.063169 | 0.287705 | 0.746611882 | MPHOSPH8 | protein_coding | 43.3 | -0.035713705 | 0.035713705 | ENSG00000196199 | n.s. |
| ENSG00000269940 | 87.83832 | -0.283121732 | 0.267113 | -1.059934 | 0.289174 | 0.748034104 | | sense_intronic | 51.23 | -0.035695592 | 0.035695592 | ENSG00000269940 | n.s. |
| ENSG00000164190 | 2428.907 | -0.266940666 | 0.240355 | -1.110609 | 0.266737 | 0.734987182 | NIPBL | protein_coding | 35.04 | -0.035695369 | 0.035695369 | ENSG00000164190 | n.s. |
| ENSG00000145882 | 304.5648 | -0.299592208 | 0.289581 | -1.034572 | 0.300869 | 0.760115318 | PCYOX1L | protein_coding | 50.16 | -0.035687578 | 0.035687578 | ENSG00000145882 | n.s. |
| ENSG00000167566 | 432.2959 | -0.261048664 | 0.231924 | -1.125579 | 0.260344 | 0.730043745 | NCKAP5L | protein_coding | 52.89 | -0.035672591 | 0.035672591 | ENSG00000167566 | n.s. |
| ENSG00000230714 | 7.782278 | 0.568939652 | 0.797049 | 0.7138073 | 0.475346 | 0.865616576 | | lincRNA | 32.75 | 0.035657972 | 0.035657972 | ENSG00000230714 | n.s. |
| ENSG00000184009 | 32781.88 | -0.290572283 | 0.276993 | -1.049023 | 0.294167 | 0.753915388 | ACTG1 | protein_coding | 61.09 | -0.03564665 | 0.03564665 | ENSG00000184009 | n.s. |
| ENSG00000117868 | 578.1839 | -0.291139917 | 0.277984 | -1.047327 | 0.294949 | 0.754466497 | ESYT2 | protein_coding | 43.39 | -0.035623892 | 0.035623892 | ENSG00000117868 | n.s. |
| ENSG00000276575 | 3.239408 | -0.545764593 | 0.747373 | -0.730244 | 0.465241 | 0.860461298 | MIR6895 | miRNA | 53.85 | -0.035621323 | 0.035621323 | ENSG00000276575 | n.s. |

| | | | | | | | | | | | | | |
|---|---|---|---|---|---|---|---|---|---|---|---|---|---|
| ENSG00000117036 | 870.742 | -0.261334944 | 0.232483 | -1.124105 | 0.260968 | 0.730784841 | ETV3 | protein_coding | 43.29 | -0.035596555 | 0.035596555 | ENSG00000117036 | n.s. |
| ENSG00000272288 | 17.17413 | 0.482755456 | 0.622167 | 0.7759258 | 0.437793 | 0.843950276 | | antisense | 62.48 | 0.035570938 | 0.035570938 | ENSG00000272288 | n.s. |
| ENSG00000148019 | 2724.236 | -0.278449733 | 0.259859 | -1.071543 | 0.283925 | 0.745300357 | CEP78 | protein_coding | 37.2 | -0.035549307 | 0.035549307 | ENSG00000148019 | n.s. |
| ENSG00000130725 | 474.6731 | -0.303495386 | 0.298698 | -1.01606 | 0.309601 | 0.763743892 | UBE2M | protein_coding | 61.71 | -0.035524818 | 0.035524818 | ENSG00000130725 | n.s. |
| ENSG00000261226 | 154.9277 | -0.294538899 | 0.283106 | -1.040384 | 0.298162 | 0.757537602 | | sense_intronic | 61.68 | -0.035520156 | 0.035520156 | ENSG00000261226 | n.s. |
| ENSG00000201318 | 4.21448 | -0.47778163 | 0.612399 | -0.78018 | 0.435285 | 0.842717361 | SNORD114-30 | snoRNA | 45.83 | -0.035507803 | 0.035507803 | ENSG00000201318 | n.s. |
| ENSG00000221500 | 143.1367 | 0.265525035 | 0.239213 | 1.1099955 | 0.267001 | 0.734987182 | SNORD100 | snoRNA | 50 | 0.03550607 | 0.03550607 | ENSG00000221500 | n.s. |
| ENSG00000037042 | 264.6594 | -0.25979263 | 0.230855 | -1.125348 | 0.260442 | 0.730043745 | TUBG2 | protein_coding | 51.9 | -0.035500953 | 0.035500953 | ENSG00000037042 | n.s. |
| ENSG00000128915 | 3091.876 | -0.278493933 | 0.260319 | -1.069818 | 0.284701 | 0.745682152 | ICE2 | protein_coding | 36.91 | -0.035493008 | 0.035493008 | ENSG00000128915 | n.s. |
| ENSG00000221788 | 27.60878 | -0.333360041 | 0.345618 | -0.964533 | 0.334779 | 0.782724681 | MIR1252 | miRNA | 26.15 | -0.035466499 | 0.035466499 | ENSG00000221788 | n.s. |
| ENSG00000166471 | 967.2862 | -0.276972039 | 0.257886 | -1.074011 | 0.282818 | 0.744776963 | TMEM41B | protein_coding | 42.5 | -0.035445155 | 0.035445155 | ENSG00000166471 | n.s. |
| ENSG00000279901 | 112.903 | 0.274666279 | 0.25327 | 1.08448 | 0.278152 | 0.742965563 | | TEC | 47.26 | 0.035440551 | 0.035440551 | ENSG00000279901 | n.s. |
| ENSG00000121417 | 270.7132 | -0.277895512 | 0.259794 | -1.069675 | 0.284766 | 0.745682152 | ZNF211 | protein_coding | 41.9 | -0.035416741 | 0.035416741 | ENSG00000121417 | n.s. |
| ENSG00000267160 | 175.8179 | -0.267184755 | 0.242688 | -1.100938 | 0.270924 | 0.737020734 | | antisense | 29.78 | -0.035407402 | 0.035407402 | ENSG00000267160 | n.s. |
| ENSG00000225156 | 120.187 | -0.300286173 | 0.294764 | -1.018733 | 0.30833 | 0.762364419 | | lincRNA | 50.97 | -0.035384936 | 0.035384936 | ENSG00000225156 | n.s. |
| ENSG00000198399 | 207.3283 | -0.279819608 | 0.263807 | -1.060698 | 0.288827 | 0.747460922 | ITSN2 | protein_coding | 39.9 | -0.035372419 | 0.035372419 | ENSG00000198399 | n.s. |
| ENSG00000185900 | 541.338 | -0.25707033 | 0.227752 | -1.12873 | 0.259012 | 0.728529796 | POMK | protein_coding | 45.29 | -0.035360713 | 0.035360713 | ENSG00000185900 | n.s. |
| ENSG00000224080 | 38.48445 | -0.345651786 | 0.369481 | -0.935506 | 0.349528 | 0.790140298 | UBE2FP1 | processed_pseudogene | 40.72 | -0.035358722 | 0.035358722 | ENSG00000224080 | n.s. |
| ENSG00000110066 | 1191.562 | -0.26958389 | 0.246938 | -1.091706 | 0.274963 | 0.739442303 | KMT5B | protein_coding | 42.48 | -0.035341291 | 0.035341291 | ENSG00000110066 | n.s. |
| ENSG00000020181 | 518.1767 | -0.276726725 | 0.258126 | -1.072059 | 0.283694 | 0.745300357 | ADGRA2 | protein_coding | 55.25 | -0.035329333 | 0.035329333 | ENSG00000020181 | n.s. |
| ENSG00000201367 | 12.30934 | 0.386719722 | 0.442364 | 0.874211 | 0.382003 | 0.81032471 | RNU6-522P | snRNA | 35.51 | 0.035323334 | 0.035323334 | ENSG00000201367 | n.s. |
| ENSG00000091009 | 687.4123 | -0.276014462 | 0.257078 | -1.073659 | 0.282976 | 0.744776963 | RBM27 | protein_coding | 38.7 | -0.03532261 | 0.03532261 | ENSG00000091009 | n.s. |
| ENSG00000232184 | 76.5003 | -0.327438386 | 0.336401 | -0.973357 | 0.330376 | 0.780141891 | | sense_intronic | 35.88 | -0.035306503 | 0.035306503 | ENSG00000232184 | n.s. |
| ENSG00000198453 | 214.2233 | -0.274577067 | 0.254203 | -1.080151 | 0.280075 | 0.743892631 | ZNF568 | protein_coding | 40.05 | -0.035280337 | 0.035280337 | ENSG00000198453 | n.s. |
| ENSG00000112249 | 4033.952 | -0.287509111 | 0.273835 | -1.049934 | 0.293748 | 0.753890684 | ASCC3 | protein_coding | 34.11 | -0.03527496 | 0.03527496 | ENSG00000112249 | n.s. |
| ENSG00000164008 | 93.94879 | -0.309797903 | 0.310627 | -0.997331 | 0.318604 | 0.769456789 | C1orf50 | protein_coding | 43.06 | -0.035259885 | 0.035259885 | ENSG00000164008 | n.s. |
| ENSG00000125257 | 907.0824 | -0.277348285 | 0.259968 | -1.066856 | 0.286037 | 0.746239075 | ABCC4 | protein_coding | 42.33 | -0.035257073 | 0.035257073 | ENSG00000125257 | n.s. |
| ENSG00000185046 | 17.30157 | -0.385760094 | 0.440318 | -0.876094 | 0.380979 | 0.810319533 | ANKS1B | protein_coding | 37.84 | -0.035236751 | 0.035236751 | ENSG00000185046 | n.s. |
| ENSG00000112701 | 4231.72 | -0.275290848 | 0.256397 | -1.07369 | 0.282962 | 0.744776963 | SENP6 | protein_coding | 37.44 | -0.035230006 | 0.035230006 | ENSG00000112701 | n.s. |
| ENSG00000109576 | 833.837 | -0.27346454 | 0.252763 | -1.081902 | 0.279296 | 0.743390786 | AADAT | protein_coding | 38.78 | -0.035217536 | 0.035217536 | ENSG00000109576 | n.s. |
| ENSG00000154743 | 596.3455 | -0.263270209 | 0.237142 | -1.110178 | 0.266923 | 0.734987182 | TSEN2 | protein_coding | 43.16 | -0.035204554 | 0.035204554 | ENSG00000154743 | n.s. |
| ENSG00000182872 | 1271.303 | -0.277273953 | 0.260523 | -1.064296 | 0.287195 | 0.746611882 | RBM10 | protein_coding | 50.32 | -0.03518748 | 0.03518748 | ENSG00000182872 | n.s. |

| | | | | | | | | | | | | | |
|---|---|---|---|---|---|---|---|---|---|---|---|---|---|
| ENSG00000163257 | 2459.382 | -0.263028838 | 0.237203 | -1.108875 | 0.267484 | 0.734987182 | DCAF16 | protein_coding | 39.37 | -0.035172278 | 0.035172278 | ENSG00000163257 | n.s. |
| ENSG00000156599 | 842.4285 | -0.264049283 | 0.238838 | -1.105558 | 0.268918 | 0.735983077 | ZDHHC5 | protein_coding | 45.38 | -0.035153455 | 0.035153455 | ENSG00000156599 | n.s. |
| ENSG00000255717 | 14102.47 | 0.229438368 | 0.190632 | 1.2035697 | 0.228756 | 0.702737638 | SNHG1 | processed_transcript | 41.79 | 0.035151515 | 0.035151515 | ENSG00000255717 | n.s. |
| ENSG00000055208 | 1459.201 | -0.27697343 | 0.259946 | -1.065503 | 0.286648 | 0.746611882 | TAB2 | protein_coding | 38.87 | -0.035149342 | 0.035149342 | ENSG00000055208 | n.s. |
| ENSG00000169682 | 315.4336 | -0.266522953 | 0.242907 | -1.097221 | 0.272545 | 0.738113065 | SPNS1 | protein_coding | 55.79 | -0.035148276 | 0.035148276 | ENSG00000169682 | n.s. |
| ENSG00000112592 | 699.4949 | -0.265463132 | 0.241258 | -1.10033 | 0.271188 | 0.737359702 | TBP | protein_coding | 39.46 | -0.035126241 | 0.035126241 | ENSG00000112592 | n.s. |
| ENSG00000132824 | 667.7415 | -0.286984543 | 0.274077 | -1.047094 | 0.295056 | 0.754466497 | SERINC3 | protein_coding | 42.56 | -0.03511544 | 0.03511544 | ENSG00000132824 | n.s. |
| ENSG00000088812 | 745.3898 | -0.276127821 | 0.258612 | -1.067728 | 0.285643 | 0.74621359 | ATRN | protein_coding | 40.34 | -0.03510602 | 0.03510602 | ENSG00000088812 | n.s. |
| ENSG00000130227 | 1616.596 | -0.276336118 | 0.259586 | -1.064527 | 0.28709 | 0.746611882 | XPO7 | protein_coding | 39.92 | -0.035068463 | 0.035068463 | ENSG00000130227 | n.s. |
| ENSG00000252361 | 6.131893 | 0.448727658 | 0.561608 | 0.7990054 | 0.424287 | 0.835315601 | RNU6-118P | snRNA | 36.27 | 0.0350678 | 0.0350678 | ENSG00000252361 | n.s. |
| ENSG00000143801 | 401.0931 | -0.308847696 | 0.310038 | -0.99616 | 0.319172 | 0.769994426 | PSEN2 | protein_coding | 52.42 | -0.035058049 | 0.035058049 | ENSG00000143801 | n.s. |
| ENSG00000169740 | 593.0422 | 0.315421309 | 0.319023 | 0.9887117 | 0.322804 | 0.774229646 | ZNF32 | protein_coding | 46.34 | 0.035052834 | 0.035052834 | ENSG00000169740 | n.s. |
| ENSG00000074603 | 1931.739 | -0.260349612 | 0.232883 | -1.117942 | 0.263592 | 0.733524057 | DPP8 | protein_coding | 40.97 | -0.035039319 | 0.035039319 | ENSG00000074603 | n.s. |
| ENSG00000108578 | 1878.26 | -0.275595724 | 0.258361 | -1.066708 | 0.286104 | 0.746239075 | BLMH | protein_coding | 40.18 | -0.035034284 | 0.035034284 | ENSG00000108578 | n.s. |
| ENSG00000154839 | 400.9266 | -0.270069178 | 0.248299 | -1.087678 | 0.276737 | 0.741896779 | SKA1 | protein_coding | 39.41 | -0.035016228 | 0.035016228 | ENSG00000154839 | n.s. |
| ENSG00000100395 | 587.6487 | -0.294475219 | 0.285887 | -1.030041 | 0.302991 | 0.760781413 | L3MBTL2 | protein_coding | 52.42 | -0.034966019 | 0.034966019 | ENSG00000100395 | n.s. |
| ENSG00000009413 | 3440.158 | -0.264225612 | 0.240332 | -1.099418 | 0.271586 | 0.73753588 | REV3L | protein_coding | 36.9 | -0.034935077 | 0.034935077 | ENSG00000009413 | n.s. |
| ENSG00000114904 | 798.3008 | -0.274920671 | 0.258405 | -1.063915 | 0.287367 | 0.746611882 | NEK4 | protein_coding | 42.79 | -0.034888836 | 0.034888836 | ENSG00000114904 | n.s. |
| ENSG00000252050 | 7.112342 | -0.446897128 | 0.559738 | -0.798404 | 0.424636 | 0.835529186 | | snoRNA | 40 | -0.034875126 | 0.034875126 | ENSG00000252050 | n.s. |
| ENSG00000253395 | 13.14381 | 0.40878871 | 0.485442 | 0.8420953 | 0.399735 | 0.821737289 | | lincRNA | 46.39 | 0.034856189 | 0.034856189 | ENSG00000253395 | n.s. |
| ENSG00000188739 | 1506.497 | -0.282026114 | 0.267673 | -1.053622 | 0.292056 | 0.75237733 | RBM34 | protein_coding | 44.48 | -0.034848359 | 0.034848359 | ENSG00000188739 | n.s. |
| ENSG00000153560 | 1443.336 | -0.27454824 | 0.258171 | -1.063437 | 0.287584 | 0.746611882 | UBP1 | protein_coding | 40.23 | -0.034841573 | 0.034841573 | ENSG00000153560 | n.s. |
| ENSG00000116406 | 1334.54 | -0.296432126 | 0.291349 | -1.017445 | 0.308942 | 0.762996547 | EDEM3 | protein_coding | 34.39 | -0.034824084 | 0.034824084 | ENSG00000116406 | n.s. |
| ENSG00000186105 | 158.5446 | -0.295701992 | 0.290457 | -1.018056 | 0.308651 | 0.762601855 | LRRC70 | protein_coding | 32.72 | -0.034804758 | 0.034804758 | ENSG00000186105 | n.s. |
| ENSG00000135655 | 3518.511 | -0.272522495 | 0.254172 | -1.072196 | 0.283632 | 0.745300357 | USP15 | protein_coding | 34.93 | -0.034792585 | 0.034792585 | ENSG00000135655 | n.s. |
| ENSG00000161513 | 210.6638 | -0.286215867 | 0.274285 | -1.0435 | 0.296717 | 0.755935669 | FDXR | protein_coding | 58.11 | -0.034779567 | 0.034779567 | ENSG00000161513 | n.s. |
| ENSG00000117899 | 1101.507 | -0.286142326 | 0.274453 | -1.04259 | 0.297138 | 0.756138711 | MESDC2 | protein_coding | 45.39 | -0.034737257 | 0.034737257 | ENSG00000117899 | n.s. |
| ENSG00000111885 | 615.4598 | -0.273062543 | 0.255782 | -1.06756 | 0.285719 | 0.74621359 | MAN1A1 | protein_coding | 37.6 | -0.034716311 | 0.034716311 | ENSG00000111885 | n.s. |
| ENSG00000134874 | 1497.698 | -0.269567722 | 0.249439 | -1.080695 | 0.279833 | 0.743390786 | DZIP1 | protein_coding | 41.42 | -0.034715693 | 0.034715693 | ENSG00000134874 | n.s. |
| ENSG00000100401 | 1338.357 | -0.273517465 | 0.256681 | -1.065593 | 0.286608 | 0.746611882 | RANGAP1 | protein_coding | 53.26 | -0.034710762 | 0.034710762 | ENSG00000100401 | n.s. |
| ENSG00000141959 | 290.0256 | -0.28449762 | 0.272186 | -1.045231 | 0.295916 | 0.755277852 | PFKL | protein_coding | 63.48 | -0.03467834 | 0.03467834 | ENSG00000141959 | n.s. |
| ENSG00000108061 | 813.8594 | -0.288945727 | 0.278657 | -1.036924 | 0.299771 | 0.758581944 | SHOC2 | protein_coding | 35.08 | -0.034672765 | 0.034672765 | ENSG00000108061 | n.s. |

| | | | | | | | | | | | | | |
|---|---|---|---|---|---|---|---|---|---|---|---|---|---|
| ENSG00000116793 | 772.4114 | -0.271559418 | 0.253583 | -1.070891 | 0.284219 | 0.745309826 | PHTF1 | protein_coding | 39.1 | -0.034668132 | 0.034668132 | ENSG00000116793 | n.s. |
| ENSG00000198826 | 2568.047 | -0.270334033 | 0.251433 | -1.075171 | 0.282298 | 0.744567264 | ARHGAP11A | protein_coding | 37.89 | -0.034628725 | 0.034628725 | ENSG00000198826 | n.s. |
| ENSG00000198363 | 1964.479 | -0.291626906 | 0.283038 | -1.030346 | 0.302847 | 0.760781413 | ASPH | protein_coding | 38.17 | -0.03462781 | 0.03462781 | ENSG00000198363 | n.s. |
| ENSG00000138686 | 474.8562 | -0.293158929 | 0.287158 | -1.020898 | 0.307303 | 0.761997697 | BBS7 | protein_coding | 35.63 | -0.034606339 | 0.034606339 | ENSG00000138686 | n.s. |
| ENSG00000167657 | 283.0015 | -0.281086464 | 0.267246 | -1.05179 | 0.292896 | 0.753230348 | DAPK3 | protein_coding | 58.44 | -0.034593927 | 0.034593927 | ENSG00000167657 | n.s. |
| ENSG00000070371 | 314.2377 | -0.292864768 | 0.285551 | -1.025612 | 0.305074 | 0.761997697 | CLTCL1 | protein_coding | 46.41 | -0.034571614 | 0.034571614 | ENSG00000070371 | n.s. |
| ENSG00000264706 | 26.53934 | -0.327607198 | 0.340885 | -0.961048 | 0.336528 | 0.784443005 | RN7SL217P | misc_RNA | 54.55 | -0.034542446 | 0.034542446 | ENSG00000264706 | n.s. |
| ENSG00000275342 | 243.684 | -0.2577952 | 0.231242 | -1.114827 | 0.264924 | 0.734601921 | PRAG1 | protein_coding | 43.93 | -0.034531136 | 0.034531136 | ENSG00000275342 | n.s. |
| ENSG00000258297 | 214.8234 | 0.31219682 | 0.316296 | 0.9870397 | 0.323623 | 0.775183815 | | antisense | 39.85 | 0.034527502 | 0.034527502 | ENSG00000258297 | n.s. |
| ENSG00000123575 | 439.1967 | -0.289882108 | 0.280482 | -1.033514 | 0.301363 | 0.760398818 | FAM199X | protein_coding | 36.81 | -0.03448396 | 0.03448396 | ENSG00000123575 | n.s. |
| ENSG00000114867 | 7555.634 | -0.257407006 | 0.231294 | -1.1129 | 0.265751 | 0.734612586 | EIF4G1 | protein_coding | 52.77 | -0.034477515 | 0.034477515 | ENSG00000114867 | n.s. |
| ENSG00000162729 | 111.7294 | -0.294980875 | 0.290688 | -1.014769 | 0.310216 | 0.764194956 | IGSF8 | protein_coding | 58.9 | -0.034452537 | 0.034452537 | ENSG00000162729 | n.s. |
| ENSG00000152102 | 3362.454 | -0.280765045 | 0.267492 | -1.04962 | 0.293893 | 0.753915388 | FAM168B | protein_coding | 45.79 | -0.034443524 | 0.034443524 | ENSG00000152102 | n.s. |
| ENSG00000207155 | 5.957032 | -0.460202545 | 0.58568 | -0.785758 | 0.43201 | 0.841828575 | RNY1P14 | misc_RNA | 47.22 | -0.03441226 | 0.03441226 | ENSG00000207155 | n.s. |
| ENSG00000155254 | 759.1714 | -0.258339173 | 0.233922 | -1.10438 | 0.269428 | 0.735983077 | MARVELD1 | protein_coding | 57.63 | -0.034393255 | 0.034393255 | ENSG00000155254 | n.s. |
| ENSG00000166377 | 187.1767 | -0.298987204 | 0.298008 | -1.003285 | 0.315723 | 0.767425784 | ATP9B | protein_coding | 41.82 | -0.034372649 | 0.034372649 | ENSG00000166377 | n.s. |
| ENSG00000031698 | 4932.17 | -0.289472443 | 0.281413 | -1.028639 | 0.303649 | 0.760781413 | SARS | protein_coding | 43.12 | -0.034371989 | 0.034371989 | ENSG00000031698 | n.s. |
| ENSG00000163596 | 32.94644 | -0.318635543 | 0.327548 | -0.972791 | 0.330657 | 0.780141891 | ICA1L | protein_coding | 39.94 | -0.034357324 | 0.034357324 | ENSG00000163596 | n.s. |
| ENSG00000005483 | 1648.446 | -0.261240331 | 0.239017 | -1.092979 | 0.274403 | 0.739386355 | KMT2E | protein_coding | 37.46 | -0.034256071 | 0.034256071 | ENSG00000005483 | n.s. |
| ENSG00000198208 | 202.4143 | -0.279956329 | 0.267344 | -1.047177 | 0.295018 | 0.754466497 | RPS6KL1 | protein_coding | 55.36 | -0.034255468 | 0.034255468 | ENSG00000198208 | n.s. |
| ENSG00000234773 | 97.30622 | -0.290993314 | 0.285781 | -1.01824 | 0.308564 | 0.762601855 | | transcribed_unprocessed_pseudc | 42.34 | -0.034250537 | 0.034250537 | ENSG00000234773 | n.s. |
| ENSG00000212232 | 482.5866 | 0.255063514 | 0.228612 | 1.1157049 | 0.264548 | 0.734165567 | SNORD17 | snoRNA | 49.79 | 0.034231051 | 0.034231051 | ENSG00000212232 | n.s. |
| ENSG00000204366 | 115.6853 | -0.289835162 | 0.28328 | -1.023139 | 0.306242 | 0.761997697 | ZBTB12 | protein_coding | 64.33 | -0.03421398 | 0.03421398 | ENSG00000204366 | n.s. |
| ENSG00000073111 | 4985.516 | -0.267017624 | 0.248354 | -1.07515 | 0.282308 | 0.744567264 | MCM2 | protein_coding | 50.73 | -0.034203906 | 0.034203906 | ENSG00000073111 | n.s. |
| ENSG00000160284 | 40.18973 | -0.326929562 | 0.342881 | -0.953477 | 0.340349 | 0.785979841 | SPATC1L | protein_coding | 50.77 | -0.034193103 | 0.034193103 | ENSG00000160284 | n.s. |
| ENSG00000167074 | 175.6113 | -0.293754169 | 0.289886 | -1.013342 | 0.310897 | 0.765100725 | TEF | protein_coding | 51.78 | -0.034158142 | 0.034158142 | ENSG00000167074 | n.s. |
| ENSG00000170525 | 642.9982 | -0.262146364 | 0.240647 | -1.089342 | 0.276003 | 0.740850388 | PFKFB3 | protein_coding | 50.91 | -0.034149672 | 0.034149672 | ENSG00000170525 | n.s. |
| ENSG00000127452 | 118.2328 | 0.268757872 | 0.252451 | 1.0645928 | 0.28706 | 0.746611882 | FBXL12 | protein_coding | 50.56 | 0.034106745 | 0.034106745 | ENSG00000127452 | n.s. |
| ENSG00000143702 | 6466.655 | -0.275799869 | 0.261598 | -1.054288 | 0.291751 | 0.752231212 | CEP170 | protein_coding | 36.28 | -0.034102282 | 0.034102282 | ENSG00000143702 | n.s. |
| ENSG00000141458 | 64.61049 | -0.296608178 | 0.295392 | -1.004118 | 0.315322 | 0.767425784 | NPC1 | protein_coding | 44.75 | -0.034099148 | 0.034099148 | ENSG00000141458 | n.s. |
| ENSG00000133111 | 330.4955 | -0.26897752 | 0.25334 | -1.061725 | 0.288361 | 0.747018696 | RFXAP | protein_coding | 38.45 | -0.034070987 | 0.034070987 | ENSG00000133111 | n.s. |
| ENSG00000105778 | 987.7352 | -0.288572443 | 0.28128 | -1.025927 | 0.304926 | 0.761997697 | AVL9 | protein_coding | 40.32 | -0.034064921 | 0.034064921 | ENSG00000105778 | n.s. |

| | | | | | | | | | | | | | |
|---|---|---|---|---|---|---|---|---|---|---|---|---|---|
| ENSG00000130508 | 1345.964 | -0.256484108 | 0.232366 | -1.103795 | 0.269682 | 0.73652118 | PXDN | protein_coding | 49.9 | -0.034064875 | 0.034064875 | ENSG00000130508 | n.s. |
| ENSG00000136141 | 2582.21 | -0.259760892 | 0.237767 | -1.092504 | 0.274612 | 0.739386355 | LRCH1 | protein_coding | 40.02 | -0.034062074 | 0.034062074 | ENSG00000136141 | n.s. |
| ENSG00000031003 | 1092.948 | -0.267768509 | 0.250484 | -1.069006 | 0.285067 | 0.746113205 | FAM13B | protein_coding | 40.05 | -0.034058889 | 0.034058889 | ENSG00000031003 | n.s. |
| ENSG00000128923 | 767.0124 | -0.276316913 | 0.262616 | -1.052171 | 0.292721 | 0.752972779 | MINDY2 | protein_coding | 39.89 | -0.03404797 | 0.03404797 | ENSG00000128923 | n.s. |
| ENSG00000145734 | 3713.837 | -0.26560211 | 0.246905 | -1.075724 | 0.282051 | 0.744567264 | BDP1 | protein_coding | 39.84 | -0.034022584 | 0.034022584 | ENSG00000145734 | n.s. |
| ENSG00000148835 | 454.1715 | -0.288078633 | 0.280669 | -1.0264 | 0.304703 | 0.761997697 | TAF5 | protein_coding | 43.36 | -0.034006629 | 0.034006629 | ENSG00000148835 | n.s. |
| ENSG00000162032 | 228.3492 | -0.282965021 | 0.272623 | -1.037936 | 0.2993 | 0.758335061 | SPSB3 | protein_coding | 64.01 | -0.033995096 | 0.033995096 | ENSG00000162032 | n.s. |
| ENSG00000072135 | 288.0572 | -0.26516713 | 0.246511 | -1.075679 | 0.282071 | 0.744567264 | PTPN18 | protein_coding | 47.65 | -0.033966865 | 0.033966865 | ENSG00000072135 | n.s. |
| ENSG00000100320 | 4651.679 | -0.273920509 | 0.259576 | -1.055261 | 0.291306 | 0.751641109 | RBFOX2 | protein_coding | 39.31 | -0.03396326 | 0.03396326 | ENSG00000100320 | n.s. |
| ENSG00000186205 | 767.3814 | -0.293572461 | 0.290682 | -1.009943 | 0.312522 | 0.766200005 | 1-Mar | protein_coding | 45.28 | -0.03395396 | 0.03395396 | ENSG00000186205 | n.s. |
| ENSG00000130816 | 3045.587 | -0.26150741 | 0.240272 | -1.088383 | 0.276426 | 0.741700023 | DNMT1 | protein_coding | 50.43 | -0.033936263 | 0.033936263 | ENSG00000130816 | n.s. |
| ENSG00000264757 | 9.506886 | -0.430835608 | 0.536575 | -0.802936 | 0.422012 | 0.834256463 | MIR3198-1 | miRNA | 56.25 | -0.033906943 | 0.033906943 | ENSG00000264757 | n.s. |
| ENSG00000168268 | 2061.843 | -0.276289581 | 0.26336 | -1.049094 | 0.294135 | 0.753915388 | NT5DC2 | protein_coding | 61.51 | -0.033894485 | 0.033894485 | ENSG00000168268 | n.s. |
| ENSG00000119669 | 405.5868 | -0.264162365 | 0.244748 | -1.079325 | 0.280443 | 0.744242186 | IRF2BPL | protein_coding | 62.81 | -0.033888258 | 0.033888258 | ENSG00000119669 | n.s. |
| ENSG00000234779 | 18.27813 | 0.384946023 | 0.451544 | 0.8525103 | 0.393931 | 0.81657325 | | antisense | 61.04 | 0.033877118 | 0.033877118 | ENSG00000234779 | n.s. |
| ENSG00000138380 | 218.4663 | -0.263011508 | 0.243044 | -1.082157 | 0.279183 | 0.743390786 | CARF | protein_coding | 38.49 | -0.033871365 | 0.033871365 | ENSG00000138380 | n.s. |
| ENSG00000240328 | 43.37408 | -0.327773514 | 0.346657 | -0.945526 | 0.34439 | 0.788289085 | | processed_pseudogene | 44.15 | -0.033863752 | 0.033863752 | ENSG00000240328 | n.s. |
| ENSG00000149782 | 482.1675 | -0.262942746 | 0.242824 | -1.082855 | 0.278873 | 0.743390786 | PLCB3 | protein_coding | 60.32 | -0.03386251 | 0.03386251 | ENSG00000149782 | n.s. |
| ENSG00000160014 | 1423.545 | -0.281760589 | 0.271424 | -1.038082 | 0.299232 | 0.758335061 | CALM3 | protein_coding | 57.51 | -0.033850397 | 0.033850397 | ENSG00000160014 | n.s. |
| ENSG00000255568 | 105.734 | 0.270510598 | 0.255837 | 1.0573539 | 0.29035 | 0.750286611 | BRWD1-AS2 | antisense | 59.24 | 0.033752366 | 0.033752366 | ENSG00000255568 | n.s. |
| ENSG00000243667 | 409.4056 | -0.281340528 | 0.271401 | -1.036622 | 0.299912 | 0.758800358 | WDR92 | protein_coding | 39.84 | -0.033724985 | 0.033724985 | ENSG00000243667 | n.s. |
| ENSG00000144229 | 4338.22 | -0.280715714 | 0.270635 | -1.03725 | 0.299619 | 0.758335061 | THSD7B | protein_coding | 37.67 | -0.033724867 | 0.033724867 | ENSG00000144229 | n.s. |
| ENSG00000188295 | 1000.904 | 0.241933942 | 0.21278 | 1.137012 | 0.255533 | 0.725521442 | ZNF669 | protein_coding | 43.07 | 0.033713434 | 0.033713434 | ENSG00000188295 | n.s. |
| ENSG00000184436 | 489.2341 | -0.240468709 | 0.210646 | -1.141578 | 0.25363 | 0.72418384 | THAP7 | protein_coding | 62.2 | -0.033701971 | 0.033701971 | ENSG00000184436 | n.s. |
| ENSG00000103855 | 521.9577 | -0.264812036 | 0.247907 | -1.068192 | 0.285434 | 0.746113205 | CD276 | protein_coding | 52.51 | -0.033682839 | 0.033682839 | ENSG00000103855 | n.s. |
| ENSG00000119725 | 294.0588 | 0.263780992 | 0.246121 | 1.0717514 | 0.283832 | 0.745300357 | ZNF410 | protein_coding | 41.75 | 0.033676569 | 0.033676569 | ENSG00000119725 | n.s. |
| ENSG00000240230 | 443.7707 | -0.261453599 | 0.24184 | -1.081102 | 0.279652 | 0.743390786 | COX19 | protein_coding | 58.98 | -0.033670733 | 0.033670733 | ENSG00000240230 | n.s. |
| ENSG00000162931 | 115.8084 | -0.274598065 | 0.262045 | -1.047905 | 0.294682 | 0.754039228 | TRIM17 | protein_coding | 56.75 | -0.033667387 | 0.033667387 | ENSG00000162931 | n.s. |
| ENSG00000162062 | 535.9664 | -0.265008463 | 0.249424 | -1.062483 | 0.288016 | 0.746611882 | C16orf59 | protein_coding | 62.04 | -0.033630926 | 0.033630926 | ENSG00000162062 | n.s. |
| ENSG00000102125 | 105.2581 | 0.292511633 | 0.291216 | 1.0044498 | 0.315162 | 0.767425784 | TAZ | protein_coding | 56.12 | 0.033628194 | 0.033628194 | ENSG00000102125 | n.s. |
| ENSG00000116791 | 611.13 | 0.304554467 | 0.308958 | 0.9857476 | 0.324257 | 0.775538613 | CRYZ | protein_coding | 39.61 | 0.03362177 | 0.03362177 | ENSG00000116791 | n.s. |
| ENSG00000277369 | 112.6247 | -0.266845694 | 0.251835 | -1.059606 | 0.289324 | 0.748190417 | | lincRNA | 58.79 | -0.033619318 | 0.033619318 | ENSG00000277369 | n.s. |

| | | | | | | | | | | | | | |
|---|---|---|---|---|---|---|---|---|---|---|---|---|---|
| ENSG00000151923 | 1828.933 | -0.27354102 | 0.260257 | -1.051043 | 0.293239 | 0.753555483 | TIAL1 | protein_coding | 38.16 | -0.033614024 | 0.033614024 | ENSG00000151923 | n.s. |
| ENSG00000276584 | 2.829088 | -0.640419659 | 1.011736 | -0.632991 | 0.52674 | 0.886259518 | MIR6737 | miRNA | 60 | -0.033583023 | 0.033583023 | ENSG00000276584 | n.s. |
| ENSG00000175206 | 29.99062 | -0.366992238 | 0.419559 | -0.874709 | 0.381732 | 0.81032471 | NPPA | protein_coding | 54.04 | -0.033521408 | 0.033521408 | ENSG00000175206 | n.s. |
| ENSG00000101412 | 1895.228 | -0.288024728 | 0.284425 | -1.012656 | 0.311224 | 0.765173417 | E2F1 | protein_coding | 55.58 | -0.033480031 | 0.033480031 | ENSG00000101412 | n.s. |
| ENSG00000138032 | 1060.384 | -0.278378833 | 0.268336 | -1.037428 | 0.299537 | 0.758335061 | PPM1B | protein_coding | 38.72 | -0.033444117 | 0.033444117 | ENSG00000138032 | n.s. |
| ENSG00000150990 | 756.0053 | -0.263474777 | 0.247954 | -1.062597 | 0.287965 | 0.746611882 | DHX37 | protein_coding | 54.02 | -0.033436294 | 0.033436294 | ENSG00000150990 | n.s. |
| ENSG00000106624 | 310.81 | -0.259558248 | 0.240159 | -1.080777 | 0.279796 | 0.743390786 | AEBP1 | protein_coding | 62.7 | -0.033426645 | 0.033426645 | ENSG00000106624 | n.s. |
| ENSG00000161654 | 402.5908 | -0.269605001 | 0.2556 | -1.054792 | 0.291521 | 0.751776696 | LSM12 | protein_coding | 46.77 | -0.033407063 | 0.033407063 | ENSG00000161654 | n.s. |
| ENSG00000114098 | 1808.763 | -0.281284823 | 0.273322 | -1.029134 | 0.303417 | 0.760781413 | ARMC8 | protein_coding | 39.28 | -0.03339979 | 0.03339979 | ENSG00000114098 | n.s. |
| ENSG00000171159 | 217.4435 | -0.290152611 | 0.289171 | -1.003393 | 0.315671 | 0.767425784 | C9orf16 | protein_coding | 56.88 | -0.033356993 | 0.033356993 | ENSG00000171159 | n.s. |
| ENSG00000198727 | 2356.448 | -0.259870177 | 0.241092 | -1.077888 | 0.281084 | 0.744242186 | MT-CYB | protein_coding | 46.28 | -0.033337632 | 0.033337632 | ENSG00000198727 | n.s. |
| ENSG00000131051 | 13672.53 | 0.249266103 | 0.22464 | 1.1096267 | 0.26716 | 0.734987182 | RBM39 | protein_coding | 40.28 | 0.033331922 | 0.033331922 | ENSG00000131051 | n.s. |
| ENSG00000171757 | 405.6383 | -0.290908333 | 0.290542 | -1.001261 | 0.316701 | 0.768261864 | LRRC34 | protein_coding | 38.99 | -0.033306306 | 0.033306306 | ENSG00000171757 | n.s. |
| ENSG00000198728 | 2168.204 | -0.271219114 | 0.258315 | -1.049956 | 0.293738 | 0.753890684 | LDB1 | protein_coding | 55.59 | -0.033276313 | 0.033276313 | ENSG00000198728 | n.s. |
| ENSG00000104419 | 142.9498 | 0.280150898 | 0.271861 | 1.030495 | 0.302778 | 0.760781413 | NDRG1 | protein_coding | 50.32 | 0.033265147 | 0.033265147 | ENSG00000104419 | n.s. |
| ENSG00000143398 | 1543.792 | -0.281735344 | 0.275371 | -1.023111 | 0.306255 | 0.761997697 | PIP5K1A | protein_coding | 45.21 | -0.033257827 | 0.033257827 | ENSG00000143398 | n.s. |
| ENSG00000277687 | 86.79093 | -0.270677694 | 0.257692 | -1.050391 | 0.293538 | 0.753629384 | | sense_intronic | 34.21 | -0.033250637 | 0.033250637 | ENSG00000277687 | n.s. |
| ENSG00000040633 | 471.8219 | -0.288169019 | 0.286171 | -1.006981 | 0.313944 | 0.766827179 | PHF23 | protein_coding | 59.51 | -0.033226609 | 0.033226609 | ENSG00000040633 | n.s. |
| ENSG00000152926 | 968.1266 | -0.281431617 | 0.275037 | -1.023252 | 0.306189 | 0.761997697 | ZNF117 | protein_coding | 39.14 | -0.033221973 | 0.033221973 | ENSG00000152926 | n.s. |
| ENSG00000125772 | 1188.378 | -0.276504568 | 0.26633 | -1.038201 | 0.299176 | 0.758335061 | GPCPD1 | protein_coding | 39.24 | -0.033218945 | 0.033218945 | ENSG00000125772 | n.s. |
| ENSG00000141429 | 1730.325 | -0.286192337 | 0.283018 | -1.011215 | 0.311914 | 0.765516281 | GALNT1 | protein_coding | 38.5 | -0.033211352 | 0.033211352 | ENSG00000141429 | n.s. |
| ENSG00000212017 | 15.59787 | -0.356181685 | 0.398757 | -0.89323 | 0.371734 | 0.806899249 | MIR548U | miRNA | 33.33 | -0.033189255 | 0.033189255 | ENSG00000212017 | n.s. |
| ENSG00000105251 | 825.908 | -0.272348428 | 0.260622 | -1.044993 | 0.296026 | 0.755420408 | SHD | protein_coding | 54.76 | -0.033175112 | 0.033175112 | ENSG00000105251 | n.s. |
| ENSG00000181019 | 140.538 | -0.307559748 | 0.315688 | -0.974253 | 0.329931 | 0.780141891 | NQO1 | protein_coding | 47.09 | -0.033163061 | 0.033163061 | ENSG00000181019 | n.s. |
| ENSG00000245849 | 124.1557 | -0.279273167 | 0.270837 | -1.031148 | 0.302471 | 0.760781413 | RAD51-AS1 | processed_transcript | 45.04 | -0.033160926 | 0.033160926 | ENSG00000245849 | n.s. |
| ENSG00000206751 | 8.601389 | -0.409255642 | 0.50265 | -0.814197 | 0.415532 | 0.829813075 | | misc_RNA | 42.59 | -0.03315778 | 0.03315778 | ENSG00000206751 | n.s. |
| ENSG00000200832 | 6.564184 | -0.418564435 | 0.519484 | -0.805731 | 0.420398 | 0.833311153 | SNORD114-4 | snoRNA | 42.67 | -0.033147292 | 0.033147292 | ENSG00000200832 | n.s. |
| ENSG00000200974 | 8.422266 | 0.443237004 | 0.564027 | 0.7858432 | 0.431959 | 0.841828575 | RNU4-87P | snRNA | 35.37 | 0.033143639 | 0.033143639 | ENSG00000200974 | n.s. |
| ENSG00000134574 | 589.3052 | -0.240292956 | 0.212415 | -1.131244 | 0.257952 | 0.728049978 | DDB2 | protein_coding | 47.15 | -0.033121694 | 0.033121694 | ENSG00000134574 | n.s. |
| ENSG00000131165 | 400.8924 | -0.284834208 | 0.281194 | -1.012947 | 0.311085 | 0.76510176 | CHMP1A | protein_coding | 58.66 | -0.033120749 | 0.033120749 | ENSG00000131165 | n.s. |
| ENSG00000188760 | 243.9465 | -0.291702413 | 0.292919 | -0.995848 | 0.319324 | 0.769994426 | TMEM198 | protein_coding | 56.18 | -0.033111847 | 0.033111847 | ENSG00000188760 | n.s. |
| ENSG00000239382 | 169.1936 | -0.290859133 | 0.291508 | -0.997775 | 0.318388 | 0.769456789 | ALKBH6 | protein_coding | 56.33 | -0.033104355 | 0.033104355 | ENSG00000239382 | n.s. |

| | | | | | | | | | | | | | |
|---|---|---|---|---|---|---|---|---|---|---|---|---|---|
| ENSG00000005302 | 88.04381 | 0.28908351 | 0.288737 | 1.0012016 | 0.316729 | 0.768261864 | MSL3 | protein_coding | 42.85 | 0.03309738 | 0.03309738 | ENSG00000005302 | n.s. |
| ENSG00000105708 | 359.5356 | -0.285113689 | 0.281752 | -1.011932 | 0.311571 | 0.765483724 | ZNF14 | protein_coding | 42.02 | -0.033091446 | 0.033091446 | ENSG00000105708 | n.s. |
| ENSG00000273117 | 82.4527 | 0.280273409 | 0.274611 | 1.0206194 | 0.307435 | 0.761997697 | | lincRNA | 56.34 | 0.033085251 | 0.033085251 | ENSG00000273117 | n.s. |
| ENSG00000148481 | 764.2782 | -0.280674332 | 0.275488 | -1.018825 | 0.308286 | 0.762364419 | MINDY3 | protein_coding | 35.88 | -0.033073928 | 0.033073928 | ENSG00000148481 | n.s. |
| ENSG00000214655 | 45.5675 | -0.313270366 | 0.325592 | -0.962157 | 0.335971 | 0.784195004 | ZSWIM8 | protein_coding | 54.46 | -0.03307381 | 0.03307381 | ENSG00000214655 | n.s. |
| ENSG00000103343 | 148.2736 | -0.277447953 | 0.268173 | -1.034586 | 0.300862 | 0.760115318 | ZNF174 | protein_coding | 46.45 | -0.033049743 | 0.033049743 | ENSG00000103343 | n.s. |
| ENSG00000147650 | 1424.316 | -0.288582424 | 0.288334 | -1.000862 | 0.316893 | 0.768392151 | LRP12 | protein_coding | 35.69 | -0.033018758 | 0.033018758 | ENSG00000147650 | n.s. |
| ENSG00000106524 | 201.9923 | 0.290034335 | 0.290614 | 0.9980056 | 0.318277 | 0.769456789 | ANKMY2 | protein_coding | 38.01 | 0.03301048 | 0.03301048 | ENSG00000106524 | n.s. |
| ENSG00000169249 | 184.0608 | -0.277962042 | 0.270289 | -1.028387 | 0.303768 | 0.760781413 | ZRSR2 | protein_coding | 41.5 | -0.033005242 | 0.033005242 | ENSG00000169249 | n.s. |
| ENSG00000201847 | 5.811284 | -0.449372737 | 0.58035 | -0.774314 | 0.438745 | 0.84443611 | SNORD31B | snoRNA | 44.93 | -0.03299888 | 0.03299888 | ENSG00000201847 | n.s. |
| ENSG00000201710 | 7.708726 | 0.405129793 | 0.494807 | 0.8187638 | 0.412921 | 0.829007487 | | snoRNA | 36 | 0.032994397 | 0.032994397 | ENSG00000201710 | n.s. |
| ENSG00000144366 | 2750.723 | -0.27458883 | 0.264249 | -1.039127 | 0.298745 | 0.758335061 | GULP1 | protein_coding | 35.36 | -0.03298879 | 0.03298879 | ENSG00000144366 | n.s. |
| ENSG00000263750 | 7.078636 | 0.526452159 | 0.742691 | 0.7088439 | 0.478421 | 0.865660698 | MIR548AV | miRNA | 38.71 | 0.032983438 | 0.032983438 | ENSG00000263750 | n.s. |
| ENSG00000143793 | 381.8797 | -0.283863703 | 0.280446 | -1.012188 | 0.311448 | 0.765353944 | C1orf35 | protein_coding | 60.86 | -0.032967271 | 0.032967271 | ENSG00000143793 | n.s. |
| ENSG00000196562 | 2372.158 | -0.27749383 | 0.269555 | -1.02945 | 0.303268 | 0.760781413 | SULF2 | protein_coding | 50.48 | -0.032949647 | 0.032949647 | ENSG00000196562 | n.s. |
| ENSG00000114062 | 4258.516 | -0.283289414 | 0.279702 | -1.012824 | 0.311144 | 0.765111329 | UBE3A | protein_coding | 36.21 | -0.032939581 | 0.032939581 | ENSG00000114062 | n.s. |
| ENSG00000138111 | 118.8895 | -0.279014682 | 0.273727 | -1.019316 | 0.308053 | 0.761997697 | MFSD13A | protein_coding | 53.46 | -0.032936662 | 0.032936662 | ENSG00000138111 | n.s. |
| ENSG00000224817 | 11.58954 | -0.390580927 | 0.465764 | -0.838582 | 0.401704 | 0.823540971 | | lincRNA | 39.76 | -0.032931749 | 0.032931749 | ENSG00000224817 | n.s. |
| ENSG00000199875 | 6.450872 | -0.61233149 | 0.943634 | -0.648908 | 0.516398 | 0.883546124 | | misc_RNA | 37.17 | -0.032925538 | 0.032925538 | ENSG00000199875 | n.s. |
| ENSG00000136717 | 604.9785 | -0.278852364 | 0.273516 | -1.019509 | 0.307961 | 0.761997697 | BIN1 | protein_coding | 59.66 | -0.032917501 | 0.032917501 | ENSG00000136717 | n.s. |
| ENSG00000270808 | 15.99091 | -0.366321085 | 0.423036 | -0.865934 | 0.386527 | 0.813136725 | | processed_pseudogene | 58.04 | -0.032908976 | 0.032908976 | ENSG00000270808 | n.s. |
| ENSG00000273727 | 7.845249 | -0.50614798 | 0.695675 | -0.727563 | 0.466881 | 0.861088114 | | snRNA | 45.68 | -0.032875529 | 0.032875529 | ENSG00000273727 | n.s. |
| ENSG00000100578 | 893.4439 | -0.256151419 | 0.237747 | -1.077413 | 0.281296 | 0.744242186 | KIAA0586 | protein_coding | 37.04 | -0.032860568 | 0.032860568 | ENSG00000100578 | n.s. |
| ENSG00000125977 | 2855.103 | -0.288391532 | 0.288713 | -0.998887 | 0.31785 | 0.769236954 | EIF2S2 | protein_coding | 42.27 | -0.032859291 | 0.032859291 | ENSG00000125977 | n.s. |
| ENSG00000101473 | 178.9342 | -0.277941887 | 0.271947 | -1.022044 | 0.30676 | 0.761997697 | ACOT8 | protein_coding | 49.06 | -0.032810023 | 0.032810023 | ENSG00000101473 | n.s. |
| ENSG00000006740 | 53.81614 | -0.289850187 | 0.291474 | -0.99443 | 0.320014 | 0.770589543 | ARHGAP44 | protein_coding | 42.68 | -0.032804342 | 0.032804342 | ENSG00000006740 | n.s. |
| ENSG00000176890 | 1908.45 | -0.267381488 | 0.254953 | -1.048746 | 0.294295 | 0.753915388 | TYMS | protein_coding | 45.77 | -0.032801664 | 0.032801664 | ENSG00000176890 | n.s. |
| ENSG00000081386 | 1030.2 | -0.277810892 | 0.271739 | -1.022346 | 0.306617 | 0.761997697 | ZNF510 | protein_coding | 38.98 | -0.032794559 | 0.032794559 | ENSG00000081386 | n.s. |
| ENSG00000262074 | 492.0616 | -0.243608793 | 0.21781 | -1.118447 | 0.263376 | 0.733524057 | SNORD3B-2 | snoRNA | 56.13 | -0.032786245 | 0.032786245 | ENSG00000262074 | n.s. |
| ENSG00000134684 | 1295.145 | -0.277577663 | 0.271829 | -1.021149 | 0.307184 | 0.761997697 | YARS | protein_coding | 46.35 | -0.032767028 | 0.032767028 | ENSG00000134684 | n.s. |
| ENSG00000181666 | 382.6086 | -0.25609798 | 0.238636 | -1.073175 | 0.283193 | 0.745049657 | HKR1 | protein_coding | 43.83 | -0.032733107 | 0.032733107 | ENSG00000181666 | n.s. |
| ENSG00000078177 | 1845.697 | -0.275115236 | 0.266162 | -1.03364 | 0.301304 | 0.760388234 | N4BP2 | protein_coding | 40.52 | -0.032728977 | 0.032728977 | ENSG00000078177 | n.s. |

| | | | | | | | | | | | | | |
|---|---|---|---|---|---|---|---|---|---|---|---|---|---|
| ENSG00000159202 | 1205.237 | -0.276989641 | 0.271646 | -1.01967 | 0.307885 | 0.761997697 | UBE2Z | protein_coding | 47.44 | -0.032697614 | 0.032697614 | ENSG00000159202 | n.s. |
| ENSG00000197601 | 2767.053 | -0.276961558 | 0.270281 | -1.024717 | 0.305497 | 0.761997697 | FAR1 | protein_coding | 36.03 | -0.032694299 | 0.032694299 | ENSG00000197601 | n.s. |
| ENSG00000135736 | 137.6557 | 0.283517548 | 0.281279 | 1.00796 | 0.313474 | 0.766827179 | CCDC102A | protein_coding | 56.06 | 0.032690283 | 0.032690283 | ENSG00000135736 | n.s. |
| ENSG00000245281 | 42.97901 | -0.311984488 | 0.326634 | -0.955151 | 0.339501 | 0.785717908 | | antisense | 40.7 | -0.03267518 | 0.03267518 | ENSG00000245281 | n.s. |
| ENSG00000180425 | 239.7859 | 0.253581068 | 0.234349 | 1.0820649 | 0.279224 | 0.743390786 | C11orf71 | protein_coding | 39.34 | 0.032656887 | 0.032656887 | ENSG00000180425 | n.s. |
| ENSG00000272931 | 40.86291 | 0.304379922 | 0.314769 | 0.9669953 | 0.333546 | 0.781172958 | | lincRNA | 49.93 | 0.032645599 | 0.032645599 | ENSG00000272931 | n.s. |
| ENSG00000173950 | 761.1496 | -0.283882957 | 0.28293 | -1.003368 | 0.315683 | 0.767425784 | XXYLT1 | protein_coding | 48.89 | -0.032636211 | 0.032636211 | ENSG00000173950 | n.s. |
| ENSG00000161904 | 631.693 | -0.276453624 | 0.271077 | -1.019832 | 0.307808 | 0.761997697 | LEMD2 | protein_coding | 53.18 | -0.032634339 | 0.032634339 | ENSG00000161904 | n.s. |
| ENSG00000106261 | 1805.477 | -0.263334851 | 0.249597 | -1.055039 | 0.291408 | 0.751763371 | ZKSCAN1 | protein_coding | 44.83 | -0.032632149 | 0.032632149 | ENSG00000106261 | n.s. |
| ENSG00000089916 | 2394.997 | -0.264267185 | 0.250916 | -1.053209 | 0.292245 | 0.752608927 | GPATCH2L | protein_coding | 39.76 | -0.032618666 | 0.032618666 | ENSG00000089916 | n.s. |
| ENSG00000110844 | 76.34576 | -0.282652348 | 0.280322 | -1.008311 | 0.313305 | 0.766768387 | PRPF40B | protein_coding | 47.42 | -0.032599935 | 0.032599935 | ENSG00000110844 | n.s. |
| ENSG00000107862 | 1052.056 | -0.248411872 | 0.22745 | -1.092159 | 0.274763 | 0.739386355 | GBF1 | protein_coding | 43.61 | -0.032573893 | 0.032573893 | ENSG00000107862 | n.s. |
| ENSG00000257524 | 66.41518 | -0.332064126 | 0.36341 | -0.913745 | 0.360851 | 0.797857548 | | processed_transcript | 56.77 | -0.03256707 | 0.03256707 | ENSG00000257524 | n.s. |
| ENSG00000136811 | 588.7296 | -0.275684901 | 0.27004 | -1.020904 | 0.3073 | 0.761997697 | ODF2 | protein_coding | 49.1 | -0.032543594 | 0.032543594 | ENSG00000136811 | n.s. |
| ENSG00000201944 | 80.30487 | -0.275657973 | 0.269868 | -1.021457 | 0.307038 | 0.761997697 | | snoRNA | 35.61 | -0.032540415 | 0.032540415 | ENSG00000201944 | n.s. |
| ENSG00000276645 | 16.45278 | -0.355973572 | 0.406846 | -0.87496 | 0.381596 | 0.81032471 | | misc_RNA | 49.86 | -0.032514953 | 0.032514953 | ENSG00000276645 | n.s. |
| ENSG00000216809 | 235.8831 | -0.280825466 | 0.27798 | -1.010238 | 0.312381 | 0.766004018 | | processed_pseudogene | 44.93 | -0.03251087 | 0.03251087 | ENSG00000216809 | n.s. |
| ENSG00000163214 | 1479.995 | -0.275321125 | 0.269922 | -1.020002 | 0.307728 | 0.761997697 | DHX57 | protein_coding | 41.54 | -0.032500651 | 0.032500651 | ENSG00000163214 | n.s. |
| ENSG00000109686 | 217.4407 | -0.276844437 | 0.272191 | -1.017096 | 0.309108 | 0.763204386 | SH3D19 | protein_coding | 40.79 | -0.032490226 | 0.032490226 | ENSG00000109686 | n.s. |
| ENSG00000072195 | 600.7532 | -0.279446261 | 0.27609 | -1.012158 | 0.311463 | 0.765353944 | SPEG | protein_coding | 57.68 | -0.032454239 | 0.032454239 | ENSG00000072195 | n.s. |
| ENSG00000277443 | 9436.813 | -0.255676547 | 0.240468 | -1.063246 | 0.28767 | 0.746611882 | MARCKS | protein_coding | 48.49 | -0.032446658 | 0.032446658 | ENSG00000277443 | n.s. |
| ENSG00000011523 | 1726.898 | -0.284404215 | 0.284639 | -0.999176 | 0.31771 | 0.769031698 | CEP68 | protein_coding | 47.81 | -0.032437939 | 0.032437939 | ENSG00000011523 | n.s. |
| ENSG00000242588 | 76.79593 | -0.273108781 | 0.265291 | -1.029468 | 0.30326 | 0.760781413 | | processed_transcript | 41.32 | -0.032428966 | 0.032428966 | ENSG00000242588 | n.s. |
| ENSG00000219395 | 64.99493 | -0.316773968 | 0.337967 | -0.937293 | 0.348608 | 0.790017511 | HSPA8P15 | processed_pseudogene | 47.04 | -0.032426023 | 0.032426023 | ENSG00000219395 | n.s. |
| ENSG00000086015 | 282.6483 | -0.25546923 | 0.239677 | -1.065888 | 0.286474 | 0.746611882 | MAST2 | protein_coding | 41.32 | -0.032420349 | 0.032420349 | ENSG00000086015 | n.s. |
| ENSG00000235244 | 109.7463 | -0.274549453 | 0.269357 | -1.019276 | 0.308072 | 0.761997697 | DANT2 | lincRNA | 42.01 | -0.032409558 | 0.032409558 | ENSG00000235244 | n.s. |
| ENSG00000259719 | 29.40513 | -0.308258344 | 0.32215 | -0.956879 | 0.338628 | 0.784993146 | LINC02284 | lincRNA | 42.62 | -0.032408474 | 0.032408474 | ENSG00000259719 | n.s. |
| ENSG00000259232 | 27.9208 | -0.310960117 | 0.327651 | -0.94906 | 0.34259 | 0.786719908 | | processed_pseudogene | 53.87 | -0.032395781 | 0.032395781 | ENSG00000259232 | n.s. |
| ENSG00000267575 | 401.4727 | -0.245439964 | 0.22344 | -1.098461 | 0.272003 | 0.737938074 | | lincRNA | 43.58 | -0.032393186 | 0.032393186 | ENSG00000267575 | n.s. |
| ENSG00000169564 | 1294.496 | -0.245354544 | 0.223529 | -1.097642 | 0.272361 | 0.738113065 | PCBP1 | protein_coding | 55.2 | -0.032356648 | 0.032356648 | ENSG00000169564 | n.s. |
| ENSG00000276823 | 7.214556 | -0.502458667 | 0.694348 | -0.723641 | 0.469286 | 0.862205928 | | misc_RNA | 47.4 | -0.032352809 | 0.032352809 | ENSG00000276823 | n.s. |
| ENSG00000137770 | 2872.191 | -0.27246276 | 0.264464 | -1.030245 | 0.302895 | 0.760781413 | CTDSPL2 | protein_coding | 40.06 | -0.032352257 | 0.032352257 | ENSG00000137770 | n.s. |

| | | | | | | | | | | | | | |
|---|---|---|---|---|---|---|---|---|---|---|---|---|---|
| ENSG00000142949 | 693.505 | -0.262361795 | 0.249311 | -1.052348 | 0.29264 | 0.752972779 | PTPRF | protein_coding | 55.69 | -0.03232841 | 0.03232841 | ENSG00000142949 | n.s. |
| ENSG00000143469 | 1974.026 | -0.283690711 | 0.284043 | -0.998758 | 0.317912 | 0.769254053 | SYT14 | protein_coding | 36.49 | -0.032320941 | 0.032320941 | ENSG00000143469 | n.s. |
| ENSG00000102098 | 631.9806 | -0.278470726 | 0.275346 | -1.01135 | 0.311849 | 0.765516281 | SCML2 | protein_coding | 39.57 | -0.032315294 | 0.032315294 | ENSG00000102098 | n.s. |
| ENSG00000136603 | 2635.609 | -0.273744808 | 0.267378 | -1.02381 | 0.305925 | 0.761997697 | SKIL | protein_coding | 39.63 | -0.032314573 | 0.032314573 | ENSG00000136603 | n.s. |
| ENSG00000125458 | 935.051 | -0.263334488 | 0.251097 | -1.048737 | 0.294299 | 0.753915388 | NT5C | protein_coding | 65.69 | -0.032305188 | 0.032305188 | ENSG00000125458 | n.s. |
| ENSG00000078369 | 1984.025 | -0.271903744 | 0.263946 | -1.030148 | 0.30294 | 0.760781413 | GNB1 | protein_coding | 46.7 | -0.032285879 | 0.032285879 | ENSG00000078369 | n.s. |
| ENSG00000254293 | 257.7542 | 0.239885348 | 0.214637 | 1.1176351 | 0.263723 | 0.733524057 | | lincRNA | 47.45 | 0.032285123 | 0.032285123 | ENSG00000254293 | n.s. |
| ENSG00000249020 | 42.93444 | -0.306500951 | 0.319555 | -0.959149 | 0.337483 | 0.78475552 | SNORA58 | snoRNA | 48.91 | -0.032264013 | 0.032264013 | ENSG00000249020 | n.s. |
| ENSG00000149150 | 117.8448 | -0.281929785 | 0.281753 | -1.000628 | 0.317007 | 0.768398703 | SLC43A1 | protein_coding | 50.69 | -0.032256539 | 0.032256539 | ENSG00000149150 | n.s. |
| ENSG00000082497 | 600.1741 | -0.265216778 | 0.25404 | -1.043997 | 0.296487 | 0.755765348 | SERTAD4 | protein_coding | 43.15 | -0.032253815 | 0.032253815 | ENSG00000082497 | n.s. |
| ENSG00000134815 | 326.1368 | -0.261999142 | 0.24914 | -1.051613 | 0.292977 | 0.753300243 | DHX34 | protein_coding | 54.23 | -0.03223425 | 0.03223425 | ENSG00000134815 | n.s. |
| ENSG00000267904 | 50.18375 | -0.308313129 | 0.323839 | -0.952057 | 0.341068 | 0.786054116 | | sense_intronic | 47.01 | -0.032233383 | 0.032233383 | ENSG00000267904 | n.s. |
| ENSG00000070495 | 503.0947 | -0.263490878 | 0.251826 | -1.046319 | 0.295414 | 0.75457253 | JMJD6 | protein_coding | 48.83 | -0.032224673 | 0.032224673 | ENSG00000070495 | n.s. |
| ENSG00000198952 | 2373.595 | -0.264765471 | 0.253635 | -1.043883 | 0.29654 | 0.755765348 | SMG5 | protein_coding | 49.82 | -0.03219893 | 0.03219893 | ENSG00000198952 | n.s. |
| ENSG00000198780 | 908.8759 | -0.270586411 | 0.261874 | -1.033268 | 0.301479 | 0.76041019 | FAM169A | protein_coding | 38.67 | -0.032186814 | 0.032186814 | ENSG00000198780 | n.s. |
| ENSG00000114383 | 234.0251 | -0.295247029 | 0.300958 | -0.981024 | 0.326581 | 0.778045194 | TUSC2 | protein_coding | 59.39 | -0.032180502 | 0.032180502 | ENSG00000114383 | n.s. |
| ENSG00000280440 | 13.55817 | 0.456398786 | 0.60061 | 0.7598924 | 0.447319 | 0.850164607 | | TEC | 43.35 | 0.032174736 | 0.032174736 | ENSG00000280440 | n.s. |
| ENSG00000230815 | 22.51201 | -0.346173577 | 0.389134 | -0.889601 | 0.37368 | 0.807377464 | | sense_intronic | 39.44 | -0.032167618 | 0.032167618 | ENSG00000230815 | n.s. |
| ENSG00000180096 | 17.41596 | -0.38625607 | 0.465484 | -0.829795 | 0.406655 | 0.825583527 | 1-Sep | protein_coding | 52.75 | -0.032151562 | 0.032151562 | ENSG00000180096 | n.s. |
| ENSG00000148484 | 678.8321 | -0.279539217 | 0.278442 | -1.003942 | 0.315406 | 0.767425784 | RSU1 | protein_coding | 41.15 | -0.032136838 | 0.032136838 | ENSG00000148484 | n.s. |
| ENSG00000196381 | 321.0551 | 0.318870549 | 0.343582 | 0.9280763 | 0.353368 | 0.792950339 | ZNF781 | protein_coding | 41.97 | 0.032127487 | 0.032127487 | ENSG00000196381 | n.s. |
| ENSG00000124198 | 1160.779 | -0.270559282 | 0.26304 | -1.028587 | 0.303674 | 0.760781413 | ARFGEF2 | protein_coding | 43.19 | -0.032126238 | 0.032126238 | ENSG00000124198 | n.s. |
| ENSG00000157837 | 441.2306 | -0.26569276 | 0.255153 | -1.041307 | 0.297733 | 0.757001215 | SPPL3 | protein_coding | 40.88 | -0.032123164 | 0.032123164 | ENSG00000157837 | n.s. |
| ENSG00000182796 | 23.4547 | -0.351421054 | 0.401 | -0.876361 | 0.380834 | 0.810319533 | TMEM198B | anscribed_unitary_pseudoge | 54.51 | -0.032100097 | 0.032100097 | ENSG00000182796 | n.s. |
| ENSG00000171302 | 604.1001 | -0.278365382 | 0.276462 | -1.006885 | 0.31399 | 0.766827179 | CANT1 | protein_coding | 57.12 | -0.032096225 | 0.032096225 | ENSG00000171302 | n.s. |
| ENSG00000044574 | 1699.158 | 0.252754962 | 0.237108 | 1.0659896 | 0.286428 | 0.746611882 | HSPA5 | protein_coding | 43.98 | 0.032075894 | 0.032075894 | ENSG00000044574 | n.s. |
| ENSG00000263916 | 31.3965 | -0.320983639 | 0.347065 | -0.924851 | 0.355043 | 0.794524781 | | sense_intronic | 46.34 | -0.032063875 | 0.032063875 | ENSG00000263916 | n.s. |
| ENSG00000064547 | 151.2396 | -0.26998775 | 0.262052 | -1.030283 | 0.302877 | 0.760781413 | LPAR2 | protein_coding | 58.96 | -0.032058374 | 0.032058374 | ENSG00000064547 | n.s. |
| ENSG00000231908 | 20.92662 | 0.362229358 | 0.423992 | 0.8543312 | 0.392921 | 0.815710819 | IDH1-AS1 | antisense | 52.09 | 0.032044177 | 0.032044177 | ENSG00000231908 | n.s. |
| ENSG00000108861 | 390.6195 | -0.279471882 | 0.278841 | -1.002263 | 0.316216 | 0.768128006 | DUSP3 | protein_coding | 50.91 | -0.032018087 | 0.032018087 | ENSG00000108861 | n.s. |
| ENSG00000143549 | 3267.271 | -0.277967834 | 0.276432 | -1.005557 | 0.314629 | 0.767310429 | TPM3 | protein_coding | 45.14 | -0.031974334 | 0.031974334 | ENSG00000143549 | n.s. |
| ENSG00000141965 | 1127.087 | -0.230300773 | 0.202988 | -1.134553 | 0.256562 | 0.726483516 | FEM1A | protein_coding | 55.19 | -0.031959814 | 0.031959814 | ENSG00000141965 | n.s. |

| | | | | | | | | | | | | | |
|---|---|---|---|---|---|---|---|---|---|---|---|---|---|
| ENSG00000196455 | 1152.513 | -0.268415882 | 0.259536 | -1.034214 | 0.301036 | 0.760228309 | PIK3R4 | protein_coding | 36.67 | -0.031956511 | 0.031956511 | ENSG00000196455 | n.s. |
| ENSG00000261202 | 50.49613 | 0.314435866 | 0.337289 | 0.9322433 | 0.351211 | 0.791580243 | | antisense | 46.62 | 0.031916829 | 0.031916829 | ENSG00000261202 | n.s. |
| ENSG00000071189 | 1398.602 | -0.275328259 | 0.272363 | -1.010886 | 0.312071 | 0.765767796 | SNX13 | protein_coding | 35.09 | -0.031911344 | 0.031911344 | ENSG00000071189 | n.s. |
| ENSG00000163743 | 684.9655 | -0.280590609 | 0.281483 | -0.996831 | 0.318846 | 0.769632078 | RCHY1 | protein_coding | 34.99 | -0.031907877 | 0.031907877 | ENSG00000163743 | n.s. |
| ENSG00000265112 | 167.4964 | 0.253061478 | 0.23877 | 1.0598568 | 0.28921 | 0.748034104 | MIR3153 | miRNA | 40.24 | 0.031905637 | 0.031905637 | ENSG00000265112 | n.s. |
| ENSG00000254708 | 30.34511 | -0.346967767 | 0.393569 | -0.881593 | 0.377997 | 0.809223062 | | processed_pseudogene | 39.73 | -0.031897354 | 0.031897354 | ENSG00000254708 | n.s. |
| ENSG00000119139 | 377.4626 | -0.266682425 | 0.257488 | -1.035707 | 0.300339 | 0.759327693 | TJP2 | protein_coding | 43.45 | -0.03188742 | 0.03188742 | ENSG00000119139 | n.s. |
| ENSG00000014919 | 692.8269 | -0.277152787 | 0.275539 | -1.005855 | 0.314485 | 0.767300529 | COX15 | protein_coding | 42.42 | -0.031882133 | 0.031882133 | ENSG00000014919 | n.s. |
| ENSG00000279192 | 628.7774 | -0.268227035 | 0.260537 | -1.029518 | 0.303236 | 0.760781413 | PWAR5 | TEC | 35.82 | -0.031849307 | 0.031849307 | ENSG00000279192 | n.s. |
| ENSG00000198795 | 729.4442 | -0.268196956 | 0.260278 | -1.030426 | 0.30281 | 0.760781413 | ZNF521 | protein_coding | 38.07 | -0.031845735 | 0.031845735 | ENSG00000198795 | n.s. |
| ENSG00000236609 | 179.9902 | -0.277909845 | 0.277387 | -1.001887 | 0.316398 | 0.768128006 | ZNF853 | protein_coding | 57.53 | -0.03183913 | 0.03183913 | ENSG00000236609 | n.s. |
| ENSG00000237125 | 12160.96 | -0.247230494 | 0.228257 | -1.083121 | 0.278755 | 0.743390786 | HAND2-AS1 | antisense | 41 | -0.031839042 | 0.031839042 | ENSG00000237125 | n.s. |
| ENSG00000197879 | 277.4447 | -0.268135464 | 0.260526 | -1.029207 | 0.303382 | 0.760781413 | MYO1C | protein_coding | 60.07 | -0.031838433 | 0.031838433 | ENSG00000197879 | n.s. |
| ENSG00000172992 | 687.0701 | -0.25606013 | 0.24243 | -1.056223 | 0.290866 | 0.751040718 | DCAKD | protein_coding | 51.25 | -0.031837625 | 0.031837625 | ENSG00000172992 | n.s. |
| ENSG00000142867 | 473.8101 | 0.26768683 | 0.25965 | 1.0309544 | 0.302562 | 0.760781413 | BCL10 | protein_coding | 40.57 | 0.031785163 | 0.031785163 | ENSG00000142867 | n.s. |
| ENSG00000130772 | 102.967 | -0.269226675 | 0.262509 | -1.02559 | 0.305085 | 0.761997697 | MED18 | protein_coding | 46.05 | -0.031781224 | 0.031781224 | ENSG00000130772 | n.s. |
| ENSG00000175376 | 375.7347 | -0.239981548 | 0.218117 | -1.100243 | 0.271226 | 0.737359702 | EIF1AD | protein_coding | 48.54 | -0.031754503 | 0.031754503 | ENSG00000175376 | n.s. |
| ENSG00000003393 | 1258.944 | -0.264167529 | 0.254429 | -1.038276 | 0.299142 | 0.758335061 | ALS2 | protein_coding | 38.06 | -0.031736787 | 0.031736787 | ENSG00000003393 | n.s. |
| ENSG00000158079 | 594.9783 | -0.293287976 | 0.299959 | -0.97776 | 0.328193 | 0.779459027 | PTPDC1 | protein_coding | 39.39 | -0.031735727 | 0.031735727 | ENSG00000158079 | n.s. |
| ENSG00000221676 | 9.594481 | 0.388720315 | 0.473764 | 0.8204927 | 0.411935 | 0.828691651 | RNU6ATAC | snRNA | 50 | 0.031722313 | 0.031722313 | ENSG00000221676 | n.s. |
| ENSG00000173039 | 260.3292 | -0.264011365 | 0.254198 | -1.038605 | 0.298988 | 0.758335061 | RELA | protein_coding | 55.62 | -0.031718026 | 0.031718026 | ENSG00000173039 | n.s. |
| ENSG00000233276 | 1171.756 | -0.302819418 | 0.317491 | -0.953789 | 0.340191 | 0.785979841 | GPX1 | protein_coding | 63.79 | -0.031671457 | 0.031671457 | ENSG00000233276 | n.s. |
| ENSG00000133789 | 551.1467 | -0.268248233 | 0.262797 | -1.020742 | 0.307376 | 0.761997697 | SWAP70 | protein_coding | 39.97 | -0.031665723 | 0.031665723 | ENSG00000133789 | n.s. |
| ENSG00000176623 | 619.759 | -0.275008215 | 0.27362 | -1.005074 | 0.314861 | 0.767364665 | RMDN1 | protein_coding | 38.38 | -0.03162545 | 0.03162545 | ENSG00000176623 | n.s. |
| ENSG00000067369 | 3052.149 | -0.267862072 | 0.261979 | -1.022456 | 0.306565 | 0.761997697 | TP53BP1 | protein_coding | 41.72 | -0.031620138 | 0.031620138 | ENSG00000067369 | n.s. |
| ENSG00000140575 | 2080.839 | -0.246784257 | 0.22934 | -1.076065 | 0.281898 | 0.744567264 | IQGAP1 | protein_coding | 40.77 | -0.031612092 | 0.031612092 | ENSG00000140575 | n.s. |
| ENSG00000171303 | 277.3276 | -0.28422787 | 0.287397 | -0.988973 | 0.322676 | 0.774129273 | KCNK3 | protein_coding | 53.76 | -0.031602305 | 0.031602305 | ENSG00000171303 | n.s. |
| ENSG00000139675 | 240.5322 | -0.298937913 | 0.310555 | -0.962592 | 0.335752 | 0.784079552 | HNRNPA1L2 | protein_coding | 41.98 | -0.031579763 | 0.031579763 | ENSG00000139675 | n.s. |
| ENSG00000177873 | 216.7731 | -0.238526356 | 0.216874 | -1.099836 | 0.271403 | 0.737359702 | ZNF619 | protein_coding | 45.23 | -0.031561951 | 0.031561951 | ENSG00000177873 | n.s. |
| ENSG00000178252 | 2387.289 | -0.271948034 | 0.268889 | -1.011376 | 0.311836 | 0.765516281 | WDR6 | protein_coding | 56.28 | -0.031558364 | 0.031558364 | ENSG00000178252 | n.s. |
| ENSG00000144057 | 662.2928 | -0.263837686 | 0.254741 | -1.035708 | 0.300338 | 0.759327693 | ST6GAL2 | protein_coding | 41.34 | -0.031547273 | 0.031547273 | ENSG00000144057 | n.s. |
| ENSG00000145220 | 664.3217 | -0.267008701 | 0.261071 | -1.022742 | 0.30643 | 0.761997697 | LYAR | protein_coding | 44.02 | -0.0315194 | 0.0315194 | ENSG00000145220 | n.s. |

| | | | | | | | | | | | | | |
|---|---|---|---|---|---|---|---|---|---|---|---|---|---|
| ENSG00000153933 | 1969.174 | -0.265200746 | 0.257961 | -1.028065 | 0.303919 | 0.760781413 | DGKE | protein_coding | 38.37 | -0.031489965 | 0.031489965 | ENSG00000153933 | n.s. |
| ENSG00000171763 | 900.768 | -0.256824035 | 0.245106 | -1.047809 | 0.294727 | 0.754039228 | SPATA5L1 | protein_coding | 41.47 | -0.031488183 | 0.031488183 | ENSG00000171763 | n.s. |
| ENSG00000197102 | 7663.815 | -0.256653413 | 0.244677 | -1.048948 | 0.294202 | 0.753915388 | DYNC1H1 | protein_coding | 46.94 | -0.031485571 | 0.031485571 | ENSG00000197102 | n.s. |
| ENSG00000076108 | 77.86525 | -0.275107461 | 0.274919 | -1.000684 | 0.31698 | 0.768398703 | BAZ2A | protein_coding | 48.64 | -0.031475974 | 0.031475974 | ENSG00000076108 | n.s. |
| ENSG00000124356 | 1128.342 | -0.265025256 | 0.257748 | -1.028234 | 0.30384 | 0.760781413 | STAMBP | protein_coding | 42.1 | -0.031469127 | 0.031469127 | ENSG00000124356 | n.s. |
| ENSG00000131153 | 1222.394 | -0.266505164 | 0.261331 | -1.019797 | 0.307825 | 0.761997697 | GINS2 | protein_coding | 49.77 | -0.03145996 | 0.03145996 | ENSG00000131153 | n.s. |
| ENSG00000199879 | 5.898558 | 0.498952302 | 0.695112 | 0.7178013 | 0.47288 | 0.864937704 | RNU1-120P | snRNA | 42.68 | 0.031441565 | 0.031441565 | ENSG00000199879 | n.s. |
| ENSG00000168522 | 1834 | -0.27443399 | 0.273898 | -1.001958 | 0.316364 | 0.768128006 | FNTA | protein_coding | 42.58 | -0.031440914 | 0.031440914 | ENSG00000168522 | n.s. |
| ENSG00000144712 | 258.4278 | -0.264741824 | 0.257367 | -1.028655 | 0.303642 | 0.760781413 | CAND2 | protein_coding | 50.87 | -0.031435472 | 0.031435472 | ENSG00000144712 | n.s. |
| ENSG00000138794 | 79.92621 | -0.304499037 | 0.322247 | -0.944925 | 0.344697 | 0.788437824 | CASP6 | protein_coding | 44.61 | -0.031434212 | 0.031434212 | ENSG00000138794 | n.s. |
| ENSG00000283563 | 566.644 | -0.27422282 | 0.273963 | -1.000947 | 0.316853 | 0.768392151 | | protein_coding | 37.06 | -0.031375774 | 0.031375774 | ENSG00000283563 | n.s. |
| ENSG00000198841 | 551.9778 | 0.244559632 | 0.22662 | 1.0791614 | 0.280516 | 0.744242186 | KTI12 | protein_coding | 55.37 | 0.031373508 | 0.031373508 | ENSG00000198841 | n.s. |
| ENSG00000129657 | 159.7959 | -0.260588766 | 0.250641 | -1.039691 | 0.298484 | 0.757940321 | SEC14L1 | protein_coding | 46.75 | -0.031365764 | 0.031365764 | ENSG00000129657 | n.s. |
| ENSG00000111785 | 129.4069 | -0.276020621 | 0.277144 | -0.995945 | 0.319277 | 0.769994426 | RIC8B | protein_coding | 38.48 | -0.031331768 | 0.031331768 | ENSG00000111785 | n.s. |
| ENSG00000162980 | 1779.753 | -0.283206984 | 0.287105 | -0.986422 | 0.323926 | 0.775183815 | ARL5A | protein_coding | 37.35 | -0.031321362 | 0.031321362 | ENSG00000162980 | n.s. |
| ENSG00000117528 | 2033.528 | -0.269271687 | 0.265607 | -1.013798 | 0.310679 | 0.765048922 | ABCD3 | protein_coding | 37.04 | -0.031319203 | 0.031319203 | ENSG00000117528 | n.s. |
| ENSG00000242474 | 159.8662 | 0.249237925 | 0.235404 | 1.058768 | 0.289705 | 0.748759774 | | lincRNA | 48.7 | 0.031318613 | 0.031318613 | ENSG00000242474 | n.s. |
| ENSG00000236514 | 39.48281 | 0.352207036 | 0.408944 | 0.8612605 | 0.389095 | 0.814886242 | | antisense | 49.23 | 0.031312267 | 0.031312267 | ENSG00000236514 | n.s. |
| ENSG00000145780 | 616.2756 | -0.26342252 | 0.255059 | -1.03279 | 0.301702 | 0.760565055 | FEM1C | protein_coding | 35.62 | -0.031311358 | 0.031311358 | ENSG00000145780 | n.s. |
| ENSG00000198105 | 858.8942 | -0.268053457 | 0.264128 | -1.014863 | 0.310171 | 0.764194956 | ZNF248 | protein_coding | 39.14 | -0.031307527 | 0.031307527 | ENSG00000198105 | n.s. |
| ENSG00000152465 | 464.9426 | -0.271484276 | 0.269606 | -1.006966 | 0.313951 | 0.766827179 | NMT2 | protein_coding | 43.09 | -0.031302816 | 0.031302816 | ENSG00000152465 | n.s. |
| ENSG00000116120 | 1689.572 | -0.276560207 | 0.278069 | -0.994574 | 0.319944 | 0.770589543 | FARSB | protein_coding | 38.05 | -0.031300224 | 0.031300224 | ENSG00000116120 | n.s. |
| ENSG00000162227 | 321.0552 | -0.283627232 | 0.287945 | -0.985005 | 0.324621 | 0.775648559 | TAF6L | protein_coding | 51.76 | -0.031294014 | 0.031294014 | ENSG00000162227 | n.s. |
| ENSG00000196961 | 264.2854 | -0.269520957 | 0.26643 | -1.0116 | 0.311729 | 0.765516281 | AP2A1 | protein_coding | 54.57 | -0.031276713 | 0.031276713 | ENSG00000196961 | n.s. |
| ENSG00000225264 | 45.17783 | -0.297751759 | 0.311534 | -0.955761 | 0.339193 | 0.785318029 | ZNRF2P2 | ıscribed_processed_pseudog | 40.97 | -0.031250367 | 0.031250367 | ENSG00000225264 | n.s. |
| ENSG00000253764 | 26.95764 | -0.352936368 | 0.411802 | -0.857053 | 0.391415 | 0.815710819 | | lincRNA | 65.14 | -0.031222084 | 0.031222084 | ENSG00000253764 | n.s. |
| ENSG00000102910 | 1408.298 | -0.264396554 | 0.257777 | -1.025679 | 0.305043 | 0.761997697 | LONP2 | protein_coding | 39.56 | -0.031211046 | 0.031211046 | ENSG00000102910 | n.s. |
| ENSG00000085733 | 1695.331 | -0.262503535 | 0.25533 | -1.028096 | 0.303905 | 0.760781413 | CTTN | protein_coding | 52.81 | -0.031169698 | 0.031169698 | ENSG00000085733 | n.s. |
| ENSG00000207217 | 27.7363 | 0.308966425 | 0.332407 | 0.9294819 | 0.352639 | 0.79272638 | SNORA80D | snoRNA | 58.52 | 0.03116751 | 0.03116751 | ENSG00000207217 | n.s. |
| ENSG00000163939 | 3745.506 | -0.264017972 | 0.258616 | -1.020888 | 0.307307 | 0.761997697 | PBRM1 | protein_coding | 40.81 | -0.031166356 | 0.031166356 | ENSG00000163939 | n.s. |
| ENSG00000100629 | 1194.762 | -0.263839956 | 0.257274 | -1.025522 | 0.305117 | 0.761997697 | CEP128 | protein_coding | 36.62 | -0.031145342 | 0.031145342 | ENSG00000100629 | n.s. |
| ENSG00000214338 | 842.1677 | -0.222221573 | 0.195011 | -1.139536 | 0.25448 | 0.72418384 | SOGA3 | protein_coding | 39.44 | -0.031144614 | 0.031144614 | ENSG00000214338 | n.s. |

| | | | | | | | | | | | | | |
|---|---|---|---|---|---|---|---|---|---|---|---|---|---|
| ENSG00000104695 | 782.0405 | -0.266880374 | 0.263101 | -1.014366 | 0.310408 | 0.764517074 | PPP2CB | protein_coding | 41.95 | -0.031121671 | 0.031121671 | ENSG00000104695 | n.s. |
| ENSG00000129625 | 737.0932 | -0.270541388 | 0.269013 | -1.00568 | 0.31457 | 0.767300529 | REEP5 | protein_coding | 41.72 | -0.031121594 | 0.031121594 | ENSG00000129625 | n.s. |
| ENSG00000154920 | 703.8318 | -0.245224272 | 0.230523 | -1.063774 | 0.287431 | 0.746611882 | EME1 | protein_coding | 47.88 | -0.031120212 | 0.031120212 | ENSG00000154920 | n.s. |
| ENSG00000218690 | 28.1227 | -0.346281154 | 0.399519 | -0.866746 | 0.386081 | 0.813115862 | HIST1H2APS4 | cribed_unprocessed_pseudc | 51.4 | -0.031112519 | 0.031112519 | ENSG00000218690 | n.s. |
| ENSG00000108774 | 595.7008 | -0.279524386 | 0.282659 | -0.988911 | 0.322707 | 0.774129273 | RAB5C | protein_coding | 49.38 | -0.031079341 | 0.031079341 | ENSG00000108774 | n.s. |
| ENSG00000126003 | 320.4931 | -0.253452243 | 0.241884 | -1.047827 | 0.294718 | 0.754039228 | PLAGL2 | protein_coding | 48.56 | -0.031074781 | 0.031074781 | ENSG00000126003 | n.s. |
| ENSG00000139428 | 524.2809 | -0.289369589 | 0.299085 | -0.967517 | 0.333285 | 0.781097236 | MMAB | protein_coding | 52.27 | -0.031047881 | 0.031047881 | ENSG00000139428 | n.s. |
| ENSG00000148396 | 938.54 | -0.253178719 | 0.241483 | -1.048433 | 0.294439 | 0.754039228 | SEC16A | protein_coding | 52.76 | -0.031041245 | 0.031041245 | ENSG00000148396 | n.s. |
| ENSG00000138434 | 6400.809 | 0.255215 | 0.244515 | 1.0437616 | 0.296596 | 0.755765348 | SSFA2 | protein_coding | 35.71 | 0.031037468 | 0.031037468 | ENSG00000138434 | n.s. |
| ENSG00000213600 | 15.42105 | -0.406402292 | 0.513406 | -0.791581 | 0.428605 | 0.838757697 | | sense_intronic | 57.49 | -0.031034294 | 0.031034294 | ENSG00000213600 | n.s. |
| ENSG00000130177 | 872.7283 | -0.272568472 | 0.273258 | -0.997475 | 0.318534 | 0.769456789 | CDC16 | protein_coding | 41.32 | -0.031022589 | 0.031022589 | ENSG00000130177 | n.s. |
| ENSG00000197858 | 333.1626 | -0.265553186 | 0.26156 | -1.015268 | 0.309978 | 0.764194956 | GPAA1 | protein_coding | 64.3 | -0.031015506 | 0.031015506 | ENSG00000197858 | n.s. |
| ENSG00000167693 | 453.0906 | -0.268707607 | 0.266282 | -1.009108 | 0.312923 | 0.766641623 | NXN | protein_coding | 50.99 | -0.031010902 | 0.031010902 | ENSG00000167693 | n.s. |
| ENSG00000138346 | 1340.64 | -0.261140127 | 0.253823 | -1.028828 | 0.30356 | 0.760781413 | DNA2 | protein_coding | 42.26 | -0.031007806 | 0.031007806 | ENSG00000138346 | n.s. |
| ENSG00000272606 | 82.70925 | -0.287530387 | 0.294648 | -0.975843 | 0.329142 | 0.780141891 | | antisense | 46.45 | -0.031003367 | 0.031003367 | ENSG00000272606 | n.s. |
| ENSG00000102781 | 881.2909 | -0.260383871 | 0.251747 | -1.034306 | 0.300993 | 0.760228309 | KATNAL1 | protein_coding | 38.41 | -0.031000252 | 0.031000252 | ENSG00000102781 | n.s. |
| ENSG00000135451 | 957.1805 | -0.254971608 | 0.244464 | -1.04298 | 0.296957 | 0.756014066 | TROAP | protein_coding | 53.1 | -0.030971433 | 0.030971433 | ENSG00000135451 | n.s. |
| ENSG00000206989 | 32.31009 | -0.317995008 | 0.349644 | -0.909483 | 0.363095 | 0.7992181 | SNORD63 | snoRNA | 35.29 | -0.030951945 | 0.030951945 | ENSG00000206989 | n.s. |
| ENSG00000071243 | 673.2069 | 0.268712559 | 0.267652 | 1.0039622 | 0.315397 | 0.767425784 | ING3 | protein_coding | 35.55 | 0.030892167 | 0.030892167 | ENSG00000071243 | n.s. |
| ENSG00000010810 | 4178.599 | -0.26445777 | 0.260547 | -1.015011 | 0.310101 | 0.764194956 | FYN | protein_coding | 42.42 | -0.030887566 | 0.030887566 | ENSG00000010810 | n.s. |
| ENSG00000264546 | 149.6412 | -0.268626757 | 0.26759 | -1.003873 | 0.31544 | 0.767425784 | | sense_intronic | 42.53 | -0.030882303 | 0.030882303 | ENSG00000264546 | n.s. |
| ENSG00000140718 | 111.9823 | -0.271287975 | 0.272102 | -0.997007 | 0.318761 | 0.769569607 | FTO | protein_coding | 40.2 | -0.030859575 | 0.030859575 | ENSG00000140718 | n.s. |
| ENSG00000140365 | 1328.739 | -0.269189945 | 0.268761 | -1.001597 | 0.316538 | 0.768209163 | COMMD4 | protein_coding | 53.48 | -0.030827772 | 0.030827772 | ENSG00000140365 | n.s. |
| ENSG00000104825 | 197.7134 | -0.295712689 | 0.311386 | -0.949667 | 0.342281 | 0.786682723 | NFKBIB | protein_coding | 50.76 | -0.030813377 | 0.030813377 | ENSG00000104825 | n.s. |
| ENSG00000199593 | 13.59268 | 0.491420788 | 0.691579 | 0.7105776 | 0.477346 | 0.865616576 | SNORD114-14 | snoRNA | 46.67 | 0.030799521 | 0.030799521 | ENSG00000199593 | n.s. |
| ENSG00000091428 | 206.8503 | -0.269692427 | 0.269799 | -0.999605 | 0.317502 | 0.768883445 | RAPGEF4 | protein_coding | 40.56 | -0.030782556 | 0.030782556 | ENSG00000091428 | n.s. |
| ENSG00000253320 | 142.3369 | 0.25021281 | 0.237982 | 1.0513932 | 0.293078 | 0.753420469 | AZIN1-AS1 | antisense | 44.04 | 0.030766814 | 0.030766814 | ENSG00000253320 | n.s. |
| ENSG00000140400 | 1154.323 | -0.241226251 | 0.225547 | -1.069516 | 0.284837 | 0.745729255 | MAN2C1 | protein_coding | 57.24 | -0.030736764 | 0.030736764 | ENSG00000140400 | n.s. |
| ENSG00000111961 | 66.09303 | -0.285029118 | 0.294085 | -0.969208 | 0.332441 | 0.780141891 | SASH1 | protein_coding | 43.29 | -0.030733664 | 0.030733664 | ENSG00000111961 | n.s. |
| ENSG00000249771 | 62.34749 | 0.291833566 | 0.304171 | 0.95944 | 0.337337 | 0.78475552 | | lincRNA | 41.45 | 0.030720042 | 0.030720042 | ENSG00000249771 | n.s. |
| ENSG00000123552 | 744.3404 | -0.260207448 | 0.254035 | -1.024298 | 0.305694 | 0.761997697 | USP45 | protein_coding | 37.98 | -0.030716537 | 0.030716537 | ENSG00000123552 | n.s. |
| ENSG00000120251 | 1699.113 | 0.277906389 | 0.281826 | 0.9860923 | 0.324088 | 0.775304974 | GRIA2 | protein_coding | 34.59 | 0.030716279 | 0.030716279 | ENSG00000120251 | n.s. |

| | | | | | | | | | | | | | |
|---|---|---|---|---|---|---|---|---|---|---|---|---|---|
| ENSG00000198843 | 1211.632 | -0.284818335 | 0.293698 | -0.969766 | 0.332163 | 0.780141891 | SELENOT | protein_coding | 37.86 | -0.030710936 | 0.030710936 | ENSG00000198843 | n.s. |
| ENSG00000248275 | 162.4874 | -0.276825369 | 0.280134 | -0.988189 | 0.32306 | 0.774575942 | TRIM52-AS1 | processed_transcript | 49.04 | -0.030709898 | 0.030709898 | ENSG00000248275 | n.s. |
| ENSG00000145715 | 823.028 | -0.258461283 | 0.251382 | -1.028162 | 0.303874 | 0.760781413 | RASA1 | protein_coding | 35.77 | -0.03068972 | 0.03068972 | ENSG00000145715 | n.s. |
| ENSG00000166167 | 424.3459 | -0.271032152 | 0.272524 | -0.994525 | 0.319967 | 0.770589543 | BTRC | protein_coding | 38.96 | -0.030674576 | 0.030674576 | ENSG00000166167 | n.s. |
| ENSG00000184164 | 157.3539 | -0.276437018 | 0.279719 | -0.988269 | 0.323021 | 0.774575942 | CRELD2 | protein_coding | 59.36 | -0.030666816 | 0.030666816 | ENSG00000184164 | n.s. |
| ENSG00000109220 | 207.9961 | -0.272497943 | 0.274464 | -0.992839 | 0.320789 | 0.77178858 | CHIC2 | protein_coding | 36.75 | -0.030656469 | 0.030656469 | ENSG00000109220 | n.s. |
| ENSG00000155545 | 1239.226 | -0.277110791 | 0.280885 | -0.986562 | 0.323857 | 0.775183815 | MIER3 | protein_coding | 38.91 | -0.030647152 | 0.030647152 | ENSG00000155545 | n.s. |
| ENSG00000100014 | 2027.562 | -0.258070324 | 0.250604 | -1.029794 | 0.303107 | 0.760781413 | SPECC1L | protein_coding | 42.29 | -0.030643298 | 0.030643298 | ENSG00000100014 | n.s. |
| ENSG00000125912 | 577.6239 | -0.26021425 | 0.255559 | -1.018216 | 0.308575 | 0.762601855 | NCLN | protein_coding | 60.96 | -0.030627775 | 0.030627775 | ENSG00000125912 | n.s. |
| ENSG00000259519 | 7.74725 | -0.461719366 | 0.628234 | -0.734948 | 0.462371 | 0.858395846 | | antisense | 50.21 | -0.030617716 | 0.030617716 | ENSG00000259519 | n.s. |
| ENSG00000072401 | 811.6381 | -0.26625262 | 0.265326 | -1.003493 | 0.315623 | 0.767425784 | UBE2D1 | protein_coding | 36.05 | -0.030609363 | 0.030609363 | ENSG00000072401 | n.s. |
| ENSG00000137070 | 753.5704 | -0.267093429 | 0.266541 | -1.002074 | 0.316308 | 0.768128006 | IL11RA | protein_coding | 54.2 | -0.030599933 | 0.030599933 | ENSG00000137070 | n.s. |
| ENSG00000274884 | 5.595756 | 0.494362552 | 0.700701 | 0.7055255 | 0.480483 | 0.867184458 | | misc_RNA | 68.97 | 0.030595364 | 0.030595364 | ENSG00000274884 | n.s. |
| ENSG00000119820 | 2116.17 | -0.290881979 | 0.303708 | -0.957769 | 0.338179 | 0.784993146 | YIPF4 | protein_coding | 41.2 | -0.030581625 | 0.030581625 | ENSG00000119820 | n.s. |
| ENSG00000102225 | 813.4158 | -0.256763238 | 0.24834 | -1.033919 | 0.301174 | 0.760334613 | CDK16 | protein_coding | 54.42 | -0.030553603 | 0.030553603 | ENSG00000102225 | n.s. |
| ENSG00000268750 | 89.90478 | -0.264936033 | 0.262829 | -1.008015 | 0.313447 | 0.766827179 | | protein_coding | 47.11 | -0.030547788 | 0.030547788 | ENSG00000268750 | n.s. |
| ENSG00000168924 | 658.3242 | -0.247813471 | 0.235536 | -1.052126 | 0.292742 | 0.752972779 | LETM1 | protein_coding | 53.83 | -0.030535755 | 0.030535755 | ENSG00000168924 | n.s. |
| ENSG00000241973 | 1598.479 | -0.247087825 | 0.234475 | -1.05379 | 0.291979 | 0.75237733 | PI4KA | protein_coding | 47.81 | -0.030531234 | 0.030531234 | ENSG00000241973 | n.s. |
| ENSG00000284024 | 211.4299 | 0.260188067 | 0.255977 | 1.0164522 | 0.309414 | 0.763554114 | | protein_coding | 46.24 | 0.03048368 | 0.03048368 | ENSG00000284024 | n.s. |
| ENSG00000273199 | 19.00863 | 0.409874675 | 0.525029 | 0.7806707 | 0.434996 | 0.842611157 | | antisense | 67.35 | 0.030483525 | 0.030483525 | ENSG00000273199 | n.s. |
| ENSG00000283103 | 194.9382 | -0.291481277 | 0.305872 | -0.952953 | 0.340614 | 0.786054116 | | antisense | 48.25 | -0.030473654 | 0.030473654 | ENSG00000283103 | n.s. |
| ENSG00000215908 | 278.0657 | -0.26477717 | 0.263203 | -1.005982 | 0.314424 | 0.767300529 | CROCCP2 | cribed_unprocessed_pseudc | 52.04 | -0.03045851 | 0.03045851 | ENSG00000215908 | n.s. |
| ENSG00000204463 | 1124.668 | -0.257942083 | 0.252895 | -1.019956 | 0.307749 | 0.761997697 | BAG6 | protein_coding | 52.36 | -0.030449119 | 0.030449119 | ENSG00000204463 | n.s. |
| ENSG00000164978 | 126.228 | -0.28227655 | 0.290664 | -0.971144 | 0.331477 | 0.780141891 | NUDT2 | protein_coding | 46.38 | -0.030436865 | 0.030436865 | ENSG00000164978 | n.s. |
| ENSG00000228109 | 117.0127 | -0.279928097 | 0.285587 | -0.980184 | 0.326995 | 0.778648187 | MELTF-AS1 | antisense | 61.61 | -0.03041663 | 0.03041663 | ENSG00000228109 | n.s. |
| ENSG00000272661 | 20.52846 | -0.351841609 | 0.415403 | -0.846988 | 0.397002 | 0.819511304 | | antisense | 51.39 | -0.030414966 | 0.030414966 | ENSG00000272661 | n.s. |
| ENSG00000244462 | 3805.942 | -0.282053437 | 0.290467 | -0.971033 | 0.331532 | 0.780141891 | RBM12 | protein_coding | 39.8 | -0.030412807 | 0.030412807 | ENSG00000244462 | n.s. |
| ENSG00000119718 | 891.2287 | -0.257622448 | 0.252101 | -1.0219 | 0.306828 | 0.761997697 | EIF2B2 | protein_coding | 44.08 | -0.030411387 | 0.030411387 | ENSG00000119718 | n.s. |
| ENSG00000065613 | 2219.244 | -0.25806206 | 0.25351 | -1.017955 | 0.3087 | 0.762601855 | SLK | protein_coding | 36.55 | -0.030374457 | 0.030374457 | ENSG00000065613 | n.s. |
| ENSG00000047932 | 1919.262 | -0.28239782 | 0.291594 | -0.968461 | 0.332814 | 0.780753624 | GOPC | protein_coding | 36.66 | -0.03035381 | 0.03035381 | ENSG00000047932 | n.s. |
| ENSG00000119689 | 1464.528 | -0.255574623 | 0.247854 | -1.031151 | 0.30247 | 0.760781413 | DLST | protein_coding | 44.92 | -0.030346958 | 0.030346958 | ENSG00000119689 | n.s. |
| ENSG00000135617 | 165.6393 | -0.304164661 | 0.330015 | -0.92167 | 0.3567 | 0.794806874 | PRADC1 | protein_coding | 52.11 | -0.030336892 | 0.030336892 | ENSG00000135617 | n.s. |

| | | | | | | | | | | | | | |
|---|---|---|---|---|---|---|---|---|---|---|---|---|---|
| ENSG00000140265 | 1281.026 | -0.256684895 | 0.250718 | -1.0238 | 0.30593 | 0.761997697 | ZSCAN29 | protein_coding | 44.13 | -0.030300713 | 0.030300713 | ENSG00000140265 | n.s. |
| ENSG00000187446 | 371.5064 | -0.256655501 | 0.250473 | -1.024683 | 0.305513 | 0.761997697 | CHP1 | protein_coding | 44.02 | -0.030297243 | 0.030297243 | ENSG00000187446 | n.s. |
| ENSG00000006451 | 511.6949 | -0.273720998 | 0.277468 | -0.986494 | 0.323891 | 0.775183815 | RALA | protein_coding | 39.69 | -0.030272257 | 0.030272257 | ENSG00000006451 | n.s. |
| ENSG00000272872 | 24.17519 | 0.321283937 | 0.35795 | 0.8975671 | 0.369416 | 0.805037468 | | sense_intronic | 48.56 | 0.030259776 | 0.030259776 | ENSG00000272872 | n.s. |
| ENSG00000269918 | 177.9955 | -0.294798445 | 0.313477 | -0.940415 | 0.347005 | 0.789590741 | | sense_intronic | 41.16 | -0.030245717 | 0.030245717 | ENSG00000269918 | n.s. |
| ENSG00000277283 | 164.9568 | -0.253906876 | 0.245372 | -1.034785 | 0.300769 | 0.760115318 | | antisense | 52.48 | -0.030245518 | 0.030245518 | ENSG00000277283 | n.s. |
| ENSG00000233822 | 606.3572 | -0.230576932 | 0.211127 | -1.092127 | 0.274777 | 0.739386355 | HIST1H2BN | protein_coding | 40.18 | -0.030235223 | 0.030235223 | ENSG00000233822 | n.s. |
| ENSG00000007392 | 2803.444 | 0.248886788 | 0.238561 | 1.043282 | 0.296818 | 0.756014066 | LUC7L | protein_coding | 47.33 | 0.030232309 | 0.030232309 | ENSG00000007392 | n.s. |
| ENSG00000112079 | 698.4173 | -0.278467164 | 0.284481 | -0.97886 | 0.327649 | 0.779005431 | STK38 | protein_coding | 41.52 | -0.030202413 | 0.030202413 | ENSG00000112079 | n.s. |
| ENSG00000105568 | 2480.365 | -0.264384078 | 0.264466 | -0.999692 | 0.31746 | 0.768883445 | PPP2R1A | protein_coding | 48.38 | -0.030176664 | 0.030176664 | ENSG00000105568 | n.s. |
| ENSG00000177189 | 859.2035 | -0.268596578 | 0.270665 | -0.992358 | 0.321023 | 0.772218859 | RPS6KA3 | protein_coding | 37.57 | -0.030152543 | 0.030152543 | ENSG00000177189 | n.s. |
| ENSG00000161813 | 2320.922 | -0.258063038 | 0.254253 | -1.014983 | 0.310114 | 0.764194956 | LARP4 | protein_coding | 41.45 | -0.030140687 | 0.030140687 | ENSG00000161813 | n.s. |
| ENSG00000198695 | 724.7055 | -0.261942024 | 0.260682 | -1.004835 | 0.314976 | 0.767364665 | MT-ND6 | protein_coding | 42.67 | -0.030122862 | 0.030122862 | ENSG00000198695 | n.s. |
| ENSG00000182685 | 109.6297 | -0.257607183 | 0.253643 | -1.015629 | 0.309806 | 0.763980297 | BRICD5 | protein_coding | 63.38 | -0.030118876 | 0.030118876 | ENSG00000182685 | n.s. |
| ENSG00000167670 | 1897.763 | -0.250391602 | 0.241354 | -1.037446 | 0.299528 | 0.758335061 | CHAF1A | protein_coding | 51.15 | -0.030081763 | 0.030081763 | ENSG00000167670 | n.s. |
| ENSG00000272344 | 12.91812 | 0.362549742 | 0.43806 | 0.827625 | 0.407883 | 0.826149803 | SNORD114-21 | snoRNA | 47.22 | 0.030070309 | 0.030070309 | ENSG00000272344 | n.s. |
| ENSG00000108439 | 392.5421 | -0.246091828 | 0.235293 | -1.045895 | 0.29561 | 0.754772091 | PNPO | protein_coding | 50.57 | -0.030068525 | 0.030068525 | ENSG00000108439 | n.s. |
| ENSG00000204564 | 91.25494 | -0.265594395 | 0.267157 | -0.99415 | 0.32015 | 0.77078424 | C6orf136 | protein_coding | 52.14 | -0.030030008 | 0.030030008 | ENSG00000204564 | n.s. |
| ENSG00000183735 | 971.8927 | -0.254211253 | 0.248129 | -1.024513 | 0.305593 | 0.761997697 | TBK1 | protein_coding | 37.87 | -0.030008708 | 0.030008708 | ENSG00000183735 | n.s. |
| ENSG00000198689 | 77.65928 | -0.278224817 | 0.287047 | -0.969264 | 0.332413 | 0.780141891 | SLC9A6 | protein_coding | 40.97 | -0.029999981 | 0.029999981 | ENSG00000198689 | n.s. |
| ENSG00000213079 | 3150.679 | -0.260127283 | 0.258229 | -1.00735 | 0.313767 | 0.766827179 | SCAF8 | protein_coding | 37.66 | -0.029993327 | 0.029993327 | ENSG00000213079 | n.s. |
| ENSG00000198858 | 201.7622 | -0.290938512 | 0.308663 | -0.942577 | 0.345897 | 0.788921338 | R3HDM4 | protein_coding | 58.93 | -0.029956861 | 0.029956861 | ENSG00000198858 | n.s. |
| ENSG00000103994 | 1428.509 | -0.263900747 | 0.264958 | -0.996009 | 0.319246 | 0.769994426 | ZNF106 | protein_coding | 41.35 | -0.029956012 | 0.029956012 | ENSG00000103994 | n.s. |
| ENSG00000167380 | 869.7866 | -0.240652474 | 0.227755 | -1.056627 | 0.290682 | 0.750798148 | ZNF226 | protein_coding | 40.39 | -0.029955652 | 0.029955652 | ENSG00000167380 | n.s. |
| ENSG00000168056 | 110.7365 | -0.288246874 | 0.304197 | -0.947567 | 0.34335 | 0.787295611 | LTBP3 | protein_coding | 60.22 | -0.029937946 | 0.029937946 | ENSG00000168056 | n.s. |
| ENSG00000256073 | 64.93838 | -0.294630844 | 0.315872 | -0.932755 | 0.350947 | 0.791420351 | URB1-AS1 | lincRNA | 54.63 | -0.029932368 | 0.029932368 | ENSG00000256073 | n.s. |
| ENSG00000274322 | 17.76349 | 0.352944345 | 0.420123 | 0.8400986 | 0.400853 | 0.822682497 | | protein_coding | 45.09 | 0.029918295 | 0.029918295 | ENSG00000274322 | n.s. |
| ENSG00000226210 | 232.7735 | -0.269801873 | 0.273117 | -0.987862 | 0.32322 | 0.774693381 | | unprocessed_pseudogene | 54.31 | -0.029912975 | 0.029912975 | ENSG00000226210 | n.s. |
| ENSG00000031823 | 646.4424 | -0.261073034 | 0.260494 | -1.002222 | 0.316237 | 0.768128006 | RANBP3 | protein_coding | 52.74 | -0.029910198 | 0.029910198 | ENSG00000031823 | n.s. |
| ENSG00000118985 | 48.84284 | 0.321873015 | 0.361508 | 0.8903625 | 0.373271 | 0.807377464 | ELL2 | protein_coding | 38.17 | 0.029909527 | 0.029909527 | ENSG00000118985 | n.s. |
| ENSG00000163933 | 480.6419 | -0.272421966 | 0.276978 | -0.983553 | 0.325336 | 0.776688313 | RFT1 | protein_coding | 45.77 | -0.029899191 | 0.029899191 | ENSG00000163933 | n.s. |
| ENSG00000262879 | 257.3974 | -0.259295394 | 0.257297 | -1.007769 | 0.313566 | 0.766827179 | | processed_transcript | 43.56 | -0.029897408 | 0.029897408 | ENSG00000262879 | n.s. |

| | | | | | | | | | | | | | |
|---|---|---|---|---|---|---|---|---|---|---|---|---|---|
| ENSG00000129355 | 82.64183 | -0.306298982 | 0.336147 | -0.911205 | 0.362187 | 0.798938342 | CDKN2D | protein_coding | 58.55 | -0.029860088 | 0.029860088 | ENSG00000129355 | n.s. |
| ENSG00000213722 | 514.8133 | -0.283979869 | 0.296888 | -0.956523 | 0.338808 | 0.784993146 | DDAH2 | protein_coding | 59.86 | -0.029855978 | 0.029855978 | ENSG00000213722 | n.s. |
| ENSG00000157350 | 425.7989 | -0.242902633 | 0.231098 | -1.051082 | 0.293221 | 0.753555483 | ST3GAL2 | protein_coding | 51.87 | -0.029849033 | 0.029849033 | ENSG00000157350 | n.s. |
| ENSG00000152778 | 1370.441 | 0.241485285 | 0.229206 | 1.0535744 | 0.292078 | 0.75237733 | IFIT5 | protein_coding | 41.44 | 0.02983896 | 0.02983896 | ENSG00000152778 | n.s. |
| ENSG00000176401 | 187.9651 | -0.242361301 | 0.230646 | -1.050792 | 0.293354 | 0.753623766 | EID2B | protein_coding | 48.95 | -0.029772974 | 0.029772974 | ENSG00000176401 | n.s. |
| ENSG00000267682 | 43.95426 | -0.284643632 | 0.298405 | -0.953882 | 0.340143 | 0.785979841 | | sense_intronic | 51.38 | -0.029770477 | 0.029770477 | ENSG00000267682 | n.s. |
| ENSG00000103852 | 271.2349 | -0.233150794 | 0.217739 | -1.070779 | 0.284269 | 0.745309826 | TTC23 | protein_coding | 41.8 | -0.029764765 | 0.029764765 | ENSG00000103852 | n.s. |
| ENSG00000133083 | 429.2362 | -0.27751426 | 0.286991 | -0.96698 | 0.333554 | 0.781172958 | DCLK1 | protein_coding | 40.16 | -0.029764181 | 0.029764181 | ENSG00000133083 | n.s. |
| ENSG00000137806 | 235.0432 | -0.278129969 | 0.28809 | -0.965428 | 0.33433 | 0.781938745 | NDUFAF1 | protein_coding | 46.09 | -0.029711865 | 0.029711865 | ENSG00000137806 | n.s. |
| ENSG00000244748 | 31.3595 | -0.301932367 | 0.329891 | -0.91525 | 0.360061 | 0.797273365 | RN7SL153P | misc_RNA | 49.66 | -0.029707954 | 0.029707954 | ENSG00000244748 | n.s. |
| ENSG00000270659 | 32.95164 | 0.302092554 | 0.330282 | 0.9146495 | 0.360376 | 0.797431699 | | lincRNA | 30 | 0.029697663 | 0.029697663 | ENSG00000270659 | n.s. |
| ENSG00000153989 | 1055.257 | -0.273228757 | 0.278943 | -0.979514 | 0.327326 | 0.778659927 | NUS1 | protein_coding | 37.38 | -0.029686899 | 0.029686899 | ENSG00000153989 | n.s. |
| ENSG00000283928 | 4.646612 | -0.473631782 | 0.66172 | -0.715758 | 0.474141 | 0.865616576 | MIR637 | miRNA | 61.62 | -0.029684605 | 0.029684605 | ENSG00000283928 | n.s. |
| ENSG00000100030 | 1783.619 | -0.256258731 | 0.253581 | -1.01056 | 0.312227 | 0.766004018 | MAPK1 | protein_coding | 43.8 | -0.029666805 | 0.029666805 | ENSG00000100030 | n.s. |
| ENSG00000131016 | 9784.255 | -0.249813074 | 0.242627 | -1.029618 | 0.303189 | 0.760781413 | AKAP12 | protein_coding | 43.87 | -0.029662831 | 0.029662831 | ENSG00000131016 | n.s. |
| ENSG00000187961 | 255.8715 | -0.257006788 | 0.254671 | -1.009171 | 0.312893 | 0.766641623 | KLHL17 | protein_coding | 67.87 | -0.029660538 | 0.029660538 | ENSG00000187961 | n.s. |
| ENSG00000152620 | 413.8923 | -0.28438571 | 0.299189 | -0.950522 | 0.341847 | 0.786560769 | NADK2 | protein_coding | 35.98 | -0.029652249 | 0.029652249 | ENSG00000152620 | n.s. |
| ENSG00000185414 | 1419.632 | -0.290065618 | 0.310333 | -0.934691 | 0.349947 | 0.790401174 | MRPL30 | protein_coding | 40.26 | -0.029630906 | 0.029630906 | ENSG00000185414 | n.s. |
| ENSG00000173276 | 188.1501 | 0.246633403 | 0.237641 | 1.037842 | 0.299344 | 0.758335061 | ZBTB21 | protein_coding | 42.8 | 0.029630257 | 0.029630257 | ENSG00000173276 | n.s. |
| ENSG00000174839 | 1087.009 | -0.286730389 | 0.303259 | -0.945496 | 0.344406 | 0.788289085 | DENND6A | protein_coding | 39.72 | -0.029623403 | 0.029623403 | ENSG00000174839 | n.s. |
| ENSG00000114302 | 979.2371 | -0.262612124 | 0.264415 | -0.993182 | 0.320621 | 0.771519169 | PRKAR2A | protein_coding | 43.41 | -0.029584117 | 0.029584117 | ENSG00000114302 | n.s. |
| ENSG00000164330 | 570.3562 | -0.258024427 | 0.257636 | -1.001507 | 0.316582 | 0.768209163 | EBF1 | protein_coding | 39.97 | -0.029549091 | 0.029549091 | ENSG00000164330 | n.s. |
| ENSG00000130600 | 441.9231 | -0.256962406 | 0.256055 | -1.003542 | 0.315599 | 0.767425784 | H19 | processed_transcript | 62.11 | -0.029541327 | 0.029541327 | ENSG00000130600 | n.s. |
| ENSG00000074071 | 1064.412 | -0.297716677 | 0.323989 | -0.91891 | 0.358143 | 0.795806072 | MRPS34 | protein_coding | 65.4 | -0.029531336 | 0.029531336 | ENSG00000074071 | n.s. |
| ENSG00000185236 | 173.7396 | -0.273842639 | 0.281273 | -0.973583 | 0.330264 | 0.780141891 | RAB11B | protein_coding | 59.05 | -0.029527466 | 0.029527466 | ENSG00000185236 | n.s. |
| ENSG00000169504 | 2152.76 | -0.288369055 | 0.307327 | -0.938314 | 0.348083 | 0.790017511 | CLIC4 | protein_coding | 41.67 | -0.029518403 | 0.029518403 | ENSG00000169504 | n.s. |
| ENSG00000100697 | 4126.489 | -0.250017467 | 0.245184 | -1.019713 | 0.307865 | 0.761997697 | DICER1 | protein_coding | 38 | -0.029513647 | 0.029513647 | ENSG00000100697 | n.s. |
| ENSG00000132670 | 793.2567 | -0.266196718 | 0.269442 | -0.987956 | 0.323174 | 0.774693381 | PTPRA | protein_coding | 43.04 | -0.029513271 | 0.029513271 | ENSG00000132670 | n.s. |
| ENSG00000187098 | 20.75074 | 0.355702988 | 0.430011 | 0.827196 | 0.408126 | 0.826171518 | MITF | protein_coding | 38.57 | 0.029498371 | 0.029498371 | ENSG00000187098 | n.s. |
| ENSG00000158987 | 1112.541 | -0.273557382 | 0.281728 | -0.970998 | 0.331549 | 0.780141891 | RAPGEF6 | protein_coding | 37.88 | -0.029496708 | 0.029496708 | ENSG00000158987 | n.s. |
| ENSG00000159640 | 77.26607 | -0.280805237 | 0.293817 | -0.955716 | 0.339216 | 0.785318029 | ACE | protein_coding | 58.35 | -0.029471754 | 0.029471754 | ENSG00000159640 | n.s. |
| ENSG00000023608 | 300.8687 | -0.279727776 | 0.291827 | -0.95854 | 0.337791 | 0.78475552 | SNAPC1 | protein_coding | 39.61 | -0.029445718 | 0.029445718 | ENSG00000023608 | n.s. |

| | | | | | | | | | | | | | |
|---|---|---|---|---|---|---|---|---|---|---|---|---|---|
| ENSG00000011304 | 3832.441 | -0.253130253 | 0.249724 | -1.013641 | 0.310754 | 0.765067945 | PTBP1 | protein_coding | 61.17 | -0.029439047 | 0.029439047 | ENSG00000011304 | n.s. |
| ENSG00000213064 | 1704.136 | -0.255057072 | 0.252717 | -1.009259 | 0.31285 | 0.766641623 | SFT2D2 | protein_coding | 43.37 | -0.029435526 | 0.029435526 | ENSG00000213064 | n.s. |
| ENSG00000023228 | 3431.835 | -0.272809476 | 0.280581 | -0.972302 | 0.3309 | 0.780141891 | NDUFS1 | protein_coding | 40.81 | -0.029416064 | 0.029416064 | ENSG00000023228 | n.s. |
| ENSG00000146872 | 2840.736 | -0.240347134 | 0.229756 | -1.046097 | 0.295516 | 0.754672555 | TLK2 | protein_coding | 42.17 | -0.029380381 | 0.029380381 | ENSG00000146872 | n.s. |
| ENSG00000168116 | 2093.084 | -0.259389447 | 0.260747 | -0.994793 | 0.319837 | 0.770564009 | KIAA1586 | protein_coding | 37.98 | -0.029360623 | 0.029360623 | ENSG00000168116 | n.s. |
| ENSG00000132953 | 1863.076 | -0.254511237 | 0.252699 | -1.00717 | 0.313853 | 0.766827179 | XPO4 | protein_coding | 38.5 | -0.029345782 | 0.029345782 | ENSG00000132953 | n.s. |
| ENSG00000151414 | 1649.066 | 0.270057304 | 0.275584 | 0.9799456 | 0.327113 | 0.778659927 | NEK7 | protein_coding | 35.84 | 0.029342314 | 0.029342314 | ENSG00000151414 | n.s. |
| ENSG00000167220 | 663.1794 | -0.27206759 | 0.280621 | -0.969521 | 0.332285 | 0.780141891 | HDHD2 | protein_coding | 38.68 | -0.029336069 | 0.029336069 | ENSG00000167220 | n.s. |
| ENSG00000165533 | 1993.069 | 0.254101969 | 0.251919 | 1.0086644 | 0.313136 | 0.766650888 | TTC8 | protein_coding | 36.61 | 0.029323966 | 0.029323966 | ENSG00000165533 | n.s. |
| ENSG00000175701 | 300.5666 | -0.252127776 | 0.248896 | -1.012985 | 0.311067 | 0.76510176 | LINC00116 | protein_coding | 40.57 | -0.029317619 | 0.029317619 | ENSG00000175701 | n.s. |
| ENSG00000085415 | 1069.981 | -0.255899995 | 0.255334 | -1.002218 | 0.316238 | 0.768128006 | SEH1L | protein_coding | 43.55 | -0.029317541 | 0.029317541 | ENSG00000085415 | n.s. |
| ENSG00000189043 | 4192.668 | 0.271685587 | 0.27959 | 0.9717283 | 0.331186 | 0.780141891 | NDUFA4 | protein_coding | 37.82 | 0.029294879 | 0.029294879 | ENSG00000189043 | n.s. |
| ENSG00000203809 | 231.5068 | -0.238781574 | 0.227609 | -1.049088 | 0.294138 | 0.753915388 | LIN28B-AS1 | lincRNA | 40.64 | -0.029293101 | 0.029293101 | ENSG00000203809 | n.s. |
| ENSG00000102471 | 629.3518 | -0.271549124 | 0.27925 | -0.972425 | 0.330839 | 0.780141891 | NDFIP2 | protein_coding | 35.47 | -0.029280165 | 0.029280165 | ENSG00000102471 | n.s. |
| ENSG00000139405 | 351.664 | -0.254337437 | 0.253317 | -1.004028 | 0.315365 | 0.767425784 | RITA1 | protein_coding | 51.56 | -0.029239551 | 0.029239551 | ENSG00000139405 | n.s. |
| ENSG00000227740 | 27.00476 | -0.303190026 | 0.335015 | -0.905004 | 0.365463 | 0.801033089 | | lincRNA | 38.64 | -0.029212221 | 0.029212221 | ENSG00000227740 | n.s. |
| ENSG00000264961 | 3.58273 | -0.668216061 | 1.219004 | -0.548166 | 0.583578 | 0.904504088 | MIR4730 | miRNA | 65.79 | -0.029127181 | 0.029127181 | ENSG00000264961 | n.s. |
| ENSG00000172301 | 772.3105 | -0.270039819 | 0.278476 | -0.969707 | 0.332193 | 0.780141891 | COPRS | protein_coding | 49.04 | -0.029117422 | 0.029117422 | ENSG00000172301 | n.s. |
| ENSG00000148123 | 194.8695 | -0.274228406 | 0.284513 | -0.963853 | 0.335119 | 0.783258007 | PLPPR1 | protein_coding | 38.59 | -0.029094306 | 0.029094306 | ENSG00000148123 | n.s. |
| ENSG00000134508 | 45.8829 | -0.291393864 | 0.315843 | -0.922592 | 0.35622 | 0.794806874 | CABLES1 | protein_coding | 48.32 | -0.029063154 | 0.029063154 | ENSG00000134508 | n.s. |
| ENSG00000156299 | 1940.379 | -0.246087598 | 0.241029 | -1.020988 | 0.30726 | 0.761997697 | TIAM1 | protein_coding | 43.86 | -0.029049741 | 0.029049741 | ENSG00000156299 | n.s. |
| ENSG00000168904 | 36.22726 | -0.312117236 | 0.349779 | -0.892327 | 0.372217 | 0.807099083 | LRRC28 | protein_coding | 38.12 | -0.029049733 | 0.029049733 | ENSG00000168904 | n.s. |
| ENSG00000267595 | 16.10376 | -0.344168553 | 0.410686 | -0.838033 | 0.402012 | 0.823540971 | | unprocessed_pseudogene | 35 | -0.029018499 | 0.029018499 | ENSG00000267595 | n.s. |
| ENSG00000171109 | 1212.385 | -0.27162001 | 0.281297 | -0.9656 | 0.334244 | 0.781938745 | MFN1 | protein_coding | 38.56 | -0.029016423 | 0.029016423 | ENSG00000171109 | n.s. |
| ENSG00000277590 | 57.80643 | -0.299610617 | 0.330308 | -0.907065 | 0.364372 | 0.800144301 | MIR7845 | miRNA | 72.73 | -0.0290118 | 0.0290118 | ENSG00000277590 | n.s. |
| ENSG00000184887 | 345.278 | -0.264498294 | 0.268983 | -0.983328 | 0.325446 | 0.776818484 | BTBD6 | protein_coding | 64.36 | -0.029010292 | 0.029010292 | ENSG00000184887 | n.s. |
| ENSG00000132471 | 314.105 | -0.270299094 | 0.279407 | -0.967401 | 0.333344 | 0.781097236 | WBP2 | protein_coding | 55.06 | -0.029001714 | 0.029001714 | ENSG00000132471 | n.s. |
| ENSG00000126581 | 1348.366 | -0.260802977 | 0.263717 | -0.988949 | 0.322688 | 0.774129273 | BECN1 | protein_coding | 44.97 | -0.028997773 | 0.028997773 | ENSG00000126581 | n.s. |
| ENSG00000234616 | 1610.711 | -0.238715487 | 0.228963 | -1.042596 | 0.297135 | 0.756138711 | JRK | protein_coding | 56.58 | -0.028979709 | 0.028979709 | ENSG00000234616 | n.s. |
| ENSG00000113194 | 1430.326 | -0.275258633 | 0.287067 | -0.958867 | 0.337626 | 0.78475552 | FAF2 | protein_coding | 42.96 | -0.028975271 | 0.028975271 | ENSG00000113194 | n.s. |
| ENSG00000071575 | 1021.603 | -0.236828076 | 0.226334 | -1.046365 | 0.295392 | 0.75457253 | TRIB2 | protein_coding | 47.68 | -0.028963839 | 0.028963839 | ENSG00000071575 | n.s. |
| ENSG00000168538 | 1272.1 | -0.268606142 | 0.275156 | -0.976194 | 0.328968 | 0.780141891 | TRAPPC11 | protein_coding | 38.51 | -0.028962834 | 0.028962834 | ENSG00000168538 | n.s. |

| | | | | | | | | | | | | | |
|---|---|---|---|---|---|---|---|---|---|---|---|---|---|
| ENSG00000124209 | 817.1448 | -0.268505281 | 0.27618 | -0.972212 | 0.330945 | 0.780141891 | RAB22A | protein_coding | 41.46 | -0.028951958 | 0.028951958 | ENSG00000124209 | n.s. |
| ENSG00000148358 | 808.7903 | -0.254325873 | 0.254837 | -0.997993 | 0.318283 | 0.769456789 | GPR107 | protein_coding | 46.73 | -0.028946294 | 0.028946294 | ENSG00000148358 | n.s. |
| ENSG00000103005 | 51.14147 | -0.280901991 | 0.297737 | -0.943457 | 0.345447 | 0.788921338 | USB1 | protein_coding | 49.63 | -0.028923438 | 0.028923438 | ENSG00000103005 | n.s. |
| ENSG00000184005 | 697.1242 | 0.278912912 | 0.294621 | 0.9466824 | 0.343801 | 0.787678531 | ST6GALNAC3 | protein_coding | 38.78 | 0.0289096 | 0.0289096 | ENSG00000184005 | n.s. |
| ENSG00000235919 | 172.8641 | 0.257678597 | 0.259774 | 0.991935 | 0.321229 | 0.772448139 | ASH1L-AS1 | antisense | 62.06 | 0.028893673 | 0.028893673 | ENSG00000235919 | n.s. |
| ENSG00000166685 | 670.5244 | -0.2443566 | 0.23869 | -1.023739 | 0.305959 | 0.761997697 | COG1 | protein_coding | 49.15 | -0.028845403 | 0.028845403 | ENSG00000166685 | n.s. |
| ENSG00000156642 | 978.5384 | -0.266417433 | 0.272351 | -0.978215 | 0.327968 | 0.779364976 | NPTN | protein_coding | 41.2 | -0.028842115 | 0.028842115 | ENSG00000156642 | n.s. |
| ENSG00000109929 | 491.4314 | -0.276777911 | 0.291319 | -0.950085 | 0.342069 | 0.786682723 | SC5D | protein_coding | 38.78 | -0.028840366 | 0.028840366 | ENSG00000109929 | n.s. |
| ENSG00000047644 | 286.799 | -0.253946117 | 0.255048 | -0.995681 | 0.319405 | 0.770056658 | WWC3 | protein_coding | 46.21 | -0.028817125 | 0.028817125 | ENSG00000047644 | n.s. |
| ENSG00000176732 | 84.57575 | 0.265944518 | 0.271783 | 0.9785176 | 0.327818 | 0.779275344 | PFN4 | protein_coding | 44.06 | 0.028804202 | 0.028804202 | ENSG00000176732 | n.s. |
| ENSG00000167685 | 37.32013 | -0.309265852 | 0.346636 | -0.892192 | 0.37229 | 0.807126641 | ZNF444 | protein_coding | 54.02 | -0.02877976 | 0.02877976 | ENSG00000167685 | n.s. |
| ENSG00000198265 | 3802.959 | -0.243761655 | 0.237986 | -1.02427 | 0.305708 | 0.761997697 | HELZ | protein_coding | 38.17 | -0.028775172 | 0.028775172 | ENSG00000198265 | n.s. |
| ENSG00000115091 | 3039.249 | -0.266466265 | 0.27447 | -0.970838 | 0.331629 | 0.780141891 | ACTR3 | protein_coding | 36.61 | -0.028732098 | 0.028732098 | ENSG00000115091 | n.s. |
| ENSG00000268129 | 56.22267 | 0.278863663 | 0.295682 | 0.943119 | 0.34562 | 0.788921338 | | lincRNA | 48.66 | 0.028713559 | 0.028713559 | ENSG00000268129 | n.s. |
| ENSG00000095066 | 7.957369 | -0.410082246 | 0.542667 | -0.75568 | 0.449841 | 0.85118703 | HOOK2 | protein_coding | 53.64 | -0.028695506 | 0.028695506 | ENSG00000095066 | n.s. |
| ENSG00000196365 | 1430.219 | -0.249450085 | 0.248253 | -1.004824 | 0.314982 | 0.767364665 | LONP1 | protein_coding | 57.82 | -0.028686311 | 0.028686311 | ENSG00000196365 | n.s. |
| ENSG00000183723 | 504.6251 | -0.266037246 | 0.273531 | -0.972604 | 0.33075 | 0.780141891 | CMTM4 | protein_coding | 45.26 | -0.028685839 | 0.028685839 | ENSG00000183723 | n.s. |
| ENSG00000141027 | 2889.208 | -0.242928645 | 0.237187 | -1.024209 | 0.305736 | 0.761997697 | NCOR1 | protein_coding | 39.72 | -0.028676838 | 0.028676838 | ENSG00000141027 | n.s. |
| ENSG00000082482 | 253.8435 | -0.251948239 | 0.252599 | -0.997422 | 0.31856 | 0.769456789 | KCNK2 | protein_coding | 36.37 | -0.028675681 | 0.028675681 | ENSG00000082482 | n.s. |
| ENSG00000107779 | 1375.246 | -0.27397707 | 0.287609 | -0.952602 | 0.340792 | 0.786054116 | BMPR1A | protein_coding | 39.9 | -0.028643632 | 0.028643632 | ENSG00000107779 | n.s. |
| ENSG00000159322 | 1423.548 | -0.265630278 | 0.273975 | -0.969542 | 0.332275 | 0.780141891 | ADPGK | protein_coding | 43.53 | -0.028641957 | 0.028641957 | ENSG00000159322 | n.s. |
| ENSG00000244363 | 28.718 | 0.311809111 | 0.35432 | 0.8800202 | 0.378848 | 0.809376313 | RPL7P23 | processed_pseudogene | 43.24 | 0.028639514 | 0.028639514 | ENSG00000244363 | n.s. |
| ENSG00000058272 | 3215.478 | -0.258774202 | 0.262203 | -0.986922 | 0.323681 | 0.775183815 | PPP1R12A | protein_coding | 36.07 | -0.028619211 | 0.028619211 | ENSG00000058272 | n.s. |
| ENSG00000107263 | 553.664 | -0.243845863 | 0.239821 | -1.016785 | 0.309256 | 0.763298957 | RAPGEF1 | protein_coding | 48.49 | -0.02860442 | 0.02860442 | ENSG00000107263 | n.s. |
| ENSG00000162630 | 350.2222 | 0.235463624 | 0.225764 | 1.0429644 | 0.296965 | 0.756014066 | B3GALT2 | protein_coding | 32.61 | 0.028601796 | 0.028601796 | ENSG00000162630 | n.s. |
| ENSG00000117122 | 126.7631 | -0.306905773 | 0.343454 | -0.893585 | 0.371544 | 0.806899249 | MFAP2 | protein_coding | 62.61 | -0.028597691 | 0.028597691 | ENSG00000117122 | n.s. |
| ENSG00000168556 | 140.1244 | -0.279280256 | 0.297666 | -0.938233 | 0.348125 | 0.790017511 | ING2 | protein_coding | 46.91 | -0.028588044 | 0.028588044 | ENSG00000168556 | n.s. |
| ENSG00000198176 | 1746.026 | -0.264991236 | 0.271711 | -0.975268 | 0.329427 | 0.780141891 | TFDP1 | protein_coding | 53.33 | -0.028573051 | 0.028573051 | ENSG00000198176 | n.s. |
| ENSG00000181798 | 62.22726 | -0.278040783 | 0.2952 | -0.941872 | 0.346258 | 0.789327092 | LINC00471 | lincRNA | 42.78 | -0.028566741 | 0.028566741 | ENSG00000181798 | n.s. |
| ENSG00000157343 | 36.82788 | 0.299193729 | 0.331618 | 0.9022232 | 0.366938 | 0.802652289 | ARMC12 | protein_coding | 46.92 | 0.028564789 | 0.028564789 | ENSG00000157343 | n.s. |
| ENSG00000270742 | 96.40405 | 0.271031283 | 0.282287 | 0.9601265 | 0.336992 | 0.784604238 | | sense_intronic | 41.28 | 0.02855297 | 0.02855297 | ENSG00000270742 | n.s. |
| ENSG00000171867 | 692.005 | -0.278873609 | 0.297911 | -0.936098 | 0.349222 | 0.790028419 | PRNP | protein_coding | 41.96 | -0.028544746 | 0.028544746 | ENSG00000171867 | n.s. |

| | | | | | | | | | | | | | |
|---|---|---|---|---|---|---|---|---|---|---|---|---|---|
| ENSG00000064419 | 1631.966 | -0.248210187 | 0.247 | -1.004899 | 0.314946 | 0.767364665 | TNPO3 | protein_coding | 39.8 | -0.028543725 | 0.028543725 | ENSG00000064419 | n.s. |
| ENSG00000129219 | 34.24701 | -0.306310351 | 0.343081 | -0.892822 | 0.371952 | 0.806901511 | PLD2 | protein_coding | 54.27 | -0.028541837 | 0.028541837 | ENSG00000129219 | n.s. |
| ENSG00000250251 | 132.7542 | -0.264629174 | 0.272631 | -0.970649 | 0.331723 | 0.780141891 | PKD1P6 | cribed_unprocessed_pseudc | 61.72 | -0.028534012 | 0.028534012 | ENSG00000250251 | n.s. |
| ENSG00000200823 | 11.20065 | 0.357854198 | 0.442563 | 0.8085954 | 0.418748 | 0.832304654 | SNORD114-2 | snoRNA | 42.31 | 0.028527305 | 0.028527305 | ENSG00000200823 | n.s. |
| ENSG00000183049 | 433.1339 | -0.251724493 | 0.252968 | -0.995085 | 0.319695 | 0.770355344 | CAMK1D | protein_coding | 45.61 | -0.028522626 | 0.028522626 | ENSG00000183049 | n.s. |
| ENSG00000198908 | 302.3895 | -0.264440819 | 0.27162 | -0.97357 | 0.33027 | 0.780141891 | BHLHB9 | protein_coding | 42.71 | -0.028513702 | 0.028513702 | ENSG00000198908 | n.s. |
| ENSG00000267655 | 38.15674 | -0.309356508 | 0.349904 | -0.884118 | 0.376633 | 0.808862777 | | sense_intronic | 40.34 | -0.028499515 | 0.028499515 | ENSG00000267655 | n.s. |
| ENSG00000171861 | 406.0489 | -0.241213945 | 0.236111 | -1.021612 | 0.306964 | 0.761997697 | MRM3 | protein_coding | 48.63 | -0.028474424 | 0.028474424 | ENSG00000171861 | n.s. |
| ENSG00000138738 | 130.1957 | -0.276381378 | 0.292765 | -0.94404 | 0.345149 | 0.788822231 | PRDM5 | protein_coding | 37.44 | -0.028473047 | 0.028473047 | ENSG00000138738 | n.s. |
| ENSG00000067064 | 2456.611 | -0.261099641 | 0.266208 | -0.98081 | 0.326686 | 0.778045194 | IDI1 | protein_coding | 43.32 | -0.028458601 | 0.028458601 | ENSG00000067064 | n.s. |
| ENSG00000108671 | 1488.149 | -0.237869433 | 0.229606 | -1.035989 | 0.300207 | 0.759327693 | PSMD11 | protein_coding | 43.11 | -0.028442229 | 0.028442229 | ENSG00000108671 | n.s. |
| ENSG00000106400 | 345.3887 | -0.277601139 | 0.296054 | -0.937672 | 0.348413 | 0.790017511 | ZNHIT1 | protein_coding | 54.13 | -0.028416164 | 0.028416164 | ENSG00000106400 | n.s. |
| ENSG00000153107 | 1879.514 | -0.263431907 | 0.27022 | -0.97488 | 0.32962 | 0.780141891 | ANAPC1 | protein_coding | 39.07 | -0.028404914 | 0.028404914 | ENSG00000153107 | n.s. |
| ENSG00000139291 | 618.7089 | -0.261417378 | 0.266965 | -0.979219 | 0.327472 | 0.778716704 | TMEM19 | protein_coding | 39.6 | -0.028395289 | 0.028395289 | ENSG00000139291 | n.s. |
| ENSG00000166532 | 1787.034 | -0.263325428 | 0.271194 | -0.970987 | 0.331555 | 0.780141891 | RIMKLB | protein_coding | 41.05 | -0.028393433 | 0.028393433 | ENSG00000166532 | n.s. |
| ENSG00000157593 | 368.3 | -0.275407202 | 0.291567 | -0.944575 | 0.344876 | 0.78871706 | SLC35B2 | protein_coding | 58.08 | -0.028388634 | 0.028388634 | ENSG00000157593 | n.s. |
| ENSG00000280227 | 81.98855 | -0.275182878 | 0.291442 | -0.944211 | 0.345062 | 0.788822231 | | TEC | 28.6 | -0.028349576 | 0.028349576 | ENSG00000280227 | n.s. |
| ENSG00000228502 | 78.52612 | 0.309305674 | 0.351714 | 0.8794236 | 0.379172 | 0.809746094 | EEF1A1P11 | processed_pseudogene | 46.72 | 0.028348217 | 0.028348217 | ENSG00000228502 | n.s. |
| ENSG00000146282 | 1409.949 | -0.262690187 | 0.269307 | -0.975431 | 0.329346 | 0.780141891 | RARS2 | protein_coding | 38.27 | -0.028324938 | 0.028324938 | ENSG00000146282 | n.s. |
| ENSG00000185658 | 4493.748 | -0.262284067 | 0.26872 | -0.976051 | 0.329039 | 0.780141891 | BRWD1 | protein_coding | 38.65 | -0.028281147 | 0.028281147 | ENSG00000185658 | n.s. |
| ENSG00000146063 | 606.4382 | -0.247400426 | 0.247311 | -1.000361 | 0.317136 | 0.768578012 | TRIM41 | protein_coding | 52.09 | -0.028280848 | 0.028280848 | ENSG00000146063 | n.s. |
| ENSG00000196396 | 937.3939 | -0.264650216 | 0.274007 | -0.965852 | 0.334118 | 0.781938745 | PTPN1 | protein_coding | 43.56 | -0.028271859 | 0.028271859 | ENSG00000196396 | n.s. |
| ENSG00000213588 | 237.6644 | -0.239434967 | 0.234012 | -1.023174 | 0.306225 | 0.761997697 | ZBTB9 | protein_coding | 53.65 | -0.028264422 | 0.028264422 | ENSG00000213588 | n.s. |
| ENSG00000184349 | 712.5072 | -0.240130149 | 0.235887 | -1.017986 | 0.308684 | 0.762601855 | EFNA5 | protein_coding | 38.27 | -0.028263833 | 0.028263833 | ENSG00000184349 | n.s. |
| ENSG00000058063 | 896.5854 | -0.248246261 | 0.248842 | -0.997607 | 0.31847 | 0.769456789 | ATP11B | protein_coding | 36.02 | -0.028254338 | 0.028254338 | ENSG00000058063 | n.s. |
| ENSG00000128596 | 209.0149 | -0.270106013 | 0.283325 | -0.953342 | 0.340417 | 0.786007479 | CCDC136 | protein_coding | 48.94 | -0.028245883 | 0.028245883 | ENSG00000128596 | n.s. |
| ENSG00000136068 | 2280.553 | -0.241192907 | 0.237363 | -1.016135 | 0.309565 | 0.763743892 | FLNB | protein_coding | 46.61 | -0.028232172 | 0.028232172 | ENSG00000136068 | n.s. |
| ENSG00000214199 | 60.36139 | 0.30899389 | 0.352296 | 0.8770872 | 0.380439 | 0.810288517 | EEF1A1P12 | processed_pseudogene | 46.58 | 0.028229779 | 0.028229779 | ENSG00000214199 | n.s. |
| ENSG00000083720 | 1049.41 | -0.261973742 | 0.270418 | -0.968774 | 0.332658 | 0.780517967 | OXCT1 | protein_coding | 37.48 | -0.028192853 | 0.028192853 | ENSG00000083720 | n.s. |
| ENSG00000157870 | 99.33995 | -0.284909326 | 0.310606 | -0.917268 | 0.359002 | 0.79624783 | FAM213B | protein_coding | 64.59 | -0.028192273 | 0.028192273 | ENSG00000157870 | n.s. |
| ENSG00000120910 | 163.7399 | -0.281289483 | 0.303823 | -0.925832 | 0.354533 | 0.793978768 | PPP3CC | protein_coding | 39.2 | -0.028182706 | 0.028182706 | ENSG00000120910 | n.s. |
| ENSG00000173846 | 67.71516 | -0.275125807 | 0.293304 | -0.938021 | 0.348234 | 0.790017511 | PLK3 | protein_coding | 59.69 | -0.028162781 | 0.028162781 | ENSG00000173846 | n.s. |

| | | | | | | | | | | | | | |
|---|---|---|---|---|---|---|---|---|---|---|---|---|---|
| ENSG00000250957 | 460.1336 | -0.238521806 | 0.233515 | -1.021442 | 0.307045 | 0.761997697 | | lincRNA | 34.31 | -0.028156627 | 0.028156627 | ENSG00000250957 | n.s. |
| ENSG00000177182 | 92.22447 | -0.261016147 | 0.267752 | -0.974843 | 0.329638 | 0.780141891 | CLVS1 | protein_coding | 39.85 | -0.028144432 | 0.028144432 | ENSG00000177182 | n.s. |
| ENSG00000226253 | 31.80467 | -0.296215666 | 0.329128 | -0.900003 | 0.368119 | 0.803511983 | MRPL35P3 | processed_pseudogene | 45.6 | -0.028142752 | 0.028142752 | ENSG00000226253 | n.s. |
| ENSG00000116815 | 138.3944 | 0.285514198 | 0.3116 | 0.9162842 | 0.359518 | 0.797029976 | CD58 | protein_coding | 41.62 | 0.028130385 | 0.028130385 | ENSG00000116815 | n.s. |
| ENSG00000169862 | 136.1704 | -0.260761957 | 0.267002 | -0.976629 | 0.328753 | 0.780141891 | CTNND2 | protein_coding | 39.43 | -0.028117023 | 0.028117023 | ENSG00000169862 | n.s. |
| ENSG00000116704 | 925.7637 | -0.241693222 | 0.238462 | -1.013551 | 0.310797 | 0.765067945 | SLC35D1 | protein_coding | 37.83 | -0.02810892 | 0.02810892 | ENSG00000116704 | n.s. |
| ENSG00000206727 | 4.464365 | 0.423738319 | 0.576479 | 0.7350461 | 0.462311 | 0.858395846 | SNORD116-9 | snoRNA | 45.26 | 0.028099102 | 0.028099102 | ENSG00000206727 | n.s. |
| ENSG00000116830 | 2895.494 | -0.260536368 | 0.26807 | -0.971896 | 0.331102 | 0.780141891 | TTF2 | protein_coding | 41.16 | -0.028092699 | 0.028092699 | ENSG00000116830 | n.s. |
| ENSG00000158435 | 646.8753 | -0.260299541 | 0.267424 | -0.973361 | 0.330374 | 0.780141891 | CNOT11 | protein_coding | 43.29 | -0.028067163 | 0.028067163 | ENSG00000158435 | n.s. |
| ENSG00000274475 | 23.75906 | -0.332800664 | 0.397127 | -0.838021 | 0.402019 | 0.823540971 | RN7SL650P | misc_RNA | 54.85 | -0.028060018 | 0.028060018 | ENSG00000274475 | n.s. |
| ENSG00000197977 | 487.979 | -0.274105714 | 0.29239 | -0.937467 | 0.348518 | 0.790017511 | ELOVL2 | protein_coding | 39.64 | -0.028058361 | 0.028058361 | ENSG00000197977 | n.s. |
| ENSG00000207547 | 73.79535 | -0.28123315 | 0.304574 | -0.923366 | 0.355817 | 0.794802373 | MIR25 | miRNA | 63.1 | -0.028050432 | 0.028050432 | ENSG00000207547 | n.s. |
| ENSG00000102172 | 1778.988 | -0.262532995 | 0.271784 | -0.96596 | 0.334064 | 0.781938745 | SMS | protein_coding | 43.48 | -0.028045682 | 0.028045682 | ENSG00000102172 | n.s. |
| ENSG00000136878 | 229.4305 | -0.266633239 | 0.278718 | -0.956642 | 0.338748 | 0.784993146 | USP20 | protein_coding | 52.14 | -0.028032255 | 0.028032255 | ENSG00000136878 | n.s. |
| ENSG00000264235 | 53.9587 | 0.283202377 | 0.308767 | 0.9172055 | 0.359035 | 0.79624783 | | antisense | 40.41 | 0.028023367 | 0.028023367 | ENSG00000264235 | n.s. |
| ENSG00000221265 | 27.576 | 0.307308975 | 0.352324 | 0.8722337 | 0.383081 | 0.810719585 | MIR1255A | miRNA | 33.63 | 0.028004863 | 0.028004863 | ENSG00000221265 | n.s. |
| ENSG00000242498 | 448.087 | -0.242829104 | 0.241019 | -1.007512 | 0.313689 | 0.766827179 | ARPIN | protein_coding | 49.81 | -0.027998803 | 0.027998803 | ENSG00000242498 | n.s. |
| ENSG00000137815 | 1308.931 | -0.253142353 | 0.25656 | -0.986678 | 0.3238 | 0.775183815 | RTF1 | protein_coding | 45.65 | -0.027996355 | 0.027996355 | ENSG00000137815 | n.s. |
| ENSG00000138942 | 351.3085 | -0.277649872 | 0.299126 | -0.928204 | 0.353302 | 0.792950339 | RNF185 | protein_coding | 44.98 | -0.027974338 | 0.027974338 | ENSG00000138942 | n.s. |
| ENSG00000196792 | 915.1175 | -0.257344764 | 0.262739 | -0.979468 | 0.327349 | 0.778659927 | STRN3 | protein_coding | 38.51 | -0.027961068 | 0.027961068 | ENSG00000196792 | n.s. |
| ENSG00000100209 | 130.5065 | -0.280337737 | 0.304278 | -0.921322 | 0.356882 | 0.794806874 | HSCB | protein_coding | 46.7 | -0.027960433 | 0.027960433 | ENSG00000100209 | n.s. |
| ENSG00000205903 | 471.0803 | -0.247387978 | 0.248895 | -0.993943 | 0.32025 | 0.770876263 | ZNF316 | protein_coding | 61.59 | -0.027958633 | 0.027958633 | ENSG00000205903 | n.s. |
| ENSG00000172432 | 701.2705 | -0.258912432 | 0.266362 | -0.972032 | 0.331035 | 0.780141891 | GTPBP2 | protein_coding | 46.56 | -0.027917596 | 0.027917596 | ENSG00000172432 | n.s. |
| ENSG00000166165 | 2866.113 | -0.270996573 | 0.28741 | -0.942892 | 0.345736 | 0.788921338 | CKB | protein_coding | 70.4 | -0.027903514 | 0.027903514 | ENSG00000166165 | n.s. |
| ENSG00000023318 | 732.2315 | -0.258728338 | 0.265653 | -0.973935 | 0.330089 | 0.780141891 | ERP44 | protein_coding | 37.86 | -0.027897746 | 0.027897746 | ENSG00000023318 | n.s. |
| ENSG00000062598 | 425.7548 | -0.264999569 | 0.276453 | -0.95857 | 0.337775 | 0.78475552 | ELMO2 | protein_coding | 44.15 | -0.027895344 | 0.027895344 | ENSG00000062598 | n.s. |
| ENSG00000210195 | 116.3651 | 0.271683941 | 0.288792 | 0.9407598 | 0.346828 | 0.789549679 | MT-TT | Mt_tRNA | 37.88 | 0.027880352 | 0.027880352 | ENSG00000210195 | n.s. |
| ENSG00000113300 | 1607.604 | -0.258364548 | 0.26549 | -0.973162 | 0.330473 | 0.780141891 | CNOT6 | protein_coding | 42.07 | -0.027858519 | 0.027858519 | ENSG00000113300 | n.s. |
| ENSG00000170456 | 1035.834 | -0.258207448 | 0.264532 | -0.976091 | 0.329019 | 0.780141891 | DENND5B | protein_coding | 40.95 | -0.02784158 | 0.02784158 | ENSG00000170456 | n.s. |
| ENSG00000152700 | 728.2495 | -0.268496569 | 0.28365 | -0.946578 | 0.343854 | 0.787678531 | SAR1B | protein_coding | 43.44 | -0.027829936 | 0.027829936 | ENSG00000152700 | n.s. |
| ENSG00000144406 | 243.1466 | -0.258078089 | 0.265565 | -0.971808 | 0.331146 | 0.780141891 | UNC80 | protein_coding | 37.57 | -0.027827632 | 0.027827632 | ENSG00000144406 | n.s. |
| ENSG00000004777 | 56.0108 | -0.287401207 | 0.31707 | -0.906429 | 0.364709 | 0.800289612 | ARHGAP33 | protein_coding | 60.93 | -0.027806877 | 0.027806877 | ENSG00000004777 | n.s. |

| | | | | | | | | | | | | | |
|---|---|---|---|---|---|---|---|---|---|---|---|---|---|
| ENSG00000214029 | 1603.51 | -0.257875794 | 0.2659 | -0.969823 | 0.332135 | 0.780141891 | ZNF891 | protein_coding | 39.65 | -0.027805819 | 0.027805819 | ENSG00000214029 | n.s. |
| ENSG00000179406 | 47.47977 | 0.282527722 | 0.308699 | 0.9152193 | 0.360077 | 0.797273365 | LINC00174 | processed_transcript | 44.08 | 0.027798678 | 0.027798678 | ENSG00000179406 | n.s. |
| ENSG00000101442 | 302.7489 | -0.257802102 | 0.264456 | -0.97484 | 0.32964 | 0.780141891 | ACTR5 | protein_coding | 44.84 | -0.027797873 | 0.027797873 | ENSG00000101442 | n.s. |
| ENSG00000183444 | 104.2697 | -0.263607832 | 0.274279 | -0.961095 | 0.336504 | 0.784443005 | OR7E38P | unprocessed_pseudogene | 49.34 | -0.027794442 | 0.027794442 | ENSG00000183444 | n.s. |
| ENSG00000175106 | 29.99445 | -0.309268221 | 0.356745 | -0.866917 | 0.385988 | 0.813115862 | TVP23C | protein_coding | 42.63 | -0.027786997 | 0.027786997 | ENSG00000175106 | n.s. |
| ENSG00000088882 | 1276.844 | -0.251721221 | 0.255434 | -0.985465 | 0.324396 | 0.775604549 | CPXM1 | protein_coding | 59.72 | -0.027779866 | 0.027779866 | ENSG00000088882 | n.s. |
| ENSG00000076924 | 485.7773 | -0.253277814 | 0.257785 | -0.982514 | 0.325846 | 0.777167724 | XAB2 | protein_coding | 60.82 | -0.027730183 | 0.027730183 | ENSG00000076924 | n.s. |
| ENSG00000219902 | 53.00422 | -0.27521573 | 0.296469 | -0.928312 | 0.353246 | 0.792950339 | RPL35P3 | processed_pseudogene | 56.6 | -0.027729089 | 0.027729089 | ENSG00000219902 | n.s. |
| ENSG00000144426 | 484.3617 | -0.277979761 | 0.30132 | -0.922541 | 0.356246 | 0.794806874 | NBEAL1 | protein_coding | 37.7 | -0.027725253 | 0.027725253 | ENSG00000144426 | n.s. |
| ENSG00000169607 | 1265.564 | -0.242647536 | 0.24261 | -1.000157 | 0.317235 | 0.768684142 | CKAP2L | protein_coding | 39.66 | -0.027722985 | 0.027722985 | ENSG00000169607 | n.s. |
| ENSG00000174485 | 902.3492 | -0.268476062 | 0.284037 | -0.945215 | 0.344549 | 0.788387641 | DENND4A | protein_coding | 37.31 | -0.02772289 | 0.02772289 | ENSG00000174485 | n.s. |
| ENSG00000106348 | 617.2683 | -0.257050968 | 0.263389 | -0.975937 | 0.329096 | 0.780141891 | IMPDH1 | protein_coding | 57.06 | -0.027716881 | 0.027716881 | ENSG00000106348 | n.s. |
| ENSG00000100083 | 139.0414 | -0.257009652 | 0.263859 | -0.974041 | 0.330036 | 0.780141891 | GGA1 | protein_coding | 56.62 | -0.027712426 | 0.027712426 | ENSG00000100083 | n.s. |
| ENSG00000265982 | 14.30637 | -0.353839282 | 0.441643 | -0.801188 | 0.423023 | 0.834993891 | | unprocessed_pseudogene | 72.92 | -0.027711526 | 0.027711526 | ENSG00000265982 | n.s. |
| ENSG00000162066 | 148.9407 | -0.262204577 | 0.272283 | -0.962986 | 0.335555 | 0.784011932 | AMDHD2 | protein_coding | 59.17 | -0.027709079 | 0.027709079 | ENSG00000162066 | n.s. |
| ENSG00000184056 | 365.2679 | -0.25020676 | 0.25336 | -0.987553 | 0.323372 | 0.774923125 | VPS33B | protein_coding | 46.83 | -0.027708242 | 0.027708242 | ENSG00000184056 | n.s. |
| ENSG00000112308 | 1833.482 | -0.233358975 | 0.22703 | -1.027875 | 0.304008 | 0.760868221 | C6orf62 | protein_coding | 40.31 | -0.027697506 | 0.027697506 | ENSG00000112308 | n.s. |
| ENSG00000138658 | 1744.477 | -0.26479254 | 0.277516 | -0.954153 | 0.340006 | 0.785979841 | ZGRF1 | protein_coding | 37.99 | -0.027694279 | 0.027694279 | ENSG00000138658 | n.s. |
| ENSG00000273759 | 155.1241 | -0.25406308 | 0.258947 | -0.981138 | 0.326525 | 0.778045194 | | lincRNA | 47.71 | -0.02769165 | 0.02769165 | ENSG00000273759 | n.s. |
| ENSG00000201377 | 5.112008 | -0.426318235 | 0.585133 | -0.728584 | 0.466256 | 0.861088114 | RNY4P23 | misc_RNA | 39.18 | -0.027690395 | 0.027690395 | ENSG00000201377 | n.s. |
| ENSG00000180257 | 291.9075 | -0.273447424 | 0.293817 | -0.930672 | 0.352023 | 0.792117552 | ZNF816 | protein_coding | 43.93 | -0.027675713 | 0.027675713 | ENSG00000180257 | n.s. |
| ENSG00000188917 | 174.2832 | -0.277468546 | 0.301474 | -0.920373 | 0.357378 | 0.79488131 | TRMT2B | protein_coding | 44.9 | -0.02766298 | 0.02766298 | ENSG00000188917 | n.s. |
| ENSG00000067225 | 18592.46 | -0.270241772 | 0.288775 | -0.935822 | 0.349365 | 0.790028419 | PKM | protein_coding | 48.38 | -0.027661214 | 0.027661214 | ENSG00000067225 | n.s. |
| ENSG00000214078 | 578.7384 | 0.253697928 | 0.258579 | 0.9811253 | 0.326531 | 0.778045194 | CPNE1 | protein_coding | 43.12 | 0.02765185 | 0.02765185 | ENSG00000214078 | n.s. |
| ENSG00000274015 | 55.33406 | 0.276973107 | 0.300667 | 0.9211963 | 0.356948 | 0.794806874 | | lincRNA | 43.32 | 0.02762485 | 0.02762485 | ENSG00000274015 | n.s. |
| ENSG00000201229 | 6.061438 | -0.45649485 | 0.657196 | -0.69461 | 0.4873 | 0.870164918 | SNORA63D | snoRNA | 43.18 | -0.027571572 | 0.027571572 | ENSG00000201229 | n.s. |
| ENSG00000242853 | 36.76849 | 0.296676489 | 0.332747 | 0.8915967 | 0.372609 | 0.807357957 | RN7SL749P | misc_RNA | 55.86 | 0.027571294 | 0.027571294 | ENSG00000242853 | n.s. |
| ENSG00000236489 | 17.63133 | -0.329404123 | 0.394631 | -0.834714 | 0.403879 | 0.824799944 | | processed_pseudogene | 45.15 | -0.027555109 | 0.027555109 | ENSG00000236489 | n.s. |
| ENSG00000120662 | 195.3984 | -0.248985436 | 0.252227 | -0.98715 | 0.323569 | 0.775183815 | MTRF1 | protein_coding | 39.67 | -0.02753662 | 0.02753662 | ENSG00000120662 | n.s. |
| ENSG00000232466 | 40.35138 | 0.288743528 | 0.320222 | 0.901698 | 0.367217 | 0.802884215 | | processed_pseudogene | 48.66 | 0.027530852 | 0.027530852 | ENSG00000232466 | n.s. |
| ENSG00000188211 | 831.9167 | -0.255259224 | 0.263215 | -0.969775 | 0.332159 | 0.780141891 | NCR3LG1 | protein_coding | 44.5 | -0.027523684 | 0.027523684 | ENSG00000188211 | n.s. |
| ENSG00000282988 | 150.0545 | -0.274302802 | 0.296008 | -0.926674 | 0.354096 | 0.79373747 | HIST1H3D | protein_coding | 47.53 | -0.027518912 | 0.027518912 | ENSG00000282988 | n.s. |

| | | | | | | | | | | | | | |
|---|---|---|---|---|---|---|---|---|---|---|---|---|---|
| ENSG00000112294 | 234.6477 | -0.260781798 | 0.271836 | -0.959336 | 0.337389 | 0.78475552 | ALDH5A1 | protein_coding | 43.11 | -0.027451358 | 0.027451358 | ENSG00000112294 | n.s. |
| ENSG00000278482 | 9.295557 | -0.376152039 | 0.487528 | -0.77155 | 0.440381 | 0.845502102 | | misc_RNA | 41.74 | -0.027415957 | 0.027415957 | ENSG00000278482 | n.s. |
| ENSG00000162413 | 311.9568 | -0.238374764 | 0.237116 | -1.00531 | 0.314748 | 0.767364082 | KLHL21 | protein_coding | 54.46 | -0.027412748 | 0.027412748 | ENSG00000162413 | n.s. |
| ENSG00000144535 | 200.0046 | -0.259464469 | 0.269557 | -0.96256 | 0.335769 | 0.784079552 | DIS3L2 | protein_coding | 42.48 | -0.027409793 | 0.027409793 | ENSG00000144535 | n.s. |
| ENSG00000139180 | 1426.417 | -0.255082672 | 0.263673 | -0.967421 | 0.333334 | 0.781097236 | NDUFA9 | protein_coding | 39.57 | -0.02736907 | 0.02736907 | ENSG00000139180 | n.s. |
| ENSG00000170161 | 191.6357 | 0.245274695 | 0.247568 | 0.9907361 | 0.321814 | 0.773444264 | | lincRNA | 48.19 | 0.027365537 | 0.027365537 | ENSG00000170161 | n.s. |
| ENSG00000138376 | 1039.049 | -0.253629394 | 0.259841 | -0.976095 | 0.329017 | 0.780141891 | BARD1 | protein_coding | 36.39 | -0.027347945 | 0.027347945 | ENSG00000138376 | n.s. |
| ENSG00000117525 | 580.7501 | 0.235180633 | 0.232095 | 1.0132934 | 0.31092 | 0.765100725 | F3 | protein_coding | 43.52 | 0.02734713 | 0.02734713 | ENSG00000117525 | n.s. |
| ENSG00000228981 | 62.92685 | 0.309106997 | 0.361445 | 0.8551974 | 0.392442 | 0.815710819 | | processed_pseudogene | 59.55 | 0.027344773 | 0.027344773 | ENSG00000228981 | n.s. |
| ENSG00000214226 | 120.8491 | -0.248596708 | 0.252627 | -0.984047 | 0.325092 | 0.776506863 | C17orf67 | protein_coding | 44.82 | -0.027309517 | 0.027309517 | ENSG00000214226 | n.s. |
| ENSG00000167100 | 36.56757 | -0.298969402 | 0.342176 | -0.87373 | 0.382265 | 0.81032471 | SAMD14 | protein_coding | 55.35 | -0.02730814 | 0.02730814 | ENSG00000167100 | n.s. |
| ENSG00000147679 | 955.3595 | 0.236599539 | 0.234599 | 1.0085262 | 0.313202 | 0.766650888 | UTP23 | protein_coding | 40.04 | 0.027304144 | 0.027304144 | ENSG00000147679 | n.s. |
| ENSG00000123213 | 1487.813 | -0.266734987 | 0.28439 | -0.937918 | 0.348286 | 0.790017511 | NLN | protein_coding | 40.37 | -0.027303869 | 0.027303869 | ENSG00000123213 | n.s. |
| ENSG00000228293 | 30.44471 | 0.316947265 | 0.374584 | 0.8461305 | 0.39748 | 0.820111643 | | antisense | 45.57 | 0.027297724 | 0.027297724 | ENSG00000228293 | n.s. |
| ENSG00000204574 | 2263.534 | -0.249326119 | 0.253776 | -0.982464 | 0.325871 | 0.777167724 | ABCF1 | protein_coding | 47.97 | -0.027297531 | 0.027297531 | ENSG00000204574 | n.s. |
| ENSG00000207996 | 92.86272 | 0.259031497 | 0.269602 | 0.9607936 | 0.336656 | 0.784602889 | MIR301A | miRNA | 40.7 | 0.027288993 | 0.027288993 | ENSG00000207996 | n.s. |
| ENSG00000186130 | 996.3702 | 0.235139735 | 0.232517 | 1.0112779 | 0.311883 | 0.765516281 | ZBTB6 | protein_coding | 36.76 | 0.027286924 | 0.027286924 | ENSG00000186130 | n.s. |
| ENSG00000183684 | 3689.922 | -0.249010925 | 0.253432 | -0.982554 | 0.325827 | 0.777167724 | ALYREF | protein_coding | 57.57 | -0.027263022 | 0.027263022 | ENSG00000183684 | n.s. |
| ENSG00000174744 | 510.5792 | -0.279696831 | 0.307545 | -0.909449 | 0.363113 | 0.7992181 | BRMS1 | protein_coding | 55.88 | -0.027224204 | 0.027224204 | ENSG00000174744 | n.s. |
| ENSG00000149926 | 146.7658 | -0.256210913 | 0.265864 | -0.963693 | 0.3352 | 0.783314118 | FAM57B | protein_coding | 52 | -0.027174766 | 0.027174766 | ENSG00000149926 | n.s. |
| ENSG00000111058 | 112.3651 | -0.252014171 | 0.259863 | -0.969796 | 0.332148 | 0.780141891 | ACSS3 | protein_coding | 35.63 | -0.027173781 | 0.027173781 | ENSG00000111058 | n.s. |
| ENSG00000169641 | 459.4374 | -0.23544631 | 0.233367 | -1.008912 | 0.313017 | 0.766650888 | LUZP1 | protein_coding | 43.28 | -0.027171059 | 0.027171059 | ENSG00000169641 | n.s. |
| ENSG00000198700 | 6431.545 | -0.251916888 | 0.259867 | -0.969407 | 0.332342 | 0.780141891 | IPO9 | protein_coding | 41.91 | -0.027163291 | 0.027163291 | ENSG00000198700 | n.s. |
| ENSG00000196230 | 19398.58 | -0.248004059 | 0.252328 | -0.982862 | 0.325675 | 0.777099712 | TUBB | protein_coding | 53.06 | -0.027162211 | 0.027162211 | ENSG00000196230 | n.s. |
| ENSG00000160710 | 3973.998 | -0.237163975 | 0.236815 | -1.001475 | 0.316597 | 0.768209163 | ADAR | protein_coding | 45.35 | -0.027160141 | 0.027160141 | ENSG00000160710 | n.s. |
| ENSG00000132970 | 913.6072 | -0.257780788 | 0.268427 | -0.960339 | 0.336885 | 0.784604238 | WASF3 | protein_coding | 39.78 | -0.027157039 | 0.027157039 | ENSG00000132970 | n.s. |
| ENSG00000183354 | 593.8784 | -0.260446321 | 0.274061 | -0.950321 | 0.341949 | 0.786560769 | KIAA2026 | protein_coding | 37.58 | -0.027156144 | 0.027156144 | ENSG00000183354 | n.s. |
| ENSG00000143314 | 607.873 | -0.257442824 | 0.267731 | -0.961573 | 0.336264 | 0.784379351 | MRPL24 | protein_coding | 48.93 | -0.027153485 | 0.027153485 | ENSG00000143314 | n.s. |
| ENSG00000272892 | 57.74784 | -0.27219248 | 0.29575 | -0.920348 | 0.357391 | 0.79488131 | | lincRNA | 50.33 | -0.027136968 | 0.027136968 | ENSG00000272892 | n.s. |
| ENSG00000185917 | 495.9645 | -0.234369167 | 0.231987 | -1.010269 | 0.312366 | 0.766004018 | SETD4 | protein_coding | 44.84 | -0.027132673 | 0.027132673 | ENSG00000185917 | n.s. |
| ENSG00000180370 | 1349.089 | -0.257179427 | 0.267703 | -0.960688 | 0.336709 | 0.784602889 | PAK2 | protein_coding | 41.7 | -0.027093878 | 0.027093878 | ENSG00000180370 | n.s. |
| ENSG00000160271 | 2860.964 | 0.245404003 | 0.248973 | 0.9856663 | 0.324297 | 0.775538613 | RALGDS | protein_coding | 58.92 | 0.027091761 | 0.027091761 | ENSG00000160271 | n.s. |

| | | | | | | | | | | | | | |
|---|---|---|---|---|---|---|---|---|---|---|---|---|---|
| ENSG00000099284 | 1690.978 | -0.263013959 | 0.279008 | -0.942677 | 0.345846 | 0.788921338 | H2AFY2 | protein_coding | 45.79 | -0.027081573 | 0.027081573 | ENSG00000099284 | n.s. |
| ENSG00000158711 | 1571.323 | -0.259006103 | 0.272093 | -0.951903 | 0.341146 | 0.786054116 | ELK4 | protein_coding | 41.85 | -0.027078454 | 0.027078454 | ENSG00000158711 | n.s. |
| ENSG00000275485 | 30.10821 | -0.29334422 | 0.331497 | -0.884908 | 0.376206 | 0.808694966 | | sense_intronic | 43.4 | -0.027050814 | 0.027050814 | ENSG00000275485 | n.s. |
| ENSG00000284415 | 4.187824 | 0.437067032 | 0.620039 | 0.7049021 | 0.480871 | 0.867184458 | MIR1287 | miRNA | 51.11 | 0.027049429 | 0.027049429 | ENSG00000284415 | n.s. |
| ENSG00000184207 | 426.8679 | -0.248951624 | 0.254149 | -0.979548 | 0.327309 | 0.778659927 | PGP | protein_coding | 61.69 | -0.027049135 | 0.027049135 | ENSG00000184207 | n.s. |
| ENSG00000131373 | 136.1628 | -0.256873582 | 0.268046 | -0.958319 | 0.337902 | 0.78475552 | HACL1 | protein_coding | 39.06 | -0.027039957 | 0.027039957 | ENSG00000131373 | n.s. |
| ENSG00000201990 | 5.264649 | -0.432730625 | 0.611295 | -0.707892 | 0.479012 | 0.866029009 | | misc_RNA | 44.66 | -0.027031626 | 0.027031626 | ENSG00000201990 | n.s. |
| ENSG00000025770 | 1022.691 | -0.257101508 | 0.268804 | -0.956465 | 0.338837 | 0.784993146 | NCAPH2 | protein_coding | 55.08 | -0.027030145 | 0.027030145 | ENSG00000025770 | n.s. |
| ENSG00000222071 | 6.754873 | -0.489774057 | 0.746551 | -0.656049 | 0.511793 | 0.880693287 | MIR1915 | miRNA | 78.75 | -0.027023437 | 0.027023437 | ENSG00000222071 | n.s. |
| ENSG00000122483 | 780.9495 | -0.255968529 | 0.266081 | -0.961993 | 0.336053 | 0.784254968 | CCDC18 | protein_coding | 36.98 | -0.027015615 | 0.027015615 | ENSG00000122483 | n.s. |
| ENSG00000172216 | 236.4983 | -0.263228908 | 0.279914 | -0.940393 | 0.347016 | 0.789590741 | CEBPB | protein_coding | 66.16 | -0.027006747 | 0.027006747 | ENSG00000172216 | n.s. |
| ENSG00000283515 | 58.54915 | 0.266814471 | 0.286606 | 0.9309462 | 0.351881 | 0.792117552 | | protein_coding | 47.26 | 0.02700439 | 0.02700439 | ENSG00000283515 | n.s. |
| ENSG00000261575 | 39.5238 | 0.295869697 | 0.339892 | 0.8704815 | 0.384037 | 0.810719585 | | transcribed_unprocessed_pseudc | 52.83 | 0.026962409 | 0.026962409 | ENSG00000261575 | n.s. |
| ENSG00000163681 | 843.2187 | -0.233691338 | 0.232004 | -1.007274 | 0.313803 | 0.766827179 | SLMAP | protein_coding | 38.95 | -0.026945196 | 0.026945196 | ENSG00000163681 | n.s. |
| ENSG00000269893 | 2149.949 | 0.210260642 | 0.195386 | 1.0761278 | 0.28187 | 0.744567264 | SNHG8 | lincRNA | 45.83 | 0.02693356 | 0.02693356 | ENSG00000269893 | n.s. |
| ENSG00000153207 | 4024.949 | -0.252061799 | 0.261005 | -0.965736 | 0.334176 | 0.781938745 | AHCTF1 | protein_coding | 38.75 | -0.026927073 | 0.026927073 | ENSG00000153207 | n.s. |
| ENSG00000148335 | 617.3796 | -0.255623373 | 0.266626 | -0.958732 | 0.337694 | 0.78475552 | NTMT1 | protein_coding | 55.42 | -0.026908353 | 0.026908353 | ENSG00000148335 | n.s. |
| ENSG00000230673 | 47.85241 | -0.262861529 | 0.279963 | -0.938917 | 0.347774 | 0.790017511 | PABPC1P3 | processed_pseudogene | 46.62 | -0.026907369 | 0.026907369 | ENSG00000230673 | n.s. |
| ENSG00000224152 | 125.9429 | 0.257040965 | 0.270037 | 0.951872 | 0.341162 | 0.786054116 | | antisense | 51.32 | 0.026873004 | 0.026873004 | ENSG00000224152 | n.s. |
| ENSG00000169032 | 657.0996 | -0.248908274 | 0.254975 | -0.976208 | 0.328961 | 0.780141891 | MAP2K1 | protein_coding | 44.72 | -0.026838883 | 0.026838883 | ENSG00000169032 | n.s. |
| ENSG00000188227 | 726.1901 | -0.256658005 | 0.269457 | -0.952501 | 0.340843 | 0.786054116 | ZNF793 | protein_coding | 42.72 | -0.026832967 | 0.026832967 | ENSG00000188227 | n.s. |
| ENSG00000102921 | 611.4354 | -0.243267841 | 0.247015 | -0.984829 | 0.324708 | 0.775722631 | N4BP1 | protein_coding | 42.34 | -0.026830871 | 0.026830871 | ENSG00000102921 | n.s. |
| ENSG00000023330 | 464.5214 | -0.266007647 | 0.28647 | -0.928572 | 0.353111 | 0.792950339 | ALAS1 | protein_coding | 45.47 | -0.026801337 | 0.026801337 | ENSG00000023330 | n.s. |
| ENSG00000105552 | 200.9241 | -0.280965858 | 0.311561 | -0.901799 | 0.367163 | 0.802884215 | BCAT2 | protein_coding | 54.54 | -0.026789274 | 0.026789274 | ENSG00000105552 | n.s. |
| ENSG00000269967 | 34.06552 | -0.302791904 | 0.35262 | -0.858691 | 0.390511 | 0.815710819 | | lincRNA | 43 | -0.026786116 | 0.026786116 | ENSG00000269967 | n.s. |
| ENSG00000163884 | 166.2272 | 0.274901893 | 0.30198 | 0.9103311 | 0.362648 | 0.799058295 | KLF15 | protein_coding | 57.83 | 0.026781365 | 0.026781365 | ENSG00000163884 | n.s. |
| ENSG00000091947 | 476.5033 | -0.240715933 | 0.243284 | -0.989446 | 0.322445 | 0.774034559 | TMEM101 | protein_coding | 49.94 | -0.026777156 | 0.026777156 | ENSG00000091947 | n.s. |
| ENSG00000128606 | 129.599 | 0.274857411 | 0.301728 | 0.9109452 | 0.362324 | 0.799058295 | LRRC17 | protein_coding | 39.14 | 0.026777031 | 0.026777031 | ENSG00000128606 | n.s. |
| ENSG00000065665 | 220.2844 | -0.26105653 | 0.278116 | -0.93866 | 0.347905 | 0.790017511 | SEC61A2 | protein_coding | 43.14 | -0.026722603 | 0.026722603 | ENSG00000065665 | n.s. |
| ENSG00000179241 | 589.419 | -0.243911016 | 0.248153 | -0.982905 | 0.325654 | 0.777099712 | LDLRAD3 | protein_coding | 44.12 | -0.026713928 | 0.026713928 | ENSG00000179241 | n.s. |
| ENSG00000204228 | 98.36018 | -0.260921982 | 0.278595 | -0.936563 | 0.348983 | 0.790028419 | HSD17B8 | protein_coding | 56.62 | -0.026707266 | 0.026707266 | ENSG00000204228 | n.s. |
| ENSG00000124783 | 1881.486 | -0.255405481 | 0.267954 | -0.95317 | 0.340504 | 0.786054116 | SSR1 | protein_coding | 43.56 | -0.026702018 | 0.026702018 | ENSG00000124783 | n.s. |

| | | | | | | | | | | | | | |
|---|---|---|---|---|---|---|---|---|---|---|---|---|---|
| ENSG00000182606 | 550.6853 | -0.247497236 | 0.254393 | -0.972894 | 0.330606 | 0.780141891 | TRAK1 | protein_coding | 44.94 | -0.026686736 | 0.026686736 | ENSG00000182606 | n.s. |
| ENSG00000165416 | 2466.485 | -0.239142561 | 0.241527 | -0.990127 | 0.322112 | 0.773444264 | SUGT1 | protein_coding | 36.99 | -0.026681369 | 0.026681369 | ENSG00000165416 | n.s. |
| ENSG00000155189 | 1883.883 | -0.260546968 | 0.27737 | -0.939348 | 0.347552 | 0.790017511 | AGPAT5 | protein_coding | 38.61 | -0.026670443 | 0.026670443 | ENSG00000155189 | n.s. |
| ENSG00000132294 | 913.3894 | -0.255894127 | 0.269588 | -0.949203 | 0.342517 | 0.786682723 | EFR3A | protein_coding | 35.32 | -0.026664267 | 0.026664267 | ENSG00000132294 | n.s. |
| ENSG00000259804 | 93.99077 | -0.253574712 | 0.264937 | -0.957112 | 0.338511 | 0.784993146 | | antisense | 59.84 | -0.026659358 | 0.026659358 | ENSG00000259804 | n.s. |
| ENSG00000183060 | 506.7635 | -0.255762399 | 0.269405 | -0.949359 | 0.342438 | 0.786682723 | LYSMD4 | protein_coding | 47.75 | -0.026650541 | 0.026650541 | ENSG00000183060 | n.s. |
| ENSG00000115306 | 7194.541 | -0.233431333 | 0.233541 | -0.99953 | 0.317538 | 0.768883445 | SPTBN1 | protein_coding | 42.78 | -0.026643733 | 0.026643733 | ENSG00000115306 | n.s. |
| ENSG00000150768 | 885.8158 | -0.267056785 | 0.28975 | -0.921679 | 0.356696 | 0.794806874 | DLAT | protein_coding | 41.27 | -0.026635813 | 0.026635813 | ENSG00000150768 | n.s. |
| ENSG00000254027 | 25.87957 | -0.297431949 | 0.344333 | -0.863792 | 0.387702 | 0.81373794 | | antisense | 64.09 | -0.02662475 | 0.02662475 | ENSG00000254027 | n.s. |
| ENSG00000136450 | 14939.96 | -0.23137302 | 0.230049 | -1.005755 | 0.314533 | 0.767300529 | SRSF1 | protein_coding | 43.01 | -0.02661588 | 0.02661588 | ENSG00000136450 | n.s. |
| ENSG00000280092 | 36.82196 | 0.317717101 | 0.379939 | 0.8362325 | 0.403024 | 0.824575506 | | TEC | 41.59 | 0.026615025 | 0.026615025 | ENSG00000280092 | n.s. |
| ENSG00000124257 | 55.58303 | -0.270093983 | 0.294832 | -0.916096 | 0.359617 | 0.797029976 | NEURL2 | protein_coding | 56.82 | -0.026611103 | 0.026611103 | ENSG00000124257 | n.s. |
| ENSG00000164975 | 2451.857 | -0.246752385 | 0.253997 | -0.971478 | 0.33131 | 0.780141891 | SNAPC3 | protein_coding | 39.78 | -0.026606422 | 0.026606422 | ENSG00000164975 | n.s. |
| ENSG00000276071 | 96.47563 | 0.24480609 | 0.24996 | 0.979381 | 0.327392 | 0.778659927 | | lincRNA | 45.43 | 0.026598714 | 0.026598714 | ENSG00000276071 | n.s. |
| ENSG00000075218 | 677.2118 | -0.254255884 | 0.267099 | -0.951918 | 0.341139 | 0.786054116 | GTSE1 | protein_coding | 50.13 | -0.026581831 | 0.026581831 | ENSG00000075218 | n.s. |
| ENSG00000281508 | 268.5752 | -0.228988468 | 0.226329 | -1.011749 | 0.311658 | 0.765516281 | CDR1 | antisense | 44.31 | -0.026573097 | 0.026573097 | ENSG00000281508 | n.s. |
| ENSG00000177666 | 164.3143 | -0.240792845 | 0.244412 | -0.985192 | 0.32453 | 0.775648559 | PNPLA2 | protein_coding | 64.55 | -0.026567881 | 0.026567881 | ENSG00000177666 | n.s. |
| ENSG00000101608 | 1296.989 | 0.260079733 | 0.278408 | 0.9341688 | 0.350217 | 0.790467136 | MYL12A | protein_coding | 38.81 | 0.026558349 | 0.026558349 | ENSG00000101608 | n.s. |
| ENSG00000090097 | 1386.64 | -0.254621935 | 0.268483 | -0.948373 | 0.34294 | 0.786873561 | PCBP4 | protein_coding | 58.75 | -0.026504882 | 0.026504882 | ENSG00000090097 | n.s. |
| ENSG00000262050 | 22.58646 | -0.325368697 | 0.398503 | -0.816477 | 0.414227 | 0.829007487 | | antisense | 57.9 | -0.02649853 | 0.02649853 | ENSG00000262050 | n.s. |
| ENSG00000184508 | 240.7059 | 0.253437286 | 0.266389 | 0.9513803 | 0.341411 | 0.786054116 | HDDC3 | protein_coding | 56.14 | 0.026496248 | 0.026496248 | ENSG00000184508 | n.s. |
| ENSG00000058729 | 1070.361 | -0.253101964 | 0.265992 | -0.95154 | 0.34133 | 0.786054116 | RIOK2 | protein_coding | 36.71 | -0.026461191 | 0.026461191 | ENSG00000058729 | n.s. |
| ENSG00000164305 | 750.0602 | -0.264876138 | 0.287506 | -0.921289 | 0.3569 | 0.794806874 | CASP3 | protein_coding | 40.48 | -0.026418318 | 0.026418318 | ENSG00000164305 | n.s. |
| ENSG00000259366 | 42.66028 | -0.289170354 | 0.330071 | -0.876086 | 0.380983 | 0.810319533 | | sense_intronic | 40.61 | -0.026413888 | 0.026413888 | ENSG00000259366 | n.s. |
| ENSG00000160972 | 27.43925 | 0.319659335 | 0.387509 | 0.8249077 | 0.409424 | 0.826802059 | PPP1R16A | protein_coding | 57.4 | 0.026403366 | 0.026403366 | ENSG00000160972 | n.s. |
| ENSG00000099203 | 198.8753 | -0.269106653 | 0.294586 | -0.913508 | 0.360975 | 0.797882401 | TMED1 | protein_coding | 56.25 | -0.026388898 | 0.026388898 | ENSG00000099203 | n.s. |
| ENSG00000160404 | 166.3176 | -0.23858215 | 0.241902 | -0.986277 | 0.323997 | 0.775220833 | TOR2A | protein_coding | 59.86 | -0.026381117 | 0.026381117 | ENSG00000160404 | n.s. |
| ENSG00000160563 | 351.9 | -0.252215302 | 0.264777 | -0.952558 | 0.340814 | 0.786054116 | MED27 | protein_coding | 44 | -0.026368493 | 0.026368493 | ENSG00000160563 | n.s. |
| ENSG00000005486 | 50.35244 | -0.273595741 | 0.302415 | -0.904704 | 0.365622 | 0.801033089 | RHBDD2 | protein_coding | 47.79 | -0.026360825 | 0.026360825 | ENSG00000005486 | n.s. |
| ENSG00000108424 | 12762.98 | -0.244445302 | 0.25072 | -0.974975 | 0.329573 | 0.780141891 | KPNB1 | protein_coding | 43.15 | -0.026357657 | 0.026357657 | ENSG00000108424 | n.s. |
| ENSG00000268205 | 762.7363 | -0.261055807 | 0.281324 | -0.927956 | 0.353431 | 0.792950339 | | lincRNA | 36.36 | -0.02630242 | 0.02630242 | ENSG00000268205 | n.s. |
| ENSG00000115084 | 742.9284 | -0.256723247 | 0.274386 | -0.935628 | 0.349465 | 0.790125901 | SLC35F5 | protein_coding | 36.74 | -0.026263738 | 0.026263738 | ENSG00000115084 | n.s. |

| | | | | | | | | | | | | | |
|---|---|---|---|---|---|---|---|---|---|---|---|---|---|
| ENSG00000026036 | 376.8775 | -0.248529354 | 0.258182 | -0.962612 | 0.335742 | 0.784079552 | RTEL1-TNFRSF6E | protein_coding | 59.83 | -0.026254609 | 0.026254609 | ENSG00000026036 | n.s. |
| ENSG00000126653 | 1011.041 | -0.226027719 | 0.223271 | -1.012345 | 0.311373 | 0.765353944 | NSRP1 | protein_coding | 38.57 | -0.026250334 | 0.026250334 | ENSG00000126653 | n.s. |
| ENSG00000155729 | 231.2848 | -0.25627646 | 0.273707 | -0.936316 | 0.34911 | 0.790028419 | KCTD18 | protein_coding | 40.26 | -0.026231763 | 0.026231763 | ENSG00000155729 | n.s. |
| ENSG00000183617 | 417.1261 | -0.297863819 | 0.349408 | -0.852482 | 0.393947 | 0.81657325 | MRPL54 | protein_coding | 56.43 | -0.026213461 | 0.026213461 | ENSG00000183617 | n.s. |
| ENSG00000196363 | 1012.131 | -0.255430528 | 0.27147 | -0.940916 | 0.346748 | 0.789549679 | WDR5 | protein_coding | 55.32 | -0.026212418 | 0.026212418 | ENSG00000196363 | n.s. |
| ENSG00000135002 | 672.8635 | 0.264763255 | 0.288623 | 0.9173336 | 0.358968 | 0.79624783 | RFK | protein_coding | 40.38 | 0.026198784 | 0.026198784 | ENSG00000135002 | n.s. |
| ENSG00000134186 | 5142.148 | 0.221654502 | 0.216909 | 1.0218759 | 0.30684 | 0.761997697 | PRPF38B | protein_coding | 37.93 | 0.026165503 | 0.026165503 | ENSG00000134186 | n.s. |
| ENSG00000253797 | 806.3283 | -0.268386877 | 0.294476 | -0.911405 | 0.362082 | 0.798938342 | UTP14C | protein_coding | 41.96 | -0.026164161 | 0.026164161 | ENSG00000253797 | n.s. |
| ENSG00000272472 | 69.64068 | 0.290549907 | 0.334931 | 0.8674926 | 0.385672 | 0.812844991 | | sense_intronic | 31.94 | 0.026147244 | 0.026147244 | ENSG00000272472 | n.s. |
| ENSG00000179361 | 243.9491 | -0.229712972 | 0.230305 | -0.99743 | 0.318556 | 0.769456789 | ARID3B | protein_coding | 46.67 | -0.026144957 | 0.026144957 | ENSG00000179361 | n.s. |
| ENSG00000132581 | 444.5277 | 0.249764778 | 0.262582 | 0.9511861 | 0.34151 | 0.786054339 | SDF2 | protein_coding | 47.11 | 0.026112266 | 0.026112266 | ENSG00000132581 | n.s. |
| ENSG00000185149 | 268.2047 | -0.255657576 | 0.273687 | -0.934124 | 0.35024 | 0.790467136 | NPY2R | protein_coding | 39.63 | -0.026106775 | 0.026106775 | ENSG00000185149 | n.s. |
| ENSG00000277342 | 379.9302 | 0.226911788 | 0.226053 | 1.0038008 | 0.315475 | 0.767425784 | | sense_intronic | 30.36 | 0.026086599 | 0.026086599 | ENSG00000277342 | n.s. |
| ENSG00000114737 | 187.0135 | -0.233738504 | 0.236048 | -0.990214 | 0.32207 | 0.773444264 | CISH | protein_coding | 57 | -0.026078433 | 0.026078433 | ENSG00000114737 | n.s. |
| ENSG00000207751 | 136.8983 | -0.23370091 | 0.236017 | -0.990187 | 0.322083 | 0.773444264 | MIR130B | processed_transcript | 53.58 | -0.026074239 | 0.026074239 | ENSG00000207751 | n.s. |
| ENSG00000185947 | 721.7094 | -0.254600591 | 0.271857 | -0.936523 | 0.349004 | 0.790028419 | ZNF267 | protein_coding | 38.82 | -0.026060226 | 0.026060226 | ENSG00000185947 | n.s. |
| ENSG00000111450 | 1103.055 | -0.254580134 | 0.272012 | -0.935916 | 0.349316 | 0.790028419 | STX2 | protein_coding | 45.87 | -0.026058132 | 0.026058132 | ENSG00000111450 | n.s. |
| ENSG00000204316 | 136.0819 | -0.247497911 | 0.258052 | -0.959103 | 0.337507 | 0.78475552 | MRPL38 | protein_coding | 57.66 | -0.026053021 | 0.026053021 | ENSG00000204316 | n.s. |
| ENSG00000123358 | 231.3748 | -0.251311368 | 0.265558 | -0.946351 | 0.343969 | 0.787678531 | NR4A1 | protein_coding | 56.01 | -0.026048672 | 0.026048672 | ENSG00000123358 | n.s. |
| ENSG00000262049 | 403.6864 | -0.217705437 | 0.210168 | -1.035865 | 0.300265 | 0.759327693 | | antisense | 61.96 | -0.026031205 | 0.026031205 | ENSG00000262049 | n.s. |
| ENSG00000201863 | 13.12207 | -0.362303284 | 0.473624 | -0.76496 | 0.444295 | 0.847626065 | | snoRNA | 53.23 | -0.026011817 | 0.026011817 | ENSG00000201863 | n.s. |
| ENSG00000276855 | 24.75419 | 0.316404262 | 0.38446 | 0.8229836 | 0.410517 | 0.827563884 | | lincRNA | 33.62 | 0.026007946 | 0.026007946 | ENSG00000276855 | n.s. |
| ENSG00000128641 | 6801.469 | -0.239993264 | 0.245329 | -0.97825 | 0.327951 | 0.779364976 | MYO1B | protein_coding | 39.67 | -0.025981458 | 0.025981458 | ENSG00000128641 | n.s. |
| ENSG00000239857 | 375.4283 | -0.269644711 | 0.298084 | -0.904594 | 0.36568 | 0.801033089 | GET4 | protein_coding | 59.75 | -0.025980145 | 0.025980145 | ENSG00000239857 | n.s. |
| ENSG00000128607 | 772.7787 | -0.263904409 | 0.288438 | -0.914943 | 0.360222 | 0.797273365 | KLHDC10 | protein_coding | 40.14 | -0.025966279 | 0.025966279 | ENSG00000128607 | n.s. |
| ENSG00000269439 | 51.69158 | -0.288957256 | 0.333462 | -0.866538 | 0.386195 | 0.813115862 | | lincRNA | 52.93 | -0.025962106 | 0.025962106 | ENSG00000269439 | n.s. |
| ENSG00000118482 | 4465.042 | -0.246371208 | 0.256588 | -0.960182 | 0.336964 | 0.784604238 | PHF3 | protein_coding | 36.07 | -0.025955047 | 0.025955047 | ENSG00000118482 | n.s. |
| ENSG00000220205 | 232.7742 | -0.232002495 | 0.23407 | -0.991166 | 0.321604 | 0.773083314 | VAMP2 | protein_coding | 58.14 | -0.025931777 | 0.025931777 | ENSG00000220205 | n.s. |
| ENSG00000240964 | 31.12915 | -0.30974538 | 0.372698 | -0.83109 | 0.405923 | 0.824799944 | RN7SL751P | misc_RNA | 50.35 | -0.025910628 | 0.025910628 | ENSG00000240964 | n.s. |
| ENSG00000143643 | 434.76 | -0.267571586 | 0.295053 | -0.90686 | 0.364481 | 0.800144301 | TTC13 | protein_coding | 39.73 | -0.025909407 | 0.025909407 | ENSG00000143643 | n.s. |
| ENSG00000266618 | 16.69512 | -0.312385055 | 0.377646 | -0.827191 | 0.408129 | 0.826171518 | MIR4742 | miRNA | 34.12 | -0.025906024 | 0.025906024 | ENSG00000266618 | n.s. |
| ENSG00000137502 | 1270.039 | -0.253470461 | 0.271091 | -0.935001 | 0.349788 | 0.790343315 | RAB30 | protein_coding | 41.21 | -0.02590068 | 0.02590068 | ENSG00000137502 | n.s. |

| | | | | | | | | | | | | | |
|---|---|---|---|---|---|---|---|---|---|---|---|---|---|
| ENSG00000177646 | 701.2994 | -0.248431971 | 0.261624 | -0.949576 | 0.342328 | 0.786682723 | ACAD9 | protein_coding | 49.2 | -0.025886708 | 0.025886708 | ENSG00000177646 | n.s. |
| ENSG00000264268 | 5.454097 | 0.437219825 | 0.634963 | 0.6885749 | 0.491091 | 0.872562973 | MIR4767 | miRNA | 82.05 | 0.025884821 | 0.025884821 | ENSG00000264268 | n.s. |
| ENSG00000166887 | 830.983 | -0.245737345 | 0.256246 | -0.95899 | 0.337564 | 0.78475552 | VPS39 | protein_coding | 42.61 | -0.025867694 | 0.025867694 | ENSG00000166887 | n.s. |
| ENSG00000141556 | 2415.363 | -0.242136832 | 0.2508 | -0.965458 | 0.334315 | 0.781938745 | TBCD | protein_coding | 52.84 | -0.025866816 | 0.025866816 | ENSG00000141556 | n.s. |
| ENSG00000263272 | 88.7677 | -0.252688746 | 0.269178 | -0.938741 | 0.347864 | 0.790017511 | | antisense | 45.52 | -0.025866049 | 0.025866049 | ENSG00000263272 | n.s. |
| ENSG00000137812 | 2199.676 | -0.2510433 | 0.265981 | -0.943838 | 0.345252 | 0.788822231 | KNL1 | protein_coding | 41.23 | -0.025862696 | 0.025862696 | ENSG00000137812 | n.s. |
| ENSG00000203644 | 154.2704 | -0.283961744 | 0.326582 | -0.869496 | 0.384576 | 0.810911249 | | sense_intronic | 33.96 | -0.025848093 | 0.025848093 | ENSG00000203644 | n.s. |
| ENSG00000122417 | 1332.582 | -0.250319787 | 0.264889 | -0.945 | 0.344659 | 0.788437824 | ODF2L | protein_coding | 34.88 | -0.02584115 | 0.02584115 | ENSG00000122417 | n.s. |
| ENSG00000072736 | 228.8297 | -0.231453853 | 0.23374 | -0.990219 | 0.322067 | 0.773444264 | NFATC3 | protein_coding | 41.53 | -0.025823532 | 0.025823532 | ENSG00000072736 | n.s. |
| ENSG00000276131 | 25.5919 | -0.308330659 | 0.369044 | -0.835485 | 0.403445 | 0.824699011 | | antisense | 56.64 | -0.025808672 | 0.025808672 | ENSG00000276131 | n.s. |
| ENSG00000260296 | 48.65462 | -0.378012773 | 0.506273 | -0.746658 | 0.45527 | 0.85453422 | | sense_overlapping | 32.58 | -0.025807137 | 0.025807137 | ENSG00000260296 | n.s. |
| ENSG00000198680 | 233.6 | -0.244185906 | 0.253731 | -0.962383 | 0.335857 | 0.784079552 | TUSC1 | protein_coding | 52.44 | -0.025795768 | 0.025795768 | ENSG00000198680 | n.s. |
| ENSG00000234685 | 43.62672 | 0.282194537 | 0.322446 | 0.8751675 | 0.381483 | 0.81032471 | NUS1P2 | processed_pseudogene | 54.41 | 0.025775908 | 0.025775908 | ENSG00000234685 | n.s. |
| ENSG00000259396 | 42.32334 | -0.279768866 | 0.316402 | -0.88422 | 0.376578 | 0.808862777 | | sense_intronic | 45.55 | -0.025773749 | 0.025773749 | ENSG00000259396 | n.s. |
| ENSG00000260519 | 19.61693 | 0.345929617 | 0.441476 | 0.7835755 | 0.433289 | 0.842381919 | | antisense | 54.97 | 0.02576863 | 0.02576863 | ENSG00000260519 | n.s. |
| ENSG00000147535 | 110.9169 | 0.254525069 | 0.273365 | 0.9310806 | 0.351812 | 0.792117552 | PLPP5 | protein_coding | 45.35 | 0.025760575 | 0.025760575 | ENSG00000147535 | n.s. |
| ENSG00000103126 | 385.1545 | -0.240077428 | 0.248125 | -0.967566 | 0.333261 | 0.781097236 | AXIN1 | protein_coding | 54.36 | -0.025759083 | 0.025759083 | ENSG00000103126 | n.s. |
| ENSG00000254165 | 174.7482 | 0.238814596 | 0.245146 | 0.9741747 | 0.32997 | 0.780141891 | | antisense | 31.96 | 0.025750519 | 0.025750519 | ENSG00000254165 | n.s. |
| ENSG00000113712 | 3362.564 | -0.238641896 | 0.246105 | -0.969676 | 0.332208 | 0.780141891 | CSNK1A1 | protein_coding | 39.26 | -0.025731897 | 0.025731897 | ENSG00000113712 | n.s. |
| ENSG00000182827 | 1329.341 | -0.238578487 | 0.245075 | -0.973492 | 0.330309 | 0.780141891 | ACBD3 | protein_coding | 41.45 | -0.02572506 | 0.02572506 | ENSG00000182827 | n.s. |
| ENSG00000272631 | 747.5073 | 0.217715125 | 0.21206 | 1.0266674 | 0.304577 | 0.761997697 | | antisense | 44.98 | 0.025700474 | 0.025700474 | ENSG00000272631 | n.s. |
| ENSG00000123154 | 225.5647 | -0.251027889 | 0.268181 | -0.936039 | 0.349253 | 0.790028419 | WDR83 | protein_coding | 55.04 | -0.025694534 | 0.025694534 | ENSG00000123154 | n.s. |
| ENSG00000167232 | 1891.027 | -0.245741569 | 0.258323 | -0.951298 | 0.341453 | 0.786054116 | ZNF91 | protein_coding | 40.56 | -0.02569168 | 0.02569168 | ENSG00000167232 | n.s. |
| ENSG00000186446 | 506.0435 | 0.215939215 | 0.20901 | 1.0331545 | 0.301532 | 0.76041019 | ZNF501 | protein_coding | 39.42 | 0.025686417 | 0.025686417 | ENSG00000186446 | n.s. |
| ENSG00000230828 | 279.9693 | 0.23820797 | 0.244979 | 0.9723594 | 0.330872 | 0.780141891 | | processed_pseudogene | 42.49 | 0.025685108 | 0.025685108 | ENSG00000230828 | n.s. |
| ENSG00000013275 | 1236.811 | -0.254830029 | 0.27437 | -0.928783 | 0.353001 | 0.792950339 | PSMC4 | protein_coding | 50.25 | -0.025675148 | 0.025675148 | ENSG00000013275 | n.s. |
| ENSG00000158793 | 319.5863 | -0.250774503 | 0.266928 | -0.939483 | 0.347483 | 0.790017511 | NIT1 | protein_coding | 49.13 | -0.025670101 | 0.025670101 | ENSG00000158793 | n.s. |
| ENSG00000130558 | 2750.269 | -0.261735657 | 0.286476 | -0.913638 | 0.360907 | 0.797857548 | OLFM1 | protein_coding | 55.24 | -0.025669631 | 0.025669631 | ENSG00000130558 | n.s. |
| ENSG00000119396 | 1031.743 | -0.249065962 | 0.263902 | -0.943782 | 0.345281 | 0.788822231 | RAB14 | protein_coding | 39.61 | -0.025658989 | 0.025658989 | ENSG00000119396 | n.s. |
| ENSG00000143799 | 8160.764 | -0.249158691 | 0.264355 | -0.942516 | 0.345929 | 0.788921338 | PARP1 | protein_coding | 46.8 | -0.025654948 | 0.025654948 | ENSG00000143799 | n.s. |
| ENSG00000011332 | 77.53484 | -0.275872409 | 0.309829 | -0.890403 | 0.37325 | 0.807377464 | DPF1 | protein_coding | 57.15 | -0.025634996 | 0.025634996 | ENSG00000011332 | n.s. |
| ENSG00000103507 | 167.6785 | -0.243292385 | 0.25334 | -0.960338 | 0.336885 | 0.784604238 | BCKDK | protein_coding | 57.82 | -0.025630695 | 0.025630695 | ENSG00000103507 | n.s. |

| | | | | | | | | | | | | | |
|---|---|---|---|---|---|---|---|---|---|---|---|---|---|
| ENSG00000146143 | 1090.253 | -0.250872073 | 0.268425 | -0.934608 | 0.34999 | 0.790401174 | PRIM2 | protein_coding | 37.46 | -0.02562719 | 0.02562719 | ENSG00000146143 | n.s. |
| ENSG00000141646 | 3116.697 | -0.248477027 | 0.263105 | -0.944401 | 0.344965 | 0.788790309 | SMAD4 | protein_coding | 37.94 | -0.025602684 | 0.025602684 | ENSG00000141646 | n.s. |
| ENSG00000121940 | 1272.155 | -0.250402586 | 0.267744 | -0.93523 | 0.34967 | 0.7903292 | CLCC1 | protein_coding | 41.67 | -0.025589134 | 0.025589134 | ENSG00000121940 | n.s. |
| ENSG00000140474 | 876.9907 | -0.244616537 | 0.256327 | -0.954315 | 0.339924 | 0.785979841 | ULK3 | protein_coding | 59.87 | -0.025584099 | 0.025584099 | ENSG00000140474 | n.s. |
| ENSG00000168283 | 1944.202 | 0.256427077 | 0.278383 | 0.9211306 | 0.356982 | 0.794806874 | BMI1 | protein_coding | 41.6 | 0.025575623 | 0.025575623 | ENSG00000168283 | n.s. |
| ENSG00000072518 | 71.08239 | -0.250827371 | 0.268656 | -0.933637 | 0.350491 | 0.790777278 | MARK2 | protein_coding | 49.65 | -0.025570802 | 0.025570802 | ENSG00000072518 | n.s. |
| ENSG00000136098 | 319.9557 | -0.246591693 | 0.26037 | -0.947081 | 0.343598 | 0.787624904 | NEK3 | protein_coding | 39.71 | -0.025566765 | 0.025566765 | ENSG00000136098 | n.s. |
| ENSG00000272047 | 583.0203 | -0.274334511 | 0.306879 | -0.893951 | 0.371348 | 0.806899249 | GTF2H5 | protein_coding | 40.89 | -0.025562679 | 0.025562679 | ENSG00000272047 | n.s. |
| ENSG00000083937 | 457.4579 | -0.263954407 | 0.290956 | -0.907196 | 0.364303 | 0.800144301 | CHMP2B | protein_coding | 34.78 | -0.02555915 | 0.02555915 | ENSG00000083937 | n.s. |
| ENSG00000260708 | 167.3141 | 0.237006747 | 0.244356 | 0.9699228 | 0.332085 | 0.780141891 | | antisense | 51.64 | 0.025555585 | 0.025555585 | ENSG00000260708 | n.s. |
| ENSG00000119231 | 1711.096 | -0.244322206 | 0.256103 | -0.953999 | 0.340084 | 0.785979841 | SENP5 | protein_coding | 42.03 | -0.025553316 | 0.025553316 | ENSG00000119231 | n.s. |
| ENSG00000279329 | 82.35578 | 0.256004576 | 0.277946 | 0.9210586 | 0.35702 | 0.794806874 | | TEC | 49.92 | 0.025533483 | 0.025533483 | ENSG00000279329 | n.s. |
| ENSG00000272711 | 229.9739 | -0.248501499 | 0.263916 | -0.941593 | 0.346401 | 0.789327092 | | lincRNA | 51.84 | -0.025531787 | 0.025531787 | ENSG00000272711 | n.s. |
| ENSG00000164118 | 6825.42 | -0.248700333 | 0.264341 | -0.94083 | 0.346792 | 0.789549679 | CEP44 | protein_coding | 35.66 | -0.025521762 | 0.025521762 | ENSG00000164118 | n.s. |
| ENSG00000089199 | 1703.238 | -0.236691638 | 0.242907 | -0.974411 | 0.329852 | 0.780141891 | CHGB | protein_coding | 42.87 | -0.025521608 | 0.025521608 | ENSG00000089199 | n.s. |
| ENSG00000089682 | 442.1547 | -0.243997411 | 0.255889 | -0.953529 | 0.340322 | 0.785979841 | RBM41 | protein_coding | 36.02 | -0.025519346 | 0.025519346 | ENSG00000089682 | n.s. |
| ENSG00000149503 | 1127.588 | -0.223573024 | 0.223722 | -0.999335 | 0.317632 | 0.768978814 | INCENP | protein_coding | 53.88 | -0.025506468 | 0.025506468 | ENSG00000149503 | n.s. |
| ENSG00000087995 | 959.502 | -0.244818505 | 0.258238 | -0.948034 | 0.343112 | 0.787036737 | METTL2A | protein_coding | 45.2 | -0.025462347 | 0.025462347 | ENSG00000087995 | n.s. |
| ENSG00000099901 | 3979.431 | -0.262108716 | 0.288631 | -0.908109 | 0.363821 | 0.799601391 | RANBP1 | protein_coding | 57.69 | -0.025457692 | 0.025457692 | ENSG00000099901 | n.s. |
| ENSG00000264940 | 51.37796 | -0.264082048 | 0.291671 | -0.905411 | 0.365248 | 0.801033089 | SNORD3C | snoRNA | 54.91 | -0.025444185 | 0.025444185 | ENSG00000264940 | n.s. |
| ENSG00000083290 | 705.9676 | -0.244007958 | 0.25693 | -0.949706 | 0.342261 | 0.786682723 | ULK2 | protein_coding | 41.45 | -0.025425724 | 0.025425724 | ENSG00000083290 | n.s. |
| ENSG00000162194 | 195.9675 | -0.223869453 | 0.224777 | -0.995961 | 0.319269 | 0.769994426 | LBHD1 | protein_coding | 52.45 | -0.025411963 | 0.025411963 | ENSG00000162194 | n.s. |
| ENSG00000100319 | 172.3455 | 0.380129576 | 0.513945 | 0.7396311 | 0.459524 | 0.857420077 | ZMAT5 | protein_coding | 52.77 | 0.02539507 | 0.02539507 | ENSG00000100319 | n.s. |
| ENSG00000187953 | 385.699 | -0.235219454 | 0.242653 | -0.969365 | 0.332363 | 0.780141891 | PMS2CL | cribed_unprocessed_pseudc | 48.28 | -0.025362867 | 0.025362867 | ENSG00000187953 | n.s. |
| ENSG00000165661 | 776.3429 | -0.2517038 | 0.271105 | -0.928437 | 0.353181 | 0.792950339 | QSOX2 | protein_coding | 54.55 | -0.025360167 | 0.025360167 | ENSG00000165661 | n.s. |
| ENSG00000259015 | 13.93709 | 0.331071388 | 0.417916 | 0.7921968 | 0.428246 | 0.838312087 | | processed_pseudogene | 51.75 | 0.025358172 | 0.025358172 | ENSG00000259015 | n.s. |
| ENSG00000136379 | 234.17 | -0.242289161 | 0.254104 | -0.953503 | 0.340335 | 0.785979841 | ABHD17C | protein_coding | 42.95 | -0.025340682 | 0.025340682 | ENSG00000136379 | n.s. |
| ENSG00000231566 | 32.19419 | -0.293098568 | 0.345757 | -0.847701 | 0.396605 | 0.819495485 | | lincRNA | 48.32 | -0.025339378 | 0.025339378 | ENSG00000231566 | n.s. |
| ENSG00000265566 | 13.48791 | -0.343360691 | 0.442123 | -0.776618 | 0.437384 | 0.843772375 | RN7SL605P | misc_RNA | 55.16 | -0.025331331 | 0.025331331 | ENSG00000265566 | n.s. |
| ENSG00000169223 | 820.0957 | -0.262729308 | 0.290315 | -0.90498 | 0.365476 | 0.801033089 | LMAN2 | protein_coding | 50.49 | -0.025313849 | 0.025313849 | ENSG00000169223 | n.s. |
| ENSG00000106397 | 139.7138 | -0.245625916 | 0.260218 | -0.943923 | 0.345209 | 0.788822231 | PLOD3 | protein_coding | 58.5 | -0.025304593 | 0.025304593 | ENSG00000106397 | n.s. |
| ENSG00000107164 | 1459.136 | -0.246767118 | 0.263282 | -0.937273 | 0.348618 | 0.790017511 | FUBP3 | protein_coding | 47.12 | -0.025259892 | 0.025259892 | ENSG00000107164 | n.s. |

| | | | | | | | | | | | | | |
|---|---|---|---|---|---|---|---|---|---|---|---|---|---|
| ENSG00000153179 | 489.8855 | -0.245795214 | 0.261091 | -0.941416 | 0.346492 | 0.789327092 | RASSF3 | protein_coding | 43.87 | -0.025253735 | 0.025253735 | ENSG00000153179 | n.s. |
| ENSG00000100403 | 1583.31 | -0.242538562 | 0.255707 | -0.948504 | 0.342873 | 0.786850558 | ZC3H7B | protein_coding | 54.29 | -0.025250142 | 0.025250142 | ENSG00000100403 | n.s. |
| ENSG00000207309 | 19.97186 | 0.305584723 | 0.370373 | 0.8250732 | 0.40933 | 0.826802059 | RNU4-70P | snRNA | 41.01 | 0.025240825 | 0.025240825 | ENSG00000207309 | n.s. |
| ENSG00000240024 | 197.0379 | -0.262516937 | 0.290371 | -0.904074 | 0.365956 | 0.801511738 | LINC00888 | cribed_unprocessed_pseudc | 39.85 | -0.025225282 | 0.025225282 | ENSG00000240024 | n.s. |
| ENSG00000198315 | 1417.06 | -0.240297359 | 0.25137 | -0.95595 | 0.339098 | 0.785305607 | ZKSCAN8 | protein_coding | 38.6 | -0.025221924 | 0.025221924 | ENSG00000198315 | n.s. |
| ENSG00000013810 | 861.731 | -0.244809378 | 0.259412 | -0.943708 | 0.345319 | 0.788822231 | TACC3 | protein_coding | 56.07 | -0.025220472 | 0.025220472 | ENSG00000013810 | n.s. |
| ENSG00000157184 | 366.1571 | -0.265407869 | 0.294841 | -0.900173 | 0.368028 | 0.803511983 | CPT2 | protein_coding | 44.68 | -0.025215776 | 0.025215776 | ENSG00000157184 | n.s. |
| ENSG00000121931 | 556.2855 | -0.258798916 | 0.284268 | -0.910406 | 0.362608 | 0.799058295 | LRIF1 | protein_coding | 36.58 | -0.025212588 | 0.025212588 | ENSG00000121931 | n.s. |
| ENSG00000234345 | 41.0088 | -0.285839934 | 0.335078 | -0.853054 | 0.393629 | 0.816295599 | | processed_pseudogene | 44.18 | -0.025197518 | 0.025197518 | ENSG00000234345 | n.s. |
| ENSG00000188613 | 377.5816 | 0.239406484 | 0.25031 | 0.9564399 | 0.33885 | 0.784993146 | NANOS1 | protein_coding | 50.86 | 0.025169794 | 0.025169794 | ENSG00000188613 | n.s. |
| ENSG00000283154 | 723.2233 | 0.218050188 | 0.216206 | 1.0085293 | 0.3132 | 0.766650888 | IQCJ-SCHIP1 | protein_coding | 39 | 0.025163506 | 0.025163506 | ENSG00000283154 | n.s. |
| ENSG00000188994 | 4376.875 | -0.244901977 | 0.26013 | -0.941461 | 0.346469 | 0.789327092 | ZNF292 | protein_coding | 37.67 | -0.025161962 | 0.025161962 | ENSG00000188994 | n.s. |
| ENSG00000014164 | 173.62 | -0.231461369 | 0.23631 | -0.979481 | 0.327342 | 0.778659927 | ZC3H3 | protein_coding | 58.38 | -0.025148781 | 0.025148781 | ENSG00000014164 | n.s. |
| ENSG00000177410 | 3139.469 | 0.211206681 | 0.204403 | 1.033286 | 0.30147 | 0.76041019 | ZFAS1 | antisense | 45.74 | 0.025123472 | 0.025123472 | ENSG00000177410 | n.s. |
| ENSG00000101928 | 194.7554 | -0.291610341 | 0.344849 | -0.845617 | 0.397767 | 0.820246021 | MOSPD1 | protein_coding | 40.84 | -0.025094781 | 0.025094781 | ENSG00000101928 | n.s. |
| ENSG00000204334 | 24.69949 | 0.302489842 | 0.365259 | 0.828152 | 0.407584 | 0.826146204 | ERICH2 | protein_coding | 37.22 | 0.025089442 | 0.025089442 | ENSG00000204334 | n.s. |
| ENSG00000172167 | 1018.673 | -0.240423006 | 0.253422 | -0.948705 | 0.342771 | 0.786744925 | MTBP | protein_coding | 35.31 | -0.025043915 | 0.025043915 | ENSG00000172167 | n.s. |
| ENSG00000272518 | 62.36921 | -0.282978801 | 0.32933 | -0.859255 | 0.3902 | 0.815710819 | | sense_intronic | 40.02 | -0.025033374 | 0.025033374 | ENSG00000272518 | n.s. |
| ENSG00000101752 | 3383.376 | -0.25089186 | 0.272473 | -0.920795 | 0.357157 | 0.794806874 | MIB1 | protein_coding | 38.64 | -0.025023549 | 0.025023549 | ENSG00000101752 | n.s. |
| ENSG00000173852 | 590.9237 | -0.250861096 | 0.271705 | -0.923283 | 0.35586 | 0.794802373 | DPY19L1 | protein_coding | 37.49 | -0.025021098 | 0.025021098 | ENSG00000173852 | n.s. |
| ENSG00000189079 | 3389.283 | -0.244419913 | 0.260498 | -0.938279 | 0.348101 | 0.790017511 | ARID2 | protein_coding | 36.1 | -0.025019625 | 0.025019625 | ENSG00000189079 | n.s. |
| ENSG00000259931 | 11.1753 | -0.335910448 | 0.428999 | -0.783009 | 0.433622 | 0.84244096 | | processed_pseudogene | 39.11 | -0.025012068 | 0.025012068 | ENSG00000259931 | n.s. |
| ENSG00000122741 | 626.9196 | -0.259382723 | 0.286716 | -0.904667 | 0.365642 | 0.801033089 | DCAF10 | protein_coding | 38.56 | -0.024991407 | 0.024991407 | ENSG00000122741 | n.s. |
| ENSG00000129911 | 232.2521 | -0.237817552 | 0.248706 | -0.956219 | 0.338962 | 0.785120835 | KLF16 | protein_coding | 65.43 | -0.024985944 | 0.024985944 | ENSG00000129911 | n.s. |
| ENSG00000166860 | 1554.717 | 0.217193289 | 0.215902 | 1.0059824 | 0.314424 | 0.767300529 | ZBTB39 | protein_coding | 49.81 | 0.024984722 | 0.024984722 | ENSG00000166860 | n.s. |
| ENSG00000196923 | 172.1518 | -0.246776041 | 0.265159 | -0.930671 | 0.352024 | 0.792117552 | PDLIM7 | protein_coding | 60.73 | -0.024976293 | 0.024976293 | ENSG00000196923 | n.s. |
| ENSG00000168175 | 2155.137 | -0.244564352 | 0.261749 | -0.934346 | 0.350126 | 0.79046507 | MAPK1IP1L | protein_coding | 40.17 | -0.024974255 | 0.024974255 | ENSG00000168175 | n.s. |
| ENSG00000151746 | 3514.666 | 0.231547318 | 0.23786 | 0.9734607 | 0.330324 | 0.780141891 | BICD1 | protein_coding | 40.45 | 0.024966914 | 0.024966914 | ENSG00000151746 | n.s. |
| ENSG00000188517 | 136.5381 | 0.250194978 | 0.271055 | 0.9230414 | 0.355986 | 0.794802462 | COL25A1 | protein_coding | 36.92 | 0.024954646 | 0.024954646 | ENSG00000188517 | n.s. |
| ENSG00000140830 | 169.2256 | -0.246837219 | 0.265424 | -0.929975 | 0.352384 | 0.792352298 | TXNL4B | protein_coding | 44.45 | -0.02495072 | 0.02495072 | ENSG00000140830 | n.s. |
| ENSG00000134982 | 1875.7 | -0.243611944 | 0.259907 | -0.937306 | 0.348601 | 0.790017511 | APC | protein_coding | 37.63 | -0.024936918 | 0.024936918 | ENSG00000134982 | n.s. |
| ENSG00000271793 | 149.0392 | -0.267711513 | 0.300096 | -0.892087 | 0.372346 | 0.807126641 | | protein_coding | 38.3 | -0.024912783 | 0.024912783 | ENSG00000271793 | n.s. |

| | | | | | | | | | | | | | |
|---|---|---|---|---|---|---|---|---|---|---|---|---|---|
| ENSG00000134153 | 465.4485 | -0.261508077 | 0.290119 | -0.901381 | 0.367386 | 0.803125887 | EMC7 | protein_coding | 39.72 | -0.024899851 | 0.024899851 | ENSG00000134153 | n.s. |
| ENSG00000271347 | 75.04064 | -0.272365106 | 0.310863 | -0.876157 | 0.380945 | 0.810319533 | | sense_intronic | 26.49 | -0.024878834 | 0.024878834 | ENSG00000271347 | n.s. |
| ENSG00000053900 | 752.5076 | -0.253628382 | 0.277564 | -0.913767 | 0.360839 | 0.797857548 | ANAPC4 | protein_coding | 37.97 | -0.024874513 | 0.024874513 | ENSG00000053900 | n.s. |
| ENSG00000180979 | 451.5013 | -0.244781711 | 0.26249 | -0.932537 | 0.351059 | 0.791500843 | LRRC57 | protein_coding | 43.17 | -0.024857245 | 0.024857245 | ENSG00000180979 | n.s. |
| ENSG00000234031 | 57.65205 | -0.270219034 | 0.306915 | -0.880437 | 0.378623 | 0.809266943 | RPS3AP44 | processed_pseudogene | 41.01 | -0.024835345 | 0.024835345 | ENSG00000234031 | n.s. |
| ENSG00000107341 | 1102.867 | -0.246485068 | 0.265631 | -0.927923 | 0.353448 | 0.792950339 | UBE2R2 | protein_coding | 42.11 | -0.024834359 | 0.024834359 | ENSG00000107341 | n.s. |
| ENSG00000111802 | 552.7125 | -0.244063451 | 0.261534 | -0.933199 | 0.350717 | 0.791159551 | TDP2 | protein_coding | 39.02 | -0.024830021 | 0.024830021 | ENSG00000111802 | n.s. |
| ENSG00000265432 | 6.101273 | -0.405354203 | 0.578683 | -0.700477 | 0.48363 | 0.868492686 | MIR4308 | miRNA | 40.74 | -0.024821392 | 0.024821392 | ENSG00000265432 | n.s. |
| ENSG00000022976 | 223.269 | -0.259826125 | 0.287925 | -0.902408 | 0.36684 | 0.802563919 | ZNF839 | protein_coding | 49.04 | -0.024818687 | 0.024818687 | ENSG00000022976 | n.s. |
| ENSG00000111206 | 1708.362 | -0.231285111 | 0.239111 | -0.967269 | 0.33341 | 0.781097236 | FOXM1 | protein_coding | 48.07 | -0.024815713 | 0.024815713 | ENSG00000111206 | n.s. |
| ENSG00000272034 | 156.7274 | 0.226100597 | 0.22986 | 0.9836466 | 0.325289 | 0.776688313 | SNORD14A | snoRNA | 42.86 | 0.024815271 | 0.024815271 | ENSG00000272034 | n.s. |
| ENSG00000147526 | 147.7428 | 0.23716844 | 0.248601 | 0.9540106 | 0.340078 | 0.785979841 | TACC1 | protein_coding | 42.76 | 0.024805113 | 0.024805113 | ENSG00000147526 | n.s. |
| ENSG00000196376 | 210.6184 | -0.235549817 | 0.245475 | -0.959567 | 0.337273 | 0.78475552 | SLC35F1 | protein_coding | 38.46 | -0.024795298 | 0.024795298 | ENSG00000196376 | n.s. |
| ENSG00000252622 | 35.15164 | 0.280013908 | 0.326312 | 0.8581172 | 0.390828 | 0.815710819 | RNU6-881P | snRNA | 40.2 | 0.024771088 | 0.024771088 | ENSG00000252622 | n.s. |
| ENSG00000266236 | 39.46702 | -0.269418295 | 0.305196 | -0.88277 | 0.37736 | 0.809223062 | NARF-IT1 | sense_intronic | 49.03 | -0.024768095 | 0.024768095 | ENSG00000266236 | n.s. |
| ENSG00000126870 | 299.569 | -0.248118684 | 0.269336 | -0.921225 | 0.356933 | 0.794806874 | WDR60 | protein_coding | 46.03 | -0.024746957 | 0.024746957 | ENSG00000126870 | n.s. |
| ENSG00000270181 | 782.8807 | -0.248034273 | 0.268887 | -0.922449 | 0.356295 | 0.794806874 | BIVM-ERCC5 | protein_coding | 40.01 | -0.024738538 | 0.024738538 | ENSG00000270181 | n.s. |
| ENSG00000108528 | 543.9018 | -0.244688641 | 0.263113 | -0.929976 | 0.352384 | 0.792352298 | SLC25A11 | protein_coding | 58.68 | -0.024733538 | 0.024733538 | ENSG00000108528 | n.s. |
| ENSG00000102882 | 192.8682 | -0.263795587 | 0.294326 | -0.896269 | 0.370109 | 0.805916031 | MAPK3 | protein_coding | 56.36 | -0.024720339 | 0.024720339 | ENSG00000102882 | n.s. |
| ENSG00000139132 | 2030.973 | -0.23423483 | 0.243665 | -0.961297 | 0.336403 | 0.78441387 | FGD4 | protein_coding | 40.83 | -0.024701172 | 0.024701172 | ENSG00000139132 | n.s. |
| ENSG00000089685 | 3859.676 | -0.247327791 | 0.267496 | -0.924602 | 0.355173 | 0.794644486 | BIRC5 | protein_coding | 49.27 | -0.024690022 | 0.024690022 | ENSG00000089685 | n.s. |
| ENSG00000120029 | 388.3582 | -0.247504656 | 0.268014 | -0.923476 | 0.355759 | 0.794802373 | C10orf76 | protein_coding | 42.35 | -0.024686324 | 0.024686324 | ENSG00000120029 | n.s. |
| ENSG00000143951 | 188.4421 | -0.256153664 | 0.283137 | -0.904698 | 0.365626 | 0.801033089 | WDPCP | protein_coding | 37.72 | -0.024680289 | 0.024680289 | ENSG00000143951 | n.s. |
| ENSG00000183943 | 840.4992 | -0.240348878 | 0.255369 | -0.941184 | 0.34661 | 0.789441767 | PRKX | protein_coding | 45.74 | -0.024678999 | 0.024678999 | ENSG00000183943 | n.s. |
| ENSG00000086848 | 256.3088 | -0.25310817 | 0.27778 | -0.911183 | 0.362199 | 0.798938342 | ALG9 | protein_coding | 40.69 | -0.024674689 | 0.024674689 | ENSG00000086848 | n.s. |
| ENSG00000198887 | 2246.017 | -0.236486912 | 0.248853 | -0.950308 | 0.341956 | 0.786560769 | SMC5 | protein_coding | 36.37 | -0.024657951 | 0.024657951 | ENSG00000198887 | n.s. |
| ENSG00000160305 | 145.876 | 0.277116853 | 0.321631 | 0.8615974 | 0.388909 | 0.814823372 | DIP2A | protein_coding | 44.91 | 0.024645809 | 0.024645809 | ENSG00000160305 | n.s. |
| ENSG00000222831 | 2.304953 | -0.493572496 | 0.816867 | -0.604226 | 0.545693 | 0.891406027 | MIR1537 | miRNA | 40.98 | -0.024641328 | 0.024641328 | ENSG00000222831 | n.s. |
| ENSG00000103540 | 1434.426 | -0.246942187 | 0.267963 | -0.921553 | 0.356762 | 0.794806874 | CCP110 | protein_coding | 37.85 | -0.024629615 | 0.024629615 | ENSG00000103540 | n.s. |
| ENSG00000165724 | 739.8862 | -0.243262003 | 0.261429 | -0.930507 | 0.352108 | 0.792117552 | ZMYND19 | protein_coding | 57.96 | -0.024620636 | 0.024620636 | ENSG00000165724 | n.s. |
| ENSG00000123607 | 1860.738 | -0.250189077 | 0.273459 | -0.914904 | 0.360242 | 0.797273365 | TTC21B | protein_coding | 36.17 | -0.02461679 | 0.02461679 | ENSG00000123607 | n.s. |
| ENSG00000138279 | 530.7931 | -0.258927202 | 0.28776 | -0.899802 | 0.368226 | 0.803511983 | ANXA7 | protein_coding | 40.79 | -0.024600063 | 0.024600063 | ENSG00000138279 | n.s. |

| | | | | | | | | | | | | | |
|---|---|---|---|---|---|---|---|---|---|---|---|---|---|
| ENSG00000106682 | 3070.749 | -0.248227878 | 0.270446 | -0.917847 | 0.358699 | 0.796009826 | EIF4H | protein_coding | 45.4 | -0.024594808 | 0.024594808 | ENSG00000106682 | n.s. |
| ENSG00000105379 | 295.9145 | -0.263875654 | 0.295391 | -0.893311 | 0.371691 | 0.806899249 | ETFB | protein_coding | 51.5 | -0.024588115 | 0.024588115 | ENSG00000105379 | n.s. |
| ENSG00000105723 | 518.3137 | -0.237655466 | 0.251308 | -0.945675 | 0.344314 | 0.788289085 | GSK3A | protein_coding | 53.29 | -0.024553253 | 0.024553253 | ENSG00000105723 | n.s. |
| ENSG00000101255 | 202.4226 | -0.269412029 | 0.309154 | -0.871449 | 0.383509 | 0.810719585 | TRIB3 | protein_coding | 50.47 | -0.024551339 | 0.024551339 | ENSG00000101255 | n.s. |
| ENSG00000110367 | 3212.907 | -0.227653086 | 0.234792 | -0.969596 | 0.332248 | 0.780141891 | DDX6 | protein_coding | 41.56 | -0.024547013 | 0.024547013 | ENSG00000110367 | n.s. |
| ENSG00000078053 | 259.3889 | -0.266308972 | 0.301189 | -0.884192 | 0.376592 | 0.808862777 | AMPH | protein_coding | 39.15 | -0.024533754 | 0.024533754 | ENSG00000078053 | n.s. |
| ENSG00000135045 | 353.8783 | -0.243245348 | 0.262184 | -0.927767 | 0.353529 | 0.792959293 | C9orf40 | protein_coding | 42.71 | -0.024506752 | 0.024506752 | ENSG00000135045 | n.s. |
| ENSG00000064313 | 1184.563 | -0.242136399 | 0.26022 | -0.930506 | 0.352109 | 0.792117552 | TAF2 | protein_coding | 37.5 | -0.024506713 | 0.024506713 | ENSG00000064313 | n.s. |
| ENSG00000169718 | 1776.44 | -0.244543123 | 0.264097 | -0.925958 | 0.354468 | 0.793978768 | DUS1L | protein_coding | 64.45 | -0.024501047 | 0.024501047 | ENSG00000169718 | n.s. |
| ENSG00000181915 | 1033.69 | 0.215424418 | 0.216132 | 0.9967264 | 0.318897 | 0.769632078 | ADO | protein_coding | 47.48 | 0.024497384 | 0.024497384 | ENSG00000181915 | n.s. |
| ENSG00000005812 | 1019.975 | -0.248924439 | 0.271963 | -0.915289 | 0.36004 | 0.797273365 | FBXL3 | protein_coding | 38.36 | -0.024492359 | 0.024492359 | ENSG00000005812 | n.s. |
| ENSG00000278175 | 186.7957 | 0.225923863 | 0.231081 | 0.9776821 | 0.328232 | 0.779459027 | GLIDR | lincRNA | 40.28 | 0.024446478 | 0.024446478 | ENSG00000278175 | n.s. |
| ENSG00000169925 | 722.18 | -0.231371087 | 0.240419 | -0.962367 | 0.335865 | 0.784079552 | BRD3 | protein_coding | 57.61 | -0.024442012 | 0.024442012 | ENSG00000169925 | n.s. |
| ENSG00000100591 | 2250.591 | 0.248627281 | 0.27189 | 0.914441 | 0.360485 | 0.797431699 | AHSA1 | protein_coding | 46.04 | 0.024441679 | 0.024441679 | ENSG00000100591 | n.s. |
| ENSG00000210127 | 25.19876 | -0.306338566 | 0.379349 | -0.807538 | 0.419357 | 0.832304654 | MT-TA | Mt_tRNA | 43.48 | -0.024420599 | 0.024420599 | ENSG00000210127 | n.s. |
| ENSG00000247516 | 802.1471 | 0.226373981 | 0.233057 | 0.9713225 | 0.331388 | 0.780141891 | MIR4458HG | lincRNA | 40.34 | 0.024409092 | 0.024409092 | ENSG00000247516 | n.s. |
| ENSG00000136100 | 1939.14 | -0.248054754 | 0.270893 | -0.915694 | 0.359828 | 0.797273365 | VPS36 | protein_coding | 39.35 | -0.024406788 | 0.024406788 | ENSG00000136100 | n.s. |
| ENSG00000104522 | 332.2998 | -0.262471549 | 0.29506 | -0.889552 | 0.373706 | 0.807377464 | TSTA3 | protein_coding | 65.11 | -0.024389743 | 0.024389743 | ENSG00000104522 | n.s. |
| ENSG00000118263 | 3447.908 | -0.233165139 | 0.244473 | -0.953745 | 0.340213 | 0.785979841 | KLF7 | protein_coding | 41.5 | -0.024386414 | 0.024386414 | ENSG00000118263 | n.s. |
| ENSG00000187164 | 277.4704 | -0.243205798 | 0.262604 | -0.926131 | 0.354378 | 0.793978768 | SHTN1 | protein_coding | 39.09 | -0.024367059 | 0.024367059 | ENSG00000187164 | n.s. |
| ENSG00000126767 | 271.0568 | -0.262915243 | 0.296097 | -0.887936 | 0.374575 | 0.808040232 | ELK1 | protein_coding | 49.22 | -0.024337279 | 0.024337279 | ENSG00000126767 | n.s. |
| ENSG00000169895 | 940.5891 | -0.249749701 | 0.274269 | -0.910601 | 0.362506 | 0.799058295 | SYAP1 | protein_coding | 43.74 | -0.024330999 | 0.024330999 | ENSG00000169895 | n.s. |
| ENSG00000198720 | 733.7215 | -0.223602834 | 0.228212 | -0.979805 | 0.327183 | 0.778659927 | ANKRD13B | protein_coding | 55.54 | -0.024294934 | 0.024294934 | ENSG00000198720 | n.s. |
| ENSG00000134686 | 614.2624 | -0.261257005 | 0.293424 | -0.890375 | 0.373265 | 0.807377464 | PHC2 | protein_coding | 45.03 | -0.024276883 | 0.024276883 | ENSG00000134686 | n.s. |
| ENSG00000225697 | 165.9377 | 0.232191902 | 0.243985 | 0.9516647 | 0.341267 | 0.786054116 | SLC26A6 | protein_coding | 59.37 | 0.024275095 | 0.024275095 | ENSG00000225697 | n.s. |
| ENSG00000143578 | 902.9228 | 0.237087482 | 0.252957 | 0.937265 | 0.348622 | 0.790017511 | CREB3L4 | protein_coding | 52.02 | 0.024269053 | 0.024269053 | ENSG00000143578 | n.s. |
| ENSG00000108239 | 423.6682 | -0.242148619 | 0.26154 | -0.925857 | 0.35452 | 0.793978768 | TBC1D12 | protein_coding | 38.55 | -0.024261139 | 0.024261139 | ENSG00000108239 | n.s. |
| ENSG00000125651 | 1030.262 | -0.243091971 | 0.263984 | -0.920858 | 0.357125 | 0.794806874 | GTF2F1 | protein_coding | 54.75 | -0.024245601 | 0.024245601 | ENSG00000125651 | n.s. |
| ENSG00000115760 | 6485.973 | -0.240635071 | 0.25939 | -0.927696 | 0.353565 | 0.792959293 | BIRC6 | protein_coding | 37.56 | -0.024243769 | 0.024243769 | ENSG00000115760 | n.s. |
| ENSG00000163864 | 38.72316 | 0.289626994 | 0.348571 | 0.8308973 | 0.406032 | 0.824799944 | NMNAT3 | protein_coding | 41.66 | 0.024227697 | 0.024227697 | ENSG00000163864 | n.s. |
| ENSG00000208025 | 8.810996 | -0.360257122 | 0.485199 | -0.742493 | 0.457789 | 0.856577235 | MIR591 | miRNA | 41.05 | -0.024221337 | 0.024221337 | ENSG00000208025 | n.s. |
| ENSG00000145908 | 2346.3 | 0.250864708 | 0.277222 | 0.9049223 | 0.365507 | 0.801033089 | ZNF300 | protein_coding | 36.58 | 0.0241707 | 0.0241707 | ENSG00000145908 | n.s. |

| | | | | | | | | | | | | | |
|---|---|---|---|---|---|---|---|---|---|---|---|---|---|
| ENSG00000055917 | 4159.021 | -0.244622933 | 0.266803 | -0.916869 | 0.359211 | 0.796512236 | PUM2 | protein_coding | 37.42 | -0.024170592 | 0.024170592 | ENSG00000055917 | n.s. |
| ENSG00000051009 | 255.1395 | -0.243279588 | 0.264787 | -0.918776 | 0.358213 | 0.79581885 | FAM160A2 | protein_coding | 44.55 | -0.024129875 | 0.024129875 | ENSG00000051009 | n.s. |
| ENSG00000254685 | 333.5366 | -0.243426328 | 0.265092 | -0.91827 | 0.358477 | 0.796009826 | FPGT | protein_coding | 34.35 | -0.024119063 | 0.024119063 | ENSG00000254685 | n.s. |
| ENSG00000070018 | 3805.064 | -0.23858618 | 0.256543 | -0.930003 | 0.35237 | 0.792352298 | LRP6 | protein_coding | 38.53 | -0.024116691 | 0.024116691 | ENSG00000070018 | n.s. |
| ENSG00000140993 | 621.9361 | 0.240547923 | 0.259759 | 0.9260442 | 0.354423 | 0.793978768 | TIGD7 | protein_coding | 44.07 | 0.024100764 | 0.024100764 | ENSG00000140993 | n.s. |
| ENSG00000116922 | 316.968 | -0.262926151 | 0.29947 | -0.877972 | 0.379959 | 0.809760709 | C1orf109 | protein_coding | 48.39 | -0.02409542 | 0.02409542 | ENSG00000116922 | n.s. |
| ENSG00000164209 | 1740.331 | -0.263552931 | 0.301482 | -0.874191 | 0.382014 | 0.81032471 | SLC25A46 | protein_coding | 35.23 | -0.024073167 | 0.024073167 | ENSG00000164209 | n.s. |
| ENSG00000134480 | 1371.464 | -0.241360555 | 0.26169 | -0.922314 | 0.356365 | 0.794806874 | CCNH | protein_coding | 36.69 | -0.024072912 | 0.024072912 | ENSG00000134480 | n.s. |
| ENSG00000232638 | 69.12842 | -0.253664583 | 0.282159 | -0.899012 | 0.368646 | 0.80373631 | | lincRNA | 57.97 | -0.024069322 | 0.024069322 | ENSG00000232638 | n.s. |
| ENSG00000180543 | 185.2261 | 0.236980106 | 0.254144 | 0.9324656 | 0.351096 | 0.791500843 | TSPYL5 | protein_coding | 49.7 | 0.024065002 | 0.024065002 | ENSG00000180543 | n.s. |
| ENSG00000174473 | 117.1299 | 0.241092134 | 0.260885 | 0.9241334 | 0.355417 | 0.794802373 | GALNTL6 | protein_coding | 36.53 | 0.024046733 | 0.024046733 | ENSG00000174473 | n.s. |
| ENSG00000174939 | 274.6143 | -0.263224114 | 0.300452 | -0.876092 | 0.38098 | 0.810319533 | ASPHD1 | protein_coding | 50.19 | -0.024043862 | 0.024043862 | ENSG00000174939 | n.s. |
| ENSG00000164638 | 1001.951 | -0.234826311 | 0.250885 | -0.93599 | 0.349278 | 0.790028419 | SLC29A4 | protein_coding | 57.52 | -0.024036184 | 0.024036184 | ENSG00000164638 | n.s. |
| ENSG00000130066 | 196.2983 | -0.231655384 | 0.244608 | -0.947047 | 0.343615 | 0.787624904 | SAT1 | protein_coding | 43.75 | -0.02401816 | 0.02401816 | ENSG00000130066 | n.s. |
| ENSG00000164151 | 2999.085 | -0.233736098 | 0.248537 | -0.940448 | 0.346988 | 0.789590741 | ICE1 | protein_coding | 38.9 | -0.023980845 | 0.023980845 | ENSG00000164151 | n.s. |
| ENSG00000232303 | 81.9185 | -0.259057403 | 0.291876 | -0.887559 | 0.374778 | 0.808040232 | DFFBP1 | processed_pseudogene | 56.69 | -0.023980171 | 0.023980171 | ENSG00000232303 | n.s. |
| ENSG00000184675 | 485.5909 | -0.236340245 | 0.253617 | -0.931877 | 0.3514 | 0.791673976 | AMER1 | protein_coding | 50.95 | -0.023977576 | 0.023977576 | ENSG00000184675 | n.s. |
| ENSG00000133030 | 3328.563 | -0.227705211 | 0.237364 | -0.959307 | 0.337404 | 0.78475552 | MPRIP | protein_coding | 51.29 | -0.02396953 | 0.02396953 | ENSG00000133030 | n.s. |
| ENSG00000108256 | 4044.67 | -0.236132893 | 0.253343 | -0.932067 | 0.351302 | 0.791580243 | NUFIP2 | protein_coding | 41.64 | -0.023968682 | 0.023968682 | ENSG00000108256 | n.s. |
| ENSG00000200942 | 4.562809 | -0.411989246 | 0.608555 | -0.676995 | 0.498409 | 0.874675553 | RNU6-501P | snRNA | 39 | -0.023958415 | 0.023958415 | ENSG00000200942 | n.s. |
| ENSG00000182985 | 851.7022 | -0.252905033 | 0.281491 | -0.898448 | 0.368947 | 0.804160729 | CADM1 | protein_coding | 38.54 | -0.023939267 | 0.023939267 | ENSG00000182985 | n.s. |
| ENSG00000171150 | 877.7738 | -0.234328055 | 0.250749 | -0.934511 | 0.35004 | 0.790401174 | SOCS5 | protein_coding | 37.21 | -0.023937179 | 0.023937179 | ENSG00000171150 | n.s. |
| ENSG00000049759 | 139.275 | -0.239874822 | 0.260524 | -0.92074 | 0.357186 | 0.794806874 | NEDD4L | protein_coding | 43.36 | -0.023924727 | 0.023924727 | ENSG00000049759 | n.s. |
| ENSG00000132964 | 639.228 | -0.245112564 | 0.268676 | -0.912299 | 0.361611 | 0.798780417 | CDK8 | protein_coding | 36.99 | -0.023916268 | 0.023916268 | ENSG00000132964 | n.s. |
| ENSG00000163481 | 321.0318 | -0.260218088 | 0.295606 | -0.880288 | 0.378703 | 0.809314697 | RNF25 | protein_coding | 49.77 | -0.023909507 | 0.023909507 | ENSG00000163481 | n.s. |
| ENSG00000186230 | 439.1043 | -0.238256159 | 0.257133 | -0.926588 | 0.35414 | 0.79373747 | ZNF749 | protein_coding | 48.04 | -0.023902601 | 0.023902601 | ENSG00000186230 | n.s. |
| ENSG00000143373 | 685.875 | -0.233318169 | 0.249142 | -0.936487 | 0.349022 | 0.790028419 | ZNF687 | protein_coding | 57.99 | -0.023881815 | 0.023881815 | ENSG00000143373 | n.s. |
| ENSG00000166405 | 243.2068 | -0.246554606 | 0.271927 | -0.906696 | 0.364568 | 0.800144301 | RIC3 | protein_coding | 39.93 | -0.023874298 | 0.023874298 | ENSG00000166405 | n.s. |
| ENSG00000264099 | 37.53422 | -0.283793197 | 0.33912 | -0.836852 | 0.402676 | 0.823993439 | MIR4803 | miRNA | 35.14 | -0.023860267 | 0.023860267 | ENSG00000264099 | n.s. |
| ENSG00000277767 | 70.61612 | -0.244371345 | 0.267836 | -0.91239 | 0.361563 | 0.798780417 | | lincRNA | 38.32 | -0.023843945 | 0.023843945 | ENSG00000277767 | n.s. |
| ENSG00000120088 | 523.7276 | -0.255747853 | 0.286413 | -0.892932 | 0.371893 | 0.806899249 | CRHR1 | protein_coding | 54.73 | -0.023830761 | 0.023830761 | ENSG00000120088 | n.s. |
| ENSG00000056736 | 154.9954 | -0.250649336 | 0.278654 | -0.8995 | 0.368386 | 0.803511983 | IL17RB | protein_coding | 45.37 | -0.023813602 | 0.023813602 | ENSG00000056736 | n.s. |

| | | | | | | | | | | | | | |
|---|---|---|---|---|---|---|---|---|---|---|---|---|---|
| ENSG00000268670 | 91.43508 | -0.238716916 | 0.259163 | -0.921107 | 0.356995 | 0.794806874 | | sense_intronic | 63.52 | -0.02380924 | 0.02380924 | ENSG00000268670 | n.s. |
| ENSG00000149930 | 586.1756 | -0.221845441 | 0.229184 | -0.96798 | 0.333055 | 0.781054224 | TAOK2 | protein_coding | 55.16 | -0.02380819 | 0.02380819 | ENSG00000149930 | n.s. |
| ENSG00000052749 | 711.0345 | -0.238637972 | 0.258407 | -0.923497 | 0.355748 | 0.794802373 | RRP12 | protein_coding | 50.49 | -0.023801953 | 0.023801953 | ENSG00000052749 | n.s. |
| ENSG00000071051 | 1224.966 | -0.244319104 | 0.268283 | -0.910675 | 0.362467 | 0.799058295 | NCK2 | protein_coding | 44.54 | -0.023801942 | 0.023801942 | ENSG00000071051 | n.s. |
| ENSG00000106976 | 30.81778 | -0.293265939 | 0.359902 | -0.814849 | 0.415159 | 0.829591076 | DNM1 | protein_coding | 54.26 | -0.023794404 | 0.023794404 | ENSG00000106976 | n.s. |
| ENSG00000173914 | 674.9779 | -0.231962276 | 0.247736 | -0.936328 | 0.349104 | 0.790028419 | RBM4B | protein_coding | 43.9 | -0.023743029 | 0.023743029 | ENSG00000173914 | n.s. |
| ENSG00000182158 | 403.6875 | -0.231893875 | 0.247539 | -0.936796 | 0.348863 | 0.790028419 | CREB3L2 | protein_coding | 43.66 | -0.023736028 | 0.023736028 | ENSG00000182158 | n.s. |
| ENSG00000227383 | 33.13304 | 0.285504199 | 0.344303 | 0.8292241 | 0.406978 | 0.825802394 | | processed_pseudogene | 42.81 | 0.023732212 | 0.023732212 | ENSG00000227383 | n.s. |
| ENSG00000270988 | 54.31787 | 0.265135521 | 0.307048 | 0.8634999 | 0.387863 | 0.813897772 | | lincRNA | 46.28 | 0.023711107 | 0.023711107 | ENSG00000270988 | n.s. |
| ENSG00000149084 | 1669.275 | -0.256556088 | 0.289754 | -0.885429 | 0.375925 | 0.808341321 | HSD17B12 | protein_coding | 38.16 | -0.023707122 | 0.023707122 | ENSG00000149084 | n.s. |
| ENSG00000100522 | 800.1323 | -0.254308486 | 0.284301 | -0.894505 | 0.371052 | 0.80683384 | GNPNAT1 | protein_coding | 37.88 | -0.023705593 | 0.023705593 | ENSG00000100522 | n.s. |
| ENSG00000177173 | 48.2554 | -0.267865074 | 0.311853 | -0.858947 | 0.390369 | 0.815710819 | NAP1L4P1 | processed_pseudogene | 42.35 | -0.023696356 | 0.023696356 | ENSG00000177173 | n.s. |
| ENSG00000257167 | 678.9381 | 0.205966752 | 0.204884 | 1.0052823 | 0.314761 | 0.767364082 | TMPO-AS1 | antisense | 49.57 | 0.023685874 | 0.023685874 | ENSG00000257167 | n.s. |
| ENSG00000158941 | 1795.011 | -0.231304172 | 0.247141 | -0.93592 | 0.349314 | 0.790028419 | CCAR2 | protein_coding | 50.83 | -0.023675667 | 0.023675667 | ENSG00000158941 | n.s. |
| ENSG00000087263 | 984.5838 | -0.253632164 | 0.283585 | -0.894379 | 0.371119 | 0.806853443 | OGFOD1 | protein_coding | 40.39 | -0.023639873 | 0.023639873 | ENSG00000087263 | n.s. |
| ENSG00000148843 | 2677.37 | -0.226097728 | 0.237639 | -0.951434 | 0.341384 | 0.786054116 | PDCD11 | protein_coding | 46.81 | -0.023637964 | 0.023637964 | ENSG00000148843 | n.s. |
| ENSG00000158406 | 1338.088 | -0.20903455 | 0.210329 | -0.993844 | 0.320299 | 0.770876263 | HIST1H4H | protein_coding | 41.76 | -0.023624108 | 0.023624108 | ENSG00000158406 | n.s. |
| ENSG00000176973 | 484.3353 | -0.255193235 | 0.287673 | -0.887096 | 0.375027 | 0.808040232 | FAM89B | protein_coding | 64.22 | -0.023622476 | 0.023622476 | ENSG00000176973 | n.s. |
| ENSG00000124422 | 5135.884 | -0.230684617 | 0.245559 | -0.939428 | 0.347511 | 0.790017511 | USP22 | protein_coding | 47.95 | -0.023613635 | 0.023613635 | ENSG00000124422 | n.s. |
| ENSG00000265683 | 110.7144 | 0.236189653 | 0.255241 | 0.9253595 | 0.354779 | 0.79440131 | | processed_pseudogene | 41.68 | 0.023609529 | 0.023609529 | ENSG00000265683 | n.s. |
| ENSG00000271886 | 13.43137 | -0.321425356 | 0.41509 | -0.774351 | 0.438723 | 0.84443611 | MIR98 | miRNA | 42.86 | -0.023603294 | 0.023603294 | ENSG00000271886 | n.s. |
| ENSG00000266897 | 16.76927 | -0.306823195 | 0.386733 | -0.793372 | 0.427561 | 0.837692206 | | lincRNA | 51.19 | -0.023599467 | 0.023599467 | ENSG00000266897 | n.s. |
| ENSG00000201264 | 168.0417 | 0.224381493 | 0.234468 | 0.9569814 | 0.338577 | 0.784993146 | SNORD73B | snoRNA | 27.78 | 0.023590154 | 0.023590154 | ENSG00000201264 | n.s. |
| ENSG00000260751 | 49.14117 | -0.266841979 | 0.312536 | -0.853796 | 0.393218 | 0.815931883 | | sense_intronic | 43.65 | -0.023574447 | 0.023574447 | ENSG00000260751 | n.s. |
| ENSG00000137076 | 1248.947 | -0.23222529 | 0.249141 | -0.932103 | 0.351283 | 0.791580243 | TLN1 | protein_coding | 51.19 | -0.023572041 | 0.023572041 | ENSG00000137076 | n.s. |
| ENSG00000027644 | 520.7028 | -0.232900789 | 0.250212 | -0.930813 | 0.35195 | 0.792117552 | INSRR | protein_coding | 55.51 | -0.023571974 | 0.023571974 | ENSG00000027644 | n.s. |
| ENSG00000267254 | 249.926 | 0.227200807 | 0.239928 | 0.9469523 | 0.343663 | 0.787624904 | | antisense | 40.52 | 0.023556307 | 0.023556307 | ENSG00000267254 | n.s. |
| ENSG00000125814 | 599.4233 | -0.234265046 | 0.252678 | -0.927129 | 0.35386 | 0.79336388 | NAPB | protein_coding | 40.72 | -0.023550097 | 0.023550097 | ENSG00000125814 | n.s. |
| ENSG00000252496 | 6.528066 | -0.47170751 | 0.777741 | -0.60651 | 0.544176 | 0.891406027 | RNA5SP33 | rRNA | 44.9 | -0.02354973 | 0.02354973 | ENSG00000252496 | n.s. |
| ENSG00000088970 | 184.34 | 0.245750927 | 0.272105 | 0.9031483 | 0.366447 | 0.802009039 | KIZ | protein_coding | 41.17 | 0.023548035 | 0.023548035 | ENSG00000088970 | n.s. |
| ENSG00000167815 | 3175.014 | -0.253914452 | 0.285658 | -0.888875 | 0.37407 | 0.807719995 | PRDX2 | protein_coding | 54.5 | -0.023547815 | 0.023547815 | ENSG00000167815 | n.s. |
| ENSG00000152291 | 1089.285 | -0.219426667 | 0.226839 | -0.967324 | 0.333382 | 0.781097236 | TGOLN2 | protein_coding | 47.66 | -0.023543362 | 0.023543362 | ENSG00000152291 | n.s. |

| | | | | | | | | | | | | | |
|---|---|---|---|---|---|---|---|---|---|---|---|---|---|
| ENSG00000059769 | 284.4785 | -0.243050428 | 0.268056 | -0.906716 | 0.364557 | 0.800144301 | DNAJC25 | protein_coding | 42.74 | -0.023534982 | 0.023534982 | ENSG00000059769 | n.s. |
| ENSG00000099385 | 260.5715 | -0.265897084 | 0.31096 | -0.855085 | 0.392504 | 0.815710819 | BCL7C | protein_coding | 48.81 | -0.02352226 | 0.02352226 | ENSG00000099385 | n.s. |
| ENSG00000207967 | 6.733797 | -0.407123911 | 0.605323 | -0.672573 | 0.501219 | 0.875518559 | MIR620 | miRNA | 21.05 | -0.023505154 | 0.023505154 | ENSG00000207967 | n.s. |
| ENSG00000265008 | 19.16722 | 0.321427842 | 0.415637 | 0.7733376 | 0.439323 | 0.845079983 | | antisense | 50.91 | 0.023497078 | 0.023497078 | ENSG00000265008 | n.s. |
| ENSG00000173020 | 701.9703 | -0.238620088 | 0.26075 | -0.915129 | 0.360124 | 0.797273365 | GRK2 | protein_coding | 61.02 | -0.023478485 | 0.023478485 | ENSG00000173020 | n.s. |
| ENSG00000230082 | 56.33946 | -0.25532019 | 0.288985 | -0.883505 | 0.376963 | 0.809223062 | PRRT3-AS1 | antisense | 54.05 | -0.023472032 | 0.023472032 | ENSG00000230082 | n.s. |
| ENSG00000180182 | 966.3016 | -0.229151922 | 0.24406 | -0.938914 | 0.347775 | 0.790017511 | MED14 | protein_coding | 38.86 | -0.023456743 | 0.023456743 | ENSG00000180182 | n.s. |
| ENSG00000200403 | 5.010836 | -0.416287886 | 0.628268 | -0.662596 | 0.507589 | 0.878345106 | RNU6-1099P | snRNA | 40.19 | -0.023451501 | 0.023451501 | ENSG00000200403 | n.s. |
| ENSG00000263740 | 128.2439 | -0.234827852 | 0.254934 | -0.921131 | 0.356982 | 0.794806874 | RN7SL4P | misc_RNA | 58.64 | -0.023421351 | 0.023421351 | ENSG00000263740 | n.s. |
| ENSG00000152582 | 49.13079 | -0.274912582 | 0.32674 | -0.84138 | 0.400135 | 0.821966929 | SPEF2 | protein_coding | 37.81 | -0.023407612 | 0.023407612 | ENSG00000152582 | n.s. |
| ENSG00000169371 | 240.0254 | -0.256263104 | 0.293575 | -0.872904 | 0.382715 | 0.81032471 | SNUPN | protein_coding | 47.36 | -0.023407307 | 0.023407307 | ENSG00000169371 | n.s. |
| ENSG00000149187 | 2547.495 | -0.2256189 | 0.238409 | -0.946353 | 0.343968 | 0.787678531 | CELF1 | protein_coding | 43.94 | -0.023385623 | 0.023385623 | ENSG00000149187 | n.s. |
| ENSG00000125944 | 5117.897 | -0.23923241 | 0.262097 | -0.912764 | 0.361367 | 0.7984941 | HNRNPR | protein_coding | 38.85 | -0.023379774 | 0.023379774 | ENSG00000125944 | n.s. |
| ENSG00000223803 | 26.27918 | 0.449867022 | 0.717485 | 0.6270056 | 0.530656 | 0.887229181 | RPS20P14 | processed_pseudogene | 46.11 | 0.023376972 | 0.023376972 | ENSG00000223803 | n.s. |
| ENSG00000196810 | 241.6357 | -0.218654884 | 0.2263 | -0.966215 | 0.333937 | 0.781805976 | CTBP1-AS2 | antisense | 46.78 | -0.023374429 | 0.023374429 | ENSG00000196810 | n.s. |
| ENSG00000161547 | 18423.38 | -0.21437417 | 0.218512 | -0.981064 | 0.326561 | 0.778045194 | SRSF2 | protein_coding | 49.23 | -0.02336575 | 0.02336575 | ENSG00000161547 | n.s. |
| ENSG00000137802 | 149.6103 | -0.2267149 | 0.240515 | -0.942622 | 0.345874 | 0.788921338 | MAPKBP1 | protein_coding | 49.09 | -0.023343994 | 0.023343994 | ENSG00000137802 | n.s. |
| ENSG00000178694 | 402.9021 | -0.227917575 | 0.242768 | -0.93883 | 0.347818 | 0.790017511 | NSUN3 | protein_coding | 37.97 | -0.023330391 | 0.023330391 | ENSG00000178694 | n.s. |
| ENSG00000270964 | 82.88188 | 0.253613813 | 0.288026 | 0.8805249 | 0.378575 | 0.809266943 | | lincRNA | 46.77 | 0.023309189 | 0.023309189 | ENSG00000270964 | n.s. |
| ENSG00000099910 | 390.3906 | -0.255181619 | 0.292379 | -0.872778 | 0.382784 | 0.81032471 | KLHL22 | protein_coding | 49.11 | -0.023308523 | 0.023308523 | ENSG00000099910 | n.s. |
| ENSG00000257354 | 662.0977 | -0.231301907 | 0.249073 | -0.928651 | 0.35307 | 0.792950339 | | lincRNA | 40.48 | -0.023304595 | 0.023304595 | ENSG00000257354 | n.s. |
| ENSG00000269894 | 21.47135 | 0.302902575 | 0.381835 | 0.7932808 | 0.427614 | 0.837692206 | | sense_intronic | 41.41 | 0.023297911 | 0.023297911 | ENSG00000269894 | n.s. |
| ENSG00000196284 | 137.2106 | 0.254127371 | 0.289316 | 0.8783738 | 0.379741 | 0.809760709 | SUPT3H | protein_coding | 36.3 | 0.023289071 | 0.023289071 | ENSG00000196284 | n.s. |
| ENSG00000124299 | 420.9538 | -0.254715292 | 0.291229 | -0.874623 | 0.381779 | 0.81032471 | PEPD | protein_coding | 53.89 | -0.023265928 | 0.023265928 | ENSG00000124299 | n.s. |
| ENSG00000265806 | 16.07416 | -0.394065057 | 0.573821 | -0.686739 | 0.492247 | 0.872953302 | MIR4292 | miRNA | 70.15 | -0.02325338 | 0.02325338 | ENSG00000265806 | n.s. |
| ENSG00000176915 | 3435.12 | -0.235956226 | 0.257792 | -0.915295 | 0.360037 | 0.797273365 | ANKLE2 | protein_coding | 51.03 | -0.023216381 | 0.023216381 | ENSG00000176915 | n.s. |
| ENSG00000109881 | 447.1344 | -0.247452742 | 0.275979 | -0.896637 | 0.369913 | 0.805866739 | CCDC34 | protein_coding | 37.62 | -0.02319542 | 0.02319542 | ENSG00000109881 | n.s. |
| ENSG00000120697 | 960.8894 | -0.244054434 | 0.271067 | -0.900349 | 0.367935 | 0.803511983 | ALG5 | protein_coding | 39.26 | -0.023187036 | 0.023187036 | ENSG00000120697 | n.s. |
| ENSG00000160447 | 254.6045 | 0.252203724 | 0.285753 | 0.8825943 | 0.377456 | 0.809223062 | PKN3 | protein_coding | 57.66 | 0.023185529 | 0.023185529 | ENSG00000160447 | n.s. |
| ENSG00000103932 | 166.7855 | -0.225525806 | 0.239507 | -0.941624 | 0.346385 | 0.789327092 | RPAP1 | protein_coding | 50.42 | -0.023171196 | 0.023171196 | ENSG00000103932 | n.s. |
| ENSG00000044459 | 1173.499 | -0.252746158 | 0.287772 | -0.878286 | 0.379789 | 0.809760709 | CNTLN | protein_coding | 36.43 | -0.023162492 | 0.023162492 | ENSG00000044459 | n.s. |
| ENSG00000271895 | 33.13436 | -0.276864528 | 0.333197 | -0.830934 | 0.406011 | 0.824799944 | | antisense | 42.99 | -0.023160099 | 0.023160099 | ENSG00000271895 | n.s. |

| | | | | | | | | | | | | | |
|---|---|---|---|---|---|---|---|---|---|---|---|---|---|
| ENSG00000160785 | 772.4116 | -0.22009553 | 0.229947 | -0.957157 | 0.338488 | 0.784993146 | SLC25A44 | protein_coding | 48.29 | -0.023139553 | 0.023139553 | ENSG00000160785 | n.s. |
| ENSG00000153006 | 773.9118 | -0.251720678 | 0.285891 | -0.880477 | 0.378601 | 0.809266943 | SREK1IP1 | protein_coding | 36.21 | -0.023135194 | 0.023135194 | ENSG00000153006 | n.s. |
| ENSG00000077684 | 1511.799 | -0.225612655 | 0.240004 | -0.940036 | 0.347199 | 0.789748319 | JADE1 | protein_coding | 44.21 | -0.023127844 | 0.023127844 | ENSG00000077684 | n.s. |
| ENSG00000115904 | 250.8402 | -0.243412949 | 0.270373 | -0.900286 | 0.367968 | 0.803511983 | SOS1 | protein_coding | 37.8 | -0.02312609 | 0.02312609 | ENSG00000115904 | n.s. |
| ENSG00000100888 | 1921.548 | -0.219878435 | 0.229619 | -0.95758 | 0.338274 | 0.784993146 | CHD8 | protein_coding | 42.52 | -0.023116729 | 0.023116729 | ENSG00000100888 | n.s. |
| ENSG00000117616 | 528.598 | -0.219873129 | 0.229786 | -0.956862 | 0.338637 | 0.784993146 | RSRP1 | protein_coding | 46.4 | -0.023116171 | 0.023116171 | ENSG00000117616 | n.s. |
| ENSG00000113318 | 962.0568 | -0.220725843 | 0.231684 | -0.952703 | 0.34074 | 0.786054116 | MSH3 | protein_coding | 38.03 | -0.023076347 | 0.023076347 | ENSG00000113318 | n.s. |
| ENSG00000068400 | 548.6454 | -0.224372837 | 0.238341 | -0.941393 | 0.346503 | 0.789327092 | GRIPAP1 | protein_coding | 50.22 | -0.023052736 | 0.023052736 | ENSG00000068400 | n.s. |
| ENSG00000099991 | 238.1419 | -0.232282706 | 0.252643 | -0.919411 | 0.357881 | 0.795715832 | CABIN1 | protein_coding | 50.24 | -0.023052201 | 0.023052201 | ENSG00000099991 | n.s. |
| ENSG00000215447 | 76.58307 | -0.250627619 | 0.284163 | -0.881987 | 0.377784 | 0.809223062 | | processed_transcript | 53.84 | -0.023040635 | 0.023040635 | ENSG00000215447 | n.s. |
| ENSG00000178074 | 953.9947 | -0.232674198 | 0.253624 | -0.917399 | 0.358934 | 0.79624783 | C2orf69 | protein_coding | 38 | -0.023023516 | 0.023023516 | ENSG00000178074 | n.s. |
| ENSG00000145819 | 66.73168 | -0.251991407 | 0.287399 | -0.876799 | 0.380596 | 0.810319533 | ARHGAP26 | protein_coding | 42.17 | -0.023017825 | 0.023017825 | ENSG00000145819 | n.s. |
| ENSG00000083642 | 2199.117 | -0.230641196 | 0.249517 | -0.92435 | 0.355304 | 0.794802373 | PDS5B | protein_coding | 36.62 | -0.023004348 | 0.023004348 | ENSG00000083642 | n.s. |
| ENSG00000100376 | 98.01541 | -0.251624391 | 0.288025 | -0.873619 | 0.382326 | 0.81032471 | FAM118A | protein_coding | 49.58 | -0.022983603 | 0.022983603 | ENSG00000100376 | n.s. |
| ENSG00000071054 | 10711.84 | -0.224237299 | 0.238586 | -0.939861 | 0.347289 | 0.789823702 | MAP4K4 | protein_coding | 41.04 | -0.02297756 | 0.02297756 | ENSG00000071054 | n.s. |
| ENSG00000156509 | 272.3555 | -0.230337925 | 0.250204 | -0.920599 | 0.35726 | 0.794843454 | FBXO43 | protein_coding | 38.91 | -0.022968929 | 0.022968929 | ENSG00000156509 | n.s. |
| ENSG00000197050 | 543.2812 | -0.231524783 | 0.252182 | -0.918087 | 0.358573 | 0.796009826 | ZNF420 | protein_coding | 41.84 | -0.022939839 | 0.022939839 | ENSG00000197050 | n.s. |
| ENSG00000113621 | 657.2518 | -0.24779449 | 0.279467 | -0.886668 | 0.375258 | 0.808091443 | TXNDC15 | protein_coding | 46.17 | -0.022930776 | 0.022930776 | ENSG00000113621 | n.s. |
| ENSG00000175344 | 852.684 | -0.22377792 | 0.23873 | -0.93737 | 0.348568 | 0.790017511 | CHRNA7 | protein_coding | 41.21 | -0.022906642 | 0.022906642 | ENSG00000175344 | n.s. |
| ENSG00000147256 | 131.3971 | 0.248992793 | 0.28176 | 0.8837039 | 0.376856 | 0.809217666 | ARHGAP36 | protein_coding | 46.18 | 0.022891064 | 0.022891064 | ENSG00000147256 | n.s. |
| ENSG00000067048 | 3868.333 | 0.222297774 | 0.235831 | 0.9426141 | 0.345878 | 0.788921338 | DDX3Y | protein_coding | 37.7 | 0.022889179 | 0.022889179 | ENSG00000067048 | n.s. |
| ENSG00000164924 | 5500.024 | -0.240716234 | 0.267242 | -0.900743 | 0.367725 | 0.803384223 | YWHAZ | protein_coding | 40.87 | -0.022886506 | 0.022886506 | ENSG00000164924 | n.s. |
| ENSG00000111144 | 2264.23 | 0.250232562 | 0.286091 | 0.8746614 | 0.381758 | 0.81032471 | LTA4H | protein_coding | 39.1 | 0.022856472 | 0.022856472 | ENSG00000111144 | n.s. |
| ENSG00000204962 | 90.63491 | -0.244373247 | 0.272972 | -0.895232 | 0.370663 | 0.806303209 | PCDHA8 | protein_coding | 40.02 | -0.022849292 | 0.022849292 | ENSG00000204962 | n.s. |
| ENSG00000144935 | 369.1446 | -0.244257755 | 0.272781 | -0.895437 | 0.370554 | 0.806253911 | TRPC1 | protein_coding | 35.58 | -0.02284498 | 0.02284498 | ENSG00000144935 | n.s. |
| ENSG00000143190 | 323.9938 | -0.21697748 | 0.226415 | -0.958319 | 0.337902 | 0.78475552 | POU2F1 | protein_coding | 37.96 | -0.022840269 | 0.022840269 | ENSG00000143190 | n.s. |
| ENSG00000206053 | 1902.069 | -0.245400144 | 0.275683 | -0.890154 | 0.373383 | 0.807377464 | JPT2 | protein_coding | 47.78 | -0.02280341 | 0.02280341 | ENSG00000206053 | n.s. |
| ENSG00000174886 | 1008.275 | -0.272306339 | 0.327069 | -0.832565 | 0.40509 | 0.824799944 | NDUFA11 | protein_coding | 59.05 | -0.0227788 | 0.0227788 | ENSG00000174886 | n.s. |
| ENSG00000178440 | 36.08815 | -0.37132511 | 0.529259 | -0.701594 | 0.482932 | 0.868307709 | LINC00843 | lincRNA | 43.38 | -0.022772011 | 0.022772011 | ENSG00000178440 | n.s. |
| ENSG00000164040 | 872.4823 | -0.242941256 | 0.27104 | -0.896328 | 0.370077 | 0.805916031 | PGRMC2 | protein_coding | 38.64 | -0.022766075 | 0.022766075 | ENSG00000164040 | n.s. |
| ENSG00000072958 | 429.1129 | -0.249723348 | 0.286454 | -0.871775 | 0.383331 | 0.810719585 | AP1M1 | protein_coding | 51.59 | -0.022757123 | 0.022757123 | ENSG00000072958 | n.s. |
| ENSG00000120899 | 24.60541 | -0.290438286 | 0.362713 | -0.800739 | 0.423282 | 0.834993891 | PTK2B | protein_coding | 46.91 | -0.022746169 | 0.022746169 | ENSG00000120899 | n.s. |

| | | | | | | | | | | | | | |
|---|---|---|---|---|---|---|---|---|---|---|---|---|---|
| ENSG00000275101 | 5.463845 | -0.400570181 | 0.602959 | -0.66434 | 0.506472 | 0.877502299 | MIR6766 | miRNA | 54.17 | -0.022733053 | 0.022733053 | ENSG00000275101 | n.s. |
| ENSG00000166250 | 561.2018 | -0.247251718 | 0.280432 | -0.881683 | 0.377948 | 0.809223062 | CLMP | protein_coding | 45.02 | -0.022730283 | 0.022730283 | ENSG00000166250 | n.s. |
| ENSG00000184465 | 690.9115 | -0.230704463 | 0.251813 | -0.916174 | 0.359576 | 0.797029976 | WDR27 | protein_coding | 42.01 | -0.022730237 | 0.022730237 | ENSG00000184465 | n.s. |
| ENSG00000095485 | 613.356 | -0.252298795 | 0.290762 | -0.867717 | 0.385549 | 0.812717858 | CWF19L1 | protein_coding | 42.57 | -0.022722079 | 0.022722079 | ENSG00000095485 | n.s. |
| ENSG00000135452 | 152.0396 | -0.244297874 | 0.274495 | -0.889989 | 0.373472 | 0.807377464 | TSPAN31 | protein_coding | 51.32 | -0.022700983 | 0.022700983 | ENSG00000135452 | n.s. |
| ENSG00000169764 | 682.6772 | -0.245225161 | 0.276925 | -0.885529 | 0.375871 | 0.808341321 | UGP2 | protein_coding | 38.65 | -0.022660085 | 0.022660085 | ENSG00000169764 | n.s. |
| ENSG00000133704 | 1287.843 | -0.237524992 | 0.263366 | -0.901882 | 0.36712 | 0.802884215 | IPO8 | protein_coding | 35.93 | -0.022647314 | 0.022647314 | ENSG00000133704 | n.s. |
| ENSG00000117000 | 664.0756 | -0.226679784 | 0.245108 | -0.924815 | 0.355062 | 0.794524781 | RLF | protein_coding | 39.69 | -0.022643622 | 0.022643622 | ENSG00000117000 | n.s. |
| ENSG00000259494 | 1095.236 | 0.210530037 | 0.217423 | 0.9682975 | 0.332896 | 0.780813442 | MRPL46 | protein_coding | 41.02 | 0.022622025 | 0.022622025 | ENSG00000259494 | n.s. |
| ENSG00000169118 | 553.5627 | -0.22677812 | 0.245815 | -0.922555 | 0.356239 | 0.794806874 | CSNK1G1 | protein_coding | 40.28 | -0.022618484 | 0.022618484 | ENSG00000169118 | n.s. |
| ENSG00000141456 | 545.6838 | -0.223415975 | 0.239883 | -0.931355 | 0.35167 | 0.792117552 | PELP1 | protein_coding | 48.03 | -0.022612012 | 0.022612012 | ENSG00000141456 | n.s. |
| ENSG00000172954 | 689.2008 | -0.232357547 | 0.255575 | -0.909155 | 0.363268 | 0.799270441 | LCLAT1 | protein_coding | 37.54 | -0.022609841 | 0.022609841 | ENSG00000172954 | n.s. |
| ENSG00000165322 | 874.837 | -0.241190473 | 0.269029 | -0.896521 | 0.369974 | 0.805875275 | ARHGAP12 | protein_coding | 36.94 | -0.022607306 | 0.022607306 | ENSG00000165322 | n.s. |
| ENSG00000162869 | 84.49314 | -0.255507925 | 0.299088 | -0.854289 | 0.392945 | 0.815710819 | PPP1R21 | protein_coding | 39.73 | -0.022603196 | 0.022603196 | ENSG00000162869 | n.s. |
| ENSG00000118564 | 1113.63 | -0.244034363 | 0.27497 | -0.887496 | 0.374812 | 0.808040232 | FBXL5 | protein_coding | 37.87 | -0.022589533 | 0.022589533 | ENSG00000118564 | n.s. |
| ENSG00000207626 | 6.03739 | -0.369672453 | 0.528358 | -0.699663 | 0.484138 | 0.868752293 | MIR562 | miRNA | 37.89 | -0.022588479 | 0.022588479 | ENSG00000207626 | n.s. |
| ENSG00000136709 | 3913.272 | -0.247140675 | 0.283115 | -0.872934 | 0.382699 | 0.81032471 | WDR33 | protein_coding | 41.82 | -0.022574056 | 0.022574056 | ENSG00000136709 | n.s. |
| ENSG00000105467 | 61.62382 | -0.269836594 | 0.323844 | -0.83323 | 0.404715 | 0.824799944 | SYNGR4 | protein_coding | 54.82 | -0.022572203 | 0.022572203 | ENSG00000105467 | n.s. |
| ENSG00000279561 | 244.2617 | -0.226083555 | 0.244684 | -0.923982 | 0.355496 | 0.794802373 | | TEC | 42.28 | -0.022549765 | 0.022549765 | ENSG00000279561 | n.s. |
| ENSG00000232818 | 32.37018 | 0.276833853 | 0.338133 | 0.8187133 | 0.41295 | 0.829007487 | RPS2P32 | processed_pseudogene | 61.32 | 0.022545777 | 0.022545777 | ENSG00000232818 | n.s. |
| ENSG00000137171 | 203.5688 | 0.247211139 | 0.284211 | 0.8698169 | 0.3844 | 0.810719585 | KLC4 | protein_coding | 52 | 0.022528187 | 0.022528187 | ENSG00000137171 | n.s. |
| ENSG00000118231 | 120.8894 | 0.261962589 | 0.309942 | 0.8451998 | 0.397999 | 0.820375622 | CRYGD | protein_coding | 37.37 | 0.022525443 | 0.022525443 | ENSG00000118231 | n.s. |
| ENSG00000105948 | 316.0885 | -0.244986447 | 0.278196 | -0.880626 | 0.37852 | 0.809266943 | TTC26 | protein_coding | 38.13 | -0.022516263 | 0.022516263 | ENSG00000105948 | n.s. |
| ENSG00000121152 | 1975.837 | -0.246947992 | 0.283458 | -0.871199 | 0.383645 | 0.810719585 | NCAPH | protein_coding | 45.2 | -0.022504206 | 0.022504206 | ENSG00000121152 | n.s. |
| ENSG00000264482 | 5.08683 | -0.390745518 | 0.581919 | -0.671478 | 0.501916 | 0.875813709 | MIR4705 | miRNA | 38.03 | -0.022502355 | 0.022502355 | ENSG00000264482 | n.s. |
| ENSG00000149532 | 1523.289 | -0.232203719 | 0.255841 | -0.907609 | 0.364085 | 0.800056047 | CPSF7 | protein_coding | 44.18 | -0.022495801 | 0.022495801 | ENSG00000149532 | n.s. |
| ENSG00000159685 | 268.7098 | -0.246764515 | 0.283482 | -0.870476 | 0.384041 | 0.810719585 | CHCHD6 | protein_coding | 46.24 | -0.022487486 | 0.022487486 | ENSG00000159685 | n.s. |
| ENSG00000114450 | 1414.487 | -0.242817844 | 0.273729 | -0.887073 | 0.375039 | 0.808040232 | GNB4 | protein_coding | 39.16 | -0.022476923 | 0.022476923 | ENSG00000114450 | n.s. |
| ENSG00000112149 | 159.4531 | -0.24901091 | 0.286896 | -0.867949 | 0.385422 | 0.812573251 | CD83 | protein_coding | 45.18 | -0.022445215 | 0.022445215 | ENSG00000112149 | n.s. |
| ENSG00000208772 | 89.54418 | -0.242552238 | 0.273677 | -0.886273 | 0.37547 | 0.808144397 | SNORD94 | snoRNA | 54.01 | -0.022438758 | 0.022438758 | ENSG00000208772 | n.s. |
| ENSG00000263606 | 72.00377 | -0.275458181 | 0.337089 | -0.817166 | 0.413833 | 0.829007487 | | transcribed_processed_pseudog | 40.49 | -0.02243374 | 0.02243374 | ENSG00000263606 | n.s. |
| ENSG00000007541 | 7.169218 | 0.422554113 | 0.654939 | 0.6451814 | 0.51881 | 0.88497967 | PIGQ | protein_coding | 64.36 | 0.022423555 | 0.022423555 | ENSG00000007541 | n.s. |

| | | | | | | | | | | | | | |
|---|---|---|---|---|---|---|---|---|---|---|---|---|---|
| ENSG00000176840 | 10.05508 | -0.448872594 | 0.747067 | -0.600846 | 0.547942 | 0.891406027 | MIR7-3HG | lincRNA | 51.53 | -0.02240971 | 0.02240971 | ENSG00000176840 | n.s. |
| ENSG00000112584 | 1138.96 | -0.243761957 | 0.276801 | -0.880639 | 0.378513 | 0.809266943 | FAM120B | protein_coding | 44.49 | -0.022403723 | 0.022403723 | ENSG00000112584 | n.s. |
| ENSG00000186312 | 150.9466 | -0.260509874 | 0.308311 | -0.844959 | 0.398134 | 0.820375622 | CA5BP1 | cribed_unprocessed_pseudc | 43 | -0.022400529 | 0.022400529 | ENSG00000186312 | n.s. |
| ENSG00000267197 | 32.4071 | -0.285664929 | 0.356799 | -0.800633 | 0.423344 | 0.834993891 | | sense_intronic | 49.4 | -0.022372335 | 0.022372335 | ENSG00000267197 | n.s. |
| ENSG00000151376 | 216.7843 | -0.229435428 | 0.251682 | -0.911609 | 0.361975 | 0.798938342 | ME3 | protein_coding | 42.88 | -0.022366911 | 0.022366911 | ENSG00000151376 | n.s. |
| ENSG00000070950 | 1165.439 | -0.232072095 | 0.256438 | -0.904983 | 0.365474 | 0.801033089 | RAD18 | protein_coding | 40.44 | -0.02236004 | 0.02236004 | ENSG00000070950 | n.s. |
| ENSG00000134899 | 86.01972 | -0.239880077 | 0.268324 | -0.893993 | 0.371326 | 0.806899249 | ERCC5 | protein_coding | 39.83 | -0.022352191 | 0.022352191 | ENSG00000134899 | n.s. |
| ENSG00000277583 | 7.082332 | 0.391463846 | 0.587408 | 0.6664255 | 0.505139 | 0.876821729 | | misc_RNA | 52.25 | 0.022348161 | 0.022348161 | ENSG00000277583 | n.s. |
| ENSG00000115318 | 459.4731 | -0.245188743 | 0.281187 | -0.871977 | 0.383221 | 0.810719585 | LOXL3 | protein_coding | 50.8 | -0.022343887 | 0.022343887 | ENSG00000115318 | n.s. |
| ENSG00000213145 | 18.03147 | -0.407066304 | 0.621425 | -0.655053 | 0.512433 | 0.881306695 | CRIP1 | protein_coding | 68.07 | -0.022336921 | 0.022336921 | ENSG00000213145 | n.s. |
| ENSG00000163507 | 2555.621 | -0.23303033 | 0.257952 | -0.903386 | 0.366321 | 0.801976953 | KIAA1524 | protein_coding | 34.15 | -0.022333187 | 0.022333187 | ENSG00000163507 | n.s. |
| ENSG00000134594 | 112.7211 | 0.252331831 | 0.294863 | 0.8557594 | 0.392131 | 0.815710819 | RAB33A | protein_coding | 47.75 | 0.022322227 | 0.022322227 | ENSG00000134594 | n.s. |
| ENSG00000067533 | 3152.227 | -0.243957167 | 0.278253 | -0.876747 | 0.380624 | 0.810319533 | RRP15 | protein_coding | 37.64 | -0.022283948 | 0.022283948 | ENSG00000067533 | n.s. |
| ENSG00000136936 | 555.3737 | -0.243910776 | 0.279249 | -0.873454 | 0.382416 | 0.81032471 | XPA | protein_coding | 38.79 | -0.022279034 | 0.022279034 | ENSG00000136936 | n.s. |
| ENSG00000101898 | 191.266 | -0.214431301 | 0.226264 | -0.947703 | 0.34328 | 0.787266337 | MCTS2P | processed_pseudogene | 40.48 | -0.022274763 | 0.022274763 | ENSG00000101898 | n.s. |
| ENSG00000213062 | 42.57076 | 0.272790443 | 0.33237 | 0.8207444 | 0.411792 | 0.828642587 | | lincRNA | 46.98 | 0.022268634 | 0.022268634 | ENSG00000213062 | n.s. |
| ENSG00000172757 | 4959.514 | -0.238929149 | 0.267481 | -0.893258 | 0.371719 | 0.806899249 | CFL1 | protein_coding | 50.18 | -0.022263583 | 0.022263583 | ENSG00000172757 | n.s. |
| ENSG00000175130 | 2135.026 | -0.217748805 | 0.233007 | -0.934514 | 0.350039 | 0.790401174 | MARCKSL1 | protein_coding | 64.21 | -0.022243568 | 0.022243568 | ENSG00000175130 | n.s. |
| ENSG00000197323 | 4096.023 | -0.242686266 | 0.276228 | -0.878571 | 0.379634 | 0.809760709 | TRIM33 | protein_coding | 38.2 | -0.02224057 | 0.02224057 | ENSG00000197323 | n.s. |
| ENSG00000137221 | 271.9672 | -0.240508399 | 0.27154 | -0.885719 | 0.375769 | 0.808254742 | TJAP1 | protein_coding | 51.18 | -0.02223542 | 0.02223542 | ENSG00000137221 | n.s. |
| ENSG00000283193 | 25.51761 | -0.284390741 | 0.355835 | -0.799222 | 0.424162 | 0.835315601 | MIR1243 | miRNA | 30.11 | -0.022224968 | 0.022224968 | ENSG00000283193 | n.s. |
| ENSG00000214770 | 230.9952 | 0.217050863 | 0.231167 | 0.938935 | 0.347764 | 0.790017511 | | antisense | 41.85 | 0.022218039 | 0.022218039 | ENSG00000214770 | n.s. |
| ENSG00000279520 | 39.87101 | 0.268630429 | 0.325283 | 0.8258349 | 0.408898 | 0.826692059 | | TEC | 54.69 | 0.02220398 | 0.02220398 | ENSG00000279520 | n.s. |
| ENSG00000141279 | 1658.519 | -0.227420595 | 0.249534 | -0.91138 | 0.362095 | 0.798938342 | NPEPPS | protein_coding | 41.25 | -0.022170491 | 0.022170491 | ENSG00000141279 | n.s. |
| ENSG00000256673 | 48.07383 | -0.264782277 | 0.318471 | -0.831417 | 0.405738 | 0.824799944 | | unprocessed_pseudogene | 51.28 | -0.022149402 | 0.022149402 | ENSG00000256673 | n.s. |
| ENSG00000149657 | 841.2967 | -0.238331918 | 0.267683 | -0.890353 | 0.373276 | 0.807377464 | LSM14B | protein_coding | 54.14 | -0.022146607 | 0.022146607 | ENSG00000149657 | n.s. |
| ENSG00000187607 | 1788.741 | -0.222028397 | 0.240299 | -0.923969 | 0.355503 | 0.794802373 | ZNF286A | protein_coding | 39.97 | -0.0221453 | 0.0221453 | ENSG00000187607 | n.s. |
| ENSG00000136699 | 615.3779 | -0.207619987 | 0.215145 | -0.965022 | 0.334534 | 0.782282887 | SMPD4 | protein_coding | 53.96 | -0.0221398 | 0.0221398 | ENSG00000136699 | n.s. |
| ENSG00000271852 | 66.5428 | 0.261176315 | 0.310884 | 0.8401074 | 0.400848 | 0.822682497 | SNORD11B | snoRNA | 36.67 | 0.022139327 | 0.022139327 | ENSG00000271852 | n.s. |
| ENSG00000038210 | 673.3179 | -0.233017752 | 0.258949 | -0.899859 | 0.368195 | 0.803511983 | PI4K2B | protein_coding | 41.82 | -0.022138467 | 0.022138467 | ENSG00000038210 | n.s. |
| ENSG00000091157 | 373.3288 | -0.238233976 | 0.267711 | -0.889891 | 0.373524 | 0.807377464 | WDR7 | protein_coding | 36.37 | -0.022137506 | 0.022137506 | ENSG00000091157 | n.s. |
| ENSG00000088832 | 839.5682 | -0.240927806 | 0.273742 | -0.880128 | 0.37879 | 0.809376313 | FKBP1A | protein_coding | 45.47 | -0.022129101 | 0.022129101 | ENSG00000088832 | n.s. |

| | | | | | | | | | | | | | |
|---|---|---|---|---|---|---|---|---|---|---|---|---|---|
| ENSG00000160688 | 1745.884 | -0.226987392 | 0.248917 | -0.911898 | 0.361822 | 0.798938342 | FLAD1 | protein_coding | 51.25 | -0.02212826 | 0.02212826 | ENSG00000160688 | n.s. |
| ENSG00000202058 | 30.31164 | -0.271415388 | 0.33208 | -0.817319 | 0.413746 | 0.829007487 | RN7SKP80 | misc_RNA | 56.89 | -0.022104489 | 0.022104489 | ENSG00000202058 | n.s. |
| ENSG00000168411 | 1740.594 | -0.237110166 | 0.265404 | -0.893393 | 0.371647 | 0.806899249 | RFWD3 | protein_coding | 43.53 | -0.022094089 | 0.022094089 | ENSG00000168411 | n.s. |
| ENSG00000212190 | 5.872306 | -0.389307145 | 0.585793 | -0.664582 | 0.506318 | 0.877502299 | RNU6-298P | snRNA | 34.38 | -0.022093857 | 0.022093857 | ENSG00000212190 | n.s. |
| ENSG00000102007 | 262.5504 | -0.239749431 | 0.271162 | -0.884156 | 0.376612 | 0.808862777 | PLP2 | protein_coding | 53.95 | -0.022086952 | 0.022086952 | ENSG00000102007 | n.s. |
| ENSG00000143155 | 1550.437 | -0.263932397 | 0.316194 | -0.834716 | 0.403877 | 0.824799944 | TIPRL | protein_coding | 38.59 | -0.022078309 | 0.022078309 | ENSG00000143155 | n.s. |
| ENSG00000181789 | 1418.689 | -0.23831785 | 0.268616 | -0.887206 | 0.374968 | 0.808040232 | COPG1 | protein_coding | 48.18 | -0.022060372 | 0.022060372 | ENSG00000181789 | n.s. |
| ENSG00000175054 | 1839.436 | -0.249927314 | 0.292815 | -0.853534 | 0.393363 | 0.816092708 | ATR | protein_coding | 37.14 | -0.02205871 | 0.02205871 | ENSG00000175054 | n.s. |
| ENSG00000137177 | 108.7768 | -0.239840214 | 0.272067 | -0.881548 | 0.378021 | 0.809223062 | KIF13A | protein_coding | 40.89 | -0.02204893 | 0.02204893 | ENSG00000137177 | n.s. |
| ENSG00000156011 | 2676.3 | -0.229609943 | 0.254056 | -0.903776 | 0.366114 | 0.801731143 | PSD3 | protein_coding | 39.93 | -0.022035952 | 0.022035952 | ENSG00000156011 | n.s. |
| ENSG00000064309 | 635.4834 | -0.239799876 | 0.272542 | -0.879864 | 0.378933 | 0.809433174 | CDON | protein_coding | 39.95 | -0.022018185 | 0.022018185 | ENSG00000064309 | n.s. |
| ENSG00000197557 | 336.2868 | -0.204172257 | 0.210459 | -0.970129 | 0.331982 | 0.780141891 | TTC30A | protein_coding | 39.41 | -0.02201516 | 0.02201516 | ENSG00000197557 | n.s. |
| ENSG00000143641 | 1887.202 | -0.229721251 | 0.254369 | -0.903103 | 0.366471 | 0.802009039 | GALNT2 | protein_coding | 45.94 | -0.02201206 | 0.02201206 | ENSG00000143641 | n.s. |
| ENSG00000183474 | 187.9417 | -0.241274926 | 0.276751 | -0.871813 | 0.38331 | 0.810719585 | GTF2H2C | protein_coding | 38.56 | -0.021987224 | 0.021987224 | ENSG00000183474 | n.s. |
| ENSG00000118007 | 1578.384 | -0.241061684 | 0.276869 | -0.870671 | 0.383934 | 0.810719585 | STAG1 | protein_coding | 38.94 | -0.021967791 | 0.021967791 | ENSG00000118007 | n.s. |
| ENSG00000104064 | 2332.336 | -0.208216149 | 0.216542 | -0.961551 | 0.336275 | 0.784379351 | GABPB1 | protein_coding | 39.38 | -0.021961358 | 0.021961358 | ENSG00000104064 | n.s. |
| ENSG00000168172 | 984.906 | -0.23541282 | 0.263599 | -0.893072 | 0.371819 | 0.806899249 | HOOK3 | protein_coding | 40.46 | -0.021935929 | 0.021935929 | ENSG00000168172 | n.s. |
| ENSG00000153922 | 4487.072 | -0.225139071 | 0.247319 | -0.910317 | 0.362655 | 0.799058295 | CHD1 | protein_coding | 35.16 | -0.021933394 | 0.021933394 | ENSG00000153922 | n.s. |
| ENSG00000008869 | 906.5262 | -0.2398952 | 0.273923 | -0.875775 | 0.381152 | 0.810319533 | HEATR5B | protein_coding | 38.59 | -0.021912913 | 0.021912913 | ENSG00000008869 | n.s. |
| ENSG00000265145 | 58.98761 | -0.24040066 | 0.275924 | -0.871255 | 0.383615 | 0.810719585 | SNORD53 | snoRNA | 46.15 | -0.021907552 | 0.021907552 | ENSG00000265145 | n.s. |
| ENSG00000088179 | 1896.431 | -0.238293865 | 0.269817 | -0.88317 | 0.377145 | 0.809223062 | PTPN4 | protein_coding | 37.04 | -0.021906772 | 0.021906772 | ENSG00000088179 | n.s. |
| ENSG00000101639 | 2166.471 | -0.237775173 | 0.269529 | -0.882187 | 0.377676 | 0.809223062 | CEP192 | protein_coding | 40.44 | -0.021859088 | 0.021859088 | ENSG00000101639 | n.s. |
| ENSG00000144559 | 66.72124 | -0.239287617 | 0.274039 | -0.873189 | 0.38256 | 0.81032471 | TAMM41 | protein_coding | 42.11 | -0.02185675 | 0.02185675 | ENSG00000144559 | n.s. |
| ENSG00000241878 | 303.7109 | -0.246944324 | 0.28701 | -0.860404 | 0.389567 | 0.815629346 | PISD | protein_coding | 51.17 | -0.02185634 | 0.02185634 | ENSG00000241878 | n.s. |
| ENSG00000174437 | 5073.592 | -0.219793325 | 0.23898 | -0.919713 | 0.357723 | 0.795491836 | ATP2A2 | protein_coding | 43.42 | -0.021839604 | 0.021839604 | ENSG00000174437 | n.s. |
| ENSG00000099940 | 382.96 | -0.234986756 | 0.264088 | -0.889805 | 0.373571 | 0.807377464 | SNAP29 | protein_coding | 48.86 | -0.021835763 | 0.021835763 | ENSG00000099940 | n.s. |
| ENSG00000223756 | 470.0408 | -0.216970878 | 0.234023 | -0.927135 | 0.353857 | 0.79336388 | TSSC2 | cribed_unprocessed_pseudc | 48.64 | -0.021811556 | 0.021811556 | ENSG00000223756 | n.s. |
| ENSG00000167384 | 651.8817 | -0.246496783 | 0.287999 | -0.855894 | 0.392056 | 0.815710819 | ZNF180 | protein_coding | 44.11 | -0.021806037 | 0.021806037 | ENSG00000167384 | n.s. |
| ENSG00000148362 | 537.1697 | -0.246452254 | 0.288588 | -0.853994 | 0.393108 | 0.815927924 | C9orf142 | protein_coding | 64.9 | -0.021773614 | 0.021773614 | ENSG00000148362 | n.s. |
| ENSG00000087111 | 576.4546 | -0.246121185 | 0.287225 | -0.856894 | 0.391504 | 0.815710819 | PIGS | protein_coding | 47.27 | -0.02177281 | 0.02177281 | ENSG00000087111 | n.s. |
| ENSG00000229994 | 69.07297 | 0.260195326 | 0.312861 | 0.8316642 | 0.405598 | 0.824799944 | RPL5P4 | processed_pseudogene | 42.78 | 0.021765697 | 0.021765697 | ENSG00000229994 | n.s. |
| ENSG00000137996 | 1030.469 | -0.223581282 | 0.245726 | -0.909879 | 0.362886 | 0.7992181 | RTCA | protein_coding | 39.07 | -0.021762215 | 0.021762215 | ENSG00000137996 | n.s. |

| | | | | | | | | | | | | | |
|---|---|---|---|---|---|---|---|---|---|---|---|---|---|
| ENSG00000132330 | 127.4939 | -0.242156405 | 0.279581 | -0.866139 | 0.386414 | 0.813136725 | SCLY | protein_coding | 46.25 | -0.021754465 | 0.021754465 | ENSG00000132330 | n.s. |
| ENSG00000078140 | 1670.765 | -0.242447202 | 0.28049 | -0.864371 | 0.387384 | 0.813399117 | UBE2K | protein_coding | 42.02 | -0.021746618 | 0.021746618 | ENSG00000078140 | n.s. |
| ENSG00000167555 | 1568.978 | -0.234877189 | 0.264582 | -0.88773 | 0.374686 | 0.808040232 | ZNF528 | protein_coding | 43.66 | -0.02174188 | 0.02174188 | ENSG00000167555 | n.s. |
| ENSG00000183530 | 1455.947 | -0.234766052 | 0.26468 | -0.886981 | 0.375089 | 0.808040232 | PRR14L | protein_coding | 45.25 | -0.021731593 | 0.021731593 | ENSG00000183530 | n.s. |
| ENSG00000243678 | 924.6271 | 0.211086469 | 0.223994 | 0.9423769 | 0.346 | 0.788954079 | NME2 | protein_coding | 52.1 | 0.021730988 | 0.021730988 | ENSG00000243678 | n.s. |
| ENSG00000103056 | 323.1762 | -0.223387463 | 0.24575 | -0.909002 | 0.363349 | 0.799322182 | SMPD3 | protein_coding | 51.06 | -0.021730716 | 0.021730716 | ENSG00000103056 | n.s. |
| ENSG00000103064 | 97.02993 | 0.233829083 | 0.262649 | 0.8902737 | 0.373319 | 0.807377464 | SLC7A6 | protein_coding | 47.81 | 0.021728188 | 0.021728188 | ENSG00000103064 | n.s. |
| ENSG00000176894 | 976.904 | -0.262818325 | 0.318174 | -0.826021 | 0.408792 | 0.826692059 | PXMP2 | protein_coding | 52.39 | -0.021723573 | 0.021723573 | ENSG00000176894 | n.s. |
| ENSG00000165699 | 75.10922 | -0.245543934 | 0.285781 | -0.859204 | 0.390228 | 0.815710819 | TSC1 | protein_coding | 42.78 | -0.021721744 | 0.021721744 | ENSG00000165699 | n.s. |
| ENSG00000243005 | 114.7989 | 0.232996848 | 0.260642 | 0.8939358 | 0.371356 | 0.806899249 | RN7SL16P | misc_RNA | 53.16 | 0.021710807 | 0.021710807 | ENSG00000243005 | n.s. |
| ENSG00000154328 | 185.3261 | -0.235973638 | 0.267648 | -0.881658 | 0.377962 | 0.809223062 | NEIL2 | protein_coding | 48.93 | -0.021693469 | 0.021693469 | ENSG00000154328 | n.s. |
| ENSG00000166847 | 2164.529 | -0.23657629 | 0.269027 | -0.879377 | 0.379197 | 0.809746094 | DCTN5 | protein_coding | 42.94 | -0.021682486 | 0.021682486 | ENSG00000166847 | n.s. |
| ENSG00000154305 | 3235.57 | -0.222652741 | 0.244803 | -0.909519 | 0.363076 | 0.7992181 | MIA3 | protein_coding | 39.16 | -0.021671835 | 0.021671835 | ENSG00000154305 | n.s. |
| ENSG00000163602 | 1775.033 | -0.237428797 | 0.272402 | -0.871611 | 0.383421 | 0.810719585 | RYBP | protein_coding | 38.33 | -0.021636728 | 0.021636728 | ENSG00000163602 | n.s. |
| ENSG00000166206 | 3131.916 | -0.232782607 | 0.261385 | -0.890572 | 0.373159 | 0.807377464 | GABRB3 | protein_coding | 41.34 | -0.021630946 | 0.021630946 | ENSG00000166206 | n.s. |
| ENSG00000110171 | 118.2692 | -0.235228925 | 0.26664 | -0.882197 | 0.37767 | 0.809223062 | TRIM3 | protein_coding | 48.68 | -0.021625006 | 0.021625006 | ENSG00000110171 | n.s. |
| ENSG00000123684 | 3436.756 | -0.232679086 | 0.261541 | -0.889645 | 0.373656 | 0.807377464 | LPGAT1 | protein_coding | 38.3 | -0.021621326 | 0.021621326 | ENSG00000123684 | n.s. |
| ENSG00000108666 | 834.324 | -0.247808521 | 0.291376 | -0.850476 | 0.395061 | 0.818022109 | C17orf75 | protein_coding | 44.91 | -0.021617566 | 0.021617566 | ENSG00000108666 | n.s. |
| ENSG00000176788 | 2277.613 | -0.24426803 | 0.28564 | -0.855159 | 0.392463 | 0.815710819 | BASP1 | protein_coding | 43.15 | -0.021608873 | 0.021608873 | ENSG00000176788 | n.s. |
| ENSG00000034053 | 1712.481 | -0.235779583 | 0.26819 | -0.87915 | 0.37932 | 0.809760709 | APBA2 | protein_coding | 48.63 | -0.021607619 | 0.021607619 | ENSG00000034053 | n.s. |
| ENSG00000265725 | 46.47836 | -0.279938248 | 0.352277 | -0.794653 | 0.426815 | 0.837228298 | MIR3144 | miRNA | 35.44 | -0.021598944 | 0.021598944 | ENSG00000265725 | n.s. |
| ENSG00000053371 | 209.4635 | -0.26270816 | 0.319526 | -0.82218 | 0.410974 | 0.82771506 | AKR7A2 | protein_coding | 52.87 | -0.021573369 | 0.021573369 | ENSG00000053371 | n.s. |
| ENSG00000200118 | 13.99583 | -0.323642022 | 0.438904 | -0.737387 | 0.460887 | 0.85776362 | | misc_RNA | 47 | -0.021565037 | 0.021565037 | ENSG00000200118 | n.s. |
| ENSG00000179526 | 223.038 | -0.233994846 | 0.264519 | -0.884604 | 0.37637 | 0.808862777 | SHARPIN | protein_coding | 60.75 | -0.02155681 | 0.02155681 | ENSG00000179526 | n.s. |
| ENSG00000233461 | 42.229 | 0.287183087 | 0.365052 | 0.786691 | 0.431463 | 0.841297658 | | antisense | 41.53 | 0.021553183 | 0.021553183 | ENSG00000233461 | n.s. |
| ENSG00000272994 | 665.03 | 0.206036308 | 0.216525 | 0.9515572 | 0.341322 | 0.786054116 | | lincRNA | 41.14 | 0.021540592 | 0.021540592 | ENSG00000272994 | n.s. |
| ENSG00000215114 | 615.5595 | -0.231701365 | 0.260156 | -0.890624 | 0.373131 | 0.807377464 | UBXN2B | protein_coding | 38.03 | -0.021530473 | 0.021530473 | ENSG00000215114 | n.s. |
| ENSG00000109436 | 828.6586 | -0.231645526 | 0.260349 | -0.889751 | 0.373599 | 0.807377464 | TBC1D9 | protein_coding | 41.09 | -0.021525285 | 0.021525285 | ENSG00000109436 | n.s. |
| ENSG00000187815 | 129.1998 | -0.252827326 | 0.300621 | -0.841017 | 0.400339 | 0.821989968 | ZFP69 | protein_coding | 41.16 | -0.021524071 | 0.021524071 | ENSG00000187815 | n.s. |
| ENSG00000167881 | 1207.629 | -0.231617247 | 0.260417 | -0.889409 | 0.373783 | 0.807377464 | SRP68 | protein_coding | 46.03 | -0.021522657 | 0.021522657 | ENSG00000167881 | n.s. |
| ENSG00000178719 | 215.4494 | -0.235452374 | 0.268787 | -0.875981 | 0.381041 | 0.810319533 | GRINA | protein_coding | 66.68 | -0.021507089 | 0.021507089 | ENSG00000178719 | n.s. |
| ENSG00000122694 | 283.5803 | -0.256791475 | 0.307907 | -0.833989 | 0.404287 | 0.824799944 | GLIPR2 | protein_coding | 51.06 | -0.021480961 | 0.021480961 | ENSG00000122694 | n.s. |

| | | | | | | | | | | | | | |
|---|---|---|---|---|---|---|---|---|---|---|---|---|---|
| ENSG00000241357 | 24.07695 | -0.287883175 | 0.367104 | -0.7842 | 0.432923 | 0.842286016 | | antisense | 66.67 | -0.02145893 | 0.02145893 | ENSG00000241357 | n.s. |
| ENSG00000182674 | 1164.193 | -0.220895579 | 0.243114 | -0.908608 | 0.363557 | 0.799601391 | KCNB2 | protein_coding | 38.88 | -0.021454805 | 0.021454805 | ENSG00000182674 | n.s. |
| ENSG00000179965 | 253.6645 | -0.238916736 | 0.27625 | -0.864857 | 0.387117 | 0.813399117 | ZNF771 | protein_coding | 51.85 | -0.021429948 | 0.021429948 | ENSG00000179965 | n.s. |
| ENSG00000181788 | 533.0797 | -0.229860662 | 0.257422 | -0.892932 | 0.371893 | 0.806899249 | SIAH2 | protein_coding | 44.98 | -0.021418575 | 0.021418575 | ENSG00000181788 | n.s. |
| ENSG00000264712 | 7.920418 | 0.372069952 | 0.555557 | 0.6697246 | 0.503033 | 0.875883784 | MIR4504 | miRNA | 38.04 | 0.021413933 | 0.021413933 | ENSG00000264712 | n.s. |
| ENSG00000143252 | 1431.67 | -0.242027672 | 0.283283 | -0.854367 | 0.392902 | 0.815710819 | SDHC | protein_coding | 42.28 | -0.021410682 | 0.021410682 | ENSG00000143252 | n.s. |
| ENSG00000170571 | 426.3392 | -0.241771296 | 0.282601 | -0.855523 | 0.392262 | 0.815710819 | EMB | protein_coding | 36.85 | -0.021388002 | 0.021388002 | ENSG00000170571 | n.s. |
| ENSG00000273888 | 278.7856 | 0.206219793 | 0.217899 | 0.9464021 | 0.343944 | 0.787678531 | FRMD6-AS1 | antisense | 52.67 | 0.021374886 | 0.021374886 | ENSG00000273888 | n.s. |
| ENSG00000266751 | 7.852972 | -0.383879515 | 0.582986 | -0.658472 | 0.510235 | 0.879714521 | MIR3661 | miRNA | 58.33 | -0.021366058 | 0.021366058 | ENSG00000266751 | n.s. |
| ENSG00000207008 | 35.47152 | -0.265237228 | 0.326513 | -0.812333 | 0.416601 | 0.83075513 | SNORA54 | snoRNA | 49.59 | -0.02135875 | 0.02135875 | ENSG00000207008 | n.s. |
| ENSG00000256663 | 59.62836 | -0.243507697 | 0.285909 | -0.851697 | 0.394382 | 0.817232331 | | processed_pseudogene | 62.26 | -0.021344536 | 0.021344536 | ENSG00000256663 | n.s. |
| ENSG00000139269 | 90.12322 | -0.237922643 | 0.275246 | -0.864401 | 0.387368 | 0.813399117 | INHBE | protein_coding | 53.48 | -0.021340782 | 0.021340782 | ENSG00000139269 | n.s. |
| ENSG00000162783 | 3803.088 | 0.187533868 | 0.188075 | 0.997121 | 0.318706 | 0.769569607 | IER5 | protein_coding | 49.76 | 0.02133237 | 0.02133237 | ENSG00000162783 | n.s. |
| ENSG00000207697 | 59.66854 | -0.261832099 | 0.320764 | -0.816277 | 0.414342 | 0.829007487 | MIR573 | miRNA | 43.43 | -0.021324011 | 0.021324011 | ENSG00000207697 | n.s. |
| ENSG00000131437 | 751.0437 | -0.233401114 | 0.266958 | -0.8743 | 0.381955 | 0.81032471 | KIF3A | protein_coding | 36.68 | -0.021319072 | 0.021319072 | ENSG00000131437 | n.s. |
| ENSG00000091656 | 1042.648 | -0.219437299 | 0.24161 | -0.908228 | 0.363758 | 0.799601391 | ZFHX4 | protein_coding | 36.38 | -0.021313168 | 0.021313168 | ENSG00000091656 | n.s. |
| ENSG00000165476 | 788.0065 | -0.240900449 | 0.281692 | -0.85519 | 0.392446 | 0.815710819 | REEP3 | protein_coding | 38.34 | -0.021310964 | 0.021310964 | ENSG00000165476 | n.s. |
| ENSG00000186871 | 578.0393 | -0.233839671 | 0.268853 | -0.869769 | 0.384427 | 0.810719585 | ERCC6L | protein_coding | 43.97 | -0.021309654 | 0.021309654 | ENSG00000186871 | n.s. |
| ENSG00000173258 | 443.4508 | -0.223930628 | 0.248489 | -0.90117 | 0.367498 | 0.80324549 | ZNF483 | protein_coding | 42.33 | -0.021307381 | 0.021307381 | ENSG00000173258 | n.s. |
| ENSG00000214253 | 527.8329 | -0.257875783 | 0.312653 | -0.824798 | 0.409486 | 0.826802059 | FIS1 | protein_coding | 51.92 | -0.02130014 | 0.02130014 | ENSG00000214253 | n.s. |
| ENSG00000149292 | 67.75933 | -0.233070806 | 0.266296 | -0.875231 | 0.381448 | 0.81032471 | TTC12 | protein_coding | 43.07 | -0.021288901 | 0.021288901 | ENSG00000149292 | n.s. |
| ENSG00000207357 | 25.05589 | 0.285782915 | 0.365025 | 0.7829131 | 0.433678 | 0.84244096 | RNU6-2 | snRNA | 45.79 | 0.021279546 | 0.021279546 | ENSG00000207357 | n.s. |
| ENSG00000164053 | 383.6095 | -0.210227324 | 0.225834 | -0.930894 | 0.351909 | 0.792117552 | ATRIP | protein_coding | 47.47 | -0.021277184 | 0.021277184 | ENSG00000164053 | n.s. |
| ENSG00000157259 | 662.0326 | -0.229906125 | 0.259413 | -0.886255 | 0.37548 | 0.808144397 | GATAD1 | protein_coding | 42.23 | -0.021268853 | 0.021268853 | ENSG00000157259 | n.s. |
| ENSG00000265808 | 1451.989 | -0.24020423 | 0.280315 | -0.856909 | 0.391495 | 0.815710819 | SEC22B | protein_coding | 36.13 | -0.021249374 | 0.021249374 | ENSG00000265808 | n.s. |
| ENSG00000130347 | 350.3086 | -0.232619275 | 0.266312 | -0.873483 | 0.3824 | 0.81032471 | RTN4IP1 | protein_coding | 41.03 | -0.021247658 | 0.021247658 | ENSG00000130347 | n.s. |
| ENSG00000122870 | 260.0176 | -0.236483289 | 0.273187 | -0.865646 | 0.386684 | 0.813136725 | BICC1 | protein_coding | 37.96 | -0.021244813 | 0.021244813 | ENSG00000122870 | n.s. |
| ENSG00000134851 | 961.1187 | -0.23109285 | 0.262431 | -0.880584 | 0.378543 | 0.809266943 | TMEM165 | protein_coding | 38.82 | -0.021239328 | 0.021239328 | ENSG00000134851 | n.s. |
| ENSG00000185837 | 25.20303 | -0.27763213 | 0.350789 | -0.791451 | 0.428681 | 0.838757697 | HDHD5-AS1 | antisense | 49.27 | -0.021200956 | 0.021200956 | ENSG00000185837 | n.s. |
| ENSG00000100243 | 416.4276 | -0.230648922 | 0.261712 | -0.881307 | 0.378152 | 0.809266943 | CYB5R3 | protein_coding | 55.99 | -0.021198527 | 0.021198527 | ENSG00000100243 | n.s. |
| ENSG00000270194 | 104.1611 | 0.235926751 | 0.272269 | 0.8665197 | 0.386205 | 0.813115862 | | antisense | 53.41 | 0.021197444 | 0.021197444 | ENSG00000270194 | n.s. |
| ENSG00000142556 | 749.7384 | -0.228070669 | 0.25644 | -0.889373 | 0.373802 | 0.807377464 | ZNF614 | protein_coding | 41.38 | -0.021193097 | 0.021193097 | ENSG00000142556 | n.s. |

| | | | | | | | | | | | | | |
|---|---|---|---|---|---|---|---|---|---|---|---|---|---|
| ENSG00000217128 | 462.1075 | -0.215443367 | 0.235579 | -0.914526 | 0.360441 | 0.797431699 | FNIP1 | protein_coding | 37.77 | -0.021179484 | 0.021179484 | ENSG00000217128 | n.s. |
| ENSG00000236404 | 194.5454 | -0.226277784 | 0.252625 | -0.895708 | 0.370409 | 0.806190999 | VLDLR-AS1 | antisense | 39.93 | -0.021171014 | 0.021171014 | ENSG00000236404 | n.s. |
| ENSG00000136231 | 2322.941 | -0.228734973 | 0.257987 | -0.886614 | 0.375287 | 0.808091443 | IGF2BP3 | protein_coding | 41.63 | -0.021167018 | 0.021167018 | ENSG00000136231 | n.s. |
| ENSG00000148334 | 440.8889 | -0.239189064 | 0.278052 | -0.860231 | 0.389662 | 0.81570591 | PTGES2 | protein_coding | 59.55 | -0.021160193 | 0.021160193 | ENSG00000148334 | n.s. |
| ENSG00000171365 | 515.2941 | -0.238953215 | 0.278806 | -0.857061 | 0.391411 | 0.815710819 | CLCN5 | protein_coding | 39.57 | -0.021138704 | 0.021138704 | ENSG00000171365 | n.s. |
| ENSG00000198478 | 531.3225 | -0.238839023 | 0.278449 | -0.857748 | 0.391032 | 0.815710819 | SH3BGRL2 | protein_coding | 38.84 | -0.021128602 | 0.021128602 | ENSG00000198478 | n.s. |
| ENSG00000105613 | 371.5453 | -0.231276878 | 0.264564 | -0.874182 | 0.382019 | 0.81032471 | MAST1 | protein_coding | 52.4 | -0.021125042 | 0.021125042 | ENSG00000105613 | n.s. |
| ENSG00000131477 | 320.9687 | -0.229189973 | 0.259119 | -0.884497 | 0.376428 | 0.808862777 | RAMP2 | protein_coding | 55.13 | -0.021114161 | 0.021114161 | ENSG00000131477 | n.s. |
| ENSG00000282381 | 192.3436 | -0.230310163 | 0.262248 | -0.878215 | 0.379827 | 0.809760709 | | lincRNA | 39.38 | -0.021106383 | 0.021106383 | ENSG00000282381 | n.s. |
| ENSG00000189091 | 3662.582 | -0.216754516 | 0.238194 | -0.909993 | 0.362826 | 0.7992181 | SF3B3 | protein_coding | 43.72 | -0.021097733 | 0.021097733 | ENSG00000189091 | n.s. |
| ENSG00000104980 | 674.0986 | -0.229532629 | 0.260611 | -0.88075 | 0.378453 | 0.809266943 | TIMM44 | protein_coding | 56.41 | -0.021095931 | 0.021095931 | ENSG00000104980 | n.s. |
| ENSG00000128908 | 1154.815 | -0.216640635 | 0.238219 | -0.909418 | 0.36313 | 0.7992181 | INO80 | protein_coding | 42.26 | -0.021086649 | 0.021086649 | ENSG00000128908 | n.s. |
| ENSG00000023516 | 2366.569 | -0.231384242 | 0.265826 | -0.870435 | 0.384063 | 0.810719585 | AKAP11 | protein_coding | 36.45 | -0.021085892 | 0.021085892 | ENSG00000023516 | n.s. |
| ENSG00000106459 | 249.4351 | -0.221897722 | 0.246704 | -0.899449 | 0.368414 | 0.803511983 | NRF1 | protein_coding | 41.15 | -0.021081979 | 0.021081979 | ENSG00000106459 | n.s. |
| ENSG00000229589 | 89.21193 | -0.229374261 | 0.260298 | -0.881198 | 0.378211 | 0.809266943 | ACVR2B-AS1 | antisense | 56.48 | -0.021081375 | 0.021081375 | ENSG00000229589 | n.s. |
| ENSG00000107518 | 323.5245 | -0.240837412 | 0.283022 | -0.85095 | 0.394797 | 0.817840002 | ATRNL1 | protein_coding | 35.41 | -0.021032729 | 0.021032729 | ENSG00000107518 | n.s. |
| ENSG00000115310 | 4999.449 | -0.226212688 | 0.253726 | -0.891564 | 0.372627 | 0.807357957 | RTN4 | protein_coding | 38.6 | -0.021022821 | 0.021022821 | ENSG00000115310 | n.s. |
| ENSG00000183665 | 455.6632 | 0.211831957 | 0.230529 | 0.9188962 | 0.35815 | 0.795806072 | TRMT12 | protein_coding | 43.63 | 0.021012195 | 0.021012195 | ENSG00000183665 | n.s. |
| ENSG00000147533 | 776.1426 | -0.236206109 | 0.27403 | -0.861972 | 0.388703 | 0.814823372 | GOLGA7 | protein_coding | 39.89 | -0.02100735 | 0.02100735 | ENSG00000147533 | n.s. |
| ENSG00000113569 | 3127.658 | -0.226918137 | 0.255504 | -0.88812 | 0.374476 | 0.808040232 | NUP155 | protein_coding | 41.76 | -0.021005135 | 0.021005135 | ENSG00000113569 | n.s. |
| ENSG00000099995 | 1049.954 | -0.220643512 | 0.245324 | -0.899396 | 0.368442 | 0.803511983 | SF3A1 | protein_coding | 49.04 | -0.02096282 | 0.02096282 | ENSG00000099995 | n.s. |
| ENSG00000275857 | 16.21662 | 0.334547671 | 0.471201 | 0.7099889 | 0.477711 | 0.865660698 | | antisense | 50.27 | 0.020960181 | 0.020960181 | ENSG00000275857 | n.s. |
| ENSG00000099875 | 309.2377 | -0.211420823 | 0.230169 | -0.918545 | 0.358334 | 0.795960392 | MKNK2 | protein_coding | 64.23 | -0.02095361 | 0.02095361 | ENSG00000099875 | n.s. |
| ENSG00000198598 | 137.8675 | -0.233167704 | 0.269142 | -0.866336 | 0.386306 | 0.813136725 | MMP17 | protein_coding | 61.93 | -0.020946953 | 0.020946953 | ENSG00000198598 | n.s. |
| ENSG00000139746 | 2808.914 | -0.235318743 | 0.273028 | -0.861884 | 0.388751 | 0.814823372 | RBM26 | protein_coding | 34.26 | -0.020928431 | 0.020928431 | ENSG00000139746 | n.s. |
| ENSG00000277149 | 303.0604 | -0.225941246 | 0.254594 | -0.887459 | 0.374832 | 0.808040232 | TYW1B | protein_coding | 41.66 | -0.020914707 | 0.020914707 | ENSG00000277149 | n.s. |
| ENSG00000205726 | 1042.044 | -0.217030551 | 0.239658 | -0.905586 | 0.365155 | 0.801016568 | ITSN1 | protein_coding | 42.47 | -0.020912738 | 0.020912738 | ENSG00000205726 | n.s. |
| ENSG00000279319 | 157.4212 | -0.228874303 | 0.26131 | -0.875873 | 0.381099 | 0.810319533 | | TEC | 34.93 | -0.020906224 | 0.020906224 | ENSG00000279319 | n.s. |
| ENSG00000127990 | 699.7417 | 0.239648132 | 0.281851 | 0.8502651 | 0.395178 | 0.818022109 | SGCE | protein_coding | 35.25 | 0.020905695 | 0.020905695 | ENSG00000127990 | n.s. |
| ENSG00000196428 | 2390.664 | -0.209579017 | 0.227286 | -0.922095 | 0.356479 | 0.794806874 | TSC22D2 | protein_coding | 36.52 | -0.020903073 | 0.020903073 | ENSG00000196428 | n.s. |
| ENSG00000008083 | 1009.919 | 0.215012679 | 0.23671 | 0.9083393 | 0.363699 | 0.799601391 | JARID2 | protein_coding | 43.92 | 0.02088342 | 0.02088342 | ENSG00000008083 | n.s. |
| ENSG00000076513 | 762.0647 | -0.236050822 | 0.275269 | -0.857527 | 0.391154 | 0.815710819 | ANKRD13A | protein_coding | 44 | -0.020881947 | 0.020881947 | ENSG00000076513 | n.s. |

| | | | | | | | | | | | | | |
|---|---|---|---|---|---|---|---|---|---|---|---|---|---|
| ENSG00000185344 | 1317.03 | -0.231999361 | 0.267467 | -0.867394 | 0.385726 | 0.812844991 | ATP6V0A2 | protein_coding | 44.41 | -0.020878148 | 0.020878148 | ENSG00000185344 | n.s. |
| ENSG00000173692 | 3290.987 | -0.228371966 | 0.261494 | -0.873334 | 0.382481 | 0.81032471 | PSMD1 | protein_coding | 37.87 | -0.020859705 | 0.020859705 | ENSG00000173692 | n.s. |
| ENSG00000197905 | 157.9396 | -0.22690062 | 0.257282 | -0.881913 | 0.377824 | 0.809223062 | TEAD4 | protein_coding | 51.6 | -0.020859371 | 0.020859371 | ENSG00000197905 | n.s. |
| ENSG00000118707 | 474.7039 | -0.232160063 | 0.267803 | -0.866907 | 0.385993 | 0.813115862 | TGIF2 | protein_coding | 51.95 | -0.020859017 | 0.020859017 | ENSG00000118707 | n.s. |
| ENSG00000242338 | 81.50492 | -0.256097759 | 0.31359 | -0.816663 | 0.414121 | 0.829007487 | BMS1P4 | cribed_unprocessed_pseudc | 39.67 | -0.020856998 | 0.020856998 | ENSG00000242338 | n.s. |
| ENSG00000048140 | 528.9046 | -0.248029144 | 0.296297 | -0.837096 | 0.402539 | 0.823993439 | TSPAN17 | protein_coding | 58.7 | -0.020853359 | 0.020853359 | ENSG00000048140 | n.s. |
| ENSG00000058262 | 2638.44 | -0.224350958 | 0.251921 | -0.89056 | 0.373165 | 0.807377464 | SEC61A1 | protein_coding | 47.3 | -0.020847449 | 0.020847449 | ENSG00000058262 | n.s. |
| ENSG00000269001 | 148.7122 | -0.238635746 | 0.280693 | -0.850165 | 0.395233 | 0.818022109 | | transcribed_unprocessed_pseudc | 44.81 | -0.020817379 | 0.020817379 | ENSG00000269001 | n.s. |
| ENSG00000140525 | 4134.541 | -0.234987444 | 0.275063 | -0.854303 | 0.392937 | 0.815710819 | FANCI | protein_coding | 40.11 | -0.020787877 | 0.020787877 | ENSG00000140525 | n.s. |
| ENSG00000163867 | 46.5928 | -0.252960332 | 0.307612 | -0.822335 | 0.410886 | 0.827684308 | ZMYM6 | protein_coding | 38.8 | -0.020776967 | 0.020776967 | ENSG00000163867 | n.s. |
| ENSG00000178662 | 817.3447 | -0.221393758 | 0.246854 | -0.89686 | 0.369794 | 0.805733297 | CSRNP3 | protein_coding | 36.48 | -0.020768658 | 0.020768658 | ENSG00000178662 | n.s. |
| ENSG00000112365 | 1207.407 | -0.212895188 | 0.233577 | -0.911457 | 0.362055 | 0.798938342 | ZBTB24 | protein_coding | 43.86 | -0.020754457 | 0.020754457 | ENSG00000112365 | n.s. |
| ENSG00000125885 | 2791.655 | -0.219015537 | 0.243776 | -0.89843 | 0.368956 | 0.804160729 | MCM8 | protein_coding | 39.4 | -0.020731384 | 0.020731384 | ENSG00000125885 | n.s. |
| ENSG00000214796 | 103.6645 | -0.245718623 | 0.292963 | -0.838735 | 0.401618 | 0.823540971 | | transcribed_unprocessed_pseudc | 51.87 | -0.020717714 | 0.020717714 | ENSG00000214796 | n.s. |
| ENSG00000165821 | 216.034 | -0.210509188 | 0.230068 | -0.914986 | 0.360199 | 0.797273365 | SALL2 | protein_coding | 48.58 | -0.020712577 | 0.020712577 | ENSG00000165821 | n.s. |
| ENSG00000249509 | 122.2973 | -0.225154081 | 0.254873 | -0.883397 | 0.377022 | 0.809223062 | | antisense | 40.94 | -0.020698808 | 0.020698808 | ENSG00000249509 | n.s. |
| ENSG00000109133 | 3788.049 | -0.233834097 | 0.273547 | -0.854823 | 0.392649 | 0.815710819 | TMEM33 | protein_coding | 35.29 | -0.020685848 | 0.020685848 | ENSG00000109133 | n.s. |
| ENSG00000237457 | 25.72295 | 0.284070039 | 0.369349 | 0.7691094 | 0.441828 | 0.845693795 | LINC01351 | lincRNA | 37.76 | 0.020676565 | 0.020676565 | ENSG00000237457 | n.s. |
| ENSG00000110583 | 717.9069 | -0.23238848 | 0.269769 | -0.861435 | 0.388999 | 0.814823372 | NAA40 | protein_coding | 50.29 | -0.020667823 | 0.020667823 | ENSG00000110583 | n.s. |
| ENSG00000107223 | 2114.574 | -0.249070927 | 0.301078 | -0.827264 | 0.408087 | 0.826171518 | EDF1 | protein_coding | 57.77 | -0.020655398 | 0.020655398 | ENSG00000107223 | n.s. |
| ENSG00000112367 | 355.8426 | -0.24892471 | 0.300769 | -0.827628 | 0.407881 | 0.826149803 | FIG4 | protein_coding | 38.91 | -0.020646113 | 0.020646113 | ENSG00000112367 | n.s. |
| ENSG00000188313 | 587.9551 | -0.220668202 | 0.246403 | -0.89556 | 0.370488 | 0.806236866 | PLSCR1 | protein_coding | 36.31 | -0.020640718 | 0.020640718 | ENSG00000188313 | n.s. |
| ENSG00000145833 | 2959.363 | -0.225933072 | 0.2577 | -0.876728 | 0.380634 | 0.810319533 | DDX46 | protein_coding | 42.1 | -0.020637561 | 0.020637561 | ENSG00000145833 | n.s. |
| ENSG00000137822 | 752.456 | -0.220246466 | 0.245774 | -0.896132 | 0.370182 | 0.805949078 | TUBGCP4 | protein_coding | 42.65 | -0.020635419 | 0.020635419 | ENSG00000137822 | n.s. |
| ENSG00000169062 | 1346.103 | -0.213044382 | 0.234801 | -0.907339 | 0.364228 | 0.800117344 | UPF3A | protein_coding | 47.58 | -0.020632563 | 0.020632563 | ENSG00000169062 | n.s. |
| ENSG00000167173 | 289.9128 | -0.206778756 | 0.223966 | -0.923261 | 0.355871 | 0.794802373 | C15orf39 | protein_coding | 58.35 | -0.020624288 | 0.020624288 | ENSG00000167173 | n.s. |
| ENSG00000112335 | 890.9113 | 0.260369972 | 0.323219 | 0.805552 | 0.420501 | 0.833311153 | SNX3 | protein_coding | 41.16 | 0.020619429 | 0.020619429 | ENSG00000112335 | n.s. |
| ENSG00000198185 | 442.7568 | -0.225733045 | 0.258433 | -0.873468 | 0.382408 | 0.81032471 | ZNF334 | protein_coding | 40.56 | -0.020618663 | 0.020618663 | ENSG00000198185 | n.s. |
| ENSG00000151726 | 968.5963 | -0.23304881 | 0.271925 | -0.857034 | 0.391426 | 0.815710819 | ACSL1 | protein_coding | 44.71 | -0.020616378 | 0.020616378 | ENSG00000151726 | n.s. |
| ENSG00000232838 | 139.6056 | -0.229444386 | 0.265087 | -0.865543 | 0.386741 | 0.813136725 | PET117 | protein_coding | 45.08 | -0.020612463 | 0.020612463 | ENSG00000232838 | n.s. |
| ENSG00000138587 | 179.7516 | -0.246563296 | 0.296915 | -0.830418 | 0.406302 | 0.825006384 | MNS1 | protein_coding | 35.19 | -0.020598561 | 0.020598561 | ENSG00000138587 | n.s. |
| ENSG00000132359 | 913.9404 | -0.225464889 | 0.257031 | -0.877189 | 0.380384 | 0.810288517 | RAP1GAP2 | protein_coding | 51.49 | -0.020598543 | 0.020598543 | ENSG00000132359 | n.s. |

| | | | | | | | | | | | | | |
|---|---|---|---|---|---|---|---|---|---|---|---|---|---|
| ENSG00000136378 | 373.9087 | -0.222399505 | 0.250639 | -0.887332 | 0.3749 | 0.808040232 | ADAMTS7 | protein_coding | 56.03 | -0.020586858 | 0.020586858 | ENSG00000136378 | n.s. |
| ENSG00000279432 | 303.1114 | -0.222899131 | 0.251811 | -0.885183 | 0.376058 | 0.808501396 | | TEC | 39.69 | -0.020577875 | 0.020577875 | ENSG00000279432 | n.s. |
| ENSG00000029993 | 3186.268 | -0.222162317 | 0.250326 | -0.887492 | 0.374814 | 0.808040232 | HMGB3 | protein_coding | 51.16 | -0.020564903 | 0.020564903 | ENSG00000029993 | n.s. |
| ENSG00000163710 | 153.1624 | 0.211319083 | 0.232416 | 0.9092266 | 0.363231 | 0.799270441 | PCOLCE2 | protein_coding | 40.92 | 0.020562667 | 0.020562667 | ENSG00000163710 | n.s. |
| ENSG00000213523 | 506.8416 | -0.2355603 | 0.27718 | -0.849845 | 0.395411 | 0.818083342 | SRA1 | protein_coding | 45.85 | -0.020541435 | 0.020541435 | ENSG00000213523 | n.s. |
| ENSG00000105701 | 1132.86 | -0.232100477 | 0.270463 | -0.858161 | 0.390804 | 0.815710819 | FKBP8 | protein_coding | 56.64 | -0.020532485 | 0.020532485 | ENSG00000105701 | n.s. |
| ENSG00000130175 | 811.2565 | -0.22164165 | 0.249852 | -0.887093 | 0.375029 | 0.808040232 | PRKCSH | protein_coding | 56.85 | -0.020516706 | 0.020516706 | ENSG00000130175 | n.s. |
| ENSG00000267673 | 134.0928 | -0.257174507 | 0.318227 | -0.808149 | 0.419005 | 0.832304654 | FDX2 | protein_coding | 52.77 | -0.020501354 | 0.020501354 | ENSG00000267673 | n.s. |
| ENSG00000174529 | 73.18625 | -0.243126044 | 0.289914 | -0.838614 | 0.401686 | 0.823540971 | TMEM81 | protein_coding | 47.52 | -0.020499121 | 0.020499121 | ENSG00000174529 | n.s. |
| ENSG00000115295 | 646.6573 | -0.234695632 | 0.275836 | -0.850852 | 0.394852 | 0.817840002 | CLIP4 | protein_coding | 38.44 | -0.020496357 | 0.020496357 | ENSG00000115295 | n.s. |
| ENSG00000204120 | 2620.325 | -0.220570772 | 0.247585 | -0.89089 | 0.372988 | 0.807377464 | GIGYF2 | protein_coding | 39.44 | -0.020496181 | 0.020496181 | ENSG00000204120 | n.s. |
| ENSG00000101407 | 923.1425 | -0.229974389 | 0.266582 | -0.862678 | 0.388315 | 0.814600671 | TTI1 | protein_coding | 43.33 | -0.020480424 | 0.020480424 | ENSG00000101407 | n.s. |
| ENSG00000189164 | 431.7514 | -0.224132276 | 0.256384 | -0.874206 | 0.382006 | 0.81032471 | ZNF527 | protein_coding | 40.73 | -0.020472448 | 0.020472448 | ENSG00000189164 | n.s. |
| ENSG00000269946 | 44.4096 | 0.261565278 | 0.326926 | 0.8000746 | 0.423668 | 0.835158421 | | lincRNA | 40.89 | 0.020462549 | 0.020462549 | ENSG00000269946 | n.s. |
| ENSG00000065029 | 196.9573 | -0.244355795 | 0.292884 | -0.834308 | 0.404107 | 0.824799944 | ZNF76 | protein_coding | 48.23 | -0.020440699 | 0.020440699 | ENSG00000065029 | n.s. |
| ENSG00000244687 | 992.2854 | -0.210326881 | 0.231528 | -0.908428 | 0.363652 | 0.799601391 | UBE2V1 | protein_coding | 44.49 | -0.020428305 | 0.020428305 | ENSG00000244687 | n.s. |
| ENSG00000183160 | 129.1925 | 0.23771601 | 0.281507 | 0.8444418 | 0.398423 | 0.820475937 | TMEM119 | protein_coding | 56.18 | 0.020427924 | 0.020427924 | ENSG00000183160 | n.s. |
| ENSG00000188229 | 3272.681 | -0.237350988 | 0.28068 | -0.845628 | 0.39776 | 0.820246021 | TUBB4B | protein_coding | 63.25 | -0.020425445 | 0.020425445 | ENSG00000188229 | n.s. |
| ENSG00000132423 | 359.4094 | -0.260715913 | 0.325601 | -0.800721 | 0.423293 | 0.834993891 | COQ3 | protein_coding | 39.36 | -0.02041841 | 0.02041841 | ENSG00000132423 | n.s. |
| ENSG00000140950 | 20.40285 | -0.36988688 | 0.563716 | -0.656158 | 0.511723 | 0.880681667 | TLDC1 | protein_coding | 49.64 | -0.020410744 | 0.020410744 | ENSG00000140950 | n.s. |
| ENSG00000229422 | 56.8144 | -0.262943503 | 0.330423 | -0.795778 | 0.426161 | 0.836464856 | | processed_pseudogene | 69.6 | -0.020391874 | 0.020391874 | ENSG00000229422 | n.s. |
| ENSG00000242028 | 328.7643 | -0.198029659 | 0.210094 | -0.942578 | 0.345897 | 0.788921338 | HYPK | protein_coding | 49.58 | -0.020390381 | 0.020390381 | ENSG00000242028 | n.s. |
| ENSG00000068489 | 1491.451 | -0.230370117 | 0.269362 | -0.855242 | 0.392417 | 0.815710819 | PRR11 | protein_coding | 44.47 | -0.020379411 | 0.020379411 | ENSG00000068489 | n.s. |
| ENSG00000101290 | 625.9737 | -0.230055842 | 0.268999 | -0.85523 | 0.392424 | 0.815710819 | CDS2 | protein_coding | 44.84 | -0.020351609 | 0.020351609 | ENSG00000101290 | n.s. |
| ENSG00000134779 | 1276.601 | -0.232945844 | 0.273832 | -0.850688 | 0.394942 | 0.817906338 | TPGS2 | protein_coding | 42.82 | -0.02033534 | 0.02033534 | ENSG00000134779 | n.s. |
| ENSG00000182504 | 1295.388 | -0.221037138 | 0.250352 | -0.882905 | 0.377288 | 0.809223062 | CEP97 | protein_coding | 40.61 | -0.020320331 | 0.020320331 | ENSG00000182504 | n.s. |
| ENSG00000091651 | 2142.674 | -0.22106276 | 0.251018 | -0.880665 | 0.378499 | 0.809266943 | ORC6 | protein_coding | 44.16 | -0.02031748 | 0.02031748 | ENSG00000091651 | n.s. |
| ENSG00000107614 | 46.20229 | 0.253224581 | 0.312055 | 0.8114735 | 0.417094 | 0.831381057 | TRDMT1 | protein_coding | 36.72 | 0.02030858 | 0.02030858 | ENSG00000107614 | n.s. |
| ENSG00000171806 | 287.131 | 0.208301091 | 0.228565 | 0.9113427 | 0.362115 | 0.798938342 | METTL18 | protein_coding | 40.77 | 0.020306593 | 0.020306593 | ENSG00000171806 | n.s. |
| ENSG00000156787 | 527.7309 | -0.238059209 | 0.282704 | -0.84208 | 0.399743 | 0.821737289 | TBC1D31 | protein_coding | 39.82 | -0.020298596 | 0.020298596 | ENSG00000156787 | n.s. |
| ENSG00000119616 | 649.4141 | -0.229435475 | 0.26768 | -0.857127 | 0.391375 | 0.815710819 | FCF1 | protein_coding | 40.97 | -0.020296729 | 0.020296729 | ENSG00000119616 | n.s. |
| ENSG00000252859 | 6.264454 | -0.362888935 | 0.548942 | -0.66107 | 0.508567 | 0.879193676 | RNU6-375P | snRNA | 33 | -0.020291096 | 0.020291096 | ENSG00000252859 | n.s. |

| | | | | | | | | | | | | | |
|---|---|---|---|---|---|---|---|---|---|---|---|---|---|
| ENSG00000040531 | 148.0246 | -0.249126535 | 0.304053 | -0.819353 | 0.412585 | 0.829007487 | CTNS | protein_coding | 54.27 | -0.02028925 | 0.02028925 | ENSG00000040531 | n.s. |
| ENSG00000064961 | 367.37 | -0.240948548 | 0.287694 | -0.837516 | 0.402303 | 0.823848555 | HMG20B | protein_coding | 65.34 | -0.020276451 | 0.020276451 | ENSG00000064961 | n.s. |
| ENSG00000106266 | 104.4508 | -0.254347778 | 0.31461 | -0.808454 | 0.418829 | 0.832304654 | SNX8 | protein_coding | 51.46 | -0.020276014 | 0.020276014 | ENSG00000106266 | n.s. |
| ENSG00000160124 | 634.8771 | -0.24568672 | 0.298052 | -0.82431 | 0.409764 | 0.827015612 | CCDC58 | protein_coding | 39.16 | -0.020265787 | 0.020265787 | ENSG00000160124 | n.s. |
| ENSG00000174132 | 113.1705 | -0.275950818 | 0.356247 | -0.774605 | 0.438573 | 0.84443611 | FAM174A | protein_coding | 35.6 | -0.020263953 | 0.020263953 | ENSG00000174132 | n.s. |
| ENSG00000145332 | 885.2393 | -0.222064834 | 0.255193 | -0.870185 | 0.384199 | 0.810719585 | KLHL8 | protein_coding | 38.23 | -0.020236621 | 0.020236621 | ENSG00000145332 | n.s. |
| ENSG00000067365 | 429.4831 | -0.217029418 | 0.242838 | -0.893721 | 0.371471 | 0.806899249 | METTL22 | protein_coding | 49.56 | -0.020222951 | 0.020222951 | ENSG00000067365 | n.s. |
| ENSG00000162624 | 55.33423 | 0.250312229 | 0.307818 | 0.8131815 | 0.416114 | 0.830260147 | LHX8 | protein_coding | 39.78 | 0.020221676 | 0.020221676 | ENSG00000162624 | n.s. |
| ENSG00000274964 | 187.6171 | 0.248073935 | 0.303846 | 0.8164466 | 0.414245 | 0.829007487 | | sense_intronic | 34.71 | 0.020203525 | 0.020203525 | ENSG00000274964 | n.s. |
| ENSG00000183576 | 918.7835 | -0.236717071 | 0.281077 | -0.84218 | 0.399687 | 0.821737289 | SETD3 | protein_coding | 42.67 | -0.020184156 | 0.020184156 | ENSG00000183576 | n.s. |
| ENSG00000149485 | 4696.472 | -0.220862218 | 0.253039 | -0.87284 | 0.38275 | 0.81032471 | FADS1 | protein_coding | 50.81 | -0.020173758 | 0.020173758 | ENSG00000149485 | n.s. |
| ENSG00000235859 | 122.9263 | -0.253011738 | 0.313 | -0.808345 | 0.418892 | 0.832304654 | | processed_pseudogene | 47.57 | -0.020169508 | 0.020169508 | ENSG00000235859 | n.s. |
| ENSG00000100285 | 1745.272 | -0.203559772 | 0.22176 | -0.917928 | 0.358656 | 0.796009826 | NEFH | protein_coding | 52.82 | -0.020169022 | 0.020169022 | ENSG00000100285 | n.s. |
| ENSG00000158163 | 125.3201 | -0.227976704 | 0.265715 | -0.857976 | 0.390906 | 0.815710819 | DZIP1L | protein_coding | 45.71 | -0.02016768 | 0.02016768 | ENSG00000158163 | n.s. |
| ENSG00000021776 | 2460.905 | -0.220028355 | 0.250239 | -0.879272 | 0.379254 | 0.809746094 | AQR | protein_coding | 37.54 | -0.020165849 | 0.020165849 | ENSG00000021776 | n.s. |
| ENSG00000160094 | 57.86464 | -0.247606089 | 0.302206 | -0.81933 | 0.412598 | 0.829007487 | ZNF362 | protein_coding | 50.56 | -0.020165423 | 0.020165423 | ENSG00000160094 | n.s. |
| ENSG00000108506 | 1424.132 | -0.215270694 | 0.240272 | -0.895945 | 0.370282 | 0.806041139 | INTS2 | protein_coding | 39.68 | -0.020158549 | 0.020158549 | ENSG00000108506 | n.s. |
| ENSG00000092208 | 315.0974 | -0.253123577 | 0.313728 | -0.806824 | 0.419768 | 0.832539843 | GEMIN2 | protein_coding | 39.65 | -0.020147364 | 0.020147364 | ENSG00000092208 | n.s. |
| ENSG00000200419 | 24.40898 | 0.276762387 | 0.360037 | 0.7687048 | 0.442069 | 0.845745639 | | misc_RNA | 43.14 | 0.020137296 | 0.020137296 | ENSG00000200419 | n.s. |
| ENSG00000173812 | 5438.356 | 0.186750999 | 0.192389 | 0.970696 | 0.3317 | 0.780141891 | EIF1 | protein_coding | 49.15 | 0.020136688 | 0.020136688 | ENSG00000173812 | n.s. |
| ENSG00000196951 | 43.86433 | 0.258136571 | 0.323576 | 0.7977629 | 0.425008 | 0.835658686 | SCOC-AS1 | antisense | 40.85 | 0.020127184 | 0.020127184 | ENSG00000196951 | n.s. |
| ENSG00000115825 | 3002.208 | -0.229216868 | 0.269023 | -0.852035 | 0.394195 | 0.816965091 | PRKD3 | protein_coding | 36.9 | -0.020124439 | 0.020124439 | ENSG00000115825 | n.s. |
| ENSG00000028839 | 1031.296 | -0.238678805 | 0.284732 | -0.838258 | 0.401886 | 0.823540971 | TBPL1 | protein_coding | 37.6 | -0.020124153 | 0.020124153 | ENSG00000028839 | n.s. |
| ENSG00000157764 | 979.4259 | -0.232690503 | 0.274548 | -0.84754 | 0.396694 | 0.819495485 | BRAF | protein_coding | 37.97 | -0.020116893 | 0.020116893 | ENSG00000157764 | n.s. |
| ENSG00000132305 | 2974.803 | -0.227381042 | 0.264719 | -0.858952 | 0.390367 | 0.815710819 | IMMT | protein_coding | 41.41 | -0.020114986 | 0.020114986 | ENSG00000132305 | n.s. |
| ENSG00000167904 | 65.72215 | -0.26069872 | 0.32809 | -0.794594 | 0.426849 | 0.837228298 | TMEM68 | protein_coding | 38.91 | -0.020114497 | 0.020114497 | ENSG00000167904 | n.s. |
| ENSG00000064999 | 366.2244 | -0.219606488 | 0.250192 | -0.877751 | 0.380079 | 0.809893031 | ANKS1A | protein_coding | 44.93 | -0.02010988 | 0.02010988 | ENSG00000064999 | n.s. |
| ENSG00000138190 | 587.1493 | -0.246897366 | 0.301625 | -0.818558 | 0.413039 | 0.829007487 | EXOC6 | protein_coding | 37.36 | -0.020107703 | 0.020107703 | ENSG00000138190 | n.s. |
| ENSG00000136861 | 2056.789 | -0.219399682 | 0.24959 | -0.879041 | 0.379379 | 0.809760709 | CDK5RAP2 | protein_coding | 43.21 | -0.020106511 | 0.020106511 | ENSG00000136861 | n.s. |
| ENSG00000096093 | 55.30142 | -0.276136797 | 0.358349 | -0.770582 | 0.440955 | 0.845649551 | EFHC1 | protein_coding | 41.58 | -0.020105403 | 0.020105403 | ENSG00000096093 | n.s. |
| ENSG00000214517 | 560.8808 | -0.23268812 | 0.274825 | -0.846676 | 0.397176 | 0.819605152 | PPME1 | protein_coding | 38.65 | -0.020103165 | 0.020103165 | ENSG00000214517 | n.s. |
| ENSG00000199035 | 21.03848 | -0.284959752 | 0.374937 | -0.760021 | 0.447242 | 0.850164607 | MIR103A1 | miRNA | 48.72 | -0.020088802 | 0.020088802 | ENSG00000199035 | n.s. |

| | | | | | | | | | | | | | |
|---|---|---|---|---|---|---|---|---|---|---|---|---|---|
| ENSG00000252128 | 18.92004 | 0.309016384 | 0.425205 | 0.7267466 | 0.467381 | 0.861088114 | | snoRNA | 38.89 | 0.020071358 | 0.020071358 | ENSG00000252128 | n.s. |
| ENSG00000241120 | 49.95266 | -0.2753811 | 0.357105 | -0.771149 | 0.440618 | 0.845502102 | HMGN1P8 | processed_pseudogene | 43.06 | -0.020071236 | 0.020071236 | ENSG00000241120 | n.s. |
| ENSG00000126883 | 425.4345 | 0.20085747 | 0.217112 | 0.925133 | 0.354897 | 0.794524781 | NUP214 | protein_coding | 42.09 | 0.020064165 | 0.020064165 | ENSG00000126883 | n.s. |
| ENSG00000108852 | 368.3543 | -0.226670907 | 0.26522 | -0.854652 | 0.392744 | 0.815710819 | MPP2 | protein_coding | 52.29 | -0.020052165 | 0.020052165 | ENSG00000108852 | n.s. |
| ENSG00000106780 | 566.3623 | -0.220034285 | 0.252485 | -0.871475 | 0.383495 | 0.810719585 | MEGF9 | protein_coding | 38.18 | -0.020051578 | 0.020051578 | ENSG00000106780 | n.s. |
| ENSG00000253729 | 10908.3 | -0.217997839 | 0.247204 | -0.881853 | 0.377856 | 0.809223062 | PRKDC | protein_coding | 41.93 | -0.020040923 | 0.020040923 | ENSG00000253729 | n.s. |
| ENSG00000120798 | 916.6991 | -0.223018351 | 0.257515 | -0.866039 | 0.386469 | 0.813136725 | NR2C1 | protein_coding | 39.56 | -0.020035171 | 0.020035171 | ENSG00000120798 | n.s. |
| ENSG00000179918 | 302.8257 | -0.21547493 | 0.242213 | -0.889608 | 0.373676 | 0.807377464 | SEPHS2 | protein_coding | 51.31 | -0.020022658 | 0.020022658 | ENSG00000179918 | n.s. |
| ENSG00000025039 | 803.8711 | -0.226267826 | 0.263821 | -0.857657 | 0.391082 | 0.815710819 | RRAGD | protein_coding | 42.25 | -0.020016507 | 0.020016507 | ENSG00000025039 | n.s. |
| ENSG00000206448 | 20.54824 | -0.301773085 | 0.410162 | -0.735742 | 0.461888 | 0.858395846 | PPIAP30 | processed_pseudogene | 48.55 | -0.020011295 | 0.020011295 | ENSG00000206448 | n.s. |
| ENSG00000160256 | 279.9202 | -0.232517833 | 0.275277 | -0.844668 | 0.398296 | 0.820375622 | FAM207A | protein_coding | 56.7 | -0.01999357 | 0.01999357 | ENSG00000160256 | n.s. |
| ENSG00000111271 | 76.8471 | -0.225973417 | 0.264311 | -0.854952 | 0.392578 | 0.815710819 | ACAD10 | protein_coding | 47.12 | -0.019990462 | 0.019990462 | ENSG00000111271 | n.s. |
| ENSG00000187535 | 274.9821 | -0.238823451 | 0.287341 | -0.83115 | 0.405889 | 0.824799944 | IFT140 | protein_coding | 54.44 | -0.019977911 | 0.019977911 | ENSG00000187535 | n.s. |
| ENSG00000176490 | 205.0633 | -0.238703226 | 0.286529 | -0.833085 | 0.404797 | 0.824799944 | DIRAS1 | protein_coding | 61.18 | -0.019967854 | 0.019967854 | ENSG00000176490 | n.s. |
| ENSG00000124181 | 2887.97 | -0.216909427 | 0.245794 | -0.882484 | 0.377515 | 0.809223062 | PLCG1 | protein_coding | 49.84 | -0.019940863 | 0.019940863 | ENSG00000124181 | n.s. |
| ENSG00000141219 | 1208.863 | -0.240413 | 0.290457 | -0.827707 | 0.407837 | 0.826149803 | C17orf80 | protein_coding | 44.12 | -0.019940142 | 0.019940142 | ENSG00000141219 | n.s. |
| ENSG00000225470 | 981.4574 | -0.225384834 | 0.262318 | -0.859205 | 0.390227 | 0.815710819 | JPX | lincRNA | 38.97 | -0.019938394 | 0.019938394 | ENSG00000225470 | n.s. |
| ENSG00000204524 | 278.3185 | -0.223030458 | 0.258339 | -0.863325 | 0.387959 | 0.813977289 | ZNF805 | protein_coding | 46.41 | -0.019936182 | 0.019936182 | ENSG00000204524 | n.s. |
| ENSG00000171451 | 446.8904 | 0.19464802 | 0.207536 | 0.9378987 | 0.348296 | 0.790017511 | DSEL | protein_coding | 41.51 | 0.01992481 | 0.01992481 | ENSG00000171451 | n.s. |
| ENSG00000179119 | 667.1441 | -0.231667437 | 0.274112 | -0.845156 | 0.398024 | 0.820375622 | SPTY2D1 | protein_coding | 41.92 | -0.019920447 | 0.019920447 | ENSG00000179119 | n.s. |
| ENSG00000101746 | 336.5452 | -0.213816834 | 0.239586 | -0.892443 | 0.372156 | 0.807090616 | NOL4 | protein_coding | 34.88 | -0.01990158 | 0.01990158 | ENSG00000101746 | n.s. |
| ENSG00000274578 | 122.654 | 0.231723204 | 0.274558 | 0.8439864 | 0.398677 | 0.820678855 | | sense_intronic | 32.7 | 0.019888051 | 0.019888051 | ENSG00000274578 | n.s. |
| ENSG00000224707 | 73.62384 | -0.241565354 | 0.293598 | -0.822776 | 0.410635 | 0.827563884 | E2F3-IT1 | sense_intronic | 39.27 | -0.019856303 | 0.019856303 | ENSG00000224707 | n.s. |
| ENSG00000214050 | 18.1261 | -0.296186313 | 0.399705 | -0.741013 | 0.458686 | 0.856995387 | FBXO16 | protein_coding | 46.01 | -0.019850856 | 0.019850856 | ENSG00000214050 | n.s. |
| ENSG00000018610 | 459.482 | -0.245421483 | 0.301726 | -0.813392 | 0.415993 | 0.830138034 | CXorf56 | protein_coding | 44.29 | -0.01984225 | 0.01984225 | ENSG00000018610 | n.s. |
| ENSG00000228015 | 6.130731 | -0.427939111 | 0.753101 | -0.568236 | 0.569875 | 0.898818733 | | processed_pseudogene | 65.38 | -0.019825514 | 0.019825514 | ENSG00000228015 | n.s. |
| ENSG00000201499 | 15.08454 | -0.309246414 | 0.428411 | -0.721844 | 0.47039 | 0.86276398 | RNU6-312P | snRNA | 38.32 | -0.019825168 | 0.019825168 | ENSG00000201499 | n.s. |
| ENSG00000166136 | 1768.866 | -0.244567125 | 0.300419 | -0.814086 | 0.415596 | 0.829820723 | NDUFB8 | protein_coding | 47.71 | -0.019813783 | 0.019813783 | ENSG00000166136 | n.s. |
| ENSG00000112576 | 150.0784 | -0.235534292 | 0.281423 | -0.836942 | 0.402625 | 0.823993439 | CCND3 | protein_coding | 47.06 | -0.019802839 | 0.019802839 | ENSG00000112576 | n.s. |
| ENSG00000054983 | 124.8708 | -0.255303335 | 0.320844 | -0.795724 | 0.426193 | 0.836464856 | GALC | protein_coding | 39.12 | -0.019799361 | 0.019799361 | ENSG00000054983 | n.s. |
| ENSG00000276770 | 3.471647 | 0.396530882 | 0.661371 | 0.5995587 | 0.5488 | 0.891406027 | MIR6780B | miRNA | 59.49 | 0.01979658 | 0.01979658 | ENSG00000276770 | n.s. |
| ENSG00000160753 | 231.2435 | -0.226923877 | 0.266853 | -0.850369 | 0.39512 | 0.818022109 | RUSC1 | protein_coding | 58.03 | -0.019795695 | 0.019795695 | ENSG00000160753 | n.s. |

| | | | | | | | | | | | | | |
|---|---|---|---|---|---|---|---|---|---|---|---|---|---|
| ENSG00000236047 | 33.25955 | -0.344956273 | 0.516095 | -0.668397 | 0.50388 | 0.876248111 | | processed_pseudogene | 50.08 | -0.019791144 | 0.019791144 | ENSG00000236047 | n.s. |
| ENSG00000198642 | 1697.035 | 0.213336168 | 0.240595 | 0.886702 | 0.375239 | 0.808091443 | KLHL9 | protein_coding | 39.32 | 0.019742021 | 0.019742021 | ENSG00000198642 | n.s. |
| ENSG00000134243 | 59.45728 | -0.235996289 | 0.283317 | -0.832975 | 0.404859 | 0.824799944 | SORT1 | protein_coding | 42.27 | -0.019741415 | 0.019741415 | ENSG00000134243 | n.s. |
| ENSG00000226137 | 194.0733 | -0.222944332 | 0.260097 | -0.857158 | 0.391358 | 0.815710819 | BAIAP2-AS1 | lincRNA | 59.08 | -0.019722498 | 0.019722498 | ENSG00000226137 | n.s. |
| ENSG00000169131 | 512.3817 | -0.239825187 | 0.29126 | -0.823405 | 0.410278 | 0.827509807 | ZNF354A | protein_coding | 46.36 | -0.01972007 | 0.01972007 | ENSG00000169131 | n.s. |
| ENSG00000152409 | 1151.789 | 0.20751236 | 0.230745 | 0.8993138 | 0.368486 | 0.803511983 | JMY | protein_coding | 40.2 | 0.01971526 | 0.01971526 | ENSG00000152409 | n.s. |
| ENSG00000063244 | 2679.987 | -0.228774723 | 0.27055 | -0.845591 | 0.397781 | 0.820246021 | U2AF2 | protein_coding | 56.03 | -0.019687407 | 0.019687407 | ENSG00000063244 | n.s. |
| ENSG00000135506 | 1921.127 | -0.235147037 | 0.281484 | -0.835383 | 0.403502 | 0.824699011 | OS9 | protein_coding | 43.99 | -0.019682871 | 0.019682871 | ENSG00000135506 | n.s. |
| ENSG00000273018 | 138.881 | -0.250659585 | 0.312761 | -0.801442 | 0.422876 | 0.834993891 | | processed_transcript | 39.96 | -0.019630832 | 0.019630832 | ENSG00000273018 | n.s. |
| ENSG00000232671 | 23.94554 | -0.29803629 | 0.406483 | -0.733207 | 0.463432 | 0.859283826 | | antisense | 54.93 | -0.019629672 | 0.019629672 | ENSG00000232671 | n.s. |
| ENSG00000213780 | 211.0439 | -0.221846098 | 0.25967 | -0.85434 | 0.392917 | 0.815710819 | GTF2H4 | protein_coding | 51.16 | -0.019625344 | 0.019625344 | ENSG00000213780 | n.s. |
| ENSG00000018699 | 474.0953 | -0.22167553 | 0.25898 | -0.855955 | 0.392023 | 0.815710819 | TTC27 | protein_coding | 39.46 | -0.019610255 | 0.019610255 | ENSG00000018699 | n.s. |
| ENSG00000111726 | 797.9664 | -0.24065507 | 0.294972 | -0.815857 | 0.414582 | 0.829007487 | CMAS | protein_coding | 37.51 | -0.019599321 | 0.019599321 | ENSG00000111726 | n.s. |
| ENSG00000072756 | 1683.689 | -0.202885621 | 0.223941 | -0.905979 | 0.364947 | 0.800686169 | TRNT1 | protein_coding | 38.89 | -0.019586106 | 0.019586106 | ENSG00000072756 | n.s. |
| ENSG00000112144 | 1067.679 | -0.225447042 | 0.26558 | -0.848885 | 0.395945 | 0.818701982 | ICK | protein_coding | 40.86 | -0.019585522 | 0.019585522 | ENSG00000112144 | n.s. |
| ENSG00000115183 | 945.7776 | 0.221291252 | 0.258454 | 0.8562126 | 0.39188 | 0.815710819 | TANC1 | protein_coding | 43.42 | 0.01957626 | 0.01957626 | ENSG00000115183 | n.s. |
| ENSG00000135316 | 11986.81 | -0.221249274 | 0.258007 | -0.857533 | 0.39115 | 0.815710819 | SYNCRIP | protein_coding | 40.05 | -0.019572547 | 0.019572547 | ENSG00000135316 | n.s. |
| ENSG00000210154 | 9.900471 | -0.332924886 | 0.487494 | -0.682931 | 0.494651 | 0.873443478 | MT-TD | Mt_tRNA | 23.53 | -0.019564395 | 0.019564395 | ENSG00000210154 | n.s. |
| ENSG00000165660 | 531.628 | -0.240150644 | 0.293558 | -0.81807 | 0.413317 | 0.829007487 | FAM175B | protein_coding | 42.62 | -0.01955824 | 0.01955824 | ENSG00000165660 | n.s. |
| ENSG00000107789 | 353.9175 | -0.233702395 | 0.281262 | -0.830905 | 0.406027 | 0.824799944 | MINPP1 | protein_coding | 38.05 | -0.019549527 | 0.019549527 | ENSG00000107789 | n.s. |
| ENSG00000139697 | 2717.978 | -0.211086638 | 0.237819 | -0.887593 | 0.37476 | 0.808040232 | SBNO1 | protein_coding | 43.26 | -0.01953966 | 0.01953966 | ENSG00000139697 | n.s. |
| ENSG00000168005 | 914.2682 | -0.233202334 | 0.278905 | -0.836136 | 0.403078 | 0.824575506 | C11orf84 | protein_coding | 52.9 | -0.019535259 | 0.019535259 | ENSG00000168005 | n.s. |
| ENSG00000209042 | 270.546 | 0.200347667 | 0.219725 | 0.9118117 | 0.361868 | 0.798938342 | SNORD12C | snoRNA | 41.57 | 0.01953124 | 0.01953124 | ENSG00000209042 | n.s. |
| ENSG00000131899 | 495.7077 | -0.2247337 | 0.264782 | -0.848748 | 0.396021 | 0.818701982 | LLGL1 | protein_coding | 59.89 | -0.019523551 | 0.019523551 | ENSG00000131899 | n.s. |
| ENSG00000137807 | 2687.528 | -0.20942998 | 0.234106 | -0.894593 | 0.371005 | 0.80683384 | KIF23 | protein_coding | 39.36 | -0.019522203 | 0.019522203 | ENSG00000137807 | n.s. |
| ENSG00000115361 | 173.4567 | -0.235397488 | 0.28472 | -0.826768 | 0.408368 | 0.826171518 | ACADL | protein_coding | 36.42 | -0.019521462 | 0.019521462 | ENSG00000115361 | n.s. |
| ENSG00000232412 | 13.92159 | -0.322939687 | 0.464748 | -0.69487 | 0.487137 | 0.870096539 | | lincRNA | 57.66 | -0.019516072 | 0.019516072 | ENSG00000232412 | n.s. |
| ENSG00000251869 | 19.29105 | 0.278875857 | 0.369011 | 0.755739 | 0.449806 | 0.85118703 | SCARNA23 | scaRNA | 34.62 | 0.019514339 | 0.019514339 | ENSG00000251869 | n.s. |
| ENSG00000147912 | 278.6603 | -0.213635535 | 0.244496 | -0.873778 | 0.382239 | 0.81032471 | FBXO10 | protein_coding | 46.45 | -0.019513666 | 0.019513666 | ENSG00000147912 | n.s. |
| ENSG00000198894 | 488.3945 | -0.213603382 | 0.24435 | -0.87417 | 0.382026 | 0.81032471 | CIPC | protein_coding | 43.02 | -0.019510729 | 0.019510729 | ENSG00000198894 | n.s. |
| ENSG00000198945 | 192.1708 | -0.212565831 | 0.242037 | -0.878236 | 0.379816 | 0.809760709 | L3MBTL3 | protein_coding | 38.41 | -0.019480234 | 0.019480234 | ENSG00000198945 | n.s. |
| ENSG00000091436 | 900.7494 | -0.232824973 | 0.279701 | -0.832408 | 0.405179 | 0.824799944 | MAP3K20 | protein_coding | 38.91 | -0.01947613 | 0.01947613 | ENSG00000091436 | n.s. |

| | | | | | | | | | | | | | |
|---|---|---|---|---|---|---|---|---|---|---|---|---|---|
| ENSG00000005156 | 2107.863 | -0.213139931 | 0.244207 | -0.872783 | 0.382781 | 0.81032471 | LIG3 | protein_coding | 47.23 | -0.019468397 | 0.019468397 | ENSG00000005156 | n.s. |
| ENSG00000115561 | 1467.801 | -0.234689518 | 0.283812 | -0.826918 | 0.408283 | 0.826171518 | CHMP3 | protein_coding | 40.96 | -0.01946275 | 0.01946275 | ENSG00000115561 | n.s. |
| ENSG00000272267 | 37.94971 | 0.261360799 | 0.334254 | 0.7819231 | 0.43426 | 0.84244096 | | antisense | 55.56 | 0.019461062 | 0.019461062 | ENSG00000272267 | n.s. |
| ENSG00000263276 | 235.4266 | -0.220737025 | 0.258757 | -0.853068 | 0.393622 | 0.816295599 | | sense_overlapping | 41.07 | -0.019458531 | 0.019458531 | ENSG00000263276 | n.s. |
| ENSG00000261716 | 450.6375 | 0.193091716 | 0.207968 | 0.9284699 | 0.353164 | 0.792950339 | | sense_overlapping | 38.15 | 0.019454765 | 0.019454765 | ENSG00000261716 | n.s. |
| ENSG00000109586 | 739.7957 | -0.228130413 | 0.270909 | -0.842092 | 0.399736 | 0.821737289 | GALNT7 | protein_coding | 38.98 | -0.019451997 | 0.019451997 | ENSG00000109586 | n.s. |
| ENSG00000124541 | 765.7469 | -0.232371796 | 0.279513 | -0.831345 | 0.405779 | 0.824799944 | RRP36 | protein_coding | 48.91 | -0.019438221 | 0.019438221 | ENSG00000124541 | n.s. |
| ENSG00000182831 | 5387.179 | -0.210064424 | 0.237067 | -0.886096 | 0.375566 | 0.808144397 | C16orf72 | protein_coding | 42.03 | -0.019433277 | 0.019433277 | ENSG00000182831 | n.s. |
| ENSG00000071539 | 997.9164 | -0.232270107 | 0.27919 | -0.831943 | 0.405441 | 0.824799944 | TRIP13 | protein_coding | 50.49 | -0.019429714 | 0.019429714 | ENSG00000071539 | n.s. |
| ENSG00000125971 | 422.3195 | -0.247085269 | 0.307823 | -0.802687 | 0.422156 | 0.83442324 | DYNLRB1 | protein_coding | 48.33 | -0.019424265 | 0.019424265 | ENSG00000125971 | n.s. |
| ENSG00000113763 | 192.4346 | -0.219539131 | 0.256953 | -0.854395 | 0.392886 | 0.815710819 | UNC5A | protein_coding | 58.17 | -0.019421261 | 0.019421261 | ENSG00000113763 | n.s. |
| ENSG00000120159 | 658.5129 | 0.229773136 | 0.27368 | 0.8395685 | 0.40115 | 0.823171216 | CAAP1 | protein_coding | 37.23 | 0.019418087 | 0.019418087 | ENSG00000120159 | n.s. |
| ENSG00000174136 | 1643.909 | 0.199295329 | 0.219061 | 0.9097693 | 0.362944 | 0.7992181 | RGMB | protein_coding | 43.29 | 0.019398349 | 0.019398349 | ENSG00000174136 | n.s. |
| ENSG00000132153 | 2083.41 | -0.219233345 | 0.256211 | -0.855675 | 0.392177 | 0.815710819 | DHX30 | protein_coding | 50.03 | -0.01939421 | 0.01939421 | ENSG00000132153 | n.s. |
| ENSG00000100580 | 870.6798 | -0.217856864 | 0.252897 | -0.861445 | 0.388993 | 0.814823372 | TMED8 | protein_coding | 41.16 | -0.019375432 | 0.019375432 | ENSG00000100580 | n.s. |
| ENSG00000212296 | 118.4783 | 0.21784333 | 0.252888 | 0.8614213 | 0.389006 | 0.814823372 | SNORD72 | snoRNA | 31.25 | 0.019374228 | 0.019374228 | ENSG00000212296 | n.s. |
| ENSG00000159409 | 603.3816 | -0.211212663 | 0.240564 | -0.877988 | 0.37995 | 0.809760709 | CELF3 | protein_coding | 57.78 | -0.019356225 | 0.019356225 | ENSG00000159409 | n.s. |
| ENSG00000198804 | 38049.14 | -0.212168847 | 0.243416 | -0.871629 | 0.383411 | 0.810719585 | MT-CO1 | protein_coding | 46.24 | -0.019334806 | 0.019334806 | ENSG00000198804 | n.s. |
| ENSG00000212443 | 58.1678 | 0.237311622 | 0.290877 | 0.8158485 | 0.414587 | 0.829007487 | SNORA53 | snoRNA | 39.36 | 0.019327026 | 0.019327026 | ENSG00000212443 | n.s. |
| ENSG00000213801 | 91.34299 | -0.261874519 | 0.336897 | -0.777313 | 0.436974 | 0.843772375 | ZNF321P | ıscribed_processed_pseudog | 46.09 | -0.019319713 | 0.019319713 | ENSG00000213801 | n.s. |
| ENSG00000171793 | 2236.38 | -0.211191662 | 0.241966 | -0.872815 | 0.382764 | 0.81032471 | CTPS1 | protein_coding | 45.27 | -0.01929044 | 0.01929044 | ENSG00000171793 | n.s. |
| ENSG00000171813 | 188.1658 | -0.233386773 | 0.282721 | -0.825503 | 0.409086 | 0.82678302 | PWWP2B | protein_coding | 62.25 | -0.01927972 | 0.01927972 | ENSG00000171813 | n.s. |
| ENSG00000087365 | 2991.569 | -0.217769871 | 0.254631 | -0.855236 | 0.39242 | 0.815710819 | SF3B2 | protein_coding | 48.41 | -0.019264745 | 0.019264745 | ENSG00000087365 | n.s. |
| ENSG00000138433 | 743.1326 | -0.22582992 | 0.268264 | -0.841819 | 0.399889 | 0.821869704 | CIR1 | protein_coding | 37.69 | -0.019240038 | 0.019240038 | ENSG00000138433 | n.s. |
| ENSG00000258099 | 57.5818 | 0.245668924 | 0.306549 | 0.8014029 | 0.422898 | 0.834993891 | ATXN2-AS | antisense | 61.79 | 0.01923998 | 0.01923998 | ENSG00000258099 | n.s. |
| ENSG00000070476 | 704.4253 | -0.214918124 | 0.248821 | -0.863745 | 0.387728 | 0.81373794 | ZXDC | protein_coding | 48.68 | -0.019238489 | 0.019238489 | ENSG00000070476 | n.s. |
| ENSG00000221520 | 11.08235 | -0.306942341 | 0.432096 | -0.710357 | 0.477483 | 0.865616576 | MIR1285-1 | miRNA | 47.62 | -0.019237438 | 0.019237438 | ENSG00000221520 | n.s. |
| ENSG00000260838 | 324.7803 | 0.235707405 | 0.287389 | 0.8201677 | 0.41212 | 0.828824745 | | lincRNA | 33.98 | 0.019218943 | 0.019218943 | ENSG00000260838 | n.s. |
| ENSG00000139192 | 109.5734 | -0.215816181 | 0.250501 | -0.861538 | 0.388942 | 0.814823372 | TAPBPL | protein_coding | 50.43 | -0.019193941 | 0.019193941 | ENSG00000139192 | n.s. |
| ENSG00000125398 | 118.0717 | -0.231130504 | 0.279231 | -0.827739 | 0.407818 | 0.826149803 | SOX9 | protein_coding | 53.34 | -0.019170241 | 0.019170241 | ENSG00000125398 | n.s. |
| ENSG00000172458 | 60.57561 | -0.260027528 | 0.334937 | -0.776347 | 0.437544 | 0.843873725 | IL17D | protein_coding | 52.71 | -0.019169888 | 0.019169888 | ENSG00000172458 | n.s. |
| ENSG00000134490 | 161.4708 | -0.229039873 | 0.274684 | -0.83383 | 0.404377 | 0.824799944 | TMEM241 | protein_coding | 45.61 | -0.019159501 | 0.019159501 | ENSG00000134490 | n.s. |

| | | | | | | | | | | | | | |
|---|---|---|---|---|---|---|---|---|---|---|---|---|---|
| ENSG00000143499 | 1205.217 | -0.22256027 | 0.263499 | -0.844634 | 0.398315 | 0.820375622 | SMYD2 | protein_coding | 43.58 | -0.019137346 | 0.019137346 | ENSG00000143499 | n.s. |
| ENSG00000180035 | 395.1135 | -0.209866717 | 0.240923 | -0.871096 | 0.383702 | 0.810719585 | ZNF48 | protein_coding | 53.07 | -0.019125014 | 0.019125014 | ENSG00000180035 | n.s. |
| ENSG00000214135 | 605.6382 | -0.208985336 | 0.238237 | -0.877216 | 0.380369 | 0.810288517 | | transcribed_unprocessed_pseudc | 45.89 | -0.019092966 | 0.019092966 | ENSG00000214135 | n.s. |
| ENSG00000174928 | 121.6711 | -0.239503245 | 0.295759 | -0.809792 | 0.41806 | 0.832304654 | C3orf33 | protein_coding | 40.75 | -0.019092642 | 0.019092642 | ENSG00000174928 | n.s. |
| ENSG00000179021 | 942.6667 | 0.215776229 | 0.25149 | 0.857993 | 0.390896 | 0.815710819 | C3orf38 | protein_coding | 38.06 | 0.01908838 | 0.01908838 | ENSG00000179021 | n.s. |
| ENSG00000167258 | 1789.918 | -0.198949318 | 0.220236 | -0.903347 | 0.366342 | 0.801976953 | CDK12 | protein_coding | 44.39 | -0.019066927 | 0.019066927 | ENSG00000167258 | n.s. |
| ENSG00000246067 | 587.2128 | -0.185085288 | 0.196201 | -0.943346 | 0.345504 | 0.788921338 | RAB30-AS1 | lincRNA | 40.04 | -0.019057547 | 0.019057547 | ENSG00000246067 | n.s. |
| ENSG00000238578 | 62.13305 | 0.227710506 | 0.273554 | 0.8324146 | 0.405175 | 0.824799944 | SNORD4A | snoRNA | 47.22 | 0.019048297 | 0.019048297 | ENSG00000238578 | n.s. |
| ENSG00000105429 | 491.5559 | -0.207751385 | 0.236599 | -0.878074 | 0.379903 | 0.809760709 | MEGF8 | protein_coding | 54.75 | -0.019039022 | 0.019039022 | ENSG00000105429 | n.s. |
| ENSG00000171681 | 3117.619 | -0.218327352 | 0.256924 | -0.849775 | 0.39545 | 0.818083342 | ATF7IP | protein_coding | 36.1 | -0.01903868 | 0.01903868 | ENSG00000171681 | n.s. |
| ENSG00000163006 | 1498.534 | -0.227822401 | 0.274293 | -0.83058 | 0.406211 | 0.825006384 | CCDC138 | protein_coding | 37.69 | -0.019032896 | 0.019032896 | ENSG00000163006 | n.s. |
| ENSG00000141449 | 204.2528 | -0.221697145 | 0.262638 | -0.844118 | 0.398604 | 0.820678855 | GREB1L | protein_coding | 40.54 | -0.019027547 | 0.019027547 | ENSG00000141449 | n.s. |
| ENSG00000135454 | 436.5553 | -0.220099756 | 0.259776 | -0.847267 | 0.396846 | 0.819511304 | B4GALNT1 | protein_coding | 56.73 | -0.019026535 | 0.019026535 | ENSG00000135454 | n.s. |
| ENSG00000222937 | 45.75681 | -0.255424416 | 0.326327 | -0.782726 | 0.433788 | 0.84244096 | SNORD63B | snoRNA | 42.86 | -0.019019036 | 0.019019036 | ENSG00000222937 | n.s. |
| ENSG00000104112 | 2092.946 | -0.225464395 | 0.268913 | -0.838429 | 0.40179 | 0.823540971 | SCG3 | protein_coding | 40.12 | -0.019009983 | 0.019009983 | ENSG00000104112 | n.s. |
| ENSG00000099800 | 1175.21 | -0.227116902 | 0.273279 | -0.831081 | 0.405928 | 0.824799944 | TIMM13 | protein_coding | 64.99 | -0.018998642 | 0.018998642 | ENSG00000099800 | n.s. |
| ENSG00000198840 | 1751.666 | 0.205145369 | 0.231518 | 0.8860893 | 0.375569 | 0.808144397 | MT-ND3 | protein_coding | 40.17 | 0.01897821 | 0.01897821 | ENSG00000198840 | n.s. |
| ENSG00000279500 | 215.1899 | 0.20774508 | 0.237472 | 0.8748189 | 0.381672 | 0.81032471 | | TEC | 46.58 | 0.018975626 | 0.018975626 | ENSG00000279500 | n.s. |
| ENSG00000198874 | 623.4475 | -0.211319438 | 0.244208 | -0.865324 | 0.386861 | 0.813266385 | TYW1 | protein_coding | 41.67 | -0.01896955 | 0.01896955 | ENSG00000198874 | n.s. |
| ENSG00000174953 | 3220.04 | -0.226630123 | 0.272547 | -0.831527 | 0.405676 | 0.824799944 | DHX36 | protein_coding | 35.49 | -0.018957922 | 0.018957922 | ENSG00000174953 | n.s. |
| ENSG00000132386 | 1756.21 | -0.232569724 | 0.284344 | -0.817917 | 0.413405 | 0.829007487 | SERPINF1 | protein_coding | 53.48 | -0.018940838 | 0.018940838 | ENSG00000132386 | n.s. |
| ENSG00000228782 | 286.9744 | -0.203804538 | 0.22918 | -0.889278 | 0.373854 | 0.807377464 | MRPL45P2 | cribed_unprocessed_pseudc | 42.96 | -0.018938206 | 0.018938206 | ENSG00000228782 | n.s. |
| ENSG00000172775 | 1667.1 | -0.226262495 | 0.272257 | -0.831062 | 0.405938 | 0.824799944 | FAM192A | protein_coding | 41.93 | -0.018927169 | 0.018927169 | ENSG00000172775 | n.s. |
| ENSG00000169230 | 2007.692 | -0.232334102 | 0.284082 | -0.81784 | 0.413448 | 0.829007487 | PRELID1 | protein_coding | 54.24 | -0.018921649 | 0.018921649 | ENSG00000169230 | n.s. |
| ENSG00000091972 | 447.7865 | -0.222201469 | 0.264169 | -0.841134 | 0.400273 | 0.821975757 | CD200 | protein_coding | 38.61 | -0.018918453 | 0.018918453 | ENSG00000091972 | n.s. |
| ENSG00000146830 | 944.4528 | -0.210581802 | 0.243265 | -0.865649 | 0.386682 | 0.813136725 | GIGYF1 | protein_coding | 62.01 | -0.018917916 | 0.018917916 | ENSG00000146830 | n.s. |
| ENSG00000238793 | 3.233417 | 0.520659372 | 1.101009 | 0.4728927 | 0.63629 | 0.919753089 | SNORD124 | snoRNA | 49.04 | 0.018914902 | 0.018914902 | ENSG00000238793 | n.s. |
| ENSG00000220771 | 9.928333 | -0.318106602 | 0.460644 | -0.69057 | 0.489836 | 0.872064188 | BOLA2P3 | processed_pseudogene | 60.94 | -0.01891193 | 0.01891193 | ENSG00000220771 | n.s. |
| ENSG00000086827 | 510.0415 | -0.210802756 | 0.243884 | -0.864358 | 0.387391 | 0.813399117 | ZW10 | protein_coding | 39.15 | -0.018908228 | 0.018908228 | ENSG00000086827 | n.s. |
| ENSG00000186020 | 923.9952 | -0.220320003 | 0.26119 | -0.843525 | 0.398935 | 0.820802709 | ZNF529 | protein_coding | 40.96 | -0.018894912 | 0.018894912 | ENSG00000186020 | n.s. |
| ENSG00000169972 | 107.579 | -0.226116561 | 0.272297 | -0.830403 | 0.406311 | 0.825006384 | PUSL1 | protein_coding | 66.76 | -0.018890386 | 0.018890386 | ENSG00000169972 | n.s. |
| ENSG00000152061 | 606.1127 | -0.210515645 | 0.243506 | -0.864521 | 0.387302 | 0.813399117 | RABGAP1L | protein_coding | 37.91 | -0.018882475 | 0.018882475 | ENSG00000152061 | n.s. |

| | | | | | | | | | | | | | |
|---|---|---|---|---|---|---|---|---|---|---|---|---|---|
| ENSG00000125459 | 439.3484 | -0.210877844 | 0.244088 | -0.863943 | 0.387619 | 0.81373794 | MSTO1 | protein_coding | 52.49 | -0.018876822 | 0.018876822 | ENSG00000125459 | n.s. |
| ENSG00000127081 | 425.2634 | -0.219780833 | 0.26042 | -0.843949 | 0.398698 | 0.820678855 | ZNF484 | protein_coding | 39.28 | -0.018863076 | 0.018863076 | ENSG00000127081 | n.s. |
| ENSG00000061676 | 5119.673 | -0.225472148 | 0.270508 | -0.833515 | 0.404555 | 0.824799944 | NCKAP1 | protein_coding | 35.82 | -0.018861056 | 0.018861056 | ENSG00000061676 | n.s. |
| ENSG00000170779 | 745.9214 | -0.226641174 | 0.273178 | -0.829645 | 0.406739 | 0.825635104 | CDCA4 | protein_coding | 52.06 | -0.018859231 | 0.018859231 | ENSG00000170779 | n.s. |
| ENSG00000234684 | 66.37181 | -0.23053155 | 0.280735 | -0.821172 | 0.411548 | 0.828379538 | SDCBP2-AS1 | antisense | 43.98 | -0.018850712 | 0.018850712 | ENSG00000234684 | n.s. |
| ENSG00000267030 | 31.80939 | -0.250818547 | 0.318718 | -0.78696 | 0.431305 | 0.84116282 | | antisense | 63.89 | -0.018841472 | 0.018841472 | ENSG00000267030 | n.s. |
| ENSG00000144747 | 866.8018 | -0.217655785 | 0.25693 | -0.84714 | 0.396917 | 0.819511304 | TMF1 | protein_coding | 36.59 | -0.018815266 | 0.018815266 | ENSG00000144747 | n.s. |
| ENSG00000254207 | 19.40288 | -0.293140761 | 0.405937 | -0.722134 | 0.470212 | 0.862665074 | | sense_intronic | 61.74 | -0.018807262 | 0.018807262 | ENSG00000254207 | n.s. |
| ENSG00000166484 | 678.946 | -0.192120476 | 0.210415 | -0.913056 | 0.361213 | 0.798281166 | MAPK7 | protein_coding | 59.22 | -0.018797858 | 0.018797858 | ENSG00000166484 | n.s. |
| ENSG00000149743 | 110.0913 | -0.241039589 | 0.302363 | -0.797187 | 0.425343 | 0.835738942 | TRPT1 | protein_coding | 60.14 | -0.018784061 | 0.018784061 | ENSG00000149743 | n.s. |
| ENSG00000126821 | 286.0866 | -0.240300054 | 0.300459 | -0.799777 | 0.42384 | 0.835315601 | SGPP1 | protein_coding | 39.66 | -0.018779307 | 0.018779307 | ENSG00000126821 | n.s. |
| ENSG00000153707 | 114.0006 | 0.218353571 | 0.25846 | 0.8448253 | 0.398208 | 0.820375622 | PTPRD | protein_coding | 35.53 | 0.018775624 | 0.018775624 | ENSG00000153707 | n.s. |
| ENSG00000013364 | 31.99044 | -0.26628359 | 0.349987 | -0.760839 | 0.446753 | 0.850164607 | MVP | protein_coding | 52.54 | -0.018772189 | 0.018772189 | ENSG00000013364 | n.s. |
| ENSG00000130340 | 1087.043 | -0.210897384 | 0.244725 | -0.861774 | 0.388812 | 0.814823372 | SNX9 | protein_coding | 42.45 | -0.01875648 | 0.01875648 | ENSG00000130340 | n.s. |
| ENSG00000186660 | 1383.887 | -0.224188754 | 0.269154 | -0.832939 | 0.404879 | 0.824799944 | ZFP91 | protein_coding | 36.47 | -0.018753698 | 0.018753698 | ENSG00000186660 | n.s. |
| ENSG00000166482 | 2230.056 | -0.211823432 | 0.247309 | -0.856513 | 0.391714 | 0.815710819 | MFAP4 | protein_coding | 57.09 | -0.018738701 | 0.018738701 | ENSG00000166482 | n.s. |
| ENSG00000170027 | 4742.102 | -0.225713859 | 0.272949 | -0.826944 | 0.408269 | 0.826171518 | YWHAG | protein_coding | 44.56 | -0.018718401 | 0.018718401 | ENSG00000170027 | n.s. |
| ENSG00000151466 | 940.2078 | -0.22262939 | 0.265987 | -0.836992 | 0.402597 | 0.823993439 | SCLT1 | protein_coding | 36.17 | -0.018717843 | 0.018717843 | ENSG00000151466 | n.s. |
| ENSG00000132432 | 1335.295 | 0.227375007 | 0.276041 | 0.8237013 | 0.410109 | 0.827409292 | SEC61G | protein_coding | 39.24 | 0.018708326 | 0.018708326 | ENSG00000132432 | n.s. |
| ENSG00000127993 | 602.4769 | -0.202211429 | 0.22825 | -0.885922 | 0.37566 | 0.808144397 | RBM48 | protein_coding | 39.51 | -0.018706788 | 0.018706788 | ENSG00000127993 | n.s. |
| ENSG00000135932 | 1344.751 | -0.225307055 | 0.271946 | -0.8285 | 0.407387 | 0.825999998 | CAB39 | protein_coding | 40.78 | -0.018704981 | 0.018704981 | ENSG00000135932 | n.s. |
| ENSG00000117069 | 1128.382 | 0.227702191 | 0.276905 | 0.8223117 | 0.4109 | 0.827684308 | ST6GALNAC5 | protein_coding | 39.93 | 0.018702383 | 0.018702383 | ENSG00000117069 | n.s. |
| ENSG00000256361 | 38.70646 | 0.265271452 | 0.349107 | 0.7598582 | 0.447339 | 0.850164607 | | processed_pseudogene | 37.64 | 0.018700836 | 0.018700836 | ENSG00000256361 | n.s. |
| ENSG00000120756 | 39.74333 | 0.256575017 | 0.332661 | 0.7712803 | 0.440541 | 0.845502102 | PLS1 | protein_coding | 38.52 | 0.018700549 | 0.018700549 | ENSG00000120756 | n.s. |
| ENSG00000174306 | 37.84107 | -0.251067969 | 0.320552 | -0.783235 | 0.433489 | 0.84244096 | ZHX3 | protein_coding | 41.13 | -0.018694653 | 0.018694653 | ENSG00000174306 | n.s. |
| ENSG00000181191 | 555.541 | -0.238481629 | 0.297578 | -0.801408 | 0.422896 | 0.834993891 | PJA1 | protein_coding | 51.45 | -0.018677095 | 0.018677095 | ENSG00000181191 | n.s. |
| ENSG00000108379 | 230.6508 | -0.213661578 | 0.251448 | -0.849726 | 0.395477 | 0.818083342 | WNT3 | protein_coding | 53.82 | -0.018631813 | 0.018631813 | ENSG00000108379 | n.s. |
| ENSG00000079332 | 1514.047 | -0.223837688 | 0.27001 | -0.828998 | 0.407106 | 0.825802394 | SAR1A | protein_coding | 40.56 | -0.018606253 | 0.018606253 | ENSG00000079332 | n.s. |
| ENSG00000062650 | 1893.403 | -0.224346247 | 0.271343 | -0.826798 | 0.408351 | 0.826171518 | WAPL | protein_coding | 39.87 | -0.018604985 | 0.018604985 | ENSG00000062650 | n.s. |
| ENSG00000274070 | 96.24327 | -0.221031391 | 0.263922 | -0.837488 | 0.402318 | 0.823848555 | CASTOR2 | protein_coding | 52.51 | -0.01860037 | 0.01860037 | ENSG00000274070 | n.s. |
| ENSG00000197044 | 488.8338 | -0.255302238 | 0.331717 | -0.769638 | 0.441515 | 0.84569031 | ZNF441 | protein_coding | 41.78 | -0.018583104 | 0.018583104 | ENSG00000197044 | n.s. |
| ENSG00000235885 | 6831.928 | -0.187156417 | 0.20362 | -0.919146 | 0.358019 | 0.795806072 | LINC01828 | lincRNA | 36.94 | -0.01856456 | 0.01856456 | ENSG00000235885 | n.s. |

| | | | | | | | | | | | | | |
|---|---|---|---|---|---|---|---|---|---|---|---|---|---|
| ENSG00000164099 | 2162.726 | -0.21636991 | 0.256547 | -0.843394 | 0.399008 | 0.82083191 | PRSS12 | protein_coding | 36.51 | -0.018552804 | 0.018552804 | ENSG00000164099 | n.s. |
| ENSG00000203808 | 40.01269 | -0.253633364 | 0.328298 | -0.772571 | 0.439777 | 0.845174857 | BVES-AS1 | antisense | 39.74 | -0.018528788 | 0.018528788 | ENSG00000203808 | n.s. |
| ENSG00000101782 | 943.6742 | -0.221399842 | 0.266261 | -0.831515 | 0.405683 | 0.824799944 | RIOK3 | protein_coding | 41.88 | -0.018520402 | 0.018520402 | ENSG00000101782 | n.s. |
| ENSG00000232969 | 5.698626 | -0.389728912 | 0.677383 | -0.575345 | 0.565058 | 0.896450345 | | antisense | 49.68 | -0.018501897 | 0.018501897 | ENSG00000232969 | n.s. |
| ENSG00000134138 | 247.7231 | 0.199950807 | 0.225687 | 0.885967 | 0.375635 | 0.808144397 | MEIS2 | protein_coding | 38.26 | 0.018497656 | 0.018497656 | ENSG00000134138 | n.s. |
| ENSG00000128512 | 603.6557 | -0.202915023 | 0.233183 | -0.870195 | 0.384194 | 0.810719585 | DOCK4 | protein_coding | 39.18 | -0.018491511 | 0.018491511 | ENSG00000128512 | n.s. |
| ENSG00000205208 | 2014.972 | 0.201130989 | 0.228 | 0.8821521 | 0.377695 | 0.809223062 | C4orf46 | protein_coding | 41.13 | 0.018490324 | 0.018490324 | ENSG00000205208 | n.s. |
| ENSG00000165813 | 457.4841 | -0.221026979 | 0.264732 | -0.834907 | 0.40377 | 0.824799944 | CCDC186 | protein_coding | 37.11 | -0.018489211 | 0.018489211 | ENSG00000165813 | n.s. |
| ENSG00000113597 | 383.7398 | -0.262265702 | 0.344488 | -0.76132 | 0.446466 | 0.850164607 | TRAPPC13 | protein_coding | 36.8 | -0.01848894 | 0.01848894 | ENSG00000113597 | n.s. |
| ENSG00000170653 | 45.64221 | -0.250906125 | 0.323631 | -0.775285 | 0.438172 | 0.843950276 | ATF7 | protein_coding | 42.93 | -0.018487551 | 0.018487551 | ENSG00000170653 | n.s. |
| ENSG00000168488 | 808.857 | -0.208788732 | 0.24322 | -0.858436 | 0.390652 | 0.815710819 | ATXN2L | protein_coding | 52.64 | -0.01847024 | 0.01847024 | ENSG00000168488 | n.s. |
| ENSG00000184470 | 98.95612 | -0.246945136 | 0.314171 | -0.786022 | 0.431855 | 0.841828071 | TXNRD2 | protein_coding | 54.14 | -0.018465717 | 0.018465717 | ENSG00000184470 | n.s. |
| ENSG00000120942 | 162.0195 | -0.220648114 | 0.265501 | -0.831064 | 0.405937 | 0.824799944 | UBIAD1 | protein_coding | 48.21 | -0.018457519 | 0.018457519 | ENSG00000120942 | n.s. |
| ENSG00000121775 | 130.3543 | -0.224542851 | 0.272956 | -0.822634 | 0.410716 | 0.827563884 | TMEM39B | protein_coding | 48.85 | -0.018457079 | 0.018457079 | ENSG00000121775 | n.s. |
| ENSG00000273353 | 31.03746 | -0.270343479 | 0.362659 | -0.745448 | 0.456001 | 0.85459484 | | sense_intronic | 33.02 | -0.018448167 | 0.018448167 | ENSG00000273353 | n.s. |
| ENSG00000183648 | 694.1999 | -0.237148705 | 0.297651 | -0.796735 | 0.425605 | 0.836005799 | NDUFB1 | protein_coding | 44.2 | -0.018447966 | 0.018447966 | ENSG00000183648 | n.s. |
| ENSG00000084090 | 4403.756 | -0.220520636 | 0.264233 | -0.834568 | 0.403961 | 0.824799944 | STARD7 | protein_coding | 42.97 | -0.018446855 | 0.018446855 | ENSG00000084090 | n.s. |
| ENSG00000135269 | 463.1113 | 0.228511485 | 0.281132 | 0.8128254 | 0.416318 | 0.830429454 | TES | protein_coding | 37.73 | 0.01844025 | 0.01844025 | ENSG00000135269 | n.s. |
| ENSG00000173933 | 1976.551 | 0.194063678 | 0.215702 | 0.8996828 | 0.368289 | 0.803511983 | RBM4 | protein_coding | 45.16 | 0.018437532 | 0.018437532 | ENSG00000173933 | n.s. |
| ENSG00000116353 | 128.2583 | -0.23402499 | 0.291457 | -0.802948 | 0.422005 | 0.834256463 | MECR | protein_coding | 48.21 | -0.018417865 | 0.018417865 | ENSG00000116353 | n.s. |
| ENSG00000004975 | 595.6829 | -0.226073364 | 0.276007 | -0.819085 | 0.412738 | 0.829007487 | DVL2 | protein_coding | 55.14 | -0.018411765 | 0.018411765 | ENSG00000004975 | n.s. |
| ENSG00000072501 | 3374.108 | -0.21395037 | 0.253159 | -0.845122 | 0.398043 | 0.820375622 | SMC1A | protein_coding | 45.54 | -0.018397005 | 0.018397005 | ENSG00000072501 | n.s. |
| ENSG00000214367 | 1051.249 | -0.207955341 | 0.242594 | -0.857214 | 0.391326 | 0.815710819 | HAUS3 | protein_coding | 37.25 | -0.018396515 | 0.018396515 | ENSG00000214367 | n.s. |
| ENSG00000157107 | 429.712 | -0.239561596 | 0.302218 | -0.792679 | 0.427965 | 0.838039772 | FCHO2 | protein_coding | 37.04 | -0.018382848 | 0.018382848 | ENSG00000157107 | n.s. |
| ENSG00000157110 | 5.569658 | -0.368176842 | 0.606592 | -0.60696 | 0.543878 | 0.891406027 | RBPMS | protein_coding | 42.19 | -0.01838102 | 0.01838102 | ENSG00000157110 | n.s. |
| ENSG00000112234 | 934.8782 | -0.225507312 | 0.275133 | -0.819632 | 0.412426 | 0.829007487 | FBXL4 | protein_coding | 36.71 | -0.018365664 | 0.018365664 | ENSG00000112234 | n.s. |
| ENSG00000165832 | 953.4008 | -0.225210686 | 0.274935 | -0.819142 | 0.412706 | 0.829007487 | TRUB1 | protein_coding | 36.89 | -0.018341507 | 0.018341507 | ENSG00000165832 | n.s. |
| ENSG00000111906 | 2067.7 | -0.234627493 | 0.293415 | -0.799644 | 0.423917 | 0.835315601 | HDDC2 | protein_coding | 40.11 | -0.018335999 | 0.018335999 | ENSG00000111906 | n.s. |
| ENSG00000121057 | 2486.853 | -0.206070776 | 0.238947 | -0.862413 | 0.38846 | 0.814783636 | AKAP1 | protein_coding | 49.96 | -0.018331583 | 0.018331583 | ENSG00000121057 | n.s. |
| ENSG00000165219 | 57.88156 | -0.234535316 | 0.293367 | -0.799461 | 0.424023 | 0.835315601 | GAPVD1 | protein_coding | 42.11 | -0.018328796 | 0.018328796 | ENSG00000165219 | n.s. |
| ENSG00000180098 | 151.9249 | -0.229135011 | 0.282728 | -0.810444 | 0.417685 | 0.831844959 | TRNAU1AP | protein_coding | 47.61 | -0.018321088 | 0.018321088 | ENSG00000180098 | n.s. |
| ENSG00000140691 | 185.4759 | -0.215154399 | 0.255748 | -0.841276 | 0.400193 | 0.821966929 | ARMC5 | protein_coding | 58.18 | -0.018319462 | 0.018319462 | ENSG00000140691 | n.s. |

| | | | | | | | | | | | | | |
|---|---|---|---|---|---|---|---|---|---|---|---|---|---|
| ENSG00000196123 | 663.151 | -0.215101277 | 0.255693 | -0.841248 | 0.400209 | 0.821966929 | KIAA0895L | protein_coding | 56.58 | -0.018314939 | 0.018314939 | ENSG00000196123 | n.s. |
| ENSG00000177599 | 204.8267 | -0.206966257 | 0.241113 | -0.85838 | 0.390683 | 0.815710819 | ZNF491 | protein_coding | 44.18 | -0.018309017 | 0.018309017 | ENSG00000177599 | n.s. |
| ENSG00000221978 | 1770.015 | -0.21855278 | 0.261595 | -0.835462 | 0.403457 | 0.824699011 | CCNL2 | protein_coding | 51.37 | -0.018293857 | 0.018293857 | ENSG00000221978 | n.s. |
| ENSG00000275611 | 14.90597 | -0.295846083 | 0.420661 | -0.703289 | 0.481876 | 0.867402024 | | sense_intronic | 36.59 | -0.018277243 | 0.018277243 | ENSG00000275611 | n.s. |
| ENSG00000199133 | 71.04617 | -0.245335721 | 0.313475 | -0.782632 | 0.433843 | 0.84244096 | MIRLET7D | miRNA | 48.28 | -0.018267827 | 0.018267827 | ENSG00000199133 | n.s. |
| ENSG00000258738 | 160.6454 | 0.207509383 | 0.243369 | 0.8526521 | 0.393852 | 0.81657325 | | antisense | 52.29 | 0.018261832 | 0.018261832 | ENSG00000258738 | n.s. |
| ENSG00000119844 | 795.9402 | -0.218083601 | 0.262389 | -0.831148 | 0.40589 | 0.824799944 | AFTPH | protein_coding | 37.29 | -0.018242994 | 0.018242994 | ENSG00000119844 | n.s. |
| ENSG00000185049 | 401.3485 | -0.217766326 | 0.261352 | -0.83323 | 0.404715 | 0.824799944 | NELFA | protein_coding | 48.3 | -0.018216453 | 0.018216453 | ENSG00000185049 | n.s. |
| ENSG00000207168 | 11.3449 | 0.308270956 | 0.44875 | 0.6869546 | 0.492111 | 0.872823758 | SNORA15 | snoRNA | 36.84 | 0.018210626 | 0.018210626 | ENSG00000207168 | n.s. |
| ENSG00000146247 | 7157.284 | -0.220429167 | 0.267008 | -0.825554 | 0.409057 | 0.82678302 | PHIP | protein_coding | 34.29 | -0.018209312 | 0.018209312 | ENSG00000146247 | n.s. |
| ENSG00000140396 | 1044.278 | -0.211549221 | 0.250143 | -0.845712 | 0.397714 | 0.820246021 | NCOA2 | protein_coding | 38.24 | -0.018205052 | 0.018205052 | ENSG00000140396 | n.s. |
| ENSG00000177119 | 3021.093 | -0.225024958 | 0.27661 | -0.81351 | 0.415926 | 0.830122927 | ANO6 | protein_coding | 39.44 | -0.018194976 | 0.018194976 | ENSG00000177119 | n.s. |
| ENSG00000060688 | 685.3327 | -0.222064577 | 0.270273 | -0.821632 | 0.411287 | 0.828104587 | SNRNP40 | protein_coding | 42.01 | -0.018190378 | 0.018190378 | ENSG00000060688 | n.s. |
| ENSG00000059588 | 1686.114 | -0.21882216 | 0.264 | -0.828871 | 0.407178 | 0.825802394 | TARBP1 | protein_coding | 40.27 | -0.018189343 | 0.018189343 | ENSG00000059588 | n.s. |
| ENSG00000102974 | 1697.247 | -0.217298213 | 0.260179 | -0.835187 | 0.403613 | 0.824699011 | CTCF | protein_coding | 45.32 | -0.018188844 | 0.018188844 | ENSG00000102974 | n.s. |
| ENSG00000223572 | 259.604 | -0.210295389 | 0.248123 | -0.847544 | 0.396692 | 0.819495485 | CKMT1A | protein_coding | 50.24 | -0.018180759 | 0.018180759 | ENSG00000223572 | n.s. |
| ENSG00000277558 | 32.90761 | -0.259720132 | 0.342805 | -0.757633 | 0.448671 | 0.851184545 | | sense_intronic | 50.6 | -0.018174246 | 0.018174246 | ENSG00000277558 | n.s. |
| ENSG00000204852 | 251.5522 | -0.233075716 | 0.292193 | -0.797679 | 0.425057 | 0.835658686 | TCTN1 | protein_coding | 44.42 | -0.018173163 | 0.018173163 | ENSG00000204852 | n.s. |
| ENSG00000183722 | 82.6785 | -0.226634191 | 0.279447 | -0.811009 | 0.41736 | 0.831423508 | LHFP | protein_coding | 39.43 | -0.018171008 | 0.018171008 | ENSG00000183722 | n.s. |
| ENSG00000110713 | 2448.255 | -0.219828823 | 0.266204 | -0.825791 | 0.408923 | 0.826692059 | NUP98 | protein_coding | 41.69 | -0.018170223 | 0.018170223 | ENSG00000110713 | n.s. |
| ENSG00000272779 | 348.3378 | -0.218571998 | 0.263667 | -0.828969 | 0.407122 | 0.825802394 | | unprocessed_pseudogene | 42.89 | -0.018168549 | 0.018168549 | ENSG00000272779 | n.s. |
| ENSG00000116641 | 2329.75 | -0.223021874 | 0.273327 | -0.815951 | 0.414528 | 0.829007487 | DOCK7 | protein_coding | 36.06 | -0.018163246 | 0.018163246 | ENSG00000116641 | n.s. |
| ENSG00000001629 | 1023.312 | -0.227823788 | 0.28202 | -0.807827 | 0.41919 | 0.832304654 | ANKIB1 | protein_coding | 36.77 | -0.018161583 | 0.018161583 | ENSG00000001629 | n.s. |
| ENSG00000172113 | 252.3119 | -0.208922237 | 0.246137 | -0.848805 | 0.39599 | 0.818701982 | NME6 | protein_coding | 47.93 | -0.018149943 | 0.018149943 | ENSG00000172113 | n.s. |
| ENSG00000279544 | 213.3078 | -0.24597499 | 0.316328 | -0.777594 | 0.436809 | 0.843772375 | | TEC | 37.73 | -0.01814673 | 0.01814673 | ENSG00000279544 | n.s. |
| ENSG00000186318 | 300.5333 | -0.225908024 | 0.278307 | -0.811723 | 0.416951 | 0.831214631 | BACE1 | protein_coding | 49.51 | -0.018137437 | 0.018137437 | ENSG00000186318 | n.s. |
| ENSG00000033867 | 1005.254 | -0.216760729 | 0.260545 | -0.831951 | 0.405437 | 0.824799944 | SLC4A7 | protein_coding | 36.85 | -0.018132334 | 0.018132334 | ENSG00000033867 | n.s. |
| ENSG00000274054 | 8.478865 | -0.371802336 | 0.635301 | -0.585238 | 0.558388 | 0.893837232 | MIR4727 | miRNA | 50.91 | -0.018122226 | 0.018122226 | ENSG00000274054 | n.s. |
| ENSG00000119777 | 299.9842 | -0.210739639 | 0.249475 | -0.844731 | 0.398261 | 0.820375622 | TMEM214 | protein_coding | 55.01 | -0.018120923 | 0.018120923 | ENSG00000119777 | n.s. |
| ENSG00000066422 | 953.9228 | -0.21618524 | 0.258839 | -0.835211 | 0.403599 | 0.824699011 | ZBTB11 | protein_coding | 40.45 | -0.018095683 | 0.018095683 | ENSG00000066422 | n.s. |
| ENSG00000173114 | 963.2966 | 0.219929427 | 0.267043 | 0.8235716 | 0.410183 | 0.827438178 | LRRN3 | protein_coding | 34.94 | 0.018092372 | 0.018092372 | ENSG00000173114 | n.s. |
| ENSG00000143258 | 429.8337 | -0.198309016 | 0.227962 | -0.869922 | 0.384343 | 0.810719585 | USP21 | protein_coding | 53.43 | -0.018071769 | 0.018071769 | ENSG00000143258 | n.s. |

| | | | | | | | | | | | | | |
|---|---|---|---|---|---|---|---|---|---|---|---|---|---|
| ENSG00000149927 | 45.63312 | 0.265599301 | 0.356733 | 0.7445322 | 0.456555 | 0.855072523 | DOC2A | protein_coding | 55.16 | 0.018059969 | 0.018059969 | ENSG00000149927 | n.s. |
| ENSG00000070961 | 2162.304 | -0.208747746 | 0.246459 | -0.846988 | 0.397002 | 0.819511304 | ATP2B1 | protein_coding | 36.37 | -0.01804521 | 0.01804521 | ENSG00000070961 | n.s. |
| ENSG00000089289 | 486.961 | 0.228502355 | 0.284046 | 0.8044557 | 0.421134 | 0.833823415 | IGBP1 | protein_coding | 42.89 | 0.018034757 | 0.018034757 | ENSG00000089289 | n.s. |
| ENSG00000172354 | 689.2441 | -0.222278795 | 0.272945 | -0.814373 | 0.415431 | 0.829730134 | GNB2 | protein_coding | 65.91 | -0.018018616 | 0.018018616 | ENSG00000172354 | n.s. |
| ENSG00000163156 | 484.324 | -0.221073174 | 0.269607 | -0.819983 | 0.412226 | 0.828916856 | SCNM1 | protein_coding | 48.32 | -0.018015038 | 0.018015038 | ENSG00000163156 | n.s. |
| ENSG00000139579 | 928.5211 | -0.236739861 | 0.29952 | -0.790397 | 0.429296 | 0.839274918 | NABP2 | protein_coding | 51.22 | -0.018014899 | 0.018014899 | ENSG00000139579 | n.s. |
| ENSG00000230185 | 68.36253 | -0.22108578 | 0.270875 | -0.816193 | 0.41439 | 0.829007487 | C9orf147 | antisense | 40.99 | -0.018005568 | 0.018005568 | ENSG00000230185 | n.s. |
| ENSG00000176887 | 8902.069 | -0.195570269 | 0.221673 | -0.882245 | 0.377644 | 0.809223062 | SOX11 | protein_coding | 47.75 | -0.017979117 | 0.017979117 | ENSG00000176887 | n.s. |
| ENSG00000266967 | 1295.375 | -0.21629066 | 0.260896 | -0.829029 | 0.407088 | 0.825802394 | AARSD1 | protein_coding | 48.21 | -0.017978915 | 0.017978915 | ENSG00000266967 | n.s. |
| ENSG00000148498 | 1297.173 | -0.216534605 | 0.261371 | -0.828458 | 0.407411 | 0.825999998 | PARD3 | protein_coding | 40.01 | -0.017976693 | 0.017976693 | ENSG00000148498 | n.s. |
| ENSG00000049541 | 1018.777 | -0.222503015 | 0.273663 | -0.813056 | 0.416186 | 0.830284692 | RFC2 | protein_coding | 48.96 | -0.017972229 | 0.017972229 | ENSG00000049541 | n.s. |
| ENSG00000260625 | 41.07702 | -0.254666545 | 0.335041 | -0.760106 | 0.447191 | 0.850164607 | | antisense | 48.85 | -0.017953222 | 0.017953222 | ENSG00000260625 | n.s. |
| ENSG00000101457 | 300.004 | -0.238838828 | 0.30334 | -0.787362 | 0.43107 | 0.84115455 | DNTTIP1 | protein_coding | 46 | -0.017942576 | 0.017942576 | ENSG00000101457 | n.s. |
| ENSG00000273983 | 3307.321 | -0.203091872 | 0.237874 | -0.853779 | 0.393227 | 0.815931883 | HIST1H3G | protein_coding | 42.68 | -0.017942374 | 0.017942374 | ENSG00000273983 | n.s. |
| ENSG00000207031 | 25.34328 | 0.276220365 | 0.380341 | 0.7262445 | 0.467689 | 0.861099275 | SNORD59A | snoRNA | 37.33 | 0.017939622 | 0.017939622 | ENSG00000207031 | n.s. |
| ENSG00000089597 | 1558.022 | -0.222574587 | 0.273932 | -0.812516 | 0.416495 | 0.830664142 | GANAB | protein_coding | 50.02 | -0.017933844 | 0.017933844 | ENSG00000089597 | n.s. |
| ENSG00000199636 | 9.797505 | -0.37220434 | 0.641958 | -0.579795 | 0.562053 | 0.895116382 | | misc_RNA | 36.19 | -0.017910657 | 0.017910657 | ENSG00000199636 | n.s. |
| ENSG00000215421 | 1125.428 | -0.210333567 | 0.250014 | -0.841288 | 0.400187 | 0.821966929 | ZNF407 | protein_coding | 37.97 | -0.017908989 | 0.017908989 | ENSG00000215421 | n.s. |
| ENSG00000211575 | 4.718052 | -0.514797791 | 1.13362 | -0.454119 | 0.649743 | 0.923184844 | MIR760 | miRNA | 63.75 | -0.017869318 | 0.017869318 | ENSG00000211575 | n.s. |
| ENSG00000148120 | 4157.853 | -0.200941599 | 0.233345 | -0.861136 | 0.389163 | 0.814907606 | C9orf3 | protein_coding | 42.89 | -0.017862026 | 0.017862026 | ENSG00000148120 | n.s. |
| ENSG00000103485 | 408.1792 | 0.237891787 | 0.302417 | 0.7866358 | 0.431495 | 0.841297658 | QPRT | protein_coding | 51.92 | 0.017853855 | 0.017853855 | ENSG00000103485 | n.s. |
| ENSG00000102606 | 1373.509 | -0.21712229 | 0.263938 | -0.822626 | 0.410721 | 0.827563884 | ARHGEF7 | protein_coding | 44.55 | -0.01784712 | 0.01784712 | ENSG00000102606 | n.s. |
| ENSG00000140451 | 572.245 | 0.211661811 | 0.252391 | 0.8386256 | 0.401679 | 0.823540971 | PIF1 | protein_coding | 55.24 | 0.017846221 | 0.017846221 | ENSG00000140451 | n.s. |
| ENSG00000124688 | 665.3842 | -0.204635923 | 0.240836 | -0.849689 | 0.395498 | 0.818083342 | MAD2L1BP | protein_coding | 49.25 | -0.017844754 | 0.017844754 | ENSG00000124688 | n.s. |
| ENSG00000170322 | 629.5301 | -0.213095487 | 0.256333 | -0.831323 | 0.405791 | 0.824799944 | NFRKB | protein_coding | 45.4 | -0.017825731 | 0.017825731 | ENSG00000170322 | n.s. |
| ENSG00000269044 | 92.46432 | -0.236975535 | 0.301006 | -0.787277 | 0.43112 | 0.84115455 | | sense_intronic | 46.78 | -0.017802598 | 0.017802598 | ENSG00000269044 | n.s. |
| ENSG00000223546 | 116.7937 | -0.225057666 | 0.279651 | -0.804781 | 0.420946 | 0.833570156 | LINC00630 | lincRNA | 39.11 | -0.017792574 | 0.017792574 | ENSG00000223546 | n.s. |
| ENSG00000177732 | 831.2962 | -0.201016323 | 0.23463 | -0.856737 | 0.39159 | 0.815710819 | SOX12 | protein_coding | 61.99 | -0.017782663 | 0.017782663 | ENSG00000177732 | n.s. |
| ENSG00000077454 | 156.7409 | -0.223067676 | 0.276266 | -0.807439 | 0.419414 | 0.832304654 | LRCH4 | protein_coding | 60.36 | -0.017782437 | 0.017782437 | ENSG00000077454 | n.s. |
| ENSG00000135541 | 1185.439 | -0.219050334 | 0.268908 | -0.814593 | 0.415305 | 0.829668953 | AHI1 | protein_coding | 36.31 | -0.017763922 | 0.017763922 | ENSG00000135541 | n.s. |
| ENSG00000212588 | 299.2198 | 0.194809519 | 0.223842 | 0.8702975 | 0.384138 | 0.810719585 | SNORA26 | snoRNA | 45.9 | 0.017752862 | 0.017752862 | ENSG00000212588 | n.s. |
| ENSG00000197329 | 1018.211 | -0.215908437 | 0.262356 | -0.822961 | 0.41053 | 0.827563884 | PELI1 | protein_coding | 36.9 | -0.017747343 | 0.017747343 | ENSG00000197329 | n.s. |

| | | | | | | | | | | | | | |
|---|---|---|---|---|---|---|---|---|---|---|---|---|---|
| ENSG00000043093 | 1305.872 | -0.213948017 | 0.258712 | -0.826972 | 0.408253 | 0.826171518 | DCUN1D1 | protein_coding | 39.56 | -0.017742662 | 0.017742662 | ENSG00000043093 | n.s. |
| ENSG00000172943 | 389.4879 | -0.205149627 | 0.242061 | -0.847511 | 0.39671 | 0.819495485 | PHF8 | protein_coding | 43.07 | -0.01773589 | 0.01773589 | ENSG00000172943 | n.s. |
| ENSG00000266965 | 142.037 | 0.22741176 | 0.285104 | 0.7976438 | 0.425077 | 0.835658686 | | sense_intronic | 40.97 | 0.017731538 | 0.017731538 | ENSG00000266965 | n.s. |
| ENSG00000179195 | 4090.398 | -0.217668904 | 0.266594 | -0.816481 | 0.414225 | 0.829007487 | ZNF664 | protein_coding | 44.11 | -0.017727292 | 0.017727292 | ENSG00000179195 | n.s. |
| ENSG00000125817 | 860.354 | -0.211599649 | 0.253211 | -0.835664 | 0.403344 | 0.824699011 | CENPB | protein_coding | 66.2 | -0.017711848 | 0.017711848 | ENSG00000125817 | n.s. |
| ENSG00000077585 | 158.2034 | -0.225584683 | 0.281658 | -0.800916 | 0.42318 | 0.834993891 | GPR137B | protein_coding | 43.63 | -0.017667048 | 0.017667048 | ENSG00000077585 | n.s. |
| ENSG00000157911 | 130.7968 | -0.239256516 | 0.307657 | -0.777674 | 0.436761 | 0.843772375 | PEX10 | protein_coding | 61.52 | -0.017651077 | 0.017651077 | ENSG00000157911 | n.s. |
| ENSG00000136451 | 3256.747 | -0.205539218 | 0.243666 | -0.843529 | 0.398932 | 0.820802709 | VEZF1 | protein_coding | 40.94 | -0.017627294 | 0.017627294 | ENSG00000136451 | n.s. |
| ENSG00000204435 | 836.097 | -0.224403466 | 0.279652 | -0.802437 | 0.4223 | 0.834590104 | CSNK2B | protein_coding | 52.7 | -0.017621679 | 0.017621679 | ENSG00000204435 | n.s. |
| ENSG00000115827 | 365.6704 | -0.196413928 | 0.227076 | -0.864972 | 0.387054 | 0.813399117 | DCAF17 | protein_coding | 37.36 | -0.017617603 | 0.017617603 | ENSG00000115827 | n.s. |
| ENSG00000111554 | 1056.085 | 0.215918391 | 0.263229 | 0.820269 | 0.412063 | 0.828824745 | MDM1 | protein_coding | 38.66 | 0.017605401 | 0.017605401 | ENSG00000111554 | n.s. |
| ENSG00000112685 | 642.6873 | -0.224713685 | 0.280517 | -0.801071 | 0.423091 | 0.834993891 | EXOC2 | protein_coding | 40.51 | -0.017598835 | 0.017598835 | ENSG00000112685 | n.s. |
| ENSG00000107959 | 770.0456 | -0.220685563 | 0.273037 | -0.808261 | 0.41894 | 0.832304654 | PITRM1 | protein_coding | 46.17 | -0.01759254 | 0.01759254 | ENSG00000107959 | n.s. |
| ENSG00000251323 | 78.99361 | -0.237494944 | 0.304849 | -0.779058 | 0.435945 | 0.843243823 | | lincRNA | 39.08 | -0.017585748 | 0.017585748 | ENSG00000251323 | n.s. |
| ENSG00000264607 | 6.005124 | -0.35806827 | 0.608303 | -0.588634 | 0.556107 | 0.893144718 | MIR3173 | miRNA | 48.53 | -0.017573334 | 0.017573334 | ENSG00000264607 | n.s. |
| ENSG00000130584 | 263.8861 | -0.22042522 | 0.272747 | -0.808167 | 0.418994 | 0.832304654 | ZBTB46 | protein_coding | 53.89 | -0.017571786 | 0.017571786 | ENSG00000130584 | n.s. |
| ENSG00000239577 | 18.05262 | -0.265375302 | 0.361481 | -0.734133 | 0.462868 | 0.858623382 | RN7SL388P | misc_RNA | 52.8 | -0.017567126 | 0.017567126 | ENSG00000239577 | n.s. |
| ENSG00000082014 | 114.6517 | 0.251259441 | 0.332825 | 0.7549299 | 0.450291 | 0.851309042 | SMARCD3 | protein_coding | 56.14 | 0.01756624 | 0.01756624 | ENSG00000082014 | n.s. |
| ENSG00000197056 | 277.0861 | -0.250900054 | 0.331398 | -0.757097 | 0.448992 | 0.85118703 | ZMYM1 | protein_coding | 41.67 | -0.017556732 | 0.017556732 | ENSG00000197056 | n.s. |
| ENSG00000140157 | 1182.958 | -0.224903727 | 0.281586 | -0.798704 | 0.424462 | 0.835527278 | NIPA2 | protein_coding | 43.73 | -0.017551344 | 0.017551344 | ENSG00000140157 | n.s. |
| ENSG00000099953 | 115.0931 | 0.236718259 | 0.303711 | 0.7794189 | 0.435733 | 0.843067025 | MMP11 | protein_coding | 57.83 | 0.017549794 | 0.017549794 | ENSG00000099953 | n.s. |
| ENSG00000146802 | 806.515 | -0.230476117 | 0.29189 | -0.789599 | 0.429762 | 0.839380003 | TMEM168 | protein_coding | 35.46 | -0.017525722 | 0.017525722 | ENSG00000146802 | n.s. |
| ENSG00000087460 | 13232.98 | -0.215162318 | 0.263428 | -0.816778 | 0.414055 | 0.829007487 | GNAS | protein_coding | 47.09 | -0.017523152 | 0.017523152 | ENSG00000087460 | n.s. |
| ENSG00000100354 | 274.8759 | -0.201653745 | 0.237595 | -0.848729 | 0.396032 | 0.818701982 | TNRC6B | protein_coding | 42.15 | -0.017518499 | 0.017518499 | ENSG00000100354 | n.s. |
| ENSG00000269425 | 12.41662 | -0.31619844 | 0.481183 | -0.657128 | 0.511099 | 0.880260889 | | antisense | 51.49 | -0.017513787 | 0.017513787 | ENSG00000269425 | n.s. |
| ENSG00000129493 | 19.65903 | -0.300826411 | 0.443956 | -0.677604 | 0.498023 | 0.874675553 | HEATR5A | protein_coding | 39.35 | -0.017493962 | 0.017493962 | ENSG00000129493 | n.s. |
| ENSG00000224870 | 148.9952 | -0.217621747 | 0.268032 | -0.811926 | 0.416834 | 0.831101564 | | processed_transcript | 59.37 | -0.017485015 | 0.017485015 | ENSG00000224870 | n.s. |
| ENSG00000199672 | 5.119013 | -0.364568094 | 0.631323 | -0.577467 | 0.563624 | 0.895460914 | RNU4-21P | snRNA | 42.86 | -0.017482267 | 0.017482267 | ENSG00000199672 | n.s. |
| ENSG00000239249 | 14.30264 | -0.295404404 | 0.429372 | -0.687992 | 0.491458 | 0.872664906 | RN7SL757P | misc_RNA | 54.49 | -0.017473906 | 0.017473906 | ENSG00000239249 | n.s. |
| ENSG00000233515 | 43.61496 | 0.249879577 | 0.330951 | 0.755036 | 0.450227 | 0.851309042 | LINC01518 | lincRNA | 41.67 | 0.017469769 | 0.017469769 | ENSG00000233515 | n.s. |
| ENSG00000109685 | 4247.296 | -0.211755128 | 0.256905 | -0.824254 | 0.409795 | 0.827015612 | NSD2 | protein_coding | 47.14 | -0.017466895 | 0.017466895 | ENSG00000109685 | n.s. |
| ENSG00000111224 | 59.99493 | -0.234840348 | 0.300839 | -0.780617 | 0.435028 | 0.842611157 | PARP11 | protein_coding | 39.15 | -0.017465733 | 0.017465733 | ENSG00000111224 | n.s. |

| | | | | | | | | | | | | | |
|---|---|---|---|---|---|---|---|---|---|---|---|---|---|
| ENSG00000234134 | 53.82324 | -0.246170749 | 0.322866 | -0.762454 | 0.445789 | 0.849421269 | | sense_intronic | 57.31 | -0.017447812 | 0.017447812 | ENSG00000234134 | n.s. |
| ENSG00000128791 | 515.9463 | -0.223389676 | 0.279758 | -0.79851 | 0.424575 | 0.835527278 | TWSG1 | protein_coding | 38.45 | -0.017433189 | 0.017433189 | ENSG00000128791 | n.s. |
| ENSG00000055483 | 1605.051 | -0.188329582 | 0.212182 | -0.887584 | 0.374764 | 0.808040232 | USP36 | protein_coding | 50.03 | -0.017433107 | 0.017433107 | ENSG00000055483 | n.s. |
| ENSG00000201176 | 5.559611 | -0.355731482 | 0.605011 | -0.587975 | 0.556549 | 0.893442909 | RNU6-853P | snRNA | 39.25 | -0.017407077 | 0.017407077 | ENSG00000201176 | n.s. |
| ENSG00000101489 | 267.5223 | -0.218351881 | 0.269838 | -0.809197 | 0.418402 | 0.832304654 | CELF4 | protein_coding | 52.35 | -0.017406505 | 0.017406505 | ENSG00000101489 | n.s. |
| ENSG00000173581 | 85.54137 | -0.23614095 | 0.304535 | -0.775414 | 0.438095 | 0.843950276 | CCDC106 | protein_coding | 61.17 | -0.017399607 | 0.017399607 | ENSG00000173581 | n.s. |
| ENSG00000100632 | 3055.265 | -0.235780355 | 0.303474 | -0.776938 | 0.437195 | 0.843772375 | ERH | protein_coding | 40.33 | -0.017394624 | 0.017394624 | ENSG00000100632 | n.s. |
| ENSG00000048405 | 923.7191 | -0.223040731 | 0.279573 | -0.79779 | 0.424992 | 0.835658686 | ZNF800 | protein_coding | 37.8 | -0.017390724 | 0.017390724 | ENSG00000048405 | n.s. |
| ENSG00000118513 | 183.1759 | -0.228360939 | 0.288749 | -0.790862 | 0.429024 | 0.839183171 | MYB | protein_coding | 39.74 | -0.01738814 | 0.01738814 | ENSG00000118513 | n.s. |
| ENSG00000276259 | 49.74202 | -0.259929826 | 0.351349 | -0.739804 | 0.459419 | 0.857420077 | | antisense | 42.84 | -0.017364963 | 0.017364963 | ENSG00000276259 | n.s. |
| ENSG00000170919 | 37.36153 | -0.238174662 | 0.309922 | -0.768499 | 0.442191 | 0.845745639 | TPT1-AS1 | processed_transcript | 48.88 | -0.017329644 | 0.017329644 | ENSG00000170919 | n.s. |
| ENSG00000196214 | 749.5081 | -0.220018575 | 0.273739 | -0.803754 | 0.421539 | 0.834151325 | ZNF766 | protein_coding | 44.77 | -0.017327597 | 0.017327597 | ENSG00000196214 | n.s. |
| ENSG00000105220 | 2049.891 | -0.209655842 | 0.254052 | -0.825247 | 0.409231 | 0.826802059 | GPI | protein_coding | 52.45 | -0.017317248 | 0.017317248 | ENSG00000105220 | n.s. |
| ENSG00000111371 | 12086.45 | -0.212280227 | 0.25981 | -0.817058 | 0.413895 | 0.829007487 | SLC38A1 | protein_coding | 39.96 | -0.01728843 | 0.01728843 | ENSG00000111371 | n.s. |
| ENSG00000198825 | 682.3437 | -0.210047196 | 0.254997 | -0.823723 | 0.410097 | 0.827409292 | INPP5F | protein_coding | 41.2 | -0.017282601 | 0.017282601 | ENSG00000198825 | n.s. |
| ENSG00000143248 | 46.31625 | 0.241787165 | 0.316503 | 0.7639325 | 0.444907 | 0.848321918 | RGS5 | protein_coding | 37.67 | 0.017273109 | 0.017273109 | ENSG00000143248 | n.s. |
| ENSG00000151693 | 809.4779 | -0.21116926 | 0.257185 | -0.821078 | 0.411602 | 0.828379538 | ASAP2 | protein_coding | 46.56 | -0.017267445 | 0.017267445 | ENSG00000151693 | n.s. |
| ENSG00000237310 | 33.89524 | 0.258398331 | 0.349416 | 0.7395157 | 0.459594 | 0.857420077 | | lincRNA | 54.44 | 0.01726265 | 0.01726265 | ENSG00000237310 | n.s. |
| ENSG00000166526 | 1631.255 | -0.204733214 | 0.244249 | -0.838216 | 0.401909 | 0.823540971 | ZNF3 | protein_coding | 48.95 | -0.017262038 | 0.017262038 | ENSG00000166526 | n.s. |
| ENSG00000180917 | 380.9427 | -0.208144161 | 0.251812 | -0.826586 | 0.408472 | 0.82626099 | CMTR2 | protein_coding | 39.62 | -0.017251561 | 0.017251561 | ENSG00000180917 | n.s. |
| ENSG00000100814 | 3091.268 | 0.215025653 | 0.265093 | 0.8111323 | 0.41729 | 0.831423508 | CCNB1IP1 | protein_coding | 39.52 | 0.017240262 | 0.017240262 | ENSG00000100814 | n.s. |
| ENSG00000164543 | 257.1211 | -0.223802668 | 0.282188 | -0.793098 | 0.427721 | 0.837692206 | STK17A | protein_coding | 39.76 | -0.0172139 | 0.0172139 | ENSG00000164543 | n.s. |
| ENSG00000165490 | 753.1456 | -0.219763515 | 0.274286 | -0.801219 | 0.423005 | 0.834993891 | DDIAS | protein_coding | 39.78 | -0.017211154 | 0.017211154 | ENSG00000165490 | n.s. |
| ENSG00000183155 | 284.8366 | -0.231645879 | 0.297184 | -0.77947 | 0.435703 | 0.843067025 | RABIF | protein_coding | 45.67 | -0.017173738 | 0.017173738 | ENSG00000183155 | n.s. |
| ENSG00000148384 | 273.7386 | -0.207882561 | 0.252054 | -0.824754 | 0.409511 | 0.826802059 | INPP5E | protein_coding | 63.05 | -0.017170777 | 0.017170777 | ENSG00000148384 | n.s. |
| ENSG00000105993 | 1639.128 | -0.216752501 | 0.269082 | -0.805526 | 0.420516 | 0.833311153 | DNAJB6 | protein_coding | 48.15 | -0.017165239 | 0.017165239 | ENSG00000105993 | n.s. |
| ENSG00000007202 | 2945.532 | -0.210759141 | 0.258294 | -0.815965 | 0.41452 | 0.829007487 | KIAA0100 | protein_coding | 46.75 | -0.017164551 | 0.017164551 | ENSG00000007202 | n.s. |
| ENSG00000111684 | 117.5278 | -0.230385862 | 0.294308 | -0.782806 | 0.433741 | 0.84244096 | LPCAT3 | protein_coding | 44.19 | -0.017154652 | 0.017154652 | ENSG00000111684 | n.s. |
| ENSG00000177981 | 291.2551 | -0.213706431 | 0.263537 | -0.810916 | 0.417414 | 0.831423508 | ASB8 | protein_coding | 42.11 | -0.01713449 | 0.01713449 | ENSG00000177981 | n.s. |
| ENSG00000156587 | 214.7192 | -0.235214205 | 0.305562 | -0.769775 | 0.441433 | 0.84569031 | UBE2L6 | protein_coding | 48.66 | -0.017120923 | 0.017120923 | ENSG00000156587 | n.s. |
| ENSG00000130803 | 766.5046 | -0.20842792 | 0.253543 | -0.822063 | 0.411041 | 0.827730385 | ZNF317 | protein_coding | 44.48 | -0.017114246 | 0.017114246 | ENSG00000130803 | n.s. |
| ENSG00000177707 | 552.6729 | -0.204482579 | 0.245617 | -0.832525 | 0.405112 | 0.824799944 | NECTIN3 | protein_coding | 36.67 | -0.017105249 | 0.017105249 | ENSG00000177707 | n.s. |

| | | | | | | | | | | | | | |
|---|---|---|---|---|---|---|---|---|---|---|---|---|---|
| ENSG00000176022 | 238.5792 | -0.193323277 | 0.224925 | -0.859501 | 0.390064 | 0.815710819 | B3GALT6 | protein_coding | 64.03 | -0.017102107 | 0.017102107 | ENSG00000176022 | n.s. |
| ENSG00000277462 | 557.2766 | -0.229880139 | 0.294407 | -0.780823 | 0.434906 | 0.842611157 | ZNF670 | protein_coding | 41.24 | -0.017096828 | 0.017096828 | ENSG00000277462 | n.s. |
| ENSG00000258311 | 134.187 | -0.219492335 | 0.275544 | -0.796579 | 0.425696 | 0.836005799 | | protein_coding | 53.8 | -0.017074465 | 0.017074465 | ENSG00000258311 | n.s. |
| ENSG00000090060 | 4356.264 | -0.206537325 | 0.250051 | -0.825981 | 0.408815 | 0.826692059 | PAPOLA | protein_coding | 36.57 | -0.017071597 | 0.017071597 | ENSG00000090060 | n.s. |
| ENSG00000076928 | 51.81829 | -0.232635733 | 0.300601 | -0.773901 | 0.438989 | 0.844555159 | ARHGEF1 | protein_coding | 58.25 | -0.017068944 | 0.017068944 | ENSG00000076928 | n.s. |
| ENSG00000165244 | 966.9144 | -0.213962685 | 0.264219 | -0.809794 | 0.418059 | 0.832304654 | ZNF367 | protein_coding | 42.86 | -0.017056608 | 0.017056608 | ENSG00000165244 | n.s. |
| ENSG00000165389 | 616.7301 | -0.242175762 | 0.319015 | -0.759136 | 0.447771 | 0.850405175 | SPTSSA | protein_coding | 41.92 | -0.017042903 | 0.017042903 | ENSG00000165389 | n.s. |
| ENSG00000117748 | 578.0008 | -0.24337425 | 0.321857 | -0.756156 | 0.449556 | 0.85118703 | RPA2 | protein_coding | 44.15 | -0.017030114 | 0.017030114 | ENSG00000117748 | n.s. |
| ENSG00000104835 | 346.2405 | -0.21236949 | 0.261856 | -0.811017 | 0.417356 | 0.831423508 | SARS2 | protein_coding | 53.66 | -0.017027297 | 0.017027297 | ENSG00000104835 | n.s. |
| ENSG00000174177 | 236.6446 | -0.221282608 | 0.278834 | -0.793599 | 0.427429 | 0.837692206 | CTU2 | protein_coding | 60.96 | -0.017020068 | 0.017020068 | ENSG00000174177 | n.s. |
| ENSG00000155980 | 445.6122 | -0.208756504 | 0.255347 | -0.81754 | 0.41362 | 0.829007487 | KIF5A | protein_coding | 47.7 | -0.017001453 | 0.017001453 | ENSG00000155980 | n.s. |
| ENSG00000010072 | 812.4651 | -0.213082979 | 0.263931 | -0.807344 | 0.419468 | 0.832304654 | SPRTN | protein_coding | 41.57 | -0.01698648 | 0.01698648 | ENSG00000010072 | n.s. |
| ENSG00000170264 | 459.6972 | -0.217017508 | 0.271238 | -0.8001 | 0.423653 | 0.835158421 | FAM161A | protein_coding | 41.32 | -0.016977526 | 0.016977526 | ENSG00000170264 | n.s. |
| ENSG00000186416 | 503.4443 | -0.215490963 | 0.268354 | -0.803011 | 0.421968 | 0.834256463 | NKRF | protein_coding | 42.08 | -0.016959229 | 0.016959229 | ENSG00000186416 | n.s. |
| ENSG00000088727 | 64.92745 | 0.227753485 | 0.291241 | 0.78201 | 0.434209 | 0.84244096 | KIF9 | protein_coding | 48.45 | 0.016958644 | 0.016958644 | ENSG00000088727 | n.s. |
| ENSG00000100532 | 303.7787 | -0.23868867 | 0.312906 | -0.762813 | 0.445575 | 0.849129133 | CGRRF1 | protein_coding | 37.44 | -0.016953163 | 0.016953163 | ENSG00000100532 | n.s. |
| ENSG00000149269 | 131.3223 | -0.227312527 | 0.289801 | -0.784373 | 0.432821 | 0.842286016 | PAK1 | protein_coding | 39.88 | -0.016943969 | 0.016943969 | ENSG00000149269 | n.s. |
| ENSG00000089486 | 398.1331 | -0.196071694 | 0.231566 | -0.846721 | 0.397151 | 0.819605152 | CDIP1 | protein_coding | 51.97 | -0.016939677 | 0.016939677 | ENSG00000089486 | n.s. |
| ENSG00000183580 | 667.845 | -0.191302537 | 0.223845 | -0.854622 | 0.392761 | 0.815710819 | FBXL7 | protein_coding | 39.82 | -0.016923345 | 0.016923345 | ENSG00000183580 | n.s. |
| ENSG00000207093 | 4.538609 | -0.344578084 | 0.584757 | -0.589267 | 0.555682 | 0.893144718 | SNORD116-8 | snoRNA | 44.21 | -0.01691126 | 0.01691126 | ENSG00000207093 | n.s. |
| ENSG00000026652 | 960.4576 | -0.225025416 | 0.285912 | -0.787044 | 0.431256 | 0.84116282 | AGPAT4 | protein_coding | 45.36 | -0.016903894 | 0.016903894 | ENSG00000026652 | n.s. |
| ENSG00000222561 | 8.395749 | -0.333865229 | 0.542984 | -0.614871 | 0.53864 | 0.889966281 | RNU6-1025P | snRNA | 36.54 | -0.016902411 | 0.016902411 | ENSG00000222561 | n.s. |
| ENSG00000222790 | 5.007735 | -0.344260587 | 0.582167 | -0.591343 | 0.55429 | 0.893144718 | RNU4-14P | snRNA | 37.7 | -0.016895678 | 0.016895678 | ENSG00000222790 | n.s. |
| ENSG00000265681 | 2102.723 | 0.19092085 | 0.223128 | 0.8556552 | 0.392189 | 0.815710819 | RPL17 | protein_coding | 42.91 | 0.01688958 | 0.01688958 | ENSG00000265681 | n.s. |
| ENSG00000104915 | 375.5912 | 0.244527575 | 0.325711 | 0.7507507 | 0.452803 | 0.853163076 | STX10 | protein_coding | 55.36 | 0.016864565 | 0.016864565 | ENSG00000104915 | n.s. |
| ENSG00000087269 | 1509.38 | -0.206638689 | 0.251824 | -0.820567 | 0.411893 | 0.828691651 | NOP14 | protein_coding | 49.99 | -0.016863171 | 0.016863171 | ENSG00000087269 | n.s. |
| ENSG00000145348 | 820.5492 | -0.228181769 | 0.293845 | -0.776539 | 0.437431 | 0.843772375 | TBCK | protein_coding | 35.98 | -0.016834041 | 0.016834041 | ENSG00000145348 | n.s. |
| ENSG00000139629 | 224.4955 | -0.221161965 | 0.279759 | -0.790546 | 0.429209 | 0.839274918 | GALNT6 | protein_coding | 49.89 | -0.016829487 | 0.016829487 | ENSG00000139629 | n.s. |
| ENSG00000110435 | 872.5173 | -0.225773247 | 0.28797 | -0.784016 | 0.43303 | 0.842286016 | PDHX | protein_coding | 36.68 | -0.01682923 | 0.01682923 | ENSG00000110435 | n.s. |
| ENSG00000164168 | 921.4753 | -0.216164857 | 0.271394 | -0.796499 | 0.425742 | 0.836005799 | TMEM184C | protein_coding | 38.71 | -0.016815618 | 0.016815618 | ENSG00000164168 | n.s. |
| ENSG00000148935 | 66.52356 | -0.24231839 | 0.322119 | -0.752263 | 0.451893 | 0.852336584 | GAS2 | protein_coding | 36.04 | -0.016814199 | 0.016814199 | ENSG00000148935 | n.s. |
| ENSG00000001631 | 955.7896 | -0.206302438 | 0.252767 | -0.816178 | 0.414399 | 0.829007487 | KRIT1 | protein_coding | 36.48 | -0.01680159 | 0.01680159 | ENSG00000001631 | n.s. |

| | | | | | | | | | | | | | |
|---|---|---|---|---|---|---|---|---|---|---|---|---|---|
| ENSG00000124570 | 125.0544 | 0.243644548 | 0.325026 | 0.7496166 | 0.453486 | 0.853227086 | SERPINB6 | protein_coding | 47.23 | 0.016795726 | 0.016795726 | ENSG00000124570 | n.s. |
| ENSG00000104691 | 403.8153 | -0.227907379 | 0.2939 | -0.775458 | 0.438069 | 0.843950276 | UBXN8 | protein_coding | 44.01 | -0.016792931 | 0.016792931 | ENSG00000104691 | n.s. |
| ENSG00000076604 | 886.5886 | 0.225206125 | 0.287228 | 0.7840681 | 0.433 | 0.842286016 | TRAF4 | protein_coding | 59.64 | 0.016786957 | 0.016786957 | ENSG00000076604 | n.s. |
| ENSG00000166411 | 617.8796 | -0.213136939 | 0.265223 | -0.803615 | 0.421619 | 0.83419139 | IDH3A | protein_coding | 44.46 | -0.016781187 | 0.016781187 | ENSG00000166411 | n.s. |
| ENSG00000165655 | 311.0763 | 0.191884095 | 0.225473 | 0.8510286 | 0.394753 | 0.817840002 | ZNF503 | protein_coding | 61.2 | 0.016757555 | 0.016757555 | ENSG00000165655 | n.s. |
| ENSG00000133028 | 1286.247 | -0.214620332 | 0.268968 | -0.79794 | 0.424905 | 0.835658686 | SCO1 | protein_coding | 41.24 | -0.016734177 | 0.016734177 | ENSG00000133028 | n.s. |
| ENSG00000213390 | 485.2015 | -0.229856227 | 0.298461 | -0.770138 | 0.441218 | 0.84569031 | ARHGAP19 | protein_coding | 42.77 | -0.016730923 | 0.016730923 | ENSG00000213390 | n.s. |
| ENSG00000151893 | 1211.369 | -0.205094668 | 0.250817 | -0.817706 | 0.413525 | 0.829007487 | CACUL1 | protein_coding | 39.56 | -0.016703227 | 0.016703227 | ENSG00000151893 | n.s. |
| ENSG00000007341 | 856.7846 | -0.186140097 | 0.215308 | -0.864531 | 0.387296 | 0.813399117 | ST7L | protein_coding | 39.05 | -0.016696079 | 0.016696079 | ENSG00000007341 | n.s. |
| ENSG00000275966 | 22.01165 | -0.282794201 | 0.412301 | -0.685893 | 0.492781 | 0.873000006 | | lincRNA | 72.3 | -0.016680829 | 0.016680829 | ENSG00000275966 | n.s. |
| ENSG00000162714 | 1050.88 | -0.209226194 | 0.259166 | -0.807307 | 0.41949 | 0.832304654 | ZNF496 | protein_coding | 48.34 | -0.016679026 | 0.016679026 | ENSG00000162714 | n.s. |
| ENSG00000125520 | 99.90752 | -0.223585158 | 0.285737 | -0.782486 | 0.433929 | 0.84244096 | SLC2A4RG | protein_coding | 69.47 | -0.016648268 | 0.016648268 | ENSG00000125520 | n.s. |
| ENSG00000170390 | 219.1868 | -0.213027425 | 0.266541 | -0.799228 | 0.424158 | 0.835315601 | DCLK2 | protein_coding | 40.72 | -0.016647967 | 0.016647967 | ENSG00000170390 | n.s. |
| ENSG00000279074 | 17.18465 | -0.286222811 | 0.422497 | -0.677455 | 0.498117 | 0.874675553 | | TEC | 44.73 | -0.016644719 | 0.016644719 | ENSG00000279074 | n.s. |
| ENSG00000133424 | 181.3172 | -0.19855573 | 0.237523 | -0.835944 | 0.403187 | 0.824675468 | LARGE1 | protein_coding | 43.98 | -0.016622475 | 0.016622475 | ENSG00000133424 | n.s. |
| ENSG00000199674 | 9.135618 | -0.313118912 | 0.485452 | -0.645005 | 0.518924 | 0.88497967 | RNU6-862P | snRNA | 40.95 | -0.016616189 | 0.016616189 | ENSG00000199674 | n.s. |
| ENSG00000144451 | 525.9348 | -0.222562438 | 0.283496 | -0.785065 | 0.432416 | 0.842149099 | SPAG16 | protein_coding | 35.89 | -0.016605609 | 0.016605609 | ENSG00000144451 | n.s. |
| ENSG00000104714 | 296.7677 | -0.229222754 | 0.298736 | -0.76731 | 0.442898 | 0.846576336 | ERICH1 | protein_coding | 51.17 | -0.01658057 | 0.01658057 | ENSG00000104714 | n.s. |
| ENSG00000243224 | 85.46034 | -0.217910116 | 0.275917 | -0.789768 | 0.429663 | 0.839380003 | | antisense | 54.35 | -0.016570186 | 0.016570186 | ENSG00000243224 | n.s. |
| ENSG00000011376 | 867.0925 | -0.203423936 | 0.248598 | -0.818285 | 0.413195 | 0.829007487 | LARS2 | protein_coding | 42.15 | -0.01656716 | 0.01656716 | ENSG00000011376 | n.s. |
| ENSG00000147364 | 688.9284 | -0.211636967 | 0.264507 | -0.80012 | 0.423641 | 0.835158421 | FBXO25 | protein_coding | 41.96 | -0.0165566 | 0.0165566 | ENSG00000147364 | n.s. |
| ENSG00000123178 | 231.9595 | -0.227378293 | 0.295379 | -0.769785 | 0.441427 | 0.84569031 | SPRYD7 | protein_coding | 40.33 | -0.016550558 | 0.016550558 | ENSG00000123178 | n.s. |
| ENSG00000055070 | 575.6552 | -0.224322983 | 0.288841 | -0.77663 | 0.437377 | 0.843772375 | SZRD1 | protein_coding | 48.1 | -0.01654936 | 0.01654936 | ENSG00000055070 | n.s. |
| ENSG00000105321 | 84.84295 | -0.224550981 | 0.289432 | -0.775834 | 0.437847 | 0.843950276 | CCDC9 | protein_coding | 56.02 | -0.016545621 | 0.016545621 | ENSG00000105321 | n.s. |
| ENSG00000244005 | 58.21999 | 0.238051233 | 0.316072 | 0.7531549 | 0.451357 | 0.852169954 | NFS1 | protein_coding | 46.45 | 0.01653832 | 0.01653832 | ENSG00000244005 | n.s. |
| ENSG00000263597 | 6.518337 | -0.336947726 | 0.570642 | -0.590471 | 0.554875 | 0.893144718 | MIR3936 | miRNA | 51.82 | -0.016536776 | 0.016536776 | ENSG00000263597 | n.s. |
| ENSG00000238966 | 48.86619 | 0.239846225 | 0.31981 | 0.7499647 | 0.453276 | 0.853222234 | | snoRNA | 36.11 | 0.01653448 | 0.01653448 | ENSG00000238966 | n.s. |
| ENSG00000099783 | 4909.978 | -0.201117178 | 0.244427 | -0.822812 | 0.410615 | 0.827563884 | HNRNPM | protein_coding | 46.68 | -0.016531524 | 0.016531524 | ENSG00000099783 | n.s. |
| ENSG00000196588 | 99.92362 | -0.213811192 | 0.269012 | -0.794803 | 0.426728 | 0.837228298 | MKL1 | protein_coding | 43.58 | -0.016496838 | 0.016496838 | ENSG00000196588 | n.s. |
| ENSG00000144231 | 1694.809 | -0.207117829 | 0.256751 | -0.806686 | 0.419847 | 0.832578684 | POLR2D | protein_coding | 45.35 | -0.016481342 | 0.016481342 | ENSG00000144231 | n.s. |
| ENSG00000136463 | 606.084 | -0.221425127 | 0.28352 | -0.780986 | 0.434811 | 0.842611157 | TACO1 | protein_coding | 49.33 | -0.016468006 | 0.016468006 | ENSG00000136463 | n.s. |
| ENSG00000227811 | 271.6792 | -0.195267727 | 0.233014 | -0.838008 | 0.402026 | 0.823540971 | FAM212B-AS1 | antisense | 49.47 | -0.016463957 | 0.016463957 | ENSG00000227811 | n.s. |

| | | | | | | | | | | | | | |
|---|---|---|---|---|---|---|---|---|---|---|---|---|---|
| ENSG00000133740 | 434.7147 | -0.223163808 | 0.287315 | -0.776721 | 0.437324 | 0.843772375 | E2F5 | protein_coding | 39.49 | -0.016463842 | 0.016463842 | ENSG00000133740 | n.s. |
| ENSG00000127252 | 148.2486 | -0.221920063 | 0.284677 | -0.77955 | 0.435656 | 0.843067025 | HRASLS | protein_coding | 37.84 | -0.016452687 | 0.016452687 | ENSG00000127252 | n.s. |
| ENSG00000164929 | 44.68897 | -0.262506952 | 0.369517 | -0.710405 | 0.477453 | 0.865616576 | BAALC | protein_coding | 42.75 | -0.016452475 | 0.016452475 | ENSG00000164929 | n.s. |
| ENSG00000279584 | 16.57588 | 0.315536353 | 0.500727 | 0.6301568 | 0.528592 | 0.886954686 | | TEC | 56.12 | 0.016438992 | 0.016438992 | ENSG00000279584 | n.s. |
| ENSG00000257803 | 111.1026 | -0.219674937 | 0.279706 | -0.785378 | 0.432232 | 0.841908613 | | processed_pseudogene | 41.61 | -0.016417417 | 0.016417417 | ENSG00000257803 | n.s. |
| ENSG00000100201 | 9667.448 | -0.199687065 | 0.242678 | -0.822846 | 0.410595 | 0.827563884 | DDX17 | protein_coding | 43.04 | -0.016413971 | 0.016413971 | ENSG00000100201 | n.s. |
| ENSG00000117984 | 314.4502 | -0.207570827 | 0.257909 | -0.804823 | 0.420922 | 0.833570156 | CTSD | protein_coding | 64.73 | -0.016410102 | 0.016410102 | ENSG00000117984 | n.s. |
| ENSG00000240857 | 451.6042 | -0.210164806 | 0.263166 | -0.798603 | 0.424521 | 0.835527278 | RDH14 | protein_coding | 41.88 | -0.016401128 | 0.016401128 | ENSG00000240857 | n.s. |
| ENSG00000081791 | 952.3149 | -0.20122594 | 0.245859 | -0.818461 | 0.413094 | 0.829007487 | KIAA0141 | protein_coding | 48.21 | -0.016388152 | 0.016388152 | ENSG00000081791 | n.s. |
| ENSG00000066382 | 50.45928 | -0.234359956 | 0.310589 | -0.754566 | 0.45051 | 0.851309042 | MPPED2 | protein_coding | 40.01 | -0.01638475 | 0.01638475 | ENSG00000066382 | n.s. |
| ENSG00000123983 | 2895.626 | -0.205370492 | 0.254279 | -0.807658 | 0.419288 | 0.832304654 | ACSL3 | protein_coding | 39.46 | -0.016371659 | 0.016371659 | ENSG00000123983 | n.s. |
| ENSG00000128581 | 171.0167 | -0.235908311 | 0.313504 | -0.752489 | 0.451757 | 0.852320382 | IFT22 | protein_coding | 49.56 | -0.016371359 | 0.016371359 | ENSG00000128581 | n.s. |
| ENSG00000207609 | 10.30956 | -0.298091702 | 0.455158 | -0.654919 | 0.51252 | 0.881306695 | MIR491 | miRNA | 51.19 | -0.016357165 | 0.016357165 | ENSG00000207609 | n.s. |
| ENSG00000148408 | 805.0771 | -0.199969448 | 0.243537 | -0.821105 | 0.411587 | 0.828379538 | CACNA1B | protein_coding | 49.94 | -0.016351629 | 0.016351629 | ENSG00000148408 | n.s. |
| ENSG00000110104 | 287.4562 | -0.208624797 | 0.260692 | -0.800273 | 0.423553 | 0.835158421 | CCDC86 | protein_coding | 54.76 | -0.016320955 | 0.016320955 | ENSG00000110104 | n.s. |
| ENSG00000186162 | 135.3652 | -0.204439255 | 0.252487 | -0.809701 | 0.418112 | 0.832304654 | CIDECP | cribed_unprocessed_pseudc | 47.2 | -0.016297422 | 0.016297422 | ENSG00000186162 | n.s. |
| ENSG00000204560 | 662.5083 | -0.205377751 | 0.254748 | -0.8062 | 0.420127 | 0.83301531 | DHX16 | protein_coding | 49.99 | -0.016296112 | 0.016296112 | ENSG00000204560 | n.s. |
| ENSG00000125249 | 1163.115 | -0.225835922 | 0.294756 | -0.766179 | 0.44357 | 0.846931399 | RAP2A | protein_coding | 36.06 | -0.016294461 | 0.016294461 | ENSG00000125249 | n.s. |
| ENSG00000265470 | 2.557143 | -0.479783261 | 1.091997 | -0.439363 | 0.660398 | 0.924786541 | MIR548AQ | miRNA | 34.48 | -0.01629272 | 0.01629272 | ENSG00000265470 | n.s. |
| ENSG00000156709 | 443.7757 | -0.223521125 | 0.28997 | -0.770841 | 0.440801 | 0.845502102 | AIFM1 | protein_coding | 42.98 | -0.016291406 | 0.016291406 | ENSG00000156709 | n.s. |
| ENSG00000174684 | 265.3227 | 0.22675114 | 0.296376 | 0.7650803 | 0.444224 | 0.847626065 | B4GAT1 | protein_coding | 57.6 | 0.016279756 | 0.016279756 | ENSG00000174684 | n.s. |
| ENSG00000162104 | 563.8786 | -0.194613777 | 0.234078 | -0.831405 | 0.405745 | 0.824799944 | ADCY9 | protein_coding | 47.22 | -0.016279711 | 0.016279711 | ENSG00000162104 | n.s. |
| ENSG00000085433 | 844.0754 | -0.194570981 | 0.233331 | -0.833884 | 0.404346 | 0.824799944 | WDR47 | protein_coding | 41.17 | -0.01627613 | 0.01627613 | ENSG00000085433 | n.s. |
| ENSG00000272455 | 83.096 | -0.212453286 | 0.268201 | -0.792142 | 0.428278 | 0.838312087 | | lincRNA | 49.44 | -0.016272705 | 0.016272705 | ENSG00000272455 | n.s. |
| ENSG00000068120 | 728.166 | -0.208544901 | 0.261278 | -0.798172 | 0.424771 | 0.835658686 | COASY | protein_coding | 57.56 | -0.016260469 | 0.016260469 | ENSG00000068120 | n.s. |
| ENSG00000119979 | 331.5436 | -0.214612162 | 0.272047 | -0.78888 | 0.430182 | 0.839914098 | FAM45A | protein_coding | 45.15 | -0.016260118 | 0.016260118 | ENSG00000119979 | n.s. |
| ENSG00000163661 | 327.8432 | -0.211377929 | 0.266303 | -0.79375 | 0.427341 | 0.837692206 | PTX3 | protein_coding | 42.75 | -0.016258244 | 0.016258244 | ENSG00000163661 | n.s. |
| ENSG00000163041 | 2609.777 | -0.195431941 | 0.236079 | -0.827824 | 0.40777 | 0.826149803 | H3F3A | protein_coding | 44.56 | -0.016209359 | 0.016209359 | ENSG00000163041 | n.s. |
| ENSG00000115289 | 2320.137 | -0.192199662 | 0.22952 | -0.8374 | 0.402368 | 0.823848555 | PCGF1 | protein_coding | 56.33 | -0.016174104 | 0.016174104 | ENSG00000115289 | n.s. |
| ENSG00000099817 | 872.9417 | -0.222172932 | 0.288693 | -0.769583 | 0.441548 | 0.84569031 | POLR2E | protein_coding | 61.47 | -0.016171667 | 0.016171667 | ENSG00000099817 | n.s. |
| ENSG00000188002 | 153.3983 | -0.202815375 | 0.251235 | -0.807272 | 0.41951 | 0.832304654 | | transcribed_unprocessed_pseudc | 50.36 | -0.016167971 | 0.016167971 | ENSG00000188002 | n.s. |
| ENSG00000244615 | 16.28053 | -0.280167692 | 0.417004 | -0.671858 | 0.501674 | 0.875728424 | PSPC1P2 | cribed_unprocessed_pseudc | 33.35 | -0.016146219 | 0.016146219 | ENSG00000244615 | n.s. |

| | | | | | | | | | | | | | |
|---|---|---|---|---|---|---|---|---|---|---|---|---|---|
| ENSG00000155313 | 1532.608 | -0.205078276 | 0.255344 | -0.803144 | 0.421892 | 0.834256463 | USP25 | protein_coding | 35.41 | -0.016139746 | 0.016139746 | ENSG00000155313 | n.s. |
| ENSG00000101439 | 242.5748 | -0.218899039 | 0.282278 | -0.775474 | 0.43806 | 0.843950276 | CST3 | protein_coding | 51.65 | -0.016129169 | 0.016129169 | ENSG00000101439 | n.s. |
| ENSG00000136059 | 234.5944 | -0.216572331 | 0.276661 | -0.782806 | 0.433741 | 0.84244096 | VILL | protein_coding | 55.98 | -0.01612609 | 0.01612609 | ENSG00000136059 | n.s. |
| ENSG00000242294 | 239.5693 | 0.194185637 | 0.234415 | 0.8283841 | 0.407453 | 0.825999998 | STAG3L5P | cribed_unprocessed_pseudc | 47.29 | 0.016121283 | 0.016121283 | ENSG00000242294 | n.s. |
| ENSG00000128159 | 565.3194 | -0.201889572 | 0.249513 | -0.809136 | 0.418437 | 0.832304654 | TUBGCP6 | protein_coding | 56.93 | -0.016094168 | 0.016094168 | ENSG00000128159 | n.s. |
| ENSG00000157005 | 25.02151 | -0.256599554 | 0.360812 | -0.711173 | 0.476977 | 0.865616576 | SST | protein_coding | 48.46 | -0.016082232 | 0.016082232 | ENSG00000157005 | n.s. |
| ENSG00000148840 | 2129.725 | -0.181793726 | 0.211837 | -0.858176 | 0.390795 | 0.815710819 | PPRC1 | protein_coding | 51.99 | -0.01608216 | 0.01608216 | ENSG00000148840 | n.s. |
| ENSG00000164329 | 1034.645 | -0.206220667 | 0.258629 | -0.79736 | 0.425242 | 0.835658686 | PAPD4 | protein_coding | 36.28 | -0.016079246 | 0.016079246 | ENSG00000164329 | n.s. |
| ENSG00000258337 | 42.26701 | -0.256413293 | 0.357935 | -0.716368 | 0.473764 | 0.865616576 | | sense_intronic | 37.7 | -0.016070558 | 0.016070558 | ENSG00000258337 | n.s. |
| ENSG00000197885 | 342.0902 | -0.204110168 | 0.254112 | -0.80323 | 0.421842 | 0.834256463 | NKIRAS1 | protein_coding | 42.91 | -0.016063556 | 0.016063556 | ENSG00000197885 | n.s. |
| ENSG00000137845 | 3569.423 | -0.201611257 | 0.249878 | -0.806839 | 0.419759 | 0.832539843 | ADAM10 | protein_coding | 37.38 | -0.016047243 | 0.016047243 | ENSG00000137845 | n.s. |
| ENSG00000125753 | 136.7989 | -0.220308314 | 0.28663 | -0.768614 | 0.442122 | 0.845745639 | VASP | protein_coding | 54.67 | -0.016029684 | 0.016029684 | ENSG00000125753 | n.s. |
| ENSG00000170946 | 330.8515 | -0.213367824 | 0.271099 | -0.787049 | 0.431253 | 0.84116282 | DNAJC24 | protein_coding | 36.59 | -0.016028176 | 0.016028176 | ENSG00000170946 | n.s. |
| ENSG00000141542 | 311.09 | -0.23474333 | 0.314617 | -0.746125 | 0.455592 | 0.85453422 | RAB40B | protein_coding | 53.03 | -0.016026054 | 0.016026054 | ENSG00000141542 | n.s. |
| ENSG00000078674 | 5537.076 | -0.20319214 | 0.252706 | -0.804066 | 0.421359 | 0.834032319 | PCM1 | protein_coding | 36.54 | -0.016015019 | 0.016015019 | ENSG00000078674 | n.s. |
| ENSG00000086200 | 1321.807 | -0.21482954 | 0.273911 | -0.784304 | 0.432862 | 0.842286016 | IPO11 | protein_coding | 38.3 | -0.016013482 | 0.016013482 | ENSG00000086200 | n.s. |
| ENSG00000137817 | 2428.363 | -0.202375727 | 0.251331 | -0.805216 | 0.420695 | 0.833547859 | PARP6 | protein_coding | 43.94 | -0.016001739 | 0.016001739 | ENSG00000137817 | n.s. |
| ENSG00000120832 | 765.267 | -0.200633938 | 0.247925 | -0.809252 | 0.41837 | 0.832304654 | MTERF2 | protein_coding | 41.69 | -0.015994071 | 0.015994071 | ENSG00000120832 | n.s. |
| ENSG00000144895 | 3148.681 | 0.21865699 | 0.28286 | 0.7730217 | 0.43951 | 0.845174857 | EIF2A | protein_coding | 37.58 | 0.015973643 | 0.015973643 | ENSG00000144895 | n.s. |
| ENSG00000107185 | 443.8521 | -0.196080136 | 0.239397 | -0.819059 | 0.412752 | 0.829007487 | RGP1 | protein_coding | 52.62 | -0.01596907 | 0.01596907 | ENSG00000107185 | n.s. |
| ENSG00000131669 | 78.64763 | -0.226418606 | 0.297914 | -0.760013 | 0.447247 | 0.850164607 | NINJ1 | protein_coding | 59.68 | -0.015961828 | 0.015961828 | ENSG00000131669 | n.s. |
| ENSG00000064225 | 76.96557 | -0.219338679 | 0.285468 | -0.768348 | 0.442281 | 0.845745639 | ST3GAL6 | protein_coding | 38.88 | -0.015959133 | 0.015959133 | ENSG00000064225 | n.s. |
| ENSG00000180329 | 1802.025 | -0.206905004 | 0.260861 | -0.793163 | 0.427683 | 0.837692206 | CCDC43 | protein_coding | 39.85 | -0.015914207 | 0.015914207 | ENSG00000180329 | n.s. |
| ENSG00000231663 | 70.25327 | -0.227354209 | 0.300558 | -0.756441 | 0.449385 | 0.85118703 | | antisense | 48.54 | -0.015909111 | 0.015909111 | ENSG00000231663 | n.s. |
| ENSG00000154429 | 1361.849 | -0.215614527 | 0.277345 | -0.777424 | 0.436909 | 0.843772375 | CCSAP | protein_coding | 43.58 | -0.015906896 | 0.015906896 | ENSG00000154429 | n.s. |
| ENSG00000025796 | 2698.093 | -0.201969158 | 0.251253 | -0.803848 | 0.421485 | 0.834151325 | SEC63 | protein_coding | 39.82 | -0.015906113 | 0.015906113 | ENSG00000025796 | n.s. |
| ENSG00000130299 | 208.9977 | -0.213349387 | 0.272097 | -0.784094 | 0.432985 | 0.842286016 | GTPBP3 | protein_coding | 56.97 | -0.015903151 | 0.015903151 | ENSG00000130299 | n.s. |
| ENSG00000159479 | 214.6291 | -0.217602601 | 0.28186 | -0.772024 | 0.4401 | 0.845174857 | MED8 | protein_coding | 46.78 | -0.015896617 | 0.015896617 | ENSG00000159479 | n.s. |
| ENSG00000187741 | 458.4743 | -0.195181488 | 0.238614 | -0.81798 | 0.413368 | 0.829007487 | FANCA | protein_coding | 50.45 | -0.015895882 | 0.015895882 | ENSG00000187741 | n.s. |
| ENSG00000171444 | 4615.464 | 0.195770379 | 0.240347 | 0.8145307 | 0.415341 | 0.829668953 | MCC | protein_coding | 41.55 | 0.01587603 | 0.01587603 | ENSG00000171444 | n.s. |
| ENSG00000200406 | 6.633851 | 0.311872321 | 0.504952 | 0.6176271 | 0.536821 | 0.889412093 | SNORD114-23 | snoRNA | 45.83 | 0.015873356 | 0.015873356 | ENSG00000200406 | n.s. |
| ENSG00000143776 | 3176.298 | -0.205701843 | 0.258895 | -0.794538 | 0.426882 | 0.837228298 | CDC42BPA | protein_coding | 36.66 | -0.015871152 | 0.015871152 | ENSG00000143776 | n.s. |

| | | | | | | | | | | | | | |
|---|---|---|---|---|---|---|---|---|---|---|---|---|---|
| ENSG00000110700 | 10176.88 | 0.229382651 | 0.305327 | 0.7512692 | 0.452491 | 0.852853265 | RPS13 | protein_coding | 46.54 | 0.015856233 | 0.015856233 | ENSG00000110700 | n.s. |
| ENSG00000082213 | 706.5529 | -0.19886483 | 0.246352 | -0.807237 | 0.41953 | 0.832304654 | C5orf22 | protein_coding | 39.88 | -0.015853042 | 0.015853042 | ENSG00000082213 | n.s. |
| ENSG00000170222 | 214.5425 | 0.212975405 | 0.272504 | 0.7815498 | 0.434479 | 0.842514477 | ADPRM | protein_coding | 40.57 | 0.015850189 | 0.015850189 | ENSG00000170222 | n.s. |
| ENSG00000042429 | 2769.087 | 0.194389922 | 0.238111 | 0.8163851 | 0.41428 | 0.829007487 | MED17 | protein_coding | 39.79 | 0.015831416 | 0.015831416 | ENSG00000042429 | n.s. |
| ENSG00000119335 | 14035.75 | -0.203007987 | 0.254579 | -0.797427 | 0.425203 | 0.835658686 | SET | protein_coding | 49.47 | -0.01582875 | 0.01582875 | ENSG00000119335 | n.s. |
| ENSG00000222343 | 36.83463 | 0.25255805 | 0.356358 | 0.7087207 | 0.478498 | 0.865660698 | RN7SKP139 | misc_RNA | 48.81 | 0.015823343 | 0.015823343 | ENSG00000222343 | n.s. |
| ENSG00000228350 | 9.504819 | 0.299412671 | 0.467053 | 0.6410682 | 0.521478 | 0.885523197 | CELSR3-AS1 | antisense | 48.95 | 0.015809006 | 0.015809006 | ENSG00000228350 | n.s. |
| ENSG00000094804 | 2725.058 | -0.194093345 | 0.237907 | -0.815838 | 0.414593 | 0.829007487 | CDC6 | protein_coding | 43.75 | -0.015807262 | 0.015807262 | ENSG00000094804 | n.s. |
| ENSG00000161671 | 735.4438 | -0.227954374 | 0.303254 | -0.751694 | 0.452235 | 0.852602588 | EMC10 | protein_coding | 57.11 | -0.015786605 | 0.015786605 | ENSG00000161671 | n.s. |
| ENSG00000237732 | 243.429 | 0.188470206 | 0.225498 | 0.8357947 | 0.40327 | 0.824699011 | | transcribed_unprocessed_pseudc | 39.28 | 0.015775809 | 0.015775809 | ENSG00000237732 | n.s. |
| ENSG00000161914 | 45.04619 | -0.230698963 | 0.309054 | -0.746467 | 0.455385 | 0.85453422 | ZNF653 | protein_coding | 55.67 | -0.015749943 | 0.015749943 | ENSG00000161914 | n.s. |
| ENSG00000238961 | 35.42775 | 0.229357204 | 0.306328 | 0.7487309 | 0.454019 | 0.853769906 | SNORA47 | snoRNA | 60.87 | 0.015747472 | 0.015747472 | ENSG00000238961 | n.s. |
| ENSG00000200480 | 5.380005 | -0.418290148 | 0.844872 | -0.495093 | 0.620535 | 0.916981464 | SNORD114-28 | snoRNA | 47.22 | -0.015744208 | 0.015744208 | ENSG00000200480 | n.s. |
| ENSG00000103876 | 137.0783 | -0.224153274 | 0.29543 | -0.758737 | 0.44801 | 0.850715881 | FAH | protein_coding | 49.96 | -0.015739025 | 0.015739025 | ENSG00000103876 | n.s. |
| ENSG00000146574 | 367.293 | -0.211002628 | 0.26907 | -0.784193 | 0.432927 | 0.842286016 | CCZ1B | protein_coding | 46.69 | -0.015728222 | 0.015728222 | ENSG00000146574 | n.s. |
| ENSG00000176994 | 808.7906 | -0.181843496 | 0.214477 | -0.847845 | 0.396524 | 0.819495485 | SMCR8 | protein_coding | 52.68 | -0.015720995 | 0.015720995 | ENSG00000176994 | n.s. |
| ENSG00000178935 | 175.312 | -0.210852495 | 0.269649 | -0.781952 | 0.434243 | 0.84244096 | ZNF552 | protein_coding | 47.58 | -0.015700188 | 0.015700188 | ENSG00000178935 | n.s. |
| ENSG00000265932 | 28.39434 | 0.249178255 | 0.347363 | 0.7173423 | 0.473163 | 0.865034671 | MIR3146 | miRNA | 37.97 | 0.015689879 | 0.015689879 | ENSG00000265932 | n.s. |
| ENSG00000230068 | 23.08309 | 0.270171802 | 0.399911 | 0.675579 | 0.499308 | 0.874970095 | CDC42-IT1 | sense_intronic | 41.02 | 0.015671799 | 0.015671799 | ENSG00000230068 | n.s. |
| ENSG00000007968 | 445.6351 | -0.18981406 | 0.230242 | -0.824411 | 0.409706 | 0.827015612 | E2F2 | protein_coding | 51.66 | -0.015657058 | 0.015657058 | ENSG00000007968 | n.s. |
| ENSG00000262814 | 862.8079 | -0.212066396 | 0.272639 | -0.777828 | 0.436671 | 0.843772375 | MRPL12 | protein_coding | 57.67 | -0.015645134 | 0.015645134 | ENSG00000262814 | n.s. |
| ENSG00000141696 | 393.3499 | -0.214111312 | 0.277298 | -0.772136 | 0.440034 | 0.845174857 | P3H4 | protein_coding | 56.65 | -0.015641566 | 0.015641566 | ENSG00000141696 | n.s. |
| ENSG00000120647 | 987.4088 | -0.216288073 | 0.2821 | -0.766706 | 0.443256 | 0.846745122 | CCDC77 | protein_coding | 43.8 | -0.015626229 | 0.015626229 | ENSG00000120647 | n.s. |
| ENSG00000100425 | 1701.353 | -0.185302838 | 0.22113 | -0.837983 | 0.40204 | 0.823540971 | BRD1 | protein_coding | 54 | -0.01562377 | 0.01562377 | ENSG00000100425 | n.s. |
| ENSG00000178917 | 268.2959 | -0.195929843 | 0.242346 | -0.808471 | 0.418819 | 0.832304654 | ZNF852 | protein_coding | 42.79 | -0.015619072 | 0.015619072 | ENSG00000178917 | n.s. |
| ENSG00000126067 | 763.8375 | -0.211467522 | 0.271781 | -0.778082 | 0.436521 | 0.843772375 | PSMB2 | protein_coding | 41.1 | -0.015600952 | 0.015600952 | ENSG00000126067 | n.s. |
| ENSG00000259802 | 193.743 | 0.191810901 | 0.235265 | 0.8152977 | 0.414902 | 0.829267837 | | antisense | 54.16 | 0.01559522 | 0.01559522 | ENSG00000259802 | n.s. |
| ENSG00000271797 | 23.15097 | 0.281357615 | 0.428035 | 0.6573245 | 0.510972 | 0.880260889 | | antisense | 53.6 | 0.015584003 | 0.015584003 | ENSG00000271797 | n.s. |
| ENSG00000111832 | 1525.916 | -0.209463748 | 0.268443 | -0.780292 | 0.435219 | 0.842717361 | RWDD1 | protein_coding | 36.42 | -0.015566939 | 0.015566939 | ENSG00000111832 | n.s. |
| ENSG00000174173 | 1339.067 | 0.195105585 | 0.241434 | 0.8081116 | 0.419026 | 0.832304654 | TRMT10C | protein_coding | 41.24 | 0.015553364 | 0.015553364 | ENSG00000174173 | n.s. |
| ENSG00000219626 | 120.3536 | -0.197120455 | 0.245091 | -0.804273 | 0.421239 | 0.833913628 | FAM228B | protein_coding | 43.18 | -0.01554865 | 0.01554865 | ENSG00000219626 | n.s. |
| ENSG00000134809 | 422.1082 | -0.213305274 | 0.277094 | -0.769795 | 0.441422 | 0.84569031 | TIMM10 | protein_coding | 48.01 | -0.015526202 | 0.015526202 | ENSG00000134809 | n.s. |

| | | | | | | | | | | | | | |
|---|---|---|---|---|---|---|---|---|---|---|---|---|---|
| ENSG00000115548 | 2212.673 | 0.199112583 | 0.249696 | 0.7974197 | 0.425207 | 0.835658686 | KDM3A | protein_coding | 40.15 | 0.015525021 | 0.015525021 | ENSG00000115548 | n.s. |
| ENSG00000234022 | 29.28359 | 0.238889314 | 0.328196 | 0.7278862 | 0.466683 | 0.861088114 | | antisense | 39.59 | 0.015516436 | 0.015516436 | ENSG00000234022 | n.s. |
| ENSG00000148908 | 232.5452 | 0.246407772 | 0.343555 | 0.7172298 | 0.473232 | 0.865034671 | RGS10 | protein_coding | 49.42 | 0.015515431 | 0.015515431 | ENSG00000148908 | n.s. |
| ENSG00000130348 | 826.9495 | -0.220033877 | 0.289343 | -0.760461 | 0.446979 | 0.850164607 | QRSL1 | protein_coding | 39.53 | -0.015511724 | 0.015511724 | ENSG00000130348 | n.s. |
| ENSG00000180616 | 12533.46 | 0.207969887 | 0.265121 | 0.7844339 | 0.432786 | 0.842286016 | SSTR2 | protein_coding | 47.17 | 0.015502161 | 0.015502161 | ENSG00000180616 | n.s. |
| ENSG00000188167 | 199.5973 | 0.187560184 | 0.227345 | 0.825003 | 0.40937 | 0.826802059 | TMPPE | protein_coding | 49.95 | 0.015492181 | 0.015492181 | ENSG00000188167 | n.s. |
| ENSG00000132155 | 3021.346 | -0.194148995 | 0.239952 | -0.809116 | 0.418448 | 0.832304654 | RAF1 | protein_coding | 42.69 | -0.015477107 | 0.015477107 | ENSG00000132155 | n.s. |
| ENSG00000089280 | 3950.282 | -0.190036512 | 0.231808 | -0.819801 | 0.412329 | 0.829005481 | FUS | protein_coding | 46.87 | -0.015477067 | 0.015477067 | ENSG00000089280 | n.s. |
| ENSG00000164535 | 280.4898 | -0.201191866 | 0.253515 | -0.793609 | 0.427423 | 0.837692206 | DAGLB | protein_coding | 48.19 | -0.015474778 | 0.015474778 | ENSG00000164535 | n.s. |
| ENSG00000110011 | 78.47421 | 0.21929409 | 0.288149 | 0.7610444 | 0.446631 | 0.850164607 | DNAJC4 | protein_coding | 58.22 | 0.015459571 | 0.015459571 | ENSG00000110011 | n.s. |
| ENSG00000233864 | 578.4095 | -0.215231515 | 0.281365 | -0.764954 | 0.444299 | 0.847626065 | TTTY15 | lincRNA | 39.24 | -0.015452697 | 0.015452697 | ENSG00000233864 | n.s. |
| ENSG00000116918 | 1753.916 | -0.212322027 | 0.276329 | -0.768366 | 0.44227 | 0.845745639 | TSNAX | protein_coding | 37.66 | -0.0154486 | 0.0154486 | ENSG00000116918 | n.s. |
| ENSG00000283108 | 56.09454 | -0.225054218 | 0.300728 | -0.748364 | 0.454241 | 0.853840253 | | processed_transcript | 42.44 | -0.01544398 | 0.01544398 | ENSG00000283108 | n.s. |
| ENSG00000152944 | 1205.214 | 0.203064568 | 0.257134 | 0.7897232 | 0.429689 | 0.839380003 | MED21 | protein_coding | 39.33 | 0.01544131 | 0.01544131 | ENSG00000152944 | n.s. |
| ENSG00000131462 | 984.7017 | -0.195307338 | 0.242671 | -0.804824 | 0.420922 | 0.833570156 | TUBG1 | protein_coding | 53.1 | -0.015440577 | 0.015440577 | ENSG00000131462 | n.s. |
| ENSG00000087152 | 1330.391 | -0.203009323 | 0.257119 | -0.789554 | 0.429788 | 0.839380003 | ATXN7L3 | protein_coding | 59.15 | -0.015437109 | 0.015437109 | ENSG00000087152 | n.s. |
| ENSG00000146530 | 254.2366 | -0.211155015 | 0.27335 | -0.772471 | 0.439836 | 0.845174857 | VWDE | protein_coding | 33.94 | -0.015425598 | 0.015425598 | ENSG00000146530 | n.s. |
| ENSG00000171105 | 422.5486 | -0.198056782 | 0.24853 | -0.796912 | 0.425502 | 0.83593429 | INSR | protein_coding | 47.45 | -0.015414335 | 0.015414335 | ENSG00000171105 | n.s. |
| ENSG00000012817 | 368.612 | 0.189244543 | 0.231334 | 0.8180573 | 0.413324 | 0.829007487 | KDM5D | protein_coding | 38.66 | 0.015412368 | 0.015412368 | ENSG00000012817 | n.s. |
| ENSG00000161010 | 321.9558 | -0.21120907 | 0.273806 | -0.771383 | 0.44048 | 0.845502102 | MRNIP | protein_coding | 49.4 | -0.015394038 | 0.015394038 | ENSG00000161010 | n.s. |
| ENSG00000249115 | 424.2689 | -0.208255603 | 0.267488 | -0.778559 | 0.436239 | 0.84357804 | HAUS5 | protein_coding | 54.43 | -0.015384827 | 0.015384827 | ENSG00000249115 | n.s. |
| ENSG00000074696 | 7635.454 | -0.205560837 | 0.26169 | -0.785514 | 0.432152 | 0.84187161 | HACD3 | protein_coding | 42.55 | -0.015366523 | 0.015366523 | ENSG00000074696 | n.s. |
| ENSG00000070367 | 2696.141 | -0.214835014 | 0.281207 | -0.763976 | 0.444882 | 0.848321918 | EXOC5 | protein_coding | 35.97 | -0.015347666 | 0.015347666 | ENSG00000070367 | n.s. |
| ENSG00000125351 | 611.1577 | -0.210065016 | 0.271942 | -0.772462 | 0.439841 | 0.845174857 | UPF3B | protein_coding | 42.87 | -0.01534597 | 0.01534597 | ENSG00000125351 | n.s. |
| ENSG00000148300 | 434.6447 | -0.210313416 | 0.272819 | -0.77089 | 0.440772 | 0.845502102 | REXO4 | protein_coding | 51.72 | -0.015328758 | 0.015328758 | ENSG00000148300 | n.s. |
| ENSG00000080298 | 111.9592 | -0.220693623 | 0.293193 | -0.752724 | 0.451616 | 0.852242943 | RFX3 | protein_coding | 35 | -0.015324212 | 0.015324212 | ENSG00000080298 | n.s. |
| ENSG00000179743 | 115.4741 | 0.201374749 | 0.254726 | 0.7905546 | 0.429204 | 0.839274918 | | antisense | 50.48 | 0.015323764 | 0.015323764 | ENSG00000179743 | n.s. |
| ENSG00000170677 | 1235.247 | -0.210763424 | 0.274418 | -0.768038 | 0.442464 | 0.84586464 | SOCS6 | protein_coding | 39.2 | -0.015322318 | 0.015322318 | ENSG00000170677 | n.s. |
| ENSG00000186866 | 27.79222 | -0.252887905 | 0.362871 | -0.696909 | 0.48586 | 0.869936401 | POFUT2 | protein_coding | 52.95 | -0.015302878 | 0.015302878 | ENSG00000186866 | n.s. |
| ENSG00000275835 | 928.4275 | -0.206857845 | 0.265631 | -0.778743 | 0.436131 | 0.843485989 | TUBGCP5 | protein_coding | 43.36 | -0.015291372 | 0.015291372 | ENSG00000275835 | n.s. |
| ENSG00000149531 | 134.9773 | -0.225313108 | 0.302878 | -0.743907 | 0.456933 | 0.855435357 | FRG1BP | unprocessed_pseudogene | 35.24 | -0.015279113 | 0.015279113 | ENSG00000149531 | n.s. |
| ENSG00000112655 | 1893.507 | -0.191642802 | 0.237332 | -0.807489 | 0.419385 | 0.832304654 | PTK7 | protein_coding | 49.42 | -0.015277319 | 0.015277319 | ENSG00000112655 | n.s. |

| | | | | | | | | | | | | | |
|---|---|---|---|---|---|---|---|---|---|---|---|---|---|
| ENSG00000233560 | 135.4717 | -0.230362553 | 0.312984 | -0.73602 | 0.461718 | 0.858393198 | KRT8P39 | processed_pseudogene | 55.13 | -0.015276201 | 0.015276201 | ENSG00000233560 | n.s. |
| ENSG00000125484 | 2253.659 | -0.195373405 | 0.244518 | -0.799015 | 0.424282 | 0.835315601 | GTF3C4 | protein_coding | 42.28 | -0.015268316 | 0.015268316 | ENSG00000125484 | n.s. |
| ENSG00000153044 | 1201.202 | -0.227548587 | 0.307154 | -0.740829 | 0.458797 | 0.856995387 | CENPH | protein_coding | 42.65 | -0.015250651 | 0.015250651 | ENSG00000153044 | n.s. |
| ENSG00000108651 | 1274.128 | -0.206890163 | 0.266765 | -0.775551 | 0.438014 | 0.843950276 | UTP6 | protein_coding | 44.12 | -0.015244317 | 0.015244317 | ENSG00000108651 | n.s. |
| ENSG00000154222 | 101.4789 | -0.22467317 | 0.302096 | -0.743715 | 0.457049 | 0.855538367 | CC2D1B | protein_coding | 51.22 | -0.015223968 | 0.015223968 | ENSG00000154222 | n.s. |
| ENSG00000227910 | 39.70786 | 0.232800629 | 0.318275 | 0.731444 | 0.464508 | 0.860217499 | | transcribed_processed_pseudog | 51.45 | 0.015223235 | 0.015223235 | ENSG00000227910 | n.s. |
| ENSG00000188342 | 1291.498 | -0.217553573 | 0.287996 | -0.755405 | 0.450006 | 0.851201408 | GTF2F2 | protein_coding | 39.12 | -0.015221716 | 0.015221716 | ENSG00000188342 | n.s. |
| ENSG00000222588 | 25.71475 | -0.258721453 | 0.378247 | -0.684001 | 0.493974 | 0.873348456 | | snoRNA | 36.43 | -0.015216041 | 0.015216041 | ENSG00000222588 | n.s. |
| ENSG00000163904 | 993.1946 | -0.208150218 | 0.269581 | -0.772126 | 0.44004 | 0.845174857 | SENP2 | protein_coding | 42.86 | -0.015206088 | 0.015206088 | ENSG00000163904 | n.s. |
| ENSG00000174775 | 347.4406 | -0.208797144 | 0.271443 | -0.769213 | 0.441767 | 0.845692528 | HRAS | protein_coding | 69.01 | -0.015197823 | 0.015197823 | ENSG00000174775 | n.s. |
| ENSG00000179603 | 389.2622 | 0.20401915 | 0.260338 | 0.7836716 | 0.433233 | 0.842381919 | GRM8 | protein_coding | 37.51 | 0.015197583 | 0.015197583 | ENSG00000179603 | n.s. |
| ENSG00000143368 | 652.1259 | -0.190447088 | 0.235868 | -0.807431 | 0.419418 | 0.832304654 | SF3B4 | protein_coding | 48.41 | -0.015181999 | 0.015181999 | ENSG00000143368 | n.s. |
| ENSG00000132549 | 1102.949 | -0.197318835 | 0.248634 | -0.793612 | 0.427421 | 0.837692206 | VPS13B | protein_coding | 36.65 | -0.015176882 | 0.015176882 | ENSG00000132549 | n.s. |
| ENSG00000189339 | 407.9541 | -0.203942282 | 0.261284 | -0.780539 | 0.435074 | 0.842611157 | SLC35E2B | protein_coding | 53.52 | -0.015167758 | 0.015167758 | ENSG00000189339 | n.s. |
| ENSG00000130684 | 677.102 | -0.207971561 | 0.269808 | -0.770814 | 0.440817 | 0.845502102 | ZNF337 | protein_coding | 42.68 | -0.015158071 | 0.015158071 | ENSG00000130684 | n.s. |
| ENSG00000146072 | 344.2017 | -0.195268436 | 0.245325 | -0.795959 | 0.426056 | 0.836432113 | TNFRSF21 | protein_coding | 41.38 | -0.015146836 | 0.015146836 | ENSG00000146072 | n.s. |
| ENSG00000107262 | 1882.241 | -0.193081509 | 0.240691 | -0.802196 | 0.42244 | 0.834747118 | BAG1 | protein_coding | 46.54 | -0.015146293 | 0.015146293 | ENSG00000107262 | n.s. |
| ENSG00000113282 | 1967.28 | -0.203350254 | 0.26004 | -0.781997 | 0.434216 | 0.84244096 | CLINT1 | protein_coding | 37.77 | -0.015141567 | 0.015141567 | ENSG00000113282 | n.s. |
| ENSG00000138802 | 1008.27 | -0.19911074 | 0.252056 | -0.789946 | 0.42956 | 0.839380003 | SEC24B | protein_coding | 37.89 | -0.015140656 | 0.015140656 | ENSG00000138802 | n.s. |
| ENSG00000142453 | 472.4956 | -0.198231709 | 0.250469 | -0.791442 | 0.428686 | 0.838757697 | CARM1 | protein_coding | 54.32 | -0.015137663 | 0.015137663 | ENSG00000142453 | n.s. |
| ENSG00000232065 | 20.9134 | 0.267181193 | 0.402584 | 0.6636655 | 0.506904 | 0.877705509 | LINC01063 | antisense | 50.92 | 0.015136129 | 0.015136129 | ENSG00000232065 | n.s. |
| ENSG00000138442 | 2409.109 | -0.203493664 | 0.260613 | -0.780828 | 0.434904 | 0.842611157 | WDR12 | protein_coding | 39.21 | -0.015134393 | 0.015134393 | ENSG00000138442 | n.s. |
| ENSG00000153443 | 254.7085 | 0.19714618 | 0.248738 | 0.7925871 | 0.428018 | 0.838039772 | UBALD1 | protein_coding | 64.7 | 0.015128085 | 0.015128085 | ENSG00000153443 | n.s. |
| ENSG00000141367 | 8818.767 | -0.203688852 | 0.261311 | -0.779489 | 0.435692 | 0.843067025 | CLTC | protein_coding | 38.54 | -0.015101063 | 0.015101063 | ENSG00000141367 | n.s. |
| ENSG00000260671 | 10.48164 | -0.302464496 | 0.504659 | -0.599344 | 0.548943 | 0.891406027 | | sense_intronic | 50.67 | -0.015100369 | 0.015100369 | ENSG00000260671 | n.s. |
| ENSG00000102543 | 157.6506 | -0.206573249 | 0.267473 | -0.772314 | 0.439929 | 0.845174857 | CDADC1 | protein_coding | 38.43 | -0.015090885 | 0.015090885 | ENSG00000102543 | n.s. |
| ENSG00000144647 | 216.4455 | -0.219696945 | 0.293386 | -0.748834 | 0.453958 | 0.853769906 | POMGNT2 | protein_coding | 47.68 | -0.015084207 | 0.015084207 | ENSG00000144647 | n.s. |
| ENSG00000142669 | 146.0643 | -0.218738457 | 0.291575 | -0.750196 | 0.453136 | 0.853222234 | SH3BGRL3 | protein_coding | 61.74 | -0.015079356 | 0.015079356 | ENSG00000142669 | n.s. |
| ENSG00000248333 | 218.3256 | -0.20164809 | 0.256662 | -0.785655 | 0.43207 | 0.841828575 | CDK11B | protein_coding | 51.91 | -0.015078505 | 0.015078505 | ENSG00000248333 | n.s. |
| ENSG00000170445 | 1543.575 | -0.206882879 | 0.268859 | -0.769484 | 0.441606 | 0.84569031 | HARS | protein_coding | 46.01 | -0.015058725 | 0.015058725 | ENSG00000170445 | n.s. |
| ENSG00000116809 | 250.9888 | -0.202121701 | 0.258603 | -0.781589 | 0.434456 | 0.842514477 | ZBTB17 | protein_coding | 51.22 | -0.015042428 | 0.015042428 | ENSG00000116809 | n.s. |
| ENSG00000189136 | 181.094 | 0.194909392 | 0.245306 | 0.7945552 | 0.426872 | 0.837228298 | UBE2Q2P1 | cribed_unprocessed_pseudc | 39.14 | 0.015038449 | 0.015038449 | ENSG00000189136 | n.s. |

| | | | | | | | | | | | | | |
|---|---|---|---|---|---|---|---|---|---|---|---|---|---|
| ENSG00000183762 | 1786.551 | -0.207756776 | 0.270862 | -0.76702 | 0.44307 | 0.846672751 | KREMEN1 | protein_coding | 45.49 | -0.015017577 | 0.015017577 | ENSG00000183762 | n.s. |
| ENSG00000206676 | 9.368629 | -0.285925785 | 0.448762 | -0.637144 | 0.524031 | 0.886111278 | | misc_RNA | 38.83 | -0.015014459 | 0.015014459 | ENSG00000206676 | n.s. |
| ENSG00000151366 | 1690.37 | -0.232715363 | 0.321402 | -0.724064 | 0.469026 | 0.861966035 | NDUFC2 | protein_coding | 44.63 | -0.015012432 | 0.015012432 | ENSG00000151366 | n.s. |
| ENSG00000158615 | 4514.037 | 0.181663643 | 0.220114 | 0.8253179 | 0.409191 | 0.826802059 | PPP1R15B | protein_coding | 40.44 | 0.015005135 | 0.015005135 | ENSG00000158615 | n.s. |
| ENSG00000122882 | 891.9168 | -0.200606634 | 0.255236 | -0.785965 | 0.431888 | 0.841828071 | ECD | protein_coding | 39.47 | -0.015000681 | 0.015000681 | ENSG00000122882 | n.s. |
| ENSG00000109079 | 269.8593 | -0.219284817 | 0.293549 | -0.747012 | 0.455056 | 0.854433778 | TNFAIP1 | protein_coding | 52.61 | -0.014981888 | 0.014981888 | ENSG00000109079 | n.s. |
| ENSG00000175198 | 267.0217 | -0.219038952 | 0.293015 | -0.747535 | 0.454741 | 0.854318522 | PCCA | protein_coding | 39.84 | -0.014977922 | 0.014977922 | ENSG00000175198 | n.s. |
| ENSG00000116874 | 435.1429 | -0.203248458 | 0.26217 | -0.775254 | 0.43819 | 0.843950276 | WARS2 | protein_coding | 38.1 | -0.014975985 | 0.014975985 | ENSG00000116874 | n.s. |
| ENSG00000121964 | 115.745 | -0.21392317 | 0.282813 | -0.756412 | 0.449402 | 0.85118703 | GTDC1 | protein_coding | 36.87 | -0.014969274 | 0.014969274 | ENSG00000121964 | n.s. |
| ENSG00000258450 | 54.69882 | -0.215320122 | 0.286023 | -0.752808 | 0.451565 | 0.852242943 | | sense_intronic | 50.17 | -0.014951094 | 0.014951094 | ENSG00000258450 | n.s. |
| ENSG00000149923 | 654.369 | -0.224459066 | 0.304478 | -0.737192 | 0.461006 | 0.857869611 | PPP4C | protein_coding | 54.46 | -0.014944196 | 0.014944196 | ENSG00000149923 | n.s. |
| ENSG00000196470 | 1237.305 | 0.207885661 | 0.271612 | 0.7653776 | 0.444047 | 0.847609304 | SIAH1 | protein_coding | 45.2 | 0.014927081 | 0.014927081 | ENSG00000196470 | n.s. |
| ENSG00000103197 | 46.26604 | -0.238048648 | 0.332641 | -0.715632 | 0.474218 | 0.865616576 | TSC2 | protein_coding | 60.06 | -0.014919565 | 0.014919565 | ENSG00000103197 | n.s. |
| ENSG00000030066 | 2271.672 | -0.213011571 | 0.281896 | -0.755639 | 0.449865 | 0.85118703 | NUP160 | protein_coding | 41.86 | -0.014905485 | 0.014905485 | ENSG00000030066 | n.s. |
| ENSG00000167641 | 365.6304 | -0.229757197 | 0.316607 | -0.725687 | 0.468031 | 0.861253285 | PPP1R14A | protein_coding | 58.39 | -0.014904145 | 0.014904145 | ENSG00000167641 | n.s. |
| ENSG00000234741 | 19172.01 | 0.173769313 | 0.205991 | 0.8435769 | 0.398906 | 0.820802709 | GAS5 | processed_transcript | 46.16 | 0.014902668 | 0.014902668 | ENSG00000234741 | n.s. |
| ENSG00000075240 | 355.7184 | -0.204100735 | 0.26458 | -0.771414 | 0.440462 | 0.845502102 | GRAMD4 | protein_coding | 57.68 | -0.014875945 | 0.014875945 | ENSG00000075240 | n.s. |
| ENSG00000277634 | 17.05059 | -0.266765337 | 0.404317 | -0.659793 | 0.509387 | 0.87951488 | | misc_RNA | 38.71 | -0.014873984 | 0.014873984 | ENSG00000277634 | n.s. |
| ENSG00000261766 | 21.95112 | -0.261882392 | 0.394308 | -0.664157 | 0.50659 | 0.877502299 | | antisense | 53.53 | -0.014862281 | 0.014862281 | ENSG00000261766 | n.s. |
| ENSG00000109084 | 2220.349 | -0.193170729 | 0.243532 | -0.793206 | 0.427658 | 0.837692206 | TMEM97 | protein_coding | 44.52 | -0.014857828 | 0.014857828 | ENSG00000109084 | n.s. |
| ENSG00000240694 | 1326.471 | -0.210701571 | 0.277066 | -0.760475 | 0.446971 | 0.850164607 | PNMA2 | protein_coding | 48.66 | -0.014853824 | 0.014853824 | ENSG00000240694 | n.s. |
| ENSG00000277452 | 15.50062 | 0.261727605 | 0.394172 | 0.663993 | 0.506695 | 0.877560859 | RN7SL473P | misc_RNA | 52.84 | 0.014845911 | 0.014845911 | ENSG00000277452 | n.s. |
| ENSG00000156482 | 12647.95 | 0.22980293 | 0.317275 | 0.7243014 | 0.468881 | 0.861812061 | RPL30 | protein_coding | 42.56 | 0.014842381 | 0.014842381 | ENSG00000156482 | n.s. |
| ENSG00000131778 | 1974.552 | -0.199094013 | 0.253721 | -0.784697 | 0.432631 | 0.842286016 | CHD1L | protein_coding | 40.27 | -0.014840549 | 0.014840549 | ENSG00000131778 | n.s. |
| ENSG00000124787 | 201.8764 | -0.210304549 | 0.276676 | -0.760112 | 0.447188 | 0.850164607 | RPP40 | protein_coding | 43.41 | -0.014825835 | 0.014825835 | ENSG00000124787 | n.s. |
| ENSG00000261002 | 22.1112 | 0.298617611 | 0.500121 | 0.5970908 | 0.550447 | 0.89197749 | | antisense | 38.94 | 0.014825201 | 0.014825201 | ENSG00000261002 | n.s. |
| ENSG00000090612 | 2562.928 | -0.21678825 | 0.290241 | -0.746925 | 0.455109 | 0.854433778 | ZNF268 | protein_coding | 41.98 | -0.014811318 | 0.014811318 | ENSG00000090612 | n.s. |
| ENSG00000280280 | 22.36909 | 0.257744973 | 0.386269 | 0.6672678 | 0.504601 | 0.876368641 | | TEC | 41.07 | 0.014772182 | 0.014772182 | ENSG00000280280 | n.s. |
| ENSG00000083123 | 682.9839 | -0.216358052 | 0.290087 | -0.74584 | 0.455764 | 0.85453422 | BCKDHB | protein_coding | 36.82 | -0.014770881 | 0.014770881 | ENSG00000083123 | n.s. |
| ENSG00000175449 | 126.1309 | 0.195857461 | 0.248496 | 0.7881718 | 0.430596 | 0.840604437 | RFESD | protein_coding | 37.57 | 0.014769282 | 0.014769282 | ENSG00000175449 | n.s. |
| ENSG00000139625 | 77.1996 | -0.216362039 | 0.290278 | -0.74536 | 0.456054 | 0.85459484 | MAP3K12 | protein_coding | 52.61 | -0.014764488 | 0.014764488 | ENSG00000139625 | n.s. |
| ENSG00000145375 | 1189.406 | -0.189790569 | 0.2383 | -0.796436 | 0.425779 | 0.836005799 | SPATA5 | protein_coding | 37.39 | -0.014763943 | 0.014763943 | ENSG00000145375 | n.s. |

| | | | | | | | | | | | | | |
|---|---|---|---|---|---|---|---|---|---|---|---|---|---|
| ENSG00000258796 | 32.96589 | -0.251218161 | 0.36777 | -0.683085 | 0.494553 | 0.873443478 | | processed_pseudogene | 37.17 | -0.014762884 | 0.014762884 | ENSG00000258796 | n.s. |
| ENSG00000077522 | 39.69017 | -0.251922061 | 0.370299 | -0.680321 | 0.496302 | 0.873799463 | ACTN2 | protein_coding | 43.06 | -0.014759667 | 0.014759667 | ENSG00000077522 | n.s. |
| ENSG00000227583 | 37.20641 | -0.235285734 | 0.328598 | -0.716029 | 0.473974 | 0.865616576 | RPS3AP37 | processed_pseudogene | 39.6 | -0.014746401 | 0.014746401 | ENSG00000227583 | n.s. |
| ENSG00000090372 | 1090.389 | -0.193273323 | 0.244329 | -0.791037 | 0.428922 | 0.839101451 | STRN4 | protein_coding | 57.35 | -0.01472463 | 0.01472463 | ENSG00000090372 | n.s. |
| ENSG00000176597 | 500.2188 | -0.219389402 | 0.29619 | -0.740705 | 0.458872 | 0.856995387 | B3GNT5 | protein_coding | 42.15 | -0.01470381 | 0.01470381 | ENSG00000176597 | n.s. |
| ENSG00000138074 | 1078.502 | -0.197845251 | 0.253603 | -0.780137 | 0.43531 | 0.842717361 | SLC5A6 | protein_coding | 53.48 | -0.014703475 | 0.014703475 | ENSG00000138074 | n.s. |
| ENSG00000120868 | 235.4408 | -0.201900817 | 0.262435 | -0.769337 | 0.441693 | 0.84569031 | APAF1 | protein_coding | 39.48 | -0.014696087 | 0.014696087 | ENSG00000120868 | n.s. |
| ENSG00000110422 | 832.535 | -0.197563765 | 0.253093 | -0.780598 | 0.435039 | 0.842611157 | HIPK3 | protein_coding | 38.83 | -0.014693369 | 0.014693369 | ENSG00000110422 | n.s. |
| ENSG00000170089 | 295.4243 | -0.226190488 | 0.311727 | -0.725604 | 0.468082 | 0.861253285 | | transcribed_unprocessed_pseudc | 44.8 | -0.014672775 | 0.014672775 | ENSG00000170089 | n.s. |
| ENSG00000182195 | 64.00324 | -0.220668183 | 0.300127 | -0.73525 | 0.462187 | 0.858395846 | LDOC1 | protein_coding | 57.35 | -0.014633035 | 0.014633035 | ENSG00000182195 | n.s. |
| ENSG00000181555 | 2519.445 | -0.190166447 | 0.239747 | -0.793195 | 0.427664 | 0.837692206 | SETD2 | protein_coding | 40.23 | -0.014626752 | 0.014626752 | ENSG00000181555 | n.s. |
| ENSG00000188559 | 511.892 | -0.198259261 | 0.255211 | -0.776844 | 0.437251 | 0.843772375 | RALGAPA2 | protein_coding | 40.42 | -0.014626517 | 0.014626517 | ENSG00000188559 | n.s. |
| ENSG00000105755 | 104.9416 | 0.248636055 | 0.363636 | 0.683749 | 0.494134 | 0.873348456 | ETHE1 | protein_coding | 48.58 | 0.014622894 | 0.014622894 | ENSG00000105755 | n.s. |
| ENSG00000215252 | 950.6304 | -0.202283274 | 0.263957 | -0.766351 | 0.443468 | 0.846852465 | GOLGA8B | protein_coding | 47.09 | -0.014603283 | 0.014603283 | ENSG00000215252 | n.s. |
| ENSG00000111445 | 1853.557 | -0.202005267 | 0.263325 | -0.767134 | 0.443002 | 0.846659901 | RFC5 | protein_coding | 45.74 | -0.014603164 | 0.014603164 | ENSG00000111445 | n.s. |
| ENSG00000173409 | 356.1448 | -0.233080003 | 0.328771 | -0.708944 | 0.478359 | 0.865660698 | ARV1 | protein_coding | 41.75 | -0.014602998 | 0.014602998 | ENSG00000173409 | n.s. |
| ENSG00000199942 | 5.234043 | 0.315134437 | 0.555222 | 0.5675823 | 0.570319 | 0.898818733 | SNORD114-19 | snoRNA | 40 | 0.014599512 | 0.014599512 | ENSG00000199942 | n.s. |
| ENSG00000229204 | 15.93104 | -0.285946178 | 0.461168 | -0.620048 | 0.535226 | 0.88909364 | PTGES3P3 | processed_pseudogene | 37.76 | -0.014598266 | 0.014598266 | ENSG00000229204 | n.s. |
| ENSG00000240823 | 10.66451 | -0.277882382 | 0.435395 | -0.63823 | 0.523324 | 0.886079926 | RN7SL23P | misc_RNA | 57.09 | -0.014596356 | 0.014596356 | ENSG00000240823 | n.s. |
| ENSG00000224348 | 11.88629 | -0.276412853 | 0.431105 | -0.641173 | 0.52141 | 0.885523197 | | processed_pseudogene | 59.41 | -0.014594615 | 0.014594615 | ENSG00000224348 | n.s. |
| ENSG00000121579 | 3308.987 | -0.211594729 | 0.282215 | -0.749764 | 0.453397 | 0.853222234 | NAA50 | protein_coding | 37.41 | -0.014586883 | 0.014586883 | ENSG00000121579 | n.s. |
| ENSG00000116062 | 6407.19 | -0.201839016 | 0.263281 | -0.766629 | 0.443302 | 0.846745122 | MSH6 | protein_coding | 44.65 | -0.014582323 | 0.014582323 | ENSG00000116062 | n.s. |
| ENSG00000198561 | 1468.104 | -0.197641828 | 0.25428 | -0.77726 | 0.437005 | 0.843772375 | CTNND1 | protein_coding | 42.81 | -0.014580966 | 0.014580966 | ENSG00000198561 | n.s. |
| ENSG00000075945 | 500.7375 | -0.214802762 | 0.288627 | -0.744223 | 0.456741 | 0.855307411 | KIFAP3 | protein_coding | 34.27 | -0.014580331 | 0.014580331 | ENSG00000075945 | n.s. |
| ENSG00000043355 | 1831.423 | -0.164666698 | 0.191612 | -0.859374 | 0.390134 | 0.815710819 | ZIC2 | protein_coding | 56.6 | -0.014567038 | 0.014567038 | ENSG00000043355 | n.s. |
| ENSG00000273355 | 19.23173 | -0.275495702 | 0.428757 | -0.642545 | 0.520519 | 0.885417361 | | lincRNA | 38.92 | -0.01456049 | 0.01456049 | ENSG00000273355 | n.s. |
| ENSG00000110436 | 612.869 | -0.217957206 | 0.295121 | -0.738536 | 0.460189 | 0.857496121 | SLC1A2 | protein_coding | 42.2 | -0.014552531 | 0.014552531 | ENSG00000110436 | n.s. |
| ENSG00000174080 | 348.8762 | 0.219387182 | 0.298643 | 0.7346136 | 0.462575 | 0.858395846 | CTSF | protein_coding | 58.67 | 0.014548089 | 0.014548089 | ENSG00000174080 | n.s. |
| ENSG00000153956 | 7151.99 | -0.20632765 | 0.271764 | -0.759216 | 0.447724 | 0.850405175 | CACNA2D1 | protein_coding | 34.98 | -0.014520124 | 0.014520124 | ENSG00000153956 | n.s. |
| ENSG00000168795 | 800.8738 | -0.206722258 | 0.272484 | -0.758658 | 0.448057 | 0.850715881 | ZBTB5 | protein_coding | 44.23 | -0.014515098 | 0.014515098 | ENSG00000168795 | n.s. |
| ENSG00000102554 | 29.39268 | 0.262404577 | 0.399638 | 0.6566057 | 0.511434 | 0.880578023 | KLF5 | protein_coding | 38.99 | 0.014493171 | 0.014493171 | ENSG00000102554 | n.s. |
| ENSG00000135931 | 96.24149 | -0.228198938 | 0.317442 | -0.718868 | 0.472222 | 0.863962245 | ARMC9 | protein_coding | 45.69 | -0.014491827 | 0.014491827 | ENSG00000135931 | n.s. |

| | | | | | | | | | | | | | |
|---|---|---|---|---|---|---|---|---|---|---|---|---|---|
| ENSG00000198218 | 1150.975 | -0.194592626 | 0.248765 | -0.782236 | 0.434076 | 0.84244096 | QRICH1 | protein_coding | 46.08 | -0.014489469 | 0.014489469 | ENSG00000198218 | n.s. |
| ENSG00000169826 | 529.9196 | -0.198931773 | 0.258506 | -0.769545 | 0.44157 | 0.84569031 | CSGALNACT2 | protein_coding | 38.88 | -0.014479974 | 0.014479974 | ENSG00000169826 | n.s. |
| ENSG00000130731 | 356.5538 | -0.222515659 | 0.305052 | -0.729436 | 0.465735 | 0.860954429 | METTL26 | protein_coding | 67.84 | -0.014467932 | 0.014467932 | ENSG00000130731 | n.s. |
| ENSG00000213380 | 283.4245 | -0.18501189 | 0.231399 | -0.799535 | 0.42398 | 0.835315601 | COG8 | protein_coding | 52.29 | -0.01445857 | 0.01445857 | ENSG00000213380 | n.s. |
| ENSG00000173272 | 1133.161 | -0.215462395 | 0.2904 | -0.74195 | 0.458118 | 0.856847578 | MZT2A | protein_coding | 51.73 | -0.014456757 | 0.014456757 | ENSG00000173272 | n.s. |
| ENSG00000252282 | 3.69299 | -0.348283797 | 0.666968 | -0.52219 | 0.601538 | 0.908868121 | RNU6-1089P | snRNA | 43.27 | -0.014453474 | 0.014453474 | ENSG00000252282 | n.s. |
| ENSG00000095002 | 3046.961 | -0.207900938 | 0.276081 | -0.753043 | 0.451424 | 0.852169954 | MSH2 | protein_coding | 44.66 | -0.014443664 | 0.014443664 | ENSG00000095002 | n.s. |
| ENSG00000197114 | 98.39978 | -0.209040248 | 0.278309 | -0.751109 | 0.452587 | 0.852919747 | ZGPAT | protein_coding | 49.61 | -0.014442973 | 0.014442973 | ENSG00000197114 | n.s. |
| ENSG00000082898 | 9472.186 | 0.204485945 | 0.268976 | 0.76024 | 0.447111 | 0.850164607 | XPO1 | protein_coding | 40.9 | 0.014415641 | 0.014415641 | ENSG00000082898 | n.s. |
| ENSG00000115539 | 555.3728 | -0.197640161 | 0.256678 | -0.769993 | 0.441304 | 0.84569031 | PDCL3 | protein_coding | 46.99 | -0.01438596 | 0.01438596 | ENSG00000115539 | n.s. |
| ENSG00000054356 | 594.7873 | -0.194892671 | 0.250964 | -0.776577 | 0.437408 | 0.843772375 | PTPRN | protein_coding | 53.59 | -0.014378148 | 0.014378148 | ENSG00000054356 | n.s. |
| ENSG00000125375 | 954.1515 | -0.205390979 | 0.271157 | -0.757463 | 0.448773 | 0.85118703 | ATP5S | protein_coding | 40.64 | -0.014372234 | 0.014372234 | ENSG00000125375 | n.s. |
| ENSG00000145604 | 3369.861 | -0.197459552 | 0.256916 | -0.768576 | 0.442145 | 0.845745639 | SKP2 | protein_coding | 39.76 | -0.014367203 | 0.014367203 | ENSG00000145604 | n.s. |
| ENSG00000101596 | 4704.431 | -0.195548839 | 0.252621 | -0.77408 | 0.438883 | 0.844468612 | SMCHD1 | protein_coding | 36.8 | -0.014356509 | 0.014356509 | ENSG00000101596 | n.s. |
| ENSG00000147592 | 184.9132 | -0.220990615 | 0.304183 | -0.726505 | 0.467529 | 0.861088114 | LACTB2 | protein_coding | 38.38 | -0.014353872 | 0.014353872 | ENSG00000147592 | n.s. |
| ENSG00000162604 | 989.5254 | -0.208133862 | 0.277318 | -0.750524 | 0.452939 | 0.853222234 | TM2D1 | protein_coding | 39.63 | -0.014348298 | 0.014348298 | ENSG00000162604 | n.s. |
| ENSG00000277972 | 142.9539 | -0.213973199 | 0.288747 | -0.74104 | 0.458669 | 0.856995387 | CISD3 | protein_coding | 56.94 | -0.014340808 | 0.014340808 | ENSG00000277972 | n.s. |
| ENSG00000056998 | 202.2078 | -0.21177736 | 0.28489 | -0.743364 | 0.457261 | 0.85570529 | GYG2 | protein_coding | 43.67 | -0.014332199 | 0.014332199 | ENSG00000056998 | n.s. |
| ENSG00000150760 | 1367.874 | -0.195165057 | 0.252082 | -0.774211 | 0.438806 | 0.84443611 | DOCK1 | protein_coding | 45.16 | -0.014331595 | 0.014331595 | ENSG00000150760 | n.s. |
| ENSG00000115137 | 137.7679 | 0.192451958 | 0.246466 | 0.7808465 | 0.434893 | 0.842611157 | DNAJC27 | protein_coding | 40.71 | 0.01431319 | 0.01431319 | ENSG00000115137 | n.s. |
| ENSG00000273329 | 430.0515 | -0.171053512 | 0.205287 | -0.833242 | 0.404708 | 0.824799944 | | lincRNA | 46.42 | -0.014308862 | 0.014308862 | ENSG00000273329 | n.s. |
| ENSG00000174469 | 14.05621 | -0.286464975 | 0.47602 | -0.601792 | 0.547312 | 0.891406027 | CNTNAP2 | protein_coding | 37.52 | -0.014301602 | 0.014301602 | ENSG00000174469 | n.s. |
| ENSG00000221914 | 1976.939 | -0.192139739 | 0.245805 | -0.781676 | 0.434405 | 0.842514477 | PPP2R2A | protein_coding | 37.61 | -0.014299544 | 0.014299544 | ENSG00000221914 | n.s. |
| ENSG00000108588 | 3131.234 | -0.198764086 | 0.259594 | -0.765674 | 0.44387 | 0.847389065 | CCDC47 | protein_coding | 42.02 | -0.014294546 | 0.014294546 | ENSG00000108588 | n.s. |
| ENSG00000237380 | 175.1906 | -0.224242607 | 0.31173 | -0.719349 | 0.471926 | 0.863532918 | HOXD-AS2 | antisense | 51.9 | -0.014288986 | 0.014288986 | ENSG00000237380 | n.s. |
| ENSG00000164169 | 382.7221 | -0.194339277 | 0.250974 | -0.774341 | 0.438729 | 0.84443611 | PRMT9 | protein_coding | 39.69 | -0.014270956 | 0.014270956 | ENSG00000164169 | n.s. |
| ENSG00000168612 | 168.707 | -0.196039627 | 0.254607 | -0.769969 | 0.441318 | 0.84569031 | ZSWIM1 | protein_coding | 50.32 | -0.014269459 | 0.014269459 | ENSG00000168612 | n.s. |
| ENSG00000113196 | 1583.786 | -0.176193822 | 0.216508 | -0.813796 | 0.415762 | 0.830032752 | HAND1 | protein_coding | 56.03 | -0.014254921 | 0.014254921 | ENSG00000113196 | n.s. |
| ENSG00000162434 | 1438.944 | -0.178807677 | 0.221439 | -0.807479 | 0.41939 | 0.832304654 | JAK1 | protein_coding | 42.69 | -0.014254133 | 0.014254133 | ENSG00000162434 | n.s. |
| ENSG00000252892 | 26.21161 | -0.244925515 | 0.36205 | -0.676497 | 0.498725 | 0.874768485 | RNU6-548P | snRNA | 46.96 | -0.014231855 | 0.014231855 | ENSG00000252892 | n.s. |
| ENSG00000162873 | 141.1353 | -0.214593205 | 0.292134 | -0.734571 | 0.462601 | 0.858395846 | KLHDC8A | protein_coding | 50.88 | -0.014230189 | 0.014230189 | ENSG00000162873 | n.s. |
| ENSG00000118939 | 107.4537 | -0.227006264 | 0.317374 | -0.715264 | 0.474446 | 0.865616576 | UCHL3 | protein_coding | 35.78 | -0.01422749 | 0.01422749 | ENSG00000118939 | n.s. |

| | | | | | | | | | | | | | |
|---|---|---|---|---|---|---|---|---|---|---|---|---|---|
| ENSG00000120733 | 1777.896 | -0.188805743 | 0.239647 | -0.78785 | 0.430785 | 0.840854265 | KDM3B | protein_coding | 43.18 | -0.014213158 | 0.014213158 | ENSG00000120733 | n.s. |
| ENSG00000221823 | 1297.487 | -0.206147779 | 0.275051 | -0.749488 | 0.453563 | 0.853257558 | PPP3R1 | protein_coding | 37.41 | -0.014207675 | 0.014207675 | ENSG00000221823 | n.s. |
| ENSG00000135999 | 2361.666 | -0.195280777 | 0.254214 | -0.768174 | 0.442384 | 0.845827195 | EPC2 | protein_coding | 37.09 | -0.014200497 | 0.014200497 | ENSG00000135999 | n.s. |
| ENSG00000199053 | 3.559982 | 0.434022524 | 1.002693 | 0.4328568 | 0.665119 | 0.927447429 | MIR324 | miRNA | 61.45 | 0.01419718 | 0.01419718 | ENSG00000199053 | n.s. |
| ENSG00000122490 | 296.2246 | -0.212644236 | 0.288082 | -0.738139 | 0.46043 | 0.857551846 | PQLC1 | protein_coding | 52.94 | -0.014191794 | 0.014191794 | ENSG00000122490 | n.s. |
| ENSG00000159335 | 1809.908 | -0.207486789 | 0.277529 | -0.747621 | 0.454689 | 0.854318522 | PTMS | protein_coding | 61.84 | -0.014187984 | 0.014187984 | ENSG00000159335 | n.s. |
| ENSG00000102900 | 1493.554 | -0.206437063 | 0.275705 | -0.748761 | 0.454001 | 0.853769906 | NUP93 | protein_coding | 42.43 | -0.014173795 | 0.014173795 | ENSG00000102900 | n.s. |
| ENSG00000120314 | 362.8475 | -0.196097434 | 0.255743 | -0.766775 | 0.443215 | 0.846745122 | WDR55 | protein_coding | 53.17 | -0.014167509 | 0.014167509 | ENSG00000120314 | n.s. |
| ENSG00000139218 | 4455.616 | -0.20214595 | 0.267433 | -0.755876 | 0.449724 | 0.85118703 | SCAF11 | protein_coding | 36.52 | -0.014145163 | 0.014145163 | ENSG00000139218 | n.s. |
| ENSG00000048540 | 21.87366 | -0.245893201 | 0.367774 | -0.668598 | 0.503752 | 0.876169148 | LMO3 | protein_coding | 35.63 | -0.014117231 | 0.014117231 | ENSG00000048540 | n.s. |
| ENSG00000152359 | 111.218 | 0.211203639 | 0.285643 | 0.7393978 | 0.459665 | 0.857438653 | POC5 | protein_coding | 37.27 | 0.014107757 | 0.014107757 | ENSG00000152359 | n.s. |
| ENSG00000221838 | 44.64137 | -0.229002143 | 0.325988 | -0.702487 | 0.482375 | 0.867933207 | AP4M1 | protein_coding | 56.13 | -0.014086768 | 0.014086768 | ENSG00000221838 | n.s. |
| ENSG00000129103 | 767.6988 | -0.201133516 | 0.265389 | -0.757883 | 0.448521 | 0.851088293 | SUMF2 | protein_coding | 49.29 | -0.014084452 | 0.014084452 | ENSG00000129103 | n.s. |
| ENSG00000107020 | 322.0601 | -0.224033104 | 0.312544 | -0.716804 | 0.473495 | 0.865380616 | PLGRKT | protein_coding | 39.88 | -0.014067674 | 0.014067674 | ENSG00000107020 | n.s. |
| ENSG00000207631 | 9.975403 | 0.286170005 | 0.48338 | 0.5920184 | 0.553838 | 0.893144718 | MIR641 | miRNA | 47.47 | 0.014044699 | 0.014044699 | ENSG00000207631 | n.s. |
| ENSG00000125247 | 1152.29 | -0.216207818 | 0.29727 | -0.727311 | 0.467036 | 0.861088114 | TMTC4 | protein_coding | 44.69 | -0.014043218 | 0.014043218 | ENSG00000125247 | n.s. |
| ENSG00000047849 | 1300.584 | -0.198873467 | 0.261721 | -0.759868 | 0.447333 | 0.850164607 | MAP4 | protein_coding | 43.43 | -0.014019979 | 0.014019979 | ENSG00000047849 | n.s. |
| ENSG00000197978 | 381.2112 | -0.190004157 | 0.244453 | -0.777264 | 0.437003 | 0.843772375 | GOLGA6L9 | protein_coding | 47.43 | -0.014017499 | 0.014017499 | ENSG00000197978 | n.s. |
| ENSG00000131943 | 287.4841 | -0.226659009 | 0.321962 | -0.703993 | 0.481437 | 0.867349802 | C19orf12 | protein_coding | 50.38 | -0.014008822 | 0.014008822 | ENSG00000131943 | n.s. |
| ENSG00000166262 | 55.02783 | -0.223421088 | 0.313932 | -0.711686 | 0.476659 | 0.865616576 | FAM227B | protein_coding | 36.14 | -0.014002791 | 0.014002791 | ENSG00000166262 | n.s. |
| ENSG00000235090 | 22.71927 | -0.269355823 | 0.42983 | -0.626656 | 0.530885 | 0.887229181 | RPL7L1P3 | processed_pseudogene | 46.3 | -0.013996855 | 0.013996855 | ENSG00000235090 | n.s. |
| ENSG00000197959 | 128.5615 | -0.196887656 | 0.258008 | -0.763108 | 0.445399 | 0.849026825 | DNM3 | protein_coding | 37.28 | -0.013994497 | 0.013994497 | ENSG00000197959 | n.s. |
| ENSG00000141446 | 706.8746 | -0.208467958 | 0.280932 | -0.742059 | 0.458051 | 0.856847578 | ESCO1 | protein_coding | 41.11 | -0.013987455 | 0.013987455 | ENSG00000141446 | n.s. |
| ENSG00000115459 | 323.5425 | -0.18762267 | 0.2398 | -0.782413 | 0.433972 | 0.84244096 | ELMOD3 | protein_coding | 46.14 | -0.013970483 | 0.013970483 | ENSG00000115459 | n.s. |
| ENSG00000171320 | 1309.376 | -0.178649908 | 0.223582 | -0.799035 | 0.42427 | 0.835315601 | ESCO2 | protein_coding | 37.71 | -0.013961384 | 0.013961384 | ENSG00000171320 | n.s. |
| ENSG00000228988 | 11.86993 | 0.334195968 | 0.62498 | 0.5347311 | 0.592836 | 0.908297635 | | antisense | 36.76 | 0.013959973 | 0.013959973 | ENSG00000228988 | n.s. |
| ENSG00000141580 | 2817.144 | -0.208904425 | 0.282775 | -0.738766 | 0.460049 | 0.857465447 | WDR45B | protein_coding | 49.34 | -0.013951342 | 0.013951342 | ENSG00000141580 | n.s. |
| ENSG00000097046 | 2890.063 | -0.208067605 | 0.280738 | -0.741146 | 0.458605 | 0.856995387 | CDC7 | protein_coding | 36.61 | -0.013945006 | 0.013945006 | ENSG00000097046 | n.s. |
| ENSG00000196233 | 1882.794 | -0.189827633 | 0.245035 | -0.774696 | 0.438519 | 0.84443611 | LCOR | protein_coding | 39.58 | -0.013939651 | 0.013939651 | ENSG00000196233 | n.s. |
| ENSG00000112200 | 5075.277 | -0.189160832 | 0.243864 | -0.775681 | 0.437937 | 0.843950276 | ZNF451 | protein_coding | 36.73 | -0.013937964 | 0.013937964 | ENSG00000112200 | n.s. |
| ENSG00000165338 | 928.3367 | -0.210519104 | 0.286781 | -0.734077 | 0.462902 | 0.858623382 | HECTD2 | protein_coding | 36.7 | -0.013935794 | 0.013935794 | ENSG00000165338 | n.s. |
| ENSG00000011275 | 1305.232 | -0.1950351 | 0.255202 | -0.764239 | 0.444725 | 0.848321661 | RNF216 | protein_coding | 45.52 | -0.013933199 | 0.013933199 | ENSG00000011275 | n.s. |

| | | | | | | | | | | | | | |
|---|---|---|---|---|---|---|---|---|---|---|---|---|---|
| ENSG00000159459 | 1830.327 | -0.200082456 | 0.265475 | -0.753678 | 0.451043 | 0.851854263 | UBR1 | protein_coding | 39.81 | -0.013932682 | 0.013932682 | ENSG00000159459 | n.s. |
| ENSG00000114770 | 1446.898 | -0.198574183 | 0.261859 | -0.758325 | 0.448257 | 0.850978686 | ABCC5 | protein_coding | 44.7 | -0.01391634 | 0.01391634 | ENSG00000114770 | n.s. |
| ENSG00000111790 | 1350.913 | -0.201788167 | 0.26914 | -0.749752 | 0.453404 | 0.853222234 | FGFR1OP2 | protein_coding | 35.97 | -0.013910839 | 0.013910839 | ENSG00000111790 | n.s. |
| ENSG00000108947 | 116.8808 | -0.230168231 | 0.331233 | -0.694884 | 0.487128 | 0.870096539 | EFNB3 | protein_coding | 58.09 | -0.013909655 | 0.013909655 | ENSG00000108947 | n.s. |
| ENSG00000184786 | 590.5854 | 0.170703061 | 0.208651 | 0.818126 | 0.413285 | 0.829007487 | TCTE3 | protein_coding | 38.54 | 0.013902321 | 0.013902321 | ENSG00000184786 | n.s. |
| ENSG00000168818 | 408.8476 | -0.200239034 | 0.266279 | -0.75199 | 0.452057 | 0.852382105 | STX18 | protein_coding | 41.43 | -0.013889716 | 0.013889716 | ENSG00000168818 | n.s. |
| ENSG00000066629 | 132.1655 | -0.221602217 | 0.310116 | -0.714579 | 0.474869 | 0.865616576 | EML1 | protein_coding | 45.19 | -0.013888794 | 0.013888794 | ENSG00000066629 | n.s. |
| ENSG00000036257 | 2486.37 | -0.198073411 | 0.262144 | -0.75559 | 0.449895 | 0.85118703 | CUL3 | protein_coding | 37.23 | -0.013860187 | 0.013860187 | ENSG00000036257 | n.s. |
| ENSG00000265100 | 63.56335 | 0.214426087 | 0.296036 | 0.7243235 | 0.468867 | 0.861812061 | | antisense | 57.86 | 0.01384923 | 0.01384923 | ENSG00000265100 | n.s. |
| ENSG00000131089 | 128.1245 | -0.196421984 | 0.258298 | -0.760446 | 0.446988 | 0.850164607 | ARHGEF9 | protein_coding | 40.8 | -0.013847156 | 0.013847156 | ENSG00000131089 | n.s. |
| ENSG00000185596 | 118.3964 | -0.197999878 | 0.262337 | -0.754755 | 0.450396 | 0.851309042 | WASH3P | cribed_unprocessed_pseudc | 54.04 | -0.013842717 | 0.013842717 | ENSG00000185596 | n.s. |
| ENSG00000008256 | 966.5365 | -0.207681738 | 0.281643 | -0.737393 | 0.460884 | 0.85776362 | CYTH3 | protein_coding | 44.78 | -0.013838328 | 0.013838328 | ENSG00000008256 | n.s. |
| ENSG00000078124 | 496.7278 | -0.202544316 | 0.271078 | -0.747181 | 0.454954 | 0.854433778 | ACER3 | protein_coding | 38.74 | -0.01383815 | 0.01383815 | ENSG00000078124 | n.s. |
| ENSG00000179981 | 299.8042 | -0.196251265 | 0.25778 | -0.761312 | 0.446471 | 0.850164607 | TSHZ1 | protein_coding | 44.58 | -0.013835121 | 0.013835121 | ENSG00000179981 | n.s. |
| ENSG00000100714 | 2403.282 | -0.200509358 | 0.267217 | -0.750362 | 0.453037 | 0.853222234 | MTHFD1 | protein_coding | 43.75 | -0.013822681 | 0.013822681 | ENSG00000100714 | n.s. |
| ENSG00000104722 | 710.693 | -0.169974605 | 0.208464 | -0.815365 | 0.414864 | 0.829267837 | NEFM | protein_coding | 49.83 | -0.013819816 | 0.013819816 | ENSG00000104722 | n.s. |
| ENSG00000003756 | 2684.876 | -0.196269601 | 0.258452 | -0.759404 | 0.447611 | 0.850405175 | RBM5 | protein_coding | 44.45 | -0.013812298 | 0.013812298 | ENSG00000003756 | n.s. |
| ENSG00000238446 | 6.690066 | 0.319856032 | 0.590676 | 0.5415083 | 0.588157 | 0.905352521 | RNU7-38P | snRNA | 38.33 | 0.013812114 | 0.013812114 | ENSG00000238446 | n.s. |
| ENSG00000020256 | 146.0614 | -0.202175747 | 0.271074 | -0.745832 | 0.455769 | 0.85453422 | ZFP64 | protein_coding | 44.05 | -0.013802648 | 0.013802648 | ENSG00000020256 | n.s. |
| ENSG00000234072 | 374.1932 | 0.16942765 | 0.207504 | 0.8165018 | 0.414213 | 0.829007487 | | antisense | 49.35 | 0.01379845 | 0.01379845 | ENSG00000234072 | n.s. |
| ENSG00000184613 | 4181.9 | 0.202056218 | 0.2709 | 0.7458698 | 0.455746 | 0.85453422 | NELL2 | protein_coding | 36.23 | 0.013794487 | 0.013794487 | ENSG00000184613 | n.s. |
| ENSG00000071553 | 500.0601 | -0.211966076 | 0.291093 | -0.728173 | 0.466508 | 0.861088114 | ATP6AP1 | protein_coding | 56.78 | -0.013767707 | 0.013767707 | ENSG00000071553 | n.s. |
| ENSG00000274428 | 394.5016 | 0.191755893 | 0.250601 | 0.7651836 | 0.444162 | 0.847626065 | | snRNA | 45.62 | 0.013767248 | 0.013767248 | ENSG00000274428 | n.s. |
| ENSG00000122386 | 138.8149 | -0.211917296 | 0.291754 | -0.726355 | 0.467621 | 0.861088114 | ZNF205 | protein_coding | 58.51 | -0.013764538 | 0.013764538 | ENSG00000122386 | n.s. |
| ENSG00000087470 | 4320.72 | -0.196552694 | 0.259366 | -0.75782 | 0.448559 | 0.851088293 | DNM1L | protein_coding | 38.21 | -0.013763678 | 0.013763678 | ENSG00000087470 | n.s. |
| ENSG00000050405 | 2323.199 | 0.201541381 | 0.269943 | 0.746606 | 0.455301 | 0.85453422 | LIMA1 | protein_coding | 42.32 | 0.013759339 | 0.013759339 | ENSG00000050405 | n.s. |
| ENSG00000171606 | 132.4999 | 0.184969108 | 0.236896 | 0.7808018 | 0.434919 | 0.842611157 | ZNF274 | protein_coding | 47.52 | 0.01375667 | 0.01375667 | ENSG00000171606 | n.s. |
| ENSG00000255153 | 23.20147 | 0.263805558 | 0.41737 | 0.6320662 | 0.527344 | 0.886936263 | TOLLIP-AS1 | antisense | 60.38 | 0.013746271 | 0.013746271 | ENSG00000255153 | n.s. |
| ENSG00000136875 | 1066.795 | -0.201251927 | 0.269989 | -0.745407 | 0.456026 | 0.85459484 | PRPF4 | protein_coding | 42.73 | -0.013733378 | 0.013733378 | ENSG00000136875 | n.s. |
| ENSG00000081870 | 856.898 | -0.215393688 | 0.299282 | -0.719701 | 0.471709 | 0.863532918 | HSPB11 | protein_coding | 40.17 | -0.013725123 | 0.013725123 | ENSG00000081870 | n.s. |
| ENSG00000171729 | 100.5404 | -0.238460376 | 0.35608 | -0.669681 | 0.503061 | 0.875883784 | TMEM51 | protein_coding | 48.41 | -0.013724233 | 0.013724233 | ENSG00000171729 | n.s. |
| ENSG00000100412 | 815.0141 | -0.205565613 | 0.278391 | -0.738406 | 0.460268 | 0.857528492 | ACO2 | protein_coding | 50.27 | -0.013721801 | 0.013721801 | ENSG00000100412 | n.s. |

| | | | | | | | | | | | | | |
|---|---|---|---|---|---|---|---|---|---|---|---|---|---|
| ENSG00000159346 | 1216.245 | -0.211101235 | 0.289701 | -0.728685 | 0.466194 | 0.861088114 | ADIPOR1 | protein_coding | 44.28 | -0.013711533 | 0.013711533 | ENSG00000159346 | n.s. |
| ENSG00000177054 | 688.9488 | -0.205656721 | 0.278798 | -0.737656 | 0.460724 | 0.857689162 | ZDHHC13 | protein_coding | 37.99 | -0.01371115 | 0.01371115 | ENSG00000177054 | n.s. |
| ENSG00000210151 | 33.29225 | -0.235674491 | 0.347645 | -0.677917 | 0.497824 | 0.874675553 | MT-TS1 | Mt_tRNA | 49.28 | -0.013705181 | 0.013705181 | ENSG00000210151 | n.s. |
| ENSG00000136631 | 1495.236 | -0.204786616 | 0.276912 | -0.739538 | 0.45958 | 0.857420077 | VPS45 | protein_coding | 39.17 | -0.013681047 | 0.013681047 | ENSG00000136631 | n.s. |
| ENSG00000175029 | 1618.325 | -0.205432097 | 0.278813 | -0.736809 | 0.461238 | 0.857958626 | CTBP2 | protein_coding | 50.11 | -0.013668148 | 0.013668148 | ENSG00000175029 | n.s. |
| ENSG00000165916 | 2030.163 | -0.205978949 | 0.280349 | -0.734722 | 0.462509 | 0.858395846 | PSMC3 | protein_coding | 54.93 | -0.013658957 | 0.013658957 | ENSG00000165916 | n.s. |
| ENSG00000084628 | 220.408 | -0.194960722 | 0.257162 | -0.758125 | 0.448376 | 0.851088293 | NKAIN1 | protein_coding | 50.46 | -0.013652199 | 0.013652199 | ENSG00000084628 | n.s. |
| ENSG00000113368 | 3136.48 | -0.184180198 | 0.236362 | -0.779231 | 0.435844 | 0.843164453 | LMNB1 | protein_coding | 42.33 | -0.013645489 | 0.013645489 | ENSG00000113368 | n.s. |
| ENSG00000135018 | 2674.236 | -0.194866248 | 0.257521 | -0.756701 | 0.449229 | 0.85118703 | UBQLN1 | protein_coding | 38.75 | -0.013635766 | 0.013635766 | ENSG00000135018 | n.s. |
| ENSG00000151413 | 162.2863 | 0.193384178 | 0.254175 | 0.7608321 | 0.446757 | 0.850164607 | NUBPL | protein_coding | 38.49 | 0.013633 | 0.013633 | ENSG00000151413 | n.s. |
| ENSG00000207344 | 6.558923 | 0.304218109 | 0.547488 | 0.5556617 | 0.578442 | 0.901989749 | SNORA22C | snoRNA | 42.54 | 0.013628484 | 0.013628484 | ENSG00000207344 | n.s. |
| ENSG00000182247 | 321.6978 | 0.203289455 | 0.274481 | 0.7406327 | 0.458916 | 0.856995387 | UBE2E2 | protein_coding | 37.68 | 0.013624767 | 0.013624767 | ENSG00000182247 | n.s. |
| ENSG00000163960 | 1314.946 | -0.18470543 | 0.238095 | -0.775763 | 0.437889 | 0.843950276 | UBXN7 | protein_coding | 42.28 | -0.013609676 | 0.013609676 | ENSG00000163960 | n.s. |
| ENSG00000178397 | 186.9509 | -0.206276576 | 0.281385 | -0.733075 | 0.463513 | 0.859318855 | FAM220A | protein_coding | 48.26 | -0.013582417 | 0.013582417 | ENSG00000178397 | n.s. |
| ENSG00000120727 | 2347.211 | 0.199262717 | 0.267497 | 0.7449158 | 0.456323 | 0.854868125 | PAIP2 | protein_coding | 43.58 | 0.013569966 | 0.013569966 | ENSG00000120727 | n.s. |
| ENSG00000135549 | 815.8774 | 0.227176338 | 0.328139 | 0.6923171 | 0.488738 | 0.871503806 | PKIB | protein_coding | 37.28 | 0.013569404 | 0.013569404 | ENSG00000135549 | n.s. |
| ENSG00000189060 | 1988.166 | 0.196635621 | 0.262191 | 0.7499711 | 0.453272 | 0.853222234 | H1F0 | protein_coding | 53.2 | 0.013555634 | 0.013555634 | ENSG00000189060 | n.s. |
| ENSG00000239119 | 29.62833 | 0.230452845 | 0.336611 | 0.6846265 | 0.49358 | 0.873348456 | RNU7-119P | snRNA | 36.51 | 0.013553495 | 0.013553495 | ENSG00000239119 | n.s. |
| ENSG00000231889 | 705.1773 | 0.178104706 | 0.225344 | 0.7903689 | 0.429312 | 0.839274918 | TRAF3IP2-AS1 | antisense | 41.83 | 0.013553012 | 0.013553012 | ENSG00000231889 | n.s. |
| ENSG00000255135 | 353.5519 | -0.172998597 | 0.21587 | -0.801401 | 0.422899 | 0.834993891 | | lincRNA | 43.63 | -0.01354868 | 0.01354868 | ENSG00000255135 | n.s. |
| ENSG00000095906 | 632.4005 | -0.20623439 | 0.281524 | -0.732563 | 0.463825 | 0.859669362 | NUBP2 | protein_coding | 62.18 | -0.013543114 | 0.013543114 | ENSG00000095906 | n.s. |
| ENSG00000106799 | 2937.959 | -0.202725124 | 0.274374 | -0.738863 | 0.45999 | 0.857465447 | TGFBR1 | protein_coding | 39.03 | -0.013538667 | 0.013538667 | ENSG00000106799 | n.s. |
| ENSG00000173960 | 617.191 | -0.206751051 | 0.282508 | -0.731842 | 0.464265 | 0.860105883 | UBXN2A | protein_coding | 42.94 | -0.01353146 | 0.01353146 | ENSG00000173960 | n.s. |
| ENSG00000122691 | 3420.512 | -0.172748519 | 0.21567 | -0.800986 | 0.42314 | 0.834993891 | TWIST1 | protein_coding | 37.36 | -0.013529094 | 0.013529094 | ENSG00000122691 | n.s. |
| ENSG00000104320 | 3901.995 | 0.194004677 | 0.257324 | 0.7539322 | 0.45089 | 0.851782482 | NBN | protein_coding | 37.15 | 0.013516557 | 0.013516557 | ENSG00000104320 | n.s. |
| ENSG00000279527 | 72.8209 | -0.215442902 | 0.302497 | -0.712214 | 0.476332 | 0.865616576 | | TEC | 37.48 | -0.013502762 | 0.013502762 | ENSG00000279527 | n.s. |
| ENSG00000276293 | 920.4907 | -0.203079728 | 0.275758 | -0.736441 | 0.461462 | 0.858145726 | PIP4K2B | protein_coding | 49.36 | -0.013492405 | 0.013492405 | ENSG00000276293 | n.s. |
| ENSG00000233797 | 30.37188 | -0.228068847 | 0.331886 | -0.687191 | 0.491963 | 0.872671134 | UFL1-AS1 | antisense | 37.42 | -0.013490133 | 0.013490133 | ENSG00000233797 | n.s. |
| ENSG00000180694 | 646.757 | -0.198057571 | 0.265937 | -0.744754 | 0.456421 | 0.854936721 | TMEM64 | protein_coding | 38.54 | -0.013480992 | 0.013480992 | ENSG00000180694 | n.s. |
| ENSG00000198964 | 114.4413 | -0.2072371 | 0.28504 | -0.727045 | 0.467199 | 0.861088114 | SGMS1 | protein_coding | 39.37 | -0.013460548 | 0.013460548 | ENSG00000198964 | n.s. |
| ENSG00000177885 | 1956.069 | -0.201375068 | 0.27229 | -0.73956 | 0.459567 | 0.857420077 | GRB2 | protein_coding | 44.22 | -0.013453134 | 0.013453134 | ENSG00000177885 | n.s. |
| ENSG00000122515 | 77.84854 | 0.198197539 | 0.266389 | 0.744015 | 0.456867 | 0.85542832 | ZMIZ2 | protein_coding | 56.51 | 0.013441039 | 0.013441039 | ENSG00000122515 | n.s. |

| | | | | | | | | | | | | | |
|---|---|---|---|---|---|---|---|---|---|---|---|---|---|
| ENSG00000178567 | 3133.627 | -0.165627392 | 0.203272 | -0.814807 | 0.415183 | 0.829591076 | EPM2AIP1 | protein_coding | 38.52 | -0.013438332 | 0.013438332 | ENSG00000178567 | n.s. |
| ENSG00000124193 | 7753.407 | 0.184330161 | 0.239073 | 0.771022 | 0.440694 | 0.845502102 | SRSF6 | protein_coding | 41.69 | 0.013434961 | 0.013434961 | ENSG00000124193 | n.s. |
| ENSG00000111667 | 2257.574 | -0.200454021 | 0.270435 | -0.741229 | 0.458555 | 0.856995387 | USP5 | protein_coding | 52.77 | -0.013434732 | 0.013434732 | ENSG00000111667 | n.s. |
| ENSG00000144028 | 5892.857 | -0.18452162 | 0.240031 | -0.76874 | 0.442048 | 0.845745639 | SNRNP200 | protein_coding | 47.58 | -0.013425836 | 0.013425836 | ENSG00000144028 | n.s. |
| ENSG00000050393 | 874.896 | -0.202075651 | 0.274902 | -0.735084 | 0.462289 | 0.858395846 | MCUR1 | protein_coding | 41.24 | -0.01340012 | 0.01340012 | ENSG00000050393 | n.s. |
| ENSG00000082258 | 4150.337 | -0.189918879 | 0.249674 | -0.760667 | 0.446856 | 0.850164607 | CCNT2 | protein_coding | 37.48 | -0.013388707 | 0.013388707 | ENSG00000082258 | n.s. |
| ENSG00000122085 | 865.8713 | 0.1814441 | 0.233626 | 0.7766442 | 0.437369 | 0.843772375 | MTERF4 | protein_coding | 47.67 | 0.013385984 | 0.013385984 | ENSG00000122085 | n.s. |
| ENSG00000100216 | 1045.078 | -0.204954947 | 0.280619 | -0.730366 | 0.465166 | 0.86043697 | TOMM22 | protein_coding | 48.7 | -0.013379651 | 0.013379651 | ENSG00000100216 | n.s. |
| ENSG00000257800 | 441.9803 | 0.16890586 | 0.209651 | 0.805651 | 0.420444 | 0.833311153 | FNBP1P1 | processed_pseudogene | 41.55 | 0.013376129 | 0.013376129 | ENSG00000257800 | n.s. |
| ENSG00000148308 | 1058.547 | -0.204517139 | 0.279741 | -0.731096 | 0.464721 | 0.860217499 | GTF3C5 | protein_coding | 51.81 | -0.013373729 | 0.013373729 | ENSG00000148308 | n.s. |
| ENSG00000120324 | 197.0814 | -0.19098077 | 0.252568 | -0.756156 | 0.449556 | 0.85118703 | PCDHB10 | protein_coding | 48.48 | -0.01336388 | 0.01336388 | ENSG00000120324 | n.s. |
| ENSG00000130522 | 670.5291 | 0.181005558 | 0.233057 | 0.7766585 | 0.43736 | 0.843772375 | JUND | protein_coding | 69.67 | 0.01335363 | 0.01335363 | ENSG00000130522 | n.s. |
| ENSG00000248152 | 45.25657 | 0.220865014 | 0.317577 | 0.6954699 | 0.486761 | 0.870096539 | | antisense | 37.21 | 0.013347438 | 0.013347438 | ENSG00000248152 | n.s. |
| ENSG00000168096 | 63.28604 | -0.20547362 | 0.282348 | -0.727731 | 0.466779 | 0.861088114 | ANKS3 | protein_coding | 49.79 | -0.013346006 | 0.013346006 | ENSG00000168096 | n.s. |
| ENSG00000003096 | 1087.74 | -0.20942056 | 0.291122 | -0.719356 | 0.471921 | 0.863532918 | KLHL13 | protein_coding | 36.56 | -0.013344509 | 0.013344509 | ENSG00000003096 | n.s. |
| ENSG00000250568 | 126.8646 | 0.19028255 | 0.251462 | 0.756706 | 0.449226 | 0.85118703 | | processed_pseudogene | 46.98 | 0.013315022 | 0.013315022 | ENSG00000250568 | n.s. |
| ENSG00000198087 | 988.1911 | -0.182140614 | 0.235587 | -0.773136 | 0.439442 | 0.845174857 | CD2AP | protein_coding | 36.38 | -0.013305997 | 0.013305997 | ENSG00000198087 | n.s. |
| ENSG00000167965 | 380.1211 | -0.20601085 | 0.284338 | -0.724528 | 0.468742 | 0.861812061 | MLST8 | protein_coding | 61.98 | -0.013305712 | 0.013305712 | ENSG00000167965 | n.s. |
| ENSG00000207174 | 6.747883 | -0.286840906 | 0.505905 | -0.566985 | 0.570724 | 0.898818733 | SNORD116-15 | snoRNA | 41.3 | -0.013288732 | 0.013288732 | ENSG00000207174 | n.s. |
| ENSG00000279372 | 23.22669 | 0.270751619 | 0.457758 | 0.5914734 | 0.554203 | 0.893144718 | | TEC | 51.01 | 0.013287993 | 0.013287993 | ENSG00000279372 | n.s. |
| ENSG00000091140 | 1689.606 | -0.20056913 | 0.273318 | -0.733832 | 0.463051 | 0.858692716 | DLD | protein_coding | 36.67 | -0.013270099 | 0.013270099 | ENSG00000091140 | n.s. |
| ENSG00000266758 | 12.2203 | -0.285222001 | 0.500043 | -0.570395 | 0.56841 | 0.898419074 | MIR3680-2 | miRNA | 43.68 | -0.013268823 | 0.013268823 | ENSG00000266758 | n.s. |
| ENSG00000254860 | 40.93642 | 0.219258658 | 0.314553 | 0.6970475 | 0.485773 | 0.869936401 | TMEM9B-AS1 | antisense | 43.8 | 0.013267889 | 0.013267889 | ENSG00000254860 | n.s. |
| ENSG00000110900 | 951.9084 | -0.194407463 | 0.260753 | -0.745563 | 0.455931 | 0.85459484 | TSPAN11 | protein_coding | 51.34 | -0.013266314 | 0.013266314 | ENSG00000110900 | n.s. |
| ENSG00000160714 | 1135.571 | -0.199993057 | 0.272231 | -0.734644 | 0.462557 | 0.858395846 | UBE2Q1 | protein_coding | 50.99 | -0.013262018 | 0.013262018 | ENSG00000160714 | n.s. |
| ENSG00000154640 | 1925.628 | 0.192204737 | 0.256037 | 0.7506907 | 0.452839 | 0.853163076 | BTG3 | protein_coding | 37.74 | 0.013255966 | 0.013255966 | ENSG00000154640 | n.s. |
| ENSG00000200742 | 18.01883 | -0.25100356 | 0.391174 | -0.641667 | 0.521089 | 0.885523197 | | misc_RNA | 42.57 | -0.013253002 | 0.013253002 | ENSG00000200742 | n.s. |
| ENSG00000244184 | 40.94845 | 0.228507589 | 0.338454 | 0.6751508 | 0.49958 | 0.875034827 | | antisense | 50.54 | 0.01324765 | 0.01324765 | ENSG00000244184 | n.s. |
| ENSG00000109814 | 733.4713 | -0.189246616 | 0.250534 | -0.755374 | 0.450025 | 0.851201408 | UGDH | protein_coding | 41.86 | -0.013241144 | 0.013241144 | ENSG00000109814 | n.s. |
| ENSG00000166946 | 531.6035 | 0.215908124 | 0.307909 | 0.7012077 | 0.483173 | 0.868366486 | CCNDBP1 | protein_coding | 40.91 | 0.013234509 | 0.013234509 | ENSG00000166946 | n.s. |
| ENSG00000138018 | 2701.911 | -0.198268794 | 0.268558 | -0.738271 | 0.46035 | 0.857551846 | SELENOI | protein_coding | 42.63 | -0.013232383 | 0.013232383 | ENSG00000138018 | n.s. |
| ENSG00000266441 | 40.45514 | -0.228658348 | 0.339248 | -0.674015 | 0.500302 | 0.875342302 | | antisense | 59.4 | -0.013221502 | 0.013221502 | ENSG00000266441 | n.s. |

| | | | | | | | | | | | | | |
|---|---|---|---|---|---|---|---|---|---|---|---|---|---|
| ENSG00000065243 | 2148.193 | -0.210857649 | 0.295587 | -0.713352 | 0.475628 | 0.865616576 | PKN2 | protein_coding | 36.25 | -0.013215384 | 0.013215384 | ENSG00000065243 | n.s. |
| ENSG00000116750 | 2263.97 | -0.203127186 | 0.278381 | -0.729674 | 0.46559 | 0.86087715 | UCHL5 | protein_coding | 35 | -0.013215215 | 0.013215215 | ENSG00000116750 | n.s. |
| ENSG00000101079 | 459.5623 | 0.211325301 | 0.298351 | 0.7083109 | 0.478752 | 0.86589592 | NDRG3 | protein_coding | 44.49 | 0.013215081 | 0.013215081 | ENSG00000101079 | n.s. |
| ENSG00000092758 | 157.2643 | -0.197913975 | 0.267979 | -0.738544 | 0.460184 | 0.857496121 | COL9A3 | protein_coding | 61.84 | -0.013214288 | 0.013214288 | ENSG00000092758 | n.s. |
| ENSG00000070718 | 1204.699 | -0.197797428 | 0.268003 | -0.738043 | 0.460488 | 0.857551846 | AP3M2 | protein_coding | 41.4 | -0.013200924 | 0.013200924 | ENSG00000070718 | n.s. |
| ENSG00000165275 | 19.7461 | -0.317823089 | 0.598158 | -0.531337 | 0.595186 | 0.908868121 | TRMT10B | protein_coding | 42.56 | -0.013189382 | 0.013189382 | ENSG00000165275 | n.s. |
| ENSG00000280063 | 90.00041 | -0.194711704 | 0.261895 | -0.743472 | 0.457196 | 0.855698283 | | TEC | 53.91 | -0.01317796 | 0.01317796 | ENSG00000280063 | n.s. |
| ENSG00000163322 | 126.6067 | 0.212738716 | 0.301419 | 0.7057918 | 0.480318 | 0.867184458 | FAM175A | protein_coding | 42.23 | 0.013166083 | 0.013166083 | ENSG00000163322 | n.s. |
| ENSG00000146540 | 363.5561 | -0.216878441 | 0.310647 | -0.698151 | 0.485083 | 0.869663535 | C7orf50 | protein_coding | 57.69 | -0.013153404 | 0.013153404 | ENSG00000146540 | n.s. |
| ENSG00000201435 | 39.82967 | -0.226469734 | 0.33526 | -0.675506 | 0.499354 | 0.874970095 | RNU4-24P | snRNA | 38.85 | -0.013136782 | 0.013136782 | ENSG00000201435 | n.s. |
| ENSG00000166925 | 130.6941 | 0.209610203 | 0.295328 | 0.7097533 | 0.477857 | 0.865660698 | TSC22D4 | protein_coding | 58.03 | 0.013132561 | 0.013132561 | ENSG00000166925 | n.s. |
| ENSG00000114573 | 935.4016 | -0.197985101 | 0.269346 | -0.735059 | 0.462304 | 0.858395846 | ATP6V1A | protein_coding | 38.22 | -0.013128866 | 0.013128866 | ENSG00000114573 | n.s. |
| ENSG00000199933 | 31.08857 | -0.24988705 | 0.390909 | -0.639246 | 0.522663 | 0.886079926 | RNY1P16 | misc_RNA | 45.05 | -0.013125843 | 0.013125843 | ENSG00000199933 | n.s. |
| ENSG00000182087 | 1239.493 | -0.192213194 | 0.257721 | -0.74582 | 0.455776 | 0.85453422 | TMEM259 | protein_coding | 65.7 | -0.013122499 | 0.013122499 | ENSG00000182087 | n.s. |
| ENSG00000181218 | 766.9935 | 0.164441078 | 0.20326 | 0.8090168 | 0.418506 | 0.832304654 | HIST3H2A | protein_coding | 57.65 | 0.013108861 | 0.013108861 | ENSG00000181218 | n.s. |
| ENSG00000111328 | 2260.601 | -0.209132754 | 0.292124 | -0.715903 | 0.474051 | 0.865616576 | CDK2AP1 | protein_coding | 58.95 | -0.013107277 | 0.013107277 | ENSG00000111328 | n.s. |
| ENSG00000127948 | 283.1878 | -0.209097192 | 0.293505 | -0.712414 | 0.476208 | 0.865616576 | POR | protein_coding | 50.98 | -0.013105049 | 0.013105049 | ENSG00000127948 | n.s. |
| ENSG00000178425 | 798.0702 | -0.203997434 | 0.282397 | -0.722378 | 0.470062 | 0.862503771 | NT5DC1 | protein_coding | 36.29 | -0.013104591 | 0.013104591 | ENSG00000178425 | n.s. |
| ENSG00000115159 | 1279.304 | -0.197423766 | 0.268378 | -0.735618 | 0.461963 | 0.858395846 | GPD2 | protein_coding | 37.9 | -0.013091642 | 0.013091642 | ENSG00000115159 | n.s. |
| ENSG00000196177 | 398.9217 | -0.215362262 | 0.308328 | -0.698483 | 0.484875 | 0.869415694 | ACADSB | protein_coding | 40.72 | -0.013088108 | 0.013088108 | ENSG00000196177 | n.s. |
| ENSG00000105705 | 491.7913 | -0.20014783 | 0.273903 | -0.730725 | 0.464947 | 0.860217499 | SUGP1 | protein_coding | 51.37 | -0.013088012 | 0.013088012 | ENSG00000105705 | n.s. |
| ENSG00000175087 | 240.8112 | -0.201472607 | 0.277365 | -0.726382 | 0.467605 | 0.861088114 | PDIK1L | protein_coding | 43.1 | -0.01308613 | 0.01308613 | ENSG00000175087 | n.s. |
| ENSG00000136149 | 26.70587 | 0.31321892 | 0.586442 | 0.5341001 | 0.593272 | 0.908297635 | RPL13AP25 | processed_pseudogene | 57.19 | 0.013083723 | 0.013083723 | ENSG00000136149 | n.s. |
| ENSG00000164048 | 371.3302 | -0.186946446 | 0.247413 | -0.755606 | 0.449886 | 0.85118703 | ZNF589 | protein_coding | 45.4 | -0.013081578 | 0.013081578 | ENSG00000164048 | n.s. |
| ENSG00000267322 | 306.3469 | -0.171869504 | 0.21751 | -0.790169 | 0.429429 | 0.839380003 | SNHG22 | antisense | 43.78 | -0.013069194 | 0.013069194 | ENSG00000267322 | n.s. |
| ENSG00000137504 | 4351.922 | 0.173557896 | 0.220366 | 0.7875904 | 0.430936 | 0.841032617 | CREBZF | protein_coding | 40.72 | 0.013049325 | 0.013049325 | ENSG00000137504 | n.s. |
| ENSG00000207342 | 18.63689 | -0.248437154 | 0.3911 | -0.635227 | 0.52528 | 0.886111278 | | misc_RNA | 37.37 | -0.013045866 | 0.013045866 | ENSG00000207342 | n.s. |
| ENSG00000254469 | 76.49337 | -0.199363583 | 0.272832 | -0.730719 | 0.464951 | 0.860217499 | | protein_coding | 42.82 | -0.013036729 | 0.013036729 | ENSG00000254469 | n.s. |
| ENSG00000198093 | 755.1593 | -0.196452789 | 0.267067 | -0.735594 | 0.461978 | 0.858395846 | ZNF649 | protein_coding | 43.38 | -0.013027254 | 0.013027254 | ENSG00000198093 | n.s. |
| ENSG00000131584 | 568.0261 | -0.194267718 | 0.262378 | -0.740411 | 0.45905 | 0.856995387 | ACAP3 | protein_coding | 67.3 | -0.013020117 | 0.013020117 | ENSG00000131584 | n.s. |
| ENSG00000124201 | 445.5741 | -0.185921433 | 0.24533 | -0.757842 | 0.448546 | 0.851088293 | ZNFX1 | protein_coding | 45.85 | -0.01301922 | 0.01301922 | ENSG00000124201 | n.s. |
| ENSG00000159792 | 204.8519 | -0.199002671 | 0.272287 | -0.730857 | 0.464866 | 0.860217499 | PSKH1 | protein_coding | 52.29 | -0.013013128 | 0.013013128 | ENSG00000159792 | n.s. |

| | | | | | | | | | | | | | |
|---|---|---|---|---|---|---|---|---|---|---|---|---|---|
| ENSG00000197608 | 667.2177 | -0.200474579 | 0.27628 | -0.725622 | 0.468071 | 0.861253285 | ZNF841 | protein_coding | 41.11 | -0.013004607 | 0.013004607 | ENSG00000197608 | n.s. |
| ENSG00000010165 | 1155.533 | -0.185992724 | 0.2464 | -0.754842 | 0.450344 | 0.851309042 | METTL13 | protein_coding | 43.61 | -0.013003264 | 0.013003264 | ENSG00000010165 | n.s. |
| ENSG00000178057 | 336.5386 | -0.185583329 | 0.245099 | -0.757177 | 0.448944 | 0.85118703 | NDUFAF3 | protein_coding | 64.14 | -0.012986194 | 0.012986194 | ENSG00000178057 | n.s. |
| ENSG00000027697 | 607.8978 | -0.213523171 | 0.305505 | -0.698918 | 0.484603 | 0.869379601 | IFNGR1 | protein_coding | 38.81 | -0.012980192 | 0.012980192 | ENSG00000027697 | n.s. |
| ENSG00000054392 | 186.4239 | -0.199771721 | 0.274824 | -0.726908 | 0.467282 | 0.861088114 | HHAT | protein_coding | 42.26 | -0.012975654 | 0.012975654 | ENSG00000054392 | n.s. |
| ENSG00000278462 | 71.77315 | 0.199721559 | 0.274938 | 0.726424 | 0.467579 | 0.861088114 | | sense_intronic | 32.33 | 0.012972396 | 0.012972396 | ENSG00000278462 | n.s. |
| ENSG00000256269 | 147.4354 | -0.207580431 | 0.293365 | -0.707585 | 0.479203 | 0.866036769 | HMBS | protein_coding | 50.91 | -0.012966235 | 0.012966235 | ENSG00000256269 | n.s. |
| ENSG00000230199 | 2.672953 | 0.373520902 | 0.822866 | 0.4539269 | 0.649881 | 0.923184844 | | processed_pseudogene | 53.75 | 0.012965409 | 0.012965409 | ENSG00000230199 | n.s. |
| ENSG00000170396 | 693.1626 | 0.178084375 | 0.231484 | 0.7693171 | 0.441705 | 0.84569031 | ZNF804A | protein_coding | 34.04 | 0.01296252 | 0.01296252 | ENSG00000170396 | n.s. |
| ENSG00000183963 | 26.80495 | -0.246818697 | 0.389112 | -0.634313 | 0.525876 | 0.886111278 | SMTN | protein_coding | 54.79 | -0.012960878 | 0.012960878 | ENSG00000183963 | n.s. |
| ENSG00000053438 | 2821.335 | -0.20323009 | 0.282177 | -0.720221 | 0.471389 | 0.863532918 | NNAT | protein_coding | 54.05 | -0.012950045 | 0.012950045 | ENSG00000053438 | n.s. |
| ENSG00000231327 | 52.48863 | -0.215772077 | 0.311228 | -0.693293 | 0.488126 | 0.870959298 | LINC01816 | lincRNA | 57.49 | -0.012946786 | 0.012946786 | ENSG00000231327 | n.s. |
| ENSG00000165671 | 2710.591 | -0.175474797 | 0.225927 | -0.776687 | 0.437343 | 0.843772375 | NSD1 | protein_coding | 42.34 | -0.0129456 | 0.0129456 | ENSG00000165671 | n.s. |
| ENSG00000185274 | 367.6073 | -0.186450977 | 0.247791 | -0.752452 | 0.451779 | 0.852320382 | WBSCR17 | protein_coding | 44.5 | -0.012939162 | 0.012939162 | ENSG00000185274 | n.s. |
| ENSG00000032742 | 397.2696 | -0.193718018 | 0.262098 | -0.739106 | 0.459843 | 0.857465447 | IFT88 | protein_coding | 38.74 | -0.012937142 | 0.012937142 | ENSG00000032742 | n.s. |
| ENSG00000177239 | 282.5512 | -0.199125889 | 0.273496 | -0.728076 | 0.466567 | 0.861088114 | MAN1B1 | protein_coding | 54.87 | -0.012933705 | 0.012933705 | ENSG00000177239 | n.s. |
| ENSG00000153201 | 10812.62 | -0.183657361 | 0.241919 | -0.759168 | 0.447752 | 0.850405175 | RANBP2 | protein_coding | 39.01 | -0.012924723 | 0.012924723 | ENSG00000153201 | n.s. |
| ENSG00000221740 | 32.18038 | -0.223839786 | 0.332755 | -0.672687 | 0.501146 | 0.875518559 | SNORD93 | snoRNA | 41.89 | -0.01292331 | 0.01292331 | ENSG00000221740 | n.s. |
| ENSG00000185324 | 563.1453 | -0.187325711 | 0.249832 | -0.749807 | 0.453371 | 0.853222234 | CDK10 | protein_coding | 55.36 | -0.012913829 | 0.012913829 | ENSG00000185324 | n.s. |
| ENSG00000269938 | 104.4405 | 0.213331177 | 0.306252 | 0.696586 | 0.486062 | 0.869962727 | | sense_intronic | 34.11 | 0.012906398 | 0.012906398 | ENSG00000269938 | n.s. |
| ENSG00000165480 | 638.6774 | -0.197356572 | 0.269895 | -0.731236 | 0.464635 | 0.860217499 | SKA3 | protein_coding | 39.84 | -0.012905487 | 0.012905487 | ENSG00000165480 | n.s. |
| ENSG00000102893 | 72.84912 | -0.200866096 | 0.277977 | -0.7226 | 0.469926 | 0.862503771 | PHKB | protein_coding | 38.19 | -0.012903437 | 0.012903437 | ENSG00000102893 | n.s. |
| ENSG00000182952 | 933.2138 | 0.217231906 | 0.314702 | 0.6902791 | 0.490019 | 0.872216858 | HMGN4 | protein_coding | 42.28 | 0.012898258 | 0.012898258 | ENSG00000182952 | n.s. |
| ENSG00000258056 | 154.7042 | -0.1855372 | 0.2464 | -0.752992 | 0.451455 | 0.852169954 | | antisense | 56.95 | -0.012889971 | 0.012889971 | ENSG00000258056 | n.s. |
| ENSG00000126934 | 607.6353 | -0.205377773 | 0.286997 | -0.715609 | 0.474233 | 0.865616576 | MAP2K2 | protein_coding | 57.42 | -0.012871936 | 0.012871936 | ENSG00000126934 | n.s. |
| ENSG00000089195 | 826.9347 | -0.198022891 | 0.272125 | -0.72769 | 0.466803 | 0.861088114 | TRMT6 | protein_coding | 42.56 | -0.012862063 | 0.012862063 | ENSG00000089195 | n.s. |
| ENSG00000114353 | 526.9668 | -0.205107306 | 0.287391 | -0.713687 | 0.475421 | 0.865616576 | GNAI2 | protein_coding | 57.36 | -0.012854985 | 0.012854985 | ENSG00000114353 | n.s. |
| ENSG00000207333 | 9.344559 | 0.267103175 | 0.460682 | 0.5797989 | 0.56205 | 0.895116382 | RNU6-680P | snRNA | 40.95 | 0.012853137 | 0.012853137 | ENSG00000207333 | n.s. |
| ENSG00000203721 | 57.55954 | -0.207662027 | 0.294441 | -0.705277 | 0.480638 | 0.867184458 | LINC00862 | lincRNA | 42.07 | -0.012851894 | 0.012851894 | ENSG00000203721 | n.s. |
| ENSG00000122406 | 30018.65 | 0.211467916 | 0.302753 | 0.6984833 | 0.484875 | 0.869415694 | RPL5 | protein_coding | 40.34 | 0.012851439 | 0.012851439 | ENSG00000122406 | n.s. |
| ENSG00000266240 | 9.855868 | -0.298787515 | 0.552511 | -0.540781 | 0.588659 | 0.905723898 | MIR5091 | miRNA | 63.44 | -0.01284911 | 0.01284911 | ENSG00000266240 | n.s. |
| ENSG00000081087 | 419.4302 | -0.207614152 | 0.294608 | -0.704713 | 0.480989 | 0.867184458 | OSTM1 | protein_coding | 42.48 | -0.012848931 | 0.012848931 | ENSG00000081087 | n.s. |

| | | | | | | | | | | | | | |
|---|---|---|---|---|---|---|---|---|---|---|---|---|---|
| ENSG00000107554 | 437.5803 | -0.175547997 | 0.227356 | -0.772128 | 0.440038 | 0.845174857 | DNMBP | protein_coding | 43.7 | -0.012824384 | 0.012824384 | ENSG00000107554 | n.s. |
| ENSG00000271601 | 567.3196 | -0.191305869 | 0.258374 | -0.740421 | 0.459045 | 0.856995387 | LIX1L | protein_coding | 43.6 | -0.012821609 | 0.012821609 | ENSG00000271601 | n.s. |
| ENSG00000074266 | 529.5374 | -0.186652213 | 0.249371 | -0.748492 | 0.454164 | 0.853840253 | EED | protein_coding | 37.1 | -0.012808705 | 0.012808705 | ENSG00000074266 | n.s. |
| ENSG00000266640 | 9.397715 | -0.305626178 | 0.57072 | -0.53551 | 0.592297 | 0.908010607 | MIR4754 | miRNA | 69.66 | -0.012808511 | 0.012808511 | ENSG00000266640 | n.s. |
| ENSG00000240718 | 16.91124 | -0.243512019 | 0.383446 | -0.635061 | 0.525388 | 0.886111278 | RN7SL851P | misc_RNA | 54.79 | -0.012787239 | 0.012787239 | ENSG00000240718 | n.s. |
| ENSG00000119509 | 642.6116 | -0.19603519 | 0.268848 | -0.729167 | 0.465899 | 0.860954429 | INVS | protein_coding | 38.92 | -0.012746176 | 0.012746176 | ENSG00000119509 | n.s. |
| ENSG00000160799 | 561.1084 | -0.183447351 | 0.243582 | -0.753124 | 0.451376 | 0.852169954 | CCDC12 | protein_coding | 51.44 | -0.012744781 | 0.012744781 | ENSG00000160799 | n.s. |
| ENSG00000155974 | 477.829 | 0.205654838 | 0.291257 | 0.7060942 | 0.48013 | 0.867148032 | GRIP1 | protein_coding | 38.92 | 0.012731424 | 0.012731424 | ENSG00000155974 | n.s. |
| ENSG00000269958 | 107.3628 | -0.196957401 | 0.271858 | -0.724485 | 0.468768 | 0.861812061 | | sense_intronic | 49.08 | -0.012720973 | 0.012720973 | ENSG00000269958 | n.s. |
| ENSG00000232956 | 3537.541 | 0.155836508 | 0.191043 | 0.8157151 | 0.414663 | 0.829028492 | SNHG15 | lincRNA | 50.27 | 0.012689851 | 0.012689851 | ENSG00000232956 | n.s. |
| ENSG00000176619 | 2298.787 | -0.18131081 | 0.239814 | -0.756049 | 0.44962 | 0.85118703 | LMNB2 | protein_coding | 59.78 | -0.012687224 | 0.012687224 | ENSG00000176619 | n.s. |
| ENSG00000147419 | 958.1667 | -0.202077997 | 0.283554 | -0.712661 | 0.476056 | 0.865616576 | CCDC25 | protein_coding | 40.07 | -0.012665124 | 0.012665124 | ENSG00000147419 | n.s. |
| ENSG00000092108 | 1318.276 | -0.198749899 | 0.276291 | -0.719351 | 0.471925 | 0.863532918 | SCFD1 | protein_coding | 36.17 | -0.012664563 | 0.012664563 | ENSG00000092108 | n.s. |
| ENSG00000163394 | 638.9236 | -0.180951949 | 0.239185 | -0.756535 | 0.449329 | 0.85118703 | CCKAR | protein_coding | 42.88 | -0.012662113 | 0.012662113 | ENSG00000163394 | n.s. |
| ENSG00000206693 | 9.577473 | -0.282588725 | 0.508767 | -0.555439 | 0.578594 | 0.901989749 | SNORA56 | snoRNA | 48.84 | -0.012659522 | 0.012659522 | ENSG00000206693 | n.s. |
| ENSG00000104343 | 687.7735 | -0.194796203 | 0.268173 | -0.726382 | 0.467605 | 0.861088114 | UBE2W | protein_coding | 39.29 | -0.012652482 | 0.012652482 | ENSG00000104343 | n.s. |
| ENSG00000198060 | 943.6044 | -0.201768849 | 0.283418 | -0.711912 | 0.476519 | 0.865616576 | 5-Mar | protein_coding | 40.44 | -0.012645749 | 0.012645749 | ENSG00000198060 | n.s. |
| ENSG00000185942 | 58.3855 | -0.200825333 | 0.279904 | -0.717479 | 0.473079 | 0.865034671 | NKAIN3 | protein_coding | 37.47 | -0.012645265 | 0.012645265 | ENSG00000185942 | n.s. |
| ENSG00000215126 | 43.98844 | -0.225905412 | 0.341532 | -0.661448 | 0.508325 | 0.879072588 | CBWD6 | protein_coding | 36.14 | -0.012645115 | 0.012645115 | ENSG00000215126 | n.s. |
| ENSG00000201574 | 7.829738 | -0.371503544 | 0.840059 | -0.442235 | 0.658319 | 0.924786541 | RNU1-93P | snRNA | 44.97 | -0.012615703 | 0.012615703 | ENSG00000201574 | n.s. |
| ENSG00000149929 | 452.7471 | -0.172611694 | 0.223556 | -0.772118 | 0.440045 | 0.845174857 | HIRIP3 | protein_coding | 58.04 | -0.012609877 | 0.012609877 | ENSG00000149929 | n.s. |
| ENSG00000167840 | 350.7288 | -0.180915588 | 0.239991 | -0.753843 | 0.450944 | 0.851782482 | ZNF232 | protein_coding | 50.38 | -0.012604623 | 0.012604623 | ENSG00000167840 | n.s. |
| ENSG00000155876 | 721.4504 | -0.180234287 | 0.238844 | -0.754611 | 0.450482 | 0.851309042 | RRAGA | protein_coding | 52.17 | -0.012600675 | 0.012600675 | ENSG00000155876 | n.s. |
| ENSG00000181894 | 334.5662 | -0.196144437 | 0.271422 | -0.722655 | 0.469892 | 0.862503771 | ZNF329 | protein_coding | 46.41 | -0.012600122 | 0.012600122 | ENSG00000181894 | n.s. |
| ENSG00000104738 | 5136.471 | -0.184674398 | 0.247827 | -0.745176 | 0.456165 | 0.854688385 | MCM4 | protein_coding | 45.8 | -0.012593353 | 0.012593353 | ENSG00000104738 | n.s. |
| ENSG00000257303 | 35.77414 | -0.226225539 | 0.34348 | -0.658628 | 0.510135 | 0.879714521 | | antisense | 46.83 | -0.012591315 | 0.012591315 | ENSG00000257303 | n.s. |
| ENSG00000275810 | 11.03348 | -0.259156878 | 0.445013 | -0.582358 | 0.560325 | 0.894198873 | MIR8082 | miRNA | 50.62 | -0.012586182 | 0.012586182 | ENSG00000275810 | n.s. |
| ENSG00000116957 | 867.9466 | -0.200806396 | 0.280908 | -0.714846 | 0.474704 | 0.865616576 | TBCE | protein_coding | 44.4 | -0.012585428 | 0.012585428 | ENSG00000116957 | n.s. |
| ENSG00000149346 | 254.3788 | -0.209374877 | 0.301809 | -0.693734 | 0.487849 | 0.870756877 | SLX4IP | protein_coding | 40.51 | -0.012584076 | 0.012584076 | ENSG00000149346 | n.s. |
| ENSG00000138107 | 1371.777 | -0.200710131 | 0.282541 | -0.710376 | 0.477471 | 0.865616576 | ACTR1A | protein_coding | 50.48 | -0.012579394 | 0.012579394 | ENSG00000138107 | n.s. |
| ENSG00000115233 | 3603.851 | -0.203123158 | 0.288238 | -0.704707 | 0.480992 | 0.867184458 | PSMD14 | protein_coding | 35.42 | -0.012570991 | 0.012570991 | ENSG00000115233 | n.s. |
| ENSG00000138071 | 2451.707 | -0.20053193 | 0.280523 | -0.714851 | 0.474701 | 0.865616576 | ACTR2 | protein_coding | 39.48 | -0.012568226 | 0.012568226 | ENSG00000138071 | n.s. |

| | | | | | | | | | | | | | |
|---|---|---|---|---|---|---|---|---|---|---|---|---|---|
| ENSG00000101464 | 266.9515 | -0.203180127 | 0.288803 | -0.703526 | 0.481728 | 0.867402024 | PIGU | protein_coding | 44.54 | -0.012552381 | 0.012552381 | ENSG00000101464 | n.s. |
| ENSG00000222489 | 175.5053 | 0.207439525 | 0.298491 | 0.6949602 | 0.48708 | 0.870096539 | SNORA79B | snoRNA | 45.95 | 0.012536101 | 0.012536101 | ENSG00000222489 | n.s. |
| ENSG00000102178 | 513.4493 | -0.201472632 | 0.284995 | -0.706934 | 0.479607 | 0.866541775 | UBL4A | protein_coding | 61.17 | -0.012533712 | 0.012533712 | ENSG00000102178 | n.s. |
| ENSG00000125779 | 1762.224 | -0.17902446 | 0.236677 | -0.756409 | 0.449404 | 0.85118703 | PANK2 | protein_coding | 45.01 | -0.012527237 | 0.012527237 | ENSG00000125779 | n.s. |
| ENSG00000168827 | 2887.861 | 0.178962363 | 0.236359 | 0.757162 | 0.448953 | 0.85118703 | GFM1 | protein_coding | 36.9 | 0.012522892 | 0.012522892 | ENSG00000168827 | n.s. |
| ENSG00000115226 | 105.5648 | -0.215554577 | 0.319171 | -0.675358 | 0.499449 | 0.875024797 | FNDC4 | protein_coding | 57.84 | -0.012497777 | 0.012497777 | ENSG00000115226 | n.s. |
| ENSG00000200792 | 22.05108 | -0.237817923 | 0.373827 | -0.636171 | 0.524665 | 0.886111278 | SNORA80A | snoRNA | 55.88 | -0.012488232 | 0.012488232 | ENSG00000200792 | n.s. |
| ENSG00000119946 | 330.8837 | -0.182571326 | 0.244175 | -0.747706 | 0.454638 | 0.854318522 | CNNM1 | protein_coding | 44.48 | -0.01248426 | 0.01248426 | ENSG00000119946 | n.s. |
| ENSG00000168014 | 122.5665 | 0.186925922 | 0.253019 | 0.7387836 | 0.460038 | 0.857465447 | C2CD3 | protein_coding | 40.35 | 0.012483543 | 0.012483543 | ENSG00000168014 | n.s. |
| ENSG00000274898 | 19.99561 | 0.254286662 | 0.429437 | 0.5921399 | 0.553757 | 0.893144718 | | lincRNA | 43.78 | 0.012479923 | 0.012479923 | ENSG00000274898 | n.s. |
| ENSG00000201098 | 63.05455 | 0.210774812 | 0.306064 | 0.6886633 | 0.491035 | 0.872562973 | RNY1 | misc_RNA | 44.25 | 0.012478548 | 0.012478548 | ENSG00000201098 | n.s. |
| ENSG00000186019 | 314.3969 | 0.16417169 | 0.207974 | 0.7893843 | 0.429887 | 0.839456232 | | antisense | 39.85 | 0.012477367 | 0.012477367 | ENSG00000186019 | n.s. |
| ENSG00000122068 | 1673.45 | -0.195801723 | 0.271762 | -0.720489 | 0.471224 | 0.863532918 | FYTTD1 | protein_coding | 42.18 | -0.012476702 | 0.012476702 | ENSG00000122068 | n.s. |
| ENSG00000131002 | 2532.992 | -0.203534776 | 0.290471 | -0.700707 | 0.483486 | 0.868366486 | TXLNGY | cribed_unprocessed_pseudc | 36.42 | -0.01247606 | 0.01247606 | ENSG00000131002 | n.s. |
| ENSG00000101365 | 1148.999 | -0.190713189 | 0.260769 | -0.73135 | 0.464565 | 0.860217499 | IDH3B | protein_coding | 51.38 | -0.012471065 | 0.012471065 | ENSG00000101365 | n.s. |
| ENSG00000266900 | 158.2327 | -0.201470181 | 0.285751 | -0.705054 | 0.480777 | 0.867184458 | | sense_intronic | 34.37 | -0.01246869 | 0.01246869 | ENSG00000266900 | n.s. |
| ENSG00000198626 | 2614.842 | -0.194005328 | 0.268527 | -0.722479 | 0.47 | 0.862503771 | RYR2 | protein_coding | 38.53 | -0.012462708 | 0.012462708 | ENSG00000198626 | n.s. |
| ENSG00000108010 | 1685.515 | -0.201333097 | 0.285383 | -0.705484 | 0.480509 | 0.867184458 | GLRX3 | protein_coding | 42.15 | -0.012460206 | 0.012460206 | ENSG00000108010 | n.s. |
| ENSG00000198298 | 207.4136 | 0.179486139 | 0.238604 | 0.7522339 | 0.45191 | 0.852336584 | ZNF485 | protein_coding | 40.92 | 0.012454341 | 0.012454341 | ENSG00000198298 | n.s. |
| ENSG00000279591 | 41.67075 | -0.229690299 | 0.352551 | -0.651509 | 0.514718 | 0.882641141 | | TEC | 51.96 | -0.012452851 | 0.012452851 | ENSG00000279591 | n.s. |
| ENSG00000147251 | 527.3256 | -0.198670569 | 0.278778 | -0.712649 | 0.476063 | 0.865616576 | DOCK11 | protein_coding | 39.68 | -0.012451566 | 0.012451566 | ENSG00000147251 | n.s. |
| ENSG00000159023 | 1460.89 | -0.1903992 | 0.260518 | -0.730848 | 0.464872 | 0.860217499 | EPB41 | protein_coding | 41.87 | -0.012450533 | 0.012450533 | ENSG00000159023 | n.s. |
| ENSG00000078403 | 1467.461 | -0.185758273 | 0.250833 | -0.740565 | 0.458957 | 0.856995387 | MLLT10 | protein_coding | 39.77 | -0.012449801 | 0.012449801 | ENSG00000078403 | n.s. |
| ENSG00000162521 | 2757.911 | -0.191276135 | 0.262346 | -0.7291 | 0.465941 | 0.860954429 | RBBP4 | protein_coding | 42.94 | -0.012436743 | 0.012436743 | ENSG00000162521 | n.s. |
| ENSG00000188610 | 120.3515 | -0.205221475 | 0.294105 | -0.697783 | 0.485313 | 0.869852312 | FAM72B | protein_coding | 41.53 | -0.012427079 | 0.012427079 | ENSG00000188610 | n.s. |
| ENSG00000100865 | 339.0949 | -0.209798801 | 0.304937 | -0.688007 | 0.491448 | 0.872664906 | CINP | protein_coding | 46.49 | -0.012410121 | 0.012410121 | ENSG00000100865 | n.s. |
| ENSG00000166454 | 1399.641 | -0.197847232 | 0.277645 | -0.712591 | 0.476099 | 0.865616576 | ATMIN | protein_coding | 44.63 | -0.012399964 | 0.012399964 | ENSG00000166454 | n.s. |
| ENSG00000054277 | 81.06304 | 0.215937849 | 0.322917 | 0.66871 | 0.50368 | 0.876169148 | OPN3 | protein_coding | 39.62 | 0.012397433 | 0.012397433 | ENSG00000054277 | n.s. |
| ENSG00000210144 | 16.79066 | -0.249381351 | 0.417028 | -0.597997 | 0.549842 | 0.891861801 | MT-TY | Mt_tRNA | 42.42 | -0.012394861 | 0.012394861 | ENSG00000210144 | n.s. |
| ENSG00000207951 | 69.22862 | -0.204818098 | 0.294306 | -0.695935 | 0.486469 | 0.870096539 | MIR561 | miRNA | 45.36 | -0.012377682 | 0.012377682 | ENSG00000207951 | n.s. |
| ENSG00000183258 | 973.0849 | -0.185347986 | 0.251156 | -0.73798 | 0.460527 | 0.857551846 | DDX41 | protein_coding | 59.55 | -0.012370053 | 0.012370053 | ENSG00000183258 | n.s. |
| ENSG00000166747 | 992.599 | -0.188687498 | 0.257688 | -0.732231 | 0.464027 | 0.859929897 | AP1G1 | protein_coding | 40.73 | -0.012366003 | 0.012366003 | ENSG00000166747 | n.s. |

| | | | | | | | | | | | | | |
|---|---|---|---|---|---|---|---|---|---|---|---|---|---|
| ENSG00000131055 | 28.24383 | -0.24596544 | 0.401694 | -0.612321 | 0.540326 | 0.890708138 | COX4I2 | protein_coding | 52.35 | -0.012363349 | 0.012363349 | ENSG00000131055 | n.s. |
| ENSG00000125966 | 487.5512 | -0.197241225 | 0.276045 | -0.714524 | 0.474903 | 0.865616576 | MMP24 | protein_coding | 49.16 | -0.012361982 | 0.012361982 | ENSG00000125966 | n.s. |
| ENSG00000140416 | 1867.516 | -0.190172694 | 0.261694 | -0.7267 | 0.46741 | 0.861088114 | TPM1 | protein_coding | 45.06 | -0.012352174 | 0.012352174 | ENSG00000140416 | n.s. |
| ENSG00000127483 | 2103.9 | -0.19347264 | 0.26836 | -0.720943 | 0.470944 | 0.863325865 | HP1BP3 | protein_coding | 41.17 | -0.012348439 | 0.012348439 | ENSG00000127483 | n.s. |
| ENSG00000099999 | 106.3408 | -0.204292345 | 0.293499 | -0.696057 | 0.486393 | 0.870096539 | RNF215 | protein_coding | 53.32 | -0.012345909 | 0.012345909 | ENSG00000099999 | n.s. |
| ENSG00000169683 | 487.2007 | 0.190231981 | 0.262056 | 0.72592 | 0.467888 | 0.861253285 | LRRC45 | protein_coding | 64.94 | 0.012340179 | 0.012340179 | ENSG00000169683 | n.s. |
| ENSG00000092853 | 902.7066 | -0.18595739 | 0.252943 | -0.735175 | 0.462233 | 0.858395846 | CLSPN | protein_coding | 42.79 | -0.012331279 | 0.012331279 | ENSG00000092853 | n.s. |
| ENSG00000197798 | 229.4023 | -0.206325138 | 0.2983 | -0.691671 | 0.489144 | 0.871557795 | FAM118B | protein_coding | 41.9 | -0.012318397 | 0.012318397 | ENSG00000197798 | n.s. |
| ENSG00000141385 | 2081.594 | -0.189640309 | 0.260367 | -0.728359 | 0.466394 | 0.861088114 | AFG3L2 | protein_coding | 44.7 | -0.012317594 | 0.012317594 | ENSG00000141385 | n.s. |
| ENSG00000126803 | 244.5911 | 0.17464488 | 0.230011 | 0.7592893 | 0.44768 | 0.850405175 | HSPA2 | protein_coding | 45.54 | 0.012290477 | 0.012290477 | ENSG00000126803 | n.s. |
| ENSG00000005436 | 1353.28 | -0.195008768 | 0.271748 | -0.717609 | 0.472999 | 0.865034671 | GCFC2 | protein_coding | 36.39 | -0.012279017 | 0.012279017 | ENSG00000005436 | n.s. |
| ENSG00000172340 | 220.8063 | -0.208901658 | 0.306243 | -0.682143 | 0.495148 | 0.873485531 | SUCLG2 | protein_coding | 39.38 | -0.012271778 | 0.012271778 | ENSG00000172340 | n.s. |
| ENSG00000134419 | 18413.1 | 0.213437354 | 0.319151 | 0.6687661 | 0.503645 | 0.876169148 | RPS15A | protein_coding | 46.41 | 0.012253875 | 0.012253875 | ENSG00000134419 | n.s. |
| ENSG00000236756 | 16.87667 | 0.267451445 | 0.474629 | 0.5634962 | 0.573097 | 0.899879367 | DNAJC9-AS1 | antisense | 41.21 | 0.012253477 | 0.012253477 | ENSG00000236756 | n.s. |
| ENSG00000158710 | 120.2243 | 0.215252179 | 0.323379 | 0.665634 | 0.505645 | 0.877161293 | TAGLN2 | protein_coding | 56.56 | 0.01225227 | 0.01225227 | ENSG00000158710 | n.s. |
| ENSG00000256060 | 125.2144 | 0.200132844 | 0.285814 | 0.7002211 | 0.483789 | 0.868574814 | | protein_coding | 50.95 | 0.012246683 | 0.012246683 | ENSG00000256060 | n.s. |
| ENSG00000165246 | 238.4049 | -0.190485942 | 0.263685 | -0.722399 | 0.470049 | 0.862503771 | NLGN4Y | protein_coding | 38.62 | -0.012236626 | 0.012236626 | ENSG00000165246 | n.s. |
| ENSG00000196576 | 796.0938 | -0.184820854 | 0.251802 | -0.733994 | 0.462952 | 0.858623382 | PLXNB2 | protein_coding | 64.97 | -0.01223464 | 0.01223464 | ENSG00000196576 | n.s. |
| ENSG00000158373 | 1380.966 | -0.167437419 | 0.216696 | -0.772684 | 0.439709 | 0.845174857 | HIST1H2BD | protein_coding | 42.24 | -0.012231878 | 0.012231878 | ENSG00000158373 | n.s. |
| ENSG00000099821 | 408.6926 | -0.201531711 | 0.288727 | -0.698 | 0.485177 | 0.869720883 | POLRMT | protein_coding | 61.15 | -0.012216873 | 0.012216873 | ENSG00000099821 | n.s. |
| ENSG00000258366 | 66.39058 | -0.211368417 | 0.314468 | -0.672145 | 0.501491 | 0.875518559 | RTEL1 | protein_coding | 59.63 | -0.01220328 | 0.01220328 | ENSG00000258366 | n.s. |
| ENSG00000167969 | 409.2561 | -0.212074449 | 0.316892 | -0.669233 | 0.503347 | 0.876011739 | ECI1 | protein_coding | 55.22 | -0.012192176 | 0.012192176 | ENSG00000167969 | n.s. |
| ENSG00000099341 | 1546.277 | -0.212387246 | 0.317788 | -0.66833 | 0.503923 | 0.876248111 | PSMD8 | protein_coding | 53.6 | -0.012185273 | 0.012185273 | ENSG00000099341 | n.s. |
| ENSG00000204256 | 7056.854 | -0.160214329 | 0.202908 | -0.789589 | 0.429768 | 0.839380003 | BRD2 | protein_coding | 51.46 | -0.012182919 | 0.012182919 | ENSG00000204256 | n.s. |
| ENSG00000242540 | 112.8753 | -0.194257688 | 0.27147 | -0.715577 | 0.474253 | 0.865616576 | | 3prime_overlapping_ncRNA | 46.07 | -0.012174991 | 0.012174991 | ENSG00000242540 | n.s. |
| ENSG00000247400 | 135.802 | -0.181414012 | 0.244506 | -0.741961 | 0.458111 | 0.856847578 | DNAJC3-AS1 | lincRNA | 38.92 | -0.012172232 | 0.012172232 | ENSG00000247400 | n.s. |
| ENSG00000175213 | 217.7901 | -0.17581057 | 0.233929 | -0.751554 | 0.452319 | 0.852645722 | ZNF408 | protein_coding | 54.13 | -0.012171609 | 0.012171609 | ENSG00000175213 | n.s. |
| ENSG00000119242 | 92.87308 | -0.198549824 | 0.283122 | -0.701288 | 0.483124 | 0.868366486 | CCDC92 | protein_coding | 47.49 | -0.012170498 | 0.012170498 | ENSG00000119242 | n.s. |
| ENSG00000269176 | 38.24925 | -0.2169324 | 0.327814 | -0.661754 | 0.508129 | 0.878842567 | | antisense | 52.69 | -0.012167503 | 0.012167503 | ENSG00000269176 | n.s. |
| ENSG00000258890 | 3677.68 | -0.187311941 | 0.257166 | -0.72837 | 0.466387 | 0.861088114 | CEP95 | protein_coding | 40.77 | -0.012166361 | 0.012166361 | ENSG00000258890 | n.s. |
| ENSG00000136158 | 297.3785 | 0.174153576 | 0.230919 | 0.7541775 | 0.450743 | 0.851633857 | SPRY2 | protein_coding | 47.95 | 0.012146704 | 0.012146704 | ENSG00000136158 | n.s. |
| ENSG00000145147 | 62.09922 | 0.213926655 | 0.3219 | 0.6645742 | 0.506323 | 0.877502299 | SLIT2 | protein_coding | 36.01 | 0.012140709 | 0.012140709 | ENSG00000145147 | n.s. |

| | | | | | | | | | | | | | |
|---|---|---|---|---|---|---|---|---|---|---|---|---|---|
| ENSG00000119950 | 717.9014 | 0.193610847 | 0.272324 | 0.7109573 | 0.477111 | 0.865616576 | MXI1 | protein_coding | 40.13 | 0.012134451 | 0.012134451 | ENSG00000119950 | n.s. |
| ENSG00000100568 | 936.8349 | -0.19348202 | 0.271561 | -0.712481 | 0.476167 | 0.865616576 | VTI1B | protein_coding | 42.97 | -0.012126376 | 0.012126376 | ENSG00000100568 | n.s. |
| ENSG00000256248 | 14.1385 | 0.249623537 | 0.428713 | 0.5822624 | 0.56039 | 0.894198873 | | lincRNA | 38.25 | 0.012123187 | 0.012123187 | ENSG00000256248 | n.s. |
| ENSG00000227172 | 20.98624 | 0.261388349 | 0.45921 | 0.5692126 | 0.569212 | 0.898745341 | | antisense | 40.78 | 0.012118839 | 0.012118839 | ENSG00000227172 | n.s. |
| ENSG00000261420 | 109.8208 | -0.188115996 | 0.259991 | -0.723549 | 0.469343 | 0.862205928 | | antisense | 54.52 | -0.0121126 | 0.0121126 | ENSG00000261420 | n.s. |
| ENSG00000135486 | 33241.05 | -0.173104045 | 0.229354 | -0.754745 | 0.450402 | 0.851309042 | HNRNPA1 | protein_coding | 42.13 | -0.01210218 | 0.01210218 | ENSG00000135486 | n.s. |
| ENSG00000133619 | 64.93157 | -0.206010188 | 0.302028 | -0.68209 | 0.495182 | 0.873485531 | KRBA1 | protein_coding | 57.32 | -0.012101921 | 0.012101921 | ENSG00000133619 | n.s. |
| ENSG00000168003 | 2529.166 | -0.170287594 | 0.223222 | -0.762863 | 0.445545 | 0.849129133 | SLC3A2 | protein_coding | 48.27 | -0.012094891 | 0.012094891 | ENSG00000168003 | n.s. |
| ENSG00000111642 | 5643.756 | -0.177046257 | 0.237239 | -0.746278 | 0.4555 | 0.85453422 | CHD4 | protein_coding | 48.35 | -0.012087044 | 0.012087044 | ENSG00000111642 | n.s. |
| ENSG00000113391 | 538.8606 | -0.188081701 | 0.260119 | -0.723061 | 0.469642 | 0.862503771 | FAM172A | protein_coding | 36.4 | -0.01208218 | 0.01208218 | ENSG00000113391 | n.s. |
| ENSG00000106636 | 942.8095 | -0.193212703 | 0.272859 | -0.708104 | 0.478881 | 0.865903214 | YKT6 | protein_coding | 48.11 | -0.012081716 | 0.012081716 | ENSG00000106636 | n.s. |
| ENSG00000223496 | 1100.35 | -0.167121133 | 0.218018 | -0.766547 | 0.443351 | 0.846745122 | EXOSC6 | protein_coding | 45.72 | -0.01207405 | 0.01207405 | ENSG00000223496 | n.s. |
| ENSG00000115042 | 412.6565 | -0.18944071 | 0.263175 | -0.719827 | 0.471632 | 0.863532918 | FAHD2A | protein_coding | 49.55 | -0.012071371 | 0.012071371 | ENSG00000115042 | n.s. |
| ENSG00000173349 | 1453.229 | 0.154119949 | 0.19237 | 0.8011662 | 0.423035 | 0.834993891 | SFT2D3 | protein_coding | 48.54 | 0.012070166 | 0.012070166 | ENSG00000173349 | n.s. |
| ENSG00000065978 | 7345.725 | -0.187793205 | 0.259806 | -0.722821 | 0.46979 | 0.862503771 | YBX1 | protein_coding | 40.41 | -0.012063647 | 0.012063647 | ENSG00000065978 | n.s. |
| ENSG00000131116 | 1050.955 | -0.1992607 | 0.285873 | -0.697025 | 0.485787 | 0.869936401 | ZNF428 | protein_coding | 52.67 | -0.012057762 | 0.012057762 | ENSG00000131116 | n.s. |
| ENSG00000189403 | 18007.43 | -0.187252926 | 0.258741 | -0.723708 | 0.469245 | 0.862205928 | HMGB1 | protein_coding | 41.99 | -0.012057028 | 0.012057028 | ENSG00000189403 | n.s. |
| ENSG00000124279 | 451.9433 | -0.185425389 | 0.254272 | -0.72924 | 0.465855 | 0.860954429 | FASTKD3 | protein_coding | 37.73 | -0.012056328 | 0.012056328 | ENSG00000124279 | n.s. |
| ENSG00000254986 | 365.0312 | -0.192207508 | 0.270177 | -0.711414 | 0.476828 | 0.865616576 | DPP3 | protein_coding | 52.26 | -0.012046497 | 0.012046497 | ENSG00000254986 | n.s. |
| ENSG00000168303 | 843.672 | -0.185187294 | 0.254526 | -0.727577 | 0.466873 | 0.861088114 | MPLKIP | protein_coding | 40.64 | -0.01202836 | 0.01202836 | ENSG00000168303 | n.s. |
| ENSG00000183137 | 316.2055 | -0.204889092 | 0.300536 | -0.681746 | 0.495399 | 0.87358987 | CEP57L1 | protein_coding | 36.59 | -0.012025435 | 0.012025435 | ENSG00000183137 | n.s. |
| ENSG00000230910 | 298.1037 | 0.173173894 | 0.230274 | 0.7520342 | 0.45203 | 0.852382105 | | antisense | 37.68 | 0.012012324 | 0.012012324 | ENSG00000230910 | n.s. |
| ENSG00000111602 | 2225.051 | -0.191508909 | 0.269457 | -0.710721 | 0.477257 | 0.865616576 | TIMELESS | protein_coding | 46.62 | -0.012002713 | 0.012002713 | ENSG00000111602 | n.s. |
| ENSG00000161835 | 86.94983 | -0.203393829 | 0.296408 | -0.686196 | 0.492589 | 0.873000006 | GRASP | protein_coding | 58.66 | -0.011997338 | 0.011997338 | ENSG00000161835 | n.s. |
| ENSG00000266887 | 1.839163 | -0.445318232 | 1.216339 | -0.366114 | 0.71428 | 0.939885098 | MIR4535 | miRNA | 55.93 | -0.011990299 | 0.011990299 | ENSG00000266887 | n.s. |
| ENSG00000103671 | 295.4935 | -0.203831026 | 0.297972 | -0.684061 | 0.493936 | 0.873348456 | TRIP4 | protein_coding | 42.11 | -0.011987801 | 0.011987801 | ENSG00000103671 | n.s. |
| ENSG00000198301 | 1726.378 | -0.191161565 | 0.267512 | -0.714589 | 0.474863 | 0.865616576 | SDAD1 | protein_coding | 40.57 | -0.011980943 | 0.011980943 | ENSG00000198301 | n.s. |
| ENSG00000235109 | 63.08874 | 0.204261134 | 0.299886 | 0.6811292 | 0.49579 | 0.873669126 | ZSCAN31 | protein_coding | 41.29 | 0.011980531 | 0.011980531 | ENSG00000235109 | n.s. |
| ENSG00000113269 | 924.2523 | -0.202537647 | 0.294639 | -0.68741 | 0.491824 | 0.872671134 | RNF130 | protein_coding | 42.83 | -0.011979978 | 0.011979978 | ENSG00000113269 | n.s. |
| ENSG00000091592 | 132.7964 | -0.208911718 | 0.313239 | -0.66694 | 0.50481 | 0.876368641 | NLRP1 | protein_coding | 46.74 | -0.011973393 | 0.011973393 | ENSG00000091592 | n.s. |
| ENSG00000163655 | 3037.664 | -0.191028815 | 0.26949 | -0.708853 | 0.478416 | 0.865660698 | GMPS | protein_coding | 38.88 | -0.011968395 | 0.011968395 | ENSG00000163655 | n.s. |
| ENSG00000170248 | 2090.939 | -0.190946243 | 0.268298 | -0.711695 | 0.476654 | 0.865616576 | PDCD6IP | protein_coding | 36.93 | -0.011967448 | 0.011967448 | ENSG00000170248 | n.s. |

| | | | | | | | | | | | | | |
|---|---|---|---|---|---|---|---|---|---|---|---|---|---|
| ENSG00000134697 | 1541.985 | -0.184018831 | 0.252373 | -0.729153 | 0.465908 | 0.860954429 | GNL2 | protein_coding | 42.77 | -0.011964874 | 0.011964874 | ENSG00000134697 | n.s. |
| ENSG00000100991 | 1090.947 | -0.191326595 | 0.270166 | -0.708181 | 0.478833 | 0.865903214 | TRPC4AP | protein_coding | 42.95 | -0.011963776 | 0.011963776 | ENSG00000100991 | n.s. |
| ENSG00000125630 | 1802.05 | -0.182788364 | 0.249789 | -0.731772 | 0.464308 | 0.860105883 | POLR1B | protein_coding | 42.17 | -0.011963148 | 0.011963148 | ENSG00000125630 | n.s. |
| ENSG00000112339 | 2995.373 | -0.184182241 | 0.253341 | -0.727013 | 0.467218 | 0.861088114 | HBS1L | protein_coding | 36.66 | -0.01196308 | 0.01196308 | ENSG00000112339 | n.s. |
| ENSG00000128654 | 1052.583 | -0.197884322 | 0.284749 | -0.694943 | 0.487091 | 0.870096539 | MTX2 | protein_coding | 35.02 | -0.011958656 | 0.011958656 | ENSG00000128654 | n.s. |
| ENSG00000278053 | 1506.799 | -0.190844724 | 0.269082 | -0.709245 | 0.478172 | 0.865660698 | DDX52 | protein_coding | 39.57 | -0.011956861 | 0.011956861 | ENSG00000278053 | n.s. |
| ENSG00000148672 | 1917.787 | -0.193073107 | 0.273699 | -0.705423 | 0.480547 | 0.867184458 | GLUD1 | protein_coding | 42.11 | -0.011949008 | 0.011949008 | ENSG00000148672 | n.s. |
| ENSG00000186193 | 561.2524 | -0.191288954 | 0.270311 | -0.707663 | 0.479154 | 0.866036769 | SAPCD2 | protein_coding | 63.61 | -0.01194861 | 0.01194861 | ENSG00000186193 | n.s. |
| ENSG00000175575 | 484.9976 | -0.1781827 | 0.240658 | -0.740398 | 0.459059 | 0.856995387 | PAAF1 | protein_coding | 42.99 | -0.011942075 | 0.011942075 | ENSG00000175575 | n.s. |
| ENSG00000150456 | 96.59009 | -0.206518407 | 0.306536 | -0.673717 | 0.500491 | 0.875342302 | EEF1AKMT1 | protein_coding | 41.53 | -0.011941325 | 0.011941325 | ENSG00000150456 | n.s. |
| ENSG00000212384 | 8.352661 | -0.276655536 | 0.511104 | -0.54129 | 0.588308 | 0.905402821 | SNORD113-2 | snoRNA | 31.94 | -0.011939943 | 0.011939943 | ENSG00000212384 | n.s. |
| ENSG00000109265 | 1126.262 | -0.183664462 | 0.25284 | -0.726405 | 0.467591 | 0.861088114 | KIAA1211 | protein_coding | 41.29 | -0.011929449 | 0.011929449 | ENSG00000109265 | n.s. |
| ENSG00000196371 | 330.4362 | 0.173034116 | 0.230798 | 0.7497226 | 0.453422 | 0.853222234 | FUT4 | protein_coding | 49.18 | 0.011928597 | 0.011928597 | ENSG00000196371 | n.s. |
| ENSG00000150756 | 282.3172 | -0.200553613 | 0.290438 | -0.690521 | 0.489867 | 0.872064188 | FAM173B | protein_coding | 42.65 | -0.011923223 | 0.011923223 | ENSG00000150756 | n.s. |
| ENSG00000196756 | 1492.809 | -0.16006007 | 0.204283 | -0.78352 | 0.433322 | 0.842381919 | SNHG17 | processed_transcript | 51.31 | -0.011923029 | 0.011923029 | ENSG00000196756 | n.s. |
| ENSG00000137872 | 1053.307 | -0.179130796 | 0.243037 | -0.737051 | 0.461091 | 0.857914774 | SEMA6D | protein_coding | 38.82 | -0.011922203 | 0.011922203 | ENSG00000137872 | n.s. |
| ENSG00000221639 | 21.12149 | -0.226993777 | 0.358076 | -0.633927 | 0.526129 | 0.88615396 | | snoRNA | 52.8 | -0.011915089 | 0.011915089 | ENSG00000221639 | n.s. |
| ENSG00000279259 | 55.0495 | -0.204942565 | 0.302955 | -0.676479 | 0.498737 | 0.874768485 | | TEC | 57.68 | -0.011908571 | 0.011908571 | ENSG00000279259 | n.s. |
| ENSG00000143179 | 2993.91 | -0.184371229 | 0.254526 | -0.72437 | 0.468839 | 0.861812061 | UCK2 | protein_coding | 44.22 | -0.011908064 | 0.011908064 | ENSG00000143179 | n.s. |
| ENSG00000182263 | 656.7556 | -0.178252019 | 0.241276 | -0.73879 | 0.460034 | 0.857465447 | FIGN | protein_coding | 36.3 | -0.011904271 | 0.011904271 | ENSG00000182263 | n.s. |
| ENSG00000156976 | 19331.3 | 0.167092286 | 0.218868 | 0.7634381 | 0.445202 | 0.848767576 | EIF4A2 | protein_coding | 41.94 | 0.011898845 | 0.011898845 | ENSG00000156976 | n.s. |
| ENSG00000132535 | 94.14 | 0.18551766 | 0.257085 | 0.7216203 | 0.470528 | 0.862789535 | DLG4 | protein_coding | 55.08 | 0.011890779 | 0.011890779 | ENSG00000132535 | n.s. |
| ENSG00000176834 | 32.18391 | 0.227942617 | 0.36096 | 0.6314896 | 0.52772 | 0.886936263 | VSIG10 | protein_coding | 47.55 | 0.01187754 | 0.01187754 | ENSG00000176834 | n.s. |
| ENSG00000185619 | 1471.475 | -0.191842748 | 0.271685 | -0.706121 | 0.480113 | 0.867148032 | PCGF3 | protein_coding | 56.09 | -0.011876362 | 0.011876362 | ENSG00000185619 | n.s. |
| ENSG00000102384 | 419.8609 | -0.192105639 | 0.27295 | -0.703813 | 0.48155 | 0.867349802 | CENPI | protein_coding | 40.62 | -0.011873227 | 0.011873227 | ENSG00000102384 | n.s. |
| ENSG00000108179 | 1891.767 | -0.173525146 | 0.232085 | -0.74768 | 0.454653 | 0.854318522 | PPIF | protein_coding | 55.59 | -0.01186568 | 0.01186568 | ENSG00000108179 | n.s. |
| ENSG00000184182 | 356.8307 | -0.195920417 | 0.280938 | -0.69738 | 0.485565 | 0.869934871 | UBE2F | protein_coding | 44.85 | -0.011855783 | 0.011855783 | ENSG00000184182 | n.s. |
| ENSG00000174197 | 2546.584 | -0.180867646 | 0.247539 | -0.730662 | 0.464986 | 0.860217499 | MGA | protein_coding | 40.41 | -0.011827248 | 0.011827248 | ENSG00000174197 | n.s. |
| ENSG00000127837 | 1245.733 | -0.189994188 | 0.268714 | -0.707051 | 0.479535 | 0.866523947 | AAMP | protein_coding | 56.53 | -0.01182133 | 0.01182133 | ENSG00000127837 | n.s. |
| ENSG00000145029 | 90.87264 | -0.207067125 | 0.310869 | -0.666092 | 0.505352 | 0.876821729 | NICN1 | protein_coding | 52.25 | -0.011821192 | 0.011821192 | ENSG00000145029 | n.s. |
| ENSG00000129696 | 330.1427 | -0.185405334 | 0.257498 | -0.720026 | 0.471509 | 0.863532918 | TTI2 | protein_coding | 42.98 | -0.011814232 | 0.011814232 | ENSG00000129696 | n.s. |
| ENSG00000164743 | 58.37295 | -0.195396927 | 0.281127 | -0.695048 | 0.487025 | 0.870096539 | C8orf48 | protein_coding | 42.28 | -0.011808336 | 0.011808336 | ENSG00000164743 | n.s. |

| | | | | | | | | | | | | | |
|---|---|---|---|---|---|---|---|---|---|---|---|---|---|
| ENSG00000100258 | 294.2476 | -0.18833471 | 0.265225 | -0.710095 | 0.477645 | 0.865660698 | LMF2 | protein_coding | 67.15 | -0.011799603 | 0.011799603 | ENSG00000100258 | n.s. |
| ENSG00000261934 | 100.0594 | -0.188273012 | 0.265638 | -0.708757 | 0.478475 | 0.865660698 | PCDHGA9 | protein_coding | 45.5 | -0.011795737 | 0.011795737 | ENSG00000261934 | n.s. |
| ENSG00000150627 | 695.2389 | -0.19274434 | 0.275361 | -0.69997 | 0.483946 | 0.868580821 | WDR17 | protein_coding | 34.28 | -0.011793981 | 0.011793981 | ENSG00000150627 | n.s. |
| ENSG00000207561 | 6.422563 | 0.284171382 | 0.538758 | 0.5274564 | 0.597877 | 0.908868121 | MIR635 | miRNA | 43.88 | 0.011792865 | 0.011792865 | ENSG00000207561 | n.s. |
| ENSG00000138286 | 237.3763 | -0.196199418 | 0.282849 | -0.693655 | 0.487898 | 0.870756877 | FAM149B1 | protein_coding | 39.89 | -0.011792191 | 0.011792191 | ENSG00000138286 | n.s. |
| ENSG00000141013 | 254.1046 | -0.188093517 | 0.263951 | -0.712608 | 0.476088 | 0.865616576 | GAS8 | protein_coding | 52.63 | -0.011788655 | 0.011788655 | ENSG00000141013 | n.s. |
| ENSG00000123219 | 1125.789 | -0.187835433 | 0.264266 | -0.710782 | 0.477219 | 0.865616576 | CENPK | protein_coding | 37.27 | -0.01177248 | 0.01177248 | ENSG00000123219 | n.s. |
| ENSG00000050030 | 533.5133 | -0.181213258 | 0.249279 | -0.726949 | 0.467257 | 0.861088114 | KIAA2022 | protein_coding | 39.49 | -0.011770237 | 0.011770237 | ENSG00000050030 | n.s. |
| ENSG00000223189 | 8.173592 | 0.281627506 | 0.527752 | 0.5336361 | 0.593593 | 0.908297635 | RNU6-177P | snRNA | 40.38 | 0.011764093 | 0.011764093 | ENSG00000223189 | n.s. |
| ENSG00000174720 | 1794.456 | -0.194247588 | 0.278714 | -0.696942 | 0.485839 | 0.869936401 | LARP7 | protein_coding | 37.92 | -0.011754406 | 0.011754406 | ENSG00000174720 | n.s. |
| ENSG00000205765 | 662.7237 | -0.187304653 | 0.264244 | -0.708833 | 0.478428 | 0.865660698 | C5orf51 | protein_coding | 36.01 | -0.011735067 | 0.011735067 | ENSG00000205765 | n.s. |
| ENSG00000105750 | 266.0114 | -0.197990843 | 0.287225 | -0.689323 | 0.49062 | 0.872495697 | ZNF85 | protein_coding | 37.87 | -0.011728325 | 0.011728325 | ENSG00000105750 | n.s. |
| ENSG00000187079 | 3235.507 | -0.182093569 | 0.25193 | -0.722794 | 0.469806 | 0.862503771 | TEAD1 | protein_coding | 42.75 | -0.011697508 | 0.011697508 | ENSG00000187079 | n.s. |
| ENSG00000116574 | 1168.08 | -0.192410173 | 0.275303 | -0.698904 | 0.484612 | 0.869379601 | RHOU | protein_coding | 46.54 | -0.011696721 | 0.011696721 | ENSG00000116574 | n.s. |
| ENSG00000131013 | 1145.098 | -0.189246533 | 0.268746 | -0.704184 | 0.481318 | 0.867349802 | PPIL4 | protein_coding | 38.44 | -0.011696517 | 0.011696517 | ENSG00000131013 | n.s. |
| ENSG00000272155 | 25.36214 | -0.326597297 | 0.697646 | -0.468142 | 0.639683 | 0.920848926 | | antisense | 39.55 | -0.011695976 | 0.011695976 | ENSG00000272155 | n.s. |
| ENSG00000130695 | 648.3452 | -0.190572254 | 0.271501 | -0.70192 | 0.482729 | 0.868238739 | CEP85 | protein_coding | 44.77 | -0.011693674 | 0.011693674 | ENSG00000130695 | n.s. |
| ENSG00000188529 | 6815.245 | 0.174450927 | 0.23548 | 0.7408315 | 0.458796 | 0.856995387 | SRSF10 | protein_coding | 39.02 | 0.011691966 | 0.011691966 | ENSG00000188529 | n.s. |
| ENSG00000275056 | 99.04226 | 0.186416564 | 0.26204 | 0.7114037 | 0.476834 | 0.865616576 | | sense_intronic | 45.74 | 0.011683553 | 0.011683553 | ENSG00000275056 | n.s. |
| ENSG00000258947 | 1440.876 | -0.200366603 | 0.295021 | -0.679159 | 0.497037 | 0.874430631 | TUBB3 | protein_coding | 58.5 | -0.011676291 | 0.011676291 | ENSG00000258947 | n.s. |
| ENSG00000279702 | 19.49519 | -0.291729539 | 0.577345 | -0.505295 | 0.613352 | 0.911981962 | | TEC | 32.92 | -0.011673193 | 0.011673193 | ENSG00000279702 | n.s. |
| ENSG00000092203 | 625.0668 | -0.197304433 | 0.287051 | -0.687351 | 0.491862 | 0.872671134 | TOX4 | protein_coding | 43.81 | -0.011670437 | 0.011670437 | ENSG00000092203 | n.s. |
| ENSG00000171914 | 42.81208 | -0.204130239 | 0.306407 | -0.666206 | 0.50528 | 0.876821729 | TLN2 | protein_coding | 43.6 | -0.011653529 | 0.011653529 | ENSG00000171914 | n.s. |
| ENSG00000006125 | 1374.014 | -0.185894137 | 0.26025 | -0.714292 | 0.475047 | 0.865616576 | AP2B1 | protein_coding | 39.93 | -0.01165081 | 0.01165081 | ENSG00000006125 | n.s. |
| ENSG00000077458 | 1076.091 | 0.186117553 | 0.262687 | 0.7085156 | 0.478625 | 0.865778531 | FAM76B | protein_coding | 34.6 | 0.011649691 | 0.011649691 | ENSG00000077458 | n.s. |
| ENSG00000152642 | 513.1672 | 0.195654083 | 0.283199 | 0.6908715 | 0.489646 | 0.871894847 | GPD1L | protein_coding | 44.49 | 0.01164844 | 0.01164844 | ENSG00000152642 | n.s. |
| ENSG00000167553 | 173.9422 | 0.221731477 | 0.347159 | 0.6387026 | 0.523016 | 0.886079926 | TUBA1C | protein_coding | 45.2 | 0.011646912 | 0.011646912 | ENSG00000167553 | n.s. |
| ENSG00000196357 | 91.31217 | -0.188115075 | 0.266949 | -0.704684 | 0.481007 | 0.867184458 | ZNF565 | protein_coding | 45.12 | -0.011642163 | 0.011642163 | ENSG00000196357 | n.s. |
| ENSG00000122643 | 1106.902 | -0.189242322 | 0.269399 | -0.702461 | 0.482392 | 0.867933207 | NT5C3A | protein_coding | 37.98 | -0.011640994 | 0.011640994 | ENSG00000122643 | n.s. |
| ENSG00000154310 | 624.1822 | 0.189885735 | 0.270832 | 0.7011202 | 0.483228 | 0.868366486 | TNIK | protein_coding | 39.55 | 0.011639416 | 0.011639416 | ENSG00000154310 | n.s. |
| ENSG00000135632 | 611.4747 | -0.175187719 | 0.237866 | -0.736497 | 0.461429 | 0.858145726 | SMYD5 | protein_coding | 50.97 | -0.011639289 | 0.011639289 | ENSG00000135632 | n.s. |
| ENSG00000163950 | 1692.435 | -0.196376476 | 0.284904 | -0.689272 | 0.490652 | 0.872495697 | SLBP | protein_coding | 47.01 | -0.011632696 | 0.011632696 | ENSG00000163950 | n.s. |

| | | | | | | | | | | | | | |
|---|---|---|---|---|---|---|---|---|---|---|---|---|---|
| ENSG00000110218 | 969.8579 | -0.185590155 | 0.261136 | -0.710704 | 0.477268 | 0.865616576 | PANX1 | protein_coding | 43.1 | -0.011631758 | 0.011631758 | ENSG00000110218 | n.s. |
| ENSG00000272028 | 7.227492 | 0.280269661 | 0.533098 | 0.525738 | 0.59907 | 0.908868121 | RNU6-87P | snRNA | 41.67 | 0.011630947 | 0.011630947 | ENSG00000272028 | n.s. |
| ENSG00000274308 | 32.27142 | -0.20693291 | 0.312539 | -0.662103 | 0.507905 | 0.878673133 | | processed_pseudogene | 55.77 | -0.01162397 | 0.01162397 | ENSG00000274308 | n.s. |
| ENSG00000138180 | 1098.306 | -0.197722073 | 0.289738 | -0.682416 | 0.494976 | 0.873453717 | CEP55 | protein_coding | 40.68 | -0.011618169 | 0.011618169 | ENSG00000138180 | n.s. |
| ENSG00000120694 | 5461.966 | -0.179741892 | 0.247928 | -0.724977 | 0.468466 | 0.861732907 | HSPH1 | protein_coding | 36.82 | -0.011616237 | 0.011616237 | ENSG00000120694 | n.s. |
| ENSG00000164002 | 281.5232 | -0.188597347 | 0.268507 | -0.702392 | 0.482434 | 0.867933207 | EXO5 | protein_coding | 43.67 | -0.01160132 | 0.01160132 | ENSG00000164002 | n.s. |
| ENSG00000165802 | 843.8152 | -0.189203416 | 0.269989 | -0.700782 | 0.483439 | 0.868366486 | NSMF | protein_coding | 65.22 | -0.011597592 | 0.011597592 | ENSG00000165802 | n.s. |
| ENSG00000145494 | 775.4919 | -0.200523172 | 0.297861 | -0.67321 | 0.500814 | 0.875342302 | NDUFS6 | protein_coding | 48.37 | -0.011594668 | 0.011594668 | ENSG00000145494 | n.s. |
| ENSG00000101935 | 669.2915 | 0.199369728 | 0.294305 | 0.677426 | 0.498136 | 0.874675553 | AMMECR1 | protein_coding | 38.78 | 0.01159395 | 0.01159395 | ENSG00000101935 | n.s. |
| ENSG00000179841 | 354.8597 | -0.164197943 | 0.215522 | -0.761863 | 0.446142 | 0.849976609 | AKAP5 | protein_coding | 37.55 | -0.01159123 | 0.01159123 | ENSG00000179841 | n.s. |
| ENSG00000123473 | 104.6295 | -0.187551342 | 0.2667 | -0.70323 | 0.481912 | 0.867402024 | STIL | protein_coding | 40.02 | -0.011586841 | 0.011586841 | ENSG00000123473 | n.s. |
| ENSG00000131791 | 937.6371 | -0.184439966 | 0.258805 | -0.712659 | 0.476057 | 0.865616576 | PRKAB2 | protein_coding | 39.83 | -0.011559671 | 0.011559671 | ENSG00000131791 | n.s. |
| ENSG00000146223 | 4533.881 | -0.184503636 | 0.259976 | -0.709696 | 0.477893 | 0.865660698 | RPL7L1 | protein_coding | 46.45 | -0.011559577 | 0.011559577 | ENSG00000146223 | n.s. |
| ENSG00000170270 | 357.0798 | -0.203631 | 0.306607 | -0.664144 | 0.506598 | 0.877502299 | GON7 | protein_coding | 43.35 | -0.011556413 | 0.011556413 | ENSG00000170270 | n.s. |
| ENSG00000112029 | 2698.22 | -0.183029483 | 0.254884 | -0.71809 | 0.472702 | 0.864725667 | FBXO5 | protein_coding | 38.53 | -0.011553123 | 0.011553123 | ENSG00000112029 | n.s. |
| ENSG00000112118 | 7438.996 | -0.184329607 | 0.25984 | -0.709396 | 0.478078 | 0.865660698 | MCM3 | protein_coding | 43.17 | -0.011548674 | 0.011548674 | ENSG00000112118 | n.s. |
| ENSG00000132716 | 2020.634 | -0.184133517 | 0.258263 | -0.712968 | 0.475865 | 0.865616576 | DCAF8 | protein_coding | 42.25 | -0.011540464 | 0.011540464 | ENSG00000132716 | n.s. |
| ENSG00000070785 | 455.3314 | -0.196679268 | 0.288863 | -0.680873 | 0.495952 | 0.873669126 | EIF2B3 | protein_coding | 42.55 | -0.011535831 | 0.011535831 | ENSG00000070785 | n.s. |
| ENSG00000178449 | 316.7574 | 0.222238679 | 0.355232 | 0.6256159 | 0.531567 | 0.887366671 | COX14 | protein_coding | 49.49 | 0.011533496 | 0.011533496 | ENSG00000178449 | n.s. |
| ENSG00000136051 | 2727.136 | -0.190582943 | 0.273594 | -0.69659 | 0.48606 | 0.869962727 | WASHC4 | protein_coding | 35.27 | -0.011530145 | 0.011530145 | ENSG00000136051 | n.s. |
| ENSG00000145687 | 41.78125 | 0.230716363 | 0.378724 | 0.6091942 | 0.542396 | 0.891406027 | SSBP2 | protein_coding | 35.97 | 0.011518384 | 0.011518384 | ENSG00000145687 | n.s. |
| ENSG00000103356 | 846.4497 | -0.183765153 | 0.257591 | -0.713399 | 0.475599 | 0.865616576 | EARS2 | protein_coding | 47.93 | -0.011517377 | 0.011517377 | ENSG00000103356 | n.s. |
| ENSG00000064601 | 86.04746 | 0.199851808 | 0.297694 | 0.6713341 | 0.502008 | 0.875813709 | CTSA | protein_coding | 56.04 | 0.011509118 | 0.011509118 | ENSG00000064601 | n.s. |
| ENSG00000147649 | 2481.48 | -0.190183411 | 0.272727 | -0.697339 | 0.485591 | 0.869934871 | MTDH | protein_coding | 40.6 | -0.011508618 | 0.011508618 | ENSG00000147649 | n.s. |
| ENSG00000262904 | 79.65042 | 0.196418337 | 0.288781 | 0.6801644 | 0.4964 | 0.873863041 | TMPOP2 | processed_pseudogene | 43.63 | 0.011501596 | 0.011501596 | ENSG00000262904 | n.s. |
| ENSG00000188725 | 1424.461 | 0.178828624 | 0.247235 | 0.7233137 | 0.469487 | 0.862357563 | SMIM15 | protein_coding | 36.57 | 0.011500938 | 0.011500938 | ENSG00000188725 | n.s. |
| ENSG00000182134 | 251.3527 | -0.2023258 | 0.304145 | -0.665228 | 0.505905 | 0.87732845 | TDRKH | protein_coding | 43.72 | -0.011499751 | 0.011499751 | ENSG00000182134 | n.s. |
| ENSG00000099942 | 2379.824 | -0.177037567 | 0.242993 | -0.728571 | 0.466264 | 0.861088114 | CRKL | protein_coding | 45.76 | -0.011499016 | 0.011499016 | ENSG00000099942 | n.s. |
| ENSG00000148158 | 798.1654 | -0.180353077 | 0.250691 | -0.719423 | 0.47188 | 0.863532918 | SNX30 | protein_coding | 44.26 | -0.011492297 | 0.011492297 | ENSG00000148158 | n.s. |
| ENSG00000130726 | 4461.402 | -0.179044508 | 0.248086 | -0.721703 | 0.470477 | 0.862789535 | TRIM28 | protein_coding | 59.3 | -0.011475881 | 0.011475881 | ENSG00000130726 | n.s. |
| ENSG00000148798 | 2126.59 | -0.182997662 | 0.25719 | -0.711527 | 0.476758 | 0.865616576 | INA | protein_coding | 45.88 | -0.011469275 | 0.011469275 | ENSG00000148798 | n.s. |
| ENSG00000253457 | 136.3948 | 0.220087724 | 0.350056 | 0.6287212 | 0.529532 | 0.886954686 | SMIM18 | protein_coding | 36.33 | 0.011466255 | 0.011466255 | ENSG00000253457 | n.s. |

| | | | | | | | | | | | | | |
|---|---|---|---|---|---|---|---|---|---|---|---|---|---|
| ENSG00000137965 | 2631.059 | -0.185491266 | 0.263794 | -0.703166 | 0.481952 | 0.867402024 | IFI44 | protein_coding | 35.31 | -0.01145957 | 0.01145957 | ENSG00000137965 | n.s. |
| ENSG00000076248 | 2663.663 | -0.182796308 | 0.256979 | -0.711327 | 0.476881 | 0.865616576 | UNG | protein_coding | 47.32 | -0.011456655 | 0.011456655 | ENSG00000076248 | n.s. |
| ENSG00000157916 | 416.9475 | -0.196995677 | 0.290872 | -0.677259 | 0.498241 | 0.874675553 | RER1 | protein_coding | 53.95 | -0.011455892 | 0.011455892 | ENSG00000157916 | n.s. |
| ENSG00000160408 | 568.7553 | -0.189527243 | 0.272703 | -0.694996 | 0.487058 | 0.870096539 | ST6GALNAC6 | protein_coding | 53.52 | -0.011453616 | 0.011453616 | ENSG00000160408 | n.s. |
| ENSG00000119862 | 500.0468 | 0.18270223 | 0.256333 | 0.7127548 | 0.475998 | 0.865616576 | LGALSL | protein_coding | 44.8 | 0.011450759 | 0.011450759 | ENSG00000119862 | n.s. |
| ENSG00000122958 | 843.0669 | -0.194849152 | 0.285359 | -0.682822 | 0.494719 | 0.873443478 | VPS26A | protein_coding | 39.68 | -0.011450348 | 0.011450348 | ENSG00000122958 | n.s. |
| ENSG00000170619 | 291.9861 | -0.197384648 | 0.291978 | -0.676025 | 0.499024 | 0.874967757 | COMMD5 | protein_coding | 53.13 | -0.011449879 | 0.011449879 | ENSG00000170619 | n.s. |
| ENSG00000111731 | 2028.16 | -0.193889348 | 0.282499 | -0.686336 | 0.492502 | 0.873000006 | C2CD5 | protein_coding | 34.95 | -0.011436709 | 0.011436709 | ENSG00000111731 | n.s. |
| ENSG00000101150 | 1081.744 | 0.205477669 | 0.312048 | 0.6584805 | 0.510229 | 0.879714521 | TPD52L2 | protein_coding | 54.36 | 0.011436525 | 0.011436525 | ENSG00000101150 | n.s. |
| ENSG00000264947 | 4.589186 | 0.329453748 | 0.718004 | 0.4588469 | 0.646344 | 0.923184844 | MIR3181 | miRNA | 83.56 | 0.011435779 | 0.011435779 | ENSG00000264947 | n.s. |
| ENSG00000128881 | 1192.114 | 0.194420911 | 0.284087 | 0.6843718 | 0.49374 | 0.873348456 | TTBK2 | protein_coding | 39.87 | 0.011434369 | 0.011434369 | ENSG00000128881 | n.s. |
| ENSG00000127870 | 1243.492 | 0.176310819 | 0.24304 | 0.7254391 | 0.468183 | 0.861324991 | RNF6 | protein_coding | 40.18 | 0.011430751 | 0.011430751 | ENSG00000127870 | n.s. |
| ENSG00000266019 | 144.7042 | 0.175619568 | 0.240662 | 0.7297343 | 0.465553 | 0.86087715 | MIR3609 | miRNA | 38.75 | 0.011425602 | 0.011425602 | ENSG00000266019 | n.s. |
| ENSG00000198885 | 622.7041 | -0.172139531 | 0.23382 | -0.736204 | 0.461606 | 0.858299252 | ITPRIPL1 | protein_coding | 50.7 | -0.011423397 | 0.011423397 | ENSG00000198885 | n.s. |
| ENSG00000132570 | 139.2053 | -0.20109542 | 0.302401 | -0.664996 | 0.506053 | 0.87743153 | PCBD2 | protein_coding | 42.21 | -0.011419558 | 0.011419558 | ENSG00000132570 | n.s. |
| ENSG00000007923 | 597.8575 | -0.192700692 | 0.279625 | -0.689139 | 0.490736 | 0.872495697 | DNAJC11 | protein_coding | 46.6 | -0.011414954 | 0.011414954 | ENSG00000007923 | n.s. |
| ENSG00000178031 | 740.4398 | -0.182124311 | 0.255558 | -0.712654 | 0.47606 | 0.865616576 | ADAMTSL1 | protein_coding | 38.59 | -0.011414538 | 0.011414538 | ENSG00000178031 | n.s. |
| ENSG00000105655 | 145.4485 | -0.198575075 | 0.296994 | -0.668616 | 0.50374 | 0.876169148 | ISYNA1 | protein_coding | 66.58 | -0.011400601 | 0.011400601 | ENSG00000105655 | n.s. |
| ENSG00000247708 | 63.36037 | 0.192508609 | 0.279616 | 0.6884752 | 0.491154 | 0.872562973 | STX18-AS1 | antisense | 45.62 | 0.01139713 | 0.01139713 | ENSG00000247708 | n.s. |
| ENSG00000198354 | 94.38714 | 0.185718806 | 0.265027 | 0.7007536 | 0.483457 | 0.868366486 | DCAF12L2 | protein_coding | 54.87 | 0.011383996 | 0.011383996 | ENSG00000198354 | n.s. |
| ENSG00000129003 | 2634.976 | -0.183869811 | 0.260818 | -0.704975 | 0.480826 | 0.867184458 | VPS13C | protein_coding | 34.88 | -0.011379429 | 0.011379429 | ENSG00000129003 | n.s. |
| ENSG00000221464 | 4.783429 | -0.310194147 | 0.639178 | -0.485302 | 0.627462 | 0.919033489 | MIR1271 | miRNA | 51.16 | -0.011374405 | 0.011374405 | ENSG00000221464 | n.s. |
| ENSG00000180834 | 87.94785 | -0.203198568 | 0.307418 | -0.660985 | 0.508622 | 0.879193676 | MAP6D1 | protein_coding | 54.58 | -0.011361938 | 0.011361938 | ENSG00000180834 | n.s. |
| ENSG00000119729 | 774.7118 | -0.193516017 | 0.283792 | -0.681893 | 0.495306 | 0.87358987 | RHOQ | protein_coding | 44.36 | -0.011357922 | 0.011357922 | ENSG00000119729 | n.s. |
| ENSG00000244754 | 3298.075 | -0.188499547 | 0.271582 | -0.694079 | 0.487632 | 0.870535462 | N4BP2L2 | protein_coding | 39.02 | -0.011350224 | 0.011350224 | ENSG00000244754 | n.s. |
| ENSG00000117151 | 335.9884 | -0.188841966 | 0.27227 | -0.693583 | 0.487944 | 0.870756877 | CTBS | protein_coding | 36.59 | -0.011349985 | 0.011349985 | ENSG00000117151 | n.s. |
| ENSG00000106028 | 3060.997 | 0.181031866 | 0.253726 | 0.7134949 | 0.47554 | 0.865616576 | SSBP1 | protein_coding | 40.78 | 0.01134607 | 0.01134607 | ENSG00000106028 | n.s. |
| ENSG00000160957 | 780.5301 | -0.176873792 | 0.244973 | -0.722012 | 0.470287 | 0.862688564 | RECQL4 | protein_coding | 65.51 | -0.011345739 | 0.011345739 | ENSG00000160957 | n.s. |
| ENSG00000137700 | 307.4779 | -0.196184284 | 0.29124 | -0.673616 | 0.500555 | 0.875342302 | SLC37A4 | protein_coding | 55.91 | -0.011343784 | 0.011343784 | ENSG00000137700 | n.s. |
| ENSG00000112972 | 3915.77 | -0.180911015 | 0.255028 | -0.709378 | 0.47809 | 0.865660698 | HMGCS1 | protein_coding | 38.81 | -0.011334491 | 0.011334491 | ENSG00000112972 | n.s. |
| ENSG00000137880 | 250.5268 | -0.217410907 | 0.345098 | -0.629997 | 0.528696 | 0.886954686 | GCHFR | protein_coding | 57.52 | -0.011326797 | 0.011326797 | ENSG00000137880 | n.s. |
| ENSG00000144642 | 4100.976 | -0.1806503 | 0.252691 | -0.714905 | 0.474668 | 0.865616576 | RBMS3 | protein_coding | 36.91 | -0.011322156 | 0.011322156 | ENSG00000144642 | n.s. |

| | | | | | | | | | | | | | |
|---|---|---|---|---|---|---|---|---|---|---|---|---|---|
| ENSG00000100764 | 1635.723 | -0.179794979 | 0.250686 | -0.717212 | 0.473243 | 0.865034671 | PSMC1 | protein_coding | 44.79 | -0.011321058 | 0.011321058 | ENSG00000100764 | n.s. |
| ENSG00000164068 | 323.2824 | -0.180618173 | 0.252429 | -0.715519 | 0.474288 | 0.865616576 | RNF123 | protein_coding | 56.42 | -0.011320142 | 0.011320142 | ENSG00000164068 | n.s. |
| ENSG00000103591 | 1386.014 | -0.19246838 | 0.281481 | -0.683771 | 0.49412 | 0.873348456 | AAGAB | protein_coding | 39.49 | -0.011319536 | 0.011319536 | ENSG00000103591 | n.s. |
| ENSG00000172175 | 655.2859 | -0.18439496 | 0.263151 | -0.700719 | 0.483479 | 0.868366486 | MALT1 | protein_coding | 38.97 | -0.011302848 | 0.011302848 | ENSG00000172175 | n.s. |
| ENSG00000166173 | 196.3873 | -0.180263505 | 0.252923 | -0.712721 | 0.476019 | 0.865616576 | LARP6 | protein_coding | 44.62 | -0.011297913 | 0.011297913 | ENSG00000166173 | n.s. |
| ENSG00000210049 | 6.892245 | -0.279473357 | 0.550649 | -0.507534 | 0.61178 | 0.911134742 | MT-TF | Mt_tRNA | 40.85 | -0.011295585 | 0.011295585 | ENSG00000210049 | n.s. |
| ENSG00000138031 | 589.1375 | -0.180204089 | 0.251639 | -0.716121 | 0.473916 | 0.865616576 | ADCY3 | protein_coding | 50.99 | -0.01129419 | 0.01129419 | ENSG00000138031 | n.s. |
| ENSG00000164889 | 461.7772 | 0.197011916 | 0.29527 | 0.6672263 | 0.504628 | 0.876368641 | SLC4A2 | protein_coding | 59.83 | 0.011291378 | 0.011291378 | ENSG00000164889 | n.s. |
| ENSG00000269386 | 69.71969 | -0.192954415 | 0.28382 | -0.679848 | 0.496601 | 0.873971799 | RAB11B-AS1 | antisense | 48.55 | -0.011288331 | 0.011288331 | ENSG00000269386 | n.s. |
| ENSG00000064607 | 2528.192 | -0.182654541 | 0.259678 | -0.703389 | 0.481814 | 0.867402024 | SUGP2 | protein_coding | 50 | -0.011284319 | 0.011284319 | ENSG00000064607 | n.s. |
| ENSG00000105738 | 66.20202 | -0.196854283 | 0.294894 | -0.667543 | 0.504426 | 0.876355973 | SIPA1L3 | protein_coding | 49.76 | -0.011283579 | 0.011283579 | ENSG00000105738 | n.s. |
| ENSG00000131507 | 1144.184 | -0.198731831 | 0.299022 | -0.664605 | 0.506303 | 0.877502299 | NDFIP1 | protein_coding | 39.29 | -0.011278377 | 0.011278377 | ENSG00000131507 | n.s. |
| ENSG00000231188 | 34.41658 | 0.220835331 | 0.356185 | 0.6200012 | 0.535257 | 0.88909364 | | antisense | 48.9 | 0.011274195 | 0.011274195 | ENSG00000231188 | n.s. |
| ENSG00000241939 | 26.89382 | 0.224197107 | 0.365744 | 0.612989 | 0.539884 | 0.890684245 | RN7SL517P | misc_RNA | 51.48 | 0.011271785 | 0.011271785 | ENSG00000241939 | n.s. |
| ENSG00000251728 | 4.848104 | -0.302608566 | 0.614617 | -0.492353 | 0.62247 | 0.917861492 | | misc_RNA | 36.46 | -0.011263953 | 0.011263953 | ENSG00000251728 | n.s. |
| ENSG00000108107 | 2407.45 | -0.176597318 | 0.245218 | -0.720163 | 0.471424 | 0.863532918 | RPL28 | protein_coding | 52.88 | -0.011252976 | 0.011252976 | ENSG00000108107 | n.s. |
| ENSG00000272163 | 172.7735 | -0.169644353 | 0.230664 | -0.73546 | 0.462059 | 0.858395846 | | antisense | 56.17 | -0.011249523 | 0.011249523 | ENSG00000272163 | n.s. |
| ENSG00000100239 | 501.3304 | -0.19126008 | 0.280063 | -0.682918 | 0.494658 | 0.873443478 | PPP6R2 | protein_coding | 51.13 | -0.011239436 | 0.011239436 | ENSG00000100239 | n.s. |
| ENSG00000264249 | 8.467498 | -0.323590601 | 0.707177 | -0.457581 | 0.647254 | 0.923184844 | MIR3912 | miRNA | 31.43 | -0.011232261 | 0.011232261 | ENSG00000264249 | n.s. |
| ENSG00000155592 | 977.7282 | -0.17310919 | 0.238538 | -0.725709 | 0.468017 | 0.861253285 | ZKSCAN2 | protein_coding | 43.2 | -0.011229439 | 0.011229439 | ENSG00000155592 | n.s. |
| ENSG00000166803 | 1719.476 | -0.2044252 | 0.312145 | -0.654905 | 0.512529 | 0.881306695 | PCLAF | protein_coding | 43.34 | -0.01121741 | 0.01121741 | ENSG00000166803 | n.s. |
| ENSG00000201586 | 10.83659 | -0.25415462 | 0.460701 | -0.551669 | 0.581175 | 0.903393748 | RNU6-593P | snRNA | 36.61 | -0.011214044 | 0.011214044 | ENSG00000201586 | n.s. |
| ENSG00000141756 | 778.9608 | 0.195209965 | 0.292299 | 0.667843 | 0.504234 | 0.876255897 | FKBP10 | protein_coding | 56.18 | 0.011199009 | 0.011199009 | ENSG00000141756 | n.s. |
| ENSG00000258359 | 47.7846 | -0.213081274 | 0.333821 | -0.63831 | 0.523272 | 0.886079926 | PCNPP1 | processed_pseudogene | 41.29 | -0.011192542 | 0.011192542 | ENSG00000258359 | n.s. |
| ENSG00000118181 | 7860.673 | 0.214699864 | 0.339739 | 0.6319556 | 0.527416 | 0.886936263 | RPS25 | protein_coding | 47.62 | 0.011187492 | 0.011187492 | ENSG00000118181 | n.s. |
| ENSG00000198984 | 8.82928 | -0.304920969 | 0.635217 | -0.480027 | 0.631208 | 0.919033489 | MIR345 | miRNA | 60.2 | -0.011181044 | 0.011181044 | ENSG00000198984 | n.s. |
| ENSG00000147383 | 359.2595 | -0.196641219 | 0.295618 | -0.665188 | 0.50593 | 0.87732845 | NSDHL | protein_coding | 45.97 | -0.011176652 | 0.011176652 | ENSG00000147383 | n.s. |
| ENSG00000276023 | 260.9472 | -0.211328269 | 0.32835 | -0.643606 | 0.519831 | 0.885364478 | DUSP14 | protein_coding | 46.28 | -0.011174596 | 0.011174596 | ENSG00000276023 | n.s. |
| ENSG00000159720 | 439.5768 | -0.194890909 | 0.292145 | -0.667104 | 0.504706 | 0.876368641 | ATP6V0D1 | protein_coding | 53.78 | -0.011169816 | 0.011169816 | ENSG00000159720 | n.s. |
| ENSG00000112159 | 6977.692 | -0.174399908 | 0.241781 | -0.721314 | 0.470716 | 0.863021191 | MDN1 | protein_coding | 41.54 | -0.011157852 | 0.011157852 | ENSG00000112159 | n.s. |
| ENSG00000207701 | 6.756248 | -0.268725603 | 0.517773 | -0.519003 | 0.603759 | 0.908868121 | MIR597 | miRNA | 44.33 | -0.011151879 | 0.011151879 | ENSG00000207701 | n.s. |
| ENSG00000174652 | 879.2104 | -0.18131566 | 0.258216 | -0.702187 | 0.482563 | 0.868051445 | ZNF266 | protein_coding | 45 | -0.01114267 | 0.01114267 | ENSG00000174652 | n.s. |

| | | | | | | | | | | | | | |
|---|---|---|---|---|---|---|---|---|---|---|---|---|---|
| ENSG00000166971 | 100.1502 | -0.18748641 | 0.271647 | -0.690185 | 0.490078 | 0.872216858 | AKTIP | protein_coding | 45.45 | -0.011132104 | 0.011132104 | ENSG00000166971 | n.s. |
| ENSG00000011021 | 49.86863 | -0.203249728 | 0.310576 | -0.654427 | 0.512837 | 0.881523829 | CLCN6 | protein_coding | 47.79 | -0.011131163 | 0.011131163 | ENSG00000011021 | n.s. |
| ENSG00000275939 | 11.85582 | 0.236693147 | 0.413337 | 0.5726395 | 0.566889 | 0.897394878 | | misc_RNA | 57.23 | 0.011128463 | 0.011128463 | ENSG00000275939 | n.s. |
| ENSG00000171316 | 4528.395 | -0.171135353 | 0.234884 | -0.728596 | 0.466249 | 0.861088114 | CHD7 | protein_coding | 39.98 | -0.011115653 | 0.011115653 | ENSG00000171316 | n.s. |
| ENSG00000253347 | 18.37735 | -0.308261173 | 0.654265 | -0.471156 | 0.637529 | 0.920373167 | | antisense | 38.81 | -0.011108515 | 0.011108515 | ENSG00000253347 | n.s. |
| ENSG00000160932 | 987.5664 | -0.189353503 | 0.277977 | -0.681183 | 0.495756 | 0.873669126 | LY6E | protein_coding | 63.58 | -0.011106153 | 0.011106153 | ENSG00000160932 | n.s. |
| ENSG00000132768 | 351.8426 | -0.185897067 | 0.268577 | -0.692156 | 0.488839 | 0.871557795 | DPH2 | protein_coding | 56.05 | -0.011098763 | 0.011098763 | ENSG00000132768 | n.s. |
| ENSG00000165943 | 956.6717 | 0.183721554 | 0.264452 | 0.6947261 | 0.487227 | 0.870146312 | MOAP1 | protein_coding | 51.56 | 0.0110982 | 0.0110982 | ENSG00000165943 | n.s. |
| ENSG00000180354 | 982.8251 | -0.181201094 | 0.258713 | -0.700394 | 0.483681 | 0.868492686 | MTURN | protein_coding | 46.82 | -0.011095638 | 0.011095638 | ENSG00000180354 | n.s. |
| ENSG00000211455 | 1466.44 | -0.179133605 | 0.253859 | -0.705642 | 0.480411 | 0.867184458 | STK38L | protein_coding | 37.77 | -0.011086313 | 0.011086313 | ENSG00000211455 | n.s. |
| ENSG00000273162 | 102.6711 | -0.183303862 | 0.263273 | -0.696251 | 0.486272 | 0.870096539 | | lincRNA | 51.28 | -0.011077521 | 0.011077521 | ENSG00000273162 | n.s. |
| ENSG00000055118 | 682.7714 | -0.17922356 | 0.25472 | -0.703611 | 0.481675 | 0.867402024 | KCNH2 | protein_coding | 59.57 | -0.011072354 | 0.011072354 | ENSG00000055118 | n.s. |
| ENSG00000264653 | 11.57022 | -0.266793144 | 0.511363 | -0.52173 | 0.601859 | 0.908868121 | MIR5194 | miRNA | 45 | -0.011071683 | 0.011071683 | ENSG00000264653 | n.s. |
| ENSG00000162997 | 53.4812 | -0.209640764 | 0.326424 | -0.642235 | 0.520721 | 0.885523197 | PRORSD1P | anscribed_unitary_pseudoge | 44.56 | -0.011069044 | 0.011069044 | ENSG00000162997 | n.s. |
| ENSG00000138336 | 1196.822 | -0.17870827 | 0.253003 | -0.706349 | 0.479971 | 0.867086691 | TET1 | protein_coding | 42.09 | -0.01106874 | 0.01106874 | ENSG00000138336 | n.s. |
| ENSG00000138081 | 2038.192 | -0.179086403 | 0.254307 | -0.704214 | 0.481299 | 0.867349802 | FBXO11 | protein_coding | 39.46 | -0.011068563 | 0.011068563 | ENSG00000138081 | n.s. |
| ENSG00000066279 | 8671.592 | -0.182090531 | 0.260689 | -0.698496 | 0.484867 | 0.869415694 | ASPM | protein_coding | 34.66 | -0.011066101 | 0.011066101 | ENSG00000066279 | n.s. |
| ENSG00000113593 | 983.1663 | -0.191350937 | 0.284168 | -0.673372 | 0.50071 | 0.875342302 | PPWD1 | protein_coding | 36.46 | -0.01106431 | 0.01106431 | ENSG00000113593 | n.s. |
| ENSG00000204178 | 493.9807 | -0.191609149 | 0.284957 | -0.672414 | 0.50132 | 0.875518559 | TMEM57 | protein_coding | 40.69 | -0.011062486 | 0.011062486 | ENSG00000204178 | n.s. |
| ENSG00000198000 | 1601.062 | -0.17890383 | 0.253965 | -0.704442 | 0.481157 | 0.867316901 | NOL8 | protein_coding | 38.87 | -0.011060227 | 0.011060227 | ENSG00000198000 | n.s. |
| ENSG00000246308 | 83.64524 | 0.191606839 | 0.285787 | 0.6704534 | 0.502569 | 0.875813709 | | antisense | 39.66 | 0.011034305 | 0.011034305 | ENSG00000246308 | n.s. |
| ENSG00000250938 | 27.94699 | 0.220876011 | 0.367972 | 0.6002517 | 0.548338 | 0.891406027 | | antisense | 36.15 | 0.01102711 | 0.01102711 | ENSG00000250938 | n.s. |
| ENSG00000136877 | 531.1921 | -0.186414956 | 0.271027 | -0.68781 | 0.491573 | 0.872671134 | FPGS | protein_coding | 53.5 | -0.011026331 | 0.011026331 | ENSG00000136877 | n.s. |
| ENSG00000132792 | 906.47 | -0.175919121 | 0.2471 | -0.711936 | 0.476504 | 0.865616576 | CTNNBL1 | protein_coding | 42.04 | -0.011025632 | 0.011025632 | ENSG00000132792 | n.s. |
| ENSG00000073792 | 45.30875 | -0.207899486 | 0.322657 | -0.644335 | 0.519358 | 0.885178473 | IGF2BP2 | protein_coding | 42.54 | -0.01101226 | 0.01101226 | ENSG00000073792 | n.s. |
| ENSG00000248932 | 113.683 | -0.175643365 | 0.245398 | -0.71575 | 0.474146 | 0.865616576 | | antisense | 42.28 | -0.011008349 | 0.011008349 | ENSG00000248932 | n.s. |
| ENSG00000144445 | 709.7196 | -0.189235197 | 0.27968 | -0.676613 | 0.498652 | 0.874768485 | KANSL1L | protein_coding | 36.36 | -0.010995865 | 0.010995865 | ENSG00000144445 | n.s. |
| ENSG00000212464 | 96.4577 | 0.266916305 | 0.516661 | 0.5166179 | 0.605423 | 0.909522325 | SNORA12 | snoRNA | 42.18 | 0.010993385 | 0.010993385 | ENSG00000212464 | n.s. |
| ENSG00000169813 | 3007.227 | -0.190061872 | 0.28187 | -0.674289 | 0.500128 | 0.875342302 | HNRNPF | protein_coding | 47.66 | -0.010989773 | 0.010989773 | ENSG00000169813 | n.s. |
| ENSG00000033050 | 1363.755 | -0.188933353 | 0.278648 | -0.678036 | 0.497749 | 0.874675553 | ABCF2 | protein_coding | 49.19 | -0.010987044 | 0.010987044 | ENSG00000033050 | n.s. |
| ENSG00000212482 | 3.461167 | 0.32350508 | 0.72555 | 0.4458759 | 0.655687 | 0.924786541 | RNU6-530P | snRNA | 34.58 | 0.010985747 | 0.010985747 | ENSG00000212482 | n.s. |
| ENSG00000010818 | 31.00926 | -0.211663587 | 0.338188 | -0.625875 | 0.531397 | 0.887366671 | HIVEP2 | protein_coding | 38.72 | -0.010984681 | 0.010984681 | ENSG00000010818 | n.s. |

| | | | | | | | | | | | | | |
|---|---|---|---|---|---|---|---|---|---|---|---|---|---|
| ENSG00000123352 | 741.7792 | -0.181748309 | 0.261303 | -0.695546 | 0.486713 | 0.870096539 | SPATS2 | protein_coding | 41.68 | -0.010983515 | 0.010983515 | ENSG00000123352 | n.s. |
| ENSG00000187772 | 2063.793 | -0.183923179 | 0.26591 | -0.691675 | 0.489142 | 0.871557795 | LIN28B | protein_coding | 36.8 | -0.010980914 | 0.010980914 | ENSG00000187772 | n.s. |
| ENSG00000177042 | 128.6997 | -0.190641666 | 0.283862 | -0.6716 | 0.501838 | 0.875813709 | TMEM80 | protein_coding | 55.94 | -0.010978722 | 0.010978722 | ENSG00000177042 | n.s. |
| ENSG00000169299 | 846.2355 | -0.189063286 | 0.279862 | -0.675559 | 0.49932 | 0.874970095 | PGM2 | protein_coding | 39.92 | -0.010966954 | 0.010966954 | ENSG00000169299 | n.s. |
| ENSG00000206605 | 6.513372 | -0.287036075 | 0.576917 | -0.497535 | 0.618812 | 0.915786649 | RNU6-946P | snRNA | 42.99 | -0.010966412 | 0.010966412 | ENSG00000206605 | n.s. |
| ENSG00000252473 | 42.58368 | -0.208837262 | 0.32933 | -0.634127 | 0.525998 | 0.886111278 | | snoRNA | 45.19 | -0.010966407 | 0.010966407 | ENSG00000252473 | n.s. |
| ENSG00000213366 | 168.4661 | -0.189565304 | 0.281383 | -0.673692 | 0.500507 | 0.875342302 | GSTM2 | protein_coding | 46.7 | -0.010961061 | 0.010961061 | ENSG00000213366 | n.s. |
| ENSG00000275084 | 25.58802 | -0.219504405 | 0.364303 | -0.602533 | 0.546819 | 0.891406027 | SNORD91B | snoRNA | 40.99 | -0.010958633 | 0.010958633 | ENSG00000275084 | n.s. |
| ENSG00000227374 | 16.06528 | -0.298832281 | 0.626264 | -0.477167 | 0.633243 | 0.919033489 | | lincRNA | 60.47 | -0.01095778 | 0.01095778 | ENSG00000227374 | n.s. |
| ENSG00000202474 | 4.466186 | -0.311596463 | 0.67033 | -0.464841 | 0.642046 | 0.922225673 | RNA5SP283 | rRNA | 58.82 | -0.010956602 | 0.010956602 | ENSG00000202474 | n.s. |
| ENSG00000278129 | 856.6944 | -0.186209471 | 0.272344 | -0.683728 | 0.494147 | 0.873348456 | ZNF8 | protein_coding | 48.3 | -0.010951434 | 0.010951434 | ENSG00000278129 | n.s. |
| ENSG00000186073 | 44.42252 | 0.210104074 | 0.33265 | 0.6316066 | 0.527644 | 0.886936263 | C15orf41 | protein_coding | 37.44 | 0.010948017 | 0.010948017 | ENSG00000186073 | n.s. |
| ENSG00000127952 | 184.4498 | -0.207314748 | 0.323425 | -0.640999 | 0.521523 | 0.885523197 | STYXL1 | protein_coding | 48.42 | -0.010946231 | 0.010946231 | ENSG00000127952 | n.s. |
| ENSG00000278022 | 34.9708 | -0.217690301 | 0.355334 | -0.612636 | 0.540117 | 0.890684245 | | sense_intronic | 45.1 | -0.010944648 | 0.010944648 | ENSG00000278022 | n.s. |
| ENSG00000183891 | 816.2504 | 0.164915061 | 0.224527 | 0.7344989 | 0.462645 | 0.858395846 | TTC32 | protein_coding | 39.39 | 0.010935912 | 0.010935912 | ENSG00000183891 | n.s. |
| ENSG00000258017 | 34.64201 | -0.218704808 | 0.358846 | -0.609467 | 0.542215 | 0.891404544 | | antisense | 46.25 | -0.010918872 | 0.010918872 | ENSG00000258017 | n.s. |
| ENSG00000221990 | 110.8212 | -0.180562362 | 0.25927 | -0.696427 | 0.486162 | 0.870029373 | EXOC3-AS1 | antisense | 55.32 | -0.010917899 | 0.010917899 | ENSG00000221990 | n.s. |
| ENSG00000138688 | 2534.552 | -0.18547556 | 0.270584 | -0.685465 | 0.493051 | 0.873266128 | KIAA1109 | protein_coding | 34.76 | -0.010915865 | 0.010915865 | ENSG00000138688 | n.s. |
| ENSG00000178761 | 1367.685 | -0.17957457 | 0.257096 | -0.698472 | 0.484882 | 0.869415694 | FAM219B | protein_coding | 51.13 | -0.0109132 | 0.0109132 | ENSG00000178761 | n.s. |
| ENSG00000143393 | 692.9883 | -0.184322539 | 0.267762 | -0.688382 | 0.491212 | 0.872562973 | PI4KB | protein_coding | 48.46 | -0.010912488 | 0.010912488 | ENSG00000143393 | n.s. |
| ENSG00000165359 | 547.7967 | -0.185829705 | 0.272751 | -0.681317 | 0.495671 | 0.873669126 | INTS6L | protein_coding | 38.59 | -0.010899472 | 0.010899472 | ENSG00000165359 | n.s. |
| ENSG00000043514 | 402.0084 | -0.188431384 | 0.279774 | -0.673513 | 0.500621 | 0.875342302 | TRIT1 | protein_coding | 42.46 | -0.010895495 | 0.010895495 | ENSG00000043514 | n.s. |
| ENSG00000075131 | 608.9643 | -0.18517483 | 0.270924 | -0.683493 | 0.494295 | 0.873359609 | TIPIN | protein_coding | 44.05 | -0.010889557 | 0.010889557 | ENSG00000075131 | n.s. |
| ENSG00000136319 | 490.6237 | -0.184568529 | 0.2691 | -0.685873 | 0.492793 | 0.873000006 | TTC5 | protein_coding | 39.84 | -0.010886914 | 0.010886914 | ENSG00000136319 | n.s. |
| ENSG00000183765 | 110.9923 | 0.206188564 | 0.321633 | 0.6410671 | 0.521479 | 0.885523197 | CHEK2 | protein_coding | 43.89 | 0.010886768 | 0.010886768 | ENSG00000183765 | n.s. |
| ENSG00000145907 | 6993.119 | -0.182290755 | 0.263544 | -0.69169 | 0.489132 | 0.871557795 | G3BP1 | protein_coding | 39.44 | -0.010883452 | 0.010883452 | ENSG00000145907 | n.s. |
| ENSG00000092621 | 4440.482 | -0.185546848 | 0.272548 | -0.680786 | 0.496007 | 0.873669126 | PHGDH | protein_coding | 44.18 | -0.010882881 | 0.010882881 | ENSG00000092621 | n.s. |
| ENSG00000204347 | 121.0012 | -0.20062658 | 0.307932 | -0.651529 | 0.514705 | 0.882641141 | BTBD17 | protein_coding | 58.45 | -0.010877137 | 0.010877137 | ENSG00000204347 | n.s. |
| ENSG00000224142 | 24.93481 | 0.208722918 | 0.331585 | 0.6294704 | 0.529041 | 0.886954686 | AMMECR1-IT1 | sense_intronic | 43.78 | 0.010874165 | 0.010874165 | ENSG00000224142 | n.s. |
| ENSG00000110237 | 433.4143 | -0.173510396 | 0.244806 | -0.708768 | 0.478468 | 0.865660698 | ARHGEF17 | protein_coding | 55.27 | -0.010870825 | 0.010870825 | ENSG00000110237 | n.s. |
| ENSG00000169609 | 782.1798 | -0.173434267 | 0.242439 | -0.715372 | 0.474379 | 0.865616576 | C15orf40 | protein_coding | 44.6 | -0.010869895 | 0.010869895 | ENSG00000169609 | n.s. |
| ENSG00000143458 | 412.773 | -0.184987204 | 0.270998 | -0.682615 | 0.49485 | 0.873453717 | GABPB2 | protein_coding | 44.85 | -0.010869867 | 0.010869867 | ENSG00000143458 | n.s. |

| | | | | | | | | | | | | | |
|---|---|---|---|---|---|---|---|---|---|---|---|---|---|
| ENSG00000252652 | 42.52149 | 0.213470968 | 0.345987 | 0.6169906 | 0.537241 | 0.889417954 | | misc_RNA | 39.58 | 0.010864414 | 0.010864414 | ENSG00000252652 | n.s. |
| ENSG00000137492 | 2378.176 | -0.188576099 | 0.280976 | -0.671147 | 0.502127 | 0.875813709 | THAP12 | protein_coding | 40.11 | -0.01085977 | 0.01085977 | ENSG00000137492 | n.s. |
| ENSG00000161526 | 2589.092 | -0.173180221 | 0.243232 | -0.711995 | 0.476468 | 0.865616576 | SAP30BP | protein_coding | 48.61 | -0.010853973 | 0.010853973 | ENSG00000161526 | n.s. |
| ENSG00000200105 | 4.449399 | -0.318849591 | 0.714023 | -0.446554 | 0.655197 | 0.924621046 | RNU6-251P | snRNA | 39.42 | -0.010852437 | 0.010852437 | ENSG00000200105 | n.s. |
| ENSG00000252148 | 6.585626 | -0.264348449 | 0.515362 | -0.512938 | 0.607995 | 0.909933765 | RNU6-1206P | snRNA | 30.1 | -0.010835701 | 0.010835701 | ENSG00000252148 | n.s. |
| ENSG00000175970 | 921.3176 | -0.179163996 | 0.25772 | -0.695189 | 0.486937 | 0.870096539 | UNC119B | protein_coding | 48.18 | -0.010827339 | 0.010827339 | ENSG00000175970 | n.s. |
| ENSG00000117020 | 4040.619 | -0.179072485 | 0.257281 | -0.69602 | 0.486416 | 0.870096539 | AKT3 | protein_coding | 37.34 | -0.010821808 | 0.010821808 | ENSG00000117020 | n.s. |
| ENSG00000141503 | 243.332 | -0.183998766 | 0.268836 | -0.684429 | 0.493705 | 0.873348456 | MINK1 | protein_coding | 51.93 | -0.010821417 | 0.010821417 | ENSG00000141503 | n.s. |
| ENSG00000135951 | 31.94447 | -0.212899109 | 0.345202 | -0.616739 | 0.537407 | 0.88958741 | TSGA10 | protein_coding | 37.63 | -0.010817696 | 0.010817696 | ENSG00000135951 | n.s. |
| ENSG00000103978 | 1442.641 | 0.172569531 | 0.242279 | 0.7122749 | 0.476295 | 0.865616576 | TMEM87A | protein_coding | 38.93 | 0.010815698 | 0.010815698 | ENSG00000103978 | n.s. |
| ENSG00000232593 | 365.4426 | -0.194153937 | 0.294615 | -0.659008 | 0.50989 | 0.879714521 | KANTR | protein_coding | 42.45 | -0.010806266 | 0.010806266 | ENSG00000232593 | n.s. |
| ENSG00000269399 | 130.9158 | -0.176265542 | 0.251545 | -0.700731 | 0.483471 | 0.868366486 | | lincRNA | 47.66 | -0.010804539 | 0.010804539 | ENSG00000269399 | n.s. |
| ENSG00000172893 | 314.2126 | -0.178497753 | 0.255864 | -0.697627 | 0.485411 | 0.869915127 | DHCR7 | protein_coding | 51.71 | -0.01080324 | 0.01080324 | ENSG00000172893 | n.s. |
| ENSG00000249605 | 28.24466 | -0.21146381 | 0.341057 | -0.620025 | 0.535241 | 0.88909364 | | sense_intronic | 37.3 | -0.010795755 | 0.010795755 | ENSG00000249605 | n.s. |
| ENSG00000132383 | 1041.044 | -0.186704833 | 0.277068 | -0.673858 | 0.500402 | 0.875342302 | RPA1 | protein_coding | 44.49 | -0.010795662 | 0.010795662 | ENSG00000132383 | n.s. |
| ENSG00000273702 | 229.5423 | 0.16930843 | 0.235009 | 0.7204348 | 0.471257 | 0.863532918 | | lincRNA | 49.57 | 0.01078852 | 0.01078852 | ENSG00000273702 | n.s. |
| ENSG00000139641 | 942.8954 | -0.185500198 | 0.27402 | -0.676958 | 0.498433 | 0.874675553 | ESYT1 | protein_coding | 50.49 | -0.010787395 | 0.010787395 | ENSG00000139641 | n.s. |
| ENSG00000164611 | 2305.155 | 0.1879759 | 0.281345 | 0.6681334 | 0.504048 | 0.876255897 | PTTG1 | protein_coding | 39.4 | 0.010783998 | 0.010783998 | ENSG00000164611 | n.s. |
| ENSG00000005075 | 567.4966 | -0.200655117 | 0.30987 | -0.647547 | 0.517278 | 0.88372671 | POLR2J | protein_coding | 55.04 | -0.010771572 | 0.010771572 | ENSG00000005075 | n.s. |
| ENSG00000198625 | 3842.719 | -0.18010119 | 0.260489 | -0.691395 | 0.489317 | 0.871754603 | MDM4 | protein_coding | 42.52 | -0.010735067 | 0.010735067 | ENSG00000198625 | n.s. |
| ENSG00000157881 | 191.1436 | -0.192443771 | 0.291733 | -0.659658 | 0.509473 | 0.87951488 | PANK4 | protein_coding | 60.69 | -0.010730051 | 0.010730051 | ENSG00000157881 | n.s. |
| ENSG00000181404 | 107.0487 | -0.197908654 | 0.303858 | -0.651319 | 0.514841 | 0.882656329 | WASHC1 | protein_coding | 54.03 | -0.010728304 | 0.010728304 | ENSG00000181404 | n.s. |
| ENSG00000231181 | 26.75883 | -0.211793547 | 0.344739 | -0.614359 | 0.538978 | 0.889966281 | | processed_pseudogene | 44.54 | -0.010722355 | 0.010722355 | ENSG00000231181 | n.s. |
| ENSG00000206888 | 8.429003 | -0.267946576 | 0.530289 | -0.505284 | 0.613359 | 0.911981962 | RNU6-48P | snRNA | 45.79 | -0.010721548 | 0.010721548 | ENSG00000206888 | n.s. |
| ENSG00000172733 | 197.1225 | 0.180883461 | 0.262267 | 0.6896927 | 0.490387 | 0.872495697 | PURG | protein_coding | 36.48 | 0.01071494 | 0.01071494 | ENSG00000172733 | n.s. |
| ENSG00000196507 | 267.9832 | -0.186015699 | 0.277456 | -0.670433 | 0.502582 | 0.875813709 | TCEAL3 | protein_coding | 41.11 | -0.010712321 | 0.010712321 | ENSG00000196507 | n.s. |
| ENSG00000185798 | 222.5951 | -0.17850766 | 0.257583 | -0.693011 | 0.488303 | 0.87106191 | WDR53 | protein_coding | 43.07 | -0.010701709 | 0.010701709 | ENSG00000185798 | n.s. |
| ENSG00000135945 | 2031.244 | -0.177337465 | 0.255401 | -0.694348 | 0.487464 | 0.870346032 | REV1 | protein_coding | 39.33 | -0.010694876 | 0.010694876 | ENSG00000135945 | n.s. |
| ENSG00000224790 | 89.14436 | -0.183828483 | 0.271511 | -0.677057 | 0.49837 | 0.874675553 | | lincRNA | 47.75 | -0.01069018 | 0.01069018 | ENSG00000224790 | n.s. |
| ENSG00000116005 | 451.7016 | 0.183955775 | 0.271861 | 0.676654 | 0.498625 | 0.874768485 | PCYOX1 | protein_coding | 45.23 | 0.010689095 | 0.010689095 | ENSG00000116005 | n.s. |
| ENSG00000131067 | 370.0785 | -0.172941496 | 0.245687 | -0.703911 | 0.481488 | 0.867349802 | GGT7 | protein_coding | 49.05 | -0.010688773 | 0.010688773 | ENSG00000131067 | n.s. |
| ENSG00000139505 | 1132.528 | -0.191056246 | 0.289109 | -0.660846 | 0.508711 | 0.879193676 | MTMR6 | protein_coding | 37.84 | -0.010682995 | 0.010682995 | ENSG00000139505 | n.s. |

| | | | | | | | | | | | | | |
|---|---|---|---|---|---|---|---|---|---|---|---|---|---|
| ENSG00000115282 | 483.9315 | -0.18032337 | 0.261888 | -0.68855 | 0.491106 | 0.872562973 | TTC31 | protein_coding | 50.85 | -0.010675724 | 0.010675724 | ENSG00000115282 | n.s. |
| ENSG00000035115 | 490.401 | -0.182477599 | 0.26844 | -0.679769 | 0.49665 | 0.873971799 | SH3YL1 | protein_coding | 40.78 | -0.01067541 | 0.01067541 | ENSG00000035115 | n.s. |
| ENSG00000278447 | 6.933407 | -0.291025975 | 0.610339 | -0.476827 | 0.633485 | 0.919033489 | MIR6781 | miRNA | 62.5 | -0.010671533 | 0.010671533 | ENSG00000278447 | n.s. |
| ENSG00000252274 | 12.77572 | 0.246701473 | 0.455158 | 0.5420131 | 0.58781 | 0.905217543 | | scaRNA | 31.39 | 0.010669108 | 0.010669108 | ENSG00000252274 | n.s. |
| ENSG00000176208 | 2025.933 | -0.17023027 | 0.23831 | -0.714322 | 0.475028 | 0.865616576 | ATAD5 | protein_coding | 41.02 | -0.010669086 | 0.010669086 | ENSG00000176208 | n.s. |
| ENSG00000188811 | 643.4617 | -0.170222911 | 0.238591 | -0.71345 | 0.475567 | 0.865616576 | NHLRC3 | protein_coding | 37.72 | -0.010668625 | 0.010668625 | ENSG00000188811 | n.s. |
| ENSG00000154845 | 1523.004 | -0.180780313 | 0.263604 | -0.685802 | 0.492838 | 0.873000006 | PPP4R1 | protein_coding | 39.14 | -0.010663463 | 0.010663463 | ENSG00000154845 | n.s. |
| ENSG00000229152 | 445.499 | -0.177906368 | 0.256776 | -0.692847 | 0.488406 | 0.87113391 | ANKRD10-IT1 | sense_intronic | 38.6 | -0.010659274 | 0.010659274 | ENSG00000229152 | n.s. |
| ENSG00000151502 | 412.3384 | -0.179761782 | 0.260773 | -0.689341 | 0.490608 | 0.872495697 | VPS26B | protein_coding | 48.41 | -0.010648496 | 0.010648496 | ENSG00000151502 | n.s. |
| ENSG00000023191 | 498.9985 | -0.181012041 | 0.264653 | -0.683959 | 0.494001 | 0.873348456 | RNH1 | protein_coding | 60.36 | -0.01064576 | 0.01064576 | ENSG00000023191 | n.s. |
| ENSG00000079462 | 665.0394 | -0.191146291 | 0.290077 | -0.658949 | 0.509928 | 0.879714521 | PAFAH1B3 | protein_coding | 53.73 | -0.010638866 | 0.010638866 | ENSG00000079462 | n.s. |
| ENSG00000166292 | 62.82774 | 0.196228119 | 0.30132 | 0.6512289 | 0.514899 | 0.882656329 | TMEM100 | protein_coding | 40.18 | 0.010637205 | 0.010637205 | ENSG00000166292 | n.s. |
| ENSG00000162623 | 1948.273 | -0.170287155 | 0.240638 | -0.707649 | 0.479163 | 0.866036769 | TYW3 | protein_coding | 40.26 | -0.010636761 | 0.010636761 | ENSG00000162623 | n.s. |
| ENSG00000201217 | 22.23227 | -0.216640795 | 0.365799 | -0.592239 | 0.55369 | 0.893144718 | | misc_RNA | 41.59 | -0.010632333 | 0.010632333 | ENSG00000201217 | n.s. |
| ENSG00000122642 | 558.894 | -0.179619474 | 0.260979 | -0.688253 | 0.491293 | 0.872595457 | FKBP9 | protein_coding | 45.26 | -0.010631147 | 0.010631147 | ENSG00000122642 | n.s. |
| ENSG00000213626 | 150.1242 | -0.184524439 | 0.275076 | -0.670813 | 0.50234 | 0.875813709 | LBH | protein_coding | 46.97 | -0.010626442 | 0.010626442 | ENSG00000213626 | n.s. |
| ENSG00000125901 | 720.5682 | -0.182712522 | 0.269375 | -0.678282 | 0.497593 | 0.874675553 | MRPS26 | protein_coding | 61.08 | -0.010625283 | 0.010625283 | ENSG00000125901 | n.s. |
| ENSG00000141198 | 1435.976 | 0.183728941 | 0.272842 | 0.6733896 | 0.500699 | 0.875342302 | TOM1L1 | protein_coding | 40.86 | 0.01062359 | 0.01062359 | ENSG00000141198 | n.s. |
| ENSG00000105419 | 759.6513 | -0.197783684 | 0.305452 | -0.647512 | 0.5173 | 0.88372671 | MEIS3 | protein_coding | 56.51 | -0.010617427 | 0.010617427 | ENSG00000105419 | n.s. |
| ENSG00000105483 | 86.17192 | -0.212609108 | 0.348905 | -0.609361 | 0.542285 | 0.891406027 | CARD8 | protein_coding | 44.39 | -0.010614389 | 0.010614389 | ENSG00000105483 | n.s. |
| ENSG00000046647 | 173.7626 | -0.175394374 | 0.252322 | -0.695122 | 0.486979 | 0.870096539 | GEMIN8 | protein_coding | 40.79 | -0.010599531 | 0.010599531 | ENSG00000046647 | n.s. |
| ENSG00000110492 | 4615.568 | -0.182153747 | 0.26904 | -0.677051 | 0.498374 | 0.874675553 | MDK | protein_coding | 66.26 | -0.010592789 | 0.010592789 | ENSG00000110492 | n.s. |
| ENSG00000077616 | 564.4225 | 0.180262392 | 0.264124 | 0.6824917 | 0.494928 | 0.873453717 | NAALAD2 | protein_coding | 34.98 | 0.010592237 | 0.010592237 | ENSG00000077616 | n.s. |
| ENSG00000110756 | 153.2434 | 0.172628255 | 0.246064 | 0.7015586 | 0.482954 | 0.868307709 | HPS5 | protein_coding | 39.75 | 0.01058666 | 0.01058666 | ENSG00000110756 | n.s. |
| ENSG00000154945 | 1475.563 | -0.179440904 | 0.261348 | -0.686598 | 0.492336 | 0.873000006 | ANKRD40 | protein_coding | 44.45 | -0.010584457 | 0.010584457 | ENSG00000154945 | n.s. |
| ENSG00000233922 | 271.9039 | -0.180222923 | 0.264412 | -0.681599 | 0.495492 | 0.87358987 | LINC01694 | lincRNA | 52.25 | -0.010577718 | 0.010577718 | ENSG00000233922 | n.s. |
| ENSG00000157827 | 2652.564 | 0.178723947 | 0.25993 | 0.6875851 | 0.491714 | 0.872671134 | FMNL2 | protein_coding | 40.57 | 0.010571412 | 0.010571412 | ENSG00000157827 | n.s. |
| ENSG00000151348 | 634.1021 | -0.183440798 | 0.273349 | -0.671086 | 0.502166 | 0.875813709 | EXT2 | protein_coding | 40.36 | -0.010564037 | 0.010564037 | ENSG00000151348 | n.s. |
| ENSG00000222346 | 4.436351 | -0.310798567 | 0.70702 | -0.43959 | 0.660234 | 0.924786541 | RNA5SP237 | rRNA | 45.13 | -0.010554253 | 0.010554253 | ENSG00000222346 | n.s. |
| ENSG00000242493 | 30.54622 | 0.214765438 | 0.364228 | 0.5896463 | 0.555428 | 0.893144718 | RN7SL37P | misc_RNA | 50.49 | 0.010540294 | 0.010540294 | ENSG00000242493 | n.s. |
| ENSG00000106290 | 288.4236 | -0.173916677 | 0.249593 | -0.696801 | 0.485927 | 0.869945355 | TAF6 | protein_coding | 53.95 | -0.010523355 | 0.010523355 | ENSG00000106290 | n.s. |
| ENSG00000173402 | 28.95524 | 0.210785509 | 0.348128 | 0.605483 | 0.544858 | 0.891406027 | DAG1 | protein_coding | 47.51 | 0.010523347 | 0.010523347 | ENSG00000173402 | n.s. |

| | | | | | | | | | | | | | |
|---|---|---|---|---|---|---|---|---|---|---|---|---|---|
| ENSG00000207988 | 15.43785 | -0.230098773 | 0.409989 | -0.561232 | 0.57464 | 0.900202572 | MIR576 | miRNA | 36.73 | -0.010506253 | 0.010506253 | ENSG00000207988 | n.s. |
| ENSG00000007168 | 2859.889 | -0.182404505 | 0.271843 | -0.670993 | 0.502225 | 0.875813709 | PAFAH1B1 | protein_coding | 42.4 | -0.010504358 | 0.010504358 | ENSG00000007168 | n.s. |
| ENSG00000162599 | 2009.113 | -0.167541045 | 0.235013 | -0.712901 | 0.475907 | 0.865616576 | NFIA | protein_coding | 40.05 | -0.01050054 | 0.01050054 | ENSG00000162599 | n.s. |
| ENSG00000175324 | 623.8434 | -0.195559369 | 0.302067 | -0.647405 | 0.51737 | 0.88373742 | LSM1 | protein_coding | 42.96 | -0.010496992 | 0.010496992 | ENSG00000175324 | n.s. |
| ENSG00000262919 | 163.0217 | -0.199868696 | 0.315082 | -0.634338 | 0.52586 | 0.886111278 | FAM58A | protein_coding | 56.23 | -0.010495452 | 0.010495452 | ENSG00000262919 | n.s. |
| ENSG00000148688 | 1297.662 | -0.182356068 | 0.272307 | -0.66967 | 0.503068 | 0.875883784 | RPP30 | protein_coding | 37.93 | -0.010495233 | 0.010495233 | ENSG00000148688 | n.s. |
| ENSG00000198933 | 1055.394 | -0.175787635 | 0.254067 | -0.691895 | 0.489004 | 0.871557795 | TBKBP1 | protein_coding | 58.66 | -0.010495191 | 0.010495191 | ENSG00000198933 | n.s. |
| ENSG00000114948 | 576.3179 | -0.177892814 | 0.259274 | -0.686118 | 0.492639 | 0.873000006 | ADAM23 | protein_coding | 38.27 | -0.010493142 | 0.010493142 | ENSG00000114948 | n.s. |
| ENSG00000215041 | 404.2312 | -0.18031439 | 0.265844 | -0.67827 | 0.4976 | 0.874675553 | NEURL4 | protein_coding | 57.23 | -0.010485825 | 0.010485825 | ENSG00000215041 | n.s. |
| ENSG00000237605 | 17.04987 | 0.254527389 | 0.494354 | 0.514869 | 0.606645 | 0.909522325 | | antisense | 56.26 | 0.010483127 | 0.010483127 | ENSG00000237605 | n.s. |
| ENSG00000115207 | 1636.742 | -0.176884705 | 0.256643 | -0.689223 | 0.490683 | 0.872495697 | GTF3C2 | protein_coding | 46.39 | -0.010478067 | 0.010478067 | ENSG00000115207 | n.s. |
| ENSG00000119522 | 223.2206 | -0.181188218 | 0.268642 | -0.674461 | 0.500019 | 0.875342302 | DENND1A | protein_coding | 42.52 | -0.01047668 | 0.01047668 | ENSG00000119522 | n.s. |
| ENSG00000162517 | 274.4196 | -0.18191481 | 0.271175 | -0.670839 | 0.502323 | 0.875813709 | PEF1 | protein_coding | 48.36 | -0.010476158 | 0.010476158 | ENSG00000162517 | n.s. |
| ENSG00000130827 | 381.6641 | -0.167042671 | 0.23368 | -0.714834 | 0.474712 | 0.865616576 | PLXNA3 | protein_coding | 60.44 | -0.010469305 | 0.010469305 | ENSG00000130827 | n.s. |
| ENSG00000207719 | 8.175494 | 0.250589665 | 0.469119 | 0.534171 | 0.593223 | 0.908297635 | MIR623 | miRNA | 52.04 | 0.010467586 | 0.010467586 | ENSG00000207719 | n.s. |
| ENSG00000188243 | 859.6077 | 0.251898255 | 0.474303 | 0.5310913 | 0.595356 | 0.908868121 | COMMD6 | protein_coding | 40 | 0.010453558 | 0.010453558 | ENSG00000188243 | n.s. |
| ENSG00000059145 | 37.70344 | -0.207746308 | 0.33947 | -0.611973 | 0.540556 | 0.890708138 | UNKL | protein_coding | 57.82 | -0.010442281 | 0.010442281 | ENSG00000059145 | n.s. |
| ENSG00000143319 | 1044.82 | -0.159350677 | 0.218037 | -0.730842 | 0.464875 | 0.860217499 | ISG20L2 | protein_coding | 50.3 | -0.010420216 | 0.010420216 | ENSG00000143319 | n.s. |
| ENSG00000247271 | 94.47021 | -0.179612362 | 0.265737 | -0.675902 | 0.499103 | 0.874967757 | ZBED5-AS1 | antisense | 40.5 | -0.010418945 | 0.010418945 | ENSG00000247271 | n.s. |
| ENSG00000274978 | 355.7649 | -0.163275755 | 0.226843 | -0.719775 | 0.471664 | 0.863532918 | RNU11 | snRNA | 51.49 | -0.010404111 | 0.010404111 | ENSG00000274978 | n.s. |
| ENSG00000135372 | 1125.57 | -0.174546741 | 0.252501 | -0.691272 | 0.489394 | 0.871780808 | NAT10 | protein_coding | 45.79 | -0.010401711 | 0.010401711 | ENSG00000135372 | n.s. |
| ENSG00000260293 | 477.9604 | -0.16596134 | 0.233542 | -0.710629 | 0.477314 | 0.865616576 | | sense_intronic | 51.91 | -0.010401533 | 0.010401533 | ENSG00000260293 | n.s. |
| ENSG00000239040 | 13.63377 | -0.239852778 | 0.439415 | -0.545846 | 0.585172 | 0.905026039 | | misc_RNA | 42.72 | -0.010394962 | 0.010394962 | ENSG00000239040 | n.s. |
| ENSG00000205078 | 65.29957 | 0.196569119 | 0.305777 | 0.6428512 | 0.520321 | 0.885364478 | SYCE1L | protein_coding | 46.1 | 0.010394164 | 0.010394164 | ENSG00000205078 | n.s. |
| ENSG00000169905 | 3667.797 | -0.165873113 | 0.233723 | -0.7097 | 0.47789 | 0.865660698 | TOR1AIP2 | protein_coding | 38.85 | -0.010392332 | 0.010392332 | ENSG00000169905 | n.s. |
| ENSG00000005700 | 1823.553 | -0.19782544 | 0.311308 | -0.635465 | 0.525125 | 0.886111278 | IBTK | protein_coding | 36.07 | -0.010388157 | 0.010388157 | ENSG00000005700 | n.s. |
| ENSG00000237550 | 965.5055 | 0.250302946 | 0.480856 | 0.5205361 | 0.60269 | 0.908868121 | RPL9P9 | nscribed_processed_pseudog | 41.84 | 0.010387354 | 0.010387354 | ENSG00000237550 | n.s. |
| ENSG00000103381 | 226.2734 | -0.196400729 | 0.305586 | -0.642701 | 0.520418 | 0.885364478 | CPPED1 | protein_coding | 44.42 | -0.010385259 | 0.010385259 | ENSG00000103381 | n.s. |
| ENSG00000136104 | 4587.947 | -0.185488369 | 0.280729 | -0.660737 | 0.508781 | 0.879205366 | RNASEH2B | protein_coding | 38.33 | -0.010370593 | 0.010370593 | ENSG00000136104 | n.s. |
| ENSG00000142892 | 651.3619 | -0.19598447 | 0.304887 | -0.642809 | 0.520348 | 0.885364478 | PIGK | protein_coding | 35.84 | -0.010363248 | 0.010363248 | ENSG00000142892 | n.s. |
| ENSG00000044115 | 2229.196 | 0.183251543 | 0.276337 | 0.663146 | 0.507237 | 0.877953916 | CTNNA1 | protein_coding | 44.15 | 0.010358894 | 0.010358894 | ENSG00000044115 | n.s. |
| ENSG00000132846 | 272.4609 | -0.178016754 | 0.262508 | -0.678138 | 0.497684 | 0.874675553 | ZBED3 | protein_coding | 43.7 | -0.01035221 | 0.01035221 | ENSG00000132846 | n.s. |

| | | | | | | | | | | | | | |
|---|---|---|---|---|---|---|---|---|---|---|---|---|---|
| ENSG00000184402 | 798.7165 | -0.185647248 | 0.281466 | -0.659572 | 0.509529 | 0.87951488 | SS18L1 | protein_coding | 55.02 | -0.010351098 | 0.010351098 | ENSG00000184402 | n.s. |
| ENSG00000125868 | 1670.825 | -0.197099446 | 0.311209 | -0.633335 | 0.526515 | 0.88615396 | DSTN | protein_coding | 39.56 | -0.010345911 | 0.010345911 | ENSG00000125868 | n.s. |
| ENSG00000100028 | 1927.175 | -0.180331722 | 0.270066 | -0.667732 | 0.504305 | 0.876255897 | SNRPD3 | protein_coding | 50.71 | -0.010345459 | 0.010345459 | ENSG00000100028 | n.s. |
| ENSG00000092051 | 267.8782 | 0.180420483 | 0.270342 | 0.667379 | 0.50453 | 0.876368641 | JPH4 | protein_coding | 56.69 | 0.01034047 | 0.01034047 | ENSG00000092051 | n.s. |
| ENSG00000270381 | 18.11527 | -0.223168343 | 0.393803 | -0.5667 | 0.570918 | 0.898818733 | | processed_pseudogene | 42.48 | -0.010338917 | 0.010338917 | ENSG00000270381 | n.s. |
| ENSG00000132128 | 428.9991 | -0.168510174 | 0.240127 | -0.701754 | 0.482833 | 0.868307709 | LRRC41 | protein_coding | 47.41 | -0.010334113 | 0.010334113 | ENSG00000132128 | n.s. |
| ENSG00000077463 | 142.2116 | -0.195691843 | 0.30505 | -0.641508 | 0.521192 | 0.885523197 | SIRT6 | protein_coding | 57.91 | -0.010332541 | 0.010332541 | ENSG00000077463 | n.s. |
| ENSG00000065457 | 576.5648 | -0.159972855 | 0.220803 | -0.724506 | 0.468755 | 0.861812061 | ADAT1 | protein_coding | 44.65 | -0.010332236 | 0.010332236 | ENSG00000065457 | n.s. |
| ENSG00000185946 | 1201.325 | -0.176452089 | 0.259504 | -0.679959 | 0.496531 | 0.873971799 | RNPC3 | protein_coding | 35.03 | -0.010322903 | 0.010322903 | ENSG00000185946 | n.s. |
| ENSG00000148343 | 43.42229 | -0.200314899 | 0.321775 | -0.62253 | 0.533593 | 0.888237452 | MIGA2 | protein_coding | 53.65 | -0.010310392 | 0.010310392 | ENSG00000148343 | n.s. |
| ENSG00000229117 | 9615.553 | 0.193039263 | 0.298769 | 0.6461151 | 0.518205 | 0.884295326 | RPL41 | protein_coding | 46.1 | 0.010308812 | 0.010308812 | ENSG00000229117 | n.s. |
| ENSG00000152894 | 92.47284 | 0.181121849 | 0.272186 | 0.6654348 | 0.505772 | 0.877272953 | PTPRK | protein_coding | 35.41 | 0.010299541 | 0.010299541 | ENSG00000152894 | n.s. |
| ENSG00000232519 | 41.7429 | 0.206255608 | 0.341545 | 0.60389 | 0.545917 | 0.891406027 | | antisense | 64.13 | 0.010297194 | 0.010297194 | ENSG00000232519 | n.s. |
| ENSG00000142330 | 94.6045 | -0.191444743 | 0.295057 | -0.64884 | 0.516442 | 0.883546124 | CAPN10 | protein_coding | 54.59 | -0.010294132 | 0.010294132 | ENSG00000142330 | n.s. |
| ENSG00000271427 | 25.8263 | -0.209706819 | 0.353359 | -0.593467 | 0.552868 | 0.893144718 | | lincRNA | 29.39 | -0.010292026 | 0.010292026 | ENSG00000271427 | n.s. |
| ENSG00000142039 | 162.5976 | -0.185792709 | 0.282697 | -0.657214 | 0.511043 | 0.880260889 | CCDC97 | protein_coding | 53.51 | -0.010290797 | 0.010290797 | ENSG00000142039 | n.s. |
| ENSG00000223599 | 158.3428 | -0.184870846 | 0.280573 | -0.658904 | 0.509958 | 0.879714521 | | processed_pseudogene | 49.2 | -0.010289586 | 0.010289586 | ENSG00000223599 | n.s. |
| ENSG00000247092 | 620.3561 | -0.150651493 | 0.201933 | -0.746047 | 0.455639 | 0.85453422 | SNHG10 | antisense | 47.24 | -0.010285059 | 0.010285059 | ENSG00000247092 | n.s. |
| ENSG00000166669 | 415.3082 | 0.163932348 | 0.229632 | 0.7138917 | 0.475294 | 0.865616576 | ATF7IP2 | protein_coding | 41.4 | 0.010274367 | 0.010274367 | ENSG00000166669 | n.s. |
| ENSG00000160194 | 601.675 | -0.18440492 | 0.279925 | -0.658765 | 0.510047 | 0.879714521 | NDUFV3 | protein_coding | 46.83 | -0.010263653 | 0.010263653 | ENSG00000160194 | n.s. |
| ENSG00000112245 | 3437.878 | 0.1529812 | 0.206223 | 0.7418256 | 0.458193 | 0.856873236 | PTP4A1 | protein_coding | 39.41 | 0.010262502 | 0.010262502 | ENSG00000112245 | n.s. |
| ENSG00000270696 | 262.9271 | -0.174345094 | 0.254759 | -0.684354 | 0.493752 | 0.873348456 | | antisense | 39.27 | -0.010253661 | 0.010253661 | ENSG00000270696 | n.s. |
| ENSG00000183207 | 1225.805 | -0.195190577 | 0.306561 | -0.636711 | 0.524313 | 0.886111278 | RUVBL2 | protein_coding | 53.06 | -0.010249796 | 0.010249796 | ENSG00000183207 | n.s. |
| ENSG00000272301 | 54.44817 | -0.196406888 | 0.311029 | -0.631474 | 0.527731 | 0.886936263 | | lincRNA | 33.68 | -0.010234289 | 0.010234289 | ENSG00000272301 | n.s. |
| ENSG00000035681 | 910.4177 | -0.18365629 | 0.27894 | -0.658408 | 0.510276 | 0.879714521 | NSMAF | protein_coding | 38.41 | -0.010221986 | 0.010221986 | ENSG00000035681 | n.s. |
| ENSG00000174796 | 181.9361 | 0.172504284 | 0.25016 | 0.6895744 | 0.490462 | 0.872495697 | THAP6 | protein_coding | 40.52 | 0.010218586 | 0.010218586 | ENSG00000174796 | n.s. |
| ENSG00000197928 | 565.1924 | -0.17610365 | 0.26051 | -0.675995 | 0.499044 | 0.874967757 | ZNF677 | protein_coding | 38.84 | -0.010215412 | 0.010215412 | ENSG00000197928 | n.s. |
| ENSG00000165898 | 431.7884 | 0.175285951 | 0.258077 | 0.6791992 | 0.497012 | 0.874430631 | ISCA2 | protein_coding | 45.59 | 0.010214725 | 0.010214725 | ENSG00000165898 | n.s. |
| ENSG00000178502 | 463.6995 | -0.166815415 | 0.238306 | -0.700006 | 0.483923 | 0.868580821 | KLHL11 | protein_coding | 47.67 | -0.010207396 | 0.010207396 | ENSG00000178502 | n.s. |
| ENSG00000176842 | 173.2105 | -0.172320642 | 0.250664 | -0.687455 | 0.491796 | 0.872671134 | IRX5 | protein_coding | 59.38 | -0.010192661 | 0.010192661 | ENSG00000176842 | n.s. |
| ENSG00000151239 | 1855.639 | 0.177053447 | 0.264311 | 0.6698677 | 0.502942 | 0.875883784 | TWF1 | protein_coding | 34.16 | 0.010190048 | 0.010190048 | ENSG00000151239 | n.s. |
| ENSG00000057468 | 130.2112 | -0.176426664 | 0.26248 | -0.672152 | 0.501487 | 0.875518559 | MSH4 | protein_coding | 37.01 | -0.01018593 | 0.01018593 | ENSG00000057468 | n.s. |

| | | | | | | | | | | | | | |
|---|---|---|---|---|---|---|---|---|---|---|---|---|---|
| ENSG00000181904 | 1900.118 | -0.18386145 | 0.279751 | -0.657233 | 0.511031 | 0.880260889 | C5orf24 | protein_coding | 41.93 | -0.010183827 | 0.010183827 | ENSG00000181904 | n.s. |
| ENSG00000113966 | 350.4646 | -0.187437211 | 0.287244 | -0.652537 | 0.514055 | 0.882409358 | ARL6 | protein_coding | 34.36 | -0.010183444 | 0.010183444 | ENSG00000113966 | n.s. |
| ENSG00000183605 | 463.0036 | 0.195369581 | 0.309666 | 0.6309041 | 0.528103 | 0.886954686 | SFXN4 | protein_coding | 47.78 | 0.010178475 | 0.010178475 | ENSG00000183605 | n.s. |
| ENSG00000092148 | 3774.365 | -0.175962573 | 0.261415 | -0.673117 | 0.500873 | 0.875342302 | HECTD1 | protein_coding | 39.35 | -0.010174523 | 0.010174523 | ENSG00000092148 | n.s. |
| ENSG00000147894 | 1147.105 | 0.189520519 | 0.29239 | 0.6481774 | 0.51687 | 0.88372671 | C9orf72 | protein_coding | 35.84 | 0.010173844 | 0.010173844 | ENSG00000147894 | n.s. |
| ENSG00000166997 | 142.5847 | 0.195768349 | 0.311954 | 0.6275562 | 0.530295 | 0.887229181 | CNPY4 | protein_coding | 50.62 | 0.010172942 | 0.010172942 | ENSG00000166997 | n.s. |
| ENSG00000108604 | 3035.288 | -0.185691314 | 0.283722 | -0.654483 | 0.512801 | 0.881523829 | SMARCD2 | protein_coding | 55.99 | -0.01016956 | 0.01016956 | ENSG00000108604 | n.s. |
| ENSG00000173786 | 434.9852 | -0.174858498 | 0.257953 | -0.67787 | 0.497854 | 0.874675553 | CNP | protein_coding | 54.63 | -0.010168548 | 0.010168548 | ENSG00000173786 | n.s. |
| ENSG00000207133 | 23.06024 | 0.211240043 | 0.364133 | 0.5801179 | 0.561835 | 0.895116382 | SNORD116-7 | snoRNA | 44.21 | 0.010164975 | 0.010164975 | ENSG00000207133 | n.s. |
| ENSG00000125388 | 259.0791 | -0.171836355 | 0.249945 | -0.687495 | 0.491771 | 0.872671134 | GRK4 | protein_coding | 46.38 | -0.010164015 | 0.010164015 | ENSG00000125388 | n.s. |
| ENSG00000234215 | 19.13718 | -0.244901621 | 0.462472 | -0.529549 | 0.596425 | 0.908868121 | | lincRNA | 44.2 | -0.010163204 | 0.010163204 | ENSG00000234215 | n.s. |
| ENSG00000102241 | 2969.899 | -0.174715385 | 0.257563 | -0.67834 | 0.497556 | 0.874675553 | HTATSF1 | protein_coding | 37.88 | -0.010160226 | 0.010160226 | ENSG00000102241 | n.s. |
| ENSG00000185608 | 377.8035 | 0.179688433 | 0.270936 | 0.6632132 | 0.507194 | 0.877953916 | MRPL40 | protein_coding | 51.63 | 0.010157478 | 0.010157478 | ENSG00000185608 | n.s. |
| ENSG00000214223 | 70.38728 | -0.195360883 | 0.311521 | -0.62712 | 0.530581 | 0.887229181 | HNRNPA1P10 | processed_pseudogene | 47.51 | -0.010151769 | 0.010151769 | ENSG00000214223 | n.s. |
| ENSG00000171067 | 279.2476 | -0.189230935 | 0.292504 | -0.646936 | 0.517674 | 0.883822248 | C11orf24 | protein_coding | 52.05 | -0.010149415 | 0.010149415 | ENSG00000171067 | n.s. |
| ENSG00000106443 | 5528.465 | -0.172456423 | 0.252276 | -0.683602 | 0.494226 | 0.873348456 | PHF14 | protein_coding | 34.81 | -0.010142584 | 0.010142584 | ENSG00000106443 | n.s. |
| ENSG00000185624 | 4051.294 | -0.183458704 | 0.27944 | -0.656522 | 0.511488 | 0.880578023 | P4HB | protein_coding | 55.79 | -0.01013282 | 0.01013282 | ENSG00000185624 | n.s. |
| ENSG00000153936 | 1773.365 | -0.186977819 | 0.287195 | -0.651048 | 0.515015 | 0.882747369 | HS2ST1 | protein_coding | 36.82 | -0.010127386 | 0.010127386 | ENSG00000153936 | n.s. |
| ENSG00000229325 | 82.49358 | -0.192670279 | 0.301299 | -0.639465 | 0.522521 | 0.886079926 | ACAP2-IT1 | sense_intronic | 33.33 | -0.010120411 | 0.010120411 | ENSG00000229325 | n.s. |
| ENSG00000166965 | 641.9412 | -0.167231146 | 0.239823 | -0.69731 | 0.485609 | 0.869934871 | RCCD1 | protein_coding | 54.47 | -0.010119702 | 0.010119702 | ENSG00000166965 | n.s. |
| ENSG00000120253 | 1638.824 | -0.175499091 | 0.261759 | -0.670459 | 0.502565 | 0.875813709 | NUP43 | protein_coding | 43.95 | -0.010106687 | 0.010106687 | ENSG00000120253 | n.s. |
| ENSG00000184702 | 395.3448 | -0.165149665 | 0.235956 | -0.699916 | 0.48398 | 0.868580821 | 5-Sep | protein_coding | 66.81 | -0.010105469 | 0.010105469 | ENSG00000184702 | n.s. |
| ENSG00000000419 | 1158.39 | -0.190980191 | 0.297055 | -0.642912 | 0.520281 | 0.885364478 | DPM1 | protein_coding | 39.85 | -0.010098633 | 0.010098633 | ENSG00000000419 | n.s. |
| ENSG00000182150 | 3123.783 | -0.172151073 | 0.25289 | -0.680735 | 0.496039 | 0.873669126 | ERCC6L2 | protein_coding | 37.1 | -0.010097179 | 0.010097179 | ENSG00000182150 | n.s. |
| ENSG00000084092 | 479.1337 | 0.185639362 | 0.28464 | 0.65219 | 0.514279 | 0.882409358 | NOA1 | protein_coding | 45.66 | 0.010085767 | 0.010085767 | ENSG00000084092 | n.s. |
| ENSG00000156990 | 320.1223 | -0.190525577 | 0.295767 | -0.644175 | 0.519462 | 0.885247561 | RPUSD3 | protein_coding | 52.21 | -0.010085521 | 0.010085521 | ENSG00000156990 | n.s. |
| ENSG00000062716 | 1592.963 | -0.175898421 | 0.26374 | -0.66694 | 0.50481 | 0.876368641 | VMP1 | protein_coding | 39.75 | -0.010081296 | 0.010081296 | ENSG00000062716 | n.s. |
| ENSG00000171004 | 189.2732 | -0.197426058 | 0.318296 | -0.620259 | 0.535087 | 0.88909364 | HS6ST2 | protein_coding | 41.21 | -0.010079093 | 0.010079093 | ENSG00000171004 | n.s. |
| ENSG00000180287 | 45.38398 | -0.197410029 | 0.318319 | -0.620165 | 0.535149 | 0.88909364 | PLD5 | protein_coding | 40.38 | -0.010078275 | 0.010078275 | ENSG00000180287 | n.s. |
| ENSG00000199865 | 4.371711 | -0.334969716 | 0.816346 | -0.410328 | 0.681565 | 0.933075339 | RNU6-495P | snRNA | 44.86 | -0.010076991 | 0.010076991 | ENSG00000199865 | n.s. |
| ENSG00000144554 | 1616.056 | -0.1717477 | 0.252342 | -0.680615 | 0.496115 | 0.873692584 | FANCD2 | protein_coding | 42.77 | -0.010071517 | 0.010071517 | ENSG00000144554 | n.s. |
| ENSG00000137145 | 850.5588 | -0.171038244 | 0.249765 | -0.684796 | 0.493473 | 0.873348456 | DENND4C | protein_coding | 38.75 | -0.010059177 | 0.010059177 | ENSG00000137145 | n.s. |

| | | | | | | | | | | | | | |
|---|---|---|---|---|---|---|---|---|---|---|---|---|---|
| ENSG00000197170 | 3244.597 | -0.172952682 | 0.255367 | -0.677272 | 0.498233 | 0.874675553 | PSMD12 | protein_coding | 40.05 | -0.010057719 | 0.010057719 | ENSG00000197170 | n.s. |
| ENSG00000150776 | 643.6502 | -0.180353909 | 0.273239 | -0.66006 | 0.509215 | 0.87951488 | C11orf57 | protein_coding | 40.81 | -0.010055958 | 0.010055958 | ENSG00000150776 | n.s. |
| ENSG00000182307 | 1523.336 | -0.170451853 | 0.248524 | -0.685856 | 0.492804 | 0.873000006 | C8orf33 | protein_coding | 51.6 | -0.010054231 | 0.010054231 | ENSG00000182307 | n.s. |
| ENSG00000101868 | 2080.017 | -0.177763941 | 0.268115 | -0.663015 | 0.507321 | 0.877989988 | POLA1 | protein_coding | 38.8 | -0.010045518 | 0.010045518 | ENSG00000101868 | n.s. |
| ENSG00000108828 | 384.854 | -0.186756932 | 0.287656 | -0.649238 | 0.516185 | 0.883546124 | VAT1 | protein_coding | 54.7 | -0.010042065 | 0.010042065 | ENSG00000108828 | n.s. |
| ENSG00000275066 | 77.38316 | -0.193859068 | 0.310522 | -0.624301 | 0.53243 | 0.887576332 | SYNRG | protein_coding | 40.78 | -0.010040792 | 0.010040792 | ENSG00000275066 | n.s. |
| ENSG00000200785 | 11.94972 | 0.232162359 | 0.428045 | 0.5423789 | 0.587558 | 0.905217543 | SNORD8 | snoRNA | 50 | 0.010040334 | 0.010040334 | ENSG00000200785 | n.s. |
| ENSG00000200882 | 40.58458 | 0.201100578 | 0.335583 | 0.5992582 | 0.549001 | 0.891406027 | RNU6-681P | snRNA | 35.58 | 0.010039833 | 0.010039833 | ENSG00000200882 | n.s. |
| ENSG00000196235 | 861.8109 | -0.162020703 | 0.230142 | -0.704005 | 0.48143 | 0.867349802 | SUPT5H | protein_coding | 51.42 | -0.010013806 | 0.010013806 | ENSG00000196235 | n.s. |
| ENSG00000160208 | 1809.6 | -0.17300465 | 0.257029 | -0.673094 | 0.500887 | 0.875342302 | RRP1B | protein_coding | 47.51 | -0.010003489 | 0.010003489 | ENSG00000160208 | n.s. |
| ENSG00000114115 | 126.9311 | 0.196133101 | 0.317145 | 0.6184329 | 0.53629 | 0.889201649 | RBP1 | protein_coding | 48.01 | 0.010002737 | 0.010002737 | ENSG00000114115 | n.s. |
| ENSG00000166295 | 929.9348 | -0.19038823 | 0.300135 | -0.634341 | 0.525858 | 0.886111278 | ANAPC16 | protein_coding | 43.46 | -0.009997616 | 0.009997616 | ENSG00000166295 | n.s. |
| ENSG00000033178 | 1284.454 | -0.179290031 | 0.271748 | -0.659765 | 0.509405 | 0.87951488 | UBA6 | protein_coding | 35.52 | -0.00999664 | 0.00999664 | ENSG00000033178 | n.s. |
| ENSG00000149091 | 384.8661 | -0.176039436 | 0.264926 | -0.664485 | 0.50638 | 0.877502299 | DGKZ | protein_coding | 56.98 | -0.009990544 | 0.009990544 | ENSG00000149091 | n.s. |
| ENSG00000170906 | 410.121 | -0.199806378 | 0.327709 | -0.609707 | 0.542056 | 0.891404544 | NDUFA3 | protein_coding | 54.99 | -0.009975365 | 0.009975365 | ENSG00000170906 | n.s. |
| ENSG00000179912 | 111.7171 | -0.185408974 | 0.285765 | -0.648816 | 0.516457 | 0.883546124 | R3HDM2 | protein_coding | 43.16 | -0.009969584 | 0.009969584 | ENSG00000179912 | n.s. |
| ENSG00000175643 | 332.6761 | -0.185674153 | 0.286698 | -0.647629 | 0.517225 | 0.88372671 | RMI2 | protein_coding | 50.87 | -0.009967363 | 0.009967363 | ENSG00000175643 | n.s. |
| ENSG00000130638 | 1980.89 | -0.183194414 | 0.280448 | -0.653221 | 0.513614 | 0.882409358 | ATXN10 | protein_coding | 40.57 | -0.009952933 | 0.009952933 | ENSG00000130638 | n.s. |
| ENSG00000104973 | 196.173 | -0.169307215 | 0.248056 | -0.682535 | 0.494901 | 0.873453717 | MED25 | protein_coding | 56.01 | -0.009948509 | 0.009948509 | ENSG00000104973 | n.s. |
| ENSG00000132967 | 583.5178 | 0.195059782 | 0.31555 | 0.6181574 | 0.536472 | 0.889201649 | HMGB1P5 | ıscribed_processed_pseudog | 39.42 | 0.009947998 | 0.009947998 | ENSG00000132967 | n.s. |
| ENSG00000005249 | 561.5449 | -0.173378415 | 0.25954 | -0.668021 | 0.50412 | 0.876255897 | PRKAR2B | protein_coding | 38.63 | -0.009946554 | 0.009946554 | ENSG00000005249 | n.s. |
| ENSG00000106617 | 106.1257 | -0.180623006 | 0.275506 | -0.655606 | 0.512078 | 0.88091768 | PRKAG2 | protein_coding | 48.91 | -0.009945947 | 0.009945947 | ENSG00000106617 | n.s. |
| ENSG00000181852 | 586.1873 | -0.190858215 | 0.303072 | -0.629746 | 0.528861 | 0.886954686 | RNF41 | protein_coding | 47.86 | -0.00994344 | 0.00994344 | ENSG00000181852 | n.s. |
| ENSG00000181038 | 634.0564 | 0.189307032 | 0.298415 | 0.6343755 | 0.525836 | 0.886111278 | METTL23 | protein_coding | 47.85 | 0.009940841 | 0.009940841 | ENSG00000181038 | n.s. |
| ENSG00000254402 | 63.40291 | 0.195202543 | 0.316156 | 0.6174253 | 0.536954 | 0.889412093 | LRRC24 | protein_coding | 63.08 | 0.009935218 | 0.009935218 | ENSG00000254402 | n.s. |
| ENSG00000271122 | 247.3113 | -0.155893133 | 0.21654 | -0.719927 | 0.47157 | 0.863532918 | | antisense | 46 | -0.009933682 | 0.009933682 | ENSG00000271122 | n.s. |
| ENSG00000145335 | 176.8788 | 0.189061685 | 0.297924 | 0.634597 | 0.525691 | 0.886111278 | SNCA | protein_coding | 36.12 | 0.009927957 | 0.009927957 | ENSG00000145335 | n.s. |
| ENSG00000141295 | 378.2584 | -0.182662644 | 0.280042 | -0.65227 | 0.514227 | 0.882409358 | SCRN2 | protein_coding | 59.28 | -0.009924042 | 0.009924042 | ENSG00000141295 | n.s. |
| ENSG00000155099 | 240.9516 | -0.190287552 | 0.302355 | -0.629351 | 0.529119 | 0.886954686 | TMEM55A | protein_coding | 35.6 | -0.009913709 | 0.009913709 | ENSG00000155099 | n.s. |
| ENSG00000175895 | 137.9922 | 0.198573525 | 0.329332 | 0.6029589 | 0.546536 | 0.891406027 | PLEKHF2 | protein_coding | 37.16 | 0.009913671 | 0.009913671 | ENSG00000175895 | n.s. |
| ENSG00000058799 | 392.4043 | -0.187633508 | 0.292793 | -0.640839 | 0.521627 | 0.885576517 | YIPF1 | protein_coding | 43.46 | -0.009902153 | 0.009902153 | ENSG00000058799 | n.s. |
| ENSG00000104388 | 992.5476 | -0.184097163 | 0.283593 | -0.64916 | 0.516235 | 0.883546124 | RAB2A | protein_coding | 39.18 | -0.009899047 | 0.009899047 | ENSG00000104388 | n.s. |

| | | | | | | | | | | | | | |
|---|---|---|---|---|---|---|---|---|---|---|---|---|---|
| ENSG00000164116 | 464.5338 | 0.172138003 | 0.257258 | 0.6691272 | 0.503414 | 0.876020205 | GUCY1A3 | protein_coding | 36.88 | 0.009895504 | 0.009895504 | ENSG00000164116 | n.s. |
| ENSG00000087842 | 198.84 | 0.196800699 | 0.321196 | 0.6127117 | 0.540067 | 0.890684245 | PIR | protein_coding | 38.99 | 0.009894397 | 0.009894397 | ENSG00000087842 | n.s. |
| ENSG00000109618 | 414.4414 | -0.171769054 | 0.255978 | -0.671031 | 0.502201 | 0.875813709 | SEPSECS | protein_coding | 36.27 | -0.009891881 | 0.009891881 | ENSG00000109618 | n.s. |
| ENSG00000158470 | 894.2722 | -0.168696955 | 0.247933 | -0.680414 | 0.496242 | 0.873799463 | B4GALT5 | protein_coding | 42.32 | -0.009883655 | 0.009883655 | ENSG00000158470 | n.s. |
| ENSG00000156931 | 309.5074 | -0.172177253 | 0.257854 | -0.667731 | 0.504305 | 0.876255897 | VPS8 | protein_coding | 38.45 | -0.009877645 | 0.009877645 | ENSG00000156931 | n.s. |
| ENSG00000153904 | 582.7287 | -0.169765511 | 0.250549 | -0.677575 | 0.498041 | 0.874675553 | DDAH1 | protein_coding | 39.19 | -0.009872376 | 0.009872376 | ENSG00000153904 | n.s. |
| ENSG00000004866 | 574.0669 | -0.16776458 | 0.245206 | -0.684178 | 0.493863 | 0.873348456 | ST7 | protein_coding | 39.15 | -0.009866645 | 0.009866645 | ENSG00000004866 | n.s. |
| ENSG00000164494 | 210.4916 | 0.186467496 | 0.290133 | 0.6426964 | 0.520421 | 0.885364478 | PDSS2 | protein_coding | 39.82 | 0.009860011 | 0.009860011 | ENSG00000164494 | n.s. |
| ENSG00000204843 | 3992.267 | -0.167778645 | 0.245627 | -0.683061 | 0.494568 | 0.873443478 | DCTN1 | protein_coding | 49.96 | -0.009859545 | 0.009859545 | ENSG00000204843 | n.s. |
| ENSG00000160446 | 88.79112 | -0.187732826 | 0.295002 | -0.636379 | 0.52453 | 0.886111278 | ZDHHC12 | protein_coding | 61.4 | -0.009858177 | 0.009858177 | ENSG00000160446 | n.s. |
| ENSG00000066379 | 245.0297 | 0.187531173 | 0.294231 | 0.6373598 | 0.52389 | 0.886111278 | ZNRD1 | protein_coding | 41.39 | 0.009847587 | 0.009847587 | ENSG00000066379 | n.s. |
| ENSG00000097033 | 2078.765 | -0.183513902 | 0.28362 | -0.647042 | 0.517605 | 0.883822248 | SH3GLB1 | protein_coding | 36.11 | -0.009842781 | 0.009842781 | ENSG00000097033 | n.s. |
| ENSG00000204272 | 210.223 | 0.18732632 | 0.293503 | 0.6382425 | 0.523316 | 0.886079926 | NBDY | protein_coding | 45.2 | 0.009839709 | 0.009839709 | ENSG00000204272 | n.s. |
| ENSG00000112996 | 2664.601 | 0.165256084 | 0.239173 | 0.6909491 | 0.489598 | 0.871894847 | MRPS30 | protein_coding | 36.6 | 0.009838668 | 0.009838668 | ENSG00000112996 | n.s. |
| ENSG00000231312 | 59.76127 | -0.199418314 | 0.334819 | -0.595601 | 0.551442 | 0.892641916 | | antisense | 41.25 | -0.009835854 | 0.009835854 | ENSG00000231312 | n.s. |
| ENSG00000198648 | 871.6993 | -0.183173968 | 0.282739 | -0.647854 | 0.517079 | 0.88372671 | STK39 | protein_coding | 40.16 | -0.009833148 | 0.009833148 | ENSG00000198648 | n.s. |
| ENSG00000148291 | 192.3083 | -0.19683678 | 0.327984 | -0.600142 | 0.548412 | 0.891406027 | SURF2 | protein_coding | 59.53 | -0.009826965 | 0.009826965 | ENSG00000148291 | n.s. |
| ENSG00000177951 | 234.5373 | -0.166913303 | 0.244028 | -0.683993 | 0.49398 | 0.873348456 | BET1L | protein_coding | 48.36 | -0.009816579 | 0.009816579 | ENSG00000177951 | n.s. |
| ENSG00000143418 | 1486.27 | -0.175925988 | 0.266671 | -0.659711 | 0.509439 | 0.87951488 | CERS2 | protein_coding | 52.38 | -0.009809072 | 0.009809072 | ENSG00000143418 | n.s. |
| ENSG00000206573 | 2654.048 | 0.158413861 | 0.224809 | 0.7046602 | 0.481022 | 0.867184458 | THUMPD3-AS1 | antisense | 41.7 | 0.009803999 | 0.009803999 | ENSG00000206573 | n.s. |
| ENSG00000143633 | 814.3344 | -0.165608365 | 0.240969 | -0.687261 | 0.491918 | 0.872671134 | C1orf131 | protein_coding | 43.5 | -0.009795634 | 0.009795634 | ENSG00000143633 | n.s. |
| ENSG00000270060 | 64.12153 | -0.18654713 | 0.294332 | -0.633798 | 0.526212 | 0.88615396 | | sense_intronic | 48.71 | -0.009792011 | 0.009792011 | ENSG00000270060 | n.s. |
| ENSG00000209482 | 67.60943 | 0.180214173 | 0.276051 | 0.6528283 | 0.513867 | 0.882409358 | SNORD83A | snoRNA | 54.74 | 0.009791017 | 0.009791017 | ENSG00000209482 | n.s. |
| ENSG00000171612 | 340.6065 | -0.188257257 | 0.300235 | -0.627033 | 0.530638 | 0.887229181 | SLC25A33 | protein_coding | 45.5 | -0.009782634 | 0.009782634 | ENSG00000171612 | n.s. |
| ENSG00000149541 | 315.6188 | -0.177598931 | 0.270915 | -0.655551 | 0.512113 | 0.88091768 | B3GAT3 | protein_coding | 53.95 | -0.009779427 | 0.009779427 | ENSG00000149541 | n.s. |
| ENSG00000099381 | 300.9234 | -0.164119265 | 0.237493 | -0.691048 | 0.489535 | 0.871894847 | SETD1A | protein_coding | 55.71 | -0.009770986 | 0.009770986 | ENSG00000099381 | n.s. |
| ENSG00000178202 | 438.391 | -0.184992531 | 0.288423 | -0.641393 | 0.521267 | 0.885523197 | KDELC2 | protein_coding | 41.21 | -0.009767616 | 0.009767616 | ENSG00000178202 | n.s. |
| ENSG00000104969 | 787.4718 | -0.18344505 | 0.284167 | -0.645553 | 0.518569 | 0.884700249 | SGTA | protein_coding | 57.8 | -0.009759983 | 0.009759983 | ENSG00000104969 | n.s. |
| ENSG00000252291 | 4.726799 | -0.280971138 | 0.61761 | -0.454933 | 0.649157 | 0.923184844 | | snoRNA | 44.34 | -0.009752883 | 0.009752883 | ENSG00000252291 | n.s. |
| ENSG00000088543 | 121.2718 | 0.190201037 | 0.305863 | 0.6218507 | 0.53404 | 0.888662588 | C3orf18 | protein_coding | 55.98 | 0.009750295 | 0.009750295 | ENSG00000088543 | n.s. |
| ENSG00000184677 | 692.4183 | -0.169361094 | 0.252933 | -0.669587 | 0.503121 | 0.875883784 | ZBTB40 | protein_coding | 42.7 | -0.009747326 | 0.009747326 | ENSG00000184677 | n.s. |
| ENSG00000213593 | 1371.857 | -0.193040228 | 0.314625 | -0.613556 | 0.539509 | 0.890328323 | TMX2 | protein_coding | 45.19 | -0.009738843 | 0.009738843 | ENSG00000213593 | n.s. |

| | | | | | | | | | | | | | |
|---|---|---|---|---|---|---|---|---|---|---|---|---|---|
| ENSG00000167325 | 3806.774 | -0.181091446 | 0.279179 | -0.648657 | 0.51656 | 0.883546124 | RRM1 | protein_coding | 40.39 | -0.009737427 | 0.009737427 | ENSG00000167325 | n.s. |
| ENSG00000176371 | 772.6612 | -0.165472151 | 0.242054 | -0.683617 | 0.494217 | 0.873348456 | ZSCAN2 | protein_coding | 48.03 | -0.009731821 | 0.009731821 | ENSG00000176371 | n.s. |
| ENSG00000186591 | 506.8187 | 0.1696256 | 0.25401 | 0.6677914 | 0.504267 | 0.876255897 | UBE2H | protein_coding | 41.85 | 0.009731259 | 0.009731259 | ENSG00000186591 | n.s. |
| ENSG00000133641 | 1424.932 | -0.17403114 | 0.263196 | -0.661223 | 0.508469 | 0.879193676 | C12orf29 | protein_coding | 36.21 | -0.009731029 | 0.009731029 | ENSG00000133641 | n.s. |
| ENSG00000128594 | 188.1166 | -0.167057818 | 0.246045 | -0.678972 | 0.497156 | 0.874529412 | LRRC4 | protein_coding | 56.07 | -0.009727038 | 0.009727038 | ENSG00000128594 | n.s. |
| ENSG00000104889 | 940.6215 | -0.187585775 | 0.300474 | -0.624299 | 0.532431 | 0.887576332 | RNASEH2A | protein_coding | 52.97 | -0.009715871 | 0.009715871 | ENSG00000104889 | n.s. |
| ENSG00000185246 | 1668.725 | -0.174779505 | 0.26561 | -0.658031 | 0.510518 | 0.879913979 | PRPF39 | protein_coding | 36.34 | -0.009710711 | 0.009710711 | ENSG00000185246 | n.s. |
| ENSG00000180011 | 871.9709 | -0.165538719 | 0.24297 | -0.681313 | 0.495674 | 0.873669126 | ZADH2 | protein_coding | 46.75 | -0.009709344 | 0.009709344 | ENSG00000180011 | n.s. |
| ENSG00000255529 | 1691.826 | 0.1700086 | 0.255166 | 0.666267 | 0.50524 | 0.876821729 | POLR2M | protein_coding | 42.66 | 0.009705569 | 0.009705569 | ENSG00000255529 | n.s. |
| ENSG00000113790 | 30.87845 | -0.19439945 | 0.322319 | -0.603127 | 0.546424 | 0.891406027 | EHHADH | protein_coding | 39.78 | -0.009705283 | 0.009705283 | ENSG00000113790 | n.s. |
| ENSG00000077721 | 507.2272 | -0.186175378 | 0.296258 | -0.628424 | 0.529726 | 0.886954686 | UBE2A | protein_coding | 40.14 | -0.00969947 | 0.00969947 | ENSG00000077721 | n.s. |
| ENSG00000166171 | 138.2373 | -0.186073227 | 0.295904 | -0.62883 | 0.529461 | 0.886954686 | DPCD | protein_coding | 50.19 | -0.009694149 | 0.009694149 | ENSG00000166171 | n.s. |
| ENSG00000082701 | 1808.54 | -0.163737452 | 0.237716 | -0.688794 | 0.490953 | 0.872562973 | GSK3B | protein_coding | 37.15 | -0.009693784 | 0.009693784 | ENSG00000082701 | n.s. |
| ENSG00000123136 | 1918.959 | -0.184509151 | 0.291354 | -0.633282 | 0.52655 | 0.88615396 | DDX39A | protein_coding | 55.52 | -0.009685036 | 0.009685036 | ENSG00000123136 | n.s. |
| ENSG00000263652 | 2.813463 | -0.32917079 | 0.818117 | -0.402352 | 0.687425 | 0.934506008 | MIR548AX | miRNA | 39.73 | -0.009683514 | 0.009683514 | ENSG00000263652 | n.s. |
| ENSG00000272888 | 10145 | 0.156399511 | 0.221774 | 0.7052208 | 0.480673 | 0.867184458 | LINC01578 | processed_transcript | 36.16 | 0.009679333 | 0.009679333 | ENSG00000272888 | n.s. |
| ENSG00000165233 | 135.9251 | -0.193650014 | 0.322683 | -0.600125 | 0.548423 | 0.891406027 | CARD19 | protein_coding | 49.96 | -0.009667867 | 0.009667867 | ENSG00000165233 | n.s. |
| ENSG00000116747 | 4005.582 | -0.18409446 | 0.28884 | -0.637358 | 0.523891 | 0.886111278 | TROVE2 | protein_coding | 35.99 | -0.00966712 | 0.00966712 | ENSG00000116747 | n.s. |
| ENSG00000102390 | 384.6166 | 0.166116211 | 0.24515 | 0.6776107 | 0.498019 | 0.874675553 | PBDC1 | protein_coding | 42.82 | 0.009660158 | 0.009660158 | ENSG00000102390 | n.s. |
| ENSG00000239523 | 52.13771 | -0.183649271 | 0.286816 | -0.640304 | 0.521975 | 0.885950923 | MYLK-AS1 | antisense | 44.17 | -0.009658177 | 0.009658177 | ENSG00000239523 | n.s. |
| ENSG00000274023 | 53.07377 | -0.185194302 | 0.293584 | -0.630806 | 0.528168 | 0.886954686 | | sense_overlapping | 41.8 | -0.009648358 | 0.009648358 | ENSG00000274023 | n.s. |
| ENSG00000146350 | 1038.655 | -0.178856244 | 0.275099 | -0.650152 | 0.515594 | 0.883195516 | TBC1D32 | protein_coding | 35.75 | -0.009648068 | 0.009648068 | ENSG00000146350 | n.s. |
| ENSG00000075151 | 181.0625 | 0.164324328 | 0.241057 | 0.6816816 | 0.49544 | 0.87358987 | EIF4G3 | protein_coding | 39.36 | 0.009644591 | 0.009644591 | ENSG00000075151 | n.s. |
| ENSG00000279968 | 21.68483 | 0.214084704 | 0.384618 | 0.5566159 | 0.57779 | 0.901500147 | GVQW2 | protein_coding | 41.79 | 0.009641133 | 0.009641133 | ENSG00000279968 | n.s. |
| ENSG00000149295 | 55.11029 | 0.191085869 | 0.311447 | 0.613543 | 0.539517 | 0.890328323 | DRD2 | protein_coding | 48.35 | 0.009640246 | 0.009640246 | ENSG00000149295 | n.s. |
| ENSG00000103266 | 492.7156 | -0.183523613 | 0.287118 | -0.639191 | 0.522699 | 0.886079926 | STUB1 | protein_coding | 64.94 | -0.009639964 | 0.009639964 | ENSG00000103266 | n.s. |
| ENSG00000182154 | 549.4482 | -0.193942223 | 0.324517 | -0.597633 | 0.550085 | 0.891877082 | MRPL41 | protein_coding | 72.44 | -0.009637958 | 0.009637958 | ENSG00000182154 | n.s. |
| ENSG00000168802 | 734.9865 | -0.185623855 | 0.29657 | -0.625903 | 0.531378 | 0.887366671 | CHTF8 | protein_coding | 48.17 | -0.0096333 | 0.0096333 | ENSG00000168802 | n.s. |
| ENSG00000025800 | 661.2614 | -0.163671968 | 0.238929 | -0.685024 | 0.493329 | 0.873348456 | KPNA6 | protein_coding | 44.21 | -0.009625948 | 0.009625948 | ENSG00000025800 | n.s. |
| ENSG00000258289 | 1162.432 | -0.185028024 | 0.294727 | -0.627795 | 0.530138 | 0.88715322 | CHURC1 | protein_coding | 41.09 | -0.00962171 | 0.00962171 | ENSG00000258289 | n.s. |
| ENSG00000126822 | 73.27704 | -0.181364367 | 0.281268 | -0.64481 | 0.51905 | 0.88508678 | PLEKHG3 | protein_coding | 52.94 | -0.009614877 | 0.009614877 | ENSG00000126822 | n.s. |
| ENSG00000176101 | 645.6085 | -0.192581976 | 0.317872 | -0.605848 | 0.544616 | 0.891406027 | SSNA1 | protein_coding | 65.89 | -0.009614546 | 0.009614546 | ENSG00000176101 | n.s. |

| | | | | | | | | | | | | | |
|---|---|---|---|---|---|---|---|---|---|---|---|---|---|
| ENSG00000171155 | 185.9785 | 0.192482667 | 0.320149 | 0.6012285 | 0.547688 | 0.891406027 | C1GALT1C1 | protein_coding | 40.73 | 0.009609588 | 0.009609588 | ENSG00000171155 | n.s. |
| ENSG00000267034 | 294.7809 | 0.162784276 | 0.237187 | 0.6863116 | 0.492517 | 0.873000006 | | lincRNA | 41.22 | 0.009601953 | 0.009601953 | ENSG00000267034 | n.s. |
| ENSG00000172469 | 762.4115 | -0.18953908 | 0.308296 | -0.614797 | 0.538689 | 0.889966281 | MANEA | protein_coding | 33.8 | -0.00959569 | 0.00959569 | ENSG00000172469 | n.s. |
| ENSG00000165280 | 5737.099 | -0.172695325 | 0.26243 | -0.658062 | 0.510498 | 0.879913979 | VCP | protein_coding | 47.81 | -0.009594915 | 0.009594915 | ENSG00000165280 | n.s. |
| ENSG00000140263 | 1034.831 | -0.182546022 | 0.287882 | -0.634101 | 0.526015 | 0.886111278 | SORD | protein_coding | 45.28 | -0.009585809 | 0.009585809 | ENSG00000140263 | n.s. |
| ENSG00000104964 | 1574.405 | -0.181253676 | 0.281471 | -0.643952 | 0.519607 | 0.885353782 | AES | protein_coding | 60.37 | -0.009585266 | 0.009585266 | ENSG00000104964 | n.s. |
| ENSG00000136448 | 2838.259 | -0.171905334 | 0.260592 | -0.659671 | 0.509465 | 0.87951488 | NMT1 | protein_coding | 48.75 | -0.009584893 | 0.009584893 | ENSG00000136448 | n.s. |
| ENSG00000204130 | 326.6858 | -0.178531031 | 0.275465 | -0.648109 | 0.516914 | 0.88372671 | RUFY2 | protein_coding | 40.87 | -0.009583906 | 0.009583906 | ENSG00000204130 | n.s. |
| ENSG00000132196 | 428.909 | -0.16040233 | 0.231871 | -0.691774 | 0.489079 | 0.871557795 | HSD17B7 | protein_coding | 39.34 | -0.00957663 | 0.00957663 | ENSG00000132196 | n.s. |
| ENSG00000202031 | 5.665235 | -0.299467289 | 0.706192 | -0.424059 | 0.671522 | 0.92901502 | SNORD38A | snoRNA | 47.14 | -0.009576145 | 0.009576145 | ENSG00000202031 | n.s. |
| ENSG00000200164 | 10.39025 | -0.265846928 | 0.565035 | -0.470496 | 0.638001 | 0.920410519 | | misc_RNA | 41.58 | -0.009575388 | 0.009575388 | ENSG00000200164 | n.s. |
| ENSG00000071246 | 329.9981 | -0.165530084 | 0.245657 | -0.673826 | 0.500422 | 0.875342302 | VASH1 | protein_coding | 54.83 | -0.009571294 | 0.009571294 | ENSG00000071246 | n.s. |
| ENSG00000198563 | 2428.428 | -0.150181041 | 0.20863 | -0.719844 | 0.471621 | 0.863532918 | DDX39B | protein_coding | 47.24 | -0.009569702 | 0.009569702 | ENSG00000198563 | n.s. |
| ENSG00000066468 | 323.4348 | -0.166224111 | 0.248273 | -0.669522 | 0.503163 | 0.875883784 | FGFR2 | protein_coding | 45.41 | -0.009566782 | 0.009566782 | ENSG00000066468 | n.s. |
| ENSG00000179598 | 632.2993 | 0.162664023 | 0.237946 | 0.6836174 | 0.494217 | 0.873348456 | PLD6 | protein_coding | 55.87 | 0.009566669 | 0.009566669 | ENSG00000179598 | n.s. |
| ENSG00000255624 | 75.58638 | 0.180931182 | 0.282025 | 0.641543 | 0.52117 | 0.885523197 | | transcribed_processed_pseudog | 37.42 | 0.009553177 | 0.009553177 | ENSG00000255624 | n.s. |
| ENSG00000109323 | 159.8888 | -0.180860717 | 0.28178 | -0.64185 | 0.520971 | 0.885523197 | MANBA | protein_coding | 39.25 | -0.009549456 | 0.009549456 | ENSG00000109323 | n.s. |
| ENSG00000149554 | 1401.245 | -0.169485368 | 0.255838 | -0.662471 | 0.507669 | 0.878374335 | CHEK1 | protein_coding | 37.72 | -0.009545477 | 0.009545477 | ENSG00000149554 | n.s. |
| ENSG00000115657 | 981.67 | -0.165203323 | 0.245639 | -0.672546 | 0.501236 | 0.875518559 | ABCB6 | protein_coding | 54.46 | -0.009537955 | 0.009537955 | ENSG00000115657 | n.s. |
| ENSG00000114416 | 5407.7 | -0.160997125 | 0.233526 | -0.689418 | 0.49056 | 0.872495697 | FXR1 | protein_coding | 38.94 | -0.009536939 | 0.009536939 | ENSG00000114416 | n.s. |
| ENSG00000237441 | 292.561 | 0.181378558 | 0.284991 | 0.6364367 | 0.524492 | 0.886111278 | RGL2 | protein_coding | 57.12 | 0.009524503 | 0.009524503 | ENSG00000237441 | n.s. |
| ENSG00000163788 | 918.2895 | -0.170307719 | 0.257685 | -0.660913 | 0.508668 | 0.879193676 | SNRK | protein_coding | 41.92 | -0.009522832 | 0.009522832 | ENSG00000163788 | n.s. |
| ENSG00000176853 | 1202.944 | -0.180296768 | 0.281089 | -0.641422 | 0.521248 | 0.885523197 | FAM91A1 | protein_coding | 36.91 | -0.00951968 | 0.00951968 | ENSG00000176853 | n.s. |
| ENSG00000175279 | 64.62038 | -0.190624729 | 0.313362 | -0.608321 | 0.542974 | 0.891406027 | CENPS | protein_coding | 46.89 | -0.009516832 | 0.009516832 | ENSG00000175279 | n.s. |
| ENSG00000198918 | 5588.448 | 0.186430935 | 0.301243 | 0.6188729 | 0.536 | 0.889201649 | RPL39 | protein_coding | 46.11 | 0.009507929 | 0.009507929 | ENSG00000198918 | n.s. |
| ENSG00000121361 | 332.9049 | -0.15140299 | 0.213165 | -0.710263 | 0.477541 | 0.865616576 | KCNJ8 | protein_coding | 40.62 | -0.009489097 | 0.009489097 | ENSG00000121361 | n.s. |
| ENSG00000188976 | 744.6555 | -0.179409559 | 0.278894 | -0.643289 | 0.520036 | 0.885364478 | NOC2L | protein_coding | 59.55 | -0.009486802 | 0.009486802 | ENSG00000188976 | n.s. |
| ENSG00000202071 | 4.682623 | -0.289994274 | 0.669582 | -0.433097 | 0.664944 | 0.927447429 | | misc_RNA | 39.22 | -0.009485915 | 0.009485915 | ENSG00000202071 | n.s. |
| ENSG00000131236 | 893.2626 | -0.180274786 | 0.281616 | -0.640144 | 0.522079 | 0.885953729 | CAP1 | protein_coding | 43.27 | -0.009480463 | 0.009480463 | ENSG00000131236 | n.s. |
| ENSG00000145817 | 901.7506 | -0.183003531 | 0.292912 | -0.624772 | 0.532121 | 0.887559389 | YIPF5 | protein_coding | 35.8 | -0.009480055 | 0.009480055 | ENSG00000145817 | n.s. |
| ENSG00000210194 | 61.70287 | -0.194749732 | 0.33314 | -0.584589 | 0.558824 | 0.894036264 | MT-TE | Mt_tRNA | 39.13 | -0.009473574 | 0.009473574 | ENSG00000210194 | n.s. |
| ENSG00000154582 | 1031.912 | -0.18038484 | 0.283365 | -0.63658 | 0.524398 | 0.886111278 | ELOC | protein_coding | 41.19 | -0.009472321 | 0.009472321 | ENSG00000154582 | n.s. |

| | | | | | | | | | | | | | |
|---|---|---|---|---|---|---|---|---|---|---|---|---|---|
| ENSG00000266910 | 42.41546 | -0.189716276 | 0.312617 | -0.606866 | 0.54394 | 0.891406027 | | processed_pseudogene | 42.23 | -0.009471478 | 0.009471478 | ENSG00000266910 | n.s. |
| ENSG00000134590 | 385.229 | -0.170096722 | 0.25831 | -0.658499 | 0.510218 | 0.879714521 | RTL8C | protein_coding | 60.29 | -0.009467284 | 0.009467284 | ENSG00000134590 | n.s. |
| ENSG00000182552 | 519.6929 | -0.189085875 | 0.309617 | -0.610708 | 0.541393 | 0.891123173 | RWDD4 | protein_coding | 40.27 | -0.009466067 | 0.009466067 | ENSG00000182552 | n.s. |
| ENSG00000212389 | 20.30823 | -0.222928098 | 0.414365 | -0.538 | 0.590577 | 0.906900529 | RNU6-1275P | snRNA | 39.25 | -0.009461145 | 0.009461145 | ENSG00000212389 | n.s. |
| ENSG00000095380 | 418.2081 | -0.164253106 | 0.245008 | -0.670399 | 0.502604 | 0.875813709 | NANS | protein_coding | 46.23 | -0.009459051 | 0.009459051 | ENSG00000095380 | n.s. |
| ENSG00000144161 | 1375.157 | -0.180099752 | 0.282697 | -0.637078 | 0.524074 | 0.886111278 | ZC3H8 | protein_coding | 38.5 | -0.009457351 | 0.009457351 | ENSG00000144161 | n.s. |
| ENSG00000029725 | 2046.456 | -0.175218651 | 0.269572 | -0.649988 | 0.5157 | 0.883268772 | RABEP1 | protein_coding | 41.37 | -0.009445533 | 0.009445533 | ENSG00000029725 | n.s. |
| ENSG00000184575 | 6094.204 | -0.179785435 | 0.282388 | -0.63666 | 0.524346 | 0.886111278 | XPOT | protein_coding | 39.76 | -0.009440845 | 0.009440845 | ENSG00000184575 | n.s. |
| ENSG00000127603 | 52.05217 | -0.182169432 | 0.291423 | -0.625102 | 0.531904 | 0.88755367 | MACF1 | protein_coding | 42.57 | -0.009437356 | 0.009437356 | ENSG00000127603 | n.s. |
| ENSG00000184939 | 701.9552 | -0.181372977 | 0.289276 | -0.62699 | 0.530666 | 0.887229181 | ZFP90 | protein_coding | 44.64 | -0.009424898 | 0.009424898 | ENSG00000184939 | n.s. |
| ENSG00000252568 | 6.457466 | 0.255559038 | 0.522422 | 0.4891815 | 0.624713 | 0.918660999 | RNU7-28P | snRNA | 40.32 | 0.009416001 | 0.009416001 | ENSG00000252568 | n.s. |
| ENSG00000172586 | 745.5987 | 0.180676113 | 0.286358 | 0.6309449 | 0.528077 | 0.886954686 | CHCHD1 | protein_coding | 45.52 | 0.009412967 | 0.009412967 | ENSG00000172586 | n.s. |
| ENSG00000173064 | 30.60885 | -0.192303455 | 0.326995 | -0.588092 | 0.55647 | 0.893419675 | HECTD4 | protein_coding | 43.29 | -0.009412194 | 0.009412194 | ENSG00000173064 | n.s. |
| ENSG00000164292 | 3349.115 | -0.180222457 | 0.285669 | -0.630878 | 0.52812 | 0.886954686 | RHOBTB3 | protein_coding | 40.56 | -0.009389332 | 0.009389332 | ENSG00000164292 | n.s. |
| ENSG00000244165 | 163.1435 | -0.174847018 | 0.269786 | -0.648096 | 0.516923 | 0.88372671 | P2RY11 | protein_coding | 62.43 | -0.009386141 | 0.009386141 | ENSG00000244165 | n.s. |
| ENSG00000132475 | 37912.76 | 0.161780011 | 0.239389 | 0.6758048 | 0.499165 | 0.874967757 | H3F3B | protein_coding | 47.38 | 0.009384527 | 0.009384527 | ENSG00000132475 | n.s. |
| ENSG00000115541 | 3630.734 | 0.178672575 | 0.280024 | 0.638062 | 0.523433 | 0.886111278 | HSPE1 | protein_coding | 44.77 | 0.009382407 | 0.009382407 | ENSG00000115541 | n.s. |
| ENSG00000140382 | 1524.678 | -0.174425632 | 0.268869 | -0.648738 | 0.516508 | 0.883546124 | HMG20A | protein_coding | 38.22 | -0.009379001 | 0.009379001 | ENSG00000140382 | n.s. |
| ENSG00000133104 | 1645.763 | 0.179795709 | 0.286133 | 0.6283651 | 0.529765 | 0.886954686 | SPG20 | protein_coding | 39.04 | 0.009367099 | 0.009367099 | ENSG00000133104 | n.s. |
| ENSG00000167972 | 386.2445 | -0.168263882 | 0.255391 | -0.658848 | 0.509993 | 0.879714521 | ABCA3 | protein_coding | 53.03 | -0.009365271 | 0.009365271 | ENSG00000167972 | n.s. |
| ENSG00000274605 | 99.85069 | -0.18012546 | 0.287427 | -0.626682 | 0.530868 | 0.887229181 | | lincRNA | 45.07 | -0.009360072 | 0.009360072 | ENSG00000274605 | n.s. |
| ENSG00000068383 | 276.994 | -0.180068146 | 0.288291 | -0.624606 | 0.53223 | 0.887559389 | INPP5A | protein_coding | 54.13 | -0.009327994 | 0.009327994 | ENSG00000068383 | n.s. |
| ENSG00000281780 | 17.52278 | -0.224635087 | 0.427948 | -0.524912 | 0.599644 | 0.908868121 | | snoRNA | 35.96 | -0.009322161 | 0.009322161 | ENSG00000281780 | n.s. |
| ENSG00000141510 | 367.042 | -0.167078042 | 0.253222 | -0.659809 | 0.509377 | 0.87951488 | TP53 | protein_coding | 48.86 | -0.009315738 | 0.009315738 | ENSG00000141510 | n.s. |
| ENSG00000083223 | 819.145 | -0.168537802 | 0.256742 | -0.656448 | 0.511536 | 0.880578023 | ZCCHC6 | protein_coding | 37.99 | -0.009308706 | 0.009308706 | ENSG00000083223 | n.s. |
| ENSG00000161057 | 1547.924 | -0.179476361 | 0.287464 | -0.624343 | 0.532402 | 0.887576332 | PSMC2 | protein_coding | 38.99 | -0.00929585 | 0.00929585 | ENSG00000161057 | n.s. |
| ENSG00000148572 | 388.6657 | -0.188291424 | 0.316368 | -0.595167 | 0.551732 | 0.892641916 | NRBF2 | protein_coding | 42.23 | -0.009287046 | 0.009287046 | ENSG00000148572 | n.s. |
| ENSG00000125450 | 2361.05 | -0.172949178 | 0.266909 | -0.64797 | 0.517004 | 0.88372671 | NUP85 | protein_coding | 48.14 | -0.009284261 | 0.009284261 | ENSG00000125450 | n.s. |
| ENSG00000094880 | 1237 | -0.178821638 | 0.285461 | -0.626432 | 0.531032 | 0.887366671 | CDC23 | protein_coding | 41.21 | -0.009280286 | 0.009280286 | ENSG00000094880 | n.s. |
| ENSG00000165304 | 864.0906 | -0.17542835 | 0.272764 | -0.643151 | 0.520126 | 0.885364478 | MELK | protein_coding | 41.98 | -0.009276284 | 0.009276284 | ENSG00000165304 | n.s. |
| ENSG00000230081 | 11.15791 | -0.239383331 | 0.47888 | -0.499881 | 0.617159 | 0.914690044 | HSPE1P28 | processed_pseudogene | 39.11 | -0.00927037 | 0.00927037 | ENSG00000230081 | n.s. |
| ENSG00000177889 | 3279.314 | -0.177102089 | 0.279997 | -0.632514 | 0.527051 | 0.886660976 | UBE2N | protein_coding | 40.11 | -0.009252239 | 0.009252239 | ENSG00000177889 | n.s. |

| | | | | | | | | | | | | | |
|---|---|---|---|---|---|---|---|---|---|---|---|---|---|
| ENSG00000268584 | 82.16534 | -0.172781689 | 0.267224 | -0.646581 | 0.517903 | 0.884019009 | | antisense | 49.31 | -0.009250455 | 0.009250455 | ENSG00000268584 | n.s. |
| ENSG00000198408 | 4293.081 | -0.166688974 | 0.253444 | -0.657697 | 0.510733 | 0.880066439 | MGEA5 | protein_coding | 41.92 | -0.009248661 | 0.009248661 | ENSG00000198408 | n.s. |
| ENSG00000202191 | 11.33025 | 0.222839547 | 0.430495 | 0.5176358 | 0.604712 | 0.908868121 | SNORD113-1 | snoRNA | 33.33 | 0.009247647 | 0.009247647 | ENSG00000202191 | n.s. |
| ENSG00000149136 | 4664.475 | -0.167537799 | 0.255317 | -0.656196 | 0.511698 | 0.880681667 | SSRP1 | protein_coding | 50.54 | -0.009244911 | 0.009244911 | ENSG00000149136 | n.s. |
| ENSG00000066455 | 542.9322 | -0.177714159 | 0.283472 | -0.62692 | 0.530712 | 0.887229181 | GOLGA5 | protein_coding | 39.61 | -0.009234771 | 0.009234771 | ENSG00000066455 | n.s. |
| ENSG00000101193 | 840.7659 | -0.161690447 | 0.242764 | -0.666039 | 0.505386 | 0.876821729 | GID8 | protein_coding | 48.86 | -0.009230697 | 0.009230697 | ENSG00000101193 | n.s. |
| ENSG00000180318 | 295.814 | -0.169231251 | 0.258825 | -0.653844 | 0.513212 | 0.882046073 | ALX1 | protein_coding | 31.86 | -0.00922458 | 0.00922458 | ENSG00000180318 | n.s. |
| ENSG00000241370 | 220.1972 | -0.187080385 | 0.314479 | -0.59489 | 0.551917 | 0.892697967 | RPP21 | protein_coding | 53.76 | -0.009222213 | 0.009222213 | ENSG00000241370 | n.s. |
| ENSG00000115761 | 1490.642 | -0.170930689 | 0.262871 | -0.650245 | 0.515534 | 0.883195516 | NOL10 | protein_coding | 40.73 | -0.009220539 | 0.009220539 | ENSG00000115761 | n.s. |
| ENSG00000196683 | 1318.9 | 0.185374185 | 0.309882 | 0.5982083 | 0.549701 | 0.891861801 | TOMM7 | protein_coding | 41.61 | 0.009213549 | 0.009213549 | ENSG00000196683 | n.s. |
| ENSG00000120885 | 263.9775 | 0.184529286 | 0.303527 | 0.6079509 | 0.54322 | 0.891406027 | CLU | protein_coding | 49.9 | 0.00921252 | 0.00921252 | ENSG00000120885 | n.s. |
| ENSG00000204217 | 1327.935 | -0.175496107 | 0.27692 | -0.633744 | 0.526248 | 0.88615396 | BMPR2 | protein_coding | 41.04 | -0.009211934 | 0.009211934 | ENSG00000204217 | n.s. |
| ENSG00000238113 | 41.4712 | -0.189517721 | 0.324689 | -0.58369 | 0.559429 | 0.894140829 | LINC01410 | lincRNA | 40.99 | -0.009209438 | 0.009209438 | ENSG00000238113 | n.s. |
| ENSG00000182013 | 504.9857 | 0.176614443 | 0.281061 | 0.6283841 | 0.529752 | 0.886954686 | PNMA8A | protein_coding | 52.55 | 0.009201359 | 0.009201359 | ENSG00000182013 | n.s. |
| ENSG00000115942 | 1357.475 | -0.175116229 | 0.274214 | -0.63861 | 0.523076 | 0.886079926 | ORC2 | protein_coding | 39.05 | -0.009198348 | 0.009198348 | ENSG00000115942 | n.s. |
| ENSG00000121749 | 1351.007 | -0.169216296 | 0.259176 | -0.6529 | 0.513821 | 0.882409358 | TBC1D15 | protein_coding | 37.01 | -0.009193504 | 0.009193504 | ENSG00000121749 | n.s. |
| ENSG00000089220 | 7283.419 | -0.183925396 | 0.301764 | -0.6095 | 0.542193 | 0.891404544 | PEBP1 | protein_coding | 49.19 | -0.009182504 | 0.009182504 | ENSG00000089220 | n.s. |
| ENSG00000272086 | 79.85843 | 0.179933279 | 0.290604 | 0.6191702 | 0.535804 | 0.889201649 | | lincRNA | 52.49 | 0.009176551 | 0.009176551 | ENSG00000272086 | n.s. |
| ENSG00000144021 | 1733.771 | -0.176092433 | 0.279605 | -0.62979 | 0.528832 | 0.886954686 | CIAO1 | protein_coding | 49.71 | -0.009174163 | 0.009174163 | ENSG00000144021 | n.s. |
| ENSG00000244380 | 48.34525 | -0.186800294 | 0.316159 | -0.590843 | 0.554626 | 0.893144718 | | antisense | 48.45 | -0.009167816 | 0.009167816 | ENSG00000244380 | n.s. |
| ENSG00000171135 | 575.4468 | 0.182312755 | 0.297927 | 0.6119374 | 0.540579 | 0.890708138 | JAGN1 | protein_coding | 47.58 | 0.009163874 | 0.009163874 | ENSG00000171135 | n.s. |
| ENSG00000174428 | 84.77213 | -0.177404447 | 0.284657 | -0.623222 | 0.533138 | 0.887904939 | GTF2IRD2B | protein_coding | 46.64 | -0.009160017 | 0.009160017 | ENSG00000174428 | n.s. |
| ENSG00000155755 | 3835.902 | -0.173467593 | 0.270626 | -0.640986 | 0.521532 | 0.885523197 | TMEM237 | protein_coding | 38.83 | -0.009159099 | 0.009159099 | ENSG00000155755 | n.s. |
| ENSG00000239002 | 303.2901 | 0.165782197 | 0.252438 | 0.6567251 | 0.511358 | 0.880578023 | SCARNA10 | snoRNA | 45.45 | 0.009156508 | 0.009156508 | ENSG00000239002 | n.s. |
| ENSG00000087095 | 680.445 | -0.158352776 | 0.234925 | -0.674057 | 0.500275 | 0.875342302 | NLK | protein_coding | 37.3 | -0.009156287 | 0.009156287 | ENSG00000087095 | n.s. |
| ENSG00000144560 | 763.1804 | -0.172966406 | 0.268931 | -0.643163 | 0.520119 | 0.885364478 | VGLL4 | protein_coding | 44.93 | -0.009146101 | 0.009146101 | ENSG00000144560 | n.s. |
| ENSG00000166266 | 1219.465 | -0.175508534 | 0.277967 | -0.6314 | 0.527779 | 0.886936263 | CUL5 | protein_coding | 39.41 | -0.009145326 | 0.009145326 | ENSG00000166266 | n.s. |
| ENSG00000124575 | 3097.557 | 0.154390532 | 0.224212 | 0.6885907 | 0.491081 | 0.872562973 | HIST1H1D | protein_coding | 54.35 | 0.009140417 | 0.009140417 | ENSG00000124575 | n.s. |
| ENSG00000258325 | 46.23909 | 0.185268796 | 0.310992 | 0.5957343 | 0.551353 | 0.892641916 | ITFG2-AS1 | antisense | 47.69 | 0.009137962 | 0.009137962 | ENSG00000258325 | n.s. |
| ENSG00000138629 | 616.8563 | -0.183021759 | 0.30466 | -0.60074 | 0.548013 | 0.891406027 | UBL7 | protein_coding | 47.95 | -0.009137258 | 0.009137258 | ENSG00000138629 | n.s. |
| ENSG00000008277 | 1961.175 | -0.169165901 | 0.260194 | -0.650154 | 0.515593 | 0.883195516 | ADAM22 | protein_coding | 37.29 | -0.009125341 | 0.009125341 | ENSG00000008277 | n.s. |
| ENSG00000162849 | 1418.084 | -0.15841273 | 0.236632 | -0.669447 | 0.50321 | 0.875883784 | KIF26B | protein_coding | 44.8 | -0.009117209 | 0.009117209 | ENSG00000162849 | n.s. |

| | | | | | | | | | | | | | |
|---|---|---|---|---|---|---|---|---|---|---|---|---|---|
| ENSG00000148824 | 492.6488 | -0.175671089 | 0.280766 | -0.625685 | 0.531522 | 0.887366671 | MTG1 | protein_coding | 47.71 | -0.009116783 | 0.009116783 | ENSG00000148824 | n.s. |
| ENSG00000171792 | 928.6544 | -0.175899165 | 0.281847 | -0.624095 | 0.532565 | 0.887587464 | RHNO1 | protein_coding | 47.54 | -0.009109614 | 0.009109614 | ENSG00000171792 | n.s. |
| ENSG00000122376 | 1041.442 | -0.182467315 | 0.301683 | -0.604831 | 0.545292 | 0.891406027 | FAM35A | protein_coding | 40.46 | -0.009109577 | 0.009109577 | ENSG00000122376 | n.s. |
| ENSG00000120256 | 689.0703 | -0.175454966 | 0.280254 | -0.626056 | 0.531278 | 0.887366671 | LRP11 | protein_coding | 43.94 | -0.009105567 | 0.009105567 | ENSG00000120256 | n.s. |
| ENSG00000100603 | 1636.934 | -0.173199345 | 0.273471 | -0.633337 | 0.526514 | 0.88615396 | SNW1 | protein_coding | 41.12 | -0.009091375 | 0.009091375 | ENSG00000100603 | n.s. |
| ENSG00000263628 | 5.503665 | 0.26762267 | 0.605627 | 0.4418932 | 0.658566 | 0.924786541 | MIR3155A | miRNA | 65.85 | 0.009088064 | 0.009088064 | ENSG00000263628 | n.s. |
| ENSG00000177788 | 17.37374 | -0.218967247 | 0.41853 | -0.523182 | 0.600848 | 0.908868121 | | lincRNA | 44.3 | -0.00908695 | 0.00908695 | ENSG00000177788 | n.s. |
| ENSG00000251602 | 67.73044 | -0.180691749 | 0.295074 | -0.612362 | 0.540299 | 0.890708138 | | antisense | 59.49 | -0.009082395 | 0.009082395 | ENSG00000251602 | n.s. |
| ENSG00000128585 | 2301.906 | -0.171348915 | 0.265884 | -0.644449 | 0.519285 | 0.885161398 | MKLN1 | protein_coding | 39.28 | -0.009077643 | 0.009077643 | ENSG00000128585 | n.s. |
| ENSG00000000003 | 790.6962 | 0.17930201 | 0.291761 | 0.61455 | 0.538852 | 0.889966281 | TSPAN6 | protein_coding | 40.4 | 0.009077424 | 0.009077424 | ENSG00000000003 | n.s. |
| ENSG00000114735 | 202.1259 | -0.168681053 | 0.259949 | -0.648902 | 0.516402 | 0.883546124 | HEMK1 | protein_coding | 53.84 | -0.009070111 | 0.009070111 | ENSG00000114735 | n.s. |
| ENSG00000180340 | 218.8001 | -0.157028037 | 0.233532 | -0.672405 | 0.501326 | 0.875518559 | FZD2 | protein_coding | 64.91 | -0.009065958 | 0.009065958 | ENSG00000180340 | n.s. |
| ENSG00000130702 | 39.75099 | 0.209602265 | 0.38639 | 0.5424627 | 0.5875 | 0.905217543 | LAMA5 | protein_coding | 61.89 | 0.009064677 | 0.009064677 | ENSG00000130702 | n.s. |
| ENSG00000187051 | 614.6035 | -0.184486576 | 0.312111 | -0.591093 | 0.554458 | 0.893144718 | RPS19BP1 | protein_coding | 54.82 | -0.009054263 | 0.009054263 | ENSG00000187051 | n.s. |
| ENSG00000249122 | 288.5927 | -0.156174452 | 0.231369 | -0.675002 | 0.499675 | 0.875090797 | | lincRNA | 45.08 | -0.009049823 | 0.009049823 | ENSG00000249122 | n.s. |
| ENSG00000100422 | 3394.748 | -0.173557244 | 0.276121 | -0.628556 | 0.52964 | 0.886954686 | CERK | protein_coding | 51.44 | -0.009042084 | 0.009042084 | ENSG00000100422 | n.s. |
| ENSG00000130939 | 298.6323 | -0.172182721 | 0.271386 | -0.634457 | 0.525783 | 0.886111278 | UBE4B | protein_coding | 44.12 | -0.009041614 | 0.009041614 | ENSG00000130939 | n.s. |
| ENSG00000141543 | 3679.498 | -0.166232182 | 0.254903 | -0.65214 | 0.514311 | 0.882409358 | EIF4A3 | protein_coding | 46.23 | -0.009031377 | 0.009031377 | ENSG00000141543 | n.s. |
| ENSG00000173674 | 2588.277 | -0.173314324 | 0.275507 | -0.629074 | 0.5293 | 0.886954686 | EIF1AX | protein_coding | 38.58 | -0.009029428 | 0.009029428 | ENSG00000173674 | n.s. |
| ENSG00000268364 | 73.16634 | -0.171933648 | 0.27094 | -0.634582 | 0.525701 | 0.886111278 | SMC5-AS1 | antisense | 39.57 | -0.009028534 | 0.009028534 | ENSG00000268364 | n.s. |
| ENSG00000140829 | 777.0434 | -0.156340691 | 0.232541 | -0.672316 | 0.501383 | 0.875518559 | DHX38 | protein_coding | 51.51 | -0.009026274 | 0.009026274 | ENSG00000140829 | n.s. |
| ENSG00000152348 | 192.3846 | -0.155636552 | 0.230508 | -0.675188 | 0.499556 | 0.875034827 | ATG10 | protein_coding | 38.03 | -0.009022976 | 0.009022976 | ENSG00000152348 | n.s. |
| ENSG00000223486 | 72.77883 | -0.181501819 | 0.303576 | -0.597879 | 0.549921 | 0.891861801 | | lincRNA | 56.45 | -0.009021083 | 0.009021083 | ENSG00000223486 | n.s. |
| ENSG00000170043 | 189.8698 | -0.180645337 | 0.300534 | -0.601082 | 0.547786 | 0.891406027 | TRAPPC1 | protein_coding | 57.62 | -0.009018616 | 0.009018616 | ENSG00000170043 | n.s. |
| ENSG00000202343 | 7.21809 | -0.258715168 | 0.559099 | -0.462736 | 0.643553 | 0.92287513 | | snoRNA | 41.48 | -0.009018049 | 0.009018049 | ENSG00000202343 | n.s. |
| ENSG00000121621 | 1107.614 | -0.178056159 | 0.289246 | -0.615588 | 0.538166 | 0.889966281 | KIF18A | protein_coding | 34.02 | -0.009014351 | 0.009014351 | ENSG00000121621 | n.s. |
| ENSG00000273237 | 55.40533 | 0.187299242 | 0.323175 | 0.5795593 | 0.562212 | 0.895116382 | | antisense | 39.99 | 0.009012932 | 0.009012932 | ENSG00000273237 | n.s. |
| ENSG00000034533 | 442.1489 | -0.149087028 | 0.214259 | -0.695825 | 0.486538 | 0.870096539 | ASTE1 | protein_coding | 39.7 | -0.009009711 | 0.009009711 | ENSG00000034533 | n.s. |
| ENSG00000121644 | 1829.591 | -0.165820103 | 0.254103 | -0.65257 | 0.514033 | 0.882409358 | DESI2 | protein_coding | 40.41 | -0.009008989 | 0.009008989 | ENSG00000121644 | n.s. |
| ENSG00000164615 | 635.3757 | 0.183058106 | 0.307864 | 0.5946064 | 0.552107 | 0.892900483 | CAMLG | protein_coding | 43.3 | 0.009005899 | 0.009005899 | ENSG00000164615 | n.s. |
| ENSG00000079950 | 207.9935 | -0.176436531 | 0.285163 | -0.618722 | 0.5361 | 0.889201649 | STX7 | protein_coding | 37.41 | -0.008998217 | 0.008998217 | ENSG00000079950 | n.s. |
| ENSG00000172530 | 58.81259 | -0.180084431 | 0.299419 | -0.601446 | 0.547543 | 0.891406027 | BANP | protein_coding | 51.17 | -0.008990613 | 0.008990613 | ENSG00000172530 | n.s. |

| | | | | | | | | | | | | | |
|---|---|---|---|---|---|---|---|---|---|---|---|---|---|
| ENSG00000105202 | 1441.861 | -0.175999841 | 0.283535 | -0.620734 | 0.534774 | 0.88909364 | FBL | protein_coding | 48.35 | -0.008985231 | 0.008985231 | ENSG00000105202 | n.s. |
| ENSG00000173209 | 2746.388 | -0.161099919 | 0.244225 | -0.659637 | 0.509487 | 0.87951488 | AHSA2 | protein_coding | 44.61 | -0.008982417 | 0.008982417 | ENSG00000173209 | n.s. |
| ENSG00000266619 | 13.44463 | -0.229809277 | 0.458554 | -0.501161 | 0.616258 | 0.914046563 | MIR4642 | miRNA | 58.54 | -0.008969842 | 0.008969842 | ENSG00000266619 | n.s. |
| ENSG00000262119 | 76.38321 | 0.185655593 | 0.319484 | 0.5811103 | 0.561166 | 0.894765412 | | sense_intronic | 31.17 | 0.008965459 | 0.008965459 | ENSG00000262119 | n.s. |
| ENSG00000169964 | 189.5042 | -0.154143838 | 0.227294 | -0.678169 | 0.497665 | 0.874675553 | TMEM42 | protein_coding | 52.21 | -0.008963928 | 0.008963928 | ENSG00000169964 | n.s. |
| ENSG00000156170 | 393.7834 | -0.170693407 | 0.269119 | -0.634269 | 0.525906 | 0.886111278 | NDUFAF6 | protein_coding | 42.88 | -0.008963407 | 0.008963407 | ENSG00000156170 | n.s. |
| ENSG00000112874 | 434.1511 | -0.183524523 | 0.312527 | -0.587228 | 0.557051 | 0.893643698 | NUDT12 | protein_coding | 34.05 | -0.008962531 | 0.008962531 | ENSG00000112874 | n.s. |
| ENSG00000111266 | 498.5832 | 0.174061301 | 0.279574 | 0.6225944 | 0.533551 | 0.888237452 | DUSP16 | protein_coding | 40.28 | 0.008959095 | 0.008959095 | ENSG00000111266 | n.s. |
| ENSG00000197111 | 5415.545 | -0.155546703 | 0.231981 | -0.670515 | 0.502529 | 0.875813709 | PCBP2 | protein_coding | 43.79 | -0.008957664 | 0.008957664 | ENSG00000197111 | n.s. |
| ENSG00000144224 | 3288.962 | -0.173069025 | 0.277517 | -0.623634 | 0.532868 | 0.887866798 | UBXN4 | protein_coding | 39.83 | -0.008939393 | 0.008939393 | ENSG00000144224 | n.s. |
| ENSG00000108639 | 238.0344 | 0.177825647 | 0.290445 | 0.6122515 | 0.540371 | 0.890708138 | SYNGR2 | protein_coding | 62.13 | 0.008938331 | 0.008938331 | ENSG00000108639 | n.s. |
| ENSG00000133065 | 411.8973 | -0.154522645 | 0.229198 | -0.674188 | 0.500191 | 0.875342302 | SLC41A1 | protein_coding | 50.84 | -0.008934821 | 0.008934821 | ENSG00000133065 | n.s. |
| ENSG00000185043 | 151.5164 | -0.18376187 | 0.314451 | -0.58439 | 0.558958 | 0.89411481 | CIB1 | protein_coding | 56.35 | -0.00893206 | 0.00893206 | ENSG00000185043 | n.s. |
| ENSG00000108106 | 3411.961 | -0.178902797 | 0.296347 | -0.603695 | 0.546047 | 0.891406027 | UBE2S | protein_coding | 54.62 | -0.008931621 | 0.008931621 | ENSG00000108106 | n.s. |
| ENSG00000279384 | 37.24726 | 0.181893941 | 0.308999 | 0.5886548 | 0.556093 | 0.893144718 | | TEC | 38.1 | 0.008927021 | 0.008927021 | ENSG00000279384 | n.s. |
| ENSG00000142444 | 240.9549 | -0.168774256 | 0.262413 | -0.643162 | 0.520119 | 0.885364478 | TIMM29 | protein_coding | 52.55 | -0.008924429 | 0.008924429 | ENSG00000142444 | n.s. |
| ENSG00000118579 | 2068.124 | -0.169945122 | 0.267323 | -0.63573 | 0.524952 | 0.886111278 | MED28 | protein_coding | 44.87 | -0.008924113 | 0.008924113 | ENSG00000118579 | n.s. |
| ENSG00000111912 | 87.43031 | -0.178646695 | 0.29504 | -0.6055 | 0.544847 | 0.891406027 | NCOA7 | protein_coding | 39.05 | -0.008918835 | 0.008918835 | ENSG00000111912 | n.s. |
| ENSG00000265565 | 38.39255 | -0.214845994 | 0.407387 | -0.527375 | 0.597933 | 0.908868121 | MIR3143 | miRNA | 41.27 | -0.008915922 | 0.008915922 | ENSG00000265565 | n.s. |
| ENSG00000167543 | 150.9027 | 0.181657688 | 0.307672 | 0.590426 | 0.554905 | 0.893144718 | TP53I13 | protein_coding | 63.48 | 0.008915426 | 0.008915426 | ENSG00000167543 | n.s. |
| ENSG00000118816 | 2033.688 | -0.163439424 | 0.250378 | -0.652771 | 0.513904 | 0.882409358 | CCNI | protein_coding | 39.09 | -0.008879647 | 0.008879647 | ENSG00000118816 | n.s. |
| ENSG00000088035 | 466.6515 | -0.178673704 | 0.299107 | -0.597358 | 0.550269 | 0.89197749 | ALG6 | protein_coding | 37.93 | -0.008870454 | 0.008870454 | ENSG00000088035 | n.s. |
| ENSG00000126970 | 347.6152 | -0.180741106 | 0.306413 | -0.589862 | 0.555283 | 0.893144718 | ZC4H2 | protein_coding | 39.24 | -0.008870442 | 0.008870442 | ENSG00000126970 | n.s. |
| ENSG00000112282 | 1903.724 | -0.166097392 | 0.257124 | -0.645981 | 0.518292 | 0.884335626 | MED23 | protein_coding | 36.17 | -0.008866757 | 0.008866757 | ENSG00000112282 | n.s. |
| ENSG00000130287 | 280.9502 | -0.168778115 | 0.264662 | -0.637713 | 0.523661 | 0.886111278 | NCAN | protein_coding | 51.55 | -0.008862832 | 0.008862832 | ENSG00000130287 | n.s. |
| ENSG00000272253 | 4.978032 | -0.328716282 | 0.893038 | -0.368088 | 0.712808 | 0.939817855 | RNA5SP386 | rRNA | 53.27 | -0.008860977 | 0.008860977 | ENSG00000272253 | n.s. |
| ENSG00000075239 | 697.6067 | 0.177464752 | 0.293721 | 0.6041947 | 0.545714 | 0.891406027 | ACAT1 | protein_coding | 42.49 | 0.008859827 | 0.008859827 | ENSG00000075239 | n.s. |
| ENSG00000157800 | 948.1778 | -0.173439894 | 0.279283 | -0.621019 | 0.534587 | 0.88909364 | SLC37A3 | protein_coding | 45.22 | -0.00885454 | 0.00885454 | ENSG00000157800 | n.s. |
| ENSG00000110917 | 2284.333 | -0.155923938 | 0.234733 | -0.66426 | 0.506524 | 0.877502299 | MLEC | protein_coding | 47.71 | -0.008848954 | 0.008848954 | ENSG00000110917 | n.s. |
| ENSG00000162702 | 3012.422 | 0.150869042 | 0.221618 | 0.6807603 | 0.496023 | 0.873669126 | ZNF281 | protein_coding | 43.03 | 0.008848924 | 0.008848924 | ENSG00000162702 | n.s. |
| ENSG00000198862 | 1733.247 | -0.166874437 | 0.258836 | -0.64471 | 0.519115 | 0.885089424 | LTN1 | protein_coding | 36.74 | -0.008846489 | 0.008846489 | ENSG00000198862 | n.s. |
| ENSG00000248098 | 367.1031 | -0.180237277 | 0.305662 | -0.589662 | 0.555417 | 0.893144718 | BCKDHA | protein_coding | 52.32 | -0.008845715 | 0.008845715 | ENSG00000248098 | n.s. |

| | | | | | | | | | | | | | |
|---|---|---|---|---|---|---|---|---|---|---|---|---|---|
| ENSG00000128268 | 285.4884 | -0.161365025 | 0.246579 | -0.654414 | 0.512845 | 0.881523829 | MGAT3 | protein_coding | 56.74 | -0.008837308 | 0.008837308 | ENSG00000128268 | n.s. |
| ENSG00000118246 | 2107.444 | -0.168355669 | 0.265775 | -0.633452 | 0.526439 | 0.88615396 | FASTKD2 | protein_coding | 37.72 | -0.008837127 | 0.008837127 | ENSG00000118246 | n.s. |
| ENSG00000224597 | 310.3167 | -0.167097566 | 0.25988 | -0.642981 | 0.520237 | 0.885364478 | SVIL-AS1 | antisense | 42.95 | -0.008835769 | 0.008835769 | ENSG00000224597 | n.s. |
| ENSG00000077380 | 2870.286 | -0.172294381 | 0.277052 | -0.621884 | 0.534018 | 0.888662588 | DYNC1I2 | protein_coding | 36.49 | -0.008832345 | 0.008832345 | ENSG00000077380 | n.s. |
| ENSG00000041988 | 104.8207 | -0.194326052 | 0.348 | -0.558408 | 0.576566 | 0.900751128 | THAP3 | protein_coding | 56.77 | -0.008821467 | 0.008821467 | ENSG00000041988 | n.s. |
| ENSG00000164134 | 3754.177 | -0.162308998 | 0.24888 | -0.652158 | 0.514299 | 0.882409358 | NAA15 | protein_coding | 39.06 | -0.008818231 | 0.008818231 | ENSG00000164134 | n.s. |
| ENSG00000197372 | 1058.08 | -0.162036901 | 0.247995 | -0.653389 | 0.513506 | 0.882332213 | ZNF675 | protein_coding | 41.44 | -0.0088096 | 0.0088096 | ENSG00000197372 | n.s. |
| ENSG00000253250 | 480.0628 | 0.179208771 | 0.301644 | 0.5941078 | 0.55244 | 0.893099715 | C8orf88 | protein_coding | 32.98 | 0.008799159 | 0.008799159 | ENSG00000253250 | n.s. |
| ENSG00000136682 | 426.7824 | -0.167511748 | 0.262187 | -0.638903 | 0.522886 | 0.886079926 | CBWD2 | protein_coding | 36.27 | -0.008798907 | 0.008798907 | ENSG00000136682 | n.s. |
| ENSG00000139644 | 4244.505 | -0.16886817 | 0.268268 | -0.629476 | 0.529038 | 0.886954686 | TMBIM6 | protein_coding | 43.42 | -0.00879779 | 0.00879779 | ENSG00000139644 | n.s. |
| ENSG00000279845 | 106.9024 | 0.175900801 | 0.288186 | 0.6103716 | 0.541616 | 0.891255806 | | TEC | 28.76 | 0.008794623 | 0.008794623 | ENSG00000279845 | n.s. |
| ENSG00000111786 | 3226.908 | -0.16724074 | 0.262985 | -0.635933 | 0.52482 | 0.886111278 | SRSF9 | protein_coding | 46.53 | -0.008782102 | 0.008782102 | ENSG00000111786 | n.s. |
| ENSG00000146535 | 394.1153 | -0.172508033 | 0.279499 | -0.617205 | 0.5371 | 0.889417954 | GNA12 | protein_coding | 45.87 | -0.008779642 | 0.008779642 | ENSG00000146535 | n.s. |
| ENSG00000142409 | 399.7544 | -0.173357942 | 0.281552 | -0.615722 | 0.538078 | 0.889966281 | ZNF787 | protein_coding | 58.61 | -0.008776497 | 0.008776497 | ENSG00000142409 | n.s. |
| ENSG00000178537 | 188.4691 | -0.177914862 | 0.298864 | -0.595303 | 0.551641 | 0.892641916 | SLC25A20 | protein_coding | 47.4 | -0.008775246 | 0.008775246 | ENSG00000178537 | n.s. |
| ENSG00000202184 | 3.733229 | -0.291563213 | 0.709157 | -0.411141 | 0.680969 | 0.933075339 | RNU6-1283P | snRNA | 35.24 | -0.00877118 | 0.00877118 | ENSG00000202184 | n.s. |
| ENSG00000132275 | 887.548 | -0.151663139 | 0.224876 | -0.67443 | 0.500038 | 0.875342302 | RRP8 | protein_coding | 47.4 | -0.008769479 | 0.008769479 | ENSG00000132275 | n.s. |
| ENSG00000177932 | 1297.427 | -0.16858893 | 0.268989 | -0.62675 | 0.530823 | 0.887229181 | ZNF354C | protein_coding | 42.43 | -0.008760586 | 0.008760586 | ENSG00000177932 | n.s. |
| ENSG00000159210 | 1818.756 | -0.162840342 | 0.250855 | -0.649141 | 0.516247 | 0.883546124 | SNF8 | protein_coding | 47.13 | -0.008756051 | 0.008756051 | ENSG00000159210 | n.s. |
| ENSG00000185104 | 1622.305 | -0.166681881 | 0.260482 | -0.639898 | 0.522239 | 0.886074796 | FAF1 | protein_coding | 38.27 | -0.008755735 | 0.008755735 | ENSG00000185104 | n.s. |
| ENSG00000151304 | 899.4548 | -0.161784156 | 0.248568 | -0.650865 | 0.515133 | 0.882841153 | SRFBP1 | protein_coding | 36.16 | -0.008755343 | 0.008755343 | ENSG00000151304 | n.s. |
| ENSG00000116717 | 2155.841 | 0.139683185 | 0.195838 | 0.7132592 | 0.475685 | 0.865616576 | GADD45A | protein_coding | 52.9 | 0.008754565 | 0.008754565 | ENSG00000116717 | n.s. |
| ENSG00000103274 | 370.5138 | -0.175187003 | 0.292106 | -0.599739 | 0.54868 | 0.891406027 | NUBP1 | protein_coding | 50.62 | -0.008746112 | 0.008746112 | ENSG00000103274 | n.s. |
| ENSG00000100150 | 128.5706 | 0.162987095 | 0.251864 | 0.6471223 | 0.517553 | 0.883822248 | DEPDC5 | protein_coding | 45.23 | 0.008741824 | 0.008741824 | ENSG00000100150 | n.s. |
| ENSG00000261236 | 1043.431 | -0.173899057 | 0.284158 | -0.61198 | 0.540551 | 0.890708138 | BOP1 | protein_coding | 62.27 | -0.008740963 | 0.008740963 | ENSG00000261236 | n.s. |
| ENSG00000112305 | 761.6362 | -0.168706144 | 0.270304 | -0.624135 | 0.532539 | 0.887587464 | SMAP1 | protein_coding | 37.73 | -0.008737096 | 0.008737096 | ENSG00000112305 | n.s. |
| ENSG00000257218 | 1105.701 | -0.177944114 | 0.299971 | -0.593205 | 0.553044 | 0.893144718 | GATC | protein_coding | 46.68 | -0.008733171 | 0.008733171 | ENSG00000257218 | n.s. |
| ENSG00000169733 | 211.0557 | 0.172970982 | 0.281796 | 0.6138153 | 0.539337 | 0.890328323 | RFNG | protein_coding | 67.12 | 0.008726353 | 0.008726353 | ENSG00000169733 | n.s. |
| ENSG00000164306 | 138.648 | 0.166170637 | 0.261635 | 0.635124 | 0.525348 | 0.886111278 | PRIMPOL | protein_coding | 41.31 | 0.008725909 | 0.008725909 | ENSG00000164306 | n.s. |
| ENSG00000119900 | 665.239 | -0.164425856 | 0.254902 | -0.645054 | 0.518892 | 0.88497967 | OGFRL1 | protein_coding | 36.22 | -0.008725538 | 0.008725538 | ENSG00000119900 | n.s. |
| ENSG00000137142 | 2372.127 | -0.167470858 | 0.265636 | -0.630452 | 0.528399 | 0.886954686 | IGFBPL1 | protein_coding | 49.74 | -0.008724992 | 0.008724992 | ENSG00000137142 | n.s. |
| ENSG00000047188 | 1876.109 | -0.167388781 | 0.265023 | -0.6316 | 0.527648 | 0.886936263 | YTHDC2 | protein_coding | 34.67 | -0.008722226 | 0.008722226 | ENSG00000047188 | n.s. |

| | | | | | | | | | | | | | |
|---|---|---|---|---|---|---|---|---|---|---|---|---|---|
| ENSG00000104687 | 982.9245 | -0.167343325 | 0.265869 | -0.629421 | 0.529073 | 0.886954686 | GSR | protein_coding | 44.72 | -0.008718347 | 0.008718347 | ENSG00000104687 | n.s. |
| ENSG00000108039 | 975.5858 | -0.170710675 | 0.27501 | -0.620743 | 0.534769 | 0.88909364 | XPNPEP1 | protein_coding | 44.62 | -0.008715206 | 0.008715206 | ENSG00000108039 | n.s. |
| ENSG00000183023 | 465.7344 | 0.155079209 | 0.234267 | 0.661976 | 0.507987 | 0.878705084 | SLC8A1 | protein_coding | 37.43 | 0.008708762 | 0.008708762 | ENSG00000183023 | n.s. |
| ENSG00000208002 | 12.08057 | 0.258823083 | 0.592082 | 0.4371406 | 0.662009 | 0.925519646 | MIR643 | miRNA | 38.14 | 0.008700172 | 0.008700172 | ENSG00000208002 | n.s. |
| ENSG00000125611 | 165.2934 | -0.193170126 | 0.347006 | -0.556677 | 0.577748 | 0.901500147 | CHCHD5 | protein_coding | 53.69 | -0.008699262 | 0.008699262 | ENSG00000125611 | n.s. |
| ENSG00000117155 | 1416.627 | -0.172415564 | 0.280985 | -0.613612 | 0.539472 | 0.890328323 | SSX2IP | protein_coding | 37.23 | -0.008698333 | 0.008698333 | ENSG00000117155 | n.s. |
| ENSG00000104899 | 567.9986 | -0.162354622 | 0.251137 | -0.646478 | 0.51797 | 0.884019009 | AMH | protein_coding | 71.68 | -0.008692206 | 0.008692206 | ENSG00000104899 | n.s. |
| ENSG00000262728 | 42.91469 | -0.186698194 | 0.327714 | -0.569699 | 0.568882 | 0.898419074 | | antisense | 45.14 | -0.008685393 | 0.008685393 | ENSG00000262728 | n.s. |
| ENSG00000214046 | 1214.291 | -0.173929857 | 0.287366 | -0.605256 | 0.545009 | 0.891406027 | SMIM7 | protein_coding | 47.85 | -0.00868335 | 0.00868335 | ENSG00000214046 | n.s. |
| ENSG00000263634 | 27.49167 | 0.200588995 | 0.368877 | 0.5437832 | 0.586591 | 0.905217543 | MIR3919 | miRNA | 43.82 | 0.00867488 | 0.00867488 | ENSG00000263634 | n.s. |
| ENSG00000138674 | 2162.702 | -0.16421314 | 0.255783 | -0.642003 | 0.520872 | 0.885523197 | SEC31A | protein_coding | 38.78 | -0.008670463 | 0.008670463 | ENSG00000138674 | n.s. |
| ENSG00000184857 | 107.7052 | -0.16986307 | 0.27464 | -0.618494 | 0.536249 | 0.889201649 | TMEM186 | protein_coding | 47.7 | -0.008662972 | 0.008662972 | ENSG00000184857 | n.s. |
| ENSG00000168300 | 283.708 | -0.167711901 | 0.269088 | -0.623261 | 0.533113 | 0.887904939 | PCMTD1 | protein_coding | 36.74 | -0.008659557 | 0.008659557 | ENSG00000168300 | n.s. |
| ENSG00000113460 | 2827.726 | 0.164904122 | 0.258737 | 0.6373429 | 0.523902 | 0.886111278 | BRIX1 | protein_coding | 38.42 | 0.008659402 | 0.008659402 | ENSG00000113460 | n.s. |
| ENSG00000228201 | 21.97897 | -0.198912388 | 0.36324 | -0.547606 | 0.583963 | 0.904712915 | | antisense | 66.35 | -0.008650543 | 0.008650543 | ENSG00000228201 | n.s. |
| ENSG00000124486 | 4060.794 | -0.164715133 | 0.258486 | -0.63723 | 0.523975 | 0.886111278 | USP9X | protein_coding | 38.5 | -0.008649478 | 0.008649478 | ENSG00000124486 | n.s. |
| ENSG00000121578 | 339.9041 | -0.18004906 | 0.311175 | -0.578609 | 0.562853 | 0.895317554 | B4GALT4 | protein_coding | 41.78 | -0.008646478 | 0.008646478 | ENSG00000121578 | n.s. |
| ENSG00000147669 | 1553.204 | 0.173186317 | 0.288166 | 0.6009943 | 0.547844 | 0.891406027 | POLR2K | protein_coding | 37.38 | 0.008646229 | 0.008646229 | ENSG00000147669 | n.s. |
| ENSG00000080503 | 125.8283 | -0.169762413 | 0.27513 | -0.617027 | 0.537217 | 0.889417954 | SMARCA2 | protein_coding | 40.37 | -0.008639907 | 0.008639907 | ENSG00000080503 | n.s. |
| ENSG00000033800 | 2031.489 | -0.155664063 | 0.236636 | -0.657821 | 0.510653 | 0.880037893 | PIAS1 | protein_coding | 38.09 | -0.008639142 | 0.008639142 | ENSG00000033800 | n.s. |
| ENSG00000117143 | 1162.549 | -0.166223759 | 0.265152 | -0.626899 | 0.530725 | 0.887229181 | UAP1 | protein_coding | 39.68 | -0.008637682 | 0.008637682 | ENSG00000117143 | n.s. |
| ENSG00000109103 | 159.6448 | -0.172690867 | 0.284191 | -0.607659 | 0.543414 | 0.891406027 | UNC119 | protein_coding | 57.56 | -0.008621494 | 0.008621494 | ENSG00000109103 | n.s. |
| ENSG00000161011 | 850.2728 | -0.164001698 | 0.258029 | -0.635595 | 0.52504 | 0.886111278 | SQSTM1 | protein_coding | 54.99 | -0.008612014 | 0.008612014 | ENSG00000161011 | n.s. |
| ENSG00000135587 | 139.2596 | -0.163977059 | 0.258577 | -0.634151 | 0.525983 | 0.886111278 | SMPD2 | protein_coding | 58.66 | -0.00861072 | 0.00861072 | ENSG00000135587 | n.s. |
| ENSG00000107404 | 278.5365 | 0.172471557 | 0.286813 | 0.6013377 | 0.547615 | 0.891406027 | DVL1 | protein_coding | 66.25 | 0.008610545 | 0.008610545 | ENSG00000107404 | n.s. |
| ENSG00000139725 | 579.3112 | 0.143417046 | 0.206891 | 0.6932023 | 0.488183 | 0.870959298 | RHOF | protein_coding | 56.89 | 0.00860533 | 0.00860533 | ENSG00000139725 | n.s. |
| ENSG00000198954 | 186.7592 | -0.169969488 | 0.276415 | -0.614907 | 0.538616 | 0.889966281 | KIF1BP | protein_coding | 43.19 | -0.008604951 | 0.008604951 | ENSG00000198954 | n.s. |
| ENSG00000170412 | 119.1946 | -0.176115369 | 0.299972 | -0.587106 | 0.557133 | 0.893658638 | GPRC5C | protein_coding | 54.5 | -0.008599422 | 0.008599422 | ENSG00000170412 | n.s. |
| ENSG00000163964 | 535.3442 | -0.170982324 | 0.27897 | -0.612907 | 0.539938 | 0.890684245 | PIGX | protein_coding | 45.36 | -0.008596347 | 0.008596347 | ENSG00000163964 | n.s. |
| ENSG00000259826 | 75.11985 | 0.175066323 | 0.295547 | 0.5923462 | 0.553619 | 0.893144718 | | antisense | 47.18 | 0.008591934 | 0.008591934 | ENSG00000259826 | n.s. |
| ENSG00000018510 | 3358.979 | -0.166203272 | 0.266541 | -0.623555 | 0.53292 | 0.887866798 | AGPS | protein_coding | 36.51 | -0.008584762 | 0.008584762 | ENSG00000018510 | n.s. |
| ENSG00000261220 | 99.89221 | 0.169454479 | 0.275399 | 0.6153051 | 0.538353 | 0.889966281 | | lincRNA | 46.54 | 0.008578878 | 0.008578878 | ENSG00000261220 | n.s. |

| | | | | | | | | | | | | | |
|---|---|---|---|---|---|---|---|---|---|---|---|---|---|
| ENSG00000135862 | 1143.22 | -0.165129771 | 0.263882 | -0.625771 | 0.531465 | 0.887366671 | LAMC1 | protein_coding | 40.08 | -0.008569721 | 0.008569721 | ENSG00000135862 | n.s. |
| ENSG00000146918 | 1740.298 | -0.163099345 | 0.255449 | -0.63848 | 0.523161 | 0.886079926 | NCAPG2 | protein_coding | 41.79 | -0.008567136 | 0.008567136 | ENSG00000146918 | n.s. |
| ENSG00000273568 | 62.24602 | -0.177941786 | 0.306992 | -0.579629 | 0.562165 | 0.895116382 | | sense_intronic | 53.66 | -0.008562647 | 0.008562647 | ENSG00000273568 | n.s. |
| ENSG00000011143 | 419.9138 | -0.169101251 | 0.275217 | -0.614429 | 0.538932 | 0.889966281 | MKS1 | protein_coding | 47.46 | -0.008560996 | 0.008560996 | ENSG00000011143 | n.s. |
| ENSG00000173744 | 1747.673 | -0.163825926 | 0.259042 | -0.63243 | 0.527106 | 0.886660976 | AGFG1 | protein_coding | 37.09 | -0.00855866 | 0.00855866 | ENSG00000173744 | n.s. |
| ENSG00000145916 | 174.688 | -0.169042172 | 0.274823 | -0.615095 | 0.538492 | 0.889966281 | RMND5B | protein_coding | 49.48 | -0.008558005 | 0.008558005 | ENSG00000145916 | n.s. |
| ENSG00000068745 | 2576.316 | 0.159070626 | 0.244976 | 0.6493309 | 0.516125 | 0.883546124 | IP6K2 | protein_coding | 47.61 | 0.008553351 | 0.008553351 | ENSG00000068745 | n.s. |
| ENSG00000236269 | 199.88 | -0.157468134 | 0.241517 | -0.651997 | 0.514403 | 0.88245884 | ENO1-IT1 | sense_intronic | 48.92 | -0.008551392 | 0.008551392 | ENSG00000236269 | n.s. |
| ENSG00000273035 | 91.3823 | -0.171252382 | 0.285286 | -0.600284 | 0.548317 | 0.891406027 | | sense_intronic | 31.45 | -0.008549678 | 0.008549678 | ENSG00000273035 | n.s. |
| ENSG00000268093 | 20.08865 | 0.210544156 | 0.414 | 0.5085603 | 0.61106 | 0.910755785 | | antisense | 50.5 | 0.008547685 | 0.008547685 | ENSG00000268093 | n.s. |
| ENSG00000211456 | 1614.64 | -0.159654796 | 0.246967 | -0.646463 | 0.517979 | 0.884019009 | SACM1L | protein_coding | 37.99 | -0.008547662 | 0.008547662 | ENSG00000211456 | n.s. |
| ENSG00000147124 | 253.6078 | -0.17114296 | 0.284734 | -0.601063 | 0.547798 | 0.891406027 | ZNF41 | protein_coding | 43.28 | -0.008544215 | 0.008544215 | ENSG00000147124 | n.s. |
| ENSG00000166439 | 1271.799 | -0.163976388 | 0.260872 | -0.62857 | 0.529631 | 0.886954686 | RNF169 | protein_coding | 39.49 | -0.008542935 | 0.008542935 | ENSG00000166439 | n.s. |
| ENSG00000129255 | 797.3694 | -0.162632318 | 0.254786 | -0.63831 | 0.523272 | 0.886079926 | MPDU1 | protein_coding | 56.21 | -0.008542604 | 0.008542604 | ENSG00000129255 | n.s. |
| ENSG00000184990 | 964.4151 | -0.177865213 | 0.30726 | -0.578875 | 0.562674 | 0.895317554 | SIVA1 | protein_coding | 52.32 | -0.008541603 | 0.008541603 | ENSG00000184990 | n.s. |
| ENSG00000087586 | 739.6026 | -0.171060233 | 0.281272 | -0.608166 | 0.543078 | 0.891406027 | AURKA | protein_coding | 42.93 | -0.008540085 | 0.008540085 | ENSG00000087586 | n.s. |
| ENSG00000212125 | 64.6232 | -0.173108978 | 0.290452 | -0.595999 | 0.551176 | 0.892639336 | TAS2R15P | processed_pseudogene | 38.47 | -0.008538424 | 0.008538424 | ENSG00000212125 | n.s. |
| ENSG00000156467 | 2796.009 | 0.178007299 | 0.308133 | 0.5776962 | 0.563469 | 0.895460914 | UQCRB | protein_coding | 38.18 | 0.008536049 | 0.008536049 | ENSG00000156467 | n.s. |
| ENSG00000152558 | 2179.452 | 0.165183339 | 0.26495 | 0.6234514 | 0.532988 | 0.887866798 | TMEM123 | protein_coding | 40.62 | 0.00853208 | 0.00853208 | ENSG00000152558 | n.s. |
| ENSG00000134375 | 2456.739 | 0.16977892 | 0.277503 | 0.6118102 | 0.540663 | 0.890741378 | TIMM17A | protein_coding | 42.32 | 0.008531114 | 0.008531114 | ENSG00000134375 | n.s. |
| ENSG00000168393 | 1293 | -0.173746948 | 0.292886 | -0.593223 | 0.553032 | 0.893144718 | DTYMK | protein_coding | 51.17 | -0.008527182 | 0.008527182 | ENSG00000168393 | n.s. |
| ENSG00000138685 | 586.6046 | -0.167169363 | 0.270185 | -0.618722 | 0.536099 | 0.889201649 | FGF2 | protein_coding | 38.74 | -0.008525594 | 0.008525594 | ENSG00000138685 | n.s. |
| ENSG00000148690 | 339.3254 | -0.164995392 | 0.264644 | -0.623461 | 0.532982 | 0.887866798 | FRA10AC1 | protein_coding | 34.82 | -0.008522372 | 0.008522372 | ENSG00000148690 | n.s. |
| ENSG00000184205 | 334.9968 | 0.156990286 | 0.240835 | 0.6518571 | 0.514493 | 0.882504745 | TSPYL2 | protein_coding | 52.46 | 0.008521896 | 0.008521896 | ENSG00000184205 | n.s. |
| ENSG00000163166 | 2439.262 | -0.162224972 | 0.253928 | -0.638861 | 0.522913 | 0.886079926 | IWS1 | protein_coding | 41.3 | -0.008521208 | 0.008521208 | ENSG00000163166 | n.s. |
| ENSG00000008735 | 543.1206 | -0.162207067 | 0.253572 | -0.639688 | 0.522375 | 0.886079926 | MAPK8IP2 | protein_coding | 63.07 | -0.008520267 | 0.008520267 | ENSG00000008735 | n.s. |
| ENSG00000206560 | 1741.729 | -0.162186979 | 0.254463 | -0.63737 | 0.523884 | 0.886111278 | ANKRD28 | protein_coding | 37.11 | -0.00851672 | 0.00851672 | ENSG00000206560 | n.s. |
| ENSG00000144118 | 252.9205 | -0.172052984 | 0.288417 | -0.596542 | 0.550813 | 0.89235881 | RALB | protein_coding | 46.37 | -0.008509824 | 0.008509824 | ENSG00000144118 | n.s. |
| ENSG00000137267 | 779.0121 | -0.170446221 | 0.282649 | -0.603032 | 0.546487 | 0.891406027 | TUBB2A | protein_coding | 56.17 | -0.008509431 | 0.008509431 | ENSG00000137267 | n.s. |
| ENSG00000142864 | 19828.12 | -0.161805257 | 0.252674 | -0.640373 | 0.52193 | 0.885950923 | SERBP1 | protein_coding | 38.87 | -0.008509393 | 0.008509393 | ENSG00000142864 | n.s. |
| ENSG00000132329 | 206.7712 | -0.177354709 | 0.306887 | -0.577915 | 0.563321 | 0.895435774 | RAMP1 | protein_coding | 52.41 | -0.008506918 | 0.008506918 | ENSG00000132329 | n.s. |
| ENSG00000134709 | 253.6252 | 0.167988297 | 0.273483 | 0.6142541 | 0.539047 | 0.889974736 | HOOK1 | protein_coding | 34.37 | 0.008503958 | 0.008503958 | ENSG00000134709 | n.s. |

| | | | | | | | | | | | | | |
|---|---|---|---|---|---|---|---|---|---|---|---|---|---|
| ENSG00000163900 | 819.6178 | -0.147667323 | 0.220382 | -0.67005 | 0.502826 | 0.875883784 | TMEM41A | protein_coding | 44.99 | -0.008498773 | 0.008498773 | ENSG00000163900 | n.s. |
| ENSG00000116221 | 1409.379 | -0.173140064 | 0.292727 | -0.591473 | 0.554204 | 0.893144718 | MRPL37 | protein_coding | 48.9 | -0.008497397 | 0.008497397 | ENSG00000116221 | n.s. |
| ENSG00000142173 | 531.3196 | -0.163701248 | 0.26143 | -0.626176 | 0.5312 | 0.887366671 | COL6A2 | protein_coding | 55.88 | -0.008495585 | 0.008495585 | ENSG00000142173 | n.s. |
| ENSG00000187954 | 121.8102 | -0.177142497 | 0.306836 | -0.57732 | 0.563723 | 0.895460914 | CYHR1 | protein_coding | 61.94 | -0.008494579 | 0.008494579 | ENSG00000187954 | n.s. |
| ENSG00000125166 | 1469.886 | -0.17012495 | 0.281722 | -0.603875 | 0.545927 | 0.891406027 | GOT2 | protein_coding | 44.18 | -0.008493392 | 0.008493392 | ENSG00000125166 | n.s. |
| ENSG00000197265 | 625.0079 | 0.174778515 | 0.299685 | 0.583207 | 0.559754 | 0.894140829 | GTF2E2 | protein_coding | 39.44 | 0.0084932 | 0.0084932 | ENSG00000197265 | n.s. |
| ENSG00000012983 | 1918.514 | -0.161640638 | 0.254084 | -0.636169 | 0.524666 | 0.886111278 | MAP4K5 | protein_coding | 37.27 | -0.008488031 | 0.008488031 | ENSG00000012983 | n.s. |
| ENSG00000174521 | 149.0841 | 0.178996166 | 0.311482 | 0.5746604 | 0.565521 | 0.896578051 | TTC9B | protein_coding | 63.92 | 0.008486548 | 0.008486548 | ENSG00000174521 | n.s. |
| ENSG00000006118 | 2416.218 | -0.156723882 | 0.240854 | -0.650701 | 0.515239 | 0.882914384 | TMEM132A | protein_coding | 58.71 | -0.008475848 | 0.008475848 | ENSG00000006118 | n.s. |
| ENSG00000008282 | 383.4929 | -0.169298452 | 0.277126 | -0.610907 | 0.541261 | 0.891123173 | SYPL1 | protein_coding | 38.18 | -0.008475464 | 0.008475464 | ENSG00000008282 | n.s. |
| ENSG00000119383 | 544.7987 | -0.1690678 | 0.276845 | -0.610696 | 0.541401 | 0.891123173 | PTPA | protein_coding | 52.59 | -0.008463917 | 0.008463917 | ENSG00000119383 | n.s. |
| ENSG00000276859 | 10.26277 | 0.243774057 | 0.532157 | 0.4580869 | 0.64689 | 0.923184844 | | misc_RNA | 65.05 | 0.008461723 | 0.008461723 | ENSG00000276859 | n.s. |
| ENSG00000149182 | 1483.75 | -0.159959302 | 0.248982 | -0.642453 | 0.520579 | 0.885417361 | ARFGAP2 | protein_coding | 53.19 | -0.008454164 | 0.008454164 | ENSG00000149182 | n.s. |
| ENSG00000142082 | 131.1208 | -0.173072028 | 0.294624 | -0.587433 | 0.556913 | 0.893643698 | SIRT3 | protein_coding | 48.61 | -0.008452077 | 0.008452077 | ENSG00000142082 | n.s. |
| ENSG00000140577 | 44.75899 | -0.178162258 | 0.310243 | -0.574268 | 0.565787 | 0.896600563 | CRTC3 | protein_coding | 44.02 | -0.008445068 | 0.008445068 | ENSG00000140577 | n.s. |
| ENSG00000204356 | 1020.829 | -0.172051064 | 0.291495 | -0.590237 | 0.555032 | 0.893144718 | NELFE | protein_coding | 49.59 | -0.008443951 | 0.008443951 | ENSG00000204356 | n.s. |
| ENSG00000206663 | 3.207671 | 0.317025313 | 0.873009 | 0.3631409 | 0.7165 | 0.94052839 | | misc_RNA | 44.12 | 0.008441779 | 0.008441779 | ENSG00000206663 | n.s. |
| ENSG00000260369 | 206.5061 | 0.160535549 | 0.25272 | 0.6352314 | 0.525277 | 0.886111278 | | lincRNA | 61.81 | 0.00843 | 0.00843 | ENSG00000260369 | n.s. |
| ENSG00000164944 | 1773.664 | -0.165250089 | 0.267071 | -0.618749 | 0.536082 | 0.889201649 | KIAA1429 | protein_coding | 38.74 | -0.008427712 | 0.008427712 | ENSG00000164944 | n.s. |
| ENSG00000066117 | 2195.897 | -0.155396556 | 0.238543 | -0.651439 | 0.514763 | 0.882641141 | SMARCD1 | protein_coding | 48.68 | -0.008424954 | 0.008424954 | ENSG00000066117 | n.s. |
| ENSG00000235363 | 127.9405 | -0.242577496 | 0.52908 | -0.458489 | 0.646601 | 0.923184844 | SNRPGP10 | processed_pseudogene | 39.11 | -0.008420188 | 0.008420188 | ENSG00000235363 | n.s. |
| ENSG00000251648 | 33.83209 | 0.182884648 | 0.323149 | 0.565945 | 0.571431 | 0.899418628 | | processed_pseudogene | 51.61 | 0.008419666 | 0.008419666 | ENSG00000251648 | n.s. |
| ENSG00000009954 | 5229.758 | -0.159215468 | 0.247635 | -0.642944 | 0.52026 | 0.885364478 | BAZ1B | protein_coding | 43.19 | -0.008418981 | 0.008418981 | ENSG00000009954 | n.s. |
| ENSG00000236617 | 23.51939 | 0.202842211 | 0.389876 | 0.5202742 | 0.602872 | 0.908868121 | | antisense | 52.43 | 0.008417775 | 0.008417775 | ENSG00000236617 | n.s. |
| ENSG00000224451 | 90.45182 | -0.171485039 | 0.289137 | -0.593092 | 0.55312 | 0.893144718 | ATP5F1P1 | processed_pseudogene | 44.31 | -0.008416171 | 0.008416171 | ENSG00000224451 | n.s. |
| ENSG00000108518 | 3039.732 | -0.168554662 | 0.277953 | -0.606415 | 0.544239 | 0.891406027 | PFN1 | protein_coding | 57.98 | -0.008414996 | 0.008414996 | ENSG00000108518 | n.s. |
| ENSG00000100749 | 1856.356 | -0.174817598 | 0.301677 | -0.579486 | 0.562261 | 0.895116382 | VRK1 | protein_coding | 38.99 | -0.008412309 | 0.008412309 | ENSG00000100749 | n.s. |
| ENSG00000144455 | 280.9039 | -0.16482491 | 0.266547 | -0.61837 | 0.536331 | 0.889201649 | SUMF1 | protein_coding | 39.53 | -0.008406028 | 0.008406028 | ENSG00000144455 | n.s. |
| ENSG00000167257 | 301.8014 | -0.164697491 | 0.26598 | -0.61921 | 0.535778 | 0.889201649 | RNF214 | protein_coding | 42.99 | -0.008399529 | 0.008399529 | ENSG00000167257 | n.s. |
| ENSG00000132507 | 3611.258 | -0.168038514 | 0.278891 | -0.602524 | 0.546826 | 0.891406027 | EIF5A | protein_coding | 55.3 | -0.008389228 | 0.008389228 | ENSG00000132507 | n.s. |
| ENSG00000123444 | 466.382 | -0.161644138 | 0.258147 | -0.62617 | 0.531203 | 0.887366671 | KBTBD4 | protein_coding | 48.95 | -0.008388828 | 0.008388828 | ENSG00000123444 | n.s. |
| ENSG00000197008 | 629.8402 | -0.172573647 | 0.295713 | -0.583584 | 0.5595 | 0.894140829 | ZNF138 | protein_coding | 39.99 | -0.008386056 | 0.008386056 | ENSG00000197008 | n.s. |

| | | | | | | | | | | | | | |
|---|---|---|---|---|---|---|---|---|---|---|---|---|---|
| ENSG00000051596 | 445.4937 | -0.168608911 | 0.281999 | -0.597906 | 0.549902 | 0.891861801 | THOC3 | protein_coding | 43.01 | -0.008380274 | 0.008380274 | ENSG00000051596 | n.s. |
| ENSG00000090565 | 453.3837 | -0.161730337 | 0.258915 | -0.624647 | 0.532203 | 0.887559389 | RAB11FIP3 | protein_coding | 52.85 | -0.008378048 | 0.008378048 | ENSG00000090565 | n.s. |
| ENSG00000222858 | 9.007513 | -0.241109233 | 0.532341 | -0.452922 | 0.650605 | 0.923184844 | RNU6-920P | snRNA | 41.3 | -0.008369223 | 0.008369223 | ENSG00000222858 | n.s. |
| ENSG00000138468 | 24.63624 | 0.197197548 | 0.366621 | 0.5378783 | 0.590661 | 0.906900529 | SENP7 | protein_coding | 37.56 | 0.008369132 | 0.008369132 | ENSG00000138468 | n.s. |
| ENSG00000147180 | 1903.462 | 0.167463731 | 0.277389 | 0.6037139 | 0.546034 | 0.891406027 | ZNF711 | protein_coding | 34.34 | 0.008360532 | 0.008360532 | ENSG00000147180 | n.s. |
| ENSG00000134294 | 10359.44 | -0.155692249 | 0.240239 | -0.648074 | 0.516937 | 0.88372671 | SLC38A2 | protein_coding | 38.54 | -0.008357874 | 0.008357874 | ENSG00000134294 | n.s. |
| ENSG00000250197 | 50.95132 | 0.186840972 | 0.336743 | 0.5548467 | 0.579 | 0.902167117 | HMGN1P15 | processed_pseudogene | 43.05 | 0.008354222 | 0.008354222 | ENSG00000250197 | n.s. |
| ENSG00000150593 | 1570.037 | 0.167045145 | 0.273677 | 0.610374 | 0.541614 | 0.891255806 | PDCD4 | protein_coding | 34.69 | 0.008351861 | 0.008351861 | ENSG00000150593 | n.s. |
| ENSG00000252712 | 21.15397 | -0.204864098 | 0.400649 | -0.51133 | 0.60912 | 0.910431619 | SCARNA14 | scaRNA | 42.65 | -0.008348759 | 0.008348759 | ENSG00000252712 | n.s. |
| ENSG00000105617 | 124.2317 | -0.16705581 | 0.278582 | -0.599665 | 0.548729 | 0.891406027 | LENG1 | protein_coding | 56.01 | -0.008340167 | 0.008340167 | ENSG00000105617 | n.s. |
| ENSG00000137713 | 632.1719 | -0.16189305 | 0.260048 | -0.622552 | 0.533579 | 0.888237452 | PPP2R1B | protein_coding | 44.57 | -0.008332784 | 0.008332784 | ENSG00000137713 | n.s. |
| ENSG00000101182 | 3283.387 | -0.170101518 | 0.289114 | -0.588355 | 0.556294 | 0.893342237 | PSMA7 | protein_coding | 52.28 | -0.008331935 | 0.008331935 | ENSG00000101182 | n.s. |
| ENSG00000198015 | 4030.544 | -0.169710073 | 0.287471 | -0.590356 | 0.554952 | 0.893144718 | MRPL42 | protein_coding | 39.33 | -0.008329059 | 0.008329059 | ENSG00000198015 | n.s. |
| ENSG00000257365 | 379.7747 | -0.164459806 | 0.267693 | -0.614361 | 0.538977 | 0.889966281 | FNTB | protein_coding | 43.84 | -0.008326016 | 0.008326016 | ENSG00000257365 | n.s. |
| ENSG00000135048 | 52.01083 | -0.181766017 | 0.323309 | -0.562204 | 0.573977 | 0.899911687 | TMEM2 | protein_coding | 40.46 | -0.008324903 | 0.008324903 | ENSG00000135048 | n.s. |
| ENSG00000257267 | 1185.115 | -0.163213695 | 0.263492 | -0.619427 | 0.535635 | 0.889201649 | ZNF271P | anscribed_unitary_pseudoge | 38.12 | -0.008323856 | 0.008323856 | ENSG00000257267 | n.s. |
| ENSG00000230124 | 2195.463 | -0.159766425 | 0.253692 | -0.629765 | 0.528848 | 0.886954686 | ACBD6 | protein_coding | 37.67 | -0.008323602 | 0.008323602 | ENSG00000230124 | n.s. |
| ENSG00000273345 | 71.56956 | 0.166627143 | 0.277989 | 0.5994024 | 0.548905 | 0.891406027 | | processed_transcript | 41.3 | 0.008318766 | 0.008318766 | ENSG00000273345 | n.s. |
| ENSG00000143669 | 33.07267 | -0.200410396 | 0.383925 | -0.522005 | 0.601667 | 0.908868121 | LYST | protein_coding | 37.77 | -0.008316857 | 0.008316857 | ENSG00000143669 | n.s. |
| ENSG00000239305 | 1021.53 | 0.159886578 | 0.254627 | 0.6279251 | 0.530053 | 0.88715322 | RNF103 | protein_coding | 39.04 | 0.008314321 | 0.008314321 | ENSG00000239305 | n.s. |
| ENSG00000051825 | 2852.799 | -0.159578728 | 0.253455 | -0.629614 | 0.528947 | 0.886954686 | MPHOSPH9 | protein_coding | 42.99 | -0.008313823 | 0.008313823 | ENSG00000051825 | n.s. |
| ENSG00000259935 | 23.77686 | -0.192223856 | 0.35452 | -0.542208 | 0.587675 | 0.905217543 | | sense_intronic | 30.82 | -0.008313112 | 0.008313112 | ENSG00000259935 | n.s. |
| ENSG00000077147 | 3598.292 | -0.162958946 | 0.263485 | -0.618476 | 0.536262 | 0.889201649 | TM9SF3 | protein_coding | 37.03 | -0.008310864 | 0.008310864 | ENSG00000077147 | n.s. |
| ENSG00000267194 | 153.1558 | -0.165889562 | 0.271345 | -0.611361 | 0.540961 | 0.891073457 | | lincRNA | 36.86 | -0.008308826 | 0.008308826 | ENSG00000267194 | n.s. |
| ENSG00000151292 | 2194.164 | 0.166406774 | 0.27671 | 0.6013753 | 0.54759 | 0.891406027 | CSNK1G3 | protein_coding | 34.48 | 0.008307764 | 0.008307764 | ENSG00000151292 | n.s. |
| ENSG00000133961 | 80.86514 | -0.169170924 | 0.286735 | -0.58999 | 0.555198 | 0.893144718 | NUMB | protein_coding | 40.56 | -0.008302599 | 0.008302599 | ENSG00000133961 | n.s. |
| ENSG00000197965 | 1446.664 | -0.160259729 | 0.256376 | -0.625097 | 0.531907 | 0.88755367 | MPZL1 | protein_coding | 41.24 | -0.008302315 | 0.008302315 | ENSG00000197965 | n.s. |
| ENSG00000110801 | 620.148 | -0.170762646 | 0.292672 | -0.58346 | 0.559584 | 0.894140829 | PSMD9 | protein_coding | 47.48 | -0.008298053 | 0.008298053 | ENSG00000110801 | n.s. |
| ENSG00000119402 | 1455.45 | -0.157944694 | 0.248407 | -0.63583 | 0.524887 | 0.886111278 | FBXW2 | protein_coding | 41.5 | -0.00829395 | 0.00829395 | ENSG00000119402 | n.s. |
| ENSG00000227345 | 1027.24 | -0.166084282 | 0.272763 | -0.608896 | 0.542593 | 0.891406027 | PARG | protein_coding | 38.39 | -0.008291664 | 0.008291664 | ENSG00000227345 | n.s. |
| ENSG00000185973 | 190.6257 | -0.168107457 | 0.282443 | -0.595192 | 0.551715 | 0.892641916 | TMLHE | protein_coding | 37.38 | -0.008291518 | 0.008291518 | ENSG00000185973 | n.s. |
| ENSG00000228623 | 193.5527 | -0.173240907 | 0.300332 | -0.576831 | 0.564054 | 0.895679463 | ZNF883 | protein_coding | 38.73 | -0.008289124 | 0.008289124 | ENSG00000228623 | n.s. |

| | | | | | | | | | | | | | |
|---|---|---|---|---|---|---|---|---|---|---|---|---|---|
| ENSG00000264529 | 49.93732 | 0.178827581 | 0.314442 | 0.5687149 | 0.56955 | 0.898810685 | | transcribed_processed_pseudog | 39.88 | 0.008285399 | 0.008285399 | ENSG00000264529 | n.s. |
| ENSG00000101544 | 1315.004 | -0.162490287 | 0.262934 | -0.61799 | 0.536582 | 0.889279067 | ADNP2 | protein_coding | 41.81 | -0.008280819 | 0.008280819 | ENSG00000101544 | n.s. |
| ENSG00000272663 | 39.40417 | 0.187637154 | 0.340666 | 0.5507951 | 0.581774 | 0.90349457 | | lincRNA | 47.03 | 0.008270005 | 0.008270005 | ENSG00000272663 | n.s. |
| ENSG00000272172 | 12.26009 | -0.229039867 | 0.485871 | -0.4714 | 0.637355 | 0.920241952 | | antisense | 66.37 | -0.008267875 | 0.008267875 | ENSG00000272172 | n.s. |
| ENSG00000129473 | 338.7427 | -0.155984339 | 0.242256 | -0.643882 | 0.519652 | 0.885353782 | BCL2L2 | protein_coding | 51.4 | -0.008248944 | 0.008248944 | ENSG00000129473 | n.s. |
| ENSG00000148719 | 580.5431 | -0.158215325 | 0.251389 | -0.629366 | 0.52911 | 0.886954686 | DNAJB12 | protein_coding | 53.03 | -0.008242792 | 0.008242792 | ENSG00000148719 | n.s. |
| ENSG00000200795 | 695.0087 | 0.13788958 | 0.199155 | 0.6923738 | 0.488703 | 0.871503806 | RNU4-1 | snRNA | 48.23 | 0.008236243 | 0.008236243 | ENSG00000200795 | n.s. |
| ENSG00000178149 | 140.6644 | -0.16484034 | 0.272166 | -0.605662 | 0.544739 | 0.891406027 | DALRD3 | protein_coding | 60.06 | -0.008229561 | 0.008229561 | ENSG00000178149 | n.s. |
| ENSG00000169919 | 619.7681 | -0.153011888 | 0.235861 | -0.648739 | 0.516507 | 0.883546124 | GUSB | protein_coding | 52.03 | -0.008227568 | 0.008227568 | ENSG00000169919 | n.s. |
| ENSG00000198663 | 656.5783 | -0.166558381 | 0.279363 | -0.596208 | 0.551036 | 0.89251749 | C6orf89 | protein_coding | 42.84 | -0.008225197 | 0.008225197 | ENSG00000198663 | n.s. |
| ENSG00000158805 | 268.95 | -0.164744152 | 0.273501 | -0.602354 | 0.546939 | 0.891406027 | ZNF276 | protein_coding | 56.58 | -0.008224759 | 0.008224759 | ENSG00000158805 | n.s. |
| ENSG00000166575 | 95.37152 | -0.164630268 | 0.274592 | -0.599545 | 0.54881 | 0.891406027 | TMEM135 | protein_coding | 36.32 | -0.008219073 | 0.008219073 | ENSG00000166575 | n.s. |
| ENSG00000188290 | 105.784 | -0.183165471 | 0.329453 | -0.555969 | 0.578232 | 0.901839975 | HES4 | protein_coding | 74.32 | -0.00821873 | 0.00821873 | ENSG00000188290 | n.s. |
| ENSG00000169744 | 61.04879 | 0.165306046 | 0.276492 | 0.5978695 | 0.549927 | 0.891861801 | LDB2 | protein_coding | 40.21 | 0.008216113 | 0.008216113 | ENSG00000169744 | n.s. |
| ENSG00000225349 | 24.12409 | -0.254798153 | 0.597133 | -0.426702 | 0.669596 | 0.928471963 | | processed_pseudogene | 47.09 | -0.008212452 | 0.008212452 | ENSG00000225349 | n.s. |
| ENSG00000273899 | 346.832 | -0.164474543 | 0.274078 | -0.6001 | 0.54844 | 0.891406027 | NOL12 | protein_coding | 54.42 | -0.008211298 | 0.008211298 | ENSG00000273899 | n.s. |
| ENSG00000056097 | 3100.535 | -0.164453986 | 0.272277 | -0.603995 | 0.545847 | 0.891406027 | ZFR | protein_coding | 38.33 | -0.008210272 | 0.008210272 | ENSG00000056097 | n.s. |
| ENSG00000213397 | 286.4546 | -0.160781163 | 0.259129 | -0.620468 | 0.534949 | 0.88909364 | HAUS7 | protein_coding | 58.62 | -0.00820828 | 0.00820828 | ENSG00000213397 | n.s. |
| ENSG00000185842 | 695.8468 | -0.167036933 | 0.282929 | -0.590385 | 0.554933 | 0.893144718 | DNAH14 | protein_coding | 37.4 | -0.008197866 | 0.008197866 | ENSG00000185842 | n.s. |
| ENSG00000270894 | 51.68271 | -0.167002404 | 0.283374 | -0.589336 | 0.555636 | 0.893144718 | | sense_overlapping | 35.62 | -0.008196172 | 0.008196172 | ENSG00000270894 | n.s. |
| ENSG00000123143 | 740.357 | -0.166982762 | 0.281824 | -0.592507 | 0.553511 | 0.893144718 | PKN1 | protein_coding | 53.14 | -0.008195208 | 0.008195208 | ENSG00000123143 | n.s. |
| ENSG00000104299 | 311.1137 | -0.1641241 | 0.272356 | -0.602609 | 0.546769 | 0.891406027 | INTS9 | protein_coding | 42.01 | -0.008193803 | 0.008193803 | ENSG00000104299 | n.s. |
| ENSG00000143437 | 911.3286 | -0.155944789 | 0.244292 | -0.638353 | 0.523244 | 0.886079926 | ARNT | protein_coding | 41.21 | -0.008191328 | 0.008191328 | ENSG00000143437 | n.s. |
| ENSG00000125812 | 979.9549 | 0.160386523 | 0.258611 | 0.6201846 | 0.535136 | 0.88909364 | GZF1 | protein_coding | 44.84 | 0.008188132 | 0.008188132 | ENSG00000125812 | n.s. |
| ENSG00000283172 | 34.0946 | -0.186657865 | 0.340034 | -0.548939 | 0.583047 | 0.903983136 | | miRNA | 32.99 | -0.008183019 | 0.008183019 | ENSG00000283172 | n.s. |
| ENSG00000200972 | 195.3838 | 0.155832851 | 0.246131 | 0.6331288 | 0.52665 | 0.886214816 | RNU5A-8P | snRNA | 39.47 | 0.008175146 | 0.008175146 | ENSG00000200972 | n.s. |
| ENSG00000213639 | 3130.082 | -0.166512907 | 0.282278 | -0.589891 | 0.555264 | 0.893144718 | PPP1CB | protein_coding | 38.98 | -0.008172148 | 0.008172148 | ENSG00000213639 | n.s. |
| ENSG00000167112 | 741.1698 | -0.175051144 | 0.306354 | -0.571402 | 0.567727 | 0.898133129 | TRUB2 | protein_coding | 51.14 | -0.008167761 | 0.008167761 | ENSG00000167112 | n.s. |
| ENSG00000090674 | 83.39035 | -0.163588665 | 0.273036 | -0.599147 | 0.549075 | 0.891414853 | MCOLN1 | protein_coding | 57.01 | -0.008166368 | 0.008166368 | ENSG00000090674 | n.s. |
| ENSG00000124194 | 766.5634 | -0.170287568 | 0.294886 | -0.577468 | 0.563623 | 0.895460914 | GDAP1L1 | protein_coding | 50.31 | -0.008165862 | 0.008165862 | ENSG00000124194 | n.s. |
| ENSG00000048392 | 458.7042 | -0.171940102 | 0.299009 | -0.575033 | 0.565269 | 0.896450345 | RRM2B | protein_coding | 37.99 | -0.008162643 | 0.008162643 | ENSG00000048392 | n.s. |
| ENSG00000207221 | 6.463069 | -0.292980071 | 0.767141 | -0.381912 | 0.702527 | 0.937867182 | SNORA70E | snoRNA | 49.63 | -0.008162032 | 0.008162032 | ENSG00000207221 | n.s. |

| | | | | | | | | | | | | | |
|---|---|---|---|---|---|---|---|---|---|---|---|---|---|
| ENSG00000196911 | 1377.493 | -0.163481659 | 0.271427 | -0.602304 | 0.546972 | 0.891406027 | KPNA5 | protein_coding | 35.56 | -0.008161729 | 0.008161729 | ENSG00000196911 | n.s. |
| ENSG00000143162 | 364.9399 | -0.176156926 | 0.310315 | -0.567672 | 0.570258 | 0.898818733 | CREG1 | protein_coding | 44.46 | -0.008160978 | 0.008160978 | ENSG00000143162 | n.s. |
| ENSG00000122674 | 473.1037 | -0.163444809 | 0.272689 | -0.599382 | 0.548918 | 0.891406027 | CCZ1 | protein_coding | 46.24 | -0.00815989 | 0.00815989 | ENSG00000122674 | n.s. |
| ENSG00000280406 | 114.8418 | -0.161072641 | 0.261997 | -0.614788 | 0.538695 | 0.889966281 | | TEC | 30.49 | -0.008154536 | 0.008154536 | ENSG00000280406 | n.s. |
| ENSG00000180104 | 333.5561 | -0.161051493 | 0.262033 | -0.614623 | 0.538803 | 0.889966281 | EXOC3 | protein_coding | 51.24 | -0.008153465 | 0.008153465 | ENSG00000180104 | n.s. |
| ENSG00000163069 | 1023.797 | -0.167581227 | 0.287352 | -0.583192 | 0.559764 | 0.894140829 | SGCB | protein_coding | 38.68 | -0.008143454 | 0.008143454 | ENSG00000163069 | n.s. |
| ENSG00000144320 | 835.6431 | -0.169316561 | 0.29232 | -0.579216 | 0.562444 | 0.895173193 | LNPK | protein_coding | 35.26 | -0.00814293 | 0.00814293 | ENSG00000144320 | n.s. |
| ENSG00000237373 | 23.35287 | -0.182914021 | 0.330327 | -0.553737 | 0.579759 | 0.902603782 | BRWD1-IT1 | sense_overlapping | 37.38 | -0.008140195 | 0.008140195 | ENSG00000237373 | n.s. |
| ENSG00000205220 | 130.6145 | -0.187089385 | 0.341535 | -0.547789 | 0.583837 | 0.904703099 | PSMB10 | protein_coding | 61.37 | -0.008137251 | 0.008137251 | ENSG00000205220 | n.s. |
| ENSG00000172661 | 556.7591 | -0.162957309 | 0.270476 | -0.602484 | 0.546852 | 0.891406027 | WASHC2C | protein_coding | 42.53 | -0.008135551 | 0.008135551 | ENSG00000172661 | n.s. |
| ENSG00000123268 | 838.7626 | -0.162911488 | 0.270821 | -0.601547 | 0.547476 | 0.891406027 | ATF1 | protein_coding | 41.93 | -0.008133264 | 0.008133264 | ENSG00000123268 | n.s. |
| ENSG00000169592 | 498.573 | -0.16477202 | 0.276682 | -0.595529 | 0.55149 | 0.892641916 | INO80E | protein_coding | 57.4 | -0.008127005 | 0.008127005 | ENSG00000169592 | n.s. |
| ENSG00000174669 | 309.8464 | -0.153292175 | 0.237822 | -0.644567 | 0.519208 | 0.885138943 | SLC29A2 | protein_coding | 58.27 | -0.00812273 | 0.00812273 | ENSG00000174669 | n.s. |
| ENSG00000152455 | 1662.52 | -0.162651433 | 0.269532 | -0.603459 | 0.546203 | 0.891406027 | SUV39H2 | protein_coding | 39.96 | -0.008120281 | 0.008120281 | ENSG00000152455 | n.s. |
| ENSG00000082516 | 1597.666 | -0.154581396 | 0.24184 | -0.639188 | 0.522701 | 0.886079926 | GEMIN5 | protein_coding | 41.33 | -0.008119713 | 0.008119713 | ENSG00000082516 | n.s. |
| ENSG00000180398 | 1771.996 | -0.162574827 | 0.270268 | -0.601532 | 0.547486 | 0.891406027 | MCFD2 | protein_coding | 45.38 | -0.008116456 | 0.008116456 | ENSG00000180398 | n.s. |
| ENSG00000149480 | 1710.664 | -0.158958879 | 0.255931 | -0.621101 | 0.534533 | 0.88909364 | MTA2 | protein_coding | 52.38 | -0.008115248 | 0.008115248 | ENSG00000149480 | n.s. |
| ENSG00000133835 | 1594.692 | 0.165227223 | 0.280101 | 0.5898834 | 0.555269 | 0.893144718 | HSD17B4 | protein_coding | 37.34 | 0.008109049 | 0.008109049 | ENSG00000133835 | n.s. |
| ENSG00000111605 | 8412.36 | -0.161238303 | 0.263166 | -0.612687 | 0.540083 | 0.890684245 | CPSF6 | protein_coding | 35.78 | -0.008106454 | 0.008106454 | ENSG00000111605 | n.s. |
| ENSG00000123728 | 347.9597 | 0.165993221 | 0.282594 | 0.5873902 | 0.556942 | 0.893643698 | RAP2C | protein_coding | 37.77 | 0.008106379 | 0.008106379 | ENSG00000123728 | n.s. |
| ENSG00000205054 | 17.55295 | -0.220996435 | 0.461372 | -0.478998 | 0.63194 | 0.919033489 | LINC01121 | processed_transcript | 44.5 | -0.008103644 | 0.008103644 | ENSG00000205054 | n.s. |
| ENSG00000125656 | 317.6181 | -0.170692197 | 0.296637 | -0.575425 | 0.565004 | 0.896450345 | CLPP | protein_coding | 54.14 | -0.008103401 | 0.008103401 | ENSG00000125656 | n.s. |
| ENSG00000166598 | 10802.8 | 0.16003805 | 0.259934 | 0.6156868 | 0.538101 | 0.889966281 | HSP90B1 | protein_coding | 39.78 | 0.008102158 | 0.008102158 | ENSG00000166598 | n.s. |
| ENSG00000105171 | 365.2499 | -0.181177552 | 0.326558 | -0.554809 | 0.579025 | 0.902167117 | POP4 | protein_coding | 49.76 | -0.008100993 | 0.008100993 | ENSG00000105171 | n.s. |
| ENSG00000133884 | 468.9386 | -0.162249803 | 0.268518 | -0.604242 | 0.545683 | 0.891406027 | DPF2 | protein_coding | 48.63 | -0.00810023 | 0.00810023 | ENSG00000133884 | n.s. |
| ENSG00000107562 | 190.8897 | -0.166132718 | 0.283661 | -0.585672 | 0.558096 | 0.893837232 | CXCL12 | protein_coding | 52.37 | -0.008097568 | 0.008097568 | ENSG00000107562 | n.s. |
| ENSG00000165915 | 157.5884 | 0.164951059 | 0.279625 | 0.5899011 | 0.555257 | 0.893144718 | SLC39A13 | protein_coding | 60.6 | 0.008095495 | 0.008095495 | ENSG00000165915 | n.s. |
| ENSG00000249853 | 209.3288 | -0.158568278 | 0.255481 | -0.620667 | 0.534819 | 0.88909364 | HS3ST5 | protein_coding | 37.56 | -0.008095306 | 0.008095306 | ENSG00000249853 | n.s. |
| ENSG00000196476 | 89.28348 | 0.16417398 | 0.275946 | 0.5949497 | 0.551877 | 0.892697967 | C20orf96 | protein_coding | 48.38 | 0.008093031 | 0.008093031 | ENSG00000196476 | n.s. |
| ENSG00000169016 | 1025.255 | -0.164844736 | 0.279224 | -0.590367 | 0.554945 | 0.893144718 | E2F6 | protein_coding | 41.96 | -0.008090277 | 0.008090277 | ENSG00000169016 | n.s. |
| ENSG00000071626 | 1697.376 | -0.168621813 | 0.292051 | -0.577371 | 0.563689 | 0.895460914 | DAZAP1 | protein_coding | 58.75 | -0.008085983 | 0.008085983 | ENSG00000071626 | n.s. |
| ENSG00000075975 | 966.7105 | -0.161945946 | 0.267669 | -0.605024 | 0.545163 | 0.891406027 | MKRN2 | protein_coding | 45.18 | -0.00808506 | 0.00808506 | ENSG00000075975 | n.s. |

| | | | | | | | | | | | | | |
|---|---|---|---|---|---|---|---|---|---|---|---|---|---|
| ENSG00000183010 | 2041.043 | -0.167187008 | 0.287501 | -0.581518 | 0.560892 | 0.894637751 | PYCR1 | protein_coding | 62.3 | -0.008083957 | 0.008083957 | ENSG00000183010 | n.s. |
| ENSG00000224745 | 153.6413 | -0.162766444 | 0.272561 | -0.597174 | 0.550391 | 0.89197749 | | sense_intronic | 34.91 | -0.00808072 | 0.00808072 | ENSG00000224745 | n.s. |
| ENSG00000272886 | 885.0064 | -0.158433483 | 0.256134 | -0.618556 | 0.536209 | 0.889201649 | DCP1A | protein_coding | 41.19 | -0.008080067 | 0.008080067 | ENSG00000272886 | n.s. |
| ENSG00000211450 | 720.7092 | -0.161805306 | 0.268002 | -0.603747 | 0.546012 | 0.891406027 | SELENOH | protein_coding | 51.67 | -0.008078038 | 0.008078038 | ENSG00000211450 | n.s. |
| ENSG00000120451 | 991.8372 | -0.153595094 | 0.239953 | -0.640106 | 0.522104 | 0.885953729 | SNX19 | protein_coding | 42.46 | -0.008077406 | 0.008077406 | ENSG00000120451 | n.s. |
| ENSG00000107897 | 816.6332 | -0.164527351 | 0.277214 | -0.593502 | 0.552845 | 0.893144718 | ACBD5 | protein_coding | 40.95 | -0.008074701 | 0.008074701 | ENSG00000107897 | n.s. |
| ENSG00000204438 | 338.3963 | -0.154979389 | 0.245778 | -0.630567 | 0.528324 | 0.886954686 | GPANK1 | protein_coding | 53.99 | -0.008074204 | 0.008074204 | ENSG00000204438 | n.s. |
| ENSG00000261013 | 28.7677 | -0.191345602 | 0.356497 | -0.536738 | 0.591449 | 0.907409399 | | lincRNA | 36.34 | -0.008074158 | 0.008074158 | ENSG00000261013 | n.s. |
| ENSG00000153721 | 1234.147 | -0.161615532 | 0.268643 | -0.6016 | 0.547441 | 0.891406027 | CNKSR3 | protein_coding | 42.73 | -0.008068564 | 0.008068564 | ENSG00000153721 | n.s. |
| ENSG00000102226 | 932.8801 | -0.165419009 | 0.282609 | -0.585328 | 0.558327 | 0.893837232 | USP11 | protein_coding | 48.33 | -0.00806278 | 0.00806278 | ENSG00000102226 | n.s. |
| ENSG00000223745 | 378.0745 | -0.160396723 | 0.261996 | -0.612211 | 0.540398 | 0.890708138 | CCDC18-AS1 | processed_transcript | 39.07 | -0.008062274 | 0.008062274 | ENSG00000223745 | n.s. |
| ENSG00000158985 | 534.0548 | -0.159075415 | 0.258083 | -0.616372 | 0.537649 | 0.889881485 | CDC42SE2 | protein_coding | 39.71 | -0.008060006 | 0.008060006 | ENSG00000158985 | n.s. |
| ENSG00000024048 | 954.2386 | -0.155286607 | 0.248237 | -0.625559 | 0.531604 | 0.887366671 | UBR2 | protein_coding | 39.1 | -0.008058891 | 0.008058891 | ENSG00000024048 | n.s. |
| ENSG00000155393 | 665.3249 | -0.159161348 | 0.258978 | -0.614576 | 0.538835 | 0.889966281 | HEATR3 | protein_coding | 42.6 | -0.008057774 | 0.008057774 | ENSG00000155393 | n.s. |
| ENSG00000228906 | 95.93158 | -0.161311997 | 0.2662 | -0.60598 | 0.544528 | 0.891406027 | | lincRNA | 45.9 | -0.00805341 | 0.00805341 | ENSG00000228906 | n.s. |
| ENSG00000166228 | 321.1945 | 0.165464689 | 0.283465 | 0.5837226 | 0.559407 | 0.894140829 | PCBD1 | protein_coding | 51.19 | 0.008040603 | 0.008040603 | ENSG00000166228 | n.s. |
| ENSG00000134769 | 20.51667 | -0.197644374 | 0.387987 | -0.50941 | 0.610465 | 0.910595741 | DTNA | protein_coding | 37.99 | -0.008039063 | 0.008039063 | ENSG00000134769 | n.s. |
| ENSG00000179262 | 862.4151 | -0.157550925 | 0.254684 | -0.618614 | 0.53617 | 0.889201649 | RAD23A | protein_coding | 54.44 | -0.008035056 | 0.008035056 | ENSG00000179262 | n.s. |
| ENSG00000213949 | 2288.071 | -0.163714836 | 0.276505 | -0.592085 | 0.553793 | 0.893144718 | ITGA1 | protein_coding | 36.8 | -0.008034824 | 0.008034824 | ENSG00000213949 | n.s. |
| ENSG00000152332 | 2733.612 | -0.15728044 | 0.253595 | -0.620203 | 0.535124 | 0.88909364 | UHMK1 | protein_coding | 38.32 | -0.008029559 | 0.008029559 | ENSG00000152332 | n.s. |
| ENSG00000224699 | 732.0303 | -0.141968191 | 0.214076 | -0.663168 | 0.507223 | 0.877953916 | LAMTOR5-AS1 | processed_transcript | 43.14 | -0.008025218 | 0.008025218 | ENSG00000224699 | n.s. |
| ENSG00000087074 | 511.7706 | 0.152496238 | 0.240186 | 0.6349098 | 0.525487 | 0.886111278 | PPP1R15A | protein_coding | 56.68 | 0.008007842 | 0.008007842 | ENSG00000087074 | n.s. |
| ENSG00000174574 | 952.8138 | -0.174764794 | 0.310242 | -0.563317 | 0.573219 | 0.899883575 | AKIRIN1 | protein_coding | 43.9 | -0.008006617 | 0.008006617 | ENSG00000174574 | n.s. |
| ENSG00000180806 | 63.60834 | -0.185052243 | 0.341386 | -0.542061 | 0.587776 | 0.905217543 | HOXC9 | protein_coding | 56.89 | -0.008002961 | 0.008002961 | ENSG00000180806 | n.s. |
| ENSG00000129197 | 2476.987 | -0.16023884 | 0.262702 | -0.609964 | 0.541886 | 0.891404544 | RPAIN | protein_coding | 45.35 | -0.007999949 | 0.007999949 | ENSG00000129197 | n.s. |
| ENSG00000265690 | 108.7839 | -0.162940876 | 0.275261 | -0.59195 | 0.553884 | 0.893144718 | | protein_coding | 67.54 | -0.007996839 | 0.007996839 | ENSG00000265690 | n.s. |
| ENSG00000143771 | 1437.64 | -0.168428081 | 0.292978 | -0.574883 | 0.565371 | 0.896450345 | CNIH4 | protein_coding | 43.22 | -0.007995915 | 0.007995915 | ENSG00000143771 | n.s. |
| ENSG00000178035 | 2989.237 | 0.160460863 | 0.26797 | 0.5988009 | 0.549306 | 0.891685023 | IMPDH2 | protein_coding | 53.38 | 0.00798911 | 0.00798911 | ENSG00000178035 | n.s. |
| ENSG00000171224 | 135.3782 | 0.159961581 | 0.265723 | 0.6019856 | 0.547184 | 0.891406027 | C10orf35 | protein_coding | 56.17 | 0.007985991 | 0.007985991 | ENSG00000171224 | n.s. |
| ENSG00000105879 | 809.592 | -0.156496377 | 0.252807 | -0.619035 | 0.535893 | 0.889201649 | CBLL1 | protein_coding | 36.56 | -0.007981275 | 0.007981275 | ENSG00000105879 | n.s. |
| ENSG00000126858 | 926.9805 | 0.175211348 | 0.313002 | 0.5597778 | 0.575631 | 0.900453394 | RHOT1 | protein_coding | 42.18 | 0.007978908 | 0.007978908 | ENSG00000126858 | n.s. |
| ENSG00000117308 | 76.46786 | -0.184092786 | 0.337888 | -0.544834 | 0.585868 | 0.905026039 | GALE | protein_coding | 60.1 | -0.007978383 | 0.007978383 | ENSG00000117308 | n.s. |

| | | | | | | | | | | | | | |
|---|---|---|---|---|---|---|---|---|---|---|---|---|---|
| ENSG00000114125 | 1215.561 | 0.172149256 | 0.303305 | 0.5675774 | 0.570322 | 0.898818733 | RNF7 | protein_coding | 39.34 | 0.007975311 | 0.007975311 | ENSG00000114125 | n.s. |
| ENSG00000143515 | 789.0237 | -0.159685858 | 0.262385 | -0.608594 | 0.542793 | 0.891406027 | ATP8B2 | protein_coding | 52.37 | -0.007972226 | 0.007972226 | ENSG00000143515 | n.s. |
| ENSG00000139687 | 1150.343 | -0.163939209 | 0.28067 | -0.584099 | 0.559154 | 0.894140829 | RB1 | protein_coding | 36.91 | -0.007966474 | 0.007966474 | ENSG00000139687 | n.s. |
| ENSG00000151876 | 105.0331 | 0.159509292 | 0.263009 | 0.6064792 | 0.544197 | 0.891406027 | FBXO4 | protein_coding | 38.88 | 0.007963411 | 0.007963411 | ENSG00000151876 | n.s. |
| ENSG00000144681 | 1064.511 | -0.159508231 | 0.264786 | -0.602405 | 0.546904 | 0.891406027 | STAC | protein_coding | 39.86 | -0.007963358 | 0.007963358 | ENSG00000144681 | n.s. |
| ENSG00000143093 | 1543.539 | -0.153240742 | 0.244408 | -0.626987 | 0.530668 | 0.887229181 | STRIP1 | protein_coding | 51.41 | -0.00796303 | 0.00796303 | ENSG00000143093 | n.s. |
| ENSG00000104131 | 2102.522 | -0.163370433 | 0.278545 | -0.586513 | 0.557531 | 0.893837232 | EIF3J | protein_coding | 42.09 | -0.00796293 | 0.00796293 | ENSG00000104131 | n.s. |
| ENSG00000152377 | 1027.86 | -0.152822519 | 0.243013 | -0.628865 | 0.529437 | 0.886954686 | SPOCK1 | protein_coding | 42.8 | -0.007961834 | 0.007961834 | ENSG00000152377 | n.s. |
| ENSG00000149600 | 202.997 | -0.175134273 | 0.313405 | -0.558811 | 0.576291 | 0.900628466 | COMMD7 | protein_coding | 48.22 | -0.007960611 | 0.007960611 | ENSG00000149600 | n.s. |
| ENSG00000072571 | 1119.433 | 0.173015257 | 0.306551 | 0.5643924 | 0.572487 | 0.899492275 | HMMR | protein_coding | 36.97 | 0.007959145 | 0.007959145 | ENSG00000072571 | n.s. |
| ENSG00000117479 | 650.9152 | -0.159392889 | 0.265359 | -0.600669 | 0.548061 | 0.891406027 | SLC19A2 | protein_coding | 38.72 | -0.0079576 | 0.0079576 | ENSG00000117479 | n.s. |
| ENSG00000110344 | 670.2939 | -0.159343788 | 0.264823 | -0.601699 | 0.547375 | 0.891406027 | UBE4A | protein_coding | 40.54 | -0.007955148 | 0.007955148 | ENSG00000110344 | n.s. |
| ENSG00000085274 | 1413.025 | -0.152686017 | 0.242195 | -0.630427 | 0.528416 | 0.886954686 | MYNN | protein_coding | 37.29 | -0.007954723 | 0.007954723 | ENSG00000085274 | n.s. |
| ENSG00000197217 | 1343.376 | -0.159270485 | 0.262271 | -0.607275 | 0.543669 | 0.891406027 | ENTPD4 | protein_coding | 44.43 | -0.007951489 | 0.007951489 | ENSG00000197217 | n.s. |
| ENSG00000183255 | 1479.392 | -0.163556193 | 0.280386 | -0.583325 | 0.559674 | 0.894140829 | PTTG1IP | protein_coding | 51 | -0.007947862 | 0.007947862 | ENSG00000183255 | n.s. |
| ENSG00000177963 | 753.5654 | -0.161907143 | 0.273833 | -0.591262 | 0.554345 | 0.893144718 | RIC8A | protein_coding | 57.49 | -0.007946106 | 0.007946106 | ENSG00000177963 | n.s. |
| ENSG00000103549 | 1530.88 | -0.155662344 | 0.251778 | -0.618253 | 0.536409 | 0.889201649 | RNF40 | protein_coding | 55.81 | -0.007938739 | 0.007938739 | ENSG00000103549 | n.s. |
| ENSG00000146463 | 1096.036 | -0.167188269 | 0.290718 | -0.575087 | 0.565232 | 0.896450345 | ZMYM4 | protein_coding | 38.64 | -0.007937056 | 0.007937056 | ENSG00000146463 | n.s. |
| ENSG00000172977 | 599.4539 | -0.163129107 | 0.279064 | -0.584558 | 0.558845 | 0.894036264 | KAT5 | protein_coding | 55.16 | -0.007935393 | 0.007935393 | ENSG00000172977 | n.s. |
| ENSG00000050426 | 1251.858 | -0.151021363 | 0.236331 | -0.639026 | 0.522806 | 0.886079926 | LETMD1 | protein_coding | 45.34 | -0.007932715 | 0.007932715 | ENSG00000050426 | n.s. |
| ENSG00000234183 | 104.8861 | -0.15883272 | 0.265052 | -0.599251 | 0.549006 | 0.891406027 | LINC01952 | antisense | 49.87 | -0.007929634 | 0.007929634 | ENSG00000234183 | n.s. |
| ENSG00000188641 | 818.1064 | -0.162665065 | 0.277829 | -0.585487 | 0.55822 | 0.893837232 | DPYD | protein_coding | 35.09 | -0.007928549 | 0.007928549 | ENSG00000188641 | n.s. |
| ENSG00000265917 | 25.08322 | -0.190870129 | 0.363974 | -0.524406 | 0.599996 | 0.908868121 | MIR3685 | miRNA | 45.16 | -0.007920944 | 0.007920944 | ENSG00000265917 | n.s. |
| ENSG00000175768 | 1412.932 | 0.150789081 | 0.235776 | 0.6395427 | 0.52247 | 0.886079926 | TOMM5 | protein_coding | 40.98 | 0.007920513 | 0.007920513 | ENSG00000175768 | n.s. |
| ENSG00000233597 | 23.16181 | 0.190299065 | 0.357151 | 0.5328254 | 0.594154 | 0.908656857 | | processed_pseudogene | 52.26 | 0.007916459 | 0.007916459 | ENSG00000233597 | n.s. |
| ENSG00000165512 | 2384.69 | 0.158564436 | 0.261283 | 0.6068688 | 0.543938 | 0.891406027 | ZNF22 | protein_coding | 41.86 | 0.00791624 | 0.00791624 | ENSG00000165512 | n.s. |
| ENSG00000134184 | 377.4124 | -0.15846471 | 0.261717 | -0.60548 | 0.54486 | 0.891406027 | GSTM1 | protein_coding | 46.17 | -0.007911261 | 0.007911261 | ENSG00000134184 | n.s. |
| ENSG00000276791 | 265.9265 | -0.151732026 | 0.240911 | -0.629826 | 0.528808 | 0.886954686 | | lincRNA | 58.86 | -0.007905021 | 0.007905021 | ENSG00000276791 | n.s. |
| ENSG00000120948 | 2133.964 | -0.155986315 | 0.253463 | -0.615421 | 0.538277 | 0.889966281 | TARDBP | protein_coding | 43.58 | -0.007897033 | 0.007897033 | ENSG00000120948 | n.s. |
| ENSG00000162878 | 599.9871 | 0.158156173 | 0.260797 | 0.6064348 | 0.544226 | 0.891406027 | PKDCC | protein_coding | 56.22 | 0.007895857 | 0.007895857 | ENSG00000162878 | n.s. |
| ENSG00000135249 | 629.1537 | -0.157508553 | 0.257697 | -0.611216 | 0.541057 | 0.891073457 | RINT1 | protein_coding | 41.55 | -0.007889051 | 0.007889051 | ENSG00000135249 | n.s. |
| ENSG00000129691 | 741.9651 | -0.157764878 | 0.2616 | -0.603076 | 0.546458 | 0.891406027 | ASH2L | protein_coding | 42.99 | -0.007876322 | 0.007876322 | ENSG00000129691 | n.s. |

| | | | | | | | | | | | | | |
|---|---|---|---|---|---|---|---|---|---|---|---|---|---|
| ENSG00000100220 | 2115.177 | -0.163749215 | 0.282712 | -0.579208 | 0.562449 | 0.895173193 | RTCB | protein_coding | 41.29 | -0.00787518 | 0.00787518 | ENSG00000100220 | n.s. |
| ENSG00000197852 | 1215.088 | -0.158403888 | 0.264728 | -0.598365 | 0.549596 | 0.891861801 | FAM212B | protein_coding | 43.96 | -0.007873059 | 0.007873059 | ENSG00000197852 | n.s. |
| ENSG00000138386 | 1406.827 | -0.162036458 | 0.278213 | -0.582419 | 0.560284 | 0.894198873 | NAB1 | protein_coding | 37.7 | -0.007869443 | 0.007869443 | ENSG00000138386 | n.s. |
| ENSG00000235262 | 23.59225 | -0.190899149 | 0.370722 | -0.514939 | 0.606596 | 0.909522325 | KDM5C-IT1 | sense_intronic | 45.91 | -0.007862494 | 0.007862494 | ENSG00000235262 | n.s. |
| ENSG00000180008 | 2648.837 | -0.16160076 | 0.276871 | -0.583669 | 0.559443 | 0.894140829 | SOCS4 | protein_coding | 37.83 | -0.007852839 | 0.007852839 | ENSG00000180008 | n.s. |
| ENSG00000181274 | 202.5176 | 0.150994993 | 0.240507 | 0.6278193 | 0.530122 | 0.88715322 | FRAT2 | protein_coding | 60.23 | 0.007851946 | 0.007851946 | ENSG00000181274 | n.s. |
| ENSG00000170004 | 6405.468 | -0.149436576 | 0.235065 | -0.635725 | 0.524955 | 0.886111278 | CHD3 | protein_coding | 53.17 | -0.007847174 | 0.007847174 | ENSG00000170004 | n.s. |
| ENSG00000144048 | 584.3107 | -0.166687153 | 0.291013 | -0.572782 | 0.566792 | 0.897367655 | DUSP11 | protein_coding | 40.29 | -0.007839228 | 0.007839228 | ENSG00000144048 | n.s. |
| ENSG00000272841 | 144.2174 | -0.156992865 | 0.258457 | -0.607423 | 0.54357 | 0.891406027 | | antisense | 53.38 | -0.00783778 | 0.00783778 | ENSG00000272841 | n.s. |
| ENSG00000180530 | 3750.798 | 0.160448341 | 0.273202 | 0.587289 | 0.55701 | 0.893643698 | NRIP1 | protein_coding | 36.59 | 0.007835591 | 0.007835591 | ENSG00000180530 | n.s. |
| ENSG00000106263 | 4791.545 | -0.157433011 | 0.262987 | -0.598633 | 0.549417 | 0.891762473 | EIF3B | protein_coding | 51.61 | -0.007832419 | 0.007832419 | ENSG00000106263 | n.s. |
| ENSG00000183856 | 668.0692 | -0.159581946 | 0.269787 | -0.59151 | 0.554179 | 0.893144718 | IQGAP3 | protein_coding | 50.86 | -0.007831989 | 0.007831989 | ENSG00000183856 | n.s. |
| ENSG00000197808 | 110.5944 | -0.165150437 | 0.287551 | -0.574335 | 0.565741 | 0.896600563 | ZNF461 | protein_coding | 40.3 | -0.007828295 | 0.007828295 | ENSG00000197808 | n.s. |
| ENSG00000141378 | 2033.55 | 0.156192634 | 0.255585 | 0.6111189 | 0.541121 | 0.891073457 | PTRH2 | protein_coding | 39.24 | 0.007823141 | 0.007823141 | ENSG00000141378 | n.s. |
| ENSG00000136122 | 690.1851 | -0.143981952 | 0.220755 | -0.652226 | 0.514255 | 0.882409358 | BORA | protein_coding | 36.79 | -0.007822524 | 0.007822524 | ENSG00000136122 | n.s. |
| ENSG00000141384 | 800.9074 | 0.150955826 | 0.241462 | 0.6251737 | 0.531857 | 0.88755367 | TAF4B | protein_coding | 38.65 | 0.007820323 | 0.007820323 | ENSG00000141384 | n.s. |
| ENSG00000119812 | 800.3223 | -0.156636483 | 0.260722 | -0.60078 | 0.547986 | 0.891406027 | FAM98A | protein_coding | 37.55 | -0.007819988 | 0.007819988 | ENSG00000119812 | n.s. |
| ENSG00000145860 | 1860.487 | -0.156625767 | 0.257371 | -0.608561 | 0.542816 | 0.891406027 | RNF145 | protein_coding | 36.97 | -0.007819453 | 0.007819453 | ENSG00000145860 | n.s. |
| ENSG00000261407 | 13.12879 | -0.230227334 | 0.519407 | -0.44325 | 0.657585 | 0.924786541 | | antisense | 52.27 | -0.007818175 | 0.007818175 | ENSG00000261407 | n.s. |
| ENSG00000165525 | 2385.962 | -0.150334941 | 0.239373 | -0.628036 | 0.52998 | 0.88715322 | NEMF | protein_coding | 40.35 | -0.007817622 | 0.007817622 | ENSG00000165525 | n.s. |
| ENSG00000208004 | 3.089536 | 0.265144336 | 0.65627 | 0.4040172 | 0.6862 | 0.934368276 | MIR323B | miRNA | 48.78 | 0.007816963 | 0.007816963 | ENSG00000208004 | n.s. |
| ENSG00000119559 | 68.86648 | -0.163138579 | 0.282694 | -0.577085 | 0.563882 | 0.895611086 | C19orf25 | protein_coding | 62.65 | -0.007811163 | 0.007811163 | ENSG00000119559 | n.s. |
| ENSG00000170275 | 1337.409 | -0.160687594 | 0.275707 | -0.582821 | 0.560014 | 0.894163182 | CRTAP | protein_coding | 44.28 | -0.00780672 | 0.00780672 | ENSG00000170275 | n.s. |
| ENSG00000175334 | 727.7068 | -0.175144969 | 0.316047 | -0.554174 | 0.579459 | 0.902507682 | BANF1 | protein_coding | 58.14 | -0.007802549 | 0.007802549 | ENSG00000175334 | n.s. |
| ENSG00000021762 | 103.6906 | -0.156287006 | 0.259608 | -0.602012 | 0.547166 | 0.891406027 | OSBPL5 | protein_coding | 57 | -0.00780254 | 0.00780254 | ENSG00000021762 | n.s. |
| ENSG00000213024 | 1359.062 | -0.153265806 | 0.248251 | -0.617383 | 0.536982 | 0.889412093 | NUP62 | protein_coding | 54.29 | -0.007800765 | 0.007800765 | ENSG00000213024 | n.s. |
| ENSG00000265623 | 6.228939 | 0.252526927 | 0.602716 | 0.4189817 | 0.67523 | 0.931353024 | MIR3139 | miRNA | 46.05 | 0.007799464 | 0.007799464 | ENSG00000265623 | n.s. |
| ENSG00000087191 | 2366.882 | -0.151128037 | 0.242557 | -0.623062 | 0.533244 | 0.88797399 | PSMC5 | protein_coding | 50.95 | -0.00779817 | 0.00779817 | ENSG00000087191 | n.s. |
| ENSG00000134352 | 1260.149 | -0.152515275 | 0.245485 | -0.621281 | 0.534415 | 0.88909364 | IL6ST | protein_coding | 37.41 | -0.007786286 | 0.007786286 | ENSG00000134352 | n.s. |
| ENSG00000002919 | 401.8431 | -0.158647995 | 0.26867 | -0.590495 | 0.554859 | 0.893144718 | SNX11 | protein_coding | 48 | -0.007786153 | 0.007786153 | ENSG00000002919 | n.s. |
| ENSG00000136807 | 499.5758 | -0.147151768 | 0.229272 | -0.641822 | 0.520989 | 0.885523197 | CDK9 | protein_coding | 57.43 | -0.007769622 | 0.007769622 | ENSG00000136807 | n.s. |
| ENSG00000154978 | 365.2743 | -0.167641265 | 0.295124 | -0.568036 | 0.57001 | 0.898818733 | VOPP1 | protein_coding | 46.42 | -0.007766465 | 0.007766465 | ENSG00000154978 | n.s. |

| | | | | | | | | | | | | | |
|---|---|---|---|---|---|---|---|---|---|---|---|---|---|
| ENSG00000134057 | 2867.177 | 0.16763484 | 0.295823 | 0.5666724 | 0.570937 | 0.898818733 | CCNB1 | protein_coding | 43.63 | 0.007766168 | 0.007766168 | ENSG00000134057 | n.s. |
| ENSG00000150433 | 327.9328 | -0.167606078 | 0.295707 | -0.566798 | 0.570851 | 0.898818733 | TMEM218 | protein_coding | 41.01 | -0.007764835 | 0.007764835 | ENSG00000150433 | n.s. |
| ENSG00000152256 | 922.8508 | -0.155494527 | 0.257597 | -0.603636 | 0.546086 | 0.891406027 | PDK1 | protein_coding | 39.52 | -0.007762976 | 0.007762976 | ENSG00000152256 | n.s. |
| ENSG00000132004 | 76.71212 | -0.16997503 | 0.302929 | -0.561106 | 0.574725 | 0.900202572 | FBXW9 | protein_coding | 56.55 | -0.007761018 | 0.007761018 | ENSG00000132004 | n.s. |
| ENSG00000236535 | 120.7642 | 0.163462269 | 0.284344 | 0.5748747 | 0.565376 | 0.896450345 | RC3H1-IT1 | sense_intronic | 39.39 | 0.007760169 | 0.007760169 | ENSG00000236535 | n.s. |
| ENSG00000136273 | 540.3487 | -0.148940859 | 0.236194 | -0.630588 | 0.52831 | 0.886954686 | HUS1 | protein_coding | 44.35 | -0.007759605 | 0.007759605 | ENSG00000136273 | n.s. |
| ENSG00000185808 | 160.4758 | 0.160665955 | 0.276571 | 0.5809222 | 0.561293 | 0.894765412 | PIGP | protein_coding | 42.2 | 0.00775869 | 0.00775869 | ENSG00000185808 | n.s. |
| ENSG00000225808 | 61.14599 | 0.228393605 | 0.517154 | 0.4416352 | 0.658753 | 0.924786541 | DNAJC19P5 | processed_pseudogene | 41.14 | 0.007755904 | 0.007755904 | ENSG00000225808 | n.s. |
| ENSG00000171202 | 294.9323 | 0.167151874 | 0.294819 | 0.5669648 | 0.570738 | 0.898818733 | TMEM126A | protein_coding | 37.69 | 0.007743793 | 0.007743793 | ENSG00000171202 | n.s. |
| ENSG00000260806 | 104.536 | 0.155107128 | 0.257704 | 0.6018813 | 0.547253 | 0.891406027 | | antisense | 52.38 | 0.007743636 | 0.007743636 | ENSG00000260806 | n.s. |
| ENSG00000168734 | 156.4242 | -0.183147768 | 0.341114 | -0.536911 | 0.591329 | 0.907325858 | PKIG | protein_coding | 44.32 | -0.00773556 | 0.00773556 | ENSG00000168734 | n.s. |
| ENSG00000100353 | 4209.246 | 0.158668671 | 0.270591 | 0.5863772 | 0.557622 | 0.893837232 | EIF3D | protein_coding | 47.04 | 0.007733758 | 0.007733758 | ENSG00000100353 | n.s. |
| ENSG00000182768 | 6157.16 | -0.167938429 | 0.29728 | -0.564916 | 0.572131 | 0.899491841 | NGRN | protein_coding | 45.25 | -0.007725633 | 0.007725633 | ENSG00000182768 | n.s. |
| ENSG00000176946 | 598.9106 | -0.158467374 | 0.270119 | -0.586657 | 0.557434 | 0.893837232 | THAP4 | protein_coding | 49.81 | -0.007723947 | 0.007723947 | ENSG00000176946 | n.s. |
| ENSG00000115271 | 322.5412 | 0.162688662 | 0.282884 | 0.5751077 | 0.565218 | 0.896450345 | GCA | protein_coding | 36.25 | 0.007723443 | 0.007723443 | ENSG00000115271 | n.s. |
| ENSG00000117362 | 1453.597 | -0.155385875 | 0.259988 | -0.597665 | 0.550063 | 0.891877082 | APH1A | protein_coding | 53.83 | -0.0077219 | 0.0077219 | ENSG00000117362 | n.s. |
| ENSG00000189042 | 544.9392 | -0.160694743 | 0.277578 | -0.578917 | 0.562645 | 0.895317554 | ZNF567 | protein_coding | 40.54 | -0.007717028 | 0.007717028 | ENSG00000189042 | n.s. |
| ENSG00000197930 | 958.0949 | -0.153134611 | 0.249783 | -0.61307 | 0.53983 | 0.890684245 | ERO1A | protein_coding | 37.59 | -0.007699031 | 0.007699031 | ENSG00000197930 | n.s. |
| ENSG00000146386 | 402.7556 | -0.177622598 | 0.324879 | -0.546735 | 0.584561 | 0.905026039 | ABRACL | protein_coding | 44.2 | -0.007697972 | 0.007697972 | ENSG00000146386 | n.s. |
| ENSG00000166263 | 913.7719 | -0.158350042 | 0.271554 | -0.583125 | 0.559809 | 0.894140829 | STXBP4 | protein_coding | 36.22 | -0.007694874 | 0.007694874 | ENSG00000166263 | n.s. |
| ENSG00000090889 | 1365.083 | -0.157860927 | 0.269751 | -0.585209 | 0.558407 | 0.893837232 | KIF4A | protein_coding | 40.48 | -0.007694388 | 0.007694388 | ENSG00000090889 | n.s. |
| ENSG00000162300 | 277.4303 | -0.154120644 | 0.253717 | -0.60745 | 0.543552 | 0.891406027 | ZFPL1 | protein_coding | 56.15 | -0.007694386 | 0.007694386 | ENSG00000162300 | n.s. |
| ENSG00000185008 | 684.4412 | 0.158356111 | 0.271708 | 0.5828177 | 0.560016 | 0.894163182 | ROBO2 | protein_coding | 35.65 | 0.007693449 | 0.007693449 | ENSG00000185008 | n.s. |
| ENSG00000250299 | 96.52115 | -0.157755834 | 0.269623 | -0.585099 | 0.558481 | 0.893837232 | MRPS31P4 | cribed_unprocessed_pseudc | 41.83 | -0.007689265 | 0.007689265 | ENSG00000250299 | n.s. |
| ENSG00000170191 | 429.9734 | -0.156631704 | 0.263972 | -0.593365 | 0.552937 | 0.893144718 | NANP | protein_coding | 43.71 | -0.007687197 | 0.007687197 | ENSG00000170191 | n.s. |
| ENSG00000111641 | 1259.307 | -0.153960496 | 0.254885 | -0.60404 | 0.545817 | 0.891406027 | NOP2 | protein_coding | 49.45 | -0.007686391 | 0.007686391 | ENSG00000111641 | n.s. |
| ENSG00000182405 | 286.2695 | 0.141466017 | 0.216891 | 0.6522461 | 0.514242 | 0.882409358 | PGBD4 | protein_coding | 42.18 | 0.007685834 | 0.007685834 | ENSG00000182405 | n.s. |
| ENSG00000200279 | 13.44253 | 0.209513862 | 0.438766 | 0.4775069 | 0.633001 | 0.919033489 | SNORD114-10 | snoRNA | 36.11 | 0.007682593 | 0.007682593 | ENSG00000200279 | n.s. |
| ENSG00000090061 | 220.4184 | -0.153797014 | 0.256564 | -0.599448 | 0.548874 | 0.891406027 | CCNK | protein_coding | 50.14 | -0.007678229 | 0.007678229 | ENSG00000090061 | n.s. |
| ENSG00000252614 | 42.26446 | -0.172492651 | 0.311428 | -0.553877 | 0.579663 | 0.902582314 | RNU6-807P | snRNA | 35.51 | -0.007678197 | 0.007678197 | ENSG00000252614 | n.s. |
| ENSG00000143061 | 2445.476 | -0.153785771 | 0.255989 | -0.600751 | 0.548006 | 0.891406027 | IGSF3 | protein_coding | 46.14 | -0.007677668 | 0.007677668 | ENSG00000143061 | n.s. |
| ENSG00000251247 | 30.67422 | -0.184971157 | 0.348465 | -0.530816 | 0.595546 | 0.908868121 | ZNF345 | protein_coding | 39.29 | -0.007676142 | 0.007676142 | ENSG00000251247 | n.s. |

| | | | | | | | | | | | | | |
|---|---|---|---|---|---|---|---|---|---|---|---|---|---|
| ENSG00000055332 | 1491.482 | -0.157463672 | 0.268606 | -0.586224 | 0.557725 | 0.893837232 | EIF2AK2 | protein_coding | 41.14 | -0.007675025 | 0.007675025 | ENSG00000055332 | n.s. |
| ENSG00000170802 | 1178.457 | -0.151566332 | 0.246188 | -0.615652 | 0.538124 | 0.889966281 | FOXN2 | protein_coding | 36.31 | -0.007673265 | 0.007673265 | ENSG00000170802 | n.s. |
| ENSG00000135052 | 1899.577 | -0.157385691 | 0.268887 | -0.585323 | 0.55833 | 0.893837232 | GOLM1 | protein_coding | 45.43 | -0.007671224 | 0.007671224 | ENSG00000135052 | n.s. |
| ENSG00000280365 | 54.26002 | -0.162776596 | 0.284029 | -0.573098 | 0.566579 | 0.897233219 | | TEC | 46.56 | -0.007665908 | 0.007665908 | ENSG00000280365 | n.s. |
| ENSG00000186222 | 301.4428 | 0.147012056 | 0.233711 | 0.629034 | 0.529327 | 0.886954686 | BLOC1S4 | protein_coding | 54.05 | 0.007659118 | 0.007659118 | ENSG00000186222 | n.s. |
| ENSG00000153975 | 815.098 | -0.15525963 | 0.26084 | -0.595228 | 0.551691 | 0.892641916 | ZUFSP | protein_coding | 36.54 | -0.007657828 | 0.007657828 | ENSG00000153975 | n.s. |
| ENSG00000243927 | 361.606 | 0.176649002 | 0.324336 | 0.5446488 | 0.585995 | 0.905026039 | MRPS6 | protein_coding | 40.3 | 0.007655778 | 0.007655778 | ENSG00000243927 | n.s. |
| ENSG00000134440 | 4916.569 | -0.161175009 | 0.279988 | -0.57565 | 0.564852 | 0.896450345 | NARS | protein_coding | 42.81 | -0.007651584 | 0.007651584 | ENSG00000134440 | n.s. |
| ENSG00000167523 | 161.8962 | -0.149837136 | 0.241261 | -0.621058 | 0.534562 | 0.88909364 | SPATA33 | protein_coding | 52.08 | -0.00764956 | 0.00764956 | ENSG00000167523 | n.s. |
| ENSG00000138660 | 781.6231 | -0.166932315 | 0.296698 | -0.562633 | 0.573685 | 0.899911687 | AP1AR | protein_coding | 36.49 | -0.007645518 | 0.007645518 | ENSG00000138660 | n.s. |
| ENSG00000120306 | 76.81926 | -0.184176119 | 0.353624 | -0.520824 | 0.602489 | 0.908868121 | CYSTM1 | protein_coding | 44.09 | -0.007643149 | 0.007643149 | ENSG00000120306 | n.s. |
| ENSG00000168575 | 400.1438 | 0.14553521 | 0.228349 | 0.6373375 | 0.523905 | 0.886111278 | SLC20A2 | protein_coding | 44.17 | 0.007642307 | 0.007642307 | ENSG00000168575 | n.s. |
| ENSG00000111011 | 7780.406 | 0.142453643 | 0.220166 | 0.6470285 | 0.517613 | 0.883822248 | RSRC2 | protein_coding | 39.85 | 0.007640511 | 0.007640511 | ENSG00000111011 | n.s. |
| ENSG00000100802 | 95.36222 | 0.16161224 | 0.281686 | 0.5737318 | 0.566149 | 0.896859308 | C14orf93 | protein_coding | 45.67 | 0.007640329 | 0.007640329 | ENSG00000100802 | n.s. |
| ENSG00000196132 | 422.7683 | 0.166795351 | 0.296716 | 0.5621372 | 0.574023 | 0.899911687 | MYT1 | protein_coding | 54.58 | 0.007639245 | 0.007639245 | ENSG00000196132 | n.s. |
| ENSG00000136111 | 458.5957 | -0.157202935 | 0.27001 | -0.582212 | 0.560424 | 0.894198873 | TBC1D4 | protein_coding | 38.08 | -0.007634699 | 0.007634699 | ENSG00000136111 | n.s. |
| ENSG00000168615 | 847.7141 | -0.154341823 | 0.258767 | -0.59645 | 0.550874 | 0.89235881 | ADAM9 | protein_coding | 37.5 | -0.007633821 | 0.007633821 | ENSG00000168615 | n.s. |
| ENSG00000252690 | 431.6255 | -0.146380101 | 0.231806 | -0.631478 | 0.527728 | 0.886936263 | SCARNA15 | processed_transcript | 38.13 | -0.007627514 | 0.007627514 | ENSG00000252690 | n.s. |
| ENSG00000163597 | 11417.99 | -0.138230179 | 0.210741 | -0.655924 | 0.511873 | 0.880722549 | SNHG16 | processed_transcript | 45.22 | -0.007624899 | 0.007624899 | ENSG00000163597 | n.s. |
| ENSG00000154511 | 494.415 | 0.175838317 | 0.322477 | 0.5452744 | 0.585565 | 0.905026039 | FAM69A | protein_coding | 39.43 | 0.007620644 | 0.007620644 | ENSG00000154511 | n.s. |
| ENSG00000183098 | 931.7912 | -0.152627296 | 0.251852 | -0.606019 | 0.544502 | 0.891406027 | GPC6 | protein_coding | 37.15 | -0.007619831 | 0.007619831 | ENSG00000183098 | n.s. |
| ENSG00000266415 | 10.93632 | -0.207529212 | 0.432842 | -0.479457 | 0.631613 | 0.919033489 | MIR4636 | miRNA | 45 | -0.007609819 | 0.007609819 | ENSG00000266415 | n.s. |
| ENSG00000124444 | 254.9221 | -0.148132826 | 0.238079 | -0.6222 | 0.53381 | 0.888492562 | ZNF576 | protein_coding | 50.29 | -0.007606058 | 0.007606058 | ENSG00000124444 | n.s. |
| ENSG00000227627 | 243.0139 | -0.145965092 | 0.231956 | -0.629279 | 0.529166 | 0.886954686 | | antisense | 42.78 | -0.007604572 | 0.007604572 | ENSG00000227627 | n.s. |
| ENSG00000131435 | 723.7803 | -0.165274503 | 0.292897 | -0.564275 | 0.572567 | 0.899492275 | PDLIM4 | protein_coding | 56.99 | -0.007603051 | 0.007603051 | ENSG00000131435 | n.s. |
| ENSG00000172687 | 1128.463 | -0.150150073 | 0.244022 | -0.615314 | 0.538347 | 0.889966281 | ZNF738 | protein_coding | 40.08 | -0.007601565 | 0.007601565 | ENSG00000172687 | n.s. |
| ENSG00000198937 | 297.4964 | -0.179105053 | 0.332959 | -0.53792 | 0.590632 | 0.906900529 | CCDC167 | protein_coding | 52.31 | -0.00760128 | 0.00760128 | ENSG00000198937 | n.s. |
| ENSG00000205423 | 336.8298 | -0.149251815 | 0.241882 | -0.617043 | 0.537206 | 0.889417954 | CNEP1R1 | protein_coding | 40.53 | -0.007596038 | 0.007596038 | ENSG00000205423 | n.s. |
| ENSG00000154767 | 941.762 | -0.16391814 | 0.288162 | -0.56884 | 0.569465 | 0.898810685 | XPC | protein_coding | 45.2 | -0.007594618 | 0.007594618 | ENSG00000154767 | n.s. |
| ENSG00000077782 | 192.5599 | 0.154723595 | 0.261564 | 0.5915321 | 0.554164 | 0.893144718 | FGFR1 | protein_coding | 50.55 | 0.00759355 | 0.00759355 | ENSG00000077782 | n.s. |
| ENSG00000241553 | 430.9944 | -0.155712368 | 0.265728 | -0.585984 | 0.557886 | 0.893837232 | ARPC4 | protein_coding | 48.44 | -0.007589664 | 0.007589664 | ENSG00000241553 | n.s. |
| ENSG00000267871 | 29.48553 | 0.18287566 | 0.344275 | 0.5311903 | 0.595287 | 0.908868121 | | antisense | 45.15 | 0.007589181 | 0.007589181 | ENSG00000267871 | n.s. |

| | | | | | | | | | | | | | |
|---|---|---|---|---|---|---|---|---|---|---|---|---|---|
| ENSG00000207223 | 12.31285 | -0.257844194 | 0.641343 | -0.402038 | 0.687656 | 0.934506008 | | misc_RNA | 39.22 | -0.007585235 | 0.007585235 | ENSG00000207223 | n.s. |
| ENSG00000143157 | 2820.97 | -0.154552636 | 0.261175 | -0.591758 | 0.554013 | 0.893144718 | POGK | protein_coding | 47.02 | -0.00758516 | 0.00758516 | ENSG00000143157 | n.s. |
| ENSG00000017483 | 475.3619 | 0.169193842 | 0.304659 | 0.5553547 | 0.578652 | 0.901989749 | SLC38A5 | protein_coding | 54.89 | 0.007579613 | 0.007579613 | ENSG00000017483 | n.s. |
| ENSG00000198794 | 1023.42 | -0.151815813 | 0.249776 | -0.607808 | 0.543315 | 0.891406027 | SCAMP5 | protein_coding | 48.31 | -0.007579319 | 0.007579319 | ENSG00000198794 | n.s. |
| ENSG00000205978 | 467.8468 | -0.151313171 | 0.247775 | -0.610687 | 0.541407 | 0.891123173 | NYNRIN | protein_coding | 52.95 | -0.00757508 | 0.00757508 | ENSG00000205978 | n.s. |
| ENSG00000122378 | 2606.265 | -0.162770949 | 0.285639 | -0.569849 | 0.56878 | 0.898419074 | FAM213A | protein_coding | 47.5 | -0.007572273 | 0.007572273 | ENSG00000122378 | n.s. |
| ENSG00000243176 | 31.85165 | 0.182463772 | 0.346694 | 0.5262959 | 0.598683 | 0.908868121 | | processed_transcript | 39.16 | 0.007572088 | 0.007572088 | ENSG00000243176 | n.s. |
| ENSG00000117153 | 1066.89 | -0.15733235 | 0.271261 | -0.580003 | 0.561913 | 0.895116382 | KLHL12 | protein_coding | 41.92 | -0.00757091 | 0.00757091 | ENSG00000117153 | n.s. |
| ENSG00000177728 | 62.38429 | -0.162658489 | 0.284924 | -0.570885 | 0.568078 | 0.898381247 | TMEM94 | protein_coding | 51.96 | -0.007570016 | 0.007570016 | ENSG00000177728 | n.s. |
| ENSG00000115275 | 428.5312 | -0.154229426 | 0.261969 | -0.588732 | 0.556041 | 0.893144718 | MOGS | protein_coding | 57.9 | -0.007569297 | 0.007569297 | ENSG00000115275 | n.s. |
| ENSG00000149131 | 187.2723 | -0.155200656 | 0.264826 | -0.586048 | 0.557843 | 0.893837232 | SERPING1 | protein_coding | 47.51 | -0.007564722 | 0.007564722 | ENSG00000149131 | n.s. |
| ENSG00000107021 | 225.6423 | -0.151027504 | 0.247096 | -0.611209 | 0.541061 | 0.891073457 | TBC1D13 | protein_coding | 52.07 | -0.007564438 | 0.007564438 | ENSG00000107021 | n.s. |
| ENSG00000135968 | 1630.059 | -0.156441835 | 0.269251 | -0.581026 | 0.561223 | 0.894765412 | GCC2 | protein_coding | 38.51 | -0.007554703 | 0.007554703 | ENSG00000135968 | n.s. |
| ENSG00000139737 | 2640.071 | 0.1512568 | 0.248101 | 0.6096579 | 0.542088 | 0.891404544 | SLAIN1 | protein_coding | 36.78 | 0.007551519 | 0.007551519 | ENSG00000139737 | n.s. |
| ENSG00000164751 | 586.4193 | -0.16465667 | 0.292847 | -0.562261 | 0.573938 | 0.899911687 | PEX2 | protein_coding | 38.37 | -0.007541293 | 0.007541293 | ENSG00000164751 | n.s. |
| ENSG00000101811 | 668.7545 | -0.154666803 | 0.264169 | -0.585485 | 0.558222 | 0.893837232 | CSTF2 | protein_coding | 38.87 | -0.007538701 | 0.007538701 | ENSG00000101811 | n.s. |
| ENSG00000260267 | 337.1684 | -0.138641034 | 0.212175 | -0.653427 | 0.513481 | 0.882332213 | | antisense | 59.88 | -0.007537617 | 0.007537617 | ENSG00000260267 | n.s. |
| ENSG00000064655 | 804.1317 | 0.157817163 | 0.273751 | 0.5764989 | 0.564278 | 0.89593377 | EYA2 | protein_coding | 45.44 | 0.007531681 | 0.007531681 | ENSG00000064655 | n.s. |
| ENSG00000160285 | 1062.783 | -0.150738096 | 0.248813 | -0.60583 | 0.544628 | 0.891406027 | LSS | protein_coding | 53.33 | -0.007525514 | 0.007525514 | ENSG00000160285 | n.s. |
| ENSG00000104356 | 1071.636 | -0.15688701 | 0.271432 | -0.577997 | 0.563266 | 0.895435774 | POP1 | protein_coding | 41.8 | -0.007525173 | 0.007525173 | ENSG00000104356 | n.s. |
| ENSG00000196510 | 1922.821 | -0.164287667 | 0.291903 | -0.562815 | 0.573561 | 0.899911687 | ANAPC7 | protein_coding | 45.87 | -0.007524393 | 0.007524393 | ENSG00000196510 | n.s. |
| ENSG00000243370 | 17.8457 | -0.184603405 | 0.361185 | -0.511104 | 0.609278 | 0.910431619 | RN7SL775P | misc_RNA | 53.57 | -0.007523081 | 0.007523081 | ENSG00000243370 | n.s. |
| ENSG00000183808 | 2589.542 | -0.150687794 | 0.248113 | -0.607336 | 0.543628 | 0.891406027 | RBM12B | protein_coding | 39 | -0.007523003 | 0.007523003 | ENSG00000183808 | n.s. |
| ENSG00000164051 | 244.8193 | -0.170169032 | 0.308388 | -0.551802 | 0.581084 | 0.903352939 | CCDC51 | protein_coding | 51.92 | -0.007511693 | 0.007511693 | ENSG00000164051 | n.s. |
| ENSG00000105364 | 346.8902 | -0.167937879 | 0.302622 | -0.554942 | 0.578934 | 0.902167117 | MRPL4 | protein_coding | 56.13 | -0.007509007 | 0.007509007 | ENSG00000105364 | n.s. |
| ENSG00000146670 | 1348.479 | -0.152786628 | 0.259456 | -0.588874 | 0.555946 | 0.893144718 | CDCA5 | protein_coding | 52.67 | -0.007498487 | 0.007498487 | ENSG00000146670 | n.s. |
| ENSG00000114388 | 167.2174 | 0.174782084 | 0.323968 | 0.5395048 | 0.589539 | 0.906089161 | NPRL2 | protein_coding | 56.73 | 0.007485753 | 0.007485753 | ENSG00000114388 | n.s. |
| ENSG00000259623 | 548.0957 | -0.152409856 | 0.258866 | -0.58876 | 0.556022 | 0.893144718 | | sense_overlapping | 34.51 | -0.007479996 | 0.007479996 | ENSG00000259623 | n.s. |
| ENSG00000144362 | 397.0741 | 0.149610041 | 0.249361 | 0.5999736 | 0.548524 | 0.891406027 | PHOSPHO2 | protein_coding | 37.24 | 0.007469197 | 0.007469197 | ENSG00000144362 | n.s. |
| ENSG00000164241 | 108.4163 | -0.149606436 | 0.24876 | -0.601409 | 0.547568 | 0.891406027 | C5orf63 | protein_coding | 39.03 | -0.007469017 | 0.007469017 | ENSG00000164241 | n.s. |
| ENSG00000163832 | 952.1662 | -0.146681787 | 0.237546 | -0.617488 | 0.536913 | 0.889412093 | ELP6 | protein_coding | 48.73 | -0.007465658 | 0.007465658 | ENSG00000163832 | n.s. |
| ENSG00000186298 | 5053.759 | 0.155602175 | 0.269151 | 0.578123 | 0.563181 | 0.895435774 | PPP1CC | protein_coding | 41.52 | 0.007463545 | 0.007463545 | ENSG00000186298 | n.s. |

| | | | | | | | | | | | | | |
|---|---|---|---|---|---|---|---|---|---|---|---|---|---|
| ENSG00000008838 | 828.9734 | -0.149407475 | 0.245038 | -0.609731 | 0.54204 | 0.891404544 | MED24 | protein_coding | 50.63 | -0.007459192 | 0.007459192 | ENSG00000008838 | n.s. |
| ENSG00000004455 | 1354.472 | -0.163662114 | 0.292604 | -0.55933 | 0.575937 | 0.900580379 | AK2 | protein_coding | 46.32 | -0.007442947 | 0.007442947 | ENSG00000004455 | n.s. |
| ENSG00000206756 | 8.284586 | -0.214422878 | 0.466977 | -0.459172 | 0.646111 | 0.923184844 | | misc_RNA | 41.18 | -0.007442904 | 0.007442904 | ENSG00000206756 | n.s. |
| ENSG00000137509 | 414.1493 | 0.17150852 | 0.31396 | 0.5462758 | 0.584876 | 0.905026039 | PRCP | protein_coding | 39.37 | 0.007432995 | 0.007432995 | ENSG00000137509 | n.s. |
| ENSG00000144746 | 791.3884 | 0.177022468 | 0.330334 | 0.5358897 | 0.592035 | 0.907847371 | ARL6IP5 | protein_coding | 42.5 | 0.007432671 | 0.007432671 | ENSG00000144746 | n.s. |
| ENSG00000244346 | 46.68412 | -0.169980368 | 0.309898 | -0.548504 | 0.583346 | 0.90424479 | | processed_pseudogene | 39.53 | -0.007430519 | 0.007430519 | ENSG00000244346 | n.s. |
| ENSG00000099331 | 431.061 | -0.148708767 | 0.247381 | -0.601133 | 0.547751 | 0.891406027 | MYO9B | protein_coding | 52.36 | -0.007424201 | 0.007424201 | ENSG00000099331 | n.s. |
| ENSG00000024526 | 2754.424 | -0.171223798 | 0.313479 | -0.546206 | 0.584924 | 0.905026039 | DEPDC1 | protein_coding | 34.94 | -0.007420655 | 0.007420655 | ENSG00000024526 | n.s. |
| ENSG00000106554 | 1551.235 | -0.162004458 | 0.287958 | -0.562597 | 0.573709 | 0.899911687 | CHCHD3 | protein_coding | 39.02 | -0.007419822 | 0.007419822 | ENSG00000106554 | n.s. |
| ENSG00000134248 | 1692.605 | 0.156114688 | 0.271366 | 0.5752915 | 0.565094 | 0.896450345 | LAMTOR5 | protein_coding | 42.55 | 0.007411351 | 0.007411351 | ENSG00000134248 | n.s. |
| ENSG00000123505 | 6142.792 | -0.148317423 | 0.247449 | -0.599386 | 0.548915 | 0.891406027 | AMD1 | protein_coding | 41.53 | -0.007404663 | 0.007404663 | ENSG00000123505 | n.s. |
| ENSG00000111653 | 386.9035 | 0.178316289 | 0.335478 | 0.5315295 | 0.595052 | 0.908868121 | ING4 | protein_coding | 47.73 | 0.007399971 | 0.007399971 | ENSG00000111653 | n.s. |
| ENSG00000270885 | 293.8424 | -0.152308865 | 0.261265 | -0.582967 | 0.559916 | 0.894163182 | RASL10B | protein_coding | 54.98 | -0.007399655 | 0.007399655 | ENSG00000270885 | n.s. |
| ENSG00000182670 | 7901.75 | -0.151496941 | 0.257992 | -0.587215 | 0.557059 | 0.893643698 | TTC3 | protein_coding | 38.7 | -0.007398444 | 0.007398444 | ENSG00000182670 | n.s. |
| ENSG00000142156 | 376.6112 | -0.151692663 | 0.259317 | -0.584969 | 0.558569 | 0.893837232 | COL6A1 | protein_coding | 64.67 | -0.007393737 | 0.007393737 | ENSG00000142156 | n.s. |
| ENSG00000102393 | 256.3698 | 0.160674361 | 0.284319 | 0.5651192 | 0.571993 | 0.899491841 | GLA | protein_coding | 43.7 | 0.007391466 | 0.007391466 | ENSG00000102393 | n.s. |
| ENSG00000204152 | 566.9179 | -0.152981698 | 0.263304 | -0.581008 | 0.561235 | 0.894765412 | TIMM23B | protein_coding | 40.65 | -0.00738761 | 0.00738761 | ENSG00000204152 | n.s. |
| ENSG00000266270 | 2.124265 | -0.491106099 | 2.036471 | -0.241155 | 0.809435 | 0.965963425 | MIR5096 | miRNA | 54.29 | -0.0073859 | 0.0073859 | ENSG00000266270 | n.s. |
| ENSG00000181652 | 116.347 | 0.151989354 | 0.260316 | 0.5838652 | 0.559311 | 0.894140829 | ATG9B | protein_coding | 58.12 | 0.007385782 | 0.007385782 | ENSG00000181652 | n.s. |
| ENSG00000151503 | 1728.182 | -0.150485013 | 0.25383 | -0.592857 | 0.553277 | 0.893144718 | NCAPD3 | protein_coding | 43.33 | -0.007385528 | 0.007385528 | ENSG00000151503 | n.s. |
| ENSG00000182165 | 71.5777 | -0.170863907 | 0.315394 | -0.541748 | 0.587992 | 0.905341301 | TP53TG1 | lincRNA | 38.94 | -0.007379214 | 0.007379214 | ENSG00000182165 | n.s. |
| ENSG00000164164 | 1705.267 | -0.145728989 | 0.237197 | -0.61438 | 0.538964 | 0.889966281 | OTUD4 | protein_coding | 37.41 | -0.007377741 | 0.007377741 | ENSG00000164164 | n.s. |
| ENSG00000091732 | 458.6322 | -0.153753108 | 0.265999 | -0.578022 | 0.563249 | 0.895435774 | ZC3HC1 | protein_coding | 44.46 | -0.007374854 | 0.007374854 | ENSG00000091732 | n.s. |
| ENSG00000166164 | 2273.285 | -0.147623049 | 0.242219 | -0.609462 | 0.542218 | 0.891404544 | BRD7 | protein_coding | 39.72 | -0.007370104 | 0.007370104 | ENSG00000166164 | n.s. |
| ENSG00000122566 | 20932.42 | -0.140294943 | 0.221236 | -0.634141 | 0.525989 | 0.886111278 | HNRNPA2B1 | protein_coding | 40.07 | -0.007367131 | 0.007367131 | ENSG00000122566 | n.s. |
| ENSG00000145050 | 474.958 | -0.151857043 | 0.260945 | -0.581951 | 0.5606 | 0.894377272 | MANF | protein_coding | 52.06 | -0.007361915 | 0.007361915 | ENSG00000145050 | n.s. |
| ENSG00000073803 | 469.4485 | -0.167016902 | 0.303581 | -0.550156 | 0.582212 | 0.90349457 | MAP3K13 | protein_coding | 40.07 | -0.007361179 | 0.007361179 | ENSG00000073803 | n.s. |
| ENSG00000158669 | 1861.015 | -0.150941261 | 0.25735 | -0.586521 | 0.557525 | 0.893837232 | GPAT4 | protein_coding | 45.7 | -0.007357112 | 0.007357112 | ENSG00000158669 | n.s. |
| ENSG00000185565 | 1335.549 | -0.149902679 | 0.254642 | -0.588681 | 0.556075 | 0.893144718 | LSAMP | protein_coding | 37.2 | -0.007356949 | 0.007356949 | ENSG00000185565 | n.s. |
| ENSG00000126777 | 5218.033 | -0.151373256 | 0.259985 | -0.582239 | 0.560406 | 0.894198873 | KTN1 | protein_coding | 37.04 | -0.007351576 | 0.007351576 | ENSG00000126777 | n.s. |
| ENSG00000158186 | 156.9193 | -0.147199424 | 0.244916 | -0.60102 | 0.547826 | 0.891406027 | MRAS | protein_coding | 48.52 | -0.007348848 | 0.007348848 | ENSG00000158186 | n.s. |
| ENSG00000135164 | 3376.079 | -0.149717292 | 0.253434 | -0.590755 | 0.554684 | 0.893144718 | DMTF1 | protein_coding | 35.84 | -0.00734785 | 0.00734785 | ENSG00000135164 | n.s. |

| | | | | | | | | | | | | | |
|---|---|---|---|---|---|---|---|---|---|---|---|---|---|
| ENSG00000207445 | 126.3153 | 0.146953951 | 0.241955 | 0.6073618 | 0.543611 | 0.891406027 | SNORD15B | snoRNA | 47.95 | 0.007336593 | 0.007336593 | ENSG00000207445 | n.s. |
| ENSG00000122257 | 2569.497 | -0.150455263 | 0.257111 | -0.585177 | 0.558429 | 0.893837232 | RBBP6 | protein_coding | 37.61 | -0.007333424 | 0.007333424 | ENSG00000122257 | n.s. |
| ENSG00000106460 | 1152.59 | -0.158227946 | 0.278316 | -0.568518 | 0.569683 | 0.898818733 | TMEM106B | protein_coding | 34.61 | -0.007330366 | 0.007330366 | ENSG00000106460 | n.s. |
| ENSG00000278356 | 33.84736 | -0.176607967 | 0.333433 | -0.529666 | 0.596343 | 0.908868121 | | sense_intronic | 44.42 | -0.007329077 | 0.007329077 | ENSG00000278356 | n.s. |
| ENSG00000198899 | 1859.593 | -0.139479786 | 0.220226 | -0.633349 | 0.526506 | 0.88615396 | MT-ATP6 | protein_coding | 44.2 | -0.007321408 | 0.007321408 | ENSG00000198899 | n.s. |
| ENSG00000142494 | 181.343 | -0.152594534 | 0.264004 | -0.578 | 0.563264 | 0.895435774 | SLC47A1 | protein_coding | 47.07 | -0.007319282 | 0.007319282 | ENSG00000142494 | n.s. |
| ENSG00000178403 | 1439.425 | 0.129061288 | 0.194407 | 0.6638721 | 0.506772 | 0.877585745 | NEUROG2 | protein_coding | 56.57 | 0.007319122 | 0.007319122 | ENSG00000178403 | n.s. |
| ENSG00000115694 | 1671.23 | -0.157258641 | 0.275865 | -0.570057 | 0.568639 | 0.898419074 | STK25 | protein_coding | 59.31 | -0.007315835 | 0.007315835 | ENSG00000115694 | n.s. |
| ENSG00000065057 | 309.1555 | 0.176215501 | 0.333093 | 0.5290278 | 0.596786 | 0.908868121 | NTHL1 | protein_coding | 61.44 | 0.00731279 | 0.00731279 | ENSG00000065057 | n.s. |
| ENSG00000078808 | 312.6976 | -0.160564285 | 0.286846 | -0.559757 | 0.575645 | 0.900453394 | SDF4 | protein_coding | 61.35 | -0.007311899 | 0.007311899 | ENSG00000078808 | n.s. |
| ENSG00000107186 | 2079.536 | -0.157676991 | 0.277069 | -0.56909 | 0.569295 | 0.898775285 | MPDZ | protein_coding | 35.81 | -0.007308152 | 0.007308152 | ENSG00000107186 | n.s. |
| ENSG00000131828 | 1566.85 | -0.157699978 | 0.277589 | -0.568105 | 0.569964 | 0.898818733 | PDHA1 | protein_coding | 44.45 | -0.007305906 | 0.007305906 | ENSG00000131828 | n.s. |
| ENSG00000204590 | 343.8619 | -0.15984944 | 0.284701 | -0.561465 | 0.574481 | 0.900202572 | GNL1 | protein_coding | 49.56 | -0.007298686 | 0.007298686 | ENSG00000204590 | n.s. |
| ENSG00000160803 | 646.9803 | -0.153877699 | 0.268019 | -0.57413 | 0.56588 | 0.896600563 | UBQLN4 | protein_coding | 50.71 | -0.007293956 | 0.007293956 | ENSG00000160803 | n.s. |
| ENSG00000101216 | 365.8829 | -0.148471594 | 0.249895 | -0.594136 | 0.552421 | 0.893099715 | GMEB2 | protein_coding | 53.42 | -0.007289963 | 0.007289963 | ENSG00000101216 | n.s. |
| ENSG00000175265 | 940.2379 | -0.158441591 | 0.280772 | -0.564307 | 0.572545 | 0.899492275 | GOLGA8A | protein_coding | 47.08 | -0.007288719 | 0.007288719 | ENSG00000175265 | n.s. |
| ENSG00000186480 | 1354.337 | -0.148233739 | 0.249364 | -0.594446 | 0.552214 | 0.892970078 | INSIG1 | protein_coding | 44.41 | -0.007287629 | 0.007287629 | ENSG00000186480 | n.s. |
| ENSG00000206937 | 9.608562 | -0.214536195 | 0.480948 | -0.446069 | 0.655547 | 0.924786541 | SNORA70B | snoRNA | 49.63 | -0.007285327 | 0.007285327 | ENSG00000206937 | n.s. |
| ENSG00000177359 | 155.0118 | 0.170052663 | 0.315319 | 0.5393041 | 0.589677 | 0.906089161 | | protein_coding | 38.21 | 0.007283197 | 0.007283197 | ENSG00000177359 | n.s. |
| ENSG00000056050 | 980.4272 | -0.160362942 | 0.287167 | -0.558431 | 0.57655 | 0.900751128 | HPF1 | protein_coding | 38.6 | -0.007279705 | 0.007279705 | ENSG00000056050 | n.s. |
| ENSG00000276291 | 445.0796 | -0.148175812 | 0.249478 | -0.593943 | 0.552551 | 0.893099715 | FRG1HP | cribed_unprocessed_pseudc | 39.23 | -0.00727544 | 0.00727544 | ENSG00000276291 | n.s. |
| ENSG00000182704 | 741.3468 | 0.142928858 | 0.23141 | 0.6176442 | 0.53681 | 0.889412093 | TSKU | protein_coding | 55.35 | 0.007274646 | 0.007274646 | ENSG00000182704 | n.s. |
| ENSG00000137210 | 1227.527 | -0.167396101 | 0.30593 | -0.547171 | 0.584261 | 0.904857859 | TMEM14B | protein_coding | 42.53 | -0.007268278 | 0.007268278 | ENSG00000137210 | n.s. |
| ENSG00000156531 | 1564.535 | -0.153074395 | 0.265902 | -0.575679 | 0.564832 | 0.896450345 | PHF6 | protein_coding | 37.47 | -0.007267017 | 0.007267017 | ENSG00000156531 | n.s. |
| ENSG00000171453 | 671.3161 | -0.149083566 | 0.254357 | -0.58612 | 0.557795 | 0.893837232 | POLR1C | protein_coding | 50.11 | -0.007266565 | 0.007266565 | ENSG00000171453 | n.s. |
| ENSG00000133935 | 290.196 | -0.160482791 | 0.287667 | -0.557877 | 0.576928 | 0.901017607 | C14orf1 | protein_coding | 44.19 | -0.00726453 | 0.00726453 | ENSG00000133935 | n.s. |
| ENSG00000236654 | 25.65192 | -0.176326304 | 0.34269 | -0.514535 | 0.606878 | 0.909522325 | | processed_pseudogene | 46.86 | -0.007262287 | 0.007262287 | ENSG00000236654 | n.s. |
| ENSG00000184178 | 692.1742 | -0.147091358 | 0.246992 | -0.595531 | 0.551489 | 0.892641916 | SCFD2 | protein_coding | 39.78 | -0.007254946 | 0.007254946 | ENSG00000184178 | n.s. |
| ENSG00000280789 | 547.5214 | -0.160783305 | 0.2884 | -0.5575 | 0.577186 | 0.901318367 | PAGR1 | protein_coding | 54.75 | -0.007254829 | 0.007254829 | ENSG00000280789 | n.s. |
| ENSG00000254505 | 191.413 | -0.145304155 | 0.239605 | -0.606432 | 0.544228 | 0.891406027 | CHMP4A | protein_coding | 50.64 | -0.007254228 | 0.007254228 | ENSG00000254505 | n.s. |
| ENSG00000164896 | 547.9042 | -0.147009829 | 0.24702 | -0.595133 | 0.551754 | 0.892641916 | FASTK | protein_coding | 61.75 | -0.007250925 | 0.007250925 | ENSG00000164896 | n.s. |
| ENSG00000113638 | 307.4012 | 0.172637106 | 0.322173 | 0.5358529 | 0.59206 | 0.907847371 | TTC33 | protein_coding | 38.63 | 0.007248542 | 0.007248542 | ENSG00000113638 | n.s. |

| | | | | | | | | | | | | | |
|---|---|---|---|---|---|---|---|---|---|---|---|---|---|
| ENSG00000224568 | 55.89351 | 0.15748842 | 0.27883 | 0.5648192 | 0.572197 | 0.899491841 | LINC01886 | lincRNA | 38.69 | 0.007244904 | 0.007244904 | ENSG00000224568 | n.s. |
| ENSG00000069956 | 3159.18 | -0.155643028 | 0.273147 | -0.569815 | 0.568803 | 0.898419074 | MAPK6 | protein_coding | 43.14 | -0.007240675 | 0.007240675 | ENSG00000069956 | n.s. |
| ENSG00000086065 | 1098.238 | 0.174338621 | 0.332387 | 0.5245046 | 0.599928 | 0.908868121 | CHMP5 | protein_coding | 41.65 | 0.007234901 | 0.007234901 | ENSG00000086065 | n.s. |
| ENSG00000172732 | 362.2895 | -0.155371531 | 0.272122 | -0.570963 | 0.568025 | 0.898381247 | MUS81 | protein_coding | 60.2 | -0.007230886 | 0.007230886 | ENSG00000172732 | n.s. |
| ENSG00000025772 | 424.0572 | -0.157485304 | 0.279858 | -0.562733 | 0.573617 | 0.899911687 | TOMM34 | protein_coding | 45.11 | -0.007212844 | 0.007212844 | ENSG00000025772 | n.s. |
| ENSG00000122140 | 573.11 | -0.156688438 | 0.27755 | -0.564542 | 0.572385 | 0.899492275 | MRPS2 | protein_coding | 63.07 | -0.00720807 | 0.00720807 | ENSG00000122140 | n.s. |
| ENSG00000128463 | 1712.072 | 0.155573207 | 0.274057 | 0.5676668 | 0.570261 | 0.898818733 | EMC4 | protein_coding | 43.89 | 0.007207378 | 0.007207378 | ENSG00000128463 | n.s. |
| ENSG00000133612 | 860.2974 | -0.15181611 | 0.264028 | -0.575 | 0.565291 | 0.896450345 | AGAP3 | protein_coding | 57.73 | -0.007207282 | 0.007207282 | ENSG00000133612 | n.s. |
| ENSG00000248643 | 163.007 | 0.145099933 | 0.243009 | 0.5970964 | 0.550443 | 0.89197749 | RBM14-RBM4 | protein_coding | 47.43 | 0.007203647 | 0.007203647 | ENSG00000248643 | n.s. |
| ENSG00000173511 | 135.3821 | -0.158065529 | 0.282284 | -0.559953 | 0.575511 | 0.900453394 | VEGFB | protein_coding | 61.25 | -0.007198108 | 0.007198108 | ENSG00000173511 | n.s. |
| ENSG00000213341 | 1372.703 | -0.15175391 | 0.264247 | -0.574288 | 0.565773 | 0.896600563 | CHUK | protein_coding | 38.39 | -0.007193286 | 0.007193286 | ENSG00000213341 | n.s. |
| ENSG00000227946 | 99.7844 | -0.158109015 | 0.282808 | -0.559068 | 0.576115 | 0.900614489 | | lincRNA | 63.47 | -0.007187805 | 0.007187805 | ENSG00000227946 | n.s. |
| ENSG00000114107 | 746.1911 | 0.161373427 | 0.291366 | 0.5538515 | 0.57968 | 0.902582314 | CEP70 | protein_coding | 39.55 | 0.007183245 | 0.007183245 | ENSG00000114107 | n.s. |
| ENSG00000197771 | 3775.476 | -0.151432284 | 0.263758 | -0.574134 | 0.565877 | 0.896600563 | MCMBP | protein_coding | 41.74 | -0.007178041 | 0.007178041 | ENSG00000197771 | n.s. |
| ENSG00000103544 | 648.2615 | -0.156008078 | 0.275807 | -0.565642 | 0.571637 | 0.899491841 | C16orf62 | protein_coding | 45.1 | -0.007176804 | 0.007176804 | ENSG00000103544 | n.s. |
| ENSG00000169188 | 241.6005 | -0.149256266 | 0.257962 | -0.578597 | 0.562861 | 0.895317554 | APEX2 | protein_coding | 49.09 | -0.007167719 | 0.007167719 | ENSG00000169188 | n.s. |
| ENSG00000182544 | 225.8779 | -0.145965412 | 0.247723 | -0.589228 | 0.555708 | 0.893144718 | MFSD5 | protein_coding | 57.75 | -0.007163715 | 0.007163715 | ENSG00000182544 | n.s. |
| ENSG00000100906 | 953.0196 | -0.144079663 | 0.240812 | -0.598308 | 0.549635 | 0.891861801 | NFKBIA | protein_coding | 56.53 | -0.00716111 | 0.00716111 | ENSG00000100906 | n.s. |
| ENSG00000068878 | 3934.783 | -0.153481161 | 0.268707 | -0.571183 | 0.567876 | 0.898265279 | PSME4 | protein_coding | 37.98 | -0.007151514 | 0.007151514 | ENSG00000068878 | n.s. |
| ENSG00000138231 | 1005.393 | -0.14393435 | 0.241105 | -0.596977 | 0.550523 | 0.891997058 | DBR1 | protein_coding | 39.44 | -0.007144409 | 0.007144409 | ENSG00000138231 | n.s. |
| ENSG00000177380 | 337.4497 | -0.152570281 | 0.266804 | -0.571844 | 0.567427 | 0.897862583 | PPFIA3 | protein_coding | 55.2 | -0.007138783 | 0.007138783 | ENSG00000177380 | n.s. |
| ENSG00000204311 | 102.3938 | 0.151776426 | 0.265081 | 0.5725661 | 0.566939 | 0.897394878 | DFNB59 | protein_coding | 39.52 | 0.007135983 | 0.007135983 | ENSG00000204311 | n.s. |
| ENSG00000187498 | 732.7316 | -0.14588778 | 0.248252 | -0.587659 | 0.556761 | 0.893643698 | COL4A1 | protein_coding | 44.9 | -0.007124518 | 0.007124518 | ENSG00000187498 | n.s. |
| ENSG00000179454 | 633.401 | 0.142687951 | 0.235644 | 0.6055226 | 0.544832 | 0.891406027 | KLHL28 | protein_coding | 37.29 | 0.007123615 | 0.007123615 | ENSG00000179454 | n.s. |
| ENSG00000214654 | 65.65956 | -0.156364109 | 0.279355 | -0.559733 | 0.575662 | 0.900453394 | | protein_coding | 41.27 | -0.007120628 | 0.007120628 | ENSG00000214654 | n.s. |
| ENSG00000067596 | 3187.187 | -0.145008176 | 0.244583 | -0.592879 | 0.553262 | 0.893144718 | DHX8 | protein_coding | 46.36 | -0.007116735 | 0.007116735 | ENSG00000067596 | n.s. |
| ENSG00000052841 | 1814.008 | -0.153574262 | 0.270056 | -0.568676 | 0.569576 | 0.898810685 | TTC17 | protein_coding | 37.89 | -0.007115368 | 0.007115368 | ENSG00000052841 | n.s. |
| ENSG00000213347 | 353.6598 | 0.151029262 | 0.263522 | 0.5731193 | 0.566564 | 0.897233219 | MXD3 | protein_coding | 58.63 | 0.007112671 | 0.007112671 | ENSG00000213347 | n.s. |
| ENSG00000279799 | 59.53696 | 0.161362236 | 0.2935 | 0.5497865 | 0.582466 | 0.90349457 | | TEC | 63.46 | 0.007111952 | 0.007111952 | ENSG00000279799 | n.s. |
| ENSG00000241547 | 46.45871 | 0.164429662 | 0.302355 | 0.5438297 | 0.586559 | 0.905217543 | ACTG1P20 | processed_pseudogene | 55.56 | 0.007111096 | 0.007111096 | ENSG00000241547 | n.s. |
| ENSG00000198554 | 1821.946 | -0.155983974 | 0.278204 | -0.560683 | 0.575014 | 0.900452118 | WDHD1 | protein_coding | 39.43 | -0.007103413 | 0.007103413 | ENSG00000198554 | n.s. |
| ENSG00000168216 | 410.3158 | -0.161108335 | 0.292737 | -0.550351 | 0.582078 | 0.90349457 | LMBRD1 | protein_coding | 35 | -0.007100762 | 0.007100762 | ENSG00000168216 | n.s. |

| | | | | | | | | | | | | | |
|---|---|---|---|---|---|---|---|---|---|---|---|---|---|
| ENSG00000207117 | 19.25558 | -0.191425956 | 0.390122 | -0.490682 | 0.623651 | 0.918167932 | | misc_RNA | 42.72 | -0.007097669 | 0.007097669 | ENSG00000207117 | n.s. |
| ENSG00000257815 | 160.6102 | -0.145871524 | 0.250473 | -0.582383 | 0.560309 | 0.894198873 | | antisense | 38.83 | -0.007084379 | 0.007084379 | ENSG00000257815 | n.s. |
| ENSG00000128245 | 1747.338 | -0.152201766 | 0.26697 | -0.570108 | 0.568604 | 0.898419074 | YWHAH | protein_coding | 45.04 | -0.007080584 | 0.007080584 | ENSG00000128245 | n.s. |
| ENSG00000102309 | 608.8853 | -0.162470363 | 0.296803 | -0.547402 | 0.584103 | 0.904712915 | PIN4 | protein_coding | 44.57 | -0.007065708 | 0.007065708 | ENSG00000102309 | n.s. |
| ENSG00000185122 | 452.0912 | -0.144834294 | 0.246887 | -0.586643 | 0.557444 | 0.893837232 | HSF1 | protein_coding | 56.87 | -0.007059449 | 0.007059449 | ENSG00000185122 | n.s. |
| ENSG00000161970 | 17021.12 | 0.171220391 | 0.33128 | 0.5168453 | 0.605264 | 0.909463932 | RPL26 | protein_coding | 43.78 | 0.007056766 | 0.007056766 | ENSG00000161970 | n.s. |
| ENSG00000125482 | 953.7536 | -0.148841393 | 0.259275 | -0.574068 | 0.565922 | 0.896600563 | TTF1 | protein_coding | 44.83 | -0.00705523 | 0.00705523 | ENSG00000125482 | n.s. |
| ENSG00000103811 | 67.48031 | -0.180571714 | 0.359564 | -0.502197 | 0.615529 | 0.91396346 | CTSH | protein_coding | 52.42 | -0.007055148 | 0.007055148 | ENSG00000103811 | n.s. |
| ENSG00000136932 | 272.9807 | 0.162436187 | 0.297647 | 0.5457343 | 0.585249 | 0.905026039 | TRMO | protein_coding | 41.69 | 0.00703981 | 0.00703981 | ENSG00000136932 | n.s. |
| ENSG00000182944 | 12000.52 | -0.140903539 | 0.231142 | -0.609598 | 0.542128 | 0.891404544 | EWSR1 | protein_coding | 44.02 | -0.007034631 | 0.007034631 | ENSG00000182944 | n.s. |
| ENSG00000134575 | 231.7012 | -0.140881197 | 0.233178 | -0.604179 | 0.545725 | 0.891406027 | ACP2 | protein_coding | 52.75 | -0.007033414 | 0.007033414 | ENSG00000134575 | n.s. |
| ENSG00000142684 | 184.4093 | 0.168643939 | 0.316293 | 0.5331885 | 0.593903 | 0.908531819 | ZNF593 | protein_coding | 62.91 | 0.007025683 | 0.007025683 | ENSG00000142684 | n.s. |
| ENSG00000146963 | 1724.461 | -0.15155239 | 0.267339 | -0.566892 | 0.570788 | 0.898818733 | LUC7L2 | protein_coding | 40.36 | -0.007021102 | 0.007021102 | ENSG00000146963 | n.s. |
| ENSG00000085760 | 1258.608 | -0.152865705 | 0.271148 | -0.563772 | 0.572909 | 0.899777025 | MTIF2 | protein_coding | 39.6 | -0.007011201 | 0.007011201 | ENSG00000085760 | n.s. |
| ENSG00000242622 | 411.1259 | -0.133469837 | 0.210033 | -0.63547 | 0.525122 | 0.886111278 | | lincRNA | 41.62 | -0.007008733 | 0.007008733 | ENSG00000242622 | n.s. |
| ENSG00000143355 | 773.697 | -0.143191932 | 0.243472 | -0.588126 | 0.556448 | 0.893419675 | LHX9 | protein_coding | 43.13 | -0.007008456 | 0.007008456 | ENSG00000143355 | n.s. |
| ENSG00000146243 | 1329.454 | -0.15197486 | 0.2686 | -0.565803 | 0.571528 | 0.899443932 | IRAK1BP1 | protein_coding | 37.25 | -0.00699478 | 0.00699478 | ENSG00000146243 | n.s. |
| ENSG00000156398 | 465.1423 | -0.140678393 | 0.235418 | -0.597567 | 0.550129 | 0.891877082 | SFXN2 | protein_coding | 48.44 | -0.006991012 | 0.006991012 | ENSG00000156398 | n.s. |
| ENSG00000198677 | 4243.021 | -0.151936862 | 0.269006 | -0.564808 | 0.572204 | 0.899491841 | TTC37 | protein_coding | 35.56 | -0.006989517 | 0.006989517 | ENSG00000198677 | n.s. |
| ENSG00000113272 | 382.9616 | -0.158529319 | 0.288046 | -0.55036 | 0.582072 | 0.90349457 | THG1L | protein_coding | 43.11 | -0.006987093 | 0.006987093 | ENSG00000113272 | n.s. |
| ENSG00000115419 | 3357.653 | -0.150802332 | 0.265115 | -0.568819 | 0.569479 | 0.898810685 | GLS | protein_coding | 35.11 | -0.006986939 | 0.006986939 | ENSG00000115419 | n.s. |
| ENSG00000214026 | 655.004 | -0.168302301 | 0.317641 | -0.529851 | 0.596215 | 0.908868121 | MRPL23 | protein_coding | 62.94 | -0.006984399 | 0.006984399 | ENSG00000214026 | n.s. |
| ENSG00000102078 | 212.6222 | -0.16101164 | 0.295491 | -0.544896 | 0.585825 | 0.905026039 | SLC25A14 | protein_coding | 37.86 | -0.006978071 | 0.006978071 | ENSG00000102078 | n.s. |
| ENSG00000132313 | 1392.55 | -0.15333919 | 0.274338 | -0.558943 | 0.576201 | 0.900614489 | MRPL35 | protein_coding | 40.87 | -0.006970964 | 0.006970964 | ENSG00000132313 | n.s. |
| ENSG00000184517 | 684.8268 | -0.150454052 | 0.265343 | -0.567017 | 0.570703 | 0.898818733 | ZFP1 | protein_coding | 42.26 | -0.006970218 | 0.006970218 | ENSG00000184517 | n.s. |
| ENSG00000112096 | 1813.337 | -0.139597019 | 0.230296 | -0.606164 | 0.544406 | 0.891406027 | SOD2 | protein_coding | 42.3 | -0.006969302 | 0.006969302 | ENSG00000112096 | n.s. |
| ENSG00000185187 | 85.06321 | -0.16943697 | 0.329587 | -0.514089 | 0.60719 | 0.90967308 | SIGIRR | protein_coding | 65.22 | -0.006966343 | 0.006966343 | ENSG00000185187 | n.s. |
| ENSG00000165406 | 106.8321 | 0.152826476 | 0.27269 | 0.5604406 | 0.575179 | 0.900453394 | 8-Mar | protein_coding | 40.34 | 0.006959528 | 0.006959528 | ENSG00000165406 | n.s. |
| ENSG00000121864 | 2217.504 | 0.145404691 | 0.25202 | 0.5769564 | 0.563969 | 0.895647177 | ZNF639 | protein_coding | 41.32 | 0.00695951 | 0.00695951 | ENSG00000121864 | n.s. |
| ENSG00000049449 | 2060.749 | -0.157826313 | 0.28675 | -0.550397 | 0.582047 | 0.90349457 | RCN1 | protein_coding | 43.38 | -0.006956108 | 0.006956108 | ENSG00000049449 | n.s. |
| ENSG00000142686 | 364.1099 | -0.152863201 | 0.273498 | -0.558919 | 0.576217 | 0.900614489 | C1orf216 | protein_coding | 53.73 | -0.006949325 | 0.006949325 | ENSG00000142686 | n.s. |
| ENSG00000158863 | 72.96931 | -0.16855598 | 0.327048 | -0.515385 | 0.606284 | 0.909522325 | FAM160B2 | protein_coding | 57.17 | -0.006942254 | 0.006942254 | ENSG00000158863 | n.s. |

| | | | | | | | | | | | | | |
|---|---|---|---|---|---|---|---|---|---|---|---|---|---|
| ENSG00000261135 | 30.89038 | 0.186445566 | 0.37882 | 0.4921747 | 0.622596 | 0.917861492 | | lincRNA | 41.26 | 0.006940035 | 0.006940035 | ENSG00000261135 | n.s. |
| ENSG00000212535 | 15.83706 | -0.189096275 | 0.393392 | -0.480682 | 0.630743 | 0.919033489 | RNU6-808P | snRNA | 32.41 | -0.006933908 | 0.006933908 | ENSG00000212535 | n.s. |
| ENSG00000260852 | 158.2476 | -0.14893269 | 0.261292 | -0.569986 | 0.568688 | 0.898419074 | FBXL19-AS1 | antisense | 52.62 | -0.006928503 | 0.006928503 | ENSG00000260852 | n.s. |
| ENSG00000228659 | 22.1285 | 0.188878988 | 0.388856 | 0.4857294 | 0.627159 | 0.919033489 | LINC01201 | antisense | 41.35 | 0.00692594 | 0.00692594 | ENSG00000228659 | n.s. |
| ENSG00000158796 | 401.0906 | -0.151041877 | 0.268568 | -0.562397 | 0.573846 | 0.899911687 | DEDD | protein_coding | 46.36 | -0.006917734 | 0.006917734 | ENSG00000158796 | n.s. |
| ENSG00000029364 | 1235.692 | -0.146819184 | 0.256085 | -0.573322 | 0.566426 | 0.897196154 | SLC39A9 | protein_coding | 40.2 | -0.006917033 | 0.006917033 | ENSG00000029364 | n.s. |
| ENSG00000278333 | 4.820748 | -0.232827119 | 0.571072 | -0.407702 | 0.683493 | 0.933907782 | | misc_RNA | 55.29 | -0.006914035 | 0.006914035 | ENSG00000278333 | n.s. |
| ENSG00000155330 | 372.972 | -0.159729489 | 0.293476 | -0.544268 | 0.586257 | 0.905188828 | C16orf87 | protein_coding | 37.09 | -0.006910028 | 0.006910028 | ENSG00000155330 | n.s. |
| ENSG00000200534 | 255.8318 | 0.133377549 | 0.213503 | 0.6247108 | 0.532161 | 0.887559389 | SNORA33 | snoRNA | 40.77 | 0.006909301 | 0.006909301 | ENSG00000200534 | n.s. |
| ENSG00000213453 | 42.78983 | -0.166451638 | 0.314104 | -0.529925 | 0.596164 | 0.908868121 | FTH1P3 | processed_pseudogene | 47.95 | -0.006907598 | 0.006907598 | ENSG00000213453 | n.s. |
| ENSG00000272523 | 39.13214 | -0.16763615 | 0.324996 | -0.515809 | 0.605988 | 0.909522325 | LINC01023 | lincRNA | 63.53 | -0.006904369 | 0.006904369 | ENSG00000272523 | n.s. |
| ENSG00000255690 | 176.7755 | -0.142064581 | 0.243467 | -0.583508 | 0.559552 | 0.894140829 | TRIL | protein_coding | 56.8 | -0.006903497 | 0.006903497 | ENSG00000255690 | n.s. |
| ENSG00000136628 | 5613.473 | -0.152006841 | 0.272259 | -0.558317 | 0.576628 | 0.900751128 | EPRS | protein_coding | 38.15 | -0.006900379 | 0.006900379 | ENSG00000136628 | n.s. |
| ENSG00000145439 | 901.7523 | -0.148313728 | 0.260377 | -0.569612 | 0.568941 | 0.898419074 | CBR4 | protein_coding | 38.87 | -0.006899708 | 0.006899708 | ENSG00000145439 | n.s. |
| ENSG00000270195 | 69.93537 | -0.150856131 | 0.268692 | -0.561446 | 0.574494 | 0.900202572 | | lincRNA | 60.47 | -0.006888053 | 0.006888053 | ENSG00000270195 | n.s. |
| ENSG00000270108 | 45.20602 | -0.164869771 | 0.308212 | -0.534923 | 0.592703 | 0.908297635 | | sense_intronic | 35.33 | -0.00688691 | 0.00688691 | ENSG00000270108 | n.s. |
| ENSG00000054267 | 3087.071 | -0.140164734 | 0.235989 | -0.593947 | 0.552548 | 0.893099715 | ARID4B | protein_coding | 39.95 | -0.006882095 | 0.006882095 | ENSG00000054267 | n.s. |
| ENSG00000156928 | 550.2147 | -0.149546395 | 0.26456 | -0.565265 | 0.571893 | 0.899491841 | MALSU1 | protein_coding | 41.44 | -0.006879549 | 0.006879549 | ENSG00000156928 | n.s. |
| ENSG00000169231 | 365.2479 | 0.152752527 | 0.274342 | 0.5567969 | 0.577666 | 0.901500147 | THBS3 | protein_coding | 55.3 | 0.006879088 | 0.006879088 | ENSG00000169231 | n.s. |
| ENSG00000230126 | 22.05037 | 0.189594521 | 0.401389 | 0.4723465 | 0.636679 | 0.919866014 | FGF12-AS2 | antisense | 52.64 | 0.006877622 | 0.006877622 | ENSG00000230126 | n.s. |
| ENSG00000106462 | 2758.393 | -0.142897912 | 0.246244 | -0.580309 | 0.561706 | 0.895116382 | EZH2 | protein_coding | 39.21 | -0.006876318 | 0.006876318 | ENSG00000106462 | n.s. |
| ENSG00000164898 | 125.2352 | -0.180342508 | 0.362729 | -0.497183 | 0.61906 | 0.916050756 | FMC1 | protein_coding | 46.51 | -0.006867526 | 0.006867526 | ENSG00000164898 | n.s. |
| ENSG00000165282 | 340.2622 | -0.137538852 | 0.227881 | -0.603556 | 0.546139 | 0.891406027 | PIGO | protein_coding | 53.46 | -0.006866549 | 0.006866549 | ENSG00000165282 | n.s. |
| ENSG00000116752 | 1760.465 | -0.155728703 | 0.282451 | -0.551347 | 0.581396 | 0.90349457 | BCAS2 | protein_coding | 40.33 | -0.006863657 | 0.006863657 | ENSG00000116752 | n.s. |
| ENSG00000266370 | 19.26731 | 0.187073998 | 0.392378 | 0.4767692 | 0.633526 | 0.919033489 | MIR3657 | miRNA | 29.91 | 0.006859753 | 0.006859753 | ENSG00000266370 | n.s. |
| ENSG00000213625 | 1345.788 | -0.14908615 | 0.264276 | -0.56413 | 0.572666 | 0.899506677 | LEPROT | protein_coding | 40.78 | -0.006857308 | 0.006857308 | ENSG00000213625 | n.s. |
| ENSG00000247240 | 253.6727 | -0.137335683 | 0.227369 | -0.604021 | 0.54583 | 0.891406027 | UBL7-AS1 | antisense | 43.8 | -0.006856406 | 0.006856406 | ENSG00000247240 | n.s. |
| ENSG00000074842 | 412.3629 | -0.165193558 | 0.312826 | -0.528069 | 0.597451 | 0.908868121 | MYDGF | protein_coding | 54.74 | -0.006855389 | 0.006855389 | ENSG00000074842 | n.s. |
| ENSG00000203705 | 608.1863 | -0.154863293 | 0.280221 | -0.552646 | 0.580506 | 0.903133423 | TATDN3 | protein_coding | 40.66 | -0.006852404 | 0.006852404 | ENSG00000203705 | n.s. |
| ENSG00000170577 | 325.9605 | -0.139584048 | 0.235058 | -0.593828 | 0.552627 | 0.89311943 | SIX2 | protein_coding | 59.16 | -0.006852245 | 0.006852245 | ENSG00000170577 | n.s. |
| ENSG00000104852 | 3454.073 | -0.1427868 | 0.247027 | -0.57802 | 0.56325 | 0.895435774 | SNRNP70 | protein_coding | 53.38 | -0.006848849 | 0.006848849 | ENSG00000104852 | n.s. |
| ENSG00000125744 | 353.162 | -0.151984163 | 0.273118 | -0.556477 | 0.577885 | 0.901500147 | RTN2 | protein_coding | 54.15 | -0.006844485 | 0.006844485 | ENSG00000125744 | n.s. |

| | | | | | | | | | | | | | |
|---|---|---|---|---|---|---|---|---|---|---|---|---|---|
| ENSG00000196812 | 192.6545 | 0.13923367 | 0.23633 | 0.5891489 | 0.555761 | 0.893144718 | ZSCAN16 | protein_coding | 41.5 | 0.006833333 | 0.006833333 | ENSG00000196812 | n.s. |
| ENSG00000113119 | 112.1315 | -0.149657283 | 0.266584 | -0.561389 | 0.574532 | 0.900202572 | TMCO6 | protein_coding | 51.35 | -0.006833314 | 0.006833314 | ENSG00000113119 | n.s. |
| ENSG00000069998 | 790.4968 | -0.154646295 | 0.280236 | -0.551843 | 0.581056 | 0.903352939 | HDHD5 | protein_coding | 50.12 | -0.00682648 | 0.00682648 | ENSG00000069998 | n.s. |
| ENSG00000140740 | 2483.024 | 0.157765015 | 0.289947 | 0.5441176 | 0.586361 | 0.905188828 | UQCRC2 | protein_coding | 40.33 | 0.006825043 | 0.006825043 | ENSG00000140740 | n.s. |
| ENSG00000178381 | 233.7877 | 0.148287821 | 0.262479 | 0.5649519 | 0.572106 | 0.899491841 | ZFAND2A | protein_coding | 54.85 | 0.006821651 | 0.006821651 | ENSG00000178381 | n.s. |
| ENSG00000175224 | 224.985 | -0.148244891 | 0.262374 | -0.565014 | 0.572064 | 0.899491841 | ATG13 | protein_coding | 44.43 | -0.006819676 | 0.006819676 | ENSG00000175224 | n.s. |
| ENSG00000240106 | 21.15571 | 0.185779758 | 0.389221 | 0.4773114 | 0.63314 | 0.919033489 | RN7SL146P | misc_RNA | 56.9 | 0.006812295 | 0.006812295 | ENSG00000240106 | n.s. |
| ENSG00000154358 | 213.0773 | -0.14641796 | 0.257046 | -0.569618 | 0.568937 | 0.898419074 | OBSCN | protein_coding | 53.87 | -0.006811515 | 0.006811515 | ENSG00000154358 | n.s. |
| ENSG00000280129 | 172.2149 | 0.140163382 | 0.240149 | 0.5836511 | 0.559455 | 0.894140829 | | TEC | 42.77 | 0.00681111 | 0.00681111 | ENSG00000280129 | n.s. |
| ENSG00000106330 | 67.96024 | -0.163955575 | 0.31049 | -0.528055 | 0.597461 | 0.908868121 | MOSPD3 | protein_coding | 57.23 | -0.006804014 | 0.006804014 | ENSG00000106330 | n.s. |
| ENSG00000145284 | 431.0906 | -0.156926763 | 0.287352 | -0.546113 | 0.584988 | 0.905026039 | SCD5 | protein_coding | 43.57 | -0.006801037 | 0.006801037 | ENSG00000145284 | n.s. |
| ENSG00000261430 | 13.31525 | 0.197540539 | 0.438687 | 0.4502996 | 0.652494 | 0.923800272 | | sense_intronic | 53.35 | 0.006799723 | 0.006799723 | ENSG00000261430 | n.s. |
| ENSG00000212102 | 25.95377 | 0.182311183 | 0.369878 | 0.4928954 | 0.622087 | 0.917706135 | MIR301B | miRNA | 52.56 | 0.006799544 | 0.006799544 | ENSG00000212102 | n.s. |
| ENSG00000169288 | 686.9939 | -0.163823305 | 0.309746 | -0.528895 | 0.596878 | 0.908868121 | MRPL1 | protein_coding | 37.82 | -0.006798525 | 0.006798525 | ENSG00000169288 | n.s. |
| ENSG00000225630 | 42.91138 | -0.163678085 | 0.315953 | -0.518045 | 0.604427 | 0.908868121 | MTND2P28 | unprocessed_pseudogene | 43.1 | -0.006792498 | 0.006792498 | ENSG00000225630 | n.s. |
| ENSG00000175104 | 533.091 | -0.154345589 | 0.281001 | -0.54927 | 0.58282 | 0.90373147 | TRAF6 | protein_coding | 38.03 | -0.006785124 | 0.006785124 | ENSG00000175104 | n.s. |
| ENSG00000151725 | 3729.087 | -0.153714232 | 0.279545 | -0.549874 | 0.582406 | 0.90349457 | CENPU | protein_coding | 40.02 | -0.006774871 | 0.006774871 | ENSG00000151725 | n.s. |
| ENSG00000010017 | 823.6439 | -0.147242195 | 0.260887 | -0.56439 | 0.572489 | 0.899492275 | RANBP9 | protein_coding | 36.94 | -0.006773518 | 0.006773518 | ENSG00000010017 | n.s. |
| ENSG00000114446 | 957.9198 | -0.152247203 | 0.275026 | -0.553573 | 0.579871 | 0.902677303 | IFT57 | protein_coding | 35.29 | -0.006770049 | 0.006770049 | ENSG00000114446 | n.s. |
| ENSG00000205544 | 187.7231 | -0.171475825 | 0.340381 | -0.503776 | 0.614419 | 0.913167298 | TMEM256 | protein_coding | 57.95 | -0.006764661 | 0.006764661 | ENSG00000205544 | n.s. |
| ENSG00000095574 | 468.7492 | -0.142461519 | 0.247758 | -0.575002 | 0.56529 | 0.896450345 | IKZF5 | protein_coding | 38.33 | -0.006763184 | 0.006763184 | ENSG00000095574 | n.s. |
| ENSG00000007520 | 501.716 | -0.151221536 | 0.272516 | -0.554909 | 0.578957 | 0.902167117 | TSR3 | protein_coding | 62.57 | -0.006761569 | 0.006761569 | ENSG00000007520 | n.s. |
| ENSG00000239466 | 11.49019 | 0.190549798 | 0.40843 | 0.4665423 | 0.640827 | 0.921718971 | RN7SL552P | misc_RNA | 52.54 | 0.006745744 | 0.006745744 | ENSG00000239466 | n.s. |
| ENSG00000134146 | 161.3458 | -0.162502714 | 0.305823 | -0.531361 | 0.595169 | 0.908868121 | DPH6 | protein_coding | 36.45 | -0.006743721 | 0.006743721 | ENSG00000134146 | n.s. |
| ENSG00000167716 | 384.1069 | -0.132217712 | 0.213555 | -0.619127 | 0.535833 | 0.889201649 | WDR81 | protein_coding | 57.66 | -0.006743069 | 0.006743069 | ENSG00000167716 | n.s. |
| ENSG00000135766 | 567.1427 | 0.148217697 | 0.264922 | 0.5594762 | 0.575837 | 0.900580379 | EGLN1 | protein_coding | 39.65 | 0.006740573 | 0.006740573 | ENSG00000135766 | n.s. |
| ENSG00000259956 | 2560.282 | -0.128322673 | 0.201302 | -0.637463 | 0.523823 | 0.886111278 | RBM15B | protein_coding | 55.91 | -0.006738446 | 0.006738446 | ENSG00000259956 | n.s. |
| ENSG00000085491 | 3057.975 | -0.147793918 | 0.263821 | -0.560204 | 0.57534 | 0.900453394 | SLC25A24 | protein_coding | 39.44 | -0.006730352 | 0.006730352 | ENSG00000085491 | n.s. |
| ENSG00000101132 | 886.5122 | 0.168196174 | 0.332807 | 0.5053862 | 0.613288 | 0.911981962 | PFDN4 | protein_coding | 41.84 | 0.00673016 | 0.00673016 | ENSG00000101132 | n.s. |
| ENSG00000107643 | 2490.685 | -0.145264913 | 0.255733 | -0.568033 | 0.570013 | 0.898818733 | MAPK8 | protein_coding | 37.34 | -0.006729816 | 0.006729816 | ENSG00000107643 | n.s. |
| ENSG00000259554 | 72.14959 | 0.237870696 | 0.60591 | 0.3925845 | 0.694626 | 0.936958261 | | sense_intronic | 39.21 | 0.006726925 | 0.006726925 | ENSG00000259554 | n.s. |
| ENSG00000261270 | 77.87386 | 0.151776267 | 0.274644 | 0.5526294 | 0.580517 | 0.903133423 | | sense_intronic | 47.7 | 0.006715809 | 0.006715809 | ENSG00000261270 | n.s. |

| | | | | | | | | | | | | | |
|---|---|---|---|---|---|---|---|---|---|---|---|---|---|
| ENSG00000139546 | 827.2522 | -0.139734942 | 0.241869 | -0.577729 | 0.563447 | 0.895460914 | TARBP2 | protein_coding | 57.56 | -0.006700761 | 0.006700761 | ENSG00000139546 | n.s. |
| ENSG00000272033 | 152.9023 | -0.152026582 | 0.276125 | -0.550572 | 0.581927 | 0.90349457 | | sense_intronic | 36.08 | -0.006700488 | 0.006700488 | ENSG00000272033 | n.s. |
| ENSG00000230715 | 40.49726 | -0.161434982 | 0.307153 | -0.525585 | 0.599176 | 0.908868121 | | processed_pseudogene | 65.26 | -0.006699411 | 0.006699411 | ENSG00000230715 | n.s. |
| ENSG00000205189 | 3823.473 | -0.138379809 | 0.237888 | -0.581702 | 0.560767 | 0.894542201 | ZBTB10 | protein_coding | 38.89 | -0.006697467 | 0.006697467 | ENSG00000205189 | n.s. |
| ENSG00000167186 | 609.2364 | -0.148460403 | 0.266434 | -0.557213 | 0.577382 | 0.901408004 | COQ7 | protein_coding | 46.2 | -0.006692386 | 0.006692386 | ENSG00000167186 | n.s. |
| ENSG00000278259 | 241.7535 | 0.131212142 | 0.21223 | 0.6182542 | 0.536408 | 0.889201649 | MYO19 | protein_coding | 47.58 | 0.006691785 | 0.006691785 | ENSG00000278259 | n.s. |
| ENSG00000186187 | 249.4445 | -0.138822921 | 0.239449 | -0.579759 | 0.562077 | 0.895116382 | ZNRF1 | protein_coding | 45.85 | -0.006680228 | 0.006680228 | ENSG00000186187 | n.s. |
| ENSG00000133997 | 1209.456 | 0.145678141 | 0.258614 | 0.5633024 | 0.573229 | 0.899883575 | MED6 | protein_coding | 38.94 | 0.006674051 | 0.006674051 | ENSG00000133997 | n.s. |
| ENSG00000124608 | 914.9536 | -0.135915565 | 0.229904 | -0.591183 | 0.554398 | 0.893144718 | AARS2 | protein_coding | 54.66 | -0.006670487 | 0.006670487 | ENSG00000124608 | n.s. |
| ENSG00000118363 | 653.5543 | -0.145608253 | 0.25856 | -0.563151 | 0.573332 | 0.899911687 | SPCS2 | protein_coding | 39.25 | -0.006668874 | 0.006668874 | ENSG00000118363 | n.s. |
| ENSG00000180071 | 12.01339 | 0.238581704 | 0.610921 | 0.3905279 | 0.696146 | 0.937752951 | ANKRD18A | protein_coding | 36.89 | 0.006659188 | 0.006659188 | ENSG00000180071 | n.s. |
| ENSG00000137656 | 499.9879 | -0.147670093 | 0.265121 | -0.556991 | 0.577533 | 0.901457225 | BUD13 | protein_coding | 41.56 | -0.006653258 | 0.006653258 | ENSG00000137656 | n.s. |
| ENSG00000197863 | 257.0519 | -0.14284552 | 0.25035 | -0.570584 | 0.568281 | 0.898419074 | ZNF790 | protein_coding | 42.57 | -0.006645322 | 0.006645322 | ENSG00000197863 | n.s. |
| ENSG00000143919 | 143.6289 | -0.150441811 | 0.272545 | -0.551989 | 0.580956 | 0.903352939 | CAMKMT | protein_coding | 37.93 | -0.006640883 | 0.006640883 | ENSG00000143919 | n.s. |
| ENSG00000184209 | 628.746 | -0.132992589 | 0.220155 | -0.604085 | 0.545787 | 0.891406027 | SNRNP35 | protein_coding | 47.6 | -0.00663958 | 0.00663958 | ENSG00000184209 | n.s. |
| ENSG00000139651 | 365.0288 | -0.13822421 | 0.238847 | -0.578715 | 0.562782 | 0.895317554 | ZNF740 | protein_coding | 50.62 | -0.006637927 | 0.006637927 | ENSG00000139651 | n.s. |
| ENSG00000147687 | 565.8442 | -0.15336011 | 0.282513 | -0.542843 | 0.587238 | 0.905217543 | TATDN1 | protein_coding | 40.76 | -0.006632371 | 0.006632371 | ENSG00000147687 | n.s. |
| ENSG00000109065 | 655.7816 | -0.149172286 | 0.269546 | -0.553421 | 0.579975 | 0.902738584 | NAT9 | protein_coding | 54.06 | -0.006628917 | 0.006628917 | ENSG00000109065 | n.s. |
| ENSG00000200888 | 44.29979 | -0.159605687 | 0.302148 | -0.528236 | 0.597335 | 0.908868121 | | misc_RNA | 43.14 | -0.006623497 | 0.006623497 | ENSG00000200888 | n.s. |
| ENSG00000182718 | 7595.172 | -0.15266179 | 0.279313 | -0.546562 | 0.58468 | 0.905026039 | ANXA2 | protein_coding | 44.95 | -0.006616198 | 0.006616198 | ENSG00000182718 | n.s. |
| ENSG00000182473 | 2758.558 | -0.149918831 | 0.272144 | -0.550881 | 0.581715 | 0.90349457 | EXOC7 | protein_coding | 51.59 | -0.006607591 | 0.006607591 | ENSG00000182473 | n.s. |
| ENSG00000168566 | 712.0534 | -0.145039675 | 0.258972 | -0.56006 | 0.575439 | 0.900453394 | SNRNP48 | protein_coding | 37.33 | -0.006604927 | 0.006604927 | ENSG00000168566 | n.s. |
| ENSG00000170873 | 1048.134 | -0.140664208 | 0.245774 | -0.57233 | 0.567098 | 0.8975378 | MTSS1 | protein_coding | 45.65 | -0.006603798 | 0.006603798 | ENSG00000170873 | n.s. |
| ENSG00000270820 | 54.33159 | 0.157185014 | 0.293328 | 0.5358671 | 0.59205 | 0.907847371 | | antisense | 44.62 | 0.006599752 | 0.006599752 | ENSG00000270820 | n.s. |
| ENSG00000112320 | 240.1328 | -0.149730056 | 0.272092 | -0.550292 | 0.582119 | 0.90349457 | SOBP | protein_coding | 41.63 | -0.00659927 | 0.00659927 | ENSG00000112320 | n.s. |
| ENSG00000124614 | 2578.714 | 0.158711383 | 0.306234 | 0.5182679 | 0.604271 | 0.908868121 | RPS10 | protein_coding | 47.56 | 0.006586384 | 0.006586384 | ENSG00000124614 | n.s. |
| ENSG00000223006 | 22.44256 | 0.174870253 | 0.353548 | 0.4946158 | 0.620871 | 0.917065125 | RN7SKP137 | misc_RNA | 52.12 | 0.00657509 | 0.00657509 | ENSG00000223006 | n.s. |
| ENSG00000151461 | 1259.288 | -0.144321451 | 0.257676 | -0.560089 | 0.575419 | 0.900453394 | UPF2 | protein_coding | 40.23 | -0.00657222 | 0.00657222 | ENSG00000151461 | n.s. |
| ENSG00000185418 | 762.0692 | -0.149041921 | 0.270535 | -0.550916 | 0.581691 | 0.90349457 | TARSL2 | protein_coding | 41.13 | -0.006568941 | 0.006568941 | ENSG00000185418 | n.s. |
| ENSG00000035499 | 749.0549 | -0.151883269 | 0.279343 | -0.543715 | 0.586637 | 0.905217543 | DEPDC1B | protein_coding | 38.63 | -0.006568502 | 0.006568502 | ENSG00000035499 | n.s. |
| ENSG00000182287 | 382.0612 | 0.154453503 | 0.286853 | 0.5384411 | 0.590273 | 0.906814644 | AP1S2 | protein_coding | 36.77 | 0.006561413 | 0.006561413 | ENSG00000182287 | n.s. |
| ENSG00000007866 | 200.9468 | -0.143799851 | 0.25639 | -0.560863 | 0.574891 | 0.900360771 | TEAD3 | protein_coding | 56.75 | -0.006554891 | 0.006554891 | ENSG00000007866 | n.s. |

| | | | | | | | | | | | | | |
|---|---|---|---|---|---|---|---|---|---|---|---|---|---|
| ENSG00000100379 | 64.90162 | 0.157887597 | 0.301843 | 0.5230779 | 0.60092 | 0.908868121 | KCTD17 | protein_coding | 60.2 | 0.006552198 | 0.006552198 | ENSG00000100379 | n.s. |
| ENSG00000278986 | 123.5962 | 0.140825249 | 0.247046 | 0.5700361 | 0.568653 | 0.898419074 | | TEC | 54.02 | 0.006551337 | 0.006551337 | ENSG00000278986 | n.s. |
| ENSG00000173480 | 625.4937 | -0.145664805 | 0.261864 | -0.556261 | 0.578033 | 0.901629862 | ZNF417 | protein_coding | 45.26 | -0.006550796 | 0.006550796 | ENSG00000173480 | n.s. |
| ENSG00000160949 | 690.3852 | -0.131192817 | 0.21665 | -0.605552 | 0.544812 | 0.891406027 | TONSL | protein_coding | 62.9 | -0.006549727 | 0.006549727 | ENSG00000160949 | n.s. |
| ENSG00000137288 | 1057.469 | -0.157801668 | 0.301698 | -0.523045 | 0.600943 | 0.908868121 | UQCC2 | protein_coding | 49.22 | -0.006548632 | 0.006548632 | ENSG00000137288 | n.s. |
| ENSG00000139233 | 1074.738 | -0.148126228 | 0.268289 | -0.552115 | 0.58087 | 0.903322646 | LLPH | protein_coding | 39.72 | -0.006540825 | 0.006540825 | ENSG00000139233 | n.s. |
| ENSG00000186297 | 310.558 | 0.148393853 | 0.26945 | 0.5507281 | 0.58182 | 0.90349457 | GABRA5 | protein_coding | 43.55 | 0.006540378 | 0.006540378 | ENSG00000186297 | n.s. |
| ENSG00000252349 | 5.262498 | -0.235029042 | 0.622352 | -0.377647 | 0.705693 | 0.937943012 | | snoRNA | 41.76 | -0.006539342 | 0.006539342 | ENSG00000252349 | n.s. |
| ENSG00000252577 | 15.51188 | -0.188332969 | 0.412234 | -0.45686 | 0.647772 | 0.923184844 | SCARNA20 | scaRNA | 61.07 | -0.006537289 | 0.006537289 | ENSG00000252577 | n.s. |
| ENSG00000119707 | 5678.014 | -0.141069199 | 0.248739 | -0.567139 | 0.57062 | 0.898818733 | RBM25 | protein_coding | 40.7 | -0.006535438 | 0.006535438 | ENSG00000119707 | n.s. |
| ENSG00000170638 | 348.2056 | 0.158612078 | 0.307753 | 0.5153875 | 0.606282 | 0.909522325 | TRABD | protein_coding | 62.8 | 0.006532698 | 0.006532698 | ENSG00000170638 | n.s. |
| ENSG00000157637 | 415.9221 | -0.156255885 | 0.292483 | -0.534239 | 0.593176 | 0.908297635 | SLC38A10 | protein_coding | 56.73 | -0.006527092 | 0.006527092 | ENSG00000157637 | n.s. |
| ENSG00000251216 | 34.21502 | 0.177857358 | 0.371294 | 0.4790206 | 0.631924 | 0.919033489 | | lincRNA | 36.87 | 0.006521792 | 0.006521792 | ENSG00000251216 | n.s. |
| ENSG00000005893 | 666.4885 | -0.157152296 | 0.296286 | -0.530407 | 0.59583 | 0.908868121 | LAMP2 | protein_coding | 38.79 | -0.006521683 | 0.006521683 | ENSG00000005893 | n.s. |
| ENSG00000115286 | 484.0975 | -0.160503048 | 0.315308 | -0.509035 | 0.610727 | 0.910725807 | NDUFS7 | protein_coding | 59.57 | -0.006518407 | 0.006518407 | ENSG00000115286 | n.s. |
| ENSG00000201228 | 11.98295 | -0.233386331 | 0.609892 | -0.382668 | 0.701966 | 0.937752951 | | misc_RNA | 44.12 | -0.006514177 | 0.006514177 | ENSG00000201228 | n.s. |
| ENSG00000273711 | 14.39789 | -0.198134638 | 0.456495 | -0.434035 | 0.664263 | 0.927094972 | | antisense | 32.32 | -0.00651383 | 0.00651383 | ENSG00000273711 | n.s. |
| ENSG00000170745 | 152.7543 | -0.156920966 | 0.300957 | -0.521407 | 0.602083 | 0.908868121 | KCNS3 | protein_coding | 39.79 | -0.006512083 | 0.006512083 | ENSG00000170745 | n.s. |
| ENSG00000104472 | 249.4842 | -0.141936842 | 0.251801 | -0.563687 | 0.572967 | 0.899777025 | CHRAC1 | protein_coding | 47.65 | -0.006509947 | 0.006509947 | ENSG00000104472 | n.s. |
| ENSG00000277368 | 16.48261 | -0.19017288 | 0.424221 | -0.448288 | 0.653946 | 0.924250964 | | antisense | 61.85 | -0.006505831 | 0.006505831 | ENSG00000277368 | n.s. |
| ENSG00000224043 | 31.08077 | -0.180594757 | 0.383756 | -0.470598 | 0.637928 | 0.920401057 | CCNT2-AS1 | antisense | 41.05 | -0.006505545 | 0.006505545 | ENSG00000224043 | n.s. |
| ENSG00000119048 | 943.8242 | -0.153402766 | 0.285523 | -0.53727 | 0.591081 | 0.907045444 | UBE2B | protein_coding | 41.55 | -0.006499822 | 0.006499822 | ENSG00000119048 | n.s. |
| ENSG00000123091 | 1463.563 | -0.156527446 | 0.295657 | -0.529423 | 0.596512 | 0.908868121 | RNF11 | protein_coding | 41.25 | -0.006495753 | 0.006495753 | ENSG00000123091 | n.s. |
| ENSG00000272006 | 9.268926 | 0.233106593 | 0.614124 | 0.379576 | 0.70426 | 0.937896121 | | processed_pseudogene | 34.82 | 0.006490914 | 0.006490914 | ENSG00000272006 | n.s. |
| ENSG00000130826 | 2993.243 | -0.140936444 | 0.249782 | -0.564238 | 0.572592 | 0.899492275 | DKC1 | protein_coding | 45.62 | -0.006483437 | 0.006483437 | ENSG00000130826 | n.s. |
| ENSG00000111850 | 716.1654 | -0.149591748 | 0.274208 | -0.545542 | 0.585381 | 0.905026039 | SMIM8 | protein_coding | 40.32 | -0.006483146 | 0.006483146 | ENSG00000111850 | n.s. |
| ENSG00000088766 | 824.503 | -0.150490307 | 0.278349 | -0.540653 | 0.588747 | 0.905738346 | CRLS1 | protein_coding | 39.36 | -0.006470669 | 0.006470669 | ENSG00000088766 | n.s. |
| ENSG00000163104 | 2943.819 | -0.149561299 | 0.275448 | -0.542974 | 0.587147 | 0.905217543 | SMARCAD1 | protein_coding | 36.02 | -0.006468083 | 0.006468083 | ENSG00000163104 | n.s. |
| ENSG00000163344 | 184.809 | -0.155788472 | 0.295446 | -0.527299 | 0.597986 | 0.908868121 | PMVK | protein_coding | 49.94 | -0.006465086 | 0.006465086 | ENSG00000163344 | n.s. |
| ENSG00000084693 | 444.7751 | -0.149130385 | 0.27303 | -0.546206 | 0.584924 | 0.905026039 | AGBL5 | protein_coding | 48.47 | -0.006463151 | 0.006463151 | ENSG00000084693 | n.s. |
| ENSG00000203950 | 216.5351 | -0.142336899 | 0.254876 | -0.558456 | 0.576533 | 0.900751128 | RTL8A | protein_coding | 60.16 | -0.00646141 | 0.00646141 | ENSG00000203950 | n.s. |
| ENSG00000260942 | 383.8816 | -0.129415716 | 0.21434 | -0.603787 | 0.545985 | 0.891406027 | CAPN10-AS1 | antisense | 54.57 | -0.006461006 | 0.006461006 | ENSG00000260942 | n.s. |

| | | | | | | | | | | | | | |
|---|---|---|---|---|---|---|---|---|---|---|---|---|---|
| ENSG00000166710 | 3861.239 | 0.1571452 | 0.305698 | 0.514053 | 0.607215 | 0.90967308 | B2M | protein_coding | 42.63 | 0.006460971 | 0.006460971 | ENSG00000166710 | n.s. |
| ENSG00000123353 | 248.5756 | -0.155608482 | 0.295547 | -0.526511 | 0.598533 | 0.908868121 | ORMDL2 | protein_coding | 46.4 | -0.006457617 | 0.006457617 | ENSG00000123353 | n.s. |
| ENSG00000122176 | 253.9203 | 0.148943753 | 0.273461 | 0.5446608 | 0.585987 | 0.905026039 | FMOD | protein_coding | 50.17 | 0.006455062 | 0.006455062 | ENSG00000122176 | n.s. |
| ENSG00000189266 | 1397.136 | 0.131512668 | 0.223172 | 0.5892875 | 0.555668 | 0.893144718 | PNRC2 | protein_coding | 43.71 | 0.006454401 | 0.006454401 | ENSG00000189266 | n.s. |
| ENSG00000198836 | 4060.458 | -0.145866097 | 0.263878 | -0.552779 | 0.580415 | 0.903133423 | OPA1 | protein_coding | 36.98 | -0.006454295 | 0.006454295 | ENSG00000198836 | n.s. |
| ENSG00000172572 | 392.7331 | 0.149239679 | 0.274799 | 0.543087 | 0.58707 | 0.905217543 | PDE3A | protein_coding | 35.95 | 0.006454174 | 0.006454174 | ENSG00000172572 | n.s. |
| ENSG00000170579 | 63.07383 | 0.165113684 | 0.3291 | 0.5017132 | 0.615869 | 0.91396346 | DLGAP1 | protein_coding | 39.67 | 0.006451184 | 0.006451184 | ENSG00000170579 | n.s. |
| ENSG00000114544 | 322.4319 | -0.155375372 | 0.298528 | -0.520472 | 0.602735 | 0.908868121 | SLC41A3 | protein_coding | 45.99 | -0.006447943 | 0.006447943 | ENSG00000114544 | n.s. |
| ENSG00000133226 | 1395.181 | 0.131353547 | 0.222174 | 0.5912204 | 0.554373 | 0.893144718 | SRRM1 | protein_coding | 41.76 | 0.006446591 | 0.006446591 | ENSG00000133226 | n.s. |
| ENSG00000174799 | 447.5309 | -0.138970213 | 0.244975 | -0.567283 | 0.570522 | 0.898818733 | CEP135 | protein_coding | 37.81 | -0.006438196 | 0.006438196 | ENSG00000174799 | n.s. |
| ENSG00000233906 | 25.60226 | 0.175458883 | 0.367442 | 0.477515 | 0.632995 | 0.919033489 | | lincRNA | 40.52 | 0.006433843 | 0.006433843 | ENSG00000233906 | n.s. |
| ENSG00000110429 | 965.6646 | 0.150207053 | 0.278463 | 0.5394143 | 0.589601 | 0.906089161 | FBXO3 | protein_coding | 37.03 | 0.006433228 | 0.006433228 | ENSG00000110429 | n.s. |
| ENSG00000079246 | 19637.03 | -0.148706938 | 0.274139 | -0.542451 | 0.587508 | 0.905217543 | XRCC5 | protein_coding | 39.46 | -0.006431135 | 0.006431135 | ENSG00000079246 | n.s. |
| ENSG00000272894 | 69.98321 | 0.15388006 | 0.288164 | 0.5340025 | 0.59334 | 0.908297635 | | lincRNA | 41.47 | 0.00642785 | 0.00642785 | ENSG00000272894 | n.s. |
| ENSG00000011454 | 975.3444 | -0.146618097 | 0.267185 | -0.548752 | 0.583176 | 0.904081462 | RABGAP1 | protein_coding | 38.49 | -0.006420764 | 0.006420764 | ENSG00000011454 | n.s. |
| ENSG00000120053 | 694.0033 | -0.154660864 | 0.296983 | -0.520773 | 0.602525 | 0.908868121 | GOT1 | protein_coding | 42.9 | -0.006418291 | 0.006418291 | ENSG00000120053 | n.s. |
| ENSG00000223734 | 26.16251 | -0.184894569 | 0.402983 | -0.458815 | 0.646367 | 0.923184844 | | lincRNA | 44.27 | -0.006417937 | 0.006417937 | ENSG00000223734 | n.s. |
| ENSG00000128951 | 3680.825 | -0.152330385 | 0.283961 | -0.536448 | 0.591649 | 0.907616853 | DUT | protein_coding | 39.96 | -0.006412721 | 0.006412721 | ENSG00000128951 | n.s. |
| ENSG00000109680 | 347.3621 | -0.148148996 | 0.273016 | -0.542638 | 0.587379 | 0.905217543 | TBC1D19 | protein_coding | 36.57 | -0.006407005 | 0.006407005 | ENSG00000109680 | n.s. |
| ENSG00000149792 | 523.6487 | -0.15589642 | 0.303498 | -0.513666 | 0.607486 | 0.909710915 | MRPL49 | protein_coding | 50.97 | -0.006406812 | 0.006406812 | ENSG00000149792 | n.s. |
| ENSG00000168906 | 16848.62 | -0.128309633 | 0.213304 | -0.601535 | 0.547483 | 0.891406027 | MAT2A | protein_coding | 44.46 | -0.006405786 | 0.006405786 | ENSG00000168906 | n.s. |
| ENSG00000104774 | 283.6011 | -0.14182516 | 0.254563 | -0.557132 | 0.577437 | 0.901408004 | MAN2B1 | protein_coding | 53.98 | -0.006393278 | 0.006393278 | ENSG00000104774 | n.s. |
| ENSG00000173273 | 1570.793 | -0.139968721 | 0.249226 | -0.561614 | 0.574379 | 0.900202572 | TNKS | protein_coding | 37.08 | -0.006390937 | 0.006390937 | ENSG00000173273 | n.s. |
| ENSG00000128513 | 253.8638 | -0.135864926 | 0.23717 | -0.572859 | 0.56674 | 0.897367655 | POT1 | protein_coding | 34.37 | -0.006389672 | 0.006389672 | ENSG00000128513 | n.s. |
| ENSG00000181704 | 820.4726 | -0.153851505 | 0.293296 | -0.52456 | 0.599889 | 0.908868121 | YIPF6 | protein_coding | 41.22 | -0.006384704 | 0.006384704 | ENSG00000181704 | n.s. |
| ENSG00000199400 | 7.899872 | -0.22575511 | 0.575504 | -0.392273 | 0.694856 | 0.936995862 | RNY4P19 | misc_RNA | 37.5 | -0.006380365 | 0.006380365 | ENSG00000199400 | n.s. |
| ENSG00000200608 | 8.208469 | 0.197934493 | 0.463854 | 0.426717 | 0.669585 | 0.928471963 | SNORD114-11 | snoRNA | 49.33 | 0.006379667 | 0.006379667 | ENSG00000200608 | n.s. |
| ENSG00000208005 | 3.932599 | 0.259014523 | 0.758781 | 0.3413562 | 0.732835 | 0.944876691 | MIR503 | miRNA | 59.15 | 0.006378198 | 0.006378198 | ENSG00000208005 | n.s. |
| ENSG00000276234 | 206.4882 | -0.142989557 | 0.258052 | -0.554111 | 0.579503 | 0.902507682 | TADA2A | protein_coding | 42.79 | -0.006370055 | 0.006370055 | ENSG00000276234 | n.s. |
| ENSG00000125352 | 193.4996 | -0.150238038 | 0.279487 | -0.53755 | 0.590888 | 0.907027342 | RNF113A | protein_coding | 54.55 | -0.006367031 | 0.006367031 | ENSG00000125352 | n.s. |
| ENSG00000115461 | 6923.47 | 0.13977175 | 0.249611 | 0.559958 | 0.575508 | 0.900453394 | IGFBP5 | protein_coding | 50.77 | 0.006365032 | 0.006365032 | ENSG00000115461 | n.s. |
| ENSG00000163050 | 1376.251 | 0.1471415 | 0.271465 | 0.5420282 | 0.587799 | 0.905217543 | COQ8A | protein_coding | 49.11 | 0.006363434 | 0.006363434 | ENSG00000163050 | n.s. |

| | | | | | | | | | | | | | |
|---|---|---|---|---|---|---|---|---|---|---|---|---|---|
| ENSG00000176386 | 125.6969 | -0.15643832 | 0.306692 | -0.510083 | 0.609993 | 0.910595741 | CDC26 | protein_coding | 43.07 | -0.006363032 | 0.006363032 | ENSG00000176386 | n.s. |
| ENSG00000091317 | 606.4991 | -0.154812022 | 0.301393 | -0.513655 | 0.607493 | 0.909710915 | CMTM6 | protein_coding | 38.68 | -0.006362247 | 0.006362247 | ENSG00000091317 | n.s. |
| ENSG00000241360 | 1061.68 | -0.153247892 | 0.28895 | -0.530362 | 0.595861 | 0.908868121 | PDXP | protein_coding | 55.3 | -0.006359654 | 0.006359654 | ENSG00000241360 | n.s. |
| ENSG00000134283 | 2311.757 | -0.148346739 | 0.274843 | -0.539751 | 0.589369 | 0.906089161 | PPHLN1 | protein_coding | 39.77 | -0.006353552 | 0.006353552 | ENSG00000134283 | n.s. |
| ENSG00000078070 | 571.1048 | -0.146750824 | 0.270247 | -0.543025 | 0.587113 | 0.905217543 | MCCC1 | protein_coding | 41.62 | -0.006346538 | 0.006346538 | ENSG00000078070 | n.s. |
| ENSG00000116521 | 487.7061 | -0.141623781 | 0.25499 | -0.55541 | 0.578615 | 0.901989749 | SCAMP3 | protein_coding | 51.95 | -0.006344519 | 0.006344519 | ENSG00000116521 | n.s. |
| ENSG00000196345 | 773.2838 | -0.131346645 | 0.226054 | -0.581041 | 0.561213 | 0.894765412 | ZKSCAN7 | protein_coding | 45.82 | -0.006342836 | 0.006342836 | ENSG00000196345 | n.s. |
| ENSG00000172809 | 18595.22 | 0.152841264 | 0.295151 | 0.5178409 | 0.604569 | 0.908868121 | RPL38 | protein_coding | 48.98 | 0.006342779 | 0.006342779 | ENSG00000172809 | n.s. |
| ENSG00000103275 | 2507.99 | -0.137745714 | 0.243735 | -0.565145 | 0.571975 | 0.899491841 | UBE2I | protein_coding | 59.46 | -0.006336685 | 0.006336685 | ENSG00000103275 | n.s. |
| ENSG00000204536 | 189.7471 | -0.146173333 | 0.268193 | -0.545031 | 0.585732 | 0.905026039 | CCHCR1 | protein_coding | 51.86 | -0.006334995 | 0.006334995 | ENSG00000204536 | n.s. |
| ENSG00000223305 | 26.75992 | -0.170848411 | 0.347996 | -0.49095 | 0.623462 | 0.918167932 | RN7SKP30 | misc_RNA | 50.6 | -0.006334697 | 0.006334697 | ENSG00000223305 | n.s. |
| ENSG00000125503 | 402.3352 | -0.152128126 | 0.286456 | -0.531071 | 0.59537 | 0.908868121 | PPP1R12C | protein_coding | 57.6 | -0.006313185 | 0.006313185 | ENSG00000125503 | n.s. |
| ENSG00000136802 | 617.4123 | -0.138785788 | 0.248102 | -0.559389 | 0.575896 | 0.900580379 | LRRC8A | protein_coding | 53.83 | -0.006311633 | 0.006311633 | ENSG00000136802 | n.s. |
| ENSG00000212747 | 474.5096 | 0.131136152 | 0.226088 | 0.5800218 | 0.5619 | 0.895116382 | RTL8B | protein_coding | 51.18 | 0.006310336 | 0.006310336 | ENSG00000212747 | n.s. |
| ENSG00000006715 | 1223.174 | -0.152044493 | 0.291574 | -0.52146 | 0.602046 | 0.908868121 | VPS41 | protein_coding | 38.35 | -0.006309714 | 0.006309714 | ENSG00000006715 | n.s. |
| ENSG00000155660 | 775.6954 | -0.145806572 | 0.267896 | -0.544266 | 0.586258 | 0.905188828 | PDIA4 | protein_coding | 50.87 | -0.006307711 | 0.006307711 | ENSG00000155660 | n.s. |
| ENSG00000273107 | 27.46129 | -0.177080089 | 0.379119 | -0.467084 | 0.64044 | 0.921257134 | | lincRNA | 49.08 | -0.00630744 | 0.00630744 | ENSG00000273107 | n.s. |
| ENSG00000116478 | 414.8516 | 0.151949808 | 0.292105 | 0.5201892 | 0.602932 | 0.908868121 | HDAC1 | protein_coding | 46.79 | 0.006305785 | 0.006305785 | ENSG00000116478 | n.s. |
| ENSG00000120539 | 1498.722 | -0.151906894 | 0.286595 | -0.53004 | 0.596084 | 0.908868121 | MASTL | protein_coding | 41.49 | -0.006304004 | 0.006304004 | ENSG00000120539 | n.s. |
| ENSG00000174238 | 508.765 | -0.151793421 | 0.288966 | -0.525299 | 0.599376 | 0.908868121 | PITPNA | protein_coding | 49.88 | -0.006299295 | 0.006299295 | ENSG00000174238 | n.s. |
| ENSG00000170502 | 354.9749 | -0.14483719 | 0.264524 | -0.547539 | 0.584008 | 0.904712915 | NUDT9 | protein_coding | 38.92 | -0.006298855 | 0.006298855 | ENSG00000170502 | n.s. |
| ENSG00000136824 | 3457.218 | -0.145330111 | 0.26653 | -0.545268 | 0.585569 | 0.905026039 | SMC2 | protein_coding | 35.98 | -0.006298451 | 0.006298451 | ENSG00000136824 | n.s. |
| ENSG00000100982 | 483.927 | -0.138304857 | 0.246893 | -0.56018 | 0.575357 | 0.900453394 | PCIF1 | protein_coding | 53.89 | -0.006298232 | 0.006298232 | ENSG00000100982 | n.s. |
| ENSG00000129038 | 609.2724 | -0.137499612 | 0.244545 | -0.562267 | 0.573934 | 0.899911687 | LOXL1 | protein_coding | 55.02 | -0.006297497 | 0.006297497 | ENSG00000129038 | n.s. |
| ENSG00000061936 | 2688.132 | -0.128251616 | 0.216643 | -0.591994 | 0.553855 | 0.893144718 | SFSWAP | protein_coding | 46.45 | -0.006294354 | 0.006294354 | ENSG00000061936 | n.s. |
| ENSG00000158552 | 182.6193 | -0.146530575 | 0.271202 | -0.540301 | 0.58899 | 0.90583249 | ZFAND2B | protein_coding | 50.41 | -0.006293797 | 0.006293797 | ENSG00000158552 | n.s. |
| ENSG00000169696 | 778.7479 | -0.145208825 | 0.266587 | -0.544697 | 0.585962 | 0.905026039 | ASPSCR1 | protein_coding | 60.76 | -0.006293194 | 0.006293194 | ENSG00000169696 | n.s. |
| ENSG00000173456 | 205.9109 | -0.139718233 | 0.251045 | -0.556546 | 0.577838 | 0.901500147 | RNF26 | protein_coding | 59.17 | -0.006292099 | 0.006292099 | ENSG00000173456 | n.s. |
| ENSG00000229588 | 18.72797 | 0.225221576 | 0.588849 | 0.3824776 | 0.702107 | 0.937752951 | | lincRNA | 44.6 | 0.006286286 | 0.006286286 | ENSG00000229588 | n.s. |
| ENSG00000277324 | 39.97219 | -0.177905278 | 0.382039 | -0.465673 | 0.64145 | 0.921891013 | | antisense | 51.28 | -0.006283689 | 0.006283689 | ENSG00000277324 | n.s. |
| ENSG00000048649 | 2211.465 | -0.144858997 | 0.265535 | -0.545536 | 0.585385 | 0.905026039 | RSF1 | protein_coding | 39.04 | -0.006278033 | 0.006278033 | ENSG00000048649 | n.s. |
| ENSG00000139289 | 1605.553 | 0.123981644 | 0.201359 | 0.6157241 | 0.538077 | 0.889966281 | PHLDA1 | protein_coding | 45.13 | 0.00627675 | 0.00627675 | ENSG00000139289 | n.s. |

| | | | | | | | | | | | | | |
|---|---|---|---|---|---|---|---|---|---|---|---|---|---|
| ENSG00000221539 | 27.5799 | -0.180806802 | 0.392168 | -0.461045 | 0.644767 | 0.923184844 | SNORD99 | snoRNA | 47.3 | -0.006276045 | 0.006276045 | ENSG00000221539 | n.s. |
| ENSG00000198040 | 4843.552 | -0.139152225 | 0.249752 | -0.557162 | 0.577416 | 0.901408004 | ZNF84 | protein_coding | 39.16 | -0.006272786 | 0.006272786 | ENSG00000198040 | n.s. |
| ENSG00000145996 | 1032.508 | -0.136081234 | 0.240459 | -0.565922 | 0.571447 | 0.899418628 | CDKAL1 | protein_coding | 38.04 | -0.006264924 | 0.006264924 | ENSG00000145996 | n.s. |
| ENSG00000143622 | 433.8698 | -0.141676042 | 0.256487 | -0.552371 | 0.580694 | 0.90320682 | RIT1 | protein_coding | 43.8 | -0.006263893 | 0.006263893 | ENSG00000143622 | n.s. |
| ENSG00000158286 | 127.5083 | -0.150757334 | 0.2834 | -0.53196 | 0.594754 | 0.908868121 | RNF207 | protein_coding | 57.32 | -0.006256298 | 0.006256298 | ENSG00000158286 | n.s. |
| ENSG00000106588 | 653.9728 | 0.144545119 | 0.265925 | 0.543557 | 0.586746 | 0.905217543 | PSMA2 | protein_coding | 38.3 | 0.006251148 | 0.006251148 | ENSG00000106588 | n.s. |
| ENSG00000160049 | 929.4328 | -0.150609003 | 0.286298 | -0.526056 | 0.598849 | 0.908868121 | DFFA | protein_coding | 48.52 | -0.006250143 | 0.006250143 | ENSG00000160049 | n.s. |
| ENSG00000231770 | 146.9462 | -0.134169271 | 0.235161 | -0.570543 | 0.56831 | 0.898419074 | TMEM44-AS1 | antisense | 48.38 | -0.006241694 | 0.006241694 | ENSG00000231770 | n.s. |
| ENSG00000129673 | 68.09277 | 0.150377662 | 0.286977 | 0.5240053 | 0.600275 | 0.908868121 | AANAT | protein_coding | 53.01 | 0.006240542 | 0.006240542 | ENSG00000129673 | n.s. |
| ENSG00000112699 | 153.1245 | -0.155008407 | 0.305927 | -0.506684 | 0.612376 | 0.911493921 | GMDS | protein_coding | 40.78 | -0.006238503 | 0.006238503 | ENSG00000112699 | n.s. |
| ENSG00000112130 | 615.8245 | -0.146965786 | 0.273226 | -0.53789 | 0.590653 | 0.906900529 | RNF8 | protein_coding | 43.66 | -0.006237279 | 0.006237279 | ENSG00000112130 | n.s. |
| ENSG00000256771 | 1144.335 | -0.143839961 | 0.26387 | -0.545116 | 0.585674 | 0.905026039 | ZNF253 | protein_coding | 38.27 | -0.006233869 | 0.006233869 | ENSG00000256771 | n.s. |
| ENSG00000172336 | 649.2651 | -0.152095372 | 0.296585 | -0.512822 | 0.608076 | 0.909957073 | POP7 | protein_coding | 59.81 | -0.006232731 | 0.006232731 | ENSG00000172336 | n.s. |
| ENSG00000066027 | 213.2534 | -0.139888743 | 0.252419 | -0.554193 | 0.579447 | 0.902507682 | PPP2R5A | protein_coding | 39.28 | -0.006231917 | 0.006231917 | ENSG00000066027 | n.s. |
| ENSG00000162972 | 589.8445 | 0.143968372 | 0.265063 | 0.5431486 | 0.587027 | 0.905217543 | MAIP1 | protein_coding | 39.93 | 0.006226206 | 0.006226206 | ENSG00000162972 | n.s. |
| ENSG00000004897 | 3993.35 | -0.145335453 | 0.269479 | -0.539321 | 0.589665 | 0.906089161 | CDC27 | protein_coding | 38.66 | -0.006224581 | 0.006224581 | ENSG00000004897 | n.s. |
| ENSG00000168214 | 2374.721 | -0.1411692 | 0.256092 | -0.551245 | 0.581466 | 0.90349457 | RBPJ | protein_coding | 40.79 | -0.006221955 | 0.006221955 | ENSG00000168214 | n.s. |
| ENSG00000173465 | 256.622 | -0.14991987 | 0.284748 | -0.526499 | 0.598541 | 0.908868121 | SSSCA1 | protein_coding | 61.32 | -0.006221544 | 0.006221544 | ENSG00000173465 | n.s. |
| ENSG00000115241 | 2311.641 | -0.148915265 | 0.278985 | -0.533775 | 0.593498 | 0.908297635 | PPM1G | protein_coding | 45.48 | -0.006220461 | 0.006220461 | ENSG00000115241 | n.s. |
| ENSG00000124228 | 986.0865 | -0.1404759 | 0.25422 | -0.552577 | 0.580553 | 0.903133423 | DDX27 | protein_coding | 49.05 | -0.00621579 | 0.00621579 | ENSG00000124228 | n.s. |
| ENSG00000122203 | 1437.043 | -0.143970361 | 0.265985 | -0.541272 | 0.58832 | 0.905402821 | KIAA1191 | protein_coding | 47.1 | -0.006213495 | 0.006213495 | ENSG00000122203 | n.s. |
| ENSG00000146425 | 4368.702 | -0.149715802 | 0.287827 | -0.520159 | 0.602953 | 0.908868121 | DYNLT1 | protein_coding | 45.31 | -0.006213075 | 0.006213075 | ENSG00000146425 | n.s. |
| ENSG00000267526 | 151.1888 | -0.155236339 | 0.307184 | -0.505354 | 0.61331 | 0.911981962 | | processed_pseudogene | 38.22 | -0.006211588 | 0.006211588 | ENSG00000267526 | n.s. |
| ENSG00000156876 | 978.3161 | -0.143271784 | 0.262934 | -0.544896 | 0.585825 | 0.905026039 | SASS6 | protein_coding | 36.48 | -0.006209245 | 0.006209245 | ENSG00000156876 | n.s. |
| ENSG00000198034 | 19575.83 | 0.152561362 | 0.299319 | 0.5096941 | 0.610266 | 0.910595741 | RPS4X | protein_coding | 46.44 | 0.00620534 | 0.00620534 | ENSG00000198034 | n.s. |
| ENSG00000131788 | 184.028 | -0.14078445 | 0.256076 | -0.549776 | 0.582473 | 0.90349457 | PIAS3 | protein_coding | 52.28 | -0.006204998 | 0.006204998 | ENSG00000131788 | n.s. |
| ENSG00000083635 | 514.4909 | -0.140676744 | 0.25531 | -0.551003 | 0.581632 | 0.90349457 | NUFIP1 | protein_coding | 38.71 | -0.006200251 | 0.006200251 | ENSG00000083635 | n.s. |
| ENSG00000186314 | 91.50354 | -0.146163012 | 0.271991 | -0.537381 | 0.591004 | 0.907027342 | PRELID2 | protein_coding | 38.53 | -0.006194333 | 0.006194333 | ENSG00000186314 | n.s. |
| ENSG00000104723 | 2143.105 | -0.149189976 | 0.280233 | -0.532379 | 0.594464 | 0.908830158 | TUSC3 | protein_coding | 36.98 | -0.006193961 | 0.006193961 | ENSG00000104723 | n.s. |
| ENSG00000228638 | 46.44208 | -0.168902361 | 0.34717 | -0.486512 | 0.626604 | 0.919033489 | FCF1P2 | processed_pseudogene | 40.48 | -0.006193424 | 0.006193424 | ENSG00000228638 | n.s. |
| ENSG00000114738 | 247.1468 | -0.149210827 | 0.287871 | -0.518326 | 0.604231 | 0.908868121 | MAPKAPK3 | protein_coding | 53.69 | -0.006192119 | 0.006192119 | ENSG00000114738 | n.s. |
| ENSG00000183579 | 1314.051 | -0.143088618 | 0.26419 | -0.541613 | 0.588085 | 0.905341301 | ZNRF3 | protein_coding | 45.29 | -0.006179664 | 0.006179664 | ENSG00000183579 | n.s. |

| | | | | | | | | | | | | | |
|---|---|---|---|---|---|---|---|---|---|---|---|---|---|
| ENSG00000090432 | 160.1588 | -0.148908486 | 0.285808 | -0.521009 | 0.60236 | 0.908868121 | MUL1 | protein_coding | 47.52 | -0.006179573 | 0.006179573 | ENSG00000090432 | n.s. |
| ENSG00000174013 | 1044.465 | -0.148762926 | 0.282891 | -0.525866 | 0.598981 | 0.908868121 | FBXO45 | protein_coding | 41.25 | -0.006173532 | 0.006173532 | ENSG00000174013 | n.s. |
| ENSG00000100129 | 3391.411 | 0.148711577 | 0.286324 | 0.5193817 | 0.603495 | 0.908868121 | EIF3L | protein_coding | 46 | 0.006171401 | 0.006171401 | ENSG00000100129 | n.s. |
| ENSG00000048544 | 640.7611 | -0.151334444 | 0.295798 | -0.511614 | 0.608921 | 0.910431619 | MRPS10 | protein_coding | 41.04 | -0.006167282 | 0.006167282 | ENSG00000048544 | n.s. |
| ENSG00000023697 | 1069.163 | -0.148541603 | 0.28266 | -0.525513 | 0.599227 | 0.908868121 | DERA | protein_coding | 39.16 | -0.006164347 | 0.006164347 | ENSG00000023697 | n.s. |
| ENSG00000075407 | 1569.937 | -0.148336812 | 0.283355 | -0.523501 | 0.600625 | 0.908868121 | ZNF37A | protein_coding | 39.11 | -0.006155849 | 0.006155849 | ENSG00000075407 | n.s. |
| ENSG00000094914 | 814.5634 | -0.148271765 | 0.283191 | -0.523575 | 0.600574 | 0.908868121 | AAAS | protein_coding | 49.39 | -0.006153149 | 0.006153149 | ENSG00000094914 | n.s. |
| ENSG00000172531 | 1100.525 | -0.151275279 | 0.296828 | -0.50964 | 0.610304 | 0.910595741 | PPP1CA | protein_coding | 61.16 | -0.006153029 | 0.006153029 | ENSG00000172531 | n.s. |
| ENSG00000136021 | 1529.16 | -0.145184625 | 0.270166 | -0.53739 | 0.590999 | 0.907027342 | SCYL2 | protein_coding | 37.97 | -0.006152869 | 0.006152869 | ENSG00000136021 | n.s. |
| ENSG00000174227 | 337.966 | 0.131040843 | 0.228995 | 0.5722424 | 0.567158 | 0.8975378 | PIGG | protein_coding | 49.98 | 0.006152007 | 0.006152007 | ENSG00000174227 | n.s. |
| ENSG00000136842 | 409.2555 | -0.151162097 | 0.296658 | -0.509551 | 0.610366 | 0.910595741 | TMOD1 | protein_coding | 46.76 | -0.006148425 | 0.006148425 | ENSG00000136842 | n.s. |
| ENSG00000115841 | 377.6754 | -0.148130464 | 0.281452 | -0.526308 | 0.598675 | 0.908868121 | RMDN2 | protein_coding | 39.02 | -0.006147285 | 0.006147285 | ENSG00000115841 | n.s. |
| ENSG00000181817 | 56.41125 | -0.157662516 | 0.314729 | -0.500947 | 0.616408 | 0.914172888 | LSM10 | protein_coding | 50.65 | -0.00614437 | 0.00614437 | ENSG00000181817 | n.s. |
| ENSG00000102753 | 1330.35 | -0.143450659 | 0.265737 | -0.539821 | 0.58932 | 0.906089161 | KPNA3 | protein_coding | 38.71 | -0.006143858 | 0.006143858 | ENSG00000102753 | n.s. |
| ENSG00000107362 | 777.2271 | -0.147999779 | 0.283283 | -0.522446 | 0.60136 | 0.908868121 | ABHD17B | protein_coding | 37.98 | -0.006141862 | 0.006141862 | ENSG00000107362 | n.s. |
| ENSG00000036549 | 3045.133 | -0.142734348 | 0.263939 | -0.540784 | 0.588656 | 0.905723898 | ZZZ3 | protein_coding | 37.17 | -0.006138173 | 0.006138173 | ENSG00000036549 | n.s. |
| ENSG00000129757 | 148.6811 | 0.147860181 | 0.281812 | 0.5246776 | 0.599807 | 0.908868121 | CDKN1C | protein_coding | 67.44 | 0.006136069 | 0.006136069 | ENSG00000129757 | n.s. |
| ENSG00000196821 | 584.6808 | -0.141074367 | 0.257672 | -0.547496 | 0.584038 | 0.904712915 | C6orf106 | protein_coding | 41.83 | -0.006135213 | 0.006135213 | ENSG00000196821 | n.s. |
| ENSG00000277130 | 117.4271 | -0.149020435 | 0.290244 | -0.513431 | 0.60765 | 0.909710915 | | sense_intronic | 33.28 | -0.006124232 | 0.006124232 | ENSG00000277130 | n.s. |
| ENSG00000100852 | 2666.192 | -0.147539141 | 0.278909 | -0.528987 | 0.596815 | 0.908868121 | ARHGAP5 | protein_coding | 36.95 | -0.006122746 | 0.006122746 | ENSG00000100852 | n.s. |
| ENSG00000136830 | 400.4156 | -0.147421444 | 0.280043 | -0.526425 | 0.598593 | 0.908868121 | FAM129B | protein_coding | 54.31 | -0.006117862 | 0.006117862 | ENSG00000136830 | n.s. |
| ENSG00000251050 | 13.26263 | -0.227888277 | 0.623972 | -0.365222 | 0.714946 | 0.940060506 | | processed_pseudogene | 47.44 | -0.006117477 | 0.006117477 | ENSG00000251050 | n.s. |
| ENSG00000173208 | 466.4949 | 0.125495391 | 0.214546 | 0.584934 | 0.558592 | 0.893837232 | ABCD2 | protein_coding | 34.95 | 0.006116841 | 0.006116841 | ENSG00000173208 | n.s. |
| ENSG00000010292 | 2980.775 | -0.14113383 | 0.258251 | -0.546499 | 0.584723 | 0.905026039 | NCAPD2 | protein_coding | 46.34 | -0.006116588 | 0.006116588 | ENSG00000010292 | n.s. |
| ENSG00000119684 | 254.5461 | -0.146383639 | 0.2742 | -0.533857 | 0.593441 | 0.908297635 | MLH3 | protein_coding | 41.37 | -0.006114711 | 0.006114711 | ENSG00000119684 | n.s. |
| ENSG00000212402 | 18.74047 | -0.176137071 | 0.386883 | -0.455272 | 0.648914 | 0.923184844 | SNORA74B | snoRNA | 54.73 | -0.006113953 | 0.006113953 | ENSG00000212402 | n.s. |
| ENSG00000204856 | 915.5144 | -0.151904596 | 0.299914 | -0.506493 | 0.61251 | 0.911498898 | FAM216A | protein_coding | 43.77 | -0.006113226 | 0.006113226 | ENSG00000204856 | n.s. |
| ENSG00000171130 | 315.885 | -0.147207677 | 0.282679 | -0.520759 | 0.602534 | 0.908868121 | ATP6V0E2 | protein_coding | 54.93 | -0.00610899 | 0.00610899 | ENSG00000171130 | n.s. |
| ENSG00000179431 | 824.9531 | 0.124308008 | 0.209646 | 0.5929413 | 0.55322 | 0.893144718 | FJX1 | protein_coding | 63.8 | 0.006100809 | 0.006100809 | ENSG00000179431 | n.s. |
| ENSG00000123080 | 2674.789 | -0.13798614 | 0.249831 | -0.552318 | 0.58073 | 0.90320682 | CDKN2C | protein_coding | 40.97 | -0.006100752 | 0.006100752 | ENSG00000123080 | n.s. |
| ENSG00000145808 | 761.8078 | 0.14569024 | 0.272988 | 0.5336882 | 0.593557 | 0.908297635 | ADAMTS19 | protein_coding | 34.94 | 0.006085746 | 0.006085746 | ENSG00000145808 | n.s. |
| ENSG00000182923 | 549.083 | -0.131351111 | 0.231291 | -0.567905 | 0.5701 | 0.898818733 | CEP63 | protein_coding | 38.49 | -0.006085219 | 0.006085219 | ENSG00000182923 | n.s. |

| | | | | | | | | | | | | | |
|---|---|---|---|---|---|---|---|---|---|---|---|---|---|
| ENSG00000278540 | 3147.426 | -0.14034085 | 0.256739 | -0.546628 | 0.584634 | 0.905026039 | ACACA | protein_coding | 41.27 | -0.006082222 | 0.006082222 | ENSG00000278540 | n.s. |
| ENSG00000258315 | 295.5238 | -0.155635417 | 0.310024 | -0.50201 | 0.61566 | 0.91396346 | C17orf49 | protein_coding | 58.07 | -0.006080857 | 0.006080857 | ENSG00000258315 | n.s. |
| ENSG00000174628 | 29.29412 | 0.164000376 | 0.334231 | 0.4906804 | 0.623653 | 0.918167932 | IQCK | protein_coding | 43.85 | 0.006080786 | 0.006080786 | ENSG00000174628 | n.s. |
| ENSG00000233325 | 29.34804 | -0.165655525 | 0.340012 | -0.487206 | 0.626113 | 0.918973809 | MIPEPP3 | cribed_unprocessed_pseudc | 39.33 | -0.006079039 | 0.006079039 | ENSG00000233325 | n.s. |
| ENSG00000105173 | 106.2322 | -0.140484799 | 0.258648 | -0.543151 | 0.587026 | 0.905217543 | CCNE1 | protein_coding | 47.46 | -0.006075551 | 0.006075551 | ENSG00000105173 | n.s. |
| ENSG00000143862 | 462.5332 | -0.146326371 | 0.275743 | -0.530663 | 0.595652 | 0.908868121 | ARL8A | protein_coding | 53.33 | -0.006072417 | 0.006072417 | ENSG00000143862 | n.s. |
| ENSG00000252464 | 25.2357 | 0.165601956 | 0.34721 | 0.4769503 | 0.633397 | 0.919033489 | RN7SKP70 | misc_RNA | 51.29 | 0.006072402 | 0.006072402 | ENSG00000252464 | n.s. |
| ENSG00000181072 | 1530.508 | -0.126141776 | 0.217692 | -0.579452 | 0.562285 | 0.895116382 | CHRM2 | protein_coding | 37.16 | -0.006070005 | 0.006070005 | ENSG00000181072 | n.s. |
| ENSG00000102218 | 208.4454 | -0.149205434 | 0.292616 | -0.509902 | 0.61012 | 0.910595741 | RP2 | protein_coding | 39.34 | -0.006068839 | 0.006068839 | ENSG00000102218 | n.s. |
| ENSG00000182621 | 751.8877 | -0.142797846 | 0.26528 | -0.53829 | 0.590377 | 0.906863989 | PLCB1 | protein_coding | 38.19 | -0.006062889 | 0.006062889 | ENSG00000182621 | n.s. |
| ENSG00000107863 | 1573.111 | -0.137556281 | 0.250078 | -0.550053 | 0.582283 | 0.90349457 | ARHGAP21 | protein_coding | 38.96 | -0.006062718 | 0.006062718 | ENSG00000107863 | n.s. |
| ENSG00000012822 | 349.7081 | 0.148139764 | 0.289095 | 0.5124256 | 0.608353 | 0.910176404 | CALCOCO1 | protein_coding | 49.2 | 0.006055128 | 0.006055128 | ENSG00000012822 | n.s. |
| ENSG00000179562 | 646.9037 | 0.125717964 | 0.216783 | 0.579925 | 0.561965 | 0.895116382 | GCC1 | protein_coding | 52.07 | 0.006049611 | 0.006049611 | ENSG00000179562 | n.s. |
| ENSG00000127804 | 951.5035 | -0.144662951 | 0.270292 | -0.535209 | 0.592505 | 0.908229748 | METTL16 | protein_coding | 44.67 | -0.00604753 | 0.00604753 | ENSG00000127804 | n.s. |
| ENSG00000120526 | 1627.31 | -0.148902902 | 0.292891 | -0.50839 | 0.61118 | 0.910755785 | NUDCD1 | protein_coding | 36.6 | -0.006045169 | 0.006045169 | ENSG00000120526 | n.s. |
| ENSG00000198604 | 4097.359 | -0.146486505 | 0.284235 | -0.515371 | 0.606294 | 0.909522325 | BAZ1A | protein_coding | 39.75 | -0.006033286 | 0.006033286 | ENSG00000198604 | n.s. |
| ENSG00000086232 | 1597.641 | -0.145367402 | 0.277283 | -0.524257 | 0.6001 | 0.908868121 | EIF2AK1 | protein_coding | 44.7 | -0.006032621 | 0.006032621 | ENSG00000086232 | n.s. |
| ENSG00000200632 | 3.089427 | 0.236134057 | 0.678535 | 0.3480059 | 0.727836 | 0.942883666 | SNORD113-7 | snoRNA | 41.56 | 0.00603131 | 0.00603131 | ENSG00000200632 | n.s. |
| ENSG00000276390 | 64.27354 | -0.164261997 | 0.340175 | -0.482875 | 0.629184 | 0.919033489 | | sense_intronic | 32.88 | -0.006023268 | 0.006023268 | ENSG00000276390 | n.s. |
| ENSG00000219200 | 222.5595 | -0.145716677 | 0.281761 | -0.517163 | 0.605042 | 0.909228554 | RNASEK | protein_coding | 57.04 | -0.006022023 | 0.006022023 | ENSG00000219200 | n.s. |
| ENSG00000108515 | 54.60893 | -0.168018077 | 0.358473 | -0.468706 | 0.63928 | 0.920825292 | ENO3 | protein_coding | 58.22 | -0.006018871 | 0.006018871 | ENSG00000108515 | n.s. |
| ENSG00000105223 | 570.7453 | -0.144982755 | 0.274129 | -0.528885 | 0.596885 | 0.908868121 | PLD3 | protein_coding | 52.33 | -0.006016658 | 0.006016658 | ENSG00000105223 | n.s. |
| ENSG00000151849 | 1699.895 | -0.14497422 | 0.280031 | -0.517708 | 0.604662 | 0.908868121 | CENPJ | protein_coding | 41.37 | -0.006016304 | 0.006016304 | ENSG00000151849 | n.s. |
| ENSG00000110931 | 863.4392 | -0.14493886 | 0.279727 | -0.518145 | 0.604357 | 0.908868121 | CAMKK2 | protein_coding | 50.83 | -0.006014837 | 0.006014837 | ENSG00000110931 | n.s. |
| ENSG00000184076 | 1143.399 | 0.153680524 | 0.306429 | 0.5015208 | 0.616005 | 0.91396346 | UQCR10 | protein_coding | 47.62 | 0.006004477 | 0.006004477 | ENSG00000184076 | n.s. |
| ENSG00000175110 | 1137.676 | 0.144221154 | 0.27075 | 0.5326727 | 0.59426 | 0.908718767 | MRPS22 | protein_coding | 42.84 | 0.005995346 | 0.005995346 | ENSG00000175110 | n.s. |
| ENSG00000267606 | 40.70402 | 0.154637334 | 0.30926 | 0.5000236 | 0.617058 | 0.9146497 | | lincRNA | 40.14 | 0.005991455 | 0.005991455 | ENSG00000267606 | n.s. |
| ENSG00000197157 | 2748.612 | -0.14437023 | 0.273939 | -0.527017 | 0.598182 | 0.908868121 | SND1 | protein_coding | 42.44 | -0.005991239 | 0.005991239 | ENSG00000197157 | n.s. |
| ENSG00000261120 | 21.85997 | 0.177539703 | 0.405676 | 0.4376386 | 0.661648 | 0.925277395 | | antisense | 59.54 | 0.005988068 | 0.005988068 | ENSG00000261120 | n.s. |
| ENSG00000152492 | 2490.447 | -0.143061003 | 0.267882 | -0.534045 | 0.59331 | 0.908297635 | CCDC50 | protein_coding | 36.59 | -0.005975918 | 0.005975918 | ENSG00000152492 | n.s. |
| ENSG00000003249 | 132.6729 | -0.143901855 | 0.27172 | -0.529596 | 0.596392 | 0.908868121 | DBNDD1 | protein_coding | 59.39 | -0.005971802 | 0.005971802 | ENSG00000003249 | n.s. |
| ENSG00000136982 | 780.3502 | -0.144983907 | 0.280867 | -0.516201 | 0.605714 | 0.909522325 | DSCC1 | protein_coding | 42.08 | -0.005971399 | 0.005971399 | ENSG00000136982 | n.s. |

| | | | | | | | | | | | | | |
|---|---|---|---|---|---|---|---|---|---|---|---|---|---|
| ENSG00000105438 | 707.0427 | -0.142839547 | 0.267309 | -0.534362 | 0.593091 | 0.908297635 | KDELR1 | protein_coding | 53.35 | -0.005966667 | 0.005966667 | ENSG00000105438 | n.s. |
| ENSG00000135775 | 1986.575 | -0.146398665 | 0.286456 | -0.511069 | 0.609303 | 0.910431619 | COG2 | protein_coding | 39.42 | -0.005966136 | 0.005966136 | ENSG00000135775 | n.s. |
| ENSG00000198556 | 492.0858 | -0.137927872 | 0.253968 | -0.543091 | 0.587067 | 0.905217543 | ZNF789 | protein_coding | 47.78 | -0.005964972 | 0.005964972 | ENSG00000198556 | n.s. |
| ENSG00000170515 | 6061.373 | -0.144683848 | 0.280607 | -0.515611 | 0.606126 | 0.909522325 | PA2G4 | protein_coding | 46.6 | -0.005959041 | 0.005959041 | ENSG00000170515 | n.s. |
| ENSG00000265083 | 2.701851 | -0.244292164 | 0.72272 | -0.338018 | 0.73535 | 0.945434013 | MIR3691 | miRNA | 57.78 | -0.005953101 | 0.005953101 | ENSG00000265083 | n.s. |
| ENSG00000184271 | 46.40803 | -0.161295873 | 0.329539 | -0.489458 | 0.624517 | 0.918566713 | POU6F1 | protein_coding | 50.08 | -0.005950091 | 0.005950091 | ENSG00000184271 | n.s. |
| ENSG00000162819 | 953.737 | -0.142985808 | 0.27095 | -0.527721 | 0.597693 | 0.908868121 | BROX | protein_coding | 36.81 | -0.005933787 | 0.005933787 | ENSG00000162819 | n.s. |
| ENSG00000103037 | 1111.801 | 0.128041268 | 0.225373 | 0.5681306 | 0.569946 | 0.898818733 | SETD6 | protein_coding | 48.83 | 0.005931881 | 0.005931881 | ENSG00000103037 | n.s. |
| ENSG00000107281 | 570.4775 | -0.137958995 | 0.255242 | -0.540502 | 0.588851 | 0.905738346 | NPDC1 | protein_coding | 65.86 | -0.005931857 | 0.005931857 | ENSG00000107281 | n.s. |
| ENSG00000094841 | 175.2189 | -0.151720444 | 0.302458 | -0.501625 | 0.615931 | 0.91396346 | UPRT | protein_coding | 37.52 | -0.005927895 | 0.005927895 | ENSG00000094841 | n.s. |
| ENSG00000163781 | 3768.375 | -0.142761272 | 0.270441 | -0.527882 | 0.597581 | 0.908868121 | TOPBP1 | protein_coding | 37.82 | -0.005924469 | 0.005924469 | ENSG00000163781 | n.s. |
| ENSG00000207523 | 123.0351 | -0.136661766 | 0.250454 | -0.545657 | 0.585302 | 0.905026039 | SNORA66 | snoRNA | 47.37 | -0.005922774 | 0.005922774 | ENSG00000207523 | n.s. |
| ENSG00000140521 | 1544.019 | -0.129682694 | 0.231097 | -0.561162 | 0.574687 | 0.900202572 | POLG | protein_coding | 49.14 | -0.005921279 | 0.005921279 | ENSG00000140521 | n.s. |
| ENSG00000138439 | 1304.783 | -0.141236614 | 0.26373 | -0.535536 | 0.592279 | 0.908010607 | FAM117B | protein_coding | 39.53 | -0.005919096 | 0.005919096 | ENSG00000138439 | n.s. |
| ENSG00000114857 | 5406.584 | -0.137772603 | 0.255043 | -0.540193 | 0.589064 | 0.905847072 | NKTR | protein_coding | 37.54 | -0.00591666 | 0.00591666 | ENSG00000114857 | n.s. |
| ENSG00000075292 | 6917.935 | -0.14250776 | 0.273219 | -0.521588 | 0.601957 | 0.908868121 | ZNF638 | protein_coding | 38.16 | -0.005913948 | 0.005913948 | ENSG00000075292 | n.s. |
| ENSG00000037241 | 441.5751 | -0.161103902 | 0.330748 | -0.487089 | 0.626195 | 0.918973809 | RPL26L1 | protein_coding | 46.12 | -0.005912008 | 0.005912008 | ENSG00000037241 | n.s. |
| ENSG00000115840 | 699.1771 | -0.145320987 | 0.284663 | -0.510502 | 0.6097 | 0.910595741 | SLC25A12 | protein_coding | 40.22 | -0.005910842 | 0.005910842 | ENSG00000115840 | n.s. |
| ENSG00000170364 | 973.8584 | 0.136382581 | 0.249978 | 0.5455791 | 0.585355 | 0.905026039 | SETMAR | protein_coding | 40.2 | 0.005910674 | 0.005910674 | ENSG00000170364 | n.s. |
| ENSG00000234884 | 39.20792 | -0.161042585 | 0.332387 | -0.484503 | 0.628029 | 0.919033489 | | antisense | 45.01 | -0.005905216 | 0.005905216 | ENSG00000234884 | n.s. |
| ENSG00000103047 | 418.476 | -0.142249094 | 0.27291 | -0.521231 | 0.602206 | 0.908868121 | TANGO6 | protein_coding | 43.08 | -0.005903214 | 0.005903214 | ENSG00000103047 | n.s. |
| ENSG00000111596 | 3474.75 | -0.142175939 | 0.274683 | -0.5176 | 0.604737 | 0.908868121 | CNOT2 | protein_coding | 35.15 | -0.005900178 | 0.005900178 | ENSG00000111596 | n.s. |
| ENSG00000158164 | 1359.294 | 0.16080635 | 0.333365 | 0.4823734 | 0.629541 | 0.919033489 | TMSB15A | protein_coding | 43.49 | 0.005896554 | 0.005896554 | ENSG00000158164 | n.s. |
| ENSG00000152952 | 874.3619 | 0.144963787 | 0.284023 | 0.5103951 | 0.609775 | 0.910595741 | PLOD2 | protein_coding | 36.22 | 0.005896313 | 0.005896313 | ENSG00000152952 | n.s. |
| ENSG00000135535 | 2673.948 | 0.137616399 | 0.255001 | 0.5396692 | 0.589425 | 0.906089161 | CD164 | protein_coding | 38.11 | 0.005893982 | 0.005893982 | ENSG00000135535 | n.s. |
| ENSG00000067248 | 1353.418 | -0.144599576 | 0.282969 | -0.511008 | 0.609346 | 0.910431619 | DHX29 | protein_coding | 37.08 | -0.005892818 | 0.005892818 | ENSG00000067248 | n.s. |
| ENSG00000227354 | 247.453 | 0.128966744 | 0.229798 | 0.5612183 | 0.574649 | 0.900202572 | RBM26-AS1 | antisense | 38.6 | 0.005888589 | 0.005888589 | ENSG00000227354 | n.s. |
| ENSG00000254485 | 35.76372 | 0.161887698 | 0.342259 | 0.472997 | 0.636215 | 0.919741018 | | antisense | 43.09 | 0.0058821 | 0.0058821 | ENSG00000254485 | n.s. |
| ENSG00000104047 | 661.6667 | -0.141726857 | 0.267377 | -0.530063 | 0.596068 | 0.908868121 | DTWD1 | protein_coding | 36.31 | -0.005881541 | 0.005881541 | ENSG00000104047 | n.s. |
| ENSG00000188042 | 492.9505 | 0.131254575 | 0.236296 | 0.5554669 | 0.578575 | 0.901989749 | ARL4C | protein_coding | 50.16 | 0.005879995 | 0.005879995 | ENSG00000188042 | n.s. |
| ENSG00000146757 | 1600.214 | -0.142992874 | 0.278496 | -0.513446 | 0.607639 | 0.909710915 | ZNF92 | protein_coding | 39.16 | -0.00587652 | 0.00587652 | ENSG00000146757 | n.s. |
| ENSG00000101210 | 793.2046 | -0.141543692 | 0.273018 | -0.518442 | 0.60415 | 0.908868121 | EEF1A2 | protein_coding | 63.5 | -0.00587394 | 0.00587394 | ENSG00000101210 | n.s. |

| | | | | | | | | | | | | | |
|---|---|---|---|---|---|---|---|---|---|---|---|---|---|
| ENSG00000197037 | 259.0718 | -0.129258827 | 0.231564 | -0.558199 | 0.576708 | 0.900775165 | ZSCAN25 | protein_coding | 48.45 | -0.00586623 | 0.00586623 | ENSG00000197037 | n.s. |
| ENSG00000165669 | 731.6117 | -0.14129905 | 0.269511 | -0.52428 | 0.600084 | 0.908868121 | FAM204A | protein_coding | 38.05 | -0.005863788 | 0.005863788 | ENSG00000165669 | n.s. |
| ENSG00000250479 | 185.8061 | 0.141234034 | 0.268317 | 0.5263699 | 0.598631 | 0.908868121 | CHCHD10 | protein_coding | 64.1 | 0.005861089 | 0.005861089 | ENSG00000250479 | n.s. |
| ENSG00000151572 | 104.0042 | -0.141214785 | 0.271702 | -0.519741 | 0.603244 | 0.908868121 | ANO4 | protein_coding | 39.06 | -0.005860291 | 0.005860291 | ENSG00000151572 | n.s. |
| ENSG00000101888 | 137.765 | 0.154695883 | 0.311612 | 0.4964372 | 0.619586 | 0.916487226 | NXT2 | protein_coding | 37.4 | 0.005858888 | 0.005858888 | ENSG00000101888 | n.s. |
| ENSG00000160695 | 461.6457 | 0.135402446 | 0.249529 | 0.5426319 | 0.587383 | 0.905217543 | VPS11 | protein_coding | 48.94 | 0.005855755 | 0.005855755 | ENSG00000160695 | n.s. |
| ENSG00000137309 | 2647.248 | -0.132311551 | 0.239316 | -0.552874 | 0.58035 | 0.903133423 | HMGA1 | protein_coding | 57.83 | -0.005854533 | 0.005854533 | ENSG00000137309 | n.s. |
| ENSG00000044090 | 391.209 | 0.140104597 | 0.262532 | 0.5336669 | 0.593572 | 0.908297635 | CUL7 | protein_coding | 54.7 | 0.005852424 | 0.005852424 | ENSG00000044090 | n.s. |
| ENSG00000162736 | 620.2451 | -0.140891415 | 0.269333 | -0.523111 | 0.600897 | 0.908868121 | NCSTN | protein_coding | 46.85 | -0.005846871 | 0.005846871 | ENSG00000162736 | n.s. |
| ENSG00000123965 | 33.31244 | -0.162252504 | 0.344639 | -0.470789 | 0.637791 | 0.920401057 | PMS2P5 | unprocessed_pseudogene | 38.92 | -0.005844804 | 0.005844804 | ENSG00000123965 | n.s. |
| ENSG00000214432 | 19.09596 | -0.194250082 | 0.472308 | -0.411279 | 0.680868 | 0.933075339 | | antisense | 47.04 | -0.005843681 | 0.005843681 | ENSG00000214432 | n.s. |
| ENSG00000228606 | 11.04542 | -0.219811875 | 0.608335 | -0.361334 | 0.71785 | 0.940629569 | | antisense | 58.74 | -0.005842901 | 0.005842901 | ENSG00000228606 | n.s. |
| ENSG00000154473 | 7945.576 | -0.137517189 | 0.255404 | -0.538431 | 0.590279 | 0.906814644 | BUB3 | protein_coding | 39.33 | -0.005841933 | 0.005841933 | ENSG00000154473 | n.s. |
| ENSG00000140688 | 449.843 | -0.148062654 | 0.293896 | -0.503793 | 0.614407 | 0.913167298 | C16orf58 | protein_coding | 51.18 | -0.00584102 | 0.00584102 | ENSG00000140688 | n.s. |
| ENSG00000013306 | 2244.963 | -0.140727403 | 0.270692 | -0.51988 | 0.603147 | 0.908868121 | SLC25A39 | protein_coding | 61.11 | -0.005840065 | 0.005840065 | ENSG00000013306 | n.s. |
| ENSG00000145041 | 1112.994 | -0.140692249 | 0.271585 | -0.518042 | 0.604429 | 0.908868121 | DCAF1 | protein_coding | 43.34 | -0.005838606 | 0.005838606 | ENSG00000145041 | n.s. |
| ENSG00000133142 | 806.1132 | -0.141750586 | 0.27455 | -0.516302 | 0.605644 | 0.909522325 | TCEAL4 | protein_coding | 39.32 | -0.00583823 | 0.00583823 | ENSG00000133142 | n.s. |
| ENSG00000276102 | 11.86177 | 0.192827213 | 0.46749 | 0.4124735 | 0.679992 | 0.932670115 | MIR6747 | miRNA | 52.46 | 0.005837254 | 0.005837254 | ENSG00000276102 | n.s. |
| ENSG00000213995 | 848.5857 | -0.140317915 | 0.26332 | -0.532879 | 0.594117 | 0.908656857 | NAXD | protein_coding | 50.13 | -0.005837238 | 0.005837238 | ENSG00000213995 | n.s. |
| ENSG00000172766 | 2198.131 | -0.13510943 | 0.249419 | -0.541697 | 0.588027 | 0.905341301 | NAA16 | protein_coding | 37.34 | -0.005835061 | 0.005835061 | ENSG00000172766 | n.s. |
| ENSG00000071894 | 714.3626 | -0.134864407 | 0.248194 | -0.543384 | 0.586866 | 0.905217543 | CPSF1 | protein_coding | 59.98 | -0.005832486 | 0.005832486 | ENSG00000071894 | n.s. |
| ENSG00000171466 | 1632.386 | -0.139584007 | 0.261477 | -0.533828 | 0.593461 | 0.908297635 | ZNF562 | protein_coding | 43.08 | -0.005830678 | 0.005830678 | ENSG00000171466 | n.s. |
| ENSG00000167721 | 2627.435 | -0.132525201 | 0.241224 | -0.549385 | 0.582741 | 0.903709421 | TSR1 | protein_coding | 45.82 | -0.005827291 | 0.005827291 | ENSG00000167721 | n.s. |
| ENSG00000103326 | 651.4784 | -0.1304028 | 0.235103 | -0.554661 | 0.579126 | 0.902224015 | CAPN15 | protein_coding | 65.01 | -0.00582713 | 0.00582713 | ENSG00000103326 | n.s. |
| ENSG00000188735 | 986.8369 | -0.134637705 | 0.247406 | -0.544198 | 0.586305 | 0.905188828 | TMEM120B | protein_coding | 51.94 | -0.005824537 | 0.005824537 | ENSG00000188735 | n.s. |
| ENSG00000092096 | 275.0481 | -0.141574178 | 0.275792 | -0.513337 | 0.607716 | 0.909711346 | SLC22A17 | protein_coding | 57.86 | -0.005818187 | 0.005818187 | ENSG00000092096 | n.s. |
| ENSG00000175197 | 254.3962 | 0.140158807 | 0.270166 | 0.518787 | 0.603909 | 0.908868121 | DDIT3 | protein_coding | 49.44 | 0.005816468 | 0.005816468 | ENSG00000175197 | n.s. |
| ENSG00000140367 | 1564.911 | -0.141149871 | 0.273885 | -0.515362 | 0.6063 | 0.909522325 | UBE2Q2 | protein_coding | 38.58 | -0.005813488 | 0.005813488 | ENSG00000140367 | n.s. |
| ENSG00000213160 | 2992.887 | -0.14007465 | 0.267453 | -0.523735 | 0.600463 | 0.908868121 | KLHL23 | protein_coding | 42.29 | -0.005812976 | 0.005812976 | ENSG00000213160 | n.s. |
| ENSG00000104381 | 2132.059 | -0.139904202 | 0.269924 | -0.51831 | 0.604242 | 0.908868121 | GDAP1 | protein_coding | 38 | -0.005805903 | 0.005805903 | ENSG00000104381 | n.s. |
| ENSG00000123636 | 2319.48 | -0.14243785 | 0.278527 | -0.511397 | 0.609073 | 0.910431619 | BAZ2B | protein_coding | 36.16 | -0.005804722 | 0.005804722 | ENSG00000123636 | n.s. |
| ENSG00000089048 | 2581.461 | -0.139858857 | 0.26713 | -0.523561 | 0.600584 | 0.908868121 | ESF1 | protein_coding | 37.97 | -0.005804021 | 0.005804021 | ENSG00000089048 | n.s. |

| | | | | | | | | | | | | | |
|---|---|---|---|---|---|---|---|---|---|---|---|---|---|
| ENSG00000152413 | 883.4475 | -0.139830181 | 0.267245 | -0.523228 | 0.600816 | 0.908868121 | HOMER1 | protein_coding | 37.23 | -0.005802831 | 0.005802831 | ENSG00000152413 | n.s. |
| ENSG00000134278 | 939.7979 | -0.139683509 | 0.265889 | -0.525345 | 0.599343 | 0.908868121 | SPIRE1 | protein_coding | 40.66 | -0.005796744 | 0.005796744 | ENSG00000134278 | n.s. |
| ENSG00000112659 | 33.47759 | 0.16681311 | 0.368318 | 0.4529047 | 0.650617 | 0.923184844 | CUL9 | protein_coding | 49.09 | 0.005790306 | 0.005790306 | ENSG00000112659 | n.s. |
| ENSG00000177426 | 622.129 | -0.124408798 | 0.218314 | -0.569861 | 0.568772 | 0.898419074 | TGIF1 | protein_coding | 43.85 | -0.005787626 | 0.005787626 | ENSG00000177426 | n.s. |
| ENSG00000019995 | 1120.442 | -0.139454958 | 0.267965 | -0.520421 | 0.60277 | 0.908868121 | ZRANB1 | protein_coding | 39.68 | -0.005787259 | 0.005787259 | ENSG00000019995 | n.s. |
| ENSG00000255150 | 247.5214 | 0.138522745 | 0.259383 | 0.5340475 | 0.593309 | 0.908297635 | EID3 | protein_coding | 44.72 | 0.005786347 | 0.005786347 | ENSG00000255150 | n.s. |
| ENSG00000108018 | 1153.419 | -0.139381224 | 0.267245 | -0.521548 | 0.601985 | 0.908868121 | SORCS1 | protein_coding | 40.11 | -0.005784199 | 0.005784199 | ENSG00000108018 | n.s. |
| ENSG00000003509 | 702.7511 | -0.142160658 | 0.278657 | -0.510164 | 0.609936 | 0.910595741 | NDUFAF7 | protein_coding | 38.97 | -0.005782297 | 0.005782297 | ENSG00000003509 | n.s. |
| ENSG00000104679 | 452.3392 | -0.141172115 | 0.27537 | -0.512663 | 0.608187 | 0.91002515 | R3HCC1 | protein_coding | 47.98 | -0.005780519 | 0.005780519 | ENSG00000104679 | n.s. |
| ENSG00000198420 | 1135.126 | -0.139247774 | 0.263395 | -0.528666 | 0.597037 | 0.908868121 | TCAF1 | protein_coding | 39.07 | -0.005778661 | 0.005778661 | ENSG00000198420 | n.s. |
| ENSG00000131495 | 812.9488 | 0.155664979 | 0.316776 | 0.4914033 | 0.623141 | 0.918094734 | NDUFA2 | protein_coding | 50.9 | 0.005777117 | 0.005777117 | ENSG00000131495 | n.s. |
| ENSG00000156136 | 817.3779 | -0.147928426 | 0.295056 | -0.501357 | 0.61612 | 0.914037491 | DCK | protein_coding | 38.98 | -0.005774533 | 0.005774533 | ENSG00000156136 | n.s. |
| ENSG00000273402 | 63.11102 | 0.139084599 | 0.267465 | 0.5200097 | 0.603057 | 0.908868121 | | antisense | 27.52 | 0.00577189 | 0.00577189 | ENSG00000273402 | n.s. |
| ENSG00000067829 | 386.8845 | -0.1390075 | 0.262718 | -0.529112 | 0.596727 | 0.908868121 | IDH3G | protein_coding | 58.77 | -0.00576869 | 0.00576869 | ENSG00000067829 | n.s. |
| ENSG00000169057 | 193.2634 | -0.134153697 | 0.24821 | -0.540484 | 0.588863 | 0.905738346 | MECP2 | protein_coding | 46.41 | -0.005768239 | 0.005768239 | ENSG00000169057 | n.s. |
| ENSG00000272143 | 258.534 | 0.123344055 | 0.215751 | 0.5716958 | 0.567528 | 0.897919764 | FGF14-AS2 | lincRNA | 51.49 | 0.005767873 | 0.005767873 | ENSG00000272143 | n.s. |
| ENSG00000075413 | 1499.612 | -0.138898577 | 0.261932 | -0.530284 | 0.595915 | 0.908868121 | MARK3 | protein_coding | 42.15 | -0.00576417 | 0.00576417 | ENSG00000075413 | n.s. |
| ENSG00000006712 | 534.0014 | -0.138815642 | 0.261334 | -0.531181 | 0.595293 | 0.908868121 | PAF1 | protein_coding | 52.73 | -0.005760728 | 0.005760728 | ENSG00000006712 | n.s. |
| ENSG00000279366 | 36.50783 | 0.157027737 | 0.32739 | 0.4796349 | 0.631487 | 0.919033489 | | TEC | 38.59 | 0.005757997 | 0.005757997 | ENSG00000279366 | n.s. |
| ENSG00000163527 | 2795.418 | -0.139455745 | 0.270759 | -0.515054 | 0.606515 | 0.909522325 | STT3B | protein_coding | 35.96 | -0.005743713 | 0.005743713 | ENSG00000163527 | n.s. |
| ENSG00000277450 | 42.88513 | -0.154293595 | 0.313541 | -0.4921 | 0.622649 | 0.917861492 | | sense_intronic | 45.6 | -0.005743247 | 0.005743247 | ENSG00000277450 | n.s. |
| ENSG00000171206 | 276.6451 | -0.138329074 | 0.266692 | -0.518685 | 0.60398 | 0.908868121 | TRIM8 | protein_coding | 57.48 | -0.005740536 | 0.005740536 | ENSG00000171206 | n.s. |
| ENSG00000083312 | 17178.59 | -0.141124107 | 0.276696 | -0.510032 | 0.610029 | 0.910595741 | TNPO1 | protein_coding | 37.25 | -0.005740136 | 0.005740136 | ENSG00000083312 | n.s. |
| ENSG00000011295 | 1877.248 | -0.139879431 | 0.272608 | -0.513116 | 0.60787 | 0.909844575 | TTC19 | protein_coding | 41.07 | -0.005739643 | 0.005739643 | ENSG00000011295 | n.s. |
| ENSG00000173041 | 866.1025 | -0.138273179 | 0.267115 | -0.517653 | 0.6047 | 0.908868121 | ZNF680 | protein_coding | 38.98 | -0.005738217 | 0.005738217 | ENSG00000173041 | n.s. |
| ENSG00000201569 | 7.304026 | 0.198113075 | 0.499206 | 0.396856 | 0.691474 | 0.935515296 | SNORD114-17 | snoRNA | 42.67 | 0.005735197 | 0.005735197 | ENSG00000201569 | n.s. |
| ENSG00000248266 | 60.48531 | -0.154665072 | 0.315142 | -0.490778 | 0.623583 | 0.918167932 | | sense_intronic | 36.47 | -0.005734653 | 0.005734653 | ENSG00000248266 | n.s. |
| ENSG00000158528 | 513.4306 | 0.137921901 | 0.263593 | 0.5232381 | 0.600809 | 0.908868121 | PPP1R9A | protein_coding | 36.45 | 0.005723639 | 0.005723639 | ENSG00000158528 | n.s. |
| ENSG00000100038 | 516.7057 | -0.138944245 | 0.270082 | -0.514452 | 0.606936 | 0.909522325 | TOP3B | protein_coding | 54.98 | -0.005722646 | 0.005722646 | ENSG00000100038 | n.s. |
| ENSG00000068796 | 1776.934 | -0.137869287 | 0.263778 | -0.522671 | 0.601203 | 0.908868121 | KIF2A | protein_coding | 38.86 | -0.005721455 | 0.005721455 | ENSG00000068796 | n.s. |
| ENSG00000111269 | 667.7214 | -0.151748071 | 0.306227 | -0.49554 | 0.620219 | 0.916893708 | CREBL2 | protein_coding | 40.66 | -0.00571802 | 0.00571802 | ENSG00000111269 | n.s. |
| ENSG00000116237 | 679.1466 | -0.13762424 | 0.265118 | -0.519106 | 0.603687 | 0.908868121 | ICMT | protein_coding | 49.59 | -0.005711286 | 0.005711286 | ENSG00000116237 | n.s. |

| | | | | | | | | | | | | | |
|---|---|---|---|---|---|---|---|---|---|---|---|---|---|
| ENSG00000280138 | 893.0385 | 0.124698325 | 0.221579 | 0.5627717 | 0.57359 | 0.899911687 | | TEC | 44.97 | 0.005711197 | 0.005711197 | ENSG00000280138 | n.s. |
| ENSG00000153140 | 454.0409 | -0.154062026 | 0.314058 | -0.490553 | 0.623743 | 0.918203974 | CETN3 | protein_coding | 33.64 | -0.005709667 | 0.005709667 | ENSG00000153140 | n.s. |
| ENSG00000202270 | 4.770021 | -0.214358881 | 0.590574 | -0.362967 | 0.716629 | 0.94052839 | SNORD114-12 | snoRNA | 44 | -0.005707968 | 0.005707968 | ENSG00000202270 | n.s. |
| ENSG00000188938 | 461.9741 | -0.137504407 | 0.264631 | -0.519607 | 0.603337 | 0.908868121 | FAM120AOS | protein_coding | 50.78 | -0.005706313 | 0.005706313 | ENSG00000188938 | n.s. |
| ENSG00000107105 | 3083.121 | -0.137382448 | 0.258611 | -0.531232 | 0.595258 | 0.908868121 | ELAVL2 | protein_coding | 38.63 | -0.005701252 | 0.005701252 | ENSG00000107105 | n.s. |
| ENSG00000260563 | 45.25846 | -0.155386746 | 0.320825 | -0.484335 | 0.628148 | 0.919033489 | | lincRNA | 47.36 | -0.005697824 | 0.005697824 | ENSG00000260563 | n.s. |
| ENSG00000107669 | 741.9557 | -0.139932721 | 0.274665 | -0.509467 | 0.610425 | 0.910595741 | ATE1 | protein_coding | 39.1 | -0.005691677 | 0.005691677 | ENSG00000107669 | n.s. |
| ENSG00000241127 | 864.1386 | 0.134190333 | 0.249555 | 0.5377186 | 0.590771 | 0.906969769 | YAE1D1 | protein_coding | 40.07 | 0.005690635 | 0.005690635 | ENSG00000241127 | n.s. |
| ENSG00000196227 | 735.8285 | -0.122748272 | 0.216483 | -0.567011 | 0.570707 | 0.898818733 | FAM217B | protein_coding | 42.7 | -0.005686668 | 0.005686668 | ENSG00000196227 | n.s. |
| ENSG00000134900 | 3011.515 | -0.137002483 | 0.262249 | -0.522414 | 0.601382 | 0.908868121 | TPP2 | protein_coding | 37.69 | -0.005685484 | 0.005685484 | ENSG00000134900 | n.s. |
| ENSG00000266555 | 6.138488 | -0.209683085 | 0.566404 | -0.3702 | 0.711233 | 0.939526932 | MIR3145 | miRNA | 31.71 | -0.005680475 | 0.005680475 | ENSG00000266555 | n.s. |
| ENSG00000116977 | 37.61226 | 0.154677562 | 0.322543 | 0.4795571 | 0.631542 | 0.919033489 | LGALS8 | protein_coding | 43.48 | 0.005671819 | 0.005671819 | ENSG00000116977 | n.s. |
| ENSG00000277846 | 68.30827 | -0.14509529 | 0.289256 | -0.501616 | 0.615937 | 0.91396346 | SNORD30 | snoRNA | 41.43 | -0.005669042 | 0.005669042 | ENSG00000277846 | n.s. |
| ENSG00000145246 | 360.0008 | -0.14415807 | 0.286471 | -0.503221 | 0.614809 | 0.913454511 | ATP10D | protein_coding | 37.6 | -0.005667297 | 0.005667297 | ENSG00000145246 | n.s. |
| ENSG00000188986 | 556.7531 | -0.139269995 | 0.273163 | -0.509841 | 0.610163 | 0.910595741 | NELFB | protein_coding | 56.65 | -0.005664721 | 0.005664721 | ENSG00000188986 | n.s. |
| ENSG00000199121 | 7.031031 | -0.212981575 | 0.589525 | -0.361277 | 0.717893 | 0.940629569 | MIR26B | miRNA | 55.84 | -0.005661343 | 0.005661343 | ENSG00000199121 | n.s. |
| ENSG00000124596 | 824.2985 | -0.134819005 | 0.251482 | -0.536098 | 0.591891 | 0.907847371 | OARD1 | protein_coding | 44.19 | -0.005660667 | 0.005660667 | ENSG00000124596 | n.s. |
| ENSG00000196387 | 1488.532 | 0.12851954 | 0.233929 | 0.5493958 | 0.582734 | 0.903709421 | ZNF140 | protein_coding | 41.08 | 0.005651157 | 0.005651157 | ENSG00000196387 | n.s. |
| ENSG00000185896 | 898.9198 | -0.136103577 | 0.26211 | -0.519261 | 0.603579 | 0.908868121 | LAMP1 | protein_coding | 50.64 | -0.00564818 | 0.00564818 | ENSG00000185896 | n.s. |
| ENSG00000239082 | 3.848215 | -0.238549235 | 0.717997 | -0.332243 | 0.739706 | 0.946996213 | RNU7-170P | snRNA | 42.62 | -0.005642109 | 0.005642109 | ENSG00000239082 | n.s. |
| ENSG00000214756 | 578.3744 | 0.122554396 | 0.216827 | 0.5652178 | 0.571926 | 0.899491841 | METTL12 | protein_coding | 52.54 | 0.005637842 | 0.005637842 | ENSG00000214756 | n.s. |
| ENSG00000173575 | 6487.181 | -0.122809791 | 0.218432 | -0.562235 | 0.573956 | 0.899911687 | CHD2 | protein_coding | 39.49 | -0.005624702 | 0.005624702 | ENSG00000173575 | n.s. |
| ENSG00000159348 | 394.8278 | -0.138211475 | 0.27142 | -0.509216 | 0.610601 | 0.910701627 | CYB5R1 | protein_coding | 50.33 | -0.005614687 | 0.005614687 | ENSG00000159348 | n.s. |
| ENSG00000144867 | 1251.062 | -0.13944599 | 0.275155 | -0.50679 | 0.612302 | 0.911480916 | SRPRB | protein_coding | 41.15 | -0.005613039 | 0.005613039 | ENSG00000144867 | n.s. |
| ENSG00000217555 | 129.6315 | 0.158800998 | 0.340597 | 0.4662424 | 0.641042 | 0.921891013 | CKLF | protein_coding | 41.69 | 0.005608918 | 0.005608918 | ENSG00000217555 | n.s. |
| ENSG00000202048 | 23.08263 | -0.242808729 | 0.743279 | -0.326672 | 0.743916 | 0.948261154 | SNORD114-20 | snoRNA | 47.22 | -0.005602093 | 0.005602093 | ENSG00000202048 | n.s. |
| ENSG00000243989 | 263.4983 | -0.152771172 | 0.317098 | -0.481778 | 0.629963 | 0.919033489 | ACY1 | protein_coding | 57.5 | -0.005601915 | 0.005601915 | ENSG00000243989 | n.s. |
| ENSG00000185432 | 334.263 | -0.134939496 | 0.255395 | -0.528357 | 0.597252 | 0.908868121 | METTL7A | protein_coding | 42.85 | -0.005599872 | 0.005599872 | ENSG00000185432 | n.s. |
| ENSG00000247735 | 86.11181 | -0.143166288 | 0.28546 | -0.501529 | 0.615999 | 0.91396346 | | antisense | 47.76 | -0.005593674 | 0.005593674 | ENSG00000247735 | n.s. |
| ENSG00000179933 | 566.8505 | 0.151859413 | 0.310805 | 0.4886002 | 0.625125 | 0.918928785 | C14orf119 | protein_coding | 47.09 | 0.005575996 | 0.005575996 | ENSG00000179933 | n.s. |
| ENSG00000135698 | 435.7781 | -0.149791237 | 0.304449 | -0.492008 | 0.622714 | 0.917861492 | MPHOSPH6 | protein_coding | 40.61 | -0.005575657 | 0.005575657 | ENSG00000135698 | n.s. |
| ENSG00000146938 | 836.4262 | 0.135295555 | 0.26235 | 0.5157067 | 0.606059 | 0.909522325 | NLGN4X | protein_coding | 38.89 | 0.005572369 | 0.005572369 | ENSG00000146938 | n.s. |

| | | | | | | | | | | | | | |
|---|---|---|---|---|---|---|---|---|---|---|---|---|---|
| ENSG00000163682 | 17572.99 | 0.151947103 | 0.314441 | 0.4832295 | 0.628933 | 0.919033489 | RPL9 | protein_coding | 41.12 | 0.005571697 | 0.005571697 | ENSG00000163682 | n.s. |
| ENSG00000232503 | 28.99602 | -0.162050955 | 0.360363 | -0.449688 | 0.652935 | 0.923905149 | CYCSP8 | processed_pseudogene | 46.47 | -0.005570115 | 0.005570115 | ENSG00000232503 | n.s. |
| ENSG00000112739 | 3358.3 | -0.134220049 | 0.257643 | -0.520954 | 0.602399 | 0.908868121 | PRPF4B | protein_coding | 38.36 | -0.005570015 | 0.005570015 | ENSG00000112739 | n.s. |
| ENSG00000144655 | 408.9888 | -0.138757898 | 0.274171 | -0.5061 | 0.612786 | 0.911714626 | CSRNP1 | protein_coding | 55.85 | -0.005569892 | 0.005569892 | ENSG00000144655 | n.s. |
| ENSG00000132323 | 1101.47 | -0.134177669 | 0.257339 | -0.521405 | 0.602085 | 0.908868121 | ILKAP | protein_coding | 44.85 | -0.005568256 | 0.005568256 | ENSG00000132323 | n.s. |
| ENSG00000162076 | 224.6137 | -0.15752619 | 0.338116 | -0.465893 | 0.641292 | 0.921891013 | FLYWCH2 | protein_coding | 51.49 | -0.005563891 | 0.005563891 | ENSG00000162076 | n.s. |
| ENSG00000013375 | 1851.222 | -0.138956977 | 0.27486 | -0.505555 | 0.613169 | 0.911981962 | PGM3 | protein_coding | 38.66 | -0.00556019 | 0.00556019 | ENSG00000013375 | n.s. |
| ENSG00000101546 | 1079.71 | -0.126038617 | 0.22865 | -0.55123 | 0.581476 | 0.90349457 | RBFA | protein_coding | 47.61 | -0.005555083 | 0.005555083 | ENSG00000101546 | n.s. |
| ENSG00000137074 | 89.22432 | -0.133857583 | 0.25416 | -0.526666 | 0.598425 | 0.908868121 | APTX | protein_coding | 42.68 | -0.005554973 | 0.005554973 | ENSG00000137074 | n.s. |
| ENSG00000225733 | 2870.772 | -0.133606461 | 0.253582 | -0.526877 | 0.598279 | 0.908868121 | FGD5-AS1 | antisense | 45.41 | -0.005544552 | 0.005544552 | ENSG00000225733 | n.s. |
| ENSG00000149483 | 927.3167 | -0.113708664 | 0.194266 | -0.585323 | 0.55833 | 0.893837232 | TMEM138 | protein_coding | 48.08 | -0.005542338 | 0.005542338 | ENSG00000149483 | n.s. |
| ENSG00000234841 | 181.113 | 0.133469988 | 0.253092 | 0.5273584 | 0.597945 | 0.908868121 | | processed_pseudogene | 41.2 | 0.005538888 | 0.005538888 | ENSG00000234841 | n.s. |
| ENSG00000165775 | 858.9525 | 0.150006581 | 0.306187 | 0.4899187 | 0.624191 | 0.91849357 | FUNDC2 | protein_coding | 39.84 | 0.005538825 | 0.005538825 | ENSG00000165775 | n.s. |
| ENSG00000014824 | 2779.965 | -0.144583516 | 0.290283 | -0.498078 | 0.618429 | 0.915570902 | SLC30A9 | protein_coding | 36.98 | -0.005538708 | 0.005538708 | ENSG00000014824 | n.s. |
| ENSG00000112473 | 775.1494 | -0.132944166 | 0.249281 | -0.533311 | 0.593818 | 0.908531819 | SLC39A7 | protein_coding | 52.92 | -0.005538435 | 0.005538435 | ENSG00000112473 | n.s. |
| ENSG00000139197 | 1220.842 | -0.133454696 | 0.256868 | -0.519546 | 0.60338 | 0.908868121 | PEX5 | protein_coding | 42.58 | -0.005538254 | 0.005538254 | ENSG00000139197 | n.s. |
| ENSG00000206585 | 99.60075 | 0.142550555 | 0.284884 | 0.5003812 | 0.616807 | 0.91447122 | RNVU1-7 | snRNA | 54.88 | 0.005535232 | 0.005535232 | ENSG00000206585 | n.s. |
| ENSG00000167792 | 1879.797 | -0.135901336 | 0.266387 | -0.510166 | 0.609935 | 0.910595741 | NDUFV1 | protein_coding | 55.96 | -0.005527703 | 0.005527703 | ENSG00000167792 | n.s. |
| ENSG00000111254 | 14.13208 | -0.198343877 | 0.520631 | -0.380968 | 0.703227 | 0.937867182 | AKAP3 | protein_coding | 38.99 | -0.005525595 | 0.005525595 | ENSG00000111254 | n.s. |
| ENSG00000106244 | 1377.3 | -0.135754593 | 0.266226 | -0.509923 | 0.610105 | 0.910595741 | PDAP1 | protein_coding | 54.05 | -0.005521735 | 0.005521735 | ENSG00000106244 | n.s. |
| ENSG00000145349 | 854.9265 | 0.132991162 | 0.253731 | 0.5241419 | 0.60018 | 0.908868121 | CAMK2D | protein_coding | 35.99 | 0.005519017 | 0.005519017 | ENSG00000145349 | n.s. |
| ENSG00000090273 | 826.468 | -0.132986649 | 0.25267 | -0.526324 | 0.598663 | 0.908868121 | NUDC | protein_coding | 48.43 | -0.00551883 | 0.00551883 | ENSG00000090273 | n.s. |
| ENSG00000160783 | 345.3274 | -0.150471264 | 0.315767 | -0.476527 | 0.633699 | 0.919033489 | PMF1 | protein_coding | 48.91 | -0.00551758 | 0.00551758 | ENSG00000160783 | n.s. |
| ENSG00000059728 | 364.2003 | -0.121275035 | 0.216881 | -0.559177 | 0.576041 | 0.900614489 | MXD1 | protein_coding | 40.74 | -0.005513293 | 0.005513293 | ENSG00000059728 | n.s. |
| ENSG00000105397 | 204.0081 | 0.134091408 | 0.260864 | 0.5140275 | 0.607233 | 0.90967308 | TYK2 | protein_coding | 54.34 | 0.005513122 | 0.005513122 | ENSG00000105397 | n.s. |
| ENSG00000144136 | 4003.059 | -0.125019502 | 0.227321 | -0.549968 | 0.582341 | 0.90349457 | SLC20A1 | protein_coding | 42.45 | -0.005510166 | 0.005510166 | ENSG00000144136 | n.s. |
| ENSG00000115677 | 6233.15 | -0.132546745 | 0.255736 | -0.518296 | 0.604252 | 0.908868121 | HDLBP | protein_coding | 46.99 | -0.005500575 | 0.005500575 | ENSG00000115677 | n.s. |
| ENSG00000062485 | 2717.328 | -0.133807851 | 0.260434 | -0.513788 | 0.6074 | 0.909710915 | CS | protein_coding | 46.16 | -0.005499047 | 0.005499047 | ENSG00000062485 | n.s. |
| ENSG00000076242 | 1611.302 | -0.133383193 | 0.258742 | -0.515506 | 0.606199 | 0.909522325 | MLH1 | protein_coding | 41.89 | -0.005493605 | 0.005493605 | ENSG00000076242 | n.s. |
| ENSG00000214765 | 814.9021 | -0.133642315 | 0.260256 | -0.513504 | 0.607599 | 0.909710915 | SEPT7P2 | cribed_unprocessed_pseudc | 37.6 | -0.005492244 | 0.005492244 | ENSG00000214765 | n.s. |
| ENSG00000166347 | 467.4959 | -0.149645435 | 0.310621 | -0.481762 | 0.629975 | 0.919033489 | CYB5A | protein_coding | 44.31 | -0.005487298 | 0.005487298 | ENSG00000166347 | n.s. |
| ENSG00000133318 | 935.4588 | -0.141580671 | 0.283386 | -0.499604 | 0.617354 | 0.914690044 | RTN3 | protein_coding | 42.12 | -0.00548286 | 0.00548286 | ENSG00000133318 | n.s. |

| | | | | | | | | | | | | | |
|---|---|---|---|---|---|---|---|---|---|---|---|---|---|
| ENSG00000122873 | 716.4337 | -0.149360678 | 0.309122 | -0.483178 | 0.62897 | 0.919033489 | CISD1 | protein_coding | 36.84 | -0.005476856 | 0.005476856 | ENSG00000122873 | n.s. |
| ENSG00000119487 | 1161.639 | -0.145455187 | 0.294062 | -0.49464 | 0.620854 | 0.917065125 | MAPKAP1 | protein_coding | 41.82 | -0.005469089 | 0.005469089 | ENSG00000119487 | n.s. |
| ENSG00000177954 | 22332.38 | 0.149816031 | 0.314983 | 0.4756317 | 0.634337 | 0.919380266 | RPS27 | protein_coding | 45.91 | 0.005469008 | 0.005469008 | ENSG00000177954 | n.s. |
| ENSG00000240032 | 45.08851 | -0.148065324 | 0.302384 | -0.489659 | 0.624375 | 0.918517547 | LNCSRLR | antisense | 34.2 | -0.005465468 | 0.005465468 | ENSG00000240032 | n.s. |
| ENSG00000138160 | 3196.059 | -0.134085112 | 0.262414 | -0.510967 | 0.609374 | 0.910431619 | KIF11 | protein_coding | 39.18 | -0.005464326 | 0.005464326 | ENSG00000138160 | n.s. |
| ENSG00000166823 | 93.33546 | -0.148973093 | 0.30875 | -0.482503 | 0.629448 | 0.919033489 | MESP1 | protein_coding | 60.98 | -0.005462644 | 0.005462644 | ENSG00000166823 | n.s. |
| ENSG00000201988 | 5.720836 | -0.210209362 | 0.5909 | -0.355744 | 0.722032 | 0.941942395 | | misc_RNA | 37.84 | -0.005460326 | 0.005460326 | ENSG00000201988 | n.s. |
| ENSG00000171824 | 997.8248 | -0.131547919 | 0.251818 | -0.522393 | 0.601397 | 0.908868121 | EXOSC10 | protein_coding | 46.38 | -0.005459124 | 0.005459124 | ENSG00000171824 | n.s. |
| ENSG00000139168 | 1199.952 | -0.149679767 | 0.315076 | -0.475059 | 0.634745 | 0.919524756 | ZCRB1 | protein_coding | 37.28 | -0.005453818 | 0.005453818 | ENSG00000139168 | n.s. |
| ENSG00000134202 | 1094.489 | 0.133794282 | 0.261751 | 0.5111518 | 0.609245 | 0.910431619 | GSTM3 | protein_coding | 47.16 | 0.005452474 | 0.005452474 | ENSG00000134202 | n.s. |
| ENSG00000060982 | 11131.86 | -0.139471534 | 0.277888 | -0.501899 | 0.615739 | 0.91396346 | BCAT1 | protein_coding | 40.06 | -0.005449316 | 0.005449316 | ENSG00000060982 | n.s. |
| ENSG00000248521 | 136.3565 | 0.144878885 | 0.293069 | 0.4943515 | 0.621058 | 0.917065125 | | processed_pseudogene | 55.01 | 0.00544742 | 0.00544742 | ENSG00000248521 | n.s. |
| ENSG00000234771 | 255.0496 | -0.131222202 | 0.250279 | -0.524305 | 0.600067 | 0.908868121 | SLC25A25-AS1 | antisense | 53.53 | -0.005445607 | 0.005445607 | ENSG00000234771 | n.s. |
| ENSG00000170160 | 542.0538 | -0.148422563 | 0.306731 | -0.483885 | 0.628468 | 0.919033489 | CCDC144A | protein_coding | 38.37 | -0.005442457 | 0.005442457 | ENSG00000170160 | n.s. |
| ENSG00000123739 | 426.4775 | -0.148174031 | 0.304304 | -0.486928 | 0.626309 | 0.918973809 | PLA2G12A | protein_coding | 39 | -0.005437523 | 0.005437523 | ENSG00000123739 | n.s. |
| ENSG00000234222 | 53.31342 | -0.145479163 | 0.294866 | -0.493373 | 0.621749 | 0.917630258 | LIX1L-AS1 | antisense | 44.35 | -0.005431068 | 0.005431068 | ENSG00000234222 | n.s. |
| ENSG00000272853 | 69.92447 | 0.138890577 | 0.276731 | 0.501898 | 0.615739 | 0.91396346 | | lincRNA | 56.86 | 0.005426617 | 0.005426617 | ENSG00000272853 | n.s. |
| ENSG00000123810 | 64.97521 | -0.147870842 | 0.308024 | -0.480063 | 0.631183 | 0.919033489 | B9D2 | protein_coding | 52.34 | -0.005422226 | 0.005422226 | ENSG00000123810 | n.s. |
| ENSG00000165632 | 401.7569 | -0.130559285 | 0.247939 | -0.526579 | 0.598486 | 0.908868121 | TAF3 | protein_coding | 39.51 | -0.005418097 | 0.005418097 | ENSG00000165632 | n.s. |
| ENSG00000260038 | 24.948 | 0.166679482 | 0.387708 | 0.4299097 | 0.667261 | 0.927899924 | | sense_intronic | 40.86 | 0.005416893 | 0.005416893 | ENSG00000260038 | n.s. |
| ENSG00000246705 | 92.70054 | -0.148502486 | 0.313273 | -0.474036 | 0.635474 | 0.919532207 | H2AFJ | protein_coding | 45.61 | -0.005410399 | 0.005410399 | ENSG00000246705 | n.s. |
| ENSG00000114391 | 13658.4 | 0.147169151 | 0.30406 | 0.4840136 | 0.628376 | 0.919033489 | RPL24 | protein_coding | 42.52 | 0.005396496 | 0.005396496 | ENSG00000114391 | n.s. |
| ENSG00000250432 | 28.44259 | 0.156346086 | 0.34616 | 0.4516589 | 0.651515 | 0.923637036 | | lincRNA | 48.26 | 0.005393731 | 0.005393731 | ENSG00000250432 | n.s. |
| ENSG00000145592 | 22321.16 | 0.143281319 | 0.289518 | 0.4948964 | 0.620673 | 0.916981464 | RPL37 | protein_coding | 42.83 | 0.005393029 | 0.005393029 | ENSG00000145592 | n.s. |
| ENSG00000151553 | 721.4349 | -0.130899469 | 0.25429 | -0.514764 | 0.606718 | 0.909522325 | FAM160B1 | protein_coding | 39.9 | -0.005391309 | 0.005391309 | ENSG00000151553 | n.s. |
| ENSG00000106603 | 1876.985 | -0.137820338 | 0.274142 | -0.502733 | 0.615152 | 0.913866731 | COA1 | protein_coding | 40.39 | -0.005391136 | 0.005391136 | ENSG00000106603 | n.s. |
| ENSG00000207965 | 6.547924 | -0.200701612 | 0.550396 | -0.364649 | 0.715373 | 0.940060506 | MIR629 | miRNA | 59.79 | -0.005387673 | 0.005387673 | ENSG00000207965 | n.s. |
| ENSG00000137941 | 475.4038 | -0.130788117 | 0.253616 | -0.515694 | 0.606068 | 0.909522325 | TTLL7 | protein_coding | 35.53 | -0.005386723 | 0.005386723 | ENSG00000137941 | n.s. |
| ENSG00000210100 | 53.03902 | 0.147808921 | 0.311814 | 0.4740296 | 0.635479 | 0.919532207 | MT-TI | Mt_tRNA | 28.99 | 0.005385131 | 0.005385131 | ENSG00000210100 | n.s. |
| ENSG00000164616 | 217.1493 | 0.146703299 | 0.30047 | 0.4882463 | 0.625375 | 0.918973809 | FBXL21 | anscribed_unitary_pseudoge | 37.89 | 0.005383551 | 0.005383551 | ENSG00000164616 | n.s. |
| ENSG00000156873 | 221.3114 | -0.146696619 | 0.303934 | -0.482659 | 0.629338 | 0.919033489 | PHKG2 | protein_coding | 52.63 | -0.005379169 | 0.005379169 | ENSG00000156873 | n.s. |
| ENSG00000279416 | 46.40022 | 0.146619045 | 0.302266 | 0.4850659 | 0.62763 | 0.919033489 | | TEC | 40.09 | 0.005376324 | 0.005376324 | ENSG00000279416 | n.s. |

| | | | | | | | | | | | | | |
|---|---|---|---|---|---|---|---|---|---|---|---|---|---|
| ENSG00000168159 | 1049.738 | -0.143954553 | 0.291777 | -0.493371 | 0.62175 | 0.917630258 | RNF187 | protein_coding | 56.59 | -0.005374151 | 0.005374151 | ENSG00000168159 | n.s. |
| ENSG00000207475 | 47.28016 | -0.146512667 | 0.307174 | -0.476969 | 0.633384 | 0.919033489 | SNORA80E | snoRNA | 54.74 | -0.005372424 | 0.005372424 | ENSG00000207475 | n.s. |
| ENSG00000080608 | 2702.296 | -0.129457208 | 0.248619 | -0.520706 | 0.602572 | 0.908868121 | PUM3 | protein_coding | 39.61 | -0.005372361 | 0.005372361 | ENSG00000080608 | n.s. |
| ENSG00000185252 | 516.4403 | -0.129442391 | 0.248715 | -0.520445 | 0.602753 | 0.908868121 | ZNF74 | protein_coding | 50.81 | -0.005371747 | 0.005371747 | ENSG00000185252 | n.s. |
| ENSG00000105647 | 426.6645 | -0.129422402 | 0.247633 | -0.522638 | 0.601226 | 0.908868121 | PIK3R2 | protein_coding | 57.77 | -0.005370917 | 0.005370917 | ENSG00000105647 | n.s. |
| ENSG00000183840 | 125.7163 | -0.129175304 | 0.246814 | -0.523371 | 0.600716 | 0.908868121 | GPR39 | protein_coding | 42.93 | -0.005360663 | 0.005360663 | ENSG00000183840 | n.s. |
| ENSG00000140006 | 583.0925 | -0.13631359 | 0.270818 | -0.50334 | 0.614725 | 0.913454511 | WDR89 | protein_coding | 41.2 | -0.005358906 | 0.005358906 | ENSG00000140006 | n.s. |
| ENSG00000108641 | 94.53177 | -0.153948038 | 0.333259 | -0.461947 | 0.644119 | 0.923072262 | B9D1 | protein_coding | 51.57 | -0.005351896 | 0.005351896 | ENSG00000108641 | n.s. |
| ENSG00000076555 | 55.63549 | -0.145857559 | 0.302732 | -0.481804 | 0.629945 | 0.919033489 | ACACB | protein_coding | 46.78 | -0.005348402 | 0.005348402 | ENSG00000076555 | n.s. |
| ENSG00000152784 | 971.2281 | 0.123346532 | 0.226267 | 0.5451371 | 0.585659 | 0.905026039 | PRDM8 | protein_coding | 47.9 | 0.005345706 | 0.005345706 | ENSG00000152784 | n.s. |
| ENSG00000185880 | 202.5756 | 0.149035021 | 0.317737 | 0.4690514 | 0.639033 | 0.920750614 | TRIM69 | protein_coding | 39.92 | 0.005344095 | 0.005344095 | ENSG00000185880 | n.s. |
| ENSG00000271737 | 52.88921 | -0.14888115 | 0.317184 | -0.469384 | 0.638795 | 0.920750614 | | lincRNA | 62.26 | -0.005338578 | 0.005338578 | ENSG00000271737 | n.s. |
| ENSG00000182108 | 373.9525 | -0.129546454 | 0.251382 | -0.515337 | 0.606317 | 0.909522325 | DEXI | protein_coding | 51.35 | -0.005335583 | 0.005335583 | ENSG00000182108 | n.s. |
| ENSG00000241837 | 2907.239 | 0.146294351 | 0.308521 | 0.4741799 | 0.635372 | 0.919532207 | ATP5O | protein_coding | 43.51 | 0.00532995 | 0.00532995 | ENSG00000241837 | n.s. |
| ENSG00000162851 | 767.9601 | 0.137646183 | 0.275626 | 0.4993947 | 0.617501 | 0.914721963 | TFB2M | protein_coding | 41.95 | 0.005328407 | 0.005328407 | ENSG00000162851 | n.s. |
| ENSG00000109654 | 884.9638 | -0.142664731 | 0.289345 | -0.493061 | 0.621969 | 0.917630258 | TRIM2 | protein_coding | 42.05 | -0.005325999 | 0.005325999 | ENSG00000109654 | n.s. |
| ENSG00000136238 | 1444.811 | -0.130660738 | 0.255195 | -0.512004 | 0.608648 | 0.91042178 | RAC1 | protein_coding | 45.86 | -0.005325387 | 0.005325387 | ENSG00000136238 | n.s. |
| ENSG00000153113 | 959.3613 | -0.131097952 | 0.257628 | -0.508864 | 0.610847 | 0.910725807 | CAST | protein_coding | 39.41 | -0.005324197 | 0.005324197 | ENSG00000153113 | n.s. |
| ENSG00000206120 | 37.54193 | -0.15139333 | 0.325751 | -0.464752 | 0.642109 | 0.922225673 | EGFEM1P | anscribed_unitary_pseudoge | 36.09 | -0.005323412 | 0.005323412 | ENSG00000206120 | n.s. |
| ENSG00000076554 | 841.4174 | 0.145001928 | 0.297138 | 0.4879949 | 0.625553 | 0.918973809 | TPD52 | protein_coding | 41.15 | 0.005321116 | 0.005321116 | ENSG00000076554 | n.s. |
| ENSG00000171222 | 439.1735 | -0.130154734 | 0.254062 | -0.512295 | 0.608445 | 0.910215508 | SCAND1 | protein_coding | 47.71 | -0.005317572 | 0.005317572 | ENSG00000171222 | n.s. |
| ENSG00000208892 | 60.35434 | 0.153167729 | 0.334564 | 0.4578127 | 0.647087 | 0.923184844 | SNORA49 | snoRNA | 48.53 | 0.005316656 | 0.005316656 | ENSG00000208892 | n.s. |
| ENSG00000197982 | 165.5539 | -0.146558809 | 0.310295 | -0.47232 | 0.636698 | 0.919866014 | C1orf122 | protein_coding | 63.25 | -0.005316483 | 0.005316483 | ENSG00000197982 | n.s. |
| ENSG00000092010 | 285.8122 | 0.153111823 | 0.337354 | 0.4538612 | 0.649929 | 0.923184844 | PSME1 | protein_coding | 57.4 | 0.005314716 | 0.005314716 | ENSG00000092010 | n.s. |
| ENSG00000015676 | 1218.273 | -0.127831761 | 0.244044 | -0.523807 | 0.600413 | 0.908868121 | NUDCD3 | protein_coding | 44.52 | -0.005304907 | 0.005304907 | ENSG00000015676 | n.s. |
| ENSG00000121957 | 1296.265 | 0.14551922 | 0.307053 | 0.4739219 | 0.635556 | 0.919532207 | GPSM2 | protein_coding | 38.78 | 0.00530171 | 0.00530171 | ENSG00000121957 | n.s. |
| ENSG00000235354 | 29.93789 | 0.155648543 | 0.348336 | 0.4468349 | 0.654994 | 0.924621046 | RPS29P16 | processed_pseudogene | 50.29 | 0.005297689 | 0.005297689 | ENSG00000235354 | n.s. |
| ENSG00000201689 | 4.324645 | 0.207347017 | 0.595815 | 0.3480055 | 0.727836 | 0.942883666 | SNORD114-29 | snoRNA | 42.86 | 0.005296034 | 0.005296034 | ENSG00000201689 | n.s. |
| ENSG00000108395 | 2263.657 | -0.127566211 | 0.242889 | -0.525205 | 0.599441 | 0.908868121 | TRIM37 | protein_coding | 39.71 | -0.005293887 | 0.005293887 | ENSG00000108395 | n.s. |
| ENSG00000204822 | 433.3593 | -0.147580592 | 0.314828 | -0.468766 | 0.639237 | 0.920825292 | MRPL53 | protein_coding | 53.92 | -0.005286744 | 0.005286744 | ENSG00000204822 | n.s. |
| ENSG00000125347 | 144.0357 | -0.13079101 | 0.257732 | -0.50747 | 0.611825 | 0.911134742 | IRF1 | protein_coding | 53.85 | -0.005286232 | 0.005286232 | ENSG00000125347 | n.s. |
| ENSG00000176986 | 1083.176 | -0.127198665 | 0.243679 | -0.521992 | 0.601676 | 0.908868121 | SEC24C | protein_coding | 47.03 | -0.005278634 | 0.005278634 | ENSG00000176986 | n.s. |

| | | | | | | | | | | | | | |
|---|---|---|---|---|---|---|---|---|---|---|---|---|---|
| ENSG00000160213 | 267.0983 | -0.128119718 | 0.24891 | -0.514722 | 0.606747 | 0.909522325 | CSTB | protein_coding | 54.21 | -0.00527682 | 0.00527682 | ENSG00000160213 | n.s. |
| ENSG00000125648 | 197.8163 | -0.126518703 | 0.237312 | -0.533132 | 0.593942 | 0.908531819 | SLC25A23 | protein_coding | 52.6 | -0.005270751 | 0.005270751 | ENSG00000125648 | n.s. |
| ENSG00000261447 | 22.11057 | 0.183392766 | 0.464326 | 0.3949653 | 0.692868 | 0.936038047 | | lincRNA | 53.89 | 0.005264564 | 0.005264564 | ENSG00000261447 | n.s. |
| ENSG00000100426 | 1269.126 | -0.130245753 | 0.256786 | -0.507215 | 0.612004 | 0.911134742 | ZBED4 | protein_coding | 47.49 | -0.005264194 | 0.005264194 | ENSG00000100426 | n.s. |
| ENSG00000239719 | 9.198934 | 0.189073429 | 0.501476 | 0.3770339 | 0.706148 | 0.937943012 | | processed_pseudogene | 50.83 | 0.005260694 | 0.005260694 | ENSG00000239719 | n.s. |
| ENSG00000059915 | 268.1618 | 0.126735763 | 0.239244 | 0.5297333 | 0.596297 | 0.908868121 | PSD | protein_coding | 59.67 | 0.005259424 | 0.005259424 | ENSG00000059915 | n.s. |
| ENSG00000035928 | 3180.177 | -0.129338096 | 0.254161 | -0.508883 | 0.610834 | 0.910725807 | RFC1 | protein_coding | 39.26 | -0.005252725 | 0.005252725 | ENSG00000035928 | n.s. |
| ENSG00000102081 | 1484.774 | -0.128843216 | 0.252116 | -0.511047 | 0.609318 | 0.910431619 | FMR1 | protein_coding | 36.35 | -0.005250705 | 0.005250705 | ENSG00000102081 | n.s. |
| ENSG00000123600 | 113.8681 | -0.127408146 | 0.246724 | -0.5164 | 0.605575 | 0.909522325 | METTL8 | protein_coding | 39.39 | -0.005247513 | 0.005247513 | ENSG00000123600 | n.s. |
| ENSG00000237436 | 31.43253 | 0.15451827 | 0.350201 | 0.4412279 | 0.659048 | 0.924786541 | | antisense | 59.77 | 0.005247209 | 0.005247209 | ENSG00000237436 | n.s. |
| ENSG00000168887 | 830.5978 | -0.126416611 | 0.239369 | -0.528125 | 0.597412 | 0.908868121 | C2orf68 | protein_coding | 51.15 | -0.005246179 | 0.005246179 | ENSG00000168887 | n.s. |
| ENSG00000261431 | 34.03872 | 0.157160274 | 0.360443 | 0.4360199 | 0.662822 | 0.926088818 | | antisense | 55.08 | 0.00524088 | 0.00524088 | ENSG00000261431 | n.s. |
| ENSG00000132780 | 4327.581 | -0.126270344 | 0.243765 | -0.518001 | 0.604458 | 0.908868121 | NASP | protein_coding | 42.9 | -0.005240109 | 0.005240109 | ENSG00000132780 | n.s. |
| ENSG00000178974 | 473.9227 | -0.135251553 | 0.270716 | -0.499607 | 0.617352 | 0.914690044 | FBXO34 | protein_coding | 40.98 | -0.005237758 | 0.005237758 | ENSG00000178974 | n.s. |
| ENSG00000137547 | 1037.297 | -0.142830888 | 0.297069 | -0.4808 | 0.630659 | 0.919033489 | MRPL15 | protein_coding | 43.26 | -0.005237418 | 0.005237418 | ENSG00000137547 | n.s. |
| ENSG00000136874 | 227.2778 | -0.129538437 | 0.255382 | -0.507234 | 0.611991 | 0.911134742 | STX17 | protein_coding | 36.32 | -0.005235606 | 0.005235606 | ENSG00000136874 | n.s. |
| ENSG00000171421 | 300.5282 | 0.15137336 | 0.334963 | 0.4519108 | 0.651333 | 0.923473769 | MRPL36 | protein_coding | 55.79 | 0.0052338 | 0.0052338 | ENSG00000171421 | n.s. |
| ENSG00000099795 | 813.3327 | -0.149804337 | 0.323185 | -0.463524 | 0.642989 | 0.922767027 | NDUFB7 | protein_coding | 57.48 | -0.00522936 | 0.00522936 | ENSG00000099795 | n.s. |
| ENSG00000149428 | 1020.669 | -0.128675075 | 0.253136 | -0.508324 | 0.611226 | 0.910755785 | HYOU1 | protein_coding | 53.98 | -0.005223959 | 0.005223959 | ENSG00000149428 | n.s. |
| ENSG00000104824 | 4118.873 | -0.125878302 | 0.241564 | -0.521096 | 0.6023 | 0.908868121 | HNRNPL | protein_coding | 53.73 | -0.00522384 | 0.00522384 | ENSG00000104824 | n.s. |
| ENSG00000167842 | 1453.405 | 0.125739868 | 0.236505 | 0.5316588 | 0.594962 | 0.908868121 | MIS12 | protein_coding | 44.08 | 0.005218095 | 0.005218095 | ENSG00000167842 | n.s. |
| ENSG00000115457 | 923.0309 | -0.141047691 | 0.28791 | -0.489902 | 0.624204 | 0.91849357 | IGFBP2 | protein_coding | 50.33 | -0.005208028 | 0.005208028 | ENSG00000115457 | n.s. |
| ENSG00000006576 | 1081.691 | -0.142015843 | 0.293724 | -0.483501 | 0.62874 | 0.919033489 | PHTF2 | protein_coding | 37.65 | -0.005207531 | 0.005207531 | ENSG00000006576 | n.s. |
| ENSG00000140943 | 1042.932 | -0.133046504 | 0.264715 | -0.502603 | 0.615243 | 0.91390508 | MBTPS1 | protein_coding | 45.14 | -0.005201973 | 0.005201973 | ENSG00000140943 | n.s. |
| ENSG00000115520 | 262.6178 | 0.14260899 | 0.300088 | 0.4752244 | 0.634627 | 0.919524756 | COQ10B | protein_coding | 41.73 | 0.005196183 | 0.005196183 | ENSG00000115520 | n.s. |
| ENSG00000138035 | 2742.669 | -0.13565487 | 0.272503 | -0.497811 | 0.618617 | 0.915595985 | PNPT1 | protein_coding | 39.78 | -0.005195055 | 0.005195055 | ENSG00000138035 | n.s. |
| ENSG00000048342 | 183.6903 | 0.127335922 | 0.249148 | 0.5110855 | 0.609291 | 0.910431619 | CC2D2A | protein_coding | 40.21 | 0.005189279 | 0.005189279 | ENSG00000048342 | n.s. |
| ENSG00000139146 | 6302.099 | -0.140096243 | 0.285774 | -0.490235 | 0.623968 | 0.91834044 | FAM60A | protein_coding | 41.46 | -0.005183041 | 0.005183041 | ENSG00000139146 | n.s. |
| ENSG00000099968 | 607.8515 | -0.14206603 | 0.299003 | -0.475133 | 0.634692 | 0.919524756 | BCL2L13 | protein_coding | 42.8 | -0.005176399 | 0.005176399 | ENSG00000099968 | n.s. |
| ENSG00000013583 | 87.27834 | -0.149024313 | 0.324919 | -0.45865 | 0.646485 | 0.923184844 | HEBP1 | protein_coding | 43.52 | -0.005172833 | 0.005172833 | ENSG00000013583 | n.s. |
| ENSG00000139637 | 525.0409 | -0.143682381 | 0.305624 | -0.470128 | 0.638263 | 0.920457187 | C12orf10 | protein_coding | 51.21 | -0.005172049 | 0.005172049 | ENSG00000139637 | n.s. |
| ENSG00000136897 | 1493.797 | -0.141033927 | 0.295922 | -0.476591 | 0.633653 | 0.919033489 | MRPL50 | protein_coding | 35.67 | -0.005171526 | 0.005171526 | ENSG00000136897 | n.s. |

| | | | | | | | | | | | | | |
|---|---|---|---|---|---|---|---|---|---|---|---|---|---|
| ENSG00000107833 | 449.3751 | 0.152266123 | 0.342074 | 0.4451266 | 0.656228 | 0.924786541 | NPM3 | protein_coding | 57.44 | 0.005170729 | 0.005170729 | ENSG00000107833 | n.s. |
| ENSG00000034677 | 616.7393 | -0.133306943 | 0.266565 | -0.500092 | 0.61701 | 0.9146497 | RNF19A | protein_coding | 38.61 | -0.005165005 | 0.005165005 | ENSG00000034677 | n.s. |
| ENSG00000225206 | 1621.376 | 0.124258732 | 0.238972 | 0.5199727 | 0.603083 | 0.908868121 | MIR137HG | lincRNA | 34.52 | 0.005156629 | 0.005156629 | ENSG00000225206 | n.s. |
| ENSG00000171617 | 3378.062 | -0.124245869 | 0.239587 | -0.518583 | 0.604052 | 0.908868121 | ENC1 | protein_coding | 45.65 | -0.005156095 | 0.005156095 | ENSG00000171617 | n.s. |
| ENSG00000241217 | 46.42509 | -0.14058617 | 0.293185 | -0.479514 | 0.631573 | 0.919033489 | RN7SL809P | misc_RNA | 50.84 | -0.005155107 | 0.005155107 | ENSG00000241217 | n.s. |
| ENSG00000105245 | 339.5861 | -0.138438707 | 0.281368 | -0.492019 | 0.622706 | 0.917861492 | NUMBL | protein_coding | 52.55 | -0.005153083 | 0.005153083 | ENSG00000105245 | n.s. |
| ENSG00000259398 | 34.53891 | -0.157011587 | 0.36211 | -0.433602 | 0.664578 | 0.927238553 | | sense_intronic | 47.21 | -0.005151318 | 0.005151318 | ENSG00000259398 | n.s. |
| ENSG00000177485 | 1334.735 | -0.12412215 | 0.239036 | -0.519261 | 0.603579 | 0.908868121 | ZBTB33 | protein_coding | 39.37 | -0.005150961 | 0.005150961 | ENSG00000177485 | n.s. |
| ENSG00000008988 | 11975.96 | 0.1495105 | 0.331418 | 0.451124 | 0.6519 | 0.923748122 | RPS20 | protein_coding | 41.97 | 0.005150103 | 0.005150103 | ENSG00000008988 | n.s. |
| ENSG00000132286 | 678.6839 | -0.124095961 | 0.239004 | -0.519222 | 0.603606 | 0.908868121 | TIMM10B | protein_coding | 47.6 | -0.005149874 | 0.005149874 | ENSG00000132286 | n.s. |
| ENSG00000199168 | 13.66161 | 0.174608386 | 0.432298 | 0.4039076 | 0.686281 | 0.934368276 | MIR374A | miRNA | 31.94 | 0.00514779 | 0.00514779 | ENSG00000199168 | n.s. |
| ENSG00000108561 | 3786.926 | 0.148251682 | 0.326901 | 0.4535069 | 0.650184 | 0.923184844 | C1QBP | protein_coding | 46.67 | 0.005146014 | 0.005146014 | ENSG00000108561 | n.s. |
| ENSG00000177556 | 230.8336 | -0.143204114 | 0.304806 | -0.46982 | 0.638483 | 0.920630217 | ATOX1 | protein_coding | 46.3 | -0.005143143 | 0.005143143 | ENSG00000177556 | n.s. |
| ENSG00000163875 | 422.3786 | -0.14194928 | 0.300818 | -0.471878 | 0.637014 | 0.92003534 | MEAF6 | protein_coding | 42.74 | -0.005137924 | 0.005137924 | ENSG00000163875 | n.s. |
| ENSG00000232821 | 362.2045 | -0.118117079 | 0.215587 | -0.547887 | 0.583769 | 0.904699627 | | antisense | 39.99 | -0.005137572 | 0.005137572 | ENSG00000232821 | n.s. |
| ENSG00000170144 | 9595.368 | -0.1237924 | 0.235021 | -0.526729 | 0.598382 | 0.908868121 | HNRNPA3 | protein_coding | 40 | -0.005137277 | 0.005137277 | ENSG00000170144 | n.s. |
| ENSG00000176142 | 501.1394 | 0.123737765 | 0.238986 | 0.5177626 | 0.604624 | 0.908868121 | TMEM39A | protein_coding | 39.09 | 0.00513501 | 0.00513501 | ENSG00000176142 | n.s. |
| ENSG00000121741 | 5124.915 | -0.138234016 | 0.281303 | -0.491406 | 0.623139 | 0.918094734 | ZMYM2 | protein_coding | 37.62 | -0.00513021 | 0.00513021 | ENSG00000121741 | n.s. |
| ENSG00000074054 | 2553.044 | -0.123501473 | 0.237031 | -0.521036 | 0.602342 | 0.908868121 | CLASP1 | protein_coding | 41.05 | -0.005125204 | 0.005125204 | ENSG00000074054 | n.s. |
| ENSG00000170759 | 6124.86 | -0.124354364 | 0.241488 | -0.51495 | 0.606588 | 0.909522325 | KIF5B | protein_coding | 37.58 | -0.005121738 | 0.005121738 | ENSG00000170759 | n.s. |
| ENSG00000006744 | 1215.248 | -0.136474307 | 0.276223 | -0.494072 | 0.621255 | 0.917256964 | ELAC2 | protein_coding | 48.08 | -0.005119012 | 0.005119012 | ENSG00000006744 | n.s. |
| ENSG00000143977 | 2259.991 | 0.13939386 | 0.287043 | 0.4856199 | 0.627237 | 0.919033489 | SNRPG | protein_coding | 41.09 | 0.005111387 | 0.005111387 | ENSG00000143977 | n.s. |
| ENSG00000207524 | 2.932437 | 0.328869159 | 1.306582 | 0.2517019 | 0.801272 | 0.964862231 | RNU6-33P | snRNA | 44.86 | 0.005108882 | 0.005108882 | ENSG00000207524 | n.s. |
| ENSG00000241058 | 657.8967 | -0.118128522 | 0.21788 | -0.542172 | 0.5877 | 0.905217543 | NSUN6 | protein_coding | 40.26 | -0.005108709 | 0.005108709 | ENSG00000241058 | n.s. |
| ENSG00000101844 | 115.286 | -0.13927094 | 0.292318 | -0.476437 | 0.633763 | 0.919033489 | ATG4A | protein_coding | 39.97 | -0.005106879 | 0.005106879 | ENSG00000101844 | n.s. |
| ENSG00000188191 | 358.5869 | -0.139231506 | 0.288068 | -0.483329 | 0.628862 | 0.919033489 | PRKAR1B | protein_coding | 54.13 | -0.005105433 | 0.005105433 | ENSG00000188191 | n.s. |
| ENSG00000187049 | 184.2968 | -0.139218289 | 0.286326 | -0.486223 | 0.626809 | 0.919033489 | TMEM216 | protein_coding | 48.57 | -0.005104949 | 0.005104949 | ENSG00000187049 | n.s. |
| ENSG00000197714 | 1712.167 | 0.125264137 | 0.244834 | 0.5116293 | 0.60891 | 0.910431619 | ZNF460 | protein_coding | 46.11 | 0.005104848 | 0.005104848 | ENSG00000197714 | n.s. |
| ENSG00000176209 | 504.0894 | 0.144301417 | 0.309814 | 0.4657673 | 0.641382 | 0.921891013 | SMIM19 | protein_coding | 44.11 | 0.005096786 | 0.005096786 | ENSG00000176209 | n.s. |
| ENSG00000167118 | 487.3857 | -0.139542462 | 0.293422 | -0.475569 | 0.634381 | 0.919380266 | URM1 | protein_coding | 52.12 | -0.005093973 | 0.005093973 | ENSG00000167118 | n.s. |
| ENSG00000263742 | 5.021131 | -0.220754359 | 0.675858 | -0.326628 | 0.743949 | 0.948261154 | MIR3165 | miRNA | 53.33 | -0.005093253 | 0.005093253 | ENSG00000263742 | n.s. |
| ENSG00000065911 | 7845.305 | -0.138651083 | 0.287951 | -0.481509 | 0.630155 | 0.919033489 | MTHFD2 | protein_coding | 43.3 | -0.00508415 | 0.00508415 | ENSG00000065911 | n.s. |

| | | | | | | | | | | | | | |
|---|---|---|---|---|---|---|---|---|---|---|---|---|---|
| ENSG00000157020 | 1267.927 | -0.138845707 | 0.291706 | -0.475978 | 0.63409 | 0.919149817 | SEC13 | protein_coding | 46.85 | -0.005083654 | 0.005083654 | ENSG00000157020 | n.s. |
| ENSG00000038219 | 4099.543 | -0.123412234 | 0.239834 | -0.514573 | 0.606851 | 0.909522325 | BOD1L1 | protein_coding | 38.88 | -0.005082935 | 0.005082935 | ENSG00000038219 | n.s. |
| ENSG00000166133 | 396.2747 | -0.122467989 | 0.23298 | -0.525659 | 0.599125 | 0.908868121 | RPUSD2 | protein_coding | 48.69 | -0.005082315 | 0.005082315 | ENSG00000166133 | n.s. |
| ENSG00000255198 | 73.91436 | -0.146399194 | 0.317975 | -0.460412 | 0.645221 | 0.923184844 | SNHG9 | lincRNA | 66.42 | -0.005081711 | 0.005081711 | ENSG00000255198 | n.s. |
| ENSG00000118193 | 2310.564 | -0.132171812 | 0.265095 | -0.498583 | 0.618073 | 0.915276425 | KIF14 | protein_coding | 37.83 | -0.005081705 | 0.005081705 | ENSG00000118193 | n.s. |
| ENSG00000144827 | 935.6499 | -0.134729871 | 0.271806 | -0.495685 | 0.620117 | 0.916893708 | ABHD10 | protein_coding | 35.84 | -0.005076757 | 0.005076757 | ENSG00000144827 | n.s. |
| ENSG00000107951 | 1437.334 | -0.131841544 | 0.264404 | -0.498636 | 0.618036 | 0.915276425 | MTPAP | protein_coding | 43.36 | -0.005069007 | 0.005069007 | ENSG00000107951 | n.s. |
| ENSG00000182986 | 754.4525 | -0.138230931 | 0.288742 | -0.478736 | 0.632127 | 0.919033489 | ZNF320 | protein_coding | 43.93 | -0.005068743 | 0.005068743 | ENSG00000182986 | n.s. |
| ENSG00000234420 | 1725.259 | -0.134752209 | 0.272585 | -0.49435 | 0.621059 | 0.917065125 | ZNF37BP | ıscribed_processed_pseudog | 39.07 | -0.005066659 | 0.005066659 | ENSG00000234420 | n.s. |
| ENSG00000159164 | 657.239 | -0.124751099 | 0.245342 | -0.508479 | 0.611117 | 0.910755785 | SV2A | protein_coding | 52.21 | -0.005064653 | 0.005064653 | ENSG00000159164 | n.s. |
| ENSG00000153234 | 143.9302 | 0.124508366 | 0.244313 | 0.5096273 | 0.610313 | 0.910595741 | NR4A2 | protein_coding | 45.15 | 0.005064301 | 0.005064301 | ENSG00000153234 | n.s. |
| ENSG00000199202 | 8.631197 | -0.181270958 | 0.470462 | -0.385304 | 0.700012 | 0.937752951 | RNA5SP289 | rRNA | 47.45 | -0.005059555 | 0.005059555 | ENSG00000199202 | n.s. |
| ENSG00000263697 | 3.377201 | -0.239199373 | 0.773076 | -0.309412 | 0.757008 | 0.95246603 | MIR3621 | miRNA | 72.94 | -0.005059187 | 0.005059187 | ENSG00000263697 | n.s. |
| ENSG00000081138 | 816.4418 | 0.137820404 | 0.283982 | 0.4853145 | 0.627453 | 0.919033489 | CDH7 | protein_coding | 35.27 | 0.00505369 | 0.00505369 | ENSG00000081138 | n.s. |
| ENSG00000170293 | 52.31569 | 0.148696451 | 0.33615 | 0.442352 | 0.658235 | 0.924786541 | CMTM8 | protein_coding | 43.06 | 0.005049508 | 0.005049508 | ENSG00000170293 | n.s. |
| ENSG00000272516 | 44.83035 | 0.138525651 | 0.292061 | 0.4743037 | 0.635283 | 0.919532207 | | lincRNA | 33.22 | 0.005046913 | 0.005046913 | ENSG00000272516 | n.s. |
| ENSG00000139826 | 953.8329 | 0.137517799 | 0.281982 | 0.4876838 | 0.625774 | 0.918973809 | ABHD13 | protein_coding | 35.43 | 0.005046472 | 0.005046472 | ENSG00000139826 | n.s. |
| ENSG00000186625 | 474.8537 | -0.124343742 | 0.244681 | -0.508188 | 0.611322 | 0.910800656 | KATNA1 | protein_coding | 42.14 | -0.005045454 | 0.005045454 | ENSG00000186625 | n.s. |
| ENSG00000180817 | 3685.33 | 0.144895466 | 0.31337 | 0.462378 | 0.64381 | 0.922958206 | PPA1 | protein_coding | 39.81 | 0.005044965 | 0.005044965 | ENSG00000180817 | n.s. |
| ENSG00000113761 | 265.5607 | -0.133947598 | 0.270466 | -0.495248 | 0.620425 | 0.916981464 | ZNF346 | protein_coding | 46.38 | -0.005041713 | 0.005041713 | ENSG00000113761 | n.s. |
| ENSG00000276704 | 78.67386 | 0.137427229 | 0.288489 | 0.4763688 | 0.633812 | 0.919033489 | | sense_intronic | 33.88 | 0.005039273 | 0.005039273 | ENSG00000276704 | n.s. |
| ENSG00000136940 | 769.4881 | -0.137229339 | 0.281562 | -0.487386 | 0.625985 | 0.918973809 | PDCL | protein_coding | 45.28 | -0.005035887 | 0.005035887 | ENSG00000136940 | n.s. |
| ENSG00000070831 | 1013.63 | -0.137322919 | 0.285746 | -0.480576 | 0.630818 | 0.919033489 | CDC42 | protein_coding | 41.82 | -0.005035448 | 0.005035448 | ENSG00000070831 | n.s. |
| ENSG00000204116 | 326.3523 | -0.135792049 | 0.276538 | -0.491043 | 0.623396 | 0.918167932 | CHIC1 | protein_coding | 37.63 | -0.005034881 | 0.005034881 | ENSG00000204116 | n.s. |
| ENSG00000087338 | 799.053 | -0.129171071 | 0.257905 | -0.500847 | 0.616479 | 0.914179911 | GMCL1 | protein_coding | 37.77 | -0.00503358 | 0.00503358 | ENSG00000087338 | n.s. |
| ENSG00000173915 | 1761.278 | 0.149691982 | 0.342777 | 0.4367033 | 0.662327 | 0.925519646 | USMG5 | protein_coding | 43.23 | 0.0050318 | 0.0050318 | ENSG00000173915 | n.s. |
| ENSG00000152795 | 19932.2 | 0.121248229 | 0.230961 | 0.5249716 | 0.599603 | 0.908868121 | HNRNPDL | protein_coding | 42.24 | 0.005031696 | 0.005031696 | ENSG00000152795 | n.s. |
| ENSG00000264007 | 30.2916 | 0.14815661 | 0.337618 | 0.4388296 | 0.660785 | 0.924786541 | | sense_intronic | 53.29 | 0.005031176 | 0.005031176 | ENSG00000264007 | n.s. |
| ENSG00000134970 | 1153.511 | 0.13713903 | 0.285744 | 0.4799373 | 0.631272 | 0.919033489 | TMED7 | protein_coding | 37.91 | 0.005028705 | 0.005028705 | ENSG00000134970 | n.s. |
| ENSG00000202392 | 15.56793 | -0.165192144 | 0.398153 | -0.414896 | 0.678218 | 0.932323161 | RN7SKP292 | misc_RNA | 53.63 | -0.00502738 | 0.00502738 | ENSG00000202392 | n.s. |
| ENSG00000236618 | 134.0438 | 0.144757881 | 0.314114 | 0.4608448 | 0.64491 | 0.923184844 | PITPNA-AS1 | antisense | 51.63 | 0.005024739 | 0.005024739 | ENSG00000236618 | n.s. |
| ENSG00000100221 | 622.4846 | -0.13194979 | 0.265492 | -0.497002 | 0.619188 | 0.916050756 | JOSD1 | protein_coding | 47.81 | -0.005024709 | 0.005024709 | ENSG00000100221 | n.s. |

| | | | | | | | | | | | | | |
|---|---|---|---|---|---|---|---|---|---|---|---|---|---|
| ENSG00000277202 | 5.421914 | 0.193463549 | 0.547324 | 0.3534717 | 0.723735 | 0.942009991 | MIR8063 | miRNA | 38.27 | 0.005019313 | 0.005019313 | ENSG00000277202 | n.s. |
| ENSG00000141380 | 2684.418 | -0.129585156 | 0.25942 | -0.499519 | 0.617414 | 0.914690044 | SS18 | protein_coding | 35.21 | -0.005018321 | 0.005018321 | ENSG00000141380 | n.s. |
| ENSG00000240682 | 646.3176 | -0.135596918 | 0.276582 | -0.490259 | 0.623951 | 0.91834044 | ISY1 | protein_coding | 45.22 | -0.005016583 | 0.005016583 | ENSG00000240682 | n.s. |
| ENSG00000225973 | 141.2491 | 0.137690474 | 0.290838 | 0.4734261 | 0.635909 | 0.919565577 | PIGBOS1 | protein_coding | 47.12 | 0.005014315 | 0.005014315 | ENSG00000225973 | n.s. |
| ENSG00000112081 | 14183.72 | -0.120623717 | 0.227584 | -0.530018 | 0.596099 | 0.908868121 | SRSF3 | protein_coding | 40.22 | -0.005005779 | 0.005005779 | ENSG00000112081 | n.s. |
| ENSG00000215973 | 53.42463 | 0.144108604 | 0.317887 | 0.4533329 | 0.650309 | 0.923184844 | MIR933 | miRNA | 66.23 | 0.005002202 | 0.005002202 | ENSG00000215973 | n.s. |
| ENSG00000225945 | 15.70121 | -0.173753094 | 0.43913 | -0.395676 | 0.692344 | 0.935906213 | | antisense | 43.87 | -0.004998472 | 0.004998472 | ENSG00000225945 | n.s. |
| ENSG00000184672 | 101.7116 | -0.130384248 | 0.261875 | -0.497888 | 0.618563 | 0.915595985 | RALYL | protein_coding | 35.32 | -0.004993211 | 0.004993211 | ENSG00000184672 | n.s. |
| ENSG00000183431 | 1978.675 | -0.133635463 | 0.270948 | -0.493215 | 0.621861 | 0.917630258 | SF3A3 | protein_coding | 44.59 | -0.004988916 | 0.004988916 | ENSG00000183431 | n.s. |
| ENSG00000139719 | 861.2603 | -0.135930423 | 0.279083 | -0.487062 | 0.626215 | 0.918973809 | VPS33A | protein_coding | 44.63 | -0.004988221 | 0.004988221 | ENSG00000139719 | n.s. |
| ENSG00000198010 | 46.27496 | -0.138531064 | 0.294696 | -0.470081 | 0.638297 | 0.920457187 | DLGAP2 | protein_coding | 47.91 | -0.004986621 | 0.004986621 | ENSG00000198010 | n.s. |
| ENSG00000162437 | 1147.59 | -0.134451466 | 0.273836 | -0.490992 | 0.623432 | 0.918167932 | RAVER2 | protein_coding | 37.2 | -0.004985175 | 0.004985175 | ENSG00000162437 | n.s. |
| ENSG00000150977 | 157.7525 | -0.122743823 | 0.241233 | -0.508819 | 0.610879 | 0.910725807 | RILPL2 | protein_coding | 49.36 | -0.004984916 | 0.004984916 | ENSG00000150977 | n.s. |
| ENSG00000118495 | 321.1696 | -0.135800346 | 0.282077 | -0.481431 | 0.63021 | 0.919033489 | PLAGL1 | protein_coding | 40.32 | -0.004979617 | 0.004979617 | ENSG00000118495 | n.s. |
| ENSG00000254093 | 551.3526 | -0.135788614 | 0.279492 | -0.485841 | 0.62708 | 0.919033489 | PINX1 | protein_coding | 43.17 | -0.004979187 | 0.004979187 | ENSG00000254093 | n.s. |
| ENSG00000186106 | 660.8444 | -0.143428721 | 0.312443 | -0.459056 | 0.646194 | 0.923184844 | ANKRD46 | protein_coding | 39.74 | -0.004978602 | 0.004978602 | ENSG00000186106 | n.s. |
| ENSG00000121281 | 295.2575 | -0.137208607 | 0.290292 | -0.472657 | 0.636458 | 0.919866014 | ADCY7 | protein_coding | 54.43 | -0.004977301 | 0.004977301 | ENSG00000121281 | n.s. |
| ENSG00000115128 | 2484.403 | 0.146495302 | 0.328592 | 0.4458273 | 0.655722 | 0.924786541 | SF3B6 | protein_coding | 43.19 | 0.004974761 | 0.004974761 | ENSG00000115128 | n.s. |
| ENSG00000221923 | 593.1856 | -0.119869395 | 0.231567 | -0.517644 | 0.604707 | 0.908868121 | ZNF880 | protein_coding | 41.86 | -0.004974476 | 0.004974476 | ENSG00000221923 | n.s. |
| ENSG00000276174 | 76.238 | -0.135622421 | 0.280828 | -0.482937 | 0.62914 | 0.919033489 | | sense_intronic | 31.9 | -0.004973093 | 0.004973093 | ENSG00000276174 | n.s. |
| ENSG00000252186 | 75.02451 | -0.146388032 | 0.329544 | -0.444214 | 0.656888 | 0.924786541 | RNU6-781P | snRNA | 48.51 | -0.004971118 | 0.004971118 | ENSG00000252186 | n.s. |
| ENSG00000038358 | 895.2954 | -0.124689015 | 0.246988 | -0.504839 | 0.613672 | 0.912313064 | EDC4 | protein_coding | 57.32 | -0.004969619 | 0.004969619 | ENSG00000038358 | n.s. |
| ENSG00000205791 | 55.94898 | -0.138753546 | 0.29625 | -0.468367 | 0.639522 | 0.920848926 | LOH12CR2 | lincRNA | 45.19 | -0.004968988 | 0.004968988 | ENSG00000205791 | n.s. |
| ENSG00000226950 | 4899.258 | 0.119731535 | 0.228567 | 0.5238363 | 0.600392 | 0.908868121 | DANCR | processed_transcript | 44.16 | 0.004968754 | 0.004968754 | ENSG00000226950 | n.s. |
| ENSG00000228808 | 18.79466 | 0.153941909 | 0.360061 | 0.427544 | 0.668983 | 0.928390061 | HMGB3P4 | processed_pseudogene | 51.56 | 0.004967631 | 0.004967631 | ENSG00000228808 | n.s. |
| ENSG00000126814 | 977.2189 | 0.124137444 | 0.245521 | 0.5056081 | 0.613132 | 0.911981962 | TRMT5 | protein_coding | 37.97 | 0.004967205 | 0.004967205 | ENSG00000126814 | n.s. |
| ENSG00000164933 | 992.0989 | -0.138590824 | 0.296338 | -0.467678 | 0.640015 | 0.920914184 | SLC25A32 | protein_coding | 36.96 | -0.004958896 | 0.004958896 | ENSG00000164933 | n.s. |
| ENSG00000206754 | 197.4335 | 0.119421089 | 0.224315 | 0.5323815 | 0.594462 | 0.908830158 | SNORD101 | snoRNA | 36.99 | 0.004958038 | 0.004958038 | ENSG00000206754 | n.s. |
| ENSG00000188848 | 1078.84 | -0.135166087 | 0.280475 | -0.481918 | 0.629864 | 0.919033489 | BEND4 | protein_coding | 42.3 | -0.00495636 | 0.00495636 | ENSG00000188848 | n.s. |
| ENSG00000212493 | 64.15111 | 0.135139918 | 0.283649 | 0.4764339 | 0.633765 | 0.919033489 | SNORD19 | snoRNA | 45.59 | 0.0049554 | 0.0049554 | ENSG00000212493 | n.s. |
| ENSG00000147099 | 305.376 | -0.136934194 | 0.290299 | -0.4717 | 0.637141 | 0.920123463 | HDAC8 | protein_coding | 40.05 | -0.004950704 | 0.004950704 | ENSG00000147099 | n.s. |
| ENSG00000259637 | 40.76538 | -0.137928946 | 0.294043 | -0.469077 | 0.639015 | 0.920750614 | DNM1P38 | unprocessed_pseudogene | 58.18 | -0.004945854 | 0.004945854 | ENSG00000259637 | n.s. |

| | | | | | | | | | | | | | |
|---|---|---|---|---|---|---|---|---|---|---|---|---|---|
| ENSG00000122481 | 424.0241 | -0.142465202 | 0.311945 | -0.4567 | 0.647887 | 0.923184844 | RWDD3 | protein_coding | 40.77 | -0.004945157 | 0.004945157 | ENSG00000122481 | n.s. |
| ENSG00000127540 | 651.3791 | 0.143425555 | 0.31811 | 0.4508675 | 0.652085 | 0.923748122 | UQCR11 | protein_coding | 56.38 | 0.004940499 | 0.004940499 | ENSG00000127540 | n.s. |
| ENSG00000182362 | 210.7061 | 0.142313562 | 0.309349 | 0.4600415 | 0.645486 | 0.923184844 | YBEY | protein_coding | 53.1 | 0.004939894 | 0.004939894 | ENSG00000182362 | n.s. |
| ENSG00000111540 | 528.7449 | -0.134703103 | 0.280747 | -0.479802 | 0.631368 | 0.919033489 | RAB5B | protein_coding | 46.01 | -0.004939383 | 0.004939383 | ENSG00000111540 | n.s. |
| ENSG00000256571 | 23.45788 | -0.160772379 | 0.384619 | -0.418004 | 0.675944 | 0.931706455 | | sense_intronic | 40.99 | -0.004939071 | 0.004939071 | ENSG00000256571 | n.s. |
| ENSG00000106268 | 631.6454 | -0.134690486 | 0.280201 | -0.480692 | 0.630735 | 0.919033489 | NUDT1 | protein_coding | 56.77 | -0.00493892 | 0.00493892 | ENSG00000106268 | n.s. |
| ENSG00000174840 | 1462.642 | 0.118982419 | 0.22686 | 0.524475 | 0.599948 | 0.908868121 | PDE12 | protein_coding | 41.24 | 0.004937667 | 0.004937667 | ENSG00000174840 | n.s. |
| ENSG00000105819 | 2149.48 | 0.128655926 | 0.258164 | 0.4983501 | 0.618237 | 0.915422518 | PMPCB | protein_coding | 38.52 | 0.00493761 | 0.00493761 | ENSG00000105819 | n.s. |
| ENSG00000105583 | 609.8913 | -0.14201169 | 0.307338 | -0.46207 | 0.644031 | 0.923072262 | WDR83OS | protein_coding | 57.12 | -0.004936937 | 0.004936937 | ENSG00000105583 | n.s. |
| ENSG00000133169 | 4262.654 | -0.134616012 | 0.276712 | -0.486484 | 0.626624 | 0.919033489 | BEX1 | protein_coding | 56.79 | -0.004936189 | 0.004936189 | ENSG00000133169 | n.s. |
| ENSG00000101452 | 668.7079 | -0.134513984 | 0.280908 | -0.478854 | 0.632042 | 0.919033489 | DHX35 | protein_coding | 41.33 | -0.004932448 | 0.004932448 | ENSG00000101452 | n.s. |
| ENSG00000213096 | 1011.672 | -0.135847953 | 0.287495 | -0.472524 | 0.636553 | 0.919866014 | ZNF254 | protein_coding | 41.43 | -0.004927942 | 0.004927942 | ENSG00000213096 | n.s. |
| ENSG00000115762 | 1556.454 | -0.130079519 | 0.262048 | -0.496397 | 0.619615 | 0.916487226 | PLEKHB2 | protein_coding | 42.42 | -0.004926578 | 0.004926578 | ENSG00000115762 | n.s. |
| ENSG00000259943 | 226.3584 | -0.118616325 | 0.223072 | -0.531741 | 0.594906 | 0.908868121 | | antisense | 40.43 | -0.004922474 | 0.004922474 | ENSG00000259943 | n.s. |
| ENSG00000026559 | 2636.773 | -0.136134357 | 0.288761 | -0.471442 | 0.637325 | 0.920241952 | KCNG1 | protein_coding | 52.21 | -0.004914174 | 0.004914174 | ENSG00000026559 | n.s. |
| ENSG00000276757 | 28.38763 | -0.151934844 | 0.354941 | -0.428057 | 0.66861 | 0.928267143 | RN7SL192P | misc_RNA | 56.19 | -0.004911601 | 0.004911601 | ENSG00000276757 | n.s. |
| ENSG00000119953 | 1865.962 | 0.119245 | 0.231563 | 0.5149571 | 0.606583 | 0.909522325 | SMNDC1 | protein_coding | 37.32 | 0.0049113 | 0.0049113 | ENSG00000119953 | n.s. |
| ENSG00000156521 | 285.7106 | -0.118301624 | 0.225696 | -0.524163 | 0.600165 | 0.908868121 | TYSND1 | protein_coding | 53.24 | -0.004909414 | 0.004909414 | ENSG00000156521 | n.s. |
| ENSG00000165030 | 314.8683 | 0.131426343 | 0.266246 | 0.4936277 | 0.621569 | 0.917623498 | NFIL3 | protein_coding | 41.93 | 0.004906865 | 0.004906865 | ENSG00000165030 | n.s. |
| ENSG00000271918 | 175.27 | -0.120727963 | 0.237397 | -0.508548 | 0.611069 | 0.910755785 | | antisense | 52.67 | -0.004901321 | 0.004901321 | ENSG00000271918 | n.s. |
| ENSG00000164414 | 380.5833 | 0.141137661 | 0.311582 | 0.4529718 | 0.650569 | 0.923184844 | SLC35A1 | protein_coding | 39.25 | 0.004899076 | 0.004899076 | ENSG00000164414 | n.s. |
| ENSG00000148248 | 1342.217 | -0.133505893 | 0.278392 | -0.479561 | 0.63154 | 0.919033489 | SURF4 | protein_coding | 52.38 | -0.004895483 | 0.004895483 | ENSG00000148248 | n.s. |
| ENSG00000121073 | 1667.223 | -0.133359169 | 0.273904 | -0.486884 | 0.626341 | 0.918973809 | SLC35B1 | protein_coding | 47.79 | -0.004893864 | 0.004893864 | ENSG00000121073 | n.s. |
| ENSG00000136827 | 544.9519 | -0.140982769 | 0.311589 | -0.452464 | 0.650934 | 0.923184844 | TOR1A | protein_coding | 52.06 | -0.0048937 | 0.0048937 | ENSG00000136827 | n.s. |
| ENSG00000280206 | 75.89499 | -0.143352536 | 0.320412 | -0.447401 | 0.654586 | 0.924413646 | | lincRNA | 40.82 | -0.004893146 | 0.004893146 | ENSG00000280206 | n.s. |
| ENSG00000130005 | 237.4477 | -0.140952458 | 0.31122 | -0.452903 | 0.650619 | 0.923184844 | GAMT | protein_coding | 64.25 | -0.004892648 | 0.004892648 | ENSG00000130005 | n.s. |
| ENSG00000131475 | 1421.342 | -0.138396757 | 0.297281 | -0.465543 | 0.641543 | 0.921891013 | VPS25 | protein_coding | 49.25 | -0.004888231 | 0.004888231 | ENSG00000131475 | n.s. |
| ENSG00000165259 | 338.7466 | -0.133196566 | 0.273468 | -0.487065 | 0.626212 | 0.918973809 | HDX | protein_coding | 34.93 | -0.004887897 | 0.004887897 | ENSG00000165259 | n.s. |
| ENSG00000129245 | 786.3015 | -0.134147163 | 0.283102 | -0.473848 | 0.635609 | 0.919532207 | FXR2 | protein_coding | 48.07 | -0.004887391 | 0.004887391 | ENSG00000129245 | n.s. |
| ENSG00000215939 | 11.05433 | 0.174993155 | 0.452528 | 0.3867012 | 0.698977 | 0.937752951 | MIR873 | miRNA | 46.75 | 0.004884332 | 0.004884332 | ENSG00000215939 | n.s. |
| ENSG00000148341 | 382.6948 | -0.133086648 | 0.279292 | -0.476514 | 0.633709 | 0.919033489 | SH3GLB2 | protein_coding | 55.75 | -0.004880109 | 0.004880109 | ENSG00000148341 | n.s. |
| ENSG00000217527 | 88.11018 | -0.134707024 | 0.28538 | -0.472027 | 0.636908 | 0.919977793 | RPS16P5 | ıscribed_processed_pseudog | 50 | -0.004879446 | 0.004879446 | ENSG00000217527 | n.s. |

| | | | | | | | | | | | | | |
|---|---|---|---|---|---|---|---|---|---|---|---|---|---|
| ENSG00000100410 | 917.7389 | -0.132942412 | 0.276711 | -0.480439 | 0.630916 | 0.919033489 | PHF5A | protein_coding | 46.26 | -0.004874821 | 0.004874821 | ENSG00000100410 | n.s. |
| ENSG00000041880 | 123.6046 | -0.125870784 | 0.251875 | -0.499736 | 0.617261 | 0.914690044 | PARP3 | protein_coding | 59.21 | -0.004874478 | 0.004874478 | ENSG00000041880 | n.s. |
| ENSG00000115514 | 1155.887 | -0.141665087 | 0.315679 | -0.448764 | 0.653602 | 0.923962103 | TXNDC9 | protein_coding | 42.12 | -0.004865607 | 0.004865607 | ENSG00000115514 | n.s. |
| ENSG00000201899 | 6.148629 | 0.184929357 | 0.516822 | 0.35782 | 0.720478 | 0.941256551 | SNORD114-24 | snoRNA | 44.44 | 0.00486216 | 0.00486216 | ENSG00000201899 | n.s. |
| ENSG00000115484 | 9346.274 | 0.13317427 | 0.280086 | 0.475477 | 0.634447 | 0.919380266 | CCT4 | protein_coding | 41.18 | 0.004861503 | 0.004861503 | ENSG00000115484 | n.s. |
| ENSG00000154813 | 653.064 | -0.140033852 | 0.304229 | -0.460292 | 0.645307 | 0.923184844 | DPH3 | protein_coding | 40.8 | -0.004860762 | 0.004860762 | ENSG00000154813 | n.s. |
| ENSG00000145919 | 2079.221 | 0.131676655 | 0.268857 | 0.4897648 | 0.6243 | 0.918517547 | BOD1 | protein_coding | 44.51 | 0.00486052 | 0.00486052 | ENSG00000145919 | n.s. |
| ENSG00000108469 | 406.6649 | -0.120191709 | 0.236926 | -0.507297 | 0.611946 | 0.911134742 | RECQL5 | protein_coding | 53.18 | -0.004857836 | 0.004857836 | ENSG00000108469 | n.s. |
| ENSG00000146083 | 598.1196 | -0.117055554 | 0.223876 | -0.522859 | 0.601073 | 0.908868121 | RNF44 | protein_coding | 60.11 | -0.004857704 | 0.004857704 | ENSG00000146083 | n.s. |
| ENSG00000178053 | 271.972 | -0.142914375 | 0.323797 | -0.441371 | 0.658945 | 0.924786541 | MLF1 | protein_coding | 37.51 | -0.004853158 | 0.004853158 | ENSG00000178053 | n.s. |
| ENSG00000117139 | 1995.431 | 0.120520376 | 0.237937 | 0.5065212 | 0.612491 | 0.911498898 | KDM5B | protein_coding | 41.09 | 0.004850204 | 0.004850204 | ENSG00000117139 | n.s. |
| ENSG00000120327 | 322.765 | -0.121666357 | 0.241028 | -0.504781 | 0.613713 | 0.912313064 | PCDHB14 | protein_coding | 44.93 | -0.004849147 | 0.004849147 | ENSG00000120327 | n.s. |
| ENSG00000092931 | 108.6084 | 0.123329454 | 0.245076 | 0.5032304 | 0.614802 | 0.913454511 | MFSD11 | protein_coding | 44.16 | 0.00484846 | 0.00484846 | ENSG00000092931 | n.s. |
| ENSG00000173457 | 1706.171 | -0.133052575 | 0.280962 | -0.473561 | 0.635813 | 0.919565577 | PPP1R14B | protein_coding | 65.56 | -0.004845415 | 0.004845415 | ENSG00000173457 | n.s. |
| ENSG00000189050 | 257.493 | -0.132064203 | 0.276995 | -0.476775 | 0.633522 | 0.919033489 | RNFT1 | protein_coding | 38.09 | -0.004842618 | 0.004842618 | ENSG00000189050 | n.s. |
| ENSG00000198960 | 151.0902 | -0.132054253 | 0.275713 | -0.478955 | 0.63197 | 0.919033489 | ARMCX6 | protein_coding | 51.56 | -0.004842253 | 0.004842253 | ENSG00000198960 | n.s. |
| ENSG00000266570 | 24.68686 | -0.149779651 | 0.350024 | -0.427912 | 0.668715 | 0.928267143 | MIR5579 | miRNA | 46.55 | -0.00484193 | 0.00484193 | ENSG00000266570 | n.s. |
| ENSG00000113141 | 2024.141 | -0.12863178 | 0.259838 | -0.495047 | 0.620567 | 0.916981464 | IK | protein_coding | 45.21 | -0.004841629 | 0.004841629 | ENSG00000113141 | n.s. |
| ENSG00000273181 | 16.30962 | 0.177092953 | 0.475324 | 0.3725731 | 0.709466 | 0.939005324 | | lincRNA | 33.06 | 0.004840295 | 0.004840295 | ENSG00000273181 | n.s. |
| ENSG00000085982 | 643.6496 | -0.131962548 | 0.275126 | -0.479645 | 0.63148 | 0.919033489 | USP40 | protein_coding | 38.86 | -0.00483889 | 0.00483889 | ENSG00000085982 | n.s. |
| ENSG00000158717 | 211.8017 | -0.14210997 | 0.317915 | -0.447007 | 0.65487 | 0.924621046 | RNF166 | protein_coding | 62.53 | -0.004836887 | 0.004836887 | ENSG00000158717 | n.s. |
| ENSG00000150459 | 2114.996 | 0.13189416 | 0.276002 | 0.4778741 | 0.63274 | 0.919033489 | SAP18 | protein_coding | 42.74 | 0.004836383 | 0.004836383 | ENSG00000150459 | n.s. |
| ENSG00000122547 | 921.258 | -0.117396977 | 0.227908 | -0.515107 | 0.606478 | 0.909522325 | EEPD1 | protein_coding | 45.46 | -0.004835187 | 0.004835187 | ENSG00000122547 | n.s. |
| ENSG00000128872 | 3520.356 | -0.131750419 | 0.270498 | -0.487067 | 0.626211 | 0.918973809 | TMOD2 | protein_coding | 41.52 | -0.004834827 | 0.004834827 | ENSG00000128872 | n.s. |
| ENSG00000126247 | 663.0494 | -0.139113713 | 0.305999 | -0.454622 | 0.649381 | 0.923184844 | CAPNS1 | protein_coding | 54.14 | -0.004828823 | 0.004828823 | ENSG00000126247 | n.s. |
| ENSG00000176783 | 503.6568 | -0.131431738 | 0.270557 | -0.485782 | 0.627121 | 0.919033489 | RUFY1 | protein_coding | 47.71 | -0.004819426 | 0.004819426 | ENSG00000176783 | n.s. |
| ENSG00000107815 | 721.6748 | -0.116116802 | 0.220233 | -0.527246 | 0.598023 | 0.908868121 | TWNK | protein_coding | 52.35 | -0.004818746 | 0.004818746 | ENSG00000107815 | n.s. |
| ENSG00000130713 | 1686.864 | -0.118736286 | 0.2337 | -0.508071 | 0.611403 | 0.91082469 | EXOSC2 | protein_coding | 47.74 | -0.004816562 | 0.004816562 | ENSG00000130713 | n.s. |
| ENSG00000145354 | 973.1408 | -0.138711252 | 0.302268 | -0.458902 | 0.646305 | 0.923184844 | CISD2 | protein_coding | 39.45 | -0.004814853 | 0.004814853 | ENSG00000145354 | n.s. |
| ENSG00000135677 | 1037.034 | -0.131183282 | 0.274675 | -0.477594 | 0.632939 | 0.919033489 | GNS | protein_coding | 42.2 | -0.004810316 | 0.004810316 | ENSG00000135677 | n.s. |
| ENSG00000198995 | 5.31378 | -0.203357811 | 0.612592 | -0.331963 | 0.739917 | 0.946996213 | MIR340 | miRNA | 38.95 | -0.00480977 | 0.00480977 | ENSG00000198995 | n.s. |
| ENSG00000163629 | 2525.046 | 0.131959175 | 0.278388 | 0.4740117 | 0.635492 | 0.919532207 | PTPN13 | protein_coding | 36.16 | 0.004807676 | 0.004807676 | ENSG00000163629 | n.s. |

| | | | | | | | | | | | | | |
|---|---|---|---|---|---|---|---|---|---|---|---|---|---|
| ENSG00000040341 | 535.9423 | -0.134190776 | 0.286588 | -0.468235 | 0.639616 | 0.920848926 | STAU2 | protein_coding | 38.25 | -0.004805588 | 0.004805588 | ENSG00000040341 | n.s. |
| ENSG00000243926 | 25.22881 | 0.159505918 | 0.3873 | 0.4118404 | 0.680456 | 0.932999624 | TIPARP-AS1 | antisense | 43.54 | 0.004804084 | 0.004804084 | ENSG00000243926 | n.s. |
| ENSG00000103351 | 456.8789 | -0.130992465 | 0.27366 | -0.478669 | 0.632174 | 0.919033489 | CLUAP1 | protein_coding | 45.59 | -0.004803319 | 0.004803319 | ENSG00000103351 | n.s. |
| ENSG00000104907 | 1203.699 | -0.130268493 | 0.266598 | -0.488632 | 0.625102 | 0.918928785 | TRMT1 | protein_coding | 53.66 | -0.004783217 | 0.004783217 | ENSG00000104907 | n.s. |
| ENSG00000124571 | 2879.258 | -0.130341891 | 0.267592 | -0.487091 | 0.626194 | 0.918973809 | XPO5 | protein_coding | 43.8 | -0.004783139 | 0.004783139 | ENSG00000124571 | n.s. |
| ENSG00000117505 | 3110.123 | -0.12692562 | 0.255997 | -0.495809 | 0.620029 | 0.916893708 | DR1 | protein_coding | 36.47 | -0.004782685 | 0.004782685 | ENSG00000117505 | n.s. |
| ENSG00000213676 | 351.1629 | -0.130388961 | 0.268973 | -0.484767 | 0.627842 | 0.919033489 | ATF6B | protein_coding | 50.77 | -0.004781189 | 0.004781189 | ENSG00000213676 | n.s. |
| ENSG00000124523 | 64.24041 | -0.140770103 | 0.317299 | -0.443652 | 0.657294 | 0.924786541 | SIRT5 | protein_coding | 42.7 | -0.004780341 | 0.004780341 | ENSG00000124523 | n.s. |
| ENSG00000197713 | 2073.362 | -0.128806195 | 0.262054 | -0.491526 | 0.623055 | 0.918094734 | RPE | protein_coding | 38.52 | -0.00478032 | 0.00478032 | ENSG00000197713 | n.s. |
| ENSG00000133112 | 18099.62 | 0.137579494 | 0.303332 | 0.4535613 | 0.650145 | 0.923184844 | TPT1 | protein_coding | 44.18 | 0.004775568 | 0.004775568 | ENSG00000133112 | n.s. |
| ENSG00000198513 | 615.8999 | -0.126721546 | 0.256298 | -0.49443 | 0.621002 | 0.917065125 | ATL1 | protein_coding | 37.9 | -0.004764708 | 0.004764708 | ENSG00000198513 | n.s. |
| ENSG00000149925 | 3573.196 | -0.129932347 | 0.270743 | -0.47991 | 0.631291 | 0.919033489 | ALDOA | protein_coding | 54.43 | -0.004764445 | 0.004764445 | ENSG00000149925 | n.s. |
| ENSG00000124177 | 3272.767 | -0.117880124 | 0.232243 | -0.507573 | 0.611753 | 0.911134742 | CHD6 | protein_coding | 41.1 | -0.004764407 | 0.004764407 | ENSG00000124177 | n.s. |
| ENSG00000162817 | 127.5891 | 0.137248544 | 0.30227 | 0.4540599 | 0.649786 | 0.923184844 | C1orf115 | protein_coding | 48.31 | 0.00476408 | 0.00476408 | ENSG00000162817 | n.s. |
| ENSG00000089693 | 1409.413 | -0.129894387 | 0.272612 | -0.47648 | 0.633732 | 0.919033489 | MLF2 | protein_coding | 52.27 | -0.004763054 | 0.004763054 | ENSG00000089693 | n.s. |
| ENSG00000120333 | 982.6109 | -0.138299102 | 0.307157 | -0.450256 | 0.652526 | 0.923800272 | MRPS14 | protein_coding | 40.16 | -0.00476052 | 0.00476052 | ENSG00000120333 | n.s. |
| ENSG00000159873 | 780.0751 | -0.1264361 | 0.255404 | -0.495043 | 0.62057 | 0.916981464 | CCDC117 | protein_coding | 44.33 | -0.004758984 | 0.004758984 | ENSG00000159873 | n.s. |
| ENSG00000240429 | 222.6268 | 0.114615269 | 0.219595 | 0.5219387 | 0.601713 | 0.908868121 | LRRFIP1P1 | processed_pseudogene | 47.15 | 0.004756434 | 0.004756434 | ENSG00000240429 | n.s. |
| ENSG00000100473 | 1100.44 | -0.129561327 | 0.269954 | -0.479938 | 0.631272 | 0.919033489 | COCH | protein_coding | 39.92 | -0.004750841 | 0.004750841 | ENSG00000100473 | n.s. |
| ENSG00000108946 | 4869.883 | -0.128033575 | 0.260647 | -0.491214 | 0.623275 | 0.918167932 | PRKAR1A | protein_coding | 41.97 | -0.004747213 | 0.004747213 | ENSG00000108946 | n.s. |
| ENSG00000134597 | 545.4843 | -0.139774321 | 0.316003 | -0.442319 | 0.658258 | 0.924786541 | RBMX2 | protein_coding | 43.77 | -0.004746526 | 0.004746526 | ENSG00000134597 | n.s. |
| ENSG00000229618 | 40.13079 | -0.14093598 | 0.322606 | -0.436867 | 0.662208 | 0.925519646 | | lincRNA | 35.74 | -0.004737472 | 0.004737472 | ENSG00000229618 | n.s. |
| ENSG00000173928 | 54.11921 | -0.136355761 | 0.299051 | -0.455962 | 0.648418 | 0.923184844 | SWSAP1 | protein_coding | 53.77 | -0.00473309 | 0.00473309 | ENSG00000173928 | n.s. |
| ENSG00000155561 | 5021.158 | -0.127920897 | 0.261277 | -0.489599 | 0.624418 | 0.918517547 | NUP205 | protein_coding | 38.99 | -0.004721886 | 0.004721886 | ENSG00000155561 | n.s. |
| ENSG00000181104 | 833.4424 | 0.136014373 | 0.29682 | 0.4582379 | 0.646782 | 0.923184844 | F2R | protein_coding | 42.79 | 0.00472124 | 0.00472124 | ENSG00000181104 | n.s. |
| ENSG00000161036 | 198.7495 | -0.131642061 | 0.280527 | -0.469267 | 0.638879 | 0.920750614 | LRWD1 | protein_coding | 61.06 | -0.004720419 | 0.004720419 | ENSG00000161036 | n.s. |
| ENSG00000168944 | 1934.118 | -0.125335583 | 0.253239 | -0.49493 | 0.62065 | 0.916981464 | CEP120 | protein_coding | 36.19 | -0.004717562 | 0.004717562 | ENSG00000168944 | n.s. |
| ENSG00000034510 | 15652.37 | -0.144777781 | 0.335903 | -0.431011 | 0.66646 | 0.927727183 | TMSB10 | protein_coding | 63.37 | -0.004716819 | 0.004716819 | ENSG00000034510 | n.s. |
| ENSG00000275709 | 160.7552 | 0.123127669 | 0.247234 | 0.4980213 | 0.618469 | 0.915570902 | | lincRNA | 40.69 | 0.004716777 | 0.004716777 | ENSG00000275709 | n.s. |
| ENSG00000252396 | 21.91928 | 0.156515836 | 0.381811 | 0.4099302 | 0.681857 | 0.933075339 | RN7SKP195 | misc_RNA | 53.47 | 0.004708511 | 0.004708511 | ENSG00000252396 | n.s. |
| ENSG00000164253 | 959.8359 | -0.129277851 | 0.27305 | -0.473459 | 0.635886 | 0.919565577 | WDR41 | protein_coding | 37.31 | -0.00470795 | 0.00470795 | ENSG00000164253 | n.s. |
| ENSG00000230898 | 96.27906 | -0.128330291 | 0.267815 | -0.479175 | 0.631814 | 0.919033489 | | sense_intronic | 46.04 | -0.0047057 | 0.0047057 | ENSG00000230898 | n.s. |

| | | | | | | | | | | | | | |
|---|---|---|---|---|---|---|---|---|---|---|---|---|---|
| ENSG00000188283 | 26.22775 | 0.151533316 | 0.361215 | 0.4195097 | 0.674844 | 0.931111765 | ZNF383 | protein_coding | 41.4 | 0.004697258 | 0.004697258 | ENSG00000188283 | n.s. |
| ENSG00000207279 | 11.84131 | -0.168288189 | 0.433711 | -0.388019 | 0.698002 | 0.937752951 | SNORD116-24 | snoRNA | 40.22 | -0.004697186 | 0.004697186 | ENSG00000207279 | n.s. |
| ENSG00000198042 | 1380.527 | -0.12033848 | 0.240087 | -0.501228 | 0.616211 | 0.914046563 | MAK16 | protein_coding | 40.72 | -0.004697013 | 0.004697013 | ENSG00000198042 | n.s. |
| ENSG00000083520 | 3423.192 | 0.128053814 | 0.266041 | 0.481331 | 0.630281 | 0.919033489 | DIS3 | protein_coding | 36.46 | 0.004695562 | 0.004695562 | ENSG00000083520 | n.s. |
| ENSG00000101343 | 1278.194 | -0.128026396 | 0.265189 | -0.482774 | 0.629256 | 0.919033489 | CRNKL1 | protein_coding | 39.08 | -0.004694557 | 0.004694557 | ENSG00000101343 | n.s. |
| ENSG00000154240 | 284.5734 | -0.12787418 | 0.262358 | -0.487402 | 0.625973 | 0.918973809 | CEP112 | protein_coding | 37.96 | -0.004692582 | 0.004692582 | ENSG00000154240 | n.s. |
| ENSG00000110851 | 1125.413 | -0.130081963 | 0.276367 | -0.470685 | 0.637866 | 0.920401057 | PRDM4 | protein_coding | 41.21 | -0.004685928 | 0.004685928 | ENSG00000110851 | n.s. |
| ENSG00000143315 | 390.3601 | -0.116451096 | 0.229975 | -0.506364 | 0.612601 | 0.911535838 | PIGM | protein_coding | 41.28 | -0.004684391 | 0.004684391 | ENSG00000143315 | n.s. |
| ENSG00000255423 | 276.7061 | -0.134793819 | 0.295757 | -0.455759 | 0.648563 | 0.923184844 | EBLN2 | protein_coding | 41.75 | -0.004678873 | 0.004678873 | ENSG00000255423 | n.s. |
| ENSG00000127399 | 177.2495 | -0.127394492 | 0.261117 | -0.487883 | 0.625632 | 0.918973809 | LRRC61 | protein_coding | 54.13 | -0.004674979 | 0.004674979 | ENSG00000127399 | n.s. |
| ENSG00000123395 | 432.2219 | -0.128303782 | 0.270484 | -0.474349 | 0.635251 | 0.919532207 | ATG101 | protein_coding | 51.45 | -0.004674499 | 0.004674499 | ENSG00000123395 | n.s. |
| ENSG00000151835 | 6293.615 | -0.130564341 | 0.279128 | -0.467757 | 0.639958 | 0.920914184 | SACS | protein_coding | 40.65 | -0.004671702 | 0.004671702 | ENSG00000151835 | n.s. |
| ENSG00000205629 | 285.9902 | -0.13673149 | 0.30533 | -0.447815 | 0.654287 | 0.92436641 | LCMT1 | protein_coding | 45.35 | -0.004670179 | 0.004670179 | ENSG00000205629 | n.s. |
| ENSG00000114993 | 214.8319 | -0.12734 | 0.263161 | -0.483886 | 0.628467 | 0.919033489 | RTKN | protein_coding | 53.87 | -0.004669388 | 0.004669388 | ENSG00000114993 | n.s. |
| ENSG00000151422 | 308.8081 | -0.128154667 | 0.270394 | -0.473955 | 0.635532 | 0.919532207 | FER | protein_coding | 35.73 | -0.004669066 | 0.004669066 | ENSG00000151422 | n.s. |
| ENSG00000065548 | 5145.259 | 0.127174409 | 0.262662 | 0.4841757 | 0.628261 | 0.919033489 | ZC3H15 | protein_coding | 37.22 | 0.004663316 | 0.004663316 | ENSG00000065548 | n.s. |
| ENSG00000175155 | 365.8247 | 0.11236447 | 0.216205 | 0.5197133 | 0.603263 | 0.908868121 | YPEL2 | protein_coding | 46.62 | 0.004663028 | 0.004663028 | ENSG00000175155 | n.s. |
| ENSG00000115464 | 4021.142 | -0.127072418 | 0.262456 | -0.484166 | 0.628268 | 0.919033489 | USP34 | protein_coding | 38.76 | -0.004659576 | 0.004659576 | ENSG00000115464 | n.s. |
| ENSG00000223508 | 105.6399 | -0.134025431 | 0.291558 | -0.459688 | 0.64574 | 0.923184844 | RPL23AP53 | ıscribed_processed_pseudog | 43.7 | -0.004652202 | 0.004652202 | ENSG00000223508 | n.s. |
| ENSG00000198373 | 42.07971 | -0.138382027 | 0.31663 | -0.437047 | 0.662077 | 0.925519646 | WWP2 | protein_coding | 45.6 | -0.004651623 | 0.004651623 | ENSG00000198373 | n.s. |
| ENSG00000226754 | 51.36102 | 0.142758059 | 0.331013 | 0.4312761 | 0.666268 | 0.927727183 | | antisense | 44.9 | 0.004651017 | 0.004651017 | ENSG00000226754 | n.s. |
| ENSG00000110315 | 633.6044 | -0.131714554 | 0.283463 | -0.464662 | 0.642173 | 0.922225673 | RNF141 | protein_coding | 36.1 | -0.004631451 | 0.004631451 | ENSG00000110315 | n.s. |
| ENSG00000207820 | 21.78405 | -0.157333296 | 0.390049 | -0.403368 | 0.686677 | 0.934474326 | MIR545 | miRNA | 41.51 | -0.004630732 | 0.004630732 | ENSG00000207820 | n.s. |
| ENSG00000108479 | 339.5055 | -0.142477757 | 0.332301 | -0.428761 | 0.668097 | 0.927917746 | GALK1 | protein_coding | 60.24 | -0.004629176 | 0.004629176 | ENSG00000108479 | n.s. |
| ENSG00000053254 | 1952.973 | -0.125605511 | 0.256753 | -0.489207 | 0.624695 | 0.918660999 | FOXN3 | protein_coding | 44.31 | -0.0046279 | 0.0046279 | ENSG00000053254 | n.s. |
| ENSG00000213753 | 337.3818 | -0.133130986 | 0.29349 | -0.453614 | 0.650107 | 0.923184844 | CENPBD1P1 | ıscribed_processed_pseudog | 45.66 | -0.004621154 | 0.004621154 | ENSG00000213753 | n.s. |
| ENSG00000196535 | 273.401 | -0.125846816 | 0.263277 | -0.478001 | 0.63265 | 0.919033489 | MYO18A | protein_coding | 53.74 | -0.004614635 | 0.004614635 | ENSG00000196535 | n.s. |
| ENSG00000132002 | 931.1193 | -0.123867108 | 0.25181 | -0.491907 | 0.622785 | 0.917861492 | DNAJB1 | protein_coding | 52.02 | -0.004610687 | 0.004610687 | ENSG00000132002 | n.s. |
| ENSG00000105443 | 477.939 | -0.126577009 | 0.267362 | -0.47343 | 0.635906 | 0.919565577 | CYTH2 | protein_coding | 58.29 | -0.004609592 | 0.004609592 | ENSG00000105443 | n.s. |
| ENSG00000279098 | 18.57479 | -0.156595876 | 0.389524 | -0.402018 | 0.68767 | 0.934506008 | | TEC | 27.38 | -0.004606722 | 0.004606722 | ENSG00000279098 | n.s. |
| ENSG00000223768 | 71.38366 | -0.135601269 | 0.307739 | -0.440638 | 0.659475 | 0.924786541 | LINC00205 | lincRNA | 51.62 | -0.004604816 | 0.004604816 | ENSG00000223768 | n.s. |
| ENSG00000251972 | 16.80562 | 0.159138653 | 0.401431 | 0.3964281 | 0.691789 | 0.935578594 | RNU6-123P | snRNA | 33.33 | 0.004602246 | 0.004602246 | ENSG00000251972 | n.s. |

| | | | | | | | | | | | | | |
|---|---|---|---|---|---|---|---|---|---|---|---|---|---|
| ENSG00000245937 | 95.98992 | -0.125407068 | 0.260189 | -0.481985 | 0.629816 | 0.919033489 | LINC01184 | lincRNA | 37.74 | -0.00459851 | 0.00459851 | ENSG00000245937 | n.s. |
| ENSG00000106305 | 865.6774 | 0.132429596 | 0.290699 | 0.4555552 | 0.64871 | 0.923184844 | AIMP2 | protein_coding | 46.93 | 0.004596808 | 0.004596808 | ENSG00000106305 | n.s. |
| ENSG00000114354 | 1695.407 | -0.125338444 | 0.262032 | -0.478333 | 0.632413 | 0.919033489 | TFG | protein_coding | 36.38 | -0.004595993 | 0.004595993 | ENSG00000114354 | n.s. |
| ENSG00000133597 | 164.1103 | -0.128428757 | 0.274649 | -0.46761 | 0.640063 | 0.920914184 | ADCK2 | protein_coding | 50.51 | -0.004595289 | 0.004595289 | ENSG00000133597 | n.s. |
| ENSG00000111817 | 1057.345 | -0.132381373 | 0.287745 | -0.460066 | 0.645469 | 0.923184844 | DSE | protein_coding | 38.2 | -0.004595134 | 0.004595134 | ENSG00000111817 | n.s. |
| ENSG00000117114 | 5150.317 | -0.126387577 | 0.267156 | -0.473085 | 0.636152 | 0.919741018 | ADGRL2 | protein_coding | 36.78 | -0.004592222 | 0.004592222 | ENSG00000117114 | n.s. |
| ENSG00000201512 | 100.2728 | -0.12499726 | 0.260869 | -0.479158 | 0.631826 | 0.919033489 | SNORA71C | snoRNA | 57.46 | -0.004583482 | 0.004583482 | ENSG00000201512 | n.s. |
| ENSG00000130520 | 2065.048 | -0.140916453 | 0.327577 | -0.430178 | 0.667066 | 0.927899924 | LSM4 | protein_coding | 56.37 | -0.004579624 | 0.004579624 | ENSG00000130520 | n.s. |
| ENSG00000161980 | 360.6342 | -0.131913142 | 0.288667 | -0.456973 | 0.64769 | 0.923184844 | POLR3K | protein_coding | 48.85 | -0.004578881 | 0.004578881 | ENSG00000161980 | n.s. |
| ENSG00000128000 | 130.9648 | -0.124813057 | 0.260715 | -0.478733 | 0.632129 | 0.919033489 | ZNF780B | protein_coding | 39.65 | -0.004576728 | 0.004576728 | ENSG00000128000 | n.s. |
| ENSG00000176055 | 324.8299 | -0.126991945 | 0.269572 | -0.471088 | 0.637578 | 0.920373167 | MBLAC2 | protein_coding | 36.86 | -0.004576288 | 0.004576288 | ENSG00000176055 | n.s. |
| ENSG00000202273 | 10.69732 | -0.179143689 | 0.514631 | -0.348102 | 0.727764 | 0.942883666 | | misc_RNA | 40.78 | -0.004575668 | 0.004575668 | ENSG00000202273 | n.s. |
| ENSG00000198612 | 2246.438 | 0.12469811 | 0.259377 | 0.4807594 | 0.630687 | 0.919033489 | COPS8 | protein_coding | 39.13 | 0.004572513 | 0.004572513 | ENSG00000198612 | n.s. |
| ENSG00000164332 | 724.0595 | -0.131684102 | 0.28817 | -0.456967 | 0.647695 | 0.923184844 | UBLCP1 | protein_coding | 36.18 | -0.004570931 | 0.004570931 | ENSG00000164332 | n.s. |
| ENSG00000182919 | 161.2994 | 0.138936307 | 0.320184 | 0.433927 | 0.664341 | 0.927094972 | C11orf54 | protein_coding | 41.12 | 0.004567639 | 0.004567639 | ENSG00000182919 | n.s. |
| ENSG00000196776 | 2289.846 | 0.131493556 | 0.286727 | 0.458602 | 0.64652 | 0.923184844 | CD47 | protein_coding | 36.88 | 0.004564317 | 0.004564317 | ENSG00000196776 | n.s. |
| ENSG00000006634 | 2032.594 | -0.124447634 | 0.258071 | -0.482223 | 0.629648 | 0.919033489 | DBF4 | protein_coding | 36.72 | -0.004563328 | 0.004563328 | ENSG00000006634 | n.s. |
| ENSG00000138028 | 241.4265 | -0.134352539 | 0.306101 | -0.438916 | 0.660722 | 0.924786541 | CGREF1 | protein_coding | 50.24 | -0.004562411 | 0.004562411 | ENSG00000138028 | n.s. |
| ENSG00000228485 | 72.42812 | 0.131432465 | 0.288332 | 0.4558375 | 0.648507 | 0.923184844 | GRK5-IT1 | sense_intronic | 39.16 | 0.004562196 | 0.004562196 | ENSG00000228485 | n.s. |
| ENSG00000178700 | 476.2914 | 0.109868841 | 0.21118 | 0.5202605 | 0.602882 | 0.908868121 | DHFR2 | protein_coding | 35.09 | 0.004559461 | 0.004559461 | ENSG00000178700 | n.s. |
| ENSG00000166913 | 2918.402 | -0.124316153 | 0.259634 | -0.478813 | 0.632072 | 0.919033489 | YWHAB | protein_coding | 40.42 | -0.004558507 | 0.004558507 | ENSG00000166913 | n.s. |
| ENSG00000108984 | 396.5264 | 0.131072732 | 0.287256 | 0.4562916 | 0.64818 | 0.923184844 | MAP2K6 | protein_coding | 41 | 0.004549709 | 0.004549709 | ENSG00000108984 | n.s. |
| ENSG00000159200 | 262.8302 | -0.133666166 | 0.299277 | -0.44663 | 0.655142 | 0.924621046 | RCAN1 | protein_coding | 44.24 | -0.004549492 | 0.004549492 | ENSG00000159200 | n.s. |
| ENSG00000109606 | 9621.992 | 0.127215565 | 0.272144 | 0.4674567 | 0.640173 | 0.920968304 | DHX15 | protein_coding | 37.26 | 0.004548633 | 0.004548633 | ENSG00000109606 | n.s. |
| ENSG00000221916 | 27.65209 | -0.153247332 | 0.37758 | -0.405867 | 0.68484 | 0.934117877 | C19orf73 | protein_coding | 63.04 | -0.004535863 | 0.004535863 | ENSG00000221916 | n.s. |
| ENSG00000184216 | 1096.008 | -0.123685787 | 0.258659 | -0.47818 | 0.632522 | 0.919033489 | IRAK1 | protein_coding | 61.78 | -0.004535392 | 0.004535392 | ENSG00000184216 | n.s. |
| ENSG00000163155 | 196.5233 | -0.123585457 | 0.253335 | -0.487833 | 0.625668 | 0.918973809 | LYSMD1 | protein_coding | 46.07 | -0.004535199 | 0.004535199 | ENSG00000163155 | n.s. |
| ENSG00000129596 | 151.6889 | 0.123657741 | 0.256873 | 0.481397 | 0.630234 | 0.919033489 | CDO1 | protein_coding | 40.48 | 0.004534364 | 0.004534364 | ENSG00000129596 | n.s. |
| ENSG00000177565 | 5917.413 | -0.124426989 | 0.262085 | -0.474758 | 0.634959 | 0.919532207 | TBL1XR1 | protein_coding | 39.31 | -0.004533255 | 0.004533255 | ENSG00000177565 | n.s. |
| ENSG00000228300 | 107.4686 | -0.131660067 | 0.2925 | -0.450119 | 0.652625 | 0.923800272 | C19orf24 | protein_coding | 63.69 | -0.004531992 | 0.004531992 | ENSG00000228300 | n.s. |
| ENSG00000122692 | 1689.724 | -0.123481683 | 0.256678 | -0.481076 | 0.630463 | 0.919033489 | SMU1 | protein_coding | 41.17 | -0.004527908 | 0.004527908 | ENSG00000122692 | n.s. |
| ENSG00000163644 | 550.9143 | -0.130358436 | 0.283933 | -0.459118 | 0.64615 | 0.923184844 | PPM1K | protein_coding | 39.9 | -0.004524915 | 0.004524915 | ENSG00000163644 | n.s. |

| | | | | | | | | | | | | | |
|---|---|---|---|---|---|---|---|---|---|---|---|---|---|
| ENSG00000178922 | 75.89205 | -0.132263573 | 0.295097 | -0.448204 | 0.654006 | 0.924250964 | HYI | protein_coding | 59.36 | -0.004524748 | 0.004524748 | ENSG00000178922 | n.s. |
| ENSG00000253771 | 33.78226 | -0.144021169 | 0.342102 | -0.420989 | 0.673763 | 0.930265705 | TPTE2P1 | cribed_unprocessed_pseudc | 41.61 | -0.004521255 | 0.004521255 | ENSG00000253771 | n.s. |
| ENSG00000126522 | 137.3232 | -0.131507302 | 0.292659 | -0.449354 | 0.653177 | 0.923905149 | ASL | protein_coding | 54.6 | -0.004520249 | 0.004520249 | ENSG00000126522 | n.s. |
| ENSG00000239672 | 5616.514 | 0.132779389 | 0.297288 | 0.4466358 | 0.655138 | 0.924621046 | NME1 | protein_coding | 46.45 | 0.004519309 | 0.004519309 | ENSG00000239672 | n.s. |
| ENSG00000163931 | 2784.241 | -0.126015078 | 0.26871 | -0.468963 | 0.639096 | 0.920750614 | TKT | protein_coding | 54.36 | -0.004518646 | 0.004518646 | ENSG00000163931 | n.s. |
| ENSG00000265321 | 7.613229 | 0.173939711 | 0.488352 | 0.3561767 | 0.721708 | 0.941942395 | MIR4263 | miRNA | 45.78 | 0.004518198 | 0.004518198 | ENSG00000265321 | n.s. |
| ENSG00000272617 | 434.7508 | -0.123215946 | 0.256776 | -0.479857 | 0.631329 | 0.919033489 | COG8 | protein_coding | 51.52 | -0.004518164 | 0.004518164 | ENSG00000272617 | n.s. |
| ENSG00000185591 | 2105.306 | -0.122970748 | 0.251705 | -0.488551 | 0.625159 | 0.918928785 | SP1 | protein_coding | 45.65 | -0.004515258 | 0.004515258 | ENSG00000185591 | n.s. |
| ENSG00000207508 | 6.958669 | 0.183361273 | 0.537834 | 0.3409252 | 0.73316 | 0.944876691 | RNU6-1237P | snRNA | 38.32 | 0.004515247 | 0.004515247 | ENSG00000207508 | n.s. |
| ENSG00000196236 | 508.5031 | -0.120937373 | 0.245259 | -0.493101 | 0.621941 | 0.917630258 | XPNPEP3 | protein_coding | 41.48 | -0.004514867 | 0.004514867 | ENSG00000196236 | n.s. |
| ENSG00000164107 | 9541.918 | -0.108719294 | 0.210012 | -0.517682 | 0.60468 | 0.908868121 | HAND2 | protein_coding | 55.84 | -0.004511756 | 0.004511756 | ENSG00000164107 | n.s. |
| ENSG00000111775 | 3019.37 | -0.13976141 | 0.327404 | -0.426877 | 0.669469 | 0.928471963 | COX6A1 | protein_coding | 48.25 | -0.004504679 | 0.004504679 | ENSG00000111775 | n.s. |
| ENSG00000115966 | 1991.199 | -0.122816104 | 0.257104 | -0.47769 | 0.632871 | 0.919033489 | ATF2 | protein_coding | 36.05 | -0.004503502 | 0.004503502 | ENSG00000115966 | n.s. |
| ENSG00000139970 | 1072.019 | -0.132540953 | 0.300829 | -0.440585 | 0.659513 | 0.924786541 | RTN1 | protein_coding | 39.2 | -0.004500892 | 0.004500892 | ENSG00000139970 | n.s. |
| ENSG00000092841 | 4441.355 | -0.132477573 | 0.300084 | -0.441469 | 0.658874 | 0.924786541 | MYL6 | protein_coding | 51.56 | -0.00449874 | 0.00449874 | ENSG00000092841 | n.s. |
| ENSG00000264357 | 4.280913 | -0.226389965 | 0.768285 | -0.294669 | 0.768247 | 0.955275572 | MIR4648 | miRNA | 63.89 | -0.004498669 | 0.004498669 | ENSG00000264357 | n.s. |
| ENSG00000202314 | 58.2585 | -0.12885252 | 0.278022 | -0.463462 | 0.643034 | 0.922767027 | SNORD6 | snoRNA | 45.21 | -0.004497975 | 0.004497975 | ENSG00000202314 | n.s. |
| ENSG00000249502 | 18.74085 | -0.16089803 | 0.418449 | -0.384511 | 0.7006 | 0.937752951 | | antisense | 43.81 | -0.004490915 | 0.004490915 | ENSG00000249502 | n.s. |
| ENSG00000078018 | 3974.381 | -0.122581785 | 0.257478 | -0.476086 | 0.634013 | 0.919134382 | MAP2 | protein_coding | 35.9 | -0.004489066 | 0.004489066 | ENSG00000078018 | n.s. |
| ENSG00000104517 | 4472.164 | -0.122413957 | 0.252354 | -0.485089 | 0.627613 | 0.919033489 | UBR5 | protein_coding | 38.38 | -0.004488756 | 0.004488756 | ENSG00000104517 | n.s. |
| ENSG00000165675 | 562.2141 | -0.132162167 | 0.301139 | -0.438874 | 0.660753 | 0.924786541 | ENOX2 | protein_coding | 38.87 | -0.004488029 | 0.004488029 | ENSG00000165675 | n.s. |
| ENSG00000049245 | 353.7913 | 0.122316063 | 0.253548 | 0.4824183 | 0.629509 | 0.919033489 | VAMP3 | protein_coding | 42.77 | 0.004485167 | 0.004485167 | ENSG00000049245 | n.s. |
| ENSG00000083828 | 522.5491 | -0.122309573 | 0.254761 | -0.480096 | 0.631159 | 0.919033489 | ZNF586 | protein_coding | 46.13 | -0.004484929 | 0.004484929 | ENSG00000083828 | n.s. |
| ENSG00000159840 | 263.0726 | -0.114774825 | 0.228691 | -0.501877 | 0.615754 | 0.91396346 | ZYX | protein_coding | 58.32 | -0.004484386 | 0.004484386 | ENSG00000159840 | n.s. |
| ENSG00000204442 | 475.6152 | -0.120387822 | 0.24468 | -0.492022 | 0.622704 | 0.917861492 | FAM155A | protein_coding | 37.4 | -0.004481178 | 0.004481178 | ENSG00000204442 | n.s. |
| ENSG00000140262 | 6211.225 | 0.122059131 | 0.252284 | 0.4838156 | 0.628517 | 0.919033489 | TCF12 | protein_coding | 37.17 | 0.004475745 | 0.004475745 | ENSG00000140262 | n.s. |
| ENSG00000135336 | 1366.934 | -0.131744296 | 0.298286 | -0.441671 | 0.658727 | 0.924786541 | ORC3 | protein_coding | 38.95 | -0.004473839 | 0.004473839 | ENSG00000135336 | n.s. |
| ENSG00000201240 | 6.207298 | 0.178987823 | 0.517785 | 0.3456796 | 0.729584 | 0.944145838 | SNORD114-9 | snoRNA | 47.22 | 0.0044677 | 0.0044677 | ENSG00000201240 | n.s. |
| ENSG00000115540 | 1165.281 | -0.131519754 | 0.299329 | -0.439382 | 0.660385 | 0.924786541 | MOB4 | protein_coding | 40.5 | -0.004466213 | 0.004466213 | ENSG00000115540 | n.s. |
| ENSG00000178193 | 47.85753 | -0.146555314 | 0.353593 | -0.414475 | 0.678526 | 0.932323161 | | antisense | 39.54 | -0.004460195 | 0.004460195 | ENSG00000178193 | n.s. |
| ENSG00000248593 | 38.20943 | -0.13900047 | 0.327443 | -0.424503 | 0.671199 | 0.92884421 | DSTNP2 | cribed_unprocessed_pseudc | 46.24 | -0.004455955 | 0.004455955 | ENSG00000248593 | n.s. |
| ENSG00000080189 | 320.9022 | -0.13119906 | 0.29495 | -0.444819 | 0.656451 | 0.924786541 | SLC35C2 | protein_coding | 53.97 | -0.004455323 | 0.004455323 | ENSG00000080189 | n.s. |

| | | | | | | | | | | | | | |
|---|---|---|---|---|---|---|---|---|---|---|---|---|---|
| ENSG00000137714 | 258.1315 | -0.130941875 | 0.29577 | -0.442715 | 0.657972 | 0.924786541 | FDX1 | protein_coding | 39.82 | -0.00444659 | 0.00444659 | ENSG00000137714 | n.s. |
| ENSG00000169019 | 287.2573 | -0.136451294 | 0.31634 | -0.431344 | 0.666218 | 0.927727183 | COMMD8 | protein_coding | 36.41 | -0.004445545 | 0.004445545 | ENSG00000169019 | n.s. |
| ENSG00000176268 | 12.28787 | -0.163365428 | 0.439345 | -0.371839 | 0.710013 | 0.939338716 | CYCSP34 | processed_pseudogene | 46.71 | -0.004439909 | 0.004439909 | ENSG00000176268 | n.s. |
| ENSG00000116991 | 1328.123 | -0.117713992 | 0.237516 | -0.495605 | 0.620173 | 0.916893708 | SIPA1L2 | protein_coding | 41.11 | -0.004435582 | 0.004435582 | ENSG00000116991 | n.s. |
| ENSG00000233927 | 2113.175 | -0.131929486 | 0.302124 | -0.436673 | 0.662348 | 0.925519646 | RPS28 | protein_coding | 57.12 | -0.004434725 | 0.004434725 | ENSG00000233927 | n.s. |
| ENSG00000164091 | 3268.508 | -0.123869679 | 0.264712 | -0.467941 | 0.639827 | 0.920914184 | WDR82 | protein_coding | 44.88 | -0.004432161 | 0.004432161 | ENSG00000164091 | n.s. |
| ENSG00000060762 | 882.054 | -0.138170486 | 0.32567 | -0.424265 | 0.671373 | 0.928900122 | MPC1 | protein_coding | 38.93 | -0.004425736 | 0.004425736 | ENSG00000060762 | n.s. |
| ENSG00000223705 | 175.415 | -0.120690851 | 0.252516 | -0.477953 | 0.632684 | 0.919033489 | NSUN5P1 | cribed_unprocessed_pseudc | 53.54 | -0.004425572 | 0.004425572 | ENSG00000223705 | n.s. |
| ENSG00000146007 | 1440.597 | -0.126706016 | 0.273038 | -0.46406 | 0.642605 | 0.922750294 | ZMAT2 | protein_coding | 43.7 | -0.004424043 | 0.004424043 | ENSG00000146007 | n.s. |
| ENSG00000158882 | 325.7851 | -0.127042874 | 0.2747 | -0.462479 | 0.643738 | 0.922949131 | TOMM40L | protein_coding | 52.77 | -0.004423917 | 0.004423917 | ENSG00000158882 | n.s. |
| ENSG00000205981 | 940.2994 | 0.12647892 | 0.273056 | 0.4631972 | 0.643223 | 0.922767027 | DNAJC19 | protein_coding | 39.17 | 0.004415118 | 0.004415118 | ENSG00000205981 | n.s. |
| ENSG00000102796 | 77.62827 | 0.127165897 | 0.278269 | 0.4569895 | 0.647679 | 0.923184844 | DHRS12 | protein_coding | 47.24 | 0.004414098 | 0.004414098 | ENSG00000102796 | n.s. |
| ENSG00000114796 | 1380.663 | 0.129929004 | 0.295405 | 0.4398341 | 0.660057 | 0.924786541 | KLHL24 | protein_coding | 39.02 | 0.004412194 | 0.004412194 | ENSG00000114796 | n.s. |
| ENSG00000205323 | 1705.725 | -0.12494713 | 0.268555 | -0.465257 | 0.641748 | 0.921994846 | SARNP | protein_coding | 42.48 | -0.004407074 | 0.004407074 | ENSG00000205323 | n.s. |
| ENSG00000199471 | 9.733284 | -0.163381122 | 0.443499 | -0.368391 | 0.712582 | 0.939817855 | | misc_RNA | 43.56 | -0.004404152 | 0.004404152 | ENSG00000199471 | n.s. |
| ENSG00000103642 | 209.5522 | 0.126851762 | 0.280405 | 0.4523879 | 0.65099 | 0.923184844 | LACTB | protein_coding | 40 | 0.004403194 | 0.004403194 | ENSG00000103642 | n.s. |
| ENSG00000002834 | 230.732 | -0.119969024 | 0.249856 | -0.480153 | 0.631119 | 0.919033489 | LASP1 | protein_coding | 53.69 | -0.004399104 | 0.004399104 | ENSG00000002834 | n.s. |
| ENSG00000196458 | 1409.71 | -0.126727236 | 0.277513 | -0.456653 | 0.647921 | 0.923184844 | ZNF605 | protein_coding | 46.54 | -0.004398871 | 0.004398871 | ENSG00000196458 | n.s. |
| ENSG00000273008 | 118.8044 | 0.119881549 | 0.249359 | 0.4807597 | 0.630687 | 0.919033489 | | lincRNA | 54.4 | 0.004395896 | 0.004395896 | ENSG00000273008 | n.s. |
| ENSG00000119912 | 966.6671 | -0.12662608 | 0.278833 | -0.454129 | 0.649736 | 0.923184844 | IDE | protein_coding | 40.71 | -0.00439536 | 0.00439536 | ENSG00000119912 | n.s. |
| ENSG00000131469 | 23358.69 | 0.137414297 | 0.324117 | 0.4239656 | 0.671591 | 0.929017422 | RPL27 | protein_coding | 49.84 | 0.004393979 | 0.004393979 | ENSG00000131469 | n.s. |
| ENSG00000232732 | 49.50404 | -0.133149053 | 0.306363 | -0.434612 | 0.663844 | 0.926901501 | | processed_transcript | 38.64 | -0.004389447 | 0.004389447 | ENSG00000232732 | n.s. |
| ENSG00000100104 | 405.2737 | -0.117568693 | 0.238441 | -0.493073 | 0.621961 | 0.917630258 | SRRD | protein_coding | 45.15 | -0.004389107 | 0.004389107 | ENSG00000100104 | n.s. |
| ENSG00000134077 | 1033.278 | -0.124591778 | 0.267942 | -0.464995 | 0.641935 | 0.922169321 | THUMPD3 | protein_coding | 40.09 | -0.004384301 | 0.004384301 | ENSG00000134077 | n.s. |
| ENSG00000136943 | 244.6944 | 0.121790536 | 0.258957 | 0.4703111 | 0.638133 | 0.920457187 | CTSV | protein_coding | 44.69 | 0.004384022 | 0.004384022 | ENSG00000136943 | n.s. |
| ENSG00000250988 | 79.49832 | -0.126285357 | 0.273909 | -0.461049 | 0.644764 | 0.923184844 | SNHG21 | antisense | 40.87 | -0.004383533 | 0.004383533 | ENSG00000250988 | n.s. |
| ENSG00000183775 | 1134.229 | 0.119530261 | 0.248758 | 0.4805084 | 0.630866 | 0.919033489 | KCTD16 | protein_coding | 37.85 | 0.004383015 | 0.004383015 | ENSG00000183775 | n.s. |
| ENSG00000118690 | 63.65916 | -0.137240787 | 0.324028 | -0.423546 | 0.671897 | 0.929142491 | ARMC2 | protein_coding | 41.62 | -0.004380407 | 0.004380407 | ENSG00000118690 | n.s. |
| ENSG00000251682 | 211.1617 | 0.126076531 | 0.27446 | 0.4593619 | 0.645974 | 0.923184844 | | processed_pseudogene | 44.33 | 0.004376285 | 0.004376285 | ENSG00000251682 | n.s. |
| ENSG00000120860 | 247.3566 | 0.132492094 | 0.304659 | 0.434886 | 0.663645 | 0.926866481 | WASHC3 | protein_coding | 36.36 | 0.004369963 | 0.004369963 | ENSG00000120860 | n.s. |
| ENSG00000113658 | 2910.191 | -0.120437892 | 0.254907 | -0.472478 | 0.636586 | 0.919866014 | SMAD5 | protein_coding | 37.17 | -0.004368936 | 0.004368936 | ENSG00000113658 | n.s. |
| ENSG00000224616 | 57.51546 | -0.128840314 | 0.293959 | -0.438294 | 0.661173 | 0.924898514 | RTCA-AS1 | antisense | 42.8 | -0.004368449 | 0.004368449 | ENSG00000224616 | n.s. |

| | | | | | | | | | | | | | |
|---|---|---|---|---|---|---|---|---|---|---|---|---|---|
| ENSG00000067715 | 9115.342 | 0.120518359 | 0.255252 | 0.4721551 | 0.636816 | 0.919940854 | SYT1 | protein_coding | 35.61 | 0.004367597 | 0.004367597 | ENSG00000067715 | n.s. |
| ENSG00000186577 | 158.2213 | 0.125745136 | 0.277724 | 0.4527701 | 0.650714 | 0.923184844 | SMIM29 | protein_coding | 61.63 | 0.004364781 | 0.004364781 | ENSG00000186577 | n.s. |
| ENSG00000107960 | 263.4898 | -0.124840817 | 0.269305 | -0.463566 | 0.642958 | 0.922767027 | STN1 | protein_coding | 41.91 | -0.004357935 | 0.004357935 | ENSG00000107960 | n.s. |
| ENSG00000163374 | 1066.567 | -0.118828214 | 0.244435 | -0.486134 | 0.626872 | 0.919033489 | YY1AP1 | protein_coding | 44.63 | -0.004357272 | 0.004357272 | ENSG00000163374 | n.s. |
| ENSG00000266338 | 334.2516 | 0.128228962 | 0.28914 | 0.4434836 | 0.657416 | 0.924786541 | NBPF15 | protein_coding | 42.75 | 0.004354463 | 0.004354463 | ENSG00000266338 | n.s. |
| ENSG00000136159 | 1294.794 | -0.127918949 | 0.286333 | -0.446748 | 0.655057 | 0.924621046 | NUDT15 | protein_coding | 41.59 | -0.004353878 | 0.004353878 | ENSG00000136159 | n.s. |
| ENSG00000235034 | 26.16679 | 0.15077307 | 0.38076 | 0.3959793 | 0.69212 | 0.935753589 | C19orf81 | protein_coding | 50.87 | 0.004348069 | 0.004348069 | ENSG00000235034 | n.s. |
| ENSG00000109919 | 1674.529 | -0.132859376 | 0.306678 | -0.433221 | 0.664855 | 0.927447429 | MTCH2 | protein_coding | 43.47 | -0.004345923 | 0.004345923 | ENSG00000109919 | n.s. |
| ENSG00000169045 | 25957.54 | -0.1111976 | 0.221555 | -0.501897 | 0.61574 | 0.91396346 | HNRNPH1 | protein_coding | 45.97 | -0.00434462 | 0.00434462 | ENSG00000169045 | n.s. |
| ENSG00000275765 | 107.4924 | -0.125002967 | 0.272658 | -0.458461 | 0.646621 | 0.923184844 | | lincRNA | 61.96 | -0.00433902 | 0.00433902 | ENSG00000275765 | n.s. |
| ENSG00000165973 | 421.2499 | 0.127739247 | 0.288481 | 0.4428003 | 0.65791 | 0.924786541 | NELL1 | protein_coding | 38.29 | 0.004337833 | 0.004337833 | ENSG00000165973 | n.s. |
| ENSG00000099804 | 437.4177 | -0.127914914 | 0.291707 | -0.438505 | 0.66102 | 0.924898514 | CDC34 | protein_coding | 64.75 | -0.004337073 | 0.004337073 | ENSG00000099804 | n.s. |
| ENSG00000146281 | 1160.035 | 0.124861871 | 0.27433 | 0.4551522 | 0.649 | 0.923184844 | PM20D2 | protein_coding | 39.61 | 0.004334122 | 0.004334122 | ENSG00000146281 | n.s. |
| ENSG00000178802 | 83.22639 | -0.124854856 | 0.273624 | -0.4563 | 0.648174 | 0.923184844 | MPI | protein_coding | 51.86 | -0.004333879 | 0.004333879 | ENSG00000178802 | n.s. |
| ENSG00000102103 | 572.4643 | -0.127443247 | 0.287181 | -0.443774 | 0.657206 | 0.924786541 | PQBP1 | protein_coding | 52.66 | -0.004327781 | 0.004327781 | ENSG00000102103 | n.s. |
| ENSG00000163521 | 17.22662 | -0.166420152 | 0.471192 | -0.353189 | 0.723946 | 0.942009991 | GLB1L | protein_coding | 49.58 | -0.004317686 | 0.004317686 | ENSG00000163521 | n.s. |
| ENSG00000155868 | 396.519 | -0.117720045 | 0.243745 | -0.482964 | 0.629121 | 0.919033489 | MED7 | protein_coding | 41.05 | -0.004316637 | 0.004316637 | ENSG00000155868 | n.s. |
| ENSG00000160867 | 96.92069 | -0.124324501 | 0.269962 | -0.460526 | 0.645139 | 0.923184844 | FGFR4 | protein_coding | 60.18 | -0.004315469 | 0.004315469 | ENSG00000160867 | n.s. |
| ENSG00000020922 | 1010.133 | -0.122157124 | 0.262468 | -0.465417 | 0.641633 | 0.921925013 | MRE11 | protein_coding | 37.31 | -0.004312684 | 0.004312684 | ENSG00000020922 | n.s. |
| ENSG00000197448 | 275.0819 | -0.123378869 | 0.266011 | -0.463811 | 0.642783 | 0.922767027 | GSTK1 | protein_coding | 41.49 | -0.004306901 | 0.004306901 | ENSG00000197448 | n.s. |
| ENSG00000116903 | 989.2772 | 0.117340602 | 0.242936 | 0.4830111 | 0.629088 | 0.919033489 | EXOC8 | protein_coding | 43.25 | 0.004302723 | 0.004302723 | ENSG00000116903 | n.s. |
| ENSG00000133983 | 753.8498 | -0.126644373 | 0.286618 | -0.441858 | 0.658592 | 0.924786541 | COX16 | protein_coding | 39.38 | -0.004300653 | 0.004300653 | ENSG00000133983 | n.s. |
| ENSG00000111652 | 491.1103 | -0.123781015 | 0.270765 | -0.457153 | 0.647561 | 0.923184844 | COPS7A | protein_coding | 50.55 | -0.004296604 | 0.004296604 | ENSG00000111652 | n.s. |
| ENSG00000067167 | 970.6023 | -0.123745043 | 0.272659 | -0.453846 | 0.64994 | 0.923184844 | TRAM1 | protein_coding | 38.27 | -0.004295356 | 0.004295356 | ENSG00000067167 | n.s. |
| ENSG00000171649 | 1062.121 | -0.121544251 | 0.260911 | -0.465845 | 0.641326 | 0.921891013 | ZIK1 | protein_coding | 47.45 | -0.004292994 | 0.004292994 | ENSG00000171649 | n.s. |
| ENSG00000251733 | 19.19703 | -0.147484281 | 0.370971 | -0.397563 | 0.690953 | 0.935225433 | SCARNA8 | scaRNA | 47.33 | -0.004289387 | 0.004289387 | ENSG00000251733 | n.s. |
| ENSG00000069943 | 120.5533 | -0.119576288 | 0.255317 | -0.468344 | 0.639538 | 0.920848926 | PIGB | protein_coding | 39.99 | -0.00428222 | 0.00428222 | ENSG00000069943 | n.s. |
| ENSG00000143797 | 1300.111 | -0.123311805 | 0.269206 | -0.458058 | 0.646911 | 0.923184844 | MBOAT2 | protein_coding | 40.17 | -0.004280317 | 0.004280317 | ENSG00000143797 | n.s. |
| ENSG00000111231 | 649.3923 | -0.12601049 | 0.286669 | -0.439568 | 0.66025 | 0.924786541 | GPN3 | protein_coding | 45.03 | -0.004279127 | 0.004279127 | ENSG00000111231 | n.s. |
| ENSG00000180385 | 578.4584 | -0.116600237 | 0.238898 | -0.488074 | 0.625497 | 0.918973809 | EMC3-AS1 | cribed_unprocessed_pseudc | 43.98 | -0.004278863 | 0.004278863 | ENSG00000180385 | n.s. |
| ENSG00000261114 | 43.44784 | 0.131694926 | 0.307185 | 0.4287157 | 0.66813 | 0.927917746 | | sense_intronic | 38.63 | 0.004278837 | 0.004278837 | ENSG00000261114 | n.s. |
| ENSG00000267213 | 134.1179 | -0.116499794 | 0.244691 | -0.476109 | 0.633997 | 0.919134382 | | antisense | 44.48 | -0.004266338 | 0.004266338 | ENSG00000267213 | n.s. |

| | | | | | | | | | | | | | |
|---|---|---|---|---|---|---|---|---|---|---|---|---|---|
| ENSG00000112992 | 1747.367 | -0.123837773 | 0.274604 | -0.450969 | 0.652012 | 0.923748122 | NNT | protein_coding | 37.88 | -0.004265769 | 0.004265769 | ENSG00000112992 | n.s. |
| ENSG00000163798 | 1285.451 | -0.122689996 | 0.268287 | -0.457309 | 0.647449 | 0.923184844 | SLC4A1AP | protein_coding | 40.08 | -0.004258733 | 0.004258733 | ENSG00000163798 | n.s. |
| ENSG00000012660 | 3446.494 | -0.123623354 | 0.274279 | -0.450722 | 0.65219 | 0.923748122 | ELOVL5 | protein_coding | 41.88 | -0.004258383 | 0.004258383 | ENSG00000012660 | n.s. |
| ENSG00000125821 | 870.8817 | -0.129733777 | 0.299171 | -0.433644 | 0.664547 | 0.927238553 | DTD1 | protein_coding | 43.21 | -0.004256373 | 0.004256373 | ENSG00000125821 | n.s. |
| ENSG00000151552 | 290.5771 | -0.122605566 | 0.27066 | -0.452988 | 0.650558 | 0.923184844 | QDPR | protein_coding | 45.09 | -0.004255803 | 0.004255803 | ENSG00000151552 | n.s. |
| ENSG00000186132 | 144.5715 | -0.130574586 | 0.302734 | -0.431318 | 0.666237 | 0.927727183 | C2orf76 | protein_coding | 39.21 | -0.004254083 | 0.004254083 | ENSG00000186132 | n.s. |
| ENSG00000201501 | 3.668107 | -0.212950377 | 0.717263 | -0.296893 | 0.766548 | 0.955053865 | | misc_RNA | 42.16 | -0.004253073 | 0.004253073 | ENSG00000201501 | n.s. |
| ENSG00000164104 | 12201.27 | -0.123732014 | 0.275457 | -0.449187 | 0.653296 | 0.923905149 | HMGB2 | protein_coding | 50.22 | -0.004252993 | 0.004252993 | ENSG00000164104 | n.s. |
| ENSG00000115274 | 160.1074 | -0.123178808 | 0.272796 | -0.451542 | 0.651599 | 0.923661977 | INO80B | protein_coding | 53.1 | -0.004248059 | 0.004248059 | ENSG00000115274 | n.s. |
| ENSG00000113719 | 770.3807 | -0.125080763 | 0.281789 | -0.443881 | 0.657129 | 0.924786541 | ERGIC1 | protein_coding | 50.91 | -0.004247555 | 0.004247555 | ENSG00000113719 | n.s. |
| ENSG00000008952 | 3208.582 | -0.122264215 | 0.270058 | -0.452733 | 0.650741 | 0.923184844 | SEC62 | protein_coding | 36.6 | -0.004243954 | 0.004243954 | ENSG00000008952 | n.s. |
| ENSG00000175175 | 823.412 | 0.12872969 | 0.296219 | 0.4345763 | 0.66387 | 0.926901501 | PPM1E | protein_coding | 39.61 | 0.004243756 | 0.004243756 | ENSG00000175175 | n.s. |
| ENSG00000235560 | 185.8904 | 0.112549893 | 0.226941 | 0.4959437 | 0.619934 | 0.916862388 | | antisense | 49.9 | 0.004242663 | 0.004242663 | ENSG00000235560 | n.s. |
| ENSG00000204237 | 470.5465 | -0.115684017 | 0.23994 | -0.482137 | 0.629708 | 0.919033489 | OXLD1 | protein_coding | 62.12 | -0.004241978 | 0.004241978 | ENSG00000204237 | n.s. |
| ENSG00000106100 | 48.24415 | 0.131985583 | 0.309936 | 0.4258485 | 0.670218 | 0.928687131 | NOD1 | protein_coding | 46.77 | 0.004240772 | 0.004240772 | ENSG00000106100 | n.s. |
| ENSG00000135148 | 576.5424 | -0.130145165 | 0.30107 | -0.432275 | 0.665541 | 0.927727183 | TRAFD1 | protein_coding | 44.88 | -0.004240093 | 0.004240093 | ENSG00000135148 | n.s. |
| ENSG00000196323 | 1033.277 | -0.122147028 | 0.266104 | -0.45902 | 0.64622 | 0.923184844 | ZBTB44 | protein_coding | 39.13 | -0.004239886 | 0.004239886 | ENSG00000196323 | n.s. |
| ENSG00000119574 | 109.186 | -0.115614433 | 0.240757 | -0.480213 | 0.631076 | 0.919033489 | ZBTB45 | protein_coding | 53.64 | -0.004239427 | 0.004239427 | ENSG00000119574 | n.s. |
| ENSG00000134086 | 1334.132 | -0.12205243 | 0.266047 | -0.458763 | 0.646405 | 0.923184844 | VHL | protein_coding | 47.55 | -0.004236603 | 0.004236603 | ENSG00000134086 | n.s. |
| ENSG00000172795 | 2238.422 | -0.116279644 | 0.24534 | -0.473953 | 0.635534 | 0.919532207 | DCP2 | protein_coding | 37.79 | -0.004236423 | 0.004236423 | ENSG00000172795 | n.s. |
| ENSG00000135093 | 350.1378 | -0.123295129 | 0.274648 | -0.448921 | 0.653488 | 0.923962103 | USP30 | protein_coding | 42.28 | -0.004234675 | 0.004234675 | ENSG00000135093 | n.s. |
| ENSG00000145495 | 4239.907 | -0.121520891 | 0.262689 | -0.462603 | 0.643649 | 0.922916579 | 6-Mar | protein_coding | 39.32 | -0.00423349 | 0.00423349 | ENSG00000145495 | n.s. |
| ENSG00000162688 | 1685.147 | -0.123941855 | 0.276703 | -0.447924 | 0.654208 | 0.92436641 | AGL | protein_coding | 36.47 | -0.004233339 | 0.004233339 | ENSG00000162688 | n.s. |
| ENSG00000105197 | 1412.621 | -0.121877975 | 0.264196 | -0.461316 | 0.644572 | 0.923184844 | TIMM50 | protein_coding | 53.07 | -0.004230547 | 0.004230547 | ENSG00000105197 | n.s. |
| ENSG00000138768 | 1235.548 | 0.124469967 | 0.281844 | 0.4416273 | 0.658759 | 0.924786541 | USO1 | protein_coding | 37.52 | 0.004226813 | 0.004226813 | ENSG00000138768 | n.s. |
| ENSG00000155906 | 1036.453 | -0.122924495 | 0.273916 | -0.448768 | 0.653599 | 0.923962103 | RMND1 | protein_coding | 42.3 | -0.004221945 | 0.004221945 | ENSG00000155906 | n.s. |
| ENSG00000134824 | 2414.949 | -0.121605315 | 0.268123 | -0.453544 | 0.650157 | 0.923184844 | FADS2 | protein_coding | 51.23 | -0.004221083 | 0.004221083 | ENSG00000134824 | n.s. |
| ENSG00000112319 | 3236.915 | 0.119488871 | 0.256445 | 0.4659434 | 0.641256 | 0.921891013 | EYA4 | protein_coding | 36.95 | 0.004220397 | 0.004220397 | ENSG00000112319 | n.s. |
| ENSG00000127922 | 1676.131 | 0.12157936 | 0.268654 | 0.4525492 | 0.650873 | 0.923184844 | SEM1 | protein_coding | 37.66 | 0.004220182 | 0.004220182 | ENSG00000127922 | n.s. |
| ENSG00000063046 | 10534.89 | 0.123600049 | 0.276211 | 0.4474838 | 0.654526 | 0.924413646 | EIF4B | protein_coding | 42.8 | 0.004218921 | 0.004218921 | ENSG00000063046 | n.s. |
| ENSG00000114127 | 1773.623 | -0.124178319 | 0.280377 | -0.442897 | 0.65784 | 0.924786541 | XRN1 | protein_coding | 36.98 | -0.004216909 | 0.004216909 | ENSG00000114127 | n.s. |
| ENSG00000104133 | 1566.706 | -0.124154249 | 0.281685 | -0.440755 | 0.65939 | 0.924786541 | SPG11 | protein_coding | 41.28 | -0.004216092 | 0.004216092 | ENSG00000104133 | n.s. |

| | | | | | | | | | | | | | |
|---|---|---|---|---|---|---|---|---|---|---|---|---|---|
| ENSG00000164167 | 944.1832 | -0.121433355 | 0.267212 | -0.454445 | 0.649508 | 0.923184844 | LSM6 | protein_coding | 39.71 | -0.004215114 | 0.004215114 | ENSG00000164167 | n.s. |
| ENSG00000185651 | 1176.404 | -0.123885239 | 0.277765 | -0.446007 | 0.655592 | 0.924786541 | UBE2L3 | protein_coding | 49.73 | -0.004206957 | 0.004206957 | ENSG00000185651 | n.s. |
| ENSG00000232046 | 22795.16 | -0.100695532 | 0.188408 | -0.534454 | 0.593027 | 0.908297635 | LINC01798 | lincRNA | 39.07 | -0.004206235 | 0.004206235 | ENSG00000232046 | n.s. |
| ENSG00000189221 | 208.8569 | 0.121138876 | 0.266337 | 0.4548332 | 0.649229 | 0.923184844 | MAOA | protein_coding | 40.27 | 0.004204892 | 0.004204892 | ENSG00000189221 | n.s. |
| ENSG00000165410 | 1412.749 | 0.117225215 | 0.249778 | 0.469317 | 0.638843 | 0.920750614 | CFL2 | protein_coding | 39.68 | 0.00420346 | 0.00420346 | ENSG00000165410 | n.s. |
| ENSG00000218739 | 748.807 | -0.12371846 | 0.281723 | -0.439149 | 0.660554 | 0.924786541 | CEBPZOS | protein_coding | 38.06 | -0.004201293 | 0.004201293 | ENSG00000218739 | n.s. |
| ENSG00000172915 | 2914.334 | -0.12363254 | 0.280468 | -0.440808 | 0.659352 | 0.924786541 | NBEA | protein_coding | 35.01 | -0.004198376 | 0.004198376 | ENSG00000172915 | n.s. |
| ENSG00000172262 | 1509.848 | -0.11718314 | 0.250323 | -0.468128 | 0.639693 | 0.920848926 | ZNF131 | protein_coding | 44.17 | -0.004196517 | 0.004196517 | ENSG00000172262 | n.s. |
| ENSG00000207088 | 20.10802 | 0.148199727 | 0.376504 | 0.3936204 | 0.693861 | 0.936924725 | SNORA7B | snoRNA | 56.83 | 0.004193356 | 0.004193356 | ENSG00000207088 | n.s. |
| ENSG00000213190 | 4154.388 | -0.118703352 | 0.254973 | -0.465552 | 0.641536 | 0.921891013 | MLLT11 | protein_coding | 46.4 | -0.004192652 | 0.004192652 | ENSG00000213190 | n.s. |
| ENSG00000115866 | 2683.069 | -0.128655284 | 0.297985 | -0.431751 | 0.665922 | 0.927727183 | DARS | protein_coding | 37.04 | -0.004191553 | 0.004191553 | ENSG00000115866 | n.s. |
| ENSG00000273148 | 104.483 | 0.117030544 | 0.24995 | 0.4682162 | 0.63963 | 0.920848926 | | lincRNA | 41.13 | 0.004191053 | 0.004191053 | ENSG00000273148 | n.s. |
| ENSG00000160469 | 391.5238 | -0.123398768 | 0.280568 | -0.439817 | 0.66007 | 0.924786541 | BRSK1 | protein_coding | 54.48 | -0.004190437 | 0.004190437 | ENSG00000160469 | n.s. |
| ENSG00000198924 | 592.6523 | -0.112474195 | 0.228631 | -0.491947 | 0.622757 | 0.917861492 | DCLRE1A | protein_coding | 36.75 | -0.00418661 | 0.00418661 | ENSG00000198924 | n.s. |
| ENSG00000122218 | 3704.999 | -0.119902399 | 0.258806 | -0.463291 | 0.643156 | 0.922767027 | COPA | protein_coding | 40.67 | -0.004185545 | 0.004185545 | ENSG00000122218 | n.s. |
| ENSG00000185129 | 560.4572 | -0.113974563 | 0.237788 | -0.479313 | 0.631716 | 0.919033489 | PURA | protein_coding | 41.49 | -0.004179295 | 0.004179295 | ENSG00000185129 | n.s. |
| ENSG00000117859 | 2425.838 | -0.12039613 | 0.265505 | -0.453461 | 0.650217 | 0.923184844 | OSBPL9 | protein_coding | 40.08 | -0.00417911 | 0.00417911 | ENSG00000117859 | n.s. |
| ENSG00000213183 | 34.75742 | 0.13534275 | 0.323184 | 0.4187794 | 0.675377 | 0.931386586 | RPS7P8 | processed_pseudogene | 46.13 | 0.004178034 | 0.004178034 | ENSG00000213183 | n.s. |
| ENSG00000164284 | 1336.437 | -0.114670581 | 0.241887 | -0.474066 | 0.635453 | 0.919532207 | GRPEL2 | protein_coding | 41.28 | -0.0041778 | 0.0041778 | ENSG00000164284 | n.s. |
| ENSG00000233484 | 6.592909 | -0.16956481 | 0.496717 | -0.341371 | 0.732824 | 0.944876691 | | transcribed_processed_pseudog | 46.46 | -0.00417551 | 0.00417551 | ENSG00000233484 | n.s. |
| ENSG00000079819 | 2452.062 | -0.114573366 | 0.242047 | -0.473352 | 0.635962 | 0.919565577 | EPB41L2 | protein_coding | 38.6 | -0.004172452 | 0.004172452 | ENSG00000079819 | n.s. |
| ENSG00000003147 | 1273.559 | -0.121339421 | 0.270128 | -0.449192 | 0.653293 | 0.923905149 | ICA1 | protein_coding | 40.3 | -0.004170753 | 0.004170753 | ENSG00000003147 | n.s. |
| ENSG00000236287 | 2581.662 | 0.113717779 | 0.235361 | 0.483164 | 0.628979 | 0.919033489 | ZBED5 | protein_coding | 40.77 | 0.004169879 | 0.004169879 | ENSG00000236287 | n.s. |
| ENSG00000179889 | 1168.826 | -0.120075074 | 0.263466 | -0.455752 | 0.648569 | 0.923184844 | PDXDC1 | protein_coding | 46.49 | -0.004167966 | 0.004167966 | ENSG00000179889 | n.s. |
| ENSG00000153551 | 212.6988 | -0.122719215 | 0.27913 | -0.439649 | 0.660192 | 0.924786541 | CMTM7 | protein_coding | 45.15 | -0.00416736 | 0.00416736 | ENSG00000153551 | n.s. |
| ENSG00000116560 | 8567.424 | -0.113517448 | 0.233176 | -0.486831 | 0.626378 | 0.918973809 | SFPQ | protein_coding | 41.83 | -0.004165735 | 0.004165735 | ENSG00000116560 | n.s. |
| ENSG00000245870 | 7128.43 | -0.119222265 | 0.257116 | -0.46369 | 0.64287 | 0.922767027 | LINC00682 | lincRNA | 51.34 | -0.004161803 | 0.004161803 | ENSG00000245870 | n.s. |
| ENSG00000140259 | 1072.242 | -0.122553987 | 0.278999 | -0.439264 | 0.66047 | 0.924786541 | MFAP1 | protein_coding | 43.98 | -0.004161749 | 0.004161749 | ENSG00000140259 | n.s. |
| ENSG00000011485 | 1169.908 | -0.12241531 | 0.275666 | -0.444071 | 0.656991 | 0.924786541 | PPP5C | protein_coding | 50.75 | -0.00415704 | 0.00415704 | ENSG00000011485 | n.s. |
| ENSG00000006459 | 533.0878 | -0.119729614 | 0.261339 | -0.458139 | 0.646853 | 0.923184844 | KDM7A | protein_coding | 39.18 | -0.004155975 | 0.004155975 | ENSG00000006459 | n.s. |
| ENSG00000178996 | 2086.509 | 0.113321451 | 0.236917 | 0.4783168 | 0.632425 | 0.919033489 | SNX18 | protein_coding | 42.68 | 0.004155346 | 0.004155346 | ENSG00000178996 | n.s. |
| ENSG00000119929 | 365.9187 | -0.136503316 | 0.328913 | -0.415013 | 0.678132 | 0.932323161 | CUTC | protein_coding | 40.02 | -0.004154277 | 0.004154277 | ENSG00000119929 | n.s. |

| | | | | | | | | | | | | | |
|---|---|---|---|---|---|---|---|---|---|---|---|---|---|
| ENSG00000165637 | 2281.123 | 0.122293773 | 0.276227 | 0.4427295 | 0.657961 | 0.924786541 | VDAC2 | protein_coding | 44.04 | 0.004152913 | 0.004152913 | ENSG00000165637 | n.s. |
| ENSG00000275097 | 75.36765 | -0.122223115 | 0.276389 | -0.442215 | 0.658334 | 0.924786541 | | 3prime_overlapping_ncRNA | 31.22 | -0.004150514 | 0.004150514 | ENSG00000275097 | n.s. |
| ENSG00000179950 | 2130.832 | -0.122133568 | 0.277077 | -0.440792 | 0.659364 | 0.924786541 | PUF60 | protein_coding | 62.04 | -0.004147473 | 0.004147473 | ENSG00000179950 | n.s. |
| ENSG00000273084 | 32.03335 | 0.140929589 | 0.352202 | 0.400138 | 0.689055 | 0.934506008 | | lincRNA | 59.24 | 0.004145853 | 0.004145853 | ENSG00000273084 | n.s. |
| ENSG00000127774 | 312.2139 | -0.127550339 | 0.297269 | -0.429074 | 0.66787 | 0.927899924 | EMC6 | protein_coding | 59.6 | -0.004145241 | 0.004145241 | ENSG00000127774 | n.s. |
| ENSG00000112695 | 3390.409 | 0.134187683 | 0.320293 | 0.4189528 | 0.675251 | 0.931353024 | COX7A2 | protein_coding | 40.34 | 0.004144477 | 0.004144477 | ENSG00000112695 | n.s. |
| ENSG00000203879 | 1402.737 | -0.120554553 | 0.268483 | -0.449022 | 0.653416 | 0.923962103 | GDI1 | protein_coding | 59.79 | -0.004140548 | 0.004140548 | ENSG00000203879 | n.s. |
| ENSG00000240303 | 514.0582 | 0.123160557 | 0.281918 | 0.436867 | 0.662208 | 0.925519646 | ACAD11 | protein_coding | 37.61 | 0.004139963 | 0.004139963 | ENSG00000240303 | n.s. |
| ENSG00000134825 | 1225.848 | 0.121849309 | 0.276269 | 0.4410529 | 0.659175 | 0.924786541 | TMEM258 | protein_coding | 58.49 | 0.00413782 | 0.00413782 | ENSG00000134825 | n.s. |
| ENSG00000135930 | 1128.554 | 0.121724766 | 0.273278 | 0.445425 | 0.656013 | 0.924786541 | EIF4E2 | protein_coding | 44.37 | 0.00413359 | 0.00413359 | ENSG00000135930 | n.s. |
| ENSG00000175193 | 681.6022 | -0.126870618 | 0.293801 | -0.431825 | 0.665868 | 0.927727183 | PARL | protein_coding | 41.83 | -0.004133409 | 0.004133409 | ENSG00000175193 | n.s. |
| ENSG00000126945 | 1826.975 | -0.119050888 | 0.262471 | -0.453577 | 0.650133 | 0.923184844 | HNRNPH2 | protein_coding | 39.49 | -0.004132415 | 0.004132415 | ENSG00000126945 | n.s. |
| ENSG00000172053 | 1264.933 | 0.121622928 | 0.273818 | 0.4441737 | 0.656917 | 0.924786541 | QARS | protein_coding | 53.42 | 0.004130132 | 0.004130132 | ENSG00000172053 | n.s. |
| ENSG00000212207 | 5.788144 | -0.175708972 | 0.531799 | -0.330405 | 0.741094 | 0.94732386 | RNU6-1321P | snRNA | 33.64 | -0.004129429 | 0.004129429 | ENSG00000212207 | n.s. |
| ENSG00000187736 | 587.4084 | -0.118920967 | 0.261659 | -0.454489 | 0.649477 | 0.923184844 | NHEJ1 | protein_coding | 43.12 | -0.004127905 | 0.004127905 | ENSG00000187736 | n.s. |
| ENSG00000214160 | 280.9377 | -0.118150228 | 0.255025 | -0.463289 | 0.643157 | 0.922767027 | ALG3 | protein_coding | 51.41 | -0.00412438 | 0.00412438 | ENSG00000214160 | n.s. |
| ENSG00000167377 | 370.0872 | -0.114544945 | 0.243639 | -0.470142 | 0.638254 | 0.920457187 | ZNF23 | protein_coding | 44.44 | -0.004123206 | 0.004123206 | ENSG00000167377 | n.s. |
| ENSG00000244086 | 9.538214 | -0.216584595 | 0.76134 | -0.284478 | 0.776044 | 0.957114084 | | processed_pseudogene | 44.54 | -0.004122968 | 0.004122968 | ENSG00000244086 | n.s. |
| ENSG00000103353 | 1631.834 | -0.118118875 | 0.255196 | -0.462855 | 0.643468 | 0.922847992 | UBFD1 | protein_coding | 47.48 | -0.004118785 | 0.004118785 | ENSG00000103353 | n.s. |
| ENSG00000213585 | 3591.975 | -0.128319412 | 0.3022 | -0.424618 | 0.671115 | 0.92884421 | VDAC1 | protein_coding | 47.42 | -0.004113551 | 0.004113551 | ENSG00000213585 | n.s. |
| ENSG00000171262 | 1074.126 | -0.11846158 | 0.260691 | -0.454414 | 0.649531 | 0.923184844 | FAM98B | protein_coding | 36.34 | -0.004111959 | 0.004111959 | ENSG00000171262 | n.s. |
| ENSG00000182173 | 610.6163 | -0.120990434 | 0.275583 | -0.439034 | 0.660637 | 0.924786541 | TSEN54 | protein_coding | 54.69 | -0.004108654 | 0.004108654 | ENSG00000182173 | n.s. |
| ENSG00000162636 | 697.8729 | -0.118198831 | 0.255956 | -0.461794 | 0.644229 | 0.923083372 | FAM102B | protein_coding | 39.64 | -0.004108482 | 0.004108482 | ENSG00000162636 | n.s. |
| ENSG00000224985 | 21.87672 | -0.15025425 | 0.403334 | -0.372531 | 0.709498 | 0.939005324 | | antisense | 46.94 | -0.004106741 | 0.004106741 | ENSG00000224985 | n.s. |
| ENSG00000157869 | 365.2062 | -0.129373843 | 0.305928 | -0.42289 | 0.672376 | 0.929537806 | RAB28 | protein_coding | 36.22 | -0.004105413 | 0.004105413 | ENSG00000157869 | n.s. |
| ENSG00000108587 | 975.3417 | -0.111921118 | 0.233886 | -0.478528 | 0.632275 | 0.919033489 | GOSR1 | protein_coding | 39.72 | -0.004103998 | 0.004103998 | ENSG00000108587 | n.s. |
| ENSG00000257103 | 1916.386 | -0.121479452 | 0.277483 | -0.43779 | 0.661538 | 0.925277395 | LSM14A | protein_coding | 39.83 | -0.004097265 | 0.004097265 | ENSG00000257103 | n.s. |
| ENSG00000142534 | 25853.16 | 0.130280375 | 0.309177 | 0.4213777 | 0.673479 | 0.930145515 | RPS11 | protein_coding | 53.05 | 0.004097201 | 0.004097201 | ENSG00000142534 | n.s. |
| ENSG00000135047 | 586.5587 | 0.117906235 | 0.25933 | 0.454657 | 0.649356 | 0.923184844 | CTSL | protein_coding | 46.28 | 0.004092683 | 0.004092683 | ENSG00000135047 | n.s. |
| ENSG00000114021 | 494.9827 | -0.130122251 | 0.308932 | -0.4212 | 0.673609 | 0.930145515 | NIT2 | protein_coding | 43.01 | -0.004092228 | 0.004092228 | ENSG00000114021 | n.s. |
| ENSG00000111679 | 35.41108 | 0.15055063 | 0.405839 | 0.3709612 | 0.710666 | 0.939480526 | PTPN6 | protein_coding | 57.71 | 0.004081761 | 0.004081761 | ENSG00000111679 | n.s. |
| ENSG00000079459 | 2477.985 | -0.120158365 | 0.273852 | -0.438771 | 0.660828 | 0.924786541 | FDFT1 | protein_coding | 44.36 | -0.004080398 | 0.004080398 | ENSG00000079459 | n.s. |

| | | | | | | | | | | | | | |
|---|---|---|---|---|---|---|---|---|---|---|---|---|---|
| ENSG00000114631 | 2074.261 | -0.119936269 | 0.270569 | -0.443274 | 0.657568 | 0.924786541 | PODXL2 | protein_coding | 48.64 | -0.004072856 | 0.004072856 | ENSG00000114631 | n.s. |
| ENSG00000269556 | 70.59663 | -0.129780695 | 0.308662 | -0.420462 | 0.674148 | 0.930520512 | TMEM185A | protein_coding | 42.64 | -0.004058768 | 0.004058768 | ENSG00000269556 | n.s. |
| ENSG00000185787 | 6690.447 | -0.116844684 | 0.253719 | -0.460527 | 0.645138 | 0.923184844 | MORF4L1 | protein_coding | 45.4 | -0.004055835 | 0.004055835 | ENSG00000185787 | n.s. |
| ENSG00000067208 | 289.489 | -0.1158988 | 0.250286 | -0.463066 | 0.643317 | 0.922767027 | EVI5 | protein_coding | 38.19 | -0.004045787 | 0.004045787 | ENSG00000067208 | n.s. |
| ENSG00000157224 | 721.1181 | -0.119123945 | 0.267615 | -0.445131 | 0.656225 | 0.924786541 | CLDN12 | protein_coding | 39.02 | -0.00404527 | 0.00404527 | ENSG00000157224 | n.s. |
| ENSG00000100124 | 361.5052 | -0.110248396 | 0.229861 | -0.47963 | 0.63149 | 0.919033489 | ANKRD54 | protein_coding | 50.74 | -0.004042661 | 0.004042661 | ENSG00000100124 | n.s. |
| ENSG00000147654 | 405.8726 | -0.119784601 | 0.27376 | -0.437554 | 0.66171 | 0.925277395 | EBAG9 | protein_coding | 34.99 | -0.004040101 | 0.004040101 | ENSG00000147654 | n.s. |
| ENSG00000134222 | 525.2886 | -0.117239628 | 0.260092 | -0.450762 | 0.652161 | 0.923748122 | PSRC1 | protein_coding | 53.65 | -0.004038487 | 0.004038487 | ENSG00000134222 | n.s. |
| ENSG00000241525 | 35.98028 | -0.149808536 | 0.407517 | -0.367613 | 0.713162 | 0.93983441 | | antisense | 50.98 | -0.004037139 | 0.004037139 | ENSG00000241525 | n.s. |
| ENSG00000156802 | 3791.983 | -0.118870598 | 0.270311 | -0.439754 | 0.660115 | 0.924786541 | ATAD2 | protein_coding | 40.58 | -0.004036667 | 0.004036667 | ENSG00000156802 | n.s. |
| ENSG00000198865 | 141.2609 | 0.130987142 | 0.313014 | 0.4184712 | 0.675603 | 0.931512729 | CCDC152 | protein_coding | 35.74 | 0.004035872 | 0.004035872 | ENSG00000198865 | n.s. |
| ENSG00000110958 | 4817.042 | 0.122684632 | 0.282528 | 0.4342395 | 0.664115 | 0.927094972 | PTGES3 | protein_coding | 44.13 | 0.004033352 | 0.004033352 | ENSG00000110958 | n.s. |
| ENSG00000204650 | 983.198 | -0.11872474 | 0.269176 | -0.441067 | 0.659164 | 0.924786541 | LINC02210 | anscribed_unitary_pseudoge | 46.27 | -0.004031714 | 0.004031714 | ENSG00000204650 | n.s. |
| ENSG00000144120 | 374.0961 | -0.115918815 | 0.251917 | -0.460146 | 0.645411 | 0.923184844 | TMEM177 | protein_coding | 52.47 | -0.004023697 | 0.004023697 | ENSG00000144120 | n.s. |
| ENSG00000105058 | 1550.33 | -0.125396754 | 0.294922 | -0.425186 | 0.670701 | 0.928780923 | FAM32A | protein_coding | 52.93 | -0.00402357 | 0.00402357 | ENSG00000105058 | n.s. |
| ENSG00000221926 | 266.3752 | -0.109717724 | 0.229262 | -0.478569 | 0.632246 | 0.919033489 | TRIM16 | protein_coding | 45.66 | -0.004023202 | 0.004023202 | ENSG00000221926 | n.s. |
| ENSG00000115073 | 896.0682 | -0.123327239 | 0.285352 | -0.432193 | 0.665601 | 0.927727183 | ACTR1B | protein_coding | 58.67 | -0.004017967 | 0.004017967 | ENSG00000115073 | n.s. |
| ENSG00000186648 | 206.3322 | -0.117847834 | 0.263633 | -0.447015 | 0.654864 | 0.924621046 | CARMIL3 | protein_coding | 57.83 | -0.004011095 | 0.004011095 | ENSG00000186648 | n.s. |
| ENSG00000115685 | 633.5665 | -0.123409823 | 0.287428 | -0.429359 | 0.667662 | 0.927899924 | PPP1R7 | protein_coding | 48.38 | -0.004010679 | 0.004010679 | ENSG00000115685 | n.s. |
| ENSG00000171951 | 2267.565 | 0.11554172 | 0.2543 | 0.4543517 | 0.649576 | 0.923184844 | SCG2 | protein_coding | 36.25 | 0.004010607 | 0.004010607 | ENSG00000171951 | n.s. |
| ENSG00000065154 | 1071.159 | -0.118072511 | 0.267024 | -0.44218 | 0.658359 | 0.924786541 | OAT | protein_coding | 42.49 | -0.004009565 | 0.004009565 | ENSG00000065154 | n.s. |
| ENSG00000099290 | 535.3435 | -0.116474214 | 0.258707 | -0.450216 | 0.652554 | 0.923800272 | WASHC2A | protein_coding | 42.59 | -0.004009265 | 0.004009265 | ENSG00000099290 | n.s. |
| ENSG00000203667 | 1081.639 | 0.113390033 | 0.243265 | 0.466117 | 0.641132 | 0.921891013 | COX20 | protein_coding | 39.53 | 0.004004983 | 0.004004983 | ENSG00000203667 | n.s. |
| ENSG00000010295 | 246.1436 | -0.115322923 | 0.254772 | -0.452652 | 0.650799 | 0.923184844 | IFFO1 | protein_coding | 54.32 | -0.004003012 | 0.004003012 | ENSG00000010295 | n.s. |
| ENSG00000151729 | 751.7373 | -0.111076607 | 0.235976 | -0.470711 | 0.637847 | 0.920401057 | SLC25A4 | protein_coding | 46.96 | -0.004001301 | 0.004001301 | ENSG00000151729 | n.s. |
| ENSG00000206527 | 628.6567 | -0.115223274 | 0.254243 | -0.453201 | 0.650404 | 0.923184844 | HACD2 | protein_coding | 40.56 | -0.003999554 | 0.003999554 | ENSG00000206527 | n.s. |
| ENSG00000266538 | 19.29532 | -0.147559644 | 0.398752 | -0.370054 | 0.711342 | 0.939526932 | | lincRNA | 50.16 | -0.003997504 | 0.003997504 | ENSG00000266538 | n.s. |
| ENSG00000001497 | 1087.978 | -0.117662289 | 0.265565 | -0.443064 | 0.657719 | 0.924786541 | LAS1L | protein_coding | 50.45 | -0.003995635 | 0.003995635 | ENSG00000001497 | n.s. |
| ENSG00000124587 | 270.4338 | 0.117550893 | 0.266623 | 0.4408874 | 0.659295 | 0.924786541 | PEX6 | protein_coding | 52.03 | 0.003991852 | 0.003991852 | ENSG00000124587 | n.s. |
| ENSG00000132424 | 9207.209 | 0.114898375 | 0.2522 | 0.455585 | 0.648688 | 0.923184844 | PNISR | protein_coding | 36 | 0.003988276 | 0.003988276 | ENSG00000132424 | n.s. |
| ENSG00000133731 | 892.357 | -0.122729849 | 0.286162 | -0.428882 | 0.668009 | 0.927917746 | IMPA1 | protein_coding | 37.59 | -0.003987557 | 0.003987557 | ENSG00000133731 | n.s. |
| ENSG00000152147 | 432.6875 | -0.114597844 | 0.253121 | -0.452739 | 0.650737 | 0.923184844 | GEMIN6 | protein_coding | 45.01 | -0.003977844 | 0.003977844 | ENSG00000152147 | n.s. |

| | | | | | | | | | | | | | |
|---|---|---|---|---|---|---|---|---|---|---|---|---|---|
| ENSG00000079739 | 659.339 | -0.124610896 | 0.294321 | -0.423384 | 0.672015 | 0.929142491 | PGM1 | protein_coding | 42.42 | -0.003977291 | 0.003977291 | ENSG00000079739 | n.s. |
| ENSG00000132463 | 2756.163 | -0.116991891 | 0.264569 | -0.442197 | 0.658346 | 0.924786541 | GRSF1 | protein_coding | 43.53 | -0.003972869 | 0.003972869 | ENSG00000132463 | n.s. |
| ENSG00000156671 | 652.0142 | -0.120662618 | 0.27801 | -0.434022 | 0.664272 | 0.927094972 | SAMD8 | protein_coding | 43.24 | -0.003966877 | 0.003966877 | ENSG00000156671 | n.s. |
| ENSG00000147155 | 492.7834 | -0.129093547 | 0.308769 | -0.418091 | 0.675881 | 0.931706455 | EBP | protein_coding | 49.69 | -0.003965869 | 0.003965869 | ENSG00000147155 | n.s. |
| ENSG00000133398 | 512.2474 | 0.113516957 | 0.245163 | 0.4630263 | 0.643346 | 0.922767027 | MED10 | protein_coding | 45.03 | 0.003962642 | 0.003962642 | ENSG00000133398 | n.s. |
| ENSG00000184903 | 234.1542 | 0.110493693 | 0.235595 | 0.4689985 | 0.639071 | 0.920750614 | IMMP2L | protein_coding | 36.11 | 0.003962081 | 0.003962081 | ENSG00000184903 | n.s. |
| ENSG00000214578 | 55.54875 | 0.134349445 | 0.332429 | 0.4041442 | 0.686107 | 0.934368276 | HMGN2P15 | processed_pseudogene | 50 | 0.003960879 | 0.003960879 | ENSG00000214578 | n.s. |
| ENSG00000202354 | 3.000721 | -0.225765354 | 0.833114 | -0.27099 | 0.786399 | 0.960409479 | RNY3 | misc_RNA | 45.1 | -0.003960729 | 0.003960729 | ENSG00000202354 | n.s. |
| ENSG00000186310 | 733.9792 | 0.116787221 | 0.266458 | 0.438295 | 0.661172 | 0.924898514 | NAP1L3 | protein_coding | 42.29 | 0.003959778 | 0.003959778 | ENSG00000186310 | n.s. |
| ENSG00000232040 | 346.2363 | -0.115026592 | 0.255547 | -0.450118 | 0.652625 | 0.923800272 | ZBED9 | protein_coding | 39.71 | -0.003959435 | 0.003959435 | ENSG00000232040 | n.s. |
| ENSG00000225663 | 1182.062 | -0.124007377 | 0.292652 | -0.423736 | 0.671758 | 0.929142491 | MCRIP1 | protein_coding | 59.8 | -0.003958028 | 0.003958028 | ENSG00000225663 | n.s. |
| ENSG00000109736 | 369.4297 | -0.113862056 | 0.24932 | -0.456691 | 0.647893 | 0.923184844 | MFSD10 | protein_coding | 67.55 | -0.003952304 | 0.003952304 | ENSG00000109736 | n.s. |
| ENSG00000152990 | 2833.242 | -0.116545766 | 0.265836 | -0.438412 | 0.661088 | 0.924898514 | ADGRA3 | protein_coding | 39.05 | -0.003951591 | 0.003951591 | ENSG00000152990 | n.s. |
| ENSG00000186468 | 17696.47 | 0.129822736 | 0.313779 | 0.4137394 | 0.679065 | 0.932323161 | RPS23 | protein_coding | 44.77 | 0.003950964 | 0.003950964 | ENSG00000186468 | n.s. |
| ENSG00000102805 | 163.7909 | 0.116298749 | 0.264912 | 0.4390094 | 0.660655 | 0.924786541 | CLN5 | protein_coding | 38.81 | 0.003949331 | 0.003949331 | ENSG00000102805 | n.s. |
| ENSG00000108389 | 3782.873 | -0.113647265 | 0.249156 | -0.45613 | 0.648297 | 0.923184844 | MTMR4 | protein_coding | 47.85 | -0.003944848 | 0.003944848 | ENSG00000108389 | n.s. |
| ENSG00000157600 | 640.5602 | -0.114509741 | 0.25401 | -0.450808 | 0.652128 | 0.923748122 | TMEM164 | protein_coding | 42.06 | -0.003944452 | 0.003944452 | ENSG00000157600 | n.s. |
| ENSG00000280432 | 100.09 | 0.180055384 | 0.569799 | 0.3159981 | 0.752004 | 0.950815756 | | TEC | 41.42 | 0.003943867 | 0.003943867 | ENSG00000280432 | n.s. |
| ENSG00000184220 | 1115.224 | -0.122988351 | 0.289409 | -0.424964 | 0.670863 | 0.928840915 | CMSS1 | protein_coding | 38.77 | -0.003942842 | 0.003942842 | ENSG00000184220 | n.s. |
| ENSG00000124641 | 312.5737 | -0.124245566 | 0.293876 | -0.422783 | 0.672454 | 0.929537806 | MED20 | protein_coding | 46.57 | -0.003942677 | 0.003942677 | ENSG00000124641 | n.s. |
| ENSG00000122705 | 2102.03 | -0.130115847 | 0.315292 | -0.412683 | 0.679839 | 0.932670115 | CLTA | protein_coding | 44.48 | -0.003938859 | 0.003938859 | ENSG00000122705 | n.s. |
| ENSG00000018869 | 189.9688 | 0.115877355 | 0.262678 | 0.4411386 | 0.659113 | 0.924786541 | ZNF582 | protein_coding | 40.84 | 0.003935021 | 0.003935021 | ENSG00000018869 | n.s. |
| ENSG00000168936 | 366.6357 | -0.120769992 | 0.279556 | -0.432007 | 0.665736 | 0.927727183 | TMEM129 | protein_coding | 62.46 | -0.003934652 | 0.003934652 | ENSG00000168936 | n.s. |
| ENSG00000142856 | 1145.749 | -0.129237605 | 0.310319 | -0.416467 | 0.677068 | 0.932323161 | ITGB3BP | protein_coding | 36.68 | -0.003933156 | 0.003933156 | ENSG00000142856 | n.s. |
| ENSG00000261186 | 41.22419 | -0.133407102 | 0.330153 | -0.404076 | 0.686157 | 0.934368276 | LINC01238 | antisense | 48.42 | -0.003933097 | 0.003933097 | ENSG00000261186 | n.s. |
| ENSG00000083750 | 334.0229 | -0.12094784 | 0.281057 | -0.430331 | 0.666955 | 0.927899924 | RRAGB | protein_coding | 40.87 | -0.003930667 | 0.003930667 | ENSG00000083750 | n.s. |
| ENSG00000173473 | 5159.916 | -0.113103069 | 0.24849 | -0.455162 | 0.648993 | 0.923184844 | SMARCC1 | protein_coding | 41.6 | -0.003925958 | 0.003925958 | ENSG00000173473 | n.s. |
| ENSG00000004478 | 5720.256 | 0.120788049 | 0.28139 | 0.4292546 | 0.667738 | 0.927899924 | FKBP4 | protein_coding | 51.56 | 0.003925474 | 0.003925474 | ENSG00000004478 | n.s. |
| ENSG00000157077 | 1631.617 | -0.112812023 | 0.247848 | -0.455165 | 0.64899 | 0.923184844 | ZFYVE9 | protein_coding | 40.84 | -0.003915856 | 0.003915856 | ENSG00000157077 | n.s. |
| ENSG00000244879 | 5500.799 | -0.104956989 | 0.212992 | -0.492773 | 0.622173 | 0.917736263 | GABPB1-AS1 | antisense | 42.62 | -0.003913018 | 0.003913018 | ENSG00000244879 | n.s. |
| ENSG00000230454 | 169.394 | 0.108974554 | 0.232116 | 0.4694837 | 0.638724 | 0.920750614 | | lincRNA | 58.51 | 0.003907608 | 0.003907608 | ENSG00000230454 | n.s. |
| ENSG00000136783 | 222.1467 | -0.132827287 | 0.332326 | -0.399689 | 0.689385 | 0.934506008 | NIPSNAP3A | protein_coding | 36.49 | -0.0039075 | 0.0039075 | ENSG00000136783 | n.s. |

| | | | | | | | | | | | | | |
|---|---|---|---|---|---|---|---|---|---|---|---|---|---|
| ENSG00000212241 | 3.521642 | -0.259732531 | 1.074252 | -0.24178 | 0.808951 | 0.965963425 | | misc_RNA | 40.43 | -0.0039062 | 0.0039062 | ENSG00000212241 | n.s. |
| ENSG00000131653 | 771.8057 | -0.112489844 | 0.248663 | -0.452378 | 0.650997 | 0.923184844 | TRAF7 | protein_coding | 63.76 | -0.003904673 | 0.003904673 | ENSG00000131653 | n.s. |
| ENSG00000122545 | 5485.865 | -0.119623248 | 0.276616 | -0.432452 | 0.665413 | 0.927668764 | 7-Sep | protein_coding | 36.13 | -0.003900563 | 0.003900563 | ENSG00000122545 | n.s. |
| ENSG00000140836 | 836.6328 | -0.112360704 | 0.247389 | -0.454186 | 0.649695 | 0.923184844 | ZFHX3 | protein_coding | 46.45 | -0.00390019 | 0.00390019 | ENSG00000140836 | n.s. |
| ENSG00000206763 | 22.38823 | -0.139691585 | 0.358545 | -0.389606 | 0.696828 | 0.937752951 | RNU6-10P | snRNA | 44.86 | -0.00389901 | 0.00389901 | ENSG00000206763 | n.s. |
| ENSG00000134884 | 3154.859 | -0.114780751 | 0.261044 | -0.439698 | 0.660156 | 0.924786541 | ARGLU1 | protein_coding | 37.66 | -0.003897782 | 0.003897782 | ENSG00000134884 | n.s. |
| ENSG00000119537 | 109.453 | 0.113392721 | 0.252401 | 0.449257 | 0.653246 | 0.923905149 | KDSR | protein_coding | 42.29 | 0.003897604 | 0.003897604 | ENSG00000119537 | n.s. |
| ENSG00000063854 | 440.6261 | -0.131120524 | 0.321393 | -0.407975 | 0.683292 | 0.933907782 | HAGH | protein_coding | 53.33 | -0.003893756 | 0.003893756 | ENSG00000063854 | n.s. |
| ENSG00000237819 | 353.2828 | 0.132340271 | 0.328471 | 0.4028975 | 0.687024 | 0.934506008 | | antisense | 37.79 | 0.003893173 | 0.003893173 | ENSG00000237819 | n.s. |
| ENSG00000163297 | 2513.365 | 0.114602521 | 0.258922 | 0.4426148 | 0.658044 | 0.924786541 | ANTXR2 | protein_coding | 35.81 | 0.00389173 | 0.00389173 | ENSG00000163297 | n.s. |
| ENSG00000255152 | 191.7792 | -0.120516007 | 0.282015 | -0.427338 | 0.669133 | 0.928390061 | MSH5-SAPCD1 | protein_coding | 48.78 | -0.003888993 | 0.003888993 | ENSG00000255152 | n.s. |
| ENSG00000199032 | 4.159485 | -0.237159679 | 0.921174 | -0.257454 | 0.796829 | 0.962958487 | MIR425 | miRNA | 57.47 | -0.003887625 | 0.003887625 | ENSG00000199032 | n.s. |
| ENSG00000181061 | 1050.56 | -0.132122845 | 0.329466 | -0.401022 | 0.688404 | 0.934506008 | HIGD1A | protein_coding | 42.66 | -0.003886777 | 0.003886777 | ENSG00000181061 | n.s. |
| ENSG00000117543 | 1156.147 | 0.111298294 | 0.240104 | 0.4635429 | 0.642975 | 0.922767027 | DPH5 | protein_coding | 37.71 | 0.003885193 | 0.003885193 | ENSG00000117543 | n.s. |
| ENSG00000213742 | 74.44564 | 0.119421391 | 0.278283 | 0.4291358 | 0.667824 | 0.927899924 | ZNF337-AS1 | antisense | 41.18 | 0.003881059 | 0.003881059 | ENSG00000213742 | n.s. |
| ENSG00000188186 | 339.9001 | -0.128987125 | 0.313954 | -0.410847 | 0.681185 | 0.933075339 | LAMTOR4 | protein_coding | 53.24 | -0.003880357 | 0.003880357 | ENSG00000188186 | n.s. |
| ENSG00000238317 | 122.7272 | 0.114246081 | 0.259334 | 0.4405367 | 0.659548 | 0.924786541 | SNORD11 | snoRNA | 36.9 | 0.003879625 | 0.003879625 | ENSG00000238317 | n.s. |
| ENSG00000164932 | 72.97588 | -0.124971941 | 0.297685 | -0.419813 | 0.674622 | 0.931014301 | CTHRC1 | protein_coding | 39.94 | -0.003879585 | 0.003879585 | ENSG00000164932 | n.s. |
| ENSG00000143761 | 3039.772 | -0.120340238 | 0.282105 | -0.42658 | 0.669685 | 0.928503293 | ARF1 | protein_coding | 52.33 | -0.003876948 | 0.003876948 | ENSG00000143761 | n.s. |
| ENSG00000149474 | 522.7063 | -0.114159662 | 0.256446 | -0.44516 | 0.656204 | 0.924786541 | KAT14 | protein_coding | 42.25 | -0.003876691 | 0.003876691 | ENSG00000149474 | n.s. |
| ENSG00000188312 | 233.2981 | -0.119254558 | 0.27758 | -0.429622 | 0.667471 | 0.927899924 | CENPP | protein_coding | 40.03 | -0.003875637 | 0.003875637 | ENSG00000188312 | n.s. |
| ENSG00000180776 | 1039.882 | -0.119173733 | 0.277789 | -0.429008 | 0.667917 | 0.927899924 | ZDHHC20 | protein_coding | 39.65 | -0.003873011 | 0.003873011 | ENSG00000180776 | n.s. |
| ENSG00000166197 | 5477.115 | -0.112872829 | 0.251632 | -0.448563 | 0.653747 | 0.924073065 | NOLC1 | protein_coding | 47.22 | -0.003870826 | 0.003870826 | ENSG00000166197 | n.s. |
| ENSG00000170113 | 421.5653 | -0.119068307 | 0.27653 | -0.43058 | 0.666774 | 0.927885071 | NIPA1 | protein_coding | 46.41 | -0.003870412 | 0.003870412 | ENSG00000170113 | n.s. |
| ENSG00000178980 | 405.9177 | 0.131053597 | 0.323715 | 0.4048424 | 0.685593 | 0.934328291 | SELENOW | protein_coding | 54.44 | 0.003866147 | 0.003866147 | ENSG00000178980 | n.s. |
| ENSG00000036054 | 745.8434 | -0.119790586 | 0.280383 | -0.427238 | 0.669206 | 0.928390061 | TBC1D23 | protein_coding | 36.46 | -0.003865584 | 0.003865584 | ENSG00000036054 | n.s. |
| ENSG00000088451 | 338.7142 | -0.118867578 | 0.27687 | -0.429326 | 0.667686 | 0.927899924 | TGDS | protein_coding | 36.98 | -0.003863061 | 0.003863061 | ENSG00000088451 | n.s. |
| ENSG00000145216 | 2099.683 | -0.111098784 | 0.243745 | -0.4558 | 0.648534 | 0.923184844 | FIP1L1 | protein_coding | 35.71 | -0.003856387 | 0.003856387 | ENSG00000145216 | n.s. |
| ENSG00000186834 | 1220.131 | -0.10506368 | 0.215789 | -0.486883 | 0.626342 | 0.918973809 | HEXIM1 | protein_coding | 48.22 | -0.003855508 | 0.003855508 | ENSG00000186834 | n.s. |
| ENSG00000079134 | 2224.77 | 0.113460817 | 0.257884 | 0.4399683 | 0.65996 | 0.924786541 | THOC1 | protein_coding | 37.31 | 0.003852959 | 0.003852959 | ENSG00000079134 | n.s. |
| ENSG00000139613 | 1329.031 | -0.1133225 | 0.254608 | -0.445086 | 0.656258 | 0.924786541 | SMARCC2 | protein_coding | 48.99 | -0.003848262 | 0.003848262 | ENSG00000139613 | n.s. |
| ENSG00000215386 | 1214.402 | -0.110606445 | 0.239461 | -0.461897 | 0.644155 | 0.923072262 | MIR99AHG | lincRNA | 35.47 | -0.003845156 | 0.003845156 | ENSG00000215386 | n.s. |

| | | | | | | | | | | | | | |
|---|---|---|---|---|---|---|---|---|---|---|---|---|---|
| ENSG00000171766 | 219.1832 | 0.118009259 | 0.273481 | 0.431508 | 0.666099 | 0.927727183 | GATM | protein_coding | 40.25 | 0.003844708 | 0.003844708 | ENSG00000171766 | n.s. |
| ENSG00000248206 | 42.50204 | -0.126281598 | 0.303807 | -0.415663 | 0.677656 | 0.932323161 | | sense_intronic | 48.09 | -0.003843194 | 0.003843194 | ENSG00000248206 | n.s. |
| ENSG00000105290 | 83.56274 | -0.126199931 | 0.304993 | -0.41378 | 0.679035 | 0.932323161 | APLP1 | protein_coding | 57.02 | -0.003840709 | 0.003840709 | ENSG00000105290 | n.s. |
| ENSG00000163807 | 1092.675 | -0.125552606 | 0.300969 | -0.417162 | 0.67656 | 0.932022724 | KIAA1143 | protein_coding | 41.53 | -0.003838582 | 0.003838582 | ENSG00000163807 | n.s. |
| ENSG00000100813 | 3259.844 | -0.110549114 | 0.239577 | -0.461435 | 0.644487 | 0.923184844 | ACIN1 | protein_coding | 45.36 | -0.003837307 | 0.003837307 | ENSG00000100813 | n.s. |
| ENSG00000056277 | 812.1144 | -0.112973386 | 0.256239 | -0.440891 | 0.659292 | 0.924786541 | ZNF280C | protein_coding | 38.03 | -0.003836407 | 0.003836407 | ENSG00000056277 | n.s. |
| ENSG00000100034 | 331.3107 | -0.112868236 | 0.25383 | -0.44466 | 0.656565 | 0.924786541 | PPM1F | protein_coding | 55.32 | -0.003832836 | 0.003832836 | ENSG00000100034 | n.s. |
| ENSG00000070444 | 388.9203 | 0.106212936 | 0.225652 | 0.4706934 | 0.63786 | 0.920401057 | MNT | protein_coding | 61.67 | 0.003826097 | 0.003826097 | ENSG00000070444 | n.s. |
| ENSG00000171033 | 1193.76 | -0.129502921 | 0.320088 | -0.404585 | 0.685783 | 0.934328291 | PKIA | protein_coding | 34.75 | -0.003820401 | 0.003820401 | ENSG00000171033 | n.s. |
| ENSG00000147140 | 9184.222 | -0.112483239 | 0.255915 | -0.439534 | 0.660275 | 0.924786541 | NONO | protein_coding | 44.11 | -0.003819762 | 0.003819762 | ENSG00000147140 | n.s. |
| ENSG00000247137 | 494.9572 | -0.098212715 | 0.196207 | -0.500557 | 0.616683 | 0.914385618 | | processed_transcript | 40.91 | -0.003817588 | 0.003817588 | ENSG00000247137 | n.s. |
| ENSG00000171097 | 92.9994 | 0.13677105 | 0.352169 | 0.3883671 | 0.697744 | 0.937752951 | KYAT1 | protein_coding | 49.05 | 0.003817493 | 0.003817493 | ENSG00000171097 | n.s. |
| ENSG00000200175 | 2.676832 | -0.247019808 | 0.99243 | -0.248904 | 0.803435 | 0.965047325 | RNU6-1309P | snRNA | 37.96 | -0.003816799 | 0.003816799 | ENSG00000200175 | n.s. |
| ENSG00000100441 | 743.0506 | -0.112392165 | 0.254542 | -0.441547 | 0.658817 | 0.924786541 | KHNYN | protein_coding | 52.2 | -0.003816669 | 0.003816669 | ENSG00000100441 | n.s. |
| ENSG00000130821 | 208.1466 | -0.110708706 | 0.2453 | -0.451319 | 0.65176 | 0.923702385 | SLC6A8 | protein_coding | 64.7 | -0.003815901 | 0.003815901 | ENSG00000130821 | n.s. |
| ENSG00000170846 | 507.4958 | -0.100191213 | 0.201583 | -0.497022 | 0.619174 | 0.916050756 | | lincRNA | 53.75 | -0.003815328 | 0.003815328 | ENSG00000170846 | n.s. |
| ENSG00000159063 | 1044.73 | -0.109889371 | 0.239392 | -0.459035 | 0.646209 | 0.923184844 | ALG8 | protein_coding | 42.15 | -0.003814407 | 0.003814407 | ENSG00000159063 | n.s. |
| ENSG00000185825 | 637.8017 | -0.128791225 | 0.316713 | -0.406649 | 0.684266 | 0.934117877 | BCAP31 | protein_coding | 54.34 | -0.003812003 | 0.003812003 | ENSG00000185825 | n.s. |
| ENSG00000283122 | 201.2251 | 0.109477189 | 0.236822 | 0.4622763 | 0.643883 | 0.922967763 | HYMAI | non_coding | 40.45 | 0.003811281 | 0.003811281 | ENSG00000283122 | n.s. |
| ENSG00000197223 | 1022.545 | 0.112028398 | 0.254604 | 0.4400098 | 0.65993 | 0.924786541 | C1D | protein_coding | 38.27 | 0.003804316 | 0.003804316 | ENSG00000197223 | n.s. |
| ENSG00000105135 | 435.2957 | -0.124969418 | 0.301041 | -0.415125 | 0.678051 | 0.932323161 | ILVBL | protein_coding | 57.25 | -0.00380326 | 0.00380326 | ENSG00000105135 | n.s. |
| ENSG00000242251 | 27.01721 | -0.136493913 | 0.35959 | -0.379582 | 0.704256 | 0.937896121 | RN7SL20P | misc_RNA | 52.19 | -0.003800709 | 0.003800709 | ENSG00000242251 | n.s. |
| ENSG00000072134 | 662.0847 | -0.119689838 | 0.283144 | -0.422718 | 0.672501 | 0.929537806 | EPN2 | protein_coding | 48.72 | -0.003798111 | 0.003798111 | ENSG00000072134 | n.s. |
| ENSG00000198270 | 329.2089 | -0.103557813 | 0.217318 | -0.476527 | 0.633699 | 0.919033489 | TMEM116 | protein_coding | 40.27 | -0.003797327 | 0.003797327 | ENSG00000198270 | n.s. |
| ENSG00000166896 | 140.9868 | -0.12876825 | 0.319362 | -0.403205 | 0.686798 | 0.934506008 | ATP23 | protein_coding | 40.87 | -0.003788092 | 0.003788092 | ENSG00000166896 | n.s. |
| ENSG00000116489 | 2587.109 | -0.112288859 | 0.256593 | -0.437615 | 0.661665 | 0.925277395 | CAPZA1 | protein_coding | 40.34 | -0.003787284 | 0.003787284 | ENSG00000116489 | n.s. |
| ENSG00000230397 | 65.76462 | -0.125095847 | 0.303252 | -0.412515 | 0.679962 | 0.932670115 | SPTLC1P1 | ıscribed_processed_pseudog | 44.83 | -0.003786894 | 0.003786894 | ENSG00000230397 | n.s. |
| ENSG00000014123 | 1319.253 | -0.109941366 | 0.243986 | -0.450605 | 0.652275 | 0.923773611 | UFL1 | protein_coding | 33.38 | -0.00378577 | 0.00378577 | ENSG00000014123 | n.s. |
| ENSG00000261188 | 309.3318 | -0.103201508 | 0.215432 | -0.479044 | 0.631907 | 0.919033489 | | antisense | 42.72 | -0.003784261 | 0.003784261 | ENSG00000261188 | n.s. |
| ENSG00000146476 | 652.673 | -0.11603769 | 0.269151 | -0.431125 | 0.666377 | 0.927727183 | ARMT1 | protein_coding | 39.28 | -0.003780475 | 0.003780475 | ENSG00000146476 | n.s. |
| ENSG00000248124 | 622.707 | -0.108888233 | 0.238864 | -0.455859 | 0.648491 | 0.923184844 | RRN3P1 | cribed_unprocessed_pseudc | 41.35 | -0.003779656 | 0.003779656 | ENSG00000248124 | n.s. |
| ENSG00000120158 | 882.5384 | -0.116251968 | 0.2702 | -0.430244 | 0.667018 | 0.927899924 | RCL1 | protein_coding | 42.83 | -0.003778057 | 0.003778057 | ENSG00000120158 | n.s. |

| | | | | | | | | | | | | | |
|---|---|---|---|---|---|---|---|---|---|---|---|---|---|
| ENSG00000227962 | 21.83885 | -0.135785328 | 0.360029 | -0.377151 | 0.706061 | 0.937943012 | | processed_pseudogene | 40.43 | -0.00377803 | 0.00377803 | ENSG00000227962 | n.s. |
| ENSG00000165996 | 167.4384 | -0.127636899 | 0.314522 | -0.405813 | 0.68488 | 0.934117877 | HACD1 | protein_coding | 41.97 | -0.003777837 | 0.003777837 | ENSG00000165996 | n.s. |
| ENSG00000204227 | 522.7637 | -0.120748748 | 0.287157 | -0.420498 | 0.674122 | 0.930520512 | RING1 | protein_coding | 53.03 | -0.003776302 | 0.003776302 | ENSG00000204227 | n.s. |
| ENSG00000231365 | 368.8977 | -0.103612459 | 0.218387 | -0.474445 | 0.635183 | 0.919532207 | | antisense | 40.62 | -0.003774918 | 0.003774918 | ENSG00000231365 | n.s. |
| ENSG00000167468 | 1203.432 | -0.130359099 | 0.328327 | -0.39704 | 0.691338 | 0.935503831 | GPX4 | protein_coding | 69.47 | -0.003774473 | 0.003774473 | ENSG00000167468 | n.s. |
| ENSG00000169727 | 1864.056 | -0.119806479 | 0.284406 | -0.421251 | 0.673572 | 0.930145515 | GPS1 | protein_coding | 66.61 | -0.003767806 | 0.003767806 | ENSG00000169727 | n.s. |
| ENSG00000131467 | 2480.613 | -0.109310972 | 0.242156 | -0.451408 | 0.651695 | 0.923702385 | PSME3 | protein_coding | 46.04 | -0.003767724 | 0.003767724 | ENSG00000131467 | n.s. |
| ENSG00000265241 | 4143.587 | -0.109543909 | 0.243705 | -0.449494 | 0.653076 | 0.923905149 | RBM8A | protein_coding | 46.78 | -0.00376531 | 0.00376531 | ENSG00000265241 | n.s. |
| ENSG00000247596 | 323.8302 | -0.127558918 | 0.315335 | -0.404518 | 0.685832 | 0.934328291 | TWF2 | protein_coding | 59.9 | -0.003763052 | 0.003763052 | ENSG00000247596 | n.s. |
| ENSG00000120519 | 125.8489 | 0.116608158 | 0.27305 | 0.4270581 | 0.669337 | 0.928390061 | SLC10A7 | protein_coding | 36.95 | 0.003762889 | 0.003762889 | ENSG00000120519 | n.s. |
| ENSG00000115947 | 1757.55 | -0.116311046 | 0.271659 | -0.428151 | 0.668541 | 0.928267143 | ORC4 | protein_coding | 36.15 | -0.00375999 | 0.00375999 | ENSG00000115947 | n.s. |
| ENSG00000214293 | 147.8864 | -0.126894943 | 0.311661 | -0.407157 | 0.683892 | 0.93406267 | APTR | lincRNA | 43.28 | -0.003759134 | 0.003759134 | ENSG00000214293 | n.s. |
| ENSG00000180660 | 6881.775 | -0.097606468 | 0.195746 | -0.498639 | 0.618034 | 0.915276425 | MAB21L1 | protein_coding | 46.24 | -0.003752746 | 0.003752746 | ENSG00000180660 | n.s. |
| ENSG00000221491 | 17.40221 | 0.13882949 | 0.37601 | 0.3692173 | 0.711966 | 0.939770546 | SNORA2C | snoRNA | 50.36 | 0.003745366 | 0.003745366 | ENSG00000221491 | n.s. |
| ENSG00000166348 | 29.27455 | -0.134355924 | 0.352455 | -0.381201 | 0.703054 | 0.937867182 | USP54 | protein_coding | 42.16 | -0.003742976 | 0.003742976 | ENSG00000166348 | n.s. |
| ENSG00000140543 | 49.74027 | 0.12336404 | 0.299025 | 0.4125538 | 0.679934 | 0.932670115 | DET1 | protein_coding | 43.27 | 0.003734469 | 0.003734469 | ENSG00000140543 | n.s. |
| ENSG00000146729 | 986.9326 | -0.124023153 | 0.302206 | -0.410393 | 0.681518 | 0.933075339 | NIPSNAP2 | protein_coding | 45.28 | -0.003731024 | 0.003731024 | ENSG00000146729 | n.s. |
| ENSG00000259151 | 74.60766 | -0.115846525 | 0.271703 | -0.426372 | 0.669837 | 0.928528381 | CAP2P1 | ıscribed_processed_pseudog | 48.92 | -0.003730816 | 0.003730816 | ENSG00000259151 | n.s. |
| ENSG00000155666 | 37.98767 | -0.133647206 | 0.347874 | -0.384183 | 0.700843 | 0.937752951 | KDM8 | protein_coding | 51.92 | -0.003730302 | 0.003730302 | ENSG00000155666 | n.s. |
| ENSG00000152240 | 759.8934 | -0.123978465 | 0.302187 | -0.410271 | 0.681607 | 0.933075339 | HAUS1 | protein_coding | 41.32 | -0.00372968 | 0.00372968 | ENSG00000152240 | n.s. |
| ENSG00000075914 | 1005.429 | -0.116843532 | 0.27592 | -0.423468 | 0.671953 | 0.929142491 | EXOSC7 | protein_coding | 46.87 | -0.003729374 | 0.003729374 | ENSG00000075914 | n.s. |
| ENSG00000112658 | 743.3726 | -0.115391273 | 0.269732 | -0.427799 | 0.668797 | 0.928288938 | SRF | protein_coding | 55.7 | -0.00372908 | 0.00372908 | ENSG00000112658 | n.s. |
| ENSG00000231842 | 23.67442 | -0.1340218 | 0.355886 | -0.376586 | 0.706481 | 0.937943012 | | lincRNA | 35.46 | -0.003728962 | 0.003728962 | ENSG00000231842 | n.s. |
| ENSG00000238344 | 60.97355 | 0.118719611 | 0.282066 | 0.4208924 | 0.673834 | 0.930271283 | SNORD126 | snoRNA | 42.86 | 0.003726654 | 0.003726654 | ENSG00000238344 | n.s. |
| ENSG00000277478 | 4.34855 | -0.219208901 | 0.831327 | -0.263686 | 0.792022 | 0.961629391 | MIR6165 | miRNA | 66.67 | -0.003724857 | 0.003724857 | ENSG00000277478 | n.s. |
| ENSG00000255769 | 544.9363 | -0.111760035 | 0.256594 | -0.435552 | 0.663162 | 0.926284538 | GOLGA2P10 | cribed_unprocessed_pseudc | 42.97 | -0.003716645 | 0.003716645 | ENSG00000255769 | n.s. |
| ENSG00000273456 | 117.3771 | -0.114282793 | 0.266263 | -0.42921 | 0.66777 | 0.927899924 | | lincRNA | 51.56 | -0.003714061 | 0.003714061 | ENSG00000273456 | n.s. |
| ENSG00000213463 | 1077.937 | -0.113459547 | 0.261972 | -0.433097 | 0.664944 | 0.927447429 | SYNJ2BP | protein_coding | 39.54 | -0.003711341 | 0.003711341 | ENSG00000213463 | n.s. |
| ENSG00000225439 | 229.2959 | -0.107871081 | 0.239721 | -0.449985 | 0.652721 | 0.923842041 | BOLA3-AS1 | antisense | 49.2 | -0.003711011 | 0.003711011 | ENSG00000225439 | n.s. |
| ENSG00000110107 | 1520.758 | -0.106883309 | 0.231827 | -0.461047 | 0.644765 | 0.923184844 | PRPF19 | protein_coding | 46.52 | -0.003710062 | 0.003710062 | ENSG00000110107 | n.s. |
| ENSG00000113594 | 1288.648 | 0.112436857 | 0.258642 | 0.4347208 | 0.663765 | 0.926901501 | LIFR | protein_coding | 39.45 | 0.00370664 | 0.00370664 | ENSG00000113594 | n.s. |
| ENSG00000152503 | 215.3516 | 0.108463476 | 0.2422 | 0.447827 | 0.654278 | 0.92436641 | TRIM36 | protein_coding | 40.18 | 0.003704662 | 0.003704662 | ENSG00000152503 | n.s. |

| | | | | | | | | | | | | | |
|---|---|---|---|---|---|---|---|---|---|---|---|---|---|
| ENSG00000116030 | 3594.432 | -0.121696594 | 0.294096 | -0.413799 | 0.679021 | 0.932323161 | SUMO1 | protein_coding | 41.66 | -0.003703657 | 0.003703657 | ENSG00000116030 | n.s. |
| ENSG00000177602 | 368.5599 | -0.103454712 | 0.221245 | -0.467602 | 0.640069 | 0.920914184 | GSG2 | protein_coding | 53.91 | -0.003701696 | 0.003701696 | ENSG00000177602 | n.s. |
| ENSG00000170860 | 1681.485 | -0.125344549 | 0.311951 | -0.401808 | 0.687825 | 0.934506008 | LSM3 | protein_coding | 41.66 | -0.003687374 | 0.003687374 | ENSG00000170860 | n.s. |
| ENSG00000251141 | 37.97968 | 0.132061466 | 0.344125 | 0.3837599 | 0.701156 | 0.937752951 | | antisense | 36.75 | 0.003686042 | 0.003686042 | ENSG00000251141 | n.s. |
| ENSG00000198855 | 139.7554 | -0.108503111 | 0.245168 | -0.442566 | 0.65808 | 0.924786541 | FICD | protein_coding | 50.41 | -0.003684603 | 0.003684603 | ENSG00000198855 | n.s. |
| ENSG00000207245 | 9.271686 | 0.150361513 | 0.442602 | 0.3397218 | 0.734066 | 0.945151747 | SNORD116-29 | snoRNA | 40.96 | 0.003683625 | 0.003683625 | ENSG00000207245 | n.s. |
| ENSG00000275023 | 1385.256 | -0.107162915 | 0.23835 | -0.449603 | 0.652997 | 0.923905149 | MLLT6 | protein_coding | 56.24 | -0.003683469 | 0.003683469 | ENSG00000275023 | n.s. |
| ENSG00000136718 | 1431.316 | -0.11299838 | 0.261808 | -0.431608 | 0.666026 | 0.927727183 | IMP4 | protein_coding | 53.58 | -0.003681455 | 0.003681455 | ENSG00000136718 | n.s. |
| ENSG00000186184 | 1249.801 | 0.121593651 | 0.294431 | 0.4129784 | 0.679622 | 0.932670115 | POLR1D | protein_coding | 40.03 | 0.003680876 | 0.003680876 | ENSG00000186184 | n.s. |
| ENSG00000210156 | 7.548762 | -0.175876062 | 0.572691 | -0.307105 | 0.758764 | 0.95300702 | MT-TK | Mt_tRNA | 32.86 | -0.003676496 | 0.003676496 | ENSG00000210156 | n.s. |
| ENSG00000167447 | 1726.262 | 0.099944325 | 0.204443 | 0.4888627 | 0.624939 | 0.918895805 | SMG8 | protein_coding | 44.25 | 0.003671328 | 0.003671328 | ENSG00000167447 | n.s. |
| ENSG00000118096 | 223.1104 | 0.124765691 | 0.311755 | 0.4002047 | 0.689006 | 0.934506008 | IFT46 | protein_coding | 45 | 0.003670345 | 0.003670345 | ENSG00000118096 | n.s. |
| ENSG00000115425 | 249.3547 | -0.114401893 | 0.269477 | -0.424533 | 0.671177 | 0.92884421 | PECR | protein_coding | 42.56 | -0.003667395 | 0.003667395 | ENSG00000115425 | n.s. |
| ENSG00000120696 | 637.5585 | 0.099965035 | 0.206868 | 0.4832313 | 0.628932 | 0.919033489 | KBTBD7 | protein_coding | 42.21 | 0.003665584 | 0.003665584 | ENSG00000120696 | n.s. |
| ENSG00000260032 | 5501.568 | -0.105568668 | 0.233176 | -0.452743 | 0.650734 | 0.923184844 | NORAD | lincRNA | 41.56 | -0.003664429 | 0.003664429 | ENSG00000260032 | n.s. |
| ENSG00000108021 | 3615.115 | -0.11412092 | 0.268306 | -0.425338 | 0.67059 | 0.928780923 | FAM208B | protein_coding | 38.83 | -0.003661765 | 0.003661765 | ENSG00000108021 | n.s. |
| ENSG00000186432 | 2129.457 | -0.111912446 | 0.258547 | -0.432851 | 0.665123 | 0.927447429 | KPNA4 | protein_coding | 37.2 | -0.003660734 | 0.003660734 | ENSG00000186432 | n.s. |
| ENSG00000181544 | 436.153 | -0.114079377 | 0.268124 | -0.425473 | 0.670492 | 0.928780923 | FANCB | protein_coding | 34.57 | -0.003660432 | 0.003660432 | ENSG00000181544 | n.s. |
| ENSG00000084623 | 1335.217 | -0.124312972 | 0.310967 | -0.399762 | 0.689332 | 0.934506008 | EIF3I | protein_coding | 50.5 | -0.003657027 | 0.003657027 | ENSG00000084623 | n.s. |
| ENSG00000139631 | 27.97878 | -0.13571905 | 0.36969 | -0.367116 | 0.713532 | 0.939884276 | CSAD | protein_coding | 47.44 | -0.003654319 | 0.003654319 | ENSG00000139631 | n.s. |
| ENSG00000204103 | 162.2599 | 0.112313537 | 0.260711 | 0.4307966 | 0.666616 | 0.927851489 | MAFB | protein_coding | 56.44 | 0.003652608 | 0.003652608 | ENSG00000204103 | n.s. |
| ENSG00000200959 | 21.83317 | 0.137411176 | 0.38036 | 0.3612664 | 0.7179 | 0.940629569 | SNORA74A | snoRNA | 48.48 | 0.003652578 | 0.003652578 | ENSG00000200959 | n.s. |
| ENSG00000115738 | 4943.691 | -0.11594395 | 0.275165 | -0.421362 | 0.673491 | 0.930145515 | ID2 | protein_coding | 49.26 | -0.003646333 | 0.003646333 | ENSG00000115738 | n.s. |
| ENSG00000196976 | 173.8518 | -0.13063326 | 0.33564 | -0.389206 | 0.697124 | 0.937752951 | LAGE3 | protein_coding | 63.33 | -0.003646178 | 0.003646178 | ENSG00000196976 | n.s. |
| ENSG00000239953 | 13.00518 | -0.148816654 | 0.438166 | -0.339635 | 0.734131 | 0.945151747 | RN7SL273P | misc_RNA | 52.84 | -0.003645779 | 0.003645779 | ENSG00000239953 | n.s. |
| ENSG00000140043 | 136.3343 | -0.124002556 | 0.310776 | -0.399009 | 0.689886 | 0.934548254 | PTGR2 | protein_coding | 42.1 | -0.003645461 | 0.003645461 | ENSG00000140043 | n.s. |
| ENSG00000105176 | 2415.765 | -0.119767713 | 0.288666 | -0.414901 | 0.678214 | 0.932323161 | URI1 | protein_coding | 37.88 | -0.003644954 | 0.003644954 | ENSG00000105176 | n.s. |
| ENSG00000112031 | 1088.914 | 0.107326243 | 0.242674 | 0.4422654 | 0.658297 | 0.924786541 | MTRF1L | protein_coding | 37.68 | 0.003644638 | 0.003644638 | ENSG00000112031 | n.s. |
| ENSG00000141562 | 1622.013 | -0.111705062 | 0.258311 | -0.432444 | 0.665419 | 0.927668764 | NARF | protein_coding | 51.72 | -0.003642374 | 0.003642374 | ENSG00000141562 | n.s. |
| ENSG00000249085 | 32.69756 | -0.136762541 | 0.377487 | -0.362298 | 0.71713 | 0.94052839 | | antisense | 38.46 | -0.003641725 | 0.003641725 | ENSG00000249085 | n.s. |
| ENSG00000165644 | 64.80504 | 0.130371121 | 0.339045 | 0.3845249 | 0.700589 | 0.937752951 | COMTD1 | protein_coding | 70.61 | 0.003638861 | 0.003638861 | ENSG00000165644 | n.s. |
| ENSG00000129636 | 555.5766 | -0.119541015 | 0.287975 | -0.415108 | 0.678063 | 0.932323161 | ITFG1 | protein_coding | 37.51 | -0.003638055 | 0.003638055 | ENSG00000129636 | n.s. |

| | | | | | | | | | | | | | |
|---|---|---|---|---|---|---|---|---|---|---|---|---|---|
| ENSG00000104897 | 729.7251 | -0.104735521 | 0.22935 | -0.456662 | 0.647914 | 0.923184844 | SF3A2 | protein_coding | 58.67 | -0.00363551 | 0.00363551 | ENSG00000104897 | n.s. |
| ENSG00000179222 | 817.2289 | -0.114627552 | 0.271253 | -0.422586 | 0.672597 | 0.929578611 | MAGED1 | protein_coding | 39 | -0.003635284 | 0.003635284 | ENSG00000179222 | n.s. |
| ENSG00000271818 | 40.1159 | 0.130130367 | 0.334966 | 0.3884884 | 0.697655 | 0.937752951 | RN7SKP4 | misc_RNA | 52.8 | 0.003632142 | 0.003632142 | ENSG00000271818 | n.s. |
| ENSG00000006007 | 665.4488 | 0.126524383 | 0.320309 | 0.3950073 | 0.692838 | 0.936038047 | GDE1 | protein_coding | 40.17 | 0.003632072 | 0.003632072 | ENSG00000006007 | n.s. |
| ENSG00000249592 | 294.5542 | -0.10462365 | 0.228387 | -0.458098 | 0.646882 | 0.923184844 | | antisense | 50.65 | -0.003631626 | 0.003631626 | ENSG00000249592 | n.s. |
| ENSG00000151014 | 289.7545 | -0.106888112 | 0.241417 | -0.442753 | 0.657945 | 0.924786541 | NOCT | protein_coding | 44.81 | -0.00362976 | 0.00362976 | ENSG00000151014 | n.s. |
| ENSG00000250742 | 154.2995 | 0.123213307 | 0.307003 | 0.4013423 | 0.688168 | 0.934506008 | LINC02381 | lincRNA | 51.73 | 0.003624677 | 0.003624677 | ENSG00000250742 | n.s. |
| ENSG00000119705 | 1738.912 | 0.12328257 | 0.308853 | 0.3991631 | 0.689773 | 0.934548254 | SLIRP | protein_coding | 41.29 | 0.003624294 | 0.003624294 | ENSG00000119705 | n.s. |
| ENSG00000105926 | 400.7524 | -0.123482729 | 0.309629 | -0.398809 | 0.690034 | 0.934657569 | MPP6 | protein_coding | 35.88 | -0.003623906 | 0.003623906 | ENSG00000105926 | n.s. |
| ENSG00000173598 | 1558.952 | 0.105429759 | 0.234525 | 0.4495451 | 0.653038 | 0.923905149 | NUDT4 | protein_coding | 41.97 | 0.003623896 | 0.003623896 | ENSG00000173598 | n.s. |
| ENSG00000261692 | 27.50064 | -0.136772496 | 0.379846 | -0.360074 | 0.718792 | 0.940842655 | | sense_intronic | 52.6 | -0.003622146 | 0.003622146 | ENSG00000261692 | n.s. |
| ENSG00000175879 | 411.1655 | -0.104638964 | 0.231533 | -0.45194 | 0.651312 | 0.923473769 | HOXD8 | protein_coding | 52.17 | -0.003617938 | 0.003617938 | ENSG00000175879 | n.s. |
| ENSG00000236088 | 194.7608 | -0.118859189 | 0.287085 | -0.41402 | 0.678859 | 0.932323161 | COX10-AS1 | processed_transcript | 40.99 | -0.003617304 | 0.003617304 | ENSG00000236088 | n.s. |
| ENSG00000112763 | 634.5351 | -0.106289681 | 0.239731 | -0.44337 | 0.657498 | 0.924786541 | BTN2A1 | protein_coding | 45.31 | -0.003609438 | 0.003609438 | ENSG00000112763 | n.s. |
| ENSG00000168288 | 2369.881 | 0.123930743 | 0.311767 | 0.3975109 | 0.690991 | 0.935225433 | MMADHC | protein_coding | 35.15 | 0.003604364 | 0.003604364 | ENSG00000168288 | n.s. |
| ENSG00000158201 | 249.0347 | 0.116586617 | 0.278274 | 0.4189631 | 0.675243 | 0.931353024 | ABHD3 | protein_coding | 42.45 | 0.003600856 | 0.003600856 | ENSG00000158201 | n.s. |
| ENSG00000122884 | 337.6954 | 0.118881777 | 0.288081 | 0.4126679 | 0.67985 | 0.932670115 | P4HA1 | protein_coding | 38.79 | 0.003598782 | 0.003598782 | ENSG00000122884 | n.s. |
| ENSG00000151914 | 12031.35 | 0.1093824 | 0.252049 | 0.4339722 | 0.664309 | 0.927094972 | DST | protein_coding | 38.07 | 0.003596031 | 0.003596031 | ENSG00000151914 | n.s. |
| ENSG00000216775 | 1004.518 | -0.103568338 | 0.228361 | -0.453529 | 0.650168 | 0.923184844 | | transcribed_unprocessed_pseudc | 41.39 | -0.003594995 | 0.003594995 | ENSG00000216775 | n.s. |
| ENSG00000168385 | 7386.476 | -0.118071854 | 0.283684 | -0.416209 | 0.677257 | 0.932323161 | 2-Sep | protein_coding | 41.89 | -0.003593343 | 0.003593343 | ENSG00000168385 | n.s. |
| ENSG00000270362 | 147.4674 | 0.10524135 | 0.235164 | 0.4475239 | 0.654497 | 0.924413646 | HMGN3-AS1 | antisense | 42.24 | 0.003592272 | 0.003592272 | ENSG00000270362 | n.s. |
| ENSG00000100372 | 504.5543 | -0.122110898 | 0.303194 | -0.402748 | 0.687133 | 0.934506008 | SLC25A17 | protein_coding | 41.77 | -0.003592246 | 0.003592246 | ENSG00000100372 | n.s. |
| ENSG00000202293 | 3.536594 | 0.18583509 | 0.64639 | 0.2874967 | 0.773732 | 0.956474979 | SNORD114-22 | snoRNA | 45.83 | 0.003591521 | 0.003591521 | ENSG00000202293 | n.s. |
| ENSG00000002822 | 533.1199 | -0.105703254 | 0.238678 | -0.44287 | 0.65786 | 0.924786541 | MAD1L1 | protein_coding | 55.65 | -0.003589524 | 0.003589524 | ENSG00000002822 | n.s. |
| ENSG00000177868 | 158.3617 | 0.119254036 | 0.29088 | 0.4099773 | 0.681823 | 0.933075339 | SVBP | protein_coding | 41.39 | 0.003587554 | 0.003587554 | ENSG00000177868 | n.s. |
| ENSG00000281649 | 723.3459 | -0.110822762 | 0.258951 | -0.427969 | 0.668674 | 0.928267143 | EBLN3P | lincRNA | 38.61 | -0.00358257 | 0.00358257 | ENSG00000281649 | n.s. |
| ENSG00000163918 | 1924.647 | -0.116766459 | 0.279695 | -0.417478 | 0.676329 | 0.931868404 | RFC4 | protein_coding | 40.5 | -0.003578356 | 0.003578356 | ENSG00000163918 | n.s. |
| ENSG00000242110 | 235.2465 | -0.11004257 | 0.255855 | -0.430097 | 0.667125 | 0.927899924 | AMACR | protein_coding | 41.36 | -0.003576258 | 0.003576258 | ENSG00000242110 | n.s. |
| ENSG00000116679 | 13744.99 | -0.110003783 | 0.255478 | -0.43058 | 0.666774 | 0.927885071 | IVNS1ABP | protein_coding | 37.35 | -0.003575762 | 0.003575762 | ENSG00000116679 | n.s. |
| ENSG00000137216 | 137.4237 | 0.117412964 | 0.283729 | 0.4138208 | 0.679005 | 0.932323161 | TMEM63B | protein_coding | 54.28 | 0.003573291 | 0.003573291 | ENSG00000137216 | n.s. |
| ENSG00000001630 | 3677.893 | -0.110638374 | 0.259035 | -0.427118 | 0.669293 | 0.928390061 | CYP51A1 | protein_coding | 39.91 | -0.003570247 | 0.003570247 | ENSG00000001630 | n.s. |
| ENSG00000131844 | 762.1941 | -0.10952978 | 0.254094 | -0.43106 | 0.666425 | 0.927727183 | MCCC2 | protein_coding | 41.2 | -0.003568449 | 0.003568449 | ENSG00000131844 | n.s. |

| | | | | | | | | | | | | | |
|---|---|---|---|---|---|---|---|---|---|---|---|---|---|
| ENSG00000143942 | 279.626 | -0.117094292 | 0.281427 | -0.416073 | 0.677356 | 0.932323161 | CHAC2 | protein_coding | 39.11 | -0.003563592 | 0.003563592 | ENSG00000143942 | n.s. |
| ENSG00000237854 | 221.6731 | -0.104888418 | 0.236597 | -0.443321 | 0.657534 | 0.924786541 | LINC00674 | cribed_unprocessed_pseudc | 40.03 | -0.003561853 | 0.003561853 | ENSG00000237854 | n.s. |
| ENSG00000064703 | 2254.492 | -0.109555289 | 0.255337 | -0.429062 | 0.667878 | 0.927899924 | DDX20 | protein_coding | 40.24 | -0.003560422 | 0.003560422 | ENSG00000064703 | n.s. |
| ENSG00000196305 | 7762.747 | -0.113133208 | 0.268284 | -0.421692 | 0.67325 | 0.930111256 | IARS | protein_coding | 41.23 | -0.003559748 | 0.003559748 | ENSG00000196305 | n.s. |
| ENSG00000103657 | 3883.264 | -0.106885472 | 0.24534 | -0.435663 | 0.663082 | 0.926265201 | HERC1 | protein_coding | 38.81 | -0.003555508 | 0.003555508 | ENSG00000103657 | n.s. |
| ENSG00000188352 | 860.7995 | -0.110953731 | 0.261456 | -0.424369 | 0.671297 | 0.928887382 | FOCAD | protein_coding | 36.92 | -0.003554618 | 0.003554618 | ENSG00000188352 | n.s. |
| ENSG00000175161 | 36.68167 | -0.127634138 | 0.335273 | -0.380687 | 0.703435 | 0.937896121 | CADM2 | protein_coding | 34 | -0.003554006 | 0.003554006 | ENSG00000175161 | n.s. |
| ENSG00000141447 | 1084.695 | -0.116592229 | 0.281806 | -0.413733 | 0.67907 | 0.932323161 | OSBPL1A | protein_coding | 40.87 | -0.003548313 | 0.003548313 | ENSG00000141447 | n.s. |
| ENSG00000156502 | 922.3023 | -0.11265959 | 0.267441 | -0.42125 | 0.673572 | 0.930145515 | SUPV3L1 | protein_coding | 41.8 | -0.003543043 | 0.003543043 | ENSG00000156502 | n.s. |
| ENSG00000266237 | 40.06134 | 0.126920181 | 0.327049 | 0.3880773 | 0.697959 | 0.937752951 | | sense_intronic | 40.4 | 0.00354254 | 0.00354254 | ENSG00000266237 | n.s. |
| ENSG00000129484 | 902.9797 | -0.110164418 | 0.258675 | -0.425879 | 0.670196 | 0.928687131 | PARP2 | protein_coding | 42.36 | -0.003539646 | 0.003539646 | ENSG00000129484 | n.s. |
| ENSG00000163389 | 663.2953 | -0.110016743 | 0.25845 | -0.425679 | 0.670342 | 0.928765958 | POGLUT1 | protein_coding | 41.18 | -0.003530846 | 0.003530846 | ENSG00000163389 | n.s. |
| ENSG00000177853 | 195.5921 | 0.103900388 | 0.23614 | 0.4399944 | 0.659941 | 0.924786541 | ZNF518A | protein_coding | 38.02 | 0.003528301 | 0.003528301 | ENSG00000177853 | n.s. |
| ENSG00000239827 | 122.2797 | 0.117105575 | 0.284458 | 0.4116803 | 0.680574 | 0.933007979 | SUGT1P3 | cribed_unprocessed_pseudc | 40.63 | 0.003526593 | 0.003526593 | ENSG00000239827 | n.s. |
| ENSG00000182899 | 14432.89 | 0.126343429 | 0.323326 | 0.3907613 | 0.695974 | 0.937752951 | RPL35A | protein_coding | 44.91 | 0.003526442 | 0.003526442 | ENSG00000182899 | n.s. |
| ENSG00000176903 | 1088.368 | -0.108215677 | 0.250872 | -0.431358 | 0.666208 | 0.927727183 | PNMA1 | protein_coding | 53.75 | -0.003525636 | 0.003525636 | ENSG00000176903 | n.s. |
| ENSG00000198837 | 564.7003 | -0.105958966 | 0.243179 | -0.435724 | 0.663037 | 0.926265201 | DENND4B | protein_coding | 56.48 | -0.003524688 | 0.003524688 | ENSG00000198837 | n.s. |
| ENSG00000132429 | 1089.065 | 0.119804478 | 0.298824 | 0.4009197 | 0.688479 | 0.934506008 | POPDC3 | protein_coding | 39.7 | 0.003524396 | 0.003524396 | ENSG00000132429 | n.s. |
| ENSG00000176171 | 767.4271 | -0.1190705 | 0.293301 | -0.405967 | 0.684767 | 0.934117877 | BNIP3 | protein_coding | 49.42 | -0.003524286 | 0.003524286 | ENSG00000176171 | n.s. |
| ENSG00000115368 | 2807.577 | -0.11571525 | 0.278652 | -0.415267 | 0.677946 | 0.932323161 | WDR75 | protein_coding | 37.03 | -0.003521623 | 0.003521623 | ENSG00000115368 | n.s. |
| ENSG00000100918 | 951.2022 | -0.109076396 | 0.255187 | -0.427438 | 0.669061 | 0.928390061 | REC8 | protein_coding | 55.21 | -0.003519843 | 0.003519843 | ENSG00000100918 | n.s. |
| ENSG00000239335 | 32.08947 | -0.126469361 | 0.333966 | -0.37869 | 0.704918 | 0.93793739 | LLPH-AS1 | antisense | 44.42 | -0.003519156 | 0.003519156 | ENSG00000239335 | n.s. |
| ENSG00000180801 | 53.30905 | 0.124335163 | 0.316529 | 0.3928087 | 0.694461 | 0.936958261 | ARSJ | protein_coding | 36.4 | 0.003516168 | 0.003516168 | ENSG00000180801 | n.s. |
| ENSG00000004059 | 687.7239 | -0.119516 | 0.29878 | -0.400013 | 0.689147 | 0.934506008 | ARF5 | protein_coding | 56.5 | -0.00351591 | 0.00351591 | ENSG00000004059 | n.s. |
| ENSG00000007255 | 130.5875 | 0.125921241 | 0.327026 | 0.3850497 | 0.700201 | 0.937752951 | TRAPPC6A | protein_coding | 56.86 | 0.003514658 | 0.003514658 | ENSG00000007255 | n.s. |
| ENSG00000091527 | 3402.132 | -0.107704651 | 0.249508 | -0.431668 | 0.665983 | 0.927727183 | CDV3 | protein_coding | 42.4 | -0.003508987 | 0.003508987 | ENSG00000091527 | n.s. |
| ENSG00000177839 | 265.3449 | -0.104386867 | 0.239005 | -0.436757 | 0.662288 | 0.925519646 | PCDHB9 | protein_coding | 46.34 | -0.003508898 | 0.003508898 | ENSG00000177839 | n.s. |
| ENSG00000271147 | 170.2438 | -0.119232612 | 0.296037 | -0.402763 | 0.687123 | 0.934506008 | \RMCX5-GPRASP | processed_transcript | 39.81 | -0.003507573 | 0.003507573 | ENSG00000271147 | n.s. |
| ENSG00000174695 | 2276.813 | -0.121185078 | 0.305486 | -0.396696 | 0.691591 | 0.935578594 | TMEM167A | protein_coding | 35.95 | -0.003504639 | 0.003504639 | ENSG00000174695 | n.s. |
| ENSG00000132639 | 1146.111 | -0.116471121 | 0.283604 | -0.410683 | 0.681305 | 0.933075339 | SNAP25 | protein_coding | 40.89 | -0.003503834 | 0.003503834 | ENSG00000132639 | n.s. |
| ENSG00000178796 | 47.97045 | -0.133209244 | 0.372493 | -0.357616 | 0.720631 | 0.941256551 | RIIAD1 | protein_coding | 51.01 | -0.003502336 | 0.003502336 | ENSG00000178796 | n.s. |
| ENSG00000204344 | 135.4663 | -0.114280874 | 0.273688 | -0.417559 | 0.67627 | 0.931868404 | STK19 | protein_coding | 53.1 | -0.003502185 | 0.003502185 | ENSG00000204344 | n.s. |

| | | | | | | | | | | | | | |
|---|---|---|---|---|---|---|---|---|---|---|---|---|---|
| ENSG00000112357 | 253.489 | -0.115067836 | 0.278042 | -0.41385 | 0.678984 | 0.932323161 | PEX7 | protein_coding | 38.37 | -0.00350192 | 0.00350192 | ENSG00000112357 | n.s. |
| ENSG00000165997 | 4662.424 | 0.113743753 | 0.271973 | 0.4182175 | 0.675788 | 0.93167622 | ARL5B | protein_coding | 37.23 | 0.003495913 | 0.003495913 | ENSG00000165997 | n.s. |
| ENSG00000104164 | 2466.814 | -0.11461975 | 0.276029 | -0.415245 | 0.677962 | 0.932323161 | BLOC1S6 | protein_coding | 39.68 | -0.003488283 | 0.003488283 | ENSG00000104164 | n.s. |
| ENSG00000206897 | 23.65853 | -0.130924751 | 0.361174 | -0.362498 | 0.71698 | 0.94052839 | SNORA9B | snoRNA | 50.38 | -0.003486276 | 0.003486276 | ENSG00000206897 | n.s. |
| ENSG00000072832 | 4528.155 | -0.114452243 | 0.27639 | -0.414098 | 0.678803 | 0.932323161 | CRMP1 | protein_coding | 48.98 | -0.003483185 | 0.003483185 | ENSG00000072832 | n.s. |
| ENSG00000163728 | 2686.559 | 0.102556796 | 0.231117 | 0.4437433 | 0.657228 | 0.924786541 | TTC14 | protein_coding | 35.13 | 0.003482675 | 0.003482675 | ENSG00000163728 | n.s. |
| ENSG00000137154 | 30073.52 | 0.120288733 | 0.303405 | 0.3964621 | 0.691764 | 0.935578594 | RPS6 | protein_coding | 40.73 | 0.003478717 | 0.003478717 | ENSG00000137154 | n.s. |
| ENSG00000101773 | 1541.048 | -0.107992513 | 0.253228 | -0.426464 | 0.66977 | 0.92852823 | RBBP8 | protein_coding | 37.83 | -0.003477887 | 0.003477887 | ENSG00000101773 | n.s. |
| ENSG00000269246 | 9.416285 | 0.161283067 | 0.516653 | 0.3121691 | 0.754912 | 0.951579479 | | lincRNA | 56.07 | 0.003476446 | 0.003476446 | ENSG00000269246 | n.s. |
| ENSG00000162384 | 1384.391 | -0.100071403 | 0.218827 | -0.457309 | 0.647449 | 0.923184844 | C1orf123 | protein_coding | 46.92 | -0.003473612 | 0.003473612 | ENSG00000162384 | n.s. |
| ENSG00000107372 | 7894.033 | 0.100036732 | 0.221008 | 0.4526395 | 0.650808 | 0.923184844 | ZFAND5 | protein_coding | 39.89 | 0.003472408 | 0.003472408 | ENSG00000107372 | n.s. |
| ENSG00000105401 | 1708.969 | -0.116907466 | 0.286424 | -0.408162 | 0.683155 | 0.933907782 | CDC37 | protein_coding | 54.04 | -0.003471685 | 0.003471685 | ENSG00000105401 | n.s. |
| ENSG00000057757 | 407.0984 | -0.108168489 | 0.254438 | -0.425126 | 0.670745 | 0.928780923 | PITHD1 | protein_coding | 44.59 | -0.003470771 | 0.003470771 | ENSG00000057757 | n.s. |
| ENSG00000073921 | 952.9898 | -0.11403207 | 0.274035 | -0.416123 | 0.67732 | 0.932323161 | PICALM | protein_coding | 37.33 | -0.003470398 | 0.003470398 | ENSG00000073921 | n.s. |
| ENSG00000274303 | 15.12846 | 0.146657433 | 0.441393 | 0.3322606 | 0.739693 | 0.946996213 | | misc_RNA | 38.26 | 0.003468706 | 0.003468706 | ENSG00000274303 | n.s. |
| ENSG00000166548 | 22.34549 | -0.135445091 | 0.388976 | -0.34821 | 0.727683 | 0.942883666 | TK2 | protein_coding | 49.54 | -0.003459523 | 0.003459523 | ENSG00000166548 | n.s. |
| ENSG00000137312 | 923.7234 | -0.119938611 | 0.30286 | -0.39602 | 0.69209 | 0.935753589 | FLOT1 | protein_coding | 49.72 | -0.003458849 | 0.003458849 | ENSG00000137312 | n.s. |
| ENSG00000115307 | 2054.047 | -0.11114535 | 0.264765 | -0.419789 | 0.674639 | 0.931014301 | AUP1 | protein_coding | 55.6 | -0.003450357 | 0.003450357 | ENSG00000115307 | n.s. |
| ENSG00000165264 | 824.1095 | 0.123891283 | 0.326549 | 0.379396 | 0.704394 | 0.937896121 | NDUFB6 | protein_coding | 41.42 | 0.003449785 | 0.003449785 | ENSG00000165264 | n.s. |
| ENSG00000113811 | 789.5349 | 0.117260935 | 0.292565 | 0.4008037 | 0.688565 | 0.934506008 | SELENOK | protein_coding | 40.23 | 0.003449571 | 0.003449571 | ENSG00000113811 | n.s. |
| ENSG00000021574 | 1108.361 | -0.117228284 | 0.293718 | -0.399118 | 0.689806 | 0.934548254 | SPAST | protein_coding | 39.98 | -0.003446309 | 0.003446309 | ENSG00000021574 | n.s. |
| ENSG00000126267 | 1716.104 | 0.125601325 | 0.336 | 0.3738135 | 0.708543 | 0.938902143 | COX6B1 | protein_coding | 49.06 | 0.003438923 | 0.003438923 | ENSG00000126267 | n.s. |
| ENSG00000131174 | 2162.643 | 0.124547797 | 0.331821 | 0.3753464 | 0.707403 | 0.938471423 | COX7B | protein_coding | 40.81 | 0.003434897 | 0.003434897 | ENSG00000131174 | n.s. |
| ENSG00000089154 | 3304.723 | -0.105622528 | 0.24588 | -0.42957 | 0.667509 | 0.927899924 | GCN1 | protein_coding | 49.67 | -0.003432612 | 0.003432612 | ENSG00000089154 | n.s. |
| ENSG00000127054 | 351.8134 | -0.115931761 | 0.28522 | -0.406464 | 0.684401 | 0.934117877 | INTS11 | protein_coding | 60.46 | -0.003431385 | 0.003431385 | ENSG00000127054 | n.s. |
| ENSG00000180233 | 422.1148 | -0.115454038 | 0.283151 | -0.407747 | 0.68346 | 0.933907782 | ZNRF2 | protein_coding | 36.75 | -0.003428524 | 0.003428524 | ENSG00000180233 | n.s. |
| ENSG00000166582 | 3567.023 | -0.116444071 | 0.289845 | -0.401746 | 0.687871 | 0.934506008 | CENPV | protein_coding | 48.01 | -0.00342554 | 0.00342554 | ENSG00000166582 | n.s. |
| ENSG00000101460 | 64.50861 | -0.127412122 | 0.349336 | -0.364727 | 0.715315 | 0.940060506 | MAP1LC3A | protein_coding | 52.56 | -0.003420276 | 0.003420276 | ENSG00000101460 | n.s. |
| ENSG00000143815 | 3691.06 | -0.112732679 | 0.272744 | -0.413328 | 0.679366 | 0.932637768 | LBR | protein_coding | 41.69 | -0.003414335 | 0.003414335 | ENSG00000143815 | n.s. |
| ENSG00000260018 | 22.6319 | -0.139831662 | 0.412584 | -0.338917 | 0.734672 | 0.945347052 | | antisense | 65.64 | -0.003413113 | 0.003413113 | ENSG00000260018 | n.s. |
| ENSG00000173141 | 1023.361 | -0.106050395 | 0.248918 | -0.426046 | 0.670074 | 0.928672635 | MRPL57 | protein_coding | 50 | -0.003408179 | 0.003408179 | ENSG00000173141 | n.s. |
| ENSG00000151576 | 1209.897 | -0.108212123 | 0.256505 | -0.421871 | 0.673119 | 0.930111256 | QTRT2 | protein_coding | 38.31 | -0.003404905 | 0.003404905 | ENSG00000151576 | n.s. |

| | | | | | | | | | | | | | |
|---|---|---|---|---|---|---|---|---|---|---|---|---|---|
| ENSG00000150991 | 5471.998 | -0.115624374 | 0.28847 | -0.40082 | 0.688553 | 0.934506008 | UBC | protein_coding | 50.7 | -0.003401427 | 0.003401427 | ENSG00000150991 | n.s. |
| ENSG00000100711 | 204.0363 | -0.114859129 | 0.282567 | -0.406485 | 0.684387 | 0.934117877 | ZFYVE21 | protein_coding | 56.98 | -0.003399637 | 0.003399637 | ENSG00000100711 | n.s. |
| ENSG00000163577 | 405.1907 | -0.111639786 | 0.269625 | -0.414056 | 0.678833 | 0.932323161 | EIF5A2 | protein_coding | 38.29 | -0.003397592 | 0.003397592 | ENSG00000163577 | n.s. |
| ENSG00000142230 | 2551.806 | -0.115490206 | 0.288277 | -0.400623 | 0.688698 | 0.934506008 | SAE1 | protein_coding | 47.18 | -0.00339748 | 0.00339748 | ENSG00000142230 | n.s. |
| ENSG00000256747 | 23.65863 | 0.13297044 | 0.378561 | 0.3512519 | 0.725399 | 0.942883666 | | antisense | 73.54 | 0.003396316 | 0.003396316 | ENSG00000256747 | n.s. |
| ENSG00000198712 | 10975.16 | -0.099976222 | 0.225672 | -0.443016 | 0.657754 | 0.924786541 | MT-CO2 | protein_coding | 46.2 | -0.003395042 | 0.003395042 | ENSG00000198712 | n.s. |
| ENSG00000132388 | 1133.672 | -0.111435528 | 0.269088 | -0.414124 | 0.678784 | 0.932323161 | UBE2G1 | protein_coding | 41.77 | -0.003391376 | 0.003391376 | ENSG00000132388 | n.s. |
| ENSG00000143321 | 6324.031 | -0.115274746 | 0.286923 | -0.401762 | 0.687859 | 0.934506008 | HDGF | protein_coding | 51.35 | -0.003391141 | 0.003391141 | ENSG00000143321 | n.s. |
| ENSG00000175756 | 420.7783 | -0.11527201 | 0.28666 | -0.402121 | 0.687595 | 0.934506008 | AURKAIP1 | protein_coding | 66.87 | -0.003391061 | 0.003391061 | ENSG00000175756 | n.s. |
| ENSG00000161956 | 373.6365 | -0.119900822 | 0.305318 | -0.392707 | 0.694536 | 0.936958261 | SENP3 | protein_coding | 51.55 | -0.003390766 | 0.003390766 | ENSG00000161956 | n.s. |
| ENSG00000197696 | 46.21664 | -0.121411245 | 0.317118 | -0.382858 | 0.701825 | 0.937752951 | NMB | protein_coding | 56.33 | -0.003388777 | 0.003388777 | ENSG00000197696 | n.s. |
| ENSG00000144369 | 1327.512 | -0.109213971 | 0.260294 | -0.419579 | 0.674793 | 0.931111765 | FAM171B | protein_coding | 37.17 | -0.003385435 | 0.003385435 | ENSG00000144369 | n.s. |
| ENSG00000182628 | 4133.454 | -0.114341113 | 0.281714 | -0.405877 | 0.684833 | 0.934117877 | SKA2 | protein_coding | 41.24 | -0.003384304 | 0.003384304 | ENSG00000182628 | n.s. |
| ENSG00000151694 | 906.8162 | -0.110964264 | 0.266267 | -0.416741 | 0.676868 | 0.932241845 | ADAM17 | protein_coding | 41 | -0.003381237 | 0.003381237 | ENSG00000151694 | n.s. |
| ENSG00000123416 | 20787.39 | -0.114918371 | 0.287076 | -0.400306 | 0.688931 | 0.934506008 | TUBA1B | protein_coding | 53.12 | -0.003380657 | 0.003380657 | ENSG00000123416 | n.s. |
| ENSG00000164024 | 1428.885 | -0.112305014 | 0.273707 | -0.410311 | 0.681578 | 0.933075339 | METAP1 | protein_coding | 39.1 | -0.003378504 | 0.003378504 | ENSG00000164024 | n.s. |
| ENSG00000176978 | 1089.768 | -0.114423621 | 0.282677 | -0.404785 | 0.685635 | 0.934328291 | DPP7 | protein_coding | 68.7 | -0.003375554 | 0.003375554 | ENSG00000176978 | n.s. |
| ENSG00000140854 | 664.729 | -0.111778411 | 0.271223 | -0.412127 | 0.680246 | 0.932926007 | KATNB1 | protein_coding | 56.59 | -0.003370432 | 0.003370432 | ENSG00000140854 | n.s. |
| ENSG00000115392 | 1479.995 | -0.113283052 | 0.277999 | -0.407494 | 0.683645 | 0.933907782 | FANCL | protein_coding | 34.76 | -0.003364054 | 0.003364054 | ENSG00000115392 | n.s. |
| ENSG00000143353 | 603.7101 | -0.104537854 | 0.245277 | -0.426203 | 0.66996 | 0.92860681 | LYPLAL1 | protein_coding | 33.96 | -0.003362788 | 0.003362788 | ENSG00000143353 | n.s. |
| ENSG00000119403 | 1179.753 | -0.103437774 | 0.240843 | -0.429483 | 0.667572 | 0.927899924 | PHF19 | protein_coding | 51.61 | -0.00336161 | 0.00336161 | ENSG00000119403 | n.s. |
| ENSG00000136240 | 1257.601 | -0.114309312 | 0.286393 | -0.399134 | 0.689794 | 0.934548254 | KDELR2 | protein_coding | 46.8 | -0.003360496 | 0.003360496 | ENSG00000136240 | n.s. |
| ENSG00000174373 | 84.97365 | -0.11165078 | 0.271615 | -0.411063 | 0.681027 | 0.933075339 | RALGAPA1 | protein_coding | 36.27 | -0.003358823 | 0.003358823 | ENSG00000174373 | n.s. |
| ENSG00000153774 | 773.6128 | -0.110257873 | 0.265462 | -0.415343 | 0.677891 | 0.932323161 | CFDP1 | protein_coding | 42.3 | -0.003355536 | 0.003355536 | ENSG00000153774 | n.s. |
| ENSG00000243406 | 744.3515 | -0.098214271 | 0.219473 | -0.447501 | 0.654513 | 0.924413646 | MRPS31P5 | cribed_unprocessed_pseudc | 39.42 | -0.003352412 | 0.003352412 | ENSG00000243406 | n.s. |
| ENSG00000156831 | 208.8853 | -0.120089749 | 0.307669 | -0.390321 | 0.696299 | 0.937752951 | NSMCE2 | protein_coding | 39.89 | -0.003351892 | 0.003351892 | ENSG00000156831 | n.s. |
| ENSG00000198075 | 185.6382 | 0.098609928 | 0.221822 | 0.4445459 | 0.656648 | 0.924786541 | SULT1C4 | protein_coding | 40.61 | 0.003348645 | 0.003348645 | ENSG00000198075 | n.s. |
| ENSG00000100461 | 960.6063 | -0.113031832 | 0.27812 | -0.406413 | 0.684439 | 0.934117877 | RBM23 | protein_coding | 45.96 | -0.003345552 | 0.003345552 | ENSG00000100461 | n.s. |
| ENSG00000136044 | 888.7898 | -0.10991825 | 0.264696 | -0.415263 | 0.67795 | 0.932323161 | APPL2 | protein_coding | 43.65 | -0.0033452 | 0.0033452 | ENSG00000136044 | n.s. |
| ENSG00000184924 | 523.1264 | 0.115493576 | 0.290831 | 0.3971159 | 0.691282 | 0.935503831 | PTRHD1 | protein_coding | 45.88 | 0.003344051 | 0.003344051 | ENSG00000184924 | n.s. |
| ENSG00000175606 | 734.6785 | -0.110438506 | 0.267326 | -0.413123 | 0.679517 | 0.932670115 | TMEM70 | protein_coding | 43.8 | -0.003343188 | 0.003343188 | ENSG00000175606 | n.s. |
| ENSG00000120685 | 935.0447 | -0.104188325 | 0.245083 | -0.425115 | 0.670753 | 0.928780923 | PROSER1 | protein_coding | 37.33 | -0.003343061 | 0.003343061 | ENSG00000120685 | n.s. |

| | | | | | | | | | | | | | |
|---|---|---|---|---|---|---|---|---|---|---|---|---|---|
| ENSG00000225138 | 199.0088 | -0.113588917 | 0.283331 | -0.400905 | 0.68849 | 0.934506008 | SLC9A3-AS1 | processed_transcript | 65.93 | -0.003341548 | 0.003341548 | ENSG00000225138 | n.s. |
| ENSG00000066654 | 2362.093 | 0.104196058 | 0.245374 | 0.4246415 | 0.671098 | 0.92884421 | THUMPD1 | protein_coding | 40.19 | 0.003340226 | 0.003340226 | ENSG00000066654 | n.s. |
| ENSG00000186364 | 63.28923 | 0.119613842 | 0.312638 | 0.3825949 | 0.70202 | 0.937752951 | NUDT17 | protein_coding | 54.02 | 0.003338609 | 0.003338609 | ENSG00000186364 | n.s. |
| ENSG00000130985 | 1808.608 | -0.113154409 | 0.27936 | -0.405049 | 0.685441 | 0.934328291 | UBA1 | protein_coding | 51.31 | -0.003338112 | 0.003338112 | ENSG00000130985 | n.s. |
| ENSG00000102978 | 1407.492 | -0.110890915 | 0.270088 | -0.410573 | 0.681386 | 0.933075339 | POLR2C | protein_coding | 48.07 | -0.003335963 | 0.003335963 | ENSG00000102978 | n.s. |
| ENSG00000172009 | 659.8214 | -0.110824265 | 0.270284 | -0.410028 | 0.681785 | 0.933075339 | THOP1 | protein_coding | 59.11 | -0.003333958 | 0.003333958 | ENSG00000172009 | n.s. |
| ENSG00000100883 | 867.9466 | -0.113082597 | 0.280019 | -0.403839 | 0.686331 | 0.934368276 | SRP54 | protein_coding | 39.06 | -0.003333892 | 0.003333892 | ENSG00000100883 | n.s. |
| ENSG00000111716 | 28524.43 | 0.113265489 | 0.28328 | 0.3998356 | 0.689278 | 0.934506008 | LDHB | protein_coding | 38.01 | 0.003332033 | 0.003332033 | ENSG00000111716 | n.s. |
| ENSG00000184867 | 381.8282 | 0.119281175 | 0.306655 | 0.388975 | 0.697295 | 0.937752951 | ARMCX2 | protein_coding | 51.79 | 0.003329324 | 0.003329324 | ENSG00000184867 | n.s. |
| ENSG00000119720 | 350.0452 | 0.11237942 | 0.276157 | 0.4069397 | 0.684052 | 0.934117877 | NRDE2 | protein_coding | 44.18 | 0.003326242 | 0.003326242 | ENSG00000119720 | n.s. |
| ENSG00000071082 | 22791.6 | 0.119163417 | 0.307519 | 0.3874991 | 0.698387 | 0.937752951 | RPL31 | protein_coding | 42.84 | 0.003326037 | 0.003326037 | ENSG00000071082 | n.s. |
| ENSG00000256712 | 224.3627 | 0.102789947 | 0.240142 | 0.4280389 | 0.668623 | 0.928267143 | | sense_intronic | 41.28 | 0.003322893 | 0.003322893 | ENSG00000256712 | n.s. |
| ENSG00000117906 | 4979.434 | -0.119049765 | 0.305996 | -0.389056 | 0.697235 | 0.937752951 | RCN2 | protein_coding | 38.97 | -0.003322865 | 0.003322865 | ENSG00000117906 | n.s. |
| ENSG00000169519 | 136.2569 | -0.11292817 | 0.282518 | -0.39972 | 0.689363 | 0.934506008 | METTL15 | protein_coding | 36.8 | -0.00332211 | 0.00332211 | ENSG00000169519 | n.s. |
| ENSG00000105136 | 520.0233 | -0.102027103 | 0.237516 | -0.429558 | 0.667517 | 0.927899924 | ZNF419 | protein_coding | 46.6 | -0.003315765 | 0.003315765 | ENSG00000105136 | n.s. |
| ENSG00000102572 | 895.0037 | -0.104864365 | 0.248344 | -0.422255 | 0.672839 | 0.929819879 | STK24 | protein_coding | 47.42 | -0.003313837 | 0.003313837 | ENSG00000102572 | n.s. |
| ENSG00000072778 | 775.2088 | -0.110124517 | 0.268664 | -0.409897 | 0.681882 | 0.933075339 | ACADVL | protein_coding | 58.5 | -0.003312908 | 0.003312908 | ENSG00000072778 | n.s. |
| ENSG00000143575 | 1603.946 | -0.118806773 | 0.313124 | -0.379425 | 0.704373 | 0.937896121 | HAX1 | protein_coding | 49.78 | -0.003308206 | 0.003308206 | ENSG00000143575 | n.s. |
| ENSG00000168282 | 678.7124 | 0.097294097 | 0.21847 | 0.4453423 | 0.656072 | 0.924786541 | MGAT2 | protein_coding | 47.6 | 0.003303962 | 0.003303962 | ENSG00000168282 | n.s. |
| ENSG00000170892 | 427.5531 | -0.118624661 | 0.312331 | -0.379805 | 0.70409 | 0.937896121 | TSEN34 | protein_coding | 55.95 | -0.003303135 | 0.003303135 | ENSG00000170892 | n.s. |
| ENSG00000130413 | 195.6088 | -0.111868875 | 0.276339 | -0.404825 | 0.685606 | 0.934328291 | STK33 | protein_coding | 37.65 | -0.003300188 | 0.003300188 | ENSG00000130413 | n.s. |
| ENSG00000122390 | 80.84568 | 0.109685345 | 0.267671 | 0.4097774 | 0.681969 | 0.933083508 | NAA60 | protein_coding | 50.69 | 0.003299279 | 0.003299279 | ENSG00000122390 | n.s. |
| ENSG00000092201 | 6973.033 | -0.10950169 | 0.265867 | -0.411866 | 0.680438 | 0.932999624 | SUPT16H | protein_coding | 40.41 | -0.00329803 | 0.00329803 | ENSG00000092201 | n.s. |
| ENSG00000184445 | 5011.034 | -0.11309525 | 0.284156 | -0.398004 | 0.690627 | 0.935187772 | KNTC1 | protein_coding | 41.24 | -0.003291205 | 0.003291205 | ENSG00000184445 | n.s. |
| ENSG00000140463 | 420.9572 | -0.116669922 | 0.297681 | -0.391929 | 0.695111 | 0.937157772 | BBS4 | protein_coding | 40.76 | -0.003288609 | 0.003288609 | ENSG00000140463 | n.s. |
| ENSG00000275569 | 73.89956 | -0.111516484 | 0.27631 | -0.403593 | 0.686512 | 0.934438006 | | antisense | 32.5 | -0.003284105 | 0.003284105 | ENSG00000275569 | n.s. |
| ENSG00000125356 | 1001.543 | 0.123235065 | 0.339564 | 0.3629214 | 0.716664 | 0.94052839 | NDUFA1 | protein_coding | 43.57 | 0.003281515 | 0.003281515 | ENSG00000125356 | n.s. |
| ENSG00000160813 | 77.44221 | -0.117471597 | 0.306764 | -0.382938 | 0.701766 | 0.937752951 | PPP1R35 | protein_coding | 67.13 | -0.003278816 | 0.003278816 | ENSG00000160813 | n.s. |
| ENSG00000166794 | 3775.45 | -0.11743261 | 0.300828 | -0.390364 | 0.696267 | 0.937752951 | PPIB | protein_coding | 52.92 | -0.003277727 | 0.003277727 | ENSG00000166794 | n.s. |
| ENSG00000250600 | 4.344747 | -0.237036273 | 1.058021 | -0.224037 | 0.822728 | 0.968673589 | ROPN1L-AS1 | antisense | 57.06 | -0.003276444 | 0.003276444 | ENSG00000250600 | n.s. |
| ENSG00000137601 | 778.9234 | -0.108818993 | 0.265076 | -0.410521 | 0.681424 | 0.933075339 | NEK1 | protein_coding | 37.07 | -0.003273633 | 0.003273633 | ENSG00000137601 | n.s. |
| ENSG00000063660 | 810.9884 | -0.108921776 | 0.265962 | -0.409539 | 0.682144 | 0.933138975 | GPC1 | protein_coding | 64.63 | -0.003273499 | 0.003273499 | ENSG00000063660 | n.s. |

| | | | | | | | | | | | | | |
|---|---|---|---|---|---|---|---|---|---|---|---|---|---|
| ENSG00000108591 | 367.2899 | -0.117245772 | 0.300801 | -0.389778 | 0.696701 | 0.937752951 | DRG2 | protein_coding | 53.55 | -0.003272512 | 0.003272512 | ENSG00000108591 | n.s. |
| ENSG00000205339 | 6887.466 | 0.110520194 | 0.272159 | 0.4060872 | 0.684679 | 0.934117877 | IPO7 | protein_coding | 40.4 | 0.003271212 | 0.003271212 | ENSG00000205339 | n.s. |
| ENSG00000110811 | 397.8181 | -0.106736064 | 0.25564 | -0.417525 | 0.676295 | 0.931868404 | P3H3 | protein_coding | 56.26 | -0.003270971 | 0.003270971 | ENSG00000110811 | n.s. |
| ENSG00000139880 | 864.5678 | -0.10189268 | 0.239584 | -0.425291 | 0.670625 | 0.928780923 | CDH24 | protein_coding | 57.26 | -0.003269401 | 0.003269401 | ENSG00000139880 | n.s. |
| ENSG00000138326 | 27540.06 | 0.117317087 | 0.309172 | 0.3794562 | 0.704349 | 0.937896121 | RPS24 | protein_coding | 47.43 | 0.003266725 | 0.003266725 | ENSG00000138326 | n.s. |
| ENSG00000166780 | 459.9526 | -0.117396997 | 0.311337 | -0.377074 | 0.706119 | 0.937943012 | C16orf45 | protein_coding | 45.91 | -0.003266401 | 0.003266401 | ENSG00000166780 | n.s. |
| ENSG00000118705 | 3240.012 | -0.110308463 | 0.271571 | -0.406186 | 0.684606 | 0.934117877 | RPN2 | protein_coding | 44.36 | -0.003264945 | 0.003264945 | ENSG00000118705 | n.s. |
| ENSG00000103174 | 76.51713 | 0.115411003 | 0.293947 | 0.3926247 | 0.694597 | 0.936958261 | NAGPA | protein_coding | 56.14 | 0.003263795 | 0.003263795 | ENSG00000103174 | n.s. |
| ENSG00000225975 | 51.52818 | 0.116903437 | 0.306088 | 0.3819274 | 0.702515 | 0.937867182 | LINC01534 | lincRNA | 47.16 | 0.003256773 | 0.003256773 | ENSG00000225975 | n.s. |
| ENSG00000173221 | 267.1988 | 0.109961075 | 0.270884 | 0.4059349 | 0.68479 | 0.934117877 | GLRX | protein_coding | 41.1 | 0.003254663 | 0.003254663 | ENSG00000173221 | n.s. |
| ENSG00000172007 | 354.6407 | -0.109509807 | 0.268466 | -0.407909 | 0.683341 | 0.933907782 | RAB33B | protein_coding | 41.25 | -0.003252004 | 0.003252004 | ENSG00000172007 | n.s. |
| ENSG00000106333 | 1425.242 | -0.106851765 | 0.257675 | -0.414677 | 0.678378 | 0.932323161 | PCOLCE | protein_coding | 60.91 | -0.003251876 | 0.003251876 | ENSG00000106333 | n.s. |
| ENSG00000138785 | 413.5686 | 0.106793424 | 0.257549 | 0.4146521 | 0.678397 | 0.932323161 | INTS12 | protein_coding | 36.56 | 0.003250101 | 0.003250101 | ENSG00000138785 | n.s. |
| ENSG00000136536 | 3810.873 | -0.11041341 | 0.275263 | -0.401119 | 0.688332 | 0.934506008 | 7-Mar | protein_coding | 38.09 | -0.003248131 | 0.003248131 | ENSG00000136536 | n.s. |
| ENSG00000253739 | 15.93546 | 0.131759532 | 0.386725 | 0.3407064 | 0.733325 | 0.944876691 | RNU6-323P | snRNA | 42.06 | 0.003244561 | 0.003244561 | ENSG00000253739 | n.s. |
| ENSG00000017260 | 2643.53 | -0.106474382 | 0.256703 | -0.414777 | 0.678305 | 0.932323161 | ATP2C1 | protein_coding | 37.56 | -0.003240391 | 0.003240391 | ENSG00000017260 | n.s. |
| ENSG00000101084 | 331.2418 | -0.109282706 | 0.268381 | -0.407192 | 0.683867 | 0.93406267 | C20orf24 | protein_coding | 49.21 | -0.003237389 | 0.003237389 | ENSG00000101084 | n.s. |
| ENSG00000228624 | 40.97089 | 0.16436872 | 0.566304 | 0.2902481 | 0.771626 | 0.955688505 | HDAC2-AS2 | antisense | 37.94 | 0.003235374 | 0.003235374 | ENSG00000228624 | n.s. |
| ENSG00000154079 | 302.6896 | 0.108896552 | 0.267182 | 0.4075751 | 0.683586 | 0.933907782 | SDHAF4 | protein_coding | 40.87 | 0.003233793 | 0.003233793 | ENSG00000154079 | n.s. |
| ENSG00000085719 | 1158.767 | -0.116042882 | 0.30785 | -0.376946 | 0.706213 | 0.937943012 | CPNE3 | protein_coding | 37.62 | -0.003228725 | 0.003228725 | ENSG00000085719 | n.s. |
| ENSG00000101391 | 523.303 | -0.106041495 | 0.255373 | -0.415242 | 0.677965 | 0.932323161 | CDK5RAP1 | protein_coding | 44.4 | -0.003227217 | 0.003227217 | ENSG00000101391 | n.s. |
| ENSG00000187778 | 1509.62 | -0.109013978 | 0.268353 | -0.406233 | 0.684571 | 0.934117877 | MCRS1 | protein_coding | 55.12 | -0.00322663 | 0.00322663 | ENSG00000187778 | n.s. |
| ENSG00000102383 | 47.34984 | -0.115911682 | 0.306927 | -0.377653 | 0.705689 | 0.937943012 | ZDHHC15 | protein_coding | 38.32 | -0.003225074 | 0.003225074 | ENSG00000102383 | n.s. |
| ENSG00000159256 | 930.4084 | -0.106523541 | 0.258204 | -0.412557 | 0.679932 | 0.932670115 | MORC3 | protein_coding | 42.3 | -0.003224674 | 0.003224674 | ENSG00000159256 | n.s. |
| ENSG00000077348 | 478.1647 | 0.115284652 | 0.294724 | 0.3911613 | 0.695678 | 0.937650061 | EXOSC5 | protein_coding | 52.53 | 0.003223268 | 0.003223268 | ENSG00000077348 | n.s. |
| ENSG00000224885 | 63.89032 | -0.115428462 | 0.298848 | -0.386245 | 0.699315 | 0.937752951 | TSSC1-IT1 | sense_intronic | 47.97 | -0.003221788 | 0.003221788 | ENSG00000224885 | n.s. |
| ENSG00000174444 | 48680.04 | 0.115257312 | 0.295981 | 0.3894082 | 0.696974 | 0.937752951 | RPL4 | protein_coding | 42.06 | 0.003217011 | 0.003217011 | ENSG00000174444 | n.s. |
| ENSG00000164323 | 1672.422 | -0.113744316 | 0.289683 | -0.392651 | 0.694577 | 0.936958261 | CFAP97 | protein_coding | 41.18 | -0.003216661 | 0.003216661 | ENSG00000164323 | n.s. |
| ENSG00000170091 | 1011.554 | -0.115422636 | 0.30301 | -0.38092 | 0.703263 | 0.937867182 | | protein_coding | 45.73 | -0.00321552 | 0.00321552 | ENSG00000170091 | n.s. |
| ENSG00000139842 | 1353.327 | -0.106873657 | 0.260498 | -0.410267 | 0.68161 | 0.933075339 | CUL4A | protein_coding | 45.97 | -0.003215111 | 0.003215111 | ENSG00000139842 | n.s. |
| ENSG00000207375 | 3.996597 | 0.170083192 | 0.599888 | 0.2835249 | 0.776775 | 0.957426384 | SNORD116-23 | snoRNA | 44.57 | 0.003213655 | 0.003213655 | ENSG00000207375 | n.s. |
| ENSG00000122034 | 1680.976 | 0.117505345 | 0.31504 | 0.3729859 | 0.709159 | 0.938989999 | GTF3A | protein_coding | 44.92 | 0.003212483 | 0.003212483 | ENSG00000122034 | n.s. |

| | | | | | | | | | | | | | |
|---|---|---|---|---|---|---|---|---|---|---|---|---|---|
| ENSG00000169139 | 1980.468 | -0.118506977 | 0.319982 | -0.370355 | 0.711118 | 0.939526932 | UBE2V2 | protein_coding | 42.29 | -0.003210445 | 0.003210445 | ENSG00000169139 | n.s. |
| ENSG00000113742 | 68.19825 | -0.115183231 | 0.302375 | -0.380928 | 0.703256 | 0.937867182 | CPEB4 | protein_coding | 37.31 | -0.003208851 | 0.003208851 | ENSG00000113742 | n.s. |
| ENSG00000135245 | 894.2981 | -0.092344554 | 0.203227 | -0.454392 | 0.649547 | 0.923184844 | HILPDA | protein_coding | 51.52 | -0.003205403 | 0.003205403 | ENSG00000135245 | n.s. |
| ENSG00000259740 | 38.01889 | -0.118790659 | 0.321943 | -0.36898 | 0.712142 | 0.939817855 | | sense_intronic | 39.95 | -0.003202158 | 0.003202158 | ENSG00000259740 | n.s. |
| ENSG00000176731 | 793.615 | 0.113208454 | 0.288138 | 0.3928965 | 0.694396 | 0.936958261 | C8orf59 | protein_coding | 38.73 | 0.003201507 | 0.003201507 | ENSG00000176731 | n.s. |
| ENSG00000079616 | 2317.59 | -0.108512929 | 0.268158 | -0.40466 | 0.685728 | 0.934328291 | KIF22 | protein_coding | 50.19 | -0.003201186 | 0.003201186 | ENSG00000079616 | n.s. |
| ENSG00000138050 | 1636.279 | 0.108813725 | 0.271831 | 0.4002986 | 0.688937 | 0.934506008 | THUMPD2 | protein_coding | 36.6 | 0.003201071 | 0.003201071 | ENSG00000138050 | n.s. |
| ENSG00000153982 | 286.3723 | 0.114878566 | 0.301176 | 0.3814333 | 0.702882 | 0.937867182 | GDPD1 | protein_coding | 40.89 | 0.003200363 | 0.003200363 | ENSG00000153982 | n.s. |
| ENSG00000147421 | 488.6268 | -0.104587834 | 0.250774 | -0.41706 | 0.676635 | 0.932022724 | HMBOX1 | protein_coding | 37.72 | -0.003197616 | 0.003197616 | ENSG00000147421 | n.s. |
| ENSG00000227540 | 130.2338 | -0.106279427 | 0.259419 | -0.409682 | 0.682039 | 0.933087478 | | antisense | 42.27 | -0.003196634 | 0.003196634 | ENSG00000227540 | n.s. |
| ENSG00000121413 | 463.1354 | -0.114507159 | 0.297519 | -0.384874 | 0.700331 | 0.937752951 | ZSCAN18 | protein_coding | 50.18 | -0.003196073 | 0.003196073 | ENSG00000121413 | n.s. |
| ENSG00000104375 | 67.64132 | -0.125111035 | 0.356842 | -0.350606 | 0.725884 | 0.942883666 | STK3 | protein_coding | 39.36 | -0.003195572 | 0.003195572 | ENSG00000104375 | n.s. |
| ENSG00000133878 | 630.7501 | 0.114697798 | 0.30351 | 0.3779043 | 0.705502 | 0.937943012 | DUSP26 | protein_coding | 52.92 | 0.0031913 | 0.0031913 | ENSG00000133878 | n.s. |
| ENSG00000118508 | 88.44881 | -0.114219185 | 0.294006 | -0.388493 | 0.697652 | 0.937752951 | RAB32 | protein_coding | 39.47 | -0.003188036 | 0.003188036 | ENSG00000118508 | n.s. |
| ENSG00000199024 | 15.21335 | -0.141806559 | 0.439995 | -0.322292 | 0.747232 | 0.949642637 | MIR103A2 | miRNA | 44.87 | -0.00318211 | 0.00318211 | ENSG00000199024 | n.s. |
| ENSG00000128272 | 9918.057 | -0.107807953 | 0.265967 | -0.405344 | 0.685225 | 0.934324562 | ATF4 | protein_coding | 58.94 | -0.003180575 | 0.003180575 | ENSG00000128272 | n.s. |
| ENSG00000105523 | 35.19802 | -0.122374879 | 0.343692 | -0.35606 | 0.721796 | 0.941942395 | FAM83E | protein_coding | 56.4 | -0.003178768 | 0.003178768 | ENSG00000105523 | n.s. |
| ENSG00000105829 | 165.6825 | -0.11377334 | 0.293979 | -0.387012 | 0.698747 | 0.937752951 | BET1 | protein_coding | 35.67 | -0.003175591 | 0.003175591 | ENSG00000105829 | n.s. |
| ENSG00000182534 | 1398.602 | -0.113623569 | 0.295763 | -0.38417 | 0.700852 | 0.937752951 | MXRA7 | protein_coding | 52.39 | -0.003171411 | 0.003171411 | ENSG00000182534 | n.s. |
| ENSG00000100347 | 652.8125 | -0.11389858 | 0.302479 | -0.37655 | 0.706508 | 0.937943012 | SAMM50 | protein_coding | 47.18 | -0.003169063 | 0.003169063 | ENSG00000100347 | n.s. |
| ENSG00000116212 | 788.5131 | 0.113875356 | 0.300575 | 0.3788581 | 0.704793 | 0.93793739 | LRRC42 | protein_coding | 43.06 | 0.003168713 | 0.003168713 | ENSG00000116212 | n.s. |
| ENSG00000119414 | 964.463 | -0.110374965 | 0.2796 | -0.39476 | 0.69302 | 0.936061063 | PPP6C | protein_coding | 42.35 | -0.0031673 | 0.0031673 | ENSG00000119414 | n.s. |
| ENSG00000159592 | 167.263 | -0.105522002 | 0.257771 | -0.409364 | 0.682273 | 0.933223457 | GPBP1L1 | protein_coding | 41.19 | -0.003167175 | 0.003167175 | ENSG00000159592 | n.s. |
| ENSG00000115365 | 3363.015 | -0.113314607 | 0.292306 | -0.387658 | 0.698269 | 0.937752951 | LANCL1 | protein_coding | 38.12 | -0.003162788 | 0.003162788 | ENSG00000115365 | n.s. |
| ENSG00000198961 | 2420.026 | 0.107441606 | 0.267681 | 0.4013792 | 0.688141 | 0.934506008 | PJA2 | protein_coding | 37.66 | 0.003160707 | 0.003160707 | ENSG00000198961 | n.s. |
| ENSG00000136153 | 278.0617 | 0.113230172 | 0.292926 | 0.3865491 | 0.69909 | 0.937752951 | LMO7 | protein_coding | 38.67 | 0.003160431 | 0.003160431 | ENSG00000136153 | n.s. |
| ENSG00000237976 | 63.1626 | -0.115574298 | 0.309514 | -0.373406 | 0.708846 | 0.938989999 | | antisense | 51.56 | -0.00315969 | 0.00315969 | ENSG00000237976 | n.s. |
| ENSG00000188706 | 330.5647 | -0.103810016 | 0.250817 | -0.413887 | 0.678957 | 0.932323161 | ZDHHC9 | protein_coding | 43.4 | -0.003159305 | 0.003159305 | ENSG00000188706 | n.s. |
| ENSG00000099246 | 1295.585 | -0.107389427 | 0.267424 | -0.40157 | 0.688001 | 0.934506008 | RAB18 | protein_coding | 36.66 | -0.003159172 | 0.003159172 | ENSG00000099246 | n.s. |
| ENSG00000263389 | 11.54334 | -0.144228591 | 0.456771 | -0.315757 | 0.752187 | 0.950815756 | MIR3973 | miRNA | 46.73 | -0.00315913 | 0.00315913 | ENSG00000263389 | n.s. |
| ENSG00000154174 | 2366.902 | -0.10904781 | 0.274708 | -0.396959 | 0.691398 | 0.935503831 | TOMM70 | protein_coding | 38.61 | -0.003157417 | 0.003157417 | ENSG00000154174 | n.s. |
| ENSG00000184368 | 515.022 | -0.109384558 | 0.276071 | -0.396219 | 0.691944 | 0.935696596 | MAP7D2 | protein_coding | 44.44 | -0.00315738 | 0.00315738 | ENSG00000184368 | n.s. |

| | | | | | | | | | | | | | |
|---|---|---|---|---|---|---|---|---|---|---|---|---|---|
| ENSG00000132589 | 306.263 | -0.113001767 | 0.294798 | -0.383319 | 0.701483 | 0.937752951 | FLOT2 | protein_coding | 54.35 | -0.003154056 | 0.003154056 | ENSG00000132589 | n.s. |
| ENSG00000145730 | 282.4071 | 0.112880435 | 0.290157 | 0.3890321 | 0.697252 | 0.937752951 | PAM | protein_coding | 35.92 | 0.003150669 | 0.003150669 | ENSG00000145730 | n.s. |
| ENSG00000260947 | 191.495 | -0.11438716 | 0.305423 | -0.374521 | 0.708017 | 0.938580742 | | lincRNA | 38.24 | -0.003148891 | 0.003148891 | ENSG00000260947 | n.s. |
| ENSG00000065600 | 722.7492 | 0.115848065 | 0.311415 | 0.3720053 | 0.709889 | 0.939338716 | TMEM206 | protein_coding | 44.57 | 0.003148493 | 0.003148493 | ENSG00000065600 | n.s. |
| ENSG00000196652 | 606.7167 | -0.106269597 | 0.26192 | -0.405732 | 0.684939 | 0.934117877 | ZKSCAN5 | protein_coding | 46.16 | -0.003145401 | 0.003145401 | ENSG00000196652 | n.s. |
| ENSG00000168028 | 16926 | 0.11267089 | 0.2919 | 0.3859915 | 0.699503 | 0.937752951 | RPSA | protein_coding | 45.15 | 0.00314482 | 0.00314482 | ENSG00000168028 | n.s. |
| ENSG00000114902 | 671.2864 | -0.113006867 | 0.300144 | -0.376509 | 0.706538 | 0.937943012 | SPCS1 | protein_coding | 47.37 | -0.003144252 | 0.003144252 | ENSG00000114902 | n.s. |
| ENSG00000170633 | 999.8676 | -0.108106381 | 0.27189 | -0.39761 | 0.690918 | 0.935225433 | RNF34 | protein_coding | 43.68 | -0.003144133 | 0.003144133 | ENSG00000170633 | n.s. |
| ENSG00000104671 | 782.2519 | -0.11259309 | 0.291972 | -0.385629 | 0.699771 | 0.937752951 | DCTN6 | protein_coding | 41.56 | -0.003142649 | 0.003142649 | ENSG00000104671 | n.s. |
| ENSG00000278013 | 19.82423 | -0.136194963 | 0.415509 | -0.327778 | 0.743079 | 0.948261154 | | lincRNA | 45.68 | -0.003142296 | 0.003142296 | ENSG00000278013 | n.s. |
| ENSG00000198585 | 1053.147 | 0.104415716 | 0.254212 | 0.410742 | 0.681262 | 0.933075339 | NUDT16 | protein_coding | 48.83 | 0.003141168 | 0.003141168 | ENSG00000198585 | n.s. |
| ENSG00000100321 | 317.1256 | 0.112502275 | 0.290038 | 0.387888 | 0.698099 | 0.937752951 | SYNGR1 | protein_coding | 55.76 | 0.003140114 | 0.003140114 | ENSG00000100321 | n.s. |
| ENSG00000133247 | 241.6937 | -0.111247403 | 0.283821 | -0.391963 | 0.695086 | 0.937157772 | KMT5C | protein_coding | 64.88 | -0.003135763 | 0.003135763 | ENSG00000133247 | n.s. |
| ENSG00000184752 | 1385.645 | 0.114627536 | 0.306803 | 0.373619 | 0.708688 | 0.938989999 | NDUFA12 | protein_coding | 43.14 | 0.003133806 | 0.003133806 | ENSG00000184752 | n.s. |
| ENSG00000167525 | 63.81134 | -0.112138715 | 0.288727 | -0.388391 | 0.697727 | 0.937752951 | PROCA1 | protein_coding | 55.73 | -0.003129967 | 0.003129967 | ENSG00000167525 | n.s. |
| ENSG00000170153 | 1696.326 | -0.102104494 | 0.244826 | -0.41705 | 0.676642 | 0.932022724 | RNF150 | protein_coding | 38.46 | -0.003121692 | 0.003121692 | ENSG00000170153 | n.s. |
| ENSG00000188599 | 80.81837 | -0.111681691 | 0.292042 | -0.382416 | 0.702153 | 0.937752951 | NPIPP1 | cribed_unprocessed_pseudc | 45.29 | -0.00311721 | 0.00311721 | ENSG00000188599 | n.s. |
| ENSG00000175634 | 452.0735 | -0.111676083 | 0.289914 | -0.385205 | 0.700086 | 0.937752951 | RPS6KB2 | protein_coding | 58.79 | -0.003117054 | 0.003117054 | ENSG00000175634 | n.s. |
| ENSG00000112559 | 139.2821 | -0.121992184 | 0.349759 | -0.348789 | 0.727247 | 0.942883666 | MDFI | protein_coding | 58.13 | -0.003115911 | 0.003115911 | ENSG00000112559 | n.s. |
| ENSG00000117394 | 618.1162 | -0.114798364 | 0.309254 | -0.37121 | 0.710481 | 0.939449151 | SLC2A1 | protein_coding | 52.89 | -0.003114103 | 0.003114103 | ENSG00000117394 | n.s. |
| ENSG00000106829 | 2452.434 | -0.095162315 | 0.219853 | -0.432846 | 0.665127 | 0.927447429 | TLE4 | protein_coding | 39.28 | -0.003112826 | 0.003112826 | ENSG00000106829 | n.s. |
| ENSG00000140600 | 282.6251 | 0.111818552 | 0.295661 | 0.3781983 | 0.705283 | 0.937943012 | SH3GL3 | protein_coding | 40.8 | 0.003111189 | 0.003111189 | ENSG00000140600 | n.s. |
| ENSG00000104960 | 1377.482 | -0.104023946 | 0.254376 | -0.408937 | 0.682586 | 0.933468077 | PTOV1 | protein_coding | 61.87 | -0.003110371 | 0.003110371 | ENSG00000104960 | n.s. |
| ENSG00000161653 | 92.63926 | -0.105679987 | 0.26355 | -0.400987 | 0.68843 | 0.934506008 | NAGS | protein_coding | 63.61 | -0.003108884 | 0.003108884 | ENSG00000161653 | n.s. |
| ENSG00000213551 | 2817.913 | -0.100827928 | 0.240938 | -0.418481 | 0.675595 | 0.931512729 | DNAJC9 | protein_coding | 40 | -0.00310663 | 0.00310663 | ENSG00000213551 | n.s. |
| ENSG00000100614 | 2201.613 | -0.107950457 | 0.272948 | -0.395499 | 0.692475 | 0.935906213 | PPM1A | protein_coding | 36.08 | -0.003105483 | 0.003105483 | ENSG00000100614 | n.s. |
| ENSG00000171311 | 585.502 | -0.104557918 | 0.256444 | -0.407722 | 0.683478 | 0.933907782 | EXOSC1 | protein_coding | 46.3 | -0.003104953 | 0.003104953 | ENSG00000171311 | n.s. |
| ENSG00000117691 | 873.027 | 0.115282017 | 0.314439 | 0.3666271 | 0.713897 | 0.939884276 | NENF | protein_coding | 47.72 | 0.003104039 | 0.003104039 | ENSG00000117691 | n.s. |
| ENSG00000233844 | 324.4675 | 0.107887195 | 0.272732 | 0.39558 | 0.692415 | 0.935906213 | KCNQ5-IT1 | sense_intronic | 36.35 | 0.003103663 | 0.003103663 | ENSG00000233844 | n.s. |
| ENSG00000164347 | 1211.37 | -0.112636663 | 0.300875 | -0.374364 | 0.708133 | 0.938580742 | GFM2 | protein_coding | 36.42 | -0.003100703 | 0.003100703 | ENSG00000164347 | n.s. |
| ENSG00000120802 | 15018.38 | -0.111042859 | 0.286448 | -0.387654 | 0.698272 | 0.937752951 | TMPO | protein_coding | 39.75 | -0.003099379 | 0.003099379 | ENSG00000120802 | n.s. |
| ENSG00000163607 | 569.225 | 0.102963542 | 0.250246 | 0.4114492 | 0.680743 | 0.933075339 | GTPBP8 | protein_coding | 37.14 | 0.003097482 | 0.003097482 | ENSG00000163607 | n.s. |

| | | | | | | | | | | | | | |
|---|---|---|---|---|---|---|---|---|---|---|---|---|---|
| ENSG00000185000 | 209.8552 | 0.107789046 | 0.272739 | 0.395209 | 0.692689 | 0.935976757 | DGAT1 | protein_coding | 63.02 | 0.003097311 | 0.003097311 | ENSG00000185000 | n.s. |
| ENSG00000089737 | 5094.143 | -0.101653721 | 0.24554 | -0.414001 | 0.678874 | 0.932323161 | DDX24 | protein_coding | 43.37 | -0.003093681 | 0.003093681 | ENSG00000089737 | n.s. |
| ENSG00000197756 | 35729.28 | 0.110831857 | 0.286988 | 0.3861902 | 0.699356 | 0.937752951 | RPL37A | protein_coding | 44.46 | 0.00309349 | 0.00309349 | ENSG00000197756 | n.s. |
| ENSG00000128652 | 237.0881 | -0.098237847 | 0.232957 | -0.421699 | 0.673245 | 0.930111256 | HOXD3 | protein_coding | 51.11 | -0.003091064 | 0.003091064 | ENSG00000128652 | n.s. |
| ENSG00000272764 | 207.889 | 0.099966365 | 0.238638 | 0.4189042 | 0.675286 | 0.931353024 | | lincRNA | 46.83 | 0.003087528 | 0.003087528 | ENSG00000272764 | n.s. |
| ENSG00000145191 | 1814.111 | 0.101975787 | 0.246919 | 0.4129937 | 0.679611 | 0.932670115 | EIF2B5 | protein_coding | 48.78 | 0.003087005 | 0.003087005 | ENSG00000145191 | n.s. |
| ENSG00000238998 | 21.62708 | -0.127173712 | 0.377409 | -0.336965 | 0.736143 | 0.945680734 | RNU7-187P | snRNA | 29.41 | -0.003084657 | 0.003084657 | ENSG00000238998 | n.s. |
| ENSG00000239883 | 502.5049 | -0.105441085 | 0.264647 | -0.398421 | 0.69032 | 0.934862275 | PARGP1 | cribed_unprocessed_pseudc | 38.95 | -0.003084401 | 0.003084401 | ENSG00000239883 | n.s. |
| ENSG00000089876 | 327.2623 | 0.117056687 | 0.32688 | 0.3581034 | 0.720266 | 0.941256551 | DHX32 | protein_coding | 41.45 | 0.003077653 | 0.003077653 | ENSG00000089876 | n.s. |
| ENSG00000117682 | 58.57044 | -0.111764125 | 0.298124 | -0.374891 | 0.707741 | 0.938571345 | DHDDS | protein_coding | 45.47 | -0.003077169 | 0.003077169 | ENSG00000117682 | n.s. |
| ENSG00000184634 | 536.5962 | -0.103550029 | 0.254113 | -0.407497 | 0.683643 | 0.933907782 | MED12 | protein_coding | 51.08 | -0.003075022 | 0.003075022 | ENSG00000184634 | n.s. |
| ENSG00000253738 | 1177.088 | 0.096191081 | 0.227102 | 0.4235589 | 0.671888 | 0.929142491 | OTUD6B-AS1 | antisense | 35.41 | 0.003070196 | 0.003070196 | ENSG00000253738 | n.s. |
| ENSG00000103051 | 747.0666 | 0.109927286 | 0.283937 | 0.3871534 | 0.698643 | 0.937752951 | COG4 | protein_coding | 47.24 | 0.003068242 | 0.003068242 | ENSG00000103051 | n.s. |
| ENSG00000169689 | 627.5693 | 0.118248532 | 0.334092 | 0.35394 | 0.723384 | 0.942009991 | CENPX | protein_coding | 65.43 | 0.003067898 | 0.003067898 | ENSG00000169689 | n.s. |
| ENSG00000181610 | 2285.984 | -0.11013192 | 0.290166 | -0.379548 | 0.704281 | 0.937896121 | MRPS23 | protein_coding | 44.44 | -0.003066652 | 0.003066652 | ENSG00000181610 | n.s. |
| ENSG00000187109 | 17505.23 | -0.104762644 | 0.262876 | -0.398525 | 0.690243 | 0.934849816 | NAP1L1 | protein_coding | 37.2 | -0.003065161 | 0.003065161 | ENSG00000187109 | n.s. |
| ENSG00000112304 | 166.7588 | 0.112086461 | 0.300865 | 0.372548 | 0.709485 | 0.939005324 | ACOT13 | protein_coding | 41.43 | 0.003063541 | 0.003063541 | ENSG00000112304 | n.s. |
| ENSG00000136271 | 1372.178 | -0.10397964 | 0.259056 | -0.401379 | 0.688141 | 0.934506008 | DDX56 | protein_coding | 53.74 | -0.003058863 | 0.003058863 | ENSG00000136271 | n.s. |
| ENSG00000256658 | 66.99326 | -0.116127749 | 0.323531 | -0.358938 | 0.719641 | 0.941153871 | | antisense | 51.61 | -0.003058731 | 0.003058731 | ENSG00000256658 | n.s. |
| ENSG00000118855 | 673.494 | -0.109742577 | 0.288547 | -0.380329 | 0.703701 | 0.937896121 | MFSD1 | protein_coding | 40.83 | -0.003055811 | 0.003055811 | ENSG00000118855 | n.s. |
| ENSG00000269971 | 11.07324 | 0.153451682 | 0.517825 | 0.2963391 | 0.766971 | 0.955198262 | | sense_intronic | 41.85 | 0.003054682 | 0.003054682 | ENSG00000269971 | n.s. |
| ENSG00000156411 | 1787.275 | 0.114887219 | 0.318101 | 0.3611658 | 0.717975 | 0.940629569 | C14orf2 | protein_coding | 47.77 | 0.00305386 | 0.00305386 | ENSG00000156411 | n.s. |
| ENSG00000107175 | 217.0164 | 0.109188885 | 0.284881 | 0.3832786 | 0.701513 | 0.937752951 | CREB3 | protein_coding | 49.01 | 0.003047632 | 0.003047632 | ENSG00000107175 | n.s. |
| ENSG00000176049 | 1061.61 | -0.102574164 | 0.251268 | -0.408225 | 0.683108 | 0.933907782 | JAKMIP2 | protein_coding | 38.29 | -0.003046043 | 0.003046043 | ENSG00000176049 | n.s. |
| ENSG00000147854 | 2376.124 | -0.099996201 | 0.240563 | -0.415676 | 0.677647 | 0.932323161 | UHRF2 | protein_coding | 38.99 | -0.003043237 | 0.003043237 | ENSG00000147854 | n.s. |
| ENSG00000135250 | 966.2506 | -0.10325316 | 0.255839 | -0.403587 | 0.686516 | 0.934438006 | SRPK2 | protein_coding | 41.06 | -0.003040755 | 0.003040755 | ENSG00000135250 | n.s. |
| ENSG00000164938 | 513.7976 | -0.10746725 | 0.273256 | -0.393284 | 0.69411 | 0.936958261 | TP53INP1 | protein_coding | 42.53 | -0.003039148 | 0.003039148 | ENSG00000164938 | n.s. |
| ENSG00000163754 | 243.6144 | 0.108853725 | 0.284141 | 0.3830982 | 0.701647 | 0.937752951 | GYG1 | protein_coding | 42.09 | 0.003038277 | 0.003038277 | ENSG00000163754 | n.s. |
| ENSG00000270061 | 78.64173 | -0.109093551 | 0.287554 | -0.379385 | 0.704402 | 0.937896121 | | sense_intronic | 45.63 | -0.003037738 | 0.003037738 | ENSG00000270061 | n.s. |
| ENSG00000164038 | 282.0129 | -0.103161423 | 0.256653 | -0.401949 | 0.687722 | 0.934506008 | SLC9B2 | protein_coding | 37.26 | -0.003034793 | 0.003034793 | ENSG00000164038 | n.s. |
| ENSG00000100479 | 699.527 | -0.108706908 | 0.284196 | -0.382506 | 0.702086 | 0.937752951 | POLE2 | protein_coding | 41.33 | -0.003034179 | 0.003034179 | ENSG00000100479 | n.s. |
| ENSG00000155438 | 2004.522 | -0.108779779 | 0.285868 | -0.380525 | 0.703556 | 0.937896121 | NIFK | protein_coding | 43.01 | -0.003029001 | 0.003029001 | ENSG00000155438 | n.s. |

| | | | | | | | | | | | | | |
|---|---|---|---|---|---|---|---|---|---|---|---|---|---|
| ENSG00000105372 | 14656.29 | 0.114619478 | 0.318544 | 0.3598232 | 0.718979 | 0.940969272 | RPS19 | protein_coding | 53.52 | 0.003028769 | 0.003028769 | ENSG00000105372 | n.s. |
| ENSG00000138413 | 2019.164 | -0.108425026 | 0.282987 | -0.383145 | 0.701612 | 0.937752951 | IDH1 | protein_coding | 41.27 | -0.003026312 | 0.003026312 | ENSG00000138413 | n.s. |
| ENSG00000042753 | 487.0165 | -0.111621573 | 0.301673 | -0.370008 | 0.711376 | 0.939526932 | AP2S1 | protein_coding | 52.87 | -0.003023914 | 0.003023914 | ENSG00000042753 | n.s. |
| ENSG00000100596 | 793.4814 | -0.10235538 | 0.252767 | -0.404939 | 0.685522 | 0.934328291 | SPTLC2 | protein_coding | 41.35 | -0.003019535 | 0.003019535 | ENSG00000100596 | n.s. |
| ENSG00000143753 | 1985.366 | 0.108152508 | 0.277509 | 0.3897257 | 0.696739 | 0.937752951 | DEGS1 | protein_coding | 46.06 | 0.003018705 | 0.003018705 | ENSG00000143753 | n.s. |
| ENSG00000281842 | 3.277568 | 0.197704414 | 0.803937 | 0.2459203 | 0.805744 | 0.9654664 | MIR1291 | miRNA | 51.72 | 0.00301753 | 0.00301753 | ENSG00000281842 | n.s. |
| ENSG00000102409 | 1342.143 | -0.106695863 | 0.271684 | -0.392721 | 0.694526 | 0.936958261 | BEX4 | protein_coding | 52.02 | -0.003017333 | 0.003017333 | ENSG00000102409 | n.s. |
| ENSG00000175137 | 614.0476 | -0.108086923 | 0.2816 | -0.383832 | 0.701103 | 0.937752951 | SH3BP5L | protein_coding | 52.31 | -0.003016875 | 0.003016875 | ENSG00000175137 | n.s. |
| ENSG00000137500 | 775.4294 | -0.108386972 | 0.285843 | -0.379183 | 0.704552 | 0.93793739 | CCDC90B | protein_coding | 37.59 | -0.003015992 | 0.003015992 | ENSG00000137500 | n.s. |
| ENSG00000131100 | 972.7552 | -0.113437722 | 0.314038 | -0.361223 | 0.717933 | 0.940629569 | ATP6V1E1 | protein_coding | 45.05 | -0.00301533 | 0.00301533 | ENSG00000131100 | n.s. |
| ENSG00000100162 | 608.0678 | -0.093396082 | 0.218603 | -0.42724 | 0.669205 | 0.928390061 | CENPM | protein_coding | 53.29 | -0.003013847 | 0.003013847 | ENSG00000100162 | n.s. |
| ENSG00000155111 | 1201.496 | -0.102036926 | 0.252028 | -0.404863 | 0.685578 | 0.934328291 | CDK19 | protein_coding | 39.87 | -0.00301014 | 0.00301014 | ENSG00000155111 | n.s. |
| ENSG00000158623 | 466.9056 | -0.107802327 | 0.28028 | -0.384623 | 0.700516 | 0.937752951 | COPG2 | protein_coding | 40.91 | -0.003008931 | 0.003008931 | ENSG00000158623 | n.s. |
| ENSG00000100029 | 1823.449 | -0.106312967 | 0.270764 | -0.392641 | 0.694584 | 0.936958261 | PES1 | protein_coding | 50.48 | -0.003006505 | 0.003006505 | ENSG00000100029 | n.s. |
| ENSG00000163376 | 398.7942 | -0.107461749 | 0.275885 | -0.389517 | 0.696894 | 0.937752951 | KBTBD8 | protein_coding | 36.69 | -0.002999425 | 0.002999425 | ENSG00000163376 | n.s. |
| ENSG00000083457 | 274.1007 | -0.113126776 | 0.313948 | -0.360336 | 0.718596 | 0.940842655 | ITGAE | protein_coding | 50.88 | -0.002995937 | 0.002995937 | ENSG00000083457 | n.s. |
| ENSG00000068903 | 65.22548 | -0.115274222 | 0.32441 | -0.355335 | 0.722339 | 0.941942395 | SIRT2 | protein_coding | 53.55 | -0.002994324 | 0.002994324 | ENSG00000068903 | n.s. |
| ENSG00000275453 | 7.870379 | -0.14900505 | 0.500089 | -0.297957 | 0.765736 | 0.954823343 | MIR548O2 | miRNA | 41.43 | -0.00299157 | 0.00299157 | ENSG00000275453 | n.s. |
| ENSG00000115504 | 149.0317 | -0.106743895 | 0.272686 | -0.391454 | 0.695462 | 0.93752693 | EHBP1 | protein_coding | 36.7 | -0.002990564 | 0.002990564 | ENSG00000115504 | n.s. |
| ENSG00000139726 | 1902.596 | -0.107017447 | 0.277824 | -0.385199 | 0.70009 | 0.937752951 | DENR | protein_coding | 42.12 | -0.002987024 | 0.002987024 | ENSG00000139726 | n.s. |
| ENSG00000111358 | 872.0624 | -0.110917958 | 0.302892 | -0.366196 | 0.714218 | 0.939885098 | GTF2H3 | protein_coding | 42.84 | -0.002986492 | 0.002986492 | ENSG00000111358 | n.s. |
| ENSG00000137760 | 500.7899 | -0.104731707 | 0.265643 | -0.394257 | 0.693391 | 0.936471375 | ALKBH8 | protein_coding | 36.18 | -0.002985429 | 0.002985429 | ENSG00000137760 | n.s. |
| ENSG00000114439 | 1644.632 | 0.106953241 | 0.280829 | 0.3808478 | 0.703316 | 0.937867182 | BBX | protein_coding | 36.95 | 0.002979574 | 0.002979574 | ENSG00000114439 | n.s. |
| ENSG00000105556 | 67.96688 | 0.106750517 | 0.275833 | 0.3870108 | 0.698748 | 0.937752951 | MIER2 | protein_coding | 53.11 | 0.002979574 | 0.002979574 | ENSG00000105556 | n.s. |
| ENSG00000132522 | 392.8501 | -0.100634429 | 0.247999 | -0.405786 | 0.6849 | 0.934117877 | GPS2 | protein_coding | 56.28 | -0.00297861 | 0.00297861 | ENSG00000132522 | n.s. |
| ENSG00000011009 | 194.565 | -0.106660951 | 0.276663 | -0.385526 | 0.699848 | 0.937752951 | LYPLA2 | protein_coding | 62.39 | -0.002977074 | 0.002977074 | ENSG00000011009 | n.s. |
| ENSG00000200354 | 101.1622 | 0.100734207 | 0.248396 | 0.4055393 | 0.685081 | 0.934219886 | SNORA71D | snoRNA | 53.73 | 0.002976786 | 0.002976786 | ENSG00000200354 | n.s. |
| ENSG00000148700 | 824.8692 | -0.106939929 | 0.282338 | -0.378765 | 0.704862 | 0.93793739 | ADD3 | protein_coding | 38.65 | -0.002975727 | 0.002975727 | ENSG00000148700 | n.s. |
| ENSG00000151491 | 122.0755 | -0.106447278 | 0.274034 | -0.388445 | 0.697686 | 0.937752951 | EPS8 | protein_coding | 37.25 | -0.00297111 | 0.00297111 | ENSG00000151491 | n.s. |
| ENSG00000166326 | 2697.624 | -0.103456237 | 0.262044 | -0.394804 | 0.692987 | 0.936061063 | TRIM44 | protein_coding | 39.96 | -0.002968762 | 0.002968762 | ENSG00000166326 | n.s. |
| ENSG00000204920 | 93.4486 | -0.113083555 | 0.316502 | -0.357291 | 0.720874 | 0.941354863 | ZNF155 | protein_coding | 40.41 | -0.002968062 | 0.002968062 | ENSG00000204920 | n.s. |
| ENSG00000161203 | 2086.356 | -0.106374931 | 0.27886 | -0.381463 | 0.70286 | 0.937867182 | AP2M1 | protein_coding | 54.13 | -0.002963463 | 0.002963463 | ENSG00000161203 | n.s. |

| | | | | | | | | | | | | | |
|---|---|---|---|---|---|---|---|---|---|---|---|---|---|
| ENSG00000233369 | 252.0387 | 0.111892878 | 0.310677 | 0.3601585 | 0.718729 | 0.940842655 | GTF2IP4 | ıscribed_processed_pseudog | 46.79 | 0.002963259 | 0.002963259 | ENSG00000233369 | n.s. |
| ENSG00000011677 | 289.5918 | 0.108221589 | 0.289922 | 0.373278 | 0.708942 | 0.938989999 | GABRA3 | protein_coding | 37.36 | 0.002958674 | 0.002958674 | ENSG00000011677 | n.s. |
| ENSG00000136436 | 104.705 | 0.110199183 | 0.302477 | 0.3643227 | 0.715617 | 0.940060506 | CALCOCO2 | protein_coding | 43.98 | 0.002958208 | 0.002958208 | ENSG00000136436 | n.s. |
| ENSG00000177683 | 890.5395 | -0.099600676 | 0.244075 | -0.408074 | 0.68322 | 0.933907782 | THAP5 | protein_coding | 39.2 | -0.002957742 | 0.002957742 | ENSG00000177683 | n.s. |
| ENSG00000159228 | 509.5588 | -0.10983532 | 0.29944 | -0.366803 | 0.713766 | 0.939884276 | CBR1 | protein_coding | 50.62 | -0.002957384 | 0.002957384 | ENSG00000159228 | n.s. |
| ENSG00000167766 | 713.2714 | -0.105855836 | 0.275535 | -0.384182 | 0.700843 | 0.937752951 | ZNF83 | protein_coding | 44.66 | -0.002954602 | 0.002954602 | ENSG00000167766 | n.s. |
| ENSG00000070081 | 297.5591 | -0.109052278 | 0.294356 | -0.370477 | 0.711027 | 0.939526932 | NUCB2 | protein_coding | 41.44 | -0.00295431 | 0.00295431 | ENSG00000070081 | n.s. |
| ENSG00000133706 | 5953.556 | -0.105815103 | 0.274389 | -0.38564 | 0.699764 | 0.937752951 | LARS | protein_coding | 38.69 | -0.002953465 | 0.002953465 | ENSG00000133706 | n.s. |
| ENSG00000157456 | 2217.635 | 0.109137024 | 0.295265 | 0.3696236 | 0.711663 | 0.939638057 | CCNB2 | protein_coding | 42.64 | 0.002951 | 0.002951 | ENSG00000157456 | n.s. |
| ENSG00000145241 | 1652.01 | -0.101409745 | 0.255054 | -0.3976 | 0.690925 | 0.935225433 | CENPC | protein_coding | 36.85 | -0.00294937 | 0.00294937 | ENSG00000145241 | n.s. |
| ENSG00000117222 | 926.4673 | -0.09796063 | 0.239 | -0.409877 | 0.681896 | 0.933075339 | RBBP5 | protein_coding | 40.77 | -0.002946978 | 0.002946978 | ENSG00000117222 | n.s. |
| ENSG00000055163 | 48.4368 | 0.109400647 | 0.298806 | 0.3661255 | 0.714271 | 0.939885098 | CYFIP2 | protein_coding | 44.29 | 0.002945638 | 0.002945638 | ENSG00000055163 | n.s. |
| ENSG00000111666 | 956.0766 | -0.105716579 | 0.2766 | -0.3822 | 0.702313 | 0.937867182 | CHPT1 | protein_coding | 38.42 | -0.002945122 | 0.002945122 | ENSG00000111666 | n.s. |
| ENSG00000086289 | 270.7214 | -0.108493295 | 0.29302 | -0.370258 | 0.71119 | 0.939526932 | EPDR1 | protein_coding | 38.37 | -0.002939166 | 0.002939166 | ENSG00000086289 | n.s. |
| ENSG00000164610 | 365.0625 | -0.106028838 | 0.281967 | -0.376033 | 0.706892 | 0.938232612 | RP9 | protein_coding | 42.71 | -0.002935883 | 0.002935883 | ENSG00000164610 | n.s. |
| ENSG00000145391 | 1643.239 | -0.108943814 | 0.296875 | -0.366968 | 0.713643 | 0.939884276 | SETD7 | protein_coding | 42.84 | -0.002933379 | 0.002933379 | ENSG00000145391 | n.s. |
| ENSG00000141076 | 192.4134 | 0.099617538 | 0.249543 | 0.3992 | 0.689746 | 0.934548254 | UTP4 | protein_coding | 43.98 | 0.002928583 | 0.002928583 | ENSG00000141076 | n.s. |
| ENSG00000172845 | 5027.827 | -0.099543873 | 0.248681 | -0.400287 | 0.688945 | 0.934506008 | SP3 | protein_coding | 37.7 | -0.002928372 | 0.002928372 | ENSG00000172845 | n.s. |
| ENSG00000199325 | 4.539565 | -0.226775267 | 1.056859 | -0.214575 | 0.830099 | 0.970726167 | RNU4-39P | snRNA | 45.67 | -0.002926141 | 0.002926141 | ENSG00000199325 | n.s. |
| ENSG00000158019 | 376.9087 | -0.105121524 | 0.277741 | -0.378488 | 0.705068 | 0.937939118 | BABAM2 | protein_coding | 39.38 | -0.002925043 | 0.002925043 | ENSG00000158019 | n.s. |
| ENSG00000165716 | 422.2839 | -0.10502025 | 0.276287 | -0.380113 | 0.703861 | 0.937896121 | FAM69B | protein_coding | 63.98 | -0.002924316 | 0.002924316 | ENSG00000165716 | n.s. |
| ENSG00000165804 | 236.4095 | 0.105005003 | 0.2763 | 0.3800395 | 0.703916 | 0.937896121 | ZNF219 | protein_coding | 55.48 | 0.002923892 | 0.002923892 | ENSG00000165804 | n.s. |
| ENSG00000188130 | 381.0234 | 0.108529904 | 0.296071 | 0.3665669 | 0.713942 | 0.939884276 | MAPK12 | protein_coding | 64.68 | 0.002922235 | 0.002922235 | ENSG00000188130 | n.s. |
| ENSG00000074201 | 2744.363 | 0.108840825 | 0.298167 | 0.3650327 | 0.715087 | 0.940060506 | CLNS1A | protein_coding | 41.76 | 0.002921744 | 0.002921744 | ENSG00000074201 | n.s. |
| ENSG00000222112 | 26.1176 | -0.11892506 | 0.349613 | -0.340162 | 0.733734 | 0.945030973 | RN7SKP16 | misc_RNA | 54.18 | -0.002920081 | 0.002920081 | ENSG00000222112 | n.s. |
| ENSG00000276334 | 194.6451 | -0.105985004 | 0.282722 | -0.374874 | 0.707754 | 0.938571345 | | sense_intronic | 32.88 | -0.002918054 | 0.002918054 | ENSG00000276334 | n.s. |
| ENSG00000183850 | 219.6893 | -0.104500985 | 0.26975 | -0.387399 | 0.698461 | 0.937752951 | ZNF730 | protein_coding | 41.52 | -0.002916786 | 0.002916786 | ENSG00000183850 | n.s. |
| ENSG00000136810 | 4205.338 | 0.112375284 | 0.316978 | 0.3545208 | 0.722949 | 0.942009991 | TXN | protein_coding | 42.28 | 0.00291552 | 0.00291552 | ENSG00000136810 | n.s. |
| ENSG00000073464 | 281.6772 | -0.108141599 | 0.293662 | -0.368253 | 0.712685 | 0.939817855 | CLCN4 | protein_coding | 45.24 | -0.002915098 | 0.002915098 | ENSG00000073464 | n.s. |
| ENSG00000118922 | 882.5358 | -0.108528028 | 0.296932 | -0.365498 | 0.714739 | 0.940053625 | KLF12 | protein_coding | 37.64 | -0.002913692 | 0.002913692 | ENSG00000118922 | n.s. |
| ENSG00000149636 | 746.0355 | -0.104356423 | 0.268125 | -0.389209 | 0.697122 | 0.937752951 | DSN1 | protein_coding | 44.84 | -0.002912751 | 0.002912751 | ENSG00000149636 | n.s. |
| ENSG00000109738 | 130.4978 | 0.112261663 | 0.317739 | 0.3533141 | 0.723853 | 0.942009991 | GLRB | protein_coding | 35.21 | 0.002912572 | 0.002912572 | ENSG00000109738 | n.s. |

| | | | | | | | | | | | | | |
|---|---|---|---|---|---|---|---|---|---|---|---|---|---|
| ENSG00000136045 | 2249.461 | -0.107158743 | 0.288006 | -0.372071 | 0.70984 | 0.939338716 | PWP1 | protein_coding | 40.59 | -0.002912336 | 0.002912336 | ENSG00000136045 | n.s. |
| ENSG00000147677 | 4064.703 | 0.105252153 | 0.280138 | 0.375716 | 0.707128 | 0.938366719 | EIF3H | protein_coding | 37.79 | 0.002907844 | 0.002907844 | ENSG00000147677 | n.s. |
| ENSG00000198498 | 1371.078 | -0.104867331 | 0.279045 | -0.375808 | 0.70706 | 0.938365427 | TMA16 | protein_coding | 36.88 | -0.002897275 | 0.002897275 | ENSG00000198498 | n.s. |
| ENSG00000054965 | 1119.692 | -0.099312274 | 0.249795 | -0.397576 | 0.690943 | 0.935225433 | FAM168A | protein_coding | 41.55 | -0.002888368 | 0.002888368 | ENSG00000054965 | n.s. |
| ENSG00000140598 | 710.1923 | -0.103387688 | 0.265791 | -0.38898 | 0.697291 | 0.937752951 | EFL1 | protein_coding | 38.04 | -0.002885712 | 0.002885712 | ENSG00000140598 | n.s. |
| ENSG00000144713 | 18368.87 | 0.10975447 | 0.306693 | 0.3578637 | 0.720445 | 0.941256551 | RPL32 | protein_coding | 47.4 | 0.002885663 | 0.002885663 | ENSG00000144713 | n.s. |
| ENSG00000188582 | 104.8105 | 0.103351806 | 0.264644 | 0.3905319 | 0.696143 | 0.937752951 | PAQR9 | protein_coding | 41.7 | 0.00288471 | 0.00288471 | ENSG00000188582 | n.s. |
| ENSG00000137038 | 1370.277 | -0.103338666 | 0.266276 | -0.388089 | 0.69795 | 0.937752951 | TMEM261 | protein_coding | 38.15 | -0.002884343 | 0.002884343 | ENSG00000137038 | n.s. |
| ENSG00000101266 | 2612.399 | -0.101973643 | 0.259812 | -0.39249 | 0.694696 | 0.936961766 | CSNK2A1 | protein_coding | 41.23 | -0.002883624 | 0.002883624 | ENSG00000101266 | n.s. |
| ENSG00000229180 | 317.5186 | -0.103245365 | 0.269482 | -0.383126 | 0.701627 | 0.937752951 | | transcribed_unprocessed_pseudc | 47.61 | -0.002881739 | 0.002881739 | ENSG00000229180 | n.s. |
| ENSG00000115317 | 485.7574 | -0.10530829 | 0.282466 | -0.372818 | 0.709284 | 0.938989999 | HTRA2 | protein_coding | 54.73 | -0.002879027 | 0.002879027 | ENSG00000115317 | n.s. |
| ENSG00000206737 | 61.31508 | 0.111714207 | 0.316833 | 0.3525966 | 0.724391 | 0.942412114 | RNVU1-18 | snRNA | 54.27 | 0.002877662 | 0.002877662 | ENSG00000206737 | n.s. |
| ENSG00000239569 | 70.64418 | 0.103293627 | 0.273128 | 0.3781879 | 0.705291 | 0.937943012 | KMT2E-AS1 | antisense | 64.44 | 0.002873995 | 0.002873995 | ENSG00000239569 | n.s. |
| ENSG00000112697 | 2486.242 | -0.102945059 | 0.267776 | -0.384445 | 0.700649 | 0.937752951 | TMEM30A | protein_coding | 37.37 | -0.002873357 | 0.002873357 | ENSG00000112697 | n.s. |
| ENSG00000260526 | 108.7532 | 0.102846264 | 0.266801 | 0.3854789 | 0.699883 | 0.937752951 | | antisense | 35.9 | 0.0028706 | 0.0028706 | ENSG00000260526 | n.s. |
| ENSG00000131188 | 67.64467 | -0.110493097 | 0.310799 | -0.355513 | 0.722205 | 0.941942395 | PRR7 | protein_coding | 62.31 | -0.002870131 | 0.002870131 | ENSG00000131188 | n.s. |
| ENSG00000100281 | 1052.621 | -0.097533207 | 0.243812 | -0.400035 | 0.689131 | 0.934506008 | HMGXB4 | protein_coding | 41.49 | -0.002869222 | 0.002869222 | ENSG00000100281 | n.s. |
| ENSG00000133812 | 1097.82 | 0.103056585 | 0.273717 | 0.376508 | 0.706539 | 0.937943012 | SBF2 | protein_coding | 38.31 | 0.0028674 | 0.0028674 | ENSG00000133812 | n.s. |
| ENSG00000101417 | 255.651 | -0.10953763 | 0.306944 | -0.356865 | 0.721193 | 0.94150619 | PXMP4 | protein_coding | 49.45 | -0.002867347 | 0.002867347 | ENSG00000101417 | n.s. |
| ENSG00000105928 | 168.7266 | -0.10742485 | 0.295426 | -0.363627 | 0.716136 | 0.940399775 | DFNA5 | protein_coding | 45.73 | -0.002866899 | 0.002866899 | ENSG00000105928 | n.s. |
| ENSG00000128311 | 158.6425 | 0.106218917 | 0.288189 | 0.3685734 | 0.712446 | 0.939817855 | TST | protein_coding | 57.04 | 0.00286327 | 0.00286327 | ENSG00000128311 | n.s. |
| ENSG00000113312 | 630.548 | 0.108741663 | 0.30356 | 0.3582214 | 0.720178 | 0.941240331 | TTC1 | protein_coding | 41.36 | 0.002859848 | 0.002859848 | ENSG00000113312 | n.s. |
| ENSG00000102984 | 80.63413 | 0.107347498 | 0.296345 | 0.3622385 | 0.717174 | 0.94052839 | ZNF821 | protein_coding | 45.9 | 0.002858459 | 0.002858459 | ENSG00000102984 | n.s. |
| ENSG00000117632 | 8917.981 | 0.10792958 | 0.299403 | 0.3604829 | 0.718486 | 0.940842655 | STMN1 | protein_coding | 48.24 | 0.002858299 | 0.002858299 | ENSG00000117632 | n.s. |
| ENSG00000094631 | 683.4878 | -0.102703797 | 0.271302 | -0.378559 | 0.705015 | 0.937939118 | HDAC6 | protein_coding | 50.55 | -0.002857769 | 0.002857769 | ENSG00000094631 | n.s. |
| ENSG00000131508 | 1278.183 | -0.10550536 | 0.285238 | -0.369885 | 0.711468 | 0.939559221 | UBE2D2 | protein_coding | 44.35 | -0.002856646 | 0.002856646 | ENSG00000131508 | n.s. |
| ENSG00000235162 | 659.4858 | -0.109983129 | 0.309775 | -0.355041 | 0.722559 | 0.941964628 | C12orf75 | protein_coding | 41.95 | -0.002855757 | 0.002855757 | ENSG00000235162 | n.s. |
| ENSG00000185065 | 92.91518 | -0.1022721 | 0.266037 | -0.384428 | 0.700661 | 0.937752951 | | antisense | 49.84 | -0.002854574 | 0.002854574 | ENSG00000185065 | n.s. |
| ENSG00000173545 | 753.4685 | -0.097033065 | 0.241513 | -0.401771 | 0.687852 | 0.934506008 | ZNF622 | protein_coding | 40.67 | -0.002854509 | 0.002854509 | ENSG00000173545 | n.s. |
| ENSG00000067445 | 857.8688 | 0.105278356 | 0.283881 | 0.3708537 | 0.710747 | 0.939497193 | TRO | protein_coding | 51.41 | 0.002853519 | 0.002853519 | ENSG00000067445 | n.s. |
| ENSG00000174748 | 16116.95 | 0.106259874 | 0.291577 | 0.3644321 | 0.715535 | 0.940060506 | RPL15 | protein_coding | 46.46 | 0.002852461 | 0.002852461 | ENSG00000174748 | n.s. |
| ENSG00000149823 | 876.2617 | -0.102167367 | 0.262037 | -0.389896 | 0.696613 | 0.937752951 | VPS51 | protein_coding | 50.69 | -0.002851651 | 0.002851651 | ENSG00000149823 | n.s. |

| | | | | | | | | | | | | | |
|---|---|---|---|---|---|---|---|---|---|---|---|---|---|
| ENSG00000166529 | 283.2619 | -0.106160542 | 0.290748 | -0.365129 | 0.715015 | 0.940060506 | ZSCAN21 | protein_coding | 47.87 | -0.002849794 | 0.002849794 | ENSG00000166529 | n.s. |
| ENSG00000221184 | 41.47934 | -0.111573017 | 0.319711 | -0.348981 | 0.727104 | 0.942883666 | MIR1254-1 | miRNA | 50.52 | -0.002849785 | 0.002849785 | ENSG00000221184 | n.s. |
| ENSG00000163291 | 1632.646 | 0.106988121 | 0.294999 | 0.3626728 | 0.716849 | 0.94052839 | PAQR3 | protein_coding | 35.92 | 0.002848889 | 0.002848889 | ENSG00000163291 | n.s. |
| ENSG00000204386 | 222.8699 | -0.102185525 | 0.267773 | -0.381612 | 0.702749 | 0.937867182 | NEU1 | protein_coding | 52.86 | -0.002846752 | 0.002846752 | ENSG00000204386 | n.s. |
| ENSG00000119041 | 2837.441 | -0.105706398 | 0.288009 | -0.367025 | 0.7136 | 0.939884276 | GTF3C3 | protein_coding | 36.72 | -0.00284621 | 0.00284621 | ENSG00000119041 | n.s. |
| ENSG00000221983 | 6474.565 | 0.107422452 | 0.298266 | 0.3601561 | 0.71873 | 0.940842655 | UBA52 | protein_coding | 53.24 | 0.002844869 | 0.002844869 | ENSG00000221983 | n.s. |
| ENSG00000100242 | 497.6431 | -0.101917328 | 0.262069 | -0.388895 | 0.697354 | 0.937752951 | SUN2 | protein_coding | 54.74 | -0.002844672 | 0.002844672 | ENSG00000100242 | n.s. |
| ENSG00000118418 | 2595.866 | 0.107985361 | 0.301298 | 0.3584001 | 0.720044 | 0.941153871 | HMGN3 | protein_coding | 38.62 | 0.002844266 | 0.002844266 | ENSG00000118418 | n.s. |
| ENSG00000159131 | 1986.048 | -0.101870852 | 0.266209 | -0.382672 | 0.701963 | 0.937752951 | GART | protein_coding | 41.63 | -0.002843374 | 0.002843374 | ENSG00000159131 | n.s. |
| ENSG00000100528 | 1959.056 | 0.10213134 | 0.270512 | 0.3775484 | 0.705766 | 0.937943012 | CNIH1 | protein_coding | 38.26 | 0.002841656 | 0.002841656 | ENSG00000100528 | n.s. |
| ENSG00000073050 | 526.6198 | -0.101640121 | 0.261999 | -0.38794 | 0.69806 | 0.937752951 | XRCC1 | protein_coding | 50.94 | -0.002836934 | 0.002836934 | ENSG00000073050 | n.s. |
| ENSG00000168872 | 600.0557 | -0.101514235 | 0.265452 | -0.38242 | 0.70215 | 0.937752951 | DDX19A | protein_coding | 47.09 | -0.002833421 | 0.002833421 | ENSG00000168872 | n.s. |
| ENSG00000151532 | 622.4713 | -0.096295507 | 0.240296 | -0.400737 | 0.688614 | 0.934506008 | VTI1A | protein_coding | 39.1 | -0.002832812 | 0.002832812 | ENSG00000151532 | n.s. |
| ENSG00000207468 | 71.80866 | 0.103528267 | 0.277569 | 0.3729827 | 0.709161 | 0.938989999 | SNORA19 | snoRNA | 32.81 | 0.002830363 | 0.002830363 | ENSG00000207468 | n.s. |
| ENSG00000124767 | 4512.862 | 0.110765585 | 0.3165 | 0.3499702 | 0.726361 | 0.942883666 | GLO1 | protein_coding | 39.8 | 0.002829162 | 0.002829162 | ENSG00000124767 | n.s. |
| ENSG00000174718 | 2299.573 | -0.099958017 | 0.253879 | -0.393722 | 0.693786 | 0.936913963 | KIAA1551 | protein_coding | 41.61 | -0.002828841 | 0.002828841 | ENSG00000174718 | n.s. |
| ENSG00000149231 | 672.1252 | -0.102654488 | 0.274242 | -0.374321 | 0.708166 | 0.938580742 | CCDC82 | protein_coding | 34.71 | -0.00282591 | 0.00282591 | ENSG00000149231 | n.s. |
| ENSG00000102908 | 787.8294 | -0.101515123 | 0.268045 | -0.378725 | 0.704892 | 0.93793739 | NFAT5 | protein_coding | 38.64 | -0.002824775 | 0.002824775 | ENSG00000102908 | n.s. |
| ENSG00000175928 | 993.5166 | -0.09599152 | 0.23963 | -0.400582 | 0.688728 | 0.934506008 | LRRN1 | protein_coding | 40.88 | -0.002823869 | 0.002823869 | ENSG00000175928 | n.s. |
| ENSG00000214706 | 617.269 | -0.101320033 | 0.265881 | -0.381073 | 0.703149 | 0.937867182 | IFRD2 | protein_coding | 59.19 | -0.00282264 | 0.00282264 | ENSG00000214706 | n.s. |
| ENSG00000129472 | 401.5144 | 0.102645152 | 0.274342 | 0.3741501 | 0.708293 | 0.938659694 | RAB2B | protein_coding | 41.19 | 0.002821903 | 0.002821903 | ENSG00000129472 | n.s. |
| ENSG00000197694 | 3326.798 | -0.09970383 | 0.253896 | -0.392696 | 0.694544 | 0.936958261 | SPTAN1 | protein_coding | 46.82 | -0.0028196 | 0.0028196 | ENSG00000197694 | n.s. |
| ENSG00000075711 | 1938.689 | -0.101235604 | 0.266514 | -0.379851 | 0.704056 | 0.937896121 | DLG1 | protein_coding | 36.59 | -0.002818932 | 0.002818932 | ENSG00000075711 | n.s. |
| ENSG00000227057 | 852.0658 | -0.103898772 | 0.279998 | -0.37107 | 0.710585 | 0.939462329 | WDR46 | protein_coding | 51.88 | -0.0028178 | 0.0028178 | ENSG00000227057 | n.s. |
| ENSG00000013374 | 864.3866 | -0.104592321 | 0.284954 | -0.36705 | 0.713581 | 0.939884276 | NUB1 | protein_coding | 44.96 | -0.002816213 | 0.002816213 | ENSG00000013374 | n.s. |
| ENSG00000105677 | 566.185 | 0.106861768 | 0.29757 | 0.3591151 | 0.719509 | 0.941153871 | TMEM147 | protein_coding | 58.59 | 0.002814671 | 0.002814671 | ENSG00000105677 | n.s. |
| ENSG00000104872 | 572.2538 | 0.107506817 | 0.301199 | 0.3569297 | 0.721144 | 0.94150619 | PIH1D1 | protein_coding | 55.81 | 0.002814187 | 0.002814187 | ENSG00000104872 | n.s. |
| ENSG00000136986 | 977.8609 | -0.104340676 | 0.283419 | -0.36815 | 0.712762 | 0.939817855 | DERL1 | protein_coding | 41.26 | -0.002812639 | 0.002812639 | ENSG00000136986 | n.s. |
| ENSG00000136888 | 2051.794 | -0.110001476 | 0.315387 | -0.348783 | 0.727252 | 0.942883666 | ATP6V1G1 | protein_coding | 44.34 | -0.002809645 | 0.002809645 | ENSG00000136888 | n.s. |
| ENSG00000104290 | 1580.081 | -0.102018805 | 0.272133 | -0.374886 | 0.707746 | 0.938571345 | FZD3 | protein_coding | 36.28 | -0.002808854 | 0.002808854 | ENSG00000104290 | n.s. |
| ENSG00000233223 | 92.30181 | -0.103528385 | 0.278733 | -0.371425 | 0.710321 | 0.939449151 | | antisense | 61.02 | -0.002808386 | 0.002808386 | ENSG00000233223 | n.s. |
| ENSG00000263816 | 4.680539 | 0.189658588 | 0.804936 | 0.2356193 | 0.813728 | 0.966492561 | MIR548AW | miRNA | 41.54 | 0.002807229 | 0.002807229 | ENSG00000263816 | n.s. |

| | | | | | | | | | | | | | |
|---|---|---|---|---|---|---|---|---|---|---|---|---|---|
| ENSG00000185650 | 134.3458 | 0.097289422 | 0.246015 | 0.3954619 | 0.692502 | 0.935906213 | ZFP36L1 | protein_coding | 53.26 | 0.00279879 | 0.00279879 | ENSG00000185650 | n.s. |
| ENSG00000272092 | 53.08494 | 0.109245922 | 0.310508 | 0.3518292 | 0.724966 | 0.942808406 | | lincRNA | 65.03 | 0.002794134 | 0.002794134 | ENSG00000272092 | n.s. |
| ENSG00000153485 | 214.3993 | 0.101324508 | 0.270658 | 0.3743642 | 0.708133 | 0.938580742 | TMEM251 | protein_coding | 48.11 | 0.002789298 | 0.002789298 | ENSG00000153485 | n.s. |
| ENSG00000240322 | 55.90542 | -0.103818371 | 0.284053 | -0.36549 | 0.714746 | 0.940053625 | RN7SL481P | misc_RNA | 50.5 | -0.00278725 | 0.00278725 | ENSG00000240322 | n.s. |
| ENSG00000101974 | 1807.712 | -0.101197087 | 0.27017 | -0.374569 | 0.707981 | 0.938580742 | ATP11C | protein_coding | 39.51 | -0.00278579 | 0.00278579 | ENSG00000101974 | n.s. |
| ENSG00000155115 | 1368.39 | -0.101790364 | 0.272903 | -0.372991 | 0.709155 | 0.938989999 | GTF3C6 | protein_coding | 45.43 | -0.00278285 | 0.00278285 | ENSG00000155115 | n.s. |
| ENSG00000047579 | 157.4571 | -0.103625265 | 0.28395 | -0.364943 | 0.715154 | 0.940060506 | DTNBP1 | protein_coding | 39.93 | -0.002781737 | 0.002781737 | ENSG00000047579 | n.s. |
| ENSG00000166797 | 1150.021 | 0.113848225 | 0.336004 | 0.33883 | 0.734738 | 0.945347052 | FAM96A | protein_coding | 41.73 | 0.00277889 | 0.00277889 | ENSG00000166797 | n.s. |
| ENSG00000231165 | 23.12686 | 0.119416046 | 0.363013 | 0.3289579 | 0.742188 | 0.947851467 | TRBV26OR9-2 | TR_V_pseudogene | 44.37 | 0.002777583 | 0.002777583 | ENSG00000231165 | n.s. |
| ENSG00000039123 | 3072.18 | -0.102349634 | 0.275655 | -0.371296 | 0.710417 | 0.939449151 | SKIV2L2 | protein_coding | 36.85 | -0.00277641 | 0.00277641 | ENSG00000039123 | n.s. |
| ENSG00000255046 | 52.06608 | 0.113830084 | 0.33664 | 0.3381362 | 0.735261 | 0.945422299 | | lincRNA | 47.88 | 0.002774513 | 0.002774513 | ENSG00000255046 | n.s. |
| ENSG00000109113 | 930.4121 | -0.099540074 | 0.26123 | -0.381044 | 0.703171 | 0.937867182 | RAB34 | protein_coding | 58.35 | -0.002773053 | 0.002773053 | ENSG00000109113 | n.s. |
| ENSG00000066855 | 484.1795 | 0.106459612 | 0.29973 | 0.3551846 | 0.722451 | 0.941964628 | MTFR1 | protein_coding | 40.22 | 0.002764267 | 0.002764267 | ENSG00000066855 | n.s. |
| ENSG00000109390 | 512.1754 | 0.108048808 | 0.309861 | 0.3487006 | 0.727314 | 0.942883666 | NDUFC1 | protein_coding | 42.39 | 0.002759771 | 0.002759771 | ENSG00000109390 | n.s. |
| ENSG00000049860 | 342.1469 | 0.104766142 | 0.292105 | 0.3586586 | 0.719851 | 0.941153871 | HEXB | protein_coding | 42.11 | 0.002759474 | 0.002759474 | ENSG00000049860 | n.s. |
| ENSG00000130935 | 5037.231 | -0.099168609 | 0.263288 | -0.376655 | 0.70643 | 0.937943012 | NOL11 | protein_coding | 40.57 | -0.002759223 | 0.002759223 | ENSG00000130935 | n.s. |
| ENSG00000094975 | 2387.368 | -0.099929211 | 0.266238 | -0.375338 | 0.707409 | 0.938471423 | SUCO | protein_coding | 36.42 | -0.002755942 | 0.002755942 | ENSG00000094975 | n.s. |
| ENSG00000152484 | 1169.408 | -0.098661708 | 0.253005 | -0.389959 | 0.696567 | 0.937752951 | USP12 | protein_coding | 38.86 | -0.002753802 | 0.002753802 | ENSG00000152484 | n.s. |
| ENSG00000271387 | 78.78026 | 0.103555398 | 0.286842 | 0.3610191 | 0.718085 | 0.940629569 | | lincRNA | 56.72 | 0.002752645 | 0.002752645 | ENSG00000271387 | n.s. |
| ENSG00000116209 | 2053.209 | 0.107759047 | 0.30899 | 0.3487457 | 0.72728 | 0.942883666 | TMEM59 | protein_coding | 38.88 | 0.002752369 | 0.002752369 | ENSG00000116209 | n.s. |
| ENSG00000171988 | 4270.3 | -0.098861225 | 0.261933 | -0.377429 | 0.705855 | 0.937943012 | JMJD1C | protein_coding | 38.11 | -0.00275067 | 0.00275067 | ENSG00000171988 | n.s. |
| ENSG00000155961 | 387.2561 | 0.102168872 | 0.279209 | 0.3659225 | 0.714423 | 0.939975765 | RAB39B | protein_coding | 41.08 | 0.002746641 | 0.002746641 | ENSG00000155961 | n.s. |
| ENSG00000242173 | 110.4605 | -0.105675679 | 0.297115 | -0.355673 | 0.722086 | 0.941942395 | ARHGDIG | protein_coding | 63.5 | -0.002744995 | 0.002744995 | ENSG00000242173 | n.s. |
| ENSG00000072415 | 482.4727 | -0.098631588 | 0.260255 | -0.378981 | 0.704702 | 0.93793739 | MPP5 | protein_coding | 37.63 | -0.002744537 | 0.002744537 | ENSG00000072415 | n.s. |
| ENSG00000106484 | 217.111 | 0.107428667 | 0.307614 | 0.3492317 | 0.726915 | 0.942883666 | MEST | protein_coding | 44.94 | 0.002743931 | 0.002743931 | ENSG00000106484 | n.s. |
| ENSG00000105186 | 498.9187 | -0.09825889 | 0.253471 | -0.387653 | 0.698273 | 0.937752951 | ANKRD27 | protein_coding | 48.98 | -0.002742559 | 0.002742559 | ENSG00000105186 | n.s. |
| ENSG00000253230 | 1030.976 | -0.092986174 | 0.229958 | -0.404361 | 0.685947 | 0.934368276 | LINC00599 | lincRNA | 53.37 | -0.002741411 | 0.002741411 | ENSG00000253230 | n.s. |
| ENSG00000136866 | 464.1345 | -0.103419933 | 0.28718 | -0.360122 | 0.718756 | 0.940842655 | ZFP37 | protein_coding | 39.08 | -0.00273887 | 0.00273887 | ENSG00000136866 | n.s. |
| ENSG00000255112 | 1186.261 | 0.098339972 | 0.258717 | 0.3801066 | 0.703866 | 0.937896121 | CHMP1B | protein_coding | 41.52 | 0.002738302 | 0.002738302 | ENSG00000255112 | n.s. |
| ENSG00000164587 | 15317.94 | 0.105439992 | 0.297561 | 0.3543474 | 0.723079 | 0.942009991 | RPS14 | protein_coding | 48.97 | 0.002735587 | 0.002735587 | ENSG00000164587 | n.s. |
| ENSG00000114423 | 216.6178 | 0.10097447 | 0.272754 | 0.3702035 | 0.711231 | 0.939526932 | CBLB | protein_coding | 35.4 | 0.002735476 | 0.002735476 | ENSG00000114423 | n.s. |
| ENSG00000135972 | 735.5621 | -0.101439052 | 0.275684 | -0.367954 | 0.712907 | 0.939817855 | MRPS9 | protein_coding | 36.63 | -0.002734422 | 0.002734422 | ENSG00000135972 | n.s. |

| | | | | | | | | | | | | | |
|---|---|---|---|---|---|---|---|---|---|---|---|---|---|
| ENSG00000269997 | 61.39593 | -0.106465114 | 0.302323 | -0.352156 | 0.724721 | 0.94257778 | | sense_intronic | 41.59 | -0.002734323 | 0.002734323 | ENSG00000269997 | n.s. |
| ENSG00000124659 | 989.5241 | -0.097961155 | 0.251472 | -0.389551 | 0.696868 | 0.937752951 | TBCC | protein_coding | 54.33 | -0.002734249 | 0.002734249 | ENSG00000124659 | n.s. |
| ENSG00000166681 | 3547.161 | -0.105259448 | 0.297509 | -0.353802 | 0.723487 | 0.942009991 | BEX3 | protein_coding | 57.48 | -0.002730903 | 0.002730903 | ENSG00000166681 | n.s. |
| ENSG00000117758 | 251.9168 | -0.106813402 | 0.305894 | -0.349185 | 0.726951 | 0.942883666 | STX12 | protein_coding | 41.41 | -0.002728216 | 0.002728216 | ENSG00000117758 | n.s. |
| ENSG00000037749 | 901.7195 | -0.09798755 | 0.258941 | -0.378416 | 0.705122 | 0.937939118 | MFAP3 | protein_coding | 43.54 | -0.002726538 | 0.002726538 | ENSG00000037749 | n.s. |
| ENSG00000234676 | 17.82794 | 0.134752592 | 0.450419 | 0.2991718 | 0.764809 | 0.954536941 | IFT74-AS1 | antisense | 39.53 | 0.002722981 | 0.002722981 | ENSG00000234676 | n.s. |
| ENSG00000185359 | 1986.395 | -0.097526889 | 0.254537 | -0.383154 | 0.701605 | 0.937752951 | HGS | protein_coding | 60.6 | -0.002722128 | 0.002722128 | ENSG00000185359 | n.s. |
| ENSG00000170037 | 667.142 | -0.094687495 | 0.23957 | -0.395239 | 0.692667 | 0.935976757 | CNTROB | protein_coding | 48.77 | -0.002720839 | 0.002720839 | ENSG00000170037 | n.s. |
| ENSG00000169618 | 201.0436 | -0.102315373 | 0.283158 | -0.361337 | 0.717847 | 0.940629569 | PROKR1 | protein_coding | 46.47 | -0.002719683 | 0.002719683 | ENSG00000169618 | n.s. |
| ENSG00000157064 | 1127.14 | -0.094039455 | 0.237208 | -0.396443 | 0.691778 | 0.935578594 | NMNAT2 | protein_coding | 44.41 | -0.002719595 | 0.002719595 | ENSG00000157064 | n.s. |
| ENSG00000255248 | 392.7815 | -0.097268003 | 0.251956 | -0.386051 | 0.699459 | 0.937752951 | MIR100HG | processed_transcript | 39.21 | -0.002714902 | 0.002714902 | ENSG00000255248 | n.s. |
| ENSG00000133874 | 225.1683 | 0.103136294 | 0.288435 | 0.3575725 | 0.720663 | 0.941256551 | RNF122 | protein_coding | 49.07 | 0.002711658 | 0.002711658 | ENSG00000133874 | n.s. |
| ENSG00000112578 | 658.6022 | -0.09706215 | 0.249449 | -0.389106 | 0.697198 | 0.937752951 | BYSL | protein_coding | 48.76 | -0.002709156 | 0.002709156 | ENSG00000112578 | n.s. |
| ENSG00000108848 | 18186.49 | -0.097019167 | 0.252304 | -0.384532 | 0.700584 | 0.937752951 | LUC7L3 | protein_coding | 39.68 | -0.002707956 | 0.002707956 | ENSG00000108848 | n.s. |
| ENSG00000100246 | 103.7315 | -0.111575221 | 0.331825 | -0.336248 | 0.736684 | 0.945680734 | DNAL4 | protein_coding | 51.3 | -0.002706308 | 0.002706308 | ENSG00000100246 | n.s. |
| ENSG00000106367 | 339.176 | 0.10921037 | 0.317056 | 0.3444513 | 0.730507 | 0.944543782 | AP1S1 | protein_coding | 54.82 | 0.002706004 | 0.002706004 | ENSG00000106367 | n.s. |
| ENSG00000164941 | 881.8748 | 0.10285764 | 0.287639 | 0.3575931 | 0.720648 | 0.941256551 | INTS8 | protein_coding | 40.21 | 0.002704332 | 0.002704332 | ENSG00000164941 | n.s. |
| ENSG00000174807 | 159.761 | -0.100003284 | 0.270645 | -0.3695 | 0.711755 | 0.939670252 | CD248 | protein_coding | 65.4 | -0.002702541 | 0.002702541 | ENSG00000174807 | n.s. |
| ENSG00000187742 | 2792.676 | 0.095577064 | 0.243633 | 0.3922992 | 0.694837 | 0.936995862 | SECISBP2 | protein_coding | 42.36 | 0.00270123 | 0.00270123 | ENSG00000187742 | n.s. |
| ENSG00000240972 | 4845.322 | -0.111996675 | 0.334165 | -0.335154 | 0.737509 | 0.946129051 | MIF | protein_coding | 69.2 | -0.002693478 | 0.002693478 | ENSG00000240972 | n.s. |
| ENSG00000133265 | 293.4474 | -0.106391136 | 0.306842 | -0.346729 | 0.728795 | 0.943381216 | HSPBP1 | protein_coding | 51.91 | -0.002693055 | 0.002693055 | ENSG00000133265 | n.s. |
| ENSG00000124802 | 1101.477 | 0.103792288 | 0.293188 | 0.3540124 | 0.72333 | 0.942009991 | EEF1E1 | protein_coding | 39.63 | 0.002692838 | 0.002692838 | ENSG00000124802 | n.s. |
| ENSG00000087087 | 1826.469 | -0.091473602 | 0.22702 | -0.402933 | 0.686998 | 0.934506008 | SRRT | protein_coding | 53.17 | -0.002690961 | 0.002690961 | ENSG00000087087 | n.s. |
| ENSG00000156261 | 6017.969 | 0.10370278 | 0.292904 | 0.3540509 | 0.723301 | 0.942009991 | CCT8 | protein_coding | 38.32 | 0.002690516 | 0.002690516 | ENSG00000156261 | n.s. |
| ENSG00000075188 | 851.513 | -0.100201554 | 0.274098 | -0.365568 | 0.714687 | 0.940053625 | NUP37 | protein_coding | 36.52 | -0.002690148 | 0.002690148 | ENSG00000075188 | n.s. |
| ENSG00000213186 | 811.9192 | 0.098954292 | 0.266128 | 0.37183 | 0.710019 | 0.939338716 | TRIM59 | protein_coding | 42.27 | 0.002689358 | 0.002689358 | ENSG00000213186 | n.s. |
| ENSG00000164182 | 476.0759 | 0.109073467 | 0.319285 | 0.3416182 | 0.732638 | 0.944876691 | NDUFAF2 | protein_coding | 36.3 | 0.002685919 | 0.002685919 | ENSG00000164182 | n.s. |
| ENSG00000187118 | 157.5017 | 0.114453326 | 0.346618 | 0.3301999 | 0.741249 | 0.94743491 | CMC1 | protein_coding | 35.93 | 0.002684001 | 0.002684001 | ENSG00000187118 | n.s. |
| ENSG00000260167 | 44.84368 | 0.108974841 | 0.318895 | 0.3417263 | 0.732557 | 0.944876691 | | antisense | 46.69 | 0.002683491 | 0.002683491 | ENSG00000260167 | n.s. |
| ENSG00000235316 | 62.20622 | -0.101840517 | 0.283735 | -0.358928 | 0.719649 | 0.941153871 | DUSP8P5 | processed_pseudogene | 67.6 | -0.002682415 | 0.002682415 | ENSG00000235316 | n.s. |
| ENSG00000165704 | 316.2934 | 0.110575395 | 0.328031 | 0.3370887 | 0.73605 | 0.945680734 | HPRT1 | protein_coding | 40.13 | 0.002682057 | 0.002682057 | ENSG00000165704 | n.s. |
| ENSG00000258815 | 79.35201 | -0.108765375 | 0.31712 | -0.342978 | 0.731615 | 0.944807198 | | lincRNA | 35.13 | -0.002681807 | 0.002681807 | ENSG00000258815 | n.s. |

| | | | | | | | | | | | | | |
|---|---|---|---|---|---|---|---|---|---|---|---|---|---|
| ENSG00000055044 | 7272.387 | -0.096067192 | 0.249559 | -0.384948 | 0.700276 | 0.937752951 | NOP58 | protein_coding | 41.15 | -0.002681385 | 0.002681385 | ENSG00000055044 | n.s. |
| ENSG00000082068 | 834.8549 | -0.101774923 | 0.283829 | -0.358579 | 0.71991 | 0.941153871 | WDR70 | protein_coding | 39 | -0.002680687 | 0.002680687 | ENSG00000082068 | n.s. |
| ENSG00000198873 | 325.8163 | 0.097352255 | 0.259573 | 0.3750481 | 0.707625 | 0.938571345 | GRK5 | protein_coding | 50.47 | 0.002680371 | 0.002680371 | ENSG00000198873 | n.s. |
| ENSG00000134775 | 355.5626 | -0.10314581 | 0.291829 | -0.353446 | 0.723754 | 0.942009991 | FHOD3 | protein_coding | 43.06 | -0.002676066 | 0.002676066 | ENSG00000134775 | n.s. |
| ENSG00000255358 | 29.20333 | -0.11920808 | 0.370151 | -0.322052 | 0.747413 | 0.949642637 | | processed_pseudogene | 44.64 | -0.002675005 | 0.002675005 | ENSG00000255358 | n.s. |
| ENSG00000260572 | 117.3166 | 0.094515016 | 0.240315 | 0.3932964 | 0.694101 | 0.936958261 | | antisense | 45.71 | 0.002672862 | 0.002672862 | ENSG00000260572 | n.s. |
| ENSG00000127463 | 481.4726 | -0.097718025 | 0.261842 | -0.373194 | 0.709004 | 0.938989999 | EMC1 | protein_coding | 46.63 | -0.002671516 | 0.002671516 | ENSG00000127463 | n.s. |
| ENSG00000111247 | 2031.975 | 0.099152447 | 0.270265 | 0.3668718 | 0.713715 | 0.939884276 | RAD51AP1 | protein_coding | 36.4 | 0.002669741 | 0.002669741 | ENSG00000111247 | n.s. |
| ENSG00000111325 | 86.17331 | -0.100730466 | 0.279652 | -0.360199 | 0.718698 | 0.940842655 | OGFOD2 | protein_coding | 59.54 | -0.002667645 | 0.002667645 | ENSG00000111325 | n.s. |
| ENSG00000041802 | 1966.443 | -0.095546509 | 0.245153 | -0.389743 | 0.696727 | 0.937752951 | LSG1 | protein_coding | 41.69 | -0.002666852 | 0.002666852 | ENSG00000041802 | n.s. |
| ENSG00000177156 | 725.1289 | -0.10825397 | 0.317042 | -0.34145 | 0.732765 | 0.944876691 | TALDO1 | protein_coding | 52.03 | -0.002665739 | 0.002665739 | ENSG00000177156 | n.s. |
| ENSG00000215256 | 172.1033 | -0.095454644 | 0.246947 | -0.386539 | 0.699098 | 0.937752951 | DHRS4-AS1 | antisense | 43.99 | -0.002664288 | 0.002664288 | ENSG00000215256 | n.s. |
| ENSG00000003056 | 1251.754 | -0.100031408 | 0.276118 | -0.362277 | 0.717145 | 0.94052839 | M6PR | protein_coding | 42.71 | -0.002663645 | 0.002663645 | ENSG00000003056 | n.s. |
| ENSG00000017797 | 1095.924 | -0.095395114 | 0.24796 | -0.38472 | 0.700445 | 0.937752951 | RALBP1 | protein_coding | 41.38 | -0.002662626 | 0.002662626 | ENSG00000017797 | n.s. |
| ENSG00000110514 | 123.0853 | 0.096701004 | 0.258154 | 0.374586 | 0.707968 | 0.938580742 | MADD | protein_coding | 47.1 | 0.00266202 | 0.00266202 | ENSG00000110514 | n.s. |
| ENSG00000205336 | 12.4894 | -0.133812103 | 0.458905 | -0.29159 | 0.7706 | 0.955275572 | ADGRG1 | protein_coding | 55.51 | -0.002659024 | 0.002659024 | ENSG00000205336 | n.s. |
| ENSG00000130830 | 178.3183 | 0.100282606 | 0.278441 | 0.3601569 | 0.71873 | 0.940842655 | MPP1 | protein_coding | 41.68 | 0.002655784 | 0.002655784 | ENSG00000130830 | n.s. |
| ENSG00000162378 | 1847.177 | -0.098615119 | 0.268647 | -0.367081 | 0.713559 | 0.939884276 | ZYG11B | protein_coding | 41.29 | -0.002655273 | 0.002655273 | ENSG00000162378 | n.s. |
| ENSG00000167283 | 1656.077 | 0.109143164 | 0.323187 | 0.3377093 | 0.735582 | 0.945536652 | ATP5L | protein_coding | 42.83 | 0.00265454 | 0.00265454 | ENSG00000167283 | n.s. |
| ENSG00000133422 | 954.6936 | -0.095030751 | 0.246796 | -0.385058 | 0.700194 | 0.937752951 | MORC2 | protein_coding | 45.11 | -0.002652457 | 0.002652457 | ENSG00000133422 | n.s. |
| ENSG00000005194 | 1059.701 | -0.108208376 | 0.318663 | -0.33957 | 0.734181 | 0.945151747 | CIAPIN1 | protein_coding | 44.41 | -0.002650938 | 0.002650938 | ENSG00000005194 | n.s. |
| ENSG00000198721 | 553.6155 | 0.106566564 | 0.308749 | 0.3451564 | 0.729977 | 0.944354757 | ECI2 | protein_coding | 41.7 | 0.002649759 | 0.002649759 | ENSG00000198721 | n.s. |
| ENSG00000272831 | 123.5379 | 0.099320636 | 0.274213 | 0.3622021 | 0.717201 | 0.94052839 | | antisense | 49.55 | 0.002644719 | 0.002644719 | ENSG00000272831 | n.s. |
| ENSG00000283749 | 2.00854 | -0.231476298 | 1.280737 | -0.180737 | 0.856574 | 0.974056274 | MIR4781 | miRNA | 53.95 | -0.002642522 | 0.002642522 | ENSG00000283749 | n.s. |
| ENSG00000112855 | 904.7657 | -0.094527042 | 0.245789 | -0.384586 | 0.700544 | 0.937752951 | HARS2 | protein_coding | 45.91 | -0.002638397 | 0.002638397 | ENSG00000112855 | n.s. |
| ENSG00000093144 | 2147.47 | -0.094495923 | 0.246458 | -0.383416 | 0.701411 | 0.937752951 | ECHDC1 | protein_coding | 36 | -0.002637529 | 0.002637529 | ENSG00000093144 | n.s. |
| ENSG00000178896 | 298.1592 | -0.097951828 | 0.267323 | -0.366418 | 0.714053 | 0.939885098 | EXOSC4 | protein_coding | 58.65 | -0.002637376 | 0.002637376 | ENSG00000178896 | n.s. |
| ENSG00000163449 | 402.4441 | 0.103230119 | 0.294334 | 0.350725 | 0.725795 | 0.942883666 | TMEM169 | protein_coding | 43.95 | 0.002636692 | 0.002636692 | ENSG00000163449 | n.s. |
| ENSG00000122335 | 330.7868 | -0.103165938 | 0.294131 | -0.350748 | 0.725778 | 0.942883666 | SERAC1 | protein_coding | 39.81 | -0.002635053 | 0.002635053 | ENSG00000122335 | n.s. |
| ENSG00000105221 | 368.2381 | -0.094837613 | 0.252007 | -0.376329 | 0.706672 | 0.938030367 | AKT2 | protein_coding | 53.61 | -0.002634883 | 0.002634883 | ENSG00000105221 | n.s. |
| ENSG00000115839 | 1962.858 | -0.097129993 | 0.261683 | -0.371174 | 0.710508 | 0.939449151 | RAB3GAP1 | protein_coding | 37.96 | -0.002634818 | 0.002634818 | ENSG00000115839 | n.s. |
| ENSG00000013441 | 3926.62 | 0.089515242 | 0.22391 | 0.3997819 | 0.689317 | 0.934506008 | CLK1 | protein_coding | 41.85 | 0.002633351 | 0.002633351 | ENSG00000013441 | n.s. |

| | | | | | | | | | | | | | |
|---|---|---|---|---|---|---|---|---|---|---|---|---|---|
| ENSG00000219665 | 246.2555 | 0.09451574 | 0.247628 | 0.3816838 | 0.702696 | 0.937867182 | | processed_transcript | 44.43 | 0.002633082 | 0.002633082 | ENSG00000219665 | n.s. |
| ENSG00000184787 | 1853.528 | -0.094308878 | 0.244707 | -0.385394 | 0.699945 | 0.937752951 | UBE2G2 | protein_coding | 45 | -0.002632308 | 0.002632308 | ENSG00000184787 | n.s. |
| ENSG00000105355 | 109.5653 | -0.107654415 | 0.317757 | -0.338795 | 0.734764 | 0.945347052 | PLIN3 | protein_coding | 52.37 | -0.002627707 | 0.002627707 | ENSG00000105355 | n.s. |
| ENSG00000135720 | 3318.986 | -0.094019704 | 0.24319 | -0.38661 | 0.699045 | 0.937752951 | DYNC1LI2 | protein_coding | 45.09 | -0.002624237 | 0.002624237 | ENSG00000135720 | n.s. |
| ENSG00000169660 | 519.9887 | 0.096865793 | 0.261414 | 0.3705452 | 0.710976 | 0.939526932 | HEXDC | protein_coding | 55.13 | 0.002624169 | 0.002624169 | ENSG00000169660 | n.s. |
| ENSG00000112306 | 16677.84 | 0.109064021 | 0.325336 | 0.3352355 | 0.737447 | 0.946129051 | RPS12 | protein_coding | 42.96 | 0.002622949 | 0.002622949 | ENSG00000112306 | n.s. |
| ENSG00000266916 | 112.6432 | 0.095933847 | 0.25699 | 0.3732974 | 0.708927 | 0.938989999 | ZNF793-AS1 | antisense | 44.46 | 0.002622739 | 0.002622739 | ENSG00000266916 | n.s. |
| ENSG00000168806 | 1437.271 | 0.081775766 | 0.192472 | 0.424871 | 0.670931 | 0.928842458 | LCMT2 | protein_coding | 46.08 | 0.002621563 | 0.002621563 | ENSG00000168806 | n.s. |
| ENSG00000206651 | 8.349454 | -0.14642067 | 0.531017 | -0.275736 | 0.78275 | 0.959619148 | | misc_RNA | 41.59 | -0.00262109 | 0.00262109 | ENSG00000206651 | n.s. |
| ENSG00000198730 | 1198.119 | -0.093812193 | 0.242062 | -0.387554 | 0.698346 | 0.937752951 | CTR9 | protein_coding | 39.07 | -0.002618445 | 0.002618445 | ENSG00000198730 | n.s. |
| ENSG00000196781 | 816.3228 | 0.093811396 | 0.241192 | 0.3889485 | 0.697314 | 0.937752951 | TLE1 | protein_coding | 43.43 | 0.002618422 | 0.002618422 | ENSG00000196781 | n.s. |
| ENSG00000136381 | 3172.516 | -0.09847777 | 0.272024 | -0.362019 | 0.717338 | 0.940619258 | IREB2 | protein_coding | 39.42 | -0.002618143 | 0.002618143 | ENSG00000136381 | n.s. |
| ENSG00000213281 | 2426.132 | -0.100705742 | 0.283248 | -0.355539 | 0.722186 | 0.941942395 | NRAS | protein_coding | 39 | -0.002615898 | 0.002615898 | ENSG00000213281 | n.s. |
| ENSG00000141391 | 137.6545 | -0.093644939 | 0.244049 | -0.383714 | 0.701191 | 0.937752951 | PRELID3A | protein_coding | 50.59 | -0.002613776 | 0.002613776 | ENSG00000141391 | n.s. |
| ENSG00000179409 | 721.3527 | -0.093502843 | 0.244314 | -0.382716 | 0.701931 | 0.937752951 | GEMIN4 | protein_coding | 56.12 | -0.00260981 | 0.00260981 | ENSG00000179409 | n.s. |
| ENSG00000207653 | 13.71806 | -0.160777798 | 0.627692 | -0.256141 | 0.797842 | 0.963346831 | MIR558 | miRNA | 42.55 | -0.002607386 | 0.002607386 | ENSG00000207653 | n.s. |
| ENSG00000108349 | 1465.068 | -0.093685146 | 0.247296 | -0.378838 | 0.704808 | 0.93793739 | CASC3 | protein_coding | 45.51 | -0.002606897 | 0.002606897 | ENSG00000108349 | n.s. |
| ENSG00000178773 | 100.3285 | -0.100353525 | 0.282378 | -0.355387 | 0.7223 | 0.941942395 | CPNE7 | protein_coding | 60.32 | -0.002606749 | 0.002606749 | ENSG00000178773 | n.s. |
| ENSG00000088930 | 4854.807 | 0.100446009 | 0.284196 | 0.3534389 | 0.723759 | 0.942009991 | XRN2 | protein_coding | 37.22 | 0.002606021 | 0.002606021 | ENSG00000088930 | n.s. |
| ENSG00000112742 | 2443.316 | -0.097024149 | 0.266411 | -0.364189 | 0.715717 | 0.940060506 | TTK | protein_coding | 34.98 | -0.002604535 | 0.002604535 | ENSG00000112742 | n.s. |
| ENSG00000155304 | 1627.384 | -0.105741176 | 0.310133 | -0.340955 | 0.733138 | 0.944876691 | HSPA13 | protein_coding | 36.7 | -0.002603862 | 0.002603862 | ENSG00000155304 | n.s. |
| ENSG00000057608 | 3918.487 | -0.101938353 | 0.293108 | -0.347784 | 0.728002 | 0.942883666 | GDI2 | protein_coding | 43.74 | -0.002603698 | 0.002603698 | ENSG00000057608 | n.s. |
| ENSG00000225195 | 18.10576 | -0.14945172 | 0.558323 | -0.26768 | 0.788946 | 0.960716322 | | sense_intronic | 45.19 | -0.002601182 | 0.002601182 | ENSG00000225195 | n.s. |
| ENSG00000118507 | 91.8555 | -0.101704714 | 0.289914 | -0.35081 | 0.725731 | 0.942883666 | AKAP7 | protein_coding | 38.22 | -0.00259773 | 0.00259773 | ENSG00000118507 | n.s. |
| ENSG00000169398 | 1950.657 | -0.096578574 | 0.264732 | -0.364817 | 0.715248 | 0.940060506 | PTK2 | protein_coding | 40.07 | -0.002592574 | 0.002592574 | ENSG00000169398 | n.s. |
| ENSG00000106615 | 1541.271 | -0.10144397 | 0.290776 | -0.348873 | 0.727185 | 0.942883666 | RHEB | protein_coding | 42 | -0.00259107 | 0.00259107 | ENSG00000106615 | n.s. |
| ENSG00000104904 | 4851.544 | -0.105026283 | 0.308446 | -0.340501 | 0.733479 | 0.944876691 | OAZ1 | protein_coding | 64.06 | -0.002586258 | 0.002586258 | ENSG00000104904 | n.s. |
| ENSG00000182810 | 435.9965 | 0.084387356 | 0.202075 | 0.4176034 | 0.676237 | 0.931868404 | DDX28 | protein_coding | 58.41 | 0.002586085 | 0.002586085 | ENSG00000182810 | n.s. |
| ENSG00000087206 | 952.9035 | -0.092645618 | 0.240357 | -0.38545 | 0.699904 | 0.937752951 | UIMC1 | protein_coding | 42.38 | -0.002585884 | 0.002585884 | ENSG00000087206 | n.s. |
| ENSG00000163001 | 605.856 | -0.101187547 | 0.289574 | -0.349436 | 0.726762 | 0.942883666 | CFAP36 | protein_coding | 40.44 | -0.002584521 | 0.002584521 | ENSG00000163001 | n.s. |
| ENSG00000204351 | 700.7309 | 0.09369752 | 0.249545 | 0.3754738 | 0.707308 | 0.938471423 | SKIV2L | protein_coding | 54.18 | 0.002584079 | 0.002584079 | ENSG00000204351 | n.s. |
| ENSG00000160214 | 742.7678 | -0.092557455 | 0.239476 | -0.3865 | 0.699127 | 0.937752951 | RRP1 | protein_coding | 57.27 | -0.002583423 | 0.002583423 | ENSG00000160214 | n.s. |

| | | | | | | | | | | | | | |
|---|---|---|---|---|---|---|---|---|---|---|---|---|---|
| ENSG00000159111 | 628.8632 | -0.099428689 | 0.280599 | -0.354344 | 0.723081 | 0.942009991 | MRPL10 | protein_coding | 49.17 | -0.002579627 | 0.002579627 | ENSG00000159111 | n.s. |
| ENSG00000067334 | 5254.591 | 0.092592919 | 0.242454 | 0.3818985 | 0.702537 | 0.937867182 | DNTTIP2 | protein_coding | 39.28 | 0.002579515 | 0.002579515 | ENSG00000067334 | n.s. |
| ENSG00000237649 | 1253.252 | -0.095980118 | 0.263442 | -0.364331 | 0.715611 | 0.940060506 | KIFC1 | protein_coding | 47.68 | -0.002576509 | 0.002576509 | ENSG00000237649 | n.s. |
| ENSG00000141499 | 325.849 | 0.099497885 | 0.282003 | 0.3528259 | 0.724219 | 0.942276487 | WRAP53 | protein_coding | 50.13 | 0.002569199 | 0.002569199 | ENSG00000141499 | n.s. |
| ENSG00000112902 | 2814.471 | 0.095690808 | 0.261935 | 0.3653232 | 0.71487 | 0.940060506 | SEMA5A | protein_coding | 40.18 | 0.002568742 | 0.002568742 | ENSG00000112902 | n.s. |
| ENSG00000221949 | 158.733 | 0.095375779 | 0.260117 | 0.3666649 | 0.713869 | 0.939884276 | LINC01465 | lincRNA | 58.2 | 0.002568052 | 0.002568052 | ENSG00000221949 | n.s. |
| ENSG00000085365 | 1669.923 | 0.102478224 | 0.296538 | 0.3455823 | 0.729657 | 0.944145838 | SCAMP1 | protein_coding | 37.59 | 0.00255795 | 0.00255795 | ENSG00000085365 | n.s. |
| ENSG00000144036 | 784.0115 | -0.095220513 | 0.261491 | -0.364145 | 0.71575 | 0.940060506 | EXOC6B | protein_coding | 37.12 | -0.002556118 | 0.002556118 | ENSG00000144036 | n.s. |
| ENSG00000207554 | 2.861228 | 0.20438142 | 0.96557 | 0.2116692 | 0.832365 | 0.971662543 | MIR647 | miRNA | 66.67 | 0.002551608 | 0.002551608 | ENSG00000207554 | n.s. |
| ENSG00000173905 | 3385.796 | -0.091537527 | 0.242446 | -0.377558 | 0.705759 | 0.937943012 | GOLIM4 | protein_coding | 38.33 | -0.002546899 | 0.002546899 | ENSG00000173905 | n.s. |
| ENSG00000158042 | 1158.696 | -0.084523209 | 0.205263 | -0.41178 | 0.680501 | 0.932999624 | MRPL17 | protein_coding | 48.63 | -0.002545715 | 0.002545715 | ENSG00000158042 | n.s. |
| ENSG00000256400 | 8.765552 | -0.133005651 | 0.46672 | -0.28498 | 0.77566 | 0.95689442 | | processed_pseudogene | 36.46 | -0.002545193 | 0.002545193 | ENSG00000256400 | n.s. |
| ENSG00000128626 | 339.1661 | -0.099618373 | 0.285965 | -0.348359 | 0.727571 | 0.942883666 | MRPS12 | protein_coding | 55.6 | -0.002544441 | 0.002544441 | ENSG00000128626 | n.s. |
| ENSG00000221630 | 3.564729 | -0.212288944 | 1.038153 | -0.204487 | 0.837973 | 0.97278132 | MIR1179 | miRNA | 41.76 | -0.002544236 | 0.002544236 | ENSG00000221630 | n.s. |
| ENSG00000198026 | 420.7368 | -0.08641894 | 0.214239 | -0.403376 | 0.686672 | 0.934474326 | ZNF335 | protein_coding | 53.47 | -0.002543536 | 0.002543536 | ENSG00000198026 | n.s. |
| ENSG00000179091 | 1091.687 | -0.099485831 | 0.284705 | -0.349435 | 0.726763 | 0.942883666 | CYC1 | protein_coding | 63.31 | -0.002541056 | 0.002541056 | ENSG00000179091 | n.s. |
| ENSG00000201573 | 27.79029 | -0.116681434 | 0.372602 | -0.313153 | 0.754165 | 0.951113891 | | misc_RNA | 39.22 | -0.00253986 | 0.00253986 | ENSG00000201573 | n.s. |
| ENSG00000232034 | 21.04573 | 0.122621749 | 0.404281 | 0.3033082 | 0.761655 | 0.95344167 | | sense_intronic | 43.86 | 0.00253899 | 0.00253899 | ENSG00000232034 | n.s. |
| ENSG00000138757 | 3959.077 | -0.097799036 | 0.2759 | -0.354472 | 0.722985 | 0.942009991 | G3BP2 | protein_coding | 41.77 | -0.002537346 | 0.002537346 | ENSG00000138757 | n.s. |
| ENSG00000176105 | 1596.099 | -0.096244625 | 0.267805 | -0.359383 | 0.719308 | 0.941113204 | YES1 | protein_coding | 40.01 | -0.002536829 | 0.002536829 | ENSG00000176105 | n.s. |
| ENSG00000121211 | 398.4811 | 0.09112824 | 0.241882 | 0.3767461 | 0.706362 | 0.937943012 | MND1 | protein_coding | 39.03 | 0.002535511 | 0.002535511 | ENSG00000121211 | n.s. |
| ENSG00000105518 | 233.4676 | -0.103873897 | 0.306797 | -0.338575 | 0.73493 | 0.945347052 | TMEM205 | protein_coding | 54.21 | -0.002535429 | 0.002535429 | ENSG00000105518 | n.s. |
| ENSG00000246090 | 64.25555 | -0.10195307 | 0.29551 | -0.345007 | 0.730089 | 0.944354757 | | antisense | 38.9 | -0.002535046 | 0.002535046 | ENSG00000246090 | n.s. |
| ENSG00000235449 | 33.42427 | 0.112960146 | 0.351176 | 0.3216626 | 0.747708 | 0.949642637 | | processed_pseudogene | 47.37 | 0.002534802 | 0.002534802 | ENSG00000235449 | n.s. |
| ENSG00000198707 | 2152.786 | -0.100121391 | 0.288739 | -0.346753 | 0.728777 | 0.943381216 | CEP290 | protein_coding | 33.9 | -0.00253435 | 0.00253435 | ENSG00000198707 | n.s. |
| ENSG00000132950 | 355.199 | -0.097323584 | 0.27553 | -0.353223 | 0.723921 | 0.942009991 | ZMYM5 | protein_coding | 42.61 | -0.002525011 | 0.002525011 | ENSG00000132950 | n.s. |
| ENSG00000105255 | 478.1282 | 0.102372389 | 0.298397 | 0.3430741 | 0.731543 | 0.944807198 | FSD1 | protein_coding | 54.71 | 0.002524176 | 0.002524176 | ENSG00000105255 | n.s. |
| ENSG00000152127 | 2097.157 | 0.095638788 | 0.266635 | 0.3586882 | 0.719828 | 0.941153871 | MGAT5 | protein_coding | 42.88 | 0.002519065 | 0.002519065 | ENSG00000152127 | n.s. |
| ENSG00000163866 | 317.6834 | -0.094535473 | 0.260278 | -0.36321 | 0.716448 | 0.94052839 | SMIM12 | protein_coding | 47.95 | -0.002517299 | 0.002517299 | ENSG00000163866 | n.s. |
| ENSG00000047410 | 14805.56 | -0.093605083 | 0.255863 | -0.36584 | 0.714484 | 0.939975765 | TPR | protein_coding | 34.16 | -0.002516418 | 0.002516418 | ENSG00000047410 | n.s. |
| ENSG00000153253 | 531.0724 | -0.098445897 | 0.281584 | -0.349614 | 0.726628 | 0.942883666 | SCN3A | protein_coding | 35.07 | -0.002514494 | 0.002514494 | ENSG00000153253 | n.s. |
| ENSG00000101310 | 1117.713 | -0.098438599 | 0.280196 | -0.351321 | 0.725348 | 0.942883666 | SEC23B | protein_coding | 42.98 | -0.002514308 | 0.002514308 | ENSG00000101310 | n.s. |

| | | | | | | | | | | | | | |
|---|---|---|---|---|---|---|---|---|---|---|---|---|---|
| ENSG00000182004 | 4538.934 | 0.103639415 | 0.308065 | 0.3364208 | 0.736554 | 0.945680734 | SNRPE | protein_coding | 42.59 | 0.002513822 | 0.002513822 | ENSG00000182004 | n.s. |
| ENSG00000118960 | 120.7243 | -0.105318158 | 0.315469 | -0.333846 | 0.738496 | 0.94653895 | HS1BP3 | protein_coding | 52.46 | -0.00251305 | 0.00251305 | ENSG00000118960 | n.s. |
| ENSG00000155957 | 157.3257 | 0.09535053 | 0.265489 | 0.3591511 | 0.719482 | 0.941153871 | TMBIM4 | protein_coding | 38.11 | 0.002511473 | 0.002511473 | ENSG00000155957 | n.s. |
| ENSG00000207442 | 20.82141 | 0.114670447 | 0.364536 | 0.3145658 | 0.753091 | 0.950880711 | SNORD116-6 | snoRNA | 41.67 | 0.002508297 | 0.002508297 | ENSG00000207442 | n.s. |
| ENSG00000133872 | 2311.429 | 0.098834979 | 0.284656 | 0.347208 | 0.728435 | 0.943266521 | SARAF | protein_coding | 40.19 | 0.002507007 | 0.002507007 | ENSG00000133872 | n.s. |
| ENSG00000269906 | 73.25574 | 0.103319055 | 0.307079 | 0.3364579 | 0.736526 | 0.945680734 | | sense_intronic | 35.18 | 0.002506051 | 0.002506051 | ENSG00000269906 | n.s. |
| ENSG00000144749 | 31.30532 | -0.112360725 | 0.351684 | -0.319493 | 0.749353 | 0.949944274 | LRIG1 | protein_coding | 46.07 | -0.002505854 | 0.002505854 | ENSG00000144749 | n.s. |
| ENSG00000197927 | 81.11522 | -0.104974562 | 0.314208 | -0.334093 | 0.73831 | 0.94653895 | C2orf27A | cribed_unprocessed_pseudc | 40.76 | -0.002504852 | 0.002504852 | ENSG00000197927 | n.s. |
| ENSG00000124172 | 4817.024 | 0.101634734 | 0.297385 | 0.3417613 | 0.732531 | 0.944876691 | ATP5E | protein_coding | 39.46 | 0.002502742 | 0.002502742 | ENSG00000124172 | n.s. |
| ENSG00000276213 | 8.108284 | 0.161887103 | 0.648397 | 0.2496728 | 0.80284 | 0.965030363 | | misc_RNA | 66.31 | 0.002502616 | 0.002502616 | ENSG00000276213 | n.s. |
| ENSG00000115568 | 414.8667 | 0.085038301 | 0.213105 | 0.3990448 | 0.68986 | 0.934548254 | ZNF142 | protein_coding | 49.52 | 0.002499979 | 0.002499979 | ENSG00000115568 | n.s. |
| ENSG00000033327 | 414.7436 | 0.092733812 | 0.252124 | 0.367811 | 0.713014 | 0.939817855 | GAB2 | protein_coding | 41.71 | 0.002499761 | 0.002499761 | ENSG00000033327 | n.s. |
| ENSG00000177551 | 3391.462 | -0.083517051 | 0.204174 | -0.409047 | 0.682505 | 0.933449397 | NHLH2 | protein_coding | 46.33 | -0.00249793 | 0.00249793 | ENSG00000177551 | n.s. |
| ENSG00000202242 | 2.42037 | -0.225003653 | 1.275203 | -0.176445 | 0.859944 | 0.974768375 | RNU6-1276P | snRNA | 33.65 | -0.002497219 | 0.002497219 | ENSG00000202242 | n.s. |
| ENSG00000107937 | 3146.16 | -0.096082567 | 0.269896 | -0.355998 | 0.721842 | 0.941942395 | GTPBP4 | protein_coding | 47.14 | -0.002495808 | 0.002495808 | ENSG00000107937 | n.s. |
| ENSG00000274737 | 50.70065 | 0.102677744 | 0.304427 | 0.3372824 | 0.735904 | 0.945638419 | | sense_intronic | 56.98 | 0.002492491 | 0.002492491 | ENSG00000274737 | n.s. |
| ENSG00000083845 | 9066.757 | 0.101641496 | 0.299026 | 0.3399081 | 0.733926 | 0.94510246 | RPS5 | protein_coding | 51.15 | 0.002492362 | 0.002492362 | ENSG00000083845 | n.s. |
| ENSG00000268996 | 64.45144 | 0.100995821 | 0.294029 | 0.3434888 | 0.731231 | 0.944807198 | MAN1B1-AS1 | antisense | 60.58 | 0.002490234 | 0.002490234 | ENSG00000268996 | n.s. |
| ENSG00000162664 | 4477.241 | -0.093262872 | 0.256563 | -0.363509 | 0.716225 | 0.940399775 | ZNF326 | protein_coding | 35.67 | -0.002488951 | 0.002488951 | ENSG00000162664 | n.s. |
| ENSG00000089157 | 37956.06 | 0.102062678 | 0.301814 | 0.3381644 | 0.735239 | 0.945422299 | RPLP0 | protein_coding | 48.92 | 0.002487692 | 0.002487692 | ENSG00000089157 | n.s. |
| ENSG00000152601 | 1858.712 | -0.093561168 | 0.259088 | -0.361117 | 0.718012 | 0.940629569 | MBNL1 | protein_coding | 35.71 | -0.002486984 | 0.002486984 | ENSG00000152601 | n.s. |
| ENSG00000178695 | 2656.211 | -0.084294071 | 0.208507 | -0.404275 | 0.686011 | 0.934368276 | KCTD12 | protein_coding | 43.57 | -0.002485151 | 0.002485151 | ENSG00000178695 | n.s. |
| ENSG00000273139 | 12.93191 | 0.129557611 | 0.453869 | 0.2854515 | 0.775298 | 0.95679591 | | lincRNA | 56.13 | 0.002485004 | 0.002485004 | ENSG00000273139 | n.s. |
| ENSG00000167193 | 961.4486 | -0.10177928 | 0.300691 | -0.338484 | 0.734998 | 0.945347052 | CRK | protein_coding | 49.13 | -0.002484303 | 0.002484303 | ENSG00000167193 | n.s. |
| ENSG00000196422 | 234.4291 | -0.092094814 | 0.250228 | -0.368044 | 0.71284 | 0.939817855 | PPP1R26 | protein_coding | 59.96 | -0.002482536 | 0.002482536 | ENSG00000196422 | n.s. |
| ENSG00000173660 | 1861.902 | 0.104917285 | 0.315993 | 0.3320245 | 0.739871 | 0.946996213 | UQCRH | protein_coding | 46.3 | 0.002481478 | 0.002481478 | ENSG00000173660 | n.s. |
| ENSG00000203819 | 71.02146 | 0.099386878 | 0.287543 | 0.3456418 | 0.729612 | 0.944145838 | | processed_pseudogene | 60 | 0.002480788 | 0.002480788 | ENSG00000203819 | n.s. |
| ENSG00000105258 | 534.0351 | -0.104862798 | 0.316104 | -0.331735 | 0.740089 | 0.946996213 | POLR2I | protein_coding | 60.72 | -0.00248019 | 0.00248019 | ENSG00000105258 | n.s. |
| ENSG00000263409 | 1.993499 | -0.212120737 | 1.053778 | -0.201295 | 0.840468 | 0.973446151 | MIR4747 | miRNA | 62.96 | -0.002479282 | 0.002479282 | ENSG00000263409 | n.s. |
| ENSG00000173230 | 3006.413 | -0.091820234 | 0.250023 | -0.367247 | 0.713435 | 0.939884276 | GOLGB1 | protein_coding | 38.07 | -0.002472316 | 0.002472316 | ENSG00000173230 | n.s. |
| ENSG00000137497 | 4692.43 | -0.088736194 | 0.235129 | -0.377393 | 0.705882 | 0.937943012 | NUMA1 | protein_coding | 46.62 | -0.002468956 | 0.002468956 | ENSG00000137497 | n.s. |
| ENSG00000202479 | 12.93604 | 0.136204635 | 0.490118 | 0.2779015 | 0.781088 | 0.959134639 | | snoRNA | 40 | 0.002468086 | 0.002468086 | ENSG00000202479 | n.s. |

| | | | | | | | | | | | | | |
|---|---|---|---|---|---|---|---|---|---|---|---|---|---|
| ENSG00000204498 | 155.2736 | -0.091501484 | 0.248717 | -0.367894 | 0.712952 | 0.939817855 | NFKBIL1 | protein_coding | 48.1 | -0.002466542 | 0.002466542 | ENSG00000204498 | n.s. |
| ENSG00000129824 | 4861.807 | 0.099829122 | 0.290892 | 0.3431827 | 0.731461 | 0.944807198 | RPS4Y1 | protein_coding | 40.56 | 0.002461467 | 0.002461467 | ENSG00000129824 | n.s. |
| ENSG00000182901 | 396.2621 | 0.096358879 | 0.274414 | 0.351144 | 0.72548 | 0.942883666 | RGS7 | protein_coding | 39.15 | 0.002461188 | 0.002461188 | ENSG00000182901 | n.s. |
| ENSG00000170629 | 257.2369 | -0.091140881 | 0.247211 | -0.368677 | 0.712369 | 0.939817855 | DPY19L2P2 | cribed_unprocessed_pseudc | 36.36 | -0.002456822 | 0.002456822 | ENSG00000170629 | n.s. |
| ENSG00000112312 | 2594.098 | 0.0904721 | 0.243457 | 0.3716143 | 0.71018 | 0.939449151 | GMNN | protein_coding | 36.82 | 0.002454211 | 0.002454211 | ENSG00000112312 | n.s. |
| ENSG00000171148 | 538.8378 | -0.091390273 | 0.250479 | -0.364862 | 0.715215 | 0.940060506 | TADA3 | protein_coding | 49.41 | -0.002453298 | 0.002453298 | ENSG00000171148 | n.s. |
| ENSG00000169902 | 230.6096 | 0.099537553 | 0.292432 | 0.3403789 | 0.733571 | 0.944908144 | TPST1 | protein_coding | 42.23 | 0.00244966 | 0.00244966 | ENSG00000169902 | n.s. |
| ENSG00000143379 | 1633.639 | -0.091922768 | 0.253473 | -0.362654 | 0.716864 | 0.94052839 | SETDB1 | protein_coding | 44.16 | -0.002447728 | 0.002447728 | ENSG00000143379 | n.s. |
| ENSG00000142192 | 4129.763 | -0.094344478 | 0.266724 | -0.353715 | 0.723552 | 0.942009991 | APP | protein_coding | 39.73 | -0.002447719 | 0.002447719 | ENSG00000142192 | n.s. |
| ENSG00000183837 | 140.4122 | -0.096323089 | 0.277304 | -0.347355 | 0.728324 | 0.943210846 | PNMA3 | protein_coding | 60.7 | -0.00244576 | 0.00244576 | ENSG00000183837 | n.s. |
| ENSG00000137106 | 740.9264 | -0.099234302 | 0.289786 | -0.34244 | 0.73202 | 0.944876691 | GRHPR | protein_coding | 53.09 | -0.002443631 | 0.002443631 | ENSG00000137106 | n.s. |
| ENSG00000235875 | 92.03229 | -0.091697012 | 0.253591 | -0.361594 | 0.717655 | 0.940629569 | ARHGEF7-AS2 | antisense | 60.85 | -0.002437432 | 0.002437432 | ENSG00000235875 | n.s. |
| ENSG00000091640 | 662.3752 | -0.097578971 | 0.28211 | -0.345889 | 0.729426 | 0.944110343 | SPAG7 | protein_coding | 51.65 | -0.002437254 | 0.002437254 | ENSG00000091640 | n.s. |
| ENSG00000243749 | 41.19605 | -0.105613331 | 0.324294 | -0.325671 | 0.744673 | 0.948261154 | TMEM35B | protein_coding | 51.77 | -0.002436715 | 0.002436715 | ENSG00000243749 | n.s. |
| ENSG00000164039 | 371.8371 | 0.104516365 | 0.317342 | 0.3293491 | 0.741892 | 0.94773468 | BDH2 | protein_coding | 38.56 | 0.002436614 | 0.002436614 | ENSG00000164039 | n.s. |
| ENSG00000105767 | 127.3382 | -0.098753051 | 0.287205 | -0.343842 | 0.730965 | 0.944807198 | CADM4 | protein_coding | 53.28 | -0.002434935 | 0.002434935 | ENSG00000105767 | n.s. |
| ENSG00000151500 | 338.9285 | -0.095312879 | 0.272812 | -0.349372 | 0.72681 | 0.942883666 | THYN1 | protein_coding | 43.95 | -0.002434471 | 0.002434471 | ENSG00000151500 | n.s. |
| ENSG00000144792 | 1200.966 | -0.087495596 | 0.231593 | -0.3778 | 0.705579 | 0.937943012 | ZNF660 | protein_coding | 45.57 | -0.002434438 | 0.002434438 | ENSG00000144792 | n.s. |
| ENSG00000182774 | 28459.53 | 0.105512513 | 0.322675 | 0.3269935 | 0.743673 | 0.948261154 | RPS17 | protein_coding | 49.9 | 0.002434389 | 0.002434389 | ENSG00000182774 | n.s. |
| ENSG00000160131 | 2027.586 | 0.093104927 | 0.261035 | 0.3566761 | 0.721334 | 0.941602997 | VMA21 | protein_coding | 41.5 | 0.002433034 | 0.002433034 | ENSG00000160131 | n.s. |
| ENSG00000106609 | 1225.21 | -0.088976672 | 0.238641 | -0.372847 | 0.709263 | 0.938989999 | TMEM248 | protein_coding | 44.8 | -0.002432536 | 0.002432536 | ENSG00000106609 | n.s. |
| ENSG00000221829 | 692.0818 | -0.090982789 | 0.250241 | -0.363581 | 0.716171 | 0.940399775 | FANCG | protein_coding | 53.25 | -0.002428102 | 0.002428102 | ENSG00000221829 | n.s. |
| ENSG00000283409 | 7.92878 | 0.135676884 | 0.493845 | 0.2747359 | 0.783519 | 0.959653909 | MIR3662 | miRNA | 37.89 | 0.00242663 | 0.00242663 | ENSG00000283409 | n.s. |
| ENSG00000166938 | 1022.044 | -0.092170404 | 0.257843 | -0.357467 | 0.720742 | 0.94127135 | DIS3L | protein_coding | 42.92 | -0.002422714 | 0.002422714 | ENSG00000166938 | n.s. |
| ENSG00000151743 | 329.4685 | 0.103973754 | 0.315957 | 0.3290752 | 0.742099 | 0.947825184 | AMN1 | protein_coding | 40.55 | 0.002419652 | 0.002419652 | ENSG00000151743 | n.s. |
| ENSG00000152082 | 2208.249 | -0.104839811 | 0.320139 | -0.327482 | 0.743303 | 0.948261154 | MZT2B | protein_coding | 57.61 | -0.002418868 | 0.002418868 | ENSG00000152082 | n.s. |
| ENSG00000069667 | 48.77042 | 0.112080718 | 0.359362 | 0.3118885 | 0.755125 | 0.951579479 | RORA | protein_coding | 41.64 | 0.002415892 | 0.002415892 | ENSG00000069667 | n.s. |
| ENSG00000223959 | 123.5688 | 0.090874121 | 0.251443 | 0.3614109 | 0.717792 | 0.940629569 | AFG3L1P | anscribed_unitary_pseudoge | 50.98 | 0.002415559 | 0.002415559 | ENSG00000223959 | n.s. |
| ENSG00000213614 | 870.5878 | -0.094489894 | 0.270565 | -0.349231 | 0.726916 | 0.942883666 | HEXA | protein_coding | 45.73 | -0.00241345 | 0.00241345 | ENSG00000213614 | n.s. |
| ENSG00000117054 | 2071.115 | -0.098806737 | 0.291789 | -0.338624 | 0.734893 | 0.945347052 | ACADM | protein_coding | 40.45 | -0.002411747 | 0.002411747 | ENSG00000117054 | n.s. |
| ENSG00000115414 | 89.26406 | -0.097841291 | 0.285799 | -0.342343 | 0.732093 | 0.944876691 | FN1 | protein_coding | 39.44 | -0.002409329 | 0.002409329 | ENSG00000115414 | n.s. |
| ENSG00000140848 | 233.9938 | -0.09427492 | 0.27028 | -0.348804 | 0.727236 | 0.942883666 | CPNE2 | protein_coding | 54.06 | -0.002407959 | 0.002407959 | ENSG00000140848 | n.s. |

| | | | | | | | | | | | | | |
|---|---|---|---|---|---|---|---|---|---|---|---|---|---|
| ENSG00000085998 | 334.7217 | -0.097657413 | 0.284532 | -0.343221 | 0.731432 | 0.944807198 | POMGNT1 | protein_coding | 49.39 | -0.00240792 | 0.00240792 | ENSG00000085998 | n.s. |
| ENSG00000244045 | 962.6797 | -0.089238601 | 0.241662 | -0.369271 | 0.711926 | 0.939770546 | TMEM199 | protein_coding | 47.27 | -0.002407494 | 0.002407494 | ENSG00000244045 | n.s. |
| ENSG00000234327 | 83.44118 | -0.103428372 | 0.314233 | -0.329146 | 0.742046 | 0.947825184 | | processed_transcript | 48.26 | -0.00240696 | 0.00240696 | ENSG00000234327 | n.s. |
| ENSG00000276026 | 10.83958 | 0.137139492 | 0.50769 | 0.2701244 | 0.787065 | 0.960409479 | | antisense | 44.87 | 0.002405915 | 0.002405915 | ENSG00000276026 | n.s. |
| ENSG00000146707 | 141.9316 | 0.091273366 | 0.253999 | 0.3593452 | 0.719337 | 0.941113204 | POMZP3 | protein_coding | 49.24 | 0.002405796 | 0.002405796 | ENSG00000146707 | n.s. |
| ENSG00000275710 | 13.0602 | -0.15974741 | 0.66402 | -0.240576 | 0.809884 | 0.965963425 | | antisense | 25.73 | -0.002402492 | 0.002402492 | ENSG00000275710 | n.s. |
| ENSG00000207638 | 69.02604 | 0.099857862 | 0.297792 | 0.3353275 | 0.737378 | 0.946129051 | MIR99A | miRNA | 53.09 | 0.002401544 | 0.002401544 | ENSG00000207638 | n.s. |
| ENSG00000099977 | 338.2408 | -0.105624378 | 0.326225 | -0.323778 | 0.746106 | 0.949025905 | DDT | protein_coding | 54.22 | -0.00239999 | 0.00239999 | ENSG00000099977 | n.s. |
| ENSG00000132478 | 2158.763 | -0.097437268 | 0.285146 | -0.34171 | 0.732569 | 0.944876691 | UNK | protein_coding | 50.87 | -0.00239938 | 0.00239938 | ENSG00000132478 | n.s. |
| ENSG00000136243 | 1139.505 | 0.088420187 | 0.238042 | 0.3714486 | 0.710303 | 0.939449151 | NUPL2 | protein_coding | 40.52 | 0.00239855 | 0.00239855 | ENSG00000136243 | n.s. |
| ENSG00000270640 | 57.86285 | 0.100462755 | 0.300953 | 0.3338159 | 0.738518 | 0.94653895 | | sense_intronic | 39.86 | 0.002397193 | 0.002397193 | ENSG00000270640 | n.s. |
| ENSG00000164654 | 1570.469 | -0.090666831 | 0.252051 | -0.359717 | 0.719059 | 0.940969272 | MIOS | protein_coding | 36.45 | -0.002395831 | 0.002395831 | ENSG00000164654 | n.s. |
| ENSG00000164418 | 482.87 | 0.090640091 | 0.252007 | 0.359673 | 0.719092 | 0.940969272 | GRIK2 | protein_coding | 33.94 | 0.002395124 | 0.002395124 | ENSG00000164418 | n.s. |
| ENSG00000132254 | 547.2595 | 0.096160968 | 0.278481 | 0.3453057 | 0.729865 | 0.944327121 | ARFIP2 | protein_coding | 52.8 | 0.002392248 | 0.002392248 | ENSG00000132254 | n.s. |
| ENSG00000167699 | 1237.159 | -0.10203651 | 0.309188 | -0.330014 | 0.741389 | 0.947527535 | GLOD4 | protein_coding | 45.54 | -0.002388487 | 0.002388487 | ENSG00000167699 | n.s. |
| ENSG00000168092 | 1585.167 | -0.093406395 | 0.26694 | -0.349916 | 0.726402 | 0.942883666 | PAFAH1B2 | protein_coding | 43.38 | -0.002385776 | 0.002385776 | ENSG00000168092 | n.s. |
| ENSG00000169612 | 409.8309 | -0.096811049 | 0.284135 | -0.340722 | 0.733313 | 0.944876691 | FAM103A1 | protein_coding | 41.1 | -0.002383959 | 0.002383959 | ENSG00000169612 | n.s. |
| ENSG00000067601 | 56.74129 | 0.108973157 | 0.34653 | 0.3144692 | 0.753165 | 0.950880711 | PMS2P4 | cribed_unprocessed_pseudc | 43.32 | 0.002383675 | 0.002383675 | ENSG00000067601 | n.s. |
| ENSG00000170142 | 1345.595 | 0.101244532 | 0.306378 | 0.3304567 | 0.741055 | 0.94732386 | UBE2E1 | protein_coding | 41.45 | 0.002379401 | 0.002379401 | ENSG00000170142 | n.s. |
| ENSG00000123933 | 644.7463 | 0.093054329 | 0.267064 | 0.3484344 | 0.727514 | 0.942883666 | MXD4 | protein_coding | 62.91 | 0.002376783 | 0.002376783 | ENSG00000123933 | n.s. |
| ENSG00000110660 | 402.7808 | -0.092657243 | 0.265405 | -0.349117 | 0.727002 | 0.942883666 | SLC35F2 | protein_coding | 43.51 | -0.002366641 | 0.002366641 | ENSG00000110660 | n.s. |
| ENSG00000175581 | 738.0829 | 0.100499832 | 0.303401 | 0.3312446 | 0.74046 | 0.947244602 | MRPL48 | protein_coding | 43.35 | 0.002365551 | 0.002365551 | ENSG00000175581 | n.s. |
| ENSG00000240225 | 949.038 | -0.088965132 | 0.246431 | -0.361014 | 0.718089 | 0.940629569 | ZNF542P | cribed_unprocessed_pseudc | 40.44 | -0.002364815 | 0.002364815 | ENSG00000240225 | n.s. |
| ENSG00000229512 | 8.869072 | 0.134659584 | 0.497842 | 0.2704864 | 0.786786 | 0.960409479 | | sense_intronic | 67.61 | 0.002362409 | 0.002362409 | ENSG00000229512 | n.s. |
| ENSG00000096063 | 2468.062 | -0.092467179 | 0.264908 | -0.349053 | 0.727049 | 0.942883666 | SRPK1 | protein_coding | 40.48 | -0.002361786 | 0.002361786 | ENSG00000096063 | n.s. |
| ENSG00000170891 | 72.17909 | -0.106856867 | 0.335808 | -0.318208 | 0.750327 | 0.950388734 | CYTL1 | protein_coding | 43.07 | -0.0023614 | 0.0023614 | ENSG00000170891 | n.s. |
| ENSG00000112715 | 984.0231 | -0.087602981 | 0.238289 | -0.367633 | 0.713147 | 0.93983441 | VEGFA | protein_coding | 56.4 | -0.002360783 | 0.002360783 | ENSG00000112715 | n.s. |
| ENSG00000170832 | 1632.998 | -0.092326612 | 0.264968 | -0.348444 | 0.727507 | 0.942883666 | USP32 | protein_coding | 40.19 | -0.002358196 | 0.002358196 | ENSG00000170832 | n.s. |
| ENSG00000258559 | 48.63245 | -0.105234625 | 0.327724 | -0.321107 | 0.748129 | 0.949712939 | | sense_overlapping | 52.46 | -0.00235806 | 0.00235806 | ENSG00000258559 | n.s. |
| ENSG00000272986 | 22.12957 | 0.118549038 | 0.401573 | 0.2952118 | 0.767832 | 0.955275572 | | antisense | 41.48 | 0.002355727 | 0.002355727 | ENSG00000272986 | n.s. |
| ENSG00000116754 | 7614.299 | -0.095447252 | 0.277489 | -0.343968 | 0.730871 | 0.944807198 | SRSF11 | protein_coding | 37.01 | -0.002353424 | 0.002353424 | ENSG00000116754 | n.s. |
| ENSG00000166949 | 243.2042 | -0.089840468 | 0.25171 | -0.35692 | 0.721151 | 0.94150619 | SMAD3 | protein_coding | 48.34 | -0.002351738 | 0.002351738 | ENSG00000166949 | n.s. |

| | | | | | | | | | | | | | |
|---|---|---|---|---|---|---|---|---|---|---|---|---|---|
| ENSG00000147162 | 4509.99 | -0.089253639 | 0.248826 | -0.358698 | 0.719821 | 0.941153871 | OGT | protein_coding | 40.6 | -0.002350884 | 0.002350884 | ENSG00000147162 | n.s. |
| ENSG00000009830 | 70.44854 | -0.098165555 | 0.293419 | -0.334558 | 0.737959 | 0.946357308 | POMT2 | protein_coding | 47.08 | -0.00235056 | 0.00235056 | ENSG00000009830 | n.s. |
| ENSG00000121297 | 633.2919 | -0.099304689 | 0.299278 | -0.331815 | 0.740029 | 0.946996213 | TSHZ3 | protein_coding | 46.88 | -0.00234873 | 0.00234873 | ENSG00000121297 | n.s. |
| ENSG00000148337 | 644.0885 | -0.091939312 | 0.261816 | -0.35116 | 0.725468 | 0.942883666 | CIZ1 | protein_coding | 52.19 | -0.002348304 | 0.002348304 | ENSG00000148337 | n.s. |
| ENSG00000207005 | 191.0697 | 0.087089411 | 0.236691 | 0.3679463 | 0.712913 | 0.939817855 | RNU1-2 | snRNA | 54.27 | 0.002347609 | 0.002347609 | ENSG00000207005 | n.s. |
| ENSG00000238197 | 122.0139 | -0.09043756 | 0.25536 | -0.354157 | 0.723222 | 0.942009991 | PAXBP1-AS1 | antisense | 39.6 | -0.002346356 | 0.002346356 | ENSG00000238197 | n.s. |
| ENSG00000118242 | 403.1274 | 0.095118187 | 0.277652 | 0.342581 | 0.731914 | 0.944876691 | MREG | protein_coding | 42.57 | 0.002342272 | 0.002342272 | ENSG00000118242 | n.s. |
| ENSG00000166595 | 628.3174 | -0.106955706 | 0.339767 | -0.314791 | 0.75292 | 0.950880711 | FAM96B | protein_coding | 55.72 | -0.002339545 | 0.002339545 | ENSG00000166595 | n.s. |
| ENSG00000175611 | 471.2463 | -0.083445709 | 0.213211 | -0.391376 | 0.695519 | 0.93752693 | LINC00476 | processed_transcript | 42.23 | -0.002337836 | 0.002337836 | ENSG00000175611 | n.s. |
| ENSG00000225091 | 106.7869 | -0.091412022 | 0.262585 | -0.348124 | 0.727747 | 0.942883666 | SNORA71A | snoRNA | 53.73 | -0.002334836 | 0.002334836 | ENSG00000225091 | n.s. |
| ENSG00000145741 | 8433.532 | 0.097065939 | 0.289694 | 0.3350637 | 0.737577 | 0.946129051 | BTF3 | protein_coding | 38.63 | 0.002334399 | 0.002334399 | ENSG00000145741 | n.s. |
| ENSG00000172922 | 812.1013 | -0.090831875 | 0.257931 | -0.352156 | 0.724721 | 0.94257778 | RNASEH2C | protein_coding | 55.83 | -0.002332817 | 0.002332817 | ENSG00000172922 | n.s. |
| ENSG00000100902 | 844.8786 | 0.089319144 | 0.251343 | 0.3553679 | 0.722314 | 0.941942395 | PSMA6 | protein_coding | 42.27 | 0.002320123 | 0.002320123 | ENSG00000100902 | n.s. |
| ENSG00000160318 | 14.12429 | -0.132236073 | 0.488961 | -0.270443 | 0.78682 | 0.960409479 | CLDND2 | protein_coding | 66.05 | -0.002319892 | 0.002319892 | ENSG00000160318 | n.s. |
| ENSG00000106245 | 812.3491 | -0.095344244 | 0.282537 | -0.337457 | 0.735772 | 0.945556068 | BUD31 | protein_coding | 48.46 | -0.002318077 | 0.002318077 | ENSG00000106245 | n.s. |
| ENSG00000166734 | 1274.948 | -0.095097306 | 0.2814 | -0.337944 | 0.735405 | 0.945434013 | CASC4 | protein_coding | 40.59 | -0.002317405 | 0.002317405 | ENSG00000166734 | n.s. |
| ENSG00000273841 | 1352.962 | 0.089295102 | 0.251767 | 0.3546729 | 0.722835 | 0.942009991 | TAF9 | protein_coding | 44.77 | 0.002316716 | 0.002316716 | ENSG00000273841 | n.s. |
| ENSG00000113387 | 5518.698 | 0.096534926 | 0.288316 | 0.3348228 | 0.737759 | 0.946275003 | SUB1 | protein_coding | 42.87 | 0.002315162 | 0.002315162 | ENSG00000113387 | n.s. |
| ENSG00000102977 | 220.7947 | -0.096617546 | 0.288682 | -0.334684 | 0.737863 | 0.946321655 | ACD | protein_coding | 62.02 | -0.002315074 | 0.002315074 | ENSG00000102977 | n.s. |
| ENSG00000137563 | 794.4806 | -0.100315935 | 0.305766 | -0.328081 | 0.742851 | 0.948261154 | GGH | protein_coding | 38.86 | -0.002314493 | 0.002314493 | ENSG00000137563 | n.s. |
| ENSG00000186141 | 995.7722 | -0.090114327 | 0.255788 | -0.352301 | 0.724613 | 0.94257778 | POLR3C | protein_coding | 43.11 | -0.002314388 | 0.002314388 | ENSG00000186141 | n.s. |
| ENSG00000108784 | 602.6202 | 0.083045847 | 0.22037 | 0.3768476 | 0.706287 | 0.937943012 | NAGLU | protein_coding | 57.01 | 0.00231063 | 0.00231063 | ENSG00000108784 | n.s. |
| ENSG00000170473 | 395.3857 | -0.102908618 | 0.320151 | -0.321437 | 0.747879 | 0.949642637 | PYM1 | protein_coding | 45.5 | -0.002309248 | 0.002309248 | ENSG00000170473 | n.s. |
| ENSG00000165006 | 351.758 | -0.095870004 | 0.285824 | -0.335416 | 0.737312 | 0.946129051 | UBAP1 | protein_coding | 42.8 | -0.002305637 | 0.002305637 | ENSG00000165006 | n.s. |
| ENSG00000167747 | 2161.385 | -0.085085108 | 0.229879 | -0.37013 | 0.711285 | 0.939526932 | C19orf48 | protein_coding | 54.79 | -0.002305021 | 0.002305021 | ENSG00000167747 | n.s. |
| ENSG00000116774 | 78.33909 | -0.093574485 | 0.274631 | -0.340728 | 0.733308 | 0.944876691 | OLFML3 | protein_coding | 45.3 | -0.002304259 | 0.002304259 | ENSG00000116774 | n.s. |
| ENSG00000125835 | 5279.677 | -0.09960093 | 0.306097 | -0.325391 | 0.744885 | 0.948261154 | SNRPB | protein_coding | 46.19 | -0.002297997 | 0.002297997 | ENSG00000125835 | n.s. |
| ENSG00000139370 | 643.5692 | 0.082517254 | 0.217513 | 0.3793665 | 0.704416 | 0.937896121 | SLC15A4 | protein_coding | 45.87 | 0.002297714 | 0.002297714 | ENSG00000139370 | n.s. |
| ENSG00000251136 | 85.87497 | 0.093280633 | 0.273432 | 0.3411475 | 0.732993 | 0.944876691 | | lincRNA | 38.48 | 0.002297023 | 0.002297023 | ENSG00000251136 | n.s. |
| ENSG00000151353 | 1258.155 | -0.096156438 | 0.287606 | -0.334334 | 0.738128 | 0.946486879 | TMEM18 | protein_coding | 53.39 | -0.002296735 | 0.002296735 | ENSG00000151353 | n.s. |
| ENSG00000137876 | 6646.842 | 0.099480233 | 0.305685 | 0.3254337 | 0.744853 | 0.948261154 | RSL24D1 | protein_coding | 36.26 | 0.002295212 | 0.002295212 | ENSG00000137876 | n.s. |
| ENSG00000123106 | 1152.198 | -0.096104519 | 0.288194 | -0.333471 | 0.738778 | 0.94653895 | CCDC91 | protein_coding | 35.52 | -0.002293199 | 0.002293199 | ENSG00000123106 | n.s. |

| | | | | | | | | | | | | | |
|---|---|---|---|---|---|---|---|---|---|---|---|---|---|
| ENSG00000260830 | 24.3987 | -0.115384733 | 0.395236 | -0.291939 | 0.770333 | 0.955275572 | | antisense | 54.69 | -0.002292848 | 0.002292848 | ENSG00000260830 | n.s. |
| ENSG00000164022 | 1330.25 | -0.096623402 | 0.290203 | -0.332952 | 0.739171 | 0.946864924 | AIMP1 | protein_coding | 36.71 | -0.002291131 | 0.002291131 | ENSG00000164022 | n.s. |
| ENSG00000164331 | 275.1352 | 0.094395983 | 0.2801 | 0.337008 | 0.736111 | 0.945680734 | ANKRA2 | protein_coding | 37.28 | 0.002289618 | 0.002289618 | ENSG00000164331 | n.s. |
| ENSG00000138796 | 566.4149 | -0.096691125 | 0.291429 | -0.331783 | 0.740053 | 0.946996213 | HADH | protein_coding | 43.13 | -0.002286915 | 0.002286915 | ENSG00000138796 | n.s. |
| ENSG00000065809 | 2399.993 | -0.099091802 | 0.303834 | -0.326138 | 0.74432 | 0.948261154 | FAM107B | protein_coding | 43.19 | -0.00228625 | 0.00228625 | ENSG00000065809 | n.s. |
| ENSG00000143889 | 2047.077 | -0.0894961 | 0.256275 | -0.349219 | 0.726925 | 0.942883666 | HNRNPLL | protein_coding | 37.07 | -0.002285899 | 0.002285899 | ENSG00000143889 | n.s. |
| ENSG00000100519 | 3168.996 | 0.088081415 | 0.249281 | 0.3533416 | 0.723832 | 0.942009991 | PSMC6 | protein_coding | 37.9 | 0.002285227 | 0.002285227 | ENSG00000100519 | n.s. |
| ENSG00000278847 | 30.26016 | 0.109708502 | 0.359123 | 0.3054902 | 0.759993 | 0.953277452 | | lincRNA | 32.79 | 0.002279817 | 0.002279817 | ENSG00000278847 | n.s. |
| ENSG00000272674 | 396.705 | -0.081823623 | 0.214819 | -0.380895 | 0.703281 | 0.937867182 | PCDHB16 | protein_coding | 44.75 | -0.002279497 | 0.002279497 | ENSG00000272674 | n.s. |
| ENSG00000206932 | 2.713336 | 0.194812894 | 0.967633 | 0.2013293 | 0.840441 | 0.973446151 | RNU6-4P | snRNA | 45.79 | 0.002276986 | 0.002276986 | ENSG00000206932 | n.s. |
| ENSG00000166478 | 640.8671 | -0.083987385 | 0.227184 | -0.369689 | 0.711614 | 0.939638057 | ZNF143 | protein_coding | 42.79 | -0.002270969 | 0.002270969 | ENSG00000166478 | n.s. |
| ENSG00000104660 | 1330.535 | -0.092980142 | 0.274495 | -0.338732 | 0.734812 | 0.945347052 | LEPROTL1 | protein_coding | 42.61 | -0.002269527 | 0.002269527 | ENSG00000104660 | n.s. |
| ENSG00000110651 | 939.4422 | -0.09209033 | 0.270214 | -0.340805 | 0.73325 | 0.944876691 | CD81 | protein_coding | 63.44 | -0.002267712 | 0.002267712 | ENSG00000110651 | n.s. |
| ENSG00000138764 | 536.8936 | -0.092010053 | 0.26906 | -0.341969 | 0.732374 | 0.944876691 | CCNG2 | protein_coding | 43.19 | -0.002265735 | 0.002265735 | ENSG00000138764 | n.s. |
| ENSG00000032389 | 605.4323 | -0.097827077 | 0.299756 | -0.326356 | 0.744155 | 0.948261154 | TSSC1 | protein_coding | 48.27 | -0.00225707 | 0.00225707 | ENSG00000032389 | n.s. |
| ENSG00000258789 | 26.48445 | -0.117637841 | 0.411718 | -0.285724 | 0.775089 | 0.95679591 | | antisense | 54.07 | -0.002256375 | 0.002256375 | ENSG00000258789 | n.s. |
| ENSG00000275964 | 145.3698 | -0.084705844 | 0.233418 | -0.362893 | 0.716685 | 0.94052839 | | lincRNA | 65.99 | -0.002255555 | 0.002255555 | ENSG00000275964 | n.s. |
| ENSG00000213762 | 1424.405 | 0.09168117 | 0.269595 | 0.3400701 | 0.733804 | 0.945032749 | ZNF134 | protein_coding | 46.33 | 0.002251061 | 0.002251061 | ENSG00000213762 | n.s. |
| ENSG00000197226 | 1006.632 | -0.09136584 | 0.267415 | -0.341663 | 0.732605 | 0.944876691 | TBC1D9B | protein_coding | 54.22 | -0.002249871 | 0.002249871 | ENSG00000197226 | n.s. |
| ENSG00000163634 | 717.1296 | -0.100635529 | 0.31426 | -0.32023 | 0.748794 | 0.949841385 | THOC7 | protein_coding | 39.89 | -0.002249095 | 0.002249095 | ENSG00000163634 | n.s. |
| ENSG00000126012 | 371.1021 | -0.085384932 | 0.238239 | -0.3584 | 0.720044 | 0.941153871 | KDM5C | protein_coding | 48.11 | -0.002248985 | 0.002248985 | ENSG00000126012 | n.s. |
| ENSG00000177432 | 420.6349 | -0.092111561 | 0.272065 | -0.338564 | 0.734938 | 0.945347052 | NAP1L5 | protein_coding | 45.8 | -0.002248326 | 0.002248326 | ENSG00000177432 | n.s. |
| ENSG00000124380 | 789.2778 | -0.092219012 | 0.272583 | -0.338315 | 0.735126 | 0.945422299 | SNRNP27 | protein_coding | 40.55 | -0.002247761 | 0.002247761 | ENSG00000124380 | n.s. |
| ENSG00000147853 | 1093.624 | -0.097010318 | 0.297489 | -0.326098 | 0.744351 | 0.948261154 | AK3 | protein_coding | 41.47 | -0.002238226 | 0.002238226 | ENSG00000147853 | n.s. |
| ENSG00000151779 | 49123.5 | -0.08762868 | 0.251964 | -0.347782 | 0.728004 | 0.942883666 | NBAS | protein_coding | 38.12 | -0.002238202 | 0.002238202 | ENSG00000151779 | n.s. |
| ENSG00000011243 | 1072.634 | -0.084034649 | 0.231814 | -0.362509 | 0.716972 | 0.94052839 | AKAP8L | protein_coding | 48.86 | -0.002237682 | 0.002237682 | ENSG00000011243 | n.s. |
| ENSG00000100823 | 2061.662 | 0.091940379 | 0.272289 | 0.3376577 | 0.735621 | 0.945536652 | APEX1 | protein_coding | 47.06 | 0.00223614 | 0.00223614 | ENSG00000100823 | n.s. |
| ENSG00000160741 | 188.8052 | 0.09190256 | 0.272262 | 0.3375515 | 0.735701 | 0.945552245 | CRTC2 | protein_coding | 53.2 | 0.002234562 | 0.002234562 | ENSG00000160741 | n.s. |
| ENSG00000143498 | 754.756 | -0.08316728 | 0.228402 | -0.364126 | 0.715764 | 0.940060506 | TAF1A | protein_coding | 36.34 | -0.002232558 | 0.002232558 | ENSG00000143498 | n.s. |
| ENSG00000226696 | 70.42514 | 0.092185396 | 0.274337 | 0.3360296 | 0.736849 | 0.945804562 | LENG8-AS1 | antisense | 48.1 | 0.002230757 | 0.002230757 | ENSG00000226696 | n.s. |
| ENSG00000242125 | 3746.829 | -0.075879997 | 0.19012 | -0.399116 | 0.689808 | 0.934548254 | SNHG3 | sense_intronic | 46.41 | -0.002230741 | 0.002230741 | ENSG00000242125 | n.s. |
| ENSG00000055147 | 87.06519 | -0.094600717 | 0.285957 | -0.330821 | 0.74078 | 0.947269752 | FAM114A2 | protein_coding | 37.35 | -0.002225608 | 0.002225608 | ENSG00000055147 | n.s. |

| | | | | | | | | | | | | | |
|---|---|---|---|---|---|---|---|---|---|---|---|---|---|
| ENSG00000152556 | 2615.391 | 0.093491345 | 0.280555 | 0.3332373 | 0.738955 | 0.94667583 | PFKM | protein_coding | 44.84 | 0.002224973 | 0.002224973 | ENSG00000152556 | n.s. |
| ENSG00000100288 | 248.2228 | 0.094470681 | 0.285736 | 0.3306222 | 0.74093 | 0.94732386 | CHKB | protein_coding | 50.66 | 0.002220205 | 0.002220205 | ENSG00000100288 | n.s. |
| ENSG00000090863 | 807.2058 | -0.089271448 | 0.258732 | -0.345034 | 0.730069 | 0.944354757 | GLG1 | protein_coding | 42.38 | -0.002219719 | 0.002219719 | ENSG00000090863 | n.s. |
| ENSG00000263001 | 2565.324 | -0.090105095 | 0.264283 | -0.340942 | 0.733147 | 0.944876691 | GTF2I | protein_coding | 42.79 | -0.002218826 | 0.002218826 | ENSG00000263001 | n.s. |
| ENSG00000183340 | 209.808 | -0.085417613 | 0.240549 | -0.355094 | 0.722519 | 0.941964628 | JRKL | protein_coding | 38.8 | -0.002217903 | 0.002217903 | ENSG00000183340 | n.s. |
| ENSG00000072506 | 683.076 | -0.101329402 | 0.321402 | -0.315273 | 0.752555 | 0.950880711 | HSD17B10 | protein_coding | 52.02 | -0.002216476 | 0.002216476 | ENSG00000072506 | n.s. |
| ENSG00000135334 | 1035.168 | -0.082212408 | 0.224539 | -0.366139 | 0.714261 | 0.939885098 | AKIRIN2 | protein_coding | 40.89 | -0.002213588 | 0.002213588 | ENSG00000135334 | n.s. |
| ENSG00000146416 | 289.7895 | 0.09360888 | 0.282307 | 0.331586 | 0.740202 | 0.947053038 | AIG1 | protein_coding | 39.67 | 0.002211575 | 0.002211575 | ENSG00000146416 | n.s. |
| ENSG00000137075 | 887.2533 | -0.089056944 | 0.258274 | -0.344816 | 0.730233 | 0.944452523 | RNF38 | protein_coding | 41.31 | -0.002210382 | 0.002210382 | ENSG00000137075 | n.s. |
| ENSG00000137692 | 658.8712 | -0.092557862 | 0.276998 | -0.334147 | 0.738269 | 0.94653895 | DCUN1D5 | protein_coding | 36.06 | -0.00220857 | 0.00220857 | ENSG00000137692 | n.s. |
| ENSG00000163382 | 439.8536 | -0.098074754 | 0.30509 | -0.321461 | 0.747861 | 0.949642637 | NAXE | protein_coding | 56.38 | -0.002200777 | 0.002200777 | ENSG00000163382 | n.s. |
| ENSG00000163171 | 2876.813 | 0.086160425 | 0.247313 | 0.3483866 | 0.72755 | 0.942883666 | CDC42EP3 | protein_coding | 41.38 | 0.0022007 | 0.0022007 | ENSG00000163171 | n.s. |
| ENSG00000206503 | 559.7403 | -0.092573314 | 0.278778 | -0.332068 | 0.739838 | 0.946996213 | HLA-A | protein_coding | 55.18 | -0.002189522 | 0.002189522 | ENSG00000206503 | n.s. |
| ENSG00000135821 | 8552.047 | -0.083958743 | 0.235909 | -0.355894 | 0.72192 | 0.941942395 | GLUL | protein_coding | 47.77 | -0.002180883 | 0.002180883 | ENSG00000135821 | n.s. |
| ENSG00000214022 | 1453.301 | 0.087976658 | 0.256415 | 0.3431025 | 0.731521 | 0.944807198 | REPIN1 | protein_coding | 62.28 | 0.002169223 | 0.002169223 | ENSG00000214022 | n.s. |
| ENSG00000084073 | 682.6298 | -0.092663844 | 0.280889 | -0.329895 | 0.741479 | 0.947527948 | ZMPSTE24 | protein_coding | 40.78 | -0.002169073 | 0.002169073 | ENSG00000084073 | n.s. |
| ENSG00000085224 | 4947.311 | -0.089879633 | 0.267707 | -0.335739 | 0.737068 | 0.945999074 | ATRX | protein_coding | 37.73 | -0.002166934 | 0.002166934 | ENSG00000085224 | n.s. |
| ENSG00000226853 | 61.49374 | 0.099380486 | 0.31694 | 0.3135623 | 0.753853 | 0.951060328 | | lincRNA | 45.09 | 0.002165693 | 0.002165693 | ENSG00000226853 | n.s. |
| ENSG00000136247 | 170.706 | -0.099177805 | 0.316036 | -0.313818 | 0.753659 | 0.951060328 | ZDHHC4 | protein_coding | 47.01 | -0.002161276 | 0.002161276 | ENSG00000136247 | n.s. |
| ENSG00000067646 | 424.0207 | -0.097487769 | 0.306683 | -0.317878 | 0.750577 | 0.950388734 | ZFY | protein_coding | 37.27 | -0.002154355 | 0.002154355 | ENSG00000067646 | n.s. |
| ENSG00000143774 | 1593.112 | -0.096577962 | 0.302025 | -0.319768 | 0.749144 | 0.94993942 | GUK1 | protein_coding | 61.76 | -0.002154084 | 0.002154084 | ENSG00000143774 | n.s. |
| ENSG00000131263 | 3807.878 | 0.087332386 | 0.255887 | 0.3412926 | 0.732883 | 0.944876691 | RLIM | protein_coding | 38.47 | 0.002150548 | 0.002150548 | ENSG00000131263 | n.s. |
| ENSG00000205707 | 485.115 | 0.08715143 | 0.25353 | 0.3437514 | 0.731033 | 0.944807198 | ETFRF1 | protein_coding | 38.05 | 0.002148876 | 0.002148876 | ENSG00000205707 | n.s. |
| ENSG00000198783 | 870.8833 | -0.079560903 | 0.216103 | -0.368162 | 0.712752 | 0.939817855 | ZNF830 | protein_coding | 42.79 | -0.002144668 | 0.002144668 | ENSG00000198783 | n.s. |
| ENSG00000197647 | 111.4687 | -0.092883349 | 0.285458 | -0.325383 | 0.744891 | 0.948261154 | ZNF433 | protein_coding | 43.27 | -0.002143008 | 0.002143008 | ENSG00000197647 | n.s. |
| ENSG00000131747 | 12862.28 | 0.092733673 | 0.282543 | 0.3282107 | 0.742752 | 0.948261154 | TOP2A | protein_coding | 39.83 | 0.002139555 | 0.002139555 | ENSG00000131747 | n.s. |
| ENSG00000116001 | 4385.755 | -0.095614953 | 0.298046 | -0.320806 | 0.748358 | 0.949807185 | TIA1 | protein_coding | 39.67 | -0.002138385 | 0.002138385 | ENSG00000116001 | n.s. |
| ENSG00000175166 | 3188.788 | -0.086782294 | 0.254801 | -0.340588 | 0.733414 | 0.944876691 | PSMD2 | protein_coding | 49 | -0.002137002 | 0.002137002 | ENSG00000175166 | n.s. |
| ENSG00000173757 | 835.5825 | -0.092572321 | 0.282829 | -0.327308 | 0.743435 | 0.948261154 | STAT5B | protein_coding | 43.63 | -0.002135832 | 0.002135832 | ENSG00000173757 | n.s. |
| ENSG00000152904 | 845.196 | -0.091216981 | 0.276713 | -0.329645 | 0.741668 | 0.947536205 | GGPS1 | protein_coding | 43.08 | -0.002134859 | 0.002134859 | ENSG00000152904 | n.s. |
| ENSG00000235701 | 43.32528 | -0.096367607 | 0.30248 | -0.318592 | 0.750036 | 0.95029168 | PCBP2P1 | processed_pseudogene | 49.54 | -0.002133875 | 0.002133875 | ENSG00000235701 | n.s. |
| ENSG00000075856 | 3435.233 | -0.084169195 | 0.242612 | -0.346929 | 0.728645 | 0.943381216 | SART3 | protein_coding | 42.65 | -0.002130556 | 0.002130556 | ENSG00000075856 | n.s. |

| | | | | | | | | | | | | | |
|---|---|---|---|---|---|---|---|---|---|---|---|---|---|
| ENSG00000182979 | 1117.749 | -0.086362694 | 0.251783 | -0.343004 | 0.731595 | 0.944807198 | MTA1 | protein_coding | 59.42 | -0.002129428 | 0.002129428 | ENSG00000182979 | n.s. |
| ENSG00000139116 | 15678.41 | -0.090025435 | 0.270574 | -0.33272 | 0.739346 | 0.946996213 | KIF21A | protein_coding | 36.22 | -0.00212926 | 0.00212926 | ENSG00000139116 | n.s. |
| ENSG00000176393 | 283.8766 | -0.090954787 | 0.275903 | -0.329662 | 0.741655 | 0.947536205 | RNPEP | protein_coding | 48.75 | -0.002128723 | 0.002128723 | ENSG00000176393 | n.s. |
| ENSG00000167088 | 4175.51 | -0.092089633 | 0.28288 | -0.325543 | 0.74477 | 0.948261154 | SNRPD1 | protein_coding | 42.32 | -0.002124696 | 0.002124696 | ENSG00000167088 | n.s. |
| ENSG00000167536 | 277.4201 | -0.0919808 | 0.281288 | -0.326998 | 0.743669 | 0.948261154 | DHRS13 | protein_coding | 54.96 | -0.002122185 | 0.002122185 | ENSG00000167536 | n.s. |
| ENSG00000168259 | 2375.66 | 0.078663956 | 0.213246 | 0.3688886 | 0.712211 | 0.939817855 | DNAJC7 | protein_coding | 45.79 | 0.00212049 | 0.00212049 | ENSG00000168259 | n.s. |
| ENSG00000210107 | 83.98321 | -0.096289979 | 0.303389 | -0.317381 | 0.750954 | 0.950589782 | MT-TQ | Mt_tRNA | 48.61 | -0.00211904 | 0.00211904 | ENSG00000210107 | n.s. |
| ENSG00000161800 | 2032.434 | -0.095012409 | 0.297273 | -0.319614 | 0.749261 | 0.949944274 | RACGAP1 | protein_coding | 43.65 | -0.002118954 | 0.002118954 | ENSG00000161800 | n.s. |
| ENSG00000129084 | 2544.407 | 0.094979029 | 0.296779 | 0.320033 | 0.748943 | 0.94993942 | PSMA1 | protein_coding | 40.21 | 0.002118421 | 0.002118421 | ENSG00000129084 | n.s. |
| ENSG00000210164 | 7.299293 | -0.142649507 | 0.604606 | -0.235938 | 0.813481 | 0.966390136 | MT-TG | Mt_tRNA | 26.47 | -0.00211799 | 0.00211799 | ENSG00000210164 | n.s. |
| ENSG00000110330 | 630.5017 | 0.087899226 | 0.262332 | 0.3350683 | 0.737574 | 0.946129051 | BIRC2 | protein_coding | 38.86 | 0.002113943 | 0.002113943 | ENSG00000110330 | n.s. |
| ENSG00000274104 | 25.75388 | 0.106562845 | 0.366233 | 0.2909702 | 0.771074 | 0.955385292 | | lincRNA | 49.81 | 0.00211223 | 0.00211223 | ENSG00000274104 | n.s. |
| ENSG00000196704 | 1289.823 | -0.089828157 | 0.271783 | -0.330514 | 0.741011 | 0.94732386 | AMZ2 | protein_coding | 43.73 | -0.002111099 | 0.002111099 | ENSG00000196704 | n.s. |
| ENSG00000120533 | 3013.853 | 0.087026697 | 0.258444 | 0.3367333 | 0.736318 | 0.945680734 | ENY2 | protein_coding | 37.11 | 0.002110872 | 0.002110872 | ENSG00000120533 | n.s. |
| ENSG00000261485 | 77.82065 | -0.089103411 | 0.268219 | -0.332204 | 0.739735 | 0.946996213 | PAN3-AS1 | antisense | 43.15 | -0.002107452 | 0.002107452 | ENSG00000261485 | n.s. |
| ENSG00000023287 | 1918.247 | -0.091278214 | 0.278918 | -0.327258 | 0.743473 | 0.948261154 | RB1CC1 | protein_coding | 37.73 | -0.002105975 | 0.002105975 | ENSG00000023287 | n.s. |
| ENSG00000189159 | 5966.744 | 0.085366451 | 0.248825 | 0.3430784 | 0.731539 | 0.944807198 | JPT1 | protein_coding | 46.27 | 0.002104864 | 0.002104864 | ENSG00000189159 | n.s. |
| ENSG00000104313 | 2157.294 | 0.082230607 | 0.234724 | 0.3503295 | 0.726091 | 0.942883666 | EYA1 | protein_coding | 37.53 | 0.002100325 | 0.002100325 | ENSG00000104313 | n.s. |
| ENSG00000166012 | 4553.45 | -0.079675584 | 0.222806 | -0.3576 | 0.720643 | 0.941256551 | TAF1D | protein_coding | 41.79 | -0.00209483 | 0.00209483 | ENSG00000166012 | n.s. |
| ENSG00000220868 | 22.89479 | -0.107429591 | 0.372223 | -0.288616 | 0.772875 | 0.956170275 | MRPL35P1 | processed_pseudogene | 41.96 | -0.002091091 | 0.002091091 | ENSG00000220868 | n.s. |
| ENSG00000105254 | 982.6871 | -0.095590505 | 0.303219 | -0.315252 | 0.75257 | 0.950880711 | TBCB | protein_coding | 55.29 | -0.002090943 | 0.002090943 | ENSG00000105254 | n.s. |
| ENSG00000199279 | 13.66198 | 0.114743815 | 0.411863 | 0.2785967 | 0.780554 | 0.958939185 | RNU6-661P | snRNA | 39.25 | 0.002089363 | 0.002089363 | ENSG00000199279 | n.s. |
| ENSG00000154518 | 3651.32 | 0.089254316 | 0.270604 | 0.3298334 | 0.741526 | 0.947527948 | ATP5G3 | protein_coding | 40.96 | 0.002089262 | 0.002089262 | ENSG00000154518 | n.s. |
| ENSG00000247556 | 3702.603 | -0.088311935 | 0.266166 | -0.331793 | 0.740046 | 0.946996213 | OIP5-AS1 | processed_transcript | 41.46 | -0.002088733 | 0.002088733 | ENSG00000247556 | n.s. |
| ENSG00000274471 | 113.9192 | -0.092685529 | 0.28697 | -0.322979 | 0.746711 | 0.949448038 | | unprocessed_pseudogene | 45.2 | -0.002088094 | 0.002088094 | ENSG00000274471 | n.s. |
| ENSG00000238835 | 35.42935 | 0.099879243 | 0.324188 | 0.3080907 | 0.758013 | 0.95300702 | SCARNA18 | snoRNA | 40.3 | 0.002087866 | 0.002087866 | ENSG00000238835 | n.s. |
| ENSG00000137842 | 161.3725 | 0.086050338 | 0.255773 | 0.336432 | 0.736545 | 0.945680734 | TMEM62 | protein_coding | 40.55 | 0.00208719 | 0.00208719 | ENSG00000137842 | n.s. |
| ENSG00000272991 | 27.24707 | -0.118886147 | 0.440078 | -0.270148 | 0.787047 | 0.960409479 | | antisense | 34.57 | -0.002085687 | 0.002085687 | ENSG00000272991 | n.s. |
| ENSG00000136193 | 1860.888 | -0.090366638 | 0.277218 | -0.325976 | 0.744442 | 0.948261154 | SCRN1 | protein_coding | 43.91 | -0.002084943 | 0.002084943 | ENSG00000136193 | n.s. |
| ENSG00000247315 | 973.1771 | -0.080356124 | 0.226965 | -0.354047 | 0.723304 | 0.942009991 | ZCCHC3 | protein_coding | 55.16 | -0.002084799 | 0.002084799 | ENSG00000247315 | n.s. |
| ENSG00000119326 | 1221.189 | -0.092951054 | 0.289614 | -0.320949 | 0.748249 | 0.949756199 | CTNNAL1 | protein_coding | 39.11 | -0.002080975 | 0.002080975 | ENSG00000119326 | n.s. |
| ENSG00000006042 | 1000.579 | -0.095586318 | 0.305244 | -0.313147 | 0.754169 | 0.951113891 | TMEM98 | protein_coding | 48.47 | -0.002080673 | 0.002080673 | ENSG00000006042 | n.s. |

| | | | | | | | | | | | | | |
|---|---|---|---|---|---|---|---|---|---|---|---|---|---|
| ENSG00000174547 | 1183.807 | 0.099251606 | 0.323108 | 0.3071774 | 0.758708 | 0.95300702 | MRPL11 | protein_coding | 48.04 | 0.002074746 | 0.002074746 | ENSG00000174547 | n.s. |
| ENSG00000087274 | 1453.888 | -0.08989904 | 0.275753 | -0.326013 | 0.744414 | 0.948261154 | ADD1 | protein_coding | 44.84 | -0.002074154 | 0.002074154 | ENSG00000087274 | n.s. |
| ENSG00000106066 | 238.4834 | 0.104085728 | 0.351535 | 0.296089 | 0.767162 | 0.955198262 | CPVL | protein_coding | 41 | 0.00207198 | 0.00207198 | ENSG00000106066 | n.s. |
| ENSG00000106526 | 29.3824 | 0.102456128 | 0.342459 | 0.2991778 | 0.764804 | 0.954536941 | ACTR3C | protein_coding | 43.31 | 0.002070358 | 0.002070358 | ENSG00000106526 | n.s. |
| ENSG00000207137 | 14.27871 | 0.115698889 | 0.42077 | 0.2749691 | 0.78334 | 0.959653909 | SNORD116-13 | snoRNA | 42.39 | 0.002069317 | 0.002069317 | ENSG00000207137 | n.s. |
| ENSG00000155827 | 2286.769 | -0.084551439 | 0.249163 | -0.339342 | 0.734352 | 0.945214194 | RNF20 | protein_coding | 38.17 | -0.002068954 | 0.002068954 | ENSG00000155827 | n.s. |
| ENSG00000187672 | 104.6701 | -0.092446299 | 0.288547 | -0.320385 | 0.748676 | 0.949841385 | ERC2 | protein_coding | 40.68 | -0.002066074 | 0.002066074 | ENSG00000187672 | n.s. |
| ENSG00000250021 | 237.1032 | -0.094439436 | 0.299946 | -0.314855 | 0.752872 | 0.950880711 | C15orf38-AP3S2 | protein_coding | 45.41 | -0.002065765 | 0.002065765 | ENSG00000250021 | n.s. |
| ENSG00000163463 | 516.4769 | 0.094221499 | 0.298138 | 0.3160336 | 0.751977 | 0.950815756 | KRTCAP2 | protein_coding | 50.64 | 0.002063793 | 0.002063793 | ENSG00000163463 | n.s. |
| ENSG00000158864 | 1811.557 | -0.089309644 | 0.273126 | -0.326991 | 0.743675 | 0.948261154 | NDUFS2 | protein_coding | 49.73 | -0.002060556 | 0.002060556 | ENSG00000158864 | n.s. |
| ENSG00000274929 | 71.16167 | 0.093103841 | 0.292367 | 0.3184483 | 0.750145 | 0.950319058 | | sense_intronic | 49.57 | 0.00206044 | 0.00206044 | ENSG00000274929 | n.s. |
| ENSG00000139352 | 1477.052 | -0.076385765 | 0.207473 | -0.368172 | 0.712745 | 0.939817855 | ASCL1 | protein_coding | 51.29 | -0.002059078 | 0.002059078 | ENSG00000139352 | n.s. |
| ENSG00000077254 | 3405.525 | -0.089141116 | 0.271887 | -0.327861 | 0.743017 | 0.948261154 | USP33 | protein_coding | 38.35 | -0.002056667 | 0.002056667 | ENSG00000077254 | n.s. |
| ENSG00000178982 | 1653.708 | 0.098873561 | 0.323349 | 0.3057801 | 0.759772 | 0.953277452 | EIF3K | protein_coding | 50.37 | 0.00205466 | 0.00205466 | ENSG00000178982 | n.s. |
| ENSG00000185875 | 375.1794 | -0.093803707 | 0.297932 | -0.31485 | 0.752876 | 0.950880711 | THNSL1 | protein_coding | 37.6 | -0.002051859 | 0.002051859 | ENSG00000185875 | n.s. |
| ENSG00000123130 | 411.3512 | 0.095868121 | 0.308147 | 0.3111113 | 0.755716 | 0.951943706 | ACOT9 | protein_coding | 44.61 | 0.002050498 | 0.002050498 | ENSG00000123130 | n.s. |
| ENSG00000167770 | 901.5521 | -0.085923331 | 0.257665 | -0.333469 | 0.73878 | 0.94653895 | OTUB1 | protein_coding | 59.12 | -0.00205026 | 0.00205026 | ENSG00000167770 | n.s. |
| ENSG00000129562 | 829.5617 | 0.101430475 | 0.338406 | 0.2997302 | 0.764383 | 0.954536941 | DAD1 | protein_coding | 41.95 | 0.002049632 | 0.002049632 | ENSG00000129562 | n.s. |
| ENSG00000137409 | 1662.438 | -0.09170723 | 0.286378 | -0.320231 | 0.748793 | 0.949841385 | MTCH1 | protein_coding | 52.18 | -0.002049557 | 0.002049557 | ENSG00000137409 | n.s. |
| ENSG00000143543 | 1518.203 | 0.098040141 | 0.319237 | 0.3071074 | 0.758762 | 0.95300702 | JTB | protein_coding | 50.96 | 0.002049421 | 0.002049421 | ENSG00000143543 | n.s. |
| ENSG00000156795 | 397.0408 | 0.093970411 | 0.29934 | 0.3139254 | 0.753578 | 0.9510571 | WDYHV1 | protein_coding | 45.17 | 0.002047935 | 0.002047935 | ENSG00000156795 | n.s. |
| ENSG00000143514 | 3233.699 | 0.0844102 | 0.250621 | 0.3368037 | 0.736265 | 0.945680734 | TP53BP2 | protein_coding | 39.79 | 0.002047408 | 0.002047408 | ENSG00000143514 | n.s. |
| ENSG00000167315 | 1607.365 | -0.088660236 | 0.272534 | -0.325318 | 0.744941 | 0.948261154 | ACAA2 | protein_coding | 39.54 | -0.002045573 | 0.002045573 | ENSG00000167315 | n.s. |
| ENSG00000197343 | 2038.935 | -0.085604674 | 0.256676 | -0.333513 | 0.738747 | 0.94653895 | ZNF655 | protein_coding | 42.73 | -0.002042657 | 0.002042657 | ENSG00000197343 | n.s. |
| ENSG00000163002 | 1923.429 | -0.085544681 | 0.256516 | -0.333486 | 0.738767 | 0.94653895 | NUP35 | protein_coding | 38.01 | -0.002041225 | 0.002041225 | ENSG00000163002 | n.s. |
| ENSG00000206838 | 10.16184 | -0.119964865 | 0.456366 | -0.26287 | 0.792651 | 0.961629391 | SNORA5A | snoRNA | 52.24 | -0.002038476 | 0.002038476 | ENSG00000206838 | n.s. |
| ENSG00000136630 | 297.0642 | -0.078340353 | 0.221656 | -0.353431 | 0.723765 | 0.942009991 | HLX | protein_coding | 54.42 | -0.002032501 | 0.002032501 | ENSG00000136630 | n.s. |
| ENSG00000134001 | 2998.942 | -0.091244363 | 0.285801 | -0.319258 | 0.749531 | 0.950083507 | EIF2S1 | protein_coding | 37.7 | -0.002029112 | 0.002029112 | ENSG00000134001 | n.s. |
| ENSG00000179152 | 437.9297 | 0.082389382 | 0.241918 | 0.3405681 | 0.733429 | 0.944876691 | TCAIM | protein_coding | 39.47 | 0.002028827 | 0.002028827 | ENSG00000179152 | n.s. |
| ENSG00000188010 | 212.0386 | 0.101686105 | 0.343618 | 0.2959278 | 0.767285 | 0.955198262 | MORN2 | protein_coding | 39.59 | 0.002024212 | 0.002024212 | ENSG00000188010 | n.s. |
| ENSG00000170348 | 1576.322 | -0.090482504 | 0.282503 | -0.320289 | 0.748749 | 0.949841385 | TMED10 | protein_coding | 42.21 | -0.002022186 | 0.002022186 | ENSG00000170348 | n.s. |
| ENSG00000120063 | 2970.281 | -0.087340244 | 0.265784 | -0.328613 | 0.742448 | 0.948097149 | GNA13 | protein_coding | 40.35 | -0.002021679 | 0.002021679 | ENSG00000120063 | n.s. |

| | | | | | | | | | | | | | |
|---|---|---|---|---|---|---|---|---|---|---|---|---|---|
| ENSG00000114650 | 738.3806 | -0.081526788 | 0.236596 | -0.344583 | 0.730408 | 0.944503674 | SCAP | protein_coding | 48.04 | -0.002021567 | 0.002021567 | ENSG00000114650 | n.s. |
| ENSG00000100505 | 359.5078 | -0.087549546 | 0.269059 | -0.325392 | 0.744884 | 0.948261154 | TRIM9 | protein_coding | 40.47 | -0.002019947 | 0.002019947 | ENSG00000100505 | n.s. |
| ENSG00000127022 | 9675.761 | -0.087537145 | 0.269103 | -0.325292 | 0.74496 | 0.948261154 | CANX | protein_coding | 45.8 | -0.002019661 | 0.002019661 | ENSG00000127022 | n.s. |
| ENSG00000107679 | 987.4927 | -0.094313899 | 0.303014 | -0.311252 | 0.755609 | 0.95189487 | PLEKHA1 | protein_coding | 37.12 | -0.002019356 | 0.002019356 | ENSG00000107679 | n.s. |
| ENSG00000111860 | 3071.779 | 0.082956172 | 0.246663 | 0.3363137 | 0.736634 | 0.945680734 | CEP85L | protein_coding | 38.17 | 0.00201214 | 0.00201214 | ENSG00000111860 | n.s. |
| ENSG00000207642 | 28.69828 | -0.099531698 | 0.331963 | -0.299828 | 0.764308 | 0.954536941 | MIR571 | miRNA | 50 | -0.002011263 | 0.002011263 | ENSG00000207642 | n.s. |
| ENSG00000117408 | 187.1784 | -0.0891301 | 0.275784 | -0.323188 | 0.746553 | 0.949420457 | IPO13 | protein_coding | 51.66 | -0.002009119 | 0.002009119 | ENSG00000117408 | n.s. |
| ENSG00000163558 | 1499.32 | -0.088345416 | 0.272856 | -0.32378 | 0.746104 | 0.949025905 | PRKCI | protein_coding | 39.82 | -0.002007379 | 0.002007379 | ENSG00000163558 | n.s. |
| ENSG00000196591 | 9455.88 | -0.086782707 | 0.265725 | -0.326588 | 0.743979 | 0.948261154 | HDAC2 | protein_coding | 38.55 | -0.002002254 | 0.002002254 | ENSG00000196591 | n.s. |
| ENSG00000086666 | 1385.115 | -0.092854409 | 0.297502 | -0.312114 | 0.754954 | 0.951579479 | ZFAND6 | protein_coding | 38.17 | -0.00200147 | 0.00200147 | ENSG00000086666 | n.s. |
| ENSG00000113456 | 1254.303 | -0.083744897 | 0.250946 | -0.333717 | 0.738593 | 0.94653895 | RAD1 | protein_coding | 40.32 | -0.00199828 | 0.00199828 | ENSG00000113456 | n.s. |
| ENSG00000224223 | 178.6205 | 0.094987828 | 0.307454 | 0.3089497 | 0.75736 | 0.952721388 | VSTM2A-OT1 | sense_overlapping | 35.79 | 0.001997982 | 0.001997982 | ENSG00000224223 | n.s. |
| ENSG00000179335 | 1059.991 | -0.077888791 | 0.221622 | -0.351449 | 0.725251 | 0.942883666 | CLK3 | protein_coding | 54.25 | -0.001989427 | 0.001989427 | ENSG00000179335 | n.s. |
| ENSG00000145247 | 709.1496 | 0.093820147 | 0.302622 | 0.3100244 | 0.756542 | 0.95246603 | OCIAD2 | protein_coding | 41.87 | 0.001984343 | 0.001984343 | ENSG00000145247 | n.s. |
| ENSG00000225791 | 263.6884 | 0.077621899 | 0.221266 | 0.3508081 | 0.725732 | 0.942883666 | TRAM2-AS1 | lincRNA | 44.52 | 0.00198261 | 0.00198261 | ENSG00000225791 | n.s. |
| ENSG00000142065 | 397.4113 | -0.088879849 | 0.277832 | -0.319905 | 0.74904 | 0.94993942 | ZFP14 | protein_coding | 42.69 | -0.001982384 | 0.001982384 | ENSG00000142065 | n.s. |
| ENSG00000138073 | 503.9048 | -0.095695507 | 0.315566 | -0.303251 | 0.761699 | 0.95344167 | PREB | protein_coding | 57.47 | -0.001981459 | 0.001981459 | ENSG00000138073 | n.s. |
| ENSG00000115524 | 15870.93 | 0.086849286 | 0.267637 | 0.3245036 | 0.745557 | 0.948847093 | SF3B1 | protein_coding | 39.41 | 0.001980491 | 0.001980491 | ENSG00000115524 | n.s. |
| ENSG00000136504 | 4074.854 | 0.078205046 | 0.22552 | 0.346776 | 0.72876 | 0.943381216 | KAT7 | protein_coding | 42.42 | 0.001979587 | 0.001979587 | ENSG00000136504 | n.s. |
| ENSG00000074800 | 10622.47 | -0.094552127 | 0.30708 | -0.307907 | 0.758153 | 0.95300702 | ENO1 | protein_coding | 50.23 | -0.001976508 | 0.001976508 | ENSG00000074800 | n.s. |
| ENSG00000130748 | 220.5958 | -0.099083891 | 0.334931 | -0.295834 | 0.767357 | 0.955198262 | TMEM160 | protein_coding | 61.6 | -0.001972411 | 0.001972411 | ENSG00000130748 | n.s. |
| ENSG00000134453 | 1693.319 | -0.083768383 | 0.2531 | -0.330969 | 0.740668 | 0.947269752 | RBM17 | protein_coding | 41.27 | -0.001970763 | 0.001970763 | ENSG00000134453 | n.s. |
| ENSG00000123131 | 1551.192 | 0.097426448 | 0.323634 | 0.3010391 | 0.763385 | 0.954522181 | PRDX4 | protein_coding | 40.65 | 0.001969376 | 0.001969376 | ENSG00000123131 | n.s. |
| ENSG00000171094 | 1885.629 | -0.077023347 | 0.221374 | -0.347933 | 0.72789 | 0.942883666 | ALK | protein_coding | 43.51 | -0.001967322 | 0.001967322 | ENSG00000171094 | n.s. |
| ENSG00000106605 | 495.3147 | -0.087904944 | 0.274154 | -0.320641 | 0.748483 | 0.949841385 | BLVRA | protein_coding | 44.31 | -0.00196458 | 0.00196458 | ENSG00000106605 | n.s. |
| ENSG00000156384 | 288.1461 | -0.079520769 | 0.231895 | -0.342917 | 0.731661 | 0.944807198 | SFR1 | protein_coding | 37.96 | -0.001960728 | 0.001960728 | ENSG00000156384 | n.s. |
| ENSG00000168118 | 1385.841 | -0.092582017 | 0.29916 | -0.309473 | 0.756962 | 0.95246603 | RAB4A | protein_coding | 41.32 | -0.001958156 | 0.001958156 | ENSG00000168118 | n.s. |
| ENSG00000207233 | 45.73727 | 0.096860551 | 0.321712 | 0.3010785 | 0.763355 | 0.954522181 | SNORA37 | snoRNA | 54.26 | 0.001957937 | 0.001957937 | ENSG00000207233 | n.s. |
| ENSG00000100612 | 559.461 | -0.091092258 | 0.29232 | -0.311618 | 0.755331 | 0.951717043 | DHRS7 | protein_coding | 39.65 | -0.001957769 | 0.001957769 | ENSG00000100612 | n.s. |
| ENSG00000165912 | 137.8757 | 0.089306116 | 0.281773 | 0.3169431 | 0.751287 | 0.950815756 | PACSIN3 | protein_coding | 59.66 | 0.001956128 | 0.001956128 | ENSG00000165912 | n.s. |
| ENSG00000167797 | 111.3792 | -0.084708234 | 0.257977 | -0.328356 | 0.742643 | 0.948258939 | CDK2AP2 | protein_coding | 62.33 | -0.001954478 | 0.001954478 | ENSG00000167797 | n.s. |
| ENSG00000254510 | 60.29195 | 0.09819375 | 0.336632 | 0.291695 | 0.77052 | 0.955275572 | | processed_transcript | 53.5 | 0.00195124 | 0.00195124 | ENSG00000254510 | n.s. |

| | | | | | | | | | | | | | |
|---|---|---|---|---|---|---|---|---|---|---|---|---|---|
| ENSG00000011426 | 2708.073 | -0.078955288 | 0.229732 | -0.343684 | 0.731084 | 0.944807198 | ANLN | protein_coding | 37.88 | -0.001946785 | 0.001946785 | ENSG00000011426 | n.s. |
| ENSG00000223509 | 90.6142 | -0.096925426 | 0.32549 | -0.297783 | 0.765868 | 0.954823343 | | transcribed_processed_pseudog | 39.96 | -0.001945969 | 0.001945969 | ENSG00000223509 | n.s. |
| ENSG00000197579 | 1355.344 | 0.084148405 | 0.257024 | 0.327395 | 0.743369 | 0.948261154 | TOPORS | protein_coding | 42.02 | 0.001941475 | 0.001941475 | ENSG00000197579 | n.s. |
| ENSG00000156650 | 1268.264 | -0.082030337 | 0.246772 | -0.332413 | 0.739577 | 0.946996213 | KAT6B | protein_coding | 40.6 | -0.001940162 | 0.001940162 | ENSG00000156650 | n.s. |
| ENSG00000137871 | 1685.895 | -0.087674323 | 0.275628 | -0.318089 | 0.750418 | 0.950388734 | ZNF280D | protein_coding | 37.52 | -0.00193749 | 0.00193749 | ENSG00000137871 | n.s. |
| ENSG00000180304 | 650.3665 | 0.093148765 | 0.304833 | 0.3055732 | 0.75993 | 0.953277452 | OAZ2 | protein_coding | 47.83 | 0.001935695 | 0.001935695 | ENSG00000180304 | n.s. |
| ENSG00000230989 | 3206.403 | 0.093165855 | 0.306519 | 0.3039476 | 0.761168 | 0.953332446 | HSBP1 | protein_coding | 44.98 | 0.001933716 | 0.001933716 | ENSG00000230989 | n.s. |
| ENSG00000205542 | 5684.749 | -0.093044197 | 0.305987 | -0.304079 | 0.761067 | 0.953332446 | TMSB4X | protein_coding | 50.47 | -0.001931191 | 0.001931191 | ENSG00000205542 | n.s. |
| ENSG00000111530 | 9888.151 | -0.085668632 | 0.265153 | -0.323092 | 0.746626 | 0.949426449 | CAND1 | protein_coding | 36.92 | -0.001930857 | 0.001930857 | ENSG00000111530 | n.s. |
| ENSG00000235652 | 63.97792 | -0.088233628 | 0.280244 | -0.314846 | 0.752879 | 0.950880711 | | antisense | 36.99 | -0.001930019 | 0.001930019 | ENSG00000235652 | n.s. |
| ENSG00000187240 | 694.5717 | -0.08822316 | 0.280325 | -0.314718 | 0.752976 | 0.950880711 | DYNC2H1 | protein_coding | 35.1 | -0.00192979 | 0.00192979 | ENSG00000187240 | n.s. |
| ENSG00000104731 | 77.55218 | -0.088173807 | 0.280058 | -0.314841 | 0.752882 | 0.950880711 | KLHDC4 | protein_coding | 53.01 | -0.001928711 | 0.001928711 | ENSG00000104731 | n.s. |
| ENSG00000198146 | 2074.621 | 0.081823978 | 0.247244 | 0.3309442 | 0.740687 | 0.947269752 | ZNF770 | protein_coding | 37.91 | 0.001925018 | 0.001925018 | ENSG00000198146 | n.s. |
| ENSG00000083535 | 1178.511 | -0.089185311 | 0.285987 | -0.311851 | 0.755153 | 0.951579479 | PIBF1 | protein_coding | 36.99 | -0.001922383 | 0.001922383 | ENSG00000083535 | n.s. |
| ENSG00000114988 | 344.3773 | 0.094265286 | 0.312534 | 0.301616 | 0.762945 | 0.954143629 | LMAN2L | protein_coding | 44.97 | 0.001921715 | 0.001921715 | ENSG00000114988 | n.s. |
| ENSG00000149932 | 231.5781 | 0.096620742 | 0.329059 | 0.2936272 | 0.769043 | 0.955275572 | TMEM219 | protein_coding | 47.42 | 0.001919982 | 0.001919982 | ENSG00000149932 | n.s. |
| ENSG00000204576 | 582.6084 | -0.078286854 | 0.230597 | -0.339497 | 0.734236 | 0.945151747 | PRR3 | protein_coding | 44.95 | -0.001917907 | 0.001917907 | ENSG00000204576 | n.s. |
| ENSG00000155366 | 173.6144 | 0.094805491 | 0.315611 | 0.3003875 | 0.763882 | 0.954536941 | RHOC | protein_coding | 57.18 | 0.001915759 | 0.001915759 | ENSG00000155366 | n.s. |
| ENSG00000135829 | 16772.16 | 0.08401097 | 0.259172 | 0.3241513 | 0.745823 | 0.948877395 | DHX9 | protein_coding | 38.24 | 0.001914602 | 0.001914602 | ENSG00000135829 | n.s. |
| ENSG00000141569 | 778.0535 | 0.090480522 | 0.292036 | 0.3098266 | 0.756693 | 0.95246603 | TRIM65 | protein_coding | 54.93 | 0.001913709 | 0.001913709 | ENSG00000141569 | n.s. |
| ENSG00000156110 | 735.3233 | 0.090447834 | 0.29209 | 0.3096576 | 0.756821 | 0.95246603 | ADK | protein_coding | 37.42 | 0.001913017 | 0.001913017 | ENSG00000156110 | n.s. |
| ENSG00000135446 | 5140.17 | -0.086283242 | 0.270816 | -0.318604 | 0.750027 | 0.95029168 | CDK4 | protein_coding | 53.24 | -0.001910576 | 0.001910576 | ENSG00000135446 | n.s. |
| ENSG00000201302 | 109.2181 | 0.08493355 | 0.264108 | 0.3215862 | 0.747766 | 0.949642637 | SNORA65 | snoRNA | 54.62 | 0.001905891 | 0.001905891 | ENSG00000201302 | n.s. |
| ENSG00000149100 | 4168.549 | 0.086907335 | 0.274151 | 0.3170052 | 0.75124 | 0.950815756 | EIF3M | protein_coding | 37.78 | 0.001903586 | 0.001903586 | ENSG00000149100 | n.s. |
| ENSG00000228223 | 635.5603 | -0.077296483 | 0.226732 | -0.340915 | 0.733167 | 0.944876691 | HCG11 | lincRNA | 39.3 | -0.001903415 | 0.001903415 | ENSG00000228223 | n.s. |
| ENSG00000144848 | 603.429 | -0.084490016 | 0.262824 | -0.32147 | 0.747855 | 0.949642637 | ATG3 | protein_coding | 36.32 | -0.001895939 | 0.001895939 | ENSG00000144848 | n.s. |
| ENSG00000268403 | 537.3318 | 0.074211069 | 0.212683 | 0.3489278 | 0.727144 | 0.942883666 | | antisense | 49.35 | 0.001895491 | 0.001895491 | ENSG00000268403 | n.s. |
| ENSG00000238741 | 1413.288 | -0.076838144 | 0.223721 | -0.343456 | 0.731256 | 0.944807198 | SCARNA7 | scaRNA | 37.27 | -0.001894583 | 0.001894583 | ENSG00000238741 | n.s. |
| ENSG00000188542 | 330.9474 | -0.074173893 | 0.211609 | -0.350523 | 0.725946 | 0.942883666 | DUSP28 | protein_coding | 53.35 | -0.001894541 | 0.001894541 | ENSG00000188542 | n.s. |
| ENSG00000171469 | 939.215 | 0.076380037 | 0.22165 | 0.3445976 | 0.730397 | 0.944503674 | ZNF561 | protein_coding | 44.62 | 0.001893946 | 0.001893946 | ENSG00000171469 | n.s. |
| ENSG00000079999 | 972.3175 | -0.084369211 | 0.262557 | -0.321337 | 0.747955 | 0.949642637 | KEAP1 | protein_coding | 54.01 | -0.001893228 | 0.001893228 | ENSG00000079999 | n.s. |
| ENSG00000118873 | 2852.281 | -0.084359203 | 0.262238 | -0.321689 | 0.747688 | 0.949642637 | RAB3GAP2 | protein_coding | 37.8 | -0.001893003 | 0.001893003 | ENSG00000118873 | n.s. |

| | | | | | | | | | | | | | |
|---|---|---|---|---|---|---|---|---|---|---|---|---|---|
| ENSG00000130312 | 446.6906 | 0.095173725 | 0.326524 | 0.2914757 | 0.770688 | 0.955275572 | MRPL34 | protein_coding | 54.25 | 0.001891228 | 0.001891228 | ENSG00000130312 | n.s. |
| ENSG00000101367 | 2236.211 | -0.08678927 | 0.277059 | -0.313252 | 0.754089 | 0.951113891 | MAPRE1 | protein_coding | 44.44 | -0.001889183 | 0.001889183 | ENSG00000101367 | n.s. |
| ENSG00000065882 | 631.5736 | -0.08583738 | 0.270329 | -0.317529 | 0.750842 | 0.950585745 | TBC1D1 | protein_coding | 42.84 | -0.001889169 | 0.001889169 | ENSG00000065882 | n.s. |
| ENSG00000072041 | 3530.713 | 0.087476475 | 0.280128 | 0.3122737 | 0.754832 | 0.951579479 | SLC6A15 | protein_coding | 34.54 | 0.001885549 | 0.001885549 | ENSG00000072041 | n.s. |
| ENSG00000258984 | 280.559 | 0.094811539 | 0.324443 | 0.2922285 | 0.770112 | 0.955275572 | UBE2F-SCLY | protein_coding | 45.35 | 0.001884031 | 0.001884031 | ENSG00000258984 | n.s. |
| ENSG00000170365 | 495.9602 | -0.082667117 | 0.255004 | -0.32418 | 0.745802 | 0.948877395 | SMAD1 | protein_coding | 38.45 | -0.001883975 | 0.001883975 | ENSG00000170365 | n.s. |
| ENSG00000266472 | 2189.14 | -0.090059332 | 0.292941 | -0.307431 | 0.758515 | 0.95300702 | MRPS21 | protein_coding | 46.58 | -0.001882591 | 0.001882591 | ENSG00000266472 | n.s. |
| ENSG00000182325 | 213.7147 | -0.085908893 | 0.27209 | -0.315737 | 0.752202 | 0.950815756 | FBXL6 | protein_coding | 66.45 | -0.001881717 | 0.001881717 | ENSG00000182325 | n.s. |
| ENSG00000089006 | 4467.923 | 0.077568299 | 0.230453 | 0.3365912 | 0.736425 | 0.945680734 | SNX5 | protein_coding | 42.13 | 0.001881455 | 0.001881455 | ENSG00000089006 | n.s. |
| ENSG00000260260 | 361.3258 | 0.076380457 | 0.222836 | 0.3427654 | 0.731775 | 0.944867156 | SNHG19 | lincRNA | 54.09 | 0.001881193 | 0.001881193 | ENSG00000260260 | n.s. |
| ENSG00000126001 | 82.90343 | 0.084086561 | 0.262317 | 0.3205538 | 0.748549 | 0.949841385 | CEP250 | protein_coding | 47.9 | 0.001879243 | 0.001879243 | ENSG00000126001 | n.s. |
| ENSG00000168785 | 648.3564 | -0.084839606 | 0.266215 | -0.318689 | 0.749963 | 0.95029168 | TSPAN5 | protein_coding | 41.93 | -0.00187861 | 0.00187861 | ENSG00000168785 | n.s. |
| ENSG00000178229 | 421.5444 | 0.079811053 | 0.241249 | 0.3308249 | 0.740777 | 0.947269752 | ZNF543 | protein_coding | 47.36 | 0.001877661 | 0.001877661 | ENSG00000178229 | n.s. |
| ENSG00000119471 | 400.7249 | 0.090462636 | 0.297756 | 0.3038146 | 0.761269 | 0.953332446 | HSDL2 | protein_coding | 40.81 | 0.001877609 | 0.001877609 | ENSG00000119471 | n.s. |
| ENSG00000159579 | 1599.904 | -0.083627026 | 0.259494 | -0.322269 | 0.747249 | 0.949642637 | RSPRY1 | protein_coding | 41.63 | -0.001876573 | 0.001876573 | ENSG00000159579 | n.s. |
| ENSG00000166260 | 2640.431 | -0.086026813 | 0.273919 | -0.314059 | 0.753476 | 0.951015428 | COX11 | protein_coding | 41.57 | -0.001876454 | 0.001876454 | ENSG00000166260 | n.s. |
| ENSG00000188647 | 3588.622 | 0.085560665 | 0.270431 | 0.3163862 | 0.751709 | 0.950815756 | PTAR1 | protein_coding | 36.03 | 0.001874089 | 0.001874089 | ENSG00000188647 | n.s. |
| ENSG00000144034 | 946.4027 | -0.093915646 | 0.317588 | -0.295715 | 0.767447 | 0.955198262 | TPRKB | protein_coding | 40.52 | -0.00186953 | 0.00186953 | ENSG00000144034 | n.s. |
| ENSG00000135404 | 1804.276 | -0.093897704 | 0.322368 | -0.291275 | 0.770841 | 0.955275572 | CD63 | protein_coding | 59.18 | -0.001865872 | 0.001865872 | ENSG00000135404 | n.s. |
| ENSG00000137776 | 4843.944 | -0.082768172 | 0.256099 | -0.323188 | 0.746553 | 0.949420457 | SLTM | protein_coding | 38.73 | -0.001865712 | 0.001865712 | ENSG00000137776 | n.s. |
| ENSG00000092330 | 545.1405 | -0.083095932 | 0.258377 | -0.321608 | 0.74775 | 0.949642637 | TINF2 | protein_coding | 53.3 | -0.001864656 | 0.001864656 | ENSG00000092330 | n.s. |
| ENSG00000213965 | 281.0089 | -0.080801418 | 0.246728 | -0.327492 | 0.743296 | 0.948261154 | NUDT19 | protein_coding | 46.13 | -0.001864254 | 0.001864254 | ENSG00000213965 | n.s. |
| ENSG00000170464 | 453.1132 | -0.089811413 | 0.294825 | -0.304626 | 0.760651 | 0.953332446 | DNAJC18 | protein_coding | 44.33 | -0.001864092 | 0.001864092 | ENSG00000170464 | n.s. |
| ENSG00000185760 | 1394.554 | -0.083063569 | 0.258469 | -0.321367 | 0.747932 | 0.949642637 | KCNQ5 | protein_coding | 37.35 | -0.001863929 | 0.001863929 | ENSG00000185760 | n.s. |
| ENSG00000064102 | 1080.153 | 0.081774951 | 0.252305 | 0.324112 | 0.745853 | 0.948877395 | INTS13 | protein_coding | 36.38 | 0.001863643 | 0.001863643 | ENSG00000064102 | n.s. |
| ENSG00000151135 | 820.2156 | 0.08703116 | 0.279579 | 0.3112942 | 0.755577 | 0.95189487 | TMEM263 | protein_coding | 38.27 | 0.001863425 | 0.001863425 | ENSG00000151135 | n.s. |
| ENSG00000188690 | 615.7422 | -0.089602009 | 0.292901 | -0.305913 | 0.759671 | 0.953277452 | UROS | protein_coding | 46.1 | -0.001861991 | 0.001861991 | ENSG00000188690 | n.s. |
| ENSG00000113649 | 4898.568 | -0.083089087 | 0.258777 | -0.321084 | 0.748147 | 0.949712939 | TCERG1 | protein_coding | 36.35 | -0.001861831 | 0.001861831 | ENSG00000113649 | n.s. |
| ENSG00000213760 | 163.1925 | -0.08423922 | 0.264938 | -0.317958 | 0.750517 | 0.950388734 | ATP6V1G2 | protein_coding | 49.67 | -0.001861579 | 0.001861579 | ENSG00000213760 | n.s. |
| ENSG00000116161 | 4340.594 | 0.085026194 | 0.270028 | 0.3148791 | 0.752853 | 0.950880711 | CACYBP | protein_coding | 40.31 | 0.00185986 | 0.00185986 | ENSG00000116161 | n.s. |
| ENSG00000109917 | 775.6694 | 0.08060589 | 0.246673 | 0.3267727 | 0.74384 | 0.948261154 | ZPR1 | protein_coding | 44.53 | 0.001859742 | 0.001859742 | ENSG00000109917 | n.s. |
| ENSG00000168496 | 3292.13 | -0.080474088 | 0.245925 | -0.32723 | 0.743494 | 0.948261154 | FEN1 | protein_coding | 49.78 | -0.001856701 | 0.001856701 | ENSG00000168496 | n.s. |

| | | | | | | | | | | | | | |
|---|---|---|---|---|---|---|---|---|---|---|---|---|---|
| ENSG00000278126 | 32.34111 | -0.098242428 | 0.347798 | -0.28247 | 0.777583 | 0.957426384 | | sense_intronic | 43.43 | -0.001856252 | 0.001856252 | ENSG00000278126 | n.s. |
| ENSG00000256185 | 8.081095 | 0.158557366 | 0.851602 | 0.1861873 | 0.852298 | 0.973446151 | | 3prime_overlapping_ncRNA | 36.43 | 0.001853229 | 0.001853229 | ENSG00000256185 | n.s. |
| ENSG00000273064 | 98.10764 | 0.083684232 | 0.262596 | 0.3186802 | 0.749969 | 0.95029168 | | antisense | 46.27 | 0.001853026 | 0.001853026 | ENSG00000273064 | n.s. |
| ENSG00000178927 | 833.8854 | -0.080307357 | 0.246411 | -0.325909 | 0.744493 | 0.948261154 | C17orf62 | protein_coding | 60.43 | -0.001852855 | 0.001852855 | ENSG00000178927 | n.s. |
| ENSG00000233170 | 44.44386 | 0.093136813 | 0.317193 | 0.2936284 | 0.769042 | 0.955275572 | | processed_pseudogene | 49.33 | 0.001850752 | 0.001850752 | ENSG00000233170 | n.s. |
| ENSG00000278834 | 83.87565 | -0.084357682 | 0.267065 | -0.31587 | 0.752102 | 0.950815756 | | antisense | 50.53 | -0.001847739 | 0.001847739 | ENSG00000278834 | n.s. |
| ENSG00000235908 | 46.69639 | -0.092799454 | 0.317131 | -0.292622 | 0.769811 | 0.955275572 | RHOA-IT1 | sense_intronic | 47.91 | -0.001844048 | 0.001844048 | ENSG00000235908 | n.s. |
| ENSG00000157538 | 562.0556 | -0.086509341 | 0.278412 | -0.310724 | 0.75601 | 0.952142297 | DSCR3 | protein_coding | 44.22 | -0.001842488 | 0.001842488 | ENSG00000157538 | n.s. |
| ENSG00000272969 | 32.64932 | -0.103897168 | 0.381604 | -0.272264 | 0.785419 | 0.960002107 | | antisense | 50.77 | -0.00184187 | 0.00184187 | ENSG00000272969 | n.s. |
| ENSG00000116604 | 131.0319 | -0.082058804 | 0.254792 | -0.322062 | 0.747406 | 0.949642637 | MEF2D | protein_coding | 54.23 | -0.001841383 | 0.001841383 | ENSG00000116604 | n.s. |
| ENSG00000129187 | 925.3143 | -0.083055897 | 0.261297 | -0.317861 | 0.750591 | 0.950388734 | DCTD | protein_coding | 46.1 | -0.001835429 | 0.001835429 | ENSG00000129187 | n.s. |
| ENSG00000129128 | 3216.118 | 0.081635317 | 0.253888 | 0.3215402 | 0.747801 | 0.949642637 | SPCS3 | protein_coding | 36.77 | 0.00183188 | 0.00183188 | ENSG00000129128 | n.s. |
| ENSG00000186665 | 205.2628 | 0.084070101 | 0.268254 | 0.3133974 | 0.753979 | 0.951113891 | C17orf58 | protein_coding | 52.41 | 0.001829994 | 0.001829994 | ENSG00000186665 | n.s. |
| ENSG00000080824 | 23790.7 | 0.086377185 | 0.278506 | 0.3101448 | 0.756451 | 0.952438506 | HSP90AA1 | protein_coding | 47.25 | 0.001828005 | 0.001828005 | ENSG00000080824 | n.s. |
| ENSG00000205531 | 3063.587 | -0.079150334 | 0.243214 | -0.325434 | 0.744852 | 0.948261154 | NAP1L4 | protein_coding | 44.27 | -0.00182616 | 0.00182616 | ENSG00000205531 | n.s. |
| ENSG00000201772 | 28.77428 | 0.096695415 | 0.343042 | 0.2818759 | 0.778039 | 0.957539143 | SNORA5C | snoRNA | 59.12 | 0.001822077 | 0.001822077 | ENSG00000201772 | n.s. |
| ENSG00000272768 | 77.41145 | 0.088203753 | 0.291602 | 0.3024797 | 0.762286 | 0.953662831 | | antisense | 63.71 | 0.001817451 | 0.001817451 | ENSG00000272768 | n.s. |
| ENSG00000174851 | 445.0771 | -0.088118491 | 0.290942 | -0.302874 | 0.761986 | 0.953630085 | YIF1A | protein_coding | 58.9 | -0.001817008 | 0.001817008 | ENSG00000174851 | n.s. |
| ENSG00000143363 | 322.3539 | -0.089774596 | 0.300438 | -0.298813 | 0.765083 | 0.954536941 | PRUNE1 | protein_coding | 46.61 | -0.001814099 | 0.001814099 | ENSG00000143363 | n.s. |
| ENSG00000116761 | 485.9984 | -0.082717412 | 0.261051 | -0.316862 | 0.751348 | 0.950815756 | CTH | protein_coding | 39.51 | -0.001811812 | 0.001811812 | ENSG00000116761 | n.s. |
| ENSG00000092094 | 639.0626 | -0.08055335 | 0.25011 | -0.322072 | 0.747398 | 0.949642637 | OSGEP | protein_coding | 45.15 | -0.001807601 | 0.001807601 | ENSG00000092094 | n.s. |
| ENSG00000239779 | 517.7449 | -0.090211401 | 0.303271 | -0.297461 | 0.766114 | 0.954933771 | WBP1 | protein_coding | 56.07 | -0.001806641 | 0.001806641 | ENSG00000239779 | n.s. |
| ENSG00000101189 | 343.5353 | -0.087657229 | 0.289607 | -0.302677 | 0.762136 | 0.953662831 | MRGBP | protein_coding | 59.23 | -0.00180619 | 0.00180619 | ENSG00000101189 | n.s. |
| ENSG00000116584 | 1138.992 | -0.081539479 | 0.255699 | -0.318888 | 0.749811 | 0.95029168 | ARHGEF2 | protein_coding | 50.95 | -0.001805535 | 0.001805535 | ENSG00000116584 | n.s. |
| ENSG00000077097 | 8130.779 | -0.081447357 | 0.255816 | -0.318383 | 0.750194 | 0.950319058 | TOP2B | protein_coding | 35.14 | -0.001802476 | 0.001802476 | ENSG00000077097 | n.s. |
| ENSG00000105700 | 481.7716 | 0.086211542 | 0.280884 | 0.3069292 | 0.758897 | 0.95300702 | KXD1 | protein_coding | 53.42 | 0.001802157 | 0.001802157 | ENSG00000105700 | n.s. |
| ENSG00000075415 | 9326.224 | 0.082244815 | 0.260012 | 0.3163112 | 0.751766 | 0.950815756 | SLC25A3 | protein_coding | 42.39 | 0.00180146 | 0.00180146 | ENSG00000075415 | n.s. |
| ENSG00000055211 | 439.6314 | -0.086171393 | 0.280558 | -0.307143 | 0.758735 | 0.95300702 | GINM1 | protein_coding | 42.05 | -0.001801318 | 0.001801318 | ENSG00000055211 | n.s. |
| ENSG00000171720 | 733.8903 | -0.085131271 | 0.274912 | -0.309667 | 0.756814 | 0.95246603 | HDAC3 | protein_coding | 45.27 | -0.001800569 | 0.001800569 | ENSG00000171720 | n.s. |
| ENSG00000036448 | 56.25736 | 0.090568368 | 0.309014 | 0.2930878 | 0.769455 | 0.955275572 | MYOM2 | protein_coding | 46.64 | 0.001799714 | 0.001799714 | ENSG00000036448 | n.s. |
| ENSG00000172115 | 6844.76 | 0.080542055 | 0.251866 | 0.319781 | 0.749134 | 0.94993942 | CYCS | protein_coding | 43.01 | 0.001796417 | 0.001796417 | ENSG00000172115 | n.s. |
| ENSG00000109832 | 124.2096 | 0.084929622 | 0.274397 | 0.3095142 | 0.75693 | 0.95246603 | DDX25 | protein_coding | 44.92 | 0.001796304 | 0.001796304 | ENSG00000109832 | n.s. |

| | | | | | | | | | | | | | |
|---|---|---|---|---|---|---|---|---|---|---|---|---|---|
| ENSG00000198597 | 541.2495 | -0.082119727 | 0.261133 | -0.314475 | 0.75316 | 0.950880711 | ZNF536 | protein_coding | 46.3 | -0.001796284 | 0.001796284 | ENSG00000198597 | n.s. |
| ENSG00000149809 | 107.221 | -0.092173989 | 0.319683 | -0.28833 | 0.773094 | 0.956186787 | TM7SF2 | protein_coding | 60.18 | -0.001793453 | 0.001793453 | ENSG00000149809 | n.s. |
| ENSG00000046653 | 514.9448 | 0.086400097 | 0.283464 | 0.3048009 | 0.760518 | 0.953332446 | GPM6B | protein_coding | 41.5 | 0.001793288 | 0.001793288 | ENSG00000046653 | n.s. |
| ENSG00000261334 | 61.57961 | 0.093481137 | 0.327038 | 0.2858419 | 0.774999 | 0.95679591 | | lincRNA | 59.15 | 0.001793033 | 0.001793033 | ENSG00000261334 | n.s. |
| ENSG00000166275 | 633.6052 | -0.089873724 | 0.307627 | -0.292151 | 0.770171 | 0.955275572 | BORCS7 | protein_coding | 41.69 | -0.00178591 | 0.00178591 | ENSG00000166275 | n.s. |
| ENSG00000276644 | 1645.172 | -0.078130766 | 0.240644 | -0.324674 | 0.745428 | 0.948769378 | DACH1 | protein_coding | 33.93 | -0.001784455 | 0.001784455 | ENSG00000276644 | n.s. |
| ENSG00000249669 | 531.0198 | 0.085843888 | 0.280584 | 0.3059473 | 0.759645 | 0.953277452 | CARMN | lincRNA | 51.97 | 0.001783894 | 0.001783894 | ENSG00000249669 | n.s. |
| ENSG00000148090 | 321.1945 | -0.077285157 | 0.235612 | -0.328019 | 0.742897 | 0.948261154 | AUH | protein_coding | 38.91 | -0.001783126 | 0.001783126 | ENSG00000148090 | n.s. |
| ENSG00000121905 | 323.2288 | 0.092673422 | 0.323162 | 0.2867705 | 0.774288 | 0.956728167 | HPCA | protein_coding | 57.12 | 0.00178039 | 0.00178039 | ENSG00000121905 | n.s. |
| ENSG00000197961 | 4084.146 | 0.079771121 | 0.249664 | 0.3195133 | 0.749337 | 0.949944274 | ZNF121 | protein_coding | 41.98 | 0.001779045 | 0.001779045 | ENSG00000197961 | n.s. |
| ENSG00000100138 | 2026.119 | 0.087979788 | 0.2938 | 0.2994545 | 0.764593 | 0.954536941 | SNU13 | protein_coding | 46.79 | 0.00177783 | 0.00177783 | ENSG00000100138 | n.s. |
| ENSG00000131269 | 540.0254 | -0.08499496 | 0.277152 | -0.306672 | 0.759093 | 0.95300702 | ABCB7 | protein_coding | 35.98 | -0.001776726 | 0.001776726 | ENSG00000131269 | n.s. |
| ENSG00000143222 | 1057.347 | -0.085541726 | 0.280791 | -0.304645 | 0.760636 | 0.953332446 | UFC1 | protein_coding | 46.11 | -0.001775472 | 0.001775472 | ENSG00000143222 | n.s. |
| ENSG00000090263 | 591.7911 | 0.089178223 | 0.306269 | 0.2911757 | 0.770917 | 0.955275572 | MRPS33 | protein_coding | 41.83 | 0.00177209 | 0.00177209 | ENSG00000090263 | n.s. |
| ENSG00000231584 | 100.2649 | -0.089078163 | 0.301734 | -0.295221 | 0.767825 | 0.955275572 | FAHD2CP | cribed_unprocessed_pseudc | 49.42 | -0.001770101 | 0.001770101 | ENSG00000231584 | n.s. |
| ENSG00000183527 | 1348.753 | 0.081175688 | 0.258776 | 0.3136911 | 0.753756 | 0.951060328 | PSMG1 | protein_coding | 41.56 | 0.001768975 | 0.001768975 | ENSG00000183527 | n.s. |
| ENSG00000168502 | 108.2435 | -0.085782941 | 0.283468 | -0.30262 | 0.76218 | 0.953662831 | MTCL1 | protein_coding | 45.45 | -0.00176757 | 0.00176757 | ENSG00000168502 | n.s. |
| ENSG00000066084 | 2140.142 | -0.08067357 | 0.255163 | -0.316165 | 0.751877 | 0.950815756 | DIP2B | protein_coding | 41.84 | -0.001767044 | 0.001767044 | ENSG00000066084 | n.s. |
| ENSG00000143033 | 4218.83 | -0.080626596 | 0.255461 | -0.315612 | 0.752297 | 0.950819931 | MTF2 | protein_coding | 37.37 | -0.001765861 | 0.001765861 | ENSG00000143033 | n.s. |
| ENSG00000137207 | 544.3559 | 0.084443486 | 0.274096 | 0.3080797 | 0.758022 | 0.95300702 | YIPF3 | protein_coding | 52.46 | 0.001765198 | 0.001765198 | ENSG00000137207 | n.s. |
| ENSG00000061918 | 169.8048 | 0.091882507 | 0.321484 | 0.2858072 | 0.775026 | 0.95679591 | GUCY1B3 | protein_coding | 36.52 | 0.00176237 | 0.00176237 | ENSG00000061918 | n.s. |
| ENSG00000115946 | 1272.394 | 0.084754913 | 0.277302 | 0.3056414 | 0.759878 | 0.953277452 | PNO1 | protein_coding | 41.05 | 0.001761265 | 0.001761265 | ENSG00000115946 | n.s. |
| ENSG00000164162 | 537.7516 | 0.080397909 | 0.254542 | 0.3158536 | 0.752114 | 0.950815756 | ANAPC10 | protein_coding | 37.19 | 0.001761006 | 0.001761006 | ENSG00000164162 | n.s. |
| ENSG00000138138 | 1652.241 | -0.084804384 | 0.278105 | -0.304937 | 0.760414 | 0.953332446 | ATAD1 | protein_coding | 36.12 | -0.001760168 | 0.001760168 | ENSG00000138138 | n.s. |
| ENSG00000266063 | 4.227674 | 0.130487205 | 0.589941 | 0.2211869 | 0.824947 | 0.969452399 | MIR4771-2 | miRNA | 44.59 | 0.001758121 | 0.001758121 | ENSG00000266063 | n.s. |
| ENSG00000156983 | 273.6583 | 0.074646793 | 0.225378 | 0.3312076 | 0.740488 | 0.947244602 | BRPF1 | protein_coding | 51.62 | 0.001757026 | 0.001757026 | ENSG00000156983 | n.s. |
| ENSG00000137434 | 29.65332 | 0.096906463 | 0.349196 | 0.2775129 | 0.781386 | 0.959134639 | C6orf52 | protein_coding | 41.8 | 0.001755986 | 0.001755986 | ENSG00000137434 | n.s. |
| ENSG00000111229 | 2137.898 | -0.093107545 | 0.330315 | -0.281875 | 0.778039 | 0.957539143 | ARPC3 | protein_coding | 45.85 | -0.001754469 | 0.001754469 | ENSG00000111229 | n.s. |
| ENSG00000142252 | 204.626 | 0.089080654 | 0.306785 | 0.2903684 | 0.771534 | 0.955688505 | GEMIN7 | protein_coding | 50.45 | 0.001753431 | 0.001753431 | ENSG00000142252 | n.s. |
| ENSG00000242259 | 948.0687 | -0.080003221 | 0.25341 | -0.315706 | 0.752225 | 0.950815756 | C22orf39 | protein_coding | 47.03 | -0.001752361 | 0.001752361 | ENSG00000242259 | n.s. |
| ENSG00000138448 | 1553.701 | -0.087966249 | 0.297107 | -0.296076 | 0.767172 | 0.955198262 | ITGAV | protein_coding | 35.78 | -0.001751098 | 0.001751098 | ENSG00000138448 | n.s. |
| ENSG00000176390 | 357.7605 | -0.084434976 | 0.278312 | -0.303382 | 0.761599 | 0.95344167 | CRLF3 | protein_coding | 42.46 | -0.0017483 | 0.0017483 | ENSG00000176390 | n.s. |

| | | | | | | | | | | | | | |
|---|---|---|---|---|---|---|---|---|---|---|---|---|---|
| ENSG00000142657 | 1031.22 | -0.084122082 | 0.275607 | -0.305225 | 0.760195 | 0.953332446 | PGD | protein_coding | 47.69 | -0.001746006 | 0.001746006 | ENSG00000142657 | n.s. |
| ENSG00000197958 | 17002.35 | -0.089325644 | 0.308826 | -0.289243 | 0.772395 | 0.956001592 | RPL12 | protein_coding | 51.23 | -0.001745547 | 0.001745547 | ENSG00000197958 | n.s. |
| ENSG00000185437 | 51.53958 | -0.089590816 | 0.310573 | -0.28847 | 0.772987 | 0.956186787 | SH3BGR | protein_coding | 40.49 | -0.001743192 | 0.001743192 | ENSG00000185437 | n.s. |
| ENSG00000188636 | 1073.787 | -0.075519698 | 0.231297 | -0.326506 | 0.744042 | 0.948261154 | RTL6 | protein_coding | 56.28 | -0.001742394 | 0.001742394 | ENSG00000188636 | n.s. |
| ENSG00000182916 | 617.4825 | -0.090650952 | 0.317569 | -0.285453 | 0.775297 | 0.95679591 | TCEAL7 | protein_coding | 44.3 | -0.001738748 | 0.001738748 | ENSG00000182916 | n.s. |
| ENSG00000110172 | 2447.671 | -0.076177916 | 0.234979 | -0.32419 | 0.745794 | 0.948877395 | CHORDC1 | protein_coding | 35.9 | -0.001736087 | 0.001736087 | ENSG00000110172 | n.s. |
| ENSG00000160908 | 383.4626 | 0.080249287 | 0.2566 | 0.3127404 | 0.754478 | 0.951417039 | ZNF394 | protein_coding | 47.6 | 0.001735718 | 0.001735718 | ENSG00000160908 | n.s. |
| ENSG00000116337 | 88.97368 | 0.087336521 | 0.299711 | 0.2914026 | 0.770743 | 0.955275572 | AMPD2 | protein_coding | 56.74 | 0.001735493 | 0.001735493 | ENSG00000116337 | n.s. |
| ENSG00000185386 | 166.6844 | -0.085801205 | 0.287348 | -0.298597 | 0.765248 | 0.954536941 | MAPK11 | protein_coding | 65.39 | -0.001733807 | 0.001733807 | ENSG00000185386 | n.s. |
| ENSG00000174243 | 2727.291 | -0.085658313 | 0.28526 | -0.300282 | 0.763962 | 0.954536941 | DDX23 | protein_coding | 49.66 | -0.00173092 | 0.00173092 | ENSG00000174243 | n.s. |
| ENSG00000109536 | 1146.669 | -0.082778838 | 0.269857 | -0.306751 | 0.759033 | 0.95300702 | FRG1 | protein_coding | 35.99 | -0.001730401 | 0.001730401 | ENSG00000109536 | n.s. |
| ENSG00000014216 | 283.1251 | -0.085589788 | 0.285109 | -0.3002 | 0.764025 | 0.954536941 | CAPN1 | protein_coding | 51.18 | -0.001729535 | 0.001729535 | ENSG00000014216 | n.s. |
| ENSG00000200150 | 11.76533 | 0.14636841 | 0.718668 | 0.2036661 | 0.838614 | 0.973164573 | | snoRNA | 35.62 | 0.001729154 | 0.001729154 | ENSG00000200150 | n.s. |
| ENSG00000184983 | 1740.737 | -0.086811854 | 0.296031 | -0.293253 | 0.769329 | 0.955275572 | NDUFA6 | protein_coding | 48.79 | -0.001725067 | 0.001725067 | ENSG00000184983 | n.s. |
| ENSG00000213930 | 151.1445 | 0.082997335 | 0.270969 | 0.3062983 | 0.759378 | 0.953277452 | GALT | protein_coding | 50.16 | 0.001724741 | 0.001724741 | ENSG00000213930 | n.s. |
| ENSG00000101474 | 953.1897 | -0.085765186 | 0.287533 | -0.298279 | 0.76549 | 0.954754096 | APMAP | protein_coding | 46.54 | -0.001724607 | 0.001724607 | ENSG00000101474 | n.s. |
| ENSG00000162409 | 859.592 | -0.085265714 | 0.285419 | -0.298739 | 0.765139 | 0.954536941 | PRKAA2 | protein_coding | 36.59 | -0.001722986 | 0.001722986 | ENSG00000162409 | n.s. |
| ENSG00000147224 | 933.2471 | -0.086626572 | 0.29602 | -0.292637 | 0.769799 | 0.955275572 | PRPS1 | protein_coding | 41.83 | -0.001721385 | 0.001721385 | ENSG00000147224 | n.s. |
| ENSG00000136813 | 3037.32 | -0.078542966 | 0.248717 | -0.315792 | 0.75216 | 0.950815756 | KIAA0368 | protein_coding | 39.33 | -0.001720376 | 0.001720376 | ENSG00000136813 | n.s. |
| ENSG00000101911 | 531.6513 | -0.087239463 | 0.300252 | -0.290555 | 0.771392 | 0.955609187 | PRPS2 | protein_coding | 42.76 | -0.001720335 | 0.001720335 | ENSG00000101911 | n.s. |
| ENSG00000115204 | 485.8038 | -0.082182861 | 0.267748 | -0.306941 | 0.758888 | 0.95300702 | MPV17 | protein_coding | 50.33 | -0.001717942 | 0.001717942 | ENSG00000115204 | n.s. |
| ENSG00000224578 | 395.2139 | -0.082140784 | 0.266425 | -0.308307 | 0.757849 | 0.95300702 | HNRNPA1P48 | nscribed_processed_pseudog | 45.03 | -0.001717063 | 0.001717063 | ENSG00000224578 | n.s. |
| ENSG00000066926 | 578.7219 | -0.086318861 | 0.292877 | -0.294727 | 0.768202 | 0.955275572 | FECH | protein_coding | 42.66 | -0.00171527 | 0.00171527 | ENSG00000066926 | n.s. |
| ENSG00000177600 | 5744.427 | -0.08631359 | 0.296416 | -0.291191 | 0.770906 | 0.955275572 | RPLP2 | protein_coding | 57.05 | -0.001715166 | 0.001715166 | ENSG00000177600 | n.s. |
| ENSG00000207633 | 10.0939 | -0.117763099 | 0.508785 | -0.23146 | 0.816958 | 0.967029262 | MIR505 | miRNA | 52.38 | -0.001714676 | 0.001714676 | ENSG00000207633 | n.s. |
| ENSG00000276081 | 4.21576 | -0.129544907 | 0.596325 | -0.217239 | 0.828022 | 0.970009871 | MIR7156 | miRNA | 55 | -0.001713082 | 0.001713082 | ENSG00000276081 | n.s. |
| ENSG00000272247 | 27.94748 | 0.100664841 | 0.3833 | 0.2626264 | 0.792839 | 0.961629391 | | antisense | 51.84 | 0.001710524 | 0.001710524 | ENSG00000272247 | n.s. |
| ENSG00000151276 | 170.5832 | -0.078378736 | 0.249506 | -0.314135 | 0.753418 | 0.951015428 | MAGI1 | protein_coding | 40.3 | -0.001709631 | 0.001709631 | ENSG00000151276 | n.s. |
| ENSG00000141933 | 173.7688 | -0.086976576 | 0.299903 | -0.290016 | 0.771804 | 0.955779792 | TPGS1 | protein_coding | 61.72 | -0.001708407 | 0.001708407 | ENSG00000141933 | n.s. |
| ENSG00000146409 | 179.5405 | 0.0858109 | 0.294418 | 0.2914599 | 0.7707 | 0.955275572 | SLC18B1 | protein_coding | 39.58 | 0.001705177 | 0.001705177 | ENSG00000146409 | n.s. |
| ENSG00000112514 | 1365.208 | -0.084223507 | 0.281585 | -0.299105 | 0.76486 | 0.954536941 | CUTA | protein_coding | 54.9 | -0.001701926 | 0.001701926 | ENSG00000112514 | n.s. |
| ENSG00000114520 | 1362.181 | -0.085606871 | 0.292252 | -0.292921 | 0.769582 | 0.955275572 | SNX4 | protein_coding | 40.09 | -0.001701122 | 0.001701122 | ENSG00000114520 | n.s. |

| | | | | | | | | | | | | | |
|---|---|---|---|---|---|---|---|---|---|---|---|---|---|
| ENSG00000198722 | 980.6869 | -0.081948011 | 0.269018 | -0.304619 | 0.760657 | 0.953332446 | UNC13B | protein_coding | 41.63 | -0.001700882 | 0.001700882 | ENSG00000198722 | n.s. |
| ENSG00000121680 | 177.9254 | -0.085336833 | 0.288636 | -0.295656 | 0.767493 | 0.955198262 | PEX16 | protein_coding | 58.04 | -0.001698756 | 0.001698756 | ENSG00000121680 | n.s. |
| ENSG00000169189 | 401.0103 | 0.088484342 | 0.310118 | 0.2853247 | 0.775395 | 0.956822701 | NSMCE1 | protein_coding | 47.29 | 0.001696115 | 0.001696115 | ENSG00000169189 | n.s. |
| ENSG00000101146 | 1432.371 | 0.081066615 | 0.263061 | 0.3081669 | 0.757955 | 0.95300702 | RAE1 | protein_coding | 44.61 | 0.001694608 | 0.001694608 | ENSG00000101146 | n.s. |
| ENSG00000113575 | 4360.107 | -0.081809086 | 0.269856 | -0.303159 | 0.761769 | 0.953443822 | PPP2CA | protein_coding | 40.38 | -0.001693848 | 0.001693848 | ENSG00000113575 | n.s. |
| ENSG00000113068 | 892.9621 | -0.083753334 | 0.280278 | -0.298822 | 0.765076 | 0.954536941 | PFDN1 | protein_coding | 40.78 | -0.001692425 | 0.001692425 | ENSG00000113068 | n.s. |
| ENSG00000235499 | 116.2991 | 0.083732633 | 0.279653 | 0.2994166 | 0.764622 | 0.954536941 | | lincRNA | 69.31 | 0.001692007 | 0.001692007 | ENSG00000235499 | n.s. |
| ENSG00000264792 | 8.401279 | 0.136705044 | 0.655641 | 0.208506 | 0.834834 | 0.971936785 | MIR4637 | miRNA | 30.95 | 0.001689945 | 0.001689945 | ENSG00000264792 | n.s. |
| ENSG00000080986 | 1347.954 | -0.081406704 | 0.267856 | -0.303919 | 0.761189 | 0.953332446 | NDC80 | protein_coding | 36.36 | -0.001689647 | 0.001689647 | ENSG00000080986 | n.s. |
| ENSG00000169684 | 854.0808 | -0.080640559 | 0.261843 | -0.307973 | 0.758103 | 0.95300702 | CHRNA5 | protein_coding | 40.43 | -0.001685702 | 0.001685702 | ENSG00000169684 | n.s. |
| ENSG00000207692 | 3.65699 | -0.13623879 | 0.647906 | -0.210276 | 0.833453 | 0.971936785 | MIR592 | miRNA | 45.36 | -0.001684181 | 0.001684181 | ENSG00000207692 | n.s. |
| ENSG00000204392 | 513.0504 | -0.089100749 | 0.314803 | -0.283036 | 0.777149 | 0.957426384 | LSM2 | protein_coding | 48.13 | -0.001683524 | 0.001683524 | ENSG00000204392 | n.s. |
| ENSG00000130396 | 4940.186 | -0.079180554 | 0.255056 | -0.310443 | 0.756224 | 0.952324948 | AFDN | protein_coding | 38.79 | -0.001679803 | 0.001679803 | ENSG00000130396 | n.s. |
| ENSG00000234707 | 28.93764 | 0.098826209 | 0.376217 | 0.262684 | 0.792794 | 0.961629391 | | antisense | 42.96 | 0.001679282 | 0.001679282 | ENSG00000234707 | n.s. |
| ENSG00000218596 | 105.1692 | -0.084479751 | 0.287407 | -0.293937 | 0.768806 | 0.955275572 | | processed_pseudogene | 47.63 | -0.001678725 | 0.001678725 | ENSG00000218596 | n.s. |
| ENSG00000100219 | 1243.863 | 0.079318323 | 0.255688 | 0.3102152 | 0.756397 | 0.952438506 | XBP1 | protein_coding | 44.93 | 0.001678618 | 0.001678618 | ENSG00000100219 | n.s. |
| ENSG00000167371 | 126.4632 | -0.083534861 | 0.280417 | -0.297895 | 0.765783 | 0.954823343 | PRRT2 | protein_coding | 61.52 | -0.001677127 | 0.001677127 | ENSG00000167371 | n.s. |
| ENSG00000162641 | 179.1996 | -0.076890997 | 0.245212 | -0.313569 | 0.753848 | 0.951060328 | AKNAD1 | protein_coding | 43.48 | -0.001675604 | 0.001675604 | ENSG00000162641 | n.s. |
| ENSG00000198242 | 6899.568 | 0.082903877 | 0.276908 | 0.2993914 | 0.764641 | 0.954536941 | RPL23A | protein_coding | 49.99 | 0.00167526 | 0.00167526 | ENSG00000198242 | n.s. |
| ENSG00000253586 | 19.56076 | 0.102141837 | 0.396575 | 0.2575597 | 0.796747 | 0.962958487 | | sense_intronic | 38.09 | 0.001674353 | 0.001674353 | ENSG00000253586 | n.s. |
| ENSG00000075790 | 258.0341 | -0.084108893 | 0.283899 | -0.296264 | 0.767029 | 0.955198262 | BCAP29 | protein_coding | 37.42 | -0.001674312 | 0.001674312 | ENSG00000075790 | n.s. |
| ENSG00000196656 | 12.63239 | -0.11498878 | 0.49603 | -0.231818 | 0.816679 | 0.967029262 | | transcribed_processed_pseudog | 35.46 | -0.001674281 | 0.001674281 | ENSG00000196656 | n.s. |
| ENSG00000227671 | 573.7506 | -0.082749718 | 0.276519 | -0.299255 | 0.764745 | 0.954536941 | | transcribed_unprocessed_pseudc | 40.91 | -0.001672145 | 0.001672145 | ENSG00000227671 | n.s. |
| ENSG00000092036 | 292.1574 | 0.08309529 | 0.279071 | 0.297757 | 0.765889 | 0.954823343 | HAUS4 | protein_coding | 45.59 | 0.001668302 | 0.001668302 | ENSG00000092036 | n.s. |
| ENSG00000265396 | 27.17056 | -0.093921643 | 0.344094 | -0.272953 | 0.784889 | 0.959966858 | MIR3128 | miRNA | 30.3 | -0.001666523 | 0.001666523 | ENSG00000265396 | n.s. |
| ENSG00000188933 | 600.1424 | 0.083812057 | 0.284866 | 0.2942154 | 0.768593 | 0.955275572 | USP32P1 | cribed_unprocessed_pseudc | 42.86 | 0.001665457 | 0.001665457 | ENSG00000188933 | n.s. |
| ENSG00000106052 | 1367.531 | -0.083348112 | 0.280578 | -0.297059 | 0.766422 | 0.955053865 | TAX1BP1 | protein_coding | 36.81 | -0.00166464 | 0.00166464 | ENSG00000106052 | n.s. |
| ENSG00000138614 | 1099.495 | -0.083751063 | 0.287297 | -0.291514 | 0.770658 | 0.955275572 | INTS14 | protein_coding | 40.67 | -0.001664245 | 0.001664245 | ENSG00000138614 | n.s. |
| ENSG00000102030 | 509.6869 | -0.087798176 | 0.30987 | -0.283339 | 0.776917 | 0.957426384 | NAA10 | protein_coding | 58.17 | -0.001658912 | 0.001658912 | ENSG00000102030 | n.s. |
| ENSG00000115216 | 816.1556 | -0.079288243 | 0.257777 | -0.307585 | 0.758398 | 0.95300702 | NRBP1 | protein_coding | 48.61 | -0.001657434 | 0.001657434 | ENSG00000115216 | n.s. |
| ENSG00000126947 | 276.0583 | -0.084598951 | 0.292331 | -0.289395 | 0.772279 | 0.955942907 | ARMCX1 | protein_coding | 48.99 | -0.001655436 | 0.001655436 | ENSG00000126947 | n.s. |
| ENSG00000263843 | 114.3721 | -0.079704167 | 0.262203 | -0.303979 | 0.761144 | 0.953332446 | | antisense | 60.78 | -0.00165431 | 0.00165431 | ENSG00000263843 | n.s. |

| | | | | | | | | | | | | | |
|---|---|---|---|---|---|---|---|---|---|---|---|---|---|
| ENSG00000203965 | 412.1696 | 0.086814876 | 0.305036 | 0.2846051 | 0.775947 | 0.957078818 | EFCAB7 | protein_coding | 35.15 | 0.001654023 | 0.001654023 | ENSG00000203965 | n.s. |
| ENSG00000147044 | 221.3744 | -0.076498203 | 0.244693 | -0.312629 | 0.754563 | 0.951437806 | CASK | protein_coding | 38.73 | -0.00165386 | 0.00165386 | ENSG00000147044 | n.s. |
| ENSG00000188554 | 1918.699 | 0.084368902 | 0.291523 | 0.289407 | 0.77227 | 0.955942907 | NBR1 | protein_coding | 44.16 | 0.001650934 | 0.001650934 | ENSG00000188554 | n.s. |
| ENSG00000185189 | 150.6833 | -0.078907975 | 0.257229 | -0.306761 | 0.759025 | 0.95300702 | NRBP2 | protein_coding | 61.1 | -0.001649484 | 0.001649484 | ENSG00000185189 | n.s. |
| ENSG00000198492 | 1786.399 | 0.075592792 | 0.240692 | 0.3140639 | 0.753472 | 0.951015428 | YTHDF2 | protein_coding | 43.57 | 0.001648863 | 0.001648863 | ENSG00000198492 | n.s. |
| ENSG00000251072 | 36.61296 | -0.094161523 | 0.349618 | -0.269327 | 0.787678 | 0.960530983 | | lincRNA | 44.31 | -0.001646755 | 0.001646755 | ENSG00000251072 | n.s. |
| ENSG00000272821 | 109.5489 | 0.08284557 | 0.280806 | 0.295028 | 0.767973 | 0.955275572 | | antisense | 61.4 | 0.001646251 | 0.001646251 | ENSG00000272821 | n.s. |
| ENSG00000169976 | 1329.8 | -0.088883108 | 0.317352 | -0.280078 | 0.779418 | 0.958292132 | SF3B5 | protein_coding | 56.57 | -0.001644522 | 0.001644522 | ENSG00000169976 | n.s. |
| ENSG00000154917 | 345.4185 | -0.08186113 | 0.2748 | -0.297893 | 0.765785 | 0.954823343 | RAB6B | protein_coding | 49.59 | -0.001643524 | 0.001643524 | ENSG00000154917 | n.s. |
| ENSG00000183828 | 81.27414 | -0.092685326 | 0.340772 | -0.271986 | 0.785633 | 0.960059735 | NUDT14 | protein_coding | 64.58 | -0.001640692 | 0.001640692 | ENSG00000183828 | n.s. |
| ENSG00000144566 | 753.2762 | -0.081162087 | 0.270931 | -0.299568 | 0.764507 | 0.954536941 | RAB5A | protein_coding | 39.8 | -0.001640063 | 0.001640063 | ENSG00000144566 | n.s. |
| ENSG00000143947 | 18638.53 | 0.088693405 | 0.317243 | 0.2795756 | 0.779803 | 0.958327464 | RPS27A | protein_coding | 45.81 | 0.001639592 | 0.001639592 | ENSG00000143947 | n.s. |
| ENSG00000219747 | 33.54213 | 0.123653911 | 0.570963 | 0.2165708 | 0.828543 | 0.970009871 | | processed_pseudogene | 51.11 | 0.00163518 | 0.00163518 | ENSG00000219747 | n.s. |
| ENSG00000103145 | 98.50253 | 0.08523875 | 0.297397 | 0.2866157 | 0.774407 | 0.956789714 | HCFC1R1 | protein_coding | 63.71 | 0.001635178 | 0.001635178 | ENSG00000103145 | n.s. |
| ENSG00000164953 | 173.8697 | 0.079315641 | 0.262147 | 0.3025621 | 0.762224 | 0.953662831 | TMEM67 | protein_coding | 37.38 | 0.00163431 | 0.00163431 | ENSG00000164953 | n.s. |
| ENSG00000133119 | 3529.418 | -0.078700818 | 0.258868 | -0.304019 | 0.761113 | 0.953332446 | RFC3 | protein_coding | 37.82 | -0.001633485 | 0.001633485 | ENSG00000133119 | n.s. |
| ENSG00000035141 | 1452.758 | -0.082149031 | 0.281743 | -0.291575 | 0.770612 | 0.955275572 | FAM136A | protein_coding | 48.46 | -0.00163241 | 0.00163241 | ENSG00000035141 | n.s. |
| ENSG00000266498 | 29.21826 | 0.099546841 | 0.386424 | 0.2576104 | 0.796708 | 0.962958487 | | lincRNA | 48.69 | 0.001631815 | 0.001631815 | ENSG00000266498 | n.s. |
| ENSG00000163468 | 11580.38 | 0.08628616 | 0.305351 | 0.2825806 | 0.777498 | 0.957426384 | CCT3 | protein_coding | 45.93 | 0.001630343 | 0.001630343 | ENSG00000163468 | n.s. |
| ENSG00000163946 | 814.986 | -0.08202982 | 0.279777 | -0.293197 | 0.769372 | 0.955275572 | FAM208A | protein_coding | 38.7 | -0.001630041 | 0.001630041 | ENSG00000163946 | n.s. |
| ENSG00000148399 | 522.5081 | 0.081590505 | 0.274904 | 0.296796 | 0.766622 | 0.955053865 | DPH7 | protein_coding | 52.97 | 0.001629536 | 0.001629536 | ENSG00000148399 | n.s. |
| ENSG00000205133 | 100.3546 | 0.081218901 | 0.272896 | 0.2976185 | 0.765994 | 0.95486968 | TRIQK | protein_coding | 34.5 | 0.001628918 | 0.001628918 | ENSG00000205133 | n.s. |
| ENSG00000147604 | 18745.18 | 0.081863854 | 0.280583 | 0.2917629 | 0.770468 | 0.955275572 | RPL7 | protein_coding | 50.72 | 0.001626743 | 0.001626743 | ENSG00000147604 | n.s. |
| ENSG00000163930 | 1197.84 | -0.078332374 | 0.256904 | -0.304909 | 0.760436 | 0.953332446 | BAP1 | protein_coding | 56.55 | -0.001625837 | 0.001625837 | ENSG00000163930 | n.s. |
| ENSG00000214941 | 193.4273 | -0.092899976 | 0.344796 | -0.269434 | 0.787595 | 0.960530983 | ZSWIM7 | protein_coding | 44.09 | -0.001624693 | 0.001624693 | ENSG00000214941 | n.s. |
| ENSG00000066557 | 2099.279 | -0.08120456 | 0.273582 | -0.296819 | 0.766604 | 0.955053865 | LRRC40 | protein_coding | 35.14 | -0.001621828 | 0.001621828 | ENSG00000066557 | n.s. |
| ENSG00000124786 | 356.7759 | -0.080173327 | 0.266743 | -0.300564 | 0.763747 | 0.954536941 | SLC35B3 | protein_coding | 34.49 | -0.001620083 | 0.001620083 | ENSG00000124786 | n.s. |
| ENSG00000141519 | 100.7592 | -0.077756361 | 0.255644 | -0.304158 | 0.761007 | 0.953332446 | CCDC40 | protein_coding | 50.67 | -0.001613882 | 0.001613882 | ENSG00000141519 | n.s. |
| ENSG00000187325 | 650.4532 | -0.081208117 | 0.275385 | -0.29489 | 0.768078 | 0.955275572 | TAF9B | protein_coding | 38.54 | -0.001613713 | 0.001613713 | ENSG00000187325 | n.s. |
| ENSG00000199150 | 12.633 | -0.107353361 | 0.449384 | -0.23889 | 0.811191 | 0.966064268 | MIRLET7G | miRNA | 50 | -0.001609654 | 0.001609654 | ENSG00000199150 | n.s. |
| ENSG00000150316 | 1105.949 | 0.080988031 | 0.277184 | 0.2921812 | 0.770148 | 0.955275572 | CWC15 | protein_coding | 37.4 | 0.00160934 | 0.00160934 | ENSG00000150316 | n.s. |
| ENSG00000141627 | 832.0384 | -0.08079283 | 0.273007 | -0.295937 | 0.767278 | 0.955198262 | DYM | protein_coding | 38.58 | -0.001608301 | 0.001608301 | ENSG00000141627 | n.s. |

| | | | | | | | | | | | | | |
|---|---|---|---|---|---|---|---|---|---|---|---|---|---|
| ENSG00000142541 | 24774.62 | 0.089917812 | 0.327679 | 0.2744081 | 0.783771 | 0.959653909 | RPL13A | protein_coding | 53.4 | 0.001608213 | 0.001608213 | ENSG00000142541 | n.s. |
| ENSG00000089818 | 1114.67 | -0.079550132 | 0.266066 | -0.298986 | 0.76495 | 0.954536941 | NECAP1 | protein_coding | 41.14 | -0.00160749 | 0.00160749 | ENSG00000089818 | n.s. |
| ENSG00000226756 | 32.74056 | 0.094508897 | 0.360217 | 0.2623664 | 0.793039 | 0.961633009 | | lincRNA | 40.49 | 0.001605766 | 0.001605766 | ENSG00000226756 | n.s. |
| ENSG00000178498 | 1166.323 | 0.077258761 | 0.254277 | 0.3038374 | 0.761252 | 0.953332446 | DTX3 | protein_coding | 55.6 | 0.001603554 | 0.001603554 | ENSG00000178498 | n.s. |
| ENSG00000047634 | 650.9912 | 0.079347849 | 0.264475 | 0.3000197 | 0.764162 | 0.954536941 | SCML1 | protein_coding | 38.89 | 0.001603403 | 0.001603403 | ENSG00000047634 | n.s. |
| ENSG00000164548 | 1767.353 | -0.073189995 | 0.23138 | -0.316319 | 0.75176 | 0.950815756 | TRA2A | protein_coding | 41.09 | -0.001603127 | 0.001603127 | ENSG00000164548 | n.s. |
| ENSG00000067177 | 292.6833 | -0.077053812 | 0.251867 | -0.305931 | 0.759657 | 0.953277452 | PHKA1 | protein_coding | 37.87 | -0.001601231 | 0.001601231 | ENSG00000067177 | n.s. |
| ENSG00000127445 | 1073.197 | -0.080554919 | 0.27512 | -0.292799 | 0.769676 | 0.955275572 | PIN1 | protein_coding | 56.31 | -0.001600733 | 0.001600733 | ENSG00000127445 | n.s. |
| ENSG00000207952 | 7.117194 | -0.136702109 | 0.709634 | -0.192638 | 0.847243 | 0.973446151 | MIR624 | miRNA | 40.21 | -0.001597783 | 0.001597783 | ENSG00000207952 | n.s. |
| ENSG00000117751 | 757.7343 | 0.076396172 | 0.248673 | 0.3072153 | 0.758679 | 0.95300702 | PPP1R8 | protein_coding | 44.44 | 0.001596978 | 0.001596978 | ENSG00000117751 | n.s. |
| ENSG00000199867 | 4.192266 | -0.150957682 | 0.948069 | -0.159226 | 0.87349 | 0.976022242 | | misc_RNA | 39.22 | -0.001591137 | 0.001591137 | ENSG00000199867 | n.s. |
| ENSG00000171863 | 24354.38 | -0.087605156 | 0.314996 | -0.278115 | 0.780924 | 0.959054979 | RPS7 | protein_coding | 44.76 | -0.001590603 | 0.001590603 | ENSG00000171863 | n.s. |
| ENSG00000135298 | 1244.721 | 0.07558514 | 0.244568 | 0.3090559 | 0.757279 | 0.952705822 | ADGRB3 | protein_coding | 35.14 | 0.001590401 | 0.001590401 | ENSG00000135298 | n.s. |
| ENSG00000108590 | 277.2545 | -0.07651498 | 0.250283 | -0.305714 | 0.759823 | 0.953277452 | MED31 | protein_coding | 40.22 | -0.001590033 | 0.001590033 | ENSG00000108590 | n.s. |
| ENSG00000085662 | 1749.615 | 0.086469305 | 0.309806 | 0.2791078 | 0.780162 | 0.958565181 | AKR1B1 | protein_coding | 50.04 | 0.001589163 | 0.001589163 | ENSG00000085662 | n.s. |
| ENSG00000233203 | 25.46294 | -0.100986452 | 0.398699 | -0.25329 | 0.800044 | 0.964417137 | | antisense | 47.36 | -0.00158903 | 0.00158903 | ENSG00000233203 | n.s. |
| ENSG00000128708 | 1770.467 | -0.079498485 | 0.267825 | -0.29683 | 0.766596 | 0.955053865 | HAT1 | protein_coding | 39.93 | -0.001587754 | 0.001587754 | ENSG00000128708 | n.s. |
| ENSG00000272205 | 36.76518 | -0.095908844 | 0.369361 | -0.259662 | 0.795125 | 0.962626178 | | lincRNA | 58.21 | -0.001586556 | 0.001586556 | ENSG00000272205 | n.s. |
| ENSG00000123297 | 836.392 | -0.08483275 | 0.301508 | -0.281361 | 0.778433 | 0.957854669 | TSFM | protein_coding | 44.22 | -0.001586405 | 0.001586405 | ENSG00000123297 | n.s. |
| ENSG00000145337 | 1132.817 | 0.082084239 | 0.285613 | 0.287397 | 0.773808 | 0.956474979 | PYURF | protein_coding | 44.72 | 0.001586392 | 0.001586392 | ENSG00000145337 | n.s. |
| ENSG00000176396 | 293.6416 | -0.079824592 | 0.270954 | -0.294605 | 0.768295 | 0.955275572 | EID2 | protein_coding | 51.5 | -0.001586221 | 0.001586221 | ENSG00000176396 | n.s. |
| ENSG00000122035 | 109.8549 | 0.077104768 | 0.25525 | 0.302076 | 0.762594 | 0.953844492 | RASL11A | protein_coding | 53.83 | 0.001582377 | 0.001582377 | ENSG00000122035 | n.s. |
| ENSG00000059758 | 2060.306 | -0.076411554 | 0.251765 | -0.303504 | 0.761506 | 0.95344167 | CDK17 | protein_coding | 37.82 | -0.001582168 | 0.001582168 | ENSG00000059758 | n.s. |
| ENSG00000144381 | 23304.93 | 0.078287737 | 0.262176 | 0.2986072 | 0.76524 | 0.954536941 | HSPD1 | protein_coding | 42.78 | 0.001581981 | 0.001581981 | ENSG00000144381 | n.s. |
| ENSG00000135905 | 734.455 | 0.079546041 | 0.271221 | 0.2932891 | 0.769301 | 0.955275572 | DOCK10 | protein_coding | 39.37 | 0.001580685 | 0.001580685 | ENSG00000135905 | n.s. |
| ENSG00000244187 | 124.2354 | 0.088287026 | 0.320117 | 0.2757959 | 0.782705 | 0.959619148 | TMEM141 | protein_coding | 62.59 | 0.001580434 | 0.001580434 | ENSG00000244187 | n.s. |
| ENSG00000259706 | 87.63531 | -0.081760931 | 0.284433 | -0.287452 | 0.773766 | 0.956474979 | HSP90B2P | processed_pseudogene | 40.31 | -0.001580143 | 0.001580143 | ENSG00000259706 | n.s. |
| ENSG00000140307 | 1613.557 | 0.08718314 | 0.314023 | 0.2776329 | 0.781294 | 0.959134639 | GTF2A2 | protein_coding | 39.73 | 0.001579795 | 0.001579795 | ENSG00000140307 | n.s. |
| ENSG00000115163 | 512.2806 | 0.085177784 | 0.303735 | 0.2804349 | 0.779144 | 0.958210479 | CENPA | protein_coding | 48.97 | 0.001579118 | 0.001579118 | ENSG00000115163 | n.s. |
| ENSG00000068697 | 1015.152 | -0.083551379 | 0.295076 | -0.283152 | 0.77706 | 0.957426384 | LAPTM4A | protein_coding | 41.61 | -0.001578671 | 0.001578671 | ENSG00000068697 | n.s. |
| ENSG00000135940 | 2055.965 | 0.08816595 | 0.320728 | 0.2748933 | 0.783398 | 0.959653909 | COX5B | protein_coding | 50.3 | 0.00157688 | 0.00157688 | ENSG00000135940 | n.s. |
| ENSG00000124795 | 8619.258 | -0.080515171 | 0.277872 | -0.289756 | 0.772003 | 0.955940517 | DEK | protein_coding | 39.43 | -0.001575612 | 0.001575612 | ENSG00000124795 | n.s. |

| | | | | | | | | | | | | | |
|---|---|---|---|---|---|---|---|---|---|---|---|---|---|
| ENSG00000109270 | 1220.665 | 0.079160448 | 0.268609 | 0.2947056 | 0.768219 | 0.955275572 | LAMTOR3 | protein_coding | 34.97 | 0.001573023 | 0.001573023 | ENSG00000109270 | n.s. |
| ENSG00000170899 | 1613.985 | 0.084990108 | 0.303734 | 0.2798178 | 0.779617 | 0.958295408 | GSTA4 | protein_coding | 41.01 | 0.001572367 | 0.001572367 | ENSG00000170899 | n.s. |
| ENSG00000136925 | 833.1762 | -0.081888955 | 0.286306 | -0.286019 | 0.774864 | 0.95679591 | TSTD2 | protein_coding | 42.44 | -0.001570687 | 0.001570687 | ENSG00000136925 | n.s. |
| ENSG00000103415 | 344.1691 | -0.081741848 | 0.286177 | -0.285634 | 0.775159 | 0.95679591 | HMOX2 | protein_coding | 47.62 | -0.001567865 | 0.001567865 | ENSG00000103415 | n.s. |
| ENSG00000263828 | 5.551453 | 0.124842942 | 0.589099 | 0.2119218 | 0.832168 | 0.971513984 | MIR4439 | miRNA | 42.5 | 0.001566897 | 0.001566897 | ENSG00000263828 | n.s. |
| ENSG00000104490 | 71.11031 | -0.081723115 | 0.286898 | -0.284851 | 0.775758 | 0.95693134 | NCALD | protein_coding | 40.58 | -0.001562482 | 0.001562482 | ENSG00000104490 | n.s. |
| ENSG00000204348 | 208.8521 | -0.07530366 | 0.248222 | -0.303372 | 0.761606 | 0.95344167 | DXO | protein_coding | 58.72 | -0.001559228 | 0.001559228 | ENSG00000204348 | n.s. |
| ENSG00000196943 | 970.2194 | -0.074537733 | 0.242916 | -0.306846 | 0.75896 | 0.95300702 | NOP9 | protein_coding | 50.61 | -0.001558129 | 0.001558129 | ENSG00000196943 | n.s. |
| ENSG00000145782 | 2002.2 | -0.078355329 | 0.266447 | -0.294074 | 0.768701 | 0.955275572 | ATG12 | protein_coding | 36.61 | -0.001557024 | 0.001557024 | ENSG00000145782 | n.s. |
| ENSG00000197444 | 754.5495 | -0.080003347 | 0.277456 | -0.288346 | 0.773082 | 0.956186787 | OGDHL | protein_coding | 55.46 | -0.001556646 | 0.001556646 | ENSG00000197444 | n.s. |
| ENSG00000099899 | 440.1731 | 0.077878435 | 0.262141 | 0.2970856 | 0.766401 | 0.955053865 | TRMT2A | protein_coding | 63.27 | 0.001555399 | 0.001555399 | ENSG00000099899 | n.s. |
| ENSG00000256683 | 355.1775 | -0.075748691 | 0.250807 | -0.30202 | 0.762637 | 0.953844492 | ZNF350 | protein_coding | 42.15 | -0.001554547 | 0.001554547 | ENSG00000256683 | n.s. |
| ENSG00000121988 | 286.8181 | -0.083730222 | 0.298368 | -0.280628 | 0.778996 | 0.958210479 | ZRANB3 | protein_coding | 38.97 | -0.001552282 | 0.001552282 | ENSG00000121988 | n.s. |
| ENSG00000157823 | 220.515 | 0.074246846 | 0.240715 | 0.3084435 | 0.757745 | 0.95300702 | AP3S2 | protein_coding | 44.42 | 0.001552049 | 0.001552049 | ENSG00000157823 | n.s. |
| ENSG00000198586 | 1704.221 | -0.076681586 | 0.255387 | -0.300257 | 0.763981 | 0.954536941 | TLK1 | protein_coding | 38.08 | -0.001549525 | 0.001549525 | ENSG00000198586 | n.s. |
| ENSG00000086589 | 1330.461 | 0.072346797 | 0.232625 | 0.3110017 | 0.755799 | 0.951962583 | RBM22 | protein_coding | 43.27 | 0.001546783 | 0.001546783 | ENSG00000086589 | n.s. |
| ENSG00000198668 | 7099.749 | -0.073981012 | 0.239912 | -0.308368 | 0.757803 | 0.95300702 | CALM1 | protein_coding | 47.09 | -0.001546492 | 0.001546492 | ENSG00000198668 | n.s. |
| ENSG00000111728 | 166.4324 | 0.078874595 | 0.27247 | 0.2894804 | 0.772214 | 0.955942907 | ST8SIA1 | protein_coding | 38.38 | 0.001543422 | 0.001543422 | ENSG00000111728 | n.s. |
| ENSG00000135090 | 317.8005 | -0.074311918 | 0.243963 | -0.304603 | 0.760668 | 0.953332446 | TAOK3 | protein_coding | 39.71 | -0.00154239 | 0.00154239 | ENSG00000135090 | n.s. |
| ENSG00000100027 | 289.5968 | -0.078748248 | 0.272082 | -0.289428 | 0.772254 | 0.955942907 | YPEL1 | protein_coding | 49.29 | -0.001540949 | 0.001540949 | ENSG00000100027 | n.s. |
| ENSG00000240137 | 35.05721 | -0.116360872 | 0.531554 | -0.218907 | 0.826722 | 0.970009871 | ERICH6-AS1 | antisense | 42.97 | -0.001538738 | 0.001538738 | ENSG00000240137 | n.s. |
| ENSG00000083844 | 245.3513 | 0.069914095 | 0.220219 | 0.3174755 | 0.750883 | 0.950585745 | ZNF264 | protein_coding | 44.47 | 0.001538719 | 0.001538719 | ENSG00000083844 | n.s. |
| ENSG00000149273 | 18455.51 | 0.08142759 | 0.287628 | 0.2831006 | 0.7771 | 0.957426384 | RPS3 | protein_coding | 48.52 | 0.001538542 | 0.001538542 | ENSG00000149273 | n.s. |
| ENSG00000134987 | 3246.532 | -0.076039711 | 0.25353 | -0.299923 | 0.764236 | 0.954536941 | WDR36 | protein_coding | 33.98 | -0.001536554 | 0.001536554 | ENSG00000134987 | n.s. |
| ENSG00000138078 | 2172.737 | -0.07959741 | 0.277192 | -0.287156 | 0.773993 | 0.956541519 | PREPL | protein_coding | 37.28 | -0.001535926 | 0.001535926 | ENSG00000138078 | n.s. |
| ENSG00000100445 | 470.772 | -0.07125413 | 0.228443 | -0.311913 | 0.755107 | 0.951579479 | SDR39U1 | protein_coding | 53.54 | -0.001535878 | 0.001535878 | ENSG00000100445 | n.s. |
| ENSG00000090971 | 156.6708 | -0.081173652 | 0.287448 | -0.282395 | 0.777641 | 0.957426384 | NAT14 | protein_coding | 66.08 | -0.001533744 | 0.001533744 | ENSG00000090971 | n.s. |
| ENSG00000158402 | 353.1792 | -0.077169242 | 0.263647 | -0.292699 | 0.769752 | 0.955275572 | CDC25C | protein_coding | 44.09 | -0.001533455 | 0.001533455 | ENSG00000158402 | n.s. |
| ENSG00000124217 | 404.6974 | -0.068324741 | 0.212159 | -0.322045 | 0.747419 | 0.949642637 | MOCS3 | protein_coding | 45.61 | -0.001533193 | 0.001533193 | ENSG00000124217 | n.s. |
| ENSG00000238531 | 4.668553 | 0.134214701 | 0.74864 | 0.1792779 | 0.857719 | 0.974056274 | SNORD105B | snoRNA | 55.7 | 0.001532189 | 0.001532189 | ENSG00000238531 | n.s. |
| ENSG00000124074 | 158.2699 | -0.086260311 | 0.316193 | -0.272809 | 0.785 | 0.959966858 | ENKD1 | protein_coding | 60.38 | -0.001530582 | 0.001530582 | ENSG00000124074 | n.s. |
| ENSG00000274828 | 178.9555 | 0.070748241 | 0.226369 | 0.312535 | 0.754634 | 0.951441537 | | lincRNA | 49.81 | 0.001529428 | 0.001529428 | ENSG00000274828 | n.s. |

| | | | | | | | | | | | | | |
|---|---|---|---|---|---|---|---|---|---|---|---|---|---|
| ENSG00000090520 | 1183.723 | 0.078506469 | 0.271825 | 0.2888127 | 0.772725 | 0.956156154 | DNAJB11 | protein_coding | 40.07 | 0.001528613 | 0.001528613 | ENSG00000090520 | n.s. |
| ENSG00000129460 | 1459.811 | 0.075638318 | 0.251787 | 0.3004054 | 0.763868 | 0.954536941 | NGDN | protein_coding | 42.99 | 0.001528443 | 0.001528443 | ENSG00000129460 | n.s. |
| ENSG00000136950 | 611.5288 | -0.076826565 | 0.260854 | -0.29452 | 0.768361 | 0.955275572 | ARPC5L | protein_coding | 52.55 | -0.001526646 | 0.001526646 | ENSG00000136950 | n.s. |
| ENSG00000213246 | 1691.646 | 0.084992116 | 0.3083 | 0.2756799 | 0.782794 | 0.959619148 | SUPT4H1 | protein_coding | 47.01 | 0.001521452 | 0.001521452 | ENSG00000213246 | n.s. |
| ENSG00000165288 | 548.044 | -0.076379899 | 0.259701 | -0.294107 | 0.768676 | 0.955275572 | BRWD3 | protein_coding | 36.15 | -0.00151777 | 0.00151777 | ENSG00000165288 | n.s. |
| ENSG00000059691 | 329.5675 | -0.078683753 | 0.274962 | -0.286162 | 0.774754 | 0.95679591 | GATB | protein_coding | 43.81 | -0.001509209 | 0.001509209 | ENSG00000059691 | n.s. |
| ENSG00000126746 | 373.3169 | -0.072708415 | 0.238808 | -0.304464 | 0.760774 | 0.953332446 | ZNF384 | protein_coding | 46.35 | -0.001509109 | 0.001509109 | ENSG00000126746 | n.s. |
| ENSG00000141504 | 343.793 | 0.084365132 | 0.307481 | 0.2743747 | 0.783797 | 0.959653909 | SAT2 | protein_coding | 56.73 | 0.001508901 | 0.001508901 | ENSG00000141504 | n.s. |
| ENSG00000123562 | 4626.145 | 0.085018264 | 0.31181 | 0.2726607 | 0.785114 | 0.959966858 | MORF4L2 | protein_coding | 39.39 | 0.001508544 | 0.001508544 | ENSG00000123562 | n.s. |
| ENSG00000025293 | 1378.159 | 0.077292471 | 0.267707 | 0.2887208 | 0.772795 | 0.956156154 | PHF20 | protein_coding | 44.08 | 0.001504975 | 0.001504975 | ENSG00000025293 | n.s. |
| ENSG00000257475 | 149.4694 | -0.07559644 | 0.259455 | -0.291367 | 0.770771 | 0.955275572 | | sense_intronic | 44.99 | -0.001502202 | 0.001502202 | ENSG00000257475 | n.s. |
| ENSG00000217930 | 432.4143 | -0.086149281 | 0.321369 | -0.26807 | 0.788646 | 0.960646942 | PAM16 | protein_coding | 56.77 | -0.001502116 | 0.001502116 | ENSG00000217930 | n.s. |
| ENSG00000143401 | 5383.719 | -0.075284663 | 0.254476 | -0.295842 | 0.767351 | 0.955198262 | ANP32E | protein_coding | 42.62 | -0.001498652 | 0.001498652 | ENSG00000143401 | n.s. |
| ENSG00000160752 | 3503.508 | -0.080741616 | 0.287933 | -0.280418 | 0.779157 | 0.958210479 | FDPS | protein_coding | 50.48 | -0.001496876 | 0.001496876 | ENSG00000160752 | n.s. |
| ENSG00000064726 | 1872.767 | -0.078009849 | 0.272785 | -0.285976 | 0.774897 | 0.95679591 | BTBD1 | protein_coding | 39.53 | -0.001496283 | 0.001496283 | ENSG00000064726 | n.s. |
| ENSG00000222345 | 66.8705 | -0.082372222 | 0.296144 | -0.278149 | 0.780898 | 0.959054979 | | snoRNA | 35.53 | -0.001495591 | 0.001495591 | ENSG00000222345 | n.s. |
| ENSG00000109184 | 1719.393 | -0.073975688 | 0.247446 | -0.298957 | 0.764973 | 0.954536941 | DCUN1D4 | protein_coding | 40.07 | -0.001494846 | 0.001494846 | ENSG00000109184 | n.s. |
| ENSG00000142168 | 3698.378 | 0.083422825 | 0.304863 | 0.2736403 | 0.784361 | 0.959720432 | SOD1 | protein_coding | 44.51 | 0.001489536 | 0.001489536 | ENSG00000142168 | n.s. |
| ENSG00000197062 | 315.7203 | -0.075617582 | 0.260569 | -0.290202 | 0.771662 | 0.955688505 | ZSCAN26 | protein_coding | 41.65 | -0.001488429 | 0.001488429 | ENSG00000197062 | n.s. |
| ENSG00000128789 | 746.5674 | -0.083183256 | 0.303314 | -0.274248 | 0.783894 | 0.959653909 | PSMG2 | protein_coding | 42.01 | -0.001487763 | 0.001487763 | ENSG00000128789 | n.s. |
| ENSG00000101138 | 890.2041 | -0.073579704 | 0.244906 | -0.300441 | 0.763841 | 0.954536941 | CSTF1 | protein_coding | 41.14 | -0.001486844 | 0.001486844 | ENSG00000101138 | n.s. |
| ENSG00000105887 | 2560.099 | -0.078650753 | 0.277959 | -0.282958 | 0.777209 | 0.957426384 | MTPN | protein_coding | 36.4 | -0.001486075 | 0.001486075 | ENSG00000105887 | n.s. |
| ENSG00000182973 | 962.4363 | -0.08313665 | 0.303621 | -0.273817 | 0.784225 | 0.959720432 | CNOT10 | protein_coding | 41.97 | -0.001484426 | 0.001484426 | ENSG00000182973 | n.s. |
| ENSG00000115649 | 235.963 | -0.083589029 | 0.306042 | -0.273129 | 0.784754 | 0.959948286 | CNPPD1 | protein_coding | 54.28 | -0.001483886 | 0.001483886 | ENSG00000115649 | n.s. |
| ENSG00000264796 | 13.2304 | -0.099865503 | 0.423051 | -0.23606 | 0.813386 | 0.966390136 | MIR5009 | miRNA | 34 | -0.001482754 | 0.001482754 | ENSG00000264796 | n.s. |
| ENSG00000180190 | 189.3441 | -0.082854587 | 0.301952 | -0.274396 | 0.78378 | 0.959653909 | TDRP | protein_coding | 42.86 | -0.001481884 | 0.001481884 | ENSG00000180190 | n.s. |
| ENSG00000132842 | 919.3617 | -0.078418543 | 0.277582 | -0.282506 | 0.777556 | 0.957426384 | AP3B1 | protein_coding | 36.53 | -0.001481688 | 0.001481688 | ENSG00000132842 | n.s. |
| ENSG00000150527 | 42.43588 | -0.085391839 | 0.31942 | -0.267334 | 0.789212 | 0.96084271 | CTAGE5 | protein_coding | 38.28 | -0.001481352 | 0.001481352 | ENSG00000150527 | n.s. |
| ENSG00000169554 | 355.8124 | -0.070019389 | 0.22631 | -0.309396 | 0.75702 | 0.95246603 | ZEB2 | protein_coding | 38.76 | -0.001480945 | 0.001480945 | ENSG00000169554 | n.s. |
| ENSG00000263813 | 18.18278 | 0.096968112 | 0.393996 | 0.2461146 | 0.805593 | 0.9654664 | MIR3679 | miRNA | 52.94 | 0.001480008 | 0.001480008 | ENSG00000263813 | n.s. |
| ENSG00000236977 | 72.61117 | -0.084606489 | 0.314859 | -0.268712 | 0.788151 | 0.960530983 | ANKRD44-IT1 | sense_intronic | 40.66 | -0.001479651 | 0.001479651 | ENSG00000236977 | n.s. |
| ENSG00000263884 | 68.00308 | -0.07992166 | 0.285525 | -0.279911 | 0.779546 | 0.958292132 | | lincRNA | 47.4 | -0.001478717 | 0.001478717 | ENSG00000263884 | n.s. |

| | | | | | | | | | | | | | |
|---|---|---|---|---|---|---|---|---|---|---|---|---|---|
| ENSG00000169359 | 700.8317 | -0.070718771 | 0.230068 | -0.307382 | 0.758552 | 0.95300702 | SLC33A1 | protein_coding | 39.65 | -0.001478298 | 0.001478298 | ENSG00000169359 | n.s. |
| ENSG00000090316 | 929.8734 | -0.086772931 | 0.327148 | -0.265241 | 0.790824 | 0.961602736 | MAEA | protein_coding | 55.03 | -0.001475514 | 0.001475514 | ENSG00000090316 | n.s. |
| ENSG00000112276 | 1718.647 | 0.074120122 | 0.250231 | 0.2962065 | 0.767072 | 0.955198262 | BVES | protein_coding | 40.62 | 0.00147547 | 0.00147547 | ENSG00000112276 | n.s. |
| ENSG00000232388 | 518.6254 | -0.085044681 | 0.318465 | -0.267046 | 0.789434 | 0.960872447 | SMIM26 | protein_coding | 44.25 | -0.001474187 | 0.001474187 | ENSG00000232388 | n.s. |
| ENSG00000162642 | 836.3892 | -0.076722616 | 0.268992 | -0.285223 | 0.775473 | 0.956833928 | C1orf52 | protein_coding | 39.66 | -0.001470269 | 0.001470269 | ENSG00000162642 | n.s. |
| ENSG00000137720 | 222.6505 | 0.083969534 | 0.312492 | 0.2687091 | 0.788154 | 0.960530983 | C11orf1 | protein_coding | 42.92 | 0.001468511 | 0.001468511 | ENSG00000137720 | n.s. |
| ENSG00000133393 | 978.1193 | -0.086374594 | 0.327753 | -0.263535 | 0.792138 | 0.961629391 | FOPNL | protein_coding | 41.7 | -0.001467701 | 0.001467701 | ENSG00000133393 | n.s. |
| ENSG00000164830 | 567.9011 | -0.082038461 | 0.298615 | -0.27473 | 0.783524 | 0.959653909 | OXR1 | protein_coding | 36.58 | -0.001467288 | 0.001467288 | ENSG00000164830 | n.s. |
| ENSG00000106399 | 1045.327 | 0.073815937 | 0.253329 | 0.291384 | 0.770758 | 0.955275572 | RPA3 | protein_coding | 37.19 | 0.001466821 | 0.001466821 | ENSG00000106399 | n.s. |
| ENSG00000167280 | 558.2138 | -0.076211699 | 0.265672 | -0.286864 | 0.774216 | 0.956724376 | ENGASE | protein_coding | 58.4 | -0.001464268 | 0.001464268 | ENSG00000167280 | n.s. |
| ENSG00000271155 | 81.72672 | -0.077529609 | 0.275002 | -0.281924 | 0.778002 | 0.957539143 | | antisense | 62.05 | -0.001460926 | 0.001460926 | ENSG00000271155 | n.s. |
| ENSG00000148218 | 241.1987 | -0.08353322 | 0.31092 | -0.268665 | 0.788188 | 0.960530983 | ALAD | protein_coding | 51.45 | -0.001460881 | 0.001460881 | ENSG00000148218 | n.s. |
| ENSG00000067836 | 82.69618 | 0.082341577 | 0.3028 | 0.2719341 | 0.785673 | 0.960059735 | ROGDI | protein_coding | 63.26 | 0.00145759 | 0.00145759 | ENSG00000067836 | n.s. |
| ENSG00000229273 | 57.59245 | -0.083010764 | 0.307525 | -0.269931 | 0.787213 | 0.960409479 | | processed_pseudogene | 62.66 | -0.001456305 | 0.001456305 | ENSG00000229273 | n.s. |
| ENSG00000265094 | 11.40382 | 0.099958466 | 0.431845 | 0.2314682 | 0.816951 | 0.967029262 | | sense_intronic | 38.9 | 0.001455434 | 0.001455434 | ENSG00000265094 | n.s. |
| ENSG00000167658 | 23069.13 | -0.077131702 | 0.273725 | -0.281785 | 0.778108 | 0.957539143 | EEF2 | protein_coding | 59.53 | -0.001453428 | 0.001453428 | ENSG00000167658 | n.s. |
| ENSG00000109475 | 11303.78 | 0.081878575 | 0.300636 | 0.2723513 | 0.785352 | 0.960002107 | RPL34 | protein_coding | 41.51 | 0.001451528 | 0.001451528 | ENSG00000109475 | n.s. |
| ENSG00000164294 | 207.1555 | -0.072942004 | 0.249139 | -0.292777 | 0.769693 | 0.955275572 | GPX8 | protein_coding | 39.03 | -0.001449454 | 0.001449454 | ENSG00000164294 | n.s. |
| ENSG00000136908 | 280.695 | -0.07808144 | 0.278028 | -0.28084 | 0.778833 | 0.958177474 | DPM2 | protein_coding | 56.79 | -0.001448726 | 0.001448726 | ENSG00000136908 | n.s. |
| ENSG00000165175 | 832.8314 | -0.070274466 | 0.23243 | -0.302346 | 0.762388 | 0.953704425 | MID1IP1 | protein_coding | 53.8 | -0.001446684 | 0.001446684 | ENSG00000165175 | n.s. |
| ENSG00000149196 | 433.5205 | 0.082415557 | 0.304727 | 0.2704574 | 0.786808 | 0.960409479 | HIKESHI | protein_coding | 37.85 | 0.001445862 | 0.001445862 | ENSG00000149196 | n.s. |
| ENSG00000185633 | 27.66128 | 0.094712543 | 0.384238 | 0.2464945 | 0.805299 | 0.9654664 | NDUFA4L2 | protein_coding | 62.05 | 0.001445582 | 0.001445582 | ENSG00000185633 | n.s. |
| ENSG00000005007 | 108.677 | -0.07646164 | 0.27002 | -0.28317 | 0.777046 | 0.957426384 | UPF1 | protein_coding | 55.16 | -0.001444713 | 0.001444713 | ENSG00000005007 | n.s. |
| ENSG00000104613 | 1485.673 | 0.078497893 | 0.281271 | 0.2790831 | 0.780181 | 0.958565181 | INTS10 | protein_coding | 41.11 | 0.001442662 | 0.001442662 | ENSG00000104613 | n.s. |
| ENSG00000010361 | 280.6588 | 0.079422061 | 0.286329 | 0.2773805 | 0.781488 | 0.959134639 | FUZ | protein_coding | 52.29 | 0.001439161 | 0.001439161 | ENSG00000010361 | n.s. |
| ENSG00000198815 | 383.4151 | -0.072228346 | 0.246424 | -0.293106 | 0.769441 | 0.955275572 | FOXJ3 | protein_coding | 38.57 | -0.001435273 | 0.001435273 | ENSG00000198815 | n.s. |
| ENSG00000138399 | 848.764 | 0.07884068 | 0.283431 | 0.2781649 | 0.780886 | 0.959054979 | FASTKD1 | protein_coding | 37.97 | 0.00143147 | 0.00143147 | ENSG00000138399 | n.s. |
| ENSG00000143409 | 130.6575 | -0.075902539 | 0.269042 | -0.282121 | 0.777851 | 0.957539143 | MINDY1 | protein_coding | 49.22 | -0.001430267 | 0.001430267 | ENSG00000143409 | n.s. |
| ENSG00000277544 | 60.28954 | 0.087235475 | 0.337083 | 0.2587951 | 0.795793 | 0.962958487 | | unprocessed_pseudogene | 40.59 | 0.001430002 | 0.001430002 | ENSG00000277544 | n.s. |
| ENSG00000122550 | 804.2282 | -0.078940081 | 0.284925 | -0.277056 | 0.781737 | 0.959207618 | KLHL7 | protein_coding | 38.72 | -0.001427819 | 0.001427819 | ENSG00000122550 | n.s. |
| ENSG00000125898 | 161.5573 | 0.071729738 | 0.243503 | 0.2945747 | 0.768319 | 0.955275572 | FAM110A | protein_coding | 51.88 | 0.001425365 | 0.001425365 | ENSG00000125898 | n.s. |
| ENSG00000173653 | 63.27411 | 0.078429098 | 0.281809 | 0.2783064 | 0.780777 | 0.959054979 | RCE1 | protein_coding | 60.51 | 0.001423998 | 0.001423998 | ENSG00000173653 | n.s. |

| | | | | | | | | | | | | | |
|---|---|---|---|---|---|---|---|---|---|---|---|---|---|
| ENSG00000261970 | 44.41329 | -0.088142585 | 0.34547 | -0.255138 | 0.798616 | 0.963515749 | | processed_pseudogene | 37.06 | -0.001422726 | 0.001422726 | ENSG00000261970 | n.s. |
| ENSG00000177917 | 1174.76 | -0.078415908 | 0.282657 | -0.277424 | 0.781455 | 0.959134639 | ARL6IP6 | protein_coding | 36.91 | -0.00142093 | 0.00142093 | ENSG00000177917 | n.s. |
| ENSG00000172765 | 261.9573 | -0.073382926 | 0.255214 | -0.287535 | 0.773703 | 0.956474979 | TMCC1 | protein_coding | 39.37 | -0.001418227 | 0.001418227 | ENSG00000172765 | n.s. |
| ENSG00000196678 | 182.4753 | 0.068229295 | 0.223209 | 0.3056745 | 0.759853 | 0.953277452 | ERI2 | protein_coding | 40.84 | 0.001417851 | 0.001417851 | ENSG00000196678 | n.s. |
| ENSG00000199204 | 8.177527 | 0.102912983 | 0.460197 | 0.2236282 | 0.823047 | 0.968826328 | | misc_RNA | 40.2 | 0.001415472 | 0.001415472 | ENSG00000199204 | n.s. |
| ENSG00000105726 | 140.6158 | -0.071211213 | 0.24401 | -0.291838 | 0.770411 | 0.955275572 | ATP13A1 | protein_coding | 57.98 | -0.001415061 | 0.001415061 | ENSG00000105726 | n.s. |
| ENSG00000204186 | 1645.484 | 0.073662806 | 0.258065 | 0.2854427 | 0.775305 | 0.95679591 | ZDBF2 | protein_coding | 37.53 | 0.001412903 | 0.001412903 | ENSG00000204186 | n.s. |
| ENSG00000008405 | 1495.56 | -0.078812947 | 0.286355 | -0.275228 | 0.783141 | 0.959653909 | CRY1 | protein_coding | 37.62 | -0.001409598 | 0.001409598 | ENSG00000008405 | n.s. |
| ENSG00000167733 | 89.16534 | 0.079993243 | 0.296527 | 0.269767 | 0.78734 | 0.960410738 | HSD11B1L | protein_coding | 58.24 | 0.001403321 | 0.001403321 | ENSG00000167733 | n.s. |
| ENSG00000207931 | 4.343466 | -0.133091616 | 0.822549 | -0.161804 | 0.87146 | 0.976022242 | MIR577 | miRNA | 39.58 | -0.001402824 | 0.001402824 | ENSG00000207931 | n.s. |
| ENSG00000198189 | 461.3563 | 0.084836873 | 0.326958 | 0.2594735 | 0.79527 | 0.962662389 | HSD17B11 | protein_coding | 39.92 | 0.001402014 | 0.001402014 | ENSG00000198189 | n.s. |
| ENSG00000125304 | 1733.209 | -0.078975733 | 0.289769 | -0.272547 | 0.785201 | 0.959989405 | TM9SF2 | protein_coding | 38.93 | -0.001400521 | 0.001400521 | ENSG00000125304 | n.s. |
| ENSG00000185728 | 2187.784 | -0.075445269 | 0.269078 | -0.280385 | 0.779182 | 0.958210479 | YTHDF3 | protein_coding | 37.56 | -0.001398686 | 0.001398686 | ENSG00000185728 | n.s. |
| ENSG00000164096 | 1079.194 | -0.07965048 | 0.294088 | -0.270839 | 0.786515 | 0.960409479 | C4orf3 | protein_coding | 42.89 | -0.001397353 | 0.001397353 | ENSG00000164096 | n.s. |
| ENSG00000143971 | 9221.097 | 0.063780708 | 0.201462 | 0.3165893 | 0.751555 | 0.950815756 | ETAA1 | protein_coding | 33.26 | 0.001397029 | 0.001397029 | ENSG00000143971 | n.s. |
| ENSG00000273604 | 289.7739 | -0.070152366 | 0.237362 | -0.295551 | 0.767573 | 0.955213071 | EPOP | protein_coding | 63.89 | -0.001396014 | 0.001396014 | ENSG00000273604 | n.s. |
| ENSG00000077312 | 1291.451 | -0.07545007 | 0.269538 | -0.279924 | 0.779536 | 0.958292132 | SNRPA | protein_coding | 50.77 | -0.001395983 | 0.001395983 | ENSG00000077312 | n.s. |
| ENSG00000169679 | 2100.842 | -0.07860753 | 0.288092 | -0.272856 | 0.784964 | 0.959966858 | BUB1 | protein_coding | 38.24 | -0.001394793 | 0.001394793 | ENSG00000169679 | n.s. |
| ENSG00000205309 | 209.6178 | -0.073696574 | 0.260448 | -0.282961 | 0.777207 | 0.957426384 | NT5M | protein_coding | 49.17 | -0.001392468 | 0.001392468 | ENSG00000205309 | n.s. |
| ENSG00000279035 | 66.28742 | -0.079363042 | 0.2934 | -0.270494 | 0.78678 | 0.960409479 | | TEC | 60.02 | -0.00139231 | 0.00139231 | ENSG00000279035 | n.s. |
| ENSG00000108055 | 6987.156 | -0.072570677 | 0.253969 | -0.285746 | 0.775073 | 0.95679591 | SMC3 | protein_coding | 36.86 | -0.001391956 | 0.001391956 | ENSG00000108055 | n.s. |
| ENSG00000150779 | 871.6893 | 0.077739845 | 0.283557 | 0.2741595 | 0.783962 | 0.959653909 | TIMM8B | protein_coding | 46.22 | 0.001390405 | 0.001390405 | ENSG00000150779 | n.s. |
| ENSG00000139163 | 2975.79 | -0.077565124 | 0.282022 | -0.275032 | 0.783291 | 0.959653909 | ETNK1 | protein_coding | 34.4 | -0.00138728 | 0.00138728 | ENSG00000139163 | n.s. |
| ENSG00000170921 | 2505.626 | 0.066833602 | 0.219001 | 0.3051752 | 0.760233 | 0.953332446 | TANC2 | protein_coding | 37.68 | 0.001387173 | 0.001387173 | ENSG00000170921 | n.s. |
| ENSG00000173171 | 468.2275 | -0.071410823 | 0.24803 | -0.287912 | 0.773414 | 0.956326929 | MTX1 | protein_coding | 55.87 | -0.001384914 | 0.001384914 | ENSG00000173171 | n.s. |
| ENSG00000159128 | 551.409 | 0.078850697 | 0.290802 | 0.2711492 | 0.786276 | 0.960409479 | IFNGR2 | protein_coding | 44.83 | 0.001383322 | 0.001383322 | ENSG00000159128 | n.s. |
| ENSG00000250950 | 86.10412 | -0.074599691 | 0.266089 | -0.280356 | 0.779204 | 0.958210479 | | lincRNA | 44.72 | -0.00138301 | 0.00138301 | ENSG00000250950 | n.s. |
| ENSG00000259746 | 22.9401 | 0.091944583 | 0.378461 | 0.242943 | 0.80805 | 0.965963425 | HSPE1P3 | processed_pseudogene | 41.42 | 0.001382784 | 0.001382784 | ENSG00000259746 | n.s. |
| ENSG00000133056 | 272.2372 | -0.06839653 | 0.228533 | -0.299286 | 0.764722 | 0.954536941 | PIK3C2B | protein_coding | 49.35 | -0.001382106 | 0.001382106 | ENSG00000133056 | n.s. |
| ENSG00000112237 | 2130.305 | 0.077213417 | 0.281488 | 0.2743042 | 0.783851 | 0.959653909 | CCNC | protein_coding | 35.29 | 0.00138099 | 0.00138099 | ENSG00000112237 | n.s. |
| ENSG00000174405 | 328.8019 | 0.07885261 | 0.293168 | 0.2689671 | 0.787955 | 0.960530983 | LIG4 | protein_coding | 40.58 | 0.001379023 | 0.001379023 | ENSG00000174405 | n.s. |
| ENSG00000135390 | 3188.53 | -0.076964844 | 0.278959 | -0.2759 | 0.782625 | 0.959619148 | ATP5G2 | protein_coding | 44.44 | -0.001377755 | 0.001377755 | ENSG00000135390 | n.s. |

| | | | | | | | | | | | | | |
|---|---|---|---|---|---|---|---|---|---|---|---|---|---|
| ENSG00000164076 | 2318.558 | -0.077041697 | 0.281473 | -0.273709 | 0.784308 | 0.959720432 | CAMKV | protein_coding | 56.12 | -0.001375599 | 0.001375599 | ENSG00000164076 | n.s. |
| ENSG00000151468 | 568.6269 | -0.072689356 | 0.257347 | -0.282456 | 0.777594 | 0.957426384 | CCDC3 | protein_coding | 43.73 | -0.001373437 | 0.001373437 | ENSG00000151468 | n.s. |
| ENSG00000066651 | 1803.828 | -0.072425695 | 0.256507 | -0.282353 | 0.777673 | 0.957426384 | TRMT11 | protein_coding | 36.02 | -0.001368455 | 0.001368455 | ENSG00000066651 | n.s. |
| ENSG00000151498 | 164.0141 | 0.077965968 | 0.287967 | 0.270746 | 0.786586 | 0.960409479 | ACAD8 | protein_coding | 47.88 | 0.001367801 | 0.001367801 | ENSG00000151498 | n.s. |
| ENSG00000278238 | 29.82166 | -0.09079709 | 0.377124 | -0.240762 | 0.80974 | 0.965963425 | | lincRNA | 45.06 | -0.001365526 | 0.001365526 | ENSG00000278238 | n.s. |
| ENSG00000107872 | 255.1609 | -0.072222429 | 0.254795 | -0.283453 | 0.77683 | 0.957426384 | FBXL15 | protein_coding | 61.93 | -0.001364615 | 0.001364615 | ENSG00000107872 | n.s. |
| ENSG00000184305 | 618.5793 | 0.078256078 | 0.292108 | 0.2679015 | 0.788775 | 0.960646942 | CCSER1 | protein_coding | 35.16 | 0.001364488 | 0.001364488 | ENSG00000184305 | n.s. |
| ENSG00000130159 | 504.2687 | 0.079194399 | 0.297273 | 0.2664033 | 0.789929 | 0.961126954 | ECSIT | protein_coding | 50.87 | 0.001363668 | 0.001363668 | ENSG00000130159 | n.s. |
| ENSG00000259775 | 45.2015 | 0.083697592 | 0.326007 | 0.2567358 | 0.797383 | 0.963287146 | | antisense | 35.16 | 0.001359603 | 0.001359603 | ENSG00000259775 | n.s. |
| ENSG00000132963 | 1694.302 | 0.079690138 | 0.299907 | 0.2657165 | 0.790458 | 0.961563689 | POMP | protein_coding | 39.25 | 0.001356481 | 0.001356481 | ENSG00000132963 | n.s. |
| ENSG00000241343 | 9610.613 | 0.075595464 | 0.274906 | 0.2749862 | 0.783327 | 0.959653909 | RPL36A | protein_coding | 41.83 | 0.001352052 | 0.001352052 | ENSG00000241343 | n.s. |
| ENSG00000177946 | 114.2634 | 0.069356537 | 0.240195 | 0.2887505 | 0.772772 | 0.956156154 | CENPBD1 | protein_coding | 52.11 | 0.001350453 | 0.001350453 | ENSG00000177946 | n.s. |
| ENSG00000068885 | 1212.97 | 0.079372696 | 0.301099 | 0.2636102 | 0.79208 | 0.961629391 | IFT80 | protein_coding | 36.52 | 0.001348722 | 0.001348722 | ENSG00000068885 | n.s. |
| ENSG00000085063 | 504.2295 | -0.079182453 | 0.30125 | -0.262846 | 0.792669 | 0.961629391 | CD59 | protein_coding | 44.03 | -0.00134549 | 0.00134549 | ENSG00000085063 | n.s. |
| ENSG00000039650 | 83.27513 | 0.078652358 | 0.296602 | 0.2651783 | 0.790872 | 0.961602736 | PNKP | protein_coding | 59.53 | 0.001337429 | 0.001337429 | ENSG00000039650 | n.s. |
| ENSG00000145088 | 72.14402 | -0.081510526 | 0.316109 | -0.257856 | 0.796518 | 0.962958487 | EAF2 | protein_coding | 36.14 | -0.001336156 | 0.001336156 | ENSG00000145088 | n.s. |
| ENSG00000166477 | 2191.503 | -0.068737869 | 0.238609 | -0.288077 | 0.773288 | 0.956292618 | LEO1 | protein_coding | 43.13 | -0.001334147 | 0.001334147 | ENSG00000166477 | n.s. |
| ENSG00000160410 | 240.2264 | -0.075976944 | 0.281544 | -0.269858 | 0.78727 | 0.960409479 | SHKBP1 | protein_coding | 53.53 | -0.001332906 | 0.001332906 | ENSG00000160410 | n.s. |
| ENSG00000165097 | 434.6969 | -0.07679697 | 0.287883 | -0.266765 | 0.78965 | 0.960872447 | KDM1B | protein_coding | 42.05 | -0.001331219 | 0.001331219 | ENSG00000165097 | n.s. |
| ENSG00000242779 | 74.01172 | 0.075868441 | 0.28112 | 0.2698796 | 0.787253 | 0.960409479 | ZNF702P | ıscribed_processed_pseudog | 45.63 | 0.001331003 | 0.001331003 | ENSG00000242779 | n.s. |
| ENSG00000116213 | 127.1673 | -0.078235459 | 0.296979 | -0.263438 | 0.792213 | 0.961629391 | WRAP73 | protein_coding | 55.25 | -0.001329398 | 0.001329398 | ENSG00000116213 | n.s. |
| ENSG00000258466 | 23.22149 | 0.088319975 | 0.366931 | 0.240699 | 0.809788 | 0.965963425 | | protein_coding | 43.56 | 0.001328272 | 0.001328272 | ENSG00000258466 | n.s. |
| ENSG00000075089 | 979.766 | 0.081870283 | 0.319817 | 0.2559911 | 0.797958 | 0.963403129 | ACTR6 | protein_coding | 41 | 0.001325639 | 0.001325639 | ENSG00000075089 | n.s. |
| ENSG00000134758 | 1639.141 | -0.078009493 | 0.294322 | -0.265048 | 0.790972 | 0.961629391 | RNF138 | protein_coding | 40.11 | -0.001325558 | 0.001325558 | ENSG00000134758 | n.s. |
| ENSG00000227666 | 25.88723 | -0.087999487 | 0.363214 | -0.24228 | 0.808563 | 0.965963425 | CYCSP24 | processed_pseudogene | 43.67 | -0.001323452 | 0.001323452 | ENSG00000227666 | n.s. |
| ENSG00000143158 | 527.9543 | 0.077861623 | 0.295479 | 0.2635102 | 0.792157 | 0.961629391 | MPC2 | protein_coding | 37.87 | 0.001323046 | 0.001323046 | ENSG00000143158 | n.s. |
| ENSG00000205476 | 589.861 | -0.075703354 | 0.282511 | -0.267966 | 0.788726 | 0.960646942 | CCDC85C | protein_coding | 56.04 | -0.001319978 | 0.001319978 | ENSG00000205476 | n.s. |
| ENSG00000167771 | 1323.185 | -0.068775056 | 0.240603 | -0.285845 | 0.774997 | 0.95679591 | RCOR2 | protein_coding | 60.76 | -0.001319153 | 0.001319153 | ENSG00000167771 | n.s. |
| ENSG00000124216 | 1107.162 | -0.067689833 | 0.234244 | -0.288972 | 0.772603 | 0.956156154 | SNAI1 | protein_coding | 55.16 | -0.001318001 | 0.001318001 | ENSG00000124216 | n.s. |
| ENSG00000215712 | 486.9256 | -0.07367647 | 0.268943 | -0.273948 | 0.784125 | 0.959684247 | TMEM242 | protein_coding | 43.34 | -0.001316719 | 0.001316719 | ENSG00000215712 | n.s. |
| ENSG00000163412 | 701.388 | -0.075199026 | 0.279434 | -0.269112 | 0.787843 | 0.960530983 | EIF4E3 | protein_coding | 41.79 | -0.001315127 | 0.001315127 | ENSG00000163412 | n.s. |
| ENSG00000156504 | 521.0845 | -0.077400131 | 0.295208 | -0.262188 | 0.793176 | 0.961633009 | FAM122B | protein_coding | 40.17 | -0.001315078 | 0.001315078 | ENSG00000156504 | n.s. |

| | | | | | | | | | | | | | |
|---|---|---|---|---|---|---|---|---|---|---|---|---|---|
| ENSG00000277161 | 655.3769 | -0.065998889 | 0.226192 | -0.291782 | 0.770453 | 0.955275572 | PIGW | protein_coding | 44.58 | -0.001311486 | 0.001311486 | ENSG00000277161 | n.s. |
| ENSG00000173848 | 2391.498 | -0.070857071 | 0.2535 | -0.279515 | 0.77985 | 0.958327464 | NET1 | protein_coding | 37.02 | -0.001309868 | 0.001309868 | ENSG00000173848 | n.s. |
| ENSG00000207001 | 8.459848 | -0.101421017 | 0.472391 | -0.214697 | 0.830004 | 0.970726167 | SNORD116-2 | snoRNA | 42.11 | -0.001308662 | 0.001308662 | ENSG00000207001 | n.s. |
| ENSG00000134371 | 1708.027 | -0.067416697 | 0.234055 | -0.288038 | 0.773317 | 0.956292618 | CDC73 | protein_coding | 35.71 | -0.001308504 | 0.001308504 | ENSG00000134371 | n.s. |
| ENSG00000202337 | 53.11347 | -0.087548483 | 0.368616 | -0.237506 | 0.812264 | 0.966187324 | RNU6-8 | snRNA | 46.73 | -0.001307857 | 0.001307857 | ENSG00000202337 | n.s. |
| ENSG00000135924 | 183.884 | -0.074746771 | 0.277711 | -0.269153 | 0.787812 | 0.960530983 | DNAJB2 | protein_coding | 56.92 | -0.001307218 | 0.001307218 | ENSG00000135924 | n.s. |
| ENSG00000131652 | 502.6317 | -0.076907404 | 0.292824 | -0.262641 | 0.792828 | 0.961629391 | THOC6 | protein_coding | 57.25 | -0.001306831 | 0.001306831 | ENSG00000131652 | n.s. |
| ENSG00000185909 | 106.628 | -0.073967214 | 0.272302 | -0.271636 | 0.785902 | 0.960171079 | KLHDC8B | protein_coding | 58.06 | -0.001305623 | 0.001305623 | ENSG00000185909 | n.s. |
| ENSG00000165733 | 2736.126 | -0.068189299 | 0.239188 | -0.285087 | 0.775578 | 0.95687778 | BMS1 | protein_coding | 40.62 | -0.001305384 | 0.001305384 | ENSG00000165733 | n.s. |
| ENSG00000173599 | 355.446 | -0.075298953 | 0.282091 | -0.266931 | 0.789522 | 0.960872447 | PC | protein_coding | 51.92 | -0.001305252 | 0.001305252 | ENSG00000173599 | n.s. |
| ENSG00000117640 | 252.2434 | -0.078796444 | 0.303277 | -0.259817 | 0.795005 | 0.962626178 | MTFR1L | protein_coding | 49.43 | -0.001303477 | 0.001303477 | ENSG00000117640 | n.s. |
| ENSG00000125454 | 321.3258 | -0.065325207 | 0.224178 | -0.291399 | 0.770746 | 0.955275572 | SLC25A19 | protein_coding | 50.68 | -0.001298099 | 0.001298099 | ENSG00000125454 | n.s. |
| ENSG00000166342 | 80.77941 | 0.074202395 | 0.276386 | 0.2684734 | 0.788335 | 0.960530983 | NETO1 | protein_coding | 36.85 | 0.001297698 | 0.001297698 | ENSG00000166342 | n.s. |
| ENSG00000125826 | 171.8288 | 0.076342758 | 0.289057 | 0.2641096 | 0.791695 | 0.961629391 | RBCK1 | protein_coding | 51.12 | 0.001297237 | 0.001297237 | ENSG00000125826 | n.s. |
| ENSG00000169241 | 168.423 | -0.081207181 | 0.319629 | -0.254067 | 0.799444 | 0.963961956 | SLC50A1 | protein_coding | 55.69 | -0.001294451 | 0.001294451 | ENSG00000169241 | n.s. |
| ENSG00000169957 | 703.2653 | -0.072282317 | 0.262242 | -0.275632 | 0.782831 | 0.959619148 | ZNF768 | protein_coding | 64.52 | -0.001293933 | 0.001293933 | ENSG00000169957 | n.s. |
| ENSG00000137414 | 750.08 | -0.0761405 | 0.289114 | -0.263358 | 0.792274 | 0.961629391 | FAM8A1 | protein_coding | 40.23 | -0.0012938 | 0.0012938 | ENSG00000137414 | n.s. |
| ENSG00000172840 | 370.358 | -0.071675393 | 0.259171 | -0.276556 | 0.782121 | 0.95928878 | PDP2 | protein_coding | 48.25 | -0.001293786 | 0.001293786 | ENSG00000172840 | n.s. |
| ENSG00000144741 | 309.4769 | -0.076035594 | 0.286725 | -0.265187 | 0.790866 | 0.961602736 | SLC25A26 | protein_coding | 40.15 | -0.001292933 | 0.001292933 | ENSG00000144741 | n.s. |
| ENSG00000128050 | 7855.833 | -0.074228306 | 0.277335 | -0.267648 | 0.78897 | 0.960716322 | PAICS | protein_coding | 39.99 | -0.001291931 | 0.001291931 | ENSG00000128050 | n.s. |
| ENSG00000100116 | 373.7192 | -0.07601 | 0.289506 | -0.262551 | 0.792897 | 0.961629391 | GCAT | protein_coding | 54.21 | -0.001291583 | 0.001291583 | ENSG00000100116 | n.s. |
| ENSG00000263307 | 22.8941 | -0.088673345 | 0.383032 | -0.231503 | 0.816924 | 0.967029262 | | antisense | 41.96 | -0.001291118 | 0.001291118 | ENSG00000263307 | n.s. |
| ENSG00000102401 | 727.7772 | -0.074353924 | 0.278675 | -0.266813 | 0.789613 | 0.960872447 | ARMCX3 | protein_coding | 45.41 | -0.00128887 | 0.00128887 | ENSG00000102401 | n.s. |
| ENSG00000271882 | 14.11914 | 0.095694769 | 0.433958 | 0.2205163 | 0.825469 | 0.969464905 | | lincRNA | 33.33 | 0.001288808 | 0.001288808 | ENSG00000271882 | n.s. |
| ENSG00000103152 | 206.7106 | 0.08443377 | 0.343482 | 0.245817 | 0.805824 | 0.9654664 | MPG | protein_coding | 61.91 | 0.001288699 | 0.001288699 | ENSG00000103152 | n.s. |
| ENSG00000064545 | 211.4242 | 0.078576392 | 0.305132 | 0.2575159 | 0.796781 | 0.962958487 | TMEM161A | protein_coding | 51.6 | 0.001288058 | 0.001288058 | ENSG00000064545 | n.s. |
| ENSG00000263895 | 16.96367 | -0.109866409 | 0.586842 | -0.187216 | 0.851491 | 0.973446151 | | sense_intronic | 37.13 | -0.001284126 | 0.001284126 | ENSG00000263895 | n.s. |
| ENSG00000135392 | 903.5202 | -0.067850305 | 0.24 | -0.28271 | 0.777399 | 0.957426384 | DNAJC14 | protein_coding | 49.07 | -0.001282005 | 0.001282005 | ENSG00000135392 | n.s. |
| ENSG00000258199 | 44.53827 | -0.087124096 | 0.371838 | -0.234307 | 0.814747 | 0.966779245 | | sense_overlapping | 49.39 | -0.001278344 | 0.001278344 | ENSG00000258199 | n.s. |
| ENSG00000164576 | 401.1739 | -0.075636836 | 0.28883 | -0.261874 | 0.793419 | 0.961843346 | SAP30L | protein_coding | 45.76 | -0.001277934 | 0.001277934 | ENSG00000164576 | n.s. |
| ENSG00000123159 | 322.0359 | -0.082182356 | 0.325882 | -0.252184 | 0.800899 | 0.964862231 | GIPC1 | protein_coding | 53.7 | -0.001276678 | 0.001276678 | ENSG00000123159 | n.s. |
| ENSG00000128891 | 203.0538 | -0.076761104 | 0.295661 | -0.259625 | 0.795153 | 0.962626178 | C15orf57 | protein_coding | 45.99 | -0.001269808 | 0.001269808 | ENSG00000128891 | n.s. |

| | | | | | | | | | | | | | |
|---|---|---|---|---|---|---|---|---|---|---|---|---|---|
| ENSG00000180992 | 702.5686 | -0.083088002 | 0.340264 | -0.244187 | 0.807086 | 0.9654664 | MRPL14 | protein_coding | 46.16 | -0.001268159 | 0.001268159 | ENSG00000180992 | n.s. |
| ENSG00000073670 | 293.103 | 0.071833845 | 0.26439 | 0.2716965 | 0.785855 | 0.960171079 | ADAM11 | protein_coding | 59.28 | 0.001267966 | 0.001267966 | ENSG00000073670 | n.s. |
| ENSG00000166073 | 264.2967 | -0.072394575 | 0.269608 | -0.268518 | 0.788301 | 0.960530983 | GPR176 | protein_coding | 40.8 | -0.001266081 | 0.001266081 | ENSG00000166073 | n.s. |
| ENSG00000219529 | 54.13251 | -0.082733766 | 0.336873 | -0.245593 | 0.805997 | 0.9654664 | | processed_pseudogene | 46.09 | -0.001262752 | 0.001262752 | ENSG00000219529 | n.s. |
| ENSG00000166707 | 185.717 | 0.072389027 | 0.26986 | 0.2682469 | 0.788509 | 0.960646942 | ZCCHC18 | protein_coding | 52.61 | 0.001262189 | 0.001262189 | ENSG00000166707 | n.s. |
| ENSG00000137135 | 227.1041 | 0.065357745 | 0.227611 | 0.2871472 | 0.774 | 0.956541519 | ARHGEF39 | protein_coding | 49.95 | 0.001261154 | 0.001261154 | ENSG00000137135 | n.s. |
| ENSG00000224856 | 29.21491 | 0.083805069 | 0.346743 | 0.2416924 | 0.809019 | 0.965963425 | | processed_pseudogene | 45.34 | 0.001260371 | 0.001260371 | ENSG00000224856 | n.s. |
| ENSG00000053770 | 751.7395 | 0.071642648 | 0.264035 | 0.2713373 | 0.786132 | 0.96036766 | AP5M1 | protein_coding | 35.17 | 0.001258222 | 0.001258222 | ENSG00000053770 | n.s. |
| ENSG00000162735 | 1101.112 | -0.076744172 | 0.296465 | -0.258864 | 0.79574 | 0.962958487 | PEX19 | protein_coding | 46.35 | -0.001258024 | 0.001258024 | ENSG00000162735 | n.s. |
| ENSG00000174456 | 151.7004 | -0.076044368 | 0.292377 | -0.26009 | 0.794794 | 0.962626178 | C12orf76 | protein_coding | 45.73 | -0.001257951 | 0.001257951 | ENSG00000174456 | n.s. |
| ENSG00000256092 | 67.77068 | 0.073937257 | 0.28198 | 0.2622078 | 0.793161 | 0.961633009 | | lincRNA | 59.97 | 0.001256241 | 0.001256241 | ENSG00000256092 | n.s. |
| ENSG00000171530 | 2018.629 | -0.081584427 | 0.32879 | -0.248135 | 0.80403 | 0.965293316 | TBCA | protein_coding | 39.32 | -0.001251562 | 0.001251562 | ENSG00000171530 | n.s. |
| ENSG00000228393 | 163.6981 | -0.069854345 | 0.253304 | -0.275773 | 0.782723 | 0.959619148 | LINC01004 | antisense | 42 | -0.001250469 | 0.001250469 | ENSG00000228393 | n.s. |
| ENSG00000213741 | 7744.335 | 0.080034059 | 0.316873 | 0.2525744 | 0.800597 | 0.964665853 | RPS29 | protein_coding | 43.43 | 0.00125038 | 0.00125038 | ENSG00000213741 | n.s. |
| ENSG00000266383 | 9.492976 | 0.106977181 | 0.537955 | 0.1988589 | 0.842373 | 0.973446151 | MIR5002 | miRNA | 48.45 | 0.001250357 | 0.001250357 | ENSG00000266383 | n.s. |
| ENSG00000137968 | 1007.679 | 0.073523152 | 0.279558 | 0.2629982 | 0.792552 | 0.961629391 | SLC44A5 | protein_coding | 36.99 | 0.001249325 | 0.001249325 | ENSG00000137968 | n.s. |
| ENSG00000070087 | 5755.996 | -0.06961222 | 0.25272 | -0.275452 | 0.782969 | 0.959653909 | PFN2 | protein_coding | 39.59 | -0.00124504 | 0.00124504 | ENSG00000070087 | n.s. |
| ENSG00000204673 | 309.3824 | 0.065890207 | 0.233118 | 0.2826479 | 0.777447 | 0.957426384 | AKT1S1 | protein_coding | 59.67 | 0.00124497 | 0.00124497 | ENSG00000204673 | n.s. |
| ENSG00000220848 | 32.57721 | -0.080127938 | 0.320201 | -0.250242 | 0.8024 | 0.964862231 | RPS18P9 | processed_pseudogene | 51.74 | -0.001244763 | 0.001244763 | ENSG00000220848 | n.s. |
| ENSG00000173085 | 275.3967 | -0.073209431 | 0.276627 | -0.26465 | 0.791279 | 0.961629391 | COQ2 | protein_coding | 39.35 | -0.001243995 | 0.001243995 | ENSG00000173085 | n.s. |
| ENSG00000204065 | 205.8996 | -0.073029593 | 0.27489 | -0.265668 | 0.790495 | 0.961563689 | TCEAL5 | protein_coding | 50.63 | -0.001243106 | 0.001243106 | ENSG00000204065 | n.s. |
| ENSG00000182512 | 939.8501 | 0.076922694 | 0.301436 | 0.2551878 | 0.798578 | 0.963515749 | GLRX5 | protein_coding | 45.13 | 0.001241623 | 0.001241623 | ENSG00000182512 | n.s. |
| ENSG00000128228 | 105.2127 | 0.079356521 | 0.314188 | 0.2525767 | 0.800595 | 0.964665853 | SDF2L1 | protein_coding | 62.46 | 0.001239794 | 0.001239794 | ENSG00000128228 | n.s. |
| ENSG00000233061 | 231.386 | 0.068410126 | 0.24704 | 0.276919 | 0.781842 | 0.959251841 | TTLL7-IT1 | sense_intronic | 37.82 | 0.00123599 | 0.00123599 | ENSG00000233061 | n.s. |
| ENSG00000067840 | 903.1741 | -0.067916123 | 0.244477 | -0.277802 | 0.781164 | 0.959134639 | PDZD4 | protein_coding | 56.35 | -0.001230669 | 0.001230669 | ENSG00000067840 | n.s. |
| ENSG00000011405 | 2889.38 | -0.069425662 | 0.255259 | -0.271981 | 0.785637 | 0.960059735 | PIK3C2A | protein_coding | 39.84 | -0.001228955 | 0.001228955 | ENSG00000011405 | n.s. |
| ENSG00000178096 | 275.0009 | 0.067767256 | 0.24445 | 0.2772229 | 0.781609 | 0.959134639 | BOLA1 | protein_coding | 45.46 | 0.001227971 | 0.001227971 | ENSG00000178096 | n.s. |
| ENSG00000231607 | 2214.819 | -0.067767004 | 0.244449 | -0.277224 | 0.781608 | 0.959134639 | DLEU2 | antisense | 38.45 | -0.001227967 | 0.001227967 | ENSG00000231607 | n.s. |
| ENSG00000130021 | 160.4863 | -0.079450069 | 0.31847 | -0.249474 | 0.802994 | 0.965047325 | PUDP | protein_coding | 43.76 | -0.001227614 | 0.001227614 | ENSG00000130021 | n.s. |
| ENSG00000198168 | 547.2575 | -0.073012185 | 0.279326 | -0.261387 | 0.793794 | 0.962046743 | SVIP | protein_coding | 36.62 | -0.001226884 | 0.001226884 | ENSG00000198168 | n.s. |
| ENSG00000123104 | 1961.573 | -0.072172767 | 0.274191 | -0.263221 | 0.79238 | 0.961629391 | ITPR2 | protein_coding | 38.25 | -0.001226379 | 0.001226379 | ENSG00000123104 | n.s. |
| ENSG00000144635 | 1361.952 | -0.074089676 | 0.285386 | -0.259612 | 0.795163 | 0.962626178 | DYNC1LI1 | protein_coding | 39.16 | -0.001225616 | 0.001225616 | ENSG00000144635 | n.s. |

| | | | | | | | | | | | | | |
|---|---|---|---|---|---|---|---|---|---|---|---|---|---|
| ENSG00000120709 | 1034.739 | 0.064820265 | 0.228695 | 0.2834351 | 0.776843 | 0.957426384 | FAM53C | protein_coding | 48.45 | 0.001224754 | 0.001224754 | ENSG00000120709 | n.s. |
| ENSG00000156313 | 47.82396 | -0.078830031 | 0.31449 | -0.25066 | 0.802077 | 0.964862231 | RPGR | protein_coding | 37.62 | -0.0012246 | 0.0012246 | ENSG00000156313 | n.s. |
| ENSG00000164117 | 1627.391 | -0.06162621 | 0.210305 | -0.293033 | 0.769497 | 0.955275572 | FBXO8 | protein_coding | 33.92 | -0.001224595 | 0.001224595 | ENSG00000164117 | n.s. |
| ENSG00000163808 | 2565.971 | -0.070337283 | 0.262894 | -0.26755 | 0.789046 | 0.960724353 | KIF15 | protein_coding | 41.48 | -0.001223953 | 0.001223953 | ENSG00000163808 | n.s. |
| ENSG00000111669 | 10675.78 | -0.074625038 | 0.289056 | -0.258168 | 0.796277 | 0.962958487 | TPI1 | protein_coding | 59.16 | -0.001223286 | 0.001223286 | ENSG00000111669 | n.s. |
| ENSG00000162910 | 414.2715 | -0.0745162 | 0.289101 | -0.257751 | 0.796599 | 0.962958487 | MRPL55 | protein_coding | 56.68 | -0.001221502 | 0.001221502 | ENSG00000162910 | n.s. |
| ENSG00000264864 | 18.02574 | 0.104502238 | 0.520319 | 0.2008425 | 0.840822 | 0.973446151 | MIR3613 | miRNA | 35.63 | 0.001221429 | 0.001221429 | ENSG00000264864 | n.s. |
| ENSG00000115350 | 480.9269 | 0.071830021 | 0.273359 | 0.2627679 | 0.792729 | 0.961629391 | POLE4 | protein_coding | 39.19 | 0.001220555 | 0.001220555 | ENSG00000115350 | n.s. |
| ENSG00000129071 | 895.5851 | -0.065349157 | 0.23242 | -0.281169 | 0.778581 | 0.957951797 | MBD4 | protein_coding | 39.9 | -0.001219176 | 0.001219176 | ENSG00000129071 | n.s. |
| ENSG00000131143 | 2162.94 | -0.076148156 | 0.299377 | -0.254356 | 0.799221 | 0.96384161 | COX4I1 | protein_coding | 50.84 | -0.001217939 | 0.001217939 | ENSG00000131143 | n.s. |
| ENSG00000183087 | 200.7803 | -0.075696964 | 0.297025 | -0.254851 | 0.798838 | 0.963630706 | GAS6 | protein_coding | 57.59 | -0.001217916 | 0.001217916 | ENSG00000183087 | n.s. |
| ENSG00000110074 | 349.6874 | -0.07635944 | 0.300613 | -0.254012 | 0.799486 | 0.963961956 | FOXRED1 | protein_coding | 51.05 | -0.001217178 | 0.001217178 | ENSG00000110074 | n.s. |
| ENSG00000130734 | 329.2351 | -0.069748822 | 0.260352 | -0.267902 | 0.788774 | 0.960646942 | ATG4D | protein_coding | 55.3 | -0.001216154 | 0.001216154 | ENSG00000130734 | n.s. |
| ENSG00000073849 | 650.871 | -0.072207644 | 0.276144 | -0.261485 | 0.793718 | 0.962039026 | ST6GAL1 | protein_coding | 44.38 | -0.001213616 | 0.001213616 | ENSG00000073849 | n.s. |
| ENSG00000165061 | 321.009 | -0.073906693 | 0.286744 | -0.257745 | 0.796604 | 0.962958487 | ZMAT4 | protein_coding | 40.92 | -0.001211511 | 0.001211511 | ENSG00000165061 | n.s. |
| ENSG00000159086 | 4361.82 | -0.068031381 | 0.248881 | -0.273349 | 0.784585 | 0.959826078 | PAXBP1 | protein_coding | 38.15 | -0.001211466 | 0.001211466 | ENSG00000159086 | n.s. |
| ENSG00000162144 | 364.9202 | -0.06134951 | 0.211049 | -0.290688 | 0.77129 | 0.955567674 | CYB561A3 | protein_coding | 52.14 | -0.00121095 | 0.00121095 | ENSG00000162144 | n.s. |
| ENSG00000111348 | 90.00097 | -0.071252186 | 0.270042 | -0.263855 | 0.791891 | 0.961629391 | ARHGDIB | protein_coding | 41.12 | -0.001210736 | 0.001210736 | ENSG00000111348 | n.s. |
| ENSG00000168246 | 881.9584 | -0.074648959 | 0.29113 | -0.256411 | 0.797634 | 0.963346062 | UBTD2 | protein_coding | 41.41 | -0.001210632 | 0.001210632 | ENSG00000168246 | n.s. |
| ENSG00000016864 | 466.2762 | -0.074321336 | 0.289465 | -0.256754 | 0.797369 | 0.963287146 | GLT8D1 | protein_coding | 44.49 | -0.001207293 | 0.001207293 | ENSG00000016864 | n.s. |
| ENSG00000255302 | 7891.409 | -0.072296291 | 0.27706 | -0.260941 | 0.794138 | 0.962379789 | EID1 | protein_coding | 41.64 | -0.001203987 | 0.001203987 | ENSG00000255302 | n.s. |
| ENSG00000171456 | 1755.09 | -0.067509602 | 0.246846 | -0.273489 | 0.784478 | 0.959778695 | ASXL1 | protein_coding | 42.17 | -0.001203622 | 0.001203622 | ENSG00000171456 | n.s. |
| ENSG00000272655 | 159.2102 | -0.066630136 | 0.240712 | -0.276804 | 0.781931 | 0.959275817 | POLR2J4 | cribed_unprocessed_pseudc | 54.06 | -0.001203107 | 0.001203107 | ENSG00000272655 | n.s. |
| ENSG00000155893 | 880.5258 | -0.072655538 | 0.279666 | -0.259794 | 0.795023 | 0.962626178 | PXYLP1 | protein_coding | 44.59 | -0.001201892 | 0.001201892 | ENSG00000155893 | n.s. |
| ENSG00000119333 | 601.5555 | -0.0760539 | 0.299998 | -0.253515 | 0.79987 | 0.964290713 | WDR34 | protein_coding | 53.82 | -0.001201044 | 0.001201044 | ENSG00000119333 | n.s. |
| ENSG00000128309 | 873.7561 | 0.077300966 | 0.308127 | 0.2508741 | 0.801911 | 0.964862231 | MPST | protein_coding | 55.28 | 0.001200847 | 0.001200847 | ENSG00000128309 | n.s. |
| ENSG00000095787 | 3555.84 | -0.069126712 | 0.259086 | -0.26681 | 0.789616 | 0.960872447 | WAC | protein_coding | 37.79 | -0.00119826 | 0.00119826 | ENSG00000095787 | n.s. |
| ENSG00000070761 | 1004.564 | -0.076120883 | 0.300773 | -0.253084 | 0.800203 | 0.964435904 | CFAP20 | protein_coding | 44.93 | -0.001197125 | 0.001197125 | ENSG00000070761 | n.s. |
| ENSG00000144659 | 673.0885 | 0.071993896 | 0.276153 | 0.260703 | 0.794322 | 0.962493005 | SLC25A38 | protein_coding | 46.41 | 0.001195273 | 0.001195273 | ENSG00000144659 | n.s. |
| ENSG00000181873 | 584.1518 | -0.068147745 | 0.25377 | -0.268542 | 0.788282 | 0.960530983 | IBA57 | protein_coding | 54.59 | -0.00119181 | 0.00119181 | ENSG00000181873 | n.s. |
| ENSG00000164609 | 1248.779 | -0.070046393 | 0.266747 | -0.262595 | 0.792863 | 0.961629391 | SLU7 | protein_coding | 36.94 | -0.001190247 | 0.001190247 | ENSG00000164609 | n.s. |
| ENSG00000253719 | 7400.631 | 0.061958197 | 0.216998 | 0.285524 | 0.775243 | 0.95679591 | ATXN7L3B | protein_coding | 38.7 | 0.001188401 | 0.001188401 | ENSG00000253719 | n.s. |

| | | | | | | | | | | | | | |
|---|---|---|---|---|---|---|---|---|---|---|---|---|---|
| ENSG00000111237 | 1225.125 | -0.073149313 | 0.285135 | -0.256543 | 0.797532 | 0.963306785 | VPS29 | protein_coding | 40.72 | -0.001187607 | 0.001187607 | ENSG00000111237 | n.s. |
| ENSG00000229992 | 28.94682 | -0.08091837 | 0.346409 | -0.233592 | 0.815302 | 0.966779245 | HMGB3P9 | processed_pseudogene | 44.08 | -0.001187289 | 0.001187289 | ENSG00000229992 | n.s. |
| ENSG00000213462 | 652.654 | 0.062694614 | 0.22152 | 0.2830199 | 0.777162 | 0.957426384 | ERV3-1 | protein_coding | 45.75 | 0.00118459 | 0.00118459 | ENSG00000213462 | n.s. |
| ENSG00000203877 | 775.4771 | 0.066148675 | 0.241025 | 0.2744475 | 0.783741 | 0.959653909 | RIPPLY2 | protein_coding | 42.89 | 0.001183093 | 0.001183093 | ENSG00000203877 | n.s. |
| ENSG00000068308 | 348.0279 | -0.067276401 | 0.248214 | -0.271042 | 0.786359 | 0.960409479 | OTUD5 | protein_coding | 49.93 | -0.001180268 | 0.001180268 | ENSG00000068308 | n.s. |
| ENSG00000135211 | 216.0823 | -0.078365434 | 0.322969 | -0.24264 | 0.808284 | 0.965963425 | TMEM60 | protein_coding | 42.55 | -0.001178563 | 0.001178563 | ENSG00000135211 | n.s. |
| ENSG00000126756 | 607.4076 | 0.078599309 | 0.329146 | 0.238798 | 0.811262 | 0.966064268 | UXT | protein_coding | 44.96 | 0.001178517 | 0.001178517 | ENSG00000126756 | n.s. |
| ENSG00000206535 | 323.0266 | 0.063670683 | 0.227243 | 0.2801873 | 0.779334 | 0.958285301 | LNP1 | protein_coding | 39.31 | 0.001178237 | 0.001178237 | ENSG00000206535 | n.s. |
| ENSG00000074582 | 577.0858 | -0.071841937 | 0.278528 | -0.257934 | 0.796458 | 0.962958487 | BCS1L | protein_coding | 54.42 | -0.001177664 | 0.001177664 | ENSG00000074582 | n.s. |
| ENSG00000119446 | 931.0353 | -0.071838344 | 0.277633 | -0.258753 | 0.795826 | 0.962958487 | RBM18 | protein_coding | 40.59 | -0.001177605 | 0.001177605 | ENSG00000119446 | n.s. |
| ENSG00000198356 | 573.8877 | -0.078268557 | 0.322709 | -0.242536 | 0.808365 | 0.965963425 | ASNA1 | protein_coding | 53.33 | -0.001177106 | 0.001177106 | ENSG00000198356 | n.s. |
| ENSG00000137944 | 928.7058 | 0.077092831 | 0.313127 | 0.2462028 | 0.805525 | 0.9654664 | KYAT3 | protein_coding | 38.59 | 0.001176655 | 0.001176655 | ENSG00000137944 | n.s. |
| ENSG00000186395 | 85.80359 | -0.078222263 | 0.322903 | -0.242247 | 0.808589 | 0.965963425 | KRT10 | protein_coding | 42.64 | -0.001176409 | 0.001176409 | ENSG00000186395 | n.s. |
| ENSG00000224470 | 609.0656 | -0.064947241 | 0.234873 | -0.276521 | 0.782148 | 0.95928878 | ATXN1L | protein_coding | 47.18 | -0.001172339 | 0.001172339 | ENSG00000224470 | n.s. |
| ENSG00000104442 | 873.1633 | -0.075397555 | 0.300678 | -0.250759 | 0.802001 | 0.964862231 | ARMC1 | protein_coding | 42.6 | -0.001171278 | 0.001171278 | ENSG00000104442 | n.s. |
| ENSG00000161920 | 112.1194 | -0.075323594 | 0.300902 | -0.250326 | 0.802335 | 0.964862231 | MED11 | protein_coding | 52.08 | -0.001170129 | 0.001170129 | ENSG00000161920 | n.s. |
| ENSG00000174109 | 247.5966 | -0.0687473 | 0.259781 | -0.264635 | 0.79129 | 0.961629391 | C16orf91 | protein_coding | 56.95 | -0.001168173 | 0.001168173 | ENSG00000174109 | n.s. |
| ENSG00000182858 | 208.11 | -0.071167043 | 0.275212 | -0.25859 | 0.795951 | 0.962958487 | ALG12 | protein_coding | 55.69 | -0.001166601 | 0.001166601 | ENSG00000182858 | n.s. |
| ENSG00000185513 | 34.37633 | 0.078526034 | 0.332253 | 0.2363444 | 0.813165 | 0.966390136 | L3MBTL1 | protein_coding | 49.19 | 0.001165916 | 0.001165916 | ENSG00000185513 | n.s. |
| ENSG00000103429 | 990.7576 | -0.074905933 | 0.298482 | -0.250956 | 0.801848 | 0.964862231 | BFAR | protein_coding | 46.07 | -0.001163641 | 0.001163641 | ENSG00000103429 | n.s. |
| ENSG00000047230 | 773.9886 | 0.069145903 | 0.264419 | 0.2615009 | 0.793706 | 0.962039026 | CTPS2 | protein_coding | 43.51 | 0.001162157 | 0.001162157 | ENSG00000047230 | n.s. |
| ENSG00000108582 | 1832.824 | -0.068269355 | 0.257158 | -0.265476 | 0.790643 | 0.961575527 | CPD | protein_coding | 38.53 | -0.001161712 | 0.001161712 | ENSG00000108582 | n.s. |
| ENSG00000100418 | 492.0578 | -0.075095729 | 0.301366 | -0.249184 | 0.803218 | 0.965047325 | DESI1 | protein_coding | 46.95 | -0.001160333 | 0.001160333 | ENSG00000100418 | n.s. |
| ENSG00000111911 | 583.0149 | -0.076012669 | 0.308923 | -0.246057 | 0.805638 | 0.9654664 | HINT3 | protein_coding | 38.08 | -0.001160169 | 0.001160169 | ENSG00000111911 | n.s. |
| ENSG00000126561 | 359.7897 | -0.066855242 | 0.250574 | -0.266808 | 0.789617 | 0.960872447 | STAT5A | protein_coding | 54.63 | -0.001158886 | 0.001158886 | ENSG00000126561 | n.s. |
| ENSG00000130150 | 110.9165 | 0.06778546 | 0.257517 | 0.2632275 | 0.792375 | 0.961629391 | MOSPD2 | protein_coding | 36.08 | 0.001151829 | 0.001151829 | ENSG00000130150 | n.s. |
| ENSG00000119397 | 1696.224 | -0.070187727 | 0.272621 | -0.257455 | 0.796828 | 0.962958487 | CNTRL | protein_coding | 38.88 | -0.001150548 | 0.001150548 | ENSG00000119397 | n.s. |
| ENSG00000063177 | 10134.62 | 0.076454499 | 0.317336 | 0.2409258 | 0.809613 | 0.965963425 | RPL18 | protein_coding | 56.83 | 0.001149823 | 0.001149823 | ENSG00000063177 | n.s. |
| ENSG00000222095 | 4.159561 | 0.108319252 | 0.63543 | 0.1704661 | 0.864644 | 0.975860117 | | snoRNA | 47.3 | 0.001149531 | 0.001149531 | ENSG00000222095 | n.s. |
| ENSG00000070756 | 6205.589 | -0.067639501 | 0.255658 | -0.26457 | 0.791341 | 0.961629391 | PABPC1 | protein_coding | 40.75 | -0.001149349 | 0.001149349 | ENSG00000070756 | n.s. |
| ENSG00000173113 | 1196.976 | 0.078130529 | 0.334178 | 0.2337988 | 0.815141 | 0.966779245 | TRMT112 | protein_coding | 56.49 | 0.001146384 | 0.001146384 | ENSG00000173113 | n.s. |
| ENSG00000204387 | 1741.84 | -0.064411012 | 0.236175 | -0.272726 | 0.785064 | 0.959966858 | C6orf48 | protein_coding | 48.85 | -0.001142894 | 0.001142894 | ENSG00000204387 | n.s. |

| | | | | | | | | | | | | | |
|---|---|---|---|---|---|---|---|---|---|---|---|---|---|
| ENSG00000204519 | 1736.367 | 0.063851164 | 0.232008 | 0.2752107 | 0.783154 | 0.959653909 | ZNF551 | protein_coding | 45.44 | 0.001142001 | 0.001142001 | ENSG00000204519 | n.s. |
| ENSG00000162368 | 537.6133 | -0.075754897 | 0.31233 | -0.242548 | 0.808356 | 0.965963425 | CMPK1 | protein_coding | 42.2 | -0.001139302 | 0.001139302 | ENSG00000162368 | n.s. |
| ENSG00000147996 | 205.8178 | -0.068409295 | 0.262509 | -0.260598 | 0.794403 | 0.962493005 | CBWD5 | protein_coding | 36.05 | -0.00113576 | 0.00113576 | ENSG00000147996 | n.s. |
| ENSG00000123144 | 2028.233 | -0.075434393 | 0.31274 | -0.241205 | 0.809396 | 0.965963425 | C19orf43 | protein_coding | 54.86 | -0.001134482 | 0.001134482 | ENSG00000123144 | n.s. |
| ENSG00000072274 | 6404.726 | 0.072897022 | 0.28966 | 0.251664 | 0.801301 | 0.964862231 | TFRC | protein_coding | 42.47 | 0.001132433 | 0.001132433 | ENSG00000072274 | n.s. |
| ENSG00000147123 | 1692.172 | -0.077472862 | 0.33274 | -0.232833 | 0.815891 | 0.96692204 | NDUFB11 | protein_coding | 52.3 | -0.001131765 | 0.001131765 | ENSG00000147123 | n.s. |
| ENSG00000260682 | 9.928181 | 0.096770161 | 0.496557 | 0.1948824 | 0.845485 | 0.973446151 | RN7SKP176 | misc_RNA | 55.66 | 0.001131056 | 0.001131056 | ENSG00000260682 | n.s. |
| ENSG00000162244 | 10745.73 | 0.075128266 | 0.312454 | 0.2404461 | 0.809984 | 0.965963425 | RPL29 | protein_coding | 53.82 | 0.001129878 | 0.001129878 | ENSG00000162244 | n.s. |
| ENSG00000225880 | 53.35085 | 0.075593183 | 0.31734 | 0.2382085 | 0.811719 | 0.966187324 | LINC00115 | lincRNA | 52.7 | 0.001129261 | 0.001129261 | ENSG00000225880 | n.s. |
| ENSG00000182934 | 529.985 | 0.06247208 | 0.225915 | 0.276529 | 0.782142 | 0.95928878 | SRPRA | protein_coding | 49.55 | 0.001127661 | 0.001127661 | ENSG00000182934 | n.s. |
| ENSG00000082556 | 218.206 | -0.07247552 | 0.288145 | -0.251525 | 0.801408 | 0.964862231 | OPRK1 | protein_coding | 42.12 | -0.001125885 | 0.001125885 | ENSG00000082556 | n.s. |
| ENSG00000108406 | 4187.242 | -0.072374564 | 0.288071 | -0.251239 | 0.80163 | 0.964862231 | DHX40 | protein_coding | 39.64 | -0.001124317 | 0.001124317 | ENSG00000108406 | n.s. |
| ENSG00000164109 | 4912.871 | -0.072249873 | 0.287183 | -0.251581 | 0.801365 | 0.964862231 | MAD2L1 | protein_coding | 38.16 | -0.00112238 | 0.00112238 | ENSG00000164109 | n.s. |
| ENSG00000155959 | 2054.683 | -0.07462537 | 0.308631 | -0.241795 | 0.808939 | 0.965963425 | VBP1 | protein_coding | 40.41 | -0.001122315 | 0.001122315 | ENSG00000155959 | n.s. |
| ENSG00000123349 | 3338.735 | 0.077042045 | 0.332231 | 0.2318931 | 0.816621 | 0.967029262 | PFDN5 | protein_coding | 47.28 | 0.001121762 | 0.001121762 | ENSG00000123349 | n.s. |
| ENSG00000143153 | 4523.214 | 0.067674487 | 0.259893 | 0.2603932 | 0.79456 | 0.962556833 | ATP1B1 | protein_coding | 43.48 | 0.001121611 | 0.001121611 | ENSG00000143153 | n.s. |
| ENSG00000155275 | 137.5325 | 0.065990271 | 0.24919 | 0.2648187 | 0.791149 | 0.961629391 | TRMT44 | protein_coding | 49.1 | 0.001121325 | 0.001121325 | ENSG00000155275 | n.s. |
| ENSG00000165443 | 1004.271 | -0.06590643 | 0.250626 | -0.262967 | 0.792576 | 0.961629391 | PHYHIPL | protein_coding | 35.18 | -0.0011199 | 0.0011199 | ENSG00000165443 | n.s. |
| ENSG00000117133 | 1809.005 | 0.069373477 | 0.271804 | 0.2552334 | 0.798543 | 0.963515749 | RPF1 | protein_coding | 37.23 | 0.00111977 | 0.00111977 | ENSG00000117133 | n.s. |
| ENSG00000172057 | 311.2136 | -0.072009603 | 0.286765 | -0.25111 | 0.801729 | 0.964862231 | ORMDL3 | protein_coding | 57.09 | -0.001118647 | 0.001118647 | ENSG00000172057 | n.s. |
| ENSG00000184602 | 724.8733 | -0.073278257 | 0.298679 | -0.245341 | 0.806192 | 0.9654664 | SNN | protein_coding | 59.43 | -0.001118434 | 0.001118434 | ENSG00000184602 | n.s. |
| ENSG00000114395 | 140.5643 | -0.072263777 | 0.290379 | -0.24886 | 0.803469 | 0.965047325 | CYB561D2 | protein_coding | 55 | -0.001116576 | 0.001116576 | ENSG00000114395 | n.s. |
| ENSG00000196262 | 12356.81 | -0.073121596 | 0.297571 | -0.245728 | 0.805893 | 0.9654664 | PPIA | protein_coding | 47.06 | -0.001116043 | 0.001116043 | ENSG00000196262 | n.s. |
| ENSG00000198276 | 450.3201 | 0.074705875 | 0.314641 | 0.2374319 | 0.812322 | 0.966187324 | UCKL1 | protein_coding | 62.54 | 0.001116006 | 0.001116006 | ENSG00000198276 | n.s. |
| ENSG00000207352 | 7.870608 | 0.105789083 | 0.64773 | 0.1633228 | 0.870264 | 0.976022242 | RNU6-540P | snRNA | 40.59 | 0.001115047 | 0.001115047 | ENSG00000207352 | n.s. |
| ENSG00000279342 | 394.2628 | 0.067077404 | 0.257444 | 0.2605511 | 0.794439 | 0.962493005 | | sense_intronic | 39.09 | 0.001113647 | 0.001113647 | ENSG00000279342 | n.s. |
| ENSG00000054116 | 218.1674 | 0.095188816 | 0.480802 | 0.1979791 | 0.843061 | 0.973446151 | TRAPPC3 | protein_coding | 49.8 | 0.001112573 | 0.001112573 | ENSG00000054116 | n.s. |
| ENSG00000122026 | 9652.414 | -0.07476828 | 0.316082 | -0.236547 | 0.813008 | 0.966390136 | RPL21 | protein_coding | 41.95 | -0.001110123 | 0.001110123 | ENSG00000122026 | n.s. |
| ENSG00000271931 | 15.69109 | -0.094956976 | 0.470703 | -0.201734 | 0.840124 | 0.973446151 | | antisense | 55.95 | -0.001109864 | 0.001109864 | ENSG00000271931 | n.s. |
| ENSG00000198833 | 1394.771 | -0.070566913 | 0.278861 | -0.253054 | 0.800227 | 0.964435904 | UBE2J1 | protein_coding | 40.13 | -0.00110978 | 0.00110978 | ENSG00000198833 | n.s. |
| ENSG00000137100 | 402.3702 | -0.07424106 | 0.313172 | -0.237062 | 0.812609 | 0.966187324 | DCTN3 | protein_coding | 49.85 | -0.001109062 | 0.001109062 | ENSG00000137100 | n.s. |
| ENSG00000113643 | 2034.519 | -0.06854343 | 0.268255 | -0.255516 | 0.798325 | 0.963515749 | RARS | protein_coding | 39.43 | -0.001106372 | 0.001106372 | ENSG00000113643 | n.s. |

| | | | | | | | | | | | | | |
|---|---|---|---|---|---|---|---|---|---|---|---|---|---|
| ENSG00000276115 | 228.3066 | 0.070173358 | 0.277364 | 0.2530006 | 0.800268 | 0.964435904 | | sense_intronic | 36.19 | 0.00110359 | 0.00110359 | ENSG00000276115 | n.s. |
| ENSG00000075043 | 632.9982 | -0.06836963 | 0.267691 | -0.255405 | 0.79841 | 0.963515749 | KCNQ2 | protein_coding | 62.3 | -0.001103567 | 0.001103567 | ENSG00000075043 | n.s. |
| ENSG00000168066 | 2991.573 | -0.059636099 | 0.213219 | -0.279695 | 0.779712 | 0.958326852 | SF1 | protein_coding | 49.33 | -0.001102453 | 0.001102453 | ENSG00000168066 | n.s. |
| ENSG00000178127 | 2474.95 | 0.066611646 | 0.256591 | 0.2596019 | 0.795171 | 0.962626178 | NDUFV2 | protein_coding | 37.35 | 0.001101912 | 0.001101912 | ENSG00000178127 | n.s. |
| ENSG00000104765 | 1295.231 | 0.070928936 | 0.282921 | 0.2507021 | 0.802044 | 0.964862231 | BNIP3L | protein_coding | 43.64 | 0.001101859 | 0.001101859 | ENSG00000104765 | n.s. |
| ENSG00000171044 | 184.8358 | -0.064760733 | 0.244843 | -0.264499 | 0.791395 | 0.961629391 | XKR6 | protein_coding | 46.08 | -0.001100432 | 0.001100432 | ENSG00000171044 | n.s. |
| ENSG00000108344 | 1468.195 | -0.073157539 | 0.305852 | -0.239193 | 0.810956 | 0.965963425 | PSMD3 | protein_coding | 49.75 | -0.001100239 | 0.001100239 | ENSG00000108344 | n.s. |
| ENSG00000203880 | 1425.294 | 0.070868417 | 0.283555 | 0.2499285 | 0.802643 | 0.964959246 | PCMTD2 | protein_coding | 45.7 | 0.001097825 | 0.001097825 | ENSG00000203880 | n.s. |
| ENSG00000138069 | 1674.251 | -0.067686053 | 0.264208 | -0.256185 | 0.797808 | 0.963346831 | RAB1A | protein_coding | 42.71 | -0.001097687 | 0.001097687 | ENSG00000138069 | n.s. |
| ENSG00000162961 | 835.016 | 0.072979399 | 0.303805 | 0.2402177 | 0.810161 | 0.965963425 | DPY30 | protein_coding | 39 | 0.00109756 | 0.00109756 | ENSG00000162961 | n.s. |
| ENSG00000276931 | 26.34184 | 0.07852989 | 0.34849 | 0.2253432 | 0.821712 | 0.968355922 | | sense_intronic | 44.83 | 0.001096669 | 0.001096669 | ENSG00000276931 | n.s. |
| ENSG00000175061 | 9416.061 | 0.071250847 | 0.286695 | 0.2485246 | 0.803729 | 0.965181308 | LRRC75A-AS1 | processed_transcript | 52.34 | 0.001096629 | 0.001096629 | ENSG00000175061 | n.s. |
| ENSG00000162086 | 1243.548 | -0.064143494 | 0.241125 | -0.266018 | 0.790225 | 0.96140387 | ZNF75A | protein_coding | 45.02 | -0.001096477 | 0.001096477 | ENSG00000162086 | n.s. |
| ENSG00000126602 | 2418.29 | -0.072036268 | 0.295247 | -0.243987 | 0.807241 | 0.965568426 | TRAP1 | protein_coding | 50.88 | -0.001096172 | 0.001096172 | ENSG00000126602 | n.s. |
| ENSG00000144504 | 145.2562 | 0.062460695 | 0.231216 | 0.2701401 | 0.787052 | 0.960409479 | ANKMY1 | protein_coding | 49.38 | 0.001095783 | 0.001095783 | ENSG00000144504 | n.s. |
| ENSG00000270095 | 28.69049 | -0.081251497 | 0.368334 | -0.220592 | 0.82541 | 0.969464905 | | sense_intronic | 35.27 | -0.001094288 | 0.001094288 | ENSG00000270095 | n.s. |
| ENSG00000164754 | 6868.939 | -0.067520484 | 0.263929 | -0.255829 | 0.798083 | 0.963471105 | RAD21 | protein_coding | 36.02 | -0.001091219 | 0.001091219 | ENSG00000164754 | n.s. |
| ENSG00000184900 | 1399.833 | -0.074728693 | 0.324069 | -0.230595 | 0.81763 | 0.967029262 | SUMO3 | protein_coding | 54.29 | -0.001088078 | 0.001088078 | ENSG00000184900 | n.s. |
| ENSG00000256061 | 73.36746 | -0.072777601 | 0.306968 | -0.237086 | 0.81259 | 0.966187324 | DNAAF4 | protein_coding | 42.67 | -0.0010872 | 0.0010872 | ENSG00000256061 | n.s. |
| ENSG00000197841 | 486.6092 | -0.06632158 | 0.256202 | -0.258864 | 0.79574 | 0.962958487 | ZNF181 | protein_coding | 39.74 | -0.001087172 | 0.001087172 | ENSG00000197841 | n.s. |
| ENSG00000171700 | 103.2898 | -0.074042098 | 0.315516 | -0.23467 | 0.814465 | 0.966779245 | RGS19 | protein_coding | 63.53 | -0.001086396 | 0.001086396 | ENSG00000171700 | n.s. |
| ENSG00000171160 | 151.2935 | -0.074313125 | 0.319075 | -0.232902 | 0.815838 | 0.96692204 | MORN4 | protein_coding | 46.51 | -0.001085606 | 0.001085606 | ENSG00000171160 | n.s. |
| ENSG00000108298 | 20222.46 | 0.073855821 | 0.314642 | 0.2347298 | 0.814418 | 0.966779245 | RPL19 | protein_coding | 49.65 | 0.001083663 | 0.001083663 | ENSG00000108298 | n.s. |
| ENSG00000196459 | 332.0278 | -0.067090819 | 0.262802 | -0.25529 | 0.798499 | 0.963515749 | TRAPPC2 | protein_coding | 42.18 | -0.001082925 | 0.001082925 | ENSG00000196459 | n.s. |
| ENSG00000183161 | 761.7096 | 0.059757432 | 0.215223 | 0.2776541 | 0.781278 | 0.959134639 | FANCF | protein_coding | 42.42 | 0.00108283 | 0.00108283 | ENSG00000183161 | n.s. |
| ENSG00000104763 | 236.2735 | -0.071906104 | 0.300339 | -0.239417 | 0.810783 | 0.965963425 | ASAH1 | protein_coding | 40.73 | -0.001081419 | 0.001081419 | ENSG00000104763 | n.s. |
| ENSG00000093167 | 427.0475 | 0.063454537 | 0.240272 | 0.2640949 | 0.791707 | 0.961629391 | LRRFIP2 | protein_coding | 38.6 | 0.001078237 | 0.001078237 | ENSG00000093167 | n.s. |
| ENSG00000233382 | 85.94856 | -0.074856683 | 0.32707 | -0.228871 | 0.81897 | 0.96738044 | NKAPP1 | ıscribed_processed_pseudog | 42.11 | -0.001078138 | 0.001078138 | ENSG00000233382 | n.s. |
| ENSG00000188629 | 170.4317 | -0.069317769 | 0.275634 | -0.251485 | 0.801439 | 0.964862231 | ZNF177 | protein_coding | 38.88 | -0.00107683 | 0.00107683 | ENSG00000188629 | n.s. |
| ENSG00000185420 | 479.4138 | 0.071407265 | 0.295715 | 0.2414734 | 0.809188 | 0.965963425 | SMYD3 | protein_coding | 40.6 | 0.001073917 | 0.001073917 | ENSG00000185420 | n.s. |
| ENSG00000116691 | 141.7518 | -0.071346915 | 0.297004 | -0.240222 | 0.810158 | 0.965963425 | MIIP | protein_coding | 55.62 | -0.001073009 | 0.001073009 | ENSG00000116691 | n.s. |
| ENSG00000145632 | 644.0971 | -0.061143198 | 0.226224 | -0.270277 | 0.786947 | 0.960409479 | PLK2 | protein_coding | 42.35 | -0.00107267 | 0.00107267 | ENSG00000145632 | n.s. |

| | | | | | | | | | | | | | |
|---|---|---|---|---|---|---|---|---|---|---|---|---|---|
| ENSG00000162757 | 236.3819 | -0.06119449 | 0.227852 | -0.268572 | 0.788259 | 0.960530983 | C1orf74 | protein_coding | 44.56 | -0.001070207 | 0.001070207 | ENSG00000162757 | n.s. |
| ENSG00000198355 | 585.863 | -0.068784854 | 0.274165 | -0.250889 | 0.8019 | 0.964862231 | PIM3 | protein_coding | 69.51 | -0.001068552 | 0.001068552 | ENSG00000198355 | n.s. |
| ENSG00000160113 | 312.3769 | -0.069145015 | 0.27786 | -0.248848 | 0.803478 | 0.965047325 | NR2F6 | protein_coding | 58.96 | -0.001068386 | 0.001068386 | ENSG00000160113 | n.s. |
| ENSG00000156030 | 118.3526 | -0.0628654 | 0.239673 | -0.262296 | 0.793093 | 0.961633009 | ELMSAN1 | protein_coding | 51.27 | -0.001068123 | 0.001068123 | ENSG00000156030 | n.s. |
| ENSG00000083099 | 1938.802 | -0.062800602 | 0.238898 | -0.262876 | 0.792646 | 0.961629391 | LYRM2 | protein_coding | 44.37 | -0.001067125 | 0.001067125 | ENSG00000083099 | n.s. |
| ENSG00000007080 | 452.2509 | -0.06900652 | 0.277117 | -0.249016 | 0.803349 | 0.965047325 | CCDC124 | protein_coding | 55.84 | -0.001066247 | 0.001066247 | ENSG00000007080 | n.s. |
| ENSG00000274885 | 116.9889 | -0.07174619 | 0.303924 | -0.236067 | 0.813381 | 0.966390136 | | sense_intronic | 33.33 | -0.001065252 | 0.001065252 | ENSG00000274885 | n.s. |
| ENSG00000123545 | 1139.387 | 0.068917652 | 0.276273 | 0.2494545 | 0.803009 | 0.965047325 | NDUFAF4 | protein_coding | 38.11 | 0.001064873 | 0.001064873 | ENSG00000123545 | n.s. |
| ENSG00000165113 | 471.7378 | -0.069372219 | 0.280938 | -0.246931 | 0.804962 | 0.9654664 | GKAP1 | protein_coding | 38.75 | -0.001058817 | 0.001058817 | ENSG00000165113 | n.s. |
| ENSG00000136937 | 2453.39 | 0.065508846 | 0.256786 | 0.2551111 | 0.798637 | 0.963515749 | NCBP1 | protein_coding | 37.78 | 0.00105739 | 0.00105739 | ENSG00000136937 | n.s. |
| ENSG00000172878 | 262.855 | -0.069274109 | 0.28364 | -0.244233 | 0.807051 | 0.9654664 | METAP1D | protein_coding | 40.05 | -0.001057319 | 0.001057319 | ENSG00000172878 | n.s. |
| ENSG00000162704 | 1417.078 | -0.073068169 | 0.318833 | -0.229174 | 0.818734 | 0.967247655 | ARPC5 | protein_coding | 38.38 | -0.001056735 | 0.001056735 | ENSG00000162704 | n.s. |
| ENSG00000117360 | 2525.123 | 0.06471558 | 0.251611 | 0.2572051 | 0.79702 | 0.963106735 | PRPF3 | protein_coding | 44.24 | 0.001056519 | 0.001056519 | ENSG00000117360 | n.s. |
| ENSG00000120658 | 1142.501 | -0.066213347 | 0.260726 | -0.253958 | 0.799528 | 0.963961956 | ENOX1 | protein_coding | 39.04 | -0.001055448 | 0.001055448 | ENSG00000120658 | n.s. |
| ENSG00000232677 | 580.7948 | -0.065823142 | 0.258714 | -0.254424 | 0.799168 | 0.96384161 | LINC00665 | lincRNA | 45.72 | -0.001052797 | 0.001052797 | ENSG00000232677 | n.s. |
| ENSG00000134897 | 619.8159 | -0.067962881 | 0.272028 | -0.249838 | 0.802713 | 0.96496019 | BIVM | protein_coding | 40.15 | -0.001052786 | 0.001052786 | ENSG00000134897 | n.s. |
| ENSG00000156052 | 1206.2 | -0.0686107 | 0.276318 | -0.248303 | 0.8039 | 0.965293316 | GNAQ | protein_coding | 39.13 | -0.001052536 | 0.001052536 | ENSG00000156052 | n.s. |
| ENSG00000167964 | 193.5099 | 0.069974795 | 0.289743 | 0.2415068 | 0.809162 | 0.965963425 | RAB26 | protein_coding | 58.26 | 0.001052373 | 0.001052373 | ENSG00000167964 | n.s. |
| ENSG00000143653 | 1334.419 | -0.070384198 | 0.296287 | -0.237554 | 0.812227 | 0.966187324 | SCCPDH | protein_coding | 41.94 | -0.001051446 | 0.001051446 | ENSG00000143653 | n.s. |
| ENSG00000144744 | 1679.473 | 0.06880872 | 0.279775 | 0.2459433 | 0.805726 | 0.9654664 | UBA3 | protein_coding | 37.06 | 0.001050216 | 0.001050216 | ENSG00000144744 | n.s. |
| ENSG00000259363 | 52.23601 | 0.072046204 | 0.312753 | 0.2303613 | 0.817811 | 0.967029262 | | lincRNA | 36.24 | 0.00104902 | 0.00104902 | ENSG00000259363 | n.s. |
| ENSG00000114405 | 950.3536 | -0.06973842 | 0.29055 | -0.240022 | 0.810313 | 0.965963425 | C3orf14 | protein_coding | 35.7 | -0.001048818 | 0.001048818 | ENSG00000114405 | n.s. |
| ENSG00000149499 | 215.8195 | -0.065533958 | 0.257458 | -0.254542 | 0.799077 | 0.963834903 | EML3 | protein_coding | 57.83 | -0.00104837 | 0.00104837 | ENSG00000149499 | n.s. |
| ENSG00000030419 | 27.1902 | -0.079273934 | 0.365245 | -0.217043 | 0.828174 | 0.970009871 | IKZF2 | protein_coding | 34.84 | -0.001048306 | 0.001048306 | ENSG00000030419 | n.s. |
| ENSG00000158411 | 358.3281 | -0.071312055 | 0.305315 | -0.233568 | 0.81532 | 0.966779245 | MITD1 | protein_coding | 39.09 | -0.001046339 | 0.001046339 | ENSG00000158411 | n.s. |
| ENSG00000143256 | 910.2576 | -0.067409331 | 0.269571 | -0.250062 | 0.802539 | 0.964918516 | PFDN2 | protein_coding | 44.14 | -0.001045476 | 0.001045476 | ENSG00000143256 | n.s. |
| ENSG00000180185 | 605.1121 | 0.063748946 | 0.247565 | 0.2575039 | 0.79679 | 0.962958487 | FAHD1 | protein_coding | 45.47 | 0.001045 | 0.001045 | ENSG00000180185 | n.s. |
| ENSG00000198856 | 1413.267 | 0.071745675 | 0.310402 | 0.2311377 | 0.817208 | 0.967029262 | OSTC | protein_coding | 39.57 | 0.001044645 | 0.001044645 | ENSG00000198856 | n.s. |
| ENSG00000183291 | 2388.574 | 0.074274488 | 0.327082 | 0.2270822 | 0.82036 | 0.96813703 | SELENOF | protein_coding | 37.87 | 0.001044535 | 0.001044535 | ENSG00000183291 | n.s. |
| ENSG00000270419 | 90.64389 | 0.067199878 | 0.267519 | 0.2511968 | 0.801662 | 0.964862231 | CAHM | lincRNA | 60.94 | 0.00104393 | 0.00104393 | ENSG00000270419 | n.s. |
| ENSG00000172409 | 371.2644 | -0.068386778 | 0.279995 | -0.244242 | 0.807043 | 0.9654664 | CLP1 | protein_coding | 47.28 | -0.001043776 | 0.001043776 | ENSG00000172409 | n.s. |
| ENSG00000155621 | 191.3947 | -0.06833017 | 0.278417 | -0.245424 | 0.806128 | 0.9654664 | C9orf85 | protein_coding | 38.63 | -0.001042912 | 0.001042912 | ENSG00000155621 | n.s. |

| | | | | | | | | | | | | | |
|---|---|---|---|---|---|---|---|---|---|---|---|---|---|
| ENSG00000105810 | 12557.67 | -0.067090068 | 0.267868 | -0.250459 | 0.802232 | 0.964862231 | CDK6 | protein_coding | 38.2 | -0.001042224 | 0.001042224 | ENSG00000105810 | n.s. |
| ENSG00000163686 | 162.4522 | 0.071579251 | 0.310812 | 0.2302974 | 0.817861 | 0.967029262 | ABHD6 | protein_coding | 45.11 | 0.001042221 | 0.001042221 | ENSG00000163686 | n.s. |
| ENSG00000140455 | 2678.056 | -0.068194639 | 0.278587 | -0.244787 | 0.806621 | 0.9654664 | USP3 | protein_coding | 38.44 | -0.001040844 | 0.001040844 | ENSG00000140455 | n.s. |
| ENSG00000272106 | 62.73998 | -0.068123094 | 0.2779 | -0.245136 | 0.806351 | 0.9654664 | | antisense | 57.36 | -0.001039752 | 0.001039752 | ENSG00000272106 | n.s. |
| ENSG00000141698 | 1119.954 | 0.069886202 | 0.295268 | 0.236687 | 0.8129 | 0.966390136 | NT5C3B | protein_coding | 48.83 | 0.001037636 | 0.001037636 | ENSG00000141698 | n.s. |
| ENSG00000136521 | 1014.873 | 0.088761828 | 0.468782 | 0.1893456 | 0.849822 | 0.973446151 | NDUFB5 | protein_coding | 39.72 | 0.001037454 | 0.001037454 | ENSG00000136521 | n.s. |
| ENSG00000115053 | 22239.15 | -0.060931632 | 0.229432 | -0.265576 | 0.790566 | 0.96156597 | NCL | protein_coding | 47.89 | -0.001037112 | 0.001037112 | ENSG00000115053 | n.s. |
| ENSG00000176387 | 81.75776 | 0.068802251 | 0.285759 | 0.2407699 | 0.809733 | 0.965963425 | HSD11B2 | protein_coding | 59.84 | 0.001034739 | 0.001034739 | ENSG00000176387 | n.s. |
| ENSG00000139263 | 1957.361 | -0.05832623 | 0.214092 | -0.272435 | 0.785287 | 0.960002107 | LRIG3 | protein_coding | 39.12 | -0.001033997 | 0.001033997 | ENSG00000139263 | n.s. |
| ENSG00000233016 | 1581.32 | -0.057851285 | 0.211127 | -0.274011 | 0.784076 | 0.959684247 | SNHG7 | antisense | 58.94 | -0.001033897 | 0.001033897 | ENSG00000233016 | n.s. |
| ENSG00000163539 | 2642.617 | 0.067706914 | 0.274389 | 0.2467553 | 0.805098 | 0.9654664 | CLASP2 | protein_coding | 36.82 | 0.0010334 | 0.0010334 | ENSG00000163539 | n.s. |
| ENSG00000206417 | 194.1388 | -0.067678726 | 0.275159 | -0.245962 | 0.805712 | 0.9654664 | H1FX-AS1 | antisense | 57.61 | -0.001032969 | 0.001032969 | ENSG00000206417 | n.s. |
| ENSG00000141030 | 1847.251 | -0.066834903 | 0.268028 | -0.249358 | 0.803084 | 0.965047325 | COPS3 | protein_coding | 44.79 | -0.001032692 | 0.001032692 | ENSG00000141030 | n.s. |
| ENSG00000102181 | 294.8468 | -0.069047406 | 0.291275 | -0.237052 | 0.812616 | 0.966187324 | CD99L2 | protein_coding | 43.02 | -0.001031476 | 0.001031476 | ENSG00000102181 | n.s. |
| ENSG00000160948 | 285.4256 | -0.070200773 | 0.300251 | -0.233807 | 0.815135 | 0.966779245 | VPS28 | protein_coding | 63.23 | -0.001030034 | 0.001030034 | ENSG00000160948 | n.s. |
| ENSG00000240731 | 15.27064 | -0.088116995 | 0.466899 | -0.188728 | 0.850306 | 0.973446151 | | sense_intronic | 47.97 | -0.001029917 | 0.001029917 | ENSG00000240731 | n.s. |
| ENSG00000092020 | 716.0266 | -0.072124234 | 0.316388 | -0.227962 | 0.819676 | 0.967663544 | PPP2R3C | protein_coding | 42.2 | -0.001029618 | 0.001029618 | ENSG00000092020 | n.s. |
| ENSG00000203668 | 5687.029 | -0.067385867 | 0.274062 | -0.245878 | 0.805776 | 0.9654664 | CHML | protein_coding | 37.83 | -0.001028499 | 0.001028499 | ENSG00000203668 | n.s. |
| ENSG00000245317 | 63.763 | 0.070622709 | 0.306339 | 0.2305374 | 0.817674 | 0.967029262 | | lincRNA | 46.28 | 0.001028294 | 0.001028294 | ENSG00000245317 | n.s. |
| ENSG00000255097 | 33.35466 | 0.073905357 | 0.329128 | 0.2245491 | 0.82233 | 0.968500556 | | sense_intronic | 45.92 | 0.001027294 | 0.001027294 | ENSG00000255097 | n.s. |
| ENSG00000159140 | 6366.071 | -0.062655628 | 0.242508 | -0.258365 | 0.796125 | 0.962958487 | SON | protein_coding | 38.94 | -0.001027078 | 0.001027078 | ENSG00000159140 | n.s. |
| ENSG00000138778 | 3861.282 | -0.066027677 | 0.263887 | -0.250212 | 0.802423 | 0.964862231 | CENPE | protein_coding | 34.67 | -0.00102572 | 0.00102572 | ENSG00000138778 | n.s. |
| ENSG00000164808 | 338.4458 | -0.06319979 | 0.24665 | -0.256233 | 0.797771 | 0.963346831 | SPIDR | protein_coding | 40.78 | -0.001024932 | 0.001024932 | ENSG00000164808 | n.s. |
| ENSG00000158290 | 711.8074 | -0.068122415 | 0.284188 | -0.239709 | 0.810556 | 0.965963425 | CUL4B | protein_coding | 39.25 | -0.001024515 | 0.001024515 | ENSG00000158290 | n.s. |
| ENSG00000170632 | 343.948 | 0.07004357 | 0.300833 | 0.2328321 | 0.815892 | 0.96692204 | ARMC10 | protein_coding | 40.09 | 0.001023234 | 0.001023234 | ENSG00000170632 | n.s. |
| ENSG00000205581 | 7565.532 | 0.066149582 | 0.265964 | 0.2487167 | 0.80358 | 0.96508612 | HMGN1 | protein_coding | 42.26 | 0.001020948 | 0.001020948 | ENSG00000205581 | n.s. |
| ENSG00000174780 | 2748.523 | -0.066850238 | 0.272132 | -0.245654 | 0.80595 | 0.9654664 | SRP72 | protein_coding | 38.04 | -0.001020324 | 0.001020324 | ENSG00000174780 | n.s. |
| ENSG00000275901 | 10.83446 | -0.089239177 | 0.493069 | -0.180987 | 0.856378 | 0.974056274 | | misc_RNA | 60.63 | -0.00101875 | 0.00101875 | ENSG00000275901 | n.s. |
| ENSG00000140395 | 1399.367 | -0.069286024 | 0.296781 | -0.233459 | 0.815405 | 0.966779245 | WDR61 | protein_coding | 43.39 | -0.001016612 | 0.001016612 | ENSG00000140395 | n.s. |
| ENSG00000108883 | 4687.647 | -0.065381015 | 0.260086 | -0.251382 | 0.801519 | 0.964862231 | EFTUD2 | protein_coding | 46.25 | -0.001015674 | 0.001015674 | ENSG00000108883 | n.s. |
| ENSG00000171862 | 2968.979 | -0.061333049 | 0.235671 | -0.260249 | 0.794672 | 0.96260812 | PTEN | protein_coding | 35.77 | -0.001015091 | 0.001015091 | ENSG00000171862 | n.s. |
| ENSG00000145388 | 1328.932 | -0.062204742 | 0.242369 | -0.256653 | 0.797447 | 0.963287146 | METTL14 | protein_coding | 37 | -0.001010468 | 0.001010468 | ENSG00000145388 | n.s. |

| | | | | | | | | | | | | | |
|---|---|---|---|---|---|---|---|---|---|---|---|---|---|
| ENSG00000120925 | 314.1913 | -0.072322654 | 0.321397 | -0.225026 | 0.821959 | 0.968355922 | RNF170 | protein_coding | 42.83 | -0.001009985 | 0.001009985 | ENSG00000120925 | n.s. |
| ENSG00000101843 | 408.3649 | 0.072313325 | 0.320265 | 0.2257924 | 0.821363 | 0.968355922 | PSMD10 | protein_coding | 37.94 | 0.001009855 | 0.001009855 | ENSG00000101843 | n.s. |
| ENSG00000006282 | 328.1325 | 0.067259914 | 0.281662 | 0.2387962 | 0.811264 | 0.966064268 | SPATA20 | protein_coding | 57.05 | 0.001008494 | 0.001008494 | ENSG00000006282 | n.s. |
| ENSG00000144401 | 440.1494 | 0.070197966 | 0.306959 | 0.2286884 | 0.819111 | 0.967465568 | METTL21A | protein_coding | 40.9 | 0.001008357 | 0.001008357 | ENSG00000144401 | n.s. |
| ENSG00000171103 | 723.3077 | 0.065860242 | 0.267748 | 0.2459783 | 0.805699 | 0.9654664 | TRMT61B | protein_coding | 41.37 | 0.001005214 | 0.001005214 | ENSG00000171103 | n.s. |
| ENSG00000109472 | 1551.846 | 0.068398724 | 0.29176 | 0.2344352 | 0.814647 | 0.966779245 | CPE | protein_coding | 38.23 | 0.001003593 | 0.001003593 | ENSG00000109472 | n.s. |
| ENSG00000233757 | 467.1939 | 0.05897471 | 0.224283 | 0.2629472 | 0.792591 | 0.961629391 | | protein_coding | 41.52 | 0.001002114 | 0.001002114 | ENSG00000233757 | n.s. |
| ENSG00000135437 | 45.5914 | -0.068803189 | 0.298954 | -0.230147 | 0.817978 | 0.967029262 | RDH5 | protein_coding | 52.34 | -0.001001801 | 0.001001801 | ENSG00000135437 | n.s. |
| ENSG00000145743 | 220.4055 | -0.065629003 | 0.267808 | -0.24506 | 0.80641 | 0.9654664 | FBXL17 | protein_coding | 35.81 | -0.001001685 | 0.001001685 | ENSG00000145743 | n.s. |
| ENSG00000163159 | 1030.959 | -0.075720174 | 0.349166 | -0.21686 | 0.828317 | 0.970009871 | VPS72 | protein_coding | 47.36 | -0.001001312 | 0.001001312 | ENSG00000163159 | n.s. |
| ENSG00000177830 | 278.3138 | -0.066913064 | 0.28096 | -0.238159 | 0.811758 | 0.966187324 | CHID1 | protein_coding | 59.37 | -0.000999592 | 0.000999592 | ENSG00000177830 | n.s. |
| ENSG00000143933 | 16341.26 | 0.067187041 | 0.284436 | 0.2362117 | 0.813268 | 0.966390136 | CALM2 | protein_coding | 39.71 | 0.00099756 | 0.00099756 | ENSG00000143933 | n.s. |
| ENSG00000129235 | 583.0725 | 0.067640522 | 0.28754 | 0.2352385 | 0.814024 | 0.96661324 | TXNDC17 | protein_coding | 42.73 | 0.000997512 | 0.000997512 | ENSG00000129235 | n.s. |
| ENSG00000140391 | 7964.737 | -0.068354002 | 0.293914 | -0.232565 | 0.816099 | 0.966962096 | TSPAN3 | protein_coding | 41.92 | -0.000997322 | 0.000997322 | ENSG00000140391 | n.s. |
| ENSG00000282826 | 188.8725 | -0.066660263 | 0.280587 | -0.237574 | 0.812212 | 0.966187324 | FRG1CP | unprocessed_pseudogene | 35.24 | -0.000995815 | 0.000995815 | ENSG00000282826 | n.s. |
| ENSG00000172671 | 368.693 | -0.065231338 | 0.265567 | -0.245631 | 0.805968 | 0.9654664 | ZFAND4 | protein_coding | 37.95 | -0.000995615 | 0.000995615 | ENSG00000172671 | n.s. |
| ENSG00000124226 | 1183.364 | 0.068331898 | 0.296306 | 0.2306126 | 0.817616 | 0.967029262 | RNF114 | protein_coding | 46.63 | 0.000994939 | 0.000994939 | ENSG00000124226 | n.s. |
| ENSG00000136003 | 1591.137 | -0.065123708 | 0.265061 | -0.245693 | 0.80592 | 0.9654664 | ISCU | protein_coding | 45.76 | -0.000993972 | 0.000993972 | ENSG00000136003 | n.s. |
| ENSG00000283891 | 6.759405 | -0.094272197 | 0.581462 | -0.16213 | 0.871204 | 0.976022242 | MIR628 | miRNA | 41.05 | -0.000993656 | 0.000993656 | ENSG00000283891 | n.s. |
| ENSG00000148634 | 557.501 | -0.065031944 | 0.264628 | -0.245748 | 0.805877 | 0.9654664 | HERC4 | protein_coding | 38 | -0.000992572 | 0.000992572 | ENSG00000148634 | n.s. |
| ENSG00000103769 | 2547.889 | -0.068150275 | 0.296087 | -0.23017 | 0.81796 | 0.967029262 | RAB11A | protein_coding | 40.1 | -0.000992294 | 0.000992294 | ENSG00000103769 | n.s. |
| ENSG00000087884 | 114.931 | 0.067912579 | 0.292041 | 0.2325444 | 0.816115 | 0.966962096 | AAMDC | protein_coding | 41.82 | 0.000990882 | 0.000990882 | ENSG00000087884 | n.s. |
| ENSG00000162174 | 580.991 | -0.066555879 | 0.281579 | -0.236367 | 0.813148 | 0.966390136 | ASRGL1 | protein_coding | 44.11 | -0.000988189 | 0.000988189 | ENSG00000162174 | n.s. |
| ENSG00000009844 | 1861.962 | -0.069159657 | 0.303188 | -0.228108 | 0.819562 | 0.967663544 | VTA1 | protein_coding | 35.53 | -0.000987297 | 0.000987297 | ENSG00000009844 | n.s. |
| ENSG00000136146 | 1319.049 | -0.065495845 | 0.27292 | -0.239982 | 0.810345 | 0.965963425 | MED4 | protein_coding | 40.77 | -0.000985013 | 0.000985013 | ENSG00000136146 | n.s. |
| ENSG00000259488 | 711.8109 | 0.056271544 | 0.208998 | 0.2692444 | 0.787742 | 0.960530983 | | antisense | 43.49 | 0.000984112 | 0.000984112 | ENSG00000259488 | n.s. |
| ENSG00000164081 | 183.5752 | -0.065804302 | 0.277222 | -0.237371 | 0.812369 | 0.966187324 | TEX264 | protein_coding | 54.18 | -0.000983028 | 0.000983028 | ENSG00000164081 | n.s. |
| ENSG00000160097 | 732.0136 | -0.067443913 | 0.290307 | -0.232319 | 0.81629 | 0.967004949 | FNDC5 | protein_coding | 54.01 | -0.000982746 | 0.000982746 | ENSG00000160097 | n.s. |
| ENSG00000068912 | 739.778 | -0.068284435 | 0.298839 | -0.228499 | 0.819258 | 0.967475307 | ERLEC1 | protein_coding | 38.1 | -0.000980572 | 0.000980572 | ENSG00000068912 | n.s. |
| ENSG00000141314 | 251.3251 | 0.065568014 | 0.275958 | 0.2376017 | 0.81219 | 0.966187324 | RHBDL3 | protein_coding | 51.16 | 0.000979499 | 0.000979499 | ENSG00000141314 | n.s. |
| ENSG00000122507 | 39.32332 | 0.079224204 | 0.379677 | 0.208662 | 0.834712 | 0.971936785 | BBS9 | protein_coding | 37.38 | 0.000979368 | 0.000979368 | ENSG00000122507 | n.s. |
| ENSG00000200376 | 7.457585 | -0.091651518 | 0.532198 | -0.172213 | 0.86327 | 0.975741574 | RNU5E-10P | snRNA | 36.44 | -0.000977481 | 0.000977481 | ENSG00000200376 | n.s. |

| | | | | | | | | | | | | | |
|---|---|---|---|---|---|---|---|---|---|---|---|---|---|
| ENSG00000054148 | 530.8602 | -0.073367603 | 0.333921 | -0.219715 | 0.826093 | 0.969833104 | PHPT1 | protein_coding | 66.23 | -0.000976009 | 0.000976009 | ENSG00000054148 | n.s. |
| ENSG00000162191 | 1178.688 | -0.066425487 | 0.283655 | -0.234177 | 0.814847 | 0.966779245 | UBXN1 | protein_coding | 52.86 | -0.00097464 | 0.00097464 | ENSG00000162191 | n.s. |
| ENSG00000232437 | 16.46579 | -0.083046886 | 0.420187 | -0.197643 | 0.843325 | 0.973446151 | RPS26P42 | processed_pseudogene | 47.21 | -0.000970658 | 0.000970658 | ENSG00000232437 | n.s. |
| ENSG00000106803 | 1152.939 | 0.072008539 | 0.325379 | 0.2213065 | 0.824854 | 0.969452399 | SEC61B | protein_coding | 42.24 | 0.000970208 | 0.000970208 | ENSG00000106803 | n.s. |
| ENSG00000059573 | 1726.071 | -0.065776157 | 0.279638 | -0.235219 | 0.814039 | 0.96661324 | ALDH18A1 | protein_coding | 43.16 | -0.000970018 | 0.000970018 | ENSG00000059573 | n.s. |
| ENSG00000167635 | 4868.431 | -0.065323962 | 0.276869 | -0.235938 | 0.813481 | 0.966390136 | ZNF146 | protein_coding | 42.84 | -0.000969898 | 0.000969898 | ENSG00000167635 | n.s. |
| ENSG00000213782 | 2124.858 | 0.06447883 | 0.26959 | 0.2391738 | 0.810971 | 0.965963425 | DDX47 | protein_coding | 40.84 | 0.000969718 | 0.000969718 | ENSG00000213782 | n.s. |
| ENSG00000092531 | 320.6136 | 0.064909708 | 0.273489 | 0.2373394 | 0.812394 | 0.966187324 | SNAP23 | protein_coding | 42.25 | 0.000969664 | 0.000969664 | ENSG00000092531 | n.s. |
| ENSG00000245694 | 1236.575 | 0.064909677 | 0.273575 | 0.2372643 | 0.812452 | 0.966187324 | CRNDE | lincRNA | 39.73 | 0.000969664 | 0.000969664 | ENSG00000245694 | n.s. |
| ENSG00000137513 | 873.7318 | 0.063187945 | 0.254565 | 0.2482194 | 0.803965 | 0.965293316 | NARS2 | protein_coding | 38.77 | 0.000969347 | 0.000969347 | ENSG00000137513 | n.s. |
| ENSG00000129480 | 428.5953 | -0.069386703 | 0.308115 | -0.225197 | 0.821826 | 0.968355922 | DTD2 | protein_coding | 38.52 | -0.000968984 | 0.000968984 | ENSG00000129480 | n.s. |
| ENSG00000113558 | 5333.094 | -0.064417925 | 0.269006 | -0.239466 | 0.810744 | 0.965963425 | SKP1 | protein_coding | 41.26 | -0.000968802 | 0.000968802 | ENSG00000113558 | n.s. |
| ENSG00000140326 | 520.9055 | 0.062327608 | 0.248554 | 0.2507607 | 0.801999 | 0.964862231 | CDAN1 | protein_coding | 54.46 | 0.00096824 | 0.00096824 | ENSG00000140326 | n.s. |
| ENSG00000137193 | 383.5272 | -0.059027414 | 0.229037 | -0.25772 | 0.796623 | 0.962958487 | PIM1 | protein_coding | 53.6 | -0.000967603 | 0.000967603 | ENSG00000137193 | n.s. |
| ENSG00000171497 | 1247.525 | 0.066663645 | 0.290975 | 0.2291045 | 0.818788 | 0.967247655 | PPID | protein_coding | 38.97 | 0.000964111 | 0.000964111 | ENSG00000171497 | n.s. |
| ENSG00000135083 | 207.6269 | -0.061952136 | 0.247301 | -0.250513 | 0.80219 | 0.964862231 | CCNJL | protein_coding | 46.79 | -0.000962407 | 0.000962407 | ENSG00000135083 | n.s. |
| ENSG00000277146 | 14.26196 | -0.082249827 | 0.427457 | -0.192417 | 0.847416 | 0.973446151 | | processed_pseudogene | 43.24 | -0.000961342 | 0.000961342 | ENSG00000277146 | n.s. |
| ENSG00000270083 | 134.3644 | -0.062952119 | 0.255253 | -0.246626 | 0.805198 | 0.9654664 | | sense_intronic | 45.11 | -0.000960828 | 0.000960828 | ENSG00000270083 | n.s. |
| ENSG00000153132 | 759.5118 | 0.068750146 | 0.304387 | 0.2258641 | 0.821307 | 0.968355922 | CLGN | protein_coding | 36.07 | 0.000960095 | 0.000960095 | ENSG00000153132 | n.s. |
| ENSG00000101966 | 44.58941 | -0.071224351 | 0.321902 | -0.221261 | 0.824889 | 0.969452399 | XIAP | protein_coding | 41.98 | -0.000959642 | 0.000959642 | ENSG00000101966 | n.s. |
| ENSG00000172594 | 260.0074 | -0.064532674 | 0.272719 | -0.236627 | 0.812946 | 0.966390136 | SMPDL3A | protein_coding | 40.47 | -0.00095815 | 0.00095815 | ENSG00000172594 | n.s. |
| ENSG00000182272 | 1342.651 | -0.063546767 | 0.262175 | -0.242383 | 0.808483 | 0.965963425 | B4GALNT4 | protein_coding | 65.98 | -0.0009557 | 0.0009557 | ENSG00000182272 | n.s. |
| ENSG00000070540 | 230.256 | 0.068979104 | 0.307528 | 0.2243017 | 0.822523 | 0.968618894 | WIPI1 | protein_coding | 46.58 | 0.000955158 | 0.000955158 | ENSG00000070540 | n.s. |
| ENSG00000005801 | 3170.156 | -0.063508585 | 0.264344 | -0.24025 | 0.810136 | 0.965963425 | ZNF195 | protein_coding | 41.43 | -0.000955126 | 0.000955126 | ENSG00000005801 | n.s. |
| ENSG00000179295 | 3470.308 | -0.063834364 | 0.268873 | -0.237414 | 0.812335 | 0.966187324 | PTPN11 | protein_coding | 43.16 | -0.0009536 | 0.0009536 | ENSG00000179295 | n.s. |
| ENSG00000270170 | 611.4608 | 0.064815871 | 0.27762 | 0.2334699 | 0.815397 | 0.966779245 | NCBP2-AS2 | protein_coding | 60.89 | 0.000951023 | 0.000951023 | ENSG00000270170 | n.s. |
| ENSG00000271971 | 37.64256 | -0.07394971 | 0.345711 | -0.213906 | 0.83062 | 0.970884671 | | antisense | 31.39 | -0.000948949 | 0.000948949 | ENSG00000271971 | n.s. |
| ENSG00000100949 | 126.2334 | -0.064589817 | 0.275123 | -0.234767 | 0.814389 | 0.966779245 | RABGGTA | protein_coding | 57.92 | -0.000947706 | 0.000947706 | ENSG00000100949 | n.s. |
| ENSG00000166189 | 212.5071 | -0.062082375 | 0.253917 | -0.244498 | 0.806845 | 0.9654664 | HPS6 | protein_coding | 64.21 | -0.000947553 | 0.000947553 | ENSG00000166189 | n.s. |
| ENSG00000139946 | 2875.203 | 0.061996752 | 0.253554 | 0.2445113 | 0.806835 | 0.9654664 | PELI2 | protein_coding | 39.91 | 0.000946246 | 0.000946246 | ENSG00000139946 | n.s. |
| ENSG00000147121 | 89.09846 | 0.062904035 | 0.262627 | 0.2395182 | 0.810704 | 0.965963425 | KRBOX4 | protein_coding | 41.38 | 0.000946034 | 0.000946034 | ENSG00000147121 | n.s. |
| ENSG00000020426 | 686.44 | -0.065285715 | 0.284243 | -0.229683 | 0.818338 | 0.96724455 | MNAT1 | protein_coding | 36.62 | -0.000944274 | 0.000944274 | ENSG00000020426 | n.s. |

| | | | | | | | | | | | | | |
|---|---|---|---|---|---|---|---|---|---|---|---|---|---|
| ENSG00000163608 | 1281.919 | -0.061773888 | 0.249932 | -0.247163 | 0.804782 | 0.9654664 | NEPRO | protein_coding | 39.07 | -0.000942845 | 0.000942845 | ENSG00000163608 | n.s. |
| ENSG00000064995 | 968.0054 | 0.064734676 | 0.280769 | 0.2305621 | 0.817655 | 0.967029262 | TAF11 | protein_coding | 41.49 | 0.000942562 | 0.000942562 | ENSG00000064995 | n.s. |
| ENSG00000186952 | 35.39067 | 0.071206161 | 0.325919 | 0.2184779 | 0.827057 | 0.970009871 | TMEM232 | protein_coding | 36.01 | 0.000941619 | 0.000941619 | ENSG00000186952 | n.s. |
| ENSG00000119328 | 237.9366 | -0.06466659 | 0.280047 | -0.230913 | 0.817382 | 0.967029262 | FAM206A | protein_coding | 39.01 | -0.00094157 | 0.00094157 | ENSG00000119328 | n.s. |
| ENSG00000163638 | 2211.681 | 0.061644681 | 0.250483 | 0.2461029 | 0.805603 | 0.9654664 | ADAMTS9 | protein_coding | 40.09 | 0.000940873 | 0.000940873 | ENSG00000163638 | n.s. |
| ENSG00000130511 | 570.83 | -0.064528136 | 0.279171 | -0.231142 | 0.817204 | 0.967029262 | SSBP4 | protein_coding | 64.53 | -0.000939554 | 0.000939554 | ENSG00000130511 | n.s. |
| ENSG00000119650 | 200.7821 | 0.073707856 | 0.345374 | 0.2134146 | 0.831004 | 0.971092758 | IFT43 | protein_coding | 43.54 | 0.000938985 | 0.000938985 | ENSG00000119650 | n.s. |
| ENSG00000214113 | 560.4801 | 0.067211415 | 0.298789 | 0.2249463 | 0.822021 | 0.968355922 | LYRM4 | protein_coding | 43.41 | 0.000938607 | 0.000938607 | ENSG00000214113 | n.s. |
| ENSG00000167302 | 523.1134 | -0.061287902 | 0.2489 | -0.246235 | 0.8055 | 0.9654664 | TEPSIN | protein_coding | 62.93 | -0.000935427 | 0.000935427 | ENSG00000167302 | n.s. |
| ENSG00000180964 | 915.7535 | 0.070605169 | 0.32225 | 0.2191005 | 0.826572 | 0.969965817 | TCEAL8 | protein_coding | 46.4 | 0.000935065 | 0.000935065 | ENSG00000180964 | n.s. |
| ENSG00000122042 | 434.811 | -0.064555741 | 0.281573 | -0.229268 | 0.818661 | 0.967247655 | UBL3 | protein_coding | 36.43 | -0.000933625 | 0.000933625 | ENSG00000122042 | n.s. |
| ENSG00000140450 | 278.6306 | 0.061123661 | 0.248964 | 0.2455117 | 0.80606 | 0.9654664 | ARRDC4 | protein_coding | 39.4 | 0.00093292 | 0.00093292 | ENSG00000140450 | n.s. |
| ENSG00000101493 | 801.107 | -0.062026763 | 0.257561 | -0.240824 | 0.809692 | 0.965963425 | ZNF516 | protein_coding | 47.61 | -0.00093284 | 0.00093284 | ENSG00000101493 | n.s. |
| ENSG00000205659 | 406.0641 | -0.063000995 | 0.267454 | -0.235559 | 0.813775 | 0.966492561 | LIN52 | protein_coding | 40.2 | -0.000932508 | 0.000932508 | ENSG00000205659 | n.s. |
| ENSG00000280069 | 360.7277 | -0.061061788 | 0.24964 | -0.2446 | 0.806766 | 0.9654664 | | TEC | 37.97 | -0.000931976 | 0.000931976 | ENSG00000280069 | n.s. |
| ENSG00000126261 | 6006.94 | -0.066234965 | 0.291752 | -0.227025 | 0.820405 | 0.96813703 | UBA2 | protein_coding | 42.17 | -0.000931473 | 0.000931473 | ENSG00000126261 | n.s. |
| ENSG00000230513 | 113.5999 | -0.061857763 | 0.257789 | -0.239955 | 0.810365 | 0.965963425 | THAP7-AS1 | antisense | 50.16 | -0.000930299 | 0.000930299 | ENSG00000230513 | n.s. |
| ENSG00000054282 | 575.0473 | -0.0612878 | 0.251498 | -0.243691 | 0.80747 | 0.965676593 | SDCCAG8 | protein_coding | 39.81 | -0.000929631 | 0.000929631 | ENSG00000054282 | n.s. |
| ENSG00000156925 | 886.8767 | 0.057167436 | 0.222647 | 0.2567623 | 0.797362 | 0.963287146 | ZIC3 | protein_coding | 48.02 | 0.000928641 | 0.000928641 | ENSG00000156925 | n.s. |
| ENSG00000141337 | 94.88667 | 0.06597526 | 0.291018 | 0.226705 | 0.820653 | 0.96813703 | ARSG | protein_coding | 45.96 | 0.000927821 | 0.000927821 | ENSG00000141337 | n.s. |
| ENSG00000274620 | 15.25246 | -0.079293288 | 0.423591 | -0.187193 | 0.851509 | 0.973446151 | MIR378J | miRNA | 42.2 | -0.000926785 | 0.000926785 | ENSG00000274620 | n.s. |
| ENSG00000163872 | 2220.913 | -0.061618551 | 0.254684 | -0.241941 | 0.808826 | 0.965963425 | YEATS2 | protein_coding | 42.58 | -0.000926701 | 0.000926701 | ENSG00000163872 | n.s. |
| ENSG00000136444 | 955.4652 | 0.059646352 | 0.238003 | 0.2506116 | 0.802114 | 0.964862231 | RSAD1 | protein_coding | 50.31 | 0.000926588 | 0.000926588 | ENSG00000136444 | n.s. |
| ENSG00000172239 | 2047.751 | -0.064057149 | 0.279049 | -0.229555 | 0.818438 | 0.96724455 | PAIP1 | protein_coding | 38.36 | -0.000926504 | 0.000926504 | ENSG00000172239 | n.s. |
| ENSG00000126456 | 132.5599 | 0.068743319 | 0.310184 | 0.2216211 | 0.824609 | 0.969452399 | IRF3 | protein_coding | 57.17 | 0.000926214 | 0.000926214 | ENSG00000126456 | n.s. |
| ENSG00000197045 | 3448.384 | 0.062880494 | 0.268363 | 0.2343112 | 0.814743 | 0.966779245 | GMFB | protein_coding | 36.79 | 0.000922625 | 0.000922625 | ENSG00000197045 | n.s. |
| ENSG00000268105 | 52.19339 | -0.065487634 | 0.288787 | -0.226768 | 0.820604 | 0.96813703 | | processed_pseudogene | 36.78 | -0.000920964 | 0.000920964 | ENSG00000268105 | n.s. |
| ENSG00000252620 | 21.47903 | -0.078693264 | 0.399968 | -0.196749 | 0.844024 | 0.973446151 | RNU6ATAC24P | snRNA | 51.11 | -0.000919772 | 0.000919772 | ENSG00000252620 | n.s. |
| ENSG00000100968 | 56.44702 | -0.071101734 | 0.330947 | -0.214843 | 0.82989 | 0.970726167 | NFATC4 | protein_coding | 56.84 | -0.000917444 | 0.000917444 | ENSG00000100968 | n.s. |
| ENSG00000100227 | 1199.094 | -0.062966398 | 0.272164 | -0.231355 | 0.817039 | 0.967029262 | POLDIP3 | protein_coding | 49.1 | -0.000916815 | 0.000916815 | ENSG00000100227 | n.s. |
| ENSG00000224505 | 115.2516 | 0.059935332 | 0.243108 | 0.2465384 | 0.805266 | 0.9654664 | | antisense | 46.18 | 0.000914783 | 0.000914783 | ENSG00000224505 | n.s. |
| ENSG00000053108 | 559.2336 | 0.060732621 | 0.253541 | 0.2395381 | 0.810688 | 0.965963425 | FSTL4 | protein_coding | 46.69 | 0.000913377 | 0.000913377 | ENSG00000053108 | n.s. |

| | | | | | | | | | | | | | |
|---|---|---|---|---|---|---|---|---|---|---|---|---|---|
| ENSG00000128694 | 507.3396 | 0.059729302 | 0.241579 | 0.247245 | 0.804719 | 0.9654664 | OSGEPL1 | protein_coding | 39.12 | 0.000911639 | 0.000911639 | ENSG00000128694 | n.s. |
| ENSG00000133134 | 729.0681 | -0.068846074 | 0.317186 | -0.217053 | 0.828167 | 0.970009871 | BEX2 | protein_coding | 57.03 | -0.00091041 | 0.00091041 | ENSG00000133134 | n.s. |
| ENSG00000249850 | 220.1806 | -0.060942139 | 0.256373 | -0.237709 | 0.812107 | 0.966187324 | KRT18P31 | processed_pseudogene | 55.88 | -0.000910394 | 0.000910394 | ENSG00000249850 | n.s. |
| ENSG00000133256 | 42.44727 | -0.068727773 | 0.313719 | -0.219074 | 0.826592 | 0.969965817 | PDE6B | protein_coding | 56.84 | -0.000910201 | 0.000910201 | ENSG00000133256 | n.s. |
| ENSG00000276900 | 69.36232 | -0.065164576 | 0.289465 | -0.22512 | 0.821886 | 0.968355922 | | antisense | 46.74 | -0.000910022 | 0.000910022 | ENSG00000276900 | n.s. |
| ENSG00000095059 | 845.5308 | 0.065135969 | 0.288667 | 0.2256443 | 0.821478 | 0.968355922 | DHPS | protein_coding | 54.74 | 0.000909623 | 0.000909623 | ENSG00000095059 | n.s. |
| ENSG00000245910 | 2436.869 | 0.062269281 | 0.270472 | 0.2302247 | 0.817917 | 0.967029262 | SNHG6 | processed_transcript | 44.54 | 0.000906665 | 0.000906665 | ENSG00000245910 | n.s. |
| ENSG00000111361 | 1869.503 | -0.062023329 | 0.27009 | -0.22964 | 0.818372 | 0.96724455 | EIF2B1 | protein_coding | 45.42 | -0.000897087 | 0.000897087 | ENSG00000111361 | n.s. |
| ENSG00000120265 | 1654.015 | -0.067810085 | 0.311648 | -0.217586 | 0.827752 | 0.970009871 | PCMT1 | protein_coding | 41.51 | -0.00089671 | 0.00089671 | ENSG00000120265 | n.s. |
| ENSG00000196981 | 398.245 | 0.054544779 | 0.211416 | 0.2579976 | 0.796409 | 0.962958487 | WDR5B | protein_coding | 40.54 | 0.000894122 | 0.000894122 | ENSG00000196981 | n.s. |
| ENSG00000081181 | 129.7526 | 0.067487041 | 0.311477 | 0.2166675 | 0.828467 | 0.970009871 | ARG2 | protein_coding | 42.2 | 0.000892438 | 0.000892438 | ENSG00000081181 | n.s. |
| ENSG00000107371 | 1190.533 | 0.055274678 | 0.216709 | 0.2550638 | 0.798674 | 0.963515749 | EXOSC3 | protein_coding | 43.49 | 0.000892199 | 0.000892199 | ENSG00000107371 | n.s. |
| ENSG00000131370 | 189.3604 | -0.063874838 | 0.282419 | -0.226171 | 0.821069 | 0.968355922 | SH3BP5 | protein_coding | 46.74 | -0.000892011 | 0.000892011 | ENSG00000131370 | n.s. |
| ENSG00000278600 | 471.8683 | 0.057725791 | 0.231828 | 0.2490026 | 0.803359 | 0.965047325 | | sense_intronic | 41.18 | 0.000891944 | 0.000891944 | ENSG00000278600 | n.s. |
| ENSG00000167996 | 2508.461 | -0.058326262 | 0.237576 | -0.245506 | 0.806065 | 0.9654664 | FTH1 | protein_coding | 50.42 | -0.000890224 | 0.000890224 | ENSG00000167996 | n.s. |
| ENSG00000228878 | 30.38844 | 0.076100859 | 0.376348 | 0.2022085 | 0.839754 | 0.973446151 | SEPT7-AS1 | lincRNA | 39.03 | 0.000889472 | 0.000889472 | ENSG00000228878 | n.s. |
| ENSG00000230498 | 40.45739 | 0.067197425 | 0.309647 | 0.2170133 | 0.828198 | 0.970009871 | | antisense | 34.89 | 0.000888608 | 0.000888608 | ENSG00000230498 | n.s. |
| ENSG00000175283 | 240.9182 | -0.059076707 | 0.246967 | -0.239209 | 0.810944 | 0.965963425 | DOLK | protein_coding | 56.89 | -0.000888473 | 0.000888473 | ENSG00000175283 | n.s. |
| ENSG00000156103 | 1667.911 | -0.063916865 | 0.284602 | -0.224583 | 0.822304 | 0.968500556 | MMP16 | protein_coding | 35.15 | -0.000888452 | 0.000888452 | ENSG00000156103 | n.s. |
| ENSG00000231961 | 28.26671 | -0.071989839 | 0.346441 | -0.207799 | 0.835386 | 0.971988837 | | processed_pseudogene | 44.88 | -0.000888263 | 0.000888263 | ENSG00000231961 | n.s. |
| ENSG00000103365 | 1631.92 | -0.058539538 | 0.240498 | -0.24341 | 0.807688 | 0.965694801 | GGA2 | protein_coding | 47.76 | -0.000887466 | 0.000887466 | ENSG00000103365 | n.s. |
| ENSG00000279744 | 121.804 | -0.059369961 | 0.248921 | -0.23851 | 0.811486 | 0.966187324 | | TEC | 54.26 | -0.000886908 | 0.000886908 | ENSG00000279744 | n.s. |
| ENSG00000168397 | 1013.102 | -0.061297641 | 0.267496 | -0.229154 | 0.81875 | 0.967247655 | ATG4B | protein_coding | 52.33 | -0.000886506 | 0.000886506 | ENSG00000168397 | n.s. |
| ENSG00000163013 | 375.7202 | -0.059283519 | 0.249509 | -0.237601 | 0.812191 | 0.966187324 | FBXO41 | protein_coding | 52.56 | -0.000885617 | 0.000885617 | ENSG00000163013 | n.s. |
| ENSG00000135763 | 2056.183 | -0.057003966 | 0.227052 | -0.251061 | 0.801767 | 0.964862231 | URB2 | protein_coding | 43.59 | -0.000885539 | 0.000885539 | ENSG00000135763 | n.s. |
| ENSG00000276728 | 139.7003 | -0.05820569 | 0.238767 | -0.243776 | 0.807404 | 0.965676593 | | lincRNA | 54.45 | -0.000882881 | 0.000882881 | ENSG00000276728 | n.s. |
| ENSG00000149476 | 198.4129 | -0.057836114 | 0.236753 | -0.244288 | 0.807007 | 0.9654664 | TKFC | protein_coding | 54.64 | -0.000882743 | 0.000882743 | ENSG00000149476 | n.s. |
| ENSG00000033030 | 1130.023 | -0.058507682 | 0.241694 | -0.242073 | 0.808724 | 0.965963425 | ZCCHC8 | protein_coding | 42.68 | -0.000879916 | 0.000879916 | ENSG00000033030 | n.s. |
| ENSG00000237651 | 333.5539 | 0.066513291 | 0.304592 | 0.2183688 | 0.827142 | 0.970009871 | C2orf74 | protein_coding | 44.26 | 0.000879562 | 0.000879562 | ENSG00000237651 | n.s. |
| ENSG00000164346 | 2044.778 | 0.062836177 | 0.278923 | 0.2252813 | 0.82176 | 0.968355922 | NSA2 | protein_coding | 37.93 | 0.000877506 | 0.000877506 | ENSG00000164346 | n.s. |
| ENSG00000137343 | 180.3172 | -0.060182834 | 0.260643 | -0.230901 | 0.817391 | 0.967029262 | ATAT1 | protein_coding | 48.87 | -0.000876285 | 0.000876285 | ENSG00000137343 | n.s. |
| ENSG00000148187 | 513.7644 | -0.060103232 | 0.258629 | -0.232392 | 0.816234 | 0.967004949 | MRRF | protein_coding | 41.15 | -0.000875782 | 0.000875782 | ENSG00000148187 | n.s. |

| | | | | | | | | | | | | | |
|---|---|---|---|---|---|---|---|---|---|---|---|---|---|
| ENSG00000112511 | 238.85 | -0.059867571 | 0.2572 | -0.232767 | 0.815942 | 0.96692204 | PHF1 | protein_coding | 57.49 | -0.000874578 | 0.000874578 | ENSG00000112511 | n.s. |
| ENSG00000072849 | 838.8568 | -0.060038247 | 0.258768 | -0.232015 | 0.816526 | 0.967029262 | DERL2 | protein_coding | 41.38 | -0.00087418 | 0.00087418 | ENSG00000072849 | n.s. |
| ENSG00000100316 | 18670.1 | 0.062508202 | 0.277501 | 0.225254 | 0.821782 | 0.968355922 | RPL3 | protein_coding | 53.41 | 0.000872926 | 0.000872926 | ENSG00000100316 | n.s. |
| ENSG00000236073 | 6.960768 | -0.082610876 | 0.494046 | -0.167213 | 0.867202 | 0.975964429 | | processed_pseudogene | 41.74 | -0.000872867 | 0.000872867 | ENSG00000236073 | n.s. |
| ENSG00000111674 | 632.1706 | 0.062454027 | 0.276535 | 0.2258446 | 0.821322 | 0.968355922 | ENO2 | protein_coding | 55.33 | 0.00087217 | 0.00087217 | ENSG00000111674 | n.s. |
| ENSG00000172500 | 581.8053 | -0.069435364 | 0.326926 | -0.212389 | 0.831804 | 0.97149614 | FIBP | protein_coding | 55.12 | -0.000872033 | 0.000872033 | ENSG00000172500 | n.s. |
| ENSG00000100934 | 1204.289 | -0.065847365 | 0.302006 | -0.218033 | 0.827403 | 0.970009871 | SEC23A | protein_coding | 38.8 | -0.000870755 | 0.000870755 | ENSG00000100934 | n.s. |
| ENSG00000118257 | 3188.045 | 0.059206864 | 0.252464 | 0.2345163 | 0.814584 | 0.966779245 | NRP2 | protein_coding | 45.88 | 0.000868723 | 0.000868723 | ENSG00000118257 | n.s. |
| ENSG00000207789 | 3.841351 | -0.097881691 | 0.724142 | -0.135169 | 0.892478 | 0.979788048 | MIR26A2 | miRNA | 47.62 | -0.000868001 | 0.000868001 | ENSG00000207789 | n.s. |
| ENSG00000200120 | 74.84397 | 0.072401811 | 0.353034 | 0.2050846 | 0.837506 | 0.97278132 | | misc_RNA | 42.16 | 0.00086772 | 0.00086772 | ENSG00000200120 | n.s. |
| ENSG00000173567 | 25.98474 | -0.074187056 | 0.370834 | -0.200055 | 0.841438 | 0.973446151 | ADGRF3 | protein_coding | 45.62 | -0.000867103 | 0.000867103 | ENSG00000173567 | n.s. |
| ENSG00000142875 | 855.9475 | 0.06429803 | 0.290862 | 0.2210606 | 0.825045 | 0.969452399 | PRKACB | protein_coding | 34.7 | 0.00086632 | 0.00086632 | ENSG00000142875 | n.s. |
| ENSG00000163541 | 1361.002 | -0.065419298 | 0.302031 | -0.216598 | 0.828522 | 0.970009871 | SUCLG1 | protein_coding | 39.25 | -0.000865095 | 0.000865095 | ENSG00000163541 | n.s. |
| ENSG00000161542 | 509.1942 | -0.061490636 | 0.270846 | -0.227032 | 0.820399 | 0.96813703 | PRPSAP1 | protein_coding | 48.2 | -0.000864753 | 0.000864753 | ENSG00000161542 | n.s. |
| ENSG00000118600 | 828.2683 | 0.056959269 | 0.234012 | 0.2434027 | 0.807693 | 0.965694801 | TMEM5 | protein_coding | 38.5 | 0.000863509 | 0.000863509 | ENSG00000118600 | n.s. |
| ENSG00000127527 | 60.50779 | -0.069357304 | 0.328516 | -0.211123 | 0.832791 | 0.971819596 | EPS15L1 | protein_coding | 50.79 | -0.000861026 | 0.000861026 | ENSG00000127527 | n.s. |
| ENSG00000141540 | 318.9028 | -0.058781793 | 0.252104 | -0.233165 | 0.815634 | 0.96692204 | TTYH2 | protein_coding | 54.47 | -0.000858716 | 0.000858716 | ENSG00000141540 | n.s. |
| ENSG00000151470 | 225.089 | -0.061669981 | 0.274674 | -0.22452 | 0.822352 | 0.968500556 | C4orf33 | protein_coding | 35.63 | -0.00085722 | 0.00085722 | ENSG00000151470 | n.s. |
| ENSG00000166352 | 323.0756 | 0.070180724 | 0.33911 | 0.2069558 | 0.836044 | 0.972288085 | C11orf74 | protein_coding | 36.82 | 0.000856558 | 0.000856558 | ENSG00000166352 | n.s. |
| ENSG00000121104 | 362.0449 | -0.058353541 | 0.249886 | -0.23352 | 0.815357 | 0.966779245 | FAM117A | protein_coding | 46.7 | -0.000856203 | 0.000856203 | ENSG00000121104 | n.s. |
| ENSG00000278311 | 1613.281 | -0.0560503 | 0.229202 | -0.244546 | 0.806808 | 0.9654664 | GGNBP2 | protein_coding | 42.1 | -0.000855487 | 0.000855487 | ENSG00000278311 | n.s. |
| ENSG00000196628 | 16621.75 | -0.057623446 | 0.244351 | -0.235822 | 0.813571 | 0.966414697 | TCF4 | protein_coding | 37.78 | -0.000854929 | 0.000854929 | ENSG00000196628 | n.s. |
| ENSG00000197381 | 17.07739 | -0.074675454 | 0.41799 | -0.178653 | 0.85821 | 0.974056274 | ADARB1 | protein_coding | 45.43 | -0.000852491 | 0.000852491 | ENSG00000197381 | n.s. |
| ENSG00000100941 | 8023.896 | -0.056656226 | 0.234176 | -0.241939 | 0.808827 | 0.965963425 | PNN | protein_coding | 40.65 | -0.000852071 | 0.000852071 | ENSG00000100941 | n.s. |
| ENSG00000111640 | 48156.04 | -0.063222263 | 0.286217 | -0.220889 | 0.825178 | 0.969452399 | GAPDH | protein_coding | 60.61 | -0.000851826 | 0.000851826 | ENSG00000111640 | n.s. |
| ENSG00000101856 | 2228.158 | -0.064200473 | 0.292947 | -0.219154 | 0.82653 | 0.969965817 | PGRMC1 | protein_coding | 42.58 | -0.000850244 | 0.000850244 | ENSG00000101856 | n.s. |
| ENSG00000154723 | 1294.563 | 0.065740512 | 0.305217 | 0.2153894 | 0.829464 | 0.970726167 | ATP5J | protein_coding | 41.96 | 0.000848267 | 0.000848267 | ENSG00000154723 | n.s. |
| ENSG00000177842 | 437.0908 | -0.056360078 | 0.234116 | -0.240736 | 0.80976 | 0.965963425 | ZNF620 | protein_coding | 43.94 | -0.000847617 | 0.000847617 | ENSG00000177842 | n.s. |
| ENSG00000186010 | 440.4286 | 0.069827362 | 0.33939 | 0.2057434 | 0.836991 | 0.972496895 | NDUFA13 | protein_coding | 53.15 | 0.000845733 | 0.000845733 | ENSG00000186010 | n.s. |
| ENSG00000233230 | 60.62241 | -0.065827801 | 0.307172 | -0.214303 | 0.830311 | 0.970878832 | | antisense | 56.26 | -0.000844898 | 0.000844898 | ENSG00000233230 | n.s. |
| ENSG00000141552 | 1089.174 | -0.068219821 | 0.325629 | -0.209502 | 0.834057 | 0.971936785 | ANAPC11 | protein_coding | 55.3 | -0.000843332 | 0.000843332 | ENSG00000141552 | n.s. |
| ENSG00000125968 | 4353.292 | 0.060345383 | 0.267939 | 0.2252209 | 0.821807 | 0.968355922 | ID1 | protein_coding | 58.56 | 0.000842722 | 0.000842722 | ENSG00000125968 | n.s. |

| | | | | | | | | | | | | | |
|---|---|---|---|---|---|---|---|---|---|---|---|---|---|
| ENSG00000119203 | 2120.809 | -0.063270709 | 0.288031 | -0.219666 | 0.826131 | 0.969833104 | CPSF3 | protein_coding | 42.1 | -0.00084169 | 0.00084169 | ENSG00000119203 | n.s. |
| ENSG00000141428 | 426.4701 | 0.063638069 | 0.292434 | 0.217615 | 0.827729 | 0.970009871 | C18orf21 | protein_coding | 42.7 | 0.00084154 | 0.00084154 | ENSG00000141428 | n.s. |
| ENSG00000254999 | 1182.171 | 0.066023128 | 0.309481 | 0.213335 | 0.831066 | 0.971092758 | BRK1 | protein_coding | 45.07 | 0.000841087 | 0.000841087 | ENSG00000254999 | n.s. |
| ENSG00000185800 | 128.9161 | -0.057300343 | 0.24521 | -0.233679 | 0.815235 | 0.966779245 | DMWD | protein_coding | 57.55 | -0.00084075 | 0.00084075 | ENSG00000185800 | n.s. |
| ENSG00000222259 | 22.19866 | -0.073614674 | 0.411986 | -0.178683 | 0.858187 | 0.974056274 | RN7SKP114 | misc_RNA | 54.12 | -0.000840382 | 0.000840382 | ENSG00000222259 | n.s. |
| ENSG00000181929 | 346.1722 | -0.06351154 | 0.292272 | -0.217303 | 0.827973 | 0.970009871 | PRKAG1 | protein_coding | 45.41 | -0.000839867 | 0.000839867 | ENSG00000181929 | n.s. |
| ENSG00000047315 | 4940.806 | -0.062109383 | 0.279953 | -0.221856 | 0.824426 | 0.969452399 | POLR2B | protein_coding | 40.19 | -0.000836831 | 0.000836831 | ENSG00000047315 | n.s. |
| ENSG00000278514 | 20.04021 | 0.07828943 | 0.454483 | 0.1722606 | 0.863233 | 0.975741574 | | lincRNA | 37.7 | 0.000834972 | 0.000834972 | ENSG00000278514 | n.s. |
| ENSG00000142676 | 10053.27 | 0.06194909 | 0.279475 | 0.2216621 | 0.824577 | 0.969452399 | RPL11 | protein_coding | 46.18 | 0.000834672 | 0.000834672 | ENSG00000142676 | n.s. |
| ENSG00000170558 | 2548.707 | -0.059754523 | 0.264385 | -0.226013 | 0.821191 | 0.968355922 | CDH2 | protein_coding | 37.22 | -0.000834471 | 0.000834471 | ENSG00000170558 | n.s. |
| ENSG00000223224 | 8.879321 | -0.094500333 | 0.707046 | -0.133655 | 0.893675 | 0.979889821 | SNORD71 | snoRNA | 48.84 | -0.000833753 | 0.000833753 | ENSG00000223224 | n.s. |
| ENSG00000114209 | 1191.496 | 0.062224766 | 0.282558 | 0.2202196 | 0.8257 | 0.969654445 | PDCD10 | protein_coding | 35.73 | 0.000832755 | 0.000832755 | ENSG00000114209 | n.s. |
| ENSG00000166140 | 147.8325 | 0.060240532 | 0.26907 | 0.2238844 | 0.822847 | 0.968673589 | ZFYVE19 | protein_coding | 53.53 | 0.000832677 | 0.000832677 | ENSG00000166140 | n.s. |
| ENSG00000138495 | 1179.647 | -0.061776362 | 0.279938 | -0.220679 | 0.825342 | 0.969464905 | COX17 | protein_coding | 41.28 | -0.000831998 | 0.000831998 | ENSG00000138495 | n.s. |
| ENSG00000198130 | 590.5631 | 0.061662193 | 0.278959 | 0.2210442 | 0.825058 | 0.969452399 | HIBCH | protein_coding | 38.94 | 0.000830806 | 0.000830806 | ENSG00000198130 | n.s. |
| ENSG00000171298 | 452.2804 | -0.056616694 | 0.242463 | -0.233506 | 0.815368 | 0.966779245 | GAA | protein_coding | 58.33 | -0.000830719 | 0.000830719 | ENSG00000171298 | n.s. |
| ENSG00000151778 | 86.87427 | 0.071056188 | 0.367587 | 0.1933042 | 0.846721 | 0.973446151 | SERP2 | protein_coding | 44.49 | 0.00083051 | 0.00083051 | ENSG00000151778 | n.s. |
| ENSG00000156697 | 831.4888 | -0.058144038 | 0.25515 | -0.227882 | 0.819738 | 0.967663544 | UTP14A | protein_coding | 45.09 | -0.000830042 | 0.000830042 | ENSG00000156697 | n.s. |
| ENSG00000260589 | 279.3426 | -0.05437131 | 0.22207 | -0.244838 | 0.806582 | 0.9654664 | STAM-AS1 | antisense | 38.61 | -0.00082986 | 0.00082986 | ENSG00000260589 | n.s. |
| ENSG00000274695 | 26.9063 | 0.072681552 | 0.395767 | 0.1836475 | 0.85429 | 0.974056274 | | lincRNA | 38.31 | 0.000829729 | 0.000829729 | ENSG00000274695 | n.s. |
| ENSG00000112208 | 911.24 | -0.057328484 | 0.249907 | -0.229399 | 0.818559 | 0.967247655 | BAG2 | protein_coding | 38.77 | -0.000829103 | 0.000829103 | ENSG00000112208 | n.s. |
| ENSG00000166181 | 3373.881 | -0.059935155 | 0.267647 | -0.223933 | 0.822809 | 0.968673589 | API5 | protein_coding | 37.75 | -0.000828456 | 0.000828456 | ENSG00000166181 | n.s. |
| ENSG00000051620 | 665.8481 | -0.066963654 | 0.320928 | -0.208656 | 0.834717 | 0.971936785 | HEBP2 | protein_coding | 47.12 | -0.000827803 | 0.000827803 | ENSG00000051620 | n.s. |
| ENSG00000064490 | 117.6624 | -0.070041323 | 0.343808 | -0.203722 | 0.838571 | 0.973164573 | RFXANK | protein_coding | 54.66 | -0.000827448 | 0.000827448 | ENSG00000064490 | n.s. |
| ENSG00000229320 | 114.83 | 0.057559448 | 0.251844 | 0.2285517 | 0.819217 | 0.967475307 | KRT8P12 | ıscribed_processed_pseudog | 50.42 | 0.00082656 | 0.00082656 | ENSG00000229320 | n.s. |
| ENSG00000171204 | 508.4191 | 0.065136133 | 0.305888 | 0.2129409 | 0.831373 | 0.971210458 | TMEM126B | protein_coding | 37.32 | 0.000826359 | 0.000826359 | ENSG00000171204 | n.s. |
| ENSG00000213516 | 2012.801 | 0.062474844 | 0.2863 | 0.2182145 | 0.827262 | 0.970009871 | RBMXL1 | protein_coding | 41.18 | 0.000826158 | 0.000826158 | ENSG00000213516 | n.s. |
| ENSG00000207816 | 25.63992 | 0.070667786 | 0.369818 | 0.1910878 | 0.848457 | 0.973446151 | MIR124-2 | miRNA | 49.54 | 0.00082597 | 0.00082597 | ENSG00000207816 | n.s. |
| ENSG00000179008 | 54.57758 | 0.064798907 | 0.303823 | 0.2132785 | 0.83111 | 0.971092758 | C14orf39 | protein_coding | 37.01 | 0.000825492 | 0.000825492 | ENSG00000179008 | n.s. |
| ENSG00000113140 | 999.3947 | 0.061260138 | 0.277242 | 0.2209628 | 0.825121 | 0.969452399 | SPARC | protein_coding | 47.91 | 0.000825389 | 0.000825389 | ENSG00000113140 | n.s. |
| ENSG00000140990 | 891.815 | 0.066517122 | 0.31643 | 0.210211 | 0.833503 | 0.971936785 | NDUFB10 | protein_coding | 58.95 | 0.000822283 | 0.000822283 | ENSG00000140990 | n.s. |
| ENSG00000079785 | 174414.6 | -0.059479943 | 0.265603 | -0.223943 | 0.822802 | 0.968673589 | DDX1 | protein_coding | 39.49 | -0.000822164 | 0.000822164 | ENSG00000079785 | n.s. |

| | | | | | | | | | | | | | |
|---|---|---|---|---|---|---|---|---|---|---|---|---|---|
| ENSG00000170876 | 977.9996 | -0.061997181 | 0.282602 | -0.21938 | 0.826354 | 0.969931403 | TMEM43 | protein_coding | 52.73 | -0.000822019 | 0.000822019 | ENSG00000170876 | n.s. |
| ENSG00000171448 | 442.8712 | -0.062101445 | 0.284764 | -0.218081 | 0.827366 | 0.970009871 | ZBTB26 | protein_coding | 39.97 | -0.00082122 | 0.00082122 | ENSG00000171448 | n.s. |
| ENSG00000251022 | 3470.251 | 0.054520982 | 0.226853 | 0.2403362 | 0.81007 | 0.965963425 | THAP9-AS1 | antisense | 39.31 | 0.000819958 | 0.000819958 | ENSG00000251022 | n.s. |
| ENSG00000013523 | 423.8713 | 0.054806651 | 0.229885 | 0.2384086 | 0.811564 | 0.966187324 | ANGEL1 | protein_coding | 44.52 | 0.000818738 | 0.000818738 | ENSG00000013523 | n.s. |
| ENSG00000189077 | 82.01317 | -0.070028265 | 0.347964 | -0.201251 | 0.840502 | 0.973446151 | TMEM120A | protein_coding | 61.59 | -0.000818495 | 0.000818495 | ENSG00000189077 | n.s. |
| ENSG00000177700 | 531.809 | -0.065913715 | 0.312002 | -0.211261 | 0.832684 | 0.971819596 | POLR2L | protein_coding | 61.14 | -0.000818276 | 0.000818276 | ENSG00000177700 | n.s. |
| ENSG00000234882 | 64.11477 | 0.069978606 | 0.367957 | 0.1901817 | 0.849167 | 0.973446151 | EIF3EP1 | processed_pseudogene | 39.97 | 0.000817915 | 0.000817915 | ENSG00000234882 | n.s. |
| ENSG00000131732 | 736.8901 | 0.056078234 | 0.243399 | 0.2303967 | 0.817784 | 0.967029262 | ZCCHC9 | protein_coding | 38.15 | 0.000816521 | 0.000816521 | ENSG00000131732 | n.s. |
| ENSG00000222145 | 12.54413 | -0.085428547 | 0.573353 | -0.148998 | 0.881555 | 0.978242302 | | snoRNA | 47.62 | -0.000816147 | 0.000816147 | ENSG00000222145 | n.s. |
| ENSG00000140280 | 328.5119 | -0.065962968 | 0.313235 | -0.210587 | 0.83321 | 0.971936785 | LYSMD2 | protein_coding | 41.49 | -0.000815433 | 0.000815433 | ENSG00000140280 | n.s. |
| ENSG00000136237 | 1184.978 | 0.053262901 | 0.215021 | 0.2477102 | 0.804359 | 0.9654664 | RAPGEF5 | protein_coding | 39.42 | 0.000812943 | 0.000812943 | ENSG00000136237 | n.s. |
| ENSG00000143126 | 579.0815 | -0.054012482 | 0.223245 | -0.241943 | 0.808824 | 0.965963425 | CELSR2 | protein_coding | 59.75 | -0.000812311 | 0.000812311 | ENSG00000143126 | n.s. |
| ENSG00000124702 | 1217.207 | 0.06582406 | 0.316309 | 0.2081004 | 0.835151 | 0.971988837 | KLHDC3 | protein_coding | 52.39 | 0.000812185 | 0.000812185 | ENSG00000124702 | n.s. |
| ENSG00000171865 | 1167.248 | -0.05351592 | 0.219748 | -0.243533 | 0.807592 | 0.965694801 | RNASEH1 | protein_coding | 45.37 | -0.000811307 | 0.000811307 | ENSG00000171865 | n.s. |
| ENSG00000239789 | 491.1576 | -0.065720379 | 0.316029 | -0.207957 | 0.835263 | 0.971988837 | MRPS17 | protein_coding | 49.9 | -0.000810906 | 0.000810906 | ENSG00000239789 | n.s. |
| ENSG00000198846 | 298.7534 | 0.057839711 | 0.256536 | 0.2254641 | 0.821618 | 0.968355922 | TOX | protein_coding | 37.57 | 0.000807731 | 0.000807731 | ENSG00000198846 | n.s. |
| ENSG00000159884 | 123.74 | 0.062580432 | 0.291281 | 0.2148453 | 0.829888 | 0.970726167 | CCDC107 | protein_coding | 52.82 | 0.000807492 | 0.000807492 | ENSG00000159884 | n.s. |
| ENSG00000177479 | 1691.645 | -0.059902929 | 0.271363 | -0.220749 | 0.825288 | 0.969464905 | ARIH2 | protein_coding | 44.51 | -0.000806767 | 0.000806767 | ENSG00000177479 | n.s. |
| ENSG00000151287 | 496.1933 | -0.062462793 | 0.290795 | -0.2148 | 0.829923 | 0.970726167 | TEX30 | protein_coding | 41.14 | -0.000805974 | 0.000805974 | ENSG00000151287 | n.s. |
| ENSG00000145979 | 434.9535 | -0.055289891 | 0.239481 | -0.230874 | 0.817413 | 0.967029262 | TBC1D7 | protein_coding | 44.56 | -0.000805042 | 0.000805042 | ENSG00000145979 | n.s. |
| ENSG00000143612 | 2749.299 | 0.065015479 | 0.310313 | 0.2095161 | 0.834045 | 0.971936785 | C1orf43 | protein_coding | 45.88 | 0.00080372 | 0.00080372 | ENSG00000143612 | n.s. |
| ENSG00000212694 | 387.7381 | -0.052609017 | 0.215011 | -0.244681 | 0.806704 | 0.9654664 | LINC01089 | lincRNA | 58.94 | -0.000802963 | 0.000802963 | ENSG00000212694 | n.s. |
| ENSG00000267470 | 22.77135 | -0.068674308 | 0.36714 | -0.187052 | 0.85162 | 0.973446151 | ZNF571-AS1 | antisense | 41.03 | -0.00080267 | 0.00080267 | ENSG00000267470 | n.s. |
| ENSG00000227210 | 15.38367 | 0.076124969 | 0.47545 | 0.1601113 | 0.872793 | 0.976022242 | | antisense | 39.4 | 0.000802379 | 0.000802379 | ENSG00000227210 | n.s. |
| ENSG00000121766 | 292.5417 | -0.064991072 | 0.31269 | -0.207845 | 0.83535 | 0.971988837 | ZCCHC17 | protein_coding | 41.86 | -0.000801907 | 0.000801907 | ENSG00000121766 | n.s. |
| ENSG00000102897 | 398.1088 | 0.060622715 | 0.277221 | 0.2186804 | 0.826899 | 0.970009871 | LYRM1 | protein_coding | 44.36 | 0.000801665 | 0.000801665 | ENSG00000102897 | n.s. |
| ENSG00000278916 | 157.4185 | 0.056896 | 0.250803 | 0.226855 | 0.820536 | 0.96813703 | CEP83-AS1 | lincRNA | 44.48 | 0.000800138 | 0.000800138 | ENSG00000278916 | n.s. |
| ENSG00000139173 | 131.003 | 0.064507924 | 0.307146 | 0.2100238 | 0.833649 | 0.971936785 | TMEM117 | protein_coding | 37.35 | 0.000797446 | 0.000797446 | ENSG00000139173 | n.s. |
| ENSG00000175854 | 82.22002 | -0.077315002 | 0.494514 | -0.156345 | 0.875761 | 0.97660852 | SWI5 | protein_coding | 52.59 | -0.000794759 | 0.000794759 | ENSG00000175854 | n.s. |
| ENSG00000234740 | 96.5037 | -0.05898114 | 0.265427 | -0.222213 | 0.824148 | 0.969452399 | | sense_intronic | 43.07 | -0.000794683 | 0.000794683 | ENSG00000234740 | n.s. |
| ENSG00000108309 | 878.092 | -0.063035186 | 0.296513 | -0.212588 | 0.831648 | 0.971395614 | RUNDC3A | protein_coding | 59.03 | -0.000794487 | 0.000794487 | ENSG00000108309 | n.s. |
| ENSG00000268266 | 56.10773 | 0.065186306 | 0.315862 | 0.2063761 | 0.836497 | 0.972328444 | | lincRNA | 42.65 | 0.000794426 | 0.000794426 | ENSG00000268266 | n.s. |

| | | | | | | | | | | | | | |
|---|---|---|---|---|---|---|---|---|---|---|---|---|---|
| ENSG00000146731 | 8410.065 | 0.058911015 | 0.266599 | 0.2209726 | 0.825114 | 0.969452399 | CCT6A | protein_coding | 41.33 | 0.000793738 | 0.000793738 | ENSG00000146731 | n.s. |
| ENSG00000116898 | 580.2459 | 0.065267823 | 0.316765 | 0.2060446 | 0.836756 | 0.972412121 | MRPS15 | protein_coding | 49.28 | 0.00079298 | 0.00079298 | ENSG00000116898 | n.s. |
| ENSG00000085382 | 151.8119 | -0.056467634 | 0.249238 | -0.226561 | 0.820765 | 0.968186961 | HACE1 | protein_coding | 37.39 | -0.000792849 | 0.000792849 | ENSG00000085382 | n.s. |
| ENSG00000187555 | 3875.849 | -0.054304133 | 0.235191 | -0.230894 | 0.817397 | 0.967029262 | USP7 | protein_coding | 46.24 | -0.000790689 | 0.000790689 | ENSG00000187555 | n.s. |
| ENSG00000198169 | 278.3494 | 0.059693238 | 0.275691 | 0.2165224 | 0.828581 | 0.970009871 | ZNF251 | protein_coding | 47.51 | 0.000789374 | 0.000789374 | ENSG00000198169 | n.s. |
| ENSG00000200428 | 10.06266 | 0.090789242 | 0.685479 | 0.1324464 | 0.894631 | 0.980194243 | | misc_RNA | 34.19 | 0.000788764 | 0.000788764 | ENSG00000200428 | n.s. |
| ENSG00000113240 | 487.817 | 0.054315769 | 0.236399 | 0.2297631 | 0.818276 | 0.96724455 | CLK4 | protein_coding | 39.49 | 0.000785607 | 0.000785607 | ENSG00000113240 | n.s. |
| ENSG00000170419 | 619.0121 | 0.059398615 | 0.272929 | 0.2176337 | 0.827715 | 0.970009871 | VSTM2A | protein_coding | 39.92 | 0.000785478 | 0.000785478 | ENSG00000170419 | n.s. |
| ENSG00000175183 | 688.4746 | 0.063469369 | 0.303371 | 0.2092138 | 0.834281 | 0.971936785 | CSRP2 | protein_coding | 42.39 | 0.000784607 | 0.000784607 | ENSG00000175183 | n.s. |
| ENSG00000225479 | 48.93571 | 0.067085867 | 0.34245 | 0.1958996 | 0.844689 | 0.973446151 | PLCB1-IT1 | sense_intronic | 37.7 | 0.000784104 | 0.000784104 | ENSG00000225479 | n.s. |
| ENSG00000198162 | 2935.913 | -0.059206329 | 0.271465 | -0.2181 | 0.827352 | 0.970009871 | MAN1A2 | protein_coding | 35.34 | -0.000782935 | 0.000782935 | ENSG00000198162 | n.s. |
| ENSG00000203618 | 99.69304 | -0.060614407 | 0.28202 | -0.21493 | 0.829822 | 0.970726167 | GP1BB | protein_coding | 70.83 | -0.000782124 | 0.000782124 | ENSG00000203618 | n.s. |
| ENSG00000257511 | 38.13316 | 0.063149104 | 0.29947 | 0.2108696 | 0.832989 | 0.971901974 | | processed_pseudogene | 39.4 | 0.00078163 | 0.00078163 | ENSG00000257511 | n.s. |
| ENSG00000145365 | 635.1957 | -0.057102078 | 0.255742 | -0.22328 | 0.823317 | 0.969007976 | TIFA | protein_coding | 40.42 | -0.000780737 | 0.000780737 | ENSG00000145365 | n.s. |
| ENSG00000263072 | 355.1352 | -0.053442853 | 0.229369 | -0.232999 | 0.815762 | 0.96692204 | ZNF213-AS1 | antisense | 53.29 | -0.000780722 | 0.000780722 | ENSG00000263072 | n.s. |
| ENSG00000273682 | 29.10573 | -0.066594112 | 0.343917 | -0.193634 | 0.846462 | 0.973446151 | | processed_pseudogene | 56.59 | -0.000778356 | 0.000778356 | ENSG00000273682 | n.s. |
| ENSG00000112984 | 2494.513 | 0.054498982 | 0.239191 | 0.2278469 | 0.819765 | 0.967663544 | KIF20A | protein_coding | 45.27 | 0.000778007 | 0.000778007 | ENSG00000112984 | n.s. |
| ENSG00000156471 | 1030.575 | -0.057619138 | 0.260612 | -0.221092 | 0.825021 | 0.969452399 | PTDSS1 | protein_coding | 43.71 | -0.000776332 | 0.000776332 | ENSG00000156471 | n.s. |
| ENSG00000147872 | 1132.362 | 0.058604427 | 0.270048 | 0.2170145 | 0.828197 | 0.970009871 | PLIN2 | protein_coding | 44.99 | 0.000774976 | 0.000774976 | ENSG00000147872 | n.s. |
| ENSG00000101052 | 627.087 | -0.062646489 | 0.30039 | -0.20855 | 0.834799 | 0.971936785 | IFT52 | protein_coding | 44.51 | -0.000774435 | 0.000774435 | ENSG00000101052 | n.s. |
| ENSG00000168286 | 563.0333 | -0.05988512 | 0.278376 | -0.215123 | 0.829671 | 0.970726167 | THAP11 | protein_coding | 60.31 | -0.000772713 | 0.000772713 | ENSG00000168286 | n.s. |
| ENSG00000201263 | 8.291572 | 0.077346238 | 0.504883 | 0.1531963 | 0.878243 | 0.977264479 | SNORD114-6 | snoRNA | 40.28 | 0.000772525 | 0.000772525 | ENSG00000201263 | n.s. |
| ENSG00000197956 | 451.8101 | 0.06607894 | 0.345385 | 0.1913198 | 0.848275 | 0.973446151 | S100A6 | protein_coding | 58.51 | 0.000772335 | 0.000772335 | ENSG00000197956 | n.s. |
| ENSG00000254377 | 771.7138 | 0.054050986 | 0.237012 | 0.228052 | 0.819606 | 0.967663544 | MIR124-2HG | lincRNA | 44.12 | 0.000771611 | 0.000771611 | ENSG00000254377 | n.s. |
| ENSG00000263753 | 1567.018 | 0.052947639 | 0.229734 | 0.2304734 | 0.817724 | 0.967029262 | LINC00667 | lincRNA | 41.91 | 0.000770938 | 0.000770938 | ENSG00000263753 | n.s. |
| ENSG00000226471 | 102.2815 | 0.056941844 | 0.255703 | 0.2226872 | 0.823779 | 0.969312504 | | antisense | 45.98 | 0.000770775 | 0.000770775 | ENSG00000226471 | n.s. |
| ENSG00000090932 | 322.3624 | 0.059631689 | 0.277791 | 0.2146638 | 0.830029 | 0.970726167 | DLL3 | protein_coding | 56.84 | 0.000769443 | 0.000769443 | ENSG00000090932 | n.s. |
| ENSG00000197746 | 2620.817 | 0.061966228 | 0.29361 | 0.2110495 | 0.832849 | 0.971819596 | PSAP | protein_coding | 47.72 | 0.00076927 | 0.00076927 | ENSG00000197746 | n.s. |
| ENSG00000250001 | 68.11348 | -0.059879599 | 0.279643 | -0.214128 | 0.830447 | 0.970884671 | | sense_intronic | 46.18 | -0.000768396 | 0.000768396 | ENSG00000250001 | n.s. |
| ENSG00000171552 | 672.7376 | 0.052074683 | 0.221444 | 0.2351593 | 0.814085 | 0.96661324 | BCL2L1 | protein_coding | 48.2 | 0.000767959 | 0.000767959 | ENSG00000171552 | n.s. |
| ENSG00000135049 | 2058.596 | -0.061167552 | 0.288156 | -0.212272 | 0.831895 | 0.971513984 | AGTPBP1 | protein_coding | 36.66 | -0.000767711 | 0.000767711 | ENSG00000135049 | n.s. |
| ENSG00000023734 | 3676.637 | 0.059699392 | 0.279053 | 0.2139353 | 0.830597 | 0.970884671 | STRAP | protein_coding | 36.48 | 0.000766084 | 0.000766084 | ENSG00000023734 | n.s. |

| | | | | | | | | | | | | | |
|---|---|---|---|---|---|---|---|---|---|---|---|---|---|
| ENSG00000159352 | 2824.1 | -0.062938136 | 0.305349 | -0.206119 | 0.836698 | 0.972412121 | PSMD4 | protein_coding | 47.67 | -0.000764675 | 0.000764675 | ENSG00000159352 | n.s. |
| ENSG00000282855 | 15.901 | 0.072041115 | 0.423054 | 0.1702881 | 0.864784 | 0.975860117 | | processed_pseudogene | 44.48 | 0.000764531 | 0.000764531 | ENSG00000282855 | n.s. |
| ENSG00000186603 | 270.9339 | 0.055909603 | 0.250434 | 0.2232511 | 0.82334 | 0.969007976 | HPDL | protein_coding | 62.8 | 0.000764432 | 0.000764432 | ENSG00000186603 | n.s. |
| ENSG00000126768 | 371.5458 | -0.065278726 | 0.336356 | -0.194077 | 0.846116 | 0.973446151 | TIMM17B | protein_coding | 52.95 | -0.000762982 | 0.000762982 | ENSG00000126768 | n.s. |
| ENSG00000118496 | 2031.249 | -0.056582499 | 0.256001 | -0.221025 | 0.825073 | 0.969452399 | FBXO30 | protein_coding | 35.2 | -0.000762365 | 0.000762365 | ENSG00000118496 | n.s. |
| ENSG00000158270 | 405.269 | 0.061638244 | 0.292602 | 0.2106556 | 0.833156 | 0.971936785 | COLEC12 | protein_coding | 43.13 | 0.000761971 | 0.000761971 | ENSG00000158270 | n.s. |
| ENSG00000014641 | 3568.997 | 0.06159778 | 0.294426 | 0.2092132 | 0.834282 | 0.971936785 | MDH1 | protein_coding | 38.05 | 0.000761471 | 0.000761471 | ENSG00000014641 | n.s. |
| ENSG00000091039 | 4320.528 | -0.058886937 | 0.27417 | -0.214783 | 0.829937 | 0.970726167 | OSBPL8 | protein_coding | 36.48 | -0.000759834 | 0.000759834 | ENSG00000091039 | n.s. |
| ENSG00000273254 | 25.56987 | -0.066464659 | 0.369082 | -0.180081 | 0.857089 | 0.974056274 | | antisense | 53.46 | -0.000758757 | 0.000758757 | ENSG00000273254 | n.s. |
| ENSG00000103253 | 69.27967 | 0.064803883 | 0.326089 | 0.1987305 | 0.842474 | 0.973446151 | HAGHL | protein_coding | 65.72 | 0.000757432 | 0.000757432 | ENSG00000103253 | n.s. |
| ENSG00000272146 | 59.87033 | 0.061365826 | 0.295218 | 0.207866 | 0.835334 | 0.971988837 | ARF4-AS1 | antisense | 43.98 | 0.000757176 | 0.000757176 | ENSG00000272146 | n.s. |
| ENSG00000127337 | 1537.75 | 0.064753478 | 0.331904 | 0.1950973 | 0.845317 | 0.973446151 | YEATS4 | protein_coding | 39.36 | 0.000756843 | 0.000756843 | ENSG00000127337 | n.s. |
| ENSG00000231128 | 54.81095 | 0.064664212 | 0.326614 | 0.1979835 | 0.843058 | 0.973446151 | | antisense | 38.96 | 0.0007558 | 0.0007558 | ENSG00000231128 | n.s. |
| ENSG00000164070 | 534.2709 | 0.05606518 | 0.25281 | 0.2217684 | 0.824494 | 0.969452399 | HSPA4L | protein_coding | 35.4 | 0.000755395 | 0.000755395 | ENSG00000164070 | n.s. |
| ENSG00000207933 | 7.645949 | -0.084599444 | 0.597674 | -0.141548 | 0.887437 | 0.979788048 | MIR9-1 | miRNA | 40.45 | -0.000750216 | 0.000750216 | ENSG00000207933 | n.s. |
| ENSG00000208883 | 4.309899 | -0.083530915 | 0.584304 | -0.142958 | 0.886323 | 0.979542794 | SNORD96B | snoRNA | 50.63 | -0.000749822 | 0.000749822 | ENSG00000208883 | n.s. |
| ENSG00000270157 | 59.61674 | -0.06398953 | 0.337096 | -0.189826 | 0.849446 | 0.973446151 | | sense_intronic | 42.92 | -0.000747914 | 0.000747914 | ENSG00000270157 | n.s. |
| ENSG00000115806 | 1194.243 | -0.056545921 | 0.258706 | -0.218572 | 0.826984 | 0.970009871 | GORASP2 | protein_coding | 41.04 | -0.000747755 | 0.000747755 | ENSG00000115806 | n.s. |
| ENSG00000230251 | 55.08896 | -0.063918457 | 0.331896 | -0.192586 | 0.847283 | 0.973446151 | PHBP4 | processed_pseudogene | 50.66 | -0.000747083 | 0.000747083 | ENSG00000230251 | n.s. |
| ENSG00000113356 | 197.606 | -0.060368609 | 0.287726 | -0.209813 | 0.833814 | 0.971936785 | POLR3G | protein_coding | 37.84 | -0.000746276 | 0.000746276 | ENSG00000113356 | n.s. |
| ENSG00000223501 | 574.9566 | -0.055376684 | 0.249938 | -0.221562 | 0.824655 | 0.969452399 | VPS52 | protein_coding | 45.57 | -0.000746118 | 0.000746118 | ENSG00000223501 | n.s. |
| ENSG00000184611 | 879.009 | 0.053419352 | 0.23647 | 0.2259037 | 0.821276 | 0.968355922 | KCNH7 | protein_coding | 35.03 | 0.000746001 | 0.000746001 | ENSG00000184611 | n.s. |
| ENSG00000119760 | 1717.978 | 0.056781408 | 0.262822 | 0.2160451 | 0.828953 | 0.970282176 | SUPT7L | protein_coding | 41.29 | 0.000743947 | 0.000743947 | ENSG00000119760 | n.s. |
| ENSG00000126432 | 1201.277 | -0.063584583 | 0.31565 | -0.20144 | 0.840354 | 0.973446151 | PRDX5 | protein_coding | 57.55 | -0.000743181 | 0.000743181 | ENSG00000126432 | n.s. |
| ENSG00000203875 | 4425.003 | 0.049409297 | 0.203484 | 0.2428166 | 0.808147 | 0.965963425 | SNHG5 | processed_transcript | 40.9 | 0.000743082 | 0.000743082 | ENSG00000203875 | n.s. |
| ENSG00000129515 | 1107.628 | 0.063538284 | 0.322322 | 0.1971267 | 0.843728 | 0.973446151 | SNX6 | protein_coding | 44.34 | 0.00074264 | 0.00074264 | ENSG00000129515 | n.s. |
| ENSG00000169018 | 3319.357 | -0.057740373 | 0.269527 | -0.214229 | 0.830369 | 0.970878832 | FEM1B | protein_coding | 39.07 | -0.000741096 | 0.000741096 | ENSG00000169018 | n.s. |
| ENSG00000109255 | 132.8083 | 0.063345338 | 0.328383 | 0.1929007 | 0.847037 | 0.973446151 | NMU | protein_coding | 38.09 | 0.000740385 | 0.000740385 | ENSG00000109255 | n.s. |
| ENSG00000169814 | 80.37495 | -0.05898727 | 0.27826 | -0.211986 | 0.832118 | 0.971513984 | BTD | protein_coding | 41.69 | -0.000740346 | 0.000740346 | ENSG00000169814 | n.s. |
| ENSG00000197894 | 3070.487 | -0.060808253 | 0.295233 | -0.205967 | 0.836817 | 0.972412121 | ADH5 | protein_coding | 40.37 | -0.000738798 | 0.000738798 | ENSG00000197894 | n.s. |
| ENSG00000111843 | 1068.605 | 0.063124416 | 0.317969 | 0.1985239 | 0.842635 | 0.973446151 | TMEM14C | protein_coding | 47.49 | 0.000737802 | 0.000737802 | ENSG00000111843 | n.s. |
| ENSG00000242265 | 3993.711 | -0.055665248 | 0.255106 | -0.218204 | 0.82727 | 0.970009871 | PEG10 | protein_coding | 43.22 | -0.000736109 | 0.000736109 | ENSG00000242265 | n.s. |

| | | | | | | | | | | | | | |
|---|---|---|---|---|---|---|---|---|---|---|---|---|---|
| ENSG00000183971 | 181.8551 | -0.060250288 | 0.291291 | -0.206839 | 0.836136 | 0.972288085 | NPW | protein_coding | 55.12 | -0.000735357 | 0.000735357 | ENSG00000183971 | n.s. |
| ENSG00000129195 | 1270.544 | -0.060249684 | 0.291025 | -0.207026 | 0.83599 | 0.972288085 | PIMREG | protein_coding | 53.2 | -0.00073535 | 0.00073535 | ENSG00000129195 | n.s. |
| ENSG00000260083 | 205.3433 | 0.0536171 | 0.240355 | 0.2230743 | 0.823478 | 0.969088012 | MIR762HG | antisense | 52.7 | 0.000731165 | 0.000731165 | ENSG00000260083 | n.s. |
| ENSG00000270115 | 29.83244 | -0.062535588 | 0.319327 | -0.195836 | 0.844739 | 0.973446151 | | lincRNA | 48.19 | -0.00073092 | 0.00073092 | ENSG00000270115 | n.s. |
| ENSG00000273343 | 37.42671 | 0.062345823 | 0.320397 | 0.1945892 | 0.845715 | 0.973446151 | | sense_intronic | 45.32 | 0.000728702 | 0.000728702 | ENSG00000273343 | n.s. |
| ENSG00000274066 | 3.474433 | 0.103264941 | 0.867055 | 0.1190985 | 0.905197 | 0.983983433 | MIR6514 | miRNA | 48.57 | 0.000724116 | 0.000724116 | ENSG00000274066 | n.s. |
| ENSG00000138663 | 893.2742 | 0.059299871 | 0.286788 | 0.2067726 | 0.836187 | 0.972288085 | COPS4 | protein_coding | 40.06 | 0.000723757 | 0.000723757 | ENSG00000138663 | n.s. |
| ENSG00000205155 | 248.2279 | 0.06190797 | 0.306332 | 0.2020943 | 0.839843 | 0.973446151 | PSENEN | protein_coding | 52 | 0.000723585 | 0.000723585 | ENSG00000205155 | n.s. |
| ENSG00000260400 | 147.5879 | -0.06188267 | 0.323486 | -0.1913 | 0.848291 | 0.973446151 | | sense_overlapping | 33.03 | -0.000723289 | 0.000723289 | ENSG00000260400 | n.s. |
| ENSG00000112667 | 370.7458 | -0.061863726 | 0.308216 | -0.200716 | 0.840921 | 0.973446151 | DNPH1 | protein_coding | 55.32 | -0.000723067 | 0.000723067 | ENSG00000112667 | n.s. |
| ENSG00000132823 | 503.3618 | 0.058460829 | 0.278607 | 0.2098327 | 0.833798 | 0.971936785 | OSER1 | protein_coding | 40.61 | 0.000722692 | 0.000722692 | ENSG00000132823 | n.s. |
| ENSG00000198898 | 1744.762 | -0.06168625 | 0.304492 | -0.202588 | 0.839457 | 0.973446151 | CAPZA2 | protein_coding | 38.99 | -0.000720993 | 0.000720993 | ENSG00000198898 | n.s. |
| ENSG00000117600 | 1091.078 | -0.060110219 | 0.292963 | -0.20518 | 0.837431 | 0.97278132 | PLPPR4 | protein_coding | 36.26 | -0.000720408 | 0.000720408 | ENSG00000117600 | n.s. |
| ENSG00000273749 | 1476.152 | -0.05733243 | 0.270269 | -0.212131 | 0.832005 | 0.971513984 | CYFIP1 | protein_coding | 48.83 | -0.000719576 | 0.000719576 | ENSG00000273749 | n.s. |
| ENSG00000275560 | 615.9304 | -0.047811282 | 0.199864 | -0.239219 | 0.810936 | 0.965963425 | | sense_intronic | 43.17 | -0.000719049 | 0.000719049 | ENSG00000275560 | n.s. |
| ENSG00000108960 | 3254.015 | -0.05815406 | 0.276727 | -0.21015 | 0.833551 | 0.971936785 | MMD | protein_coding | 41.06 | -0.000718899 | 0.000718899 | ENSG00000108960 | n.s. |
| ENSG00000034713 | 1033.445 | -0.061487913 | 0.310462 | -0.198053 | 0.843004 | 0.973446151 | GABARAPL2 | protein_coding | 47.18 | -0.000718675 | 0.000718675 | ENSG00000034713 | n.s. |
| ENSG00000203805 | 120.5873 | -0.061452521 | 0.316156 | -0.194374 | 0.845883 | 0.973446151 | PLPP4 | protein_coding | 42.31 | -0.000718261 | 0.000718261 | ENSG00000203805 | n.s. |
| ENSG00000119801 | 476.1292 | -0.061447952 | 0.305077 | -0.201418 | 0.840372 | 0.973446151 | YPEL5 | protein_coding | 41.9 | -0.000718208 | 0.000718208 | ENSG00000119801 | n.s. |
| ENSG00000243317 | 1239.53 | -0.061388809 | 0.311893 | -0.196826 | 0.843963 | 0.973446151 | C7orf73 | protein_coding | 43.1 | -0.000717517 | 0.000717517 | ENSG00000243317 | n.s. |
| ENSG00000101901 | 489.9225 | 0.05984972 | 0.292435 | 0.20466 | 0.837838 | 0.97278132 | ALG13 | protein_coding | 37.51 | 0.000717286 | 0.000717286 | ENSG00000101901 | n.s. |
| ENSG00000264585 | 41.05065 | 0.061238157 | 0.311919 | 0.1963269 | 0.844354 | 0.973446151 | MIR4449 | miRNA | 86.36 | 0.000715756 | 0.000715756 | ENSG00000264585 | n.s. |
| ENSG00000278291 | 159.4825 | -0.053072666 | 0.24027 | -0.220888 | 0.82518 | 0.969452399 | | antisense | 45.9 | -0.000715075 | 0.000715075 | ENSG00000278291 | n.s. |
| ENSG00000107036 | 1147.387 | -0.053961806 | 0.24917 | -0.216567 | 0.828546 | 0.970009871 | RIC1 | protein_coding | 37.47 | -0.000713583 | 0.000713583 | ENSG00000107036 | n.s. |
| ENSG00000114742 | 1512.267 | -0.057516983 | 0.275605 | -0.208693 | 0.834688 | 0.971936785 | WDR48 | protein_coding | 39.95 | -0.000711024 | 0.000711024 | ENSG00000114742 | n.s. |
| ENSG00000164542 | 129.9703 | 0.060778275 | 0.305146 | 0.1991777 | 0.842124 | 0.973446151 | KIAA0895 | protein_coding | 39.56 | 0.000710381 | 0.000710381 | ENSG00000164542 | n.s. |
| ENSG00000102024 | 976.6647 | 0.060575762 | 0.316472 | 0.1914095 | 0.848205 | 0.973446151 | PLS3 | protein_coding | 38.9 | 0.000708014 | 0.000708014 | ENSG00000102024 | n.s. |
| ENSG00000172172 | 888.701 | -0.06054078 | 0.325512 | -0.185986 | 0.852456 | 0.973446151 | MRPL13 | protein_coding | 35.86 | -0.000707605 | 0.000707605 | ENSG00000172172 | n.s. |
| ENSG00000277586 | 774.838 | 0.05247874 | 0.23611 | 0.2222641 | 0.824108 | 0.969452399 | NEFL | protein_coding | 44.93 | 0.000707073 | 0.000707073 | ENSG00000277586 | n.s. |
| ENSG00000253043 | 8.404502 | 0.079574908 | 0.569887 | 0.1396327 | 0.88895 | 0.979788048 | RNU7-181P | snRNA | 36.49 | 0.000705659 | 0.000705659 | ENSG00000253043 | n.s. |
| ENSG00000169251 | 2341.005 | -0.057045744 | 0.27246 | -0.209373 | 0.834157 | 0.971936785 | NMD3 | protein_coding | 37.99 | -0.000705198 | 0.000705198 | ENSG00000169251 | n.s. |
| ENSG00000167005 | 4928.827 | -0.057669801 | 0.278633 | -0.206974 | 0.83603 | 0.972288085 | NUDT21 | protein_coding | 39.54 | -0.000703862 | 0.000703862 | ENSG00000167005 | n.s. |

| | | | | | | | | | | | | | |
|---|---|---|---|---|---|---|---|---|---|---|---|---|---|
| ENSG00000038274 | 1749.671 | -0.058726477 | 0.2874 | -0.204337 | 0.83809 | 0.97278132 | MAT2B | protein_coding | 38.25 | -0.000703824 | 0.000703824 | ENSG00000038274 | n.s. |
| ENSG00000126950 | 223.6242 | -0.060077616 | 0.305295 | -0.196786 | 0.843995 | 0.973446151 | TMEM35A | protein_coding | 44.18 | -0.000702191 | 0.000702191 | ENSG00000126950 | n.s. |
| ENSG00000151665 | 411.5792 | 0.060103458 | 0.323772 | 0.1856353 | 0.852731 | 0.973483743 | PIGF | protein_coding | 36.13 | 0.000701485 | 0.000701485 | ENSG00000151665 | n.s. |
| ENSG00000165782 | 325.5942 | -0.052560715 | 0.23898 | -0.219938 | 0.82592 | 0.969830562 | TMEM55B | protein_coding | 51.18 | -0.000699275 | 0.000699275 | ENSG00000165782 | n.s. |
| ENSG00000276533 | 24.08069 | -0.065830278 | 0.387347 | -0.169952 | 0.865048 | 0.975860117 | | sense_intronic | 50.8 | -0.000698619 | 0.000698619 | ENSG00000276533 | n.s. |
| ENSG00000225648 | 418.7478 | 0.055047748 | 0.258585 | 0.212881 | 0.83142 | 0.971210458 | SBDSP1 | cribed_unprocessed_pseudc | 43.81 | 0.000698371 | 0.000698371 | ENSG00000225648 | n.s. |
| ENSG00000127980 | 718.1997 | -0.059632135 | 0.316403 | -0.188469 | 0.850509 | 0.973446151 | PEX1 | protein_coding | 37.11 | -0.000696984 | 0.000696984 | ENSG00000127980 | n.s. |
| ENSG00000201839 | 9.635496 | -0.070954442 | 0.467197 | -0.151873 | 0.879287 | 0.977664771 | SNORD114-3 | snoRNA | 48 | -0.000696065 | 0.000696065 | ENSG00000201839 | n.s. |
| ENSG00000145390 | 202.5771 | 0.055968115 | 0.265105 | 0.2111167 | 0.832796 | 0.971819596 | USP53 | protein_coding | 36.03 | 0.000694808 | 0.000694808 | ENSG00000145390 | n.s. |
| ENSG00000229797 | 38.64985 | -0.059476551 | 0.3209 | -0.185343 | 0.85296 | 0.973483743 | | antisense | 45.12 | -0.000694168 | 0.000694168 | ENSG00000229797 | n.s. |
| ENSG00000263934 | 10909.3 | -0.046099388 | 0.192308 | -0.239716 | 0.810551 | 0.965963425 | SNORD3A | snoRNA | 54.22 | -0.000693303 | 0.000693303 | ENSG00000263934 | n.s. |
| ENSG00000041357 | 4396.903 | 0.059282259 | 0.311936 | 0.1900465 | 0.849273 | 0.973446151 | PSMA4 | protein_coding | 42.32 | 0.000692895 | 0.000692895 | ENSG00000041357 | n.s. |
| ENSG00000119431 | 87.64929 | 0.059262954 | 0.293363 | 0.2020126 | 0.839907 | 0.973446151 | HDHD3 | protein_coding | 54.87 | 0.000692669 | 0.000692669 | ENSG00000119431 | n.s. |
| ENSG00000283045 | 156.2263 | -0.055992131 | 0.267995 | -0.208929 | 0.834503 | 0.971936785 | | sense_intronic | 33.63 | -0.000692174 | 0.000692174 | ENSG00000283045 | n.s. |
| ENSG00000113583 | 909.8287 | -0.060611296 | 0.339771 | -0.178389 | 0.858418 | 0.974056274 | C5orf15 | protein_coding | 39.91 | -0.000691936 | 0.000691936 | ENSG00000113583 | n.s. |
| ENSG00000156469 | 653.1311 | -0.052286835 | 0.239467 | -0.218347 | 0.827159 | 0.970009871 | MTERF3 | protein_coding | 38.81 | -0.000691433 | 0.000691433 | ENSG00000156469 | n.s. |
| ENSG00000239486 | 14.5908 | -0.072843969 | 0.492418 | -0.147931 | 0.882397 | 0.978387771 | | unprocessed_pseudogene | 43.29 | -0.000691215 | 0.000691215 | ENSG00000239486 | n.s. |
| ENSG00000170469 | 33.44403 | 0.064780208 | 0.375092 | 0.1727049 | 0.862883 | 0.975738908 | SPATA24 | protein_coding | 51.17 | 0.00069097 | 0.00069097 | ENSG00000170469 | n.s. |
| ENSG00000070610 | 735.5679 | -0.055724935 | 0.266732 | -0.208917 | 0.834513 | 0.971936785 | GBA2 | protein_coding | 50.7 | -0.000688871 | 0.000688871 | ENSG00000070610 | n.s. |
| ENSG00000137947 | 1170.369 | 0.057446867 | 0.280999 | 0.204438 | 0.838011 | 0.97278132 | GTF2B | protein_coding | 39.88 | 0.000688488 | 0.000688488 | ENSG00000137947 | n.s. |
| ENSG00000227704 | 126.079 | 0.053292286 | 0.247587 | 0.2152463 | 0.829575 | 0.970726167 | | antisense | 40 | 0.000687644 | 0.000687644 | ENSG00000227704 | n.s. |
| ENSG00000117450 | 3259.074 | -0.058786667 | 0.303817 | -0.193494 | 0.846572 | 0.973446151 | PRDX1 | protein_coding | 47.46 | -0.000687103 | 0.000687103 | ENSG00000117450 | n.s. |
| ENSG00000161395 | 53.89036 | -0.058761857 | 0.315726 | -0.186116 | 0.852354 | 0.973446151 | PGAP3 | protein_coding | 51.61 | -0.000686813 | 0.000686813 | ENSG00000161395 | n.s. |
| ENSG00000165819 | 795.2732 | -0.055528218 | 0.265925 | -0.208812 | 0.834595 | 0.971936785 | METTL3 | protein_coding | 42.49 | -0.000686439 | 0.000686439 | ENSG00000165819 | n.s. |
| ENSG00000166902 | 391.8322 | -0.058713175 | 0.306856 | -0.191338 | 0.848261 | 0.973446151 | MRPL16 | protein_coding | 44.6 | -0.000686244 | 0.000686244 | ENSG00000166902 | n.s. |
| ENSG00000179115 | 834.2658 | -0.060051058 | 0.331584 | -0.181104 | 0.856286 | 0.974056274 | FARSA | protein_coding | 53.91 | -0.00068554 | 0.00068554 | ENSG00000179115 | n.s. |
| ENSG00000177733 | 5879.336 | -0.048682418 | 0.214645 | -0.226804 | 0.820576 | 0.96813703 | HNRNPA0 | protein_coding | 41.07 | -0.000684629 | 0.000684629 | ENSG00000177733 | n.s. |
| ENSG00000254004 | 1783.993 | -0.056091734 | 0.271383 | -0.206689 | 0.836253 | 0.972288085 | ZNF260 | protein_coding | 40.16 | -0.000684602 | 0.000684602 | ENSG00000254004 | n.s. |
| ENSG00000134330 | 740.6598 | -0.058197385 | 0.298889 | -0.194713 | 0.845618 | 0.973446151 | IAH1 | protein_coding | 45.86 | -0.000680215 | 0.000680215 | ENSG00000134330 | n.s. |
| ENSG00000185869 | 671.9601 | -0.05667514 | 0.277205 | -0.204452 | 0.838 | 0.97278132 | ZNF829 | protein_coding | 39.33 | -0.000679239 | 0.000679239 | ENSG00000185869 | n.s. |
| ENSG00000207197 | 4.817994 | -0.076570784 | 0.571244 | -0.134042 | 0.893369 | 0.979788048 | SNORD116-12 | snoRNA | 35.87 | -0.000679019 | 0.000679019 | ENSG00000207197 | n.s. |
| ENSG00000173157 | 791.4104 | 0.055698225 | 0.269857 | 0.2063989 | 0.836479 | 0.972328444 | ADAMTS20 | protein_coding | 35.33 | 0.000678795 | 0.000678795 | ENSG00000173157 | n.s. |

| | | | | | | | | | | | | | |
|---|---|---|---|---|---|---|---|---|---|---|---|---|---|
| ENSG00000275055 | 64.88662 | -0.058044295 | 0.295189 | -0.196634 | 0.844114 | 0.973446151 | | antisense | 45.72 | -0.000678426 | 0.000678426 | ENSG00000275055 | n.s. |
| ENSG00000103254 | 205.0911 | -0.057945291 | 0.297802 | -0.194576 | 0.845725 | 0.973446151 | FAM173A | protein_coding | 71.94 | -0.000677268 | 0.000677268 | ENSG00000103254 | n.s. |
| ENSG00000145982 | 272.7597 | 0.057927886 | 0.286038 | 0.2025183 | 0.839512 | 0.973446151 | FARS2 | protein_coding | 41.44 | 0.000677065 | 0.000677065 | ENSG00000145982 | n.s. |
| ENSG00000259788 | 38.12782 | 0.063668749 | 0.37374 | 0.1703556 | 0.864731 | 0.975860117 | | antisense | 47.6 | 0.00067568 | 0.00067568 | ENSG00000259788 | n.s. |
| ENSG00000116157 | 758.389 | -0.057645631 | 0.294039 | -0.196047 | 0.844573 | 0.973446151 | GPX7 | protein_coding | 49.67 | -0.000673766 | 0.000673766 | ENSG00000116157 | n.s. |
| ENSG00000128989 | 6595.245 | -0.054437544 | 0.260037 | -0.209346 | 0.834178 | 0.971936785 | ARPP19 | protein_coding | 38.46 | -0.000672956 | 0.000672956 | ENSG00000128989 | n.s. |
| ENSG00000198860 | 1354.336 | -0.057566291 | 0.290504 | -0.19816 | 0.84292 | 0.973446151 | TSEN15 | protein_coding | 35.04 | -0.000672839 | 0.000672839 | ENSG00000198860 | n.s. |
| ENSG00000242247 | 442.8158 | -0.057551186 | 0.290644 | -0.198013 | 0.843035 | 0.973446151 | ARFGAP3 | protein_coding | 45 | -0.000672662 | 0.000672662 | ENSG00000242247 | n.s. |
| ENSG00000067560 | 3063.549 | -0.057632638 | 0.312034 | -0.1847 | 0.853464 | 0.97351991 | RHOA | protein_coding | 46.57 | -0.000671718 | 0.000671718 | ENSG00000067560 | n.s. |
| ENSG00000101574 | 244.1366 | -0.070714756 | 0.477801 | -0.148001 | 0.882342 | 0.978387771 | METTL4 | protein_coding | 36.78 | -0.000671011 | 0.000671011 | ENSG00000101574 | n.s. |
| ENSG00000078804 | 163.8063 | 0.050699018 | 0.23372 | 0.2169217 | 0.828269 | 0.970009871 | TP53INP2 | protein_coding | 55.18 | 0.000670436 | 0.000670436 | ENSG00000078804 | n.s. |
| ENSG00000111678 | 3138.722 | 0.054325656 | 0.261582 | 0.2076811 | 0.835478 | 0.971988837 | C12orf57 | protein_coding | 52.58 | 0.000670309 | 0.000670309 | ENSG00000111678 | n.s. |
| ENSG00000164318 | 451.752 | 0.055908765 | 0.273254 | 0.2046039 | 0.837882 | 0.97278132 | EGFLAM | protein_coding | 42.6 | 0.000670054 | 0.000670054 | ENSG00000164318 | n.s. |
| ENSG00000184840 | 338.9581 | -0.055836959 | 0.271911 | -0.20535 | 0.837298 | 0.972772376 | TMED9 | protein_coding | 53.16 | -0.000669416 | 0.000669416 | ENSG00000184840 | n.s. |
| ENSG00000132676 | 1691.98 | -0.057269304 | 0.284871 | -0.201036 | 0.84067 | 0.973446151 | DAP3 | protein_coding | 44.89 | -0.000669367 | 0.000669367 | ENSG00000132676 | n.s. |
| ENSG00000197043 | 2515.595 | -0.054897439 | 0.265768 | -0.206561 | 0.836352 | 0.972322592 | ANXA6 | protein_coding | 49.02 | -0.000669179 | 0.000669179 | ENSG00000197043 | n.s. |
| ENSG00000004534 | 2227.303 | -0.052110795 | 0.243663 | -0.213864 | 0.830653 | 0.970884671 | RBM6 | protein_coding | 44.14 | -0.000668704 | 0.000668704 | ENSG00000004534 | n.s. |
| ENSG00000258355 | 91.77017 | 0.054460965 | 0.262679 | 0.2073286 | 0.835753 | 0.972194165 | | sense_intronic | 52.08 | 0.000666983 | 0.000666983 | ENSG00000258355 | n.s. |
| ENSG00000137575 | 907.1777 | -0.057026074 | 0.282938 | -0.20155 | 0.840269 | 0.973446151 | SDCBP | protein_coding | 35.86 | -0.000666525 | 0.000666525 | ENSG00000137575 | n.s. |
| ENSG00000134463 | 111.161 | -0.056940909 | 0.293518 | -0.193995 | 0.84618 | 0.973446151 | ECHDC3 | protein_coding | 47.51 | -0.000665529 | 0.000665529 | ENSG00000134463 | n.s. |
| ENSG00000112146 | 282.9681 | -0.055530946 | 0.271854 | -0.204268 | 0.838144 | 0.97278132 | FBXO9 | protein_coding | 39.22 | -0.000665526 | 0.000665526 | ENSG00000112146 | n.s. |
| ENSG00000171503 | 288.6935 | 0.056924308 | 0.284884 | 0.1998158 | 0.841625 | 0.973446151 | ETFDH | protein_coding | 38.19 | 0.000665335 | 0.000665335 | ENSG00000171503 | n.s. |
| ENSG00000096872 | 409.3166 | -0.056901855 | 0.288513 | -0.197225 | 0.843652 | 0.973446151 | IFT74 | protein_coding | 37.56 | -0.000665073 | 0.000665073 | ENSG00000096872 | n.s. |
| ENSG00000170266 | 62.752 | -0.056888141 | 0.301515 | -0.188674 | 0.850348 | 0.973446151 | GLB1 | protein_coding | 44.03 | -0.000664912 | 0.000664912 | ENSG00000170266 | n.s. |
| ENSG00000273611 | 725.6445 | 0.057032504 | 0.308704 | 0.1847484 | 0.853426 | 0.97351991 | ZNHIT3 | protein_coding | 45.77 | 0.000664723 | 0.000664723 | ENSG00000273611 | n.s. |
| ENSG00000147586 | 508.451 | -0.056773148 | 0.287094 | -0.197751 | 0.84324 | 0.973446151 | MRPS28 | protein_coding | 37.19 | -0.000663568 | 0.000663568 | ENSG00000147586 | n.s. |
| ENSG00000137200 | 1184.896 | -0.053776672 | 0.258985 | -0.207644 | 0.835507 | 0.971988837 | CMTR1 | protein_coding | 45.88 | -0.000663535 | 0.000663535 | ENSG00000137200 | n.s. |
| ENSG00000182197 | 498.9766 | -0.049930082 | 0.227445 | -0.219526 | 0.82624 | 0.969879438 | EXT1 | protein_coding | 41.16 | -0.000663184 | 0.000663184 | ENSG00000182197 | n.s. |
| ENSG00000273061 | 51.43868 | -0.057969877 | 0.319182 | -0.18162 | 0.855881 | 0.974056274 | CDC37L1-AS1 | lincRNA | 41.23 | -0.000661781 | 0.000661781 | ENSG00000273061 | n.s. |
| ENSG00000167863 | 2687.737 | 0.057969679 | 0.320092 | 0.1811032 | 0.856287 | 0.974056274 | ATP5H | protein_coding | 47.61 | 0.000661779 | 0.000661779 | ENSG00000167863 | n.s. |
| ENSG00000268987 | 27.09762 | -0.062573398 | 0.373676 | -0.167454 | 0.867013 | 0.975964429 | | sense_intronic | 47.59 | -0.000661151 | 0.000661151 | ENSG00000268987 | n.s. |
| ENSG00000271749 | 13.89113 | -0.062707834 | 0.395123 | -0.158704 | 0.873902 | 0.976022242 | | unprocessed_pseudogene | 43.53 | -0.000660958 | 0.000660958 | ENSG00000271749 | n.s. |

| | | | | | | | | | | | | | |
|---|---|---|---|---|---|---|---|---|---|---|---|---|---|
| ENSG00000213967 | 162.7532 | -0.051852229 | 0.243117 | -0.213281 | 0.831108 | 0.971092758 | ZNF726 | protein_coding | 37.37 | -0.00066056 | 0.00066056 | ENSG00000213967 | n.s. |
| ENSG00000169490 | 460.8874 | 0.055026247 | 0.268367 | 0.2050413 | 0.83754 | 0.97278132 | TM2D2 | protein_coding | 42.34 | 0.000659477 | 0.000659477 | ENSG00000169490 | n.s. |
| ENSG00000101166 | 1812.686 | 0.056335684 | 0.289906 | 0.1943237 | 0.845922 | 0.973446151 | PRELID3B | protein_coding | 40.89 | 0.000658455 | 0.000658455 | ENSG00000101166 | n.s. |
| ENSG00000159377 | 3002.928 | 0.056247749 | 0.302131 | 0.1861704 | 0.852311 | 0.973446151 | PSMB4 | protein_coding | 49.27 | 0.000657427 | 0.000657427 | ENSG00000159377 | n.s. |
| ENSG00000232112 | 2398.974 | -0.056209133 | 0.301295 | -0.186559 | 0.852007 | 0.973446151 | TMA7 | protein_coding | 47.75 | -0.000656976 | 0.000656976 | ENSG00000232112 | n.s. |
| ENSG00000198919 | 1314.827 | 0.056183086 | 0.29381 | 0.1912226 | 0.848351 | 0.973446151 | DZIP3 | protein_coding | 35.7 | 0.000656672 | 0.000656672 | ENSG00000198919 | n.s. |
| ENSG00000100271 | 115.516 | 0.057471643 | 0.315395 | 0.1822212 | 0.855409 | 0.974056274 | TTLL1 | protein_coding | 51.32 | 0.000656094 | 0.000656094 | ENSG00000100271 | n.s. |
| ENSG00000270133 | 62.47303 | 0.056078803 | 0.277019 | 0.2024368 | 0.839575 | 0.973446151 | | sense_intronic | 36.21 | 0.000655453 | 0.000655453 | ENSG00000270133 | n.s. |
| ENSG00000119787 | 1986.935 | -0.056041044 | 0.276858 | -0.202418 | 0.83959 | 0.973446151 | ATL2 | protein_coding | 39.46 | -0.000655012 | 0.000655012 | ENSG00000119787 | n.s. |
| ENSG00000078304 | 1310.228 | 0.055975118 | 0.281146 | 0.1990962 | 0.842187 | 0.973446151 | PPP2R5C | protein_coding | 44.63 | 0.000654241 | 0.000654241 | ENSG00000078304 | n.s. |
| ENSG00000106537 | 485.7181 | -0.057284356 | 0.318462 | -0.179878 | 0.857248 | 0.974056274 | TSPAN13 | protein_coding | 39.23 | -0.000653955 | 0.000653955 | ENSG00000106537 | n.s. |
| ENSG00000158850 | 626.8116 | 0.055836325 | 0.280055 | 0.1993761 | 0.841969 | 0.973446151 | B4GALT3 | protein_coding | 52.28 | 0.000652619 | 0.000652619 | ENSG00000158850 | n.s. |
| ENSG00000139684 | 2621.299 | 0.055754878 | 0.298496 | 0.1867861 | 0.851828 | 0.973446151 | ESD | protein_coding | 34.63 | 0.000651667 | 0.000651667 | ENSG00000139684 | n.s. |
| ENSG00000249421 | 34.19531 | -0.061370423 | 0.359505 | -0.170708 | 0.864453 | 0.975860117 | ADAMTS19-AS1 | antisense | 63.75 | -0.000651289 | 0.000651289 | ENSG00000249421 | n.s. |
| ENSG00000099864 | 71.20119 | -0.055698915 | 0.292485 | -0.190433 | 0.84897 | 0.973446151 | PALM | protein_coding | 59.66 | -0.000651013 | 0.000651013 | ENSG00000099864 | n.s. |
| ENSG00000105193 | 3226.54 | 0.055657897 | 0.280247 | 0.1986029 | 0.842573 | 0.973446151 | RPS16 | protein_coding | 50.95 | 0.000650533 | 0.000650533 | ENSG00000105193 | n.s. |
| ENSG00000157181 | 1568.781 | -0.054236982 | 0.264673 | -0.204921 | 0.837634 | 0.97278132 | C1orf27 | protein_coding | 35.17 | -0.000650018 | 0.000650018 | ENSG00000157181 | n.s. |
| ENSG00000106049 | 248.7825 | -0.055613166 | 0.289661 | -0.191994 | 0.847747 | 0.973446151 | HIBADH | protein_coding | 36.88 | -0.00065001 | 0.00065001 | ENSG00000106049 | n.s. |
| ENSG00000091483 | 2676.948 | -0.055592259 | 0.289836 | -0.191806 | 0.847894 | 0.973446151 | FH | protein_coding | 36.87 | -0.000649766 | 0.000649766 | ENSG00000091483 | n.s. |
| ENSG00000207280 | 109.7976 | 0.052509318 | 0.250609 | 0.2095272 | 0.834037 | 0.971936785 | SNORD20 | snoRNA | 33.75 | 0.000649119 | 0.000649119 | ENSG00000207280 | n.s. |
| ENSG00000233654 | 68.08397 | 0.055520081 | 0.287264 | 0.1932718 | 0.846746 | 0.973446151 | | antisense | 39.04 | 0.000648922 | 0.000648922 | ENSG00000233654 | n.s. |
| ENSG00000087302 | 3737.587 | 0.055483375 | 0.277434 | 0.1999877 | 0.84149 | 0.973446151 | C14orf166 | protein_coding | 36.73 | 0.000648493 | 0.000648493 | ENSG00000087302 | n.s. |
| ENSG00000166441 | 14591.36 | 0.055482812 | 0.297326 | 0.1866061 | 0.851969 | 0.973446151 | RPL27A | protein_coding | 50.57 | 0.000648487 | 0.000648487 | ENSG00000166441 | n.s. |
| ENSG00000143156 | 704.1846 | -0.055472379 | 0.292265 | -0.189801 | 0.849465 | 0.973446151 | NME7 | protein_coding | 36.61 | -0.000648365 | 0.000648365 | ENSG00000143156 | n.s. |
| ENSG00000143545 | 277.5322 | -0.055449172 | 0.276917 | -0.200237 | 0.841295 | 0.973446151 | RAB13 | protein_coding | 48.71 | -0.000648094 | 0.000648094 | ENSG00000143545 | n.s. |
| ENSG00000119421 | 399.8634 | 0.056738155 | 0.318205 | 0.1783069 | 0.858482 | 0.974056274 | NDUFA8 | protein_coding | 41.5 | 0.00064772 | 0.00064772 | ENSG00000119421 | n.s. |
| ENSG00000204859 | 109.512 | 0.055364609 | 0.276846 | 0.1999832 | 0.841494 | 0.973446151 | ZBTB48 | protein_coding | 57.78 | 0.000647105 | 0.000647105 | ENSG00000204859 | n.s. |
| ENSG00000099797 | 287.104 | -0.055297314 | 0.29164 | -0.189608 | 0.849616 | 0.973446151 | TECR | protein_coding | 53.54 | -0.000646319 | 0.000646319 | ENSG00000099797 | n.s. |
| ENSG00000267191 | 27.56041 | -0.061128375 | 0.363371 | -0.168226 | 0.866406 | 0.975964429 | | antisense | 40.01 | -0.000645883 | 0.000645883 | ENSG00000267191 | n.s. |
| ENSG00000248866 | 232.2445 | 0.047630748 | 0.213926 | 0.2226503 | 0.823808 | 0.969312504 | USP46-AS1 | lincRNA | 49.93 | 0.000644739 | 0.000644739 | ENSG00000248866 | n.s. |
| ENSG00000134291 | 876.266 | -0.055240475 | 0.298157 | -0.185273 | 0.853015 | 0.973483743 | TMEM106C | protein_coding | 46.67 | -0.000644728 | 0.000644728 | ENSG00000134291 | n.s. |
| ENSG00000100225 | 2201.465 | -0.052243046 | 0.251369 | -0.207834 | 0.835359 | 0.971988837 | FBXO7 | protein_coding | 38.81 | -0.000644612 | 0.000644612 | ENSG00000100225 | n.s. |

| | | | | | | | | | | | | | |
|---|---|---|---|---|---|---|---|---|---|---|---|---|---|
| ENSG00000065150 | 5995.232 | 0.055141539 | 0.279867 | 0.1970277 | 0.843806 | 0.973446151 | IPO5 | protein_coding | 40.27 | 0.000644498 | 0.000644498 | ENSG00000065150 | n.s. |
| ENSG00000108679 | 652.0147 | -0.055139673 | 0.280763 | -0.196393 | 0.844303 | 0.973446151 | LGALS3BP | protein_coding | 58.85 | -0.000644476 | 0.000644476 | ENSG00000108679 | n.s. |
| ENSG00000234456 | 2312.534 | -0.055115383 | 0.277255 | -0.19879 | 0.842427 | 0.973446151 | MAGI2-AS3 | processed_transcript | 38.5 | -0.000644192 | 0.000644192 | ENSG00000234456 | n.s. |
| ENSG00000112893 | 933.0446 | -0.050765228 | 0.238325 | -0.213009 | 0.83132 | 0.971210458 | MAN2A1 | protein_coding | 36.58 | -0.000644041 | 0.000644041 | ENSG00000112893 | n.s. |
| ENSG00000174917 | 255.3414 | -0.059902064 | 0.345459 | -0.173398 | 0.862338 | 0.975554135 | C19orf70 | protein_coding | 61.19 | -0.000643865 | 0.000643865 | ENSG00000174917 | n.s. |
| ENSG00000198546 | 433.2179 | 0.055022877 | 0.292865 | 0.1878777 | 0.850973 | 0.973446151 | ZNF511 | protein_coding | 57.36 | 0.000643111 | 0.000643111 | ENSG00000198546 | n.s. |
| ENSG00000207863 | 14.94454 | 0.062487533 | 0.401816 | 0.1555129 | 0.876417 | 0.976610381 | MIR125B2 | miRNA | 49.44 | 0.000642288 | 0.000642288 | ENSG00000207863 | n.s. |
| ENSG00000126524 | 1086.9 | -0.054894211 | 0.282864 | -0.194066 | 0.846124 | 0.973446151 | SBDS | protein_coding | 42.81 | -0.000641607 | 0.000641607 | ENSG00000126524 | n.s. |
| ENSG00000174233 | 369.178 | 0.054861446 | 0.278105 | 0.1972687 | 0.843617 | 0.973446151 | ADCY6 | protein_coding | 54.36 | 0.000641224 | 0.000641224 | ENSG00000174233 | n.s. |
| ENSG00000178234 | 794.5196 | -0.054690159 | 0.270058 | -0.202512 | 0.839516 | 0.973446151 | GALNT11 | protein_coding | 41.23 | -0.000639222 | 0.000639222 | ENSG00000178234 | n.s. |
| ENSG00000232119 | 656.968 | 0.054625035 | 0.290337 | 0.1881437 | 0.850764 | 0.973446151 | MCTS1 | protein_coding | 39.24 | 0.000638461 | 0.000638461 | ENSG00000232119 | n.s. |
| ENSG00000149089 | 478.2665 | -0.056697468 | 0.319309 | -0.177563 | 0.859066 | 0.974414762 | APIP | protein_coding | 40.49 | -0.000638195 | 0.000638195 | ENSG00000149089 | n.s. |
| ENSG00000269846 | 37.53045 | 0.060362445 | 0.358957 | 0.1681609 | 0.866457 | 0.975964429 | | antisense | 55.52 | 0.00063779 | 0.00063779 | ENSG00000269846 | n.s. |
| ENSG00000240344 | 558.3665 | 0.054415684 | 0.267834 | 0.2031695 | 0.839003 | 0.973382833 | PPIL3 | protein_coding | 41.39 | 0.000637551 | 0.000637551 | ENSG00000240344 | n.s. |
| ENSG00000196368 | 816.4057 | 0.054472719 | 0.286295 | 0.1902677 | 0.849099 | 0.973446151 | NUDT11 | protein_coding | 39.05 | 0.000636681 | 0.000636681 | ENSG00000196368 | n.s. |
| ENSG00000123595 | 241.6148 | -0.054370924 | 0.273122 | -0.199072 | 0.842207 | 0.973446151 | RAB9A | protein_coding | 41.6 | -0.000635491 | 0.000635491 | ENSG00000123595 | n.s. |
| ENSG00000272870 | 75.96886 | -0.05434442 | 0.289699 | -0.187589 | 0.851199 | 0.973446151 | | antisense | 38.2 | -0.000635181 | 0.000635181 | ENSG00000272870 | n.s. |
| ENSG00000155463 | 808.0062 | -0.05128535 | 0.245508 | -0.208895 | 0.83453 | 0.971936785 | OXA1L | protein_coding | 46.66 | -0.000633989 | 0.000633989 | ENSG00000155463 | n.s. |
| ENSG00000156959 | 455.327 | -0.052166498 | 0.253326 | -0.205926 | 0.836849 | 0.972412121 | LHFPL4 | protein_coding | 46.67 | -0.000633804 | 0.000633804 | ENSG00000156959 | n.s. |
| ENSG00000196782 | 1223.346 | -0.048028259 | 0.222005 | -0.216338 | 0.828724 | 0.970096103 | MAML3 | protein_coding | 41.6 | -0.000633264 | 0.000633264 | ENSG00000196782 | n.s. |
| ENSG00000196605 | 18.37252 | -0.061505148 | 0.391719 | -0.157013 | 0.875234 | 0.976582458 | ZNF846 | protein_coding | 45.78 | -0.000632954 | 0.000632954 | ENSG00000196605 | n.s. |
| ENSG00000149646 | 84.54228 | -0.054197872 | 0.292388 | -0.185363 | 0.852944 | 0.973483743 | CNBD2 | protein_coding | 45.62 | -0.000632559 | 0.000632559 | ENSG00000149646 | n.s. |
| ENSG00000114346 | 3805.115 | 0.054046354 | 0.270575 | 0.1997461 | 0.841679 | 0.973446151 | ECT2 | protein_coding | 34.96 | 0.000631697 | 0.000631697 | ENSG00000114346 | n.s. |
| ENSG00000184203 | 844.1459 | -0.053836324 | 0.27573 | -0.19525 | 0.845197 | 0.973446151 | PPP1R2 | protein_coding | 39 | -0.000629243 | 0.000629243 | ENSG00000184203 | n.s. |
| ENSG00000143436 | 1413.673 | -0.053761879 | 0.287676 | -0.186883 | 0.851752 | 0.973446151 | MRPL9 | protein_coding | 48.42 | -0.000628372 | 0.000628372 | ENSG00000143436 | n.s. |
| ENSG00000130332 | 1175.414 | -0.055018574 | 0.306321 | -0.179611 | 0.857458 | 0.974056274 | LSM7 | protein_coding | 57.71 | -0.000628089 | 0.000628089 | ENSG00000130332 | n.s. |
| ENSG00000170310 | 214.8436 | 0.058813148 | 0.342034 | 0.171951 | 0.863476 | 0.975741574 | STX8 | protein_coding | 43.43 | 0.000627253 | 0.000627253 | ENSG00000170310 | n.s. |
| ENSG00000143457 | 275.2541 | -0.053649699 | 0.27982 | -0.191729 | 0.847954 | 0.973446151 | GOLPH3L | protein_coding | 40.09 | -0.000627061 | 0.000627061 | ENSG00000143457 | n.s. |
| ENSG00000166407 | 72.27455 | 0.05490051 | 0.306801 | 0.1789452 | 0.857981 | 0.974056274 | LMO1 | protein_coding | 52.88 | 0.000626742 | 0.000626742 | ENSG00000166407 | n.s. |
| ENSG00000213420 | 477.1442 | -0.05362031 | 0.271943 | -0.197175 | 0.843691 | 0.973446151 | GPC2 | protein_coding | 55.45 | -0.000626718 | 0.000626718 | ENSG00000213420 | n.s. |
| ENSG00000142188 | 208.2627 | -0.053596258 | 0.287726 | -0.186275 | 0.852229 | 0.973446151 | TMEM50B | protein_coding | 43.36 | -0.000626437 | 0.000626437 | ENSG00000142188 | n.s. |
| ENSG00000189241 | 1750.92 | 0.050673025 | 0.241756 | 0.2096042 | 0.833977 | 0.971936785 | TSPYL1 | protein_coding | 48.02 | 0.000626419 | 0.000626419 | ENSG00000189241 | n.s. |

| | | | | | | | | | | | | | |
|---|---|---|---|---|---|---|---|---|---|---|---|---|---|
| ENSG00000125863 | 617.1109 | -0.054818174 | 0.300638 | -0.18234 | 0.855316 | 0.974056274 | MKKS | protein_coding | 38.18 | -0.000625802 | 0.000625802 | ENSG00000125863 | n.s. |
| ENSG00000201343 | 56.08963 | 0.054579425 | 0.297782 | 0.1832863 | 0.854573 | 0.974056274 | | misc_RNA | 40.91 | 0.000623076 | 0.000623076 | ENSG00000201343 | n.s. |
| ENSG00000134308 | 9242.309 | -0.053295951 | 0.284298 | -0.187465 | 0.851296 | 0.973446151 | YWHAQ | protein_coding | 40.33 | -0.000622927 | 0.000622927 | ENSG00000134308 | n.s. |
| ENSG00000102054 | 2407.743 | 0.053265169 | 0.276116 | 0.1929084 | 0.847031 | 0.973446151 | RBBP7 | protein_coding | 42.25 | 0.000622567 | 0.000622567 | ENSG00000102054 | n.s. |
| ENSG00000144834 | 1143.51 | 0.056014121 | 0.317705 | 0.1763084 | 0.860052 | 0.974768375 | TAGLN3 | protein_coding | 45.75 | 0.000621677 | 0.000621677 | ENSG00000144834 | n.s. |
| ENSG00000256591 | 295.8268 | 0.046731079 | 0.212565 | 0.2198436 | 0.825993 | 0.969833104 | | protein_coding | 48.75 | 0.000621663 | 0.000621663 | ENSG00000256591 | n.s. |
| ENSG00000131848 | 85.40494 | 0.053130332 | 0.272084 | 0.1952719 | 0.84518 | 0.973446151 | ZSCAN5A | protein_coding | 44.09 | 0.000620991 | 0.000620991 | ENSG00000131848 | n.s. |
| ENSG00000141376 | 245.0122 | -0.053063012 | 0.279471 | -0.189869 | 0.849411 | 0.973446151 | BCAS3 | protein_coding | 39.85 | -0.000620204 | 0.000620204 | ENSG00000141376 | n.s. |
| ENSG00000159199 | 1561.708 | 0.053015038 | 0.28137 | 0.1884172 | 0.85055 | 0.973446151 | ATP5G1 | protein_coding | 50.08 | 0.000619643 | 0.000619643 | ENSG00000159199 | n.s. |
| ENSG00000018236 | 2831.276 | 0.054202506 | 0.300135 | 0.180594 | 0.856686 | 0.974056274 | CNTN1 | protein_coding | 35.21 | 0.000618773 | 0.000618773 | ENSG00000018236 | n.s. |
| ENSG00000196531 | 17813.39 | 0.053075067 | 0.287271 | 0.1847559 | 0.853421 | 0.97351991 | NACA | protein_coding | 47.57 | 0.000618599 | 0.000618599 | ENSG00000196531 | n.s. |
| ENSG00000178999 | 979.6302 | -0.050010915 | 0.238645 | -0.209562 | 0.83401 | 0.971936785 | AURKB | protein_coding | 52.69 | -0.000618234 | 0.000618234 | ENSG00000178999 | n.s. |
| ENSG00000100300 | 346.9711 | -0.058356786 | 0.346118 | -0.168604 | 0.866108 | 0.975964429 | TSPO | protein_coding | 57.81 | -0.000616598 | 0.000616598 | ENSG00000100300 | n.s. |
| ENSG00000039560 | 5144.416 | -0.052733732 | 0.27183 | -0.193995 | 0.84618 | 0.973446151 | RAI14 | protein_coding | 42.13 | -0.000616355 | 0.000616355 | ENSG00000039560 | n.s. |
| ENSG00000198089 | 22.07699 | 0.061690031 | 0.40341 | 0.1529214 | 0.87846 | 0.977264479 | SFI1 | protein_coding | 46.58 | 0.000616153 | 0.000616153 | ENSG00000198089 | n.s. |
| ENSG00000221066 | 25.25743 | 0.0583031 | 0.352057 | 0.1656068 | 0.868466 | 0.975964429 | SNORD111 | snoRNA | 37.23 | 0.000616031 | 0.000616031 | ENSG00000221066 | n.s. |
| ENSG00000117500 | 1492.442 | 0.052702876 | 0.280409 | 0.1879499 | 0.850916 | 0.973446151 | TMED5 | protein_coding | 38.24 | 0.000615995 | 0.000615995 | ENSG00000117500 | n.s. |
| ENSG00000183624 | 410.2718 | -0.05265891 | 0.269729 | -0.195229 | 0.845214 | 0.973446151 | HMCES | protein_coding | 44.53 | -0.000615481 | 0.000615481 | ENSG00000183624 | n.s. |
| ENSG00000237004 | 29.09676 | -0.058392527 | 0.366157 | -0.159474 | 0.873295 | 0.976022242 | ZNRF2P1 | processed_pseudogene | 75.62 | -0.000615474 | 0.000615474 | ENSG00000237004 | n.s. |
| ENSG00000105640 | 8395.446 | -0.052580626 | 0.279384 | -0.188202 | 0.850719 | 0.973446151 | RPL18A | protein_coding | 58.32 | -0.000614566 | 0.000614566 | ENSG00000105640 | n.s. |
| ENSG00000204922 | 302.5373 | -0.052510772 | 0.276698 | -0.189776 | 0.849484 | 0.973446151 | UQCC3 | protein_coding | 52.86 | -0.000613749 | 0.000613749 | ENSG00000204922 | n.s. |
| ENSG00000101363 | 275.2647 | -0.053697044 | 0.292383 | -0.183653 | 0.854286 | 0.974056274 | MANBAL | protein_coding | 47.37 | -0.000613003 | 0.000613003 | ENSG00000101363 | n.s. |
| ENSG00000231500 | 18399.84 | 0.053651867 | 0.300507 | 0.178538 | 0.858301 | 0.974056274 | RPS18 | protein_coding | 46.95 | 0.000612487 | 0.000612487 | ENSG00000231500 | n.s. |
| ENSG00000149311 | 2471.436 | -0.052305021 | 0.25925 | -0.201755 | 0.840108 | 0.973446151 | ATM | protein_coding | 37.52 | -0.000611345 | 0.000611345 | ENSG00000149311 | n.s. |
| ENSG00000160691 | 808.9691 | 0.052262393 | 0.270688 | 0.1930727 | 0.846902 | 0.973446151 | SHC1 | protein_coding | 53.79 | 0.000610846 | 0.000610846 | ENSG00000160691 | n.s. |
| ENSG00000118217 | 1293.274 | -0.052234777 | 0.26243 | -0.199043 | 0.842229 | 0.973446151 | ATF6 | protein_coding | 37.21 | -0.000610524 | 0.000610524 | ENSG00000118217 | n.s. |
| ENSG00000167004 | 3682.408 | -0.052206291 | 0.270396 | -0.193073 | 0.846902 | 0.973446151 | PDIA3 | protein_coding | 43.02 | -0.000610191 | 0.000610191 | ENSG00000167004 | n.s. |
| ENSG00000120509 | 317.873 | 0.05342941 | 0.293157 | 0.1822551 | 0.855383 | 0.974056274 | PDZD11 | protein_coding | 49.29 | 0.000609948 | 0.000609948 | ENSG00000120509 | n.s. |
| ENSG00000267430 | 89.44193 | 0.052178421 | 0.268069 | 0.1946454 | 0.845671 | 0.973446151 | | processed_pseudogene | 48.83 | 0.000609865 | 0.000609865 | ENSG00000267430 | n.s. |
| ENSG00000111875 | 1520.757 | 0.053399638 | 0.297778 | 0.1793269 | 0.857681 | 0.974056274 | ASF1A | protein_coding | 39.13 | 0.000609608 | 0.000609608 | ENSG00000111875 | n.s. |
| ENSG00000185904 | 919.1317 | 0.052081544 | 0.277258 | 0.187845 | 0.850998 | 0.973446151 | LINC00839 | lincRNA | 43.47 | 0.000608733 | 0.000608733 | ENSG00000185904 | n.s. |
| ENSG00000198746 | 142.5008 | -0.052066784 | 0.26735 | -0.194752 | 0.845587 | 0.973446151 | GPATCH3 | protein_coding | 50.12 | -0.00060856 | 0.00060856 | ENSG00000198746 | n.s. |

| | | | | | | | | | | | | | |
|---|---|---|---|---|---|---|---|---|---|---|---|---|---|
| ENSG00000164221 | 1151.382 | 0.052028816 | 0.265699 | 0.1958185 | 0.844752 | 0.973446151 | CCDC112 | protein_coding | 36.27 | 0.000608116 | 0.000608116 | ENSG00000164221 | n.s. |
| ENSG00000105618 | 1017.943 | 0.053220441 | 0.291064 | 0.1828477 | 0.854918 | 0.974056274 | PRPF31 | protein_coding | 57.08 | 0.000607562 | 0.000607562 | ENSG00000105618 | n.s. |
| ENSG00000143486 | 697.5674 | -0.051923866 | 0.268146 | -0.19364 | 0.846458 | 0.973446151 | EIF2D | protein_coding | 46.17 | -0.00060689 | 0.00060689 | ENSG00000143486 | n.s. |
| ENSG00000100462 | 1616.08 | 0.051911448 | 0.275821 | 0.188207 | 0.850714 | 0.973446151 | PRMT5 | protein_coding | 47.75 | 0.000606745 | 0.000606745 | ENSG00000100462 | n.s. |
| ENSG00000119318 | 4439.681 | -0.051888681 | 0.272108 | -0.190692 | 0.848767 | 0.973446151 | RAD23B | protein_coding | 38 | -0.000606478 | 0.000606478 | ENSG00000119318 | n.s. |
| ENSG00000142937 | 12578.26 | 0.055113491 | 0.315568 | 0.1746484 | 0.861356 | 0.97501288 | RPS8 | protein_coding | 55.11 | 0.000605678 | 0.000605678 | ENSG00000142937 | n.s. |
| ENSG00000108797 | 435.9342 | -0.051805098 | 0.268492 | -0.192948 | 0.847 | 0.973446151 | CNTNAP1 | protein_coding | 54.63 | -0.000605501 | 0.000605501 | ENSG00000108797 | n.s. |
| ENSG00000164985 | 5511.07 | -0.051798213 | 0.263359 | -0.196683 | 0.844075 | 0.973446151 | PSIP1 | protein_coding | 37.88 | -0.000605421 | 0.000605421 | ENSG00000164985 | n.s. |
| ENSG00000198160 | 1914.37 | -0.051656106 | 0.254267 | -0.203157 | 0.839012 | 0.973382833 | MIER1 | protein_coding | 36.15 | -0.000605219 | 0.000605219 | ENSG00000198160 | n.s. |
| ENSG00000272654 | 86.34256 | -0.054313712 | 0.307028 | -0.176901 | 0.859586 | 0.974673271 | | lincRNA | 44.22 | -0.000605106 | 0.000605106 | ENSG00000272654 | n.s. |
| ENSG00000119632 | 166.9681 | -0.057302794 | 0.348934 | -0.164223 | 0.869556 | 0.975997758 | IFI27L2 | protein_coding | 54.59 | -0.000604612 | 0.000604612 | ENSG00000119632 | n.s. |
| ENSG00000253352 | 4283.978 | -0.050406696 | 0.246691 | -0.204331 | 0.838095 | 0.97278132 | TUG1 | antisense | 39.53 | -0.000604113 | 0.000604113 | ENSG00000253352 | n.s. |
| ENSG00000127989 | 727.1083 | 0.051651378 | 0.268337 | 0.1924872 | 0.847361 | 0.973446151 | MTERF1 | protein_coding | 38.41 | 0.000603705 | 0.000603705 | ENSG00000127989 | n.s. |
| ENSG00000109756 | 1070.313 | -0.051147792 | 0.251352 | -0.203491 | 0.838751 | 0.973242264 | RAPGEF2 | protein_coding | 36.57 | -0.000602472 | 0.000602472 | ENSG00000109756 | n.s. |
| ENSG00000104177 | 4631.538 | 0.051496604 | 0.27301 | 0.1886255 | 0.850386 | 0.973446151 | MYEF2 | protein_coding | 33.92 | 0.000601896 | 0.000601896 | ENSG00000104177 | n.s. |
| ENSG00000197989 | 636.6972 | -0.045425053 | 0.209782 | -0.216534 | 0.828571 | 0.970009871 | SNHG12 | antisense | 51.03 | -0.000600694 | 0.000600694 | ENSG00000197989 | n.s. |
| ENSG00000253730 | 25.93725 | 0.0605124 | 0.397235 | 0.1523339 | 0.878924 | 0.97741985 | | antisense | 42.46 | 0.000600213 | 0.000600213 | ENSG00000253730 | n.s. |
| ENSG00000169981 | 940.7063 | 0.051341703 | 0.254354 | 0.2018513 | 0.840033 | 0.973446151 | ZNF35 | protein_coding | 42.6 | 0.000600085 | 0.000600085 | ENSG00000169981 | n.s. |
| ENSG00000100938 | 137.4597 | 0.052562696 | 0.293626 | 0.1790126 | 0.857928 | 0.974056274 | GMPR2 | protein_coding | 46.62 | 0.000600053 | 0.000600053 | ENSG00000100938 | n.s. |
| ENSG00000004142 | 892.7067 | -0.051366445 | 0.277295 | -0.185241 | 0.85304 | 0.973483743 | POLDIP2 | protein_coding | 50.16 | -0.000599513 | 0.000599513 | ENSG00000004142 | n.s. |
| ENSG00000173436 | 310.4228 | 0.05128921 | 0.273342 | 0.1876375 | 0.851161 | 0.973446151 | MINOS1 | protein_coding | 42.96 | 0.000599472 | 0.000599472 | ENSG00000173436 | n.s. |
| ENSG00000183978 | 1108.91 | 0.05458391 | 0.312981 | 0.1743999 | 0.861551 | 0.975075415 | COA3 | protein_coding | 54.78 | 0.000598338 | 0.000598338 | ENSG00000183978 | n.s. |
| ENSG00000175893 | 1002.415 | -0.051144329 | 0.268115 | -0.190755 | 0.848717 | 0.973446151 | ZDHHC21 | protein_coding | 35.18 | -0.000597778 | 0.000597778 | ENSG00000175893 | n.s. |
| ENSG00000113407 | 7065.919 | -0.051142062 | 0.269097 | -0.19005 | 0.84927 | 0.973446151 | TARS | protein_coding | 38.63 | -0.000597752 | 0.000597752 | ENSG00000113407 | n.s. |
| ENSG00000212158 | 4.105864 | -0.10465419 | 1.026513 | -0.101951 | 0.918795 | 0.986959954 | SNORD66 | snoRNA | 51.32 | -0.000596578 | 0.000596578 | ENSG00000212158 | n.s. |
| ENSG00000039523 | 38.49223 | 0.055929227 | 0.325112 | 0.1720305 | 0.863414 | 0.975741574 | RIPOR1 | protein_coding | 57.05 | 0.000596496 | 0.000596496 | ENSG00000039523 | n.s. |
| ENSG00000172046 | 939.5167 | -0.051014077 | 0.262971 | -0.193991 | 0.846183 | 0.973446151 | USP19 | protein_coding | 56.09 | -0.000596256 | 0.000596256 | ENSG00000172046 | n.s. |
| ENSG00000196155 | 499.2679 | -0.051012617 | 0.270363 | -0.188682 | 0.850342 | 0.973446151 | PLEKHG4 | protein_coding | 59.77 | -0.000596239 | 0.000596239 | ENSG00000196155 | n.s. |
| ENSG00000214174 | 405.4614 | -0.050993605 | 0.262747 | -0.194079 | 0.846114 | 0.973446151 | AMZ2P1 | cribed_unprocessed_pseudc | 41.33 | -0.000596017 | 0.000596017 | ENSG00000214174 | n.s. |
| ENSG00000265996 | 54.77016 | 0.056529828 | 0.350146 | 0.1614466 | 0.871742 | 0.976022242 | MIR3671 | miRNA | 27.27 | 0.000595841 | 0.000595841 | ENSG00000265996 | n.s. |
| ENSG00000230180 | 4.145056 | -0.104515237 | 1.035554 | -0.100927 | 0.919608 | 0.986968832 | RPL12P49 | processed_pseudogene | 46.28 | -0.000595378 | 0.000595378 | ENSG00000230180 | n.s. |
| ENSG00000152404 | 640.9673 | -0.050869951 | 0.25518 | -0.199349 | 0.84199 | 0.973446151 | CWF19L2 | protein_coding | 36.3 | -0.000594571 | 0.000594571 | ENSG00000152404 | n.s. |

| | | | | | | | | | | | | | |
|---|---|---|---|---|---|---|---|---|---|---|---|---|---|
| ENSG00000115944 | 1560.001 | 0.056002227 | 0.329399 | 0.1700134 | 0.865 | 0.975860117 | COX7A2L | protein_coding | 44 | 0.00059432 | 0.00059432 | ENSG00000115944 | n.s. |
| ENSG00000119599 | 333.4755 | 0.050825395 | 0.256789 | 0.197927 | 0.843102 | 0.973446151 | DCAF4 | protein_coding | 48.94 | 0.000594051 | 0.000594051 | ENSG00000119599 | n.s. |
| ENSG00000075391 | 469.7712 | -0.050797301 | 0.265802 | -0.191109 | 0.84844 | 0.973446151 | RASAL2 | protein_coding | 36.99 | -0.000593722 | 0.000593722 | ENSG00000075391 | n.s. |
| ENSG00000273004 | 41.31354 | 0.055221555 | 0.318361 | 0.1734556 | 0.862293 | 0.975554135 | | lincRNA | 46.54 | 0.000593556 | 0.000593556 | ENSG00000273004 | n.s. |
| ENSG00000221614 | 2.916551 | 0.083995783 | 0.697185 | 0.1204785 | 0.904104 | 0.983886286 | MIR1185-2 | miRNA | 38.37 | 0.000592598 | 0.000592598 | ENSG00000221614 | n.s. |
| ENSG00000258568 | 84.67342 | -0.050682215 | 0.260325 | -0.194688 | 0.845637 | 0.973446151 | RHOQP1 | processed_pseudogene | 48.88 | -0.000592377 | 0.000592377 | ENSG00000258568 | n.s. |
| ENSG00000117475 | 839.5885 | -0.050661744 | 0.266704 | -0.189955 | 0.849344 | 0.973446151 | BLZF1 | protein_coding | 35.78 | -0.000592138 | 0.000592138 | ENSG00000117475 | n.s. |
| ENSG00000068366 | 801.6098 | -0.053164655 | 0.302096 | -0.175986 | 0.860305 | 0.974774428 | ACSL4 | protein_coding | 39.16 | -0.000589908 | 0.000589908 | ENSG00000068366 | n.s. |
| ENSG00000167671 | 371.2056 | -0.053516568 | 0.305639 | -0.175097 | 0.861003 | 0.974949369 | UBXN6 | protein_coding | 59.86 | -0.000589642 | 0.000589642 | ENSG00000167671 | n.s. |
| ENSG00000198431 | 5802.493 | 0.050432148 | 0.260504 | 0.1935948 | 0.846493 | 0.973446151 | TXNRD1 | protein_coding | 42.37 | 0.000589454 | 0.000589454 | ENSG00000198431 | n.s. |
| ENSG00000133103 | 192.2442 | -0.051634213 | 0.285619 | -0.18078 | 0.85654 | 0.974056274 | COG6 | protein_coding | 35.35 | -0.000589454 | 0.000589454 | ENSG00000133103 | n.s. |
| ENSG00000130560 | 590.4561 | 0.05160525 | 0.282748 | 0.1825131 | 0.85518 | 0.974056274 | UBAC1 | protein_coding | 51.82 | 0.000589123 | 0.000589123 | ENSG00000130560 | n.s. |
| ENSG00000188785 | 470.3029 | -0.050177182 | 0.259877 | -0.19308 | 0.846896 | 0.973446151 | ZNF548 | protein_coding | 45.35 | -0.000586474 | 0.000586474 | ENSG00000188785 | n.s. |
| ENSG00000246089 | 153.8967 | -0.050122785 | 0.250665 | -0.199959 | 0.841513 | 0.973446151 | | antisense | 49.69 | -0.000585839 | 0.000585839 | ENSG00000246089 | n.s. |
| ENSG00000256981 | 296.1948 | -0.050116109 | 0.256804 | -0.195153 | 0.845273 | 0.973446151 | | processed_pseudogene | 39.01 | -0.00058576 | 0.00058576 | ENSG00000256981 | n.s. |
| ENSG00000121064 | 612.7794 | 0.05127582 | 0.283104 | 0.18112 | 0.856273 | 0.974056274 | SCPEP1 | protein_coding | 45.31 | 0.000585362 | 0.000585362 | ENSG00000121064 | n.s. |
| ENSG00000204619 | 230.1133 | 0.049987334 | 0.253357 | 0.1972999 | 0.843593 | 0.973446151 | PPP1R11 | protein_coding | 49.16 | 0.000584255 | 0.000584255 | ENSG00000204619 | n.s. |
| ENSG00000252010 | 190.0366 | -0.049961889 | 0.262249 | -0.190513 | 0.848907 | 0.973446151 | SCARNA5 | scaRNA | 47.46 | -0.000583958 | 0.000583958 | ENSG00000252010 | n.s. |
| ENSG00000132664 | 693.7473 | -0.049958103 | 0.260738 | -0.191602 | 0.848054 | 0.973446151 | POLR3F | protein_coding | 40.58 | -0.000583914 | 0.000583914 | ENSG00000132664 | n.s. |
| ENSG00000167785 | 447.2068 | -0.051095068 | 0.282727 | -0.180722 | 0.856586 | 0.974056274 | ZNF558 | protein_coding | 44.39 | -0.000583299 | 0.000583299 | ENSG00000167785 | n.s. |
| ENSG00000073584 | 5634.223 | -0.049768981 | 0.262782 | -0.189392 | 0.849785 | 0.973446151 | SMARCE1 | protein_coding | 37.72 | -0.000581703 | 0.000581703 | ENSG00000073584 | n.s. |
| ENSG00000135469 | 322.9555 | 0.049777221 | 0.268533 | 0.1853669 | 0.852941 | 0.973483743 | COQ10A | protein_coding | 50.99 | 0.000580965 | 0.000580965 | ENSG00000135469 | n.s. |
| ENSG00000137942 | 5459.925 | -0.049683186 | 0.25912 | -0.191738 | 0.847947 | 0.973446151 | FNBP1L | protein_coding | 37.34 | -0.0005807 | 0.0005807 | ENSG00000137942 | n.s. |
| ENSG00000153147 | 6401.128 | -0.049680466 | 0.25479 | -0.194986 | 0.845404 | 0.973446151 | SMARCA5 | protein_coding | 35.51 | -0.000580669 | 0.000580669 | ENSG00000153147 | n.s. |
| ENSG00000009307 | 20393.8 | -0.049653853 | 0.266466 | -0.186342 | 0.852176 | 0.973446151 | CSDE1 | protein_coding | 38.28 | -0.000580358 | 0.000580358 | ENSG00000009307 | n.s. |
| ENSG00000201592 | 103.9282 | 0.049645989 | 0.265559 | 0.1869492 | 0.8517 | 0.973446151 | | snoRNA | 43.75 | 0.000580266 | 0.000580266 | ENSG00000201592 | n.s. |
| ENSG00000123737 | 2378.327 | 0.049589845 | 0.251316 | 0.1973208 | 0.843577 | 0.973446151 | EXOSC9 | protein_coding | 37.71 | 0.000579609 | 0.000579609 | ENSG00000123737 | n.s. |
| ENSG00000267298 | 48.92388 | -0.05199786 | 0.294373 | -0.176639 | 0.859792 | 0.974768375 | | sense_intronic | 31.13 | -0.000577102 | 0.000577102 | ENSG00000267298 | n.s. |
| ENSG00000104635 | 967.4042 | 0.051959067 | 0.295417 | 0.1758839 | 0.860385 | 0.974785744 | SLC39A14 | protein_coding | 48.28 | 0.000576269 | 0.000576269 | ENSG00000104635 | n.s. |
| ENSG00000119865 | 1507.685 | 0.049274899 | 0.26128 | 0.1885901 | 0.850414 | 0.973446151 | CNRIP1 | protein_coding | 40.04 | 0.000575928 | 0.000575928 | ENSG00000119865 | n.s. |
| ENSG00000133606 | 1122.495 | -0.049269604 | 0.256625 | -0.19199 | 0.84775 | 0.973446151 | MKRN1 | protein_coding | 45.78 | -0.000575866 | 0.000575866 | ENSG00000133606 | n.s. |
| ENSG00000115641 | 507.0369 | -0.049193572 | 0.262828 | -0.18717 | 0.851527 | 0.973446151 | FHL2 | protein_coding | 44.54 | -0.000574978 | 0.000574978 | ENSG00000115641 | n.s. |

| | | | | | | | | | | | | | |
|---|---|---|---|---|---|---|---|---|---|---|---|---|---|
| ENSG00000128059 | 2229.616 | -0.049156203 | 0.263896 | -0.186271 | 0.852232 | 0.973446151 | PPAT | protein_coding | 37.44 | -0.000574541 | 0.000574541 | ENSG00000128059 | n.s. |
| ENSG00000118162 | 88.4677 | 0.054329058 | 0.32367 | 0.167853 | 0.866699 | 0.975964429 | KPTN | protein_coding | 53.48 | 0.000574041 | 0.000574041 | ENSG00000118162 | n.s. |
| ENSG00000089335 | 1652.308 | -0.049073573 | 0.257644 | -0.190471 | 0.84894 | 0.973446151 | ZNF302 | protein_coding | 39.83 | -0.000573575 | 0.000573575 | ENSG00000089335 | n.s. |
| ENSG00000168090 | 658.4862 | 0.051159019 | 0.288712 | 0.1771975 | 0.859353 | 0.974568335 | COPS6 | protein_coding | 52.76 | 0.000572352 | 0.000572352 | ENSG00000168090 | n.s. |
| ENSG00000120008 | 2059.631 | -0.048965823 | 0.26067 | -0.187846 | 0.850997 | 0.973446151 | WDR11 | protein_coding | 36.82 | -0.000572316 | 0.000572316 | ENSG00000120008 | n.s. |
| ENSG00000084463 | 3126.613 | 0.048950959 | 0.2432 | 0.2012788 | 0.840481 | 0.973446151 | WBP11 | protein_coding | 37.17 | 0.000572142 | 0.000572142 | ENSG00000084463 | n.s. |
| ENSG00000110200 | 589.8349 | -0.045981857 | 0.217436 | -0.211473 | 0.832518 | 0.971759608 | ANAPC15 | protein_coding | 52.32 | -0.000572067 | 0.000572067 | ENSG00000110200 | n.s. |
| ENSG00000102898 | 1148.601 | -0.050066525 | 0.280398 | -0.178555 | 0.858287 | 0.974056274 | NUTF2 | protein_coding | 51.22 | -0.000571557 | 0.000571557 | ENSG00000102898 | n.s. |
| ENSG00000105968 | 2702.069 | -0.049283876 | 0.2672 | -0.184446 | 0.853664 | 0.973667644 | H2AFV | protein_coding | 41.41 | -0.000571164 | 0.000571164 | ENSG00000105968 | n.s. |
| ENSG00000167136 | 373.7858 | -0.053984242 | 0.319569 | -0.168929 | 0.865853 | 0.975964429 | ENDOG | protein_coding | 61.54 | -0.000570398 | 0.000570398 | ENSG00000167136 | n.s. |
| ENSG00000147130 | 438.0793 | 0.0487886 | 0.253223 | 0.1926707 | 0.847217 | 0.973446151 | ZMYM3 | protein_coding | 53.76 | 0.000570244 | 0.000570244 | ENSG00000147130 | n.s. |
| ENSG00000108829 | 4241.55 | -0.049595246 | 0.26951 | -0.18402 | 0.853998 | 0.973888905 | LRRC59 | protein_coding | 47.65 | -0.000569878 | 0.000569878 | ENSG00000108829 | n.s. |
| ENSG00000168002 | 850.7381 | 0.053855237 | 0.32615 | 0.1651243 | 0.868846 | 0.975964429 | POLR2G | protein_coding | 50.57 | 0.000569035 | 0.000569035 | ENSG00000168002 | n.s. |
| ENSG00000169992 | 192.7393 | -0.048621019 | 0.261611 | -0.185852 | 0.852561 | 0.973446151 | NLGN2 | protein_coding | 59.54 | -0.000568286 | 0.000568286 | ENSG00000169992 | n.s. |
| ENSG00000111364 | 2194.236 | 0.048619631 | 0.244152 | 0.199137 | 0.842156 | 0.973446151 | DDX55 | protein_coding | 45.83 | 0.00056827 | 0.00056827 | ENSG00000111364 | n.s. |
| ENSG00000131115 | 870.297 | -0.048616139 | 0.252236 | -0.192741 | 0.847162 | 0.973446151 | ZNF227 | protein_coding | 41.09 | -0.000568229 | 0.000568229 | ENSG00000131115 | n.s. |
| ENSG00000186635 | 318.1718 | -0.048563273 | 0.261199 | -0.185925 | 0.852504 | 0.973446151 | ARAP1 | protein_coding | 54.91 | -0.000567611 | 0.000567611 | ENSG00000186635 | n.s. |
| ENSG00000128595 | 2423.895 | -0.049535632 | 0.275072 | -0.180083 | 0.857088 | 0.974056274 | CALU | protein_coding | 39.99 | -0.000565496 | 0.000565496 | ENSG00000128595 | n.s. |
| ENSG00000134056 | 494.4598 | 0.053585387 | 0.332642 | 0.1610901 | 0.872022 | 0.976022242 | MRPS36 | protein_coding | 42.53 | 0.000564805 | 0.000564805 | ENSG00000134056 | n.s. |
| ENSG00000278475 | 67.07082 | -0.053156544 | 0.311589 | -0.170598 | 0.86454 | 0.975860117 | | sense_intronic | 33.05 | -0.00056412 | 0.00056412 | ENSG00000278475 | n.s. |
| ENSG00000272636 | 662.4376 | 0.048193451 | 0.253691 | 0.1899692 | 0.849333 | 0.973446151 | DOC2B | protein_coding | 56.44 | 0.000563288 | 0.000563288 | ENSG00000272636 | n.s. |
| ENSG00000243836 | 39.02482 | 0.052975463 | 0.311609 | 0.1700062 | 0.865005 | 0.975860117 | WDR86-AS1 | processed_transcript | 60.18 | 0.000562198 | 0.000562198 | ENSG00000243836 | n.s. |
| ENSG00000179915 | 664.5713 | 0.049162309 | 0.274692 | 0.1789727 | 0.857959 | 0.974056274 | NRXN1 | protein_coding | 35.81 | 0.000561235 | 0.000561235 | ENSG00000179915 | n.s. |
| ENSG00000198805 | 1297.018 | 0.048017463 | 0.247282 | 0.1941809 | 0.846034 | 0.973446151 | PNP | protein_coding | 47.11 | 0.000561231 | 0.000561231 | ENSG00000198805 | n.s. |
| ENSG00000160310 | 340.7924 | 0.053044898 | 0.31352 | 0.1691913 | 0.865646 | 0.975964429 | PRMT2 | protein_coding | 45.52 | 0.000560473 | 0.000560473 | ENSG00000160310 | n.s. |
| ENSG00000166169 | 183.4929 | -0.047893672 | 0.256356 | -0.186825 | 0.851798 | 0.973446151 | POLL | protein_coding | 52.96 | -0.000559784 | 0.000559784 | ENSG00000166169 | n.s. |
| ENSG00000070501 | 557.7823 | -0.052511485 | 0.306035 | -0.171587 | 0.863763 | 0.975753891 | POLB | protein_coding | 40.44 | -0.000559757 | 0.000559757 | ENSG00000070501 | n.s. |
| ENSG00000187866 | 384.7564 | 0.04536047 | 0.218285 | 0.2078039 | 0.835382 | 0.971988837 | FAM122A | protein_coding | 40.85 | 0.00055969 | 0.00055969 | ENSG00000187866 | n.s. |
| ENSG00000226688 | 376.2476 | -0.047702356 | 0.244584 | -0.195035 | 0.845366 | 0.973446151 | ENTPD1-AS1 | antisense | 39.46 | -0.000557548 | 0.000557548 | ENSG00000226688 | n.s. |
| ENSG00000163870 | 353.0636 | -0.048771618 | 0.2677 | -0.182188 | 0.855436 | 0.974056274 | TPRA1 | protein_coding | 55.15 | -0.000556774 | 0.000556774 | ENSG00000163870 | n.s. |
| ENSG00000282978 | 36.4486 | 0.054021113 | 0.345014 | 0.1565767 | 0.875578 | 0.976582458 | | processed_pseudogene | 43.09 | 0.000555936 | 0.000555936 | ENSG00000282978 | n.s. |
| ENSG00000149179 | 46.36202 | 0.052599547 | 0.31881 | 0.164987 | 0.868954 | 0.975964429 | C11orf49 | protein_coding | 41.37 | 0.000555767 | 0.000555767 | ENSG00000149179 | n.s. |

| | | | | | | | | | | | | | |
|---|---|---|---|---|---|---|---|---|---|---|---|---|---|
| ENSG00000280828 | 95.45753 | 0.051695266 | 0.298157 | 0.1733826 | 0.862351 | 0.975554135 | | unprocessed_pseudogene | 41.25 | 0.000555653 | 0.000555653 | ENSG00000280828 | n.s. |
| ENSG00000274001 | 49.26272 | -0.052042003 | 0.301542 | -0.172586 | 0.862977 | 0.975738908 | | sense_intronic | 34.3 | -0.0005551 | 0.0005551 | ENSG00000274001 | n.s. |
| ENSG00000066427 | 156.5315 | 0.048488312 | 0.27003 | 0.1795665 | 0.857493 | 0.974056274 | ATXN3 | protein_coding | 42.78 | 0.00055354 | 0.00055354 | ENSG00000066427 | n.s. |
| ENSG00000165672 | 3180.718 | 0.052377378 | 0.316594 | 0.1654403 | 0.868597 | 0.975964429 | PRDX3 | protein_coding | 46.03 | 0.00055342 | 0.00055342 | ENSG00000165672 | n.s. |
| ENSG00000128944 | 1595.977 | -0.048452921 | 0.264906 | -0.182906 | 0.854872 | 0.974056274 | KNSTRN | protein_coding | 44.95 | -0.000553136 | 0.000553136 | ENSG00000128944 | n.s. |
| ENSG00000105821 | 1935.849 | -0.047449142 | 0.256562 | -0.184942 | 0.853275 | 0.97351991 | DNAJC2 | protein_coding | 40.56 | -0.000553027 | 0.000553027 | ENSG00000105821 | n.s. |
| ENSG00000149547 | 872.6845 | -0.051809124 | 0.300507 | -0.172406 | 0.863119 | 0.975738908 | EI24 | protein_coding | 43.49 | -0.000552616 | 0.000552616 | ENSG00000149547 | n.s. |
| ENSG00000127884 | 620.7048 | -0.052418085 | 0.327281 | -0.160162 | 0.872753 | 0.976022242 | ECHS1 | protein_coding | 56.3 | -0.000552502 | 0.000552502 | ENSG00000127884 | n.s. |
| ENSG00000146776 | 66.78949 | -0.048368018 | 0.270299 | -0.178943 | 0.857983 | 0.974056274 | ATXN7L1 | protein_coding | 45.57 | -0.000552167 | 0.000552167 | ENSG00000146776 | n.s. |
| ENSG00000239467 | 66.58545 | 0.052276697 | 0.318644 | 0.1640601 | 0.869684 | 0.975997758 | | lincRNA | 40.11 | 0.000551581 | 0.000551581 | ENSG00000239467 | n.s. |
| ENSG00000154727 | 1120.408 | -0.049934074 | 0.285055 | -0.175173 | 0.860943 | 0.974949369 | GABPA | protein_coding | 36.67 | -0.000550171 | 0.000550171 | ENSG00000154727 | n.s. |
| ENSG00000100664 | 7816.294 | 0.046983004 | 0.251877 | 0.1865318 | 0.852028 | 0.973446151 | EIF5 | protein_coding | 44.94 | 0.000549141 | 0.000549141 | ENSG00000100664 | n.s. |
| ENSG00000133316 | 1948.678 | -0.043667879 | 0.205943 | -0.212039 | 0.832077 | 0.971513984 | WDR74 | protein_coding | 51.65 | -0.000548073 | 0.000548073 | ENSG00000133316 | n.s. |
| ENSG00000167702 | 146.5949 | -0.05049449 | 0.290326 | -0.173923 | 0.861926 | 0.975340686 | KIFC2 | protein_coding | 63.6 | -0.000547545 | 0.000547545 | ENSG00000167702 | n.s. |
| ENSG00000161981 | 516.9939 | -0.05182039 | 0.313991 | -0.165038 | 0.868914 | 0.975964429 | SNRNP25 | protein_coding | 56.7 | -0.000547535 | 0.000547535 | ENSG00000161981 | n.s. |
| ENSG00000151465 | 1548.068 | -0.048617425 | 0.273563 | -0.177719 | 0.858943 | 0.974414762 | CDC123 | protein_coding | 42.91 | -0.000547245 | 0.000547245 | ENSG00000151465 | n.s. |
| ENSG00000125962 | 639.9947 | 0.046729232 | 0.250852 | 0.1862822 | 0.852224 | 0.973446151 | ARMCX5 | protein_coding | 45.27 | 0.000546174 | 0.000546174 | ENSG00000125962 | n.s. |
| ENSG00000173226 | 1171.488 | 0.047828796 | 0.265464 | 0.1801703 | 0.857019 | 0.974056274 | IQCB1 | protein_coding | 36.52 | 0.000546011 | 0.000546011 | ENSG00000173226 | n.s. |
| ENSG00000164338 | 1381.087 | -0.046701979 | 0.233861 | -0.1997 | 0.841716 | 0.973446151 | UTP15 | protein_coding | 37.9 | -0.000545856 | 0.000545856 | ENSG00000164338 | n.s. |
| ENSG00000127955 | 807.5481 | -0.047800621 | 0.264089 | -0.181002 | 0.856366 | 0.974056274 | GNAI1 | protein_coding | 36.74 | -0.00054569 | 0.00054569 | ENSG00000127955 | n.s. |
| ENSG00000128185 | 296.0514 | -0.051632319 | 0.312734 | -0.1651 | 0.868866 | 0.975964429 | DGCR6L | protein_coding | 57.73 | -0.000545548 | 0.000545548 | ENSG00000128185 | n.s. |
| ENSG00000163428 | 2668.32 | -0.046673207 | 0.252034 | -0.185186 | 0.853083 | 0.973483743 | LRRC58 | protein_coding | 38.94 | -0.000544737 | 0.000544737 | ENSG00000163428 | n.s. |
| ENSG00000144306 | 647.3362 | -0.04928052 | 0.281736 | -0.174918 | 0.861144 | 0.974949369 | SCRN3 | protein_coding | 37.9 | -0.00054297 | 0.00054297 | ENSG00000144306 | n.s. |
| ENSG00000145425 | 18896.37 | 0.051507378 | 0.323099 | 0.1594168 | 0.87334 | 0.976022242 | RPS3A | protein_coding | 41.56 | 0.000542902 | 0.000542902 | ENSG00000145425 | n.s. |
| ENSG00000108064 | 2727.456 | -0.047549702 | 0.260402 | -0.182601 | 0.855111 | 0.974056274 | TFAM | protein_coding | 38.21 | -0.000542825 | 0.000542825 | ENSG00000108064 | n.s. |
| ENSG00000198912 | 210.7237 | 0.047433618 | 0.261135 | 0.1816443 | 0.855862 | 0.974056274 | C1orf174 | protein_coding | 49.32 | 0.0005415 | 0.0005415 | ENSG00000198912 | n.s. |
| ENSG00000145414 | 737.9361 | 0.046236081 | 0.239109 | 0.1933679 | 0.846671 | 0.973446151 | NAF1 | protein_coding | 36.36 | 0.00054041 | 0.00054041 | ENSG00000145414 | n.s. |
| ENSG00000104408 | 8634.372 | 0.051206037 | 0.322663 | 0.158698 | 0.873907 | 0.976022242 | EIF3E | protein_coding | 37.25 | 0.000539726 | 0.000539726 | ENSG00000104408 | n.s. |
| ENSG00000053501 | 155.2022 | -0.046146455 | 0.245182 | -0.188213 | 0.850709 | 0.973446151 | USE1 | protein_coding | 55.26 | -0.000539363 | 0.000539363 | ENSG00000053501 | n.s. |
| ENSG00000228889 | 163.891 | 0.04600258 | 0.238763 | 0.1926703 | 0.847217 | 0.973446151 | UBAC2-AS1 | lincRNA | 44.85 | 0.000537681 | 0.000537681 | ENSG00000228889 | n.s. |
| ENSG00000170584 | 965.3793 | -0.045876505 | 0.245866 | -0.186591 | 0.851981 | 0.973446151 | NUDCD2 | protein_coding | 37.65 | -0.000536208 | 0.000536208 | ENSG00000170584 | n.s. |
| ENSG00000131871 | 888.9014 | 0.046952201 | 0.263136 | 0.178433 | 0.858383 | 0.974056274 | SELENOS | protein_coding | 45.06 | 0.000536004 | 0.000536004 | ENSG00000131871 | n.s. |

| | | | | | | | | | | | | | |
|---|---|---|---|---|---|---|---|---|---|---|---|---|---|
| ENSG00000154957 | 262.3877 | -0.046881742 | 0.26184 | -0.179047 | 0.857901 | 0.974056274 | ZNF18 | protein_coding | 41.67 | -0.0005352 | 0.0005352 | ENSG00000154957 | n.s. |
| ENSG00000108559 | 1619.802 | -0.047486502 | 0.267457 | -0.177548 | 0.859078 | 0.974414762 | NUP88 | protein_coding | 42.72 | -0.000534515 | 0.000534515 | ENSG00000108559 | n.s. |
| ENSG00000137133 | 335.6982 | -0.050652066 | 0.309495 | -0.163661 | 0.869998 | 0.975997758 | HINT2 | protein_coding | 54.66 | -0.000534439 | 0.000534439 | ENSG00000137133 | n.s. |
| ENSG00000162068 | 95.4711 | -0.046481663 | 0.259844 | -0.178883 | 0.85803 | 0.974056274 | NTN3 | protein_coding | 70.83 | -0.000530632 | 0.000530632 | ENSG00000162068 | n.s. |
| ENSG00000161677 | 49.50618 | -0.053106094 | 0.347433 | -0.152853 | 0.878514 | 0.977264479 | JOSD2 | protein_coding | 61.76 | -0.000530418 | 0.000530418 | ENSG00000161677 | n.s. |
| ENSG00000135597 | 1972.586 | -0.046462402 | 0.259767 | -0.178862 | 0.858046 | 0.974056274 | REPS1 | protein_coding | 37.32 | -0.000530413 | 0.000530413 | ENSG00000135597 | n.s. |
| ENSG00000146457 | 3358.719 | -0.047847826 | 0.272651 | -0.175491 | 0.860694 | 0.97481838 | WTAP | protein_coding | 38.63 | -0.000529976 | 0.000529976 | ENSG00000146457 | n.s. |
| ENSG00000169193 | 147.2697 | -0.049925925 | 0.291694 | -0.171159 | 0.864099 | 0.975860117 | CCDC126 | protein_coding | 41.98 | -0.000529835 | 0.000529835 | ENSG00000169193 | n.s. |
| ENSG00000147202 | 573.1803 | 0.047741248 | 0.271251 | 0.1760038 | 0.860291 | 0.974774428 | DIAPH2 | protein_coding | 37 | 0.000529731 | 0.000529731 | ENSG00000147202 | n.s. |
| ENSG00000178105 | 1384.756 | 0.046385016 | 0.259138 | 0.1789975 | 0.85794 | 0.974056274 | DDX10 | protein_coding | 37.41 | 0.000529529 | 0.000529529 | ENSG00000178105 | n.s. |
| ENSG00000110696 | 4018.52 | -0.050172371 | 0.306751 | -0.16356 | 0.870077 | 0.975997758 | C11orf58 | protein_coding | 38.23 | -0.000529378 | 0.000529378 | ENSG00000110696 | n.s. |
| ENSG00000144524 | 152.485 | -0.047162518 | 0.265944 | -0.17734 | 0.859241 | 0.974520785 | COPS7B | protein_coding | 44.9 | -0.00052864 | 0.00052864 | ENSG00000144524 | n.s. |
| ENSG00000128829 | 870.9991 | -0.049506805 | 0.287623 | -0.172124 | 0.86334 | 0.975741574 | EIF2AK4 | protein_coding | 40.17 | -0.000527999 | 0.000527999 | ENSG00000128829 | n.s. |
| ENSG00000074935 | 2028.431 | 0.045278529 | 0.244731 | 0.1850137 | 0.853218 | 0.97351991 | TUBE1 | protein_coding | 34.64 | 0.000527729 | 0.000527729 | ENSG00000074935 | n.s. |
| ENSG00000219545 | 177.4224 | 0.049962792 | 0.30679 | 0.1628564 | 0.870632 | 0.976022242 | UMAD1 | protein_coding | 37.6 | 0.000526622 | 0.000526622 | ENSG00000219545 | n.s. |
| ENSG00000188603 | 47.00493 | -0.051315093 | 0.331242 | -0.154917 | 0.876887 | 0.97668357 | CLN3 | protein_coding | 50.61 | -0.00052578 | 0.00052578 | ENSG00000188603 | n.s. |
| ENSG00000196419 | 8743.541 | -0.049792004 | 0.304307 | -0.163624 | 0.870027 | 0.975997758 | XRCC6 | protein_coding | 46.22 | -0.000525364 | 0.000525364 | ENSG00000196419 | n.s. |
| ENSG00000152219 | 1208.173 | -0.049549246 | 0.295665 | -0.167586 | 0.866909 | 0.975964429 | ARL14EP | protein_coding | 36.48 | -0.000523538 | 0.000523538 | ENSG00000152219 | n.s. |
| ENSG00000133627 | 327.7712 | -0.049529218 | 0.293355 | -0.168837 | 0.865925 | 0.975964429 | ACTR3B | protein_coding | 43.36 | -0.000523326 | 0.000523326 | ENSG00000133627 | n.s. |
| ENSG00000172292 | 2417.132 | 0.047161723 | 0.268737 | 0.1754936 | 0.860692 | 0.97481838 | CERS6 | protein_coding | 39.32 | 0.000522377 | 0.000522377 | ENSG00000172292 | n.s. |
| ENSG00000163923 | 427.6998 | -0.049418551 | 0.299785 | -0.164847 | 0.869065 | 0.975964429 | RPL39L | protein_coding | 41.84 | -0.000522157 | 0.000522157 | ENSG00000163923 | n.s. |
| ENSG00000008324 | 380.5289 | 0.049338945 | 0.293309 | 0.1682152 | 0.866414 | 0.975964429 | SS18L2 | protein_coding | 47.77 | 0.000521316 | 0.000521316 | ENSG00000008324 | n.s. |
| ENSG00000111196 | 2043.432 | 0.046784291 | 0.265562 | 0.1761708 | 0.86016 | 0.974768375 | MAGOHB | protein_coding | 36.42 | 0.000519239 | 0.000519239 | ENSG00000111196 | n.s. |
| ENSG00000159596 | 179.2822 | -0.045369024 | 0.248293 | -0.182724 | 0.855015 | 0.974056274 | TMEM69 | protein_coding | 44.47 | -0.000517931 | 0.000517931 | ENSG00000159596 | n.s. |
| ENSG00000152382 | 1143.92 | 0.04901496 | 0.292609 | 0.1675098 | 0.866969 | 0.975964429 | TADA1 | protein_coding | 40.95 | 0.000517893 | 0.000517893 | ENSG00000152382 | n.s. |
| ENSG00000078319 | 191.4775 | -0.049124462 | 0.305166 | -0.160976 | 0.872112 | 0.976022242 | PMS2P1 | unprocessed_pseudogene | 44.3 | -0.000517786 | 0.000517786 | ENSG00000078319 | n.s. |
| ENSG00000065518 | 1367.379 | 0.049011488 | 0.29837 | 0.164264 | 0.869523 | 0.975997758 | NDUFB4 | protein_coding | 39.39 | 0.000517129 | 0.000517129 | ENSG00000065518 | n.s. |
| ENSG00000092421 | 155.3371 | 0.048477585 | 0.28097 | 0.1725363 | 0.863016 | 0.975738908 | SEMA6A | protein_coding | 40.58 | 0.00051708 | 0.00051708 | ENSG00000092421 | n.s. |
| ENSG00000145293 | 1450.023 | 0.049050103 | 0.300904 | 0.1630089 | 0.870511 | 0.976022242 | ENOPH1 | protein_coding | 43.17 | 0.000517002 | 0.000517002 | ENSG00000145293 | n.s. |
| ENSG00000248487 | 154.9806 | -0.04904651 | 0.303583 | -0.161559 | 0.871653 | 0.976022242 | ABHD14A | protein_coding | 54.78 | -0.000516964 | 0.000516964 | ENSG00000248487 | n.s. |
| ENSG00000155329 | 465.4127 | -0.048756707 | 0.291616 | -0.167195 | 0.867217 | 0.975964429 | ZCCHC10 | protein_coding | 42.73 | -0.000515164 | 0.000515164 | ENSG00000155329 | n.s. |
| ENSG00000141682 | 2778.817 | 0.044031198 | 0.236658 | 0.1860543 | 0.852402 | 0.973446151 | PMAIP1 | protein_coding | 39.83 | 0.00051464 | 0.00051464 | ENSG00000141682 | n.s. |

| | | | | | | | | | | | | | |
|---|---|---|---|---|---|---|---|---|---|---|---|---|---|
| ENSG00000110955 | 13389.09 | 0.048676787 | 0.287744 | 0.1691673 | 0.865665 | 0.975964429 | ATP5B | protein_coding | 45.43 | 0.000514319 | 0.000514319 | ENSG00000110955 | n.s. |
| ENSG00000186908 | 2645.71 | -0.048194631 | 0.278858 | -0.172829 | 0.862786 | 0.975738908 | ZDHHC17 | protein_coding | 36.11 | -0.000514062 | 0.000514062 | ENSG00000186908 | n.s. |
| ENSG00000263727 | 26.5427 | -0.050652916 | 0.329038 | -0.153943 | 0.877655 | 0.976942004 | | sense_intronic | 47.33 | -0.000513176 | 0.000513176 | ENSG00000263727 | n.s. |
| ENSG00000187189 | 2447.429 | -0.043811971 | 0.225001 | -0.194719 | 0.845613 | 0.973446151 | TSPYL4 | protein_coding | 47.75 | -0.000512077 | 0.000512077 | ENSG00000187189 | n.s. |
| ENSG00000153310 | 1581.918 | -0.048575453 | 0.299367 | -0.16226 | 0.871101 | 0.976022242 | FAM49B | protein_coding | 42.04 | -0.000511999 | 0.000511999 | ENSG00000153310 | n.s. |
| ENSG00000049656 | 1035.735 | -0.04844908 | 0.288463 | -0.167956 | 0.866618 | 0.975964429 | CLPTM1L | protein_coding | 56.17 | -0.000511914 | 0.000511914 | ENSG00000049656 | n.s. |
| ENSG00000104412 | 668.1343 | 0.048501654 | 0.296287 | 0.1636983 | 0.869969 | 0.975997758 | EMC2 | protein_coding | 34.69 | 0.00051175 | 0.00051175 | ENSG00000104412 | n.s. |
| ENSG00000184428 | 282.1659 | 0.048157797 | 0.281389 | 0.1711432 | 0.864111 | 0.975860117 | TOP1MT | protein_coding | 54.3 | 0.000511071 | 0.000511071 | ENSG00000184428 | n.s. |
| ENSG00000165506 | 646.615 | -0.044707261 | 0.248683 | -0.179776 | 0.857328 | 0.974056274 | DNAAF2 | protein_coding | 46.14 | -0.000510376 | 0.000510376 | ENSG00000165506 | n.s. |
| ENSG00000108826 | 2362.995 | -0.046008069 | 0.261847 | -0.175706 | 0.860525 | 0.974785806 | MRPL27 | protein_coding | 47.14 | -0.000510266 | 0.000510266 | ENSG00000108826 | n.s. |
| ENSG00000120137 | 8890.412 | -0.045916366 | 0.260359 | -0.176358 | 0.860013 | 0.974768375 | PANK3 | protein_coding | 37.86 | -0.000509606 | 0.000509606 | ENSG00000120137 | n.s. |
| ENSG00000196655 | 366.7035 | 0.0482112 | 0.28514 | 0.1690791 | 0.865734 | 0.975964429 | TRAPPC4 | protein_coding | 49.51 | 0.0005094 | 0.0005094 | ENSG00000196655 | n.s. |
| ENSG00000273391 | 112.2931 | 0.04798087 | 0.282258 | 0.1699894 | 0.865018 | 0.975860117 | | antisense | 62.06 | 0.000509194 | 0.000509194 | ENSG00000273391 | n.s. |
| ENSG00000106211 | 270.6144 | -0.048186576 | 0.290878 | -0.165659 | 0.868425 | 0.975964429 | HSPB1 | protein_coding | 63.41 | -0.00050914 | 0.00050914 | ENSG00000106211 | n.s. |
| ENSG00000125871 | 947.9296 | 0.043481421 | 0.228889 | 0.1899671 | 0.849335 | 0.973446151 | MGME1 | protein_coding | 42.82 | 0.000508214 | 0.000508214 | ENSG00000125871 | n.s. |
| ENSG00000182180 | 1425.155 | -0.044483439 | 0.249138 | -0.178549 | 0.858292 | 0.974056274 | MRPS16 | protein_coding | 47.47 | -0.000507821 | 0.000507821 | ENSG00000182180 | n.s. |
| ENSG00000125991 | 1194.435 | 0.0481172 | 0.29448 | 0.1633971 | 0.870206 | 0.976022242 | ERGIC3 | protein_coding | 50.94 | 0.000507169 | 0.000507169 | ENSG00000125991 | n.s. |
| ENSG00000160606 | 42.70197 | -0.048092678 | 0.302992 | -0.158726 | 0.873885 | 0.976022242 | TLCD1 | protein_coding | 60.34 | -0.000506911 | 0.000506911 | ENSG00000160606 | n.s. |
| ENSG00000228439 | 42.15317 | -0.051068863 | 0.334822 | -0.152526 | 0.878772 | 0.977404381 | TSTD3 | protein_coding | 40.48 | -0.000506895 | 0.000506895 | ENSG00000228439 | n.s. |
| ENSG00000248773 | 114.3449 | 0.048035553 | 0.292882 | 0.1640099 | 0.869723 | 0.975997758 | | sense_intronic | 33.4 | 0.000506832 | 0.000506832 | ENSG00000248773 | n.s. |
| ENSG00000156603 | 345.4717 | -0.047434501 | 0.276032 | -0.171844 | 0.86356 | 0.975741574 | MED19 | protein_coding | 45.7 | -0.000505898 | 0.000505898 | ENSG00000156603 | n.s. |
| ENSG00000252615 | 9.219076 | -0.061779372 | 0.480352 | -0.128613 | 0.897664 | 0.981326694 | | misc_RNA | 41 | -0.00050575 | 0.00050575 | ENSG00000252615 | n.s. |
| ENSG00000114026 | 175.329 | 0.044228618 | 0.241612 | 0.1830566 | 0.854754 | 0.974056274 | OGG1 | protein_coding | 49.74 | 0.000504912 | 0.000504912 | ENSG00000114026 | n.s. |
| ENSG00000072121 | 413.6518 | -0.043177874 | 0.232193 | -0.185957 | 0.852478 | 0.973446151 | ZFYVE26 | protein_coding | 43 | -0.000504666 | 0.000504666 | ENSG00000072121 | n.s. |
| ENSG00000161265 | 112.4409 | -0.047744375 | 0.286492 | -0.166652 | 0.867644 | 0.975964429 | U2AF1L4 | protein_coding | 55.84 | -0.000504468 | 0.000504468 | ENSG00000161265 | n.s. |
| ENSG00000216031 | 3.440343 | 0.074012666 | 0.640958 | 0.1154719 | 0.908071 | 0.984433066 | MIR298 | miRNA | 53.41 | 0.000504308 | 0.000504308 | ENSG00000216031 | n.s. |
| ENSG00000177030 | 403.3031 | 0.047819878 | 0.296321 | 0.1613785 | 0.871795 | 0.976022242 | DEAF1 | protein_coding | 53.21 | 0.000504035 | 0.000504035 | ENSG00000177030 | n.s. |
| ENSG00000108465 | 4443.921 | -0.04471379 | 0.251501 | -0.177787 | 0.85889 | 0.974414762 | CDK5RAP3 | protein_coding | 51.51 | -0.000503305 | 0.000503305 | ENSG00000108465 | n.s. |
| ENSG00000181541 | 7426.555 | -0.043051131 | 0.229792 | -0.187348 | 0.851388 | 0.973446151 | MAB21L2 | protein_coding | 52.09 | -0.000503185 | 0.000503185 | ENSG00000181541 | n.s. |
| ENSG00000260608 | 36.72837 | -0.049051553 | 0.31693 | -0.154771 | 0.877002 | 0.97668357 | | antisense | 39.41 | -0.000502588 | 0.000502588 | ENSG00000260608 | n.s. |
| ENSG00000178467 | 186.4707 | 0.047424185 | 0.27931 | 0.1697907 | 0.865175 | 0.975894269 | P4HTM | protein_coding | 54.55 | 0.000502565 | 0.000502565 | ENSG00000178467 | n.s. |
| ENSG00000198056 | 1092.762 | -0.047299295 | 0.278282 | -0.169969 | 0.865035 | 0.975860117 | PRIM1 | protein_coding | 42.82 | -0.000501961 | 0.000501961 | ENSG00000198056 | n.s. |

| | | | | | | | | | | | | | |
|---|---|---|---|---|---|---|---|---|---|---|---|---|---|
| ENSG00000258101 | 41.06413 | -0.052430253 | 0.351682 | -0.149084 | 0.881487 | 0.978242302 | | antisense | 46.26 | -0.000500896 | 0.000500896 | ENSG00000258101 | n.s. |
| ENSG00000099904 | 188.4188 | -0.042786821 | 0.227622 | -0.187973 | 0.850898 | 0.973446151 | ZDHHC8 | protein_coding | 63.57 | -0.000500095 | 0.000500095 | ENSG00000099904 | n.s. |
| ENSG00000162613 | 13240 | 0.04503282 | 0.255539 | 0.1762267 | 0.860116 | 0.974768375 | FUBP1 | protein_coding | 36.32 | 0.0004998 | 0.0004998 | ENSG00000162613 | n.s. |
| ENSG00000272195 | 79.22153 | 0.047294414 | 0.281068 | 0.1682667 | 0.866373 | 0.975964429 | | antisense | 62.79 | 0.000499713 | 0.000499713 | ENSG00000272195 | n.s. |
| ENSG00000185115 | 506.4845 | 0.046777582 | 0.271246 | 0.1724546 | 0.86308 | 0.975738908 | NSMCE3 | protein_coding | 41.84 | 0.000498947 | 0.000498947 | ENSG00000185115 | n.s. |
| ENSG00000172315 | 1486.073 | -0.042654822 | 0.224613 | -0.189904 | 0.849385 | 0.973446151 | TP53RK | protein_coding | 45.1 | -0.000498552 | 0.000498552 | ENSG00000172315 | n.s. |
| ENSG00000123472 | 457.6067 | -0.047309684 | 0.299109 | -0.158169 | 0.874324 | 0.976043167 | ATPAF1 | protein_coding | 41.86 | -0.000498217 | 0.000498217 | ENSG00000123472 | n.s. |
| ENSG00000138085 | 1030.654 | -0.04721672 | 0.288228 | -0.163817 | 0.869875 | 0.975997758 | ATRAID | protein_coding | 45.56 | -0.000498192 | 0.000498192 | ENSG00000138085 | n.s. |
| ENSG00000148303 | 20635.52 | 0.048434622 | 0.311768 | 0.1553545 | 0.876542 | 0.976610381 | RPL7A | protein_coding | 50.42 | 0.000497843 | 0.000497843 | ENSG00000148303 | n.s. |
| ENSG00000284062 | 2.996847 | 0.087241333 | 0.863215 | 0.1010656 | 0.919498 | 0.986968832 | MIR4469 | miRNA | 72.15 | 0.000496976 | 0.000496976 | ENSG00000284062 | n.s. |
| ENSG00000166451 | 732.1362 | -0.047654836 | 0.302476 | -0.157549 | 0.874812 | 0.976274971 | CENPN | protein_coding | 43.26 | -0.000496937 | 0.000496937 | ENSG00000166451 | n.s. |
| ENSG00000126804 | 1070.94 | -0.046992193 | 0.2792 | -0.16831 | 0.866339 | 0.975964429 | ZBTB1 | protein_coding | 39.55 | -0.00049652 | 0.00049652 | ENSG00000126804 | n.s. |
| ENSG00000057663 | 1089.828 | -0.047052952 | 0.292611 | -0.160804 | 0.872248 | 0.976022242 | ATG5 | protein_coding | 39.61 | -0.000495952 | 0.000495952 | ENSG00000057663 | n.s. |
| ENSG00000100380 | 4603.335 | -0.047014282 | 0.293729 | -0.16006 | 0.872834 | 0.976022242 | ST13 | protein_coding | 38.45 | -0.000495544 | 0.000495544 | ENSG00000100380 | n.s. |
| ENSG00000130640 | 229.0906 | -0.04687361 | 0.278397 | -0.16837 | 0.866292 | 0.975964429 | TUBGCP2 | protein_coding | 55.72 | -0.000495267 | 0.000495267 | ENSG00000130640 | n.s. |
| ENSG00000117410 | 529.6188 | -0.048798347 | 0.31728 | -0.153802 | 0.877766 | 0.976949771 | ATP6V0B | protein_coding | 57.08 | -0.000494218 | 0.000494218 | ENSG00000117410 | n.s. |
| ENSG00000125652 | 144.0345 | -0.050615235 | 0.333841 | -0.151615 | 0.879491 | 0.977776254 | ALKBH7 | protein_coding | 64.35 | -0.000494031 | 0.000494031 | ENSG00000125652 | n.s. |
| ENSG00000101190 | 264.8003 | 0.046107256 | 0.266437 | 0.1730515 | 0.862611 | 0.975663031 | TCFL5 | protein_coding | 48.88 | 0.000493355 | 0.000493355 | ENSG00000101190 | n.s. |
| ENSG00000100567 | 1824.656 | -0.046667676 | 0.279965 | -0.166691 | 0.867613 | 0.975964429 | PSMA3 | protein_coding | 40.29 | -0.000493091 | 0.000493091 | ENSG00000100567 | n.s. |
| ENSG00000131043 | 759.4172 | 0.044745309 | 0.25595 | 0.1748207 | 0.86122 | 0.974949369 | AAR2 | protein_coding | 47.41 | 0.000493001 | 0.000493001 | ENSG00000131043 | n.s. |
| ENSG00000077080 | 338.8295 | -0.046748091 | 0.289707 | -0.161363 | 0.871807 | 0.976022242 | ACTL6B | protein_coding | 53.08 | -0.000492738 | 0.000492738 | ENSG00000077080 | n.s. |
| ENSG00000163053 | 290.4328 | -0.046722496 | 0.295131 | -0.158311 | 0.874212 | 0.976022242 | SLC16A14 | protein_coding | 42.86 | -0.000492468 | 0.000492468 | ENSG00000163053 | n.s. |
| ENSG00000103042 | 59.16104 | -0.046602347 | 0.278733 | -0.167193 | 0.867218 | 0.975964429 | SLC38A7 | protein_coding | 52.78 | -0.000492401 | 0.000492401 | ENSG00000103042 | n.s. |
| ENSG00000174915 | 324.0403 | -0.046502134 | 0.277414 | -0.167627 | 0.866877 | 0.975964429 | PTDSS2 | protein_coding | 56.05 | -0.000491342 | 0.000491342 | ENSG00000174915 | n.s. |
| ENSG00000203761 | 193.4867 | 0.043032331 | 0.239769 | 0.1794738 | 0.857566 | 0.974056274 | MSTO2P | unprocessed_pseudogene | 52.8 | 0.000491255 | 0.000491255 | ENSG00000203761 | n.s. |
| ENSG00000090054 | 854.1672 | -0.04636902 | 0.281097 | -0.164957 | 0.868978 | 0.975964429 | SPTLC1 | protein_coding | 41.6 | -0.000489936 | 0.000489936 | ENSG00000090054 | n.s. |
| ENSG00000120833 | 1755.547 | -0.042911137 | 0.236712 | -0.18128 | 0.856148 | 0.974056274 | SOCS2 | protein_coding | 47.34 | -0.000489871 | 0.000489871 | ENSG00000120833 | n.s. |
| ENSG00000109861 | 2239.272 | -0.046424748 | 0.282033 | -0.164607 | 0.869253 | 0.975997758 | CTSC | protein_coding | 38.6 | -0.000489836 | 0.000489836 | ENSG00000109861 | n.s. |
| ENSG00000165501 | 642.266 | -0.042884812 | 0.239781 | -0.17885 | 0.858055 | 0.974056274 | LRR1 | protein_coding | 45.06 | -0.000489571 | 0.000489571 | ENSG00000165501 | n.s. |
| ENSG00000207392 | 28.26365 | 0.054474188 | 0.379776 | 0.1434376 | 0.885945 | 0.979542794 | SNORA20 | snoRNA | 40.15 | 0.000488992 | 0.000488992 | ENSG00000207392 | n.s. |
| ENSG00000140553 | 840.4469 | -0.046113432 | 0.273576 | -0.168558 | 0.866144 | 0.975964429 | UNC45A | protein_coding | 53.82 | -0.000487235 | 0.000487235 | ENSG00000140553 | n.s. |
| ENSG00000122565 | 10861.78 | -0.046000559 | 0.278842 | -0.16497 | 0.868968 | 0.975964429 | CBX3 | protein_coding | 41.57 | -0.000486042 | 0.000486042 | ENSG00000122565 | n.s. |

| | | | | | | | | | | | | | |
|---|---|---|---|---|---|---|---|---|---|---|---|---|---|
| ENSG00000071994 | 2936.81 | 0.043785981 | 0.249196 | 0.1757091 | 0.860523 | 0.974785806 | PDCD2 | protein_coding | 41.64 | 0.000485622 | 0.000485622 | ENSG00000071994 | n.s. |
| ENSG00000174738 | 1419.978 | 0.045809704 | 0.276931 | 0.1654194 | 0.868614 | 0.975964429 | NR1D2 | protein_coding | 40.69 | 0.000484026 | 0.000484026 | ENSG00000174738 | n.s. |
| ENSG00000166200 | 4424.904 | -0.045894001 | 0.28746 | -0.159654 | 0.873154 | 0.976022242 | COPS2 | protein_coding | 35.44 | -0.000483736 | 0.000483736 | ENSG00000166200 | n.s. |
| ENSG00000144045 | 8.656254 | 0.06686081 | 0.548106 | 0.1219851 | 0.902911 | 0.983510168 | DQX1 | protein_coding | 49.43 | 0.000482812 | 0.000482812 | ENSG00000144045 | n.s. |
| ENSG00000108773 | 588.9925 | 0.045651851 | 0.276938 | 0.1648449 | 0.869066 | 0.975964429 | KAT2A | protein_coding | 58.67 | 0.000482358 | 0.000482358 | ENSG00000108773 | n.s. |
| ENSG00000015475 | 1084.725 | -0.046463125 | 0.295434 | -0.157271 | 0.875032 | 0.976441858 | BID | protein_coding | 52.51 | -0.000481061 | 0.000481061 | ENSG00000015475 | n.s. |
| ENSG00000075702 | 295.1191 | -0.04112426 | 0.209823 | -0.195995 | 0.844614 | 0.973446151 | WDR62 | protein_coding | 51.45 | -0.000480663 | 0.000480663 | ENSG00000075702 | n.s. |
| ENSG00000082153 | 5692.892 | -0.044953376 | 0.261694 | -0.171779 | 0.863612 | 0.975741574 | BZW1 | protein_coding | 41.12 | -0.000479436 | 0.000479436 | ENSG00000082153 | n.s. |
| ENSG00000100979 | 124.8871 | 0.045412874 | 0.283722 | 0.1600613 | 0.872833 | 0.976022242 | PLTP | protein_coding | 51.86 | 0.000478665 | 0.000478665 | ENSG00000100979 | n.s. |
| ENSG00000188021 | 805.4372 | 0.041887357 | 0.230367 | 0.1818285 | 0.855717 | 0.974056274 | UBQLN2 | protein_coding | 44.3 | 0.000478184 | 0.000478184 | ENSG00000188021 | n.s. |
| ENSG00000170606 | 4483.916 | 0.045349511 | 0.28403 | 0.1596642 | 0.873146 | 0.976022242 | HSPA4 | protein_coding | 41.31 | 0.000477997 | 0.000477997 | ENSG00000170606 | n.s. |
| ENSG00000155508 | 1055.021 | 0.045091499 | 0.272095 | 0.1657197 | 0.868378 | 0.975964429 | CNOT8 | protein_coding | 44.08 | 0.000476437 | 0.000476437 | ENSG00000155508 | n.s. |
| ENSG00000116857 | 812.4327 | 0.045089657 | 0.272304 | 0.1655857 | 0.868483 | 0.975964429 | TMEM9 | protein_coding | 48.53 | 0.000476418 | 0.000476418 | ENSG00000116857 | n.s. |
| ENSG00000006194 | 1535.229 | -0.041706002 | 0.231683 | -0.180013 | 0.857142 | 0.974056274 | ZNF263 | protein_coding | 47.09 | -0.000476114 | 0.000476114 | ENSG00000006194 | n.s. |
| ENSG00000275052 | 2960.633 | -0.045047958 | 0.266412 | -0.169091 | 0.865725 | 0.975964429 | PPP4R3B | protein_coding | 38.14 | -0.000475977 | 0.000475977 | ENSG00000275052 | n.s. |
| ENSG00000129083 | 2242.197 | 0.046245382 | 0.295231 | 0.1566415 | 0.875527 | 0.976582458 | COPB1 | protein_coding | 38.27 | 0.000475915 | 0.000475915 | ENSG00000129083 | n.s. |
| ENSG00000110075 | 3376.785 | -0.044892817 | 0.264488 | -0.169735 | 0.865219 | 0.975894269 | PPP6R3 | protein_coding | 41.89 | -0.00047574 | 0.00047574 | ENSG00000110075 | n.s. |
| ENSG00000179240 | 360.8951 | -0.040665451 | 0.218027 | -0.186515 | 0.852041 | 0.973446151 | | protein_coding | 43.61 | -0.000475301 | 0.000475301 | ENSG00000179240 | n.s. |
| ENSG00000015479 | 2376.633 | 0.041625487 | 0.230196 | 0.1808265 | 0.856504 | 0.974056274 | MATR3 | protein_coding | 38.8 | 0.000475195 | 0.000475195 | ENSG00000015479 | n.s. |
| ENSG00000088205 | 4838.201 | -0.044875477 | 0.273498 | -0.16408 | 0.869668 | 0.975997758 | DDX18 | protein_coding | 39.26 | -0.000473489 | 0.000473489 | ENSG00000088205 | n.s. |
| ENSG00000125676 | 2637.491 | -0.044916191 | 0.276114 | -0.162673 | 0.870776 | 0.976022242 | THOC2 | protein_coding | 38.09 | -0.000473429 | 0.000473429 | ENSG00000125676 | n.s. |
| ENSG00000153037 | 1091.434 | 0.044736981 | 0.269983 | 0.165703 | 0.868391 | 0.975964429 | SRP19 | protein_coding | 41.11 | 0.000472691 | 0.000472691 | ENSG00000153037 | n.s. |
| ENSG00000116729 | 1093.655 | -0.044671024 | 0.272988 | -0.163637 | 0.870017 | 0.975997758 | WLS | protein_coding | 41.57 | -0.000471332 | 0.000471332 | ENSG00000116729 | n.s. |
| ENSG00000064115 | 829.1766 | -0.044596187 | 0.26906 | -0.165748 | 0.868355 | 0.975964429 | TM7SF3 | protein_coding | 40.51 | -0.000471204 | 0.000471204 | ENSG00000064115 | n.s. |
| ENSG00000087301 | 616.146 | 0.04464987 | 0.272899 | 0.1636132 | 0.870036 | 0.975997758 | TXNDC16 | protein_coding | 36.83 | 0.000471109 | 0.000471109 | ENSG00000087301 | n.s. |
| ENSG00000115211 | 600.3791 | -0.04124415 | 0.228346 | -0.180621 | 0.856665 | 0.974056274 | EIF2B4 | protein_coding | 50.64 | -0.000470841 | 0.000470841 | ENSG00000115211 | n.s. |
| ENSG00000134744 | 2482.163 | -0.044517103 | 0.269257 | -0.165333 | 0.868682 | 0.975964429 | ZCCHC11 | protein_coding | 38.59 | -0.000470368 | 0.000470368 | ENSG00000134744 | n.s. |
| ENSG00000172731 | 449.2183 | -0.044513615 | 0.265684 | -0.167544 | 0.866942 | 0.975964429 | LRRC20 | protein_coding | 49.41 | -0.000470331 | 0.000470331 | ENSG00000172731 | n.s. |
| ENSG00000166557 | 939.8353 | -0.045743402 | 0.29239 | -0.156447 | 0.875681 | 0.97660852 | TMED3 | protein_coding | 42.58 | -0.000470219 | 0.000470219 | ENSG00000166557 | n.s. |
| ENSG00000229833 | 744.587 | -0.046392571 | 0.301351 | -0.153948 | 0.87765 | 0.976942004 | PET100 | protein_coding | 55.95 | -0.000470013 | 0.000470013 | ENSG00000229833 | n.s. |
| ENSG00000114331 | 1667.744 | -0.044470208 | 0.280372 | -0.158612 | 0.873975 | 0.976022242 | ACAP2 | protein_coding | 38.17 | -0.000468729 | 0.000468729 | ENSG00000114331 | n.s. |
| ENSG00000133863 | 742.0105 | -0.044431777 | 0.279527 | -0.158954 | 0.873705 | 0.976022242 | TEX15 | protein_coding | 35.93 | -0.000468324 | 0.000468324 | ENSG00000133863 | n.s. |

| | | | | | | | | | | | | | |
|---|---|---|---|---|---|---|---|---|---|---|---|---|---|
| ENSG00000140995 | 177.3752 | -0.044374704 | 0.277292 | -0.160029 | 0.872858 | 0.976022242 | DEF8 | protein_coding | 57.08 | -0.000467722 | 0.000467722 | ENSG00000140995 | n.s. |
| ENSG00000109775 | 459.639 | 0.048363363 | 0.321041 | 0.1506456 | 0.880255 | 0.97803802 | UFSP2 | protein_coding | 38.42 | 0.000466429 | 0.000466429 | ENSG00000109775 | n.s. |
| ENSG00000164934 | 1997.884 | -0.04424883 | 0.277872 | -0.159242 | 0.873478 | 0.976022242 | DCAF13 | protein_coding | 37.09 | -0.000466395 | 0.000466395 | ENSG00000164934 | n.s. |
| ENSG00000170468 | 369.4619 | 0.039867846 | 0.212091 | 0.1879749 | 0.850896 | 0.973446151 | RIOX1 | protein_coding | 58.62 | 0.000465978 | 0.000465978 | ENSG00000170468 | n.s. |
| ENSG00000174365 | 290.7054 | 0.040817006 | 0.227514 | 0.179404 | 0.85762 | 0.974056274 | SNHG11 | processed_transcript | 53.2 | 0.000465965 | 0.000465965 | ENSG00000174365 | n.s. |
| ENSG00000183496 | 2475.17 | -0.039866648 | 0.20058 | -0.198757 | 0.842453 | 0.973446151 | MEX3B | protein_coding | 55.84 | -0.000465964 | 0.000465964 | ENSG00000183496 | n.s. |
| ENSG00000144597 | 875.4398 | -0.043874613 | 0.256882 | -0.170797 | 0.864384 | 0.975860117 | EAF1 | protein_coding | 43.63 | -0.000465616 | 0.000465616 | ENSG00000144597 | n.s. |
| ENSG00000129055 | 1238.766 | -0.04398036 | 0.266101 | -0.165277 | 0.868726 | 0.975964429 | ANAPC13 | protein_coding | 45.33 | -0.000464697 | 0.000464697 | ENSG00000129055 | n.s. |
| ENSG00000151093 | 213.6117 | -0.040641898 | 0.224239 | -0.181244 | 0.856176 | 0.974056274 | OXSM | protein_coding | 42 | -0.000463966 | 0.000463966 | ENSG00000151093 | n.s. |
| ENSG00000109320 | 77.62483 | -0.043962592 | 0.273173 | -0.160933 | 0.872146 | 0.976022242 | NFKB1 | protein_coding | 37.86 | -0.000463378 | 0.000463378 | ENSG00000109320 | n.s. |
| ENSG00000179941 | 592.1878 | -0.04173485 | 0.236604 | -0.176391 | 0.859986 | 0.974768375 | BBS10 | protein_coding | 37.34 | -0.000463197 | 0.000463197 | ENSG00000179941 | n.s. |
| ENSG00000168036 | 10905.18 | 0.043886443 | 0.267869 | 0.1638352 | 0.869861 | 0.975997758 | CTNNB1 | protein_coding | 39.91 | 0.000463054 | 0.000463054 | ENSG00000168036 | n.s. |
| ENSG00000247626 | 682.7853 | -0.040206647 | 0.218354 | -0.184135 | 0.853908 | 0.973865848 | MARS2 | protein_coding | 48.39 | -0.000462411 | 0.000462411 | ENSG00000247626 | n.s. |
| ENSG00000073969 | 1019.096 | -0.04497882 | 0.288087 | -0.156129 | 0.875931 | 0.976610381 | NSF | protein_coding | 38.62 | -0.000462322 | 0.000462322 | ENSG00000073969 | n.s. |
| ENSG00000273066 | 336.1729 | 0.040469757 | 0.221425 | 0.1827699 | 0.854979 | 0.974056274 | | processed_transcript | 60.91 | 0.000462001 | 0.000462001 | ENSG00000273066 | n.s. |
| ENSG00000207380 | 17.23265 | -0.052057598 | 0.376731 | -0.138182 | 0.890096 | 0.979788048 | | misc_RNA | 38.05 | -0.00046164 | 0.00046164 | ENSG00000207380 | n.s. |
| ENSG00000136738 | 1040.823 | 0.043464041 | 0.25441 | 0.1708428 | 0.864347 | 0.975860117 | STAM | protein_coding | 37.21 | 0.000461259 | 0.000461259 | ENSG00000136738 | n.s. |
| ENSG00000096384 | 29179.82 | 0.043703312 | 0.270573 | 0.1615212 | 0.871683 | 0.976022242 | HSP90AB1 | protein_coding | 49.1 | 0.000460645 | 0.000460645 | ENSG00000096384 | n.s. |
| ENSG00000136930 | 2655.823 | -0.043656484 | 0.274797 | -0.158868 | 0.873773 | 0.976022242 | PSMB7 | protein_coding | 42.8 | -0.000460152 | 0.000460152 | ENSG00000136930 | n.s. |
| ENSG00000226709 | 18.47833 | 0.051879348 | 0.374302 | 0.1386029 | 0.889764 | 0.979788048 | FGF12-AS3 | antisense | 34.08 | 0.000460059 | 0.000460059 | ENSG00000226709 | n.s. |
| ENSG00000052723 | 1879.393 | -0.043635294 | 0.268903 | -0.162272 | 0.871092 | 0.976022242 | SIKE1 | protein_coding | 36.9 | -0.000459928 | 0.000459928 | ENSG00000052723 | n.s. |
| ENSG00000163482 | 388.9639 | -0.042926631 | 0.248094 | -0.173026 | 0.862631 | 0.975663031 | STK36 | protein_coding | 46.05 | -0.000459321 | 0.000459321 | ENSG00000163482 | n.s. |
| ENSG00000104823 | 268.5115 | -0.047960333 | 0.31951 | -0.150106 | 0.880681 | 0.978189412 | ECH1 | protein_coding | 51.47 | -0.000459318 | 0.000459318 | ENSG00000104823 | n.s. |
| ENSG00000106080 | 404.6821 | 0.043518027 | 0.270979 | 0.1605956 | 0.872412 | 0.976022242 | FKBP14 | protein_coding | 38.7 | 0.000458692 | 0.000458692 | ENSG00000106080 | n.s. |
| ENSG00000176422 | 990.6589 | 0.039230746 | 0.195147 | 0.2010323 | 0.840673 | 0.973446151 | SPRYD4 | protein_coding | 48.14 | 0.000458532 | 0.000458532 | ENSG00000176422 | n.s. |
| ENSG00000107443 | 1289.137 | -0.043363331 | 0.261765 | -0.165657 | 0.868427 | 0.975964429 | CCNJ | protein_coding | 39.76 | -0.000458177 | 0.000458177 | ENSG00000107443 | n.s. |
| ENSG00000184014 | 1298.848 | -0.043380844 | 0.27143 | -0.159823 | 0.87302 | 0.976022242 | DENND5A | protein_coding | 43.01 | -0.000457246 | 0.000457246 | ENSG00000184014 | n.s. |
| ENSG00000131876 | 1699.629 | 0.04090449 | 0.231025 | 0.1770569 | 0.859464 | 0.974614188 | SNRPA1 | protein_coding | 42.28 | 0.000456792 | 0.000456792 | ENSG00000131876 | n.s. |
| ENSG00000182518 | 186.4592 | -0.046974283 | 0.311237 | -0.150928 | 0.880033 | 0.977868793 | FAM104B | protein_coding | 36.9 | -0.000456562 | 0.000456562 | ENSG00000182518 | n.s. |
| ENSG00000163811 | 5259.185 | -0.041404168 | 0.236789 | -0.174857 | 0.861192 | 0.974949369 | WDR43 | protein_coding | 39.29 | -0.000456189 | 0.000456189 | ENSG00000163811 | n.s. |
| ENSG00000198467 | 156.9215 | -0.043176372 | 0.271397 | -0.159089 | 0.873599 | 0.976022242 | TPM2 | protein_coding | 56.38 | -0.000455091 | 0.000455091 | ENSG00000198467 | n.s. |
| ENSG00000150093 | 4163.214 | -0.043127975 | 0.272345 | -0.158358 | 0.874175 | 0.976022242 | ITGB1 | protein_coding | 39.15 | -0.000454581 | 0.000454581 | ENSG00000150093 | n.s. |

| | | | | | | | | | | | | | |
|---|---|---|---|---|---|---|---|---|---|---|---|---|---|
| ENSG00000125703 | 437.7996 | -0.044819265 | 0.291496 | -0.153756 | 0.877802 | 0.976949771 | ATG4C | protein_coding | 35.44 | -0.000453919 | 0.000453919 | ENSG00000125703 | n.s. |
| ENSG00000124784 | 1503.856 | 0.042949927 | 0.253795 | 0.169231 | 0.865615 | 0.975964429 | RIOK1 | protein_coding | 40.11 | 0.000453809 | 0.000453809 | ENSG00000124784 | n.s. |
| ENSG00000112983 | 1439.47 | -0.042949455 | 0.271348 | -0.158282 | 0.874235 | 0.976022242 | BRD8 | protein_coding | 43.02 | -0.0004527 | 0.0004527 | ENSG00000112983 | n.s. |
| ENSG00000140931 | 188.0523 | 0.042844744 | 0.264944 | 0.1617124 | 0.871532 | 0.976022242 | CMTM3 | protein_coding | 58.84 | 0.000451596 | 0.000451596 | ENSG00000140931 | n.s. |
| ENSG00000245958 | 745.5115 | -0.041458842 | 0.238084 | -0.174135 | 0.861759 | 0.975231621 | | transcribed_unprocessed_pseudc | 36.55 | -0.000451579 | 0.000451579 | ENSG00000245958 | n.s. |
| ENSG00000156875 | 229.5673 | -0.042778464 | 0.262853 | -0.162747 | 0.870718 | 0.976022242 | MFSD14A | protein_coding | 39.5 | -0.000450897 | 0.000450897 | ENSG00000156875 | n.s. |
| ENSG00000117461 | 1245.307 | -0.041939072 | 0.241965 | -0.173327 | 0.862395 | 0.975554135 | PIK3R3 | protein_coding | 40.27 | -0.000450787 | 0.000450787 | ENSG00000117461 | n.s. |
| ENSG00000196659 | 294.3274 | -0.039479517 | 0.215523 | -0.18318 | 0.854657 | 0.974056274 | TTC30B | protein_coding | 43.32 | -0.000450696 | 0.000450696 | ENSG00000196659 | n.s. |
| ENSG00000266412 | 2203.01 | -0.043836517 | 0.281977 | -0.155461 | 0.876458 | 0.976610381 | NCOA4 | protein_coding | 41.24 | -0.000450581 | 0.000450581 | ENSG00000266412 | n.s. |
| ENSG00000105472 | 52.29323 | 0.046980149 | 0.313501 | 0.1498565 | 0.880878 | 0.978189412 | CLEC11A | protein_coding | 66.4 | 0.000449931 | 0.000449931 | ENSG00000105472 | n.s. |
| ENSG00000277170 | 113.265 | 0.042324765 | 0.247919 | 0.1707203 | 0.864444 | 0.975860117 | | sense_intronic | 45.3 | 0.000449169 | 0.000449169 | ENSG00000277170 | n.s. |
| ENSG00000072364 | 3077.914 | -0.042475523 | 0.256357 | -0.165689 | 0.868402 | 0.975964429 | AFF4 | protein_coding | 40.42 | -0.000448797 | 0.000448797 | ENSG00000072364 | n.s. |
| ENSG00000155229 | 1140.794 | 0.042450338 | 0.252662 | 0.1680125 | 0.866573 | 0.975964429 | MMS19 | protein_coding | 44.91 | 0.000448531 | 0.000448531 | ENSG00000155229 | n.s. |
| ENSG00000104341 | 3219.065 | -0.04671586 | 0.310734 | -0.15034 | 0.880496 | 0.978149421 | LAPTM4B | protein_coding | 44 | -0.000448229 | 0.000448229 | ENSG00000104341 | n.s. |
| ENSG00000123643 | 37.34105 | 0.049266818 | 0.339454 | 0.1451355 | 0.884604 | 0.979274867 | SLC36A1 | protein_coding | 42.72 | 0.000448101 | 0.000448101 | ENSG00000123643 | n.s. |
| ENSG00000210117 | 151.691 | 0.044170948 | 0.286841 | 0.1539909 | 0.877617 | 0.976942004 | MT-TW | Mt_tRNA | 35.29 | 0.000447505 | 0.000447505 | ENSG00000210117 | n.s. |
| ENSG00000164105 | 738.2379 | 0.043479771 | 0.277659 | 0.1565939 | 0.875565 | 0.976582458 | SAP30 | protein_coding | 42.24 | 0.000447454 | 0.000447454 | ENSG00000164105 | n.s. |
| ENSG00000143751 | 1212.877 | 0.042317685 | 0.25591 | 0.1653613 | 0.86866 | 0.975964429 | SDE2 | protein_coding | 40.61 | 0.000447129 | 0.000447129 | ENSG00000143751 | n.s. |
| ENSG00000130224 | 543.8784 | 0.043462754 | 0.279992 | 0.1552285 | 0.876641 | 0.976610381 | LRCH2 | protein_coding | 37.13 | 0.000446739 | 0.000446739 | ENSG00000130224 | n.s. |
| ENSG00000165240 | 451.9772 | -0.044080035 | 0.286211 | -0.154013 | 0.8776 | 0.976942004 | ATP7A | protein_coding | 38.61 | -0.000446584 | 0.000446584 | ENSG00000165240 | n.s. |
| ENSG00000168876 | 402.2455 | -0.04236782 | 0.264066 | -0.160444 | 0.872531 | 0.976022242 | ANKRD49 | protein_coding | 37.68 | -0.000446569 | 0.000446569 | ENSG00000168876 | n.s. |
| ENSG00000164402 | 815.8678 | -0.04225225 | 0.264854 | -0.15953 | 0.873251 | 0.976022242 | 8-Sep | protein_coding | 48.12 | -0.000445351 | 0.000445351 | ENSG00000164402 | n.s. |
| ENSG00000123179 | 785.5323 | -0.048303515 | 0.331546 | -0.145692 | 0.884165 | 0.979008503 | EBPL | protein_coding | 45.56 | -0.000445046 | 0.000445046 | ENSG00000123179 | n.s. |
| ENSG00000108953 | 5283.693 | -0.043169889 | 0.276998 | -0.155849 | 0.876152 | 0.976610381 | YWHAE | protein_coding | 44.81 | -0.000443729 | 0.000443729 | ENSG00000108953 | n.s. |
| ENSG00000279632 | 130.1816 | 0.041967943 | 0.252659 | 0.1661049 | 0.868074 | 0.975964429 | | TEC | 42.26 | 0.000443434 | 0.000443434 | ENSG00000279632 | n.s. |
| ENSG00000106144 | 971.1855 | 0.040145529 | 0.229654 | 0.1748089 | 0.86123 | 0.974949369 | CASP2 | protein_coding | 43.43 | 0.000442321 | 0.000442321 | ENSG00000106144 | n.s. |
| ENSG00000132300 | 5458.868 | -0.041796311 | 0.252819 | -0.165321 | 0.868691 | 0.975964429 | PTCD3 | protein_coding | 41.22 | -0.00044162 | 0.00044162 | ENSG00000132300 | n.s. |
| ENSG00000198246 | 107.5378 | -0.041883056 | 0.261554 | -0.160132 | 0.872777 | 0.976022242 | SLC29A3 | protein_coding | 51.86 | -0.000441459 | 0.000441459 | ENSG00000198246 | n.s. |
| ENSG00000117650 | 1125.403 | 0.041752318 | 0.257157 | 0.1623611 | 0.871021 | 0.976022242 | NEK2 | protein_coding | 39.5 | 0.000440081 | 0.000440081 | ENSG00000117650 | n.s. |
| ENSG00000100348 | 307.6182 | -0.042724848 | 0.275143 | -0.155282 | 0.876599 | 0.976610381 | TXN2 | protein_coding | 51.41 | -0.000439154 | 0.000439154 | ENSG00000100348 | n.s. |
| ENSG00000128534 | 1734.205 | 0.041341982 | 0.245241 | 0.1685769 | 0.866129 | 0.975964429 | LSM8 | protein_coding | 36.89 | 0.00043682 | 0.00043682 | ENSG00000128534 | n.s. |
| ENSG00000166704 | 594.4896 | 0.037957658 | 0.207074 | 0.1833045 | 0.854559 | 0.974056274 | ZNF606 | protein_coding | 45 | 0.000433323 | 0.000433323 | ENSG00000166704 | n.s. |

| | | | | | | | | | | | | | |
|---|---|---|---|---|---|---|---|---|---|---|---|---|---|
| ENSG00000149806 | 4315.544 | 0.047590752 | 0.328738 | 0.144768 | 0.884894 | 0.97927567 | FAU | protein_coding | 55.53 | 0.00043284 | 0.00043284 | ENSG00000149806 | n.s. |
| ENSG00000165119 | 17780.57 | -0.040962255 | 0.245866 | -0.166604 | 0.867681 | 0.975964429 | HNRNPK | protein_coding | 39.12 | -0.000432808 | 0.000432808 | ENSG00000165119 | n.s. |
| ENSG00000250317 | 218.0263 | 0.048732945 | 0.346573 | 0.1406137 | 0.888175 | 0.979788048 | SMIM20 | protein_coding | 45.19 | 0.000432157 | 0.000432157 | ENSG00000250317 | n.s. |
| ENSG00000155096 | 3595.195 | -0.042033936 | 0.270971 | -0.155124 | 0.876724 | 0.976610381 | AZIN1 | protein_coding | 40.82 | -0.000432053 | 0.000432053 | ENSG00000155096 | n.s. |
| ENSG00000117395 | 750.7837 | -0.043503453 | 0.285587 | -0.15233 | 0.878927 | 0.97741985 | EBNA1BP2 | protein_coding | 44.38 | -0.000431504 | 0.000431504 | ENSG00000117395 | n.s. |
| ENSG00000138835 | 210.194 | 0.04091417 | 0.251839 | 0.1624616 | 0.870942 | 0.976022242 | RGS3 | protein_coding | 48.79 | 0.000431247 | 0.000431247 | ENSG00000138835 | n.s. |
| ENSG00000134108 | 1047.53 | -0.044930653 | 0.301934 | -0.14881 | 0.881704 | 0.978242302 | ARL8B | protein_coding | 40.82 | -0.000429248 | 0.000429248 | ENSG00000134108 | n.s. |
| ENSG00000132773 | 518.286 | 0.037356481 | 0.20785 | 0.1797279 | 0.857366 | 0.974056274 | TOE1 | protein_coding | 53.53 | 0.00042646 | 0.00042646 | ENSG00000132773 | n.s. |
| ENSG00000162971 | 1004.662 | -0.041481305 | 0.267002 | -0.155359 | 0.876538 | 0.976610381 | TYW5 | protein_coding | 37.41 | -0.000426372 | 0.000426372 | ENSG00000162971 | n.s. |
| ENSG00000207014 | 4.28006 | -0.077066666 | 0.795302 | -0.096902 | 0.922804 | 0.987349084 | SNORD116-3 | snoRNA | 45.26 | -0.000426123 | 0.000426123 | ENSG00000207014 | n.s. |
| ENSG00000100644 | 4194.944 | 0.040260659 | 0.254351 | 0.1582881 | 0.87423 | 0.976022242 | HIF1A | protein_coding | 37.25 | 0.000424359 | 0.000424359 | ENSG00000100644 | n.s. |
| ENSG00000196418 | 2179.409 | 0.044493227 | 0.299698 | 0.1484604 | 0.881979 | 0.978288882 | ZNF124 | protein_coding | 42.19 | 0.000424149 | 0.000424149 | ENSG00000196418 | n.s. |
| ENSG00000198830 | 3199.507 | 0.038637182 | 0.221537 | 0.174405 | 0.861547 | 0.975075415 | HMGN2 | protein_coding | 48.34 | 0.000423533 | 0.000423533 | ENSG00000198830 | n.s. |
| ENSG00000119185 | 659.1663 | -0.04570733 | 0.311846 | -0.14657 | 0.883471 | 0.978903461 | ITGB1BP1 | protein_coding | 45.09 | -0.000423256 | 0.000423256 | ENSG00000119185 | n.s. |
| ENSG00000196653 | 509.2042 | 0.044092863 | 0.293252 | 0.1503583 | 0.880482 | 0.978149421 | ZNF502 | protein_coding | 40.62 | 0.000423062 | 0.000423062 | ENSG00000196653 | n.s. |
| ENSG00000241420 | 29.6761 | 0.047678397 | 0.355469 | 0.134128 | 0.893301 | 0.979788048 | RN7SL505P | misc_RNA | 54.61 | 0.000422805 | 0.000422805 | ENSG00000241420 | n.s. |
| ENSG00000141252 | 80.38367 | 0.041622806 | 0.269974 | 0.1541731 | 0.877473 | 0.976942004 | VPS53 | protein_coding | 45.86 | 0.00042169 | 0.00042169 | ENSG00000141252 | n.s. |
| ENSG00000138698 | 1095.017 | -0.044085058 | 0.294811 | -0.149537 | 0.88113 | 0.978242302 | RAP1GDS1 | protein_coding | 36.01 | -0.000421169 | 0.000421169 | ENSG00000138698 | n.s. |
| ENSG00000002330 | 220.7473 | 0.04563418 | 0.312477 | 0.1460403 | 0.88389 | 0.979008503 | BAD | protein_coding | 53.92 | 0.000420452 | 0.000420452 | ENSG00000002330 | n.s. |
| ENSG00000145194 | 138.8693 | 0.045374906 | 0.310018 | 0.146362 | 0.883636 | 0.978903461 | ECE2 | protein_coding | 51.09 | 0.000420178 | 0.000420178 | ENSG00000145194 | n.s. |
| ENSG00000102038 | 1698.44 | -0.04202584 | 0.275085 | -0.152774 | 0.878576 | 0.977264479 | SMARCA1 | protein_coding | 36.87 | -0.000419749 | 0.000419749 | ENSG00000102038 | n.s. |
| ENSG00000110711 | 446.4785 | -0.047186712 | 0.335031 | -0.140843 | 0.887994 | 0.979788048 | AIP | protein_coding | 56.24 | -0.000418445 | 0.000418445 | ENSG00000110711 | n.s. |
| ENSG00000175536 | 88.49219 | 0.040632558 | 0.259998 | 0.1562801 | 0.875812 | 0.97660852 | LIPT2 | protein_coding | 55.44 | 0.000417682 | 0.000417682 | ENSG00000175536 | n.s. |
| ENSG00000143337 | 1230.62 | 0.039532472 | 0.24945 | 0.1584788 | 0.87408 | 0.976022242 | TOR1AIP1 | protein_coding | 38.76 | 0.000416684 | 0.000416684 | ENSG00000143337 | n.s. |
| ENSG00000243725 | 977.3636 | -0.040439312 | 0.259923 | -0.155582 | 0.876363 | 0.976610381 | TTC4 | protein_coding | 42.06 | -0.000415662 | 0.000415662 | ENSG00000243725 | n.s. |
| ENSG00000084207 | 3949.79 | -0.046276004 | 0.321765 | -0.143819 | 0.885643 | 0.979542794 | GSTP1 | protein_coding | 63.05 | -0.000415401 | 0.000415401 | ENSG00000084207 | n.s. |
| ENSG00000164211 | 2896.803 | 0.040313256 | 0.257417 | 0.1566066 | 0.875555 | 0.976582458 | STARD4 | protein_coding | 36.15 | 0.000414867 | 0.000414867 | ENSG00000164211 | n.s. |
| ENSG00000198839 | 496.315 | -0.040301321 | 0.256817 | -0.156926 | 0.875303 | 0.976582458 | ZNF277 | protein_coding | 37.36 | -0.000414744 | 0.000414744 | ENSG00000198839 | n.s. |
| ENSG00000140941 | 502.4308 | -0.04030596 | 0.25891 | -0.155676 | 0.876289 | 0.976610381 | MAP1LC3B | protein_coding | 47.47 | -0.000414291 | 0.000414291 | ENSG00000140941 | n.s. |
| ENSG00000235159 | 194.09 | -0.039323552 | 0.249034 | -0.157904 | 0.874532 | 0.976119374 | | lincRNA | 41.72 | -0.000412782 | 0.000412782 | ENSG00000235159 | n.s. |
| ENSG00000184897 | 2783.996 | 0.039230454 | 0.248215 | 0.1580506 | 0.874417 | 0.976068897 | H1FX | protein_coding | 68.61 | 0.000412686 | 0.000412686 | ENSG00000184897 | n.s. |
| ENSG00000258655 | 119.4484 | 0.040096617 | 0.258486 | 0.1551212 | 0.876726 | 0.976610381 | ARHGAP5-AS1 | antisense | 46 | 0.00041214 | 0.00041214 | ENSG00000258655 | n.s. |

| | | | | | | | | | | | | | |
|---|---|---|---|---|---|---|---|---|---|---|---|---|---|
| ENSG00000114023 | 270.296 | -0.046294106 | 0.344518 | -0.134374 | 0.893107 | 0.979788048 | FAM162A | protein_coding | 38.78 | -0.00041053 | 0.00041053 | ENSG00000114023 | n.s. |
| ENSG00000163719 | 416.7577 | -0.041924749 | 0.276608 | -0.151567 | 0.879528 | 0.977776254 | MTMR14 | protein_coding | 49.36 | -0.000409207 | 0.000409207 | ENSG00000163719 | n.s. |
| ENSG00000231473 | 52.93801 | -0.045567632 | 0.318234 | -0.143189 | 0.886141 | 0.979542794 | LINC00441 | lincRNA | 39.88 | -0.000409042 | 0.000409042 | ENSG00000231473 | n.s. |
| ENSG00000124535 | 1040.949 | -0.039758385 | 0.255926 | -0.155351 | 0.876544 | 0.976610381 | WRNIP1 | protein_coding | 44.69 | -0.000408663 | 0.000408663 | ENSG00000124535 | n.s. |
| ENSG00000050165 | 130.9959 | -0.042745825 | 0.286245 | -0.149333 | 0.881291 | 0.978242302 | DKK3 | protein_coding | 49.18 | -0.000408375 | 0.000408375 | ENSG00000050165 | n.s. |
| ENSG00000181264 | 315.0603 | 0.04188984 | 0.277493 | 0.1509582 | 0.880009 | 0.977868793 | TMEM136 | protein_coding | 43.45 | 0.000407145 | 0.000407145 | ENSG00000181264 | n.s. |
| ENSG00000132436 | 1872.031 | -0.038535335 | 0.238588 | -0.161514 | 0.871689 | 0.976022242 | FIGNL1 | protein_coding | 43.84 | -0.000406173 | 0.000406173 | ENSG00000132436 | n.s. |
| ENSG00000112078 | 2324.649 | -0.042279457 | 0.283102 | -0.149343 | 0.881283 | 0.978242302 | KCTD20 | protein_coding | 42.39 | -0.000403919 | 0.000403919 | ENSG00000112078 | n.s. |
| ENSG00000174498 | 189.5186 | 0.038301363 | 0.238176 | 0.1608112 | 0.872242 | 0.976022242 | IGDCC3 | protein_coding | 51.86 | 0.000403707 | 0.000403707 | ENSG00000174498 | n.s. |
| ENSG00000266594 | 9.526758 | 0.057518179 | 0.485789 | 0.1184015 | 0.90575 | 0.983983433 | MIR4766 | miRNA | 36.84 | 0.00040333 | 0.00040333 | ENSG00000266594 | n.s. |
| ENSG00000171723 | 361.8468 | 0.041400791 | 0.274068 | 0.1510605 | 0.879928 | 0.977868793 | GPHN | protein_coding | 36.43 | 0.000402391 | 0.000402391 | ENSG00000171723 | n.s. |
| ENSG00000138750 | 1700.939 | -0.041930189 | 0.279489 | -0.150025 | 0.880745 | 0.978189412 | NUP54 | protein_coding | 36.8 | -0.000401567 | 0.000401567 | ENSG00000138750 | n.s. |
| ENSG00000116096 | 252.7437 | -0.042639501 | 0.288949 | -0.147568 | 0.882684 | 0.978550055 | SPR | protein_coding | 54.78 | -0.000401534 | 0.000401534 | ENSG00000116096 | n.s. |
| ENSG00000153391 | 269.8323 | -0.037953248 | 0.235502 | -0.161159 | 0.871968 | 0.976022242 | INO80C | protein_coding | 42.29 | -0.000400038 | 0.000400038 | ENSG00000153391 | n.s. |
| ENSG00000166333 | 381.5539 | -0.04249225 | 0.288646 | -0.147212 | 0.882964 | 0.978576629 | ILK | protein_coding | 49.78 | -0.000399646 | 0.000399646 | ENSG00000166333 | n.s. |
| ENSG00000167775 | 407.2298 | -0.043361027 | 0.297188 | -0.145904 | 0.883997 | 0.979008503 | CD320 | protein_coding | 58.35 | -0.000399508 | 0.000399508 | ENSG00000167775 | n.s. |
| ENSG00000106853 | 143.0451 | -0.042474863 | 0.288589 | -0.147181 | 0.882989 | 0.978576629 | PTGR1 | protein_coding | 43.23 | -0.000399483 | 0.000399483 | ENSG00000106853 | n.s. |
| ENSG00000273587 | 6.937935 | -0.069851195 | 0.690878 | -0.101105 | 0.919467 | 0.986968832 | SNORA78 | snoRNA | 72.44 | -0.000397912 | 0.000397912 | ENSG00000273587 | n.s. |
| ENSG00000099326 | 300.3457 | -0.03973807 | 0.260072 | -0.152797 | 0.878559 | 0.977264479 | MZF1 | protein_coding | 55.33 | -0.000396899 | 0.000396899 | ENSG00000099326 | n.s. |
| ENSG00000137413 | 209.8486 | -0.037541969 | 0.230021 | -0.163211 | 0.870352 | 0.976022242 | TAF8 | protein_coding | 45.88 | -0.000395703 | 0.000395703 | ENSG00000137413 | n.s. |
| ENSG00000077152 | 2433.78 | -0.04460539 | 0.325271 | -0.137133 | 0.890926 | 0.979788048 | UBE2T | protein_coding | 40.71 | -0.000395554 | 0.000395554 | ENSG00000077152 | n.s. |
| ENSG00000131966 | 1046.343 | 0.044602374 | 0.31772 | 0.1403826 | 0.888358 | 0.979788048 | ACTR10 | protein_coding | 38.67 | 0.000395528 | 0.000395528 | ENSG00000131966 | n.s. |
| ENSG00000115875 | 11696.32 | 0.037167033 | 0.217937 | 0.1705402 | 0.864585 | 0.975860117 | SRSF7 | protein_coding | 41.45 | 0.000394433 | 0.000394433 | ENSG00000115875 | n.s. |
| ENSG00000200413 | 7.794823 | -0.056994141 | 0.488766 | -0.116608 | 0.907171 | 0.984198171 | SNORD114-26 | snoRNA | 47.22 | -0.000394254 | 0.000394254 | ENSG00000200413 | n.s. |
| ENSG00000114942 | 12877.28 | 0.044323606 | 0.313946 | 0.1411822 | 0.887726 | 0.979788048 | EEF1B2 | protein_coding | 45.25 | 0.000393056 | 0.000393056 | ENSG00000114942 | n.s. |
| ENSG00000276934 | 31.44409 | -0.044287104 | 0.318588 | -0.139011 | 0.889442 | 0.979788048 | | sense_intronic | 38.72 | -0.000392732 | 0.000392732 | ENSG00000276934 | n.s. |
| ENSG00000134313 | 3513.563 | -0.040228723 | 0.266322 | -0.151053 | 0.879934 | 0.977868793 | KIDINS220 | protein_coding | 38.39 | -0.000391 | 0.000391 | ENSG00000134313 | n.s. |
| ENSG00000173120 | 707.8614 | -0.036906382 | 0.221358 | -0.166727 | 0.867585 | 0.975964429 | KDM2A | protein_coding | 42.99 | -0.000389953 | 0.000389953 | ENSG00000173120 | n.s. |
| ENSG00000198901 | 5948.082 | 0.040705719 | 0.27372 | 0.1487132 | 0.88178 | 0.978248817 | PRC1 | protein_coding | 45.15 | 0.000388767 | 0.000388767 | ENSG00000198901 | n.s. |
| ENSG00000058804 | 2651.326 | -0.041971881 | 0.286679 | -0.146407 | 0.8836 | 0.978903461 | NDC1 | protein_coding | 40.26 | -0.000388665 | 0.000388665 | ENSG00000058804 | n.s. |
| ENSG00000149313 | 2113.866 | 0.039117849 | 0.256947 | 0.1522407 | 0.878997 | 0.977420056 | AASDHPPT | protein_coding | 36.57 | 0.000388 | 0.000388 | ENSG00000149313 | n.s. |
| ENSG00000179958 | 859.1023 | 0.043736909 | 0.308478 | 0.141783 | 0.887251 | 0.979788048 | DCTPP1 | protein_coding | 53.23 | 0.000387853 | 0.000387853 | ENSG00000179958 | n.s. |

| | | | | | | | | | | | | | |
|---|---|---|---|---|---|---|---|---|---|---|---|---|---|
| ENSG00000165169 | 423.8066 | 0.043696599 | 0.309634 | 0.1411234 | 0.887772 | 0.979788048 | DYNLT3 | protein_coding | 37.62 | 0.000387495 | 0.000387495 | ENSG00000165169 | n.s. |
| ENSG00000048052 | 1948.249 | 0.036736737 | 0.231602 | 0.1586202 | 0.873968 | 0.976022242 | HDAC9 | protein_coding | 36.13 | 0.000387216 | 0.000387216 | ENSG00000048052 | n.s. |
| ENSG00000277182 | 60.39638 | -0.043412838 | 0.307783 | -0.14105 | 0.88783 | 0.979788048 | | antisense | 56.2 | -0.000384979 | 0.000384979 | ENSG00000277182 | n.s. |
| ENSG00000104093 | 3205.12 | -0.040174029 | 0.268714 | -0.149505 | 0.881155 | 0.978242302 | DMXL2 | protein_coding | 36.95 | -0.000383805 | 0.000383805 | ENSG00000104093 | n.s. |
| ENSG00000111845 | 982.505 | -0.040153689 | 0.268945 | -0.149301 | 0.881316 | 0.978242302 | PAK1IP1 | protein_coding | 40.83 | -0.000383611 | 0.000383611 | ENSG00000111845 | n.s. |
| ENSG00000216866 | 57.64735 | 0.046220736 | 0.357134 | 0.1294212 | 0.897024 | 0.981075379 | RPS2P55 | processed_pseudogene | 60.82 | 0.000383522 | 0.000383522 | ENSG00000216866 | n.s. |
| ENSG00000182149 | 1379.477 | 0.041368251 | 0.282513 | 0.1464296 | 0.883582 | 0.978903461 | IST1 | protein_coding | 45.18 | 0.000383076 | 0.000383076 | ENSG00000182149 | n.s. |
| ENSG00000100911 | 474.7942 | -0.042532799 | 0.295161 | -0.144101 | 0.885421 | 0.979542794 | PSME2 | protein_coding | 52.29 | -0.000381799 | 0.000381799 | ENSG00000100911 | n.s. |
| ENSG00000074319 | 746.0397 | 0.043008937 | 0.314324 | 0.13683 | 0.891165 | 0.979788048 | TSG101 | protein_coding | 39.7 | 0.000381397 | 0.000381397 | ENSG00000074319 | n.s. |
| ENSG00000183718 | 794.7241 | -0.036149665 | 0.222336 | -0.16259 | 0.870841 | 0.976022242 | TRIM52 | protein_coding | 43.45 | -0.000381028 | 0.000381028 | ENSG00000183718 | n.s. |
| ENSG00000166770 | 511.7945 | 0.041773472 | 0.288336 | 0.1448777 | 0.884807 | 0.97927567 | ZNF667-AS1 | lincRNA | 43.34 | 0.000379931 | 0.000379931 | ENSG00000166770 | n.s. |
| ENSG00000187630 | 82.70991 | -0.042708571 | 0.313353 | -0.136295 | 0.891588 | 0.979788048 | DHRS4L2 | protein_coding | 43.65 | -0.000378734 | 0.000378734 | ENSG00000187630 | n.s. |
| ENSG00000135070 | 1296.219 | 0.042099421 | 0.293642 | 0.1433697 | 0.885998 | 0.979542794 | ISCA1 | protein_coding | 41.36 | 0.000377909 | 0.000377909 | ENSG00000135070 | n.s. |
| ENSG00000172613 | 160.0523 | -0.039518605 | 0.265461 | -0.148868 | 0.881658 | 0.978242302 | RAD9A | protein_coding | 52.03 | -0.000377543 | 0.000377543 | ENSG00000172613 | n.s. |
| ENSG00000153130 | 1217.485 | 0.042546515 | 0.307793 | 0.1382311 | 0.890058 | 0.979788048 | SCOC | protein_coding | 40.94 | 0.000377297 | 0.000377297 | ENSG00000153130 | n.s. |
| ENSG00000085231 | 407.4629 | -0.042319792 | 0.311291 | -0.135949 | 0.891861 | 0.979788048 | AK6 | protein_coding | 41.64 | -0.000375286 | 0.000375286 | ENSG00000085231 | n.s. |
| ENSG00000158301 | 709.6163 | 0.040707262 | 0.278833 | 0.1459915 | 0.883928 | 0.979008503 | GPRASP2 | protein_coding | 51.49 | 0.000375058 | 0.000375058 | ENSG00000158301 | n.s. |
| ENSG00000101158 | 1618.194 | -0.039488033 | 0.266877 | -0.147964 | 0.882372 | 0.978387771 | NELFCD | protein_coding | 48.57 | -0.000374701 | 0.000374701 | ENSG00000101158 | n.s. |
| ENSG00000213719 | 1040.907 | -0.042768262 | 0.321087 | -0.133198 | 0.894037 | 0.980054355 | CLIC1 | protein_coding | 51 | -0.000374215 | 0.000374215 | ENSG00000213719 | n.s. |
| ENSG00000154059 | 1102.424 | 0.039046457 | 0.260571 | 0.1498495 | 0.880883 | 0.978189412 | IMPACT | protein_coding | 38.35 | 0.00037395 | 0.00037395 | ENSG00000154059 | n.s. |
| ENSG00000107581 | 9702.904 | -0.036659419 | 0.237197 | -0.154552 | 0.877174 | 0.976797347 | EIF3A | protein_coding | 41.68 | -0.000373762 | 0.000373762 | ENSG00000107581 | n.s. |
| ENSG00000163605 | 2749.89 | -0.040520203 | 0.278345 | -0.145575 | 0.884257 | 0.979008503 | PPP4R2 | protein_coding | 38.82 | -0.000373334 | 0.000373334 | ENSG00000163605 | n.s. |
| ENSG00000156298 | 473.2437 | 0.042077129 | 0.304286 | 0.1382816 | 0.890018 | 0.979788048 | TSPAN7 | protein_coding | 39.6 | 0.000373134 | 0.000373134 | ENSG00000156298 | n.s. |
| ENSG00000272277 | 62.08253 | 0.041873293 | 0.298249 | 0.140397 | 0.888346 | 0.979788048 | | antisense | 65.06 | 0.000371327 | 0.000371327 | ENSG00000272277 | n.s. |
| ENSG00000168894 | 528.2551 | -0.041809586 | 0.307509 | -0.135962 | 0.891851 | 0.979788048 | RNF181 | protein_coding | 54.37 | -0.000370762 | 0.000370762 | ENSG00000168894 | n.s. |
| ENSG00000116191 | 553.6803 | -0.041756192 | 0.301011 | -0.13872 | 0.889672 | 0.979788048 | RALGPS2 | protein_coding | 36.82 | -0.000370288 | 0.000370288 | ENSG00000116191 | n.s. |
| ENSG00000228649 | 501.6132 | -0.034809202 | 0.203097 | -0.171392 | 0.863916 | 0.975847571 | SNHG26 | processed_transcript | 43.38 | -0.000369605 | 0.000369605 | ENSG00000228649 | n.s. |
| ENSG00000010270 | 806.5723 | -0.041652585 | 0.310749 | -0.134039 | 0.893371 | 0.979788048 | STARD3NL | protein_coding | 38.4 | -0.000369369 | 0.000369369 | ENSG00000010270 | n.s. |
| ENSG00000181751 | 1085.241 | -0.04008124 | 0.275417 | -0.145529 | 0.884293 | 0.979008503 | C5orf30 | protein_coding | 37.54 | -0.00036929 | 0.00036929 | ENSG00000181751 | n.s. |
| ENSG00000167085 | 5195.186 | -0.041637429 | 0.297466 | -0.139974 | 0.888681 | 0.979788048 | PHB | protein_coding | 48.42 | -0.000369235 | 0.000369235 | ENSG00000167085 | n.s. |
| ENSG00000198718 | 556.0105 | -0.035006788 | 0.216846 | -0.161436 | 0.87175 | 0.976022242 | TOGARAM1 | protein_coding | 37.85 | -0.000368982 | 0.000368982 | ENSG00000198718 | n.s. |
| ENSG00000108961 | 195.3512 | 0.04159674 | 0.296491 | 0.1402966 | 0.888426 | 0.979788048 | RANGRF | protein_coding | 59.59 | 0.000368874 | 0.000368874 | ENSG00000108961 | n.s. |

| | | | | | | | | | | | | | |
|---|---|---|---|---|---|---|---|---|---|---|---|---|---|
| ENSG00000100387 | 1180.024 | -0.04257711 | 0.322517 | -0.132015 | 0.894972 | 0.980308504 | RBX1 | protein_coding | 44.92 | -0.000367748 | 0.000367748 | ENSG00000100387 | n.s. |
| ENSG00000108654 | 42540.12 | 0.035847164 | 0.231538 | 0.1548219 | 0.876962 | 0.97668357 | DDX5 | protein_coding | 45.11 | 0.000367294 | 0.000367294 | ENSG00000108654 | n.s. |
| ENSG00000261324 | 249.8686 | 0.034646917 | 0.218715 | 0.1584111 | 0.874133 | 0.976022242 | | sense_overlapping | 36.81 | 0.000365188 | 0.000365188 | ENSG00000261324 | n.s. |
| ENSG00000068001 | 829.9464 | -0.03847214 | 0.260275 | -0.147813 | 0.88249 | 0.978412868 | HYAL2 | protein_coding | 61.05 | -0.000364633 | 0.000364633 | ENSG00000068001 | n.s. |
| ENSG00000183011 | 536.913 | -0.041718003 | 0.314467 | -0.132662 | 0.89446 | 0.980194243 | NAA38 | protein_coding | 44.28 | -0.00036244 | 0.00036244 | ENSG00000183011 | n.s. |
| ENSG00000251485 | 58.32061 | 0.041057849 | 0.307098 | 0.1336963 | 0.893643 | 0.979889821 | | processed_pseudogene | 49.11 | 0.000362243 | 0.000362243 | ENSG00000251485 | n.s. |
| ENSG00000269889 | 85.41812 | 0.040751572 | 0.301885 | 0.1349904 | 0.892619 | 0.979788048 | | antisense | 21.04 | 0.000361379 | 0.000361379 | ENSG00000269889 | n.s. |
| ENSG00000179010 | 3285.68 | -0.040685785 | 0.288423 | -0.141063 | 0.88782 | 0.979788048 | MRFAP1 | protein_coding | 53.11 | -0.000360796 | 0.000360796 | ENSG00000179010 | n.s. |
| ENSG00000064300 | 915.2438 | 0.039104161 | 0.268522 | 0.1456272 | 0.884216 | 0.979008503 | NGFR | protein_coding | 56.74 | 0.000360288 | 0.000360288 | ENSG00000064300 | n.s. |
| ENSG00000029363 | 15172.73 | 0.037035692 | 0.244902 | 0.1512264 | 0.879797 | 0.977868793 | BCLAF1 | protein_coding | 35.59 | 0.000359965 | 0.000359965 | ENSG00000029363 | n.s. |
| ENSG00000162775 | 4068.175 | -0.033766996 | 0.196723 | -0.171647 | 0.863715 | 0.975753891 | RBM15 | protein_coding | 44.01 | -0.000359946 | 0.000359946 | ENSG00000162775 | n.s. |
| ENSG00000128567 | 263.3101 | -0.037876553 | 0.255582 | -0.148197 | 0.882187 | 0.978387771 | PODXL | protein_coding | 50.47 | -0.00035941 | 0.00035941 | ENSG00000128567 | n.s. |
| ENSG00000224349 | 88.98875 | 0.040472762 | 0.285258 | 0.1418812 | 0.887174 | 0.979788048 | | antisense | 64.38 | 0.000358907 | 0.000358907 | ENSG00000224349 | n.s. |
| ENSG00000142945 | 767.587 | 0.040324967 | 0.287287 | 0.1403649 | 0.888372 | 0.979788048 | KIF2C | protein_coding | 49.08 | 0.000357596 | 0.000357596 | ENSG00000142945 | n.s. |
| ENSG00000136152 | 923.6574 | -0.037459019 | 0.252419 | -0.1484 | 0.882027 | 0.978288882 | COG3 | protein_coding | 39.15 | -0.000357092 | 0.000357092 | ENSG00000136152 | n.s. |
| ENSG00000065427 | 2885.676 | 0.040238521 | 0.294191 | 0.1367771 | 0.891207 | 0.979788048 | KARS | protein_coding | 45.57 | 0.00035683 | 0.00035683 | ENSG00000065427 | n.s. |
| ENSG00000258445 | 39.94128 | 0.043552115 | 0.338586 | 0.1286296 | 0.897651 | 0.981326694 | | processed_pseudogene | 41.59 | 0.000356534 | 0.000356534 | ENSG00000258445 | n.s. |
| ENSG00000113845 | 1109.861 | 0.040178872 | 0.295355 | 0.1360357 | 0.891793 | 0.979788048 | TIMMDC1 | protein_coding | 38.48 | 0.000356301 | 0.000356301 | ENSG00000113845 | n.s. |
| ENSG00000113851 | 1077.936 | 0.040154125 | 0.29107 | 0.1379534 | 0.890277 | 0.979788048 | CRBN | protein_coding | 37.25 | 0.000356081 | 0.000356081 | ENSG00000113851 | n.s. |
| ENSG00000100324 | 265.422 | -0.039944426 | 0.294286 | -0.135733 | 0.892032 | 0.979788048 | TAB1 | protein_coding | 51.38 | -0.000354222 | 0.000354222 | ENSG00000100324 | n.s. |
| ENSG00000117862 | 1088.445 | 0.039872815 | 0.294084 | 0.1355832 | 0.892151 | 0.979788048 | TXNDC12 | protein_coding | 43.03 | 0.000353587 | 0.000353587 | ENSG00000117862 | n.s. |
| ENSG00000132603 | 989.1623 | 0.036902799 | 0.245997 | 0.1500132 | 0.880754 | 0.978189412 | NIP7 | protein_coding | 45.71 | 0.00035342 | 0.00035342 | ENSG00000132603 | n.s. |
| ENSG00000100897 | 827.0143 | -0.038758737 | 0.267876 | -0.144689 | 0.884956 | 0.97927567 | DCAF11 | protein_coding | 50.99 | -0.000352512 | 0.000352512 | ENSG00000100897 | n.s. |
| ENSG00000143870 | 4842.024 | 0.039741871 | 0.28827 | 0.1378636 | 0.890348 | 0.979788048 | PDIA6 | protein_coding | 46.27 | 0.000352425 | 0.000352425 | ENSG00000143870 | n.s. |
| ENSG00000131558 | 1795.859 | -0.038688735 | 0.267086 | -0.144855 | 0.884825 | 0.97927567 | EXOC4 | protein_coding | 38.3 | -0.000351875 | 0.000351875 | ENSG00000131558 | n.s. |
| ENSG00000179818 | 1188.707 | 0.033536642 | 0.212541 | 0.1577888 | 0.874623 | 0.976142659 | PCBP1-AS1 | processed_transcript | 45.84 | 0.000351689 | 0.000351689 | ENSG00000179818 | n.s. |
| ENSG00000253559 | 56.88599 | 0.039616461 | 0.288858 | 0.1371486 | 0.890913 | 0.979788048 | OSGEPL1-AS1 | antisense | 40.66 | 0.000351313 | 0.000351313 | ENSG00000253559 | n.s. |
| ENSG00000144724 | 816.2103 | 0.035170498 | 0.229842 | 0.1530203 | 0.878382 | 0.977264479 | PTPRG | protein_coding | 40.79 | 0.000351279 | 0.000351279 | ENSG00000144724 | n.s. |
| ENSG00000259318 | 51.32492 | -0.041417407 | 0.31731 | -0.130526 | 0.89615 | 0.980688401 | | processed_pseudogene | 45.92 | -0.000350762 | 0.000350762 | ENSG00000259318 | n.s. |
| ENSG00000221988 | 216.3972 | 0.03954856 | 0.29189 | 0.1354912 | 0.892224 | 0.979788048 | PPT2 | protein_coding | 50.59 | 0.000350711 | 0.000350711 | ENSG00000221988 | n.s. |
| ENSG00000202538 | 610.1381 | 0.033256873 | 0.204524 | 0.1626066 | 0.870828 | 0.976022242 | RNU4-2 | snRNA | 45.39 | 0.000350537 | 0.000350537 | ENSG00000202538 | n.s. |
| ENSG00000143553 | 365.7081 | -0.041286018 | 0.314995 | -0.131069 | 0.895721 | 0.98066548 | SNAPIN | protein_coding | 51.31 | -0.000350069 | 0.000350069 | ENSG00000143553 | n.s. |

| | | | | | | | | | | | | | |
|---|---|---|---|---|---|---|---|---|---|---|---|---|---|
| ENSG00000143621 | 10992.64 | 0.039405004 | 0.283422 | 0.1390329 | 0.889424 | 0.979788048 | ILF2 | protein_coding | 44.37 | 0.000349438 | 0.000349438 | ENSG00000143621 | n.s. |
| ENSG00000115234 | 1142.223 | 0.038394974 | 0.265208 | 0.1447729 | 0.88489 | 0.97927567 | SNX17 | protein_coding | 53.11 | 0.000349204 | 0.000349204 | ENSG00000115234 | n.s. |
| ENSG00000105607 | 375.5629 | -0.039375245 | 0.281519 | -0.139867 | 0.888765 | 0.979788048 | GCDH | protein_coding | 51.4 | -0.000349174 | 0.000349174 | ENSG00000105607 | n.s. |
| ENSG00000134452 | 719.1426 | 0.03927318 | 0.282019 | 0.1392573 | 0.889247 | 0.979788048 | FBXO18 | protein_coding | 45.82 | 0.000348269 | 0.000348269 | ENSG00000134452 | n.s. |
| ENSG00000115239 | 251.2233 | -0.034628351 | 0.225613 | -0.153486 | 0.878015 | 0.977108686 | ASB3 | protein_coding | 39.31 | -0.000348262 | 0.000348262 | ENSG00000115239 | n.s. |
| ENSG00000179271 | 1493.692 | -0.039248113 | 0.292422 | -0.134217 | 0.893231 | 0.979788048 | GADD45GIP1 | protein_coding | 55.03 | -0.000348047 | 0.000348047 | ENSG00000179271 | n.s. |
| ENSG00000102158 | 640.0326 | -0.039157351 | 0.288338 | -0.135804 | 0.891977 | 0.979788048 | MAGT1 | protein_coding | 40.93 | -0.000347242 | 0.000347242 | ENSG00000102158 | n.s. |
| ENSG00000166839 | 190.5854 | -0.039150178 | 0.290147 | -0.134932 | 0.892666 | 0.979788048 | ANKDD1A | protein_coding | 46.75 | -0.000347178 | 0.000347178 | ENSG00000166839 | n.s. |
| ENSG00000132406 | 519.8299 | 0.038604555 | 0.2681 | 0.1439929 | 0.885506 | 0.979542794 | TMEM128 | protein_coding | 41.69 | 0.000346537 | 0.000346537 | ENSG00000132406 | n.s. |
| ENSG00000149357 | 446.6381 | -0.039006304 | 0.290014 | -0.134498 | 0.893009 | 0.979788048 | LAMTOR1 | protein_coding | 48.06 | -0.000345903 | 0.000345903 | ENSG00000149357 | n.s. |
| ENSG00000129518 | 468.732 | -0.040797591 | 0.312621 | -0.130502 | 0.896169 | 0.980688401 | EAPP | protein_coding | 44.09 | -0.000345513 | 0.000345513 | ENSG00000129518 | n.s. |
| ENSG00000164237 | 289.1349 | -0.038948861 | 0.287893 | -0.135289 | 0.892383 | 0.979788048 | CMBL | protein_coding | 46.43 | -0.000345393 | 0.000345393 | ENSG00000164237 | n.s. |
| ENSG00000159921 | 863.6781 | -0.03893763 | 0.278943 | -0.13959 | 0.888984 | 0.979788048 | GNE | protein_coding | 43.82 | -0.000345294 | 0.000345294 | ENSG00000159921 | n.s. |
| ENSG00000167081 | 1179.022 | -0.036635721 | 0.248814 | -0.147241 | 0.882942 | 0.978576629 | PBX3 | protein_coding | 37.29 | -0.000344565 | 0.000344565 | ENSG00000167081 | n.s. |
| ENSG00000119013 | 1191.81 | -0.04269879 | 0.334031 | -0.127829 | 0.898284 | 0.981590948 | NDUFB3 | protein_coding | 41.49 | -0.000344556 | 0.000344556 | ENSG00000119013 | n.s. |
| ENSG00000186376 | 350.8685 | 0.038366299 | 0.267842 | 0.1432422 | 0.886099 | 0.979542794 | ZNF75D | protein_coding | 41.67 | 0.000344398 | 0.000344398 | ENSG00000186376 | n.s. |
| ENSG00000114784 | 465.2603 | -0.03717965 | 0.253583 | -0.146617 | 0.883434 | 0.978903461 | EIF1B | protein_coding | 41.77 | -0.000344289 | 0.000344289 | ENSG00000114784 | n.s. |
| ENSG00000140299 | 2150.93 | -0.038756907 | 0.276086 | -0.14038 | 0.88836 | 0.979788048 | BNIP2 | protein_coding | 38.55 | -0.000343691 | 0.000343691 | ENSG00000140299 | n.s. |
| ENSG00000181045 | 139.9949 | -0.038751084 | 0.274963 | -0.140932 | 0.887924 | 0.979788048 | SLC26A11 | protein_coding | 53.49 | -0.000343639 | 0.000343639 | ENSG00000181045 | n.s. |
| ENSG00000163714 | 9783.257 | -0.038215194 | 0.265578 | -0.143895 | 0.885584 | 0.979542794 | U2SURP | protein_coding | 38.45 | -0.000343042 | 0.000343042 | ENSG00000163714 | n.s. |
| ENSG00000264895 | 770.0446 | 0.035931719 | 0.241899 | 0.1485399 | 0.881917 | 0.978288882 | | sense_intronic | 36.41 | 0.000342533 | 0.000342533 | ENSG00000264895 | n.s. |
| ENSG00000216813 | 29.75374 | -0.045123018 | 0.361949 | -0.124667 | 0.900787 | 0.982691813 | | processed_pseudogene | 44.58 | -0.000342153 | 0.000342153 | ENSG00000216813 | n.s. |
| ENSG00000113971 | 122.354 | -0.038563497 | 0.271978 | -0.141789 | 0.887247 | 0.979788048 | NPHP3 | protein_coding | 36.77 | -0.000341976 | 0.000341976 | ENSG00000113971 | n.s. |
| ENSG00000072803 | 1459.536 | -0.038554277 | 0.274581 | -0.140411 | 0.888335 | 0.979788048 | FBXW11 | protein_coding | 39.72 | -0.000341894 | 0.000341894 | ENSG00000072803 | n.s. |
| ENSG00000157514 | 410.5122 | 0.038375567 | 0.282883 | 0.1356588 | 0.892091 | 0.979788048 | TSC22D3 | protein_coding | 48.75 | 0.000340309 | 0.000340309 | ENSG00000157514 | n.s. |
| ENSG00000022277 | 1200.805 | 0.038323423 | 0.280262 | 0.1367416 | 0.891235 | 0.979788048 | RTFDC1 | protein_coding | 43.71 | 0.000339847 | 0.000339847 | ENSG00000022277 | n.s. |
| ENSG00000160058 | 387.9715 | 0.038727102 | 0.290377 | 0.1333684 | 0.893902 | 0.980032055 | BSDC1 | protein_coding | 47.99 | 0.000339239 | 0.000339239 | ENSG00000160058 | n.s. |
| ENSG00000141127 | 1132.382 | -0.039033982 | 0.294586 | -0.132505 | 0.894585 | 0.980194243 | PRPSAP2 | protein_coding | 44.41 | -0.000339121 | 0.000339121 | ENSG00000141127 | n.s. |
| ENSG00000165516 | 1153.614 | 0.038234968 | 0.284649 | 0.1343231 | 0.893147 | 0.979788048 | KLHDC2 | protein_coding | 41.17 | 0.000339062 | 0.000339062 | ENSG00000165516 | n.s. |
| ENSG00000082146 | 549.879 | 0.038142865 | 0.283718 | 0.1344393 | 0.893055 | 0.979788048 | STRADB | protein_coding | 38.23 | 0.000338246 | 0.000338246 | ENSG00000082146 | n.s. |
| ENSG00000144824 | 2159.29 | 0.038094486 | 0.281992 | 0.1350905 | 0.89254 | 0.979788048 | PHLDB2 | protein_coding | 38.78 | 0.000337817 | 0.000337817 | ENSG00000144824 | n.s. |
| ENSG00000127334 | 7715.711 | 0.035890397 | 0.243423 | 0.1474402 | 0.882785 | 0.978576629 | DYRK2 | protein_coding | 43.53 | 0.000337555 | 0.000337555 | ENSG00000127334 | n.s. |

| | | | | | | | | | | | | | |
|---|---|---|---|---|---|---|---|---|---|---|---|---|---|
| ENSG00000008382 | 117.5672 | 0.038032776 | 0.281396 | 0.1351577 | 0.892487 | 0.979788048 | MPND | protein_coding | 55.95 | 0.000337269 | 0.000337269 | ENSG00000008382 | n.s. |
| ENSG00000109332 | 7264.371 | -0.037564003 | 0.262716 | -0.142983 | 0.886303 | 0.979542794 | UBE2D3 | protein_coding | 38.48 | -0.000337197 | 0.000337197 | ENSG00000109332 | n.s. |
| ENSG00000107438 | 456.198 | -0.037883604 | 0.273016 | -0.13876 | 0.88964 | 0.979788048 | PDLIM1 | protein_coding | 43.74 | -0.000335947 | 0.000335947 | ENSG00000107438 | n.s. |
| ENSG00000136859 | 359.3471 | -0.037748612 | 0.276614 | -0.136467 | 0.891452 | 0.979788048 | ANGPTL2 | protein_coding | 49.92 | -0.00033475 | 0.00033475 | ENSG00000136859 | n.s. |
| ENSG00000183077 | 705.4949 | 0.03773338 | 0.269196 | 0.1401709 | 0.888525 | 0.979788048 | AFMID | protein_coding | 51.08 | 0.000334614 | 0.000334614 | ENSG00000183077 | n.s. |
| ENSG00000120889 | 1378.07 | -0.037730636 | 0.26521 | -0.142267 | 0.886869 | 0.979788048 | TNFRSF10B | protein_coding | 43.23 | -0.00033459 | 0.00033459 | ENSG00000120889 | n.s. |
| ENSG00000111897 | 4857.618 | -0.03768795 | 0.281072 | -0.134086 | 0.893334 | 0.979788048 | SERINC1 | protein_coding | 35.93 | -0.000334212 | 0.000334212 | ENSG00000111897 | n.s. |
| ENSG00000147471 | 367.3575 | -0.037591162 | 0.266571 | -0.141018 | 0.887856 | 0.979788048 | PLPBP | protein_coding | 43.59 | -0.000333353 | 0.000333353 | ENSG00000147471 | n.s. |
| ENSG00000277563 | 5.084603 | -0.070319463 | 0.838003 | -0.083913 | 0.933125 | 0.98914507 | | misc_RNA | 43.18 | -0.000333315 | 0.000333315 | ENSG00000277563 | n.s. |
| ENSG00000165572 | 1400.63 | -0.034276118 | 0.226441 | -0.151369 | 0.879685 | 0.977868793 | KBTBD6 | protein_coding | 42.89 | -0.000333144 | 0.000333144 | ENSG00000165572 | n.s. |
| ENSG00000138780 | 347.2633 | 0.036702151 | 0.254103 | 0.1444382 | 0.885154 | 0.979372226 | GSTCD | protein_coding | 35.68 | 0.000332236 | 0.000332236 | ENSG00000138780 | n.s. |
| ENSG00000174021 | 675.5339 | 0.037433131 | 0.272764 | 0.1372365 | 0.890844 | 0.979788048 | GNG5 | protein_coding | 40.66 | 0.000331952 | 0.000331952 | ENSG00000174021 | n.s. |
| ENSG00000267374 | 144.003 | -0.037406544 | 0.276817 | -0.135131 | 0.892509 | 0.979788048 | | lincRNA | 37.1 | -0.000331716 | 0.000331716 | ENSG00000267374 | n.s. |
| ENSG00000165185 | 936.1843 | -0.037390639 | 0.267293 | -0.139886 | 0.88875 | 0.979788048 | KIAA1958 | protein_coding | 38.95 | -0.000331575 | 0.000331575 | ENSG00000165185 | n.s. |
| ENSG00000201076 | 8.17522 | 0.063973629 | 0.712978 | 0.0897274 | 0.928504 | 0.988157204 | RNU4-51P | snRNA | 37.14 | 0.000330997 | 0.000330997 | ENSG00000201076 | n.s. |
| ENSG00000139354 | 767.7321 | -0.03728006 | 0.274479 | -0.135821 | 0.891963 | 0.979788048 | GAS2L3 | protein_coding | 36.61 | -0.000330594 | 0.000330594 | ENSG00000139354 | n.s. |
| ENSG00000265527 | 20.97767 | 0.047080026 | 0.40108 | 0.1173831 | 0.906556 | 0.983983433 | MIR5690 | miRNA | 30.14 | 0.000330135 | 0.000330135 | ENSG00000265527 | n.s. |
| ENSG00000260774 | 91.45644 | 0.037196642 | 0.261315 | 0.1423443 | 0.886808 | 0.979788048 | | processed_transcript | 46.87 | 0.000329855 | 0.000329855 | ENSG00000260774 | n.s. |
| ENSG00000114054 | 922.3505 | -0.037653206 | 0.282442 | -0.133313 | 0.893946 | 0.980032055 | PCCB | protein_coding | 41.76 | -0.000329832 | 0.000329832 | ENSG00000114054 | n.s. |
| ENSG00000085721 | 2382.202 | -0.037326615 | 0.279172 | -0.133705 | 0.893636 | 0.979889821 | RRN3 | protein_coding | 42.09 | -0.000329324 | 0.000329324 | ENSG00000085721 | n.s. |
| ENSG00000147400 | 317.9246 | -0.042674496 | 0.339667 | -0.125636 | 0.90002 | 0.98243649 | CETN2 | protein_coding | 43.47 | -0.000328402 | 0.000328402 | ENSG00000147400 | n.s. |
| ENSG00000154265 | 80.64891 | 0.037013479 | 0.274884 | 0.1346514 | 0.892887 | 0.979788048 | ABCA5 | protein_coding | 34.61 | 0.00032823 | 0.00032823 | ENSG00000154265 | n.s. |
| ENSG00000185009 | 1326.187 | -0.03687395 | 0.271488 | -0.135822 | 0.891962 | 0.979788048 | AP3M1 | protein_coding | 41.25 | -0.000326993 | 0.000326993 | ENSG00000185009 | n.s. |
| ENSG00000125995 | 635.3224 | -0.039177693 | 0.302056 | -0.129704 | 0.896801 | 0.980999371 | ROMO1 | protein_coding | 50.26 | -0.0003264 | 0.0003264 | ENSG00000125995 | n.s. |
| ENSG00000134905 | 744.4125 | -0.0366953 | 0.262489 | -0.139797 | 0.88882 | 0.979788048 | CARS2 | protein_coding | 48.05 | -0.000325409 | 0.000325409 | ENSG00000134905 | n.s. |
| ENSG00000164649 | 951.2089 | -0.036692635 | 0.263414 | -0.139297 | 0.889216 | 0.979788048 | CDCA7L | protein_coding | 41.49 | -0.000325385 | 0.000325385 | ENSG00000164649 | n.s. |
| ENSG00000234545 | 1521.487 | 0.035299414 | 0.242077 | 0.1458188 | 0.884064 | 0.979008503 | FAM133B | protein_coding | 37.43 | 0.000325232 | 0.000325232 | ENSG00000234545 | n.s. |
| ENSG00000162694 | 1099.453 | 0.038178087 | 0.291562 | 0.1309435 | 0.89582 | 0.980688401 | EXTL2 | protein_coding | 38.7 | 0.000323329 | 0.000323329 | ENSG00000162694 | n.s. |
| ENSG00000004487 | 3614.859 | -0.036314051 | 0.268185 | -0.135407 | 0.89229 | 0.979788048 | KDM1A | protein_coding | 39.16 | -0.000322028 | 0.000322028 | ENSG00000004487 | n.s. |
| ENSG00000214455 | 44.5733 | -0.042375913 | 0.338982 | -0.125009 | 0.900516 | 0.982670048 | RCN1P2 | processed_pseudogene | 48.63 | -0.00032173 | 0.00032173 | ENSG00000214455 | n.s. |
| ENSG00000164902 | 1443.075 | -0.035799329 | 0.250163 | -0.143104 | 0.886208 | 0.979542794 | PHAX | protein_coding | 43.51 | -0.000321356 | 0.000321356 | ENSG00000164902 | n.s. |
| ENSG00000198736 | 56.12556 | -0.043157186 | 0.348077 | -0.123987 | 0.901325 | 0.983013503 | MSRB1 | protein_coding | 58.61 | -0.000321112 | 0.000321112 | ENSG00000198736 | n.s. |

| | | | | | | | | | | | | | |
|---|---|---|---|---|---|---|---|---|---|---|---|---|---|
| ENSG00000152804 | 246.4788 | 0.036146259 | 0.264864 | 0.1364711 | 0.891449 | 0.979788048 | HHEX | protein_coding | 46.38 | 0.00032054 | 0.00032054 | ENSG00000152804 | n.s. |
| ENSG00000205937 | 3252.69 | -0.033511284 | 0.224833 | -0.14905 | 0.881515 | 0.978242302 | RNPS1 | protein_coding | 49.19 | -0.000320152 | 0.000320152 | ENSG00000205937 | n.s. |
| ENSG00000197555 | 23.41658 | -0.045227004 | 0.374671 | -0.120711 | 0.90392 | 0.983886286 | SIPA1L1 | protein_coding | 39.56 | -0.000319081 | 0.000319081 | ENSG00000197555 | n.s. |
| ENSG00000006695 | 494.1688 | 0.035470168 | 0.248094 | 0.1429706 | 0.886313 | 0.979542794 | COX10 | protein_coding | 38.47 | 0.000318401 | 0.000318401 | ENSG00000006695 | n.s. |
| ENSG00000114779 | 111.0481 | 0.03581316 | 0.261285 | 0.1370656 | 0.890979 | 0.979788048 | ABHD14B | protein_coding | 55.05 | 0.000317586 | 0.000317586 | ENSG00000114779 | n.s. |
| ENSG00000186111 | 187.1344 | -0.035348052 | 0.246935 | -0.143147 | 0.886174 | 0.979542794 | PIP5K1C | protein_coding | 60.55 | -0.000317305 | 0.000317305 | ENSG00000186111 | n.s. |
| ENSG00000204427 | 355.6617 | -0.037456594 | 0.287309 | -0.130371 | 0.896273 | 0.980688401 | ABHD16A | protein_coding | 50.35 | -0.000317218 | 0.000317218 | ENSG00000204427 | n.s. |
| ENSG00000160209 | 15.96389 | 0.047878656 | 0.427918 | 0.1118874 | 0.910913 | 0.984879111 | PDXK | protein_coding | 57.26 | 0.000316817 | 0.000316817 | ENSG00000160209 | n.s. |
| ENSG00000263786 | 551.72 | -0.03313919 | 0.222675 | -0.148823 | 0.881693 | 0.978242302 | | sense_intronic | 44.17 | -0.000316597 | 0.000316597 | ENSG00000263786 | n.s. |
| ENSG00000162441 | 205.0123 | -0.03810267 | 0.294213 | -0.129507 | 0.896957 | 0.981075379 | LZIC | protein_coding | 43.89 | -0.000316162 | 0.000316162 | ENSG00000162441 | n.s. |
| ENSG00000139131 | 901.5174 | -0.035629252 | 0.255032 | -0.139705 | 0.888893 | 0.979788048 | YARS2 | protein_coding | 39.51 | -0.000315955 | 0.000315955 | ENSG00000139131 | n.s. |
| ENSG00000198522 | 1242.635 | 0.036158744 | 0.271746 | 0.1330607 | 0.894145 | 0.980096525 | GPN1 | protein_coding | 41.23 | 0.000315707 | 0.000315707 | ENSG00000198522 | n.s. |
| ENSG00000068394 | 311.4288 | -0.037234118 | 0.285691 | -0.13033 | 0.896305 | 0.980688401 | GPKOW | protein_coding | 50.87 | -0.000315334 | 0.000315334 | ENSG00000068394 | n.s. |
| ENSG00000141101 | 576.9836 | 0.03555284 | 0.257495 | 0.1380721 | 0.890183 | 0.979788048 | NOB1 | protein_coding | 48.27 | 0.000315278 | 0.000315278 | ENSG00000141101 | n.s. |
| ENSG00000101654 | 1700.061 | -0.035409048 | 0.252639 | -0.140157 | 0.888536 | 0.979788048 | RNMT | protein_coding | 39.02 | -0.000314003 | 0.000314003 | ENSG00000101654 | n.s. |
| ENSG00000121690 | 386.021 | -0.035398558 | 0.260213 | -0.136037 | 0.891792 | 0.979788048 | DEPDC7 | protein_coding | 38.52 | -0.00031391 | 0.00031391 | ENSG00000121690 | n.s. |
| ENSG00000167552 | 13711.69 | -0.040654971 | 0.321841 | -0.12632 | 0.899479 | 0.982393201 | TUBA1A | protein_coding | 48.62 | -0.000313639 | 0.000313639 | ENSG00000167552 | n.s. |
| ENSG00000137411 | 462.3888 | 0.034885134 | 0.244591 | 0.1426266 | 0.886585 | 0.979754256 | VARS2 | protein_coding | 53.81 | 0.000309879 | 0.000309879 | ENSG00000137411 | n.s. |
| ENSG00000136144 | 998.5936 | -0.038894199 | 0.30571 | -0.127226 | 0.898762 | 0.981832619 | RCBTB1 | protein_coding | 42.24 | -0.000309697 | 0.000309697 | ENSG00000136144 | n.s. |
| ENSG00000204370 | 601.7999 | 0.037269021 | 0.288129 | 0.1293486 | 0.897082 | 0.981075379 | SDHD | protein_coding | 41.79 | 0.000309244 | 0.000309244 | ENSG00000204370 | n.s. |
| ENSG00000114541 | 755.7641 | 0.035793368 | 0.271383 | 0.1318923 | 0.895069 | 0.980337632 | FRMD4B | protein_coding | 41.42 | 0.000308694 | 0.000308694 | ENSG00000114541 | n.s. |
| ENSG00000205250 | 1063.779 | -0.034747766 | 0.251219 | -0.138316 | 0.88999 | 0.979788048 | E2F4 | protein_coding | 56.55 | -0.000308138 | 0.000308138 | ENSG00000205250 | n.s. |
| ENSG00000129315 | 3021.809 | -0.034657291 | 0.246327 | -0.140696 | 0.88811 | 0.979788048 | CCNT1 | protein_coding | 40.95 | -0.000307336 | 0.000307336 | ENSG00000129315 | n.s. |
| ENSG00000134717 | 2483.642 | -0.036292262 | 0.27901 | -0.130075 | 0.896507 | 0.980755 | BTF3L4 | protein_coding | 41.19 | -0.000306287 | 0.000306287 | ENSG00000134717 | n.s. |
| ENSG00000099219 | 179.3557 | -0.034430905 | 0.256279 | -0.134349 | 0.893126 | 0.979788048 | ERMP1 | protein_coding | 39 | -0.000305329 | 0.000305329 | ENSG00000099219 | n.s. |
| ENSG00000274292 | 176.7236 | -0.03437129 | 0.253944 | -0.13535 | 0.892335 | 0.979788048 | | lincRNA | 50.9 | -0.0003048 | 0.0003048 | ENSG00000274292 | n.s. |
| ENSG00000068323 | 140.5672 | 0.034253419 | 0.255188 | 0.1342283 | 0.893222 | 0.979788048 | TFE3 | protein_coding | 52.58 | 0.000303755 | 0.000303755 | ENSG00000068323 | n.s. |
| ENSG00000146410 | 95.53794 | 0.034247452 | 0.251963 | 0.1359226 | 0.891882 | 0.979788048 | MTFR2 | protein_coding | 39.81 | 0.000303702 | 0.000303702 | ENSG00000146410 | n.s. |
| ENSG00000108592 | 1870.088 | -0.034205589 | 0.246307 | -0.138874 | 0.88955 | 0.979788048 | FTSJ3 | protein_coding | 50.46 | -0.00030333 | 0.00030333 | ENSG00000108592 | n.s. |
| ENSG00000177548 | 27.58127 | -0.044046817 | 0.379934 | -0.115933 | 0.907706 | 0.984343751 | RABEP2 | protein_coding | 52.63 | -0.000301862 | 0.000301862 | ENSG00000177548 | n.s. |
| ENSG00000215039 | 620.7842 | 0.035448179 | 0.270221 | 0.1311821 | 0.895631 | 0.980644403 | CD27-AS1 | processed_transcript | 49.5 | 0.0003009 | 0.0003009 | ENSG00000215039 | n.s. |
| ENSG00000124383 | 2573.049 | -0.033911107 | 0.241821 | -0.140232 | 0.888477 | 0.979788048 | MPHOSPH10 | protein_coding | 40.5 | -0.000300719 | 0.000300719 | ENSG00000124383 | n.s. |

| | | | | | | | | | | | | | |
|---|---|---|---|---|---|---|---|---|---|---|---|---|---|
| ENSG00000115526 | 1103.833 | 0.03319168 | 0.229859 | 0.1444005 | 0.885184 | 0.979372226 | CHST10 | protein_coding | 48.91 | 0.000300458 | 0.000300458 | ENSG00000115526 | n.s. |
| ENSG00000163938 | 5876.463 | -0.033753652 | 0.250953 | -0.134502 | 0.893006 | 0.979788048 | GNL3 | protein_coding | 43.2 | -0.000299323 | 0.000299323 | ENSG00000163938 | n.s. |
| ENSG00000199436 | 7.259912 | -0.053660192 | 0.542681 | -0.09888 | 0.921234 | 0.987238446 | SNORD9 | snoRNA | 47.12 | -0.000299313 | 0.000299313 | ENSG00000199436 | n.s. |
| ENSG00000251013 | 78.72777 | -0.035343831 | 0.271522 | -0.130169 | 0.896433 | 0.980750424 | GAPDHP62 | processed_pseudogene | 50.81 | -0.000298355 | 0.000298355 | ENSG00000251013 | n.s. |
| ENSG00000175455 | 6162.236 | -0.036589683 | 0.285562 | -0.128132 | 0.898044 | 0.981476624 | CCDC14 | protein_coding | 36.3 | -0.00029711 | 0.00029711 | ENSG00000175455 | n.s. |
| ENSG00000132467 | 886.279 | 0.03349941 | 0.241208 | 0.1388818 | 0.889544 | 0.979788048 | UTP3 | protein_coding | 44.81 | 0.000297068 | 0.000297068 | ENSG00000132467 | n.s. |
| ENSG00000197345 | 796.6097 | -0.039003186 | 0.313232 | -0.124518 | 0.900905 | 0.982691813 | MRPL21 | protein_coding | 50.69 | -0.000295748 | 0.000295748 | ENSG00000197345 | n.s. |
| ENSG00000010256 | 1487.36 | -0.036190398 | 0.282554 | -0.128083 | 0.898083 | 0.981476624 | UQCRC1 | protein_coding | 53.04 | -0.000293867 | 0.000293867 | ENSG00000010256 | n.s. |
| ENSG00000135387 | 8793.636 | -0.034652505 | 0.265562 | -0.130487 | 0.896181 | 0.980688401 | CAPRIN1 | protein_coding | 40.22 | -0.000293471 | 0.000293471 | ENSG00000135387 | n.s. |
| ENSG00000166788 | 1085.103 | 0.034521035 | 0.263908 | 0.130807 | 0.895928 | 0.980688401 | SAAL1 | protein_coding | 39.77 | 0.000292357 | 0.000292357 | ENSG00000166788 | n.s. |
| ENSG00000008018 | 4153.312 | 0.03783446 | 0.299519 | 0.1263174 | 0.899481 | 0.982393201 | PSMB1 | protein_coding | 39.81 | 0.00029188 | 0.00029188 | ENSG00000008018 | n.s. |
| ENSG00000171714 | 531.1371 | 0.037828315 | 0.299487 | 0.1263102 | 0.899486 | 0.982393201 | ANO5 | protein_coding | 34.49 | 0.000291832 | 0.000291832 | ENSG00000171714 | n.s. |
| ENSG00000083896 | 2898.183 | -0.033850304 | 0.256894 | -0.131768 | 0.895168 | 0.980368399 | YTHDC1 | protein_coding | 35.4 | -0.000291475 | 0.000291475 | ENSG00000083896 | n.s. |
| ENSG00000139133 | 690.7462 | -0.032366981 | 0.225401 | -0.143597 | 0.885819 | 0.979542794 | ALG10 | protein_coding | 36.5 | -0.000290545 | 0.000290545 | ENSG00000139133 | n.s. |
| ENSG00000234465 | 160.2266 | -0.032742497 | 0.241537 | -0.135559 | 0.89217 | 0.979788048 | PINLYP | protein_coding | 51.61 | -0.000290356 | 0.000290356 | ENSG00000234465 | n.s. |
| ENSG00000227542 | 101.0489 | -0.033373 | 0.251131 | -0.132891 | 0.89428 | 0.980166821 | | lincRNA | 42.43 | -0.000290345 | 0.000290345 | ENSG00000227542 | n.s. |
| ENSG00000113108 | 127.8252 | 0.034075303 | 0.260858 | 0.1306278 | 0.89607 | 0.980688401 | APBB3 | protein_coding | 53.75 | 0.000288582 | 0.000288582 | ENSG00000113108 | n.s. |
| ENSG00000257433 | 35.54263 | 0.041440663 | 0.355646 | 0.1165224 | 0.907239 | 0.984198171 | | antisense | 50.18 | 0.000286664 | 0.000286664 | ENSG00000257433 | n.s. |
| ENSG00000221200 | 10.18511 | -0.050014657 | 0.492423 | -0.101569 | 0.919099 | 0.986968832 | MIR1253 | miRNA | 45.71 | -0.000284912 | 0.000284912 | ENSG00000221200 | n.s. |
| ENSG00000180376 | 996.1036 | -0.031916159 | 0.231364 | -0.137948 | 0.890282 | 0.979788048 | CCDC66 | protein_coding | 39.8 | -0.000283028 | 0.000283028 | ENSG00000180376 | n.s. |
| ENSG00000156970 | 2290.666 | 0.034922512 | 0.273317 | 0.1277728 | 0.898329 | 0.981590948 | BUB1B | protein_coding | 38.74 | 0.000281806 | 0.000281806 | ENSG00000156970 | n.s. |
| ENSG00000130255 | 8479.654 | 0.039028369 | 0.321118 | 0.121539 | 0.903264 | 0.98351261 | RPL36 | protein_coding | 58.81 | 0.000281787 | 0.000281787 | ENSG00000130255 | n.s. |
| ENSG00000124151 | 392.9637 | -0.036599686 | 0.290326 | -0.126064 | 0.899681 | 0.98243649 | NCOA3 | protein_coding | 41.15 | -0.000281653 | 0.000281653 | ENSG00000124151 | n.s. |
| ENSG00000125457 | 86.742 | -0.038933834 | 0.319981 | -0.121675 | 0.903156 | 0.98351261 | MIF4GD | protein_coding | 55.74 | -0.000281105 | 0.000281105 | ENSG00000125457 | n.s. |
| ENSG00000168495 | 1406.681 | 0.031612251 | 0.230057 | 0.1374106 | 0.890706 | 0.979788048 | POLR3D | protein_coding | 49.5 | 0.000280333 | 0.000280333 | ENSG00000168495 | n.s. |
| ENSG00000125691 | 18597.75 | -0.037679162 | 0.304628 | -0.123689 | 0.901561 | 0.98302057 | RPL23 | protein_coding | 44.44 | -0.000280235 | 0.000280235 | ENSG00000125691 | n.s. |
| ENSG00000135974 | 926.7159 | -0.032548093 | 0.247277 | -0.131626 | 0.89528 | 0.980413918 | C2orf49 | protein_coding | 38.53 | -0.000279605 | 0.000279605 | ENSG00000135974 | n.s. |
| ENSG00000143228 | 3025.55 | 0.036769498 | 0.294411 | 0.1248919 | 0.900609 | 0.982691813 | NUF2 | protein_coding | 36.44 | 0.000278811 | 0.000278811 | ENSG00000143228 | n.s. |
| ENSG00000201793 | 25.84818 | 0.041332295 | 0.364262 | 0.1134686 | 0.909659 | 0.984599969 | RN7SKP9 | misc_RNA | 54.52 | 0.000278587 | 0.000278587 | ENSG00000201793 | n.s. |
| ENSG00000100577 | 227.4549 | -0.039129515 | 0.324748 | -0.120492 | 0.904093 | 0.983886286 | GSTZ1 | protein_coding | 54.52 | -0.000276062 | 0.000276062 | ENSG00000100577 | n.s. |
| ENSG00000107938 | 1838.466 | 0.031773528 | 0.239465 | 0.1326857 | 0.894442 | 0.980194243 | EDRF1 | protein_coding | 39.09 | 0.000276044 | 0.000276044 | ENSG00000107938 | n.s. |
| ENSG00000237594 | 138.796 | 0.031767954 | 0.239914 | 0.1324139 | 0.894657 | 0.980194243 | | antisense | 58.57 | 0.000275995 | 0.000275995 | ENSG00000237594 | n.s. |

| | | | | | | | | | | | | | |
|---|---|---|---|---|---|---|---|---|---|---|---|---|---|
| ENSG00000165462 | 2438.954 | 0.031816384 | 0.240759 | 0.1321505 | 0.894865 | 0.980308504 | PHOX2A | protein_coding | 61.28 | 0.000274805 | 0.000274805 | ENSG00000165462 | n.s. |
| ENSG00000257621 | 2195.374 | 0.031714915 | 0.240135 | 0.1320711 | 0.894928 | 0.980308504 | PSMA3-AS1 | antisense | 38.57 | 0.000273929 | 0.000273929 | ENSG00000257621 | n.s. |
| ENSG00000102316 | 1387.202 | -0.037091388 | 0.301425 | -0.123053 | 0.902065 | 0.9832041 | MAGED2 | protein_coding | 52.37 | -0.000272856 | 0.000272856 | ENSG00000102316 | n.s. |
| ENSG00000187735 | 1580.51 | -0.035431202 | 0.281622 | -0.125811 | 0.899881 | 0.98243649 | TCEA1 | protein_coding | 39.44 | -0.000272661 | 0.000272661 | ENSG00000187735 | n.s. |
| ENSG00000168883 | 2195.135 | 0.033720775 | 0.264199 | 0.1276339 | 0.898439 | 0.981633983 | USP39 | protein_coding | 46.55 | 0.000271466 | 0.000271466 | ENSG00000168883 | n.s. |
| ENSG00000176261 | 234.7084 | 0.037440256 | 0.308284 | 0.1214472 | 0.903337 | 0.98351261 | ZBTB8OS | protein_coding | 44.37 | 0.000270321 | 0.000270321 | ENSG00000176261 | n.s. |
| ENSG00000106610 | 350.5966 | -0.03627169 | 0.294043 | -0.123355 | 0.901826 | 0.98302057 | STAG3L4 | cribed_unprocessed_pseudc | 44.75 | -0.000269767 | 0.000269767 | ENSG00000106610 | n.s. |
| ENSG00000230695 | 25.83421 | 0.043633223 | 0.401868 | 0.1085761 | 0.913539 | 0.985890439 | | antisense | 51.89 | 0.000269276 | 0.000269276 | ENSG00000230695 | n.s. |
| ENSG00000279513 | 62.48357 | 0.038383867 | 0.327208 | 0.1173071 | 0.906617 | 0.983983433 | | TEC | 33.18 | 0.000269156 | 0.000269156 | ENSG00000279513 | n.s. |
| ENSG00000207721 | 83.34343 | 0.036782997 | 0.299975 | 0.12262 | 0.902408 | 0.983313374 | MIR186 | miRNA | 36.05 | 0.000268812 | 0.000268812 | ENSG00000207721 | n.s. |
| ENSG00000198890 | 1473.736 | -0.03028842 | 0.224433 | -0.134955 | 0.892647 | 0.979788048 | PRMT6 | protein_coding | 50.31 | -0.000268594 | 0.000268594 | ENSG00000198890 | n.s. |
| ENSG00000266668 | 12.15645 | -0.045183804 | 0.431157 | -0.104797 | 0.916537 | 0.986622242 | MIR5692C2 | miRNA | 32.47 | -0.000264285 | 0.000264285 | ENSG00000266668 | n.s. |
| ENSG00000279873 | 332.2153 | 0.02973925 | 0.219578 | 0.1354385 | 0.892265 | 0.979788048 | | lincRNA | 56.62 | 0.000263724 | 0.000263724 | ENSG00000279873 | n.s. |
| ENSG00000135441 | 289.8417 | -0.037599242 | 0.313382 | -0.119979 | 0.9045 | 0.983983433 | BLOC1S1 | protein_coding | 55.05 | -0.000263654 | 0.000263654 | ENSG00000135441 | n.s. |
| ENSG00000213339 | 580.3774 | 0.03425369 | 0.271957 | 0.1259525 | 0.899769 | 0.98243649 | QTRT1 | protein_coding | 52.81 | 0.0002636 | 0.0002636 | ENSG00000213339 | n.s. |
| ENSG00000162627 | 579.6865 | 0.034660813 | 0.278047 | 0.1246581 | 0.900794 | 0.982691813 | SNX7 | protein_coding | 36.05 | 0.000262821 | 0.000262821 | ENSG00000162627 | n.s. |
| ENSG00000180198 | 676.5385 | -0.034565234 | 0.277746 | -0.124449 | 0.90096 | 0.982691813 | RCC1 | protein_coding | 50 | -0.000262097 | 0.000262097 | ENSG00000180198 | n.s. |
| ENSG00000100575 | 393.5939 | -0.035887274 | 0.293034 | -0.122468 | 0.902528 | 0.983324419 | TIMM9 | protein_coding | 40.88 | -0.000262091 | 0.000262091 | ENSG00000100575 | n.s. |
| ENSG00000116586 | 391.169 | 0.037365716 | 0.31743 | 0.1177134 | 0.906295 | 0.983983433 | LAMTOR2 | protein_coding | 50.94 | 0.000262016 | 0.000262016 | ENSG00000116586 | n.s. |
| ENSG00000170231 | 151.6694 | -0.038380602 | 0.332933 | -0.11528 | 0.908223 | 0.984436653 | FABP6 | protein_coding | 48.54 | -0.000261457 | 0.000261457 | ENSG00000170231 | n.s. |
| ENSG00000099204 | 1625.082 | -0.03182025 | 0.247238 | -0.128703 | 0.897593 | 0.981326694 | ABLIM1 | protein_coding | 42.59 | -0.000260493 | 0.000260493 | ENSG00000099204 | n.s. |
| ENSG00000069329 | 2149.74 | -0.033837768 | 0.269148 | -0.125722 | 0.899952 | 0.98243649 | VPS35 | protein_coding | 38.86 | -0.000260399 | 0.000260399 | ENSG00000069329 | n.s. |
| ENSG00000243943 | 2170.774 | -0.033794475 | 0.268763 | -0.125741 | 0.899937 | 0.98243649 | ZNF512 | protein_coding | 41.21 | -0.000260066 | 0.000260066 | ENSG00000243943 | n.s. |
| ENSG00000168078 | 1308.887 | 0.036949804 | 0.312992 | 0.1180534 | 0.906025 | 0.983983433 | PBK | protein_coding | 37.25 | 0.0002591 | 0.0002591 | ENSG00000168078 | n.s. |
| ENSG00000088682 | 532.1262 | -0.036882548 | 0.315018 | -0.117081 | 0.906796 | 0.983983433 | COQ9 | protein_coding | 45.83 | -0.000258628 | 0.000258628 | ENSG00000088682 | n.s. |
| ENSG00000114767 | 763.3018 | -0.035565728 | 0.2911 | -0.122177 | 0.902759 | 0.983421408 | RRP9 | protein_coding | 55.15 | -0.000258219 | 0.000258219 | ENSG00000114767 | n.s. |
| ENSG00000134882 | 472.1529 | -0.036587772 | 0.303321 | -0.120624 | 0.903989 | 0.983886286 | UBAC2 | protein_coding | 41.75 | -0.00025813 | 0.00025813 | ENSG00000134882 | n.s. |
| ENSG00000093010 | 595.5891 | 0.037865982 | 0.331521 | 0.1142189 | 0.909064 | 0.984436653 | COMT | protein_coding | 53.41 | 0.000257952 | 0.000257952 | ENSG00000093010 | n.s. |
| ENSG00000099282 | 72.63772 | 0.03755726 | 0.324457 | 0.1157542 | 0.907847 | 0.984343751 | TSPAN15 | protein_coding | 51.44 | 0.000257388 | 0.000257388 | ENSG00000099282 | n.s. |
| ENSG00000160796 | 342.3731 | 0.031378157 | 0.243466 | 0.128881 | 0.897452 | 0.981325779 | NBEAL2 | protein_coding | 59.9 | 0.000256886 | 0.000256886 | ENSG00000160796 | n.s. |
| ENSG00000037474 | 2256.422 | -0.031549601 | 0.245943 | -0.12828 | 0.897927 | 0.981460132 | NSUN2 | protein_coding | 43.29 | -0.000256414 | 0.000256414 | ENSG00000037474 | n.s. |
| ENSG00000168653 | 1568.308 | -0.036475512 | 0.309717 | -0.117771 | 0.906249 | 0.983983433 | NDUFS5 | protein_coding | 46.54 | -0.000255774 | 0.000255774 | ENSG00000168653 | n.s. |

| | | | | | | | | | | | | | |
|---|---|---|---|---|---|---|---|---|---|---|---|---|---|
| ENSG00000216179 | 3.331122 | -0.053921876 | 0.654973 | -0.082327 | 0.934387 | 0.98914507 | MIR541 | miRNA | 53.57 | -0.00025559 | 0.00025559 | ENSG00000216179 | n.s. |
| ENSG00000212452 | 37.81933 | -0.037302476 | 0.3242 | -0.11506 | 0.908397 | 0.984436653 | SNORD69 | snoRNA | 41.56 | -0.000254113 | 0.000254113 | ENSG00000212452 | n.s. |
| ENSG00000156162 | 923.3277 | -0.036217177 | 0.30177 | -0.120016 | 0.904471 | 0.983983433 | DPY19L4 | protein_coding | 40.06 | -0.000253963 | 0.000253963 | ENSG00000156162 | n.s. |
| ENSG00000184117 | 2169.321 | 0.036176423 | 0.302194 | 0.1197124 | 0.904711 | 0.983983433 | NIPSNAP1 | protein_coding | 50.23 | 0.000253677 | 0.000253677 | ENSG00000184117 | n.s. |
| ENSG00000120992 | 1583.279 | 0.036190783 | 0.309484 | 0.116939 | 0.906908 | 0.984015436 | LYPLA1 | protein_coding | 41.54 | 0.000253266 | 0.000253266 | ENSG00000120992 | n.s. |
| ENSG00000168439 | 3083.816 | 0.03501962 | 0.287835 | 0.1216658 | 0.903164 | 0.98351261 | STIP1 | protein_coding | 50.06 | 0.000252844 | 0.000252844 | ENSG00000168439 | n.s. |
| ENSG00000204231 | 415.0831 | -0.033992503 | 0.275054 | -0.123585 | 0.901644 | 0.98302057 | RXRB | protein_coding | 53.95 | -0.000252816 | 0.000252816 | ENSG00000204231 | n.s. |
| ENSG00000253710 | 954.5983 | 0.029468251 | 0.224334 | 0.1313588 | 0.895491 | 0.980568428 | ALG11 | protein_coding | 41.48 | 0.000251131 | 0.000251131 | ENSG00000253710 | n.s. |
| ENSG00000121350 | 570.6293 | -0.030470906 | 0.236004 | -0.129112 | 0.897269 | 0.981202978 | PYROXD1 | protein_coding | 36.31 | -0.000251115 | 0.000251115 | ENSG00000121350 | n.s. |
| ENSG00000089351 | 330.7407 | -0.030765107 | 0.239576 | -0.128415 | 0.897821 | 0.981420652 | GRAMD1A | protein_coding | 55.68 | -0.000250576 | 0.000250576 | ENSG00000089351 | n.s. |
| ENSG00000064933 | 1573.191 | 0.033482225 | 0.270684 | 0.123695 | 0.901557 | 0.98302057 | PMS1 | protein_coding | 36.81 | 0.000249021 | 0.000249021 | ENSG00000064933 | n.s. |
| ENSG00000126249 | 153.1048 | -0.036555731 | 0.320315 | -0.114124 | 0.909139 | 0.984441204 | PDCD2L | protein_coding | 45.07 | -0.000248952 | 0.000248952 | ENSG00000126249 | n.s. |
| ENSG00000173418 | 712.608 | -0.035430295 | 0.301867 | -0.11737 | 0.906566 | 0.983983433 | NAA20 | protein_coding | 41.08 | -0.000248445 | 0.000248445 | ENSG00000173418 | n.s. |
| ENSG00000140612 | 1505.687 | 0.035376063 | 0.299337 | 0.1181813 | 0.905924 | 0.983983433 | SEC11A | protein_coding | 42.41 | 0.000248065 | 0.000248065 | ENSG00000140612 | n.s. |
| ENSG00000125124 | 329.4304 | -0.035369039 | 0.300087 | -0.117863 | 0.906176 | 0.983983433 | BBS2 | protein_coding | 40.21 | -0.000248015 | 0.000248015 | ENSG00000125124 | n.s. |
| ENSG00000173264 | 18.4928 | 0.042007008 | 0.397063 | 0.1057942 | 0.915746 | 0.986554184 | GPR137 | protein_coding | 56.27 | 0.000246962 | 0.000246962 | ENSG00000173264 | n.s. |
| ENSG00000177971 | 1753.38 | 0.032077371 | 0.255975 | 0.1253144 | 0.900275 | 0.982560436 | IMP3 | protein_coding | 52.2 | 0.000245094 | 0.000245094 | ENSG00000177971 | n.s. |
| ENSG00000164885 | 233.0706 | 0.035688748 | 0.308071 | 0.1158457 | 0.907775 | 0.984343751 | CDK5 | protein_coding | 58.08 | 0.000244583 | 0.000244583 | ENSG00000164885 | n.s. |
| ENSG00000055950 | 409.0143 | 0.035901782 | 0.311956 | 0.1150861 | 0.908377 | 0.984436653 | MRPL43 | protein_coding | 53.18 | 0.000244571 | 0.000244571 | ENSG00000055950 | n.s. |
| ENSG00000107625 | 1493.861 | -0.034825281 | 0.296432 | -0.117482 | 0.906478 | 0.983983433 | DDX50 | protein_coding | 40.32 | -0.000244202 | 0.000244202 | ENSG00000107625 | n.s. |
| ENSG00000109180 | 3142.653 | -0.034732338 | 0.294292 | -0.11802 | 0.906052 | 0.983983433 | OCIAD1 | protein_coding | 39.5 | -0.000243551 | 0.000243551 | ENSG00000109180 | n.s. |
| ENSG00000107537 | 226.6577 | -0.036109422 | 0.317835 | -0.113611 | 0.909547 | 0.984599969 | PHYH | protein_coding | 46.07 | -0.000243384 | 0.000243384 | ENSG00000107537 | n.s. |
| ENSG00000081154 | 2166.245 | -0.035243374 | 0.303185 | -0.116244 | 0.907459 | 0.984229696 | PCNP | protein_coding | 39.82 | -0.000243304 | 0.000243304 | ENSG00000081154 | n.s. |
| ENSG00000271780 | 40.11201 | 0.036723828 | 0.32839 | 0.1118298 | 0.910958 | 0.984879111 | | lincRNA | 58.94 | 0.000243004 | 0.000243004 | ENSG00000271780 | n.s. |
| ENSG00000170889 | 6783.652 | 0.035083791 | 0.301486 | 0.1163696 | 0.90736 | 0.984198171 | RPS9 | protein_coding | 51.79 | 0.00024269 | 0.00024269 | ENSG00000170889 | n.s. |
| ENSG00000174721 | 161.2096 | 0.030164808 | 0.236546 | 0.127522 | 0.898527 | 0.981653704 | FGFBP3 | protein_coding | 52.5 | 0.000242576 | 0.000242576 | ENSG00000174721 | n.s. |
| ENSG00000152954 | 210.5608 | 0.034457241 | 0.289403 | 0.119063 | 0.905225 | 0.983983433 | NRSN1 | protein_coding | 40.3 | 0.000241622 | 0.000241622 | ENSG00000152954 | n.s. |
| ENSG00000022267 | 1315.204 | 0.034445959 | 0.286943 | 0.1200445 | 0.904448 | 0.983983433 | FHL1 | protein_coding | 42.07 | 0.000241542 | 0.000241542 | ENSG00000022267 | n.s. |
| ENSG00000082515 | 450.9974 | -0.035456131 | 0.309827 | -0.114439 | 0.90889 | 0.984436653 | MRPL22 | protein_coding | 38.83 | -0.000241535 | 0.000241535 | ENSG00000082515 | n.s. |
| ENSG00000183726 | 353.2627 | -0.034363125 | 0.287897 | -0.119359 | 0.904991 | 0.983983433 | TMEM50A | protein_coding | 42.04 | -0.000240962 | 0.000240962 | ENSG00000183726 | n.s. |
| ENSG00000117724 | 27259.65 | 0.032200092 | 0.260914 | 0.1234126 | 0.90178 | 0.98302057 | CENPF | protein_coding | 40.92 | 0.000239485 | 0.000239485 | ENSG00000117724 | n.s. |
| ENSG00000144674 | 2311.3 | 0.032191525 | 0.260675 | 0.1234929 | 0.901717 | 0.98302057 | GOLGA4 | protein_coding | 37.82 | 0.000239421 | 0.000239421 | ENSG00000144674 | n.s. |

| | | | | | | | | | | | | | |
|---|---|---|---|---|---|---|---|---|---|---|---|---|---|
| ENSG00000165792 | 1028.824 | 0.031101004 | 0.246871 | 0.125981 | 0.899747 | 0.98243649 | METTL17 | protein_coding | 45.05 | 0.000239338 | 0.000239338 | ENSG00000165792 | n.s. |
| ENSG00000121691 | 999.754 | 0.033931356 | 0.287011 | 0.118223 | 0.905891 | 0.983983433 | CAT | protein_coding | 40.6 | 0.000237934 | 0.000237934 | ENSG00000121691 | n.s. |
| ENSG00000122033 | 291.2081 | -0.03483566 | 0.305707 | -0.113951 | 0.909276 | 0.984513298 | MTIF3 | protein_coding | 42.6 | -0.000236131 | 0.000236131 | ENSG00000122033 | n.s. |
| ENSG00000112110 | 1153.89 | 0.034557558 | 0.301869 | 0.1144788 | 0.908858 | 0.984436653 | MRPL18 | protein_coding | 43.85 | 0.000235414 | 0.000235414 | ENSG00000112110 | n.s. |
| ENSG00000274210 | 6.903121 | -0.06037416 | 0.842624 | -0.07165 | 0.94288 | 0.991070672 | | snRNA | 51.52 | -0.00023518 | 0.00023518 | ENSG00000274210 | n.s. |
| ENSG00000249915 | 1153.662 | -0.034507533 | 0.299509 | -0.115214 | 0.908276 | 0.984436653 | PDCD6 | protein_coding | 50.84 | -0.000235073 | 0.000235073 | ENSG00000249915 | n.s. |
| ENSG00000162607 | 6739.693 | -0.032145453 | 0.262732 | -0.122351 | 0.902621 | 0.983348618 | USP1 | protein_coding | 39.48 | -0.00023442 | 0.00023442 | ENSG00000162607 | n.s. |
| ENSG00000169855 | 1359.007 | 0.030679311 | 0.244819 | 0.1253145 | 0.900275 | 0.982560436 | ROBO1 | protein_coding | 35.1 | 0.000234412 | 0.000234412 | ENSG00000169855 | n.s. |
| ENSG00000105953 | 509.4567 | -0.033328944 | 0.280069 | -0.119003 | 0.905273 | 0.983983433 | OGDH | protein_coding | 46.23 | -0.00023371 | 0.00023371 | ENSG00000105953 | n.s. |
| ENSG00000211460 | 3494.28 | -0.03271244 | 0.2702 | -0.121068 | 0.903637 | 0.983762962 | TSN | protein_coding | 44 | -0.00023257 | 0.00023257 | ENSG00000211460 | n.s. |
| ENSG00000088325 | 2630.232 | -0.031796667 | 0.259416 | -0.12257 | 0.902448 | 0.983313374 | TPX2 | protein_coding | 41.52 | -0.000232372 | 0.000232372 | ENSG00000088325 | n.s. |
| ENSG00000168291 | 1811.515 | 0.03057189 | 0.244407 | 0.125086 | 0.900455 | 0.982670048 | PDHB | protein_coding | 43.56 | 0.00023211 | 0.00023211 | ENSG00000168291 | n.s. |
| ENSG00000078618 | 2788.065 | -0.032785942 | 0.272027 | -0.120525 | 0.904068 | 0.983886286 | NRDC | protein_coding | 39.35 | -0.000231308 | 0.000231308 | ENSG00000078618 | n.s. |
| ENSG00000185238 | 1104.614 | 0.033394805 | 0.286865 | 0.1164129 | 0.907325 | 0.984198171 | PRMT3 | protein_coding | 36.02 | 0.000231007 | 0.000231007 | ENSG00000185238 | n.s. |
| ENSG00000272993 | 61.21793 | 0.032895829 | 0.280384 | 0.1173242 | 0.906603 | 0.983983433 | | lincRNA | 33.98 | 0.000230673 | 0.000230673 | ENSG00000272993 | n.s. |
| ENSG00000237753 | 66.8919 | 0.03284848 | 0.277099 | 0.1185442 | 0.905637 | 0.983983433 | | lincRNA | 48.77 | 0.000230341 | 0.000230341 | ENSG00000237753 | n.s. |
| ENSG00000150867 | 415.8741 | -0.03070871 | 0.248584 | -0.123534 | 0.901684 | 0.98302057 | PIP4K2A | protein_coding | 41.89 | -0.000228393 | 0.000228393 | ENSG00000150867 | n.s. |
| ENSG00000067191 | 19.91189 | 0.038759103 | 0.367388 | 0.1054992 | 0.91598 | 0.986554184 | CACNB1 | protein_coding | 54.39 | 0.000227867 | 0.000227867 | ENSG00000067191 | n.s. |
| ENSG00000180900 | 759.6084 | -0.031157574 | 0.253743 | -0.122792 | 0.902272 | 0.983313374 | SCRIB | protein_coding | 62.5 | -0.000227701 | 0.000227701 | ENSG00000180900 | n.s. |
| ENSG00000076382 | 2602.27 | -0.032438608 | 0.271633 | -0.119421 | 0.904942 | 0.983983433 | SPAG5 | protein_coding | 46.5 | -0.000227466 | 0.000227466 | ENSG00000076382 | n.s. |
| ENSG00000174442 | 1569.543 | -0.033332323 | 0.291618 | -0.114301 | 0.908999 | 0.984436653 | ZWILCH | protein_coding | 40.27 | -0.000227067 | 0.000227067 | ENSG00000174442 | n.s. |
| ENSG00000253190 | 49.36738 | 0.033257724 | 0.290467 | 0.1144975 | 0.908843 | 0.984436653 | | lincRNA | 53.21 | 0.000226559 | 0.000226559 | ENSG00000253190 | n.s. |
| ENSG00000113734 | 200.05 | -0.036578626 | 0.335237 | -0.109113 | 0.913113 | 0.985890439 | BNIP1 | protein_coding | 45.87 | -0.000225739 | 0.000225739 | ENSG00000113734 | n.s. |
| ENSG00000207559 | 19.27901 | -0.039929033 | 0.399274 | -0.100004 | 0.920341 | 0.987068378 | MIR578 | miRNA | 34.38 | -0.000225709 | 0.000225709 | ENSG00000207559 | n.s. |
| ENSG00000112759 | 556.1156 | -0.033112581 | 0.288536 | -0.114761 | 0.908635 | 0.984436653 | SLC29A1 | protein_coding | 56.86 | -0.00022557 | 0.00022557 | ENSG00000112759 | n.s. |
| ENSG00000136891 | 1461.727 | 0.029731562 | 0.238816 | 0.1244959 | 0.900923 | 0.982691813 | TEX10 | protein_coding | 37.95 | 0.000225444 | 0.000225444 | ENSG00000136891 | n.s. |
| ENSG00000153157 | 216.899 | -0.032135296 | 0.272693 | -0.117844 | 0.906191 | 0.983983433 | SYCP2L | protein_coding | 40.47 | -0.00022534 | 0.00022534 | ENSG00000153157 | n.s. |
| ENSG00000139921 | 1831.326 | -0.031933236 | 0.269565 | -0.118462 | 0.905702 | 0.983983433 | TMX1 | protein_coding | 36.96 | -0.000223923 | 0.000223923 | ENSG00000139921 | n.s. |
| ENSG00000157653 | 93.19273 | -0.031907817 | 0.266016 | -0.119947 | 0.904525 | 0.983983433 | C9orf43 | protein_coding | 43.34 | -0.000223744 | 0.000223744 | ENSG00000157653 | n.s. |
| ENSG00000164713 | 218.3761 | 0.034011307 | 0.305326 | 0.1113935 | 0.911304 | 0.98498594 | BRI3 | protein_coding | 51.97 | 0.000223453 | 0.000223453 | ENSG00000164713 | n.s. |
| ENSG00000158457 | 101.1993 | 0.032772461 | 0.285547 | 0.1147706 | 0.908627 | 0.984436653 | TSPAN33 | protein_coding | 48.14 | 0.000223253 | 0.000223253 | ENSG00000158457 | n.s. |
| ENSG00000183878 | 224.0262 | -0.030538359 | 0.248922 | -0.122683 | 0.902358 | 0.983313374 | UTY | protein_coding | 37.47 | -0.000223176 | 0.000223176 | ENSG00000183878 | n.s. |

| | | | | | | | | | | | | | |
|---|---|---|---|---|---|---|---|---|---|---|---|---|---|
| ENSG00000183520 | 798.2309 | 0.031782743 | 0.268994 | 0.1181539 | 0.905946 | 0.983983433 | UTP11 | protein_coding | 44.58 | 0.000222867 | 0.000222867 | ENSG00000183520 | n.s. |
| ENSG00000117592 | 2497.821 | 0.033198046 | 0.295422 | 0.112375 | 0.910526 | 0.984676239 | PRDX6 | protein_coding | 40.72 | 0.000222644 | 0.000222644 | ENSG00000117592 | n.s. |
| ENSG00000164603 | 349.4851 | -0.031746964 | 0.267786 | -0.118553 | 0.905629 | 0.983983433 | BMT2 | protein_coding | 36.97 | -0.000222616 | 0.000222616 | ENSG00000164603 | n.s. |
| ENSG00000175354 | 567.6423 | -0.031689477 | 0.265886 | -0.119185 | 0.905129 | 0.983983433 | PTPN2 | protein_coding | 42.19 | -0.000222213 | 0.000222213 | ENSG00000175354 | n.s. |
| ENSG00000175792 | 2123.109 | -0.033669478 | 0.301479 | -0.111681 | 0.911076 | 0.98493031 | RUVBL1 | protein_coding | 45.86 | -0.000222033 | 0.000222033 | ENSG00000175792 | n.s. |
| ENSG00000128739 | 1137.716 | -0.032914917 | 0.292011 | -0.112718 | 0.910254 | 0.984599969 | SNRPN | protein_coding | 41.92 | -0.000221853 | 0.000221853 | ENSG00000128739 | n.s. |
| ENSG00000153879 | 3375.34 | 0.030726357 | 0.252574 | 0.121653 | 0.903174 | 0.98351261 | CEBPG | protein_coding | 44.81 | 0.000221846 | 0.000221846 | ENSG00000153879 | n.s. |
| ENSG00000166086 | 1043.015 | 0.031934969 | 0.279297 | 0.1143407 | 0.908968 | 0.984436653 | JAM3 | protein_coding | 41.98 | 0.000217548 | 0.000217548 | ENSG00000166086 | n.s. |
| ENSG00000183513 | 930.9822 | 0.031883924 | 0.278823 | 0.1143518 | 0.908959 | 0.984436653 | COA5 | protein_coding | 42.89 | 0.0002172 | 0.0002172 | ENSG00000183513 | n.s. |
| ENSG00000150477 | 145.6886 | -0.030926761 | 0.262078 | -0.118006 | 0.906063 | 0.983983433 | KIAA1328 | protein_coding | 37.13 | -0.000216865 | 0.000216865 | ENSG00000150477 | n.s. |
| ENSG00000140374 | 2386.508 | -0.03208892 | 0.284283 | -0.112877 | 0.910128 | 0.984599969 | ETFA | protein_coding | 40.14 | -0.000216285 | 0.000216285 | ENSG00000140374 | n.s. |
| ENSG00000106069 | 57.24403 | -0.032222945 | 0.287077 | -0.112245 | 0.910629 | 0.984676239 | CHN2 | protein_coding | 42.02 | -0.000216105 | 0.000216105 | ENSG00000106069 | n.s. |
| ENSG00000049167 | 445.2944 | 0.033055084 | 0.298321 | 0.1108037 | 0.911772 | 0.985070348 | ERCC8 | protein_coding | 37.16 | 0.000215941 | 0.000215941 | ENSG00000049167 | n.s. |
| ENSG00000163320 | 3449.918 | 0.030657845 | 0.258699 | 0.1185079 | 0.905665 | 0.983983433 | CGGBP1 | protein_coding | 35.99 | 0.000214979 | 0.000214979 | ENSG00000163320 | n.s. |
| ENSG00000230679 | 18.68555 | 0.038484771 | 0.393777 | 0.0977325 | 0.922145 | 0.987238446 | ENO1-AS1 | antisense | 59.53 | 0.000214666 | 0.000214666 | ENSG00000230679 | n.s. |
| ENSG00000185222 | 992.0851 | 0.030927284 | 0.267176 | 0.1157563 | 0.907846 | 0.984343751 | TCEAL9 | protein_coding | 44.15 | 0.000211951 | 0.000211951 | ENSG00000185222 | n.s. |
| ENSG00000144840 | 493.0228 | -0.031416878 | 0.278456 | -0.112825 | 0.910169 | 0.984599969 | RABL3 | protein_coding | 37.93 | -0.000211755 | 0.000211755 | ENSG00000144840 | n.s. |
| ENSG00000164841 | 113.4755 | -0.030012054 | 0.254066 | -0.118127 | 0.905967 | 0.983983433 | TMEM74 | protein_coding | 37.41 | -0.000210451 | 0.000210451 | ENSG00000164841 | n.s. |
| ENSG00000128709 | 105.3767 | -0.031153876 | 0.274906 | -0.113326 | 0.909772 | 0.984599969 | HOXD9 | protein_coding | 60.19 | -0.000209983 | 0.000209983 | ENSG00000128709 | n.s. |
| ENSG00000127184 | 3079.969 | -0.034847751 | 0.325136 | -0.107179 | 0.914647 | 0.986282937 | COX7C | protein_coding | 42.27 | -0.000209034 | 0.000209034 | ENSG00000127184 | n.s. |
| ENSG00000103502 | 325.2608 | -0.031889269 | 0.28702 | -0.111105 | 0.911533 | 0.985068187 | CDIPT | protein_coding | 56.47 | -0.000208355 | 0.000208355 | ENSG00000103502 | n.s. |
| ENSG00000106638 | 178.9688 | -0.030483256 | 0.265365 | -0.114873 | 0.908546 | 0.984436653 | TBL2 | protein_coding | 52.83 | -0.000207659 | 0.000207659 | ENSG00000106638 | n.s. |
| ENSG00000237440 | 828.3591 | -0.030386509 | 0.263051 | -0.115516 | 0.908036 | 0.984433066 | ZNF737 | protein_coding | 39.46 | -0.000207048 | 0.000207048 | ENSG00000237440 | n.s. |
| ENSG00000196636 | 258.4816 | -0.034385679 | 0.320943 | -0.10714 | 0.914678 | 0.986282937 | SDHAF3 | protein_coding | 38.69 | -0.000206262 | 0.000206262 | ENSG00000196636 | n.s. |
| ENSG00000150787 | 548.0616 | 0.031212182 | 0.280148 | 0.1114133 | 0.911289 | 0.98498594 | PTS | protein_coding | 41.9 | 0.000205063 | 0.000205063 | ENSG00000150787 | n.s. |
| ENSG00000120837 | 2417.737 | -0.030419272 | 0.269041 | -0.113066 | 0.909979 | 0.984599969 | NFYB | protein_coding | 39.19 | -0.000205031 | 0.000205031 | ENSG00000120837 | n.s. |
| ENSG00000264112 | 1737.862 | 0.030241408 | 0.267302 | 0.1131358 | 0.909923 | 0.984599969 | | lincRNA | 40.9 | 0.000203833 | 0.000203833 | ENSG00000264112 | n.s. |
| ENSG00000171566 | 1625.888 | 0.031066815 | 0.280056 | 0.1109306 | 0.911671 | 0.985070348 | PLRG1 | protein_coding | 34.71 | 0.000202952 | 0.000202952 | ENSG00000171566 | n.s. |
| ENSG00000125870 | 3028.569 | -0.032865539 | 0.301057 | -0.109167 | 0.91307 | 0.985890439 | SNRPB2 | protein_coding | 38.18 | -0.000202825 | 0.000202825 | ENSG00000125870 | n.s. |
| ENSG00000004779 | 1400.009 | 0.035287259 | 0.345205 | 0.1022211 | 0.918581 | 0.986959954 | NDUFAB1 | protein_coding | 48.8 | 0.000201154 | 0.000201154 | ENSG00000004779 | n.s. |
| ENSG00000214194 | 571.3965 | -0.034310247 | 0.327981 | -0.104611 | 0.916685 | 0.986622242 | LINC00998 | protein_coding | 37.39 | -0.000200684 | 0.000200684 | ENSG00000214194 | n.s. |
| ENSG00000182224 | 228.9188 | -0.028618777 | 0.244412 | -0.117092 | 0.906787 | 0.983983433 | CYB5D1 | protein_coding | 48.18 | -0.000200681 | 0.000200681 | ENSG00000182224 | n.s. |

| | | | | | | | | | | | | | |
|---|---|---|---|---|---|---|---|---|---|---|---|---|---|
| ENSG00000207982 | 8.385281 | -0.047786226 | 0.647916 | -0.073754 | 0.941206 | 0.990390549 | MIR548B | miRNA | 37.11 | -0.000200392 | 0.000200392 | ENSG00000207982 | n.s. |
| ENSG00000167363 | 96.6294 | 0.029759774 | 0.265017 | 0.1122937 | 0.910591 | 0.984676239 | FN3K | protein_coding | 51.33 | 0.000199585 | 0.000199585 | ENSG00000167363 | n.s. |
| ENSG00000147274 | 11161.93 | 0.030355844 | 0.272621 | 0.1113483 | 0.91134 | 0.98498594 | RBMX | protein_coding | 43.01 | 0.000199437 | 0.000199437 | ENSG00000147274 | n.s. |
| ENSG00000146085 | 934.5417 | -0.031709713 | 0.288907 | -0.109757 | 0.912602 | 0.98566081 | MUT | protein_coding | 34.4 | -0.0001989 | 0.0001989 | ENSG00000146085 | n.s. |
| ENSG00000110321 | 21501 | -0.02834007 | 0.242088 | -0.117065 | 0.906808 | 0.983983433 | EIF4G2 | protein_coding | 41.78 | -0.000198727 | 0.000198727 | ENSG00000110321 | n.s. |
| ENSG00000181029 | 306.6225 | -0.033785924 | 0.318712 | -0.106008 | 0.915576 | 0.986554184 | TRAPPC5 | protein_coding | 58.05 | -0.000198629 | 0.000198629 | ENSG00000181029 | n.s. |
| ENSG00000230383 | 51.792 | -0.035906299 | 0.371179 | -0.096736 | 0.922936 | 0.987349084 | | processed_pseudogene | 46.64 | -0.000198536 | 0.000198536 | ENSG00000230383 | n.s. |
| ENSG00000044446 | 22.27604 | -0.035789104 | 0.370567 | -0.096579 | 0.92306 | 0.987349084 | PHKA2 | protein_coding | 45.75 | -0.000197888 | 0.000197888 | ENSG00000044446 | n.s. |
| ENSG00000203778 | 316.2423 | -0.0336753 | 0.32367 | -0.104042 | 0.917136 | 0.986622242 | FAM229B | protein_coding | 36.12 | -0.00019697 | 0.00019697 | ENSG00000203778 | n.s. |
| ENSG00000198690 | 1271.722 | -0.028073386 | 0.236331 | -0.118789 | 0.905443 | 0.983983433 | FAN1 | protein_coding | 42.55 | -0.000196857 | 0.000196857 | ENSG00000198690 | n.s. |
| ENSG00000281896 | 59.24689 | -0.032660102 | 0.302974 | -0.107798 | 0.914156 | 0.986223353 | | transcribed_unprocessed_pseudc | 48.09 | -0.000196768 | 0.000196768 | ENSG00000281896 | n.s. |
| ENSG00000113552 | 1019.41 | 0.033372065 | 0.31521 | 0.1058726 | 0.915683 | 0.986554184 | GNPDA1 | protein_coding | 44.88 | 0.000196196 | 0.000196196 | ENSG00000113552 | n.s. |
| ENSG00000234358 | 37.83505 | -0.034252172 | 0.339749 | -0.100816 | 0.919696 | 0.986968832 | | processed_pseudogene | 48.76 | -0.00019512 | 0.00019512 | ENSG00000234358 | n.s. |
| ENSG00000144591 | 455.1286 | 0.02979707 | 0.268263 | 0.1110739 | 0.911558 | 0.985068187 | GMPPA | protein_coding | 54.38 | 0.000194685 | 0.000194685 | ENSG00000144591 | n.s. |
| ENSG00000117593 | 1318.142 | -0.029789844 | 0.268683 | -0.110874 | 0.911717 | 0.985070348 | DARS2 | protein_coding | 40.56 | -0.00019461 | 0.00019461 | ENSG00000117593 | n.s. |
| ENSG00000138002 | 606.716 | 0.028858795 | 0.256089 | 0.1126906 | 0.910276 | 0.984599969 | IFT172 | protein_coding | 45.51 | 0.000194514 | 0.000194514 | ENSG00000138002 | n.s. |
| ENSG00000105373 | 1955.008 | -0.028846583 | 0.255945 | -0.112706 | 0.910263 | 0.984599969 | NOP53 | protein_coding | 57.2 | -0.000194431 | 0.000194431 | ENSG00000105373 | n.s. |
| ENSG00000100099 | 225.0249 | 0.026925834 | 0.221592 | 0.1215109 | 0.903286 | 0.98351261 | HPS4 | protein_coding | 46.81 | 0.000194406 | 0.000194406 | ENSG00000100099 | n.s. |
| ENSG00000198331 | 213.2047 | 0.03193694 | 0.298539 | 0.1069773 | 0.914807 | 0.986282937 | HYLS1 | protein_coding | 41.34 | 0.000191573 | 0.000191573 | ENSG00000198331 | n.s. |
| ENSG00000148834 | 863.4739 | -0.033521406 | 0.333194 | -0.100606 | 0.919863 | 0.986968832 | GSTO1 | protein_coding | 41.43 | -0.000190957 | 0.000190957 | ENSG00000148834 | n.s. |
| ENSG00000114739 | 2959.212 | -0.028118403 | 0.247237 | -0.113731 | 0.909451 | 0.984599969 | ACVR2B | protein_coding | 47.64 | -0.000189523 | 0.000189523 | ENSG00000114739 | n.s. |
| ENSG00000160360 | 786.2245 | 0.030709624 | 0.283019 | 0.1085074 | 0.913593 | 0.985890439 | GPSM1 | protein_coding | 66.05 | 0.00018952 | 0.00018952 | ENSG00000160360 | n.s. |
| ENSG00000163636 | 1642.008 | 0.029646928 | 0.269355 | 0.1100662 | 0.912357 | 0.985472788 | PSMD6 | protein_coding | 39.12 | 0.000188417 | 0.000188417 | ENSG00000163636 | n.s. |
| ENSG00000169228 | 289.4476 | 0.029192656 | 0.264389 | 0.1104156 | 0.91208 | 0.985326378 | RAB24 | protein_coding | 57.83 | 0.000187414 | 0.000187414 | ENSG00000169228 | n.s. |
| ENSG00000105854 | 228.3629 | -0.032691038 | 0.317383 | -0.103002 | 0.917961 | 0.986959954 | PON2 | protein_coding | 38.28 | -0.000186354 | 0.000186354 | ENSG00000105854 | n.s. |
| ENSG00000146109 | 854.496 | -0.027610796 | 0.244619 | -0.112873 | 0.910132 | 0.984599969 | ABT1 | protein_coding | 47.69 | -0.000186102 | 0.000186102 | ENSG00000146109 | n.s. |
| ENSG00000134058 | 480.5197 | -0.030029289 | 0.2757 | -0.10892 | 0.913266 | 0.985890439 | CDK7 | protein_coding | 40.47 | -0.000185321 | 0.000185321 | ENSG00000134058 | n.s. |
| ENSG00000242485 | 485.4001 | -0.031500001 | 0.300957 | -0.104666 | 0.916641 | 0.986622242 | MRPL20 | protein_coding | 53.09 | -0.000184247 | 0.000184247 | ENSG00000242485 | n.s. |
| ENSG00000111880 | 1547.858 | -0.029796521 | 0.274253 | -0.108646 | 0.913483 | 0.985890439 | RNGTT | protein_coding | 36.64 | -0.000183885 | 0.000183885 | ENSG00000111880 | n.s. |
| ENSG00000166562 | 667.4921 | -0.032272828 | 0.319368 | -0.101052 | 0.919509 | 0.986968832 | SEC11C | protein_coding | 43.41 | -0.000183844 | 0.000183844 | ENSG00000166562 | n.s. |
| ENSG00000254328 | 46.48952 | 0.031288849 | 0.297374 | 0.1052173 | 0.916203 | 0.986606571 | | processed_pseudogene | 50.1 | 0.000183227 | 0.000183227 | ENSG00000254328 | n.s. |
| ENSG00000261526 | 211.3404 | 0.028674443 | 0.260414 | 0.1101111 | 0.912321 | 0.985472788 | | lincRNA | 55.35 | 0.000182237 | 0.000182237 | ENSG00000261526 | n.s. |

| | | | | | | | | | | | | | |
|---|---|---|---|---|---|---|---|---|---|---|---|---|---|
| ENSG00000260329 | 280.0167 | -0.025957605 | 0.218584 | -0.118754 | 0.90547 | 0.983983433 | | antisense | 45.75 | -0.00018202 | 0.00018202 | ENSG00000260329 | n.s. |
| ENSG00000100526 | 592.529 | -0.030940594 | 0.293124 | -0.105555 | 0.915936 | 0.986554184 | CDKN3 | protein_coding | 40.35 | -0.000181902 | 0.000181902 | ENSG00000100526 | n.s. |
| ENSG00000248121 | 52.04255 | -0.031925069 | 0.319429 | -0.099944 | 0.920389 | 0.987068378 | SMURF2P1 | cribed_unprocessed_pseudc | 37.46 | -0.000180465 | 0.000180465 | ENSG00000248121 | n.s. |
| ENSG00000187479 | 454.1288 | -0.030621263 | 0.288385 | -0.106182 | 0.915438 | 0.986554184 | C11orf96 | protein_coding | 53.45 | -0.000180024 | 0.000180024 | ENSG00000187479 | n.s. |
| ENSG00000124562 | 1838.23 | -0.031565942 | 0.311156 | -0.101447 | 0.919195 | 0.986968832 | SNRPC | protein_coding | 42.71 | -0.000179817 | 0.000179817 | ENSG00000124562 | n.s. |
| ENSG00000132341 | 17364.34 | 0.028988404 | 0.266535 | 0.1087603 | 0.913393 | 0.985890439 | RAN | protein_coding | 45.79 | 0.000178897 | 0.000178897 | ENSG00000132341 | n.s. |
| ENSG00000257169 | 67.56022 | -0.031374064 | 0.311518 | -0.100713 | 0.919778 | 0.986968832 | | processed_pseudogene | 41.64 | -0.000178724 | 0.000178724 | ENSG00000257169 | n.s. |
| ENSG00000103202 | 1479.97 | -0.031315691 | 0.304807 | -0.102739 | 0.91817 | 0.986959954 | NME4 | protein_coding | 58.71 | -0.000178514 | 0.000178514 | ENSG00000103202 | n.s. |
| ENSG00000078668 | 2113.455 | 0.030363465 | 0.288383 | 0.1052886 | 0.916147 | 0.986606571 | VDAC3 | protein_coding | 40.42 | 0.000177808 | 0.000177808 | ENSG00000078668 | n.s. |
| ENSG00000165152 | 714.9174 | 0.030192927 | 0.285413 | 0.1057868 | 0.915751 | 0.986554184 | TMEM246 | protein_coding | 39.12 | 0.000177506 | 0.000177506 | ENSG00000165152 | n.s. |
| ENSG00000143224 | 223.0434 | 0.030125343 | 0.285331 | 0.1055804 | 0.915915 | 0.986554184 | PPOX | protein_coding | 53.01 | 0.000177109 | 0.000177109 | ENSG00000143224 | n.s. |
| ENSG00000122482 | 5205.167 | -0.02862843 | 0.262008 | -0.109265 | 0.912992 | 0.985890439 | ZNF644 | protein_coding | 36.39 | -0.000176676 | 0.000176676 | ENSG00000122482 | n.s. |
| ENSG00000148229 | 2286.112 | 0.030165297 | 0.28695 | 0.1051238 | 0.916278 | 0.986606571 | POLE3 | protein_coding | 49.33 | 0.000176648 | 0.000176648 | ENSG00000148229 | n.s. |
| ENSG00000153339 | 185.7761 | -0.02978655 | 0.28168 | -0.105746 | 0.915784 | 0.986554184 | TRAPPC8 | protein_coding | 38.41 | -0.000175117 | 0.000175117 | ENSG00000153339 | n.s. |
| ENSG00000083544 | 462.1611 | -0.028948859 | 0.269284 | -0.107503 | 0.91439 | 0.986223353 | TDRD3 | protein_coding | 36.41 | -0.000174409 | 0.000174409 | ENSG00000083544 | n.s. |
| ENSG00000114686 | 3553.012 | -0.029591286 | 0.278335 | -0.106315 | 0.915332 | 0.986554184 | MRPL3 | protein_coding | 35.94 | -0.000173969 | 0.000173969 | ENSG00000114686 | n.s. |
| ENSG00000088298 | 98.80924 | -0.030932702 | 0.3104 | -0.099654 | 0.920619 | 0.987197667 | EDEM2 | protein_coding | 45.82 | -0.000173096 | 0.000173096 | ENSG00000088298 | n.s. |
| ENSG00000111615 | 4392.401 | -0.027956904 | 0.256519 | -0.108986 | 0.913214 | 0.985890439 | KRR1 | protein_coding | 35.47 | -0.000172532 | 0.000172532 | ENSG00000111615 | n.s. |
| ENSG00000130643 | 65.90469 | 0.033341566 | 0.370769 | 0.0899255 | 0.928346 | 0.988157204 | CALY | protein_coding | 60.02 | 0.000172508 | 0.000172508 | ENSG00000130643 | n.s. |
| ENSG00000109445 | 1372.806 | -0.027947951 | 0.256649 | -0.108896 | 0.913285 | 0.985890439 | ZNF330 | protein_coding | 34.91 | -0.000172476 | 0.000172476 | ENSG00000109445 | n.s. |
| ENSG00000095139 | 1097.323 | -0.029329692 | 0.276126 | -0.106219 | 0.915409 | 0.986554184 | ARCN1 | protein_coding | 42.33 | -0.000172431 | 0.000172431 | ENSG00000095139 | n.s. |
| ENSG00000196990 | 73.76869 | -0.031086861 | 0.324167 | -0.095898 | 0.923602 | 0.987349084 | FAM163B | protein_coding | 57.82 | -0.000171888 | 0.000171888 | ENSG00000196990 | n.s. |
| ENSG00000134152 | 915.9452 | 0.029183557 | 0.27411 | 0.1064665 | 0.915212 | 0.986554184 | KATNBL1 | protein_coding | 40.54 | 0.000171572 | 0.000171572 | ENSG00000134152 | n.s. |
| ENSG00000162600 | 212.6316 | -0.030708581 | 0.314214 | -0.097732 | 0.922145 | 0.987238446 | OMA1 | protein_coding | 39.98 | -0.000171291 | 0.000171291 | ENSG00000162600 | n.s. |
| ENSG00000260230 | 378.2809 | 0.030063297 | 0.299371 | 0.1004217 | 0.92001 | 0.986968832 | FRRS1L | protein_coding | 42.27 | 0.000171257 | 0.000171257 | ENSG00000260230 | n.s. |
| ENSG00000117614 | 548.7485 | 0.028534681 | 0.266547 | 0.1070533 | 0.914747 | 0.986282937 | SYF2 | protein_coding | 43.61 | 0.000171165 | 0.000171165 | ENSG00000117614 | n.s. |
| ENSG00000100554 | 1151.094 | 0.029186136 | 0.280588 | 0.1040177 | 0.917155 | 0.986622242 | ATP6V1D | protein_coding | 38.54 | 0.000170713 | 0.000170713 | ENSG00000100554 | n.s. |
| ENSG00000107745 | 736.7486 | -0.030463764 | 0.312654 | -0.097436 | 0.92238 | 0.987238446 | MICU1 | protein_coding | 40.17 | -0.000169925 | 0.000169925 | ENSG00000107745 | n.s. |
| ENSG00000198692 | 707.0186 | -0.02977973 | 0.290195 | -0.10262 | 0.918265 | 0.986959954 | EIF1AY | protein_coding | 35.64 | -0.000169758 | 0.000169758 | ENSG00000198692 | n.s. |
| ENSG00000069849 | 1365.615 | -0.029754421 | 0.29496 | -0.100876 | 0.919649 | 0.986968832 | ATP1B3 | protein_coding | 40.61 | -0.000169498 | 0.000169498 | ENSG00000069849 | n.s. |
| ENSG00000196730 | 191.5321 | -0.027442715 | 0.252852 | -0.108533 | 0.913573 | 0.985890439 | DAPK1 | protein_coding | 45.13 | -0.000169358 | 0.000169358 | ENSG00000196730 | n.s. |
| ENSG00000128524 | 409.6328 | 0.030619894 | 0.323183 | 0.0947447 | 0.924518 | 0.987349084 | ATP6V1F | protein_coding | 52.57 | 0.000169306 | 0.000169306 | ENSG00000128524 | n.s. |

| | | | | | | | | | | | | | |
|---|---|---|---|---|---|---|---|---|---|---|---|---|---|
| ENSG00000023041 | 1005.316 | -0.02773779 | 0.256499 | -0.10814 | 0.913885 | 0.986051968 | ZDHHC6 | protein_coding | 38.61 | -0.000169206 | 0.000169206 | ENSG00000023041 | n.s. |
| ENSG00000171703 | 140.2636 | -0.030280511 | 0.311256 | -0.097285 | 0.9225 | 0.987238446 | TCEA2 | protein_coding | 61.58 | -0.000168903 | 0.000168903 | ENSG00000171703 | n.s. |
| ENSG00000245614 | 45.61521 | -0.030526841 | 0.317201 | -0.096238 | 0.923331 | 0.987349084 | DDX11-AS1 | antisense | 44.82 | -0.000168791 | 0.000168791 | ENSG00000245614 | n.s. |
| ENSG00000184047 | 1411.782 | 0.02800645 | 0.260541 | 0.1074934 | 0.914398 | 0.986223353 | DIABLO | protein_coding | 47.05 | 0.000168731 | 0.000168731 | ENSG00000184047 | n.s. |
| ENSG00000165156 | 1910.365 | -0.028832214 | 0.277504 | -0.103898 | 0.91725 | 0.986622242 | ZHX1 | protein_coding | 38.83 | -0.000168642 | 0.000168642 | ENSG00000165156 | n.s. |
| ENSG00000070061 | 1964.466 | -0.027898778 | 0.260017 | -0.107296 | 0.914554 | 0.986282937 | IKBKAP | protein_coding | 39.91 | -0.00016735 | 0.00016735 | ENSG00000070061 | n.s. |
| ENSG00000069248 | 3099.359 | -0.028417149 | 0.271666 | -0.104603 | 0.916691 | 0.986622242 | NUP133 | protein_coding | 40.25 | -0.000166215 | 0.000166215 | ENSG00000069248 | n.s. |
| ENSG00000135540 | 93.54781 | -0.028306951 | 0.26944 | -0.105059 | 0.916329 | 0.986606571 | NHSL1 | protein_coding | 40.97 | -0.000165765 | 0.000165765 | ENSG00000135540 | n.s. |
| ENSG00000235272 | 15.7398 | -0.034903117 | 0.409159 | -0.085305 | 0.932019 | 0.989130548 | FAM103A2P | processed_pseudogene | 47.74 | -0.000165664 | 0.000165664 | ENSG00000235272 | n.s. |
| ENSG00000173726 | 11235.52 | 0.028311931 | 0.272262 | 0.1039878 | 0.917179 | 0.986622242 | TOMM20 | protein_coding | 41.67 | 0.000165599 | 0.000165599 | ENSG00000173726 | n.s. |
| ENSG00000163785 | 1399.642 | -0.028308876 | 0.271306 | -0.104343 | 0.916897 | 0.986622242 | RYK | protein_coding | 40.6 | -0.000165581 | 0.000165581 | ENSG00000163785 | n.s. |
| ENSG00000171960 | 437.1945 | 0.029556695 | 0.298374 | 0.0990594 | 0.921091 | 0.987238446 | PPIH | protein_coding | 43.8 | 0.000164865 | 0.000164865 | ENSG00000171960 | n.s. |
| ENSG00000138029 | 977.8878 | 0.028911823 | 0.285295 | 0.10134 | 0.919281 | 0.986968832 | HADHB | protein_coding | 41.22 | 0.000164698 | 0.000164698 | ENSG00000138029 | n.s. |
| ENSG00000204220 | 562.6711 | -0.029464251 | 0.298113 | -0.098836 | 0.921269 | 0.987238446 | PFDN6 | protein_coding | 54.13 | -0.00016435 | 0.00016435 | ENSG00000204220 | n.s. |
| ENSG00000164163 | 5300.447 | -0.029061069 | 0.291949 | -0.099542 | 0.920708 | 0.987206598 | ABCE1 | protein_coding | 35.08 | -0.000162508 | 0.000162508 | ENSG00000164163 | n.s. |
| ENSG00000228474 | 1203.579 | -0.029121829 | 0.295629 | -0.098508 | 0.921529 | 0.987238446 | OST4 | protein_coding | 56.45 | -0.00016244 | 0.00016244 | ENSG00000228474 | n.s. |
| ENSG00000136518 | 2166.934 | -0.029111815 | 0.297494 | -0.097857 | 0.922046 | 0.987238446 | ACTL6A | protein_coding | 39.25 | -0.000162384 | 0.000162384 | ENSG00000136518 | n.s. |
| ENSG00000153395 | 673.7513 | -0.028484046 | 0.27978 | -0.101809 | 0.918908 | 0.986959954 | LPCAT1 | protein_coding | 56.14 | -0.000162372 | 0.000162372 | ENSG00000153395 | n.s. |
| ENSG00000175600 | 110.7805 | 0.031240473 | 0.349183 | 0.0894674 | 0.92871 | 0.988157204 | SUGCT | protein_coding | 38.71 | 0.000161637 | 0.000161637 | ENSG00000175600 | n.s. |
| ENSG00000066583 | 370.0925 | -0.028816403 | 0.289743 | -0.099455 | 0.920777 | 0.987206598 | ISOC1 | protein_coding | 38.93 | -0.00016114 | 0.00016114 | ENSG00000066583 | n.s. |
| ENSG00000187522 | 789.12 | -0.028229984 | 0.281518 | -0.100278 | 0.920124 | 0.986971016 | HSPA14 | protein_coding | 39.55 | -0.000160787 | 0.000160787 | ENSG00000187522 | n.s. |
| ENSG00000165487 | 606.4305 | 0.028664301 | 0.288443 | 0.099376 | 0.92084 | 0.987206598 | MICU2 | protein_coding | 38.3 | 0.000160289 | 0.000160289 | ENSG00000165487 | n.s. |
| ENSG00000179104 | 335.692 | -0.026156289 | 0.241524 | -0.108297 | 0.91376 | 0.985994396 | TMTC2 | protein_coding | 37.42 | -0.000160222 | 0.000160222 | ENSG00000179104 | n.s. |
| ENSG00000151892 | 1218.395 | 0.026331702 | 0.244694 | 0.107611 | 0.914304 | 0.986223353 | GFRA1 | protein_coding | 44 | 0.000158641 | 0.000158641 | ENSG00000151892 | n.s. |
| ENSG00000213123 | 221.3937 | -0.030624317 | 0.334613 | -0.091522 | 0.927078 | 0.988157204 | TCTEX1D2 | protein_coding | 45.15 | -0.000158449 | 0.000158449 | ENSG00000213123 | n.s. |
| ENSG00000117399 | 438.0703 | 0.027079382 | 0.260715 | 0.1038657 | 0.917276 | 0.986622242 | CDC20 | protein_coding | 52.2 | 0.00015839 | 0.00015839 | ENSG00000117399 | n.s. |
| ENSG00000135241 | 515.4256 | -0.027764642 | 0.272569 | -0.101863 | 0.918866 | 0.986959954 | PNPLA8 | protein_coding | 37.75 | -0.000158271 | 0.000158271 | ENSG00000135241 | n.s. |
| ENSG00000165526 | 772.7433 | 0.026907774 | 0.254394 | 0.1057721 | 0.915763 | 0.986554184 | RPUSD4 | protein_coding | 43.35 | 0.000158192 | 0.000158192 | ENSG00000165526 | n.s. |
| ENSG00000072210 | 558.9353 | 0.027079302 | 0.261376 | 0.1036027 | 0.917485 | 0.986678534 | ALDH3A2 | protein_coding | 44.22 | 0.000157719 | 0.000157719 | ENSG00000072210 | n.s. |
| ENSG00000162650 | 716.9425 | 0.023499478 | 0.209261 | 0.1122973 | 0.910588 | 0.984676239 | ATXN7L2 | protein_coding | 56.87 | 0.0001576 | 0.0001576 | ENSG00000162650 | n.s. |
| ENSG00000132912 | 2703.631 | 0.026931869 | 0.259404 | 0.1038221 | 0.917311 | 0.986622242 | DCTN4 | protein_coding | 38.73 | 0.000157527 | 0.000157527 | ENSG00000132912 | n.s. |
| ENSG00000259673 | 69.32964 | 0.028135683 | 0.287696 | 0.0977964 | 0.922094 | 0.987238446 | IQCH-AS1 | lincRNA | 41.68 | 0.000156939 | 0.000156939 | ENSG00000259673 | n.s. |

| | | | | | | | | | | | | | |
|---|---|---|---|---|---|---|---|---|---|---|---|---|---|
| ENSG00000055130 | 1595.789 | -0.02832126 | 0.294681 | -0.096108 | 0.923435 | 0.987349084 | CUL1 | protein_coding | 41.22 | -0.000156596 | 0.000156596 | ENSG00000055130 | n.s. |
| ENSG00000164897 | 148.9701 | 0.027381256 | 0.26809 | 0.1021346 | 0.91865 | 0.986959954 | TMUB1 | protein_coding | 67.61 | 0.000156086 | 0.000156086 | ENSG00000164897 | n.s. |
| ENSG00000076706 | 185.2698 | 0.027132018 | 0.263673 | 0.1029002 | 0.918042 | 0.986959954 | MCAM | protein_coding | 59.59 | 0.000154665 | 0.000154665 | ENSG00000076706 | n.s. |
| ENSG00000083444 | 214.6826 | -0.027676554 | 0.283942 | -0.097473 | 0.922351 | 0.987238446 | PLOD1 | protein_coding | 53.54 | -0.000154378 | 0.000154378 | ENSG00000083444 | n.s. |
| ENSG00000111581 | 3851.39 | -0.027074498 | 0.265555 | -0.101954 | 0.918793 | 0.986959954 | NUP107 | protein_coding | 39.32 | -0.000154337 | 0.000154337 | ENSG00000111581 | n.s. |
| ENSG00000200520 | 21.80767 | -0.032488452 | 0.37862 | -0.085807 | 0.931619 | 0.989130548 | RNU6-1214P | snRNA | 34.86 | -0.000154203 | 0.000154203 | ENSG00000200520 | n.s. |
| ENSG00000118197 | 544.5478 | 0.027052162 | 0.269525 | 0.1003696 | 0.920051 | 0.986968832 | DDX59 | protein_coding | 42 | 0.000154104 | 0.000154104 | ENSG00000118197 | n.s. |
| ENSG00000198931 | 497.5488 | -0.029060821 | 0.312197 | -0.093085 | 0.925836 | 0.987870825 | APRT | protein_coding | 64.35 | -0.000154018 | 0.000154018 | ENSG00000198931 | n.s. |
| ENSG00000082996 | 1066.196 | 0.027026159 | 0.267049 | 0.101203 | 0.919389 | 0.986968832 | RNF13 | protein_coding | 36.39 | 0.000153956 | 0.000153956 | ENSG00000082996 | n.s. |
| ENSG00000272525 | 95.18493 | 0.027006712 | 0.265116 | 0.1018674 | 0.918862 | 0.986959954 | | lincRNA | 48.25 | 0.000153951 | 0.000153951 | ENSG00000272525 | n.s. |
| ENSG00000114374 | 869.7334 | -0.027599315 | 0.282299 | -0.097766 | 0.922118 | 0.987238446 | USP9Y | protein_coding | 36.81 | -0.000153947 | 0.000153947 | ENSG00000114374 | n.s. |
| ENSG00000105248 | 315.2231 | 0.026974001 | 0.268402 | 0.1004987 | 0.919948 | 0.986968832 | CCDC94 | protein_coding | 52.8 | 0.000153659 | 0.000153659 | ENSG00000105248 | n.s. |
| ENSG00000167862 | 866.4924 | -0.029611634 | 0.326085 | -0.090809 | 0.927644 | 0.988157204 | MRPL58 | protein_coding | 48.66 | -0.000153209 | 0.000153209 | ENSG00000167862 | n.s. |
| ENSG00000272562 | 119.4787 | 0.026183102 | 0.252216 | 0.1038121 | 0.917318 | 0.986622242 | | antisense | 41.12 | 0.000153148 | 0.000153148 | ENSG00000272562 | n.s. |
| ENSG00000196950 | 3092.599 | -0.026819412 | 0.262781 | -0.10206 | 0.918709 | 0.986959954 | SLC39A10 | protein_coding | 38.48 | -0.000152883 | 0.000152883 | ENSG00000196950 | n.s. |
| ENSG00000166128 | 819.2285 | 0.026760036 | 0.260446 | 0.1027468 | 0.918164 | 0.986959954 | RAB8B | protein_coding | 39.83 | 0.000152545 | 0.000152545 | ENSG00000166128 | n.s. |
| ENSG00000155380 | 7640.977 | -0.026035242 | 0.247975 | -0.104991 | 0.916383 | 0.986606571 | SLC16A1 | protein_coding | 40.76 | -0.000152462 | 0.000152462 | ENSG00000155380 | n.s. |
| ENSG00000152234 | 8300.915 | 0.027513743 | 0.28862 | 0.0953285 | 0.924054 | 0.987349084 | ATP5A1 | protein_coding | 44.44 | 0.000152131 | 0.000152131 | ENSG00000152234 | n.s. |
| ENSG00000141425 | 3193.495 | 0.027433541 | 0.28802 | 0.0952488 | 0.924117 | 0.987349084 | RPRD1A | protein_coding | 38.63 | 0.000151688 | 0.000151688 | ENSG00000141425 | n.s. |
| ENSG00000257698 | 210.0928 | 0.029463971 | 0.333453 | 0.0883601 | 0.92959 | 0.988216325 | | lincRNA | 40.3 | 0.00015168 | 0.00015168 | ENSG00000257698 | n.s. |
| ENSG00000280904 | 5.868511 | 0.050989102 | 0.784063 | 0.0650319 | 0.948149 | 0.993188446 | | snoRNA | 43.75 | 0.000151353 | 0.000151353 | ENSG00000280904 | n.s. |
| ENSG00000120725 | 57.19187 | 0.027327999 | 0.285366 | 0.0957646 | 0.923708 | 0.987349084 | SIL1 | protein_coding | 44.78 | 0.000151104 | 0.000151104 | ENSG00000120725 | n.s. |
| ENSG00000141759 | 1576.335 | -0.027300417 | 0.288422 | -0.094654 | 0.924589 | 0.987349084 | TXNL4A | protein_coding | 45.6 | -0.000150951 | 0.000150951 | ENSG00000141759 | n.s. |
| ENSG00000142546 | 479.3845 | 0.027632899 | 0.293104 | 0.0942767 | 0.924889 | 0.98751796 | NOSIP | protein_coding | 50.75 | 0.000150737 | 0.000150737 | ENSG00000142546 | n.s. |
| ENSG00000249725 | 14.63786 | -0.042829395 | 0.620452 | -0.069029 | 0.944966 | 0.991932047 | | lincRNA | 38.54 | -0.000150677 | 0.000150677 | ENSG00000249725 | n.s. |
| ENSG00000275180 | 942.982 | -0.022215741 | 0.197021 | -0.112758 | 0.910222 | 0.984599969 | | lincRNA | 36.25 | -0.000149738 | 0.000149738 | ENSG00000275180 | n.s. |
| ENSG00000134333 | 9052.848 | 0.026797087 | 0.27077 | 0.0989663 | 0.921165 | 0.987238446 | LDHA | protein_coding | 44.3 | 0.000149473 | 0.000149473 | ENSG00000134333 | n.s. |
| ENSG00000158773 | 915.0096 | -0.02701164 | 0.28002 | -0.096463 | 0.923153 | 0.987349084 | USF1 | protein_coding | 50.71 | -0.000149355 | 0.000149355 | ENSG00000158773 | n.s. |
| ENSG00000090266 | 1709.749 | 0.028659953 | 0.312099 | 0.0918296 | 0.926833 | 0.988157204 | NDUFB2 | protein_coding | 43.57 | 0.000148285 | 0.000148285 | ENSG00000090266 | n.s. |
| ENSG00000272008 | 310.1901 | -0.024541975 | 0.227751 | -0.107758 | 0.914188 | 0.986223353 | | antisense | 45.53 | -0.000147858 | 0.000147858 | ENSG00000272008 | n.s. |
| ENSG00000168374 | 1276.119 | -0.026485262 | 0.27065 | -0.097858 | 0.922045 | 0.987238446 | ARF4 | protein_coding | 41.71 | -0.000147733 | 0.000147733 | ENSG00000168374 | n.s. |
| ENSG00000186469 | 383.8054 | 0.027388884 | 0.29214 | 0.0937525 | 0.925306 | 0.987659658 | GNG2 | protein_coding | 40.23 | 0.0001477 | 0.0001477 | ENSG00000186469 | n.s. |

| | | | | | | | | | | | | | |
|---|---|---|---|---|---|---|---|---|---|---|---|---|---|
| ENSG00000015153 | 2185.568 | 0.025211829 | 0.241882 | 0.1042318 | 0.916985 | 0.986622242 | YAF2 | protein_coding | 39.02 | 0.000147466 | 0.000147466 | ENSG00000015153 | n.s. |
| ENSG00000170903 | 1022.721 | 0.026535836 | 0.273968 | 0.0968574 | 0.92284 | 0.987349084 | MSANTD4 | protein_coding | 38.77 | 0.000146724 | 0.000146724 | ENSG00000170903 | n.s. |
| ENSG00000152942 | 1075.81 | 0.026525494 | 0.278212 | 0.0953428 | 0.924043 | 0.987349084 | RAD17 | protein_coding | 39.7 | 0.000146667 | 0.000146667 | ENSG00000152942 | n.s. |
| ENSG00000225921 | 1150.15 | -0.026525129 | 0.278343 | -0.095297 | 0.924079 | 0.987349084 | NOL7 | protein_coding | 38.69 | -0.000146665 | 0.000146665 | ENSG00000225921 | n.s. |
| ENSG00000213672 | 262.6061 | -0.026266631 | 0.266142 | -0.098694 | 0.921381 | 0.987238446 | NCKIPSD | protein_coding | 56.62 | -0.000146514 | 0.000146514 | ENSG00000213672 | n.s. |
| ENSG00000090581 | 324.837 | -0.026469873 | 0.27916 | -0.09482 | 0.924458 | 0.987349084 | GNPTG | protein_coding | 56.48 | -0.000146359 | 0.000146359 | ENSG00000090581 | n.s. |
| ENSG00000162385 | 1317.729 | 0.02645727 | 0.279119 | 0.0947886 | 0.924483 | 0.987349084 | MAGOH | protein_coding | 41.63 | 0.000146289 | 0.000146289 | ENSG00000162385 | n.s. |
| ENSG00000099364 | 421.8017 | 0.026419897 | 0.278273 | 0.0949424 | 0.924361 | 0.987349084 | FBXL19 | protein_coding | 55.33 | 0.000146083 | 0.000146083 | ENSG00000099364 | n.s. |
| ENSG00000137285 | 2874.248 | -0.026381708 | 0.273888 | -0.096323 | 0.923264 | 0.987349084 | TUBB2B | protein_coding | 56.75 | -0.000145872 | 0.000145872 | ENSG00000137285 | n.s. |
| ENSG00000150753 | 9434.084 | 0.027036228 | 0.288034 | 0.0938646 | 0.925217 | 0.987659658 | CCT5 | protein_coding | 43.56 | 0.000145798 | 0.000145798 | ENSG00000150753 | n.s. |
| ENSG00000212978 | 109.0979 | 0.026097539 | 0.263609 | 0.0990009 | 0.921138 | 0.987238446 | | processed_transcript | 45.77 | 0.000145571 | 0.000145571 | ENSG00000212978 | n.s. |
| ENSG00000247728 | 40.55658 | 0.028202379 | 0.317832 | 0.0887336 | 0.929294 | 0.988186394 | | antisense | 41.21 | 0.000145556 | 0.000145556 | ENSG00000247728 | n.s. |
| ENSG00000246465 | 227.3688 | -0.025528924 | 0.247669 | -0.103077 | 0.917902 | 0.986959954 | | lincRNA | 49.16 | -0.000145527 | 0.000145527 | ENSG00000246465 | n.s. |
| ENSG00000125462 | 164.5919 | 0.025287989 | 0.246595 | 0.1025485 | 0.918321 | 0.986959954 | C1orf61 | protein_coding | 51.53 | 0.000144153 | 0.000144153 | ENSG00000125462 | n.s. |
| ENSG00000170540 | 6499.84 | -0.027021924 | 0.291219 | -0.092789 | 0.926071 | 0.987870825 | ARL6IP1 | protein_coding | 40.91 | -0.000143212 | 0.000143212 | ENSG00000170540 | n.s. |
| ENSG00000170854 | 2195.448 | 0.025581908 | 0.260536 | 0.0981896 | 0.921782 | 0.987238446 | RIOX2 | protein_coding | 41.54 | 0.000142694 | 0.000142694 | ENSG00000170854 | n.s. |
| ENSG00000103245 | 189.5381 | -0.027544592 | 0.303877 | -0.090644 | 0.927775 | 0.988157204 | NARFL | protein_coding | 60.82 | -0.000142515 | 0.000142515 | ENSG00000103245 | n.s. |
| ENSG00000118680 | 1816.067 | 0.02742942 | 0.307776 | 0.0891214 | 0.928985 | 0.988157204 | MYL12B | protein_coding | 40.39 | 0.000141919 | 0.000141919 | ENSG00000118680 | n.s. |
| ENSG00000156256 | 1918.532 | -0.02560359 | 0.27027 | -0.094733 | 0.924527 | 0.987349084 | USP16 | protein_coding | 38.76 | -0.000141569 | 0.000141569 | ENSG00000156256 | n.s. |
| ENSG00000228409 | 611.4434 | 0.024850279 | 0.246714 | 0.1007253 | 0.919769 | 0.986968832 | CCT6P1 | cribed_unprocessed_pseudc | 40.94 | 0.000141561 | 0.000141561 | ENSG00000228409 | n.s. |
| ENSG00000164588 | 11.44902 | 0.033646876 | 0.45639 | 0.0737239 | 0.94123 | 0.990390549 | HCN1 | protein_coding | 34.76 | 0.000141099 | 0.000141099 | ENSG00000164588 | n.s. |
| ENSG00000063176 | 219.0647 | -0.025491284 | 0.264448 | -0.096394 | 0.923207 | 0.987349084 | SPHK2 | protein_coding | 56.78 | -0.000140948 | 0.000140948 | ENSG00000063176 | n.s. |
| ENSG00000252759 | 123.4354 | 0.025245415 | 0.2595 | 0.0972847 | 0.9225 | 0.987238446 | | misc_RNA | 39.58 | 0.000140817 | 0.000140817 | ENSG00000252759 | n.s. |
| ENSG00000081177 | 45.83829 | 0.02960121 | 0.350279 | 0.0845075 | 0.932653 | 0.989132128 | EXD2 | protein_coding | 41.37 | 0.000140478 | 0.000140478 | ENSG00000081177 | n.s. |
| ENSG00000008300 | 968.0096 | 0.024076853 | 0.232475 | 0.1035676 | 0.917513 | 0.986678534 | CELSR3 | protein_coding | 59.89 | 0.000140231 | 0.000140231 | ENSG00000008300 | n.s. |
| ENSG00000107560 | 1173.549 | 0.025332824 | 0.263518 | 0.0961331 | 0.923415 | 0.987349084 | RAB11FIP2 | protein_coding | 35.43 | 0.000140072 | 0.000140072 | ENSG00000107560 | n.s. |
| ENSG00000166848 | 1791.715 | 0.026314163 | 0.283208 | 0.0929145 | 0.925971 | 0.987870825 | TERF2IP | protein_coding | 42.42 | 0.000139461 | 0.000139461 | ENSG00000166848 | n.s. |
| ENSG00000204209 | 1083.066 | -0.024911809 | 0.255468 | -0.097514 | 0.922318 | 0.987238446 | DAXX | protein_coding | 51.32 | -0.000138957 | 0.000138957 | ENSG00000204209 | n.s. |
| ENSG00000120438 | 7728.527 | -0.024848547 | 0.254893 | -0.097486 | 0.92234 | 0.987238446 | TCP1 | protein_coding | 40.93 | -0.000138604 | 0.000138604 | ENSG00000120438 | n.s. |
| ENSG00000101224 | 573.8524 | -0.024990759 | 0.260736 | -0.095847 | 0.923642 | 0.987349084 | CDC25B | protein_coding | 53.03 | -0.000138181 | 0.000138181 | ENSG00000101224 | n.s. |
| ENSG00000158887 | 72.71997 | 0.02592342 | 0.279239 | 0.0928359 | 0.926034 | 0.987870825 | MPZ | protein_coding | 50.52 | 0.00013739 | 0.00013739 | ENSG00000158887 | n.s. |
| ENSG00000233593 | 31.16911 | -0.030238296 | 0.378627 | -0.079863 | 0.936346 | 0.98960211 | | processed_transcript | 41.8 | -0.000137263 | 0.000137263 | ENSG00000233593 | n.s. |

| | | | | | | | | | | | | | |
|---|---|---|---|---|---|---|---|---|---|---|---|---|---|
| ENSG00000121058 | 1874.306 | 0.025727314 | 0.275335 | 0.09344 | 0.925554 | 0.98779046 | COIL | protein_coding | 43 | 0.00013726 | 0.00013726 | ENSG00000121058 | n.s. |
| ENSG00000119986 | 83.09005 | 0.026470177 | 0.297825 | 0.0888783 | 0.929179 | 0.988157204 | AVPI1 | protein_coding | 49.03 | 0.000136956 | 0.000136956 | ENSG00000119986 | n.s. |
| ENSG00000088833 | 417.2499 | -0.0245262 | 0.250539 | -0.097894 | 0.922017 | 0.987238446 | NSFL1C | protein_coding | 45.86 | -0.000136806 | 0.000136806 | ENSG00000088833 | n.s. |
| ENSG00000165732 | 7726.151 | -0.02534811 | 0.270347 | -0.093761 | 0.925299 | 0.987659658 | DDX21 | protein_coding | 41.17 | -0.000136694 | 0.000136694 | ENSG00000165732 | n.s. |
| ENSG00000259818 | 88.1907 | 0.026407303 | 0.29535 | 0.0894102 | 0.928756 | 0.988157204 | | antisense | 47.12 | 0.00013663 | 0.00013663 | ENSG00000259818 | n.s. |
| ENSG00000136758 | 4039.001 | 0.024657015 | 0.258906 | 0.0952355 | 0.924128 | 0.987349084 | YME1L1 | protein_coding | 38.36 | 0.000136335 | 0.000136335 | ENSG00000136758 | n.s. |
| ENSG00000142507 | 1313.726 | -0.026294377 | 0.292593 | -0.089867 | 0.928393 | 0.988157204 | PSMB6 | protein_coding | 52.71 | -0.000136046 | 0.000136046 | ENSG00000142507 | n.s. |
| ENSG00000276744 | 98.58931 | -0.024580935 | 0.256116 | -0.095976 | 0.92354 | 0.987349084 | | sense_intronic | 57.72 | -0.000135915 | 0.000135915 | ENSG00000276744 | n.s. |
| ENSG00000115661 | 406.4186 | -0.023831124 | 0.232637 | -0.102439 | 0.918408 | 0.986959954 | STK16 | protein_coding | 53.12 | -0.000135849 | 0.000135849 | ENSG00000115661 | n.s. |
| ENSG00000180228 | 1785.566 | 0.025400338 | 0.271895 | 0.0934196 | 0.92557 | 0.98779046 | PRKRA | protein_coding | 39.65 | 0.000135515 | 0.000135515 | ENSG00000180228 | n.s. |
| ENSG00000111276 | 3768.63 | -0.023951165 | 0.239762 | -0.099896 | 0.920427 | 0.987068378 | CDKN1B | protein_coding | 45.43 | -0.00013539 | 0.00013539 | ENSG00000111276 | n.s. |
| ENSG00000126226 | 1229.562 | -0.026057492 | 0.287008 | -0.09079 | 0.927659 | 0.988157204 | PCID2 | protein_coding | 43.4 | -0.00013482 | 0.00013482 | ENSG00000126226 | n.s. |
| ENSG00000136261 | 2509.933 | -0.025736248 | 0.278495 | -0.092412 | 0.926371 | 0.988038912 | BZW2 | protein_coding | 37.77 | -0.000134496 | 0.000134496 | ENSG00000136261 | n.s. |
| ENSG00000062582 | 688.4179 | 0.028288711 | 0.336873 | 0.0839744 | 0.933077 | 0.98914507 | MRPS24 | protein_coding | 50.93 | 0.000134089 | 0.000134089 | ENSG00000062582 | n.s. |
| ENSG00000185721 | 1135.69 | -0.025854493 | 0.284276 | -0.090948 | 0.927534 | 0.988157204 | DRG1 | protein_coding | 44.41 | -0.00013377 | 0.00013377 | ENSG00000185721 | n.s. |
| ENSG00000183172 | 276.3538 | 0.025908697 | 0.292951 | 0.0884404 | 0.929527 | 0.988216325 | SMDT1 | protein_coding | 48.41 | 0.000133377 | 0.000133377 | ENSG00000183172 | n.s. |
| ENSG00000025156 | 2289.213 | 0.025713767 | 0.287931 | 0.0893053 | 0.928839 | 0.988157204 | HSF2 | protein_coding | 35.55 | 0.000133042 | 0.000133042 | ENSG00000025156 | n.s. |
| ENSG00000176102 | 1639.676 | 0.025682415 | 0.28868 | 0.0889649 | 0.92911 | 0.988157204 | CSTF3 | protein_coding | 37.22 | 0.00013288 | 0.00013288 | ENSG00000176102 | n.s. |
| ENSG00000136942 | 13339.75 | 0.027603423 | 0.327821 | 0.0842028 | 0.932895 | 0.98914507 | RPL35 | protein_coding | 56.22 | 0.00013084 | 0.00013084 | ENSG00000136942 | n.s. |
| ENSG00000176714 | 132.6262 | -0.023982319 | 0.254643 | -0.09418 | 0.924966 | 0.987524058 | CCDC121 | protein_coding | 42.38 | -0.000130759 | 0.000130759 | ENSG00000176714 | n.s. |
| ENSG00000199459 | 9.222994 | 0.046648298 | 0.772899 | 0.060355 | 0.951873 | 0.99357526 | | misc_RNA | 43.64 | 0.000130579 | 0.000130579 | ENSG00000199459 | n.s. |
| ENSG00000235954 | 150.172 | -0.023911038 | 0.253409 | -0.094357 | 0.924825 | 0.98751796 | TTC28-AS1 | processed_transcript | 43.33 | -0.000130435 | 0.000130435 | ENSG00000235954 | n.s. |
| ENSG00000163689 | 38.53806 | 0.027408449 | 0.31752 | 0.0863204 | 0.931212 | 0.989110489 | C3orf67 | protein_coding | 38.65 | 0.000130332 | 0.000130332 | ENSG00000163689 | n.s. |
| ENSG00000168301 | 600.3747 | -0.023482195 | 0.244276 | -0.09613 | 0.923418 | 0.987349084 | KCTD6 | protein_coding | 40.78 | -0.000129839 | 0.000129839 | ENSG00000168301 | n.s. |
| ENSG00000109534 | 372.2427 | -0.023415688 | 0.245263 | -0.095472 | 0.92394 | 0.987349084 | GAR1 | protein_coding | 39.37 | -0.000129472 | 0.000129472 | ENSG00000109534 | n.s. |
| ENSG00000151929 | 103.7911 | 0.025020906 | 0.278661 | 0.0897897 | 0.928454 | 0.988157204 | BAG3 | protein_coding | 49.77 | 0.000129457 | 0.000129457 | ENSG00000151929 | n.s. |
| ENSG00000117519 | 5256.236 | 0.025017887 | 0.279928 | 0.0893727 | 0.928786 | 0.988157204 | CNN3 | protein_coding | 43.03 | 0.000129442 | 0.000129442 | ENSG00000117519 | n.s. |
| ENSG00000276529 | 16.4232 | -0.030794688 | 0.42225 | -0.07293 | 0.941862 | 0.990390549 | | lincRNA | 55.76 | -0.000129138 | 0.000129138 | ENSG00000276529 | n.s. |
| ENSG00000165502 | 1775.62 | 0.024948412 | 0.274038 | 0.0910399 | 0.927461 | 0.988157204 | RPL36AL | protein_coding | 45.59 | 0.000129082 | 0.000129082 | ENSG00000165502 | n.s. |
| ENSG00000199753 | 97.84252 | -0.024403518 | 0.263398 | -0.092649 | 0.926183 | 0.987913954 | SNORD104 | snoRNA | 51.43 | -0.000128872 | 0.000128872 | ENSG00000199753 | n.s. |
| ENSG00000164951 | 1141.693 | 0.023292526 | 0.244833 | 0.0951364 | 0.924207 | 0.987349084 | PDP1 | protein_coding | 42.9 | 0.000128791 | 0.000128791 | ENSG00000164951 | n.s. |
| ENSG00000272391 | 154.2955 | 0.024745471 | 0.270337 | 0.0915357 | 0.927067 | 0.988157204 | POM121C | protein_coding | 45.74 | 0.000128032 | 0.000128032 | ENSG00000272391 | n.s. |

| | | | | | | | | | | | | | |
|---|---|---|---|---|---|---|---|---|---|---|---|---|---|
| ENSG00000155256 | 421.6652 | -0.024696584 | 0.275691 | -0.089581 | 0.92862 | 0.988157204 | ZFYVE27 | protein_coding | 48.53 | -0.000127779 | 0.000127779 | ENSG00000155256 | n.s. |
| ENSG00000166313 | 692.4155 | 0.025727045 | 0.294337 | 0.0874066 | 0.930348 | 0.988645945 | APBB1 | protein_coding | 51.87 | 0.000127586 | 0.000127586 | ENSG00000166313 | n.s. |
| ENSG00000063241 | 223.3773 | 0.027757592 | 0.344414 | 0.0805937 | 0.935765 | 0.989493204 | ISOC2 | protein_coding | 56.69 | 0.000127329 | 0.000127329 | ENSG00000063241 | n.s. |
| ENSG00000166900 | 515.6568 | -0.024458542 | 0.269432 | -0.090778 | 0.927669 | 0.988157204 | STX3 | protein_coding | 44.87 | -0.000126547 | 0.000126547 | ENSG00000166900 | n.s. |
| ENSG00000114120 | 3273.248 | 0.024456966 | 0.269374 | 0.090792 | 0.927658 | 0.988157204 | SLC25A36 | protein_coding | 35.47 | 0.000126539 | 0.000126539 | ENSG00000114120 | n.s. |
| ENSG00000199977 | 28.14683 | -0.02656718 | 0.319435 | -0.083169 | 0.933717 | 0.98914507 | | snoRNA | 46.83 | -0.000125929 | 0.000125929 | ENSG00000199977 | n.s. |
| ENSG00000104529 | 3972.956 | -0.024327562 | 0.271625 | -0.089563 | 0.928634 | 0.988157204 | EEF1D | protein_coding | 61.99 | -0.00012587 | 0.00012587 | ENSG00000104529 | n.s. |
| ENSG00000074695 | 2965.46 | 0.024232216 | 0.270829 | 0.0894741 | 0.928705 | 0.988157204 | LMAN1 | protein_coding | 37.33 | 0.000125376 | 0.000125376 | ENSG00000074695 | n.s. |
| ENSG00000172939 | 1256.523 | -0.024075249 | 0.262063 | -0.091868 | 0.926803 | 0.988157204 | OXSR1 | protein_coding | 38.67 | -0.000124564 | 0.000124564 | ENSG00000172939 | n.s. |
| ENSG00000164919 | 2388.829 | 0.026933931 | 0.333145 | 0.0808475 | 0.935563 | 0.98943003 | COX6C | protein_coding | 40.79 | 0.000124298 | 0.000124298 | ENSG00000164919 | n.s. |
| ENSG00000126698 | 1125.325 | -0.026932645 | 0.332916 | -0.080899 | 0.935522 | 0.98943003 | DNAJC8 | protein_coding | 43.16 | -0.000124292 | 0.000124292 | ENSG00000126698 | n.s. |
| ENSG00000231508 | 15.44397 | -0.029637594 | 0.404385 | -0.073291 | 0.941575 | 0.990390549 | RPL34P20 | processed_pseudogene | 44.73 | -0.000124286 | 0.000124286 | ENSG00000231508 | n.s. |
| ENSG00000228506 | 257.4688 | 0.022177098 | 0.225773 | 0.0982273 | 0.921752 | 0.987238446 | | antisense | 42.95 | 0.000123703 | 0.000123703 | ENSG00000228506 | n.s. |
| ENSG00000115486 | 117.2552 | -0.023904318 | 0.268166 | -0.08914 | 0.928971 | 0.988157204 | GGCX | protein_coding | 46.19 | -0.00012368 | 0.00012368 | ENSG00000115486 | n.s. |
| ENSG00000120805 | 1002.638 | -0.026034239 | 0.302939 | -0.085939 | 0.931515 | 0.989130548 | ARL1 | protein_coding | 39.51 | -0.000123569 | 0.000123569 | ENSG00000120805 | n.s. |
| ENSG00000081307 | 1698.521 | -0.023275223 | 0.249919 | -0.093131 | 0.9258 | 0.987870825 | UBA5 | protein_coding | 36.76 | -0.000123355 | 0.000123355 | ENSG00000081307 | n.s. |
| ENSG00000165678 | 2421.785 | 0.025218124 | 0.290365 | 0.0868496 | 0.930791 | 0.988828751 | GHITM | protein_coding | 37.88 | 0.000123037 | 0.000123037 | ENSG00000165678 | n.s. |
| ENSG00000111711 | 1505.343 | -0.025179796 | 0.28979 | -0.08689 | 0.930759 | 0.988828751 | GOLT1B | protein_coding | 35.99 | -0.00012285 | 0.00012285 | ENSG00000111711 | n.s. |
| ENSG00000196465 | 894.7445 | -0.027035415 | 0.339385 | -0.07966 | 0.936508 | 0.98960211 | MYL6B | protein_coding | 53.91 | -0.000122724 | 0.000122724 | ENSG00000196465 | n.s. |
| ENSG00000155090 | 1022.52 | -0.023662107 | 0.259565 | -0.091161 | 0.927365 | 0.988157204 | KLF10 | protein_coding | 48.47 | -0.000122427 | 0.000122427 | ENSG00000155090 | n.s. |
| ENSG00000136108 | 3182.934 | -0.023646743 | 0.262925 | -0.089937 | 0.928337 | 0.988157204 | CKAP2 | protein_coding | 38.86 | -0.000122347 | 0.000122347 | ENSG00000136108 | n.s. |
| ENSG00000121440 | 767.1896 | -0.023567606 | 0.257358 | -0.091575 | 0.927036 | 0.988157204 | PDZRN3 | protein_coding | 41.51 | -0.000121938 | 0.000121938 | ENSG00000121440 | n.s. |
| ENSG00000169762 | 237.0906 | 0.02566551 | 0.302083 | 0.0849619 | 0.932292 | 0.989130548 | TAPT1 | protein_coding | 38.85 | 0.000121818 | 0.000121818 | ENSG00000169762 | n.s. |
| ENSG00000167397 | 418.6391 | -0.025580331 | 0.301856 | -0.084743 | 0.932465 | 0.989130548 | VKORC1 | protein_coding | 55.01 | -0.000121414 | 0.000121414 | ENSG00000167397 | n.s. |
| ENSG00000148943 | 1921.691 | 0.023405332 | 0.261533 | 0.0894928 | 0.92869 | 0.988157204 | LIN7C | protein_coding | 37.11 | 0.000121098 | 0.000121098 | ENSG00000148943 | n.s. |
| ENSG00000081026 | 2930.849 | 0.023454164 | 0.264525 | 0.0886651 | 0.929348 | 0.988186394 | MAGI3 | protein_coding | 37.11 | 0.00012105 | 0.00012105 | ENSG00000081026 | n.s. |
| ENSG00000228043 | 83.82641 | 0.023696609 | 0.269921 | 0.087791 | 0.930043 | 0.98839672 | | lincRNA | 38.27 | 0.000120111 | 0.000120111 | ENSG00000228043 | n.s. |
| ENSG00000163399 | 8857.826 | 0.02313377 | 0.255844 | 0.0904213 | 0.927952 | 0.988157204 | ATP1A1 | protein_coding | 44.46 | 0.000119693 | 0.000119693 | ENSG00000163399 | n.s. |
| ENSG00000225377 | 53.79765 | 0.025207689 | 0.298903 | 0.0843341 | 0.932791 | 0.98914507 | NRSN2-AS1 | antisense | 52 | 0.000119485 | 0.000119485 | ENSG00000225377 | n.s. |
| ENSG00000037897 | 417.1413 | -0.021387562 | 0.218326 | -0.097961 | 0.921963 | 0.987238446 | METTL1 | protein_coding | 54.22 | -0.000119299 | 0.000119299 | ENSG00000037897 | n.s. |
| ENSG00000229989 | 1007.605 | 0.022373892 | 0.240451 | 0.0930496 | 0.925864 | 0.987870825 | MIR181A1HG | lincRNA | 35.03 | 0.000118578 | 0.000118578 | ENSG00000229989 | n.s. |
| ENSG00000205758 | 473.7314 | 0.024973244 | 0.295001 | 0.0846547 | 0.932536 | 0.989130548 | CRYZL1 | protein_coding | 41.59 | 0.000118533 | 0.000118533 | ENSG00000205758 | n.s. |

| | | | | | | | | | | | | | |
|---|---|---|---|---|---|---|---|---|---|---|---|---|---|
| ENSG00000134759 | 2592.383 | -0.022985816 | 0.261151 | -0.088017 | 0.929863 | 0.988281026 | ELP2 | protein_coding | 40.55 | -0.000117677 | 0.000117677 | ENSG00000134759 | n.s. |
| ENSG00000230844 | 192.1991 | 0.022472021 | 0.250112 | 0.0898479 | 0.928408 | 0.988157204 | ZNF674-AS1 | lincRNA | 47.02 | 0.000116269 | 0.000116269 | ENSG00000230844 | n.s. |
| ENSG00000116667 | 221.2544 | -0.024465976 | 0.293883 | -0.083251 | 0.933652 | 0.98914507 | C1orf21 | protein_coding | 40.26 | -0.000115969 | 0.000115969 | ENSG00000116667 | n.s. |
| ENSG00000169567 | 4172.243 | 0.024342482 | 0.295759 | 0.0823052 | 0.934404 | 0.98914507 | HINT1 | protein_coding | 43.5 | 0.000115384 | 0.000115384 | ENSG00000169567 | n.s. |
| ENSG00000252391 | 21.2795 | -0.027411318 | 0.374477 | -0.073199 | 0.941648 | 0.990390549 | RNU6-638P | snRNA | 36.54 | -0.00011495 | 0.00011495 | ENSG00000252391 | n.s. |
| ENSG00000120616 | 3276.379 | 0.022216375 | 0.242791 | 0.091504 | 0.927092 | 0.988157204 | EPC1 | protein_coding | 38.9 | 0.000114947 | 0.000114947 | ENSG00000120616 | n.s. |
| ENSG00000242114 | 238.8153 | -0.024071225 | 0.293045 | -0.082142 | 0.934534 | 0.98914507 | MTFP1 | protein_coding | 56.98 | -0.000114098 | 0.000114098 | ENSG00000242114 | n.s. |
| ENSG00000170166 | 56.94528 | -0.025954924 | 0.32932 | -0.078814 | 0.937181 | 0.989947183 | HOXD4 | protein_coding | 59.55 | -0.00011389 | 0.00011389 | ENSG00000170166 | n.s. |
| ENSG00000131019 | 65.8829 | -0.023934338 | 0.286983 | -0.0834 | 0.933534 | 0.98914507 | ULBP3 | protein_coding | 54.66 | -0.000113449 | 0.000113449 | ENSG00000131019 | n.s. |
| ENSG00000197622 | 1376.741 | -0.02388715 | 0.280096 | -0.085282 | 0.932037 | 0.989130548 | CDC42SE1 | protein_coding | 46.96 | -0.000113378 | 0.000113378 | ENSG00000197622 | n.s. |
| ENSG00000163535 | 1208.447 | -0.023849597 | 0.278632 | -0.085595 | 0.931788 | 0.989130548 | SGO2 | protein_coding | 37.75 | -0.000113199 | 0.000113199 | ENSG00000163535 | n.s. |
| ENSG00000134644 | 2144.14 | -0.021835252 | 0.245599 | -0.088906 | 0.929157 | 0.988157204 | PUM1 | protein_coding | 41.05 | -0.000112975 | 0.000112975 | ENSG00000134644 | n.s. |
| ENSG00000263793 | 4.354091 | 0.040307 | 0.670785 | 0.0600893 | 0.952085 | 0.99357526 | MIR3115 | miRNA | 41.18 | 0.000112829 | 0.000112829 | ENSG00000263793 | n.s. |
| ENSG00000172171 | 168.1836 | 0.023758277 | 0.286696 | 0.0828693 | 0.933955 | 0.98914507 | TEFM | protein_coding | 43.17 | 0.000112614 | 0.000112614 | ENSG00000172171 | n.s. |
| ENSG00000143106 | 2292.872 | -0.024381142 | 0.301223 | -0.080941 | 0.935489 | 0.98943003 | PSMA5 | protein_coding | 40.74 | -0.000112517 | 0.000112517 | ENSG00000143106 | n.s. |
| ENSG00000105185 | 1692.571 | -0.023635725 | 0.276088 | -0.085609 | 0.931777 | 0.989130548 | PDCD5 | protein_coding | 45.84 | -0.000112184 | 0.000112184 | ENSG00000105185 | n.s. |
| ENSG00000136143 | 1050.982 | -0.023480239 | 0.277548 | -0.084599 | 0.93258 | 0.989130548 | SUCLA2 | protein_coding | 38.08 | -0.000111446 | 0.000111446 | ENSG00000136143 | n.s. |
| ENSG00000182168 | 1219.011 | 0.021616873 | 0.244375 | 0.0884577 | 0.929513 | 0.988216325 | UNC5C | protein_coding | 37.41 | 0.000111283 | 0.000111283 | ENSG00000182168 | n.s. |
| ENSG00000166450 | 228.01 | 0.021381926 | 0.238882 | 0.0895084 | 0.928678 | 0.988157204 | PRTG | protein_coding | 38.58 | 0.000110629 | 0.000110629 | ENSG00000166450 | n.s. |
| ENSG00000147905 | 514.0923 | 0.023327662 | 0.281881 | 0.0827573 | 0.934045 | 0.98914507 | ZCCHC7 | protein_coding | 36.62 | 0.000110573 | 0.000110573 | ENSG00000147905 | n.s. |
| ENSG00000201047 | 3.477277 | -0.039356038 | 0.662716 | -0.059386 | 0.952645 | 0.99357526 | | misc_RNA | 38.24 | -0.000110167 | 0.000110167 | ENSG00000201047 | n.s. |
| ENSG00000137185 | 904.052 | -0.021263373 | 0.232805 | -0.091336 | 0.927226 | 0.988157204 | ZSCAN9 | protein_coding | 43.42 | -0.000110016 | 0.000110016 | ENSG00000137185 | n.s. |
| ENSG00000106355 | 1710.452 | 0.02305757 | 0.266822 | 0.0864155 | 0.931136 | 0.989105712 | LSM5 | protein_coding | 39.73 | 0.000109692 | 0.000109692 | ENSG00000106355 | n.s. |
| ENSG00000184432 | 2717.92 | 0.023090304 | 0.268538 | 0.0859851 | 0.931478 | 0.989130548 | COPB2 | protein_coding | 37.63 | 0.000109595 | 0.000109595 | ENSG00000184432 | n.s. |
| ENSG00000127914 | 4384.095 | 0.023035502 | 0.274044 | 0.0840575 | 0.933011 | 0.98914507 | AKAP9 | protein_coding | 36.72 | 0.000109188 | 0.000109188 | ENSG00000127914 | n.s. |
| ENSG00000156508 | 123363.2 | 0.024044667 | 0.300514 | 0.0800119 | 0.936228 | 0.98960211 | EEF1A1 | protein_coding | 47.14 | 0.000109148 | 0.000109148 | ENSG00000156508 | n.s. |
| ENSG00000132591 | 441.4836 | -0.022163998 | 0.25412 | -0.087218 | 0.930498 | 0.988729417 | ERAL1 | protein_coding | 50.7 | -0.000109103 | 0.000109103 | ENSG00000132591 | n.s. |
| ENSG00000252835 | 497.7312 | -0.020859165 | 0.226035 | -0.092283 | 0.926473 | 0.988072469 | SCARNA21 | scaRNA | 64.03 | -0.000108701 | 0.000108701 | ENSG00000252835 | n.s. |
| ENSG00000182551 | 1352.039 | -0.022925213 | 0.278198 | -0.082406 | 0.934324 | 0.98914507 | ADI1 | protein_coding | 44.71 | -0.000108666 | 0.000108666 | ENSG00000182551 | n.s. |
| ENSG00000214114 | 205.5246 | -0.02574348 | 0.331431 | -0.077674 | 0.938088 | 0.990343731 | MYCBP | protein_coding | 44.31 | -0.000108484 | 0.000108484 | ENSG00000214114 | n.s. |
| ENSG00000102119 | 286.7335 | -0.022758886 | 0.268894 | -0.084639 | 0.932549 | 0.989130548 | EMD | protein_coding | 62.14 | -0.000108022 | 0.000108022 | ENSG00000102119 | n.s. |
| ENSG00000089053 | 3568.286 | -0.022725329 | 0.268606 | -0.084605 | 0.932576 | 0.989130548 | ANAPC5 | protein_coding | 44.55 | -0.000107863 | 0.000107863 | ENSG00000089053 | n.s. |

| | | | | | | | | | | | | | |
|---|---|---|---|---|---|---|---|---|---|---|---|---|---|
| ENSG00000278845 | 557.1666 | 0.024567195 | 0.312466 | 0.0786236 | 0.937332 | 0.989947183 | MRPL45 | protein_coding | 45.16 | 0.0001078 | 0.0001078 | ENSG00000278845 | n.s. |
| ENSG00000155100 | 943.6381 | -0.023286304 | 0.287701 | -0.080939 | 0.93549 | 0.98943003 | OTUD6B | protein_coding | 34.94 | -0.000107464 | 0.000107464 | ENSG00000155100 | n.s. |
| ENSG00000129116 | 388.3899 | -0.022615843 | 0.26576 | -0.085099 | 0.932183 | 0.989130548 | PALLD | protein_coding | 39.87 | -0.000107344 | 0.000107344 | ENSG00000129116 | n.s. |
| ENSG00000107798 | 506.0284 | -0.024737428 | 0.316096 | -0.078259 | 0.937622 | 0.990063608 | LIPA | protein_coding | 41.43 | -0.000107284 | 0.000107284 | ENSG00000107798 | n.s. |
| ENSG00000121892 | 4136.898 | -0.022594449 | 0.272715 | -0.08285 | 0.933971 | 0.98914507 | PDS5A | protein_coding | 40.35 | -0.000107098 | 0.000107098 | ENSG00000121892 | n.s. |
| ENSG00000210174 | 3.800498 | -0.038059613 | 0.658159 | -0.057827 | 0.953886 | 0.99357526 | MT-TR | Mt_tRNA | 23.08 | -0.000106538 | 0.000106538 | ENSG00000210174 | n.s. |
| ENSG00000144199 | 146.125 | -0.025221451 | 0.325226 | -0.07755 | 0.938186 | 0.990343731 | FAHD2B | protein_coding | 48.77 | -0.000106284 | 0.000106284 | ENSG00000144199 | n.s. |
| ENSG00000263069 | 19.33563 | -0.028432285 | 0.404213 | -0.07034 | 0.943923 | 0.991514406 | | antisense | 48.14 | -0.000105227 | 0.000105227 | ENSG00000263069 | n.s. |
| ENSG00000099949 | 573.9916 | 0.022153741 | 0.268027 | 0.082655 | 0.934126 | 0.98914507 | LZTR1 | protein_coding | 58.44 | 0.000105009 | 0.000105009 | ENSG00000099949 | n.s. |
| ENSG00000136270 | 1363.428 | 0.020257563 | 0.227127 | 0.0891904 | 0.928931 | 0.988157204 | TBRG4 | protein_coding | 53.25 | 0.000104812 | 0.000104812 | ENSG00000136270 | n.s. |
| ENSG00000176244 | 157.7594 | -0.024936384 | 0.325036 | -0.076719 | 0.938847 | 0.990390549 | ACBD7 | protein_coding | 46.04 | -0.000104571 | 0.000104571 | ENSG00000176244 | n.s. |
| ENSG00000182481 | 11945.38 | 0.021402184 | 0.246476 | 0.0868328 | 0.930804 | 0.988828751 | KPNA2 | protein_coding | 43.84 | 0.000104419 | 0.000104419 | ENSG00000182481 | n.s. |
| ENSG00000196967 | 120.0575 | -0.024827859 | 0.333591 | -0.074426 | 0.940671 | 0.990390549 | ZNF585A | protein_coding | 40.77 | -0.000104116 | 0.000104116 | ENSG00000196967 | n.s. |
| ENSG00000011258 | 2550.357 | -0.021924554 | 0.265098 | -0.082703 | 0.934087 | 0.98914507 | MBTD1 | protein_coding | 39.5 | -0.000103923 | 0.000103923 | ENSG00000011258 | n.s. |
| ENSG00000196839 | 114.5135 | 0.024780256 | 0.327324 | 0.0757055 | 0.939653 | 0.990390549 | ADA | protein_coding | 53.46 | 0.000103916 | 0.000103916 | ENSG00000196839 | n.s. |
| ENSG00000071794 | 3321.16 | 0.022513436 | 0.278289 | 0.0808996 | 0.935522 | 0.98943003 | HLTF | protein_coding | 35.95 | 0.000103898 | 0.000103898 | ENSG00000071794 | n.s. |
| ENSG00000101444 | 2847.256 | -0.023325835 | 0.294467 | -0.079214 | 0.936863 | 0.989819451 | AHCY | protein_coding | 50.38 | -0.00010366 | 0.00010366 | ENSG00000101444 | n.s. |
| ENSG00000138430 | 2897.719 | -0.022453318 | 0.276196 | -0.081295 | 0.935207 | 0.98943003 | OLA1 | protein_coding | 36.5 | -0.00010362 | 0.00010362 | ENSG00000138430 | n.s. |
| ENSG00000128656 | 1601.932 | 0.021849282 | 0.266196 | 0.0820797 | 0.934583 | 0.98914507 | CHN1 | protein_coding | 37.22 | 0.000103566 | 0.000103566 | ENSG00000128656 | n.s. |
| ENSG00000170035 | 2152.757 | 0.021658536 | 0.262092 | 0.0826372 | 0.93414 | 0.98914507 | UBE2E3 | protein_coding | 35.61 | 0.000102662 | 0.000102662 | ENSG00000170035 | n.s. |
| ENSG00000121060 | 655.4066 | 0.021654165 | 0.260614 | 0.083089 | 0.933781 | 0.98914507 | TRIM25 | protein_coding | 49.18 | 0.000102641 | 0.000102641 | ENSG00000121060 | n.s. |
| ENSG00000198521 | 2129.17 | -0.021961493 | 0.268743 | -0.081719 | 0.93487 | 0.98931345 | ZNF43 | protein_coding | 41.78 | -0.000102474 | 0.000102474 | ENSG00000198521 | n.s. |
| ENSG00000198755 | 12121.98 | 0.023700052 | 0.303493 | 0.078091 | 0.937756 | 0.990094174 | RPL10A | protein_coding | 57.02 | 0.000102467 | 0.000102467 | ENSG00000198755 | n.s. |
| ENSG00000272398 | 12776.23 | 0.021489771 | 0.261104 | 0.0823036 | 0.934405 | 0.98914507 | CD24 | protein_coding | 50.84 | 0.000101862 | 0.000101862 | ENSG00000272398 | n.s. |
| ENSG00000154719 | 729.4076 | 0.022158944 | 0.274638 | 0.0806843 | 0.935693 | 0.989492194 | MRPL39 | protein_coding | 37.94 | 0.000101657 | 0.000101657 | ENSG00000154719 | n.s. |
| ENSG00000221716 | 5.481836 | 0.036304287 | 0.589224 | 0.0616137 | 0.95087 | 0.99357526 | SNORA11 | snoRNA | 53.91 | 0.000101624 | 0.000101624 | ENSG00000221716 | n.s. |
| ENSG00000274618 | 278.3223 | -0.019690917 | 0.223296 | -0.088183 | 0.929731 | 0.988216325 | HIST1H4F | protein_coding | 57.37 | -0.000101368 | 0.000101368 | ENSG00000274618 | n.s. |
| ENSG00000203724 | 103.9164 | -0.024167532 | 0.31804 | -0.075989 | 0.939428 | 0.990390549 | C1orf53 | protein_coding | 40.35 | -0.000101347 | 0.000101347 | ENSG00000203724 | n.s. |
| ENSG00000172428 | 846.6295 | -0.024161569 | 0.330014 | -0.073214 | 0.941636 | 0.990390549 | COPS9 | protein_coding | 54.66 | -0.000101322 | 0.000101322 | ENSG00000172428 | n.s. |
| ENSG00000212607 | 84.96062 | -0.021339456 | 0.251696 | -0.084783 | 0.932434 | 0.989130548 | SNORA3B | snoRNA | 52.67 | -0.000101285 | 0.000101285 | ENSG00000212607 | n.s. |
| ENSG00000109519 | 1014.359 | -0.02169706 | 0.265567 | -0.081701 | 0.934885 | 0.98931345 | GRPEL1 | protein_coding | 47.5 | -0.00010124 | 0.00010124 | ENSG00000109519 | n.s. |
| ENSG00000105671 | 708.5416 | 0.022300781 | 0.278673 | 0.0800249 | 0.936217 | 0.98960211 | DDX49 | protein_coding | 57.12 | 0.000101232 | 0.000101232 | ENSG00000105671 | n.s. |

| | | | | | | | | | | | | | |
|---|---|---|---|---|---|---|---|---|---|---|---|---|---|
| ENSG00000110328 | 151.2042 | 0.021933883 | 0.269479 | 0.0813938 | 0.935129 | 0.98943003 | GALNT18 | protein_coding | 46.35 | 0.000101223 | 0.000101223 | ENSG00000110328 | n.s. |
| ENSG00000140905 | 218.3835 | 0.02126921 | 0.252926 | 0.0840925 | 0.932983 | 0.98914507 | GCSH | protein_coding | 43.92 | 0.000100816 | 0.000100816 | ENSG00000140905 | n.s. |
| ENSG00000273080 | 85.6184 | -0.022068847 | 0.276804 | -0.079727 | 0.936454 | 0.98960211 | | antisense | 51.74 | -0.000100179 | 0.000100179 | ENSG00000273080 | n.s. |
| ENSG00000168237 | 171.7632 | 0.021109055 | 0.25292 | 0.0834613 | 0.933485 | 0.98914507 | GLYCTK | protein_coding | 56.59 | 0.000100057 | 0.000100057 | ENSG00000168237 | n.s. |
| ENSG00000128805 | 43.62879 | 0.023763951 | 0.315533 | 0.0753137 | 0.939965 | 0.990390549 | ARHGAP22 | protein_coding | 47.75 | 9.97E-05 | 9.97E-05 | ENSG00000128805 | n.s. |
| ENSG00000006377 | 118.0098 | 0.020986913 | 0.254336 | 0.0825165 | 0.934236 | 0.98914507 | DLX6 | protein_coding | 49.51 | 9.95E-05 | 9.95E-05 | ENSG00000006377 | n.s. |
| ENSG00000103024 | 169.1853 | 0.02358672 | 0.306231 | 0.0770227 | 0.938605 | 0.990390549 | NME3 | protein_coding | 69.2 | 9.89E-05 | 9.89E-05 | ENSG00000103024 | n.s. |
| ENSG00000146352 | 5112.247 | 0.020838822 | 0.243566 | 0.0855572 | 0.931818 | 0.989130548 | CLVS2 | protein_coding | 36.42 | 9.89E-05 | 9.89E-05 | ENSG00000146352 | n.s. |
| ENSG00000100916 | 637.1141 | 0.021781257 | 0.271389 | 0.0802584 | 0.936032 | 0.98960211 | BRMS1L | protein_coding | 38.94 | 9.89E-05 | 9.89E-05 | ENSG00000100916 | n.s. |
| ENSG00000197860 | 665.2589 | 0.021321784 | 0.26355 | 0.0809023 | 0.93552 | 0.98943003 | SGTB | protein_coding | 39.79 | 9.84E-05 | 9.84E-05 | ENSG00000197860 | n.s. |
| ENSG00000263412 | 448.6999 | -0.019049299 | 0.215861 | -0.088248 | 0.929679 | 0.988216325 | | processed_transcript | 49.4 | -9.81E-05 | 9.81E-05 | ENSG00000263412 | n.s. |
| ENSG00000129353 | 137.9018 | -0.022503387 | 0.287705 | -0.078217 | 0.937656 | 0.990063608 | SLC44A2 | protein_coding | 51.73 | -9.76E-05 | 9.76E-05 | ENSG00000129353 | n.s. |
| ENSG00000140319 | 2573.45 | 0.022239219 | 0.282737 | 0.078657 | 0.937305 | 0.989947183 | SRP14 | protein_coding | 43.51 | 9.76E-05 | 9.76E-05 | ENSG00000140319 | n.s. |
| ENSG00000160294 | 1336.528 | 0.020575963 | 0.248442 | 0.0828201 | 0.933995 | 0.98914507 | MCM3AP | protein_coding | 47.4 | 9.75E-05 | 9.75E-05 | ENSG00000160294 | n.s. |
| ENSG00000120699 | 2134.069 | 0.02050145 | 0.238318 | 0.0860255 | 0.931446 | 0.989130548 | EXOSC8 | protein_coding | 40.36 | 9.73E-05 | 9.73E-05 | ENSG00000120699 | n.s. |
| ENSG00000122786 | 2167.113 | 0.021423862 | 0.268446 | 0.0798068 | 0.936391 | 0.98960211 | CALD1 | protein_coding | 40.05 | 9.73E-05 | 9.73E-05 | ENSG00000122786 | n.s. |
| ENSG00000175203 | 1662.048 | 0.023148954 | 0.300124 | 0.0771314 | 0.938519 | 0.990390549 | DCTN2 | protein_coding | 48.08 | 9.71E-05 | 9.71E-05 | ENSG00000175203 | n.s. |
| ENSG00000002549 | 995.4812 | -0.021301741 | 0.266518 | -0.079926 | 0.936296 | 0.98960211 | LAP3 | protein_coding | 44.17 | -9.67E-05 | 9.67E-05 | ENSG00000002549 | n.s. |
| ENSG00000182220 | 736.6939 | -0.023034264 | 0.3091 | -0.07452 | 0.940596 | 0.990390549 | ATP6AP2 | protein_coding | 41.77 | -9.66E-05 | 9.66E-05 | ENSG00000182220 | n.s. |
| ENSG00000255864 | 39.11006 | -0.022921468 | 0.310009 | -0.073938 | 0.94106 | 0.990390549 | | lincRNA | 37.59 | -9.61E-05 | 9.61E-05 | ENSG00000255864 | n.s. |
| ENSG00000111859 | 288.1615 | -0.020276201 | 0.24702 | -0.082083 | 0.934581 | 0.98914507 | NEDD9 | protein_coding | 43.09 | -9.61E-05 | 9.61E-05 | ENSG00000111859 | n.s. |
| ENSG00000101940 | 328.0357 | 0.02153568 | 0.272145 | 0.0791332 | 0.936927 | 0.989819451 | WDR13 | protein_coding | 51.72 | 9.57E-05 | 9.57E-05 | ENSG00000101940 | n.s. |
| ENSG00000171045 | 171.931 | -0.021033882 | 0.263683 | -0.07977 | 0.936421 | 0.98960211 | TSNARE1 | protein_coding | 55.7 | -9.55E-05 | 9.55E-05 | ENSG00000171045 | n.s. |
| ENSG00000089063 | 1618.103 | 0.022635968 | 0.297282 | 0.0761431 | 0.939305 | 0.990390549 | TMEM230 | protein_coding | 44.98 | 9.49E-05 | 9.49E-05 | ENSG00000089063 | n.s. |
| ENSG00000163378 | 129.6507 | 0.020023276 | 0.238839 | 0.0838359 | 0.933187 | 0.98914507 | EOGT | protein_coding | 41.82 | 9.49E-05 | 9.49E-05 | ENSG00000163378 | n.s. |
| ENSG00000111142 | 4813.678 | -0.021346672 | 0.269474 | -0.079216 | 0.936861 | 0.989819451 | METAP2 | protein_coding | 39.23 | -9.49E-05 | 9.49E-05 | ENSG00000111142 | n.s. |
| ENSG00000164062 | 859.1786 | 0.022569233 | 0.294019 | 0.0767611 | 0.938814 | 0.990390549 | APEH | protein_coding | 58.08 | 9.46E-05 | 9.46E-05 | ENSG00000164062 | n.s. |
| ENSG00000196998 | 279.4075 | -0.022351572 | 0.303828 | -0.073567 | 0.941355 | 0.990390549 | WDR45 | protein_coding | 50.92 | -9.37E-05 | 9.37E-05 | ENSG00000196998 | n.s. |
| ENSG00000141371 | 9.939708 | -0.033409634 | 0.56276 | -0.059367 | 0.952659 | 0.99357526 | C17orf64 | protein_coding | 45.51 | -9.35E-05 | 9.35E-05 | ENSG00000141371 | n.s. |
| ENSG00000104231 | 1275.701 | 0.022226561 | 0.288429 | 0.0770607 | 0.938575 | 0.990390549 | ZFAND1 | protein_coding | 37.45 | 9.32E-05 | 9.32E-05 | ENSG00000104231 | n.s. |
| ENSG00000106591 | 2393.587 | -0.021084455 | 0.268149 | -0.07863 | 0.937327 | 0.989947183 | MRPL32 | protein_coding | 40.85 | -9.25E-05 | 9.25E-05 | ENSG00000106591 | n.s. |
| ENSG00000084733 | 1832.158 | 0.021849308 | 0.292784 | 0.0746261 | 0.940512 | 0.990390549 | RAB10 | protein_coding | 40.62 | 9.16E-05 | 9.16E-05 | ENSG00000084733 | n.s. |

| | | | | | | | | | | | | | |
|---|---|---|---|---|---|---|---|---|---|---|---|---|---|
| ENSG00000105393 | 734.557 | -0.021795392 | 0.291821 | -0.074687 | 0.940463 | 0.990390549 | BABAM1 | protein_coding | 52.45 | -9.14E-05 | 9.14E-05 | ENSG00000105393 | n.s. |
| ENSG00000008513 | 74.11793 | -0.021662706 | 0.27953 | -0.077497 | 0.938228 | 0.990343731 | ST3GAL1 | protein_coding | 49.4 | -9.13E-05 | 9.13E-05 | ENSG00000008513 | n.s. |
| ENSG00000163479 | 2719.107 | -0.021608719 | 0.286019 | -0.07555 | 0.939777 | 0.990390549 | SSR2 | protein_coding | 46.99 | -9.06E-05 | 9.06E-05 | ENSG00000163479 | n.s. |
| ENSG00000127995 | 1759.589 | -0.021519069 | 0.292073 | -0.073677 | 0.941267 | 0.990390549 | CASD1 | protein_coding | 34.66 | -9.02E-05 | 9.02E-05 | ENSG00000127995 | n.s. |
| ENSG00000117335 | 3183.402 | -0.021479511 | 0.277833 | -0.077311 | 0.938376 | 0.990373886 | CD46 | protein_coding | 38.21 | -9.02E-05 | 9.02E-05 | ENSG00000117335 | n.s. |
| ENSG00000061794 | 1644.115 | -0.021415519 | 0.287232 | -0.074558 | 0.940566 | 0.990390549 | MRPS35 | protein_coding | 38.64 | -8.98E-05 | 8.98E-05 | ENSG00000061794 | n.s. |
| ENSG00000197837 | 857.5759 | -0.018864375 | 0.22028 | -0.085638 | 0.931754 | 0.989130548 | HIST4H4 | protein_coding | 49.57 | -8.95E-05 | 8.95E-05 | ENSG00000197837 | n.s. |
| ENSG00000215717 | 758.5573 | 0.021217106 | 0.278572 | 0.0761639 | 0.939289 | 0.990390549 | TMEM167B | protein_coding | 43.73 | 8.90E-05 | 8.90E-05 | ENSG00000215717 | n.s. |
| ENSG00000126088 | 416.2199 | -0.021145705 | 0.285213 | -0.07414 | 0.940899 | 0.990390549 | UROD | protein_coding | 52.29 | -8.87E-05 | 8.87E-05 | ENSG00000126088 | n.s. |
| ENSG00000138801 | 1436.729 | -0.021139128 | 0.286887 | -0.073684 | 0.941261 | 0.990390549 | PAPSS1 | protein_coding | 38.61 | -8.86E-05 | 8.86E-05 | ENSG00000138801 | n.s. |
| ENSG00000139668 | 2474.92 | -0.018671973 | 0.218867 | -0.085312 | 0.932013 | 0.989130548 | WDFY2 | protein_coding | 39.68 | -8.86E-05 | 8.86E-05 | ENSG00000139668 | n.s. |
| ENSG00000230269 | 188.6753 | 0.021077258 | 0.284155 | 0.0741753 | 0.940871 | 0.990390549 | | lincRNA | 45.16 | 8.84E-05 | 8.84E-05 | ENSG00000230269 | n.s. |
| ENSG00000067704 | 2467.814 | 0.02103048 | 0.280468 | 0.0749836 | 0.940228 | 0.990390549 | IARS2 | protein_coding | 39.21 | 8.82E-05 | 8.82E-05 | ENSG00000067704 | n.s. |
| ENSG00000170634 | 98.97995 | -0.020959895 | 0.274567 | -0.076338 | 0.93915 | 0.990390549 | ACYP2 | protein_coding | 40.31 | -8.79E-05 | 8.79E-05 | ENSG00000170634 | n.s. |
| ENSG00000158604 | 1098.364 | 0.020748226 | 0.273247 | 0.0759322 | 0.939473 | 0.990390549 | TMED4 | protein_coding | 48.93 | 8.70E-05 | 8.70E-05 | ENSG00000158604 | n.s. |
| ENSG00000100564 | 298.8961 | -0.022288528 | 0.311031 | -0.07166 | 0.942872 | 0.991070672 | PIGH | protein_coding | 43.92 | -8.68E-05 | 8.68E-05 | ENSG00000100564 | n.s. |
| ENSG00000163170 | 499.4336 | 0.022933577 | 0.324049 | 0.070772 | 0.943579 | 0.991373218 | BOLA3 | protein_coding | 50.59 | 8.63E-05 | 8.63E-05 | ENSG00000163170 | n.s. |
| ENSG00000222767 | 3.644835 | -0.035196436 | 0.655885 | -0.053663 | 0.957204 | 0.994376566 | | misc_RNA | 44.21 | -8.62E-05 | 8.62E-05 | ENSG00000222767 | n.s. |
| ENSG00000223749 | 210.3033 | -0.020142782 | 0.260121 | -0.077436 | 0.938277 | 0.990343731 | MIR503HG | lincRNA | 53.09 | -8.49E-05 | 8.49E-05 | ENSG00000223749 | n.s. |
| ENSG00000113328 | 2132.625 | 0.020212338 | 0.266718 | 0.0757818 | 0.939593 | 0.990390549 | CCNG1 | protein_coding | 37.5 | 8.48E-05 | 8.48E-05 | ENSG00000113328 | n.s. |
| ENSG00000178971 | 312.7114 | -0.02019297 | 0.266145 | -0.075872 | 0.939521 | 0.990390549 | CTC1 | protein_coding | 48.84 | -8.47E-05 | 8.47E-05 | ENSG00000178971 | n.s. |
| ENSG00000075142 | 1098.661 | 0.020188804 | 0.276087 | 0.0731247 | 0.941707 | 0.990390549 | SRI | protein_coding | 39.53 | 8.47E-05 | 8.47E-05 | ENSG00000075142 | n.s. |
| ENSG00000114978 | 2705.45 | -0.020032282 | 0.271633 | -0.073748 | 0.941211 | 0.990390549 | MOB1A | protein_coding | 41.02 | -8.40E-05 | 8.40E-05 | ENSG00000114978 | n.s. |
| ENSG00000163659 | 927.561 | -0.018998877 | 0.241931 | -0.07853 | 0.937406 | 0.989950514 | TIPARP | protein_coding | 38.27 | -8.33E-05 | 8.33E-05 | ENSG00000163659 | n.s. |
| ENSG00000132718 | 1572.084 | 0.020876634 | 0.289597 | 0.0720885 | 0.942532 | 0.990853681 | SYT11 | protein_coding | 46.5 | 8.33E-05 | 8.33E-05 | ENSG00000132718 | n.s. |
| ENSG00000134986 | 6719.603 | 0.019824224 | 0.267704 | 0.0740527 | 0.940968 | 0.990390549 | NREP | protein_coding | 38.48 | 8.31E-05 | 8.31E-05 | ENSG00000134986 | n.s. |
| ENSG00000183283 | 1686.199 | 0.019813646 | 0.266251 | 0.0744172 | 0.940678 | 0.990390549 | DAZAP2 | protein_coding | 46.84 | 8.31E-05 | 8.31E-05 | ENSG00000183283 | n.s. |
| ENSG00000197780 | 535.7327 | -0.019539672 | 0.258991 | -0.075445 | 0.93986 | 0.990390549 | TAF13 | protein_coding | 43.54 | -8.19E-05 | 8.19E-05 | ENSG00000197780 | n.s. |
| ENSG00000012061 | 557.5684 | -0.020985309 | 0.293819 | -0.071423 | 0.943061 | 0.991162611 | ERCC1 | protein_coding | 52.1 | -8.09E-05 | 8.09E-05 | ENSG00000012061 | n.s. |
| ENSG00000005810 | 8968.627 | 0.019278052 | 0.260878 | 0.0738969 | 0.941092 | 0.990390549 | MYCBP2 | protein_coding | 35.72 | 8.08E-05 | 8.08E-05 | ENSG00000005810 | n.s. |
| ENSG00000086061 | 7833.862 | 0.01908051 | 0.2615 | 0.0729656 | 0.941834 | 0.990390549 | DNAJA1 | protein_coding | 41.22 | 8.00E-05 | 8.00E-05 | ENSG00000086061 | n.s. |
| ENSG00000102531 | 2590.257 | -0.018944081 | 0.260487 | -0.072726 | 0.942024 | 0.990470225 | FNDC3A | protein_coding | 37.08 | -7.88E-05 | 7.88E-05 | ENSG00000102531 | n.s. |

| | | | | | | | | | | | | | |
|---|---|---|---|---|---|---|---|---|---|---|---|---|---|
| ENSG00000171126 | 158.7261 | -0.019284801 | 0.266491 | -0.072366 | 0.942311 | 0.990696552 | KCNG3 | protein_coding | 42.42 | -7.83E-05 | 7.83E-05 | ENSG00000171126 | n.s. |
| ENSG00000157796 | 732.0958 | 0.020864619 | 0.296694 | 0.0703237 | 0.943936 | 0.991514406 | WDR19 | protein_coding | 39.3 | 7.72E-05 | 7.72E-05 | ENSG00000157796 | n.s. |
| ENSG00000115446 | 855.887 | -0.018220604 | 0.242439 | -0.075155 | 0.940091 | 0.990390549 | UNC50 | protein_coding | 42.81 | -7.64E-05 | 7.64E-05 | ENSG00000115446 | n.s. |
| ENSG00000102931 | 651.4005 | -0.021309242 | 0.308736 | -0.069021 | 0.944973 | 0.991932047 | ARL2BP | protein_coding | 45.49 | -7.50E-05 | 7.50E-05 | ENSG00000102931 | n.s. |
| ENSG00000145386 | 3548.796 | -0.017846094 | 0.237685 | -0.075083 | 0.940149 | 0.990390549 | CCNA2 | protein_coding | 39.44 | -7.48E-05 | 7.48E-05 | ENSG00000145386 | n.s. |
| ENSG00000107874 | 847.1365 | 0.021267082 | 0.30746 | 0.0691702 | 0.944854 | 0.991932047 | CUEDC2 | protein_coding | 52.94 | 7.48E-05 | 7.48E-05 | ENSG00000107874 | n.s. |
| ENSG00000221669 | 4.099828 | 0.037864283 | 0.849067 | 0.0445951 | 0.96443 | 0.995533737 | MIR548N | miRNA | 36 | 7.36E-05 | 7.36E-05 | ENSG00000221669 | n.s. |
| ENSG00000162928 | 914.7862 | -0.019471169 | 0.274342 | -0.070974 | 0.943418 | 0.991337013 | PEX13 | protein_coding | 39.79 | -7.36E-05 | 7.36E-05 | ENSG00000162928 | n.s. |
| ENSG00000177879 | 992.6952 | -0.02064521 | 0.295675 | -0.069824 | 0.944334 | 0.991835548 | AP3S1 | protein_coding | 36.62 | -7.35E-05 | 7.35E-05 | ENSG00000177879 | n.s. |
| ENSG00000160766 | 184.2117 | -0.017462518 | 0.235361 | -0.074195 | 0.940855 | 0.990390549 | GBAP1 | cribed_unprocessed_pseudo | 49.29 | -7.32E-05 | 7.32E-05 | ENSG00000160766 | n.s. |
| ENSG00000188242 | 553.0937 | 0.017452908 | 0.23853 | 0.0731685 | 0.941672 | 0.990390549 | | antisense | 58.18 | 7.32E-05 | 7.32E-05 | ENSG00000188242 | n.s. |
| ENSG00000189319 | 711.7329 | 0.017428455 | 0.231166 | 0.0753936 | 0.939902 | 0.990390549 | FAM53B | protein_coding | 52.68 | 7.31E-05 | 7.31E-05 | ENSG00000189319 | n.s. |
| ENSG00000226498 | 50.29128 | -0.020524193 | 0.294248 | -0.069751 | 0.944392 | 0.991835548 | RPSAP21 | scribed_processed_pseudog | 51.75 | -7.31E-05 | 7.31E-05 | ENSG00000226498 | n.s. |
| ENSG00000122952 | 2383.772 | -0.017419952 | 0.237882 | -0.07323 | 0.941623 | 0.990390549 | ZWINT | protein_coding | 48.07 | -7.31E-05 | 7.31E-05 | ENSG00000122952 | n.s. |
| ENSG00000263764 | 2.694358 | 0.041874082 | 1.049718 | 0.0398908 | 0.96818 | 0.995991293 | SNORD43 | snoRNA | 45.31 | 7.30E-05 | 7.30E-05 | ENSG00000263764 | n.s. |
| ENSG00000267260 | 203.2288 | -0.01865422 | 0.261405 | -0.071361 | 0.94311 | 0.991162611 | | lincRNA | 45.82 | -7.19E-05 | 7.19E-05 | ENSG00000267260 | n.s. |
| ENSG00000120656 | 145.568 | 0.02038475 | 0.294063 | 0.069321 | 0.944734 | 0.991932047 | TAF12 | protein_coding | 47.17 | 7.17E-05 | 7.17E-05 | ENSG00000120656 | n.s. |
| ENSG00000273344 | 292.9872 | 0.016656404 | 0.228451 | 0.0729102 | 0.941878 | 0.990390549 | PAXIP1-AS1 | lincRNA | 48.32 | 6.98E-05 | 6.98E-05 | ENSG00000273344 | n.s. |
| ENSG00000165689 | 782.5981 | -0.020523466 | 0.300338 | -0.068334 | 0.945519 | 0.992271538 | SDCCAG3 | protein_coding | 56.51 | -6.92E-05 | 6.92E-05 | ENSG00000165689 | n.s. |
| ENSG00000177613 | 1834.243 | -0.016448996 | 0.213868 | -0.076912 | 0.938694 | 0.990390549 | CSTF2T | protein_coding | 43.36 | -6.90E-05 | 6.90E-05 | ENSG00000177613 | n.s. |
| ENSG00000081721 | 1578.82 | -0.018184294 | 0.258614 | -0.070314 | 0.943943 | 0.991514406 | DUSP12 | protein_coding | 39.87 | -6.73E-05 | 6.73E-05 | ENSG00000081721 | n.s. |
| ENSG00000165209 | 1030.927 | -0.018860111 | 0.270851 | -0.069633 | 0.944486 | 0.99185978 | STRBP | protein_coding | 39.12 | -6.69E-05 | 6.69E-05 | ENSG00000165209 | n.s. |
| ENSG00000235904 | 54.46181 | -0.020055223 | 0.294267 | -0.068153 | 0.945664 | 0.992348211 | RBMS3-AS3 | antisense | 37.6 | -6.69E-05 | 6.69E-05 | ENSG00000235904 | n.s. |
| ENSG00000177352 | 258.2528 | -0.015952087 | 0.216634 | -0.073636 | 0.9413 | 0.990390549 | CCDC71 | protein_coding | 56.27 | -6.69E-05 | 6.69E-05 | ENSG00000177352 | n.s. |
| ENSG00000140264 | 4658.907 | -0.020826101 | 0.310897 | -0.066987 | 0.946592 | 0.992723592 | SERF2 | protein_coding | 49.15 | -6.61E-05 | 6.61E-05 | ENSG00000140264 | n.s. |
| ENSG00000100650 | 7513.277 | 0.017027953 | 0.238945 | 0.071263 | 0.943188 | 0.991170089 | SRSF5 | protein_coding | 43.68 | 6.56E-05 | 6.56E-05 | ENSG00000100650 | n.s. |
| ENSG00000147576 | 27.75155 | -0.023394785 | 0.399255 | -0.058596 | 0.953274 | 0.99357526 | ADHFE1 | protein_coding | 43.01 | -6.55E-05 | 6.55E-05 | ENSG00000147576 | n.s. |
| ENSG00000090487 | 910.3988 | 0.018906133 | 0.274946 | 0.0687632 | 0.945178 | 0.99206296 | SPG21 | protein_coding | 44.04 | 6.54E-05 | 6.54E-05 | ENSG00000090487 | n.s. |
| ENSG00000120686 | 1626.42 | -0.019571282 | 0.288312 | -0.067882 | 0.945879 | 0.992391383 | UFM1 | protein_coding | 33.9 | -6.49E-05 | 6.49E-05 | ENSG00000120686 | n.s. |
| ENSG00000252130 | 40.69975 | -0.021571741 | 0.326041 | -0.066163 | 0.947248 | 0.993112862 | RNU6-1045P | snRNA | 31.78 | -6.47E-05 | 6.47E-05 | ENSG00000252130 | n.s. |
| ENSG00000163590 | 557.3787 | 0.018201022 | 0.2619 | 0.0694961 | 0.944595 | 0.991899272 | PPM1L | protein_coding | 38.44 | 6.43E-05 | 6.43E-05 | ENSG00000163590 | n.s. |
| ENSG00000170631 | 343.7608 | -0.016984525 | 0.240058 | -0.070752 | 0.943595 | 0.991373218 | ZNF16 | protein_coding | 47.4 | -6.39E-05 | 6.39E-05 | ENSG00000170631 | n.s. |

| | | | | | | | | | | | | | |
|---|---|---|---|---|---|---|---|---|---|---|---|---|---|
| ENSG00000038002 | 135.3463 | 0.019132732 | 0.282009 | 0.0678445 | 0.945909 | 0.992391383 | AGA | protein_coding | 39.89 | 6.35E-05 | 6.35E-05 | ENSG00000038002 | n.s. |
| ENSG00000143420 | 3794.828 | -0.018947427 | 0.279814 | -0.067714 | 0.946013 | 0.992391383 | ENSA | protein_coding | 45.32 | -6.28E-05 | 6.28E-05 | ENSG00000143420 | n.s. |
| ENSG00000177383 | 1819.919 | -0.018930672 | 0.279984 | -0.067613 | 0.946093 | 0.992391383 | MAGEF1 | protein_coding | 52.87 | -6.28E-05 | 6.28E-05 | ENSG00000177383 | n.s. |
| ENSG00000270487 | 3.659162 | -0.034996632 | 0.94334 | -0.037099 | 0.970406 | 0.995991293 | | antisense | 42.64 | -6.11E-05 | 6.11E-05 | ENSG00000270487 | n.s. |
| ENSG00000186767 | 389.3036 | -0.017305823 | 0.250775 | -0.069009 | 0.944982 | 0.991932047 | SPIN4 | protein_coding | 37.43 | -6.09E-05 | 6.09E-05 | ENSG00000186767 | n.s. |
| ENSG00000164405 | 2356.91 | 0.019823994 | 0.30566 | 0.0648564 | 0.948288 | 0.993188446 | UQCRQ | protein_coding | 51.29 | 5.88E-05 | 5.88E-05 | ENSG00000164405 | n.s. |
| ENSG00000170315 | 11400.01 | 0.020568989 | 0.324393 | 0.0634076 | 0.949442 | 0.993482322 | UBB | protein_coding | 51.44 | 5.84E-05 | 5.84E-05 | ENSG00000170315 | n.s. |
| ENSG00000127415 | 13.70831 | -0.023666883 | 0.44122 | -0.05364 | 0.957222 | 0.994376566 | IDUA | protein_coding | 65.45 | -5.80E-05 | 5.80E-05 | ENSG00000127415 | n.s. |
| ENSG00000121486 | 392.0229 | -0.017141015 | 0.253368 | -0.067653 | 0.946062 | 0.992391383 | TRMT1L | protein_coding | 35.55 | -5.69E-05 | 5.69E-05 | ENSG00000121486 | n.s. |
| ENSG00000234329 | 38.15428 | -0.020010533 | 0.317569 | -0.063012 | 0.949757 | 0.993482322 | | processed_pseudogene | 50.13 | -5.68E-05 | 5.68E-05 | ENSG00000234329 | n.s. |
| ENSG00000252695 | 3.837129 | 0.032515132 | 0.789139 | 0.0412033 | 0.967134 | 0.995991293 | MIR2276 | miRNA | 57.3 | 5.67E-05 | 5.67E-05 | ENSG00000252695 | n.s. |
| ENSG00000110442 | 559.3899 | -0.018166379 | 0.272273 | -0.066721 | 0.946804 | 0.992870867 | COMMD9 | protein_coding | 42.17 | -5.64E-05 | 5.64E-05 | ENSG00000110442 | n.s. |
| ENSG00000006015 | 327.7048 | -0.020135123 | 0.328438 | -0.061306 | 0.951116 | 0.99357526 | C19orf60 | protein_coding | 60.24 | -5.64E-05 | 5.64E-05 | ENSG00000006015 | n.s. |
| ENSG00000162909 | 1402.201 | -0.018667819 | 0.282629 | -0.066051 | 0.947338 | 0.993131614 | CAPN2 | protein_coding | 48.73 | -5.59E-05 | 5.59E-05 | ENSG00000162909 | n.s. |
| ENSG00000164258 | 982.1257 | 0.019890059 | 0.319905 | 0.062175 | 0.950423 | 0.99357526 | NDUFS4 | protein_coding | 36.57 | 5.57E-05 | 5.57E-05 | ENSG00000164258 | n.s. |
| ENSG00000254330 | 68.26165 | -0.01986595 | 0.331989 | -0.059839 | 0.952284 | 0.99357526 | | sense_intronic | 36.98 | -5.56E-05 | 5.56E-05 | ENSG00000254330 | n.s. |
| ENSG00000284154 | 5.651086 | -0.026866848 | 0.56181 | -0.047822 | 0.961858 | 0.995247333 | MIR3605 | miRNA | 55 | -5.56E-05 | 5.56E-05 | ENSG00000284154 | n.s. |
| ENSG00000109805 | 4943.458 | -0.016119584 | 0.234883 | -0.068628 | 0.945286 | 0.992101029 | NCAPG | protein_coding | 35.41 | -5.55E-05 | 5.55E-05 | ENSG00000109805 | n.s. |
| ENSG00000168152 | 402.1579 | 0.019738225 | 0.316852 | 0.0622947 | 0.950328 | 0.99357526 | THAP9 | protein_coding | 37.02 | 5.53E-05 | 5.53E-05 | ENSG00000168152 | n.s. |
| ENSG00000132405 | 1668.056 | -0.018403804 | 0.277559 | -0.066306 | 0.947134 | 0.993112742 | TBC1D14 | protein_coding | 49.5 | -5.52E-05 | 5.52E-05 | ENSG00000132405 | n.s. |
| ENSG00000126787 | 1462.419 | 0.01858287 | 0.2862 | 0.0649298 | 0.94823 | 0.993188446 | DLGAP5 | protein_coding | 39.11 | 5.52E-05 | 5.52E-05 | ENSG00000126787 | n.s. |
| ENSG00000201223 | 98.37176 | -0.016535289 | 0.244734 | -0.067564 | 0.946133 | 0.992391383 | RNU6-1305P | snRNA | 46.15 | -5.48E-05 | 5.48E-05 | ENSG00000201223 | n.s. |
| ENSG00000226328 | 99.8014 | -0.019442029 | 0.335016 | -0.058033 | 0.953722 | 0.99357526 | NUP50-AS1 | lincRNA | 41.42 | -5.44E-05 | 5.44E-05 | ENSG00000226328 | n.s. |
| ENSG00000072042 | 1705.836 | 0.018299658 | 0.280557 | 0.0652261 | 0.947994 | 0.993188446 | RDH11 | protein_coding | 42.66 | 5.43E-05 | 5.43E-05 | ENSG00000072042 | n.s. |
| ENSG00000214736 | 1604.889 | 0.019074153 | 0.302673 | 0.0630191 | 0.949751 | 0.993482322 | TOMM6 | protein_coding | 50.87 | 5.42E-05 | 5.42E-05 | ENSG00000214736 | n.s. |
| ENSG00000136156 | 1923.803 | -0.019309939 | 0.314595 | -0.06138 | 0.951056 | 0.99357526 | ITM2B | protein_coding | 37.31 | -5.41E-05 | 5.41E-05 | ENSG00000136156 | n.s. |
| ENSG00000181938 | 1646.305 | 0.018334304 | 0.284171 | 0.0645185 | 0.948557 | 0.993288546 | GINS3 | protein_coding | 41.93 | 5.36E-05 | 5.36E-05 | ENSG00000181938 | n.s. |
| ENSG00000187713 | 427.4194 | -0.018056788 | 0.276844 | -0.065224 | 0.947996 | 0.993188446 | TMEM203 | protein_coding | 56.71 | -5.36E-05 | 5.36E-05 | ENSG00000187713 | n.s. |
| ENSG00000215021 | 6075.385 | -0.019145359 | 0.31677 | -0.060439 | 0.951806 | 0.99357526 | PHB2 | protein_coding | 53.17 | -5.36E-05 | 5.36E-05 | ENSG00000215021 | n.s. |
| ENSG00000106153 | 4861.199 | 0.019012337 | 0.312917 | 0.0607585 | 0.951552 | 0.99357526 | CHCHD2 | protein_coding | 45.29 | 5.32E-05 | 5.32E-05 | ENSG00000106153 | n.s. |
| ENSG00000162998 | 787.071 | -0.018110869 | 0.280274 | -0.064618 | 0.948478 | 0.99327997 | FRZB | protein_coding | 36.85 | -5.30E-05 | 5.30E-05 | ENSG00000162998 | n.s. |
| ENSG00000238390 | 39.74777 | -0.018656837 | 0.295202 | -0.0632 | 0.949607 | 0.993482322 | | snoRNA | 41.24 | -5.30E-05 | 5.30E-05 | ENSG00000238390 | n.s. |

| | | | | | | | | | | | | | |
|---|---|---|---|---|---|---|---|---|---|---|---|---|---|
| ENSG00000184831 | 259.3098 | 0.018892695 | 0.314006 | 0.0601666 | 0.952023 | 0.99357526 | APOO | protein_coding | 43.27 | 5.29E-05 | 5.29E-05 | ENSG00000184831 | n.s. |
| ENSG00000215301 | 11496.86 | 0.015857168 | 0.2351 | 0.0674487 | 0.946224 | 0.992413077 | DDX3X | protein_coding | 42.68 | 5.24E-05 | 5.24E-05 | ENSG00000215301 | n.s. |
| ENSG00000247077 | 823.3669 | 0.018303745 | 0.28607 | 0.0639835 | 0.948983 | 0.993435766 | PGAM5 | protein_coding | 55.09 | 5.24E-05 | 5.24E-05 | ENSG00000247077 | n.s. |
| ENSG00000138821 | 272.6259 | 0.017597623 | 0.269232 | 0.0653624 | 0.947885 | 0.993188446 | SLC39A8 | protein_coding | 38.15 | 5.22E-05 | 5.22E-05 | ENSG00000138821 | n.s. |
| ENSG00000116171 | 1142.086 | -0.018366056 | 0.289178 | -0.063511 | 0.949359 | 0.993482322 | SCP2 | protein_coding | 40.01 | -5.22E-05 | 5.22E-05 | ENSG00000116171 | n.s. |
| ENSG00000110871 | 616.6176 | -0.018595816 | 0.309305 | -0.060121 | 0.952059 | 0.99357526 | COQ5 | protein_coding | 45.87 | -5.21E-05 | 5.21E-05 | ENSG00000110871 | n.s. |
| ENSG00000265706 | 37.19751 | -0.019229466 | 0.338922 | -0.056737 | 0.954755 | 0.993798482 | SNORD53B | snoRNA | 46.15 | -5.20E-05 | 5.20E-05 | ENSG00000265706 | n.s. |
| ENSG00000086712 | 950.5985 | -0.017485112 | 0.265998 | -0.065734 | 0.94759 | 0.993188446 | TXLNG | protein_coding | 42.93 | -5.19E-05 | 5.19E-05 | ENSG00000086712 | n.s. |
| ENSG00000115268 | 6226.373 | -0.017722174 | 0.275087 | -0.064424 | 0.948633 | 0.993292641 | RPS15 | protein_coding | 62.7 | -5.18E-05 | 5.18E-05 | ENSG00000115268 | n.s. |
| ENSG00000161016 | 11693.97 | -0.018338907 | 0.319059 | -0.057478 | 0.954164 | 0.99357526 | RPL8 | protein_coding | 60.89 | -5.13E-05 | 5.13E-05 | ENSG00000161016 | n.s. |
| ENSG00000118961 | 1305.272 | -0.018301619 | 0.304897 | -0.060026 | 0.952135 | 0.99357526 | LDAH | protein_coding | 38.21 | -5.12E-05 | 5.12E-05 | ENSG00000118961 | n.s. |
| ENSG00000010244 | 9037.056 | 0.016863479 | 0.254534 | 0.0662524 | 0.947177 | 0.993112742 | ZNF207 | protein_coding | 38.47 | 5.06E-05 | 5.06E-05 | ENSG00000010244 | n.s. |
| ENSG00000113384 | 948.9072 | -0.017984771 | 0.296764 | -0.060603 | 0.951675 | 0.99357526 | GOLPH3 | protein_coding | 41.12 | -5.03E-05 | 5.03E-05 | ENSG00000113384 | n.s. |
| ENSG00000207546 | 9.59027 | -0.024314836 | 0.504219 | -0.048223 | 0.961539 | 0.995247333 | MIR548C | miRNA | 34.02 | -5.03E-05 | 5.03E-05 | ENSG00000207546 | n.s. |
| ENSG00000182208 | 79.69694 | 0.017892211 | 0.303801 | 0.0588946 | 0.953036 | 0.99357526 | MOB2 | protein_coding | 54.17 | 5.01E-05 | 5.01E-05 | ENSG00000182208 | n.s. |
| ENSG00000003989 | 660.362 | 0.017565769 | 0.27579 | 0.0636926 | 0.949215 | 0.993482322 | SLC7A2 | protein_coding | 40.13 | 4.99E-05 | 4.99E-05 | ENSG00000003989 | n.s. |
| ENSG00000105341 | 346.1459 | -0.016785238 | 0.258961 | -0.064818 | 0.948319 | 0.993188446 | ATP5SL | protein_coding | 51.01 | -4.98E-05 | 4.98E-05 | ENSG00000105341 | n.s. |
| ENSG00000279691 | 68.70345 | -0.017544586 | 0.277363 | -0.063255 | 0.949563 | 0.993482322 | | TEC | 58.13 | -4.98E-05 | 4.98E-05 | ENSG00000279691 | n.s. |
| ENSG00000150764 | 673.1838 | 0.017783984 | 0.303792 | 0.0585401 | 0.953318 | 0.99357526 | DIXDC1 | protein_coding | 43.3 | 4.98E-05 | 4.98E-05 | ENSG00000150764 | n.s. |
| ENSG00000235284 | 10.86087 | -0.023546145 | 0.481359 | -0.048916 | 0.960986 | 0.995151118 | SNORD62A | snoRNA | 46.51 | -4.97E-05 | 4.97E-05 | ENSG00000235284 | n.s. |
| ENSG00000120896 | 98.50779 | -0.017750144 | 0.2877 | -0.061697 | 0.950804 | 0.99357526 | SORBS3 | protein_coding | 55.84 | -4.97E-05 | 4.97E-05 | ENSG00000120896 | n.s. |
| ENSG00000116675 | 991.9792 | 0.017451537 | 0.275363 | 0.0633765 | 0.949467 | 0.993482322 | DNAJC6 | protein_coding | 40.01 | 4.96E-05 | 4.96E-05 | ENSG00000116675 | n.s. |
| ENSG00000175573 | 155.193 | -0.016603838 | 0.25415 | -0.065331 | 0.947911 | 0.993188446 | C11orf68 | protein_coding | 61.82 | -4.93E-05 | 4.93E-05 | ENSG00000175573 | n.s. |
| ENSG00000092199 | 15017.96 | -0.017591223 | 0.304916 | -0.057692 | 0.953994 | 0.99357526 | HNRNPC | protein_coding | 39.78 | -4.92E-05 | 4.92E-05 | ENSG00000092199 | n.s. |
| ENSG00000166226 | 11211.82 | 0.016857438 | 0.26249 | 0.0642212 | 0.948794 | 0.993312256 | CCT2 | protein_coding | 39.53 | 4.91E-05 | 4.91E-05 | ENSG00000166226 | n.s. |
| ENSG00000175390 | 2827.8 | 0.016511881 | 0.254056 | 0.0649931 | 0.948179 | 0.993188446 | EIF3F | protein_coding | 41.38 | 4.90E-05 | 4.90E-05 | ENSG00000175390 | n.s. |
| ENSG00000154639 | 903.1272 | -0.017227548 | 0.274096 | -0.062852 | 0.949884 | 0.993482322 | CXADR | protein_coding | 40.2 | -4.89E-05 | 4.89E-05 | ENSG00000154639 | n.s. |
| ENSG00000169738 | 1058.869 | -0.017466421 | 0.29223 | -0.059769 | 0.952339 | 0.99357526 | DCXR | protein_coding | 61.99 | -4.89E-05 | 4.89E-05 | ENSG00000169738 | n.s. |
| ENSG00000254682 | 65.06488 | -0.017296724 | 0.293166 | -0.059 | 0.952952 | 0.99357526 | | antisense | 45.01 | -4.84E-05 | 4.84E-05 | ENSG00000254682 | n.s. |
| ENSG00000139372 | 2753.289 | 0.017271289 | 0.289003 | 0.0597617 | 0.952345 | 0.99357526 | TDG | protein_coding | 40.65 | 4.83E-05 | 4.83E-05 | ENSG00000139372 | n.s. |
| ENSG00000178952 | 1814.493 | -0.017216998 | 0.290171 | -0.059334 | 0.952686 | 0.99357526 | TUFM | protein_coding | 55.48 | -4.82E-05 | 4.82E-05 | ENSG00000178952 | n.s. |
| ENSG00000163512 | 1221.089 | -0.017173999 | 0.276978 | -0.062005 | 0.950559 | 0.99357526 | AZI2 | protein_coding | 34.24 | -4.81E-05 | 4.81E-05 | ENSG00000163512 | n.s. |

| | | | | | | | | | | | | | |
|---|---|---|---|---|---|---|---|---|---|---|---|---|---|
| ENSG00000199961 | 52.56924 | 0.017127306 | 0.296971 | 0.0576734 | 0.954009 | 0.99357526 | SNORD1B | snoRNA | 40.48 | 4.79E-05 | 4.79E-05 | ENSG00000199961 | n.s. |
| ENSG00000164828 | 1863.614 | -0.01711165 | 0.282252 | -0.060625 | 0.951658 | 0.99357526 | SUN1 | protein_coding | 52.69 | -4.79E-05 | 4.79E-05 | ENSG00000164828 | n.s. |
| ENSG00000239911 | 140.7863 | -0.01708471 | 0.286836 | -0.059563 | 0.952504 | 0.99357526 | PRKAG2-AS1 | antisense | 60.13 | -4.78E-05 | 4.78E-05 | ENSG00000239911 | n.s. |
| ENSG00000205302 | 1153.946 | -0.016233793 | 0.25265 | -0.064254 | 0.948768 | 0.993312256 | SNX2 | protein_coding | 36.08 | -4.73E-05 | 4.73E-05 | ENSG00000205302 | n.s. |
| ENSG00000106392 | 199.1118 | -0.016843561 | 0.283963 | -0.059316 | 0.9527 | 0.99357526 | C1GALT1 | protein_coding | 37.33 | -4.71E-05 | 4.71E-05 | ENSG00000106392 | n.s. |
| ENSG00000265282 | 34.34852 | 0.019078135 | 0.357886 | 0.0533079 | 0.957487 | 0.994376566 | | lincRNA | 48.05 | 4.67E-05 | 4.67E-05 | ENSG00000265282 | n.s. |
| ENSG00000252213 | 64.63864 | 0.016673254 | 0.271532 | 0.0614043 | 0.951037 | 0.99357526 | SNORA74D | snoRNA | 53.19 | 4.67E-05 | 4.67E-05 | ENSG00000252213 | n.s. |
| ENSG00000138760 | 2679.533 | -0.016640376 | 0.276943 | -0.060086 | 0.952087 | 0.99357526 | SCARB2 | protein_coding | 41.79 | -4.66E-05 | 4.66E-05 | ENSG00000138760 | n.s. |
| ENSG00000169714 | 8906.439 | -0.017142025 | 0.300008 | -0.057138 | 0.954435 | 0.993762848 | CNBP | protein_coding | 42.41 | -4.66E-05 | 4.66E-05 | ENSG00000169714 | n.s. |
| ENSG00000226508 | 144.8671 | 0.015683032 | 0.240486 | 0.0652139 | 0.948004 | 0.993188446 | LINC01918 | lincRNA | 57.47 | 4.66E-05 | 4.66E-05 | ENSG00000226508 | n.s. |
| ENSG00000204371 | 1033.25 | 0.015548089 | 0.237132 | 0.0655673 | 0.947722 | 0.993188446 | EHMT2 | protein_coding | 53.82 | 4.62E-05 | 4.62E-05 | ENSG00000204371 | n.s. |
| ENSG00000137486 | 201.9539 | 0.016477345 | 0.271966 | 0.0605861 | 0.951689 | 0.99357526 | ARRB1 | protein_coding | 51.96 | 4.61E-05 | 4.61E-05 | ENSG00000137486 | n.s. |
| ENSG00000210176 | 4.128175 | -0.026168613 | 0.688888 | -0.037987 | 0.969698 | 0.995991293 | MT-TH | Mt_tRNA | 36.23 | -4.57E-05 | 4.57E-05 | ENSG00000210176 | n.s. |
| ENSG00000205213 | 641.9581 | 0.015975677 | 0.254178 | 0.0628523 | 0.949884 | 0.993482322 | LGR4 | protein_coding | 39.08 | 4.54E-05 | 4.54E-05 | ENSG00000205213 | n.s. |
| ENSG00000137818 | 13134.42 | 0.016713741 | 0.293584 | 0.0569299 | 0.954601 | 0.993787187 | RPLP1 | protein_coding | 47.38 | 4.52E-05 | 4.52E-05 | ENSG00000137818 | n.s. |
| ENSG00000199179 | 86.61306 | 0.015994203 | 0.265753 | 0.0601845 | 0.952009 | 0.99357526 | MIRLET7I | miRNA | 53.57 | 4.48E-05 | 4.48E-05 | ENSG00000199179 | n.s. |
| ENSG00000274380 | 7.74533 | -0.022963159 | 0.492734 | -0.046604 | 0.962829 | 0.995533737 | MIR6801 | miRNA | 59.49 | -4.46E-05 | 4.46E-05 | ENSG00000274380 | n.s. |
| ENSG00000134853 | 2474.66 | 0.015944858 | 0.277523 | 0.0574542 | 0.954183 | 0.99357526 | PDGFRA | protein_coding | 43.65 | 4.46E-05 | 4.46E-05 | ENSG00000134853 | n.s. |
| ENSG00000196290 | 1041.088 | -0.015932557 | 0.263094 | -0.060558 | 0.951711 | 0.99357526 | NIF3L1 | protein_coding | 41.02 | -4.46E-05 | 4.46E-05 | ENSG00000196290 | n.s. |
| ENSG00000196504 | 8182.123 | -0.015858124 | 0.262218 | -0.060477 | 0.951776 | 0.99357526 | PRPF40A | protein_coding | 36.31 | -4.44E-05 | 4.44E-05 | ENSG00000196504 | n.s. |
| ENSG00000135473 | 135.556 | -0.015767719 | 0.252581 | -0.062426 | 0.950223 | 0.99357526 | PAN2 | protein_coding | 47.67 | -4.41E-05 | 4.41E-05 | ENSG00000135473 | n.s. |
| ENSG00000161179 | 307.1717 | 0.015756415 | 0.263504 | 0.0597958 | 0.952318 | 0.99357526 | YDJC | protein_coding | 67.56 | 4.41E-05 | 4.41E-05 | ENSG00000161179 | n.s. |
| ENSG00000128266 | 217.8033 | -0.015678854 | 0.26297 | -0.059622 | 0.952457 | 0.99357526 | GNAZ | protein_coding | 57.44 | -4.39E-05 | 4.39E-05 | ENSG00000128266 | n.s. |
| ENSG00000141258 | 876.8481 | -0.015672969 | 0.264583 | -0.059236 | 0.952764 | 0.99357526 | SGSM2 | protein_coding | 54.99 | -4.39E-05 | 4.39E-05 | ENSG00000141258 | n.s. |
| ENSG00000276550 | 774.5862 | -0.015429898 | 0.242941 | -0.063513 | 0.949358 | 0.993482322 | HERC2P2 | cribed_unprocessed_pseudc | 42.65 | -4.38E-05 | 4.38E-05 | ENSG00000276550 | n.s. |
| ENSG00000233098 | 114.0214 | 0.01778111 | 0.333119 | 0.0533777 | 0.957431 | 0.994376566 | CCDC144NL-AS1 | antisense | 43.23 | 4.35E-05 | 4.35E-05 | ENSG00000233098 | n.s. |
| ENSG00000202141 | 53.55549 | -0.01773356 | 0.326498 | -0.054314 | 0.956685 | 0.994376566 | | misc_RNA | 42.48 | -4.34E-05 | 4.34E-05 | ENSG00000202141 | n.s. |
| ENSG00000226823 | 54.11708 | 0.017667283 | 0.322229 | 0.0548284 | 0.956275 | 0.994376566 | SUGT1P1 | unprocessed_pseudogene | 38.61 | 4.33E-05 | 4.33E-05 | ENSG00000226823 | n.s. |
| ENSG00000278467 | 107.2408 | 0.015358075 | 0.261363 | 0.0587616 | 0.953142 | 0.99357526 | | lincRNA | 44.92 | 4.30E-05 | 4.30E-05 | ENSG00000278467 | n.s. |
| ENSG00000116455 | 1182.615 | -0.015172697 | 0.262496 | -0.057802 | 0.953907 | 0.99357526 | WDR77 | protein_coding | 44.57 | -4.25E-05 | 4.25E-05 | ENSG00000116455 | n.s. |
| ENSG00000132635 | 642.3328 | -0.015138261 | 0.246118 | -0.061508 | 0.950954 | 0.99357526 | PCED1A | protein_coding | 57.7 | -4.24E-05 | 4.24E-05 | ENSG00000132635 | n.s. |
| ENSG00000124207 | 5787.583 | -0.015634343 | 0.274224 | -0.057013 | 0.954535 | 0.993787187 | CSE1L | protein_coding | 41.33 | -4.23E-05 | 4.23E-05 | ENSG00000124207 | n.s. |

| | | | | | | | | | | | | | |
|---|---|---|---|---|---|---|---|---|---|---|---|---|---|
| ENSG00000200922 | 19.1802 | 0.019962524 | 0.406372 | 0.0491237 | 0.960821 | 0.995135871 | | misc_RNA | 42.71 | 4.23E-05 | 4.23E-05 | ENSG00000200922 | n.s. |
| ENSG00000256537 | 2093.169 | 0.014956139 | 0.238593 | 0.0626846 | 0.950018 | 0.993547345 | SMIM10L1 | protein_coding | 38.73 | 4.20E-05 | 4.20E-05 | ENSG00000256537 | n.s. |
| ENSG00000237289 | 347.8655 | -0.014982316 | 0.254129 | -0.058956 | 0.952987 | 0.99357526 | CKMT1B | protein_coding | 49.85 | -4.19E-05 | 4.19E-05 | ENSG00000237289 | n.s. |
| ENSG00000113048 | 1463.639 | 0.014977621 | 0.258782 | 0.0578773 | 0.953846 | 0.99357526 | MRPS27 | protein_coding | 38.95 | 4.19E-05 | 4.19E-05 | ENSG00000113048 | n.s. |
| ENSG00000197128 | 114.0868 | -0.014904098 | 0.25566 | -0.058296 | 0.953512 | 0.99357526 | ZNF772 | protein_coding | 42.84 | -4.17E-05 | 4.17E-05 | ENSG00000197128 | n.s. |
| ENSG00000115355 | 7057.932 | 0.015803779 | 0.28035 | 0.0563716 | 0.955046 | 0.993954679 | CCDC88A | protein_coding | 36.45 | 4.16E-05 | 4.16E-05 | ENSG00000115355 | n.s. |
| ENSG00000131408 | 199.6118 | 0.015787836 | 0.280078 | 0.0563695 | 0.955047 | 0.993954679 | NR1H2 | protein_coding | 51.27 | 4.16E-05 | 4.16E-05 | ENSG00000131408 | n.s. |
| ENSG00000204308 | 542.8811 | 0.016926615 | 0.315356 | 0.0536746 | 0.957194 | 0.994376566 | RNF5 | protein_coding | 51.43 | 4.15E-05 | 4.15E-05 | ENSG00000204308 | n.s. |
| ENSG00000138777 | 1244.407 | 0.017171205 | 0.326829 | 0.0525387 | 0.958099 | 0.994479281 | PPA2 | protein_coding | 36.65 | 4.13E-05 | 4.13E-05 | ENSG00000138777 | n.s. |
| ENSG00000221303 | 26.70835 | 0.02352754 | 0.602596 | 0.0390437 | 0.968856 | 0.995991293 | SNORA79 | snoRNA | 34.29 | 4.10E-05 | 4.10E-05 | ENSG00000221303 | n.s. |
| ENSG00000102738 | 557.4445 | 0.015152317 | 0.266917 | 0.0567679 | 0.95473 | 0.993798482 | MRPS31 | protein_coding | 39.19 | 4.09E-05 | 4.09E-05 | ENSG00000102738 | n.s. |
| ENSG00000105676 | 678.4022 | 0.016699365 | 0.31295 | 0.0533611 | 0.957444 | 0.994376566 | ARMC6 | protein_coding | 51.45 | 4.09E-05 | 4.09E-05 | ENSG00000105676 | n.s. |
| ENSG00000138668 | 5647.178 | -0.014603112 | 0.247157 | -0.059084 | 0.952885 | 0.99357526 | HNRNPD | protein_coding | 38.75 | -4.09E-05 | 4.09E-05 | ENSG00000138668 | n.s. |
| ENSG00000143740 | 544.0662 | -0.016632756 | 0.304706 | -0.054586 | 0.956468 | 0.994376566 | SNAP47 | protein_coding | 53.99 | -4.07E-05 | 4.07E-05 | ENSG00000143740 | n.s. |
| ENSG00000221763 | 35.21072 | -0.018323271 | 0.363684 | -0.050382 | 0.959818 | 0.994902161 | MIR1289-1 | miRNA | 41.67 | -4.07E-05 | 4.07E-05 | ENSG00000221763 | n.s. |
| ENSG00000163818 | 363.9444 | -0.016234619 | 0.291441 | -0.055705 | 0.955577 | 0.994312186 | LZTFL1 | protein_coding | 42.03 | -4.02E-05 | 4.02E-05 | ENSG00000163818 | n.s. |
| ENSG00000142396 | 862.8722 | 0.014361093 | 0.247636 | 0.0579928 | 0.953754 | 0.99357526 | | protein_coding | 45.5 | 4.02E-05 | 4.02E-05 | ENSG00000142396 | n.s. |
| ENSG00000126953 | 338.5704 | -0.014098357 | 0.220848 | -0.063837 | 0.9491 | 0.993482322 | TIMM8A | protein_coding | 45.87 | -4.00E-05 | 4.00E-05 | ENSG00000126953 | n.s. |
| ENSG00000103363 | 2134.873 | -0.016709968 | 0.320493 | -0.052138 | 0.958419 | 0.994524382 | ELOB | protein_coding | 55.52 | -3.98E-05 | 3.98E-05 | ENSG00000103363 | n.s. |
| ENSG00000247934 | 133.7562 | -0.014212816 | 0.242975 | -0.058495 | 0.953354 | 0.99357526 | | antisense | 37.35 | -3.98E-05 | 3.98E-05 | ENSG00000247934 | n.s. |
| ENSG00000270647 | 2285.668 | -0.013341478 | 0.203232 | -0.065647 | 0.947659 | 0.993188446 | TAF15 | protein_coding | 46.76 | -3.96E-05 | 3.96E-05 | ENSG00000270647 | n.s. |
| ENSG00000175548 | 1368.335 | 0.014131926 | 0.2395 | 0.0590059 | 0.952947 | 0.99357526 | ALG10B | protein_coding | 34.07 | 3.96E-05 | 3.96E-05 | ENSG00000175548 | n.s. |
| ENSG00000147403 | 13101.09 | -0.015977915 | 0.289827 | -0.055129 | 0.956036 | 0.994376566 | RPL10 | protein_coding | 54.28 | -3.91E-05 | 3.91E-05 | ENSG00000147403 | n.s. |
| ENSG00000207756 | 5.05177 | -0.022350033 | 0.587706 | -0.038029 | 0.969664 | 0.995991293 | MIR580 | miRNA | 29.9 | -3.90E-05 | 3.90E-05 | ENSG00000207756 | n.s. |
| ENSG00000171858 | 5397.606 | 0.016829346 | 0.328394 | 0.0512474 | 0.959128 | 0.994707196 | RPS21 | protein_coding | 59.5 | 3.88E-05 | 3.88E-05 | ENSG00000171858 | n.s. |
| ENSG00000176399 | 273.0421 | -0.013843753 | 0.23797 | -0.058174 | 0.95361 | 0.99357526 | DMRTA1 | protein_coding | 37.55 | -3.88E-05 | 3.88E-05 | ENSG00000176399 | n.s. |
| ENSG00000006625 | 1705.851 | -0.015770294 | 0.292956 | -0.053832 | 0.957069 | 0.994376566 | GGCT | protein_coding | 40.55 | -3.86E-05 | 3.86E-05 | ENSG00000006625 | n.s. |
| ENSG00000101347 | 446.8886 | -0.016063887 | 0.305088 | -0.052653 | 0.958008 | 0.994479281 | SAMHD1 | protein_coding | 43.82 | -3.86E-05 | 3.86E-05 | ENSG00000101347 | n.s. |
| ENSG00000096092 | 447.3248 | 0.016739888 | 0.32866 | 0.0509337 | 0.959378 | 0.994744177 | TMEM14A | protein_coding | 41.73 | 3.83E-05 | 3.83E-05 | ENSG00000096092 | n.s. |
| ENSG00000060339 | 3787.939 | 0.015262173 | 0.272892 | 0.0559276 | 0.9554 | 0.994246758 | CCAR1 | protein_coding | 41.75 | 3.82E-05 | 3.82E-05 | ENSG00000060339 | n.s. |
| ENSG00000102003 | 933.2264 | -0.015341532 | 0.288045 | -0.053261 | 0.957524 | 0.994376566 | SYP | protein_coding | 52.69 | -3.76E-05 | 3.76E-05 | ENSG00000102003 | n.s. |
| ENSG00000185480 | 1116.571 | 0.015729658 | 0.303809 | 0.0517749 | 0.958708 | 0.994524382 | PARPBP | protein_coding | 36.45 | 3.75E-05 | 3.75E-05 | ENSG00000185480 | n.s. |

| | | | | | | | | | | | | | |
|---|---|---|---|---|---|---|---|---|---|---|---|---|---|
| ENSG00000121851 | 182.6204 | 0.015300358 | 0.280621 | 0.0545233 | 0.956518 | 0.994376566 | POLR3GL | protein_coding | 44.9 | 3.75E-05 | 3.75E-05 | ENSG00000121851 | n.s. |
| ENSG00000167986 | 5334.719 | -0.015102022 | 0.271281 | -0.055669 | 0.955605 | 0.994312186 | DDB1 | protein_coding | 47.67 | -3.74E-05 | 3.74E-05 | ENSG00000167986 | n.s. |
| ENSG00000277534 | 429.3138 | -0.013254183 | 0.230383 | -0.057531 | 0.954122 | 0.99357526 | | sense_intronic | 37.54 | -3.71E-05 | 3.71E-05 | ENSG00000277534 | n.s. |
| ENSG00000163902 | 2170.383 | 0.015094676 | 0.276746 | 0.0545434 | 0.956502 | 0.994376566 | RPN1 | protein_coding | 46.98 | 3.70E-05 | 3.70E-05 | ENSG00000163902 | n.s. |
| ENSG00000169446 | 584.8936 | 0.015048437 | 0.278692 | 0.0539967 | 0.956938 | 0.994376566 | MMGT1 | protein_coding | 39.98 | 3.69E-05 | 3.69E-05 | ENSG00000169446 | n.s. |
| ENSG00000284329 | 27.28594 | -0.017327347 | 0.34847 | -0.049724 | 0.960342 | 0.995125117 | MIR9-3 | miRNA | 44.44 | -3.68E-05 | 3.68E-05 | ENSG00000284329 | n.s. |
| ENSG00000170837 | 371.1644 | -0.014921979 | 0.26868 | -0.055538 | 0.95571 | 0.994346619 | GPR27 | protein_coding | 54.84 | -3.67E-05 | 3.67E-05 | ENSG00000170837 | n.s. |
| ENSG00000111639 | 2628.216 | -0.015261481 | 0.292822 | -0.052119 | 0.958434 | 0.994524382 | MRPL51 | protein_coding | 48.72 | -3.64E-05 | 3.64E-05 | ENSG00000111639 | n.s. |
| ENSG00000144736 | 687.1917 | -0.01481336 | 0.27388 | -0.054087 | 0.956866 | 0.994376566 | SHQ1 | protein_coding | 39.56 | -3.63E-05 | 3.63E-05 | ENSG00000144736 | n.s. |
| ENSG00000145781 | 455.0636 | 0.014684228 | 0.275405 | 0.0533186 | 0.957478 | 0.994376566 | COMMD10 | protein_coding | 37.46 | 3.60E-05 | 3.60E-05 | ENSG00000145781 | n.s. |
| ENSG00000167272 | 514.0399 | 0.014672155 | 0.269138 | 0.0545153 | 0.956525 | 0.994376566 | POP5 | protein_coding | 48.65 | 3.59E-05 | 3.59E-05 | ENSG00000167272 | n.s. |
| ENSG00000152464 | 276.5722 | -0.014899744 | 0.281324 | -0.052963 | 0.957761 | 0.994474733 | RPP38 | protein_coding | 43.41 | -3.59E-05 | 3.59E-05 | ENSG00000152464 | n.s. |
| ENSG00000189184 | 333.4974 | -0.012807182 | 0.213014 | -0.060124 | 0.952057 | 0.99357526 | PCDH18 | protein_coding | 36.59 | -3.59E-05 | 3.59E-05 | ENSG00000189184 | n.s. |
| ENSG00000158169 | 206.6351 | 0.014433984 | 0.263963 | 0.0546818 | 0.956392 | 0.994376566 | FANCC | protein_coding | 41.78 | 3.54E-05 | 3.54E-05 | ENSG00000158169 | n.s. |
| ENSG00000163430 | 1021.212 | -0.014675573 | 0.279349 | -0.052535 | 0.958103 | 0.994479281 | FSTL1 | protein_coding | 43.92 | -3.53E-05 | 3.53E-05 | ENSG00000163430 | n.s. |
| ENSG00000134910 | 966.2875 | -0.01435753 | 0.264209 | -0.054342 | 0.956663 | 0.994376566 | STT3A | protein_coding | 41.03 | -3.52E-05 | 3.52E-05 | ENSG00000134910 | n.s. |
| ENSG00000105669 | 1086.535 | 0.015789774 | 0.314569 | 0.050195 | 0.959967 | 0.994909955 | COPE | protein_coding | 57.06 | 3.50E-05 | 3.50E-05 | ENSG00000105669 | n.s. |
| ENSG00000133243 | 433.7481 | 0.014265357 | 0.260034 | 0.0548595 | 0.95625 | 0.994376566 | BTBD2 | protein_coding | 55.3 | 3.49E-05 | 3.49E-05 | ENSG00000133243 | n.s. |
| ENSG00000070669 | 399.0522 | -0.014175066 | 0.265464 | -0.053397 | 0.957415 | 0.994376566 | ASNS | protein_coding | 38.42 | -3.47E-05 | 3.47E-05 | ENSG00000070669 | n.s. |
| ENSG00000252080 | 4.600444 | -0.026332811 | 0.919173 | -0.028648 | 0.977145 | 0.9969708 | RNU7-99P | snRNA | 35.48 | -3.47E-05 | 3.47E-05 | ENSG00000252080 | n.s. |
| ENSG00000168298 | 2845.833 | 0.012261338 | 0.208512 | 0.0588039 | 0.953108 | 0.99357526 | HIST1H1E | protein_coding | 60.08 | 3.43E-05 | 3.43E-05 | ENSG00000168298 | n.s. |
| ENSG00000120688 | 511.3868 | -0.014004104 | 0.259917 | -0.053879 | 0.957032 | 0.994376566 | WBP4 | protein_coding | 38.92 | -3.43E-05 | 3.43E-05 | ENSG00000120688 | n.s. |
| ENSG00000110536 | 673.7058 | -0.015142673 | 0.299085 | -0.05063 | 0.95962 | 0.994828899 | PTPMT1 | protein_coding | 47.83 | -3.41E-05 | 3.41E-05 | ENSG00000110536 | n.s. |
| ENSG00000242372 | 993.8491 | 0.015880287 | 0.323589 | 0.0490756 | 0.960859 | 0.995135871 | EIF6 | protein_coding | 52.05 | 3.36E-05 | 3.36E-05 | ENSG00000242372 | n.s. |
| ENSG00000215251 | 815.7427 | -0.014081286 | 0.272156 | -0.05174 | 0.958736 | 0.994524382 | FASTKD5 | protein_coding | 44.53 | -3.36E-05 | 3.36E-05 | ENSG00000215251 | n.s. |
| ENSG00000211574 | 10.44671 | -0.019208138 | 0.474015 | -0.040522 | 0.967677 | 0.995991293 | MIR770 | miRNA | 65.31 | -3.35E-05 | 3.35E-05 | ENSG00000211574 | n.s. |
| ENSG00000154144 | 571.243 | 0.013902565 | 0.262368 | 0.0529888 | 0.957741 | 0.994474733 | TBRG1 | protein_coding | 40.05 | 3.35E-05 | 3.35E-05 | ENSG00000154144 | n.s. |
| ENSG00000156273 | 2023.442 | 0.013633588 | 0.254242 | 0.0536244 | 0.957234 | 0.994376566 | BACH1 | protein_coding | 39.61 | 3.34E-05 | 3.34E-05 | ENSG00000156273 | n.s. |
| ENSG00000114503 | 3072.304 | 0.013948387 | 0.269599 | 0.0517376 | 0.958738 | 0.994524382 | NCBP2 | protein_coding | 44 | 3.33E-05 | 3.33E-05 | ENSG00000114503 | n.s. |
| ENSG00000140386 | 771.8498 | -0.014349155 | 0.281332 | -0.051004 | 0.959322 | 0.994744177 | SCAPER | protein_coding | 37.7 | -3.28E-05 | 3.28E-05 | ENSG00000140386 | n.s. |
| ENSG00000079150 | 900.3009 | 0.013382215 | 0.248829 | 0.0537807 | 0.95711 | 0.994376566 | FKBP7 | protein_coding | 37.58 | 3.28E-05 | 3.28E-05 | ENSG00000079150 | n.s. |
| ENSG00000086598 | 4258.914 | 0.014122409 | 0.276696 | 0.0510394 | 0.959294 | 0.994744177 | TMED2 | protein_coding | 42.27 | 3.23E-05 | 3.23E-05 | ENSG00000086598 | n.s. |

| | | | | | | | | | | | | | |
|---|---|---|---|---|---|---|---|---|---|---|---|---|---|
| ENSG00000116350 | 1494.353 | -0.013443327 | 0.259557 | -0.051793 | 0.958693 | 0.994524382 | SRSF4 | protein_coding | 43 | -3.21E-05 | 3.21E-05 | ENSG00000116350 | n.s. |
| ENSG00000129534 | 2296.407 | -0.013193441 | 0.25139 | -0.052482 | 0.958145 | 0.994479281 | MIS18BP1 | protein_coding | 37.65 | -3.17E-05 | 3.17E-05 | ENSG00000129534 | n.s. |
| ENSG00000113810 | 12544.88 | -0.013192794 | 0.249969 | -0.052778 | 0.957909 | 0.994479281 | SMC4 | protein_coding | 36.23 | -3.17E-05 | 3.17E-05 | ENSG00000113810 | n.s. |
| ENSG00000165138 | 594.511 | -0.013682091 | 0.26694 | -0.051255 | 0.959122 | 0.994707196 | ANKS6 | protein_coding | 46.95 | -3.15E-05 | 3.15E-05 | ENSG00000165138 | n.s. |
| ENSG00000107672 | 1369.299 | 0.014340255 | 0.286877 | 0.0499875 | 0.960132 | 0.995007255 | NSMCE4A | protein_coding | 43.84 | 3.12E-05 | 3.12E-05 | ENSG00000107672 | n.s. |
| ENSG00000071859 | 599.3401 | 0.013478566 | 0.262618 | 0.0513238 | 0.959068 | 0.994707196 | FAM50A | protein_coding | 60.48 | 3.11E-05 | 3.11E-05 | ENSG00000071859 | n.s. |
| ENSG00000163882 | 847.7045 | -0.014599805 | 0.295574 | -0.049395 | 0.960605 | 0.995135871 | POLR2H | protein_coding | 52.77 | -3.09E-05 | 3.09E-05 | ENSG00000163882 | n.s. |
| ENSG00000109466 | 194.1827 | -0.012902836 | 0.248308 | -0.051963 | 0.958558 | 0.994524382 | KLHL2 | protein_coding | 38.15 | -3.08E-05 | 3.08E-05 | ENSG00000109466 | n.s. |
| ENSG00000144468 | 68.37105 | -0.014858312 | 0.311342 | -0.047723 | 0.961937 | 0.995247333 | RHBDD1 | protein_coding | 39.57 | -3.07E-05 | 3.07E-05 | ENSG00000144468 | n.s. |
| ENSG00000272477 | 569.1528 | -0.012443935 | 0.237391 | -0.05242 | 0.958194 | 0.994479281 | | sense_intronic | 31.69 | -2.99E-05 | 2.99E-05 | ENSG00000272477 | n.s. |
| ENSG00000123815 | 196.6767 | 0.013175591 | 0.260582 | 0.0505622 | 0.959674 | 0.994828899 | COQ8B | protein_coding | 52.66 | 2.97E-05 | 2.97E-05 | ENSG00000123815 | n.s. |
| ENSG00000271040 | 18.08224 | -0.016991201 | 0.445423 | -0.038146 | 0.969571 | 0.995991293 | | lincRNA | 52.72 | -2.96E-05 | 2.96E-05 | ENSG00000271040 | n.s. |
| ENSG00000165995 | 44.66303 | -0.015335166 | 0.344818 | -0.044473 | 0.964527 | 0.995560243 | CACNB2 | protein_coding | 40.47 | -2.96E-05 | 2.96E-05 | ENSG00000165995 | n.s. |
| ENSG00000116266 | 641.2644 | -0.015110171 | 0.328996 | -0.045928 | 0.963368 | 0.995533737 | STXBP3 | protein_coding | 36.03 | -2.94E-05 | 2.94E-05 | ENSG00000116266 | n.s. |
| ENSG00000130703 | 306.5667 | -0.014190395 | 0.295269 | -0.048059 | 0.961669 | 0.995247333 | OSBPL2 | protein_coding | 52.75 | -2.94E-05 | 2.94E-05 | ENSG00000130703 | n.s. |
| ENSG00000252277 | 11.55384 | -0.016822512 | 0.422095 | -0.039855 | 0.968209 | 0.995991293 | SNORD116-30 | snoRNA | 35.29 | -2.93E-05 | 2.93E-05 | ENSG00000252277 | n.s. |
| ENSG00000102699 | 1199.131 | -0.013795876 | 0.27818 | -0.049593 | 0.960446 | 0.995125117 | PARP4 | protein_coding | 42.88 | -2.93E-05 | 2.93E-05 | ENSG00000102699 | n.s. |
| ENSG00000283210 | 9.096824 | 0.016655042 | 0.453373 | 0.0367359 | 0.970696 | 0.995991293 | MIR1322 | miRNA | 33.8 | 2.91E-05 | 2.91E-05 | ENSG00000283210 | n.s. |
| ENSG00000163281 | 1280.816 | -0.012876963 | 0.253804 | -0.050736 | 0.959536 | 0.994828899 | GNPDA2 | protein_coding | 33.84 | -2.90E-05 | 2.90E-05 | ENSG00000163281 | n.s. |
| ENSG00000144278 | 1568.539 | 0.013719768 | 0.282155 | 0.0486249 | 0.961218 | 0.995243634 | GALNT13 | protein_coding | 35.16 | 2.84E-05 | 2.84E-05 | ENSG00000144278 | n.s. |
| ENSG00000206704 | 3.017466 | -0.021435534 | 0.739105 | -0.029002 | 0.976863 | 0.9969708 | | misc_RNA | 45.54 | -2.82E-05 | 2.82E-05 | ENSG00000206704 | n.s. |
| ENSG00000103249 | 72.07875 | -0.013559367 | 0.28362 | -0.047808 | 0.961869 | 0.995247333 | CLCN7 | protein_coding | 60.03 | -2.81E-05 | 2.81E-05 | ENSG00000103249 | n.s. |
| ENSG00000205269 | 729.3514 | -0.014390178 | 0.310196 | -0.046391 | 0.962999 | 0.995533737 | TMEM170B | protein_coding | 36.55 | -2.80E-05 | 2.80E-05 | ENSG00000205269 | n.s. |
| ENSG00000086504 | 483.3507 | 0.014316313 | 0.318997 | 0.0448791 | 0.964204 | 0.995533737 | MRPL28 | protein_coding | 64.85 | 2.78E-05 | 2.78E-05 | ENSG00000086504 | n.s. |
| ENSG00000185615 | 83.6758 | -0.013406823 | 0.277179 | -0.048369 | 0.961422 | 0.995247333 | PDIA2 | protein_coding | 64.25 | -2.77E-05 | 2.77E-05 | ENSG00000185615 | n.s. |
| ENSG00000120915 | 35.4519 | -0.014907907 | 0.337493 | -0.044172 | 0.964767 | 0.99573389 | EPHX2 | protein_coding | 47.54 | -2.77E-05 | 2.77E-05 | ENSG00000120915 | n.s. |
| ENSG00000172590 | 348.8557 | 0.014230185 | 0.313331 | 0.0454159 | 0.963776 | 0.995533737 | MRPL52 | protein_coding | 47.12 | 2.77E-05 | 2.77E-05 | ENSG00000172590 | n.s. |
| ENSG00000066032 | 593.12 | -0.01327054 | 0.276839 | -0.047936 | 0.961767 | 0.995247333 | CTNNA2 | protein_coding | 38.65 | -2.75E-05 | 2.75E-05 | ENSG00000066032 | n.s. |
| ENSG00000113448 | 67.00574 | 0.014088189 | 0.306516 | 0.0459623 | 0.96334 | 0.995533737 | PDE4D | protein_coding | 37.37 | 2.74E-05 | 2.74E-05 | ENSG00000113448 | n.s. |
| ENSG00000004961 | 272.5817 | 0.01290924 | 0.263128 | 0.0490608 | 0.960871 | 0.995135871 | HCCS | protein_coding | 42.99 | 2.73E-05 | 2.73E-05 | ENSG00000004961 | n.s. |
| ENSG00000086189 | 959.2 | -0.012817704 | 0.258548 | -0.049576 | 0.96046 | 0.995125117 | DIMT1 | protein_coding | 39.95 | -2.72E-05 | 2.72E-05 | ENSG00000086189 | n.s. |
| ENSG00000117448 | 511.3847 | 0.013929587 | 0.3105 | 0.0448618 | 0.964217 | 0.995533737 | AKR1A1 | protein_coding | 49.1 | 2.71E-05 | 2.71E-05 | ENSG00000117448 | n.s. |

| | | | | | | | | | | | | | |
|---|---|---|---|---|---|---|---|---|---|---|---|---|---|
| ENSG00000112651 | 293.8822 | -0.012979379 | 0.269395 | -0.04818 | 0.961573 | 0.995247333 | MRPL2 | protein_coding | 49.64 | -2.69E-05 | 2.69E-05 | ENSG00000112651 | n.s. |
| ENSG00000260219 | 27.33105 | -0.015330245 | 0.370941 | -0.041328 | 0.967034 | 0.995991293 | | lincRNA | 58.47 | -2.67E-05 | 2.67E-05 | ENSG00000260219 | n.s. |
| ENSG00000116906 | 1597.77 | 0.013705828 | 0.304872 | 0.044956 | 0.964142 | 0.995533737 | GNPAT | protein_coding | 41.78 | 2.66E-05 | 2.66E-05 | ENSG00000116906 | n.s. |
| ENSG00000243314 | 19.46824 | 0.015121582 | 0.378248 | 0.039978 | 0.968111 | 0.995991293 | | processed_pseudogene | 42.78 | 2.64E-05 | 2.64E-05 | ENSG00000243314 | n.s. |
| ENSG00000147601 | 1611.701 | -0.012439086 | 0.252461 | -0.049271 | 0.960703 | 0.995135871 | TERF1 | protein_coding | 36.98 | -2.63E-05 | 2.63E-05 | ENSG00000147601 | n.s. |
| ENSG00000075539 | 2523.803 | 0.012729821 | 0.264533 | 0.0481219 | 0.961619 | 0.995247333 | FRYL | protein_coding | 36.94 | 2.63E-05 | 2.63E-05 | ENSG00000075539 | n.s. |
| ENSG00000227161 | 25.5149 | -0.015095934 | 0.35696 | -0.04229 | 0.966267 | 0.995991293 | | sense_overlapping | 38.3 | -2.63E-05 | 2.63E-05 | ENSG00000227161 | n.s. |
| ENSG00000119673 | 349.9706 | 0.01347355 | 0.290292 | 0.0464138 | 0.96298 | 0.995533737 | ACOT2 | protein_coding | 48.61 | 2.62E-05 | 2.62E-05 | ENSG00000119673 | n.s. |
| ENSG00000170312 | 4740.016 | 0.012806826 | 0.269303 | 0.0475555 | 0.96207 | 0.99531187 | CDK1 | protein_coding | 35.07 | 2.61E-05 | 2.61E-05 | ENSG00000170312 | n.s. |
| ENSG00000174891 | 1050.09 | -0.013318513 | 0.298369 | -0.044638 | 0.964396 | 0.995533737 | RSRC1 | protein_coding | 35.48 | -2.59E-05 | 2.59E-05 | ENSG00000174891 | n.s. |
| ENSG00000104805 | 337.6649 | 0.013264531 | 0.297435 | 0.0445965 | 0.964429 | 0.995533737 | NUCB1 | protein_coding | 52.81 | 2.58E-05 | 2.58E-05 | ENSG00000104805 | n.s. |
| ENSG00000237973 | 141.8985 | -0.013247021 | 0.286986 | -0.046159 | 0.963183 | 0.995533737 | MTCO1P12 | unprocessed_pseudogene | 45.76 | -2.58E-05 | 2.58E-05 | ENSG00000237973 | n.s. |
| ENSG00000135365 | 244.0466 | 0.01201064 | 0.244992 | 0.0490246 | 0.9609 | 0.995135871 | PHF21A | protein_coding | 38.79 | 2.54E-05 | 2.54E-05 | ENSG00000135365 | n.s. |
| ENSG00000067113 | 219.9973 | 0.013004999 | 0.286386 | 0.0454108 | 0.96378 | 0.995533737 | PLPP1 | protein_coding | 39.62 | 2.53E-05 | 2.53E-05 | ENSG00000067113 | n.s. |
| ENSG00000253982 | 122.1519 | 0.012841986 | 0.280726 | 0.0457456 | 0.963513 | 0.995533737 | | antisense | 64.54 | 2.50E-05 | 2.50E-05 | ENSG00000253982 | n.s. |
| ENSG00000250838 | 275.5619 | 0.011245871 | 0.223602 | 0.0502941 | 0.959888 | 0.994902161 | | antisense | 61.43 | 2.50E-05 | 2.50E-05 | ENSG00000250838 | n.s. |
| ENSG00000168591 | 463.0528 | 0.012036597 | 0.251383 | 0.0478814 | 0.961811 | 0.995247333 | TMUB2 | protein_coding | 52.96 | 2.49E-05 | 2.49E-05 | ENSG00000168591 | n.s. |
| ENSG00000272758 | 40.72704 | 0.01409219 | 0.324387 | 0.0434425 | 0.965349 | 0.995965295 | | antisense | 39.37 | 2.47E-05 | 2.47E-05 | ENSG00000272758 | n.s. |
| ENSG00000011478 | 85.99764 | -0.012715002 | 0.270021 | -0.047089 | 0.962442 | 0.995533737 | QPCTL | protein_coding | 52.56 | -2.47E-05 | 2.47E-05 | ENSG00000011478 | n.s. |
| ENSG00000141424 | 3527.729 | -0.012655716 | 0.276085 | -0.04584 | 0.963438 | 0.995533737 | SLC39A6 | protein_coding | 39.45 | -2.46E-05 | 2.46E-05 | ENSG00000141424 | n.s. |
| ENSG00000107719 | 317.3678 | -0.012636836 | 0.275277 | -0.045906 | 0.963385 | 0.995533737 | PALD1 | protein_coding | 52.2 | -2.46E-05 | 2.46E-05 | ENSG00000107719 | n.s. |
| ENSG00000185495 | 181.012 | -0.012594496 | 0.282105 | -0.044645 | 0.964391 | 0.995533737 | | transcribed_unprocessed_pseudc | 36.59 | -2.45E-05 | 2.45E-05 | ENSG00000185495 | n.s. |
| ENSG00000150471 | 131.558 | 0.012510656 | 0.274549 | 0.045568 | 0.963655 | 0.995533737 | ADGRL3 | protein_coding | 35.25 | 2.43E-05 | 2.43E-05 | ENSG00000150471 | n.s. |
| ENSG00000101361 | 6262.954 | -0.011514042 | 0.235814 | -0.048827 | 0.961057 | 0.995151118 | NOP56 | protein_coding | 54.33 | -2.43E-05 | 2.43E-05 | ENSG00000101361 | n.s. |
| ENSG00000143198 | 685.0591 | -0.013909934 | 0.348904 | -0.039868 | 0.968199 | 0.995991293 | MGST3 | protein_coding | 43.51 | -2.43E-05 | 2.43E-05 | ENSG00000143198 | n.s. |
| ENSG00000137161 | 538.3287 | -0.012337042 | 0.274586 | -0.04493 | 0.964163 | 0.995533737 | CNPY3 | protein_coding | 50.57 | -2.40E-05 | 2.40E-05 | ENSG00000137161 | n.s. |
| ENSG00000108799 | 785.8231 | -0.012277653 | 0.26856 | -0.045717 | 0.963536 | 0.995533737 | EZH1 | protein_coding | 45.44 | -2.39E-05 | 2.39E-05 | ENSG00000108799 | n.s. |
| ENSG00000102362 | 2084.864 | 0.012258595 | 0.260065 | 0.0471366 | 0.962404 | 0.995533737 | SYTL4 | protein_coding | 41.39 | 2.38E-05 | 2.38E-05 | ENSG00000102362 | n.s. |
| ENSG00000117477 | 206.8347 | 0.012082878 | 0.269221 | 0.044881 | 0.964202 | 0.995533737 | CCDC181 | protein_coding | 36.8 | 2.35E-05 | 2.35E-05 | ENSG00000117477 | n.s. |
| ENSG00000138385 | 5618.672 | 0.012070633 | 0.269025 | 0.0448681 | 0.964212 | 0.995533737 | SSB | protein_coding | 39.88 | 2.35E-05 | 2.35E-05 | ENSG00000138385 | n.s. |
| ENSG00000114850 | 3498.299 | 0.011987959 | 0.261349 | 0.0458696 | 0.963414 | 0.995533737 | SSR3 | protein_coding | 38.72 | 2.33E-05 | 2.33E-05 | ENSG00000114850 | n.s. |
| ENSG00000145824 | 351.7386 | 0.013275657 | 0.311826 | 0.0425739 | 0.966041 | 0.995991293 | CXCL14 | protein_coding | 52.05 | 2.32E-05 | 2.32E-05 | ENSG00000145824 | n.s. |

| | | | | | | | | | | | | | |
|---|---|---|---|---|---|---|---|---|---|---|---|---|---|
| ENSG00000100804 | 2255.297 | -0.013270783 | 0.311488 | -0.042604 | 0.966017 | 0.995991293 | PSMB5 | protein_coding | 46.64 | -2.32E-05 | 2.32E-05 | ENSG00000100804 | n.s. |
| ENSG00000277791 | 1246.283 | 0.013039518 | 0.326046 | 0.0399929 | 0.968099 | 0.995991293 | PSMB3 | protein_coding | 49.05 | 2.27E-05 | 2.27E-05 | ENSG00000277791 | n.s. |
| ENSG00000197283 | 37.86495 | 0.012938797 | 0.330221 | 0.0391822 | 0.968745 | 0.995991293 | SYNGAP1 | protein_coding | 53.86 | 2.26E-05 | 2.26E-05 | ENSG00000197283 | n.s. |
| ENSG00000096401 | 3468.066 | -0.011592958 | 0.252406 | -0.04593 | 0.963366 | 0.995533737 | CDC5L | protein_coding | 38.5 | -2.25E-05 | 2.25E-05 | ENSG00000096401 | n.s. |
| ENSG00000145912 | 1860.104 | -0.012915618 | 0.312459 | -0.041335 | 0.967029 | 0.995991293 | NHP2 | protein_coding | 52.15 | -2.25E-05 | 2.25E-05 | ENSG00000145912 | n.s. |
| ENSG00000264346 | 40.6698 | 0.012883539 | 0.362764 | 0.0355149 | 0.971669 | 0.995991293 | SNORA77B | snoRNA | 55.2 | 2.25E-05 | 2.25E-05 | ENSG00000264346 | n.s. |
| ENSG00000129158 | 52.80252 | -0.012706767 | 0.318106 | -0.039945 | 0.968137 | 0.995991293 | SERGEF | protein_coding | 42.67 | -2.22E-05 | 2.22E-05 | ENSG00000129158 | n.s. |
| ENSG00000270557 | 31.14223 | -0.012705524 | 0.34217 | -0.037132 | 0.97038 | 0.995991293 | | lincRNA | 40.43 | -2.22E-05 | 2.22E-05 | ENSG00000270557 | n.s. |
| ENSG00000197006 | 1762.025 | 0.012582585 | 0.307974 | 0.040856 | 0.967411 | 0.995991293 | METTL9 | protein_coding | 42.83 | 2.19E-05 | 2.19E-05 | ENSG00000197006 | n.s. |
| ENSG00000134748 | 2263.419 | -0.011285149 | 0.244459 | -0.046164 | 0.96318 | 0.995533737 | PRPF38A | protein_coding | 41.39 | -2.19E-05 | 2.19E-05 | ENSG00000134748 | n.s. |
| ENSG00000103510 | 579.8766 | 0.012272231 | 0.290212 | 0.0422871 | 0.96627 | 0.995991293 | KAT8 | protein_coding | 52.56 | 2.14E-05 | 2.14E-05 | ENSG00000103510 | n.s. |
| ENSG00000117419 | 311.251 | 0.012247916 | 0.310795 | 0.0394083 | 0.968565 | 0.995991293 | ERI3 | protein_coding | 47.73 | 2.14E-05 | 2.14E-05 | ENSG00000117419 | n.s. |
| ENSG00000106404 | 123.771 | -0.012223533 | 0.288371 | -0.042388 | 0.966189 | 0.995991293 | CLDN15 | protein_coding | 59.59 | -2.13E-05 | 2.13E-05 | ENSG00000106404 | n.s. |
| ENSG00000226769 | 33.63761 | -0.012069529 | 0.323999 | -0.037252 | 0.970284 | 0.995991293 | GAPDHP54 | processed_pseudogene | 51.18 | -2.11E-05 | 2.11E-05 | ENSG00000226769 | n.s. |
| ENSG00000253327 | 53.53913 | -0.012045905 | 0.330795 | -0.036415 | 0.970951 | 0.995991293 | RAD21-AS1 | antisense | 42.45 | -2.10E-05 | 2.10E-05 | ENSG00000253327 | n.s. |
| ENSG00000137269 | 214.4387 | -0.011391817 | 0.259931 | -0.043826 | 0.965043 | 0.995792945 | LRRC1 | protein_coding | 39.72 | -2.09E-05 | 2.09E-05 | ENSG00000137269 | n.s. |
| ENSG00000170855 | 1009.739 | -0.010687463 | 0.2341 | -0.045654 | 0.963586 | 0.995533737 | TRIAP1 | protein_coding | 44.54 | -2.08E-05 | 2.08E-05 | ENSG00000170855 | n.s. |
| ENSG00000269082 | 57.5685 | -0.011894366 | 0.317845 | -0.037422 | 0.970149 | 0.995991293 | | processed_transcript | 44.76 | -2.07E-05 | 2.07E-05 | ENSG00000269082 | n.s. |
| ENSG00000171517 | 342.8236 | -0.011793979 | 0.303064 | -0.038916 | 0.968958 | 0.995991293 | LPAR3 | protein_coding | 41.6 | -2.06E-05 | 2.06E-05 | ENSG00000171517 | n.s. |
| ENSG00000115364 | 1745.825 | -0.011094207 | 0.251874 | -0.044047 | 0.964867 | 0.995763554 | MRPL19 | protein_coding | 36.97 | -2.05E-05 | 2.05E-05 | ENSG00000115364 | n.s. |
| ENSG00000063180 | 632.7343 | -0.011719809 | 0.291788 | -0.040165 | 0.967961 | 0.995991293 | CA11 | protein_coding | 53.83 | -2.04E-05 | 2.04E-05 | ENSG00000063180 | n.s. |
| ENSG00000157426 | 561.5184 | -0.011718322 | 0.299317 | -0.03915 | 0.968771 | 0.995991293 | AASDH | protein_coding | 38.98 | -2.04E-05 | 2.04E-05 | ENSG00000157426 | n.s. |
| ENSG00000064763 | 534.9892 | 0.011656769 | 0.289699 | 0.0402375 | 0.967904 | 0.995991293 | FAR2 | protein_coding | 38.9 | 2.03E-05 | 2.03E-05 | ENSG00000064763 | n.s. |
| ENSG00000179085 | 128.9379 | -0.011605087 | 0.325367 | -0.035668 | 0.971547 | 0.995991293 | DPM3 | protein_coding | 60.71 | -2.02E-05 | 2.02E-05 | ENSG00000179085 | n.s. |
| ENSG00000267321 | 233.142 | 0.011538354 | 0.286413 | 0.0402857 | 0.967865 | 0.995991293 | LINC02001 | lincRNA | 44.42 | 2.01E-05 | 2.01E-05 | ENSG00000267321 | n.s. |
| ENSG00000100075 | 409.9624 | 0.011481597 | 0.29231 | 0.0392788 | 0.968668 | 0.995991293 | SLC25A1 | protein_coding | 67.54 | 2.00E-05 | 2.00E-05 | ENSG00000100075 | n.s. |
| ENSG00000279117 | 113.5817 | -0.011446389 | 0.26557 | -0.043101 | 0.965621 | 0.995991293 | | TEC | 51.75 | -2.00E-05 | 2.00E-05 | ENSG00000279117 | n.s. |
| ENSG00000013293 | 1128.646 | -0.011331254 | 0.291277 | -0.038902 | 0.968969 | 0.995991293 | SLC7A14 | protein_coding | 42.15 | -1.98E-05 | 1.98E-05 | ENSG00000013293 | n.s. |
| ENSG00000131374 | 1314.256 | -0.010734557 | 0.244334 | -0.043934 | 0.964957 | 0.99578251 | TBC1D5 | protein_coding | 37.62 | -1.97E-05 | 1.97E-05 | ENSG00000131374 | n.s. |
| ENSG00000147684 | 2460.534 | -0.011239056 | 0.314413 | -0.035746 | 0.971485 | 0.995991293 | NDUFB9 | protein_coding | 45.62 | -1.96E-05 | 1.96E-05 | ENSG00000147684 | n.s. |
| ENSG00000167619 | 137.808 | -0.011217915 | 0.269498 | -0.041625 | 0.966797 | 0.995991293 | TMEM145 | protein_coding | 57.14 | -1.96E-05 | 1.96E-05 | ENSG00000167619 | n.s. |
| ENSG00000145087 | 382.7925 | 0.011155183 | 0.27148 | 0.0410902 | 0.967224 | 0.995991293 | STXBP5L | protein_coding | 36.35 | 1.95E-05 | 1.95E-05 | ENSG00000145087 | n.s. |

| | | | | | | | | | | | | | |
|---|---|---|---|---|---|---|---|---|---|---|---|---|---|
| ENSG00000156253 | 316.432 | 0.011282433 | 0.32076 | 0.035174 | 0.971941 | 0.996041878 | RWDD2B | protein_coding | 45.42 | 1.94E-05 | 1.94E-05 | ENSG00000156253 | n.s. |
| ENSG00000124785 | 296.3359 | 0.011038172 | 0.27795 | 0.0397128 | 0.968322 | 0.995991293 | NRN1 | protein_coding | 52.74 | 1.93E-05 | 1.93E-05 | ENSG00000124785 | n.s. |
| ENSG00000111052 | 1111.919 | 0.011002661 | 0.303706 | 0.0362279 | 0.971101 | 0.995991293 | LIN7A | protein_coding | 36.73 | 1.92E-05 | 1.92E-05 | ENSG00000111052 | n.s. |
| ENSG00000198589 | 6181.29 | 0.010980343 | 0.262802 | 0.0417818 | 0.966673 | 0.995991293 | LRBA | protein_coding | 36.23 | 1.92E-05 | 1.92E-05 | ENSG00000198589 | n.s. |
| ENSG00000089009 | 30092.74 | 0.010911895 | 0.297277 | 0.0367061 | 0.970719 | 0.995991293 | RPL6 | protein_coding | 45.67 | 1.90E-05 | 1.90E-05 | ENSG00000089009 | n.s. |
| ENSG00000164111 | 1268.955 | -0.010903311 | 0.296848 | -0.03673 | 0.9707 | 0.995991293 | ANXA5 | protein_coding | 39.13 | -1.90E-05 | 1.90E-05 | ENSG00000164111 | n.s. |
| ENSG00000127838 | 408.8759 | 0.011041838 | 0.315869 | 0.034957 | 0.972114 | 0.996041878 | PNKD | protein_coding | 51.32 | 1.90E-05 | 1.90E-05 | ENSG00000127838 | n.s. |
| ENSG00000278231 | 15.76569 | -0.014391065 | 0.515216 | -0.027932 | 0.977716 | 0.9969708 | | lincRNA | 47.64 | -1.90E-05 | 1.90E-05 | ENSG00000278231 | n.s. |
| ENSG00000170291 | 475.4514 | -0.010861005 | 0.310581 | -0.03497 | 0.972104 | 0.996041878 | ELP5 | protein_coding | 51.3 | -1.87E-05 | 1.87E-05 | ENSG00000170291 | n.s. |
| ENSG00000050130 | 1293.738 | 0.010719067 | 0.260552 | 0.0411399 | 0.967184 | 0.995991293 | JKAMP | protein_coding | 36.78 | 1.87E-05 | 1.87E-05 | ENSG00000050130 | n.s. |
| ENSG00000102144 | 4746.194 | 0.010712998 | 0.294647 | 0.0363588 | 0.970996 | 0.995991293 | PGK1 | protein_coding | 41.35 | 1.87E-05 | 1.87E-05 | ENSG00000102144 | n.s. |
| ENSG00000135521 | 1984.225 | 0.010674919 | 0.269438 | 0.0396192 | 0.968397 | 0.995991293 | LTV1 | protein_coding | 39.74 | 1.86E-05 | 1.86E-05 | ENSG00000135521 | n.s. |
| ENSG00000128699 | 2531.598 | -0.010674428 | 0.273233 | -0.039067 | 0.968837 | 0.995991293 | ORMDL1 | protein_coding | 37.58 | -1.86E-05 | 1.86E-05 | ENSG00000128699 | n.s. |
| ENSG00000130193 | 162.2171 | -0.010663911 | 0.295217 | -0.036122 | 0.971185 | 0.995991293 | THEM6 | protein_coding | 52.84 | -1.86E-05 | 1.86E-05 | ENSG00000130193 | n.s. |
| ENSG00000103035 | 1568.65 | 0.010581462 | 0.268139 | 0.0394626 | 0.968522 | 0.995991293 | PSMD7 | protein_coding | 43.55 | 1.85E-05 | 1.85E-05 | ENSG00000103035 | n.s. |
| ENSG00000222624 | 8.456494 | -0.013912537 | 0.462919 | -0.030054 | 0.976024 | 0.9969708 | RNU2-15P | snRNA | 38.74 | -1.83E-05 | 1.83E-05 | ENSG00000222624 | n.s. |
| ENSG00000280120 | 279.2869 | 0.010499891 | 0.284057 | 0.036964 | 0.970514 | 0.995991293 | | TEC | 39.37 | 1.83E-05 | 1.83E-05 | ENSG00000280120 | n.s. |
| ENSG00000179523 | 167.9514 | 0.010464737 | 0.255784 | 0.0409125 | 0.967366 | 0.995991293 | EIF3J-AS1 | lincRNA | 44.04 | 1.83E-05 | 1.83E-05 | ENSG00000179523 | n.s. |
| ENSG00000279310 | 9.417952 | -0.013769975 | 0.53275 | -0.025847 | 0.979379 | 0.9969708 | | TEC | 37.93 | -1.81E-05 | 1.81E-05 | ENSG00000279310 | n.s. |
| ENSG00000237137 | 6.761093 | -0.013739339 | 0.579057 | -0.023727 | 0.98107 | 0.9969708 | | processed_pseudogene | 44.88 | -1.81E-05 | 1.81E-05 | ENSG00000237137 | n.s. |
| ENSG00000179456 | 7305.984 | 0.0093064 | 0.205653 | 0.0452529 | 0.963906 | 0.995533737 | ZBTB18 | protein_coding | 44.35 | 1.81E-05 | 1.81E-05 | ENSG00000179456 | n.s. |
| ENSG00000116863 | 125.3401 | -0.010343398 | 0.281052 | -0.036802 | 0.970643 | 0.995991293 | ADPRHL2 | protein_coding | 55.16 | -1.80E-05 | 1.80E-05 | ENSG00000116863 | n.s. |
| ENSG00000233327 | 334.5033 | 0.01046682 | 0.302005 | 0.0346578 | 0.972353 | 0.996041878 | USP32P2 | cribed_unprocessed_pseudc | 42.09 | 1.80E-05 | 1.80E-05 | ENSG00000233327 | n.s. |
| ENSG00000189376 | 193.3194 | 0.01029139 | 0.273669 | 0.0376053 | 0.970002 | 0.995991293 | C8orf76 | protein_coding | 43.75 | 1.80E-05 | 1.80E-05 | ENSG00000189376 | n.s. |
| ENSG00000024862 | 251.2971 | 0.010272581 | 0.28098 | 0.0365598 | 0.970836 | 0.995991293 | CCDC28A | protein_coding | 39.18 | 1.79E-05 | 1.79E-05 | ENSG00000024862 | n.s. |
| ENSG00000119977 | 549.9626 | -0.010220325 | 0.26599 | -0.038424 | 0.96935 | 0.995991293 | TCTN3 | protein_coding | 39.54 | -1.78E-05 | 1.78E-05 | ENSG00000119977 | n.s. |
| ENSG00000179387 | 2362.224 | -0.010178354 | 0.274407 | -0.037092 | 0.970411 | 0.995991293 | ELMOD2 | protein_coding | 34.81 | -1.78E-05 | 1.78E-05 | ENSG00000179387 | n.s. |
| ENSG00000246379 | 154.5231 | -0.010171982 | 0.235284 | -0.043233 | 0.965516 | 0.995991293 | | lincRNA | 38.46 | -1.77E-05 | 1.77E-05 | ENSG00000246379 | n.s. |
| ENSG00000166170 | 1487.488 | 0.010156546 | 0.240962 | 0.04215 | 0.966379 | 0.995991293 | BAG5 | protein_coding | 46.41 | 1.77E-05 | 1.77E-05 | ENSG00000166170 | n.s. |
| ENSG00000170581 | 301.4812 | 0.010153161 | 0.271324 | 0.0374209 | 0.970149 | 0.995991293 | STAT2 | protein_coding | 47.77 | 1.77E-05 | 1.77E-05 | ENSG00000170581 | n.s. |
| ENSG00000177034 | 3342.242 | 0.010082266 | 0.247173 | 0.0407903 | 0.967463 | 0.995991293 | MTX3 | protein_coding | 37.49 | 1.76E-05 | 1.76E-05 | ENSG00000177034 | n.s. |
| ENSG00000181924 | 948.4529 | 0.01003587 | 0.266981 | 0.0375902 | 0.970014 | 0.995991293 | COA4 | protein_coding | 46.92 | 1.75E-05 | 1.75E-05 | ENSG00000181924 | n.s. |

| | | | | | | | | | | | | | |
|---|---|---|---|---|---|---|---|---|---|---|---|---|---|
| ENSG00000165609 | 1666.019 | 0.010152988 | 0.292178 | 0.0347493 | 0.97228 | 0.996041878 | NUDT5 | protein_coding | 44.24 | 1.75E-05 | 1.75E-05 | ENSG00000165609 | n.s. |
| ENSG00000062194 | 2635.259 | 0.009981284 | 0.254063 | 0.0392866 | 0.968662 | 0.995991293 | GPBP1 | protein_coding | 40.02 | 1.74E-05 | 1.74E-05 | ENSG00000062194 | n.s. |
| ENSG00000108384 | 2116.831 | -0.009469314 | 0.216482 | -0.043742 | 0.96511 | 0.995792945 | RAD51C | protein_coding | 41.71 | -1.73E-05 | 1.73E-05 | ENSG00000108384 | n.s. |
| ENSG00000069345 | 1722.51 | 0.009927846 | 0.279467 | 0.0355243 | 0.971662 | 0.995991293 | DNAJA2 | protein_coding | 42.72 | 1.73E-05 | 1.73E-05 | ENSG00000069345 | n.s. |
| ENSG00000047249 | 701.271 | -0.009888447 | 0.272896 | -0.036235 | 0.971095 | 0.995991293 | ATP6V1H | protein_coding | 39.2 | -1.72E-05 | 1.72E-05 | ENSG00000047249 | n.s. |
| ENSG00000179218 | 3770.906 | -0.009883083 | 0.261078 | -0.037855 | 0.969803 | 0.995991293 | CALR | protein_coding | 53.01 | -1.72E-05 | 1.72E-05 | ENSG00000179218 | n.s. |
| ENSG00000253910 | 88.47907 | 0.009869668 | 0.262241 | 0.0376359 | 0.969978 | 0.995991293 | PCDHGB2 | protein_coding | 45.22 | 1.72E-05 | 1.72E-05 | ENSG00000253910 | n.s. |
| ENSG00000201654 | 7.273816 | 0.013019036 | 0.568183 | 0.0229135 | 0.981719 | 0.9969708 | RNU6-7 | snRNA | 46.73 | 1.72E-05 | 1.72E-05 | ENSG00000201654 | n.s. |
| ENSG00000153561 | 2734.095 | -0.009821695 | 0.258828 | -0.037947 | 0.96973 | 0.995991293 | RMND5A | protein_coding | 40.38 | -1.71E-05 | 1.71E-05 | ENSG00000153561 | n.s. |
| ENSG00000212529 | 13.23479 | 0.013668917 | 0.731953 | 0.0186746 | 0.985101 | 0.997132464 | | snoRNA | 53.1 | 1.70E-05 | 1.70E-05 | ENSG00000212529 | n.s. |
| ENSG00000096717 | 1171.557 | 0.00976477 | 0.242477 | 0.0402709 | 0.967877 | 0.995991293 | SIRT1 | protein_coding | 40.87 | 1.70E-05 | 1.70E-05 | ENSG00000096717 | n.s. |
| ENSG00000132485 | 5433.575 | 0.009759503 | 0.272879 | 0.035765 | 0.97147 | 0.995991293 | ZRANB2 | protein_coding | 34.21 | 1.70E-05 | 1.70E-05 | ENSG00000132485 | n.s. |
| ENSG00000170915 | 251.6811 | -0.009737819 | 0.238496 | -0.04083 | 0.967431 | 0.995991293 | PAQR8 | protein_coding | 42.53 | -1.70E-05 | 1.70E-05 | ENSG00000170915 | n.s. |
| ENSG00000114853 | 186.7656 | -0.009681879 | 0.243345 | -0.039787 | 0.968263 | 0.995991293 | ZBTB47 | protein_coding | 62.11 | -1.69E-05 | 1.69E-05 | ENSG00000114853 | n.s. |
| ENSG00000180447 | 233.9305 | -0.009663367 | 0.225391 | -0.042874 | 0.965802 | 0.995991293 | GAS1 | protein_coding | 57.73 | -1.69E-05 | 1.69E-05 | ENSG00000180447 | n.s. |
| ENSG00000167654 | 737.5587 | -0.009594556 | 0.269993 | -0.035536 | 0.971652 | 0.995991293 | ATCAY | protein_coding | 51.6 | -1.67E-05 | 1.67E-05 | ENSG00000167654 | n.s. |
| ENSG00000165629 | 2306.805 | 0.010884331 | 0.333125 | 0.0326734 | 0.973935 | 0.99646746 | ATP5C1 | protein_coding | 40.13 | 1.67E-05 | 1.67E-05 | ENSG00000165629 | n.s. |
| ENSG00000148450 | 172.9881 | 0.010099089 | 0.296431 | 0.0340689 | 0.972822 | 0.996219704 | MSRB2 | protein_coding | 43.06 | 1.66E-05 | 1.66E-05 | ENSG00000148450 | n.s. |
| ENSG00000196268 | 1246.567 | -0.009509248 | 0.234838 | -0.040493 | 0.9677 | 0.995991293 | ZNF493 | protein_coding | 38.43 | -1.66E-05 | 1.66E-05 | ENSG00000196268 | n.s. |
| ENSG00000154122 | 242.9037 | -0.009405058 | 0.220951 | -0.042566 | 0.966047 | 0.995991293 | ANKH | protein_coding | 43.86 | -1.64E-05 | 1.64E-05 | ENSG00000154122 | n.s. |
| ENSG00000273015 | 2721.126 | 0.00925589 | 0.226302 | 0.0409007 | 0.967375 | 0.995991293 | | lincRNA | 38.04 | 1.61E-05 | 1.61E-05 | ENSG00000273015 | n.s. |
| ENSG00000108448 | 31.19221 | 0.010800288 | 0.338478 | 0.0319084 | 0.974545 | 0.996591368 | TRIM16L | protein_coding | 44.38 | 1.60E-05 | 1.60E-05 | ENSG00000108448 | n.s. |
| ENSG00000162129 | 1025.96 | -0.009168323 | 0.255792 | -0.035843 | 0.971408 | 0.995991293 | CLPB | protein_coding | 45.92 | -1.60E-05 | 1.60E-05 | ENSG00000162129 | n.s. |
| ENSG00000147416 | 2474.713 | -0.009249969 | 0.267766 | -0.034545 | 0.972443 | 0.996041878 | ATP6V1B2 | protein_coding | 39.51 | -1.59E-05 | 1.59E-05 | ENSG00000147416 | n.s. |
| ENSG00000131748 | 226.7752 | 0.010262165 | 0.308054 | 0.0333129 | 0.973425 | 0.99646746 | STARD3 | protein_coding | 54.9 | 1.58E-05 | 1.58E-05 | ENSG00000131748 | n.s. |
| ENSG00000267909 | 217.1137 | -0.009018876 | 0.21449 | -0.042048 | 0.96646 | 0.995991293 | CCDC177 | protein_coding | 62.35 | -1.57E-05 | 1.57E-05 | ENSG00000267909 | n.s. |
| ENSG00000198155 | 135.2945 | 0.010192936 | 0.306668 | 0.0332377 | 0.973485 | 0.99646746 | ZNF876P | cribed_unprocessed_pseudc | 38.73 | 1.57E-05 | 1.57E-05 | ENSG00000198155 | n.s. |
| ENSG00000178307 | 465.347 | 0.008964956 | 0.233097 | 0.0384603 | 0.969321 | 0.995991293 | TMEM11 | protein_coding | 51.96 | 1.56E-05 | 1.56E-05 | ENSG00000178307 | n.s. |
| ENSG00000130758 | 134.1138 | 0.010160524 | 0.310776 | 0.032694 | 0.973919 | 0.99646746 | MAP3K10 | protein_coding | 52.96 | 1.56E-05 | 1.56E-05 | ENSG00000130758 | n.s. |
| ENSG00000167985 | 311.5245 | 0.010142657 | 0.314194 | 0.0322815 | 0.974248 | 0.99646746 | SDHAF2 | protein_coding | 45.09 | 1.56E-05 | 1.56E-05 | ENSG00000167985 | n.s. |
| ENSG00000102580 | 663.3298 | 0.008921433 | 0.240516 | 0.0370929 | 0.970411 | 0.995991293 | DNAJC3 | protein_coding | 39.79 | 1.56E-05 | 1.56E-05 | ENSG00000102580 | n.s. |
| ENSG00000068438 | 556.7912 | -0.010062396 | 0.311702 | -0.032282 | 0.974247 | 0.99646746 | FTSJ1 | protein_coding | 50.6 | -1.55E-05 | 1.55E-05 | ENSG00000068438 | n.s. |

| | | | | | | | | | | | | | |
|---|---|---|---|---|---|---|---|---|---|---|---|---|---|
| ENSG00000175063 | 1648.024 | -0.008899037 | 0.258243 | -0.03446 | 0.97251 | 0.996041878 | UBE2C | protein_coding | 51.32 | -1.53E-05 | 1.53E-05 | ENSG00000175063 | n.s. |
| ENSG00000168765 | 94.54287 | 0.009845264 | 0.303336 | 0.0324566 | 0.974108 | 0.99646746 | GSTM4 | protein_coding | 49.98 | 1.51E-05 | 1.51E-05 | ENSG00000168765 | n.s. |
| ENSG00000103034 | 1410.417 | 0.008766733 | 0.254084 | 0.0345033 | 0.972476 | 0.996041878 | NDRG4 | protein_coding | 53.85 | 1.51E-05 | 1.51E-05 | ENSG00000103034 | n.s. |
| ENSG00000235145 | 65.09689 | -0.00975057 | 0.293485 | -0.033223 | 0.973496 | 0.99646746 | RPSAP16 | processed_pseudogene | 49.82 | -1.50E-05 | 1.50E-05 | ENSG00000235145 | n.s. |
| ENSG00000171314 | 1206.619 | 0.008669792 | 0.252202 | 0.0343764 | 0.972577 | 0.996041878 | PGAM1 | protein_coding | 47.47 | 1.49E-05 | 1.49E-05 | ENSG00000171314 | n.s. |
| ENSG00000126457 | 3425.066 | -0.009703863 | 0.297883 | -0.032576 | 0.974013 | 0.99646746 | PRMT1 | protein_coding | 58.23 | -1.49E-05 | 1.49E-05 | ENSG00000126457 | n.s. |
| ENSG00000160679 | 3553.971 | 0.008539095 | 0.234667 | 0.0363881 | 0.970973 | 0.995991293 | CHTOP | protein_coding | 44.38 | 1.49E-05 | 1.49E-05 | ENSG00000160679 | n.s. |
| ENSG00000134323 | 48939.42 | 0.008508108 | 0.20981 | 0.0405514 | 0.967653 | 0.995991293 | MYCN | protein_coding | 53.76 | 1.48E-05 | 1.48E-05 | ENSG00000134323 | n.s. |
| ENSG00000248283 | 67.38392 | -0.009494583 | 0.291133 | -0.032613 | 0.973984 | 0.99646746 | CCNL2P1 | processed_pseudogene | 48.61 | -1.46E-05 | 1.46E-05 | ENSG00000248283 | n.s. |
| ENSG00000151150 | 1250.552 | 0.008392693 | 0.241077 | 0.0348134 | 0.972229 | 0.996041878 | ANK3 | protein_coding | 37.75 | 1.45E-05 | 1.45E-05 | ENSG00000151150 | n.s. |
| ENSG00000119638 | 381.5669 | -0.008215215 | 0.218342 | -0.037625 | 0.969986 | 0.995991293 | NEK9 | protein_coding | 42.1 | -1.43E-05 | 1.43E-05 | ENSG00000119638 | n.s. |
| ENSG00000125037 | 557.9287 | -0.009299253 | 0.288292 | -0.032256 | 0.974268 | 0.99646746 | EMC3 | protein_coding | 44.9 | -1.43E-05 | 1.43E-05 | ENSG00000125037 | n.s. |
| ENSG00000096746 | 3674.465 | 0.008097813 | 0.231631 | 0.0349599 | 0.972112 | 0.996041878 | HNRNPH3 | protein_coding | 39.82 | 1.39E-05 | 1.39E-05 | ENSG00000096746 | n.s. |
| ENSG00000147164 | 353.3879 | -0.009378589 | 0.293734 | -0.031929 | 0.974529 | 0.996591368 | SNX12 | protein_coding | 42.55 | -1.39E-05 | 1.39E-05 | ENSG00000147164 | n.s. |
| ENSG00000159593 | 2851.49 | 0.009027535 | 0.273203 | 0.0330433 | 0.97364 | 0.99646746 | NAE1 | protein_coding | 46.31 | 1.39E-05 | 1.39E-05 | ENSG00000159593 | n.s. |
| ENSG00000141971 | 169.801 | -0.010388507 | 0.336397 | -0.030882 | 0.975364 | 0.996940486 | MVB12A | protein_coding | 50.72 | -1.38E-05 | 1.38E-05 | ENSG00000141971 | n.s. |
| ENSG00000153094 | 283.2747 | 0.007859024 | 0.227982 | 0.0344722 | 0.972501 | 0.996041878 | BCL2L11 | protein_coding | 44.42 | 1.35E-05 | 1.35E-05 | ENSG00000153094 | n.s. |
| ENSG00000105404 | 282.4802 | -0.010090591 | 0.324285 | -0.031116 | 0.975177 | 0.996919824 | RABAC1 | protein_coding | 61.62 | -1.35E-05 | 1.35E-05 | ENSG00000105404 | n.s. |
| ENSG00000221808 | 5.159661 | 0.010764189 | 0.54896 | 0.0196083 | 0.984356 | 0.997115786 | MIR1256 | miRNA | 42.02 | 1.35E-05 | 1.35E-05 | ENSG00000221808 | n.s. |
| ENSG00000235245 | 34.53565 | -0.010226962 | 0.360205 | -0.028392 | 0.977349 | 0.9969708 | | antisense | 58.53 | -1.35E-05 | 1.35E-05 | ENSG00000235245 | n.s. |
| ENSG00000165527 | 1554.329 | 0.007691719 | 0.203263 | 0.0378413 | 0.969814 | 0.995991293 | ARF6 | protein_coding | 43.17 | 1.34E-05 | 1.34E-05 | ENSG00000165527 | n.s. |
| ENSG00000137821 | 405.3484 | 0.008589529 | 0.25552 | 0.0336159 | 0.973183 | 0.99646746 | LRRC49 | protein_coding | 38.17 | 1.32E-05 | 1.32E-05 | ENSG00000137821 | n.s. |
| ENSG00000129933 | 723.8252 | 0.008558937 | 0.26019 | 0.032895 | 0.973758 | 0.99646746 | MAU2 | protein_coding | 54.38 | 1.32E-05 | 1.32E-05 | ENSG00000129933 | n.s. |
| ENSG00000263004 | 37.87235 | -0.00996983 | 0.343974 | -0.028984 | 0.976877 | 0.9969708 | | lincRNA | 45.5 | -1.31E-05 | 1.31E-05 | ENSG00000263004 | n.s. |
| ENSG00000162236 | 184.3235 | -0.008443927 | 0.26191 | -0.03224 | 0.974281 | 0.99646746 | STX5 | protein_coding | 47.05 | -1.30E-05 | 1.30E-05 | ENSG00000162236 | n.s. |
| ENSG00000140988 | 22817.53 | -0.009592983 | 0.308886 | -0.031057 | 0.975224 | 0.996919824 | RPS2 | protein_coding | 57.46 | -1.29E-05 | 1.29E-05 | ENSG00000140988 | n.s. |
| ENSG00000048162 | 1167.996 | -0.007185904 | 0.201705 | -0.035626 | 0.971581 | 0.995991293 | NOP16 | protein_coding | 49.33 | -1.25E-05 | 1.25E-05 | ENSG00000048162 | n.s. |
| ENSG00000243789 | 37.04413 | -0.009487222 | 0.336574 | -0.028188 | 0.977512 | 0.9969708 | JMJD7 | protein_coding | 54.81 | -1.25E-05 | 1.25E-05 | ENSG00000243789 | n.s. |
| ENSG00000261864 | 101.8942 | -0.008725455 | 0.275565 | -0.031664 | 0.97474 | 0.996717549 | | processed_transcript | 43.16 | -1.25E-05 | 1.25E-05 | ENSG00000261864 | n.s. |
| ENSG00000088448 | 2916.286 | -0.008072234 | 0.241086 | -0.033483 | 0.97329 | 0.99646746 | ANKRD10 | protein_coding | 40 | -1.24E-05 | 1.24E-05 | ENSG00000088448 | n.s. |
| ENSG00000134602 | 527.0058 | 0.0093721 | 0.311091 | 0.0301265 | 0.975966 | 0.9969708 | STK26 | protein_coding | 36.62 | 1.23E-05 | 1.23E-05 | ENSG00000134602 | n.s. |
| ENSG00000130724 | 372.9188 | -0.009159586 | 0.311996 | -0.029358 | 0.976579 | 0.9969708 | CHMP2A | protein_coding | 54.26 | -1.21E-05 | 1.21E-05 | ENSG00000130724 | n.s. |

| | | | | | | | | | | | | | |
|---|---|---|---|---|---|---|---|---|---|---|---|---|---|
| ENSG00000174010 | 1157.655 | 0.008708616 | 0.277813 | 0.0313471 | 0.974993 | 0.996829487 | KLHL15 | protein_coding | 41.85 | 1.20E-05 | 1.20E-05 | ENSG00000174010 | n.s. |
| ENSG00000109794 | 50.90756 | -0.009058297 | 0.322292 | -0.028106 | 0.977578 | 0.9969708 | FAM149A | protein_coding | 44.16 | -1.19E-05 | 1.19E-05 | ENSG00000109794 | n.s. |
| ENSG00000130818 | 1005.58 | 0.007660361 | 0.231324 | 0.0331153 | 0.973583 | 0.99646746 | ZNF426 | protein_coding | 43.51 | 1.18E-05 | 1.18E-05 | ENSG00000130818 | n.s. |
| ENSG00000254662 | 13.16949 | 0.00937819 | 0.474184 | 0.0197775 | 0.984221 | 0.997115786 | | antisense | 50.38 | 1.18E-05 | 1.18E-05 | ENSG00000254662 | n.s. |
| ENSG00000130733 | 128.5746 | -0.008691086 | 0.332972 | -0.026102 | 0.979176 | 0.9969708 | YIPF2 | protein_coding | 58.8 | -1.15E-05 | 1.15E-05 | ENSG00000130733 | n.s. |
| ENSG00000167110 | 166.2414 | -0.008669543 | 0.299955 | -0.028903 | 0.976942 | 0.9969708 | GOLGA2 | protein_coding | 52.34 | -1.14E-05 | 1.14E-05 | ENSG00000167110 | n.s. |
| ENSG00000114744 | 1231.952 | -0.008595929 | 0.285942 | -0.030062 | 0.976018 | 0.9969708 | COMMD2 | protein_coding | 37.91 | -1.13E-05 | 1.13E-05 | ENSG00000114744 | n.s. |
| ENSG00000202297 | 19.15213 | 0.008553392 | 0.388945 | 0.0219913 | 0.982455 | 0.9969708 | RNY3P12 | misc_RNA | 39.22 | 1.13E-05 | 1.13E-05 | ENSG00000202297 | n.s. |
| ENSG00000005448 | 393.4137 | 0.008528462 | 0.324501 | 0.0262817 | 0.979033 | 0.9969708 | WDR54 | protein_coding | 54.68 | 1.12E-05 | 1.12E-05 | ENSG00000005448 | n.s. |
| ENSG00000206601 | 21.39215 | 0.008426817 | 0.385572 | 0.0218554 | 0.982563 | 0.9969708 | RNU6-431P | snRNA | 43.93 | 1.11E-05 | 1.11E-05 | ENSG00000206601 | n.s. |
| ENSG00000156171 | 745.3813 | -0.008417516 | 0.287144 | -0.029315 | 0.976614 | 0.9969708 | DRAM2 | protein_coding | 37.94 | -1.11E-05 | 1.11E-05 | ENSG00000156171 | n.s. |
| ENSG00000169020 | 1428.303 | 0.008413474 | 0.333774 | 0.0252071 | 0.97989 | 0.9969708 | ATP5I | protein_coding | 57.33 | 1.11E-05 | 1.11E-05 | ENSG00000169020 | n.s. |
| ENSG00000206728 | 11.88999 | -0.008807325 | 0.468603 | -0.018795 | 0.985005 | 0.997132464 | | misc_RNA | 44.23 | -1.10E-05 | 1.10E-05 | ENSG00000206728 | n.s. |
| ENSG00000172331 | 380.1447 | -0.00830785 | 0.303336 | -0.027388 | 0.97815 | 0.9969708 | BPGM | protein_coding | 39.66 | -1.09E-05 | 1.09E-05 | ENSG00000172331 | n.s. |
| ENSG00000120705 | 3072.911 | 0.008297819 | 0.280734 | 0.0295575 | 0.97642 | 0.9969708 | ETF1 | protein_coding | 43.04 | 1.09E-05 | 1.09E-05 | ENSG00000120705 | n.s. |
| ENSG00000121410 | 151.4216 | -0.008280456 | 0.324549 | -0.025514 | 0.979645 | 0.9969708 | A1BG | protein_coding | 55.83 | -1.09E-05 | 1.09E-05 | ENSG00000121410 | n.s. |
| ENSG00000201672 | 17.49168 | -0.00862576 | 0.425483 | -0.020273 | 0.983826 | 0.997095274 | SNORD113-4 | snoRNA | 40 | -1.09E-05 | 1.09E-05 | ENSG00000201672 | n.s. |
| ENSG00000254531 | 47.88428 | 0.008263488 | 0.320819 | 0.0257575 | 0.979451 | 0.9969708 | | lincRNA | 62.32 | 1.09E-05 | 1.09E-05 | ENSG00000254531 | n.s. |
| ENSG00000188343 | 995.2558 | -0.007809215 | 0.248134 | -0.031472 | 0.974893 | 0.996800984 | FAM92A | protein_coding | 38.54 | -1.09E-05 | 1.09E-05 | ENSG00000188343 | n.s. |
| ENSG00000265354 | 1299.658 | 0.00820261 | 0.282891 | 0.0289956 | 0.976868 | 0.9969708 | TIMM23 | protein_coding | 40.16 | 1.08E-05 | 1.08E-05 | ENSG00000265354 | n.s. |
| ENSG00000111481 | 1376.208 | 0.008178562 | 0.299366 | 0.0273196 | 0.978205 | 0.9969708 | COPZ1 | protein_coding | 46.59 | 1.08E-05 | 1.08E-05 | ENSG00000111481 | n.s. |
| ENSG00000072133 | 428.674 | -0.008177969 | 0.273769 | -0.029872 | 0.976169 | 0.9969708 | RPS6KA6 | protein_coding | 35.17 | -1.08E-05 | 1.08E-05 | ENSG00000072133 | n.s. |
| ENSG00000171735 | 208.8552 | -0.008141988 | 0.3369 | -0.024167 | 0.980719 | 0.9969708 | CAMTA1 | protein_coding | 48 | -1.07E-05 | 1.07E-05 | ENSG00000171735 | n.s. |
| ENSG00000113732 | 556.795 | -0.008089959 | 0.318261 | -0.025419 | 0.979721 | 0.9969708 | ATP6V0E1 | protein_coding | 44.09 | -1.07E-05 | 1.07E-05 | ENSG00000113732 | n.s. |
| ENSG00000147457 | 666.0552 | -0.008089711 | 0.295843 | -0.027345 | 0.978185 | 0.9969708 | CHMP7 | protein_coding | 46.73 | -1.07E-05 | 1.07E-05 | ENSG00000147457 | n.s. |
| ENSG00000138640 | 1369.079 | 0.008078417 | 0.277027 | 0.0291611 | 0.976736 | 0.9969708 | FAM13A | protein_coding | 38.27 | 1.06E-05 | 1.06E-05 | ENSG00000138640 | n.s. |
| ENSG00000171928 | 655.7427 | -0.007885379 | 0.26029 | -0.030295 | 0.975832 | 0.9969708 | TVP23B | protein_coding | 41.13 | -1.04E-05 | 1.04E-05 | ENSG00000171928 | n.s. |
| ENSG00000198791 | 3252.935 | 0.007859665 | 0.264816 | 0.0296797 | 0.976322 | 0.9969708 | CNOT7 | protein_coding | 36.74 | 1.04E-05 | 1.04E-05 | ENSG00000198791 | n.s. |
| ENSG00000165283 | 1590.52 | 0.007823759 | 0.297819 | 0.0262702 | 0.979042 | 0.9969708 | STOML2 | protein_coding | 53.1 | 1.03E-05 | 1.03E-05 | ENSG00000165283 | n.s. |
| ENSG00000276712 | 73.47483 | 0.007768648 | 0.285621 | 0.0271991 | 0.978301 | 0.9969708 | MIR7111 | miRNA | 58.33 | 1.02E-05 | 1.02E-05 | ENSG00000276712 | n.s. |
| ENSG00000102901 | 574.9286 | -0.007768495 | 0.256593 | -0.030276 | 0.975847 | 0.9969708 | CENPT | protein_coding | 53.77 | -1.02E-05 | 1.02E-05 | ENSG00000102901 | n.s. |
| ENSG00000120549 | 84.24679 | 0.007749175 | 0.275474 | 0.0281303 | 0.977558 | 0.9969708 | KIAA1217 | protein_coding | 40.56 | 1.02E-05 | 1.02E-05 | ENSG00000120549 | n.s. |

| | | | | | | | | | | | | | |
|---|---|---|---|---|---|---|---|---|---|---|---|---|---|
| ENSG00000138381 | 2300.193 | 0.007697791 | 0.265864 | 0.0289539 | 0.976901 | 0.9969708 | ASNSD1 | protein_coding | 37.75 | 1.01E-05 | 1.01E-05 | ENSG00000138381 | n.s. |
| ENSG00000105849 | 1190.63 | -0.007622046 | 0.26092 | -0.029212 | 0.976695 | 0.9969708 | TWISTNB | protein_coding | 36.3 | -1.00E-05 | 1.00E-05 | ENSG00000105849 | n.s. |
| ENSG00000028203 | 3164.771 | 0.007608719 | 0.272266 | 0.0279459 | 0.977705 | 0.9969708 | VEZT | protein_coding | 38.6 | 1.00E-05 | 1.00E-05 | ENSG00000028203 | n.s. |
| ENSG00000256377 | 106.8612 | 0.007585426 | 0.304391 | 0.02492 | 0.980119 | 0.9969708 | | antisense | 42.77 | 9.99E-06 | 9.99E-06 | ENSG00000256377 | n.s. |
| ENSG00000228203 | 264.7535 | 0.00754467 | 0.254592 | 0.0296343 | 0.976359 | 0.9969708 | RNF144A-AS1 | processed_transcript | 51.22 | 9.94E-06 | 9.94E-06 | ENSG00000228203 | n.s. |
| ENSG00000117118 | 442.4608 | 0.007456857 | 0.299469 | 0.0249003 | 0.980134 | 0.9969708 | SDHB | protein_coding | 43.51 | 9.82E-06 | 9.82E-06 | ENSG00000117118 | n.s. |
| ENSG00000165806 | 168.3578 | -0.007433119 | 0.289998 | -0.025632 | 0.979551 | 0.9969708 | CASP7 | protein_coding | 42.96 | -9.79E-06 | 9.79E-06 | ENSG00000165806 | n.s. |
| ENSG00000167645 | 286.6747 | -0.007415313 | 0.314318 | -0.023592 | 0.981178 | 0.9969708 | YIF1B | protein_coding | 55.37 | -9.77E-06 | 9.77E-06 | ENSG00000167645 | n.s. |
| ENSG00000166337 | 361.6552 | -0.007255765 | 0.252157 | -0.028775 | 0.977044 | 0.9969708 | TAF10 | protein_coding | 52.56 | -9.56E-06 | 9.56E-06 | ENSG00000166337 | n.s. |
| ENSG00000136933 | 1021.157 | 0.007231848 | 0.31863 | 0.0226967 | 0.981892 | 0.9969708 | RABEPK | protein_coding | 46.49 | 9.53E-06 | 9.53E-06 | ENSG00000136933 | n.s. |
| ENSG00000135624 | 8665.711 | -0.00720935 | 0.282748 | -0.025497 | 0.979658 | 0.9969708 | CCT7 | protein_coding | 46.07 | -9.50E-06 | 9.50E-06 | ENSG00000135624 | n.s. |
| ENSG00000084754 | 2585.217 | -0.007199746 | 0.286938 | -0.025092 | 0.979982 | 0.9969708 | HADHA | protein_coding | 41.93 | -9.49E-06 | 9.49E-06 | ENSG00000084754 | n.s. |
| ENSG00000104976 | 78.09649 | 0.007173674 | 0.314397 | 0.0228173 | 0.981796 | 0.9969708 | SNAPC2 | protein_coding | 64.16 | 9.45E-06 | 9.45E-06 | ENSG00000104976 | n.s. |
| ENSG00000100097 | 69.08947 | 0.007524458 | 0.375817 | 0.0200216 | 0.984026 | 0.997115786 | LGALS1 | protein_coding | 56.78 | 9.44E-06 | 9.44E-06 | ENSG00000100097 | n.s. |
| ENSG00000214548 | 1748.725 | -0.007142638 | 0.243306 | -0.029357 | 0.97658 | 0.9969708 | MEG3 | lincRNA | 51.68 | -9.41E-06 | 9.41E-06 | ENSG00000214548 | n.s. |
| ENSG00000157404 | 585.6777 | 0.007098174 | 0.270311 | 0.0262593 | 0.979051 | 0.9969708 | KIT | protein_coding | 40.87 | 9.35E-06 | 9.35E-06 | ENSG00000157404 | n.s. |
| ENSG00000171425 | 195.5878 | 0.007085332 | 0.27868 | 0.0254247 | 0.979716 | 0.9969708 | ZNF581 | protein_coding | 58.42 | 9.34E-06 | 9.34E-06 | ENSG00000171425 | n.s. |
| ENSG00000028116 | 344.7158 | 0.007071143 | 0.249543 | 0.0283364 | 0.977394 | 0.9969708 | VRK2 | protein_coding | 35.24 | 9.32E-06 | 9.32E-06 | ENSG00000028116 | n.s. |
| ENSG00000133895 | 238.4258 | -0.00705304 | 0.285262 | -0.024725 | 0.980275 | 0.9969708 | MEN1 | protein_coding | 57.78 | -9.29E-06 | 9.29E-06 | ENSG00000133895 | n.s. |
| ENSG00000005882 | 90.90712 | 0.006992658 | 0.316414 | 0.0220997 | 0.982368 | 0.9969708 | PDK2 | protein_coding | 56.67 | 9.21E-06 | 9.21E-06 | ENSG00000005882 | n.s. |
| ENSG00000179632 | 549.5082 | -0.006944197 | 0.29505 | -0.023536 | 0.981223 | 0.9969708 | MAF1 | protein_coding | 63.19 | -9.15E-06 | 9.15E-06 | ENSG00000179632 | n.s. |
| ENSG00000163528 | 250.757 | -0.006915164 | 0.304126 | -0.022738 | 0.981859 | 0.9969708 | CHCHD4 | protein_coding | 48.18 | -9.11E-06 | 9.11E-06 | ENSG00000163528 | n.s. |
| ENSG00000182117 | 985.0143 | 0.006908146 | 0.328988 | 0.0209981 | 0.983247 | 0.9969708 | NOP10 | protein_coding | 49.56 | 9.10E-06 | 9.10E-06 | ENSG00000182117 | n.s. |
| ENSG00000234604 | 34.52353 | 0.006877837 | 0.323746 | 0.0212446 | 0.983051 | 0.9969708 | | processed_pseudogene | 35 | 9.06E-06 | 9.06E-06 | ENSG00000234604 | n.s. |
| ENSG00000221241 | 24.15347 | -0.007222903 | 0.37926 | -0.019045 | 0.984805 | 0.997115786 | SNORD88A | snoRNA | 70.33 | -9.06E-06 | 9.06E-06 | ENSG00000221241 | n.s. |
| ENSG00000212027 | 9.656539 | -0.008458544 | 0.475183 | -0.017801 | 0.985798 | 0.997542208 | MIR374B | miRNA | 41.67 | -9.04E-06 | 9.04E-06 | ENSG00000212027 | n.s. |
| ENSG00000146066 | 487.4497 | 0.006851119 | 0.280529 | 0.0244222 | 0.980516 | 0.9969708 | HIGD2A | protein_coding | 57.37 | 9.03E-06 | 9.03E-06 | ENSG00000146066 | n.s. |
| ENSG00000116288 | 1201.166 | -0.006787508 | 0.317569 | -0.021373 | 0.982948 | 0.9969708 | PARK7 | protein_coding | 45.41 | -8.94E-06 | 8.94E-06 | ENSG00000116288 | n.s. |
| ENSG00000270019 | 418.6187 | 0.006748503 | 0.225633 | 0.0299092 | 0.976139 | 0.9969708 | | lincRNA | 44.12 | 8.89E-06 | 8.89E-06 | ENSG00000270019 | n.s. |
| ENSG00000176884 | 240.8626 | 0.0067207 | 0.2525 | 0.0266167 | 0.978765 | 0.9969708 | GRIN1 | protein_coding | 62.1 | 8.85E-06 | 8.85E-06 | ENSG00000176884 | n.s. |
| ENSG00000148290 | 307.5833 | 0.006642717 | 0.301384 | 0.0220407 | 0.982416 | 0.9969708 | SURF1 | protein_coding | 53.96 | 8.75E-06 | 8.75E-06 | ENSG00000148290 | n.s. |
| ENSG00000164967 | 148.366 | -0.006627598 | 0.277631 | -0.023872 | 0.980955 | 0.9969708 | RPP25L | protein_coding | 58.74 | -8.73E-06 | 8.73E-06 | ENSG00000164967 | n.s. |

| | | | | | | | | | | | | | |
|---|---|---|---|---|---|---|---|---|---|---|---|---|---|
| ENSG00000143727 | 4876.402 | 0.006625943 | 0.290148 | 0.0228364 | 0.981781 | 0.9969708 | ACP1 | protein_coding | 42.67 | 8.73E-06 | 8.73E-06 | ENSG00000143727 | n.s. |
| ENSG00000278637 | 414.5621 | -0.00651523 | 0.240579 | -0.027081 | 0.978395 | 0.9969708 | HIST1H4A | protein_coding | 60.58 | -8.58E-06 | 8.58E-06 | ENSG00000278637 | n.s. |
| ENSG00000138303 | 267.4757 | 0.006493561 | 0.291268 | 0.0222941 | 0.982213 | 0.9969708 | ASCC1 | protein_coding | 41.33 | 8.56E-06 | 8.56E-06 | ENSG00000138303 | n.s. |
| ENSG00000167515 | 445.8837 | -0.006487876 | 0.289965 | -0.022375 | 0.982149 | 0.9969708 | TRAPPC2L | protein_coding | 58.39 | -8.55E-06 | 8.55E-06 | ENSG00000167515 | n.s. |
| ENSG00000154040 | 83.74536 | 0.006464089 | 0.268049 | 0.0241153 | 0.980761 | 0.9969708 | CABYR | protein_coding | 43.36 | 8.52E-06 | 8.52E-06 | ENSG00000154040 | n.s. |
| ENSG00000167700 | 144.1484 | -0.006458978 | 0.282771 | -0.022842 | 0.981777 | 0.9969708 | MFSD3 | protein_coding | 66.96 | -8.51E-06 | 8.51E-06 | ENSG00000167700 | n.s. |
| ENSG00000103121 | 559.1133 | -0.006414276 | 0.278693 | -0.023016 | 0.981638 | 0.9969708 | CMC2 | protein_coding | 40.64 | -8.45E-06 | 8.45E-06 | ENSG00000103121 | n.s. |
| ENSG00000213619 | 1243.664 | 0.00641123 | 0.29152 | 0.0219924 | 0.982454 | 0.9969708 | NDUFS3 | protein_coding | 48.67 | 8.45E-06 | 8.45E-06 | ENSG00000213619 | n.s. |
| ENSG00000078142 | 5802.834 | 0.00639557 | 0.255263 | 0.0250548 | 0.980011 | 0.9969708 | PIK3C3 | protein_coding | 35.33 | 8.43E-06 | 8.43E-06 | ENSG00000078142 | n.s. |
| ENSG00000108100 | 1059.577 | -0.006348502 | 0.259496 | -0.024465 | 0.980482 | 0.9969708 | CCNY | protein_coding | 43.24 | -8.36E-06 | 8.36E-06 | ENSG00000108100 | n.s. |
| ENSG00000164597 | 738.5895 | -0.006337752 | 0.28118 | -0.02254 | 0.982017 | 0.9969708 | COG5 | protein_coding | 37.1 | -8.35E-06 | 8.35E-06 | ENSG00000164597 | n.s. |
| ENSG00000241859 | 203.615 | 0.00631828 | 0.258461 | 0.0244458 | 0.980497 | 0.9969708 | ANOS2P | cribed_unprocessed_pseudc | 40.21 | 8.32E-06 | 8.32E-06 | ENSG00000241859 | n.s. |
| ENSG00000166483 | 3203.53 | 0.006236516 | 0.202144 | 0.0308518 | 0.975388 | 0.996940486 | WEE1 | protein_coding | 40.54 | 8.30E-06 | 8.30E-06 | ENSG00000166483 | n.s. |
| ENSG00000176593 | 257.8435 | -0.006287993 | 0.214034 | -0.029379 | 0.976563 | 0.9969708 | | processed_transcript | 50.45 | -8.28E-06 | 8.28E-06 | ENSG00000176593 | n.s. |
| ENSG00000182093 | 391.1945 | 0.006586631 | 0.332858 | 0.0197881 | 0.984212 | 0.997115786 | WRB | protein_coding | 44.24 | 8.26E-06 | 8.26E-06 | ENSG00000182093 | n.s. |
| ENSG00000276517 | 360.9312 | -0.006234173 | 0.279283 | -0.022322 | 0.982191 | 0.9969708 | | sense_intronic | 38.95 | -8.21E-06 | 8.21E-06 | ENSG00000276517 | n.s. |
| ENSG00000206770 | 4.98951 | 0.009594603 | 0.737237 | 0.0130143 | 0.989616 | 0.998039352 | | misc_RNA | 37.27 | 8.18E-06 | 8.18E-06 | ENSG00000206770 | n.s. |
| ENSG00000154832 | 551.3814 | 0.006177878 | 0.264898 | 0.0233217 | 0.981394 | 0.9969708 | CXXC1 | protein_coding | 59.29 | 8.14E-06 | 8.14E-06 | ENSG00000154832 | n.s. |
| ENSG00000108669 | 344.4214 | 0.00612398 | 0.267715 | 0.022875 | 0.98175 | 0.9969708 | CYTH1 | protein_coding | 45.26 | 8.07E-06 | 8.07E-06 | ENSG00000108669 | n.s. |
| ENSG00000100360 | 208.4662 | 0.006376802 | 0.315247 | 0.0202279 | 0.983862 | 0.997095274 | IFT27 | protein_coding | 51.18 | 8.06E-06 | 8.06E-06 | ENSG00000100360 | n.s. |
| ENSG00000100983 | 507.4993 | 0.006107952 | 0.293862 | 0.0207851 | 0.983417 | 0.9969708 | GSS | protein_coding | 46.29 | 8.05E-06 | 8.05E-06 | ENSG00000100983 | n.s. |
| ENSG00000135845 | 1309.778 | -0.006079362 | 0.227091 | -0.026771 | 0.978643 | 0.9969708 | PIGC | protein_coding | 38.22 | -8.01E-06 | 8.01E-06 | ENSG00000135845 | n.s. |
| ENSG00000116251 | 3888.294 | 0.006062167 | 0.284725 | 0.0212913 | 0.983013 | 0.9969708 | RPL22 | protein_coding | 50.61 | 7.99E-06 | 7.99E-06 | ENSG00000116251 | n.s. |
| ENSG00000175216 | 6140.772 | 0.006061176 | 0.263615 | 0.0229926 | 0.981656 | 0.9969708 | CKAP5 | protein_coding | 40.42 | 7.99E-06 | 7.99E-06 | ENSG00000175216 | n.s. |
| ENSG00000169599 | 421.5423 | 0.006349316 | 0.328955 | 0.0193015 | 0.984601 | 0.997115786 | NFU1 | protein_coding | 41.47 | 7.96E-06 | 7.96E-06 | ENSG00000169599 | n.s. |
| ENSG00000121022 | 1200.603 | -0.005997389 | 0.259858 | -0.02308 | 0.981587 | 0.9969708 | COPS5 | protein_coding | 39.82 | -7.90E-06 | 7.90E-06 | ENSG00000121022 | n.s. |
| ENSG00000215305 | 546.888 | 0.005972772 | 0.287097 | 0.020804 | 0.983402 | 0.9969708 | VPS16 | protein_coding | 48.8 | 7.87E-06 | 7.87E-06 | ENSG00000215305 | n.s. |
| ENSG00000152518 | 826.6749 | 0.00595177 | 0.209613 | 0.0283941 | 0.977348 | 0.9969708 | ZFP36L2 | protein_coding | 55.96 | 7.84E-06 | 7.84E-06 | ENSG00000152518 | n.s. |
| ENSG00000216895 | 112.7322 | 0.005896505 | 0.266999 | 0.0220844 | 0.982381 | 0.9969708 | | antisense | 42.24 | 7.77E-06 | 7.77E-06 | ENSG00000216895 | n.s. |
| ENSG00000090238 | 152.1922 | 0.006134468 | 0.304334 | 0.020157 | 0.983918 | 0.997095274 | YPEL3 | protein_coding | 58.41 | 7.75E-06 | 7.75E-06 | ENSG00000090238 | n.s. |
| ENSG00000085788 | 570.5233 | -0.005839687 | 0.251732 | -0.023198 | 0.981492 | 0.9969708 | DDHD2 | protein_coding | 42.25 | -7.69E-06 | 7.69E-06 | ENSG00000085788 | n.s. |
| ENSG00000172466 | 4672.611 | -0.005812473 | 0.270234 | -0.021509 | 0.98284 | 0.9969708 | ZNF24 | protein_coding | 39.65 | -7.66E-06 | 7.66E-06 | ENSG00000172466 | n.s. |

| | | | | | | | | | | | | | |
|---|---|---|---|---|---|---|---|---|---|---|---|---|---|
| ENSG00000162377 | 2079.971 | 0.005800793 | 0.278175 | 0.020853 | 0.983363 | 0.9969708 | COA7 | protein_coding | 46.05 | 7.64E-06 | 7.64E-06 | ENSG00000162377 | n.s. |
| ENSG00000184277 | 1290.866 | 0.005793429 | 0.241513 | 0.0239881 | 0.980862 | 0.9969708 | TM2D3 | protein_coding | 40.91 | 7.63E-06 | 7.63E-06 | ENSG00000184277 | n.s. |
| ENSG00000151806 | 3234.073 | -0.00578399 | 0.24629 | -0.023484 | 0.981264 | 0.9969708 | GUF1 | protein_coding | 33.75 | -7.62E-06 | 7.62E-06 | ENSG00000151806 | n.s. |
| ENSG00000164758 | 295.1518 | 0.006057526 | 0.31766 | 0.0190692 | 0.984786 | 0.997115786 | MED30 | protein_coding | 34.58 | 7.60E-06 | 7.60E-06 | ENSG00000164758 | n.s. |
| ENSG00000100416 | 277.8319 | 0.005763095 | 0.254809 | 0.0226173 | 0.981956 | 0.9969708 | TRMU | protein_coding | 52.44 | 7.59E-06 | 7.59E-06 | ENSG00000100416 | n.s. |
| ENSG00000148541 | 563.2757 | 0.005750282 | 0.240948 | 0.0238652 | 0.98096 | 0.9969708 | FAM13C | protein_coding | 40.06 | 7.58E-06 | 7.58E-06 | ENSG00000148541 | n.s. |
| ENSG00000203760 | 784.0142 | -0.005970128 | 0.293563 | -0.020337 | 0.983775 | 0.997095274 | CENPW | protein_coding | 35.46 | -7.54E-06 | 7.54E-06 | ENSG00000203760 | n.s. |
| ENSG00000106628 | 1632.91 | 0.005638272 | 0.256448 | 0.021986 | 0.982459 | 0.9969708 | POLD2 | protein_coding | 55.2 | 7.43E-06 | 7.43E-06 | ENSG00000106628 | n.s. |
| ENSG00000060971 | 89.08875 | 0.005620531 | 0.267223 | 0.0210331 | 0.983219 | 0.9969708 | ACAA1 | protein_coding | 51.46 | 7.41E-06 | 7.41E-06 | ENSG00000060971 | n.s. |
| ENSG00000176749 | 709.0822 | -0.005612856 | 0.210252 | -0.026696 | 0.978702 | 0.9969708 | CDK5R1 | protein_coding | 58.04 | -7.40E-06 | 7.40E-06 | ENSG00000176749 | n.s. |
| ENSG00000107949 | 2511.75 | -0.005575584 | 0.267639 | -0.020832 | 0.983379 | 0.9969708 | BCCIP | protein_coding | 42.86 | -7.35E-06 | 7.35E-06 | ENSG00000107949 | n.s. |
| ENSG00000122008 | 251.8284 | -0.005564963 | 0.24499 | -0.022715 | 0.981878 | 0.9969708 | POLK | protein_coding | 36.45 | -7.33E-06 | 7.33E-06 | ENSG00000122008 | n.s. |
| ENSG00000188846 | 10844.14 | -0.0055637 | 0.267372 | -0.020809 | 0.983398 | 0.9969708 | RPL14 | protein_coding | 44.36 | -7.33E-06 | 7.33E-06 | ENSG00000188846 | n.s. |
| ENSG00000079215 | 89.06545 | 0.005555588 | 0.265168 | 0.0209512 | 0.983285 | 0.9969708 | SLC1A3 | protein_coding | 40.36 | 7.32E-06 | 7.32E-06 | ENSG00000079215 | n.s. |
| ENSG00000162231 | 995.4898 | 0.005553978 | 0.221993 | 0.0250187 | 0.98004 | 0.9969708 | NXF1 | protein_coding | 50.07 | 7.32E-06 | 7.32E-06 | ENSG00000162231 | n.s. |
| ENSG00000050344 | 176.7466 | 0.005552918 | 0.267484 | 0.0207598 | 0.983437 | 0.9969708 | NFE2L3 | protein_coding | 44.64 | 7.32E-06 | 7.32E-06 | ENSG00000050344 | n.s. |
| ENSG00000175582 | 845.2354 | -0.005491339 | 0.257963 | -0.021287 | 0.983016 | 0.9969708 | RAB6A | protein_coding | 41.4 | -7.24E-06 | 7.24E-06 | ENSG00000175582 | n.s. |
| ENSG00000231871 | 54.92375 | -0.00569298 | 0.298097 | -0.019098 | 0.984763 | 0.997115786 | IPO9-AS1 | antisense | 46.27 | -7.14E-06 | 7.14E-06 | ENSG00000231871 | n.s. |
| ENSG00000200834 | 17.90499 | -0.006638812 | 0.375026 | -0.017702 | 0.985876 | 0.997542208 | | misc_RNA | 41.58 | -7.10E-06 | 7.10E-06 | ENSG00000200834 | n.s. |
| ENSG00000188452 | 765.1624 | 0.005507 | 0.283446 | 0.0194287 | 0.984499 | 0.997115786 | CERKL | protein_coding | 37.01 | 6.91E-06 | 6.91E-06 | ENSG00000188452 | n.s. |
| ENSG00000100483 | 620.0012 | 0.005217296 | 0.232969 | 0.0223948 | 0.982133 | 0.9969708 | VCPKMT | protein_coding | 39.75 | 6.87E-06 | 6.87E-06 | ENSG00000100483 | n.s. |
| ENSG00000143742 | 7935.657 | -0.005799864 | 0.315114 | -0.018406 | 0.985315 | 0.99727715 | SRP9 | protein_coding | 40.6 | -6.87E-06 | 6.87E-06 | ENSG00000143742 | n.s. |
| ENSG00000198938 | 15083.62 | 0.005156161 | 0.23378 | 0.0220557 | 0.982404 | 0.9969708 | MT-CO3 | protein_coding | 46.56 | 6.79E-06 | 6.79E-06 | ENSG00000198938 | n.s. |
| ENSG00000243554 | 109.4128 | -0.005382892 | 0.274153 | -0.019635 | 0.984335 | 0.997115786 | | unprocessed_pseudogene | 46.12 | -6.75E-06 | 6.75E-06 | ENSG00000243554 | n.s. |
| ENSG00000171877 | 188.6526 | 0.005061225 | 0.241185 | 0.0209848 | 0.983258 | 0.9969708 | FRMD5 | protein_coding | 41.23 | 6.67E-06 | 6.67E-06 | ENSG00000171877 | n.s. |
| ENSG00000130414 | 1770.255 | 0.005148517 | 0.268049 | 0.0192074 | 0.984676 | 0.997115786 | NDUFA10 | protein_coding | 48.44 | 6.46E-06 | 6.46E-06 | ENSG00000130414 | n.s. |
| ENSG00000198081 | 625.1642 | -0.004891241 | 0.235152 | -0.0208 | 0.983405 | 0.9969708 | ZBTB14 | protein_coding | 49 | -6.44E-06 | 6.44E-06 | ENSG00000198081 | n.s. |
| ENSG00000239437 | 13.61745 | -0.006785813 | 0.441537 | -0.015369 | 0.987738 | 0.997819739 | RN7SL752P | misc_RNA | 55.4 | -6.43E-06 | 6.43E-06 | ENSG00000239437 | n.s. |
| ENSG00000112852 | 141.6612 | 0.005044829 | 0.260739 | 0.0193482 | 0.984563 | 0.997115786 | PCDHB2 | protein_coding | 48.38 | 6.33E-06 | 6.33E-06 | ENSG00000112852 | n.s. |
| ENSG00000231699 | 22.40323 | 0.005833728 | 0.352692 | 0.0165406 | 0.986803 | 0.997558623 | | processed_pseudogene | 40.49 | 6.19E-06 | 6.19E-06 | ENSG00000231699 | n.s. |
| ENSG00000266931 | 10.7751 | 0.00663369 | 0.484386 | 0.013695 | 0.989073 | 0.99786564 | | unprocessed_pseudogene | 41.96 | 6.16E-06 | 6.16E-06 | ENSG00000266931 | n.s. |
| ENSG00000213465 | 283.5381 | -0.005606432 | 0.336124 | -0.01668 | 0.986692 | 0.997558623 | ARL2 | protein_coding | 55.34 | -5.95E-06 | 5.95E-06 | ENSG00000213465 | n.s. |

| | | | | | | | | | | | | | |
|---|---|---|---|---|---|---|---|---|---|---|---|---|---|
| ENSG00000083093 | 925.4214 | -0.004670319 | 0.229767 | -0.020326 | 0.983783 | 0.997095274 | PALB2 | protein_coding | 43.53 | -5.90E-06 | 5.90E-06 | ENSG00000083093 | n.s. |
| ENSG00000151575 | 50.91372 | 0.005296095 | 0.303524 | 0.0174487 | 0.986079 | 0.997542208 | TEX9 | protein_coding | 36.87 | 5.66E-06 | 5.66E-06 | ENSG00000151575 | n.s. |
| ENSG00000100109 | 471.2166 | 0.004510731 | 0.241502 | 0.0186778 | 0.985098 | 0.997132464 | TFIP11 | protein_coding | 44.89 | 5.63E-06 | 5.63E-06 | ENSG00000100109 | n.s. |
| ENSG00000136560 | 1159.59 | -0.004438531 | 0.234043 | -0.018965 | 0.984869 | 0.997115786 | TANK | protein_coding | 35.76 | -5.57E-06 | 5.57E-06 | ENSG00000136560 | n.s. |
| ENSG00000157045 | 314.9143 | 0.005097229 | 0.290386 | 0.0175533 | 0.985995 | 0.997542208 | NTAN1 | protein_coding | 47.6 | 5.45E-06 | 5.45E-06 | ENSG00000157045 | n.s. |
| ENSG00000143164 | 798.1223 | -0.005036875 | 0.280969 | -0.017927 | 0.985697 | 0.997542208 | DCAF6 | protein_coding | 35.91 | -5.38E-06 | 5.38E-06 | ENSG00000143164 | n.s. |
| ENSG00000137804 | 3657.567 | -0.005052946 | 0.293421 | -0.017221 | 0.98626 | 0.997558623 | NUSAP1 | protein_coding | 44.52 | -5.36E-06 | 5.36E-06 | ENSG00000137804 | n.s. |
| ENSG00000131069 | 216.0638 | 0.005123917 | 0.312756 | 0.0163831 | 0.986929 | 0.997604841 | ACSS2 | protein_coding | 45.42 | 5.34E-06 | 5.34E-06 | ENSG00000131069 | n.s. |
| ENSG00000138095 | 7265.38 | 0.004841493 | 0.276238 | 0.0175265 | 0.986017 | 0.997542208 | LRPPRC | protein_coding | 37.13 | 5.17E-06 | 5.17E-06 | ENSG00000138095 | n.s. |
| ENSG00000185627 | 1213.891 | -0.005098634 | 0.320127 | -0.015927 | 0.987293 | 0.997733163 | PSMD13 | protein_coding | 44.35 | -5.03E-06 | 5.03E-06 | ENSG00000185627 | n.s. |
| ENSG00000144567 | 668.735 | -0.004698073 | 0.281068 | -0.016715 | 0.986664 | 0.997558623 | RETREG2 | protein_coding | 53.85 | -4.99E-06 | 4.99E-06 | ENSG00000144567 | n.s. |
| ENSG00000189046 | 920.4938 | -0.00464917 | 0.263616 | -0.017636 | 0.985929 | 0.997542208 | ALKBH2 | protein_coding | 51.05 | -4.97E-06 | 4.97E-06 | ENSG00000189046 | n.s. |
| ENSG00000201231 | 5.256933 | 0.006238489 | 0.572195 | 0.0109027 | 0.991301 | 0.998180522 | RNU4-50P | snRNA | 41.13 | 4.93E-06 | 4.93E-06 | ENSG00000201231 | n.s. |
| ENSG00000105880 | 70.95183 | -0.004632481 | 0.280243 | -0.01653 | 0.986811 | 0.997558623 | DLX5 | protein_coding | 55.55 | -4.92E-06 | 4.92E-06 | ENSG00000105880 | n.s. |
| ENSG00000240463 | 24.91125 | 0.005150947 | 0.361016 | 0.0142679 | 0.988616 | 0.99786564 | RPS19P3 | processed_pseudogene | 56.24 | 4.78E-06 | 4.78E-06 | ENSG00000240463 | n.s. |
| ENSG00000204304 | 346.9029 | -0.004467321 | 0.268857 | -0.016616 | 0.986743 | 0.997558623 | PBX2 | protein_coding | 54.92 | -4.74E-06 | 4.74E-06 | ENSG00000204304 | n.s. |
| ENSG00000132781 | 111.5832 | 0.004293106 | 0.253776 | 0.0169169 | 0.986503 | 0.997558623 | MUTYH | protein_coding | 51.03 | 4.56E-06 | 4.56E-06 | ENSG00000132781 | n.s. |
| ENSG00000144354 | 3334.025 | 0.004196428 | 0.243708 | 0.0172191 | 0.986262 | 0.997558623 | CDCA7 | protein_coding | 43.24 | 4.45E-06 | 4.45E-06 | ENSG00000144354 | n.s. |
| ENSG00000110717 | 641.199 | -0.004635672 | 0.304612 | -0.015218 | 0.987858 | 0.997819739 | NDUFS8 | protein_coding | 57.63 | -4.39E-06 | 4.39E-06 | ENSG00000110717 | n.s. |
| ENSG00000196526 | 2582.287 | 0.004105277 | 0.241318 | 0.0170119 | 0.986427 | 0.997558623 | AFAP1 | protein_coding | 46.39 | 4.36E-06 | 4.36E-06 | ENSG00000196526 | n.s. |
| ENSG00000234911 | 126.8651 | -0.004294178 | 0.265101 | -0.016198 | 0.987076 | 0.997681444 | TEX21P | unitary_pseudogene | 42.85 | -4.33E-06 | 4.33E-06 | ENSG00000234911 | n.s. |
| ENSG00000117598 | 1022.065 | -0.004544024 | 0.297244 | -0.015287 | 0.987803 | 0.997819739 | PLPPR5 | protein_coding | 37.21 | -4.31E-06 | 4.31E-06 | ENSG00000117598 | n.s. |
| ENSG00000125445 | 2320.477 | 0.004334928 | 0.274571 | 0.015788 | 0.987404 | 0.997733163 | MRPS7 | protein_coding | 50.49 | 4.27E-06 | 4.27E-06 | ENSG00000125445 | n.s. |
| ENSG00000180879 | 1286.438 | -0.004491452 | 0.309359 | -0.014519 | 0.988416 | 0.99786564 | SSR4 | protein_coding | 61.54 | -4.17E-06 | 4.17E-06 | ENSG00000180879 | n.s. |
| ENSG00000101751 | 545.5842 | -0.003905618 | 0.229469 | -0.01702 | 0.98642 | 0.997558623 | POLI | protein_coding | 36.47 | -4.15E-06 | 4.15E-06 | ENSG00000101751 | n.s. |
| ENSG00000206768 | 10.51883 | 0.005205199 | 0.502114 | 0.0103666 | 0.991729 | 0.998180522 | | misc_RNA | 41.18 | 4.12E-06 | 4.12E-06 | ENSG00000206768 | n.s. |
| ENSG00000153015 | 843.0463 | -0.004143681 | 0.262676 | -0.015775 | 0.987414 | 0.997733163 | CWC27 | protein_coding | 35.58 | -4.08E-06 | 4.08E-06 | ENSG00000153015 | n.s. |
| ENSG00000069509 | 139.9119 | 0.004342528 | 0.296532 | 0.0146444 | 0.988316 | 0.99786564 | FUNDC1 | protein_coding | 41.18 | 4.03E-06 | 4.03E-06 | ENSG00000069509 | n.s. |
| ENSG00000117697 | 1288.964 | 0.004267549 | 0.302114 | 0.0141256 | 0.98873 | 0.99786564 | NSL1 | protein_coding | 39.75 | 3.96E-06 | 3.96E-06 | ENSG00000117697 | n.s. |
| ENSG00000165688 | 899.4002 | 0.004116423 | 0.280066 | 0.014698 | 0.988273 | 0.99786564 | PMPCA | protein_coding | 53.93 | 3.82E-06 | 3.82E-06 | ENSG00000165688 | n.s. |
| ENSG00000198088 | 85.4802 | 0.003980821 | 0.286825 | 0.0138789 | 0.988927 | 0.99786564 | NUP62CL | protein_coding | 37.06 | 3.69E-06 | 3.69E-06 | ENSG00000198088 | n.s. |
| ENSG00000186814 | 569.4916 | 0.003965823 | 0.283118 | 0.0140077 | 0.988824 | 0.99786564 | ZSCAN30 | protein_coding | 42.35 | 3.68E-06 | 3.68E-06 | ENSG00000186814 | n.s. |

| | | | | | | | | | | | | | |
|---|---|---|---|---|---|---|---|---|---|---|---|---|---|
| ENSG00000240853 | 17.97433 | -0.00461284 | 0.405407 | -0.011378 | 0.990922 | 0.998180522 | RN7SL328P | misc_RNA | 54 | -3.65E-06 | 3.65E-06 | ENSG00000240853 | n.s. |
| ENSG00000151092 | 355.2801 | 0.003924696 | 0.287232 | 0.0136638 | 0.989098 | 0.99786564 | NGLY1 | protein_coding | 37.55 | 3.64E-06 | 3.64E-06 | ENSG00000151092 | n.s. |
| ENSG00000129347 | 683.1186 | -0.003891031 | 0.263684 | -0.014756 | 0.988227 | 0.99786564 | KRI1 | protein_coding | 55.09 | -3.61E-06 | 3.61E-06 | ENSG00000129347 | n.s. |
| ENSG00000197586 | 484.4074 | 0.003866726 | 0.273488 | 0.0141386 | 0.988719 | 0.99786564 | ENTPD6 | protein_coding | 50.79 | 3.59E-06 | 3.59E-06 | ENSG00000197586 | n.s. |
| ENSG00000080822 | 1401.324 | 0.003856492 | 0.261919 | 0.014724 | 0.988252 | 0.99786564 | CLDND1 | protein_coding | 38.91 | 3.58E-06 | 3.58E-06 | ENSG00000080822 | n.s. |
| ENSG00000214413 | 341.5685 | 0.004113247 | 0.311725 | 0.0131951 | 0.989472 | 0.998002069 | BBIP1 | protein_coding | 38.64 | 3.57E-06 | 3.57E-06 | ENSG00000214413 | n.s. |
| ENSG00000171791 | 4064.632 | -0.00352724 | 0.21959 | -0.016063 | 0.987184 | 0.997718241 | BCL2 | protein_coding | 42.74 | -3.50E-06 | 3.50E-06 | ENSG00000171791 | n.s. |
| ENSG00000130844 | 455.6479 | 0.003710497 | 0.257987 | 0.0143825 | 0.988525 | 0.99786564 | ZNF331 | protein_coding | 44.28 | 3.44E-06 | 3.44E-06 | ENSG00000130844 | n.s. |
| ENSG00000143570 | 1271.105 | 0.003679151 | 0.271638 | 0.0135443 | 0.989194 | 0.99786564 | SLC39A1 | protein_coding | 52.3 | 3.41E-06 | 3.41E-06 | ENSG00000143570 | n.s. |
| ENSG00000131773 | 710.3384 | 0.003619522 | 0.263671 | 0.0137274 | 0.989047 | 0.99786564 | KHDRBS3 | protein_coding | 38.22 | 3.36E-06 | 3.36E-06 | ENSG00000131773 | n.s. |
| ENSG00000187147 | 563.1909 | 0.003595348 | 0.265081 | 0.0135632 | 0.989178 | 0.99786564 | RNF220 | protein_coding | 47.21 | 3.34E-06 | 3.34E-06 | ENSG00000187147 | n.s. |
| ENSG00000016082 | 2921.54 | 0.00347921 | 0.225525 | 0.0154272 | 0.987691 | 0.997819739 | ISL1 | protein_coding | 44.32 | 3.30E-06 | 3.30E-06 | ENSG00000016082 | n.s. |
| ENSG00000240771 | 758.6441 | 0.003539704 | 0.255271 | 0.0138665 | 0.988937 | 0.99786564 | ARHGEF25 | protein_coding | 56.4 | 3.28E-06 | 3.28E-06 | ENSG00000240771 | n.s. |
| ENSG00000113441 | 1318.167 | 0.003385062 | 0.243195 | 0.0139191 | 0.988894 | 0.99786564 | LNPEP | protein_coding | 37.19 | 3.14E-06 | 3.14E-06 | ENSG00000113441 | n.s. |
| ENSG00000165521 | 6123.032 | -0.003599806 | 0.271545 | -0.013257 | 0.989423 | 0.998002069 | EML5 | protein_coding | 37.41 | -3.13E-06 | 3.13E-06 | ENSG00000165521 | n.s. |
| ENSG00000241745 | 41.81173 | 0.003944043 | 0.320511 | 0.0123055 | 0.990182 | 0.998180522 | RN7SL788P | misc_RNA | 55.06 | 3.12E-06 | 3.12E-06 | ENSG00000241745 | n.s. |
| ENSG00000221227 | 13.05837 | 0.004270415 | 0.438815 | 0.0097317 | 0.992235 | 0.998334008 | MIR1305 | miRNA | 36.05 | 3.09E-06 | 3.09E-06 | ENSG00000221227 | n.s. |
| ENSG00000178741 | 1407.863 | -0.003761801 | 0.296622 | -0.012682 | 0.989881 | 0.998125629 | COX5A | protein_coding | 44.93 | -3.07E-06 | 3.07E-06 | ENSG00000178741 | n.s. |
| ENSG00000173145 | 1768.869 | 0.003418576 | 0.264115 | 0.0129435 | 0.989673 | 0.998039352 | NOC3L | protein_coding | 37.01 | 2.91E-06 | 2.91E-06 | ENSG00000173145 | n.s. |
| ENSG00000109132 | 8177.958 | 0.003071862 | 0.201722 | 0.0152282 | 0.98785 | 0.997819739 | PHOX2B | protein_coding | 52.1 | 2.91E-06 | 2.91E-06 | ENSG00000109132 | n.s. |
| ENSG00000119878 | 393.3307 | 0.003665398 | 0.313002 | 0.0117105 | 0.990657 | 0.998180522 | CRIPT | protein_coding | 39.14 | 2.90E-06 | 2.90E-06 | ENSG00000119878 | n.s. |
| ENSG00000241468 | 2188.837 | 0.003656833 | 0.335551 | 0.010898 | 0.991305 | 0.998180522 | ATP5J2 | protein_coding | 53.67 | 2.89E-06 | 2.89E-06 | ENSG00000241468 | n.s. |
| ENSG00000104979 | 610.1996 | 0.003642344 | 0.32746 | 0.011123 | 0.991125 | 0.998180522 | C19orf53 | protein_coding | 54.88 | 2.88E-06 | 2.88E-06 | ENSG00000104979 | n.s. |
| ENSG00000168275 | 388.9188 | -0.003636976 | 0.297736 | -0.012215 | 0.990254 | 0.998180522 | COA6 | protein_coding | 44.99 | -2.88E-06 | 2.88E-06 | ENSG00000168275 | n.s. |
| ENSG00000197818 | 54.35464 | 0.003609273 | 0.298165 | 0.0121049 | 0.990342 | 0.998180522 | SLC9A8 | protein_coding | 45.56 | 2.85E-06 | 2.85E-06 | ENSG00000197818 | n.s. |
| ENSG00000204899 | 1178.208 | 0.00359747 | 0.311773 | 0.0115388 | 0.990794 | 0.998180522 | MZT1 | protein_coding | 38.9 | 2.85E-06 | 2.85E-06 | ENSG00000204899 | n.s. |
| ENSG00000139343 | 2316.093 | -0.003320551 | 0.257822 | -0.012879 | 0.989724 | 0.998039352 | SNRPF | protein_coding | 39.09 | -2.83E-06 | 2.83E-06 | ENSG00000139343 | n.s. |
| ENSG00000279757 | 27.24607 | 0.003505151 | 0.343998 | 0.0101894 | 0.99187 | 0.998180522 | | TEC | 49.02 | 2.77E-06 | 2.77E-06 | ENSG00000279757 | n.s. |
| ENSG00000129559 | 2076.332 | 0.003459552 | 0.294607 | 0.0117429 | 0.990631 | 0.998180522 | NEDD8 | protein_coding | 43.63 | 2.74E-06 | 2.74E-06 | ENSG00000129559 | n.s. |
| ENSG00000168958 | 1548.909 | -0.003439661 | 0.286611 | -0.012001 | 0.990425 | 0.998180522 | MFF | protein_coding | 36.79 | -2.72E-06 | 2.72E-06 | ENSG00000168958 | n.s. |
| ENSG00000104147 | 288.2446 | 0.003429821 | 0.308824 | 0.0111061 | 0.991139 | 0.998180522 | OIP5 | protein_coding | 45.46 | 2.71E-06 | 2.71E-06 | ENSG00000104147 | n.s. |
| ENSG00000185305 | 99.57921 | -0.003313145 | 0.290172 | -0.011418 | 0.99089 | 0.998180522 | ARL15 | protein_coding | 38.5 | -2.62E-06 | 2.62E-06 | ENSG00000185305 | n.s. |

| | | | | | | | | | | | | | |
|---|---|---|---|---|---|---|---|---|---|---|---|---|---|
| ENSG00000130811 | 1377.938 | -0.003304089 | 0.281063 | -0.011756 | 0.990621 | 0.998180522 | EIF3G | protein_coding | 59.24 | -2.61E-06 | 2.61E-06 | ENSG00000130811 | n.s. |
| ENSG00000165121 | 465.6394 | -0.003256813 | 0.281589 | -0.011566 | 0.990772 | 0.998180522 | | transcribed_unprocessed_pseudc | 37.75 | -2.58E-06 | 2.58E-06 | ENSG00000165121 | n.s. |
| ENSG00000111665 | 224.8625 | -0.003226676 | 0.271219 | -0.011897 | 0.990508 | 0.998180522 | CDCA3 | protein_coding | 52.08 | -2.55E-06 | 2.55E-06 | ENSG00000111665 | n.s. |
| ENSG00000108468 | 9504.244 | 0.003111012 | 0.273803 | 0.0113622 | 0.990934 | 0.998180522 | CBX1 | protein_coding | 45.96 | 2.46E-06 | 2.46E-06 | ENSG00000108468 | n.s. |
| ENSG00000164054 | 209.1697 | 0.002953101 | 0.285628 | 0.010339 | 0.991751 | 0.998180522 | SHISA5 | protein_coding | 50.15 | 2.34E-06 | 2.34E-06 | ENSG00000164054 | n.s. |
| ENSG00000137124 | 688.715 | 0.002919742 | 0.26755 | 0.0109129 | 0.991293 | 0.998180522 | ALDH1B1 | protein_coding | 51.48 | 2.31E-06 | 2.31E-06 | ENSG00000137124 | n.s. |
| ENSG00000235786 | 98.90819 | 0.002854434 | 0.280185 | 0.0101877 | 0.991872 | 0.998180522 | ZNRF3-IT1 | sense_intronic | 45.22 | 2.26E-06 | 2.26E-06 | ENSG00000235786 | n.s. |
| ENSG00000273210 | 29.51067 | -0.003365761 | 0.381155 | -0.00883 | 0.992954 | 0.998476414 | | antisense | 47.15 | -2.23E-06 | 2.23E-06 | ENSG00000273210 | n.s. |
| ENSG00000118689 | 963.0583 | 0.00280077 | 0.250031 | 0.0112017 | 0.991063 | 0.998180522 | FOXO3 | protein_coding | 41.71 | 2.22E-06 | 2.22E-06 | ENSG00000118689 | n.s. |
| ENSG00000116044 | 2953.864 | 0.00271525 | 0.263797 | 0.0102929 | 0.991788 | 0.998180522 | NFE2L2 | protein_coding | 40.63 | 2.15E-06 | 2.15E-06 | ENSG00000116044 | n.s. |
| ENSG00000157557 | 1443.121 | 0.00266812 | 0.252827 | 0.0105532 | 0.99158 | 0.998180522 | ETS2 | protein_coding | 46.04 | 2.11E-06 | 2.11E-06 | ENSG00000157557 | n.s. |
| ENSG00000130741 | 2293.492 | -0.00260977 | 0.254438 | -0.010257 | 0.991816 | 0.998180522 | EIF2S3 | protein_coding | 41.58 | -2.06E-06 | 2.06E-06 | ENSG00000130741 | n.s. |
| ENSG00000125743 | 3779.845 | 0.002978209 | 0.321263 | 0.0092703 | 0.992603 | 0.998460202 | SNRPD2 | protein_coding | 52.25 | 1.99E-06 | 1.99E-06 | ENSG00000125743 | n.s. |
| ENSG00000146676 | 1803.303 | 0.002490129 | 0.239106 | 0.0104143 | 0.991691 | 0.998180522 | PURB | protein_coding | 40.58 | 1.97E-06 | 1.97E-06 | ENSG00000146676 | n.s. |
| ENSG00000103479 | 1352.609 | 0.002673211 | 0.274826 | 0.0097269 | 0.992239 | 0.998334008 | RBL2 | protein_coding | 38.71 | 1.94E-06 | 1.94E-06 | ENSG00000103479 | n.s. |
| ENSG00000173207 | 1787.132 | 0.00277916 | 0.293259 | 0.0094768 | 0.992439 | 0.998460202 | CKS1B | protein_coding | 44.4 | 1.86E-06 | 1.86E-06 | ENSG00000173207 | n.s. |
| ENSG00000275700 | 728.0411 | -0.002532844 | 0.257076 | -0.009853 | 0.992139 | 0.998334008 | AATF | protein_coding | 40.51 | -1.83E-06 | 1.83E-06 | ENSG00000275700 | n.s. |
| ENSG00000167526 | 15130.91 | -0.002802367 | 0.325779 | -0.008602 | 0.993137 | 0.998515412 | RPL13 | protein_coding | 55.95 | -1.81E-06 | 1.81E-06 | ENSG00000167526 | n.s. |
| ENSG00000164032 | 10126.29 | 0.002694315 | 0.302099 | 0.0089186 | 0.992884 | 0.998476414 | H2AFZ | protein_coding | 49.59 | 1.78E-06 | 1.78E-06 | ENSG00000164032 | n.s. |
| ENSG00000181163 | 42805.96 | 0.002651894 | 0.291479 | 0.0090981 | 0.992741 | 0.998476414 | NPM1 | protein_coding | 43.6 | 1.76E-06 | 1.76E-06 | ENSG00000181163 | n.s. |
| ENSG00000108523 | 421.8455 | 0.002612287 | 0.279332 | 0.0093519 | 0.992538 | 0.998460202 | RNF167 | protein_coding | 52.54 | 1.75E-06 | 1.75E-06 | ENSG00000108523 | n.s. |
| ENSG00000115816 | 4487.09 | 0.002453373 | 0.266376 | 0.0092102 | 0.992651 | 0.998460202 | CEBPZ | protein_coding | 38.07 | 1.64E-06 | 1.64E-06 | ENSG00000115816 | n.s. |
| ENSG00000124098 | 279.3721 | 0.002439626 | 0.270836 | 0.0090078 | 0.992813 | 0.998476414 | FAM210B | protein_coding | 49.44 | 1.62E-06 | 1.62E-06 | ENSG00000124098 | n.s. |
| ENSG00000091164 | 1387.48 | 0.002313881 | 0.272857 | 0.0084802 | 0.993234 | 0.998541052 | TXNL1 | protein_coding | 38.58 | 1.47E-06 | 1.47E-06 | ENSG00000091164 | n.s. |
| ENSG00000136636 | 3483.393 | -0.002264681 | 0.261402 | -0.008664 | 0.993088 | 0.998515412 | KCTD3 | protein_coding | 36.04 | -1.46E-06 | 1.46E-06 | ENSG00000136636 | n.s. |
| ENSG00000244038 | 944.6902 | -0.002441076 | 0.303027 | -0.008056 | 0.993573 | 0.99880948 | DDOST | protein_coding | 50.61 | -1.26E-06 | 1.26E-06 | ENSG00000244038 | n.s. |
| ENSG00000176340 | 1981.02 | -0.002416295 | 0.339111 | -0.007125 | 0.994315 | 0.998821569 | COX8A | protein_coding | 57 | -1.24E-06 | 1.24E-06 | ENSG00000176340 | n.s. |
| ENSG00000141741 | 217.3863 | -0.00232557 | 0.310028 | -0.007501 | 0.994015 | 0.998821569 | MIEN1 | protein_coding | 57.02 | -1.19E-06 | 1.19E-06 | ENSG00000141741 | n.s. |
| ENSG00000138382 | 780.3104 | -0.002242031 | 0.310541 | -0.00722 | 0.99424 | 0.998821569 | METTL5 | protein_coding | 39.58 | -1.15E-06 | 1.15E-06 | ENSG00000138382 | n.s. |
| ENSG00000163444 | 1511.187 | -0.002208352 | 0.278176 | -0.007939 | 0.993666 | 0.998821569 | TMEM183A | protein_coding | 41.5 | -1.13E-06 | 1.13E-06 | ENSG00000163444 | n.s. |
| ENSG00000115758 | 10821.21 | -0.002192314 | 0.289099 | -0.007583 | 0.993949 | 0.998821569 | ODC1 | protein_coding | 46.16 | -1.12E-06 | 1.12E-06 | ENSG00000115758 | n.s. |
| ENSG00000213563 | 269.2895 | -0.002156385 | 0.278268 | -0.007749 | 0.993817 | 0.998821569 | C8orf82 | protein_coding | 67.88 | -1.10E-06 | 1.10E-06 | ENSG00000213563 | n.s. |

| | | | | | | | | | | | | | |
|---|---|---|---|---|---|---|---|---|---|---|---|---|---|
| ENSG00000261514 | 223.9267 | -0.002148144 | 0.302863 | -0.007093 | 0.994341 | 0.998821569 | LINC01976 | lincRNA | 63.99 | -1.10E-06 | 1.10E-06 | ENSG00000261514 | n.s. |
| ENSG00000005022 | 2068.789 | 0.002138064 | 0.299974 | 0.0071275 | 0.994313 | 0.998821569 | SLC25A5 | protein_coding | 53.25 | 1.09E-06 | 1.09E-06 | ENSG00000005022 | n.s. |
| ENSG00000178988 | 2487.679 | 0.001899907 | 0.245679 | 0.0077333 | 0.99383 | 0.998821569 | MRFAP1L1 | protein_coding | 44.91 | 9.73E-07 | 9.73E-07 | ENSG00000178988 | n.s. |
| ENSG00000008294 | 8708.03 | 0.001893177 | 0.258811 | 0.0073149 | 0.994164 | 0.998821569 | SPAG9 | protein_coding | 40 | 9.69E-07 | 9.69E-07 | ENSG00000008294 | n.s. |
| ENSG00000163161 | 1628.933 | -0.001862038 | 0.264062 | -0.007052 | 0.994374 | 0.998821569 | ERCC3 | protein_coding | 46.18 | -9.54E-07 | 9.54E-07 | ENSG00000163161 | n.s. |
| ENSG00000171490 | 5798.638 | 0.00181211 | 0.262221 | 0.0069106 | 0.994486 | 0.998862413 | RSL1D1 | protein_coding | 45.24 | 8.96E-07 | 8.96E-07 | ENSG00000171490 | n.s. |
| ENSG00000240342 | 3269.447 | -0.001957401 | 0.310902 | -0.006296 | 0.994977 | 0.99899544 | | processed_pseudogene | 61.21 | -8.54E-07 | 8.54E-07 | ENSG00000240342 | n.s. |
| ENSG00000089248 | 1150.262 | 0.001885329 | 0.294475 | 0.0064024 | 0.994892 | 0.99899544 | ERP29 | protein_coding | 47.22 | 8.23E-07 | 8.23E-07 | ENSG00000089248 | n.s. |
| ENSG00000157036 | 75.39722 | -0.00187233 | 0.282886 | -0.006619 | 0.994719 | 0.99899544 | EXOG | protein_coding | 42.1 | -8.17E-07 | 8.17E-07 | ENSG00000157036 | n.s. |
| ENSG00000160051 | 79.32617 | -0.001833421 | 0.28105 | -0.006523 | 0.994795 | 0.99899544 | IQCC | protein_coding | 55.4 | -8.00E-07 | 8.00E-07 | ENSG00000160051 | n.s. |
| ENSG00000243147 | 1485.527 | -0.00183331 | 0.303378 | -0.006043 | 0.995178 | 0.99899544 | MRPL33 | protein_coding | 39.12 | -8.00E-07 | 8.00E-07 | ENSG00000243147 | n.s. |
| ENSG00000088986 | 2499.281 | 0.001828602 | 0.324912 | 0.005628 | 0.99551 | 0.99899544 | DYNLL1 | protein_coding | 47.04 | 7.98E-07 | 7.98E-07 | ENSG00000088986 | n.s. |
| ENSG00000136710 | 221.0391 | 0.001779708 | 0.313178 | 0.0056827 | 0.995466 | 0.99899544 | CCDC115 | protein_coding | 51.35 | 7.77E-07 | 7.77E-07 | ENSG00000136710 | n.s. |
| ENSG00000104325 | 814.1776 | 0.001733923 | 0.309958 | 0.0055941 | 0.995537 | 0.99899544 | DECR1 | protein_coding | 38.15 | 7.57E-07 | 7.57E-07 | ENSG00000104325 | n.s. |
| ENSG00000155368 | 2360.501 | -0.001668764 | 0.302363 | -0.005519 | 0.995596 | 0.99899544 | DBI | protein_coding | 50.37 | -7.28E-07 | 7.28E-07 | ENSG00000155368 | n.s. |
| ENSG00000001084 | 1371.659 | 0.001651489 | 0.264633 | 0.0062407 | 0.995021 | 0.99899544 | GCLC | protein_coding | 40.16 | 7.21E-07 | 7.21E-07 | ENSG00000001084 | n.s. |
| ENSG00000180773 | 238.3435 | -0.001594061 | 0.27913 | -0.005711 | 0.995443 | 0.99899544 | SLC36A4 | protein_coding | 35.08 | -6.96E-07 | 6.96E-07 | ENSG00000180773 | n.s. |
| ENSG00000283078 | 106.6336 | -0.001576606 | 0.293737 | -0.005367 | 0.995717 | 0.999005291 | | lincRNA | 51.24 | -6.81E-07 | 6.81E-07 | ENSG00000283078 | n.s. |
| ENSG00000169750 | 614.2284 | -0.00155504 | 0.293766 | -0.005293 | 0.995776 | 0.999005291 | RAC3 | protein_coding | 68.77 | -6.72E-07 | 6.72E-07 | ENSG00000169750 | n.s. |
| ENSG00000207067 | 161.7321 | -0.001424864 | 0.247445 | -0.005758 | 0.995406 | 0.99899544 | SNORA72 | snoRNA | 40.15 | -6.22E-07 | 6.22E-07 | ENSG00000207067 | n.s. |
| ENSG00000101972 | 4453.55 | 0.001415707 | 0.257831 | 0.0054908 | 0.995619 | 0.99899544 | STAG2 | protein_coding | 39.19 | 6.18E-07 | 6.18E-07 | ENSG00000101972 | n.s. |
| ENSG00000281398 | 512.2818 | -0.00117342 | 0.213908 | -0.005486 | 0.995623 | 0.99899544 | SNHG4 | processed_transcript | 43.8 | -5.12E-07 | 5.12E-07 | ENSG00000281398 | n.s. |
| ENSG00000280198 | 63.30863 | -0.001486852 | 0.310425 | -0.00479 | 0.996178 | 0.999237607 | | TEC | 51.71 | -4.92E-07 | 4.92E-07 | ENSG00000280198 | n.s. |
| ENSG00000108733 | 158.9778 | -0.00118933 | 0.232849 | -0.005108 | 0.995925 | 0.999081968 | PEX12 | protein_coding | 42.42 | -4.74E-07 | 4.74E-07 | ENSG00000108733 | n.s. |
| ENSG00000109083 | 229.0282 | 0.001374299 | 0.299707 | 0.0045855 | 0.996341 | 0.999237607 | IFT20 | protein_coding | 47.79 | 4.55E-07 | 4.55E-07 | ENSG00000109083 | n.s. |
| ENSG00000112640 | 817.2463 | 0.001307152 | 0.287066 | 0.0045535 | 0.996367 | 0.999237607 | PPP2R5D | protein_coding | 49.89 | 4.33E-07 | 4.33E-07 | ENSG00000112640 | n.s. |
| ENSG00000105866 | 654.1735 | 0.001117214 | 0.241374 | 0.0046286 | 0.996307 | 0.999237607 | SP4 | protein_coding | 38.35 | 3.70E-07 | 3.70E-07 | ENSG00000105866 | n.s. |
| ENSG00000138175 | 447.547 | -0.001194129 | 0.308563 | -0.00387 | 0.996912 | 0.999319207 | ARL3 | protein_coding | 44.57 | -3.53E-07 | 3.53E-07 | ENSG00000138175 | n.s. |
| ENSG00000154277 | 7609.047 | -0.001144873 | 0.290277 | -0.003944 | 0.996853 | 0.999319207 | UCHL1 | protein_coding | 44.57 | -3.39E-07 | 3.39E-07 | ENSG00000154277 | n.s. |
| ENSG00000114126 | 1528.982 | 0.001101828 | 0.270959 | 0.0040664 | 0.996755 | 0.999319207 | TFDP2 | protein_coding | 39.89 | 3.26E-07 | 3.26E-07 | ENSG00000114126 | n.s. |
| ENSG00000157191 | 211.6391 | -0.001094803 | 0.264037 | -0.004146 | 0.996692 | 0.999319207 | NECAP2 | protein_coding | 49.4 | -3.24E-07 | 3.24E-07 | ENSG00000157191 | n.s. |
| ENSG00000204628 | 29735.14 | -0.001089323 | 0.292145 | -0.003729 | 0.997025 | 0.999319207 | RACK1 | protein_coding | 52.35 | -3.22E-07 | 3.22E-07 | ENSG00000204628 | n.s. |

| | | | | | | | | | | | | | |
|---|---|---|---|---|---|---|---|---|---|---|---|---|---|
| ENSG00000138036 | 328.4086 | -0.001073698 | 0.305145 | -0.003519 | 0.997193 | 0.999319207 | DYNC2LI1 | protein_coding | 37.57 | -3.18E-07 | 3.18E-07 | ENSG00000138036 | n.s. |
| ENSG00000165630 | 669.6714 | 0.001048081 | 0.270904 | 0.0038688 | 0.996913 | 0.999319207 | PRPF18 | protein_coding | 38.95 | 3.10E-07 | 3.10E-07 | ENSG00000165630 | n.s. |
| ENSG00000137168 | 1084.706 | 0.001044545 | 0.323924 | 0.0032247 | 0.997427 | 0.999319207 | PPIL1 | protein_coding | 44.26 | 3.09E-07 | 3.09E-07 | ENSG00000137168 | n.s. |
| ENSG00000170296 | 1742.749 | 0.001025612 | 0.274381 | 0.0037379 | 0.997018 | 0.999319207 | GABARAP | protein_coding | 50.38 | 3.03E-07 | 3.03E-07 | ENSG00000170296 | n.s. |
| ENSG00000136522 | 1157.373 | -0.000981576 | 0.290052 | -0.003384 | 0.9973 | 0.999319207 | MRPL47 | protein_coding | 38.56 | -2.90E-07 | 2.90E-07 | ENSG00000136522 | n.s. |
| ENSG00000128609 | 2287.756 | 0.000964352 | 0.297137 | 0.0032455 | 0.99741 | 0.999319207 | NDUFA5 | protein_coding | 35.76 | 2.85E-07 | 2.85E-07 | ENSG00000128609 | n.s. |
| ENSG00000103995 | 3030.042 | 0.000959171 | 0.238708 | 0.0040182 | 0.996794 | 0.999319207 | CEP152 | protein_coding | 36.6 | 2.84E-07 | 2.84E-07 | ENSG00000103995 | n.s. |
| ENSG00000109189 | 2340.114 | 0.000888917 | 0.273944 | 0.0032449 | 0.997411 | 0.999319207 | USP46 | protein_coding | 41.39 | 2.63E-07 | 2.63E-07 | ENSG00000109189 | n.s. |
| ENSG00000142731 | 1933.413 | 0.000828113 | 0.259424 | 0.0031921 | 0.997453 | 0.999319207 | PLK4 | protein_coding | 36.34 | 2.45E-07 | 2.45E-07 | ENSG00000142731 | n.s. |
| ENSG00000277918 | 232.9458 | -0.000708072 | 0.234974 | -0.003013 | 0.997596 | 0.999390148 | | snRNA | 53.66 | -1.88E-07 | 1.88E-07 | ENSG00000277918 | n.s. |
| ENSG00000082212 | 1818.651 | 0.000732972 | 0.289089 | 0.0025355 | 0.997977 | 0.999657391 | ME2 | protein_coding | 38.73 | 1.09E-07 | 1.09E-07 | ENSG00000082212 | n.s. |
| ENSG00000141480 | 460.4803 | -0.000667368 | 0.267044 | -0.002499 | 0.998006 | 0.999657391 | ARRB2 | protein_coding | 56.76 | -9.93E-08 | 9.93E-08 | ENSG00000141480 | n.s. |
| ENSG00000198832 | 98.15597 | 0.000547625 | 0.315087 | 0.001738 | 0.998613 | 0.999937086 | SELENOM | protein_coding | 49.48 | 1.50E-08 | 1.50E-08 | ENSG00000198832 | n.s. |
| ENSG00000087086 | 4143.99 | -0.000547027 | 0.271866 | -0.002012 | 0.998395 | 0.999937086 | FTL | protein_coding | 55.96 | -1.49E-08 | 1.49E-08 | ENSG00000087086 | n.s. |
| ENSG00000205352 | 304.6869 | -0.000480901 | 0.275748 | -0.001744 | 0.998608 | 0.999937086 | PRR13 | protein_coding | 48.6 | -1.31E-08 | 1.31E-08 | ENSG00000205352 | n.s. |
| ENSG00000058600 | 484.5443 | -0.000453723 | 0.284008 | -0.001598 | 0.998725 | 0.999937086 | POLR3E | protein_coding | 51.67 | -1.24E-08 | 1.24E-08 | ENSG00000058600 | n.s. |
| ENSG00000100442 | 1963.21 | 0.000412222 | 0.292191 | 0.0014108 | 0.998874 | 0.999937086 | FKBP3 | protein_coding | 40.63 | 1.13E-08 | 1.13E-08 | ENSG00000100442 | n.s. |
| ENSG00000143374 | 828.8253 | -0.000380014 | 0.259355 | -0.001465 | 0.998831 | 0.999937086 | TARS2 | protein_coding | 48.52 | -1.04E-08 | 1.04E-08 | ENSG00000143374 | n.s. |
| ENSG00000198258 | 1842.215 | 0.000365209 | 0.310568 | 0.0011759 | 0.999062 | 0.999937086 | UBL5 | protein_coding | 52.38 | 9.98E-09 | 9.98E-09 | ENSG00000198258 | n.s. |
| ENSG00000114956 | 1263.194 | -0.000336959 | 0.308546 | -0.001092 | 0.999129 | 0.999937086 | DGUOK | protein_coding | 42.81 | -9.21E-09 | 9.21E-09 | ENSG00000114956 | n.s. |
| ENSG00000004864 | 982.7791 | 0.000322627 | 0.292925 | 0.0011014 | 0.999121 | 0.999937086 | SLC25A13 | protein_coding | 39.76 | 8.82E-09 | 8.82E-09 | ENSG00000004864 | n.s. |
| ENSG00000116459 | 4131.471 | -0.000298478 | 0.30489 | -0.000979 | 0.999219 | 0.999937086 | ATP5F1 | protein_coding | 39.38 | -8.16E-09 | 8.16E-09 | ENSG00000116459 | n.s. |
| ENSG00000254615 | 149.7371 | -0.000295979 | 0.264825 | -0.001118 | 0.999108 | 0.999937086 | | lincRNA | 57.15 | -8.09E-09 | 8.09E-09 | ENSG00000254615 | n.s. |
| ENSG00000138613 | 276.4278 | 0.000285131 | 0.283168 | 0.0010069 | 0.999197 | 0.999937086 | APH1B | protein_coding | 40.76 | 7.79E-09 | 7.79E-09 | ENSG00000138613 | n.s. |
| ENSG00000207780 | 11.86098 | 0.00025008 | 0.486914 | 0.0005136 | 0.99959 | 0.999943647 | MIR648 | miRNA | 63.83 | 6.12E-09 | 6.12E-09 | ENSG00000207780 | n.s. |
| ENSG00000075426 | 603.2603 | 0.000210513 | 0.21365 | 0.0009853 | 0.999214 | 0.999937086 | FOSL2 | protein_coding | 51.8 | 5.75E-09 | 5.75E-09 | ENSG00000075426 | n.s. |
| ENSG00000172380 | 992.4229 | 0.000229315 | 0.281391 | 0.0008149 | 0.99935 | 0.999943647 | GNG12 | protein_coding | 41.55 | 5.61E-09 | 5.61E-09 | ENSG00000172380 | n.s. |
| ENSG00000132646 | 7651.464 | -0.00013605 | 0.274364 | -0.000496 | 0.999604 | 0.999943647 | PCNA | protein_coding | 42.54 | -3.33E-09 | 3.33E-09 | ENSG00000132646 | n.s. |
| ENSG00000132661 | 448.5102 | 0.000124142 | 0.29341 | 0.0004231 | 0.999662 | 0.999943647 | NXT1 | protein_coding | 48.94 | 3.04E-09 | 3.04E-09 | ENSG00000132661 | n.s. |
| ENSG00000187837 | 3606.958 | 9.99E-05 | 0.210223 | 0.0004754 | 0.999621 | 0.999943647 | HIST1H1C | protein_coding | 57.48 | 2.45E-09 | 2.45E-09 | ENSG00000187837 | n.s. |
| ENSG00000023572 | 265.8112 | 9.88E-05 | 0.306706 | 0.000322 | 0.999743 | 0.999943647 | GLRX2 | protein_coding | 41.92 | 2.42E-09 | 2.42E-09 | ENSG00000023572 | n.s. |
| ENSG00000113013 | 9218.984 | -9.59E-05 | 0.268618 | -0.000357 | 0.999715 | 0.999943647 | HSPA9 | protein_coding | 42.74 | -2.35E-09 | 2.35E-09 | ENSG00000113013 | n.s. |

| | | | | | | | | | | | | | |
|---|---|---|---|---|---|---|---|---|---|---|---|---|---|
| ENSG00000011260 | 2828.636 | 4.86E-05 | 0.265359 | 0.000183 | 0.999854 | 0.999943647 | UTP18 | protein_coding | 40.38 | 1.19E-09 | 1.19E-09 | ENSG00000011260 | n.s. |
| ENSG00000085377 | 1975.111 | 3.42E-05 | 0.248933 | 0.0001374 | 0.99989 | 0.999943647 | PREP | protein_coding | 40.95 | 8.37E-10 | 8.37E-10 | ENSG00000085377 | n.s. |
| ENSG00000139620 | 2470.26 | 1.63E-05 | 0.231402 | 7.06E-05 | 0.999944 | 0.999943647 | KANSL2 | protein_coding | 41.97 | 4.00E-10 | 4.00E-10 | ENSG00000139620 | n.s. |
