## Supplementary material for "CDK12 loss in cancer cells affects DNA damage response genes through premature cleavage and polyadenylation"

**Supplementary Table 3: DDR genes from http://repairtoire.genesilico.pl/proteins/ and https://www.mdanderson.org/documents/Labs/Wood-Laboratory/human-dna-repair-genes.html**

| Gene Name (synonyms) | Activity | Chromosome_location | Accession_number |
|---|---|---|---|
| Base excision repair (BER) | DNA glycosylases: major altered base released | Top of Page | |
| UNG | U | 12q24.11 | NM_080911 |
| SMUG1 | U | 12q13.13 | NM_014311 |
| MBD4 | U or T opposite G at CpG sequences | 3q21.3 | NM_003925 |
| TDG | U, T or ethenoC opposite G | 12q23.3 | NM_003211 |
| OGG1 | 8-oxoG opposite C | 3p25.3 | NM_016821 |
| MUTYH (MYH) | A opposite 8-oxoG | 1p34.1 | NM_012222 |
| NTHL1 (NTH1) | Ring-saturated or fragmented pyrimidines | 16p13.3 | NM_002528 |
| MPG | 3-meA, ethenoA, hypoxanthine | 16p13.3 | NM_002434 |
| NEIL1 | Removes thymine glycol | 15q24.2 | NM_024608 |
| NEIL2 | Removes oxidative products of pyrimidines | 8p23.1 | NM_145043 |
| NEIL3 | Removes oxidative products of pyrimidines | 4q34 | NM_018248 |
| Other BER and strand break joining factors | | Top of Page | |
| APEX1 (APE1) | AP endonuclease | 14q11.2 | NM_001641 |
| APEX2 | AP endonuclease | Xp11.21 | NM_014481 |
| LIG3 | DNA Ligase III | 17q12 | NM_013975 |
| XRCC1 | LIG3 accessory factor | 19q13.31 | NM_006297 |
| PNKP | Converts some DNA breaks to ligatable ends | 19q13.33 | NM_007254 |
| APLF (C2ORF13) | Accessory factor for DNA end-joining | 2p13.3 | NM_173545 |
| Mismatch excision repair (MMR) | Top of Page | | |
| MSH2 | Mismatch (MSH2-MSH6) and loop (MSH2-MSH3) recognition | 2p21 | NM_000251 |
| MSH3 | | 5q14.1 | NM_002439 |
| MSH6 | MSH2, MSH3, MSH6 | 2p16.3 | NM_000179 |
| MLH1 | MutL homologs, forming heterodimer | 3p22.3 | NM_000249 |
| PMS2 | MLH1, PMS2 | 7p22.1 | NM_000535 |
| MSH4 | MutS homologs specialized for meiosis | 1p31.1 | NM_002440 |
| MSH5 | MSH4, MSH5 | 6p21.33 | NM_002441 |
| MLH3 | MutL homologs of unknown function | 14q24.3 | NM_014381 |
| PMS1 | | 2q32.2 | NM_000534 |
| PMS2L3 | MLH3, PMS1, PMS2L3 | 7q11.23 | NM_005395 |
| Nucleotide excision repair (NER) | (XP = xeroderma pigmentosum) | Top of Page | |
| XPC | Binds DNA distortions | 3p25.1 | NM_004628 |
| RAD23B | | 9q31.2 | NM_002874 |
| CETN2 | XPC, RAD23B, CETN2 | Xq28 | NM_004344 |
| RAD23A | Substitutes for RAD23B | 19p13.13 | NM_005053 |
| XPA | Binds damaged DNA in preincision complex | 9q22.33 | NM_000380 |

| | | | |
|---|---|---|---|
| DDB1 | Complex defective in XP group E | 11q12.2 | NM_001923 |
| DDB2 (XPE) | DDB1, DDB2 | 11p11.2 | NM_000107 |
| RPA1 | Binds DNA in preincision complex | 17p13.3 | NM_002945 |
| RPA2 | | 1p35.3 | NM_002946 |
| RPA3 | RPA1, RPA2, RPA3 | 7p21.3 | NM_002947 |
| TFIIH | Catalyzes unwinding in preincision complex | Top of Page | |
| ERCC3 (XPB) | 3' to 5' DNA helicase | 2q14.3 | NM_000122 |
| ERCC2 (XPD) | 5' to 3' DNA helicase | 19q13.32 | NM_000400 |
| GTF2H1 | Core TFIIH subunit p62 | 11p15.1 | NM_005316 |
| GTF2H2 | Core TFIIH subunit p44 | 5q13.2 | NM_001515 |
| GTF2H3 | Core TFIIH subunit p34 | 12q24.31 | NM_001516 |
| GTF2H4 | Core TFIIH subunit p52 | 6p21.33 | NM_001517 |
| GTF2H5 (TTDA) | Core TFIIH subunit p8 | 6p25.3 | NM_207118 |
| CDK7 | Kinase subunits of TFIIH | 5q13.2 | NM_001799 |
| CCNH | | 5q14.3 | NM_001239 |
| MNAT1 | CDK7, CCNH, MNAT1 | 14q23.1 | NM_002431 |
| ERCC5 (XPG) | 3' incision | 13q33.1 | NM_000123 |
| ERCC1 | 5' incision DNA binding subunit | 19q13.32 | NM_001983 |
| ERCC4 (XPF) | 5' incision catalytic subunit | 16p13.12 | NM_005236 |
| LIG1 | DNA ligase | 19q13.32 | NM_000234 |
| NER-related | Top of Page | | |
| ERCC8 (CSA) | Cockayne syndrome and UV-Sensitive Syndrome; Needed for transcription-coupled NER | 5q12.1 | NM_000082 |
| ERCC6 (CSB) | | 10q11.23 | NM_000124 |
| UVSSA (KIAA1530) | ERCC8, ERCC6, UV-sensitive syndrome | 4p16.3 | NM_020894 |
| XAB2 (HCNP) | XAB2 | 19p13.2 | NM_020196 |
| MMS19 | Iron-sulfur cluster loading and transport | 10q24.1 | NM_022362 |
| Homologous recombination | Top of Page | | |
| RAD51 | Homologous pairing | 15q15.1 | NM_002875 |
| RAD51B | Rad51 homolog | 14q24.1 | NM_002877 |
| RAD51D | Rad51 homolog | 17q12 | NM_002878 |
| DMC1 | Rad51 homolog, meiosis | 22q13.1 | NM_007068 |
| XRCC2 | DNA break and crosslink repair | 7q36.1 | NM_005431 |
| XRCC3 | XRCC2, XRCC3 | 14q32.33 | NM_005432 |
| RAD52 | Accessory factors for recombination | 12p13.33 | NM_002879 |
| RAD54L | | 1p34.1 | NM_003579 |
| RAD54B | RAD52, RAD54L, RAD54B | 8q22.1 | NM_012415 |
| BRCA1 | Accessory factor for transcription and recombination, E3 Ubiquitin ligase | 17q21.31 | NM_007295 |
| SHFM1 (DSS1) | BRCA2 associated | 7q21.3 | NM_006304 |
| RAD50 | ATPase in complex with MRE11A, NBS1 | 5q23.3 | NM_005732 |
| MRE11A | 3' exonuclease, defective in ATLD (ataxia-telangiectasia-like disorder) | 11q21 | NM_005590 |
| NBN (NBS1) | Mutated in Nijmegen breakage syndrome | 8q21.3 | NM_002485 |

| | | | |
|---|---|---|---|
| RBBP8 (CtIP) | Promotes DNA end resection | 18q11.2 | NM_002894 |
| MUS81 | Subunits of structure-specific DNA nuclease | 11q13.1 | NM_025128 |
| EME1 (MMS4L) | | 17q21.33 | NM_152463 |
| EME2 | MUS81, EME1, EME2 | 16p13.3 | NM_001010865 |
| GIYD1 (SLX1A) | nit of SLX1-SLX4 structure-specific nuclease, two identical tandem genes in the human gen | 16p11.2 | NM_001014999 |
| GIYD2 (SLX1B) | | 16p11.2 | NM_024044 |
| GEN1 | Nuclease cleaving Holliday junctions | 2p24.2 | NM_182625 |
| Fanconi anemia | Tolerance and repair of DNA crosslinks and other adducts in DNA | | Top of Page |
| FANCA | FANCA | 16q24.3 | NM_000135 |
| FANCB | FANCB | Xp22.31 | NM_152633 |
| FANCC | FANCC | 9q22.32 | NM_000136 |
| BRCA2 (FANCD1) | Cooperation with RAD51, essential function | 13q13.1 | NM_000059 |
| FANCD2 | target for monoubiquitination | 3p25.3 | NM_033084 |
| FANCE | FANCE | 6p21.31 | NM_021922 |
| FANCF | FANCF | 11p14.3 | NM_022725 |
| FANCG (XRCC9) | FANCG | 9p13.3 | NM_004629 |
| FANCI (KIAA1794) | target for monoubiquitination | 15q26.1 | NM_018193 |
| BRIP1 (FANCJ) | DNA helicase, BRCA1-interacting | 17q23 | NM_032043 |
| FANCL | FANCL | 2p16.1 | NM_018062 |
| FANCM | helicase/translocase | 14q21.3 | NM_020937 |
| PALB2 (FANCN) | co-localizes with BRCA2 (FANCD1) | 16p12.1 | NM_024675 |
| RAD51C (FANCO) | Rad51 homolog FANCO | 17q23.2 | NM_002876 |
| BTBD12 (SLX4) (FANCP) | nuclease subunit/scaffold BTBD12 (SLX4) FANCP | 16p13.3 | NM_032444 |
| FAAP20 (C1orf86) | FANCA - associated | 1p36.33 | NM_182533.2 |
| FAAP24 (C19orf40) | FAAP24 | 19q13.11 | NM_152266 |
| Non-homologous end-joining | Top of Page | | |
| XRCC6 (Ku70) | DNA end binding subunit | 22q13.2 | NM_001469 |
| XRCC5 (Ku80) | DNA end binding subunit | 2q35 | NM_021141 |
| PRKDC | DNA-dependent protein kinase catalytic subunit | 8q11.21 | NM_006904 |
| LIG4 | Ligase | 13q33.3 | NM_002312 |
| XRCC4 | Ligase accessory factor | 5q14.2 | NM_003401 |
| DCLRE1C (Artemis) | Nuclease | 10p13 | NM_022487 |
| NHEJ1 (XLF, Cernunnos) | End-joining factor | 2q35 | NM_024782 |
| Other conserved DNA damage response genes | Top of Page | | |
| ATR | ATM- and PI-3K-like essential kinase | 3q23 | NM_001184 |
| ATRIP | ATR-interacting protein | 3p21.31 | NM_130384 |
| MDC1 | Mediator of DNA damage checkpoint | 6p21.3 | NM_014641 |
| RAD1 | subunits of PCNA-like sensor of damaged DNA | 5p13.2 | NM_002853 |
| RAD9A | | 11q13.2 | NM_004584 |
| HUS1 | RAD1, RAD9, HUS1 | 7p12.3 | NM_004507 |
| RAD17 (RAD24) | RFC-like DNA damage sensor | 5q13.2 | NM_002873 |
| CHEK1 | Effector kinases | 11q24.2 | NM_001274 |

| | | | |
|---|---|---|---|
| ATM | | | |
| CHEK2 | CHEK1, CHEK2 | 22q12.1 | NM_007194 |
| TP53 | Regulation of the cell cycle | 17p13.1 | NM_000546 |
| TP53BP1 (53BP1) | chromatin-binding checkpoint protein | 15q15-q21 | NM_001141980 |
| RIF1 | suppressor of 5'-end-resection | 2q23.3 | NM_001177665 |
| TOPBP1 | DNA damage checkpoint control | 3q22.1 | NM_007027 |
| CLK2 | S-phase check point and biological clock protein | 1q21 | NM_003993 |
| PER1 | S-phase check point and biological clock protein | 17p12 | NM_002616 |
