## Supplementary material for "CDK12 loss in cancer cells affects DNA damage response genes through premature cleavage and polyadenylation"

**Supplementary Table 4: PCPA and DDR gene info**

| gene_name | ensembl_gene_id | chr | start | stop | width | PCPA | DDR | length_groups | Oh_et_al_4h | Oh_et_al_8h |
|---|---|---|---|---|---|---|---|---|---|---|
| ABCD2 | ENSG00000173208 | chr12 | 39550033 | 39619751 | 69718 | PCPA | other | long | no | no |
| ABHD17C | ENSG00000136379 | chr15 | 80695303 | 80755621 | 60318 | PCPA | other | medium-long | no | yes |
| AC007240.1 | ENSG00000284681 | chr2 | 10122739 | 10199288 | 76549 | PCPA | other | long | no | no |
| AC009779.3 | ENSG00000258311 | chr12 | 55716036 | 55724703 | 8667 | PCPA | other | short | no | no |
| AC013394.1 | ENSG00000279765 | chr15 | 92883413 | 92949230 | 65817 | PCPA | other | long | no | no |
| AC090360.1 | ENSG00000267127 | chr18 | 80045943 | 80051545 | 5602 | PCPA | other | short | no | no |
| ACAD10 | ENSG00000111271 | chr12 | 111686077 | 111692749 | 6672 | PCPA | other | short | no | no |
| ACER2 | ENSG00000177076 | chr9 | 19409059 | 19452020 | 42961 | PCPA | other | medium-long | no | no |
| ADHFE1 | ENSG00000147576 | chr8 | 66432510 | 66444648 | 12138 | PCPA | other | medium-short | no | no |
| ADIPOR2 | ENSG00000006831 | chr12 | 1691027 | 1788678 | 97651 | PCPA | other | long | yes | no |
| AEN | ENSG00000181026 | chr15 | 88621296 | 88632282 | 10986 | PCPA | other | medium-short | yes | yes |
| AFF1 | ENSG00000172493 | chr4 | 87006988 | 87141038 | 134050 | PCPA | other | long | yes | yes |
| AGA | ENSG00000038002 | chr4 | 177430770 | 177442503 | 11733 | PCPA | other | medium-short | no | no |
| AGL | ENSG00000162688 | chr1 | 99850084 | 99924020 | 73936 | PCPA | other | long | no | yes |
| AK6 | ENSG00000085231 | chr5 | 69350984 | 69370013 | 19029 | PCPA | other | medium-short | yes | yes |
| AKIRIN2 | ENSG00000135334 | chr6 | 87675072 | 87702209 | 27137 | PCPA | other | medium-long | no | no |
| AL161911.1 | ENSG00000214654 | chr9 | 120793551 | 120798632 | 5081 | PCPA | other | short | no | no |
| AL513165.2 | ENSG00000256966 | chr9 | 37512547 | 37592469 | 79922 | PCPA | other | long | no | no |
| ALDH1L2 | ENSG00000136010 | chr12 | 105019790 | 105084577 | 64787 | PCPA | other | long | no | no |
| ALG5 | ENSG00000120697 | chr13 | 36949830 | 37000261 | 50431 | PCPA | other | medium-long | no | no |
| ALKBH1 | ENSG00000100601 | chr14 | 77672404 | 77708020 | 35616 | PCPA | DDR | medium-long | yes | yes |
| ALKBH3 | ENSG00000166199 | chr11 | 43880831 | 43920266 | 39435 | PCPA | DDR | medium-long | no | no |
| ALKBH8 | ENSG00000137760 | chr11 | 107502726 | 107565746 | 63020 | PCPA | DDR | medium-long | no | no |
| ALMS1 | ENSG00000116127 | chr2 | 73385758 | 73610793 | 225035 | PCPA | other | long | no | no |
| ALS2 | ENSG00000003393 | chr2 | 201700554 | 201781189 | 80635 | PCPA | other | long | no | no |
| AMER3 | ENSG00000178171 | chr2 | 130755435 | 130768134 | 12699 | PCPA | other | medium-short | no | no |
| AMOTL1 | ENSG00000166025 | chr11 | 94768342 | 94876753 | 108411 | PCPA | other | long | yes | yes |
| ANGEL1 | ENSG00000013523 | chr14 | 76786178 | 76812940 | 26762 | PCPA | other | medium-long | no | no |
| ANKMY1 | ENSG00000144504 | chr2 | 240554979 | 240560826 | 5847 | PCPA | other | short | no | no |
| ANKRD13C | ENSG00000118454 | chr1 | 70260588 | 70354734 | 94146 | PCPA | other | long | yes | yes |

| | | | | | | | | | | |
|---|---|---|---|---|---|---|---|---|---|---|
| ANKRD17 | ENSG00000132466 | chr4 | 73073376 | 73258785 | 185409 | PCPA | other | long | no | yes |
| ANKRD50 | ENSG00000151458 | chr4 | 124664052 | 124712732 | 48680 | PCPA | other | medium-long | yes | yes |
| ANLN | ENSG00000011426 | chr7 | 36389806 | 36453791 | 63985 | PCPA | other | medium-long | yes | yes |
| AP5S1 | ENSG00000125843 | chr20 | 3820547 | 3828837 | 8290 | PCPA | DDR | short | no | no |
| APPBP2 | ENSG00000062725 | chr17 | 60443149 | 60526219 | 83070 | PCPA | other | long | yes | no |
| APTX | ENSG00000137074 | chr9 | 33015888 | 33025110 | 9222 | PCPA | DDR | short | no | no |
| AQR | ENSG00000021776 | chr15 | 34851782 | 34969839 | 118057 | PCPA | other | long | no | yes |
| ARHGAP11B | ENSG00000187951 | chr15 | 30624548 | 30772993 | 148445 | PCPA | other | long | no | yes |
| ARHGAP5 | ENSG00000100852 | chr14 | 32076967 | 32159728 | 82761 | PCPA | other | long | no | yes |
| ARID1A | ENSG00000117713 | chr1 | 26696033 | 26782104 | 86071 | PCPA | other | long | yes | no |
| ARID1B | ENSG00000049618 | chr6 | 156777374 | 157210779 | 433405 | PCPA | other | long | no | no |
| ARID2 | ENSG00000189079 | chr12 | 45729665 | 45908040 | 178375 | PCPA | other | long | yes | yes |
| ARID4B | ENSG00000054267 | chr1 | 235166895 | 235328217 | 161322 | PCPA | other | long | no | no |
| ARIH1 | ENSG00000166233 | chr15 | 72474326 | 72602985 | 128659 | PCPA | other | long | no | no |
| ARL14EP | ENSG00000152219 | chr11 | 30323051 | 30338227 | 15176 | PCPA | other | medium-short | no | no |
| ARL5A | ENSG00000162980 | chr2 | 151789635 | 151828492 | 38857 | PCPA | other | medium-long | no | no |
| ARMC8 | ENSG00000114098 | chr3 | 138187342 | 138296541 | 109199 | PCPA | other | long | yes | yes |
| ARMT1 | ENSG00000146476 | chr6 | 151452287 | 151470101 | 17814 | PCPA | other | medium-short | yes | yes |
| ARNTL2 | ENSG00000029153 | chr12 | 27332854 | 27425289 | 92435 | PCPA | other | long | no | no |
| ASB7 | ENSG00000183475 | chr15 | 100602534 | 100651705 | 49171 | PCPA | other | medium-long | yes | yes |
| ASH1L | ENSG00000116539 | chr1 | 155335268 | 155562807 | 227539 | PCPA | other | long | no | no |
| ASTE1 | ENSG00000034533 | chr3 | 131013875 | 131026802 | 12927 | PCPA | other | medium-short | no | no |
| ASXL1 | ENSG00000171456 | chr20 | 32359565 | 32439318 | 79753 | PCPA | other | long | no | no |
| ATAD5 | ENSG00000176208 | chr17 | 30831970 | 30895869 | 63899 | PCPA | other | medium-long | yes | yes |
| ATF2 | ENSG00000115966 | chr2 | 175072250 | 175168206 | 95956 | PCPA | DDR | long | no | no |
| ATG2B | ENSG00000066739 | chr14 | 96279202 | 96363870 | 84668 | PCPA | other | long | no | no |
| ATIC | ENSG00000138363 | chr2 | 215311992 | 215349773 | 37781 | PCPA | other | medium-long | no | no |
| ATOH8 | ENSG00000168874 | chr2 | 85753894 | 85788066 | 34172 | PCPA | other | medium-long | no | no |
| ATP1A1 | ENSG00000163399 | chr1 | 116373260 | 116410261 | 37001 | PCPA | other | medium-long | no | no |
| ATRIP | ENSG00000164053 | chr3 | 48446710 | 48465716 | 19006 | PCPA | DDR | medium-short | no | yes |
| ATXN2 | ENSG00000204842 | chr12 | 111469818 | 111597807 | 127989 | PCPA | other | long | yes | no |
| AUH | ENSG00000148090 | chr9 | 91213815 | 91361913 | 148098 | PCPA | other | long | yes | yes |

| | | | | | | | | | | |
|---|---|---|---|---|---|---|---|---|---|---|
| B3GALT2 | ENSG00000162630 | chr1 | 193179045 | 193186654 | 7609 | PCPA | other | short | no | no |
| BAG3 | ENSG00000151929 | chr10 | 119651370 | 119677819 | 26449 | PCPA | other | medium-short | yes | yes |
| BAIAP2 | ENSG00000175866 | chr17 | 81035151 | 81110512 | 75361 | PCPA | other | long | no | yes |
| BARD1 | ENSG00000138376 | chr2 | 214725646 | 214809704 | 84058 | PCPA | DDR | long | yes | yes |
| BAZ1A | ENSG00000198604 | chr14 | 34752731 | 34875647 | 122916 | PCPA | DDR | long | yes | yes |
| BAZ1B | ENSG00000009954 | chr7 | 73440398 | 73522278 | 81880 | PCPA | DDR | long | yes | yes |
| BCDIN3D | ENSG00000186666 | chr12 | 49836039 | 49843129 | 7090 | PCPA | other | short | yes | yes |
| BCL11A | ENSG00000119866 | chr2 | 60451781 | 60553500 | 101719 | PCPA | other | long | no | no |
| BCL11B | ENSG00000127152 | chr14 | 99169287 | 99271524 | 102237 | PCPA | other | long | no | no |
| BCL2 | ENSG00000171791 | chr18 | 63123346 | 63320128 | 196782 | PCPA | DDR | long | no | yes |
| BCL2L11 | ENSG00000153094 | chr2 | 111120914 | 111164231 | 43317 | PCPA | other | medium-long | no | no |
| BDP1 | ENSG00000145734 | chr5 | 71455615 | 71567820 | 112205 | PCPA | other | long | yes | yes |
| BLM | ENSG00000197299 | chr15 | 90717327 | 90816165 | 98838 | PCPA | DDR | long | no | yes |
| BLZF1 | ENSG00000117475 | chr1 | 169367970 | 169396540 | 28570 | PCPA | other | medium-long | no | yes |
| BMPR2 | ENSG00000204217 | chr2 | 202376936 | 202567751 | 190815 | PCPA | other | long | no | no |
| BMS1 | ENSG00000165733 | chr10 | 42782801 | 42834937 | 52136 | PCPA | other | medium-long | no | yes |
| BNIP2 | ENSG00000140299 | chr15 | 59659146 | 59689534 | 30388 | PCPA | other | medium-long | yes | yes |
| BOC | ENSG00000144857 | chr3 | 113211534 | 113268426 | 56892 | PCPA | other | medium-long | no | no |
| BORA | ENSG00000136122 | chr13 | 72727752 | 72756198 | 28446 | PCPA | other | medium-long | yes | yes |
| BRAF | ENSG00000157764 | chr7 | 140719327 | 140924709 | 205382 | PCPA | DDR | long | no | yes |
| BRCA2 | ENSG00000139618 | chr13 | 32315474 | 32400266 | 84792 | PCPA | DDR | long | no | yes |
| BRD1 | ENSG00000100425 | chr22 | 49773283 | 49827512 | 54229 | PCPA | other | medium-long | no | no |
| BRI3BP | ENSG00000184992 | chr12 | 124993700 | 125031231 | 37531 | PCPA | other | medium-long | no | no |
| BRIP1 | ENSG00000136492 | chr17 | 61681266 | 61863521 | 182255 | PCPA | DDR | long | yes | yes |
| BTAF1 | ENSG00000095564 | chr10 | 91923769 | 92030325 | 106556 | PCPA | other | long | yes | yes |
| BTBD11 | ENSG00000151136 | chr12 | 107318413 | 107659642 | 341229 | PCPA | other | long | no | no |
| C14orf28 | ENSG00000179476 | chr14 | 44897295 | 44907257 | 9962 | PCPA | other | medium-short | no | no |
| C16orf52 | ENSG00000185716 | chr16 | 22007638 | 22084643 | 77005 | PCPA | other | long | no | no |
| C16orf72 | ENSG00000182831 | chr16 | 9091648 | 9121640 | 29992 | PCPA | other | medium-long | yes | yes |
| C18orf25 | ENSG00000152242 | chr18 | 46173553 | 46266991 | 93438 | PCPA | other | long | yes | yes |
| C19orf54 | ENSG00000188493 | chr19 | 40740856 | 40750553 | 9697 | PCPA | other | short | no | no |
| C1orf27 | ENSG00000157181 | chr1 | 186375838 | 186421378 | 45540 | PCPA | other | medium-long | no | no |

| | | | | | | | | | | |
|---|---|---|---|---|---|---|---|---|---|---|
| C21orf58 | ENSG00000160298 | chr21 | 46314871 | 46323869 | 8998 | PCPA | other | short | yes | yes |
| C21orf91 | ENSG00000154642 | chr21 | 17788967 | 17819386 | 30419 | PCPA | other | medium-long | no | no |
| C22orf46 | ENSG00000184208 | chr22 | 41688939 | 41698136 | 9197 | PCPA | other | short | yes | yes |
| C2orf42 | ENSG00000115998 | chr2 | 70149898 | 70248615 | 98717 | PCPA | other | long | no | no |
| C5orf22 | ENSG00000082213 | chr5 | 31532295 | 31554839 | 22544 | PCPA | other | medium-short | yes | yes |
| C5orf34 | ENSG00000172244 | chr5 | 43486701 | 43515145 | 28444 | PCPA | other | medium-long | yes | yes |
| C5orf42 | ENSG00000197603 | chr5 | 37106228 | 37249428 | 143200 | PCPA | other | long | no | yes |
| C6orf203 | ENSG00000130349 | chr6 | 107028213 | 107051342 | 23129 | PCPA | other | medium-short | yes | yes |
| C6orf58 | ENSG00000184530 | chr6 | 127519455 | 127578707 | 59252 | PCPA | other | medium-long | no | no |
| C9orf43 | ENSG00000157653 | chr9 | 113410742 | 113429684 | 18942 | PCPA | other | medium-short | no | no |
| CABLES1 | ENSG00000134508 | chr18 | 23135452 | 23194540 | 59088 | PCPA | other | medium-long | yes | yes |
| CARNMT1 | ENSG00000156017 | chr9 | 74981020 | 75028423 | 47403 | PCPA | other | medium-long | no | no |
| CBFA2T3 | ENSG00000129993 | chr16 | 88892285 | 88941200 | 48915 | PCPA | other | medium-long | no | no |
| CBFB | ENSG00000067955 | chr16 | 67029147 | 67101058 | 71911 | PCPA | other | long | yes | yes |
| CCDC117 | ENSG00000159873 | chr22 | 28772674 | 28789301 | 16627 | PCPA | other | medium-short | yes | yes |
| CCDC137 | ENSG00000185298 | chr17 | 81666364 | 81673904 | 7540 | PCPA | other | short | yes | no |
| CCDC138 | ENSG00000163006 | chr2 | 108786761 | 108885475 | 98714 | PCPA | other | long | no | yes |
| CCDC66 | ENSG00000180376 | chr3 | 56557219 | 56615417 | 58198 | PCPA | other | medium-long | no | no |
| CCDC92 | ENSG00000119242 | chr12 | 123937765 | 123972090 | 34325 | PCPA | other | medium-long | no | no |
| CCT8 | ENSG00000156261 | chr21 | 29055805 | 29073797 | 17992 | PCPA | other | medium-short | no | no |
| CD200 | ENSG00000091972 | chr3 | 112332841 | 112362812 | 29971 | PCPA | other | medium-long | no | no |
| CDC5L | ENSG00000096401 | chr6 | 44387525 | 44450426 | 62901 | PCPA | other | medium-long | no | no |
| CDC6 | ENSG00000094804 | chr17 | 40287633 | 40304657 | 17024 | PCPA | DDR | medium-short | yes | yes |
| CDCA2 | ENSG00000184661 | chr8 | 25458997 | 25507920 | 48923 | PCPA | other | medium-long | yes | yes |
| CDK12 | ENSG00000167258 | chr17 | 39461511 | 39535146 | 73635 | PCPA | other | long | yes | yes |
| CDK13 | ENSG00000065883 | chr7 | 39950037 | 40097134 | 147097 | PCPA | other | long | yes | yes |
| CDO1 | ENSG00000129596 | chr5 | 115804733 | 115816954 | 12221 | PCPA | other | medium-short | no | no |
| CDON | ENSG00000064309 | chr11 | 126005393 | 126062811 | 57418 | PCPA | other | medium-long | no | no |
| CDR2L | ENSG00000109089 | chr17 | 74987632 | 75005800 | 18168 | PCPA | other | medium-short | no | yes |
| CENPC | ENSG00000145241 | chr4 | 67468748 | 67545606 | 76858 | PCPA | other | long | yes | no |
| CENPL | ENSG00000120334 | chr1 | 173799550 | 173824473 | 24923 | PCPA | other | medium-short | yes | yes |
| CENPN | ENSG00000166451 | chr16 | 81006552 | 81029309 | 22757 | PCPA | other | medium-short | no | no |

| | | | | | | | | | | |
|---|---|---|---|---|---|---|---|---|---|---|
| CENPQ | ENSG00000031691 | chr6 | 49463378 | 49493107 | 29729 | PCPA | other | medium-long | yes | yes |
| CEP120 | ENSG00000168944 | chr5 | 123344885 | 123423592 | 78707 | PCPA | other | long | no | yes |
| CEP152 | ENSG00000103995 | chr15 | 48712928 | 48811146 | 98218 | PCPA | other | long | yes | yes |
| CEP250 | ENSG00000126001 | chr20 | 35455201 | 35462304 | 7103 | PCPA | other | short | no | no |
| CEP350 | ENSG00000135837 | chr1 | 179954738 | 180114880 | 160142 | PCPA | other | long | yes | yes |
| CEP44 | ENSG00000164118 | chr4 | 174283677 | 174333380 | 49703 | PCPA | other | medium-long | no | no |
| CEP68 | ENSG00000011523 | chr2 | 65056372 | 65087004 | 30632 | PCPA | other | medium-long | no | no |
| CEP95 | ENSG00000258890 | chr17 | 64506588 | 64537948 | 31360 | PCPA | other | medium-long | yes | yes |
| CEP97 | ENSG00000182504 | chr3 | 101723925 | 101770562 | 46637 | PCPA | other | medium-long | no | yes |
| CHAC2 | ENSG00000143942 | chr2 | 53767792 | 53775196 | 7404 | PCPA | other | short | no | no |
| CHAF1A | ENSG00000167670 | chr19 | 4402662 | 4443392 | 40730 | PCPA | other | medium-long | no | yes |
| CHAMP1 | ENSG00000198824 | chr13 | 114314513 | 114327328 | 12815 | PCPA | other | medium-short | no | yes |
| CHD2 | ENSG00000173575 | chr15 | 92900189 | 93027942 | 127753 | PCPA | other | long | no | no |
| CHPT1 | ENSG00000111666 | chr12 | 101696947 | 101744140 | 47193 | PCPA | other | medium-long | no | no |
| CHRM2 | ENSG00000181072 | chr7 | 136868669 | 137020255 | 151586 | PCPA | other | long | no | no |
| CHST10 | ENSG00000115526 | chr2 | 100391860 | 100417656 | 25796 | PCPA | other | medium-short | no | no |
| CHST8 | ENSG00000124302 | chr19 | 33621955 | 33773508 | 151553 | PCPA | other | long | no | no |
| CLSPN | ENSG00000092853 | chr1 | 35720218 | 35769967 | 49749 | PCPA | DDR | medium-long | no | yes |
| CLVS2 | ENSG00000146352 | chr6 | 122995971 | 123072927 | 76956 | PCPA | other | long | no | no |
| CNOT2 | ENSG00000111596 | chr12 | 70243030 | 70354993 | 111963 | PCPA | other | long | yes | yes |
| CNR1 | ENSG00000118432 | chr6 | 88139864 | 88166359 | 26495 | PCPA | other | medium-long | no | no |
| CNTN5 | ENSG00000149972 | chr11 | 99020953 | 99049844 | 28891 | PCPA | other | medium-long | no | no |
| COL4A3BP | ENSG00000113163 | chr5 | 75368487 | 75512138 | 143651 | PCPA | DDR | long | yes | no |
| COPS2 | ENSG00000166200 | chr15 | 49106068 | 49155661 | 49593 | PCPA | other | medium-long | no | no |
| CORO2B | ENSG00000103647 | chr15 | 68578969 | 68727806 | 148837 | PCPA | other | long | no | no |
| COX10 | ENSG00000006695 | chr17 | 14069496 | 14208677 | 139181 | PCPA | other | long | yes | yes |
| COX11 | ENSG00000166260 | chr17 | 54951902 | 54968785 | 16883 | PCPA | other | medium-short | yes | yes |
| COX18 | ENSG00000163626 | chr4 | 73056080 | 73069755 | 13675 | PCPA | other | medium-short | no | yes |
| CPD | ENSG00000108582 | chr17 | 30378905 | 30469989 | 91084 | PCPA | other | long | no | yes |
| CPEB4 | ENSG00000113742 | chr5 | 173890666 | 173910818 | 20152 | PCPA | other | medium-short | no | no |
| CREB1 | ENSG00000118260 | chr2 | 207529912 | 207603431 | 73519 | PCPA | DDR | long | no | no |
| CRISPLD1 | ENSG00000121005 | chr8 | 74984515 | 75034558 | 50043 | PCPA | other | medium-long | no | no |

| | | | | | | | | | | |
|---|---|---|---|---|---|---|---|---|---|---|
| CSGALNACT2 | ENSG00000169826 | chr10 | 43138486 | 43185308 | 46822 | PCPA | other | medium-long | no | no |
| CSPP1 | ENSG00000104218 | chr8 | 67062426 | 67196263 | 133837 | PCPA | other | long | yes | yes |
| CSTF1 | ENSG00000101138 | chr20 | 56392371 | 56406369 | 13998 | PCPA | other | medium-short | yes | yes |
| CTH | ENSG00000116761 | chr1 | 70411272 | 70439570 | 28298 | PCPA | other | medium-long | yes | yes |
| CTHRC1 | ENSG00000164932 | chr8 | 103371550 | 103382989 | 11439 | PCPA | other | medium-short | no | no |
| CUL1 | ENSG00000055130 | chr7 | 148698912 | 148801036 | 102124 | PCPA | other | long | yes | yes |
| CYP2U1 | ENSG00000155016 | chr4 | 107931369 | 107953457 | 22088 | PCPA | other | medium-short | no | no |
| DACH1 | ENSG00000276644 | chr13 | 71437966 | 71867192 | 429226 | PCPA | other | long | no | no |
| DBF4 | ENSG00000006634 | chr7 | 87876229 | 87909541 | 33312 | PCPA | DDR | medium-long | yes | yes |
| DCAF10 | ENSG00000122741 | chr9 | 37800502 | 37867666 | 67164 | PCPA | other | long | yes | yes |
| DCAF17 | ENSG00000115827 | chr2 | 171434664 | 171474384 | 39720 | PCPA | other | medium-long | no | no |
| DCP1A | ENSG00000272886 | chr3 | 53283428 | 53347610 | 64182 | PCPA | other | medium-long | yes | yes |
| DDB2 | ENSG00000134574 | chr11 | 47215058 | 47239216 | 24158 | PCPA | DDR | medium-short | yes | yes |
| DDX31 | ENSG00000125485 | chr9 | 132594386 | 132670401 | 76015 | PCPA | other | long | yes | yes |
| DENND5B | ENSG00000170456 | chr12 | 31382223 | 31591097 | 208874 | PCPA | other | long | yes | yes |
| DESI2 | ENSG00000121644 | chr1 | 244652935 | 244709033 | 56098 | PCPA | other | medium-long | no | yes |
| DET1 | ENSG00000140543 | chr15 | 88531868 | 88546337 | 14469 | PCPA | other | medium-short | no | no |
| DFNB59 | ENSG00000204311 | chr2 | 178451436 | 178461390 | 9954 | PCPA | other | short | no | no |
| DGKE | ENSG00000153933 | chr17 | 56834099 | 56869567 | 35468 | PCPA | other | medium-long | no | yes |
| DHFR2 | ENSG00000178700 | chr3 | 94047836 | 94062987 | 15151 | PCPA | other | medium-short | no | no |
| DIDO1 | ENSG00000101191 | chr20 | 62877738 | 62937952 | 60214 | PCPA | other | medium-long | yes | yes |
| DIEXF | ENSG00000117597 | chr1 | 209828007 | 209857565 | 29558 | PCPA | other | medium-long | yes | yes |
| DNAJC24 | ENSG00000170946 | chr11 | 31369840 | 31431849 | 62009 | PCPA | other | medium-long | no | no |
| DNTTIP2 | ENSG00000067334 | chr1 | 93866283 | 93879918 | 13635 | PCPA | other | medium-short | yes | yes |
| DOPEY1 | ENSG00000083097 | chr6 | 83067666 | 83168719 | 101053 | PCPA | other | long | no | no |
| DRGX | ENSG00000165606 | chr10 | 49364181 | 49396016 | 31835 | PCPA | other | medium-long | no | no |
| DTL | ENSG00000143476 | chr1 | 212035577 | 212107400 | 71823 | PCPA | DDR | long | no | no |
| DTWD1 | ENSG00000104047 | chr15 | 49621037 | 49643657 | 22620 | PCPA | other | medium-short | yes | yes |
| DUSP10 | ENSG00000143507 | chr1 | 221701424 | 221742176 | 40752 | PCPA | other | medium-long | no | yes |
| DYRK3 | ENSG00000143479 | chr1 | 206635536 | 206649197 | 13661 | PCPA | DDR | medium-short | no | yes |
| E2F2 | ENSG00000007968 | chr1 | 23506430 | 23531220 | 24790 | PCPA | other | medium-short | yes | yes |
| E2F3 | ENSG00000112242 | chr6 | 20401906 | 20493715 | 91809 | PCPA | other | long | yes | yes |

| | | | | | | | | | | |
|---|---|---|---|---|---|---|---|---|---|---|
| E2F5 | ENSG00000133740 | chr8 | 85177225 | 85217158 | 39933 | PCPA | other | medium-long | yes | yes |
| E2F7 | ENSG00000165891 | chr12 | 77021247 | 77065580 | 44333 | PCPA | other | medium-long | yes | yes |
| E2F8 | ENSG00000129173 | chr11 | 19224063 | 19241620 | 17557 | PCPA | other | medium-short | yes | yes |
| EDRF1 | ENSG00000107938 | chr10 | 125719515 | 125764139 | 44624 | PCPA | other | medium-long | yes | yes |
| EEPD1 | ENSG00000122547 | chr7 | 36153149 | 36301543 | 148394 | PCPA | other | long | no | no |
| EIF4ENIF1 | ENSG00000184708 | chr22 | 31439365 | 31489937 | 50572 | PCPA | other | medium-long | no | no |
| ELAVL3 | ENSG00000196361 | chr19 | 11451326 | 11481046 | 29720 | PCPA | other | medium-long | no | no |
| ELAVL4 | ENSG00000162374 | chr1 | 50105690 | 50203786 | 98096 | PCPA | other | long | no | no |
| ELK3 | ENSG00000111145 | chr12 | 96194382 | 96269835 | 75453 | PCPA | other | long | no | no |
| ELK4 | ENSG00000158711 | chr1 | 205607943 | 205631962 | 24019 | PCPA | other | medium-short | no | no |
| EME1 | ENSG00000154920 | chr17 | 50373220 | 50381483 | 8263 | PCPA | other | short | no | no |
| ENC1 | ENSG00000171617 | chr5 | 74627406 | 74641424 | 14018 | PCPA | other | medium-short | no | no |
| EP300 | ENSG00000100393 | chr22 | 41091786 | 41180079 | 88293 | PCPA | DDR | long | yes | yes |
| EP400 | ENSG00000183495 | chr12 | 131949920 | 132080466 | 130546 | PCPA | DDR | long | no | no |
| EPCAM | ENSG00000119888 | chr2 | 47345158 | 47387601 | 42443 | PCPA | other | medium-long | no | no |
| EPHA7 | ENSG00000135333 | chr6 | 93240020 | 93419547 | 179527 | PCPA | other | long | no | no |
| ERCC4 | ENSG00000175595 | chr16 | 13920157 | 13933247 | 13090 | PCPA | DDR | medium-short | no | no |
| ERCC6L2 | ENSG00000182150 | chr9 | 95875701 | 96014571 | 138870 | PCPA | other | long | yes | yes |
| ERN1 | ENSG00000178607 | chr17 | 64053967 | 64130819 | 76852 | PCPA | DDR | long | no | no |
| ETS2 | ENSG00000157557 | chr21 | 38805831 | 38824955 | 19124 | PCPA | other | medium-short | yes | yes |
| ETV5 | ENSG00000244405 | chr3 | 186046308 | 186109112 | 62804 | PCPA | other | medium-long | no | no |
| EXO1 | ENSG00000174371 | chr1 | 241848180 | 241895148 | 46968 | PCPA | DDR | medium-long | yes | yes |
| EXO5 | ENSG00000164002 | chr1 | 40508747 | 40516556 | 7809 | PCPA | other | short | no | no |
| EXTL2 | ENSG00000162694 | chr1 | 100872387 | 100895998 | 23611 | PCPA | other | medium-short | no | yes |
| EYA1 | ENSG00000104313 | chr8 | 71197433 | 71362232 | 164799 | PCPA | other | long | no | no |
| FAM120A | ENSG00000048828 | chr9 | 93451722 | 93566107 | 114385 | PCPA | other | long | no | no |
| FAM120B | ENSG00000112584 | chr6 | 170295321 | 170407065 | 111744 | PCPA | other | long | no | no |
| FAM124A | ENSG00000150510 | chr13 | 51222334 | 51284241 | 61907 | PCPA | other | medium-long | no | no |
| FAM135A | ENSG00000082269 | chr6 | 70413550 | 70504960 | 91410 | PCPA | other | long | no | no |
| FAM160B1 | ENSG00000151553 | chr10 | 114821744 | 114864717 | 42973 | PCPA | other | medium-long | no | no |
| FAM200A | ENSG00000221909 | chr7 | 99546308 | 99558536 | 12228 | PCPA | other | medium-short | yes | yes |
| FAM200B | ENSG00000237765 | chr4 | 15681729 | 15690439 | 8710 | PCPA | other | short | yes | yes |

| | | | | | | | | | | |
|---|---|---|---|---|---|---|---|---|---|---|
| FAM20C | ENSG00000177706 | chr7 | 192969 | 260745 | 67776 | PCPA | DDR | long | no | no |
| FAM222A | ENSG00000139438 | chr12 | 109714228 | 109770507 | 56279 | PCPA | other | medium-long | yes | yes |
| FAM57A | ENSG00000167695 | chr17 | 732412 | 742972 | 10560 | PCPA | other | medium-short | no | yes |
| FAM84A | ENSG00000162981 | chr2 | 14632686 | 14650814 | 18128 | PCPA | other | medium-short | no | no |
| FAN1 | ENSG00000198690 | chr15 | 30903893 | 30943108 | 39215 | PCPA | DDR | medium-long | yes | yes |
| FANCB | ENSG00000181544 | chrX | 14843407 | 14873069 | 29662 | PCPA | other | medium-long | no | no |
| FANCM | ENSG00000187790 | chr14 | 45135940 | 45167423 | 31483 | PCPA | other | medium-long | no | yes |
| FASTKD2 | ENSG00000118246 | chr2 | 206765357 | 206792509 | 27152 | PCPA | other | medium-long | no | no |
| FAXC | ENSG00000146267 | chr6 | 99271169 | 99350062 | 78893 | PCPA | other | long | no | no |
| FBXO10 | ENSG00000147912 | chr9 | 37510892 | 37576349 | 65457 | PCPA | other | long | no | no |
| FBXO28 | ENSG00000143756 | chr1 | 224114087 | 224162047 | 47960 | PCPA | other | medium-long | yes | yes |
| FBXO30 | ENSG00000118496 | chr6 | 145793502 | 145814753 | 21251 | PCPA | other | medium-short | no | no |
| FBXO8 | ENSG00000164117 | chr4 | 174236658 | 174284264 | 47606 | PCPA | other | medium-long | yes | yes |
| FBXW2 | ENSG00000119402 | chr9 | 120756974 | 120793412 | 36438 | PCPA | other | medium-long | yes | yes |
| FGD4 | ENSG00000139132 | chr12 | 32502049 | 32646050 | 144001 | PCPA | other | long | no | no |
| FGF2 | ENSG00000138685 | chr4 | 122826708 | 122898236 | 71528 | PCPA | other | long | yes | yes |
| FIGN | ENSG00000182263 | chr2 | 163593396 | 163736012 | 142616 | PCPA | other | long | no | no |
| FLRT3 | ENSG00000125848 | chr20 | 14323995 | 14337616 | 13621 | PCPA | other | medium-short | no | no |
| FNBP4 | ENSG00000109920 | chr11 | 47716517 | 47767443 | 50926 | PCPA | other | medium-long | yes | yes |
| FNIP2 | ENSG00000052795 | chr4 | 158769138 | 158908049 | 138911 | PCPA | DDR | long | no | no |
| FOSL2 | ENSG00000075426 | chr2 | 28392858 | 28417312 | 24454 | PCPA | other | medium-short | yes | yes |
| FOXK1 | ENSG00000164916 | chr7 | 4682309 | 4771443 | 89134 | PCPA | other | long | yes | yes |
| FOXK2 | ENSG00000141568 | chr17 | 82519715 | 82604607 | 84892 | PCPA | other | long | yes | yes |
| FPGT | ENSG00000254685 | chr1 | 74198212 | 74207234 | 9022 | PCPA | other | short | no | no |
| FRMD6 | ENSG00000139926 | chr14 | 51651858 | 51730727 | 78869 | PCPA | other | long | no | no |
| FSTL4 | ENSG00000053108 | chr5 | 133196455 | 133612564 | 416109 | PCPA | other | long | no | no |
| FZD3 | ENSG00000104290 | chr8 | 28494214 | 28574268 | 80054 | PCPA | other | long | no | no |
| GAN | ENSG00000261609 | chr16 | 81314952 | 81390884 | 75932 | PCPA | other | long | yes | yes |
| GART | ENSG00000159131 | chr21 | 33503931 | 33542820 | 38889 | PCPA | other | medium-long | no | yes |
| GATAD2B | ENSG00000143614 | chr1 | 153804725 | 153922975 | 118250 | PCPA | other | long | yes | yes |
| GCC2 | ENSG00000135968 | chr2 | 108449191 | 108509397 | 60206 | PCPA | other | medium-long | no | no |
| GCFC2 | ENSG00000005436 | chr2 | 75662706 | 75710899 | 48193 | PCPA | other | medium-long | yes | yes |

| | | | | | | | | | | |
|---|---|---|---|---|---|---|---|---|---|---|
| GCLM | ENSG00000023909 | chr1 | 93885205 | 93909410 | 24205 | PCPA | other | medium-short | no | no |
| GEM | ENSG00000164949 | chr8 | 94249258 | 94262345 | 13087 | PCPA | other | medium-short | no | no |
| GEN1 | ENSG00000178295 | chr2 | 17753895 | 17785365 | 31470 | PCPA | DDR | medium-long | yes | yes |
| GFOD1 | ENSG00000145990 | chr6 | 13357830 | 13487662 | 129832 | PCPA | other | long | yes | yes |
| GIN1 | ENSG00000145723 | chr5 | 103102276 | 103120138 | 17862 | PCPA | other | medium-short | no | no |
| GINS1 | ENSG00000101003 | chr20 | 25407727 | 25448555 | 40828 | PCPA | other | medium-long | no | no |
| GINS4 | ENSG00000147536 | chr8 | 41529206 | 41545046 | 15840 | PCPA | other | medium-short | no | no |
| GJC1 | ENSG00000182963 | chr17 | 44798448 | 44830813 | 32365 | PCPA | other | medium-long | no | no |
| GOLGA8B | ENSG00000215252 | chr15 | 34525282 | 34588503 | 63221 | PCPA | other | medium-long | no | no |
| GOSR1 | ENSG00000108587 | chr17 | 30477362 | 30527592 | 50230 | PCPA | other | medium-long | yes | yes |
| GPATCH8 | ENSG00000186566 | chr17 | 44395284 | 44503430 | 108146 | PCPA | other | long | yes | yes |
| GPR39 | ENSG00000183840 | chr2 | 132604441 | 132646554 | 42113 | PCPA | other | medium-long | no | no |
| GPR63 | ENSG00000112218 | chr6 | 96794126 | 96837463 | 43337 | PCPA | other | medium-long | no | no |
| GPRIN3 | ENSG00000185477 | chr4 | 89236386 | 89308010 | 71624 | PCPA | other | long | no | no |
| GTF2A1 | ENSG00000165417 | chr14 | 81175452 | 81221377 | 45925 | PCPA | other | medium-long | no | no |
| GTF2H1 | ENSG00000110768 | chr11 | 18322567 | 18367044 | 44477 | PCPA | DDR | medium-long | yes | yes |
| GTF3C4 | ENSG00000125484 | chr9 | 132670035 | 132694955 | 24920 | PCPA | other | medium-short | yes | yes |
| GUCY1A3 | ENSG00000164116 | chr4 | 155666726 | 155732349 | 65623 | PCPA | other | long | no | no |
| GULP1 | ENSG00000144366 | chr2 | 188291946 | 188595931 | 303985 | PCPA | other | long | no | no |
| HAUS3 | ENSG00000214367 | chr4 | 2227464 | 2242164 | 14700 | PCPA | other | medium-short | no | no |
| HBS1L | ENSG00000112339 | chr6 | 134960378 | 135054898 | 94520 | PCPA | other | long | no | no |
| HECTD2 | ENSG00000165338 | chr10 | 91410344 | 91514761 | 104417 | PCPA | other | long | no | no |
| HEG1 | ENSG00000173706 | chr3 | 124965710 | 125055958 | 90248 | PCPA | other | long | no | yes |
| HELB | ENSG00000127311 | chr12 | 66302545 | 66347645 | 45100 | PCPA | DDR | medium-long | no | yes |
| HELZ | ENSG00000198265 | chr17 | 67070438 | 67245187 | 174749 | PCPA | other | long | no | yes |
| HERPUD1 | ENSG00000051108 | chr16 | 56932048 | 56944863 | 12815 | PCPA | other | medium-short | yes | yes |
| HIF1AN | ENSG00000166135 | chr10 | 100535859 | 100559998 | 24139 | PCPA | other | medium-short | yes | no |
| HKR1 | ENSG00000181666 | chr19 | 37317900 | 37359776 | 41876 | PCPA | other | medium-long | no | no |
| HMGXB4 | ENSG00000100281 | chr22 | 35257489 | 35295764 | 38275 | PCPA | other | medium-long | yes | yes |
| HNRNPU | ENSG00000153187 | chr1 | 244840638 | 244864560 | 23922 | PCPA | other | medium-short | no | no |
| HRK | ENSG00000135116 | chr12 | 116856144 | 116881441 | 25297 | PCPA | other | medium-short | no | no |
| HS3ST3A1 | ENSG00000153976 | chr17 | 13495689 | 13601927 | 106238 | PCPA | other | long | no | no |

| | | | | | | | | | | |
|---|---|---|---|---|---|---|---|---|---|---|
| HS3ST5 | ENSG00000249853 | chr6 | 114055586 | 114343045 | 287459 | PCPA | other | long | no | no |
| HS6ST1 | ENSG00000136720 | chr2 | 128265480 | 128318577 | 53097 | PCPA | other | medium-long | no | yes |
| HSP90B1 | ENSG00000166598 | chr12 | 103930334 | 103953645 | 23311 | PCPA | other | medium-short | no | no |
| HTR1E | ENSG00000168830 | chr6 | 86937306 | 87016683 | 79377 | PCPA | other | long | no | no |
| HUNK | ENSG00000142149 | chr21 | 31873315 | 32004064 | 130749 | PCPA | DDR | long | no | no |
| ICE1 | ENSG00000164151 | chr5 | 5420664 | 5490234 | 69570 | PCPA | other | long | yes | yes |
| IFRD1 | ENSG00000006652 | chr7 | 112450428 | 112477200 | 26772 | PCPA | other | medium-long | no | no |
| ING3 | ENSG00000071243 | chr7 | 120950763 | 120975014 | 24251 | PCPA | DDR | medium-short | no | yes |
| INO80 | ENSG00000128908 | chr15 | 40978880 | 41116354 | 137474 | PCPA | DDR | long | yes | yes |
| INO80C | ENSG00000153391 | chr18 | 35468584 | 35497960 | 29376 | PCPA | other | medium-long | no | yes |
| INO80D | ENSG00000114933 | chr2 | 205993721 | 206086303 | 92582 | PCPA | other | long | no | no |
| INSIG1 | ENSG00000186480 | chr7 | 155297776 | 155310235 | 12459 | PCPA | other | medium-short | no | yes |
| INTS5 | ENSG00000185085 | chr11 | 62646848 | 62653302 | 6454 | PCPA | other | short | yes | yes |
| INTS6 | ENSG00000102786 | chr13 | 51354077 | 51453207 | 99130 | PCPA | other | long | yes | yes |
| INTS7 | ENSG00000143493 | chr1 | 211940399 | 212035542 | 95143 | PCPA | DDR | long | yes | yes |
| IPMK | ENSG00000151151 | chr10 | 58191517 | 58267934 | 76417 | PCPA | other | long | yes | yes |
| IRS2 | ENSG00000185950 | chr13 | 109752698 | 109786568 | 33870 | PCPA | other | medium-long | no | no |
| ITGA1 | ENSG00000213949 | chr5 | 52787896 | 52959210 | 171314 | PCPA | other | long | yes | yes |
| ITGA4 | ENSG00000115232 | chr2 | 181457202 | 181536187 | 78985 | PCPA | other | long | no | no |
| JAKMIP2 | ENSG00000176049 | chr5 | 147588566 | 147782775 | 194209 | PCPA | other | long | no | no |
| JDP2 | ENSG00000140044 | chr14 | 75427806 | 75470457 | 42651 | PCPA | other | medium-long | no | no |
| KANSL1L | ENSG00000144445 | chr2 | 210097849 | 210170654 | 72805 | PCPA | other | long | no | no |
| KAT14 | ENSG00000149474 | chr20 | 18138118 | 18188387 | 50269 | PCPA | other | medium-long | no | no |
| KAT7 | ENSG00000136504 | chr17 | 49788555 | 49835030 | 46475 | PCPA | DDR | medium-long | yes | yes |
| KATNAL1 | ENSG00000102781 | chr13 | 30202630 | 30307484 | 104854 | PCPA | other | long | yes | yes |
| KCNG3 | ENSG00000171126 | chr2 | 42442017 | 42494097 | 52080 | PCPA | other | medium-long | no | no |
| KCNH7 | ENSG00000184611 | chr2 | 162423247 | 162838730 | 415483 | PCPA | other | long | no | no |
| KCNJ8 | ENSG00000121361 | chr12 | 21764955 | 21775581 | 10626 | PCPA | other | medium-short | no | no |
| KCNMB4 | ENSG00000135643 | chr12 | 70366276 | 70434292 | 68016 | PCPA | other | long | no | no |
| KCNQ5 | ENSG00000185760 | chr6 | 72621792 | 73198848 | 577056 | PCPA | other | long | no | no |
| KCTD15 | ENSG00000153885 | chr19 | 33796416 | 33815763 | 19347 | PCPA | other | medium-short | no | no |
| KCTD3 | ENSG00000136636 | chr1 | 215567392 | 215621807 | 54415 | PCPA | other | medium-long | no | yes |

| | | | | | | | | | | |
|---|---|---|---|---|---|---|---|---|---|---|
| KDM5A | ENSG00000073614 | chr12 | 280129 | 389454 | 109325 | PCPA | other | long | yes | yes |
| KIAA0586 | ENSG00000100578 | chr14 | 58427684 | 58548045 | 120361 | PCPA | other | long | no | yes |
| KIAA1841 | ENSG00000162929 | chr2 | 61065871 | 61093364 | 27493 | PCPA | other | medium-long | no | no |
| KIF15 | ENSG00000163808 | chr3 | 44761717 | 44873376 | 111659 | PCPA | other | long | no | yes |
| KIF20B | ENSG00000138182 | chr10 | 89701610 | 89774939 | 73329 | PCPA | other | long | no | yes |
| KITLG | ENSG00000049130 | chr12 | 88492793 | 88580471 | 87678 | PCPA | other | long | yes | yes |
| KLF3 | ENSG00000109787 | chr4 | 38664196 | 38701042 | 36846 | PCPA | other | medium-long | yes | yes |
| KLF7 | ENSG00000118263 | chr2 | 207074137 | 207167246 | 93109 | PCPA | other | long | no | no |
| KLHL20 | ENSG00000076321 | chr1 | 173714941 | 173786702 | 71761 | PCPA | other | long | no | yes |
| KLHL28 | ENSG00000179454 | chr14 | 44924319 | 44962190 | 37871 | PCPA | other | medium-long | no | no |
| KLHL42 | ENSG00000087448 | chr12 | 27780020 | 27803040 | 23020 | PCPA | other | medium-short | yes | yes |
| KMT2E | ENSG00000005483 | chr7 | 105014179 | 105114085 | 99906 | PCPA | other | long | yes | yes |
| L2HGDH | ENSG00000087299 | chr14 | 50242434 | 50312548 | 70114 | PCPA | other | long | no | no |
| L3HYPDH | ENSG00000126790 | chr14 | 59474454 | 59484408 | 9954 | PCPA | other | short | yes | yes |
| LACTB | ENSG00000103642 | chr15 | 63121800 | 63142061 | 20261 | PCPA | other | medium-short | no | yes |
| LANCL2 | ENSG00000132434 | chr7 | 55365448 | 55433742 | 68294 | PCPA | other | long | no | yes |
| LARP4 | ENSG00000161813 | chr12 | 50400889 | 50480004 | 79115 | PCPA | other | long | yes | yes |
| LATS1 | ENSG00000131023 | chr6 | 149658153 | 149718256 | 60103 | PCPA | DDR | medium-long | yes | yes |
| LCORL | ENSG00000178177 | chr4 | 17880595 | 18021755 | 141160 | PCPA | other | long | yes | yes |
| LDLR | ENSG00000130164 | chr19 | 11089463 | 11133816 | 44353 | PCPA | other | medium-long | no | no |
| LEMD3 | ENSG00000174106 | chr12 | 65169571 | 65248327 | 78756 | PCPA | other | long | yes | yes |
| LIN28B | ENSG00000187772 | chr6 | 104936616 | 105083332 | 146716 | PCPA | other | long | no | no |
| LIN9 | ENSG00000183814 | chr1 | 226231157 | 226309869 | 78712 | PCPA | other | long | yes | yes |
| LMNA | ENSG00000160789 | chr1 | 156082600 | 156093811 | 11211 | PCPA | other | medium-short | no | no |
| LMO4 | ENSG00000143013 | chr1 | 87328468 | 87348923 | 20455 | PCPA | other | medium-short | yes | yes |
| LRPAP1 | ENSG00000163956 | chr4 | 3506376 | 3532559 | 26183 | PCPA | other | medium-short | no | no |
| LRRC8C | ENSG00000171488 | chr1 | 89633072 | 89719533 | 86461 | PCPA | other | long | no | no |
| LRRC8D | ENSG00000171492 | chr1 | 89821014 | 89936001 | 114987 | PCPA | other | long | yes | yes |
| LRRN1 | ENSG00000175928 | chr3 | 3799437 | 3847703 | 48266 | PCPA | other | medium-long | no | no |
| LRRN3 | ENSG00000173114 | chr7 | 111091006 | 111125451 | 34445 | PCPA | other | medium-long | no | no |
| LSM6 | ENSG00000164167 | chr4 | 146175685 | 146200000 | 24315 | PCPA | other | medium-short | no | no |
| LYPLAL1 | ENSG00000143353 | chr1 | 219173876 | 219212865 | 38989 | PCPA | other | medium-long | no | no |

| | | | | | | | | | | |
|---|---|---|---|---|---|---|---|---|---|---|
| MAK16 | ENSG00000198042 | chr8 | 33484750 | 33501260 | 16510 | PCPA | other | medium-short | no | no |
| MAML1 | ENSG00000161021 | chr5 | 179732850 | 179777286 | 44436 | PCPA | DDR | medium-long | yes | yes |
| MAN1A1 | ENSG00000111885 | chr6 | 119177209 | 119349761 | 172552 | PCPA | other | long | yes | yes |
| MAN1A2 | ENSG00000198162 | chr1 | 117367449 | 117528872 | 161423 | PCPA | other | long | no | yes |
| MAN2A1 | ENSG00000112893 | chr5 | 109689366 | 109867178 | 177812 | PCPA | other | long | yes | yes |
| MANEA | ENSG00000172469 | chr6 | 95577543 | 95609457 | 31914 | PCPA | other | medium-long | no | no |
| MAP1LC3B | ENSG00000140941 | chr16 | 87391800 | 87404779 | 12979 | PCPA | other | medium-short | no | no |
| MAP2K4 | ENSG00000065559 | chr17 | 12020824 | 12143823 | 122999 | PCPA | other | long | no | no |
| MAP3K14 | ENSG00000006062 | chr17 | 45263121 | 45317040 | 53919 | PCPA | other | medium-long | yes | yes |
| MAP3K20 | ENSG00000091436 | chr2 | 173075836 | 173227146 | 151310 | PCPA | DDR | long | no | no |
| MAP3K3 | ENSG00000198909 | chr17 | 63622442 | 63696303 | 73861 | PCPA | other | long | no | yes |
| MAPK4 | ENSG00000141639 | chr18 | 50560078 | 50731824 | 171746 | PCPA | DDR | long | no | no |
| MAPRE2 | ENSG00000166974 | chr18 | 35041360 | 35143470 | 102110 | PCPA | other | long | no | no |
| 7-Mar | ENSG00000136536 | chr2 | 159712457 | 159771027 | 58570 | PCPA | other | medium-long | no | no |
| MARK1 | ENSG00000116141 | chr1 | 220528183 | 220664456 | 136273 | PCPA | DDR | long | no | yes |
| MBNL1 | ENSG00000152601 | chr3 | 152268906 | 152465780 | 196874 | PCPA | other | long | no | no |
| MCM8 | ENSG00000125885 | chr20 | 5950652 | 5998977 | 48325 | PCPA | DDR | medium-long | no | yes |
| MED26 | ENSG00000105085 | chr19 | 16577585 | 16624661 | 47076 | PCPA | other | medium-long | no | no |
| MED9 | ENSG00000141026 | chr17 | 17476986 | 17493226 | 16240 | PCPA | other | medium-short | yes | yes |
| MEIS1 | ENSG00000143995 | chr2 | 66435133 | 66573869 | 138736 | PCPA | other | long | yes | yes |
| MEIS2 | ENSG00000134138 | chr15 | 37066297 | 37096191 | 29894 | PCPA | other | medium-long | no | no |
| METTL14 | ENSG00000145388 | chr4 | 118685368 | 118715433 | 30065 | PCPA | other | medium-long | no | no |
| MFSD14C | ENSG00000196312 | chr9 | 96897917 | 97013708 | 115791 | PCPA | other | long | no | no |
| MFSD4B | ENSG00000173214 | chr6 | 111259348 | 111271167 | 11819 | PCPA | other | medium-short | no | no |
| MGME1 | ENSG00000125871 | chr20 | 17968913 | 17991122 | 22209 | PCPA | other | medium-short | yes | yes |
| MIER1 | ENSG00000198160 | chr1 | 66924957 | 66988031 | 63074 | PCPA | other | medium-long | yes | yes |
| MIOS | ENSG00000164654 | chr7 | 7566875 | 7607223 | 40348 | PCPA | other | medium-long | no | yes |
| MLXIP | ENSG00000175727 | chr12 | 122078722 | 122147347 | 68625 | PCPA | other | long | no | yes |
| MMS22L | ENSG00000146263 | chr6 | 97142161 | 97283217 | 141056 | PCPA | DDR | long | no | no |
| MON2 | ENSG00000061987 | chr12 | 62467062 | 62593081 | 126019 | PCPA | other | long | yes | yes |
| MPP5 | ENSG00000072415 | chr14 | 67241423 | 67335819 | 94396 | PCPA | other | long | yes | yes |
| MRPL42 | ENSG00000198015 | chr12 | 93467488 | 93516213 | 48725 | PCPA | other | medium-long | no | yes |

| | | | | | | | | | | |
|---|---|---|---|---|---|---|---|---|---|---|
| MSH3 | ENSG00000113318 | chr5 | 80654648 | 80876460 | 221812 | PCPA | DDR | long | no | yes |
| MTBP | ENSG00000172167 | chr8 | 120445400 | 120523635 | 78235 | PCPA | other | long | no | no |
| MTHFD1L | ENSG00000120254 | chr6 | 150865716 | 151101885 | 236169 | PCPA | other | long | no | no |
| MTO1 | ENSG00000135297 | chr6 | 73461578 | 73509236 | 47658 | PCPA | other | medium-long | yes | yes |
| MVB12B | ENSG00000196814 | chr9 | 126326849 | 126507041 | 180192 | PCPA | other | long | no | no |
| MXD1 | ENSG00000059728 | chr2 | 69915071 | 69942945 | 27874 | PCPA | other | medium-long | no | yes |
| MYLIP | ENSG00000007944 | chr6 | 16129125 | 16148248 | 19123 | PCPA | other | medium-short | no | no |
| MYNN | ENSG00000085274 | chr3 | 169772831 | 169789181 | 16350 | PCPA | other | medium-short | yes | yes |
| MYRIP | ENSG00000170011 | chr3 | 39809605 | 40260321 | 450716 | PCPA | other | long | no | no |
| NABP1 | ENSG00000173559 | chr2 | 191678136 | 191686157 | 8021 | PCPA | DDR | short | no | no |
| NAF1 | ENSG00000145414 | chr4 | 163110073 | 163166831 | 56758 | PCPA | other | medium-long | yes | no |
| NAGLU | ENSG00000108784 | chr17 | 42536172 | 42544449 | 8277 | PCPA | other | short | yes | no |
| NAPEPLD | ENSG00000161048 | chr7 | 103126790 | 103149235 | 22445 | PCPA | other | medium-short | no | no |
| NCOA5 | ENSG00000124160 | chr20 | 46060985 | 46089952 | 28967 | PCPA | other | medium-long | yes | yes |
| NCOR2 | ENSG00000196498 | chr12 | 124354837 | 124567589 | 212752 | PCPA | other | long | no | no |
| NCR3LG1 | ENSG00000188211 | chr11 | 17351726 | 17377341 | 25615 | PCPA | other | medium-short | no | no |
| NDNF | ENSG00000173376 | chr4 | 121035613 | 121072518 | 36905 | PCPA | other | medium-long | no | no |
| NDUFAF5 | ENSG00000101247 | chr20 | 13784950 | 13821582 | 36632 | PCPA | other | medium-long | yes | yes |
| NEIL3 | ENSG00000109674 | chr4 | 177309836 | 177362943 | 53107 | PCPA | other | medium-long | yes | yes |
| NEK4 | ENSG00000114904 | chr3 | 52708449 | 52770949 | 62500 | PCPA | DDR | medium-long | no | no |
| NETO2 | ENSG00000171208 | chr16 | 47077703 | 47143997 | 66294 | PCPA | other | long | yes | yes |
| NFE2L2 | ENSG00000116044 | chr2 | 177227595 | 177265131 | 37536 | PCPA | other | medium-long | no | no |
| NFIB | ENSG00000147862 | chr9 | 14081843 | 14322338 | 240495 | PCPA | other | long | no | yes |
| NFX1 | ENSG00000086102 | chr9 | 33290511 | 33371157 | 80646 | PCPA | other | long | yes | yes |
| NHLH1 | ENSG00000171786 | chr1 | 160367067 | 160372848 | 5781 | PCPA | other | short | no | no |
| NHLRC2 | ENSG00000196865 | chr10 | 113854661 | 113917194 | 62533 | PCPA | other | medium-long | yes | yes |
| NINL | ENSG00000101004 | chr20 | 25452705 | 25585517 | 132812 | PCPA | other | long | no | no |
| NIPBL | ENSG00000164190 | chr5 | 36876790 | 37064190 | 187400 | PCPA | DDR | long | yes | yes |
| NLK | ENSG00000087095 | chr17 | 28042156 | 28196381 | 154225 | PCPA | DDR | long | no | yes |
| NOCT | ENSG00000151014 | chr4 | 139015789 | 139045939 | 30150 | PCPA | other | medium-long | no | no |
| NOL10 | ENSG00000115761 | chr2 | 10570766 | 10689975 | 119209 | PCPA | other | long | yes | yes |
| NOM1 | ENSG00000146909 | chr7 | 156949723 | 156973182 | 23459 | PCPA | other | medium-short | yes | yes |

| | | | | | | | | | | |
|---|---|---|---|---|---|---|---|---|---|---|
| NPAT | ENSG00000149308 | chr11 | 108157215 | 108222642 | 65427 | PCPA | other | long | yes | yes |
| NR2C2 | ENSG00000177463 | chr3 | 14947584 | 15049273 | 101689 | PCPA | other | long | yes | yes |
| NR3C1 | ENSG00000113580 | chr5 | 143277931 | 143403700 | 125769 | PCPA | other | long | yes | yes |
| NRF1 | ENSG00000106459 | chr7 | 129611739 | 129709097 | 97358 | PCPA | other | long | yes | yes |
| NRP1 | ENSG00000099250 | chr10 | 33195424 | 33334642 | 139218 | PCPA | other | long | no | no |
| NSRP1 | ENSG00000126653 | chr17 | 30116781 | 30186475 | 69694 | PCPA | other | long | no | no |
| NSUN3 | ENSG00000178694 | chr3 | 94062916 | 94128545 | 65629 | PCPA | other | long | yes | yes |
| NT5DC1 | ENSG00000178425 | chr6 | 116100849 | 116249497 | 148648 | PCPA | other | long | no | no |
| NT5DC3 | ENSG00000111696 | chr12 | 103772310 | 103841197 | 68887 | PCPA | other | long | yes | yes |
| NUB1 | ENSG00000013374 | chr7 | 151341699 | 151378449 | 36750 | PCPA | other | medium-long | no | yes |
| NUDCD2 | ENSG00000170584 | chr5 | 163446526 | 163460140 | 13614 | PCPA | other | medium-short | no | no |
| NUDT19 | ENSG00000213965 | chr19 | 32691961 | 32713796 | 21835 | PCPA | other | medium-short | no | no |
| NUFIP2 | ENSG00000108256 | chr17 | 29255836 | 29294118 | 38282 | PCPA | other | medium-long | no | no |
| NUP98 | ENSG00000110713 | chr11 | 3675010 | 3797792 | 122782 | PCPA | DDR | long | yes | yes |
| NUPL2 | ENSG00000136243 | chr7 | 23181841 | 23201011 | 19170 | PCPA | other | medium-short | yes | yes |
| NXPH2 | ENSG00000144227 | chr2 | 138670772 | 138780348 | 109576 | PCPA | other | long | no | no |
| OGFRL1 | ENSG00000119900 | chr6 | 71288803 | 71308950 | 20147 | PCPA | other | medium-short | no | yes |
| ORC5 | ENSG00000164815 | chr7 | 104126341 | 104208047 | 81706 | PCPA | DDR | long | yes | yes |
| OXSM | ENSG00000151093 | chr3 | 25790080 | 25794534 | 4454 | PCPA | other | short | yes | yes |
| PACRGL | ENSG00000163138 | chr4 | 20700439 | 20724889 | 24450 | PCPA | other | medium-short | no | yes |
| PAG1 | ENSG00000076641 | chr8 | 80967810 | 81112068 | 144258 | PCPA | other | long | no | no |
| PALB2 | ENSG00000083093 | chr16 | 23603170 | 23641310 | 38140 | PCPA | DDR | medium-long | yes | yes |
| PANK1 | ENSG00000152782 | chr10 | 89582988 | 89643956 | 60968 | PCPA | other | medium-long | no | no |
| PARS2 | ENSG00000162396 | chr1 | 54756898 | 54764514 | 7616 | PCPA | other | short | no | yes |
| PATL1 | ENSG00000166889 | chr11 | 59636716 | 59668980 | 32264 | PCPA | other | medium-long | yes | yes |
| PAXBP1 | ENSG00000159086 | chr21 | 32733900 | 32771792 | 37892 | PCPA | other | medium-long | yes | yes |
| PCDH18 | ENSG00000189184 | chr4 | 137519703 | 137532494 | 12791 | PCPA | other | medium-short | no | no |
| PCF11 | ENSG00000165494 | chr11 | 83156988 | 83187451 | 30463 | PCPA | other | medium-long | yes | yes |
| PCLO | ENSG00000186472 | chr7 | 82822097 | 82894454 | 72357 | PCPA | other | long | no | no |
| PCM1 | ENSG00000078674 | chr8 | 17922857 | 18029944 | 107087 | PCPA | other | long | no | yes |
| PDE4B | ENSG00000184588 | chr1 | 65992422 | 66374573 | 382151 | PCPA | other | long | no | no |
| PDGFRA | ENSG00000134853 | chr4 | 54229097 | 54298247 | 69150 | PCPA | other | long | no | no |

| | | | | | | | | | | |
|---|---|---|---|---|---|---|---|---|---|---|
| PDS5B | ENSG00000083642 | chr13 | 32586427 | 32778019 | 191592 | PCPA | DDR | long | no | no |
| PEAK1 | ENSG00000173517 | chr15 | 77155200 | 77420094 | 264894 | PCPA | other | long | no | no |
| PEX13 | ENSG00000162928 | chr2 | 61017562 | 61051990 | 34428 | PCPA | other | medium-long | yes | yes |
| PFN4 | ENSG00000176732 | chr2 | 24115371 | 24123477 | 8106 | PCPA | other | short | no | no |
| PGBD1 | ENSG00000137338 | chr6 | 28281572 | 28302549 | 20977 | PCPA | other | medium-short | no | no |
| PGBD5 | ENSG00000177614 | chr1 | 230314482 | 230426371 | 111889 | PCPA | other | long | no | no |
| PHF14 | ENSG00000106443 | chr7 | 10973872 | 11169630 | 195758 | PCPA | other | long | yes | yes |
| PHF20L1 | ENSG00000129292 | chr8 | 132775372 | 132848807 | 73435 | PCPA | other | long | no | yes |
| PHF3 | ENSG00000118482 | chr6 | 63635836 | 63715481 | 79645 | PCPA | other | long | no | no |
| PHOSPHO2 | ENSG00000144362 | chr2 | 169694454 | 169701708 | 7254 | PCPA | other | short | no | no |
| PHYHIPL | ENSG00000165443 | chr10 | 59176590 | 59247774 | 71184 | PCPA | other | long | no | no |
| PIBF1 | ENSG00000083535 | chr13 | 72782059 | 73016461 | 234402 | PCPA | other | long | no | yes |
| PIGG | ENSG00000174227 | chr4 | 499240 | 524911 | 25671 | PCPA | other | medium-short | no | no |
| PIK3C3 | ENSG00000078142 | chr18 | 41955206 | 42087830 | 132624 | PCPA | other | long | no | no |
| PIK3R3 | ENSG00000117461 | chr1 | 46040140 | 46132642 | 92502 | PCPA | other | long | no | no |
| PIKFYVE | ENSG00000115020 | chr2 | 208266267 | 208358751 | 92484 | PCPA | other | long | no | yes |
| PKD2 | ENSG00000118762 | chr4 | 88007668 | 88077777 | 70109 | PCPA | other | long | no | yes |
| PLCL2 | ENSG00000154822 | chr3 | 16884959 | 17090594 | 205635 | PCPA | other | long | no | no |
| PLEKHA5 | ENSG00000052126 | chr12 | 19129796 | 19373668 | 243872 | PCPA | other | long | no | no |
| PLPBP | ENSG00000147471 | chr8 | 37762593 | 37779767 | 17174 | PCPA | other | medium-short | no | no |
| PMEPA1 | ENSG00000124225 | chr20 | 57648392 | 57709902 | 61510 | PCPA | other | medium-long | yes | yes |
| POGZ | ENSG00000143442 | chr1 | 151402724 | 151459465 | 56741 | PCPA | other | medium-long | yes | no |
| POLH | ENSG00000170734 | chr6 | 43576150 | 43615660 | 39510 | PCPA | DDR | medium-long | yes | yes |
| POLQ | ENSG00000051341 | chr3 | 121431431 | 121546006 | 114575 | PCPA | other | long | yes | yes |
| POLR2A | ENSG00000181222 | chr17 | 7484366 | 7514616 | 30250 | PCPA | other | medium-long | yes | yes |
| POLR3B | ENSG00000013503 | chr12 | 106357658 | 106510198 | 152540 | PCPA | other | long | no | yes |
| POMZP3 | ENSG00000146707 | chr7 | 76609986 | 76627241 | 17255 | PCPA | other | medium-short | yes | yes |
| POU2F1 | ENSG00000143190 | chr1 | 167220886 | 167396825 | 175939 | PCPA | other | long | no | yes |
| PPHLN1 | ENSG00000134283 | chr12 | 42326149 | 42448623 | 122474 | PCPA | other | long | no | no |
| PPIP5K2 | ENSG00000145725 | chr5 | 103120149 | 103212799 | 92650 | PCPA | other | long | no | yes |
| PPM1D | ENSG00000170836 | chr17 | 60600183 | 60666280 | 66097 | PCPA | other | long | yes | yes |
| PPM1E | ENSG00000175175 | chr17 | 58755869 | 58985176 | 229307 | PCPA | other | long | no | no |

| | | | | | | | | | | |
|---|---|---|---|---|---|---|---|---|---|---|
| PPP1R10 | ENSG00000204569 | chr6 | 30600400 | 30617244 | 16844 | PCPA | other | medium-short | yes | no |
| PPP2CB | ENSG00000104695 | chr8 | 30774457 | 30812872 | 38415 | PCPA | other | medium-long | no | yes |
| PPP2R2A | ENSG00000221914 | chr8 | 26291491 | 26372680 | 81189 | PCPA | other | long | no | no |
| PPP2R2D | ENSG00000175470 | chr10 | 131901031 | 131956555 | 55524 | PCPA | other | medium-long | no | yes |
| PPP4R3B | ENSG00000275052 | chr2 | 55547292 | 55618880 | 71588 | PCPA | other | long | no | no |
| PRAG1 | ENSG00000275342 | chr8 | 8317737 | 8386498 | 68761 | PCPA | other | long | no | no |
| PRDM11 | ENSG00000019485 | chr11 | 45146675 | 45235110 | 88435 | PCPA | other | long | no | no |
| PRDM4 | ENSG00000110851 | chr12 | 107732866 | 107761160 | 28294 | PCPA | other | medium-long | no | yes |
| PRICKLE1 | ENSG00000139174 | chr12 | 42469450 | 42589622 | 120172 | PCPA | other | long | no | no |
| PRIMPOL | ENSG00000164306 | chr4 | 184649756 | 184657654 | 7898 | PCPA | other | short | no | no |
| PRKAA2 | ENSG00000162409 | chr1 | 56645322 | 56715335 | 70013 | PCPA | DDR | long | no | yes |
| PRKCA | ENSG00000154229 | chr17 | 66302636 | 66810743 | 508107 | PCPA | DDR | long | no | no |
| PROCA1 | ENSG00000167525 | chr17 | 28704081 | 28711299 | 7218 | PCPA | other | short | no | no |
| PROSER1 | ENSG00000120685 | chr13 | 39010153 | 39037837 | 27684 | PCPA | other | medium-long | yes | yes |
| PROX1 | ENSG00000117707 | chr1 | 213987943 | 214041502 | 53559 | PCPA | other | medium-long | no | no |
| PRR4 | ENSG00000111215 | chr12 | 11005934 | 11171608 | 165674 | PCPA | other | long | no | no |
| PSPC1 | ENSG00000121390 | chr13 | 19674756 | 19783019 | 108263 | PCPA | other | long | yes | yes |
| PTCD1 | ENSG00000106246 | chr7 | 99416739 | 99466163 | 49424 | PCPA | other | medium-long | no | no |
| PTCD2 | ENSG00000049883 | chr5 | 72320373 | 72359228 | 38855 | PCPA | other | medium-long | yes | yes |
| PTCH1 | ENSG00000185920 | chr9 | 95442980 | 95517057 | 74077 | PCPA | other | long | yes | yes |
| PTEN | ENSG00000171862 | chr10 | 87863113 | 87971930 | 108817 | PCPA | other | long | no | no |
| PTPN6 | ENSG00000111679 | chr12 | 6946468 | 6955262 | 8794 | PCPA | other | short | no | no |
| PTPRO | ENSG00000151490 | chr12 | 15322397 | 15597393 | 274996 | PCPA | other | long | no | no |
| PUM3 | ENSG00000080608 | chr9 | 2720469 | 2844241 | 123772 | PCPA | other | long | no | no |
| PWWP2A | ENSG00000170234 | chr5 | 160075882 | 160119423 | 43541 | PCPA | other | medium-long | no | no |
| PYGO1 | ENSG00000171016 | chr15 | 55538890 | 55588947 | 50057 | PCPA | other | medium-long | no | no |
| PYROXD1 | ENSG00000121350 | chr12 | 21437915 | 21470350 | 32435 | PCPA | other | medium-long | yes | yes |
| QSER1 | ENSG00000060749 | chr11 | 32893178 | 32980266 | 87088 | PCPA | other | long | yes | yes |
| QSOX1 | ENSG00000116260 | chr1 | 180154834 | 180204030 | 49196 | PCPA | other | medium-long | no | no |
| QTRT2 | ENSG00000151576 | chr3 | 114056735 | 114088349 | 31614 | PCPA | other | medium-long | no | no |
| RAB12 | ENSG00000206418 | chr18 | 8609445 | 8639381 | 29936 | PCPA | other | medium-long | no | yes |
| RAB30 | ENSG00000137502 | chr11 | 82973133 | 83071923 | 98790 | PCPA | other | long | no | no |

| | | | | | | | | | | |
|---|---|---|---|---|---|---|---|---|---|---|
| RAB5A | ENSG00000144566 | chr3 | 19947079 | 19985175 | 38096 | PCPA | other | medium-long | yes | yes |
| RABGAP1 | ENSG00000011454 | chr9 | 122941009 | 123104866 | 163857 | PCPA | other | long | no | no |
| RAD17 | ENSG00000152942 | chr5 | 69369818 | 69414801 | 44983 | PCPA | DDR | medium-long | yes | yes |
| RAD51C | ENSG00000108384 | chr17 | 58692573 | 58735611 | 43038 | PCPA | DDR | medium-long | no | no |
| RALGAPB | ENSG00000170471 | chr20 | 38472816 | 38578861 | 106045 | PCPA | other | long | no | yes |
| RANBP17 | ENSG00000204764 | chr5 | 170861870 | 171029200 | 167330 | PCPA | other | long | no | no |
| RAP1B | ENSG00000127314 | chr12 | 68610839 | 68671901 | 61062 | PCPA | other | medium-long | no | no |
| RAPH1 | ENSG00000173166 | chr2 | 203433682 | 203535410 | 101728 | PCPA | other | long | no | yes |
| RBBP5 | ENSG00000117222 | chr1 | 205086142 | 205122015 | 35873 | PCPA | DDR | medium-long | no | no |
| RBBP6 | ENSG00000122257 | chr16 | 24539585 | 24572863 | 33278 | PCPA | DDR | medium-long | yes | yes |
| RBL1 | ENSG00000080839 | chr20 | 36996349 | 37095995 | 99646 | PCPA | other | long | yes | yes |
| RBM4 | ENSG00000173933 | chr11 | 66638617 | 66649930 | 11313 | PCPA | other | medium-short | yes | yes |
| RC3H1 | ENSG00000135870 | chr1 | 173931214 | 174022297 | 91083 | PCPA | other | long | no | no |
| RC3H2 | ENSG00000056586 | chr9 | 122844556 | 122905280 | 60724 | PCPA | other | medium-long | no | no |
| REV3L | ENSG00000009413 | chr6 | 111299031 | 111482919 | 183888 | PCPA | DDR | long | yes | yes |
| REXO5 | ENSG00000005189 | chr16 | 20806708 | 20814992 | 8284 | PCPA | other | short | no | no |
| RFT1 | ENSG00000163933 | chr3 | 53088483 | 53130462 | 41979 | PCPA | other | medium-long | no | no |
| RFX7 | ENSG00000181827 | chr15 | 56087283 | 56243266 | 155983 | PCPA | other | long | yes | yes |
| RHOF | ENSG00000139725 | chr12 | 121779608 | 121802928 | 23320 | PCPA | other | medium-short | yes | yes |
| RMI1 | ENSG00000178966 | chr9 | 83980711 | 84004074 | 23363 | PCPA | DDR | medium-short | yes | yes |
| RNASEH2B | ENSG00000136104 | chr13 | 50909678 | 50973745 | 64067 | PCPA | other | medium-long | no | no |
| RNF145 | ENSG00000145860 | chr5 | 159157409 | 159209791 | 52382 | PCPA | other | medium-long | yes | no |
| RNF19A | ENSG00000034677 | chr8 | 100257060 | 100309910 | 52850 | PCPA | other | medium-long | no | no |
| RNF217 | ENSG00000146373 | chr6 | 124963249 | 125045411 | 82162 | PCPA | other | long | yes | yes |
| RNF24 | ENSG00000101236 | chr20 | 3927309 | 4015545 | 88236 | PCPA | other | long | no | no |
| RNF44 | ENSG00000146083 | chr5 | 176526697 | 176537420 | 10723 | PCPA | other | medium-short | no | no |
| RNF6 | ENSG00000127870 | chr13 | 26212762 | 26222089 | 9327 | PCPA | other | short | no | yes |
| RNMT | ENSG00000101654 | chr18 | 13726661 | 13764558 | 37897 | PCPA | other | medium-long | yes | yes |
| RPAP2 | ENSG00000122484 | chr1 | 92298965 | 92402056 | 103091 | PCPA | other | long | yes | yes |
| RPL30 | ENSG00000156482 | chr8 | 98024851 | 98045590 | 20739 | PCPA | other | medium-short | no | no |
| RPUSD2 | ENSG00000166133 | chr15 | 40569300 | 40574943 | 5643 | PCPA | other | short | no | no |
| RRP15 | ENSG00000067533 | chr1 | 218285287 | 218337983 | 52696 | PCPA | other | medium-long | no | no |

| | | | | | | | | | | |
|---|---|---|---|---|---|---|---|---|---|---|
| RSBN1 | ENSG00000081019 | chr1 | 113761833 | 113812448 | 50615 | PCPA | other | medium-long | yes | yes |
| RSPRY1 | ENSG00000159579 | chr16 | 57186137 | 57240475 | 54338 | PCPA | other | medium-long | yes | yes |
| RTKN2 | ENSG00000182010 | chr10 | 62183035 | 62268707 | 85672 | PCPA | other | long | no | no |
| RUNX1T1 | ENSG00000079102 | chr8 | 91959793 | 92103286 | 143493 | PCPA | other | long | no | no |
| SATB1 | ENSG00000182568 | chr3 | 18345387 | 18444873 | 99486 | PCPA | other | long | no | no |
| SAV1 | ENSG00000151748 | chr14 | 50633636 | 50668331 | 34695 | PCPA | other | medium-long | yes | yes |
| SCAF4 | ENSG00000156304 | chr21 | 31671033 | 31732075 | 61042 | PCPA | other | medium-long | yes | yes |
| SCLY | ENSG00000132330 | chr2 | 238060966 | 238096186 | 35220 | PCPA | other | medium-long | no | no |
| SEC22A | ENSG00000121542 | chr3 | 123201927 | 123274130 | 72203 | PCPA | other | long | no | no |
| SEC31A | ENSG00000138674 | chr4 | 82818661 | 82891184 | 72523 | PCPA | other | long | yes | no |
| SECISBP2L | ENSG00000138593 | chr15 | 48988638 | 49046563 | 57925 | PCPA | other | medium-long | yes | yes |
| SELENOT | ENSG00000198843 | chr3 | 150603288 | 150630435 | 27147 | PCPA | other | medium-long | no | no |
| SENP6 | ENSG00000112701 | chr6 | 75601509 | 75718278 | 116769 | PCPA | other | long | yes | yes |
| SEPSECS | ENSG00000109618 | chr4 | 25120014 | 25160442 | 40428 | PCPA | other | medium-long | no | no |
| SERTAD2 | ENSG00000179833 | chr2 | 64631621 | 64653913 | 22292 | PCPA | other | medium-short | no | no |
| SERTAD4 | ENSG00000082497 | chr1 | 210232799 | 210246628 | 13829 | PCPA | other | medium-short | yes | yes |
| SESN3 | ENSG00000149212 | chr11 | 95165513 | 95231197 | 65684 | PCPA | other | long | no | no |
| SETD4 | ENSG00000185917 | chr21 | 36034541 | 36060261 | 25720 | PCPA | other | medium-short | yes | yes |
| SETD5 | ENSG00000168137 | chr3 | 9397615 | 9478154 | 80539 | PCPA | other | long | yes | yes |
| SETMAR | ENSG00000170364 | chr3 | 4303304 | 4317567 | 14263 | PCPA | other | medium-short | no | yes |
| SFSWAP | ENSG00000061936 | chr12 | 131711081 | 131799737 | 88656 | PCPA | other | long | yes | yes |
| SFXN2 | ENSG00000156398 | chr10 | 102714540 | 102743492 | 28952 | PCPA | other | medium-long | yes | yes |
| SGO1 | ENSG00000129810 | chr3 | 20160593 | 20186292 | 25699 | PCPA | other | medium-short | no | no |
| SH3RF1 | ENSG00000154447 | chr4 | 169094256 | 169271105 | 176849 | PCPA | other | long | yes | yes |
| SHB | ENSG00000107338 | chr9 | 37919134 | 38069211 | 150077 | PCPA | other | long | yes | yes |
| SIRT1 | ENSG00000096717 | chr10 | 67884669 | 67918390 | 33721 | PCPA | DDR | medium-long | no | no |
| SKA1 | ENSG00000154839 | chr18 | 50374995 | 50394173 | 19178 | PCPA | other | medium-short | no | no |
| SKA3 | ENSG00000165480 | chr13 | 21153595 | 21176602 | 23007 | PCPA | other | medium-short | no | yes |
| SKI | ENSG00000157933 | chr1 | 2228695 | 2310119 | 81424 | PCPA | other | long | no | no |
| SLAIN1 | ENSG00000139737 | chr13 | 77697854 | 77764242 | 66388 | PCPA | other | long | no | no |
| SLC12A2 | ENSG00000064651 | chr5 | 128083766 | 128189688 | 105922 | PCPA | other | long | yes | no |
| SLC15A4 | ENSG00000139370 | chr12 | 128794173 | 128819867 | 25694 | PCPA | other | medium-short | no | no |

| | | | | | | | | | | |
|---|---|---|---|---|---|---|---|---|---|---|
| SLC16A10 | ENSG00000112394 | chr6 | 111087503 | 111231194 | 143691 | PCPA | other | long | no | no |
| SLC1A4 | ENSG00000115902 | chr2 | 64988477 | 65023865 | 35388 | PCPA | other | medium-long | yes | yes |
| SLC22A15 | ENSG00000163393 | chr1 | 115976498 | 116070054 | 93556 | PCPA | other | long | no | no |
| SLC25A25 | ENSG00000148339 | chr9 | 128068201 | 128109244 | 41043 | PCPA | other | medium-long | no | no |
| SLC30A6 | ENSG00000152683 | chr2 | 32165841 | 32224379 | 58538 | PCPA | other | medium-long | no | yes |
| SLC30A7 | ENSG00000162695 | chr1 | 100896076 | 100981753 | 85677 | PCPA | other | long | yes | yes |
| SLC35D1 | ENSG00000116704 | chr1 | 66999332 | 67054099 | 54767 | PCPA | other | medium-long | yes | yes |
| SLC35G2 | ENSG00000168917 | chr3 | 136818647 | 136855892 | 37245 | PCPA | other | medium-long | no | no |
| SLC5A3 | ENSG00000198743 | chr21 | 34073570 | 34106262 | 32692 | PCPA | other | medium-long | yes | yes |
| SLC6A2 | ENSG00000103546 | chr16 | 55656086 | 55703791 | 47705 | PCPA | other | medium-long | no | no |
| SLC7A1 | ENSG00000139514 | chr13 | 29509410 | 29595688 | 86278 | PCPA | other | long | yes | yes |
| SLC7A5 | ENSG00000103257 | chr16 | 87830023 | 87869488 | 39465 | PCPA | other | medium-long | no | no |
| SLC9B2 | ENSG00000164038 | chr4 | 103048591 | 103076336 | 27745 | PCPA | other | medium-long | no | no |
| SLCO5A1 | ENSG00000137571 | chr8 | 69667047 | 69835064 | 168017 | PCPA | other | long | no | no |
| SLF2 | ENSG00000119906 | chr10 | 100912963 | 100926760 | 13797 | PCPA | DDR | medium-short | no | no |
| SMAD2 | ENSG00000175387 | chr18 | 47808957 | 47931146 | 122189 | PCPA | other | long | no | yes |
| SMAD6 | ENSG00000137834 | chr15 | 66702228 | 66782848 | 80620 | PCPA | other | long | yes | yes |
| SMARCAD1 | ENSG00000163104 | chr4 | 94207611 | 94291292 | 83681 | PCPA | other | long | yes | yes |
| SMC4 | ENSG00000113810 | chr3 | 160399639 | 160434954 | 35315 | PCPA | other | medium-long | no | no |
| SMC5 | ENSG00000198887 | chr9 | 70258962 | 70354888 | 95926 | PCPA | DDR | long | yes | yes |
| SMCHD1 | ENSG00000101596 | chr18 | 2655738 | 2805017 | 149279 | PCPA | other | long | yes | yes |
| SMCR8 | ENSG00000176994 | chr17 | 18315310 | 18328055 | 12745 | PCPA | other | medium-short | no | no |
| SNX16 | ENSG00000104497 | chr8 | 81799587 | 81842866 | 43279 | PCPA | other | medium-long | no | no |
| SNX5 | ENSG00000089006 | chr20 | 17941597 | 17968980 | 27383 | PCPA | other | medium-long | no | no |
| SOBP | ENSG00000112320 | chr6 | 107490106 | 107558287 | 68181 | PCPA | other | long | no | no |
| SOGA3 | ENSG00000214338 | chr6 | 127472794 | 127519191 | 46397 | PCPA | other | medium-long | no | no |
| SOGA3 | ENSG00000255330 | chr6 | 127438406 | 127519001 | 80595 | PCPA | other | long | no | no |
| SON | ENSG00000159140 | chr21 | 33543052 | 33577481 | 34429 | PCPA | other | medium-long | yes | yes |
| SOS1 | ENSG00000115904 | chr2 | 39024121 | 39124345 | 100224 | PCPA | other | long | yes | yes |
| SOS2 | ENSG00000100485 | chr14 | 50117120 | 50231558 | 114438 | PCPA | other | long | no | no |
| SP4 | ENSG00000105866 | chr7 | 21428034 | 21514822 | 86788 | PCPA | other | long | no | no |
| SPATA5L1 | ENSG00000171763 | chr15 | 45402331 | 45421419 | 19088 | PCPA | other | medium-short | no | no |

| | | | | | | | | | | |
|---|---|---|---|---|---|---|---|---|---|---|
| SPRED1 | ENSG00000166068 | chr15 | 38252326 | 38357249 | 104923 | PCPA | other | long | yes | yes |
| SPRED2 | ENSG00000198369 | chr2 | 65310851 | 65432637 | 121786 | PCPA | other | long | yes | yes |
| SPSB4 | ENSG00000175093 | chr3 | 141051402 | 141148611 | 97209 | PCPA | other | long | no | no |
| SRCAP | ENSG00000080603 | chr16 | 30698209 | 30736263 | 38054 | PCPA | other | medium-long | no | no |
| SRFBP1 | ENSG00000151304 | chr5 | 121961961 | 122075570 | 113609 | PCPA | other | long | yes | yes |
| SRRM4 | ENSG00000139767 | chr12 | 118981495 | 119163051 | 181556 | PCPA | other | long | no | no |
| SSFA2 | ENSG00000138434 | chr2 | 181891833 | 181930738 | 38905 | PCPA | other | medium-long | yes | yes |
| ST3GAL6 | ENSG00000064225 | chr3 | 98733456 | 98756546 | 23090 | PCPA | other | medium-short | no | no |
| ST6GAL2 | ENSG00000144057 | chr2 | 106801604 | 106886646 | 85042 | PCPA | other | long | no | no |
| ST7 | ENSG00000004866 | chr7 | 117020218 | 117199144 | 178926 | PCPA | other | long | no | no |
| ST7L | ENSG00000007341 | chr1 | 112525876 | 112619672 | 93796 | PCPA | other | long | yes | yes |
| ST8SIA2 | ENSG00000140557 | chr15 | 92393828 | 92468728 | 74900 | PCPA | other | long | no | no |
| STK17B | ENSG00000081320 | chr2 | 196133566 | 196171575 | 38009 | PCPA | DDR | medium-long | yes | no |
| SUCLA2 | ENSG00000136143 | chr13 | 47936491 | 48001354 | 64863 | PCPA | other | long | yes | yes |
| SUCO | ENSG00000094975 | chr1 | 172533118 | 172611831 | 78713 | PCPA | other | long | no | yes |
| SUGT1 | ENSG00000165416 | chr13 | 52652709 | 52700909 | 48200 | PCPA | other | medium-long | no | no |
| SULT1C4 | ENSG00000198075 | chr2 | 108377911 | 108388057 | 10146 | PCPA | other | medium-short | no | no |
| SUZ12 | ENSG00000178691 | chr17 | 31937018 | 32001045 | 64027 | PCPA | other | medium-long | yes | yes |
| SYN2 | ENSG00000157152 | chr3 | 12004406 | 12191385 | 186979 | PCPA | other | long | no | no |
| SYNDIG1 | ENSG00000101463 | chr20 | 24469199 | 24666616 | 197417 | PCPA | other | long | no | no |
| SYT14 | ENSG00000143469 | chr1 | 209938174 | 210171389 | 233215 | PCPA | other | long | no | no |
| TAB2 | ENSG00000055208 | chr6 | 149318582 | 149411613 | 93031 | PCPA | other | long | no | no |
| TAF1B | ENSG00000115750 | chr2 | 9843354 | 9934416 | 91062 | PCPA | other | long | yes | yes |
| TAPBPL | ENSG00000139192 | chr12 | 6452042 | 6462318 | 10276 | PCPA | other | medium-short | no | no |
| TASP1 | ENSG00000089123 | chr20 | 13483227 | 13638903 | 155676 | PCPA | other | long | no | no |
| TBK1 | ENSG00000183735 | chr12 | 64451880 | 64502108 | 50228 | PCPA | DDR | medium-long | yes | yes |
| TBPL1 | ENSG00000028839 | chr6 | 133952170 | 133990432 | 38262 | PCPA | other | medium-long | yes | yes |
| TCERG1 | ENSG00000113649 | chr5 | 146447311 | 146511961 | 64650 | PCPA | other | long | yes | yes |
| TES | ENSG00000135269 | chr7 | 116210493 | 116258783 | 48290 | PCPA | other | medium-long | yes | yes |
| TET2 | ENSG00000168769 | chr4 | 105145875 | 105242771 | 96896 | PCPA | other | long | no | no |
| TEX14 | ENSG00000121101 | chr17 | 58556678 | 58692055 | 135377 | PCPA | other | long | no | no |
| TFAP2B | ENSG00000008196 | chr6 | 50818723 | 50847613 | 28890 | PCPA | other | medium-long | no | no |

| | | | | | | | | | | |
|---|---|---|---|---|---|---|---|---|---|---|
| THAP6 | ENSG00000174796 | chr4 | 75513946 | 75527214 | 13268 | PCPA | other | medium-short | no | yes |
| THAP9 | ENSG00000168152 | chr4 | 82900684 | 82919969 | 19285 | PCPA | other | medium-short | no | no |
| THNSL1 | ENSG00000185875 | chr10 | 25016658 | 25026664 | 10006 | PCPA | other | medium-short | no | no |
| THUMPD2 | ENSG00000138050 | chr2 | 39736060 | 39779267 | 43207 | PCPA | other | medium-long | yes | yes |
| TICRR | ENSG00000140534 | chr15 | 89575482 | 89631056 | 55574 | PCPA | DDR | medium-long | yes | yes |
| TIMM21 | ENSG00000075336 | chr18 | 74148511 | 74160530 | 12019 | PCPA | other | medium-short | no | yes |
| TIMP2 | ENSG00000035862 | chr17 | 78852977 | 78925387 | 72410 | PCPA | other | long | no | no |
| TIMP3 | ENSG00000100234 | chr22 | 32801701 | 32863043 | 61342 | PCPA | other | medium-long | no | no |
| TIPARP | ENSG00000163659 | chr3 | 156674342 | 156706770 | 32428 | PCPA | other | medium-long | no | no |
| TLE1 | ENSG00000196781 | chr9 | 81583683 | 81689305 | 105622 | PCPA | other | long | yes | yes |
| TLE3 | ENSG00000140332 | chr15 | 70048977 | 70097917 | 48940 | PCPA | other | medium-long | no | no |
| TLE4 | ENSG00000106829 | chr9 | 79571773 | 79726882 | 155109 | PCPA | other | long | yes | yes |
| TMEM245 | ENSG00000106771 | chr9 | 109015152 | 109119945 | 104793 | PCPA | other | long | no | no |
| TMEM39A | ENSG00000176142 | chr3 | 119436944 | 119468830 | 31886 | PCPA | other | medium-long | yes | yes |
| TMEM5 | ENSG00000118600 | chr12 | 63779803 | 63809558 | 29755 | PCPA | other | medium-long | no | yes |
| TMEM87A | ENSG00000103978 | chr15 | 42210452 | 42273663 | 63211 | PCPA | other | medium-long | yes | yes |
| TMX3 | ENSG00000166479 | chr18 | 68673688 | 68715298 | 41610 | PCPA | other | medium-long | yes | yes |
| TNRC18 | ENSG00000182095 | chr7 | 5306800 | 5425414 | 118614 | PCPA | other | long | no | yes |
| TNRC6A | ENSG00000090905 | chr16 | 24729695 | 24826221 | 96526 | PCPA | other | long | yes | yes |
| TNRC6C | ENSG00000078687 | chr17 | 78041047 | 78093299 | 52252 | PCPA | other | medium-long | no | no |
| TOB2 | ENSG00000183864 | chr22 | 41433492 | 41447023 | 13531 | PCPA | other | medium-short | yes | yes |
| TOP3A | ENSG00000177302 | chr17 | 18271428 | 18315007 | 43579 | PCPA | DDR | medium-long | yes | yes |
| TOPBP1 | ENSG00000163781 | chr3 | 133598175 | 133661893 | 63718 | PCPA | DDR | medium-long | yes | yes |
| TP53INP2 | ENSG00000078804 | chr20 | 34704290 | 34713439 | 9149 | PCPA | other | short | no | no |
| TPCN1 | ENSG00000186815 | chr12 | 113221429 | 113260412 | 38983 | PCPA | other | medium-long | no | no |
| TPP2 | ENSG00000134900 | chr13 | 102597003 | 102679958 | 82955 | PCPA | other | long | yes | yes |
| TRIB2 | ENSG00000071575 | chr2 | 12716889 | 12742734 | 25845 | PCPA | other | medium-short | no | no |
| TRIM13 | ENSG00000204977 | chr13 | 49996813 | 50020481 | 23668 | PCPA | other | medium-short | no | no |
| TRIM36 | ENSG00000152503 | chr5 | 115147068 | 115169730 | 22662 | PCPA | other | medium-short | no | no |
| TRIM67 | ENSG00000119283 | chr1 | 231162112 | 231221556 | 59444 | PCPA | other | medium-long | no | no |
| TRPS1 | ENSG00000104447 | chr8 | 115619747 | 115668294 | 48547 | PCPA | other | medium-long | no | yes |
| TSC22D2 | ENSG00000196428 | chr3 | 150408335 | 150466431 | 58096 | PCPA | other | medium-long | yes | yes |

| | | | | | | | | | | |
|---|---|---|---|---|---|---|---|---|---|---|
| TSEN2 | ENSG00000154743 | chr3 | 12484432 | 12539620 | 55188 | PCPA | other | medium-long | no | yes |
| TSFM | ENSG00000123297 | chr12 | 57782589 | 57802631 | 20042 | PCPA | other | medium-short | yes | yes |
| TSHZ1 | ENSG00000179981 | chr18 | 75210755 | 75289950 | 79195 | PCPA | other | long | yes | yes |
| TTC14 | ENSG00000163728 | chr3 | 180602193 | 180617828 | 15635 | PCPA | other | medium-short | yes | yes |
| TTC17 | ENSG00000052841 | chr11 | 43358932 | 43494933 | 136001 | PCPA | other | long | no | no |
| TTC21B | ENSG00000123607 | chr2 | 165873362 | 165953843 | 80481 | PCPA | other | long | no | no |
| TTLL7 | ENSG00000137941 | chr1 | 83912996 | 83999150 | 86154 | PCPA | other | long | no | no |
| TTPAL | ENSG00000124120 | chr20 | 44475886 | 44494603 | 18717 | PCPA | other | medium-short | yes | yes |
| TUBGCP3 | ENSG00000126216 | chr13 | 112485005 | 112588167 | 103162 | PCPA | other | long | yes | yes |
| TWIST1 | ENSG00000122691 | chr7 | 19020991 | 19117672 | 96681 | PCPA | other | long | no | no |
| TWNK | ENSG00000107815 | chr10 | 100987795 | 100994401 | 6606 | PCPA | other | short | no | no |
| U2SURP | ENSG00000163714 | chr3 | 143001524 | 143060546 | 59022 | PCPA | other | medium-long | yes | yes |
| UBE2E3 | ENSG00000170035 | chr2 | 180980385 | 181076585 | 96200 | PCPA | other | long | no | no |
| UBE2QL1 | ENSG00000215218 | chr5 | 6448623 | 6494909 | 46286 | PCPA | other | medium-long | no | no |
| UBE3C | ENSG00000009335 | chr7 | 157138913 | 157269372 | 130459 | PCPA | other | long | yes | yes |
| UBN1 | ENSG00000118900 | chr16 | 4846665 | 4882360 | 35695 | PCPA | other | medium-long | no | yes |
| UBN2 | ENSG00000157741 | chr7 | 139230356 | 139308236 | 77880 | PCPA | other | long | yes | yes |
| UBOX5 | ENSG00000185019 | chr20 | 3107573 | 3160196 | 52623 | PCPA | other | medium-long | no | no |
| UBR1 | ENSG00000159459 | chr15 | 42942897 | 43106113 | 163216 | PCPA | other | long | yes | yes |
| UHRF1BP1L | ENSG00000111647 | chr12 | 100037076 | 100142848 | 105772 | PCPA | other | long | yes | yes |
| UMPS | ENSG00000114491 | chr3 | 124730366 | 124749273 | 18907 | PCPA | other | medium-short | no | no |
| UNC5B | ENSG00000107731 | chr10 | 71212570 | 71302864 | 90294 | PCPA | other | long | no | no |
| UPF1 | ENSG00000005007 | chr19 | 18833130 | 18850691 | 17561 | PCPA | DDR | medium-short | yes | yes |
| USP10 | ENSG00000103194 | chr16 | 84699978 | 84779922 | 79944 | PCPA | DDR | long | yes | yes |
| USP37 | ENSG00000135913 | chr2 | 218450251 | 218568361 | 118110 | PCPA | other | long | no | yes |
| USP45 | ENSG00000123552 | chr6 | 99462464 | 99515489 | 53025 | PCPA | DDR | medium-long | yes | yes |
| USPL1 | ENSG00000132952 | chr13 | 30617693 | 30660770 | 43077 | PCPA | other | medium-long | yes | no |
| UTP15 | ENSG00000164338 | chr5 | 73565741 | 73583377 | 17636 | PCPA | other | medium-short | yes | yes |
| VANGL1 | ENSG00000173218 | chr1 | 115641953 | 115698224 | 56271 | PCPA | other | medium-long | no | no |
| VEZT | ENSG00000028203 | chr12 | 95217746 | 95302790 | 85044 | PCPA | other | long | yes | yes |
| VIRMA | ENSG00000164944 | chr8 | 94487693 | 94553529 | 65836 | PCPA | other | long | no | no |
| VMP1 | ENSG00000062716 | chr17 | 59707465 | 59842255 | 134790 | PCPA | other | long | no | no |

| | | | | | | | | | | |
|---|---|---|---|---|---|---|---|---|---|---|
| VRK1 | ENSG00000100749 | chr14 | 96797304 | 96931722 | 134418 | PCPA | DDR | long | no | no |
| VWDE | ENSG00000146530 | chr7 | 12330885 | 12403941 | 73056 | PCPA | other | long | no | yes |
| WAC | ENSG00000095787 | chr10 | 28532493 | 28623112 | 90619 | PCPA | DDR | long | no | no |
| WAPL | ENSG00000062650 | chr10 | 86435256 | 86521815 | 86559 | PCPA | other | long | no | no |
| WASHC4 | ENSG00000136051 | chr12 | 105107324 | 105169134 | 61810 | PCPA | other | medium-long | no | no |
| WDFY3 | ENSG00000163625 | chr4 | 84669610 | 84966391 | 296781 | PCPA | other | long | no | no |
| WDR20 | ENSG00000140153 | chr14 | 102139897 | 102198429 | 58532 | PCPA | other | medium-long | yes | yes |
| WDR26 | ENSG00000162923 | chr1 | 224385143 | 224434299 | 49156 | PCPA | other | medium-long | no | no |
| WDR33 | ENSG00000136709 | chr2 | 127701022 | 127811187 | 110165 | PCPA | other | long | no | no |
| WDR36 | ENSG00000134987 | chr5 | 111092172 | 111130502 | 38330 | PCPA | other | medium-long | yes | yes |
| WDR37 | ENSG00000047056 | chr10 | 1049538 | 1132297 | 82759 | PCPA | other | long | yes | yes |
| WDR53 | ENSG00000185798 | chr3 | 196554177 | 196568639 | 14462 | PCPA | other | medium-short | yes | no |
| WDR76 | ENSG00000092470 | chr15 | 43826963 | 43868419 | 41456 | PCPA | other | medium-long | yes | yes |
| WEE1 | ENSG00000166483 | chr11 | 9573681 | 9589984 | 16303 | PCPA | other | medium-short | no | no |
| WRN | ENSG00000165392 | chr8 | 31033801 | 31173769 | 139968 | PCPA | DDR | long | no | no |
| XKR7 | ENSG00000260903 | chr20 | 31968002 | 32003387 | 35385 | PCPA | other | medium-long | no | no |
| XRN1 | ENSG00000114127 | chr3 | 142306607 | 142448062 | 141455 | PCPA | other | long | no | yes |
| YAF2 | ENSG00000015153 | chr12 | 42157104 | 42238248 | 81144 | PCPA | other | long | no | no |
| YARS2 | ENSG00000139131 | chr12 | 32727490 | 32755902 | 28412 | PCPA | other | medium-long | yes | yes |
| YEATS2 | ENSG00000163872 | chr3 | 183697818 | 183812625 | 114807 | PCPA | other | long | yes | yes |
| ZBED5 | ENSG00000236287 | chr11 | 10849897 | 10858073 | 8176 | PCPA | other | short | no | no |
| ZBED8 | ENSG00000221886 | chr5 | 160393148 | 160400097 | 6949 | PCPA | other | short | yes | yes |
| ZBED9 | ENSG00000232040 | chr6 | 28571630 | 28587335 | 15705 | PCPA | other | medium-short | yes | yes |
| ZBTB21 | ENSG00000173276 | chr21 | 41993740 | 42010366 | 16626 | PCPA | other | medium-short | yes | yes |
| ZBTB24 | ENSG00000112365 | chr6 | 109462594 | 109483237 | 20643 | PCPA | other | medium-short | yes | yes |
| ZBTB7A | ENSG00000178951 | chr19 | 4044364 | 4066945 | 22581 | PCPA | other | medium-short | no | no |
| ZC3H12C | ENSG00000149289 | chr11 | 110131004 | 110171839 | 40835 | PCPA | other | medium-long | yes | yes |
| ZC3H13 | ENSG00000123200 | chr13 | 45954465 | 46052759 | 98294 | PCPA | other | long | no | yes |
| ZCCHC6 | ENSG00000083223 | chr9 | 86287959 | 86354412 | 66453 | PCPA | other | long | no | yes |
| ZCCHC7 | ENSG00000147905 | chr9 | 37120539 | 37358149 | 237610 | PCPA | other | long | yes | yes |
| ZDHHC5 | ENSG00000156599 | chr11 | 57667896 | 57701187 | 33291 | PCPA | other | medium-long | no | yes |
| ZFAND4 | ENSG00000172671 | chr10 | 45616385 | 45672780 | 56395 | PCPA | other | medium-long | no | no |

| | | | | | | | | | | |
|---|---|---|---|---|---|---|---|---|---|---|
| ZFX | ENSG00000005889 | chrX | 24149173 | 24179219 | 30046 | PCPA | other | medium-long | yes | yes |
| ZFY | ENSG00000067646 | chrY | 2935281 | 2982506 | 47225 | PCPA | other | medium-long | no | no |
| ZKSCAN1 | ENSG00000106261 | chr7 | 100015581 | 100041689 | 26108 | PCPA | other | medium-short | yes | yes |
| ZKSCAN2 | ENSG00000155592 | chr16 | 25236001 | 25257931 | 21930 | PCPA | other | medium-short | yes | yes |
| ZKSCAN7 | ENSG00000196345 | chr3 | 44555203 | 44583483 | 28280 | PCPA | other | medium-long | no | no |
| ZMAT3 | ENSG00000172667 | chr3 | 179017223 | 179072279 | 55056 | PCPA | other | medium-long | no | no |
| ZNF10 | ENSG00000256223 | chr12 | 133130592 | 133159465 | 28873 | PCPA | other | medium-long | no | no |
| ZNF100 | ENSG00000197020 | chr19 | 21722766 | 21767628 | 44862 | PCPA | other | medium-long | no | no |
| ZNF107 | ENSG00000196247 | chr7 | 64666083 | 64711582 | 45499 | PCPA | other | medium-long | no | no |
| ZNF124 | ENSG00000196418 | chr1 | 247121975 | 247172016 | 50041 | PCPA | other | medium-long | no | no |
| ZNF138 | ENSG00000197008 | chr7 | 64794388 | 64833681 | 39293 | PCPA | other | medium-long | no | no |
| ZNF140 | ENSG00000196387 | chr12 | 133080916 | 133107428 | 26512 | PCPA | other | medium-long | no | no |
| ZNF143 | ENSG00000166478 | chr11 | 9460965 | 9528524 | 67559 | PCPA | other | long | yes | yes |
| ZNF200 | ENSG00000010539 | chr16 | 3222345 | 3235167 | 12822 | PCPA | other | medium-short | yes | yes |
| ZNF223 | ENSG00000178386 | chr19 | 44052009 | 44067991 | 15982 | PCPA | other | medium-short | no | no |
| ZNF225 | ENSG00000256294 | chr19 | 44112181 | 44134816 | 22635 | PCPA | other | medium-short | yes | yes |
| ZNF234 | ENSG00000263002 | chr19 | 44141606 | 44160309 | 18703 | PCPA | other | medium-short | yes | no |
| ZNF253 | ENSG00000256771 | chr19 | 19865886 | 19894674 | 28788 | PCPA | other | medium-long | no | no |
| ZNF254 | ENSG00000213096 | chr19 | 24087193 | 24129852 | 42659 | PCPA | other | medium-long | yes | no |
| ZNF260 | ENSG00000254004 | chr19 | 36510695 | 36528660 | 17965 | PCPA | other | medium-short | yes | yes |
| ZNF273 | ENSG00000198039 | chr7 | 64903253 | 64930966 | 27713 | PCPA | other | medium-long | no | no |
| ZNF280C | ENSG00000056277 | chrX | 130202711 | 130268899 | 66188 | PCPA | other | long | no | yes |
| ZNF286A | ENSG00000187607 | chr17 | 15699577 | 15720787 | 21210 | PCPA | other | medium-short | no | yes |
| ZNF286B | ENSG00000249459 | chr17 | 18658429 | 18682262 | 23833 | PCPA | other | medium-short | no | no |
| ZNF287 | ENSG00000141040 | chr17 | 16551387 | 16569206 | 17819 | PCPA | other | medium-short | no | no |
| ZNF326 | ENSG00000162664 | chr1 | 89995112 | 90035531 | 40419 | PCPA | other | medium-long | yes | yes |
| ZNF367 | ENSG00000165244 | chr9 | 96385941 | 96418329 | 32388 | PCPA | other | medium-long | yes | yes |
| ZNF397 | ENSG00000186812 | chr18 | 35241030 | 35249803 | 8773 | PCPA | other | short | no | no |
| ZNF420 | ENSG00000197050 | chr19 | 37078418 | 37130311 | 51893 | PCPA | other | medium-long | no | no |
| ZNF426 | ENSG00000130818 | chr19 | 9523224 | 9538645 | 15421 | PCPA | other | medium-short | yes | yes |
| ZNF430 | ENSG00000118620 | chr19 | 21020620 | 21060050 | 39430 | PCPA | other | medium-long | yes | yes |
| ZNF431 | ENSG00000196705 | chr19 | 21142024 | 21196053 | 54029 | PCPA | other | medium-long | yes | yes |

| | | | | | | | | | | |
|---|---|---|---|---|---|---|---|---|---|---|
| ZNF451 | ENSG00000112200 | chr6 | 57090010 | 57170307 | 80297 | PCPA | other | long | no | no |
| ZNF462 | ENSG00000148143 | chr9 | 106863097 | 107013634 | 150537 | PCPA | other | long | no | no |
| ZNF480 | ENSG00000198464 | chr19 | 52297177 | 52325922 | 28745 | PCPA | other | medium-long | no | yes |
| ZNF483 | ENSG00000173258 | chr9 | 111525189 | 111544432 | 19243 | PCPA | other | medium-short | no | no |
| ZNF486 | ENSG00000256229 | chr19 | 20167228 | 20200490 | 33262 | PCPA | other | medium-long | no | no |
| ZNF493 | ENSG00000196268 | chr19 | 21397129 | 21427573 | 30444 | PCPA | other | medium-long | no | no |
| ZNF501 | ENSG00000186446 | chr3 | 44729596 | 44737083 | 7487 | PCPA | other | short | no | no |
| ZNF507 | ENSG00000168813 | chr19 | 32345594 | 32387667 | 42073 | PCPA | other | medium-long | yes | yes |
| ZNF526 | ENSG00000167625 | chr19 | 42220271 | 42228201 | 7930 | PCPA | other | short | no | yes |
| ZNF547 | ENSG00000152433 | chr19 | 57363477 | 57379565 | 16088 | PCPA | other | medium-short | no | no |
| ZNF557 | ENSG00000130544 | chr19 | 7069444 | 7087968 | 18524 | PCPA | other | medium-short | yes | yes |
| ZNF572 | ENSG00000180938 | chr8 | 124973298 | 124979389 | 6091 | PCPA | other | short | no | no |
| ZNF582 | ENSG00000018869 | chr19 | 56382752 | 56393545 | 10793 | PCPA | other | medium-short | no | no |
| ZNF594 | ENSG00000180626 | chr17 | 5179536 | 5191862 | 12326 | PCPA | other | medium-short | yes | yes |
| ZNF596 | ENSG00000172748 | chr8 | 232397 | 247340 | 14943 | PCPA | other | medium-short | no | no |
| ZNF608 | ENSG00000168916 | chr5 | 124636917 | 124746627 | 109710 | PCPA | other | long | no | no |
| ZNF619 | ENSG00000177873 | chr3 | 40477113 | 40490236 | 13123 | PCPA | other | medium-short | yes | no |
| ZNF623 | ENSG00000183309 | chr8 | 143636162 | 143656418 | 20256 | PCPA | other | medium-short | no | no |
| ZNF625-ZNF20 | ENSG00000213297 | chr19 | 12132117 | 12156731 | 24614 | PCPA | other | medium-short | no | no |
| ZNF644 | ENSG00000122482 | chr1 | 90915298 | 91022255 | 106957 | PCPA | other | long | yes | yes |
| ZNF654 | ENSG00000175105 | chr3 | 88059274 | 88144665 | 85391 | PCPA | other | long | no | no |
| ZNF66 | ENSG00000160229 | chr19 | 20776304 | 20807322 | 31018 | PCPA | other | medium-long | no | no |
| ZNF670-ZNF695 | ENSG00000135747 | chr1 | 246945547 | 247078811 | 133264 | PCPA | other | long | no | no |
| ZNF695 | ENSG00000197472 | chr1 | 246945547 | 247008056 | 62509 | PCPA | other | medium-long | no | no |
| ZNF711 | ENSG00000147180 | chrX | 85244032 | 85273362 | 29330 | PCPA | other | medium-long | no | no |
| ZNF726 | ENSG00000213967 | chr19 | 23914876 | 23923893 | 9017 | PCPA | other | short | no | no |
| ZNF737 | ENSG00000237440 | chr19 | 20543431 | 20565809 | 22378 | PCPA | other | medium-short | no | no |
| ZNF74 | ENSG00000185252 | chr22 | 20394188 | 20408455 | 14267 | PCPA | other | medium-short | no | no |
| ZNF749 | ENSG00000186230 | chr19 | 57435329 | 57445485 | 10156 | PCPA | other | medium-short | yes | yes |
| ZNF766 | ENSG00000196214 | chr19 | 52269571 | 52296046 | 26475 | PCPA | other | medium-long | no | no |
| ZNF776 | ENSG00000152443 | chr19 | 57746796 | 57758159 | 11363 | PCPA | other | medium-short | yes | no |
| ZNF79 | ENSG00000196152 | chr9 | 127424374 | 127445372 | 20998 | PCPA | other | medium-short | no | yes |

| | | | | | | | | | | |
|---|---|---|---|---|---|---|---|---|---|---|
| ZNF8 | ENSG00000278129 | chr19 | 58278951 | 58302805 | 23854 | PCPA | other | medium-short | yes | yes |
| ZNF804A | ENSG00000170396 | chr2 | 184598366 | 184939492 | 341126 | PCPA | other | long | no | no |
| ZNF829 | ENSG00000185869 | chr19 | 36888124 | 36916291 | 28167 | PCPA | other | medium-long | no | no |
| ZNF84 | ENSG00000198040 | chr12 | 133037390 | 133063297 | 25907 | PCPA | other | medium-short | yes | yes |
| ZNF880 | ENSG00000221923 | chr19 | 52369936 | 52385795 | 15859 | PCPA | other | medium-short | no | no |
| ZNF92 | ENSG00000146757 | chr7 | 65373799 | 65401125 | 27326 | PCPA | other | medium-long | yes | yes |
| ZNF93 | ENSG00000184635 | chr19 | 19900913 | 19935575 | 34662 | PCPA | other | medium-long | no | no |
| ZSCAN2 | ENSG00000176371 | chr15 | 84601005 | 84623716 | 22711 | PCPA | other | medium-short | no | yes |
| ZSCAN23 | ENSG00000187987 | chr6 | 28431930 | 28443502 | 11572 | PCPA | other | medium-short | no | no |
| ZXDC | ENSG00000070476 | chr3 | 126437601 | 126475919 | 38318 | PCPA | other | medium-long | yes | yes |
| ZZZ3 | ENSG00000036549 | chr1 | 77562416 | 77683403 | 120987 | PCPA | other | long | yes | yes |
